# Repeated LPS induces training and tolerance of microglial responses across brain regions

**DOI:** 10.1101/2024.04.08.588502

**Authors:** Jennifer Kim, Olivia Sullivan, Kristen Lee, Justin Jao, Juan Tamayo, Abdullah Muhammad Madany, Brandon Wong, Paul Ashwood, Annie Vogel Ciernia

## Abstract

**Background:** Neuroinflammation is involved in the pathogenesis of almost every central nervous system disorder. As the brain’s innate immune cells, microglia fine tune their activity to a dynamic brain environment. Previous studies have shown that repeated bouts of peripheral inflammation can trigger long-term changes in microglial gene expression and function, a form of innate immune memory.

**Methods and Results:** In this study, we used multiple low-dose lipopolysaccharide (LPS) injections in adult mice to study the acute cytokine, transcriptomic, and microglia morphological changes that contribute to the formation of immune memory in the frontal cortex, hippocampus, and striatum, as well as the long-term effects of these changes on behavior. Training and tolerance of gene expression was shared across regions, and we identified 3 unique clusters of DEGs (2xLPS-sensitive, 4xLPS-sensitive, LPS-decreased) with different biological functions. 2xLPS-sensitive DEG promoters were enriched for binding sites for IRF and NFkB family transcription factors, two key regulators of innate immune memory. We quantified shifts in microglia morphological populations and found that while the proportion of ramified and rod-like microglia mostly remained consistent within brain regions and sexes with LPS treatment, there was a shift from ameboid towards hypertrophic morphological states across immune memory states and a dynamic emergence and resolution of trains of rod-like microglia with repeated LPS.

**Conclusions:** Together, findings support the dynamic regulation of microglia during the formation of immune memories in the brain and support future work to exploit this model in brain disease contexts.

## Background

Innate immune cells dynamically adjust their responses to immune challenges through innate immune memory, a process influenced by their past encounters with inflammatory events. (1–4) As the brain’s resident immune cells, microglia rapidly respond to inflammation and engage in classic immune functions including cytokine and chemokine production and phagocytosis of pathogens and cellular debris. In addition to their responsibilities as immune cells, microglia play critical roles in establishing and maintaining brain homeostasis including the regulation of neuron and oligodendrocyte cell numbers (5), shaping of brain circuitry (6,7), and fine-tuning of neuronal connections (3,8,9). Many of these processes are disrupted in brain disorders, supporting a key role for microglia in health and disease. Several clinical and post-mortem studies point to a neuroinflammatory component of brain disorder etiologies, including Schizophrenia, Autism Spectrum Disorder, and Alzheimer’s Disease, driven by increased activation of microglia. (10–13) Microglia directly communicate with neurons and modulate their function by releasing and responding to various molecular substrates in the brain environment including cytokines, chemokines, and neurotransmitters. (14) Thus, dysregulated microglial responses can disrupt the homeostatic brain environment and normal communication across cell types, ultimately altering brain function and behavior.

Innate immune cells update their responses to inflammatory events based on previous exposures to immune stimuli. (4,15,16) After the resolution of an initial immune activating event, innate immune cells can remain in a ‘trained’ state, characterized by exaggerated responses to subsequent immune challenges, or in a ‘tolerized’ state characterized by blunted responses. (4) Both forms of innate immune memory serve as mechanisms that enhance an organism’s ability to respond effectively to recurring infections or damage, but can become detrimental if dysregulated. While immune training allows proactive adaptation of immune responses that have a protective benefit to the organism, this process can become maladaptive in the context of disease or injury when inflammation cannot be resolved. (4) Similarly, tolerized immune responses play a crucial role in averting chronic inflammatory conditions when encountering commensal microbes. (4,17) However, these responses can be counterproductive when facing a new pathogen. As the resident innate immune cells of the brain, microglia are unique in that they are long-lived cells that locally self-renew over the lifespan (18,19), suggesting that formation of innate immune memories in microglia could have long-lasting impacts on brain function and disease development throughout life.

Innate immune cells, including microglia, express receptors that respond to microbe-associated molecular patterns. Microglia Toll-like receptor 4 (TLR4) binds lipopolysaccharide (LPS), a major structural component of gram-negative bacteria. Repeated peripheral injections of low-dose LPS has recently been demonstrated as a robust model to study microglial immune memory responses in the brain. (4,20) Repeated LPS induces long-lasting immune training and tolerance in brain microglia that has been shown to persist for at least six months. (20) Similarly, LPS preconditioning has also been shown to reduce damage in traumatic brain injury by preventing tolerance (21). Interestingly, Wendeln et al., 2018 (20) showed that pathological hallmarks of Alzheimer’s disease and stroke in mouse models are exacerbated by LPS-induced immune training and alleviated by immune tolerance. Long-term functional changes in microglia were paralleled by long-term reprogramming of microglial enhancers and gene expression in these models. Genes involved in hypoxia-induced factor-a (HIF-1α) regulation, metabolic glycolysis, and Rap1-mediated phagocytosis were increased several months after training but not tolerance with repeated LPS. Together, findings in the literature support a model in which long-term reprogramming of gene expression underlies long-term innate immune memory in the brain relevant to disease contexts. However, the mechanisms involved in the initial formation of innate immune memories are not as well characterized in microglia and hence form the major focus of this study.

Here, we used a mouse model of repeated LPS to identify how gene expression, cytokine expression, and microglia morphology changes drive the formation of training and tolerance in the brain. Unlike prior studies using the same repeated low-dose LPS paradigm, ours focused specifically on characterizing immediate, acute microglial changes across multiple brain regions including the frontal cortex, striatum, and hippocampus, which have known differences in microglial morphology and function. (22,23) Furthermore, we assessed long-term impacts of innate immune memory induction in a comprehensive battery of anxiety-like, depressive-like, repetitive, and learning and memory behaviors and performed a systematic evaluation of LPS-induced shifts in microglia morphological states across brain regions. We describe shared patterns of gene expression across brain regions with shared and unique regulatory pathways. We also describe the dynamic emergence and resolution of “trains” of rod-like microglia that align end-to-end with the formation of immune priming and tolerance, a morphological phenotype which has not previously been described in the context of repeated LPS. Together, these findings offer new insight into microglial changes in the formation of immune memory and support future work to further explore described mechanisms in the context of brain disease.

## Methods

### Animals

All experiments were conducted in accordance with the National Institutes of Health Guidelines for the Care and Use of Laboratory Animals, with approval from the Institutional Animal Care and Use Committee at the University of California, Davis and the Canadian Council on Animal Care guidelines, with approval from the Animal Care Committee at the University of British Columbia. Mice in experiments undertaken at the University of California, Davis (RNA-seq, Luminex protein array) were housed in treatment-matched groups of two to four on a regular 12-hr light/12-hr dark cycle and all experiments were performed in regular light during the mouse’s regular light cycle. Mice in experiments undertaken at the University of British Columbia (RT-qPCR, microglia morphology, behavioral battery) were housed in treatment-matched groups of two to four on a reversed 12-hr light/12-hr dark cycle. Because mice are nocturnal animals, all behavior experiments were performed in red light during the mouse’s dark cycle when they are most active. All mice had ad libitum food and water and cages were changed biweekly.

### *In vivo* injections

Adult male and female C57BL/6J mice (Jackson Laboratories, over 10 weeks old) were given daily intraperitoneal injections of 0.5 mg/kg LPS (from *E. coli* O55:B5, Sigma-Aldrich L5418) or an equal volume of vehicle solution (1xPBS; Phosphate buffered saline, Fisher Scientific BP3991) for 4 days between 09:00 and 10:00 on each day. Animals received either four PBS vehicle injections on four consecutive days (PBS), three vehicle injections on three consecutive days followed by a single LPS injection on the fourth day (1xLPS), two vehicle injections on two consecutive days followed by two LPS injections on the following two days (2xLPS), a single vehicle injection followed by three LPS injections on the following three days (3xLPS, males only), or four LPS injections (4xLPS). PBS, 1xLPS, 2xLPS, 3xLPS, and 4xLPS were the five experimental groups analyzed in this study. (Fig. 1A) Body weight was measured right before administering injections each day.

**Figure 1:**
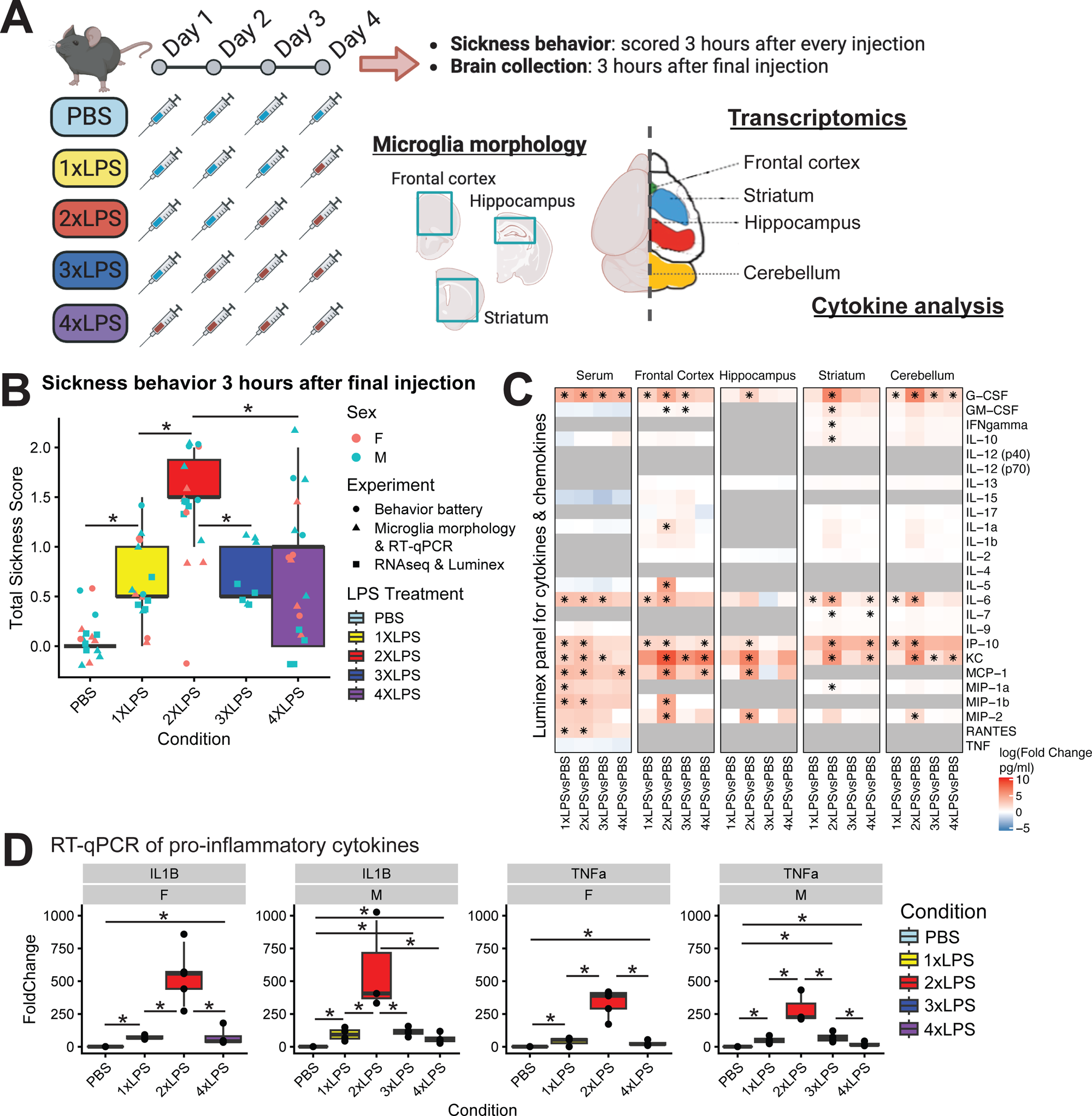
Repeated LPS produces robust training and tolerance in the mouse brain. (A) Outline of repeated LPS injection paradigm and experiments. (B) Sickness behaviors measured 3 hours after final injections for all experimental cohorts in this paper. (*p<0.05, BH-corrected) (C) Luminex protein quantification for in serum and four brain regions at 3hrs post-final injection. Log10 Fold Change between LPS treatments and PBS controls (pg/ml) are shown for each cytokine and chemokine on the Luminex mouse protein array. Columns are grouped by tissue sample and LPS treatment condition. (*p<0.05, BH-corrected). Genes without readings above background are shown as grey. (D) RT-qPCR fold change comparisons for pro-inflammatory cytokines IL1b and TNFa (*p<0.05, BH-corrected)

### Sickness behavior scoring and analysis

Changes in sickness behavior were assessed 3 hours-post injection each day. Sickness behavior was scored according to a 4-point grading scale (0 to 3 points total). Lethargy signified by diminished locomotion and curled body posture, ptosis signified by drooping eyelids, and piloerection signified by ruffled and greasy fur were each assessed individually for either 0, 0.5, or 1 point. A total cumulative score of 0 indicated no symptoms of sickness behavior and a total cumulative score of 3 points indicated that all symptoms of sickness behavior were maximally present. We assessed how cumulative scores for each mouse on the last day of assessment (3 hours after final injection) changed with LPS treatment by fitting a non-parametric Aligned Ranks Transformation (ART) model to measure scores as a factor of LPS Treatment (Total Score ∼ Treatment) using the ARTool R package. (24) The model was assessed using ART ANOVA and tests between Treatment groups were corrected for multiple comparisons using the Benjamini-Hochberg (BH) method. Corrected p-values < 0.05 were considered statistically significant.

### Behavioral battery

Animals underwent a battery of behavioral tasks measuring anxiety-like behavior, repetitive behavior, learning and memory, and depressive-like behaviors. All behavioral tests were performed during the dark cycle under red light (40-100lux) between the hours of 0900-1700. Behavioral testing began on day 9, or 5 days after the last injection, and ended on day 50. (Fig. 2B) ANY-maze tracking software was used to record and track animal movement during the task. All statistical analyses of behavior were performed using GraphPad Prism 9 software. (Supp. File S1) Behavior was analyzed using a one-way ANOVA assessing treatment as the main effect and using the BH-correction for multiple comparisons. Corrected p-values < 0.05 were considered statistically significant.

**Figure 2:**
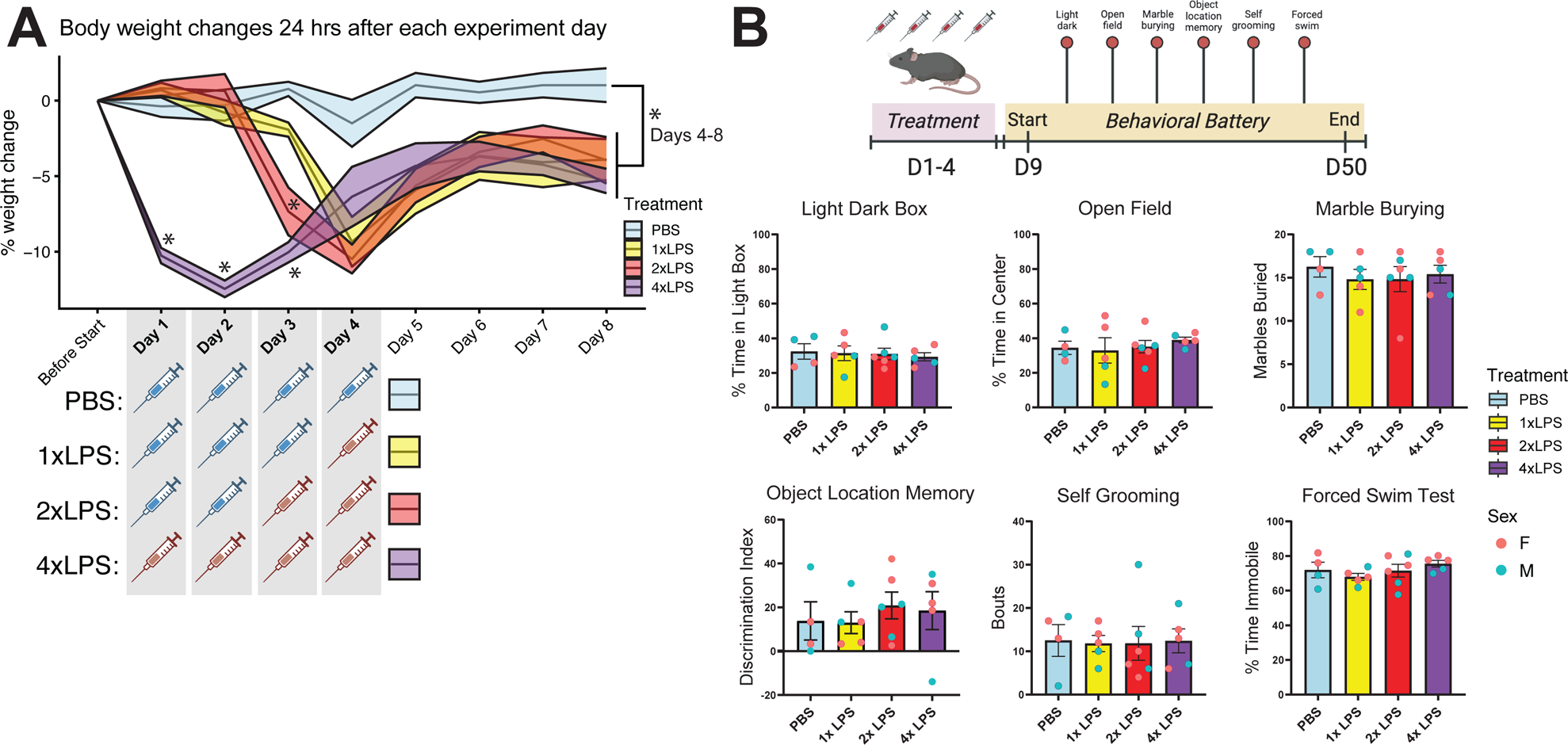
There are no long-term changes in behavior weeks after repeated LPS. (A) Percentage body weight changes for each treatment group, 24 hours after each experiment day 1-8, prior to beginning behavioral battery on day 9. Mean +/− SEM shown in line graphs. (*p<0.05, BH-corrected) (B) Timeline of behavioral assessment for anxiety-like behaviors, repetitive behaviors, learning and memory tasks, and depressive-like behaviors after 1-4xLPS paradigm. Group differences for each of the behavior tasks in behavioral battery are plotted. Mean +/− SEM shown in bar graphs. No significant differences. (*p<0.05, BH-corrected)

#### Light dark box

The light-dark box test was performed to measure anxiety-like behavior. The light chamber was an open-top 25cm x 25cm x 30cm (*lwh*) white box. The dark chamber was a closed-top 16cm x 25cm x 30cm (*lwh*) black box that was connected to the light box through a 10cm x 5cm (*wh*) passage and sliding door. Mice were individually placed into the dark chamber and were allowed to freely explore for 10 minutes once the sliding door was opened. Video footage was recorded from above and ANY-maze recorded time spent in the light chamber.

#### Open field test

The open field test was performed to measure anxiety-like behavior. The box was 65cm x 65cm x 40cm (*lwh*). One at a time, mice were placed in the center of the box and allowed to explore freely for 20 minutes. Video footage was recorded from above and ANY-maze recorded time spent in the inner and outer zones. The zone boundary was defined at 10cm from the box wall.

#### Marble burying test

The marble burying test was performed to measure repetitive behavior. To prepare for the task, a 30 cm x 19.4 cm x 39.4 cm rodent cage was filled with bedding 5cm deep and 20 marbles were equally spaced on top of the bedding in a 4×5 grid using a premade stencil. Mice were placed into the centre of the box one at a time and recorded for 20 minutes. Video footage subsequently hand scored at a later time to count for the number of marbles buried after 20 minutes. A marble was scored as buried if < 50% of the marble was visible.

#### Object Location Memory Test

The OLM task was performed in an open-topped square box 40cm x 40cm x 40 cm (*lwh*) lined with bedding ∼1cm in depth. One wall of the box was lined with duct tape as a visual cue. Objects were cleaned with 1:16 Saber solution (Rododonto, Cat # 09-12400-04) between trials and allowed to fully dry before re-use. The OLM took 8 days in total with 6 days of habituation, 1 day of training and 1 day of testing (25). Bedding was kept in the same box each day but shuffled around between mice. Mice were habituated for 10 minutes/day for 6 days, where they were allowed to freely explore the same box each day with no objects present. On the training day, mice were given 10 minutes to investigate two identical glass beakers filled with dry cement. The 5-minute testing phase took place 24 hours after training. During the test, one object was moved to a novel location and the other remained in the same spot. An experienced scorer blinded to experimental conditions hand-scored the testing videos for investigation time with the unmoved and moved objects. Investigation time was scored when the mouse’s nose was within 1cm of the object and pointing directly at the object. Looking over or past the object and climbing on top of the object were not counted as investigation. For the training and testing days, a discrimination index (DI) was calculated as [(T_M_ - T_S_) / (T_M_ + T_S_)]*100, where T_M_ is time investigating the moved object and T_S_ is time investigating the unmoved object.

#### Self-Grooming

Self-grooming was performed to measure repetitive-like behaviour. Mice were placed in a cage without bedding and filmed straight-on for 20 minutes. The last 10 minutes of video footage were hand-scored for the number of grooming bouts.

#### Forced Swim Test

Mice were placed individually into 1L glass beakers filled with 900mL of 23-25°C water. Mice were recorded for 6 minutes and closely monitored during the task. Mice were placed in a cage with dry paper towel on top of a heating pad after the task and were returned to the home cage once they were dry and exhibiting normal motor and grooming behaviours. The last four minutes of footage was hand-scored by a researcher blind to experimental conditions for time spent immobile, defined as the absence of escape-related movements.

### Weight change analysis

Body weights were measured 24 hours after each day before undergoing the behavioral battery described above. (Fig. 2A) Mice received LPS or PBS injections on Days 1-4 and were left alone to recover other than weight measurements on Days 5-8 before starting the behavioral assessments (outlined in ‘Behavioral Battery’ section) on Day 9. Each animal’s weights were normalized to their weights taken before the start of injections to calculate % weight change. We assessed how % weight differences change with LPS treatment in relation to experiment day by fitting a linear mixed effects model to measure % weight change as a factor of treatment and experiment day interactions with mouseID as a repeated measure (% weight change ∼ Treatment*Timepoint + (1|MouseID)). The model was assessed using ANOVA and two separate sets of posthoc analyses using the BH-method were run: 1) to correct across all comparisons made between timepoints within Treatment groups (∼Timepoint|Treatment) and 2) to correct across comparisons made against PBS (1-4xLPS vs. PBS) within each timepoint (∼Treatment|Timepoint). BH-corrected p-values < 0.05 were considered statistically significant.

### Brain tissue collection

In each of the separate cohorts outlined here, mice were euthanized 3 hours after the final injection immediately after the final day’s sickness behavior was scored. Mice were quickly anesthetized with isofluorane and transcardially perfused with 15ml of 1xPBS before brains were extracted for downstream experiments. <u>For the RNAseq and Luminex cohort</u>: one hemisphere from each brain was microdissected for brain regions (frontal cortex, striatum, hippocampus) for RNAseq and the other hemisphere was microdissected for brain regions (frontal cortex, striatum, hippocampus, cerebellum) for Luminex protein arrays. Dissected brain regions were immediately flash-frozen on dry ice and stored at −80°C until RNA and protein extractions for RNAseq and Luminex protein arrays, respectively. <u>For the microglia morphology & RT-qPCR cohort</u>: one hemisphere from each brain was collected for downstream histology analysis and the other hemisphere was microdissected for brain regions (frontal cortex, striatum, hippocampus) for RT-qPCR. Extracted hemispheres for histology were immersion-fixed in 4% paraformaldehyde for 48 hours before cryoprotecting in 30% sucrose for 48 hours prior to cryosectioning. Cryoprotected brains were then sectioned at 30um on the cryostat, collected in 1xPBS for long-term storage, and processed for immunohistochemistry. Dissected brain regions were immediately flash-frozen on dry ice and stored at −80°C until RNA extractions for RT-qPCR. <u>For the isolated microglia</u> <u>experiments (Fig. S4C,E)</u>: cortex from both hemispheres was collected for microglia isolations, subsequent RNA extractions, and downstream RT-qPCR.

### RNA-seq library preparation

RNA was extracted from whole tissue in the hippocampus, striatum and frontal cortex using the RNA/DNA Miniprep Plus Zymo Kit (Cat D7003) with on-column DNase digestion as described in the kit manual. RNA quality was assessed by bioanalyzer and all samples had RNA Integrity Numbers >7 (Supp. Table S3). QuantSeq 3’mRNA sequencing FWD (RNAseq) libraries (Lexogen, Cat No 015) were prepared per manufacturer’s instructions using 75ng of total input RNA per sample and 17 cycles of PCR amplification. All samples were prepared with incorporation of Unique Molecular Identifiers as part of the second strand synthesis step (Lexogen, Cat No 081) and sample unique, dual index barcodes. All samples were then pooled, exonuclease VII treated and sequenced across two lanes of a HiSeq4000 to generate Single End 100 base pair reads.

### RNA-Seq alignment

Raw fastq sequencing reads were assessed for quality control using FastQC (https://www.bioinformatics.babraham.ac.uk/projects/fastqc/) and multiqc (27) (PreTrim_multiqc_report.html). The UMI index was then added to the header of each read using custom scripts from Lexogen (umi2index). Reads were then trimmed to remove the first 11 bases, adapter contamination, polyA read through and low-quality reads using BBDuk from BBtools (<u>sourceforge.net/projects/bbmap/</u>). All samples were then re-assessed for quality using FastQC and multiqc (PostTrim_multiqc_report.html). Each sample was then aligned to the mouse genome (GRCm38.92) using STAR(28). Aligned bam files were then filtered for unique UMIs using custom Lexogen script (collapse_UMI_bam). Alignments for unfiltered and UMI filtered bam files were then compared using multiqc (STARAlignment_multiqc_report.html). All subsequent analysis was performed on UMI filtered bam files. These files were indexed using samtools(29) and then reads per ensembl gene (Mus_musculus.GRCm38.92.gtf) were counted using FeatureCounts(30) with options for forward strand (-s 1). Counts were then assess using multiqc (Counts_collapsedUMI_multiqc_report.html) and read into R for statistical analysis using EdgeR(31) and LimmaVoom(32). All scripts and R code are available on the CierniaLab github.

### Differential Expression (DE) analysis

Low expressing genes were removed by filtering for genes with more than 1 count per million (CPM) in at least 4 samples (Figure S1C). The remaining genes (n=16,788) were then normalized using Trimmed Mean of M-values (TMM) to correct for library composition using calcNormFactors, method= TMM (Figure S1E). To evaluate the contribution of each factor to the overall gene expression profile, Multi-Dimensional Scaling (MDS) plots were constructed for Brain Region, LPS Treatment, and Hemisphere (Figure S1B). CPM values were then fed into voom using a model design matrix for LPS treatment differences in gene expression which vary with brain region (∼0 + Treatment:BrainRegion). Individual contrast comparisons for each brain region were then called using contrasts.fit followed by eBayes. Differentially expressed genes (DEGs) were identified for each comparison of interest using Toptreat with a BH p-value correction (significance at adjusted p-values < 0.05 and fold change > +/−2). Heatmaps of normalized CPM values of significant DEGs (n=432) were made using the pheatmap R package. CPM values for each gene were first scaled within each brain region and then scaled across brain regions as input for the DEG heatmap (Fig. 3A), before hierarchical clustering. Breaks between each color of the heatmap scale were adjusted to fit the distribution of the normalized gene expression values such that 10% of the data was contained within each color break of the inferno color palette from the viridisLite R package (Fig. S1A). This allowed for the stark visualization of the strongest patterns of gene expression across brain regions and characterization of cluster IDs as 2xLPS-sensitive, 4xLPS-sensitive, and LPS-decreased.

**Figure 3:**
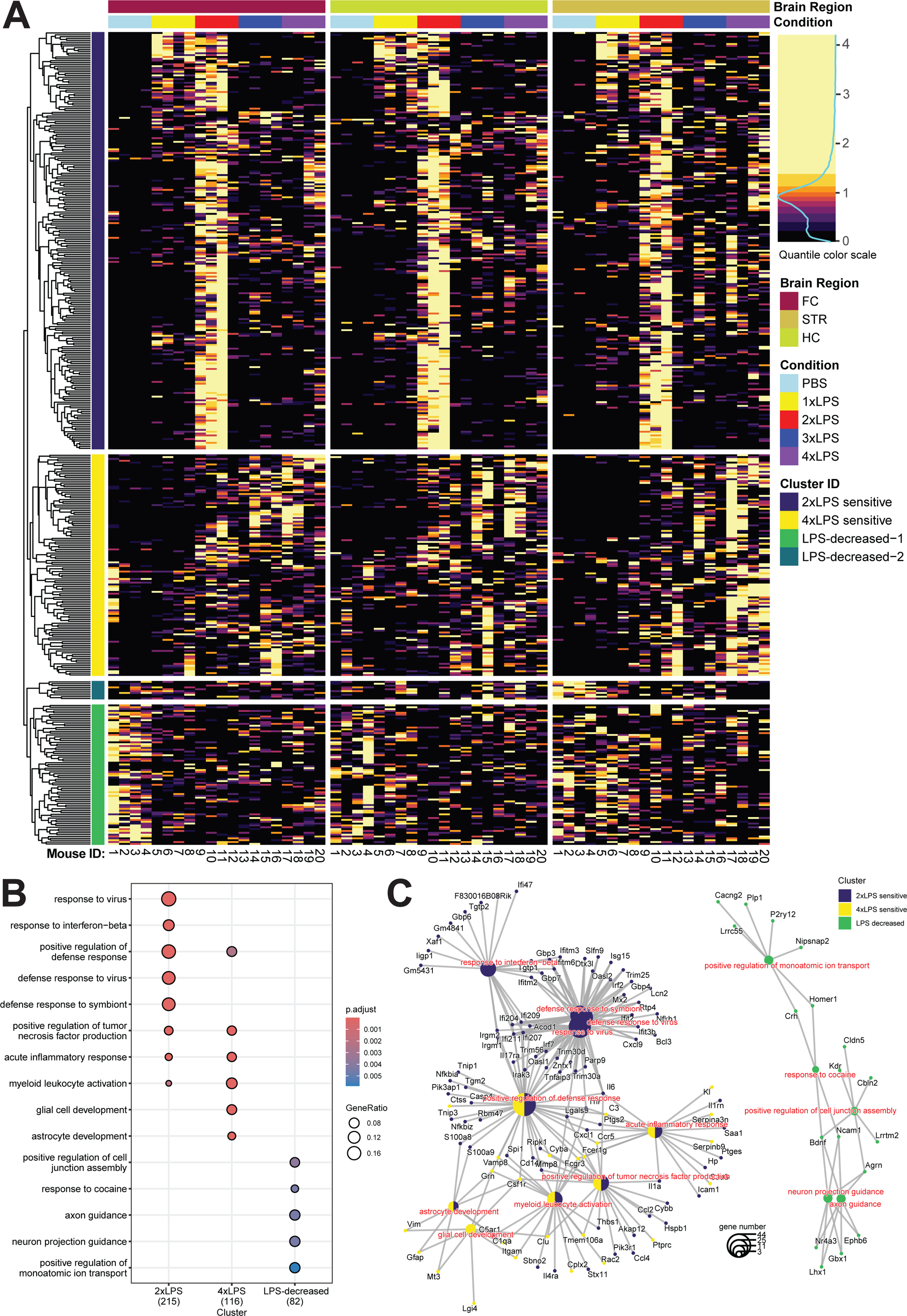
Repeated LPS induces shared gene expression signatures across brain regions. (A) Heatmap of scaled gene expression for each sample (columns) is shown for each differentially expressed gene (DEG) (rows). Only significant DEGs are shown (*p<0.05, BH-corrected). Genes are hierarchically clustered by patterns of expression: 2xLPS-sensitive cluster (226 total genes), 4xLPS-sensitive cluster (120 total genes), LPS-decreased-1 and 2 cluster (86 total genes). CPM values for each gene were first scaled within each brain region and then scaled across brain regions for plotting. Breaks between each color of the heatmap scale were adjusted to fit the distribution of the scaled gene expression values such that 10% of the data was contained within each color break of the quantile scale (Fig. S1A). (B) GO term enrichment dotplot for each cluster identified in (A). LPS-decreased-1 and LPS-decreased-2 genes were combined into one representative LPS-decreased cluster. Only significant GO terms shown (*p<0.05, BH-corrected). (C) GO terms in (B) represented as cnet plot showing specific genes from DEG clusters that were present in GO term gene lists.

### Gene Ontology (GO) enrichment analysis

Enrichment for gene ontology (GO) terms was subsequently performed on the DEG clusters using the clusterProfiler (version 4.4.4) and org.Mm.eg.db (version 3.15.0) R packages to further explore the functional role of the identified genes. GO enrichment was run on all 3 DEG cluster lists together using the compareCluster function in clusterProfiler with the “enrichGO” option and a p-value cutoff < 0.01 and BH-adjusted q-value cutoff < 0.05 against the Biological Processes (BP) ontology, with all genes interrogated in the experiment as background. Dotplot and cnetplot functions within the clusterProfiler package was used to create visualizations of the significant GO enrichments.

### MGEnrichment analysis

Gene list enrichment analysis was performed using the 2024 database of the MGEnrichmentApp (33) with all genes interrogated in the experiment as background. Significant enrichments against mouse microglial gene lists in MGEnrichment were tested by a one-tail Fischer’s exact test and BH-corrected to reach significance at an adjusted p-value < 0.01. Target % and Background % values for significant gene list enrichments were plotted using ggplot in R.

### HOMER promoter motif enrichment analysis

HOMER Motif Analysis Software, v4.11 (http://homer.ucsd.edu/homer/) (34) was used to find motif enrichments within promoters (+1000, −200 to transcription start site) for significant DEGs in each cluster identified (2xLPS-sensitive, 4xLPS-senstive, LPSDownregulated) and run against background genes (all genes interrogated in the experiment, n=16,788) using the following command: findMotifs.pl [cluster-specific genes file] mm10_promoters [output directory] -bg [background genes file] -start -1000 -end 200 -p 12. Enrichment output was interpreted according to recommendations on the HOMER website (http://homer.ucsd.edu/homer/). Top-ranked *de novo* motifs were defined as having p-values < 1e-10 and not marked as a possible false positive in the Homer *de novo* Motif Results output (Supp. File S2-S4). Top-ranked known motifs were defined as having BH-corrected q-values < 0.05 in the Homer Known Motif Enrichment Results output. (Supp. File S2-S4) Target % and Background % values for significant promoter enrichments were plotted using ggplot in R. All known motifs were concatenated into one file and used as input to determine target gene promoter locations with known motifs using the following command: findMotifs.pl [cluster-specific genes file] mm10_promoters [output directory] -find [all known motifs file] > [desired name of output file] -bg [background genes file] - start -1000 -end 200 -p 12. Identified instances of significant known motifs in each gene promoter were binarized (yes = present, no = not present) and plotted side-by-side with each gene’s normalized gene expression using the R package pheatmap.

### Published ChIP-seq promoter enrichment analysis

Publicly available ChIP-seq peaks (35–39) from relevant microglia and bone marrow-derived macrophage (BMDM) studies were downloaded from the NCBI GEO repository and processed using sort-bed functions and -intersect and -difference options within the bedops software (version 2.4.41) (40) to isolate out ChIP-seq mm10 promoter peaks and to characterize lost vs. gained promoter peaks in any studies with treatment conditions, respectively. Final ChIP-seq peaks were curated into collections for each study to use as input for analysis, according to the requirements outlined in the LOLA R package. ChIP-seq peak promoter enrichment analysis was performed using LOLA and significant enrichments were defined as BH-corrected adjusted p-values < 0.05. Target % and Background % values for significant enrichments were plotted using ggplot in R.

### Luminex protein arrays

As described previously (26), blood was collected by cardiac punch and brain tissue for each region (frontal cortex, striatum, hippocampus, and cerebellum) was collected and flash frozen. Blood was placed on ice and subsequently processed into serum by centrifugation. Brain tissue was lysed in cell lysis buffer (Cell Signaling Technologies, Danvers, MA, USA) containing protease and phosphatase inhibitors and incubated with agitation for 20 min on ice followed by sonication for 30 seconds. The cell lysate was then vortexed for 30 seconds and centrifuged at 20,000× *g* for 10 min at 4°C. Protein concentrations were measured using a Bio-Rad Benchmark Plus Spectrophotometer system and all samples were standardized to 70 µg/mL for subsequent immunoassays.

Analysis of serum cytokines was performed using a multiplex mouse 25-plex bead immunoassay (Milliplex Mouse Cytokine/Chemokine Magnetic Bead Panel #MCYTMAG70PMX25BK). The following cytokines were quantified: G-CSF, GM-CSF, IFN-γ, IL-1α, IL-1β, IL-2, IL-4, IL-5, IL-6, IL-7, IL-9, IL-10, IL-12 (p40), IL-12 (p70), IL-13, IL-15, IL-17, IP-10, KC, MCP-1, MIP-1α, MIP-1β, MIP-2, RANTES, and TNF-α. Standards and reagents were all prepared according to the manufacturers’ recommendations. Each serum and brain sample were diluted to a standardized concentration and run in duplicates. 25uL of sample, standards, or blanks was loaded into a 96-well plate with appropriate amounts of assay buffer and matrix solution. The plate was then incubated overnight with antibody-coupled magnetic beads. The following day, after a series of washes, the plate was incubated with a biotinylated detection antibody on a shaker for 1 hour. Streptavidin-phycoerythrin was added and incubated while shaking continued for 30 minutes. Plates were washed using a Bio-Plex handheld magnet (Bio-Rad Laboratories, Hercules, CA, USA). After the final wash, the plate was analyzed using a Bio-Rad Bio-Plex 200 plate reader (Bio-Rad Laboratories, Hercules, CA, USA) and analyzed using Bio-Plex Manager software (Bio-Rad Laboratories, Hercules, CA, USA). The following were the minimal amounts of detectable cytokine concentrations: G-CSF: 1.7 pg/mL; GM-CSF: 10.9 pg/mL; IFNγ: 1.1 pg/mL; IL-1α: 10.3 pg/mL; IL-1β: 5.4 pg/mL; IL-2: 1.0 pg/mL; IL-4: 0.4 pg/mL; IL-5: 1.0 pg/mL; IL-6: 1.1 pg/mL; IL-7: 1.4 pg/mL; IL-9: 17.3 pg/mL; IL-10: 2.0 pg/mL; IL-12 (p40): 3.9 pg/mL; IL-12 (p70): 4.8 pg/mL; IL-13: 7.8 pg/mL; IL-15: 7.4 pg/mL; IL-17: 0.5 pg/mL; IP-10: 0.8 pg/mL; KC: 2.3 pg/mL; MCP-1: 6.7 pg/mL; MIP-1α: 7.7 pg/mL; MIP-1β: 11.9 pg/mL; MIP-2: 30.6 pg/mL; RANTES: 2.7 pg/mL; TNF-α: 2.3 pg/mL. As per previous studies, sample concentrations that fell below minimal detection value were given a proxy value of half the limit of detection for statistical comparisons using non-parametric Mann–Whitney U analyses and BH-correction for multiple comparisons.

### Immunohistochemistry

30um brain sections were stained for immunofluorescent analysis with a commonly used microglia marker, purinergic receptor P2Y12 (P2ry12), to analyze microglial morphology. Free-floating brain sections were washed 3 times for 5 minutes each in 1xPBS, permeabilized in 1xPBS + 0.5% Triton (Fisher BioReagents BP151-500) for 5 minutes, and incubated in blocking solution made of 1xPBS + 0.03% Triton + 1% Bovine Serum Albumin (BSA; Bio-techne Tocris 5217) for 1 hour. After the blocking steps, sections were incubated overnight at 4°C in primary antibody solution containing 2% Normal Donkey Serum (NDS; Jackson Immunoresearch Laboratories Inc. 017-000-121) + 1xPBS + 0.03% Triton + primary antibody (rabbit anti-P2RY12: 1:500, Anaspec AS-55043A). After primary antibody incubation, sections were washed 3 times for 5 minutes each with 1xPBS + 0.03% Triton before incubating for 2 hours in secondary solution containing 2% NDS + DAPI (1:1000; Biolegend 422801) + secondary antibodies (Alexafluor 647 donkey anti-rabbit: 1:500, Jackson Immunoresearch Laboratories Inc. 711-606-152). Sections were washed 3 times for 5 minutes each with 1xPBS + 0.03% Triton before being transferred into 1xPBS for temporary storage before mounting. Sections were mounted onto microscope slides (Premium Superfrost Plus Microscrope Slides, VWR CA48311-703) and air dried before being coverslipped with mounting media (ProLong Glass Antifade Mountant, Invitrogen P36980; 24×60mm 1.5H High Performance Coverslips, Marienfield 0107242).

### Imaging and microglia morphology analysis

All mounted brain sections were imaged on the ZEISS Axioscan 6 microscope slide scanner at 20x magnification with a step-size of 1um using the z-stack acquisition parameters within the imaging software (ZEISS ZEN 3.7). During image acquisition, Extended Depth of Focus (EDF) images were created using maximum projection settings and saved as the outputs in the final .czi files. Maximum projection EDF images only compile the pixels of highest intensity at any given position in a z-stack to construct a new 2D image which retains the 3D information. Using ImageJ, we created .tiff images of each fluorescent channel from .czi files and selected and saved .tiffs of brain regions of interest (ROIs) to use as input for downstream morphological analysis in MicrogliaMorphology and MicrogliaMorphologyR, as detailed in Kim et al., 2023 (41). The parameters used for MicrogliaMorphology included: auto local thresholding using the Mean method and radius 100 with an area filter of 0.01613 - 0.1808 in^2 (or 153.3358 – 1718.73 um^2 with .czi conversion applied factor of .325um/pixel applied to scale). After performing principal components analysis on the 27 morphology measures output from MicrogliaMorphology, the first 3 principal components were used as input for K-means clustering in MicrogliaMorphologyR to define four distinct classes of microglia morphology (ameboid, hypertrophic, rod-like, ramified). We focused our analyses on multiple brain regions including the hippocampus, frontal cortex, and striatum. Images of coronal brain sections containing these regions were aligned to the Allen Brain Atlas (mouse.brain-map.org) (42) using the ImageJ macro FASTMAP (43). We assessed how percentages of morphological clusters for each mouse changed with LPS treatment by fitting a generalized linear mixed model to measure percentage changes as a factor of cluster and treatment interactions for each brain region and sex with mouseID as a repeated measure (percentage ∼ ClusterID*Treatment + (1|MouseID)) using MicrogliaMorphologyR, which calls to the glmmTMB R package. (41,44) We had to analyze each sex and brain region separately because the males had an additional treatment group (3xLPS) that was not present in the females and the model converged when considering ClusterID*Treatment*BrainRegion interactions together. The model was assessed using ANOVA and tests between groups were corrected across for comparisons made within each cluster (∼Treatment|Cluster) using the BH method. Corrected p-values < 0.05 were considered statistically significant.

Events of rod-like microglia trains were manually quantified within each brain region of interest (ROI) for every image using ImageJ software. Manual counts were normalized to the areas of the ROIs to calculate densities of event observations for each brain region analyzed. Microglial density was quantified by dividing the number of microglia analyzed in each ROI by the area of each ROI. We assessed how rod train event densities and microglial densities change with LPS treatment across brain regions by fitting separate linear mixed effects models to measure either rod-train density or microglial density as a factor of treatment and brain region interactions with mouseID as a repeated measure, for each sex separately (RodTrainDensity ∼ Treatment*BrainRegion + (1|MouseID) and MicroglialDensity ∼ Treatment*BrainRegion + (1|MouseID)). Both models were assessed using ANOVA and tests between groups were corrected across for comparisons made within each brain region (∼Treatment|BrainRegion) using the BH method. Corrected p-values < 0.05 were considered statistically significant.

### Microglia isolations

Isolated cortical tissue from each mouse was placed on individual petri dishes on ice with 1mL of digestion buffer, made up of 10ul DNase I (STEMCELL Technologies; Cat. 07900) + 990ul activated Papain solution (Worthington Biochemical, Cat. LS003124), added to each brain sample before it was minced into small (<1mm) pieces using a scalpel. Minced samples in digestion solution were transferred to individual wells within a 24-well cell culture plate and incubated on ice for 30 minutes. Digested brain solution was transferred into a glass dounce homogenizer on ice with 5 mL ice-cold flow cytometry (FACS) buffer comprising 2.5% bovine serum albumin (BSA; Tocris Bioscience Cat. 5217/100G) and 1 M ethylenediaminetetraacetic acid (EDTA; Invitrogen Cat. 17892) in Hanks’ Balanced Salt Solution (HBSS). Brains were slowly and gently homogenized 20 times using a loose pestle, followed by 2 times using a tight pestle. Brain homogenate was strained through a 70um nylon mesh strainer (Thermo Fisher, Cat. 08-771-1) with FACS buffer into 50ml tubes on ice. 50ml tubes were centrifuged at 300xg for 10 minutes at 4C, at the machine’s minimum brake setting. After carefully and discarding the supernatant, cell pellets were gently resuspended in 30% Percoll solution and transferred to new 15ml tubes, where 2ml of 37% Percoll solution was gently underlaid. To make the Percoll solutions, 3.6ml of Isotonic Percoll (Millipore Sigma, Cat. GE17-0891-02) was added to 0.4ml 10xHBSS per brain to make a 100% Percoll solution, which was diluted down to 30% Percoll and 37% Percoll in 1xHBSS. 15ml tubes containing the layered Percoll gradients were centrifuged at 700xg for 10 minutes at 4C, at the machine’s minimum brake setting. Myelin layers and supernatant layers were discarded using a transfer pipette, leaving 1ml of the 37% layer at the bottom of the tube. Cell pellets were gently resuspended and topped up to 10ml with ice-cold FACS buffer then centrifuged at 300xg for 10 minutes at 4C, at the machine’s minimum brake setting. Supernatant in tubes was discarded down to 200ul, 5ml of ice-cold FACS buffer added, and tubes spun again at 300xg for 10 min at 4C at the machine’s minimum brake setting. Finally, supernatant was removed and cells resuspended in 200ul remaining FACS buffer before being counted on the hemocytometer. After counting, samples were diluted to a concentration of 2.5×10^7 cells/ml in 0.05-0.2ml of FACS buffer and transferred to a round-bottom, non-adherent 96-well plate (Corning, Cat. 3788). CD11b+ microglial cells were isolated by magnetic bead enrichment using the EasySep™ Mouse CD11b Positive Selection Kit (STEMCELL Technologies, Cat. 18970), according to the manufacturer’s protocol.

### RT-qPCR

Isolated brain region tissues and isolated microglia cells were homogenized using QIAshredder columns (Qiagen 79656). RNA from homogenized tissue was isolated using the Monarch Total RNA Miniprep kit, according to the manufacturer’s protocol (New England Biolabs T2010S). For isolated microglia, cells were homogenized and RNA extracted using the Total RNA Purification from Cultured Mammalian Cells version of the protocol for the Monarch Total RNA Miniprep kit. Isolated RNA from brain regions and purified microglia was converted to cDNA and prepared for Real Time quantitative PCR (RT-qPCR) reactions using the LunaScript® RT SuperMix Kit (New England Biolabs E3010) and Luna® Universal qPCR Master Mix (New England Biolabs M3003), according to the manufacturer’s Protocol for Two-step RT-qPCR. The primers in Table 1 were used for RT-qPCR reactions and RT-qPCR reactions were loaded into MicroAmp^TM^ Fast Optical 96-Well Reaction plates (Thermo Fisher Scientific 43-469-07) sealed with optical adhesive film (Thermo Fisher Scientific AB-1170). All biological samples were tested in triplicates. Reactions were run on the QuantStudio 6 qPCR machine (Life Technologies, California, USA) and samples were analyzed by the comparative CT method (Λ1Λ1CT). The Λ1CT was calculated by subtracting the average CT value of the housekeeping gene hypoxanthine phosphoribosyltransferase 1 (*Hprt1*) from the average CT value from the gene of interest. Λ1Λ1CT values were calculated by subtracting the Λ1CT from the gene of interest by the average Λ1CT from its respective control. Fold change was calculated by using the equation 2^−ΔΔCT^. We assessed how expression of each target gene changes with LPS treatment by fitting a linear mixed-effects model to measure fold change values as a factor of LPS Treatment for each gene independently (gene fold change ∼ Treatment + (1|MouseID)) using the nlme R package. We analyzed each sex separately, as the males had an additional treatment group, 3xLPS, that was not present in the females. The model was assessed using ANOVA and tests between LPS treatments were corrected for multiple comparisons using the BH method. Corrected p-values < 0.05 were considered statistically significant.

**Table 1.**
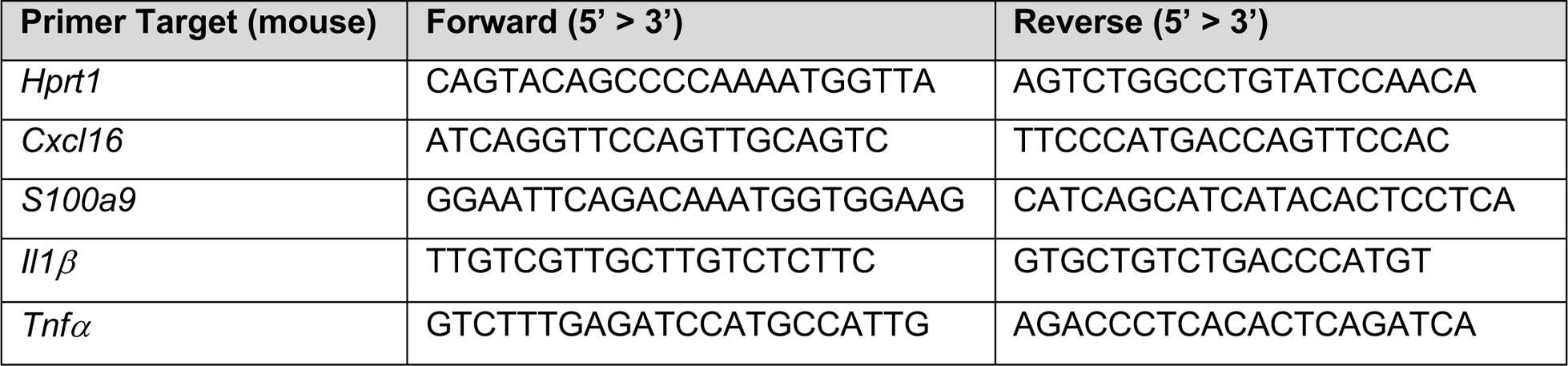
RT-qPCR primer sequences used in this study.

## Results

### Repeated LPS produces robust priming and tolerance of acute sickness behaviors, but not long-term behavioral change

In our experimental model, mice were given daily intraperitoneal injections of low-dose LPS (0.5 mg/kg) or vehicle (1xPBS) over a span of four days and sickness behavior was scored 3 hours following the final treatment of PBS, 1xLPS, 2xLPS, 3xLPS or 4xLPS. (Fig. 1A-B) Repeated LPS injections produced both training and tolerance of sickness behaviors: sickness behavior peaked and the greatest drop in body weight was observed across treatment groups after the second injection of LPS relative to PBS. (Fig. 1B, 2A; BH-adjusted p<0.05, Supp. Table S2) Sickness behavior showed recovery towards baseline levels after the third and fourth LPS injections (Fig. 1B, 2A). While body weights recovered by 2 days after peak weight loss from the second exposure to LPS across treatment groups, weights in LPS-treated groups was still significantly lower than that of the PBS group for up to 6 days after peak loss (BH-adjusted p<0.05, Supp. Table S2) and none of the mice in the LPS-treated groups fully recovered to their pre-treatment body weights prior to the start of the behavioral experiment (Fig. 2A; BH-adjusted p<0.05, Supp. Table S2). However, there were no long-term differences in anxiety-like, depressive-like, learning and memory, nor repetitive behaviors across treatment groups when behaviors were then assessed, indicating that all treatment conditions had fully recovered and did not have long-term detrimental impacts of repeated LPS. (Fig. 2B; BH-adjusted p<0.05, Supp. File S1)

### Cytokine and chemokine protein levels increase first in serum and then in brain

To examine protein levels of known inflammatory proteins, including chemokines, cytokines, and other signaling molecules, we performed a Luminex multiplex protein quantification on serum and tissue collected from four different brain regions (frontal cortex, hippocampus, striatum, cerebellum) 3 hours after final injections. We identified significant changes in protein levels of both pro- and anti-inflammatory molecules (Fig. 1C; BH-adjusted p<0.05, Supp. Table S1). In serum, we found significant increases of G-CSF, IL-6, IP-10, KC, MCP-1, MIP-1a, MIP-1b and RANTES in response to 1xLPS that was maintained in response to 2xLPS and then returned to baseline level in most cases after 3xLPS and 4xLPS. In frontal cortex we found significant increases in G-CSF, IL-6, and IP-10 to 1xLPS, but the majority of changes were in response to the 2xLPS (10 proteins), most of which again were no longer significant after 3xLPS and 4xLPS. Similar patterns were observed in hippocampus, striatum, and cerebellum with the majority of significant increases occurring after 2xLPS (Fig. 1C; BH-adjusted p<0.05, Supp. Table S1). We further explored these findings in males and females using RT-qPCR on RNA isolated from frontal cortex tissue for *Il-1β* and *Tnf-α*, two hallmark pro-inflammatory cytokines that are known to be produced by microglia during immune activation. Similar to the patterns observed in the protein analysis, we found the greatest induction of *Il-1β* and *Tnf-α* in response to 2xLPS, which returned towards baseline levels after 3xLPS and 4xLPS (Fig. 1D; BH-adjusted p<0.05, Supp. Table S5). Together, these findings suggest that 2xLPS produces the largest increase in inflammatory signaling molecules and that these responses are blunted after 3xLPS and 4xLPS. These patterns also parallel the peak in sickness behaviors and drop in body weight observed after 2xLPS that resolve towards baseline levels by 4xLPS.

### Repeated LPS produces priming and tolerance of gene expression within brain regions that is likely microglia-driven

To examine changes in gene expression in response to repeated LPS stimulation, we performed RNA-sequencing on brain tissue from the hippocampus, striatum and frontal cortex 3 hours after the last LPS or PBS injection (n=4/treatment/region; Fig. 1A, Fig. S1). Differentially expressed genes were identified across brain regions for differences in expression between the PBS controls and each LPS condition (BH-adjusted p<0.05 and fold change>+/−2, Supp. Table S3) and clustered according to their gene expression patterns across brain regions and treatments, which were the greatest explanatory variables in the data as assessed by MDS analysis (Fig. 3A, Fig. S1B). Similar to the impacts on brain cytokine protein levels, sickness behaviors, and body weight changes, changes in gene expression were most striking following 2xLPS (Fig. S2A-C). Very few genes were decreased in response to LPS across all comparisons and brain regions. Clusters that emerged from patterns of gene expression were characterized as “2xLPS-sensitive”, “4xLPS-sensitive”, and “LPS-decreased”. Across brain regions, 2xLPS-sensitive genes (n=226, 52% of DEGs) were uniquely increased in response to 2xLPS, 4xLPS-sensitive genes (n=120, 28% of DEGs) were uniquely increased in response to 4xLPS, and LPS-decreased genes (n=86, 20% of DEGs) were repressed from baseline levels across LPS treatments. (Fig. 3A)

The different DEG clusters were enriched for shared and unique GO terms. (Fig. 3B; BH-corrected p<0.05, Supp. Table S3). The majority of 2xLPS-sensitive genes were not engaged at 3xLPS and 4xLPS, suggesting the formation of tolerance for the majority of these genes. 2xLPS-senstive genes were enriched for biological processes related to immune regulation including response to virus, response to interferon-beta, positive regulation of defense response, defense response to virus, defense response to symbiont, positive regulation of tumor necrosis factor production, acute inflammatory response, and myeloid leukocyte activation. These findings are consistent with previous work examining LPS-activated gene expression in the brain and periphery and indicate that alterations in immune cell signaling underlie the enhancement in gene expression to 2xLPS and repression to 3xLPS and 4xLPS in the brain (20,45–47). The 4xLPS-sensitive genes were enriched for many of the same processes, as well as glial cell development and astrocyte development. LPS-decreased genes were enriched for processes related to neuronal regulation including positive regulation of cell junction assembly, response to cocaine, axon guidance, neuron projection guidance, and positive regulation of monoatomic ion transport. Together, findings suggest that separate gene expression signatures are engaged in response to repeated injections of LPS.

To identify whether the differential gene expression observed in our bulk RNAseq data could be driven by microglia, we performed enrichment analysis of our DEGs against curated lists in MGEnrichment, our recently developed database of published microglial gene lists (33). We identified shared and unique enrichment between 2xLPS and 4xLPS-sensitive but not LPS-decreased genes with published microglia-specific gene lists important for maintaining microglia homeostasis, disease-associated microglia (DAM) phenotypes, immune primed microglia, microglia development, and microglial changes in microbiome perturbations, among many others (Fig. S3; BH-corrected p<0.01, Supp. Table S3). Furthermore, two LPS-sensitive genes from our DEG list that are highly expressed in microglia (Fig. S4B, E), *Cxcl16* and *S100a9*, also displayed the same training and tolerance gene expression patterns when assessed by RT-qPCR in repeat experiments done in frontal cortex in both males and females, as well as in microglia isolated from cortical regions in males. (Fig. S4C, F; BH-corrected p<0.05, Supp. Table S5) Together, findings suggest microglia as a likely candidate cell type driving the differential expression observed in the 2xLPS-sensitive and 4xLPS-sensitive gene clusters.

### Repeated LPS regulates immune memory through engagement with IRF and NFkB family transcription factors

To examine which up-stream regulatory factors could be driving gene expression in response to repeated LPS, we examined promoter regions of DEGs for enrichment of transcription factor binding motifs (Fig. 4A,C). Promoters of 2xLPS-sensitive cluster DEGs were significantly enriched for interferon regulatory factor (IRF), basic helix-loop-helix/bZIP (bHLH/bZIP), Rel homology domain (RHD), C2H2 zinc finger (Zf), and signal transducer and activator of transcription (Stat) transcription family factors across *de novo* and known Homer motifs. (Supp. File S2) Different sets of genes within the 2xLPS-sensitive DEG cluster had differences in the presence of binding motifs for transcription factors across the enriched transcription factor families (Fig. 4D) in their promoters. Notably, 2xLPS-sensitive genes that increased in response to 1xLPS and stayed increase after 2xLPS did not have binding sites for bZIP family transcription factors, which were present throughout most of the other 2xLPS cluster genes. Moreover, only subsets of 2xLPS-sensitive genes had binding sites for IRF family transcription factors. Binding sites for RHD family transcription factor NFkB, another critical regulator of immune genes, were found in most 2xLPS cluster DEG promoters. DEG promoters were also significantly enriched for direct binding sites of RHD family (NFkB, p65) and IRF family (IRF8) from publicly available ChIP-seq data (35–39), further supporting the roles of these transcription factors as major regulators of LPS-induced gene expression differences in immune training and tolerance. (Fig. 4B; BH-adjusted p<0.05, Supp. Table S3) DEG promoters were also enriched for H3K27ac ChIP-seq peaks that were lost or gained with conditional IRF8 knockouts in microglia in adults (35), suggesting that IRF8 may be interacting with other transcriptional machinery such as histone acetyltransferases and deacetylases to mediate microglial gene expression at these sites.

**Figure 4:**
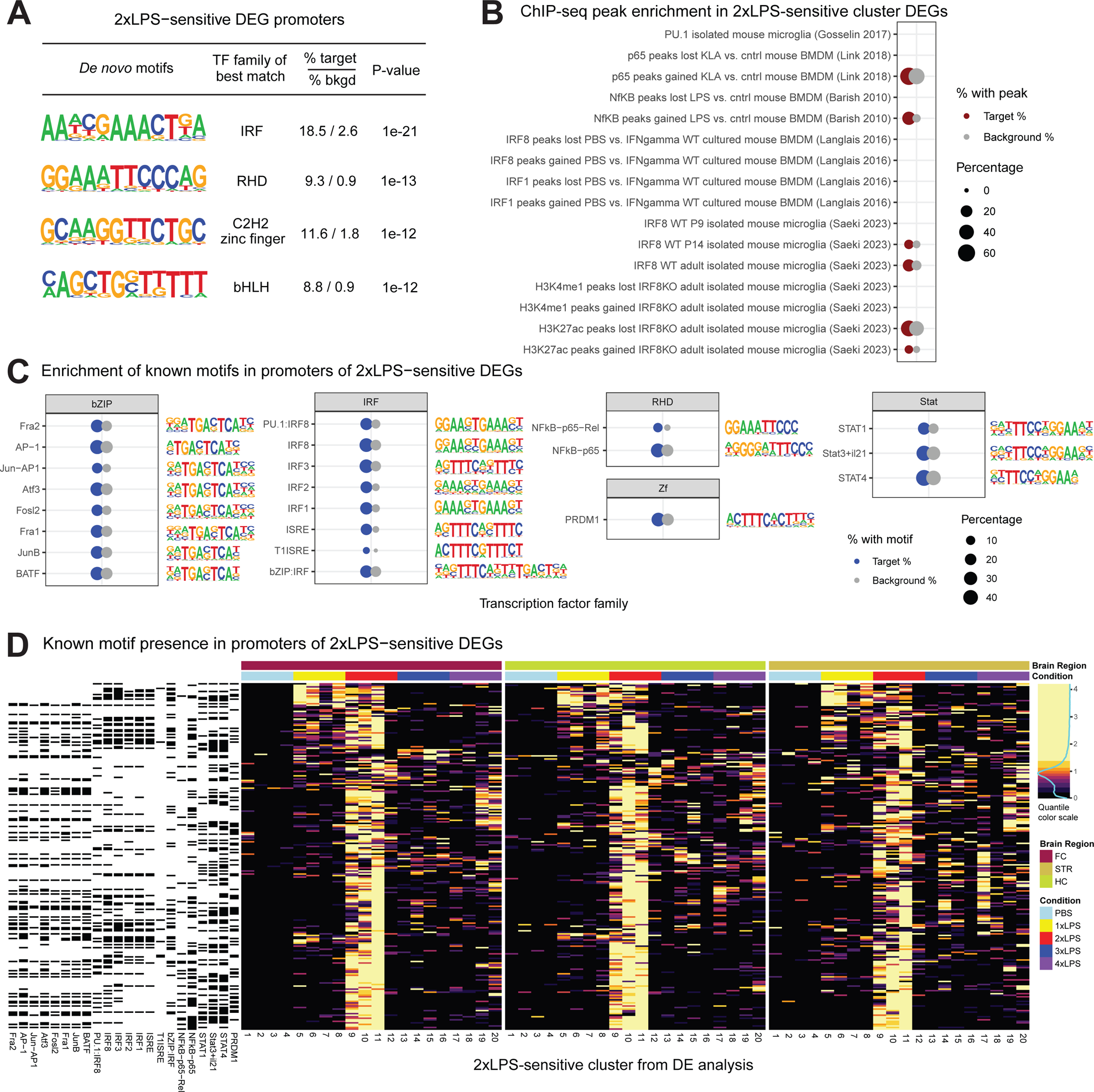
IRF and NFkB transcription factor motifs are enriched in 2xLPS-sensitive DEGs. (A) Homer *de novo* transcription factor motif enrichments within promoters (+1000, −200 to Transcription Start Site) of 2xLPS-sensitive DEGs. Top ranked *de novo* motifs are shown (p<1e-10). (B) Enrichment of 2xLPS-sensitive DEGs within ChIP-seq promoter peaks from publicly available microglial and bone marrow derived macrophage (BMDM) studies. Red circles denote a significant enrichment in the percent of DEGs containing a ChIP-seq peak compared to the background of all genes detected in the RNA-Seq experiment containing a ChIP-seq peak from a given study (grey circles). *p<0.05, BH-corrected (C) Homer known transcription factor motif enrichments within promoters of 2xLPS-sensitive DEGs. Blue circles denote a significant enrichment in the percent of DEGs containing that promoter motif compared to the background of all genes detected in the RNA-Seq experiment with that motif (grey circles). Top ranked known motifs are shown (p<0.05, BH-corrected) (D) Heatmap of scaled gene expression of 2xLPS-sensitive DEGs and depiction of whether a binding site for each known promoter motif from (C) is present in promoters of 2xLPS-sensitive DEGs (black=yes, white=no).

### Microglia change their morphology across brain regions in the formation of immune priming and tolerance

Using MicrogliaMorphology and MicrogliaMorphologyR, as previously described (41), we quantified shifts in morphological populations within the same brain regions assessed for RNA-seq (frontal cortex, hippocampus, striatum) from male and female mice given 1-4xLPS. (Fig. 5A, S5A-E) Using unbiased clustering, we identified four clusters of microglial morphologies: ramified, rod-like, hypertrophic, and ameboid. While the percentage of ramified and rod-like microglia mostly remained consistent within the different brain regions and sexes with LPS treatment, there was an LPS-mediated shift between ameboid and hypertrophic states (Fig. 5A-B; BH-corrected p<0.05, Supp. Table S4). Within males, the hippocampus and striatum showed a significant shift from ameboid to hypertrophic morphological states in the formation of immune priming (2xLPS) and back with immune tolerance (4xLPS). In males the frontal cortex maintained the 2xLPS-induced hypertrophic morphology with immune tolerance. Within females, the frontal cortex, hippocampus, and striatum showed a morphological switch to a hypertrophic morphology which was sustained with immune tolerance.

**Figure 5:**
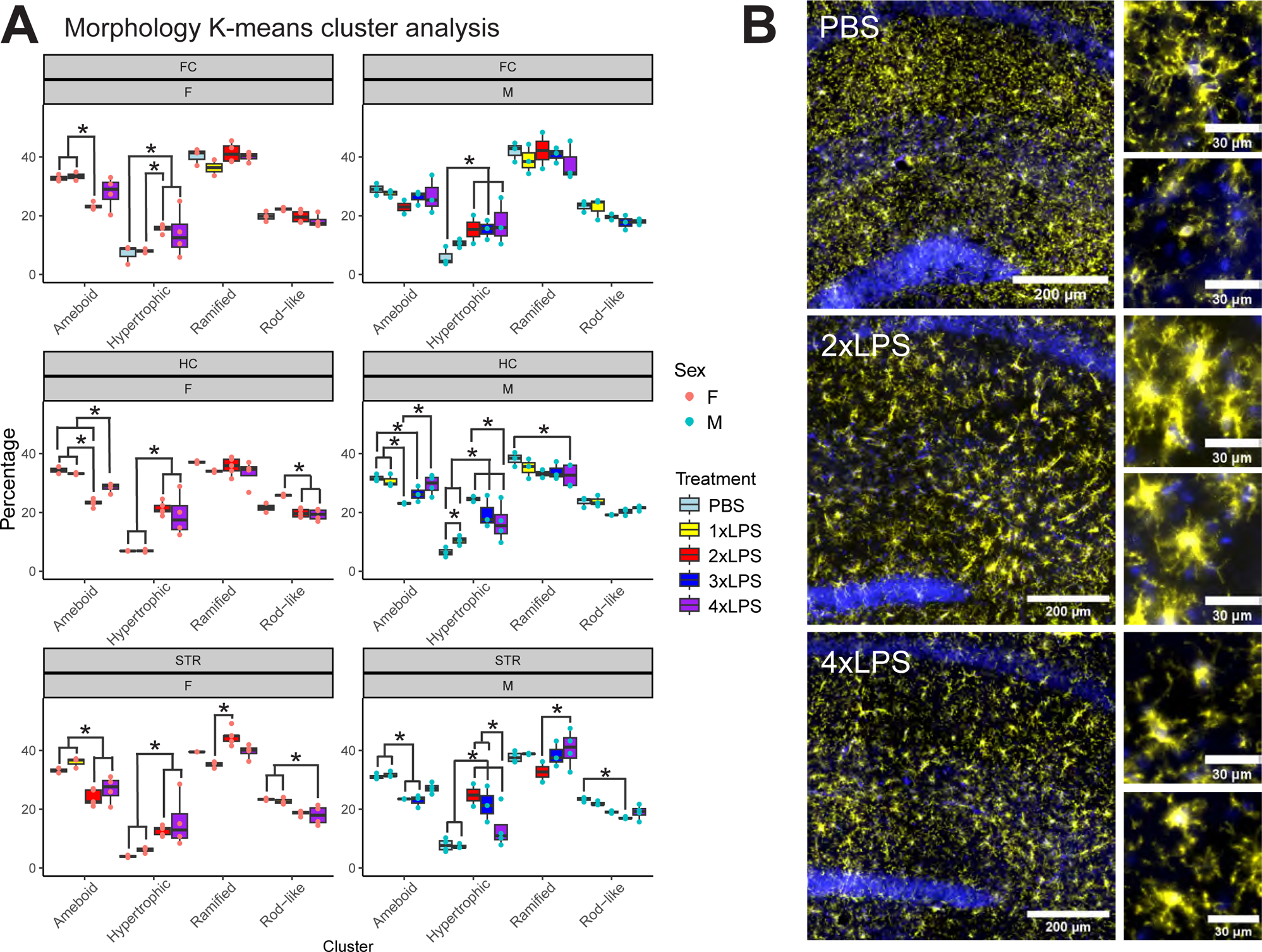
Microglia morphology shifts mainly between ameboid and hypertrophic states in immune memory. (A) LPS-induced shifts in morphological populations between ameboid and hypertrophic states within brain regions and sexes. (*p<0.05, BH-corrected) (B) Example immunofluorescent images of hippocampal microglial cells stained with P2ry12 (yellow) in PBS, 2xLPS, and 4xLPS conditions. Nuclei are stained with DAPI (blue). Scale bars are 200um and 30um.

Furthermore, we also observed the formation of trains of rod-like microglia with increasing exposures to LPS. Rod-like microglia have previously been observed to align end-to-end with one another to form ‘trains’ of microglia adjacent to neuronal processes in diffuse traumatic brain injury (TBI) in rats. (48–50) Within males, the occurrence of microglial rod-trains increased with 2xLPS, began to resolve after 3xLPS, and re-emerged with 4xLPS across the frontal cortex, hippocampus, and striatum. (Fig. 6A-B; BH-corrected p < 0.05, Supp. Table S4) Within females, microglial-rod train occurrence significantly increased with 4xLPS in the frontal cortex and hippocampus, but not in the striatum. There were no significant differences in microglial density with repeated LPS across brain regions and in both sexes, indicating that the changes observed in rod-train occurrence were not due to changes in cell numbers. (Fig. S5F; BH-corrected p < 0.05, Supp. Table S4)

**Figure 6:**
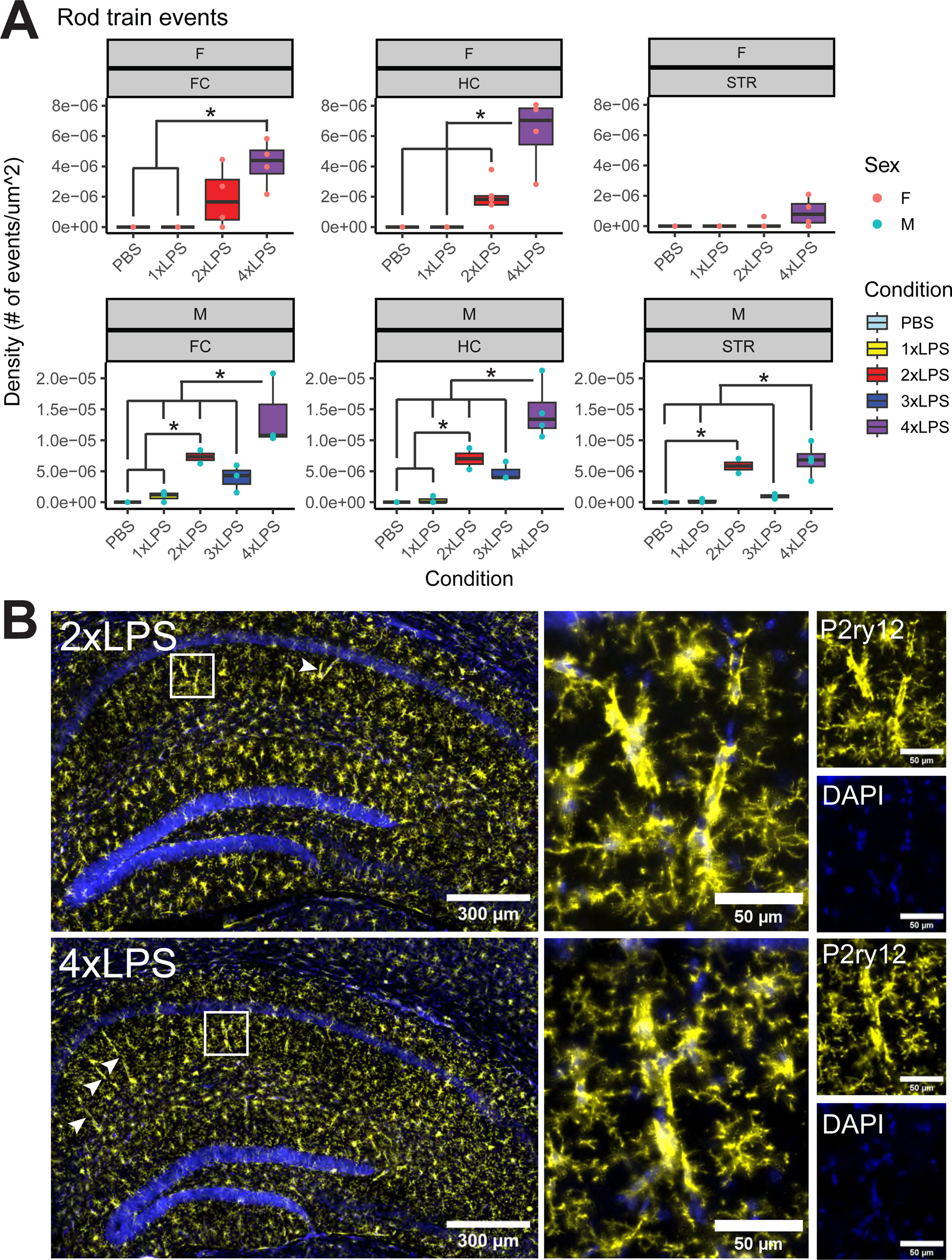
Trains of rod-like microglia dynamically emerge and resolve with repeated LPS. (A) Quantification of increasing microglial rod-train events that occurred with repeated LPS across brain regions within sexes. (*p<0.05, BH-corrected) (B) Example immunofluorescent images of microglial rod trains in hippocampus, stained with P2ry12 (yellow), in 2xLPS, and 4xLPS conditions. Nuclei are stained with DAPI (blue). Scale bars are 300um and 50um. Arrows point to examples of rod-train events in the images.

## Discussion

Disruption of microglial activity in the brain has been well-documented in the pathology of a wide range of brain disorders, but the molecular mechanisms underlying how innate immune memory directly contributes to the onset and progression of brain disorders remains understudied. In our study, we used a recently described mouse model of repeated LPS (20) to investigate the immediate transcriptomic and microglia morphological changes that contribute to the formation of immune memory in the frontal cortex, hippocampus, and striatum, as well as the long-term effects of these changes on various behaviors relevant to brain disorders. Mice showed robust changes in acute sickness behavior, body weight, and pro-inflammatory cytokine expression with LPS-induced immune training, which resolved with the formation of immune tolerance. We found transcriptomic changes in a subset of genes that showed a training response after 2xLPS and a tolerized response at 3xLPS and 4xLPS. These “2xLPS-sensitive” genes were enriched for biological processes related to immune regulation and were targeted by IRF and RHD family transcription factors, two key families whose coordinated activity has previously been linked to the formation of innate immune memories in microglia and other immune cells. (45,51–57) Together with the characterized LPS-induced shifts in morphology, our work identifies several key changes in microglial regulation that occur during the formation of innate immune memories that inform previous findings on innate immune memory in disease contexts in the brain.

Microglia are the only tissue-resident macrophages in the brain parenchyma (58), self-renew without contribution from the periphery (18), and exhibit different properties morphologically and functionally when placed outside of the brain environment (59), suggesting that brain-intrinsic signals that drive microglial identity make them distinct from macrophages in other tissues. However, the majority of what is known about regulation of immune training and tolerance comes from work in cultured peripheral macrophages. Work in *in vitro* macrophage culture models has demonstrated a clear role for epigenetic regulation of gene expression programs mediating training and tolerance (60,61). Recent work has demonstrated that training and tolerance occur in microglia in response to repeated peripheral injections of LPS (20,62). Wendeln et al. (20) demonstrated alterations in microglial gene expression and enhancer activity 6 months after LPS injection in the Alzheimer’s disease model, suggesting long-term reprogramming of microglial function after repeated LPS. Zhang et al. (45) found that enrichment of permissive epigenetic marks at accessible enhancer regions could explain training of microglia to LPS and that tolerized responses were regulated by the loss of permissive epigenetic marks at these sites. In line with both studies, our study identified LPS-sensitive DEGs that were enriched for Interferon (IRF) family and NFkB transcription factor motifs in their promoters. IRFs are induced in response to interferon (IFN), LPS, and other immune stimuli, and initiate gene expression by forming regulatory complexes with IRF family members and other transcription factors, such as PU.1, STAT1, and NF-kB, which further bind with the interferon-sensitive response elements (ISRE) to regulate IFN and TLR signaling pathways. (54,63) IRF family transcription factors have been shown to be required for *Il-1β* expression in microglia after peripheral nerve injury (64) and regulate disease-associated microglial gene expression of markers including *Apoe* and *Trem2*. (65) IRF8 is particularly well-characterized in microglia, and has been shown to bind to and regulate enhancer regions of microglia during postnatal development to contribute to the establishment of microglial identity and function (35). IRF8-deficient microglia have documented changes in morphology and function marked by reduced Iba-1 expression, less elaborated processes, decreased phagocytic capacity, reduced expression of pro-inflammatory genes, and less proliferative potential in mixed glial cultures. (56,66–68) Accumulation of p50/RelA, subunits of NFkB, in the nucleus has also been tied to increased transcription of pro-inflammatory cytokines including i*NOS*, *Tnfα, Il1β,* and *Il6* in the formation of pro-inflammatory phenotypes in microglia. (55,56,69) We identified enrichment for direct binding sites of NfkB/p65 and IRF8 from published microglia and bone marrow-derived macrophage studies (35,37,38) within our DEGs, together pointing to the coordinated activity of these transcription factors as key regulators of immune priming and tolerance in the brain.

Although LPS given peripherally reliably induces a neuroinflammatory response in the brain, LPS has been shown to be largely unable to penetrate the blood brain barrier (BBB) in healthy animals, unless at very high doses (70). Instead, impacts of peripherally administered LPS are likely conveyed to the brain parenchyma by TLR4 receptors on meningeal and endothelial cells, which then signal to other cell types in the brain including microglia. (14,71) Microglia are likely cooperating with other cell types such as astrocytes to perpetuate the expression of pro-inflammatory cytokines, as microglial depletion alone fails to alleviate LPS-induced sickness behaviors. (14,72) Once immune signaling molecules reach the brain parenchyma, they can bind to cytokine receptors expressed on both neurons and glial cells to directly or indirectly modulate neuronal firing properties, which could ultimately shape circuit function and downstream behavior. (14) A recent study using the same repeated low-dose LPS model (73) defined a temporal sequence after 2xLPS by which changes in microglia morphology, density, and expression of phagocytic markers precedes GABAergic synapse loss, followed by memory impairment on the novel object recognition task. However, this study focused on short-term changes within a range of 6 days post-2xLPS, and investigation into the prolonged effects of repeated low-dose LPS on behaviors long after recovery had yet to be thoroughly examined. We failed to identify long-term effects of repeated LPS on a battery of behaviors relevant to brain disorders. Mice showed immediate sickness behaviors including lethargy, ptosis, and piloerection 3 hours after 2xLPS exposure, but they did not exhibit any long-term deficits in learning and memory tasks or anxiety-like, depressive-like, and repetitive behaviors when assessed weeks after LPS treatment. While our data suggest that our LPS model does not contribute to any obvious observable long-term behavioral deficits in healthy mice, preconditioning with repeated peripheral injections of low-dose LPS has been demonstrated to alleviate disease pathology that develops months later in mouse models of Alzheimer’s Disease and neuroinflammatory responses in stroke and traumatic brain injury (21,57,74–76). This suggests that a secondary disease pathology or injury is necessary to reveal the long-term reprogramming effects of LPS on behavior.

Changes in pro-inflammatory gene expression with immune priming and tolerance in the brain have been well-documented to be accompanied by changes in microglial morphology. (20,77,78) 2xLPS-sensitive and 4xLPS-sensitive DEGs identified in our study were enriched for DEGs previously identified as dynamically expressed in the transition from embryonic to postnatal mouse microglia development in both sexes (Fig. S3; BH-corrected p<0.01, Supp. Table S3). (79) During embryonic development, immature microglia display an ameboid morphology which aids in phagocytosis of apoptotic debris, synaptic pruning of developing circuits, and increased motility as microglia migrate along developing white matter tracts to populate gray matter. As development progresses postnatally, microglia morphology transforms into a fully ramified, mature form that is maintained throughout life. (80–82) Enrichment of our LPS innate immune memory genes for genes with differential expression between the ameboid and mature ramified development states, indicates that the morphological adaptations in the adult brain to repeated LPS exposure involve shared developmental genes and pathways.

While the proportions of ramified and rod-like microglial forms remained mostly consistent across LPS treatments, we observed a switch between ameboid and hypertrophic morphology states in the formation of immune memory that was observed in most brain regions within both sexes. Hypertrophic morphology is described by enlarged cell soma with shorter, thicker, hyper-ramified processes (22,83), which give microglia a “bushy” appearance. This morphological form is commonly observed in regions of brain pathology in human cases of Alzheimer’s Disease and Huntington’s Disease, as well as in mouse models of Alzheimer’s Disease, stroke, accelerated aging, chronic stress, depression, and traumatic brain injury. (22) Hypertrophic microglia have been described in close vicinity to beta-amyloid plaques in Alzheimer’s Disease in both human tissue and mouse models. Hypertrophic microglia also exhibit increased surface markers for the disease-associated microglia (DAM) phenotype including CD-33 and triggering receptor expressed on myeloid cells 2 (TREM2), a key switch in the change to the DAM phenotype. (22,81,84,85) TREM2 has recently been described in the context of the same repeated LPS model used in our study to mediate decreases in microglial phagocytosis of synapses and morphology changes with the formation of immune tolerance in the hippocampus. (78) TREM2 knockout prevented acute increases in both microglial soma size and changes in process length and arborization that aligned with a hypertrophic morphology. (78) While we did not identify *Trem2* as a significant DEG in our bulk tissue analysis, we did observe enrichment of 2xLPS-sensitive DEGs with published microglia-specific gene lists for the DAM phenotype (Fig. S3; BH-corrected p<0.01, Supp. Table S3), pointing to the significance of the hypertrophic microglia morphology in regulating immune memory states in the brain.

We also observed the formation of rod-like train events with increasing exposures to LPS. Rod-like microglia have previously been observed to align end-to-end with one another to form ‘trains’ of microglia adjacent to neuronal processes in diffuse traumatic brain injury (TBI) in rats. (48–50) Rod-trains have been shown to emerge and resolve dynamically with the damage and repair cascades that evolve over the first week post-TBI, where rod trains accumulate at 2 and 6 hours post-injury, with a decrease in percentage microglia population at 1 and 2 days post-injury, followed by a subsequent re-accumulation at 7 days post-injury. (48) Similar to these documented patterns, the occurrence of microglial rod-trains in our study increased with 2xLPS, began to resolve after 3xLPS, and re-emerged with 4xLPS (Fig. 5A), suggesting that rod-like microglia may be engaged in separate, delayed waves of repair after inflammation in the brain in the switch from an immune trained to tolerized state after the resolution of cytokine responses and obvious sickness behaviors. This could explain the short-term learning and memory impairments described by Jung et al., 2023 (73), which were observed at 6 days after LPS, and why we failed to observe them in our study in which behavioral testing was performed weeks after LPS.

Together, findings from our study offer new insight into microglial changes in the formation of immune memory. Given that microglia are long-lived cells (4-20 years in humans) (86) that self-renew without contribution from the periphery (87,88) and epigenetically tune their function to the local brain environment through differential enhancer activity (89,90), the acute changes induced during the formation of innate immune memories could have long lasting impacts on the brain’s response to disease and injury. Future work can link our identified acute changes to the long-lasting impacts identified in other model systems to identify how innate immune memories are maintained in microglia and impact disease risk. Understanding these underlying mechanisms will allow for future development of therapies to either enhance or blunt inappropriate microglial activity in the context of brain diseases.

---

***Supplementary Table S1:*** Kruskal-Wallis rank sum test results, Dunn-corrected posthocs, and input data for Luminex heatmap Figure 1C.

***Supplementary Table S2:*** ANOVA results and BH-corrected posthocs from sickness behavior and weight change analysis, and 1-way ANOVA results for each behavioral task in behavioral battery. Related to Figures 1B and 2.

***Supplementary Table S3:*** RNAseq data (Figure 3): Sample information, differential expression analysis, genes in DEG clusters, and gene ontology terms for genes in DEG clusters; TF motif analysis data (Figure 4D): locations of Homer known motifs in promoters of 2xLPS-sensitive cluster DEGs and (Figure 4B): LOLA ChIPseq peak enrichment results; MGEnrichment data (Figure S3): Enrichments against microglial gene lists results.

***Supplementary Table S4:*** ANOVA results and BH-corrected posthocs from MicrogliaMorphology cluster analysis, rod train event occurrence analysis, and microglial density analysis related to Figures 5, 6, and S5F.

***Supplementary Table S5:*** ANOVA results and BH-corrected posthocs from RT-qPCR data analysis, related to Figures 1D, S4C, and S4F.

***Supplementary File S1:*** Homer software output for transcription factor motif analysis of 2xLPS-sensitive cluster gene promoters. Related to Figures 4A and 4C.

***Supplementary File S2:*** Homer software output for transcription factor motif analysis of 4xLPS-sensitive cluster gene promoters.

***Supplementary File S3:*** Homer software output for transcription factor motif analysis of LPS-decreased cluster gene promoters.

## List of Abbreviations

LPS: lipopolysaccharide
PBS: phosphate buffered saline
TLR4: toll-like receptor 4
MDS: multi-dimensional scaling
CPM: counts per million
DEG: differentially expressed gene
TMM: trimmed mean of m-values
GO: gene ontology
BMDM: bone marrow-derived macrophage
P2ry12: 
ROI: region of interest
DAM: disease associated microglia
IRF: interferon regulatory factor
bHLH/bZIP: basic helix-loop-helix/bZip
RHD: rel homology domain
Zf: C2H2 zinc finger
Stat: signal transducer and activator of transcription
BBB: blood brain barrier
TREM2: triggering receptor expressed on myeloid cells 2
TBI: traumatic brain injury

## Declarations

### Ethics approval and consent to participate

Not applicable

### Consent for publication

Not applicable

### Availability of data and materials

The datasets generated and/or analyzed during the current study are freely available to download and all relevant materials including input images, data, and supporting code used in this study are available on the Open Science Framework (OSF) website at [link will be available with final publication]. The ImageJ and R code underlying the analysis demonstrated in this paper are all available through the Ciernia Lab Github at https://github.com/ciernialab/Kim2024_1-4xLPS. The RNA sequencing data generated in this study will be made available through the NCBI GEO database with the final publication.

### Authors’ contributions

P.A. and A.V.C designed the initial experiments and A.V.C. and J.K. conceptualized the project. J.K. performed the majority of the experiments, analysis, and manuscript writing. A.V.C., J.K., and J.J. performed bioinformatic analysis. J.T. and A.M.M. assisted with injections and tissue collection for the RNAseq experiment and performed the Luminex assay. O.S. and K.L. performed the behavioral battery and analysis of the behavioral battery. B.W. assisted with RT-qPCR experiments. All authors read and edited the manuscript.

## Acknowledgements

We thank the Ciernia Lab, Ashwood Lab, and Pavlidis Lab members for their thoughtful feedback and suggestions during lab meetings throughout the progression of this project. We are grateful for the resources provided by the Neuroimaging & Neurocomputation Centre and the Dynamic Brain Circuits in Health & Disease Research Cluster at the University of British Columbia, as well as the University of California, Davis DNA Technologies & Expression Analysis Core.

---

**Supplementary Figure S1:**
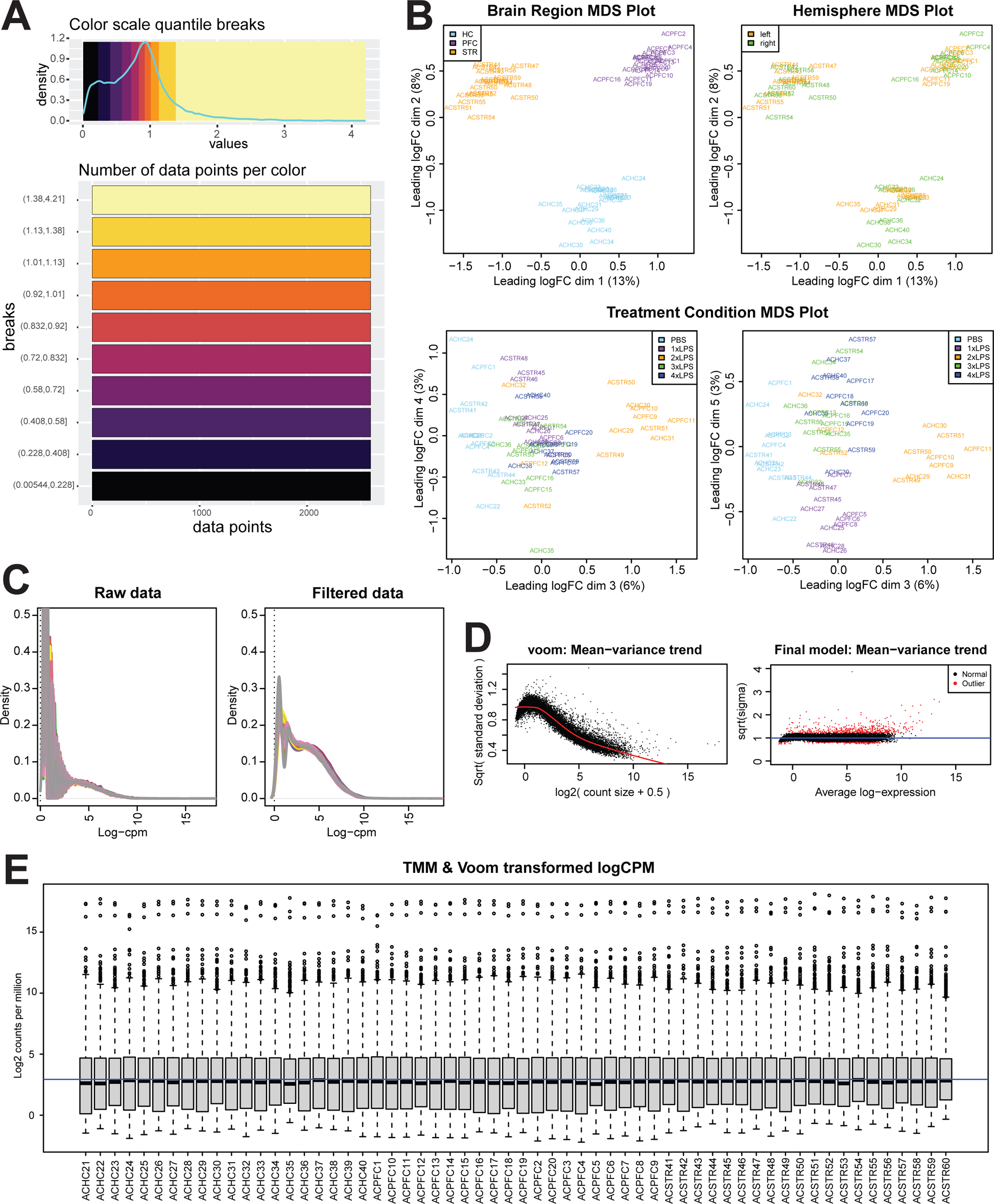
Differential gene expression analysis using lima-voom. (A) Reference for color scale for scaled gene expression in DEG heatmap in Figure 3. Counts per million (CPM) for each DEG was scaled within each brain region first, then scaled within genes across brain regions (rows) in the final heatmap (Fig. 3A). Breaks between each color of the heatmap scale were adjusted to fit the distribution of the scaled gene expression values such that 10% of the data was contained within each color break. (B) MDS plots colored by brain region and hemisphere along dimensions 1 and 2 and colored by treatment along dimensions 3, 4, and 5. (C) Distribution of log CPM for each sample before and after filtering. (D) Mean-variance trend of counts and final statistical model after limma-voom normalization. (E) Boxplots of Log2CPM values for each sample after TMM and limma-voom normalization.

**Supplementary Figure S2:**
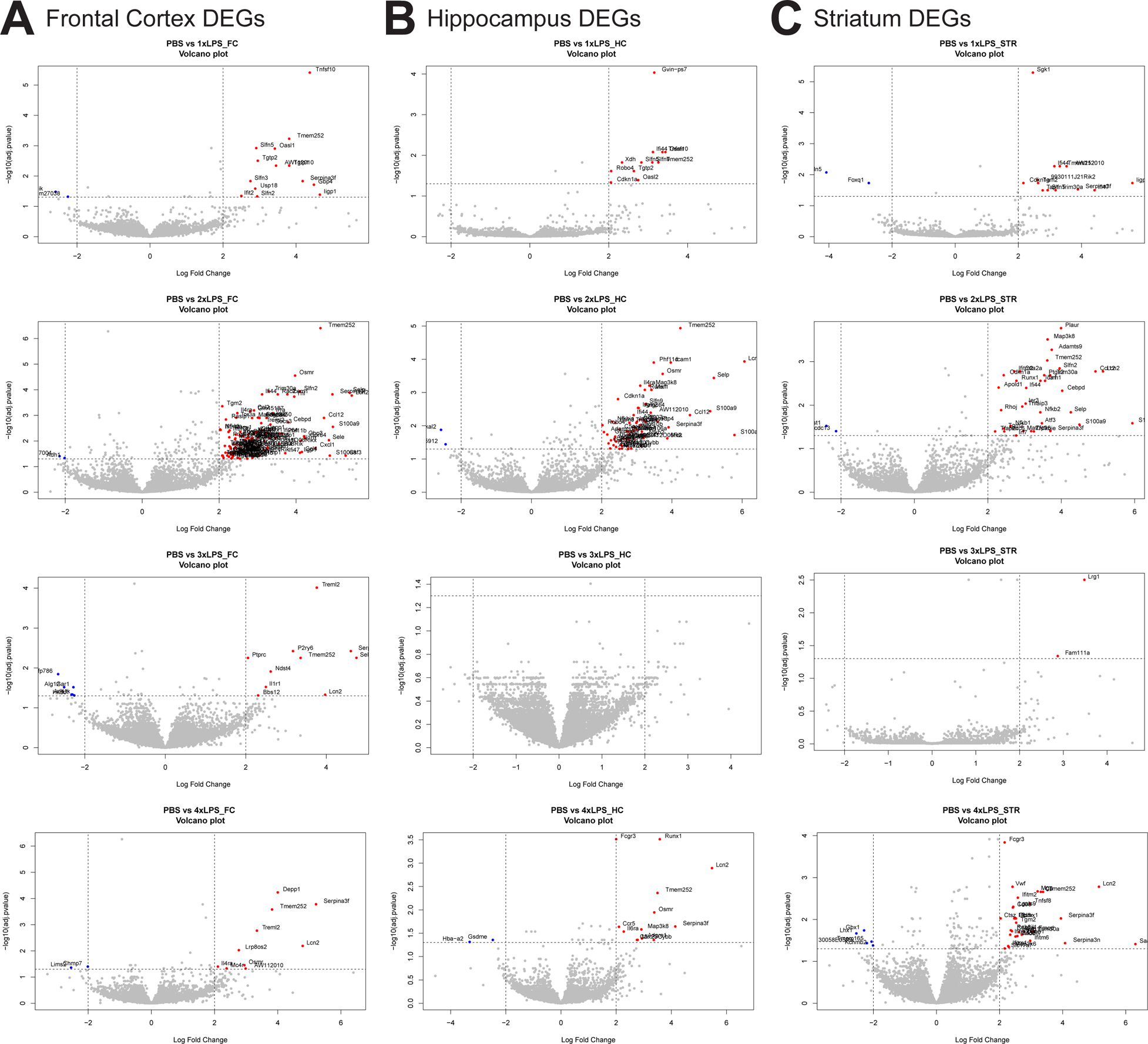
DEGs within each brain region for each LPS versus PBS comparison. Volcano plots for Log Fold Change versus −log(corrected pvalue) for PBS versus each LPS treatment condition for (A) Frontal Cortex, (B) Hippocampus, and (C) Striatum. Genes with BH-corrected p-value < 0.05 and fold change>+/− 2 for each comparison are colored red (increased) or blue (decreased) and labeled with the gene name.

**Supplementary Figure S3:**
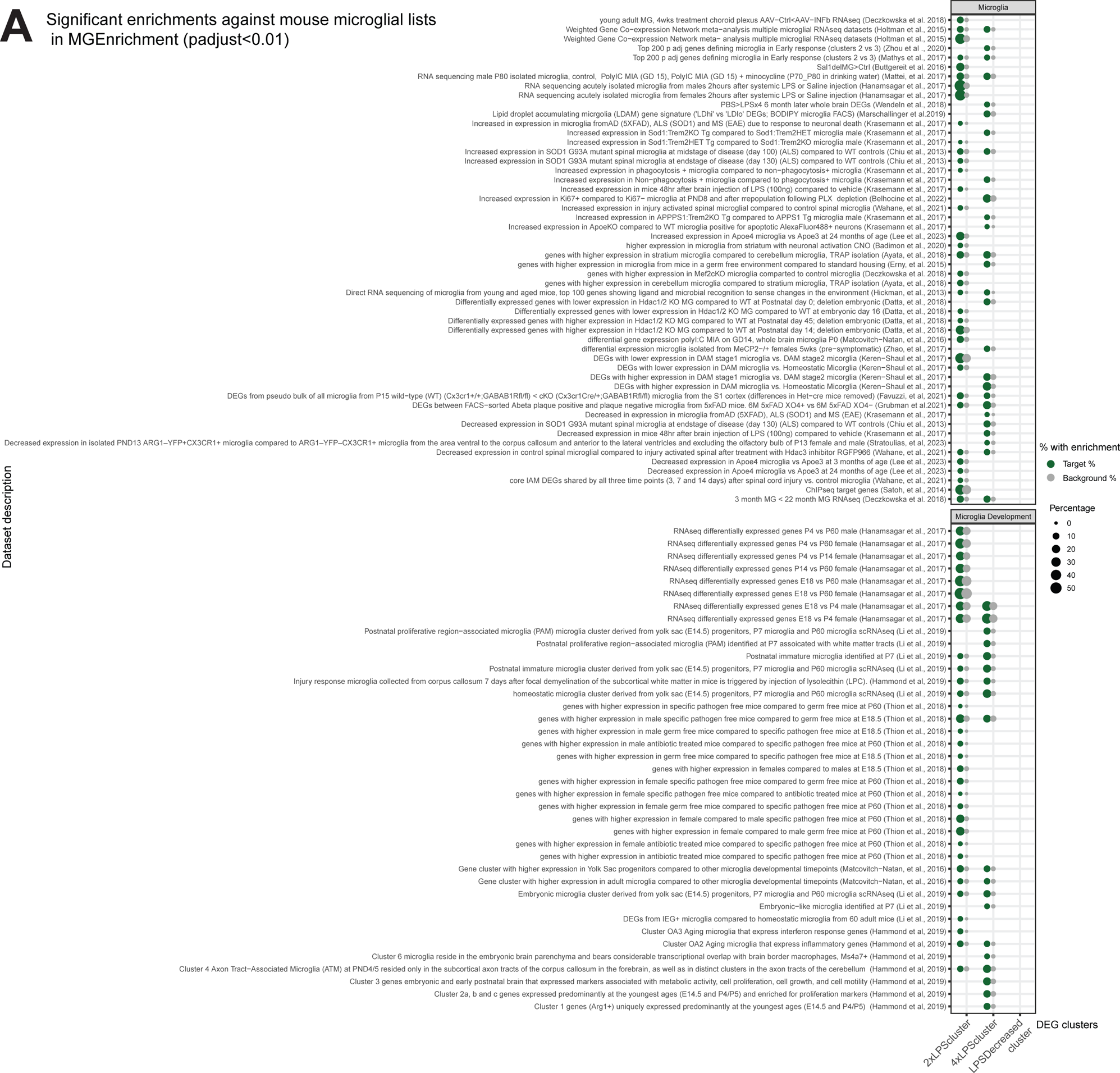
2xLPS-sensitive and 4xLPS-sensitive cluster DEGs are enriched for microglia-relevant genes. (A) Significant enrichments against mouse microglial gene lists in MGEnrichment (33). Green circles denote a significant enrichment in the percent of DEGs for a given gene list compared to that of the background of all genes detected in the RNA-Seq experiment (grey circles). Only significant enrichments are shown (*p<0.01, BH-corrected).

**Supplementary Figure S4:**
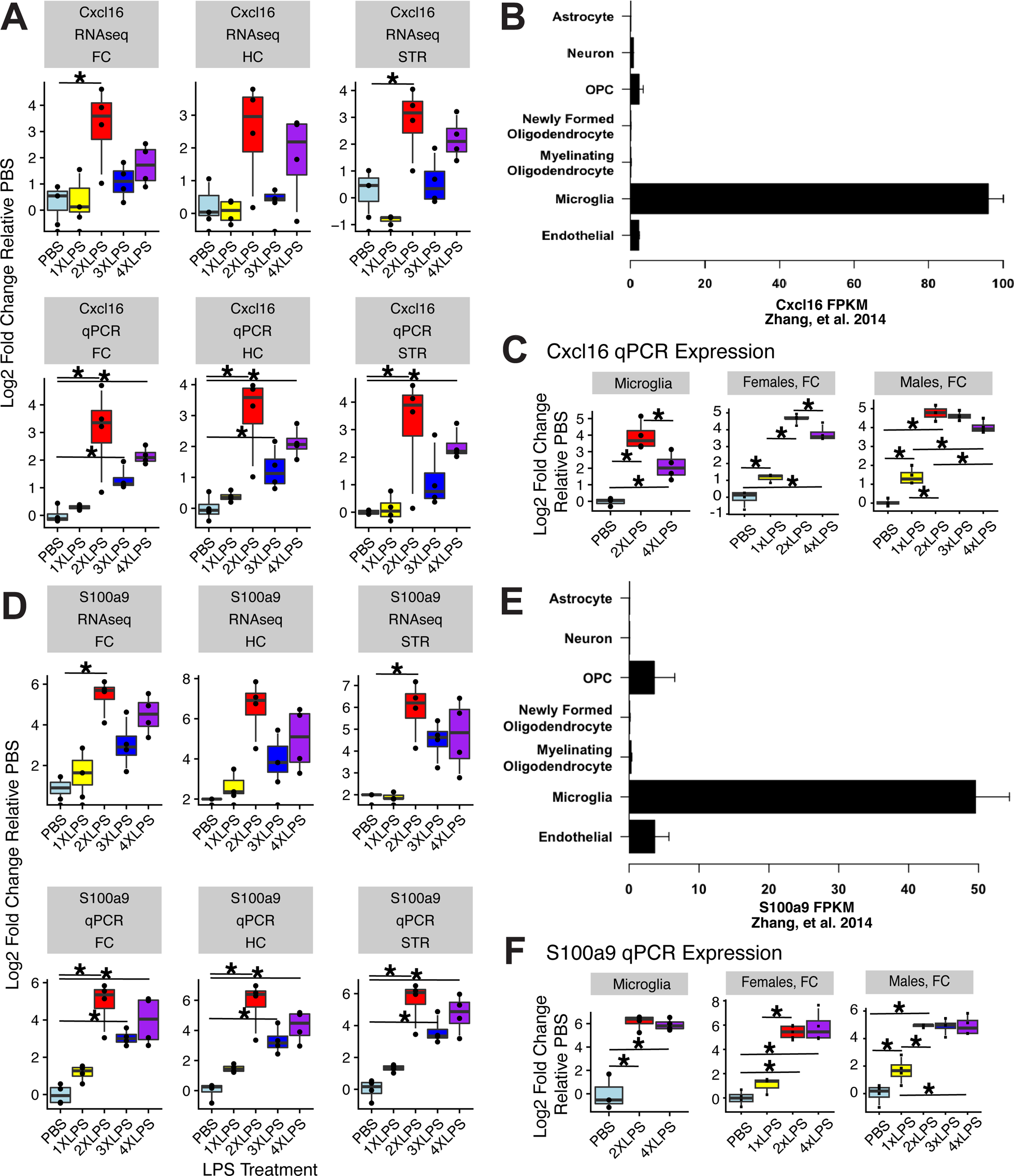
Cxcl16 and S100a9 expression in microglia is dynamic in response to repeated LPS. (A) Log2 fold change relative to PBS controls for each LPS treatment condition for *Cxcl16* in RNAseq (top row) compared to RT-qPCR expression (bottom row) from the same mice. Patterns of expression were conserved across the two techniques for each brain region (Frontal Cortex, FC; Hippocampus, HC; and Striatum, STR). For RNAseq *p<0.05, BH-corrected for multiple comparison across all genes in the RNAseq. For RT-qPCR *p<0.05, corrected for multiple comparisons within the gene of interest. (B) Expression of *Cxcl16* within different isolated brain cell types from Zhang et al., 2014 (https://web.stanford.edu/group/barres_lab/brain_rnaseq.html). (C) Expression of *Cxcl16* by RT-qPCR in isolated cortical microglia and frontal cortex tissue following LPS treatments. *p<0.05, BH-corrected for multiple comparisons for 1-4xLPS treatment versus PBS. (D) Log2 fold change relative to PBS controls for each LPS treatment condition for *S100a9* in RNAseq (top row) compared to RT-qPCR expression (bottom row) from the same mice. Patterns of expression were conserved across the two techniques for each brain region (Frontal Cortex, FC; Hippocampus, HC; and Striatum, STR). For RNAseq *p<0.05, BH-corrected for multiple comparison across all genes in the RNAseq. For RT-qPCR *p<0.05, corrected for multiple comparisons within the gene of interest. (E) Expression of *S100a9* in isolated brain cell types from Zhang et al., 2014. (F) Expression of *S100a9* by RT-qPCR in isolated cortical microglia and frontal cortex tissue following LPS treatments. *p<0.05, BH-corrected for multiple comparisons for 1-4xLPS treatment versus PBS.

**Supplementary Figure S5:**
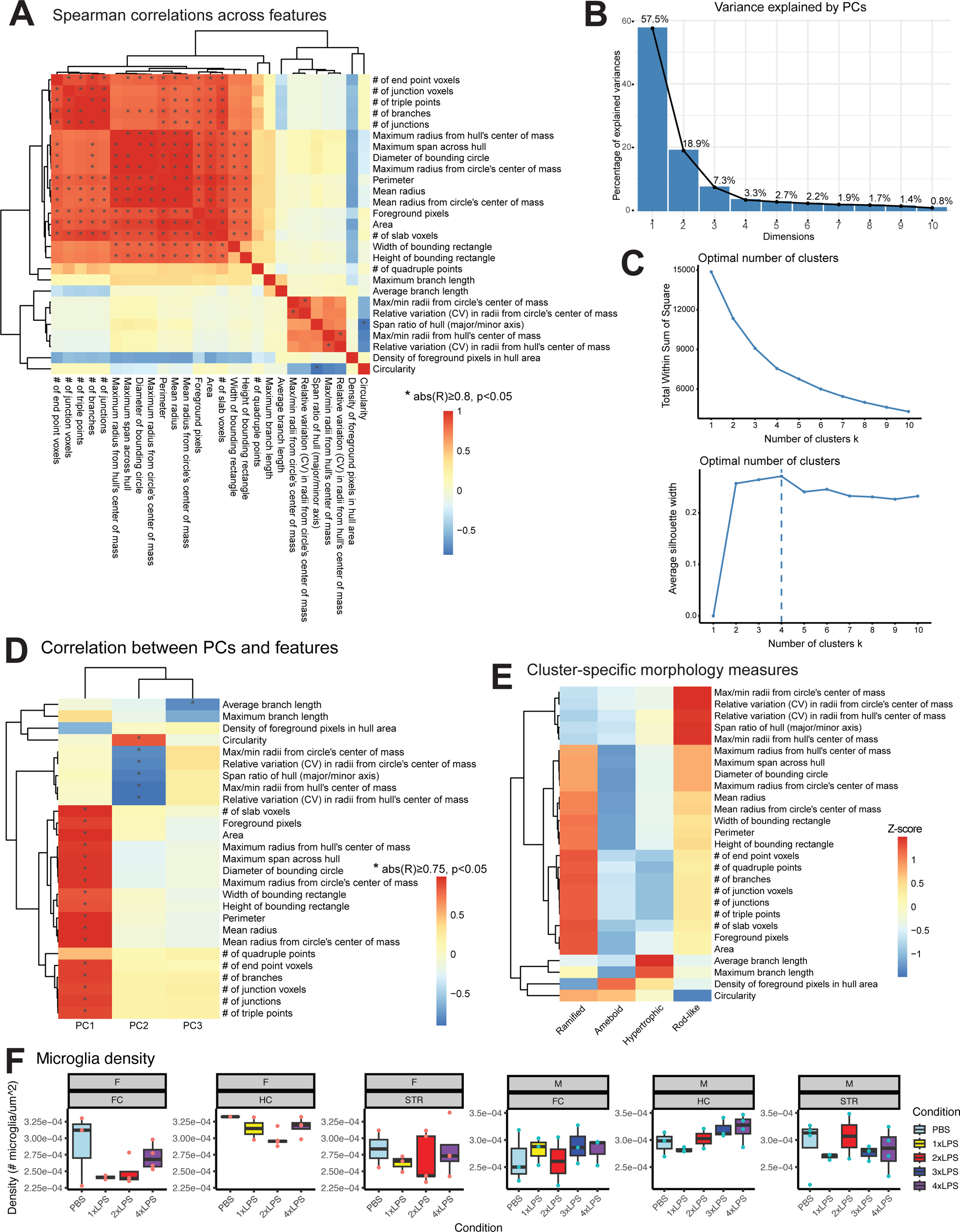
Analysis of morphology measures and clusters using MicrogliaMorphology and MicrogliaMorphologyR. (A) Spearman’s correlation matrix of 27 features measured by MicrogliaMorphology. (*abs(R)≥0.8, p<0.05) (B) Variance in dataset described by first 10 principal components. (C) Optimal k-means clustering parameters determined using within sum of squares and gap statistic techniques. (D) Spearman’s correlation of morphology measures to first 3 PCs after dimensionality reduction. (*abs(R)≥0.75, p<0.05) (E) Average values for all 27 morphology features, centered and scaled across clusters. (F) Quantification of microglia density with repeated LPS across brain regions within sexes. No significant differences between groups. (*p<0.05, BH-corrected)

