## Supplementary material for "Repeated LPS induces training and tolerance of microglial responses across brain regions": FileS1: geneOntology.html

Gene Ontology Results

### Gene Ontology Enrichment Results

Homer *de novo* Motif Enrichment Results  
Known Motif Enrichment Results  

###### Text file version of complete results (i.e. open with Excel) - biological process: Functional groupings of proteins (Gene Ontology) - molecular function: Mechanistic actions of proteins (Gene Ontology) - cellular component: Protein localization (Gene Ontology) - chromosome location: Genes with similar chromosome localization (NCBI Gene) - KEGG pathways: Groups of proteins in the same pathways (KEGG) - protein interactions: "Proteins interacting with a common protein (BIND, EcoCyc, HPRD)" (NCBI Gene) - interpro domains: Proteins with similar domains and features (Interpro) - pfam domains: Proteins with similar domains and features (Pfam) - smart domains: Proteins with similar domains and features (SMART) - gene3d domains: Proteins with similar domains and features (Gene3D) - prosite domains: Proteins with similar domains and features (Prosite) - prints domains: Proteins with similar domains and features (PRINTS) - MSigDB lists: "Genes sets for pathways, factor/miRNA target predictions, expression patterns, etc." (MSigDB) - BIOCYC pathways: Groups of proteins in the same pathways (BIOCYC) - COSMIC cancer mutations: Genes mutated in similar cancers (COSMIC) - GWAS genes: Genes mutated in similar diseases (GWAS Catalog) - Lipid Maps pathways: Groups of proteins in the same lipid pathways (Lipid Maps/Biosystems) - Pathway Interaction DB: Groups of proteins in the same pathways (Pathway Interaction Database) - REACTOME pathways: Groups of proteins in the same pathways (REACTOME) - SMPDB pathways: Groups of proteins in the same pathways (SMPDB) - WikiPathways: Groups of proteins in the same pathways (Wikipathways) Enriched Categories | | | | | | | | | | | | --- | --- | --- | --- | --- | --- | --- | --- | --- | --- | | P-value | ln(P) | Term | GO Tree | GO ID | # of Genes in Term | # of Target Genes in Term | # of Total Genes | # of Target Genes | Common Genes | | 3.985e-49 | -111.44 | ALTEMEIER\_RESPONSE\_TO\_LPS\_WITH\_MECHANICAL\_VENTILATION | MSigDB lists | ALTEMEIER\_RESPONSE\_TO\_LPS\_WITH\_MECHANICAL\_VENTILATION | 104 | 41 | 12187 | 179 | Irgm2,C5ar1,Gem,Irf7,Slfn4,Nfkbia,Casp4,Plek,Csf2rb,Plaur,Socs3,Timp1,Il1rn,Atf3,Cxcl1,Bcl3,Osmr,Cmpk2,Gbp3,Ifit2,Map3k8,Maff,Ccl4,Gpr84,Usp18,Cybb,Hcar2,Ccl12,Tnfaip3,Il6,Selp,Isg15,Cd14,Slfn3,Thbs1,Nfkbiz,Gadd45g,Il1a,Akap12,Ms4a6d,Cebpd | | 6.593e-49 | -110.94 | response to biotic stimulus | biological process | GO:0009607 | 738 | 82 | 13711 | 214 | Gbp6,Wfdc21,Irgm2,AA467197,Bcl3,Usp18,S100a9,Cxcl16,Irf2,Ccl12,S100a8,Tnip3,Ifitm2,Gbp3,Ifi204,Lcn2,Adamts9,Tnfaip8,Irak3,Il1rn,Ccl2,Cxcl9,Tnfaip3,Il17ra,Trim56,Rtp4,Irgm1,Nfkb1,Slfn4,Gbp4,Lgals9,Mx1,Iigp1,Ifitm3,Oasl1,Cybb,Ifit2,Noct,Gbp7,Ifit3b,Serpina3f,Mt2,Hp,Hspa5,C5ar1,Cxcl1,Trim30a,Cflar,Cd14,Ncf1,Zfp36,Saa1,Ifi44,Acod1,Tnf,Trim25,Resf1,Ier3,Ptgs2,Mx2,Ccl4,Herc6,Ifi211,Ifitm6,Stx11,Parp9,Sbno2,Isg15,Slfn2,Cmpk2,Tgtp1,Trib1,Parp14,Ccl7,Dtx3l,Irf7,Il6,Nfkbia,Oasl2,Il4ra,Slfn9,Casp4 | | 9.230e-49 | -110.60 | response to other organism | biological process | GO:0051707 | 718 | 81 | 13711 | 214 | Casp4,Il4ra,Oasl2,Slfn9,Nfkbia,Il6,Irf7,Dtx3l,Ccl7,Parp14,Trib1,Tgtp1,Slfn2,Cmpk2,Sbno2,Isg15,Parp9,Ifitm6,Stx11,Ifi211,Herc6,Ccl4,Mx2,Ptgs2,Ier3,Trim25,Resf1,Acod1,Tnf,Ifi44,Zfp36,Saa1,Ncf1,Cd14,Cflar,Trim30a,C5ar1,Cxcl1,Hp,Mt2,Serpina3f,Ifit3b,Gbp7,Noct,Ifit2,Cybb,Oasl1,Ifitm3,Iigp1,Mx1,Lgals9,Gbp4,Slfn4,Nfkb1,Irgm1,Rtp4,Trim56,Il17ra,Tnfaip3,Cxcl9,Ccl2,Il1rn,Irak3,Tnfaip8,Adamts9,Lcn2,Ifi204,Gbp3,Ifitm2,Tnip3,S100a8,Irf2,Ccl12,Cxcl16,Usp18,S100a9,Bcl3,AA467197,Irgm2,Wfdc21,Gbp6 | | 1.148e-48 | -110.39 | response to external biotic stimulus | biological process | GO:0043207 | 720 | 81 | 13711 | 214 | Ncf1,Cflar,Cd14,Trim30a,Acod1,Tnf,Ifi44,Saa1,Zfp36,Serpina3f,Ifit3b,Gbp7,Noct,Ifit2,Cxcl1,C5ar1,Mt2,Hp,Irf7,Dtx3l,Ccl7,Trib1,Parp14,Casp4,Slfn9,Oasl2,Il4ra,Nfkbia,Il6,Ifitm6,Stx11,Ifi211,Herc6,Ccl4,Mx2,Ptgs2,Ier3,Trim25,Resf1,Tgtp1,Slfn2,Cmpk2,Sbno2,Isg15,Parp9,Il1rn,Irak3,Adamts9,Tnfaip8,Lcn2,Ifi204,Tnfaip3,Cxcl9,Ccl2,S100a9,Usp18,Bcl3,AA467197,Irgm2,Wfdc21,Gbp6,Gbp3,Ifitm2,Tnip3,S100a8,Ccl12,Irf2,Cxcl16,Oasl1,Ifitm3,Iigp1,Mx1,Lgals9,Cybb,Irgm1,Rtp4,Trim56,Il17ra,Gbp4,Slfn4,Nfkb1 | | 2.572e-41 | -93.46 | HALLMARK\_INTERFERON\_GAMMA\_RESPONSE | MSigDB lists | HALLMARK\_INTERFERON\_GAMMA\_RESPONSE | 175 | 43 | 12187 | 179 | Nfkb1,Rnf213,Isg15,Herc6,Ifitm3,Selp,Tap1,Il6,Ccl12,Rtp4,Cmpk2,Gbp3,Ifi44,Ifit3b,Vcam1,Socs3,Ifitm2,Ptgs2,Csf2rb,Nfkbia,Xaf1,Parp12,Gbp4,Gbp6,Ptpn1,Irf2,Il4ra,Ifit2,Usp18,Ripk1,Tnfaip3,Arid5b,Cdkn1a,Samd9l,Znfx1,Trim25,Icam1,Irf9,Casp4,Parp14,Oasl1,Irf7,Tnfsf10 | | 8.508e-40 | -89.96 | immune system process | biological process | GO:0002376 | 1296 | 93 | 13711 | 214 | Ncf1,Sele,Trim30a,Pik3ap1,Acod1,Msn,Gbp7,Selp,C5ar1,Parp14,Ccl7,Hcar2,Nfkbia,Slfn9,Il4ra,Mx2,Herc6,Ifi211,Cebpd,Trim25,Runx1,Tiparp,Isg15,Csf3,Zbtb16,Il1rn,C3ar1,Retnlg,Lcn2,Cxcl9,Tnfaip3,Ccl2,S100a9,Vcam1,Ifitm2,Nfkbiz,Cxcl16,Tnip3,Iigp1,Oasl1,Lgals9,Gadd45g,Nfkb2,Fyb,Icam1,Rtp4,Irgm1,Map3k8,Cd14,Tnip1,Tnf,Ifi44,Ifit3b,Ifit2,Cxcl1,Hp,Prg4,Dtx3l,Irf7,Pik3r1,Il1a,Oasl2,Casp4,Il6,Rbm47,Ccl4,Spi1,Ifitm6,Stx11,Ifi209,Parp9,Sbno2,Tgtp1,Irak3,Ifi204,Tnfsf10,Irf9,Gbp6,Irgm2,Samd9l,Bcl3,Gbp3,Ccl12,Irf2,Tap1,S100a8,Ifitm3,Mx1,Cybb,Il17ra,Trim56,Gbp4,Nfkb1 | | 8.577e-40 | -89.95 | defense response | biological process | GO:0006952 | 731 | 73 | 13711 | 214 | Il1rn,C3ar1,Tnfaip8,Ifi47,Lcn2,Tnfaip3,Cxcl9,Ccl2,S100a9,Bcl3,Gbp6,Irgm2,Nfkbiz,Gbp3,Ifitm2,S100a8,Tap1,Cxcl16,Ccl12,Irf2,Thbs1,Ifitm3,Gm4841,Oasl1,Iigp1,Mx1,Icam1,Cybb,Irgm1,Rtp4,Trim56,Il17ra,Gbp4,F830016B08Rik,Ncf1,Sele,Cd14,Tnip1,Trim30a,Acod1,Tnf,Saa1,Nfe2l2,Ifit3b,Gbp7,Selp,Ifit2,C5ar1,Cxcl1,Hp,Dtx3l,Irf7,Parp14,Ccl7,Oasl2,Il4ra,Slfn9,Casp4,Il1a,Gm5431,Il6,Tgtp2,Ifitm6,Stx11,Mx2,Herc6,Ccl4,Ptgs2,Trim25,Timp1,Tgtp1,Parp9,Isg15 | | 2.651e-37 | -84.22 | GSE13485\_DAY3\_VS\_DAY7\_YF17D\_VACCINE\_PBMC\_DN | MSigDB lists | GSE13485\_DAY3\_VS\_DAY7\_YF17D\_VACCINE\_PBMC\_DN | 136 | 37 | 12187 | 179 | Phf11d,Ifi211,Dtx3l,Rtp4,Cmpk2,Usp18,Gpr84,Ccl4,Ifit2,Gadd45b,Ccl12,Slfn5,Isg15,Herc6,Rnf213,Ifitm3,Tap1,Parp9,Phf11b,Irf7,C3ar1,Tnfsf10,Xaf1,Parp14,Tnf,Oasl1,Parp12,Ifitm2,Trim25,Ifit3b,Ifi44,Ifi209,Znfx1,Ifi207,Samd9l,Il1rn,Ifi204 | | 3.843e-37 | -83.85 | response to cytokine | biological process | GO:0034097 | 603 | 65 | 13711 | 214 | Irgm2,Wfdc21,Gbp6,Ccl12,Cxcl16,Gbp3,Ifitm2,Ifi204,Il1rn,Arid5b,Irak3,Ifi47,Csf3,Ccl2,Cxcl9,Trim56,Il17ra,Irgm1,Nfkb1,F830016B08Rik,Gbp4,Birc3,Osmr,Gm4841,Ifitm3,Iigp1,Icam1,Gbp7,Csf2rb,Selp,Ifit2,Serpina3f,Nfe2l2,Cxcl1,Hspa5,Sele,Ifi207,Cd14,Ripk1,Zfp36,Acod1,Tnf,Ptgs2,Trim25,Ifitm6,Stx11,Ifi211,Xaf1,Spi1,Ccl4,Tgtp1,Isg15,Sbno2,Timp1,Ifi209,Fzd4,Socs3,Irf7,Ccl7,Il6,Gm5431,Tgtp2,Il4ra,Il1a,Nfkbia | | 1.321e-35 | -80.31 | response to external stimulus | biological process | GO:0009605 | 1456 | 93 | 13711 | 214 | Dtx3l,Irf7,Trib1,Oasl2,Casp4,Il6,Cdkn1a,Stx11,Ifitm6,Ccl4,Ptgs2,Ier3,Cmpk2,Tgtp1,Parp9,Sbno2,Cd14,Cflar,Fstl1,Ifi44,Tnf,Zfp36,Saa1,Ifit3b,Ifit2,Cxcl1,Rrp8,Mt2,Hp,Thbs1,Ifitm3,Mx1,Cybb,Trim56,Il17ra,Slfn4,Gbp4,Nfkb1,Irak3,Adamts9,Ifi204,Usp18,AA467197,Bcl3,Gbp6,Irgm2,Gbp3,S100a8,Ccl12,Irf2,Parp14,Ccl7,Slfn9,Il4ra,Nfkbia,Ifi211,Mx2,Herc6,Trim25,Resf1,Fam107a,Slfn2,Isg15,Ncf1,Trim30a,Acod1,Atf3,Nfe2l2,Serpina3f,Angptl4,Noct,Gbp7,C5ar1,Hspa5,Oasl1,Iigp1,Lgals9,Irgm1,Rtp4,Il1rn,C3ar1,Tnfaip8,Retnlg,Lcn2,Tnfaip3,Cxcl9,Ccl2,Vcam1,S100a9,Wfdc21,Ifitm2,Tnip3,Cxcl16 | | 4.114e-35 | -79.18 | response to stimulus | biological process | GO:0050896 | 5062 | 167 | 13711 | 214 | Oasl1,Iigp1,Gadd45g,Lgals9,Mt1,Icam1,Rgs16,Fyb,Nfkb2,Rassf4,Irgm1,Rtp4,Pik3r5,Map3k8,Birc3,Rhoj,Nfkbie,C3ar1,Il1rn,Csf3,Tnfaip8,Lcn2,Retnlg,Tnfaip3,Cxcl9,Ccl2,Vcam1,S100a9,Wfdc21,Nfkbiz,Ifitm2,Ptpn1,Tnip3,Iqgap1,Col4a1,Cxcl16,Ccl7,Parp14,Hcar2,Ccnd2,Il4ra,Slfn9,Akap12,Nfkbia,Ip6k2,Gm5431,Rasip1,Xaf1,Ifi211,Herc6,Mx2,Runx1,Resf1,Trim25,Timp1,Tiparp,Fam107a,Map3k6,Slfn2,Isg15,Sele,Ncf1,Trim30a,Cp,Acod1,Pik3ap1,Atf3,Rhou,Serpina3f,Nfe2l2,Rasd1,Gbp7,Noct,Selp,Angptl4,Msn,Tgm2,C5ar1,Clic4,Hspa5,Gm4841,Thbs1,Ifitm3,Mx1,P2ry6,Errfi1,Cybb,Bach1,Kcna5,Trim56,Plaur,Il17ra,Gbp4,Slfn4,Osmr,Nfkb1,F830016B08Rik,Pygm,Irak3,Arid5b,Ifi47,Adamts9,Ifi204,Plek,Ksr1,Tnfsf10,Xdh,Sgk3,Usp18,Bcl3,AA467197,Irgm2,Gbp6,Gbp3,S100a8,Tap1,Irf2,Ccl12,Irf7,Socs3,Dtx3l,Trib1,Pik3r1,Casp4,Oasl2,Gem,Il1a,Il6,Tgtp2,Cdkn1a,Stx11,Ifitm6,Ccl4,Spi1,Sgk1,Ptgs2,Ier3,Ifi209,Fzd4,Tgtp1,Cmpk2,Sdc4,Sbno2,Parp9,Hspb1,Slc24a4,Ifi207,Cd14,Cflar,Gpr84,Ripk1,Tnip1,Fstl1,Rhoc,Ifi44,Tnf,Saa1,Zfp36,Ifit3b,Csf2rb,Ifit2,Dnajb1,Rrp8,Cxcl1,Prg4,Hp,Mt2 | | 1.167e-34 | -78.13 | response to bacterium | biological process | GO:0009617 | 379 | 52 | 13711 | 214 | Lgals9,Iigp1,Nfkb1,Slfn4,Irgm1,Ccl2,Tnfaip3,Cxcl9,Lcn2,Ifi204,Irak3,Il1rn,Adamts9,Tnfaip8,Tnip3,Ccl12,Cxcl16,Gbp3,Bcl3,AA467197,Irgm2,Wfdc21,Gbp6,Usp18,Il6,Nfkbia,Trib1,Tgtp1,Cmpk2,Slfn2,Sbno2,Isg15,Ptgs2,Resf1,Ifi211,Herc6,Saa1,Zfp36,Tnf,Acod1,Ifi44,Trim30a,Ncf1,Cflar,Cd14,Hp,Mt2,Cxcl1,C5ar1,Gbp7,Noct,Serpina3f | | 4.823e-34 | -76.71 | GSE38681\_WT\_VS\_LYL1\_KO\_LYMPHOID\_PRIMED\_MULTIPOTENT\_PROGENITOR\_DN | MSigDB lists | GSE38681\_WT\_VS\_LYL1\_KO\_LYMPHOID\_PRIMED\_MULTIPOTENT\_PROGENITOR\_DN | 177 | 38 | 12187 | 179 | Usp18,Ifit2,Map3k8,Nfe2l2,Ripk1,Gbp7,Gbp3,Rtp4,Cmpk2,Nfkbiz,Hspa5,Gadd45g,Slfn9,Isg15,Sgk3,Tap1,Irf2,Parp9,Slfn3,Tor3a,Parp14,Lgals9,Birc3,Parp12,Oasl1,Gbp4,Irf7,Gbp6,Gem,Irgm2,Tnfsf10,Slfn4,Ifit3b,Socs3,Znfx1,Irf9,Clic4,Trim25 | | 1.394e-33 | -75.65 | GSE9316\_IL6\_KO\_VS\_IFNG\_KO\_INVIVO\_EXPANDED\_CD4\_TCELL\_DN | MSigDB lists | GSE9316\_IL6\_KO\_VS\_IFNG\_KO\_INVIVO\_EXPANDED\_CD4\_TCELL\_DN | 168 | 37 | 12187 | 179 | Rtp4,Cxcl1,Ptges,Csf3,Gadd45b,Tnfaip3,Ccl12,Gpr84,Ccl2,Ccl4,Maff,Rhou,Tap1,Cd14,Selp,Ifitm3,Ier3,Gadd45g,Nfkbiz,Tnfsf10,Xdh,Irf7,Cflar,Parp12,Oasl1,Nfkbia,Tor3a,Tnf,Clic4,Icam1,Saa1,Atf3,Col4a1,Timp1,Il1rn,Socs3,Cdkn1a | | 2.229e-33 | -75.18 | HALLMARK\_TNFA\_SIGNALING\_VIA\_NFKB | MSigDB lists | HALLMARK\_TNFA\_SIGNALING\_VIA\_NFKB | 170 | 37 | 12187 | 179 | Tap1,Sgk1,Tiparp,Nfkb1,Il1a,Ier3,Cebpd,Nfkbie,Cxcl1,Bcl3,Tnfaip8,Ifit2,Map3k8,Maff,Ccl4,Ccl12,Tnfaip3,Nfe2l2,Il6,Gadd45b,Ptgs2,Tnip1,Icam1,Zfp36,Plek,Plaur,Cdkn1a,Socs3,Atf3,Cflar,Gem,Nfkb2,Tnf,Birc3,Sdc4,Trib1,Nfkbia | | 5.590e-33 | -74.26 | GSE18281\_CORTICAL\_VS\_MEDULLARY\_THYMOCYTE\_UP | MSigDB lists | GSE18281\_CORTICAL\_VS\_MEDULLARY\_THYMOCYTE\_UP | 174 | 37 | 12187 | 179 | Gadd45g,Lcn2,Cebpd,Ifitm3,Isg15,Cd14,Slfn3,Tap1,Ifit2,Ccnd2,Tgm2,Ccl2,Usp18,Gadd45b,Cxcl1,Bcl3,Osmr,Cmpk2,Gbp3,Ifit3b,Il1rn,Znfx1,Rasd1,Atf3,Ptgs2,Saa1,Icam1,Plaur,Clic4,Irf9,Tnf,Casp4,Nfkbia,Irgm2,Irf7,Slfn4,Tnfsf10 | | 1.829e-32 | -73.08 | multi-organism process | biological process | GO:0051704 | 1450 | 89 | 13711 | 214 | Gbp3,A2m,S100a8,Irf2,Ccl12,Usp18,Bcl3,AA467197,Irgm2,Gbp6,Irak3,Adamts9,Ifi204,Gbp4,Slfn4,Nfkb1,Adamts1,Trim56,Il17ra,Cybb,Ifitm3,Mx1,Cxcl1,Hp,Mt2,Ifit3b,Ifit2,Ifi44,Tnf,Saa1,Zfp36,Cd14,Cflar,Ripk1,Tnip1,Tgtp1,Cmpk2,Sbno2,Parp9,Ifitm6,Stx11,Ccl4,Ptgs2,Ier3,Casp4,Oasl2,Il6,Irf7,Dtx3l,Trib1,Ifitm2,Tnip3,Cxcl16,Vcam1,S100a9,Wfdc21,Tnfaip3,Cxcl9,Ccl2,Il1rn,Zbtb16,Tnfaip8,Lcn2,Irgm1,Rtp4,Icam1,Oasl1,Iigp1,Lgals9,C5ar1,Clic4,Serpina3f,Noct,Gbp7,Acod1,Ncf1,Trim30a,Slfn2,Isg15,Ifi211,Herc6,Mx2,Trim25,Resf1,Slfn9,Il4ra,Nfkbia,Ccl7,Parp14 | | 2.051e-32 | -72.96 | GSE19888\_ADENOSINE\_A3R\_INH\_PRETREAT\_AND\_ACT\_BY\_A3R\_VS\_TCELL\_MEMBRANES\_ACT\_MAST\_CELL\_UP | MSigDB lists | GSE19888\_ADENOSINE\_A3R\_INH\_PRETREAT\_AND\_ACT\_BY\_A3R\_VS\_TCELL\_MEMBRANES\_ACT\_MAST\_CELL\_UP | 166 | 36 | 12187 | 179 | Tap1,Parp9,Pik3ap1,Slfn9,Slfn5,Herc6,Isg15,Rnf213,Ifitm3,Gbp3,Cmpk2,Rtp4,Gbp7,Dtx3l,Ccl12,Usp18,Ifit2,Irf9,Trim25,Znfx1,Samd9l,Timp1,Ifit3b,Ifi44,Tnfsf10,Irf7,Gbp6,Irgm2,Parp12,Oasl1,Gbp4,Tubb6,Casp4,Xaf1,Parp14,Tor3a | | 6.410e-32 | -71.82 | GSE36527\_CD69\_NEG\_VS\_POS\_TREG\_CD62L\_LOS\_KLRG1\_NEG\_UP | MSigDB lists | GSE36527\_CD69\_NEG\_VS\_POS\_TREG\_CD62L\_LOS\_KLRG1\_NEG\_UP | 171 | 36 | 12187 | 179 | Icam1,Trim25,Irf9,Clic4,Ifi44,Ifit3b,Socs3,Znfx1,Gbp6,Irgm2,Irf7,Slfn4,Tnfsf10,Casp4,Parp14,Tor3a,Parp12,Oasl1,Gbp4,Isg15,Selp,Slfn9,Slfn3,Tap1,Parp9,Gadd45g,Gbp7,Bcl3,Cmpk2,Rtp4,Gbp3,Ccnd2,Ifit2,Map3k8,Usp18,Ripk1 | | 1.247e-31 | -71.16 | GSE35825\_UNTREATED\_VS\_IFNG\_STIM\_MACROPHAGE\_UP | MSigDB lists | GSE35825\_UNTREATED\_VS\_IFNG\_STIM\_MACROPHAGE\_UP | 174 | 36 | 12187 | 179 | Slfn5,Irak3,Parp9,Il1a,Nfkbie,Csf3,Pik3r5,Ptges,Bcl3,Rtp4,Gbp3,Map3k8,Cybb,Hcar2,Ptgs2,Saa1,Plek,Clic4,Irf9,Cdkn1a,Vcam1,Ifit3b,Stx11,Il1rn,Hp,Znfx1,Cflar,Gbp6,Irf7,Igsf6,Birc3,Tor3a,Nfkbia,Casp4,Tubb6,Gbp4 | | 5.279e-31 | -69.72 | GSE13484\_UNSTIM\_VS\_YF17D\_VACCINE\_STIM\_PBMC\_DN | MSigDB lists | GSE13484\_UNSTIM\_VS\_YF17D\_VACCINE\_STIM\_PBMC\_DN | 153 | 34 | 12187 | 179 | Tnfsf10,Irf7,Cflar,Oasl1,Parp12,Xaf1,Casp4,Tnf,Irf9,Trim25,Atf3,Ifi204,Il1rn,Ifi207,Socs3,Ifit3b,Ifi209,Ifi44,Sbno2,Tnfaip8,Rtp4,Ifi211,Phf11d,Arid5b,Gadd45b,Il6,Ccl4,Usp18,Ifit2,Tap1,Ifitm3,Herc6,Isg15,Phf11b | | 2.139e-30 | -68.32 | GSE13485\_CTRL\_VS\_DAY7\_YF17D\_VACCINE\_PBMC\_DN | MSigDB lists | GSE13485\_CTRL\_VS\_DAY7\_YF17D\_VACCINE\_PBMC\_DN | 159 | 34 | 12187 | 179 | Oasl1,Parp12,Xaf1,Parp14,Tnfsf10,Irf7,C3ar1,Ifi204,Il1rn,Samd9l,Ifi207,Ifit3b,Ifi209,Ifi44,Irf9,Trim25,Ifitm2,Usp18,Ifit2,Msr1,Gbp3,Rtp4,Cmpk2,Ifi211,Phf11d,Dtx3l,Phf11b,Parp9,Tap1,Pik3ap1,Ifitm3,Herc6,Rnf213,Isg15 | | 2.328e-30 | -68.23 | GSE21360\_SECONDARY\_VS\_QUATERNARY\_MEMORY\_CD8\_TCELL\_UP | MSigDB lists | GSE21360\_SECONDARY\_VS\_QUATERNARY\_MEMORY\_CD8\_TCELL\_UP | 146 | 33 | 12187 | 179 | Ccl12,Tnfaip3,Il6,Gadd45b,Ccl4,Map3k8,Sbno2,Ptges,Cxcl1,Pik3r5,Bcl3,Lcn2,Nfkb1,Il1a,Nfkbiz,Irak3,Cd14,Nfkbia,Casp4,Tnf,Nfkb2,S100a9,Cflar,Tnip3,Il1rn,Stx11,Socs3,Vcam1,Clic4,Zfp36,Plek,Tnip1,Ptgs2 | | 2.451e-30 | -68.18 | cellular response to cytokine stimulus | biological process | GO:0071345 | 529 | 55 | 13711 | 214 | F830016B08Rik,Nfkb1,Birc3,Osmr,Gbp4,Il17ra,Irgm1,Icam1,Iigp1,Gm4841,Ifitm3,Ccl12,Ifitm2,Gbp3,Irgm2,Gbp6,Ccl2,Cxcl9,Ifi204,Ifi47,Csf3,Il1rn,Irak3,Arid5b,Sbno2,Tgtp1,Fzd4,Ifi209,Trim25,Spi1,Ccl4,Stx11,Ifitm6,Ifi211,Tgtp2,Il6,Gm5431,Il1a,Nfkbia,Il4ra,Ccl7,Irf7,Socs3,Hspa5,Cxcl1,Ifit2,Selp,Gbp7,Csf2rb,Nfe2l2,Zfp36,Tnf,Acod1,Ripk1,Ifi207 | | 2.666e-30 | -68.10 | GSE18281\_SUBCAPSULAR\_VS\_CENTRAL\_CORTICAL\_REGION\_OF\_THYMUS\_DN | MSigDB lists | GSE18281\_SUBCAPSULAR\_VS\_CENTRAL\_CORTICAL\_REGION\_OF\_THYMUS\_DN | 174 | 35 | 12187 | 179 | Cxcl16,Tap1,Slfn3,Nfkbiz,Lcn2,Mmp8,Gbp7,Nfkbie,S100a8,Cmpk2,Usp18,Ifit2,Tnfaip3,Cybb,Hcar2,Clic4,Socs3,Ifit3b,Cdkn1a,Ifi44,Znfx1,Timp1,Hp,S100a9,Irf7,Irgm2,Cflar,Slfn4,Parp14,Tor3a,Casp4,Tnf,Lgals9,Oasl1,Tubb6 | | 2.827e-30 | -68.04 | GSE18791\_CTRL\_VS\_NEWCASTLE\_VIRUS\_DC\_8H\_DN | MSigDB lists | GSE18791\_CTRL\_VS\_NEWCASTLE\_VIRUS\_DC\_8H\_DN | 134 | 32 | 12187 | 179 | Ptgs2,Errfi1,Trim25,Ifit3b,Ifi44,Znfx1,Samd9l,Trim56,Irf7,Tnfsf10,Xaf1,Parp14,Tnf,Parp12,Oasl1,Slfn5,Ifitm3,Isg15,Herc6,Rnf213,Parp9,Tap1,Phf11b,Phf11d,Bcl3,Dtx3l,Gbp3,Cmpk2,Rtp4,Usp18,Ifit2,Gadd45b | | 3.772e-30 | -67.75 | GSE2706\_UNSTIM\_VS\_2H\_R848\_DC\_DN | MSigDB lists | GSE2706\_UNSTIM\_VS\_2H\_R848\_DC\_DN | 148 | 33 | 12187 | 179 | Plek,Icam1,Plaur,Ptgs2,Stx11,Ifit3b,Socs3,Gem,Cflar,AA467197,Birc3,Zfp189,Sdc4,Nfkbia,Tap1,Ptpn1,Herc6,Isg15,Il1a,Nfkb1,Nfkbiz,Cmpk2,Tnfaip8,Bcl3,Pik3r5,Cxcl1,Il6,Gadd45b,Tnfaip3,Maff,Map3k8,Ifit2,Ccl4 | | 6.067e-30 | -67.27 | GSE2706\_UNSTIM\_VS\_2H\_LPS\_DC\_DN | MSigDB lists | GSE2706\_UNSTIM\_VS\_2H\_LPS\_DC\_DN | 150 | 33 | 12187 | 179 | Irf7,Gem,Cflar,Oasl1,AA467197,Nfkbia,Parp14,Birc3,Plaur,Icam1,Tnip1,Atf3,Znfx1,Stx11,Tnip3,Ifit3b,Socs3,Ifi44,Cdkn1a,Tnfaip8,Pik3r5,Cxcl1,Il6,Gadd45b,Tnfaip3,Map3k8,Tap1,Ptpn1,Herc6,Isg15,Il1a,Nfkb1,Nfkbiz | | 8.118e-30 | -66.98 | defense response to other organism | biological process | GO:0098542 | 473 | 52 | 13711 | 214 | Ifit2,Gbp7,Ifit3b,Hp,C5ar1,Trim30a,Cd14,Ncf1,Acod1,Tnf,Trim25,Ccl4,Herc6,Mx2,Stx11,Ifitm6,Isg15,Parp9,Tgtp1,Ccl7,Parp14,Irf7,Dtx3l,Il6,Casp4,Il4ra,Oasl2,Slfn9,Irgm2,Gbp6,Bcl3,S100a9,Irf2,Ccl12,Cxcl16,S100a8,Ifitm2,Gbp3,Lcn2,Tnfaip8,Ccl2,Cxcl9,Il17ra,Trim56,Rtp4,Irgm1,Gbp4,Mx1,Iigp1,Oasl1,Ifitm3,Cybb | | 1.558e-29 | -66.33 | GSE45365\_NK\_CELL\_VS\_CD11B\_DC\_DN | MSigDB lists | GSE45365\_NK\_CELL\_VS\_CD11B\_DC\_DN | 168 | 34 | 12187 | 179 | S100a9,Sdc4,Nfkbia,Birc3,AA467197,Tubb6,Tnip1,Saa1,Msn,Csf2rb,Clic4,Zfp36,Sele,Icam1,Il1rn,Timp1,Stx11,Ptges,Cxcl1,Csf3,S100a8,Osmr,Adamts9,Sntb2,Spi1,Ccl4,Tgm2,Maff,Il6,Tnfaip3,Cd14,Nfkbiz,Ier3,Tiparp | | 2.139e-29 | -66.01 | immune response | biological process | GO:0006955 | 625 | 58 | 13711 | 214 | Prg4,Cxcl1,Gbp7,Ifit2,Ifi44,Acod1,Tnf,Trim30a,Cd14,Tgtp1,Parp9,Isg15,Sbno2,Trim25,Stx11,Ifitm6,Mx2,Herc6,Ccl4,Il6,Il4ra,Oasl2,Casp4,Il1a,Dtx3l,Irf7,Parp14,Ccl7,S100a8,Tap1,Cxcl16,Ccl12,Gbp3,Ifitm2,Bcl3,Gbp6,Irgm2,S100a9,Tnfsf10,Ccl2,Tnfaip3,Cxcl9,Lcn2,Il1rn,C3ar1,Csf3,Gbp4,Trim56,Irgm1,Cybb,Icam1,Fyb,Nfkb2,Gadd45g,Mx1,Ifitm3,Oasl1,Iigp1 | | 4.442e-29 | -65.28 | GO\_DEFENSE\_RESPONSE | MSigDB lists | GO\_DEFENSE\_RESPONSE | 718 | 59 | 12187 | 179 | Ip6k2,Irf2,Il4ra,Ifit2,Ccl2,Tnfaip3,Bcl3,Cxcl1,Ifi211,S100a8,Ifi209,Hp,Ifi204,Saa1,Icam1,Trim25,Irf9,Casp4,Oasl1,S100a9,Irf7,Thbs1,Nfkbiz,Il1a,Nfkb1,Lcn2,Isg15,Ifitm3,Selp,Cd14,Il17ra,Tap1,Cxcl16,Ccl4,Cybb,Nfe2l2,Il6,Ccl12,Ptges,Ifit3b,Vcam1,Ifi207,Il1rn,Tnip3,Ptgs2,Ifitm2,Ncf1,Tnip1,Sele,Zfp189,Tnf,Xaf1,Gbp4,Gbp6,C3ar1,Irgm2,C5ar1,Trim56,Nfkb2 | | 6.901e-29 | -64.84 | GSE19888\_ADENOSINE\_A3R\_INH\_VS\_ACT\_WITH\_INHIBITOR\_PRETREATMENT\_IN\_MAST\_CELL\_UP | MSigDB lists | GSE19888\_ADENOSINE\_A3R\_INH\_VS\_ACT\_WITH\_INHIBITOR\_PRETREATMENT\_IN\_MAST\_CELL\_UP | 147 | 32 | 12187 | 179 | Ifitm3,Isg15,Rnf213,Herc6,Slfn5,Slfn9,Pik3ap1,Parp9,Tap1,Dtx3l,Gbp7,Cmpk2,Rtp4,Gbp3,Ifit2,Ccl2,Usp18,Clic4,Irf9,Ifi44,Ifit3b,Samd9l,Znfx1,Irgm2,Gbp6,Irf7,Tnfsf10,Parp14,Xaf1,Gbp4,Parp12,Oasl1 | | 1.364e-28 | -64.16 | GSE21546\_UNSTIM\_VS\_ANTI\_CD3\_STIM\_ELK1\_KO\_DP\_THYMOCYTES\_UP | MSigDB lists | GSE21546\_UNSTIM\_VS\_ANTI\_CD3\_STIM\_ELK1\_KO\_DP\_THYMOCYTES\_UP | 124 | 30 | 12187 | 179 | Ifit3b,Ifi44,Ifi209,Znfx1,Ifi207,Ifi204,Irf9,Csf2rb,Trim25,Xaf1,Parp14,Parp12,Oasl1,Trim56,Irf7,C5ar1,Phf11b,Rnf213,Herc6,Isg15,Ifitm3,Tap1,Parp9,Usp18,Ifit2,Ifi211,Phf11d,Dtx3l,Rtp4,Cmpk2 | | 5.749e-28 | -62.72 | NEMETH\_INFLAMMATORY\_RESPONSE\_LPS\_UP | MSigDB lists | NEMETH\_INFLAMMATORY\_RESPONSE\_LPS\_UP | 84 | 26 | 12187 | 179 | Slfn4,C3ar1,C5ar1,Cflar,Nfkbia,Casp4,Tnf,Trim25,Icam1,Tnip1,Ptgs2,Atf3,Ifi207,Il1rn,Ifi204,Cdkn1a,Ifi209,Rgs16,Ifi211,Rhoc,Msr1,Tap1,Slfn3,Ier3,Cebpd,Nfkb1 | | 8.219e-28 | -62.37 | GO\_IMMUNE\_SYSTEM\_PROCESS | MSigDB lists | GO\_IMMUNE\_SYSTEM\_PROCESS | 1290 | 75 | 12187 | 179 | C3ar1,Gem,Gbp6,C5ar1,Irgm2,Trim56,Igsf6,Nfkb2,Birc3,Tnf,Nfkbia,Xaf1,Gbp4,Ifitm2,Ncf1,Tnip1,Rbm47,Sele,Msn,Ifit3b,Vcam1,Zbtb16,Ifi207,Il1rn,Tnip3,Csf3,Sbno2,Map3k8,Ccl4,Cybb,Il6,Ccl12,Herc6,Isg15,Selp,Ifitm3,Cd14,Pik3ap1,Il17ra,Irak3,Tap1,Cxcl16,Thbs1,Il1a,Nfkb1,Tiparp,Lcn2,Irf7,S100a9,Tnfsf10,Runx1,Casp4,Oasl1,Saa1,Plek,Trim25,Icam1,Irf9,Ifi209,Hp,Ifi204,Bcl3,Cxcl1,Ifi211,S100a8,Il4ra,Pik3r1,Ifit2,Spi1,Ccl2,Hcar2,Ripk1,Tnfaip3,Ip6k2,Irf2 | | 1.308e-27 | -61.90 | GSE33424\_CD161\_INT\_VS\_NEG\_CD8\_TCELL\_UP | MSigDB lists | GSE33424\_CD161\_INT\_VS\_NEG\_CD8\_TCELL\_UP | 175 | 33 | 12187 | 179 | Dtx3l,Gbp7,Rtp4,Cmpk2,Usp18,Ccnd2,Ifit2,Cybb,Slfn9,Slfn5,Herc6,Isg15,Rnf213,Ifitm3,Parp9,Slfn3,Hspa5,Irf7,Gbp6,Irgm2,Tnfsf10,Slfn4,Casp4,Tor3a,Parp14,Xaf1,Oasl1,Parp12,Gbp4,Irf9,Ifit3b,Ifi44,Samd9l | | 1.533e-27 | -61.74 | GSE2706\_UNSTIM\_VS\_2H\_LPS\_AND\_R848\_DC\_DN | MSigDB lists | GSE2706\_UNSTIM\_VS\_2H\_LPS\_AND\_R848\_DC\_DN | 147 | 31 | 12187 | 179 | Cxcl1,Tnfaip8,Cmpk2,Rtp4,Map3k8,Ifit2,Tnfaip3,Il6,Isg15,Herc6,Nfkbiz,Nfkb1,Il1a,Cflar,Gem,Irf7,Birc3,Nfkbia,AA467197,Oasl1,Ptgs2,Tnip1,Icam1,Plaur,Cdkn1a,Ifi44,Socs3,Ifit3b,Stx11,Znfx1,Atf3 | | 1.933e-27 | -61.51 | GSE35685\_CD34POS\_CD38NEG\_VS\_CD34POS\_CD10POS\_BONE\_MARROW\_DN | MSigDB lists | GSE35685\_CD34POS\_CD38NEG\_VS\_CD34POS\_CD10POS\_BONE\_MARROW\_DN | 177 | 33 | 12187 | 179 | Tiparp,Nfkb1,Gadd45g,Ier3,Cebpd,Nfkbiz,Il17ra,Ptpn1,Tap1,Selp,Isg15,Ccl12,Ifit2,Map3k8,Il4ra,Ccl4,Ccl2,Usp18,Gbp3,Gbp7,Bcl3,Timp1,Znfx1,Aff1,Zfp36,Csf2rb,Irf9,Bach1,Lgals9,Tor3a,Nfkbia,Casp4,Irgm2 | | 2.948e-27 | -61.09 | GO\_RESPONSE\_TO\_BIOTIC\_STIMULUS | MSigDB lists | GO\_RESPONSE\_TO\_BIOTIC\_STIMULUS | 553 | 51 | 12187 | 179 | Hspa5,S100a8,Bcl3,Cxcl1,Ifi211,Tnfaip3,Il4ra,Ifit2,Irf9,Icam1,Trim25,Hp,Ifi204,Ifi209,S100a9,Irf7,Oasl1,Trib1,Cxcl16,Il17ra,Irak3,Cd14,Isg15,Ifitm3,Selp,Nfkb1,Sbno2,Cmpk2,Ptges,Csf3,Il6,Ccl12,Ccl4,Sele,Hspb1,Ifitm2,Ptgs2,Ifi207,Tnip3,Ifit3b,Vcam1,Socs3,Ifi44,Nfkb2,Trim56,Gbp6,C5ar1,Gbp4,Nfkbia,Zfp189,Tnf | | 5.658e-27 | -60.44 | GSE18791\_UNSTIM\_VS\_NEWCATSLE\_VIRUS\_DC\_10H\_DN | MSigDB lists | GSE18791\_UNSTIM\_VS\_NEWCATSLE\_VIRUS\_DC\_10H\_DN | 126 | 29 | 12187 | 179 | Oasl1,Xaf1,Parp14,Tnf,Tnfsf10,Irf7,Znfx1,Samd9l,Stx11,Ifit3b,Ifi44,Ptgs2,Gadd45b,Il6,Usp18,Map3k8,Ifit2,Gbp3,Rtp4,Cmpk2,Phf11d,Dtx3l,Nfkbiz,Phf11b,Tap1,Parp9,Slfn5,Isg15,Herc6 | | 9.982e-27 | -59.87 | GO\_IMMUNE\_RESPONSE | MSigDB lists | GO\_IMMUNE\_RESPONSE | 594 | 52 | 12187 | 179 | Irf9,Trim25,Icam1,Saa1,Ifi204,Ifi209,Tnfsf10,Irf7,S100a9,Oasl1,Casp4,Irf2,Ip6k2,S100a8,Bcl3,Ifi211,Cxcl1,Ccl2,Il4ra,Ifit2,Ncf1,Ifitm2,Ifi207,Il1rn,Ifit3b,Vcam1,Nfkb2,Igsf6,Trim56,C3ar1,Gem,Gbp6,Irgm2,C5ar1,Gbp4,Xaf1,Tnf,Tap1,Cxcl16,Cd14,Isg15,Ifitm3,Lcn2,Il1a,Nfkb1,Thbs1,Sbno2,Csf3,Il6,Ccl12,Cybb,Ccl4 | | 1.376e-26 | -59.55 | ZHANG\_RESPONSE\_TO\_IKK\_INHIBITOR\_AND\_TNF\_UP | MSigDB lists | ZHANG\_RESPONSE\_TO\_IKK\_INHIBITOR\_AND\_TNF\_UP | 172 | 32 | 12187 | 179 | Tnfsf10,Nfkb2,Oasl1,AA467197,Birc3,Nfkbia,Sdc4,Icam1,Irf9,Plaur,Tnip1,Il1rn,Atf3,Znfx1,Cdkn1a,Ifit3b,Socs3,Cmpk2,Tnfaip8,Nfkbie,Bcl3,Cxcl1,Tnfaip3,Maff,Usp18,Tap1,Cxcl16,Slfn5,Il1a,Nfkb1,Sgk1,Nfkbiz | | 2.730e-26 | -58.86 | response to organic substance | biological process | GO:0010033 | 1870 | 92 | 13711 | 214 | Isg15,Tiparp,Timp1,Fam107a,Runx1,Trim25,Xaf1,Ifi211,Ip6k2,Gm5431,Il4ra,Nfkbia,Ccl7,C5ar1,Hspa5,Gbp7,Noct,Selp,Msn,Serpina3f,Nfe2l2,Atf3,Acod1,Sele,Birc3,Irgm1,Icam1,Lgals9,Iigp1,Tnip3,Iqgap1,Col4a1,Cxcl16,Ifitm2,Ptpn1,Wfdc21,Vcam1,Ccl2,Tnfaip3,Cxcl9,Il1rn,Csf3,Tgtp1,Cmpk2,Sbno2,Ifi209,Fzd4,Sgk1,Ptgs2,Stx11,Ifitm6,Ccl4,Spi1,Il6,Tgtp2,Casp4,Il1a,Pik3r1,Irf7,Socs3,Trib1,Hp,Cxcl1,Csf2rb,Ifit2,Zfp36,Tnf,Hspb1,Ifi207,Cd14,Cflar,Ripk1,Nfkb1,F830016B08Rik,Gbp4,Osmr,Trim56,Il17ra,P2ry6,Thbs1,Ifitm3,Gm4841,Ccl12,Gbp3,Irgm2,Gbp6,Usp18,Ifi204,Arid5b,Pygm,Irak3,Ifi47 | | 4.981e-26 | -58.26 | GSE21546\_WT\_VS\_SAP1A\_KO\_DP\_THYMOCYTES\_UP | MSigDB lists | GSE21546\_WT\_VS\_SAP1A\_KO\_DP\_THYMOCYTES\_UP | 149 | 30 | 12187 | 179 | Phf11b,Ifitm3,Rnf213,Isg15,Herc6,Irf2,Parp9,Tap1,Ifit2,Usp18,Dtx3l,Ifi211,Phf11d,Cmpk2,Ifi209,Ifi44,Ifit3b,Ifi204,Samd9l,Ifi207,Znfx1,Ifitm2,Trim25,Irf9,Parp14,Xaf1,Parp12,Oasl1,Irf7,Trim56 | | 9.856e-26 | -57.58 | response to chemical | biological process | GO:0042221 | 2591 | 108 | 13711 | 214 | Ptpn1,Ifitm2,Cxcl16,Col4a1,Iqgap1,Tnip3,S100a9,Vcam1,Wfdc21,Cxcl9,Tnfaip3,Ccl2,Csf3,C3ar1,Il1rn,Retnlg,Lcn2,Birc3,Nfkbie,Rtp4,Irgm1,Icam1,Mt1,Iigp1,Lgals9,Clic4,Hspa5,C5ar1,Nfe2l2,Serpina3f,Msn,Gbp7,Noct,Selp,Acod1,Atf3,Ncf1,Sele,Cp,Fam107a,Tiparp,Timp1,Isg15,Xaf1,Ifi211,Trim25,Runx1,Nfkbia,Il4ra,Gm5431,Ip6k2,Ccl7,Gbp3,Ccl12,S100a8,Usp18,Gbp6,Irgm2,Xdh,Ifi47,Arid5b,Pygm,Irak3,Ifi204,Osmr,Gbp4,F830016B08Rik,Nfkb1,Bach1,Il17ra,Trim56,Cybb,Gm4841,Thbs1,Ifitm3,P2ry6,Cxcl1,Mt2,Hp,Ifit2,Csf2rb,Tnf,Zfp36,Saa1,Ripk1,Cflar,Cd14,Ifi207,Hspb1,Fzd4,Ifi209,Sbno2,Cmpk2,Tgtp1,Ccl4,Spi1,Stx11,Ifitm6,Sgk1,Ptgs2,Il1a,Casp4,Tgtp2,Il6,Trib1,Irf7,Socs3,Pik3r1 | | 1.717e-25 | -57.02 | cellular response to chemical stimulus | biological process | GO:0070887 | 1836 | 90 | 13711 | 214 | Iigp1,Icam1,Mt1,Irgm1,Birc3,Csf3,Il1rn,Retnlg,Lcn2,Cxcl9,Tnfaip3,Ccl2,S100a9,Vcam1,Ptpn1,Ifitm2,Cxcl16,Iqgap1,Col4a1,Tnip3,Ccl7,Nfkbia,Il4ra,Gm5431,Ip6k2,Ifi211,Trim25,Runx1,Fam107a,Tiparp,Ncf1,Acod1,Atf3,Nfe2l2,Msn,Selp,Gbp7,Hspa5,Clic4,C5ar1,Thbs1,Gm4841,Ifitm3,P2ry6,Cybb,Bach1,Il17ra,Osmr,Gbp4,F830016B08Rik,Nfkb1,Ifi47,Irak3,Arid5b,Ifi204,Gbp6,Irgm2,Gbp3,Ccl12,S100a8,Irf7,Socs3,Pik3r1,Il1a,Casp4,Tgtp2,Il6,Ccl4,Spi1,Ifitm6,Stx11,Sgk1,Fzd4,Ifi209,Sbno2,Cmpk2,Tgtp1,Ripk1,Cflar,Cd14,Ifi207,Hspb1,Tnf,Zfp36,Saa1,Ifit2,Csf2rb,Cxcl1,Mt2,Hp | | 3.503e-25 | -56.31 | GSE18791\_UNSTIM\_VS\_NEWCATSLE\_VIRUS\_DC\_6H\_DN | MSigDB lists | GSE18791\_UNSTIM\_VS\_NEWCATSLE\_VIRUS\_DC\_6H\_DN | 144 | 29 | 12187 | 179 | Tnfsf10,Trim56,Irf7,Oasl1,Xaf1,Parp14,Irf9,Trim25,Znfx1,Ifi204,Samd9l,Ifi207,Ifit3b,Ifi209,Ifi44,Rtp4,Cmpk2,Ifi211,Phf11d,Dtx3l,Ripk1,Usp18,Ifit2,Parp9,Tap1,Herc6,Isg15,Rnf213,Phf11b | | 5.015e-25 | -55.95 | response to stress | biological process | GO:0006950 | 2282 | 100 | 13711 | 214 | Gbp3,S100a8,Tap1,Ccl12,Irf2,Bcl3,Gbp6,Irgm2,Plek,Pygm,Ifi47,Ifi204,Gbp4,Nfkb1,F830016B08Rik,Bach1,Trim56,Il17ra,Cybb,Errfi1,Ifitm3,Thbs1,Gm4841,Mx1,Rrp8,Dnajb1,Cxcl1,Mt2,Hp,Ifit3b,Ifit2,Tnf,Zfp36,Saa1,Hspb1,Ripk1,Cd14,Cflar,Fstl1,Tnip1,Tgtp1,Parp9,Sdc4,Ifitm6,Stx11,Ccl4,Ptgs2,Sgk1,Ier3,Oasl2,Casp4,Il1a,Il6,Cdkn1a,Tgtp2,Dtx3l,Irf7,Trib1,Pik3r1,Nfkbiz,Ptpn1,Ifitm2,Cxcl16,S100a9,Tnfaip3,Cxcl9,Ccl2,Il1rn,C3ar1,Tnfaip8,Lcn2,Map3k8,Irgm1,Rtp4,Icam1,Mt1,Oasl1,Iigp1,C5ar1,Hspa5,Nfe2l2,Angptl4,Selp,Gbp7,Acod1,Atf3,Ncf1,Sele,Trim30a,Timp1,Isg15,Mx2,Herc6,Runx1,Trim25,Il4ra,Slfn9,Gm5431,Parp14,Ccl7 | | 5.461e-25 | -55.87 | GO\_INFLAMMATORY\_RESPONSE | MSigDB lists | GO\_INFLAMMATORY\_RESPONSE | 283 | 37 | 12187 | 179 | Il1a,Nfkb1,Nfkbiz,Thbs1,Cd14,Selp,Il6,Nfe2l2,Tnfaip3,Ccl12,Cybb,Ccl2,Ccl4,Il4ra,S100a8,Ptges,Cxcl1,Ifi211,Hp,Ifi207,Tnip3,Il1rn,Ifi204,Vcam1,Ifi209,Sele,Icam1,Tnip1,Saa1,Ptgs2,Casp4,Tnf,Nfkb2,S100a9,C3ar1,Irgm2,C5ar1 | | 5.654e-25 | -55.83 | GSE19888\_ADENOSINE\_A3R\_INH\_VS\_TCELL\_MEMBRANES\_ACT\_MAST\_CELL\_UP | MSigDB lists | GSE19888\_ADENOSINE\_A3R\_INH\_VS\_TCELL\_MEMBRANES\_ACT\_MAST\_CELL\_UP | 161 | 30 | 12187 | 179 | Gbp7,Dtx3l,Cmpk2,Rtp4,Gbp3,Ifit2,Usp18,Sgk3,Isg15,Rnf213,Herc6,Slfn5,Slfn9,Tap1,Parp9,Gbp6,Irgm2,Trim56,Tnfsf10,Casp4,Parp14,Xaf1,Parp12,Gbp4,Trim25,Irf9,Clic4,Ifi44,Ifit3b,Samd9l | | 6.564e-25 | -55.68 | GSE9988\_ANTI\_TREM1\_VS\_LOW\_LPS\_MONOCYTE\_DN | MSigDB lists | GSE9988\_ANTI\_TREM1\_VS\_LOW\_LPS\_MONOCYTE\_DN | 147 | 29 | 12187 | 179 | Nfkbiz,Nfkb1,Il1a,Ier3,Pik3ap1,Map3k8,Rhou,Ccl4,Gpr84,Tnfaip3,Il6,Nfe2l2,Csf3,Cxcl1,Tnfaip8,Socs3,Stx11,Tnip3,Ptgs2,Tnip1,Icam1,Clic4,Tnf,Birc3,Nfkbia,Sdc4,Oasl1,Cflar,Igsf6 | | 6.738e-25 | -55.66 | response to interferon-beta | biological process | GO:0035456 | 45 | 20 | 13711 | 214 | Tgtp2,Gm5431,Acod1,Ifi204,Iigp1,Ifi47,Gm4841,Ifi207,Ifitm3,F830016B08Rik,Tgtp1,Ifitm2,Gbp3,Ifi209,Gbp6,Irgm2,Ifi211,Irgm1,Xaf1,Ifitm6 | | 8.302e-25 | -55.45 | GSE26030\_TH1\_VS\_TH17\_DAY5\_POST\_POLARIZATION\_UP | MSigDB lists | GSE26030\_TH1\_VS\_TH17\_DAY5\_POST\_POLARIZATION\_UP | 163 | 30 | 12187 | 179 | Hspa5,Parp9,Slfn3,Slfn9,Slfn5,Ifitm3,Herc6,Isg15,Rnf213,Usp18,Ifit2,Cmpk2,Rtp4,Gbp7,Dtx3l,Samd9l,Ifit3b,Ifi44,Irf9,Gbp4,Parp12,Oasl1,Parp14,Tor3a,Xaf1,Tnfsf10,Slfn4,Irf7,Irgm2,Gbp6 | | 3.730e-24 | -53.95 | GSE2706\_UNSTIM\_VS\_8H\_R848\_DC\_DN | MSigDB lists | GSE2706\_UNSTIM\_VS\_8H\_R848\_DC\_DN | 141 | 28 | 12187 | 179 | Ifit3b,Socs3,Ifi44,Samd9l,Stx11,Tnfsf10,Xaf1,Parp14,Birc3,Tnf,AA467197,Slfn5,Isg15,Rnf213,Herc6,Parp9,Nfkbiz,Thbs1,Sox11,Nfkb1,Cmpk2,Tnfaip8,Usp18,Ifit2,Map3k8,Il6,Gadd45b,Tnfaip3 | | 4.377e-24 | -53.79 | GSE43863\_TFH\_VS\_LY6C\_LOW\_CXCR5NEG\_EFFECTOR\_CD4\_TCELL\_UP | MSigDB lists | GSE43863\_TFH\_VS\_LY6C\_LOW\_CXCR5NEG\_EFFECTOR\_CD4\_TCELL\_UP | 172 | 30 | 12187 | 179 | Nfkb1,Ms4a6d,Rnf213,Slfn9,Pik3ap1,Parp9,Tap1,Ifit2,Cybb,Il6,Gadd45b,Dtx3l,Bcl3,Cmpk2,Rtp4,Gbp3,Socs3,Ksr1,Znfx1,Ncf1,Irf9,Lgals9,Xaf1,Parp14,Nfkbia,Gbp4,Parp12,Irgm2,Gbp6,Irf7 | | 5.611e-24 | -53.54 | GO\_RESPONSE\_TO\_CYTOKINE | MSigDB lists | GO\_RESPONSE\_TO\_CYTOKINE | 487 | 45 | 12187 | 179 | Oasl1,Irf7,Icam1,Trim25,Irf9,Ripk1,Ifit2,Il4ra,Ccl2,Cxcl1,Hspa5,Ip6k2,Irf2,Gbp4,Tnf,Birc3,Xaf1,Nfkbia,Nfkb2,Gbp6,Trim56,Timp1,Socs3,Vcam1,Ifit3b,Sele,Csf2rb,Ptgs2,Ifitm2,Ccl12,Nfe2l2,Il6,Ccl4,Osmr,Sbno2,Csf3,Ptges,Nfkb1,Il1a,Irak3,Il17ra,Cxcl16,Ifitm3,Isg15,Cd14 | | 8.466e-24 | -53.13 | GSE42021\_TREG\_VS\_TCONV\_PLN\_UP | MSigDB lists | GSE42021\_TREG\_VS\_TCONV\_PLN\_UP | 160 | 29 | 12187 | 179 | Oasl1,Parp12,Xaf1,Tnfsf10,Irf7,Ifi207,Ifi204,Atf3,Ifi44,Ifi209,Ifit3b,Irf9,Plaur,Ifitm2,Ccl12,Maff,Ifit2,Usp18,Rtp4,Osmr,Phf11d,Cxcl1,Ifi211,Cebpd,Phf11b,Tap1,Herc6,Isg15,Ifitm3 | | 1.035e-23 | -52.93 | GSE18791\_CTRL\_VS\_NEWCASTLE\_VIRUS\_DC\_10H\_DN | MSigDB lists | GSE18791\_CTRL\_VS\_NEWCASTLE\_VIRUS\_DC\_10H\_DN | 146 | 28 | 12187 | 179 | Dtx3l,Phf11d,Rtp4,Cmpk2,Gbp3,Ifit2,Map3k8,Usp18,Il6,Tnfaip3,Herc6,Rnf213,Isg15,Slfn9,Tap1,Irf2,Phf11b,Tnfsf10,Tnf,Xaf1,Parp14,Oasl1,Ptgs2,Ifi44,Ifit3b,Samd9l,Stx11,Znfx1 | | 1.771e-23 | -52.39 | GSE22140\_GERMFREE\_VS\_SPF\_MOUSE\_CD4\_TCELL\_UP | MSigDB lists | GSE22140\_GERMFREE\_VS\_SPF\_MOUSE\_CD4\_TCELL\_UP | 164 | 29 | 12187 | 179 | Tnf,Trib1,Casp4,Xaf1,Oasl1,Gem,Irf7,Tnfsf10,Ifi44,Cdkn1a,Ifi209,Ifit3b,Socs3,Ifi207,Ifi204,Plek,Zfp36,Icam1,Trim25,Irf9,Csf2rb,Ifit2,Ccl2,Gadd45b,Bcl3,Ifi211,Sbno2,Isg15,Tap1 | | 1.872e-23 | -52.33 | GSE18791\_CTRL\_VS\_NEWCASTLE\_VIRUS\_DC\_6H\_DN | MSigDB lists | GSE18791\_CTRL\_VS\_NEWCASTLE\_VIRUS\_DC\_6H\_DN | 149 | 28 | 12187 | 179 | Ifit2,Usp18,Dtx3l,Bcl3,Phf11d,Angptl4,Cmpk2,Rtp4,Phf11b,Rnf213,Herc6,Isg15,Tap1,Parp9,Parp14,Xaf1,Parp12,Oasl1,Trim56,Irf7,Tnfsf10,Ifi44,Ifit3b,Samd9l,Stx11,Znfx1,Trim25,Errfi1 | | 2.389e-23 | -52.09 | GALINDO\_IMMUNE\_RESPONSE\_TO\_ENTEROTOXIN | MSigDB lists | GALINDO\_IMMUNE\_RESPONSE\_TO\_ENTEROTOXIN | 74 | 22 | 12187 | 179 | Tnip1,Cd14,Ptgs2,Errfi1,Csf2rb,Plaur,Icam1,Slfn3,Socs3,Ier3,Nfkb1,Il1rn,Cxcl1,Bcl3,Csf3,Nfkb2,Slfn4,Ccl4,Nfkbia,Tnf,Tnfaip3,Gadd45b | | 5.879e-23 | -51.19 | GSE9601\_NFKB\_INHIBITOR\_VS\_PI3K\_INHIBITOR\_TREATED\_HCMV\_INF\_MONOCYTE\_DN | MSigDB lists | GSE9601\_NFKB\_INHIBITOR\_VS\_PI3K\_INHIBITOR\_TREATED\_HCMV\_INF\_MONOCYTE\_DN | 155 | 28 | 12187 | 179 | Tubb6,Sdc4,Trib1,Tnfsf10,Irf7,Gem,Znfx1,Il1rn,Timp1,Ifit3b,Cdkn1a,Clic4,Saa1,Ptgs2,Ccl12,Cybb,Usp18,Ccl2,Ccl4,Maff,Ifit2,Map3k8,Rhou,Rtp4,Csf3,Il1a,Cd14,Selp | | 6.131e-23 | -51.15 | GSE26343\_UNSTIM\_VS\_LPS\_STIM\_NFAT5\_KO\_MACROPHAGE\_DN | MSigDB lists | GSE26343\_UNSTIM\_VS\_LPS\_STIM\_NFAT5\_KO\_MACROPHAGE\_DN | 171 | 29 | 12187 | 179 | Cflar,Nfkb2,Igsf6,Casp4,Tnip1,Saa1,Csf2rb,Errfi1,Icam1,Zfp36,Socs3,Cdkn1a,Il1rn,Cxcl1,Pik3r5,Bcl3,Nfkbie,Gbp3,Gpr84,Map3k8,Maff,Nfe2l2,Gadd45b,Il6,Cd14,Cxcl16,Nfkbiz,Ier3,Il1a | | 1.025e-22 | -50.63 | GSE34392\_ST2\_KO\_VS\_WT\_DAY8\_LCMV\_EFFECTOR\_CD8\_TCELL\_DN | MSigDB lists | GSE34392\_ST2\_KO\_VS\_WT\_DAY8\_LCMV\_EFFECTOR\_CD8\_TCELL\_DN | 174 | 29 | 12187 | 179 | Gadd45b,Il6,Nfe2l2,Hcar2,Tgm2,Gpr84,Ccl2,Usp18,Ifit2,Cmpk2,Cxcl1,Bcl3,Nfkbie,Nfkbiz,Irak3,Isg15,Tor3a,Parp14,Casp4,Birc3,Nfkb2,Cflar,Gem,Znfx1,Atf3,Stx11,Plek,Tagln2,Saa1 | | 1.465e-22 | -50.28 | ICHIBA\_GRAFT\_VERSUS\_HOST\_DISEASE\_D7\_UP | MSigDB lists | ICHIBA\_GRAFT\_VERSUS\_HOST\_DISEASE\_D7\_UP | 91 | 23 | 12187 | 179 | Il1rn,Ifi204,Ifi207,Col4a1,Vcam1,Ifit3b,Ifi209,Icam1,Nfkbia,Xdh,Irf7,Irgm2,Nfkbiz,Ptpn1,Tap1,Cd14,Isg15,Ccl12,Usp18,Msr1,Gbp3,Ifi211,Gbp7 | | 1.646e-22 | -50.16 | HALLMARK\_INTERFERON\_ALPHA\_RESPONSE | MSigDB lists | HALLMARK\_INTERFERON\_ALPHA\_RESPONSE | 80 | 22 | 12187 | 179 | Trim25,Parp9,Irf2,Irf9,Tap1,Ifitm3,Herc6,Ifitm2,Isg15,Samd9l,Ifi44,Ifit3b,Rtp4,Cmpk2,Gbp3,Irf7,Oasl1,Parp12,Ifit2,Il4ra,Parp14,Usp18 | | 1.796e-22 | -50.07 | GSE18791\_CTRL\_VS\_NEWCASTLE\_VIRUS\_DC\_4H\_DN | MSigDB lists | GSE18791\_CTRL\_VS\_NEWCASTLE\_VIRUS\_DC\_4H\_DN | 131 | 26 | 12187 | 179 | Dtx3l,Phf11d,Rtp4,Cmpk2,Ifit2,Usp18,Gadd45b,Rnf213,Herc6,Isg15,Tap1,Parp9,Phf11b,Irf7,Trim56,Tnfsf10,Nfkb2,Xaf1,Parp14,Oasl1,Trim25,Errfi1,Ifi44,Ifit3b,Samd9l,Znfx1 | | 1.870e-22 | -50.03 | SEKI\_INFLAMMATORY\_RESPONSE\_LPS\_UP | MSigDB lists | SEKI\_INFLAMMATORY\_RESPONSE\_LPS\_UP | 60 | 20 | 12187 | 179 | Cxcl1,Nfkbie,Nfkb2,Gbp3,Osmr,Nfkbia,Casp4,Maff,Birc3,Ccl12,Tnfaip3,Gadd45b,Tnip1,Cxcl16,Icam1,Sele,Vcam1,Nfkbiz,Ier3,Stx11 | | 2.978e-22 | -49.57 | MCLACHLAN\_DENTAL\_CARIES\_UP | MSigDB lists | MCLACHLAN\_DENTAL\_CARIES\_UP | 164 | 28 | 12187 | 179 | Msr1,Maff,Cybb,Il6,Tnfaip3,Ccl12,Bcl3,S100a8,Tnfaip8,Ms4a6d,Ier3,Cebpd,Cd14,Birc3,Trib1,C3ar1,C5ar1,S100a9,Socs3,Vcam1,Il1rn,Timp1,Ptgs2,Plek,Sele,Icam1,Csf2rb,Plaur | | 3.844e-22 | -49.31 | GSE35685\_CD34POS\_CD10NEG\_CD62LPOS\_VS\_CD34POS\_CD10POS\_BONE\_MARROW\_DN | MSigDB lists | GSE35685\_CD34POS\_CD10NEG\_CD62LPOS\_VS\_CD34POS\_CD10POS\_BONE\_MARROW\_DN | 182 | 29 | 12187 | 179 | Timp1,Csf2rb,Zfp36,Casp4,Lgals9,Slfn4,Irgm2,Ier3,Cebpd,Gadd45g,Nfkbiz,Hspa5,Ptpn1,Tap1,Slfn3,Ifitm3,Selp,Isg15,Ccl12,Ccl4,Ccl2,Map3k8,Ifit2,Pik3r1,Il4ra,Gbp3,Tnfaip8,Bcl3,Gbp7 | | 4.067e-22 | -49.25 | GSE9988\_LOW\_LPS\_VS\_VEHICLE\_TREATED\_MONOCYTE\_UP | MSigDB lists | GSE9988\_LOW\_LPS\_VS\_VEHICLE\_TREATED\_MONOCYTE\_UP | 135 | 26 | 12187 | 179 | Nfkbia,Sdc4,Birc3,Tnf,Gem,Cflar,Socs3,Tnip3,Ptgs2,Plaur,Errfi1,Plek,Icam1,Gpr84,Ccl4,Maff,Map3k8,Nfe2l2,Gadd45b,Tnfaip3,Cxcl1,Nfkbiz,Ier3,Il1a,Nfkb1,Pik3ap1 | | 4.210e-22 | -49.22 | GSE19198\_CTRL\_VS\_IL21\_TREATED\_TCELL\_24H\_UP | MSigDB lists | GSE19198\_CTRL\_VS\_IL21\_TREATED\_TCELL\_24H\_UP | 166 | 28 | 12187 | 179 | Tnf,Trib1,Xdh,Ifi44,Cdkn1a,Cp,Il1rn,Timp1,Atf3,Tnip1,Icam1,Errfi1,Maff,Ccl2,Tgm2,Ccl4,Nfe2l2,Il6,Gadd45b,Csf3,Rhoj,Cxcl1,Gbp3,Gadd45g,Akap12,Il1a,Map3k6,Ifitm3 | | 4.997e-22 | -49.05 | GSE34006\_WT\_VS\_A2AR\_KO\_TREG\_DN | MSigDB lists | GSE34006\_WT\_VS\_A2AR\_KO\_TREG\_DN | 167 | 28 | 12187 | 179 | Irf7,Gbp6,Irgm2,Tnfsf10,Slfn4,Parp14,Tor3a,Lgals9,Parp12,Oasl1,Gbp4,Irf9,Trim25,Ifit3b,Znfx1,Gbp7,Gbp3,Rtp4,Cmpk2,Usp18,Ccnd2,Ifit2,Ripk1,Slfn9,Isg15,Tap1,Parp9,Slfn3 | | 4.997e-22 | -49.05 | GSE5589\_LPS\_AND\_IL10\_VS\_LPS\_AND\_IL6\_STIM\_IL6\_KO\_MACROPHAGE\_45MIN\_UP | MSigDB lists | GSE5589\_LPS\_AND\_IL10\_VS\_LPS\_AND\_IL6\_STIM\_IL6\_KO\_MACROPHAGE\_45MIN\_UP | 167 | 28 | 12187 | 179 | Gbp6,Cflar,Tnfsf10,Xdh,Slfn4,Sdc4,Casp4,Tor3a,Parp14,Parp12,Gbp4,Irf9,Cdkn1a,Znfx1,Samd9l,Gbp7,Dtx3l,Ccl2,Msr1,Ccl12,Isg15,Ifitm3,Tap1,Cxcl16,Parp9,Pik3ap1,Slfn3,Ms4a6d | | 6.056e-22 | -48.86 | GSE21546\_ELK1\_KO\_VS\_SAP1A\_KO\_AND\_ELK1\_KO\_DP\_THYMOCYTES\_UP | MSigDB lists | GSE21546\_ELK1\_KO\_VS\_SAP1A\_KO\_AND\_ELK1\_KO\_DP\_THYMOCYTES\_UP | 137 | 26 | 12187 | 179 | Ifitm3,Herc6,Rnf213,Isg15,P2ry6,Parp9,Phf11b,Dtx3l,Phf11d,Ifi211,Adamts9,Cmpk2,Ifit2,Usp18,Ifitm2,Trim25,Irf9,Ifi209,Ifit3b,Ifi204,Ifi207,Samd9l,Irf7,Parp14,Oasl1,Parp12 | | 7.036e-22 | -48.71 | GSE13485\_PRE\_VS\_POST\_YF17D\_VACCINATION\_PBMC\_DN | MSigDB lists | GSE13485\_PRE\_VS\_POST\_YF17D\_VACCINATION\_PBMC\_DN | 153 | 27 | 12187 | 179 | Pik3ap1,Parp9,Tap1,Ifitm3,Herc6,Isg15,Ifit2,Msr1,Usp18,Rtp4,Cmpk2,Dtx3l,Ifi211,Ifi204,Samd9l,Ifi207,Ifi209,Ifi44,Ifit3b,Irf9,Ifitm2,Parp12,Oasl1,Xaf1,Parp14,C3ar1,Irf7 | | 1.034e-21 | -48.32 | MODULE\_84 | MSigDB lists | MODULE\_84 | 373 | 38 | 12187 | 179 | S100a9,A2m,C3ar1,Tnfsf10,Olfml2b,Igsf6,Parp14,Birc3,Tnf,Parp12,Plek,Icam1,Vcam1,Ifi44,Ifi209,Cdkn1a,Ifi207,Col4a1,Ifi204,Il1rn,Kcna5,Stx11,Ifi211,S100a8,Tnfaip8,Ccl4,Tgm2,Il4ra,Ccnd2,Il6,Tnfaip3,Ccl12,Cybb,Cd14,Isg15,Tap1,Cxcl16,Ier3 | | 1.316e-21 | -48.08 | GSE9988\_LOW\_LPS\_VS\_CTRL\_TREATED\_MONOCYTE\_UP | MSigDB lists | GSE9988\_LOW\_LPS\_VS\_CTRL\_TREATED\_MONOCYTE\_UP | 141 | 26 | 12187 | 179 | Nfkb1,Il1a,Ier3,Nfkbiz,Tnfaip3,Nfe2l2,Il6,Gadd45b,Map3k8,Maff,Ccl4,Gpr84,Csf3,Cxcl1,Tnip3,Socs3,Icam1,Plek,Errfi1,Plaur,Ptgs2,Tnf,Birc3,Sdc4,Nfkbia,Cflar | | 1.363e-21 | -48.04 | GSE32986\_CURDLAN\_HIGHDOSE\_VS\_GMCSF\_AND\_CURDLAN\_HIGHDOSE\_STIM\_DC\_DN | MSigDB lists | GSE32986\_CURDLAN\_HIGHDOSE\_VS\_GMCSF\_AND\_CURDLAN\_HIGHDOSE\_STIM\_DC\_DN | 173 | 28 | 12187 | 179 | Socs3,Stx11,Ptgs2,Tnip1,Zfp36,Icam1,Birc3,Tnf,Nfkbia,Sdc4,Casp4,Gem,Cflar,Slfn4,Nfkb2,Nfkbiz,Il1a,Nfkb1,Isg15,Slfn3,Maff,Gpr84,Ccl4,Rassf4,Gadd45b,Tnfaip3,Nfkbie,Bcl3 | | 1.363e-21 | -48.04 | GSE22601\_IMMATURE\_CD4\_SINGLE\_POSITIVE\_VS\_CD8\_SINGLE\_POSITIVE\_THYMOCYTE\_DN | MSigDB lists | GSE22601\_IMMATURE\_CD4\_SINGLE\_POSITIVE\_VS\_CD8\_SINGLE\_POSITIVE\_THYMOCYTE\_DN | 173 | 28 | 12187 | 179 | Tnfaip8,Rtp4,Dtx3l,Nfe2l2,Map3k8,Tgm2,Ptpn1,Selp,Ifitm3,Slfn9,Tiparp,Cebpd,Sgk1,Nfkbiz,Irgm2,Gbp6,Gem,Irf7,Gbp4,Birc3,Xaf1,Parp14,Casp4,Trim25,Zfp36,Irf9,Stx11,Ksr1 | | 1.656e-21 | -47.85 | innate immune response | biological process | GO:0045087 | 351 | 38 | 13711 | 214 | Oasl2,Casp4,Cybb,Ifitm3,Dtx3l,Oasl1,Irf7,Parp14,Ccl7,Iigp1,Mx1,Gbp4,Tgtp1,Parp9,Isg15,Irgm1,Ifitm6,Stx11,Mx2,Ccl4,Herc6,Trim56,Trim25,Acod1,Ccl2,Cd14,Lcn2,Trim30a,Gbp3,Ifitm2,S100a8,Cxcl16,Ccl12,S100a9,Gbp7,Gbp6,Irgm2,Ifit2 | | 1.934e-21 | -47.69 | cellular response to stimulus | biological process | GO:0051716 | 4061 | 130 | 13711 | 214 | Fzd4,Ifi209,Sbno2,Parp9,Tgtp1,Cmpk2,Spi1,Ccl4,Ifitm6,Stx11,Ier3,Sgk1,Ptgs2,Gem,Il1a,Casp4,Cdkn1a,Tgtp2,Il6,Trib1,Irf7,Socs3,Dtx3l,Pik3r1,Rrp8,Cxcl1,Dnajb1,Hp,Mt2,Ifit2,Csf2rb,Tnf,Zfp36,Saa1,Cflar,Cd14,Gpr84,Ripk1,Hspb1,Ifi207,Rhoc,Tnip1,Osmr,Gbp4,F830016B08Rik,Nfkb1,Kcna5,Bach1,Plaur,Il17ra,Errfi1,Cybb,Thbs1,Ifitm3,Gm4841,P2ry6,Gbp3,Ccl12,S100a8,Sgk3,Irgm2,Gbp6,Bcl3,Plek,Ksr1,Ifi47,Arid5b,Irak3,Ifi204,Fam107a,Map3k6,Tiparp,Isg15,Rasip1,Ifi211,Trim25,Runx1,Akap12,Nfkbia,Il4ra,Ip6k2,Gm5431,Ccl7,Ccnd2,Hcar2,Clic4,Hspa5,Tgm2,C5ar1,Rasd1,Nfe2l2,Gbp7,Selp,Msn,Angptl4,Pik3ap1,Acod1,Rhou,Atf3,Ncf1,Birc3,Map3k8,Rhoj,Irgm1,Pik3r5,Rgs16,Nfkb2,Fyb,Mt1,Icam1,Rassf4,Iigp1,Lgals9,Ifitm2,Ptpn1,Nfkbiz,Cxcl16,Tnip3,Col4a1,Iqgap1,S100a9,Vcam1,Cxcl9,Tnfaip3,Ccl2,Csf3,C3ar1,Il1rn,Lcn2,Retnlg | | 2.110e-21 | -47.61 | FOSTER\_TOLERANT\_MACROPHAGE\_DN | MSigDB lists | FOSTER\_TOLERANT\_MACROPHAGE\_DN | 357 | 37 | 12187 | 179 | Sgk3,Selp,Slfn5,Irf2,Nfkbiz,Gadd45g,Il1a,Nfkb1,Tiparp,Cebpd,Gbp7,Bcl3,Ifi211,Rgs16,Gbp3,Il4ra,Ccnd2,Map3k8,Gpr84,Il6,Tnfaip3,Ptgs2,Zfp36,Plaur,Errfi1,Ifi209,Cdkn1a,Aff1,Socs3,Vcam1,Cp,Ifi207,Ifi204,Tnfsf10,Sdc4,Parp14,Tubb6 | | 2.217e-21 | -47.56 | GSE14769\_UNSTIM\_VS\_40MIN\_LPS\_BMDM\_DN | MSigDB lists | GSE14769\_UNSTIM\_VS\_40MIN\_LPS\_BMDM\_DN | 176 | 28 | 12187 | 179 | Socs3,Atf3,Ptgs2,Zfp36,Errfi1,Tnf,Birc3,Sdc4,Nfkbia,Cflar,Slfn4,Sgk1,Nfkbiz,Tiparp,Ier3,Cd14,Slfn3,Map3k8,Maff,Ccl4,Ccl2,Gpr84,Usp18,Ccl12,Tnfaip3,Gadd45b,Nfkbie,Cxcl1 | | 2.217e-21 | -47.56 | GSE34006\_A2AR\_KO\_VS\_A2AR\_AGONIST\_TREATED\_TREG\_UP | MSigDB lists | GSE34006\_A2AR\_KO\_VS\_A2AR\_AGONIST\_TREATED\_TREG\_UP | 176 | 28 | 12187 | 179 | Slfn9,Isg15,Tap1,Parp9,Slfn3,Usp18,Ifit2,Ripk1,Gbp7,Gbp3,Rtp4,Cmpk2,Ifit3b,Ifi44,Znfx1,Irf9,Trim25,Parp14,Tor3a,Lgals9,Parp12,Oasl1,Gbp4,Irf7,Gbp6,Irgm2,Tnfsf10,Slfn4 | | 2.560e-21 | -47.41 | GO\_INNATE\_IMMUNE\_RESPONSE | MSigDB lists | GO\_INNATE\_IMMUNE\_RESPONSE | 336 | 36 | 12187 | 179 | Xaf1,Casp4,Gbp4,Oasl1,Irf7,S100a9,Trim56,Irgm2,Gbp6,Nfkb2,Vcam1,Ifit3b,Ifi209,Ifi204,Ifi207,Ncf1,Ifitm2,Saa1,Irf9,Trim25,Icam1,Ccl4,Ccl2,Ifit2,Ccl12,Cybb,Ifi211,S100a8,Lcn2,Nfkb1,Cd14,Ifitm3,Isg15,Irf2,Cxcl16,Ip6k2 | | 2.873e-21 | -47.30 | GSE19888\_CTRL\_VS\_TCELL\_MEMBRANES\_ACT\_MAST\_CELL\_PRETREAT\_A3R\_INH\_DN | MSigDB lists | GSE19888\_CTRL\_VS\_TCELL\_MEMBRANES\_ACT\_MAST\_CELL\_PRETREAT\_A3R\_INH\_DN | 161 | 27 | 12187 | 179 | Irf7,Parp14,Tor3a,Tubb6,AA467197,Trim25,Irf9,Clic4,Vcam1,Samd9l,Dtx3l,Gbp7,Bcl3,Pik3r5,Rtp4,Cmpk2,Rgs16,Ifit2,Usp18,Il6,Ccl12,Ifitm3,Slfn5,Slfn9,Pik3ap1,Parp9,Nfkb1 | | 2.873e-21 | -47.30 | GSE42021\_CD24HI\_VS\_CD24INT\_TREG\_THYMUS\_DN | MSigDB lists | GSE42021\_CD24HI\_VS\_CD24INT\_TREG\_THYMUS\_DN | 161 | 27 | 12187 | 179 | Ifitm2,Plaur,Irf9,Trim25,Zfp36,Ifit3b,Ifi209,Ifi44,Ifi204,Ifi207,Irf7,Tnfsf10,Nfkbia,Parp12,Oasl1,Bach1,Herc6,Isg15,Tap1,Phf11b,Cxcl1,Ifi211,Phf11d,Rtp4,Tgm2,Usp18,Ifit2 | | 2.961e-21 | -47.27 | cellular response to organic substance | biological process | GO:0071310 | 1457 | 74 | 13711 | 214 | Ptpn1,Ifitm2,Gbp3,Ccl12,Cxcl16,Tnip3,Iqgap1,Col4a1,Vcam1,Irgm2,Gbp6,Cxcl9,Tnfaip3,Ccl2,Ifi47,Csf3,Arid5b,Il1rn,Irak3,Ifi204,Birc3,Osmr,Gbp4,F830016B08Rik,Nfkb1,Irgm1,Il17ra,Icam1,Iigp1,Gm4841,Ifitm3,P2ry6,Hspa5,Cxcl1,Nfe2l2,Ifit2,Gbp7,Csf2rb,Selp,Msn,Tnf,Acod1,Atf3,Zfp36,Cd14,Ripk1,Hspb1,Ifi207,Fam107a,Fzd4,Tiparp,Ifi209,Sbno2,Tgtp1,Cmpk2,Ccl4,Spi1,Stx11,Ifitm6,Ifi211,Trim25,Sgk1,Runx1,Il1a,Nfkbia,Casp4,Il4ra,Tgtp2,Il6,Gm5431,Ccl7,Irf7,Socs3,Pik3r1 | | 4.502e-21 | -46.85 | GSE21546\_UNSTIM\_VS\_ANTI\_CD3\_STIM\_SAP1A\_KO\_AND\_ELK1\_KO\_DP\_THYMOCYTES\_UP | MSigDB lists | GSE21546\_UNSTIM\_VS\_ANTI\_CD3\_STIM\_SAP1A\_KO\_AND\_ELK1\_KO\_DP\_THYMOCYTES\_UP | 118 | 24 | 12187 | 179 | Oasl1,Parp12,Parp14,Usp18,Cmpk2,Phf11d,Ifi211,Irf7,C5ar1,Dtx3l,Ifi204,Ifi207,Ifit3b,Ifi209,Phf11b,Parp9,Irf9,Trim25,P2ry6,Ifitm3,Herc6,Ifitm2,Rnf213,Isg15 | | 4.817e-21 | -46.78 | GSE9988\_ANTI\_TREM1\_VS\_LPS\_MONOCYTE\_DN | MSigDB lists | GSE9988\_ANTI\_TREM1\_VS\_LPS\_MONOCYTE\_DN | 148 | 26 | 12187 | 179 | Socs3,Stx11,Tnip3,Ptgs2,Clic4,Icam1,Sdc4,Nfkbia,Birc3,Tnf,Oasl1,Cflar,Nfkbiz,Ier3,Il1a,Nfkb1,Pik3ap1,Ccl4,Map3k8,Rhou,Il6,Nfe2l2,Tnfaip3,Cxcl1,Csf3,Tnfaip8 | | 5.074e-21 | -46.73 | regulation of response to external stimulus | biological process | GO:0032101 | 697 | 51 | 13711 | 214 | Tap1,S100a8,A2m,Tnip3,Ccl12,Nfkbiz,Irgm2,Usp18,S100a9,Ccl2,Tnfaip3,Plek,Ifi204,Irak3,C3ar1,Nfkb1,Gbp4,Il17ra,Irgm1,Lgals9,Thbs1,Cxcl1,C5ar1,Tgm2,Selp,Nfe2l2,Mmp8,Zfp36,Acod1,Tnf,Pik3ap1,Tnip1,Trim30a,Hspb1,Cd14,Parp9,Sbno2,Ifi209,Ptgs2,Ier3,Ifi211,Ccl4,Il6,Cdkn1a,Nfkbia,Dtx3l,Irf7,Socs3,Ctla2a,Parp14,Trib1 | | 9.215e-21 | -46.13 | GSE26890\_CXCR1\_NEG\_VS\_POS\_EFFECTOR\_CD8\_TCELL\_UP | MSigDB lists | GSE26890\_CXCR1\_NEG\_VS\_POS\_EFFECTOR\_CD8\_TCELL\_UP | 168 | 27 | 12187 | 179 | Ifit2,Usp18,Cybb,Dtx3l,Gbp7,Cmpk2,Rtp4,Ifitm3,Rnf213,Isg15,Herc6,Slfn5,Slfn9,Parp9,Parp14,Tor3a,Xaf1,Gbp4,Parp12,Oasl1,Gbp6,Irf7,Xdh,Tnfsf10,Ifi44,Samd9l,Irf9 | | 1.279e-20 | -45.81 | TAKEDA\_TARGETS\_OF\_NUP98\_HOXA9\_FUSION\_3D\_UP | MSigDB lists | TAKEDA\_TARGETS\_OF\_NUP98\_HOXA9\_FUSION\_3D\_UP | 123 | 24 | 12187 | 179 | Ifi204,Samd9l,Ifi207,Ifi209,Ifi44,Ifit3b,Trim25,Irf9,Ptgs2,Oasl1,Parp12,Xaf1,Parp14,Irf7,Parp9,Isg15,Herc6,Rnf213,Ifit2,Usp18,Cmpk2,Dtx3l,Ifi211,Arid5b | | 2.285e-20 | -45.23 | GSE42021\_CD24INT\_VS\_CD24LOW\_TREG\_THYMUS\_DN | MSigDB lists | GSE42021\_CD24INT\_VS\_CD24LOW\_TREG\_THYMUS\_DN | 141 | 25 | 12187 | 179 | Ifitm3,Isg15,Herc6,Phf11b,Thbs1,Mmp8,Phf11d,Ifi211,Rtp4,Ifit2,Usp18,Ifitm2,Ifi209,Ifi44,Ifit3b,Ifi204,Ifi207,Cflar,Robo4,Irf7,Tnfsf10,Xaf1,Nfkbia,Parp12,Oasl1 | | 3.063e-20 | -44.93 | GO\_RESPONSE\_TO\_EXTERNAL\_STIMULUS | MSigDB lists | GO\_RESPONSE\_TO\_EXTERNAL\_STIMULUS | 1251 | 64 | 12187 | 179 | Hspa5,Tnfaip3,Ifit2,Il4ra,Ccl2,S100a8,Cxcl1,Ifi211,Bcl3,Ifi204,Hp,Cdkn1a,Ifi209,Fstl1,Trim25,Icam1,Irf9,Saa1,Oasl1,Trib1,Irf7,S100a9,Nfkb1,Il17ra,Irak3,Cxcl16,Ifitm3,Selp,Isg15,Cd14,Ccl12,Il6,Nfe2l2,Ccl4,Cmpk2,Sbno2,Csf3,Ptges,Rrp8,Tnip3,Kcna5,Cp,Ifi207,Atf3,Adamts1,Ifi44,Vcam1,Socs3,Ifit3b,Hspb1,Sele,Zfp36,Plaur,Ptgs2,Ifitm2,Gbp4,Tnf,Zfp189,Nfkbia,Nfkb2,C5ar1,C3ar1,Gbp6,Trim56 | | 3.620e-20 | -44.77 | REACTOME\_CYTOKINE\_SIGNALING\_IN\_IMMUNE\_SYSTEM | MSigDB lists | REACTOME\_CYTOKINE\_SIGNALING\_IN\_IMMUNE\_SYSTEM | 213 | 29 | 12187 | 179 | Ifitm2,Csf2rb,Irf9,Trim25,Icam1,Vcam1,Socs3,Ifit3b,Il1rn,Irf7,Gbp6,Nfkb2,Xaf1,Gbp4,Oasl1,Ifitm3,Isg15,Irf2,Ptpn1,Irak3,Ip6k2,Il1a,Gbp7,Gbp3,Usp18,Map3k8,Ifit2,Pik3r1,Il6 | | 3.803e-20 | -44.72 | ZWANG\_CLASS\_3\_TRANSIENTLY\_INDUCED\_BY\_EGF | MSigDB lists | ZWANG\_CLASS\_3\_TRANSIENTLY\_INDUCED\_BY\_EGF | 177 | 27 | 12187 | 179 | Tnfaip3,Il6,Map3k8,Maff,Sbno2,Pygm,Adamts9,Cxcl1,Bcl3,Cebpd,Ier3,Nfkb1,Tiparp,Il1a,Nfkbiz,Trib1,Runx1,Birc3,Nfkb2,Gem,Atf3,Il1rn,Stx11,Socs3,Aff1,Plaur,Ptgs2 | | 3.912e-20 | -44.69 | cellular response to interferon-beta | biological process | GO:0035458 | 36 | 16 | 13711 | 214 | Irgm1,Ifi211,Gbp6,Irgm2,Ifi209,Gbp3,Tgtp1,F830016B08Rik,Gm4841,Ifi207,Iigp1,Ifi47,Ifi204,Acod1,Gm5431,Tgtp2 | | 4.428e-20 | -44.56 | GSE42021\_TCONV\_PLN\_VS\_TREG\_PRECURSORS\_THYMUS\_DN | MSigDB lists | GSE42021\_TCONV\_PLN\_VS\_TREG\_PRECURSORS\_THYMUS\_DN | 178 | 27 | 12187 | 179 | Gbp7,Ptges,Angptl4,Cxcl1,Bcl3,Nfkb2,Map3k8,Ccnd2,Il4ra,Msr1,Birc3,Ccl2,Selp,Ptgs2,Cd14,Icam1,Errfi1,Plaur,Cdkn1a,Hspa5,Socs3,Nfkbiz,Il1rn,Nfkb1,Il1a,Cebpd,Lcn2 | | 5.198e-20 | -44.40 | MARKEY\_RB1\_ACUTE\_LOF\_DN | MSigDB lists | MARKEY\_RB1\_ACUTE\_LOF\_DN | 197 | 28 | 12187 | 179 | C5ar1,C3ar1,Irf7,Slfn4,Xdh,Tor3a,Casp4,Saa1,Irf9,Ifi209,Socs3,Ifit3b,Ifi204,Ifi207,Hp,Znfx1,Gbp7,Ifi211,Cxcl1,Iqgap1,Cmpk2,Gbp3,Ifit2,Ccl4,Usp18,Isg15,Slfn3,Tap1 | | 6.950e-20 | -44.11 | OSWALD\_HEMATOPOIETIC\_STEM\_CELL\_IN\_COLLAGEN\_GEL\_UP | MSigDB lists | OSWALD\_HEMATOPOIETIC\_STEM\_CELL\_IN\_COLLAGEN\_GEL\_UP | 181 | 27 | 12187 | 179 | Cflar,C3ar1,Trib1,Nfkbia,Runx1,Tnf,Birc3,Icam1,Zfp36,Ptgs2,Luc7l3,Atf3,Il1rn,Cdkn1a,Cxcl1,Ccl12,Tnfaip3,Gadd45b,Il6,Rassf4,Ccl4,Maff,Cd14,Ier3,Tiparp,Il1a,Sgk1 | | 7.884e-20 | -43.99 | GSE13485\_DAY1\_VS\_DAY7\_YF17D\_VACCINE\_PBMC\_DN | MSigDB lists | GSE13485\_DAY1\_VS\_DAY7\_YF17D\_VACCINE\_PBMC\_DN | 148 | 25 | 12187 | 179 | C3ar1,Irf7,Tnfsf10,Parp14,Xaf1,Parp12,Oasl1,Ifi44,Ifit3b,Il1rn,Samd9l,Dtx3l,Phf11d,Rtp4,Cmpk2,Gbp3,Ifit2,Usp18,Ifitm3,Herc6,Isg15,Pik3ap1,Parp9,Tap1,Phf11b | | 8.941e-20 | -43.86 | GSE9988\_ANTI\_TREM1\_VS\_ANTI\_TREM1\_AND\_LPS\_MONOCYTE\_DN | MSigDB lists | GSE9988\_ANTI\_TREM1\_VS\_ANTI\_TREM1\_AND\_LPS\_MONOCYTE\_DN | 133 | 24 | 12187 | 179 | Tnfaip8,Csf3,Cxcl1,Il6,Tnfaip3,Map3k8,Gpr84,Ccl4,Il1a,Ier3,Nfkbiz,Cflar,Birc3,Tnf,Sdc4,Nfkbia,Plek,Zfp36,Icam1,Plaur,Clic4,Ptgs2,Tnip3,Socs3 | | 9.279e-20 | -43.82 | response to virus | biological process | GO:0009615 | 171 | 27 | 13711 | 214 | Lcn2,Trim30a,Irak3,Cxcl9,Tnf,Irgm2,Ifit2,Bcl3,Ifit3b,Irf2,Ifitm2,Lgals9,Mx1,Dtx3l,Ifitm3,Irf7,Oasl1,Il6,Oasl2,Slfn9,Trim25,Trim56,Mx2,Rtp4,Ifitm6,Parp9,Isg15 | | 9.349e-20 | -43.82 | MODULE\_75 | MSigDB lists | MODULE\_75 | 240 | 30 | 12187 | 179 | Nfkbia,Oasl1,Gem,C3ar1,C5ar1,Irf7,Igsf6,Tnfsf10,Ifi44,Cdkn1a,Hp,Timp1,Ifitm2,Saa1,Csf2rb,Il4ra,Map3k8,Ccl2,Ccl4,Cybb,Il6,Ccl12,Csf3,Cxcl1,Osmr,Il1a,Isg15,Ifitm3,Selp,Cd14 | | 1.382e-19 | -43.43 | FULCHER\_INFLAMMATORY\_RESPONSE\_LECTIN\_VS\_LPS\_DN | MSigDB lists | FULCHER\_INFLAMMATORY\_RESPONSE\_LECTIN\_VS\_LPS\_DN | 354 | 35 | 12187 | 179 | Sgk1,Hspa5,Tiparp,Ms4a6d,Cebpd,Ifitm3,Herc6,Rnf213,Slfn5,Cd14,Pik3ap1,Map3k8,Ifit2,Usp18,Gadd45b,Ifi211,Arid5b,Cmpk2,Ifi209,Aff1,Ifi44,Socs3,Ifit3b,Ifi204,Ifi207,Samd9l,Ifitm2,Zfp36,Rbm47,Xaf1,Bach1,Oasl1,C3ar1,Olfml2b,Tnfsf10 | | 1.592e-19 | -43.28 | GSE8384\_CTRL\_VS\_B\_ABORTUS\_4H\_MAC\_CELL\_LINE\_DN | MSigDB lists | GSE8384\_CTRL\_VS\_B\_ABORTUS\_4H\_MAC\_CELL\_LINE\_DN | 169 | 26 | 12187 | 179 | Slfn3,Nfkbiz,Il1a,Ier3,Csf3,Bcl3,Sntb2,Ccl2,Ccl4,Cybb,Gadd45b,Ptgs2,Tnip1,Zfp36,Msn,Plaur,Cdkn1a,Aff1,Socs3,Il1rn,Znfx1,C5ar1,Slfn4,Nfkb2,Tnf,Nfkbia | | 1.761e-19 | -43.18 | GO\_CELLULAR\_RESPONSE\_TO\_CYTOKINE\_STIMULUS | MSigDB lists | GO\_CELLULAR\_RESPONSE\_TO\_CYTOKINE\_STIMULUS | 406 | 37 | 12187 | 179 | Csf3,Cxcl1,Osmr,Sbno2,Il4ra,Ifit2,Ccl2,Ccl4,Nfe2l2,Il6,Ccl12,Ripk1,Isg15,Ifitm3,Ip6k2,Il17ra,Irak3,Irf2,Hspa5,Il1a,Nfkb1,Gbp6,Irf7,Birc3,Tnf,Nfkbia,Xaf1,Oasl1,Gbp4,Ifitm2,Icam1,Trim25,Irf9,Csf2rb,Ifit3b,Socs3,Vcam1 | | 1.808e-19 | -43.16 | NUYTTEN\_EZH2\_TARGETS\_UP | MSigDB lists | NUYTTEN\_EZH2\_TARGETS\_UP | 853 | 52 | 12187 | 179 | Ifi44,Ifi207,Atf3,Ptgs2,Errfi1,Birc3,Xaf1,Nfkbia,Parp12,Gem,Robo4,Phf11b,Thbs1,Herc6,Parp9,Tap1,Tgm2,Rhoc,Il6,Nfkbie,Angptl4,Phf11d,Rtp4,Gbp3,Cdkn1a,Ifi209,Ifi204,Samd9l,Znfx1,Saa1,Trim25,Icam1,Clic4,Parp14,Bach1,AA467197,Oasl1,Cflar,Tnfsf10,Hspa5,Sgk1,Slfn5,Ptpn1,Irf2,Rhou,Usp18,Tnfaip3,Dtx3l,Cxcl1,Ifi211,Arid5b,Tnfaip8 | | 1.856e-19 | -43.13 | GSE14000\_UNSTIM\_VS\_4H\_LPS\_DC\_TRANSLATED\_RNA\_DN | MSigDB lists | GSE14000\_UNSTIM\_VS\_4H\_LPS\_DC\_TRANSLATED\_RNA\_DN | 137 | 24 | 12187 | 179 | Tnfsf10,Trim56,Irf7,Cflar,Oasl1,Xaf1,Birc3,Errfi1,Atf3,Stx11,Socs3,Ifit3b,Ifi44,Rtp4,Cmpk2,Dtx3l,Ccl4,Usp18,Ifit2,Parp9,Slfn5,Herc6,Isg15,Nfkbiz | | 1.856e-19 | -43.13 | GSE9988\_LPS\_VS\_VEHICLE\_TREATED\_MONOCYTE\_UP | MSigDB lists | GSE9988\_LPS\_VS\_VEHICLE\_TREATED\_MONOCYTE\_UP | 137 | 24 | 12187 | 179 | Nfkbiz,Nfkb1,Il1a,Ier3,Pik3ap1,Map3k8,Maff,Ccl4,Gpr84,Tnfaip3,Gadd45b,Il6,Cxcl1,Socs3,Tnip3,Ptgs2,Icam1,Plek,Plaur,Tnf,Birc3,Nfkbia,Cflar,Gem | | 2.545e-19 | -42.82 | GSE2770\_TGFB\_AND\_IL4\_ACT\_VS\_ACT\_CD4\_TCELL\_2H\_DN | MSigDB lists | GSE2770\_TGFB\_AND\_IL4\_ACT\_VS\_ACT\_CD4\_TCELL\_2H\_DN | 155 | 25 | 12187 | 179 | Tnfsf10,Irf7,Trim56,Parp12,Oasl1,Xaf1,Trim25,Irf9,Ifitm2,Ifi204,Ifi207,Samd9l,Znfx1,Ifi209,Ifi44,Dtx3l,Phf11d,Ifi211,Ifit2,Irf2,Parp9,Ifitm3,Isg15,Tiparp,Phf11b | | 2.926e-19 | -42.68 | GSE34156\_NOD2\_LIGAND\_VS\_TLR1\_TLR2\_LIGAND\_6H\_TREATED\_MONOCYTE\_UP | MSigDB lists | GSE34156\_NOD2\_LIGAND\_VS\_TLR1\_TLR2\_LIGAND\_6H\_TREATED\_MONOCYTE\_UP | 173 | 26 | 12187 | 179 | Irf7,Irgm2,Xdh,Tnfsf10,Slfn4,Tor3a,Casp4,Lgals9,Clic4,Irf9,Trim25,Zfp36,Ifit3b,Znfx1,Gbp7,Gbp3,Tnfaip8,Osmr,Usp18,Ifit2,Il6,Ifitm3,Isg15,Tap1,Slfn3,Gadd45g | | 3.942e-19 | -42.38 | GSE19198\_1H\_VS\_6H\_IL21\_TREATED\_TCELL\_DN | MSigDB lists | GSE19198\_1H\_VS\_6H\_IL21\_TREATED\_TCELL\_DN | 175 | 26 | 12187 | 179 | Tiparp,Akap12,Il1a,Thbs1,Tap1,Ifitm3,Map3k6,Il6,Hcar2,Ccl4,Ccnd2,Maff,Ptges,Rhoj,Timp1,Hp,Socs3,Ifit3b,Cdkn1a,Clic4,Irf9,Tubb6,Bach1,Parp14,Casp4,Irf7 | | 3.966e-19 | -42.37 | regulation of defense response | biological process | GO:0031347 | 460 | 40 | 13711 | 214 | Tap1,S100a8,A2m,Tnip3,Cxcl1,C5ar1,Tgm2,Nfkbiz,Irgm2,S100a9,Mmp8,Usp18,Zfp36,Tnf,Tnfaip3,Acod1,Pik3ap1,Tnip1,Ifi204,Trim30a,Irak3,Cd14,Nfkb1,Parp9,Sbno2,Ifi209,Gbp4,Ptgs2,Il17ra,Ier3,Ifi211,Irgm1,Il6,Nfkbia,Lgals9,Dtx3l,Irf7,Socs3,Ctla2a,Parp14 | | 4.836e-19 | -42.17 | GSE37533\_PPARG1\_FOXP3\_VS\_PPARG2\_FOXP3\_TRANSDUCED\_CD4\_TCELL\_PIOGLITAZONE\_TREATED\_DN | MSigDB lists | GSE37533\_PPARG1\_FOXP3\_VS\_PPARG2\_FOXP3\_TRANSDUCED\_CD4\_TCELL\_PIOGLITAZONE\_TREATED\_DN | 159 | 25 | 12187 | 179 | Trib1,Xaf1,Parp12,Bach1,Ifit3b,Socs3,Ifi44,Aff1,Atf3,Irf9,Sntb2,Usp18,Ifit2,Ripk1,Fam107a,Bcl3,Phf11d,Sbno2,Rtp4,Dnajb1,Phf11b,Herc6,Isg15,Tap1,Irf2 | | 5.302e-19 | -42.08 | GSE42724\_NAIVE\_BCELL\_VS\_PLASMABLAST\_UP | MSigDB lists | GSE42724\_NAIVE\_BCELL\_VS\_PLASMABLAST\_UP | 143 | 24 | 12187 | 179 | Dtx3l,Irf7,Rtp4,Cmpk2,Tnfsf10,Gbp3,Ifit2,Usp18,Parp14,Xaf1,Oasl1,Parp12,Ripk1,AA467197,Herc6,Isg15,Trim25,Tap1,Irf9,Parp9,Ifi44,Ifit3b,Samd9l,Znfx1 | | 7.437e-19 | -41.74 | GSE14000\_UNSTIM\_VS\_4H\_LPS\_DC\_DN | MSigDB lists | GSE14000\_UNSTIM\_VS\_4H\_LPS\_DC\_DN | 145 | 24 | 12187 | 179 | Tnfsf10,Irf7,Cflar,Oasl1,Nfkbia,Xaf1,Parp14,Zfp189,Irf9,Plek,Atf3,Znfx1,Ifit3b,Ifi44,Rtp4,Cmpk2,Dtx3l,Usp18,Ccl4,Ifit2,Parp9,Irf2,Isg15,Nfkbiz | | 1.273e-18 | -41.20 | regulation of immune system process | biological process | GO:0002682 | 953 | 56 | 13711 | 214 | Irgm2,Vcam1,Ccl12,A2m,Tnip3,Tap1,Nfkbiz,Ifi204,Zbtb16,Csf3,C3ar1,Irak3,Ccl2,Tnfaip3,Irgm1,Gbp4,Lgals9,Thbs1,Sox11,Fyb,Icam1,Selp,Msn,Mmp8,Nfe2l2,Cxcl1,C5ar1,Trim30a,Tnip1,Cd14,Ripk1,Sele,Zfp36,Pik3ap1,Tnf,Acod1,Runx1,Spi1,Ccl4,Ifi211,Isg15,Sdc4,Parp9,Ifi209,Hcar2,Pik3r1,Ctla2a,Trib1,Parp14,Irf7,Dtx3l,Cdkn1a,Il6,Il1a,Nfkbia,Il4ra | | 2.119e-18 | -40.70 | GSE9988\_LPS\_VS\_CTRL\_TREATED\_MONOCYTE\_UP | MSigDB lists | GSE9988\_LPS\_VS\_CTRL\_TREATED\_MONOCYTE\_UP | 135 | 23 | 12187 | 179 | Il1a,Ier3,Nfkbiz,Il6,Gadd45b,Tnfaip3,Maff,Map3k8,Gpr84,Ccl4,Csf3,Cxcl1,Tnip3,Cdkn1a,Socs3,Plek,Icam1,Plaur,Ptgs2,Birc3,Tnf,Nfkbia,Cflar | | 2.332e-18 | -40.60 | GSE42021\_TREG\_PLN\_VS\_TREG\_PRECURSORS\_THYMUS\_DN | MSigDB lists | GSE42021\_TREG\_PLN\_VS\_TREG\_PRECURSORS\_THYMUS\_DN | 152 | 24 | 12187 | 179 | Ifitm2,Irf9,Trim25,Ifit3b,Ifi44,Ifi209,Ifi207,Ifi204,Irf7,Tnfsf10,Xaf1,Parp12,Oasl1,Isg15,Herc6,Ifitm3,Tap1,Phf11b,Ifi211,Phf11d,Osmr,Usp18,Maff,Ifit2 | | 2.453e-18 | -40.55 | GRAESSMANN\_APOPTOSIS\_BY\_SERUM\_DEPRIVATION\_UP | MSigDB lists | GRAESSMANN\_APOPTOSIS\_BY\_SERUM\_DEPRIVATION\_UP | 439 | 37 | 12187 | 179 | Cebpd,Sgk1,Slfn3,Cxcl16,Irf2,Parp9,Isg15,Slfn9,Gadd45b,Ccl12,Ifit2,Usp18,Osmr,Cmpk2,Rtp4,Gbp7,Cxcl1,Ifi211,Cp,Ifi207,Ifi204,Ifi44,Ifi209,Ifit3b,Fstl1,Trim25,Hspb1,Irf9,Saa1,Parp12,Oasl1,Runx1,Tor3a,Slfn4,Xdh,Irgm2,Irf7 | | 2.732e-18 | -40.44 | GSE22886\_CTRL\_VS\_LPS\_24H\_DC\_DN | MSigDB lists | GSE22886\_CTRL\_VS\_LPS\_24H\_DC\_DN | 153 | 24 | 12187 | 179 | Ifitm3,Herc6,Ifitm2,Isg15,Tnip1,Icam1,Tap1,Irf9,Phf11b,Ifi44,Ifit3b,Nfkb1,Phf11d,Irf7,Tnfaip8,Rtp4,Nfkb2,Tnfsf10,Ifit2,Birc3,Xaf1,Usp18,Tnfaip3,Oasl1 | | 5.092e-18 | -39.82 | MODULE\_46 | MSigDB lists | MODULE\_46 | 233 | 28 | 12187 | 179 | Osmr,Cxcl1,Csf3,Ccl12,Il6,Cybb,Ccl4,Ccl2,Il4ra,Cd14,Selp,Ifitm3,Isg15,Il1a,Tnfsf10,Igsf6,Irf7,C5ar1,Gem,C3ar1,Oasl1,Nfkbia,Csf2rb,Ifitm2,Saa1,Timp1,Hp,Ifi44 | | 6.436e-18 | -39.58 | inflammatory response | biological process | GO:0006954 | 302 | 32 | 13711 | 214 | Il1a,Casp4,Icam1,Il4ra,Il6,Cybb,Ccl7,Thbs1,Timp1,Ccl4,Ptgs2,Cxcl9,Tnf,Acod1,Tnfaip3,Ccl2,Saa1,Cd14,Sele,Il1rn,C3ar1,Ncf1,Tnip1,Nfkbiz,Cxcl1,C5ar1,Ccl12,S100a8,Hp,S100a9,Nfe2l2,Selp | | 6.994e-18 | -39.50 | GSE42021\_TREG\_PLN\_VS\_CD24INT\_TREG\_THYMUS\_UP | MSigDB lists | GSE42021\_TREG\_PLN\_VS\_CD24INT\_TREG\_THYMUS\_UP | 177 | 25 | 12187 | 179 | Socs3,Cdkn1a,Il1rn,Ptgs2,Errfi1,Plaur,Icam1,Birc3,Nfkbiz,Cebpd,Lcn2,Il1a,Nfkb1,Cd14,Selp,Sntb2,Ccl2,Il4ra,Msr1,Map3k8,Ccnd2,Gadd45b,Cxcl1,Ptges,Gbp7 | | 7.200e-18 | -39.47 | GO\_RESPONSE\_TO\_BACTERIUM | MSigDB lists | GO\_RESPONSE\_TO\_BACTERIUM | 302 | 31 | 12187 | 179 | Nfkb1,Isg15,Selp,Cd14,Irak3,Cxcl16,Il6,Tnfaip3,Ccl12,Csf3,Bcl3,Ptges,Cxcl1,Cmpk2,S100a8,Sbno2,Vcam1,Socs3,Hp,Tnip3,Ptgs2,Sele,Icam1,Tnf,Trib1,Nfkbia,Gbp4,Gbp6,C5ar1,S100a9,Nfkb2 | | 8.611e-18 | -39.29 | regulation of response to stress | biological process | GO:0080134 | 1063 | 58 | 13711 | 214 | Tnfaip3,Plek,Xdh,Ccl2,Il1rn,Irak3,Ifi204,Nfkbiz,Ptpn1,S100a8,Tap1,A2m,Tnip3,S100a9,Usp18,Irgm2,Errfi1,Thbs1,Gadd45g,Lgals9,Gbp4,Nfkb1,Irgm1,Il17ra,Acod1,Tnf,Pik3ap1,Zfp36,Ncf1,Hspb1,Ripk1,Cd14,Tnip1,Trim30a,Cxcl1,C5ar1,Tgm2,Nfe2l2,Mmp8,Selp,Nfkbia,Il1a,Il6,Cdkn1a,Dtx3l,Socs3,Irf7,Parp14,Ctla2a,Gadd45b,Pik3r1,Ifi209,Fzd4,Parp9,Sbno2,Ifi211,Ptgs2,Ier3 | | 8.819e-18 | -39.27 | GSE36891\_POLYIC\_TLR3\_VS\_PAM\_TLR2\_STIM\_PERITONEAL\_MACROPHAGE\_UP | MSigDB lists | GSE36891\_POLYIC\_TLR3\_VS\_PAM\_TLR2\_STIM\_PERITONEAL\_MACROPHAGE\_UP | 112 | 21 | 12187 | 179 | Maff,Birc3,Rhou,Sdc4,Trib1,Il6,Ccl12,Tnfaip3,Gem,Cxcl1,Sgk1,Adamts1,Nfkbiz,Il1a,Akap12,Atf3,Ier3,Cebpd,Ptgs2,Zfp36,Apold1 | | 1.414e-17 | -38.80 | GO\_CYTOKINE\_MEDIATED\_SIGNALING\_PATHWAY | MSigDB lists | GO\_CYTOKINE\_MEDIATED\_SIGNALING\_PATHWAY | 286 | 30 | 12187 | 179 | Isg15,Ifitm3,Ip6k2,Irak3,Il17ra,Irf2,Il1a,Csf3,Cxcl1,Osmr,Il4ra,Ifit2,Ccl2,Ccl4,Il6,Ripk1,Ccl12,Ifitm2,Trim25,Icam1,Irf9,Csf2rb,Ifit3b,Vcam1,Socs3,Irf7,Birc3,Tnf,Xaf1,Oasl1 | | 1.536e-17 | -38.71 | GSE13485\_CTRL\_VS\_DAY3\_YF17D\_VACCINE\_PBMC\_DN | MSigDB lists | GSE13485\_CTRL\_VS\_DAY3\_YF17D\_VACCINE\_PBMC\_DN | 147 | 23 | 12187 | 179 | C3ar1,Irf7,Oasl1,Parp12,Xaf1,Parp14,Ifitm2,Samd9l,Ifi44,Ifit3b,Cmpk2,Rtp4,Dtx3l,Phf11d,Ifit2,Usp18,Pik3ap1,Parp9,Tap1,Ifitm3,Herc6,Isg15,Phf11b | | 2.221e-17 | -38.35 | GSE3982\_CTRL\_VS\_LPS\_4H\_MAC\_DN | MSigDB lists | GSE3982\_CTRL\_VS\_LPS\_4H\_MAC\_DN | 167 | 24 | 12187 | 179 | Oasl1,Tubb6,Bach1,Nfkb2,Cflar,Il1rn,Ifi204,Ifi207,Ifi209,Ifi44,Clic4,Trim25,Plek,Tnfaip3,Gadd45b,Usp18,Ifit2,Maff,Ifi211,Pik3r5,Arid5b,Cebpd,Tiparp,Irf2 | | 2.559e-17 | -38.20 | GSE14769\_UNSTIM\_VS\_80MIN\_LPS\_BMDM\_DN | MSigDB lists | GSE14769\_UNSTIM\_VS\_80MIN\_LPS\_BMDM\_DN | 168 | 24 | 12187 | 179 | Nfkbie,Usp18,Gpr84,Ccl2,Tgm2,Ccl4,Maff,Map3k8,Gadd45b,Isg15,Slfn3,Tiparp,Cflar,Rbm39,Slfn4,Birc3,Plaur,Plek,Sele,Icam1,Adamts1,Atf3,Stx11,Il1rn | | 2.603e-17 | -38.19 | REACTOME\_IMMUNE\_SYSTEM | MSigDB lists | REACTOME\_IMMUNE\_SYSTEM | 682 | 44 | 12187 | 179 | Tap1,Irf2,Ptpn1,Ip6k2,Pik3ap1,Irak3,Cd14,Isg15,Ifitm3,Il1a,Hspa5,Gbp3,Gbp7,Dtx3l,Nfkbie,Il6,Tnfaip3,Ripk1,Cybb,Usp18,Pik3r1,Map3k8,Ifit2,Irf9,Csf2rb,Trim25,Icam1,Saa1,Ifitm2,Il1rn,Ifit3b,Zbtb16,Socs3,Vcam1,Cdkn1a,Nfkb2,Irf7,Gbp6,Oasl1,Gbp4,Casp4,Nfkbia,Xaf1,Birc3 | | 3.271e-17 | -37.96 | REACTOME\_INTERFERON\_SIGNALING | MSigDB lists | REACTOME\_INTERFERON\_SIGNALING | 119 | 21 | 12187 | 179 | Ifitm2,Isg15,Ifitm3,Ip6k2,Trim25,Icam1,Irf9,Ptpn1,Irf2,Ifit3b,Vcam1,Socs3,Gbp7,Gbp6,Irf7,Gbp3,Ifit2,Usp18,Xaf1,Oasl1,Gbp4 | | 3.388e-17 | -37.92 | GSE14769\_UNSTIM\_VS\_60MIN\_LPS\_BMDM\_DN | MSigDB lists | GSE14769\_UNSTIM\_VS\_60MIN\_LPS\_BMDM\_DN | 170 | 24 | 12187 | 179 | Nfkbie,Gadd45b,Map3k8,Ccl4,Ccl2,Gpr84,Slfn3,Cd14,Tiparp,Ier3,Sgk1,Slfn4,Rbm39,Cflar,Gem,Birc3,Icam1,Plek,Sele,Plaur,Errfi1,Ptgs2,Stx11,Atf3 | | 3.499e-17 | -37.89 | WALLACE\_PROSTATE\_CANCER\_RACE\_UP | MSigDB lists | WALLACE\_PROSTATE\_CANCER\_RACE\_UP | 189 | 25 | 12187 | 179 | C5ar1,Xaf1,Parp12,Ifitm2,Irf9,Csf2rb,Sele,Plek,Apold1,Ifit3b,Ifi44,Ifi209,Adamts1,Ifi207,Ifi204,Ifi211,Rgs16,Ccl4,Herc6,Isg15,Selp,Ifitm3,Tap1,Cebpd,Gadd45g | | 4.307e-17 | -37.68 | HECKER\_IFNB1\_TARGETS | MSigDB lists | HECKER\_IFNB1\_TARGETS | 55 | 16 | 12187 | 179 | Cmpk2,Rtp4,Dtx3l,C3ar1,Irf7,Ccl12,Parp12,Oasl1,Ifit2,Xaf1,Parp9,Isg15,Herc6,Samd9l,Ifi44,Ifit3b | | 4.477e-17 | -37.64 | GSE43863\_TFH\_VS\_LY6C\_INT\_CXCR5POS\_MEMORY\_CD4\_TCELL\_UP | MSigDB lists | GSE43863\_TFH\_VS\_LY6C\_INT\_CXCR5POS\_MEMORY\_CD4\_TCELL\_UP | 154 | 23 | 12187 | 179 | Socs3,Hspa5,Ifi44,Atf3,Il1rn,Slfn9,Isg15,Ifitm3,Tap1,Icam1,Nfkbia,Usp18,Xaf1,Parp14,Lgals9,Il6,Ripk1,Gbp4,Ptges,Dtx3l,Gbp6,Irgm2,Rtp4 | | 4.948e-17 | -37.54 | GO\_REGULATION\_OF\_RESPONSE\_TO\_STRESS | MSigDB lists | GO\_REGULATION\_OF\_RESPONSE\_TO\_STRESS | 1121 | 56 | 12187 | 179 | Saa1,Plek,Ifi209,Hp,Ifi204,Fzd4,S100a9,Irf7,Casp4,Map3k6,Ptpn1,Hspa5,Ier3,Ifi211,S100a8,Pik3r1,Ccl2,Usp18,Gadd45b,Ripk1,Tnfaip3,Ptgs2,Tnip1,Sele,Hspb1,Plaur,Errfi1,Socs3,Ifi207,Tnip3,C3ar1,C5ar1,A2m,Xdh,Birc3,Zfp189,Tnf,Nfkbia,Selp,Cd14,Pik3ap1,Irak3,Il17ra,Tap1,Parp9,Thbs1,Dnajb1,Il1a,Gadd45g,Nfkb1,Osmr,Sbno2,Ccl4,Il6,Nfe2l2,Ccl12 | | 5.161e-17 | -37.50 | response to molecule of bacterial origin | biological process | GO:0002237 | 198 | 26 | 13711 | 214 | C5ar1,Cxcl1,Ccl12,Cxcl16,Tnip3,Irgm2,Wfdc21,Gbp6,Noct,Cxcl9,Tnfaip3,Tnf,Acod1,Ccl2,Zfp36,Cd14,Irak3,Il1rn,Sbno2,Nfkb1,Cmpk2,Ptgs2,Nfkbia,Il6,Trib1,Lgals9 | | 5.192e-17 | -37.50 | HALLMARK\_INFLAMMATORY\_RESPONSE | MSigDB lists | HALLMARK\_INFLAMMATORY\_RESPONSE | 155 | 23 | 12187 | 179 | Sele,Icam1,Plaur,Timp1,Cdkn1a,Tnfsf10,C3ar1,C5ar1,Irf7,Nfkbia,Cd14,Il1a,Nfkb1,Osmr,Rtp4,Rgs16,Csf3,Pik3r5,Cybb,Il6,Ccl12,Msr1,Il4ra | | 5.343e-17 | -37.47 | GSE18791\_CTRL\_VS\_NEWCASTLE\_VIRUS\_DC\_12H\_DN | MSigDB lists | GSE18791\_CTRL\_VS\_NEWCASTLE\_VIRUS\_DC\_12H\_DN | 138 | 22 | 12187 | 179 | Ifit2,Map3k8,Il6,Dtx3l,Cmpk2,Rtp4,Gbp3,Il1a,Isg15,Rnf213,Herc6,Slfn5,Tap1,Tnf,Parp14,Xaf1,Irf7,Ifi44,Ifit3b,Samd9l,Znfx1,Trim25 | | 5.870e-17 | -37.37 | GSE27434\_WT\_VS\_DNMT1\_KO\_TREG\_DN | MSigDB lists | GSE27434\_WT\_VS\_DNMT1\_KO\_TREG\_DN | 174 | 24 | 12187 | 179 | Tnf,Birc3,Nfkbia,Casp4,Icam1,Zfp36,Errfi1,Plaur,Ptgs2,Il1rn,Cdkn1a,Socs3,Tnfaip8,Cxcl1,Nfe2l2,Gadd45b,Map3k8,Il4ra,Ccl2,Selp,Cd14,Il1a,Sgk1,Nfkbiz | | 5.957e-17 | -37.36 | TNF signaling pathway | KEGG pathways | ko04668 | 94 | 21 | 5248 | 107 | Ifi47,Map3k8,Ccl2,Pik3r1,Cflar,Nfkbia,Sele,Cxcl1,Ccl12,Tnfaip3,Birc3,Icam1,Tnf,Vcam1,Ripk1,Socs3,Il6,Nfkb1,Gm5431,Ptgs2,Bcl3 | | 5.957e-17 | -37.36 | TNF signaling pathway | KEGG pathways | mmu04668 | 94 | 21 | 5248 | 107 | Vcam1,Ripk1,Birc3,Tnf,Icam1,Nfkb1,Socs3,Il6,Gm5431,Bcl3,Ptgs2,Cflar,Ccl2,Pik3r1,Ifi47,Map3k8,Nfkbia,Sele,Cxcl1,Ccl12,Tnfaip3 | | 8.192e-17 | -37.04 | response to lipopolysaccharide | biological process | GO:0032496 | 183 | 25 | 13711 | 214 | Nfkbia,Il6,Trib1,Lgals9,Nfkb1,Cmpk2,Sbno2,Ptgs2,Tnfaip3,Tnf,Acod1,Cxcl9,Zfp36,Ccl2,Il1rn,Irak3,Cd14,Cxcl1,Tnip3,Ccl12,Cxcl16,Noct,Wfdc21,Irgm2,Gbp6 | | 9.197e-17 | -36.93 | MOSERLE\_IFNA\_RESPONSE | MSigDB lists | MOSERLE\_IFNA\_RESPONSE | 30 | 13 | 12187 | 179 | Ifi204,Ifi207,Samd9l,Ifi209,Ifi44,Ifit3b,Cmpk2,Rtp4,Tnfsf10,Ifi211,Oasl1,Ifit2,Usp18 | | 9.459e-17 | -36.90 | ZHOU\_INFLAMMATORY\_RESPONSE\_LPS\_UP | MSigDB lists | ZHOU\_INFLAMMATORY\_RESPONSE\_LPS\_UP | 260 | 28 | 12187 | 179 | Ccl4,Usp18,Gpr84,Ifit2,Maff,Ripk1,Il6,Cxcl1,Arid5b,Csf3,Rtp4,Ier3,Nfkb1,Slfn5,Isg15,Ptpn1,Sdc4,Tnf,Oasl1,Irf7,Fzd4,Nfkb2,Ifit3b,Ifi44,Znfx1,Ptgs2,Plaur,Icam1 | | 1.562e-16 | -36.40 | GO\_REGULATION\_OF\_RESPONSE\_TO\_WOUNDING | MSigDB lists | GO\_REGULATION\_OF\_RESPONSE\_TO\_WOUNDING | 265 | 28 | 12187 | 179 | C3ar1,C5ar1,A2m,S100a9,Birc3,Tnf,Casp4,Plek,Sele,Plaur,Ptgs2,Saa1,Tnip1,Socs3,S100a8,Osmr,Sbno2,Nfe2l2,Il6,Tnfaip3,Ccl12,Ccl2,Ccl4,Pik3ap1,Il17ra,Selp,Nfkb1,Thbs1 | | 1.634e-16 | -36.35 | GSE37533\_PPARG1\_FOXP3\_VS\_FOXP3\_TRANSDUCED\_CD4\_TCELL\_DN | MSigDB lists | GSE37533\_PPARG1\_FOXP3\_VS\_FOXP3\_TRANSDUCED\_CD4\_TCELL\_DN | 163 | 23 | 12187 | 179 | Phf11b,Tap1,Irf2,Herc6,Isg15,Ripk1,Usp18,Ifit2,Rtp4,Phf11d,Ifi211,Ifi207,Ifi204,Ifit3b,Ifi44,Ifi209,Irf9,Ifitm2,Parp12,Oasl1,Xaf1,Tnfsf10,Irf7 | | 1.775e-16 | -36.27 | GO\_REGULATION\_OF\_CYTOKINE\_PRODUCTION | MSigDB lists | GO\_REGULATION\_OF\_CYTOKINE\_PRODUCTION | 389 | 33 | 12187 | 179 | Cd14,Isg15,Il17ra,Irak3,Thbs1,Il1a,Nfkb1,Bcl3,Ifi211,Iqgap1,Pik3r1,Il4ra,Il6,Tnfaip3,Ripk1,Ccl12,Ptgs2,Saa1,Errfi1,Zfp36,Hspb1,Trim25,Ifi209,Ifi207,Ifi204,Irf7,Trim56,C3ar1,C5ar1,Nfkb2,Birc3,Tnf,Runx1 | | 1.877e-16 | -36.21 | GSE41978\_ID2\_KO\_VS\_BIM\_KO\_KLRG1\_LOW\_EFFECTOR\_CD8\_TCELL\_UP | MSigDB lists | GSE41978\_ID2\_KO\_VS\_BIM\_KO\_KLRG1\_LOW\_EFFECTOR\_CD8\_TCELL\_UP | 164 | 23 | 12187 | 179 | Rtp4,Phf11d,Ifi211,Cxcl1,Arid5b,Ccl12,Maff,Tap1,Ifitm3,Isg15,Cebpd,Phf11b,Irf7,Bach1,Parp12,Oasl1,Icam1,Zfp36,Irf9,Ifi204,Ifi207,Ifi209,Ifit3b | | 2.154e-16 | -36.07 | GSE21546\_WT\_VS\_SAP1A\_KO\_AND\_ELK1\_KO\_DP\_THYMOCYTES\_UP | MSigDB lists | GSE21546\_WT\_VS\_SAP1A\_KO\_AND\_ELK1\_KO\_DP\_THYMOCYTES\_UP | 165 | 23 | 12187 | 179 | Nfkbia,Tor3a,Xaf1,Ifit3b,Tagln2,Icam1,Ifit2,Map3k8,Gpr84,Ccl2,Ccl4,Cybb,Nfe2l2,Nfkbie,Gbp7,Pik3r5,Ptges,Cmpk2,Rtp4,Nfkb1,Lcn2,Ptpn1,Parp9 | | 2.283e-16 | -36.02 | regulation of multi-organism process | biological process | GO:0043900 | 468 | 37 | 13711 | 214 | Irak3,Cd14,Tnip1,Trim30a,Ifi204,Acod1,Tnfaip3,Tnf,Pik3ap1,Zfp36,Irgm2,Cxcl1,Ifitm2,A2m,Tnip3,Tap1,Irf7,Oasl1,Ifitm3,Dtx3l,Trib1,Parp14,Lgals9,Oasl2,Nfkbia,Ifitm6,Ifi211,Irgm1,Ccl4,Trim56,Resf1,Trim25,Ifi209,Gbp4,Timp1,Isg15,Parp9 | | 2.888e-16 | -35.78 | GSE42021\_CD24HI\_VS\_CD24LOW\_TREG\_THYMUS\_DN | MSigDB lists | GSE42021\_CD24HI\_VS\_CD24LOW\_TREG\_THYMUS\_DN | 149 | 22 | 12187 | 179 | Usp18,Ifit2,Tnfaip3,Ifi211,Phf11d,Phf11b,Ifitm3,Isg15,Herc6,Xaf1,Oasl1,Parp12,Irf7,Tnfsf10,Ifit3b,Ifi209,Ifi44,Atf3,Ifi204,Ifi207,Ifitm2,Irf9 | | 3.057e-16 | -35.72 | GO\_REGULATION\_OF\_DEFENSE\_RESPONSE | MSigDB lists | GO\_REGULATION\_OF\_DEFENSE\_RESPONSE | 538 | 38 | 12187 | 179 | Nfkb1,Tap1,Parp9,Ptpn1,Pik3ap1,Il17ra,Irak3,Cd14,Il6,Ripk1,Tnfaip3,Ccl12,Usp18,Ccl2,Ccl4,Sbno2,S100a8,Osmr,Ifi211,Ifi207,Tnip3,Ifi204,Socs3,Ifi209,Sele,Tnip1,Saa1,Ptgs2,Casp4,Nfkbia,Birc3,Tnf,Zfp189,Irf7,S100a9,A2m,C3ar1,C5ar1 | | 6.958e-16 | -34.90 | RASHI\_RESPONSE\_TO\_IONIZING\_RADIATION\_2 | MSigDB lists | RASHI\_RESPONSE\_TO\_IONIZING\_RADIATION\_2 | 105 | 19 | 12187 | 179 | Cxcl1,Nfkb2,Nfkbia,Birc3,Rhou,Map3k8,Nfe2l2,Tnfaip3,Cd14,Tagln2,Tnip1,Icam1,Nfkbiz,Thbs1,Cdkn1a,Cebpd,Lcn2,Atf3,Hp | | 7.045e-16 | -34.89 | REACTOME\_INTERFERON\_ALPHA\_BETA\_SIGNALING | MSigDB lists | REACTOME\_INTERFERON\_ALPHA\_BETA\_SIGNALING | 43 | 14 | 12187 | 179 | Irf7,Usp18,Xaf1,Ifit2,Oasl1,Ifitm2,Isg15,Ifitm3,Irf9,Irf2,Ptpn1,Ip6k2,Ifit3b,Socs3 | | 9.205e-16 | -34.62 | GSE37301\_MULTIPOTENT\_PROGENITOR\_VS\_GRAN\_MONO\_PROGENITOR\_DN | MSigDB lists | GSE37301\_MULTIPOTENT\_PROGENITOR\_VS\_GRAN\_MONO\_PROGENITOR\_DN | 176 | 23 | 12187 | 179 | Tap1,Clic4,Icam1,Isg15,Lcn2,Znfx1,Gadd45g,Ifit3b,Nfkb2,Xdh,Gbp3,Osmr,Cmpk2,Irf7,Gbp7,Irgm2,Ripk1,Tubb6,Ccl2,Usp18,Casp4,Birc3,Ifit2 | | 1.232e-15 | -34.33 | MODULE\_5 | MSigDB lists | MODULE\_5 | 336 | 30 | 12187 | 179 | Tap1,Cd14,Lcn2,Ier3,Thbs1,S100a8,Cxcl1,Ifi211,Cybb,Il6,Tnfaip3,Ccl12,Maff,Il4ra,Tgm2,Icam1,Irf9,Csf2rb,Plaur,Col4a1,Hp,Ifi207,Ifi204,Il1rn,Cdkn1a,Ifi209,Socs3,Tnfsf10,S100a9,Casp4 | | 1.363e-15 | -34.23 | defense response to virus | biological process | GO:0051607 | 133 | 21 | 13711 | 214 | Trim30a,Mx1,Dtx3l,Ifitm3,Oasl1,Irf7,Il6,Cxcl9,Slfn9,Oasl2,Trim25,Ifit2,Trim56,Mx2,Ifit3b,Rtp4,Ifitm6,Parp9,Irf2,Isg15,Ifitm2 | | 1.479e-15 | -34.15 | positive regulation of immune system process | biological process | GO:0002684 | 612 | 41 | 13711 | 214 | Runx1,Ccl4,Ifi211,Irgm1,Parp9,Isg15,Ifi209,Lgals9,Hcar2,Trib1,Thbs1,Irf7,Cdkn1a,Il6,Nfkbia,Fyb,Il1a,Il4ra,Icam1,Irgm2,Selp,Mmp8,Vcam1,Ccl12,Tnip3,C5ar1,Cxcl1,Nfkbiz,Ifi204,Tnip1,Zbtb16,Ripk1,Csf3,Cd14,C3ar1,Irak3,Sele,Ccl2,Pik3ap1,Tnf,Acod1 | | 1.533e-15 | -34.11 | Immune System | REACTOME pathways | R-MMU-168256 | 1188 | 56 | 6297 | 105 | Lgals9,Serpina3f,Ksr1,Il17ra,Csf2rb,Cxcl1,Osmr,Iqgap1,Irak3,Fyb,Rnf213,Vcam1,C5ar1,Ptpn1,Lcn2,Ier3,Csf3,Birc3,Pik3ap1,Map3k8,Irf2,Ifitm3,Plaur,Mmp8,Il4ra,Nfkb2,Icam1,Socs3,Usp18,C3ar1,Hp,Isg15,Herc6,Tnf,Ncf1,Nfkbia,Nfkbie,S100a8,Tap1,Tnfaip3,Il1a,Cd14,Irf7,Nfkb1,Xdh,Ifitm2,Trim56,Irf9,Pik3r1,Gpr84,S100a9,Dtx3l,Il1rn,Cybb,Il6,Cdkn1a | | 1.715e-15 | -34.00 | positive regulation of response to external stimulus | biological process | GO:0032103 | 341 | 31 | 13711 | 214 | Ccl4,Ifi211,Irgm1,Il17ra,Ptgs2,Ifi209,Parp9,Irf7,Thbs1,Lgals9,Nfkbia,Il6,S100a9,Mmp8,Irgm2,Nfkbiz,Tgm2,Cxcl1,C5ar1,Tnip3,S100a8,Cd14,Hspb1,Irak3,C3ar1,Ifi204,Tnip1,Pik3ap1,Tnf,Acod1,Ccl2 | | 2.171e-15 | -33.76 | Immunity-related\_GTPase-like | interpro domains | IPR007743 | 11 | 9 | 13788 | 212 | Iigp1,F830016B08Rik,Ifi47,Irgm1,9930111J21Rik2,Irgm2,Tgtp1,Gm5431,Gm4841 | | 2.260e-15 | -33.72 | positive regulation of response to stimulus | biological process | GO:0048584 | 1637 | 69 | 13711 | 214 | Thbs1,Gadd45g,P2ry6,Lgals9,Icam1,Fyb,Sox11,Irgm1,Pik3r5,Il17ra,Plaur,Map3k8,Nfkb1,C3ar1,Irak3,Il1rn,Csf3,Ifi204,Tnfaip3,Plek,Xdh,Tnfsf10,Ksr1,Ccl2,S100a9,Irgm2,Nfkbiz,Ptpn1,S100a8,Iqgap1,Tnip3,Ccl12,Dtx3l,Irf7,Parp14,Ccl7,Gadd45b,Pik3r1,Il4ra,Casp4,Nfkbia,Il1a,Akap12,Il6,Ifi211,Ccl4,Runx1,Ptgs2,Trim25,Ifi209,Map3k6,Fzd4,Parp9,Ncf1,Hspb1,Ripk1,Cflar,Cd14,Tnip1,Tnf,Acod1,Pik3ap1,Atf3,Nfe2l2,Mmp8,Selp,C5ar1,Cxcl1,Tgm2 | | 2.456e-15 | -33.64 | IIGP | pfam domains | PF05049 | 11 | 9 | 12881 | 201 | Tgtp1,Irgm2,Irgm1,Gm4841,Iigp1,Ifi47,9930111J21Rik2,Gm5431,F830016B08Rik | | 2.466e-15 | -33.64 | regulation of cytokine production | biological process | GO:0001817 | 504 | 37 | 13711 | 214 | Irf7,Thbs1,Lgals9,Pik3r1,Il1a,Akap12,Casp4,Il4ra,Il6,Errfi1,Cybb,Ccl4,Ifi211,Il17ra,Ptgs2,Runx1,Gbp4,Isg15,Nfkb1,Cd14,Ripk1,Hspb1,C3ar1,Il1rn,Irak3,Trim30a,Ifi204,Tnfaip3,Tnf,Acod1,Ccl2,Zfp36,Mmp8,Bcl3,C5ar1,Prg4,Iqgap1 | | 2.747e-15 | -33.53 | GO\_CELLULAR\_RESPONSE\_TO\_ORGANIC\_SUBSTANCE | MSigDB lists | GO\_CELLULAR\_RESPONSE\_TO\_ORGANIC\_SUBSTANCE | 1393 | 60 | 12187 | 179 | Nfkb1,Tiparp,Il1a,Cxcl16,Irak3,Il17ra,Cd14,Ifitm3,Isg15,Ccl12,Il6,Nfe2l2,Ccl4,Msr1,Sbno2,Iqgap1,Cmpk2,Osmr,Csf3,Atf3,Tnip3,Vcam1,Socs3,Ifit3b,Adamts1,Errfi1,Csf2rb,Msn,Hspb1,Ifitm2,Ptgs2,Gbp4,Xaf1,Nfkbia,Tnf,Birc3,Gbp6,Sgk1,Hspa5,Irf2,Ptpn1,Ip6k2,P2ry6,Tnfaip3,Ripk1,Ccl2,Spi1,Ifit2,Pik3r1,Il4ra,Cxcl1,Col4a1,Fstl1,Irf9,Trim25,Icam1,Oasl1,Casp4,Irf7,Fzd4 | | 2.995e-15 | -33.44 | GRAESSMANN\_RESPONSE\_TO\_MC\_AND\_SERUM\_DEPRIVATION\_UP | MSigDB lists | GRAESSMANN\_RESPONSE\_TO\_MC\_AND\_SERUM\_DEPRIVATION\_UP | 166 | 22 | 12187 | 179 | Tor3a,Parp14,Usp18,Ifit2,Ccl12,Parp12,Irf7,Irgm2,Gbp7,Slfn4,Rtp4,Cmpk2,Fstl1,Ifit3b,Ifi44,Cp,Isg15,Irf2,Parp9,Irf9,Trim25,Slfn3 | | 3.090e-15 | -33.41 | ZHANG\_INTERFERON\_RESPONSE | MSigDB lists | ZHANG\_INTERFERON\_RESPONSE | 22 | 11 | 12187 | 179 | Ifi209,Ifi44,Ifi204,Ifi207,Ifitm3,Isg15,Irf9,Ifit2,Oasl1,Ifi211,Rtp4 | | 3.097e-15 | -33.41 | GSE7509\_UNSTIM\_VS\_IFNA\_STIM\_IMMATURE\_DC\_DN | MSigDB lists | GSE7509\_UNSTIM\_VS\_IFNA\_STIM\_IMMATURE\_DC\_DN | 130 | 20 | 12187 | 179 | Gbp6,Phf11d,Tnfsf10,Ifit2,Xaf1,Tgm2,Nfkbia,Usp18,Casp4,Gbp4,Parp12,Isg15,Icam1,Trim25,Phf11b,Ifi44,Vcam1,Ifit3b,Stx11,Samd9l | | 3.156e-15 | -33.39 | GO\_RESPONSE\_TO\_MOLECULE\_OF\_BACTERIAL\_ORIGIN | MSigDB lists | GO\_RESPONSE\_TO\_MOLECULE\_OF\_BACTERIAL\_ORIGIN | 228 | 25 | 12187 | 179 | Nfkb1,Cd14,Selp,Cxcl16,Irak3,Ccl12,Tnfaip3,Il6,Cxcl1,Ptges,Csf3,Sbno2,S100a8,Cmpk2,Vcam1,Socs3,Tnip3,Ptgs2,Icam1,Sele,Nfkbia,Trib1,Tnf,C5ar1,Nfkb2 | | 3.549e-15 | -33.27 | GO\_REGULATION\_OF\_INFLAMMATORY\_RESPONSE | MSigDB lists | GO\_REGULATION\_OF\_INFLAMMATORY\_RESPONSE | 187 | 23 | 12187 | 179 | Sele,Tnip1,Saa1,Ptgs2,Socs3,A2m,S100a9,C3ar1,C5ar1,Casp4,Birc3,Tnf,Pik3ap1,Il17ra,Nfkb1,Sbno2,Osmr,S100a8,Il6,Tnfaip3,Ccl12,Ccl2,Ccl4 | | 3.965e-15 | -33.16 | GO\_DEFENSE\_RESPONSE\_TO\_OTHER\_ORGANISM | MSigDB lists | GO\_DEFENSE\_RESPONSE\_TO\_OTHER\_ORGANISM | 276 | 27 | 12187 | 179 | Gbp4,Oasl1,Zfp189,Tnf,Trim56,S100a9,Irf7,C5ar1,Gbp6,Ifi204,Hp,Ifi207,Ifit3b,Ifi209,Irf9,Trim25,Ifitm2,Il6,Ifit2,Il4ra,S100a8,Ifi211,Bcl3,Il17ra,Selp,Ifitm3,Isg15 | | 5.651e-15 | -32.81 | GSE42021\_CD24HI\_TREG\_VS\_CD24HI\_TCONV\_THYMUS\_DN | MSigDB lists | GSE42021\_CD24HI\_TREG\_VS\_CD24HI\_TCONV\_THYMUS\_DN | 171 | 22 | 12187 | 179 | Gadd45g,Il1a,Ier3,Sgk1,Nfkbiz,Selp,Cd14,Gadd45b,Map3k8,Ccnd2,Il4ra,Ccl2,Cxcl1,Il1rn,Cdkn1a,Socs3,Icam1,Plaur,Errfi1,Ptgs2,Birc3,Nfkbia | | 5.848e-15 | -32.77 | TIAN\_TNF\_SIGNALING\_VIA\_NFKB | MSigDB lists | TIAN\_TNF\_SIGNALING\_VIA\_NFKB | 23 | 11 | 12187 | 179 | Nfkbia,Sdc4,Nfkb1,Il6,Tnfaip3,Nfkbie,Ptgs2,Cxcl1,Tnip1,Zfp36,Nfkb2 | | 7.702e-15 | -32.50 | LIANG\_SILENCED\_BY\_METHYLATION\_2 | MSigDB lists | LIANG\_SILENCED\_BY\_METHYLATION\_2 | 31 | 12 | 12187 | 179 | Ifi44,Ifit3b,Lcn2,Ptgs2,Icam1,Ifit2,Xaf1,Tnfaip3,Il6,Cxcl1,Irf7,Tnfsf10 | | 8.569e-15 | -32.39 | G\_IRG\_dom | interpro domains | IPR030385 | 12 | 9 | 13788 | 212 | 9930111J21Rik2,Irgm2,Tgtp1,Gm5431,Gm4841,F830016B08Rik,Iigp1,Ifi47,Irgm1 | | 8.794e-15 | -32.36 | GSE7348\_UNSTIM\_VS\_LPS\_STIM\_MACROPHAGE\_DN | MSigDB lists | GSE7348\_UNSTIM\_VS\_LPS\_STIM\_MACROPHAGE\_DN | 137 | 20 | 12187 | 179 | Plek,Icam1,Tnip1,Ptgs2,Isg15,Atf3,Il1a,Nfkb1,Stx11,Vcam1,Socs3,Igsf6,Arid5b,Ptges,Cflar,Tnfaip3,Casp4,Ccl4,Birc3,Map3k8 | | 8.794e-15 | -32.36 | GSE35825\_IFNA\_VS\_IFNG\_STIM\_MACROPHAGE\_UP | MSigDB lists | GSE35825\_IFNA\_VS\_IFNG\_STIM\_MACROPHAGE\_UP | 137 | 20 | 12187 | 179 | Zfp36,Tap1,Errfi1,Slfn5,Nfkb1,Il1rn,Atf3,Ier3,Vcam1,Rtp4,Cmpk2,Gbp3,Gbp6,Cflar,Irf7,Cxcl1,Parp12,Gbp4,Ifit2,Parp14 | | 9.655e-15 | -32.27 | GSE46606\_IRF4MID\_VS\_WT\_CD40L\_IL2\_IL5\_DAY1\_STIMULATED\_BCELL\_DN | MSigDB lists | GSE46606\_IRF4MID\_VS\_WT\_CD40L\_IL2\_IL5\_DAY1\_STIMULATED\_BCELL\_DN | 156 | 21 | 12187 | 179 | Socs3,Nfkbiz,Nfkb1,Cebpd,Tnip1,Icam1,Irak3,Plek,Ptpn1,Tnf,Birc3,Maff,Ccl4,Nfkbia,Hcar2,Tnfaip3,Cflar,Csf3,Nfkbie,Pik3r5,Nfkb2 | | 1.049e-14 | -32.19 | PHONG\_TNF\_TARGETS\_UP | MSigDB lists | PHONG\_TNF\_TARGETS\_UP | 51 | 14 | 12187 | 179 | Icam1,Zfp36,Ier3,Atf3,Cebpd,Adamts9,Nfkb2,Gem,Cxcl1,Tnfaip3,Il6,Birc3,Nfkbia,Sdc4 | | 1.080e-14 | -32.16 | MODULE\_45 | MSigDB lists | MODULE\_45 | 419 | 32 | 12187 | 179 | Ifi209,Ifi44,Ifit3b,Stx11,Kcna5,Il1rn,Ifi204,Ifi207,Plek,Csf2rb,Tnf,Birc3,Casp4,Oasl1,S100a9,Olfml2b,Igsf6,Xdh,Nfkb2,Tnfsf10,Ier3,Cd14,Tap1,Ccnd2,Il4ra,Spi1,Cybb,Tnfaip3,Ifi211,Tnfaip8,S100a8,Sbno2 | | 1.171e-14 | -32.08 | GSE35825\_UNTREATED\_VS\_IFNA\_STIM\_MACROPHAGE\_DN | MSigDB lists | GSE35825\_UNTREATED\_VS\_IFNA\_STIM\_MACROPHAGE\_DN | 139 | 20 | 12187 | 179 | Parp12,Il6,Tnfaip3,Usp18,Xaf1,Tgm2,Nfkb2,Gbp3,Cmpk2,Pik3r5,Nfkbie,Cflar,Atf3,Znfx1,Timp1,Il1rn,Nfkbiz,Ifit3b,Tap1,Isg15 | | 1.182e-14 | -32.07 | immune effector process | biological process | GO:0002252 | 316 | 29 | 13711 | 214 | Icam1,Oasl2,Slfn9,Il4ra,Il6,Oasl1,Irf7,Dtx3l,Ifitm3,Mx1,Gadd45g,Sbno2,Isg15,Parp9,Rtp4,Mx2,Stx11,Ifitm6,Trim25,Trim56,Cxcl9,Tnfaip3,Ncf1,Trim30a,Ifitm2,Irf2,Ifit3b,Ifit2,Bcl3 | | 1.201e-14 | -32.05 | BASSO\_CD40\_SIGNALING\_UP | MSigDB lists | BASSO\_CD40\_SIGNALING\_UP | 76 | 16 | 12187 | 179 | Nfkb2,Tnfaip8,Arid5b,Nfkbie,Cflar,Gadd45b,Tnfaip3,Nfkbia,Ccl4,Map3k8,Tnf,Irf9,Tap1,Plek,Icam1,Ier3 | | 2.350e-14 | -31.38 | GSE42021\_TREG\_PLN\_VS\_CD24LO\_TREG\_THYMUS\_DN | MSigDB lists | GSE42021\_TREG\_PLN\_VS\_CD24LO\_TREG\_THYMUS\_DN | 144 | 20 | 12187 | 179 | Ifi204,Il1rn,Ifi207,Ifit3b,Phf11b,Ifi209,Ifi44,Irf9,Tap1,Zfp36,Ifitm3,Herc6,Ifitm2,Isg15,Oasl1,Parp12,Usp18,Ifi211,Phf11d,Irf7 | | 2.350e-14 | -31.38 | GSE360\_CTRL\_VS\_M\_TUBERCULOSIS\_DC\_DN | MSigDB lists | GSE360\_CTRL\_VS\_M\_TUBERCULOSIS\_DC\_DN | 144 | 20 | 12187 | 179 | Tnip1,Ifitm2,Irf9,Csf2rb,Icam1,Dnajb1,Ifi44,Ier3,Il1rn,Stx11,S100a9,C3ar1,Tnfsf10,Sbno2,S100a8,Nfkbia,Trib1,Sntb2,Ccl4,Ifit2 | | 2.692e-14 | -31.25 | GSE6259\_FLT3L\_INDUCED\_DEC205\_POS\_DC\_VS\_CD8\_TCELL\_DN | MSigDB lists | GSE6259\_FLT3L\_INDUCED\_DEC205\_POS\_DC\_VS\_CD8\_TCELL\_DN | 145 | 20 | 12187 | 179 | Parp12,Parp14,Ccl4,Map3k8,Sbno2,Gbp3,Cmpk2,Rtp4,Phf11d,Irf7,Cflar,Ifit3b,Phf11b,Cdkn1a,Irf9,Trim25,Icam1,Slfn5,Isg15,Rnf213 | | 3.034e-14 | -31.13 | GSE37534\_UNTREATED\_VS\_PIOGLITAZONE\_TREATED\_CD4\_TCELL\_PPARG1\_AND\_FOXP3\_TRASDUCED\_DN | MSigDB lists | GSE37534\_UNTREATED\_VS\_PIOGLITAZONE\_TREATED\_CD4\_TCELL\_PPARG1\_AND\_FOXP3\_TRASDUCED\_DN | 165 | 21 | 12187 | 179 | Ifit3b,Ifi44,Atf3,Ifitm2,Isg15,Herc6,Ifitm3,Tap1,Irf9,Trim25,Usp18,Xaf1,Ifit2,Il6,Oasl1,Parp12,Gadd45b,Ccl12,Irf7,Cflar,Rtp4 | | 3.036e-14 | -31.13 | DER\_IFN\_BETA\_RESPONSE\_UP | MSigDB lists | DER\_IFN\_BETA\_RESPONSE\_UP | 95 | 17 | 12187 | 179 | Iqgap1,Ifi211,Il6,Oasl1,Rhoc,Ifit2,Irf9,Tap1,Plaur,Irf2,Isg15,Cebpd,Ifi207,Ifi204,Ifit3b,Ifi44,Ifi209 | | 3.752e-14 | -30.91 | HELLER\_SILENCED\_BY\_METHYLATION\_UP | MSigDB lists | HELLER\_SILENCED\_BY\_METHYLATION\_UP | 187 | 22 | 12187 | 179 | Sbno2,Olfml2b,C5ar1,Usp18,Birc3,Msr1,Map3k8,Irf9,Csf2rb,Plek,Icam1,Ncf1,Herc6,Isg15,Map3k6,Ier3,Akap12,Ifit3b,Socs3,Vcam1,Ifi44,Cdkn1a | | 3.985e-14 | -30.85 | HINATA\_NFKB\_TARGETS\_KERATINOCYTE\_UP | MSigDB lists | HINATA\_NFKB\_TARGETS\_KERATINOCYTE\_UP | 68 | 15 | 12187 | 179 | Nfkbia,Tnf,Ccl12,Tnfaip3,Il6,Cflar,Tnfaip8,Cdkn1a,Il1rn,Nfkb1,Il1a,Tnip1,Saa1,Tap1,Icam1 | | 4.363e-14 | -30.76 | G\_IRG | prosite domains | PS51716 | 12 | 9 | 8845 | 164 | Irgm2,Gm5431,F830016B08Rik,Tgtp1,9930111J21Rik2,Gm4841,Irgm1,Iigp1,Ifi47 | | 4.366e-14 | -30.76 | GO\_REGULATION\_OF\_IMMUNE\_SYSTEM\_PROCESS | MSigDB lists | GO\_REGULATION\_OF\_IMMUNE\_SYSTEM\_PROCESS | 947 | 47 | 12187 | 179 | Csf3,Ccl4,Map3k8,Nfe2l2,Il6,Ccl12,Cd14,Isg15,Selp,Tap1,Parp9,Pik3ap1,Irak3,Thbs1,Nfkb1,A2m,C3ar1,C5ar1,Nfkbia,Birc3,Zfp189,Tnf,Tnip1,Msn,Vcam1,Socs3,Zbtb16,Ifi207,Tnip3,Cxcl1,Ifi211,Spi1,Usp18,Pik3r1,Il4ra,Tnfaip3,Ripk1,Hcar2,Ptpn1,Sox11,Irf7,Trib1,Runx1,Icam1,Ifi209,Cdkn1a,Ifi204 | | 4.421e-14 | -30.75 | BOSCO\_INTERFERON\_INDUCED\_ANTIVIRAL\_MODULE | MSigDB lists | BOSCO\_INTERFERON\_INDUCED\_ANTIVIRAL\_MODULE | 56 | 14 | 12187 | 179 | Parp12,Oasl1,Ccl12,Xaf1,Tnfsf10,Gbp3,Irf7,Gbp7,Ms4a6d,Samd9l,Tnip3,Ifit3b,Ifi44,Ifitm3 | | 4.756e-14 | -30.68 | GO\_POSITIVE\_REGULATION\_OF\_IMMUNE\_SYSTEM\_PROCESS | MSigDB lists | GO\_POSITIVE\_REGULATION\_OF\_IMMUNE\_SYSTEM\_PROCESS | 596 | 37 | 12187 | 179 | Hcar2,Tnfaip3,Ripk1,Ccl12,Il6,Map3k8,Il4ra,Pik3r1,Ccl4,Csf3,Ifi211,Cxcl1,Nfkb1,Thbs1,Irak3,Pik3ap1,Tap1,Selp,Isg15,Cd14,Tnf,Runx1,Birc3,Trib1,Nfkbia,C5ar1,C3ar1,Irf7,Ifi204,Tnip3,Ifi207,Ifi209,Cdkn1a,Zbtb16,Vcam1,Icam1,Tnip1 | | 4.917e-14 | -30.64 | cellular response to lipopolysaccharide | biological process | GO:0071222 | 123 | 19 | 13711 | 214 | Cd14,Il1rn,Nfkbia,Cxcl9,Acod1,Tnf,Tnfaip3,Ccl2,Il6,Zfp36,Gbp6,Irgm2,Cxcl1,Cxcl16,Ccl12,Sbno2,Cmpk2,Tnip3,Nfkb1 | | 4.931e-14 | -30.64 | GSE37533\_PPARG1\_FOXP3\_VS\_FOXP3\_TRANSDUCED\_CD4\_TCELL\_PIOGLITAZONE\_TREATED\_UP | MSigDB lists | GSE37533\_PPARG1\_FOXP3\_VS\_FOXP3\_TRANSDUCED\_CD4\_TCELL\_PIOGLITAZONE\_TREATED\_UP | 169 | 21 | 12187 | 179 | Rtp4,Tnfsf10,Phf11d,Ifi211,Irf7,Parp12,Oasl1,Ifit2,Xaf1,Usp18,Trim25,Ifitm3,Isg15,Ifitm2,Herc6,Ifi204,Ifi207,Ifi209,Phf11b,Ifi44,Ifit3b | | 5.224e-14 | -30.58 | GSE21360\_PRIMARY\_VS\_QUATERNARY\_MEMORY\_CD8\_TCELL\_UP | MSigDB lists | GSE21360\_PRIMARY\_VS\_QUATERNARY\_MEMORY\_CD8\_TCELL\_UP | 150 | 20 | 12187 | 179 | Rnf213,Ifitm3,Clic4,Trim25,Ifit3b,Ifi44,Atf3,Znfx1,Samd9l,Tnip3,Il1rn,Mmp8,Irf7,Cmpk2,Usp18,Xaf1,Ifit2,Ccl12,Rassf4,Cybb | | 5.224e-14 | -30.58 | GSE46606\_IRF4\_KO\_VS\_WT\_UNSTIM\_BCELL\_DN | MSigDB lists | GSE46606\_IRF4\_KO\_VS\_WT\_UNSTIM\_BCELL\_DN | 150 | 20 | 12187 | 179 | Birc3,Tnf,Nfkbia,Ccl4,Hcar2,Cybb,Tnfaip3,Nfkbie,Csf3,Cflar,Irf7,Ptges,Nfkb2,Nfkbiz,Socs3,Lcn2,Cebpd,Tnip1,Plek,Irak3 | | 5.224e-14 | -30.58 | GSE16755\_CTRL\_VS\_IFNA\_TREATED\_MAC\_DN | MSigDB lists | GSE16755\_CTRL\_VS\_IFNA\_TREATED\_MAC\_DN | 150 | 20 | 12187 | 179 | Parp12,Il6,Sdc4,Nfkbia,Cmpk2,Tnfaip8,Tnfsf10,Dtx3l,Trim56,Samd9l,Stx11,Tnip3,Ifi44,Cdkn1a,Socs3,Vcam1,Icam1,Herc6,Ptgs2,Isg15 | | 5.902e-14 | -30.46 | SARRIO\_EPITHELIAL\_MESENCHYMAL\_TRANSITION\_DN | MSigDB lists | SARRIO\_EPITHELIAL\_MESENCHYMAL\_TRANSITION\_DN | 115 | 18 | 12187 | 179 | S100a8,Ifi211,Gem,Nfe2l2,Tnfaip3,Nfkbia,Birc3,Errfi1,Ifitm2,Ifitm3,Atf3,Gadd45g,Ifi207,Ifi204,Il1rn,Zbtb16,Ifi44,Ifi209 | | 6.898e-14 | -30.30 | GSE37605\_TREG\_VS\_TCONV\_NOD\_FOXP3\_FUSION\_GFP\_UP | MSigDB lists | GSE37605\_TREG\_VS\_TCONV\_NOD\_FOXP3\_FUSION\_GFP\_UP | 116 | 18 | 12187 | 179 | Adamts9,Fzd4,Slc24a4,Ccl12,Birc3,Maff,Trib1,Zfp36,Sele,Apold1,Ptgs2,Il1a,Akap12,Cebpd,Atf3,Ier3,Sgk1,Adamts1 | | 7.908e-14 | -30.17 | GO\_RESPONSE\_TO\_VIRUS | MSigDB lists | GO\_RESPONSE\_TO\_VIRUS | 173 | 21 | 12187 | 179 | Oasl1,Il6,Ccl4,Tnf,Ifit2,Irf7,Trim56,Bcl3,Ifi211,Ifi207,Ifi204,Ifit3b,Ifi44,Ifi209,Irf9,Hspb1,Irak3,Trim25,Isg15,Ifitm2,Ifitm3 | | 8.376e-14 | -30.11 | PHONG\_TNF\_RESPONSE\_NOT\_VIA\_P38 | MSigDB lists | PHONG\_TNF\_RESPONSE\_NOT\_VIA\_P38 | 287 | 26 | 12187 | 179 | Arid5b,Bcl3,Ptges,Phf11d,Sbno2,Tnfaip8,Adamts9,Irf2,Dnajb1,Hspa5,Sgk1,Phf11b,Nfkb1,Gem,Cflar,Nfkb2,Trib1,Nfkbia,Sdc4,Birc3,Parp12,Tnip1,Plaur,Rbm47,Icam1,Vcam1 | | 8.463e-14 | -30.10 | GSE41867\_NAIVE\_VS\_DAY30\_LCMV\_CLONE13\_EXHAUSTED\_CD8\_TCELL\_UP | MSigDB lists | GSE41867\_NAIVE\_VS\_DAY30\_LCMV\_CLONE13\_EXHAUSTED\_CD8\_TCELL\_UP | 135 | 19 | 12187 | 179 | Plek,Errfi1,Rnf213,Herc6,Ptgs2,Tnip3,Stx11,Samd9l,Znfx1,Cflar,Pik3r5,Ripk1,Il6,Oasl1,Map3k8,Parp14,Ccl4,Usp18,Sdc4 | | 8.497e-14 | -30.10 | MODULE\_44 | MSigDB lists | MODULE\_44 | 239 | 24 | 12187 | 179 | Ifi211,S100a9,Sbno2,Tnfaip8,S100a8,Tgm2,Casp4,Il4ra,Pik3r1,Birc3,Tnfaip3,Fam107a,Cd14,Isg15,Tap1,Irf9,Fstl1,Vcam1,Ifit3b,Ifi209,Ifi44,Ifi204,Col4a1,Ifi207 | | 8.716e-14 | -30.07 | GSE21360\_NAIVE\_VS\_QUATERNARY\_MEMORY\_CD8\_TCELL\_DN | MSigDB lists | GSE21360\_NAIVE\_VS\_QUATERNARY\_MEMORY\_CD8\_TCELL\_DN | 154 | 20 | 12187 | 179 | Ripk1,Parp12,Oasl1,Xaf1,Usp18,Casp4,Ifit2,Rtp4,Phf11d,Irf7,Nfkbie,Ifit3b,Phf11b,Ifi44,Tap1,Irf9,Ifitm3,Ifitm2,Isg15,Herc6 | | 8.716e-14 | -30.07 | GSE41176\_UNSTIM\_VS\_ANTI\_IGM\_STIM\_BCELL\_1H\_UP | MSigDB lists | GSE41176\_UNSTIM\_VS\_ANTI\_IGM\_STIM\_BCELL\_1H\_UP | 154 | 20 | 12187 | 179 | Il6,Gadd45b,Casp4,Trib1,Nfkbia,Ccl4,Tnfaip8,Nfkbie,Bcl3,Cxcl1,Il1a,Nfkb1,Tnip3,Ier3,Socs3,Icam1,Clic4,Plaur,Ptgs2,Tnip1 | | 8.881e-14 | -30.05 | GSE36527\_CD62L\_HIGH\_VS\_CD62L\_LOW\_TREG\_CD69\_NEG\_KLRG1\_NEG\_UP | MSigDB lists | GSE36527\_CD62L\_HIGH\_VS\_CD62L\_LOW\_TREG\_CD69\_NEG\_KLRG1\_NEG\_UP | 174 | 21 | 12187 | 179 | Parp12,Oasl1,Map3k8,Ifit2,Usp18,Parp14,Tor3a,Rtp4,Irgm2,Irf7,Znfx1,Hspa5,Ifit3b,Socs3,Trim25,Tap1,Irf9,Parp9,Clic4,Isg15,Slfn9 | | 1.042e-13 | -29.89 | GO\_POSITIVE\_REGULATION\_OF\_DEFENSE\_RESPONSE | MSigDB lists | GO\_POSITIVE\_REGULATION\_OF\_DEFENSE\_RESPONSE | 265 | 25 | 12187 | 179 | S100a8,Osmr,Ifi211,Il6,Ripk1,Ccl12,Tnfaip3,Ccl2,Ccl4,Pik3ap1,Il17ra,Irak3,Cd14,Nfkb1,Irf7,S100a9,Nfkbia,Birc3,Tnf,Tnip1,Ptgs2,Ifi207,Ifi204,Tnip3,Ifi209 | | 1.114e-13 | -29.83 | GSE25123\_WT\_VS\_PPARG\_KO\_MACROPHAGE\_UP | MSigDB lists | GSE25123\_WT\_VS\_PPARG\_KO\_MACROPHAGE\_UP | 137 | 19 | 12187 | 179 | Cxcl1,Ptges,S100a8,Map3k8,Oasl1,Il6,Tnfaip3,AA467197,Ifitm2,Slfn5,Irak3,Msn,Ifi44,Thbs1,Nfkbiz,Socs3,Tnip3,Lcn2,Cebpd | | 1.275e-13 | -29.69 | GSE6269\_HEALTHY\_VS\_FLU\_INF\_PBMC\_DN | MSigDB lists | GSE6269\_HEALTHY\_VS\_FLU\_INF\_PBMC\_DN | 138 | 19 | 12187 | 179 | Ifi211,Sbno2,Igsf6,Xaf1,Usp18,Spi1,Oasl1,Tagln2,Ifitm3,Isg15,Ifitm2,Herc6,Msn,Trim25,Ifit3b,Ifi209,Ifi44,Ifi204,Ifi207 | | 1.286e-13 | -29.68 | response to interferon-gamma | biological process | GO:0034341 | 97 | 17 | 13711 | 214 | Irgm1,Ifitm6,Stx11,Ccl4,Gbp7,Gbp6,Irgm2,Gbp4,Gbp3,Ifitm2,Tgtp1,Cxcl16,Ccl12,Ifitm3,Ccl7,Acod1,Ccl2 | | 1.390e-13 | -29.60 | cellular response to molecule of bacterial origin | biological process | GO:0071219 | 130 | 19 | 13711 | 214 | Cxcl1,Cxcl16,Sbno2,Ccl12,Cmpk2,Tnip3,Nfkb1,Gbp6,Irgm2,Cxcl9,Nfkbia,Tnf,Tnfaip3,Acod1,Ccl2,Il6,Zfp36,Cd14,Il1rn | | 1.618e-13 | -29.45 | GSE42021\_TREG\_PLN\_VS\_CD24INT\_TREG\_THYMUS\_DN | MSigDB lists | GSE42021\_TREG\_PLN\_VS\_CD24INT\_TREG\_THYMUS\_DN | 159 | 20 | 12187 | 179 | Cxcl1,Ifi211,Phf11d,Rtp4,Casp4,Tgm2,Parp12,Oasl1,Isg15,Zfp36,Trim25,Tap1,Irf9,Plaur,Ifi44,Ifi209,Phf11b,Ifit3b,Ifi207,Ifi204 | | 1.622e-13 | -29.45 | cellular response to biotic stimulus | biological process | GO:0071216 | 149 | 20 | 13711 | 214 | Cd14,Il1rn,Ccl2,Zfp36,Il6,Nfkbia,Cxcl9,Tnf,Tnfaip3,Acod1,Gbp6,Irgm2,Cxcl16,Ccl12,Sbno2,Cmpk2,Tnip3,Nfkb1,Hspa5,Cxcl1 | | 1.826e-13 | -29.33 | GSE46606\_UNSTIM\_VS\_CD40L\_IL2\_IL5\_1DAY\_STIMULATED\_IRF4HIGH\_SORTED\_BCELL\_DN | MSigDB lists | GSE46606\_UNSTIM\_VS\_CD40L\_IL2\_IL5\_1DAY\_STIMULATED\_IRF4HIGH\_SORTED\_BCELL\_DN | 160 | 20 | 12187 | 179 | Errfi1,Cxcl16,Plek,Zfp36,Icam1,Ptgs2,Ier3,Nfkbiz,Socs3,Cdkn1a,Gem,Il6,Tnfaip3,Bach1,Nfkbia,Trib1,Ccl4,Maff,Map3k8,Tnf | | 1.887e-13 | -29.30 | GO\_POSITIVE\_REGULATION\_OF\_RESPONSE\_TO\_STIMULUS | MSigDB lists | GO\_POSITIVE\_REGULATION\_OF\_RESPONSE\_TO\_STIMULUS | 1397 | 57 | 12187 | 179 | Hspb1,Plaur,Ptgs2,Tnip1,Ifi207,Tnip3,Atf3,Socs3,Xdh,C3ar1,C5ar1,Birc3,Tnf,Nfkbia,Pik3ap1,Irak3,Il17ra,Selp,Cd14,Gadd45g,Akap12,Il1a,Nfkb1,Thbs1,Osmr,Csf3,Pik3r5,Rhoc,Nfe2l2,Il6,Ccl12,Map3k8,Ccl4,Plek,Icam1,Trim25,Saa1,Ifi204,Ksr1,Ifi209,Tnfsf10,Cflar,Fzd4,Irf7,S100a9,Ptpn1,Map3k6,Sox11,S100a8,Cxcl1,Ifi211,Gadd45b,Ripk1,Tnfaip3,Il4ra,Pik3r1,Ccl2 | | 2.083e-13 | -29.20 | negative regulation of multi-organism process | biological process | GO:0043901 | 170 | 21 | 13711 | 214 | Ifitm2,Timp1,Isg15,Tap1,A2m,Ccl4,Ifitm6,Resf1,Trim25,Trim56,Tnf,Tnfaip3,Acod1,Oasl2,Zfp36,Trib1,Parp14,Irak3,Ifitm3,Oasl1,Lgals9 | | 2.360e-13 | -29.08 | GSE36891\_UNSTIM\_VS\_POLYIC\_TLR3\_STIM\_PERITONEAL\_MACROPHAGE\_UP | MSigDB lists | GSE36891\_UNSTIM\_VS\_POLYIC\_TLR3\_STIM\_PERITONEAL\_MACROPHAGE\_UP | 107 | 17 | 12187 | 179 | Nfkbiz,Adamts1,Rasd1,Atf3,Ier3,Cebpd,Akap12,Ptgs2,Apold1,Zfp36,Sdc4,Maff,Ccl12,Tnfaip3,Il6,Cxcl1,Adamts9 | | 2.610e-13 | -28.97 | GSE21927\_SPLEEN\_C57BL6\_VS\_4T1\_TUMOR\_BALBC\_MONOCYTES\_DN | MSigDB lists | GSE21927\_SPLEEN\_C57BL6\_VS\_4T1\_TUMOR\_BALBC\_MONOCYTES\_DN | 163 | 20 | 12187 | 179 | Herc6,Isg15,Ifitm2,Ifitm3,Irf9,Ifit3b,Ifi44,Ifi209,Phf11b,Ifi207,Ifi204,Irf7,Phf11d,Ifi211,Rtp4,Usp18,Xaf1,Ifit2,Oasl1,Parp12 | | 2.808e-13 | -28.90 | GROSS\_HYPOXIA\_VIA\_ELK3\_DN | MSigDB lists | GROSS\_HYPOXIA\_VIA\_ELK3\_DN | 144 | 19 | 12187 | 179 | Nfkbia,Ccl12,Il6,Csf3,Cxcl1,Ifi211,S100a8,Aff1,Ifi209,Hspa5,Ifi204,Mmp8,Akap12,Ifi207,Atf3,Ptgs2,Zfp36,Plek,Errfi1 | | 2.935e-13 | -28.86 | GSE37605\_C57BL6\_VS\_NOD\_FOXP3\_FUSION\_GFP\_TREG\_DN | MSigDB lists | GSE37605\_C57BL6\_VS\_NOD\_FOXP3\_FUSION\_GFP\_TREG\_DN | 164 | 20 | 12187 | 179 | Cd14,Ptgs2,Plaur,Apold1,Zfp36,Nfkbiz,Adamts1,Cebpd,Atf3,Ier3,Cp,Il1a,Akap12,C3ar1,Sdc4,Sntb2,Maff,Tnfaip3,Ccl12,Il6 | | 2.935e-13 | -28.86 | GSE10325\_MYELOID\_VS\_LUPUS\_MYELOID\_DN | MSigDB lists | GSE10325\_MYELOID\_VS\_LUPUS\_MYELOID\_DN | 164 | 20 | 12187 | 179 | Xaf1,Usp18,Ifit2,Ccl12,Oasl1,Parp12,Ifi211,Irf7,Igsf6,Rtp4,Ifit3b,Ifi209,Ifi44,Ifi204,Il1rn,Ifi207,Isg15,Herc6,Tap1,Plek | | 3.191e-13 | -28.77 | GSE10325\_CD4\_TCELL\_VS\_LUPUS\_CD4\_TCELL\_DN | MSigDB lists | GSE10325\_CD4\_TCELL\_VS\_LUPUS\_CD4\_TCELL\_DN | 145 | 19 | 12187 | 179 | Ifit3b,Ifi44,Ifi209,Phf11b,Ifi207,Ifi204,Isg15,Herc6,Selp,Tap1,Usp18,Xaf1,Oasl1,Parp12,Irf7,Arid5b,Phf11d,Ifi211,Rtp4 | | 3.298e-13 | -28.74 | GSE19401\_NAIVE\_VS\_IMMUNIZED\_MOUSE\_PLN\_FOLLICULAR\_DC\_UP | MSigDB lists | GSE19401\_NAIVE\_VS\_IMMUNIZED\_MOUSE\_PLN\_FOLLICULAR\_DC\_UP | 165 | 20 | 12187 | 179 | Ifit3b,Dnajb1,Cdkn1a,Thbs1,Ifi44,Atf3,Tiparp,Nfkb1,Trim25,Icam1,P2ry6,Zfp36,Ccl4,Tnf,Oasl1,Cxcl1,S100a9,Bcl3,Gem,S100a8 | | 3.298e-13 | -28.74 | GSE44649\_WT\_VS\_MIR155\_KO\_NAIVE\_CD8\_TCELL\_DN | MSigDB lists | GSE44649\_WT\_VS\_MIR155\_KO\_NAIVE\_CD8\_TCELL\_DN | 165 | 20 | 12187 | 179 | Irf7,Gbp6,Gbp4,Gadd45b,Oasl1,Parp12,Usp18,Sdc4,Nfkbia,Ccnd2,Pik3r1,Clic4,Slfn5,Tagln2,Ifitm2,Saa1,Isg15,Il1rn,Socs3,Cdkn1a | | 4.108e-13 | -28.52 | GSE9960\_GRAM\_NEG\_VS\_GRAM\_POS\_SEPSIS\_PBMC\_UP | MSigDB lists | GSE9960\_GRAM\_NEG\_VS\_GRAM\_POS\_SEPSIS\_PBMC\_UP | 147 | 19 | 12187 | 179 | Xaf1,Usp18,Trib1,Cflar,Dtx3l,Bcl3,Irf7,Tnfaip8,Cdkn1a,Ifi44,Ifit3b,Samd9l,Ifitm3,Ifitm2,Tagln2,Icam1,Pik3ap1,Plaur,Csf2rb | | 4.108e-13 | -28.52 | GSE8835\_CD4\_VS\_CD8\_TCELL\_CLL\_PATIENT\_UP | MSigDB lists | GSE8835\_CD4\_VS\_CD8\_TCELL\_CLL\_PATIENT\_UP | 147 | 19 | 12187 | 179 | Oasl1,Gbp4,Usp18,Ccl2,Ifit2,Tnfsf10,Cmpk2,Rtp4,Irf7,Gbp6,Irgm2,Samd9l,Stx11,Ifit3b,Hspa5,Slfn9,Herc6,Isg15,Rnf213 | | 4.150e-13 | -28.51 | positive regulation of defense response | biological process | GO:0031349 | 240 | 24 | 13711 | 214 | Nfkbia,Il6,Irf7,Lgals9,Ifi209,Parp9,Irgm1,Ifi211,Il17ra,Ptgs2,Pik3ap1,Tnf,Cd14,Irak3,Ifi204,Tnip1,Cxcl1,Tgm2,Nfkbiz,S100a8,Tnip3,S100a9,Mmp8,Irgm2 | | 4.279e-13 | -28.48 | response to oxygen-containing compound | biological process | GO:1901700 | 969 | 48 | 13711 | 214 | Pygm,Irak3,Il1rn,Lcn2,Tnfaip3,Cxcl9,Ccl2,Vcam1,Gbp6,Irgm2,Wfdc21,Ptpn1,Col4a1,Tnip3,Cxcl16,Ccl12,Thbs1,P2ry6,Lgals9,Icam1,Mt1,Nfkb1,Ncf1,Ripk1,Cflar,Cd14,Acod1,Tnf,Zfp36,Nfe2l2,Serpina3f,Msn,Noct,C5ar1,Cxcl1,Hp,Trib1,Ccl7,Pik3r1,Casp4,Nfkbia,Il6,Ip6k2,Runx1,Ptgs2,Sgk1,Cmpk2,Sbno2 | | 4.654e-13 | -28.40 | GSE3982\_CTRL\_VS\_LPS\_48H\_DC\_DN | MSigDB lists | GSE3982\_CTRL\_VS\_LPS\_48H\_DC\_DN | 148 | 19 | 12187 | 179 | Irf7,Cflar,Ccl4,Nfkbia,Trib1,Tnfaip3,Parp12,Gadd45b,Tnip1,Ifitm3,Ifitm2,Isg15,Plaur,Irf9,P2ry6,Ifit3b,Ifi44,Atf3,Gadd45g | | 5.496e-13 | -28.23 | leukocyte migration | biological process | GO:0050900 | 140 | 19 | 13711 | 214 | Vcam1,S100a9,Ccl4,Selp,Msn,Il17ra,Cxcl1,C5ar1,S100a8,Ccl12,Cxcl16,Sele,Il1rn,Ccl7,Retnlg,Icam1,Tnf,Cxcl9,Ccl2 | | 5.955e-13 | -28.15 | GSE6269\_FLU\_VS\_STREP\_PNEUMO\_INF\_PBMC\_UP | MSigDB lists | GSE6269\_FLU\_VS\_STREP\_PNEUMO\_INF\_PBMC\_UP | 150 | 19 | 12187 | 179 | Xaf1,Usp18,Parp12,Oasl1,Phf11d,Ifi211,Irf7,Rtp4,Phf11b,Ifi209,Ifi44,Ifit3b,Ifi204,Ifi207,Ifitm3,Isg15,Ifitm2,Herc6,Irf9 | | 6.447e-13 | -28.07 | GO\_POSITIVE\_REGULATION\_OF\_CYTOKINE\_PRODUCTION | MSigDB lists | GO\_POSITIVE\_REGULATION\_OF\_CYTOKINE\_PRODUCTION | 262 | 24 | 12187 | 179 | Ripk1,Ccl12,Il6,Pik3r1,Il4ra,Ifi211,Bcl3,Nfkb1,Il1a,Thbs1,Il17ra,Cd14,Runx1,Tnf,Nfkb2,Irf7,C5ar1,C3ar1,Ifi204,Ifi207,Ifi209,Hspb1,Saa1,Ptgs2 | | 6.656e-13 | -28.04 | cytokine-mediated signaling pathway | biological process | GO:0019221 | 201 | 22 | 13711 | 214 | Ccl12,Cxcl1,Ifitm2,Osmr,Fzd4,Csf2rb,Ifitm6,Spi1,Ccl4,Il6,Ccl2,Tnf,Il4ra,Il1a,Cxcl9,Nfkbia,Irf7,Il1rn,Irak3,Ifitm3,Iigp1,Ccl7 | | 6.726e-13 | -28.03 | GSE39382\_IL3\_VS\_IL3\_IL33\_TREATED\_MAST\_CELL\_DN | MSigDB lists | GSE39382\_IL3\_VS\_IL3\_IL33\_TREATED\_MAST\_CELL\_DN | 151 | 19 | 12187 | 179 | Il1rn,Stx11,Nfkb1,Gadd45g,Icam1,Ifitm3,Isg15,Herc6,Tnip1,Tagln2,Tnfaip3,Ifit2,Il4ra,Birc3,Usp18,Sbno2,Nfkb2,Cflar,Irf7 | | 6.726e-13 | -28.03 | GSE36826\_WT\_VS\_IL1R\_KO\_SKIN\_STAPH\_AUREUS\_INF\_DN | MSigDB lists | GSE36826\_WT\_VS\_IL1R\_KO\_SKIN\_STAPH\_AUREUS\_INF\_DN | 151 | 19 | 12187 | 179 | Socs3,Fstl1,Phf11b,Ptpn1,Tap1,Hspb1,Icam1,Ifitm2,Herc6,Isg15,Tnfaip3,Gadd45b,Oasl1,Tgm2,Usp18,Tnf,Rtp4,Phf11d,Cflar | | 7.591e-13 | -27.91 | GSE36888\_UNTREATED\_VS\_IL2\_TREATED\_STAT5\_AB\_KNOCKIN\_TCELL\_2H\_UP | MSigDB lists | GSE36888\_UNTREATED\_VS\_IL2\_TREATED\_STAT5\_AB\_KNOCKIN\_TCELL\_2H\_UP | 152 | 19 | 12187 | 179 | Tnf,Gpr84,Ccl4,Ccl12,AA467197,Tnfaip3,S100a9,Pik3r5,Cxcl1,Tnfaip8,Nfkb2,Nfkbiz,Socs3,Il1a,Tnip3,Tnip1,Icam1,Errfi1,Plaur | | 7.792e-13 | -27.88 | GO\_REGULATION\_OF\_INTRACELLULAR\_SIGNAL\_TRANSDUCTION | MSigDB lists | GO\_REGULATION\_OF\_INTRACELLULAR\_SIGNAL\_TRANSDUCTION | 1271 | 53 | 12187 | 179 | Selp,Pik3ap1,Irak3,Thbs1,Gadd45g,Il1a,Akap12,Csf3,Rhoj,Pik3r5,Banp,Map3k8,Ccl4,Rhoc,Nfe2l2,Il6,Ccl12,Ptgs2,Tnip1,Hspb1,Errfi1,Plaur,Socs3,Tnip3,Atf3,C5ar1,A2m,Xdh,Birc3,Tnf,Nfkbia,Map3k6,Ptpn1,Sox11,Ier3,Bcl3,S100a8,Pik3r1,Rhou,Ccl2,Gadd45b,Ripk1,Tnfaip3,Saa1,Plek,Icam1,Trim25,Ksr1,Cflar,Fzd4,S100a9,Tnfsf10,Trib1 | | 7.839e-13 | -27.87 | negative regulation of biological process | biological process | GO:0048519 | 4173 | 115 | 13711 | 214 | Fam107a,Tiparp,Timp1,Isg15,Slfn2,Xaf1,Ifi211,Rasip1,Cebpd,Trim25,Resf1,Runx1,Nfkbia,Il4ra,Slfn3,Ip6k2,Parp14,Ccnd2,Hcar2,Hspa5,Clic4,C5ar1,Rasd1,Mmp8,Nfe2l2,Serpina3f,Angptl4,Noct,Pik3ap1,Acod1,Atf3,Ncf1,Trim30a,Rnf213,Birc3,Nfkbie,Rgs16,Icam1,Mt1,Bbs12,Msr1,Oasl1,Lgals9,Gadd45g,Ifitm2,Ptpn1,Iqgap1,Tnip3,Wfdc21,Znfx1,Tnfaip3,Ccl2,Csf3,Tnfaip8,Zbtb16,Il1rn,Fzd4,Ifi209,Parp9,Sbno2,Ptges,Sdc4,Spi1,Ccl4,Ifitm6,Ier3,Sgk1,Ptgs2,Gem,Il1a,Oasl2,Cdkn1a,Il6,Trib1,Ctla2a,Dtx3l,Socs3,Gadd45b,Pik3r1,Rrp8,Dnajb1,Hp,Prg4,Tnf,Robo4,Zfp36,Ripk1,Cflar,Ifi207,Hspb1,Tnip1,Gbp4,Adamts1,Nfkb1,Kcna5,Bach1,Plaur,Trim56,Sox11,Mxd4,Errfi1,Thbs1,Ifitm3,Banp,Ccl12,Tap1,A2m,Sgk3,Bcl3,Plek,Xdh,Adamts9,Irak3,Arid5b,Ifi204 | | 8.143e-13 | -27.84 | GSE15930\_STIM\_VS\_STIM\_AND\_IFNAB\_48H\_CD8\_T\_CELL\_DN | MSigDB lists | GSE15930\_STIM\_VS\_STIM\_AND\_IFNAB\_48H\_CD8\_T\_CELL\_DN | 173 | 20 | 12187 | 179 | Ifit3b,Znfx1,Ncf1,Ifitm3,Isg15,Clic4,Tap1,Irf9,Slfn3,Tgm2,Casp4,Usp18,Ifit2,Map3k8,Lgals9,Irf7,Irgm2,Nfkb2,Slfn4,Cmpk2 | | 9.754e-13 | -27.66 | GO\_RESPONSE\_TO\_TYPE\_I\_INTERFERON | MSigDB lists | GO\_RESPONSE\_TO\_TYPE\_I\_INTERFERON | 44 | 12 | 12187 | 179 | Isg15,Ifitm2,Ifitm3,Ip6k2,Irf9,Irf2,Ifit3b,Trim56,Irf7,Ifit2,Xaf1,Oasl1 | | 1.006e-12 | -27.63 | extracellular region part | cellular component | GO:0044421 | 1048 | 49 | 13825 | 212 | Prg4,A2m,Csf3,Serpina3f,Timp1,Icam1,Wfdc21,Il1a,Gbp7,Sele,Vcam1,Olfml2b,Cp,Cxcl9,Cxcl16,Msr1,Thbs1,Tnf,Ccl2,Lcn2,Ccl4,Retnlg,Adamts1,S100a8,Gbp6,S100a9,Tnfsf10,Il6,Ccl12,Hp,Fstl1,Lgals9,Ccl7,Mmp8,Xdh,Irgm2,Runx1,Gbp3,Col4a1,Angptl4,Adamts9,Il1rn,Cd14,Cxcl1,Iigp1,Tgm2,Selp,Il4ra,Saa1 | | 1.042e-12 | -27.59 | regulation of response to biotic stimulus | biological process | GO:0002831 | 274 | 25 | 13711 | 214 | Irgm2,Cxcl1,Tnip3,A2m,Tap1,Cd14,Irak3,Trim30a,Ifi204,Tnip1,Pik3ap1,Tnfaip3,Tnf,Acod1,Ifi211,Irgm1,Ifi209,Gbp4,Parp9,Parp14,Trib1,Irf7,Dtx3l,Lgals9,Nfkbia | | 1.071e-12 | -27.56 | VERHAAK\_AML\_WITH\_NPM1\_MUTATED\_UP | MSigDB lists | VERHAAK\_AML\_WITH\_NPM1\_MUTATED\_UP | 117 | 17 | 12187 | 179 | Tnfsf10,Nfkb2,Cxcl1,C3ar1,C5ar1,Il6,Tnfaip3,Trib1,Nfkbia,Ccl4,Tnf,Runx1,Plaur,Plek,Cd14,Ier3,Thbs1 | | 1.071e-12 | -27.56 | GERY\_CEBP\_TARGETS | MSigDB lists | GERY\_CEBP\_TARGETS | 117 | 17 | 12187 | 179 | Gadd45b,Ifi211,Angptl4,Slfn4,S100a8,Xdh,Rgs16,Ifi209,Dnajb1,Ifi204,Gadd45g,Hp,Ifi207,Atf3,Lcn2,Slfn3,Plaur | | 1.071e-12 | -27.56 | GSE7218\_IGM\_VS\_IGG\_SIGNAL\_THGOUGH\_ANTIGEN\_BCELL\_DN | MSigDB lists | GSE7218\_IGM\_VS\_IGG\_SIGNAL\_THGOUGH\_ANTIGEN\_BCELL\_DN | 117 | 17 | 12187 | 179 | Parp12,Gbp4,Usp18,Parp14,Birc3,Ifit2,Rgs16,Tnfsf10,Rtp4,Nfkbie,Gbp6,Gem,Lcn2,Ifit3b,Ifi44,Icam1,Hspb1 | | 1.184e-12 | -27.46 | WONG\_ADULT\_TISSUE\_STEM\_MODULE | MSigDB lists | WONG\_ADULT\_TISSUE\_STEM\_MODULE | 594 | 35 | 12187 | 179 | Cxcl1,Pygm,Tgm2,Rhou,Il6,Hcar2,Selp,Ifitm3,Isg15,Nfkbiz,Thbs1,Ier3,Cebpd,Gadd45g,Irf7,Gem,Nfkbia,Casp4,Tubb6,Bach1,Ptgs2,Errfi1,Clic4,Msn,Icam1,Plek,Vcam1,Fstl1,Socs3,Adamts1,Cdkn1a,Ifi44,Rasd1,Atf3,Col4a1 | | 1.220e-12 | -27.43 | GSE13485\_DAY7\_VS\_DAY21\_YF17D\_VACCINE\_PBMC\_UP | MSigDB lists | GSE13485\_DAY7\_VS\_DAY21\_YF17D\_VACCINE\_PBMC\_UP | 156 | 19 | 12187 | 179 | Parp14,Xaf1,Usp18,Oasl1,Parp12,Phf11d,Irf7,Dtx3l,Ifit3b,Phf11b,Cdkn1a,Znfx1,Ksr1,Samd9l,Isg15,Herc6,Parp9,Trim25,Pik3ap1 | | 1.313e-12 | -27.36 | GAURNIER\_PSMD4\_TARGETS | MSigDB lists | GAURNIER\_PSMD4\_TARGETS | 45 | 12 | 12187 | 179 | Cxcl1,C3ar1,C5ar1,Nfkb2,Cxcl16,Il17ra,Ccl2,Tnf,Il6,Ccl12,Il1a,Nfkb1 | | 1.321e-12 | -27.35 | GSE36888\_UNTREATED\_VS\_IL2\_TREATED\_TCELL\_6H\_UP | MSigDB lists | GSE36888\_UNTREATED\_VS\_IL2\_TREATED\_TCELL\_6H\_UP | 137 | 18 | 12187 | 179 | Ptgs2,Cd14,Pik3ap1,P2ry6,Nfkbiz,Socs3,Tnip3,Ksr1,Ier3,C5ar1,S100a9,Ptges,S100a8,Nfkb2,Il4ra,Runx1,Trib1,Slc24a4 | | 1.370e-12 | -27.32 | GSE37605\_FOXP3\_FUSION\_GFP\_VS\_IRES\_GFP\_TREG\_C57BL6\_UP | MSigDB lists | GSE37605\_FOXP3\_FUSION\_GFP\_VS\_IRES\_GFP\_TREG\_C57BL6\_UP | 157 | 19 | 12187 | 179 | Cxcl1,Gem,Sdc4,Sntb2,Maff,Ccl12,Ptgs2,Plaur,Zfp36,Apold1,Trim25,Sgk1,Thbs1,Adamts1,Atf3,Ier3,Cebpd,Il1a,Akap12 | | 1.370e-12 | -27.32 | GSE43955\_TH0\_VS\_TGFB\_IL6\_TH17\_ACT\_CD4\_TCELL\_1H\_UP | MSigDB lists | GSE43955\_TH0\_VS\_TGFB\_IL6\_TH17\_ACT\_CD4\_TCELL\_1H\_UP | 157 | 19 | 12187 | 179 | Csf2rb,Errfi1,Ptpn1,Sele,Il17ra,Saa1,Selp,Cebpd,Rasd1,Gadd45g,Il1rn,Socs3,Gbp3,Il6,Bach1,Ccl4,Il4ra,Birc3,Map3k8 | | 1.424e-12 | -27.28 | STK33\_UP | MSigDB lists | STK33\_UP | 223 | 22 | 12187 | 179 | Samd9l,Il1a,Ifi207,Ifi204,Sgk1,Cdkn1a,Ifi209,Zfp36,Icam1,Irf9,Bbs12,Rhoc,AA467197,Maff,Tnf,Trib1,Ccl4,C5ar1,Bcl3,Arid5b,Cxcl1,Ifi211 | | 1.470e-12 | -27.25 | CHEN\_METABOLIC\_SYNDROM\_NETWORK | MSigDB lists | CHEN\_METABOLIC\_SYNDROM\_NETWORK | 963 | 45 | 12187 | 179 | Tagln2,Plek,Ifi209,Col4a1,Ifi204,Bach1,Oasl1,P2ry6,Ptpn1,Ms4a6d,Ifi211,Cxcl1,S100a8,Tnfaip8,Ifit2,Spi1,Gadd45b,Tnip1,Plaur,Ifit3b,Vcam1,Ifi207,Tnip3,Atf3,C3ar1,Igsf6,Slfn4,Nfkb2,Tnf,Tubb6,Cd14,Slfn3,Pik3ap1,Cxcl16,Phf11b,Rhoj,Phf11d,Msr1,Map3k8,Ccl4,Rassf4,Rhoc,Cybb,Nfe2l2,Ccl12 | | 1.500e-12 | -27.23 | GSE18791\_CTRL\_VS\_NEWCASTLE\_VIRUS\_DC\_16H\_DN | MSigDB lists | GSE18791\_CTRL\_VS\_NEWCASTLE\_VIRUS\_DC\_16H\_DN | 138 | 18 | 12187 | 179 | Oasl1,Tnf,Ifit2,Xaf1,Parp14,Cmpk2,Rtp4,Gbp3,Irf7,Phf11d,Ifi44,Phf11b,Ifit3b,Trim25,Ptgs2,Isg15,Herc6,Slfn5 | | 1.766e-12 | -27.06 | ICHIBA\_GRAFT\_VERSUS\_HOST\_DISEASE\_35D\_UP | MSigDB lists | ICHIBA\_GRAFT\_VERSUS\_HOST\_DISEASE\_35D\_UP | 103 | 16 | 12187 | 179 | Gbp3,Iqgap1,Slfn4,S100a8,S100a9,Irgm2,Ccl12,Cybb,Ccl4,Msr1,Ptpn1,Tap1,Slfn3,Cd14,Vcam1,Nfkbiz | | 1.928e-12 | -26.97 | GSE9988\_ANTI\_TREM1\_AND\_LPS\_VS\_CTRL\_TREATED\_MONOCYTES\_UP | MSigDB lists | GSE9988\_ANTI\_TREM1\_AND\_LPS\_VS\_CTRL\_TREATED\_MONOCYTES\_UP | 140 | 18 | 12187 | 179 | Ptgs2,Plaur,Errfi1,Sgk1,Cdkn1a,Nfkbiz,Socs3,Il1a,Ier3,Csf3,Gem,Cflar,Banp,Cxcl1,Map3k8,Tnf,Gadd45b,Il6 | | 2.164e-12 | -26.86 | GSE19888\_ADENOSINE\_A3R\_ACT\_VS\_A3R\_ACT\_WITH\_A3R\_INH\_PRETREATMENT\_IN\_MAST\_CELL\_DN | MSigDB lists | GSE19888\_ADENOSINE\_A3R\_ACT\_VS\_A3R\_ACT\_WITH\_A3R\_INH\_PRETREATMENT\_IN\_MAST\_CELL\_DN | 161 | 19 | 12187 | 179 | Gbp4,Usp18,Parp14,Tor3a,Xaf1,Lgals9,Tnfsf10,Cmpk2,Slfn4,Irf7,Gbp6,Irgm2,Ifit3b,Slfn3,Ip6k2,Saa1,Rnf213,Isg15,Herc6 | | 2.209e-12 | -26.84 | GO\_RESPONSE\_TO\_INTERFERON\_GAMMA | MSigDB lists | GO\_RESPONSE\_TO\_INTERFERON\_GAMMA | 88 | 15 | 12187 | 179 | Irf7,Gbp6,Ccl2,Ccl4,Oasl1,Gbp4,Ccl12,Ifitm2,Ifitm3,Irf9,Cxcl16,Irf2,Trim25,Icam1,Vcam1 | | 2.318e-12 | -26.79 | IL-17 signaling pathway | KEGG pathways | mmu04657 | 67 | 15 | 5248 | 107 | Cxcl1,Ccl12,Tnfaip3,Ccl7,S100a8,Ccl2,Nfkbia,S100a9,Il17ra,Lcn2,Csf3,Ptgs2,Tnf,Il6,Nfkb1 | | 2.318e-12 | -26.79 | IL-17 signaling pathway | KEGG pathways | ko04657 | 67 | 15 | 5248 | 107 | Il6,Nfkb1,Tnf,Ptgs2,Csf3,Lcn2,Nfkbia,S100a9,Il17ra,Ccl7,S100a8,Ccl2,Tnfaip3,Ccl12,Cxcl1 | | 2.356e-12 | -26.77 | BOYLAN\_MULTIPLE\_MYELOMA\_C\_D\_DN | MSigDB lists | BOYLAN\_MULTIPLE\_MYELOMA\_C\_D\_DN | 183 | 20 | 12187 | 179 | Ifi209,Vcam1,Zbtb16,Ifi204,Mmp8,Cp,Ifi207,Hp,Lcn2,Ifitm3,Plek,Ccnd2,Casp4,AA467197,Ifi211,S100a9,Igsf6,Rtp4,S100a8,Xdh | | 2.420e-12 | -26.75 | GSE32034\_LY6C\_HIGH\_VS\_LOW\_ROSIGLIZATONE\_TREATED\_MONOCYTE\_DN | MSigDB lists | GSE32034\_LY6C\_HIGH\_VS\_LOW\_ROSIGLIZATONE\_TREATED\_MONOCYTE\_DN | 162 | 19 | 12187 | 179 | Bcl3,Rgs16,Ccl2,Casp4,Ccl4,Map3k8,Ccl12,Bach1,Ifitm3,Ptpn1,Errfi1,Zfp36,Il17ra,Nfkbiz,Aff1,Cebpd,Gadd45g,Tiparp,Timp1 | | 2.493e-12 | -26.72 | LINDGREN\_BLADDER\_CANCER\_CLUSTER\_2B | MSigDB lists | LINDGREN\_BLADDER\_CANCER\_CLUSTER\_2B | 305 | 25 | 12187 | 179 | Nfkbie,Adamts9,Ccl2,Cybb,Ccl12,Tnfaip3,Gadd45b,Ifitm3,Nfkbiz,Il1a,Akap12,Gem,C3ar1,A2m,Birc3,Nfkbia,Ifitm2,Icam1,Sele,Plaur,Clic4,Msn,Aff1,Vcam1,Socs3 | | 2.551e-12 | -26.69 | response to lipid | biological process | GO:0033993 | 476 | 32 | 13711 | 214 | Cflar,Cd14,Irak3,Il1rn,Ccl2,Zfp36,Cxcl9,Acod1,Tnfaip3,Tnf,Wfdc21,Irgm2,Gbp6,Noct,Msn,Ccl12,Cxcl16,Tnip3,Cxcl1,Lgals9,P2ry6,Pik3r1,Trib1,Il6,Nfkbia,Sgk1,Ptgs2,Runx1,Sbno2,Nfkb1,Cmpk2,Fam107a | | 2.552e-12 | -26.69 | regulation of protein metabolic process | biological process | GO:0051246 | 2256 | 77 | 13711 | 214 | Spi1,Ccl4,Rasip1,Ier3,Ptgs2,Map3k6,Fzd4,Fam107a,Tiparp,Timp1,Parp9,Sdc4,Isg15,Trib1,Parp14,Ctla2a,Ccl7,Dtx3l,Socs3,Gadd45b,Ccnd2,Pik3r1,Nfkbia,Il1a,Akap12,Casp4,Cdkn1a,Il6,Mmp8,Serpina3f,Msn,Hspa5,C5ar1,Prg4,Ripk1,Cflar,Ncf1,Hspb1,Trim30a,Tnip1,Pik3ap1,Tnf,Atf3,Zfp36,Plaur,Pik3r5,Birc3,Map3k8,Gbp4,Nfkb1,Thbs1,P2ry6,Lgals9,Gadd45g,Banp,Icam1,Cybb,Errfi1,S100a9,Irgm2,Wfdc21,Samd9l,Bcl3,Ptpn1,Ccl12,S100a8,Iqgap1,A2m,Tnfaip8,Csf3,Il1rn,Irak3,Tnfaip3,Ccl2,Xdh,Ksr1,Tnfsf10 | | 2.631e-12 | -26.66 | GO\_RESPONSE\_TO\_LIPID | MSigDB lists | GO\_RESPONSE\_TO\_LIPID | 645 | 36 | 12187 | 179 | Tnf,Nfkbia,Trib1,C5ar1,Fzd4,Nfkb2,Adamts1,Cdkn1a,Vcam1,Socs3,Tnip3,Il1rn,Ptgs2,Sele,Icam1,Trim25,Msn,Errfi1,Il4ra,Il6,Ccl12,Tnfaip3,Csf3,Cxcl1,Ptges,Cmpk2,S100a8,Sbno2,Thbs1,Sgk1,Nfkb1,Selp,Cd14,P2ry6,Irak3,Cxcl16 | | 2.663e-12 | -26.65 | STK33\_NOMO\_UP | MSigDB lists | STK33\_NOMO\_UP | 230 | 22 | 12187 | 179 | Zfp36,Icam1,Bbs12,Samd9l,Ifi207,Il1a,Ifi204,Ier3,Sgk1,Ifi209,C3ar1,C5ar1,A2m,Arid5b,Bcl3,Cxcl1,Ifi211,AA467197,Maff,Map3k8,Sdc4,Ccl4 | | 3.040e-12 | -26.52 | STK33\_SKM\_UP | MSigDB lists | STK33\_SKM\_UP | 208 | 21 | 12187 | 179 | Tnf,Ccl4,Nfkbia,Trib1,Rhoc,Cybb,C5ar1,Ifi211,Bcl3,Arid5b,Ifi209,Cdkn1a,Sgk1,Nfkbiz,Ifi204,Il1a,Ifi207,Bbs12,Zfp36,Errfi1,Irf9 | | 3.062e-12 | -26.51 | REACTOME\_INTERFERON\_GAMMA\_SIGNALING | MSigDB lists | REACTOME\_INTERFERON\_GAMMA\_SIGNALING | 48 | 12 | 12187 | 179 | Gbp3,Irf7,Gbp6,Gbp7,Gbp4,Oasl1,Irf2,Ptpn1,Irf9,Icam1,Socs3,Vcam1 | | 3.062e-12 | -26.51 | LINDSTEDT\_DENDRITIC\_CELL\_MATURATION\_A | MSigDB lists | LINDSTEDT\_DENDRITIC\_CELL\_MATURATION\_A | 48 | 12 | 12187 | 179 | Plek,Icam1,Plaur,Isg15,Nfkbie,Cxcl1,Il1rn,Il6,Ccl12,Ifit2,Tnf,Ccl4 | | 3.145e-12 | -26.49 | JAATINEN\_HEMATOPOIETIC\_STEM\_CELL\_DN | MSigDB lists | JAATINEN\_HEMATOPOIETIC\_STEM\_CELL\_DN | 144 | 18 | 12187 | 179 | S100a9,Pik3r5,C5ar1,S100a8,Igsf6,Sntb2,Ccl4,Tnfaip3,Cybb,Cd14,Cxcl16,Thbs1,Sgk1,Cebpd,Ms4a6d,Il1rn,Stx11,Mmp8 | | 3.204e-12 | -26.47 | signal transduction | biological process | GO:0007165 | 2703 | 86 | 13711 | 214 | Pik3r1,Dtx3l,Irf7,Socs3,Trib1,Il6,Cdkn1a,Casp4,Gem,Il1a,Sgk1,Ier3,Ifitm6,Ccl4,Spi1,Fzd4,Tnip1,Rhoc,Hspb1,Ripk1,Cd14,Gpr84,Zfp36,Tnf,Csf2rb,Hp,Mt2,Cxcl1,Rrp8,P2ry6,Ifitm3,Cybb,Errfi1,Plaur,Kcna5,Nfkb1,Osmr,Ifi204,Arid5b,Irak3,Ksr1,Plek,Bcl3,Irgm2,Sgk3,Ccl12,Hcar2,Ccl7,Il4ra,Nfkbia,Akap12,Rasip1,Isg15,Tiparp,Map3k6,Atf3,Rhou,Pik3ap1,Selp,Nfe2l2,Rasd1,C5ar1,Tgm2,Hspa5,Lgals9,Iigp1,Rassf4,Mt1,Fyb,Nfkb2,Rgs16,Pik3r5,Rhoj,Map3k8,Lcn2,Il1rn,C3ar1,Ccl2,Cxcl9,Vcam1,Col4a1,Iqgap1,Tnip3,Nfkbiz,Ifitm2,Ptpn1 | | 3.352e-12 | -26.42 | MODULE\_170 | MSigDB lists | MODULE\_170 | 75 | 14 | 12187 | 179 | Cybb,Ccl12,Ccnd2,Tnfsf10,C3ar1,A2m,Timp1,Cebpd,Ier3,Thbs1,Vcam1,Ifitm2,Ifitm3,Cd14 | | 3.674e-12 | -26.33 | regulation of response to stimulus | biological process | GO:0048583 | 2962 | 91 | 13711 | 214 | Il6,Cdkn1a,Casp4,Il1a,Pik3r1,Gadd45b,Irf7,Socs3,Dtx3l,Ctla2a,Trib1,Sbno2,Parp9,Ifi209,Fzd4,Ptgs2,Ier3,Ccl4,Zfp36,Tnf,Tnip1,Fstl1,Hspb1,Cd14,Cflar,Ripk1,Cxcl1,Errfi1,Sox11,P2ry6,Thbs1,Nfkb1,Gbp4,Plaur,Il17ra,Ksr1,Tnfsf10,Xdh,Plek,Ifi204,Irak3,A2m,S100a8,Tap1,Ccl12,Bcl3,Irgm2,Sgk3,Usp18,Il4ra,Akap12,Nfkbia,Ccl7,Parp14,Timp1,Map3k6,Runx1,Trim25,Rasip1,Ifi211,Atf3,Acod1,Pik3ap1,Trim30a,Rnf213,Ncf1,Tgm2,C5ar1,Hspa5,Selp,Nfe2l2,Mmp8,Icam1,Rgs16,Fyb,Gadd45g,Lgals9,Map3k8,Birc3,Pik3r5,Irgm1,Ccl2,Tnfaip3,Il1rn,C3ar1,Csf3,Tnip3,Iqgap1,Nfkbiz,Ptpn1,S100a9 | | 3.759e-12 | -26.31 | GSE7219\_UNSTIM\_VS\_LPS\_AND\_ANTI\_CD40\_STIM\_NIK\_NFKB2\_KO\_DC\_DN | MSigDB lists | GSE7219\_UNSTIM\_VS\_LPS\_AND\_ANTI\_CD40\_STIM\_NIK\_NFKB2\_KO\_DC\_DN | 166 | 19 | 12187 | 179 | Tnfsf10,Rgs16,Nfkbie,Gadd45b,Tnf,Nfkbia,Trib1,Ccl4,Zfp36,Pik3ap1,Icam1,Clic4,Rnf213,Ifitm2,Ifitm3,Atf3,Ifi44,Cdkn1a,Dnajb1 | | 3.993e-12 | -26.25 | GSE360\_HIGH\_DOSE\_B\_MALAYI\_VS\_M\_TUBERCULOSIS\_DC\_DN | MSigDB lists | GSE360\_HIGH\_DOSE\_B\_MALAYI\_VS\_M\_TUBERCULOSIS\_DC\_DN | 146 | 18 | 12187 | 179 | Timp1,Il1rn,Nfkb1,Stx11,Ier3,Ifi44,Vcam1,Icam1,Csf2rb,Cybb,Gadd45b,Ccl12,Ifit2,Nfkbia,Ccl4,S100a8,Sbno2,S100a9 | | 3.993e-12 | -26.25 | GSE17721\_PAM3CSK4\_VS\_GADIQUIMOD\_1H\_BMDC\_UP | MSigDB lists | GSE17721\_PAM3CSK4\_VS\_GADIQUIMOD\_1H\_BMDC\_UP | 146 | 18 | 12187 | 179 | Cxcl1,Gem,Igsf6,Parp14,Nfkbia,Ccl2,Casp4,Birc3,Maff,Il6,Tnip1,Errfi1,Icam1,Plek,Zfp36,Fstl1,Nfkbiz,Atf3 | | 4.465e-12 | -26.13 | positive regulation of cytokine production | biological process | GO:0001819 | 318 | 26 | 13711 | 214 | C5ar1,Bcl3,Mmp8,Ccl2,Tnf,Ifi204,Ripk1,Cd14,Il1rn,C3ar1,Hspb1,Il17ra,Runx1,Ptgs2,Ccl4,Ifi211,Cybb,Il6,Il1a,Akap12,Il4ra,Casp4,Lgals9,Pik3r1,Thbs1,Irf7 | | 5.374e-12 | -25.95 | regulation of inflammatory response | biological process | GO:0050727 | 246 | 23 | 13711 | 214 | Tnip1,Pik3ap1,Acod1,Tnfaip3,Tnf,Zfp36,Usp18,S100a9,Mmp8,C5ar1,Nfkbiz,Tgm2,S100a8,Ctla2a,Socs3,Lgals9,Nfkbia,Il6,Il17ra,Ier3,Ptgs2,Sbno2,Nfkb1 | | 5.914e-12 | -25.85 | DAUER\_STAT3\_TARGETS\_UP | MSigDB lists | DAUER\_STAT3\_TARGETS\_UP | 39 | 11 | 12187 | 179 | Icam1,Zfp36,Bcl3,Ccl12,Atf3,Cebpd,Akap12,Socs3,Trib1,Thbs1,Maff | | 6.111e-12 | -25.82 | GO\_REGULATION\_OF\_CELL\_DEATH | MSigDB lists | GO\_REGULATION\_OF\_CELL\_DEATH | 1126 | 48 | 12187 | 179 | Nfkb1,Il1a,Gadd45g,Thbs1,Sgk3,Ccl12,Il6,Nfe2l2,Tgm2,Angptl4,Csf3,Atf3,Timp1,Zbtb16,Socs3,Ifit3b,Plaur,Hspb1,Ptgs2,Nfkbia,Tnf,Birc3,Xdh,C5ar1,Robo4,Ier3,Sox11,Hspa5,Sgk1,Ptpn1,Ip6k2,Tnfaip3,Ripk1,Gadd45b,Hcar2,Ifit2,Pik3r1,Tnfaip8,S100a8,Bcl3,Hp,Cdkn1a,Icam1,Casp4,Tnfsf10,Irf7,S100a9,Cflar | | 6.814e-12 | -25.71 | CHARAFE\_BREAST\_CANCER\_LUMINAL\_VS\_BASAL\_DN | MSigDB lists | CHARAFE\_BREAST\_CANCER\_LUMINAL\_VS\_BASAL\_DN | 347 | 26 | 12187 | 179 | Ptgs2,Clic4,Msn,Icam1,Zbtb16,Fstl1,Ifit3b,Ifi209,Ifi44,Ifi204,Col4a1,Ifi207,Cflar,Casp4,Birc3,Tubb6,Cd14,Sgk1,Cebpd,Il1a,Ifi211,Cxcl1,Tnfaip8,Osmr,Tnfaip3,Nfe2l2 | | 7.112e-12 | -25.67 | GSE25085\_FETAL\_BM\_VS\_ADULT\_BM\_SP4\_THYMIC\_IMPLANT\_UP | MSigDB lists | GSE25085\_FETAL\_BM\_VS\_ADULT\_BM\_SP4\_THYMIC\_IMPLANT\_UP | 172 | 19 | 12187 | 179 | Cmpk2,Rtp4,Gbp3,Irf7,Hcar2,Oasl1,Parp12,Nfe2l2,Ifit2,Sdc4,Ccl4,Parp14,Isg15,Herc6,Slfn9,Samd9l,Atf3,Ifit3b,Socs3 | | 7.113e-12 | -25.67 | extracellular region | cellular component | GO:0005576 | 1307 | 54 | 13825 | 212 | A2m,Prg4,Wfdc21,Il1a,Icam1,Timp1,Serpina3f,Csf3,Vcam1,Gbp7,Sele,Olfml2b,Tnf,Thbs1,Cxcl9,Cp,Cxcl16,Msr1,Ccl2,Ccl4,Lcn2,Retnlg,Adamts1,Gbp6,S100a8,Isg15,Ctla2a,Plaur,Il6,S100a9,Tnfsf10,Ccl12,Hp,Xdh,Mmp8,Ccl7,Lgals9,Fstl1,Irgm2,Angptl4,Gbp3,Col4a1,Runx1,Tgm2,Casp4,Sdc4,Cd14,Il1rn,Iigp1,Cxcl1,Adamts9,Selp,Il4ra,Saa1 | | 7.991e-12 | -25.55 | GSE17721\_CTRL\_VS\_LPS\_1H\_BMDC\_DN | MSigDB lists | GSE17721\_CTRL\_VS\_LPS\_1H\_BMDC\_DN | 152 | 18 | 12187 | 179 | Cflar,Arid5b,Igsf6,Maff,Map3k8,Ccl2,Tgm2,Nfe2l2,Oasl1,Zfp36,Icam1,Errfi1,Hspa5,Cdkn1a,Fstl1,Tiparp,Atf3,Ier3 | | 8.745e-12 | -25.46 | GSE42021\_TREG\_PLN\_VS\_CD24HI\_TREG\_THYMUS\_UP | MSigDB lists | GSE42021\_TREG\_PLN\_VS\_CD24HI\_TREG\_THYMUS\_UP | 174 | 19 | 12187 | 179 | Nfkbiz,Socs3,Sgk1,Cdkn1a,Ier3,Il1a,Il1rn,Cd14,Ptgs2,Plaur,Errfi1,Icam1,Ccl2,Nfkbia,Birc3,Tnf,Gadd45b,Nfe2l2,Cxcl1 | | 8.745e-12 | -25.46 | GSE14769\_UNSTIM\_VS\_120MIN\_LPS\_BMDM\_DN | MSigDB lists | GSE14769\_UNSTIM\_VS\_120MIN\_LPS\_BMDM\_DN | 174 | 19 | 12187 | 179 | Vcam1,Il1rn,Tiparp,Atf3,Znfx1,Isg15,Tnip1,Zfp36,Sele,Plaur,Clic4,Ccl2,Sdc4,Tgm2,Parp14,AA467197,Cxcl1,Igsf6,Gbp3 | | 8.819e-12 | -25.45 | GSE18791\_UNSTIM\_VS\_NEWCATSLE\_VIRUS\_DC\_18H\_DN | MSigDB lists | GSE18791\_UNSTIM\_VS\_NEWCATSLE\_VIRUS\_DC\_18H\_DN | 114 | 16 | 12187 | 179 | Samd9l,Nfkbiz,Ifi44,Phf11b,Herc6,Ptgs2,Gadd45b,Oasl1,Parp14,Xaf1,Ifit2,Map3k8,Tnfsf10,Cmpk2,Irf7,Phf11d | | 8.942e-12 | -25.44 | GSE21360\_PRIMARY\_VS\_TERTIARY\_MEMORY\_CD8\_TCELL\_DN | MSigDB lists | GSE21360\_PRIMARY\_VS\_TERTIARY\_MEMORY\_CD8\_TCELL\_DN | 153 | 18 | 12187 | 179 | Gem,Irf7,Rrp8,Sbno2,Tnf,Sdc4,Casp4,Trib1,Il6,Gadd45b,Zfp36,Plek,Trim25,Icam1,Tap1,Cdkn1a,Socs3,Gadd45g | | 9.419e-12 | -25.39 | SMID\_BREAST\_CANCER\_LUMINAL\_B\_DN | MSigDB lists | SMID\_BREAST\_CANCER\_LUMINAL\_B\_DN | 352 | 26 | 12187 | 179 | Ifi204,Ifi207,Adamts1,Ifi209,Vcam1,Icam1,Sele,Clic4,Msn,Ptgs2,Birc3,Gem,S100a9,Lcn2,Sox11,Tap1,Ccl12,Fam107a,Tnfaip3,Il6,Ccl2,Tnfaip8,S100a8,Cxcl1,Ifi211,Angptl4 | | 1.048e-11 | -25.28 | regulation of metabolic process | biological process | GO:0019222 | 4733 | 122 | 13711 | 214 | Icam1,Nfkb2,Oasl1,Iigp1,Msr1,Gadd45g,Lgals9,Map3k8,Birc3,Nfkbie,Irgm1,Pik3r5,Znfx1,Tnfaip3,Depp1,Ccl2,C3ar1,Il1rn,Aff1,Zbtb16,Tnfaip8,Csf3,Lcn2,Ptpn1,Iqgap1,S100a9,Wfdc21,Il4ra,Nfkbia,Akap12,Parp14,Ccl7,Ccnd2,Hcar2,Timp1,Tiparp,Map3k6,Fam107a,Isg15,Ifi211,Cebpd,Rasip1,Runx1,Trim25,Resf1,Acod1,Pik3ap1,Atf3,Ncf1,Sele,Trim30a,C5ar1,Maff,Hspa5,Nfe2l2,Serpina3f,Mmp8,Rasd1,Angptl4,Msn,Noct,Mxd4,Sox11,Cybb,Errfi1,Thbs1,Rbm39,Banp,P2ry6,Gbp4,Nfkb1,Bach1,Plaur,Plek,Xdh,Ksr1,Tnfsf10,Arid5b,Irak3,Ifi204,S100a8,A2m,Ccl12,Irf2,Irf9,Bcl3,Samd9l,Irgm2,Oasl2,Casp4,Il1a,Il6,Cdkn1a,Dtx3l,Irf7,Socs3,Trib1,Ctla2a,Gadd45b,Pik3r1,Ifi209,Fzd4,Parp9,Sdc4,Sbno2,Spi1,Ccl4,Ptgs2,Ier3,Tnf,Zfp36,Ifi207,Hspb1,Ripk1,Cflar,Fstl1,Tnip1,Cxcl1,Dnajb1,Rrp8,Hp,Prg4 | | 1.095e-11 | -25.24 | GO\_CELL\_DEATH | MSigDB lists | GO\_CELL\_DEATH | 787 | 39 | 12187 | 179 | Thbs1,Sgk1,Ier3,Lcn2,Nfkb1,Gadd45g,Il1a,Cd14,Spi1,Map3k8,Ifit2,Pik3r1,Tnfaip3,Ripk1,Il6,Gadd45b,Hcar2,Ifi211,Rrp8,Bcl3,Tnfaip8,S100a8,Zbtb16,Cdkn1a,Ifi209,Ifi204,Ifi207,Plaur,Clic4,Xaf1,Nfkbia,Casp4,Tnf,Birc3,S100a9,Cflar,C5ar1,Xdh,Tnfsf10 | | 1.098e-11 | -25.24 | BOWIE\_RESPONSE\_TO\_TAMOXIFEN | MSigDB lists | BOWIE\_RESPONSE\_TO\_TAMOXIFEN | 22 | 9 | 12187 | 179 | Irf7,Ifi211,Isg15,Irf9,Ifit3b,Ifit2,Ifi209,Ifi207,Ifi204 | | 1.098e-11 | -25.24 | BENNETT\_SYSTEMIC\_LUPUS\_ERYTHEMATOSUS | MSigDB lists | BENNETT\_SYSTEMIC\_LUPUS\_ERYTHEMATOSUS | 22 | 9 | 12187 | 179 | Ifit3b,Xaf1,Oasl1,Isg15,Ifitm3,Irf7,S100a8,Tnfsf10,Tap1 | | 1.117e-11 | -25.22 | EGFR\_UP.V1\_UP | MSigDB lists | EGFR\_UP.V1\_UP | 155 | 18 | 12187 | 179 | Rgs16,Nfkb2,S100a8,Tnfaip8,S100a9,Bcl3,Arid5b,A2m,Angptl4,Ccl12,Il4ra,Tap1,P2ry6,Icam1,Cebpd,Atf3,Akap12,Ifit3b | | 1.120e-11 | -25.22 | GO\_POSITIVE\_REGULATION\_OF\_RESPONSE\_TO\_WOUNDING | MSigDB lists | GO\_POSITIVE\_REGULATION\_OF\_RESPONSE\_TO\_WOUNDING | 98 | 15 | 12187 | 179 | Thbs1,Ptgs2,Selp,Tnip1,Plek,Il17ra,Tnf,Ccl2,Ccl4,Nfe2l2,Il6,Ccl12,S100a9,S100a8,Osmr | | 1.216e-11 | -25.13 | BOYLAN\_MULTIPLE\_MYELOMA\_PCA1\_UP | MSigDB lists | BOYLAN\_MULTIPLE\_MYELOMA\_PCA1\_UP | 67 | 13 | 12187 | 179 | S100a9,Ifi211,Xdh,S100a8,Igsf6,Zbtb16,Ifi209,Lcn2,Cp,Ifi207,Hp,Ifi204,Ifitm3 | | 1.239e-11 | -25.11 | ZHOU\_INFLAMMATORY\_RESPONSE\_LIVE\_UP | MSigDB lists | ZHOU\_INFLAMMATORY\_RESPONSE\_LIVE\_UP | 328 | 25 | 12187 | 179 | Il6,Tnfaip3,Gpr84,Ccl4,Maff,Rgs16,Cxcl1,Csf3,Ier3,Il1a,Akap12,Mmp8,Nfkb1,Ptpn1,Rnf213,Nfkbia,Sdc4,Tnf,Nfkb2,Cflar,Col4a1,Plaur,Icam1,Tnip1,Ptgs2 | | 1.247e-11 | -25.11 | GSE5679\_PPARG\_LIGAND\_ROSIGLITAZONE\_VS\_RARA\_AGONIST\_AM580\_TREATED\_DC\_DN | MSigDB lists | GSE5679\_PPARG\_LIGAND\_ROSIGLITAZONE\_VS\_RARA\_AGONIST\_AM580\_TREATED\_DC\_DN | 156 | 18 | 12187 | 179 | Samd9l,Znfx1,Sgk1,Irak3,Irf9,Rnf213,Pik3r1,Il4ra,Xaf1,Usp18,Rtp4,Cmpk2,Tnfsf10,Irgm2,Gbp7,Pik3r5,Irf7,Bcl3 | | 1.390e-11 | -25.00 | GSE7509\_DC\_VS\_MONOCYTE\_WITH\_FCGRIIB\_STIM\_DN | MSigDB lists | GSE7509\_DC\_VS\_MONOCYTE\_WITH\_FCGRIIB\_STIM\_DN | 157 | 18 | 12187 | 179 | Oasl1,Il6,Ripk1,Ccl12,Ccl2,Xaf1,Ifit2,Sbno2,Irf7,Atf3,Ifit3b,Ifi44,Irf9,Zfp36,P2ry6,Trim25,Isg15,Ifitm3 | | 1.396e-11 | -24.99 | DIRMEIER\_LMP1\_RESPONSE\_EARLY | MSigDB lists | DIRMEIER\_LMP1\_RESPONSE\_EARLY | 54 | 12 | 12187 | 179 | Nfkbiz,Icam1,Zfp36,Tnfaip3,Gadd45b,Tnf,Birc3,Ccl4,Nfkbia,Nfkb2,Nfkbie,Irf7 | | 1.472e-11 | -24.94 | YANG\_BCL3\_TARGETS\_UP | MSigDB lists | YANG\_BCL3\_TARGETS\_UP | 303 | 24 | 12187 | 179 | Herc6,Sgk3,Tap1,Rbm47,Ifit3b,Timp1,Akap12,Hp,Cp,Angptl4,Irf7,A2m,Dtx3l,Gbp3,Rtp4,Cmpk2,Parp14,Usp18,Ccnd2,Tnf,Ifit2,Ccl12,Il6,Oasl1 | | 1.638e-11 | -24.83 | NF-kappa B signaling pathway | KEGG pathways | mmu04064 | 76 | 15 | 5248 | 107 | Cflar,Gadd45b,Nfkbia,Cd14,Ccl4,Tnfaip3,Vcam1,Ripk1,Icam1,Tnf,Birc3,Nfkb1,Trim25,Nfkb2,Ptgs2 | | 1.638e-11 | -24.83 | NF-kappa B signaling pathway | KEGG pathways | ko04064 | 76 | 15 | 5248 | 107 | Cflar,Nfkbia,Gadd45b,Ccl4,Cd14,Tnfaip3,Tnf,Icam1,Birc3,Ripk1,Vcam1,Trim25,Nfkb1,Nfkb2,Ptgs2 | | 1.679e-11 | -24.81 | cell communication | biological process | GO:0007154 | 3040 | 91 | 13711 | 214 | Akap12,Nfkbia,Il4ra,Hcar2,Ccl7,Isg15,Fam107a,Map3k6,Tiparp,Rasip1,Rhou,Atf3,Pik3ap1,Hspa5,Tgm2,C5ar1,Selp,Angptl4,Rasd1,Nfe2l2,Rassf4,Rgs16,Nfkb2,Fyb,Mt1,Lgals9,Iigp1,Rhoj,Map3k8,Pik3r5,Ccl2,Cxcl9,Lcn2,Il1rn,C3ar1,Tnip3,Iqgap1,Col4a1,Ptpn1,Ifitm2,Nfkbiz,S100a9,Vcam1,Cdkn1a,Il6,Gem,Il1a,Casp4,Pik3r1,Trib1,Socs3,Irf7,Dtx3l,Fzd4,Ier3,Sgk1,Spi1,Ccl4,Ifitm6,Stx11,Zfp36,Tnf,Rhoc,Tnip1,Gpr84,Cd14,Ripk1,Hspb1,Mt2,Hp,Cxcl1,Rrp8,Csf2rb,Errfi1,Cybb,P2ry6,Ifitm3,Nfkb1,Osmr,Plaur,Kcna5,Ksr1,Plek,Ifi204,Irak3,Arid5b,Ccl12,S100a8,Irgm2,Bcl3,Sgk3 | | 1.726e-11 | -24.78 | GSE2706\_R848\_VS\_R848\_AND\_LPS\_2H\_STIM\_DC\_DN | MSigDB lists | GSE2706\_R848\_VS\_R848\_AND\_LPS\_2H\_STIM\_DC\_DN | 119 | 16 | 12187 | 179 | Ifit3b,Socs3,Ifi44,Znfx1,Il1a,Tnip3,Isg15,Ptgs2,Herc6,Tap1,Pik3ap1,Ifit2,Tnf,Oasl1,Tnfaip3,Cxcl1 | | 1.746e-11 | -24.77 | signaling | biological process | GO:0023052 | 2939 | 89 | 13711 | 214 | Sgk3,Bcl3,Irgm2,S100a8,Ccl12,Arid5b,Irak3,Ifi204,Plek,Ksr1,Kcna5,Plaur,Osmr,Nfkb1,Ifitm3,P2ry6,Errfi1,Cybb,Csf2rb,Cxcl1,Rrp8,Mt2,Hp,Hspb1,Cd14,Gpr84,Ripk1,Tnip1,Rhoc,Tnf,Zfp36,Ifitm6,Stx11,Ccl4,Spi1,Sgk1,Ier3,Fzd4,Irf7,Socs3,Dtx3l,Trib1,Pik3r1,Casp4,Il1a,Gem,Il6,Cdkn1a,Vcam1,S100a9,Nfkbiz,Ifitm2,Ptpn1,Tnip3,Col4a1,Iqgap1,C3ar1,Il1rn,Lcn2,Cxcl9,Ccl2,Pik3r5,Map3k8,Rhoj,Iigp1,Lgals9,Mt1,Rgs16,Fyb,Nfkb2,Rassf4,Nfe2l2,Rasd1,Selp,Tgm2,C5ar1,Hspa5,Pik3ap1,Atf3,Rhou,Rasip1,Tiparp,Map3k6,Isg15,Ccl7,Hcar2,Il4ra,Akap12,Nfkbia | | 1.805e-11 | -24.74 | DER\_IFN\_GAMMA\_RESPONSE\_UP | MSigDB lists | DER\_IFN\_GAMMA\_RESPONSE\_UP | 69 | 13 | 12187 | 179 | Ifit3b,Ifi209,Cebpd,Ifi204,Ifi207,Isg15,Tap1,Irf9,Icam1,Ifit2,Il6,Rhoc,Ifi211 | | 1.805e-11 | -24.74 | DER\_IFN\_ALPHA\_RESPONSE\_UP | MSigDB lists | DER\_IFN\_ALPHA\_RESPONSE\_UP | 69 | 13 | 12187 | 179 | Ifi211,Isg15,Irf2,Irf9,Tap1,Ifit3b,Ifi209,Ifi44,Oasl1,Il6,Rhoc,Ifi204,Ifi207 | | 1.854e-11 | -24.71 | GSE7768\_OVA\_ALONE\_VS\_OVA\_WITH\_LPS\_IMMUNIZED\_MOUSE\_WHOLE\_SPLEEN\_6H\_DN | MSigDB lists | GSE7768\_OVA\_ALONE\_VS\_OVA\_WITH\_LPS\_IMMUNIZED\_MOUSE\_WHOLE\_SPLEEN\_6H\_DN | 139 | 17 | 12187 | 179 | Nfkbiz,Samd9l,Nfkb1,Ptgs2,Cd14,Plek,Icam1,Plaur,Rhou,Nfkbia,Casp4,Ccl4,Cybb,Nfe2l2,Nfkbie,Pik3r5,Igsf6 | | 1.890e-11 | -24.69 | regulation of innate immune response | biological process | GO:0045088 | 214 | 21 | 13711 | 214 | Parp9,Tap1,Tnip3,A2m,Ifi209,Irgm2,Irgm1,Ifi211,Nfkbia,Pik3ap1,Tnfaip3,Tnf,Acod1,Ifi204,Trim30a,Lgals9,Tnip1,Parp14,Cd14,Irak3,Irf7 | | 1.908e-11 | -24.68 | regulation of primary metabolic process | biological process | GO:0080090 | 4246 | 113 | 13711 | 214 | Plek,Tnfsf10,Ksr1,Xdh,Irak3,Arid5b,Ifi204,Irf2,Ccl12,A2m,S100a8,Irf9,Irgm2,Bcl3,Samd9l,Sox11,Mxd4,Errfi1,Cybb,Thbs1,P2ry6,Rbm39,Banp,Gbp4,Nfkb1,Bach1,Plaur,Tnf,Zfp36,Cflar,Ripk1,Hspb1,Ifi207,Tnip1,Fstl1,Dnajb1,Rrp8,Prg4,Il1a,Casp4,Oasl2,Cdkn1a,Il6,Trib1,Ctla2a,Irf7,Socs3,Dtx3l,Pik3r1,Gadd45b,Fzd4,Ifi209,Sbno2,Sdc4,Parp9,Ccl4,Spi1,Ier3,Ptgs2,Znfx1,Tnfaip3,Ccl2,Tnfaip8,Csf3,Zbtb16,Aff1,C3ar1,Il1rn,Ptpn1,Iqgap1,S100a9,Wfdc21,Nfkb2,Icam1,Oasl1,Lgals9,Gadd45g,Birc3,Map3k8,Nfkbie,Pik3r5,Pik3ap1,Acod1,Atf3,Ncf1,Trim30a,Hspa5,C5ar1,Maff,Mmp8,Rasd1,Serpina3f,Nfe2l2,Noct,Msn,Angptl4,Akap12,Nfkbia,Ccl7,Parp14,Hcar2,Ccnd2,Fam107a,Map3k6,Tiparp,Timp1,Isg15,Cebpd,Rasip1,Ifi211,Resf1,Trim25,Runx1 | | 2.000e-11 | -24.64 | regulation of tumor necrosis factor production | biological process | GO:0032680 | 131 | 17 | 13711 | 214 | Bcl3,Mmp8,Ccl4,Pik3r1,Lgals9,Trim30a,Hspb1,Irak3,Thbs1,Cd14,Ripk1,Zfp36,Errfi1,Cybb,Ccl2,Tnfaip3,Akap12 | | 2.063e-11 | -24.60 | positive regulation of protein metabolic process | biological process | GO:0051247 | 1402 | 56 | 13711 | 214 | Bcl3,Irgm2,S100a9,S100a8,Iqgap1,Ccl12,Ptpn1,Il1rn,Csf3,Tnfsf10,Ksr1,Xdh,Ccl2,Tnfaip3,Pik3r5,Plaur,Map3k8,Birc3,Gadd45g,Lgals9,P2ry6,Thbs1,Cybb,Icam1,Msn,Mmp8,C5ar1,Hspa5,Tnip1,Trim30a,Hspb1,Ncf1,Cflar,Ripk1,Tnf,Ptgs2,Ier3,Ccl4,Sdc4,Parp9,Tiparp,Fam107a,Map3k6,Fzd4,Ccnd2,Pik3r1,Gadd45b,Ccl7,Parp14,Trib1,Il6,Cdkn1a,Casp4,Akap12,Il1a,Nfkbia | | 2.083e-11 | -24.59 | GSE13484\_3H\_UNSTIM\_VS\_YF17D\_VACCINE\_STIM\_PBMC\_DN | MSigDB lists | GSE13484\_3H\_UNSTIM\_VS\_YF17D\_VACCINE\_STIM\_PBMC\_DN | 140 | 17 | 12187 | 179 | Oasl1,Bach1,Xaf1,Tor3a,Ccl4,Igsf6,Tnfaip8,Cflar,Gem,Il1a,Aff1,Plaur,Clic4,Irf9,Trim25,Icam1,Ptgs2 | | 2.134e-11 | -24.57 | GSE5679\_CTRL\_VS\_PPARG\_LIGAND\_ROSIGLITAZONE\_AND\_RARA\_AGONIST\_AM580\_TREATED\_DC\_UP | MSigDB lists | GSE5679\_CTRL\_VS\_PPARG\_LIGAND\_ROSIGLITAZONE\_AND\_RARA\_AGONIST\_AM580\_TREATED\_DC\_UP | 161 | 18 | 12187 | 179 | Clic4,Tap1,Icam1,Slfn3,Isg15,Znfx1,Ifit3b,Aff1,Gbp3,Slfn4,S100a8,Irf7,Irgm2,Cflar,Nfkbie,Gbp7,Cybb,Usp18 | | 2.186e-11 | -24.55 | positive regulation of biological process | biological process | GO:0048518 | 4720 | 121 | 13711 | 214 | Atf3,Acod1,Pik3ap1,Trim30a,Sele,Ncf1,Tgm2,C5ar1,Maff,Hspa5,Selp,Noct,Angptl4,Msn,Nfe2l2,Mmp8,Ip6k2,Il4ra,Akap12,Nfkbia,Hcar2,Ccnd2,Ccl7,Parp14,Isg15,Tiparp,Timp1,Fam107a,Map3k6,Runx1,Resf1,Trim25,Rasip1,Cebpd,Ifi211,Ccl2,Tnfaip3,Cxcl9,Lcn2,Aff1,Il1rn,C3ar1,Tnfaip8,Zbtb16,Csf3,Tnip3,Iqgap1,Cxcl16,Nfkbiz,Ptpn1,Vcam1,S100a9,Icam1,Fyb,Nfkb2,Gadd45g,Lgals9,Msr1,Rhoj,Map3k8,Birc3,Pik3r5,Irgm1,Zfp36,Tnf,Tnip1,Rhoc,Hspb1,Cflar,Cd14,Ripk1,Hp,Rrp8,Cxcl1,Ifit2,Il6,Cdkn1a,Casp4,Il1a,Pik3r1,Gadd45b,Irf7,Socs3,Dtx3l,Trib1,Sbno2,Sdc4,Parp9,Ifi209,Fzd4,Sgk1,Ptgs2,Ier3,Spi1,Ccl4,Ksr1,Tnfsf10,Xdh,Plek,Ifi204,Arid5b,Irak3,Adamts9,S100a8,Ccl12,Irf2,Bcl3,Irgm2,Cybb,Sox11,Banp,Rbm39,P2ry6,Thbs1,Nfkb1,Adamts1,Osmr,Plaur,Il17ra,Bach1,Kcna5 | | 2.189e-11 | -24.54 | HALLMARK\_IL6\_JAK\_STAT3\_SIGNALING | MSigDB lists | HALLMARK\_IL6\_JAK\_STAT3\_SIGNALING | 70 | 13 | 12187 | 179 | Il6,Il4ra,Tnf,Map3k8,Osmr,A2m,Pik3r5,Socs3,Il17ra,Irf9,Csf2rb,Ptpn1,Cd14 | | 2.228e-11 | -24.53 | MODULE\_1 | MSigDB lists | MODULE\_1 | 309 | 24 | 12187 | 179 | Ier3,Tiparp,Akap12,Gadd45g,Sgk1,Cd14,Tnfaip3,Ccl12,Nfe2l2,Il6,Tgm2,Ccnd2,Maff,Iqgap1,Ifi211,Atf3,Ifi204,Ifi207,Col4a1,Ifi209,Cdkn1a,Irf9,Sele,Gem | | 2.264e-11 | -24.51 | regulation of tumor necrosis factor superfamily cytokine production | biological process | GO:1903555 | 132 | 17 | 13711 | 214 | Cybb,Errfi1,Zfp36,Ccl2,Tnfaip3,Akap12,Pik3r1,Trim30a,Lgals9,Thbs1,Irak3,Hspb1,Ripk1,Cd14,Bcl3,Ccl4,Mmp8 | | 2.339e-11 | -24.48 | GSE40685\_TREG\_VS\_FOXP3\_KO\_TREG\_PRECURSOR\_UP | MSigDB lists | GSE40685\_TREG\_VS\_FOXP3\_KO\_TREG\_PRECURSOR\_UP | 141 | 17 | 12187 | 179 | Znfx1,Samd9l,Ifit3b,Irf9,Tap1,Parp9,Il17ra,Parp12,Oasl1,Casp4,Parp14,Xaf1,Ifit2,Tnfsf10,Gbp3,Rtp4,Bcl3 | | 2.369e-11 | -24.47 | response to tumor necrosis factor | biological process | GO:0034612 | 114 | 16 | 13711 | 214 | Birc3,Cxcl16,Ccl12,Nfkb1,Ccl4,Nfe2l2,Wfdc21,Nfkbia,Tnf,Acod1,Ccl2,Zfp36,Il6,Ripk1,Ccl7,Cd14 | | 2.918e-11 | -24.26 | GSE43863\_DAY6\_EFF\_VS\_DAY150\_MEM\_TH1\_CD4\_TCELL\_DN | MSigDB lists | GSE43863\_DAY6\_EFF\_VS\_DAY150\_MEM\_TH1\_CD4\_TCELL\_DN | 164 | 18 | 12187 | 179 | Socs3,Nfkbiz,Nfkb1,Ksr1,Ms4a6d,Tagln2,Ncf1,Irf9,Birc3,Gbp4,Tnfaip3,Gadd45b,Il6,Bcl3,Cflar,Irgm2,Gbp6,Gbp3 | | 2.935e-11 | -24.25 | cellular response to oxygen-containing compound | biological process | GO:1901701 | 686 | 37 | 13711 | 214 | Vcam1,Nfe2l2,Gbp6,Irgm2,Msn,Ptpn1,Cxcl1,Cxcl16,Ccl12,Col4a1,Tnip3,Ripk1,Cflar,Cd14,Il1rn,Ncf1,Lcn2,Cxcl9,Acod1,Tnfaip3,Tnf,Ccl2,Zfp36,Sgk1,Sbno2,Cmpk2,Nfkb1,Ccl7,Thbs1,P2ry6,Pik3r1,Nfkbia,Icam1,Mt1,Casp4,Ip6k2,Il6 | | 2.940e-11 | -24.25 | GSE43955\_TH0\_VS\_TGFB\_IL6\_TH17\_ACT\_CD4\_TCELL\_30H\_UP | MSigDB lists | GSE43955\_TH0\_VS\_TGFB\_IL6\_TH17\_ACT\_CD4\_TCELL\_30H\_UP | 143 | 17 | 12187 | 179 | Gbp3,Gbp7,Bach1,Gadd45b,Ccl12,Il4ra,Map3k8,Casp4,Ccl4,Il17ra,Errfi1,Ptpn1,Csf2rb,Ptgs2,Selp,Tiparp,Socs3 | | 3.015e-11 | -24.22 | positive regulation of metabolic process | biological process | GO:0009893 | 2865 | 87 | 13711 | 214 | Cybb,Sox11,P2ry6,Banp,Rbm39,Thbs1,Nfkb1,Plaur,Bach1,Xdh,Ksr1,Tnfsf10,Plek,Arid5b,Ccl12,Irf2,S100a8,Irgm2,Bcl3,Cdkn1a,Il6,Il1a,Casp4,Gadd45b,Pik3r1,Trib1,Dtx3l,Irf7,Parp9,Sbno2,Sdc4,Fzd4,Ier3,Ptgs2,Spi1,Ccl4,Zfp36,Tnf,Tnip1,Ripk1,Cflar,Hspb1,Cxcl1,Nfkb2,Icam1,Lgals9,Gadd45g,Birc3,Map3k8,Pik3r5,Irgm1,Ccl2,Tnfaip3,Lcn2,Csf3,Zbtb16,Il1rn,Aff1,C3ar1,Iqgap1,Ptpn1,S100a9,Nfkbia,Akap12,Il4ra,Ccnd2,Parp14,Ccl7,Map3k6,Fam107a,Tiparp,Resf1,Runx1,Cebpd,Atf3,Acod1,Trim30a,Ncf1,Sele,Hspa5,C5ar1,Maff,Msn,Angptl4,Noct,Mmp8,Nfe2l2 | | 3.121e-11 | -24.19 | GO\_NEGATIVE\_REGULATION\_OF\_MULTI\_ORGANISM\_PROCESS | MSigDB lists | GO\_NEGATIVE\_REGULATION\_OF\_MULTI\_ORGANISM\_PROCESS | 105 | 15 | 12187 | 179 | Tnfaip3,Oasl1,Tnf,Ccl4,Trib1,Ifi211,Timp1,Ifi204,Ifi207,Ifi209,Trim25,Ifitm3,Ifitm2,Isg15,Tnip1 | | 3.141e-11 | -24.18 | response to protozoan | biological process | GO:0001562 | 23 | 9 | 13711 | 214 | Iigp1,Gbp6,Irgm2,Ier3,Gbp7,Bcl3,Il4ra,Gbp3,Il6 | | 3.188e-11 | -24.17 | MODULE\_128 | MSigDB lists | MODULE\_128 | 72 | 13 | 12187 | 179 | C3ar1,A2m,Cybb,Ccl12,Ccnd2,Ifitm3,Ifitm2,Cd14,Timp1,Ier3,Cebpd,Thbs1,Vcam1 | | 3.511e-11 | -24.07 | UZONYI\_RESPONSE\_TO\_LEUKOTRIENE\_AND\_THROMBIN | MSigDB lists | UZONYI\_RESPONSE\_TO\_LEUKOTRIENE\_AND\_THROMBIN | 34 | 10 | 12187 | 179 | Ptgs2,Gem,Arid5b,Cxcl1,Sele,Zfp36,Apold1,Map3k8,Adamts1,Atf3 | | 3.829e-11 | -23.99 | MODULE\_79 | MSigDB lists | MODULE\_79 | 73 | 13 | 12187 | 179 | Timp1,Cebpd,Ier3,Thbs1,Vcam1,Ifitm3,Ifitm2,Cd14,Cybb,Ccl12,Ccnd2,C3ar1,A2m | | 3.963e-11 | -23.95 | GSE46606\_UNSTIM\_VS\_CD40L\_IL2\_IL5\_3DAY\_STIMULATED\_IRF4MID\_SORTED\_BCELL\_DN | MSigDB lists | GSE46606\_UNSTIM\_VS\_CD40L\_IL2\_IL5\_3DAY\_STIMULATED\_IRF4MID\_SORTED\_BCELL\_DN | 167 | 18 | 12187 | 179 | Gbp4,Hcar2,Casp4,Nfkbia,Nfkb2,Rtp4,Gbp6,Gem,Znfx1,Samd9l,Tiparp,Socs3,Cdkn1a,Tnip1,Rnf213,Ptgs2,Isg15,Ifitm3 | | 3.984e-11 | -23.95 | regulation of cellular process | biological process | GO:0050794 | 7419 | 162 | 13711 | 214 | Ptpn1,Ifitm2,Nfkbiz,Cxcl16,Tnip3,Col4a1,Iqgap1,S100a9,Vcam1,Wfdc21,Cxcl9,Tnfaip3,Znfx1,Ccl2,Depp1,Tnfaip8,Zbtb16,Csf3,C3ar1,Aff1,Il1rn,Lcn2,Birc3,Map3k8,Nfkbie,Apold1,Rhoj,Irgm1,Pik3r5,Rgs16,Nfkb2,Fyb,Mt1,Icam1,Rassf4,Msr1,Iigp1,Bbs12,Oasl1,Lgals9,Gadd45g,Hspa5,Clic4,Tgm2,C5ar1,Maff,Mmp8,Rasd1,Serpina3f,Nfe2l2,Noct,Selp,Angptl4,Msn,Pik3ap1,Acod1,Rhou,Atf3,Sele,Ncf1,Trim30a,Rnf213,Fam107a,Map3k6,Tiparp,Timp1,Isg15,Slfn2,Cebpd,Rasip1,Xaf1,Ifi211,Resf1,Trim25,Runx1,Akap12,Nfkbia,Il4ra,Slfn3,Ip6k2,Ccl7,Parp14,Hcar2,Ccnd2,Ccl12,Irf2,A2m,S100a8,Sgk3,Irf9,Irgm2,Bcl3,Plek,Ksr1,Tnfsf10,Xdh,Adamts9,Arid5b,Irak3,Ifi204,Osmr,Gbp4,Adamts1,Nfkb1,Kcna5,Bach1,Plaur,Il17ra,Sox11,Mxd4,Errfi1,Cybb,Ifitm3,Thbs1,P2ry6,Banp,Rbm39,Dnajb1,Rrp8,Cxcl1,Prg4,Mt2,Hp,Ifit2,Csf2rb,Robo4,Tnf,Zfp36,Gpr84,Cd14,Cflar,Ripk1,Hspb1,Ifi207,Rhoc,Tnip1,Fstl1,Fzd4,Ifi209,Sdc4,Ptges,Sbno2,Parp9,Spi1,Ccl4,Ifitm6,Ier3,Sgk1,Ptgs2,Il1a,Gem,Casp4,Oasl2,Cdkn1a,Il6,Ctla2a,Trib1,Irf7,Socs3,Dtx3l,Pik3r1,Gadd45b | | 4.022e-11 | -23.94 | RASHI\_NFKB1\_TARGETS | MSigDB lists | RASHI\_NFKB1\_TARGETS | 17 | 8 | 12187 | 179 | Tnip1,Cxcl1,Nfkb2,Birc3,Map3k8,Nfkbia,Lcn2,Tnfaip3 | | 4.277e-11 | -23.88 | MODULE\_76 | MSigDB lists | MODULE\_76 | 59 | 12 | 12187 | 179 | Cxcl1,S100a9,C3ar1,Ccl12,Cybb,Ccl4,Ccl2,Cd14,Ptgs2,Saa1,Nfkb1,Il1a | | 4.395e-11 | -23.85 | regulation of nitrogen compound metabolic process | biological process | GO:0051171 | 4126 | 110 | 13711 | 214 | Il1rn,Aff1,C3ar1,Csf3,Tnfaip8,Zbtb16,Ccl2,Znfx1,Tnfaip3,Wfdc21,S100a9,Iqgap1,Ptpn1,Gadd45g,Lgals9,Oasl1,Icam1,Nfkb2,Pik3r5,Nfkbie,Map3k8,Birc3,Trim30a,Ncf1,Atf3,Acod1,Pik3ap1,Msn,Noct,Nfe2l2,Serpina3f,Rasd1,Mmp8,C5ar1,Maff,Hspa5,Ccnd2,Parp14,Ccl7,Nfkbia,Akap12,Runx1,Resf1,Trim25,Ifi211,Cebpd,Rasip1,Isg15,Timp1,Tiparp,Map3k6,Fam107a,Ifi204,Arid5b,Irak3,Xdh,Tnfsf10,Ksr1,Bcl3,Samd9l,Irgm2,Irf9,S100a8,A2m,Irf2,Ccl12,Banp,Rbm39,P2ry6,Thbs1,Cybb,Errfi1,Mxd4,Sox11,Plaur,Bach1,Nfkb1,Gbp4,Fstl1,Tnip1,Ifi207,Hspb1,Ripk1,Cflar,Zfp36,Tnf,Prg4,Rrp8,Dnajb1,Gadd45b,Pik3r1,Dtx3l,Socs3,Irf7,Trib1,Ctla2a,Il6,Cdkn1a,Oasl2,Casp4,Il1a,Ptgs2,Ier3,Ccl4,Spi1,Parp9,Sbno2,Sdc4,Ifi209,Fzd4 | | 4.593e-11 | -23.80 | HALLMARK\_APOPTOSIS | MSigDB lists | HALLMARK\_APOPTOSIS | 147 | 17 | 12187 | 179 | Gadd45b,Il6,Tnf,Ccnd2,Birc3,Casp4,Tnfsf10,Cflar,Timp1,Il1a,Atf3,Ier3,Cdkn1a,Hspb1,Tap1,Ifitm3,Cd14 | | 4.782e-11 | -23.76 | regulation of cellular metabolic process | biological process | GO:0031323 | 4418 | 115 | 13711 | 214 | Ifi209,Fzd4,Sdc4,Sbno2,Parp9,Ccl4,Spi1,Ptgs2,Ier3,Casp4,Oasl2,Il1a,Il6,Cdkn1a,Irf7,Socs3,Dtx3l,Trib1,Ctla2a,Pik3r1,Gadd45b,Rrp8,Dnajb1,Cxcl1,Prg4,Hp,Tnf,Zfp36,Hspb1,Ifi207,Cflar,Ripk1,Tnip1,Fstl1,Gbp4,Nfkb1,Bach1,Plaur,Mxd4,Sox11,Errfi1,Cybb,Thbs1,Banp,Rbm39,P2ry6,A2m,S100a8,Ccl12,Irf2,Irf9,Bcl3,Irgm2,Plek,Tnfsf10,Ksr1,Xdh,Irak3,Arid5b,Ifi204,Timp1,Fam107a,Map3k6,Isg15,Rasip1,Cebpd,Ifi211,Runx1,Resf1,Trim25,Akap12,Nfkbia,Ccl7,Parp14,Ccnd2,C5ar1,Maff,Hspa5,Serpina3f,Nfe2l2,Rasd1,Mmp8,Noct,Msn,Acod1,Pik3ap1,Atf3,Sele,Ncf1,Trim30a,Map3k8,Birc3,Nfkbie,Irgm1,Pik3r5,Icam1,Nfkb2,Oasl1,Iigp1,Gadd45g,Lgals9,Ptpn1,Iqgap1,S100a9,Wfdc21,Tnfaip3,Znfx1,Ccl2,Depp1,Aff1,C3ar1,Il1rn,Csf3,Tnfaip8,Zbtb16 | | 4.844e-11 | -23.75 | GSE20715\_0H\_VS\_24H\_OZONE\_TLR4\_KO\_LUNG\_DN | MSigDB lists | GSE20715\_0H\_VS\_24H\_OZONE\_TLR4\_KO\_LUNG\_DN | 169 | 18 | 12187 | 179 | Saa1,Thbs1,Hspa5,Cdkn1a,Socs3,Zbtb16,Col4a1,Hp,Gadd45g,Timp1,Lcn2,Bcl3,Cxcl1,Angptl4,Osmr,Maff,Rhou,Tubb6 | | 5.411e-11 | -23.64 | GO\_DEFENSE\_RESPONSE\_TO\_VIRUS | MSigDB lists | GO\_DEFENSE\_RESPONSE\_TO\_VIRUS | 109 | 15 | 12187 | 179 | Ifi211,Trim56,Irf7,Oasl1,Il6,Ifit2,Irf9,Trim25,Ifitm3,Ifitm2,Isg15,Ifi204,Ifi207,Ifit3b,Ifi209 | | 5.510e-11 | -23.62 | positive regulation of cellular process | biological process | GO:0048522 | 4254 | 112 | 13711 | 214 | Sele,Ncf1,Acod1,Pik3ap1,Atf3,Nfe2l2,Mmp8,Selp,Noct,Msn,Tgm2,C5ar1,Maff,Hspa5,Ccl7,Parp14,Ccnd2,Hcar2,Il4ra,Akap12,Nfkbia,Ip6k2,Rasip1,Cebpd,Runx1,Trim25,Resf1,Timp1,Fam107a,Map3k6,Isg15,Il1rn,C3ar1,Aff1,Csf3,Tnfaip8,Zbtb16,Lcn2,Tnfaip3,Cxcl9,Ccl2,Vcam1,S100a9,Nfkbiz,Ptpn1,Iqgap1,Cxcl16,Msr1,Gadd45g,Lgals9,Icam1,Nfkb2,Irgm1,Pik3r5,Map3k8,Birc3,Rhoj,Hspb1,Cd14,Cflar,Ripk1,Tnip1,Rhoc,Tnf,Zfp36,Ifit2,Rrp8,Cxcl1,Hp,Socs3,Irf7,Dtx3l,Trib1,Pik3r1,Gadd45b,Casp4,Il1a,Il6,Cdkn1a,Spi1,Ccl4,Sgk1,Ptgs2,Fzd4,Sbno2,Sdc4,Parp9,Arid5b,Adamts9,Ifi204,Plek,Ksr1,Tnfsf10,Xdh,Bcl3,Irgm2,S100a8,Ccl12,Irf2,Thbs1,Rbm39,Banp,P2ry6,Sox11,Cybb,Bach1,Kcna5,Plaur,Il17ra,Osmr,Nfkb1,Adamts1 | | 5.643e-11 | -23.60 | GO\_POSITIVE\_REGULATION\_OF\_INTRACELLULAR\_SIGNAL\_TRANSDUCTION | MSigDB lists | GO\_POSITIVE\_REGULATION\_OF\_INTRACELLULAR\_SIGNAL\_TRANSDUCTION | 645 | 34 | 12187 | 179 | Birc3,Tnf,Fzd4,S100a9,Cflar,C5ar1,Tnfsf10,Xdh,Socs3,Ksr1,Saa1,Ptgs2,Icam1,Trim25,Ccl2,Ccl4,Map3k8,Il6,Gadd45b,Ripk1,Ccl12,Rhoc,Pik3r5,Csf3,S100a8,Thbs1,Sox11,Gadd45g,Il1a,Akap12,Map3k6,Selp,Ptpn1,Pik3ap1 | | 6.360e-11 | -23.48 | GSE14000\_UNSTIM\_VS\_16H\_LPS\_DC\_TRANSLATED\_RNA\_DN | MSigDB lists | GSE14000\_UNSTIM\_VS\_16H\_LPS\_DC\_TRANSLATED\_RNA\_DN | 150 | 17 | 12187 | 179 | Parp12,Birc3,Ifit2,Usp18,Ccl4,Cmpk2,Tnfsf10,Cflar,Trim56,Irf7,Phf11d,Samd9l,Atf3,Phf11b,Ifit3b,Nfkbiz,Herc6 | | 6.360e-11 | -23.48 | GSE22886\_NAIVE\_BCELL\_VS\_NEUTROPHIL\_DN | MSigDB lists | GSE22886\_NAIVE\_BCELL\_VS\_NEUTROPHIL\_DN | 150 | 17 | 12187 | 179 | C5ar1,Cflar,Bcl3,S100a9,Igsf6,S100a8,Tnfsf10,Trib1,Casp4,Ifitm3,Ptgs2,Ifitm2,Plek,Csf2rb,Ifit3b,Ier3,Cebpd | | 6.360e-11 | -23.48 | GSE8515\_CTRL\_VS\_IL1\_4H\_STIM\_MAC\_DN | MSigDB lists | GSE8515\_CTRL\_VS\_IL1\_4H\_STIM\_MAC\_DN | 150 | 17 | 12187 | 179 | Tnip1,Icam1,Plek,Nfkb1,Tnip3,Ier3,Nfkbie,Cxcl1,Bcl3,Nfkb2,Tnf,Ccl4,Nfkbia,Sdc4,Tnfaip3,Gadd45b,Il6 | | 6.901e-11 | -23.40 | DEBIASI\_APOPTOSIS\_BY\_REOVIRUS\_INFECTION\_UP | MSigDB lists | DEBIASI\_APOPTOSIS\_BY\_REOVIRUS\_INFECTION\_UP | 271 | 22 | 12187 | 179 | Oasl1,Tnfaip3,Bach1,Nfkbia,Ifit2,Zfp189,Irf7,Cxcl1,Ifi211,Gem,Atf3,Ifi207,Ifi204,Tiparp,Ifit3b,Ifi44,Ifi209,Tap1,Irf9,Zfp36,Isg15,Ifitm3 | | 7.076e-11 | -23.37 | GSE22589\_HEALTHY\_VS\_HIV\_INFECTED\_DC\_DN | MSigDB lists | GSE22589\_HEALTHY\_VS\_HIV\_INFECTED\_DC\_DN | 151 | 17 | 12187 | 179 | Irf9,Tap1,Herc6,Ifitm3,Slfn5,Cybb,Gbp4,Tnf,Tor3a,Parp14,Rtp4,Gbp3,Gbp7,Gbp6,Gem,C5ar1,Irf7 | | 7.076e-11 | -23.37 | GSE360\_L\_DONOVANI\_VS\_M\_TUBERCULOSIS\_DC\_DN | MSigDB lists | GSE360\_L\_DONOVANI\_VS\_M\_TUBERCULOSIS\_DC\_DN | 151 | 17 | 12187 | 179 | Plek,Sele,Isg15,Cd14,Mmp8,Ifi44,Olfml2b,S100a8,Tnfsf10,Pygm,C3ar1,S100a9,Irf7,Ccl12,Oasl1,Ifit2,Ccl4 | | 7.080e-11 | -23.37 | NUYTTEN\_NIPP1\_TARGETS\_UP | MSigDB lists | NUYTTEN\_NIPP1\_TARGETS\_UP | 580 | 32 | 12187 | 179 | Sgk1,Slfn5,Rnf213,Isg15,Tap1,Parp9,Ccdc58,Ptpn1,Tgm2,Tnfaip3,Cxcl1,Ifi211,Csf3,Rtp4,Ifi44,Ifi209,Atf3,Luc7l3,Znfx1,Ifi207,Samd9l,Ifi204,Ptgs2,Casp4,Parp14,Xaf1,Birc3,Parp12,Oasl1,Gem,Robo4,Tnfsf10 | | 7.327e-11 | -23.34 | COLINA\_TARGETS\_OF\_4EBP1\_AND\_4EBP2 | MSigDB lists | COLINA\_TARGETS\_OF\_4EBP1\_AND\_4EBP2 | 299 | 23 | 12187 | 179 | Irgm2,Sdc4,Tor3a,Parp12,Trim25,Irf9,Ifi44,Adamts1,Vcam1,Cp,Il1rn,Rtp4,Gbp3,Ifit2,Usp18,Tgm2,Isg15,Ifitm3,Cd14,Thbs1,Akap12,Nfkb1,Lcn2 | | 8.738e-11 | -23.16 | GSE22886\_DAY1\_VS\_DAY7\_MONOCYTE\_IN\_CULTURE\_UP | MSigDB lists | GSE22886\_DAY1\_VS\_DAY7\_MONOCYTE\_IN\_CULTURE\_UP | 153 | 17 | 12187 | 179 | Ptgs2,Tnip1,Icam1,Irak3,Il1a,Tnip3,Ier3,Cflar,Irf7,Banp,Tnfaip8,Runx1,Nfkbia,Oasl1,Il6,Gadd45b,Tnfaip3 | | 9.698e-11 | -23.06 | GSE37301\_MULTIPOTENT\_PROGENITOR\_VS\_GRAN\_MONO\_PROGENITOR\_UP | MSigDB lists | GSE37301\_MULTIPOTENT\_PROGENITOR\_VS\_GRAN\_MONO\_PROGENITOR\_UP | 154 | 17 | 12187 | 179 | Il17ra,Ptpn1,Isg15,Cd14,Col4a1,Cebpd,Znfx1,Cmpk2,Xdh,Gbp7,Irgm2,C5ar1,Bcl3,Cxcl1,Ifit2,Usp18,Tor3a | | 9.698e-11 | -23.06 | GSE37533\_PPARG2\_FOXP3\_VS\_FOXP3\_TRANSDUCED\_CD4\_TCELL\_DN | MSigDB lists | GSE37533\_PPARG2\_FOXP3\_VS\_FOXP3\_TRANSDUCED\_CD4\_TCELL\_DN | 154 | 17 | 12187 | 179 | Rtp4,Irf7,Phf11d,Parp12,Oasl1,Ripk1,Usp18,Xaf1,Ifit2,Irf9,Tap1,Irf2,Herc6,Isg15,Ifit3b,Ifi44,Phf11b | | 9.698e-11 | -23.06 | GSE360\_LOW\_DOSE\_B\_MALAYI\_VS\_M\_TUBERCULOSIS\_DC\_DN | MSigDB lists | GSE360\_LOW\_DOSE\_B\_MALAYI\_VS\_M\_TUBERCULOSIS\_DC\_DN | 154 | 17 | 12187 | 179 | Icam1,Plaur,Tnip1,Ier3,Atf3,Ifi44,Hspa5,S100a8,Cflar,C3ar1,Gem,S100a9,Cybb,Ccl12,Oasl1,Ifit2,Birc3 | | 1.076e-10 | -22.95 | GSE8515\_IL1\_VS\_IL6\_4H\_STIM\_MAC\_UP | MSigDB lists | GSE8515\_IL1\_VS\_IL6\_4H\_STIM\_MAC\_UP | 155 | 17 | 12187 | 179 | Cxcl1,Nfkb2,Ccl4,Tor3a,Nfkbia,Sdc4,Tnf,Maff,Tnfaip3,Il6,Tnip1,Ptgs2,Icam1,Plek,Ier3,Nfkb1,Stx11 | | 1.078e-10 | -22.95 | GSE41176\_UNSTIM\_VS\_ANTI\_IGM\_STIM\_BCELL\_3H\_UP | MSigDB lists | GSE41176\_UNSTIM\_VS\_ANTI\_IGM\_STIM\_BCELL\_3H\_UP | 134 | 16 | 12187 | 179 | Irak3,Plaur,Rgs16,Cflar,Ptgs2,Tnip1,Cxcl1,Tnip3,Il1a,Tnfaip3,Ier3,Il6,Tnf,Maff,Socs3,Ccl4 | | 1.095e-10 | -22.94 | regulation of protein phosphorylation | biological process | GO:0001932 | 1172 | 49 | 13711 | 214 | Iqgap1,Ccl12,Ptpn1,Irgm2,Xdh,Ksr1,Ccl2,Tnfaip3,Il1rn,Irak3,Csf3,Map3k8,Gbp4,Pik3r5,Plaur,Errfi1,Icam1,Gadd45g,P2ry6,Lgals9,Thbs1,C5ar1,Mmp8,Atf3,Tnf,Pik3ap1,Tnip1,Ncf1,Hspb1,Ripk1,Cflar,Parp9,Sdc4,Map3k6,Fzd4,Ptgs2,Rasip1,Ccl4,Il6,Cdkn1a,Akap12,Il1a,Gadd45b,Ccnd2,Pik3r1,Socs3,Trib1,Parp14,Ccl7 | | 1.203e-10 | -22.84 | MCDOWELL\_ACUTE\_LUNG\_INJURY\_UP | MSigDB lists | MCDOWELL\_ACUTE\_LUNG\_INJURY\_UP | 38 | 10 | 12187 | 179 | Akap12,Lcn2,Il4ra,Cdkn1a,Nfkbia,Zfp36,Osmr,S100a8,Ptges,Angptl4 | | 1.203e-10 | -22.84 | RADAEVA\_RESPONSE\_TO\_IFNA1\_UP | MSigDB lists | RADAEVA\_RESPONSE\_TO\_IFNA1\_UP | 38 | 10 | 12187 | 179 | Tnfsf10,Isg15,Ifitm2,Ifitm3,Irf7,Rrp8,Mxd4,Ifi44,Ifit2,Vcam1 | | 1.225e-10 | -22.82 | regulation of macromolecule metabolic process | biological process | GO:0060255 | 4362 | 113 | 13711 | 214 | Zfp36,Tnf,Fstl1,Tnip1,Ifi207,Hspb1,Ripk1,Cflar,Prg4,Dnajb1,Rrp8,Il6,Cdkn1a,Oasl2,Casp4,Il1a,Gadd45b,Pik3r1,Dtx3l,Irf7,Socs3,Ctla2a,Trib1,Parp9,Sdc4,Sbno2,Ifi209,Fzd4,Ptgs2,Ier3,Ccl4,Spi1,Xdh,Ksr1,Tnfsf10,Ifi204,Arid5b,Irak3,S100a8,A2m,Ccl12,Irf2,Samd9l,Bcl3,Irgm2,Irf9,Cybb,Errfi1,Mxd4,Sox11,Banp,Rbm39,P2ry6,Thbs1,Nfkb1,Gbp4,Plaur,Bach1,Atf3,Acod1,Pik3ap1,Trim30a,Ncf1,Sele,C5ar1,Maff,Hspa5,Msn,Noct,Nfe2l2,Serpina3f,Rasd1,Mmp8,Nfkbia,Akap12,Ccnd2,Parp14,Ccl7,Isg15,Tiparp,Timp1,Map3k6,Fam107a,Runx1,Resf1,Trim25,Ifi211,Rasip1,Cebpd,Ccl2,Tnfaip3,Znfx1,Lcn2,C3ar1,Il1rn,Aff1,Tnfaip8,Csf3,Zbtb16,Iqgap1,Ptpn1,Wfdc21,S100a9,Icam1,Nfkb2,Gadd45g,Lgals9,Oasl1,Msr1,Nfkbie,Map3k8,Birc3,Pik3r5 | | 1.320e-10 | -22.75 | GSE19888\_ADENOSINE\_A3R\_INH\_VS\_INH\_PRETREAT\_AND\_ACT\_WITH\_TCELL\_MEMBRANES\_MAST\_CELL\_UP | MSigDB lists | GSE19888\_ADENOSINE\_A3R\_INH\_VS\_INH\_PRETREAT\_AND\_ACT\_WITH\_TCELL\_MEMBRANES\_MAST\_CELL\_UP | 157 | 17 | 12187 | 179 | Isg15,Rnf213,Samd9l,Ms4a6d,Akap12,Gbp3,Rtp4,Gbp6,Gbp7,Dtx3l,Irgm2,Parp12,Gbp4,Usp18,Casp4,Tor3a,Parp14 | | 1.413e-10 | -22.68 | ST\_TUMOR\_NECROSIS\_FACTOR\_PATHWAY | MSigDB lists | ST\_TUMOR\_NECROSIS\_FACTOR\_PATHWAY | 28 | 9 | 12187 | 179 | Nfkb2,Nfkbie,Cflar,Ripk1,Tnfaip3,Nfkb1,Nfkbia,Birc3,Tnf | | 1.460e-10 | -22.65 | GSE39864\_WT\_VS\_GATA3\_KO\_TREG\_UP | MSigDB lists | GSE39864\_WT\_VS\_GATA3\_KO\_TREG\_UP | 158 | 17 | 12187 | 179 | Timp1,Ptgs2,Selp,Ifitm3,Slfn3,Tap1,Il4ra,Ifit2,Map3k8,Ccl2,Casp4,Usp18,Ccl12,Gbp7,Irgm2,Slfn4,Gbp3 | | 1.598e-10 | -22.56 | GO\_NEGATIVE\_REGULATION\_OF\_VIRAL\_GENOME\_REPLICATION | MSigDB lists | GO\_NEGATIVE\_REGULATION\_OF\_VIRAL\_GENOME\_REPLICATION | 39 | 10 | 12187 | 179 | Tnip1,Ifi211,Isg15,Ifitm2,Ifitm3,Tnf,Ifi209,Oasl1,Ifi207,Ifi204 | | 1.614e-10 | -22.55 | GSE45365\_WT\_VS\_IFNAR\_KO\_CD8A\_DC\_DN | MSigDB lists | GSE45365\_WT\_VS\_IFNAR\_KO\_CD8A\_DC\_DN | 159 | 17 | 12187 | 179 | Tap1,Ptpn1,Tnip1,Herc6,Ifi44,Phf11b,Irf7,Cxcl1,Phf11d,Robo4,Csf3,Parp12,Tnfaip3,Rassf4,Nfkbia,Sdc4,Il4ra | | 1.719e-10 | -22.48 | GSE17974\_IL4\_AND\_ANTI\_IL12\_VS\_UNTREATED\_48H\_ACT\_CD4\_TCELL\_DN | MSigDB lists | GSE17974\_IL4\_AND\_ANTI\_IL12\_VS\_UNTREATED\_48H\_ACT\_CD4\_TCELL\_DN | 118 | 15 | 12187 | 179 | Atf3,Ksr1,Ifi204,Ifi207,Cdkn1a,Ifi209,Phf11b,Ifi44,Parp9,Parp14,Map3k8,Pik3r5,Phf11d,Ifi211,Irf7 | | 1.720e-10 | -22.48 | GO\_NEGATIVE\_REGULATION\_OF\_VIRAL\_PROCESS | MSigDB lists | GO\_NEGATIVE\_REGULATION\_OF\_VIRAL\_PROCESS | 66 | 12 | 12187 | 179 | Ifi207,Ifi204,Ifi209,Trim25,Tnip1,Ifitm2,Isg15,Ifitm3,Oasl1,Ccl4,Tnf,Ifi211 | | 1.727e-10 | -22.48 | regulation of signal transduction | biological process | GO:0009966 | 2258 | 73 | 13711 | 214 | Parp9,Timp1,Map3k6,Fzd4,Ptgs2,Runx1,Ier3,Trim25,Rasip1,Ccl4,Il6,Casp4,Il1a,Akap12,Nfkbia,Pik3r1,Gadd45b,Irf7,Socs3,Ccl7,Parp14,Trib1,Tgm2,C5ar1,Hspa5,Selp,Nfe2l2,Mmp8,Atf3,Tnf,Acod1,Pik3ap1,Tnip1,Fstl1,Trim30a,Rnf213,Hspb1,Ncf1,Cd14,Cflar,Ripk1,Nfkb1,Map3k8,Birc3,Pik3r5,Plaur,Irgm1,Errfi1,Icam1,Rgs16,Sox11,Gadd45g,Lgals9,P2ry6,Thbs1,Tnip3,S100a8,Iqgap1,Ccl12,Ptpn1,Bcl3,Irgm2,Sgk3,S100a9,Ksr1,Tnfsf10,Xdh,Ccl2,Tnfaip3,Plek,Il1rn,Irak3,Csf3 | | 1.735e-10 | -22.48 | BOQUEST\_STEM\_CELL\_CULTURED\_VS\_FRESH\_UP | MSigDB lists | BOQUEST\_STEM\_CELL\_CULTURED\_VS\_FRESH\_UP | 341 | 24 | 12187 | 179 | Nfkbia,Trib1,Oasl1,Gem,A2m,Tnfsf10,Ifi44,Vcam1,Zbtb16,Atf3,Apold1,Zfp36,Map3k8,Pik3r1,Tgm2,Tnfaip3,Ccl12,Gadd45b,Angptl4,Cxcl1,Rgs16,Thbs1,Dnajb1,Gadd45g | | 1.765e-10 | -22.46 | positive regulation of macromolecule metabolic process | biological process | GO:0010604 | 2653 | 81 | 13711 | 214 | Nfe2l2,Mmp8,Msn,Noct,C5ar1,Maff,Hspa5,Ncf1,Hspb1,Sele,Ripk1,Cflar,Tnip1,Trim30a,Tnf,Atf3,Zfp36,Cebpd,Ccl4,Spi1,Runx1,Ptgs2,Resf1,Ier3,Tiparp,Fzd4,Map3k6,Fam107a,Parp9,Sdc4,Sbno2,Dtx3l,Irf7,Parp14,Trib1,Ccl7,Gadd45b,Pik3r1,Ccnd2,Casp4,Nfkbia,Il1a,Akap12,Il6,Cdkn1a,S100a9,Bcl3,Irgm2,Ptpn1,S100a8,Iqgap1,Irf2,Ccl12,Arid5b,C3ar1,Il1rn,Aff1,Zbtb16,Csf3,Lcn2,Tnfaip3,Xdh,Tnfsf10,Ksr1,Ccl2,Bach1,Pik3r5,Plaur,Map3k8,Birc3,Nfkb1,Thbs1,Banp,Gadd45g,Rbm39,P2ry6,Lgals9,Icam1,Nfkb2,Sox11,Cybb | | 1.766e-10 | -22.46 | Type II interferon signaling (IFNG) | WikiPathways | WP1253 | 25 | 10 | 3756 | 96 | Socs3,Irf2,Ifit2,Tap1,Icam1,Spi1,Isg15,Cxcl9,Cybb,Irf9 | | 2.069e-10 | -22.30 | JACKSON\_DNMT1\_TARGETS\_UP | MSigDB lists | JACKSON\_DNMT1\_TARGETS\_UP | 67 | 12 | 12187 | 179 | Isg15,Ptgs2,Trim25,Hspb1,Zfp36,Cdkn1a,Socs3,Ifit3b,Irf7,Rgs16,Ccl12,Nfe2l2 | | 2.078e-10 | -22.29 | regulation of phosphorylation | biological process | GO:0042325 | 1276 | 51 | 13711 | 214 | Tnip1,Hspb1,Ncf1,Cflar,Ripk1,Atf3,Tnf,Pik3ap1,Mmp8,C5ar1,Pik3r1,Ccnd2,Gadd45b,Socs3,Ccl7,Parp14,Trib1,Il6,Cdkn1a,Akap12,Il1a,Ptgs2,Ier3,Rasip1,Ccl4,Sdc4,Parp9,Fzd4,Map3k6,Il1rn,Irak3,Csf3,Ksr1,Xdh,Ccl2,Tnfaip3,Irgm2,Iqgap1,Ccl12,Ptpn1,Gadd45g,Lgals9,P2ry6,Thbs1,Errfi1,Icam1,Pik3r5,Plaur,Gbp4,Map3k8,Birc3 | | 2.098e-10 | -22.28 | GSE30083\_SP1\_VS\_SP2\_THYMOCYTE\_DN | MSigDB lists | GSE30083\_SP1\_VS\_SP2\_THYMOCYTE\_DN | 140 | 16 | 12187 | 179 | Irf9,Tap1,Il17ra,Slfn5,Herc6,Samd9l,Ifi44,Sox11,Rgs16,Cmpk2,Gbp6,Dtx3l,Irgm2,Gbp4,Parp14,Ccnd2 | | 2.191e-10 | -22.24 | GSE21063\_CTRL\_VS\_ANTI\_IGM\_STIM\_BCELL\_8H\_UP | MSigDB lists | GSE21063\_CTRL\_VS\_ANTI\_IGM\_STIM\_BCELL\_8H\_UP | 120 | 15 | 12187 | 179 | Nfkb2,Tnfsf10,Parp14,Trib1,Nfkbia,Birc3,Tnfaip3,Gadd45b,Isg15,Clic4,Plaur,Icam1,Zfp36,Nfkbiz,Nfkb1 | | 2.394e-10 | -22.15 | GSE2128\_CTRL\_VS\_MIMETOPE\_NEGATIVE\_SELECTION\_DP\_THYMOCYTE\_NOD\_UP | MSigDB lists | GSE2128\_CTRL\_VS\_MIMETOPE\_NEGATIVE\_SELECTION\_DP\_THYMOCYTE\_NOD\_UP | 163 | 17 | 12187 | 179 | Ccl4,Gpr84,Sdc4,Casp4,Lgals9,Gbp3,Rtp4,Bcl3,C3ar1,Gbp7,Atf3,Ms4a6d,Irak3,Cd14,Ifitm3,Ifitm2,Ptgs2 | | 2.394e-10 | -22.15 | GSE9960\_HEALTHY\_VS\_GRAM\_NEG\_SEPSIS\_PBMC\_DN | MSigDB lists | GSE9960\_HEALTHY\_VS\_GRAM\_NEG\_SEPSIS\_PBMC\_DN | 163 | 17 | 12187 | 179 | Phf11b,Nfkb1,Tnip3,Tagln2,Ifitm3,Csf2rb,Plaur,Clic4,Irf2,Pik3ap1,Gpr84,Nfkbia,Phf11d,Bcl3,Irf7,Cflar,Tnfaip8 | | 2.395e-10 | -22.15 | CHEN\_HOXA5\_TARGETS\_9HR\_UP | MSigDB lists | CHEN\_HOXA5\_TARGETS\_9HR\_UP | 186 | 18 | 12187 | 179 | Ifi209,Ifi207,Ifi204,Tiparp,Atf3,Ptgs2,Ip6k2,Maff,Birc3,Nfkbia,Bach1,Gadd45b,Tnfaip3,Gem,Nfkbie,Irf7,Ifi211,Rbm39 | | 2.600e-10 | -22.07 | GSE360\_L\_DONOVANI\_VS\_B\_MALAYI\_HIGH\_DOSE\_DC\_UP | MSigDB lists | GSE360\_L\_DONOVANI\_VS\_B\_MALAYI\_HIGH\_DOSE\_DC\_UP | 142 | 16 | 12187 | 179 | Tnfaip3,Gadd45b,Ccl4,Nfkbia,Rgs16,Irf7,Cflar,Ier3,Stx11,Nfkb1,Il1rn,Timp1,Gadd45g,Vcam1,Csf2rb,Isg15 | | 2.780e-10 | -22.00 | FOSTER\_TOLERANT\_MACROPHAGE\_UP | MSigDB lists | FOSTER\_TOLERANT\_MACROPHAGE\_UP | 122 | 15 | 12187 | 179 | Il1rn,Hp,Cp,Lcn2,Thbs1,Sgk1,Ifi44,Irak3,Pik3ap1,Msr1,Xaf1,Igsf6,Gem,Ptges,Irf7 | | 2.903e-10 | -21.96 | GSE21360\_NAIVE\_VS\_QUATERNARY\_MEMORY\_CD8\_TCELL\_UP | MSigDB lists | GSE21360\_NAIVE\_VS\_QUATERNARY\_MEMORY\_CD8\_TCELL\_UP | 165 | 17 | 12187 | 179 | Parp12,Oasl1,Ripk1,Ifit2,Usp18,Xaf1,Rtp4,Irf7,Phf11d,Ifi44,Phf11b,Ifit3b,Trim25,Irf9,Ifitm2,Herc6,Isg15 | | 2.949e-10 | -21.94 | GO\_RESPONSE\_TO\_OXYGEN\_CONTAINING\_COMPOUND | MSigDB lists | GO\_RESPONSE\_TO\_OXYGEN\_CONTAINING\_COMPOUND | 1042 | 43 | 12187 | 179 | Csf3,Ptges,Cxcl1,Cmpk2,S100a8,Pygm,Sbno2,Pik3r1,Spi1,Cybb,Nfe2l2,Il6,Tnfaip3,Ccl12,Selp,Cd14,P2ry6,Irak3,Cxcl16,Ptpn1,Thbs1,Nfkb1,C5ar1,Fzd4,Nfkb2,Tnf,Casp4,Nfkbia,Trib1,Ptgs2,Sele,Icam1,Msn,Errfi1,Cdkn1a,Adamts1,Socs3,Vcam1,Col4a1,Hp,Timp1,Tnip3,Kcna5 | | 2.954e-10 | -21.94 | extracellular space | cellular component | GO:0005615 | 869 | 40 | 13825 | 212 | Lcn2,Ccl4,Retnlg,Selp,Il4ra,S100a8,Saa1,Angptl4,Tnf,Thbs1,Col4a1,Cp,Cxcl9,Cxcl16,Msr1,Ccl2,Cd14,Il1rn,Cxcl1,Adamts9,Xdh,Vcam1,Ccl7,Mmp8,Fstl1,Sele,Olfml2b,A2m,Il6,Prg4,S100a9,Tnfsf10,Ccl12,Wfdc21,Hp,Il1a,Icam1,Timp1,Csf3,Serpina3f | | 2.982e-10 | -21.93 | regulation of MAPK cascade | biological process | GO:0043408 | 605 | 33 | 13711 | 214 | Ccl12,Iqgap1,Ptpn1,C5ar1,Mmp8,Ccl2,Atf3,Ksr1,Xdh,Pik3ap1,Tnf,Tnip1,Cflar,Ripk1,Irak3,Il1rn,Ncf1,Fzd4,Map3k6,Map3k8,Pik3r5,Ccl4,Il6,Errfi1,Akap12,Il1a,Icam1,P2ry6,Gadd45g,Gadd45b,Ccl7,Trib1,Thbs1 | | 3.126e-10 | -21.89 | TAKEDA\_TARGETS\_OF\_NUP98\_HOXA9\_FUSION\_10D\_UP | MSigDB lists | TAKEDA\_TARGETS\_OF\_NUP98\_HOXA9\_FUSION\_10D\_UP | 123 | 15 | 12187 | 179 | Oasl1,AA467197,Usp18,Xaf1,Ifit2,Rtp4,Cmpk2,Irf7,Gem,Samd9l,Ifit3b,Ifi44,Thbs1,Ptgs2,Isg15 | | 3.193e-10 | -21.86 | GSE19923\_WT\_VS\_HEB\_AND\_E2A\_KO\_DP\_THYMOCYTE\_DN | MSigDB lists | GSE19923\_WT\_VS\_HEB\_AND\_E2A\_KO\_DP\_THYMOCYTE\_DN | 166 | 17 | 12187 | 179 | Cflar,Gem,Map3k8,Tnf,Ccl4,Ccl2,Nfkbia,Il6,Zfp36,Errfi1,Sgk1,Socs3,Nfkbiz,Dnajb1,Tiparp,Ier3,Atf3 | | 3.542e-10 | -21.76 | regulation of cellular protein metabolic process | biological process | GO:0032268 | 2101 | 69 | 13711 | 214 | P2ry6,Lgals9,Gadd45g,Thbs1,Errfi1,Icam1,Plaur,Pik3r5,Birc3,Map3k8,Gbp4,Csf3,Tnfaip8,Irak3,Il1rn,Ccl2,Xdh,Ksr1,Tnfsf10,Tnfaip3,Irgm2,Wfdc21,Bcl3,S100a9,Ccl12,S100a8,Iqgap1,A2m,Ptpn1,Gadd45b,Ccnd2,Pik3r1,Parp14,Trib1,Ctla2a,Ccl7,Dtx3l,Socs3,Cdkn1a,Il6,Nfkbia,Il1a,Akap12,Casp4,Ptgs2,Ccl4,Spi1,Rasip1,Parp9,Sdc4,Isg15,Map3k6,Fzd4,Fam107a,Timp1,Tnip1,Ripk1,Cflar,Ncf1,Hspb1,Atf3,Zfp36,Pik3ap1,Tnf,Msn,Mmp8,Serpina3f,Hspa5,C5ar1 | | 3.564e-10 | -21.76 | GSE360\_CTRL\_VS\_T\_GONDII\_DC\_DN | MSigDB lists | GSE360\_CTRL\_VS\_T\_GONDII\_DC\_DN | 145 | 16 | 12187 | 179 | Nfkb2,Cflar,Tnfaip3,Gadd45b,Nfkbia,Trib1,Csf2rb,Tap1,Ifitm3,Tagln2,Stx11,Nfkb1,Cp,Cebpd,Socs3,Ifit3b | | 3.692e-10 | -21.72 | regulation of biological process | biological process | GO:0050789 | 7871 | 166 | 13711 | 214 | Cxcl1,Dnajb1,Rrp8,Mt2,Hp,Prg4,Ifit2,Csf2rb,Tnf,Robo4,Zfp36,Ripk1,Cd14,Gpr84,Cflar,Ifi207,Hspb1,Rhoc,Fstl1,Tnip1,Fzd4,Ifi209,Parp9,Sbno2,Ptges,Sdc4,Ccl4,Spi1,Ifitm6,Ier3,Ptgs2,Sgk1,Il1a,Gem,Oasl2,Casp4,Cdkn1a,Il6,Ctla2a,Trib1,Dtx3l,Irf7,Socs3,Gadd45b,Pik3r1,Ccl12,Irf2,Tap1,S100a8,A2m,Usp18,Sgk3,Irf9,Irgm2,Bcl3,Samd9l,Plek,Xdh,Tnfsf10,Ksr1,Adamts9,Irak3,Arid5b,Ifi204,Osmr,Gbp4,Adamts1,Nfkb1,Kcna5,Bach1,Il17ra,Plaur,Trim56,Sox11,Mxd4,Cybb,Errfi1,Ifitm3,Thbs1,P2ry6,Banp,Rbm39,Clic4,Hspa5,Maff,C5ar1,Tgm2,Rasd1,Mmp8,Nfe2l2,Serpina3f,Angptl4,Msn,Noct,Selp,Pik3ap1,Acod1,Rhou,Atf3,Ncf1,Sele,Rnf213,Trim30a,Map3k6,Fam107a,Tiparp,Timp1,Isg15,Slfn2,Ifi211,Xaf1,Cebpd,Rasip1,Trim25,Resf1,Runx1,Nfkbia,Akap12,Il4ra,Slfn3,Ip6k2,Parp14,Ccl7,Ccnd2,Hcar2,Ptpn1,Ifitm2,Nfkbiz,Cxcl16,Iqgap1,Col4a1,Tnip3,S100a9,Vcam1,Wfdc21,Cxcl9,Znfx1,Tnfaip3,Depp1,Ccl2,Tnfaip8,Zbtb16,Csf3,Il1rn,C3ar1,Aff1,Lcn2,Birc3,Map3k8,Nfkbie,Rhoj,Apold1,Irgm1,Pik3r5,Fyb,Nfkb2,Rgs16,Icam1,Mt1,Rassf4,Bbs12,Iigp1,Msr1,Oasl1,Lgals9,Gadd45g | | 3.717e-10 | -21.71 | GO\_NEGATIVE\_REGULATION\_OF\_IMMUNE\_SYSTEM\_PROCESS | MSigDB lists | GO\_NEGATIVE\_REGULATION\_OF\_IMMUNE\_SYSTEM\_PROCESS | 268 | 21 | 12187 | 179 | Cd14,Irak3,Pik3ap1,Tap1,Ifi209,Sox11,Thbs1,Zbtb16,Ifi204,Ifi207,Ifi211,A2m,Runx1,Tnf,Il4ra,Pik3r1,Trib1,Nfkbia,Tnfaip3,Ccl12,Nfe2l2 | | 3.856e-10 | -21.68 | HALLMARK\_HYPOXIA | MSigDB lists | HALLMARK\_HYPOXIA | 168 | 17 | 12187 | 179 | Cdkn1a,Hspa5,Tiparp,Akap12,Cp,Ier3,Atf3,Zfp36,Errfi1,Plaur,Maff,Tgm2,Sdc4,Tnfaip3,Il6,Angptl4,Pygm | | 3.856e-10 | -21.68 | GSE7548\_NAIVE\_VS\_DAY7\_PCC\_IMMUNIZATION\_CD4\_TCELL\_DN | MSigDB lists | GSE7548\_NAIVE\_VS\_DAY7\_PCC\_IMMUNIZATION\_CD4\_TCELL\_DN | 168 | 17 | 12187 | 179 | Oasl1,Ripk1,Xaf1,Ifit2,Irf7,Ifi211,Ifi207,Ifi204,Ifit3b,Ifi44,Ifi209,Tap1,Csf2rb,Zfp36,Ifitm2,Isg15,Ifitm3 | | 3.952e-10 | -21.65 | GSE2706\_UNSTIM\_VS\_8H\_LPS\_DC\_DN | MSigDB lists | GSE2706\_UNSTIM\_VS\_8H\_LPS\_DC\_DN | 146 | 16 | 12187 | 179 | Sox11,Socs3,Nfkb1,Stx11,Rnf213,Tnip1,Icam1,Parp14,Trib1,Tnfaip3,AA467197,Il6,Dtx3l,Irf7,Tnfaip8,Rtp4 | | 4.039e-10 | -21.63 | positive regulation of signal transduction | biological process | GO:0009967 | 1217 | 49 | 13711 | 214 | C5ar1,Tgm2,Selp,Mmp8,Atf3,Pik3ap1,Tnf,Ripk1,Cd14,Cflar,Ncf1,Parp9,Fzd4,Map3k6,Trim25,Ccl4,Il6,Akap12,Il1a,Casp4,Gadd45b,Pik3r1,Parp14,Ccl7,Irf7,Ccl12,Iqgap1,S100a8,Ptpn1,Irgm2,S100a9,Ccl2,Xdh,Tnfsf10,Ksr1,Tnfaip3,Csf3,Il1rn,Nfkb1,Map3k8,Plaur,Pik3r5,Irgm1,Sox11,Icam1,P2ry6,Lgals9,Gadd45g,Thbs1 | | 4.048e-10 | -21.63 | positive regulation of response to biotic stimulus | biological process | GO:0002833 | 158 | 17 | 13711 | 214 | Irgm2,Ifi211,Irgm1,Parp9,Tnip3,Ifi209,Cxcl1,Lgals9,Ifi204,Tnip1,Cd14,Irf7,Irak3,Pik3ap1,Nfkbia,Tnf,Acod1 | | 4.127e-10 | -21.61 | DEBOSSCHER\_NFKB\_TARGETS\_REPRESSED\_BY\_GLUCOCORTICOIDS | MSigDB lists | DEBOSSCHER\_NFKB\_TARGETS\_REPRESSED\_BY\_GLUCOCORTICOIDS | 14 | 7 | 12187 | 179 | Il6,Icam1,Sele,Vcam1,Ccl2,Tnf,Ptgs2 | | 4.233e-10 | -21.58 | GSE43955\_TH0\_VS\_TGFB\_IL6\_TH17\_ACT\_CD4\_TCELL\_4H\_UP | MSigDB lists | GSE43955\_TH0\_VS\_TGFB\_IL6\_TH17\_ACT\_CD4\_TCELL\_4H\_UP | 169 | 17 | 12187 | 179 | Cebpd,Gadd45g,Socs3,Adamts1,Sox11,Errfi1,Sele,Selp,Saa1,Ptgs2,Il6,Bach1,Ccl4,Map3k8,Il4ra,Birc3,Gbp3 | | 4.233e-10 | -21.58 | GSE22140\_HEALTHY\_VS\_ARTHRITIC\_GERMFREE\_MOUSE\_CD4\_TCELL\_DN | MSigDB lists | GSE22140\_HEALTHY\_VS\_ARTHRITIC\_GERMFREE\_MOUSE\_CD4\_TCELL\_DN | 169 | 17 | 12187 | 179 | Gem,Rgs16,Nfkb2,Tnf,Ifit2,Trib1,Ccl4,Oasl1,Trim25,Icam1,Thbs1,Ifi44,Cdkn1a,Ifit3b,Nfkb1,Timp1,Ier3 | | 4.233e-10 | -21.58 | GSE22935\_WT\_VS\_MYD88\_KO\_MACROPHAGE\_12H\_MBOVIS\_BCG\_STIM\_DN | MSigDB lists | GSE22935\_WT\_VS\_MYD88\_KO\_MACROPHAGE\_12H\_MBOVIS\_BCG\_STIM\_DN | 169 | 17 | 12187 | 179 | Slfn4,Igsf6,Ptges,Nfe2l2,Tnfaip3,Gpr84,Parp9,Clic4,Slfn3,Plek,Isg15,Lcn2,Ier3,Timp1,Ksr1,Nfkbiz,Aff1 | | 4.378e-10 | -21.55 | GSE30971\_CTRL\_VS\_LPS\_STIM\_MACROPHAGE\_WBP7\_HET\_2H\_UP | MSigDB lists | GSE30971\_CTRL\_VS\_LPS\_STIM\_MACROPHAGE\_WBP7\_HET\_2H\_UP | 147 | 16 | 12187 | 179 | Tnf,Nfkbia,Ccl4,Il6,Ptges,Cxcl1,Rgs16,Socs3,Il1a,Nfkb1,Tnip3,Ier3,Ptgs2,Irak3,Clic4,Plaur | | 4.494e-10 | -21.52 | cytokine activity | molecular function | GO:0005125 | 100 | 14 | 13516 | 211 | Ccl7,Ccl12,Tnf,Tnfsf10,Cxcl1,Timp1,Il1rn,Csf3,Il1a,Cxcl16,Ccl2,Il6,Cxcl9,Ccl4 | | 4.644e-10 | -21.49 | GSE43863\_DAY6\_EFF\_VS\_DAY150\_MEM\_TFH\_CD4\_TCELL\_UP | MSigDB lists | GSE43863\_DAY6\_EFF\_VS\_DAY150\_MEM\_TFH\_CD4\_TCELL\_UP | 170 | 17 | 12187 | 179 | Xaf1,Usp18,Ifit2,Lgals9,Parp12,Bach1,Irf7,Rtp4,Cmpk2,Rbm39,Ifit3b,Ifi44,Hspa5,Znfx1,Rasd1,Saa1,Pik3ap1 | | 4.847e-10 | -21.45 | GSE42724\_MEMORY\_VS\_B1\_BCELL\_DN | MSigDB lists | GSE42724\_MEMORY\_VS\_B1\_BCELL\_DN | 148 | 16 | 12187 | 179 | AA467197,Tnfaip3,Gadd45b,Sdc4,Nfkbia,Gbp3,Cflar,Arid5b,Nfkb1,Znfx1,Cdkn1a,Socs3,Nfkbiz,Parp9,Ptpn1,Tap1 | | 4.847e-10 | -21.45 | GSE6259\_DEC205\_POS\_DC\_VS\_BCELL\_DN | MSigDB lists | GSE6259\_DEC205\_POS\_DC\_VS\_BCELL\_DN | 148 | 16 | 12187 | 179 | Znfx1,Ifi44,Phf11b,Pik3ap1,Icam1,Isg15,Tnip1,Gadd45b,Msr1,Ifit2,Nfkbia,Xaf1,Parp14,Gbp3,Irf7,Phf11d | | 4.847e-10 | -21.45 | GSE12845\_PRE\_GC\_VS\_DARKZONE\_GC\_TONSIL\_BCELL\_UP | MSigDB lists | GSE12845\_PRE\_GC\_VS\_DARKZONE\_GC\_TONSIL\_BCELL\_UP | 148 | 16 | 12187 | 179 | Cd14,Ifitm2,Ifitm3,Tap1,Zfp36,Rbm47,Il17ra,Ifi44,Ier3,Cxcl1,Nfkbia,Ccnd2,Il6,Parp12,Tubb6,Rhoc | | 4.969e-10 | -21.42 | BOWIE\_RESPONSE\_TO\_EXTRACELLULAR\_MATRIX | MSigDB lists | BOWIE\_RESPONSE\_TO\_EXTRACELLULAR\_MATRIX | 22 | 8 | 12187 | 179 | Ifi209,Ifit2,Ifit3b,Ifi204,Ifi207,Isg15,Ifi211,Irf7 | | 5.091e-10 | -21.40 | GSE17301\_CTRL\_VS\_48H\_ACD3\_ACD28\_STIM\_CD8\_TCELL\_DN | MSigDB lists | GSE17301\_CTRL\_VS\_48H\_ACD3\_ACD28\_STIM\_CD8\_TCELL\_DN | 171 | 17 | 12187 | 179 | Igsf6,Nfkbie,Pik3r5,Rassf4,Tnfaip3,Tnf,Map3k8,Spi1,Gpr84,Nfkbia,Casp4,Irak3,Icam1,Tnip1,Stx11,Nfkb1,Nfkbiz | | 5.139e-10 | -21.39 | positive regulation of cellular protein metabolic process | biological process | GO:0032270 | 1309 | 51 | 13711 | 214 | Xdh,Ksr1,Tnfsf10,Ccl2,Tnfaip3,Il1rn,Csf3,S100a8,Iqgap1,Ccl12,Ptpn1,Bcl3,Irgm2,S100a9,Icam1,Gadd45g,P2ry6,Lgals9,Thbs1,Map3k8,Birc3,Pik3r5,Plaur,Tnf,Tnip1,Ncf1,Ripk1,Cflar,C5ar1,Hspa5,Msn,Mmp8,Il6,Cdkn1a,Casp4,Nfkbia,Akap12,Il1a,Gadd45b,Ccnd2,Pik3r1,Parp14,Trib1,Ccl7,Parp9,Sdc4,Map3k6,Fzd4,Fam107a,Ptgs2,Ccl4 | | 5.222e-10 | -21.37 | GO\_REGULATION\_OF\_INNATE\_IMMUNE\_RESPONSE | MSigDB lists | GO\_REGULATION\_OF\_INNATE\_IMMUNE\_RESPONSE | 273 | 21 | 12187 | 179 | A2m,Irf7,Ifi211,Nfkbia,Usp18,Birc3,Ripk1,Tnfaip3,Cd14,Tnip1,Tap1,Ptpn1,Parp9,Pik3ap1,Irak3,Socs3,Ifi209,Ifi207,Nfkb1,Tnip3,Ifi204 | | 5.252e-10 | -21.37 | RODWELL\_AGING\_KIDNEY\_UP | MSigDB lists | RODWELL\_AGING\_KIDNEY\_UP | 360 | 24 | 12187 | 179 | Ccl12,Map3k8,Ccnd2,Msr1,Osmr,Dtx3l,Ms4a6d,Nfkbiz,Cxcl16,Tap1,Il17ra,Cd14,Rnf213,AA467197,Parp12,Parp14,Birc3,A2m,Timp1,Samd9l,Vcam1,Socs3,Irf9,Tagln2 | | 5.282e-10 | -21.36 | defense response to protozoan | biological process | GO:0042832 | 21 | 8 | 13711 | 214 | Gbp3,Il4ra,Il6,Iigp1,Bcl3,Gbp7,Irgm2,Gbp6 | | 5.793e-10 | -21.27 | positive regulation of intracellular signal transduction | biological process | GO:1902533 | 762 | 37 | 13711 | 214 | Il6,Akap12,Sox11,Il1a,Icam1,P2ry6,Gadd45b,Gadd45g,Ccl7,Thbs1,Fzd4,Map3k6,Map3k8,Trim25,Pik3r5,Ccl4,Ccl2,Xdh,Tnfsf10,Ksr1,Pik3ap1,Tnf,Csf3,Ripk1,Cd14,Cflar,Il1rn,Ncf1,Ccl12,S100a8,Iqgap1,Ptpn1,C5ar1,Tgm2,Selp,Mmp8,S100a9 | | 5.926e-10 | -21.25 | GSE1432\_CTRL\_VS\_IFNG\_6H\_MICROGLIA\_DN | MSigDB lists | GSE1432\_CTRL\_VS\_IFNG\_6H\_MICROGLIA\_DN | 150 | 16 | 12187 | 179 | Ifi44,Phf11b,Akap12,Stx11,Isg15,Herc6,Tap1,Csf2rb,Plek,Icam1,Casp4,Parp12,Irf7,Phf11d,Tnfsf10,Sbno2 | | 5.926e-10 | -21.25 | GSE21360\_NAIVE\_VS\_SECONDARY\_MEMORY\_CD8\_TCELL\_UP | MSigDB lists | GSE21360\_NAIVE\_VS\_SECONDARY\_MEMORY\_CD8\_TCELL\_UP | 150 | 16 | 12187 | 179 | Gadd45b,Cybb,Sdc4,Nfkbia,Tnf,Bcl3,Pik3r5,Csf3,Nfkbie,Ier3,Cp,Nfkb1,Il1rn,Ptpn1,Irak3,Ifitm2 | | 6.018e-10 | -21.23 | positive regulation of MAPK cascade | biological process | GO:0043410 | 427 | 27 | 13711 | 214 | Il1a,Akap12,Icam1,Il6,Ccl7,Thbs1,P2ry6,Gadd45b,Gadd45g,Fzd4,Map3k6,Map3k8,Ccl4,Pik3r5,Tnf,Ccl2,Xdh,Ksr1,Ripk1,Cflar,Ncf1,Il1rn,Ptpn1,C5ar1,Ccl12,Iqgap1,Mmp8 | | 6.096e-10 | -21.22 | positive regulation of protein phosphorylation | biological process | GO:0001934 | 800 | 38 | 13711 | 214 | Tnf,Ccl2,Xdh,Ksr1,Ripk1,Csf3,Cflar,Il1rn,Ncf1,Ptpn1,C5ar1,Ccl12,Iqgap1,Mmp8,Irgm2,Akap12,Il1a,Icam1,Cdkn1a,Il6,Parp14,Ccl7,Thbs1,P2ry6,Lgals9,Gadd45b,Pik3r1,Gadd45g,Ccnd2,Map3k6,Fzd4,Map3k8,Parp9,Sdc4,Ccl4,Plaur,Pik3r5,Ptgs2 | | 6.585e-10 | -21.14 | GO\_POSITIVE\_REGULATION\_OF\_PROTEIN\_METABOLIC\_PROCESS | MSigDB lists | GO\_POSITIVE\_REGULATION\_OF\_PROTEIN\_METABOLIC\_PROCESS | 1155 | 45 | 12187 | 179 | Map3k6,Ptpn1,Hspa5,Bcl3,Ifi211,S100a8,Ccl2,Ccnd2,Gadd45b,Ripk1,Tnfaip3,Saa1,Plek,Icam1,Ifi209,Cdkn1a,Ksr1,Ifi204,Fzd4,S100a9,Tnfsf10,Trib1,Thbs1,Gadd45g,Il1a,Tiparp,Pik3r5,Csf3,Iqgap1,Ccl4,Map3k8,Nfe2l2,Il6,Ccl12,Msn,Plaur,Hspb1,Socs3,Ifi207,C5ar1,Xdh,Sdc4,Nfkbia,Birc3,Tnf | | 6.684e-10 | -21.13 | GSE7509\_DC\_VS\_MONOCYTE\_DN | MSigDB lists | GSE7509\_DC\_VS\_MONOCYTE\_DN | 174 | 17 | 12187 | 179 | Ifit3b,Znfx1,Hp,Cd14,Saa1,Clic4,Tap1,Usp18,Spi1,Nfkbia,Casp4,Tnf,Cybb,Ptges,Irgm2,Cmpk2,S100a8 | | 6.684e-10 | -21.13 | GSE339\_EX\_VIVO\_VS\_IN\_CULTURE\_CD8POS\_DC\_DN | MSigDB lists | GSE339\_EX\_VIVO\_VS\_IN\_CULTURE\_CD8POS\_DC\_DN | 174 | 17 | 12187 | 179 | Trim25,Csf2rb,Isg15,Ifitm2,Tnip1,Tiparp,Znfx1,Cdkn1a,Hspa5,Cmpk2,Nfkb2,Irgm2,Bcl3,Gadd45b,Pik3r1,Usp18,Nfkbia | | 6.895e-10 | -21.10 | regulation of catalytic activity | biological process | GO:0050790 | 1714 | 60 | 13711 | 214 | Tnfaip3,Plek,Xdh,Tnfsf10,Ksr1,Ccl2,Il1rn,Irak3,Tnfaip8,Ptpn1,Iqgap1,S100a8,A2m,Ccl12,S100a9,Irgm2,Wfdc21,Icam1,Rgs16,Errfi1,Thbs1,Oasl1,Gadd45g,Lgals9,Map3k8,Birc3,Nfkb1,Pik3r5,Plaur,Tnf,Zfp36,Ncf1,Hspb1,Sele,Ripk1,Cflar,Rhoc,Dnajb1,C5ar1,Hp,Serpina3f,Angptl4,Oasl2,Casp4,Il6,Cdkn1a,Dtx3l,Trib1,Ccl7,Gadd45b,Ccnd2,Pik3r1,Timp1,Map3k6,Fzd4,Parp9,Sdc4,Rasip1,Ccl4,Ptgs2 | | 7.023e-10 | -21.08 | positive regulation of nitrogen compound metabolic process | biological process | GO:0051173 | 2525 | 77 | 13711 | 214 | Atf3,Zfp36,Tnf,Tnip1,Trim30a,Ncf1,Hspb1,Ripk1,Cflar,Maff,C5ar1,Hspa5,Msn,Nfe2l2,Mmp8,Il6,Cdkn1a,Casp4,Nfkbia,Akap12,Il1a,Gadd45b,Ccnd2,Pik3r1,Dtx3l,Irf7,Trib1,Parp14,Ccl7,Parp9,Sbno2,Sdc4,Tiparp,Fzd4,Map3k6,Fam107a,Runx1,Ptgs2,Ier3,Cebpd,Spi1,Ccl4,Xdh,Ksr1,Tnfsf10,Ccl2,Tnfaip3,Il1rn,Arid5b,Aff1,C3ar1,Zbtb16,Csf3,Iqgap1,S100a8,Ccl12,Irf2,Ptpn1,Bcl3,Irgm2,S100a9,Cybb,Icam1,Nfkb2,Sox11,Banp,Rbm39,Gadd45g,P2ry6,Lgals9,Thbs1,Nfkb1,Map3k8,Birc3,Pik3r5,Plaur,Bach1 | | 7.221e-10 | -21.05 | GSE2706\_UNSTIM\_VS\_8H\_LPS\_AND\_R848\_DC\_DN | MSigDB lists | GSE2706\_UNSTIM\_VS\_8H\_LPS\_AND\_R848\_DC\_DN | 152 | 16 | 12187 | 179 | Irf7,Cmpk2,Tnfaip8,Map3k8,Usp18,Parp14,Gadd45b,AA467197,Herc6,Tnip1,Icam1,Socs3,Tnip3,Nfkb1,Stx11,Znfx1 | | 7.963e-10 | -20.95 | GSE9509\_LPS\_VS\_LPS\_AND\_IL10\_STIM\_IL10\_KO\_MACROPHAGE\_10MIN\_UP | MSigDB lists | GSE9509\_LPS\_VS\_LPS\_AND\_IL10\_STIM\_IL10\_KO\_MACROPHAGE\_10MIN\_UP | 153 | 16 | 12187 | 179 | Cd14,Pik3ap1,Plaur,Ptpn1,Socs3,Gadd45g,Gem,Cxcl1,Rhoj,Cmpk2,Rhou,Ccl4,Trib1,Tubb6,Ccl12,Oasl1 | | 7.963e-10 | -20.95 | GSE9988\_LOW\_LPS\_VS\_ANTI\_TREM1\_AND\_LPS\_MONOCYTE\_UP | MSigDB lists | GSE9988\_LOW\_LPS\_VS\_ANTI\_TREM1\_AND\_LPS\_MONOCYTE\_UP | 153 | 16 | 12187 | 179 | Thbs1,Nfkbiz,Tnip3,Stx11,Tnip1,Pik3ap1,Clic4,Rhou,Map3k8,Birc3,Cybb,Tnfaip3,Nfe2l2,Oasl1,Csf3,Pik3r5 | | 8.334e-10 | -20.91 | granulocyte chemotaxis | biological process | GO:0071621 | 56 | 11 | 13711 | 214 | Il1rn,Ccl7,Cxcl9,Ccl2,Ccl4,S100a9,Il17ra,C5ar1,Cxcl1,S100a8,Ccl12 | | 8.591e-10 | -20.88 | positive regulation of protein modification process | biological process | GO:0031401 | 1001 | 43 | 13711 | 214 | Tnf,Ksr1,Xdh,Ccl2,Ncf1,Il1rn,Cflar,Ripk1,Csf3,Tnip1,C5ar1,Ptpn1,Hspa5,Iqgap1,Ccl12,Mmp8,Irgm2,Icam1,Akap12,Il1a,Il6,Cdkn1a,Thbs1,Ccl7,Trib1,Parp14,Pik3r1,Gadd45g,Ccnd2,Gadd45b,Lgals9,P2ry6,Map3k8,Fam107a,Birc3,Fzd4,Map3k6,Sdc4,Parp9,Ccl4,Ptgs2,Pik3r5,Plaur | | 8.773e-10 | -20.85 | GSE30083\_SP1\_VS\_SP4\_THYMOCYTE\_DN | MSigDB lists | GSE30083\_SP1\_VS\_SP4\_THYMOCYTE\_DN | 154 | 16 | 12187 | 179 | Tap1,Il17ra,Rnf213,Herc6,Ms4a6d,Samd9l,Zbtb16,Sgk1,Ifi44,Xdh,Irf7,Gbp7,Dtx3l,Irgm2,Oasl1,Sntb2 | | 8.773e-10 | -20.85 | GSE6092\_IFNG\_VS\_IFNG\_AND\_B\_BURGDORFERI\_INF\_ENDOTHELIAL\_CELL\_DN | MSigDB lists | GSE6092\_IFNG\_VS\_IFNG\_AND\_B\_BURGDORFERI\_INF\_ENDOTHELIAL\_CELL\_DN | 154 | 16 | 12187 | 179 | Phf11b,Socs3,Stx11,Cp,Znfx1,Ncf1,Cd14,Ifit2,Sdc4,Usp18,Parp12,Cflar,Phf11d,Igsf6,Cmpk2,Tnfsf10 | | 8.782e-10 | -20.85 | regulation of reactive oxygen species metabolic process | biological process | GO:2000377 | 166 | 17 | 13711 | 214 | Hp,Birc3,Cxcl1,Ier3,Ptgs2,Mmp8,Nfe2l2,Cdkn1a,Il6,Xdh,Icam1,Tnf,Acod1,Trim30a,Cflar,Ripk1,Thbs1 | | 8.860e-10 | -20.84 | NOD-like receptor signaling pathway | KEGG pathways | mmu04621 | 133 | 17 | 5248 | 107 | Cybb,Ifi204,Nfkb1,Il6,Irf7,Ripk1,Irf9,Birc3,Tnf,Tnfaip3,Gbp7,Cxcl1,Ccl12,Nfkbia,Ccl2,Gbp3,Casp4 | | 8.860e-10 | -20.84 | NOD-like receptor signaling pathway | KEGG pathways | ko04621 | 133 | 17 | 5248 | 107 | Nfkbia,Ccl2,Gbp3,Casp4,Tnfaip3,Gbp7,Cxcl1,Ccl12,Nfkb1,Il6,Ripk1,Irf7,Irf9,Birc3,Tnf,Ifi204,Cybb | | 9.258e-10 | -20.80 | SANA\_TNF\_SIGNALING\_UP | MSigDB lists | SANA\_TNF\_SIGNALING\_UP | 60 | 11 | 12187 | 179 | Tnfaip3,Ccl12,Samd9l,Vcam1,Parp14,Nfkbia,Birc3,Icam1,Cmpk2,Cxcl1,Tnip1 | | 9.744e-10 | -20.75 | Malaria | KEGG pathways | mmu05144 | 35 | 10 | 5248 | 107 | Thbs1,Il6,Sele,Ccl2,Vcam1,Tnf,Icam1,Selp,Ccl12,Csf3 | | 9.744e-10 | -20.75 | Malaria | KEGG pathways | ko05144 | 35 | 10 | 5248 | 107 | Il6,Sele,Thbs1,Tnf,Icam1,Ccl2,Vcam1,Selp,Csf3,Ccl12 | | 1.010e-09 | -20.71 | CHIARADONNA\_NEOPLASTIC\_TRANSFORMATION\_CDC25\_UP | MSigDB lists | CHIARADONNA\_NEOPLASTIC\_TRANSFORMATION\_CDC25\_UP | 113 | 14 | 12187 | 179 | Errfi1,Cebpd,Ifi207,Ifi204,Vcam1,Thbs1,Ifi209,Xdh,Osmr,Ifi211,Il6,Ccl12,Bach1,Casp4 | | 1.023e-09 | -20.70 | GSE45365\_CD8A\_DC\_VS\_CD11B\_DC\_IFNAR\_KO\_MCMV\_INFECTION\_DN | MSigDB lists | GSE45365\_CD8A\_DC\_VS\_CD11B\_DC\_IFNAR\_KO\_MCMV\_INFECTION\_DN | 94 | 13 | 12187 | 179 | Nfkb2,Cxcl1,Tnfaip3,Ccl12,Slc24a4,Nfkbia,Sdc4,Map3k8,Ptgs2,Atf3,Ier3,Nfkbiz,Ifi44 | | 1.115e-09 | -20.61 | MODULE\_263 | MSigDB lists | MODULE\_263 | 24 | 8 | 12187 | 179 | Plaur,Cxcl1,C5ar1,C3ar1,Ccl12,Il1a,Ccl4,Ccl2 | | 1.116e-09 | -20.61 | KEGG\_APOPTOSIS | MSigDB lists | KEGG\_APOPTOSIS | 77 | 12 | 12187 | 179 | Irak3,Csf2rb,Nfkb1,Il1a,Tnfsf10,Cflar,Pik3r5,Ripk1,Tnf,Birc3,Pik3r1,Nfkbia | | 1.116e-09 | -20.61 | WINZEN\_DEGRADED\_VIA\_KHSRP | MSigDB lists | WINZEN\_DEGRADED\_VIA\_KHSRP | 77 | 12 | 12187 | 179 | Cxcl1,Gem,Sdc4,Ifit2,Tnf,Il6,Ptgs2,Errfi1,Plaur,Zfp36,Sele,Nfkbiz | | 1.136e-09 | -20.60 | cellular response to lipid | biological process | GO:0071396 | 321 | 23 | 13711 | 214 | Gbp6,Irgm2,Msn,Cxcl1,Cxcl16,Ccl12,Tnip3,Cd14,Il1rn,Cxcl9,Acod1,Tnf,Tnfaip3,Ccl2,Zfp36,Sgk1,Fam107a,Sbno2,Cmpk2,Nfkb1,P2ry6,Nfkbia,Il6 | | 1.176e-09 | -20.56 | BROWNE\_INTERFERON\_RESPONSIVE\_GENES | MSigDB lists | BROWNE\_INTERFERON\_RESPONSIVE\_GENES | 47 | 10 | 12187 | 179 | Stx11,Oasl1,Ifi44,Ifit2,Ifit3b,Xaf1,Tnfsf10,Tap1,Isg15,Irf7 | | 1.176e-09 | -20.56 | BURTON\_ADIPOGENESIS\_PEAK\_AT\_2HR | MSigDB lists | BURTON\_ADIPOGENESIS\_PEAK\_AT\_2HR | 47 | 10 | 12187 | 179 | Errfi1,Zfp36,Rhoj,Ptgs2,Cebpd,Il6,Ier3,Gadd45g,Dnajb1,Thbs1 | | 1.176e-09 | -20.56 | LINDSTEDT\_DENDRITIC\_CELL\_MATURATION\_B | MSigDB lists | LINDSTEDT\_DENDRITIC\_CELL\_MATURATION\_B | 47 | 10 | 12187 | 179 | Tnip1,Cflar,Nfkb2,Tap1,Sdc4,Nfkbia,Maff,Birc3,Tnfaip3,Nfkb1 | | 1.177e-09 | -20.56 | GSE17974\_IL4\_AND\_ANTI\_IL12\_VS\_UNTREATED\_72H\_ACT\_CD4\_TCELL\_DN | MSigDB lists | GSE17974\_IL4\_AND\_ANTI\_IL12\_VS\_UNTREATED\_72H\_ACT\_CD4\_TCELL\_DN | 135 | 15 | 12187 | 179 | Parp14,Rhou,Birc3,Maff,Gbp3,Tnfsf10,Arid5b,Dtx3l,Stx11,Ksr1,Gadd45g,Samd9l,Parp9,Tap1,Isg15 | | 1.190e-09 | -20.55 | positive regulation of cellular metabolic process | biological process | GO:0031325 | 2654 | 79 | 13711 | 214 | Tnip1,Ncf1,Sele,Hspb1,Ripk1,Cflar,Atf3,Zfp36,Acod1,Tnf,Msn,Nfe2l2,Mmp8,C5ar1,Maff,Cxcl1,Hspa5,Gadd45b,Pik3r1,Ccnd2,Dtx3l,Irf7,Trib1,Parp14,Ccl7,Il6,Cdkn1a,Casp4,Nfkbia,Il1a,Akap12,Runx1,Ptgs2,Cebpd,Spi1,Ccl4,Parp9,Sbno2,Sdc4,Map3k6,Fzd4,Fam107a,Il1rn,Aff1,C3ar1,Arid5b,Zbtb16,Csf3,Xdh,Ksr1,Tnfsf10,Ccl2,Tnfaip3,Plek,Bcl3,Irgm2,S100a9,Iqgap1,S100a8,Irf2,Ccl12,Ptpn1,Rbm39,Gadd45g,Banp,P2ry6,Lgals9,Thbs1,Cybb,Icam1,Nfkb2,Sox11,Pik3r5,Plaur,Bach1,Irgm1,Nfkb1,Map3k8,Birc3 | | 1.302e-09 | -20.46 | GO\_CELLULAR\_RESPONSE\_TO\_INTERFERON\_GAMMA | MSigDB lists | GO\_CELLULAR\_RESPONSE\_TO\_INTERFERON\_GAMMA | 78 | 12 | 12187 | 179 | Icam1,Trim25,Irf2,Irf9,Gbp6,Irf7,Gbp4,Ccl12,Oasl1,Ccl4,Vcam1,Ccl2 | | 1.342e-09 | -20.43 | WANG\_TNF\_TARGETS | MSigDB lists | WANG\_TNF\_TARGETS | 16 | 7 | 12187 | 179 | Nfkbia,Vcam1,Csf3,Selp,Il6,Tnfaip3,Sele | | 1.345e-09 | -20.43 | GO\_LEUKOCYTE\_MIGRATION | MSigDB lists | GO\_LEUKOCYTE\_MIGRATION | 182 | 17 | 12187 | 179 | Vcam1,Selp,Saa1,Cxcl16,Msn,Icam1,Il17ra,Sele,Ccl4,Ccl2,Tnf,Pik3r1,Ccl12,Il6,S100a9,C5ar1,S100a8 | | 1.365e-09 | -20.41 | positive regulation of cell communication | biological process | GO:0010647 | 1389 | 52 | 13711 | 214 | Iqgap1,S100a8,Ccl12,Ptpn1,Irgm2,S100a9,Ksr1,Tnfsf10,Xdh,Ccl2,Tnfaip3,Il1rn,Csf3,Nfkb1,Map3k8,Pik3r5,Plaur,Irgm1,Icam1,Sox11,Gadd45g,Lgals9,P2ry6,Thbs1,Tgm2,C5ar1,Selp,Mmp8,Atf3,Tnf,Pik3ap1,Ncf1,Cd14,Cflar,Ripk1,Parp9,Map3k6,Fzd4,Ptgs2,Runx1,Trim25,Ccl4,Il6,Casp4,Il1a,Akap12,Hcar2,Pik3r1,Gadd45b,Irf7,Ccl7,Parp14 | | 1.408e-09 | -20.38 | regulation of protein modification process | biological process | GO:0031399 | 1521 | 55 | 13711 | 214 | Akap12,Il1a,Cdkn1a,Il6,Ccl7,Trib1,Parp14,Socs3,Dtx3l,Pik3r1,Ccnd2,Gadd45b,Fam107a,Map3k6,Fzd4,Sdc4,Isg15,Parp9,Spi1,Ccl4,Rasip1,Ptgs2,Pik3ap1,Tnf,Atf3,Cflar,Ripk1,Hspb1,Ncf1,Tnip1,Hspa5,C5ar1,Mmp8,Icam1,Errfi1,Thbs1,Lgals9,P2ry6,Gadd45g,Birc3,Gbp4,Map3k8,Plaur,Pik3r5,Tnfaip3,Ccl2,Ksr1,Xdh,Csf3,Irak3,Il1rn,Ptpn1,Ccl12,Iqgap1,Irgm2 | | 1.409e-09 | -20.38 | GSE29618\_PDC\_VS\_MDC\_DN | MSigDB lists | GSE29618\_PDC\_VS\_MDC\_DN | 159 | 16 | 12187 | 179 | Ifitm3,Ifitm2,Ptgs2,Irak3,Msn,Sgk1,Tiparp,Il1rn,Stx11,Timp1,Cebpd,S100a9,Iqgap1,Xaf1,Trib1,Spi1 | | 1.546e-09 | -20.29 | GSE46606\_DAY1\_VS\_DAY3\_CD40L\_IL2\_IL5\_STIMULATED\_IRF4HIGH\_BCELL\_DN | MSigDB lists | GSE46606\_DAY1\_VS\_DAY3\_CD40L\_IL2\_IL5\_STIMULATED\_IRF4HIGH\_BCELL\_DN | 160 | 16 | 12187 | 179 | Dtx3l,Tnfsf10,Ccnd2,Tnf,Trib1,Ripk1,Il6,Oasl1,Plaur,Clic4,Hspa5,Zbtb16,Nfkbiz,Tiparp,Znfx1,Atf3 | | 1.555e-09 | -20.28 | ZHOU\_INFLAMMATORY\_RESPONSE\_FIMA\_UP | MSigDB lists | ZHOU\_INFLAMMATORY\_RESPONSE\_FIMA\_UP | 349 | 23 | 12187 | 179 | Csf3,Rhoj,Angptl4,Cxcl1,Rgs16,Maff,Gpr84,Ccl4,Il6,Tnfaip3,Ptpn1,Il1a,Nfkb1,Ier3,Gem,Fzd4,Nfkb2,Tnf,Sdc4,Nfkbia,Ptgs2,Icam1,Plaur | | 1.604e-09 | -20.25 | GSE22589\_SIV\_VS\_HIV\_AND\_SIV\_INFECTED\_DC\_UP | MSigDB lists | GSE22589\_SIV\_VS\_HIV\_AND\_SIV\_INFECTED\_DC\_UP | 138 | 15 | 12187 | 179 | Cmpk2,Rtp4,Bcl3,Irf7,Tnfaip3,Oasl1,Cybb,Tor3a,Usp18,Nfkbia,Ifit2,Parp9,Irf9,Trim25,Ifit3b | | 1.669e-09 | -20.21 | positive regulation of signaling | biological process | GO:0023056 | 1397 | 52 | 13711 | 214 | Thbs1,P2ry6,Lgals9,Gadd45g,Sox11,Icam1,Irgm1,Plaur,Pik3r5,Map3k8,Nfkb1,Csf3,Il1rn,Tnfaip3,Ccl2,Xdh,Ksr1,Tnfsf10,S100a9,Irgm2,Ptpn1,Ccl12,S100a8,Iqgap1,Parp14,Ccl7,Irf7,Gadd45b,Hcar2,Pik3r1,Akap12,Il1a,Casp4,Il6,Ccl4,Trim25,Runx1,Ptgs2,Map3k6,Fzd4,Parp9,Ripk1,Cd14,Cflar,Ncf1,Pik3ap1,Tnf,Atf3,Mmp8,Selp,C5ar1,Tgm2 | | 1.712e-09 | -20.19 | GO\_NEGATIVE\_REGULATION\_OF\_CELL\_DEATH | MSigDB lists | GO\_NEGATIVE\_REGULATION\_OF\_CELL\_DEATH | 658 | 32 | 12187 | 179 | Nfkbia,Birc3,Tnf,Tnfsf10,Irf7,C5ar1,Cflar,Timp1,Ifit3b,Socs3,Cdkn1a,Plaur,Hspb1,Icam1,Nfe2l2,Il6,Ripk1,Tnfaip3,Ccl12,Pik3r1,Tnfaip8,Bcl3,Angptl4,Csf3,Ier3,Il1a,Nfkb1,Hspa5,Thbs1,Sgk1,Sox11,Ptpn1 | | 1.731e-09 | -20.17 | KIM\_WT1\_TARGETS\_UP | MSigDB lists | KIM\_WT1\_TARGETS\_UP | 185 | 17 | 12187 | 179 | Plaur,Sgk1,Thbs1,Ier3,Akap12,Nfkb1,Arid5b,Nfkbie,Gem,Nfkb2,Trib1,Sdc4,Nfkbia,Il4ra,Maff,Tnfaip3,Ccl12 | | 1.857e-09 | -20.10 | GSE14000\_UNSTIM\_VS\_16H\_LPS\_DC\_DN | MSigDB lists | GSE14000\_UNSTIM\_VS\_16H\_LPS\_DC\_DN | 162 | 16 | 12187 | 179 | Irf2,Parp9,Msn,Irf9,Ifi44,Fstl1,Nfkbiz,Cmpk2,Tnfsf10,Cflar,Dtx3l,Irf7,Rassf4,Gadd45b,Parp12,Nfkbia | | 1.858e-09 | -20.10 | GO\_REGULATION\_OF\_IMMUNE\_RESPONSE | MSigDB lists | GO\_REGULATION\_OF\_IMMUNE\_RESPONSE | 551 | 29 | 12187 | 179 | Tnf,Birc3,Nfkbia,C5ar1,C3ar1,A2m,Irf7,Tnip3,Ifi204,Ifi207,Ifi209,Socs3,Vcam1,Icam1,Tnip1,Ripk1,Tnfaip3,Il6,Pik3r1,Il4ra,Usp18,Ifi211,Nfkb1,Irak3,Pik3ap1,Parp9,Ptpn1,Tap1,Cd14 | | 1.963e-09 | -20.05 | GSE24634\_IL4\_VS\_CTRL\_TREATED\_NAIVE\_CD4\_TCELL\_DAY5\_DN | MSigDB lists | GSE24634\_IL4\_VS\_CTRL\_TREATED\_NAIVE\_CD4\_TCELL\_DAY5\_DN | 140 | 15 | 12187 | 179 | Ifit3b,Phf11b,Ifitm2,Herc6,Tap1,Casp4,Xaf1,Ccnd2,Ifit2,Parp12,Ripk1,Irf7,Arid5b,Phf11d,Nfkb2 | | 1.963e-09 | -20.05 | GSE21360\_NAIVE\_VS\_SECONDARY\_MEMORY\_CD8\_TCELL\_DN | MSigDB lists | GSE21360\_NAIVE\_VS\_SECONDARY\_MEMORY\_CD8\_TCELL\_DN | 140 | 15 | 12187 | 179 | Iqgap1,Irf7,Parp12,Xaf1,Ifit2,Clic4,Trim25,Slfn5,Rnf213,Atf3,Luc7l3,Znfx1,Samd9l,Ifit3b,Ifi44 | | 1.981e-09 | -20.04 | regulation of cytokine biosynthetic process | biological process | GO:0042035 | 76 | 12 | 13711 | 214 | Bcl3,Hspb1,Thbs1,Zfp36,Il6,Prg4,Errfi1,Nfkb1,Cybb,Tnf,Akap12,Il1a | | 1.981e-09 | -20.04 | GO\_EXTRACELLULAR\_SPACE | MSigDB lists | GO\_EXTRACELLULAR\_SPACE | 738 | 34 | 12187 | 179 | S100a8,Adamts9,Csf3,Angptl4,Cxcl1,Il6,Ccl12,Msr1,Il4ra,Ccl2,Ccl4,Cxcl16,Selp,Cd14,Il1a,Mmp8,Lcn2,Thbs1,Tnfsf10,Xdh,A2m,S100a9,Tnf,Sele,Icam1,Hspb1,Msn,Saa1,Hp,Cp,Timp1,Il1rn,Vcam1,Fstl1 | | 2.090e-09 | -19.99 | regulation of intracellular signal transduction | biological process | GO:1902531 | 1320 | 50 | 13711 | 214 | Akap12,Il1a,Il6,Ccl7,Trib1,Gadd45b,Map3k6,Fzd4,Rasip1,Ccl4,Ptgs2,Trim25,Tnf,Pik3ap1,Atf3,Hspb1,Ncf1,Cd14,Cflar,Ripk1,Tnip1,Tgm2,C5ar1,Nfe2l2,Mmp8,Selp,Icam1,Sox11,Errfi1,Thbs1,Gadd45g,P2ry6,Map3k8,Birc3,Pik3r5,Plaur,Tnfaip3,Tnfsf10,Ksr1,Xdh,Ccl2,Irak3,Il1rn,Csf3,Ptpn1,Tnip3,S100a8,Iqgap1,Ccl12,S100a9 | | 2.186e-09 | -19.94 | granulocyte migration | biological process | GO:0097530 | 61 | 11 | 13711 | 214 | C5ar1,Cxcl1,S100a8,Ccl12,S100a9,Ccl4,Il17ra,Cxcl9,Ccl2,Il1rn,Ccl7 | | 2.237e-09 | -19.92 | LENAOUR\_DENDRITIC\_CELL\_MATURATION\_DN | MSigDB lists | LENAOUR\_DENDRITIC\_CELL\_MATURATION\_DN | 100 | 13 | 12187 | 179 | S100a8,S100a9,Irf7,Cxcl1,C3ar1,C5ar1,Cybb,Nfkbia,Tnf,Plaur,Cd14,Isg15,Ier3 | | 2.243e-09 | -19.92 | leukocyte chemotaxis | biological process | GO:0030595 | 94 | 13 | 13711 | 214 | Ccl2,Cxcl9,Retnlg,Ccl7,Il1rn,Cxcl16,Ccl12,S100a8,C5ar1,Cxcl1,Il17ra,Ccl4,S100a9 | | 2.432e-09 | -19.83 | GSE20715\_0H\_VS\_48H\_OZONE\_LUNG\_DN | MSigDB lists | GSE20715\_0H\_VS\_48H\_OZONE\_LUNG\_DN | 165 | 16 | 12187 | 179 | Map3k6,Saa1,Ptgs2,Cd14,Hspb1,Rbm47,Errfi1,Cdkn1a,Hspa5,Socs3,Dnajb1,Timp1,Lcn2,Banp,Rhou,Il6 | | 2.534e-09 | -19.79 | Influenza A | KEGG pathways | mmu05164 | 124 | 16 | 5248 | 107 | Nfkbia,Ccl2,Pik3r1,Il1a,Ccl12,Socs3,Il6,Trim25,Nfkb1,Icam1,Tnf,Tnfsf10,Irf9,Irf7,Dnajb1,Mx2 | | 2.534e-09 | -19.79 | Influenza A | KEGG pathways | ko05164 | 124 | 16 | 5248 | 107 | Tnfsf10,Irf9,Irf7,Icam1,Tnf,Nfkb1,Socs3,Il6,Trim25,Mx2,Dnajb1,Pik3r1,Ccl2,Nfkbia,Ccl12,Il1a | | 2.641e-09 | -19.75 | positive regulation of phosphorylation | biological process | GO:0042327 | 843 | 38 | 13711 | 214 | Cdkn1a,Il6,Il1a,Akap12,Icam1,P2ry6,Lgals9,Gadd45b,Ccnd2,Pik3r1,Gadd45g,Parp14,Ccl7,Thbs1,Parp9,Sdc4,Fzd4,Map3k6,Map3k8,Plaur,Pik3r5,Ptgs2,Ccl4,Ccl2,Xdh,Ksr1,Tnf,Ripk1,Csf3,Cflar,Ncf1,Il1rn,Ccl12,Iqgap1,Ptpn1,C5ar1,Irgm2,Mmp8 | | 2.698e-09 | -19.73 | activation of innate immune response | biological process | GO:0002218 | 78 | 12 | 13711 | 214 | Tnip3,Ifi209,Irgm2,Ifi211,Tnf,Pik3ap1,Nfkbia,Tnip1,Lgals9,Ifi204,Irak3,Cd14 | | 2.698e-09 | -19.73 | negative regulation of viral process | biological process | GO:0048525 | 78 | 12 | 13711 | 214 | Zfp36,Isg15,Tnf,Oasl2,Ifitm2,Trim56,Resf1,Trim25,Ifitm3,Oasl1,Ifitm6,Ccl4 | | 2.713e-09 | -19.73 | GTPase activity | molecular function | GO:0003924 | 253 | 20 | 13516 | 211 | Rhoc,Gbp6,Ifi47,Gm5431,Irgm1,Gbp7,Irgm2,Tubb6,Rhoj,Mx1,Rhou,Mx2,Gbp3,Iigp1,Gem,Tgtp1,F830016B08Rik,Gm4841,Rasd1,Tgtp2 | | 2.901e-09 | -19.66 | GSE45365\_HEALTHY\_VS\_MCMV\_INFECTION\_CD8A\_DC\_IFNAR\_KO\_DN | MSigDB lists | GSE45365\_HEALTHY\_VS\_MCMV\_INFECTION\_CD8A\_DC\_IFNAR\_KO\_DN | 167 | 16 | 12187 | 179 | Casp4,Tnf,Gadd45b,Phf11d,Irf7,Cflar,C3ar1,Phf11b,Ifi44,Cebpd,Il1a,Ifitm2,Plaur,Irf9,Tap1,Rbm47 | | 2.901e-09 | -19.66 | GSE22886\_DAY0\_VS\_DAY1\_MONOCYTE\_IN\_CULTURE\_DN | MSigDB lists | GSE22886\_DAY0\_VS\_DAY1\_MONOCYTE\_IN\_CULTURE\_DN | 167 | 16 | 12187 | 179 | Cxcl1,Nfkbie,Ccl4,Sdc4,Runx1,Birc3,Maff,Ccl12,Il6,Tubb6,Tnip1,Ptgs2,Cdkn1a,Ier3,Il1rn,Gadd45g | | 2.906e-09 | -19.66 | biological regulation | biological process | GO:0065007 | 8325 | 170 | 13711 | 214 | Trim56,Il17ra,Plaur,Bach1,Kcna5,Nfkb1,Adamts1,Gbp4,Osmr,Rbm39,Banp,P2ry6,Thbs1,Ifitm3,Cybb,Errfi1,Mxd4,Sox11,Samd9l,Bcl3,Irgm2,Sgk3,Irf9,Usp18,Tap1,S100a8,A2m,Irf2,Ccl12,Ifi204,Arid5b,Irak3,Pygm,Adamts9,Xdh,Tnfsf10,Ksr1,Plek,Ptgs2,Sgk1,Ier3,Stx11,Ifitm6,Ccl4,Spi1,Parp9,Sbno2,Sdc4,Ptges,Ifi209,Fzd4,Gadd45b,Pik3r1,Dtx3l,Irf7,Socs3,Trib1,Ctla2a,Il6,Cdkn1a,Oasl2,Casp4,Il1a,Gem,Csf2rb,Ifit2,Hp,Mt2,Prg4,Rrp8,Cxcl1,Dnajb1,Fstl1,Tnip1,Rhoc,Ifi207,Slc24a4,Hspb1,Ripk1,Cflar,Cd14,Gpr84,Zfp36,Tnf,Robo4,Pik3r5,Irgm1,Rhoj,Apold1,Nfkbie,Map3k8,Birc3,Gadd45g,Lgals9,Oasl1,Bbs12,Msr1,Iigp1,Rassf4,Icam1,Mt1,Nfkb2,Fyb,Rgs16,Wfdc21,Vcam1,S100a9,Iqgap1,Col4a1,Tnip3,Cxcl16,Nfkbiz,Ptpn1,Ifitm2,Lcn2,Aff1,Il1rn,C3ar1,Zbtb16,Tnfaip8,Csf3,Depp1,Ccl2,Znfx1,Tnfaip3,Cxcl9,Runx1,Trim25,Resf1,Xaf1,Ifi211,Rasip1,Cebpd,Slfn2,Isg15,Tiparp,Timp1,Map3k6,Fam107a,Ccnd2,Hcar2,Parp14,Ccl7,Ip6k2,Slfn3,Il4ra,Nfkbia,Akap12,Angptl4,Msn,Selp,Noct,Nfe2l2,Serpina3f,Mmp8,Rasd1,C5ar1,Maff,Tgm2,Clic4,Hspa5,Cp,Rnf213,Trim30a,Ncf1,Sele,Atf3,Rhou,Acod1,Pik3ap1 | | 2.911e-09 | -19.65 | GSE1432\_CTRL\_VS\_IFNG\_24H\_MICROGLIA\_DN | MSigDB lists | GSE1432\_CTRL\_VS\_IFNG\_24H\_MICROGLIA\_DN | 144 | 15 | 12187 | 179 | Icam1,Plek,Tap1,Ifitm3,Isg15,Phf11b,Ifi44,Ifit3b,Rtp4,Sbno2,Tnfsf10,Phf11d,Irf7,Parp12,Tgm2 | | 2.925e-09 | -19.65 | cellular response to tumor necrosis factor | biological process | GO:0071356 | 96 | 13 | 13711 | 214 | Nfe2l2,Ccl4,Nfkb1,Ccl12,Birc3,Ccl7,Ripk1,Il6,Zfp36,Ccl2,Acod1,Tnf,Nfkbia | | 2.939e-09 | -19.65 | DAUER\_STAT3\_TARGETS\_DN | MSigDB lists | DAUER\_STAT3\_TARGETS\_DN | 38 | 9 | 12187 | 179 | Parp12,Oasl1,Usp18,Ifit3b,Ifi44,Tnfsf10,Irf7,Herc6,Isg15 | | 2.939e-09 | -19.65 | WORSCHECH\_TUMOR\_REJECTION\_UP | MSigDB lists | WORSCHECH\_TUMOR\_REJECTION\_UP | 38 | 9 | 12187 | 179 | Ifi211,Cxcl1,Ifi207,Il1a,Ifi204,Msr1,Ifi209,Nfkbiz,Ccl4 | | 3.089e-09 | -19.60 | GO\_REGULATION\_OF\_RESPONSE\_TO\_EXTERNAL\_STIMULUS | MSigDB lists | GO\_REGULATION\_OF\_RESPONSE\_TO\_EXTERNAL\_STIMULUS | 674 | 32 | 12187 | 179 | Socs3,Plaur,Hspb1,Plek,Sele,Tnip1,Saa1,Ptgs2,Casp4,Trib1,Tnf,Zfp189,Birc3,S100a9,A2m,C5ar1,C3ar1,Nfkb1,Thbs1,Il17ra,Pik3ap1,Selp,Ccl12,Tnfaip3,Il6,Nfe2l2,Ccl4,Ccl2,Sbno2,Osmr,S100a8,Cxcl1 | | 3.129e-09 | -19.58 | GSE34156\_UNTREATED\_VS\_6H\_NOD2\_AND\_TLR1\_TLR2\_LIGAND\_TREATED\_MONOCYTE\_DN | MSigDB lists | GSE34156\_UNTREATED\_VS\_6H\_NOD2\_AND\_TLR1\_TLR2\_LIGAND\_TREATED\_MONOCYTE\_DN | 123 | 14 | 12187 | 179 | Rnf213,Slfn5,Pik3ap1,Tap1,Ifi44,Ifit3b,Samd9l,Irf7,Rtp4,Tnfsf10,Ifit2,Usp18,Xaf1,Parp12 | | 3.129e-09 | -19.58 | GSE41867\_DAY6\_VS\_DAY15\_LCMV\_ARMSTRONG\_EFFECTOR\_CD8\_TCELL\_UP | MSigDB lists | GSE41867\_DAY6\_VS\_DAY15\_LCMV\_ARMSTRONG\_EFFECTOR\_CD8\_TCELL\_UP | 123 | 14 | 12187 | 179 | Rnf213,Ptgs2,Errfi1,Stx11,Nfkb1,Il1a,Samd9l,Gem,Map3k8,Gpr84,Usp18,Ripk1,Il6,Oasl1 | | 3.166e-09 | -19.57 | GSE19401\_PLN\_VS\_PEYERS\_PATCH\_FOLLICULAR\_DC\_DN | MSigDB lists | GSE19401\_PLN\_VS\_PEYERS\_PATCH\_FOLLICULAR\_DC\_DN | 168 | 16 | 12187 | 179 | Ifit3b,Cdkn1a,Ifi44,Stx11,Ifitm3,Plaur,Tap1,Xaf1,Tgm2,Ccl4,Ifit2,Maff,Ccl12,Il6,Oasl1,Cflar | | 3.166e-09 | -19.57 | GSE3337\_CTRL\_VS\_4H\_IFNG\_IN\_CD8POS\_DC\_DN | MSigDB lists | GSE3337\_CTRL\_VS\_4H\_IFNG\_IN\_CD8POS\_DC\_DN | 168 | 16 | 12187 | 179 | Gbp7,Slfn4,Gbp3,Ifit2,Tnf,Casp4,Usp18,Tor3a,Tnfaip3,Slfn3,Tap1,Ifit3b,Gadd45g,Ksr1,Ier3,Znfx1 | | 3.176e-09 | -19.57 | HESS\_TARGETS\_OF\_HOXA9\_AND\_MEIS1\_DN | MSigDB lists | HESS\_TARGETS\_OF\_HOXA9\_AND\_MEIS1\_DN | 67 | 11 | 12187 | 179 | C3ar1,C5ar1,Cxcl1,Plek,Zfp36,Slfn3,Slfn4,Msr1,Mmp8,Cybb,Lcn2 | | 3.205e-09 | -19.56 | GSE9037\_CTRL\_VS\_LPS\_1H\_STIM\_BMDM\_DN | MSigDB lists | GSE9037\_CTRL\_VS\_LPS\_1H\_STIM\_BMDM\_DN | 145 | 15 | 12187 | 179 | Casp4,Ccl4,Parp14,Tgm2,Msr1,Maff,Map3k8,Gadd45b,Rtp4,Rbm39,Slfn4,Vcam1,Tnip1,Errfi1,Slfn3 | | 3.210e-09 | -19.56 | HORIUCHI\_WTAP\_TARGETS\_UP | MSigDB lists | HORIUCHI\_WTAP\_TARGETS\_UP | 245 | 19 | 12187 | 179 | Rnf213,Cxcl16,Sele,Icam1,Apold1,Ifit3b,Vcam1,Ifi44,Atf3,Cebpd,Col4a1,Arid5b,Cxcl1,Osmr,Sdc4,Birc3,Ifit2,Ccl12,Tnfaip3 | | 3.240e-09 | -19.55 | SCHOEN\_NFKB\_SIGNALING | MSigDB lists | SCHOEN\_NFKB\_SIGNALING | 27 | 8 | 12187 | 179 | Il1a,Ccl12,Gadd45b,Thbs1,Birc3,Nfkbia,Ptgs2,Cxcl1 | | 3.240e-09 | -19.55 | GROSS\_ELK3\_TARGETS\_DN | MSigDB lists | GROSS\_ELK3\_TARGETS\_DN | 27 | 8 | 12187 | 179 | Socs3,Adamts1,Ifi44,Maff,Il6,Cxcl1,Ptgs2,Plek | | 3.241e-09 | -19.55 | GSE17974\_IL4\_AND\_ANTI\_IL12\_VS\_UNTREATED\_24H\_ACT\_CD4\_TCELL\_DN | MSigDB lists | GSE17974\_IL4\_AND\_ANTI\_IL12\_VS\_UNTREATED\_24H\_ACT\_CD4\_TCELL\_DN | 103 | 13 | 12187 | 179 | Tnfsf10,Rhou,Ifit2,Usp18,Ccl4,Xaf1,Oasl1,Isg15,Herc6,Ifi44,Ifit3b,Samd9l,Atf3 | | 3.316e-09 | -19.52 | Nfkb1 (nuclear factor of kappa light polypeptide gene enhancer in B cells 1, p105) | protein interactions | 18033 | 16 | 7 | 6802 | 115 | Nfkbiz,Il6,Map3k8,Nfkb2,Nfkb1,Nfkbia,Sele | | 3.482e-09 | -19.48 | BROWN\_MYELOID\_CELL\_DEVELOPMENT\_UP | MSigDB lists | BROWN\_MYELOID\_CELL\_DEVELOPMENT\_UP | 124 | 14 | 12187 | 179 | Slfn3,Plaur,Samd9l,Il1rn,Mmp8,Lcn2,Cebpd,C5ar1,Slfn4,Iqgap1,Xdh,Ccnd2,Tnf,Cybb | | 3.482e-09 | -19.48 | GSE27241\_CTRL\_VS\_DIGOXIN\_TREATED\_CD4\_TCELL\_IN\_TH17\_POLARIZING\_CONDITIONS\_UP | MSigDB lists | GSE27241\_CTRL\_VS\_DIGOXIN\_TREATED\_CD4\_TCELL\_IN\_TH17\_POLARIZING\_CONDITIONS\_UP | 124 | 14 | 12187 | 179 | Gadd45b,Tnfaip3,Trib1,Nfkbia,Parp14,Birc3,Map3k8,Nfkb2,Arid5b,Ier3,Stx11,Nfkb1,Nfkbiz,Icam1 | | 3.499e-09 | -19.47 | regulation of phosphate metabolic process | biological process | GO:0019220 | 1427 | 52 | 13711 | 214 | Pik3r1,Ccnd2,Gadd45b,Socs3,Ccl7,Parp14,Trib1,Il6,Cdkn1a,Akap12,Il1a,Ptgs2,Ier3,Rasip1,Ccl4,Sdc4,Parp9,Map3k6,Fzd4,Tnip1,Hspb1,Ncf1,Cflar,Ripk1,Atf3,Tnf,Pik3ap1,Mmp8,C5ar1,Gadd45g,Lgals9,P2ry6,Thbs1,Errfi1,Icam1,Pik3r5,Plaur,Gbp4,Map3k8,Birc3,Irak3,Il1rn,Csf3,Ksr1,Xdh,Ccl2,Tnfaip3,Plek,Irgm2,Iqgap1,Ccl12,Ptpn1 | | 3.584e-09 | -19.45 | regulation of phosphorus metabolic process | biological process | GO:0051174 | 1428 | 52 | 13711 | 214 | Socs3,Trib1,Parp14,Ccl7,Gadd45b,Ccnd2,Pik3r1,Akap12,Il1a,Il6,Cdkn1a,Rasip1,Ccl4,Ptgs2,Ier3,Map3k6,Fzd4,Parp9,Sdc4,Ncf1,Hspb1,Ripk1,Cflar,Tnip1,Tnf,Pik3ap1,Atf3,Mmp8,C5ar1,Thbs1,Gadd45g,P2ry6,Lgals9,Icam1,Errfi1,Pik3r5,Plaur,Map3k8,Gbp4,Birc3,Irak3,Il1rn,Csf3,Tnfaip3,Plek,Xdh,Ksr1,Ccl2,Irgm2,Ptpn1,Iqgap1,Ccl12 | | 3.657e-09 | -19.43 | TAKEDA\_TARGETS\_OF\_NUP98\_HOXA9\_FUSION\_16D\_UP | MSigDB lists | TAKEDA\_TARGETS\_OF\_NUP98\_HOXA9\_FUSION\_16D\_UP | 104 | 13 | 12187 | 179 | Ptgs2,Isg15,Thbs1,Ifi44,Ifit3b,Gem,Cmpk2,Tnfsf10,Ifit2,Usp18,Xaf1,Oasl1,AA467197 | | 3.763e-09 | -19.40 | GSE15930\_STIM\_VS\_STIM\_AND\_TRICHOSTATINA\_48H\_CD8\_T\_CELL\_DN | MSigDB lists | GSE15930\_STIM\_VS\_STIM\_AND\_TRICHOSTATINA\_48H\_CD8\_T\_CELL\_DN | 170 | 16 | 12187 | 179 | Irf9,Tap1,Slfn3,Trim25,Isg15,Atf3,Znfx1,Nfkb1,Cmpk2,Slfn4,Irf7,Irgm2,Usp18,Lgals9,Ccnd2,Tnf | | 3.878e-09 | -19.37 | GSE12392\_IFNAR\_KO\_VS\_IFNB\_KO\_CD8\_NEG\_SPLEEN\_DC\_DN | MSigDB lists | GSE12392\_IFNAR\_KO\_VS\_IFNB\_KO\_CD8\_NEG\_SPLEEN\_DC\_DN | 147 | 15 | 12187 | 179 | Nfkbia,Il4ra,Il6,Cxcl1,Csf3,Socs3,Nfkbiz,Adamts1,Thbs1,Rasip1,Nfkb1,Mmp8,Tnip1,Errfi1,Plaur | | 3.928e-09 | -19.36 | - | gene3d domains | 3.30.950.30 | 6 | 5 | 6647 | 99 | Slfn5,Slfn3,Slfn2,Slfn4,Slfn9 | | 4.014e-09 | -19.33 | BIOCARTA\_LAIR\_PATHWAY | MSigDB lists | BIOCARTA\_LAIR\_PATHWAY | 11 | 6 | 12187 | 179 | Selp,Tnf,Vcam1,Icam1,Il1a,Il6 | | 4.099e-09 | -19.31 | GSE37416\_0H\_VS\_6H\_F\_TULARENSIS\_LVS\_NEUTROPHIL\_DN | MSigDB lists | GSE37416\_0H\_VS\_6H\_F\_TULARENSIS\_LVS\_NEUTROPHIL\_DN | 171 | 16 | 12187 | 179 | Banp,Pik3r5,Nfkbie,Gpr84,Nfkbia,Tnf,Maff,Il4ra,Plaur,Cxcl16,Icam1,Cdkn1a,Ier3,Tiparp,Nfkb1,Gadd45g | | 4.143e-09 | -19.30 | neutrophil chemotaxis | biological process | GO:0030593 | 50 | 10 | 13711 | 214 | Il1rn,Ccl4,Ccl7,S100a9,C5ar1,Cxcl1,Cxcl9,S100a8,Ccl2,Ccl12 | | 4.216e-09 | -19.28 | positive regulation of phosphorus metabolic process | biological process | GO:0010562 | 896 | 39 | 13711 | 214 | Cdkn1a,Il6,Akap12,Il1a,Icam1,Lgals9,P2ry6,Ccnd2,Pik3r1,Gadd45g,Gadd45b,Ccl7,Parp14,Thbs1,Sdc4,Parp9,Map3k6,Fzd4,Map3k8,Plaur,Ptgs2,Pik3r5,Ccl4,Ccl2,Ksr1,Xdh,Plek,Tnf,Cflar,Ripk1,Csf3,Il1rn,Ncf1,Ccl12,Iqgap1,Ptpn1,C5ar1,Irgm2,Mmp8 | | 4.216e-09 | -19.28 | positive regulation of phosphate metabolic process | biological process | GO:0045937 | 896 | 39 | 13711 | 214 | Plaur,Pik3r5,Ptgs2,Ccl4,Parp9,Sdc4,Map3k6,Fzd4,Map3k8,P2ry6,Lgals9,Gadd45b,Ccnd2,Gadd45g,Pik3r1,Parp14,Ccl7,Thbs1,Cdkn1a,Il6,Il1a,Akap12,Icam1,Irgm2,Mmp8,Ccl12,Iqgap1,Ptpn1,C5ar1,Ripk1,Csf3,Cflar,Ncf1,Il1rn,Ccl2,Xdh,Ksr1,Plek,Tnf | | 4.217e-09 | -19.28 | myeloid leukocyte migration | biological process | GO:0097529 | 81 | 12 | 13711 | 214 | C5ar1,Cxcl1,Ccl12,S100a8,Ccl4,S100a9,Il17ra,Cxcl9,Ccl2,Ccl7,Il1rn,Retnlg | | 4.261e-09 | -19.27 | GSE17721\_LPS\_VS\_POLYIC\_1H\_BMDC\_UP | MSigDB lists | GSE17721\_LPS\_VS\_POLYIC\_1H\_BMDC\_UP | 148 | 15 | 12187 | 179 | Timp1,Tiparp,Il1a,Cebpd,Cdkn1a,Hspa5,Icam1,Slfn3,Zfp36,Errfi1,Gadd45b,Maff,Msr1,Slfn4,Tnfsf10 | | 4.261e-09 | -19.27 | GSE34156\_UNTREATED\_VS\_6H\_NOD2\_LIGAND\_TREATED\_MONOCYTE\_DN | MSigDB lists | GSE34156\_UNTREATED\_VS\_6H\_NOD2\_LIGAND\_TREATED\_MONOCYTE\_DN | 148 | 15 | 12187 | 179 | Phf11d,C3ar1,Tnfsf10,Gbp3,Cmpk2,Usp18,Xaf1,Parp12,Irf2,Ifit3b,Phf11b,Znfx1,Luc7l3,Samd9l,Gadd45g | | 4.300e-09 | -19.26 | GSE7348\_UNSTIM\_VS\_TOLERIZED\_AND\_LPS\_STIM\_MACROPHAGE\_DN | MSigDB lists | GSE7348\_UNSTIM\_VS\_TOLERIZED\_AND\_LPS\_STIM\_MACROPHAGE\_DN | 126 | 14 | 12187 | 179 | Cflar,Arid5b,Ptges,Phf11d,Parp12,Tnfaip3,Map3k8,Casp4,Isg15,Tnip1,Samd9l,Nfkb1,Stx11,Phf11b | | 4.300e-09 | -19.26 | GO\_CELLULAR\_RESPONSE\_TO\_BIOTIC\_STIMULUS | MSigDB lists | GO\_CELLULAR\_RESPONSE\_TO\_BIOTIC\_STIMULUS | 126 | 14 | 12187 | 179 | Sbno2,Cmpk2,Csf3,Il6,Ccl12,Tnfaip3,Nfkbia,Tnf,Cxcl16,Icam1,Cd14,Tnip3,Nfkb1,Hspa5 | | 4.315e-09 | -19.26 | positive regulation of innate immune response | biological process | GO:0045089 | 139 | 15 | 13711 | 214 | Irak3,Irf7,Cd14,Tnip1,Ifi204,Lgals9,Tnf,Nfkbia,Pik3ap1,Irgm1,Ifi211,Irgm2,Ifi209,Tnip3,Parp9 | | 4.394e-09 | -19.24 | GO\_POSITIVE\_REGULATION\_OF\_INFLAMMATORY\_RESPONSE | MSigDB lists | GO\_POSITIVE\_REGULATION\_OF\_INFLAMMATORY\_RESPONSE | 69 | 11 | 12187 | 179 | Ccl2,Ccl4,Tnf,Il6,Ccl12,S100a9,Tnip1,Ptgs2,Osmr,S100a8,Il17ra | | 4.394e-09 | -19.24 | GO\_REGULATION\_OF\_TUMOR\_NECROSIS\_FACTOR\_SUPERFAMILY\_CYTOKINE\_PRODUCTION | MSigDB lists | GO\_REGULATION\_OF\_TUMOR\_NECROSIS\_FACTOR\_SUPERFAMILY\_CYTOKINE\_PRODUCTION | 69 | 11 | 12187 | 179 | Zfp36,Irak3,Hspb1,Errfi1,Cd14,Bcl3,Tnfaip3,Ripk1,Ccl12,Thbs1,Pik3r1 | | 4.460e-09 | -19.23 | regulation of molecular function | biological process | GO:0065009 | 2276 | 70 | 13711 | 214 | Oasl2,Casp4,Nfkbia,Gem,Il6,Cdkn1a,Dtx3l,Trib1,Ccl7,Gadd45b,Pik3r1,Ccnd2,Timp1,Fzd4,Map3k6,Parp9,Sdc4,Rasip1,Ccl4,Ptgs2,Trim25,Tnf,Acod1,Zfp36,Ncf1,Sele,Hspb1,Ripk1,Cflar,Rhoc,Dnajb1,C5ar1,Hp,Serpina3f,Mmp8,Angptl4,Ifit2,Icam1,Rgs16,Sox11,Errfi1,Thbs1,Oasl1,Gadd45g,P2ry6,Lgals9,Map3k8,Birc3,Nfkb1,Pik3r5,Plaur,Tnfaip3,Plek,Xdh,Tnfsf10,Ksr1,Ccl2,Irak3,Arid5b,Il1rn,Tnfaip8,Csf3,Ptpn1,S100a8,Iqgap1,A2m,Ccl12,S100a9,Irgm2,Wfdc21 | | 4.480e-09 | -19.22 | BOYAULT\_LIVER\_CANCER\_SUBCLASS\_G5\_DN | MSigDB lists | BOYAULT\_LIVER\_CANCER\_SUBCLASS\_G5\_DN | 28 | 8 | 12187 | 179 | Bcl3,Ifi211,Icam1,Ifi44,Ifi209,Tgm2,Ifi207,Ifi204 | | 4.536e-09 | -19.21 | QI\_PLASMACYTOMA\_UP | MSigDB lists | QI\_PLASMACYTOMA\_UP | 197 | 17 | 12187 | 179 | Rgs16,Ptges,Irgm2,Nfkbie,Tnfaip3,Rassf4,Ccl4,Tgm2,Casp4,Nfkbia,Tnf,Il4ra,Ptpn1,Irf2,Plek,Dnajb1,Sox11 | | 4.678e-09 | -19.18 | GSE2405\_S\_AUREUS\_VS\_A\_PHAGOCYTOPHILUM\_NEUTROPHIL\_UP | MSigDB lists | GSE2405\_S\_AUREUS\_VS\_A\_PHAGOCYTOPHILUM\_NEUTROPHIL\_UP | 149 | 15 | 12187 | 179 | Bcl3,Gbp3,Adamts9,Xaf1,Tor3a,Oasl1,Slfn9,Ifitm3,Parp9,Csf2rb,Trim25,Hspb1,Plek,Timp1,Nfkb1 | | 4.778e-09 | -19.16 | SENESE\_HDAC1\_TARGETS\_UP | MSigDB lists | SENESE\_HDAC1\_TARGETS\_UP | 370 | 23 | 12187 | 179 | Gem,Robo4,Nfkb2,Birc3,Nfkbia,Trib1,Bach1,Errfi1,Ifi209,Ifi207,Ifi204,Ifi211,Osmr,Iqgap1,Maff,Ccnd2,Tgm2,Nfe2l2,Tnfaip3,Ccl12,Nfkbiz,Il1a,Cebpd | | 4.808e-09 | -19.15 | GO\_POSITIVE\_REGULATION\_OF\_MOLECULAR\_FUNCTION | MSigDB lists | GO\_POSITIVE\_REGULATION\_OF\_MOLECULAR\_FUNCTION | 1413 | 49 | 12187 | 179 | Ptpn1,Map3k6,Sgk1,Hspa5,Rgs16,S100a8,Ifi211,Arid5b,Ripk1,Gadd45b,Ccl2,Ccnd2,Trim25,Icam1,Plek,Saa1,Ifi204,Ifi209,Cdkn1a,Tnfsf10,S100a9,Fzd4,Cflar,Irak3,Sgk3,Nfkb1,Gadd45g,Dnajb1,Thbs1,Iqgap1,Pik3r5,Csf3,Ccl12,Il6,Rhoc,Ccl4,Map3k8,Plaur,Errfi1,Csf2rb,Sele,Ifi207,Nfkb2,Xdh,A2m,C5ar1,Sdc4,Nfkbia,Tnf | | 4.859e-09 | -19.14 | Schlafen | interpro domains | IPR029684 | 6 | 5 | 13788 | 212 | Slfn3,Slfn5,Slfn9,Slfn4,Slfn2 | | 4.859e-09 | -19.14 | Schlafen\_AlbA\_2\_dom | interpro domains | IPR007421 | 6 | 5 | 13788 | 212 | Slfn9,Slfn3,Slfn5,Slfn2,Slfn4 | | 4.859e-09 | -19.14 | Schlafen\_AlbA\_2\_dom\_sf | interpro domains | IPR038461 | 6 | 5 | 13788 | 212 | Slfn3,Slfn5,Slfn9,Slfn4,Slfn2 | | 4.908e-09 | -19.13 | ONDER\_CDH1\_TARGETS\_2\_DN | MSigDB lists | ONDER\_CDH1\_TARGETS\_2\_DN | 309 | 21 | 12187 | 179 | Parp12,Tnfaip3,Il4ra,Birc3,Ccnd2,Sdc4,S100a8,Tnfsf10,Xdh,Csf3,S100a9,Cxcl1,Ptges,Il1a,Il1rn,Ier3,Lcn2,Rbm47,Icam1,Ptgs2,Herc6 | | 5.115e-09 | -19.09 | leukocyte cell-cell adhesion | biological process | GO:0007159 | 38 | 9 | 13711 | 214 | S100a8,Icam1,Tnf,Msn,Tnip1,Selp,Vcam1,Sele,S100a9 | | 5.119e-09 | -19.09 | LEI\_MYB\_TARGETS | MSigDB lists | LEI\_MYB\_TARGETS | 252 | 19 | 12187 | 179 | S100a9,Arid5b,Cflar,S100a8,Tgm2,Ccnd2,Ccl12,Il6,Tnip1,Ifitm2,Saa1,Isg15,Clic4,Msn,Hspb1,Dnajb1,Cdkn1a,Lcn2,Il1rn | | 5.217e-09 | -19.07 | AlbA\_2 | pfam domains | PF04326 | 6 | 5 | 12881 | 201 | Slfn2,Slfn9,Slfn5,Slfn3,Slfn4 | | 6.014e-09 | -18.93 | HUMMERICH\_SKIN\_CANCER\_PROGRESSION\_UP | MSigDB lists | HUMMERICH\_SKIN\_CANCER\_PROGRESSION\_UP | 71 | 11 | 12187 | 179 | Il4ra,Rhoc,Gadd45b,Ifitm3,Saa1,Il17ra,Cdkn1a,Hspa5,Vcam1,Il1rn,Lcn2 | | 6.109e-09 | -18.91 | KRASNOSELSKAYA\_ILF3\_TARGETS\_UP | MSigDB lists | KRASNOSELSKAYA\_ILF3\_TARGETS\_UP | 29 | 8 | 12187 | 179 | Tap1,Ifi207,Ifi204,Ifit3b,Ifi211,Xaf1,Isg15,Ifi209 | | 6.483e-09 | -18.85 | KEGG\_CYTOKINE\_CYTOKINE\_RECEPTOR\_INTERACTION | MSigDB lists | KEGG\_CYTOKINE\_CYTOKINE\_RECEPTOR\_INTERACTION | 130 | 14 | 12187 | 179 | Osmr,Il17ra,Tnfsf10,Cxcl16,Csf2rb,Csf3,Cxcl1,Il1a,Il6,Ccl12,Il4ra,Tnf,Ccl2,Ccl4 | | 7.009e-09 | -18.78 | MODULE\_208 | MSigDB lists | MODULE\_208 | 72 | 11 | 12187 | 179 | Ifitm3,Selp,Csf3,Ifitm2,Isg15,Cd14,Tnfsf10,Ifi44,Thbs1,Vcam1,Cebpd | | 7.915e-09 | -18.65 | GSE2706\_2H\_VS\_8H\_R848\_STIM\_DC\_DN | MSigDB lists | GSE2706\_2H\_VS\_8H\_R848\_STIM\_DC\_DN | 132 | 14 | 12187 | 179 | Parp9,Ifi204,Tnip3,Samd9l,Ifi207,Ifi209,Thbs1,Ifi44,Tnfsf10,Cmpk2,Ifi211,Dtx3l,Parp14,Xaf1 | | 8.018e-09 | -18.64 | MISSIAGLIA\_REGULATED\_BY\_METHYLATION\_UP | MSigDB lists | MISSIAGLIA\_REGULATED\_BY\_METHYLATION\_UP | 91 | 12 | 12187 | 179 | Isg15,Ptgs2,Plaur,Tap1,Sdc4,Cdkn1a,Birc3,Mxd4,Tnfaip3,Atf3,Lcn2,Timp1 | | 8.079e-09 | -18.63 | SENESE\_HDAC3\_TARGETS\_UP | MSigDB lists | SENESE\_HDAC3\_TARGETS\_UP | 413 | 24 | 12187 | 179 | Slfn5,Nfkbiz,Rgs16,Iqgap1,Ifi211,Ccl12,Tnfaip3,Rhoc,Ifit2,Ccnd2,Errfi1,Atf3,Ifi207,Ifi204,Ifit3b,Ifi209,Nfkb2,Robo4,Gem,Oasl1,Bach1,Trib1,Nfkbia,Birc3 | | 8.213e-09 | -18.62 | GSE7218\_UNSTIM\_VS\_ANTIGEN\_STIM\_THROUGH\_IGG\_BCELL\_DN | MSigDB lists | GSE7218\_UNSTIM\_VS\_ANTIGEN\_STIM\_THROUGH\_IGG\_BCELL\_DN | 111 | 13 | 12187 | 179 | Casp4,Tnf,Gadd45b,S100a9,Nfkbie,Gem,Rgs16,Ifi44,Atf3,Lcn2,Gadd45g,Il1rn,Irak3 | | 8.227e-09 | -18.62 | MODULE\_108 | MSigDB lists | MODULE\_108 | 30 | 8 | 12187 | 179 | Ccl2,Ccl4,Ccl12,Il1a,Cxcl1,C3ar1,C5ar1,Plaur | | 8.515e-09 | -18.58 | cell chemotaxis | biological process | GO:0060326 | 146 | 15 | 13711 | 214 | Ccl12,Cxcl16,S100a8,Cxcl1,C5ar1,Il17ra,Ccl4,S100a9,Vcam1,Ccl2,Saa1,Cxcl9,Retnlg,Ccl7,Il1rn | | 8.811e-09 | -18.55 | LTE2\_UP.V1\_DN | MSigDB lists | LTE2\_UP.V1\_DN | 156 | 15 | 12187 | 179 | Atf3,Ifi44,Phf11b,Ifit3b,Irf9,Tap1,Isg15,Herc6,Oasl1,Parp12,Usp18,Gem,Irf7,Ptges,Phf11d | | 9.141e-09 | -18.51 | neutrophil migration | biological process | GO:1990266 | 54 | 10 | 13711 | 214 | Il1rn,S100a9,Ccl4,Ccl7,S100a8,Ccl12,Ccl2,C5ar1,Cxcl1,Cxcl9 | | 9.546e-09 | -18.47 | GO\_ACUTE\_INFLAMMATORY\_RESPONSE | MSigDB lists | GO\_ACUTE\_INFLAMMATORY\_RESPONSE | 43 | 9 | 12187 | 179 | Saa1,Ptges,S100a8,Icam1,Vcam1,Il1a,Hp,Il1rn,Il6 | | 1.005e-08 | -18.42 | defense response to bacterium | biological process | GO:0042742 | 106 | 13 | 13711 | 214 | Hp,Isg15,C5ar1,Gbp3,Gbp7,Bcl3,Gbp6,Irgm2,Tnf,Lcn2,Ncf1,Tnfaip8,Iigp1 | | 1.032e-08 | -18.39 | GO\_CYTOKINE\_ACTIVITY | MSigDB lists | GO\_CYTOKINE\_ACTIVITY | 93 | 12 | 12187 | 179 | Cxcl1,Csf3,Cxcl16,Tnfsf10,Ccl4,Ccl2,Tnf,Ccl12,Il6,Timp1,Il1rn,Il1a | | 1.093e-08 | -18.33 | KEGG\_TOLL\_LIKE\_RECEPTOR\_SIGNALING\_PATHWAY | MSigDB lists | KEGG\_TOLL\_LIKE\_RECEPTOR\_SIGNALING\_PATHWAY | 75 | 11 | 12187 | 179 | Nfkb1,Cd14,Pik3r1,Map3k8,Tnf,Nfkbia,Ccl4,Il6,Ripk1,Irf7,Pik3r5 | | 1.095e-08 | -18.33 | KIM\_LRRC3B\_TARGETS | MSigDB lists | KIM\_LRRC3B\_TARGETS | 31 | 8 | 12187 | 179 | Ifi204,Ifi207,Ifi209,Ifi44,Tap1,Isg15,Ifitm2,Ifi211 | | 1.095e-08 | -18.33 | HAN\_JNK\_SINGALING\_UP | MSigDB lists | HAN\_JNK\_SINGALING\_UP | 31 | 8 | 12187 | 179 | Isg15,Gbp3,Irf9,Ccnd2,Runx1,Sgk1,Ifit3b,Usp18 | | 1.144e-08 | -18.29 | GSE7219\_WT\_VS\_NIK\_NFKB2\_KO\_DC\_UP | MSigDB lists | GSE7219\_WT\_VS\_NIK\_NFKB2\_KO\_DC\_UP | 159 | 15 | 12187 | 179 | Rnf213,Clic4,Pik3ap1,Zfp36,Dnajb1,Socs3,Ifi44,Atf3,Irf7,Gem,Nfkbia,Usp18,Ccl4,Tnf,Gadd45b | | 1.225e-08 | -18.22 | GHANDHI\_BYSTANDER\_IRRADIATION\_UP | MSigDB lists | GHANDHI\_BYSTANDER\_IRRADIATION\_UP | 59 | 10 | 12187 | 179 | Ptges,Cxcl1,Ptgs2,Irak3,Icam1,Nfkbiz,Birc3,Tnfaip3,Gadd45g,Il1a | | 1.238e-08 | -18.21 | TLR4 (Toll-like receptor 4) signaling pathway | Lipid Maps pathways | TLR4 | 61 | 17 | 354 | 26 | Icam1,Ptgs2,Tnfaip3,Nfkb2,Ripk1,Nfkb1,Nfkbie,Map3k8,Ccl4,Vcam1,Irf7,Cd14,Il6,Nfkbia,Tnf,Il1a,Irak3 | | 1.246e-08 | -18.20 | GSE25088\_ROSIGLITAZONE\_VS\_IL4\_AND\_ROSIGLITAZONE\_STIM\_STAT6\_KO\_MACROPHAGE\_DAY10\_DN | MSigDB lists | GSE25088\_ROSIGLITAZONE\_VS\_IL4\_AND\_ROSIGLITAZONE\_STIM\_STAT6\_KO\_MACROPHAGE\_DAY10\_DN | 160 | 15 | 12187 | 179 | Ier3,Il1rn,Il1a,Ifit3b,Hspa5,Plaur,Cxcl16,Tap1,Pik3ap1,Parp12,Gadd45b,Parp14,Tgm2,Nfkbia,Dtx3l | | 1.270e-08 | -18.18 | BROCKE\_APOPTOSIS\_REVERSED\_BY\_IL6 | MSigDB lists | BROCKE\_APOPTOSIS\_REVERSED\_BY\_IL6 | 115 | 13 | 12187 | 179 | Bcl3,Irf9,Rgs16,Tnfsf10,Sbno2,Zfp36,Icam1,Socs3,Sgk1,Maff,Map3k8,Gadd45b,Ccl12 | | 1.279e-08 | -18.17 | GO\_POSITIVE\_REGULATION\_OF\_MULTICELLULAR\_ORGANISMAL\_PROCESS | MSigDB lists | GO\_POSITIVE\_REGULATION\_OF\_MULTICELLULAR\_ORGANISMAL\_PROCESS | 1046 | 40 | 12187 | 179 | Saa1,Ptgs2,Icam1,Hspb1,Plek,Adamts1,Ifi209,Zbtb16,Ifi204,Ifi207,C5ar1,C3ar1,S100a9,Irf7,Nfkb2,Runx1,Tnf,Trib1,Selp,Isg15,Cd14,Irak3,Il17ra,Sox11,Thbs1,Sgk1,Nfkb1,Il1a,Csf3,Ifi211,Angptl4,Bcl3,Pik3r1,Il4ra,Hcar2,Tnfaip3,Ripk1,Ccl12,Nfe2l2,Il6 | | 1.357e-08 | -18.12 | GSE17721\_CTRL\_VS\_GARDIQUIMOD\_12H\_BMDC\_DN | MSigDB lists | GSE17721\_CTRL\_VS\_GARDIQUIMOD\_12H\_BMDC\_DN | 161 | 15 | 12187 | 179 | Slfn4,Ccl12,Ifit2,Maff,Parp14,Nfkbia,Ccl2,Casp4,Slfn3,Map3k6,Tagln2,Timp1,Nfkb1,Tiparp,Cdkn1a | | 1.357e-08 | -18.12 | GSE37533\_UNTREATED\_VS\_PIOGLIZATONE\_TREATED\_CD4\_TCELL\_PPARG1\_AND\_FOXP3\_TRASDUCED\_UP | MSigDB lists | GSE37533\_UNTREATED\_VS\_PIOGLIZATONE\_TREATED\_CD4\_TCELL\_PPARG1\_AND\_FOXP3\_TRASDUCED\_UP | 161 | 15 | 12187 | 179 | Nfkb2,Igsf6,Tnfaip8,Ccl4,Ccl2,Sdc4,Maff,Il6,Tubb6,Tnip1,Clic4,Plaur,Tnip3,Il1rn,Il1a | | 1.357e-08 | -18.12 | GO\_RESPONSE\_TO\_TUMOR\_NECROSIS\_FACTOR | MSigDB lists | GO\_RESPONSE\_TO\_TUMOR\_NECROSIS\_FACTOR | 161 | 15 | 12187 | 179 | Vcam1,Nfkb1,Ptgs2,Cd14,Icam1,Sele,Cxcl16,Tnf,Birc3,Ccl4,Ccl2,Ripk1,Ccl12,Il6,Nfe2l2 | | 1.452e-08 | -18.05 | GO\_REGULATION\_OF\_VIRAL\_GENOME\_REPLICATION | MSigDB lists | GO\_REGULATION\_OF\_VIRAL\_GENOME\_REPLICATION | 60 | 10 | 12187 | 179 | Tnip1,Ifi211,Ifitm3,Isg15,Ifitm2,Oasl1,Ifi204,Ifi207,Ifi209,Tnf | | 1.455e-08 | -18.05 | HINATA\_NFKB\_IMMU\_INF | MSigDB lists | HINATA\_NFKB\_IMMU\_INF | 13 | 6 | 12187 | 179 | Cxcl1,Icam1,Nfkbia,Ccl12,Il6,Nfkb1 | | 1.455e-08 | -18.05 | MODULE\_537 | MSigDB lists | MODULE\_537 | 13 | 6 | 12187 | 179 | Birc3,Cflar,Il1a,Nfkb1,Ier3,Tnfaip3 | | 1.476e-08 | -18.03 | GSE22140\_GERMFREE\_VS\_SPF\_ARTHRITIC\_MOUSE\_CD4\_TCELL\_UP | MSigDB lists | GSE22140\_GERMFREE\_VS\_SPF\_ARTHRITIC\_MOUSE\_CD4\_TCELL\_UP | 162 | 15 | 12187 | 179 | Tnfsf10,Olfml2b,Irf7,Il6,Oasl1,Gadd45b,Xaf1,Ifit2,Irf9,Plek,Isg15,Ifitm2,Ifit3b,Ifi44,Thbs1 | | 1.549e-08 | -17.98 | GSE6269\_FLU\_VS\_E\_COLI\_INF\_PBMC\_UP | MSigDB lists | GSE6269\_FLU\_VS\_E\_COLI\_INF\_PBMC\_UP | 139 | 14 | 12187 | 179 | Herc6,Isg15,Ifitm2,Ifitm3,Ifi207,Ifi204,Ifi44,Ifi209,Irf7,Ifi211,Parp12,Oasl1,Usp18,Xaf1 | | 1.549e-08 | -17.98 | GSE30971\_CTRL\_VS\_LPS\_STIM\_MACROPHAGE\_WBP7\_KO\_2H\_UP | MSigDB lists | GSE30971\_CTRL\_VS\_LPS\_STIM\_MACROPHAGE\_WBP7\_KO\_2H\_UP | 139 | 14 | 12187 | 179 | Clic4,Irak3,Ptgs2,Ier3,Nfkb1,Tnip3,Il1a,Socs3,Rgs16,Cxcl1,Gadd45b,Il6,Nfkbia,Tnf | | 1.699e-08 | -17.89 | GSE24634\_IL4\_VS\_CTRL\_TREATED\_NAIVE\_CD4\_TCELL\_DAY3\_DN | MSigDB lists | GSE24634\_IL4\_VS\_CTRL\_TREATED\_NAIVE\_CD4\_TCELL\_DAY3\_DN | 140 | 14 | 12187 | 179 | Thbs1,Ifit3b,Cybb,Il1a,Ccl12,Parp12,Ifitm3,Cflar,Isg15,Ifitm2,Irf7,P2ry6,Tap1,Tnfsf10 | | 1.699e-08 | -17.89 | GSE34205\_HEALTHY\_VS\_FLU\_INF\_INFANT\_PBMC\_DN | MSigDB lists | GSE34205\_HEALTHY\_VS\_FLU\_INF\_INFANT\_PBMC\_DN | 140 | 14 | 12187 | 179 | S100a9,Irf7,S100a8,Cmpk2,Usp18,Xaf1,Parp12,Oasl1,Ifitm2,Herc6,Isg15,Ifitm3,Ifi44,Mmp8 | | 1.699e-08 | -17.89 | GSE37605\_TREG\_VS\_TCONV\_NOD\_FOXP3\_FUSION\_GFP\_DN | MSigDB lists | GSE37605\_TREG\_VS\_TCONV\_NOD\_FOXP3\_FUSION\_GFP\_DN | 140 | 14 | 12187 | 179 | Sdc4,Ccl12,Gadd45b,Cxcl1,Cflar,Adamts9,Nfkbiz,Adamts1,Thbs1,Atf3,Ier3,Il1rn,Akap12,Selp | | 1.740e-08 | -17.87 | TSAI\_RESPONSE\_TO\_IONIZING\_RADIATION | MSigDB lists | TSAI\_RESPONSE\_TO\_IONIZING\_RADIATION | 118 | 13 | 12187 | 179 | Nfkbie,Cflar,Irf9,Tnfsf10,Nfkb2,Tnfaip8,Nfkbia,Ccl4,Birc3,Sgk1,Cdkn1a,Tnfaip3,Nfkb1 | | 1.743e-08 | -17.86 | regulation of apoptotic process | biological process | GO:0042981 | 1233 | 46 | 13711 | 214 | Ccl12,S100a8,Ptpn1,Bcl3,S100a9,Sgk3,Tnfsf10,Xdh,Tnfaip3,Zbtb16,Tnfaip8,Irak3,Il1rn,Nfkb1,Birc3,Plaur,Mt1,Icam1,Gadd45g,Thbs1,Hspa5,Tgm2,C5ar1,Ifit2,Angptl4,Nfe2l2,Zfp36,Atf3,Tnf,Cflar,Ripk1,Hspb1,Timp1,Ier3,Sgk1,Ptgs2,Cdkn1a,Ip6k2,Il6,Akap12,Casp4,Ccnd2,Hcar2,Pik3r1,Gadd45b,Socs3 | | 1.770e-08 | -17.85 | negative regulation of protein metabolic process | biological process | GO:0051248 | 904 | 38 | 13711 | 214 | Tnf,Tnfaip3,Zfp36,Atf3,Xdh,Hspb1,Irak3,Cflar,Tnfaip8,Tnip1,Ptpn1,Prg4,A2m,Serpina3f,Bcl3,Wfdc21,Errfi1,Il6,Cdkn1a,Socs3,Dtx3l,Thbs1,Parp14,Trib1,Ctla2a,Gadd45g,Pik3r1,Banp,Gadd45b,Timp1,Gbp4,Birc3,Nfkb1,Isg15,Rasip1,Spi1,Ptgs2,Plaur | | 1.863e-08 | -17.80 | GSE1432\_CTRL\_VS\_IFNG\_1H\_MICROGLIA\_DN | MSigDB lists | GSE1432\_CTRL\_VS\_IFNG\_1H\_MICROGLIA\_DN | 141 | 14 | 12187 | 179 | Plek,Zfp36,Cdkn1a,Ifit3b,Socs3,Gadd45g,Angptl4,Rgs16,Pygm,Ifit2,Trib1,Bach1,Gadd45b,Ccl12 | | 1.863e-08 | -17.80 | GSE36888\_UNTREATED\_VS\_IL2\_TREATED\_TCELL\_17H\_DN | MSigDB lists | GSE36888\_UNTREATED\_VS\_IL2\_TREATED\_TCELL\_17H\_DN | 141 | 14 | 12187 | 179 | Ccl4,Il6,Tnfaip3,AA467197,Ptges,Cxcl1,Nfkb2,Nfkbiz,Tnip3,Tnip1,Ptgs2,Plaur,Pik3ap1,Icam1 | | 1.917e-08 | -17.77 | CHEN\_LVAD\_SUPPORT\_OF\_FAILING\_HEART\_UP | MSigDB lists | CHEN\_LVAD\_SUPPORT\_OF\_FAILING\_HEART\_UP | 79 | 11 | 12187 | 179 | Gadd45b,Pik3r1,Zfp189,Nfkbia,Zfp36,Gadd45g,Tiparp,Cebpd,Atf3,Cdkn1a,Zbtb16 | | 2.018e-08 | -17.72 | CASTELLANO\_NRAS\_TARGETS\_UP | MSigDB lists | CASTELLANO\_NRAS\_TARGETS\_UP | 62 | 10 | 12187 | 179 | Usp18,Ifi209,Lcn2,Ifi207,Il1rn,Ifi204,Ifi211,Isg15,Irgm2,Tap1 | | 2.024e-08 | -17.72 | GO\_IMMUNE\_EFFECTOR\_PROCESS | MSigDB lists | GO\_IMMUNE\_EFFECTOR\_PROCESS | 304 | 20 | 12187 | 179 | Ifi207,Ifi204,Ifit3b,Ifi209,Irf9,Icam1,Trim25,Ifitm2,Isg15,Ifitm3,Il6,Oasl1,Il4ra,Pik3r1,Ifit2,Sbno2,Irf7,Bcl3,Trim56,Ifi211 | | 2.040e-08 | -17.71 | GSE9960\_HEALTHY\_VS\_GRAM\_POS\_SEPSIS\_PBMC\_DN | MSigDB lists | GSE9960\_HEALTHY\_VS\_GRAM\_POS\_SEPSIS\_PBMC\_DN | 142 | 14 | 12187 | 179 | Ifi44,Samd9l,Ifitm3,Isg15,Herc6,Hspb1,Ip6k2,Ifit2,Il4ra,Birc3,AA467197,Tnfaip3,Irf7,Nfkb2 | | 2.075e-08 | -17.69 | Cytokine-cytokine receptor interaction | KEGG pathways | ko04060 | 143 | 16 | 5248 | 107 | Tnf,Cxcl9,Tnfsf10,Il6,Csf3,Cxcl16,Ccl7,Ccl2,Il4ra,Osmr,Il17ra,Il1a,Ccl4,Csf2rb,Cxcl1,Ccl12 | | 2.075e-08 | -17.69 | Cytokine-cytokine receptor interaction | KEGG pathways | mmu04060 | 143 | 16 | 5248 | 107 | Ccl4,Il1a,Csf2rb,Cxcl1,Ccl12,Ccl7,Ccl2,Osmr,Il4ra,Il17ra,Csf3,Cxcl16,Tnf,Cxcl9,Tnfsf10,Il6 | | 2.144e-08 | -17.66 | MODULE\_64 | MSigDB lists | MODULE\_64 | 336 | 21 | 12187 | 179 | Rgs16,Tnfsf10,Igsf6,Cxcl1,S100a9,C5ar1,Csf3,C3ar1,Ccl12,Il6,Ccl4,Ccl2,Map3k8,Il4ra,Csf2rb,Plaur,Icam1,Cd14,Akap12,Il1a,Cdkn1a | | 2.214e-08 | -17.63 | regulation of cell communication | biological process | GO:0010646 | 2617 | 75 | 13711 | 214 | Plaur,Pik3r5,Irgm1,Nfkb1,Birc3,Map3k8,Lgals9,P2ry6,Gadd45g,Thbs1,Errfi1,Rgs16,Sox11,Icam1,Irgm2,Bcl3,S100a9,Sgk3,Ccl12,Tnip3,Iqgap1,S100a8,Ptpn1,Csf3,Irak3,Il1rn,Ccl2,Ksr1,Tnfsf10,Xdh,Plek,Tnfaip3,Ier3,Trim25,Ptgs2,Runx1,Ccl4,Rasip1,Parp9,Fam107a,Fzd4,Map3k6,Timp1,Hcar2,Pik3r1,Gadd45b,Ccl7,Trib1,Parp14,Socs3,Irf7,Il6,Il1a,Akap12,Nfkbia,Casp4,Selp,Mmp8,Nfe2l2,Hspa5,Tgm2,C5ar1,Rnf213,Trim30a,Tnip1,Fstl1,Cd14,Cflar,Ripk1,Hspb1,Ncf1,Atf3,Pik3ap1,Acod1,Tnf | | 2.233e-08 | -17.62 | HALLMARK\_ALLOGRAFT\_REJECTION | MSigDB lists | HALLMARK\_ALLOGRAFT\_REJECTION | 143 | 14 | 12187 | 179 | Irf7,Bcl3,Igsf6,Tnf,Ccnd2,Il4ra,Ccl4,Ccl2,Spi1,Ccl12,Il6,Icam1,Tap1,Timp1 | | 2.362e-08 | -17.56 | GSE18791\_CTRL\_VS\_NEWCASTLE\_VIRUS\_DC\_14H\_DN | MSigDB lists | GSE18791\_CTRL\_VS\_NEWCASTLE\_VIRUS\_DC\_14H\_DN | 121 | 13 | 12187 | 179 | Ifit2,Tnf,Parp14,Xaf1,Tnfaip3,Irf7,Cmpk2,Gbp3,Ifi44,Ifit3b,Herc6,Isg15,Trim25 | | 2.380e-08 | -17.55 | GO\_REGULATION\_OF\_TYPE\_I\_INTERFERON\_PRODUCTION | MSigDB lists | GO\_REGULATION\_OF\_TYPE\_I\_INTERFERON\_PRODUCTION | 100 | 12 | 12187 | 179 | Tnfaip3,Nfkb2,Ifi211,Trim56,Irf7,Nfkb1,Ifi204,Ifi207,Ifi209,Trim25,Isg15,Cd14 | | 2.419e-08 | -17.54 | GSE23502\_WT\_VS\_HDC\_KO\_MYELOID\_DERIVED\_SUPPRESSOR\_CELL\_BM\_DN | MSigDB lists | GSE23502\_WT\_VS\_HDC\_KO\_MYELOID\_DERIVED\_SUPPRESSOR\_CELL\_BM\_DN | 168 | 15 | 12187 | 179 | Ptpn1,Csf2rb,Irf9,Rbm47,Ptgs2,Tiparp,Il1rn,Sbno2,Bcl3,C5ar1,Gem,Dtx3l,Ripk1,Rassf4,Tgm2 | | 2.419e-08 | -17.54 | GSE24726\_WT\_VS\_E2\_2\_KO\_PDC\_DN | MSigDB lists | GSE24726\_WT\_VS\_E2\_2\_KO\_PDC\_DN | 168 | 15 | 12187 | 179 | Parp9,Plek,Tagln2,Isg15,Cebpd,Dnajb1,Nfkb2,Pik3r5,Irf7,Bcl3,Cflar,Nfe2l2,Cybb,Hcar2,Usp18 | | 2.419e-08 | -17.54 | GSE17721\_12H\_VS\_24H\_LPS\_BMDC\_UP | MSigDB lists | GSE17721\_12H\_VS\_24H\_LPS\_BMDC\_UP | 168 | 15 | 12187 | 179 | Ifit3b,Atf3,Isg15,Sgk3,P2ry6,Plek,Casp4,Gpr84,Nfkbia,Tor3a,Birc3,Il6,Parp12,Gbp4,Gbp6 | | 2.515e-08 | -17.50 | GILMORE\_CORE\_NFKB\_PATHWAY | MSigDB lists | GILMORE\_CORE\_NFKB\_PATHWAY | 14 | 6 | 12187 | 179 | Nfkb1,Nfkbia,Nfkbiz,Nfkb2,Bcl3,Nfkbie | | 2.636e-08 | -17.45 | PID\_IL23\_PATHWAY | MSigDB lists | PID\_IL23\_PATHWAY | 23 | 7 | 12187 | 179 | Pik3r1,Tnf,Nfkbia,Socs3,Nfkb1,Il6,Ccl12 | | 2.666e-08 | -17.44 | KEGG\_NOD\_LIKE\_RECEPTOR\_SIGNALING\_PATHWAY | MSigDB lists | KEGG\_NOD\_LIKE\_RECEPTOR\_SIGNALING\_PATHWAY | 48 | 9 | 12187 | 179 | Cxcl1,Ccl2,Nfkbia,Tnf,Birc3,Ccl12,Tnfaip3,Il6,Nfkb1 | | 2.706e-08 | -17.43 | NFKB\_Q6 | MSigDB lists | NFKB\_Q6 | 195 | 16 | 12187 | 179 | Luc7l3,Il1rn,Vcam1,Ip6k2,Icam1,Tnip1,Ptgs2,Cybb,Nfkbia,Runx1,Map3k8,Nfkb2,Rrp8,Bcl3,Cxcl1,Ptges | | 2.734e-08 | -17.42 | cytokine receptor binding | molecular function | GO:0005126 | 183 | 16 | 13516 | 211 | Tnf,Ccl12,Ccl7,Il1a,Cflar,Csf3,Il1rn,Cxcl1,Tnfsf10,Cxcl16,Ccl4,Pik3r1,Il6,Cxcl9,Ripk1,Ccl2 | | 2.845e-08 | -17.38 | regulation of signaling | biological process | GO:0023051 | 2632 | 75 | 13711 | 214 | Casp4,Il1a,Akap12,Nfkbia,Il6,Socs3,Irf7,Ccl7,Trib1,Parp14,Hcar2,Pik3r1,Gadd45b,Timp1,Fam107a,Fzd4,Map3k6,Parp9,Rasip1,Ccl4,Ptgs2,Runx1,Ier3,Trim25,Acod1,Tnf,Pik3ap1,Atf3,Hspb1,Ncf1,Cflar,Cd14,Ripk1,Tnip1,Fstl1,Rnf213,Trim30a,Tgm2,C5ar1,Hspa5,Nfe2l2,Mmp8,Selp,Icam1,Rgs16,Sox11,Errfi1,Thbs1,Gadd45g,Lgals9,P2ry6,Map3k8,Birc3,Nfkb1,Irgm1,Pik3r5,Plaur,Tnfaip3,Plek,Ksr1,Tnfsf10,Xdh,Ccl2,Irak3,Il1rn,Csf3,Ptpn1,Tnip3,S100a8,Iqgap1,Ccl12,Sgk3,S100a9,Bcl3,Irgm2 | | 2.851e-08 | -17.37 | regulation of programmed cell death | biological process | GO:0043067 | 1253 | 46 | 13711 | 214 | Zfp36,Atf3,Tnf,Cflar,Ripk1,Hspb1,Hspa5,Tgm2,C5ar1,Ifit2,Angptl4,Nfe2l2,Cdkn1a,Ip6k2,Il6,Akap12,Casp4,Pik3r1,Hcar2,Ccnd2,Gadd45b,Socs3,Timp1,Ier3,Sgk1,Ptgs2,Tnfsf10,Xdh,Tnfaip3,Zbtb16,Tnfaip8,Irak3,Il1rn,Ccl12,S100a8,Ptpn1,Bcl3,S100a9,Sgk3,Mt1,Icam1,Gadd45g,Thbs1,Nfkb1,Birc3,Plaur | | 2.987e-08 | -17.33 | regulation of cell death | biological process | GO:0010941 | 1386 | 49 | 13711 | 214 | Ifit2,Angptl4,Nfe2l2,Hp,Hspa5,Tgm2,C5ar1,Cflar,Ripk1,Hspb1,Zfp36,Atf3,Tnf,Ier3,Sgk1,Ptgs2,Timp1,Hcar2,Pik3r1,Ccnd2,Gadd45b,Socs3,Cdkn1a,Il6,Ip6k2,Akap12,Casp4,Bcl3,S100a9,Sgk3,Ccl12,S100a8,Ptpn1,Csf3,Zbtb16,Tnfaip8,Il1rn,Irak3,Tnfsf10,Xdh,Tnfaip3,Plaur,Nfkb1,Birc3,Gadd45g,Thbs1,Sox11,Mt1,Icam1 | | 3.071e-08 | -17.30 | GO\_REGULATION\_OF\_CYSTEINE\_TYPE\_ENDOPEPTIDASE\_ACTIVITY | MSigDB lists | GO\_REGULATION\_OF\_CYSTEINE\_TYPE\_ENDOPEPTIDASE\_ACTIVITY | 171 | 15 | 12187 | 179 | Tnf,Birc3,Ripk1,Il6,Ifi211,S100a9,Tnfaip8,S100a8,Xdh,Tnfsf10,Ifi209,Thbs1,Ifi204,Ifi207,Plaur | | 3.071e-08 | -17.30 | GO\_ACTIVATION\_OF\_INNATE\_IMMUNE\_RESPONSE | MSigDB lists | GO\_ACTIVATION\_OF\_INNATE\_IMMUNE\_RESPONSE | 171 | 15 | 12187 | 179 | Ifi211,Irf7,Birc3,Nfkbia,Tnfaip3,Ripk1,Tnip1,Cd14,Irak3,Pik3ap1,Ifi209,Tnip3,Nfkb1,Ifi204,Ifi207 | | 3.198e-08 | -17.26 | activation of immune response | biological process | GO:0002253 | 185 | 16 | 13711 | 214 | Ifi209,Nfkbiz,C5ar1,Tnip3,Ifi211,Irgm2,Tnf,Pik3ap1,Nfkbia,Fyb,C3ar1,Irak3,Cd14,Tnip1,Lgals9,Ifi204 | | 3.224e-08 | -17.25 | HAHTOLA\_MYCOSIS\_FUNGOIDES\_CD4\_UP | MSigDB lists | HAHTOLA\_MYCOSIS\_FUNGOIDES\_CD4\_UP | 49 | 9 | 12187 | 179 | Ccl4,Trib1,Cdkn1a,Atf3,Ier3,Cebpd,Cxcl1,Ptgs2,Plaur | | 3.235e-08 | -17.25 | THEILGAARD\_NEUTROPHIL\_AT\_SKIN\_WOUND\_UP | MSigDB lists | THEILGAARD\_NEUTROPHIL\_AT\_SKIN\_WOUND\_UP | 65 | 10 | 12187 | 179 | Nfkb1,Ccl12,Ier3,Nfe2l2,Tnf,Trib1,Zfp36,Plek,Cflar,Cxcl1 | | 3.290e-08 | -17.23 | negative regulation of cellular process | biological process | GO:0048523 | 3771 | 96 | 13711 | 214 | C5ar1,Clic4,Hspa5,Serpina3f,Nfe2l2,Rasd1,Noct,Angptl4,Acod1,Pik3ap1,Atf3,Trim30a,Rnf213,Timp1,Fam107a,Slfn2,Isg15,Cebpd,Rasip1,Xaf1,Ifi211,Runx1,Il4ra,Nfkbia,Ip6k2,Slfn3,Parp14,Ccnd2,Ptpn1,Tnip3,Iqgap1,Wfdc21,Znfx1,Tnfaip3,Il1rn,Tnfaip8,Csf3,Zbtb16,Birc3,Nfkbie,Mt1,Icam1,Rgs16,Bbs12,Gadd45g,Lgals9,Rrp8,Dnajb1,Prg4,Hp,Robo4,Tnf,Zfp36,Hspb1,Ifi207,Cflar,Ripk1,Tnip1,Ifi209,Fzd4,Sbno2,Sdc4,Ptges,Parp9,Spi1,Sgk1,Ptgs2,Ier3,Il1a,Il6,Cdkn1a,Socs3,Dtx3l,Trib1,Ctla2a,Pik3r1,Gadd45b,A2m,Ccl12,Sgk3,Bcl3,Plek,Xdh,Arid5b,Irak3,Adamts9,Ifi204,Gbp4,Nfkb1,Bach1,Plaur,Mxd4,Sox11,Errfi1,Thbs1,Ifitm3 | | 3.335e-08 | -17.22 | CRX\_DN.V1\_DN | MSigDB lists | CRX\_DN.V1\_DN | 103 | 12 | 12187 | 179 | Tagln2,Cdkn1a,Socs3,Ksr1,Timp1,Nfkb1,Cebpd,Bcl3,Osmr,Map3k8,Tgm2,Tubb6 | | 3.462e-08 | -17.18 | regulation of immune response | biological process | GO:0050776 | 516 | 27 | 13711 | 214 | Tnip3,A2m,Tap1,Nfkbiz,C5ar1,Irgm2,Pik3ap1,Tnfaip3,Tnf,Acod1,Trim30a,Ifi204,Tnip1,Cd14,C3ar1,Irak3,Parp9,Ifi209,Irgm1,Ifi211,Il6,Nfkbia,Fyb,Il4ra,Lgals9,Parp14,Irf7 | | 3.468e-08 | -17.18 | GSE42021\_CD24INT\_VS\_CD24LOW\_TCONV\_THYMUS\_DN | MSigDB lists | GSE42021\_CD24INT\_VS\_CD24LOW\_TCONV\_THYMUS\_DN | 148 | 14 | 12187 | 179 | Ifi44,Phf11b,Ifit3b,Irf9,Ifitm2,Herc6,Isg15,Parp12,Oasl1,Ifit2,Usp18,Tnfsf10,Irf7,Phf11d | | 3.591e-08 | -17.14 | GSE17721\_0.5H\_VS\_24H\_CPG\_BMDC\_DN | MSigDB lists | GSE17721\_0.5H\_VS\_24H\_CPG\_BMDC\_DN | 173 | 15 | 12187 | 179 | Gbp3,Irgm2,Gbp6,Gbp4,Oasl1,Gadd45b,Parp14,Usp18,Ccnd2,Birc3,Tap1,Slfn9,Tnip1,Cp,Fstl1 | | 3.591e-08 | -17.14 | GSE44649\_WT\_VS\_MIR155\_KO\_ACTIVATED\_CD8\_TCELL\_UP | MSigDB lists | GSE44649\_WT\_VS\_MIR155\_KO\_ACTIVATED\_CD8\_TCELL\_UP | 173 | 15 | 12187 | 179 | Parp14,Casp4,Irgm2,Cflar,Gbp7,Sbno2,Ifit3b,Znfx1,Il1rn,Slfn5,Slfn9,Herc6,Clic4,Parp9,Tap1 | | 3.698e-08 | -17.11 | MODULE\_171 | MSigDB lists | MODULE\_171 | 84 | 11 | 12187 | 179 | Ifitm3,Selp,Isg15,Ifitm2,Cd14,Cebpd,Thbs1,Ifi44,Vcam1,Tnfsf10,Csf3 | | 3.722e-08 | -17.11 | GO\_POSITIVE\_REGULATION\_OF\_NF\_KAPPAB\_TRANSCRIPTION\_FACTOR\_ACTIVITY | MSigDB lists | GO\_POSITIVE\_REGULATION\_OF\_NF\_KAPPAB\_TRANSCRIPTION\_FACTOR\_ACTIVITY | 104 | 12 | 12187 | 179 | Cflar,S100a9,Icam1,Irak3,Trim25,S100a8,Nfkb2,Tnf,Nfkbia,Nfkb1,Ripk1,Il6 | | 3.722e-08 | -17.11 | GSE43863\_NAIVE\_VS\_MEMORY\_TH1\_CD4\_TCELL\_D150\_LCMV\_UP | MSigDB lists | GSE43863\_NAIVE\_VS\_MEMORY\_TH1\_CD4\_TCELL\_D150\_LCMV\_UP | 104 | 12 | 12187 | 179 | Ifit3b,Samd9l,Cebpd,Atf3,Rnf213,Isg15,Tap1,Parp14,Bach1,Ccl12,Cmpk2,Rtp4 | | 3.756e-08 | -17.10 | negative regulation of immune system process | biological process | GO:0002683 | 355 | 22 | 13711 | 214 | Pik3r1,Lgals9,Trim30a,Thbs1,Irak3,Zbtb16,Parp14,Trib1,Zfp36,Tnf,Acod1,Il4ra,Tnfaip3,Pik3ap1,Sox11,Nfkbia,Runx1,Nfe2l2,A2m,Tap1,Sdc4,Ccl12 | | 3.779e-08 | -17.09 | GSE30083\_SP2\_VS\_SP4\_THYMOCYTE\_DN | MSigDB lists | GSE30083\_SP2\_VS\_SP4\_THYMOCYTE\_DN | 149 | 14 | 12187 | 179 | Tap1,Il17ra,Plek,Zbtb16,Socs3,Ifit3b,Sgk1,Ifi44,Ms4a6d,Gem,Xdh,Sdc4,Sntb2,Cybb | | 3.880e-08 | -17.06 | GSE45365\_NK\_CELL\_VS\_BCELL\_UP | MSigDB lists | GSE45365\_NK\_CELL\_VS\_BCELL\_UP | 174 | 15 | 12187 | 179 | Thbs1,Vcam1,Atf3,Selp,Zfp36,Icam1,Birc3,Ccl2,Bach1,Il6,Tnfaip3,Gem,Arid5b,Ptges,S100a8 | | 3.880e-08 | -17.06 | GSE25085\_FETAL\_BM\_VS\_ADULT\_BM\_SP4\_THYMIC\_IMPLANT\_DN | MSigDB lists | GSE25085\_FETAL\_BM\_VS\_ADULT\_BM\_SP4\_THYMIC\_IMPLANT\_DN | 174 | 15 | 12187 | 179 | Tnip1,Saa1,Icam1,Plek,Hspa5,Atf3,Il1rn,Hp,Nfkbie,Nfkbia,Gpr84,Casp4,Birc3,Tnfaip3,Rassf4 | | 4.116e-08 | -17.01 | GSE14699\_NAIVE\_VS\_DELETIONAL\_TOLERANCE\_CD8\_TCELL\_DN | MSigDB lists | GSE14699\_NAIVE\_VS\_DELETIONAL\_TOLERANCE\_CD8\_TCELL\_DN | 150 | 14 | 12187 | 179 | Irgm2,Rbm39,Rtp4,Cmpk2,Ifit2,Nfe2l2,Rnf213,Slfn9,Slfn5,Trim25,Clic4,Parp9,Sgk1,Ifit3b | | 4.190e-08 | -16.99 | GSE41867\_NAIVE\_VS\_DAY6\_LCMV\_EFFECTOR\_CD8\_TCELL\_UP | MSigDB lists | GSE41867\_NAIVE\_VS\_DAY6\_LCMV\_EFFECTOR\_CD8\_TCELL\_UP | 175 | 15 | 12187 | 179 | Cflar,Gbp6,Dtx3l,Xdh,S100a8,Nfkbia,Ifit2,Msr1,Gbp4,Rnf213,Trim25,Nfkbiz,Thbs1,Ksr1,Samd9l | | 4.226e-08 | -16.98 | cellular response to interferon-gamma | biological process | GO:0071346 | 80 | 11 | 13711 | 214 | Gbp4,Gbp3,Ccl12,Ccl4,Irgm1,Stx11,Gbp6,Gbp7,Acod1,Ccl2,Ccl7 | | 4.370e-08 | -16.95 | HINATA\_NFKB\_TARGETS\_FIBROBLAST\_UP | MSigDB lists | HINATA\_NFKB\_TARGETS\_FIBROBLAST\_UP | 67 | 10 | 12187 | 179 | Tnfaip3,Ccl12,Ier3,Il6,Nfkb1,Nfkbia,Nfkb2,Icam1,Tnfaip8,Sele | | 4.478e-08 | -16.92 | SCHUETZ\_BREAST\_CANCER\_DUCTAL\_INVASIVE\_UP | MSigDB lists | SCHUETZ\_BREAST\_CANCER\_DUCTAL\_INVASIVE\_UP | 288 | 19 | 12187 | 179 | Msr1,Ccnd2,Tgm2,Tubb6,C3ar1,Gem,Ifi211,Olfml2b,Sgk1,Ifi209,Fstl1,Ifi207,Col4a1,Ms4a6d,Ifi204,Irak3,Msn,Csf2rb,Plaur | | 4.479e-08 | -16.92 | GO\_CYTOKINE\_RECEPTOR\_BINDING | MSigDB lists | GO\_CYTOKINE\_RECEPTOR\_BINDING | 151 | 14 | 12187 | 179 | Il1rn,Il1a,Cxcl16,Ripk1,Ccl12,Il6,Ccl4,Ccl2,Tnf,Pik3r1,Tnfsf10,Cxcl1,Cflar,Csf3 | | 4.523e-08 | -16.91 | GO\_REGULATION\_OF\_I\_KAPPAB\_KINASE\_NF\_KAPPAB\_SIGNALING | MSigDB lists | GO\_REGULATION\_OF\_I\_KAPPAB\_KINASE\_NF\_KAPPAB\_SIGNALING | 176 | 15 | 12187 | 179 | Cflar,Bcl3,Tnfsf10,Tnf,Birc3,Nfkbia,Rhoc,Ripk1,Tnfaip3,Ptgs2,Tnip1,Trim25,Hspb1,Tnip3,Il1a | | 4.661e-08 | -16.88 | PHONG\_TNF\_RESPONSE\_VIA\_P38\_PARTIAL | MSigDB lists | PHONG\_TNF\_RESPONSE\_VIA\_P38\_PARTIAL | 128 | 13 | 12187 | 179 | Plaur,Thbs1,Cdkn1a,Cebpd,Ier3,Tiparp,Cxcl1,Nfkbie,Nfkb2,Il4ra,Maff,Il6,Tnfaip3 | | 4.770e-08 | -16.86 | GO\_POSITIVE\_REGULATION\_OF\_RESPONSE\_TO\_EXTERNAL\_STIMULUS | MSigDB lists | GO\_POSITIVE\_REGULATION\_OF\_RESPONSE\_TO\_EXTERNAL\_STIMULUS | 203 | 16 | 12187 | 179 | Cxcl1,S100a9,C5ar1,C3ar1,Osmr,S100a8,Ccl4,Ccl2,Tnf,Ccl12,Il6,Tnip1,Ptgs2,Il17ra,Hspb1,Thbs1 | | 4.817e-08 | -16.85 | positive regulation of immune response | biological process | GO:0050778 | 360 | 22 | 13711 | 214 | Il6,Nfkbia,Fyb,Pik3ap1,Acod1,Il4ra,Tnf,Ifi204,Lgals9,Tnip1,Cd14,Irak3,C3ar1,Irf7,Parp9,Tnip3,C5ar1,Ifi209,Nfkbiz,Irgm2,Irgm1,Ifi211 | | 4.871e-08 | -16.84 | HALLMARK\_COMPLEMENT | MSigDB lists | HALLMARK\_COMPLEMENT | 152 | 14 | 12187 | 179 | Pik3r5,Irf7,S100a9,Tnfaip3,Il6,Casp4,Maff,Plaur,Irf2,Plek,Timp1,Mmp8,Cp,Hspa5 | | 5.051e-08 | -16.80 | negative regulation of viral life cycle | biological process | GO:1903901 | 64 | 10 | 13711 | 214 | Ifitm3,Oasl1,Ifitm6,Trim56,Resf1,Trim25,Oasl2,Tnf,Ifitm2,Isg15 | | 5.058e-08 | -16.80 | NABA\_MATRISOME\_ASSOCIATED | MSigDB lists | NABA\_MATRISOME\_ASSOCIATED | 386 | 22 | 12187 | 179 | S100a8,Adamts9,Tnfsf10,Csf3,A2m,S100a9,Cxcl1,Angptl4,Il6,Ccl12,Lgals9,Tnf,Ccl2,Sdc4,Ccl4,Tgm2,Il1a,Timp1,Mmp8,Il1rn,Adamts1,Fstl1 | | 5.293e-08 | -16.75 | GSE40443\_INDUCED\_VS\_TOTAL\_TREG\_DN | MSigDB lists | GSE40443\_INDUCED\_VS\_TOTAL\_TREG\_DN | 153 | 14 | 12187 | 179 | Cmpk2,Bcl3,Irgm2,Usp18,Msn,Irf9,Ptpn1,Slfn9,Tagln2,Rnf213,Ms4a6d,Socs3,Sgk1,Ifi44 | | 5.293e-08 | -16.75 | GSE37534\_UNTREATED\_VS\_ROSIGLITAZONE\_TREATED\_CD4\_TCELL\_PPARG1\_AND\_FOXP3\_TRASDUCED\_DN | MSigDB lists | GSE37534\_UNTREATED\_VS\_ROSIGLITAZONE\_TREATED\_CD4\_TCELL\_PPARG1\_AND\_FOXP3\_TRASDUCED\_DN | 153 | 14 | 12187 | 179 | Herc6,Icam1,Trim25,Zfp36,Tap1,Ifit3b,Atf3,Ifit2,Maff,Xaf1,Trib1,Usp18,Il6,Parp12 | | 5.496e-08 | -16.72 | positive regulation of tumor necrosis factor superfamily cytokine production | biological process | GO:1903557 | 82 | 11 | 13711 | 214 | Mmp8,Ccl4,Cybb,Ccl2,Akap12,Pik3r1,Lgals9,Hspb1,Thbs1,Cd14,Ripk1 | | 5.496e-08 | -16.72 | positive regulation of tumor necrosis factor production | biological process | GO:0032760 | 82 | 11 | 13711 | 214 | Mmp8,Ccl4,Ccl2,Cybb,Akap12,Lgals9,Pik3r1,Cd14,Ripk1,Hspb1,Thbs1 | | 5.560e-08 | -16.71 | REACTOME\_RIG\_I\_MDA5\_MEDIATED\_INDUCTION\_OF\_IFN\_ALPHA\_BETA\_PATHWAYS | MSigDB lists | REACTOME\_RIG\_I\_MDA5\_MEDIATED\_INDUCTION\_OF\_IFN\_ALPHA\_BETA\_PATHWAYS | 52 | 9 | 12187 | 179 | Tnfaip3,Ripk1,Nfkbia,Trim25,Nfkb2,Irf2,Isg15,Saa1,Irf7 | | 5.793e-08 | -16.66 | GO\_NEGATIVE\_REGULATION\_OF\_RESPONSE\_TO\_STIMULUS | MSigDB lists | GO\_NEGATIVE\_REGULATION\_OF\_RESPONSE\_TO\_STIMULUS | 1060 | 39 | 12187 | 179 | Thbs1,Hspa5,Nfkb1,Il1a,Rnf213,Cd14,Irak3,Pik3ap1,Ptpn1,Tap1,Il4ra,Ccl12,Tnfaip3,Ripk1,Nfe2l2,Ifi211,Rgs16,Ifi209,Socs3,Ifi204,Tnip3,Il1rn,Hp,Ifi207,Atf3,Saa1,Tnip1,Hspb1,Icam1,Plek,Plaur,Errfi1,Tnf,Trib1,Nfkbia,Cflar,A2m,Xdh,Tnfsf10 | | 6.154e-08 | -16.60 | GSE9988\_ANTI\_TREM1\_AND\_LPS\_VS\_VEHICLE\_TREATED\_MONOCYTES\_UP | MSigDB lists | GSE9988\_ANTI\_TREM1\_AND\_LPS\_VS\_VEHICLE\_TREATED\_MONOCYTES\_UP | 131 | 13 | 12187 | 179 | Il1a,Tnip3,Ier3,Nfkbiz,Plek,Plaur,Ptgs2,Map3k8,Tnf,Gem,Cflar,Banp,Cxcl1 | | 6.218e-08 | -16.59 | GO\_NEGATIVE\_REGULATION\_OF\_INNATE\_IMMUNE\_RESPONSE | MSigDB lists | GO\_NEGATIVE\_REGULATION\_OF\_INNATE\_IMMUNE\_RESPONSE | 38 | 8 | 12187 | 179 | Ifi209,Ifi207,Ifi204,Tnfaip3,A2m,Ifi211,Irak3,Tap1 | | 6.250e-08 | -16.59 | positive regulation of reactive oxygen species metabolic process | biological process | GO:2000379 | 83 | 11 | 13711 | 214 | Il6,Xdh,Cdkn1a,Icam1,Tnf,Acod1,Cxcl1,Ptgs2,Thbs1,Nfe2l2,Mmp8 | | 6.268e-08 | -16.59 | ACEVEDO\_FGFR1\_TARGETS\_IN\_PROSTATE\_CANCER\_MODEL\_UP | MSigDB lists | ACEVEDO\_FGFR1\_TARGETS\_IN\_PROSTATE\_CANCER\_MODEL\_UP | 207 | 16 | 12187 | 179 | Plaur,Cxcl16,Socs3,Ifi44,Thbs1,Lcn2,Cp,Timp1,Fzd4,Bcl3,Cxcl1,Irgm2,Rtp4,Usp18,Tnfaip3,Ccl12 | | 6.664e-08 | -16.52 | positive regulation of multicellular organismal process | biological process | GO:0051240 | 1512 | 51 | 13711 | 214 | Selp,Nfe2l2,Mmp8,Cxcl1,C5ar1,Hspa5,Hspb1,Cd14,Cflar,Ripk1,Tnf,Sgk1,Ptgs2,Runx1,Cebpd,Ifi211,Ccl4,Isg15,Fzd4,Hcar2,Pik3r1,Irf7,Trib1,Il6,Casp4,Il4ra,Akap12,Il1a,Bcl3,S100a9,Iqgap1,Nfkbiz,Lcn2,Ifi204,Irak3,C3ar1,Il1rn,Zbtb16,Csf3,Ccl2,Tnfaip3,Plek,Il17ra,Rhoj,Adamts1,Gadd45g,Lgals9,Thbs1,Cybb,Icam1,Sox11 | | 6.729e-08 | -16.51 | GO\_POSITIVE\_REGULATION\_OF\_LEUKOCYTE\_MIGRATION | MSigDB lists | GO\_POSITIVE\_REGULATION\_OF\_LEUKOCYTE\_MIGRATION | 70 | 10 | 12187 | 179 | Ccl4,Thbs1,Tnf,Il6,Ccl12,Cxcl1,C3ar1,C5ar1,Selp,Icam1 | | 6.768e-08 | -16.51 | GSE42021\_CD24HI\_VS\_CD24LOW\_TCONV\_THYMUS\_DN | MSigDB lists | GSE42021\_CD24HI\_VS\_CD24LOW\_TCONV\_THYMUS\_DN | 156 | 14 | 12187 | 179 | Tgm2,Il4ra,Maff,Ifit2,Oasl1,Tnfaip3,Bach1,Cxcl1,Tnfsf10,Rtp4,Ifit3b,Tap1,Plaur,Zfp36 | | 7.017e-08 | -16.47 | GO\_CELL\_CHEMOTAXIS | MSigDB lists | GO\_CELL\_CHEMOTAXIS | 110 | 12 | 12187 | 179 | S100a8,Il17ra,Cxcl16,Saa1,C5ar1,S100a9,Cxcl1,Il6,Ccl12,Ccl2,Ccl4,Vcam1 | | 7.376e-08 | -16.42 | RPS14\_DN.V1\_UP | MSigDB lists | RPS14\_DN.V1\_UP | 133 | 13 | 12187 | 179 | Ifi204,Ifi207,Ms4a6d,Ifi209,Cdkn1a,Rbm47,Cd14,Ccl12,Sdc4,Birc3,Igsf6,Ifi211,C3ar1 | | 7.674e-08 | -16.38 | negative regulation of metabolic process | biological process | GO:0009892 | 2285 | 67 | 13711 | 214 | Lgals9,Gadd45g,Banp,Msr1,Thbs1,Errfi1,Sox11,Mxd4,Plaur,Bach1,Nfkbie,Nfkb1,Birc3,Gbp4,Ifi204,Zbtb16,Tnfaip8,Arid5b,Irak3,Xdh,Plek,Tnfaip3,Znfx1,Wfdc21,Bcl3,A2m,Iqgap1,Ptpn1,Pik3r1,Hcar2,Gadd45b,Parp14,Trib1,Ctla2a,Socs3,Dtx3l,Cdkn1a,Il6,Ier3,Ptgs2,Runx1,Spi1,Cebpd,Rasip1,Ifi211,Isg15,Sbno2,Parp9,Timp1,Tiparp,Ifi209,Trim30a,Tnip1,Cflar,Hspb1,Ifi207,Zfp36,Atf3,Tnf,Noct,Rasd1,Mmp8,Serpina3f,Prg4,Hp,Dnajb1,Rrp8 | | 7.675e-08 | -16.38 | GO\_DEFENSE\_RESPONSE\_TO\_BACTERIUM | MSigDB lists | GO\_DEFENSE\_RESPONSE\_TO\_BACTERIUM | 90 | 11 | 12187 | 179 | Il6,Gbp4,Hp,Tnf,S100a8,S100a9,Bcl3,Gbp6,Isg15,C5ar1,Selp | | 7.947e-08 | -16.35 | GSE19401\_UNSTIM\_VS\_RETINOIC\_ACID\_STIM\_FOLLICULAR\_DC\_DN | MSigDB lists | GSE19401\_UNSTIM\_VS\_RETINOIC\_ACID\_STIM\_FOLLICULAR\_DC\_DN | 158 | 14 | 12187 | 179 | Tnf,Ccl4,Nfkbia,Sdc4,Tnfaip3,Cflar,Gem,Nfkb2,Thbs1,Vcam1,Nfkb1,Gadd45g,Atf3,Tnip1 | | 8.027e-08 | -16.34 | negative regulation of catalytic activity | biological process | GO:0043086 | 573 | 28 | 13711 | 214 | Cflar,Tnfaip8,Hspb1,Irak3,Tnfaip3,Tnf,Zfp36,Serpina3f,Wfdc21,Angptl4,Ptpn1,A2m,Hp,Trib1,Dtx3l,Thbs1,Gadd45g,Gadd45b,Cdkn1a,Errfi1,Il6,Rasip1,Plaur,Ptgs2,Birc3,Timp1,Parp9,Nfkb1 | | 8.465e-08 | -16.28 | GO\_REGULATION\_OF\_MAPK\_CASCADE | MSigDB lists | GO\_REGULATION\_OF\_MAPK\_CASCADE | 503 | 25 | 12187 | 179 | Ksr1,Atf3,Saa1,Tnip1,Icam1,Errfi1,Tnf,Trib1,C5ar1,Fzd4,Xdh,Thbs1,Gadd45g,Il1a,Map3k6,Irak3,Ptpn1,Map3k8,Ccl2,Ccl4,Il6,Gadd45b,Ripk1,Ccl12,Pik3r5 | | 8.604e-08 | -16.27 | GSE11961\_GERMINAL\_CENTER\_BCELL\_DAY7\_VS\_PLASMA\_CELL\_DAY7\_UP | MSigDB lists | GSE11961\_GERMINAL\_CENTER\_BCELL\_DAY7\_VS\_PLASMA\_CELL\_DAY7\_UP | 159 | 14 | 12187 | 179 | Hspa5,Ifi44,Parp9,Clic4,Slfn3,AA467197,Oasl1,Ccnd2,Birc3,Sbno2,Tnfsf10,Slfn4,Cflar,Dtx3l | | 8.811e-08 | -16.24 | GSE6269\_HEALTHY\_VS\_STAPH\_PNEUMO\_INF\_PBMC\_DN | MSigDB lists | GSE6269\_HEALTHY\_VS\_STAPH\_PNEUMO\_INF\_PBMC\_DN | 135 | 13 | 12187 | 179 | Cd14,Plek,Trim25,Plaur,Csf2rb,Socs3,Il1rn,Lcn2,Bcl3,S100a9,S100a8,Spi1,Cybb | | 8.868e-08 | -16.24 | KIM\_GLIS2\_TARGETS\_UP | MSigDB lists | KIM\_GLIS2\_TARGETS\_UP | 72 | 10 | 12187 | 179 | Osmr,Icam1,Angptl4,Isg15,Ifitm3,Ccl12,Col4a1,Casp4,Vcam1,Socs3 | | 8.964e-08 | -16.23 | Cytokine Signaling in Immune system | REACTOME pathways | R-MMU-1280215 | 364 | 22 | 6297 | 105 | Usp18,Nfkb2,Socs3,Il1a,Il6,Il4ra,Il1rn,Irak3,Birc3,Pik3r1,Tnf,Map3k8,Irf9,Osmr,Csf3,Isg15,Csf2rb,Il17ra,Ksr1,Ptpn1,Lgals9,Nfkb1 | | 9.087e-08 | -16.21 | DUTTA\_APOPTOSIS\_VIA\_NFKB | MSigDB lists | DUTTA\_APOPTOSIS\_VIA\_NFKB | 27 | 7 | 12187 | 179 | Tnfaip3,Ier3,Gadd45b,Tnf,Birc3,Tnfsf10,Cflar | | 9.310e-08 | -16.19 | GSE20715\_0H\_VS\_24H\_OZONE\_LUNG\_DN | MSigDB lists | GSE20715\_0H\_VS\_24H\_OZONE\_LUNG\_DN | 160 | 14 | 12187 | 179 | Maff,Trib1,Cflar,Banp,Osmr,Zbtb16,Timp1,Col4a1,Hp,Cp,Lcn2,Map3k6,Saa1,Rbm47 | | 9.310e-08 | -16.19 | GSE17721\_LPS\_VS\_PAM3CSK4\_2H\_BMDC\_UP | MSigDB lists | GSE17721\_LPS\_VS\_PAM3CSK4\_2H\_BMDC\_UP | 160 | 14 | 12187 | 179 | Sdc4,Ccl2,Ccl4,Tnf,Il6,Ccl12,Arid5b,Csf3,Ifit3b,Il1a,Isg15,Plek,Zfp36,Sele | | 1.007e-07 | -16.11 | GSE17721\_POLYIC\_VS\_CPG\_1H\_BMDC\_DN | MSigDB lists | GSE17721\_POLYIC\_VS\_CPG\_1H\_BMDC\_DN | 161 | 14 | 12187 | 179 | Tgm2,Gadd45b,Hcar2,A2m,Nfkbie,Slfn4,Il1rn,Tiparp,Tnip1,Cd14,Saa1,Errfi1,Slfn3,Zfp36 | | 1.165e-07 | -15.97 | NFKAPPAB\_01 | MSigDB lists | NFKAPPAB\_01 | 189 | 15 | 12187 | 179 | Tnip1,Icam1,Msn,Cxcl16,Vcam1,Il1a,Il1rn,Bcl3,Cxcl1,Nfkb2,Birc3,Tnf,Map3k8,Sdc4,Gadd45b | | 1.175e-07 | -15.96 | GSE32986\_CURDLAN\_LOWDOSE\_VS\_CURDLAN\_HIGHDOSE\_STIM\_DC\_DN | MSigDB lists | GSE32986\_CURDLAN\_LOWDOSE\_VS\_CURDLAN\_HIGHDOSE\_STIM\_DC\_DN | 163 | 14 | 12187 | 179 | Il1rn,Ifit3b,Plaur,Cxcl16,Ccl12,Gadd45b,Ifit2,Tor3a,Tgm2,Ccl4,Casp4,Spi1,Sdc4,Cflar | | 1.175e-07 | -15.96 | GSE1432\_1H\_VS\_6H\_IFNG\_MICROGLIA\_DN | MSigDB lists | GSE1432\_1H\_VS\_6H\_IFNG\_MICROGLIA\_DN | 163 | 14 | 12187 | 179 | Birc3,Casp4,Sntb2,Parp12,Nfkbie,Irf7,Phf11d,Tnfsf10,Ifi44,Phf11b,Akap12,Herc6,Tap1,Csf2rb | | 1.175e-07 | -15.96 | GSE17721\_LPS\_VS\_PAM3CSK4\_1H\_BMDC\_DN | MSigDB lists | GSE17721\_LPS\_VS\_PAM3CSK4\_1H\_BMDC\_DN | 163 | 14 | 12187 | 179 | Tnip1,Ptgs2,Icam1,Vcam1,Fstl1,Nfkbiz,Cdkn1a,Cxcl1,Nfkbia,Ccnd2,Map3k8,Birc3,Parp12,Gadd45b | | 1.175e-07 | -15.96 | GSE17721\_CTRL\_VS\_POLYIC\_4H\_BMDC\_DN | MSigDB lists | GSE17721\_CTRL\_VS\_POLYIC\_4H\_BMDC\_DN | 163 | 14 | 12187 | 179 | Irgm2,Hcar2,Ccl12,Parp12,Casp4,Pik3ap1,Tap1,Ptgs2,Il1rn,Ms4a6d,Znfx1,Cebpd,Atf3,Cdkn1a | | 1.175e-07 | -15.96 | GSE9988\_LPS\_VS\_LPS\_AND\_ANTI\_TREM1\_MONOCYTE\_UP | MSigDB lists | GSE9988\_LPS\_VS\_LPS\_AND\_ANTI\_TREM1\_MONOCYTE\_UP | 163 | 14 | 12187 | 179 | Pik3r5,Birc3,Map3k8,Sdc4,Cybb,Nfe2l2,Oasl1,Tnfaip3,Tnip1,Pik3ap1,Clic4,Thbs1,Nfkbiz,Socs3 | | 1.175e-07 | -15.96 | GSE17721\_0.5H\_VS\_4H\_CPG\_BMDC\_DN | MSigDB lists | GSE17721\_0.5H\_VS\_4H\_CPG\_BMDC\_DN | 163 | 14 | 12187 | 179 | Aff1,Nfkb1,Cp,Ifitm2,Tap1,Icam1,Tgm2,Tor3a,Birc3,Ptges,Irgm2,Gbp7,Tnfsf10,Rtp4 | | 1.288e-07 | -15.87 | DELYS\_THYROID\_CANCER\_UP | MSigDB lists | DELYS\_THYROID\_CANCER\_UP | 340 | 20 | 12187 | 179 | C3ar1,Cxcl1,Igsf6,Runx1,Ccnd2,Msr1,Tgm2,Ccl2,Sdc4,Map3k6,Icam1,Cdkn1a,Thbs1,Timp1,Il1rn,Tiparp,Hp,Ms4a6d,Ier3,Lcn2 | | 1.352e-07 | -15.82 | GO\_REGULATION\_OF\_LEUKOCYTE\_MIGRATION | MSigDB lists | GO\_REGULATION\_OF\_LEUKOCYTE\_MIGRATION | 95 | 11 | 12187 | 179 | Cxcl1,C3ar1,C5ar1,Selp,Msn,Icam1,Ccl4,Thbs1,Tnf,Il6,Ccl12 | | 1.356e-07 | -15.81 | GSE30971\_WBP7\_HET\_VS\_KO\_MACROPHAGE\_DN | MSigDB lists | GSE30971\_WBP7\_HET\_VS\_KO\_MACROPHAGE\_DN | 140 | 13 | 12187 | 179 | Bach1,Il6,Gadd45b,Tnfaip8,Csf3,Cxcl1,Tnip3,Ier3,Dnajb1,Socs3,Pik3ap1,P2ry6,Ptgs2 | | 1.356e-07 | -15.81 | NAGASHIMA\_NRG1\_SIGNALING\_UP | MSigDB lists | NAGASHIMA\_NRG1\_SIGNALING\_UP | 140 | 13 | 12187 | 179 | Tiparp,Ier3,Atf3,Sgk1,Dnajb1,Zfp36,Plaur,Tubb6,Gadd45b,Maff,Map3k8,Trib1,Gem | | 1.393e-07 | -15.79 | chemokine activity | molecular function | GO:0008009 | 27 | 7 | 13516 | 211 | Cxcl9,Cxcl1,Ccl2,Ccl4,Ccl7,Cxcl16,Ccl12 | | 1.427e-07 | -15.76 | GO\_REGULATION\_OF\_INTERLEUKIN\_1\_PRODUCTION | MSigDB lists | GO\_REGULATION\_OF\_INTERLEUKIN\_1\_PRODUCTION | 42 | 8 | 12187 | 179 | Errfi1,Hspb1,Ifi211,Saa1,Tnfaip3,Ifi204,Ifi207,Ifi209 | | 1.476e-07 | -15.73 | GSE17721\_12H\_VS\_24H\_PAM3CSK4\_BMDC\_UP | MSigDB lists | GSE17721\_12H\_VS\_24H\_PAM3CSK4\_BMDC\_UP | 166 | 14 | 12187 | 179 | Gem,Bcl3,Rrp8,S100a8,Ccl2,Rhoc,Gadd45b,Nfe2l2,Icam1,P2ry6,Cdkn1a,Fstl1,Ifit3b,Kcna5 | | 1.478e-07 | -15.73 | negative regulation of nitrogen compound metabolic process | biological process | GO:0051172 | 1880 | 58 | 13711 | 214 | Isg15,Sbno2,Parp9,Ifi209,Timp1,Ier3,Ptgs2,Runx1,Spi1,Rasip1,Cebpd,Ifi211,Cdkn1a,Il6,Pik3r1,Gadd45b,Parp14,Trib1,Ctla2a,Socs3,Dtx3l,Prg4,Dnajb1,Rrp8,Noct,Rasd1,Serpina3f,Zfp36,Atf3,Tnf,Tnip1,Cflar,Hspb1,Ifi207,Nfkbie,Nfkb1,Birc3,Gbp4,Plaur,Bach1,Errfi1,Sox11,Mxd4,Gadd45g,Banp,Thbs1,A2m,Ptpn1,Wfdc21,Bcl3,Xdh,Tnfaip3,Znfx1,Ifi204,Tnfaip8,Zbtb16,Arid5b,Irak3 | | 1.480e-07 | -15.73 | TONKS\_TARGETS\_OF\_RUNX1\_RUNX1T1\_FUSION\_SUSTAINED\_IN\_GRANULOCYTE\_UP | MSigDB lists | TONKS\_TARGETS\_OF\_RUNX1\_RUNX1T1\_FUSION\_SUSTAINED\_IN\_GRANULOCYTE\_UP | 18 | 6 | 12187 | 179 | Ifi207,Ifi204,Hspb1,Ifi209,Arid5b,Ifi211 | | 1.501e-07 | -15.71 | MODULE\_345 | MSigDB lists | MODULE\_345 | 76 | 10 | 12187 | 179 | Cebpd,Parp14,Ifi44,Thbs1,Tap1,Tnfsf10,Cd14,Csf3,Isg15,Ifitm3 | | 1.501e-07 | -15.71 | MODULE\_292 | MSigDB lists | MODULE\_292 | 76 | 10 | 12187 | 179 | Vcam1,Thbs1,Ifi44,Cebpd,Cd14,Isg15,Csf3,Ifitm2,Ifitm3,Tnfsf10 | | 1.501e-07 | -15.71 | GO\_LEUKOCYTE\_CHEMOTAXIS | MSigDB lists | GO\_LEUKOCYTE\_CHEMOTAXIS | 76 | 10 | 12187 | 179 | Il17ra,S100a8,Cxcl16,C5ar1,Saa1,S100a9,Ccl12,Il6,Ccl4,Ccl2 | | 1.529e-07 | -15.69 | GO\_REGULATION\_OF\_SEQUENCE\_SPECIFIC\_DNA\_BINDING\_TRANSCRIPTION\_FACTOR\_ACTIVITY | MSigDB lists | GO\_REGULATION\_OF\_SEQUENCE\_SPECIFIC\_DNA\_BINDING\_TRANSCRIPTION\_FACTOR\_ACTIVITY | 280 | 18 | 12187 | 179 | Fzd4,Arid5b,S100a9,Cflar,Nfkb2,S100a8,Nfkbia,Trib1,Tnf,Tnfaip3,Ripk1,Il6,Sgk3,Icam1,Irak3,Trim25,Sgk1,Nfkb1 | | 1.535e-07 | -15.69 | DAZARD\_UV\_RESPONSE\_CLUSTER\_G28 | MSigDB lists | DAZARD\_UV\_RESPONSE\_CLUSTER\_G28 | 10 | 5 | 12187 | 179 | Cxcl1,Ptgs2,Atf3,Ier3,Gadd45b | | 1.535e-07 | -15.69 | GO\_CHRONIC\_INFLAMMATORY\_RESPONSE | MSigDB lists | GO\_CHRONIC\_INFLAMMATORY\_RESPONSE | 10 | 5 | 12187 | 179 | Ptges,Vcam1,Thbs1,Tnf,S100a8 | | 1.535e-07 | -15.69 | BIOCARTA\_GRANULOCYTES\_PATHWAY | MSigDB lists | BIOCARTA\_GRANULOCYTES\_PATHWAY | 10 | 5 | 12187 | 179 | Icam1,Il1a,Tnf,Selp,Csf3 | | 1.547e-07 | -15.68 | toll-like receptor signaling pathway | biological process | GO:0002224 | 40 | 8 | 13711 | 214 | Nfkbia,Pik3ap1,Tnf,Tnip3,Cd14,Irak3,Lgals9,Tnip1 | | 1.586e-07 | -15.66 | pattern recognition receptor signaling pathway | biological process | GO:0002221 | 55 | 9 | 13711 | 214 | Tnip3,Nfkbia,Pik3ap1,Tnf,Irgm2,Lgals9,Tnip1,Cd14,Irak3 | | 1.586e-07 | -15.66 | innate immune response-activating signal transduction | biological process | GO:0002758 | 55 | 9 | 13711 | 214 | Tnip3,Tnf,Pik3ap1,Nfkbia,Tnip1,Lgals9,Irgm2,Irak3,Cd14 | | 1.590e-07 | -15.65 | GSE19923\_HEB\_KO\_VS\_HEB\_AND\_E2A\_KO\_DP\_THYMOCYTE\_DN | MSigDB lists | GSE19923\_HEB\_KO\_VS\_HEB\_AND\_E2A\_KO\_DP\_THYMOCYTE\_DN | 167 | 14 | 12187 | 179 | Nfkb2,Slfn4,Il6,Gadd45b,Ccl4,Ccl2,Nfkbia,Ccnd2,Tnf,Ptpn1,Slfn3,Ier3,Atf3,Nfkb1 | | 1.590e-07 | -15.65 | GSE22282\_HYPOXIA\_VS\_NORMOXIA\_MYELOID\_DC\_UP | MSigDB lists | GSE22282\_HYPOXIA\_VS\_NORMOXIA\_MYELOID\_DC\_UP | 167 | 14 | 12187 | 179 | Hp,Il1rn,Stx11,Cxcl16,Sele,Icam1,Tnip1,Saa1,Tnfaip3,Nfkbia,Gpr84,Tnf,Igsf6,Nfkbie | | 1.590e-07 | -15.65 | GSE37301\_LYMPHOID\_PRIMED\_MPP\_VS\_COMMON\_LYMPHOID\_PROGENITOR\_DN | MSigDB lists | GSE37301\_LYMPHOID\_PRIMED\_MPP\_VS\_COMMON\_LYMPHOID\_PROGENITOR\_DN | 167 | 14 | 12187 | 179 | Il1rn,Cebpd,Rasd1,Znfx1,Sgk1,Ifit3b,Trim25,Map3k6,Cd14,Gadd45b,Ccl12,Usp18,Cmpk2,Xdh | | 1.590e-07 | -15.65 | NABA\_SECRETED\_FACTORS | MSigDB lists | NABA\_SECRETED\_FACTORS | 167 | 14 | 12187 | 179 | Ccl12,Il6,Ccl4,Ccl2,Tnf,Tnfsf10,S100a8,Cxcl1,Angptl4,S100a9,Csf3,Il1rn,Il1a,Fstl1 | | 1.606e-07 | -15.64 | GO\_POSITIVE\_REGULATION\_OF\_IMMUNE\_RESPONSE | MSigDB lists | GO\_POSITIVE\_REGULATION\_OF\_IMMUNE\_RESPONSE | 378 | 21 | 12187 | 179 | Ifi204,Nfkb1,Tnip3,Ifi207,Ifi209,Irak3,Pik3ap1,Tnip1,Cd14,Tnfaip3,Ripk1,Il6,Nfkbia,Tnf,Il4ra,Pik3r1,Birc3,Ifi211,Irf7,C5ar1,C3ar1 | | 1.684e-07 | -15.60 | Apoptosis | WikiPathways | WP1254 | 72 | 12 | 3756 | 96 | Tnf,Ripk1,Cflar,Irf2,Casp4,Nfkb1,Irf7,Pik3r1,Nfkbie,Nfkbia,Birc3,Tnfsf10 | | 1.713e-07 | -15.58 | GSE26343\_WT\_VS\_NFAT5\_KO\_MACROPHAGE\_DN | MSigDB lists | GSE26343\_WT\_VS\_NFAT5\_KO\_MACROPHAGE\_DN | 168 | 14 | 12187 | 179 | Gbp3,Irf7,Csf3,Nfe2l2,Oasl1,Tubb6,Casp4,Map3k8,Ptpn1,Irf9,Irak3,Saa1,Znfx1,Socs3 | | 1.732e-07 | -15.57 | BROWNE\_HCMV\_INFECTION\_4HR\_UP | MSigDB lists | BROWNE\_HCMV\_INFECTION\_4HR\_UP | 43 | 8 | 12187 | 179 | Ifit3b,Thbs1,Ifit2,Oasl1,Lcn2,Ripk1,Isg15,Ptgs2 | | 1.741e-07 | -15.56 | GSE30083\_SP3\_VS\_SP4\_THYMOCYTE\_DN | MSigDB lists | GSE30083\_SP3\_VS\_SP4\_THYMOCYTE\_DN | 143 | 13 | 12187 | 179 | Xdh,Irf7,Gem,Ripk1,Rhoc,Cybb,Tor3a,Sdc4,Sntb2,Irf9,Plek,Ifi44,Sgk1 | | 1.741e-07 | -15.56 | GSE360\_T\_GONDII\_VS\_M\_TUBERCULOSIS\_MAC\_DN | MSigDB lists | GSE360\_T\_GONDII\_VS\_M\_TUBERCULOSIS\_MAC\_DN | 143 | 13 | 12187 | 179 | Fstl1,Cdkn1a,Ptgs2,Ptpn1,Zfp36,Ccl4,Zfp189,Il6,Cxcl1,Ptges,Nfkb2,Sbno2,Pygm | | 1.756e-07 | -15.55 | GO\_POSITIVE\_REGULATION\_OF\_INNATE\_IMMUNE\_RESPONSE | MSigDB lists | GO\_POSITIVE\_REGULATION\_OF\_INNATE\_IMMUNE\_RESPONSE | 195 | 15 | 12187 | 179 | Ifi204,Nfkb1,Tnip3,Ifi207,Ifi209,Irak3,Pik3ap1,Tnip1,Cd14,Tnfaip3,Ripk1,Nfkbia,Birc3,Ifi211,Irf7 | | 1.867e-07 | -15.49 | GSE43863\_NAIVE\_VS\_LY6C\_LOW\_CXCR5NEG\_CD4\_EFF\_TCELL\_D6\_LCMV\_UP | MSigDB lists | GSE43863\_NAIVE\_VS\_LY6C\_LOW\_CXCR5NEG\_CD4\_EFF\_TCELL\_D6\_LCMV\_UP | 98 | 11 | 12187 | 179 | Vcam1,Atf3,Sele,Ccl4,Gbp4,AA467197,Ccl12,Cxcl1,Gbp6,Nfkbie,Nfkb2 | | 1.915e-07 | -15.47 | symbiont-containing vacuole | cellular component | GO:0020003 | 10 | 5 | 13825 | 212 | Irgm2,Gbp6,Gbp3,Iigp1,Gbp7 | | 1.966e-07 | -15.44 | GO\_POSITIVE\_REGULATION\_OF\_MAPK\_CASCADE | MSigDB lists | GO\_POSITIVE\_REGULATION\_OF\_MAPK\_CASCADE | 349 | 20 | 12187 | 179 | Ccl4,Ccl2,Map3k8,Tnf,Ccl12,Ripk1,Gadd45b,Il6,Pik3r5,Fzd4,C5ar1,Xdh,Thbs1,Ksr1,Il1a,Gadd45g,Map3k6,Saa1,Ptpn1,Icam1 | | 1.966e-07 | -15.44 | MITSIADES\_RESPONSE\_TO\_APLIDIN\_UP | MSigDB lists | MITSIADES\_RESPONSE\_TO\_APLIDIN\_UP | 349 | 20 | 12187 | 179 | Ifitm3,Icam1,Ip6k2,Timp1,Ksr1,Irf7,Bcl3,Arid5b,Gem,C3ar1,Tnfsf10,Tor3a,Trib1,Nfkbia,Runx1,Maff,Nfe2l2,Oasl1,Gadd45b,Bach1 | | 1.984e-07 | -15.43 | GSE22601\_IMMATURE\_CD4\_SINGLE\_POSITIVE\_VS\_DOUBLE\_POSITIVE\_THYMOCYTE\_DN | MSigDB lists | GSE22601\_IMMATURE\_CD4\_SINGLE\_POSITIVE\_VS\_DOUBLE\_POSITIVE\_THYMOCYTE\_DN | 170 | 14 | 12187 | 179 | Parp14,Tor3a,Nfe2l2,Gbp4,Gbp6,Irgm2,Tnfaip8,Mxd4,Irf9,Tap1,Plek,Il17ra,Trim25,Icam1 | | 1.986e-07 | -15.43 | LI\_INDUCED\_T\_TO\_NATURAL\_KILLER\_UP | MSigDB lists | LI\_INDUCED\_T\_TO\_NATURAL\_KILLER\_UP | 225 | 16 | 12187 | 179 | Errfi1,Ifitm3,Ifitm2,Ier3,Nfkb1,Stx11,Samd9l,Gadd45g,Cdkn1a,Sgk1,Tnfsf10,AA467197,Tubb6,Ccl4,Ccnd2,Tnf | | 2.007e-07 | -15.42 | HIRSCH\_CELLULAR\_TRANSFORMATION\_SIGNATURE\_UP | MSigDB lists | HIRSCH\_CELLULAR\_TRANSFORMATION\_SIGNATURE\_UP | 197 | 15 | 12187 | 179 | Casp4,Il6,Bcl3,Irf7,Csf3,Sbno2,Tnfaip8,Osmr,Socs3,Timp1,Il1a,Plaur,Irf2,Irf9,Tap1 | | 2.007e-07 | -15.42 | BIOCARTA\_IL1R\_PATHWAY | MSigDB lists | BIOCARTA\_IL1R\_PATHWAY | 30 | 7 | 12187 | 179 | Irak3,Tnf,Nfkbia,Il1rn,Nfkb1,Il1a,Il6 | | 2.132e-07 | -15.36 | cell surface receptor signaling pathway | biological process | GO:0007166 | 1296 | 45 | 13711 | 214 | Plek,Cxcl9,Ccl2,Irak3,Il1rn,Arid5b,C3ar1,Lcn2,Nfkbiz,Ptpn1,Ifitm2,Iqgap1,Col4a1,Ccl12,Fyb,Ifitm3,Iigp1,P2ry6,Osmr,Kcna5,Plaur,Tnf,Hspb1,Cd14,Ripk1,Cxcl1,C5ar1,Hp,Csf2rb,Il4ra,Il1a,Nfkbia,Il6,Socs3,Irf7,Dtx3l,Ccl7,Pik3r1,Hcar2,Tiparp,Fzd4,Isg15,Ifitm6,Ccl4,Spi1 | | 2.180e-07 | -15.34 | P53\_DN.V2\_UP | MSigDB lists | P53\_DN.V2\_UP | 79 | 10 | 12187 | 179 | Angptl4,Cxcl1,P2ry6,Nfkbia,Sdc4,Xaf1,Ccl4,Ier3,Tnfaip3,Il1rn | | 2.185e-07 | -15.34 | regulation of cell migration | biological process | GO:0030334 | 753 | 32 | 13711 | 214 | Rhoc,Adamts9,Sele,Hspb1,C3ar1,Il1rn,Ccl2,Robo4,Selp,Msn,Nfe2l2,Ccl12,Cxcl16,Iqgap1,Clic4,Hspa5,C5ar1,Lgals9,P2ry6,Pik3r1,Trib1,Thbs1,Il1a,Akap12,Icam1,Ptgs2,Ccl4,Sdc4,Adamts1,Rhoj,Fam107a,Timp1 | | 2.221e-07 | -15.32 | GSE360\_T\_GONDII\_VS\_B\_MALAYI\_HIGH\_DOSE\_DC\_UP | MSigDB lists | GSE360\_T\_GONDII\_VS\_B\_MALAYI\_HIGH\_DOSE\_DC\_UP | 146 | 13 | 12187 | 179 | Nfkb2,Cflar,Gadd45b,Tnfaip3,Nfkbia,Tap1,Ifitm3,Tagln2,Il1a,Nfkb1,Stx11,Ifit3b,Socs3 | | 2.221e-07 | -15.32 | GSE360\_L\_DONOVANI\_VS\_M\_TUBERCULOSIS\_MAC\_DN | MSigDB lists | GSE360\_L\_DONOVANI\_VS\_M\_TUBERCULOSIS\_MAC\_DN | 146 | 13 | 12187 | 179 | Il6,Tnfaip3,Casp4,Tnf,Zfp189,Cdkn1a,Pygm,Sbno2,Nfkb2,Zfp36,Ptges,Ptgs2,Cflar | | 2.233e-07 | -15.31 | negative regulation of cellular metabolic process | biological process | GO:0031324 | 2049 | 61 | 13711 | 214 | Ptpn1,A2m,Iqgap1,Wfdc21,Bcl3,Plek,Tnfaip3,Znfx1,Xdh,Tnfaip8,Zbtb16,Arid5b,Irak3,Ifi204,Birc3,Gbp4,Nfkbie,Nfkb1,Bach1,Plaur,Sox11,Mxd4,Errfi1,Thbs1,Gadd45g,Rrp8,Dnajb1,Prg4,Hp,Rasd1,Serpina3f,Noct,Tnf,Zfp36,Atf3,Cflar,Hspb1,Ifi207,Trim30a,Tnip1,Ifi209,Timp1,Sbno2,Isg15,Parp9,Spi1,Cebpd,Rasip1,Ifi211,Ier3,Ptgs2,Runx1,Cdkn1a,Il6,Ctla2a,Parp14,Trib1,Socs3,Dtx3l,Pik3r1,Gadd45b | | 2.298e-07 | -15.29 | GO\_REGULATION\_OF\_PEPTIDASE\_ACTIVITY | MSigDB lists | GO\_REGULATION\_OF\_PEPTIDASE\_ACTIVITY | 257 | 17 | 12187 | 179 | Plaur,Thbs1,Ifi209,Ifi207,Timp1,Ifi204,S100a9,A2m,Ifi211,Tnfsf10,Xdh,S100a8,Tnfaip8,Birc3,Tnf,Il6,Ripk1 | | 2.301e-07 | -15.28 | TAKEDA\_TARGETS\_OF\_NUP98\_HOXA9\_FUSION\_8D\_UP | MSigDB lists | TAKEDA\_TARGETS\_OF\_NUP98\_HOXA9\_FUSION\_8D\_UP | 100 | 11 | 12187 | 179 | Irf7,Xaf1,Usp18,Ifit2,Ptgs2,Herc6,Rnf213,Isg15,Ifit3b,Thbs1,Ifi44 | | 2.361e-07 | -15.26 | positive regulation of multi-organism process | biological process | GO:0043902 | 270 | 18 | 13711 | 214 | Parp9,Tnip3,Ifi209,Cxcl1,Irgm2,Irgm1,Ifi211,Pik3ap1,Nfkbia,Acod1,Tnf,Trim30a,Lgals9,Ifi204,Tnip1,Cd14,Irf7,Irak3 | | 2.380e-07 | -15.25 | response to interleukin-1 | biological process | GO:0070555 | 75 | 10 | 13711 | 214 | Nfkb1,Il6,Ccl2,Ccl12,Acod1,Il1a,Sele,Irak3,Ccl4,Ccl7 | | 2.405e-07 | -15.24 | GSE30971\_CTRL\_VS\_LPS\_STIM\_MACROPHAGE\_WBP7\_KO\_4H\_UP | MSigDB lists | GSE30971\_CTRL\_VS\_LPS\_STIM\_MACROPHAGE\_WBP7\_KO\_4H\_UP | 147 | 13 | 12187 | 179 | Csf3,Rgs16,Tnf,Nfkbia,Il6,Gadd45b,Ptgs2,Pik3ap1,Irak3,Socs3,Il1a,Tnip3,Ier3 | | 2.405e-07 | -15.24 | GSE360\_L\_MAJOR\_VS\_B\_MALAYI\_HIGH\_DOSE\_DC\_UP | MSigDB lists | GSE360\_L\_MAJOR\_VS\_B\_MALAYI\_HIGH\_DOSE\_DC\_UP | 147 | 13 | 12187 | 179 | Tnfaip3,Nfkbia,Spi1,Tnfaip8,Rgs16,Cflar,Gadd45g,Il1rn,Ier3,Cdkn1a,Vcam1,Icam1,Plaur | | 2.463e-07 | -15.22 | NFKB\_Q6\_01 | MSigDB lists | NFKB\_Q6\_01 | 173 | 14 | 12187 | 179 | Vcam1,Luc7l3,Tnip3,Il1rn,Tnip1,Icam1,Nfkbia,Birc3,Gadd45b,Rrp8,Bcl3,Ptges,Csf3,Nfkb2 | | 2.550e-07 | -15.18 | AZARE\_NEOPLASTIC\_TRANSFORMATION\_BY\_STAT3\_UP | MSigDB lists | AZARE\_NEOPLASTIC\_TRANSFORMATION\_BY\_STAT3\_UP | 101 | 11 | 12187 | 179 | Ptgs2,Csf3,Ifi211,S100a9,S100a8,Clic4,Ccnd2,Ifi209,Thbs1,Ifi204,Ifi207 | | 2.603e-07 | -15.16 | GSE17721\_LPS\_VS\_GARDIQUIMOD\_1H\_BMDC\_UP | MSigDB lists | GSE17721\_LPS\_VS\_GARDIQUIMOD\_1H\_BMDC\_UP | 148 | 13 | 12187 | 179 | Sele,Slfn3,Timp1,Atf3,Nfkbiz,Rtp4,Slfn4,Arid5b,Cxcl1,Gadd45b,Il6,Tnf,Ccl2 | | 2.603e-07 | -15.16 | GSE21774\_CD62L\_POS\_CD56\_BRIGHT\_VS\_CD62L\_NEG\_CD56\_DIM\_NK\_CELL\_UP | MSigDB lists | GSE21774\_CD62L\_POS\_CD56\_BRIGHT\_VS\_CD62L\_NEG\_CD56\_DIM\_NK\_CELL\_UP | 148 | 13 | 12187 | 179 | Arid5b,Ncf1,Cd14,Igsf6,P2ry6,Pik3ap1,Tnf,Spi1,Cybb,Tubb6,Timp1,Rassf4,Lcn2 | | 2.645e-07 | -15.15 | GSE37416\_0H\_VS\_3H\_F\_TULARENSIS\_LVS\_NEUTROPHIL\_DN | MSigDB lists | GSE37416\_0H\_VS\_3H\_F\_TULARENSIS\_LVS\_NEUTROPHIL\_DN | 174 | 14 | 12187 | 179 | Tiparp,Stx11,Cdkn1a,Plaur,Cxcl16,Nfe2l2,Nfkbia,Tnf,Map3k8,Il4ra,Maff,Ptges,Pik3r5,Nfkbie | | 2.720e-07 | -15.12 | GO\_POSITIVE\_REGULATION\_OF\_CELL\_COMMUNICATION | MSigDB lists | GO\_POSITIVE\_REGULATION\_OF\_CELL\_COMMUNICATION | 1170 | 40 | 12187 | 179 | Map3k6,Selp,Pik3ap1,Ptpn1,Thbs1,Sox11,Akap12,Il1a,Gadd45g,Nfkb1,Csf3,Pik3r5,S100a8,Pik3r1,Map3k8,Ccl2,Ccl4,Hcar2,Rhoc,Gadd45b,Il6,Ccl12,Ripk1,Ptgs2,Saa1,Trim25,Icam1,Plaur,Socs3,Ksr1,Atf3,Cflar,C5ar1,Fzd4,Irf7,S100a9,Tnfsf10,Xdh,Birc3,Tnf | | 2.816e-07 | -15.08 | GSE19888\_ADENOSINE\_A3R\_INH\_VS\_ACT\_IN\_MAST\_CELL\_DN | MSigDB lists | GSE19888\_ADENOSINE\_A3R\_INH\_VS\_ACT\_IN\_MAST\_CELL\_DN | 149 | 13 | 12187 | 179 | Ifit3b,Znfx1,Cp,Ifitm3,Rnf213,Parp14,Ifit2,Lgals9,Gbp4,Gbp7,Gbp6,Gbp3,Xdh | | 2.822e-07 | -15.08 | GO\_REGULATION\_OF\_RESPONSE\_TO\_CYTOKINE\_STIMULUS | MSigDB lists | GO\_REGULATION\_OF\_RESPONSE\_TO\_CYTOKINE\_STIMULUS | 102 | 11 | 12187 | 179 | Socs3,Il1rn,Parp9,Ptpn1,Irak3,Usp18,Birc3,Tnf,Ripk1,Tnfaip3,Irf7 | | 2.822e-07 | -15.08 | GSE2706\_R848\_VS\_LPS\_2H\_STIM\_DC\_DN | MSigDB lists | GSE2706\_R848\_VS\_LPS\_2H\_STIM\_DC\_DN | 102 | 11 | 12187 | 179 | Ifit2,Ifi44,Ifit3b,Tnip3,Il1a,Znfx1,Oasl1,Ptgs2,Herc6,Isg15,Pik3ap1 | | 2.838e-07 | -15.07 | VERHAAK\_GLIOBLASTOMA\_MESENCHYMAL | MSigDB lists | VERHAAK\_GLIOBLASTOMA\_MESENCHYMAL | 175 | 14 | 12187 | 179 | Cd14,Rbm47,Plaur,Phf11b,Thbs1,Timp1,C5ar1,Phf11d,Iqgap1,Tnfaip8,Il4ra,Msr1,Casp4,Tnfaip3 | | 2.838e-07 | -15.07 | GO\_POSITIVE\_REGULATION\_OF\_SEQUENCE\_SPECIFIC\_DNA\_BINDING\_TRANSCRIPTION\_FACTOR\_ACTIVITY | MSigDB lists | GO\_POSITIVE\_REGULATION\_OF\_SEQUENCE\_SPECIFIC\_DNA\_BINDING\_TRANSCRIPTION\_FACTOR\_ACTIVITY | 175 | 14 | 12187 | 179 | Cflar,S100a9,Fzd4,Arid5b,S100a8,Nfkb2,Tnf,Nfkbia,Il6,Ripk1,Irak3,Icam1,Trim25,Nfkb1 | | 2.911e-07 | -15.05 | Measles | KEGG pathways | mmu05162 | 93 | 12 | 5248 | 107 | Pik3r1,Ccnd2,Nfkbia,Il1a,Tnfaip3,Tnfsf10,Irf9,Irf7,Msn,Nfkb1,Il6,Mx2 | | 2.911e-07 | -15.05 | Measles | KEGG pathways | ko05162 | 93 | 12 | 5248 | 107 | Mx2,Il6,Msn,Nfkb1,Tnfsf10,Irf9,Irf7,Tnfaip3,Il1a,Nfkbia,Ccnd2,Pik3r1 | | 2.933e-07 | -15.04 | KRIGE\_RESPONSE\_TO\_TOSEDOSTAT\_24HR\_UP | MSigDB lists | KRIGE\_RESPONSE\_TO\_TOSEDOSTAT\_24HR\_UP | 614 | 27 | 12187 | 179 | Ifi211,Phf11d,Bcl3,Rgs16,S100a8,Gpr84,Maff,Ccl12,Cybb,Slfn9,Tap1,Phf11b,S100a9,Rbm39,Errfi1,Clic4,Apold1,Plek,Ifit3b,Ifi209,Cdkn1a,Atf3,Ifi204,Il1rn,Ksr1,Ifi207,Samd9l | | 2.964e-07 | -15.03 | NFKB\_C | MSigDB lists | NFKB\_C | 203 | 15 | 12187 | 179 | Icam1,Ptgs2,Tnip1,Tnip3,Il1rn,Il1a,Luc7l3,Lcn2,Vcam1,Nfkb2,Ptges,Bcl3,Gadd45b,Map3k8,Nfkbia | | 2.969e-07 | -15.03 | cellular response to interleukin-1 | biological process | GO:0071347 | 59 | 9 | 13711 | 214 | Ccl2,Ccl12,Nfkb1,Il6,Il1a,Acod1,Ccl4,Ccl7,Irak3 | | 3.002e-07 | -15.02 | DASU\_IL6\_SIGNALING\_UP | MSigDB lists | DASU\_IL6\_SIGNALING\_UP | 46 | 8 | 12187 | 179 | Ccl12,Ier3,Il6,Maff,Irf9,Zfp36,Ptges,Gem | | 3.002e-07 | -15.02 | GO\_GRANULOCYTE\_MIGRATION | MSigDB lists | GO\_GRANULOCYTE\_MIGRATION | 46 | 8 | 12187 | 179 | S100a8,Il17ra,S100a9,Saa1,C5ar1,Ccl12,Ccl2,Ccl4 | | 3.017e-07 | -15.01 | GESERICK\_TERT\_TARGETS\_DN | MSigDB lists | GESERICK\_TERT\_TARGETS\_DN | 20 | 6 | 12187 | 179 | Nfkbiz,Gadd45b,Ier3,Gadd45g,Errfi1,Zfp36 | | 3.069e-07 | -15.00 | positive regulation of cell migration | biological process | GO:0030335 | 468 | 24 | 13711 | 214 | Ccl2,C3ar1,Sele,Hspb1,Rhoc,C5ar1,Hspa5,Iqgap1,Cxcl16,Ccl12,Nfe2l2,Selp,Icam1,Akap12,Il1a,Thbs1,Pik3r1,P2ry6,Lgals9,Fam107a,Rhoj,Adamts1,Ccl4,Ptgs2 | | 3.234e-07 | -14.94 | mouse chr11\|11 B1.2 | chromosome location | mouse chr11\|11 B1.2 | 11 | 5 | 14556 | 220 | 9930111J21Rik2,Tgtp1,Irgm1,Gm5431,Ifi47 | | 3.264e-07 | -14.94 | GSE43863\_TH1\_VS\_LY6C\_INT\_CXCR5POS\_MEMORY\_CD4\_TCELL\_UP | MSigDB lists | GSE43863\_TH1\_VS\_LY6C\_INT\_CXCR5POS\_MEMORY\_CD4\_TCELL\_UP | 177 | 14 | 12187 | 179 | Socs3,Atf3,Ksr1,Ncf1,Tap1,Icam1,Birc3,AA467197,Gbp4,Il6,Cflar,Irgm2,Gbp6,Nfkb2 | | 3.276e-07 | -14.93 | Toll-like receptor signaling pathway | KEGG pathways | mmu04620 | 77 | 11 | 5248 | 107 | Cd14,Ccl4,Cxcl9,Ripk1,Irf7,Pik3r1,Map3k8,Tnf,Nfkb1,Il6,Nfkbia | | 3.276e-07 | -14.93 | Toll-like receptor signaling pathway | KEGG pathways | ko04620 | 77 | 11 | 5248 | 107 | Ccl4,Cd14,Map3k8,Pik3r1,Nfkbia,Tnf,Irf7,Cxcl9,Ripk1,Il6,Nfkb1 | | 3.289e-07 | -14.93 | GSE41867\_MEMORY\_VS\_EXHAUSTED\_CD8\_TCELL\_DAY30\_LCMV\_UP | MSigDB lists | GSE41867\_MEMORY\_VS\_EXHAUSTED\_CD8\_TCELL\_DAY30\_LCMV\_UP | 151 | 13 | 12187 | 179 | Sgk3,Icam1,Plek,Ms4a6d,Cebpd,Gbp7,Gbp3,Ifit2,Lgals9,Parp14,Tor3a,Spi1,Cybb | | 3.325e-07 | -14.92 | regulation of locomotion | biological process | GO:0040012 | 847 | 34 | 13711 | 214 | P2ry6,Lgals9,Pik3r1,Trib1,Thbs1,Akap12,Il1a,Icam1,Trim25,Trim56,Ptgs2,Ccl4,Adamts1,Sdc4,Rhoj,Fam107a,Timp1,Rhoc,Adamts9,Il1rn,C3ar1,Hspb1,Sele,Ccl2,Robo4,Msn,Selp,Nfe2l2,Cxcl16,Ccl12,Iqgap1,Clic4,Hspa5,C5ar1 | | 3.326e-07 | -14.92 | GO\_ACTIVATION\_OF\_IMMUNE\_RESPONSE | MSigDB lists | GO\_ACTIVATION\_OF\_IMMUNE\_RESPONSE | 295 | 18 | 12187 | 179 | Tnfaip3,Ripk1,Pik3r1,Birc3,Nfkbia,C5ar1,C3ar1,Ifi211,Irf7,Tnip3,Nfkb1,Ifi204,Ifi207,Ifi209,Irak3,Pik3ap1,Tnip1,Cd14 | | 3.377e-07 | -14.90 | negative regulation of macromolecule metabolic process | biological process | GO:0010605 | 2073 | 61 | 13711 | 214 | Socs3,Dtx3l,Ctla2a,Parp14,Trib1,Pik3r1,Gadd45b,Il6,Cdkn1a,Rasip1,Cebpd,Ifi211,Spi1,Ptgs2,Runx1,Timp1,Ifi209,Tiparp,Isg15,Sbno2,Parp9,Hspb1,Ifi207,Cflar,Tnip1,Tnf,Atf3,Zfp36,Serpina3f,Mmp8,Rasd1,Noct,Rrp8,Dnajb1,Prg4,Thbs1,Msr1,Banp,Gadd45g,Lgals9,Mxd4,Sox11,Errfi1,Bach1,Plaur,Gbp4,Birc3,Nfkb1,Nfkbie,Arid5b,Irak3,Zbtb16,Tnfaip8,Ifi204,Tnfaip3,Znfx1,Xdh,Bcl3,Wfdc21,Ptpn1,A2m | | 3.418e-07 | -14.89 | negative regulation of cellular protein metabolic process | biological process | GO:0032269 | 848 | 34 | 13711 | 214 | Serpina3f,Wfdc21,Ptpn1,A2m,Hspb1,Irak3,Cflar,Tnfaip8,Tnip1,Tnf,Tnfaip3,Atf3,Zfp36,Xdh,Rasip1,Spi1,Ptgs2,Plaur,Gbp4,Timp1,Birc3,Isg15,Socs3,Thbs1,Dtx3l,Trib1,Parp14,Ctla2a,Gadd45g,Pik3r1,Gadd45b,Il6,Errfi1,Cdkn1a | | 3.467e-07 | -14.87 | extracellular membrane-bounded organelle | cellular component | GO:0065010 | 11 | 5 | 13825 | 212 | Irgm2,Gbp6,Iigp1,Gbp3,Gbp7 | | 3.500e-07 | -14.87 | GO\_ACTIVATION\_OF\_CYSTEINE\_TYPE\_ENDOPEPTIDASE\_ACTIVITY | MSigDB lists | GO\_ACTIVATION\_OF\_CYSTEINE\_TYPE\_ENDOPEPTIDASE\_ACTIVITY | 83 | 10 | 12187 | 179 | Ifi211,S100a9,Xdh,Tnfsf10,S100a8,Ifi209,Tnf,Ripk1,Ifi204,Ifi207 | | 3.538e-07 | -14.85 | negative regulation of cytokine production | biological process | GO:0001818 | 193 | 15 | 13711 | 214 | Mmp8,Bcl3,Gbp4,Prg4,Nfkb1,Irak3,Thbs1,Lgals9,Trim30a,Tnf,Tnfaip3,Acod1,Zfp36,Errfi1,Il6 | | 3.551e-07 | -14.85 | GSE29617\_CTRL\_VS\_DAY7\_TIV\_FLU\_VACCINE\_PBMC\_2008\_UP | MSigDB lists | GSE29617\_CTRL\_VS\_DAY7\_TIV\_FLU\_VACCINE\_PBMC\_2008\_UP | 152 | 13 | 12187 | 179 | Bach1,Nfe2l2,Ifit2,C5ar1,Cxcl1,Bcl3,Tiparp,Stx11,Atf3,Cdkn1a,Sgk1,Socs3,Ptgs2 | | 3.572e-07 | -14.84 | NAGASHIMA\_EGF\_SIGNALING\_UP | MSigDB lists | NAGASHIMA\_EGF\_SIGNALING\_UP | 47 | 8 | 12187 | 179 | Atf3,Ier3,Tiparp,Trib1,Dnajb1,Maff,Zfp36,Gem | | 3.578e-07 | -14.84 | GO\_MYELOID\_LEUKOCYTE\_MIGRATION | MSigDB lists | GO\_MYELOID\_LEUKOCYTE\_MIGRATION | 64 | 9 | 12187 | 179 | Ccl4,Ccl2,S100a9,C5ar1,Saa1,Ccl12,Il6,Il17ra,S100a8 | | 3.642e-07 | -14.83 | immune response-activating signal transduction | biological process | GO:0002757 | 143 | 13 | 13711 | 214 | Cd14,Irak3,C3ar1,Lgals9,Tnip1,Nfkbia,Fyb,Pik3ap1,Tnf,Irgm2,C5ar1,Nfkbiz,Tnip3 | | 3.692e-07 | -14.81 | AGE-RAGE signaling pathway in diabetic complications | KEGG pathways | ko04933 | 95 | 12 | 5248 | 107 | Cybb,Icam1,Tnf,Vcam1,Il6,Nfkb1,Il1a,Ccl12,Col4a1,Ccl2,Pik3r1,Sele | | 3.692e-07 | -14.81 | AGE-RAGE signaling pathway in diabetic complications | KEGG pathways | mmu04933 | 95 | 12 | 5248 | 107 | Cybb,Vcam1,Tnf,Icam1,Nfkb1,Il6,Ccl12,Il1a,Pik3r1,Ccl2,Col4a1,Sele | | 3.747e-07 | -14.80 | GSE2770\_TGFB\_AND\_IL4\_VS\_IL12\_TREATED\_ACT\_CD4\_TCELL\_6H\_UP | MSigDB lists | GSE2770\_TGFB\_AND\_IL4\_VS\_IL12\_TREATED\_ACT\_CD4\_TCELL\_6H\_UP | 179 | 14 | 12187 | 179 | Birc3,Parp14,Usp18,Gbp7,Iqgap1,Rtp4,Sbno2,Aff1,Samd9l,Atf3,Rnf213,Trim25,Parp9,Msn | | 3.747e-07 | -14.80 | GSE32034\_LY6C\_HIGH\_VS\_LOW\_MONOCYTE\_DN | MSigDB lists | GSE32034\_LY6C\_HIGH\_VS\_LOW\_MONOCYTE\_DN | 179 | 14 | 12187 | 179 | Ifit3b,Irf9,Parp9,Slfn9,Tnip1,Parp12,Gbp4,Usp18,Parp14,Ifit2,Xdh,Cmpk2,Rtp4,Gbp6 | | 3.764e-07 | -14.79 | GSE7509\_UNSTIM\_VS\_FCGRIIB\_STIM\_DC\_DN | MSigDB lists | GSE7509\_UNSTIM\_VS\_FCGRIIB\_STIM\_DC\_DN | 128 | 12 | 12187 | 179 | Clic4,Icam1,Isg15,Nfkbia,Usp18,Casp4,Xaf1,Ifit2,Rgs16,Tnfsf10,Sbno2,Angptl4 | | 3.799e-07 | -14.78 | GGGNNTTTCC\_NFKB\_Q6\_01 | MSigDB lists | GGGNNTTTCC\_NFKB\_Q6\_01 | 105 | 11 | 12187 | 179 | Il6,Ccnd2,Birc3,Nfkbia,Nfkb2,Gem,Ptges,Bcl3,Il1rn,Csf2rb,Tnip1 | | 3.832e-07 | -14.77 | GSE41867\_DAY6\_EFFECTOR\_VS\_DAY30\_EXHAUSTED\_CD8\_TCELL\_LCMV\_CLONE13\_UP | MSigDB lists | GSE41867\_DAY6\_EFFECTOR\_VS\_DAY30\_EXHAUSTED\_CD8\_TCELL\_LCMV\_CLONE13\_UP | 153 | 13 | 12187 | 179 | Tor3a,Parp14,Lgals9,Parp12,Znfx1,Irf7,Dtx3l,Irgm2,Ifitm3,Gbp3,Xdh,Slfn3,Slfn4 | | 4.055e-07 | -14.72 | negative regulation of viral genome replication | biological process | GO:0045071 | 45 | 8 | 13711 | 214 | Ifitm2,Tnf,Oasl2,Isg15,Oasl1,Ifitm6,Ifitm3,Resf1 | | 4.058e-07 | -14.72 | apoptotic process | biological process | GO:0006915 | 696 | 30 | 13711 | 214 | C5ar1,Rrp8,S100a8,S100a9,Ifit2,Bcl3,Tnfaip3,Tnf,Tnfsf10,Tnfaip8,Ripk1,Cflar,Irak3,Ncf1,Ifi204,Lcn2,Birc3,Nfkb1,Spi1,Xaf1,Ier3,Sgk1,Il1a,Casp4,Cdkn1a,Il6,Gadd45b,Pik3r1,Hcar2,Gadd45g | | 4.101e-07 | -14.71 | GO\_POSITIVE\_REGULATION\_OF\_REACTIVE\_OXYGEN\_SPECIES\_METABOLIC\_PROCESS | MSigDB lists | GO\_POSITIVE\_REGULATION\_OF\_REACTIVE\_OXYGEN\_SPECIES\_METABOLIC\_PROCESS | 65 | 9 | 12187 | 179 | Ripk1,Nfe2l2,Il6,Cdkn1a,Tnf,Thbs1,Icam1,Xdh,Ptgs2 | | 4.132e-07 | -14.70 | GO\_REGULATION\_OF\_SYMBIOSIS\_ENCOMPASSING\_MUTUALISM\_THROUGH\_PARASITISM | MSigDB lists | GO\_REGULATION\_OF\_SYMBIOSIS\_ENCOMPASSING\_MUTUALISM\_THROUGH\_PARASITISM | 154 | 13 | 12187 | 179 | Ifi211,Tnf,Ccl4,Oasl1,Ifitm3,Ifitm2,Isg15,Tnip1,Trim25,Tap1,Ifi209,Ifi204,Ifi207 | | 4.172e-07 | -14.69 | BIOCARTA\_NFKB\_PATHWAY | MSigDB lists | BIOCARTA\_NFKB\_PATHWAY | 21 | 6 | 12187 | 179 | Ripk1,Tnfaip3,Il1a,Nfkb1,Nfkbia,Tnf | | 4.292e-07 | -14.66 | NFKAPPAB65\_01 | MSigDB lists | NFKAPPAB65\_01 | 181 | 14 | 12187 | 179 | Csf2rb,Cxcl16,Icam1,Tnip1,Ier3,Nfkb2,Bcl3,Rrp8,Ptges,Gadd45b,Nfkbia,Sdc4,Birc3,Map3k8 | | 4.352e-07 | -14.65 | response to type I interferon | biological process | GO:0034340 | 20 | 6 | 13711 | 214 | Trim56,Isg15,Ifitm6,Irf7,Ifitm3,Ifitm2 | | 4.453e-07 | -14.62 | GSE33162\_HDAC3\_KO\_VS\_HDAC3\_KO\_MACROPHAGE\_UP | MSigDB lists | GSE33162\_HDAC3\_KO\_VS\_HDAC3\_KO\_MACROPHAGE\_UP | 155 | 13 | 12187 | 179 | Gbp4,Parp12,Il4ra,Parp14,Usp18,Olfml2b,Rtp4,Gbp6,Ptges,Ifit3b,Parp9,Tap1,Irf9 | | 4.677e-07 | -14.58 | positive regulation of cell motility | biological process | GO:2000147 | 479 | 24 | 13711 | 214 | Icam1,Akap12,Il1a,Thbs1,Pik3r1,Lgals9,P2ry6,Fam107a,Rhoj,Adamts1,Ccl4,Ptgs2,Ccl2,Hspb1,Sele,C3ar1,Rhoc,C5ar1,Hspa5,Iqgap1,Ccl12,Cxcl16,Nfe2l2,Selp | | 4.689e-07 | -14.57 | GNF2\_HCK | MSigDB lists | GNF2\_HCK | 66 | 9 | 12187 | 179 | Spi1,Stx11,Cybb,Ms4a6d,C5ar1,Cd14,S100a9,Igsf6,Tnfsf10 | | 4.796e-07 | -14.55 | GSE45365\_NK\_CELL\_VS\_BCELL\_MCMV\_INFECTION\_DN | MSigDB lists | GSE45365\_NK\_CELL\_VS\_BCELL\_MCMV\_INFECTION\_DN | 156 | 13 | 12187 | 179 | Il4ra,Sdc4,Nfkbia,Rassf4,Tnfaip3,Gadd45b,Sbno2,Fstl1,Nfkb1,Tnip1,Plek,Ptpn1,Tap1 | | 4.845e-07 | -14.54 | GSE2128\_CTRL\_VS\_MIMETOPE\_NEGATIVE\_SELECTION\_DP\_THYMOCYTE\_C57BL6\_UP | MSigDB lists | GSE2128\_CTRL\_VS\_MIMETOPE\_NEGATIVE\_SELECTION\_DP\_THYMOCYTE\_C57BL6\_UP | 131 | 12 | 12187 | 179 | Ifi204,Nfkb1,Ifi207,Cdkn1a,Ifi209,Thbs1,Icam1,Cybb,Gadd45b,Tnf,S100a8,Ifi211 | | 4.905e-07 | -14.53 | GO\_PATTERN\_RECOGNITION\_RECEPTOR\_SIGNALING\_PATHWAY | MSigDB lists | GO\_PATTERN\_RECOGNITION\_RECEPTOR\_SIGNALING\_PATHWAY | 86 | 10 | 12187 | 179 | Nfkbia,Birc3,Ripk1,Tnfaip3,Tnip3,Tnip1,Irf7,Cd14,Irak3,Pik3ap1 | | 5.013e-07 | -14.51 | GO\_POSITIVE\_REGULATION\_OF\_BIOSYNTHETIC\_PROCESS | MSigDB lists | GO\_POSITIVE\_REGULATION\_OF\_BIOSYNTHETIC\_PROCESS | 1340 | 43 | 12187 | 179 | Zbtb16,Ifi207,Atf3,Ptgs2,Tnip1,Hspb1,Tnf,Nfkbia,C3ar1,Nfkb2,Thbs1,Nfkb1,Il1a,Akap12,Irak3,Ccl12,Il6,Nfe2l2,Csf3,Banp,Sbno2,Ifi209,Ifi204,Icam1,Trim25,Runx1,Bach1,Cflar,Irf7,S100a9,Fzd4,Sox11,Hspa5,Cebpd,Irf2,Maff,Pik3r1,Spi1,Ripk1,Ifi211,Bcl3,Arid5b,S100a8 | | 5.016e-07 | -14.51 | regulation of symbiosis, encompassing mutualism through parasitism | biological process | GO:0043903 | 172 | 14 | 13711 | 214 | Trim56,Trim25,Resf1,Ifitm6,Ccl4,Isg15,Cxcl1,Ifitm2,Trim30a,Oasl1,Ifitm3,Zfp36,Tnf,Oasl2 | | 5.047e-07 | -14.50 | MODULE\_312 | MSigDB lists | MODULE\_312 | 34 | 7 | 12187 | 179 | Ccnd2,Birc3,Ier3,Nfkb1,Il1a,Cflar,Tnfsf10 | | 5.047e-07 | -14.50 | DORN\_ADENOVIRUS\_INFECTION\_24HR\_DN | MSigDB lists | DORN\_ADENOVIRUS\_INFECTION\_24HR\_DN | 34 | 7 | 12187 | 179 | Nfkb2,Ier3,Il6,Nfe2l2,Ifit2,Sgk1,Nfkbia | | 5.047e-07 | -14.50 | MANTOVANI\_NFKB\_TARGETS\_UP | MSigDB lists | MANTOVANI\_NFKB\_TARGETS\_UP | 34 | 7 | 12187 | 179 | Vcam1,Casp4,Nfkbiz,Cp,Ccl12,Saa1,Osmr | | 5.047e-07 | -14.50 | GEISS\_RESPONSE\_TO\_DSRNA\_UP | MSigDB lists | GEISS\_RESPONSE\_TO\_DSRNA\_UP | 34 | 7 | 12187 | 179 | Cflar,Ccl4,Nfkbia,Tnfaip3,Plaur,Tap1,Atf3 | | 5.246e-07 | -14.46 | GO\_POSITIVE\_REGULATION\_OF\_PHOSPHORUS\_METABOLIC\_PROCESS | MSigDB lists | GO\_POSITIVE\_REGULATION\_OF\_PHOSPHORUS\_METABOLIC\_PROCESS | 755 | 30 | 12187 | 179 | Sdc4,Tnf,Xdh,Fzd4,C5ar1,Ksr1,Socs3,Cdkn1a,Plaur,Icam1,Plek,Saa1,Ccl12,Ripk1,Gadd45b,Il6,Ccl4,Ccl2,Map3k8,Ccnd2,Iqgap1,Pik3r5,Csf3,Akap12,Gadd45g,Il1a,Thbs1,Hspa5,Ptpn1,Map3k6 | | 5.348e-07 | -14.44 | GHANDHI\_DIRECT\_IRRADIATION\_UP | MSigDB lists | GHANDHI\_DIRECT\_IRRADIATION\_UP | 67 | 9 | 12187 | 179 | Il1a,Icam1,Il6,Tnfaip3,Birc3,Ptgs2,Cdkn1a,Nfkbiz,Cxcl1 | | 5.368e-07 | -14.44 | regulation of cell motility | biological process | GO:2000145 | 784 | 32 | 13711 | 214 | C3ar1,Il1rn,Hspb1,Sele,Adamts9,Rhoc,Robo4,Ccl2,Nfe2l2,Msn,Selp,C5ar1,Clic4,Hspa5,Iqgap1,Cxcl16,Ccl12,Thbs1,Trib1,Pik3r1,P2ry6,Lgals9,Icam1,Akap12,Il1a,Ccl4,Ptgs2,Timp1,Fam107a,Rhoj,Adamts1,Sdc4 | | 5.555e-07 | -14.40 | GSE360\_DC\_VS\_MAC\_B\_MALAYI\_HIGH\_DOSE\_DN | MSigDB lists | GSE360\_DC\_VS\_MAC\_B\_MALAYI\_HIGH\_DOSE\_DN | 158 | 13 | 12187 | 179 | S100a8,Rgs16,Ifi211,Rhoc,Maff,Plaur,Isg15,Cd14,Ifi204,Ifi207,Cebpd,Ifi209,Ifi44 | | 5.555e-07 | -14.40 | GSE5455\_EX\_VIVO\_VS\_POST\_24H\_INCUBATION\_MONOCYTES\_FROM\_TUMOR\_BEARING\_MOUSE\_DN | MSigDB lists | GSE5455\_EX\_VIVO\_VS\_POST\_24H\_INCUBATION\_MONOCYTES\_FROM\_TUMOR\_BEARING\_MOUSE\_DN | 158 | 13 | 12187 | 179 | Ifit3b,Sgk1,Samd9l,Slfn5,Rnf213,Irf9,Irak3,Xaf1,Usp18,Il4ra,Bcl3,Tnfsf10,Rtp4 | | 5.555e-07 | -14.40 | JISON\_SICKLE\_CELL\_DISEASE\_UP | MSigDB lists | JISON\_SICKLE\_CELL\_DISEASE\_UP | 158 | 13 | 12187 | 179 | Nfkbie,Selp,Tap1,Clic4,Plek,Ifit3b,Ifi44,Cdkn1a,Ifit2,Oasl1,Gadd45b,Atf3,Timp1 | | 5.667e-07 | -14.38 | FARMER\_BREAST\_CANCER\_CLUSTER\_1 | MSigDB lists | FARMER\_BREAST\_CANCER\_CLUSTER\_1 | 22 | 6 | 12187 | 179 | Isg15,Herc6,Ifi44,Xaf1,Ifit3b,Oasl1 | | 5.690e-07 | -14.38 | SWEET\_LUNG\_CANCER\_KRAS\_DN | MSigDB lists | SWEET\_LUNG\_CANCER\_KRAS\_DN | 339 | 19 | 12187 | 179 | Gbp7,S100a9,Rhoj,Ptges,S100a8,Gbp3,Nfkbia,Usp18,Gadd45b,Ifitm3,Clic4,Sox11,Ifit3b,Zbtb16,Ms4a6d,Akap12,Hp,Cp,Rasip1 | | 5.880e-07 | -14.35 | positive regulation of inflammatory response | biological process | GO:0050729 | 103 | 11 | 13711 | 214 | Tnip1,Ptgs2,Il17ra,Mmp8,S100a9,Il6,S100a8,Tgm2,Nfkbiz,Tnf,Nfkbia | | 5.973e-07 | -14.33 | GSE36009\_UNSTIM\_VS\_LPS\_STIM\_DC\_UP | MSigDB lists | GSE36009\_UNSTIM\_VS\_LPS\_STIM\_DC\_UP | 159 | 13 | 12187 | 179 | Atf3,Ksr1,Gadd45g,Ptpn1,Irf9,Tap1,Slfn9,Ifitm3,Hcar2,Parp14,Gbp3,Tnfsf10,Gbp7 | | 5.973e-07 | -14.33 | GSE360\_L\_DONOVANI\_VS\_B\_MALAYI\_LOW\_DOSE\_DC\_UP | MSigDB lists | GSE360\_L\_DONOVANI\_VS\_B\_MALAYI\_LOW\_DOSE\_DC\_UP | 159 | 13 | 12187 | 179 | Birc3,Gadd45b,Tnfaip3,Irf7,Gem,Nfkbie,Cflar,Ier3,Gadd45g,Nfkb1,Tnip1,Isg15,Csf2rb | | 6.014e-07 | -14.32 | regulation of tumor necrosis factor biosynthetic process | biological process | GO:0042534 | 21 | 6 | 13711 | 214 | Bcl3,Errfi1,Cybb,Akap12,Hspb1,Thbs1 | | 6.101e-07 | -14.31 | GO\_NEGATIVE\_REGULATION\_OF\_DEFENSE\_RESPONSE | MSigDB lists | GO\_NEGATIVE\_REGULATION\_OF\_DEFENSE\_RESPONSE | 110 | 11 | 12187 | 179 | Ifi211,A2m,Saa1,Tap1,Irak3,Socs3,Ifi209,Tnfaip3,Ifi204,Nfkb1,Ifi207 | | 6.232e-07 | -14.29 | GO\_REGULATION\_OF\_INTERLEUKIN\_1\_BETA\_PRODUCTION | MSigDB lists | GO\_REGULATION\_OF\_INTERLEUKIN\_1\_BETA\_PRODUCTION | 35 | 7 | 12187 | 179 | Ifi211,Hspb1,Errfi1,Ifi209,Ifi204,Ifi207,Tnfaip3 | | 6.232e-07 | -14.29 | GO\_CELLULAR\_RESPONSE\_TO\_GLUCOSE\_STARVATION | MSigDB lists | GO\_CELLULAR\_RESPONSE\_TO\_GLUCOSE\_STARVATION | 35 | 7 | 12187 | 179 | Nfe2l2,Ifi207,Ifi204,Rrp8,Ifi211,Hspa5,Ifi209 | | 6.328e-07 | -14.27 | regulation of viral process | biological process | GO:0050792 | 150 | 13 | 13711 | 214 | Resf1,Trim25,Trim56,Ccl4,Ifitm6,Isg15,Ifitm2,Trim30a,Oasl1,Ifitm3,Zfp36,Oasl2,Tnf | | 6.419e-07 | -14.26 | GSE24142\_EARLY\_THYMIC\_PROGENITOR\_VS\_DN2\_THYMOCYTE\_FETAL\_UP | MSigDB lists | GSE24142\_EARLY\_THYMIC\_PROGENITOR\_VS\_DN2\_THYMOCYTE\_FETAL\_UP | 160 | 13 | 12187 | 179 | Ier3,Mxd4,Hspa5,Fstl1,Plek,Icam1,Irf9,Birc3,Il4ra,Usp18,Tgm2,Pygm,Gem | | 6.626e-07 | -14.23 | GENTILE\_UV\_HIGH\_DOSE\_DN | MSigDB lists | GENTILE\_UV\_HIGH\_DOSE\_DN | 277 | 17 | 12187 | 179 | Zfp36,Ptpn1,Isg15,Tiparp,Cebpd,Atf3,Thbs1,Tnfaip8,Gem,Arid5b,Bach1,Ripk1,Tnfaip3,Nfe2l2,Runx1,Il4ra,Maff | | 6.781e-07 | -14.20 | KEGG\_JAK\_STAT\_SIGNALING\_PATHWAY | MSigDB lists | KEGG\_JAK\_STAT\_SIGNALING\_PATHWAY | 89 | 10 | 12187 | 179 | Il6,Socs3,Ccnd2,Pik3r1,Il4ra,Csf2rb,Irf9,Osmr,Pik3r5,Csf3 | | 6.781e-07 | -14.20 | GO\_NEGATIVE\_REGULATION\_OF\_IMMUNE\_RESPONSE | MSigDB lists | GO\_NEGATIVE\_REGULATION\_OF\_IMMUNE\_RESPONSE | 89 | 10 | 12187 | 179 | Tap1,Irak3,Ifi211,A2m,Tnfaip3,Ifi204,Ifi207,Ifi209,Tnf,Il4ra | | 6.781e-07 | -14.20 | GO\_ZYMOGEN\_ACTIVATION | MSigDB lists | GO\_ZYMOGEN\_ACTIVATION | 89 | 10 | 12187 | 179 | Ifi207,Ifi204,Ripk1,Tnf,Ifi209,S100a8,Tnfsf10,Xdh,S100a9,Ifi211 | | 6.894e-07 | -14.19 | GSE2770\_UNTREATED\_VS\_IL12\_TREATED\_ACT\_CD4\_TCELL\_6H\_UP | MSigDB lists | GSE2770\_UNTREATED\_VS\_IL12\_TREATED\_ACT\_CD4\_TCELL\_6H\_UP | 161 | 13 | 12187 | 179 | Aff1,Atf3,Tap1,Ptpn1,Zfp36,Icam1,Maff,AA467197,Gbp4,Trim56,Banp,Gbp6,Gbp7 | | 6.894e-07 | -14.19 | GSE14769\_UNSTIM\_VS\_20MIN\_LPS\_BMDM\_DN | MSigDB lists | GSE14769\_UNSTIM\_VS\_20MIN\_LPS\_BMDM\_DN | 161 | 13 | 12187 | 179 | Map3k8,Tnf,Usp18,Nfkbia,Ccl4,Sgk1,Nfkbiz,Socs3,Gadd45g,Ier3,Cd14,Zfp36,Parp9 | | 7.253e-07 | -14.14 | positive regulation of molecular function | biological process | GO:0044093 | 1399 | 46 | 13711 | 214 | Ncf1,Sele,Ripk1,Cflar,Rhoc,Tnf,Mmp8,Dnajb1,Dtx3l,Trib1,Ccl7,Gadd45b,Ccnd2,Casp4,Il6,Cdkn1a,Ccl4,Trim25,Fzd4,Map3k6,Parp9,Sdc4,Il1rn,Irak3,Arid5b,Csf3,Plek,Xdh,Tnfsf10,Ccl2,S100a9,Irgm2,Wfdc21,Ptpn1,S100a8,Iqgap1,Ccl12,Thbs1,Gadd45g,P2ry6,Lgals9,Icam1,Rgs16,Pik3r5,Plaur,Map3k8 | | 7.367e-07 | -14.12 | immune response-regulating signaling pathway | biological process | GO:0002764 | 152 | 13 | 13711 | 214 | Cd14,Irak3,C3ar1,Lgals9,Tnip1,Pik3ap1,Fyb,Nfkbia,Tnf,Irgm2,Nfkbiz,C5ar1,Tnip3 | | 7.400e-07 | -14.12 | GSE10325\_BCELL\_VS\_LUPUS\_BCELL\_DN | MSigDB lists | GSE10325\_BCELL\_VS\_LUPUS\_BCELL\_DN | 162 | 13 | 12187 | 179 | Tiparp,Ifi44,Ifit3b,Rbm47,Tap1,Herc6,Isg15,Oasl1,Xaf1,Usp18,Rtp4,Cflar,Irf7 | | 7.400e-07 | -14.12 | GSE23502\_WT\_VS\_HDC\_KO\_MYELOID\_DERIVED\_SUPPRESSOR\_CELL\_COLON\_TUMOR\_UP | MSigDB lists | GSE23502\_WT\_VS\_HDC\_KO\_MYELOID\_DERIVED\_SUPPRESSOR\_CELL\_COLON\_TUMOR\_UP | 162 | 13 | 12187 | 179 | Ifit3b,Vcam1,Ifitm2,Ifitm3,Tagln2,Tgm2,Gbp4,Gbp6,C5ar1,Fzd4,S100a9,S100a8,Igsf6 | | 7.425e-07 | -14.11 | GO\_RESPONSE\_TO\_EXTRACELLULAR\_STIMULUS | MSigDB lists | GO\_RESPONSE\_TO\_EXTRACELLULAR\_STIMULUS | 345 | 19 | 12187 | 179 | Ifi211,Rrp8,Ccl12,Il6,Nfe2l2,Ptgs2,Icam1,Zfp36,Adamts1,Cdkn1a,Ifi209,Hspa5,Vcam1,Fstl1,Socs3,Ifi204,Ifi207,Cp,Atf3 | | 7.533e-07 | -14.10 | TAKEDA\_TARGETS\_OF\_NUP98\_HOXA9\_FUSION\_10D\_DN | MSigDB lists | TAKEDA\_TARGETS\_OF\_NUP98\_HOXA9\_FUSION\_10D\_DN | 90 | 10 | 12187 | 179 | Olfml2b,A2m,Slfn9,Cxcl1,C5ar1,Rassf4,Mmp8,Ccl2,Ccl4,Sgk1 | | 7.563e-07 | -14.09 | XU\_HGF\_TARGETS\_INDUCED\_BY\_AKT1\_6HR | MSigDB lists | XU\_HGF\_TARGETS\_INDUCED\_BY\_AKT1\_6HR | 13 | 5 | 12187 | 179 | Plaur,Maff,Isg15,Thbs1,Irf7 | | 7.563e-07 | -14.09 | LEE\_LIVER\_CANCER\_HEPATOBLAST | MSigDB lists | LEE\_LIVER\_CANCER\_HEPATOBLAST | 13 | 5 | 12187 | 179 | Ptgs2,Ccnd2,Timp1,Il6,Plaur | | 7.641e-07 | -14.08 | VARELA\_ZMPSTE24\_TARGETS\_UP | MSigDB lists | VARELA\_ZMPSTE24\_TARGETS\_UP | 36 | 7 | 12187 | 179 | Cd14,Zbtb16,Cdkn1a,Gadd45b,Rgs16,Atf3,Gadd45g | | 7.833e-07 | -14.06 | GO\_TOLL\_LIKE\_RECEPTOR\_SIGNALING\_PATHWAY | MSigDB lists | GO\_TOLL\_LIKE\_RECEPTOR\_SIGNALING\_PATHWAY | 70 | 9 | 12187 | 179 | Tnip3,Ripk1,Birc3,Nfkbia,Pik3ap1,Irak3,Irf7,Cd14,Tnip1 | | 7.864e-07 | -14.06 | GSE30083\_SP1\_VS\_SP3\_THYMOCYTE\_DN | MSigDB lists | GSE30083\_SP1\_VS\_SP3\_THYMOCYTE\_DN | 137 | 12 | 12187 | 179 | Ms4a6d,Zbtb16,Ifi44,Sgk1,Il17ra,Slfn5,Herc6,Rnf213,Usp18,Iqgap1,Irf7,Irgm2 | | 7.864e-07 | -14.06 | GSE36888\_UNTREATED\_VS\_IL2\_TREATED\_STAT5\_AB\_KNOCKIN\_TCELL\_17H\_UP | MSigDB lists | GSE36888\_UNTREATED\_VS\_IL2\_TREATED\_STAT5\_AB\_KNOCKIN\_TCELL\_17H\_UP | 137 | 12 | 12187 | 179 | Tnfaip3,Tnip3,Slc24a4,Casp4,Vcam1,Runx1,Plaur,Arid5b,Cxcl1,Ptges,Ptgs2,C5ar1 | | 8.162e-07 | -14.02 | response to interferon-alpha | biological process | GO:0035455 | 22 | 6 | 13711 | 214 | Ifitm3,Ifitm6,Ifitm2,Tgtp1,Ifi204,Ifit2 | | 8.328e-07 | -14.00 | CHICAS\_RB1\_TARGETS\_CONFLUENT | MSigDB lists | CHICAS\_RB1\_TARGETS\_CONFLUENT | 454 | 22 | 12187 | 179 | Ifi209,Thbs1,Vcam1,Ifi204,Ifi207,Samd9l,Ifitm3,Ifitm2,Hspb1,Errfi1,Parp9,Ifit2,Xaf1,Tgm2,Sntb2,Sdc4,Tubb6,Rassf4,Dtx3l,Ifi211,Arid5b,Rhoj | | 8.371e-07 | -13.99 | regulation of cell population proliferation | biological process | GO:0042127 | 1270 | 43 | 13711 | 214 | Cflar,Ripk1,Zbtb16,Csf3,C3ar1,Tnfaip3,Tnf,Ccl2,Zfp36,Atf3,Ksr1,Xdh,Vcam1,Tgm2,Cxcl1,C5ar1,Ccl12,Prg4,Trib1,Ifitm3,Thbs1,Lgals9,P2ry6,Ccnd2,Pik3r1,Il1a,Sox11,Nfkbia,Il4ra,Slfn3,Cdkn1a,Il6,Errfi1,Kcna5,Rasip1,Ptgs2,Runx1,Osmr,Timp1,Ptges,Sdc4,Adamts1,Slfn2 | | 8.514e-07 | -13.98 | GSE19401\_UNSTIM\_VS\_RETINOIC\_ACID\_AND\_PAM2CSK4\_STIM\_FOLLICULAR\_DC\_DN | MSigDB lists | GSE19401\_UNSTIM\_VS\_RETINOIC\_ACID\_AND\_PAM2CSK4\_STIM\_FOLLICULAR\_DC\_DN | 164 | 13 | 12187 | 179 | Il6,Gadd45b,Sdc4,Map3k8,Tnf,Olfml2b,Cxcl1,Gem,C3ar1,Timp1,Thbs1,Icam1,Tnip1 | | 8.514e-07 | -13.98 | GSE42724\_MEMORY\_BCELL\_VS\_PLASMABLAST\_UP | MSigDB lists | GSE42724\_MEMORY\_BCELL\_VS\_PLASMABLAST\_UP | 164 | 13 | 12187 | 179 | Il6,Usp18,Parp14,Il4ra,Cmpk2,Irf7,Cflar,Nfkb1,Il1rn,Clic4,Icam1,Slfn5,Tagln2 | | 8.519e-07 | -13.98 | programmed cell death | biological process | GO:0012501 | 721 | 30 | 13711 | 214 | Nfkb1,Birc3,Sgk1,Ier3,Xaf1,Spi1,Il6,Cdkn1a,Casp4,Il1a,Gadd45b,Pik3r1,Hcar2,Gadd45g,S100a8,C5ar1,Rrp8,Bcl3,Ifit2,S100a9,Tnfsf10,Tnf,Tnfaip3,Ifi204,Lcn2,Ncf1,Irak3,Tnfaip8,Ripk1,Cflar | | 8.752e-07 | -13.95 | KOKKINAKIS\_METHIONINE\_DEPRIVATION\_48HR\_UP | MSigDB lists | KOKKINAKIS\_METHIONINE\_DEPRIVATION\_48HR\_UP | 114 | 11 | 12187 | 179 | Cflar,Plaur,Tnfsf10,Hspb1,Rtp4,Nfkbia,Cdkn1a,Birc3,Il6,Timp1,Il1a | | 8.757e-07 | -13.95 | regulation of cytokine-mediated signaling pathway | biological process | GO:0001959 | 86 | 10 | 13711 | 214 | Parp9,Il6,Casp4,Irgm2,Ripk1,Parp14,Irak3,Il1rn,Irgm1,Irf7 | | 8.783e-07 | -13.95 | ICSBP\_Q6 | MSigDB lists | ICSBP\_Q6 | 192 | 14 | 12187 | 179 | Nfkb1,Col4a1,Thbs1,Ifi44,Ifit3b,Cxcl16,Tap1,Isg15,Tubb6,Parp12,Usp18,Rbm39,Dtx3l,Angptl4 | | 8.832e-07 | -13.94 | GRAESSMANN\_APOPTOSIS\_BY\_DOXORUBICIN\_UP | MSigDB lists | GRAESSMANN\_APOPTOSIS\_BY\_DOXORUBICIN\_UP | 948 | 34 | 12187 | 179 | Tor3a,Oasl1,Parp12,Irf7,Irgm2,Ifit3b,Ifi209,Cdkn1a,Ifi44,Atf3,Ifi204,Ifi207,Zfp36,Tgm2,Usp18,Il4ra,Ccl12,Ifi211,Cxcl1,Nfkbie,Gbp7,Csf3,Rgs16,Cmpk2,Rtp4,Dnajb1,Thbs1,Mxd4,Tiparp,Cd14,Isg15,Ptpn1,Tap1,P2ry6 | | 8.858e-07 | -13.94 | FRIDMAN\_SENESCENCE\_UP | MSigDB lists | FRIDMAN\_SENESCENCE\_UP | 71 | 9 | 12187 | 179 | Cdkn1a,Ifi209,Thbs1,Ifi204,Ifi207,Tnfaip3,Isg15,Ifi211,Irf7 | | 9.055e-07 | -13.91 | positive regulation of cellular component movement | biological process | GO:0051272 | 497 | 24 | 13711 | 214 | Ccl4,Ptgs2,Fam107a,Adamts1,Rhoj,Thbs1,Lgals9,P2ry6,Pik3r1,Il1a,Akap12,Icam1,Nfe2l2,Selp,Hspa5,C5ar1,Ccl12,Cxcl16,Iqgap1,Sele,Hspb1,C3ar1,Rhoc,Ccl2 | | 9.190e-07 | -13.90 | GSE27859\_MACROPHAGE\_VS\_DC\_UP | MSigDB lists | GSE27859\_MACROPHAGE\_VS\_DC\_UP | 139 | 12 | 12187 | 179 | Sdc4,Tgm2,Parp14,Map3k8,Tnf,Igsf6,Nfkbie,Cflar,Atf3,Tnip3,Nfkb1,Pik3ap1 | | 9.190e-07 | -13.90 | GSE15330\_LYMPHOID\_MULTIPOTENT\_VS\_MEGAKARYOCYTE\_ERYTHROID\_PROGENITOR\_IKAROS\_KO\_DN | MSigDB lists | GSE15330\_LYMPHOID\_MULTIPOTENT\_VS\_MEGAKARYOCYTE\_ERYTHROID\_PROGENITOR\_IKAROS\_KO\_DN | 139 | 12 | 12187 | 179 | Ccl4,Casp4,Gem,Phf11d,S100a9,S100a8,Phf11b,Plek,Rbm47,Pik3ap1,Clic4,Errfi1 | | 9.225e-07 | -13.90 | GO\_CELLULAR\_RESPONSE\_TO\_LIPID | MSigDB lists | GO\_CELLULAR\_RESPONSE\_TO\_LIPID | 350 | 19 | 12187 | 179 | Cd14,Icam1,P2ry6,Cxcl16,Errfi1,Msn,Adamts1,Sgk1,Nfkb1,Tnip3,Csf3,Fzd4,Cmpk2,Sbno2,Tnf,Nfkbia,Tnfaip3,Ccl12,Il6 | | 9.341e-07 | -13.88 | GAJATE\_RESPONSE\_TO\_TRABECTEDIN\_UP | MSigDB lists | GAJATE\_RESPONSE\_TO\_TRABECTEDIN\_UP | 53 | 8 | 12187 | 179 | Casp4,Cdkn1a,Atf3,Oasl1,Il6,Tnfaip3,Isg15,Clic4 | | 9.419e-07 | -13.88 | host cell cytoplasm | cellular component | GO:0030430 | 13 | 5 | 13825 | 212 | Gbp7,Gbp3,Iigp1,Gbp6,Irgm2 | | 9.419e-07 | -13.88 | host cell cytoplasm part | cellular component | GO:0033655 | 13 | 5 | 13825 | 212 | Iigp1,Gbp3,Gbp7,Irgm2,Gbp6 | | 9.445e-07 | -13.87 | positive regulation of cytokine biosynthetic process | biological process | GO:0042108 | 50 | 8 | 13711 | 214 | Bcl3,Thbs1,Hspb1,Cybb,Il6,Tnf,Akap12,Il1a | | 9.489e-07 | -13.87 | lymphocyte migration | biological process | GO:0072676 | 35 | 7 | 13711 | 214 | Msn,Ccl7,Ccl4,Ccl12,Ccl2,Cxcl16,Icam1 | | 9.674e-07 | -13.85 | regulation of multicellular organismal process | biological process | GO:0051239 | 2546 | 69 | 13711 | 214 | Noct,Selp,Mmp8,Nfe2l2,Prg4,Hspa5,Cxcl1,C5ar1,Maff,Trim30a,Cd14,Cflar,Ripk1,Hspb1,Ncf1,Zfp36,Acod1,Tnf,Ptgs2,Sgk1,Runx1,Ccl4,Spi1,Cebpd,Ifi211,Isg15,Fzd4,Timp1,Hcar2,Pik3r1,Trib1,Ctla2a,Irf7,Il6,Akap12,Il1a,Nfkbia,Casp4,Il4ra,Bcl3,S100a9,Iqgap1,Nfkbiz,Lcn2,Ifi204,Zbtb16,Csf3,Adamts9,C3ar1,Il1rn,Irak3,Ccl2,Xdh,Plek,Tnfaip3,Il17ra,Kcna5,Adamts1,Rhoj,Nfkb1,Apold1,Gbp4,Lgals9,Gadd45g,Thbs1,Errfi1,Cybb,Sox11,Icam1 | | 9.776e-07 | -13.84 | GSE23308\_WT\_VS\_MINERALCORTICOID\_REC\_KO\_MACROPHAGE\_DN | MSigDB lists | GSE23308\_WT\_VS\_MINERALCORTICOID\_REC\_KO\_MACROPHAGE\_DN | 166 | 13 | 12187 | 179 | Ccl4,Gpr84,Casp4,Maff,Birc3,Ccl12,Tnfaip3,Nfe2l2,Gbp7,Nfkbiz,Znfx1,Parp9,Msn | | 9.901e-07 | -13.83 | negative regulation of molecular function | biological process | GO:0044092 | 847 | 33 | 13711 | 214 | Thbs1,Dtx3l,Trib1,Gadd45b,Gadd45g,Nfkbia,Gem,Sox11,Il6,Errfi1,Cdkn1a,Rasip1,Ptgs2,Plaur,Timp1,Birc3,Nfkb1,Parp9,Irak3,Hspb1,Tnfaip8,Cflar,Tnf,Tnfaip3,Acod1,Zfp36,Serpina3f,Angptl4,Wfdc21,Ifit2,Ptpn1,Hp,A2m | | 9.925e-07 | -13.82 | GSE39820\_IL1B\_IL6\_VS\_IL1B\_IL6\_IL23A\_TREATED\_CD4\_TCELL\_UP | MSigDB lists | GSE39820\_IL1B\_IL6\_VS\_IL1B\_IL6\_IL23A\_TREATED\_CD4\_TCELL\_UP | 140 | 12 | 12187 | 179 | Dtx3l,Gbp7,Gbp6,Irgm2,Cmpk2,Irf9,Gbp3,Sntb2,Usp18,Bach1,Parp12,Gbp4 | | 9.925e-07 | -13.82 | GSE6269\_HEALTHY\_VS\_STAPH\_AUREUS\_INF\_PBMC\_UP | MSigDB lists | GSE6269\_HEALTHY\_VS\_STAPH\_AUREUS\_INF\_PBMC\_UP | 140 | 12 | 12187 | 179 | Ifi204,Ifi207,Ifi209,Ifi44,Ifitm3,Herc6,Isg15,Parp12,Oasl1,Xaf1,Ifi211,Irf7 | | 9.976e-07 | -13.82 | KAMIKUBO\_MYELOID\_CEBPA\_NETWORK | MSigDB lists | KAMIKUBO\_MYELOID\_CEBPA\_NETWORK | 24 | 6 | 12187 | 179 | S100a9,Hp,S100a8,Mmp8,Cybb,Lcn2 | | 1.024e-06 | -13.79 | MODULE\_177 | MSigDB lists | MODULE\_177 | 93 | 10 | 12187 | 179 | Ifi211,Nfkb1,Ifi204,Ifi207,Tnfaip3,Ifi209,Il4ra,Sgk1,Nfkbia,Ifit3b | | 1.047e-06 | -13.77 | GSE23502\_BM\_VS\_COLON\_TUMOR\_MYELOID\_DERIVED\_SUPPRESSOR\_CELL\_UP | MSigDB lists | GSE23502\_BM\_VS\_COLON\_TUMOR\_MYELOID\_DERIVED\_SUPPRESSOR\_CELL\_UP | 167 | 13 | 12187 | 179 | Sbno2,Gbp7,Gem,Cxcl1,Ccnd2,Trib1,Casp4,Rbm47,Csf2rb,Tagln2,Tiparp,Stx11,Ier3 | | 1.068e-06 | -13.75 | GO\_INTRACELLULAR\_SIGNAL\_TRANSDUCTION | MSigDB lists | GO\_INTRACELLULAR\_SIGNAL\_TRANSDUCTION | 1281 | 41 | 12187 | 179 | Ripk1,Tnfaip3,Ccl12,Rhoc,Map3k8,Rhou,Pik3r1,Iqgap1,Ifi211,Pik3r5,Rrp8,Bcl3,Rhoj,Nfkb1,Sox11,Sgk1,Ptpn1,Cd14,Selp,Map3k6,Sgk3,Trib1,Nfkbia,Casp4,Tnf,Birc3,Nfkb2,Irf7,Gem,Rasd1,Ifi204,Ksr1,Ifi207,Vcam1,Socs3,Cdkn1a,Ifi209,Csf2rb,Hspb1,Sele,Plek | | 1.090e-06 | -13.73 | negative regulation of cytokine biosynthetic process | biological process | GO:0042036 | 23 | 6 | 13711 | 214 | Bcl3,Errfi1,Zfp36,Il6,Nfkb1,Prg4 | | 1.104e-06 | -13.72 | GO\_IMMUNE\_SYSTEM\_DEVELOPMENT | MSigDB lists | GO\_IMMUNE\_SYSTEM\_DEVELOPMENT | 425 | 21 | 12187 | 179 | Ifi209,Vcam1,Zbtb16,Ifi207,Tiparp,Ifi204,Herc6,Plek,Rbm47,Pik3r1,Runx1,Tnf,Spi1,Il6,Tnfaip3,C3ar1,Csf3,Bcl3,Ifi211,Sbno2,Nfkb2 | | 1.120e-06 | -13.70 | PICCALUGA\_ANGIOIMMUNOBLASTIC\_LYMPHOMA\_UP | MSigDB lists | PICCALUGA\_ANGIOIMMUNOBLASTIC\_LYMPHOMA\_UP | 168 | 13 | 12187 | 179 | Osmr,Adamts9,A2m,Rassf4,Fam107a,Ccl12,Irak3,Col4a1,Cp,Samd9l,Adamts1,Fstl1,Vcam1 | | 1.120e-06 | -13.70 | GSE26030\_TH1\_VS\_TH17\_DAY15\_POST\_POLARIZATION\_DN | MSigDB lists | GSE26030\_TH1\_VS\_TH17\_DAY15\_POST\_POLARIZATION\_DN | 168 | 13 | 12187 | 179 | P2ry6,Slfn3,Icam1,Ifitm3,Ncf1,Tagln2,Cebpd,Rtp4,Slfn4,Gbp3,Rassf4,Lgals9,Msr1 | | 1.132e-06 | -13.69 | MARKEY\_RB1\_CHRONIC\_LOF\_DN | MSigDB lists | MARKEY\_RB1\_CHRONIC\_LOF\_DN | 94 | 10 | 12187 | 179 | Ms4a6d,Cp,Ccl12,Il6,Msr1,Spi1,Slfn4,Slfn3,C3ar1,Cd14 | | 1.160e-06 | -13.67 | positive regulation of locomotion | biological process | GO:0040017 | 504 | 24 | 13711 | 214 | Ccl4,Ptgs2,Fam107a,Rhoj,Adamts1,Thbs1,Pik3r1,Lgals9,P2ry6,Icam1,Il1a,Akap12,Nfe2l2,Selp,C5ar1,Hspa5,Iqgap1,Ccl12,Cxcl16,Sele,Hspb1,C3ar1,Rhoc,Ccl2 | | 1.162e-06 | -13.66 | positive regulation of cytokine-mediated signaling pathway | biological process | GO:0001961 | 36 | 7 | 13711 | 214 | Parp9,Casp4,Irgm2,Ripk1,Parp14,Irf7,Irgm1 | | 1.162e-06 | -13.66 | GO\_REGULATION\_OF\_MONONUCLEAR\_CELL\_MIGRATION | MSigDB lists | GO\_REGULATION\_OF\_MONONUCLEAR\_CELL\_MIGRATION | 14 | 5 | 12187 | 179 | Ccl12,C3ar1,Thbs1,Tnf,C5ar1 | | 1.162e-06 | -13.66 | GO\_POSITIVE\_REGULATION\_OF\_COAGULATION | MSigDB lists | GO\_POSITIVE\_REGULATION\_OF\_COAGULATION | 14 | 5 | 12187 | 179 | S100a9,Selp,Thbs1,Nfe2l2,Plek | | 1.163e-06 | -13.66 | chemokine receptor binding | molecular function | GO:0042379 | 36 | 7 | 13516 | 211 | Ccl2,Cxcl1,Cxcl9,Ccl4,Ccl7,Cxcl16,Ccl12 | | 1.245e-06 | -13.60 | GSE24634\_TEFF\_VS\_TCONV\_DAY5\_IN\_CULTURE\_DN | MSigDB lists | GSE24634\_TEFF\_VS\_TCONV\_DAY5\_IN\_CULTURE\_DN | 143 | 12 | 12187 | 179 | Il17ra,Nfkb2,Ifitm2,Phf11d,Arid5b,Irf7,Il1rn,Runx1,Phf11b,Ifi44,Il4ra,Ifit3b | | 1.245e-06 | -13.60 | GSE34156\_NOD2\_LIGAND\_VS\_TLR1\_TLR2\_LIGAND\_6H\_TREATED\_MONOCYTE\_DN | MSigDB lists | GSE34156\_NOD2\_LIGAND\_VS\_TLR1\_TLR2\_LIGAND\_6H\_TREATED\_MONOCYTE\_DN | 143 | 12 | 12187 | 179 | Cd14,Sgk3,Clic4,Csf2rb,Cxcl16,Rbm47,Timp1,C3ar1,Igsf6,Msr1,AA467197,Cybb | | 1.245e-06 | -13.60 | GSE4984\_UNTREATED\_VS\_GALECTIN1\_TREATED\_DC\_UP | MSigDB lists | GSE4984\_UNTREATED\_VS\_GALECTIN1\_TREATED\_DC\_UP | 143 | 12 | 12187 | 179 | Spi1,Gpr84,Rhou,Bcl3,S100a9,Cflar,Sbno2,S100a8,Igsf6,Vcam1,Aff1,Zfp36 | | 1.249e-06 | -13.59 | TAKAO\_RESPONSE\_TO\_UVB\_RADIATION\_DN | MSigDB lists | TAKAO\_RESPONSE\_TO\_UVB\_RADIATION\_DN | 95 | 10 | 12187 | 179 | Ifi211,Zfp36,Nfkb2,Ifi209,Ccnd2,Dnajb1,Ifi204,Ifi207,Tnfaip3,Cebpd | | 1.251e-06 | -13.59 | molecular function regulator | molecular function | GO:0098772 | 1245 | 42 | 13516 | 211 | Tnf,Parp9,Il1a,Trib1,Cflar,Csf3,Rgs16,Tnfsf10,Hspb1,Cxcl16,Cdkn1a,Ccl4,Wfdc21,Irgm2,Socs3,Retnlg,Dtx3l,Cxcl9,Angptl4,Ccl2,Birc3,Ccnd2,A2m,Sgk1,Ccl12,Ncf1,Kcna5,Ccl7,Gem,Serpina3f,Il1rn,Timp1,Cxcl1,Dnajb1,Sgk3,Saa1,Iqgap1,Lgals9,Tnfaip8,Pik3r1,Pik3r5,Il6 | | 1.251e-06 | -13.59 | positive regulation of leukocyte migration | biological process | GO:0002687 | 111 | 11 | 13711 | 214 | C5ar1,Ccl12,Ccl4,Selp,Icam1,Il1a,Ccl2,C3ar1,Thbs1,Sele,Lgals9 | | 1.262e-06 | -13.58 | regulation of protein serine/threonine kinase activity | biological process | GO:0071900 | 401 | 21 | 13711 | 214 | Map3k8,Map3k6,Fzd4,Ptpn1,Iqgap1,Rasip1,Pik3r5,Irgm2,Tnfaip3,Tnf,Ksr1,Cdkn1a,Irak3,Il1rn,Thbs1,Trib1,Ripk1,Gadd45b,Gadd45g,Ccnd2,Lgals9 | | 1.264e-06 | -13.58 | regulation of protein kinase activity | biological process | GO:0045859 | 618 | 27 | 13711 | 214 | Sdc4,Map3k8,Fzd4,Map3k6,Pik3r5,Rasip1,Il6,Errfi1,Cdkn1a,Ccnd2,Gadd45g,Gadd45b,Lgals9,Thbs1,Trib1,Iqgap1,C5ar1,Ptpn1,Irgm2,Ksr1,Tnfaip3,Tnf,Hspb1,Il1rn,Irak3,Ncf1,Ripk1 | | 1.271e-06 | -13.58 | WINTER\_HYPOXIA\_METAGENE | MSigDB lists | WINTER\_HYPOXIA\_METAGENE | 198 | 14 | 12187 | 179 | Angptl4,Nfkb2,Tgm2,Tnfaip3,Il6,Ptgs2,Icam1,Plaur,Errfi1,Cdkn1a,Hspa5,Thbs1,Nfkb1,Cp | | 1.281e-06 | -13.57 | GSE19198\_CTRL\_VS\_IL21\_TREATED\_TCELL\_6H\_UP | MSigDB lists | GSE19198\_CTRL\_VS\_IL21\_TREATED\_TCELL\_6H\_UP | 170 | 13 | 12187 | 179 | Tnip1,Ifitm2,Ifitm3,Csf2rb,Plek,Aff1,Ifi209,Ifi207,Ifi204,Ifi211,Cxcl1,Ccl2,Birc3 | | 1.297e-06 | -13.56 | ONO\_AML1\_TARGETS\_DN | MSigDB lists | ONO\_AML1\_TARGETS\_DN | 25 | 6 | 12187 | 179 | Slfn3,Slfn4,Il17ra,Casp4,Socs3,Il4ra | | 1.297e-06 | -13.56 | TSAI\_RESPONSE\_TO\_RADIATION\_THERAPY | MSigDB lists | TSAI\_RESPONSE\_TO\_RADIATION\_THERAPY | 25 | 6 | 12187 | 179 | Col4a1,Il6,Isg15,Thbs1,Ifit2,A2m | | 1.315e-06 | -13.54 | Interleukin\_8-like\_sf | interpro domains | IPR036048 | 24 | 6 | 13788 | 212 | Ccl12,Ccl4,Ccl2,Ccl7,Cxcl9,Cxcl1 | | 1.315e-06 | -13.54 | Chemokine\_IL8-like\_dom | interpro domains | IPR001811 | 24 | 6 | 13788 | 212 | Cxcl1,Ccl2,Ccl4,Cxcl9,Ccl7,Ccl12 | | 1.317e-06 | -13.54 | regulation of I-kappaB kinase/NF-kappaB signaling | biological process | GO:0043122 | 135 | 12 | 13711 | 214 | Tnip3,Tgm2,Trim25,Tnfsf10,Il1a,Pik3ap1,Tnf,Tnfaip3,Tnip1,Ripk1,Il1rn,Hspb1 | | 1.325e-06 | -13.53 | SCY | smart domains | SM00199 | 22 | 6 | 7188 | 123 | Ccl7,Ccl12,Ccl4,Ccl2,Cxcl1,Cxcl9 | | 1.341e-06 | -13.52 | GSE36888\_UNTREATED\_VS\_IL2\_TREATED\_STAT5\_AB\_KNOCKIN\_TCELL\_6H\_UP | MSigDB lists | GSE36888\_UNTREATED\_VS\_IL2\_TREATED\_STAT5\_AB\_KNOCKIN\_TCELL\_6H\_UP | 144 | 12 | 12187 | 179 | S100a8,S100a9,Ptges,Ier3,Ksr1,Tnip3,Nfkbiz,Socs3,Thbs1,P2ry6,Pik3ap1,Cd14 | | 1.341e-06 | -13.52 | GSE41087\_WT\_VS\_FOXP3\_MUT\_ANTI\_CD3\_CD28\_STIM\_CD4\_TCELL\_UP | MSigDB lists | GSE41087\_WT\_VS\_FOXP3\_MUT\_ANTI\_CD3\_CD28\_STIM\_CD4\_TCELL\_UP | 144 | 12 | 12187 | 179 | Bach1,Tnfaip3,Birc3,Sdc4,Nfkbia,Nfkbie,Cflar,Bcl3,Nfkb1,Atf3,Csf2rb,Tnip1 | | 1.341e-06 | -13.52 | GSE21774\_CD62L\_POS\_CD56\_DIM\_VS\_CD62L\_NEG\_CD56\_DIM\_NK\_CELL\_DN | MSigDB lists | GSE21774\_CD62L\_POS\_CD56\_DIM\_VS\_CD62L\_NEG\_CD56\_DIM\_NK\_CELL\_DN | 144 | 12 | 12187 | 179 | Arid5b,Igsf6,Spi1,Tor3a,Gadd45b,Tubb6,Cd14,Ncf1,Lcn2,Il1a,Hp,Mmp8 | | 1.369e-06 | -13.50 | REACTOME\_INNATE\_IMMUNE\_SYSTEM | MSigDB lists | REACTOME\_INNATE\_IMMUNE\_SYSTEM | 171 | 13 | 12187 | 179 | Irak3,Trim25,Irf2,Isg15,Saa1,Cd14,Nfkb2,Irf7,Tnfaip3,Ripk1,Birc3,Casp4,Nfkbia | | 1.369e-06 | -13.50 | GSE43955\_1H\_VS\_20H\_ACT\_CD4\_TCELL\_UP | MSigDB lists | GSE43955\_1H\_VS\_20H\_ACT\_CD4\_TCELL\_UP | 171 | 13 | 12187 | 179 | Saa1,Tnip1,Aff1,Il1rn,Rhoj,Cxcl1,Tnfaip8,Msr1,Maff,Il4ra,Tnf,Nfe2l2,Il6 | | 1.382e-06 | -13.49 | NAKAMURA\_TUMOR\_ZONE\_PERIPHERAL\_VS\_CENTRAL\_DN | MSigDB lists | NAKAMURA\_TUMOR\_ZONE\_PERIPHERAL\_VS\_CENTRAL\_DN | 468 | 22 | 12187 | 179 | Ptgs2,Cxcl16,Dnajb1,Socs3,Hspa5,Lcn2,Cebpd,Cp,Hp,Timp1,Ksr1,Arid5b,S100a9,Bcl3,Trim56,Tnfsf10,Xdh,S100a8,Trib1,Birc3,Il6,Bach1 | | 1.416e-06 | -13.47 | BUYTAERT\_PHOTODYNAMIC\_THERAPY\_STRESS\_UP | MSigDB lists | BUYTAERT\_PHOTODYNAMIC\_THERAPY\_STRESS\_UP | 707 | 28 | 12187 | 179 | Cxcl1,Ifi211,Nfe2l2,Il6,Maff,Ip6k2,Il1a,Tiparp,Cebpd,Ier3,Mxd4,Dnajb1,Rbm39,Gem,Irf7,Bach1,Oasl1,Trib1,Nfkbia,Zfp36,Trim25,Irf9,Ptgs2,Ifi207,Ifi204,Atf3,Cdkn1a,Ifi209 | | 1.420e-06 | -13.46 | HUANG\_DASATINIB\_RESISTANCE\_UP | MSigDB lists | HUANG\_DASATINIB\_RESISTANCE\_UP | 75 | 9 | 12187 | 179 | Osmr,Msn,Ifi211,Ifi204,Samd9l,Ifi207,Ifi209,Fstl1,Ifit3b | | 1.427e-06 | -13.46 | IL8 | pfam domains | PF00048 | 24 | 6 | 12881 | 201 | Ccl7,Cxcl1,Ccl12,Cxcl9,Ccl4,Ccl2 | | 1.444e-06 | -13.45 | GSE30971\_WBP7\_HET\_VS\_KO\_MACROPHAGE\_2H\_LPS\_STIM\_DN | MSigDB lists | GSE30971\_WBP7\_HET\_VS\_KO\_MACROPHAGE\_2H\_LPS\_STIM\_DN | 145 | 12 | 12187 | 179 | Il4ra,Bach1,Il6,Gadd45b,Csf3,Cxcl1,Tnfaip8,Socs3,Tnip3,Ier3,Ptgs2,Pik3ap1 | | 1.444e-06 | -13.45 | GSE360\_CTRL\_VS\_L\_MAJOR\_DC\_DN | MSigDB lists | GSE360\_CTRL\_VS\_L\_MAJOR\_DC\_DN | 145 | 12 | 12187 | 179 | Ier3,Timp1,Il1rn,Tap1,Ptpn1,Hspb1,Ifitm3,Ccl12,Tnfaip3,Trib1,Tnf,Cflar | | 1.462e-06 | -13.44 | GSE29618\_BCELL\_VS\_MONOCYTE\_DAY7\_FLU\_VACCINE\_DN | MSigDB lists | GSE29618\_BCELL\_VS\_MONOCYTE\_DAY7\_FLU\_VACCINE\_DN | 172 | 13 | 12187 | 179 | Trib1,C5ar1,Sgk1,Atf3,Cebpd,Il1rn,Stx11,Cd14,Plaur,Icam1,Irak3,Il17ra,Rbm47 | | 1.462e-06 | -13.44 | GSE9037\_WT\_VS\_IRAK4\_KO\_LPS\_4H\_STIM\_BMDM\_UP | MSigDB lists | GSE9037\_WT\_VS\_IRAK4\_KO\_LPS\_4H\_STIM\_BMDM\_UP | 172 | 13 | 12187 | 179 | Gbp4,Map3k8,Nfkb2,Gbp7,Gbp6,Nfkbie,Csf3,Nfkb1,Il1rn,Cdkn1a,Tap1,Clic4,Herc6 | | 1.463e-06 | -13.44 | GSE40274\_CTRL\_VS\_FOXP3\_AND\_LEF1\_TRANSDUCED\_ACTIVATED\_CD4\_TCELL\_UP | MSigDB lists | GSE40274\_CTRL\_VS\_FOXP3\_AND\_LEF1\_TRANSDUCED\_ACTIVATED\_CD4\_TCELL\_UP | 120 | 11 | 12187 | 179 | C3ar1,S100a9,Apold1,Tnfaip8,Map3k8,Maff,Gpr84,Nfkbiz,Stx11,Bach1,Cebpd | | 1.516e-06 | -13.40 | Fluid shear stress and atherosclerosis | KEGG pathways | mmu05418 | 128 | 13 | 5248 | 107 | Nfe2l2,Cybb,Il1a,Ccl12,Sele,Ncf1,Nfkb1,Icam1,Tnf,Pik3r1,Ccl2,Vcam1,Sdc4 | | 1.516e-06 | -13.40 | Fluid shear stress and atherosclerosis | KEGG pathways | ko05418 | 128 | 13 | 5248 | 107 | Cybb,Icam1,Tnf,Vcam1,Sdc4,Nfkb1,Ncf1,Il1a,Ccl12,Nfe2l2,Pik3r1,Ccl2,Sele | | 1.522e-06 | -13.40 | GOUYER\_TATI\_TARGETS\_UP | MSigDB lists | GOUYER\_TATI\_TARGETS\_UP | 7 | 4 | 12187 | 179 | Lcn2,Ccl12,Zbtb16,Tnf | | 1.527e-06 | -13.39 | negative regulation of response to stimulus | biological process | GO:0048585 | 1299 | 43 | 13711 | 214 | Ptpn1,Ccl12,Tap1,A2m,Tnip3,Sgk3,Bcl3,Plek,Tnfaip3,Xdh,Il1rn,Irak3,Nfkb1,Plaur,Rgs16,Icam1,Errfi1,Thbs1,Lgals9,Hspa5,Nfe2l2,Pik3ap1,Tnf,Acod1,Zfp36,Atf3,Ripk1,Cflar,Hspb1,Rnf213,Trim30a,Tnip1,Rasip1,Ier3,Ptgs2,Nfkbia,Il4ra,Cdkn1a,Il6,Ctla2a,Trib1,Parp14,Socs3 | | 1.553e-06 | -13.38 | GSE40685\_TREG\_VS\_FOXP3\_KO\_TREG\_PRECURSOR\_DN | MSigDB lists | GSE40685\_TREG\_VS\_FOXP3\_KO\_TREG\_PRECURSOR\_DN | 146 | 12 | 12187 | 179 | Tnf,Ifit2,Socs3,Parp14,Sntb2,Casp4,Znfx1,Ifitm3,Ifitm2,P2ry6,Gbp3,Tnfsf10 | | 1.561e-06 | -13.37 | GSE15330\_LYMPHOID\_MULTIPOTENT\_VS\_PRO\_BCELL\_DN | MSigDB lists | GSE15330\_LYMPHOID\_MULTIPOTENT\_VS\_PRO\_BCELL\_DN | 173 | 13 | 12187 | 179 | Bcl3,Tnfaip8,Casp4,Maff,Ccnd2,Parp9,Clic4,Trim25,Il17ra,Socs3,Sgk1,Ms4a6d,Akap12 | | 1.586e-06 | -13.35 | negative regulation of apoptotic process | biological process | GO:0043066 | 743 | 30 | 13711 | 214 | Cflar,Tnfaip8,Ripk1,Hspb1,Il1rn,Tnf,Tnfaip3,Bcl3,Angptl4,Nfe2l2,Sgk3,Ccl12,Ptpn1,Hspa5,C5ar1,Pik3r1,Ccnd2,Socs3,Thbs1,Cdkn1a,Il6,Mt1,Icam1,Ier3,Plaur,Ptgs2,Sgk1,Nfkb1,Birc3,Timp1 | | 1.656e-06 | -13.31 | GERHOLD\_ADIPOGENESIS\_DN | MSigDB lists | GERHOLD\_ADIPOGENESIS\_DN | 57 | 8 | 12187 | 179 | Xdh,Ier3,Cebpd,Ccl12,Il4ra,Thbs1,Ccnd2,Dnajb1 | | 1.665e-06 | -13.31 | GO\_REGULATION\_OF\_VASCULAR\_ENDOTHELIAL\_GROWTH\_FACTOR\_PRODUCTION | MSigDB lists | GO\_REGULATION\_OF\_VASCULAR\_ENDOTHELIAL\_GROWTH\_FACTOR\_PRODUCTION | 26 | 6 | 12187 | 179 | C5ar1,C3ar1,Ptgs2,Il1a,Ccl12,Il6 | | 1.665e-06 | -13.31 | GSE41176\_WT\_VS\_TAK1\_KO\_ANTI\_IGM\_STIM\_BCELL\_6H\_DN | MSigDB lists | GSE41176\_WT\_VS\_TAK1\_KO\_ANTI\_IGM\_STIM\_BCELL\_6H\_DN | 174 | 13 | 12187 | 179 | Socs3,Nfkb1,Ptgs2,Tnip1,Birc3,Il4ra,Nfkbia,Ccl4,Tnfaip3,C3ar1,Cflar,Tnfaip8,Igsf6 | | 1.665e-06 | -13.31 | GSE17721\_POLYIC\_VS\_PAM3CSK4\_1H\_BMDC\_DN | MSigDB lists | GSE17721\_POLYIC\_VS\_PAM3CSK4\_1H\_BMDC\_DN | 174 | 13 | 12187 | 179 | Slfn3,Zfp36,Icam1,Slfn4,Errfi1,Tnip1,Hcar2,Tiparp,Ccl12,Birc3,Trib1,Ccl2,Tgm2 | | 1.665e-06 | -13.31 | BILD\_HRAS\_ONCOGENIC\_SIGNATURE | MSigDB lists | BILD\_HRAS\_ONCOGENIC\_SIGNATURE | 174 | 13 | 12187 | 179 | Nfkbiz,Timp1,Il1a,Akap12,Ier3,Ptgs2,Zfp36,Plaur,Runx1,Sdc4,Trib1,Cxcl1,Angptl4 | | 1.724e-06 | -13.27 | GSE18791\_CTRL\_VS\_NEWCASTLE\_VIRUS\_DC\_18H\_DN | MSigDB lists | GSE18791\_CTRL\_VS\_NEWCASTLE\_VIRUS\_DC\_18H\_DN | 122 | 11 | 12187 | 179 | Irf7,Dtx3l,Tnfsf10,Gbp3,Parp14,Tnf,Ptgs2,Herc6,Tap1,Adamts1,Znfx1 | | 1.776e-06 | -13.24 | GSE39152\_BRAIN\_VS\_SPLEEN\_CD103\_NEG\_MEMORY\_CD8\_TCELL\_DN | MSigDB lists | GSE39152\_BRAIN\_VS\_SPLEEN\_CD103\_NEG\_MEMORY\_CD8\_TCELL\_DN | 175 | 13 | 12187 | 179 | Clic4,Csf2rb,Ccdc58,Ptpn1,Il17ra,Icam1,Atf3,Gadd45g,Gbp7,Bach1,Ccl4,Casp4,Il4ra | | 1.776e-06 | -13.24 | GSE22196\_HEALTHY\_VS\_OBESE\_MOUSE\_SKIN\_GAMMADELTA\_TCELL\_DN | MSigDB lists | GSE22196\_HEALTHY\_VS\_OBESE\_MOUSE\_SKIN\_GAMMADELTA\_TCELL\_DN | 175 | 13 | 12187 | 179 | Ccl4,Tnfaip3,Cxcl1,S100a9,Gem,S100a8,Thbs1,Atf3,Ier3,Tiparp,Nfkb1,Tnip1,Plaur | | 1.776e-06 | -13.24 | GSE17721\_0.5H\_VS\_8H\_POLYIC\_BMDC\_DN | MSigDB lists | GSE17721\_0.5H\_VS\_8H\_POLYIC\_BMDC\_DN | 175 | 13 | 12187 | 179 | Slfn3,Trim25,Msn,Cdkn1a,Adamts1,Il1rn,Atf3,Luc7l3,Irgm2,Slfn4,Map3k8,Tgm2,Hcar2 | | 1.779e-06 | -13.24 | GO\_NEGATIVE\_REGULATION\_OF\_CYSTEINE\_TYPE\_ENDOPEPTIDASE\_ACTIVITY | MSigDB lists | GO\_NEGATIVE\_REGULATION\_OF\_CYSTEINE\_TYPE\_ENDOPEPTIDASE\_ACTIVITY | 77 | 9 | 12187 | 179 | Thbs1,Birc3,Ifi209,Ifi207,Ifi204,Il6,Ifi211,Tnfaip8,Plaur | | 1.794e-06 | -13.23 | GSE28737\_FOLLICULAR\_VS\_MARGINAL\_ZONE\_BCELL\_BCL6\_HET\_DN | MSigDB lists | GSE28737\_FOLLICULAR\_VS\_MARGINAL\_ZONE\_BCELL\_BCL6\_HET\_DN | 148 | 12 | 12187 | 179 | Dtx3l,Gbp6,Herc6,Angptl4,Irf7,Slfn4,Slfn3,Tap1,Msn,Ccnd2,Gpr84,Gbp4 | | 1.820e-06 | -13.22 | GRUETZMANN\_PANCREATIC\_CANCER\_UP | MSigDB lists | GRUETZMANN\_PANCREATIC\_CANCER\_UP | 298 | 17 | 12187 | 179 | Ifitm2,Ptgs2,Msn,Tap1,Plaur,Atf3,Lcn2,Akap12,Il1rn,Gem,Cflar,Rgs16,Iqgap1,Casp4,Il4ra,Birc3,Tnfaip3 | | 1.831e-06 | -13.21 | receptor ligand activity | molecular function | GO:0048018 | 249 | 16 | 13516 | 211 | Cxcl16,Saa1,Ccl4,Il6,Cxcl9,Retnlg,Ccl2,Tnf,Ccl12,Ccl7,Il1a,Il1rn,Csf3,Cxcl1,Timp1,Tnfsf10 | | 1.834e-06 | -13.21 | GBP\_C | interpro domains | IPR037684 | 7 | 4 | 13788 | 212 | Gbp3,Gbp7,Gbp6,Gbp4 | | 1.870e-06 | -13.19 | WU\_CELL\_MIGRATION | MSigDB lists | WU\_CELL\_MIGRATION | 123 | 11 | 12187 | 179 | Col4a1,Il1a,Akap12,Tubb6,Thbs1,Adamts1,Tgm2,Msn,Ptgs2,Arid5b,Ptges | | 1.924e-06 | -13.16 | CHICAS\_RB1\_TARGETS\_SENESCENT | MSigDB lists | CHICAS\_RB1\_TARGETS\_SENESCENT | 440 | 21 | 12187 | 179 | Il1a,Akap12,Rasd1,Sox11,Vcam1,Nfkbiz,Ifit3b,Irf9,Ptgs2,Isg15,Tnfaip3,AA467197,Il6,Birc3,Tnfaip8,Rbm39,Osmr,Dtx3l,Gem,Csf3,Cxcl1 | | 1.926e-06 | -13.16 | GSE28737\_WT\_VS\_BCL6\_HET\_MARGINAL\_ZONE\_BCELL\_DN | MSigDB lists | GSE28737\_WT\_VS\_BCL6\_HET\_MARGINAL\_ZONE\_BCELL\_DN | 149 | 12 | 12187 | 179 | Stx11,Cdkn1a,Pik3ap1,Rbm47,Icam1,Isg15,Rhoc,Tnfaip3,Ripk1,Casp4,Trib1,Gbp7 | | 1.934e-06 | -13.16 | AMIT\_EGF\_RESPONSE\_40\_HELA | MSigDB lists | AMIT\_EGF\_RESPONSE\_40\_HELA | 41 | 7 | 12187 | 179 | Sgk1,Il6,Atf3,Ier3,Ptgs2,Bcl3,Zfp36 | | 1.934e-06 | -13.16 | LEE\_LIVER\_CANCER\_DENA\_UP | MSigDB lists | LEE\_LIVER\_CANCER\_DENA\_UP | 41 | 7 | 12187 | 179 | S100a9,Tagln2,Plek,Thbs1,Vcam1,Col4a1,Lcn2 | | 1.934e-06 | -13.16 | MODULE\_254 | MSigDB lists | MODULE\_254 | 41 | 7 | 12187 | 179 | Ccnd2,Birc3,Nfkb1,Il1a,Ier3,Cflar,Tnfsf10 | | 2.006e-06 | -13.12 | regulation of response to cytokine stimulus | biological process | GO:0060759 | 94 | 10 | 13711 | 214 | Irgm2,Parp14,Ripk1,Il1rn,Irak3,Irgm1,Irf7,Parp9,Il6,Casp4 | | 2.018e-06 | -13.11 | SENESE\_HDAC1\_AND\_HDAC2\_TARGETS\_UP | MSigDB lists | SENESE\_HDAC1\_AND\_HDAC2\_TARGETS\_UP | 177 | 13 | 12187 | 179 | Nfkbiz,Il1a,Plaur,Tgm2,Maff,Birc3,Ccnd2,Tnfaip3,Robo4,Gem,Nfkb2,Osmr,Iqgap1 | | 2.067e-06 | -13.09 | GSE25123\_IL4\_VS\_IL4\_AND\_ROSIGLITAZONE\_STIM\_PPARG\_KO\_MACROPHAGE\_DAY10\_DN | MSigDB lists | GSE25123\_IL4\_VS\_IL4\_AND\_ROSIGLITAZONE\_STIM\_PPARG\_KO\_MACROPHAGE\_DAY10\_DN | 150 | 12 | 12187 | 179 | AA467197,Gpr84,Tnf,Cxcl1,Il1rn,Timp1,Il1a,Aff1,Errfi1,Trim25,Pik3ap1,Ptgs2 | | 2.114e-06 | -13.07 | regulation of viral life cycle | biological process | GO:1903900 | 117 | 11 | 13711 | 214 | Isg15,Ifitm2,Trim56,Resf1,Trim25,Ifitm6,Tnf,Oasl2,Trim30a,Oasl1,Ifitm3 | | 2.209e-06 | -13.02 | MODULE\_88 | MSigDB lists | MODULE\_88 | 520 | 23 | 12187 | 179 | Ccnd2,Il4ra,Tgm2,Trib1,Ccl12,Fam107a,S100a9,S100a8,Xdh,Sbno2,Tnfsf10,Cdkn1a,Il1rn,Kcna5,Col4a1,Hp,Cp,Gadd45g,Ier3,Lcn2,Isg15,Cd14,Sele | | 2.218e-06 | -13.02 | GSE9988\_ANTI\_TREM1\_VS\_CTRL\_TREATED\_MONOCYTES\_UP | MSigDB lists | GSE9988\_ANTI\_TREM1\_VS\_CTRL\_TREATED\_MONOCYTES\_UP | 151 | 12 | 12187 | 179 | Sgk1,Cdkn1a,Atf3,Rasd1,Ptgs2,Plaur,Tnf,Trib1,Gadd45b,Gem,Banp,Cxcl1 | | 2.245e-06 | -13.01 | Fadd (Fas (TNFRSF6)-associated via death domain) | protein interactions | 14082 | 14 | 5 | 6802 | 115 | Tnf,Cflar,Ripk1,Tnfaip3,Birc3 | | 2.255e-06 | -13.00 | signaling receptor activator activity | molecular function | GO:0030546 | 253 | 16 | 13516 | 211 | Saa1,Cxcl16,Ccl4,Ccl2,Retnlg,Cxcl9,Il6,Ccl12,Tnf,Ccl7,Csf3,Il1rn,Il1a,Tnfsf10,Timp1,Cxcl1 | | 2.293e-06 | -12.99 | LEE\_LIVER\_CANCER\_E2F1\_UP | MSigDB lists | LEE\_LIVER\_CANCER\_E2F1\_UP | 42 | 7 | 12187 | 179 | Xdh,Lcn2,Tagln2,Casp4,Usp18,Ifit2,Ifi44 | | 2.293e-06 | -12.99 | KHETCHOUMIAN\_TRIM24\_TARGETS\_UP | MSigDB lists | KHETCHOUMIAN\_TRIM24\_TARGETS\_UP | 42 | 7 | 12187 | 179 | Icam1,Cybb,Col4a1,Cp,Tgm2,Vcam1,Cdkn1a | | 2.320e-06 | -12.97 | cell death | biological process | GO:0008219 | 757 | 30 | 13711 | 214 | Tnf,Tnfaip3,Tnfsf10,Irak3,Ncf1,Ripk1,Tnfaip8,Cflar,Ifi204,Lcn2,C5ar1,Rrp8,S100a8,S100a9,Bcl3,Ifit2,Casp4,Il1a,Il6,Cdkn1a,Gadd45b,Pik3r1,Gadd45g,Hcar2,Birc3,Nfkb1,Xaf1,Spi1,Sgk1,Ier3 | | 2.352e-06 | -12.96 | LU\_AGING\_BRAIN\_UP | MSigDB lists | LU\_AGING\_BRAIN\_UP | 239 | 15 | 12187 | 179 | Ifi209,Aff1,Tiparp,Ifi204,Ifi207,Hp,Cebpd,Clic4,Msn,Tgm2,Nfkbia,Nfe2l2,Ifi211,Irf7,Iqgap1 | | 2.373e-06 | -12.95 | PETROVA\_ENDOTHELIUM\_LYMPHATIC\_VS\_BLOOD\_DN | MSigDB lists | PETROVA\_ENDOTHELIUM\_LYMPHATIC\_VS\_BLOOD\_DN | 126 | 11 | 12187 | 179 | Icam1,Sele,Selp,Isg15,Ccl12,Fam107a,Ier3,Il6,Cdkn1a,Il4ra,Tgm2 | | 2.377e-06 | -12.95 | GSE22935\_24H\_VS\_48H\_MBOVIS\_BCG\_STIM\_MACROPHAGE\_UP | MSigDB lists | GSE22935\_24H\_VS\_48H\_MBOVIS\_BCG\_STIM\_MACROPHAGE\_UP | 152 | 12 | 12187 | 179 | Gbp4,Ms4a6d,Parp14,Birc3,Ifit2,Rgs16,Trim25,Slfn9,Gbp6,Gbp7,Dtx3l,Irgm2 | | 2.377e-06 | -12.95 | GSE17721\_POLYIC\_VS\_PAM3CSK4\_2H\_BMDC\_DN | MSigDB lists | GSE17721\_POLYIC\_VS\_PAM3CSK4\_2H\_BMDC\_DN | 152 | 12 | 12187 | 179 | Gpr84,Pik3r1,Rhou,Map3k8,Igsf6,Nfkbiz,Thbs1,Il1rn,Nfkb1,Ptgs2,Errfi1,Il17ra | | 2.377e-06 | -12.95 | GSE42724\_NAIVE\_VS\_MEMORY\_BCELL\_UP | MSigDB lists | GSE42724\_NAIVE\_VS\_MEMORY\_BCELL\_UP | 152 | 12 | 12187 | 179 | Nfkbie,Gbp7,Cflar,Tnfsf10,Msr1,Birc3,Hcar2,Ptgs2,Tnip1,Plin4,Cxcl16,Hp | | 2.383e-06 | -12.95 | negative regulation of programmed cell death | biological process | GO:0043069 | 758 | 30 | 13711 | 214 | Ccnd2,Pik3r1,Thbs1,Socs3,Cdkn1a,Il6,Icam1,Mt1,Plaur,Ier3,Sgk1,Ptgs2,Nfkb1,Birc3,Timp1,Ripk1,Tnfaip8,Cflar,Il1rn,Hspb1,Tnfaip3,Tnf,Angptl4,Bcl3,Nfe2l2,Sgk3,Ccl12,Ptpn1,Hspa5,C5ar1 | | 2.391e-06 | -12.94 | regulation of vascular endothelial growth factor production | biological process | GO:0010574 | 26 | 6 | 13711 | 214 | Il1a,C5ar1,Ccl2,Il6,C3ar1,Ptgs2 | | 2.391e-06 | -12.94 | regulation of viral entry into host cell | biological process | GO:0046596 | 26 | 6 | 13711 | 214 | Ifitm2,Ifitm6,Ifitm3,Trim30a,Trim25,Trim56 | | 2.391e-06 | -12.94 | regulation of epithelial cell apoptotic process | biological process | GO:1904035 | 75 | 9 | 13711 | 214 | Icam1,Tnf,Tnfaip3,Zfp36,Ccl12,Thbs1,Nfe2l2,Cflar,Angptl4 | | 2.462e-06 | -12.91 | positive regulation of response to cytokine stimulus | biological process | GO:0060760 | 40 | 7 | 13711 | 214 | Casp4,Parp9,Ripk1,Parp14,Irgm1,Irf7,Irgm2 | | 2.462e-06 | -12.91 | negative regulation of innate immune response | biological process | GO:0045824 | 40 | 7 | 13711 | 214 | Acod1,Tnfaip3,A2m,Tap1,Parp14,Irak3,Lgals9 | | 2.470e-06 | -12.91 | GO\_REGULATION\_OF\_CYTOKINE\_BIOSYNTHETIC\_PROCESS | MSigDB lists | GO\_REGULATION\_OF\_CYTOKINE\_BIOSYNTHETIC\_PROCESS | 60 | 8 | 12187 | 179 | Thbs1,Tnf,Il6,Il1a,Nfkb1,Bcl3,Errfi1,Hspb1 | | 2.471e-06 | -12.91 | GO\_NEGATIVE\_REGULATION\_OF\_CATALYTIC\_ACTIVITY | MSigDB lists | GO\_NEGATIVE\_REGULATION\_OF\_CATALYTIC\_ACTIVITY | 603 | 25 | 12187 | 179 | Ptpn1,Irak3,Thbs1,Nfkb1,Angptl4,Ifi211,Tnfaip8,Iqgap1,Il6,Gadd45b,Tnfaip3,Errfi1,Plaur,Hspb1,Socs3,Ifi209,Cdkn1a,Ifi207,Hp,Ifi204,Timp1,A2m,Trib1,Birc3,Tnf | | 2.539e-06 | -12.88 | regulation of hemopoiesis | biological process | GO:1903706 | 318 | 18 | 13711 | 214 | Zfp36,Il6,Il4ra,Tnf,Nfkbia,Pik3r1,Lgals9,Csf3,Ripk1,Trib1,Ctla2a,Zbtb16,Isg15,Cxcl1,Nfkbiz,Runx1,Nfe2l2,Spi1 | | 2.547e-06 | -12.88 | GSE17721\_CTRL\_VS\_PAM3CSK4\_1H\_BMDC\_DN | MSigDB lists | GSE17721\_CTRL\_VS\_PAM3CSK4\_1H\_BMDC\_DN | 153 | 12 | 12187 | 179 | Igsf6,Slfn4,Cflar,Hcar2,Map3k8,Birc3,Ccl2,Icam1,Slfn3,Errfi1,Atf3,Cdkn1a | | 2.547e-06 | -12.88 | GSE13485\_DAY1\_VS\_DAY3\_YF17D\_VACCINE\_PBMC\_DN | MSigDB lists | GSE13485\_DAY1\_VS\_DAY3\_YF17D\_VACCINE\_PBMC\_DN | 153 | 12 | 12187 | 179 | Phf11b,Ifi44,Ifit3b,Sgk3,Herc6,Isg15,Parp9,Tap1,Xaf1,Usp18,Phf11d,Cmpk2 | | 2.547e-06 | -12.88 | GSE10325\_CD4\_TCELL\_VS\_MYELOID\_DN | MSigDB lists | GSE10325\_CD4\_TCELL\_VS\_MYELOID\_DN | 153 | 12 | 12187 | 179 | Cdkn1a,Msr1,Trib1,Cybb,Tubb6,Nfkbie,Icam1,Plek,Rbm47,Clic4,Csf2rb,Plaur | | 2.547e-06 | -12.88 | GSE17721\_0.5H\_VS\_4H\_LPS\_BMDC\_DN | MSigDB lists | GSE17721\_0.5H\_VS\_4H\_LPS\_BMDC\_DN | 153 | 12 | 12187 | 179 | Trim25,Rbm47,Timp1,Nfkb1,Iqgap1,Irgm2,Parp14,Tor3a,Ccl2,Tnf,Zfp189,Birc3 | | 2.590e-06 | -12.86 | GSE43957\_UNTREATED\_VS\_NACL\_TREATED\_ANTI\_CD3\_CD28\_STIM\_CD4\_TCELL\_DN | MSigDB lists | GSE43957\_UNTREATED\_VS\_NACL\_TREATED\_ANTI\_CD3\_CD28\_STIM\_CD4\_TCELL\_DN | 181 | 13 | 12187 | 179 | Errfi1,Clic4,Sele,Il17ra,Icam1,Saa1,Selp,Gadd45b,Bach1,Tubb6,Ccl2,Il4ra,Map3k8 | | 2.599e-06 | -12.86 | P-loop\_NTPase | interpro domains | IPR027417 | 732 | 29 | 13788 | 212 | Tap1,Gbp7,F830016B08Rik,Ifi47,Rhoc,Ifi44,Cmpk2,Rhou,Tor3a,Gem,Mx2,Slfn5,9930111J21Rik2,Gm5431,Tgtp1,Gbp3,Rasd1,Iqgap1,Sbno2,Iigp1,Slfn9,Gbp6,Mx1,Znfx1,Irgm1,Irgm2,Gbp4,Rhoj,Gm4841 | | 2.611e-06 | -12.86 | regulation of smooth muscle cell proliferation | biological process | GO:0048660 | 144 | 12 | 13711 | 214 | Tgm2,Adamts1,Ptgs2,Tnf,Tnfaip3,Cdkn1a,Il6,Trib1,Thbs1,C3ar1,P2ry6,Pik3r1 | | 2.627e-06 | -12.85 | GO\_REGULATION\_OF\_HYDROLASE\_ACTIVITY | MSigDB lists | GO\_REGULATION\_OF\_HYDROLASE\_ACTIVITY | 995 | 34 | 12187 | 179 | Hspa5,Thbs1,Dnajb1,Rasip1,Ptpn1,Ccl2,Ccl4,Rhoc,Il6,Ccl12,Ripk1,Angptl4,Ifi211,S100a8,Tnfaip8,Iqgap1,Rgs16,Ifi209,Ifi207,Timp1,Ifi204,Sele,Plek,Icam1,Plaur,Csf2rb,Errfi1,Birc3,Tnf,C5ar1,S100a9,A2m,Tnfsf10,Xdh | | 2.638e-06 | -12.85 | NABA\_MATRISOME | MSigDB lists | NABA\_MATRISOME | 565 | 24 | 12187 | 179 | Fstl1,Adamts1,Col4a1,Timp1,Il1rn,S100a9,A2m,Tnfsf10,Sdc4,Lgals9,Tnf,Thbs1,Il1a,Mmp8,Cxcl1,Angptl4,Csf3,S100a8,Adamts9,Ccl2,Ccl4,Tgm2,Il6,Ccl12 | | 2.659e-06 | -12.84 | AMIT\_SERUM\_RESPONSE\_40\_MCF10A | MSigDB lists | AMIT\_SERUM\_RESPONSE\_40\_MCF10A | 28 | 6 | 12187 | 179 | Atf3,Clic4,Zfp36,Il1a,Cxcl1,Thbs1 | | 2.662e-06 | -12.84 | regulation of kinase activity | biological process | GO:0043549 | 682 | 28 | 13711 | 214 | Sdc4,Map3k8,Map3k6,Fzd4,Pik3r5,Rasip1,Il6,Errfi1,Cdkn1a,Pik3r1,Gadd45g,Ccnd2,Gadd45b,Lgals9,Thbs1,Trib1,Iqgap1,C5ar1,Ptpn1,Irgm2,Ksr1,Tnfaip3,Tnf,Hspb1,Il1rn,Irak3,Ncf1,Ripk1 | | 2.727e-06 | -12.81 | TONKS\_TARGETS\_OF\_RUNX1\_RUNX1T1\_FUSION\_HSC\_UP | MSigDB lists | TONKS\_TARGETS\_OF\_RUNX1\_RUNX1T1\_FUSION\_HSC\_UP | 154 | 12 | 12187 | 179 | Ifi211,Irf7,Arid5b,Nfkb2,Runx1,Rhoc,Hspb1,Irf9,Ifi209,Fstl1,Ifi204,Ifi207 | | 2.727e-06 | -12.81 | GSE17974\_CTRL\_VS\_ACT\_IL4\_AND\_ANTI\_IL12\_72H\_CD4\_TCELL\_UP | MSigDB lists | GSE17974\_CTRL\_VS\_ACT\_IL4\_AND\_ANTI\_IL12\_72H\_CD4\_TCELL\_UP | 154 | 12 | 12187 | 179 | Maff,Casp4,Trib1,Dtx3l,Pik3r5,S100a8,Gbp3,Sgk1,Nfkbiz,Socs3,Herc6,Plek | | 2.841e-06 | -12.77 | GO\_CELLULAR\_HOMEOSTASIS | MSigDB lists | GO\_CELLULAR\_HOMEOSTASIS | 489 | 22 | 12187 | 179 | S100a9,C5ar1,C3ar1,Kcna5,Cp,Ncf1,Saa1,Clic4,Icam1,Tgm2,Ccl2,Pik3r1,Ccl12,Nfe2l2,Il6,Slc24a4,Cybb,Pygm,S100a8,Sgk1,Lcn2,Sgk3 | | 2.855e-06 | -12.77 | DANG\_REGULATED\_BY\_MYC\_DN | MSigDB lists | DANG\_REGULATED\_BY\_MYC\_DN | 212 | 14 | 12187 | 179 | Irf7,A2m,Vcam1,Fstl1,Thbs1,Cdkn1a,Ier3,Akap12,Gadd45g,Msn,Ptpn1,Errfi1,Plaur,Hspb1 | | 2.879e-06 | -12.76 | CHIARADONNA\_NEOPLASTIC\_TRANSFORMATION\_KRAS\_UP | MSigDB lists | CHIARADONNA\_NEOPLASTIC\_TRANSFORMATION\_KRAS\_UP | 104 | 10 | 12187 | 179 | Vcam1,Casp4,Nfkbia,Ccl12,Nfe2l2,Tubb6,Tiparp,Timp1,Bach1,Rgs16 | | 2.919e-06 | -12.74 | GSE45365\_NK\_CELL\_VS\_CD8\_TCELL\_MCMV\_INFECTION\_DN | MSigDB lists | GSE45365\_NK\_CELL\_VS\_CD8\_TCELL\_MCMV\_INFECTION\_DN | 155 | 12 | 12187 | 179 | Nfkb2,Gadd45b,Tnfaip3,Rassf4,Nfkbia,Sdc4,Birc3,Tap1,Ptpn1,Tnip1,Nfkb1,Cdkn1a | | 2.930e-06 | -12.74 | acute inflammatory response | biological process | GO:0002526 | 41 | 7 | 13711 | 214 | Cxcl1,Icam1,Il1a,Hp,Il6,Saa1,Il1rn | | 3.009e-06 | -12.71 | GO\_LEUKOCYTE\_MIGRATION\_INVOLVED\_IN\_INFLAMMATORY\_RESPONSE | MSigDB lists | GO\_LEUKOCYTE\_MIGRATION\_INVOLVED\_IN\_INFLAMMATORY\_RESPONSE | 8 | 4 | 12187 | 179 | Sele,S100a8,Ccl12,S100a9 | | 3.025e-06 | -12.71 | regulation of transferase activity | biological process | GO:0051338 | 767 | 30 | 13711 | 214 | Lgals9,Pik3r1,Ccnd2,Gadd45g,Gadd45b,Trib1,Dtx3l,Thbs1,Cdkn1a,Il6,Errfi1,Pik3r5,Rasip1,Sdc4,Map3k6,Fzd4,Map3k8,Ripk1,Hspb1,Il1rn,Irak3,Ncf1,Ksr1,Zfp36,Tnfaip3,Tnf,Irgm2,Iqgap1,Ptpn1,C5ar1 | | 3.063e-06 | -12.70 | BLALOCK\_ALZHEIMERS\_DISEASE\_UP | MSigDB lists | BLALOCK\_ALZHEIMERS\_DISEASE\_UP | 1384 | 42 | 12187 | 179 | Tnfsf10,Igsf6,Irf7,Cflar,Gem,Oasl1,Casp4,Nfkbia,Icam1,Zfp36,Tnip1,Ifitm2,Luc7l3,Ifi204,Ifi207,Cp,Vcam1,Socs3,Zbtb16,Ifi209,Aff1,Rgs16,Adamts9,Iqgap1,Tnfaip8,Banp,Cxcl1,Ifi211,Tnfaip3,Fam107a,Gadd45b,Il6,Maff,Tap1,Il17ra,Mxd4,Cebpd,Tiparp,Nfkb1,Il1a,Gadd45g,Sgk1 | | 3.079e-06 | -12.69 | host intracellular part | cellular component | GO:0033646 | 16 | 5 | 13825 | 212 | Irgm2,Gbp6,Iigp1,Gbp3,Gbp7 | | 3.079e-06 | -12.69 | intracellular region of host | cellular component | GO:0043656 | 16 | 5 | 13825 | 212 | Gbp3,Iigp1,Gbp7,Irgm2,Gbp6 | | 3.116e-06 | -12.68 | MARTINELLI\_IMMATURE\_NEUTROPHIL\_UP | MSigDB lists | MARTINELLI\_IMMATURE\_NEUTROPHIL\_UP | 3 | 3 | 12187 | 179 | Cybb,Mmp8,Lcn2 | | 3.122e-06 | -12.68 | GSE17721\_12H\_VS\_24H\_CPG\_BMDC\_UP | MSigDB lists | GSE17721\_12H\_VS\_24H\_CPG\_BMDC\_UP | 156 | 12 | 12187 | 179 | Rhoc,Gbp4,Msr1,Pik3r1,Ccl2,Trib1,Gbp6,Ptges,Il1a,Dnajb1,Zfp36,Clic4 | | 3.177e-06 | -12.66 | GO\_REGULATION\_OF\_EXTRINSIC\_APOPTOTIC\_SIGNALING\_PATHWAY\_VIA\_DEATH\_DOMAIN\_RECEPTORS | MSigDB lists | GO\_REGULATION\_OF\_EXTRINSIC\_APOPTOTIC\_SIGNALING\_PATHWAY\_VIA\_DEATH\_DOMAIN\_RECEPTORS | 44 | 7 | 12187 | 179 | Cflar,Icam1,Tnfsf10,Thbs1,Atf3,Ripk1,Tnfaip3 | | 3.177e-06 | -12.66 | FURUKAWA\_DUSP6\_TARGETS\_PCI35\_UP | MSigDB lists | FURUKAWA\_DUSP6\_TARGETS\_PCI35\_UP | 44 | 7 | 12187 | 179 | Phf11b,AA467197,Phf11d,Herc6,Saa1,Parp9,Rtp4 | | 3.317e-06 | -12.62 | GO\_REGULATION\_OF\_MULTI\_ORGANISM\_PROCESS | MSigDB lists | GO\_REGULATION\_OF\_MULTI\_ORGANISM\_PROCESS | 346 | 18 | 12187 | 179 | Ifi209,Ifi207,Ifi204,Timp1,Tnip1,Ifitm2,Isg15,Ifitm3,Tap1,Trim25,Trib1,Ccl4,Birc3,Tnf,Zfp189,Oasl1,Tnfaip3,Ifi211 | | 3.338e-06 | -12.61 | GSE29949\_MICROGLIA\_BRAIN\_VS\_MONOCYTE\_BONE\_MARROW\_DN | MSigDB lists | GSE29949\_MICROGLIA\_BRAIN\_VS\_MONOCYTE\_BONE\_MARROW\_DN | 157 | 12 | 12187 | 179 | Il6,Gem,Nfkbie,Ifi211,Cxcl1,Ifi44,Ifi209,Ifi207,Ifi204,Mmp8,Atf3,Isg15 | | 3.338e-06 | -12.61 | GO\_NEGATIVE\_REGULATION\_OF\_APOPTOTIC\_SIGNALING\_PATHWAY | MSigDB lists | GO\_NEGATIVE\_REGULATION\_OF\_APOPTOTIC\_SIGNALING\_PATHWAY | 157 | 12 | 12187 | 179 | Cflar,Tnfsf10,Tnf,Ripk1,Tnfaip3,Nfe2l2,Ptpn1,Plaur,Hspb1,Icam1,Thbs1,Il1a | | 3.338e-06 | -12.61 | GSE11961\_FOLLICULAR\_BCELL\_VS\_GERMINAL\_CENTER\_BCELL\_DAY7\_DN | MSigDB lists | GSE11961\_FOLLICULAR\_BCELL\_VS\_GERMINAL\_CENTER\_BCELL\_DAY7\_DN | 157 | 12 | 12187 | 179 | Ifi44,Cdkn1a,Znfx1,Rnf213,Slfn9,Pik3ap1,Il4ra,Ccnd2,Trib1,Ripk1,Dtx3l,Fzd4 | | 3.356e-06 | -12.60 | positive regulation of hemostasis | biological process | GO:1900048 | 16 | 5 | 13711 | 214 | Selp,Nfe2l2,Thbs1,Plek,S100a9 | | 3.356e-06 | -12.60 | positive regulation of blood coagulation | biological process | GO:0030194 | 16 | 5 | 13711 | 214 | Selp,Plek,S100a9,Nfe2l2,Thbs1 | | 3.361e-06 | -12.60 | MODULE\_436 | MSigDB lists | MODULE\_436 | 83 | 9 | 12187 | 179 | Parp14,Thbs1,Cebpd,Cd14,Ifitm3,Isg15,Csf3,Tnfsf10,Tap1 | | 3.477e-06 | -12.57 | TONKS\_TARGETS\_OF\_RUNX1\_RUNX1T1\_FUSION\_ERYTHROCYTE\_UP | MSigDB lists | TONKS\_TARGETS\_OF\_RUNX1\_RUNX1T1\_FUSION\_ERYTHROCYTE\_UP | 131 | 11 | 12187 | 179 | Tubb6,Il1rn,Ifi204,Ifi207,Ifi209,Ifi44,Socs3,Hspb1,Irak3,C3ar1,Ifi211 | | 3.509e-06 | -12.56 | GO\_REGULATION\_OF\_KINASE\_ACTIVITY | MSigDB lists | GO\_REGULATION\_OF\_KINASE\_ACTIVITY | 615 | 25 | 12187 | 179 | Map3k6,Ptpn1,Irak3,Hspa5,Thbs1,Gadd45g,Pik3r5,Iqgap1,Pik3r1,Ccnd2,Map3k8,Il6,Gadd45b,Ripk1,Tnfaip3,Saa1,Errfi1,Hspb1,Socs3,Cdkn1a,Fzd4,C5ar1,Trib1,Sdc4,Tnf | | 3.566e-06 | -12.54 | GSE16385\_ROSIGLITAZONE\_IFNG\_TNF\_VS\_IL4\_STIM\_MACROPHAGE\_UP | MSigDB lists | GSE16385\_ROSIGLITAZONE\_IFNG\_TNF\_VS\_IL4\_STIM\_MACROPHAGE\_UP | 158 | 12 | 12187 | 179 | Map3k8,Maff,Ccl4,Trib1,Tnfaip3,Gem,Adamts9,Hspa5,Nfkbiz,Dnajb1,Irak3,Zfp36 | | 3.566e-06 | -12.54 | GSE17721\_CTRL\_VS\_GARDIQUIMOD\_8H\_BMDC\_DN | MSigDB lists | GSE17721\_CTRL\_VS\_GARDIQUIMOD\_8H\_BMDC\_DN | 158 | 12 | 12187 | 179 | Gbp3,Pik3ap1,Il17ra,Igsf6,Cd14,Slfn9,Map3k6,Znfx1,Nfkb1,Nfkbia,Birc3,Ifit2 | | 3.586e-06 | -12.54 | symbiont-containing vacuole membrane | cellular component | GO:0020005 | 8 | 4 | 13825 | 212 | Gbp6,Gbp7,Gbp3,Iigp1 | | 3.602e-06 | -12.53 | BURTON\_ADIPOGENESIS\_2 | MSigDB lists | BURTON\_ADIPOGENESIS\_2 | 63 | 8 | 12187 | 179 | Rhou,Map3k6,Ptges,Rhoj,Timp1,Col4a1,Clic4,Xdh | | 3.602e-06 | -12.53 | MAHAJAN\_RESPONSE\_TO\_IL1A\_UP | MSigDB lists | MAHAJAN\_RESPONSE\_TO\_IL1A\_UP | 63 | 8 | 12187 | 179 | Cxcl1,Trim25,Thbs1,Ifit2,Il6,Oasl1,Ripk1,Il1a | | 3.624e-06 | -12.53 | HIN200/IF120x | interpro domains | IPR004021 | 8 | 4 | 13788 | 212 | Ifi209,Ifi211,Ifi204,Ifi207 | | 3.631e-06 | -12.53 | GO\_POSITIVE\_REGULATION\_OF\_CATALYTIC\_ACTIVITY | MSigDB lists | GO\_POSITIVE\_REGULATION\_OF\_CATALYTIC\_ACTIVITY | 1198 | 38 | 12187 | 179 | Gadd45b,Ripk1,Ccl12,Rhoc,Ccl2,Ccl4,Ccnd2,Map3k8,Rgs16,S100a8,Iqgap1,Ifi211,Pik3r5,Gadd45g,Dnajb1,Hspa5,Thbs1,Ptpn1,Map3k6,Sdc4,Tnf,Tnfsf10,Xdh,S100a9,A2m,Fzd4,C5ar1,Cflar,Ifi207,Ifi204,Cdkn1a,Ifi209,Errfi1,Csf2rb,Sele,Plek,Icam1,Saa1 | | 3.716e-06 | -12.50 | GO\_REGULATION\_OF\_ACUTE\_INFLAMMATORY\_RESPONSE | MSigDB lists | GO\_REGULATION\_OF\_ACUTE\_INFLAMMATORY\_RESPONSE | 45 | 7 | 12187 | 179 | Tnf,Il6,A2m,C3ar1,Ptgs2,C5ar1,Osmr | | 3.745e-06 | -12.50 | cellular response to toxic substance | biological process | GO:0097237 | 124 | 11 | 13711 | 214 | Tnf,Tnfaip3,Il6,Ccl7,Ripk1,Lcn2,Nfkb1,S100a8,Hp,Nfe2l2,S100a9 | | 3.809e-06 | -12.48 | GSE2405\_0H\_VS\_12H\_A\_PHAGOCYTOPHILUM\_STIM\_NEUTROPHIL\_UP | MSigDB lists | GSE2405\_0H\_VS\_12H\_A\_PHAGOCYTOPHILUM\_STIM\_NEUTROPHIL\_UP | 159 | 12 | 12187 | 179 | Tap1,Irf9,Slfn5,Nfkb1,Gadd45g,Ifit3b,Cmpk2,Irf7,Oasl1,Ccnd2,Parp14,Xaf1 | | 3.809e-06 | -12.48 | GSE360\_DC\_VS\_MAC\_DN | MSigDB lists | GSE360\_DC\_VS\_MAC\_DN | 159 | 12 | 12187 | 179 | S100a9,Irf7,Cd14,Ifi211,Pygm,Maff,Ifi209,Cp,Ifi207,Ifi204,Nfkb1,Atf3 | | 3.825e-06 | -12.47 | negative regulation of cell death | biological process | GO:0060548 | 859 | 32 | 13711 | 214 | Ier3,Plaur,Sgk1,Ptgs2,Nfkb1,Birc3,Timp1,Ccnd2,Pik3r1,Socs3,Thbs1,Cdkn1a,Il6,Sox11,Mt1,Icam1,Bcl3,Angptl4,Sgk3,Nfe2l2,Ccl12,Ptpn1,Hspa5,C5ar1,Cflar,Ripk1,Tnfaip8,Csf3,Hspb1,Il1rn,Tnf,Tnfaip3 | | 3.836e-06 | -12.47 | HIN | pfam domains | PF02760 | 8 | 4 | 12881 | 201 | Ifi211,Ifi204,Ifi207,Ifi209 | | 3.919e-06 | -12.45 | regulation of endopeptidase activity | biological process | GO:0052548 | 264 | 16 | 13711 | 214 | Casp4,Tnf,Il6,Tnfsf10,Xdh,Thbs1,Cflar,Tnfaip8,Timp1,Birc3,A2m,S100a8,Serpina3f,S100a9,Ptgs2,Plaur | | 4.079e-06 | -12.41 | GO\_NEGATIVE\_REGULATION\_OF\_MOLECULAR\_FUNCTION | MSigDB lists | GO\_NEGATIVE\_REGULATION\_OF\_MOLECULAR\_FUNCTION | 790 | 29 | 12187 | 179 | Ptpn1,Irak3,Nfkb1,Thbs1,Hspa5,Sox11,Iqgap1,Tnfaip8,Ifi211,Angptl4,Il6,Gadd45b,Tnfaip3,Ifit2,Plaur,Errfi1,Hspb1,Ifi207,Hp,Ifi204,Timp1,Socs3,Cdkn1a,Ifi209,A2m,Nfkbia,Trib1,Birc3,Tnf | | 4.090e-06 | -12.41 | PID\_AMB2\_NEUTROPHILS\_PATHWAY | MSigDB lists | PID\_AMB2\_NEUTROPHILS\_PATHWAY | 30 | 6 | 12187 | 179 | Icam1,Nfkb1,Il6,Plaur,Tnf,Selp | | 4.090e-06 | -12.41 | AMIT\_EGF\_RESPONSE\_60\_MCF10A | MSigDB lists | AMIT\_EGF\_RESPONSE\_60\_MCF10A | 30 | 6 | 12187 | 179 | Trib1,Cxcl1,Ptgs2,Ier3,Il6,Tnfaip3 | | 4.090e-06 | -12.41 | BIOCARTA\_DEATH\_PATHWAY | MSigDB lists | BIOCARTA\_DEATH\_PATHWAY | 30 | 6 | 12187 | 179 | Cflar,Birc3,Nfkbia,Nfkb1,Ripk1,Tnfsf10 | | 4.090e-06 | -12.41 | ONO\_FOXP3\_TARGETS\_DN | MSigDB lists | ONO\_FOXP3\_TARGETS\_DN | 30 | 6 | 12187 | 179 | Casp4,Il4ra,Cdkn1a,Slfn3,Il17ra,Slfn4 | | 4.090e-06 | -12.41 | regulation of endothelial cell apoptotic process | biological process | GO:2000351 | 43 | 7 | 13711 | 214 | Angptl4,Nfe2l2,Thbs1,Ccl12,Tnf,Tnfaip3,Icam1 | | 4.215e-06 | -12.38 | regulation of cellular component movement | biological process | GO:0051270 | 863 | 32 | 13711 | 214 | Nfe2l2,Selp,Msn,C5ar1,Clic4,Hspa5,Iqgap1,Ccl12,Cxcl16,Hspb1,Sele,Il1rn,C3ar1,Adamts9,Rhoc,Robo4,Ccl2,Ccl4,Ptgs2,Timp1,Fam107a,Rhoj,Sdc4,Adamts1,Thbs1,Trib1,Pik3r1,Lgals9,P2ry6,Icam1,Il1a,Akap12 | | 4.337e-06 | -12.35 | GSE11961\_FOLLICULAR\_BCELL\_VS\_MEMORY\_BCELL\_DAY7\_UP | MSigDB lists | GSE11961\_FOLLICULAR\_BCELL\_VS\_MEMORY\_BCELL\_DAY7\_UP | 161 | 12 | 12187 | 179 | Pik3ap1,Parp9,Irf9,Herc6,Rnf213,Samd9l,Ifit3b,Cmpk2,Pik3r5,Irf7,Xaf1,Trib1 | | 4.360e-06 | -12.34 | MODULE\_6 | MSigDB lists | MODULE\_6 | 284 | 16 | 12187 | 179 | Cd14,Lcn2,Ier3,Atf3,Cdkn1a,Tnfsf10,S100a8,S100a9,Gem,Ccl12,Fam107a,Tnfaip3,Gadd45b,Tgm2,Il4ra,Maff | | 4.413e-06 | -12.33 | positive regulation of catalytic activity | biological process | GO:0043085 | 1081 | 37 | 13711 | 214 | Wfdc21,Irgm2,S100a9,Ccl12,S100a8,Iqgap1,Ptpn1,Dnajb1,Rhoc,Ripk1,Cflar,Ncf1,Il1rn,Sele,Ccl2,Xdh,Tnfsf10,Plek,Tnf,Pik3r5,Ccl4,Sdc4,Fzd4,Map3k6,Map3k8,Lgals9,Gadd45b,Ccnd2,Gadd45g,Trib1,Ccl7,Dtx3l,Thbs1,Cdkn1a,Rgs16,Icam1,Casp4 | | 4.415e-06 | -12.33 | GSE6269\_HEALTHY\_VS\_E\_COLI\_INF\_PBMC\_DN | MSigDB lists | GSE6269\_HEALTHY\_VS\_E\_COLI\_INF\_PBMC\_DN | 109 | 10 | 12187 | 179 | Trim25,Irak3,Plek,Plaur,Msn,Tagln2,Cxcl1,Cybb,Bach1,Ccl2 | | 4.514e-06 | -12.31 | TARTE\_PLASMA\_CELL\_VS\_PLASMABLAST\_UP | MSigDB lists | TARTE\_PLASMA\_CELL\_VS\_PLASMABLAST\_UP | 252 | 15 | 12187 | 179 | S100a9,S100a8,Nfkb2,Runx1,Birc3,Il4ra,Nfkbia,Tnfaip3,Hspb1,Ifi44,Vcam1,Fstl1,Hp,Col4a1,Lcn2 | | 4.575e-06 | -12.29 | GO\_POSITIVE\_REGULATION\_OF\_GENE\_EXPRESSION | MSigDB lists | GO\_POSITIVE\_REGULATION\_OF\_GENE\_EXPRESSION | 1307 | 40 | 12187 | 179 | S100a8,Sbno2,Csf3,Banp,Ifi211,Bcl3,Arid5b,Ripk1,Nfe2l2,Il6,Maff,Pik3r1,Spi1,Irak3,Irf2,Nfkb1,Il1a,Cebpd,Sox11,Hspa5,Thbs1,Nfkb2,Cflar,S100a9,Fzd4,Irf7,Bach1,Tnf,Runx1,Nfkbia,Trim25,Icam1,Zfp36,Msn,Tnip1,Ifi204,Ifi207,Atf3,Ifi209,Zbtb16 | | 4.593e-06 | -12.29 | regulation of leukocyte migration | biological process | GO:0002685 | 152 | 12 | 13711 | 214 | C5ar1,Ccl12,Ccl4,Msn,Selp,Il1a,Icam1,Ccl2,Thbs1,C3ar1,Sele,Lgals9 | | 4.605e-06 | -12.29 | GO\_NEGATIVE\_REGULATION\_OF\_PROTEIN\_METABOLIC\_PROCESS | MSigDB lists | GO\_NEGATIVE\_REGULATION\_OF\_PROTEIN\_METABOLIC\_PROCESS | 839 | 30 | 12187 | 179 | Tnfaip3,Il6,Gadd45b,Spi1,Tnfaip8,Ifi211,Banp,Bcl3,Nfkb1,Thbs1,Irak3,Ptpn1,Isg15,Tnf,Birc3,Trib1,Xdh,A2m,Timp1,Ifi204,Ifi207,Atf3,Cdkn1a,Ifi209,Socs3,Hspb1,Zfp36,Errfi1,Plaur,Tnip1 | | 4.625e-06 | -12.28 | GSE2197\_IMMUNOSUPPRESSIVE\_DNA\_VS\_UNTREATED\_IN\_DC\_UP | MSigDB lists | GSE2197\_IMMUNOSUPPRESSIVE\_DNA\_VS\_UNTREATED\_IN\_DC\_UP | 162 | 12 | 12187 | 179 | Atf3,Socs3,Ifi44,Irf9,Zfp36,Pik3ap1,Icam1,Ptgs2,Hcar2,Birc3,Rgs16,Gbp7 | | 4.625e-06 | -12.28 | GSE19772\_CTRL\_VS\_HCMV\_INF\_MONOCYTES\_UP | MSigDB lists | GSE19772\_CTRL\_VS\_HCMV\_INF\_MONOCYTES\_UP | 162 | 12 | 12187 | 179 | Xdh,Rgs16,Irgm2,Tor3a,Usp18,Ifit2,Irf9,Tap1,Saa1,Nfkb1,Hp,Akap12 | | 4.625e-06 | -12.28 | GSE22601\_DOUBLE\_POSITIVE\_VS\_CD8\_SINGLE\_POSITIVE\_THYMOCYTE\_UP | MSigDB lists | GSE22601\_DOUBLE\_POSITIVE\_VS\_CD8\_SINGLE\_POSITIVE\_THYMOCYTE\_UP | 162 | 12 | 12187 | 179 | Nfkbie,Trim56,Ccnd2,Il4ra,Tor3a,Parp14,Irak3,Trim25,Il17ra,Irf9,Gadd45g,Ier3 | | 4.627e-06 | -12.28 | regulation of cysteine-type endopeptidase activity involved in apoptotic process | biological process | GO:0043281 | 179 | 13 | 13711 | 214 | Plaur,Ptgs2,S100a9,S100a8,Birc3,Cflar,Tnfaip8,Thbs1,Il6,Tnfsf10,Xdh,Casp4,Tnf | | 4.662e-06 | -12.28 | GO\_REGULATION\_OF\_MYELOID\_CELL\_DIFFERENTIATION | MSigDB lists | GO\_REGULATION\_OF\_MYELOID\_CELL\_DIFFERENTIATION | 135 | 11 | 12187 | 179 | Zbtb16,Isg15,Ripk1,Runx1,Tnf,Pik3r1,Trib1,Nfkbia,Spi1,Csf3,Irf7 | | 4.662e-06 | -12.28 | GO\_RESPONSE\_TO\_STARVATION | MSigDB lists | GO\_RESPONSE\_TO\_STARVATION | 135 | 11 | 12187 | 179 | Zfp36,Rrp8,Ifi211,Ifi207,Ifi204,Nfe2l2,Atf3,Hspa5,Cdkn1a,Ifi209,Fstl1 | | 4.695e-06 | -12.27 | acute-phase response | biological process | GO:0006953 | 17 | 5 | 13711 | 214 | Il6,Saa1,Hp,Il1rn,Il1a | | 4.695e-06 | -12.27 | negative regulation of viral entry into host cell | biological process | GO:0046597 | 17 | 5 | 13711 | 214 | Trim25,Trim56,Ifitm2,Ifitm6,Ifitm3 | | 4.734e-06 | -12.26 | GO\_REGULATION\_OF\_PROTEIN\_SERINE\_THREONINE\_KINASE\_ACTIVITY | MSigDB lists | GO\_REGULATION\_OF\_PROTEIN\_SERINE\_THREONINE\_KINASE\_ACTIVITY | 391 | 19 | 12187 | 179 | Trib1,Tnf,Map3k8,Ccnd2,Gadd45b,Ripk1,Tnfaip3,Fzd4,Pik3r5,C5ar1,Iqgap1,Thbs1,Cdkn1a,Gadd45g,Saa1,Map3k6,Ptpn1,Hspb1,Irak3 | | 4.929e-06 | -12.22 | GSE15930\_STIM\_VS\_STIM\_AND\_IL12\_48H\_CD8\_T\_CELL\_DN | MSigDB lists | GSE15930\_STIM\_VS\_STIM\_AND\_IL12\_48H\_CD8\_T\_CELL\_DN | 163 | 12 | 12187 | 179 | Socs3,Usp18,Casp4,Map3k8,Il6,Ksr1,Gadd45g,Bcl3,Selp,Ifitm3,Gem,Tnfsf10 | | 4.935e-06 | -12.22 | GO\_CELLULAR\_RESPONSE\_TO\_OXYGEN\_CONTAINING\_COMPOUND | MSigDB lists | GO\_CELLULAR\_RESPONSE\_TO\_OXYGEN\_CONTAINING\_COMPOUND | 627 | 25 | 12187 | 179 | Cd14,P2ry6,Cxcl16,Ptpn1,Nfkb1,Csf3,Cmpk2,Sbno2,Pik3r1,Spi1,Nfe2l2,Il6,Ccl12,Tnfaip3,Ptgs2,Icam1,Msn,Errfi1,Adamts1,Col4a1,Tnip3,Fzd4,Tnf,Nfkbia,Casp4 | | 4.973e-06 | -12.21 | GO\_REGULATION\_OF\_PHOSPHORUS\_METABOLIC\_PROCESS | MSigDB lists | GO\_REGULATION\_OF\_PHOSPHORUS\_METABOLIC\_PROCESS | 1214 | 38 | 12187 | 179 | Xdh,Fzd4,C5ar1,Trib1,Sdc4,Tnf,Plaur,Errfi1,Plek,Hspb1,Icam1,Tnip1,Saa1,Atf3,Ksr1,Socs3,Cdkn1a,Iqgap1,Pik3r5,Csf3,Il6,Gadd45b,Ripk1,Ccl12,Tnfaip3,Ccl2,Ccl4,Pik3r1,Ccnd2,Map3k8,Ptpn1,Irak3,Map3k6,Il1a,Akap12,Gadd45g,Hspa5,Thbs1 | | 4.988e-06 | -12.21 | MODULE\_92 | MSigDB lists | MODULE\_92 | 87 | 9 | 12187 | 179 | Il6,Ccl12,Il1a,Ccl2,Ccl4,Tnfsf10,S100a9,Csf3,Isg15 | | 5.009e-06 | -12.20 | GSE46606\_IRF4HIGH\_VS\_IRF4MID\_CD40L\_IL2\_IL5\_DAY1\_STIMULATED\_BCELL\_DN | MSigDB lists | GSE46606\_IRF4HIGH\_VS\_IRF4MID\_CD40L\_IL2\_IL5\_DAY1\_STIMULATED\_BCELL\_DN | 136 | 11 | 12187 | 179 | Gbp3,Cmpk2,Ccl4,Ccnd2,Gadd45b,Slfn9,Isg15,Parp9,Hspa5,Atf3,Samd9l | | 5.010e-06 | -12.20 | BURTON\_ADIPOGENESIS\_1 | MSigDB lists | BURTON\_ADIPOGENESIS\_1 | 31 | 6 | 12187 | 179 | Ier3,Il6,Cebpd,Sgk1,Maff,Errfi1 | | 5.010e-06 | -12.20 | GO\_POSITIVE\_REGULATION\_OF\_INTERLEUKIN\_1\_PRODUCTION | MSigDB lists | GO\_POSITIVE\_REGULATION\_OF\_INTERLEUKIN\_1\_PRODUCTION | 31 | 6 | 12187 | 179 | Hspb1,Ifi204,Ifi207,Ifi209,Saa1,Ifi211 | | 5.010e-06 | -12.20 | GO\_INTRINSIC\_APOPTOTIC\_SIGNALING\_PATHWAY\_IN\_RESPONSE\_TO\_DNA\_DAMAGE\_BY\_P53\_CLASS\_MEDIATOR | MSigDB lists | GO\_INTRINSIC\_APOPTOTIC\_SIGNALING\_PATHWAY\_IN\_RESPONSE\_TO\_DNA\_DAMAGE\_BY\_P53\_CLASS\_MEDIATOR | 31 | 6 | 12187 | 179 | Ifi207,Ifi204,Cdkn1a,Ifi209,Bcl3,Ifi211 | | 5.024e-06 | -12.20 | KEGG\_RIG\_I\_LIKE\_RECEPTOR\_SIGNALING\_PATHWAY | MSigDB lists | KEGG\_RIG\_I\_LIKE\_RECEPTOR\_SIGNALING\_PATHWAY | 47 | 7 | 12187 | 179 | Trim25,Irf7,Isg15,Ripk1,Nfkb1,Nfkbia,Tnf | | 5.145e-06 | -12.18 | GO\_INTRINSIC\_APOPTOTIC\_SIGNALING\_PATHWAY\_IN\_RESPONSE\_TO\_DNA\_DAMAGE | MSigDB lists | GO\_INTRINSIC\_APOPTOTIC\_SIGNALING\_PATHWAY\_IN\_RESPONSE\_TO\_DNA\_DAMAGE | 66 | 8 | 12187 | 179 | Ifi211,Bcl3,Cdkn1a,Tnf,Ifi209,Pik3r1,Ifi204,Ifi207 | | 5.266e-06 | -12.15 | IRF7\_01 | MSigDB lists | IRF7\_01 | 193 | 13 | 12187 | 179 | Irf2,Cxcl16,Tap1,Isg15,Stx11,Angptl4,Dtx3l,Parp12,Oasl1,Xaf1,Usp18,Ifit2,Map3k8 | | 5.392e-06 | -12.13 | GO\_REGULATION\_OF\_CELLULAR\_COMPONENT\_MOVEMENT | MSigDB lists | GO\_REGULATION\_OF\_CELLULAR\_COMPONENT\_MOVEMENT | 589 | 24 | 12187 | 179 | Robo4,C3ar1,C5ar1,Sdc4,Trib1,Tnf,Msn,Clic4,Hspb1,Icam1,Ptgs2,Timp1,Cxcl1,Il6,Ccl12,Ccl4,Pik3r1,Cxcl16,P2ry6,Sgk3,Selp,Hspa5,Sgk1,Thbs1 | | 5.425e-06 | -12.12 | HELLER\_HDAC\_TARGETS\_SILENCED\_BY\_METHYLATION\_DN | MSigDB lists | HELLER\_HDAC\_TARGETS\_SILENCED\_BY\_METHYLATION\_DN | 224 | 14 | 12187 | 179 | Hspb1,Ifi209,Phf11b,Dnajb1,Ifi207,Ifi204,Atf3,Cflar,Bcl3,Ifi211,Phf11d,Maff,Ccnd2,Trib1 | | 5.592e-06 | -12.09 | GSE17721\_PAM3CSK4\_VS\_GADIQUIMOD\_12H\_BMDC\_DN | MSigDB lists | GSE17721\_PAM3CSK4\_VS\_GADIQUIMOD\_12H\_BMDC\_DN | 165 | 12 | 12187 | 179 | Parp9,Irf2,S100a8,Ptges,Gbp6,Gbp4,Tiparp,Rhoc,Parp14,Ccl2,Ccnd2,Sgk1 | | 5.592e-06 | -12.09 | GSE10325\_LUPUS\_CD4\_TCELL\_VS\_LUPUS\_MYELOID\_DN | MSigDB lists | GSE10325\_LUPUS\_CD4\_TCELL\_VS\_LUPUS\_MYELOID\_DN | 165 | 12 | 12187 | 179 | Cybb,Rassf4,Igsf6,C5ar1,S100a9,Il1rn,Cebpd,Rbm47,Clic4,Plaur,Csf2rb,Cd14 | | 5.592e-06 | -12.09 | GSE2770\_TGFB\_AND\_IL4\_ACT\_VS\_ACT\_CD4\_TCELL\_48H\_UP | MSigDB lists | GSE2770\_TGFB\_AND\_IL4\_ACT\_VS\_ACT\_CD4\_TCELL\_48H\_UP | 165 | 12 | 12187 | 179 | Cdkn1a,Il1a,Tnip1,Sdc4,Rhou,Tnf,Nfe2l2,Tubb6,Banp,Pik3r5,Tnfaip8,Rtp4 | | 5.610e-06 | -12.09 | defense response to Gram-positive bacterium | biological process | GO:0050830 | 45 | 7 | 13711 | 214 | Tnf,C5ar1,Gbp3,Tnfaip8,Ncf1,Gbp6,Gbp7 | | 5.769e-06 | -12.06 | GSE6269\_FLU\_VS\_STAPH\_AUREUS\_INF\_PBMC\_DN | MSigDB lists | GSE6269\_FLU\_VS\_STAPH\_AUREUS\_INF\_PBMC\_DN | 138 | 11 | 12187 | 179 | Timp1,Il1rn,Socs3,Spi1,Cdkn1a,Plaur,Sbno2,Icam1,S100a8,Plek,S100a9 | | 5.770e-06 | -12.06 | JECHLINGER\_EPITHELIAL\_TO\_MESENCHYMAL\_TRANSITION\_UP | MSigDB lists | JECHLINGER\_EPITHELIAL\_TO\_MESENCHYMAL\_TRANSITION\_UP | 67 | 8 | 12187 | 179 | Ccl12,Ifit3b,Gbp3,S100a8,Bcl3,Irf7,Isg15,Ifitm3 | | 5.809e-06 | -12.06 | GO\_INTERFERON\_GAMMA\_MEDIATED\_SIGNALING\_PATHWAY | MSigDB lists | GO\_INTERFERON\_GAMMA\_MEDIATED\_SIGNALING\_PATHWAY | 48 | 7 | 12187 | 179 | Irf2,Irf9,Trim25,Icam1,Irf7,Oasl1,Vcam1 | | 5.890e-06 | -12.04 | host cell part | cellular component | GO:0033643 | 18 | 5 | 13825 | 212 | Irgm2,Gbp6,Gbp3,Iigp1,Gbp7 | | 5.896e-06 | -12.04 | RUTELLA\_RESPONSE\_TO\_HGF\_VS\_CSF2RB\_AND\_IL4\_DN | MSigDB lists | RUTELLA\_RESPONSE\_TO\_HGF\_VS\_CSF2RB\_AND\_IL4\_DN | 195 | 13 | 12187 | 179 | Cebpd,Gadd45b,Ms4a6d,Sdc4,Ccl2,Cdkn1a,Runx1,Birc3,Hspb1,Tnfaip8,Zfp36,Cflar,Map3k6 | | 5.951e-06 | -12.03 | GSE22886\_NAIVE\_CD8\_TCELL\_VS\_MONOCYTE\_DN | MSigDB lists | GSE22886\_NAIVE\_CD8\_TCELL\_VS\_MONOCYTE\_DN | 166 | 12 | 12187 | 179 | Ms4a6d,Bach1,Cybb,Cebpd,S100a8,Igsf6,Irak3,Csf2rb,C5ar1,Cd14,S100a9,Bcl3 | | 5.951e-06 | -12.03 | GSE15330\_HSC\_VS\_LYMPHOID\_PRIMED\_MULTIPOTENT\_PROGENITOR\_UP | MSigDB lists | GSE15330\_HSC\_VS\_LYMPHOID\_PRIMED\_MULTIPOTENT\_PROGENITOR\_UP | 166 | 12 | 12187 | 179 | Ifitm2,Isg15,Ifitm3,Zfp36,Icam1,Irf9,Clic4,Ptpn1,Sgk1,Socs3,Birc3,Parp12 | | 6.001e-06 | -12.02 | receptor regulator activity | molecular function | GO:0030545 | 273 | 16 | 13516 | 211 | Tnfsf10,Timp1,Cxcl1,Csf3,Il1rn,Il1a,Ccl7,Ccl12,Tnf,Ccl2,Retnlg,Cxcl9,Il6,Ccl4,Saa1,Cxcl16 | | 6.011e-06 | -12.02 | GO\_REGULATION\_OF\_HEMOPOIESIS | MSigDB lists | GO\_REGULATION\_OF\_HEMOPOIESIS | 226 | 14 | 12187 | 179 | Isg15,Zbtb16,Irf7,Csf3,Trib1,Spi1,Nfkbia,Tnf,Runx1,Pik3r1,Il4ra,Ripk1,Il6,Nfe2l2 | | 6.028e-06 | -12.02 | MODULE\_456 | MSigDB lists | MODULE\_456 | 89 | 9 | 12187 | 179 | Stx11,Tnfaip3,Il6,Tnf,Cdkn1a,Hspa5,Nfkbia,Sele,Nfkb2 | | 6.186e-06 | -11.99 | GSE22611\_MUTANT\_NOD2\_VS\_CTRL\_TRANSDUCED\_HEK293T\_CELL\_UP | MSigDB lists | GSE22611\_MUTANT\_NOD2\_VS\_CTRL\_TRANSDUCED\_HEK293T\_CELL\_UP | 139 | 11 | 12187 | 179 | Zbtb16,Icam1,Clic4,Ptgs2,Rhoc,Il6,Gadd45b,Map3k8,Tnf,Csf3,Cxcl1 | | 6.186e-06 | -11.99 | GSE9960\_GRAM\_POS\_VS\_GRAM\_NEG\_AND\_POS\_SEPSIS\_PBMC\_DN | MSigDB lists | GSE9960\_GRAM\_POS\_VS\_GRAM\_NEG\_AND\_POS\_SEPSIS\_PBMC\_DN | 139 | 11 | 12187 | 179 | Bcl3,Ncf1,A2m,Tnip1,Nfkbie,Nfkb2,Birc3,Lcn2,Rasip1,Hp,Il1a | | 6.246e-06 | -11.98 | regulation of peptidase activity | biological process | GO:0052547 | 306 | 17 | 13711 | 214 | Thbs1,Cflar,Tnfaip8,Casp4,Tnf,Il6,Tnfsf10,Xdh,Serpina3f,S100a9,Ptgs2,Wfdc21,Plaur,Timp1,Birc3,A2m,S100a8 | | 6.331e-06 | -11.97 | HALLMARK\_IL2\_STAT5\_SIGNALING | MSigDB lists | HALLMARK\_IL2\_STAT5\_SIGNALING | 167 | 12 | 12187 | 179 | Ifitm3,Selp,Nfkbiz,Rgs16,Tnfsf10,Gbp3,Gadd45b,Maff,Il4ra,Ccnd2,Map3k8,Tgm2 | | 6.361e-06 | -11.97 | YAN\_ESCAPE\_FROM\_ANOIKIS | MSigDB lists | YAN\_ESCAPE\_FROM\_ANOIKIS | 19 | 5 | 12187 | 179 | Vcam1,Irf7,Slfn4,Nfkb1,Slfn3 | | 6.361e-06 | -11.97 | WORSCHECH\_TUMOR\_EVASION\_AND\_TOLEROGENICITY\_UP | MSigDB lists | WORSCHECH\_TUMOR\_EVASION\_AND\_TOLEROGENICITY\_UP | 19 | 5 | 12187 | 179 | Nfkb1,Il6,Ccl12,Irf7,Ccl4 | | 6.361e-06 | -11.97 | GO\_CHEMOKINE\_ACTIVITY | MSigDB lists | GO\_CHEMOKINE\_ACTIVITY | 19 | 5 | 12187 | 179 | Ccl12,Cxcl16,Ccl2,Cxcl1,Ccl4 | | 6.444e-06 | -11.95 | Guanylate-bd/ATL\_C | interpro domains | IPR003191 | 9 | 4 | 13788 | 212 | Gbp4,Gbp3,Gbp7,Gbp6 | | 6.444e-06 | -11.95 | HIN-200 | interpro domains | IPR040205 | 9 | 4 | 13788 | 212 | Ifi209,Ifi204,Ifi211,Ifi207 | | 6.616e-06 | -11.93 | GSE18791\_CTRL\_VS\_NEWCASTLE\_VIRUS\_DC\_2H\_DN | MSigDB lists | GSE18791\_CTRL\_VS\_NEWCASTLE\_VIRUS\_DC\_2H\_DN | 114 | 10 | 12187 | 179 | Ptgs2,Isg15,Gem,Irf7,Errfi1,Ifit2,Tnf,Ifit3b,Dnajb1,Gadd45b | | 6.668e-06 | -11.92 | regulation of ERK1 and ERK2 cascade | biological process | GO:0070372 | 244 | 15 | 13711 | 214 | Tnip1,P2ry6,Cflar,Ccl7,Il6,Atf3,Errfi1,Ccl2,Icam1,Il1a,Akap12,Ccl4,Ccl12,C5ar1,Ptpn1 | | 6.694e-06 | -11.91 | SANA\_RESPONSE\_TO\_IFNG\_UP | MSigDB lists | SANA\_RESPONSE\_TO\_IFNG\_UP | 49 | 7 | 12187 | 179 | Parp14,Samd9l,Rnf213,Dtx3l,Tnfsf10,Parp9,Gbp3 | | 6.732e-06 | -11.91 | GSE43700\_UNTREATED\_VS\_IL10\_TREATED\_PBMC\_DN | MSigDB lists | GSE43700\_UNTREATED\_VS\_IL10\_TREATED\_PBMC\_DN | 168 | 12 | 12187 | 179 | Ccl4,Map3k8,Maff,Il6,Gbp7,Xdh,Socs3,Ifit3b,Cebpd,Gadd45g,Selp,Errfi1 | | 6.820e-06 | -11.90 | GBP\_C | pfam domains | PF02841 | 9 | 4 | 12881 | 201 | Gbp6,Gbp7,Gbp4,Gbp3 | | 6.964e-06 | -11.87 | ISRE\_01 | MSigDB lists | ISRE\_01 | 198 | 13 | 12187 | 179 | Isg15,Irf2,Irf9,Ifi44,Thbs1,Ifit3b,Dtx3l,Angptl4,Rtp4,Rbm39,Ifit2,Xaf1,Usp18 | | 7.155e-06 | -11.85 | HALLMARK\_EPITHELIAL\_MESENCHYMAL\_TRANSITION | MSigDB lists | HALLMARK\_EPITHELIAL\_MESENCHYMAL\_TRANSITION | 169 | 12 | 12187 | 179 | Tnfaip3,Gadd45b,Il6,Tgm2,Sdc4,Gem,Timp1,Col4a1,Thbs1,Vcam1,Fstl1,Plaur | | 7.185e-06 | -11.84 | GO\_POSITIVE\_REGULATION\_OF\_PROTEIN\_MODIFICATION\_PROCESS | MSigDB lists | GO\_POSITIVE\_REGULATION\_OF\_PROTEIN\_MODIFICATION\_PROCESS | 858 | 30 | 12187 | 179 | Ccl2,Ccl4,Ccnd2,Map3k8,Gadd45b,Il6,Ccl12,Ripk1,Pik3r5,Csf3,Iqgap1,Thbs1,Hspa5,Gadd45g,Il1a,Map3k6,Ptpn1,Sdc4,Birc3,Tnf,Fzd4,C5ar1,Xdh,Socs3,Cdkn1a,Ksr1,Saa1,Plaur,Plek,Icam1 | | 7.215e-06 | -11.84 | HARRIS\_HYPOXIA | MSigDB lists | HARRIS\_HYPOXIA | 69 | 8 | 12187 | 179 | Plaur,Ptgs2,Cp,Nfkb1,Il6,Ccl12,Cdkn1a,Tgm2 | | 7.355e-06 | -11.82 | PID\_HIV\_NEF\_PATHWAY | MSigDB lists | PID\_HIV\_NEF\_PATHWAY | 33 | 6 | 12187 | 179 | Nfkb1,Ripk1,Tnf,Cflar,Birc3,Nfkbia | | 7.528e-06 | -11.80 | HIN\_200 | prosite domains | PS50834 | 8 | 4 | 8845 | 164 | Ifi211,Ifi204,Ifi209,Ifi207 | | 7.587e-06 | -11.79 | Osteoclast differentiation | KEGG pathways | mmu04380 | 105 | 11 | 5248 | 107 | Il1a,Cybb,Nfkb2,Socs3,Nfkbia,Ncf1,Spi1,Nfkb1,Tnf,Pik3r1,Irf9 | | 7.587e-06 | -11.79 | Osteoclast differentiation | KEGG pathways | ko04380 | 105 | 11 | 5248 | 107 | Cybb,Nfkb2,Tnf,Irf9,Socs3,Ncf1,Nfkb1,Il1a,Pik3r1,Nfkbia,Spi1 | | 7.601e-06 | -11.79 | GSE22140\_GERMFREE\_VS\_SPF\_MOUSE\_CD4\_TCELL\_DN | MSigDB lists | GSE22140\_GERMFREE\_VS\_SPF\_MOUSE\_CD4\_TCELL\_DN | 170 | 12 | 12187 | 179 | Ifitm2,Timp1,Nfkb1,Ier3,Ifi44,Thbs1,Ifit3b,Olfml2b,Oasl1,Pik3r1,Ifit2,Ccl4 | | 7.601e-06 | -11.79 | GSE17721\_CTRL\_VS\_CPG\_6H\_BMDC\_DN | MSigDB lists | GSE17721\_CTRL\_VS\_CPG\_6H\_BMDC\_DN | 170 | 12 | 12187 | 179 | Slfn3,Tap1,Tiparp,Znfx1,Atf3,Irgm2,Ptges,Slfn4,Pygm,Maff,Parp14,Ccl12 | | 7.601e-06 | -11.79 | GSE9006\_TYPE\_1\_DIABETES\_AT\_DX\_VS\_4MONTH\_POST\_DX\_PBMC\_UP | MSigDB lists | GSE9006\_TYPE\_1\_DIABETES\_AT\_DX\_VS\_4MONTH\_POST\_DX\_PBMC\_UP | 170 | 12 | 12187 | 179 | Hp,Stx11,Socs3,Sgk1,Hspa5,Zfp36,Tnfaip3,Cybb,Trib1,Nfkbia,Nfkb2,Cxcl1 | | 7.626e-06 | -11.78 | MODULE\_55 | MSigDB lists | MODULE\_55 | 520 | 22 | 12187 | 179 | Gadd45b,Fam107a,Ccl12,Trib1,Tgm2,Il4ra,Ccnd2,Tnfsf10,Sbno2,Xdh,S100a8,S100a9,Ier3,Lcn2,Col4a1,Gadd45g,Cp,Hp,Il1rn,Sele,Cd14,Isg15 | | 7.688e-06 | -11.78 | GNF2\_INPP5D | MSigDB lists | GNF2\_INPP5D | 50 | 7 | 12187 | 179 | Phf11b,Ifi209,Ifi207,Ifi204,Ifi211,Phf11d,Tap1 | | 7.809e-06 | -11.76 | GO\_RESPONSE\_TO\_INORGANIC\_SUBSTANCE | MSigDB lists | GO\_RESPONSE\_TO\_INORGANIC\_SUBSTANCE | 368 | 18 | 12187 | 179 | Ptges,Iqgap1,S100a8,Tnfaip3,Il6,Nfe2l2,Ptgs2,Cd14,Icam1,Clic4,Hspa5,Thbs1,Vcam1,Kcna5,Nfkb1,Hp,Cp,Il1a | | 8.045e-06 | -11.73 | GO\_POSITIVE\_REGULATION\_OF\_TYPE\_I\_INTERFERON\_PRODUCTION | MSigDB lists | GO\_POSITIVE\_REGULATION\_OF\_TYPE\_I\_INTERFERON\_PRODUCTION | 70 | 8 | 12187 | 179 | Nfkb2,Irf7,Cd14,Ifi211,Ifi207,Nfkb1,Ifi204,Ifi209 | | 8.045e-06 | -11.73 | HERNANDEZ\_ABERRANT\_MITOSIS\_BY\_DOCETACEL\_2NM\_UP | MSigDB lists | HERNANDEZ\_ABERRANT\_MITOSIS\_BY\_DOCETACEL\_2NM\_UP | 70 | 8 | 12187 | 179 | Ifi211,Ifi207,Ifi204,Il6,Ifi44,Sgk1,Pik3r1,Ifi209 | | 8.071e-06 | -11.73 | GSE36009\_WT\_VS\_NLRP10\_KO\_DC\_UP | MSigDB lists | GSE36009\_WT\_VS\_NLRP10\_KO\_DC\_UP | 171 | 12 | 12187 | 179 | Il6,Parp12,Il4ra,Slfn4,Gbp3,Gbp7,Trim56,Tiparp,Atf3,Pik3ap1,Slfn3,Slfn5 | | 8.071e-06 | -11.73 | BOQUEST\_STEM\_CELL\_DN | MSigDB lists | BOQUEST\_STEM\_CELL\_DN | 171 | 12 | 12187 | 179 | Sele,Apold1,Csf2rb,Selp,Stx11,Rasip1,Adamts9,Csf3,Cflar,Fam107a,Xaf1,Tgm2 | | 8.128e-06 | -11.72 | GO\_NEGATIVE\_REGULATION\_OF\_PEPTIDASE\_ACTIVITY | MSigDB lists | GO\_NEGATIVE\_REGULATION\_OF\_PEPTIDASE\_ACTIVITY | 143 | 11 | 12187 | 179 | A2m,Ifi211,Plaur,Tnfaip8,Birc3,Thbs1,Ifi209,Il6,Ifi207,Ifi204,Timp1 | | 8.380e-06 | -11.69 | GO\_CELLULAR\_EXTRAVASATION | MSigDB lists | GO\_CELLULAR\_EXTRAVASATION | 20 | 5 | 12187 | 179 | Ccl12,Sele,Vcam1,Selp,Tnf | | 8.380e-06 | -11.69 | TIAN\_TNF\_SIGNALING\_NOT\_VIA\_NFKB | MSigDB lists | TIAN\_TNF\_SIGNALING\_NOT\_VIA\_NFKB | 20 | 5 | 12187 | 179 | Maff,Ptges,Tiparp,Rgs16,Ier3 | | 8.567e-06 | -11.67 | GSE18893\_TCONV\_VS\_TREG\_2H\_CULTURE\_DN | MSigDB lists | GSE18893\_TCONV\_VS\_TREG\_2H\_CULTURE\_DN | 172 | 12 | 12187 | 179 | Socs3,Cdkn1a,Sgk1,Nfe2l2,Parp12,Ncf1,Ifitm3,Irgm2,Dtx3l,Ptpn1,Tap1,Icam1 | | 8.600e-06 | -11.66 | positive regulation of coagulation | biological process | GO:0050820 | 19 | 5 | 13711 | 214 | Selp,Nfe2l2,Thbs1,S100a9,Plek | | 8.600e-06 | -11.66 | response to copper ion | biological process | GO:0046688 | 19 | 5 | 13711 | 214 | Mt1,Nfe2l2,Il1a,Mt2,Cp | | 8.666e-06 | -11.66 | negative regulation of response to external stimulus | biological process | GO:0032102 | 281 | 16 | 13711 | 214 | Ier3,Nfkb1,A2m,Tap1,Ccl12,Socs3,Irak3,Thbs1,Parp14,Ctla2a,Trib1,Lgals9,Tnfaip3,Acod1,Zfp36,Cdkn1a | | 8.689e-06 | -11.65 | GSE360\_T\_GONDII\_VS\_B\_MALAYI\_LOW\_DOSE\_DC\_UP | MSigDB lists | GSE360\_T\_GONDII\_VS\_B\_MALAYI\_LOW\_DOSE\_DC\_UP | 144 | 11 | 12187 | 179 | Nfkb2,Cflar,Tnfaip3,Nfkbia,Trim25,Tap1,Ifitm3,Tagln2,Tnip1,Nfkb1,Ifit3b | | 8.689e-06 | -11.65 | GSE34156\_TLR1\_TLR2\_LIGAND\_VS\_NOD2\_AND\_TLR1\_TLR2\_LIGAND\_24H\_TREATED\_MONOCYTE\_UP | MSigDB lists | GSE34156\_TLR1\_TLR2\_LIGAND\_VS\_NOD2\_AND\_TLR1\_TLR2\_LIGAND\_24H\_TREATED\_MONOCYTE\_UP | 144 | 11 | 12187 | 179 | Msr1,Cybb,AA467197,C3ar1,Igsf6,Timp1,Cd14,Rbm47,Csf2rb,Cxcl16,Clic4 | | 8.702e-06 | -11.65 | cellular extravasation | biological process | GO:0045123 | 32 | 6 | 13711 | 214 | Ccl2,Selp,Tnf,Icam1,Vcam1,Sele | | 8.802e-06 | -11.64 | GO\_INTRINSIC\_APOPTOTIC\_SIGNALING\_PATHWAY\_BY\_P53\_CLASS\_MEDIATOR | MSigDB lists | GO\_INTRINSIC\_APOPTOTIC\_SIGNALING\_PATHWAY\_BY\_P53\_CLASS\_MEDIATOR | 51 | 7 | 12187 | 179 | Ifi209,Cdkn1a,Ifi204,Ifi207,Ifi211,Rrp8,Bcl3 | | 8.821e-06 | -11.64 | GO\_POSITIVE\_REGULATION\_OF\_MACROPHAGE\_CHEMOTAXIS | MSigDB lists | GO\_POSITIVE\_REGULATION\_OF\_MACROPHAGE\_CHEMOTAXIS | 10 | 4 | 12187 | 179 | Ccl12,C5ar1,C3ar1,Thbs1 | | 8.821e-06 | -11.64 | GO\_LEUKOCYTE\_ADHESION\_TO\_VASCULAR\_ENDOTHELIAL\_CELL | MSigDB lists | GO\_LEUKOCYTE\_ADHESION\_TO\_VASCULAR\_ENDOTHELIAL\_CELL | 10 | 4 | 12187 | 179 | Tnf,Selp,Vcam1,Sele | | 8.821e-06 | -11.64 | GO\_PERK\_MEDIATED\_UNFOLDED\_PROTEIN\_RESPONSE | MSigDB lists | GO\_PERK\_MEDIATED\_UNFOLDED\_PROTEIN\_RESPONSE | 10 | 4 | 12187 | 179 | Hspa5,Ccl12,Nfe2l2,Atf3 | | 8.821e-06 | -11.64 | SHIN\_B\_CELL\_LYMPHOMA\_CLUSTER\_5 | MSigDB lists | SHIN\_B\_CELL\_LYMPHOMA\_CLUSTER\_5 | 10 | 4 | 12187 | 179 | Tnf,Bcl3,Socs3,Il6 | | 8.821e-06 | -11.64 | REACTOME\_EXTRINSIC\_PATHWAY\_FOR\_APOPTOSIS | MSigDB lists | REACTOME\_EXTRINSIC\_PATHWAY\_FOR\_APOPTOSIS | 10 | 4 | 12187 | 179 | Ripk1,Tnfsf10,Cflar,Tnf | | 8.821e-06 | -11.64 | LI\_WILMS\_TUMOR\_ANAPLASTIC\_DN | MSigDB lists | LI\_WILMS\_TUMOR\_ANAPLASTIC\_DN | 10 | 4 | 12187 | 179 | Ifi211,Ifi209,Ifi207,Ifi204 | | 8.823e-06 | -11.64 | GRAHAM\_CML\_QUIESCENT\_VS\_NORMAL\_DIVIDING\_UP | MSigDB lists | GRAHAM\_CML\_QUIESCENT\_VS\_NORMAL\_DIVIDING\_UP | 34 | 6 | 12187 | 179 | Maff,Il1rn,Gadd45b,Ptgs2,Bcl3,Cxcl1 | | 8.823e-06 | -11.64 | STAMBOLSKY\_TARGETS\_OF\_MUTATED\_TP53\_DN | MSigDB lists | STAMBOLSKY\_TARGETS\_OF\_MUTATED\_TP53\_DN | 34 | 6 | 12187 | 179 | Irf9,Dtx3l,Isg15,Herc6,Irf7,Xaf1 | | 8.854e-06 | -11.63 | negative regulation of defense response | biological process | GO:0031348 | 162 | 12 | 13711 | 214 | Tnfaip3,Acod1,Zfp36,Parp14,Ctla2a,Socs3,Irak3,Lgals9,A2m,Nfkb1,Tap1,Ier3 | | 9.006e-06 | -11.62 | GSE9960\_GRAM\_NEG\_VS\_GRAM\_NEG\_AND\_POS\_SEPSIS\_PBMC\_UP | MSigDB lists | GSE9960\_GRAM\_NEG\_VS\_GRAM\_NEG\_AND\_POS\_SEPSIS\_PBMC\_UP | 118 | 10 | 12187 | 179 | Ifitm2,Ifitm3,Ifi44,Il4ra,Ifit2,Ifit3b,Usp18,Parp14,Xaf1,Samd9l | | 9.089e-06 | -11.61 | GSE22935\_WT\_VS\_MYD88\_KO\_MACROPHAGE\_UP | MSigDB lists | GSE22935\_WT\_VS\_MYD88\_KO\_MACROPHAGE\_UP | 173 | 12 | 12187 | 179 | Cybb,Tnfaip3,Nfkbie,Igsf6,Xdh,Fstl1,Stx11,Ptgs2,Ncf1,Il17ra,Rbm47,Csf2rb | | 9.140e-06 | -11.60 | negative regulation of response to biotic stimulus | biological process | GO:0002832 | 67 | 8 | 13711 | 214 | Acod1,Tnfaip3,Tap1,A2m,Irak3,Parp14,Trib1,Lgals9 | | 9.176e-06 | -11.60 | positive regulation of developmental process | biological process | GO:0051094 | 1253 | 40 | 13711 | 214 | Hspa5,Nfkbiz,Cxcl1,C5ar1,Iqgap1,S100a9,Nfe2l2,Noct,Cxcl9,Tnfaip3,Tnf,Zfp36,Cflar,Ripk1,Zbtb16,Adamts9,Csf3,Hspb1,Irak3,C3ar1,Ifi204,Fzd4,Isg15,Adamts1,Rhoj,Cebpd,Sgk1,Ptgs2,Runx1,Il1a,Sox11,Il4ra,Il6,Cybb,Msr1,Trib1,Socs3,Thbs1,Lgals9,Pik3r1 | | 9.284e-06 | -11.59 | GSE21360\_TERTIARY\_VS\_QUATERNARY\_MEMORY\_CD8\_TCELL\_DN | MSigDB lists | GSE21360\_TERTIARY\_VS\_QUATERNARY\_MEMORY\_CD8\_TCELL\_DN | 145 | 11 | 12187 | 179 | Lcn2,Ifitm2,Icam1,Nfkbia,Usp18,Sdc4,Cybb,Cflar,Pik3r5,S100a9,Cmpk2 | | 9.428e-06 | -11.57 | positive regulation of hemopoiesis | biological process | GO:1903708 | 163 | 12 | 13711 | 214 | Il6,Tnf,Il4ra,Lgals9,Csf3,Zbtb16,Trib1,Ripk1,Isg15,Cxcl1,Nfkbiz,Runx1 | | 9.472e-06 | -11.57 | REACTOME\_SIGNALING\_BY\_ILS | MSigDB lists | REACTOME\_SIGNALING\_BY\_ILS | 94 | 9 | 12187 | 179 | Irak3,Nfkb2,Csf2rb,Map3k8,Pik3r1,Socs3,Il1rn,Il1a,Il6 | | 9.639e-06 | -11.55 | GSE4748\_CTRL\_VS\_LPS\_STIM\_DC\_3H\_UP | MSigDB lists | GSE4748\_CTRL\_VS\_LPS\_STIM\_DC\_3H\_UP | 174 | 12 | 12187 | 179 | Irgm2,Angptl4,Rhoj,Iqgap1,Rtp4,Errfi1,Adamts1,Ifi44,Fstl1,Ifit3b,Akap12,Ripk1 | | 9.709e-06 | -11.54 | GO\_REGULATION\_OF\_REACTIVE\_OXYGEN\_SPECIES\_METABOLIC\_PROCESS | MSigDB lists | GO\_REGULATION\_OF\_REACTIVE\_OXYGEN\_SPECIES\_METABOLIC\_PROCESS | 119 | 10 | 12187 | 179 | Thbs1,Cdkn1a,Tnf,Hp,Il6,Nfe2l2,Ripk1,Ptgs2,Icam1,Xdh | | 9.747e-06 | -11.54 | GO\_POSITIVE\_REGULATION\_OF\_TRANSCRIPTION\_FROM\_RNA\_POLYMERASE\_II\_PROMOTER | MSigDB lists | GO\_POSITIVE\_REGULATION\_OF\_TRANSCRIPTION\_FROM\_RNA\_POLYMERASE\_II\_PROMOTER | 738 | 27 | 12187 | 179 | Cebpd,Il1a,Nfkb1,Hspa5,Sox11,Irf2,Nfe2l2,Il6,Ripk1,Spi1,Maff,Pik3r1,Sbno2,Bcl3,Ifi211,Csf3,Atf3,Ifi207,Ifi204,Ifi209,Tnip1,Bach1,Nfkbia,Tnf,Runx1,Nfkb2,Irf7 | | 9.914e-06 | -11.52 | GSE5589\_LPS\_VS\_LPS\_AND\_IL10\_STIM\_IL10\_KO\_MACROPHAGE\_45MIN\_DN | MSigDB lists | GSE5589\_LPS\_VS\_LPS\_AND\_IL10\_STIM\_IL10\_KO\_MACROPHAGE\_45MIN\_DN | 146 | 11 | 12187 | 179 | Irgm2,S100a8,Clic4,Ifit2,Msr1,Xaf1,Ifit3b,Samd9l,Cp,Gadd45g,Tnfaip3 | | 9.914e-06 | -11.52 | GSE360\_DC\_VS\_MAC\_M\_TUBERCULOSIS\_UP | MSigDB lists | GSE360\_DC\_VS\_MAC\_M\_TUBERCULOSIS\_UP | 146 | 11 | 12187 | 179 | Trib1,Ccl4,Il6,Gadd45b,Oasl1,Ripk1,Ccl12,Irf9,Ifit3b,Nfkb1,Il1rn | | 1.005e-05 | -11.51 | GO\_I\_KAPPAB\_KINASE\_NF\_KAPPAB\_SIGNALING | MSigDB lists | GO\_I\_KAPPAB\_KINASE\_NF\_KAPPAB\_SIGNALING | 52 | 7 | 12187 | 179 | Nfkb1,Ripk1,Nfkb2,Birc3,Tnf,Bcl3,Cd14 | | 1.015e-05 | -11.50 | HOSHIDA\_LIVER\_CANCER\_SUBCLASS\_S1 | MSigDB lists | HOSHIDA\_LIVER\_CANCER\_SUBCLASS\_S1 | 205 | 13 | 12187 | 179 | Ier3,Col4a1,Ifi207,Ifi204,Ifi209,Msn,Plaur,Cybb,Ccnd2,Iqgap1,Ifi211,C3ar1,Gem | | 1.040e-05 | -11.47 | host | cellular component | GO:0018995 | 20 | 5 | 13825 | 212 | Iigp1,Gbp3,Gbp7,Irgm2,Gbp6 | | 1.040e-05 | -11.47 | host cell | cellular component | GO:0043657 | 20 | 5 | 13825 | 212 | Irgm2,Gbp6,Gbp3,Iigp1,Gbp7 | | 1.058e-05 | -11.46 | GSE3982\_EOSINOPHIL\_VS\_NEUTROPHIL\_DN | MSigDB lists | GSE3982\_EOSINOPHIL\_VS\_NEUTROPHIL\_DN | 147 | 11 | 12187 | 179 | Ptges,Ifitm3,Tnfsf10,Irf2,Plek,Icam1,Apold1,Trib1,Atf3,Tnfaip3,Il1rn | | 1.061e-05 | -11.45 | Guanylate-bd\_C\_sf | interpro domains | IPR036543 | 10 | 4 | 13788 | 212 | Gbp3,Gbp7,Gbp6,Gbp4 | | 1.083e-05 | -11.43 | GSE4748\_CTRL\_VS\_CYANOBACTERIUM\_LPSLIKE\_STIM\_DC\_3H\_DN | MSigDB lists | GSE4748\_CTRL\_VS\_CYANOBACTERIUM\_LPSLIKE\_STIM\_DC\_3H\_DN | 176 | 12 | 12187 | 179 | Gbp3,Tnfsf10,C3ar1,AA467197,Ccl12,Tubb6,Ccl4,Ccl2,Msr1,Cxcl16,Atf3,Ier3 | | 1.090e-05 | -11.43 | negative regulation of signal transduction | biological process | GO:0009968 | 991 | 34 | 13711 | 214 | Thbs1,Socs3,Parp14,Trib1,Icam1,Nfkbia,Rgs16,Errfi1,Il6,Rasip1,Ptgs2,Ier3,Plaur,Irak3,Il1rn,Hspb1,Ripk1,Cflar,Tnip1,Rnf213,Trim30a,Tnf,Tnfaip3,Acod1,Plek,Pik3ap1,Xdh,Atf3,Sgk3,Nfe2l2,Bcl3,Hspa5,Ptpn1,Tnip3 | | 1.100e-05 | -11.42 | negative regulation of phosphate metabolic process | biological process | GO:0045936 | 498 | 22 | 13711 | 214 | Iqgap1,Ptpn1,Birc3,Gbp4,Ier3,Rasip1,Cdkn1a,Il6,Errfi1,Atf3,Xdh,Plek,Tnfaip3,Tnf,Gadd45g,Tnip1,Gadd45b,Parp14,Trib1,Hspb1,Socs3,Irak3 | | 1.100e-05 | -11.42 | negative regulation of phosphorus metabolic process | biological process | GO:0010563 | 498 | 22 | 13711 | 214 | Errfi1,Il6,Atf3,Xdh,Cdkn1a,Tnfaip3,Tnf,Plek,Tnip1,Gadd45g,Gadd45b,Hspb1,Socs3,Irak3,Parp14,Trib1,Iqgap1,Gbp4,Birc3,Ptpn1,Ier3,Rasip1 | | 1.126e-05 | -11.39 | regulation of interferon-gamma-mediated signaling pathway | biological process | GO:0060334 | 10 | 4 | 13711 | 214 | Parp9,Irgm2,Parp14,Irgm1 | | 1.126e-05 | -11.39 | regulation of response to interferon-gamma | biological process | GO:0060330 | 10 | 4 | 13711 | 214 | Irgm1,Parp14,Irgm2,Parp9 | | 1.129e-05 | -11.39 | GSE43955\_TH0\_VS\_TGFB\_IL6\_TH17\_ACT\_CD4\_TCELL\_52H\_DN | MSigDB lists | GSE43955\_TH0\_VS\_TGFB\_IL6\_TH17\_ACT\_CD4\_TCELL\_52H\_DN | 148 | 11 | 12187 | 179 | Il17ra,Errfi1,Saa1,Il1rn,Tiparp,Bach1,Gadd45g,Cebpd,Map3k8,Socs3,Fstl1 | | 1.129e-05 | -11.39 | GSE3982\_BCELL\_VS\_CENT\_MEMORY\_CD4\_TCELL\_UP | MSigDB lists | GSE3982\_BCELL\_VS\_CENT\_MEMORY\_CD4\_TCELL\_UP | 148 | 11 | 12187 | 179 | Cdkn1a,Cd14,Csf2rb,Plek,Tor3a,Trib1,Map3k8,Tubb6,Cybb,Irf7,S100a8 | | 1.132e-05 | -11.39 | positive regulation of vascular endothelial growth factor production | biological process | GO:0010575 | 20 | 5 | 13711 | 214 | Il1a,C3ar1,C5ar1,Il6,Ptgs2 | | 1.143e-05 | -11.38 | MODULE\_291 | MSigDB lists | MODULE\_291 | 53 | 7 | 12187 | 179 | Spi1,Ier3,Cd14,S100a9,C3ar1,Saa1,C5ar1 | | 1.143e-05 | -11.38 | GO\_POSITIVE\_REGULATION\_OF\_LEUKOCYTE\_CHEMOTAXIS | MSigDB lists | GO\_POSITIVE\_REGULATION\_OF\_LEUKOCYTE\_CHEMOTAXIS | 53 | 7 | 12187 | 179 | C3ar1,C5ar1,Cxcl1,Thbs1,Ccl4,Il6,Ccl12 | | 1.147e-05 | -11.38 | GO\_REGULATION\_OF\_VASCULATURE\_DEVELOPMENT | MSigDB lists | GO\_REGULATION\_OF\_VASCULATURE\_DEVELOPMENT | 177 | 12 | 12187 | 179 | Thbs1,Runx1,Il6,Ccl12,Tnfaip3,Il1a,Angptl4,C3ar1,Ptgs2,C5ar1,Xdh,Hspb1 | | 1.203e-05 | -11.33 | GSE17721\_CTRL\_VS\_PAM3CSK4\_2H\_BMDC\_DN | MSigDB lists | GSE17721\_CTRL\_VS\_PAM3CSK4\_2H\_BMDC\_DN | 149 | 11 | 12187 | 179 | Arid5b,Cflar,Ptgs2,Map3k6,Slfn4,Slfn3,Zbtb16,Parp14,Pik3r1,Il1rn,Cp | | 1.211e-05 | -11.32 | GO\_TRANSCRIPTIONAL\_REPRESSOR\_ACTIVITY\_RNA\_POLYMERASE\_II\_TRANSCRIPTION\_REGULATORY\_REGION\_SEQUENCE\_SPECIFIC\_BINDING | MSigDB lists | GO\_TRANSCRIPTIONAL\_REPRESSOR\_ACTIVITY\_RNA\_POLYMERASE\_II\_TRANSCRIPTION\_REGULATORY\_REGION\_SEQUENCE\_SPECIFIC\_BINDING | 122 | 10 | 12187 | 179 | Ifi211,Arid5b,Atf3,Nfkb1,Ifi204,Bach1,Ifi207,Zbtb16,Spi1,Ifi209 | | 1.214e-05 | -11.32 | DAZARD\_RESPONSE\_TO\_UV\_NHEK\_UP | MSigDB lists | DAZARD\_RESPONSE\_TO\_UV\_NHEK\_UP | 178 | 12 | 12187 | 179 | S100a9,Cxcl1,Gem,Ptgs2,Zfp36,Trib1,Sgk1,Atf3,Il6,Gadd45b,Ier3,Tnfaip3 | | 1.214e-05 | -11.32 | LEE\_RECENT\_THYMIC\_EMIGRANT | MSigDB lists | LEE\_RECENT\_THYMIC\_EMIGRANT | 178 | 12 | 12187 | 179 | Tnfsf10,Cmpk2,Ifi211,Tnfaip3,Rhou,Tagln2,Isg15,Ifi207,Ifi204,Nfkbiz,Socs3,Ifi209 | | 1.232e-05 | -11.30 | ONKEN\_UVEAL\_MELANOMA\_UP | MSigDB lists | ONKEN\_UVEAL\_MELANOMA\_UP | 661 | 25 | 12187 | 179 | Ccnd2,Iqgap1,Tnfaip8,Phf11d,Ifi211,Ier3,Cebpd,Phf11b,Sgk1,Ifitm3,Isg15,Parp12,Trib1,Gem,Luc7l3,Ifi204,Col4a1,Ifi207,Cdkn1a,Ifi209,Clic4,Msn,Irf9,Tagln2,Ifitm2 | | 1.247e-05 | -11.29 | SAGIV\_CD24\_TARGETS\_DN | MSigDB lists | SAGIV\_CD24\_TARGETS\_DN | 36 | 6 | 12187 | 179 | Thbs1,Cdkn1a,Atf3,Oasl1,Gem,Icam1 | | 1.247e-05 | -11.29 | ZHU\_CMV\_8\_HR\_UP | MSigDB lists | ZHU\_CMV\_8\_HR\_UP | 36 | 6 | 12187 | 179 | Ptgs2,Isg15,Irf2,Il6,Ripk1,Tnfaip3 | | 1.278e-05 | -11.27 | positive regulation of programmed cell death | biological process | GO:0043068 | 541 | 23 | 13711 | 214 | Gadd45b,Hcar2,Gadd45g,Thbs1,Cdkn1a,Ip6k2,Il6,Akap12,Casp4,Plaur,Ptgs2,Tnfaip8,Ripk1,Zbtb16,Xdh,Tnfsf10,Atf3,Tnf,Ifit2,S100a9,Ccl12,S100a8,Tgm2 | | 1.282e-05 | -11.26 | GSE36392\_TYPE\_2\_MYELOID\_VS\_EOSINOPHIL\_IL25\_TREATED\_LUNG\_DN | MSigDB lists | GSE36392\_TYPE\_2\_MYELOID\_VS\_EOSINOPHIL\_IL25\_TREATED\_LUNG\_DN | 150 | 11 | 12187 | 179 | Irf9,Isg15,Herc6,Lcn2,Stx11,Ifit3b,S100a8,AA467197,Gadd45b,Hcar2,Tnf | | 1.297e-05 | -11.25 | YORDY\_RECIPROCAL\_REGULATION\_BY\_ETS1\_AND\_SP100\_DN | MSigDB lists | YORDY\_RECIPROCAL\_REGULATION\_BY\_ETS1\_AND\_SP100\_DN | 54 | 7 | 12187 | 179 | Irf9,Icam1,Ptges,Angptl4,Samd9l,Dnajb1,Socs3 | | 1.301e-05 | -11.25 | SMIRNOV\_CIRCULATING\_ENDOTHELIOCYTES\_IN\_CANCER\_UP | MSigDB lists | SMIRNOV\_CIRCULATING\_ENDOTHELIOCYTES\_IN\_CANCER\_UP | 123 | 10 | 12187 | 179 | Rgs16,Irak3,Ptges,Cd14,C5ar1,C3ar1,Ier3,Cdkn1a,Thbs1,Maff | | 1.304e-05 | -11.25 | GO\_REGULATION\_OF\_PROTEIN\_MODIFICATION\_PROCESS | MSigDB lists | GO\_REGULATION\_OF\_PROTEIN\_MODIFICATION\_PROCESS | 1315 | 39 | 12187 | 179 | Fzd4,C5ar1,Xdh,Trib1,Sdc4,Tnf,Birc3,Tnip1,Saa1,Errfi1,Plaur,Icam1,Hspb1,Plek,Socs3,Cdkn1a,Atf3,Ksr1,Pik3r5,Csf3,Iqgap1,Ccl4,Ccl2,Spi1,Map3k8,Ccnd2,Tnfaip3,Ccl12,Ripk1,Gadd45b,Il6,Isg15,Map3k6,Ptpn1,Irak3,Hspa5,Thbs1,Il1a,Gadd45g | | 1.313e-05 | -11.24 | GO\_APOPTOTIC\_SIGNALING\_PATHWAY | MSigDB lists | GO\_APOPTOTIC\_SIGNALING\_PATHWAY | 242 | 14 | 12187 | 179 | Cd14,Ifi204,Il1a,Ifi207,Ifi209,Cdkn1a,Tnfsf10,Ifi211,Bcl3,Rrp8,Ripk1,Tnf,Pik3r1,Casp4 | | 1.343e-05 | -11.22 | regulation of interleukin-6 production | biological process | GO:0032675 | 116 | 10 | 13711 | 214 | Il6,Prg4,Tnfaip3,Tnf,Il1a,Lgals9,Trim30a,Il1rn,Irak3,Mmp8 | | 1.351e-05 | -11.21 | GO\_REGULATION\_OF\_DNA\_BINDING | MSigDB lists | GO\_REGULATION\_OF\_DNA\_BINDING | 75 | 8 | 12187 | 179 | Ifi209,Sox11,Nfkbia,Ifi207,Ifi204,Bcl3,Ifi211,Plaur | | 1.351e-05 | -11.21 | GO\_MYELOID\_LEUKOCYTE\_DIFFERENTIATION | MSigDB lists | GO\_MYELOID\_LEUKOCYTE\_DIFFERENTIATION | 75 | 8 | 12187 | 179 | Ifi209,Tnf,Spi1,Ifi207,Ifi204,Csf3,Ifi211,Sbno2 | | 1.365e-05 | -11.20 | GSE46242\_CTRL\_VS\_EGR2\_DELETED\_ANERGIC\_TH1\_CD4\_TCELL\_DN | MSigDB lists | GSE46242\_CTRL\_VS\_EGR2\_DELETED\_ANERGIC\_TH1\_CD4\_TCELL\_DN | 151 | 11 | 12187 | 179 | Ccl4,Tor3a,Birc3,Rassf4,Cybb,Bcl3,Nfkbie,Nfkb2,Gbp3,Cdkn1a,Nfkb1 | | 1.370e-05 | -11.20 | GRANDVAUX\_IFN\_RESPONSE\_NOT\_VIA\_IRF3 | MSigDB lists | GRANDVAUX\_IFN\_RESPONSE\_NOT\_VIA\_IRF3 | 11 | 4 | 12187 | 179 | Irf7,Ifitm3,Irgm2,Irf9 | | 1.370e-05 | -11.20 | GO\_REGULATION\_OF\_TUMOR\_NECROSIS\_FACTOR\_BIOSYNTHETIC\_PROCESS | MSigDB lists | GO\_REGULATION\_OF\_TUMOR\_NECROSIS\_FACTOR\_BIOSYNTHETIC\_PROCESS | 11 | 4 | 12187 | 179 | Errfi1,Hspb1,Bcl3,Thbs1 | | 1.370e-05 | -11.20 | GO\_REGULATION\_OF\_MACROPHAGE\_CHEMOTAXIS | MSigDB lists | GO\_REGULATION\_OF\_MACROPHAGE\_CHEMOTAXIS | 11 | 4 | 12187 | 179 | Ccl12,C5ar1,Thbs1,C3ar1 | | 1.382e-05 | -11.19 | positive regulation of stress-activated MAPK cascade | biological process | GO:0032874 | 142 | 11 | 13711 | 214 | Gadd45g,Gadd45b,Mmp8,Ripk1,Ncf1,Il1rn,Xdh,Il1a,Ptpn1,Fzd4,Tnf | | 1.398e-05 | -11.18 | GSE1112\_HY\_CD8AB\_VS\_HY\_CD8AA\_THYMOCYTE\_RTOC\_CULTURE\_UP | MSigDB lists | GSE1112\_HY\_CD8AB\_VS\_HY\_CD8AA\_THYMOCYTE\_RTOC\_CULTURE\_UP | 124 | 10 | 12187 | 179 | Tiparp,Samd9l,Tnf,Ccnd2,Vcam1,Parp14,Rtp4,Gbp3,Ptgs2,Banp | | 1.419e-05 | -11.16 | Regulation of necroptotic cell death | REACTOME pathways | R-MMU-5675482 | 10 | 4 | 6297 | 105 | Cflar,Ripk1,Tnfsf10,Birc3 | | 1.419e-05 | -11.16 | RIPK1-mediated regulated necrosis | REACTOME pathways | R-MMU-5213460 | 10 | 4 | 6297 | 105 | Ripk1,Cflar,Tnfsf10,Birc3 | | 1.419e-05 | -11.16 | Regulated Necrosis | REACTOME pathways | R-MMU-5218859 | 10 | 4 | 6297 | 105 | Ripk1,Cflar,Tnfsf10,Birc3 | | 1.428e-05 | -11.16 | GO\_POSITIVE\_REGULATION\_OF\_LOCOMOTION | MSigDB lists | GO\_POSITIVE\_REGULATION\_OF\_LOCOMOTION | 312 | 16 | 12187 | 179 | Ccl12,Il6,Tnf,Pik3r1,Ccl4,C5ar1,C3ar1,Cxcl1,Thbs1,Hspa5,Hspb1,Icam1,P2ry6,Cxcl16,Selp,Ptgs2 | | 1.442e-05 | -11.15 | cell migration | biological process | GO:0016477 | 664 | 26 | 13711 | 214 | Rhoc,Retnlg,Sele,Il1rn,Arid5b,Ccl2,Saa1,Cxcl9,Tnf,Tnfaip3,Selp,Msn,S100a9,Vcam1,Ccl12,Cxcl16,S100a8,Iqgap1,C5ar1,Cxcl1,Ccl7,Thbs1,Icam1,Il17ra,Ccl4,Sdc4 | | 1.449e-05 | -11.14 | positive regulation of JNK cascade | biological process | GO:0046330 | 117 | 10 | 13711 | 214 | Ripk1,Mmp8,Il1rn,Ncf1,Gadd45b,Gadd45g,Fzd4,Il1a,Ptpn1,Tnf | | 1.453e-05 | -11.14 | GSE43955\_1H\_VS\_20H\_ACT\_CD4\_TCELL\_WITH\_TGFB\_IL6\_DN | MSigDB lists | GSE43955\_1H\_VS\_20H\_ACT\_CD4\_TCELL\_WITH\_TGFB\_IL6\_DN | 152 | 11 | 12187 | 179 | Casp4,Birc3,Osmr,Irf7,C3ar1,Csf3,Il1a,Nfkbiz,Csf2rb,Clic4,Ptgs2 | | 1.453e-05 | -11.14 | GSE21379\_WT\_VS\_SAP\_KO\_TFH\_CD4\_TCELL\_DN | MSigDB lists | GSE21379\_WT\_VS\_SAP\_KO\_TFH\_CD4\_TCELL\_DN | 152 | 11 | 12187 | 179 | Bach1,Ms4a6d,Gbp4,Oasl1,Ifi44,Zbtb16,Ccl4,Icam1,Rtp4,Gbp7,Gbp6 | | 1.453e-05 | -11.14 | GSE22589\_HIV\_VS\_HIV\_AND\_SIV\_INFECTED\_DC\_UP | MSigDB lists | GSE22589\_HIV\_VS\_HIV\_AND\_SIV\_INFECTED\_DC\_UP | 152 | 11 | 12187 | 179 | Slfn4,Slfn3,Errfi1,Gem,Bcl3,Il1a,Tnfaip3,Il6,Tnf,Nfkbia,Gpr84 | | 1.453e-05 | -11.14 | GSE41176\_WT\_VS\_TAK1\_KO\_ANTI\_IGM\_STIM\_BCELL\_6H\_UP | MSigDB lists | GSE41176\_WT\_VS\_TAK1\_KO\_ANTI\_IGM\_STIM\_BCELL\_6H\_UP | 152 | 11 | 12187 | 179 | Zfp36,Sbno2,Ptgs2,Cflar,Cxcl1,Gadd45g,Ier3,Adamts1,Map3k8,Ccl4,Socs3 | | 1.470e-05 | -11.13 | GO\_REGULATION\_OF\_TOLL\_LIKE\_RECEPTOR\_SIGNALING\_PATHWAY | MSigDB lists | GO\_REGULATION\_OF\_TOLL\_LIKE\_RECEPTOR\_SIGNALING\_PATHWAY | 37 | 6 | 12187 | 179 | Irf7,Cd14,Birc3,Tnfaip3,Irak3,Pik3ap1 | | 1.470e-05 | -11.13 | FRIDMAN\_IMMORTALIZATION\_DN | MSigDB lists | FRIDMAN\_IMMORTALIZATION\_DN | 37 | 6 | 12187 | 179 | Ifi209,Cdkn1a,Ifi211,Irf7,Ifi204,Ifi207 | | 1.477e-05 | -11.12 | positive regulation of stress-activated protein kinase signaling cascade | biological process | GO:0070304 | 143 | 11 | 13711 | 214 | Xdh,Tnf,Ptpn1,Il1a,Fzd4,Gadd45g,Gadd45b,Ncf1,Il1rn,Mmp8,Ripk1 | | 1.483e-05 | -11.12 | CCR2 chemokine receptor binding | molecular function | GO:0031727 | 4 | 3 | 13516 | 211 | Ccl12,Ccl2,Ccl7 | | 1.491e-05 | -11.11 | GO\_RESPONSE\_TO\_INTERLEUKIN\_1 | MSigDB lists | GO\_RESPONSE\_TO\_INTERLEUKIN\_1 | 76 | 8 | 12187 | 179 | Icam1,Irak3,Sele,Nfkb1,Ccl12,Il6,Ccl4,Ccl2 | | 1.500e-05 | -11.11 | GSE14386\_UNTREATED\_VS\_IFNA\_TREATED\_ACT\_PBMC\_MS\_PATIENT\_DN | MSigDB lists | GSE14386\_UNTREATED\_VS\_IFNA\_TREATED\_ACT\_PBMC\_MS\_PATIENT\_DN | 125 | 10 | 12187 | 179 | Parp12,Ccl12,Stx11,Ifit3b,Maff,Birc3,Ifi44,Rgs16,Rtp4,Irf7 | | 1.500e-05 | -11.11 | GSE1740\_UNSTIM\_VS\_IFNA\_STIMULATED\_MCSF\_DERIVED\_MACROPHAGE\_DN | MSigDB lists | GSE1740\_UNSTIM\_VS\_IFNA\_STIMULATED\_MCSF\_DERIVED\_MACROPHAGE\_DN | 125 | 10 | 12187 | 179 | Tnf,Parp14,Samd9l,Ptgs2,Cxcl1,Irf7,Rtp4,Gbp3,Plaur,Tap1 | | 1.546e-05 | -11.08 | HOXA9\_DN.V1\_UP | MSigDB lists | HOXA9\_DN.V1\_UP | 153 | 11 | 12187 | 179 | Ifi204,Ms4a6d,Ifi207,Ifi209,Rbm47,Sgk3,Isg15,Parp12,S100a8,Ifi211,S100a9 | | 1.546e-05 | -11.08 | GSE28737\_FOLLICULAR\_VS\_MARGINAL\_ZONE\_BCELL\_DN | MSigDB lists | GSE28737\_FOLLICULAR\_VS\_MARGINAL\_ZONE\_BCELL\_DN | 153 | 11 | 12187 | 179 | Parp9,Irf2,Cmpk2,Rtp4,Icam1,Slfn9,Irgm2,Znfx1,Samd9l,Cdkn1a,Ccnd2 | | 1.546e-05 | -11.08 | GSE29618\_MONOCYTE\_VS\_PDC\_UP | MSigDB lists | GSE29618\_MONOCYTE\_VS\_PDC\_UP | 153 | 11 | 12187 | 179 | C5ar1,S100a9,Spi1,Trib1,Ifitm2,Cd14,Irak3,Msn,Il1rn,Timp1,Ier3 | | 1.548e-05 | -11.08 | MODULE\_3 | MSigDB lists | MODULE\_3 | 350 | 17 | 12187 | 179 | Plaur,Fstl1,Cdkn1a,Ifi209,Sgk1,Thbs1,Ier3,Ifi204,Col4a1,Ifi207,Akap12,Ifi211,Iqgap1,Tgm2,Casp4,Maff,Ccl12 | | 1.602e-05 | -11.04 | PEDRIOLI\_MIR31\_TARGETS\_DN | MSigDB lists | PEDRIOLI\_MIR31\_TARGETS\_DN | 280 | 15 | 12187 | 179 | Vcam1,Ifit3b,Nfkbiz,Samd9l,Akap12,Rasd1,Icam1,Sele,Ifit2,Rhou,Birc3,Tnfaip3,Cxcl1,Adamts9,Nfkb2 | | 1.609e-05 | -11.04 | GO\_POSITIVE\_REGULATION\_OF\_PEPTIDASE\_ACTIVITY | MSigDB lists | GO\_POSITIVE\_REGULATION\_OF\_PEPTIDASE\_ACTIVITY | 126 | 10 | 12187 | 179 | Ripk1,Ifi204,Ifi207,Tnf,Ifi209,Xdh,Tnfsf10,S100a8,Ifi211,S100a9 | | 1.625e-05 | -11.03 | regulation of cysteine-type endopeptidase activity | biological process | GO:2000116 | 201 | 13 | 13711 | 214 | S100a8,Birc3,Ptgs2,Plaur,S100a9,Tnfsf10,Il6,Xdh,Casp4,Tnf,Thbs1,Cflar,Tnfaip8 | | 1.634e-05 | -11.02 | regulation of MAP kinase activity | biological process | GO:0043405 | 263 | 15 | 13711 | 214 | Pik3r5,Iqgap1,Ptpn1,Map3k6,Fzd4,Map3k8,Gadd45g,Gadd45b,Ripk1,Trib1,Irak3,Thbs1,Il1rn,Ksr1,Tnf | | 1.644e-05 | -11.02 | GSE8835\_HEALTHY\_VS\_CLL\_CD8\_TCELL\_DN | MSigDB lists | GSE8835\_HEALTHY\_VS\_CLL\_CD8\_TCELL\_DN | 154 | 11 | 12187 | 179 | Oasl1,Hp,Col4a1,Nfkbiz,Il4ra,Gbp3,Errfi1,S100a8,Ptges,Tnip1,Csf3 | | 1.644e-05 | -11.02 | GSE43955\_TH0\_VS\_TGFB\_IL6\_TH17\_ACT\_CD4\_TCELL\_10H\_DN | MSigDB lists | GSE43955\_TH0\_VS\_TGFB\_IL6\_TH17\_ACT\_CD4\_TCELL\_10H\_DN | 154 | 11 | 12187 | 179 | Il4ra,Ccl4,Il1rn,Gadd45g,Ripk1,Cebpd,Il6,Selp,Saa1,Il17ra,Gbp3 | | 1.647e-05 | -11.01 | G\_GB1\_RHD3\_dom | interpro domains | IPR030386 | 11 | 4 | 13788 | 212 | Gbp4,Gbp6,Gbp7,Gbp3 | | 1.647e-05 | -11.01 | Guanylate-bd\_N | interpro domains | IPR015894 | 11 | 4 | 13788 | 212 | Gbp6,Gbp7,Gbp3,Gbp4 | | 1.647e-05 | -11.01 | Chemokine\_CC\_CS | interpro domains | IPR000827 | 11 | 4 | 13788 | 212 | Ccl7,Ccl2,Ccl12,Ccl4 | | 1.656e-05 | -11.01 | GO\_POSITIVE\_REGULATION\_OF\_HYDROLASE\_ACTIVITY | MSigDB lists | GO\_POSITIVE\_REGULATION\_OF\_HYDROLASE\_ACTIVITY | 716 | 26 | 12187 | 179 | Iqgap1,S100a8,Rgs16,Ifi211,Rhoc,Ripk1,Ccl12,Ccl4,Ccl2,Ptpn1,Hspa5,Dnajb1,Xdh,Tnfsf10,C5ar1,A2m,S100a9,Tnf,Icam1,Sele,Plek,Errfi1,Csf2rb,Ifi204,Ifi207,Ifi209 | | 1.658e-05 | -11.01 | GO\_CELLULAR\_RESPONSE\_TO\_INTERLEUKIN\_1 | MSigDB lists | GO\_CELLULAR\_RESPONSE\_TO\_INTERLEUKIN\_1 | 56 | 7 | 12187 | 179 | Il6,Ccl12,Nfkb1,Ccl2,Ccl4,Icam1,Irak3 | | 1.709e-05 | -10.98 | positive regulation of protein serine/threonine kinase activity | biological process | GO:0071902 | 264 | 15 | 13711 | 214 | Iqgap1,Ptpn1,Map3k6,Fzd4,Map3k8,Irgm2,Pik3r5,Tnf,Lgals9,Gadd45g,Ccnd2,Gadd45b,Ripk1,Thbs1,Il1rn | | 1.710e-05 | -10.98 | SMID\_BREAST\_CANCER\_BASAL\_UP | MSigDB lists | SMID\_BREAST\_CANCER\_BASAL\_UP | 467 | 20 | 12187 | 179 | Fam107a,Ccl12,S100a9,Angptl4,Ifi211,S100a8,Rtp4,Sox11,Ifi209,Socs3,Cp,Ifi207,Ifi204,Lcn2,Ptgs2,Tagln2,Icam1,Msn,Tap1,Clic4 | | 1.722e-05 | -10.97 | other organism part | cellular component | GO:0044217 | 22 | 5 | 13825 | 212 | Gbp3,Iigp1,Gbp7,Irgm2,Gbp6 | | 1.722e-05 | -10.97 | other organism cell | cellular component | GO:0044216 | 22 | 5 | 13825 | 212 | Gbp7,Gbp3,Iigp1,Gbp6,Irgm2 | | 1.725e-05 | -10.97 | KEGG\_CYTOSOLIC\_DNA\_SENSING\_PATHWAY | MSigDB lists | KEGG\_CYTOSOLIC\_DNA\_SENSING\_PATHWAY | 38 | 6 | 12187 | 179 | Nfkb1,Ripk1,Il6,Ccl4,Nfkbia,Irf7 | | 1.730e-05 | -10.96 | GO\_RESPONSE\_TO\_METAL\_ION | MSigDB lists | GO\_RESPONSE\_TO\_METAL\_ION | 248 | 14 | 12187 | 179 | Hspa5,Thbs1,Vcam1,Nfkb1,Cp,Il1a,Ptgs2,Cd14,Icam1,Clic4,Il6,Ptges,Iqgap1,S100a8 | | 1.743e-05 | -10.96 | GBP | pfam domains | PF02263 | 11 | 4 | 12881 | 201 | Gbp6,Gbp7,Gbp4,Gbp3 | | 1.747e-05 | -10.95 | retina vasculature morphogenesis in camera-type eye | biological process | GO:0061299 | 11 | 4 | 13711 | 214 | Fzd4,Clic4,Col4a1,Rhoj | | 1.748e-05 | -10.95 | GSE360\_LOW\_DOSE\_B\_MALAYI\_VS\_M\_TUBERCULOSIS\_MAC\_DN | MSigDB lists | GSE360\_LOW\_DOSE\_B\_MALAYI\_VS\_M\_TUBERCULOSIS\_MAC\_DN | 155 | 11 | 12187 | 179 | Irak3,Irf2,Il1rn,Osmr,Olfml2b,Pygm,Msr1,Zfp189,Runx1,Sntb2,Ripk1 | | 1.748e-05 | -10.95 | GSE43863\_TFH\_VS\_LY6C\_INT\_CXCR5POS\_EFFECTOR\_CD4\_TCELL\_DN | MSigDB lists | GSE43863\_TFH\_VS\_LY6C\_INT\_CXCR5POS\_EFFECTOR\_CD4\_TCELL\_DN | 155 | 11 | 12187 | 179 | Il6,Gadd45b,Irgm2,Bcl3,Ptges,Il1rn,Stx11,Ksr1,Nfkbiz,Socs3,Ifitm3 | | 1.748e-05 | -10.95 | GSE25088\_IL4\_VS\_IL4\_AND\_ROSIGLITAZONE\_STIM\_STAT6\_KO\_MACROPHAGE\_DAY10\_DN | MSigDB lists | GSE25088\_IL4\_VS\_IL4\_AND\_ROSIGLITAZONE\_STIM\_STAT6\_KO\_MACROPHAGE\_DAY10\_DN | 155 | 11 | 12187 | 179 | Dtx3l,Pik3r5,Nfkb2,Pik3ap1,Cxcl16,Hspa5,Ifit3b,Nfkb1,Il1a,Lcn2,Ier3 | | 1.748e-05 | -10.95 | positive regulation of cell death | biological process | GO:0010942 | 591 | 24 | 13711 | 214 | Tgm2,Ccl12,Hp,S100a8,S100a9,Plaur,Ifit2,Ptgs2,Akap12,Casp4,Tnf,Cdkn1a,Ip6k2,Tnfsf10,Il6,Atf3,Xdh,Zbtb16,Ripk1,Tnfaip8,Thbs1,Gadd45g,Hcar2,Gadd45b | | 1.755e-05 | -10.95 | GO\_ACUTE\_PHASE\_RESPONSE | MSigDB lists | GO\_ACUTE\_PHASE\_RESPONSE | 23 | 5 | 12187 | 179 | Saa1,Hp,Il1a,Il1rn,Il6 | | 1.781e-05 | -10.94 | Metal sequestration by antimicrobial proteins | REACTOME pathways | R-MMU-6799990 | 4 | 3 | 6297 | 105 | Lcn2,S100a9,S100a8 | | 1.845e-05 | -10.90 | negative regulation of phosphorylation | biological process | GO:0042326 | 403 | 19 | 13711 | 214 | Birc3,Ptpn1,Gbp4,Ier3,Rasip1,Cdkn1a,Errfi1,Il6,Atf3,Xdh,Tnfaip3,Tnip1,Gadd45g,Gadd45b,Parp14,Trib1,Hspb1,Socs3,Irak3 | | 1.856e-05 | -10.89 | GSE12366\_GC\_VS\_NAIVE\_BCELL\_DN | MSigDB lists | GSE12366\_GC\_VS\_NAIVE\_BCELL\_DN | 156 | 11 | 12187 | 179 | Ifit2,Ifi209,Ifi207,Ifi204,Dtx3l,Herc6,Ifitm3,Ifi211,Irf9,Msn,Parp9 | | 1.856e-05 | -10.89 | GSE42021\_TCONV\_PLN\_VS\_CD24HI\_TCONV\_THYMUS\_UP | MSigDB lists | GSE42021\_TCONV\_PLN\_VS\_CD24HI\_TCONV\_THYMUS\_UP | 156 | 11 | 12187 | 179 | Maff,Tgm2,Bach1,Atf3,Arid5b,Cxcl1,Rtp4,Zfp36,Icam1,Tap1,Sbno2 | | 1.856e-05 | -10.89 | GSE17721\_CTRL\_VS\_POLYIC\_2H\_BMDC\_DN | MSigDB lists | GSE17721\_CTRL\_VS\_POLYIC\_2H\_BMDC\_DN | 156 | 11 | 12187 | 179 | Ripk1,Hcar2,Gpr84,Pygm,Arid5b,Cflar,C5ar1,Luc7l3,Cdkn1a,Icam1,Cd14 | | 1.865e-05 | -10.89 | PLASARI\_TGFB1\_TARGETS\_10HR\_DN | MSigDB lists | PLASARI\_TGFB1\_TARGETS\_10HR\_DN | 217 | 13 | 12187 | 179 | Usp18,Pik3r1,Ifit2,Rtp4,Arid5b,C3ar1,Il1rn,Ifit3b,Vcam1,Adamts1,Parp9,Plek,Herc6 | | 1.893e-05 | -10.87 | GO\_REGULATION\_OF\_APOPTOTIC\_SIGNALING\_PATHWAY | MSigDB lists | GO\_REGULATION\_OF\_APOPTOTIC\_SIGNALING\_PATHWAY | 284 | 15 | 12187 | 179 | Plaur,Ptpn1,Hspb1,Icam1,Thbs1,Atf3,Il1a,S100a9,Cflar,Tnfsf10,S100a8,Tnf,Nfe2l2,Tnfaip3,Ripk1 | | 1.971e-05 | -10.83 | GSE17721\_CTRL\_VS\_CPG\_4H\_BMDC\_DN | MSigDB lists | GSE17721\_CTRL\_VS\_CPG\_4H\_BMDC\_DN | 157 | 11 | 12187 | 179 | Aff1,Tgm2,Ms4a6d,Nfkb1,Il1rn,Atf3,Znfx1,Gbp7,Ptges,Pik3ap1,Gbp3 | | 1.971e-05 | -10.83 | GSE46606\_DAY1\_VS\_DAY3\_CD40L\_IL2\_IL5\_STIMULATED\_BCELL\_DN | MSigDB lists | GSE46606\_DAY1\_VS\_DAY3\_CD40L\_IL2\_IL5\_STIMULATED\_BCELL\_DN | 157 | 11 | 12187 | 179 | Rnf213,Ifi207,Il1a,Ifi204,Sgk1,Ifi209,Tnfaip8,Dtx3l,Ifi211,Birc3,Runx1 | | 1.987e-05 | -10.83 | MODULE\_174 | MSigDB lists | MODULE\_174 | 79 | 8 | 12187 | 179 | Cp,Ier3,Spi1,C3ar1,Saa1,C5ar1,S100a9,Cd14 | | 2.014e-05 | -10.81 | WUNDER\_INFLAMMATORY\_RESPONSE\_AND\_CHOLESTEROL\_UP | MSigDB lists | WUNDER\_INFLAMMATORY\_RESPONSE\_AND\_CHOLESTEROL\_UP | 39 | 6 | 12187 | 179 | Cmpk2,Gbp4,Gbp6,Ifi44,Cd14,Nfkbiz | | 2.014e-05 | -10.81 | TAVOR\_CEBPA\_TARGETS\_UP | MSigDB lists | TAVOR\_CEBPA\_TARGETS\_UP | 39 | 6 | 12187 | 179 | Akap12,Il1rn,S100a9,Trib1,C3ar1,Ccnd2 | | 2.032e-05 | -10.80 | ROETH\_TERT\_TARGETS\_UP | MSigDB lists | ROETH\_TERT\_TARGETS\_UP | 12 | 4 | 12187 | 179 | Ifi44,Herc6,Ifit3b,Oasl1 | | 2.092e-05 | -10.77 | GSE6092\_B\_BURGDOFERI\_VS\_B\_BURGDORFERI\_AND\_IFNG\_STIM\_ENDOTHELIAL\_CELL\_DN | MSigDB lists | GSE6092\_B\_BURGDOFERI\_VS\_B\_BURGDORFERI\_AND\_IFNG\_STIM\_ENDOTHELIAL\_CELL\_DN | 158 | 11 | 12187 | 179 | Gbp3,Pik3r5,Cd14,Nfkbie,Tnfaip3,Gadd45b,Lcn2,Stx11,Tnip3,Cp,Trib1 | | 2.092e-05 | -10.77 | GSE41867\_DAY15\_EFFECTOR\_VS\_DAY30\_EXHAUSTED\_CD8\_TCELL\_LCMV\_CLONE13\_UP | MSigDB lists | GSE41867\_DAY15\_EFFECTOR\_VS\_DAY30\_EXHAUSTED\_CD8\_TCELL\_LCMV\_CLONE13\_UP | 158 | 11 | 12187 | 179 | Ccl4,Parp14,Znfx1,Stx11,Irf7,Herc6,Dtx3l,Saa1,Irgm2,Rtp4,Cmpk2 | | 2.097e-05 | -10.77 | BOSCO\_TH1\_CYTOTOXIC\_MODULE | MSigDB lists | BOSCO\_TH1\_CYTOTOXIC\_MODULE | 58 | 7 | 12187 | 179 | Isg15,Herc6,Gbp6,Usp18,Cmpk2,Rtp4,Gbp4 | | 2.108e-05 | -10.77 | CHARAFE\_BREAST\_CANCER\_LUMINAL\_VS\_MESENCHYMAL\_DN | MSigDB lists | CHARAFE\_BREAST\_CANCER\_LUMINAL\_VS\_MESENCHYMAL\_DN | 396 | 18 | 12187 | 179 | Msn,Ptgs2,Ifi207,Akap12,Col4a1,Ifi204,Timp1,Socs3,Fstl1,Sgk1,Ifi209,Osmr,Cxcl1,Ifi211,Il6,Tnfaip3,Bach1,Tubb6 | | 2.123e-05 | -10.76 | Jak-STAT signaling pathway | KEGG pathways | mmu04630 | 96 | 10 | 5248 | 107 | Cdkn1a,Csf2rb,Csf3,Il6,Socs3,Il4ra,Osmr,Pik3r1,Irf9,Ccnd2 | | 2.123e-05 | -10.76 | Jak-STAT signaling pathway | KEGG pathways | ko04630 | 96 | 10 | 5248 | 107 | Cdkn1a,Csf2rb,Csf3,Il4ra,Osmr,Socs3,Il6,Irf9,Pik3r1,Ccnd2 | | 2.125e-05 | -10.76 | regulation of viral genome replication | biological process | GO:0045069 | 75 | 8 | 13711 | 214 | Ifitm3,Oasl1,Ifitm6,Resf1,Ifitm2,Oasl2,Tnf,Isg15 | | 2.180e-05 | -10.73 | GO\_NEGATIVE\_REGULATION\_OF\_EXTRINSIC\_APOPTOTIC\_SIGNALING\_PATHWAY | MSigDB lists | GO\_NEGATIVE\_REGULATION\_OF\_EXTRINSIC\_APOPTOTIC\_SIGNALING\_PATHWAY | 80 | 8 | 12187 | 179 | Cflar,Icam1,Tnfsf10,Tnf,Thbs1,Il1a,Tnfaip3,Ripk1 | | 2.209e-05 | -10.72 | Lung fibrosis | WikiPathways | WP3632 | 48 | 8 | 3756 | 96 | Nfe2l2,Timp1,Ccl2,Il6,Ccl4,Tnf,Csf3,Mt2 | | 2.219e-05 | -10.72 | GSE2826\_XID\_VS\_BTK\_KO\_BCELL\_DN | MSigDB lists | GSE2826\_XID\_VS\_BTK\_KO\_BCELL\_DN | 159 | 11 | 12187 | 179 | Lcn2,Ccl12,Hp,Il1a,Ccl2,Vcam1,Tap1,Plaur,Sele,S100a8,S100a9 | | 2.219e-05 | -10.72 | GSE17721\_CTRL\_VS\_PAM3CSK4\_4H\_BMDC\_DN | MSigDB lists | GSE17721\_CTRL\_VS\_PAM3CSK4\_4H\_BMDC\_DN | 159 | 11 | 12187 | 179 | Hp,Nfkb1,Ksr1,Sntb2,Socs3,Parp14,Ccnd2,Iqgap1,Igsf6,Ifitm2,Cflar | | 2.315e-05 | -10.67 | negative regulation of peptidase activity | biological process | GO:0010466 | 150 | 11 | 13711 | 214 | Wfdc21,Plaur,Ptgs2,Tnfaip8,Cflar,Thbs1,Serpina3f,Il6,A2m,Birc3,Timp1 | | 2.331e-05 | -10.67 | GO\_CELLULAR\_RESPONSE\_TO\_STARVATION | MSigDB lists | GO\_CELLULAR\_RESPONSE\_TO\_STARVATION | 105 | 9 | 12187 | 179 | Hspa5,Ifi209,Cdkn1a,Ifi207,Ifi204,Atf3,Nfe2l2,Rrp8,Ifi211 | | 2.331e-05 | -10.67 | GO\_POSITIVE\_REGULATION\_OF\_STRESS\_ACTIVATED\_PROTEIN\_KINASE\_SIGNALING\_CASCADE | MSigDB lists | GO\_POSITIVE\_REGULATION\_OF\_STRESS\_ACTIVATED\_PROTEIN\_KINASE\_SIGNALING\_CASCADE | 105 | 9 | 12187 | 179 | Tnf,Gadd45g,Il1a,Gadd45b,Ripk1,Map3k6,Fzd4,Ptpn1,Xdh | | 2.353e-05 | -10.66 | GSE20715\_0H\_VS\_6H\_OZONE\_LUNG\_DN | MSigDB lists | GSE20715\_0H\_VS\_6H\_OZONE\_LUNG\_DN | 160 | 11 | 12187 | 179 | Socs3,Fstl1,Hp,Cp,Cd14,Errfi1,Trib1,Rhoc,Cxcl1,Bcl3,Osmr | | 2.353e-05 | -10.66 | GSE17721\_CTRL\_VS\_LPS\_2H\_BMDC\_DN | MSigDB lists | GSE17721\_CTRL\_VS\_LPS\_2H\_BMDC\_DN | 160 | 11 | 12187 | 179 | Arid5b,Nfkb2,Birc3,Map3k8,Cdkn1a,Casp4,Nfkbia,Ccl2,Vcam1,Timp1,Atf3 | | 2.353e-05 | -10.66 | GSE29618\_MONOCYTE\_VS\_PDC\_DAY7\_FLU\_VACCINE\_UP | MSigDB lists | GSE29618\_MONOCYTE\_VS\_PDC\_DAY7\_FLU\_VACCINE\_UP | 160 | 11 | 12187 | 179 | Sgk1,Trib1,Timp1,Il1rn,Cebpd,Ier3,C5ar1,S100a9,Cd14,Igsf6,Msn | | 2.353e-05 | -10.66 | GSE23321\_CENTRAL\_MEMORY\_VS\_NAIVE\_CD8\_TCELL\_UP | MSigDB lists | GSE23321\_CENTRAL\_MEMORY\_VS\_NAIVE\_CD8\_TCELL\_UP | 160 | 11 | 12187 | 179 | Irf7,S100a9,S100a8,Ptgs2,Nfkbiz,Sgk1,Hspa5,Lcn2,Tiparp,Mmp8,Il1a | | 2.365e-05 | -10.65 | cellular response to interleukin-6 | biological process | GO:0071354 | 23 | 5 | 13711 | 214 | Spi1,Sbno2,Il6,Nfkb1,Selp | | 2.365e-05 | -10.65 | lymphocyte chemotaxis | biological process | GO:0048247 | 23 | 5 | 13711 | 214 | Cxcl16,Ccl2,Ccl12,Ccl7,Ccl4 | | 2.373e-05 | -10.65 | GO\_CELLULAR\_RESPONSE\_TO\_EXTERNAL\_STIMULUS | MSigDB lists | GO\_CELLULAR\_RESPONSE\_TO\_EXTERNAL\_STIMULUS | 222 | 13 | 12187 | 179 | Icam1,Ptgs2,Nfkb1,Ifi204,Ifi207,Atf3,Cdkn1a,Adamts1,Ifi209,Hspa5,Ifi211,Rrp8,Nfe2l2 | | 2.409e-05 | -10.63 | Cytokines and Inflammatory Response | WikiPathways | WP222 | 15 | 5 | 3756 | 96 | Il6,Cxcl1,Csf3,Tnf,Il1a | | 2.412e-05 | -10.63 | negative regulation of endopeptidase activity | biological process | GO:0010951 | 124 | 10 | 13711 | 214 | Timp1,Birc3,Il6,A2m,Thbs1,Serpina3f,Tnfaip8,Cflar,Ptgs2,Plaur | | 2.417e-05 | -10.63 | RIG-I/MDA5 mediated induction of IFN-alpha/beta pathways | REACTOME pathways | R-MMU-168928 | 36 | 6 | 6297 | 105 | Irf7,Nfkb1,Irf2,Nfkb2,Tnfaip3,Nfkbia | | 2.479e-05 | -10.61 | response to glucocorticoid | biological process | GO:0051384 | 56 | 7 | 13711 | 214 | Sgk1,Pik3r1,Il1rn,Il6,Zfp36,Tnf,Fam107a | | 2.494e-05 | -10.60 | GSE16522\_MEMORY\_VS\_NAIVE\_ANTI\_CD3CD28\_STIM\_CD8\_TCELL\_DN | MSigDB lists | GSE16522\_MEMORY\_VS\_NAIVE\_ANTI\_CD3CD28\_STIM\_CD8\_TCELL\_DN | 161 | 11 | 12187 | 179 | Casp4,Lgals9,Ccnd2,Gbp4,Irf7,Gbp6,Irgm2,Xdh,Ncf1,Rnf213,Ptpn1 | | 2.494e-05 | -10.60 | GSE21063\_WT\_VS\_NFATC1\_KO\_3H\_ANTI\_IGM\_STIM\_BCELL\_DN | MSigDB lists | GSE21063\_WT\_VS\_NFATC1\_KO\_3H\_ANTI\_IGM\_STIM\_BCELL\_DN | 161 | 11 | 12187 | 179 | Parp12,Gbp4,AA467197,Tnfsf10,Gbp3,Gbp6,Gbp7,Irgm2,Tap1,Parp9,Tnip1 | | 2.494e-05 | -10.60 | GSE7852\_LN\_VS\_FAT\_TCONV\_DN | MSigDB lists | GSE7852\_LN\_VS\_FAT\_TCONV\_DN | 161 | 11 | 12187 | 179 | Hspa5,Cdkn1a,Nfkbiz,Col4a1,Ier3,Atf3,Ripk1,Ifitm2,Tagln2,Plek,Iqgap1 | | 2.494e-05 | -10.60 | GSE23925\_DARK\_ZONE\_VS\_NAIVE\_BCELL\_DN | MSigDB lists | GSE23925\_DARK\_ZONE\_VS\_NAIVE\_BCELL\_DN | 161 | 11 | 12187 | 179 | Tnfaip3,Ccl4,Dnajb1,Trib1,Nfkbiz,Tnf,Maff,Hspa5,Ptpn1,Zfp36,Gem | | 2.530e-05 | -10.58 | Apoptosis | KEGG pathways | mmu04210 | 119 | 11 | 5248 | 107 | Gadd45g,Tnf,Birc3,Ripk1,Tnfsf10,Nfkb1,Csf2rb,Cflar,Pik3r1,Nfkbia,Gadd45b | | 2.530e-05 | -10.58 | Apoptosis | KEGG pathways | ko04210 | 119 | 11 | 5248 | 107 | Nfkbia,Gadd45b,Cflar,Pik3r1,Csf2rb,Nfkb1,Tnf,Birc3,Ripk1,Tnfsf10,Gadd45g | | 2.580e-05 | -10.57 | GSE36392\_MAC\_VS\_NEUTROPHIL\_IL25\_TREATED\_LUNG\_DN | MSigDB lists | GSE36392\_MAC\_VS\_NEUTROPHIL\_IL25\_TREATED\_LUNG\_DN | 133 | 10 | 12187 | 179 | Slfn3,Slfn4,Errfi1,Cd14,Il1a,Il1rn,Cdkn1a,Nfkbiz,Trib1,Socs3 | | 2.589e-05 | -10.56 | eosinophil chemotaxis | biological process | GO:0048245 | 12 | 4 | 13711 | 214 | Ccl7,Ccl4,Ccl12,Ccl2 | | 2.589e-05 | -10.56 | toll-like receptor 4 signaling pathway | biological process | GO:0034142 | 12 | 4 | 13711 | 214 | Tnip3,Nfkbia,Cd14,Pik3ap1 | | 2.589e-05 | -10.56 | eosinophil migration | biological process | GO:0072677 | 12 | 4 | 13711 | 214 | Ccl4,Ccl7,Ccl12,Ccl2 | | 2.619e-05 | -10.55 | regulation of cytokine secretion | biological process | GO:0050707 | 152 | 11 | 13711 | 214 | Mmp8,Il17ra,Casp4,Tnfaip3,Tnf,Il4ra,Il1a,Akap12,Il6,Cd14,Lgals9 | | 2.627e-05 | -10.55 | MODULE\_241 | MSigDB lists | MODULE\_241 | 60 | 7 | 12187 | 179 | Ier3,Spi1,C3ar1,Saa1,C5ar1,Cd14,S100a9 | | 2.627e-05 | -10.55 | GO\_REGULATION\_OF\_LEUKOCYTE\_CHEMOTAXIS | MSigDB lists | GO\_REGULATION\_OF\_LEUKOCYTE\_CHEMOTAXIS | 60 | 7 | 12187 | 179 | C3ar1,C5ar1,Cxcl1,Thbs1,Ccl4,Il6,Ccl12 | | 2.642e-05 | -10.54 | GO\_REGULATION\_OF\_ERK1\_AND\_ERK2\_CASCADE | MSigDB lists | GO\_REGULATION\_OF\_ERK1\_AND\_ERK2\_CASCADE | 162 | 11 | 12187 | 179 | C5ar1,Tnf,Ccl2,Ccl4,Il6,Ccl12,Tnip1,Icam1,Errfi1,Ptpn1,Atf3 | | 2.642e-05 | -10.54 | GSE2405\_0H\_VS\_3H\_A\_PHAGOCYTOPHILUM\_STIM\_NEUTROPHIL\_UP | MSigDB lists | GSE2405\_0H\_VS\_3H\_A\_PHAGOCYTOPHILUM\_STIM\_NEUTROPHIL\_UP | 162 | 11 | 12187 | 179 | Col4a1,Timp1,Csf2rb,Ifitm2,Cybb,Casp4,Nfkbia,Tgm2,Tor3a,Adamts9,Rhoj | | 2.642e-05 | -10.54 | GO\_CELLULAR\_RESPONSE\_TO\_EXTRACELLULAR\_STIMULUS | MSigDB lists | GO\_CELLULAR\_RESPONSE\_TO\_EXTRACELLULAR\_STIMULUS | 162 | 11 | 12187 | 179 | Icam1,Ifi211,Rrp8,Ifi204,Ifi207,Atf3,Nfe2l2,Cdkn1a,Ifi209,Adamts1,Hspa5 | | 2.642e-05 | -10.54 | GSE3039\_ALPHAALPHA\_VS\_ALPHABETA\_CD8\_TCELL\_UP | MSigDB lists | GSE3039\_ALPHAALPHA\_VS\_ALPHABETA\_CD8\_TCELL\_UP | 162 | 11 | 12187 | 179 | Il6,Sdc4,Gpr84,Ccl4,Il4ra,Maff,Ptges,Ier3,Socs3,Cdkn1a,Cd14 | | 2.642e-05 | -10.54 | GSE557\_CIITA\_KO\_VS\_I\_AB\_KO\_DC\_UP | MSigDB lists | GSE557\_CIITA\_KO\_VS\_I\_AB\_KO\_DC\_UP | 162 | 11 | 12187 | 179 | Slfn4,Slfn3,Sbno2,Gbp3,Gbp7,Ncf1,Samd9l,Map3k8,Lgals9,Xaf1,Socs3 | | 2.642e-05 | -10.54 | GSE40277\_EOS\_AND\_LEF1\_TRANSDUCED\_VS\_CTRL\_CD4\_TCELL\_UP | MSigDB lists | GSE40277\_EOS\_AND\_LEF1\_TRANSDUCED\_VS\_CTRL\_CD4\_TCELL\_UP | 162 | 11 | 12187 | 179 | Gbp4,Birc3,Aff1,Casp4,Parp14,Icam1,Irf9,Clic4,Irf2,Gbp6,Bcl3 | | 2.650e-05 | -10.54 | Tnfaip3 (tumor necrosis factor, alpha-induced protein 3) | protein interactions | 21929 | 22 | 5 | 6802 | 115 | Tnfaip3,Ripk1,Tnf,Tnip3,Tnip1 | | 2.706e-05 | -10.52 | DORN\_ADENOVIRUS\_INFECTION\_12HR\_DN | MSigDB lists | DORN\_ADENOVIRUS\_INFECTION\_12HR\_DN | 25 | 5 | 12187 | 179 | Sgk1,Nfkbia,Nfe2l2,Ier3,Il6 | | 2.709e-05 | -10.52 | LEE\_LIVER\_CANCER\_MYC\_E2F1\_UP | MSigDB lists | LEE\_LIVER\_CANCER\_MYC\_E2F1\_UP | 41 | 6 | 12187 | 179 | Ifi44,Ifit2,Usp18,Tagln2,Icam1,Lcn2 | | 2.709e-05 | -10.52 | LIAN\_LIPA\_TARGETS\_3M | MSigDB lists | LIAN\_LIPA\_TARGETS\_3M | 41 | 6 | 12187 | 179 | Igsf6,S100a9,Cxcl1,C3ar1,Mmp8,Msr1 | | 2.709e-05 | -10.52 | ZHENG\_IL22\_SIGNALING\_UP | MSigDB lists | ZHENG\_IL22\_SIGNALING\_UP | 41 | 6 | 12187 | 179 | S100a9,Cd14,S100a8,Sbno2,Hp,Timp1 | | 2.712e-05 | -10.52 | ZHU\_CMV\_ALL\_UP | MSigDB lists | ZHU\_CMV\_ALL\_UP | 107 | 9 | 12187 | 179 | Irf2,Tap1,Isg15,Ptgs2,Nfkb1,Ripk1,Tnfaip3,Il6,Nfkbia | | 2.801e-05 | -10.48 | GO\_RESPONSE\_TO\_ORGANIC\_CYCLIC\_COMPOUND | MSigDB lists | GO\_RESPONSE\_TO\_ORGANIC\_CYCLIC\_COMPOUND | 694 | 25 | 12187 | 179 | P2ry6,Irak3,Selp,Tiparp,Nfkb1,Sgk1,Thbs1,Pygm,Ptges,Il6,Ccl12,Msr1,Il4ra,Trim25,Icam1,Msn,Errfi1,Ptgs2,Il1rn,Cdkn1a,Adamts1,Vcam1,Socs3,Tnf,Nfkbia | | 2.825e-05 | -10.47 | regulation of hydrolase activity | biological process | GO:0051336 | 948 | 32 | 13711 | 214 | Iqgap1,S100a8,A2m,Ccl12,Dnajb1,Angptl4,Wfdc21,Serpina3f,S100a9,Xdh,Tnfsf10,Ccl2,Tnf,Plek,Rhoc,Sele,Tnfaip8,Cflar,Timp1,Birc3,Ptgs2,Plaur,Rasip1,Ccl4,Il6,Oasl2,Icam1,Casp4,Rgs16,Thbs1,Oasl1,Ccl7 | | 2.845e-05 | -10.47 | RUTELLA\_RESPONSE\_TO\_HGF\_UP | MSigDB lists | RUTELLA\_RESPONSE\_TO\_HGF\_UP | 330 | 16 | 12187 | 179 | Sgk1,Tnip3,Il1a,Gadd45g,Hspb1,Rbm47,Tgm2,Ccl2,Sdc4,Runx1,Birc3,Ccl12,Cybb,Cflar,C3ar1,Tnfaip8 | | 2.860e-05 | -10.46 | positive regulation of wound healing | biological process | GO:0090303 | 39 | 6 | 13711 | 214 | S100a9,Nfe2l2,Thbs1,Selp,Plek,Ccl2 | | 2.882e-05 | -10.45 | GO\_REGULATION\_OF\_PROTEOLYSIS | MSigDB lists | GO\_REGULATION\_OF\_PROTEOLYSIS | 525 | 21 | 12187 | 179 | Plaur,Ifi207,Ifi204,Timp1,Thbs1,Ifi209,Tnfsf10,Xdh,S100a8,Tnfaip8,A2m,S100a9,Ifi211,C3ar1,C5ar1,Il6,Nfe2l2,Ripk1,Trib1,Birc3,Tnf | | 2.892e-05 | -10.45 | GO\_NEGATIVE\_REGULATION\_OF\_GENE\_EXPRESSION | MSigDB lists | GO\_NEGATIVE\_REGULATION\_OF\_GENE\_EXPRESSION | 1113 | 34 | 12187 | 179 | Ifi204,Ifi207,Rasd1,Atf3,Cdkn1a,Ifi209,Zbtb16,Zfp36,Bach1,Zfp189,Tnf,Birc3,Trib1,Nfkbia,Nfkb2,Xdh,A2m,Irf7,Nfkb1,Tiparp,Mxd4,Sox11,Thbs1,Irak3,Irf2,Ifitm3,Tnfaip3,Ccl4,Spi1,Sbno2,Ifi211,Arid5b,Rrp8,Bcl3 | | 2.901e-05 | -10.45 | UROSEVIC\_RESPONSE\_TO\_IMIQUIMOD | MSigDB lists | UROSEVIC\_RESPONSE\_TO\_IMIQUIMOD | 13 | 4 | 12187 | 179 | Oasl1,Il6,Isg15,Irf7 | | 2.901e-05 | -10.45 | GO\_TOLL\_LIKE\_RECEPTOR\_4\_SIGNALING\_PATHWAY | MSigDB lists | GO\_TOLL\_LIKE\_RECEPTOR\_4\_SIGNALING\_PATHWAY | 13 | 4 | 12187 | 179 | Pik3ap1,Tnip3,Cd14,Nfkbia | | 2.901e-05 | -10.45 | TSAI\_DNAJB4\_TARGETS\_UP | MSigDB lists | TSAI\_DNAJB4\_TARGETS\_UP | 13 | 4 | 12187 | 179 | Irf9,Ifit3b,Cdkn1a,Isg15 | | 2.931e-05 | -10.44 | ST\_FAS\_SIGNALING\_PATHWAY | MSigDB lists | ST\_FAS\_SIGNALING\_PATHWAY | 61 | 7 | 12187 | 179 | Nfkbie,Hspb1,Nfkb2,Nfkbia,Il1a,Nfkb1,Ripk1 | | 2.931e-05 | -10.44 | PID\_REG\_GR\_PATHWAY | MSigDB lists | PID\_REG\_GR\_PATHWAY | 61 | 7 | 12187 | 179 | Cdkn1a,Sgk1,Spi1,Nfkb1,Il6,Icam1,Sele | | 2.931e-05 | -10.44 | GNF2\_CD53 | MSigDB lists | GNF2\_CD53 | 61 | 7 | 12187 | 179 | Tap1,Ifi204,Ifi207,Ifi211,Phf11d,Phf11b,Ifi209 | | 2.935e-05 | -10.44 | GO\_CELLULAR\_CHEMICAL\_HOMEOSTASIS | MSigDB lists | GO\_CELLULAR\_CHEMICAL\_HOMEOSTASIS | 406 | 18 | 12187 | 179 | Saa1,Sgk3,Clic4,Icam1,Sgk1,Lcn2,Cp,Kcna5,S100a9,C3ar1,C5ar1,Pygm,S100a8,Ccl2,Tgm2,Pik3r1,Ccl12,Slc24a4 | | 2.950e-05 | -10.43 | response to interleukin-6 | biological process | GO:0070741 | 24 | 5 | 13711 | 214 | Spi1,Nfkb1,Il6,Selp,Sbno2 | | 2.952e-05 | -10.43 | THUM\_SYSTOLIC\_HEART\_FAILURE\_UP | MSigDB lists | THUM\_SYSTOLIC\_HEART\_FAILURE\_UP | 331 | 16 | 12187 | 179 | Clic4,Selp,Cd14,Cp,Ms4a6d,Thbs1,Vcam1,Tnfsf10,C3ar1,Cxcl1,S100a9,Oasl1,Gadd45b,Msr1,Ccl4,Tgm2 | | 2.961e-05 | -10.43 | GSE46606\_IRF4HIGH\_VS\_WT\_CD40L\_IL2\_IL5\_DAY1\_STIMULATED\_BCELL\_DN | MSigDB lists | GSE46606\_IRF4HIGH\_VS\_WT\_CD40L\_IL2\_IL5\_DAY1\_STIMULATED\_BCELL\_DN | 164 | 11 | 12187 | 179 | Clic4,Parp9,Gbp7,Ptges,Slfn5,Znfx1,Maff,Aff1,Nfkbiz,Casp4,Vcam1 | | 2.961e-05 | -10.43 | HALLMARK\_KRAS\_SIGNALING\_UP | MSigDB lists | HALLMARK\_KRAS\_SIGNALING\_UP | 164 | 11 | 12187 | 179 | Trib1,Birc3,Ccnd2,Tnfaip3,Gadd45g,Akap12,Angptl4,Ptgs2,C3ar1,Rgs16,Plaur | | 2.998e-05 | -10.41 | regulation of vasculature development | biological process | GO:1901342 | 277 | 15 | 13711 | 214 | C5ar1,Adamts1,Rhoj,Nfe2l2,Ptgs2,Runx1,Il1a,Ccl2,Cybb,Xdh,Cflar,Adamts9,Hspb1,C3ar1,Thbs1 | | 3.010e-05 | -10.41 | Signaling by Interleukins | REACTOME pathways | R-MMU-449147 | 299 | 16 | 6297 | 105 | Il4ra,Il6,Il1rn,Il1a,Socs3,Nfkb2,Il17ra,Ksr1,Csf2rb,Nfkb1,Lgals9,Map3k8,Pik3r1,Irak3,Osmr,Csf3 | | 3.029e-05 | -10.40 | GO\_POSITIVE\_REGULATION\_OF\_CELL\_DEATH | MSigDB lists | GO\_POSITIVE\_REGULATION\_OF\_CELL\_DEATH | 486 | 20 | 12187 | 179 | Tgm2,Tnf,Ifit2,Gadd45b,Il6,Ripk1,Hcar2,S100a9,Tnfsf10,Xdh,S100a8,Zbtb16,Thbs1,Cdkn1a,Atf3,Hp,Gadd45g,Ptgs2,Plaur,Ip6k2 | | 3.062e-05 | -10.39 | GO\_WOUND\_HEALING | MSigDB lists | GO\_WOUND\_HEALING | 332 | 16 | 12187 | 179 | Saa1,Irf2,Plaur,Plek,Hspb1,Il1a,Timp1,A2m,Pik3r5,Cflar,S100a8,Sdc4,Maff,Pik3r1,Il6,Rhoc | | 3.122e-05 | -10.37 | GO\_NEGATIVE\_REGULATION\_OF\_DNA\_BINDING | MSigDB lists | GO\_NEGATIVE\_REGULATION\_OF\_DNA\_BINDING | 42 | 6 | 12187 | 179 | Sox11,Ifi209,Ifi211,Nfkbia,Ifi204,Ifi207 | | 3.122e-05 | -10.37 | ZHOU\_TNF\_SIGNALING\_30MIN | MSigDB lists | ZHOU\_TNF\_SIGNALING\_30MIN | 42 | 6 | 12187 | 179 | Tnfaip3,Il6,Hspb1,Tagln2,Nfkbia,Ptgs2 | | 3.129e-05 | -10.37 | GO\_NEGATIVE\_REGULATION\_OF\_CYTOKINE\_PRODUCTION | MSigDB lists | GO\_NEGATIVE\_REGULATION\_OF\_CYTOKINE\_PRODUCTION | 136 | 10 | 12187 | 179 | Isg15,Bcl3,Irak3,Trim25,Errfi1,Tnf,Thbs1,Nfkb1,Tnfaip3,Il6 | | 3.132e-05 | -10.37 | GSE20715\_0H\_VS\_48H\_OZONE\_TLR4\_KO\_LUNG\_DN | MSigDB lists | GSE20715\_0H\_VS\_48H\_OZONE\_TLR4\_KO\_LUNG\_DN | 165 | 11 | 12187 | 179 | Osmr,Ptpn1,Cd14,Cxcl1,Gadd45g,Lcn2,Thbs1,Cdkn1a,Rhou,Dnajb1,Socs3 | | 3.132e-05 | -10.37 | GSE39152\_CD103\_NEG\_VS\_POS\_MEMORY\_CD8\_TCELL\_DN | MSigDB lists | GSE39152\_CD103\_NEG\_VS\_POS\_MEMORY\_CD8\_TCELL\_DN | 165 | 11 | 12187 | 179 | C5ar1,Ccl4,Il4ra,Ptpn1,Zfp36,Ncf1,Ier3,Gadd45g,Timp1,Tiparp,Socs3 | | 3.132e-05 | -10.37 | GSE29618\_MONOCYTE\_VS\_MDC\_DAY7\_FLU\_VACCINE\_UP | MSigDB lists | GSE29618\_MONOCYTE\_VS\_MDC\_DAY7\_FLU\_VACCINE\_UP | 165 | 11 | 12187 | 179 | Cxcl1,Cd14,S100a9,C5ar1,C3ar1,Irak3,Icam1,S100a8,Nfkbia,Ier3,Cybb | | 3.132e-05 | -10.37 | GSE11961\_MARGINAL\_ZONE\_BCELL\_VS\_MEMORY\_BCELL\_DAY7\_UP | MSigDB lists | GSE11961\_MARGINAL\_ZONE\_BCELL\_VS\_MEMORY\_BCELL\_DAY7\_UP | 165 | 11 | 12187 | 179 | Slfn9,Slfn5,Herc6,Irf7,Rgs16,Cmpk2,Trib1,Usp18,Tor3a,Xaf1,Ifit2 | | 3.137e-05 | -10.37 | GO\_CELL\_MOTILITY | MSigDB lists | GO\_CELL\_MOTILITY | 612 | 23 | 12187 | 179 | Thbs1,Cxcl16,Parp9,Il17ra,Selp,Ccl12,Il6,Rhoc,Ccl4,Ccl2,Pik3r1,S100a8,Cxcl1,Arid5b,Vcam1,Msn,Icam1,Sele,Saa1,Sdc4,Tnf,S100a9,C5ar1 | | 3.226e-05 | -10.34 | Adaptive Immune System | REACTOME pathways | R-MMU-1280218 | 442 | 20 | 6297 | 105 | Tap1,Vcam1,Cdkn1a,Icam1,Dtx3l,Rnf213,Ifitm3,Nfkbie,Cybb,Pik3ap1,Pik3r1,Ncf1,Map3k8,Nfkbia,Fyb,Nfkb1,Herc6,Xdh,Ifitm2,Ier3 | | 3.261e-05 | -10.33 | Interferon alpha/beta signaling | REACTOME pathways | R-MMU-909733 | 12 | 4 | 6297 | 105 | Usp18,Ptpn1,Irf9,Socs3 | | 3.310e-05 | -10.32 | GO\_MYD88\_INDEPENDENT\_TOLL\_LIKE\_RECEPTOR\_SIGNALING\_PATHWAY | MSigDB lists | GO\_MYD88\_INDEPENDENT\_TOLL\_LIKE\_RECEPTOR\_SIGNALING\_PATHWAY | 26 | 5 | 12187 | 179 | Irf7,Cd14,Birc3,Ripk1,Tnip3 | | 3.310e-05 | -10.32 | GNF2\_SPI1 | MSigDB lists | GNF2\_SPI1 | 26 | 5 | 12187 | 179 | Spi1,C5ar1,Tnfsf10,Igsf6,Bach1 | | 3.312e-05 | -10.32 | GSE40274\_FOXP3\_VS\_FOXP3\_AND\_GATA1\_TRANSDUCED\_ACTIVATED\_CD4\_TCELL\_DN | MSigDB lists | GSE40274\_FOXP3\_VS\_FOXP3\_AND\_GATA1\_TRANSDUCED\_ACTIVATED\_CD4\_TCELL\_DN | 166 | 11 | 12187 | 179 | Irf9,Ifitm2,Herc6,Rnf213,Ncf1,Gadd45g,Ifi44,Bcl3,Irf7,Oasl1,Birc3 | | 3.312e-05 | -10.32 | GSE14769\_UNSTIM\_VS\_240MIN\_LPS\_BMDM\_DN | MSigDB lists | GSE14769\_UNSTIM\_VS\_240MIN\_LPS\_BMDM\_DN | 166 | 11 | 12187 | 179 | Il1rn,Atf3,Cdkn1a,Map3k8,Tor3a,Tgm2,Irf9,Csf2rb,Clic4,Gbp7,Slfn5 | | 3.333e-05 | -10.31 | GO\_INTRINSIC\_APOPTOTIC\_SIGNALING\_PATHWAY | MSigDB lists | GO\_INTRINSIC\_APOPTOTIC\_SIGNALING\_PATHWAY | 137 | 10 | 12187 | 179 | Casp4,Pik3r1,Cdkn1a,Tnf,Ifi209,Ifi207,Ifi204,Bcl3,Rrp8,Ifi211 | | 3.333e-05 | -10.31 | GSE2706\_2H\_VS\_8H\_R848\_AND\_LPS\_STIM\_DC\_DN | MSigDB lists | GSE2706\_2H\_VS\_8H\_R848\_AND\_LPS\_STIM\_DC\_DN | 137 | 10 | 12187 | 179 | Isg15,Herc6,Ifitm3,Phf11d,Irf9,Parp9,Birc3,Phf11b,Xaf1,AA467197 | | 3.336e-05 | -10.31 | cellular process | biological process | GO:0009987 | 10077 | 182 | 13711 | 214 | Msr1,Iigp1,Bbs12,Lgals9,Gadd45g,Rgs16,Nfkb2,Fyb,Mt1,Rtp4,Birc3,Map3k8,Rhoj,Tnfaip8,Luc7l3,Aff1,Znfx1,Tnfaip3,Ccl2,Depp1,Vcam1,Cxcl16,Tnip3,Iqgap1,Col4a1,Ccl7,Ccnd2,Akap12,Il4ra,Slfn9,Ip6k2,Gm5431,Tubb6,Mx2,Cebpd,Rasip1,Xaf1,Trim25,Runx1,Sele,Ncf1,Trim30a,Cp,Acod1,Atf3,Mmp8,Nfe2l2,Noct,Angptl4,Clic4,Tgm2,C5ar1,Maff,Gm4841,Ifitm3,Thbs1,P2ry6,Cemip2,Errfi1,Kcna5,Bach1,Plaur,Trim56,Slfn5,Ifi204,Plek,Ksr1,Xdh,Usp18,Sgk3,Irgm2,Gbp6,Bcl3,Gbp3,Ccl12,Trib1,Socs3,Gem,Il1a,Tgtp2,Cdkn1a,Ccl4,Rbm47,Ifitm6,Ier3,Sgk1,Ptges,Sbno2,Parp9,Cflar,Cd14,Slc24a4,Rhoc,Fstl1,Robo4,Tnf,Ifit2,Cxcl1,Icam1,Rassf4,Irgm1,Pik3r5,Zbtb16,Csf3,Il1rn,C3ar1,Lcn2,Retnlg,Cxcl9,S100a9,Wfdc21,Ifitm2,Ptpn1,Nfkbiz,Parp14,Hcar2,Nfkbia,Herc6,Ifi211,Fam107a,Map3k6,Tiparp,Timp1,Isg15,Rnf213,Pik3ap1,Rhou,Rasd1,Olfml2b,Gbp7,Selp,Msn,Hspa5,Mx1,Banp,Rbm39,Sox11,Cybb,Il17ra,Osmr,Gbp4,Tagln2,F830016B08Rik,Adamts1,Nfkb1,Ifi47,Adamts9,Arid5b,Pygm,Irak3,Parp12,Tnfsf10,Samd9l,A2m,S100a8,Irf7,Dtx3l,Pik3r1,Gadd45b,Casp4,Oasl2,Il6,Spi1,Stx11,Ptgs2,Fzd4,Ifi209,Sdc4,Tgtp1,Cmpk2,Gpr84,Ripk1,Hspb1,Ifi207,Tnip1,Saa1,Zfp36,Csf2rb,Rrp8,Dnajb1,Mt2,Hp | | 3.397e-05 | -10.29 | G\_GB1\_RHD3 | prosite domains | PS51715 | 11 | 4 | 8845 | 164 | Gbp4,Gbp6,Gbp3,Gbp7 | | 3.397e-05 | -10.29 | SMALL\_CYTOKINES\_CC | prosite domains | PS00472 | 11 | 4 | 8845 | 164 | Ccl2,Ccl12,Ccl7,Ccl4 | | 3.405e-05 | -10.29 | GOZGIT\_ESR1\_TARGETS\_DN | MSigDB lists | GOZGIT\_ESR1\_TARGETS\_DN | 531 | 21 | 12187 | 179 | Tnfsf10,Arid5b,Irf7,Tnfaip3,Ifit2,Birc3,Xaf1,Tgm2,Usp18,Cxcl16,Irf9,Herc6,Timp1,Samd9l,Cp,Rasd1,Lcn2,Thbs1,Sgk1,Nfkbiz,Ifit3b | | 3.405e-05 | -10.29 | positive regulation of apoptotic process | biological process | GO:0043065 | 536 | 22 | 13711 | 214 | Il6,Ip6k2,Casp4,Akap12,Gadd45b,Gadd45g,Hcar2,Thbs1,Ptgs2,Plaur,Xdh,Atf3,Tnfsf10,Tnf,Ripk1,Tnfaip8,Zbtb16,S100a8,Ccl12,Tgm2,Ifit2,S100a9 | | 3.469e-05 | -10.27 | interspecies interaction between organisms | biological process | GO:0044419 | 216 | 13 | 13711 | 214 | Gbp6,Irgm2,Gbp7,Ccl4,S100a9,Irgm1,Vcam1,Gbp3,Tnip1,Ripk1,Ncf1,Ccl2,Icam1 | | 3.484e-05 | -10.26 | DAPIN | interpro domains | IPR004020 | 13 | 4 | 13788 | 212 | Ifi209,Ifi211,Ifi204,Ifi207 | | 3.484e-05 | -10.26 | Chemokine\_b/g/d | interpro domains | IPR039809 | 13 | 4 | 13788 | 212 | Ccl12,Ccl4,Ccl2,Ccl7 | | 3.501e-05 | -10.26 | GSE18893\_TCONV\_VS\_TREG\_24H\_CULTURE\_DN | MSigDB lists | GSE18893\_TCONV\_VS\_TREG\_24H\_CULTURE\_DN | 167 | 11 | 12187 | 179 | Irf7,Bcl3,Rnf213,Cflar,Cmpk2,Ifit3b,Sdc4,Ifi44,Ifit2,Oasl1,Akap12 | | 3.501e-05 | -10.26 | GSE16385\_ROSIGLITAZONE\_IL4\_VS\_IL4\_ALONE\_STIM\_MACROPHAGE\_12H\_UP | MSigDB lists | GSE16385\_ROSIGLITAZONE\_IL4\_VS\_IL4\_ALONE\_STIM\_MACROPHAGE\_12H\_UP | 167 | 11 | 12187 | 179 | Zfp36,Rbm47,Csf2rb,Tap1,Phf11d,Tubb6,Kcna5,Ms4a6d,Parp12,Phf11b,Pik3r1 | | 3.505e-05 | -10.26 | response to corticosteroid | biological process | GO:0031960 | 59 | 7 | 13711 | 214 | Il1rn,Sgk1,Pik3r1,Fam107a,Tnf,Zfp36,Il6 | | 3.549e-05 | -10.25 | KEGG\_CHEMOKINE\_SIGNALING\_PATHWAY | MSigDB lists | KEGG\_CHEMOKINE\_SIGNALING\_PATHWAY | 138 | 10 | 12187 | 179 | Cxcl16,Ncf1,Cxcl1,Pik3r5,Nfkb1,Ccl12,Pik3r1,Ccl2,Nfkbia,Ccl4 | | 3.585e-05 | -10.24 | GO\_POSITIVE\_REGULATION\_OF\_TUMOR\_NECROSIS\_FACTOR\_SUPERFAMILY\_CYTOKINE\_PRODUCTION | MSigDB lists | GO\_POSITIVE\_REGULATION\_OF\_TUMOR\_NECROSIS\_FACTOR\_SUPERFAMILY\_CYTOKINE\_PRODUCTION | 43 | 6 | 12187 | 179 | Hspb1,Ripk1,Ccl12,Thbs1,Pik3r1,Cd14 | | 3.611e-05 | -10.23 | GO\_NEGATIVE\_REGULATION\_OF\_HYDROLASE\_ACTIVITY | MSigDB lists | GO\_NEGATIVE\_REGULATION\_OF\_HYDROLASE\_ACTIVITY | 265 | 14 | 12187 | 179 | Tnfaip8,Iqgap1,A2m,Angptl4,Ifi211,Il6,Birc3,Tnf,Plaur,Ifi207,Ifi204,Timp1,Thbs1,Ifi209 | | 3.631e-05 | -10.22 | FULCHER\_INFLAMMATORY\_RESPONSE\_LECTIN\_VS\_LPS\_UP | MSigDB lists | FULCHER\_INFLAMMATORY\_RESPONSE\_LECTIN\_VS\_LPS\_UP | 452 | 19 | 12187 | 179 | Tagln2,P2ry6,Plaur,Il1a,Gadd45g,Il1rn,Nfkbie,Cxcl1,Igsf6,Birc3,Maff,Runx1,Ccl2,Spi1,Tgm2,Tubb6,Cybb,Rhoc,Il6 | | 3.664e-05 | -10.21 | positive regulation of interferon-gamma-mediated signaling pathway | biological process | GO:0060335 | 5 | 3 | 13711 | 214 | Irgm1,Irgm2,Parp9 | | 3.664e-05 | -10.21 | positive regulation of response to interferon-gamma | biological process | GO:0060332 | 5 | 3 | 13711 | 214 | Irgm2,Parp9,Irgm1 | | 3.678e-05 | -10.21 | blood vessel morphogenesis | biological process | GO:0048514 | 351 | 17 | 13711 | 214 | Rnf213,Thbs1,Adamts9,Cemip2,Ccl2,Robo4,Angptl4,Ptgs2,Rasip1,Spi1,Col4a1,Apold1,Rhoj,Ccl12,Tiparp,Fzd4,Clic4 | | 3.685e-05 | -10.21 | PYRIN | pfam domains | PF02758 | 13 | 4 | 12881 | 201 | Ifi204,Ifi207,Ifi211,Ifi209 | | 3.694e-05 | -10.21 | adhesion of symbiont to host | biological process | GO:0044406 | 13 | 4 | 13711 | 214 | Gbp7,Gbp6,Icam1,Gbp3 | | 3.699e-05 | -10.20 | GSE17721\_12H\_VS\_24H\_POLYIC\_BMDC\_UP | MSigDB lists | GSE17721\_12H\_VS\_24H\_POLYIC\_BMDC\_UP | 168 | 11 | 12187 | 179 | Usp18,Aff1,Gadd45b,Il6,S100a9,Cxcl1,Ptges,Tnip1,Slfn9,Nfkb2,Igsf6 | | 3.699e-05 | -10.20 | GSE17721\_0.5H\_VS\_4H\_GARDIQUIMOD\_BMDC\_DN | MSigDB lists | GSE17721\_0.5H\_VS\_4H\_GARDIQUIMOD\_BMDC\_DN | 168 | 11 | 12187 | 179 | Msn,Plaur,Il17ra,Atf3,Fstl1,Arid5b,Nfkbia,Tor3a,Tgm2,Birc3,Zfp189 | | 3.699e-05 | -10.20 | GSE42724\_NAIVE\_VS\_MEMORY\_BCELL\_DN | MSigDB lists | GSE42724\_NAIVE\_VS\_MEMORY\_BCELL\_DN | 168 | 11 | 12187 | 179 | Clic4,Parp9,Slfn9,Sgk3,Luc7l3,Nfkbiz,Aff1,Gbp7,Sdc4,Gpr84,Casp4 | | 3.699e-05 | -10.20 | GSE40274\_CTRL\_VS\_FOXP3\_TRANSDUCED\_ACTIVATED\_CD4\_TCELL\_DN | MSigDB lists | GSE40274\_CTRL\_VS\_FOXP3\_TRANSDUCED\_ACTIVATED\_CD4\_TCELL\_DN | 168 | 11 | 12187 | 179 | Sgk1,Parp14,Sdc4,Samd9l,Bach1,Irgm2,Gbp7,Ncf1,Arid5b,Irak3,Trim25 | | 3.699e-05 | -10.20 | GSE17721\_CTRL\_VS\_CPG\_8H\_BMDC\_DN | MSigDB lists | GSE17721\_CTRL\_VS\_CPG\_8H\_BMDC\_DN | 168 | 11 | 12187 | 179 | Hcar2,Ccl12,Gbp4,Parp14,Dnajb1,Nfkbia,Rbm47,Gbp6,Ptges,Tagln2,Arid5b | | 3.776e-05 | -10.18 | TONKS\_TARGETS\_OF\_RUNX1\_RUNX1T1\_FUSION\_HSC\_DN | MSigDB lists | TONKS\_TARGETS\_OF\_RUNX1\_RUNX1T1\_FUSION\_HSC\_DN | 139 | 10 | 12187 | 179 | Il17ra,Csf2rb,Cflar,S100a9,Il1rn,Cybb,Ccl12,Birc3,Spi1,Zbtb16 | | 3.776e-05 | -10.18 | GSE42021\_CD24INT\_VS\_CD24LOW\_TCONV\_THYMUS\_UP | MSigDB lists | GSE42021\_CD24INT\_VS\_CD24LOW\_TCONV\_THYMUS\_UP | 139 | 10 | 12187 | 179 | Zfp36,Icam1,Pygm,Ptpn1,Bcl3,Arid5b,Cxcl1,Cebpd,Ccl12,Maff | | 3.776e-05 | -10.18 | GSE11961\_FOLLICULAR\_BCELL\_VS\_PLASMA\_CELL\_DAY7\_UP | MSigDB lists | GSE11961\_FOLLICULAR\_BCELL\_VS\_PLASMA\_CELL\_DAY7\_UP | 139 | 10 | 12187 | 179 | Nfkbiz,Ccl4,Tgm2,Socs3,Maff,Slfn9,Nfkbie,Isg15,Adamts9,Icam1 | | 3.820e-05 | -10.17 | adaptive immune response based on somatic recombination of immune receptors built from immunoglobulin superfamily domains | biological process | GO:0002460 | 105 | 9 | 13711 | 214 | Nfkb2,Il4ra,Icam1,Il6,Stx11,Irf7,C3ar1,Bcl3,Gadd45g | | 3.878e-05 | -10.16 | CREL\_01 | MSigDB lists | CREL\_01 | 200 | 12 | 12187 | 179 | Bcl3,Ptges,Nfkb2,Birc3,Map3k8,Nfkbia,Sdc4,Gadd45b,Tnip1,Icam1,Csf2rb,Ier3 | | 3.887e-05 | -10.16 | Death Receptor Signalling | REACTOME pathways | R-MMU-73887 | 39 | 6 | 6297 | 105 | Tnfsf10,Ripk1,Tnf,Birc3,Cflar,Tnfaip3 | | 3.891e-05 | -10.15 | regulation of developmental process | biological process | GO:0050793 | 2184 | 57 | 13711 | 214 | Tnf,Rhou,Zfp36,Ripk1,Cflar,Hspb1,Rhoc,Hspa5,Cxcl1,C5ar1,Maff,Nfe2l2,Msn,Noct,Nfkbia,Il1a,Il4ra,Cdkn1a,Il6,Ctla2a,Trib1,Socs3,Pik3r1,Fzd4,Timp1,Isg15,Spi1,Cebpd,Runx1,Sgk1,Ptgs2,Cxcl9,Tnfaip3,Ccl2,Xdh,Zbtb16,Adamts9,Csf3,Irak3,Il1rn,C3ar1,Ifi204,Nfkbiz,Ccl12,Iqgap1,S100a9,Sox11,Icam1,Cybb,Errfi1,Bbs12,Msr1,Thbs1,Lgals9,Adamts1,Apold1,Rhoj | | 3.907e-05 | -10.15 | GSE5589\_LPS\_VS\_LPS\_AND\_IL6\_STIM\_IL6\_KO\_MACROPHAGE\_45MIN\_DN | MSigDB lists | GSE5589\_LPS\_VS\_LPS\_AND\_IL6\_STIM\_IL6\_KO\_MACROPHAGE\_45MIN\_DN | 169 | 11 | 12187 | 179 | Birc3,Ccnd2,Tgm2,S100a8,Iqgap1,C5ar1,Angptl4,Il1a,Samd9l,Icam1,Trim25 | | 3.921e-05 | -10.15 | RUTELLA\_RESPONSE\_TO\_CSF2RB\_AND\_IL4\_UP | MSigDB lists | RUTELLA\_RESPONSE\_TO\_CSF2RB\_AND\_IL4\_UP | 267 | 14 | 12187 | 179 | Cdkn1a,Il1a,Tnip3,Map3k6,Rbm47,Hspb1,Birc3,Runx1,Ccl2,Sdc4,Tgm2,Gadd45b,Cflar,Tnfaip8 | | 3.921e-05 | -10.15 | GO\_REGULATION\_OF\_MAP\_KINASE\_ACTIVITY | MSigDB lists | GO\_REGULATION\_OF\_MAP\_KINASE\_ACTIVITY | 267 | 14 | 12187 | 179 | Map3k6,Saa1,Ptpn1,Irak3,Thbs1,Gadd45g,Pik3r5,Fzd4,C5ar1,Trib1,Tnf,Map3k8,Ripk1,Gadd45b | | 3.970e-05 | -10.13 | DEATH-like\_dom\_sf | interpro domains | IPR011029 | 61 | 7 | 13788 | 212 | Birc3,Casp4,Irak3,Cflar,Nfkb1,Nfkb2,Ripk1 | | 4.015e-05 | -10.12 | GO\_CHEMOKINE\_RECEPTOR\_BINDING | MSigDB lists | GO\_CHEMOKINE\_RECEPTOR\_BINDING | 27 | 5 | 12187 | 179 | Cxcl16,Ccl12,Ccl4,Cxcl1,Ccl2 | | 4.015e-05 | -10.12 | GO\_POSITIVE\_REGULATION\_OF\_INTERLEUKIN\_1\_BETA\_PRODUCTION | MSigDB lists | GO\_POSITIVE\_REGULATION\_OF\_INTERLEUKIN\_1\_BETA\_PRODUCTION | 27 | 5 | 12187 | 179 | Ifi211,Ifi209,Hspb1,Ifi204,Ifi207 | | 4.015e-05 | -10.12 | GO\_REGULATION\_OF\_GRANULOCYTE\_CHEMOTAXIS | MSigDB lists | GO\_REGULATION\_OF\_GRANULOCYTE\_CHEMOTAXIS | 27 | 5 | 12187 | 179 | C5ar1,Thbs1,C3ar1,Cxcl1,Ccl12 | | 4.015e-05 | -10.12 | BOYAULT\_LIVER\_CANCER\_SUBCLASS\_G56\_DN | MSigDB lists | BOYAULT\_LIVER\_CANCER\_SUBCLASS\_G56\_DN | 14 | 4 | 12187 | 179 | Trib1,Maff,Il4ra,Ier3 | | 4.015e-05 | -10.12 | GRANDVAUX\_IRF3\_TARGETS\_UP | MSigDB lists | GRANDVAUX\_IRF3\_TARGETS\_UP | 14 | 4 | 12187 | 179 | Ifit3b,Ifit2,Isg15,Ifi44 | | 4.015e-05 | -10.12 | BIOCARTA\_INFLAM\_PATHWAY | MSigDB lists | BIOCARTA\_INFLAM\_PATHWAY | 14 | 4 | 12187 | 179 | Tnf,Csf3,Il1a,Il6 | | 4.015e-05 | -10.12 | GSE18804\_SPLEEN\_MACROPHAGE\_VS\_BRAIN\_TUMORAL\_MACROPHAGE\_UP | MSigDB lists | GSE18804\_SPLEEN\_MACROPHAGE\_VS\_BRAIN\_TUMORAL\_MACROPHAGE\_UP | 140 | 10 | 12187 | 179 | Cxcl1,Ptgs2,Ier3,Atf3,Ccl12,Tnfaip3,Sdc4,Nfkbia,Rhou,Map3k8 | | 4.028e-05 | -10.12 | GO\_POSITIVE\_REGULATION\_OF\_CHEMOTAXIS | MSigDB lists | GO\_POSITIVE\_REGULATION\_OF\_CHEMOTAXIS | 87 | 8 | 12187 | 179 | Thbs1,Ccl4,Il6,Ccl12,C3ar1,C5ar1,Cxcl1,Hspb1 | | 4.028e-05 | -10.12 | KEEN\_RESPONSE\_TO\_ROSIGLITAZONE\_DN | MSigDB lists | KEEN\_RESPONSE\_TO\_ROSIGLITAZONE\_DN | 87 | 8 | 12187 | 179 | Nfkbiz,Atf3,Gadd45b,Cxcl1,Cd14,S100a9,Slfn4,Slfn3 | | 4.058e-05 | -10.11 | GO\_NEGATIVE\_REGULATION\_OF\_CELL\_COMMUNICATION | MSigDB lists | GO\_NEGATIVE\_REGULATION\_OF\_CELL\_COMMUNICATION | 939 | 30 | 12187 | 179 | Tnfsf10,Xdh,Cflar,Tnf,Nfkbia,Trib1,Plek,Icam1,Hspb1,Plaur,Errfi1,Ptgs2,Tnip1,Tnip3,Il1rn,Atf3,Socs3,Rgs16,Il6,Nfe2l2,Ripk1,Tnfaip3,Pik3ap1,Irak3,Ptpn1,Rnf213,Cd14,Il1a,Hspa5,Thbs1 | | 4.101e-05 | -10.10 | PID\_IL4\_2PATHWAY | MSigDB lists | PID\_IL4\_2PATHWAY | 44 | 6 | 12187 | 179 | Spi1,Parp14,Socs3,Il4ra,Pik3r1,Selp | | 4.101e-05 | -10.10 | PID\_IL6\_7\_PATHWAY | MSigDB lists | PID\_IL6\_7\_PATHWAY | 44 | 6 | 12187 | 179 | Pik3r1,Socs3,A2m,Timp1,Cebpd,Il6 | | 4.125e-05 | -10.10 | GSE17721\_LPS\_VS\_PAM3CSK4\_12H\_BMDC\_UP | MSigDB lists | GSE17721\_LPS\_VS\_PAM3CSK4\_12H\_BMDC\_UP | 170 | 11 | 12187 | 179 | Cflar,Nfkbie,Rhoc,Sdc4,Nfkbia,Clic4,Tagln2,Znfx1,Cdkn1a,Sgk1,Vcam1 | | 4.125e-05 | -10.10 | GO\_LEUKOCYTE\_CELL\_CELL\_ADHESION | MSigDB lists | GO\_LEUKOCYTE\_CELL\_CELL\_ADHESION | 170 | 11 | 12187 | 179 | Sele,S100a8,Icam1,Msn,Selp,S100a9,Bcl3,Tnip1,Il6,Tnf,Vcam1 | | 4.125e-05 | -10.10 | GSE20754\_WT\_VS\_TCF1\_KO\_MEMORY\_CD8\_TCELL\_UP | MSigDB lists | GSE20754\_WT\_VS\_TCF1\_KO\_MEMORY\_CD8\_TCELL\_UP | 170 | 11 | 12187 | 179 | C3ar1,Pik3r5,Birc3,Maff,Nfe2l2,Gadd45b,Tnfaip3,Tnip1,Icam1,Il1a,Atf3 | | 4.125e-05 | -10.10 | GSE43955\_TH0\_VS\_TGFB\_IL6\_IL23\_TH17\_ACT\_CD4\_TCELL\_52H\_DN | MSigDB lists | GSE43955\_TH0\_VS\_TGFB\_IL6\_IL23\_TH17\_ACT\_CD4\_TCELL\_52H\_DN | 170 | 11 | 12187 | 179 | Ifitm3,Osmr,Zfp36,Il17ra,Clic4,Xdh,Rhou,Casp4,Sntb2,Cebpd,Ier3 | | 4.252e-05 | -10.07 | binding | molecular function | GO:0005488 | 9835 | 178 | 13516 | 211 | Aff1,Xaf1,Banp,Msn,Tnfaip3,Rgs16,Il1a,Nfe2l2,Zfp36,Xdh,Stx11,Ifit2,Adamts1,Cd14,Map3k6,Luc7l3,Trim30a,Gadd45b,Akap12,Sdc4,S100a9,A2m,Nfkbie,Fzd4,Map3k8,Timp1,Rhoj,Gbp6,Isg15,Rtp4,Fam107a,Csf3,Cflar,Ifit3b,Resf1,Cemip2,Mmp8,Irak3,Parp12,Oasl1,Ifi211,Casp4,Nfkb1,Retnlg,Sbno2,Oasl2,Lcn2,Ccnd2,Rbm47,Olfml2b,Tap1,Ptges,Usp18,Thbs1,Sele,Tnip1,Ier3,Tnf,Hcar2,Gbp3,Tnfsf10,Cp,Cxcl16,Rhou,Prg4,Herc6,Ifi207,Ksr1,C5ar1,Fyb,Tor3a,Cmpk2,Cxcl9,Nfkb2,Pygm,Gbp4,Gem,Kcna5,Cxcl1,Mx2,Noct,Apold1,Bach1,Irf7,Pik3r1,Atf3,Hspa5,Errfi1,Ifi204,Clic4,Bcl3,Zbtb16,Ncf1,Sgk1,Plaur,Il4ra,Cybb,Sox11,Dnajb1,Vcam1,Ifi209,Iigp1,Csf2rb,Rnf213,Runx1,Tiparp,Cdkn1a,Rrp8,Osmr,Nfkbia,Mt2,Fstl1,C3ar1,Robo4,Phf11d,Nfkbiz,Rasip1,Bbs12,Il6,Parp9,Arid5b,Hspb1,Apol10b,Irgm2,Sntb2,Ripk1,Dtx3l,P2ry6,Il1rn,S100a8,Msr1,Sgk3,Samd9l,Trim56,Pik3ap1,Rhoc,Mt1,Tgtp1,Trib1,Slfn5,Irf9,Slfn9,Rbm39,Tubb6,Socs3,Col4a1,Ccl4,Ip6k2,Irgm1,Angptl4,Znfx1,Ccl7,Cebpd,Icam1,Ptgs2,Selp,Saa1,Iqgap1,Tgm2,Tnip3,Mxd4,Parp14,Hp,Irf2,Maff,Spi1,Birc3,Ccl2,Ccl12,Rasd1,Trim25,Ptpn1,Lgals9,Gadd45g,Il17ra,Mx1,Gbp7,Plek | | 4.286e-05 | -10.06 | Spinal Cord Injury | WikiPathways | WP2432 | 84 | 10 | 3756 | 96 | Col4a1,Ccl2,Rhoc,Il6,Tnf,Il1a,Selp,Icam1,Cxcl1,Zfp36 | | 4.300e-05 | -10.05 | GO\_POSITIVE\_REGULATION\_OF\_PROTEIN\_SERINE\_THREONINE\_KINASE\_ACTIVITY | MSigDB lists | GO\_POSITIVE\_REGULATION\_OF\_PROTEIN\_SERINE\_THREONINE\_KINASE\_ACTIVITY | 235 | 13 | 12187 | 179 | Ptpn1,Saa1,Map3k6,Gadd45g,Thbs1,Iqgap1,C5ar1,Pik3r5,Fzd4,Ripk1,Gadd45b,Ccnd2,Tnf | | 4.353e-05 | -10.04 | GSE24726\_WT\_VS\_E2\_22\_KO\_PDC\_DAY4\_POST\_DELETION\_DN | MSigDB lists | GSE24726\_WT\_VS\_E2\_22\_KO\_PDC\_DAY4\_POST\_DELETION\_DN | 171 | 11 | 12187 | 179 | Irak3,Pik3r5,Ptges,Bcl3,Irf7,Cd14,Nfkbie,Tnfaip3,Cybb,Tgm2,Birc3 | | 4.353e-05 | -10.04 | GSE22935\_WT\_VS\_MYD88\_KO\_MACROPHAGE\_48H\_MBOVIS\_BCG\_STIM\_DN | MSigDB lists | GSE22935\_WT\_VS\_MYD88\_KO\_MACROPHAGE\_48H\_MBOVIS\_BCG\_STIM\_DN | 171 | 11 | 12187 | 179 | Tnfaip8,Tnfaip3,Rhoc,Cybb,Bach1,Nfkbia,P2ry6,Ncf1,Stx11,Fstl1,Socs3 | | 4.376e-05 | -10.04 | MARZEC\_IL2\_SIGNALING\_UP | MSigDB lists | MARZEC\_IL2\_SIGNALING\_UP | 88 | 8 | 12187 | 179 | Csf2rb,Tnfsf10,Tnfaip8,Ccnd2,Birc3,Ier3,Gadd45b,Il1a | | 4.392e-05 | -10.03 | regulation of apoptotic signaling pathway | biological process | GO:2001233 | 356 | 17 | 13711 | 214 | Thbs1,Hspb1,Ripk1,Cflar,Tnf,Tnfaip3,Icam1,Atf3,Tnfsf10,Sgk3,Nfe2l2,S100a9,Ptgs2,Ier3,Plaur,Ptpn1,S100a8 | | 4.451e-05 | -10.02 | antimicrobial humoral response | biological process | GO:0019730 | 26 | 5 | 13711 | 214 | Ccl2,Bcl3,S100a9,Cxcl9,Cxcl1 | | 4.453e-05 | -10.02 | MANTOVANI\_VIRAL\_GPCR\_SIGNALING\_UP | MSigDB lists | MANTOVANI\_VIRAL\_GPCR\_SIGNALING\_UP | 65 | 7 | 12187 | 179 | Iqgap1,Osmr,Cp,Ccl12,Saa1,Vcam1,Nfkbiz | | 4.533e-05 | -10.00 | GSE360\_T\_GONDII\_VS\_M\_TUBERCULOSIS\_DC\_DN | MSigDB lists | GSE360\_T\_GONDII\_VS\_M\_TUBERCULOSIS\_DC\_DN | 142 | 10 | 12187 | 179 | Cd14,C3ar1,Tap1,S100a8,Ccl4,Ifit3b,Hspa5,Ccl12,Atf3,Mmp8 | | 4.533e-05 | -10.00 | GSE13522\_WT\_VS\_IFNAR\_KO\_SKING\_T\_CRUZI\_Y\_STRAIN\_INF\_UP | MSigDB lists | GSE13522\_WT\_VS\_IFNAR\_KO\_SKING\_T\_CRUZI\_Y\_STRAIN\_INF\_UP | 142 | 10 | 12187 | 179 | Cmpk2,Trim25,Gbp3,Parp9,Irgm2,Ncf1,Stx11,Birc3,Nfkbiz,Trib1 | | 4.565e-05 | -9.99 | positive regulation of cell differentiation | biological process | GO:0045597 | 927 | 31 | 13711 | 214 | Cxcl9,Tnf,Zfp36,Adamts9,Ripk1,Csf3,Zbtb16,Cflar,Hspb1,Ifi204,Hspa5,Nfkbiz,Iqgap1,S100a9,Nfe2l2,Noct,Sox11,Il4ra,Il6,Trib1,Msr1,Socs3,Lgals9,Pik3r1,Fzd4,Adamts1,Isg15,Cebpd,Runx1,Sgk1,Ptgs2 | | 4.591e-05 | -9.99 | GSE1925\_CTRL\_VS\_IFNG\_PRIMED\_MACROPHAGE\_24H\_IFNG\_STIM\_DN | MSigDB lists | GSE1925\_CTRL\_VS\_IFNG\_PRIMED\_MACROPHAGE\_24H\_IFNG\_STIM\_DN | 172 | 11 | 12187 | 179 | Irf9,Icam1,Ifitm3,Isg15,Nfkb1,Ifit3b,Gbp3,Tnfsf10,Irgm2,Usp18,Ccnd2 | | 4.591e-05 | -9.99 | GSE7852\_THYMUS\_VS\_FAT\_TCONV\_DN | MSigDB lists | GSE7852\_THYMUS\_VS\_FAT\_TCONV\_DN | 172 | 11 | 12187 | 179 | Maff,Gadd45b,Xdh,Hspa5,Nfkbiz,Col4a1,Atf3,Ier3,Ifitm2,Tagln2,Plek | | 4.591e-05 | -9.99 | GSE29618\_BCELL\_VS\_MONOCYTE\_DN | MSigDB lists | GSE29618\_BCELL\_VS\_MONOCYTE\_DN | 172 | 11 | 12187 | 179 | Timp1,Cebpd,Trib1,Il17ra,Rbm47,Plek,Plaur,C5ar1,Ptgs2,Cxcl1,Cd14 | | 4.594e-05 | -9.99 | Nfkbiz (nuclear factor of kappa light polypeptide gene enhancer in B cells inhibitor, zeta) | protein interactions | 80859 | 5 | 3 | 6802 | 115 | Il6,Nfkb2,Nfkb1 | | 4.594e-05 | -9.99 | Tmem173 (transmembrane protein 173) | protein interactions | 72512 | 5 | 3 | 6802 | 115 | Trim30a,Usp18,Trim56 | | 4.659e-05 | -9.97 | GO\_HOMEOSTATIC\_PROCESS | MSigDB lists | GO\_HOMEOSTATIC\_PROCESS | 946 | 30 | 12187 | 179 | Kcna5,Cp,Ncf1,Saa1,Clic4,Hspb1,Icam1,S100a9,C5ar1,C3ar1,Sgk1,Lcn2,Il1a,Bbs12,Sgk3,Tap1,Tgm2,Ccl2,Spi1,Pik3r1,Tnfaip3,Ccl12,Il6,Nfe2l2,Cybb,Slc24a4,Hcar2,Angptl4,Pygm,S100a8 | | 4.674e-05 | -9.97 | GO\_REGULATION\_OF\_EPITHELIAL\_CELL\_APOPTOTIC\_PROCESS | MSigDB lists | GO\_REGULATION\_OF\_EPITHELIAL\_CELL\_APOPTOTIC\_PROCESS | 45 | 6 | 12187 | 179 | Thbs1,Tnfaip3,Nfe2l2,Il6,Angptl4,Icam1 | | 4.674e-05 | -9.97 | CHANG\_IMMORTALIZED\_BY\_HPV31\_DN | MSigDB lists | CHANG\_IMMORTALIZED\_BY\_HPV31\_DN | 45 | 6 | 12187 | 179 | Lcn2,Il1rn,Trib1,Ifit2,Ifi44,Ptgs2 | | 4.674e-05 | -9.97 | ZHOU\_TNF\_SIGNALING\_4HR | MSigDB lists | ZHOU\_TNF\_SIGNALING\_4HR | 45 | 6 | 12187 | 179 | Ptges,Tnip1,Nfkbia,Nfkb1,Ccl12,Clic4 | | 4.813e-05 | -9.94 | SERVITJA\_ISLET\_HNF1A\_TARGETS\_UP | MSigDB lists | SERVITJA\_ISLET\_HNF1A\_TARGETS\_UP | 143 | 10 | 12187 | 179 | Ifitm3,Msn,Gbp3,Ifit2,Tgm2,Vcam1,Akap12,Rhoc,Atf3,Ccl12 | | 4.813e-05 | -9.94 | GSE360\_HIGH\_DOSE\_B\_MALAYI\_VS\_M\_TUBERCULOSIS\_MAC\_DN | MSigDB lists | GSE360\_HIGH\_DOSE\_B\_MALAYI\_VS\_M\_TUBERCULOSIS\_MAC\_DN | 143 | 10 | 12187 | 179 | Zfp36,Rgs16,Pygm,Ptgs2,Cxcl1,Bcl3,Timp1,Tnfaip3,Runx1,Zfp189 | | 4.829e-05 | -9.94 | GO\_NEGATIVE\_REGULATION\_OF\_EXTRINSIC\_APOPTOTIC\_SIGNALING\_PATHWAY\_VIA\_DEATH\_DOMAIN\_RECEPTORS | MSigDB lists | GO\_NEGATIVE\_REGULATION\_OF\_EXTRINSIC\_APOPTOTIC\_SIGNALING\_PATHWAY\_VIA\_DEATH\_DOMAIN\_RECEPTORS | 28 | 5 | 12187 | 179 | Cflar,Icam1,Ripk1,Tnfaip3,Tnfsf10 | | 4.829e-05 | -9.94 | ZHAN\_MULTIPLE\_MYELOMA\_DN | MSigDB lists | ZHAN\_MULTIPLE\_MYELOMA\_DN | 28 | 5 | 12187 | 179 | Cebpd,Lcn2,Vcam1,S100a9,A2m | | 4.842e-05 | -9.94 | GSE37416\_CTRL\_VS\_24H\_F\_TULARENSIS\_LVS\_NEUTROPHIL\_DN | MSigDB lists | GSE37416\_CTRL\_VS\_24H\_F\_TULARENSIS\_LVS\_NEUTROPHIL\_DN | 173 | 11 | 12187 | 179 | Irak3,Icam1,Nfkbie,Bcl3,Cxcl1,Gadd45b,Nfe2l2,Tnfaip3,Birc3,Trib1,Nfkbia | | 4.848e-05 | -9.93 | MODULE\_118 | MSigDB lists | MODULE\_118 | 308 | 15 | 12187 | 179 | Ccl12,Ccl4,Tgm2,Maff,Ccnd2,Tnf,Rgs16,S100a8,Olfml2b,Ier3,Col4a1,Akap12,Thbs1,Cdkn1a,Isg15 | | 4.865e-05 | -9.93 | GO\_POSITIVE\_REGULATION\_OF\_KINASE\_ACTIVITY | MSigDB lists | GO\_POSITIVE\_REGULATION\_OF\_KINASE\_ACTIVITY | 383 | 17 | 12187 | 179 | Ripk1,Gadd45b,Sdc4,Tnf,Ccnd2,Map3k8,Iqgap1,Pik3r5,Fzd4,C5ar1,Gadd45g,Cdkn1a,Thbs1,Hspa5,Ptpn1,Saa1,Map3k6 | | 4.903e-05 | -9.92 | MODULE\_27 | MSigDB lists | MODULE\_27 | 238 | 13 | 12187 | 179 | Ccl2,Il4ra,Cdkn1a,Ccl12,Cd14,C3ar1,Selp,C5ar1,Csf2rb,Plaur,P2ry6,Igsf6,Icam1 | | 5.021e-05 | -9.90 | GO\_RNA\_POLYMERASE\_II\_TRANSCRIPTION\_FACTOR\_ACTIVITY\_SEQUENCE\_SPECIFIC\_DNA\_BINDING | MSigDB lists | GO\_RNA\_POLYMERASE\_II\_TRANSCRIPTION\_FACTOR\_ACTIVITY\_SEQUENCE\_SPECIFIC\_DNA\_BINDING | 423 | 18 | 12187 | 179 | Nfe2l2,Bach1,Spi1,Runx1,Maff,Nfkb2,Ifi211,Arid5b,Irf7,Atf3,Cebpd,Ifi204,Nfkb1,Ifi207,Zbtb16,Ifi209,Sox11,Irf2 | | 5.031e-05 | -9.90 | KUMAR\_TARGETS\_OF\_MLL\_AF9\_FUSION | MSigDB lists | KUMAR\_TARGETS\_OF\_MLL\_AF9\_FUSION | 309 | 15 | 12187 | 179 | Cmpk2,Irf7,Ifi211,Nfe2l2,Usp18,Sdc4,Ifit2,Irf9,Isg15,Ifi207,Ifi204,Il1rn,Ifit3b,Vcam1,Ifi209 | | 5.074e-05 | -9.89 | chemokine-mediated signaling pathway | biological process | GO:0070098 | 43 | 6 | 13711 | 214 | Ccl7,Ccl4,Ccl2,Ccl12,Cxcl1,Cxcl9 | | 5.103e-05 | -9.88 | HALLMARK\_P53\_PATHWAY | MSigDB lists | HALLMARK\_P53\_PATHWAY | 174 | 11 | 12187 | 179 | Zbtb16,Cdkn1a,Ier3,Atf3,Mxd4,Il1a,Tap1,Ip6k2,Ccnd2,Rrp8,Rgs16 | | 5.109e-05 | -9.88 | type I interferon signaling pathway | biological process | GO:0060337 | 14 | 4 | 13711 | 214 | Ifitm2,Ifitm3,Ifitm6,Irf7 | | 5.109e-05 | -9.88 | cellular response to type I interferon | biological process | GO:0071357 | 14 | 4 | 13711 | 214 | Ifitm3,Ifitm6,Irf7,Ifitm2 | | 5.136e-05 | -9.88 | negative regulation of immune response | biological process | GO:0050777 | 109 | 9 | 13711 | 214 | Il4ra,Acod1,Tnf,Tnfaip3,Tap1,A2m,Parp14,Irak3,Lgals9 | | 5.136e-05 | -9.88 | positive regulation of cytokine secretion | biological process | GO:0050715 | 109 | 9 | 13711 | 214 | Il4ra,Tnf,Casp4,Akap12,Il1a,Il17ra,Lgals9,Cd14,Mmp8 | | 5.177e-05 | -9.87 | ENK\_UV\_RESPONSE\_KERATINOCYTE\_UP | MSigDB lists | ENK\_UV\_RESPONSE\_KERATINOCYTE\_UP | 424 | 18 | 12187 | 179 | Cdkn1a,Sgk1,Ier3,Atf3,Stx11,Tagln2,Isg15,Hspb1,Zfp36,Tgm2,Sdc4,Trib1,Maff,Tnfaip3,Gadd45b,Cxcl1,Gem,Rgs16 | | 5.181e-05 | -9.87 | PYRIN | smart domains | SM01289 | 13 | 4 | 7188 | 123 | Ifi204,Ifi209,Ifi207,Ifi211 | | 5.196e-05 | -9.86 | FEVR\_CTNNB1\_TARGETS\_UP | MSigDB lists | FEVR\_CTNNB1\_TARGETS\_UP | 505 | 20 | 12187 | 179 | Plek,Trim25,Tap1,Cxcl16,Gadd45g,Hp,Atf3,Cdkn1a,Ifit3b,Socs3,Cmpk2,Xdh,Irf7,Angptl4,Oasl1,Gadd45b,Tnfaip3,Maff,Usp18,Casp4 | | 5.220e-05 | -9.86 | SMID\_BREAST\_CANCER\_NORMAL\_LIKE\_UP | MSigDB lists | SMID\_BREAST\_CANCER\_NORMAL\_LIKE\_UP | 310 | 15 | 12187 | 179 | Gem,Ifi211,Cxcl1,Tnfaip8,Birc3,Pik3r1,Il6,Ccl12,Fam107a,Sele,Ifi209,Vcam1,Ifi207,Ms4a6d,Ifi204 | | 5.310e-05 | -9.84 | TONKS\_TARGETS\_OF\_RUNX1\_RUNX1T1\_FUSION\_SUSTAINDED\_IN\_ERYTHROCYTE\_UP | MSigDB lists | TONKS\_TARGETS\_OF\_RUNX1\_RUNX1T1\_FUSION\_SUSTAINDED\_IN\_ERYTHROCYTE\_UP | 46 | 6 | 12187 | 179 | Ifi207,Hspb1,Ifi204,Il1rn,Ifi211,Ifi209 | | 5.392e-05 | -9.83 | GO\_NEGATIVE\_REGULATION\_OF\_INTRACELLULAR\_SIGNAL\_TRANSDUCTION | MSigDB lists | GO\_NEGATIVE\_REGULATION\_OF\_INTRACELLULAR\_SIGNAL\_TRANSDUCTION | 348 | 16 | 12187 | 179 | Tnip1,Plek,Irak3,Hspb1,Plaur,Ptpn1,Errfi1,Thbs1,Socs3,Tnip3,Atf3,Xdh,Nfkbia,Nfe2l2,Tnfaip3,Ripk1 | | 5.413e-05 | -9.82 | GO\_RESPONSE\_TO\_INTERFERON\_ALPHA | MSigDB lists | GO\_RESPONSE\_TO\_INTERFERON\_ALPHA | 15 | 4 | 12187 | 179 | Ifitm2,Ifitm3,Ifit2,Ifit3b | | 5.413e-05 | -9.82 | MAHADEVAN\_RESPONSE\_TO\_MP470\_DN | MSigDB lists | MAHADEVAN\_RESPONSE\_TO\_MP470\_DN | 15 | 4 | 12187 | 179 | Gem,Maff,Atf3,Gadd45b | | 5.417e-05 | -9.82 | GO\_MYELOID\_CELL\_DIFFERENTIATION | MSigDB lists | GO\_MYELOID\_CELL\_DIFFERENTIATION | 145 | 10 | 12187 | 179 | Sbno2,Ifi211,Csf3,Ifi204,Ifi207,Zbtb16,Spi1,Tnf,Runx1,Ifi209 | | 5.574e-05 | -9.79 | LABBE\_WNT3A\_TARGETS\_UP | MSigDB lists | LABBE\_WNT3A\_TARGETS\_UP | 91 | 8 | 12187 | 179 | Gadd45g,Akap12,Cebpd,Sgk1,Ifit3b,Ccl2,Tgm2,Nfkb2 | | 5.743e-05 | -9.77 | MEK\_UP.V1\_DN | MSigDB lists | MEK\_UP.V1\_DN | 146 | 10 | 12187 | 179 | Oasl1,Atf3,Ifi44,Usp18,S100a8,Tap1,Irf9,Herc6,Isg15,Irf7 | | 5.743e-05 | -9.77 | GSE32901\_TH17\_EMRICHED\_VS\_TH17\_NEG\_CD4\_TCELL\_DN | MSigDB lists | GSE32901\_TH17\_EMRICHED\_VS\_TH17\_NEG\_CD4\_TCELL\_DN | 146 | 10 | 12187 | 179 | Dnajb1,Nfkbiz,Nfkbia,Nfkb1,Tnfaip3,Rtp4,Ip6k2,Zfp36,Il17ra,Irf9 | | 5.743e-05 | -9.77 | GSE27859\_MACROPHAGE\_VS\_CD11C\_INT\_F480\_HI\_MACROPHAGE\_UP | MSigDB lists | GSE27859\_MACROPHAGE\_VS\_CD11C\_INT\_F480\_HI\_MACROPHAGE\_UP | 146 | 10 | 12187 | 179 | Map3k8,Tnf,Parp14,Nfkbia,Nfkb1,Gadd45b,Atf3,Nfkbie,Plek,Pik3ap1 | | 5.743e-05 | -9.77 | GSE18281\_MEDULLARY\_THYMOCYTE\_VS\_WHOLE\_MEDULLA\_THYMUS\_DN | MSigDB lists | GSE18281\_MEDULLARY\_THYMOCYTE\_VS\_WHOLE\_MEDULLA\_THYMUS\_DN | 146 | 10 | 12187 | 179 | Ifi209,Parp14,Ifi207,Ifi204,Oasl1,Znfx1,Herc6,Ifitm2,Ifi211,Rtp4 | | 5.767e-05 | -9.76 | SARTIPY\_NORMAL\_AT\_INSULIN\_RESISTANCE\_UP | MSigDB lists | SARTIPY\_NORMAL\_AT\_INSULIN\_RESISTANCE\_UP | 29 | 5 | 12187 | 179 | Ccl12,Tubb6,Socs3,Ptges,Thbs1 | | 5.767e-05 | -9.76 | GO\_POSITIVE\_REGULATION\_OF\_WOUND\_HEALING | MSigDB lists | GO\_POSITIVE\_REGULATION\_OF\_WOUND\_HEALING | 29 | 5 | 12187 | 179 | Nfe2l2,Plek,S100a9,Selp,Thbs1 | | 5.767e-05 | -9.76 | GNF2\_FOS | MSigDB lists | GNF2\_FOS | 29 | 5 | 12187 | 179 | Ms4a6d,Zfp36,Stx11,Cybb,Nfkbia | | 5.767e-05 | -9.76 | DORN\_ADENOVIRUS\_INFECTION\_32HR\_DN | MSigDB lists | DORN\_ADENOVIRUS\_INFECTION\_32HR\_DN | 29 | 5 | 12187 | 179 | Nfkbia,Sgk1,Nfkb2,Ier3,Nfe2l2 | | 5.784e-05 | -9.76 | Transcriptional misregulation in cancer | KEGG pathways | mmu05202 | 130 | 11 | 5248 | 107 | Cd14,Runx1,Zbtb16,Ccnd2,Spi1,Aff1,Cdkn1a,Nfkbiz,Birc3,Nfkb1,Il6 | | 5.784e-05 | -9.76 | Transcriptional misregulation in cancer | KEGG pathways | ko05202 | 130 | 11 | 5248 | 107 | Ccnd2,Birc3,Nfkb1,Spi1,Il6,Cd14,Aff1,Runx1,Cdkn1a,Zbtb16,Nfkbiz | | 5.838e-05 | -9.75 | RIG-I-like receptor signaling pathway | KEGG pathways | ko04622 | 50 | 7 | 5248 | 107 | Tnf,Isg15,Irf7,Ripk1,Trim25,Nfkbia,Nfkb1 | | 5.838e-05 | -9.75 | Legionellosis | KEGG pathways | mmu05134 | 50 | 7 | 5248 | 107 | Cd14,Cxcl1,Nfkb2,Tnf,Nfkb1,Nfkbia,Il6 | | 5.838e-05 | -9.75 | RIG-I-like receptor signaling pathway | KEGG pathways | mmu04622 | 50 | 7 | 5248 | 107 | Isg15,Irf7,Ripk1,Tnf,Nfkb1,Trim25,Nfkbia | | 5.838e-05 | -9.75 | Legionellosis | KEGG pathways | ko05134 | 50 | 7 | 5248 | 107 | Il6,Nfkbia,Nfkb1,Tnf,Cxcl1,Nfkb2,Cd14 | | 5.972e-05 | -9.73 | JOHNSTONE\_PARVB\_TARGETS\_3\_UP | MSigDB lists | JOHNSTONE\_PARVB\_TARGETS\_3\_UP | 351 | 16 | 12187 | 179 | Cebpd,Timp1,Thbs1,Errfi1,Rbm47,Icam1,Tnip1,Ifitm3,Parp12,Trib1,Nfkbia,Tgm2,Tnfsf10,Osmr,Iqgap1,Bcl3 | | 6.014e-05 | -9.72 | DIRMEIER\_LMP1\_RESPONSE\_LATE\_UP | MSigDB lists | DIRMEIER\_LMP1\_RESPONSE\_LATE\_UP | 47 | 6 | 12187 | 179 | Nfkb2,Plek,Nfkb1,Tnip1,Il4ra,Cflar | | 6.084e-05 | -9.71 | GSE22886\_IGG\_IGA\_MEMORY\_BCELL\_VS\_BM\_PLASMA\_CELL\_DN | MSigDB lists | GSE22886\_IGG\_IGA\_MEMORY\_BCELL\_VS\_BM\_PLASMA\_CELL\_DN | 147 | 10 | 12187 | 179 | Cflar,Cd14,Rbm47,Runx1,Nfkbia,Trib1,Socs3,Bach1,Timp1,Mxd4 | | 6.197e-05 | -9.69 | positive regulation of MAP kinase activity | biological process | GO:0043406 | 197 | 12 | 13711 | 214 | Ripk1,Thbs1,Il1rn,Gadd45g,Gadd45b,Tnf,Pik3r5,Ptpn1,Fzd4,Map3k6,Map3k8,Iqgap1 | | 6.223e-05 | -9.68 | positive regulation of vasculature development | biological process | GO:1904018 | 167 | 11 | 13711 | 214 | Cybb,Rhoj,C5ar1,Il1a,Runx1,Ptgs2,Nfe2l2,C3ar1,Thbs1,Hspb1,Cflar | | 6.233e-05 | -9.68 | positive regulation of protein kinase activity | biological process | GO:0045860 | 403 | 18 | 13711 | 214 | Sdc4,Iqgap1,Map3k6,Fzd4,Ptpn1,Map3k8,Irgm2,Pik3r5,Cdkn1a,Tnf,Lgals9,Gadd45b,Gadd45g,Ccnd2,Ripk1,Ncf1,Thbs1,Il1rn | | 6.275e-05 | -9.68 | GSE9006\_HEALTHY\_VS\_TYPE\_1\_DIABETES\_PBMC\_4MONTH\_POST\_DX\_UP | MSigDB lists | GSE9006\_HEALTHY\_VS\_TYPE\_1\_DIABETES\_PBMC\_4MONTH\_POST\_DX\_UP | 178 | 11 | 12187 | 179 | Cybb,Tnfaip3,Atf3,Gadd45b,Pik3r1,Maff,Tor3a,Nfkbia,Nfkb2,Msn,Nfkbie | | 6.275e-05 | -9.68 | RIGGINS\_TAMOXIFEN\_RESISTANCE\_DN | MSigDB lists | RIGGINS\_TAMOXIFEN\_RESISTANCE\_DN | 178 | 11 | 12187 | 179 | Isg15,Ifitm2,Ifitm3,Arid5b,S100a8,Rbm47,Tnfsf10,Irf9,Pik3r1,Timp1,Cebpd | | 6.398e-05 | -9.66 | response to toxic substance | biological process | GO:0009636 | 262 | 14 | 13711 | 214 | Lcn2,Ripk1,Ccl7,Il6,Tnf,Tnfaip3,Mt1,Nfe2l2,Ccl4,S100a9,S100a8,Mt2,Hp,Nfkb1 | | 6.444e-05 | -9.65 | GSE29618\_MONOCYTE\_VS\_MDC\_UP | MSigDB lists | GSE29618\_MONOCYTE\_VS\_MDC\_UP | 148 | 10 | 12187 | 179 | Nfkbia,Bach1,Cybb,Ier3,C5ar1,Cd14,S100a9,Cxcl1,S100a8,Irak3 | | 6.444e-05 | -9.65 | STTTCRNTTT\_IRF\_Q6 | MSigDB lists | STTTCRNTTT\_IRF\_Q6 | 148 | 10 | 12187 | 179 | Col4a1,Ifi44,Ifit2,Ifit3b,Usp18,Xaf1,Tap1,Dtx3l,Isg15,Ncf1 | | 6.444e-05 | -9.65 | GSE17721\_PAM3CSK4\_VS\_CPG\_2H\_BMDC\_DN | MSigDB lists | GSE17721\_PAM3CSK4\_VS\_CPG\_2H\_BMDC\_DN | 148 | 10 | 12187 | 179 | Irf9,Icam1,Ccl12,Oasl1,Il1a,Ccl4,Ccl2,Tnf,Ifit2,Maff | | 6.741e-05 | -9.60 | regulation of localization | biological process | GO:0032879 | 2332 | 59 | 13711 | 214 | Gbp4,Adamts1,Rhoj,Kcna5,Il17ra,Sox11,Icam1,Cybb,Msr1,Thbs1,Lgals9,P2ry6,Ptpn1,Ccl12,Cxcl16,Iqgap1,S100a8,S100a9,Tnfaip3,Ccl2,Adamts9,Il1rn,C3ar1,Fam107a,Timp1,Sdc4,Parp9,Ccl4,Ier3,Sgk1,Ptgs2,Runx1,Gem,Il1a,Akap12,Nfkbia,Casp4,Il4ra,Il6,Trib1,Dtx3l,Pik3r1,Hcar2,Clic4,Hspa5,C5ar1,Cxcl1,Mmp8,Nfe2l2,Selp,Msn,Robo4,Tnf,Zfp36,Cd14,Ripk1,Sele,Hspb1,Rhoc | | 6.790e-05 | -9.60 | GNF2\_CARD15 | MSigDB lists | GNF2\_CARD15 | 48 | 6 | 12187 | 179 | Ms4a6d,Stx11,Cybb,S100a9,Cd14,Igsf6 | | 6.790e-05 | -9.60 | CROONQUIST\_STROMAL\_STIMULATION\_UP | MSigDB lists | CROONQUIST\_STROMAL\_STIMULATION\_UP | 48 | 6 | 12187 | 179 | Cxcl1,Ccl4,Tgm2,Nfkbia,Il6,Ier3 | | 6.790e-05 | -9.60 | PID\_CERAMIDE\_PATHWAY | MSigDB lists | PID\_CERAMIDE\_PATHWAY | 48 | 6 | 12187 | 179 | Ksr1,Nfkb1,Ripk1,Tnf,Birc3,Nfkbia | | 6.804e-05 | -9.60 | Hemostasis | REACTOME pathways | R-MMU-109582 | 391 | 18 | 6297 | 105 | Ptpn1,Irf7,Maff,Serpina3f,Timp1,Sdc4,Sele,Csf2rb,Pik3r1,Pik3r5,Irf2,Tagln2,Thbs1,Plaur,Plek,A2m,Hspa5,Selp | | 6.810e-05 | -9.59 | Tnfrsf1a (tumor necrosis factor receptor superfamily, member 1a) | protein interactions | 21937 | 14 | 4 | 6802 | 115 | Tnfaip3,Ripk1,Tnf,Tnip1 | | 6.821e-05 | -9.59 | GSE34156\_UNTREATED\_VS\_6H\_TLR1\_TLR2\_LIGAND\_TREATED\_MONOCYTE\_UP | MSigDB lists | GSE34156\_UNTREATED\_VS\_6H\_TLR1\_TLR2\_LIGAND\_TREATED\_MONOCYTE\_UP | 149 | 10 | 12187 | 179 | Cd14,Rbm47,Igsf6,Clic4,Csf2rb,Cxcl16,Msr1,Rhou,Cybb,AA467197 | | 6.821e-05 | -9.59 | GSE34179\_THPOK\_KO\_VS\_WT\_VA14I\_NKTCELL\_DN | MSigDB lists | GSE34179\_THPOK\_KO\_VS\_WT\_VA14I\_NKTCELL\_DN | 149 | 10 | 12187 | 179 | Trib1,Ccl4,Ccnd2,Rasip1,Gadd45b,Hcar2,Il1rn,Csf3,Clic4,Plek | | 6.821e-05 | -9.59 | GSE41867\_NAIVE\_VS\_DAY6\_LCMV\_ARMSTRONG\_EFFECTOR\_CD8\_TCELL\_DN | MSigDB lists | GSE41867\_NAIVE\_VS\_DAY6\_LCMV\_ARMSTRONG\_EFFECTOR\_CD8\_TCELL\_DN | 149 | 10 | 12187 | 179 | S100a8,Tnip1,Irf7,Stx11,Tnfaip3,Il6,Ier3,Atf3,Tnf,Nfkbia | | 6.838e-05 | -9.59 | DORN\_ADENOVIRUS\_INFECTION\_48HR\_DN | MSigDB lists | DORN\_ADENOVIRUS\_INFECTION\_48HR\_DN | 30 | 5 | 12187 | 179 | Il6,Nfe2l2,Ier3,Sgk1,Nfkbia | | 6.838e-05 | -9.59 | VALK\_AML\_WITH\_CEBPA | MSigDB lists | VALK\_AML\_WITH\_CEBPA | 30 | 5 | 12187 | 179 | Trib1,Mmp8,Tubb6,Hspb1,Lcn2 | | 6.838e-05 | -9.59 | GO\_REGULATION\_OF\_ENDOTHELIAL\_CELL\_APOPTOTIC\_PROCESS | MSigDB lists | GO\_REGULATION\_OF\_ENDOTHELIAL\_CELL\_APOPTOTIC\_PROCESS | 30 | 5 | 12187 | 179 | Thbs1,Angptl4,Icam1,Tnfaip3,Nfe2l2 | | 6.875e-05 | -9.59 | Apoptosis signaling pathway | Lipid Maps pathways | Apoptosis | 71 | 14 | 354 | 26 | Irak3,Pik3r1,Nfkbia,Tnf,Il1a,Cflar,Csf2rb,Birc3,Nfkbie,Nfkb1,Ripk1,Nfkb2,Tnfsf10,Pik3r5 | | 6.881e-05 | -9.58 | leukocyte tethering or rolling | biological process | GO:0050901 | 15 | 4 | 13711 | 214 | Selp,Sele,Vcam1,Tnf | | 6.932e-05 | -9.58 | regulation of myeloid cell differentiation | biological process | GO:0045637 | 169 | 11 | 13711 | 214 | Nfkbia,Tnf,Isg15,Zfp36,Csf3,Ripk1,Zbtb16,Trib1,Spi1,Runx1,Pik3r1 | | 7.033e-05 | -9.56 | MIKKELSEN\_MEF\_LCP\_WITH\_H3K4ME3 | MSigDB lists | MIKKELSEN\_MEF\_LCP\_WITH\_H3K4ME3 | 94 | 8 | 12187 | 179 | Xaf1,Casp4,Il1rn,Cp,Ccl12,Slfn4,Slfn3,Xdh | | 7.080e-05 | -9.56 | negative regulation of cell communication | biological process | GO:0010648 | 1086 | 34 | 13711 | 214 | Trim30a,Rnf213,Tnip1,Cflar,Ripk1,Hspb1,Irak3,Il1rn,Atf3,Xdh,Pik3ap1,Plek,Acod1,Tnfaip3,Tnf,Bcl3,Sgk3,Nfe2l2,Tnip3,Ptpn1,Hspa5,Trib1,Parp14,Socs3,Thbs1,Il6,Errfi1,Rgs16,Nfkbia,Icam1,Ier3,Plaur,Ptgs2,Rasip1 | | 7.086e-05 | -9.55 | negative regulation of protein phosphorylation | biological process | GO:0001933 | 370 | 17 | 13711 | 214 | Irak3,Hspb1,Socs3,Trib1,Parp14,Gadd45b,Tnip1,Gadd45g,Tnfaip3,Xdh,Errfi1,Atf3,Il6,Cdkn1a,Rasip1,Gbp4,Ptpn1 | | 7.094e-05 | -9.55 | KEGG\_PATHWAYS\_IN\_CANCER | MSigDB lists | KEGG\_PATHWAYS\_IN\_CANCER | 282 | 14 | 12187 | 179 | Ptgs2,Cdkn1a,Zbtb16,Nfkb1,Col4a1,Pik3r5,Fzd4,Nfkb2,Runx1,Birc3,Pik3r1,Nfkbia,Spi1,Il6 | | 7.134e-05 | -9.55 | BIOCARTA\_TNFR2\_PATHWAY | MSigDB lists | BIOCARTA\_TNFR2\_PATHWAY | 16 | 4 | 12187 | 179 | Nfkbia,Ripk1,Tnfaip3,Nfkb1 | | 7.134e-05 | -9.55 | PID\_NFKAPPAB\_ATYPICAL\_PATHWAY | MSigDB lists | PID\_NFKAPPAB\_ATYPICAL\_PATHWAY | 16 | 4 | 12187 | 179 | Pik3r1,Nfkbia,Bcl3,Nfkb1 | | 7.134e-05 | -9.55 | BIOCARTA\_RELA\_PATHWAY | MSigDB lists | BIOCARTA\_RELA\_PATHWAY | 16 | 4 | 12187 | 179 | Nfkbia,Tnf,Ripk1,Nfkb1 | | 7.134e-05 | -9.55 | GO\_RETINA\_VASCULATURE\_DEVELOPMENT\_IN\_CAMERA\_TYPE\_EYE | MSigDB lists | GO\_RETINA\_VASCULATURE\_DEVELOPMENT\_IN\_CAMERA\_TYPE\_EYE | 16 | 4 | 12187 | 179 | Fzd4,Rhoj,Clic4,Col4a1 | | 7.134e-05 | -9.55 | REACTOME\_RIP\_MEDIATED\_NFKB\_ACTIVATION\_VIA\_DAI | MSigDB lists | REACTOME\_RIP\_MEDIATED\_NFKB\_ACTIVATION\_VIA\_DAI | 16 | 4 | 12187 | 179 | Ripk1,Nfkb2,Saa1,Nfkbia | | 7.148e-05 | -9.55 | GO\_LEUKOCYTE\_DIFFERENTIATION | MSigDB lists | GO\_LEUKOCYTE\_DIFFERENTIATION | 213 | 12 | 12187 | 179 | Sbno2,Csf3,Bcl3,Ifi211,Ifi207,Ifi204,Il6,Pik3r1,Ifi209,Tnf,Spi1,Vcam1 | | 7.150e-05 | -9.55 | DAPIN | prosite domains | PS50824 | 13 | 4 | 8845 | 164 | Ifi204,Ifi211,Ifi209,Ifi207 | | 7.167e-05 | -9.54 | WNT\_UP.V1\_DN | MSigDB lists | WNT\_UP.V1\_DN | 121 | 9 | 12187 | 179 | Irf9,Csf2rb,Gbp3,Cmpk2,Irf7,Isg15,Ptgs2,Rasd1,Usp18 | | 7.169e-05 | -9.54 | negative regulation of apoptotic signaling pathway | biological process | GO:2001234 | 200 | 12 | 13711 | 214 | Ptgs2,Plaur,Ier3,Nfe2l2,Sgk3,Ptpn1,Hspb1,Ripk1,Cflar,Tnf,Tnfaip3,Icam1 | | 7.171e-05 | -9.54 | RUTELLA\_RESPONSE\_TO\_CSF2RB\_AND\_IL4\_DN | MSigDB lists | RUTELLA\_RESPONSE\_TO\_CSF2RB\_AND\_IL4\_DN | 247 | 13 | 12187 | 179 | S100a9,C3ar1,C5ar1,Ccl12,Cybb,Zfp36,Irak3,Cd14,Ifitm2,Cebpd,Ms4a6d,Gadd45g,Sgk1 | | 7.209e-05 | -9.54 | GO\_RESPONSE\_TO\_HEAT | MSigDB lists | GO\_RESPONSE\_TO\_HEAT | 70 | 7 | 12187 | 179 | Cd14,Socs3,Thbs1,Cdkn1a,Il6,Ccl12,Il1a | | 7.209e-05 | -9.54 | GO\_ENHANCER\_BINDING | MSigDB lists | GO\_ENHANCER\_BINDING | 70 | 7 | 12187 | 179 | Bach1,Tiparp,Nfkb1,Nfe2l2,Sox11,Spi1,Zbtb16 | | 7.216e-05 | -9.54 | GSE36078\_WT\_VS\_IL1R\_KO\_LUNG\_DC\_AFTER\_AD5\_INF\_UP | MSigDB lists | GSE36078\_WT\_VS\_IL1R\_KO\_LUNG\_DC\_AFTER\_AD5\_INF\_UP | 150 | 10 | 12187 | 179 | Sgk1,Cdkn1a,Ier3,AA467197,Ifitm2,Ifitm3,Rhoj,Pik3ap1,Plek,Rbm47 | | 7.381e-05 | -9.51 | enhancer binding | molecular function | GO:0035326 | 89 | 8 | 13516 | 211 | Tiparp,Bach1,Nfkb1,Irf7,Sox11,Zbtb16,Spi1,Nfe2l2 | | 7.585e-05 | -9.49 | WANG\_ESOPHAGUS\_CANCER\_VS\_NORMAL\_UP | MSigDB lists | WANG\_ESOPHAGUS\_CANCER\_VS\_NORMAL\_UP | 95 | 8 | 12187 | 179 | Ccl4,Parp14,Socs3,Sdc4,Tnfaip3,Atf3,Ifitm3,Csf2rb | | 7.612e-05 | -9.48 | negative regulation of signaling | biological process | GO:0023057 | 1090 | 34 | 13711 | 214 | Plek,Pik3ap1,Acod1,Tnfaip3,Tnf,Xdh,Atf3,Ripk1,Cflar,Irak3,Il1rn,Hspb1,Rnf213,Trim30a,Tnip1,Hspa5,Ptpn1,Tnip3,Sgk3,Nfe2l2,Bcl3,Nfkbia,Rgs16,Icam1,Errfi1,Il6,Trib1,Parp14,Thbs1,Socs3,Rasip1,Plaur,Ier3,Ptgs2 | | 7.631e-05 | -9.48 | GSE2826\_WT\_VS\_BTK\_KO\_BCELL\_DN | MSigDB lists | GSE2826\_WT\_VS\_BTK\_KO\_BCELL\_DN | 151 | 10 | 12187 | 179 | Znfx1,Ccl12,Hp,Il1a,Vcam1,Thbs1,S100a8,Cd14,S100a9,Gbp7 | | 7.631e-05 | -9.48 | GSE360\_L\_MAJOR\_VS\_B\_MALAYI\_LOW\_DOSE\_DC\_UP | MSigDB lists | GSE360\_L\_MAJOR\_VS\_B\_MALAYI\_LOW\_DOSE\_DC\_UP | 151 | 10 | 12187 | 179 | Gadd45g,Nfkb1,Il6,Ier3,Tnfaip3,Dnajb1,Icam1,Tap1,Ifitm3,Tnip1 | | 7.631e-05 | -9.48 | GSE43955\_1H\_VS\_60H\_ACT\_CD4\_TCELL\_DN | MSigDB lists | GSE43955\_1H\_VS\_60H\_ACT\_CD4\_TCELL\_DN | 151 | 10 | 12187 | 179 | A2m,Il17ra,Errfi1,Il4ra,Map3k8,Casp4,Bach1,Il1rn,Gadd45b,Ccl12 | | 7.644e-05 | -9.48 | KANG\_GIST\_WITH\_PDGFRA\_UP | MSigDB lists | KANG\_GIST\_WITH\_PDGFRA\_UP | 49 | 6 | 12187 | 179 | Ifi204,Ifi207,Ifi209,Isg15,Ifi211,Usp18 | | 7.644e-05 | -9.48 | KEGG\_HEMATOPOIETIC\_CELL\_LINEAGE | MSigDB lists | KEGG\_HEMATOPOIETIC\_CELL\_LINEAGE | 49 | 6 | 12187 | 179 | Cd14,Csf3,Il4ra,Tnf,Il6,Il1a | | 7.644e-05 | -9.48 | chr1q22 | MSigDB lists | chr1q22 | 49 | 6 | 12187 | 179 | Selp,Ifi209,Ifi211,Sele,Ifi207,Ifi204 | | 7.671e-05 | -9.48 | GSE26488\_WT\_VS\_HDAC7\_DELTAP\_TG\_OT2\_THYMOCYTE\_WITH\_PEPTIDE\_INJECTION\_DN | MSigDB lists | GSE26488\_WT\_VS\_HDAC7\_DELTAP\_TG\_OT2\_THYMOCYTE\_WITH\_PEPTIDE\_INJECTION\_DN | 182 | 11 | 12187 | 179 | S100a9,Isg15,Fstl1,Socs3,Ifit3b,Sdc4,Thbs1,Ier3,Tiparp,Tubb6,Cp | | 7.735e-05 | -9.47 | negative regulation of toll-like receptor signaling pathway | biological process | GO:0034122 | 29 | 5 | 13711 | 214 | Trim30a,Pik3ap1,Irak3,Acod1,Tnfaip3 | | 8.057e-05 | -9.43 | PLASARI\_TGFB1\_TARGETS\_1HR\_UP | MSigDB lists | PLASARI\_TGFB1\_TARGETS\_1HR\_UP | 31 | 5 | 12187 | 179 | Ptgs2,Gadd45g,Il6,Ier3,Gadd45b | | 8.057e-05 | -9.43 | WEINMANN\_ADAPTATION\_TO\_HYPOXIA\_DN | MSigDB lists | WEINMANN\_ADAPTATION\_TO\_HYPOXIA\_DN | 31 | 5 | 12187 | 179 | Il1a,Plaur,Il6,Ier3,Sdc4 | | 8.066e-05 | -9.43 | negative regulation of multicellular organismal process | biological process | GO:0051241 | 1047 | 33 | 13711 | 214 | Noct,Bcl3,Mmp8,Nfe2l2,Prg4,Trim30a,Ripk1,Adamts9,Zbtb16,Ncf1,Irak3,Ccl2,Zfp36,Xdh,Tnf,Acod1,Tnfaip3,Ptgs2,Runx1,Adamts1,Nfkb1,Timp1,Gbp4,Lgals9,Pik3r1,Trib1,Thbs1,Errfi1,Il6,Il1a,Sox11,Nfkbia,Il4ra | | 8.066e-05 | -9.43 | GSE17721\_CTRL\_VS\_GARDIQUIMOD\_1H\_BMDC\_DN | MSigDB lists | GSE17721\_CTRL\_VS\_GARDIQUIMOD\_1H\_BMDC\_DN | 152 | 10 | 12187 | 179 | Map3k8,Maff,Nfkbia,Ccl2,Il1rn,Hcar2,Tagln2,Arid5b,Errfi1,Gbp3 | | 8.125e-05 | -9.42 | positive regulation of ERK1 and ERK2 cascade | biological process | GO:0070374 | 172 | 11 | 13711 | 214 | Ccl4,C5ar1,Ccl12,Cflar,Ccl7,P2ry6,Il1a,Akap12,Icam1,Ccl2,Il6 | | 8.142e-05 | -9.42 | GO\_REGULATION\_OF\_EXTRINSIC\_APOPTOTIC\_SIGNALING\_PATHWAY | MSigDB lists | GO\_REGULATION\_OF\_EXTRINSIC\_APOPTOTIC\_SIGNALING\_PATHWAY | 123 | 9 | 12187 | 179 | Ripk1,Tnfaip3,Atf3,Il1a,Tnf,Thbs1,Tnfsf10,Icam1,Cflar | | 8.253e-05 | -9.40 | negative regulation of transferase activity | biological process | GO:0051348 | 235 | 13 | 13711 | 214 | Rasip1,Ptpn1,Trib1,Hspb1,Dtx3l,Irak3,Gadd45g,Gadd45b,Tnfaip3,Cdkn1a,Il6,Zfp36,Errfi1 | | 8.266e-05 | -9.40 | regulation of cell activation | biological process | GO:0050865 | 412 | 18 | 13711 | 214 | Plek,Sox11,Il4ra,Tnfaip3,Cdkn1a,Ccl2,Il6,Zbtb16,Ctla2a,Thbs1,Lgals9,C5ar1,Nfkbiz,Sdc4,Mmp8,Vcam1,Runx1,Selp | | 8.464e-05 | -9.38 | RUTELLA\_RESPONSE\_TO\_HGF\_DN | MSigDB lists | RUTELLA\_RESPONSE\_TO\_HGF\_DN | 184 | 11 | 12187 | 179 | Ms4a6d,Gadd45b,Cebpd,Cdkn1a,Zfp36,Irak3,Ifitm2,Map3k6,C5ar1,S100a9,Cd14 | | 8.522e-05 | -9.37 | KRAS.600.LUNG.BREAST\_UP.V1\_UP | MSigDB lists | KRAS.600.LUNG.BREAST\_UP.V1\_UP | 153 | 10 | 12187 | 179 | Birc3,Tnfaip3,Akap12,Kcna5,A2m,Cxcl1,Angptl4,Csf3,Ptgs2,Plaur | | 8.522e-05 | -9.37 | GSE41867\_DAY6\_VS\_DAY8\_LCMV\_CLONE13\_EFFECTOR\_CD8\_TCELL\_UP | MSigDB lists | GSE41867\_DAY6\_VS\_DAY8\_LCMV\_CLONE13\_EFFECTOR\_CD8\_TCELL\_UP | 153 | 10 | 12187 | 179 | Lgals9,Ccl4,Stx11,Rhoc,Ms4a6d,Ifitm3,Rnf213,Tnip1,Slfn4,Slfn3 | | 8.522e-05 | -9.37 | BERTUCCI\_MEDULLARY\_VS\_DUCTAL\_BREAST\_CANCER\_UP | MSigDB lists | BERTUCCI\_MEDULLARY\_VS\_DUCTAL\_BREAST\_CANCER\_UP | 153 | 10 | 12187 | 179 | Icam1,Tap1,Gbp3,Nfkb2,Dtx3l,Nfkbie,Cebpd,Birc3,Socs3,Vcam1 | | 8.522e-05 | -9.37 | WIERENGA\_STAT5A\_TARGETS\_DN | MSigDB lists | WIERENGA\_STAT5A\_TARGETS\_DN | 153 | 10 | 12187 | 179 | Plek,Il17ra,Cd14,Ifi211,Selp,Lcn2,Ifi207,Stx11,Ifi204,Ifi209 | | 8.553e-05 | -9.37 | Poly(ADP-ribose)pol\_cat\_dom | interpro domains | IPR012317 | 16 | 4 | 13788 | 212 | Parp9,Parp12,Parp14,Tiparp | | 8.582e-05 | -9.36 | LIAN\_LIPA\_TARGETS\_6M | MSigDB lists | LIAN\_LIPA\_TARGETS\_6M | 50 | 6 | 12187 | 179 | Mmp8,Msr1,Igsf6,C3ar1,S100a9,Cxcl1 | | 8.670e-05 | -9.35 | GO\_POSITIVE\_REGULATION\_OF\_HEMOPOIESIS | MSigDB lists | GO\_POSITIVE\_REGULATION\_OF\_HEMOPOIESIS | 124 | 9 | 12187 | 179 | Csf3,Isg15,Trib1,Zbtb16,Il4ra,Runx1,Tnf,Il6,Ripk1 | | 8.954e-05 | -9.32 | GO\_POSITIVE\_REGULATION\_OF\_TRANSPORT | MSigDB lists | GO\_POSITIVE\_REGULATION\_OF\_TRANSPORT | 700 | 24 | 12187 | 179 | Msn,Sele,Ptgs2,Saa1,Nfkbia,Sdc4,Tnf,S100a9,Il1a,Sgk1,Sox11,Il17ra,Cd14,Sgk3,Il6,Nfe2l2,Ccl12,Hcar2,Ccl4,Pik3r1,Il4ra,Rhou,S100a8,Csf3 | | 9.000e-05 | -9.32 | GSE25085\_FETAL\_LIVER\_VS\_FETAL\_BM\_SP4\_THYMIC\_IMPLANT\_DN | MSigDB lists | GSE25085\_FETAL\_LIVER\_VS\_FETAL\_BM\_SP4\_THYMIC\_IMPLANT\_DN | 154 | 10 | 12187 | 179 | Gbp4,Mmp8,Tubb6,Hp,Parp14,Msr1,Xdh,Trim25,Ifitm3,Gbp6 | | 9.063e-05 | -9.31 | leukocyte adhesion to vascular endothelial cell | biological process | GO:0061756 | 16 | 4 | 13711 | 214 | Sele,Vcam1,Tnf,Selp | | 9.063e-05 | -9.31 | T cell migration | biological process | GO:0072678 | 16 | 4 | 13711 | 214 | Msn,Ccl2,Cxcl16,Icam1 | | 9.211e-05 | -9.29 | GO\_CELLULAR\_RESPONSE\_TO\_STRESS | MSigDB lists | GO\_CELLULAR\_RESPONSE\_TO\_STRESS | 1280 | 36 | 12187 | 179 | Trib1,Casp4,Nfkbia,Zfp189,Tnf,Bach1,Irf7,Ifi209,Cdkn1a,Atf3,Ifi207,Ifi204,Ptgs2,Errfi1,Icam1,Trim25,Pik3r1,Map3k8,Nfe2l2,Il6,Ccl12,Tnfaip3,Ripk1,Bcl3,Rrp8,Ifi211,Dtx3l,Hspa5,Sgk1,Thbs1,Il1a,Nfkb1,Map3k6,Isg15,Ptpn1,Parp9 | | 9.223e-05 | -9.29 | GO\_LYMPHOCYTE\_CHEMOTAXIS | MSigDB lists | GO\_LYMPHOCYTE\_CHEMOTAXIS | 17 | 4 | 12187 | 179 | Cxcl16,Ccl12,Saa1,Ccl2 | | 9.223e-05 | -9.29 | GO\_TRANSCRIPTIONAL\_ACTIVATOR\_ACTIVITY\_RNA\_POLYMERASE\_II\_DISTAL\_ENHANCER\_SEQUENCE\_SPECIFIC\_BINDING | MSigDB lists | GO\_TRANSCRIPTIONAL\_ACTIVATOR\_ACTIVITY\_RNA\_POLYMERASE\_II\_DISTAL\_ENHANCER\_SEQUENCE\_SPECIFIC\_BINDING | 17 | 4 | 12187 | 179 | Spi1,Nfkb1,Bach1,Nfe2l2 | | 9.223e-05 | -9.29 | DAZARD\_UV\_RESPONSE\_CLUSTER\_G2 | MSigDB lists | DAZARD\_UV\_RESPONSE\_CLUSTER\_G2 | 17 | 4 | 12187 | 179 | Sgk1,Cxcl1,Atf3,Ier3 | | 9.326e-05 | -9.28 | DAVICIONI\_TARGETS\_OF\_PAX\_FOXO1\_FUSIONS\_UP | MSigDB lists | DAVICIONI\_TARGETS\_OF\_PAX\_FOXO1\_FUSIONS\_UP | 219 | 12 | 12187 | 179 | Tnfaip3,Sdc4,Il4ra,Maff,Ccnd2,Arid5b,Akap12,Timp1,Thbs1,Sgk1,Adamts1,Msn | | 9.383e-05 | -9.27 | negative regulation of proteolysis | biological process | GO:0045861 | 238 | 13 | 13711 | 214 | Il6,Cflar,Ctla2a,Tnfaip8,Thbs1,Pik3r1,Birc3,Timp1,A2m,Serpina3f,Wfdc21,Plaur,Ptgs2 | | 9.436e-05 | -9.27 | GNF2\_CD1D | MSigDB lists | GNF2\_CD1D | 32 | 5 | 12187 | 179 | Ms4a6d,Cybb,Igsf6,Stx11,Cd14 | | 9.436e-05 | -9.27 | SIG\_CD40PATHWAYMAP | MSigDB lists | SIG\_CD40PATHWAYMAP | 32 | 5 | 12187 | 179 | Nfkb1,Nfkb2,Nfkbie,Pik3r1,Nfkbia | | 9.436e-05 | -9.27 | PID\_IL1\_PATHWAY | MSigDB lists | PID\_IL1\_PATHWAY | 32 | 5 | 12187 | 179 | Irak3,Il1rn,Nfkb1,Il1a,Pik3r1 | | 9.443e-05 | -9.27 | SASSON\_RESPONSE\_TO\_GONADOTROPHINS\_DN | MSigDB lists | SASSON\_RESPONSE\_TO\_GONADOTROPHINS\_DN | 73 | 7 | 12187 | 179 | Akap12,Thbs1,Casp4,Trib1,Ptpn1,Cflar,Gem | | 9.500e-05 | -9.26 | GSE21063\_CTRL\_VS\_ANTI\_IGM\_STIM\_BCELL\_NFATC1\_KO\_16H\_UP | MSigDB lists | GSE21063\_CTRL\_VS\_ANTI\_IGM\_STIM\_BCELL\_NFATC1\_KO\_16H\_UP | 155 | 10 | 12187 | 179 | Casp4,Ccl4,Samd9l,Timp1,Ccl12,Ptges,Slfn3,Slfn4,Parp9,Xdh | | 9.500e-05 | -9.26 | GSE339\_EX\_VIVO\_VS\_IN\_CULTURE\_CD4POS\_DC\_DN | MSigDB lists | GSE339\_EX\_VIVO\_VS\_IN\_CULTURE\_CD4POS\_DC\_DN | 155 | 10 | 12187 | 179 | Il1rn,Tubb6,Rasd1,Cdkn1a,Ifit2,Pik3r1,Casp4,Irgm2,Cflar,Ptges | | 9.520e-05 | -9.26 | vasculature development | biological process | GO:0001944 | 455 | 19 | 13711 | 214 | Ccl2,Errfi1,Cemip2,Robo4,Rnf213,Adamts9,Thbs1,Socs3,Ccl12,Col4a1,Rhoj,Apold1,Fzd4,Clic4,Tiparp,Angptl4,Ptgs2,Spi1,Rasip1 | | 9.716e-05 | -9.24 | regulation of interleukin-1 production | biological process | GO:0032652 | 69 | 7 | 13711 | 214 | Ifi204,Lgals9,Ifi211,Hspb1,Errfi1,Tnfaip3,Casp4 | | 9.767e-05 | -9.23 | negative regulation of protein modification process | biological process | GO:0031400 | 535 | 21 | 13711 | 214 | Rasip1,Spi1,Isg15,Gbp4,Ptpn1,Tnip1,Gadd45g,Gadd45b,Hspb1,Socs3,Dtx3l,Irak3,Trib1,Parp14,Il6,Errfi1,Atf3,Xdh,Cdkn1a,Tnfaip3,Tnf | | 9.786e-05 | -9.23 | cell motility | biological process | GO:0048870 | 744 | 26 | 13711 | 214 | Ccl4,Il17ra,Sdc4,Ccl7,Thbs1,Icam1,S100a9,Vcam1,Msn,Selp,Cxcl1,C5ar1,Cxcl16,Ccl12,Iqgap1,S100a8,Il1rn,Arid5b,Sele,Retnlg,Rhoc,Cxcl9,Tnf,Tnfaip3,Ccl2,Saa1 | | 9.786e-05 | -9.23 | localization of cell | biological process | GO:0051674 | 744 | 26 | 13711 | 214 | S100a9,Vcam1,Msn,Selp,Cxcl1,C5ar1,Cxcl16,Ccl12,S100a8,Iqgap1,Arid5b,Il1rn,Sele,Retnlg,Rhoc,Cxcl9,Tnf,Tnfaip3,Ccl2,Saa1,Ccl4,Il17ra,Sdc4,Ccl7,Thbs1,Icam1 | | 9.812e-05 | -9.23 | GROSS\_HYPOXIA\_VIA\_ELK3\_AND\_HIF1A\_UP | MSigDB lists | GROSS\_HYPOXIA\_VIA\_ELK3\_AND\_HIF1A\_UP | 126 | 9 | 12187 | 179 | Sdc4,Nfkbia,Nfkbiz,Sgk1,Atf3,Angptl4,Rgs16,Errfi1,Zfp36 | | 9.979e-05 | -9.21 | GO\_RESPONSE\_TO\_WOUNDING | MSigDB lists | GO\_RESPONSE\_TO\_WOUNDING | 406 | 17 | 12187 | 179 | S100a8,Cflar,Pik3r5,A2m,Rhoc,Ccl12,Il6,Pik3r1,Maff,Sdc4,Hspb1,Plek,Irf2,Plaur,Saa1,Timp1,Il1a | | 1.002e-04 | -9.21 | GSE17721\_PAM3CSK4\_VS\_CPG\_4H\_BMDC\_DN | MSigDB lists | GSE17721\_PAM3CSK4\_VS\_CPG\_4H\_BMDC\_DN | 156 | 10 | 12187 | 179 | Tor3a,Ccl2,Ms4a6d,Rasd1,Ptgs2,Isg15,Gbp3,Clic4,Irf2,Irf9 | | 1.002e-04 | -9.21 | GSE23925\_LIGHT\_ZONE\_VS\_NAIVE\_BCELL\_UP | MSigDB lists | GSE23925\_LIGHT\_ZONE\_VS\_NAIVE\_BCELL\_UP | 156 | 10 | 12187 | 179 | Map3k8,Maff,Ccl4,Nfkbiz,Trib1,Tnfaip3,Atf3,Gem,Zfp36,Sbno2 | | 1.008e-04 | -9.20 | GO\_RESPONSE\_TO\_ABIOTIC\_STIMULUS | MSigDB lists | GO\_RESPONSE\_TO\_ABIOTIC\_STIMULUS | 797 | 26 | 12187 | 179 | Bach1,Tnf,Nfkbia,Ifi204,Kcna5,Ifi207,Ifi209,Cdkn1a,Vcam1,Socs3,Apold1,Icam1,Errfi1,Ptgs2,Ccl12,Il6,Pik3r1,Pygm,Angptl4,Ifi211,Bcl3,Nfkb1,Il1a,Sgk1,Thbs1,Cd14 | | 1.021e-04 | -9.19 | positive regulation of cell activation | biological process | GO:0050867 | 240 | 13 | 13711 | 214 | Vcam1,Mmp8,Selp,Runx1,Nfkbiz,Thbs1,Zbtb16,Lgals9,Il4ra,Plek,Il6,Cdkn1a,Ccl2 | | 1.036e-04 | -9.17 | GTP binding | molecular function | GO:0005525 | 309 | 15 | 13516 | 211 | Rasd1,Gem,Tgtp1,Hcar2,Gbp3,Iigp1,Rhou,Mx1,Rhoj,Gbp7,Tubb6,Tgm2,Rhoc,Gbp6,Irgm1 | | 1.044e-04 | -9.17 | ISHIKAWA\_STING\_SIGNALING | MSigDB lists | ISHIKAWA\_STING\_SIGNALING | 7 | 3 | 12187 | 179 | Irf2,Irf9,Irf7 | | 1.054e-04 | -9.16 | GO\_REGULATION\_OF\_TRANSFERASE\_ACTIVITY | MSigDB lists | GO\_REGULATION\_OF\_TRANSFERASE\_ACTIVITY | 753 | 25 | 12187 | 179 | Ccnd2,Map3k8,Pik3r1,Tnfaip3,Ripk1,Il6,Gadd45b,Pik3r5,Iqgap1,Hspa5,Thbs1,Gadd45g,Map3k6,Irak3,Ptpn1,Tnf,Sdc4,Trib1,C5ar1,Fzd4,Cdkn1a,Socs3,Saa1,Hspb1,Errfi1 | | 1.057e-04 | -9.15 | GSE43955\_TH0\_VS\_TGFB\_IL6\_TH17\_ACT\_CD4\_TCELL\_52H\_UP | MSigDB lists | GSE43955\_TH0\_VS\_TGFB\_IL6\_TH17\_ACT\_CD4\_TCELL\_52H\_UP | 157 | 10 | 12187 | 179 | Nfkbiz,Sgk1,Tnf,Nfe2l2,Il1a,Cxcl1,Tnip1,Gem,Plin4,Plaur | | 1.057e-04 | -9.15 | E2F1\_UP.V1\_DN | MSigDB lists | E2F1\_UP.V1\_DN | 157 | 10 | 12187 | 179 | Tgm2,Nfkbiz,Ifit2,Ccnd2,Adamts1,Sgk1,Ccl12,Nfkb1,Angptl4,Ptgs2 | | 1.073e-04 | -9.14 | NAKAYAMA\_SOFT\_TISSUE\_TUMORS\_PCA1\_UP | MSigDB lists | NAKAYAMA\_SOFT\_TISSUE\_TUMORS\_PCA1\_UP | 52 | 6 | 12187 | 179 | Il4ra,Thbs1,Ccl2,Timp1,Ccl12,Cd14 | | 1.073e-04 | -9.14 | TONKS\_TARGETS\_OF\_RUNX1\_RUNX1T1\_FUSION\_GRANULOCYTE\_UP | MSigDB lists | TONKS\_TARGETS\_OF\_RUNX1\_RUNX1T1\_FUSION\_GRANULOCYTE\_UP | 52 | 6 | 12187 | 179 | Ifi211,Arid5b,Ifi209,Hspb1,Ifi204,Ifi207 | | 1.073e-04 | -9.14 | SEITZ\_NEOPLASTIC\_TRANSFORMATION\_BY\_8P\_DELETION\_UP | MSigDB lists | SEITZ\_NEOPLASTIC\_TRANSFORMATION\_BY\_8P\_DELETION\_UP | 52 | 6 | 12187 | 179 | Ifit2,Herc6,Ifi44,Isg15,Ifit3b,Oasl1 | | 1.075e-04 | -9.14 | GO\_VASCULATURE\_DEVELOPMENT | MSigDB lists | GO\_VASCULATURE\_DEVELOPMENT | 369 | 16 | 12187 | 179 | Robo4,Fzd4,Rhoj,Angptl4,Spi1,Ccl12,Rnf213,Ptgs2,Apold1,Clic4,Errfi1,Thbs1,Socs3,Col4a1,Tiparp,Rasip1 | | 1.080e-04 | -9.13 | chemotaxis | biological process | GO:0006935 | 383 | 17 | 13711 | 214 | Cxcl9,Ccl2,Saa1,Ccl7,C3ar1,Il1rn,Lgals9,Retnlg,C5ar1,Cxcl1,Ccl12,Cxcl16,S100a8,Ccl4,S100a9,Vcam1,Il17ra | | 1.087e-04 | -9.13 | GO\_RECEPTOR\_BINDING | MSigDB lists | GO\_RECEPTOR\_BINDING | 991 | 30 | 12187 | 179 | S100a8,Rtp4,Csf3,Cxcl1,Ccl12,Ripk1,Il6,Pik3r1,Ccl4,Ccl2,Il17ra,Cxcl16,Ptpn1,Tap1,Il1a,Thbs1,Tnfsf10,Cflar,A2m,S100a9,Oasl1,Tnf,Icam1,Plaur,Msn,Saa1,Kcna5,Il1rn,Timp1,Vcam1 | | 1.087e-04 | -9.13 | positive regulation of transferase activity | biological process | GO:0051347 | 499 | 20 | 13711 | 214 | Iqgap1,Sdc4,Map3k8,Map3k6,Fzd4,Ptpn1,Pik3r5,Irgm2,Cdkn1a,Tnf,Gadd45b,Gadd45g,Ccnd2,Lgals9,Il1rn,Ncf1,Thbs1,Dtx3l,Trib1,Ripk1 | | 1.099e-04 | -9.12 | PEDERSEN\_METASTASIS\_BY\_ERBB2\_ISOFORM\_1 | MSigDB lists | PEDERSEN\_METASTASIS\_BY\_ERBB2\_ISOFORM\_1 | 33 | 5 | 12187 | 179 | Errfi1,Rasd1,Maff,Gem,Trib1 | | 1.103e-04 | -9.11 | Innate Immune System | REACTOME pathways | R-MMU-168249 | 860 | 29 | 6297 | 105 | Irf2,Cybb,Mmp8,Plaur,Nfkb2,Cdkn1a,C3ar1,Irf7,Nfkb1,Trim56,Ier3,Lcn2,Map3k8,S100a9,Pik3ap1,Pik3r1,Gpr84,S100a8,C5ar1,Tnfaip3,Cd14,Serpina3f,Hp,Ksr1,Csf2rb,Iqgap1,Cxcl1,Ncf1,Nfkbia | | 1.108e-04 | -9.11 | GO\_RESPONSE\_TO\_IONIZING\_RADIATION | MSigDB lists | GO\_RESPONSE\_TO\_IONIZING\_RADIATION | 128 | 9 | 12187 | 179 | Ifi207,Ifi204,Ccl12,Ifi209,Cdkn1a,Socs3,Vcam1,Icam1,Ifi211 | | 1.114e-04 | -9.10 | GSE17721\_POLYIC\_VS\_CPG\_8H\_BMDC\_UP | MSigDB lists | GSE17721\_POLYIC\_VS\_CPG\_8H\_BMDC\_UP | 158 | 10 | 12187 | 179 | Gbp6,C5ar1,Iqgap1,Tnfsf10,Sgk1,Tor3a,Ccl4,Akap12,Gbp4,Mxd4 | | 1.114e-04 | -9.10 | GSE37301\_RAG2\_KO\_VS\_RAG2\_AND\_ETS1\_KO\_NK\_CELL\_UP | MSigDB lists | GSE37301\_RAG2\_KO\_VS\_RAG2\_AND\_ETS1\_KO\_NK\_CELL\_UP | 158 | 10 | 12187 | 179 | Saa1,A2m,Cd14,Osmr,Xdh,Sntb2,Tiparp,Gadd45g,Ripk1,Lcn2 | | 1.122e-04 | -9.09 | GO\_ADAPTIVE\_IMMUNE\_RESPONSE\_BASED\_ON\_SOMATIC\_RECOMBINATION\_OF\_IMMUNE\_RECEPTORS\_BUILT\_FROM\_IMMUNOGLOBULIN\_SUPERFAMILY\_DOMAINS | MSigDB lists | GO\_ADAPTIVE\_IMMUNE\_RESPONSE\_BASED\_ON\_SOMATIC\_RECOMBINATION\_OF\_IMMUNE\_RECEPTORS\_BUILT\_FROM\_IMMUNOGLOBULIN\_SUPERFAMILY\_DOMAINS | 75 | 7 | 12187 | 179 | Nfkb2,Icam1,Irf7,Bcl3,C3ar1,Il6,Il4ra | | 1.122e-04 | -9.09 | GO\_REGULATION\_OF\_TRANSCRIPTION\_FACTOR\_IMPORT\_INTO\_NUCLEUS | MSigDB lists | GO\_REGULATION\_OF\_TRANSCRIPTION\_FACTOR\_IMPORT\_INTO\_NUCLEUS | 75 | 7 | 12187 | 179 | Csf3,Nfkbie,Ptgs2,Bcl3,Tnf,Pik3r1,Nfkbia | | 1.128e-04 | -9.09 | nucleoside-triphosphatase activity | molecular function | GO:0017111 | 665 | 24 | 13516 | 211 | Hspa5,Tgtp2,Gm4841,Rasd1,F830016B08Rik,Gem,Tgtp1,Tap1,Iigp1,Gbp3,Rnf213,Mx2,Rhou,Rhoj,Mx1,Irgm2,Gbp7,Tubb6,Tor3a,Ifi47,Gm5431,Irgm1,Rhoc,Gbp6 | | 1.171e-04 | -9.05 | positive regulation of macrophage chemotaxis | biological process | GO:0010759 | 17 | 4 | 13711 | 214 | Ccl2,C5ar1,Thbs1,C3ar1 | | 1.171e-04 | -9.05 | positive regulation of tumor necrosis factor biosynthetic process | biological process | GO:0042535 | 17 | 4 | 13711 | 214 | Hspb1,Thbs1,Akap12,Cybb | | 1.171e-04 | -9.05 | NAD+ ADP-ribosyltransferase activity | molecular function | GO:0003950 | 17 | 4 | 13516 | 211 | Parp9,Parp12,Tiparp,Parp14 | | 1.172e-04 | -9.05 | negative regulation of cell population proliferation | biological process | GO:0008285 | 542 | 21 | 13711 | 214 | Pik3r1,Lgals9,Ifitm3,Thbs1,Trib1,Zbtb16,Ripk1,Il6,Xdh,Slfn3,Cdkn1a,Tnf,Tnfaip3,Sox11,Il1a,Ptgs2,Runx1,Slfn2,Ccl12,Ptges,Sdc4 | | 1.172e-04 | -9.05 | FERRARI\_RESPONSE\_TO\_FENRETINIDE\_UP | MSigDB lists | FERRARI\_RESPONSE\_TO\_FENRETINIDE\_UP | 18 | 4 | 12187 | 179 | Cxcl1,Ptgs2,Atf3,Sele | | 1.172e-04 | -9.05 | WANG\_IMMORTALIZED\_BY\_HOXA9\_AND\_MEIS1\_UP | MSigDB lists | WANG\_IMMORTALIZED\_BY\_HOXA9\_AND\_MEIS1\_UP | 18 | 4 | 12187 | 179 | Igsf6,Rhoj,Ccnd2,Thbs1 | | 1.174e-04 | -9.05 | GSE17721\_POLYIC\_VS\_GARDIQUIMOD\_2H\_BMDC\_DN | MSigDB lists | GSE17721\_POLYIC\_VS\_GARDIQUIMOD\_2H\_BMDC\_DN | 159 | 10 | 12187 | 179 | Gpr84,Casp4,Socs3,Nfe2l2,Saa1,Csf3,C5ar1,Irf9,Icam1,Tnfaip8 | | 1.174e-04 | -9.05 | GSE31622\_WT\_VS\_KLF3\_KO\_BCELL\_DN | MSigDB lists | GSE31622\_WT\_VS\_KLF3\_KO\_BCELL\_DN | 159 | 10 | 12187 | 179 | Sele,Nfkb2,Ptpn1,Ifitm2,Pik3r5,Stx11,Znfx1,Ccl4,Socs3,Trib1 | | 1.174e-04 | -9.05 | GSE39820\_CTRL\_VS\_IL1B\_IL6\_CD4\_TCELL\_DN | MSigDB lists | GSE39820\_CTRL\_VS\_IL1B\_IL6\_CD4\_TCELL\_DN | 159 | 10 | 12187 | 179 | Gem,Plek,Zfp36,Hspa5,Socs3,Timp1,Il1rn,Nfe2l2,Tnfaip3,AA467197 | | 1.174e-04 | -9.05 | GSE15767\_MED\_VS\_SCS\_MAC\_LN\_UP | MSigDB lists | GSE15767\_MED\_VS\_SCS\_MAC\_LN\_UP | 159 | 10 | 12187 | 179 | Cybb,Thbs1,Msr1,Csf2rb,Rbm47,P2ry6,Ncf1,C5ar1,Ifitm3,Ifitm2 | | 1.174e-04 | -9.05 | GSE22886\_NAIVE\_CD4\_TCELL\_VS\_MONOCYTE\_DN | MSigDB lists | GSE22886\_NAIVE\_CD4\_TCELL\_VS\_MONOCYTE\_DN | 159 | 10 | 12187 | 179 | Spi1,Cebpd,Cybb,Cd14,S100a9,C5ar1,Csf2rb,S100a8,Igsf6,Irak3 | | 1.174e-04 | -9.05 | GSE5589\_IL6\_KO\_VS\_IL10\_KO\_LPS\_AND\_IL10\_STIM\_MACROPHAGE\_45MIN\_UP | MSigDB lists | GSE5589\_IL6\_KO\_VS\_IL10\_KO\_LPS\_AND\_IL10\_STIM\_MACROPHAGE\_45MIN\_UP | 159 | 10 | 12187 | 179 | Ccl4,Cdkn1a,Tnfaip3,Mmp8,Ncf1,Cd14,Nfkbie,Ifitm3,Rgs16,Plaur | | 1.176e-04 | -9.05 | HTLV-I infection | KEGG pathways | ko05166 | 218 | 14 | 5248 | 107 | Spi1,Zfp36,Nfkbia,Pik3r1,Ccnd2,Nfkb1,Il6,Vcam1,Icam1,Tnf,Cdkn1a,Atf3,Nfkb2,Fzd4 | | 1.176e-04 | -9.05 | HTLV-I infection | KEGG pathways | mmu05166 | 218 | 14 | 5248 | 107 | Zfp36,Spi1,Nfkbia,Pik3r1,Ccnd2,Nfkb1,Il6,Vcam1,Tnf,Icam1,Atf3,Cdkn1a,Nfkb2,Fzd4 | | 1.181e-04 | -9.04 | IRF\_Q6 | MSigDB lists | IRF\_Q6 | 191 | 11 | 12187 | 179 | Dtx3l,Rtp4,Ifit2,Parp12,Cybb,Ncf1,Isg15,Irf9,Irf2,Ifi44,Thbs1 | | 1.187e-04 | -9.04 | taxis | biological process | GO:0042330 | 386 | 17 | 13711 | 214 | Il1rn,C3ar1,Ccl7,Lgals9,Retnlg,Cxcl9,Saa1,Ccl2,Vcam1,Ccl4,S100a9,Il17ra,C5ar1,Cxcl1,S100a8,Ccl12,Cxcl16 | | 1.195e-04 | -9.03 | KEGG\_LEISHMANIA\_INFECTION | MSigDB lists | KEGG\_LEISHMANIA\_INFECTION | 53 | 6 | 12187 | 179 | Tnf,Ptgs2,Ncf1,Nfkbia,Nfkb1,Il1a | | 1.202e-04 | -9.03 | immune system development | biological process | GO:0002520 | 543 | 21 | 13711 | 214 | Nfkb2,Tnfaip3,Tnf,Acod1,Il6,Zbtb16,Csf3,C3ar1,Pik3r1,Gadd45g,Tiparp,Sbno2,Prg4,Nfkb1,Rbm47,Herc6,Spi1,Cebpd,Runx1,Bcl3,Samd9l | | 1.203e-04 | -9.03 | regulation of immune effector process | biological process | GO:0002697 | 278 | 14 | 13711 | 214 | Nfkbiz,Gbp4,Cxcl1,A2m,Tap1,Parp9,Dtx3l,Irak3,Lgals9,Tnfaip3,Tnf,Il4ra,Il6,Ccl2 | | 1.216e-04 | -9.01 | positive regulation of angiogenesis | biological process | GO:0045766 | 150 | 10 | 13711 | 214 | Cybb,Rhoj,C5ar1,Il1a,Runx1,Ptgs2,Thbs1,Nfe2l2,C3ar1,Hspb1 | | 1.221e-04 | -9.01 | KRAS.LUNG.BREAST\_UP.V1\_UP | MSigDB lists | KRAS.LUNG.BREAST\_UP.V1\_UP | 76 | 7 | 12187 | 179 | Ptgs2,Csf3,A2m,Cxcl1,Angptl4,Birc3,Tnfaip3 | | 1.221e-04 | -9.01 | GRAHAM\_CML\_DIVIDING\_VS\_NORMAL\_QUIESCENT\_DN | MSigDB lists | GRAHAM\_CML\_DIVIDING\_VS\_NORMAL\_QUIESCENT\_DN | 76 | 7 | 12187 | 179 | Tnfsf10,Ifi211,Cxcl1,Ifi204,Ifi207,Ifi209,Birc3 | | 1.230e-04 | -9.00 | response to inorganic substance | biological process | GO:0010035 | 350 | 16 | 13711 | 214 | Cp,Lcn2,Ripk1,Cflar,Thbs1,Ncf1,Xdh,Il6,Il1a,Tnfaip3,Mt1,Nfe2l2,Hp,Iqgap1,Mt2,Clic4 | | 1.234e-04 | -9.00 | GO\_LOCOMOTION | MSigDB lists | GO\_LOCOMOTION | 807 | 26 | 12187 | 179 | Sele,Icam1,Msn,Plaur,Saa1,Vcam1,C3ar1,C5ar1,S100a9,Fzd4,Tnf,Sdc4,Il17ra,Cxcl16,Parp9,Selp,Thbs1,S100a8,Arid5b,Cxcl1,Rhoc,Il6,Ccl12,Pik3r1,Ccl2,Ccl4 | | 1.237e-04 | -9.00 | GSE13485\_DAY3\_VS\_DAY21\_YF17D\_VACCINE\_PBMC\_UP | MSigDB lists | GSE13485\_DAY3\_VS\_DAY21\_YF17D\_VACCINE\_PBMC\_UP | 160 | 10 | 12187 | 179 | Ifi44,Ifit3b,Ksr1,Slc24a4,Ripk1,Tnfaip3,Parp12,Herc6,Pik3r5,Cmpk2 | | 1.237e-04 | -9.00 | GSE17721\_12H\_VS\_24H\_GARDIQUIMOD\_BMDC\_UP | MSigDB lists | GSE17721\_12H\_VS\_24H\_GARDIQUIMOD\_BMDC\_UP | 160 | 10 | 12187 | 179 | Parp9,Ptpn1,Zfp36,Iqgap1,Il17ra,Irf7,Nfkb1,Gpr84,Ccl4,Sgk1 | | 1.248e-04 | -8.99 | GSE6259\_33D1\_POS\_VS\_DEC205\_POS\_FLT3L\_INDUCED\_SPLENIC\_DC\_UP | MSigDB lists | GSE6259\_33D1\_POS\_VS\_DEC205\_POS\_FLT3L\_INDUCED\_SPLENIC\_DC\_UP | 130 | 9 | 12187 | 179 | Ptgs2,Tnip1,Cxcl1,Ptges,Ccl4,Il1a,Ier3,Lcn2,Tnfaip3 | | 1.273e-04 | -8.97 | GO\_REGULATION\_OF\_LIPID\_STORAGE | MSigDB lists | GO\_REGULATION\_OF\_LIPID\_STORAGE | 34 | 5 | 12187 | 179 | Nfkbia,Msr1,Tnf,Il6,Nfkb1 | | 1.273e-04 | -8.97 | VILIMAS\_NOTCH1\_TARGETS\_UP | MSigDB lists | VILIMAS\_NOTCH1\_TARGETS\_UP | 34 | 5 | 12187 | 179 | Nfkb2,Icam1,Nfkbia,Irf7,Birc3 | | 1.273e-04 | -8.97 | ABBUD\_LIF\_SIGNALING\_1\_UP | MSigDB lists | ABBUD\_LIF\_SIGNALING\_1\_UP | 34 | 5 | 12187 | 179 | Socs3,Bcl3,Rhou,Ptpn1,Lcn2 | | 1.276e-04 | -8.97 | TNF-alpha NF-kB Signaling Pathway | WikiPathways | WP246 | 177 | 14 | 3756 | 96 | Tnip1,Ripk1,Cflar,Bcl3,Tnf,Map3k8,Nfkb1,Nfkbie,Tnfaip3,Birc3,Nfkbia,Hspb1,Nfkbiz,Nfkb2 | | 1.282e-04 | -8.96 | purine ribonucleoside binding | molecular function | GO:0032550 | 315 | 15 | 13516 | 211 | Gbp3,Iigp1,Rasd1,Gem,Hcar2,Tgtp1,Tgm2,Tubb6,Gbp7,Gbp6,Rhoc,Irgm1,Rhou,Mx1,Rhoj | | 1.285e-04 | -8.96 | positive regulation of leukocyte cell-cell adhesion | biological process | GO:1903039 | 151 | 10 | 13711 | 214 | Nfkbiz,Icam1,Tnf,Il4ra,Ccl2,Il6,Zbtb16,Vcam1,Lgals9,Runx1 | | 1.302e-04 | -8.95 | GSE19923\_E2A\_KO\_VS\_E2A\_AND\_HEB\_KO\_DP\_THYMOCYTE\_DN | MSigDB lists | GSE19923\_E2A\_KO\_VS\_E2A\_AND\_HEB\_KO\_DP\_THYMOCYTE\_DN | 161 | 10 | 12187 | 179 | Ccl4,Nfkbia,Gadd45b,Il6,Isg15,C3ar1,Gem,Irf7,Slfn4,Slfn3 | | 1.302e-04 | -8.95 | PLASARI\_TGFB1\_TARGETS\_10HR\_UP | MSigDB lists | PLASARI\_TGFB1\_TARGETS\_10HR\_UP | 161 | 10 | 12187 | 179 | Tnfaip3,Il6,Ier3,Gadd45b,Runx1,Maff,Gpr84,Plaur,Rgs16,Ptgs2 | | 1.302e-04 | -8.95 | GSE43955\_TH0\_VS\_TGFB\_IL6\_TH17\_ACT\_CD4\_TCELL\_42H\_UP | MSigDB lists | GSE43955\_TH0\_VS\_TGFB\_IL6\_TH17\_ACT\_CD4\_TCELL\_42H\_UP | 161 | 10 | 12187 | 179 | Tor3a,Ifit3b,Plaur,Irf9,Tap1,Trim25,Icam1,Cxcl1,Irf7,Isg15 | | 1.302e-04 | -8.95 | GSE17721\_CTRL\_VS\_PAM3CSK4\_2H\_BMDC\_UP | MSigDB lists | GSE17721\_CTRL\_VS\_PAM3CSK4\_2H\_BMDC\_UP | 161 | 10 | 12187 | 179 | Angptl4,Sele,P2ry6,Trim25,Tap1,Msn,Irf2,Lgals9,Tor3a,Bach1 | | 1.302e-04 | -8.95 | GSE22935\_UNSTIM\_VS\_24H\_MBOVIS\_BCG\_STIM\_MACROPHAGE\_DN | MSigDB lists | GSE22935\_UNSTIM\_VS\_24H\_MBOVIS\_BCG\_STIM\_MACROPHAGE\_DN | 161 | 10 | 12187 | 179 | Sele,Msn,Parp9,Irf2,Cd14,Hcar2,Ccl12,Birc3,Maff,Casp4 | | 1.323e-04 | -8.93 | GSE360\_L\_DONOVANI\_VS\_L\_MAJOR\_MAC\_DN | MSigDB lists | GSE360\_L\_DONOVANI\_VS\_L\_MAJOR\_MAC\_DN | 131 | 9 | 12187 | 179 | Tnfaip3,Nfkbia,Pygm,Nfkb2,Plaur,Tnfaip8,Tnip1,Ptges,Cflar | | 1.323e-04 | -8.93 | GO\_RESPONSE\_TO\_CORTICOSTEROID | MSigDB lists | GO\_RESPONSE\_TO\_CORTICOSTEROID | 131 | 9 | 12187 | 179 | Errfi1,Ptgs2,Ccl12,Il6,Il1rn,Socs3,Tnf,Cdkn1a,Sgk1 | | 1.329e-04 | -8.93 | GO\_REGULATION\_OF\_COAGULATION | MSigDB lists | GO\_REGULATION\_OF\_COAGULATION | 54 | 6 | 12187 | 179 | S100a9,Selp,Thbs1,Plaur,Nfe2l2,Plek | | 1.329e-04 | -8.93 | KEGG\_ACUTE\_MYELOID\_LEUKEMIA | MSigDB lists | KEGG\_ACUTE\_MYELOID\_LEUKEMIA | 54 | 6 | 12187 | 179 | Nfkb1,Pik3r5,Zbtb16,Spi1,Runx1,Pik3r1 | | 1.343e-04 | -8.92 | DARWICHE\_PAPILLOMA\_RISK\_HIGH\_UP | MSigDB lists | DARWICHE\_PAPILLOMA\_RISK\_HIGH\_UP | 103 | 8 | 12187 | 179 | Ifitm3,S100a9,Icam1,Cmpk2,S100a8,Cxcl16,Nfe2l2,Rasip1 | | 1.343e-04 | -8.92 | GO\_RESPONSE\_TO\_TRANSITION\_METAL\_NANOPARTICLE | MSigDB lists | GO\_RESPONSE\_TO\_TRANSITION\_METAL\_NANOPARTICLE | 103 | 8 | 12187 | 179 | Icam1,S100a8,Ptgs2,Nfkb1,Cp,Il1a,Vcam1,Hspa5 | | 1.356e-04 | -8.91 | GO\_POSITIVE\_REGULATION\_OF\_CELL\_ACTIVATION | MSigDB lists | GO\_POSITIVE\_REGULATION\_OF\_CELL\_ACTIVATION | 194 | 11 | 12187 | 179 | Il4ra,Pik3r1,Map3k8,Il6,Ccl12,Selp,Plek,Thbs1,Cdkn1a,Vcam1,Zbtb16 | | 1.371e-04 | -8.90 | GAL\_LEUKEMIC\_STEM\_CELL\_DN | MSigDB lists | GAL\_LEUKEMIC\_STEM\_CELL\_DN | 162 | 10 | 12187 | 179 | Ms4a6d,Il1rn,Stx11,Cebpd,Ccl4,S100a8,Igsf6,C3ar1,Cd14,S100a9 | | 1.371e-04 | -8.90 | GSE369\_PRE\_VS\_POST\_IL6\_INJECTION\_SOCS3\_KO\_LIVER\_UP | MSigDB lists | GSE369\_PRE\_VS\_POST\_IL6\_INJECTION\_SOCS3\_KO\_LIVER\_UP | 162 | 10 | 12187 | 179 | Osmr,S100a8,Arid5b,Cflar,Gem,Bach1,Fstl1,Ccnd2,Cdkn1a,Maff | | 1.371e-04 | -8.90 | GSE22935\_WT\_VS\_MYD88\_KO\_MACROPHAGE\_12H\_MBOVIS\_BCG\_STIM\_UP | MSigDB lists | GSE22935\_WT\_VS\_MYD88\_KO\_MACROPHAGE\_12H\_MBOVIS\_BCG\_STIM\_UP | 162 | 10 | 12187 | 179 | Il6,Ripk1,Socs3,Thbs1,Sgk1,Adamts1,Zfp189,Msn,Sele,Apold1 | | 1.374e-04 | -8.89 | cardiovascular system development | biological process | GO:0072358 | 468 | 19 | 13711 | 214 | Rnf213,Thbs1,Socs3,Adamts9,Errfi1,Cemip2,Ccl2,Robo4,Angptl4,Ptgs2,Rasip1,Spi1,Col4a1,Apold1,Rhoj,Ccl12,Tiparp,Fzd4,Clic4 | | 1.382e-04 | -8.89 | BERENJENO\_TRANSFORMED\_BY\_RHOA\_UP | MSigDB lists | BERENJENO\_TRANSFORMED\_BY\_RHOA\_UP | 458 | 18 | 12187 | 179 | Runx1,Ifit2,Nfkbia,Tubb6,Tnfaip3,Ccl12,Irgm2,Rgs16,Thbs1,Cdkn1a,Nfkbiz,Timp1,Ier3,Ptgs2,Tagln2,Msn,Errfi1,Plaur | | 1.402e-04 | -8.87 | VART\_KSHV\_INFECTION\_ANGIOGENIC\_MARKERS\_UP | MSigDB lists | VART\_KSHV\_INFECTION\_ANGIOGENIC\_MARKERS\_UP | 132 | 9 | 12187 | 179 | Osmr,Plaur,Ptgs2,Robo4,Cxcl1,Fzd4,Timp1,Il6,Tnf | | 1.402e-04 | -8.87 | GO\_CORE\_PROMOTER\_BINDING | MSigDB lists | GO\_CORE\_PROMOTER\_BINDING | 132 | 9 | 12187 | 179 | Ifi204,Ifi207,Sox11,Ifi209,Runx1,Spi1,Irf9,Ifi211,Irf7 | | 1.416e-04 | -8.86 | blood vessel development | biological process | GO:0001568 | 430 | 18 | 13711 | 214 | Ccl12,Apold1,Rhoj,Col4a1,Clic4,Fzd4,Tiparp,Ptgs2,Angptl4,Spi1,Rasip1,Ccl2,Cemip2,Robo4,Rnf213,Adamts9,Socs3,Thbs1 | | 1.417e-04 | -8.86 | regulation of angiogenesis | biological process | GO:0045765 | 248 | 13 | 13711 | 214 | Nfe2l2,Ptgs2,Runx1,C5ar1,Adamts1,Rhoj,Adamts9,Hspb1,Thbs1,C3ar1,Il1a,Ccl2,Cybb | | 1.423e-04 | -8.86 | ribonucleoside binding | molecular function | GO:0032549 | 318 | 15 | 13516 | 211 | Rhoc,Gbp6,Irgm1,Gbp7,Tgm2,Tubb6,Mx1,Rhoj,Rhou,Gbp3,Iigp1,Gem,Tgtp1,Hcar2,Rasd1 | | 1.423e-04 | -8.86 | purine nucleoside binding | molecular function | GO:0001883 | 318 | 15 | 13516 | 211 | Rasd1,Gem,Tgtp1,Hcar2,Gbp3,Iigp1,Rhou,Mx1,Rhoj,Gbp7,Tubb6,Tgm2,Rhoc,Gbp6,Irgm1 | | 1.440e-04 | -8.85 | KEGG\_SMALL\_CELL\_LUNG\_CANCER | MSigDB lists | KEGG\_SMALL\_CELL\_LUNG\_CANCER | 78 | 7 | 12187 | 179 | Ptgs2,Pik3r5,Col4a1,Nfkb1,Birc3,Pik3r1,Nfkbia | | 1.440e-04 | -8.85 | BURTON\_ADIPOGENESIS\_9 | MSigDB lists | BURTON\_ADIPOGENESIS\_9 | 78 | 7 | 12187 | 179 | Ptgs2,Gem,Vcam1,Thbs1,Cebpd,Ccl12,Akap12 | | 1.442e-04 | -8.84 | GSE7460\_TREG\_VS\_TCONV\_ACT\_UP | MSigDB lists | GSE7460\_TREG\_VS\_TCONV\_ACT\_UP | 163 | 10 | 12187 | 179 | Sdc4,Nfkbiz,Tnfaip3,Stx11,Ncf1,Gbp7,Gem,Gbp3,Tnfsf10,Icam1 | | 1.442e-04 | -8.84 | GSE25123\_CTRL\_VS\_IL4\_STIM\_MACROPHAGE\_UP | MSigDB lists | GSE25123\_CTRL\_VS\_IL4\_STIM\_MACROPHAGE\_UP | 163 | 10 | 12187 | 179 | Ifi209,Runx1,Timp1,Ifi204,Rassf4,Ifi207,Ifi211,Pik3r5,Msn,Trim25 | | 1.459e-04 | -8.83 | positive regulation of smooth muscle cell proliferation | biological process | GO:0048661 | 98 | 8 | 13711 | 214 | P2ry6,Ptgs2,C3ar1,Thbs1,Adamts1,Il6,Tnf,Tgm2 | | 1.466e-04 | -8.83 | Chemokine signaling pathway | KEGG pathways | ko04062 | 144 | 11 | 5248 | 107 | Cxcl16,Ccl4,Ccl12,Cxcl1,Nfkbia,Ncf1,Nfkb1,Ccl7,Ccl2,Pik3r1,Cxcl9 | | 1.466e-04 | -8.83 | Chemokine signaling pathway | KEGG pathways | mmu04062 | 144 | 11 | 5248 | 107 | Cxcl16,Cxcl1,Ccl12,Ccl4,Ncf1,Nfkb1,Nfkbia,Ccl2,Pik3r1,Cxcl9,Ccl7 | | 1.468e-04 | -8.83 | GO\_POSITIVE\_REGULATION\_OF\_ACUTE\_INFLAMMATORY\_RESPONSE | MSigDB lists | GO\_POSITIVE\_REGULATION\_OF\_ACUTE\_INFLAMMATORY\_RESPONSE | 19 | 4 | 12187 | 179 | Osmr,Il6,Tnf,Ptgs2 | | 1.468e-04 | -8.83 | GO\_CELLULAR\_RESPONSE\_TO\_INTERLEUKIN\_6 | MSigDB lists | GO\_CELLULAR\_RESPONSE\_TO\_INTERLEUKIN\_6 | 19 | 4 | 12187 | 179 | Ccl12,Sbno2,Il6,Nfkb1 | | 1.468e-04 | -8.83 | GO\_NEGATIVE\_REGULATION\_OF\_ENDOTHELIAL\_CELL\_APOPTOTIC\_PROCESS | MSigDB lists | GO\_NEGATIVE\_REGULATION\_OF\_ENDOTHELIAL\_CELL\_APOPTOTIC\_PROCESS | 19 | 4 | 12187 | 179 | Tnfaip3,Nfe2l2,Icam1,Angptl4 | | 1.468e-04 | -8.83 | GO\_MONOCYTE\_CHEMOTAXIS | MSigDB lists | GO\_MONOCYTE\_CHEMOTAXIS | 19 | 4 | 12187 | 179 | Ccl4,Ccl2,Ccl12,Il6 | | 1.468e-04 | -8.83 | MODULE\_500 | MSigDB lists | MODULE\_500 | 19 | 4 | 12187 | 179 | Plaur,Nfkb2,Il6,P2ry6 | | 1.468e-04 | -8.83 | EINAV\_INTERFERON\_SIGNATURE\_IN\_CANCER | MSigDB lists | EINAV\_INTERFERON\_SIGNATURE\_IN\_CANCER | 19 | 4 | 12187 | 179 | Irf9,Ifit3b,Ifi44,Isg15 | | 1.468e-04 | -8.83 | XU\_AKT1\_TARGETS\_6HR | MSigDB lists | XU\_AKT1\_TARGETS\_6HR | 19 | 4 | 12187 | 179 | Isg15,Ifitm2,Ifit3b,Irf7 | | 1.473e-04 | -8.82 | RASHI\_RESPONSE\_TO\_IONIZING\_RADIATION\_4 | MSigDB lists | RASHI\_RESPONSE\_TO\_IONIZING\_RADIATION\_4 | 55 | 6 | 12187 | 179 | Ifi204,Ifi207,Ifi211,Ifit3b,Irgm2,Ifi209 | | 1.481e-04 | -8.82 | DNA-binding transcription repressor activity, RNA polymerase II-specific | molecular function | GO:0001227 | 184 | 11 | 13516 | 211 | Mxd4,Ifi209,Ifi204,Arid5b,Spi1,Ifi207,Ifi211,Bach1,Atf3,Zbtb16,Nfkb1 | | 1.481e-04 | -8.82 | DNA-binding transcription repressor activity | molecular function | GO:0001217 | 184 | 11 | 13516 | 211 | Ifi204,Arid5b,Ifi209,Mxd4,Zbtb16,Atf3,Nfkb1,Ifi211,Spi1,Ifi207,Bach1 | | 1.485e-04 | -8.81 | GSE34205\_HEALTHY\_VS\_RSV\_INF\_INFANT\_PBMC\_DN | MSigDB lists | GSE34205\_HEALTHY\_VS\_RSV\_INF\_INFANT\_PBMC\_DN | 133 | 9 | 12187 | 179 | Gpr84,Trib1,Hp,Mmp8,Il1rn,Lcn2,Ifitm3,S100a9,S100a8 | | 1.485e-04 | -8.81 | GO\_POSITIVE\_REGULATION\_OF\_I\_KAPPAB\_KINASE\_NF\_KAPPAB\_SIGNALING | MSigDB lists | GO\_POSITIVE\_REGULATION\_OF\_I\_KAPPAB\_KINASE\_NF\_KAPPAB\_SIGNALING | 133 | 9 | 12187 | 179 | Trim25,Tnfsf10,Cflar,Ptgs2,Rhoc,Il1a,Ripk1,Tnf,Birc3 | | 1.485e-04 | -8.81 | KATSANOU\_ELAVL1\_TARGETS\_UP | MSigDB lists | KATSANOU\_ELAVL1\_TARGETS\_UP | 133 | 9 | 12187 | 179 | Ifi44,Tgm2,Cybb,Irgm2,Nfkbie,C3ar1,Olfml2b,Slfn4,Slfn3 | | 1.500e-04 | -8.80 | African trypanosomiasis | KEGG pathways | ko05143 | 26 | 5 | 5248 | 107 | Tnf,Icam1,Vcam1,Il6,Sele | | 1.500e-04 | -8.80 | African trypanosomiasis | KEGG pathways | mmu05143 | 26 | 5 | 5248 | 107 | Tnf,Icam1,Vcam1,Il6,Sele | | 1.508e-04 | -8.80 | detoxification | biological process | GO:0098754 | 52 | 6 | 13711 | 214 | S100a9,Nfe2l2,S100a8,Hp,Mt2,Mt1 | | 1.516e-04 | -8.79 | GSE17721\_CTRL\_VS\_LPS\_6H\_BMDC\_DN | MSigDB lists | GSE17721\_CTRL\_VS\_LPS\_6H\_BMDC\_DN | 164 | 10 | 12187 | 179 | Ccl12,Ier3,Timp1,Ksr1,Map3k8,Msn,Slfn4,Slfn3,Ptges,Cd14 | | 1.516e-04 | -8.79 | GSE12392\_WT\_VS\_IFNAR\_KO\_CD8A\_NEG\_SPLEEN\_DC\_UP | MSigDB lists | GSE12392\_WT\_VS\_IFNAR\_KO\_CD8A\_NEG\_SPLEEN\_DC\_UP | 164 | 10 | 12187 | 179 | Casp4,Ifit3b,Sdc4,Cebpd,Nfkbie,Ifitm2,Cflar,Slfn3,Slfn4,Igsf6 | | 1.516e-04 | -8.79 | GSE5542\_UNTREATED\_VS\_IFNG\_TREATED\_EPITHELIAL\_CELLS\_24H\_UP | MSigDB lists | GSE5542\_UNTREATED\_VS\_IFNG\_TREATED\_EPITHELIAL\_CELLS\_24H\_UP | 164 | 10 | 12187 | 179 | Arid5b,Iqgap1,Slfn4,Slfn3,Sntb2,Thbs1,Mxd4,Oasl1,Ksr1,Tnip3 | | 1.516e-04 | -8.79 | GSE17721\_LPS\_VS\_POLYIC\_8H\_BMDC\_DN | MSigDB lists | GSE17721\_LPS\_VS\_POLYIC\_8H\_BMDC\_DN | 164 | 10 | 12187 | 179 | Birc3,Usp18,Tiparp,Hp,Mxd4,Isg15,Irf7,Arid5b,Iqgap1,S100a8 | | 1.516e-04 | -8.79 | GSE3337\_CTRL\_VS\_16H\_IFNG\_IN\_CD8POS\_DC\_UP | MSigDB lists | GSE3337\_CTRL\_VS\_16H\_IFNG\_IN\_CD8POS\_DC\_UP | 164 | 10 | 12187 | 179 | C5ar1,C3ar1,Cd14,Fzd4,Cp,Hp,Ier3,Adamts1,Birc3,Vcam1 | | 1.516e-04 | -8.79 | GSE17721\_ALL\_VS\_24H\_PAM3CSK4\_BMDC\_UP | MSigDB lists | GSE17721\_ALL\_VS\_24H\_PAM3CSK4\_BMDC\_UP | 164 | 10 | 12187 | 179 | Slfn4,Slfn3,Plaur,Sgk3,Gbp7,Tubb6,Il1rn,Ifit2,Hspa5,Ccl2 | | 1.516e-04 | -8.79 | GSE43955\_TH0\_VS\_TGFB\_IL6\_TH17\_ACT\_CD4\_TCELL\_4H\_DN | MSigDB lists | GSE43955\_TH0\_VS\_TGFB\_IL6\_TH17\_ACT\_CD4\_TCELL\_4H\_DN | 164 | 10 | 12187 | 179 | Irf7,Tap1,Irf9,Cmpk2,Trim25,Usp18,Nfkbiz,Tor3a,Vcam1,Ifit2 | | 1.520e-04 | -8.79 | intracellular signal transduction | biological process | GO:0035556 | 1177 | 35 | 13711 | 214 | Rasd1,Vcam1,Sgk3,Irgm2,Selp,Bcl3,Ptpn1,Rrp8,Ccl12,Mt2,Ripk1,Hspb1,Irak3,Rhoc,Ifi204,Plek,Tnf,Rhou,Zfp36,Ksr1,Ier3,Sgk1,Pik3r5,Map3k6,Map3k8,Rhoj,Trib1,Socs3,Pik3r1,Nfkbia,Nfkb2,Mt1,Casp4,Cdkn1a,Errfi1 | | 1.534e-04 | -8.78 | signaling receptor binding | molecular function | GO:0005102 | 1178 | 35 | 13516 | 211 | Pik3r1,Thbs1,Vcam1,Il6,Pik3ap1,Isg15,Saa1,Lgals9,Il17ra,Ptpn1,Il1rn,S100a8,Icam1,Plaur,Timp1,Cxcl1,Ccl12,A2m,S100a9,Kcna5,Ccl7,Tap1,Ccl4,Ripk1,Ccl2,Retnlg,Cxcl9,Cxcl16,Csf3,Cflar,Il1a,Tnfsf10,Msn,Rtp4,Tnf | | 1.535e-04 | -8.78 | positive regulation of peptide secretion | biological process | GO:0002793 | 250 | 13 | 13711 | 214 | Il6,Il1a,Akap12,Casp4,Tnf,Il4ra,Lgals9,Hcar2,Cd14,S100a8,Il17ra,S100a9,Mmp8 | | 1.536e-04 | -8.78 | GSE9960\_HEALTHY\_VS\_SEPSIS\_PBMC\_DN | MSigDB lists | GSE9960\_HEALTHY\_VS\_SEPSIS\_PBMC\_DN | 105 | 8 | 12187 | 179 | Zfp36,Hspb1,Ifi211,Ifi207,Samd9l,Ifi204,Ifi209,Aff1 | | 1.536e-04 | -8.78 | GO\_POSITIVE\_REGULATION\_OF\_VASCULATURE\_DEVELOPMENT | MSigDB lists | GO\_POSITIVE\_REGULATION\_OF\_VASCULATURE\_DEVELOPMENT | 105 | 8 | 12187 | 179 | Thbs1,Runx1,Il1a,Angptl4,C3ar1,Ptgs2,C5ar1,Hspb1 | | 1.547e-04 | -8.77 | GO\_REGULATORY\_REGION\_NUCLEIC\_ACID\_BINDING | MSigDB lists | GO\_REGULATORY\_REGION\_NUCLEIC\_ACID\_BINDING | 547 | 20 | 12187 | 179 | Nfkb2,Ifi211,Irf7,Arid5b,Bach1,Nfe2l2,Runx1,Tnf,Spi1,Irf2,Irf9,Ifi204,Tiparp,Nfkb1,Ifi207,Cebpd,Atf3,Sox11,Ifi209,Zbtb16 | | 1.560e-04 | -8.77 | HOEBEKE\_LYMPHOID\_STEM\_CELL\_DN | MSigDB lists | HOEBEKE\_LYMPHOID\_STEM\_CELL\_DN | 79 | 7 | 12187 | 179 | Ifi204,Ifi207,Ifi209,Maff,Tnfsf10,Hspb1,Ifi211 | | 1.594e-04 | -8.74 | GSE17721\_LPS\_VS\_CPG\_12H\_BMDC\_UP | MSigDB lists | GSE17721\_LPS\_VS\_CPG\_12H\_BMDC\_UP | 165 | 10 | 12187 | 179 | Gbp4,Il1rn,Timp1,Ifit3b,Casp4,Vcam1,Sox11,Ifit2,Tnfsf10,Gbp6 | | 1.594e-04 | -8.74 | GSE17721\_LPS\_VS\_GARDIQUIMOD\_4H\_BMDC\_UP | MSigDB lists | GSE17721\_LPS\_VS\_GARDIQUIMOD\_4H\_BMDC\_UP | 165 | 10 | 12187 | 179 | Usp18,Ccl2,Sdc4,Tor3a,Gadd45b,C3ar1,Bcl3,Irf7,Rtp4,Irf2 | | 1.594e-04 | -8.74 | GSE43955\_10H\_VS\_30H\_ACT\_CD4\_TCELL\_UP | MSigDB lists | GSE43955\_10H\_VS\_30H\_ACT\_CD4\_TCELL\_UP | 165 | 10 | 12187 | 179 | Il6,Il1a,Vcam1,Ccl2,Tnf,Ifit2,Plaur,Msn,Ncf1,C3ar1 | | 1.594e-04 | -8.74 | GSE3039\_ALPHAALPHA\_CD8\_TCELL\_VS\_B2\_BCELL\_DN | MSigDB lists | GSE3039\_ALPHAALPHA\_CD8\_TCELL\_VS\_B2\_BCELL\_DN | 165 | 10 | 12187 | 179 | Ifit3b,Ccl4,Fstl1,Maff,Tiparp,Timp1,Arid5b,Herc6,Cxcl16,Pik3ap1 | | 1.594e-04 | -8.74 | GSE12392\_IFNAR\_KO\_VS\_IFNB\_KO\_CD8\_NEG\_SPLEEN\_DC\_UP | MSigDB lists | GSE12392\_IFNAR\_KO\_VS\_IFNB\_KO\_CD8\_NEG\_SPLEEN\_DC\_UP | 165 | 10 | 12187 | 179 | Sdc4,Ccl4,Maff,Birc3,Sgk1,Nfe2l2,Atf3,Tagln2,Cflar,Plaur | | 1.594e-04 | -8.74 | GSE17721\_PAM3CSK4\_VS\_GADIQUIMOD\_8H\_BMDC\_DN | MSigDB lists | GSE17721\_PAM3CSK4\_VS\_GADIQUIMOD\_8H\_BMDC\_DN | 165 | 10 | 12187 | 179 | Zfp36,Gbp3,Slfn9,Parp12,Ripk1,Pik3r1,Map3k8,Aff1,Nfkbiz,Ccl4 | | 1.630e-04 | -8.72 | BIOCARTA\_HIVNEF\_PATHWAY | MSigDB lists | BIOCARTA\_HIVNEF\_PATHWAY | 56 | 6 | 12187 | 179 | Birc3,Tnf,Cflar,Nfkbia,Nfkb1,Ripk1 | | 1.641e-04 | -8.71 | DARWICHE\_SQUAMOUS\_CELL\_CARCINOMA\_UP | MSigDB lists | DARWICHE\_SQUAMOUS\_CELL\_CARCINOMA\_UP | 106 | 8 | 12187 | 179 | Nfe2l2,Rasip1,Ifitm3,S100a9,Cmpk2,S100a8,Icam1,Cxcl16 | | 1.641e-04 | -8.71 | GO\_NEGATIVE\_REGULATION\_OF\_BINDING | MSigDB lists | GO\_NEGATIVE\_REGULATION\_OF\_BINDING | 106 | 8 | 12187 | 179 | Ifi211,Ifi207,Ifi204,Nfkbia,Hspa5,Ifi209,Sox11,Ifit2 | | 1.653e-04 | -8.71 | BIOCARTA\_CYTOKINE\_PATHWAY | MSigDB lists | BIOCARTA\_CYTOKINE\_PATHWAY | 8 | 3 | 12187 | 179 | Tnf,Il1a,Il6 | | 1.653e-04 | -8.71 | IYENGAR\_RESPONSE\_TO\_ADIPOCYTE\_FACTORS | MSigDB lists | IYENGAR\_RESPONSE\_TO\_ADIPOCYTE\_FACTORS | 8 | 3 | 12187 | 179 | Atf3,Tnfaip3,Nfkb1 | | 1.653e-04 | -8.71 | BIOCARTA\_FREE\_PATHWAY | MSigDB lists | BIOCARTA\_FREE\_PATHWAY | 8 | 3 | 12187 | 179 | Tnf,Nfkb1,Xdh | | 1.664e-04 | -8.70 | GSE360\_CTRL\_VS\_L\_DONOVANI\_DC\_DN | MSigDB lists | GSE360\_CTRL\_VS\_L\_DONOVANI\_DC\_DN | 135 | 9 | 12187 | 179 | Ier3,Cebpd,Stx11,Nfkbia,Ccl4,Il4ra,Csf2rb,Nfkb2,P2ry6 | | 1.675e-04 | -8.69 | GSE24634\_IL4\_VS\_CTRL\_TREATED\_NAIVE\_CD4\_TCELL\_DAY10\_DN | MSigDB lists | GSE24634\_IL4\_VS\_CTRL\_TREATED\_NAIVE\_CD4\_TCELL\_DAY10\_DN | 166 | 10 | 12187 | 179 | Cybb,Tubb6,Stx11,Il1rn,Cdkn1a,Ifit3b,Plaur,Clic4,Gem,Irf7 | | 1.675e-04 | -8.69 | GSE17721\_0.5H\_VS\_12H\_CPG\_BMDC\_DN | MSigDB lists | GSE17721\_0.5H\_VS\_12H\_CPG\_BMDC\_DN | 166 | 10 | 12187 | 179 | Tnip1,Slfn9,Arid5b,Rtp4,Cmpk2,Tap1,Birc3,Parp14,Cp,Parp12 | | 1.678e-04 | -8.69 | regulation of toll-like receptor signaling pathway | biological process | GO:0034121 | 53 | 6 | 13711 | 214 | Trim30a,Pik3ap1,Irf7,Acod1,Tnfaip3,Irak3 | | 1.678e-04 | -8.69 | positive regulation of response to wounding | biological process | GO:1903036 | 53 | 6 | 13711 | 214 | Plek,S100a9,Nfe2l2,Thbs1,Ccl2,Selp | | 1.678e-04 | -8.69 | cellular response to chemokine | biological process | GO:1990869 | 53 | 6 | 13711 | 214 | Ccl4,Ccl7,Cxcl9,Cxcl1,Ccl12,Ccl2 | | 1.678e-04 | -8.69 | response to chemokine | biological process | GO:1990868 | 53 | 6 | 13711 | 214 | Ccl7,Ccl4,Ccl12,Ccl2,Cxcl1,Cxcl9 | | 1.683e-04 | -8.69 | MODULE\_12 | MSigDB lists | MODULE\_12 | 306 | 14 | 12187 | 179 | Ccnd2,Tgm2,Il6,Ccl12,Fam107a,Tnfaip3,Cxcl1,S100a8,Rgs16,Cdkn1a,Zbtb16,Col4a1,Isg15,Cd14 | | 1.689e-04 | -8.69 | guanyl ribonucleotide binding | molecular function | GO:0032561 | 323 | 15 | 13516 | 211 | Gbp7,Tubb6,Tgm2,Irgm1,Rhoc,Gbp6,Rhou,Rhoj,Mx1,Iigp1,Gbp3,Rasd1,Tgtp1,Gem,Hcar2 | | 1.689e-04 | -8.69 | guanyl nucleotide binding | molecular function | GO:0019001 | 323 | 15 | 13516 | 211 | Iigp1,Gbp3,Tgtp1,Gem,Hcar2,Rasd1,Irgm1,Gbp6,Rhoc,Tgm2,Tubb6,Gbp7,Mx1,Rhoj,Rhou | | 1.689e-04 | -8.69 | GO\_REGULATION\_OF\_WOUND\_HEALING | MSigDB lists | GO\_REGULATION\_OF\_WOUND\_HEALING | 80 | 7 | 12187 | 179 | Plek,Nfe2l2,Plaur,Tnfaip3,Thbs1,Selp,S100a9 | | 1.689e-04 | -8.69 | WIELAND\_UP\_BY\_HBV\_INFECTION | MSigDB lists | WIELAND\_UP\_BY\_HBV\_INFECTION | 80 | 7 | 12187 | 179 | Ifi209,Ifi207,Ifi204,Parp12,C3ar1,Ifi211,Igsf6 | | 1.742e-04 | -8.66 | PARP\_CATALYTIC | prosite domains | PS51059 | 16 | 4 | 8845 | 164 | Tiparp,Parp14,Parp12,Parp9 | | 1.751e-04 | -8.65 | DOUGLAS\_BMI1\_TARGETS\_DN | MSigDB lists | DOUGLAS\_BMI1\_TARGETS\_DN | 270 | 13 | 12187 | 179 | Herc6,Slfn5,Rbm47,Icam1,Irf9,Sgk1,Ifit3b,Nfkbiz,Gadd45g,Nfkb2,Birc3,Ifit2,Tnfaip3 | | 1.759e-04 | -8.65 | GSE22601\_DOUBLE\_NEGATIVE\_VS\_CD4\_SINGLE\_POSITIVE\_THYMOCYTE\_DN | MSigDB lists | GSE22601\_DOUBLE\_NEGATIVE\_VS\_CD4\_SINGLE\_POSITIVE\_THYMOCYTE\_DN | 167 | 10 | 12187 | 179 | Il17ra,Ptpn1,Ifitm2,Samd9l,Ksr1,Stx11,Tiparp,Il4ra,Map3k8,Parp14 | | 1.759e-04 | -8.65 | GSE9037\_CTRL\_VS\_LPS\_4H\_STIM\_IRAK4\_KO\_BMDM\_DN | MSigDB lists | GSE9037\_CTRL\_VS\_LPS\_4H\_STIM\_IRAK4\_KO\_BMDM\_DN | 167 | 10 | 12187 | 179 | Xaf1,Ccnd2,Adamts1,Cdkn1a,Parp12,Ms4a6d,Il1rn,Slfn9,Irgm2,Tap1 | | 1.759e-04 | -8.65 | RODWELL\_AGING\_KIDNEY\_NO\_BLOOD\_UP | MSigDB lists | RODWELL\_AGING\_KIDNEY\_NO\_BLOOD\_UP | 167 | 10 | 12187 | 179 | Rnf213,Cxcl1,A2m,Osmr,Msr1,Socs3,Vcam1,Nfkbiz,Timp1,Ccl12 | | 1.760e-04 | -8.65 | GSE360\_L\_MAJOR\_VS\_B\_MALAYI\_HIGH\_DOSE\_MAC\_UP | MSigDB lists | GSE360\_L\_MAJOR\_VS\_B\_MALAYI\_HIGH\_DOSE\_MAC\_UP | 136 | 9 | 12187 | 179 | Il1rn,Nfkb1,Tnfaip3,Map3k8,Nfkbia,Plaur,Cflar,Tnip1,Ptges | | 1.760e-04 | -8.65 | HAMAI\_APOPTOSIS\_VIA\_TRAIL\_DN | MSigDB lists | HAMAI\_APOPTOSIS\_VIA\_TRAIL\_DN | 136 | 9 | 12187 | 179 | Kcna5,Il4ra,Icam1,Plaur,Ifitm2,Isg15,Angptl4,Tagln2,Bcl3 | | 1.762e-04 | -8.64 | Hepatitis C | KEGG pathways | ko05160 | 100 | 9 | 5248 | 107 | Socs3,Nfkbia,Nfkb1,Tnf,Ripk1,Irf7,Irf9,Pik3r1,Cdkn1a | | 1.762e-04 | -8.64 | Hepatitis C | KEGG pathways | mmu05160 | 100 | 9 | 5248 | 107 | Irf7,Ripk1,Irf9,Pik3r1,Tnf,Nfkb1,Socs3,Nfkbia,Cdkn1a | | 1.770e-04 | -8.64 | Rela (v-rel reticuloendotheliosis viral oncogene homolog A (avian)) | protein interactions | 19697 | 32 | 5 | 6802 | 115 | Il6,Nfkbia,Nfkb1,Nfkb2,Sele | | 1.780e-04 | -8.63 | positive regulation of kinase activity | biological process | GO:0033674 | 438 | 18 | 13711 | 214 | Gadd45b,Gadd45g,Ccnd2,Lgals9,Il1rn,Ncf1,Thbs1,Ripk1,Cdkn1a,Tnf,Pik3r5,Irgm2,Iqgap1,Sdc4,Map3k8,Fzd4,Map3k6,Ptpn1 | | 1.814e-04 | -8.61 | GO\_NEGATIVE\_REGULATION\_OF\_CYTOKINE\_BIOSYNTHETIC\_PROCESS | MSigDB lists | GO\_NEGATIVE\_REGULATION\_OF\_CYTOKINE\_BIOSYNTHETIC\_PROCESS | 20 | 4 | 12187 | 179 | Nfkb1,Errfi1,Il6,Bcl3 | | 1.814e-04 | -8.61 | AZARE\_STAT3\_TARGETS | MSigDB lists | AZARE\_STAT3\_TARGETS | 20 | 4 | 12187 | 179 | S100a9,Thbs1,Ccnd2,Clic4 | | 1.814e-04 | -8.61 | GO\_NEGATIVE\_REGULATION\_OF\_TOLL\_LIKE\_RECEPTOR\_SIGNALING\_PATHWAY | MSigDB lists | GO\_NEGATIVE\_REGULATION\_OF\_TOLL\_LIKE\_RECEPTOR\_SIGNALING\_PATHWAY | 20 | 4 | 12187 | 179 | Cd14,Tnfaip3,Irak3,Pik3ap1 | | 1.814e-04 | -8.61 | REACTOME\_TRAF6\_MEDIATED\_NFKB\_ACTIVATION | MSigDB lists | REACTOME\_TRAF6\_MEDIATED\_NFKB\_ACTIVATION | 20 | 4 | 12187 | 179 | Trim25,Nfkb2,Saa1,Nfkbia | | 1.814e-04 | -8.61 | GO\_MONOCYTE\_DIFFERENTIATION | MSigDB lists | GO\_MONOCYTE\_DIFFERENTIATION | 20 | 4 | 12187 | 179 | Ifi211,Ifi209,Ifi204,Ifi207 | | 1.814e-04 | -8.61 | GO\_RESPONSE\_TO\_COPPER\_ION | MSigDB lists | GO\_RESPONSE\_TO\_COPPER\_ION | 20 | 4 | 12187 | 179 | Nfkb1,Icam1,Cp,Il1a | | 1.847e-04 | -8.60 | GSE32034\_LY6C\_HIGH\_VS\_LOW\_ROSIGLIZATONE\_TREATED\_MONOCYTE\_UP | MSigDB lists | GSE32034\_LY6C\_HIGH\_VS\_LOW\_ROSIGLIZATONE\_TREATED\_MONOCYTE\_UP | 168 | 10 | 12187 | 179 | Ms4a6d,Znfx1,Lgals9,Ifit2,Nfkbia,Ifit3b,S100a8,Cmpk2,Isg15,Irf7 | | 1.847e-04 | -8.60 | GSE40274\_FOXP3\_VS\_FOXP3\_AND\_EOS\_TRANSDUCED\_ACTIVATED\_CD4\_TCELL\_UP | MSigDB lists | GSE40274\_FOXP3\_VS\_FOXP3\_AND\_EOS\_TRANSDUCED\_ACTIVATED\_CD4\_TCELL\_UP | 168 | 10 | 12187 | 179 | Tiparp,Socs3,Sdc4,Nfkbia,Trim25,Rtp4,Zfp36,Errfi1,Irgm2,Arid5b | | 1.847e-04 | -8.60 | GSE17186\_NAIVE\_VS\_CD21LOW\_TRANSITIONAL\_BCELL\_UP | MSigDB lists | GSE17186\_NAIVE\_VS\_CD21LOW\_TRANSITIONAL\_BCELL\_UP | 168 | 10 | 12187 | 179 | Il4ra,Socs3,Timp1,Gadd45g,Selp,Slfn9,Arid5b,Bcl3,Icam1,Ptpn1 | | 1.860e-04 | -8.59 | GSE7768\_OVA\_WITH\_LPS\_VS\_OVA\_WITH\_MPL\_IMMUNIZED\_MOUSE\_WHOLE\_SPLEEN\_6H\_DN | MSigDB lists | GSE7768\_OVA\_WITH\_LPS\_VS\_OVA\_WITH\_MPL\_IMMUNIZED\_MOUSE\_WHOLE\_SPLEEN\_6H\_DN | 137 | 9 | 12187 | 179 | Map3k8,Tnip3,Nfkb1,Pik3r5,Cd14,Ptgs2,Plaur,Irf2,Zfp36 | | 1.861e-04 | -8.59 | retina vasculature development in camera-type eye | biological process | GO:0061298 | 19 | 4 | 13711 | 214 | Fzd4,Clic4,Col4a1,Rhoj | | 1.896e-04 | -8.57 | Interferon\_reg\_fac\_CS | interpro domains | IPR019817 | 8 | 3 | 13788 | 212 | Irf2,Irf9,Irf7 | | 1.896e-04 | -8.57 | Interferon\_reg\_fact\_DNA-bd\_dom | interpro domains | IPR001346 | 8 | 3 | 13788 | 212 | Irf7,Irf2,Irf9 | | 1.925e-04 | -8.56 | WOOD\_EBV\_EBNA1\_TARGETS\_DN | MSigDB lists | WOOD\_EBV\_EBNA1\_TARGETS\_DN | 37 | 5 | 12187 | 179 | Sgk1,Ptgs2,Iqgap1,Akap12,P2ry6 | | 1.925e-04 | -8.56 | GO\_POSITIVE\_REGULATION\_OF\_CYTOKINE\_BIOSYNTHETIC\_PROCESS | MSigDB lists | GO\_POSITIVE\_REGULATION\_OF\_CYTOKINE\_BIOSYNTHETIC\_PROCESS | 37 | 5 | 12187 | 179 | Bcl3,Thbs1,Tnf,Il1a,Hspb1 | | 1.925e-04 | -8.56 | ST\_B\_CELL\_ANTIGEN\_RECEPTOR | MSigDB lists | ST\_B\_CELL\_ANTIGEN\_RECEPTOR | 37 | 5 | 12187 | 179 | Nfkbia,Nfkbie,Pik3r1,Nfkb2,Nfkb1 | | 1.927e-04 | -8.55 | cellular response to reactive oxygen species | biological process | GO:0034614 | 102 | 8 | 13711 | 214 | Ncf1,Nfe2l2,Thbs1,Ripk1,Cflar,Lcn2,Tnfaip3,Il6 | | 1.931e-04 | -8.55 | nucleoside binding | molecular function | GO:0001882 | 327 | 15 | 13516 | 211 | Gbp3,Iigp1,Tgtp1,Hcar2,Gem,Rasd1,Gbp6,Rhoc,Irgm1,Tgm2,Tubb6,Gbp7,Rhoj,Mx1,Rhou | | 1.934e-04 | -8.55 | GO\_TRANSCRIPTIONAL\_ACTIVATOR\_ACTIVITY\_RNA\_POLYMERASE\_II\_TRANSCRIPTION\_REGULATORY\_REGION\_SEQUENCE\_SPECIFIC\_BINDING | MSigDB lists | GO\_TRANSCRIPTIONAL\_ACTIVATOR\_ACTIVITY\_RNA\_POLYMERASE\_II\_TRANSCRIPTION\_REGULATORY\_REGION\_SEQUENCE\_SPECIFIC\_BINDING | 202 | 11 | 12187 | 179 | Nfkb2,Irf2,Spi1,Maff,Runx1,Sox11,Nfe2l2,Atf3,Cebpd,Bach1,Nfkb1 | | 1.934e-04 | -8.55 | FLORIO\_NEOCORTEX\_BASAL\_RADIAL\_GLIA\_UP | MSigDB lists | FLORIO\_NEOCORTEX\_BASAL\_RADIAL\_GLIA\_UP | 202 | 11 | 12187 | 179 | Gbp4,Col4a1,Parp14,Thbs1,Tnfsf10,Apold1,Adamts9,Slfn4,Sele,Slfn3,Gbp6 | | 1.939e-04 | -8.55 | GSE17721\_PAM3CSK4\_VS\_CPG\_24H\_BMDC\_DN | MSigDB lists | GSE17721\_PAM3CSK4\_VS\_CPG\_24H\_BMDC\_DN | 169 | 10 | 12187 | 179 | Gem,Gbp6,Ifit2,Ccl2,Slfn9,Tor3a,Cmpk2,Trim25,Gadd45b,Gbp4 | | 1.939e-04 | -8.55 | GSE46606\_UNSTIM\_VS\_CD40L\_IL2\_IL5\_1DAY\_STIMULATED\_IRF4MID\_SORTED\_BCELL\_UP | MSigDB lists | GSE46606\_UNSTIM\_VS\_CD40L\_IL2\_IL5\_1DAY\_STIMULATED\_IRF4MID\_SORTED\_BCELL\_UP | 169 | 10 | 12187 | 179 | Zfp36,Icam1,Ptgs2,Gem,Il6,Tnfaip3,Socs3,Maff,Map3k8,Tnf | | 1.939e-04 | -8.55 | GSE42021\_CD24LO\_TREG\_VS\_CD24LO\_TCONV\_THYMUS\_UP | MSigDB lists | GSE42021\_CD24LO\_TREG\_VS\_CD24LO\_TCONV\_THYMUS\_UP | 169 | 10 | 12187 | 179 | Ccdc58,Tnfaip8,Banp,Bcl3,Lcn2,Ms4a6d,Il1a,Ccl4,Birc3,Lgals9 | | 1.949e-04 | -8.54 | negative regulation of cysteine-type endopeptidase activity involved in apoptotic process | biological process | GO:0043154 | 77 | 7 | 13711 | 214 | Thbs1,Tnfaip8,Cflar,Ptgs2,Plaur,Birc3,Il6 | | 1.959e-04 | -8.54 | negative regulation of interleukin-6 production | biological process | GO:0032715 | 35 | 5 | 13711 | 214 | Trim30a,Prg4,Tnf,Irak3,Tnfaip3 | | 1.965e-04 | -8.53 | GSE37301\_HEMATOPOIETIC\_STEM\_CELL\_VS\_CD4\_TCELL\_UP | MSigDB lists | GSE37301\_HEMATOPOIETIC\_STEM\_CELL\_VS\_CD4\_TCELL\_UP | 138 | 9 | 12187 | 179 | Gem,S100a9,Rbm47,Pik3ap1,Xdh,Csf2rb,Casp4,Rassf4,Lcn2 | | 1.972e-04 | -8.53 | KEGG\_T\_CELL\_RECEPTOR\_SIGNALING\_PATHWAY | MSigDB lists | KEGG\_T\_CELL\_RECEPTOR\_SIGNALING\_PATHWAY | 82 | 7 | 12187 | 179 | Map3k8,Tnf,Nfkbie,Pik3r1,Pik3r5,Nfkbia,Nfkb1 | | 1.979e-04 | -8.53 | IRF | pfam domains | PF00605 | 8 | 3 | 12881 | 201 | Irf9,Irf7,Irf2 | | 1.982e-04 | -8.53 | tolerance induction | biological process | GO:0002507 | 8 | 3 | 13711 | 214 | Acod1,C3ar1,Tnfaip3 | | 1.982e-04 | -8.53 | negative regulation of lipopolysaccharide-mediated signaling pathway | biological process | GO:0031665 | 8 | 3 | 13711 | 214 | Trib1,Acod1,Tnfaip3 | | 1.982e-04 | -8.53 | positive regulation of transport | biological process | GO:0051050 | 867 | 28 | 13711 | 214 | Il6,Il1a,Sox11,Akap12,Nfkbia,Casp4,Il4ra,Lgals9,P2ry6,Hcar2,Pik3r1,Sdc4,Il17ra,Sgk1,Ptgs2,Runx1,Ccl4,Ccl2,Tnf,Cd14,Sele,Ccl12,S100a8,Cxcl1,Msn,Mmp8,S100a9,Nfe2l2 | | 2.001e-04 | -8.52 | positive regulation of leukocyte differentiation | biological process | GO:1902107 | 130 | 9 | 13711 | 214 | Lgals9,Runx1,Zbtb16,Ripk1,Trib1,Il6,Nfkbiz,Il4ra,Tnf | | 2.028e-04 | -8.50 | regulation of proteolysis | biological process | GO:0030162 | 564 | 21 | 13711 | 214 | Casp4,Tnf,Il6,Tnfsf10,Xdh,Cflar,Ctla2a,Trib1,Tnfaip8,Thbs1,Pik3r1,Tnip1,Birc3,Timp1,A2m,S100a8,S100a9,Serpina3f,Wfdc21,Plaur,Ptgs2 | | 2.034e-04 | -8.50 | GSE15930\_STIM\_VS\_STIM\_AND\_IL12\_48H\_CD8\_T\_CELL\_UP | MSigDB lists | GSE15930\_STIM\_VS\_STIM\_AND\_IL12\_48H\_CD8\_T\_CELL\_UP | 170 | 10 | 12187 | 179 | Lgals9,Ccnd2,Vcam1,Nfkb1,Cmpk2,Slfn3,Slfn4,Irf9,Tap1,Ptpn1 | | 2.034e-04 | -8.50 | GSE43955\_TGFB\_IL6\_VS\_TGFB\_IL6\_IL23\_TH17\_ACT\_CD4\_TCELL\_52H\_DN | MSigDB lists | GSE43955\_TGFB\_IL6\_VS\_TGFB\_IL6\_IL23\_TH17\_ACT\_CD4\_TCELL\_52H\_DN | 170 | 10 | 12187 | 179 | Ptgs2,Irf7,Zfp36,Clic4,Birc3,Msr1,Tnf,Casp4,Fstl1,Zbtb16 | | 2.045e-04 | -8.50 | GO\_REGULATION\_OF\_MULTICELLULAR\_ORGANISMAL\_DEVELOPMENT | MSigDB lists | GO\_REGULATION\_OF\_MULTICELLULAR\_ORGANISMAL\_DEVELOPMENT | 1280 | 35 | 12187 | 179 | Nfkbia,Trib1,Tnf,Runx1,Irf7,Fzd4,C3ar1,C5ar1,Xdh,Zbtb16,Adamts1,Timp1,Il1rn,Ptgs2,Errfi1,Apold1,Hspb1,Spi1,Il4ra,Pik3r1,Maff,Il6,Nfe2l2,Tnfaip3,Ripk1,Ccl12,Angptl4,Csf3,Adamts9,Thbs1,Sgk1,Sox11,Il1a,Isg15,Irak3 | | 2.050e-04 | -8.49 | GO\_REGULATION\_OF\_CELL\_PROLIFERATION | MSigDB lists | GO\_REGULATION\_OF\_CELL\_PROLIFERATION | 1077 | 31 | 12187 | 179 | Il4ra,Il6,Tnfaip3,Ccl12,Ptges,Cxcl1,Csf3,Osmr,Sgk1,Thbs1,Sox11,Mxd4,Il1a,Sgk3,Nfkbia,Trib1,Tnf,C3ar1,C5ar1,Cflar,Xdh,Ifit3b,Zbtb16,Vcam1,Cdkn1a,Adamts1,Atf3,Timp1,Kcna5,Ptgs2,Errfi1 | | 2.055e-04 | -8.49 | negative regulation of protein kinase activity | biological process | GO:0006469 | 191 | 11 | 13711 | 214 | Trib1,Irak3,Hspb1,Rasip1,Gadd45b,Gadd45g,Ptpn1,Tnfaip3,Cdkn1a,Errfi1,Il6 | | 2.067e-04 | -8.48 | regulation of JNK cascade | biological process | GO:0046328 | 160 | 10 | 13711 | 214 | Ripk1,Mmp8,Il1rn,Ncf1,Gadd45b,Gadd45g,Fzd4,Il1a,Ptpn1,Tnf | | 2.075e-04 | -8.48 | GSE6259\_CD4\_TCELL\_VS\_CD8\_TCELL\_DN | MSigDB lists | GSE6259\_CD4\_TCELL\_VS\_CD8\_TCELL\_DN | 139 | 9 | 12187 | 179 | Casp4,Sdc4,Tnfsf10,Irf2,P2ry6,Pik3ap1,Icam1,Trim25,Irf7 | | 2.075e-04 | -8.48 | GSE17974\_IL4\_AND\_ANTI\_IL12\_VS\_UNTREATED\_6H\_ACT\_CD4\_TCELL\_DN | MSigDB lists | GSE17974\_IL4\_AND\_ANTI\_IL12\_VS\_UNTREATED\_6H\_ACT\_CD4\_TCELL\_DN | 139 | 9 | 12187 | 179 | Ifit3b,Sgk1,Ifit2,Ccl12,Il1a,Stx11,Ptgs2,Tnfsf10,Gbp3 | | 2.075e-04 | -8.48 | GSE13946\_CTRL\_VS\_DSS\_COLITIS\_GD\_TCELL\_FROM\_COLON\_UP | MSigDB lists | GSE13946\_CTRL\_VS\_DSS\_COLITIS\_GD\_TCELL\_FROM\_COLON\_UP | 139 | 9 | 12187 | 179 | Gem,Phf11d,S100a9,Irak3,Rbm47,S100a8,Phf11b,Ccl4,Gadd45b | | 2.089e-04 | -8.47 | MARTENS\_TRETINOIN\_RESPONSE\_UP | MSigDB lists | MARTENS\_TRETINOIN\_RESPONSE\_UP | 391 | 16 | 12187 | 179 | Rtp4,S100a8,Gem,Pik3r5,Ptges,Angptl4,S100a9,Ccl12,Tgm2,Ccl2,Icam1,Csf2rb,Ncf1,Cd14,Kcna5,Mxd4 | | 2.091e-04 | -8.47 | cell surface | cellular component | GO:0009986 | 618 | 22 | 13825 | 212 | Adamts9,Cd14,C5ar1,Sdc4,Fzd4,Icam1,Ifitm3,Il1a,Cxcl9,Kcna5,Il6,Plaur,Thbs1,Tnf,Osmr,Clic4,Robo4,Selp,Rtp4,Hspa5,Msn,Vcam1 | | 2.121e-04 | -8.46 | PROVENZANI\_METASTASIS\_DN | MSigDB lists | PROVENZANI\_METASTASIS\_DN | 110 | 8 | 12187 | 179 | Hspb1,Ifitm2,Ifitm3,Tnip1,Lcn2,Ccnd2,Sdc4,Tgm2 | | 2.127e-04 | -8.46 | SENGUPTA\_NASOPHARYNGEAL\_CARCINOMA\_WITH\_LMP1\_UP | MSigDB lists | SENGUPTA\_NASOPHARYNGEAL\_CARCINOMA\_WITH\_LMP1\_UP | 313 | 14 | 12187 | 179 | Ifit2,Usp18,Xaf1,Rbm39,Cmpk2,Iqgap1,Samd9l,Cp,Cebpd,Luc7l3,Adamts1,Ifit3b,Irak3,Ptgs2 | | 2.128e-04 | -8.46 | UEDA\_CENTRAL\_CLOCK | MSigDB lists | UEDA\_CENTRAL\_CLOCK | 83 | 7 | 12187 | 179 | Sgk1,Hspa5,Dnajb1,Gadd45g,Rasd1,Gadd45b,Rgs16 | | 2.128e-04 | -8.46 | REACTOME\_ACTIVATED\_TLR4\_SIGNALLING | MSigDB lists | REACTOME\_ACTIVATED\_TLR4\_SIGNALLING | 83 | 7 | 12187 | 179 | Nfkb2,Irak3,Cd14,Irf7,Saa1,Ripk1,Nfkbia | | 2.134e-04 | -8.45 | GSE23308\_CTRL\_VS\_CORTICOSTERONE\_TREATED\_MACROPHAGE\_DN | MSigDB lists | GSE23308\_CTRL\_VS\_CORTICOSTERONE\_TREATED\_MACROPHAGE\_DN | 171 | 10 | 12187 | 179 | Plaur,Igsf6,Tnfaip8,Ptges,Cxcl1,Rrp8,Saa1,Lcn2,Nfe2l2,Casp4 | | 2.134e-04 | -8.45 | GSE43955\_1H\_VS\_10H\_ACT\_CD4\_TCELL\_UP | MSigDB lists | GSE43955\_1H\_VS\_10H\_ACT\_CD4\_TCELL\_UP | 171 | 10 | 12187 | 179 | Cebpd,Tnf,Thbs1,Msr1,Ccl4,Xdh,Plaur,Saa1,Gem,Tnip1 | | 2.134e-04 | -8.45 | GSE20715\_WT\_VS\_TLR4\_KO\_LUNG\_UP | MSigDB lists | GSE20715\_WT\_VS\_TLR4\_KO\_LUNG\_UP | 171 | 10 | 12187 | 179 | Pik3r1,Hspa5,Lgals9,Ifit3b,Cybb,Parp12,Ptgs2,Ncf1,Hspb1,Clic4 | | 2.182e-04 | -8.43 | PID\_INTEGRIN1\_PATHWAY | MSigDB lists | PID\_INTEGRIN1\_PATHWAY | 59 | 6 | 12187 | 179 | Thbs1,Cd14,Vcam1,Tgm2,Col4a1,Plaur | | 2.187e-04 | -8.43 | Platelet activation, signaling and aggregation | REACTOME pathways | R-MMU-76002 | 215 | 12 | 6297 | 105 | Pik3r5,Pik3r1,Csf2rb,Timp1,Serpina3f,Ptpn1,Hspa5,Selp,A2m,Plek,Tagln2,Thbs1 | | 2.190e-04 | -8.43 | GSE6259\_FLT3L\_INDUCED\_VS\_WT\_SPLENIC\_DC\_33D1\_POS\_UP | MSigDB lists | GSE6259\_FLT3L\_INDUCED\_VS\_WT\_SPLENIC\_DC\_33D1\_POS\_UP | 140 | 9 | 12187 | 179 | Usp18,Nfkbiz,Parp12,C5ar1,Tnip1,Ncf1,Cd14,Irf9,Tnfsf10 | | 2.190e-04 | -8.43 | GSE14415\_INDUCED\_TREG\_VS\_FOXP3\_KO\_INDUCED\_TREG\_IL2\_CULTURE\_UP | MSigDB lists | GSE14415\_INDUCED\_TREG\_VS\_FOXP3\_KO\_INDUCED\_TREG\_IL2\_CULTURE\_UP | 140 | 9 | 12187 | 179 | Ier3,Phf11b,Ccl4,Nfkbia,Rtp4,Rbm47,Rgs16,Gem,Phf11d | | 2.190e-04 | -8.43 | TGFB\_UP.V1\_DN | MSigDB lists | TGFB\_UP.V1\_DN | 140 | 9 | 12187 | 179 | Col4a1,Ifit2,Adamts1,Birc3,Ifit3b,Rbm47,Angptl4,Cxcl1,Arid5b | | 2.190e-04 | -8.43 | GENTILE\_UV\_RESPONSE\_CLUSTER\_D2 | MSigDB lists | GENTILE\_UV\_RESPONSE\_CLUSTER\_D2 | 38 | 5 | 12187 | 179 | Thbs1,Runx1,Nfe2l2,Tnfaip3,Bach1 | | 2.190e-04 | -8.43 | KRIEG\_HYPOXIA\_VIA\_KDM3A | MSigDB lists | KRIEG\_HYPOXIA\_VIA\_KDM3A | 38 | 5 | 12187 | 179 | Tgm2,Nfkbie,Birc3,Plaur,AA467197 | | 2.190e-04 | -8.43 | GO\_NEGATIVE\_REGULATION\_OF\_I\_KAPPAB\_KINASE\_NF\_KAPPAB\_SIGNALING | MSigDB lists | GO\_NEGATIVE\_REGULATION\_OF\_I\_KAPPAB\_KINASE\_NF\_KAPPAB\_SIGNALING | 38 | 5 | 12187 | 179 | Ripk1,Tnfaip3,Tnip3,Tnip1,Nfkbia | | 2.208e-04 | -8.42 | MODULE\_38 | MSigDB lists | MODULE\_38 | 353 | 15 | 12187 | 179 | Cdkn1a,Gadd45g,Col4a1,Ier3,Atf3,Cd14,Csf2rb,Tnf,Maff,Il4ra,Tgm2,Fam107a,Tnfaip3,S100a9,S100a8 | | 2.215e-04 | -8.42 | PID\_TOLL\_ENDOGENOUS\_PATHWAY | MSigDB lists | PID\_TOLL\_ENDOGENOUS\_PATHWAY | 21 | 4 | 12187 | 179 | S100a8,Saa1,Cd14,S100a9 | | 2.215e-04 | -8.42 | GO\_POSITIVE\_REGULATION\_OF\_VASCULAR\_ENDOTHELIAL\_GROWTH\_FACTOR\_PRODUCTION | MSigDB lists | GO\_POSITIVE\_REGULATION\_OF\_VASCULAR\_ENDOTHELIAL\_GROWTH\_FACTOR\_PRODUCTION | 21 | 4 | 12187 | 179 | Il1a,C3ar1,Ptgs2,C5ar1 | | 2.215e-04 | -8.42 | BAUS\_TFF2\_TARGETS\_UP | MSigDB lists | BAUS\_TFF2\_TARGETS\_UP | 21 | 4 | 12187 | 179 | Irf7,Tap1,Slfn3,Slfn4 | | 2.238e-04 | -8.40 | locomotion | biological process | GO:0040011 | 919 | 29 | 13711 | 214 | Sdc4,Fzd4,Il17ra,Ccl4,Icam1,Lgals9,Ccl7,Thbs1,Cxcl16,Ccl12,Iqgap1,S100a8,C5ar1,Cxcl1,Msn,Selp,S100a9,Vcam1,Ccl2,Saa1,Cxcl9,Tnf,Tnfaip3,Retnlg,Rhoc,Il1rn,Arid5b,C3ar1,Sele | | 2.247e-04 | -8.40 | positive regulation of nitric oxide biosynthetic process | biological process | GO:0045429 | 36 | 5 | 13711 | 214 | Ptgs2,Il6,Mmp8,Icam1,Tnf | | 2.258e-04 | -8.40 | GSE3920\_UNTREATED\_VS\_IFNG\_TREATED\_ENDOTHELIAL\_CELL\_UP | MSigDB lists | GSE3920\_UNTREATED\_VS\_IFNG\_TREATED\_ENDOTHELIAL\_CELL\_UP | 111 | 8 | 12187 | 179 | Tnf,Cdkn1a,Nfkbiz,Ccl4,Bach1,Gadd45b,Tnfaip3,Rgs16 | | 2.277e-04 | -8.39 | MODULE\_117 | MSigDB lists | MODULE\_117 | 354 | 15 | 12187 | 179 | P2ry6,Csf2rb,Vcam1,Gadd45g,Hp,Mmp8,Stx11,S100a9,Igsf6,Pygm,Tnfsf10,Nfkb2,Ccl4,Xaf1,Cybb | | 2.283e-04 | -8.38 | regulation of interleukin-1 beta production | biological process | GO:0032651 | 56 | 6 | 13711 | 214 | Ifi204,Hspb1,Ifi211,Errfi1,Casp4,Tnfaip3 | | 2.297e-04 | -8.38 | positive regulation of macrophage activation | biological process | GO:0043032 | 20 | 4 | 13711 | 214 | Lgals9,Thbs1,Il4ra,Mmp8 | | 2.310e-04 | -8.37 | GSE15624\_CTRL\_VS\_3H\_HALOFUGINONE\_TREATED\_CD4\_TCELL\_UP | MSigDB lists | GSE15624\_CTRL\_VS\_3H\_HALOFUGINONE\_TREATED\_CD4\_TCELL\_UP | 141 | 9 | 12187 | 179 | Iqgap1,Irf7,Slfn5,Nfe2l2,Tnfaip3,Ifit3b,Ccl4,Parp14,Xaf1 | | 2.310e-04 | -8.37 | GSE15330\_MEGAKARYOCYTE\_ERYTHROID\_VS\_GRANULOCYTE\_MONOCYTE\_PROGENITOR\_UP | MSigDB lists | GSE15330\_MEGAKARYOCYTE\_ERYTHROID\_VS\_GRANULOCYTE\_MONOCYTE\_PROGENITOR\_UP | 141 | 9 | 12187 | 179 | Gem,S100a9,Phf11d,Pik3ap1,Plek,S100a8,Rbm47,Phf11b,Ccl4 | | 2.341e-04 | -8.36 | I-kappaB/NF-kappaB complex | cellular component | GO:0033256 | 2 | 2 | 13825 | 212 | Bcl3,Nfkb2 | | 2.341e-04 | -8.36 | Bcl3/NF-kappaB2 complex | cellular component | GO:0033257 | 2 | 2 | 13825 | 212 | Nfkb2,Bcl3 | | 2.344e-04 | -8.36 | GSE17721\_0.5H\_VS\_4H\_POLYIC\_BMDC\_DN | MSigDB lists | GSE17721\_0.5H\_VS\_4H\_POLYIC\_BMDC\_DN | 173 | 10 | 12187 | 179 | Nfkbia,Cdkn1a,Ccl12,Znfx1,Atf3,Nfkb1,Cybb,Ms4a6d,Irf2,Msn | | 2.344e-04 | -8.36 | CAMP\_UP.V1\_DN | MSigDB lists | CAMP\_UP.V1\_DN | 173 | 10 | 12187 | 179 | Fstl1,Ccnd2,Birc3,Il4ra,Nfkb1,Ms4a6d,Akap12,Gem,Plek,Zfp36 | | 2.347e-04 | -8.36 | GO\_REGULATION\_OF\_BODY\_FLUID\_LEVELS | MSigDB lists | GO\_REGULATION\_OF\_BODY\_FLUID\_LEVELS | 355 | 15 | 12187 | 179 | Plek,Hspb1,Irf2,Plaur,Saa1,Selp,Thbs1,Xdh,A2m,S100a9,Pik3r5,Nfe2l2,Il6,Pik3r1,Maff | | 2.347e-04 | -8.36 | Cytosolic DNA-sensing pathway | KEGG pathways | mmu04623 | 44 | 6 | 5248 | 107 | Ccl4,Ripk1,Irf7,Il6,Nfkbia,Nfkb1 | | 2.347e-04 | -8.36 | Cytosolic DNA-sensing pathway | KEGG pathways | ko04623 | 44 | 6 | 5248 | 107 | Ripk1,Irf7,Ccl4,Nfkb1,Il6,Nfkbia | | 2.353e-04 | -8.35 | IL6/GCSF/MGF\_CS | interpro domains | IPR030473 | 2 | 2 | 13788 | 212 | Il6,Csf3 | | 2.353e-04 | -8.35 | Anphylx\_rcpt | interpro domains | IPR002234 | 2 | 2 | 13788 | 212 | C3ar1,C5ar1 | | 2.353e-04 | -8.35 | IL-6/GCSF/MGF | interpro domains | IPR030474 | 2 | 2 | 13788 | 212 | Csf3,Il6 | | 2.374e-04 | -8.35 | Dimerization of procaspase-8 | REACTOME pathways | R-MMU-69416 | 8 | 3 | 6297 | 105 | Ripk1,Cflar,Tnfsf10 | | 2.374e-04 | -8.35 | Ligand-dependent caspase activation | REACTOME pathways | R-MMU-140534 | 8 | 3 | 6297 | 105 | Tnfsf10,Ripk1,Cflar | | 2.374e-04 | -8.35 | CASP8 activity is inhibited | REACTOME pathways | R-MMU-5218900 | 8 | 3 | 6297 | 105 | Ripk1,Cflar,Tnfsf10 | | 2.374e-04 | -8.35 | Regulation by c-FLIP | REACTOME pathways | R-MMU-3371378 | 8 | 3 | 6297 | 105 | Ripk1,Cflar,Tnfsf10 | | 2.374e-04 | -8.35 | TNFR1-induced proapoptotic signaling | REACTOME pathways | R-MMU-5357786 | 8 | 3 | 6297 | 105 | Tnf,Ripk1,Tnfaip3 | | 2.395e-04 | -8.34 | HELLEBREKERS\_SILENCED\_DURING\_TUMOR\_ANGIOGENESIS | MSigDB lists | HELLEBREKERS\_SILENCED\_DURING\_TUMOR\_ANGIOGENESIS | 60 | 6 | 12187 | 179 | Cdkn1a,Sdc4,Icam1,Il6,Ier3,Tnfaip3 | | 2.425e-04 | -8.32 | neutrophil aggregation | biological process | GO:0070488 | 2 | 2 | 13711 | 214 | S100a8,S100a9 | | 2.425e-04 | -8.32 | follicular dendritic cell differentiation | biological process | GO:0002268 | 2 | 2 | 13711 | 214 | Nfkb2,Bcl3 | | 2.425e-04 | -8.32 | neutrophil apoptotic process | biological process | GO:0001781 | 2 | 2 | 13711 | 214 | Il6,Hcar2 | | 2.425e-04 | -8.32 | negative regulation of tumor necrosis factor biosynthetic process | biological process | GO:0042536 | 2 | 2 | 13711 | 214 | Errfi1,Bcl3 | | 2.425e-04 | -8.32 | tolerance induction to lipopolysaccharide | biological process | GO:0072573 | 2 | 2 | 13711 | 214 | Acod1,Tnfaip3 | | 2.425e-04 | -8.32 | follicular dendritic cell activation | biological process | GO:0002266 | 2 | 2 | 13711 | 214 | Bcl3,Nfkb2 | | 2.432e-04 | -8.32 | angiogenesis | biological process | GO:0001525 | 262 | 13 | 13711 | 214 | Angptl4,Ptgs2,Rasip1,Ccl12,Col4a1,Rhoj,Apold1,Clic4,Rnf213,Thbs1,Ccl2,Cemip2,Robo4 | | 2.435e-04 | -8.32 | GSE9006\_1MONTH\_VS\_4MONTH\_POST\_TYPE\_1\_DIABETES\_DX\_PBMC\_UP | MSigDB lists | GSE9006\_1MONTH\_VS\_4MONTH\_POST\_TYPE\_1\_DIABETES\_DX\_PBMC\_UP | 142 | 9 | 12187 | 179 | Nfkbie,Ier3,Mmp8,Nfkbia,Ifit3b,Trib1,Tnf,Maff,Sgk1 | | 2.435e-04 | -8.32 | GO\_RESPONSE\_TO\_KETONE | MSigDB lists | GO\_RESPONSE\_TO\_KETONE | 142 | 9 | 12187 | 179 | Adamts1,Cdkn1a,Thbs1,Socs3,Ccl12,Il6,P2ry6,Errfi1,Msn | | 2.452e-04 | -8.31 | GO\_NEGATIVE\_REGULATION\_OF\_INTERLEUKIN\_12\_PRODUCTION | MSigDB lists | GO\_NEGATIVE\_REGULATION\_OF\_INTERLEUKIN\_12\_PRODUCTION | 9 | 3 | 12187 | 179 | Irak3,Nfkb1,Thbs1 | | 2.452e-04 | -8.31 | GO\_REGULATION\_OF\_NECROPTOTIC\_PROCESS | MSigDB lists | GO\_REGULATION\_OF\_NECROPTOTIC\_PROCESS | 9 | 3 | 12187 | 179 | Birc3,Cflar,Ripk1 | | 2.452e-04 | -8.31 | BIOCARTA\_LYM\_PATHWAY | MSigDB lists | BIOCARTA\_LYM\_PATHWAY | 9 | 3 | 12187 | 179 | Vcam1,Icam1,Il1a | | 2.452e-04 | -8.31 | IIZUKA\_LIVER\_CANCER\_EARLY\_RECURRENCE | MSigDB lists | IIZUKA\_LIVER\_CANCER\_EARLY\_RECURRENCE | 9 | 3 | 12187 | 179 | Ccnd2,Sgk1,Tnfaip3 | | 2.452e-04 | -8.31 | BIOCARTA\_EPONFKB\_PATHWAY | MSigDB lists | BIOCARTA\_EPONFKB\_PATHWAY | 9 | 3 | 12187 | 179 | Nfkb1,Nfkbia,Cdkn1a | | 2.464e-04 | -8.31 | GO\_CHEMICAL\_HOMEOSTASIS | MSigDB lists | GO\_CHEMICAL\_HOMEOSTASIS | 611 | 21 | 12187 | 179 | Sgk3,Saa1,Tap1,Clic4,Icam1,Sgk1,Lcn2,Cp,Kcna5,S100a9,Angptl4,C3ar1,C5ar1,Pygm,S100a8,Ccl2,Tgm2,Pik3r1,Il6,Ccl12,Slc24a4 | | 2.468e-04 | -8.31 | GO\_TRANSCRIPTIONAL\_REPRESSOR\_ACTIVITY\_RNA\_POLYMERASE\_II\_CORE\_PROMOTER\_PROXIMAL\_REGION\_SEQUENCE\_SPECIFIC\_BINDING | MSigDB lists | GO\_TRANSCRIPTIONAL\_REPRESSOR\_ACTIVITY\_RNA\_POLYMERASE\_II\_CORE\_PROMOTER\_PROXIMAL\_REGION\_SEQUENCE\_SPECIFIC\_BINDING | 85 | 7 | 12187 | 179 | Ifi204,Bach1,Ifi207,Atf3,Ifi209,Ifi211,Spi1 | | 2.476e-04 | -8.30 | enhancer sequence-specific DNA binding | molecular function | GO:0001158 | 80 | 7 | 13516 | 211 | Sox11,Zbtb16,Nfe2l2,Nfkb1,Irf7,Spi1,Bach1 | | 2.509e-04 | -8.29 | positive regulation of cell adhesion | biological process | GO:0045785 | 335 | 15 | 13711 | 214 | Il6,Ccl2,Tnf,Il4ra,Icam1,Lgals9,Thbs1,Zbtb16,Iqgap1,Sdc4,Nfkbiz,Tgm2,Runx1,Selp,Vcam1 | | 2.548e-04 | -8.28 | KRIGE\_RESPONSE\_TO\_TOSEDOSTAT\_6HR\_UP | MSigDB lists | KRIGE\_RESPONSE\_TO\_TOSEDOSTAT\_6HR\_UP | 750 | 24 | 12187 | 179 | Phf11b,Tap1,Slfn9,Cybb,Ccl12,Maff,Gpr84,S100a8,Rgs16,Phf11d,Bcl3,Ksr1,Il1rn,Samd9l,Atf3,Cdkn1a,Ifit3b,Apold1,Plek,Errfi1,Clic4,Trib1,Rbm39,S100a9 | | 2.565e-04 | -8.27 | positive regulation of nitric oxide metabolic process | biological process | GO:1904407 | 37 | 5 | 13711 | 214 | Ptgs2,Il6,Icam1,Tnf,Mmp8 | | 2.567e-04 | -8.27 | GSE21360\_NAIVE\_VS\_PRIMARY\_MEMORY\_CD8\_TCELL\_UP | MSigDB lists | GSE21360\_NAIVE\_VS\_PRIMARY\_MEMORY\_CD8\_TCELL\_UP | 143 | 9 | 12187 | 179 | Il17ra,Tnfaip8,Ptpn1,Sbno2,Thbs1,Ccnd2,Gpr84,Socs3,Hp | | 2.567e-04 | -8.27 | GSE7219\_UNSTIM\_VS\_LPS\_AND\_ANTI\_CD40\_STIM\_DC\_DN | MSigDB lists | GSE7219\_UNSTIM\_VS\_LPS\_AND\_ANTI\_CD40\_STIM\_DC\_DN | 143 | 9 | 12187 | 179 | Tnfsf10,Rgs16,Irak3,Ifitm3,Ifitm2,Nfkbie,Trib1,Cdkn1a,Tnf | | 2.572e-04 | -8.27 | IRF | smart domains | SM00348 | 8 | 3 | 7188 | 123 | Irf2,Irf7,Irf9 | | 2.573e-04 | -8.27 | GSE21546\_WT\_VS\_SAP1A\_KO\_ANTI\_CD3\_STIM\_DP\_THYMOCYTES\_DN | MSigDB lists | GSE21546\_WT\_VS\_SAP1A\_KO\_ANTI\_CD3\_STIM\_DP\_THYMOCYTES\_DN | 175 | 10 | 12187 | 179 | Tnfaip8,Slfn9,Ptges,Cybb,Nfkb1,Samd9l,Nfe2l2,Parp14,Tor3a,Usp18 | | 2.585e-04 | -8.26 | RUTELLA\_RESPONSE\_TO\_HGF\_VS\_CSF2RB\_AND\_IL4\_UP | MSigDB lists | RUTELLA\_RESPONSE\_TO\_HGF\_VS\_CSF2RB\_AND\_IL4\_UP | 319 | 14 | 12187 | 179 | Sgk1,Tnip3,Il1a,Gadd45g,Cd14,Ifitm2,Irak3,Rbm47,Tgm2,Ccl12,Cybb,S100a9,C5ar1,C3ar1 | | 2.600e-04 | -8.25 | GO\_NEGATIVE\_REGULATION\_OF\_PROTEOLYSIS | MSigDB lists | GO\_NEGATIVE\_REGULATION\_OF\_PROTEOLYSIS | 209 | 11 | 12187 | 179 | Tnfaip8,Plaur,Ifi211,A2m,Timp1,Ifi204,Ifi207,Il6,Ifi209,Thbs1,Birc3 | | 2.624e-04 | -8.25 | LINDSTEDT\_DENDRITIC\_CELL\_MATURATION\_D | MSigDB lists | LINDSTEDT\_DENDRITIC\_CELL\_MATURATION\_D | 61 | 6 | 12187 | 179 | Ifi209,Ifi211,Ifi207,P2ry6,Igsf6,Ifi204 | | 2.624e-04 | -8.25 | GO\_POSITIVE\_REGULATION\_OF\_CYTOKINE\_SECRETION | MSigDB lists | GO\_POSITIVE\_REGULATION\_OF\_CYTOKINE\_SECRETION | 61 | 6 | 12187 | 179 | Tnf,Saa1,Il4ra,Cd14,Il17ra,Il1a | | 2.624e-04 | -8.25 | REACTOME\_RESPONSE\_TO\_ELEVATED\_PLATELET\_CYTOSOLIC\_CA2\_ | MSigDB lists | REACTOME\_RESPONSE\_TO\_ELEVATED\_PLATELET\_CYTOSOLIC\_CA2\_ | 61 | 6 | 12187 | 179 | Plek,Timp1,Thbs1,Hspa5,Selp,A2m | | 2.624e-04 | -8.25 | MORI\_PRE\_BI\_LYMPHOCYTE\_DN | MSigDB lists | MORI\_PRE\_BI\_LYMPHOCYTE\_DN | 61 | 6 | 12187 | 179 | Il4ra,Birc3,Nfkbiz,Nfkb1,Zfp36,Rbm39 | | 2.650e-04 | -8.24 | GO\_NUCLEIC\_ACID\_BINDING\_TRANSCRIPTION\_FACTOR\_ACTIVITY | MSigDB lists | GO\_NUCLEIC\_ACID\_BINDING\_TRANSCRIPTION\_FACTOR\_ACTIVITY | 752 | 24 | 12187 | 179 | Bcl3,Arid5b,Ifi211,Spi1,Maff,Nfe2l2,Irf2,Sox11,Cebpd,Nfkb1,Irf7,Nfkb2,Runx1,Zfp189,Bach1,Irf9,Trim25,Zbtb16,Ifi209,Aff1,Atf3,Znfx1,Ifi207,Ifi204 | | 2.654e-04 | -8.23 | GO\_REGULATION\_OF\_CYTOKINE\_SECRETION | MSigDB lists | GO\_REGULATION\_OF\_CYTOKINE\_SECRETION | 86 | 7 | 12187 | 179 | Il1a,Il17ra,Il6,Il4ra,Saa1,Tnf,Cd14 | | 2.656e-04 | -8.23 | regulation of extrinsic apoptotic signaling pathway | biological process | GO:2001236 | 135 | 9 | 13711 | 214 | Atf3,Tnfsf10,Tnf,Icam1,Tnfaip3,Thbs1,Sgk3,Ripk1,Cflar | | 2.660e-04 | -8.23 | GO\_PROTEOLYSIS | MSigDB lists | GO\_PROTEOLYSIS | 847 | 26 | 12187 | 179 | Xdh,Tnfsf10,S100a9,Cflar,Casp4,Tnf,Trim25,Tnip1,Ifi204,Ifi207,Hp,Ifi209,Adamts1,Adamts9,S100a8,Ifi211,Ripk1,Tnfaip3,Nfe2l2,Usp18,Isg15,Rnf213,Herc6,Mmp8,Nfkb1,Hspa5 | | 2.677e-04 | -8.23 | GO\_RESPONSE\_TO\_INTERLEUKIN\_6 | MSigDB lists | GO\_RESPONSE\_TO\_INTERLEUKIN\_6 | 22 | 4 | 12187 | 179 | Ccl12,Sbno2,Il6,Nfkb1 | | 2.677e-04 | -8.23 | YAO\_TEMPORAL\_RESPONSE\_TO\_PROGESTERONE\_CLUSTER\_5 | MSigDB lists | YAO\_TEMPORAL\_RESPONSE\_TO\_PROGESTERONE\_CLUSTER\_5 | 22 | 4 | 12187 | 179 | Lcn2,Ier3,Atf3,Xdh | | 2.677e-04 | -8.23 | GNF2\_TNFSF10 | MSigDB lists | GNF2\_TNFSF10 | 22 | 4 | 12187 | 179 | C5ar1,Spi1,Igsf6,Tnfsf10 | | 2.677e-04 | -8.23 | GO\_NAD\_ADP\_RIBOSYLTRANSFERASE\_ACTIVITY | MSigDB lists | GO\_NAD\_ADP\_RIBOSYLTRANSFERASE\_ACTIVITY | 22 | 4 | 12187 | 179 | Parp9,Parp12,Tiparp,Parp14 | | 2.677e-04 | -8.23 | PID\_NFKAPPAB\_CANONICAL\_PATHWAY | MSigDB lists | PID\_NFKAPPAB\_CANONICAL\_PATHWAY | 22 | 4 | 12187 | 179 | Tnf,Nfkbia,Nfkb1,Tnfaip3 | | 2.693e-04 | -8.22 | GSE43955\_TH0\_VS\_TGFB\_IL6\_IL23\_TH17\_ACT\_CD4\_TCELL\_52H\_UP | MSigDB lists | GSE43955\_TH0\_VS\_TGFB\_IL6\_IL23\_TH17\_ACT\_CD4\_TCELL\_52H\_UP | 176 | 10 | 12187 | 179 | Irf9,Plaur,Ncf1,Irf7,Tagln2,C3ar1,Il1a,Ccl2,Ifit3b,Ifit2 | | 2.704e-04 | -8.22 | GSE37301\_MULTIPOTENT\_PROGENITOR\_VS\_LYMPHOID\_PRIMED\_MPP\_DN | MSigDB lists | GSE37301\_MULTIPOTENT\_PROGENITOR\_VS\_LYMPHOID\_PRIMED\_MPP\_DN | 144 | 9 | 12187 | 179 | Tubb6,Akap12,Ier3,Atf3,Cdkn1a,Vcam1,Nfkbia,Nfkbiz,Zfp36 | | 2.704e-04 | -8.22 | GSE9006\_HEALTHY\_VS\_TYPE\_1\_DIABETES\_PBMC\_AT\_DX\_DN | MSigDB lists | GSE9006\_HEALTHY\_VS\_TYPE\_1\_DIABETES\_PBMC\_AT\_DX\_DN | 144 | 9 | 12187 | 179 | S100a8,Zfp36,C3ar1,Ptgs2,Cxcl1,Ccl12,Sgk1,Socs3,Trib1 | | 2.783e-04 | -8.19 | kinase regulator activity | molecular function | GO:0019207 | 166 | 10 | 13516 | 211 | Irgm2,Socs3,Pik3r1,Trib1,Pik3r5,Hspb1,Ccnd2,Cdkn1a,Lgals9,Iqgap1 | | 2.803e-04 | -8.18 | monocyte chemotaxis | biological process | GO:0002548 | 21 | 4 | 13711 | 214 | Ccl4,Ccl7,Ccl12,Ccl2 | | 2.803e-04 | -8.18 | protein ADP-ribosylation | biological process | GO:0006471 | 21 | 4 | 13711 | 214 | Parp12,Parp9,Tiparp,Parp14 | | 2.804e-04 | -8.18 | BURTON\_ADIPOGENESIS\_PEAK\_AT\_8HR | MSigDB lists | BURTON\_ADIPOGENESIS\_PEAK\_AT\_8HR | 40 | 5 | 12187 | 179 | Ifi209,Bcl3,Ifi211,Ifi207,Ifi204 | | 2.805e-04 | -8.18 | CCR chemokine receptor binding | molecular function | GO:0048020 | 21 | 4 | 13516 | 211 | Ccl2,Ccl7,Ccl12,Ccl4 | | 2.819e-04 | -8.17 | GSE46606\_IRF4MID\_VS\_WT\_CD40L\_IL2\_IL5\_DAY3\_STIMULATED\_BCELL\_UP | MSigDB lists | GSE46606\_IRF4MID\_VS\_WT\_CD40L\_IL2\_IL5\_DAY3\_STIMULATED\_BCELL\_UP | 177 | 10 | 12187 | 179 | Clic4,Plaur,Il17ra,Zfp36,Cd14,Stx11,Hcar2,Vcam1,Rhou,Aff1 | | 2.834e-04 | -8.17 | PRDX4 (peroxiredoxin 4) | protein interactions | 10549 | 2 | 2 | 6802 | 115 | Bach1,Ksr1 | | 2.834e-04 | -8.17 | Relb (avian reticuloendotheliosis viral (v-rel) oncogene related B) | protein interactions | 19698 | 2 | 2 | 6802 | 115 | Nfkb1,Nfkb2 | | 2.834e-04 | -8.17 | Ccr2 (chemokine (C-C motif) receptor 2) | protein interactions | 12772 | 2 | 2 | 6802 | 115 | Ccl2,Ccl7 | | 2.834e-04 | -8.17 | Cbfa2t3 (CBFA2/RUNX1 translocation partner 3) | protein interactions | 12398 | 2 | 2 | 6802 | 115 | Runx1,Spi1 | | 2.837e-04 | -8.17 | regulation of cell-cell adhesion | biological process | GO:0022407 | 302 | 14 | 13711 | 214 | Vcam1,Runx1,Nfkbiz,Sdc4,Zbtb16,Il1rn,Lgals9,Pik3r1,Il4ra,Icam1,Tnfaip3,Tnf,Ccl2,Il6 | | 2.843e-04 | -8.17 | regulation of biosynthetic process | biological process | GO:0009889 | 2895 | 67 | 13711 | 214 | Pik3r1,Irf7,Dtx3l,Trib1,Il6,Cdkn1a,Akap12,Il1a,Nfkbia,Ptgs2,Runx1,Trim25,Resf1,Cebpd,Ifi211,Spi1,Sbno2,Parp9,Ifi209,Fzd4,Tnip1,Fstl1,Hspb1,Ifi207,Ncf1,Cflar,Ripk1,Zfp36,Atf3,Acod1,Tnf,Nfe2l2,Mmp8,Rasd1,Prg4,Dnajb1,Maff,Rrp8,Banp,Rbm39,P2ry6,Thbs1,Errfi1,Cybb,Mxd4,Icam1,Sox11,Nfkb2,Bach1,Nfkb1,Nfkbie,Gbp4,Ifi204,C3ar1,Arid5b,Irak3,Aff1,Zbtb16,Csf3,Ccl2,Tnfaip3,Znfx1,Plek,Bcl3,Irf9,S100a9,Irf2 | | 2.847e-04 | -8.16 | GSE2770\_TGFB\_AND\_IL4\_VS\_IL12\_TREATED\_ACT\_CD4\_TCELL\_2H\_DN | MSigDB lists | GSE2770\_TGFB\_AND\_IL4\_VS\_IL12\_TREATED\_ACT\_CD4\_TCELL\_2H\_DN | 145 | 9 | 12187 | 179 | Cmpk2,Plaur,Timp1,AA467197,Atf3,Oasl1,Ifit2,Maff,Nfkbiz | | 2.851e-04 | -8.16 | GSE29615\_CTRL\_VS\_DAY7\_LAIV\_FLU\_VACCINE\_PBMC\_DN | MSigDB lists | GSE29615\_CTRL\_VS\_DAY7\_LAIV\_FLU\_VACCINE\_PBMC\_DN | 87 | 7 | 12187 | 179 | Rtp4,Sele,Slfn9,Irgm2,Fam107a,Il6,Tnf | | 2.854e-04 | -8.16 | protein binding | molecular function | GO:0005515 | 7104 | 136 | 13516 | 211 | Rtp4,Fam107a,Parp9,Ifit3b,Csf3,Cflar,Resf1,Hspb1,Irak3,Ifi211,Nfkb1,Sntb2,Casp4,Irgm2,Dtx3l,Retnlg,Ripk1,Lcn2,Ccnd2,Sbno2,Tap1,Rbm47,Olfml2b,S100a8,Il1rn,Samd9l,Usp18,Trim56,Thbs1,Sele,Pik3ap1,Rhoc,Tnip1,Msn,Banp,Ifi209,Tnfaip3,Il1a,Rgs16,Iigp1,Nfe2l2,Zfp36,Csf2rb,Xdh,Runx1,Stx11,Rrp8,Cdkn1a,Map3k6,Gadd45b,Adamts1,Nfkbia,Akap12,Osmr,S100a9,Sdc4,A2m,Fstl1,C3ar1,Nfkbie,Fzd4,Robo4,Timp1,Nfkbiz,Rasip1,Rhoj,Isg15,Il6,Mxd4,Hspa5,Errfi1,Parp14,Hp,Clic4,Ifi204,Spi1,Bcl3,Ccl2,Birc3,Zbtb16,Rasd1,Ccl12,Ncf1,Sgk1,Il4ra,Plaur,Ptpn1,Dnajb1,Cybb,Mx1,Il17ra,Lgals9,Gadd45g,Plek,Vcam1,Tnf,Ier3,Trib1,Gbp3,Cp,Tnfsf10,Irf9,Herc6,Cxcl16,Rhou,Ksr1,Ifi207,Rbm39,Tor3a,Ccl4,Col4a1,C5ar1,Socs3,Fyb,Cxcl9,Angptl4,Nfkb2,Kcna5,Gem,Ccl7,Gbp4,Ptgs2,Icam1,Cebpd,Cxcl1,Mx2,Selp,Bach1,Noct,Saa1,Iqgap1,Irf7,Pik3r1,Tgm2,Tnip3,Atf3 | | 2.869e-04 | -8.16 | GO\_BLOOD\_VESSEL\_MORPHOGENESIS | MSigDB lists | GO\_BLOOD\_VESSEL\_MORPHOGENESIS | 284 | 13 | 12187 | 179 | Robo4,Fzd4,Angptl4,Ccl12,Spi1,Apold1,Clic4,Rnf213,Ptgs2,Col4a1,Tiparp,Rasip1,Thbs1 | | 2.870e-04 | -8.16 | GO\_POSITIVE\_REGULATION\_OF\_MYELOID\_CELL\_DIFFERENTIATION | MSigDB lists | GO\_POSITIVE\_REGULATION\_OF\_MYELOID\_CELL\_DIFFERENTIATION | 62 | 6 | 12187 | 179 | Ripk1,Trib1,Isg15,Csf3,Runx1,Tnf | | 2.870e-04 | -8.16 | GOTZMANN\_EPITHELIAL\_TO\_MESENCHYMAL\_TRANSITION\_UP | MSigDB lists | GOTZMANN\_EPITHELIAL\_TO\_MESENCHYMAL\_TRANSITION\_UP | 62 | 6 | 12187 | 179 | Il4ra,Irgm2,Gadd45b,Ccl12,Col4a1,Timp1 | | 2.885e-04 | -8.15 | enzyme inhibitor activity | molecular function | GO:0004857 | 232 | 12 | 13516 | 211 | Trib1,Timp1,A2m,Parp9,Serpina3f,Wfdc21,Tnfaip8,Birc3,Dtx3l,Angptl4,Hspb1,Cdkn1a | | 2.905e-04 | -8.14 | IL6 | smart domains | SM00126 | 2 | 2 | 7188 | 123 | Csf3,Il6 | | 2.917e-04 | -8.14 | negative regulation of epithelial cell apoptotic process | biological process | GO:1904036 | 38 | 5 | 13711 | 214 | Angptl4,Cflar,Icam1,Tnfaip3,Nfe2l2 | | 2.922e-04 | -8.14 | regulation of cellular biosynthetic process | biological process | GO:0031326 | 2842 | 66 | 13711 | 214 | Zfp36,Atf3,Tnf,Acod1,Fstl1,Tnip1,Ripk1,Cflar,Ifi207,Ncf1,Hspb1,Prg4,Rrp8,Maff,Dnajb1,Mmp8,Rasd1,Nfe2l2,Cdkn1a,Il6,Nfkbia,Akap12,Il1a,Pik3r1,Trib1,Dtx3l,Irf7,Parp9,Sbno2,Fzd4,Ifi209,Trim25,Resf1,Runx1,Ptgs2,Spi1,Ifi211,Cebpd,Plek,Znfx1,Tnfaip3,Ifi204,Zbtb16,Csf3,Aff1,C3ar1,Arid5b,Irak3,Irf2,Bcl3,S100a9,Irf9,Cybb,Errfi1,Nfkb2,Sox11,Mxd4,Icam1,P2ry6,Rbm39,Banp,Thbs1,Nfkbie,Nfkb1,Gbp4,Bach1 | | 2.938e-04 | -8.13 | type I interferon production | biological process | GO:0032606 | 9 | 3 | 13711 | 214 | Irf7,Irf9,Trim56 | | 2.996e-04 | -8.11 | GSE2706\_2H\_VS\_8H\_R848\_STIM\_DC\_UP | MSigDB lists | GSE2706\_2H\_VS\_8H\_R848\_STIM\_DC\_UP | 146 | 9 | 12187 | 179 | Sgk1,Maff,Sdc4,Ms4a6d,Ier3,Nfe2l2,Gem,Plek,Icam1 | | 3.006e-04 | -8.11 | pyrophosphatase activity | molecular function | GO:0016462 | 710 | 24 | 13516 | 211 | Tubb6,Irgm2,Tor3a,Gm5431,Irgm1,Rhou,Iigp1,Gbp3,Rnf213,Hspa5,Gm4841,Tgtp1,Gbp7,Ifi47,Gbp6,Rhoc,Rhoj,Mx1,Mx2,Tgtp2,Rasd1,F830016B08Rik,Tap1,Gem | | 3.035e-04 | -8.10 | regulation of leukocyte activation | biological process | GO:0002694 | 379 | 16 | 13711 | 214 | Runx1,Vcam1,Mmp8,Sdc4,C5ar1,Nfkbiz,Lgals9,Thbs1,Zbtb16,Ctla2a,Il6,Ccl2,Cdkn1a,Il4ra,Tnfaip3,Sox11 | | 3.070e-04 | -8.09 | GO\_RESPONSE\_TO\_ALCOHOL | MSigDB lists | GO\_RESPONSE\_TO\_ALCOHOL | 286 | 13 | 12187 | 179 | Cd14,Ptgs2,Errfi1,Msn,Icam1,S100a8,Vcam1,Socs3,Spi1,Adamts1,Cdkn1a,Ccl12,Il6 | | 3.085e-04 | -8.08 | GSE39152\_BRAIN\_VS\_SPLEEN\_CD103\_NEG\_MEMORY\_CD8\_TCELL\_UP | MSigDB lists | GSE39152\_BRAIN\_VS\_SPLEEN\_CD103\_NEG\_MEMORY\_CD8\_TCELL\_UP | 179 | 10 | 12187 | 179 | Ptgs2,Gbp7,Irgm2,Tap1,Irf9,Gbp3,Clic4,Iqgap1,Usp18,Atf3 | | 3.110e-04 | -8.08 | negative regulation of cysteine-type endopeptidase activity | biological process | GO:2000117 | 83 | 7 | 13711 | 214 | Thbs1,Tnfaip8,Cflar,Ptgs2,Plaur,Birc3,Il6 | | 3.126e-04 | -8.07 | GO\_SINGLE\_ORGANISM\_CELL\_ADHESION | MSigDB lists | GO\_SINGLE\_ORGANISM\_CELL\_ADHESION | 325 | 14 | 12187 | 179 | Icam1,Hspb1,Plek,Sele,Msn,Selp,Tnip1,Vcam1,S100a8,Bcl3,S100a9,Fzd4,Il6,Tnf | | 3.133e-04 | -8.07 | hydrolase activity, acting on acid anhydrides | molecular function | GO:0016817 | 712 | 24 | 13516 | 211 | Ifi47,Irgm1,Gm5431,Rhoc,Gbp6,Irgm2,Gbp7,Tubb6,Tor3a,Mx1,Rhoj,Rhou,Rnf213,Mx2,Iigp1,Gbp3,F830016B08Rik,Gem,Tgtp1,Tap1,Hspa5,Tgtp2,Gm4841,Rasd1 | | 3.133e-04 | -8.07 | hydrolase activity, acting on acid anhydrides, in phosphorus-containing anhydrides | molecular function | GO:0016818 | 712 | 24 | 13516 | 211 | Rasd1,Tgtp2,Tap1,Gem,F830016B08Rik,Mx2,Mx1,Rhoj,Gbp7,Gbp6,Rhoc,Ifi47,Gm4841,Hspa5,Tgtp1,Gbp3,Iigp1,Rnf213,Rhou,Tor3a,Tubb6,Irgm2,Gm5431,Irgm1 | | 3.140e-04 | -8.07 | Ikbkg (inhibitor of kappaB kinase gamma) | protein interactions | 16151 | 36 | 5 | 6802 | 115 | Nfkbiz,Nfkbia,Tnf,Ripk1,Tnfaip3 | | 3.148e-04 | -8.06 | GO\_NEGATIVE\_REGULATION\_OF\_MULTICELLULAR\_ORGANISMAL\_PROCESS | MSigDB lists | GO\_NEGATIVE\_REGULATION\_OF\_MULTICELLULAR\_ORGANISMAL\_PROCESS | 714 | 23 | 12187 | 179 | Timp1,Zbtb16,Plaur,Errfi1,Trim25,Ptgs2,Trib1,Nfkbia,Tnf,Runx1,Xdh,Nfkb1,Thbs1,Sox11,Irak3,Isg15,Il6,Nfe2l2,Tnfaip3,Ccl12,Il4ra,Pik3r1,Bcl3 | | 3.152e-04 | -8.06 | GSE28783\_ANTI\_MIR33\_VS\_UNTREATED\_ATHEROSCLEROSIS\_MACROPHAGE\_UP | MSigDB lists | GSE28783\_ANTI\_MIR33\_VS\_UNTREATED\_ATHEROSCLEROSIS\_MACROPHAGE\_UP | 147 | 9 | 12187 | 179 | Tnf,Zfp189,Gadd45b,Rhoc,Timp1,Rrp8,Cxcl1,Isg15,Plek | | 3.152e-04 | -8.06 | GSE19401\_UNSTIM\_VS\_PAM2CSK4\_STIM\_FOLLICULAR\_DC\_UP | MSigDB lists | GSE19401\_UNSTIM\_VS\_PAM2CSK4\_STIM\_FOLLICULAR\_DC\_UP | 147 | 9 | 12187 | 179 | Zfp36,Tnfaip8,Sbno2,Ptgs2,Col4a1,Stx11,Il6,Fam107a,Ccl4 | | 3.152e-04 | -8.06 | GSE22886\_DC\_VS\_MONOCYTE\_DN | MSigDB lists | GSE22886\_DC\_VS\_MONOCYTE\_DN | 147 | 9 | 12187 | 179 | Cebpd,Gadd45b,S100a9,Cd14,Irf7,Ifitm2,Tnfsf10,Zfp36,Irak3 | | 3.152e-04 | -8.06 | GSE36095\_WT\_VS\_HDAC9\_KO\_TREG\_UP | MSigDB lists | GSE36095\_WT\_VS\_HDAC9\_KO\_TREG\_UP | 147 | 9 | 12187 | 179 | Tnfaip3,Mxd4,Cybb,Nfkb1,Nfkbia,Nfkbiz,Usp18,Cdkn1a,Tnfaip8 | | 3.152e-04 | -8.06 | GSE37605\_FOXP3\_FUSION\_GFP\_VS\_IRES\_GFP\_TREG\_C57BL6\_DN | MSigDB lists | GSE37605\_FOXP3\_FUSION\_GFP\_VS\_IRES\_GFP\_TREG\_C57BL6\_DN | 147 | 9 | 12187 | 179 | Adamts9,Hspb1,Sntb2,Ifit3b,Thbs1,Sgk1,Adamts1,Atf3,Akap12 | | 3.156e-04 | -8.06 | MODULE\_154 | MSigDB lists | MODULE\_154 | 41 | 5 | 12187 | 179 | S100a9,Angptl4,Ptgs2,Saa1,S100a8 | | 3.156e-04 | -8.06 | CROMER\_TUMORIGENESIS\_UP | MSigDB lists | CROMER\_TUMORIGENESIS\_UP | 41 | 5 | 12187 | 179 | Col4a1,Il6,Oasl1,Tnfaip3,Plaur | | 3.203e-04 | -8.05 | GO\_REGULATION\_OF\_HUMORAL\_IMMUNE\_RESPONSE | MSigDB lists | GO\_REGULATION\_OF\_HUMORAL\_IMMUNE\_RESPONSE | 23 | 4 | 12187 | 179 | C3ar1,Tnf,C5ar1,A2m | | 3.203e-04 | -8.05 | PID\_TRAIL\_PATHWAY | MSigDB lists | PID\_TRAIL\_PATHWAY | 23 | 4 | 12187 | 179 | Tnfsf10,Ripk1,Pik3r1,Cflar | | 3.203e-04 | -8.05 | BERENJENO\_TRANSFORMED\_BY\_RHOA\_FOREVER\_DN | MSigDB lists | BERENJENO\_TRANSFORMED\_BY\_RHOA\_FOREVER\_DN | 23 | 4 | 12187 | 179 | Ptgs2,Akap12,Ier3,Plaur | | 3.203e-04 | -8.05 | GO\_LYMPHOCYTE\_MIGRATION | MSigDB lists | GO\_LYMPHOCYTE\_MIGRATION | 23 | 4 | 12187 | 179 | Ccl12,Cxcl16,Saa1,Ccl2 | | 3.203e-04 | -8.05 | MODULE\_19 | MSigDB lists | MODULE\_19 | 250 | 12 | 12187 | 179 | Sgk1,Fstl1,Zbtb16,Col4a1,Gadd45g,Cd14,Irf9,Ccnd2,Tgm2,Ccl12,Nfe2l2,Iqgap1 | | 3.208e-04 | -8.04 | Chemokine signaling pathway | WikiPathways | WP2292 | 148 | 12 | 3756 | 96 | Pik3r5,Cxcl9,Ccl12,Nfkbia,Cxcl1,Ccl4,Nfkb1,Cxcl16,Pik3r1,Ccl2,Ccl7,Ncf1 | | 3.225e-04 | -8.04 | GSE360\_HIGH\_VS\_LOW\_DOSE\_B\_MALAYI\_MAC\_UP | MSigDB lists | GSE360\_HIGH\_VS\_LOW\_DOSE\_B\_MALAYI\_MAC\_UP | 180 | 10 | 12187 | 179 | Ifi209,Cp,Ifi207,Ifi204,Nfkb1,Il1rn,Irf7,Ifi211,Igsf6,Rgs16 | | 3.262e-04 | -8.03 | Chagas disease (American trypanosomiasis) | KEGG pathways | ko05142 | 86 | 8 | 5248 | 107 | Tnf,Ccl2,Pik3r1,Cflar,Il6,Nfkbia,Nfkb1,Ccl12 | | 3.262e-04 | -8.03 | Chagas disease (American trypanosomiasis) | KEGG pathways | mmu05142 | 86 | 8 | 5248 | 107 | Tnf,Ccl2,Pik3r1,Cflar,Il6,Nfkbia,Nfkb1,Ccl12 | | 3.273e-04 | -8.02 | IRF\_2 | prosite domains | PS51507 | 8 | 3 | 8845 | 164 | Irf9,Irf7,Irf2 | | 3.273e-04 | -8.02 | IRF\_1 | prosite domains | PS00601 | 8 | 3 | 8845 | 164 | Irf2,Irf9,Irf7 | | 3.280e-04 | -8.02 | GO\_STRESS\_ACTIVATED\_PROTEIN\_KINASE\_SIGNALING\_CASCADE | MSigDB lists | GO\_STRESS\_ACTIVATED\_PROTEIN\_KINASE\_SIGNALING\_CASCADE | 89 | 7 | 12187 | 179 | Nfkb1,Ripk1,Map3k8,Tnf,Trib1,Ptpn1,Map3k6 | | 3.283e-04 | -8.02 | GO\_REGULATION\_OF\_IMMUNE\_EFFECTOR\_PROCESS | MSigDB lists | GO\_REGULATION\_OF\_IMMUNE\_EFFECTOR\_PROCESS | 288 | 13 | 12187 | 179 | Il4ra,Birc3,Tnf,Zfp189,Il6,Tnfaip3,Ccl12,A2m,C3ar1,C5ar1,Tap1,Irak3,Icam1 | | 3.304e-04 | -8.02 | regulation of extrinsic apoptotic signaling pathway via death domain receptors | biological process | GO:1902041 | 39 | 5 | 13711 | 214 | Icam1,Tnfaip3,Thbs1,Cflar,Atf3 | | 3.304e-04 | -8.02 | positive regulation of alpha-beta T cell differentiation | biological process | GO:0046638 | 39 | 5 | 13711 | 214 | Runx1,Il6,Zbtb16,Il4ra,Nfkbiz | | 3.304e-04 | -8.02 | T cell activation involved in immune response | biological process | GO:0002286 | 39 | 5 | 13711 | 214 | Bcl3,Gadd45g,Il6,Stx11,Icam1 | | 3.322e-04 | -8.01 | GO\_RESPONSE\_TO\_OXYGEN\_LEVELS | MSigDB lists | GO\_RESPONSE\_TO\_OXYGEN\_LEVELS | 251 | 12 | 12187 | 179 | Cdkn1a,Thbs1,Vcam1,Socs3,Kcna5,Ptgs2,Icam1,Apold1,Bach1,Ccl12,Angptl4,Pygm | | 3.332e-04 | -8.01 | regulation of stress-activated MAPK cascade | biological process | GO:0032872 | 202 | 11 | 13711 | 214 | Ptpn1,Fzd4,Mmp8,Il1a,Tnf,Xdh,Ripk1,Il1rn,Ncf1,Gadd45g,Gadd45b | | 3.348e-04 | -8.00 | Platelet degranulation | REACTOME pathways | R-MMU-114608 | 105 | 8 | 6297 | 105 | Plek,Thbs1,Tagln2,Timp1,Serpina3f,Selp,Hspa5,A2m | | 3.385e-04 | -7.99 | positive regulation of macrophage migration | biological process | GO:1905523 | 22 | 4 | 13711 | 214 | C5ar1,Thbs1,C3ar1,Ccl2 | | 3.385e-04 | -7.99 | regulation of macrophage chemotaxis | biological process | GO:0010758 | 22 | 4 | 13711 | 214 | Ccl2,C5ar1,C3ar1,Thbs1 | | 3.385e-04 | -7.99 | antimicrobial humoral immune response mediated by antimicrobial peptide | biological process | GO:0061844 | 22 | 4 | 13711 | 214 | Ccl2,Cxcl1,Cxcl9,S100a9 | | 3.395e-04 | -7.99 | MODULE\_24 | MSigDB lists | MODULE\_24 | 289 | 13 | 12187 | 179 | Il4ra,Tgm2,Trib1,Il1rn,Hp,Gadd45g,Col4a1,Fam107a,Ier3,S100a9,Cd14,S100a8,Sbno2 | | 3.396e-04 | -7.99 | Oxidative Damage | WikiPathways | WP1496 | 38 | 6 | 3756 | 96 | C3ar1,C5ar1,Nfkbie,Tnf,Nfkb1,Cdkn1a | | 3.417e-04 | -7.98 | GO\_CELLULAR\_RESPONSE\_TO\_KETONE | MSigDB lists | GO\_CELLULAR\_RESPONSE\_TO\_KETONE | 64 | 6 | 12187 | 179 | P2ry6,Errfi1,Msn,Adamts1,Ccl12,Il6 | | 3.417e-04 | -7.98 | INTERLEUKIN\_6 | prosite domains | PS00254 | 2 | 2 | 8845 | 164 | Csf3,Il6 | | 3.430e-04 | -7.98 | GO\_RESPONSE\_TO\_TEMPERATURE\_STIMULUS | MSigDB lists | GO\_RESPONSE\_TO\_TEMPERATURE\_STIMULUS | 118 | 8 | 12187 | 179 | Cd14,Il1a,Il6,Ccl12,Thbs1,Cdkn1a,Nfkbia,Socs3 | | 3.430e-04 | -7.98 | GSE37534\_GW1929\_VS\_PIOGLITAZONE\_TREATED\_CD4\_TCELL\_PPARG1\_FOXP3\_TRANSDUCED\_UP | MSigDB lists | GSE37534\_GW1929\_VS\_PIOGLITAZONE\_TREATED\_CD4\_TCELL\_PPARG1\_FOXP3\_TRANSDUCED\_UP | 118 | 8 | 12187 | 179 | Oasl1,Stx11,Col4a1,Ifit2,Hspb1,Ifitm3,Ifitm2,Isg15 | | 3.443e-04 | -7.97 | negative regulation of inflammatory response | biological process | GO:0050728 | 111 | 8 | 13711 | 214 | Nfkb1,Zfp36,Acod1,Tnfaip3,Ier3,Lgals9,Socs3,Ctla2a | | 3.466e-04 | -7.97 | XU\_HGF\_TARGETS\_INDUCED\_BY\_AKT1\_48HR\_UP | MSigDB lists | XU\_HGF\_TARGETS\_INDUCED\_BY\_AKT1\_48HR\_UP | 10 | 3 | 12187 | 179 | Plaur,Tnip1,Tgm2 | | 3.466e-04 | -7.97 | BIOCARTA\_SODD\_PATHWAY | MSigDB lists | BIOCARTA\_SODD\_PATHWAY | 10 | 3 | 12187 | 179 | Ripk1,Tnf,Birc3 | | 3.466e-04 | -7.97 | SANCHEZ\_MDM2\_TARGETS | MSigDB lists | SANCHEZ\_MDM2\_TARGETS | 10 | 3 | 12187 | 179 | Cdkn1a,Il1rn,Timp1 | | 3.466e-04 | -7.97 | REACTOME\_REGULATION\_OF\_IFNA\_SIGNALING | MSigDB lists | REACTOME\_REGULATION\_OF\_IFNA\_SIGNALING | 10 | 3 | 12187 | 179 | Usp18,Socs3,Ptpn1 | | 3.466e-04 | -7.97 | CROONQUIST\_STROMAL\_STIMULATION\_DN | MSigDB lists | CROONQUIST\_STROMAL\_STIMULATION\_DN | 10 | 3 | 12187 | 179 | Nfkb2,Trim25,Ncf1 | | 3.466e-04 | -7.97 | BIOCARTA\_MONOCYTE\_PATHWAY | MSigDB lists | BIOCARTA\_MONOCYTE\_PATHWAY | 10 | 3 | 12187 | 179 | Icam1,Sele,Selp | | 3.466e-04 | -7.97 | BIOCARTA\_ERYTH\_PATHWAY | MSigDB lists | BIOCARTA\_ERYTH\_PATHWAY | 10 | 3 | 12187 | 179 | Il6,Il1a,Csf3 | | 3.483e-04 | -7.96 | GO\_REGULATION\_OF\_PROTEIN\_IMPORT | MSigDB lists | GO\_REGULATION\_OF\_PROTEIN\_IMPORT | 149 | 9 | 12187 | 179 | Il6,Nfkbia,Cdkn1a,Tnf,Pik3r1,Bcl3,Nfkbie,Ptgs2,Csf3 | | 3.483e-04 | -7.96 | GSE3039\_CD4\_TCELL\_VS\_B1\_BCELL\_DN | MSigDB lists | GSE3039\_CD4\_TCELL\_VS\_B1\_BCELL\_DN | 149 | 9 | 12187 | 179 | Il4ra,Ccl4,Hp,Mmp8,Il1rn,Ier3,Pik3ap1,Tnfaip8,Msn | | 3.512e-04 | -7.95 | TNFR1-induced NFkappaB signaling pathway | REACTOME pathways | R-MMU-5357956 | 21 | 4 | 6297 | 105 | Birc3,Tnf,Tnfaip3,Ripk1 | | 3.513e-04 | -7.95 | GO\_NEGATIVE\_REGULATION\_OF\_HEMOPOIESIS | MSigDB lists | GO\_NEGATIVE\_REGULATION\_OF\_HEMOPOIESIS | 90 | 7 | 12187 | 179 | Nfe2l2,Zbtb16,Trib1,Nfkbia,Runx1,Pik3r1,Il4ra | | 3.518e-04 | -7.95 | Regulation of TLR by endogenous ligand | REACTOME pathways | R-MMU-5686938 | 9 | 3 | 6297 | 105 | S100a9,Cd14,S100a8 | | 3.540e-04 | -7.95 | VANHARANTA\_UTERINE\_FIBROID\_WITH\_7Q\_DELETION\_DN | MSigDB lists | VANHARANTA\_UTERINE\_FIBROID\_WITH\_7Q\_DELETION\_DN | 42 | 5 | 12187 | 179 | Ifi211,Ifi209,Clic4,Ifi207,Ifi204 | | 3.540e-04 | -7.95 | SMIRNOV\_RESPONSE\_TO\_IR\_2HR\_UP | MSigDB lists | SMIRNOV\_RESPONSE\_TO\_IR\_2HR\_UP | 42 | 5 | 12187 | 179 | Cdkn1a,Sgk1,Ier3,Atf3,Rgs16 | | 3.540e-04 | -7.95 | LEE\_LIVER\_CANCER\_CIPROFIBRATE\_UP | MSigDB lists | LEE\_LIVER\_CANCER\_CIPROFIBRATE\_UP | 42 | 5 | 12187 | 179 | Cdkn1a,S100a9,Tgm2,Col4a1,Lcn2 | | 3.624e-04 | -7.92 | symbiont process | biological process | GO:0044403 | 204 | 11 | 13711 | 214 | Gbp3,Irgm1,Vcam1,Ccl4,Gbp7,Gbp6,Irgm2,Icam1,Ncf1,Ripk1,Tnip1 | | 3.624e-04 | -7.92 | regulation of stress-activated protein kinase signaling cascade | biological process | GO:0070302 | 204 | 11 | 13711 | 214 | Ptpn1,Fzd4,Mmp8,Tnf,Il1a,Xdh,Il1rn,Ncf1,Ripk1,Gadd45g,Gadd45b | | 3.660e-04 | -7.91 | humoral immune response | biological process | GO:0006959 | 61 | 6 | 13711 | 214 | Bcl3,Ccl2,Cxcl1,Tnf,Cxcl9,S100a9 | | 3.672e-04 | -7.91 | Emx1 (empty spiracles homeobox 1) | protein interactions | 13796 | 9 | 3 | 6802 | 115 | Irf9,Irf7,Spi1 | | 3.759e-04 | -7.89 | KEGG\_LEUKOCYTE\_TRANSENDOTHELIAL\_MIGRATION | MSigDB lists | KEGG\_LEUKOCYTE\_TRANSENDOTHELIAL\_MIGRATION | 91 | 7 | 12187 | 179 | Icam1,Cybb,Msn,Pik3r1,Ncf1,Pik3r5,Vcam1 | | 3.759e-04 | -7.89 | COULOUARN\_TEMPORAL\_TGFB1\_SIGNATURE\_UP | MSigDB lists | COULOUARN\_TEMPORAL\_TGFB1\_SIGNATURE\_UP | 91 | 7 | 12187 | 179 | Bcl3,Clic4,Iqgap1,Rbm47,Sgk1,Ier3,Nfe2l2 | | 3.796e-04 | -7.88 | Interleukin-1 signaling | REACTOME pathways | R-MMU-446652 | 38 | 5 | 6297 | 105 | Irak3,Map3k8,Nfkb1,Il1rn,Il1a | | 3.800e-04 | -7.88 | ST\_GAQ\_PATHWAY | MSigDB lists | ST\_GAQ\_PATHWAY | 24 | 4 | 12187 | 179 | Nfkb2,Nfkb1,Nfkbia,Nfkbie | | 3.800e-04 | -7.88 | WILENSKY\_RESPONSE\_TO\_DARAPLADIB | MSigDB lists | WILENSKY\_RESPONSE\_TO\_DARAPLADIB | 24 | 4 | 12187 | 179 | Ncf1,Cybb,Il1a,Plaur | | 3.800e-04 | -7.88 | REACTOME\_CHEMOKINE\_RECEPTORS\_BIND\_CHEMOKINES | MSigDB lists | REACTOME\_CHEMOKINE\_RECEPTORS\_BIND\_CHEMOKINES | 24 | 4 | 12187 | 179 | Cxcl16,Ccl12,Ccl4,Cxcl1 | | 3.800e-04 | -7.88 | WESTON\_VEGFA\_TARGETS\_12HR | MSigDB lists | WESTON\_VEGFA\_TARGETS\_12HR | 24 | 4 | 12187 | 179 | Vcam1,Sdc4,Gem,Ccl12 | | 3.800e-04 | -7.88 | BIOCARTA\_STRESS\_PATHWAY | MSigDB lists | BIOCARTA\_STRESS\_PATHWAY | 24 | 4 | 12187 | 179 | Tnf,Nfkbia,Nfkb1,Ripk1 | | 3.800e-04 | -7.88 | KIM\_HYPOXIA | MSigDB lists | KIM\_HYPOXIA | 24 | 4 | 12187 | 179 | Il6,Bach1,Icam1,Angptl4 | | 3.808e-04 | -7.87 | Toll-Like Receptors Cascades | REACTOME pathways | R-MMU-168898 | 107 | 8 | 6297 | 105 | Irf7,Nfkb1,S100a8,Nfkb2,Map3k8,S100a9,Cd14,Nfkbia | | 3.838e-04 | -7.87 | TGANTCA\_AP1\_C | MSigDB lists | TGANTCA\_AP1\_C | 819 | 25 | 12187 | 179 | Cdkn1a,Znfx1,Stx11,Tnip3,Il1rn,Tagln2,Trim25,Trib1,Runx1,Bach1,C3ar1,Xdh,Rbm39,Sgk1,Thbs1,Gadd45g,Sgk3,Map3k6,Il6,Gadd45b,Slc24a4,Rhoc,Angptl4,Csf3,Osmr | | 3.843e-04 | -7.86 | IRF1\_01 | MSigDB lists | IRF1\_01 | 184 | 10 | 12187 | 179 | Dtx3l,Isg15,Irf9,Tap1,Usp18,Xaf1,Col4a1,Stx11,Il6,Atf3 | | 3.843e-04 | -7.86 | GSE10325\_BCELL\_VS\_MYELOID\_DN | MSigDB lists | GSE10325\_BCELL\_VS\_MYELOID\_DN | 151 | 9 | 12187 | 179 | Oasl1,Cebpd,Il1rn,Ms4a6d,Plaur,Icam1,Igsf6,C5ar1,C3ar1 | | 3.843e-04 | -7.86 | GSE6259\_CD4\_TCELL\_VS\_CD8\_TCELL\_UP | MSigDB lists | GSE6259\_CD4\_TCELL\_VS\_CD8\_TCELL\_UP | 151 | 9 | 12187 | 179 | Fstl1,Il6,Il1a,Timp1,Rhoj,Angptl4,Rgs16,Plaur,Hspb1 | | 3.845e-04 | -7.86 | carbohydrate derivative binding | molecular function | GO:0097367 | 1791 | 46 | 13516 | 211 | Thbs1,Sele,Gbp7,Rhoc,Mx1,Lgals9,Sgk3,Rasd1,Oasl2,Sgk1,Tap1,Ccl2,Ripk1,Oasl1,Irak3,Hspa5,Parp9,Bbs12,Tgm2,Gbp6,Saa1,Rhoj,Map3k8,Selp,Pygm,Ccl7,Gem,Fstl1,Map3k6,Tor3a,Tubb6,Adamts1,Cd14,Cmpk2,Ip6k2,Irgm1,Runx1,Rhou,Ksr1,Slfn9,Trib1,Gbp3,Iigp1,Slfn5,Hcar2,Tgtp1 | | 3.851e-04 | -7.86 | GO\_ION\_HOMEOSTASIS | MSigDB lists | GO\_ION\_HOMEOSTASIS | 413 | 16 | 12187 | 179 | Ccl2,Tgm2,Ccl12,Slc24a4,S100a9,C3ar1,C5ar1,Pygm,S100a8,Sgk1,Lcn2,Cp,Kcna5,Saa1,Sgk3,Clic4 | | 3.872e-04 | -7.86 | adaptive immune response | biological process | GO:0002250 | 173 | 10 | 13711 | 214 | Nfkb2,Icam1,Il4ra,Tap1,Il6,C3ar1,Irf7,Stx11,Bcl3,Gadd45g | | 3.873e-04 | -7.86 | GO\_MOVEMENT\_OF\_CELL\_OR\_SUBCELLULAR\_COMPONENT | MSigDB lists | GO\_MOVEMENT\_OF\_CELL\_OR\_SUBCELLULAR\_COMPONENT | 966 | 28 | 12187 | 179 | Sdc4,Tnf,S100a9,C5ar1,Kcna5,Vcam1,Plaur,Msn,Hspb1,Icam1,Sele,Ptgs2,Saa1,Ccl12,Il6,Rhoc,Ccl4,Ccl2,Pik3r1,S100a8,Cxcl1,Arid5b,Thbs1,Parp9,Cxcl16,Il17ra,Bbs12,Selp | | 3.879e-04 | -7.85 | SMID\_BREAST\_CANCER\_RELAPSE\_IN\_BONE\_DN | MSigDB lists | SMID\_BREAST\_CANCER\_RELAPSE\_IN\_BONE\_DN | 219 | 11 | 12187 | 179 | Ifi204,Ifi207,Cp,Ifi209,Sox11,Clic4,S100a8,Ifi211,Tagln2,S100a9,Ptgs2 | | 3.937e-04 | -7.84 | regulation of peptidyl-tyrosine phosphorylation | biological process | GO:0050730 | 206 | 11 | 13711 | 214 | Icam1,Tnf,Ptpn1,Iqgap1,Errfi1,Il6,Parp9,Ncf1,Socs3,Csf3,Parp14 | | 3.959e-04 | -7.83 | PID\_TNF\_PATHWAY | MSigDB lists | PID\_TNF\_PATHWAY | 43 | 5 | 12187 | 179 | Tnf,Birc3,Nfkb1,Tnfaip3,Ripk1 | | 3.959e-04 | -7.83 | CHIBA\_RESPONSE\_TO\_TSA | MSigDB lists | CHIBA\_RESPONSE\_TO\_TSA | 43 | 5 | 12187 | 179 | Dnajb1,Sdc4,Tgm2,Cdkn1a,Ifitm3 | | 3.959e-04 | -7.83 | BURTON\_ADIPOGENESIS\_7 | MSigDB lists | BURTON\_ADIPOGENESIS\_7 | 43 | 5 | 12187 | 179 | Timp1,Vcam1,Tagln2,Rhoj,Ifitm3 | | 3.959e-04 | -7.83 | HADDAD\_T\_LYMPHOCYTE\_AND\_NK\_PROGENITOR\_DN | MSigDB lists | HADDAD\_T\_LYMPHOCYTE\_AND\_NK\_PROGENITOR\_DN | 43 | 5 | 12187 | 179 | Plek,S100a8,Ms4a6d,Cebpd,Tnf | | 3.971e-04 | -7.83 | CD225/Dispanin\_fam | interpro domains | IPR007593 | 10 | 3 | 13788 | 212 | Ifitm6,Ifitm3,Ifitm2 | | 4.034e-04 | -7.82 | GSE3982\_NEUTROPHIL\_VS\_NKCELL\_UP | MSigDB lists | GSE3982\_NEUTROPHIL\_VS\_NKCELL\_UP | 152 | 9 | 12187 | 179 | Bach1,Il1rn,Spi1,Sgk1,Ifit2,Irf9,Trim25,Cflar,C5ar1 | | 4.034e-04 | -7.82 | GSE17721\_CTRL\_VS\_PAM3CSK4\_6H\_BMDC\_DN | MSigDB lists | GSE17721\_CTRL\_VS\_PAM3CSK4\_6H\_BMDC\_DN | 152 | 9 | 12187 | 179 | Ier3,Hcar2,Map3k8,Cdkn1a,Gbp3,Ptpn1,Msn,Tnfaip8,Ptges | | 4.034e-04 | -7.82 | GSE1432\_1H\_VS\_24H\_IFNG\_MICROGLIA\_DN | MSigDB lists | GSE1432\_1H\_VS\_24H\_IFNG\_MICROGLIA\_DN | 152 | 9 | 12187 | 179 | Sntb2,Casp4,Ifi44,Phf11b,Tap1,Tnfsf10,P2ry6,Irf7,Phf11d | | 4.042e-04 | -7.81 | PID\_CMYB\_PATHWAY | MSigDB lists | PID\_CMYB\_PATHWAY | 66 | 6 | 12187 | 179 | Ptgs2,Birc3,Cdkn1a,Spi1,Iqgap1,Cebpd | | 4.047e-04 | -7.81 | negative regulation of endothelial cell apoptotic process | biological process | GO:2000352 | 23 | 4 | 13711 | 214 | Nfe2l2,Tnfaip3,Icam1,Angptl4 | | 4.102e-04 | -7.80 | negative regulation of kinase activity | biological process | GO:0033673 | 207 | 11 | 13711 | 214 | Trib1,Rasip1,Hspb1,Irak3,Gadd45g,Gadd45b,Ptpn1,Tnfaip3,Cdkn1a,Il6,Errfi1 | | 4.126e-04 | -7.79 | regulation of anatomical structure morphogenesis | biological process | GO:0022603 | 907 | 28 | 13711 | 214 | Nfe2l2,Msn,C5ar1,Ccl12,Hspb1,Il1rn,C3ar1,Cflar,Adamts9,Rhoc,Tnf,Tnfaip3,Cxcl9,Xdh,Ccl2,Rhou,Ptgs2,Sgk1,Runx1,Fzd4,Rhoj,Adamts1,Thbs1,Il4ra,Icam1,Il1a,Il6,Cybb | | 4.143e-04 | -7.79 | CD225 | pfam domains | PF04505 | 10 | 3 | 12881 | 201 | Ifitm3,Ifitm2,Ifitm6 | | 4.149e-04 | -7.79 | leukocyte aggregation | biological process | GO:0070486 | 10 | 3 | 13711 | 214 | S100a8,Msn,S100a9 | | 4.154e-04 | -7.79 | regulation of leukocyte chemotaxis | biological process | GO:0002688 | 87 | 7 | 13711 | 214 | Ccl2,Ccl12,C5ar1,Lgals9,Ccl4,C3ar1,Thbs1 | | 4.233e-04 | -7.77 | GSE16450\_CTRL\_VS\_IFNA\_6H\_STIM\_IMMATURE\_NEURON\_CELL\_LINE\_UP | MSigDB lists | GSE16450\_CTRL\_VS\_IFNA\_6H\_STIM\_IMMATURE\_NEURON\_CELL\_LINE\_UP | 153 | 9 | 12187 | 179 | Aff1,Zfp189,Vcam1,Gpr84,Stx11,Cp,Gbp4,Gbp6,Plek | | 4.233e-04 | -7.77 | GSE43955\_TH0\_VS\_TGFB\_IL6\_TH17\_ACT\_CD4\_TCELL\_42H\_DN | MSigDB lists | GSE43955\_TH0\_VS\_TGFB\_IL6\_TH17\_ACT\_CD4\_TCELL\_42H\_DN | 153 | 9 | 12187 | 179 | Zbtb16,Il4ra,Rhou,Hp,Il1rn,Tnip1,Selp,Csf2rb,S100a8 | | 4.233e-04 | -7.77 | GSE42021\_TCONV\_PLN\_VS\_CD24LO\_TCONV\_THYMUS\_DN | MSigDB lists | GSE42021\_TCONV\_PLN\_VS\_CD24LO\_TCONV\_THYMUS\_DN | 153 | 9 | 12187 | 179 | Saa1,Bcl3,Plaur,Maff,Birc3,Fstl1,Tiparp,Cybb,Lcn2 | | 4.283e-04 | -7.76 | response to reactive oxygen species | biological process | GO:0000302 | 144 | 9 | 13711 | 214 | Hp,Il6,Tnfaip3,Lcn2,Ripk1,Cflar,Nfe2l2,Thbs1,Ncf1 | | 4.293e-04 | -7.75 | SENESE\_HDAC2\_TARGETS\_UP | MSigDB lists | SENESE\_HDAC2\_TARGETS\_UP | 93 | 7 | 12187 | 179 | Nfkbiz,Tgm2,Ccnd2,Tnfaip3,Osmr,Il1a,Iqgap1 | | 4.296e-04 | -7.75 | GSE25123\_CTRL\_VS\_ROSIGLITAZONE\_STIM\_PPARG\_KO\_MACROPHAGE\_UP | MSigDB lists | GSE25123\_CTRL\_VS\_ROSIGLITAZONE\_STIM\_PPARG\_KO\_MACROPHAGE\_UP | 122 | 8 | 12187 | 179 | Cxcl1,Tagln2,Plaur,Birc3,AA467197,Tnfaip3,Il1rn,Tnip3 | | 4.296e-04 | -7.75 | CHIANG\_LIVER\_CANCER\_SUBCLASS\_CTNNB1\_DN | MSigDB lists | CHIANG\_LIVER\_CANCER\_SUBCLASS\_CTNNB1\_DN | 122 | 8 | 12187 | 179 | Timp1,Cp,AA467197,Adamts1,Socs3,Tnfaip8,Cxcl16,Gem | | 4.339e-04 | -7.74 | regulation of multicellular organismal development | biological process | GO:2000026 | 1748 | 45 | 13711 | 214 | Ccl2,Xdh,Tnfaip3,Csf3,Zbtb16,Adamts9,C3ar1,Irak3,Il1rn,Iqgap1,Nfkbiz,S100a9,Cybb,Errfi1,Sox11,Lgals9,Thbs1,Adamts1,Rhoj,Apold1,Zfp36,Tnf,Ripk1,Cflar,Hspb1,Hspa5,Cxcl1,Maff,C5ar1,Noct,Nfe2l2,Il6,Nfkbia,Il1a,Il4ra,Pik3r1,Trib1,Ctla2a,Isg15,Fzd4,Timp1,Runx1,Sgk1,Ptgs2,Spi1 | | 4.364e-04 | -7.74 | regulation of interferon-gamma production | biological process | GO:0032649 | 63 | 6 | 13711 | 214 | Tnf,Cd14,Runx1,Bcl3,Lgals9,Isg15 | | 4.386e-04 | -7.73 | LIU\_VAV3\_PROSTATE\_CARCINOGENESIS\_UP | MSigDB lists | LIU\_VAV3\_PROSTATE\_CARCINOGENESIS\_UP | 67 | 6 | 12187 | 179 | Ccnd2,Tgm2,Ifit3b,Cybb,Tnfsf10,Tap1 | | 4.386e-04 | -7.73 | KEGG\_B\_CELL\_RECEPTOR\_SIGNALING\_PATHWAY | MSigDB lists | KEGG\_B\_CELL\_RECEPTOR\_SIGNALING\_PATHWAY | 67 | 6 | 12187 | 179 | Pik3r1,Nfkbie,Pik3r5,Nfkbia,Nfkb1,Pik3ap1 | | 4.414e-04 | -7.73 | ZHANG\_PROLIFERATING\_VS\_QUIESCENT | MSigDB lists | ZHANG\_PROLIFERATING\_VS\_QUIESCENT | 44 | 5 | 12187 | 179 | Hspb1,Plaur,Ccnd2,Casp4,Cxcl1 | | 4.414e-04 | -7.73 | MODULE\_297 | MSigDB lists | MODULE\_297 | 44 | 5 | 12187 | 179 | S100a8,Angptl4,S100a9,Ptgs2,Saa1 | | 4.414e-04 | -7.73 | GNF2\_PECAM1 | MSigDB lists | GNF2\_PECAM1 | 44 | 5 | 12187 | 179 | Cebpd,Igsf6,Cybb,Zfp36,Ms4a6d | | 4.414e-04 | -7.73 | WESTON\_VEGFA\_TARGETS\_6HR | MSigDB lists | WESTON\_VEGFA\_TARGETS\_6HR | 44 | 5 | 12187 | 179 | Gem,Sdc4,Vcam1,Tnfsf10,Ccl12 | | 4.414e-04 | -7.73 | MODULE\_357 | MSigDB lists | MODULE\_357 | 44 | 5 | 12187 | 179 | S100a8,Ptgs2,Saa1,S100a9,Angptl4 | | 4.440e-04 | -7.72 | GSE2585\_CTEC\_VS\_THYMIC\_MACROPHAGE\_UP | MSigDB lists | GSE2585\_CTEC\_VS\_THYMIC\_MACROPHAGE\_UP | 154 | 9 | 12187 | 179 | Gbp4,Col4a1,Xaf1,Fstl1,Ccnd2,Il4ra,Cflar,Isg15,Gbp6 | | 4.440e-04 | -7.72 | GSE17721\_CTRL\_VS\_POLYIC\_24H\_BMDC\_DN | MSigDB lists | GSE17721\_CTRL\_VS\_POLYIC\_24H\_BMDC\_DN | 154 | 9 | 12187 | 179 | Ptges,Slfn4,Slfn3,Rbm47,Tgm2,Dnajb1,Nfkbia,Tnf,Timp1 | | 4.440e-04 | -7.72 | GSE1925\_CTRL\_VS\_IFNG\_PRIMED\_MACROPHAGE\_24H\_IFNG\_STIM\_UP | MSigDB lists | GSE1925\_CTRL\_VS\_IFNG\_PRIMED\_MACROPHAGE\_24H\_IFNG\_STIM\_UP | 154 | 9 | 12187 | 179 | Icam1,Nfkb2,Gbp3,Irgm2,Ms4a6d,Nfe2l2,Gadd45b,Rasd1,Usp18 | | 4.440e-04 | -7.72 | GSE2197\_CPG\_DNA\_VS\_UNTREATED\_IN\_DC\_UP | MSigDB lists | GSE2197\_CPG\_DNA\_VS\_UNTREATED\_IN\_DC\_UP | 154 | 9 | 12187 | 179 | Tnip1,Slfn9,Icam1,Ptpn1,Hspa5,Maff,Tgm2,Rhoc,Hp | | 4.440e-04 | -7.72 | GSE3982\_MAST\_CELL\_VS\_NEUTROPHIL\_DN | MSigDB lists | GSE3982\_MAST\_CELL\_VS\_NEUTROPHIL\_DN | 154 | 9 | 12187 | 179 | Ifi209,Spi1,Ifi207,Ifi204,Tnfaip3,C5ar1,Cd14,Ifi211,Apold1 | | 4.472e-04 | -7.71 | DUNNE\_TARGETS\_OF\_AML1\_MTG8\_FUSION\_DN | MSigDB lists | DUNNE\_TARGETS\_OF\_AML1\_MTG8\_FUSION\_DN | 25 | 4 | 12187 | 179 | Ifi211,Ifi209,Ifi207,Ifi204 | | 4.472e-04 | -7.71 | GO\_RESPONSE\_TO\_FUNGUS | MSigDB lists | GO\_RESPONSE\_TO\_FUNGUS | 25 | 4 | 12187 | 179 | Il17ra,S100a8,Il6,S100a9 | | 4.472e-04 | -7.71 | GO\_NEGATIVE\_REGULATION\_OF\_TUMOR\_NECROSIS\_FACTOR\_SUPERFAMILY\_CYTOKINE\_PRODUCTION | MSigDB lists | GO\_NEGATIVE\_REGULATION\_OF\_TUMOR\_NECROSIS\_FACTOR\_SUPERFAMILY\_CYTOKINE\_PRODUCTION | 25 | 4 | 12187 | 179 | Irak3,Errfi1,Tnfaip3,Bcl3 | | 4.472e-04 | -7.71 | GO\_NEGATIVE\_REGULATION\_OF\_EPITHELIAL\_CELL\_APOPTOTIC\_PROCESS | MSigDB lists | GO\_NEGATIVE\_REGULATION\_OF\_EPITHELIAL\_CELL\_APOPTOTIC\_PROCESS | 25 | 4 | 12187 | 179 | Nfe2l2,Tnfaip3,Icam1,Angptl4 | | 4.472e-04 | -7.71 | BIOCARTA\_RACCYCD\_PATHWAY | MSigDB lists | BIOCARTA\_RACCYCD\_PATHWAY | 25 | 4 | 12187 | 179 | Nfkb1,Pik3r1,Cdkn1a,Nfkbia | | 4.503e-04 | -7.71 | GO\_TAXIS | MSigDB lists | GO\_TAXIS | 337 | 14 | 12187 | 179 | Il17ra,Plaur,Cxcl16,Saa1,Vcam1,S100a8,C3ar1,C5ar1,S100a9,Cxcl1,Il6,Ccl12,Ccl2,Ccl4 | | 4.527e-04 | -7.70 | positive regulation of secretion | biological process | GO:0051047 | 393 | 16 | 13711 | 214 | S100a8,Sdc4,Runx1,Il17ra,S100a9,Mmp8,Il6,Tnf,Il4ra,Casp4,Akap12,Sox11,Il1a,Hcar2,Lgals9,Cd14 | | 4.538e-04 | -7.70 | GSE13229\_MATURE\_VS\_INTMATURE\_NKCELL\_UP | MSigDB lists | GSE13229\_MATURE\_VS\_INTMATURE\_NKCELL\_UP | 123 | 8 | 12187 | 179 | Gbp6,Herc6,Tnf,Cxcl1,Sele,Nfkb1,Parp12,Gbp4 | | 4.538e-04 | -7.70 | PODAR\_RESPONSE\_TO\_ADAPHOSTIN\_UP | MSigDB lists | PODAR\_RESPONSE\_TO\_ADAPHOSTIN\_UP | 123 | 8 | 12187 | 179 | Atf3,Bach1,Dnajb1,Trib1,Maff,Hspb1,Plek,Arid5b | | 4.582e-04 | -7.69 | REACTOME\_PEPTIDE\_LIGAND\_BINDING\_RECEPTORS | MSigDB lists | REACTOME\_PEPTIDE\_LIGAND\_BINDING\_RECEPTORS | 94 | 7 | 12187 | 179 | Cxcl1,C5ar1,Saa1,C3ar1,Cxcl16,Ccl4,Ccl12 | | 4.594e-04 | -7.69 | Response to elevated platelet cytosolic Ca2+ | REACTOME pathways | R-MMU-76005 | 110 | 8 | 6297 | 105 | Hspa5,Selp,A2m,Timp1,Tagln2,Thbs1,Plek,Serpina3f | | 4.632e-04 | -7.68 | ANPHYLATOXNR | prints domains | PR01104 | 2 | 2 | 2951 | 64 | C5ar1,C3ar1 | | 4.632e-04 | -7.68 | IL6GCSFMGF | prints domains | PR00433 | 2 | 2 | 2951 | 64 | Csf3,Il6 | | 4.655e-04 | -7.67 | REACTOME\_CLASS\_A1\_RHODOPSIN\_LIKE\_RECEPTORS | MSigDB lists | REACTOME\_CLASS\_A1\_RHODOPSIN\_LIKE\_RECEPTORS | 155 | 9 | 12187 | 179 | Hcar2,Ccl12,Ccl4,P2ry6,Cxcl16,C5ar1,C3ar1,Saa1,Cxcl1 | | 4.655e-04 | -7.67 | GSE2585\_CTEC\_VS\_MTEC\_THYMUS\_UP | MSigDB lists | GSE2585\_CTEC\_VS\_MTEC\_THYMUS\_UP | 155 | 9 | 12187 | 179 | Fstl1,Col4a1,Hp,Rhoc,Cd14,Cxcl1,Saa1,Rgs16,S100a8 | | 4.655e-04 | -7.67 | GSE29949\_CD8\_NEG\_DC\_SPLEEN\_VS\_CD8\_POS\_DC\_SPLEEN\_DN | MSigDB lists | GSE29949\_CD8\_NEG\_DC\_SPLEEN\_VS\_CD8\_POS\_DC\_SPLEEN\_DN | 155 | 9 | 12187 | 179 | Isg15,Ifi211,Cxcl1,Icam1,Ifi209,Ifi207,Ifi204,Il6,Lcn2 | | 4.655e-04 | -7.67 | GSE2770\_IL12\_AND\_TGFB\_ACT\_VS\_ACT\_CD4\_TCELL\_2H\_UP | MSigDB lists | GSE2770\_IL12\_AND\_TGFB\_ACT\_VS\_ACT\_CD4\_TCELL\_2H\_UP | 155 | 9 | 12187 | 179 | Sdc4,Zfp189,Cdkn1a,Runx1,Tubb6,Gadd45g,Pik3r5,Tnip1,Hspb1 | | 4.699e-04 | -7.66 | regulation of macrophage activation | biological process | GO:0043030 | 42 | 5 | 13711 | 214 | Lgals9,Mmp8,Thbs1,C5ar1,Il4ra | | 4.714e-04 | -7.66 | GO\_RESPONSE\_TO\_PROTOZOAN | MSigDB lists | GO\_RESPONSE\_TO\_PROTOZOAN | 11 | 3 | 12187 | 179 | Bcl3,Il4ra,Il6 | | 4.714e-04 | -7.66 | MODULE\_262 | MSigDB lists | MODULE\_262 | 11 | 3 | 12187 | 179 | Ccl2,Cp,Ccl12 | | 4.714e-04 | -7.66 | GO\_ACTIVATION\_OF\_MAPKKK\_ACTIVITY | MSigDB lists | GO\_ACTIVATION\_OF\_MAPKKK\_ACTIVITY | 11 | 3 | 12187 | 179 | Tnf,Gadd45b,Gadd45g | | 4.714e-04 | -7.66 | GO\_NEGATIVE\_REGULATION\_OF\_VIRAL\_ENTRY\_INTO\_HOST\_CELL | MSigDB lists | GO\_NEGATIVE\_REGULATION\_OF\_VIRAL\_ENTRY\_INTO\_HOST\_CELL | 11 | 3 | 12187 | 179 | Ifitm3,Ifitm2,Trim25 | | 4.753e-04 | -7.65 | SWEET\_KRAS\_TARGETS\_UP | MSigDB lists | SWEET\_KRAS\_TARGETS\_UP | 68 | 6 | 12187 | 179 | Sgk1,Map3k8,Gadd45b,Ccl12,Col4a1,Hspb1 | | 4.771e-04 | -7.65 | positive regulation of I-kappaB kinase/NF-kappaB signaling | biological process | GO:0043123 | 89 | 7 | 13711 | 214 | Ripk1,Il1a,Tnf,Il1rn,Tgm2,Trim25,Tnfsf10 | | 4.771e-04 | -7.65 | modification of morphology or physiology of other organism | biological process | GO:0035821 | 89 | 7 | 13711 | 214 | Ccl2,Ncf1,Irgm1,S100a9,Ccl4,Tnip1,Irgm2 | | 4.815e-04 | -7.64 | hematopoietic or lymphoid organ development | biological process | GO:0048534 | 517 | 19 | 13711 | 214 | Rbm47,Herc6,Spi1,Cebpd,Runx1,Samd9l,Bcl3,Tiparp,Sbno2,Prg4,Nfkb1,Csf3,Zbtb16,Gadd45g,Pik3r1,Nfkb2,Tnfaip3,Tnf,Il6 | | 4.879e-04 | -7.63 | GSE7852\_THYMUS\_VS\_FAT\_TREG\_DN | MSigDB lists | GSE7852\_THYMUS\_VS\_FAT\_TREG\_DN | 156 | 9 | 12187 | 179 | Cdkn1a,Tnf,Maff,Ier3,Stx11,Rhoc,Tiparp,Fzd4,Nfkbie | | 4.907e-04 | -7.62 | GEORGANTAS\_HSC\_MARKERS | MSigDB lists | GEORGANTAS\_HSC\_MARKERS | 45 | 5 | 12187 | 179 | Birc3,Robo4,Tnfsf10,Tiparp,Hspb1 | | 4.980e-04 | -7.60 | positive regulation of peptidyl-tyrosine phosphorylation | biological process | GO:0050731 | 147 | 9 | 13711 | 214 | Ncf1,Csf3,Parp14,Il6,Iqgap1,Parp9,Tnf,Icam1,Ptpn1 | | 5.043e-04 | -7.59 | INTFRNREGFCT | prints domains | PR00267 | 8 | 3 | 2951 | 64 | Irf9,Irf2,Irf7 | | 5.056e-04 | -7.59 | GO\_REGULATION\_OF\_JNK\_CASCADE | MSigDB lists | GO\_REGULATION\_OF\_JNK\_CASCADE | 125 | 8 | 12187 | 179 | Ptpn1,Fzd4,Map3k6,Ripk1,Gadd45b,Il1a,Gadd45g,Tnf | | 5.056e-04 | -7.59 | GSE40274\_CTRL\_VS\_FOXP3\_AND\_XBP1\_TRANSDUCED\_ACTIVATED\_CD4\_TCELL\_DN | MSigDB lists | GSE40274\_CTRL\_VS\_FOXP3\_AND\_XBP1\_TRANSDUCED\_ACTIVATED\_CD4\_TCELL\_DN | 125 | 8 | 12187 | 179 | Mxd4,Rassf4,Il1a,Msr1,Sbno2,P2ry6,Irf7,Bcl3 | | 5.111e-04 | -7.58 | GSE11961\_MEMORY\_BCELL\_DAY7\_VS\_GERMINAL\_CENTER\_BCELL\_DAY7\_DN | MSigDB lists | GSE11961\_MEMORY\_BCELL\_DAY7\_VS\_GERMINAL\_CENTER\_BCELL\_DAY7\_DN | 157 | 9 | 12187 | 179 | Hcar2,Akap12,Cybb,Ifit2,Sdc4,Casp4,Tnfsf10,Gem,Slfn9 | | 5.111e-04 | -7.58 | GSE17721\_PAM3CSK4\_VS\_CPG\_12H\_BMDC\_DN | MSigDB lists | GSE17721\_PAM3CSK4\_VS\_CPG\_12H\_BMDC\_DN | 157 | 9 | 12187 | 179 | Ptgs2,Gbp3,Tnf,Nfkbiz,Nfkbia,Parp14,Tiparp,Rhoc,Gadd45b | | 5.111e-04 | -7.58 | GSE34156\_UNTREATED\_VS\_24H\_NOD2\_LIGAND\_TREATED\_MONOCYTE\_DN | MSigDB lists | GSE34156\_UNTREATED\_VS\_24H\_NOD2\_LIGAND\_TREATED\_MONOCYTE\_DN | 157 | 9 | 12187 | 179 | Cybb,AA467197,Rhou,Rbm47,Igsf6,Clic4,Csf2rb,Cxcl16,Cd14 | | 5.111e-04 | -7.58 | GSE3982\_DC\_VS\_NKCELL\_UP | MSigDB lists | GSE3982\_DC\_VS\_NKCELL\_UP | 157 | 9 | 12187 | 179 | Atf3,Il1rn,Tor3a,Sgk1,Cdkn1a,Rbm47,Igsf6,Tnip1,Nfkbie | | 5.111e-04 | -7.58 | GSE29618\_PDC\_VS\_MDC\_DAY7\_FLU\_VACCINE\_DN | MSigDB lists | GSE29618\_PDC\_VS\_MDC\_DAY7\_FLU\_VACCINE\_DN | 157 | 9 | 12187 | 179 | Irak3,Igsf6,Msn,Ifitm3,Ifitm2,Timp1,Il1rn,Cebpd,Sgk1 | | 5.146e-04 | -7.57 | regulation of cell adhesion | biological process | GO:0030155 | 562 | 20 | 13711 | 214 | Vcam1,Selp,Runx1,Fam107a,Fzd4,Tgm2,Nfkbiz,Sdc4,Iqgap1,Zbtb16,Il1rn,Thbs1,Lgals9,Pik3r1,Tnfaip3,Tnf,Il4ra,Icam1,Ccl2,Il6 | | 5.154e-04 | -7.57 | mouse chr11\|11 C | chromosome location | mouse chr11\|11 C | 67 | 6 | 14556 | 220 | Wfdc21,Slfn9,Slfn5,Slfn3,Trim25,Slfn4 | | 5.206e-04 | -7.56 | GROSS\_HYPOXIA\_VIA\_HIF1A\_DN | MSigDB lists | GROSS\_HYPOXIA\_VIA\_HIF1A\_DN | 96 | 7 | 12187 | 179 | Gadd45g,Zfp36,Mmp8,Atf3,Errfi1,Sgk1,Nfkbia | | 5.216e-04 | -7.56 | regulation of cell differentiation | biological process | GO:0045595 | 1557 | 41 | 13711 | 214 | Il4ra,Il1a,Sox11,Nfkbia,Errfi1,Il6,Socs3,Msr1,Ctla2a,Trib1,Bbs12,Pik3r1,Lgals9,Fzd4,Apold1,Isg15,Adamts1,Cebpd,Spi1,Sgk1,Ptgs2,Runx1,Tnf,Cxcl9,Zfp36,Xdh,Hspb1,Cflar,Zbtb16,Ripk1,Adamts9,Csf3,Ifi204,Nfkbiz,C5ar1,Maff,Hspa5,Iqgap1,Nfe2l2,S100a9,Noct | | 5.220e-04 | -7.56 | regulation of leukocyte cell-cell adhesion | biological process | GO:1903037 | 213 | 11 | 13711 | 214 | Il4ra,Tnf,Icam1,Il6,Ccl2,Zbtb16,Lgals9,Nfkbiz,Sdc4,Vcam1,Runx1 | | 5.225e-04 | -7.56 | GILDEA\_METASTASIS | MSigDB lists | GILDEA\_METASTASIS | 26 | 4 | 12187 | 179 | Ifi209,Ifi211,Ifi207,Ifi204 | | 5.233e-04 | -7.56 | JIANG\_HYPOXIA\_NORMAL | MSigDB lists | JIANG\_HYPOXIA\_NORMAL | 264 | 12 | 12187 | 179 | Ccnd2,Hspa5,Tgm2,Nfkbia,Sdc4,Timp1,Akap12,Gadd45b,Dtx3l,Angptl4,A2m,Zfp36 | | 5.353e-04 | -7.53 | GSE360\_CTRL\_VS\_M\_TUBERCULOSIS\_MAC\_DN | MSigDB lists | GSE360\_CTRL\_VS\_M\_TUBERCULOSIS\_MAC\_DN | 158 | 9 | 12187 | 179 | Irak3,Osmr,Irf9,Pygm,Il1rn,Gadd45b,Runx1,Zfp189,Trib1 | | 5.353e-04 | -7.53 | GSE17301\_IFNA2\_VS\_IFNA5\_STIM\_ACD3\_ACD28\_ACT\_CD8\_TCELL\_UP | MSigDB lists | GSE17301\_IFNA2\_VS\_IFNA5\_STIM\_ACD3\_ACD28\_ACT\_CD8\_TCELL\_UP | 158 | 9 | 12187 | 179 | Runx1,Il4ra,Ier3,Tnip1,Arid5b,Ifitm2,Irf2,Msn,Osmr | | 5.353e-04 | -7.53 | GSE2770\_IL12\_AND\_TGFB\_VS\_IL4\_TREATED\_ACT\_CD4\_TCELL\_2H\_UP | MSigDB lists | GSE2770\_IL12\_AND\_TGFB\_VS\_IL4\_TREATED\_ACT\_CD4\_TCELL\_2H\_UP | 158 | 9 | 12187 | 179 | Sdc4,Zfp189,Runx1,Cdkn1a,AA467197,Il1a,Rhoc,Trim56,Tnip1 | | 5.353e-04 | -7.53 | GSE17721\_LPS\_VS\_GARDIQUIMOD\_2H\_BMDC\_UP | MSigDB lists | GSE17721\_LPS\_VS\_GARDIQUIMOD\_2H\_BMDC\_UP | 158 | 9 | 12187 | 179 | Gadd45b,Nfkbiz,Ccl2,Cdkn1a,Tnfsf10,Icam1,Rtp4,Gbp7,Ifitm2 | | 5.353e-04 | -7.53 | GSE2770\_TGFB\_AND\_IL4\_VS\_IL4\_TREATED\_ACT\_CD4\_TCELL\_48H\_UP | MSigDB lists | GSE2770\_TGFB\_AND\_IL4\_VS\_IL4\_TREATED\_ACT\_CD4\_TCELL\_48H\_UP | 158 | 9 | 12187 | 179 | Pik3r5,Sdc4,Zfp189,Znfx1,Nfe2l2,Ier3,Tubb6,Nfkb1,Il1a | | 5.353e-04 | -7.53 | GSE43955\_TH0\_VS\_TGFB\_IL6\_TH17\_ACT\_CD4\_TCELL\_60H\_UP | MSigDB lists | GSE43955\_TH0\_VS\_TGFB\_IL6\_TH17\_ACT\_CD4\_TCELL\_60H\_UP | 158 | 9 | 12187 | 179 | C3ar1,A2m,Cxcl1,Tnip1,Ccdc58,Lgals9,Tnf,Il1a,Rasd1 | | 5.367e-04 | -7.53 | regulation of macromolecule biosynthetic process | biological process | GO:0010556 | 2735 | 63 | 13711 | 214 | Gbp4,Nfkbie,Nfkb1,Bach1,Sox11,Nfkb2,Mxd4,Icam1,Errfi1,Cybb,Thbs1,Rbm39,Banp,Irf2,S100a9,Irf9,Bcl3,Tnfaip3,Znfx1,Ccl2,Zbtb16,Csf3,Aff1,C3ar1,Arid5b,Irak3,Ifi204,Fzd4,Ifi209,Sbno2,Parp9,Spi1,Cebpd,Ifi211,Trim25,Resf1,Runx1,Il1a,Akap12,Nfkbia,Cdkn1a,Il6,Trib1,Irf7,Dtx3l,Pik3r1,Dnajb1,Rrp8,Maff,Prg4,Rasd1,Nfe2l2,Tnf,Acod1,Zfp36,Atf3,Cflar,Ripk1,Hspb1,Ifi207,Ncf1,Tnip1,Fstl1 | | 5.399e-04 | -7.52 | WWE\_dom\_sf | interpro domains | IPR037197 | 11 | 3 | 13788 | 212 | Parp12,Tiparp,Parp14 | | 5.441e-04 | -7.52 | PID\_IL12\_2PATHWAY | MSigDB lists | PID\_IL12\_2PATHWAY | 46 | 5 | 12187 | 179 | Ccl4,Nfkb2,Gadd45b,Nfkb1,Gadd45g | | 5.441e-04 | -7.52 | CORDENONSI\_YAP\_CONSERVED\_SIGNATURE | MSigDB lists | CORDENONSI\_YAP\_CONSERVED\_SIGNATURE | 46 | 5 | 12187 | 179 | Tgm2,Fstl1,Thbs1,Sgk1,Gadd45b | | 5.447e-04 | -7.52 | GO\_EXTRACELLULAR\_STRUCTURE\_ORGANIZATION | MSigDB lists | GO\_EXTRACELLULAR\_STRUCTURE\_ORGANIZATION | 228 | 11 | 12187 | 179 | Nfkb2,Icam1,Bcl3,A2m,Ripk1,Timp1,Mmp8,Col4a1,Vcam1,Tnf,Thbs1 | | 5.543e-04 | -7.50 | GO\_POSITIVE\_REGULATION\_OF\_LEUKOCYTE\_DIFFERENTIATION | MSigDB lists | GO\_POSITIVE\_REGULATION\_OF\_LEUKOCYTE\_DIFFERENTIATION | 97 | 7 | 12187 | 179 | Ripk1,Il6,Zbtb16,Trib1,Runx1,Tnf,Il4ra | | 5.597e-04 | -7.49 | GO\_POSITIVE\_REGULATION\_OF\_SECRETION | MSigDB lists | GO\_POSITIVE\_REGULATION\_OF\_SECRETION | 266 | 12 | 12187 | 179 | Il4ra,Tnf,Sdc4,Hcar2,Il6,S100a9,S100a8,Sox11,Il1a,Saa1,Cd14,Il17ra | | 5.604e-04 | -7.49 | GSE21670\_STAT3\_KO\_VS\_WT\_CD4\_TCELL\_DN | MSigDB lists | GSE21670\_STAT3\_KO\_VS\_WT\_CD4\_TCELL\_DN | 159 | 9 | 12187 | 179 | Sbno2,Ptpn1,Arid5b,Ccl12,Il6,Il1a,Socs3,Nfkbiz,Il4ra | | 5.604e-04 | -7.49 | GSE11057\_NAIVE\_VS\_EFF\_MEMORY\_CD4\_TCELL\_DN | MSigDB lists | GSE11057\_NAIVE\_VS\_EFF\_MEMORY\_CD4\_TCELL\_DN | 159 | 9 | 12187 | 179 | Ifi211,Iqgap1,Ifi209,Tnf,Rhou,Tor3a,Ifi207,Ifi204,Stx11 | | 5.604e-04 | -7.49 | GSE13484\_UNSTIM\_VS\_12H\_YF17D\_VACCINE\_STIM\_PBMC\_DN | MSigDB lists | GSE13484\_UNSTIM\_VS\_12H\_YF17D\_VACCINE\_STIM\_PBMC\_DN | 159 | 9 | 12187 | 179 | Irf2,Plaur,Irak3,Arid5b,Irf7,Ptgs2,Ifitm2,Trib1,Tnf | | 5.604e-04 | -7.49 | GO\_REGULATION\_OF\_STRESS\_ACTIVATED\_PROTEIN\_KINASE\_SIGNALING\_CASCADE | MSigDB lists | GO\_REGULATION\_OF\_STRESS\_ACTIVATED\_PROTEIN\_KINASE\_SIGNALING\_CASCADE | 159 | 9 | 12187 | 179 | Map3k6,Fzd4,Ptpn1,Xdh,Tnf,Il1a,Gadd45g,Gadd45b,Ripk1 | | 5.620e-04 | -7.48 | DEATH | smart domains | SM00005 | 23 | 4 | 7188 | 123 | Ripk1,Irak3,Nfkb2,Nfkb1 | | 5.640e-04 | -7.48 | cellular response to copper ion | biological process | GO:0071280 | 11 | 3 | 13711 | 214 | Mt2,Mt1,Nfe2l2 | | 5.640e-04 | -7.48 | cytokine biosynthetic process | biological process | GO:0042089 | 11 | 3 | 13711 | 214 | Gadd45g,Irf9,Irf7 | | 5.640e-04 | -7.48 | negative regulation of interleukin-12 production | biological process | GO:0032695 | 11 | 3 | 13711 | 214 | Irak3,Thbs1,Nfkb1 | | 5.642e-04 | -7.48 | positive regulation of p38MAPK cascade | biological process | GO:1900745 | 25 | 4 | 13711 | 214 | Gadd45g,Xdh,Gadd45b,Ncf1 | | 5.642e-04 | -7.48 | mononuclear cell migration | biological process | GO:0071674 | 25 | 4 | 13711 | 214 | Ccl2,Ccl12,Ccl4,Ccl7 | | 5.658e-04 | -7.48 | cellular response to drug | biological process | GO:0035690 | 250 | 12 | 13711 | 214 | Il6,Acod1,Tnfaip3,Tnf,Lcn2,Thbs1,Cflar,Ccl7,Ripk1,Nfkb1,Nfe2l2,Ccl4 | | 5.739e-04 | -7.46 | FoxO signaling pathway | KEGG pathways | mmu04068 | 117 | 9 | 5248 | 107 | Il6,Sgk3,Gadd45b,Ccnd2,Pik3r1,Tnfsf10,Cdkn1a,Gadd45g,Sgk1 | | 5.765e-04 | -7.46 | response to organic cyclic compound | biological process | GO:0014070 | 483 | 18 | 13711 | 214 | Usp18,Msn,Sgk1,Tiparp,Fam107a,Nfkb1,Ccl12,Il1rn,Irak3,Pygm,Cflar,Pik3r1,P2ry6,Acod1,Tnf,Nfkbia,Il6,Zfp36 | | 5.772e-04 | -7.46 | positive regulation of cell-cell adhesion | biological process | GO:0022409 | 182 | 10 | 13711 | 214 | Nfkbiz,Icam1,Il4ra,Tnf,Ccl2,Il6,Zbtb16,Vcam1,Lgals9,Runx1 | | 5.864e-04 | -7.44 | GSE17721\_LPS\_VS\_POLYIC\_2H\_BMDC\_UP | MSigDB lists | GSE17721\_LPS\_VS\_POLYIC\_2H\_BMDC\_UP | 160 | 9 | 12187 | 179 | Ccl2,Nfkbia,Vcam1,Nfkb1,Il1rn,Ifitm2,Saa1,Csf3,Nfkb2 | | 5.896e-04 | -7.44 | MODULE\_223 | MSigDB lists | MODULE\_223 | 98 | 7 | 12187 | 179 | Osmr,Csf3,Selp,Il6,Il1a,Hp,Timp1 | | 5.981e-04 | -7.42 | Insulin resistance | KEGG pathways | mmu04931 | 94 | 8 | 5248 | 107 | Pygm,Nfkbia,Il6,Socs3,Nfkb1,Tnf,Ptpn1,Pik3r1 | | 5.981e-04 | -7.42 | GO\_POSITIVE\_REGULATION\_OF\_CELL\_ADHESION | MSigDB lists | GO\_POSITIVE\_REGULATION\_OF\_CELL\_ADHESION | 268 | 12 | 12187 | 179 | Il6,Ccl12,Il4ra,Pik3r1,Tnf,Map3k8,Sdc4,Tgm2,Zbtb16,Vcam1,Saa1,Selp | | 5.997e-04 | -7.42 | KEGG\_CHRONIC\_MYELOID\_LEUKEMIA | MSigDB lists | KEGG\_CHRONIC\_MYELOID\_LEUKEMIA | 71 | 6 | 12187 | 179 | Nfkb1,Pik3r5,Nfkbia,Runx1,Cdkn1a,Pik3r1 | | 6.017e-04 | -7.42 | NAKAMURA\_ADIPOGENESIS\_EARLY\_UP | MSigDB lists | NAKAMURA\_ADIPOGENESIS\_EARLY\_UP | 47 | 5 | 12187 | 179 | Gadd45b,Cebpd,Ripk1,Zfp36,Pik3r1 | | 6.017e-04 | -7.42 | JOSEPH\_RESPONSE\_TO\_SODIUM\_BUTYRATE\_DN | MSigDB lists | JOSEPH\_RESPONSE\_TO\_SODIUM\_BUTYRATE\_DN | 47 | 5 | 12187 | 179 | Aff1,Fstl1,A2m,Irf2,Il6 | | 6.064e-04 | -7.41 | HUPER\_BREAST\_BASAL\_VS\_LUMINAL\_UP | MSigDB lists | HUPER\_BREAST\_BASAL\_VS\_LUMINAL\_UP | 27 | 4 | 12187 | 179 | Il1a,S100a8,S100a9,Ptges | | 6.064e-04 | -7.41 | PID\_CD40\_PATHWAY | MSigDB lists | PID\_CD40\_PATHWAY | 27 | 4 | 12187 | 179 | Tnfaip3,Nfkb1,Nfkbia,Birc3 | | 6.064e-04 | -7.41 | GO\_HETEROPHILIC\_CELL\_CELL\_ADHESION\_VIA\_PLASMA\_MEMBRANE\_CELL\_ADHESION\_MOLECULES | MSigDB lists | GO\_HETEROPHILIC\_CELL\_CELL\_ADHESION\_VIA\_PLASMA\_MEMBRANE\_CELL\_ADHESION\_MOLECULES | 27 | 4 | 12187 | 179 | Selp,Vcam1,Icam1,Sele | | 6.076e-04 | -7.41 | GO\_ANGIOGENESIS | MSigDB lists | GO\_ANGIOGENESIS | 231 | 11 | 12187 | 179 | Ptgs2,Rnf213,Apold1,Clic4,Thbs1,Col4a1,Rasip1,Robo4,Angptl4,Spi1,Ccl12 | | 6.087e-04 | -7.40 | intrinsic apoptotic signaling pathway in response to DNA damage | biological process | GO:0008630 | 67 | 6 | 13711 | 214 | Ier3,Ifi204,Pik3r1,Bcl3,Tnf,Cdkn1a | | 6.101e-04 | -7.40 | regulation of DNA-binding transcription factor activity | biological process | GO:0051090 | 326 | 14 | 13711 | 214 | Cflar,Csf3,Trib1,Ripk1,Arid5b,Irak3,Nfkbia,Tnfaip3,Acod1,Tnf,Icam1,Il6,Trim25,Fzd4 | | 6.134e-04 | -7.40 | GSE16522\_ANTI\_CD3CD28\_STIM\_VS\_UNSTIM\_MEMORY\_CD8\_TCELL\_UP | MSigDB lists | GSE16522\_ANTI\_CD3CD28\_STIM\_VS\_UNSTIM\_MEMORY\_CD8\_TCELL\_UP | 161 | 9 | 12187 | 179 | Tnf,Map3k8,Tiparp,Ksr1,Tubb6,Cflar,Hspb1,Cxcl16,Plaur | | 6.134e-04 | -7.40 | GSE17721\_PAM3CSK4\_VS\_GADIQUIMOD\_4H\_BMDC\_DN | MSigDB lists | GSE17721\_PAM3CSK4\_VS\_GADIQUIMOD\_4H\_BMDC\_DN | 161 | 9 | 12187 | 179 | Isg15,Ptges,Cmpk2,Rtp4,Irf2,Usp18,Hcar2,Atf3,Parp12 | | 6.134e-04 | -7.40 | GO\_REGULATION\_OF\_LEUKOCYTE\_DIFFERENTIATION | MSigDB lists | GO\_REGULATION\_OF\_LEUKOCYTE\_DIFFERENTIATION | 161 | 9 | 12187 | 179 | Il6,Ripk1,Trib1,Zbtb16,Il4ra,Pik3r1,Tnf,Runx1,Irf7 | | 6.218e-04 | -7.38 | GO\_POSITIVE\_REGULATION\_OF\_P38MAPK\_CASCADE | MSigDB lists | GO\_POSITIVE\_REGULATION\_OF\_P38MAPK\_CASCADE | 12 | 3 | 12187 | 179 | Xdh,Gadd45b,Gadd45g | | 6.218e-04 | -7.38 | HUMMEL\_BURKITTS\_LYMPHOMA\_DN | MSigDB lists | HUMMEL\_BURKITTS\_LYMPHOMA\_DN | 12 | 3 | 12187 | 179 | Nfkbia,Bcl3,Cflar | | 6.218e-04 | -7.38 | NUNODA\_RESPONSE\_TO\_DASATINIB\_IMATINIB\_DN | MSigDB lists | NUNODA\_RESPONSE\_TO\_DASATINIB\_IMATINIB\_DN | 12 | 3 | 12187 | 179 | Tnfaip3,Cdkn1a,Tnf | | 6.218e-04 | -7.38 | TIMOFEEVA\_GROWTH\_STRESS\_VIA\_STAT1\_DN | MSigDB lists | TIMOFEEVA\_GROWTH\_STRESS\_VIA\_STAT1\_DN | 12 | 3 | 12187 | 179 | Isg15,Ccnd2,Hspb1 | | 6.218e-04 | -7.38 | GO\_OVULATION | MSigDB lists | GO\_OVULATION | 12 | 3 | 12187 | 179 | Adamts1,Il4ra,Ptgs2 | | 6.218e-04 | -7.38 | MODULE\_191 | MSigDB lists | MODULE\_191 | 12 | 3 | 12187 | 179 | Ccl2,Ccl12,Cp | | 6.397e-04 | -7.35 | GO\_REGULATION\_OF\_TRANSPORT | MSigDB lists | GO\_REGULATION\_OF\_TRANSPORT | 1360 | 35 | 12187 | 179 | Il17ra,Ptpn1,Sgk3,Cd14,Nfkb1,Il1a,Sox11,Thbs1,Sgk1,S100a8,Nfkbie,Csf3,Bcl3,Cybb,Hcar2,Ccl12,Ripk1,Nfe2l2,Il6,Rhou,Pik3r1,Il4ra,Ccl4,Icam1,Sele,Clic4,Msn,Ptgs2,Saa1,Kcna5,Cdkn1a,S100a9,Tnf,Nfkbia,Sdc4 | | 6.414e-04 | -7.35 | GSE17721\_CTRL\_VS\_PAM3CSK4\_12H\_BMDC\_DN | MSigDB lists | GSE17721\_CTRL\_VS\_PAM3CSK4\_12H\_BMDC\_DN | 162 | 9 | 12187 | 179 | Il1rn,Atf3,Birc3,Map3k8,Tgm2,Pik3ap1,Map3k6,Cd14,Tagln2 | | 6.414e-04 | -7.35 | GSE21546\_WT\_VS\_SAP1A\_KO\_AND\_ELK1\_KO\_ANTI\_CD3\_STIM\_DP\_THYMOCYTES\_UP | MSigDB lists | GSE21546\_WT\_VS\_SAP1A\_KO\_AND\_ELK1\_KO\_ANTI\_CD3\_STIM\_DP\_THYMOCYTES\_UP | 162 | 9 | 12187 | 179 | Ifi211,Fstl1,Ifi209,Pik3r1,Rasd1,Cebpd,Ifi204,Samd9l,Ifi207 | | 6.499e-04 | -7.34 | negative regulation of tumor necrosis factor production | biological process | GO:0032720 | 45 | 5 | 13711 | 214 | Tnfaip3,Irak3,Errfi1,Bcl3,Trim30a | | 6.499e-04 | -7.34 | positive regulation of interleukin-1 production | biological process | GO:0032732 | 45 | 5 | 13711 | 214 | Ifi211,Hspb1,Casp4,Ifi204,Lgals9 | | 6.525e-04 | -7.33 | - | gene3d domains | 4.10.10.10 | 3 | 2 | 6647 | 99 | Mt1,Mt2 | | 6.527e-04 | -7.33 | TBK1.DF\_UP | MSigDB lists | TBK1.DF\_UP | 233 | 11 | 12187 | 179 | Nfkbie,Saa1,Cxcl1,S100a8,Tnfaip8,Plaur,Runx1,Il1a,Nfkb1,Atf3,Tnfaip3 | | 6.527e-04 | -7.33 | GO\_REGULATION\_OF\_BINDING | MSigDB lists | GO\_REGULATION\_OF\_BINDING | 233 | 11 | 12187 | 179 | Csf3,Ifi211,Bcl3,Plaur,Sox11,Ifi209,Ifit2,Hspa5,Nfkbia,Ifi204,Ifi207 | | 6.611e-04 | -7.32 | regulation of epithelial cell proliferation | biological process | GO:0050678 | 291 | 13 | 13711 | 214 | Ccl12,C5ar1,Xdh,Errfi1,Zfp36,Il6,Ccl2,Tnf,Tnfaip3,Sox11,Ccnd2,Thbs1,Cflar | | 6.637e-04 | -7.32 | AMIT\_SERUM\_RESPONSE\_60\_MCF10A | MSigDB lists | AMIT\_SERUM\_RESPONSE\_60\_MCF10A | 48 | 5 | 12187 | 179 | Ptgs2,Sgk1,Map3k8,Trib1,Tnfaip3 | | 6.647e-04 | -7.32 | DNA-binding transcription factor activity, RNA polymerase II-specific | molecular function | GO:0000981 | 531 | 19 | 13516 | 211 | Nfe2l2,Sox11,Ifi204,Arid5b,Cebpd,Ifi209,Nfkb2,Mxd4,Atf3,Zbtb16,Nfkb1,Ifi211,Irf7,Spi1,Ifi207,Bach1,Irf9,Irf2,Runx1 | | 6.657e-04 | -7.31 | REACTOME\_TOLL\_RECEPTOR\_CASCADES | MSigDB lists | REACTOME\_TOLL\_RECEPTOR\_CASCADES | 100 | 7 | 12187 | 179 | Ripk1,Nfkbia,Nfkb2,Irak3,Irf7,Cd14,Saa1 | | 6.657e-04 | -7.31 | ZHANG\_TLX\_TARGETS\_UP | MSigDB lists | ZHANG\_TLX\_TARGETS\_UP | 100 | 7 | 12187 | 179 | Mxd4,Gadd45b,Gadd45g,Cdkn1a,Nfkb2,Pygm,Tap1 | | 6.667e-04 | -7.31 | GO\_REGULATION\_OF\_TRANSCRIPTION\_FROM\_RNA\_POLYMERASE\_II\_PROMOTER | MSigDB lists | GO\_REGULATION\_OF\_TRANSCRIPTION\_FROM\_RNA\_POLYMERASE\_II\_PROMOTER | 1310 | 34 | 12187 | 179 | Nfe2l2,Il6,Ripk1,Spi1,Maff,Pik3r1,Sbno2,Arid5b,Bcl3,Ifi211,Csf3,Cebpd,Mxd4,Il1a,Nfkb1,Nfkbiz,Hspa5,Sox11,Irf2,Bach1,Nfkbia,Tnf,Runx1,Zfp189,Nfkb2,Irf7,Atf3,Ifi207,Ifi204,Zbtb16,Fstl1,Ifi209,Zfp36,Tnip1 | | 6.705e-04 | -7.31 | GSE19198\_CTRL\_VS\_IL21\_TREATED\_TCELL\_6H\_DN | MSigDB lists | GSE19198\_CTRL\_VS\_IL21\_TREATED\_TCELL\_6H\_DN | 163 | 9 | 12187 | 179 | Zbtb16,Sntb2,Map3k8,Gadd45b,Pik3r5,Arid5b,Sgk3,Dtx3l,Irak3 | | 6.705e-04 | -7.31 | GSE39556\_CD8A\_DC\_VS\_NK\_CELL\_MOUSE\_3H\_POST\_POLYIC\_INJ\_DN | MSigDB lists | GSE39556\_CD8A\_DC\_VS\_NK\_CELL\_MOUSE\_3H\_POST\_POLYIC\_INJ\_DN | 163 | 9 | 12187 | 179 | Zfp36,Rgs16,Nfkbie,Cxcl1,Gadd45g,Tnfaip3,Cebpd,Map3k8,Il4ra | | 6.705e-04 | -7.31 | GSE23925\_LIGHT\_ZONE\_VS\_NAIVE\_BCELL\_DN | MSigDB lists | GSE23925\_LIGHT\_ZONE\_VS\_NAIVE\_BCELL\_DN | 163 | 9 | 12187 | 179 | Nfkbie,Slfn4,Slfn3,Errfi1,Socs3,Nfkbia,Tiparp,Bach1,Luc7l3 | | 6.705e-04 | -7.31 | GSE17721\_0.5H\_VS\_24H\_LPS\_BMDC\_DN | MSigDB lists | GSE17721\_0.5H\_VS\_24H\_LPS\_BMDC\_DN | 163 | 9 | 12187 | 179 | Slfn4,Slfn3,Gbp3,Tap1,Irgm2,Slfn9,Tagln2,Cybb,Zfp189 | | 6.705e-04 | -7.31 | GSE26669\_CTRL\_VS\_COSTIM\_BLOCK\_MLR\_CD4\_TCELL\_UP | MSigDB lists | GSE26669\_CTRL\_VS\_COSTIM\_BLOCK\_MLR\_CD4\_TCELL\_UP | 163 | 9 | 12187 | 179 | Irf7,Slfn5,Rnf213,Tap1,Irf2,Cmpk2,Il4ra,Cdkn1a,Samd9l | | 6.745e-04 | -7.30 | DEx/H-box helicases activate type I IFN and inflammatory cytokines production | REACTOME pathways | R-MMU-3134963 | 11 | 3 | 6297 | 105 | Irf7,Nfkb1,Nfkb2 | | 6.790e-04 | -7.29 | anion binding | molecular function | GO:0043168 | 2265 | 54 | 13516 | 211 | Pygm,S100a9,Ccl7,Gem,Fstl1,Map3k8,Selp,Saa1,Rhoj,Iqgap1,Bbs12,Tgm2,Gbp6,Hcar2,Tgtp1,Gbp3,Trib1,Iigp1,Slfn5,Xdh,Runx1,Rhou,Ksr1,Slfn9,Tor3a,Map3k6,Adamts1,Tubb6,Ip6k2,Irgm1,Cmpk2,Lcn2,Oasl2,Rasd1,Sgk1,Ncf1,Tap1,S100a8,Sgk3,Cybb,Ptges,Mx1,Sele,Thbs1,Gbp7,Rhoc,Plek,Hspa5,Parp9,Parp14,Oasl1,Irak3,Ripk1,Ccl2 | | 6.861e-04 | -7.28 | Epstein-Barr virus infection | KEGG pathways | mmu05169 | 172 | 11 | 5248 | 107 | Nfkb2,Tnfaip3,Hspb1,Cdkn1a,Pik3r1,Ripk1,Nfkbie,Icam1,Spi1,Nfkb1,Nfkbia | | 6.861e-04 | -7.28 | Epstein-Barr virus infection | KEGG pathways | ko05169 | 172 | 11 | 5248 | 107 | Cdkn1a,Tnfaip3,Hspb1,Nfkb2,Nfkbia,Nfkb1,Spi1,Icam1,Nfkbie,Ripk1,Pik3r1 | | 6.941e-04 | -7.27 | GO\_NEGATIVE\_REGULATION\_OF\_PHOSPHORUS\_METABOLIC\_PROCESS | MSigDB lists | GO\_NEGATIVE\_REGULATION\_OF\_PHOSPHORUS\_METABOLIC\_PROCESS | 436 | 16 | 12187 | 179 | Atf3,Socs3,Cdkn1a,Errfi1,Ptpn1,Hspb1,Irak3,Plek,Tnip1,Tnfaip3,Gadd45b,Il6,Trib1,Tnf,Xdh,Iqgap1 | | 6.958e-04 | -7.27 | SASSON\_RESPONSE\_TO\_FORSKOLIN\_DN | MSigDB lists | SASSON\_RESPONSE\_TO\_FORSKOLIN\_DN | 73 | 6 | 12187 | 179 | Akap12,Thbs1,Casp4,Trib1,Ptpn1,Gem | | 6.958e-04 | -7.27 | LABBE\_TGFB1\_TARGETS\_DN | MSigDB lists | LABBE\_TGFB1\_TARGETS\_DN | 73 | 6 | 12187 | 179 | Ccl2,Saa1,Lcn2,Tap1,Plaur,Cp | | 6.988e-04 | -7.27 | IL-1\_CS | interpro domains | IPR020877 | 3 | 2 | 13788 | 212 | Il1a,Il1rn | | 6.988e-04 | -7.27 | Metalthion\_vert | interpro domains | IPR000006 | 3 | 2 | 13788 | 212 | Mt2,Mt1 | | 6.988e-04 | -7.27 | Metalthion\_vert\_metal\_BS | interpro domains | IPR018064 | 3 | 2 | 13788 | 212 | Mt2,Mt1 | | 6.988e-04 | -7.27 | GADD45 | interpro domains | IPR024824 | 3 | 2 | 13788 | 212 | Gadd45g,Gadd45b | | 6.988e-04 | -7.27 | Selectin\_superfamily | interpro domains | IPR002396 | 3 | 2 | 13788 | 212 | Selp,Sele | | 6.988e-04 | -7.27 | Metalthion\_dom\_sf | interpro domains | IPR017854 | 3 | 2 | 13788 | 212 | Mt2,Mt1 | | 6.988e-04 | -7.27 | 2-5-oligoadenylate\_synth\_CS | interpro domains | IPR006117 | 3 | 2 | 13788 | 212 | Oasl2,Oasl1 | | 6.988e-04 | -7.27 | Selectin\_CTLD | interpro domains | IPR033991 | 3 | 2 | 13788 | 212 | Selp,Sele | | 6.988e-04 | -7.27 | Metalthion | interpro domains | IPR003019 | 3 | 2 | 13788 | 212 | Mt1,Mt2 | | 6.988e-04 | -7.27 | Metalthion\_dom\_sf\_vert | interpro domains | IPR023587 | 3 | 2 | 13788 | 212 | Mt2,Mt1 | | 6.988e-04 | -7.27 | ApoL | interpro domains | IPR008405 | 3 | 2 | 13788 | 212 | Apold1,Apol10b | | 6.988e-04 | -7.27 | DUF2075 | interpro domains | IPR018647 | 3 | 2 | 13788 | 212 | Slfn9,Slfn5 | | 6.994e-04 | -7.27 | GO\_LIPOPOLYSACCHARIDE\_MEDIATED\_SIGNALING\_PATHWAY | MSigDB lists | GO\_LIPOPOLYSACCHARIDE\_MEDIATED\_SIGNALING\_PATHWAY | 28 | 4 | 12187 | 179 | Cd14,Nfkbia,Tnf,Ccl12 | | 6.994e-04 | -7.27 | STEARMAN\_TUMOR\_FIELD\_EFFECT\_UP | MSigDB lists | STEARMAN\_TUMOR\_FIELD\_EFFECT\_UP | 28 | 4 | 12187 | 179 | Lcn2,Csf2rb,Iqgap1,Socs3 | | 6.994e-04 | -7.27 | HUANG\_FOXA2\_TARGETS\_DN | MSigDB lists | HUANG\_FOXA2\_TARGETS\_DN | 28 | 4 | 12187 | 179 | Tgm2,Cxcl1,Cdkn1a,Plaur | | 6.994e-04 | -7.27 | KUROZUMI\_RESPONSE\_TO\_ONCOCYTIC\_VIRUS | MSigDB lists | KUROZUMI\_RESPONSE\_TO\_ONCOCYTIC\_VIRUS | 28 | 4 | 12187 | 179 | Cxcl1,Tnf,Ccl12,Il1a | | 6.994e-04 | -7.27 | MODULE\_300 | MSigDB lists | MODULE\_300 | 28 | 4 | 12187 | 179 | Nfkbia,Cd14,Ripk1,Tnfaip3 | | 6.994e-04 | -7.27 | NUNODA\_RESPONSE\_TO\_DASATINIB\_IMATINIB\_UP | MSigDB lists | NUNODA\_RESPONSE\_TO\_DASATINIB\_IMATINIB\_UP | 28 | 4 | 12187 | 179 | Nfkbia,Ccnd2,Gadd45b,Nfkb1 | | 7.000e-04 | -7.26 | GO\_REGULATION\_OF\_CELL\_ACTIVATION | MSigDB lists | GO\_REGULATION\_OF\_CELL\_ACTIVATION | 312 | 13 | 12187 | 179 | Il4ra,Pik3r1,Map3k8,Il6,Tnfaip3,Ccl12,Selp,Plek,Vcam1,Zbtb16,Thbs1,Cdkn1a,Sox11 | | 7.006e-04 | -7.26 | GSE17721\_0.5H\_VS\_12H\_POLYIC\_BMDC\_DN | MSigDB lists | GSE17721\_0.5H\_VS\_12H\_POLYIC\_BMDC\_DN | 164 | 9 | 12187 | 179 | Luc7l3,Znfx1,Ms4a6d,Hcar2,Cybb,Dnajb1,Birc3,Gbp3,Trim25 | | 7.006e-04 | -7.26 | GSE21379\_WT\_VS\_SAP\_KO\_TFH\_CD4\_TCELL\_UP | MSigDB lists | GSE21379\_WT\_VS\_SAP\_KO\_TFH\_CD4\_TCELL\_UP | 164 | 9 | 12187 | 179 | Selp,Rhoj,Cd14,Igsf6,Xdh,Sgk1,Col4a1,Akap12,Lcn2 | | 7.006e-04 | -7.26 | GSE22886\_NAIVE\_TCELL\_VS\_MONOCYTE\_DN | MSigDB lists | GSE22886\_NAIVE\_TCELL\_VS\_MONOCYTE\_DN | 164 | 9 | 12187 | 179 | Csf2rb,Igsf6,Irak3,Rbm47,S100a9,Cd14,Cebpd,Cybb,Bach1 | | 7.006e-04 | -7.26 | KAECH\_NAIVE\_VS\_MEMORY\_CD8\_TCELL\_DN | MSigDB lists | KAECH\_NAIVE\_VS\_MEMORY\_CD8\_TCELL\_DN | 164 | 9 | 12187 | 179 | Sntb2,Casp4,Ccl4,Gadd45b,Tnfaip3,Tnfsf10,Gbp3,Errfi1,Xdh | | 7.006e-04 | -7.26 | GSE25087\_FETAL\_VS\_ADULT\_TREG\_UP | MSigDB lists | GSE25087\_FETAL\_VS\_ADULT\_TREG\_UP | 164 | 9 | 12187 | 179 | Ier3,Tnip3,Rhoc,Cybb,Sdc4,Hspa5,Ccnd2,Cmpk2,Nfkbie | | 7.006e-04 | -7.26 | GSE6674\_CPG\_VS\_CPG\_AND\_ANTI\_IGM\_STIM\_BCELL\_DN | MSigDB lists | GSE6674\_CPG\_VS\_CPG\_AND\_ANTI\_IGM\_STIM\_BCELL\_DN | 164 | 9 | 12187 | 179 | Errfi1,Gem,Banp,Gadd45g,Atf3,Nfe2l2,Tnf,Cdkn1a,Birc3 | | 7.006e-04 | -7.26 | GSE17721\_CTRL\_VS\_POLYIC\_12H\_BMDC\_DN | MSigDB lists | GSE17721\_CTRL\_VS\_POLYIC\_12H\_BMDC\_DN | 164 | 9 | 12187 | 179 | Gbp3,Pik3ap1,Iqgap1,Dnajb1,Ccl2,Cdkn1a,Znfx1,Ms4a6d,Timp1 | | 7.043e-04 | -7.26 | Amoebiasis | KEGG pathways | ko05146 | 74 | 7 | 5248 | 107 | Pik3r1,Col4a1,Tnf,Nfkb1,Il6,Cd14,Hspb1 | | 7.043e-04 | -7.26 | Amoebiasis | KEGG pathways | mmu05146 | 74 | 7 | 5248 | 107 | Nfkb1,Il6,Pik3r1,Tnf,Col4a1,Hspb1,Cd14 | | 7.117e-04 | -7.25 | WWE-dom | interpro domains | IPR004170 | 12 | 3 | 13788 | 212 | Parp14,Tiparp,Parp12 | | 7.117e-04 | -7.25 | bZIP\_Maf | interpro domains | IPR004826 | 12 | 3 | 13788 | 212 | Nfe2l2,Bach1,Maff | | 7.123e-04 | -7.25 | Herpes simplex infection | KEGG pathways | ko05168 | 146 | 10 | 5248 | 107 | Ccl2,Irf7,Irf9,Tnf,Nfkb1,Il6,Nfkbia,Socs3,Tap1,Ccl12 | | 7.123e-04 | -7.25 | Herpes simplex infection | KEGG pathways | mmu05168 | 146 | 10 | 5248 | 107 | Ccl12,Tnf,Irf9,Irf7,Ccl2,Tap1,Il6,Socs3,Nfkbia,Nfkb1 | | 7.194e-04 | -7.24 | DUF2075 | pfam domains | PF09848 | 3 | 2 | 12881 | 201 | Slfn9,Slfn5 | | 7.194e-04 | -7.24 | ApoL | pfam domains | PF05461 | 3 | 2 | 12881 | 201 | Apol10b,Apold1 | | 7.194e-04 | -7.24 | Metallothio | pfam domains | PF00131 | 3 | 2 | 12881 | 201 | Mt1,Mt2 | | 7.200e-04 | -7.24 | interferon-beta production | biological process | GO:0032608 | 3 | 2 | 13711 | 214 | Irf7,Trim56 | | 7.200e-04 | -7.24 | myeloid cell apoptotic process | biological process | GO:0033028 | 3 | 2 | 13711 | 214 | Il6,Hcar2 | | 7.200e-04 | -7.24 | positive regulation of reactive oxygen species biosynthetic process | biological process | GO:1903428 | 46 | 5 | 13711 | 214 | Mmp8,Tnf,Icam1,Ptgs2,Il6 | | 7.200e-04 | -7.24 | negative regulation of tumor necrosis factor superfamily cytokine production | biological process | GO:1903556 | 46 | 5 | 13711 | 214 | Trim30a,Bcl3,Errfi1,Irak3,Tnfaip3 | | 7.223e-04 | -7.23 | Death\_domain | interpro domains | IPR000488 | 27 | 4 | 13788 | 212 | Ripk1,Irak3,Nfkb2,Nfkb1 | | 7.226e-04 | -7.23 | negative regulation of hemopoiesis | biological process | GO:1903707 | 124 | 8 | 13711 | 214 | Runx1,Pik3r1,Trib1,Zbtb16,Nfe2l2,Zfp36,Nfkbia,Il4ra | | 7.257e-04 | -7.23 | GO\_REGULATION\_OF\_PROTEIN\_TARGETING | MSigDB lists | GO\_REGULATION\_OF\_PROTEIN\_TARGETING | 236 | 11 | 12187 | 179 | Csf3,Nfkbie,Ptgs2,Bcl3,Pik3r1,Tnf,Rhou,Cdkn1a,Nfkbia,Il6,Ccl12 | | 7.257e-04 | -7.23 | GSE24081\_CONTROLLER\_VS\_PROGRESSOR\_HIV\_SPECIFIC\_CD8\_TCELL\_DN | MSigDB lists | GSE24081\_CONTROLLER\_VS\_PROGRESSOR\_HIV\_SPECIFIC\_CD8\_TCELL\_DN | 132 | 8 | 12187 | 179 | Parp12,Ifi44,Xaf1,Sntb2,Usp18,Plek,Irf9,Ptgs2 | | 7.282e-04 | -7.22 | Leishmaniasis | KEGG pathways | ko05140 | 54 | 6 | 5248 | 107 | Nfkbia,Nfkb1,Ncf1,Tnf,Ptgs2,Il1a | | 7.282e-04 | -7.22 | Leishmaniasis | KEGG pathways | mmu05140 | 54 | 6 | 5248 | 107 | Il1a,Ptgs2,Tnf,Nfkbia,Ncf1,Nfkb1 | | 7.311e-04 | -7.22 | Inflammatory bowel disease (IBD) | KEGG pathways | mmu05321 | 36 | 5 | 5248 | 107 | Il4ra,Il6,Nfkb1,Il1a,Tnf | | 7.311e-04 | -7.22 | Inflammatory bowel disease (IBD) | KEGG pathways | ko05321 | 36 | 5 | 5248 | 107 | Tnf,Il1a,Nfkb1,Il6,Il4ra | | 7.318e-04 | -7.22 | GSE43955\_1H\_VS\_42H\_ACT\_CD4\_TCELL\_UP | MSigDB lists | GSE43955\_1H\_VS\_42H\_ACT\_CD4\_TCELL\_UP | 165 | 9 | 12187 | 179 | Adamts1,Msr1,Thbs1,Cebpd,Cybb,S100a9,Ncf1,Gem,Nfkbie | | 7.318e-04 | -7.22 | GSE9650\_NAIVE\_VS\_MEMORY\_CD8\_TCELL\_DN | MSigDB lists | GSE9650\_NAIVE\_VS\_MEMORY\_CD8\_TCELL\_DN | 165 | 9 | 12187 | 179 | Tnfsf10,Xdh,Errfi1,Gbp3,Sntb2,Casp4,Ccl4,Gadd45b,Tnfaip3 | | 7.318e-04 | -7.22 | GSE21670\_IL6\_VS\_TGFB\_AND\_IL6\_TREATED\_STAT3\_KO\_CD4\_TCELL\_DN | MSigDB lists | GSE21670\_IL6\_VS\_TGFB\_AND\_IL6\_TREATED\_STAT3\_KO\_CD4\_TCELL\_DN | 165 | 9 | 12187 | 179 | Irf7,Gbp6,Isg15,Rtp4,Tor3a,Ccnd2,Gbp4,Mxd4,Gadd45g | | 7.318e-04 | -7.22 | GSE37416\_0H\_VS\_24H\_F\_TULARENSIS\_LVS\_NEUTROPHIL\_DN | MSigDB lists | GSE37416\_0H\_VS\_24H\_F\_TULARENSIS\_LVS\_NEUTROPHIL\_DN | 165 | 9 | 12187 | 179 | Gadd45g,Gadd45b,Ier3,Tnfaip3,Nfkbie,Cxcl1,Ptges,Icam1,Tnfaip8 | | 7.318e-04 | -7.22 | GSE29949\_MICROGLIA\_BRAIN\_VS\_CD8\_POS\_DC\_SPLEEN\_UP | MSigDB lists | GSE29949\_MICROGLIA\_BRAIN\_VS\_CD8\_POS\_DC\_SPLEEN\_UP | 165 | 9 | 12187 | 179 | Mmp8,Timp1,Atf3,Il6,Zfp189,Cdkn1a,Rgs16,Nfkbie,Gem | | 7.320e-04 | -7.22 | MODULE\_52 | MSigDB lists | MODULE\_52 | 354 | 14 | 12187 | 179 | Thbs1,Ifi209,Cdkn1a,Ier3,Col4a1,Ifi207,Akap12,Ifi204,Isg15,Tgm2,Maff,Tnf,Ccl12,Ifi211 | | 7.382e-04 | -7.21 | CEBP\_01 | MSigDB lists | CEBP\_01 | 200 | 10 | 12187 | 179 | Nfkbiz,Sntb2,Tnf,Thbs1,Slc24a4,S100a9,Tnfsf10,Ip6k2,Rbm39,Plek | | 7.398e-04 | -7.21 | LIM\_MAMMARY\_STEM\_CELL\_UP | MSigDB lists | LIM\_MAMMARY\_STEM\_CELL\_UP | 396 | 15 | 12187 | 179 | Icam1,Ptgs2,Col4a1,Rasip1,Sox11,Adamts1,Cdkn1a,Thbs1,Gem,Rhoj,Tubb6,Bach1,Il6,Ccnd2,Sntb2 | | 7.423e-04 | -7.21 | bZIP\_Maf | pfam domains | PF03131 | 12 | 3 | 12881 | 201 | Nfe2l2,Maff,Bach1 | | 7.433e-04 | -7.20 | nitric oxide mediated signal transduction | biological process | GO:0007263 | 12 | 3 | 13711 | 214 | Mt2,Mt1,Rasd1 | | 7.433e-04 | -7.20 | activation of MAPKKK activity | biological process | GO:0000185 | 12 | 3 | 13711 | 214 | Tnf,Gadd45b,Gadd45g | | 7.433e-04 | -7.20 | cytokine metabolic process | biological process | GO:0042107 | 12 | 3 | 13711 | 214 | Irf7,Irf9,Gadd45g | | 7.437e-04 | -7.20 | 1-phosphatidylinositol-3-kinase regulator activity | molecular function | GO:0046935 | 12 | 3 | 13516 | 211 | Pik3r1,Socs3,Pik3r5 | | 7.494e-04 | -7.20 | MIKKELSEN\_MCV6\_LCP\_WITH\_H3K4ME3 | MSigDB lists | MIKKELSEN\_MCV6\_LCP\_WITH\_H3K4ME3 | 102 | 7 | 12187 | 179 | Xaf1,Xdh,Tnfsf10,Slfn4,Slfn3,Csf3,Robo4 | | 7.494e-04 | -7.20 | GSE40666\_STAT1\_KO\_VS\_STAT4\_KO\_CD8\_TCELL\_WITH\_IFNA\_STIM\_90MIN\_DN | MSigDB lists | GSE40666\_STAT1\_KO\_VS\_STAT4\_KO\_CD8\_TCELL\_WITH\_IFNA\_STIM\_90MIN\_DN | 102 | 7 | 12187 | 179 | Angptl4,Cxcl1,Socs3,Ccl4,Tnf,Ier3,Tiparp | | 7.524e-04 | -7.19 | MODULE\_23 | MSigDB lists | MODULE\_23 | 355 | 14 | 12187 | 179 | Fam107a,Ccl12,Gadd45b,Tgm2,Trib1,Tnf,Il4ra,Sbno2,Xdh,Cp,Hp,Sele,Cd14,Isg15 | | 7.588e-04 | -7.18 | Interferon Signaling | REACTOME pathways | R-MMU-913531 | 44 | 5 | 6297 | 105 | Irf9,Socs3,Ptpn1,Usp18,Isg15 | | 7.622e-04 | -7.18 | Death | pfam domains | PF00531 | 27 | 4 | 12881 | 201 | Nfkb1,Irak3,Ripk1,Nfkb2 | | 7.627e-04 | -7.18 | GO\_ADAPTIVE\_IMMUNE\_RESPONSE | MSigDB lists | GO\_ADAPTIVE\_IMMUNE\_RESPONSE | 133 | 8 | 12187 | 179 | Il4ra,Il6,C3ar1,Irf7,Bcl3,Icam1,Nfkb2,Tap1 | | 7.642e-04 | -7.18 | GSE12392\_CD8A\_POS\_VS\_NEG\_SPLEEN\_DC\_DN | MSigDB lists | GSE12392\_CD8A\_POS\_VS\_NEG\_SPLEEN\_DC\_DN | 166 | 9 | 12187 | 179 | Slfn5,Irf7,Ifitm3,Rnf213,Xdh,Trim25,P2ry6,Ifi44,Oasl1 | | 7.642e-04 | -7.18 | GSE40274\_FOXP3\_VS\_FOXP3\_AND\_PBX1\_TRANSDUCED\_ACTIVATED\_CD4\_TCELL\_DN | MSigDB lists | GSE40274\_FOXP3\_VS\_FOXP3\_AND\_PBX1\_TRANSDUCED\_ACTIVATED\_CD4\_TCELL\_DN | 166 | 9 | 12187 | 179 | Ifi44,Usp18,Sdc4,Nfkb2,Rnf213,Ifitm2,Ifitm3,Bcl3,Ncf1 | | 7.642e-04 | -7.18 | GSE28737\_BCL6\_HET\_VS\_BCL6\_KO\_FOLLICULAR\_BCELL\_DN | MSigDB lists | GSE28737\_BCL6\_HET\_VS\_BCL6\_KO\_FOLLICULAR\_BCELL\_DN | 166 | 9 | 12187 | 179 | Il17ra,S100a8,Gbp3,Rhou,Socs3,Fstl1,Ms4a6d,Rasd1,Rasip1 | | 7.642e-04 | -7.18 | GSE13738\_RESTING\_VS\_BYSTANDER\_ACTIVATED\_CD4\_TCELL\_DN | MSigDB lists | GSE13738\_RESTING\_VS\_BYSTANDER\_ACTIVATED\_CD4\_TCELL\_DN | 166 | 9 | 12187 | 179 | Tagln2,Nfkbie,Csf2rb,Trib1,Tor3a,Birc3,Sgk1,Map3k8,Tnf | | 7.650e-04 | -7.18 | MODULE\_2 | MSigDB lists | MODULE\_2 | 315 | 13 | 12187 | 179 | Tnfaip3,Ccl12,Ccnd2,Maff,Tgm2,S100a8,Csf3,Gem,Akap12,Col4a1,Ier3,Atf3,Cdkn1a | | 7.780e-04 | -7.16 | GO\_TRANSCRIPTION\_FACTOR\_ACTIVITY\_RNA\_POLYMERASE\_II\_CORE\_PROMOTER\_PROXIMAL\_REGION\_SEQUENCE\_SPECIFIC\_BINDING | MSigDB lists | GO\_TRANSCRIPTION\_FACTOR\_ACTIVITY\_RNA\_POLYMERASE\_II\_CORE\_PROMOTER\_PROXIMAL\_REGION\_SEQUENCE\_SPECIFIC\_BINDING | 238 | 11 | 12187 | 179 | Cebpd,Atf3,Ifi204,Ifi207,Bach1,Spi1,Ifi209,Sox11,Nfkb2,Ifi211,Irf7 | | 7.780e-04 | -7.16 | TTCYNRGAA\_STAT5B\_01 | MSigDB lists | TTCYNRGAA\_STAT5B\_01 | 238 | 11 | 12187 | 179 | Irf2,Irf9,Icam1,Trim25,Apold1,Aff1,Adamts9,Ccl12,Ccnd2,Maff,Pik3r1 | | 7.843e-04 | -7.15 | DNA-binding transcription factor activity | molecular function | GO:0003700 | 714 | 23 | 13516 | 211 | Arid5b,Nfe2l2,Ifi204,Ifi209,Aff1,Mxd4,Nfkb1,Zbtb16,Irf2,Irf9,Runx1,Ifi211,Bcl3,Ifi207,Spi1,Maff,Cebpd,Sox11,Nfkb2,Znfx1,Atf3,Bach1,Irf7 | | 7.956e-04 | -7.14 | activation of MAPKK activity | biological process | GO:0000186 | 47 | 5 | 13711 | 214 | Ripk1,Map3k6,Map3k8,Gadd45g,Gadd45b | | 7.977e-04 | -7.13 | GSE22886\_NAIVE\_BCELL\_VS\_MONOCYTE\_DN | MSigDB lists | GSE22886\_NAIVE\_BCELL\_VS\_MONOCYTE\_DN | 167 | 9 | 12187 | 179 | Sgk1,Cd14,S100a9,Bcl3,C5ar1,Csf2rb,Irak3,S100a8,Rbm47 | | 7.977e-04 | -7.13 | GSE16697\_CD4\_TCELL\_VS\_TFH\_CD4\_TCELL\_UP | MSigDB lists | GSE16697\_CD4\_TCELL\_VS\_TFH\_CD4\_TCELL\_UP | 167 | 9 | 12187 | 179 | Cybb,Nfkbie,Plek,Icam1,Irak3,Tnfaip8,Gbp3,Nfkb2,Clic4 | | 7.977e-04 | -7.13 | GSE21380\_NON\_TFH\_VS\_TFH\_CD4\_TCELL\_UP | MSigDB lists | GSE21380\_NON\_TFH\_VS\_TFH\_CD4\_TCELL\_UP | 167 | 9 | 12187 | 179 | Ifi44,Map3k8,Fstl1,Parp14,Ccl4,Tiparp,Ifitm3,Slfn3,Slfn4 | | 7.977e-04 | -7.13 | GSE17721\_LPS\_VS\_CPG\_2H\_BMDC\_UP | MSigDB lists | GSE17721\_LPS\_VS\_CPG\_2H\_BMDC\_UP | 167 | 9 | 12187 | 179 | Znfx1,Sdc4,Maff,Hspa5,Cmpk2,Zfp36,Rtp4,Gem,Isg15 | | 7.977e-04 | -7.13 | GSE21546\_WT\_VS\_SAP1A\_KO\_AND\_ELK1\_KO\_DP\_THYMOCYTES\_DN | MSigDB lists | GSE21546\_WT\_VS\_SAP1A\_KO\_AND\_ELK1\_KO\_DP\_THYMOCYTES\_DN | 167 | 9 | 12187 | 179 | Col4a1,Ifi44,Ccnd2,Fstl1,Ccl4,Parp14,Rbm47,Trim25,Rnf213 | | 7.996e-04 | -7.13 | BIOCARTA\_CD40\_PATHWAY | MSigDB lists | BIOCARTA\_CD40\_PATHWAY | 13 | 3 | 12187 | 179 | Tnfaip3,Nfkb1,Nfkbia | | 7.996e-04 | -7.13 | DALESSIO\_TSA\_RESPONSE | MSigDB lists | DALESSIO\_TSA\_RESPONSE | 13 | 3 | 12187 | 179 | Maff,Atf3,Gadd45b | | 7.996e-04 | -7.13 | INGA\_TP53\_TARGETS | MSigDB lists | INGA\_TP53\_TARGETS | 13 | 3 | 12187 | 179 | Thbs1,Cdkn1a,Ier3 | | 7.996e-04 | -7.13 | MARSON\_FOXP3\_CORE\_DIRECT\_TARGETS | MSigDB lists | MARSON\_FOXP3\_CORE\_DIRECT\_TARGETS | 13 | 3 | 12187 | 179 | Gadd45b,Slfn3,Slfn4 | | 7.996e-04 | -7.13 | BIOCARTA\_HSP27\_PATHWAY | MSigDB lists | BIOCARTA\_HSP27\_PATHWAY | 13 | 3 | 12187 | 179 | Hspb1,Il1a,Tnf | | 8.011e-04 | -7.13 | GO\_REGULATION\_OF\_CHEMOTAXIS | MSigDB lists | GO\_REGULATION\_OF\_CHEMOTAXIS | 134 | 8 | 12187 | 179 | Ccl4,Thbs1,Il6,Ccl12,Cxcl1,C3ar1,C5ar1,Hspb1 | | 8.021e-04 | -7.13 | KASLER\_HDAC7\_TARGETS\_2\_DN | MSigDB lists | KASLER\_HDAC7\_TARGETS\_2\_DN | 29 | 4 | 12187 | 179 | Nfkbia,Ifitm2,Cdkn1a,Gadd45b | | 8.021e-04 | -7.13 | PID\_TXA2PATHWAY | MSigDB lists | PID\_TXA2PATHWAY | 50 | 5 | 12187 | 179 | Sele,Icam1,Pik3r5,Vcam1,Tgm2 | | 8.040e-04 | -7.13 | Rheumatoid arthritis | KEGG pathways | mmu05323 | 55 | 6 | 5248 | 107 | Ccl12,Il1a,Il6,Ccl2,Icam1,Tnf | | 8.040e-04 | -7.13 | Rheumatoid arthritis | KEGG pathways | ko05323 | 55 | 6 | 5248 | 107 | Ccl12,Il1a,Il6,Ccl2,Icam1,Tnf | | 8.053e-04 | -7.12 | GO\_MEMBRANE\_MICRODOMAIN | MSigDB lists | GO\_MEMBRANE\_MICRODOMAIN | 239 | 11 | 12187 | 179 | Sdc4,Birc3,Tnf,Ripk1,Kcna5,Cd14,Ptgs2,Cflar,Rgs16,Sele,Icam1 | | 8.141e-04 | -7.11 | GO\_RESPONSE\_TO\_ENDOGENOUS\_STIMULUS | MSigDB lists | GO\_RESPONSE\_TO\_ENDOGENOUS\_STIMULUS | 1115 | 30 | 12187 | 179 | Sgk1,Thbs1,Nfkb1,P2ry6,Irak3,Ptpn1,Il4ra,Pik3r1,Il6,Ccl12,Tnfaip3,Iqgap1,Pygm,Cdkn1a,Adamts1,Fstl1,Vcam1,Socs3,Col4a1,Il1rn,Timp1,Ptgs2,Icam1,Trim25,Msn,Errfi1,Tnf,Nfkbia,Casp4,C5ar1 | | 8.173e-04 | -7.11 | Metallothioneins bind metals | REACTOME pathways | R-MMU-5661231 | 3 | 2 | 6297 | 105 | Mt1,Mt2 | | 8.173e-04 | -7.11 | Response to metal ions | REACTOME pathways | R-MMU-5660526 | 3 | 2 | 6297 | 105 | Mt2,Mt1 | | 8.226e-04 | -7.10 | apoptotic signaling pathway | biological process | GO:0097190 | 225 | 11 | 13711 | 214 | Ifi204,Ier3,Lcn2,Pik3r1,Bcl3,Ripk1,Cdkn1a,Il1a,Tnf,Rrp8,Casp4 | | 8.226e-04 | -7.10 | regulation of leukocyte differentiation | biological process | GO:1902105 | 225 | 11 | 13711 | 214 | Trib1,Ctla2a,Zbtb16,Ripk1,Lgals9,Runx1,Pik3r1,Tnf,Il4ra,Nfkbiz,Il6 | | 8.227e-04 | -7.10 | Ovarian tumor domain proteases | REACTOME pathways | R-MMU-5689896 | 26 | 4 | 6297 | 105 | Tnfaip3,Ripk1,Tnip1,Tnip3 | | 8.227e-04 | -7.10 | Regulation of TNFR1 signaling | REACTOME pathways | R-MMU-5357905 | 26 | 4 | 6297 | 105 | Birc3,Tnf,Tnfaip3,Ripk1 | | 8.227e-04 | -7.10 | GO\_G\_PROTEIN\_COUPLED\_RECEPTOR\_SIGNALING\_PATHWAY | MSigDB lists | GO\_G\_PROTEIN\_COUPLED\_RECEPTOR\_SIGNALING\_PATHWAY | 443 | 16 | 12187 | 179 | Plek,P2ry6,Rasd1,Akap12,Pik3r5,Cxcl1,Fzd4,C5ar1,C3ar1,Rgs16,Ccl4,Tgm2,Ccl2,Gpr84,Ccl12,Hcar2 | | 8.279e-04 | -7.10 | RICKMAN\_METASTASIS\_DN | MSigDB lists | RICKMAN\_METASTASIS\_DN | 203 | 10 | 12187 | 179 | Tap1,Tnfaip8,Ifi211,Tagln2,Ptgs2,Ifi207,Ifi204,Spi1,Cdkn1a,Ifi209 | | 8.279e-04 | -7.10 | GO\_PROTEIN\_MATURATION | MSigDB lists | GO\_PROTEIN\_MATURATION | 203 | 10 | 12187 | 179 | Ifi209,Tnf,Ripk1,Ifi207,Ifi204,S100a9,Ifi211,Tnfsf10,Xdh,S100a8 | | 8.297e-04 | -7.09 | positive regulation of T cell differentiation | biological process | GO:0045582 | 71 | 6 | 13711 | 214 | Lgals9,Runx1,Il6,Zbtb16,Il4ra,Nfkbiz | | 8.324e-04 | -7.09 | GO\_POSITIVE\_REGULATION\_OF\_MAP\_KINASE\_ACTIVITY | MSigDB lists | GO\_POSITIVE\_REGULATION\_OF\_MAP\_KINASE\_ACTIVITY | 168 | 9 | 12187 | 179 | Ripk1,Tnf,Thbs1,Ptpn1,C5ar1,Map3k6,Saa1,Pik3r5,Fzd4 | | 8.408e-04 | -7.08 | Map3k8 (mitogen-activated protein kinase kinase kinase 8) | protein interactions | 26410 | 3 | 2 | 6802 | 115 | Nfkb1,Ripk1 | | 8.408e-04 | -7.08 | Kat6a (K(lysine) acetyltransferase 6A) | protein interactions | 244349 | 3 | 2 | 6802 | 115 | Runx1,Spi1 | | 8.411e-04 | -7.08 | GSE15733\_BM\_VS\_SPLEEN\_MEMORY\_CD4\_TCELL\_UP | MSigDB lists | GSE15733\_BM\_VS\_SPLEEN\_MEMORY\_CD4\_TCELL\_UP | 135 | 8 | 12187 | 179 | Gbp7,S100a8,Pik3ap1,Icam1,Zbtb16,Tnfaip3,Hp,Cybb | | 8.411e-04 | -7.08 | GSE17974\_CTRL\_VS\_ACT\_IL4\_AND\_ANTI\_IL12\_6H\_CD4\_TCELL\_UP | MSigDB lists | GSE17974\_CTRL\_VS\_ACT\_IL4\_AND\_ANTI\_IL12\_6H\_CD4\_TCELL\_UP | 135 | 8 | 12187 | 179 | Tagln2,Pik3r5,Ptgs2,Tnfaip3,Gadd45b,Socs3,Dnajb1,Maff | | 8.411e-04 | -7.08 | GSE369\_PRE\_VS\_POST\_IL6\_INJECTION\_IFNG\_WT\_LIVER\_UP | MSigDB lists | GSE369\_PRE\_VS\_POST\_IL6\_INJECTION\_IFNG\_WT\_LIVER\_UP | 135 | 8 | 12187 | 179 | Socs3,Tgm2,Casp4,Usp18,Nfkb1,Bcl3,Dtx3l,Gbp7 | | 8.412e-04 | -7.08 | BYSTRYKH\_HEMATOPOIESIS\_STEM\_CELL\_QTL\_CIS | MSigDB lists | BYSTRYKH\_HEMATOPOIESIS\_STEM\_CELL\_QTL\_CIS | 104 | 7 | 12187 | 179 | Ifi204,Tubb6,Ifi207,Ccnd2,Runx1,Ifi209,Ifi211 | | 8.412e-04 | -7.08 | GSE40274\_CTRL\_VS\_GATA1\_TRANSDUCED\_ACTIVATED\_CD4\_TCELL\_UP | MSigDB lists | GSE40274\_CTRL\_VS\_GATA1\_TRANSDUCED\_ACTIVATED\_CD4\_TCELL\_UP | 104 | 7 | 12187 | 179 | Cp,Irf2,Csf2rb,P2ry6,S100a8,Trim56,Slfn5 | | 8.569e-04 | -7.06 | regulation of gene expression | biological process | GO:0010468 | 3008 | 67 | 13711 | 214 | Nfkbie,Nfkb1,Birc3,Gbp4,Bach1,Nfkb2,Sox11,Icam1,Mxd4,Lgals9,Banp,Msr1,Thbs1,Irf2,Bcl3,S100a9,Irf9,Ccl2,Xdh,Tnfaip3,Znfx1,Ifi204,Lcn2,Csf3,Zbtb16,Aff1,Arid5b,Irak3,Parp9,Sbno2,Fzd4,Ifi209,Tiparp,Trim25,Resf1,Runx1,Spi1,Ifi211,Cebpd,Cdkn1a,Il6,Nfkbia,Il1a,Pik3r1,Parp14,Trib1,Ctla2a,Dtx3l,Irf7,Dnajb1,Rrp8,Maff,Msn,Noct,Rasd1,Mmp8,Nfe2l2,Atf3,Zfp36,Tnf,Acod1,Fstl1,Tnip1,Ripk1,Cflar,Ifi207,Ncf1 | | 8.618e-04 | -7.06 | MARTENS\_BOUND\_BY\_PML\_RARA\_FUSION | MSigDB lists | MARTENS\_BOUND\_BY\_PML\_RARA\_FUSION | 360 | 14 | 12187 | 179 | Icam1,Plek,Pik3ap1,Ifitm3,Tagln2,Ksr1,Zbtb16,C5ar1,Ptges,Banp,Bcl3,Tnfaip3,Runx1,Spi1 | | 8.618e-04 | -7.06 | GO\_REGULATION\_OF\_MYELOID\_LEUKOCYTE\_DIFFERENTIATION | MSigDB lists | GO\_REGULATION\_OF\_MYELOID\_LEUKOCYTE\_DIFFERENTIATION | 76 | 6 | 12187 | 179 | Ripk1,Irf7,Trib1,Pik3r1,Runx1,Tnf | | 8.618e-04 | -7.06 | GUO\_HEX\_TARGETS\_UP | MSigDB lists | GUO\_HEX\_TARGETS\_UP | 76 | 6 | 12187 | 179 | Akap12,Rbm39,Iqgap1,Ptpn1,Sox11,Fzd4 | | 8.683e-04 | -7.05 | GSE19401\_UNSTIM\_VS\_PAM2CSK4\_STIM\_FOLLICULAR\_DC\_DN | MSigDB lists | GSE19401\_UNSTIM\_VS\_PAM2CSK4\_STIM\_FOLLICULAR\_DC\_DN | 169 | 9 | 12187 | 179 | Ifitm2,Isg15,Ifitm3,Hspb1,Irf2,Map3k8,Tgm2,Il1rn,Oasl1 | | 8.683e-04 | -7.05 | GSE24102\_GRANULOCYSTIC\_MDSC\_VS\_NEUTROPHIL\_DN | MSigDB lists | GSE24102\_GRANULOCYSTIC\_MDSC\_VS\_NEUTROPHIL\_DN | 169 | 9 | 12187 | 179 | Pik3ap1,Ccdc58,Nfkbie,Tnip1,Cxcl1,Il1a,Rhoc,Timp1,Kcna5 | | 8.683e-04 | -7.05 | GSE17721\_CTRL\_VS\_GARDIQUIMOD\_6H\_BMDC\_DN | MSigDB lists | GSE17721\_CTRL\_VS\_GARDIQUIMOD\_6H\_BMDC\_DN | 169 | 9 | 12187 | 179 | Msn,Slfn3,Slfn4,Ptges,Irgm2,Atf3,Ier3,Birc3,Maff | | 8.683e-04 | -7.05 | REACTOME\_PLATELET\_ACTIVATION\_SIGNALING\_AND\_AGGREGATION | MSigDB lists | REACTOME\_PLATELET\_ACTIVATION\_SIGNALING\_AND\_AGGREGATION | 169 | 9 | 12187 | 179 | Timp1,Thbs1,Pik3r1,Hspa5,Plek,Ptpn1,Selp,Pik3r5,A2m | | 8.683e-04 | -7.05 | GSE17721\_LPS\_VS\_CPG\_1H\_BMDC\_DN | MSigDB lists | GSE17721\_LPS\_VS\_CPG\_1H\_BMDC\_DN | 169 | 9 | 12187 | 179 | Nfkbiz,Nfkbia,Tnf,Map3k8,Cdkn1a,Nfe2l2,Il1rn,Clic4,Plek | | 8.683e-04 | -7.05 | GSE9006\_TYPE\_1\_DIABETES\_AT\_DX\_VS\_1MONTH\_POST\_DX\_PBMC\_UP | MSigDB lists | GSE9006\_TYPE\_1\_DIABETES\_AT\_DX\_VS\_1MONTH\_POST\_DX\_PBMC\_UP | 169 | 9 | 12187 | 179 | Nfe2l2,Ier3,Ccl12,Cybb,Stx11,Ifit3b,Socs3,Hspa5,Map3k8 | | 8.769e-04 | -7.04 | regulation of tyrosine phosphorylation of STAT protein | biological process | GO:0042509 | 48 | 5 | 13711 | 214 | Socs3,Tnf,Parp14,Il6,Parp9 | | 8.788e-04 | -7.04 | GO\_REGULATION\_OF\_PROTEIN\_MATURATION | MSigDB lists | GO\_REGULATION\_OF\_PROTEIN\_MATURATION | 51 | 5 | 12187 | 179 | A2m,C3ar1,Birc3,Thbs1,C5ar1 | | 8.788e-04 | -7.04 | LIN\_SILENCED\_BY\_TUMOR\_MICROENVIRONMENT | MSigDB lists | LIN\_SILENCED\_BY\_TUMOR\_MICROENVIRONMENT | 51 | 5 | 12187 | 179 | Il1rn,Adamts9,AA467197,Lcn2,Ptgs2 | | 8.788e-04 | -7.04 | GO\_RNA\_POLYMERASE\_II\_DISTAL\_ENHANCER\_SEQUENCE\_SPECIFIC\_DNA\_BINDING | MSigDB lists | GO\_RNA\_POLYMERASE\_II\_DISTAL\_ENHANCER\_SEQUENCE\_SPECIFIC\_DNA\_BINDING | 51 | 5 | 12187 | 179 | Nfkb1,Bach1,Nfe2l2,Zbtb16,Spi1 | | 8.788e-04 | -7.04 | SHETH\_LIVER\_CANCER\_VS\_TXNIP\_LOSS\_PAM3 | MSigDB lists | SHETH\_LIVER\_CANCER\_VS\_TXNIP\_LOSS\_PAM3 | 51 | 5 | 12187 | 179 | Tubb6,Rhoc,Lcn2,Ifit2,Maff | | 8.805e-04 | -7.04 | regulation of cellular extravasation | biological process | GO:0002691 | 28 | 4 | 13711 | 214 | Icam1,Il1a,Selp,Ccl2 | | 8.845e-04 | -7.03 | positive regulation of leukocyte activation | biological process | GO:0002696 | 227 | 11 | 13711 | 214 | Il6,Ccl2,Cdkn1a,Nfkbiz,Il4ra,Runx1,Lgals9,Thbs1,Vcam1,Mmp8,Zbtb16 | | 8.890e-04 | -7.03 | positive regulation of biosynthetic process | biological process | GO:0009891 | 1599 | 41 | 13711 | 214 | Rbm39,Pik3r1,Banp,P2ry6,Thbs1,Dtx3l,Irf7,Cybb,Il6,Icam1,Nfkbia,Nfkb2,Akap12,Sox11,Il1a,Runx1,Ptgs2,Bach1,Cebpd,Spi1,Nfkb1,Parp9,Sbno2,Fzd4,Tnip1,Aff1,C3ar1,Ncf1,Arid5b,Hspb1,Ripk1,Csf3,Zbtb16,Atf3,Ccl2,Tnf,Bcl3,Nfe2l2,Mmp8,Irf2,Maff | | 8.902e-04 | -7.02 | side of membrane | cellular component | GO:0098552 | 385 | 15 | 13825 | 212 | Vcam1,Msn,Selp,Gem,Robo4,Errfi1,Osmr,Tnf,Thbs1,Il6,Cxcl9,Icam1,Iqgap1,Cd14,Ptpn1 | | 8.903e-04 | -7.02 | KOKKINAKIS\_METHIONINE\_DEPRIVATION\_96HR\_UP | MSigDB lists | KOKKINAKIS\_METHIONINE\_DEPRIVATION\_96HR\_UP | 105 | 7 | 12187 | 179 | Il6,Timp1,Il1a,Cdkn1a,Plaur,Hspb1,Cflar | | 8.966e-04 | -7.02 | GO\_POSITIVE\_REGULATION\_OF\_DEVELOPMENTAL\_PROCESS | MSigDB lists | GO\_POSITIVE\_REGULATION\_OF\_DEVELOPMENTAL\_PROCESS | 868 | 25 | 12187 | 179 | Socs3,Zbtb16,Adamts1,Ptgs2,Hspb1,Trib1,Runx1,Tnf,C3ar1,C5ar1,Sgk1,Thbs1,Sox11,Il1a,Nfkb1,Isg15,Irak3,Il4ra,Msr1,Il6,Ripk1,Tnfaip3,Angptl4,Csf3,Adamts9 | | 9.019e-04 | -7.01 | bZIP | interpro domains | IPR004827 | 49 | 5 | 13788 | 212 | Cebpd,Atf3,Bach1,Nfe2l2,Maff | | 9.042e-04 | -7.01 | negative regulation of immune effector process | biological process | GO:0002698 | 99 | 7 | 13711 | 214 | Il4ra,Tnf,Tnfaip3,Tap1,A2m,Irak3,Lgals9 | | 9.042e-04 | -7.01 | regulation of wound healing | biological process | GO:0061041 | 99 | 7 | 13711 | 214 | Selp,S100a9,Nfe2l2,Thbs1,Ccl2,Cdkn1a,Plek | | 9.054e-04 | -7.01 | GSE46606\_UNSTIM\_VS\_CD40L\_IL2\_IL5\_1DAY\_STIMULATED\_IRF4\_KO\_BCELL\_UP | MSigDB lists | GSE46606\_UNSTIM\_VS\_CD40L\_IL2\_IL5\_1DAY\_STIMULATED\_IRF4\_KO\_BCELL\_UP | 170 | 9 | 12187 | 179 | Rtp4,Parp9,Ifi211,Ifi207,Ifi204,Il1rn,Ccl12,Ifi209,Xaf1 | | 9.054e-04 | -7.01 | GSE11057\_NAIVE\_VS\_MEMORY\_CD4\_TCELL\_DN | MSigDB lists | GSE11057\_NAIVE\_VS\_MEMORY\_CD4\_TCELL\_DN | 170 | 9 | 12187 | 179 | Tagln2,Ifi211,Tnip1,Iqgap1,Ifi209,Tor3a,Ifi204,Timp1,Ifi207 | | 9.054e-04 | -7.01 | GSE41176\_UNSTIM\_VS\_ANTI\_IGM\_STIM\_BCELL\_1H\_DN | MSigDB lists | GSE41176\_UNSTIM\_VS\_ANTI\_IGM\_STIM\_BCELL\_1H\_DN | 170 | 9 | 12187 | 179 | Sntb2,Ifit2,Ifi209,Cebpd,Ifi207,Ifi204,Ifi211,Tap1,Tnfsf10 | | 9.150e-04 | -7.00 | HOUSTIS\_ROS | MSigDB lists | HOUSTIS\_ROS | 30 | 4 | 12187 | 179 | Cp,Hp,Xdh,Sdc4 | | 9.150e-04 | -7.00 | PATTERSON\_DOCETAXEL\_RESISTANCE | MSigDB lists | PATTERSON\_DOCETAXEL\_RESISTANCE | 30 | 4 | 12187 | 179 | Ifi207,Ifi204,Ifi209,Ifi211 | | 9.150e-04 | -7.00 | SCHLINGEMANN\_SKIN\_CARCINOGENESIS\_TPA\_UP | MSigDB lists | SCHLINGEMANN\_SKIN\_CARCINOGENESIS\_TPA\_UP | 30 | 4 | 12187 | 179 | S100a9,Il4ra,Saa1,S100a8 | | 9.150e-04 | -7.00 | RICKMAN\_HEAD\_AND\_NECK\_CANCER\_E | MSigDB lists | RICKMAN\_HEAD\_AND\_NECK\_CANCER\_E | 30 | 4 | 12187 | 179 | Gbp6,Gbp4,Cp,Il1rn | | 9.150e-04 | -7.00 | BROWNE\_HCMV\_INFECTION\_2HR\_UP | MSigDB lists | BROWNE\_HCMV\_INFECTION\_2HR\_UP | 30 | 4 | 12187 | 179 | Icam1,Atf3,Map3k8,Nfkbia | | 9.150e-04 | -7.00 | KEGG\_COMPLEMENT\_AND\_COAGULATION\_CASCADES | MSigDB lists | KEGG\_COMPLEMENT\_AND\_COAGULATION\_CASCADES | 30 | 4 | 12187 | 179 | Plaur,A2m,C5ar1,C3ar1 | | 9.150e-04 | -7.00 | GO\_CHEMOKINE\_MEDIATED\_SIGNALING\_PATHWAY | MSigDB lists | GO\_CHEMOKINE\_MEDIATED\_SIGNALING\_PATHWAY | 30 | 4 | 12187 | 179 | Ccl12,Ccl4,Cxcl1,Ccl2 | | 9.150e-04 | -7.00 | ZHAN\_MULTIPLE\_MYELOMA\_LB\_DN | MSigDB lists | ZHAN\_MULTIPLE\_MYELOMA\_LB\_DN | 30 | 4 | 12187 | 179 | Usp18,Casp4,Xaf1,Rtp4 | | 9.219e-04 | -6.99 | tube morphogenesis | biological process | GO:0035239 | 589 | 20 | 13711 | 214 | Rasip1,Spi1,Angptl4,Ptgs2,Tgm2,Tiparp,Fzd4,Clic4,Col4a1,Apold1,Rhoj,Sdc4,Ccl12,Thbs1,Adamts9,Rnf213,Robo4,Sox11,Cemip2,Ccl2 | | 9.226e-04 | -6.99 | GO\_DIVALENT\_INORGANIC\_CATION\_HOMEOSTASIS | MSigDB lists | GO\_DIVALENT\_INORGANIC\_CATION\_HOMEOSTASIS | 243 | 11 | 12187 | 179 | Saa1,Kcna5,C5ar1,C3ar1,S100a9,S100a8,Pygm,Tgm2,Ccl2,Slc24a4,Ccl12 | | 9.259e-04 | -6.98 | GSE29949\_MICROGLIA\_VS\_DC\_BRAIN\_DN | MSigDB lists | GSE29949\_MICROGLIA\_VS\_DC\_BRAIN\_DN | 137 | 8 | 12187 | 179 | Gadd45b,Stx11,Ccl2,Socs3,Tnfsf10,Trim25,Irf7,Selp | | 9.259e-04 | -6.98 | AKT\_UP\_MTOR\_DN.V1\_UP | MSigDB lists | AKT\_UP\_MTOR\_DN.V1\_UP | 137 | 8 | 12187 | 179 | Atf3,Rasd1,Msr1,Tor3a,S100a8,Nfkb2,Cxcl16,Saa1 | | 9.264e-04 | -6.98 | GO\_NEGATIVE\_REGULATION\_OF\_KINASE\_ACTIVITY | MSigDB lists | GO\_NEGATIVE\_REGULATION\_OF\_KINASE\_ACTIVITY | 206 | 10 | 12187 | 179 | Ptpn1,Errfi1,Irak3,Hspb1,Socs3,Trib1,Cdkn1a,Tnfaip3,Gadd45b,Il6 | | 9.264e-04 | -6.98 | BACH2\_01 | MSigDB lists | BACH2\_01 | 206 | 10 | 12187 | 179 | Tagln2,Csf3,Map3k6,Trib1,Sgk1,Cdkn1a,Gadd45b,Rasd1,Bach1,Rhoc | | 9.297e-04 | -6.98 | GO\_CELL\_ACTIVATION | MSigDB lists | GO\_CELL\_ACTIVATION | 405 | 15 | 12187 | 179 | Pik3r1,Spi1,Il6,Bcl3,Pik3r5,Sbno2,Nfkb2,Vcam1,Timp1,Saa1,Plek,Apold1,Hspb1,Il17ra,Icam1 | | 9.338e-04 | -6.98 | ZWANG\_CLASS\_1\_TRANSIENTLY\_INDUCED\_BY\_EGF | MSigDB lists | ZWANG\_CLASS\_1\_TRANSIENTLY\_INDUCED\_BY\_EGF | 363 | 14 | 12187 | 179 | Bach1,Parp12,Gadd45b,Parp14,Sdc4,Angptl4,Arid5b,Akap12,Col4a1,Adamts1,Sgk1,Thbs1,Zfp36,Ptpn1 | | 9.417e-04 | -6.97 | GO\_SIGNAL\_TRANSDUCTION\_BY\_P53\_CLASS\_MEDIATOR | MSigDB lists | GO\_SIGNAL\_TRANSDUCTION\_BY\_P53\_CLASS\_MEDIATOR | 106 | 7 | 12187 | 179 | Ifi204,Ifi207,Cdkn1a,Ifi209,Ifi211,Bcl3,Rrp8 | | 9.439e-04 | -6.97 | GSE22103\_UNSTIM\_VS\_LPS\_STIM\_NEUTROPHIL\_UP | MSigDB lists | GSE22103\_UNSTIM\_VS\_LPS\_STIM\_NEUTROPHIL\_UP | 171 | 9 | 12187 | 179 | Nfe2l2,Tubb6,Rhoc,Trib1,Il4ra,Errfi1,Icam1,Bcl3,Pik3r5 | | 9.439e-04 | -6.97 | GSE43956\_WT\_VS\_SGK1\_KO\_TH17\_DIFFERENTIATED\_CD4\_TCELL\_DN | MSigDB lists | GSE43956\_WT\_VS\_SGK1\_KO\_TH17\_DIFFERENTIATED\_CD4\_TCELL\_DN | 171 | 9 | 12187 | 179 | Il1a,Gadd45b,Nfe2l2,Sgk1,Tnf,Zbtb16,Rbm39,Cxcl1,Tnip1 | | 9.439e-04 | -6.97 | GSE3337\_4H\_VS\_16H\_IFNG\_IN\_CD8POS\_DC\_UP | MSigDB lists | GSE3337\_4H\_VS\_16H\_IFNG\_IN\_CD8POS\_DC\_UP | 171 | 9 | 12187 | 179 | Tnip1,Zfp36,Cdkn1a,Tnf,Birc3,Il1rn,Gadd45g,Tnfaip3,Gadd45b | | 9.439e-04 | -6.97 | GSE10239\_NAIVE\_VS\_KLRG1INT\_EFF\_CD8\_TCELL\_UP | MSigDB lists | GSE10239\_NAIVE\_VS\_KLRG1INT\_EFF\_CD8\_TCELL\_UP | 171 | 9 | 12187 | 179 | Ccnd2,Ifit3b,Socs3,Trim56,Tnip1,Rbm39,Slfn3,Slfn4,Irf9 | | 9.553e-04 | -6.95 | regulation of integrin activation | biological process | GO:0033623 | 13 | 3 | 13711 | 214 | Selp,Rasip1,Plek | | 9.596e-04 | -6.95 | regulation of G1/S transition of mitotic cell cycle | biological process | GO:2000045 | 100 | 7 | 13711 | 214 | Adamts1,Cdkn1a,Ccl12,Ccnd2,Ripk1,Fam107a,Kcna5 | | 9.609e-04 | -6.95 | GO\_POSITIVE\_REGULATION\_OF\_JUN\_KINASE\_ACTIVITY | MSigDB lists | GO\_POSITIVE\_REGULATION\_OF\_JUN\_KINASE\_ACTIVITY | 52 | 5 | 12187 | 179 | Fzd4,Map3k6,Tnf,Ptpn1,Ripk1 | | 9.644e-04 | -6.94 | regulation of blood coagulation | biological process | GO:0030193 | 49 | 5 | 13711 | 214 | S100a9,Plek,Nfe2l2,Thbs1,Selp | | 9.644e-04 | -6.94 | regulation of hemostasis | biological process | GO:1900046 | 49 | 5 | 13711 | 214 | Plek,S100a9,Thbs1,Nfe2l2,Selp | | 9.644e-04 | -6.94 | regulation of CD4-positive, alpha-beta T cell activation | biological process | GO:2000514 | 49 | 5 | 13711 | 214 | Lgals9,Runx1,Il6,Il4ra,Nfkbiz | | 9.707e-04 | -6.94 | GSE29618\_BCELL\_VS\_PDC\_UP | MSigDB lists | GSE29618\_BCELL\_VS\_PDC\_UP | 138 | 8 | 12187 | 179 | Bach1,Parp12,Il4ra,Birc3,Nfkbia,Clic4,Msn,Arid5b | | 9.738e-04 | -6.93 | Hematopoietic cell lineage | KEGG pathways | mmu04640 | 57 | 6 | 5248 | 107 | Cd14,Tnf,Il1a,Csf3,Il4ra,Il6 | | 9.738e-04 | -6.93 | Hematopoietic cell lineage | KEGG pathways | ko04640 | 57 | 6 | 5248 | 107 | Tnf,Il1a,Csf3,Cd14,Il4ra,Il6 | | 9.837e-04 | -6.92 | GSE17721\_0.5H\_VS\_12H\_LPS\_BMDC\_DN | MSigDB lists | GSE17721\_0.5H\_VS\_12H\_LPS\_BMDC\_DN | 172 | 9 | 12187 | 179 | Ccl2,Atf3,Nfkb1,Cybb,Arid5b,Slfn9,Cflar,Irgm2,Trim25 | | 9.837e-04 | -6.92 | GSE23502\_BM\_VS\_COLON\_TUMOR\_HDC\_KO\_MYELOID\_DERIVED\_SUPPRESSOR\_CELL\_DN | MSigDB lists | GSE23502\_BM\_VS\_COLON\_TUMOR\_HDC\_KO\_MYELOID\_DERIVED\_SUPPRESSOR\_CELL\_DN | 172 | 9 | 12187 | 179 | Sbno2,Il17ra,Tagln2,Pik3r5,Cxcl1,Mxd4,Zfp189,Ccnd2,Pik3r1 | | 9.837e-04 | -6.92 | GSE22886\_NAIVE\_TCELL\_VS\_NKCELL\_DN | MSigDB lists | GSE22886\_NAIVE\_TCELL\_VS\_NKCELL\_DN | 172 | 9 | 12187 | 179 | Ifi209,Map3k8,Ifi207,Ifi204,Lcn2,Cebpd,Cflar,S100a9,Ifi211 | | 9.837e-04 | -6.92 | GSE36891\_UNSTIM\_VS\_POLYIC\_TLR3\_STIM\_PERITONEAL\_MACROPHAGE\_DN | MSigDB lists | GSE36891\_UNSTIM\_VS\_POLYIC\_TLR3\_STIM\_PERITONEAL\_MACROPHAGE\_DN | 172 | 9 | 12187 | 179 | Cdkn1a,Ccl2,Nfkbiz,Akap12,Ms4a6d,Parp12,Ccl12,Irf9,Xdh | | 9.845e-04 | -6.92 | positive regulation of protein secretion | biological process | GO:0050714 | 230 | 11 | 13711 | 214 | Mmp8,Cd14,Il17ra,Lgals9,Hcar2,Il1a,Akap12,Il4ra,Tnf,Casp4,Il6 | | 9.860e-04 | -6.92 | negative regulation of gene expression | biological process | GO:0010629 | 1352 | 36 | 13711 | 214 | Dnajb1,Rrp8,Noct,Bcl3,Mmp8,Rasd1,Zfp36,Atf3,Xdh,Znfx1,Tnf,Ifi204,Zbtb16,Ifi207,Arid5b,Sbno2,Parp9,Nfkbie,Nfkb1,Birc3,Gbp4,Ifi209,Tiparp,Runx1,Spi1,Cebpd,Bach1,Ifi211,Cdkn1a,Sox11,Mxd4,Lgals9,Msr1,Ctla2a,Parp14,Thbs1 | | 9.884e-04 | -6.92 | LU\_IL4\_SIGNALING | MSigDB lists | LU\_IL4\_SIGNALING | 78 | 6 | 12187 | 179 | Tnfaip3,Il6,Cflar,Il4ra,Ccl4,Nfkbiz | | 1.007e-03 | -6.90 | MATTHEWS\_AP1\_TARGETS | MSigDB lists | MATTHEWS\_AP1\_TARGETS | 14 | 3 | 12187 | 179 | Ptgs2,Bcl3,Plaur | | 1.007e-03 | -6.90 | MARKS\_HDAC\_TARGETS\_DN | MSigDB lists | MARKS\_HDAC\_TARGETS\_DN | 14 | 3 | 12187 | 179 | Il6,Tnf,Cflar | | 1.009e-03 | -6.90 | positive regulation of protein complex disassembly | biological process | GO:0043243 | 29 | 4 | 13711 | 214 | Irgm2,Plek,Irgm1,Tnf | | 1.013e-03 | -6.90 | METALLOTHIONEIN\_VRT | prosite domains | PS00203 | 3 | 2 | 8845 | 164 | Mt2,Mt1 | | 1.013e-03 | -6.90 | INTERLEUKIN\_1 | prosite domains | PS00253 | 3 | 2 | 8845 | 164 | Il1rn,Il1a | | 1.013e-03 | -6.90 | 25A\_SYNTH\_2 | prosite domains | PS00833 | 3 | 2 | 8845 | 164 | Oasl2,Oasl1 | | 1.017e-03 | -6.89 | GSE9037\_WT\_VS\_IRAK4\_KO\_LPS\_1H\_STIM\_BMDM\_DN | MSigDB lists | GSE9037\_WT\_VS\_IRAK4\_KO\_LPS\_1H\_STIM\_BMDM\_DN | 139 | 8 | 12187 | 179 | A2m,Slfn5,Sgk3,S100a8,Rtp4,Parp14,Ifi44,Cdkn1a | | 1.019e-03 | -6.89 | negative regulation of hydrolase activity | biological process | GO:0051346 | 305 | 13 | 13711 | 214 | Cflar,Tnfaip8,Thbs1,Il6,Tnf,Plaur,Wfdc21,Ptgs2,Angptl4,Serpina3f,A2m,Birc3,Timp1 | | 1.019e-03 | -6.89 | GO\_RESPONSE\_TO\_RADIATION | MSigDB lists | GO\_RESPONSE\_TO\_RADIATION | 325 | 13 | 12187 | 179 | Ptgs2,Icam1,Sgk1,Ifi209,Cdkn1a,Vcam1,Socs3,Ifi207,Ifi204,Bcl3,Ifi211,Pik3r1,Ccl12 | | 1.020e-03 | -6.89 | RNA polymerase II regulatory region sequence-specific DNA binding | molecular function | GO:0000977 | 594 | 20 | 13516 | 211 | Bach1,Irf2,Irf9,Runx1,Ifi211,Irf7,Spi1,Ifi207,Maff,Nfkb1,Zbtb16,Atf3,Ifi209,Nfkb2,Mxd4,Arid5b,Cebpd,Nfe2l2,Sox11,Ifi204 | | 1.025e-03 | -6.88 | GSE34515\_CD16\_POS\_MONOCYTE\_VS\_DC\_DN | MSigDB lists | GSE34515\_CD16\_POS\_MONOCYTE\_VS\_DC\_DN | 173 | 9 | 12187 | 179 | C5ar1,Ifitm3,Dtx3l,Pik3ap1,Parp14,Ifit2,Map3k8,Cybb,Ms4a6d | | 1.032e-03 | -6.88 | positive regulation of interleukin-6 production | biological process | GO:0032755 | 74 | 6 | 13711 | 214 | Mmp8,Il1a,Il1rn,Tnf,Lgals9,Il6 | | 1.032e-03 | -6.88 | lymphocyte activation involved in immune response | biological process | GO:0002285 | 74 | 6 | 13711 | 214 | Bcl3,Gadd45g,Stx11,Il6,Icam1,Tnfaip3 | | 1.039e-03 | -6.87 | ZUCCHI\_METASTASIS\_DN | MSigDB lists | ZUCCHI\_METASTASIS\_DN | 31 | 4 | 12187 | 179 | Cxcl1,Nfkbia,Rasd1,Gadd45b | | 1.039e-03 | -6.87 | ONDER\_CDH1\_TARGETS\_3\_DN | MSigDB lists | ONDER\_CDH1\_TARGETS\_3\_DN | 31 | 4 | 12187 | 179 | Lcn2,Ptgs2,Csf3,Cxcl1 | | 1.039e-03 | -6.87 | GO\_TUMOR\_NECROSIS\_FACTOR\_RECEPTOR\_SUPERFAMILY\_BINDING | MSigDB lists | GO\_TUMOR\_NECROSIS\_FACTOR\_RECEPTOR\_SUPERFAMILY\_BINDING | 31 | 4 | 12187 | 179 | Tnfsf10,Ripk1,Cflar,Tnf | | 1.039e-03 | -6.87 | OHGUCHI\_LIVER\_HNF4A\_TARGETS\_UP | MSigDB lists | OHGUCHI\_LIVER\_HNF4A\_TARGETS\_UP | 31 | 4 | 12187 | 179 | Gadd45b,Il1rn,Usp18,Ifi44 | | 1.039e-03 | -6.87 | GO\_ER\_NUCLEUS\_SIGNALING\_PATHWAY | MSigDB lists | GO\_ER\_NUCLEUS\_SIGNALING\_PATHWAY | 31 | 4 | 12187 | 179 | Nfe2l2,Atf3,Ccl12,Hspa5 | | 1.039e-03 | -6.87 | HAN\_JNK\_SINGALING\_DN | MSigDB lists | HAN\_JNK\_SINGALING\_DN | 31 | 4 | 12187 | 179 | Cd14,Socs3,Cp,Lcn2 | | 1.039e-03 | -6.87 | GO\_REGULATION\_OF\_NF\_KAPPAB\_IMPORT\_INTO\_NUCLEUS | MSigDB lists | GO\_REGULATION\_OF\_NF\_KAPPAB\_IMPORT\_INTO\_NUCLEUS | 31 | 4 | 12187 | 179 | Tnf,Ptgs2,Nfkbia,Bcl3 | | 1.051e-03 | -6.86 | GSE40274\_CTRL\_VS\_EOS\_TRANSDUCED\_ACTIVATED\_CD4\_TCELL\_DN | MSigDB lists | GSE40274\_CTRL\_VS\_EOS\_TRANSDUCED\_ACTIVATED\_CD4\_TCELL\_DN | 108 | 7 | 12187 | 179 | Tnf,Nfkbia,Rtp4,Rnf213,Sgk3,Banp,Ncf1 | | 1.051e-03 | -6.86 | PKCA\_DN.V1\_DN | MSigDB lists | PKCA\_DN.V1\_DN | 108 | 7 | 12187 | 179 | Rhoj,Saa1,Map3k6,Sntb2,Nfkbiz,Trib1,Cp | | 1.051e-03 | -6.86 | GSE17974\_IL4\_AND\_ANTI\_IL12\_VS\_UNTREATED\_12H\_ACT\_CD4\_TCELL\_DN | MSigDB lists | GSE17974\_IL4\_AND\_ANTI\_IL12\_VS\_UNTREATED\_12H\_ACT\_CD4\_TCELL\_DN | 108 | 7 | 12187 | 179 | Sgk1,Rhou,Oasl1,Ccl12,Pygm,Rgs16,Gbp3 | | 1.058e-03 | -6.85 | BRLZ | smart domains | SM00338 | 46 | 5 | 7188 | 123 | Nfe2l2,Bach1,Atf3,Cebpd,Maff | | 1.058e-03 | -6.85 | positive regulation of chemokine production | biological process | GO:0032722 | 50 | 5 | 13711 | 214 | Lgals9,Il6,Il1a,Tnf,Il4ra | | 1.066e-03 | -6.84 | GSE22432\_MULTIPOTENT\_PROGENITOR\_VS\_CDC\_UP | MSigDB lists | GSE22432\_MULTIPOTENT\_PROGENITOR\_VS\_CDC\_UP | 140 | 8 | 12187 | 179 | Gem,Gbp7,Arid5b,Slfn5,Slfn9,Tap1,Irf9,Gbp3 | | 1.066e-03 | -6.84 | VECCHI\_GASTRIC\_CANCER\_ADVANCED\_VS\_EARLY\_UP | MSigDB lists | VECCHI\_GASTRIC\_CANCER\_ADVANCED\_VS\_EARLY\_UP | 140 | 8 | 12187 | 179 | Pik3r5,Rnf213,C3ar1,Il6,Col4a1,Fstl1,Msr1,Thbs1 | | 1.066e-03 | -6.84 | GSE46606\_UNSTIM\_VS\_CD40L\_IL2\_IL5\_3DAY\_STIMULATED\_IRF4\_KO\_BCELL\_DN | MSigDB lists | GSE46606\_UNSTIM\_VS\_CD40L\_IL2\_IL5\_3DAY\_STIMULATED\_IRF4\_KO\_BCELL\_DN | 140 | 8 | 12187 | 179 | Irf9,Irf2,Dtx3l,Herc6,Parp12,Znfx1,Ifit3b,Msr1 | | 1.066e-03 | -6.84 | ELVIDGE\_HYPOXIA\_UP | MSigDB lists | ELVIDGE\_HYPOXIA\_UP | 140 | 8 | 12187 | 179 | Tiparp,Akap12,Atf3,Gadd45b,Maff,Tnfaip8,Plaur,Angptl4 | | 1.066e-03 | -6.84 | GSE11864\_CSF1\_PAM3CYS\_VS\_CSF1\_IFNG\_PAM3CYS\_IN\_MAC\_DN | MSigDB lists | GSE11864\_CSF1\_PAM3CYS\_VS\_CSF1\_IFNG\_PAM3CYS\_IN\_MAC\_DN | 140 | 8 | 12187 | 179 | Nfkbie,Dtx3l,Csf2rb,Parp9,Gbp3,Maff,Sntb2,Oasl1 | | 1.066e-03 | -6.84 | GSE6269\_E\_COLI\_VS\_STAPH\_AUREUS\_INF\_PBMC\_DN | MSigDB lists | GSE6269\_E\_COLI\_VS\_STAPH\_AUREUS\_INF\_PBMC\_DN | 140 | 8 | 12187 | 179 | Icam1,Cd14,S100a9,Cxcl1,Mmp8,Ier3,Hspa5,Tor3a | | 1.067e-03 | -6.84 | GSE15930\_STIM\_VS\_STIM\_AND\_IFNAB\_72H\_CD8\_T\_CELL\_DN | MSigDB lists | GSE15930\_STIM\_VS\_STIM\_AND\_IFNAB\_72H\_CD8\_T\_CELL\_DN | 174 | 9 | 12187 | 179 | Hspa5,Adamts1,Dnajb1,Il1rn,Saa1,Isg15,Irf7,Tagln2,Ccdc58 | | 1.093e-03 | -6.82 | PI3K-Akt signaling pathway | KEGG pathways | ko04151 | 271 | 14 | 5248 | 107 | Nfkb1,Pik3ap1,Il6,Sgk1,Csf3,Cdkn1a,Pik3r1,Ccnd2,Col4a1,Thbs1,Sgk3,Il4ra,Osmr,Pik3r5 | | 1.093e-03 | -6.82 | PI3K-Akt signaling pathway | KEGG pathways | mmu04151 | 271 | 14 | 5248 | 107 | Cdkn1a,Csf3,Sgk1,Il6,Pik3ap1,Nfkb1,Pik3r5,Osmr,Il4ra,Sgk3,Thbs1,Col4a1,Ccnd2,Pik3r1 | | 1.105e-03 | -6.81 | GO\_POSITIVE\_REGULATION\_OF\_TRANSFERASE\_ACTIVITY | MSigDB lists | GO\_POSITIVE\_REGULATION\_OF\_TRANSFERASE\_ACTIVITY | 500 | 17 | 12187 | 179 | C5ar1,Pik3r5,Fzd4,Iqgap1,Map3k8,Ccnd2,Tnf,Sdc4,Ripk1,Gadd45b,Map3k6,Saa1,Ptpn1,Cdkn1a,Thbs1,Hspa5,Gadd45g | | 1.108e-03 | -6.81 | negative regulation of myeloid cell differentiation | biological process | GO:0045638 | 75 | 6 | 13711 | 214 | Trib1,Zbtb16,Runx1,Pik3r1,Nfkbia,Zfp36 | | 1.110e-03 | -6.80 | GO\_POSITIVE\_REGULATION\_OF\_ERK1\_AND\_ERK2\_CASCADE | MSigDB lists | GO\_POSITIVE\_REGULATION\_OF\_ERK1\_AND\_ERK2\_CASCADE | 109 | 7 | 12187 | 179 | Icam1,C5ar1,Il6,Ccl12,Tnf,Ccl2,Ccl4 | | 1.116e-03 | -6.80 | GSE19198\_6H\_VS\_24H\_IL21\_TREATED\_TCELL\_UP | MSigDB lists | GSE19198\_6H\_VS\_24H\_IL21\_TREATED\_TCELL\_UP | 141 | 8 | 12187 | 179 | Atf3,Tubb6,Tor3a,Birc3,Plaur,Plek,Banp,Nfkbie | | 1.116e-03 | -6.80 | GO\_RESPONSE\_TO\_TOPOLOGICALLY\_INCORRECT\_PROTEIN | MSigDB lists | GO\_RESPONSE\_TO\_TOPOLOGICALLY\_INCORRECT\_PROTEIN | 141 | 8 | 12187 | 179 | Ptpn1,Hspb1,Dnajb1,Hspa5,Thbs1,Ccl12,Nfe2l2,Atf3 | | 1.116e-03 | -6.80 | GSE2585\_THYMIC\_DC\_VS\_MTEC\_DN | MSigDB lists | GSE2585\_THYMIC\_DC\_VS\_MTEC\_DN | 141 | 8 | 12187 | 179 | Thbs1,Ifit3b,Ccl2,Irak3,Clic4,Tnfsf10,Ifitm3,Isg15 | | 1.116e-03 | -6.80 | GSE16450\_CTRL\_VS\_IFNA\_6H\_STIM\_MATURE\_NEURON\_CELL\_LINE\_UP | MSigDB lists | GSE16450\_CTRL\_VS\_IFNA\_6H\_STIM\_MATURE\_NEURON\_CELL\_LINE\_UP | 141 | 8 | 12187 | 179 | Rtp4,Gbp7,Dtx3l,Selp,Cybb,Pik3r1,Sgk1,Socs3 | | 1.116e-03 | -6.80 | GSE43863\_DAY6\_EFF\_VS\_DAY150\_MEM\_TFH\_CD4\_TCELL\_DN | MSigDB lists | GSE43863\_DAY6\_EFF\_VS\_DAY150\_MEM\_TFH\_CD4\_TCELL\_DN | 141 | 8 | 12187 | 179 | Cflar,Ifitm2,Nfkb1,Ksr1,Kcna5,Il6,Gadd45b,Socs3 | | 1.116e-03 | -6.80 | ATF2\_S\_UP.V1\_DN | MSigDB lists | ATF2\_S\_UP.V1\_DN | 141 | 8 | 12187 | 179 | Ptgs2,Angptl4,A2m,Osmr,Sbno2,Il4ra,Vcam1,Ccl12 | | 1.129e-03 | -6.79 | ZAMORA\_NOS2\_TARGETS\_DN | MSigDB lists | ZAMORA\_NOS2\_TARGETS\_DN | 80 | 6 | 12187 | 179 | Ripk1,Hspb1,Hp,Cp,Fzd4,Ifitm2 | | 1.138e-03 | -6.78 | positive regulation of cell population proliferation | biological process | GO:0008284 | 734 | 23 | 13711 | 214 | Thbs1,Ccnd2,P2ry6,Sox11,Il6,Cdkn1a,Rasip1,Kcna5,Ptgs2,Timp1,Osmr,Adamts1,C3ar1,Csf3,Cflar,Tnf,Tnfaip3,Atf3,Ccl2,Vcam1,C5ar1,Cxcl1,Tgm2 | | 1.142e-03 | -6.78 | intrinsic apoptotic signaling pathway | biological process | GO:0097193 | 133 | 8 | 13711 | 214 | Cdkn1a,Tnf,Rrp8,Casp4,Ifi204,Ier3,Pik3r1,Bcl3 | | 1.142e-03 | -6.77 | SAMOLS\_TARGETS\_OF\_KHSV\_MIRNAS\_DN | MSigDB lists | SAMOLS\_TARGETS\_OF\_KHSV\_MIRNAS\_DN | 54 | 5 | 12187 | 179 | Ifi211,Ifi209,Thbs1,Ifi204,Ifi207 | | 1.142e-03 | -6.77 | GO\_CELLULAR\_RESPONSE\_TO\_IONIZING\_RADIATION | MSigDB lists | GO\_CELLULAR\_RESPONSE\_TO\_IONIZING\_RADIATION | 54 | 5 | 12187 | 179 | Ifi207,Ifi204,Ifi211,Cdkn1a,Ifi209 | | 1.142e-03 | -6.77 | AMIT\_EGF\_RESPONSE\_120\_HELA | MSigDB lists | AMIT\_EGF\_RESPONSE\_120\_HELA | 54 | 5 | 12187 | 179 | Cxcl1,Cdkn1a,Gem,Maff,Plaur | | 1.157e-03 | -6.76 | GO\_REGULATION\_OF\_PROTEIN\_LOCALIZATION\_TO\_NUCLEUS | MSigDB lists | GO\_REGULATION\_OF\_PROTEIN\_LOCALIZATION\_TO\_NUCLEUS | 176 | 9 | 12187 | 179 | Csf3,Ptgs2,Nfkbie,Bcl3,Pik3r1,Cdkn1a,Tnf,Nfkbia,Il6 | | 1.157e-03 | -6.76 | GSE7852\_TREG\_VS\_TCONV\_FAT\_UP | MSigDB lists | GSE7852\_TREG\_VS\_TCONV\_FAT\_UP | 176 | 9 | 12187 | 179 | Nfkbiz,Cdkn1a,Atf3,Stx11,Bcl3,Gem,Nfkbie,Rgs16,Zfp36 | | 1.160e-03 | -6.76 | mouse chr1\|1 H3 | chromosome location | mouse chr1\|1 H3 | 31 | 4 | 14556 | 220 | Olfml2b,Ifi209,Ifi207,Ifi211 | | 1.161e-03 | -6.76 | extracellular organelle | cellular component | GO:0043230 | 77 | 6 | 13825 | 212 | Irgm2,Gbp6,Icam1,Iigp1,Gbp3,Gbp7 | | 1.171e-03 | -6.75 | MYC\_UP.V1\_DN | MSigDB lists | MYC\_UP.V1\_DN | 110 | 7 | 12187 | 179 | S100a8,Map3k6,Angptl4,Trim56,S100a9,Il6,Tgm2 | | 1.173e-03 | -6.75 | PID\_INTEGRIN3\_PATHWAY | MSigDB lists | PID\_INTEGRIN3\_PATHWAY | 32 | 4 | 12187 | 179 | Thbs1,Sdc4,Col4a1,Plaur | | 1.173e-03 | -6.75 | ST\_GA13\_PATHWAY | MSigDB lists | ST\_GA13\_PATHWAY | 32 | 4 | 12187 | 179 | Nfkb2,Nfkb1,Nfkbia,Nfkbie | | 1.187e-03 | -6.74 | positive regulation of cellular biosynthetic process | biological process | GO:0031328 | 1571 | 40 | 13711 | 214 | Thbs1,Dtx3l,Irf7,P2ry6,Banp,Rbm39,Pik3r1,Nfkbia,Nfkb2,Sox11,Il1a,Akap12,Icam1,Cybb,Il6,Spi1,Bach1,Cebpd,Runx1,Ptgs2,Fzd4,Parp9,Sbno2,Nfkb1,Ripk1,Csf3,Zbtb16,Aff1,Ncf1,C3ar1,Arid5b,Hspb1,Tnip1,Tnf,Atf3,Mmp8,Nfe2l2,Bcl3,Maff,Irf2 | | 1.194e-03 | -6.73 | MyD88-independent TLR3/TLR4 cascade | REACTOME pathways | R-MMU-166166 | 72 | 6 | 6297 | 105 | Irf7,Nfkb1,Cd14,Map3k8,Nfkbia,Nfkb2 | | 1.194e-03 | -6.73 | Toll Like Receptor 3 (TLR3) Cascade | REACTOME pathways | R-MMU-168164 | 72 | 6 | 6297 | 105 | Irf7,Nfkb1,Nfkb2,Cd14,Map3k8,Nfkbia | | 1.200e-03 | -6.73 | RNA polymerase II regulatory region DNA binding | molecular function | GO:0001012 | 602 | 20 | 13516 | 211 | Irf2,Irf9,Bach1,Runx1,Ifi211,Maff,Spi1,Ifi207,Irf7,Nfkb1,Zbtb16,Atf3,Nfkb2,Ifi209,Mxd4,Arid5b,Cebpd,Nfe2l2,Ifi204,Sox11 | | 1.202e-03 | -6.72 | protein ADP-ribosylase activity | molecular function | GO:1990404 | 14 | 3 | 13516 | 211 | Parp12,Tiparp,Parp14 | | 1.203e-03 | -6.72 | GSE339\_CD4POS\_VS\_CD8POS\_DC\_UP | MSigDB lists | GSE339\_CD4POS\_VS\_CD8POS\_DC\_UP | 177 | 9 | 12187 | 179 | Ccl4,Tgm2,Usp18,Nfkb1,Banp,Bcl3,Ptgs2,Ifitm2,Saa1 | | 1.203e-03 | -6.72 | GSE26343\_UNSTIM\_VS\_LPS\_STIM\_NFAT5\_KO\_MACROPHAGE\_UP | MSigDB lists | GSE26343\_UNSTIM\_VS\_LPS\_STIM\_NFAT5\_KO\_MACROPHAGE\_UP | 177 | 9 | 12187 | 179 | Plek,Rrp8,Bcl3,Cxcl1,Nfkbie,Gem,Cflar,Il6,Casp4 | | 1.212e-03 | -6.72 | positive regulation of protein transport | biological process | GO:0051222 | 350 | 14 | 13711 | 214 | Pik3r1,Hcar2,Lgals9,Cd14,Il6,Ccl2,Il4ra,Tnf,Casp4,Il1a,Akap12,Ptgs2,Il17ra,Mmp8 | | 1.217e-03 | -6.71 | WWE | prosite domains | PS50918 | 12 | 3 | 8845 | 164 | Tiparp,Parp14,Parp12 | | 1.234e-03 | -6.70 | GO\_ACTIVATION\_OF\_MAPK\_ACTIVITY | MSigDB lists | GO\_ACTIVATION\_OF\_MAPK\_ACTIVITY | 111 | 7 | 12187 | 179 | Ptpn1,C5ar1,Saa1,Map3k6,Ripk1,Tnf,Thbs1 | | 1.234e-03 | -6.70 | GSE40666\_UNTREATED\_VS\_IFNA\_STIM\_STAT1\_KO\_CD8\_TCELL\_90MIN\_UP | MSigDB lists | GSE40666\_UNTREATED\_VS\_IFNA\_STIM\_STAT1\_KO\_CD8\_TCELL\_90MIN\_UP | 111 | 7 | 12187 | 179 | Dtx3l,Rnf213,Parp14,Tor3a,Irf7,Parp9,Irf9 | | 1.236e-03 | -6.70 | positive regulation of DNA-binding transcription factor activity | biological process | GO:0051091 | 201 | 10 | 13711 | 214 | Irak3,Arid5b,Csf3,Ripk1,Cflar,Trim25,Icam1,Tnf,Fzd4,Il6 | | 1.237e-03 | -6.70 | WANG\_MLL\_TARGETS | MSigDB lists | WANG\_MLL\_TARGETS | 214 | 10 | 12187 | 179 | Plek,Rgs16,Cd14,Arid5b,Ptges,Cxcl1,Hcar2,Il1rn,Il6,Parp14 | | 1.241e-03 | -6.69 | VANHARANTA\_UTERINE\_FIBROID\_DN | MSigDB lists | VANHARANTA\_UTERINE\_FIBROID\_DN | 55 | 5 | 12187 | 179 | Gadd45b,Atf3,Trib1,Birc3,Maff | | 1.241e-03 | -6.69 | KEGG\_P53\_SIGNALING\_PATHWAY | MSigDB lists | KEGG\_P53\_SIGNALING\_PATHWAY | 55 | 5 | 12187 | 179 | Gadd45g,Gadd45b,Cdkn1a,Ccnd2,Thbs1 | | 1.245e-03 | -6.69 | CHEOK\_RESPONSE\_TO\_HD\_MTX\_UP | MSigDB lists | CHEOK\_RESPONSE\_TO\_HD\_MTX\_UP | 15 | 3 | 12187 | 179 | S100a9,Nfkb2,S100a8 | | 1.245e-03 | -6.69 | GO\_CCR\_CHEMOKINE\_RECEPTOR\_BINDING | MSigDB lists | GO\_CCR\_CHEMOKINE\_RECEPTOR\_BINDING | 15 | 3 | 12187 | 179 | Ccl12,Ccl4,Ccl2 | | 1.245e-03 | -6.69 | GO\_NEGATIVE\_REGULATION\_OF\_LIPID\_STORAGE | MSigDB lists | GO\_NEGATIVE\_REGULATION\_OF\_LIPID\_STORAGE | 15 | 3 | 12187 | 179 | Il6,Nfkbia,Tnf | | 1.245e-03 | -6.69 | BIOCARTA\_IL10\_PATHWAY | MSigDB lists | BIOCARTA\_IL10\_PATHWAY | 15 | 3 | 12187 | 179 | Tnf,Il1a,Il6 | | 1.245e-03 | -6.69 | BIOCARTA\_CDMAC\_PATHWAY | MSigDB lists | BIOCARTA\_CDMAC\_PATHWAY | 15 | 3 | 12187 | 179 | Nfkb1,Nfkbia,Tnf | | 1.252e-03 | -6.68 | GSE11057\_NAIVE\_VS\_CENT\_MEMORY\_CD4\_TCELL\_DN | MSigDB lists | GSE11057\_NAIVE\_VS\_CENT\_MEMORY\_CD4\_TCELL\_DN | 178 | 9 | 12187 | 179 | Iqgap1,Tagln2,Ifi211,Arid5b,Ifi204,Ifi207,Ifi209,Rhou,Trib1 | | 1.252e-03 | -6.68 | GO\_REGULATION\_OF\_NUCLEOCYTOPLASMIC\_TRANSPORT | MSigDB lists | GO\_REGULATION\_OF\_NUCLEOCYTOPLASMIC\_TRANSPORT | 178 | 9 | 12187 | 179 | Bcl3,Nfkbie,Ptgs2,Csf3,Il6,Nfkbia,Pik3r1,Cdkn1a,Tnf | | 1.252e-03 | -6.68 | regulation of lymphocyte activation | biological process | GO:0051249 | 312 | 13 | 13711 | 214 | Vcam1,Ctla2a,Zbtb16,Runx1,Lgals9,Nfkbiz,Il4ra,Tnfaip3,Sox11,Il6,Cdkn1a,Ccl2,Sdc4 | | 1.257e-03 | -6.68 | regulation of cell shape | biological process | GO:0008360 | 135 | 8 | 13711 | 214 | Rhoc,Msn,Icam1,Ccl12,Ccl2,Rhou,Rhoj,Il6 | | 1.262e-03 | -6.67 | LUDWICZEK\_TREATING\_IRON\_OVERLOAD | MSigDB lists | LUDWICZEK\_TREATING\_IRON\_OVERLOAD | 4 | 2 | 12187 | 179 | Cp,Lcn2 | | 1.262e-03 | -6.67 | SMIRNOV\_CIRCULATING\_ENDOTHELIOCYTES\_IN\_CANCER\_DN | MSigDB lists | SMIRNOV\_CIRCULATING\_ENDOTHELIOCYTES\_IN\_CANCER\_DN | 4 | 2 | 12187 | 179 | Angptl4,Sele | | 1.266e-03 | -6.67 | regulation of nitric oxide biosynthetic process | biological process | GO:0045428 | 52 | 5 | 13711 | 214 | Mmp8,Tnf,Icam1,Il6,Ptgs2 | | 1.266e-03 | -6.67 | positive regulation of blood vessel endothelial cell migration | biological process | GO:0043536 | 52 | 5 | 13711 | 214 | Hspb1,Thbs1,Nfe2l2,Ptgs2,Rhoj | | 1.266e-03 | -6.67 | positive regulation of alpha-beta T cell activation | biological process | GO:0046635 | 52 | 5 | 13711 | 214 | Runx1,Il6,Zbtb16,Il4ra,Nfkbiz | | 1.270e-03 | -6.67 | cell activation | biological process | GO:0001775 | 433 | 16 | 13711 | 214 | Gadd45g,Pik3r1,Tnfaip3,Tnf,Icam1,Fyb,Plek,Nfkb2,Il6,Stx11,Spi1,Bcl3,Msn,Il17ra,Timp1,Sbno2 | | 1.274e-03 | -6.67 | BAELDE\_DIABETIC\_NEPHROPATHY\_DN | MSigDB lists | BAELDE\_DIABETIC\_NEPHROPATHY\_DN | 375 | 14 | 12187 | 179 | Tap1,Isg15,Timp1,Cebpd,Luc7l3,Ifi44,Hspa5,Osmr,Cflar,Arid5b,Cxcl1,Birc3,Maff,Sdc4 | | 1.277e-03 | -6.66 | regulation of transport | biological process | GO:0051049 | 1577 | 40 | 13711 | 214 | Nfe2l2,S100a9,Mmp8,Msn,Cxcl1,Ptpn1,Clic4,S100a8,Ccl12,Il1rn,Sele,Ripk1,Cd14,Tnf,Tnfaip3,Ccl2,Kcna5,Ccl4,Runx1,Sgk1,Ptgs2,Il17ra,Ier3,Gbp4,Sdc4,Thbs1,Pik3r1,Hcar2,P2ry6,Lgals9,Il4ra,Icam1,Casp4,Nfkbia,Akap12,Sox11,Gem,Il1a,Cybb,Il6 | | 1.278e-03 | -6.66 | GO\_POSITIVE\_REGULATION\_OF\_PROTEIN\_SECRETION | MSigDB lists | GO\_POSITIVE\_REGULATION\_OF\_PROTEIN\_SECRETION | 144 | 8 | 12187 | 179 | Il4ra,Tnf,Il6,Il1a,Hcar2,Cd14,Saa1,Il17ra | | 1.278e-03 | -6.66 | GSE9650\_EXHAUSTED\_VS\_MEMORY\_CD8\_TCELL\_UP | MSigDB lists | GSE9650\_EXHAUSTED\_VS\_MEMORY\_CD8\_TCELL\_UP | 144 | 8 | 12187 | 179 | Fstl1,Il1a,Il1rn,Bcl3,Selp,Rgs16,Clic4,Osmr | | 1.278e-03 | -6.66 | GSE3039\_CD4\_TCELL\_VS\_ALPHABETA\_CD8\_TCELL\_UP | MSigDB lists | GSE3039\_CD4\_TCELL\_VS\_ALPHABETA\_CD8\_TCELL\_UP | 144 | 8 | 12187 | 179 | Trib1,Fstl1,Hp,Ms4a6d,Mmp8,Cybb,Ptgs2,Ncf1 | | 1.292e-03 | -6.65 | - | gene3d domains | 1.10.1410.20 | 4 | 2 | 6647 | 99 | Oasl1,Oasl2 | | 1.301e-03 | -6.64 | GSE26488\_WT\_VS\_HDAC7\_KO\_DOUBLE\_POSITIVE\_THYMOCYTE\_DN | MSigDB lists | GSE26488\_WT\_VS\_HDAC7\_KO\_DOUBLE\_POSITIVE\_THYMOCYTE\_DN | 112 | 7 | 12187 | 179 | Gbp4,Oasl1,Tnip3,Ifit2,Cxcl1,Gbp6,Ptgs2 | | 1.301e-03 | -6.64 | GSE6092\_CTRL\_VS\_BORRELIA\_BIRGDOFERI\_INF\_ENDOTHELIAL\_CELL\_DN | MSigDB lists | GSE6092\_CTRL\_VS\_BORRELIA\_BIRGDOFERI\_INF\_ENDOTHELIAL\_CELL\_DN | 112 | 7 | 12187 | 179 | Plaur,Nfkbie,Isg15,Tnfaip3,Gadd45b,Atf3,Nfkbia | | 1.304e-03 | -6.64 | lipopolysaccharide-mediated signaling pathway | biological process | GO:0031663 | 31 | 4 | 13711 | 214 | Ccl12,Cd14,Nfkbia,Tnf | | 1.304e-03 | -6.64 | regulation of macrophage migration | biological process | GO:1905521 | 31 | 4 | 13711 | 214 | C5ar1,Thbs1,C3ar1,Ccl2 | | 1.305e-03 | -6.64 | biosynthesis of prostaglandins | BIOCYC pathways | MOUSE\_PWY3DJ-35583 | 10 | 2 | 823 | 5 | Ptgs2,Ptges | | 1.308e-03 | -6.64 | GO\_REGULATION\_OF\_CELL\_ADHESION | MSigDB lists | GO\_REGULATION\_OF\_CELL\_ADHESION | 463 | 16 | 12187 | 179 | Ccl12,Il6,Tnf,Map3k8,Il4ra,Pik3r1,Tgm2,Sdc4,Fzd4,Il1rn,Thbs1,Zbtb16,Vcam1,Icam1,Selp,Saa1 | | 1.320e-03 | -6.63 | CROONQUIST\_NRAS\_SIGNALING\_UP | MSigDB lists | CROONQUIST\_NRAS\_SIGNALING\_UP | 33 | 4 | 12187 | 179 | Msr1,Ccl4,Ncf1,Cybb | | 1.320e-03 | -6.63 | GO\_PLATELET\_DERIVED\_GROWTH\_FACTOR\_RECEPTOR\_SIGNALING\_PATHWAY | MSigDB lists | GO\_PLATELET\_DERIVED\_GROWTH\_FACTOR\_RECEPTOR\_SIGNALING\_PATHWAY | 33 | 4 | 12187 | 179 | Ptpn1,Iqgap1,Tiparp,Arid5b | | 1.326e-03 | -6.63 | Signal Transduction | REACTOME pathways | R-MMU-162582 | 1547 | 40 | 6297 | 105 | Ripk1,Cxcl9,Tnfaip3,C5ar1,Rhou,Fstl1,Tnfsf10,Rhoc,Dnajb1,Sdc4,P2ry6,Csf2rb,Ksr1,Cxcl1,Iqgap1,Tnf,Nfkbia,Ncf1,Col4a1,Thbs1,Il6,Cybb,Rhoj,Cdkn1a,Cxcl16,Hcar2,A2m,Socs3,C3ar1,Ptpn1,Hspb1,Nfkb1,Ier3,Cflar,Fzd4,Gpr84,Pik3ap1,Birc3,Pik3r1,Pik3r5 | | 1.336e-03 | -6.62 | GSE28726\_NAIVE\_VS\_ACTIVATED\_NKTCELL\_UP | MSigDB lists | GSE28726\_NAIVE\_VS\_ACTIVATED\_NKTCELL\_UP | 145 | 8 | 12187 | 179 | Cybb,Stx11,Sgk1,Dnajb1,Irak3,Pygm,Saa1,Rrp8 | | 1.336e-03 | -6.62 | GSE41176\_UNSTIM\_VS\_ANTI\_IGM\_STIM\_TAK1\_KO\_BCELL\_24H\_DN | MSigDB lists | GSE41176\_UNSTIM\_VS\_ANTI\_IGM\_STIM\_TAK1\_KO\_BCELL\_24H\_DN | 145 | 8 | 12187 | 179 | S100a9,Ptges,Csf2rb,S100a8,P2ry6,Nfkbia,Ccl2,Tnfaip3 | | 1.337e-03 | -6.62 | GO\_REGULATION\_OF\_INTRACELLULAR\_PROTEIN\_TRANSPORT | MSigDB lists | GO\_REGULATION\_OF\_INTRACELLULAR\_PROTEIN\_TRANSPORT | 294 | 12 | 12187 | 179 | Pik3r1,Tnf,Rhou,Nfkbia,Il6,Ccl12,Csf3,Nfkbie,Bcl3,Cdkn1a,Ptgs2,Ptpn1 | | 1.340e-03 | -6.61 | GO\_DOUBLE\_STRANDED\_DNA\_BINDING | MSigDB lists | GO\_DOUBLE\_STRANDED\_DNA\_BINDING | 509 | 17 | 12187 | 179 | Runx1,Spi1,Bach1,Nfe2l2,Arid5b,Irf7,Ifi211,Nfkb2,Ifi209,Sox11,Zbtb16,Ifi207,Ifi204,Nfkb1,Cebpd,Atf3,Irf2 | | 1.347e-03 | -6.61 | MODULE\_375 | MSigDB lists | MODULE\_375 | 56 | 5 | 12187 | 179 | Rgs16,Akap12,Cxcl1,C5ar1,C3ar1 | | 1.347e-03 | -6.61 | PID\_MYC\_REPRESS\_PATHWAY | MSigDB lists | PID\_MYC\_REPRESS\_PATHWAY | 56 | 5 | 12187 | 179 | Cebpd,Mxd4,Cflar,Cdkn1a,Spi1 | | 1.347e-03 | -6.61 | GO\_POSITIVE\_REGULATION\_OF\_PROTEIN\_KINASE\_B\_SIGNALING | MSigDB lists | GO\_POSITIVE\_REGULATION\_OF\_PROTEIN\_KINASE\_B\_SIGNALING | 56 | 5 | 12187 | 179 | Csf3,Thbs1,Tnf,Pik3r5,Il6 | | 1.355e-03 | -6.60 | mouse chr18\|18 D3 | chromosome location | mouse chr18\|18 D3 | 15 | 3 | 14556 | 220 | F830016B08Rik,Iigp1,Smim3 | | 1.360e-03 | -6.60 | positive regulation of myeloid cell differentiation | biological process | GO:0045639 | 78 | 6 | 13711 | 214 | Ripk1,Csf3,Trib1,Tnf,Isg15,Runx1 | | 1.360e-03 | -6.60 | regulation of alpha-beta T cell activation | biological process | GO:0046634 | 78 | 6 | 13711 | 214 | Il4ra,Nfkbiz,Zbtb16,Runx1,Il6,Lgals9 | | 1.363e-03 | -6.60 | hemopoiesis | biological process | GO:0030097 | 478 | 17 | 13711 | 214 | Tiparp,Sbno2,Prg4,Rbm47,Spi1,Herc6,Cebpd,Runx1,Bcl3,Samd9l,Tnfaip3,Tnf,Il6,Zbtb16,Csf3,Pik3r1,Gadd45g | | 1.368e-03 | -6.59 | MENSE\_HYPOXIA\_UP | MSigDB lists | MENSE\_HYPOXIA\_UP | 83 | 6 | 12187 | 179 | Maff,Angptl4,Tiparp,Osmr,Atf3,Cebpd | | 1.368e-03 | -6.59 | Cytosolic sensors of pathogen-associated DNA | REACTOME pathways | R-MMU-1834949 | 50 | 5 | 6297 | 105 | Trim56,Nfkbia,Nfkb2,Irf7,Nfkb1 | | 1.370e-03 | -6.59 | SELECTIN | prints domains | PR00343 | 3 | 2 | 2951 | 64 | Selp,Sele | | 1.370e-03 | -6.59 | MTVERTEBRATE | prints domains | PR00860 | 3 | 2 | 2951 | 64 | Mt2,Mt1 | | 1.370e-03 | -6.59 | INTERLEUKIN1 | prints domains | PR00264 | 3 | 2 | 2951 | 64 | Il1a,Il1rn | | 1.380e-03 | -6.59 | regulation of alpha-beta T cell differentiation | biological process | GO:0046637 | 53 | 5 | 13711 | 214 | Il4ra,Nfkbiz,Zbtb16,Runx1,Il6 | | 1.380e-03 | -6.59 | regulation of coagulation | biological process | GO:0050818 | 53 | 5 | 13711 | 214 | Plek,S100a9,Thbs1,Nfe2l2,Selp | | 1.381e-03 | -6.58 | positive regulation of T cell activation | biological process | GO:0050870 | 137 | 8 | 13711 | 214 | Zbtb16,Vcam1,Lgals9,Runx1,Nfkbiz,Il4ra,Ccl2,Il6 | | 1.383e-03 | -6.58 | 2-5-oligoAdlate\_synth\_1\_dom2/C | interpro domains | IPR018952 | 4 | 2 | 13788 | 212 | Oasl2,Oasl1 | | 1.383e-03 | -6.58 | 2-5-oligoAdlate\_synth\_2\_sf | interpro domains | IPR038121 | 4 | 2 | 13788 | 212 | Oasl2,Oasl1 | | 1.383e-03 | -6.58 | IL-1\_fam | interpro domains | IPR000975 | 4 | 2 | 13788 | 212 | Il1rn,Il1a | | 1.383e-03 | -6.58 | 2-5A\_synthase | interpro domains | IPR026774 | 4 | 2 | 13788 | 212 | Oasl2,Oasl1 | | 1.396e-03 | -6.57 | GSE22025\_UNTREATED\_VS\_PROGESTERONE\_TREATED\_CD4\_TCELL\_UP | MSigDB lists | GSE22025\_UNTREATED\_VS\_PROGESTERONE\_TREATED\_CD4\_TCELL\_UP | 146 | 8 | 12187 | 179 | Ifi207,Ifi204,Ccl12,Ifi209,Xaf1,Hspb1,Isg15,Ifi211 | | 1.396e-03 | -6.57 | GSE17721\_CTRL\_VS\_LPS\_4H\_BMDC\_DN | MSigDB lists | GSE17721\_CTRL\_VS\_LPS\_4H\_BMDC\_DN | 146 | 8 | 12187 | 179 | Iqgap1,Rbm47,Irgm2,Ier3,Nfkb1,Timp1,Parp14,Birc3 | | 1.396e-03 | -6.57 | GSE22935\_UNSTIM\_VS\_12H\_MBOVIS\_BCG\_STIM\_MYD88\_KO\_MACROPHAGE\_UP | MSigDB lists | GSE22935\_UNSTIM\_VS\_12H\_MBOVIS\_BCG\_STIM\_MYD88\_KO\_MACROPHAGE\_UP | 146 | 8 | 12187 | 179 | Ifit3b,Ier3,Parp12,Cp,Cd14,Saa1,Slfn4,Slfn3 | | 1.396e-03 | -6.57 | GSE13484\_12H\_UNSTIM\_VS\_YF17D\_VACCINE\_STIM\_PBMC\_DN | MSigDB lists | GSE13484\_12H\_UNSTIM\_VS\_YF17D\_VACCINE\_STIM\_PBMC\_DN | 146 | 8 | 12187 | 179 | Plaur,Tnfaip8,Irak3,Banp,Nfkbie,Gadd45b,Atf3,Il1rn | | 1.406e-03 | -6.57 | enzyme regulator activity | molecular function | GO:0030234 | 746 | 23 | 13516 | 211 | Hspb1,Cdkn1a,Lgals9,Iqgap1,Irgm2,Socs3,Tnfaip8,Pik3r1,Wfdc21,Birc3,Angptl4,Dtx3l,Pik3r5,Ncf1,Ccnd2,A2m,Serpina3f,Parp9,Cflar,Rgs16,Trib1,Timp1,Dnajb1 | | 1.423e-03 | -6.56 | TF\_DNA-bd\_sf | interpro domains | IPR008917 | 15 | 3 | 13788 | 212 | Bach1,Nfe2l2,Maff | | 1.424e-03 | -6.55 | IL1 | pfam domains | PF00340 | 4 | 2 | 12881 | 201 | Il1a,Il1rn | | 1.424e-03 | -6.55 | OAS1\_C | pfam domains | PF10421 | 4 | 2 | 12881 | 201 | Oasl1,Oasl2 | | 1.425e-03 | -6.55 | regulation of natural killer cell chemotaxis | biological process | GO:2000501 | 4 | 2 | 13711 | 214 | Ccl4,Ccl12 | | 1.425e-03 | -6.55 | negative regulation of interleukin-1-mediated signaling pathway | biological process | GO:2000660 | 4 | 2 | 13711 | 214 | Il1rn,Il6 | | 1.425e-03 | -6.55 | fever generation | biological process | GO:0001660 | 4 | 2 | 13711 | 214 | Il1a,Il1rn | | 1.425e-03 | -6.55 | dynamin family protein polymerization involved in mitochondrial fission | biological process | GO:0003374 | 4 | 2 | 13711 | 214 | Mx2,Mx1 | | 1.425e-03 | -6.55 | regulation of calcidiol 1-monooxygenase activity | biological process | GO:0060558 | 4 | 2 | 13711 | 214 | Nfkb1,Tnf | | 1.425e-03 | -6.55 | inflammatory cell apoptotic process | biological process | GO:0006925 | 4 | 2 | 13711 | 214 | Il6,Hcar2 | | 1.425e-03 | -6.55 | dynamin family protein polymerization involved in membrane fission | biological process | GO:0003373 | 4 | 2 | 13711 | 214 | Mx2,Mx1 | | 1.426e-03 | -6.55 | complement receptor activity | molecular function | GO:0004875 | 4 | 2 | 13516 | 211 | C5ar1,C3ar1 | | 1.426e-03 | -6.55 | 2'-5'-oligoadenylate synthetase activity | molecular function | GO:0001730 | 4 | 2 | 13516 | 211 | Oasl1,Oasl2 | | 1.434e-03 | -6.55 | circulatory system development | biological process | GO:0072359 | 747 | 23 | 13711 | 214 | Rnf213,Adamts9,Ccl2,Robo4,Angptl4,Vcam1,Col4a1,Ccl12,Clic4,Thbs1,Socs3,Cemip2,Errfi1,Cdkn1a,Sox11,Ptgs2,Rasip1,Spi1,Rhoj,Apold1,Adamts1,Tiparp,Fzd4 | | 1.435e-03 | -6.55 | RIP-mediated NFkB activation via ZBP1 | REACTOME pathways | R-MMU-1810476 | 14 | 3 | 6297 | 105 | Nfkb1,Nfkb2,Nfkbia | | 1.442e-03 | -6.54 | GREGORY\_SYNTHETIC\_LETHAL\_WITH\_IMATINIB | MSigDB lists | GREGORY\_SYNTHETIC\_LETHAL\_WITH\_IMATINIB | 114 | 7 | 12187 | 179 | Ifi204,Ifi207,Rtp4,Gbp3,Ifi209,Ifi211,Irf7 | | 1.442e-03 | -6.54 | GSE18791\_CTRL\_VS\_NEWCASTLE\_VIRUS\_DC\_1H\_DN | MSigDB lists | GSE18791\_CTRL\_VS\_NEWCASTLE\_VIRUS\_DC\_1H\_DN | 114 | 7 | 12187 | 179 | Nfkb2,Gadd45b,Tiparp,Dnajb1,Map3k8,Gem,Ptgs2 | | 1.442e-03 | -6.54 | GSE37301\_PRO\_BCELL\_VS\_GRANULOCYTE\_MONOCYTE\_PROGENITOR\_UP | MSigDB lists | GSE37301\_PRO\_BCELL\_VS\_GRANULOCYTE\_MONOCYTE\_PROGENITOR\_UP | 114 | 7 | 12187 | 179 | Sgk1,Dnajb1,Bach1,Fam107a,Gadd45b,Ifitm3,Zfp36 | | 1.455e-03 | -6.53 | GO\_REGULATION\_OF\_SMOOTH\_MUSCLE\_CELL\_PROLIFERATION | MSigDB lists | GO\_REGULATION\_OF\_SMOOTH\_MUSCLE\_CELL\_PROLIFERATION | 84 | 6 | 12187 | 179 | Il6,Tnfaip3,Trib1,Ptgs2,C3ar1,Tnf | | 1.458e-03 | -6.53 | GSE19198\_1H\_VS\_24H\_IL21\_TREATED\_TCELL\_DN | MSigDB lists | GSE19198\_1H\_VS\_24H\_IL21\_TREATED\_TCELL\_DN | 147 | 8 | 12187 | 179 | Birc3,Xaf1,Tor3a,Tubb6,Slc24a4,Plek,Csf2rb,Tnfsf10 | | 1.459e-03 | -6.53 | GO\_BLOOD\_MICROPARTICLE | MSigDB lists | GO\_BLOOD\_MICROPARTICLE | 57 | 5 | 12187 | 179 | Hp,Cp,Msn,Angptl4,A2m | | 1.459e-03 | -6.53 | HOFFMANN\_LARGE\_TO\_SMALL\_PRE\_BII\_LYMPHOCYTE\_DN | MSigDB lists | HOFFMANN\_LARGE\_TO\_SMALL\_PRE\_BII\_LYMPHOCYTE\_DN | 57 | 5 | 12187 | 179 | Il4ra,Birc3,Nfkbiz,Bcl3,Ier3 | | 1.461e-03 | -6.53 | GO\_NEGATIVE\_REGULATION\_OF\_TRANSCRIPTION\_FROM\_RNA\_POLYMERASE\_II\_PROMOTER | MSigDB lists | GO\_NEGATIVE\_REGULATION\_OF\_TRANSCRIPTION\_FROM\_RNA\_POLYMERASE\_II\_PROMOTER | 559 | 18 | 12187 | 179 | Spi1,Tnf,Zfp189,Bach1,Ifi211,Irf7,Arid5b,Nfkb2,Zbtb16,Sox11,Ifi209,Mxd4,Atf3,Nfkb1,Ifi204,Ifi207,Irf2,Zfp36 | | 1.471e-03 | -6.52 | REACTOME\_GPCR\_LIGAND\_BINDING | MSigDB lists | REACTOME\_GPCR\_LIGAND\_BINDING | 219 | 10 | 12187 | 179 | Ccl12,Hcar2,Ccl4,Cxcl16,P2ry6,Cxcl1,Fzd4,C5ar1,C3ar1,Saa1 | | 1.478e-03 | -6.52 | Activated TLR4 signalling | REACTOME pathways | R-MMU-166054 | 75 | 6 | 6297 | 105 | Nfkb1,Irf7,Nfkb2,Nfkbia,Map3k8,Cd14 | | 1.479e-03 | -6.52 | KAAB\_FAILED\_HEART\_VENTRICLE\_DN | MSigDB lists | KAAB\_FAILED\_HEART\_VENTRICLE\_DN | 34 | 4 | 12187 | 179 | Ifitm3,Ifitm2,Isg15,Cd14 | | 1.479e-03 | -6.52 | CROONQUIST\_NRAS\_VS\_STROMAL\_STIMULATION\_UP | MSigDB lists | CROONQUIST\_NRAS\_VS\_STROMAL\_STIMULATION\_UP | 34 | 4 | 12187 | 179 | Cybb,Trim25,Ncf1,Spi1 | | 1.479e-03 | -6.52 | GNF2\_CBFB | MSigDB lists | GNF2\_CBFB | 34 | 4 | 12187 | 179 | Ifi204,Ifi207,Ifi209,Ifi211 | | 1.483e-03 | -6.51 | PARP | pfam domains | PF00644 | 15 | 3 | 12881 | 201 | Parp14,Parp12,Tiparp | | 1.485e-03 | -6.51 | modulation by symbiont of host cellular process | biological process | GO:0044068 | 15 | 3 | 13711 | 214 | Irgm1,Tnip1,Irgm2 | | 1.516e-03 | -6.49 | GO\_RESPONSE\_TO\_INTERFERON\_BETA | MSigDB lists | GO\_RESPONSE\_TO\_INTERFERON\_BETA | 16 | 3 | 12187 | 179 | Xaf1,Ifitm3,Ifitm2 | | 1.516e-03 | -6.49 | ZHENG\_RESPONSE\_TO\_ARSENITE\_DN | MSigDB lists | ZHENG\_RESPONSE\_TO\_ARSENITE\_DN | 16 | 3 | 12187 | 179 | Cebpd,Il1a,Adamts1 | | 1.516e-03 | -6.49 | GO\_DEFENSE\_RESPONSE\_TO\_FUNGUS | MSigDB lists | GO\_DEFENSE\_RESPONSE\_TO\_FUNGUS | 16 | 3 | 12187 | 179 | Il17ra,S100a8,S100a9 | | 1.516e-03 | -6.49 | KIM\_ALL\_DISORDERS\_CALB1\_CORR\_DN | MSigDB lists | KIM\_ALL\_DISORDERS\_CALB1\_CORR\_DN | 16 | 3 | 12187 | 179 | Gadd45b,Sbno2,S100a8 | | 1.516e-03 | -6.49 | SESTO\_RESPONSE\_TO\_UV\_C3 | MSigDB lists | SESTO\_RESPONSE\_TO\_UV\_C3 | 16 | 3 | 12187 | 179 | Ptgs2,Ier3,Atf3 | | 1.516e-03 | -6.49 | GO\_RESPONSE\_TO\_PLATELET\_DERIVED\_GROWTH\_FACTOR | MSigDB lists | GO\_RESPONSE\_TO\_PLATELET\_DERIVED\_GROWTH\_FACTOR | 16 | 3 | 12187 | 179 | Iqgap1,Ccl12,Errfi1 | | 1.516e-03 | -6.49 | HERNANDEZ\_ABERRANT\_MITOSIS\_BY\_DOCETACEL\_4NM\_UP | MSigDB lists | HERNANDEZ\_ABERRANT\_MITOSIS\_BY\_DOCETACEL\_4NM\_UP | 16 | 3 | 12187 | 179 | Cdkn1a,Gem,Sgk1 | | 1.516e-03 | -6.49 | GO\_NECROPTOTIC\_PROCESS | MSigDB lists | GO\_NECROPTOTIC\_PROCESS | 16 | 3 | 12187 | 179 | Cd14,Tnf,Ripk1 | | 1.516e-03 | -6.49 | PID\_INTEGRIN2\_PATHWAY | MSigDB lists | PID\_INTEGRIN2\_PATHWAY | 16 | 3 | 12187 | 179 | Vcam1,Icam1,Plaur | | 1.516e-03 | -6.49 | GO\_REGULATION\_OF\_VIRAL\_ENTRY\_INTO\_HOST\_CELL | MSigDB lists | GO\_REGULATION\_OF\_VIRAL\_ENTRY\_INTO\_HOST\_CELL | 16 | 3 | 12187 | 179 | Trim25,Ifitm3,Ifitm2 | | 1.516e-03 | -6.49 | BIOCARTA\_TID\_PATHWAY | MSigDB lists | BIOCARTA\_TID\_PATHWAY | 16 | 3 | 12187 | 179 | Nfkb1,Tnf,Nfkbia | | 1.523e-03 | -6.49 | GSE41176\_WT\_VS\_TAK1\_KO\_UNSTIM\_BCELL\_UP | MSigDB lists | GSE41176\_WT\_VS\_TAK1\_KO\_UNSTIM\_BCELL\_UP | 148 | 8 | 12187 | 179 | Stx11,Tnfaip8,Bach1,Nfkb2,Clic4,Map3k8,Maff,Tagln2 | | 1.529e-03 | -6.48 | BZIP | prosite domains | PS50217 | 46 | 5 | 8845 | 164 | Cebpd,Nfe2l2,Bach1,Maff,Atf3 | | 1.576e-03 | -6.45 | cytosol | cellular component | GO:0005829 | 2843 | 62 | 13825 | 212 | Zbtb16,Rnf213,Apold1,Trim25,Sgk1,Msn,Ptpn1,Rhoc,Rrp8,Il1a,Parp9,Dtx3l,Herc6,S100a8,Bcl3,Ifit2,Hspa5,Parp14,Lcn2,Nfkbie,Nfkb1,Tnip1,Msr1,Map3k8,Cflar,Ifit3b,Ifi209,Irf7,Dnajb1,Nfkb2,Nfe2l2,Irgm2,Akap12,Oasl2,Pik3ap1,Lgals9,Mt1,Xdh,Irf2,Ripk1,Cdkn1a,S100a9,Ccnd2,Plin4,Ifi211,Errfi1,Clic4,Oasl1,Zfp36,Ifi204,Ifi207,Pik3r1,Casp4,Birc3,Pik3r5,Tgm2,Runx1,Ncf1,Irf9,Gbp3,Usp18,Nfkbia | | 1.578e-03 | -6.45 | JECHLINGER\_EPITHELIAL\_TO\_MESENCHYMAL\_TRANSITION\_DN | MSigDB lists | JECHLINGER\_EPITHELIAL\_TO\_MESENCHYMAL\_TRANSITION\_DN | 58 | 5 | 12187 | 179 | Thbs1,Sgk1,Tgm2,Zfp36,Atf3 | | 1.578e-03 | -6.45 | KEGG\_ADIPOCYTOKINE\_SIGNALING\_PATHWAY | MSigDB lists | KEGG\_ADIPOCYTOKINE\_SIGNALING\_PATHWAY | 58 | 5 | 12187 | 179 | Nfkb1,Tnf,Nfkbie,Socs3,Nfkbia | | 1.578e-03 | -6.45 | SHAFFER\_IRF4\_TARGETS\_IN\_PLASMA\_CELL\_VS\_MATURE\_B\_LYMPHOCYTE | MSigDB lists | SHAFFER\_IRF4\_TARGETS\_IN\_PLASMA\_CELL\_VS\_MATURE\_B\_LYMPHOCYTE | 58 | 5 | 12187 | 179 | Zfp36,Hspb1,Atf3,Tnfaip3,Cflar | | 1.581e-03 | -6.45 | Neutrophil degranulation | REACTOME pathways | R-MMU-6798695 | 383 | 15 | 6297 | 105 | Iqgap1,Cxcl1,S100a9,Gpr84,Serpina3f,Nfkb1,Hp,Lcn2,C5ar1,C3ar1,Cd14,Mmp8,Cybb,S100a8,Plaur | | 1.588e-03 | -6.45 | leukocyte activation involved in immune response | biological process | GO:0002366 | 109 | 7 | 13711 | 214 | Icam1,Tnfaip3,Sbno2,Il6,Stx11,Gadd45g,Bcl3 | | 1.589e-03 | -6.44 | GSE7460\_CD8\_TCELL\_VS\_CD4\_TCELL\_ACT\_UP | MSigDB lists | GSE7460\_CD8\_TCELL\_VS\_CD4\_TCELL\_ACT\_UP | 149 | 8 | 12187 | 179 | Ncf1,Gbp6,Tnfsf10,Plaur,Gbp3,Plek,Icam1,Gbp4 | | 1.589e-03 | -6.44 | GSE17721\_CPG\_VS\_GARDIQUIMOD\_12H\_BMDC\_UP | MSigDB lists | GSE17721\_CPG\_VS\_GARDIQUIMOD\_12H\_BMDC\_UP | 149 | 8 | 12187 | 179 | Irgm2,Ptgs2,Gbp7,Ccl12,Vcam1,Tor3a,Tgm2,Sdc4 | | 1.589e-03 | -6.44 | MCBRYAN\_PUBERTAL\_TGFB1\_TARGETS\_UP | MSigDB lists | MCBRYAN\_PUBERTAL\_TGFB1\_TARGETS\_UP | 149 | 8 | 12187 | 179 | Cd14,Fzd4,Runx1,Cdkn1a,Tgm2,Sntb2,Cp,Ier3 | | 1.594e-03 | -6.44 | GO\_REGULATION\_OF\_CELL\_SHAPE | MSigDB lists | GO\_REGULATION\_OF\_CELL\_SHAPE | 116 | 7 | 12187 | 179 | Rhoj,Msn,Icam1,Ccl2,Rhou,Il6,Ccl12 | | 1.594e-03 | -6.44 | GSE9601\_UNTREATED\_VS\_NFKB\_INHIBITOR\_TREATED\_HCMV\_INF\_MONOCYTE\_UP | MSigDB lists | GSE9601\_UNTREATED\_VS\_NFKB\_INHIBITOR\_TREATED\_HCMV\_INF\_MONOCYTE\_UP | 116 | 7 | 12187 | 179 | Gbp3,Gbp6,Ifitm3,Samd9l,Gbp4,Cdkn1a,Socs3 | | 1.594e-03 | -6.44 | GO\_MAINTENANCE\_OF\_LOCATION | MSigDB lists | GO\_MAINTENANCE\_OF\_LOCATION | 116 | 7 | 12187 | 179 | Nfkbia,Tnf,Hspa5,S100a8,S100a9,Bcl3,Nfkbie | | 1.594e-03 | -6.44 | JOHNSTONE\_PARVB\_TARGETS\_2\_UP | MSigDB lists | JOHNSTONE\_PARVB\_TARGETS\_2\_UP | 116 | 7 | 12187 | 179 | Oasl1,Tnfaip3,Znfx1,Sbno2,Ifit3b,Isg15,Ifit2 | | 1.614e-03 | -6.43 | GO\_NEGATIVE\_REGULATION\_OF\_PHOSPHORYLATION | MSigDB lists | GO\_NEGATIVE\_REGULATION\_OF\_PHOSPHORYLATION | 342 | 13 | 12187 | 179 | Tnfaip3,Il6,Gadd45b,Trib1,Xdh,Atf3,Socs3,Cdkn1a,Errfi1,Ptpn1,Irak3,Hspb1,Tnip1 | | 1.617e-03 | -6.43 | TRAIL signaling | REACTOME pathways | R-MMU-75158 | 4 | 2 | 6297 | 105 | Tnfsf10,Cflar | | 1.619e-03 | -6.43 | regulation of catabolic process | biological process | GO:0009894 | 754 | 23 | 13711 | 214 | Irgm2,Msn,Noct,Samd9l,Hp,Ptpn1,Trim30a,Irak3,Depp1,Zfp36,Tnf,Tnfaip3,Ier3,Irgm1,Rasip1,Tiparp,Timp1,Hcar2,Banp,Trib1,Iigp1,Dtx3l,Il6 | | 1.622e-03 | -6.42 | regulation of nucleic acid-templated transcription | biological process | GO:1903506 | 2294 | 53 | 13711 | 214 | Dtx3l,Irf7,Trib1,Pik3r1,Nfkbia,Il1a,Il6,Ifi211,Cebpd,Spi1,Runx1,Resf1,Trim25,Ifi209,Fzd4,Parp9,Sbno2,Ncf1,Ifi207,Ripk1,Cflar,Fstl1,Tnip1,Tnf,Acod1,Atf3,Zfp36,Nfe2l2,Rasd1,Maff,Rrp8,Dnajb1,Banp,Rbm39,Icam1,Mxd4,Nfkb2,Sox11,Bach1,Gbp4,Nfkb1,Nfkbie,Irak3,Arid5b,Aff1,Zbtb16,Csf3,Ifi204,Tnfaip3,Znfx1,Irf9,Bcl3,Irf2 | | 1.627e-03 | -6.42 | GO\_HEMOSTASIS | MSigDB lists | GO\_HEMOSTASIS | 222 | 10 | 12187 | 179 | Plek,Hspb1,Irf2,Plaur,Saa1,A2m,Pik3r5,Il6,Maff,Pik3r1 | | 1.627e-03 | -6.42 | GO\_ACTIVATION\_OF\_PROTEIN\_KINASE\_ACTIVITY | MSigDB lists | GO\_ACTIVATION\_OF\_PROTEIN\_KINASE\_ACTIVITY | 222 | 10 | 12187 | 179 | Tnf,Map3k8,Thbs1,Ripk1,Gadd45b,Gadd45g,C5ar1,Map3k6,Saa1,Ptpn1 | | 1.651e-03 | -6.41 | CHEN\_LVAD\_SUPPORT\_OF\_FAILING\_HEART\_DN | MSigDB lists | CHEN\_LVAD\_SUPPORT\_OF\_FAILING\_HEART\_DN | 35 | 4 | 12187 | 179 | Cd14,S100a8,Ccl12,Tnfsf10 | | 1.651e-03 | -6.41 | RASHI\_RESPONSE\_TO\_IONIZING\_RADIATION\_1 | MSigDB lists | RASHI\_RESPONSE\_TO\_IONIZING\_RADIATION\_1 | 35 | 4 | 12187 | 179 | Sgk1,Atf3,Ier3,Zfp36 | | 1.651e-03 | -6.41 | SCHRAETS\_MLL\_TARGETS\_UP | MSigDB lists | SCHRAETS\_MLL\_TARGETS\_UP | 35 | 4 | 12187 | 179 | Ifi204,Ifi207,Ifi209,Ifi211 | | 1.651e-03 | -6.41 | REACTOME\_IL1\_SIGNALING | MSigDB lists | REACTOME\_IL1\_SIGNALING | 35 | 4 | 12187 | 179 | Map3k8,Il1a,Il1rn,Irak3 | | 1.651e-03 | -6.41 | GO\_REGULATION\_OF\_TUMOR\_NECROSIS\_FACTOR\_MEDIATED\_SIGNALING\_PATHWAY | MSigDB lists | GO\_REGULATION\_OF\_TUMOR\_NECROSIS\_FACTOR\_MEDIATED\_SIGNALING\_PATHWAY | 35 | 4 | 12187 | 179 | Tnfaip3,Ripk1,Tnf,Birc3 | | 1.653e-03 | -6.41 | regulation of reactive oxygen species biosynthetic process | biological process | GO:1903426 | 81 | 6 | 13711 | 214 | Il6,Icam1,Tnf,Ptgs2,Mmp8,Cflar | | 1.654e-03 | -6.40 | regulation of RNA biosynthetic process | biological process | GO:2001141 | 2296 | 53 | 13711 | 214 | Ifi204,Arid5b,Aff1,Irak3,Zbtb16,Csf3,Tnfaip3,Znfx1,Bcl3,Irf9,Irf2,Rbm39,Banp,Mxd4,Icam1,Sox11,Nfkb2,Bach1,Nfkb1,Nfkbie,Gbp4,Tnip1,Fstl1,Ncf1,Ifi207,Cflar,Ripk1,Atf3,Zfp36,Acod1,Tnf,Nfe2l2,Rasd1,Maff,Rrp8,Dnajb1,Pik3r1,Irf7,Dtx3l,Trib1,Il6,Il1a,Nfkbia,Runx1,Trim25,Resf1,Cebpd,Ifi211,Spi1,Sbno2,Parp9,Ifi209,Fzd4 | | 1.656e-03 | -6.40 | tumor necrosis factor receptor superfamily binding | molecular function | GO:0032813 | 33 | 4 | 13516 | 211 | Tnf,Cflar,Tnfsf10,Ripk1 | | 1.658e-03 | -6.40 | GSE3982\_DC\_VS\_MAC\_LPS\_STIM\_DN | MSigDB lists | GSE3982\_DC\_VS\_MAC\_LPS\_STIM\_DN | 150 | 8 | 12187 | 179 | Ifit3b,Ccl4,Msr1,Ifit2,Cybb,Cd14,Rgs16,Trim25 | | 1.658e-03 | -6.40 | GSE17721\_LPS\_VS\_GARDIQUIMOD\_24H\_BMDC\_UP | MSigDB lists | GSE17721\_LPS\_VS\_GARDIQUIMOD\_24H\_BMDC\_UP | 150 | 8 | 12187 | 179 | Timp1,Il1rn,Sgk1,Ifit3b,Casp4,Ptges,Slfn9,Ncf1 | | 1.658e-03 | -6.40 | GSE27241\_CTRL\_VS\_DIGOXIN\_TREATED\_CD4\_TCELL\_IN\_TH17\_POLARIZING\_CONDITIONS\_DN | MSigDB lists | GSE27241\_CTRL\_VS\_DIGOXIN\_TREATED\_CD4\_TCELL\_IN\_TH17\_POLARIZING\_CONDITIONS\_DN | 150 | 8 | 12187 | 179 | AA467197,Tnf,Ifit3b,Usp18,Nfkbia,Xaf1,Irf9,Irf7 | | 1.658e-03 | -6.40 | GSE369\_PRE\_VS\_POST\_IL6\_INJECTION\_IFNG\_KO\_LIVER\_DN | MSigDB lists | GSE369\_PRE\_VS\_POST\_IL6\_INJECTION\_IFNG\_KO\_LIVER\_DN | 150 | 8 | 12187 | 179 | Gem,Trim25,Dnajb1,Parp14,Socs3,Bach1,Oasl1,Mxd4 | | 1.663e-03 | -6.40 | Cd81 (CD81 antigen) | protein interactions | 12520 | 4 | 2 | 6802 | 115 | Ifitm3,Ifitm2 | | 1.663e-03 | -6.40 | Mafg (v-maf musculoaponeurotic fibrosarcoma oncogene family, protein G (avian)) | protein interactions | 17134 | 4 | 2 | 6802 | 115 | Bach1,Nfe2l2 | | 1.663e-03 | -6.40 | Il1b (interleukin 1 beta) | protein interactions | 16176 | 4 | 2 | 6802 | 115 | Ripk1,Tnfaip3 | | 1.663e-03 | -6.40 | Tnip1 (TNFAIP3 interacting protein 1) | protein interactions | 57783 | 4 | 2 | 6802 | 115 | Tnf,Tnfaip3 | | 1.663e-03 | -6.40 | MAPKAPK2 (MAPK activated protein kinase 2) | protein interactions | 9261 | 4 | 2 | 6802 | 115 | Zfp36,Hspb1 | | 1.675e-03 | -6.39 | GO\_CELLULAR\_RESPONSE\_TO\_RADIATION | MSigDB lists | GO\_CELLULAR\_RESPONSE\_TO\_RADIATION | 117 | 7 | 12187 | 179 | Ifi211,Ptgs2,Ifi204,Ifi207,Cdkn1a,Ifi209,Pik3r1 | | 1.675e-03 | -6.39 | GSE1791\_CTRL\_VS\_NEUROMEDINU\_IN\_T\_CELL\_LINE\_3H\_UP | MSigDB lists | GSE1791\_CTRL\_VS\_NEUROMEDINU\_IN\_T\_CELL\_LINE\_3H\_UP | 117 | 7 | 12187 | 179 | Rtp4,Samd9l,Tap1,Plaur,Tnf,Trim56,Banp | | 1.704e-03 | -6.37 | BOYLAN\_MULTIPLE\_MYELOMA\_D\_DN | MSigDB lists | BOYLAN\_MULTIPLE\_MYELOMA\_D\_DN | 59 | 5 | 12187 | 179 | Ksr1,Cybb,Cmpk2,Bcl3,Nfkbie | | 1.719e-03 | -6.37 | GO\_RESPONSE\_TO\_NITROGEN\_COMPOUND | MSigDB lists | GO\_RESPONSE\_TO\_NITROGEN\_COMPOUND | 662 | 20 | 12187 | 179 | Il6,Ccl12,Tnfaip3,Nfkbia,Casp4,Pik3r1,Tnf,Pygm,C5ar1,Col4a1,Nfkb1,Timp1,Socs3,Vcam1,Cdkn1a,Errfi1,Ptpn1,Icam1,Irak3,Ptgs2 | | 1.730e-03 | -6.36 | GSE19923\_WT\_VS\_HEB\_KO\_DP\_THYMOCYTE\_UP | MSigDB lists | GSE19923\_WT\_VS\_HEB\_KO\_DP\_THYMOCYTE\_UP | 151 | 8 | 12187 | 179 | Socs3,Ccl2,Nfkbia,Col4a1,Bcl3,Irgm2,Nfkb2,Tap1 | | 1.730e-03 | -6.36 | GSE19941\_LPS\_VS\_LPS\_AND\_IL10\_STIM\_IL10\_KO\_NFKBP50\_KO\_MACROPHAGE\_DN | MSigDB lists | GSE19941\_LPS\_VS\_LPS\_AND\_IL10\_STIM\_IL10\_KO\_NFKBP50\_KO\_MACROPHAGE\_DN | 151 | 8 | 12187 | 179 | Ifitm3,Ifitm2,Cdkn1a,Aff1,Nfkb1,Rhoc,AA467197,Il6 | | 1.730e-03 | -6.36 | GSE17721\_CPG\_VS\_GARDIQUIMOD\_6H\_BMDC\_UP | MSigDB lists | GSE17721\_CPG\_VS\_GARDIQUIMOD\_6H\_BMDC\_UP | 151 | 8 | 12187 | 179 | Sgk1,Birc3,Ifit3b,Casp4,Atf3,Il6,Ccl12,Sele | | 1.735e-03 | -6.36 | regulation of transcription, DNA-templated | biological process | GO:0006355 | 2246 | 52 | 13711 | 214 | Banp,Sox11,Nfkb2,Mxd4,Icam1,Bach1,Gbp4,Nfkbie,Nfkb1,Zbtb16,Csf3,Irak3,Arid5b,Aff1,Ifi204,Tnfaip3,Znfx1,Irf9,Bcl3,Irf2,Trib1,Irf7,Dtx3l,Pik3r1,Il1a,Nfkbia,Il6,Spi1,Cebpd,Ifi211,Resf1,Trim25,Runx1,Fzd4,Ifi209,Sbno2,Parp9,Cflar,Ripk1,Ifi207,Ncf1,Tnip1,Fstl1,Acod1,Tnf,Zfp36,Atf3,Rasd1,Nfe2l2,Maff,Dnajb1,Rrp8 | | 1.738e-03 | -6.36 | MARTINEZ\_RB1\_AND\_TP53\_TARGETS\_UP | MSigDB lists | MARTINEZ\_RB1\_AND\_TP53\_TARGETS\_UP | 476 | 16 | 12187 | 179 | Hcar2,Trib1,Ccnd2,Msr1,Xdh,S100a8,Atf3,Ier3,Timp1,Ms4a6d,Cp,Dnajb1,Cdkn1a,Thbs1,Plin4,Ptgs2 | | 1.744e-03 | -6.35 | DIAZ\_CHRONIC\_MEYLOGENOUS\_LEUKEMIA\_DN | MSigDB lists | DIAZ\_CHRONIC\_MEYLOGENOUS\_LEUKEMIA\_DN | 87 | 6 | 12187 | 179 | S100a8,S100a9,Irf7,Cebpd,Gadd45b,Ms4a6d | | 1.744e-03 | -6.35 | HALLMARK\_COAGULATION | MSigDB lists | HALLMARK\_COAGULATION | 87 | 6 | 12187 | 179 | Timp1,Mmp8,Thbs1,Maff,Plek,A2m | | 1.763e-03 | -6.34 | cytokine production | biological process | GO:0001816 | 111 | 7 | 13711 | 214 | Isg15,Trim56,Gadd45g,Ptgs2,Tnfaip8,Irf9,Irf7 | | 1.769e-03 | -6.34 | cellular response to hydrogen peroxide | biological process | GO:0070301 | 56 | 5 | 13711 | 214 | Il6,Lcn2,Nfe2l2,Tnfaip3,Ripk1 | | 1.773e-03 | -6.33 | MODULE\_53 | MSigDB lists | MODULE\_53 | 304 | 12 | 12187 | 179 | Ccl12,Ccnd2,Tnf,Ccl4,Olfml2b,S100a8,Rgs16,Cxcl1,Ier3,Atf3,Tap1,Isg15 | | 1.798e-03 | -6.32 | GO\_ENZYME\_INHIBITOR\_ACTIVITY | MSigDB lists | GO\_ENZYME\_INHIBITOR\_ACTIVITY | 225 | 10 | 12187 | 179 | Timp1,Birc3,Cdkn1a,Trib1,Socs3,Tnfaip8,Iqgap1,Hspb1,A2m,Angptl4 | | 1.803e-03 | -6.32 | GSE1460\_DP\_THYMOCYTE\_VS\_THYMIC\_STROMAL\_CELL\_UP | MSigDB lists | GSE1460\_DP\_THYMOCYTE\_VS\_THYMIC\_STROMAL\_CELL\_UP | 152 | 8 | 12187 | 179 | Ifi204,Ifi207,Phf11b,Ifi209,Irf2,Tnfaip8,Ifi211,Phf11d | | 1.803e-03 | -6.32 | GSE17721\_POLYIC\_VS\_CPG\_6H\_BMDC\_DN | MSigDB lists | GSE17721\_POLYIC\_VS\_CPG\_6H\_BMDC\_DN | 152 | 8 | 12187 | 179 | Nfkb2,Gadd45b,Slfn4,Slfn3,Sele,Ccl2,Thbs1,Hspa5 | | 1.803e-03 | -6.32 | GSE17721\_PAM3CSK4\_VS\_CPG\_16H\_BMDC\_DN | MSigDB lists | GSE17721\_PAM3CSK4\_VS\_CPG\_16H\_BMDC\_DN | 152 | 8 | 12187 | 179 | Ms4a6d,Rhoc,Znfx1,Gbp4,Sox11,Ccl2,Gbp3,Gbp6 | | 1.803e-03 | -6.32 | GSE9650\_GP33\_VS\_GP276\_LCMV\_SPECIFIC\_EXHAUSTED\_CD8\_TCELL\_UP | MSigDB lists | GSE9650\_GP33\_VS\_GP276\_LCMV\_SPECIFIC\_EXHAUSTED\_CD8\_TCELL\_UP | 152 | 8 | 12187 | 179 | Fstl1,Tgm2,Il1a,Cp,Selp,Bcl3,S100a8,Rgs16 | | 1.803e-03 | -6.32 | GSE21546\_UNSTIM\_VS\_ANTI\_CD3\_STIM\_DP\_THYMOCYTES\_UP | MSigDB lists | GSE21546\_UNSTIM\_VS\_ANTI\_CD3\_STIM\_DP\_THYMOCYTES\_UP | 152 | 8 | 12187 | 179 | Ifi211,Fstl1,Ifi209,Thbs1,Cebpd,Ifi204,Ifi207,Samd9l | | 1.803e-03 | -6.32 | GSE22589\_HEALTHY\_VS\_SIV\_INFECTED\_DC\_DN | MSigDB lists | GSE22589\_HEALTHY\_VS\_SIV\_INFECTED\_DC\_DN | 152 | 8 | 12187 | 179 | Samd9l,Akap12,Tor3a,Nfkbia,Tap1,Irf9,Gbp7,Herc6 | | 1.803e-03 | -6.32 | GSE24634\_TREG\_VS\_TCONV\_POST\_DAY5\_IL4\_CONVERSION\_DN | MSigDB lists | GSE24634\_TREG\_VS\_TCONV\_POST\_DAY5\_IL4\_CONVERSION\_DN | 152 | 8 | 12187 | 179 | Plek,S100a8,Igsf6,Il4ra,Cdkn1a,Ccl12,Ms4a6d,Cybb | | 1.803e-03 | -6.32 | GSE13547\_CTRL\_VS\_ANTI\_IGM\_STIM\_BCELL\_2H\_UP | MSigDB lists | GSE13547\_CTRL\_VS\_ANTI\_IGM\_STIM\_BCELL\_2H\_UP | 152 | 8 | 12187 | 179 | Ifit3b,Casp4,Usp18,Clic4,Xdh,Trim25,Ncf1,Rnf213 | | 1.807e-03 | -6.32 | regulation of programmed necrotic cell death | biological process | GO:0062098 | 16 | 3 | 13711 | 214 | Cflar,Birc3,Ripk1 | | 1.807e-03 | -6.32 | leukocyte migration involved in inflammatory response | biological process | GO:0002523 | 16 | 3 | 13711 | 214 | Ccl2,S100a8,S100a9 | | 1.807e-03 | -6.32 | necroptotic process | biological process | GO:0070266 | 16 | 3 | 13711 | 214 | Tnf,Ripk1,Birc3 | | 1.807e-03 | -6.32 | regulation of necroptotic process | biological process | GO:0060544 | 16 | 3 | 13711 | 214 | Cflar,Birc3,Ripk1 | | 1.821e-03 | -6.31 | REACTOME\_TRAF6\_MEDIATED\_IRF7\_ACTIVATION | MSigDB lists | REACTOME\_TRAF6\_MEDIATED\_IRF7\_ACTIVATION | 17 | 3 | 12187 | 179 | Irf7,Trim25,Irf2 | | 1.821e-03 | -6.31 | MURATA\_VIRULENCE\_OF\_H\_PILORI | MSigDB lists | MURATA\_VIRULENCE\_OF\_H\_PILORI | 17 | 3 | 12187 | 179 | S100a9,Ccnd2,Cdkn1a | | 1.821e-03 | -6.31 | ZIRN\_TRETINOIN\_RESPONSE\_UP | MSigDB lists | ZIRN\_TRETINOIN\_RESPONSE\_UP | 17 | 3 | 12187 | 179 | Akap12,Ccl12,Plaur | | 1.821e-03 | -6.31 | SCHEIDEREIT\_IKK\_TARGETS | MSigDB lists | SCHEIDEREIT\_IKK\_TARGETS | 17 | 3 | 12187 | 179 | Nfkb2,Nfkb1,Nfkbia | | 1.821e-03 | -6.31 | AMIT\_DELAYED\_EARLY\_GENES | MSigDB lists | AMIT\_DELAYED\_EARLY\_GENES | 17 | 3 | 12187 | 179 | Zfp36,Atf3,Maff | | 1.821e-03 | -6.31 | GO\_REGULATION\_OF\_CYSTEINE\_TYPE\_ENDOPEPTIDASE\_ACTIVITY\_INVOLVED\_IN\_APOPTOTIC\_SIGNALING\_PATHWAY | MSigDB lists | GO\_REGULATION\_OF\_CYSTEINE\_TYPE\_ENDOPEPTIDASE\_ACTIVITY\_INVOLVED\_IN\_APOPTOTIC\_SIGNALING\_PATHWAY | 17 | 3 | 12187 | 179 | Plaur,Ripk1,Tnfsf10 | | 1.836e-03 | -6.30 | PID\_FAS\_PATHWAY | MSigDB lists | PID\_FAS\_PATHWAY | 36 | 4 | 12187 | 179 | Pik3r1,Birc3,Cflar,Ripk1 | | 1.836e-03 | -6.30 | GNF2\_CD33 | MSigDB lists | GNF2\_CD33 | 36 | 4 | 12187 | 179 | Cd14,Ms4a6d,Cybb,Stx11 | | 1.837e-03 | -6.30 | GO\_TRANSCRIPTION\_FACTOR\_ACTIVITY\_RNA\_POLYMERASE\_II\_DISTAL\_ENHANCER\_SEQUENCE\_SPECIFIC\_BINDING | MSigDB lists | GO\_TRANSCRIPTION\_FACTOR\_ACTIVITY\_RNA\_POLYMERASE\_II\_DISTAL\_ENHANCER\_SEQUENCE\_SPECIFIC\_BINDING | 60 | 5 | 12187 | 179 | Zbtb16,Spi1,Nfkb1,Bach1,Nfe2l2 | | 1.837e-03 | -6.30 | GO\_NEGATIVE\_REGULATION\_OF\_MYELOID\_CELL\_DIFFERENTIATION | MSigDB lists | GO\_NEGATIVE\_REGULATION\_OF\_MYELOID\_CELL\_DIFFERENTIATION | 60 | 5 | 12187 | 179 | Zbtb16,Trib1,Nfkbia,Runx1,Pik3r1 | | 1.846e-03 | -6.29 | HUANG\_GATA2\_TARGETS\_UP | MSigDB lists | HUANG\_GATA2\_TARGETS\_UP | 119 | 7 | 12187 | 179 | Il17ra,Tnfaip8,Irak3,Hp,Spi1,Cdkn1a,Aff1 | | 1.849e-03 | -6.29 | GO\_REGULATION\_OF\_PROTEIN\_KINASE\_B\_SIGNALING | MSigDB lists | GO\_REGULATION\_OF\_PROTEIN\_KINASE\_B\_SIGNALING | 88 | 6 | 12187 | 179 | Pik3r5,Csf3,Thbs1,Tnf,Il6,Xdh | | 1.849e-03 | -6.29 | GO\_REGULATION\_OF\_EPITHELIAL\_CELL\_DIFFERENTIATION | MSigDB lists | GO\_REGULATION\_OF\_EPITHELIAL\_CELL\_DIFFERENTIATION | 88 | 6 | 12187 | 179 | Tnf,Maff,Apold1,Adamts9,Xdh,Errfi1 | | 1.854e-03 | -6.29 | negative regulation of endothelial cell proliferation | biological process | GO:0001937 | 34 | 4 | 13711 | 214 | Tnf,Thbs1,Ccl12,Xdh | | 1.856e-03 | -6.29 | regulation of cell cycle G1/S phase transition | biological process | GO:1902806 | 112 | 7 | 13711 | 214 | Fam107a,Cdkn1a,Ccl12,Adamts1,Kcna5,Ripk1,Ccnd2 | | 1.856e-03 | -6.29 | regulation of T cell differentiation | biological process | GO:0045580 | 112 | 7 | 13711 | 214 | Zbtb16,Ctla2a,Lgals9,Runx1,Il4ra,Nfkbiz,Il6 | | 1.874e-03 | -6.28 | negative regulation of extrinsic apoptotic signaling pathway | biological process | GO:2001237 | 83 | 6 | 13711 | 214 | Cflar,Ripk1,Sgk3,Tnfaip3,Tnf,Icam1 | | 1.874e-03 | -6.28 | regulation of receptor signaling pathway via JAK-STAT | biological process | GO:0046425 | 83 | 6 | 13711 | 214 | Parp9,Bcl3,Il6,Parp14,Tnf,Socs3 | | 1.874e-03 | -6.28 | positive regulation of lymphocyte differentiation | biological process | GO:0045621 | 83 | 6 | 13711 | 214 | Il6,Runx1,Lgals9,Nfkbiz,Il4ra,Zbtb16 | | 1.879e-03 | -6.28 | GSE1460\_INTRATHYMIC\_T\_PROGENITOR\_VS\_CD4\_THYMOCYTE\_DN | MSigDB lists | GSE1460\_INTRATHYMIC\_T\_PROGENITOR\_VS\_CD4\_THYMOCYTE\_DN | 153 | 8 | 12187 | 179 | Tnfsf10,Msn,Ifi44,Birc3,Sgk1,Il4ra,Tnf,Tnfaip3 | | 1.879e-03 | -6.28 | GSE43955\_1H\_VS\_10H\_ACT\_CD4\_TCELL\_WITH\_TGFB\_IL6\_DN | MSigDB lists | GSE43955\_1H\_VS\_10H\_ACT\_CD4\_TCELL\_WITH\_TGFB\_IL6\_DN | 153 | 8 | 12187 | 179 | Tagln2,Irf7,Clic4,Tap1,Ifit3b,Usp18,Tubb6,Cp | | 1.879e-03 | -6.28 | GSE25123\_CTRL\_VS\_IL4\_STIM\_PPARG\_KO\_MACROPHAGE\_DN | MSigDB lists | GSE25123\_CTRL\_VS\_IL4\_STIM\_PPARG\_KO\_MACROPHAGE\_DN | 153 | 8 | 12187 | 179 | Dtx3l,Gbp7,Errfi1,Xdh,Nfkbiz,Parp12,Ms4a6d,Kcna5 | | 1.879e-03 | -6.28 | GSE29949\_MICROGLIA\_VS\_DC\_BRAIN\_UP | MSigDB lists | GSE29949\_MICROGLIA\_VS\_DC\_BRAIN\_UP | 153 | 8 | 12187 | 179 | Rgs16,Tnfaip8,Gem,Atf3,Il6,Gadd45g,Cdkn1a,Ccnd2 | | 1.879e-03 | -6.28 | GSE12198\_CTRL\_VS\_HIGH\_IL2\_STIM\_NK\_CELL\_UP | MSigDB lists | GSE12198\_CTRL\_VS\_HIGH\_IL2\_STIM\_NK\_CELL\_UP | 153 | 8 | 12187 | 179 | Ccl4,Socs3,Birc3,Msr1,Rtp4,Il17ra,Bcl3,Selp | | 1.882e-03 | -6.28 | REACTOME\_HEMOSTASIS | MSigDB lists | REACTOME\_HEMOSTASIS | 348 | 13 | 12187 | 179 | Pik3r1,Maff,Pik3r5,Irf7,A2m,Timp1,Thbs1,Hspa5,Irf2,Plaur,Ptpn1,Plek,Selp | | 1.895e-03 | -6.27 | BONOME\_OVARIAN\_CANCER\_SURVIVAL\_OPTIMAL\_DEBULKING | MSigDB lists | BONOME\_OVARIAN\_CANCER\_SURVIVAL\_OPTIMAL\_DEBULKING | 189 | 9 | 12187 | 179 | Tagln2,Ifi211,Iqgap1,Runx1,Sox11,Ifi209,Ifi207,Cp,Ifi204 | | 1.895e-03 | -6.27 | MCBRYAN\_PUBERTAL\_BREAST\_4\_5WK\_UP | MSigDB lists | MCBRYAN\_PUBERTAL\_BREAST\_4\_5WK\_UP | 189 | 9 | 12187 | 179 | Lcn2,Timp1,Tgm2,Sdc4,Runx1,Thbs1,Errfi1,Nfkb2,Cd14 | | 1.915e-03 | -6.26 | positive regulation of phosphatidylinositol 3-kinase signaling | biological process | GO:0014068 | 57 | 5 | 13711 | 214 | Csf3,Pik3ap1,Ncf1,Tnf,Selp | | 1.935e-03 | -6.25 | Tnf (tumor necrosis factor) | protein interactions | 21926 | 32 | 4 | 6802 | 115 | Tnfaip3,Zfp36,Ripk1,Tnip1 | | 1.939e-03 | -6.25 | positive regulation of establishment of protein localization | biological process | GO:1904951 | 368 | 14 | 13711 | 214 | Ptgs2,Il17ra,Mmp8,Hcar2,Pik3r1,Lgals9,Cd14,Il6,Ccl2,Casp4,Tnf,Il4ra,Akap12,Il1a | | 1.953e-03 | -6.24 | cell activation involved in immune response | biological process | GO:0002263 | 113 | 7 | 13711 | 214 | Gadd45g,Bcl3,Stx11,Il6,Sbno2,Icam1,Tnfaip3 | | 1.958e-03 | -6.24 | GSE12845\_NAIVE\_VS\_DARKZONE\_GC\_TONSIL\_BCELL\_UP | MSigDB lists | GSE12845\_NAIVE\_VS\_DARKZONE\_GC\_TONSIL\_BCELL\_UP | 154 | 8 | 12187 | 179 | Tagln2,Herc6,Nfkbie,Msn,Irak3,Maff,Atf3,Timp1 | | 1.958e-03 | -6.24 | GSE22886\_DAY0\_VS\_DAY7\_MONOCYTE\_IN\_CULTURE\_UP | MSigDB lists | GSE22886\_DAY0\_VS\_DAY7\_MONOCYTE\_IN\_CULTURE\_UP | 154 | 8 | 12187 | 179 | Tnfsf10,Zfp36,Irak3,Irf7,S100a9,Ifitm2,Cebpd,Gadd45b | | 1.958e-03 | -6.24 | GSE27670\_CTRL\_VS\_LMP1\_TRANSDUCED\_GC\_BCELL\_UP | MSigDB lists | GSE27670\_CTRL\_VS\_LMP1\_TRANSDUCED\_GC\_BCELL\_UP | 154 | 8 | 12187 | 179 | Nfkb2,Errfi1,Ptgs2,Znfx1,Cdkn1a,Tnf,Casp4,Usp18 | | 1.958e-03 | -6.24 | GSE2585\_CTEC\_VS\_MTEC\_THYMUS\_DN | MSigDB lists | GSE2585\_CTEC\_VS\_MTEC\_THYMUS\_DN | 154 | 8 | 12187 | 179 | Cd14,Ptges,Herc6,Saa1,Tnf,S100a8,Stx11,Il1rn | | 1.958e-03 | -6.24 | GSE43955\_TH0\_VS\_TGFB\_IL6\_TH17\_ACT\_CD4\_TCELL\_20H\_DN | MSigDB lists | GSE43955\_TH0\_VS\_TGFB\_IL6\_TH17\_ACT\_CD4\_TCELL\_20H\_DN | 154 | 8 | 12187 | 179 | Sgk1,Tnf,Vcam1,Ccdc58,Plaur,Irf7,Tnip1,Cxcl1 | | 1.958e-03 | -6.24 | GSE5679\_RARA\_AGONIST\_AM580\_VS\_AM580\_AND\_ROSIGLITAZONE\_TREATED\_DC\_DN | MSigDB lists | GSE5679\_RARA\_AGONIST\_AM580\_VS\_AM580\_AND\_ROSIGLITAZONE\_TREATED\_DC\_DN | 154 | 8 | 12187 | 179 | Ifitm3,Cflar,Ifitm2,Slfn5,Pik3r5,Ncf1,Irak3,Pik3r1 | | 1.959e-03 | -6.24 | PTEN\_DN.V2\_UP | MSigDB lists | PTEN\_DN.V2\_UP | 89 | 6 | 12187 | 179 | Il4ra,Birc3,Casp4,Lcn2,Ptgs2,Angptl4 | | 2.031e-03 | -6.20 | KATSANOU\_ELAVL1\_TARGETS\_DN | MSigDB lists | KATSANOU\_ELAVL1\_TARGETS\_DN | 121 | 7 | 12187 | 179 | Ifi204,Ifi207,Ifi209,Sgk1,Adamts9,Osmr,Ifi211 | | 2.031e-03 | -6.20 | AR\_01 | MSigDB lists | AR\_01 | 121 | 7 | 12187 | 179 | Sgk1,Cdkn1a,Luc7l3,Mxd4,Fam107a,Ip6k2,Kcna5 | | 2.036e-03 | -6.20 | KEGG\_TYPE\_II\_DIABETES\_MELLITUS | MSigDB lists | KEGG\_TYPE\_II\_DIABETES\_MELLITUS | 37 | 4 | 12187 | 179 | Tnf,Pik3r1,Pik3r5,Socs3 | | 2.036e-03 | -6.20 | BROWNE\_HCMV\_INFECTION\_30MIN\_UP | MSigDB lists | BROWNE\_HCMV\_INFECTION\_30MIN\_UP | 37 | 4 | 12187 | 179 | Socs3,Thbs1,Lcn2,Zfp36 | | 2.036e-03 | -6.20 | GO\_POSITIVE\_REGULATION\_OF\_MYELOID\_LEUKOCYTE\_DIFFERENTIATION | MSigDB lists | GO\_POSITIVE\_REGULATION\_OF\_MYELOID\_LEUKOCYTE\_DIFFERENTIATION | 37 | 4 | 12187 | 179 | Ripk1,Trib1,Runx1,Tnf | | 2.036e-03 | -6.20 | GO\_POSITIVE\_REGULATION\_OF\_TRANSCRIPTION\_FACTOR\_IMPORT\_INTO\_NUCLEUS | MSigDB lists | GO\_POSITIVE\_REGULATION\_OF\_TRANSCRIPTION\_FACTOR\_IMPORT\_INTO\_NUCLEUS | 37 | 4 | 12187 | 179 | Tnf,Pik3r1,Ptgs2,Csf3 | | 2.036e-03 | -6.20 | BECKER\_TAMOXIFEN\_RESISTANCE\_UP | MSigDB lists | BECKER\_TAMOXIFEN\_RESISTANCE\_UP | 37 | 4 | 12187 | 179 | Irf2,Irf9,Isg15,Ifi44 | | 2.038e-03 | -6.20 | positive regulation of secretion by cell | biological process | GO:1903532 | 370 | 14 | 13711 | 214 | Runx1,Il17ra,Mmp8,Sdc4,Hcar2,Lgals9,Cd14,Il6,Casp4,Tnf,Il4ra,Akap12,Il1a,Sox11 | | 2.039e-03 | -6.20 | GSE19401\_UNSTIM\_VS\_RETINOIC\_ACID\_STIM\_FOLLICULAR\_DC\_UP | MSigDB lists | GSE19401\_UNSTIM\_VS\_RETINOIC\_ACID\_STIM\_FOLLICULAR\_DC\_UP | 155 | 8 | 12187 | 179 | Rgs16,Hspb1,Il17ra,S100a8,P2ry6,Irf7,S100a9,Tgm2 | | 2.039e-03 | -6.20 | GSE7764\_IL15\_NK\_CELL\_24H\_VS\_SPLENOCYTE\_DN | MSigDB lists | GSE7764\_IL15\_NK\_CELL\_24H\_VS\_SPLENOCYTE\_DN | 155 | 8 | 12187 | 179 | Rassf4,Xaf1,Irf9,Zfp36,Ip6k2,Sgk3,Ifitm2,Irgm2 | | 2.039e-03 | -6.20 | GSE23568\_CTRL\_TRANSDUCED\_VS\_WT\_CD8\_TCELL\_DN | MSigDB lists | GSE23568\_CTRL\_TRANSDUCED\_VS\_WT\_CD8\_TCELL\_DN | 155 | 8 | 12187 | 179 | Isg15,Irf7,Banp,Tnip1,Tap1,Xdh,Sgk1,Ms4a6d | | 2.054e-03 | -6.19 | regulation of epithelial cell differentiation | biological process | GO:0030856 | 114 | 7 | 13711 | 214 | Xdh,Apold1,Errfi1,Zfp36,Il1a,Maff,Tnf | | 2.054e-03 | -6.19 | stress-activated protein kinase signaling cascade | biological process | GO:0031098 | 114 | 7 | 13711 | 214 | Ripk1,Trib1,Errfi1,Zfp36,Map3k8,Tnf,Ptpn1 | | 2.067e-03 | -6.18 | response to steroid hormone | biological process | GO:0048545 | 146 | 8 | 13711 | 214 | Il1rn,Sgk1,Pik3r1,Tnf,Acod1,Fam107a,Zfp36,Il6 | | 2.084e-03 | -6.17 | GO\_SUPEROXIDE\_GENERATING\_NADPH\_OXIDASE\_ACTIVITY | MSigDB lists | GO\_SUPEROXIDE\_GENERATING\_NADPH\_OXIDASE\_ACTIVITY | 5 | 2 | 12187 | 179 | Cybb,Ncf1 | | 2.084e-03 | -6.17 | BUDHU\_LIVER\_CANCER\_METASTASIS\_DN | MSigDB lists | BUDHU\_LIVER\_CANCER\_METASTASIS\_DN | 5 | 2 | 12187 | 179 | Tnf,Il1a | | 2.084e-03 | -6.17 | SOUCEK\_MYC\_TARGETS | MSigDB lists | SOUCEK\_MYC\_TARGETS | 5 | 2 | 12187 | 179 | Cxcl1,Ccl12 | | 2.084e-03 | -6.17 | COLLER\_MYC\_TARGETS\_DN | MSigDB lists | COLLER\_MYC\_TARGETS\_DN | 5 | 2 | 12187 | 179 | A2m,Cdkn1a | | 2.084e-03 | -6.17 | BIOCARTA\_IL17\_PATHWAY | MSigDB lists | BIOCARTA\_IL17\_PATHWAY | 5 | 2 | 12187 | 179 | Il6,Csf3 | | 2.087e-03 | -6.17 | GO\_POSITIVE\_REGULATION\_OF\_PROTEOLYSIS | MSigDB lists | GO\_POSITIVE\_REGULATION\_OF\_PROTEOLYSIS | 310 | 12 | 12187 | 179 | Trib1,Tnf,Ifi209,Ripk1,Nfe2l2,Ifi204,Ifi207,Ifi211,S100a9,Xdh,Tnfsf10,S100a8 | | 2.104e-03 | -6.16 | transcription regulatory region DNA binding | molecular function | GO:0044212 | 723 | 22 | 13516 | 211 | Mxd4,Ifi209,Nfkb2,Tnf,Sox11,Ifi204,Nfe2l2,Cebpd,Arid5b,Irf7,Ifi207,Spi1,Maff,Ifi211,Runx1,Tiparp,Bach1,Irf9,Irf2,Atf3,Zbtb16,Nfkb1 | | 2.116e-03 | -6.16 | response to hydrogen peroxide | biological process | GO:0042542 | 85 | 6 | 13711 | 214 | Ripk1,Nfe2l2,Tnfaip3,Lcn2,Il6,Hp | | 2.119e-03 | -6.16 | GO\_REGULATION\_OF\_ANATOMICAL\_STRUCTURE\_MORPHOGENESIS | MSigDB lists | GO\_REGULATION\_OF\_ANATOMICAL\_STRUCTURE\_MORPHOGENESIS | 822 | 23 | 12187 | 179 | Il1rn,Hspb1,Icam1,Msn,Ptgs2,Tnf,Runx1,Xdh,C5ar1,Cflar,C3ar1,Fzd4,Il1a,Sgk1,Thbs1,Ccl12,Tnfaip3,Il6,Rhou,Il4ra,Ccl2,Angptl4,Rhoj | | 2.123e-03 | -6.15 | GSE22886\_CD4\_TCELL\_VS\_BCELL\_NAIVE\_UP | MSigDB lists | GSE22886\_CD4\_TCELL\_VS\_BCELL\_NAIVE\_UP | 156 | 8 | 12187 | 179 | Map3k6,Ifitm3,Il17ra,Pik3r1,Runx1,Socs3,Timp1,Tnfaip3 | | 2.123e-03 | -6.15 | GSE13484\_UNSTIM\_VS\_3H\_YF17D\_VACCINE\_STIM\_PBMC\_DN | MSigDB lists | GSE13484\_UNSTIM\_VS\_3H\_YF17D\_VACCINE\_STIM\_PBMC\_DN | 156 | 8 | 12187 | 179 | Irf9,Nfkbie,Ifitm2,Phf11d,Bcl3,Gadd45b,Phf11b,Thbs1 | | 2.123e-03 | -6.15 | GSE17721\_LPS\_VS\_CPG\_12H\_BMDC\_DN | MSigDB lists | GSE17721\_LPS\_VS\_CPG\_12H\_BMDC\_DN | 156 | 8 | 12187 | 179 | Plaur,P2ry6,C3ar1,Nfe2l2,Tgm2,Tnf,Msr1,Pik3r1 | | 2.126e-03 | -6.15 | GO\_MEMBRANE\_DOCKING | MSigDB lists | GO\_MEMBRANE\_DOCKING | 62 | 5 | 12187 | 179 | Plek,Stx11,Icam1,Msn,Vcam1 | | 2.128e-03 | -6.15 | GO\_EXTERNAL\_SIDE\_OF\_PLASMA\_MEMBRANE | MSigDB lists | GO\_EXTERNAL\_SIDE\_OF\_PLASMA\_MEMBRANE | 122 | 7 | 12187 | 179 | Icam1,Il6,Tnf,Selp,Thbs1,Vcam1,Cd14 | | 2.141e-03 | -6.15 | ODONNELL\_TFRC\_TARGETS\_UP | MSigDB lists | ODONNELL\_TFRC\_TARGETS\_UP | 270 | 11 | 12187 | 179 | Il4ra,Map3k8,Runx1,Gadd45b,Thbs1,Sgk1,Adamts1,Cdkn1a,Mxd4,Cp,Akap12 | | 2.144e-03 | -6.15 | MODULE\_60 | MSigDB lists | MODULE\_60 | 311 | 12 | 12187 | 179 | Cd14,Isg15,Tap1,Sbno2,Trib1,Tgm2,Ccnd2,Cdkn1a,Ier3,Ccl12,Fam107a,Kcna5 | | 2.154e-03 | -6.14 | TRAF6 mediated NF-kB activation | REACTOME pathways | R-MMU-933542 | 16 | 3 | 6297 | 105 | Nfkbia,Nfkb2,Nfkb1 | | 2.161e-03 | -6.14 | GO\_REGULATION\_OF\_PROTEIN\_ACTIVATION\_CASCADE | MSigDB lists | GO\_REGULATION\_OF\_PROTEIN\_ACTIVATION\_CASCADE | 18 | 3 | 12187 | 179 | C5ar1,C3ar1,A2m | | 2.161e-03 | -6.14 | GENTILE\_UV\_HIGH\_DOSE\_UP | MSigDB lists | GENTILE\_UV\_HIGH\_DOSE\_UP | 18 | 3 | 12187 | 179 | Atf3,Isg15,Maff | | 2.161e-03 | -6.14 | BIOCARTA\_ATM\_PATHWAY | MSigDB lists | BIOCARTA\_ATM\_PATHWAY | 18 | 3 | 12187 | 179 | Cdkn1a,Nfkbia,Nfkb1 | | 2.161e-03 | -6.14 | DELLA\_RESPONSE\_TO\_TSA\_AND\_BUTYRATE | MSigDB lists | DELLA\_RESPONSE\_TO\_TSA\_AND\_BUTYRATE | 18 | 3 | 12187 | 179 | Hspb1,Icam1,Cdkn1a | | 2.161e-03 | -6.14 | GO\_LEUKOCYTE\_APOPTOTIC\_PROCESS | MSigDB lists | GO\_LEUKOCYTE\_APOPTOTIC\_PROCESS | 18 | 3 | 12187 | 179 | Ripk1,Il6,Hcar2 | | 2.161e-03 | -6.14 | KEGG\_GRAFT\_VERSUS\_HOST\_DISEASE | MSigDB lists | KEGG\_GRAFT\_VERSUS\_HOST\_DISEASE | 18 | 3 | 12187 | 179 | Il1a,Il6,Tnf | | 2.169e-03 | -6.13 | regulation of lymphocyte chemotaxis | biological process | GO:1901623 | 17 | 3 | 13711 | 214 | Lgals9,Ccl12,Ccl4 | | 2.169e-03 | -6.13 | regulation of myoblast fusion | biological process | GO:1901739 | 17 | 3 | 13711 | 214 | Il4ra,Cflar,Cxcl9 | | 2.170e-03 | -6.13 | phosphatidylinositol 3-kinase regulator activity | molecular function | GO:0035014 | 17 | 3 | 13516 | 211 | Pik3r5,Pik3r1,Socs3 | | 2.173e-03 | -6.13 | transcription regulatory region sequence-specific DNA binding | molecular function | GO:0000976 | 633 | 20 | 13516 | 211 | Ifi204,Sox11,Nfe2l2,Cebpd,Arid5b,Mxd4,Nfkb2,Ifi209,Zbtb16,Atf3,Nfkb1,Spi1,Maff,Ifi207,Irf7,Ifi211,Runx1,Irf2,Irf9,Bach1 | | 2.173e-03 | -6.13 | Set (SET nuclear oncogene) | protein interactions | 56086 | 33 | 4 | 6802 | 115 | Pygm,Hspa5,Ccnd2,Runx1 | | 2.193e-03 | -6.12 | SMIRNOV\_RESPONSE\_TO\_IR\_6HR\_DN | MSigDB lists | SMIRNOV\_RESPONSE\_TO\_IR\_6HR\_DN | 91 | 6 | 12187 | 179 | Cybb,Zfp36,Ccl4,Trib1,Tnf,Maff | | 2.210e-03 | -6.11 | GSE20366\_EX\_VIVO\_VS\_DEC205\_CONVERSION\_DN | MSigDB lists | GSE20366\_EX\_VIVO\_VS\_DEC205\_CONVERSION\_DN | 157 | 8 | 12187 | 179 | Pik3ap1,Rbm39,Ifitm2,Angptl4,Atf3,Aff1,Sgk1,Casp4 | | 2.210e-03 | -6.11 | GO\_POSITIVE\_REGULATION\_OF\_CELL\_CELL\_ADHESION | MSigDB lists | GO\_POSITIVE\_REGULATION\_OF\_CELL\_CELL\_ADHESION | 157 | 8 | 12187 | 179 | Zbtb16,Vcam1,Map3k8,Tnf,Pik3r1,Il4ra,Ccl12,Il6 | | 2.229e-03 | -6.11 | SMITH\_TERT\_TARGETS\_UP | MSigDB lists | SMITH\_TERT\_TARGETS\_UP | 123 | 7 | 12187 | 179 | Zfp36,Cxcl1,Ptgs2,Cebpd,Atf3,Nfkb1,Hspa5 | | 2.240e-03 | -6.10 | IL-1 Signaling Pathway | WikiPathways | WP37 | 37 | 5 | 3756 | 96 | Nfkb1,Il1rn,Nfkbia,Irak3,Il1a | | 2.249e-03 | -6.10 | ST\_T\_CELL\_SIGNAL\_TRANSDUCTION | MSigDB lists | ST\_T\_CELL\_SIGNAL\_TRANSDUCTION | 38 | 4 | 12187 | 179 | Nfkb1,Nfkb2,Nfkbie,Nfkbia | | 2.251e-03 | -6.10 | response to starvation | biological process | GO:0042594 | 148 | 8 | 13711 | 214 | Nfe2l2,Fstl1,Angptl4,Rrp8,Hspa5,Zfp36,Atf3,Cdkn1a | | 2.252e-03 | -6.10 | regulatory region nucleic acid binding | molecular function | GO:0001067 | 727 | 22 | 13516 | 211 | Nfkb2,Cebpd,Sox11,Bach1,Irf7,Atf3,Mxd4,Tnf,Ifi209,Arid5b,Ifi204,Nfe2l2,Runx1,Tiparp,Irf2,Irf9,Spi1,Maff,Ifi207,Ifi211,Nfkb1,Zbtb16 | | 2.255e-03 | -6.09 | RYTTCCTG\_ETS2\_B | MSigDB lists | RYTTCCTG\_ETS2\_B | 826 | 23 | 12187 | 179 | Nfkbie,Robo4,S100a9,Rhoj,Bcl3,Pik3r1,Trib1,Spi1,Hcar2,Ccl12,Tagln2,Sele,Apold1,Trim25,Msn,Sgk1,Cdkn1a,Aff1,Fstl1,Stx11,Timp1,Nfkb1,Il1rn | | 2.269e-03 | -6.09 | MEISSNER\_BRAIN\_HCP\_WITH\_H3K4ME3\_AND\_H3K27ME3 | MSigDB lists | MEISSNER\_BRAIN\_HCP\_WITH\_H3K4ME3\_AND\_H3K27ME3 | 877 | 24 | 12187 | 179 | Gadd45b,Nfe2l2,Rhoc,Tubb6,Slc24a4,Parp14,Usp18,Sdc4,Runx1,Maff,Il4ra,Tnfaip8,Osmr,Pik3r5,Atf3,Cxcl16,Tap1,Il17ra,Trim25,Irak3,Zfp36,Pik3ap1,Tagln2,Map3k6 | | 2.270e-03 | -6.09 | ripoptosome | cellular component | GO:0097342 | 5 | 2 | 13825 | 212 | Cflar,Ripk1 | | 2.282e-03 | -6.08 | TNF\_CS | interpro domains | IPR021184 | 5 | 2 | 13788 | 212 | Tnfsf10,Tnf | | 2.282e-03 | -6.08 | NFkB/Dor | interpro domains | IPR000451 | 5 | 2 | 13788 | 212 | Nfkb1,Nfkb2 | | 2.282e-03 | -6.08 | ICAM\_VCAM\_N | interpro domains | IPR003987 | 5 | 2 | 13788 | 212 | Vcam1,Icam1 | | 2.282e-03 | -6.08 | IPT\_NFkappaB | interpro domains | IPR033926 | 5 | 2 | 13788 | 212 | Nfkb1,Nfkb2 | | 2.282e-03 | -6.08 | RHD\_CS | interpro domains | IPR030492 | 5 | 2 | 13788 | 212 | Nfkb1,Nfkb2 | | 2.298e-03 | -6.08 | positive regulation of tyrosine phosphorylation of STAT protein | biological process | GO:0042531 | 36 | 4 | 13711 | 214 | Tnf,Parp14,Il6,Parp9 | | 2.299e-03 | -6.08 | GSE7219\_WT\_VS\_NIK\_NFKB2\_KO\_LPS\_AND\_ANTI\_CD40\_STIM\_DC\_DN | MSigDB lists | GSE7219\_WT\_VS\_NIK\_NFKB2\_KO\_LPS\_AND\_ANTI\_CD40\_STIM\_DC\_DN | 158 | 8 | 12187 | 179 | Usp18,Pik3r5,Xaf1,Slfn9,Gbp7,Parp12,Il1a,Irak3 | | 2.299e-03 | -6.08 | GSE26343\_UNSTIM\_VS\_LPS\_STIM\_MACROPHAGE\_UP | MSigDB lists | GSE26343\_UNSTIM\_VS\_LPS\_STIM\_MACROPHAGE\_UP | 158 | 8 | 12187 | 179 | Zfp36,Il1rn,Csf2rb,Ptgs2,Map3k8,Nfkbiz,Cxcl1,Vcam1 | | 2.299e-03 | -6.08 | GSE20715\_WT\_VS\_TLR4\_KO\_48H\_OZONE\_LUNG\_UP | MSigDB lists | GSE20715\_WT\_VS\_TLR4\_KO\_48H\_OZONE\_LUNG\_UP | 158 | 8 | 12187 | 179 | Cebpd,Nfe2l2,Il1rn,Dnajb1,Clic4,Hspb1,Tagln2,Map3k6 | | 2.301e-03 | -6.07 | Iron Homeostasis | WikiPathways | WP1596 | 11 | 3 | 3756 | 96 | Il1a,Tnf,Il6 | | 2.304e-03 | -6.07 | regulation of RNA metabolic process | biological process | GO:0051252 | 2497 | 56 | 13711 | 214 | Irf7,Dtx3l,Trib1,Pik3r1,Oasl2,Il1a,Nfkbia,Il6,Cebpd,Ifi211,Spi1,Runx1,Trim25,Resf1,Ifi209,Fzd4,Sbno2,Parp9,Ifi207,Ncf1,Cflar,Ripk1,Tnip1,Fstl1,Tnf,Acod1,Atf3,Zfp36,Nfe2l2,Rasd1,Noct,Maff,Rrp8,Dnajb1,Oasl1,Rbm39,Banp,Icam1,Mxd4,Sox11,Nfkb2,Bach1,Gbp4,Nfkb1,Nfkbie,Aff1,Irak3,Arid5b,Zbtb16,Csf3,Ifi204,Znfx1,Tnfaip3,Irf9,Bcl3,Irf2 | | 2.322e-03 | -6.07 | KEGG\_MAPK\_SIGNALING\_PATHWAY | MSigDB lists | KEGG\_MAPK\_SIGNALING\_PATHWAY | 233 | 10 | 12187 | 179 | Hspb1,Nfkb2,Map3k6,Cd14,Nfkb1,Il1a,Gadd45g,Gadd45b,Map3k8,Tnf | | 2.334e-03 | -6.06 | SATO\_SILENCED\_BY\_METHYLATION\_IN\_PANCREATIC\_CANCER\_1 | MSigDB lists | SATO\_SILENCED\_BY\_METHYLATION\_IN\_PANCREATIC\_CANCER\_1 | 273 | 11 | 12187 | 179 | Icam1,Trim25,Nfkb2,Plaur,Irf7,Ksr1,Ier3,Birc3,Cdkn1a,Tnf,Ccl4 | | 2.334e-03 | -6.06 | GSE2770\_UNTREATED\_VS\_ACT\_CD4\_TCELL\_48H\_DN | MSigDB lists | GSE2770\_UNTREATED\_VS\_ACT\_CD4\_TCELL\_48H\_DN | 124 | 7 | 12187 | 179 | Isg15,Ifit2,Spi1,Cmpk2,Timp1,Atf3,Rasd1 | | 2.351e-03 | -6.05 | connective tissue replacement | biological process | GO:0097709 | 5 | 2 | 13711 | 214 | Il1a,Timp1 | | 2.351e-03 | -6.05 | necroptotic signaling pathway | biological process | GO:0097527 | 5 | 2 | 13711 | 214 | Ripk1,Tnf | | 2.351e-03 | -6.05 | connective tissue replacement involved in inflammatory response wound healing | biological process | GO:0002248 | 5 | 2 | 13711 | 214 | Il1a,Timp1 | | 2.351e-03 | -6.05 | peptidyl-cysteine S-nitrosylation | biological process | GO:0018119 | 5 | 2 | 13711 | 214 | S100a9,S100a8 | | 2.351e-03 | -6.05 | type I interferon biosynthetic process | biological process | GO:0045351 | 5 | 2 | 13711 | 214 | Irf9,Irf7 | | 2.351e-03 | -6.05 | detoxification of copper ion | biological process | GO:0010273 | 5 | 2 | 13711 | 214 | Mt2,Mt1 | | 2.351e-03 | -6.05 | positive regulation of platelet activation | biological process | GO:0010572 | 5 | 2 | 13711 | 214 | Selp,Plek | | 2.351e-03 | -6.05 | cellular response to nicotine | biological process | GO:0071316 | 5 | 2 | 13711 | 214 | Nfkb1,Tnf | | 2.351e-03 | -6.05 | retinal blood vessel morphogenesis | biological process | GO:0061304 | 5 | 2 | 13711 | 214 | Col4a1,Fzd4 | | 2.351e-03 | -6.05 | positive regulation of autophagosome maturation | biological process | GO:1901098 | 5 | 2 | 13711 | 214 | Irgm1,Irgm2 | | 2.351e-03 | -6.05 | regulation of vitamin D biosynthetic process | biological process | GO:0060556 | 5 | 2 | 13711 | 214 | Nfkb1,Tnf | | 2.351e-03 | -6.05 | stress response to copper ion | biological process | GO:1990169 | 5 | 2 | 13711 | 214 | Mt2,Mt1 | | 2.352e-03 | -6.05 | Toll-like receptor 4 binding | molecular function | GO:0035662 | 5 | 2 | 13516 | 211 | S100a9,S100a8 | | 2.352e-03 | -6.05 | superoxide-generating NADPH oxidase activity | molecular function | GO:0016175 | 5 | 2 | 13516 | 211 | Cybb,Ncf1 | | 2.391e-03 | -6.04 | GSE19923\_E2A\_KO\_VS\_HEB\_AND\_E2A\_KO\_DP\_THYMOCYTE\_UP | MSigDB lists | GSE19923\_E2A\_KO\_VS\_HEB\_AND\_E2A\_KO\_DP\_THYMOCYTE\_UP | 159 | 8 | 12187 | 179 | Il6,Tnf,Ccnd2,Slfn4,Slfn3,Tnfsf10,Cflar,Gem | | 2.391e-03 | -6.04 | GSE24634\_IL4\_VS\_CTRL\_TREATED\_NAIVE\_CD4\_TCELL\_DAY7\_DN | MSigDB lists | GSE24634\_IL4\_VS\_CTRL\_TREATED\_NAIVE\_CD4\_TCELL\_DAY7\_DN | 159 | 8 | 12187 | 179 | Ifit3b,Atf3,Il1rn,Bach1,Tagln2,Cd14,Clic4,Tnfsf10 | | 2.396e-03 | -6.03 | BHAT\_ESR1\_TARGETS\_VIA\_AKT1\_UP | MSigDB lists | BHAT\_ESR1\_TARGETS\_VIA\_AKT1\_UP | 234 | 10 | 12187 | 179 | Rgs16,Sbno2,Ptges,Gadd45g,Tiparp,Atf3,Parp12,Ier3,Gadd45b,Tgm2 | | 2.433e-03 | -6.02 | ZHENG\_BOUND\_BY\_FOXP3 | MSigDB lists | ZHENG\_BOUND\_BY\_FOXP3 | 402 | 14 | 12187 | 179 | Ms4a6d,Nfkbiz,Ifit3b,Irf2,Selp,AA467197,Gbp4,Tnfaip3,Pik3r1,Map3k8,Trib1,Gbp6,Arid5b,Banp | | 2.447e-03 | -6.01 | cellular response to inorganic substance | biological process | GO:0071241 | 150 | 8 | 13711 | 214 | Cflar,Nfe2l2,Thbs1,Ncf1,Clic4,Mt1,Iqgap1,Mt2 | | 2.447e-03 | -6.01 | GO\_REGULATION\_OF\_JUN\_KINASE\_ACTIVITY | MSigDB lists | GO\_REGULATION\_OF\_JUN\_KINASE\_ACTIVITY | 64 | 5 | 12187 | 179 | Ripk1,Ptpn1,Map3k6,Tnf,Fzd4 | | 2.451e-03 | -6.01 | Salmonella infection | KEGG pathways | mmu05132 | 68 | 6 | 5248 | 107 | Nfkb1,Il6,Cxcl1,Cd14,Ccl4,Il1a | | 2.451e-03 | -6.01 | Salmonella infection | KEGG pathways | ko05132 | 68 | 6 | 5248 | 107 | Il1a,Ccl4,Cd14,Cxcl1,Il6,Nfkb1 | | 2.477e-03 | -6.00 | GO\_RESPONSE\_TO\_HORMONE | MSigDB lists | GO\_RESPONSE\_TO\_HORMONE | 683 | 20 | 12187 | 179 | Tnf,Pik3r1,Il4ra,Ccl12,Il6,Adamts1,Cdkn1a,Thbs1,Sgk1,Socs3,Timp1,Nfkb1,Il1rn,Ptgs2,Icam1,Trim25,P2ry6,Errfi1,Ptpn1,Msn | | 2.478e-03 | -6.00 | GO\_POSITIVE\_REGULATION\_OF\_REACTIVE\_OXYGEN\_SPECIES\_BIOSYNTHETIC\_PROCESS | MSigDB lists | GO\_POSITIVE\_REGULATION\_OF\_REACTIVE\_OXYGEN\_SPECIES\_BIOSYNTHETIC\_PROCESS | 39 | 4 | 12187 | 179 | Ptgs2,Tnf,Icam1,Il6 | | 2.478e-03 | -6.00 | MODULE\_259 | MSigDB lists | MODULE\_259 | 39 | 4 | 12187 | 179 | Cdkn1a,Il4ra,Ccl12,Csf2rb | | 2.486e-03 | -6.00 | GSE5589\_LPS\_VS\_LPS\_AND\_IL10\_STIM\_IL6\_KO\_MACROPHAGE\_45MIN\_DN | MSigDB lists | GSE5589\_LPS\_VS\_LPS\_AND\_IL10\_STIM\_IL6\_KO\_MACROPHAGE\_45MIN\_DN | 160 | 8 | 12187 | 179 | Irgm2,Plaur,Gbp3,Lgals9,Thbs1,Tnf,Cp,Samd9l | | 2.486e-03 | -6.00 | GSE2770\_IL4\_ACT\_VS\_ACT\_CD4\_TCELL\_48H\_DN | MSigDB lists | GSE2770\_IL4\_ACT\_VS\_ACT\_CD4\_TCELL\_48H\_DN | 160 | 8 | 12187 | 179 | Tnfaip8,Msn,Selp,Nfkbie,Tnip1,Slfn9,Xaf1,Trim56 | | 2.486e-03 | -6.00 | GSE37416\_12H\_VS\_24H\_F\_TULARENSIS\_LVS\_NEUTROPHIL\_UP | MSigDB lists | GSE37416\_12H\_VS\_24H\_F\_TULARENSIS\_LVS\_NEUTROPHIL\_UP | 160 | 8 | 12187 | 179 | Banp,Cxcl16,Runx1,Il4ra,Spi1,Nfkb1,Stx11,Oasl1 | | 2.486e-03 | -6.00 | GSE43955\_TH0\_VS\_TGFB\_IL6\_TH17\_ACT\_CD4\_TCELL\_20H\_UP | MSigDB lists | GSE43955\_TH0\_VS\_TGFB\_IL6\_TH17\_ACT\_CD4\_TCELL\_20H\_UP | 160 | 8 | 12187 | 179 | Plin4,Gem,Cxcl1,Cybb,Il1a,Tnf,Sgk1,Nfkbiz | | 2.486e-03 | -6.00 | GSE17721\_LPS\_VS\_CPG\_16H\_BMDC\_UP | MSigDB lists | GSE17721\_LPS\_VS\_CPG\_16H\_BMDC\_UP | 160 | 8 | 12187 | 179 | Slfn3,Slfn4,Gbp7,Cflar,Irf7,Msr1,Cdkn1a,Ccl2 | | 2.523e-03 | -5.98 | hematopoietic progenitor cell differentiation | biological process | GO:0002244 | 88 | 6 | 13711 | 214 | Spi1,Herc6,Rbm47,Cebpd,Samd9l,Runx1 | | 2.523e-03 | -5.98 | regulation of receptor signaling pathway via STAT | biological process | GO:1904892 | 88 | 6 | 13711 | 214 | Parp9,Bcl3,Il6,Parp14,Socs3,Tnf | | 2.539e-03 | -5.98 | PARK\_TRETINOIN\_RESPONSE\_AND\_RARA\_PLZF\_FUSION | MSigDB lists | PARK\_TRETINOIN\_RESPONSE\_AND\_RARA\_PLZF\_FUSION | 19 | 3 | 12187 | 179 | Spi1,Ier3,Plek | | 2.539e-03 | -5.98 | GO\_REGULATION\_OF\_NECROTIC\_CELL\_DEATH | MSigDB lists | GO\_REGULATION\_OF\_NECROTIC\_CELL\_DEATH | 19 | 3 | 12187 | 179 | Ripk1,Birc3,Cflar | | 2.539e-03 | -5.98 | GAUSSMANN\_MLL\_AF4\_FUSION\_TARGETS\_B\_UP | MSigDB lists | GAUSSMANN\_MLL\_AF4\_FUSION\_TARGETS\_B\_UP | 19 | 3 | 12187 | 179 | Irgm2,Rbm47,Parp12 | | 2.539e-03 | -5.98 | GO\_REGULATION\_OF\_NEUTROPHIL\_CHEMOTAXIS | MSigDB lists | GO\_REGULATION\_OF\_NEUTROPHIL\_CHEMOTAXIS | 19 | 3 | 12187 | 179 | C5ar1,C3ar1,Cxcl1 | | 2.539e-03 | -5.98 | GO\_REGULATION\_OF\_CELLULAR\_EXTRAVASATION | MSigDB lists | GO\_REGULATION\_OF\_CELLULAR\_EXTRAVASATION | 19 | 3 | 12187 | 179 | Icam1,Ccl12,Selp | | 2.539e-03 | -5.98 | MCCABE\_HOXC6\_TARGETS\_DN | MSigDB lists | MCCABE\_HOXC6\_TARGETS\_DN | 19 | 3 | 12187 | 179 | Vcam1,Tnfaip8,Errfi1 | | 2.539e-03 | -5.98 | GO\_POSITIVE\_REGULATION\_OF\_NEUTROPHIL\_MIGRATION | MSigDB lists | GO\_POSITIVE\_REGULATION\_OF\_NEUTROPHIL\_MIGRATION | 19 | 3 | 12187 | 179 | Cxcl1,C3ar1,C5ar1 | | 2.555e-03 | -5.97 | GSE37534\_PIOGLITAZONE\_VS\_ROSIGLITAZONE\_TREATED\_CD4\_TCELL\_PPARG1\_FOXP3\_TRANSDUCED\_DN | MSigDB lists | GSE37534\_PIOGLITAZONE\_VS\_ROSIGLITAZONE\_TREATED\_CD4\_TCELL\_PPARG1\_FOXP3\_TRANSDUCED\_DN | 126 | 7 | 12187 | 179 | A2m,Csf2rb,Nfkb2,Dnajb1,Trib1,Birc3,Col4a1 | | 2.555e-03 | -5.97 | IL15\_UP.V1\_UP | MSigDB lists | IL15\_UP.V1\_UP | 126 | 7 | 12187 | 179 | Il1a,Gadd45b,Maff,Ccl4,Trib1,Tnfaip8,Rgs16 | | 2.555e-03 | -5.97 | IL2\_UP.V1\_UP | MSigDB lists | IL2\_UP.V1\_UP | 126 | 7 | 12187 | 179 | Rgs16,Tnfaip8,Gadd45b,Stx11,Il1a,Trib1,Maff | | 2.555e-03 | -5.97 | PRC2\_EED\_UP.V1\_UP | MSigDB lists | PRC2\_EED\_UP.V1\_UP | 126 | 7 | 12187 | 179 | Angptl4,Rbm47,Trib1,Ccnd2,Atf3,Bach1,Gadd45g | | 2.573e-03 | -5.96 | positive regulation of T-helper cell differentiation | biological process | GO:0045624 | 18 | 3 | 13711 | 214 | Il6,Il4ra,Nfkbiz | | 2.573e-03 | -5.96 | regulation of heterotypic cell-cell adhesion | biological process | GO:0034114 | 18 | 3 | 13711 | 214 | Il1rn,Tnfaip3,Tnf | | 2.573e-03 | -5.96 | regulation of lipopolysaccharide-mediated signaling pathway | biological process | GO:0031664 | 18 | 3 | 13711 | 214 | Trib1,Acod1,Tnfaip3 | | 2.576e-03 | -5.96 | regulation of T cell activation | biological process | GO:0050863 | 222 | 10 | 13711 | 214 | Il4ra,Nfkbiz,Sdc4,Ccl2,Il6,Zbtb16,Ctla2a,Vcam1,Lgals9,Runx1 | | 2.580e-03 | -5.96 | anatomical structure formation involved in morphogenesis | biological process | GO:0048646 | 735 | 22 | 13711 | 214 | Adamts1,Sbno2,Sdc4,Apold1,Rhoj,Ptgs2,Rasip1,Cemip2,Nfkb2,Sox11,Thbs1,Ccl12,Col4a1,Clic4,Tgm2,Angptl4,Bcl3,Ccl2,Robo4,Rnf213,Cflar,Slc24a4 | | 2.584e-03 | -5.96 | GSE3982\_DC\_VS\_CENT\_MEMORY\_CD4\_TCELL\_UP | MSigDB lists | GSE3982\_DC\_VS\_CENT\_MEMORY\_CD4\_TCELL\_UP | 161 | 8 | 12187 | 179 | Nfkbie,C5ar1,Igsf6,Cdkn1a,Sdc4,Tor3a,Ms4a6d,Cybb | | 2.584e-03 | -5.96 | BILD\_MYC\_ONCOGENIC\_SIGNATURE | MSigDB lists | BILD\_MYC\_ONCOGENIC\_SIGNATURE | 161 | 8 | 12187 | 179 | Cflar,Map3k6,Rnf213,Nfkb2,Irf9,Tgm2,Runx1,Akap12 | | 2.584e-03 | -5.96 | GSE37416\_12H\_VS\_48H\_F\_TULARENSIS\_LVS\_NEUTROPHIL\_UP | MSigDB lists | GSE37416\_12H\_VS\_48H\_F\_TULARENSIS\_LVS\_NEUTROPHIL\_UP | 161 | 8 | 12187 | 179 | Ptges,Banp,Nfkbie,Irf9,Spi1,Il4ra,Oasl1,Nfkb1 | | 2.584e-03 | -5.96 | GSE17721\_LPS\_VS\_GARDIQUIMOD\_0.5H\_BMDC\_UP | MSigDB lists | GSE17721\_LPS\_VS\_GARDIQUIMOD\_0.5H\_BMDC\_UP | 161 | 8 | 12187 | 179 | Map3k8,Cybb,Hp,Ier3,Nfkbie,Cd14,Ncf1,Ptpn1 | | 2.584e-03 | -5.96 | GSE17721\_CTRL\_VS\_LPS\_12H\_BMDC\_DN | MSigDB lists | GSE17721\_CTRL\_VS\_LPS\_12H\_BMDC\_DN | 161 | 8 | 12187 | 179 | Arid5b,Cd14,Slfn9,Tap1,Birc3,Zfp189,Tor3a,Timp1 | | 2.584e-03 | -5.96 | GSE36078\_UNTREATED\_VS\_AD5\_INF\_MOUSE\_LUNG\_DC\_DN | MSigDB lists | GSE36078\_UNTREATED\_VS\_AD5\_INF\_MOUSE\_LUNG\_DC\_DN | 161 | 8 | 12187 | 179 | Cdkn1a,Ccl4,Rhoc,Ifitm3,Gbp7,Gem,Pik3ap1,Errfi1 | | 2.584e-03 | -5.96 | GSE46606\_IRF4HIGH\_VS\_IRF4MID\_CD40L\_IL2\_IL5\_DAY3\_STIMULATED\_BCELL\_DN | MSigDB lists | GSE46606\_IRF4HIGH\_VS\_IRF4MID\_CD40L\_IL2\_IL5\_DAY3\_STIMULATED\_BCELL\_DN | 161 | 8 | 12187 | 179 | Icam1,Irf2,Ifitm3,Trim56,Slfn9,Atf3,Ccnd2,Tnf | | 2.584e-03 | -5.96 | GSE24634\_TREG\_VS\_TCONV\_POST\_DAY10\_IL4\_CONVERSION\_DN | MSigDB lists | GSE24634\_TREG\_VS\_TCONV\_POST\_DAY10\_IL4\_CONVERSION\_DN | 161 | 8 | 12187 | 179 | Cd14,S100a9,Rbm47,S100a8,Igsf6,Cdkn1a,Ccl2,Tubb6 | | 2.584e-03 | -5.96 | ZBP1(DAI) mediated induction of type I IFNs | REACTOME pathways | R-MMU-1606322 | 17 | 3 | 6297 | 105 | Nfkbia,Nfkb2,Nfkb1 | | 2.584e-03 | -5.96 | Caspase activation via extrinsic apoptotic signalling pathway | REACTOME pathways | R-MMU-5357769 | 17 | 3 | 6297 | 105 | Cflar,Ripk1,Tnfsf10 | | 2.588e-03 | -5.96 | cytokine receptor activity | molecular function | GO:0004896 | 61 | 5 | 13516 | 211 | Il4ra,Fzd4,Il17ra,Osmr,Csf2rb | | 2.598e-03 | -5.95 | CEBPB\_02 | MSigDB lists | CEBPB\_02 | 198 | 9 | 12187 | 179 | Rbm39,Ip6k2,S100a9,Cp,Nfe2l2,Ccnd2,Trib1,Nfkbia,Nfkbiz | | 2.620e-03 | -5.94 | Staphylococcus aureus infection | KEGG pathways | mmu05150 | 29 | 4 | 5248 | 107 | C3ar1,Selp,Icam1,C5ar1 | | 2.620e-03 | -5.94 | Staphylococcus aureus infection | KEGG pathways | ko05150 | 29 | 4 | 5248 | 107 | C3ar1,Selp,Icam1,C5ar1 | | 2.620e-03 | -5.94 | GO\_VESICLE\_LUMEN | MSigDB lists | GO\_VESICLE\_LUMEN | 65 | 5 | 12187 | 179 | Saa1,Thbs1,A2m,Timp1,Hp | | 2.620e-03 | -5.94 | KOKKINAKIS\_METHIONINE\_DEPRIVATION\_96HR\_DN | MSigDB lists | KOKKINAKIS\_METHIONINE\_DEPRIVATION\_96HR\_DN | 65 | 5 | 12187 | 179 | Birc3,Nfkbia,Irf7,Rtp4,Tnfsf10 | | 2.620e-03 | -5.94 | REACTOME\_NFKB\_AND\_MAP\_KINASES\_ACTIVATION\_MEDIATED\_BY\_TLR4\_SIGNALING\_REPERTOIRE | MSigDB lists | REACTOME\_NFKB\_AND\_MAP\_KINASES\_ACTIVATION\_MEDIATED\_BY\_TLR4\_SIGNALING\_REPERTOIRE | 65 | 5 | 12187 | 179 | Nfkb2,Ripk1,Cd14,Nfkbia,Saa1 | | 2.620e-03 | -5.94 | GO\_CYTOKINE\_BINDING | MSigDB lists | GO\_CYTOKINE\_BINDING | 65 | 5 | 12187 | 179 | Thbs1,A2m,Fzd4,Zfp36,Il1rn | | 2.621e-03 | -5.94 | peptidase regulator activity | molecular function | GO:0061134 | 119 | 7 | 13516 | 211 | Serpina3f,A2m,Timp1,Birc3,Tnfaip8,Wfdc21,Cflar | | 2.623e-03 | -5.94 | cellular response to stress | biological process | GO:0033554 | 1276 | 33 | 13711 | 214 | Bcl3,Angptl4,Nfe2l2,Rrp8,Dnajb1,Ptpn1,Hspa5,Lcn2,Ifi204,Ncf1,Cflar,Ripk1,Zfp36,Atf3,Tnfaip3,Tnf,Ptgs2,Sgk1,Ier3,Trim25,Bach1,Parp9,Map3k8,Pik3r1,Dtx3l,Thbs1,Trib1,Errfi1,Il6,Cybb,Cdkn1a,Casp4,Il1a | | 2.656e-03 | -5.93 | MAPK cascade | biological process | GO:0000165 | 152 | 8 | 13711 | 214 | Trib1,Ripk1,Map3k6,Ptpn1,Map3k8,Tnf,Ccl12,Zfp36 | | 2.671e-03 | -5.93 | positive regulation of ossification | biological process | GO:0045778 | 89 | 6 | 13711 | 214 | Ifi204,Isg15,Il6,Zbtb16,Sox11,Cebpd | | 2.685e-03 | -5.92 | GSE17721\_LPS\_VS\_PAM3CSK4\_4H\_BMDC\_UP | MSigDB lists | GSE17721\_LPS\_VS\_PAM3CSK4\_4H\_BMDC\_UP | 162 | 8 | 12187 | 179 | Clic4,Slfn3,Slfn4,Atf3,Tnfaip3,Casp4,Sdc4,Tor3a | | 2.685e-03 | -5.92 | GSE26290\_CTRL\_VS\_AKT\_INHIBITOR\_TREATED\_ANTI\_CD3\_AND\_IL2\_STIM\_CD8\_TCELL\_DN | MSigDB lists | GSE26290\_CTRL\_VS\_AKT\_INHIBITOR\_TREATED\_ANTI\_CD3\_AND\_IL2\_STIM\_CD8\_TCELL\_DN | 162 | 8 | 12187 | 179 | Tnfaip8,Plaur,Tap1,Rhoc,Map3k8,Birc3,Il4ra,Casp4 | | 2.685e-03 | -5.92 | GSE2770\_IL4\_ACT\_VS\_ACT\_CD4\_TCELL\_6H\_DN | MSigDB lists | GSE2770\_IL4\_ACT\_VS\_ACT\_CD4\_TCELL\_6H\_DN | 162 | 8 | 12187 | 179 | Sdc4,Birc3,Nfe2l2,Znfx1,Tubb6,Tnip1,Cflar,Hspb1 | | 2.685e-03 | -5.92 | GSE16450\_IMMATURE\_VS\_MATURE\_NEURON\_CELL\_LINE\_12H\_IFNA\_STIM\_DN | MSigDB lists | GSE16450\_IMMATURE\_VS\_MATURE\_NEURON\_CELL\_LINE\_12H\_IFNA\_STIM\_DN | 162 | 8 | 12187 | 179 | Gadd45b,Casp4,Cdkn1a,Clic4,Slfn4,Slfn3,Cflar,Nfkbie | | 2.686e-03 | -5.92 | GO\_SIDE\_OF\_MEMBRANE | MSigDB lists | GO\_SIDE\_OF\_MEMBRANE | 278 | 11 | 12187 | 179 | Cd14,Selp,Gem,Errfi1,Ptpn1,Iqgap1,Icam1,Vcam1,Tnf,Thbs1,Il6 | | 2.688e-03 | -5.92 | IVANOVA\_HEMATOPOIESIS\_STEM\_CELL | MSigDB lists | IVANOVA\_HEMATOPOIESIS\_STEM\_CELL | 199 | 9 | 12187 | 179 | Pygm,Gem,Nfkbie,Robo4,Rasd1,Col4a1,Nfkbia,Maff,Ifi44 | | 2.688e-03 | -5.92 | GO\_RESPONSE\_TO\_ENDOPLASMIC\_RETICULUM\_STRESS | MSigDB lists | GO\_RESPONSE\_TO\_ENDOPLASMIC\_RETICULUM\_STRESS | 199 | 9 | 12187 | 179 | Atf3,Nfe2l2,Ccl12,Pik3r1,Thbs1,Hspa5,Casp4,Trim25,Ptpn1 | | 2.722e-03 | -5.91 | YU\_MYC\_TARGETS\_DN | MSigDB lists | YU\_MYC\_TARGETS\_DN | 40 | 4 | 12187 | 179 | Pygm,Gbp3,Slfn3,Slfn4 | | 2.722e-03 | -5.91 | ENGELMANN\_CANCER\_PROGENITORS\_UP | MSigDB lists | ENGELMANN\_CANCER\_PROGENITORS\_UP | 40 | 4 | 12187 | 179 | Map3k8,Gem,Rasd1,Nfe2l2 | | 2.722e-03 | -5.91 | GO\_REGULATION\_OF\_NITRIC\_OXIDE\_BIOSYNTHETIC\_PROCESS | MSigDB lists | GO\_REGULATION\_OF\_NITRIC\_OXIDE\_BIOSYNTHETIC\_PROCESS | 40 | 4 | 12187 | 179 | Il6,Icam1,Ptgs2,Tnf | | 2.722e-03 | -5.91 | CHIARADONNA\_NEOPLASTIC\_TRANSFORMATION\_KRAS\_CDC25\_DN | MSigDB lists | CHIARADONNA\_NEOPLASTIC\_TRANSFORMATION\_KRAS\_CDC25\_DN | 40 | 4 | 12187 | 179 | Ptgs2,Vcam1,Zfp36,Osmr | | 2.722e-03 | -5.91 | MODULE\_418 | MSigDB lists | MODULE\_418 | 40 | 4 | 12187 | 179 | Sele,Cp,Rhou,Ifi44 | | 2.722e-03 | -5.91 | MARSON\_FOXP3\_TARGETS\_DN | MSigDB lists | MARSON\_FOXP3\_TARGETS\_DN | 40 | 4 | 12187 | 179 | Nfkbie,Slfn4,Slfn3,Gadd45b | | 2.722e-03 | -5.91 | HALMOS\_CEBPA\_TARGETS\_UP | MSigDB lists | HALMOS\_CEBPA\_TARGETS\_UP | 40 | 4 | 12187 | 179 | Tnfaip8,Cxcl1,Ptgs2,Il4ra | | 2.729e-03 | -5.90 | cellular chemical homeostasis | biological process | GO:0055082 | 555 | 18 | 13711 | 214 | Sgk1,Kcna5,Vcam1,Mt2,Clic4,C5ar1,Cxcl1,Tgm2,Cp,P2ry6,Lcn2,Pik3r1,C3ar1,Pygm,Slc24a4,Ccl2,Icam1,Mt1 | | 2.741e-03 | -5.90 | USP9X (ubiquitin specific peptidase 9 X-linked) | protein interactions | 8239 | 5 | 2 | 6802 | 115 | Msn,Ksr1 | | 2.741e-03 | -5.90 | Stat6 (signal transducer and activator of transcription 6) | protein interactions | 20852 | 5 | 2 | 6802 | 115 | Parp14,Il4ra | | 2.748e-03 | -5.90 | response to metal ion | biological process | GO:0010038 | 224 | 10 | 13711 | 214 | Nfe2l2,Ncf1,Thbs1,Cp,Il1a,Clic4,Mt1,Iqgap1,Xdh,Mt2 | | 2.767e-03 | -5.89 | GO\_ANATOMICAL\_STRUCTURE\_FORMATION\_INVOLVED\_IN\_MORPHOGENESIS | MSigDB lists | GO\_ANATOMICAL\_STRUCTURE\_FORMATION\_INVOLVED\_IN\_MORPHOGENESIS | 739 | 21 | 12187 | 179 | Slc24a4,Ccl12,Spi1,Sdc4,Tgm2,Nfkb2,Sbno2,Robo4,Cflar,Bcl3,Angptl4,Col4a1,Mmp8,Rasip1,Thbs1,Adamts1,Sox11,Apold1,Clic4,Ptgs2,Rnf213 | | 2.789e-03 | -5.88 | GSE19198\_6H\_VS\_24H\_IL21\_TREATED\_TCELL\_DN | MSigDB lists | GSE19198\_6H\_VS\_24H\_IL21\_TREATED\_TCELL\_DN | 163 | 8 | 12187 | 179 | Olfml2b,Dtx3l,Ifitm3,Arid5b,Pik3r5,Cebpd,Aff1,Sntb2 | | 2.789e-03 | -5.88 | CAIRO\_HEPATOBLASTOMA\_DN | MSigDB lists | CAIRO\_HEPATOBLASTOMA\_DN | 163 | 8 | 12187 | 179 | Cp,Hp,Gadd45g,Trib1,Il4ra,Xdh,Cd14,Angptl4 | | 2.789e-03 | -5.88 | GSE14308\_TH17\_VS\_NATURAL\_TREG\_DN | MSigDB lists | GSE14308\_TH17\_VS\_NATURAL\_TREG\_DN | 163 | 8 | 12187 | 179 | Luc7l3,Tnf,Nfkbia,Trib1,Tnfaip8,Cmpk2,Pik3r5,S100a9 | | 2.789e-03 | -5.88 | GSE17721\_LPS\_VS\_CPG\_4H\_BMDC\_UP | MSigDB lists | GSE17721\_LPS\_VS\_CPG\_4H\_BMDC\_UP | 163 | 8 | 12187 | 179 | Parp9,Gbp3,Icam1,Ptgs2,Parp12,Ccl4,Aff1,Ccnd2 | | 2.792e-03 | -5.88 | GSE22886\_NAIVE\_CD8\_TCELL\_VS\_NEUTROPHIL\_DN | MSigDB lists | GSE22886\_NAIVE\_CD8\_TCELL\_VS\_NEUTROPHIL\_DN | 128 | 7 | 12187 | 179 | Ptgs2,C5ar1,S100a9,Tnfsf10,Csf2rb,Cybb,Cebpd | | 2.792e-03 | -5.88 | KRAS.600\_UP.V1\_DN | MSigDB lists | KRAS.600\_UP.V1\_DN | 128 | 7 | 12187 | 179 | Zbtb16,Xaf1,Sox11,Gadd45g,Tnfsf10,Rtp4,Adamts9 | | 2.802e-03 | -5.88 | GARGALOVIC\_RESPONSE\_TO\_OXIDIZED\_PHOSPHOLIPIDS\_TURQUOISE\_UP | MSigDB lists | GARGALOVIC\_RESPONSE\_TO\_OXIDIZED\_PHOSPHOLIPIDS\_TURQUOISE\_UP | 66 | 5 | 12187 | 179 | Sgk1,Cflar,Zfp36,Akap12,Tiparp | | 2.811e-03 | -5.87 | negative regulation of I-kappaB kinase/NF-kappaB signaling | biological process | GO:0043124 | 38 | 4 | 13711 | 214 | Tnip3,Tnip1,Tnfaip3,Ripk1 | | 2.811e-03 | -5.87 | regulation of CD4-positive, alpha-beta T cell differentiation | biological process | GO:0043370 | 38 | 4 | 13711 | 214 | Il6,Runx1,Nfkbiz,Il4ra | | 2.824e-03 | -5.87 | GO\_REGULATION\_OF\_CATABOLIC\_PROCESS | MSigDB lists | GO\_REGULATION\_OF\_CATABOLIC\_PROCESS | 642 | 19 | 12187 | 179 | Nfe2l2,Tnfaip3,Hcar2,Trib1,Tnf,Banp,Ifi211,Rasip1,Hp,Ifi207,Timp1,Ifi204,Tiparp,Ifi209,Msn,Ptpn1,Zfp36,Hspb1,Irak3 | | 2.853e-03 | -5.86 | TNF signaling | REACTOME pathways | R-MMU-75893 | 36 | 4 | 6297 | 105 | Tnf,Birc3,Tnfaip3,Ripk1 | | 2.873e-03 | -5.85 | DAZARD\_RESPONSE\_TO\_UV\_SCC\_UP | MSigDB lists | DAZARD\_RESPONSE\_TO\_UV\_SCC\_UP | 96 | 6 | 12187 | 179 | Fstl1,Cxcl1,Ptgs2,Tnf,Il6,Atf3 | | 2.878e-03 | -5.85 | heparin binding | molecular function | GO:0008201 | 121 | 7 | 13516 | 211 | Thbs1,Adamts1,Selp,Ccl2,Saa1,Ccl7,Fstl1 | | 2.896e-03 | -5.84 | GSE10239\_NAIVE\_VS\_DAY4.5\_EFF\_CD8\_TCELL\_UP | MSigDB lists | GSE10239\_NAIVE\_VS\_DAY4.5\_EFF\_CD8\_TCELL\_UP | 164 | 8 | 12187 | 179 | Socs3,Rnf213,Nfkbie,Tnip1,Irf7,Il17ra,Zfp36,Rtp4 | | 2.896e-03 | -5.84 | GSE40666\_WT\_VS\_STAT1\_KO\_CD8\_TCELL\_UP | MSigDB lists | GSE40666\_WT\_VS\_STAT1\_KO\_CD8\_TCELL\_UP | 164 | 8 | 12187 | 179 | Bcl3,Clic4,Irf9,Il17ra,Il4ra,Birc3,Tiparp,Ms4a6d | | 2.896e-03 | -5.84 | GSE36891\_UNSTIM\_VS\_PAM\_TLR2\_STIM\_PERITONEAL\_MACROPHAGE\_UP | MSigDB lists | GSE36891\_UNSTIM\_VS\_PAM\_TLR2\_STIM\_PERITONEAL\_MACROPHAGE\_UP | 164 | 8 | 12187 | 179 | Il4ra,Maff,Dnajb1,Rassf4,Ksr1,Ripk1,Bcl3,Tnfaip8 | | 2.896e-03 | -5.84 | GSE17721\_LPS\_VS\_GARDIQUIMOD\_12H\_BMDC\_UP | MSigDB lists | GSE17721\_LPS\_VS\_GARDIQUIMOD\_12H\_BMDC\_UP | 164 | 8 | 12187 | 179 | Sdc4,Ifit2,Clic4,Slfn3,Trim25,Slfn4,Bcl3,Tagln2 | | 2.896e-03 | -5.84 | WIERENGA\_STAT5A\_TARGETS\_UP | MSigDB lists | WIERENGA\_STAT5A\_TARGETS\_UP | 164 | 8 | 12187 | 179 | Arid5b,Gbp3,Ccnd2,Il4ra,Tgm2,Trib1,Ccl12,Gadd45b | | 2.896e-03 | -5.84 | GSE11057\_NAIVE\_CD4\_VS\_PBMC\_CD4\_TCELL\_DN | MSigDB lists | GSE11057\_NAIVE\_CD4\_VS\_PBMC\_CD4\_TCELL\_DN | 164 | 8 | 12187 | 179 | Irf2,Csf2rb,C3ar1,Cebpd,Rhoc,Map3k8,Cdkn1a,Rhou | | 2.896e-03 | -5.84 | GSE19941\_IL10\_KO\_VS\_IL10\_KO\_AND\_NFKBP50\_KO\_UNSTIM\_MACROPHAGE\_DN | MSigDB lists | GSE19941\_IL10\_KO\_VS\_IL10\_KO\_AND\_NFKBP50\_KO\_UNSTIM\_MACROPHAGE\_DN | 164 | 8 | 12187 | 179 | Cebpd,Sgk1,Pik3r1,Plaur,Map3k6,Saa1,Gem,Pik3r5 | | 2.955e-03 | -5.82 | STAT3\_01 | MSigDB lists | STAT3\_01 | 20 | 3 | 12187 | 179 | Maff,Ccl12,Icam1 | | 2.955e-03 | -5.82 | MODULE\_488 | MSigDB lists | MODULE\_488 | 20 | 3 | 12187 | 179 | Cdkn1a,Il1a,Il6 | | 2.955e-03 | -5.82 | HOFMANN\_MYELODYSPLASTIC\_SYNDROM\_LOW\_RISK\_UP | MSigDB lists | HOFMANN\_MYELODYSPLASTIC\_SYNDROM\_LOW\_RISK\_UP | 20 | 3 | 12187 | 179 | Runx1,Cdkn1a,Atf3 | | 2.955e-03 | -5.82 | GO\_POSITIVE\_REGULATION\_OF\_TRANSCRIPTION\_FROM\_RNA\_POLYMERASE\_II\_PROMOTER\_IN\_RESPONSE\_TO\_STRESS | MSigDB lists | GO\_POSITIVE\_REGULATION\_OF\_TRANSCRIPTION\_FROM\_RNA\_POLYMERASE\_II\_PROMOTER\_IN\_RESPONSE\_TO\_STRESS | 20 | 3 | 12187 | 179 | Nfe2l2,Atf3,Hspa5 | | 2.955e-03 | -5.82 | BIOCARTA\_AKT\_PATHWAY | MSigDB lists | BIOCARTA\_AKT\_PATHWAY | 20 | 3 | 12187 | 179 | Nfkb1,Nfkbia,Pik3r1 | | 2.955e-03 | -5.82 | GO\_REGULATION\_OF\_LYMPHOCYTE\_MIGRATION | MSigDB lists | GO\_REGULATION\_OF\_LYMPHOCYTE\_MIGRATION | 20 | 3 | 12187 | 179 | Ccl4,Msn,Ccl12 | | 2.971e-03 | -5.82 | WANG\_CISPLATIN\_RESPONSE\_AND\_XPC\_DN | MSigDB lists | WANG\_CISPLATIN\_RESPONSE\_AND\_XPC\_DN | 202 | 9 | 12187 | 179 | Ifi204,Ifi207,Cdkn1a,Runx1,Phf11b,Ifi209,Casp4,Ifi211,Phf11d | | 2.971e-03 | -5.82 | GO\_GTPASE\_ACTIVITY | MSigDB lists | GO\_GTPASE\_ACTIVITY | 202 | 9 | 12187 | 179 | Tubb6,Gbp4,Rasd1,Rhou,Gbp3,Gbp7,Gem,Gbp6,Rhoj | | 2.977e-03 | -5.82 | Toll Like Receptor 4 (TLR4) Cascade | REACTOME pathways | R-MMU-166016 | 86 | 6 | 6297 | 105 | Map3k8,Cd14,Nfkbia,Nfkb2,Irf7,Nfkb1 | | 2.980e-03 | -5.82 | positive regulation of cellular component organization | biological process | GO:0051130 | 1084 | 29 | 13711 | 214 | Lcn2,Rhoc,Hspb1,Sele,Cd14,Cflar,Csf3,Tnfsf10,Tnf,Plek,Cxcl9,Msn,Irgm2,Nfe2l2,S100a9,Iqgap1,Hspa5,Il6,Icam1,Il4ra,Il1a,Sgk1,Plaur,Resf1,Rasip1,Irgm1,Sdc4,Adamts1,Fzd4 | | 2.982e-03 | -5.82 | RNA polymerase II distal enhancer sequence-specific DNA binding | molecular function | GO:0000980 | 63 | 5 | 13516 | 211 | Bach1,Nfkb1,Nfe2l2,Zbtb16,Spi1 | | 2.983e-03 | -5.81 | MODULE\_165 | MSigDB lists | MODULE\_165 | 41 | 4 | 12187 | 179 | Ptgs2,C5ar1,Spi1,Plek | | 2.987e-03 | -5.81 | cytokine binding | molecular function | GO:0019955 | 91 | 6 | 13516 | 211 | A2m,Zfp36,Osmr,Thbs1,Il1rn,Fzd4 | | 2.987e-03 | -5.81 | peptidase inhibitor activity | molecular function | GO:0030414 | 91 | 6 | 13516 | 211 | Serpina3f,A2m,Timp1,Birc3,Tnfaip8,Wfdc21 | | 2.992e-03 | -5.81 | REACTOME\_INTEGRIN\_CELL\_SURFACE\_INTERACTIONS | MSigDB lists | REACTOME\_INTEGRIN\_CELL\_SURFACE\_INTERACTIONS | 67 | 5 | 12187 | 179 | Col4a1,Icam1,Ptpn1,Thbs1,Vcam1 | | 3.006e-03 | -5.81 | GSE37534\_UNTREATED\_VS\_GW1929\_TREATED\_CD4\_TCELL\_PPARG1\_AND\_FOXP3\_TRASDUCED\_UP | MSigDB lists | GSE37534\_UNTREATED\_VS\_GW1929\_TREATED\_CD4\_TCELL\_PPARG1\_AND\_FOXP3\_TRASDUCED\_UP | 165 | 8 | 12187 | 179 | Nfkbie,Phf11d,Fzd4,Ifit2,Phf11b,Nfkbia,Tnfaip3,Il6 | | 3.006e-03 | -5.81 | GSE22886\_NAIVE\_CD8\_TCELL\_VS\_MEMORY\_TCELL\_DN | MSigDB lists | GSE22886\_NAIVE\_CD8\_TCELL\_VS\_MEMORY\_TCELL\_DN | 165 | 8 | 12187 | 179 | Timp1,Il4ra,Birc3,Ccnd2,Trim25,Tnfsf10,Arid5b,Tagln2 | | 3.006e-03 | -5.81 | GSE7831\_UNSTIM\_VS\_CPG\_STIM\_PDC\_4H\_DN | MSigDB lists | GSE7831\_UNSTIM\_VS\_CPG\_STIM\_PDC\_4H\_DN | 165 | 8 | 12187 | 179 | Isg15,Clic4,Csf2rb,Trim25,Usp18,Adamts1,Cebpd,Timp1 | | 3.013e-03 | -5.80 | negative regulation of catabolic process | biological process | GO:0009895 | 265 | 11 | 13711 | 214 | Timp1,Tnfaip3,Tnf,Zfp36,Hp,Rasip1,Irak3,Ier3,Noct,Hcar2,Banp | | 3.021e-03 | -5.80 | cellular response to cadmium ion | biological process | GO:0071276 | 19 | 3 | 13711 | 214 | Ncf1,Mt1,Mt2 | | 3.024e-03 | -5.80 | negative regulation of cell adhesion | biological process | GO:0007162 | 227 | 10 | 13711 | 214 | Il1rn,Thbs1,Pik3r1,Runx1,Lgals9,Tnfaip3,Il4ra,Fam107a,Fzd4,Sdc4 | | 3.025e-03 | -5.80 | GO\_CELLULAR\_RESPONSE\_TO\_ALCOHOL | MSigDB lists | GO\_CELLULAR\_RESPONSE\_TO\_ALCOHOL | 97 | 6 | 12187 | 179 | Adamts1,Spi1,Il6,Ccl12,Msn,Errfi1 | | 3.032e-03 | -5.80 | membrane raft | cellular component | GO:0045121 | 310 | 12 | 13825 | 212 | Ripk1,Cd14,Sdc4,Iqgap1,Icam1,Birc3,Kcna5,Tnf,Ptgs2,Cflar,Sele,Rgs16 | | 3.064e-03 | -5.79 | Interleukin-6 family signaling | REACTOME pathways | R-MMU-6783589 | 18 | 3 | 6297 | 105 | Osmr,Socs3,Il6 | | 3.064e-03 | -5.79 | Antimicrobial peptides | REACTOME pathways | R-MMU-6803157 | 18 | 3 | 6297 | 105 | Lcn2,S100a9,S100a8 | | 3.094e-03 | -5.78 | negative regulation of cytokine-mediated signaling pathway | biological process | GO:0001960 | 39 | 4 | 13711 | 214 | Il6,Il1rn,Irak3,Parp14 | | 3.094e-03 | -5.78 | regulation of p38MAPK cascade | biological process | GO:1900744 | 39 | 4 | 13711 | 214 | Gadd45g,Gadd45b,Xdh,Ncf1 | | 3.094e-03 | -5.78 | platelet-derived growth factor receptor signaling pathway | biological process | GO:0048008 | 39 | 4 | 13711 | 214 | Iqgap1,Ptpn1,Arid5b,Tiparp | | 3.094e-03 | -5.78 | positive regulation of interleukin-1 beta production | biological process | GO:0032731 | 39 | 4 | 13711 | 214 | Ifi204,Ifi211,Casp4,Hspb1 | | 3.096e-03 | -5.78 | GO\_NEGATIVE\_REGULATION\_OF\_INTERLEUKIN\_1\_BETA\_PRODUCTION | MSigDB lists | GO\_NEGATIVE\_REGULATION\_OF\_INTERLEUKIN\_1\_BETA\_PRODUCTION | 6 | 2 | 12187 | 179 | Tnfaip3,Errfi1 | | 3.096e-03 | -5.78 | GO\_SUPEROXIDE\_ANION\_GENERATION | MSigDB lists | GO\_SUPEROXIDE\_ANION\_GENERATION | 6 | 2 | 12187 | 179 | Cybb,Ncf1 | | 3.096e-03 | -5.78 | SEIKE\_LUNG\_CANCER\_POOR\_SURVIVAL | MSigDB lists | SEIKE\_LUNG\_CANCER\_POOR\_SURVIVAL | 6 | 2 | 12187 | 179 | Il1a,Tnf | | 3.096e-03 | -5.78 | KONDO\_HYPOXIA | MSigDB lists | KONDO\_HYPOXIA | 6 | 2 | 12187 | 179 | Timp1,Il6 | | 3.096e-03 | -5.78 | SHIN\_B\_CELL\_LYMPHOMA\_CLUSTER\_6 | MSigDB lists | SHIN\_B\_CELL\_LYMPHOMA\_CLUSTER\_6 | 6 | 2 | 12187 | 179 | Ptgs2,S100a8 | | 3.096e-03 | -5.78 | REICHERT\_G1S\_REGULATORS\_AS\_PI3K\_TARGETS | MSigDB lists | REICHERT\_G1S\_REGULATORS\_AS\_PI3K\_TARGETS | 6 | 2 | 12187 | 179 | Cdkn1a,Ccnd2 | | 3.096e-03 | -5.78 | BIOCARTA\_STEM\_PATHWAY | MSigDB lists | BIOCARTA\_STEM\_PATHWAY | 6 | 2 | 12187 | 179 | Csf3,Il6 | | 3.096e-03 | -5.78 | GO\_REGULATION\_OF\_CHRONIC\_INFLAMMATORY\_RESPONSE | MSigDB lists | GO\_REGULATION\_OF\_CHRONIC\_INFLAMMATORY\_RESPONSE | 6 | 2 | 12187 | 179 | Tnf,Tnfaip3 | | 3.096e-03 | -5.78 | SIMBULAN\_UV\_RESPONSE\_NORMAL\_UP | MSigDB lists | SIMBULAN\_UV\_RESPONSE\_NORMAL\_UP | 6 | 2 | 12187 | 179 | Il6,Cxcl1 | | 3.096e-03 | -5.78 | chr1p | MSigDB lists | chr1p | 6 | 2 | 12187 | 179 | Ifi44,Clic4 | | 3.104e-03 | -5.78 | protein kinase binding | molecular function | GO:0019901 | 653 | 20 | 13516 | 211 | Ksr1,Rhoj,Iqgap1,Cdkn1a,Rhou,Hspb1,Plek,Rhoc,Tnip1,Pik3r1,Irgm2,Kcna5,Sdc4,Ccnd2,Msn,Sgk1,Zfp36,Errfi1,Trib1,Ptpn1 | | 3.112e-03 | -5.77 | membrane microdomain | cellular component | GO:0098857 | 311 | 12 | 13825 | 212 | Sele,Rgs16,Kcna5,Tnf,Ptgs2,Cflar,Ripk1,Cd14,Birc3,Iqgap1,Sdc4,Icam1 | | 3.120e-03 | -5.77 | GSE33424\_CD161\_HIGH\_VS\_INT\_CD8\_TCELL\_UP | MSigDB lists | GSE33424\_CD161\_HIGH\_VS\_INT\_CD8\_TCELL\_UP | 166 | 8 | 12187 | 179 | Plaur,Gbp3,Clic4,Saa1,Rasd1,Tiparp,Tubb6,Map3k8 | | 3.120e-03 | -5.77 | GSE40685\_NAIVE\_CD4\_TCELL\_VS\_TREG\_DN | MSigDB lists | GSE40685\_NAIVE\_CD4\_TCELL\_VS\_TREG\_DN | 166 | 8 | 12187 | 179 | Lgals9,Tnip1,Tor3a,Xaf1,Ifit3b,Ksr1,Parp9,Gadd45b | | 3.120e-03 | -5.77 | GSE17721\_0.5H\_VS\_8H\_CPG\_BMDC\_DN | MSigDB lists | GSE17721\_0.5H\_VS\_8H\_CPG\_BMDC\_DN | 166 | 8 | 12187 | 179 | Usp18,Parp14,Ccl12,Ifitm2,Rtp4,Icam1,Tap1,Clic4 | | 3.120e-03 | -5.77 | GSE5542\_UNTREATED\_VS\_IFNA\_TREATED\_EPITHELIAL\_CELLS\_24H\_UP | MSigDB lists | GSE5542\_UNTREATED\_VS\_IFNA\_TREATED\_EPITHELIAL\_CELLS\_24H\_UP | 166 | 8 | 12187 | 179 | Slfn4,Slfn3,Cflar,Slfn9,Ksr1,Tnip3,Tnfaip3,Mxd4 | | 3.120e-03 | -5.77 | GSE23321\_CENTRAL\_VS\_EFFECTOR\_MEMORY\_CD8\_TCELL\_UP | MSigDB lists | GSE23321\_CENTRAL\_VS\_EFFECTOR\_MEMORY\_CD8\_TCELL\_UP | 166 | 8 | 12187 | 179 | Casp4,Dnajb1,Tgm2,Luc7l3,Bcl3,Cxcl16,Slfn3,Slfn4 | | 3.153e-03 | -5.76 | regulation of nucleobase-containing compound metabolic process | biological process | GO:0019219 | 2701 | 59 | 13711 | 214 | Parp9,Sbno2,Ifi209,Fzd4,Runx1,Trim25,Resf1,Ier3,Ifi211,Cebpd,Spi1,Il6,Cdkn1a,Oasl2,Nfkbia,Il1a,Pik3r1,Dtx3l,Irf7,Trib1,Maff,Rrp8,Dnajb1,Noct,Nfe2l2,Rasd1,Zfp36,Atf3,Tnf,Acod1,Fstl1,Tnip1,Ncf1,Ifi207,Ripk1,Cflar,Nfkb1,Nfkbie,Gbp4,Bach1,Mxd4,Icam1,Nfkb2,Sox11,Rbm39,Banp,Oasl1,Irf2,Bcl3,Irf9,Tnfaip3,Znfx1,Ifi204,Irak3,Arid5b,C3ar1,Aff1,Csf3,Zbtb16 | | 3.172e-03 | -5.75 | OISHI\_CHOLANGIOMA\_STEM\_CELL\_LIKE\_DN | MSigDB lists | OISHI\_CHOLANGIOMA\_STEM\_CELL\_LIKE\_DN | 204 | 9 | 12187 | 179 | Bach1,Timp1,Stx11,Ksr1,Vcam1,Irf2,Irak3,Ptgs2,Gem | | 3.176e-03 | -5.75 | positive regulation of macromolecule biosynthetic process | biological process | GO:0010557 | 1499 | 37 | 13711 | 214 | Nfkbia,Nfkb2,Il1a,Sox11,Akap12,Cybb,Il6,Thbs1,Dtx3l,Irf7,Rbm39,Banp,Pik3r1,Fzd4,Parp9,Sbno2,Nfkb1,Spi1,Bach1,Cebpd,Runx1,Tnf,Ccl2,Atf3,Ripk1,Csf3,Zbtb16,Ncf1,Aff1,C3ar1,Arid5b,Hspb1,Tnip1,Maff,Irf2,Nfe2l2,Bcl3 | | 3.177e-03 | -5.75 | GSE15324\_NAIVE\_VS\_ACTIVATED\_CD8\_TCELL\_UP | MSigDB lists | GSE15324\_NAIVE\_VS\_ACTIVATED\_CD8\_TCELL\_UP | 131 | 7 | 12187 | 179 | Znfx1,Xaf1,Ifit3b,Usp18,Cmpk2,Tnip1,Arid5b | | 3.177e-03 | -5.75 | GSE17974\_CTRL\_VS\_ACT\_IL4\_AND\_ANTI\_IL12\_48H\_CD4\_TCELL\_UP | MSigDB lists | GSE17974\_CTRL\_VS\_ACT\_IL4\_AND\_ANTI\_IL12\_48H\_CD4\_TCELL\_UP | 131 | 7 | 12187 | 179 | Plek,Tiparp,Dnajb1,Nfkbiz,Socs3,Sgk1,Maff | | 3.192e-03 | -5.75 | REACTOME\_TRIF\_MEDIATED\_TLR3\_SIGNALING | MSigDB lists | REACTOME\_TRIF\_MEDIATED\_TLR3\_SIGNALING | 68 | 5 | 12187 | 179 | Saa1,Nfkbia,Irf7,Nfkb2,Ripk1 | | 3.193e-03 | -5.75 | activation of cysteine-type endopeptidase activity involved in apoptotic process | biological process | GO:0006919 | 64 | 5 | 13711 | 214 | Xdh,S100a8,Casp4,Tnf,S100a9 | | 3.236e-03 | -5.73 | GSE24634\_TREG\_VS\_TCONV\_POST\_DAY3\_IL4\_CONVERSION\_DN | MSigDB lists | GSE24634\_TREG\_VS\_TCONV\_POST\_DAY3\_IL4\_CONVERSION\_DN | 167 | 8 | 12187 | 179 | Cdkn1a,Ifi44,Tgm2,Timp1,Plek,Bach1,Ccl12,Cebpd | | 3.236e-03 | -5.73 | GSE29618\_BCELL\_VS\_MDC\_DAY7\_FLU\_VACCINE\_DN | MSigDB lists | GSE29618\_BCELL\_VS\_MDC\_DAY7\_FLU\_VACCINE\_DN | 167 | 8 | 12187 | 179 | Il1rn,Rhoc,Tiparp,Timp1,Cebpd,Atf3,Icam1,Igsf6 | | 3.236e-03 | -5.73 | GSE17721\_PAM3CSK4\_VS\_GADIQUIMOD\_24H\_BMDC\_DN | MSigDB lists | GSE17721\_PAM3CSK4\_VS\_GADIQUIMOD\_24H\_BMDC\_DN | 167 | 8 | 12187 | 179 | Ccl2,Tor3a,Ccl4,Gbp4,Rhoc,Gbp6,Gbp7,Tap1 | | 3.236e-03 | -5.73 | GSE20715\_0H\_VS\_6H\_OZONE\_TLR4\_KO\_LUNG\_DN | MSigDB lists | GSE20715\_0H\_VS\_6H\_OZONE\_TLR4\_KO\_LUNG\_DN | 167 | 8 | 12187 | 179 | Hspb1,Ptgs2,Nfe2l2,Cebpd,Lcn2,Hp,Nfkbia,Aff1 | | 3.259e-03 | -5.73 | GO\_NEGATIVE\_REGULATION\_OF\_ERK1\_AND\_ERK2\_CASCADE | MSigDB lists | GO\_NEGATIVE\_REGULATION\_OF\_ERK1\_AND\_ERK2\_CASCADE | 42 | 4 | 12187 | 179 | Errfi1,Ptpn1,Atf3,Tnip1 | | 3.259e-03 | -5.73 | GO\_TRANSFERASE\_ACTIVITY\_TRANSFERRING\_PENTOSYL\_GROUPS | MSigDB lists | GO\_TRANSFERASE\_ACTIVITY\_TRANSFERRING\_PENTOSYL\_GROUPS | 42 | 4 | 12187 | 179 | Tiparp,Parp9,Parp12,Parp14 | | 3.259e-03 | -5.73 | BERENJENO\_ROCK\_SIGNALING\_NOT\_VIA\_RHOA\_DN | MSigDB lists | BERENJENO\_ROCK\_SIGNALING\_NOT\_VIA\_RHOA\_DN | 42 | 4 | 12187 | 179 | Col4a1,Luc7l3,Hspa5,Vcam1 | | 3.266e-03 | -5.72 | KRIEG\_HYPOXIA\_NOT\_VIA\_KDM3A | MSigDB lists | KRIEG\_HYPOXIA\_NOT\_VIA\_KDM3A | 602 | 18 | 12187 | 179 | Tiparp,Cp,Akap12,Ier3,Il17ra,Icam1,Tnip1,Tubb6,Bach1,Tnfaip3,Il6,Ifit2,Runx1,Maff,Nfkbia,Tnfaip8,Osmr,Angptl4 | | 3.294e-03 | -5.72 | TNF\_1 | prosite domains | PS00251 | 5 | 2 | 8845 | 164 | Tnf,Tnfsf10 | | 3.294e-03 | -5.72 | 25A\_SYNTH\_3 | prosite domains | PS50152 | 5 | 2 | 8845 | 164 | Oasl1,Oasl2 | | 3.294e-03 | -5.72 | REL\_1 | prosite domains | PS01204 | 5 | 2 | 8845 | 164 | Nfkb1,Nfkb2 | | 3.313e-03 | -5.71 | external side of plasma membrane | cellular component | GO:0009897 | 234 | 10 | 13825 | 212 | Icam1,Robo4,Cd14,Osmr,Tnf,Vcam1,Cxcl9,Thbs1,Selp,Il6 | | 3.315e-03 | -5.71 | REACTOME\_ADAPTIVE\_IMMUNE\_SYSTEM | MSigDB lists | REACTOME\_ADAPTIVE\_IMMUNE\_SYSTEM | 416 | 14 | 12187 | 179 | Pik3r1,Map3k8,Nfkbia,Cybb,Dtx3l,Nfkbie,Hspa5,Cdkn1a,Vcam1,Zbtb16,Socs3,Pik3ap1,Icam1,Tap1 | | 3.349e-03 | -5.70 | RELA\_DN.V1\_UP | MSigDB lists | RELA\_DN.V1\_UP | 99 | 6 | 12187 | 179 | Ptgs2,Ripk1,Ccl12,Atf3,Gadd45b,Zfp36 | | 3.357e-03 | -5.70 | GSE29164\_UNTREATED\_VS\_CD8\_TCELL\_TREATED\_MELANOMA\_DAY3\_DN | MSigDB lists | GSE29164\_UNTREATED\_VS\_CD8\_TCELL\_TREATED\_MELANOMA\_DAY3\_DN | 168 | 8 | 12187 | 179 | Cebpd,Bach1,Parp14,Usp18,Ptpn1,Msn,Igsf6,Bcl3 | | 3.357e-03 | -5.70 | GSE39820\_CTRL\_VS\_IL1B\_IL6\_IL23A\_CD4\_TCELL\_DN | MSigDB lists | GSE39820\_CTRL\_VS\_IL1B\_IL6\_IL23A\_CD4\_TCELL\_DN | 168 | 8 | 12187 | 179 | Nfe2l2,Gadd45g,Timp1,Nfkbiz,Plek,Bcl3,Tnip1,Gem | | 3.357e-03 | -5.70 | GSE24292\_WT\_VS\_PPARG\_KO\_MACROPHAGE\_UP | MSigDB lists | GSE24292\_WT\_VS\_PPARG\_KO\_MACROPHAGE\_UP | 168 | 8 | 12187 | 179 | Igsf6,Tnfaip8,Csf2rb,Ifitm2,Nfe2l2,Maff,Nfkbiz,Tgm2 | | 3.357e-03 | -5.70 | GSE360\_DC\_VS\_MAC\_L\_DONOVANI\_DN | MSigDB lists | GSE360\_DC\_VS\_MAC\_L\_DONOVANI\_DN | 168 | 8 | 12187 | 179 | Dnajb1,Ifi44,Atf3,Ccl12,Timp1,Cd14,Gem,Sele | | 3.357e-03 | -5.70 | GSE37416\_0H\_VS\_12H\_F\_TULARENSIS\_LVS\_NEUTROPHIL\_DN | MSigDB lists | GSE37416\_0H\_VS\_12H\_F\_TULARENSIS\_LVS\_NEUTROPHIL\_DN | 168 | 8 | 12187 | 179 | Plaur,Banp,Ptges,Nfkbie,Gadd45b,Ier3,Tnfaip3,Maff | | 3.357e-03 | -5.70 | GSE3039\_CD4\_TCELL\_VS\_B2\_BCELL\_DN | MSigDB lists | GSE3039\_CD4\_TCELL\_VS\_B2\_BCELL\_DN | 168 | 8 | 12187 | 179 | Gpr84,Ccl4,Ccnd2,Ier3,A2m,Tnip1,C3ar1,Plaur | | 3.357e-03 | -5.70 | KAECH\_DAY15\_EFF\_VS\_MEMORY\_CD8\_TCELL\_DN | MSigDB lists | KAECH\_DAY15\_EFF\_VS\_MEMORY\_CD8\_TCELL\_DN | 168 | 8 | 12187 | 179 | Socs3,Usp18,Nfkbia,Tnf,Il4ra,Xdh,Zfp36,Tnip1 | | 3.357e-03 | -5.70 | GSE17721\_PAM3CSK4\_VS\_GADIQUIMOD\_2H\_BMDC\_DN | MSigDB lists | GSE17721\_PAM3CSK4\_VS\_GADIQUIMOD\_2H\_BMDC\_DN | 168 | 8 | 12187 | 179 | Ccl2,Tor3a,Lgals9,Bach1,Ms4a6d,Rrp8,Irf2,Trim25 | | 3.368e-03 | -5.69 | Small\_GTPase\_Rho | interpro domains | IPR003578 | 20 | 3 | 13788 | 212 | Rhoj,Rhou,Rhoc | | 3.371e-03 | -5.69 | death-inducing signaling complex | cellular component | GO:0031264 | 6 | 2 | 13825 | 212 | Cflar,Ripk1 | | 3.389e-03 | -5.69 | Formyl\_rcpt-rel | interpro domains | IPR000826 | 6 | 2 | 13788 | 212 | C3ar1,C5ar1 | | 3.389e-03 | -5.69 | Interferon\_reg\_factor-3 | interpro domains | IPR019471 | 6 | 2 | 13788 | 212 | Irf9,Irf7 | | 3.389e-03 | -5.69 | TRIM5 | interpro domains | IPR032917 | 6 | 2 | 13788 | 212 | Trim30d,Trim30a | | 3.389e-03 | -5.69 | Hempt\_rcpt\_S\_F1\_CS | interpro domains | IPR003531 | 6 | 2 | 13788 | 212 | Il4ra,Csf2rb | | 3.389e-03 | -5.69 | Dynamin\_central | interpro domains | IPR000375 | 6 | 2 | 13788 | 212 | Mx1,Mx2 | | 3.389e-03 | -5.69 | Dynamin\_GTPase\_CS | interpro domains | IPR019762 | 6 | 2 | 13788 | 212 | Mx1,Mx2 | | 3.389e-03 | -5.69 | GED | interpro domains | IPR003130 | 6 | 2 | 13788 | 212 | Mx2,Mx1 | | 3.389e-03 | -5.69 | 2-5-oligoadenylate\_synth\_N | interpro domains | IPR006116 | 6 | 2 | 13788 | 212 | Oasl2,Oasl1 | | 3.389e-03 | -5.69 | GED\_dom | interpro domains | IPR020850 | 6 | 2 | 13788 | 212 | Mx1,Mx2 | | 3.396e-03 | -5.69 | regulation of granulocyte chemotaxis | biological process | GO:0071622 | 40 | 4 | 13711 | 214 | Ccl2,C3ar1,C5ar1,Thbs1 | | 3.400e-03 | -5.68 | GO\_TISSUE\_DEVELOPMENT | MSigDB lists | GO\_TISSUE\_DEVELOPMENT | 1062 | 27 | 12187 | 179 | Rasip1,Mmp8,Tiparp,Sox11,Sbno2,Iqgap1,Arid5b,Il6,Rhoc,Spi1,Tgm2,Maff,Msn,Errfi1,Clic4,Icam1,Tagln2,Ptgs2,Atf3,Col4a1,Timp1,Zbtb16,Socs3,Cdkn1a,Cflar,Sdc4,Runx1 | | 3.401e-03 | -5.68 | MORI\_SMALL\_PRE\_BII\_LYMPHOCYTE\_DN | MSigDB lists | MORI\_SMALL\_PRE\_BII\_LYMPHOCYTE\_DN | 69 | 5 | 12187 | 179 | Cdkn1a,Ccnd2,Gbp7,Dnajb1,S100a9 | | 3.411e-03 | -5.68 | GO\_MYD88\_DEPENDENT\_TOLL\_LIKE\_RECEPTOR\_SIGNALING\_PATHWAY | MSigDB lists | GO\_MYD88\_DEPENDENT\_TOLL\_LIKE\_RECEPTOR\_SIGNALING\_PATHWAY | 21 | 3 | 12187 | 179 | Irak3,Cd14,Tnip1 | | 3.411e-03 | -5.68 | NOJIMA\_SFRP2\_TARGETS\_UP | MSigDB lists | NOJIMA\_SFRP2\_TARGETS\_UP | 21 | 3 | 12187 | 179 | Hspa5,Gadd45b,Tnfaip3 | | 3.411e-03 | -5.68 | GENTILE\_UV\_LOW\_DOSE\_UP | MSigDB lists | GENTILE\_UV\_LOW\_DOSE\_UP | 21 | 3 | 12187 | 179 | Tap1,Isg15,Cdkn1a | | 3.411e-03 | -5.68 | GO\_REGULATION\_OF\_MACROPHAGE\_DERIVED\_FOAM\_CELL\_DIFFERENTIATION | MSigDB lists | GO\_REGULATION\_OF\_MACROPHAGE\_DERIVED\_FOAM\_CELL\_DIFFERENTIATION | 21 | 3 | 12187 | 179 | Nfkb1,Nfkbia,Msr1 | | 3.411e-03 | -5.68 | GO\_NEGATIVE\_REGULATION\_OF\_PROTEIN\_MATURATION | MSigDB lists | GO\_NEGATIVE\_REGULATION\_OF\_PROTEIN\_MATURATION | 21 | 3 | 12187 | 179 | A2m,Thbs1,Birc3 | | 3.411e-03 | -5.68 | REACTOME\_TAK1\_ACTIVATES\_NFKB\_BY\_PHOSPHORYLATION\_AND\_ACTIVATION\_OF\_IKKS\_COMPLEX | MSigDB lists | REACTOME\_TAK1\_ACTIVATES\_NFKB\_BY\_PHOSPHORYLATION\_AND\_ACTIVATION\_OF\_IKKS\_COMPLEX | 21 | 3 | 12187 | 179 | Nfkbia,Saa1,Nfkb2 | | 3.411e-03 | -5.68 | VALK\_AML\_CLUSTER\_15 | MSigDB lists | VALK\_AML\_CLUSTER\_15 | 21 | 3 | 12187 | 179 | Hspb1,Tubb6,Runx1 | | 3.411e-03 | -5.68 | MUELLER\_METHYLATED\_IN\_GLIOBLASTOMA | MSigDB lists | MUELLER\_METHYLATED\_IN\_GLIOBLASTOMA | 21 | 3 | 12187 | 179 | Cxcl1,Ier3,Lcn2 | | 3.411e-03 | -5.68 | GO\_NEGATIVE\_REGULATION\_OF\_INTERLEUKIN\_6\_PRODUCTION | MSigDB lists | GO\_NEGATIVE\_REGULATION\_OF\_INTERLEUKIN\_6\_PRODUCTION | 21 | 3 | 12187 | 179 | Tnf,Irak3,Tnfaip3 | | 3.432e-03 | -5.67 | GO\_RESPONSE\_TO\_STEROID\_HORMONE | MSigDB lists | GO\_RESPONSE\_TO\_STEROID\_HORMONE | 373 | 13 | 12187 | 179 | Ptgs2,Errfi1,Msn,Trim25,Socs3,Cdkn1a,Sgk1,Thbs1,Il1rn,Tnf,Il4ra,Ccl12,Il6 | | 3.433e-03 | -5.67 | GO\_RESPONSE\_TO\_ACID\_CHEMICAL | MSigDB lists | GO\_RESPONSE\_TO\_ACID\_CHEMICAL | 246 | 10 | 12187 | 179 | Col4a1,Ccl12,Il6,Tnf,Adamts1,Icam1,P2ry6,Ptgs2,Ptges,Fzd4 | | 3.441e-03 | -5.67 | LEE\_BMP2\_TARGETS\_UP | MSigDB lists | LEE\_BMP2\_TARGETS\_UP | 605 | 18 | 12187 | 179 | Fam107a,Pik3r1,Sdc4,Usp18,Gbp3,Gbp7,Dtx3l,Cp,Rasd1,Mxd4,Cebpd,Lcn2,Ier3,Zbtb16,Plek,Ip6k2,Irf2,Cd14 | | 3.443e-03 | -5.67 | BYSTRYKH\_HEMATOPOIESIS\_STEM\_CELL\_QTL\_TRANS | MSigDB lists | BYSTRYKH\_HEMATOPOIESIS\_STEM\_CELL\_QTL\_TRANS | 703 | 20 | 12187 | 179 | Ifitm3,Cd14,Slfn3,Vcam1,Col4a1,Cp,Il1rn,Nfkb1,Ier3,Nfkbie,Csf3,C5ar1,Slfn4,Rgs16,Pik3r1,Msr1,Tor3a,Ccl4,Tgm2,Ccl12 | | 3.445e-03 | -5.67 | movement of cell or subcellular component | biological process | GO:0006928 | 1095 | 29 | 13711 | 214 | Ccl4,Kcna5,Il17ra,Sdc4,Thbs1,Bbs12,Ccl7,Icam1,Vcam1,S100a9,Msn,Selp,Cxcl1,C5ar1,S100a8,Iqgap1,Cxcl16,Ccl12,Arid5b,Il1rn,Hspb1,Sele,Retnlg,Rhoc,Tnf,Tnfaip3,Cxcl9,Saa1,Ccl2 | | 3.479e-03 | -5.66 | RIGGI\_EWING\_SARCOMA\_PROGENITOR\_UP | MSigDB lists | RIGGI\_EWING\_SARCOMA\_PROGENITOR\_UP | 330 | 12 | 12187 | 179 | A2m,Fzd4,C3ar1,Ccnd2,Pik3r1,AA467197,Cd14,Ptgs2,Icam1,Zbtb16,Ifi44,Mmp8 | | 3.480e-03 | -5.66 | GSE17721\_0.5H\_VS\_24H\_GARDIQUIMOD\_BMDC\_DN | MSigDB lists | GSE17721\_0.5H\_VS\_24H\_GARDIQUIMOD\_BMDC\_DN | 169 | 8 | 12187 | 179 | C3ar1,Ifitm3,Banp,Cmpk2,Msn,Tnf,Socs3,Gadd45b | | 3.480e-03 | -5.66 | GSE17721\_0.5H\_VS\_8H\_GARDIQUIMOD\_BMDC\_DN | MSigDB lists | GSE17721\_0.5H\_VS\_8H\_GARDIQUIMOD\_BMDC\_DN | 169 | 8 | 12187 | 179 | Ier3,Ccl12,Thbs1,Nfkbia,Tnfsf10,Plaur,Gem,Ifitm3 | | 3.480e-03 | -5.66 | GSE339\_CD4POS\_VS\_CD4CD8DN\_DC\_UP | MSigDB lists | GSE339\_CD4POS\_VS\_CD4CD8DN\_DC\_UP | 169 | 8 | 12187 | 179 | Tnf,Ccnd2,Usp18,Mxd4,Tnip1,Slfn3,Slfn4,Csf2rb | | 3.480e-03 | -5.66 | GSE9650\_EFFECTOR\_VS\_MEMORY\_CD8\_TCELL\_DN | MSigDB lists | GSE9650\_EFFECTOR\_VS\_MEMORY\_CD8\_TCELL\_DN | 169 | 8 | 12187 | 179 | Il4ra,Ccnd2,Sntb2,Nfkbiz,Socs3,Rbm39,Zfp36,C3ar1 | | 3.487e-03 | -5.66 | GED | pfam domains | PF02212 | 6 | 2 | 12881 | 201 | Mx2,Mx1 | | 3.487e-03 | -5.66 | Dynamin\_M | pfam domains | PF01031 | 6 | 2 | 12881 | 201 | Mx1,Mx2 | | 3.487e-03 | -5.66 | IRF-3 | pfam domains | PF10401 | 6 | 2 | 12881 | 201 | Irf9,Irf7 | | 3.490e-03 | -5.66 | regulation of chronic inflammatory response | biological process | GO:0002676 | 6 | 2 | 13711 | 214 | Tnfaip3,Tnf | | 3.490e-03 | -5.66 | detoxification of inorganic compound | biological process | GO:0061687 | 6 | 2 | 13711 | 214 | Mt2,Mt1 | | 3.490e-03 | -5.66 | protein nitrosylation | biological process | GO:0017014 | 6 | 2 | 13711 | 214 | S100a9,S100a8 | | 3.490e-03 | -5.66 | IRE1-mediated unfolded protein response | biological process | GO:0036498 | 6 | 2 | 13711 | 214 | Ptpn1,Hspa5 | | 3.490e-03 | -5.66 | positive regulation of receptor catabolic process | biological process | GO:2000646 | 6 | 2 | 13711 | 214 | Ptpn1,Dtx3l | | 3.490e-03 | -5.66 | positive regulation of T-helper 2 cell differentiation | biological process | GO:0045630 | 6 | 2 | 13711 | 214 | Il6,Il4ra | | 3.490e-03 | -5.66 | hydrogen peroxide biosynthetic process | biological process | GO:0050665 | 6 | 2 | 13711 | 214 | Ncf1,Cybb | | 3.490e-03 | -5.66 | epithelial cell proliferation involved in salivary gland morphogenesis | biological process | GO:0060664 | 6 | 2 | 13711 | 214 | Il6,Tnf | | 3.490e-03 | -5.66 | T cell extravasation | biological process | GO:0072683 | 6 | 2 | 13711 | 214 | Ccl2,Icam1 | | 3.490e-03 | -5.66 | germinal center formation | biological process | GO:0002467 | 6 | 2 | 13711 | 214 | Nfkb2,Bcl3 | | 3.490e-03 | -5.66 | heat generation | biological process | GO:0031649 | 6 | 2 | 13711 | 214 | Il1rn,Il1a | | 3.490e-03 | -5.66 | regulation of interleukin-1-mediated signaling pathway | biological process | GO:2000659 | 6 | 2 | 13711 | 214 | Il1rn,Il6 | | 3.491e-03 | -5.66 | interleukin-1 binding | molecular function | GO:0019966 | 6 | 2 | 13516 | 211 | Il1rn,A2m | | 3.491e-03 | -5.66 | STAT family protein binding | molecular function | GO:0097677 | 6 | 2 | 13516 | 211 | Parp9,Dtx3l | | 3.505e-03 | -5.65 | cellular homeostasis | biological process | GO:0019725 | 660 | 20 | 13711 | 214 | Ccl2,Il6,Icam1,Mt1,Cp,P2ry6,Lcn2,Pik3r1,Slc24a4,Pygm,C3ar1,Mt2,Clic4,C5ar1,Cxcl1,Tgm2,Sgk1,Kcna5,Nfe2l2,Vcam1 | | 3.510e-03 | -5.65 | response to unfolded protein | biological process | GO:0006986 | 94 | 6 | 13711 | 214 | Hspb1,Thbs1,Nfe2l2,Ptpn1,Hspa5,Atf3 | | 3.513e-03 | -5.65 | leukocyte apoptotic process | biological process | GO:0071887 | 20 | 3 | 13711 | 214 | Hcar2,Il6,Ripk1 | | 3.553e-03 | -5.64 | MODULE\_399 | MSigDB lists | MODULE\_399 | 43 | 4 | 12187 | 179 | Nfkbia,Cd14,Tnfaip3,Ripk1 | | 3.553e-03 | -5.64 | ROSS\_AML\_WITH\_CBFB\_MYH11\_FUSION | MSigDB lists | ROSS\_AML\_WITH\_CBFB\_MYH11\_FUSION | 43 | 4 | 12187 | 179 | S100a9,Cebpd,S100a8,Cybb | | 3.553e-03 | -5.64 | NUTT\_GBM\_VS\_AO\_GLIOMA\_UP | MSigDB lists | NUTT\_GBM\_VS\_AO\_GLIOMA\_UP | 43 | 4 | 12187 | 179 | Rhoc,Clic4,Msn,Tagln2 | | 3.555e-03 | -5.64 | positive regulation of lymphocyte activation | biological process | GO:0051251 | 195 | 9 | 13711 | 214 | Vcam1,Zbtb16,Runx1,Lgals9,Nfkbiz,Il4ra,Il6,Cdkn1a,Ccl2 | | 3.603e-03 | -5.63 | GSE2770\_IL12\_ACT\_VS\_ACT\_CD4\_TCELL\_6H\_DN | MSigDB lists | GSE2770\_IL12\_ACT\_VS\_ACT\_CD4\_TCELL\_6H\_DN | 134 | 7 | 12187 | 179 | Arid5b,Cxcl1,Gbp6,Dtx3l,Fstl1,Rhou,Gbp4 | | 3.603e-03 | -5.63 | GO\_MORPHOGENESIS\_OF\_A\_BRANCHING\_STRUCTURE | MSigDB lists | GO\_MORPHOGENESIS\_OF\_A\_BRANCHING\_STRUCTURE | 134 | 7 | 12187 | 179 | Clic4,Socs3,Tgm2,Spi1,Col4a1,Rasip1,Il6 | | 3.607e-03 | -5.62 | GSE22886\_NAIVE\_CD8\_TCELL\_VS\_NKCELL\_DN | MSigDB lists | GSE22886\_NAIVE\_CD8\_TCELL\_VS\_NKCELL\_DN | 170 | 8 | 12187 | 179 | Cflar,S100a9,Ifi211,Ifi209,Ifi207,Ifi204,Cebpd,Lcn2 | | 3.620e-03 | -5.62 | MODULE\_121 | MSigDB lists | MODULE\_121 | 70 | 5 | 12187 | 179 | Csf3,Il6,Timp1,Il1a,Osmr | | 3.620e-03 | -5.62 | KRAS.BREAST\_UP.V1\_UP | MSigDB lists | KRAS.BREAST\_UP.V1\_UP | 70 | 5 | 12187 | 179 | Akap12,Tnfaip3,Birc3,Cxcl1,A2m | | 3.620e-03 | -5.62 | MODULE\_169 | MSigDB lists | MODULE\_169 | 70 | 5 | 12187 | 179 | Cybb,Ier3,Rgs16,Isg15,Ccl4 | | 3.620e-03 | -5.62 | GO\_JNK\_CASCADE | MSigDB lists | GO\_JNK\_CASCADE | 70 | 5 | 12187 | 179 | Trib1,Map3k6,Tnf,Ripk1,Ptpn1 | | 3.673e-03 | -5.61 | GO\_POSITIVE\_REGULATION\_OF\_ESTABLISHMENT\_OF\_PROTEIN\_LOCALIZATION | MSigDB lists | GO\_POSITIVE\_REGULATION\_OF\_ESTABLISHMENT\_OF\_PROTEIN\_LOCALIZATION | 376 | 13 | 12187 | 179 | Il1a,Saa1,Ptgs2,Cd14,Il17ra,Rhou,Tnf,Pik3r1,Il4ra,Hcar2,Ccl12,Il6,Csf3 | | 3.698e-03 | -5.60 | KAYO\_AGING\_MUSCLE\_DN | MSigDB lists | KAYO\_AGING\_MUSCLE\_DN | 101 | 6 | 12187 | 179 | Ifi207,Ifi204,Nfkb1,Ifi209,Casp4,Ifi211 | | 3.698e-03 | -5.60 | SCHAEFFER\_PROSTATE\_DEVELOPMENT\_12HR\_UP | MSigDB lists | SCHAEFFER\_PROSTATE\_DEVELOPMENT\_12HR\_UP | 101 | 6 | 12187 | 179 | Gadd45g,Errfi1,Cebpd,Map3k6,Maff,Vcam1 | | 3.718e-03 | -5.59 | regulation of alcohol biosynthetic process | biological process | GO:1902930 | 41 | 4 | 13711 | 214 | Plek,Tnf,P2ry6,Nfkb1 | | 3.718e-03 | -5.59 | regulation of interleukin-2 production | biological process | GO:0032663 | 41 | 4 | 13711 | 214 | Il1a,Tnfaip3,Runx1,Zfp36 | | 3.720e-03 | -5.59 | transferase activity, transferring pentosyl groups | molecular function | GO:0016763 | 41 | 4 | 13516 | 211 | Parp9,Parp14,Parp12,Tiparp | | 3.738e-03 | -5.59 | GSE17721\_CTRL\_VS\_GARDIQUIMOD\_0.5H\_BMDC\_UP | MSigDB lists | GSE17721\_CTRL\_VS\_GARDIQUIMOD\_0.5H\_BMDC\_UP | 171 | 8 | 12187 | 179 | Ptpn1,Msn,Iqgap1,Gbp7,Luc7l3,Parp12,Mmp8,Il1rn | | 3.738e-03 | -5.59 | GSE28726\_NAIVE\_VS\_ACTIVATED\_NKTCELL\_DN | MSigDB lists | GSE28726\_NAIVE\_VS\_ACTIVATED\_NKTCELL\_DN | 171 | 8 | 12187 | 179 | Cflar,Gem,Il4ra,Ccl4,Nfkb1,Atf3,Ier3,Nfe2l2 | | 3.738e-03 | -5.59 | GSE3039\_NKT\_CELL\_VS\_ALPHABETA\_CD8\_TCELL\_UP | MSigDB lists | GSE3039\_NKT\_CELL\_VS\_ALPHABETA\_CD8\_TCELL\_UP | 171 | 8 | 12187 | 179 | Plaur,Gbp3,Hspb1,Gpr84,Maff,Nfe2l2,Il1rn,Gadd45g | | 3.749e-03 | -5.59 | GO\_REGULATION\_OF\_CELL\_DIFFERENTIATION | MSigDB lists | GO\_REGULATION\_OF\_CELL\_DIFFERENTIATION | 1123 | 28 | 12187 | 179 | Il6,Nfe2l2,Ripk1,Msr1,Pik3r1,Il4ra,Maff,Spi1,Adamts9,Csf3,Nfkb1,Sgk1,Sox11,Isg15,Bbs12,Tnf,Runx1,Nfkbia,Trib1,Xdh,Cflar,Irf7,Adamts1,Socs3,Zbtb16,Apold1,Errfi1,Ptgs2 | | 3.754e-03 | -5.58 | GSE33513\_TCF7\_KO\_VS\_HET\_EARLY\_THYMIC\_PROGENITOR\_UP | MSigDB lists | GSE33513\_TCF7\_KO\_VS\_HET\_EARLY\_THYMIC\_PROGENITOR\_UP | 135 | 7 | 12187 | 179 | Rtp4,Gbp4,Plin4,Ifitm3,C5ar1,Gbp6,Sdc4 | | 3.754e-03 | -5.58 | GO\_POSITIVE\_REGULATION\_OF\_APOPTOTIC\_SIGNALING\_PATHWAY | MSigDB lists | GO\_POSITIVE\_REGULATION\_OF\_APOPTOTIC\_SIGNALING\_PATHWAY | 135 | 7 | 12187 | 179 | S100a9,Tnfsf10,Plaur,S100a8,Thbs1,Atf3,Ripk1 | | 3.754e-03 | -5.58 | GSE32034\_LY6C\_HIGH\_VS\_LOW\_MONOCYTE\_UP | MSigDB lists | GSE32034\_LY6C\_HIGH\_VS\_LOW\_MONOCYTE\_UP | 135 | 7 | 12187 | 179 | Ier3,Tnfsf10,Lcn2,Plaur,Sgk3,Casp4,Ccl4 | | 3.760e-03 | -5.58 | Small cell lung cancer | KEGG pathways | ko05222 | 74 | 6 | 5248 | 107 | Ptgs2,Nfkbia,Nfkb1,Col4a1,Birc3,Pik3r1 | | 3.760e-03 | -5.58 | Small cell lung cancer | KEGG pathways | mmu05222 | 74 | 6 | 5248 | 107 | Pik3r1,Col4a1,Birc3,Nfkb1,Ptgs2,Nfkbia | | 3.770e-03 | -5.58 | negative regulation of intracellular signal transduction | biological process | GO:1902532 | 439 | 15 | 13711 | 214 | Plaur,Ptgs2,Nfe2l2,Rasip1,Tnip3,Ptpn1,Tnip1,Ripk1,Thbs1,Irak3,Hspb1,Xdh,Atf3,Errfi1,Tnfaip3 | | 3.800e-03 | -5.57 | GO\_KINASE\_BINDING | MSigDB lists | GO\_KINASE\_BINDING | 515 | 16 | 12187 | 179 | Msn,Ptpn1,Errfi1,Zfp36,Plek,Hspb1,Tnip1,Kcna5,Ksr1,Sgk1,Iqgap1,Tnfaip3,Sdc4,Trib1,Pik3r1,Ccnd2 | | 3.816e-03 | -5.57 | membrane region | cellular component | GO:0098589 | 319 | 12 | 13825 | 212 | Sele,Rgs16,Ripk1,Cd14,Icam1,Iqgap1,Birc3,Sdc4,Kcna5,Tnf,Cflar,Ptgs2 | | 3.845e-03 | -5.56 | ENK\_UV\_RESPONSE\_EPIDERMIS\_DN | MSigDB lists | ENK\_UV\_RESPONSE\_EPIDERMIS\_DN | 423 | 14 | 12187 | 179 | Ptgs2,Bcl3,Cxcl1,Hspb1,Plaur,Il4ra,Adamts1,Map3k8,Trib1,Dnajb1,Zbtb16,Stx11,Atf3,Tnfaip3 | | 3.849e-03 | -5.56 | NELSON\_RESPONSE\_TO\_ANDROGEN\_UP | MSigDB lists | NELSON\_RESPONSE\_TO\_ANDROGEN\_UP | 71 | 5 | 12187 | 179 | Adamts1,Sgk1,Arid5b,Tnfaip8,Akap12 | | 3.849e-03 | -5.56 | GO\_VASCULAR\_ENDOTHELIAL\_GROWTH\_FACTOR\_RECEPTOR\_SIGNALING\_PATHWAY | MSigDB lists | GO\_VASCULAR\_ENDOTHELIAL\_GROWTH\_FACTOR\_RECEPTOR\_SIGNALING\_PATHWAY | 71 | 5 | 12187 | 179 | Ccl12,Hspb1,Cybb,Ncf1,Pik3r1 | | 3.864e-03 | -5.56 | GO\_POSITIVE\_REGULATION\_OF\_EXTRINSIC\_APOPTOTIC\_SIGNALING\_PATHWAY | MSigDB lists | GO\_POSITIVE\_REGULATION\_OF\_EXTRINSIC\_APOPTOTIC\_SIGNALING\_PATHWAY | 44 | 4 | 12187 | 179 | Atf3,Tnfsf10,Ripk1,Thbs1 | | 3.864e-03 | -5.56 | BIOCARTA\_KERATINOCYTE\_PATHWAY | MSigDB lists | BIOCARTA\_KERATINOCYTE\_PATHWAY | 44 | 4 | 12187 | 179 | Nfkb1,Ripk1,Tnf,Nfkbia | | 3.872e-03 | -5.55 | GO\_NEGATIVE\_REGULATION\_OF\_PROTEIN\_MODIFICATION\_PROCESS | MSigDB lists | GO\_NEGATIVE\_REGULATION\_OF\_PROTEIN\_MODIFICATION\_PROCESS | 516 | 16 | 12187 | 179 | Xdh,Il6,Gadd45b,Tnfaip3,Spi1,Trib1,Tnf,Ptpn1,Errfi1,Irak3,Hspb1,Tnip1,Isg15,Atf3,Socs3,Cdkn1a | | 3.873e-03 | -5.55 | GSE17186\_BLOOD\_VS\_CORD\_BLOOD\_CD21LOW\_TRANSITIONAL\_BCELL\_UP | MSigDB lists | GSE17186\_BLOOD\_VS\_CORD\_BLOOD\_CD21LOW\_TRANSITIONAL\_BCELL\_UP | 172 | 8 | 12187 | 179 | Ifit2,Rtp4,Il17ra,Msn,Ptpn1,Trim56,Tnip1,Slfn9 | | 3.873e-03 | -5.55 | GSE37301\_MULTIPOTENT\_PROGENITOR\_VS\_COMMON\_LYMPHOID\_PROGENITOR\_DN | MSigDB lists | GSE37301\_MULTIPOTENT\_PROGENITOR\_VS\_COMMON\_LYMPHOID\_PROGENITOR\_DN | 172 | 8 | 12187 | 179 | Irf9,Icam1,Tnfaip8,Osmr,Irf7,Isg15,Tubb6,Usp18 | | 3.883e-03 | -5.55 | DARWICHE\_SKIN\_TUMOR\_PROMOTER\_UP | MSigDB lists | DARWICHE\_SKIN\_TUMOR\_PROMOTER\_UP | 102 | 6 | 12187 | 179 | S100a9,Ifitm3,Cxcl16,Cmpk2,S100a8,Icam1 | | 3.883e-03 | -5.55 | GSE12963\_UNINF\_VS\_ENV\_AND\_NEF\_AND\_VPR\_DEFICIENT\_HIV1\_INF\_CD4\_TCELL\_DN | MSigDB lists | GSE12963\_UNINF\_VS\_ENV\_AND\_NEF\_AND\_VPR\_DEFICIENT\_HIV1\_INF\_CD4\_TCELL\_DN | 102 | 6 | 12187 | 179 | Thbs1,Sdc4,Ccl4,Pik3ap1,Tubb6,Icam1 | | 3.892e-03 | -5.55 | GO\_NEGATIVE\_REGULATION\_OF\_NITROGEN\_COMPOUND\_METABOLIC\_PROCESS | MSigDB lists | GO\_NEGATIVE\_REGULATION\_OF\_NITROGEN\_COMPOUND\_METABOLIC\_PROCESS | 1126 | 28 | 12187 | 179 | Irak3,Irf2,Ifitm3,Nfkb1,Mxd4,Sox11,Sbno2,Ifi211,Rrp8,Arid5b,Bcl3,Tnfaip3,Ccl4,Spi1,Zfp36,Ifi204,Ifi207,Rasd1,Atf3,Ifi209,Zbtb16,Nfkb2,Irf7,Bach1,Tnf,Zfp189,Nfkbia,Trib1 | | 3.896e-03 | -5.55 | regulation of chemokine production | biological process | GO:0032642 | 67 | 5 | 13711 | 214 | Il6,Lgals9,Tnf,Il4ra,Il1a | | 3.905e-03 | -5.55 | GO\_GUANYL\_NUCLEOTIDE\_BINDING | MSigDB lists | GO\_GUANYL\_NUCLEOTIDE\_BINDING | 292 | 11 | 12187 | 179 | Rasd1,Gbp3,Rhoj,Gem,Gbp6,Gbp7,Irgm2,Gbp4,Tubb6,Rhoc,Rhou | | 3.907e-03 | -5.54 | GO\_PROTEIN\_IMPORT\_INTO\_NUCLEUS\_TRANSLOCATION | MSigDB lists | GO\_PROTEIN\_IMPORT\_INTO\_NUCLEUS\_TRANSLOCATION | 22 | 3 | 12187 | 179 | Nfkbia,Bcl3,Tnf | | 3.907e-03 | -5.54 | BIOCARTA\_NTHI\_PATHWAY | MSigDB lists | BIOCARTA\_NTHI\_PATHWAY | 22 | 3 | 12187 | 179 | Nfkbia,Tnf,Nfkb1 | | 3.907e-03 | -5.54 | PAL\_PRMT5\_TARGETS\_DN | MSigDB lists | PAL\_PRMT5\_TARGETS\_DN | 22 | 3 | 12187 | 179 | Vcam1,Lcn2,Il1a | | 3.907e-03 | -5.54 | VALK\_AML\_CLUSTER\_1 | MSigDB lists | VALK\_AML\_CLUSTER\_1 | 22 | 3 | 12187 | 179 | Kcna5,Atf3,Irf7 | | 3.907e-03 | -5.54 | VALK\_AML\_CLUSTER\_4 | MSigDB lists | VALK\_AML\_CLUSTER\_4 | 22 | 3 | 12187 | 179 | Il4ra,Trib1,Tubb6 | | 3.910e-03 | -5.54 | GSE13547\_WT\_VS\_ZFX\_KO\_BCELL\_DN | MSigDB lists | GSE13547\_WT\_VS\_ZFX\_KO\_BCELL\_DN | 136 | 7 | 12187 | 179 | S100a9,Phf11d,Gem,Ifitm3,Clic4,Phf11b,Gadd45b | | 3.910e-03 | -5.54 | HALLMARK\_UV\_RESPONSE\_UP | MSigDB lists | HALLMARK\_UV\_RESPONSE\_UP | 136 | 7 | 12187 | 179 | Il6,Atf3,Dnajb1,Nfkbia,Tap1,Icam1,Cxcl1 | | 3.927e-03 | -5.54 | GO\_CELLULAR\_RESPONSE\_TO\_ORGANIC\_CYCLIC\_COMPOUND | MSigDB lists | GO\_CELLULAR\_RESPONSE\_TO\_ORGANIC\_CYCLIC\_COMPOUND | 379 | 13 | 12187 | 179 | Il6,Ccl12,Msr1,Tnf,Nfkbia,Tiparp,Nfkb1,Sgk1,Adamts1,P2ry6,Msn,Errfi1,Ptgs2 | | 3.950e-03 | -5.53 | mouse chr16\|16 B3 | chromosome location | mouse chr16\|16 B3 | 43 | 4 | 14556 | 220 | Dtx3l,Parp9,Ccdc58,Parp14 | | 3.955e-03 | -5.53 | Interleukin-1 processing | REACTOME pathways | R-MMU-448706 | 6 | 2 | 6297 | 105 | Nfkb1,Nfkb2 | | 3.958e-03 | -5.53 | TBK1.DF\_DN | MSigDB lists | TBK1.DF\_DN | 251 | 10 | 12187 | 179 | Luc7l3,Col4a1,Bach1,Il1rn,Sntb2,Thbs1,Ccnd2,Irf2,Ptges,Angptl4 | | 4.011e-03 | -5.52 | CHANG\_CORE\_SERUM\_RESPONSE\_DN | MSigDB lists | CHANG\_CORE\_SERUM\_RESPONSE\_DN | 173 | 8 | 12187 | 179 | Cdkn1a,Parp14,Rnf213,Arid5b,Irak3,Trim25,Tnfaip8,Errfi1 | | 4.041e-03 | -5.51 | regulation of peptide transport | biological process | GO:0090087 | 622 | 19 | 13711 | 214 | Il6,Ccl2,Casp4,Tnfaip3,Il4ra,Tnf,Akap12,Il1a,Hcar2,Pik3r1,Lgals9,Cd14,S100a8,Gbp4,Ptpn1,Ptgs2,Il17ra,S100a9,Mmp8 | | 4.052e-03 | -5.51 | positive regulation of cellular extravasation | biological process | GO:0002693 | 21 | 3 | 13711 | 214 | Ccl2,Icam1,Il1a | | 4.052e-03 | -5.51 | negative regulation of receptor signaling pathway via JAK-STAT | biological process | GO:0046426 | 21 | 3 | 13711 | 214 | Socs3,Parp14,Bcl3 | | 4.052e-03 | -5.51 | negative regulation of viral transcription | biological process | GO:0032897 | 21 | 3 | 13711 | 214 | Ifitm3,Ccl4,Zfp36 | | 4.060e-03 | -5.51 | regulation of mononuclear cell migration | biological process | GO:0071675 | 42 | 4 | 13711 | 214 | Ccl2,C5ar1,C3ar1,Thbs1 | | 4.060e-03 | -5.51 | regulation of lymphocyte migration | biological process | GO:2000401 | 42 | 4 | 13711 | 214 | Msn,Ccl12,Lgals9,Ccl4 | | 4.066e-03 | -5.51 | PSMD2 (proteasome 26S subunit, non-ATPase 2) | protein interactions | 5708 | 6 | 2 | 6802 | 115 | Ksr1,Msn | | 4.066e-03 | -5.51 | Il6 (interleukin 6) | protein interactions | 16193 | 6 | 2 | 6802 | 115 | Nfkb1,Nfkbiz | | 4.066e-03 | -5.51 | Fbxw11 (F-box and WD-40 domain protein 11) | protein interactions | 103583 | 6 | 2 | 6802 | 115 | Nfkbia,Map3k6 | | 4.066e-03 | -5.51 | Nfkb2 (nuclear factor of kappa light polypeptide gene enhancer in B cells 2, p49/p100) | protein interactions | 18034 | 6 | 2 | 6802 | 115 | Nfkbiz,Nfkb1 | | 4.066e-03 | -5.51 | Nos2 (nitric oxide synthase 2, inducible) | protein interactions | 18126 | 6 | 2 | 6802 | 115 | Nfkb1,Ptgs2 | | 4.066e-03 | -5.51 | Mafk (v-maf musculoaponeurotic fibrosarcoma oncogene family, protein K (avian)) | protein interactions | 17135 | 6 | 2 | 6802 | 115 | Bach1,Nfe2l2 | | 4.088e-03 | -5.50 | GO\_NEGATIVE\_REGULATION\_OF\_IMMUNE\_EFFECTOR\_PROCESS | MSigDB lists | GO\_NEGATIVE\_REGULATION\_OF\_IMMUNE\_EFFECTOR\_PROCESS | 72 | 5 | 12187 | 179 | A2m,Tnf,Il4ra,Tap1,Irak3 | | 4.088e-03 | -5.50 | HAHTOLA\_SEZARY\_SYNDROM\_UP | MSigDB lists | HAHTOLA\_SEZARY\_SYNDROM\_UP | 72 | 5 | 12187 | 179 | Trib1,C3ar1,Tnfsf10,Hp,Iqgap1 | | 4.088e-03 | -5.50 | LEE\_CALORIE\_RESTRICTION\_NEOCORTEX\_DN | MSigDB lists | LEE\_CALORIE\_RESTRICTION\_NEOCORTEX\_DN | 72 | 5 | 12187 | 179 | Hspb1,Col4a1,Clic4,Nfe2l2,Zbtb16 | | 4.088e-03 | -5.50 | GO\_CELLULAR\_RESPONSE\_TO\_ABIOTIC\_STIMULUS | MSigDB lists | GO\_CELLULAR\_RESPONSE\_TO\_ABIOTIC\_STIMULUS | 212 | 9 | 12187 | 179 | Ptgs2,Ifi211,Errfi1,Ifi209,Cdkn1a,Pik3r1,Ifi204,Nfkb1,Ifi207 | | 4.099e-03 | -5.50 | cellular response to antibiotic | biological process | GO:0071236 | 97 | 6 | 13711 | 214 | Tnfaip3,Il6,Ripk1,Ccl7,Nfe2l2,Lcn2 | | 4.099e-03 | -5.50 | negative regulation of locomotion | biological process | GO:0040013 | 276 | 11 | 13711 | 214 | Robo4,Thbs1,Il1rn,Trib1,Adamts9,Ccl12,Timp1,Clic4,Trim56,Trim25,Nfe2l2 | | 4.112e-03 | -5.49 | regulation of peptide secretion | biological process | GO:0002791 | 400 | 14 | 13711 | 214 | Il6,Tnfaip3,Tnf,Il4ra,Casp4,Akap12,Il1a,Hcar2,Lgals9,Cd14,S100a8,Il17ra,Mmp8,S100a9 | | 4.153e-03 | -5.48 | positive regulation of JUN kinase activity | biological process | GO:0043507 | 68 | 5 | 13711 | 214 | Il1rn,Tnf,Fzd4,Ripk1,Ptpn1 | | 4.153e-03 | -5.48 | GSE26030\_UNSTIM\_VS\_RESTIM\_TH1\_DAY15\_POST\_POLARIZATION\_DN | MSigDB lists | GSE26030\_UNSTIM\_VS\_RESTIM\_TH1\_DAY15\_POST\_POLARIZATION\_DN | 174 | 8 | 12187 | 179 | Nfkbia,Nfkbiz,Trib1,Tubb6,C5ar1,Angptl4,Plek,Msn | | 4.165e-03 | -5.48 | GED | smart domains | SM00302 | 6 | 2 | 7188 | 123 | Mx1,Mx2 | | 4.165e-03 | -5.48 | TNF | smart domains | SM00207 | 6 | 2 | 7188 | 123 | Tnf,Tnfsf10 | | 4.165e-03 | -5.48 | IRF-3 | smart domains | SM01243 | 6 | 2 | 7188 | 123 | Irf7,Irf9 | | 4.193e-03 | -5.47 | BURTON\_ADIPOGENESIS\_PEAK\_AT\_16HR | MSigDB lists | BURTON\_ADIPOGENESIS\_PEAK\_AT\_16HR | 45 | 4 | 12187 | 179 | Ifi209,Ifi211,Ifi204,Ifi207 | | 4.193e-03 | -5.47 | LEE\_LIVER\_CANCER\_MYC\_TGFA\_UP | MSigDB lists | LEE\_LIVER\_CANCER\_MYC\_TGFA\_UP | 45 | 4 | 12187 | 179 | Icam1,Lcn2,Usp18,Tagln2 | | 4.225e-03 | -5.47 | GO\_CIRCULATORY\_SYSTEM\_DEVELOPMENT | MSigDB lists | GO\_CIRCULATORY\_SYSTEM\_DEVELOPMENT | 617 | 18 | 12187 | 179 | Apold1,Errfi1,Clic4,Rnf213,Ptgs2,Col4a1,Tiparp,Rasip1,Thbs1,Sox11,Adamts1,Socs3,Robo4,Fzd4,Rhoj,Angptl4,Ccl12,Spi1 | | 4.231e-03 | -5.47 | Acute myeloid leukemia | KEGG pathways | ko05221 | 53 | 5 | 5248 | 107 | Runx1,Pik3r1,Nfkb1,Spi1,Zbtb16 | | 4.231e-03 | -5.47 | Acute myeloid leukemia | KEGG pathways | mmu05221 | 53 | 5 | 5248 | 107 | Runx1,Pik3r1,Spi1,Zbtb16,Nfkb1 | | 4.236e-03 | -5.46 | GSE17721\_CPG\_VS\_GARDIQUIMOD\_2H\_BMDC\_UP | MSigDB lists | GSE17721\_CPG\_VS\_GARDIQUIMOD\_2H\_BMDC\_UP | 138 | 7 | 12187 | 179 | Tgm2,Zbtb16,Cdkn1a,Tnf,Gadd45b,Isg15,Icam1 | | 4.236e-03 | -5.46 | GSE3982\_CTRL\_VS\_PMA\_STIM\_EOSINOPHIL\_DN | MSigDB lists | GSE3982\_CTRL\_VS\_PMA\_STIM\_EOSINOPHIL\_DN | 138 | 7 | 12187 | 179 | Osmr,Cp,Bach1,Rgs16,Plaur,Cdkn1a,Dnajb1 | | 4.236e-03 | -5.46 | ZHONG\_RESPONSE\_TO\_AZACITIDINE\_AND\_TSA\_UP | MSigDB lists | ZHONG\_RESPONSE\_TO\_AZACITIDINE\_AND\_TSA\_UP | 138 | 7 | 12187 | 179 | Apold1,Irf7,Akap12,Timp1,Atf3,Parp12,Cdkn1a | | 4.272e-03 | -5.46 | GARGALOVIC\_RESPONSE\_TO\_OXIDIZED\_PHOSPHOLIPIDS\_BLUE\_UP | MSigDB lists | GARGALOVIC\_RESPONSE\_TO\_OXIDIZED\_PHOSPHOLIPIDS\_BLUE\_UP | 104 | 6 | 12187 | 179 | Bach1,Plaur,Atf3,Banp,Dnajb1,Trib1 | | 4.292e-03 | -5.45 | GO\_RAGE\_RECEPTOR\_BINDING | MSigDB lists | GO\_RAGE\_RECEPTOR\_BINDING | 7 | 2 | 12187 | 179 | S100a9,S100a8 | | 4.292e-03 | -5.45 | GO\_REGULATION\_OF\_INTERLEUKIN\_8\_BIOSYNTHETIC\_PROCESS | MSigDB lists | GO\_REGULATION\_OF\_INTERLEUKIN\_8\_BIOSYNTHETIC\_PROCESS | 7 | 2 | 12187 | 179 | Bcl3,Tnf | | 4.292e-03 | -5.45 | ZERBINI\_RESPONSE\_TO\_SULINDAC\_UP | MSigDB lists | ZERBINI\_RESPONSE\_TO\_SULINDAC\_UP | 7 | 2 | 12187 | 179 | Il6,Atf3 | | 4.292e-03 | -5.45 | GO\_REGULATION\_OF\_VITAMIN\_METABOLIC\_PROCESS | MSigDB lists | GO\_REGULATION\_OF\_VITAMIN\_METABOLIC\_PROCESS | 7 | 2 | 12187 | 179 | Tnf,Nfkb1 | | 4.292e-03 | -5.45 | KUWANO\_RNA\_STABILIZED\_BY\_NO | MSigDB lists | KUWANO\_RNA\_STABILIZED\_BY\_NO | 7 | 2 | 12187 | 179 | Ptgs2,Gadd45b | | 4.292e-03 | -5.45 | RAY\_ALZHEIMERS\_DISEASE | MSigDB lists | RAY\_ALZHEIMERS\_DISEASE | 7 | 2 | 12187 | 179 | Il1a,Tnf | | 4.292e-03 | -5.45 | XU\_HGF\_TARGETS\_REPRESSED\_BY\_AKT1\_UP | MSigDB lists | XU\_HGF\_TARGETS\_REPRESSED\_BY\_AKT1\_UP | 7 | 2 | 12187 | 179 | Ccl12,Angptl4 | | 4.292e-03 | -5.45 | GO\_TOLERANCE\_INDUCTION | MSigDB lists | GO\_TOLERANCE\_INDUCTION | 7 | 2 | 12187 | 179 | C3ar1,Tnfaip3 | | 4.298e-03 | -5.45 | GSE38304\_MYC\_NEG\_VS\_POS\_GC\_BCELL\_UP | MSigDB lists | GSE38304\_MYC\_NEG\_VS\_POS\_GC\_BCELL\_UP | 175 | 8 | 12187 | 179 | Slfn5,Irf7,Trim56,Sgk3,Gbp7,Ifit3b,Il4ra,Birc3 | | 4.347e-03 | -5.44 | GO\_AGING | MSigDB lists | GO\_AGING | 214 | 9 | 12187 | 179 | Ccl12,Il6,Timp1,Cp,Socs3,Vcam1,Cdkn1a,Nfkb2,Icam1 | | 4.351e-03 | -5.44 | Ripk1 (receptor (TNFRSF)-interacting serine-threonine kinase 1) | protein interactions | 19766 | 20 | 3 | 6802 | 115 | Tnf,Tnfaip3,Map3k8 | | 4.351e-03 | -5.44 | Btrc (beta-transducin repeat containing protein) | protein interactions | 12234 | 20 | 3 | 6802 | 115 | Nfe2l2,Map3k6,Nfkbia | | 4.383e-03 | -5.43 | MODULE\_16 | MSigDB lists | MODULE\_16 | 384 | 13 | 12187 | 179 | Tnfaip8,S100a8,Sbno2,S100a9,Cybb,Ccl12,Fam107a,Tnf,Ccnd2,Ccl4,Plek,Plaur,Ier3 | | 4.407e-03 | -5.42 | GSE46606\_DAY1\_VS\_DAY3\_CD40L\_IL2\_IL5\_STIMULATED\_IRF4MID\_BCELL\_UP | MSigDB lists | GSE46606\_DAY1\_VS\_DAY3\_CD40L\_IL2\_IL5\_STIMULATED\_IRF4MID\_BCELL\_UP | 139 | 7 | 12187 | 179 | Il6,Tubb6,Nfkbiz,Socs3,Maff,Ccnd2,Tnip1 | | 4.421e-03 | -5.42 | positive regulation of leukocyte chemotaxis | biological process | GO:0002690 | 69 | 5 | 13711 | 214 | Ccl2,Ccl4,Thbs1,C3ar1,C5ar1 | | 4.423e-03 | -5.42 | negative regulation of response to cytokine stimulus | biological process | GO:0060761 | 43 | 4 | 13711 | 214 | Il6,Irak3,Il1rn,Parp14 | | 4.440e-03 | -5.42 | NFKBTNSCPFCT | prints domains | PR00057 | 5 | 2 | 2951 | 64 | Nfkb1,Nfkb2 | | 4.440e-03 | -5.42 | ICAMVCAM1 | prints domains | PR01472 | 5 | 2 | 2951 | 64 | Vcam1,Icam1 | | 4.445e-03 | -5.42 | GARGALOVIC\_RESPONSE\_TO\_OXIDIZED\_PHOSPHOLIPIDS\_MAGENTA\_UP | MSigDB lists | GARGALOVIC\_RESPONSE\_TO\_OXIDIZED\_PHOSPHOLIPIDS\_MAGENTA\_UP | 23 | 3 | 12187 | 179 | Rasd1,Maff,Ptgs2 | | 4.445e-03 | -5.42 | GO\_REGULATION\_OF\_NEUTROPHIL\_MIGRATION | MSigDB lists | GO\_REGULATION\_OF\_NEUTROPHIL\_MIGRATION | 23 | 3 | 12187 | 179 | C5ar1,C3ar1,Cxcl1 | | 4.445e-03 | -5.42 | GO\_NECROTIC\_CELL\_DEATH | MSigDB lists | GO\_NECROTIC\_CELL\_DEATH | 23 | 3 | 12187 | 179 | Tnf,Cd14,Ripk1 | | 4.445e-03 | -5.42 | GO\_CELLULAR\_RESPONSE\_TO\_DEXAMETHASONE\_STIMULUS | MSigDB lists | GO\_CELLULAR\_RESPONSE\_TO\_DEXAMETHASONE\_STIMULUS | 23 | 3 | 12187 | 179 | Il6,Ccl12,Errfi1 | | 4.445e-03 | -5.42 | GO\_REGULATION\_OF\_TYPE\_I\_INTERFERON\_MEDIATED\_SIGNALING\_PATHWAY | MSigDB lists | GO\_REGULATION\_OF\_TYPE\_I\_INTERFERON\_MEDIATED\_SIGNALING\_PATHWAY | 23 | 3 | 12187 | 179 | Irf7,Usp18,Ptpn1 | | 4.445e-03 | -5.42 | VERRECCHIA\_RESPONSE\_TO\_TGFB1\_C2 | MSigDB lists | VERRECCHIA\_RESPONSE\_TO\_TGFB1\_C2 | 23 | 3 | 12187 | 179 | Rhoc,Icam1,Timp1 | | 4.445e-03 | -5.42 | GO\_REGULATION\_OF\_ENDOTHELIAL\_CELL\_DIFFERENTIATION | MSigDB lists | GO\_REGULATION\_OF\_ENDOTHELIAL\_CELL\_DIFFERENTIATION | 23 | 3 | 12187 | 179 | Tnf,Apold1,Xdh | | 4.445e-03 | -5.42 | GO\_REGULATION\_OF\_P38MAPK\_CASCADE | MSigDB lists | GO\_REGULATION\_OF\_P38MAPK\_CASCADE | 23 | 3 | 12187 | 179 | Gadd45g,Xdh,Gadd45b | | 4.445e-03 | -5.42 | BERENJENO\_ROCK\_SIGNALING\_NOT\_VIA\_RHOA\_UP | MSigDB lists | BERENJENO\_ROCK\_SIGNALING\_NOT\_VIA\_RHOA\_UP | 23 | 3 | 12187 | 179 | Cp,Atf3,Ptges | | 4.445e-03 | -5.42 | MODULE\_134 | MSigDB lists | MODULE\_134 | 23 | 3 | 12187 | 179 | Ccl2,C5ar1,C3ar1 | | 4.445e-03 | -5.42 | GNF2\_CD14 | MSigDB lists | GNF2\_CD14 | 23 | 3 | 12187 | 179 | Ms4a6d,Stx11,Cd14 | | 4.445e-03 | -5.42 | TANG\_SENESCENCE\_TP53\_TARGETS\_UP | MSigDB lists | TANG\_SENESCENCE\_TP53\_TARGETS\_UP | 23 | 3 | 12187 | 179 | Ccnd2,Cdkn1a,A2m | | 4.448e-03 | -5.42 | GSE26488\_CTRL\_VS\_PEPTIDE\_INJECTION\_HDAC7\_DELTAP\_TG\_OT2\_THYMOCYTE\_DN | MSigDB lists | GSE26488\_CTRL\_VS\_PEPTIDE\_INJECTION\_HDAC7\_DELTAP\_TG\_OT2\_THYMOCYTE\_DN | 176 | 8 | 12187 | 179 | Dnajb1,Tiparp,Gem,Errfi1,Gbp3,Tnfsf10,Sele,Zfp36 | | 4.477e-03 | -5.41 | GO\_HEPARIN\_BINDING | MSigDB lists | GO\_HEPARIN\_BINDING | 105 | 6 | 12187 | 179 | Ccl12,Fstl1,Adamts1,Thbs1,Selp,Saa1 | | 4.541e-03 | -5.39 | ISSAEVA\_MLL2\_TARGETS | MSigDB lists | ISSAEVA\_MLL2\_TARGETS | 46 | 4 | 12187 | 179 | C5ar1,A2m,Fstl1,Tagln2 | | 4.541e-03 | -5.39 | PID\_ANGIOPOIETIN\_RECEPTOR\_PATHWAY | MSigDB lists | PID\_ANGIOPOIETIN\_RECEPTOR\_PATHWAY | 46 | 4 | 12187 | 179 | Nfkb1,Cdkn1a,Tnf,Pik3r1 | | 4.541e-03 | -5.39 | PAPASPYRIDONOS\_UNSTABLE\_ATEROSCLEROTIC\_PLAQUE\_UP | MSigDB lists | PAPASPYRIDONOS\_UNSTABLE\_ATEROSCLEROTIC\_PLAQUE\_UP | 46 | 4 | 12187 | 179 | Sgk1,Cxcl1,Ifit3b,Csf2rb | | 4.541e-03 | -5.39 | GO\_CELLULAR\_RESPONSE\_TO\_CORTICOSTEROID\_STIMULUS | MSigDB lists | GO\_CELLULAR\_RESPONSE\_TO\_CORTICOSTEROID\_STIMULUS | 46 | 4 | 12187 | 179 | Ccl12,Errfi1,Il6,Sgk1 | | 4.541e-03 | -5.39 | GRAHAM\_NORMAL\_QUIESCENT\_VS\_NORMAL\_DIVIDING\_UP | MSigDB lists | GRAHAM\_NORMAL\_QUIESCENT\_VS\_NORMAL\_DIVIDING\_UP | 46 | 4 | 12187 | 179 | Cxcl1,Maff,Ptgs2,Tnfsf10 | | 4.541e-03 | -5.39 | STEARMAN\_LUNG\_CANCER\_EARLY\_VS\_LATE\_DN | MSigDB lists | STEARMAN\_LUNG\_CANCER\_EARLY\_VS\_LATE\_DN | 46 | 4 | 12187 | 179 | Socs3,Iqgap1,Csf2rb,Lcn2 | | 4.547e-03 | -5.39 | kinase binding | molecular function | GO:0019900 | 723 | 21 | 13516 | 211 | Irgm2,Pik3r1,Tnip1,Plek,Rhoc,Rhou,Hspb1,Cdkn1a,Iqgap1,Rhoj,Ksr1,Ptpn1,Trib1,Errfi1,Zfp36,Sgk1,Sdc4,Ccnd2,Msn,Tnfaip3,Kcna5 | | 4.583e-03 | -5.39 | GSE7768\_OVA\_ALONE\_VS\_OVA\_WITH\_MPL\_IMMUNIZED\_MOUSE\_WHOLE\_SPLEEN\_6H\_DN | MSigDB lists | GSE7768\_OVA\_ALONE\_VS\_OVA\_WITH\_MPL\_IMMUNIZED\_MOUSE\_WHOLE\_SPLEEN\_6H\_DN | 140 | 7 | 12187 | 179 | Casp4,Xaf1,Ptges,Isg15,Irf2,Pik3ap1,Samd9l | | 4.583e-03 | -5.39 | GSE22501\_PERIPHERAL\_BLOOD\_VS\_CORD\_BLOOD\_TREG\_DN | MSigDB lists | GSE22501\_PERIPHERAL\_BLOOD\_VS\_CORD\_BLOOD\_TREG\_DN | 140 | 7 | 12187 | 179 | Il1rn,Ptpn1,Irf9,Selp,Il4ra,C3ar1,Ccl4 | | 4.583e-03 | -5.39 | VEGF\_A\_UP.V1\_UP | MSigDB lists | VEGF\_A\_UP.V1\_UP | 140 | 7 | 12187 | 179 | Sdc4,Ccl12,Atf3,Nfkbie,Icam1,Nfkb2,Irf9 | | 4.583e-03 | -5.39 | GSE20366\_EX\_VIVO\_VS\_DEC205\_CONVERSION\_NAIVE\_CD4\_TCELL\_DN | MSigDB lists | GSE20366\_EX\_VIVO\_VS\_DEC205\_CONVERSION\_NAIVE\_CD4\_TCELL\_DN | 140 | 7 | 12187 | 179 | Pik3ap1,Rgs16,Ifitm2,Atf3,Sgk1,Maff,Socs3 | | 4.583e-03 | -5.39 | GSE21678\_WT\_VS\_FOXO1\_FOXO3\_KO\_TREG\_UP | MSigDB lists | GSE21678\_WT\_VS\_FOXO1\_FOXO3\_KO\_TREG\_UP | 140 | 7 | 12187 | 179 | Gem,Sgk1,Thbs1,Map3k8,Nfkbiz,Sdc4,Socs3 | | 4.598e-03 | -5.38 | GO\_PLATELET\_DEGRANULATION | MSigDB lists | GO\_PLATELET\_DEGRANULATION | 74 | 5 | 12187 | 179 | Timp1,Plek,A2m,Selp,Thbs1 | | 4.601e-03 | -5.38 | GSE1432\_1H\_VS\_24H\_IFNG\_MICROGLIA\_UP | MSigDB lists | GSE1432\_1H\_VS\_24H\_IFNG\_MICROGLIA\_UP | 177 | 8 | 12187 | 179 | Gadd45g,Tiparp,Gadd45b,Trib1,Socs3,Rgs16,C5ar1,Cd14 | | 4.638e-03 | -5.37 | regulation of vascular associated smooth muscle cell migration | biological process | GO:1904752 | 22 | 3 | 13711 | 214 | Iqgap1,Adamts1,Nfe2l2 | | 4.638e-03 | -5.37 | CD4-positive, alpha-beta T cell differentiation involved in immune response | biological process | GO:0002294 | 22 | 3 | 13711 | 214 | Bcl3,Gadd45g,Il6 | | 4.638e-03 | -5.37 | negative regulation of receptor signaling pathway via STAT | biological process | GO:1904893 | 22 | 3 | 13711 | 214 | Bcl3,Parp14,Socs3 | | 4.638e-03 | -5.37 | positive regulation of gene silencing by miRNA | biological process | GO:2000637 | 22 | 3 | 13711 | 214 | Zfp36,Nfkb1,Ripk1 | | 4.638e-03 | -5.37 | T-helper cell differentiation | biological process | GO:0042093 | 22 | 3 | 13711 | 214 | Il6,Bcl3,Gadd45g | | 4.640e-03 | -5.37 | protein serine/threonine kinase activator activity | molecular function | GO:0043539 | 22 | 3 | 13516 | 211 | Irgm2,Iqgap1,Lgals9 | | 4.654e-03 | -5.37 | regulation of response to wounding | biological process | GO:1903034 | 132 | 7 | 13711 | 214 | Ccl2,Cdkn1a,Plek,Selp,S100a9,Thbs1,Nfe2l2 | | 4.672e-03 | -5.37 | NADPH oxidase complex | cellular component | GO:0043020 | 7 | 2 | 13825 | 212 | Cybb,Ncf1 | | 4.696e-03 | -5.36 | Dynamin\_GTPase | interpro domains | IPR001401 | 7 | 2 | 13788 | 212 | Mx2,Mx1 | | 4.696e-03 | -5.36 | Chemokine\_CXC | interpro domains | IPR001089 | 7 | 2 | 13788 | 212 | Cxcl1,Cxcl9 | | 4.696e-03 | -5.36 | Chemokine\_CXC\_CS | interpro domains | IPR018048 | 7 | 2 | 13788 | 212 | Cxcl1,Cxcl9 | | 4.719e-03 | -5.36 | sequence-specific double-stranded DNA binding | molecular function | GO:1990837 | 678 | 20 | 13516 | 211 | Bach1,Irf9,Irf2,Runx1,Ifi211,Irf7,Maff,Spi1,Ifi207,Nfkb1,Atf3,Zbtb16,Ifi209,Nfkb2,Mxd4,Arid5b,Cebpd,Nfe2l2,Sox11,Ifi204 | | 4.758e-03 | -5.35 | BLUM\_RESPONSE\_TO\_SALIRASIB\_UP | MSigDB lists | BLUM\_RESPONSE\_TO\_SALIRASIB\_UP | 217 | 9 | 12187 | 179 | Casp4,Nfkbia,Cdkn1a,Maff,Tnfaip3,Atf3,Il6,Timp1,Bach1 | | 4.759e-03 | -5.35 | positive regulation of cysteine-type endopeptidase activity involved in apoptotic process | biological process | GO:0043280 | 100 | 6 | 13711 | 214 | Tnf,Casp4,S100a9,S100a8,Xdh,Tnfsf10 | | 4.759e-03 | -5.35 | MODULE\_123 | MSigDB lists | MODULE\_123 | 178 | 8 | 12187 | 179 | Cdkn1a,Atf3,Nfkb1,Bcl3,Cxcl1,C5ar1,Nfkb2,Irf2 | | 4.759e-03 | -5.35 | STAT5B\_01 | MSigDB lists | STAT5B\_01 | 178 | 8 | 12187 | 179 | Maff,Nfkbia,Ccl12,Adamts9,Icam1,Trim25,Irf2,Irf9 | | 4.764e-03 | -5.35 | GSE2585\_CD80\_HIGH\_VS\_LOW\_AIRE\_KO\_MTEC\_UP | MSigDB lists | GSE2585\_CD80\_HIGH\_VS\_LOW\_AIRE\_KO\_MTEC\_UP | 141 | 7 | 12187 | 179 | Saa1,Csf3,Nfkbiz,Tagln2,Il1a,Irak3,Il6 | | 4.764e-03 | -5.35 | ESC\_V6.5\_UP\_LATE.V1\_DN | MSigDB lists | ESC\_V6.5\_UP\_LATE.V1\_DN | 141 | 7 | 12187 | 179 | Rasip1,Timp1,Socs3,Sgk1,Icam1,Bcl3,Angptl4 | | 4.764e-03 | -5.35 | ESC\_J1\_UP\_EARLY.V1\_DN | MSigDB lists | ESC\_J1\_UP\_EARLY.V1\_DN | 141 | 7 | 12187 | 179 | Socs3,Luc7l3,Cebpd,Angptl4,Bcl3,Rrp8,Zfp36 | | 4.764e-03 | -5.35 | GSE45739\_UNSTIM\_VS\_ACD3\_ACD28\_STIM\_WT\_CD4\_TCELL\_DN | MSigDB lists | GSE45739\_UNSTIM\_VS\_ACD3\_ACD28\_STIM\_WT\_CD4\_TCELL\_DN | 141 | 7 | 12187 | 179 | Map3k8,Maff,Vcam1,Ccl4,Tiparp,Iqgap1,Plek | | 4.807e-03 | -5.34 | negative regulation of smooth muscle cell proliferation | biological process | GO:0048662 | 44 | 4 | 13711 | 214 | Pik3r1,Cdkn1a,Tnfaip3,Trib1 | | 4.807e-03 | -5.34 | heterophilic cell-cell adhesion via plasma membrane cell adhesion molecules | biological process | GO:0007157 | 44 | 4 | 13711 | 214 | Vcam1,Sele,Lgals9,Selp | | 4.836e-03 | -5.33 | regulation of ribonuclease activity | biological process | GO:0060700 | 7 | 2 | 13711 | 214 | Oasl1,Oasl2 | | 4.836e-03 | -5.33 | nucleotide-binding oligomerization domain containing 2 signaling pathway | biological process | GO:0070431 | 7 | 2 | 13711 | 214 | Nfkbia,Irgm2 | | 4.836e-03 | -5.33 | positive regulation of granulocyte differentiation | biological process | GO:0030854 | 7 | 2 | 13711 | 214 | Trib1,Runx1 | | 4.836e-03 | -5.33 | positive regulation of tumor necrosis factor-mediated signaling pathway | biological process | GO:1903265 | 7 | 2 | 13711 | 214 | Casp4,Ripk1 | | 4.836e-03 | -5.33 | regulation of interleukin-8 biosynthetic process | biological process | GO:0045414 | 7 | 2 | 13711 | 214 | Tnf,Bcl3 | | 4.836e-03 | -5.33 | autocrine signaling | biological process | GO:0035425 | 7 | 2 | 13711 | 214 | S100a9,S100a8 | | 4.836e-03 | -5.33 | nucleotide-binding oligomerization domain containing signaling pathway | biological process | GO:0070423 | 7 | 2 | 13711 | 214 | Irgm2,Nfkbia | | 4.836e-03 | -5.33 | positive regulation of glomerular mesangial cell proliferation | biological process | GO:0072126 | 7 | 2 | 13711 | 214 | Cflar,C3ar1 | | 4.836e-03 | -5.33 | modulation by symbiont of host autophagy | biological process | GO:0075071 | 7 | 2 | 13711 | 214 | Irgm1,Irgm2 | | 4.836e-03 | -5.33 | stress response to metal ion | biological process | GO:0097501 | 7 | 2 | 13711 | 214 | Mt2,Mt1 | | 4.836e-03 | -5.33 | positive regulation by symbiont of host autophagy | biological process | GO:0075044 | 7 | 2 | 13711 | 214 | Irgm2,Irgm1 | | 4.837e-03 | -5.33 | arachidonic acid binding | molecular function | GO:0050544 | 7 | 2 | 13516 | 211 | S100a9,S100a8 | | 4.869e-03 | -5.32 | CAHOY\_ASTROGLIAL | MSigDB lists | CAHOY\_ASTROGLIAL | 75 | 5 | 12187 | 179 | Cp,Rassf4,Timp1,Lcn2,Msn | | 4.869e-03 | -5.32 | LABBE\_TGFB1\_TARGETS\_UP | MSigDB lists | LABBE\_TGFB1\_TARGETS\_UP | 75 | 5 | 12187 | 179 | Gadd45g,Iqgap1,Ptgs2,Sgk1,Sox11 | | 4.881e-03 | -5.32 | GED | prosite domains | PS51388 | 6 | 2 | 8845 | 164 | Mx2,Mx1 | | 4.881e-03 | -5.32 | G\_DYNAMIN\_1 | prosite domains | PS00410 | 6 | 2 | 8845 | 164 | Mx1,Mx2 | | 4.881e-03 | -5.32 | HEMATOPO\_REC\_S\_F1 | prosite domains | PS01355 | 6 | 2 | 8845 | 164 | Il4ra,Csf2rb | | 4.881e-03 | -5.32 | REACTOME\_SIGNALING\_BY\_GPCR | MSigDB lists | REACTOME\_SIGNALING\_BY\_GPCR | 389 | 13 | 12187 | 179 | Saa1,Cxcl16,P2ry6,Fzd4,Pik3r5,Cxcl1,C3ar1,C5ar1,Ccl4,Pik3r1,Ccl12,Hcar2,Rhoc | | 4.882e-03 | -5.32 | Th17 cell differentiation | KEGG pathways | ko04659 | 78 | 6 | 5248 | 107 | Runx1,Nfkbie,Il4ra,Il6,Nfkbia,Nfkb1 | | 4.882e-03 | -5.32 | Th17 cell differentiation | KEGG pathways | mmu04659 | 78 | 6 | 5248 | 107 | Runx1,Nfkbie,Il4ra,Il6,Nfkbia,Nfkb1 | | 4.907e-03 | -5.32 | PID\_CASPASE\_PATHWAY | MSigDB lists | PID\_CASPASE\_PATHWAY | 47 | 4 | 12187 | 179 | Tnf,Birc3,Casp4,Ripk1 | | 4.907e-03 | -5.32 | MODULE\_199 | MSigDB lists | MODULE\_199 | 47 | 4 | 12187 | 179 | Csf2rb,Ccl12,Il4ra,Cdkn1a | | 4.913e-03 | -5.32 | MARTINEZ\_RB1\_TARGETS\_UP | MSigDB lists | MARTINEZ\_RB1\_TARGETS\_UP | 529 | 16 | 12187 | 179 | Errfi1,Plin4,Zfp36,Ptgs2,Ier3,Timp1,Cp,Dnajb1,Cdkn1a,Thbs1,Xdh,Rtp4,Hcar2,Bach1,Usp18,Ccnd2 | | 4.927e-03 | -5.31 | GO\_REGULATION\_OF\_PROTEIN\_LOCALIZATION | MSigDB lists | GO\_REGULATION\_OF\_PROTEIN\_LOCALIZATION | 726 | 20 | 12187 | 179 | Rhou,Tnf,Il4ra,Pik3r1,Nfkbia,Hcar2,Ccl12,Il6,Csf3,Nfkbie,Bcl3,Cdkn1a,Kcna5,Il1a,Saa1,Ptgs2,Cd14,Il17ra,Ptpn1,Msn | | 4.950e-03 | -5.31 | GSE28783\_CTRL\_ANTI\_MIR\_VS\_UNTREATED\_ATHEROSCLEROSIS\_MACROPHAGE\_UP | MSigDB lists | GSE28783\_CTRL\_ANTI\_MIR\_VS\_UNTREATED\_ATHEROSCLEROSIS\_MACROPHAGE\_UP | 142 | 7 | 12187 | 179 | Nfkbie,Rrp8,Rgs16,Tap1,Ccl4,Zbtb16,Sdc4 | | 5.026e-03 | -5.29 | TAVOR\_CEBPA\_TARGETS\_DN | MSigDB lists | TAVOR\_CEBPA\_TARGETS\_DN | 24 | 3 | 12187 | 179 | A2m,Isg15,Tnfsf10 | | 5.026e-03 | -5.29 | GO\_REGULATION\_OF\_ENDOPLASMIC\_RETICULUM\_UNFOLDED\_PROTEIN\_RESPONSE | MSigDB lists | GO\_REGULATION\_OF\_ENDOPLASMIC\_RETICULUM\_UNFOLDED\_PROTEIN\_RESPONSE | 24 | 3 | 12187 | 179 | Ptpn1,Pik3r1,Hspa5 | | 5.026e-03 | -5.29 | BERENJENO\_TRANSFORMED\_BY\_RHOA\_REVERSIBLY\_DN | MSigDB lists | BERENJENO\_TRANSFORMED\_BY\_RHOA\_REVERSIBLY\_DN | 24 | 3 | 12187 | 179 | Errfi1,Thbs1,Irgm2 | | 5.026e-03 | -5.29 | REACTOME\_PERK\_REGULATED\_GENE\_EXPRESSION | MSigDB lists | REACTOME\_PERK\_REGULATED\_GENE\_EXPRESSION | 24 | 3 | 12187 | 179 | Ccl12,Atf3,Hspa5 | | 5.026e-03 | -5.29 | BIOCARTA\_GSK3\_PATHWAY | MSigDB lists | BIOCARTA\_GSK3\_PATHWAY | 24 | 3 | 12187 | 179 | Cd14,Pik3r1,Nfkb1 | | 5.026e-03 | -5.29 | GO\_NEGATIVE\_REGULATION\_OF\_SIGNAL\_TRANSDUCTION\_IN\_ABSENCE\_OF\_LIGAND | MSigDB lists | GO\_NEGATIVE\_REGULATION\_OF\_SIGNAL\_TRANSDUCTION\_IN\_ABSENCE\_OF\_LIGAND | 24 | 3 | 12187 | 179 | Ripk1,Il1a,Tnf | | 5.026e-03 | -5.29 | GO\_POSITIVE\_REGULATION\_OF\_BLOOD\_VESSEL\_ENDOTHELIAL\_CELL\_MIGRATION | MSigDB lists | GO\_POSITIVE\_REGULATION\_OF\_BLOOD\_VESSEL\_ENDOTHELIAL\_CELL\_MIGRATION | 24 | 3 | 12187 | 179 | Hspb1,Ptgs2,Thbs1 | | 5.043e-03 | -5.29 | mouse chr7\|7 F5 | chromosome location | mouse chr7\|7 F5 | 46 | 4 | 14556 | 220 | Ifitm3,Ifitm2,Ifitm6,Irf7 | | 5.119e-03 | -5.27 | response to nitrogen compound | biological process | GO:1901698 | 683 | 20 | 13711 | 214 | Pik3r1,P2ry6,Thbs1,Irak3,Pygm,Cflar,Tnf,Tnfaip3,Icam1,Casp4,Nfkbia,Sgk1,Trim25,Vcam1,Serpina3f,Col4a1,Nfkb1,C5ar1,Hspa5,Ptpn1 | | 5.135e-03 | -5.27 | GSE17974\_IL4\_AND\_ANTI\_IL12\_VS\_UNTREATED\_1H\_ACT\_CD4\_TCELL\_UP | MSigDB lists | GSE17974\_IL4\_AND\_ANTI\_IL12\_VS\_UNTREATED\_1H\_ACT\_CD4\_TCELL\_UP | 108 | 6 | 12187 | 179 | Rtp4,Rgs16,Irgm2,Sox11,Nfkbiz,Socs3 | | 5.135e-03 | -5.27 | BMI1\_DN\_MEL18\_DN.V1\_UP | MSigDB lists | BMI1\_DN\_MEL18\_DN.V1\_UP | 108 | 6 | 12187 | 179 | C3ar1,Birc3,Atf3,Plaur,Il1a,Tnip3 | | 5.135e-03 | -5.27 | ELVIDGE\_HYPOXIA\_BY\_DMOG\_UP | MSigDB lists | ELVIDGE\_HYPOXIA\_BY\_DMOG\_UP | 108 | 6 | 12187 | 179 | Maff,Angptl4,Akap12,Tnfaip8,Atf3,Plaur | | 5.135e-03 | -5.27 | GO\_REGULATION\_OF\_INTRINSIC\_APOPTOTIC\_SIGNALING\_PATHWAY | MSigDB lists | GO\_REGULATION\_OF\_INTRINSIC\_APOPTOTIC\_SIGNALING\_PATHWAY | 108 | 6 | 12187 | 179 | S100a9,S100a8,Hspb1,Ptpn1,Plaur,Nfe2l2 | | 5.142e-03 | -5.27 | GSE3039\_NKT\_CELL\_VS\_B1\_BCELL\_UP | MSigDB lists | GSE3039\_NKT\_CELL\_VS\_B1\_BCELL\_UP | 143 | 7 | 12187 | 179 | Cebpd,Samd9l,S100a8,Slfn3,Olfml2b,Slfn4,Saa1 | | 5.142e-03 | -5.27 | OKUMURA\_INFLAMMATORY\_RESPONSE\_LPS | MSigDB lists | OKUMURA\_INFLAMMATORY\_RESPONSE\_LPS | 143 | 7 | 12187 | 179 | Birc3,Thbs1,Runx1,Sdc4,Tnip1,Cebpd,Tnfaip3 | | 5.142e-03 | -5.27 | GSE9988\_ANTI\_TREM1\_VS\_VEHICLE\_TREATED\_MONOCYTES\_UP | MSigDB lists | GSE9988\_ANTI\_TREM1\_VS\_VEHICLE\_TREATED\_MONOCYTES\_UP | 143 | 7 | 12187 | 179 | Tubb6,Gem,Sgk1,Ptgs2,Cdkn1a,Trib1,Cxcl1 | | 5.142e-03 | -5.27 | GSE24634\_TEFF\_VS\_TCONV\_DAY3\_IN\_CULTURE\_DN | MSigDB lists | GSE24634\_TEFF\_VS\_TCONV\_DAY3\_IN\_CULTURE\_DN | 143 | 7 | 12187 | 179 | Cybb,S100a8,Ccl12,Cdkn1a,Thbs1,Ccl4,Cd14 | | 5.151e-03 | -5.27 | GO\_CYTOKINE\_PRODUCTION | MSigDB lists | GO\_CYTOKINE\_PRODUCTION | 76 | 5 | 12187 | 179 | S100a9,Trim56,Irf7,Ptgs2,S100a8 | | 5.201e-03 | -5.26 | GARY\_CD5\_TARGETS\_UP | MSigDB lists | GARY\_CD5\_TARGETS\_UP | 392 | 13 | 12187 | 179 | Rbm39,Nfkb2,Rgs16,Bcl3,Tnfaip3,Ccl4,Tor3a,Spi1,Nfkbia,Icam1,Plek,Irf9,Cdkn1a | | 5.212e-03 | -5.26 | regulation of interleukin-8 production | biological process | GO:0032677 | 45 | 4 | 13711 | 214 | Bcl3,Tnf,Cd14,Ripk1 | | 5.236e-03 | -5.25 | GO\_SEQUENCE\_SPECIFIC\_DNA\_BINDING | MSigDB lists | GO\_SEQUENCE\_SPECIFIC\_DNA\_BINDING | 630 | 18 | 12187 | 179 | Arid5b,Irf7,Ifi211,Nfkb2,Maff,Runx1,Spi1,Bach1,Nfe2l2,Irf2,Sox11,Ifi209,Zbtb16,Ifi207,Nfkb1,Ifi204,Cebpd,Atf3 | | 5.251e-03 | -5.25 | Traf6 (TNF receptor-associated factor 6) | protein interactions | 22034 | 95 | 6 | 6802 | 115 | Tnfaip3,Irf7,Ripk1,Irak3,Il17ra,Hspa5 | | 5.258e-03 | -5.25 | regulation of adaptive immune response | biological process | GO:0002819 | 135 | 7 | 13711 | 214 | Lgals9,Irf7,Il6,Nfkbiz,Tnf,Il4ra,Tnfaip3 | | 5.273e-03 | -5.25 | plasma membrane | cellular component | GO:0005886 | 3462 | 70 | 13825 | 212 | Thbs1,Msr1,Cp,Cxcl9,Tnf,Ptgs2,Cflar,Fzd4,Irgm1,Gem,C3ar1,Hspa5,Cemip2,S100a8,Bcl3,Ksr1,Kcna5,Rhoc,Ptpn1,Ifitm2,Il1a,Icam1,Rrp8,Sele,Msn,Vcam1,Sgk1,Zbtb16,Robo4,Rasd1,Ncf1,Fyb,Gpr84,Nfkbia,Osmr,Il17ra,Cd14,Pik3r5,Tgm2,Ifitm3,Casp4,Iqgap1,Il4ra,Sntb2,Selp,P2ry6,Rhou,Pik3r1,Slc24a4,Rhoj,Ifitm6,Il6,Plaur,S100a9,Tnfsf10,Plin4,Clic4,Errfi1,Cybb,Stx11,Ripk1,C5ar1,Plek,Hspb1,Akap12,Pik3ap1,Fam107a,Nfe2l2,Hcar2,Irak3 | | 5.273e-03 | -5.25 | negative regulation of protein processing | biological process | GO:0010955 | 23 | 3 | 13711 | 214 | Thbs1,Ctla2a,Birc3 | | 5.273e-03 | -5.25 | negative regulation of extrinsic apoptotic signaling pathway via death domain receptors | biological process | GO:1902042 | 23 | 3 | 13711 | 214 | Cflar,Icam1,Tnfaip3 | | 5.273e-03 | -5.25 | negative regulation of protein maturation | biological process | GO:1903318 | 23 | 3 | 13711 | 214 | Thbs1,Ctla2a,Birc3 | | 5.273e-03 | -5.25 | regulation of neuroinflammatory response | biological process | GO:0150077 | 23 | 3 | 13711 | 214 | Ptgs2,Mmp8,C5ar1 | | 5.273e-03 | -5.25 | alpha-beta T cell differentiation involved in immune response | biological process | GO:0002293 | 23 | 3 | 13711 | 214 | Il6,Gadd45g,Bcl3 | | 5.273e-03 | -5.25 | positive regulation of posttranscriptional gene silencing | biological process | GO:0060148 | 23 | 3 | 13711 | 214 | Nfkb1,Zfp36,Ripk1 | | 5.273e-03 | -5.25 | positive regulation of CD4-positive, alpha-beta T cell differentiation | biological process | GO:0043372 | 23 | 3 | 13711 | 214 | Il4ra,Nfkbiz,Il6 | | 5.273e-03 | -5.25 | programmed necrotic cell death | biological process | GO:0097300 | 23 | 3 | 13711 | 214 | Birc3,Ripk1,Tnf | | 5.273e-03 | -5.25 | regulation of humoral immune response | biological process | GO:0002920 | 23 | 3 | 13711 | 214 | Tnf,Acod1,A2m | | 5.273e-03 | -5.25 | regulation of syncytium formation by plasma membrane fusion | biological process | GO:0060142 | 23 | 3 | 13711 | 214 | Cxcl9,Cflar,Il4ra | | 5.293e-03 | -5.24 | PRAMOONJAGO\_SOX4\_TARGETS\_UP | MSigDB lists | PRAMOONJAGO\_SOX4\_TARGETS\_UP | 48 | 4 | 12187 | 179 | Maff,Ier3,Tnfaip3,Tiparp | | 5.293e-03 | -5.24 | CORRE\_MULTIPLE\_MYELOMA\_UP | MSigDB lists | CORRE\_MULTIPLE\_MYELOMA\_UP | 48 | 4 | 12187 | 179 | Rhoj,Angptl4,Maff,AA467197 | | 5.335e-03 | -5.23 | G protein-coupled receptor signaling pathway | biological process | GO:0007186 | 456 | 15 | 13711 | 214 | Fzd4,C5ar1,Tgm2,Ccl12,Ccl4,Pik3r5,Plek,Rgs16,Akap12,Ccl2,Ccl7,Gpr84,C3ar1,P2ry6,Hcar2 | | 5.339e-03 | -5.23 | GSE17974\_2H\_VS\_72H\_UNTREATED\_IN\_VITRO\_CD4\_TCELL\_UP | MSigDB lists | GSE17974\_2H\_VS\_72H\_UNTREATED\_IN\_VITRO\_CD4\_TCELL\_UP | 144 | 7 | 12187 | 179 | Bach1,Gadd45g,Mxd4,Ripk1,Il6,Zfp36,Sgk3 | | 5.339e-03 | -5.23 | GSE37605\_C57BL6\_VS\_NOD\_FOXP3\_IRES\_GFP\_TREG\_DN | MSigDB lists | GSE37605\_C57BL6\_VS\_NOD\_FOXP3\_IRES\_GFP\_TREG\_DN | 144 | 7 | 12187 | 179 | Nfe2l2,Usp18,Parp14,Zbtb16,Lgals9,Irf9,Irgm2 | | 5.369e-03 | -5.23 | GSE17974\_IL4\_AND\_ANTI\_IL12\_VS\_UNTREATED\_2H\_ACT\_CD4\_TCELL\_DN | MSigDB lists | GSE17974\_IL4\_AND\_ANTI\_IL12\_VS\_UNTREATED\_2H\_ACT\_CD4\_TCELL\_DN | 109 | 6 | 12187 | 179 | Plaur,Il6,Rhou,Gem,Maff,Fstl1 | | 5.369e-03 | -5.23 | GSE22886\_NEUTROPHIL\_VS\_DC\_UP | MSigDB lists | GSE22886\_NEUTROPHIL\_VS\_DC\_UP | 109 | 6 | 12187 | 179 | Ptgs2,Ifitm2,Tnfsf10,Kcna5,Cebpd,Oasl1 | | 5.402e-03 | -5.22 | G alpha (i) signalling events | REACTOME pathways | R-MMU-418594 | 97 | 6 | 6297 | 105 | C5ar1,Hcar2,Cxcl16,Cxcl1,C3ar1,Cxcl9 | | 5.421e-03 | -5.22 | signal transduction by protein phosphorylation | biological process | GO:0023014 | 171 | 8 | 13711 | 214 | Ptpn1,Map3k6,Map3k8,Tnf,Ccl12,Zfp36,Ripk1,Trib1 | | 5.431e-03 | -5.22 | CEBP\_Q2 | MSigDB lists | CEBP\_Q2 | 182 | 8 | 12187 | 179 | Nfe2l2,Nfkbiz,Nfkbia,Clic4,Ip6k2,Osmr,Csf3,C3ar1 | | 5.444e-03 | -5.21 | LABBE\_TARGETS\_OF\_TGFB1\_AND\_WNT3A\_DN | MSigDB lists | LABBE\_TARGETS\_OF\_TGFB1\_AND\_WNT3A\_DN | 77 | 5 | 12187 | 179 | Cp,Plek,Lcn2,Cd14,Vcam1 | | 5.468e-03 | -5.21 | Immunoregulatory interactions between a Lymphoid and a non-Lymphoid cell | REACTOME pathways | R-MMU-198933 | 43 | 4 | 6297 | 105 | Vcam1,Icam1,Ifitm3,Ifitm2 | | 5.510e-03 | -5.20 | G beta:gamma signalling through PI3Kgamma | REACTOME pathways | R-MMU-392451 | 22 | 3 | 6297 | 105 | Csf2rb,Pik3r5,Pik3r1 | | 5.510e-03 | -5.20 | Binding and Uptake of Ligands by Scavenger Receptors | REACTOME pathways | R-MMU-2173782 | 22 | 3 | 6297 | 105 | Msr1,Hp,Apol10b | | 5.510e-03 | -5.20 | Chemokine receptors bind chemokines | REACTOME pathways | R-MMU-380108 | 22 | 3 | 6297 | 105 | Cxcl16,Cxcl1,Cxcl9 | | 5.540e-03 | -5.20 | FLECHNER\_BIOPSY\_KIDNEY\_TRANSPLANT\_OK\_VS\_DONOR\_UP | MSigDB lists | FLECHNER\_BIOPSY\_KIDNEY\_TRANSPLANT\_OK\_VS\_DONOR\_UP | 488 | 15 | 12187 | 179 | Birc3,Casp4,Cflar,Ifi211,Arid5b,Tnfsf10,Aff1,Ifi209,Hspa5,Vcam1,Ifi204,Col4a1,Ifi207,Luc7l3,Msn | | 5.542e-03 | -5.20 | P53\_DN.V1\_DN | MSigDB lists | P53\_DN.V1\_DN | 145 | 7 | 12187 | 179 | Ifi207,Ifi204,Socs3,Sgk1,Ifi209,Cdkn1a,Ifi211 | | 5.542e-03 | -5.20 | GSE42021\_CD24HI\_VS\_CD24INT\_TCONV\_THYMUS\_DN | MSigDB lists | GSE42021\_CD24HI\_VS\_CD24INT\_TCONV\_THYMUS\_DN | 145 | 7 | 12187 | 179 | Ccl12,Atf3,Maff,Icam1,Tap1,Cxcl1,Arid5b | | 5.542e-03 | -5.20 | GSE19888\_CTRL\_VS\_A3R\_INHIBITOR\_TREATED\_MAST\_CELL\_DN | MSigDB lists | GSE19888\_CTRL\_VS\_A3R\_INHIBITOR\_TREATED\_MAST\_CELL\_DN | 145 | 7 | 12187 | 179 | AA467197,Tnfaip3,Bach1,Thbs1,Plaur,Zfp36,Arid5b | | 5.610e-03 | -5.18 | GSE39820\_TGFBETA1\_IL6\_VS\_TGFBETA1\_IL6\_IL23A\_TREATED\_CD4\_TCELL\_UP | MSigDB lists | GSE39820\_TGFBETA1\_IL6\_VS\_TGFBETA1\_IL6\_IL23A\_TREATED\_CD4\_TCELL\_UP | 183 | 8 | 12187 | 179 | Iqgap1,AA467197,Il1rn,Nfkb1,Timp1,Sgk1,Pik3r1,Birc3 | | 5.610e-03 | -5.18 | STAT5A\_01 | MSigDB lists | STAT5A\_01 | 183 | 8 | 12187 | 179 | Maff,Ccl12,Csf3,Icam1,Apold1,Adamts9,Irf2,Irf9 | | 5.612e-03 | -5.18 | LINDGREN\_BLADDER\_CANCER\_CLUSTER\_2A\_DN | MSigDB lists | LINDGREN\_BLADDER\_CANCER\_CLUSTER\_2A\_DN | 110 | 6 | 12187 | 179 | Cybb,Clic4,C3ar1,Sgk1,Socs3,Ptges | | 5.630e-03 | -5.18 | Cdk6 (cyclin-dependent kinase 6) | protein interactions | 12571 | 7 | 2 | 6802 | 115 | Runx1,Ccnd2 | | 5.630e-03 | -5.18 | Hnrnpa0 (heterogeneous nuclear ribonucleoprotein A0) | protein interactions | 77134 | 7 | 2 | 6802 | 115 | Tnf,Ptgs2 | | 5.640e-03 | -5.18 | intrinsic apoptotic signaling pathway by p53 class mediator | biological process | GO:0072332 | 46 | 4 | 13711 | 214 | Rrp8,Ifi204,Cdkn1a,Bcl3 | | 5.650e-03 | -5.18 | GNF2\_MMP1 | MSigDB lists | GNF2\_MMP1 | 25 | 3 | 12187 | 179 | Csf3,Il6,Ccl12 | | 5.650e-03 | -5.18 | WOTTON\_RUNX\_TARGETS\_DN | MSigDB lists | WOTTON\_RUNX\_TARGETS\_DN | 25 | 3 | 12187 | 179 | Errfi1,Xdh,Cebpd | | 5.650e-03 | -5.18 | MODULE\_355 | MSigDB lists | MODULE\_355 | 25 | 3 | 12187 | 179 | Hspb1,Hspa5,Dnajb1 | | 5.650e-03 | -5.18 | MODULE\_521 | MSigDB lists | MODULE\_521 | 25 | 3 | 12187 | 179 | Timp1,Il6,Csf3 | | 5.650e-03 | -5.18 | PID\_P38\_MKK3\_6PATHWAY | MSigDB lists | PID\_P38\_MKK3\_6PATHWAY | 25 | 3 | 12187 | 179 | Map3k6,Gadd45b,Gadd45g | | 5.654e-03 | -5.18 | TRIF-mediated TLR3/TLR4 signaling | REACTOME pathways | R-MMU-937061 | 69 | 5 | 6297 | 105 | Map3k8,Nfkbia,Nfkb2,Irf7,Nfkb1 | | 5.658e-03 | -5.17 | RHO | prosite domains | PS51420 | 20 | 3 | 8845 | 164 | Rhou,Rhoj,Rhoc | | 5.667e-03 | -5.17 | OSAWA\_TNF\_TARGETS | MSigDB lists | OSAWA\_TNF\_TARGETS | 8 | 2 | 12187 | 179 | Cflar,Ier3 | | 5.667e-03 | -5.17 | GO\_POSITIVE\_REGULATION\_OF\_MACROPHAGE\_ACTIVATION | MSigDB lists | GO\_POSITIVE\_REGULATION\_OF\_MACROPHAGE\_ACTIVATION | 8 | 2 | 12187 | 179 | Il4ra,Thbs1 | | 5.667e-03 | -5.17 | GO\_REGULATION\_OF\_CHEMOKINE\_BIOSYNTHETIC\_PROCESS | MSigDB lists | GO\_REGULATION\_OF\_CHEMOKINE\_BIOSYNTHETIC\_PROCESS | 8 | 2 | 12187 | 179 | Il6,Tnf | | 5.667e-03 | -5.17 | WHITESIDE\_CISPLATIN\_RESISTANCE\_UP | MSigDB lists | WHITESIDE\_CISPLATIN\_RESISTANCE\_UP | 8 | 2 | 12187 | 179 | Cdkn1a,Ptges | | 5.667e-03 | -5.17 | GO\_OXIDOREDUCTASE\_ACTIVITY\_ACTING\_ON\_NAD\_P\_H\_OXYGEN\_AS\_ACCEPTOR | MSigDB lists | GO\_OXIDOREDUCTASE\_ACTIVITY\_ACTING\_ON\_NAD\_P\_H\_OXYGEN\_AS\_ACCEPTOR | 8 | 2 | 12187 | 179 | Ncf1,Cybb | | 5.667e-03 | -5.17 | GO\_SEQUESTERING\_OF\_METAL\_ION | MSigDB lists | GO\_SEQUESTERING\_OF\_METAL\_ION | 8 | 2 | 12187 | 179 | S100a8,S100a9 | | 5.667e-03 | -5.17 | GO\_INTERLEUKIN\_1\_RECEPTOR\_BINDING | MSigDB lists | GO\_INTERLEUKIN\_1\_RECEPTOR\_BINDING | 8 | 2 | 12187 | 179 | Il1rn,Il1a | | 5.667e-03 | -5.17 | GALI\_TP53\_TARGETS\_APOPTOTIC\_UP | MSigDB lists | GALI\_TP53\_TARGETS\_APOPTOTIC\_UP | 8 | 2 | 12187 | 179 | Casp4,Birc3 | | 5.667e-03 | -5.17 | GUTIERREZ\_WALDENSTROEMS\_MACROGLOBULINEMIA\_1\_DN | MSigDB lists | GUTIERREZ\_WALDENSTROEMS\_MACROGLOBULINEMIA\_1\_DN | 8 | 2 | 12187 | 179 | S100a8,Il6 | | 5.667e-03 | -5.17 | GO\_NEGATIVE\_REGULATION\_OF\_INTERLEUKIN\_1\_PRODUCTION | MSigDB lists | GO\_NEGATIVE\_REGULATION\_OF\_INTERLEUKIN\_1\_PRODUCTION | 8 | 2 | 12187 | 179 | Tnfaip3,Errfi1 | | 5.667e-03 | -5.17 | REACTOME\_IKK\_COMPLEX\_RECRUITMENT\_MEDIATED\_BY\_RIP1 | MSigDB lists | REACTOME\_IKK\_COMPLEX\_RECRUITMENT\_MEDIATED\_BY\_RIP1 | 8 | 2 | 12187 | 179 | Ripk1,Cd14 | | 5.674e-03 | -5.17 | phosphatidylinositol 3-kinase complex | cellular component | GO:0005942 | 24 | 3 | 13825 | 212 | Socs3,Pik3r1,Pik3r5 | | 5.699e-03 | -5.17 | LAU\_APOPTOSIS\_CDKN2A\_UP | MSigDB lists | LAU\_APOPTOSIS\_CDKN2A\_UP | 49 | 4 | 12187 | 179 | Cdkn1a,Cflar,Nfkbia,Tnfaip3 | | 5.699e-03 | -5.17 | PID\_ATF2\_PATHWAY | MSigDB lists | PID\_ATF2\_PATHWAY | 49 | 4 | 12187 | 179 | Socs3,Sele,Il6,Atf3 | | 5.699e-03 | -5.17 | MCBRYAN\_PUBERTAL\_TGFB1\_TARGETS\_DN | MSigDB lists | MCBRYAN\_PUBERTAL\_TGFB1\_TARGETS\_DN | 49 | 4 | 12187 | 179 | Ifit3b,Bcl3,Lcn2,Nfkb2 | | 5.699e-03 | -5.17 | BIOCARTA\_PPARA\_PATHWAY | MSigDB lists | BIOCARTA\_PPARA\_PATHWAY | 49 | 4 | 12187 | 179 | Tnf,Pik3r1,Ptgs2,Nfkbia | | 5.699e-03 | -5.17 | PETROVA\_PROX1\_TARGETS\_DN | MSigDB lists | PETROVA\_PROX1\_TARGETS\_DN | 49 | 4 | 12187 | 179 | Il6,Ccl12,Fam107a,Selp | | 5.736e-03 | -5.16 | Pml (promyelocytic leukemia) | protein interactions | 18854 | 22 | 3 | 6802 | 115 | Spi1,Runx1,Zbtb16 | | 5.750e-03 | -5.16 | MODULE\_289 | MSigDB lists | MODULE\_289 | 78 | 5 | 12187 | 179 | Akap12,Rgs16,C3ar1,C5ar1,Cxcl1 | | 5.750e-03 | -5.16 | EIF4E\_DN | MSigDB lists | EIF4E\_DN | 78 | 5 | 12187 | 179 | Tnip1,Angptl4,Ptgs2,Saa1,Iqgap1 | | 5.751e-03 | -5.16 | HFH8\_01 | MSigDB lists | HFH8\_01 | 146 | 7 | 12187 | 179 | Hcar2,Rasd1,Runx1,Tnf,Fstl1,Ip6k2,Cxcl1 | | 5.751e-03 | -5.16 | GSE29618\_LAIV\_VS\_TIV\_FLU\_VACCINE\_DAY7\_PDC\_DN | MSigDB lists | GSE29618\_LAIV\_VS\_TIV\_FLU\_VACCINE\_DAY7\_PDC\_DN | 146 | 7 | 12187 | 179 | Ifi207,Tnip3,Ifi204,Ifi209,Ip6k2,Ifi211,Map3k6 | | 5.751e-03 | -5.16 | GSE15659\_RESTING\_TREG\_VS\_NONSUPPRESSIVE\_TCELL\_DN | MSigDB lists | GSE15659\_RESTING\_TREG\_VS\_NONSUPPRESSIVE\_TCELL\_DN | 146 | 7 | 12187 | 179 | Plin4,Xaf1,Rhou,Sox11,Rasd1,Timp1,Rhoc | | 5.766e-03 | -5.16 | DYNc | smart domains | SM00053 | 7 | 2 | 7188 | 123 | Mx1,Mx2 | | 5.817e-03 | -5.15 | regulation of cellular response to stress | biological process | GO:0080135 | 597 | 18 | 13711 | 214 | Il1a,Tnf,Xdh,Ripk1,Dtx3l,Il1rn,Ncf1,Hspb1,Gadd45b,Gadd45g,Pik3r1,Fzd4,Ptpn1,Parp9,Mmp8,Nfe2l2,Ier3,Ptgs2 | | 5.862e-03 | -5.14 | GO\_RESPONSE\_TO\_ETHANOL | MSigDB lists | GO\_RESPONSE\_TO\_ETHANOL | 111 | 6 | 12187 | 179 | Ccl12,S100a8,Icam1,Spi1,Cd14,Vcam1 | | 5.866e-03 | -5.14 | WEST\_ADRENOCORTICAL\_TUMOR\_DN | MSigDB lists | WEST\_ADRENOCORTICAL\_TUMOR\_DN | 444 | 14 | 12187 | 179 | Ifitm2,Zfp36,Icam1,Kcna5,Cebpd,Rasip1,Ier3,Mxd4,Gem,Tnfsf10,Maff,Runx1,Cybb,Ccl12 | | 5.874e-03 | -5.14 | T cell receptor signaling pathway | KEGG pathways | mmu04660 | 81 | 6 | 5248 | 107 | Nfkbie,Pik3r1,Map3k8,Tnf,Nfkb1,Nfkbia | | 5.874e-03 | -5.14 | T cell receptor signaling pathway | KEGG pathways | ko04660 | 81 | 6 | 5248 | 107 | Nfkbia,Nfkb1,Tnf,Map3k8,Pik3r1,Nfkbie | | 5.919e-03 | -5.13 | regulation of lymphocyte differentiation | biological process | GO:0045619 | 138 | 7 | 13711 | 214 | Il6,Nfkbiz,Il4ra,Lgals9,Runx1,Zbtb16,Ctla2a | | 5.933e-03 | -5.13 | G protein-coupled receptor binding | molecular function | GO:0001664 | 211 | 9 | 13516 | 211 | Saa1,Ccl12,Cxcl16,Rtp4,Ccl7,Ccl4,Ccl2,Cxcl1,Cxcl9 | | 5.953e-03 | -5.12 | stress-activated MAPK cascade | biological process | GO:0051403 | 74 | 5 | 13711 | 214 | Zfp36,Ptpn1,Trib1,Ripk1,Tnf | | 5.958e-03 | -5.12 | negative regulation of CD4-positive, alpha-beta T cell activation | biological process | GO:2000515 | 24 | 3 | 13711 | 214 | Il4ra,Lgals9,Runx1 | | 5.958e-03 | -5.12 | positive regulation of acute inflammatory response | biological process | GO:0002675 | 24 | 3 | 13711 | 214 | Ptgs2,Il6,Tnf | | 5.958e-03 | -5.12 | regulation of necrotic cell death | biological process | GO:0010939 | 24 | 3 | 13711 | 214 | Ripk1,Cflar,Birc3 | | 5.958e-03 | -5.12 | alpha-beta T cell activation involved in immune response | biological process | GO:0002287 | 24 | 3 | 13711 | 214 | Il6,Gadd45g,Bcl3 | | 5.966e-03 | -5.12 | GSE22935\_UNSTIM\_VS\_24H\_MBOVIS\_BCG\_STIM\_MACROPHAGE\_UP | MSigDB lists | GSE22935\_UNSTIM\_VS\_24H\_MBOVIS\_BCG\_STIM\_MACROPHAGE\_UP | 147 | 7 | 12187 | 179 | Gbp3,Irf2,Il17ra,Ptges,Ptgs2,Aff1,Cdkn1a | | 5.966e-03 | -5.12 | GSE1460\_INTRATHYMIC\_T\_PROGENITOR\_VS\_NAIVE\_CD4\_TCELL\_ADULT\_BLOOD\_DN | MSigDB lists | GSE1460\_INTRATHYMIC\_T\_PROGENITOR\_VS\_NAIVE\_CD4\_TCELL\_ADULT\_BLOOD\_DN | 147 | 7 | 12187 | 179 | Nfkb1,Timp1,Il4ra,Tnf,Irf9,Msn,Csf3 | | 5.966e-03 | -5.12 | CAIRO\_LIVER\_DEVELOPMENT\_DN | MSigDB lists | CAIRO\_LIVER\_DEVELOPMENT\_DN | 147 | 7 | 12187 | 179 | Cybb,Il1rn,Hp,Lcn2,S100a8,A2m,S100a9 | | 5.966e-03 | -5.12 | GO\_KINASE\_REGULATOR\_ACTIVITY | MSigDB lists | GO\_KINASE\_REGULATOR\_ACTIVITY | 147 | 7 | 12187 | 179 | Hspb1,Iqgap1,Pik3r5,Trib1,Socs3,Pik3r1,Cdkn1a | | 6.007e-03 | -5.11 | regulation of chemotaxis | biological process | GO:0050920 | 174 | 8 | 13711 | 214 | Ccl2,Ccl12,C5ar1,Lgals9,Ccl4,C3ar1,Thbs1,Hspb1 | | 6.067e-03 | -5.10 | WESTON\_VEGFA\_TARGETS | MSigDB lists | WESTON\_VEGFA\_TARGETS | 79 | 5 | 12187 | 179 | Tnfsf10,Ccl12,Sdc4,Vcam1,Gem | | 6.075e-03 | -5.10 | guanosine nucleotides degradation III | BIOCYC pathways | META\_PWY-6608 | 1 | 1 | 823 | 5 | Xdh | | 6.075e-03 | -5.10 | urate biosynthesis/inosine 5'-phosphate degradation | BIOCYC pathways | META\_PWY-5695 | 1 | 1 | 823 | 5 | Xdh | | 6.075e-03 | -5.10 | purine nucleotides degradation II (aerobic) | BIOCYC pathways | META\_PWY-6353 | 1 | 1 | 823 | 5 | Xdh | | 6.091e-03 | -5.10 | negative regulation of cytokine secretion | biological process | GO:0050710 | 47 | 4 | 13711 | 214 | Tnf,Tnfaip3,Lgals9,Il6 | | 6.091e-03 | -5.10 | regulation of immunoglobulin production | biological process | GO:0002637 | 47 | 4 | 13711 | 214 | Il4ra,Tnfaip3,Tnf,Il6 | | 6.091e-03 | -5.10 | positive regulation of interferon-gamma production | biological process | GO:0032729 | 47 | 4 | 13711 | 214 | Cd14,Tnf,Runx1,Bcl3 | | 6.120e-03 | -5.10 | GSE14415\_NATURAL\_TREG\_VS\_FOXP3\_KO\_NATURAL\_TREG\_UP | MSigDB lists | GSE14415\_NATURAL\_TREG\_VS\_FOXP3\_KO\_NATURAL\_TREG\_UP | 112 | 6 | 12187 | 179 | Ccl12,Samd9l,Ifit3b,Cflar,Ifi44,Gem | | 6.124e-03 | -5.10 | BROWNE\_HCMV\_INFECTION\_6HR\_UP | MSigDB lists | BROWNE\_HCMV\_INFECTION\_6HR\_UP | 50 | 4 | 12187 | 179 | Isg15,Irf7,Oasl1,Ripk1 | | 6.124e-03 | -5.10 | ZHENG\_GLIOBLASTOMA\_PLASTICITY\_DN | MSigDB lists | ZHENG\_GLIOBLASTOMA\_PLASTICITY\_DN | 50 | 4 | 12187 | 179 | Col4a1,Zfp36,Mxd4,Rhou | | 6.124e-03 | -5.10 | DEMAGALHAES\_AGING\_UP | MSigDB lists | DEMAGALHAES\_AGING\_UP | 50 | 4 | 12187 | 179 | Sgk1,Rnf213,Msn,Clic4 | | 6.124e-03 | -5.10 | SCHEIDEREIT\_IKK\_INTERACTING\_PROTEINS | MSigDB lists | SCHEIDEREIT\_IKK\_INTERACTING\_PROTEINS | 50 | 4 | 12187 | 179 | Ripk1,Tnfaip3,Map3k8,Irf7 | | 6.152e-03 | -5.09 | negative regulation of cell-cell adhesion | biological process | GO:0022408 | 139 | 7 | 13711 | 214 | Il4ra,Tnfaip3,Sdc4,Il1rn,Lgals9,Runx1,Pik3r1 | | 6.155e-03 | -5.09 | protein kinase regulator activity | molecular function | GO:0019887 | 139 | 7 | 13516 | 211 | Lgals9,Cdkn1a,Iqgap1,Hspb1,Ccnd2,Irgm2,Trib1 | | 6.174e-03 | -5.09 | PAX\_Q6 | MSigDB lists | PAX\_Q6 | 186 | 8 | 12187 | 179 | A2m,Tnfsf10,Icam1,Trib1,Runx1,Atf3,Ier3,Cp | | 6.186e-03 | -5.09 | GSE360\_L\_DONOVANI\_VS\_L\_MAJOR\_DC\_DN | MSigDB lists | GSE360\_L\_DONOVANI\_VS\_L\_MAJOR\_DC\_DN | 148 | 7 | 12187 | 179 | Gadd45g,Atf3,Casp4,Trib1,Il17ra,S100a8,Zfp36 | | 6.186e-03 | -5.09 | GSE39110\_UNTREATED\_VS\_IL2\_TREATED\_CD8\_TCELL\_DAY6\_POST\_IMMUNIZATION\_UP | MSigDB lists | GSE39110\_UNTREATED\_VS\_IL2\_TREATED\_CD8\_TCELL\_DAY6\_POST\_IMMUNIZATION\_UP | 148 | 7 | 12187 | 179 | AA467197,Ier3,Sgk1,Pik3ap1,Plek,Ifitm3,Ifitm2 | | 6.186e-03 | -5.09 | KRAS.DF.V1\_UP | MSigDB lists | KRAS.DF.V1\_UP | 148 | 7 | 12187 | 179 | Akap12,Rasip1,Ifi44,Irf9,Map3k6,Csf3,Bcl3 | | 6.199e-03 | -5.08 | CXC\_Chemokine\_domain | interpro domains | IPR033899 | 8 | 2 | 13788 | 212 | Cxcl1,Cxcl9 | | 6.199e-03 | -5.08 | TNF\_dom | interpro domains | IPR006052 | 8 | 2 | 13788 | 212 | Tnf,Tnfsf10 | | 6.226e-03 | -5.08 | BASAKI\_YBX1\_TARGETS\_DN | MSigDB lists | BASAKI\_YBX1\_TARGETS\_DN | 311 | 11 | 12187 | 179 | Ifi211,Rbm47,Rbm39,Tnfsf10,Maff,Pik3r1,Ifi209,Nfkbiz,Ifi207,Ifi204,Lcn2 | | 6.235e-03 | -5.08 | Pertussis | KEGG pathways | ko05133 | 58 | 5 | 5248 | 107 | Nfkb1,Il6,Cd14,Il1a,Tnf | | 6.235e-03 | -5.08 | Pertussis | KEGG pathways | mmu05133 | 58 | 5 | 5248 | 107 | Il6,Nfkb1,Il1a,Tnf,Cd14 | | 6.299e-03 | -5.07 | extrinsic apoptotic signaling pathway | biological process | GO:0097191 | 75 | 5 | 13711 | 214 | Tnf,Ripk1,Il1a,Pik3r1,Lcn2 | | 6.319e-03 | -5.06 | MODULE\_288 | MSigDB lists | MODULE\_288 | 26 | 3 | 12187 | 179 | C5ar1,C3ar1,Ccl2 | | 6.319e-03 | -5.06 | GO\_T\_CELL\_ACTIVATION\_INVOLVED\_IN\_IMMUNE\_RESPONSE | MSigDB lists | GO\_T\_CELL\_ACTIVATION\_INVOLVED\_IN\_IMMUNE\_RESPONSE | 26 | 3 | 12187 | 179 | Bcl3,Il6,Icam1 | | 6.319e-03 | -5.06 | REACTOME\_NEGATIVE\_REGULATORS\_OF\_RIG\_I\_MDA5\_SIGNALING | MSigDB lists | REACTOME\_NEGATIVE\_REGULATORS\_OF\_RIG\_I\_MDA5\_SIGNALING | 26 | 3 | 12187 | 179 | Tnfaip3,Trim25,Isg15 | | 6.319e-03 | -5.06 | PEPPER\_CHRONIC\_LYMPHOCYTIC\_LEUKEMIA\_UP | MSigDB lists | PEPPER\_CHRONIC\_LYMPHOCYTIC\_LEUKEMIA\_UP | 26 | 3 | 12187 | 179 | Cebpd,S100a8,Cxcl1 | | 6.319e-03 | -5.06 | GO\_REGULATION\_OF\_MULTICELLULAR\_ORGANISMAL\_METABOLIC\_PROCESS | MSigDB lists | GO\_REGULATION\_OF\_MULTICELLULAR\_ORGANISMAL\_METABOLIC\_PROCESS | 26 | 3 | 12187 | 179 | Ccl12,Errfi1,Il6 | | 6.371e-03 | -5.06 | response to extracellular stimulus | biological process | GO:0009991 | 293 | 11 | 13711 | 214 | Rrp8,Hspa5,Fam107a,Nfe2l2,Vcam1,Noct,Angptl4,Atf3,Zfp36,Cdkn1a,Fstl1 | | 6.377e-03 | -5.06 | Macro | pfam domains | PF01661 | 8 | 2 | 12881 | 201 | Parp14,Parp9 | | 6.377e-03 | -5.06 | TNF | pfam domains | PF00229 | 8 | 2 | 12881 | 201 | Tnfsf10,Tnf | | 6.382e-03 | -5.05 | regulation of cell-cell adhesion involved in gastrulation | biological process | GO:0070587 | 8 | 2 | 13711 | 214 | Il1rn,Tnfaip3 | | 6.382e-03 | -5.05 | inflammatory response to wounding | biological process | GO:0090594 | 8 | 2 | 13711 | 214 | Il1a,Timp1 | | 6.382e-03 | -5.05 | regulation of removal of superoxide radicals | biological process | GO:2000121 | 8 | 2 | 13711 | 214 | Tnf,Nfe2l2 | | 6.382e-03 | -5.05 | positive regulation of cell proliferation involved in kidney development | biological process | GO:1901724 | 8 | 2 | 13711 | 214 | C3ar1,Cflar | | 6.382e-03 | -5.05 | positive regulation of integrin activation | biological process | GO:0033625 | 8 | 2 | 13711 | 214 | Rasip1,Plek | | 6.382e-03 | -5.05 | positive regulation of response to reactive oxygen species | biological process | GO:1901033 | 8 | 2 | 13711 | 214 | Ripk1,Tnf | | 6.382e-03 | -5.05 | negative regulation of heterotypic cell-cell adhesion | biological process | GO:0034115 | 8 | 2 | 13711 | 214 | Il1rn,Tnfaip3 | | 6.382e-03 | -5.05 | complement receptor mediated signaling pathway | biological process | GO:0002430 | 8 | 2 | 13711 | 214 | C3ar1,C5ar1 | | 6.382e-03 | -5.05 | ectopic germ cell programmed cell death | biological process | GO:0035234 | 8 | 2 | 13711 | 214 | Il1a,Casp4 | | 6.382e-03 | -5.05 | positive regulation of humoral immune response | biological process | GO:0002922 | 8 | 2 | 13711 | 214 | Acod1,Tnf | | 6.382e-03 | -5.05 | nucleotide-binding domain, leucine rich repeat containing receptor signaling pathway | biological process | GO:0035872 | 8 | 2 | 13711 | 214 | Irgm2,Nfkbia | | 6.382e-03 | -5.05 | wound healing involved in inflammatory response | biological process | GO:0002246 | 8 | 2 | 13711 | 214 | Timp1,Il1a | | 6.382e-03 | -5.05 | positive regulation of hepatocyte proliferation | biological process | GO:2000347 | 8 | 2 | 13711 | 214 | Tnfaip3,Cflar | | 6.382e-03 | -5.05 | positive regulation of fever generation | biological process | GO:0031622 | 8 | 2 | 13711 | 214 | Ptgs2,Tnf | | 6.384e-03 | -5.05 | icosanoid binding | molecular function | GO:0050542 | 8 | 2 | 13516 | 211 | S100a8,S100a9 | | 6.384e-03 | -5.05 | oligosaccharide binding | molecular function | GO:0070492 | 8 | 2 | 13516 | 211 | Selp,Sele | | 6.384e-03 | -5.05 | RAGE receptor binding | molecular function | GO:0050786 | 8 | 2 | 13516 | 211 | S100a9,S100a8 | | 6.384e-03 | -5.05 | oxidoreductase activity, acting on NAD(P)H, oxygen as acceptor | molecular function | GO:0050664 | 8 | 2 | 13516 | 211 | Ncf1,Cybb | | 6.384e-03 | -5.05 | sialic acid binding | molecular function | GO:0033691 | 8 | 2 | 13516 | 211 | Selp,Sele | | 6.384e-03 | -5.05 | icosatetraenoic acid binding | molecular function | GO:0050543 | 8 | 2 | 13516 | 211 | S100a8,S100a9 | | 6.386e-03 | -5.05 | KEGG\_NEUROTROPHIN\_SIGNALING\_PATHWAY | MSigDB lists | KEGG\_NEUROTROPHIN\_SIGNALING\_PATHWAY | 113 | 6 | 12187 | 179 | Nfkbia,Pik3r1,Nfkb1,Pik3r5,Nfkbie,Irak3 | | 6.396e-03 | -5.05 | KEGG\_PROSTATE\_CANCER | MSigDB lists | KEGG\_PROSTATE\_CANCER | 80 | 5 | 12187 | 179 | Nfkb1,Cdkn1a,Pik3r1,Pik3r5,Nfkbia | | 6.396e-03 | -5.05 | GO\_PROTEASE\_BINDING | MSigDB lists | GO\_PROTEASE\_BINDING | 80 | 5 | 12187 | 179 | Timp1,Tnfaip3,Cflar,Tnf,A2m | | 6.396e-03 | -5.05 | GSE16385\_IFNG\_TNF\_VS\_ROSIGLITAZONE\_STIM\_MACROPHAGE\_DN | MSigDB lists | GSE16385\_IFNG\_TNF\_VS\_ROSIGLITAZONE\_STIM\_MACROPHAGE\_DN | 80 | 5 | 12187 | 179 | Il1a,Cmpk2,Oasl1,Isg15,Fzd4 | | 6.413e-03 | -5.05 | GSE17721\_PAM3CSK4\_VS\_GADIQUIMOD\_0.5H\_BMDC\_UP | MSigDB lists | GSE17721\_PAM3CSK4\_VS\_GADIQUIMOD\_0.5H\_BMDC\_UP | 149 | 7 | 12187 | 179 | Errfi1,Nfkbie,Cxcl1,Arid5b,Ier3,Tnf,Ccl2 | | 6.413e-03 | -5.05 | GSE7568\_IL4\_TGFB\_DEXAMETHASONE\_VS\_IL4\_TGFB\_TREATED\_MACROPHAGE\_DN | MSigDB lists | GSE7568\_IL4\_TGFB\_DEXAMETHASONE\_VS\_IL4\_TGFB\_TREATED\_MACROPHAGE\_DN | 149 | 7 | 12187 | 179 | Icam1,Cflar,Tnfaip3,Ripk1,Mxd4,Ifit2,Sntb2 | | 6.413e-03 | -5.05 | KIM\_WT1\_TARGETS\_8HR\_UP | MSigDB lists | KIM\_WT1\_TARGETS\_8HR\_UP | 149 | 7 | 12187 | 179 | Gem,Plaur,Dnajb1,Trib1,Aff1,Maff,Il4ra | | 6.413e-03 | -5.05 | ESC\_J1\_UP\_LATE.V1\_DN | MSigDB lists | ESC\_J1\_UP\_LATE.V1\_DN | 149 | 7 | 12187 | 179 | Socs3,Sntb2,Rasip1,Angptl4,Bcl3,Rrp8,Icam1 | | 6.413e-03 | -5.05 | GSE17721\_CTRL\_VS\_GARDIQUIMOD\_4H\_BMDC\_DN | MSigDB lists | GSE17721\_CTRL\_VS\_GARDIQUIMOD\_4H\_BMDC\_DN | 149 | 7 | 12187 | 179 | Maff,Tgm2,Il1rn,Atf3,Ier3,Cebpd,Oasl1 | | 6.413e-03 | -5.05 | GSE9650\_NAIVE\_VS\_EXHAUSTED\_CD8\_TCELL\_DN | MSigDB lists | GSE9650\_NAIVE\_VS\_EXHAUSTED\_CD8\_TCELL\_DN | 149 | 7 | 12187 | 179 | Gem,Tnfsf10,Rgs16,Clic4,Tgm2,Lcn2,Il1a | | 6.502e-03 | -5.04 | 3-phosphoinositide biosynthesis | BIOCYC pathways | MOUSE\_PWY-6352 | 22 | 2 | 823 | 5 | Pik3r1,Pik3r5 | | 6.523e-03 | -5.03 | GO\_MORPHOGENESIS\_OF\_AN\_EPITHELIUM | MSigDB lists | GO\_MORPHOGENESIS\_OF\_AN\_EPITHELIUM | 313 | 11 | 12187 | 179 | Col4a1,Rhoc,Il6,Rasip1,Sox11,Sdc4,Spi1,Socs3,Tgm2,Msn,Clic4 | | 6.565e-03 | -5.03 | regulation of vascular smooth muscle cell proliferation | biological process | GO:1904705 | 48 | 4 | 13711 | 214 | Tnf,Cdkn1a,P2ry6,Adamts1 | | 6.565e-03 | -5.03 | negative regulation of G1/S transition of mitotic cell cycle | biological process | GO:2000134 | 48 | 4 | 13711 | 214 | Fam107a,Ripk1,Cdkn1a,Ccl12 | | 6.565e-03 | -5.03 | regulation of transcription from RNA polymerase II promoter in response to stress | biological process | GO:0043618 | 48 | 4 | 13711 | 214 | Nfe2l2,Bach1,Dnajb1,Atf3 | | 6.565e-03 | -5.03 | endoplasmic reticulum unfolded protein response | biological process | GO:0030968 | 48 | 4 | 13711 | 214 | Atf3,Ptpn1,Hspa5,Nfe2l2 | | 6.571e-03 | -5.03 | TTGCWCAAY\_CEBPB\_02 | MSigDB lists | TTGCWCAAY\_CEBPB\_02 | 51 | 4 | 12187 | 179 | Ip6k2,Nfkbie,Saa1,Trib1 | | 6.571e-03 | -5.03 | GO\_REGULATION\_OF\_RESPONSE\_TO\_OXIDATIVE\_STRESS | MSigDB lists | GO\_REGULATION\_OF\_RESPONSE\_TO\_OXIDATIVE\_STRESS | 51 | 4 | 12187 | 179 | Nfe2l2,Hp,Hspb1,Tnf | | 6.573e-03 | -5.02 | VERHAAK\_AML\_WITH\_NPM1\_MUTATED\_DN | MSigDB lists | VERHAAK\_AML\_WITH\_NPM1\_MUTATED\_DN | 188 | 8 | 12187 | 179 | Hspb1,Ifi211,Ifi207,Ifi204,Lcn2,Ifi44,Ccnd2,Ifi209 | | 6.646e-03 | -5.01 | GSE3039\_CD4\_TCELL\_VS\_B2\_BCELL\_UP | MSigDB lists | GSE3039\_CD4\_TCELL\_VS\_B2\_BCELL\_UP | 150 | 7 | 12187 | 179 | Ncf1,Selp,Hp,Stx11,Trib1,Vcam1,Fstl1 | | 6.646e-03 | -5.01 | GSE18893\_TCONV\_VS\_TREG\_2H\_TNF\_STIM\_DN | MSigDB lists | GSE18893\_TCONV\_VS\_TREG\_2H\_TNF\_STIM\_DN | 150 | 7 | 12187 | 179 | Phf11d,Plek,Zfp36,Phf11b,Zfp189,Thbs1,Bach1 | | 6.646e-03 | -5.01 | GSE2128\_C57BL6\_VS\_NOD\_CD4CD8\_DP\_THYMOCYTE\_UP | MSigDB lists | GSE2128\_C57BL6\_VS\_NOD\_CD4CD8\_DP\_THYMOCYTE\_UP | 150 | 7 | 12187 | 179 | Ptgs2,Cxcl1,Banp,Plaur,Errfi1,Runx1,Il1a | | 6.646e-03 | -5.01 | GSE17721\_CTRL\_VS\_GARDIQUIMOD\_2H\_BMDC\_DN | MSigDB lists | GSE17721\_CTRL\_VS\_GARDIQUIMOD\_2H\_BMDC\_DN | 150 | 7 | 12187 | 179 | Igsf6,Il1rn,Slfn4,Slfn3,Socs3,Casp4,Maff | | 6.646e-03 | -5.01 | GSE36527\_CD62L\_HIGH\_CD69\_NEG\_VS\_CD62L\_LOW\_CD69\_POS\_TREG\_KLRG1\_NEG\_DN | MSigDB lists | GSE36527\_CD62L\_HIGH\_CD69\_NEG\_VS\_CD62L\_LOW\_CD69\_POS\_TREG\_KLRG1\_NEG\_DN | 150 | 7 | 12187 | 179 | Gadd45g,Cdkn1a,Il4ra,Socs3,Parp14,Parp9,Bcl3 | | 6.646e-03 | -5.01 | GSE17580\_UNINFECTED\_VS\_S\_MANSONI\_INF\_TEFF\_DN | MSigDB lists | GSE17580\_UNINFECTED\_VS\_S\_MANSONI\_INF\_TEFF\_DN | 150 | 7 | 12187 | 179 | Gadd45g,Ksr1,Casp4,Socs3,Gbp3,Icam1,Ifitm2 | | 6.661e-03 | -5.01 | GSE4748\_CYANOBACTERIUM\_LPSLIKE\_VS\_LPS\_AND\_CYANOBACTERIUM\_LPSLIKE\_STIM\_DC\_3H\_DN | MSigDB lists | GSE4748\_CYANOBACTERIUM\_LPSLIKE\_VS\_LPS\_AND\_CYANOBACTERIUM\_LPSLIKE\_STIM\_DC\_3H\_DN | 114 | 6 | 12187 | 179 | S100a9,Zfp36,Mmp8,Il1rn,Lcn2,Sbno2 | | 6.661e-03 | -5.01 | ACACTAC\_MIR1423P | MSigDB lists | ACACTAC\_MIR1423P | 114 | 6 | 12187 | 179 | Bach1,Tiparp,Sgk1,Sox11,Aff1,Arid5b | | 6.694e-03 | -5.01 | negative regulation of organic acid transport | biological process | GO:0032891 | 25 | 3 | 13711 | 214 | Thbs1,Il1rn,Tnf | | 6.696e-03 | -5.01 | MAP kinase kinase kinase activity | molecular function | GO:0004709 | 25 | 3 | 13516 | 211 | Map3k8,Map3k6,Ripk1 | | 6.696e-03 | -5.01 | scavenger receptor activity | molecular function | GO:0005044 | 25 | 3 | 13516 | 211 | Msr1,Prg4,Cxcl16 | | 6.710e-03 | -5.00 | multicellular organismal process | biological process | GO:0032501 | 4571 | 89 | 13711 | 214 | Msr1,Bbs12,Lgals9,Gadd45g,Nfkb2,Icam1,Rtp4,Rhoj,Apold1,Tnfaip8,Csf3,Zbtb16,Il1rn,C3ar1,Tnfaip3,Ccl2,S100a9,Vcam1,Col4a1,Iqgap1,Ccnd2,Akap12,Il4ra,Herc6,Cebpd,Rasip1,Runx1,Fam107a,Tiparp,Timp1,Isg15,Rnf213,Acod1,Atf3,Nfe2l2,Noct,Angptl4,Msn,Clic4,Hspa5,Tgm2,C5ar1,Maff,Thbs1,Banp,Sox11,Cemip2,Errfi1,Kcna5,Plaur,Trim56,Adamts1,Nfkb1,Adamts9,Arid5b,Ifi204,Plek,Xdh,Irf9,Samd9l,Bcl3,Ccl12,A2m,S100a8,Socs3,Irf7,Pik3r1,Gadd45b,Il1a,Cdkn1a,Il6,Ccl4,Spi1,Rbm47,Ier3,Ptgs2,Sgk1,Fzd4,Sdc4,Sbno2,Cflar,Hspb1,Slc24a4,Fstl1,Robo4,Tnf,Zfp36,Prg4,Hp | | 6.737e-03 | -5.00 | GNF2\_CASP1 | MSigDB lists | GNF2\_CASP1 | 81 | 5 | 12187 | 179 | Igsf6,Iqgap1,Stx11,Cybb,Ms4a6d | | 6.737e-03 | -5.00 | KEGG\_NATURAL\_KILLER\_CELL\_MEDIATED\_CYTOTOXICITY | MSigDB lists | KEGG\_NATURAL\_KILLER\_CELL\_MEDIATED\_CYTOTOXICITY | 81 | 5 | 12187 | 179 | Pik3r5,Pik3r1,Tnf,Tnfsf10,Icam1 | | 6.742e-03 | -5.00 | GO\_CELL\_SUBSTRATE\_JUNCTION | MSigDB lists | GO\_CELL\_SUBSTRATE\_JUNCTION | 359 | 12 | 12187 | 179 | Tgm2,Sntb2,Sdc4,Rhou,Iqgap1,Hspa5,Akap12,Plaur,Irf2,Msn,Icam1,Hspb1 | | 6.750e-03 | -5.00 | SMALL\_CYTOKINES\_CXC | prosite domains | PS00471 | 7 | 2 | 8845 | 164 | Cxcl1,Cxcl9 | | 6.849e-03 | -4.98 | double-stranded DNA binding | molecular function | GO:0003690 | 750 | 21 | 13516 | 211 | Nfkb1,Atf3,Zbtb16,Irf9,Irf2,Bach1,Runx1,Ifi211,Ifi207,Spi1,Maff,Irf7,Arid5b,Cebpd,Nfe2l2,Ifi204,Sox11,Nfkb2,Aff1,Ifi209,Mxd4 | | 6.859e-03 | -4.98 | response to oxidative stress | biological process | GO:0006979 | 296 | 11 | 13711 | 214 | Lcn2,Ptgs2,Cflar,Ripk1,Ncf1,Thbs1,Nfe2l2,Nfkb1,Il6,Hp,Tnfaip3 | | 6.885e-03 | -4.98 | GSE22886\_NAIVE\_BCELL\_VS\_BM\_PLASMA\_CELL\_DN | MSigDB lists | GSE22886\_NAIVE\_BCELL\_VS\_BM\_PLASMA\_CELL\_DN | 151 | 7 | 12187 | 179 | Cflar,Cd14,Runx1,Cdkn1a,Tor3a,Trib1,Timp1 | | 6.885e-03 | -4.98 | GSE11961\_GERMINAL\_CENTER\_BCELL\_DAY7\_VS\_MEMORY\_BCELL\_DAY40\_DN | MSigDB lists | GSE11961\_GERMINAL\_CENTER\_BCELL\_DAY7\_VS\_MEMORY\_BCELL\_DAY40\_DN | 151 | 7 | 12187 | 179 | Cflar,Cxcl1,Cmpk2,Parp9,Spi1,Rasip1,Il6 | | 6.885e-03 | -4.98 | GSE3720\_UNSTIM\_VS\_PMA\_STIM\_VD2\_GAMMADELTA\_TCELL\_DN | MSigDB lists | GSE3720\_UNSTIM\_VS\_PMA\_STIM\_VD2\_GAMMADELTA\_TCELL\_DN | 151 | 7 | 12187 | 179 | Ripk1,Plek,Dnajb1,Sdc4,Pik3r5,Ifitm2,Pik3r1 | | 6.894e-03 | -4.98 | negative regulation of ion transport | biological process | GO:0043271 | 142 | 7 | 13711 | 214 | Kcna5,Thbs1,Il1rn,Ptgs2,Gem,Tnf,Icam1 | | 7.010e-03 | -4.96 | mouse chr16 | chromosome location | mouse chr16 | 147 | 7 | 14556 | 220 | Retnlg,Runx1,Mx1,Fstl1,Mx2,Cebpd,Adamts1 | | 7.033e-03 | -4.96 | MCBRYAN\_PUBERTAL\_BREAST\_3\_4WK\_DN | MSigDB lists | MCBRYAN\_PUBERTAL\_BREAST\_3\_4WK\_DN | 27 | 3 | 12187 | 179 | Cdkn1a,Nfkb2,Hp | | 7.033e-03 | -4.96 | REACTOME\_NOD1\_2\_SIGNALING\_PATHWAY | MSigDB lists | REACTOME\_NOD1\_2\_SIGNALING\_PATHWAY | 27 | 3 | 12187 | 179 | Tnfaip3,Birc3,Casp4 | | 7.033e-03 | -4.96 | GO\_RESPONSE\_TO\_DEXAMETHASONE | MSigDB lists | GO\_RESPONSE\_TO\_DEXAMETHASONE | 27 | 3 | 12187 | 179 | Il6,Ccl12,Errfi1 | | 7.033e-03 | -4.96 | KEGG\_PRION\_DISEASES | MSigDB lists | KEGG\_PRION\_DISEASES | 27 | 3 | 12187 | 179 | Hspa5,Il1a,Il6 | | 7.033e-03 | -4.96 | GO\_SALIVARY\_GLAND\_DEVELOPMENT | MSigDB lists | GO\_SALIVARY\_GLAND\_DEVELOPMENT | 27 | 3 | 12187 | 179 | Tnf,Tgm2,Il6 | | 7.033e-03 | -4.96 | GRADE\_COLON\_VS\_RECTAL\_CANCER\_UP | MSigDB lists | GRADE\_COLON\_VS\_RECTAL\_CANCER\_UP | 27 | 3 | 12187 | 179 | Il1a,Gadd45b,Fstl1 | | 7.034e-03 | -4.96 | regulation of smooth muscle cell migration | biological process | GO:0014910 | 77 | 5 | 13711 | 214 | Trib1,Nfe2l2,Adamts1,P2ry6,Iqgap1 | | 7.034e-03 | -4.96 | regulation of phosphatidylinositol 3-kinase signaling | biological process | GO:0014066 | 77 | 5 | 13711 | 214 | Ncf1,Tnf,Pik3ap1,Csf3,Selp | | 7.038e-03 | -4.96 | GO\_REGULATION\_OF\_REACTIVE\_OXYGEN\_SPECIES\_BIOSYNTHETIC\_PROCESS | MSigDB lists | GO\_REGULATION\_OF\_REACTIVE\_OXYGEN\_SPECIES\_BIOSYNTHETIC\_PROCESS | 52 | 4 | 12187 | 179 | Il6,Icam1,Ptgs2,Tnf | | 7.038e-03 | -4.96 | GO\_PLATELET\_ALPHA\_GRANULE | MSigDB lists | GO\_PLATELET\_ALPHA\_GRANULE | 52 | 4 | 12187 | 179 | Selp,Thbs1,A2m,Timp1 | | 7.038e-03 | -4.96 | GO\_POSITIVE\_REGULATION\_OF\_SMOOTH\_MUSCLE\_CELL\_PROLIFERATION | MSigDB lists | GO\_POSITIVE\_REGULATION\_OF\_SMOOTH\_MUSCLE\_CELL\_PROLIFERATION | 52 | 4 | 12187 | 179 | Il6,C3ar1,Ptgs2,Tnf | | 7.062e-03 | -4.95 | regulation of leukocyte mediated cytotoxicity | biological process | GO:0001910 | 49 | 4 | 13711 | 214 | Tap1,Lgals9,Ccl2,Cxcl1 | | 7.130e-03 | -4.94 | HNF3ALPHA\_Q6 | MSigDB lists | HNF3ALPHA\_Q6 | 152 | 7 | 12187 | 179 | Cp,Trib1,Sgk1,Cdkn1a,Errfi1,Rbm47,Sgk3 | | 7.130e-03 | -4.94 | GSE4984\_LPS\_VS\_VEHICLE\_CTRL\_TREATED\_DC\_DN | MSigDB lists | GSE4984\_LPS\_VS\_VEHICLE\_CTRL\_TREATED\_DC\_DN | 152 | 7 | 12187 | 179 | Cflar,Slfn9,Iqgap1,Rhou,Runx1,Zbtb16,Tnfaip3 | | 7.130e-03 | -4.94 | GSE16522\_ANTI\_CD3CD28\_STIM\_VS\_UNSTIM\_NAIVE\_CD8\_TCELL\_UP | MSigDB lists | GSE16522\_ANTI\_CD3CD28\_STIM\_VS\_UNSTIM\_NAIVE\_CD8\_TCELL\_UP | 152 | 7 | 12187 | 179 | Cxcl1,Casp4,Cflar,Cxcl16,Slfn4,Slfn3,Il1a | | 7.130e-03 | -4.94 | GSE37532\_WT\_VS\_PPARG\_KO\_LN\_TREG\_UP | MSigDB lists | GSE37532\_WT\_VS\_PPARG\_KO\_LN\_TREG\_UP | 152 | 7 | 12187 | 179 | Runx1,Pik3r5,Arid5b,Ncf1,Ifitm3,Irf2,Zfp36 | | 7.156e-03 | -4.94 | regulation of intrinsic apoptotic signaling pathway | biological process | GO:2001242 | 143 | 7 | 13711 | 214 | Nfe2l2,Hspb1,Ptpn1,S100a9,S100a8,Ptgs2,Plaur | | 7.179e-03 | -4.94 | Butyrophylin\_SPRY | interpro domains | IPR003879 | 26 | 3 | 13788 | 212 | Trim30d,Trim30a,Trim25 | | 7.197e-03 | -4.93 | Adipocytokine signaling pathway | KEGG pathways | ko04920 | 60 | 5 | 5248 | 107 | Nfkbia,Socs3,Nfkb1,Tnf,Nfkbie | | 7.197e-03 | -4.93 | p53 signaling pathway | KEGG pathways | mmu04115 | 60 | 5 | 5248 | 107 | Gadd45g,Ccnd2,Thbs1,Gadd45b,Cdkn1a | | 7.197e-03 | -4.93 | p53 signaling pathway | KEGG pathways | ko04115 | 60 | 5 | 5248 | 107 | Thbs1,Cdkn1a,Gadd45b,Gadd45g,Ccnd2 | | 7.197e-03 | -4.93 | Adipocytokine signaling pathway | KEGG pathways | mmu04920 | 60 | 5 | 5248 | 107 | Socs3,Nfkbia,Nfkb1,Tnf,Nfkbie | | 7.206e-03 | -4.93 | positive regulation of NF-kappaB transcription factor activity | biological process | GO:0051092 | 109 | 6 | 13711 | 214 | Cflar,Ripk1,Irak3,Trim25,Tnf,Icam1 | | 7.206e-03 | -4.93 | leukocyte mediated immunity | biological process | GO:0002443 | 109 | 6 | 13711 | 214 | Icam1,Ncf1,Il4ra,Stx11,Irf7,Bcl3 | | 7.217e-03 | -4.93 | GO\_T\_CELL\_MIGRATION | MSigDB lists | GO\_T\_CELL\_MIGRATION | 9 | 2 | 12187 | 179 | Ccl12,Cxcl16 | | 7.217e-03 | -4.93 | HOWLIN\_CITED1\_TARGETS\_2\_UP | MSigDB lists | HOWLIN\_CITED1\_TARGETS\_2\_UP | 9 | 2 | 12187 | 179 | Ifi44,Iqgap1 | | 7.217e-03 | -4.93 | SCHMAHL\_PDGF\_SIGNALING | MSigDB lists | SCHMAHL\_PDGF\_SIGNALING | 9 | 2 | 12187 | 179 | Tiparp,Arid5b | | 7.217e-03 | -4.93 | GO\_REGULATION\_OF\_REMOVAL\_OF\_SUPEROXIDE\_RADICALS | MSigDB lists | GO\_REGULATION\_OF\_REMOVAL\_OF\_SUPEROXIDE\_RADICALS | 9 | 2 | 12187 | 179 | Tnf,Nfe2l2 | | 7.217e-03 | -4.93 | GO\_COMPLEMENT\_BINDING | MSigDB lists | GO\_COMPLEMENT\_BINDING | 9 | 2 | 12187 | 179 | C5ar1,C3ar1 | | 7.217e-03 | -4.93 | WONG\_ENDMETRIUM\_CANCER\_UP | MSigDB lists | WONG\_ENDMETRIUM\_CANCER\_UP | 9 | 2 | 12187 | 179 | Hp,Lcn2 | | 7.217e-03 | -4.93 | CHEN\_HOXA5\_TARGETS\_6HR\_UP | MSigDB lists | CHEN\_HOXA5\_TARGETS\_6HR\_UP | 9 | 2 | 12187 | 179 | Gadd45b,Tiparp | | 7.219e-03 | -4.93 | regulation of cellular macromolecule biosynthetic process | biological process | GO:2000112 | 2631 | 56 | 13711 | 214 | Tnf,Acod1,Atf3,Zfp36,Ifi207,Ncf1,Cflar,Ripk1,Tnip1,Fstl1,Dnajb1,Maff,Rrp8,Nfe2l2,Rasd1,Il1a,Nfkbia,Il6,Cdkn1a,Irf7,Dtx3l,Trib1,Pik3r1,Ifi209,Fzd4,Sbno2,Parp9,Cebpd,Ifi211,Spi1,Runx1,Resf1,Trim25,Tnfaip3,Znfx1,Irak3,Arid5b,Aff1,C3ar1,Zbtb16,Csf3,Ifi204,Irf2,Irf9,S100a9,Bcl3,Icam1,Mxd4,Sox11,Nfkb2,Thbs1,Banp,Gbp4,Nfkb1,Nfkbie,Bach1 | | 7.223e-03 | -4.93 | Cross-presentation of particulate exogenous antigens (phagosomes) | REACTOME pathways | R-MMU-1236973 | 8 | 2 | 6297 | 105 | Ncf1,Cybb | | 7.223e-03 | -4.93 | TRAF6 mediated IRF7 activation | REACTOME pathways | R-MMU-933541 | 8 | 2 | 6297 | 105 | Irf2,Irf7 | | 7.236e-03 | -4.93 | REACTOME\_G\_ALPHA\_I\_SIGNALLING\_EVENTS | MSigDB lists | REACTOME\_G\_ALPHA\_I\_SIGNALLING\_EVENTS | 116 | 6 | 12187 | 179 | C5ar1,C3ar1,Saa1,Cxcl1,Cxcl16,Hcar2 | | 7.236e-03 | -4.93 | MODULE\_122 | MSigDB lists | MODULE\_122 | 116 | 6 | 12187 | 179 | Ccl4,Vcam1,Selp,Ccl12,Col4a1,Icam1 | | 7.258e-03 | -4.93 | positive regulation of gene expression | biological process | GO:0010628 | 1577 | 37 | 13711 | 214 | Nfe2l2,Mmp8,Msn,Bcl3,Noct,Maff,Irf2,Ncf1,Aff1,Csf3,Zbtb16,Ripk1,Tnip1,Lcn2,Tnf,Atf3,Zfp36,Ccl2,Bach1,Cebpd,Spi1,Runx1,Fzd4,Nfkb1,Parp9,Sbno2,Thbs1,Dtx3l,Irf7,Pik3r1,Banp,Lgals9,Nfkb2,Nfkbia,Sox11,Il1a,Il6 | | 7.259e-03 | -4.93 | negative regulation of developmental process | biological process | GO:0051093 | 852 | 23 | 13711 | 214 | Noct,Runx1,Nfe2l2,Adamts1,Timp1,Pik3r1,Cflar,Zbtb16,Adamts9,Bbs12,Trib1,Ripk1,Thbs1,Cdkn1a,Ccl2,Il6,Zfp36,Xdh,Sox11,Il1a,Nfkbia,Il4ra,Tnf | | 7.267e-03 | -4.92 | tube development | biological process | GO:0035295 | 754 | 21 | 13711 | 214 | Tiparp,Tgm2,Clic4,Fzd4,Apold1,Rhoj,Col4a1,Ccl12,Sdc4,Rasip1,Spi1,Ptgs2,Angptl4,Robo4,Sox11,Cemip2,Errfi1,Ccl2,Thbs1,Adamts9,Rnf213 | | 7.298e-03 | -4.92 | Graft-versus-host disease | KEGG pathways | ko05332 | 20 | 3 | 5248 | 107 | Il6,Tnf,Il1a | | 7.298e-03 | -4.92 | Graft-versus-host disease | KEGG pathways | mmu05332 | 20 | 3 | 5248 | 107 | Il6,Il1a,Tnf | | 7.356e-03 | -4.91 | Traf2 (TNF receptor-associated factor 2) | protein interactions | 22030 | 24 | 3 | 6802 | 115 | Birc3,Tnfaip3,Ripk1 | | 7.382e-03 | -4.91 | GSE15930\_STIM\_VS\_STIM\_AND\_IFNAB\_24H\_CD8\_T\_CELL\_DN | MSigDB lists | GSE15930\_STIM\_VS\_STIM\_AND\_IFNAB\_24H\_CD8\_T\_CELL\_DN | 153 | 7 | 12187 | 179 | Ifit2,Lgals9,Hspa5,Ccl4,Ifit3b,Il1a,Ier3 | | 7.382e-03 | -4.91 | GSE43863\_LY6C\_INT\_CXCR5POS\_VS\_LY6C\_LOW\_CXCR5NEG\_EFFECTOR\_CD4\_TCELL\_UP | MSigDB lists | GSE43863\_LY6C\_INT\_CXCR5POS\_VS\_LY6C\_LOW\_CXCR5NEG\_EFFECTOR\_CD4\_TCELL\_UP | 153 | 7 | 12187 | 179 | Birc3,Ncf1,Tagln2,Cp,Ms4a6d,Rassf4,Il6 | | 7.382e-03 | -4.91 | GSE3039\_NKT\_CELL\_VS\_B2\_BCELL\_DN | MSigDB lists | GSE3039\_NKT\_CELL\_VS\_B2\_BCELL\_DN | 153 | 7 | 12187 | 179 | Ptgs2,Sdc4,Ccl4,Maff,Cdkn1a,Tnfaip3,Hcar2 | | 7.382e-03 | -4.91 | GSE26488\_CTRL\_VS\_PEPTIDE\_INJECTION\_OT2\_THYMOCYTE\_DN | MSigDB lists | GSE26488\_CTRL\_VS\_PEPTIDE\_INJECTION\_OT2\_THYMOCYTE\_DN | 153 | 7 | 12187 | 179 | Sgk1,Isg15,Iqgap1,Trim25,Sele,Irf2,Msn | | 7.382e-03 | -4.91 | GSE17721\_CPG\_VS\_GARDIQUIMOD\_4H\_BMDC\_UP | MSigDB lists | GSE17721\_CPG\_VS\_GARDIQUIMOD\_4H\_BMDC\_UP | 153 | 7 | 12187 | 179 | Atf3,Casp4,Ccl2,Tor3a,Tnfsf10,Rtp4,Plek | | 7.429e-03 | -4.90 | Complement and Coagulation Cascades | WikiPathways | WP449 | 31 | 4 | 3756 | 96 | A2m,Plaur,C5ar1,C3ar1 | | 7.429e-03 | -4.90 | CHEMNITZ\_RESPONSE\_TO\_PROSTAGLANDIN\_E2\_DN | MSigDB lists | CHEMNITZ\_RESPONSE\_TO\_PROSTAGLANDIN\_E2\_DN | 275 | 10 | 12187 | 179 | Bbs12,Tnfsf10,Clic4,Ip6k2,Rbm47,Sntb2,Cdkn1a,Ccnd2,Lcn2,Akap12 | | 7.429e-03 | -4.90 | KINSEY\_TARGETS\_OF\_EWSR1\_FLII\_FUSION\_DN | MSigDB lists | KINSEY\_TARGETS\_OF\_EWSR1\_FLII\_FUSION\_DN | 275 | 10 | 12187 | 179 | Ier3,Mxd4,Aff1,Nfkbiz,Fstl1,Isg15,Gem,Arid5b,A2m,Slfn5 | | 7.430e-03 | -4.90 | GO\_RESPONSE\_TO\_TOXIC\_SUBSTANCE | MSigDB lists | GO\_RESPONSE\_TO\_TOXIC\_SUBSTANCE | 192 | 8 | 12187 | 179 | S100a9,Ptgs2,Ccl4,Cdkn1a,Hspa5,Ccl12,Il6,Hp | | 7.459e-03 | -4.90 | BIOCARTA\_MAPK\_PATHWAY | MSigDB lists | BIOCARTA\_MAPK\_PATHWAY | 83 | 5 | 12187 | 179 | Nfkbia,Map3k6,Map3k8,Ripk1,Nfkb1 | | 7.459e-03 | -4.90 | GO\_REGULATION\_OF\_FAT\_CELL\_DIFFERENTIATION | MSigDB lists | GO\_REGULATION\_OF\_FAT\_CELL\_DIFFERENTIATION | 83 | 5 | 12187 | 179 | Il6,Bbs12,Zbtb16,Ptgs2,Tnf | | 7.459e-03 | -4.90 | GO\_POSITIVE\_REGULATION\_OF\_PROTEIN\_IMPORT | MSigDB lists | GO\_POSITIVE\_REGULATION\_OF\_PROTEIN\_IMPORT | 83 | 5 | 12187 | 179 | Il6,Tnf,Pik3r1,Csf3,Ptgs2 | | 7.481e-03 | -4.90 | T cell differentiation involved in immune response | biological process | GO:0002292 | 26 | 3 | 13711 | 214 | Bcl3,Il6,Gadd45g | | 7.481e-03 | -4.90 | salivary gland morphogenesis | biological process | GO:0007435 | 26 | 3 | 13711 | 214 | Tgm2,Tnf,Il6 | | 7.481e-03 | -4.90 | necrotic cell death | biological process | GO:0070265 | 26 | 3 | 13711 | 214 | Tnf,Ripk1,Birc3 | | 7.481e-03 | -4.90 | modification by symbiont of host morphology or physiology | biological process | GO:0044003 | 26 | 3 | 13711 | 214 | Irgm1,Tnip1,Irgm2 | | 7.504e-03 | -4.89 | cellular metal ion homeostasis | biological process | GO:0006875 | 429 | 14 | 13711 | 214 | Mt1,Ccl2,C3ar1,Pygm,Slc24a4,P2ry6,Cp,Lcn2,Cxcl1,C5ar1,Tgm2,Mt2,Kcna5,Sgk1 | | 7.527e-03 | -4.89 | AMIT\_EGF\_RESPONSE\_240\_HELA | MSigDB lists | AMIT\_EGF\_RESPONSE\_240\_HELA | 53 | 4 | 12187 | 179 | Akap12,Thbs1,Sdc4,Bcl3 | | 7.527e-03 | -4.89 | GO\_CELL\_REDOX\_HOMEOSTASIS | MSigDB lists | GO\_CELL\_REDOX\_HOMEOSTASIS | 53 | 4 | 12187 | 179 | Cybb,Nfe2l2,Il6,Ncf1 | | 7.536e-03 | -4.89 | MCBRYAN\_PUBERTAL\_BREAST\_5\_6WK\_DN | MSigDB lists | MCBRYAN\_PUBERTAL\_BREAST\_5\_6WK\_DN | 117 | 6 | 12187 | 179 | Fzd4,Gbp7,Sdc4,Sntb2,Runx1,Ccnd2 | | 7.584e-03 | -4.88 | positive regulation of myeloid leukocyte differentiation | biological process | GO:0002763 | 50 | 4 | 13711 | 214 | Runx1,Ripk1,Trib1,Tnf | | 7.584e-03 | -4.88 | positive regulation of extrinsic apoptotic signaling pathway | biological process | GO:2001238 | 50 | 4 | 13711 | 214 | Tnfsf10,Atf3,Thbs1,Ripk1 | | 7.591e-03 | -4.88 | MyD88 dependent cascade initiated on endosome | REACTOME pathways | R-MMU-975155 | 74 | 5 | 6297 | 105 | Nfkbia,Map3k8,Nfkb1,Irf7,Nfkb2 | | 7.591e-03 | -4.88 | Toll Like Receptor 7/8 (TLR7/8) Cascade | REACTOME pathways | R-MMU-168181 | 74 | 5 | 6297 | 105 | Nfkbia,Map3k8,Nfkb1,Irf7,Nfkb2 | | 7.602e-03 | -4.88 | A1pp | smart domains | SM00506 | 8 | 2 | 7188 | 123 | Parp14,Parp9 | | 7.640e-03 | -4.87 | GAUSSMANN\_MLL\_AF4\_FUSION\_TARGETS\_F\_UP | MSigDB lists | GAUSSMANN\_MLL\_AF4\_FUSION\_TARGETS\_F\_UP | 154 | 7 | 12187 | 179 | Isg15,Osmr,Nfkbiz,Il4ra,Ifi44,Map3k8,Cebpd | | 7.640e-03 | -4.87 | GSE2706\_LPS\_VS\_R848\_AND\_LPS\_8H\_STIM\_DC\_UP | MSigDB lists | GSE2706\_LPS\_VS\_R848\_AND\_LPS\_8H\_STIM\_DC\_UP | 154 | 7 | 12187 | 179 | Samd9l,Gbp3,Parp9,Tnfsf10,Ifit2,Xaf1,Sntb2 | | 7.640e-03 | -4.87 | GSE15330\_MEGAKARYOCYTE\_ERYTHROID\_VS\_GRANULOCYTE\_MONOCYTE\_PROGENITOR\_IKAROS\_KO\_DN | MSigDB lists | GSE15330\_MEGAKARYOCYTE\_ERYTHROID\_VS\_GRANULOCYTE\_MONOCYTE\_PROGENITOR\_IKAROS\_KO\_DN | 154 | 7 | 12187 | 179 | Tagln2,Csf3,Sgk1,Timp1,Tiparp,Hspb1,S100a8 | | 7.640e-03 | -4.87 | GSE20727\_CTRL\_VS\_H2O2\_TREATED\_DC\_UP | MSigDB lists | GSE20727\_CTRL\_VS\_H2O2\_TREATED\_DC\_UP | 154 | 7 | 12187 | 179 | Parp14,Ifi209,Il4ra,Ifi44,Ifi204,Ifi207,Ifi211 | | 7.640e-03 | -4.87 | GSE29949\_CD8\_NEG\_DC\_SPLEEN\_VS\_DC\_BRAIN\_DN | MSigDB lists | GSE29949\_CD8\_NEG\_DC\_SPLEEN\_VS\_DC\_BRAIN\_DN | 154 | 7 | 12187 | 179 | Rgs16,Irf9,Gem,Cxcl1,Ccl12,Atf3,Map3k8 | | 7.640e-03 | -4.87 | GSE22432\_MULTIPOTENT\_PROGENITOR\_VS\_PDC\_UP | MSigDB lists | GSE22432\_MULTIPOTENT\_PROGENITOR\_VS\_PDC\_UP | 154 | 7 | 12187 | 179 | Tap1,Gbp3,Cmpk2,Rtp4,Arid5b,Tor3a,Ccnd2 | | 7.651e-03 | -4.87 | GO\_RESPONSE\_TO\_PEPTIDE | MSigDB lists | GO\_RESPONSE\_TO\_PEPTIDE | 320 | 11 | 12187 | 179 | Socs3,Timp1,Nfkb1,Ptpn1,Errfi1,Casp4,Nfkbia,Pik3r1,Il6,Ccl12,Tnfaip3 | | 7.722e-03 | -4.86 | negative regulation of cell migration | biological process | GO:0030336 | 220 | 9 | 13711 | 214 | Adamts9,Trib1,Nfe2l2,Il1rn,Thbs1,Clic4,Robo4,Timp1,Ccl12 | | 7.722e-03 | -4.86 | regulation of protein modification by small protein conjugation or removal | biological process | GO:1903320 | 220 | 9 | 13711 | 214 | Dtx3l,Trib1,Tnip1,Tnfaip3,Gbp4,Hspa5,Fam107a,Birc3,Isg15 | | 7.763e-03 | -4.86 | Complement and coagulation cascades | KEGG pathways | ko04610 | 39 | 4 | 5248 | 107 | C3ar1,C5ar1,Plaur,A2m | | 7.763e-03 | -4.86 | Complement and coagulation cascades | KEGG pathways | mmu04610 | 39 | 4 | 5248 | 107 | A2m,Plaur,C5ar1,C3ar1 | | 7.793e-03 | -4.85 | TARTE\_PLASMA\_CELL\_VS\_B\_LYMPHOCYTE\_DN | MSigDB lists | TARTE\_PLASMA\_CELL\_VS\_B\_LYMPHOCYTE\_DN | 28 | 3 | 12187 | 179 | Nfkb1,Il4ra,Nfkbie | | 7.793e-03 | -4.85 | PID\_FRA\_PATHWAY | MSigDB lists | PID\_FRA\_PATHWAY | 28 | 3 | 12187 | 179 | Plaur,Ccl12,Il6 | | 7.793e-03 | -4.85 | GO\_ACTIVATION\_OF\_JUN\_KINASE\_ACTIVITY | MSigDB lists | GO\_ACTIVATION\_OF\_JUN\_KINASE\_ACTIVITY | 28 | 3 | 12187 | 179 | Map3k6,Ptpn1,Ripk1 | | 7.793e-03 | -4.85 | SEITZ\_NEOPLASTIC\_TRANSFORMATION\_BY\_8P\_DELETION\_DN | MSigDB lists | SEITZ\_NEOPLASTIC\_TRANSFORMATION\_BY\_8P\_DELETION\_DN | 28 | 3 | 12187 | 179 | Cdkn1a,Csf3,Il1a | | 7.793e-03 | -4.85 | PID\_AVB3\_OPN\_PATHWAY | MSigDB lists | PID\_AVB3\_OPN\_PATHWAY | 28 | 3 | 12187 | 179 | Nfkb1,Nfkbia,Pik3r1 | | 7.793e-03 | -4.85 | GO\_REGULATION\_OF\_IMMUNOGLOBULIN\_PRODUCTION | MSigDB lists | GO\_REGULATION\_OF\_IMMUNOGLOBULIN\_PRODUCTION | 28 | 3 | 12187 | 179 | Il6,Il4ra,Tnf | | 7.793e-03 | -4.85 | GO\_CYTOPLASMIC\_PATTERN\_RECOGNITION\_RECEPTOR\_SIGNALING\_PATHWAY | MSigDB lists | GO\_CYTOPLASMIC\_PATTERN\_RECOGNITION\_RECEPTOR\_SIGNALING\_PATHWAY | 28 | 3 | 12187 | 179 | Tnfaip3,Nfkbia,Irf7 | | 7.793e-03 | -4.85 | GO\_NEGATIVE\_REGULATION\_OF\_RESPONSE\_TO\_OXIDATIVE\_STRESS | MSigDB lists | GO\_NEGATIVE\_REGULATION\_OF\_RESPONSE\_TO\_OXIDATIVE\_STRESS | 28 | 3 | 12187 | 179 | Nfe2l2,Hp,Hspb1 | | 7.806e-03 | -4.85 | regulation of ossification | biological process | GO:0030278 | 182 | 8 | 13711 | 214 | Ifi204,Noct,Zbtb16,Cebpd,Isg15,Il6,Sox11,Tnf | | 7.826e-03 | -4.85 | modification of morphology or physiology of other organism involved in symbiotic interaction | biological process | GO:0051817 | 79 | 5 | 13711 | 214 | Irgm2,Tnip1,Ccl4,Irgm1,Ncf1 | | 7.838e-03 | -4.85 | LABBE\_TARGETS\_OF\_TGFB1\_AND\_WNT3A\_UP | MSigDB lists | LABBE\_TARGETS\_OF\_TGFB1\_AND\_WNT3A\_UP | 84 | 5 | 12187 | 179 | Ptgs2,Sgk1,Sox11,Pygm,Gadd45g | | 7.846e-03 | -4.85 | GSE37605\_NOD\_VS\_C57BL6\_IRES\_GFP\_TREG\_DN | MSigDB lists | GSE37605\_NOD\_VS\_C57BL6\_IRES\_GFP\_TREG\_DN | 118 | 6 | 12187 | 179 | Trim56,Pik3r1,Nfe2l2,Rasip1,Trim25,Irak3 | | 7.858e-03 | -4.85 | protease binding | molecular function | GO:0002020 | 111 | 6 | 13516 | 211 | Tnf,A2m,Lcn2,Cflar,Timp1,Tnfaip3 | | 7.863e-03 | -4.85 | negative regulation of transcription, DNA-templated | biological process | GO:0045892 | 958 | 25 | 13711 | 214 | Mxd4,Sox11,Nfkb1,Parp9,Nfkbie,Sbno2,Ifi209,Gbp4,Runx1,Bach1,Ifi211,Cebpd,Spi1,Zfp36,Atf3,Znfx1,Tnf,Ifi204,Ifi207,Arid5b,Zbtb16,Dnajb1,Rrp8,Bcl3,Rasd1 | | 7.889e-03 | -4.84 | Macro\_dom | interpro domains | IPR002589 | 9 | 2 | 13788 | 212 | Parp9,Parp14 | | 7.902e-03 | -4.84 | HELLER\_HDAC\_TARGETS\_DN | MSigDB lists | HELLER\_HDAC\_TARGETS\_DN | 235 | 9 | 12187 | 179 | Phf11d,Cflar,Dnajb1,Phf11b,Ccnd2,Sox11,Hspa5,Maff,Atf3 | | 7.905e-03 | -4.84 | GSE17301\_IFNA2\_VS\_IFNA2\_AND\_ACD3\_ACD28\_STIM\_CD8\_TCELL\_DN | MSigDB lists | GSE17301\_IFNA2\_VS\_IFNA2\_AND\_ACD3\_ACD28\_STIM\_CD8\_TCELL\_DN | 155 | 7 | 12187 | 179 | Ptgs2,Tnfsf10,Il4ra,Birc3,Trib1,Nfkbia,Ier3 | | 7.905e-03 | -4.84 | GSE360\_CTRL\_VS\_T\_GONDII\_MAC\_DN | MSigDB lists | GSE360\_CTRL\_VS\_T\_GONDII\_MAC\_DN | 155 | 7 | 12187 | 179 | Zfp36,Csf2rb,Pygm,Rgs16,Oasl1,Tnf,Trib1 | | 7.905e-03 | -4.84 | GSE1448\_CTRL\_VS\_ANTI\_VALPHA2\_DP\_THYMOCYTE\_DN | MSigDB lists | GSE1448\_CTRL\_VS\_ANTI\_VALPHA2\_DP\_THYMOCYTE\_DN | 155 | 7 | 12187 | 179 | Il17ra,Tnfsf10,Irf9,Isg15,A2m,Ptges,Ccl4 | | 7.905e-03 | -4.84 | GSE19888\_ADENOSINE\_A3R\_INH\_PRETREAT\_AND\_ACT\_BY\_A3R\_VS\_A3R\_INH\_AND\_TCELL\_MEMBRANES\_ACT\_MAST\_CELL\_UP | MSigDB lists | GSE19888\_ADENOSINE\_A3R\_INH\_PRETREAT\_AND\_ACT\_BY\_A3R\_VS\_A3R\_INH\_AND\_TCELL\_MEMBRANES\_ACT\_MAST\_CELL\_UP | 155 | 7 | 12187 | 179 | Zfp36,Plaur,Cflar,Arid5b,Bach1,Tnfaip3,Thbs1 | | 7.937e-03 | -4.84 | G-protein beta:gamma signalling | REACTOME pathways | R-MMU-397795 | 25 | 3 | 6297 | 105 | Pik3r1,Csf2rb,Pik3r5 | | 7.937e-03 | -4.84 | TAK1 activates NFkB by phosphorylation and activation of IKKs complex | REACTOME pathways | R-MMU-445989 | 25 | 3 | 6297 | 105 | Nfkb2,Nfkb1,Nfkbia | | 8.038e-03 | -4.82 | chr1p22 | MSigDB lists | chr1p22 | 54 | 4 | 12187 | 179 | Gbp6,Gbp7,Gbp4,Gbp3 | | 8.038e-03 | -4.82 | PID\_RB\_1PATHWAY | MSigDB lists | PID\_RB\_1PATHWAY | 54 | 4 | 12187 | 179 | Cdkn1a,Ccnd2,Spi1,Cebpd | | 8.102e-03 | -4.82 | cellular cation homeostasis | biological process | GO:0030003 | 478 | 15 | 13711 | 214 | Mt1,Ccl2,Slc24a4,C3ar1,Pygm,Lcn2,Cp,P2ry6,Tgm2,C5ar1,Cxcl1,Clic4,Mt2,Kcna5,Sgk1 | | 8.121e-03 | -4.81 | regulation of glomerular mesangial cell proliferation | biological process | GO:0072124 | 9 | 2 | 13711 | 214 | C3ar1,Cflar | | 8.121e-03 | -4.81 | toll-like receptor 3 signaling pathway | biological process | GO:0034138 | 9 | 2 | 13711 | 214 | Tnf,Lgals9 | | 8.121e-03 | -4.81 | regulation of autophagosome maturation | biological process | GO:1901096 | 9 | 2 | 13711 | 214 | Irgm1,Irgm2 | | 8.121e-03 | -4.81 | regulation of protein localization to early endosome | biological process | GO:1902965 | 9 | 2 | 13711 | 214 | Msn,Dtx3l | | 8.121e-03 | -4.81 | marginal zone B cell differentiation | biological process | GO:0002315 | 9 | 2 | 13711 | 214 | Bcl3,Tnfaip3 | | 8.121e-03 | -4.81 | regulation of fever generation | biological process | GO:0031620 | 9 | 2 | 13711 | 214 | Tnf,Ptgs2 | | 8.121e-03 | -4.81 | positive regulation of protein localization to early endosome | biological process | GO:1902966 | 9 | 2 | 13711 | 214 | Msn,Dtx3l | | 8.121e-03 | -4.81 | superoxide anion generation | biological process | GO:0042554 | 9 | 2 | 13711 | 214 | Ncf1,Cybb | | 8.121e-03 | -4.81 | negative regulation of granulocyte differentiation | biological process | GO:0030853 | 9 | 2 | 13711 | 214 | Runx1,Trib1 | | 8.131e-03 | -4.81 | positive regulation of receptor signaling pathway via JAK-STAT | biological process | GO:0046427 | 51 | 4 | 13711 | 214 | Parp14,Tnf,Parp9,Il6 | | 8.177e-03 | -4.81 | GSE34515\_CD16\_NEG\_MONOCYTE\_VS\_DC\_UP | MSigDB lists | GSE34515\_CD16\_NEG\_MONOCYTE\_VS\_DC\_UP | 156 | 7 | 12187 | 179 | Msn,C3ar1,Bach1,Rhoc,Pik3r1,Sntb2,Zbtb16 | | 8.177e-03 | -4.81 | GSE41176\_UNSTIM\_VS\_ANTI\_IGM\_STIM\_BCELL\_6H\_DN | MSigDB lists | GSE41176\_UNSTIM\_VS\_ANTI\_IGM\_STIM\_BCELL\_6H\_DN | 156 | 7 | 12187 | 179 | Ifitm3,Ifitm2,Il17ra,Timp1,Pik3ap1,Rtp4,Tnfsf10 | | 8.177e-03 | -4.81 | GSE14308\_TH17\_VS\_NAIVE\_CD4\_TCELL\_DN | MSigDB lists | GSE14308\_TH17\_VS\_NAIVE\_CD4\_TCELL\_DN | 156 | 7 | 12187 | 179 | Gbp4,Irf9,Zfp36,Ccl4,Aff1,Ccnd2,Gbp6 | | 8.177e-03 | -4.81 | SHEDDEN\_LUNG\_CANCER\_GOOD\_SURVIVAL\_A4 | MSigDB lists | SHEDDEN\_LUNG\_CANCER\_GOOD\_SURVIVAL\_A4 | 156 | 7 | 12187 | 179 | Map3k6,Gem,Icam1,Nfkbia,Zbtb16,Tiparp,Cebpd | | 8.177e-03 | -4.81 | GSE42088\_UNINF\_VS\_LEISHMANIA\_INF\_DC\_2H\_DN | MSigDB lists | GSE42088\_UNINF\_VS\_LEISHMANIA\_INF\_DC\_2H\_DN | 156 | 7 | 12187 | 179 | Zfp36,Ifitm3,Ifitm2,Tagln2,Oasl1,Cebpd,Pik3r1 | | 8.177e-03 | -4.81 | GSE21063\_CTRL\_VS\_ANTI\_IGM\_STIM\_BCELL\_NFATC1\_KO\_3H\_UP | MSigDB lists | GSE21063\_CTRL\_VS\_ANTI\_IGM\_STIM\_BCELL\_NFATC1\_KO\_3H\_UP | 156 | 7 | 12187 | 179 | Isg15,Cd14,S100a9,S100a8,Usp18,Parp12,Lcn2 | | 8.177e-03 | -4.81 | ATACCTC\_MIR202 | MSigDB lists | ATACCTC\_MIR202 | 156 | 7 | 12187 | 179 | Sox11,Ccnd2,Trib1,Ip6k2,Adamts9,Rgs16,Luc7l3 | | 8.177e-03 | -4.81 | GSE22033\_UNTREATED\_VS\_ROSIGLITAZONE\_TREATED\_MEF\_DN | MSigDB lists | GSE22033\_UNTREATED\_VS\_ROSIGLITAZONE\_TREATED\_MEF\_DN | 156 | 7 | 12187 | 179 | Tnfaip3,Nfkbia,Ifit2,Birc3,Ifi44,Nfkb2,Arid5b | | 8.232e-03 | -4.80 | CROONQUIST\_NRAS\_VS\_STROMAL\_STIMULATION\_DN | MSigDB lists | CROONQUIST\_NRAS\_VS\_STROMAL\_STIMULATION\_DN | 85 | 5 | 12187 | 179 | Col4a1,Sbno2,Il6,Ier3,Nfkbia | | 8.232e-03 | -4.80 | KEGG\_CELL\_ADHESION\_MOLECULES\_CAMS | MSigDB lists | KEGG\_CELL\_ADHESION\_MOLECULES\_CAMS | 85 | 5 | 12187 | 179 | Vcam1,Sdc4,Selp,Icam1,Sele | | 8.232e-03 | -4.80 | TAKEDA\_TARGETS\_OF\_NUP98\_HOXA9\_FUSION\_16D\_DN | MSigDB lists | TAKEDA\_TARGETS\_OF\_NUP98\_HOXA9\_FUSION\_16D\_DN | 85 | 5 | 12187 | 179 | Olfml2b,Mmp8,Hp,Slfn9,Cflar | | 8.244e-03 | -4.80 | regulation of JUN kinase activity | biological process | GO:0043506 | 80 | 5 | 13711 | 214 | Tnf,Il1rn,Ptpn1,Ripk1,Fzd4 | | 8.319e-03 | -4.79 | positive regulation of catabolic process | biological process | GO:0009896 | 390 | 13 | 13711 | 214 | Ptpn1,Tiparp,Ier3,Irgm2,Msn,Irgm1,Zfp36,Il6,Tnf,Tnfaip3,Trim30a,Trib1,Dtx3l | | 8.320e-03 | -4.79 | positive regulation of CD4-positive, alpha-beta T cell activation | biological process | GO:2000516 | 27 | 3 | 13711 | 214 | Il6,Il4ra,Nfkbiz | | 8.320e-03 | -4.79 | response to cadmium ion | biological process | GO:0046686 | 27 | 3 | 13711 | 214 | Mt1,Ncf1,Mt2 | | 8.320e-03 | -4.79 | CD4-positive, alpha-beta T cell differentiation | biological process | GO:0043367 | 27 | 3 | 13711 | 214 | Gadd45g,Bcl3,Il6 | | 8.320e-03 | -4.79 | negative regulation of signal transduction in absence of ligand | biological process | GO:1901099 | 27 | 3 | 13711 | 214 | Ripk1,Tnf,Sgk3 | | 8.320e-03 | -4.79 | negative regulation of extrinsic apoptotic signaling pathway in absence of ligand | biological process | GO:2001240 | 27 | 3 | 13711 | 214 | Ripk1,Tnf,Sgk3 | | 8.368e-03 | -4.78 | GO\_NEGATIVE\_REGULATION\_OF\_LOCOMOTION | MSigDB lists | GO\_NEGATIVE\_REGULATION\_OF\_LOCOMOTION | 196 | 8 | 12187 | 179 | Ifitm2,Ifitm3,Clic4,Trim25,Trib1,Thbs1,Ccl12,Timp1 | | 8.402e-03 | -4.78 | response to hormone | biological process | GO:0009725 | 480 | 15 | 13711 | 214 | Fam107a,Ptpn1,Timp1,Nfkb1,Serpina3f,Sgk1,Msn,Acod1,Tnf,Zfp36,Il6,Cflar,Il1rn,P2ry6,Pik3r1 | | 8.445e-03 | -4.77 | mouse chr9\|9 A1 | chromosome location | mouse chr9\|9 A1 | 28 | 3 | 14556 | 220 | Phxr4,Mmp8,Birc3 | | 8.455e-03 | -4.77 | GSE24634\_IL4\_VS\_CTRL\_TREATED\_NAIVE\_CD4\_TCELL\_DAY5\_UP | MSigDB lists | GSE24634\_IL4\_VS\_CTRL\_TREATED\_NAIVE\_CD4\_TCELL\_DAY5\_UP | 157 | 7 | 12187 | 179 | Tubb6,Rassf4,Trib1,Ccl2,Igsf6,Olfml2b,Nfkbie | | 8.455e-03 | -4.77 | GSE17721\_CTRL\_VS\_POLYIC\_8H\_BMDC\_DN | MSigDB lists | GSE17721\_CTRL\_VS\_POLYIC\_8H\_BMDC\_DN | 157 | 7 | 12187 | 179 | Ier3,Ms4a6d,Trim25,Timp1,Il1rn,Ccl2,Irgm2 | | 8.455e-03 | -4.77 | GSE15930\_STIM\_VS\_STIM\_AND\_IL12\_24H\_CD8\_T\_CELL\_UP | MSigDB lists | GSE15930\_STIM\_VS\_STIM\_AND\_IL12\_24H\_CD8\_T\_CELL\_UP | 157 | 7 | 12187 | 179 | Dnajb1,Ifit2,Adamts1,Saa1,Luc7l3,Ccdc58,Cybb | | 8.455e-03 | -4.77 | GSE339\_EX\_VIVO\_VS\_IN\_CULTURE\_CD4CD8DN\_DC\_DN | MSigDB lists | GSE339\_EX\_VIVO\_VS\_IN\_CULTURE\_CD4CD8DN\_DC\_DN | 157 | 7 | 12187 | 179 | Cdkn1a,Tgm2,Tubb6,Il1rn,Trim25,Osmr,Nfkb2 | | 8.483e-03 | -4.77 | Matrix Metalloproteinases | WikiPathways | WP441 | 17 | 3 | 3756 | 96 | Mmp8,Timp1,Tnf | | 8.492e-03 | -4.77 | GSE25846\_IL10\_POS\_VS\_NEG\_CD8\_TCELL\_DAY7\_POST\_CORONAVIRUS\_BRAIN\_UP | MSigDB lists | GSE25846\_IL10\_POS\_VS\_NEG\_CD8\_TCELL\_DAY7\_POST\_CORONAVIRUS\_BRAIN\_UP | 120 | 6 | 12187 | 179 | Cxcl1,Rgs16,Ccl4,Nfkbia,Tnfaip3,Atf3 | | 8.492e-03 | -4.77 | GSE26559\_TCF1\_KO\_VS\_WT\_LIN\_NEG\_CELL\_DN | MSigDB lists | GSE26559\_TCF1\_KO\_VS\_WT\_LIN\_NEG\_CELL\_DN | 120 | 6 | 12187 | 179 | Sgk1,Ccl2,Pygm,Selp,S100a9,Irf7 | | 8.544e-03 | -4.76 | positive regulation of chemotaxis | biological process | GO:0050921 | 113 | 6 | 13711 | 214 | C5ar1,Ccl2,C3ar1,Thbs1,Hspb1,Ccl4 | | 8.571e-03 | -4.76 | GO\_REGULATION\_OF\_DNA\_TEMPLATED\_TRANSCRIPTION\_IN\_RESPONSE\_TO\_STRESS | MSigDB lists | GO\_REGULATION\_OF\_DNA\_TEMPLATED\_TRANSCRIPTION\_IN\_RESPONSE\_TO\_STRESS | 55 | 4 | 12187 | 179 | Atf3,Nfe2l2,Bach1,Hspa5 | | 8.571e-03 | -4.76 | GO\_REGULATION\_OF\_INTERFERON\_GAMMA\_PRODUCTION | MSigDB lists | GO\_REGULATION\_OF\_INTERFERON\_GAMMA\_PRODUCTION | 55 | 4 | 12187 | 179 | Isg15,Tnf,Bcl3,Cd14 | | 8.571e-03 | -4.76 | PEDERSEN\_TARGETS\_OF\_611CTF\_ISOFORM\_OF\_ERBB2 | MSigDB lists | PEDERSEN\_TARGETS\_OF\_611CTF\_ISOFORM\_OF\_ERBB2 | 55 | 4 | 12187 | 179 | Angptl4,Ptges,Tnfaip8,Plaur | | 8.598e-03 | -4.76 | response to organonitrogen compound | biological process | GO:0010243 | 621 | 18 | 13711 | 214 | Pik3r1,P2ry6,Pygm,Irak3,Tnfaip3,Icam1,Tnf,Casp4,Nfkbia,Sgk1,Trim25,Vcam1,Serpina3f,Col4a1,Nfkb1,C5ar1,Ptpn1,Hspa5 | | 8.600e-03 | -4.76 | GO\_REGULATION\_OF\_RESPONSE\_TO\_REACTIVE\_OXYGEN\_SPECIES | MSigDB lists | GO\_REGULATION\_OF\_RESPONSE\_TO\_REACTIVE\_OXYGEN\_SPECIES | 29 | 3 | 12187 | 179 | Tnf,Hp,Nfe2l2 | | 8.600e-03 | -4.76 | MURAKAMI\_UV\_RESPONSE\_6HR\_UP | MSigDB lists | MURAKAMI\_UV\_RESPONSE\_6HR\_UP | 29 | 3 | 12187 | 179 | Casp4,Cdkn1a,Timp1 | | 8.600e-03 | -4.76 | GARGALOVIC\_RESPONSE\_TO\_OXIDIZED\_PHOSPHOLIPIDS\_BLACK\_UP | MSigDB lists | GARGALOVIC\_RESPONSE\_TO\_OXIDIZED\_PHOSPHOLIPIDS\_BLACK\_UP | 29 | 3 | 12187 | 179 | Cdkn1a,Errfi1,Nfe2l2 | | 8.600e-03 | -4.76 | VANASSE\_BCL2\_TARGETS\_UP | MSigDB lists | VANASSE\_BCL2\_TARGETS\_UP | 29 | 3 | 12187 | 179 | Il1rn,Socs3,Map3k6 | | 8.600e-03 | -4.76 | GO\_CELLULAR\_RESPONSE\_TO\_ALKALOID | MSigDB lists | GO\_CELLULAR\_RESPONSE\_TO\_ALKALOID | 29 | 3 | 12187 | 179 | Icam1,Nfkb1,Tnf | | 8.600e-03 | -4.76 | KORKOLA\_TERATOMA | MSigDB lists | KORKOLA\_TERATOMA | 29 | 3 | 12187 | 179 | Nfkbiz,Zfp36,Lcn2 | | 8.600e-03 | -4.76 | GO\_CELLULAR\_RESPONSE\_TO\_HEAT | MSigDB lists | GO\_CELLULAR\_RESPONSE\_TO\_HEAT | 29 | 3 | 12187 | 179 | Il1a,Thbs1,Cdkn1a | | 8.638e-03 | -4.75 | BROWNE\_HCMV\_INFECTION\_12HR\_UP | MSigDB lists | BROWNE\_HCMV\_INFECTION\_12HR\_UP | 86 | 5 | 12187 | 179 | Il17ra,Stx11,Tap1,Ifi44,Xaf1 | | 8.638e-03 | -4.75 | HELLER\_SILENCED\_BY\_METHYLATION\_DN | MSigDB lists | HELLER\_SILENCED\_BY\_METHYLATION\_DN | 86 | 5 | 12187 | 179 | Atf3,Rgs16,Maff,Thbs1,Dnajb1 | | 8.678e-03 | -4.75 | regulation of blood vessel endothelial cell migration | biological process | GO:0043535 | 81 | 5 | 13711 | 214 | Hspb1,Thbs1,Nfe2l2,Ptgs2,Rhoj | | 8.700e-03 | -4.74 | Integrin cell surface interactions | REACTOME pathways | R-MMU-216083 | 49 | 4 | 6297 | 105 | Col4a1,Thbs1,Icam1,Vcam1 | | 8.703e-03 | -4.74 | negative regulation of cell cycle G1/S phase transition | biological process | GO:1902807 | 52 | 4 | 13711 | 214 | Ripk1,Fam107a,Cdkn1a,Ccl12 | | 8.703e-03 | -4.74 | negative regulation of cell-substrate adhesion | biological process | GO:0010812 | 52 | 4 | 13711 | 214 | Fzd4,Fam107a,Thbs1,Pik3r1 | | 8.741e-03 | -4.74 | GSE44955\_MCSF\_VS\_MCSF\_AND\_IL27\_STIM\_MACROPHAGE\_UP | MSigDB lists | GSE44955\_MCSF\_VS\_MCSF\_AND\_IL27\_STIM\_MACROPHAGE\_UP | 158 | 7 | 12187 | 179 | Cdkn1a,Tgm2,Timp1,Cybb,Ccl12,Tagln2,Gbp3 | | 8.741e-03 | -4.74 | GSE36392\_EOSINOPHIL\_VS\_NEUTROPHIL\_IL25\_TREATED\_LUNG\_UP | MSigDB lists | GSE36392\_EOSINOPHIL\_VS\_NEUTROPHIL\_IL25\_TREATED\_LUNG\_UP | 158 | 7 | 12187 | 179 | Irf2,Plek,Pik3ap1,Stx11,Sntb2,Parp14,Tnf | | 8.741e-03 | -4.74 | GSE33292\_DN3\_THYMOCYTE\_VS\_TCF1\_KO\_TCELL\_LYMPHOMA\_UP | MSigDB lists | GSE33292\_DN3\_THYMOCYTE\_VS\_TCF1\_KO\_TCELL\_LYMPHOMA\_UP | 158 | 7 | 12187 | 179 | Cybb,Ccl4,Map3k8,Il4ra,Birc3,Hspb1,Plek | | 8.741e-03 | -4.74 | GSE22229\_UNTREATED\_VS\_IMMUNOSUPP\_THERAPY\_RENAL\_TRANSPLANT\_PATIENT\_PBMC\_DN | MSigDB lists | GSE22229\_UNTREATED\_VS\_IMMUNOSUPP\_THERAPY\_RENAL\_TRANSPLANT\_PATIENT\_PBMC\_DN | 158 | 7 | 12187 | 179 | Col4a1,Ms4a6d,Ifi44,Xaf1,Irf9,Rnf213,Slfn9 | | 8.741e-03 | -4.74 | GSE17974\_1H\_VS\_72H\_UNTREATED\_IN\_VITRO\_CD4\_TCELL\_UP | MSigDB lists | GSE17974\_1H\_VS\_72H\_UNTREATED\_IN\_VITRO\_CD4\_TCELL\_UP | 158 | 7 | 12187 | 179 | Gem,Cxcl16,Ip6k2,Zfp36,Trib1,Znfx1,Bach1 | | 8.741e-03 | -4.74 | GSE17721\_PAM3CSK4\_VS\_CPG\_6H\_BMDC\_DN | MSigDB lists | GSE17721\_PAM3CSK4\_VS\_CPG\_6H\_BMDC\_DN | 158 | 7 | 12187 | 179 | Nfkbiz,Sgk1,Irf2,Sele,Pik3ap1,Ifitm2,Sgk3 | | 8.741e-03 | -4.74 | GSE24634\_TREG\_VS\_TCONV\_POST\_DAY7\_IL4\_CONVERSION\_DN | MSigDB lists | GSE24634\_TREG\_VS\_TCONV\_POST\_DAY7\_IL4\_CONVERSION\_DN | 158 | 7 | 12187 | 179 | Cybb,Rbm47,Ccl12,Plaur,Cdkn1a,Maff,Cd14 | | 8.741e-03 | -4.74 | GSE24574\_BCL6\_LOW\_TFH\_VS\_NAIVE\_CD4\_TCELL\_UP | MSigDB lists | GSE24574\_BCL6\_LOW\_TFH\_VS\_NAIVE\_CD4\_TCELL\_UP | 158 | 7 | 12187 | 179 | Isg15,Irf7,Rrp8,Cxcl1,Sele,Ksr1,Tap1 | | 8.741e-03 | -4.74 | GSE22025\_PROGESTERONE\_VS\_TGFB1\_AND\_PROGESTERONE\_TREATED\_CD4\_TCELL\_UP | MSigDB lists | GSE22025\_PROGESTERONE\_VS\_TGFB1\_AND\_PROGESTERONE\_TREATED\_CD4\_TCELL\_UP | 158 | 7 | 12187 | 179 | Gem,Nfkbie,Rgs16,Aff1,Map3k8,Ifit3b,Atf3 | | 8.741e-03 | -4.74 | GSE15930\_NAIVE\_VS\_48H\_IN\_VITRO\_STIM\_CD8\_TCELL\_UP | MSigDB lists | GSE15930\_NAIVE\_VS\_48H\_IN\_VITRO\_STIM\_CD8\_TCELL\_UP | 158 | 7 | 12187 | 179 | Irf9,Cd14,Irf7,Bcl3,Tnfaip3,Usp18,Socs3 | | 8.741e-03 | -4.74 | GSE3982\_EOSINOPHIL\_VS\_EFF\_MEMORY\_CD4\_TCELL\_UP | MSigDB lists | GSE3982\_EOSINOPHIL\_VS\_EFF\_MEMORY\_CD4\_TCELL\_UP | 158 | 7 | 12187 | 179 | Cybb,Trib1,Thbs1,Bcl3,Cxcl1,C3ar1,Isg15 | | 8.790e-03 | -4.73 | MCCABE\_BOUND\_BY\_HOXC6 | MSigDB lists | MCCABE\_BOUND\_BY\_HOXC6 | 282 | 10 | 12187 | 179 | Vcam1,Runx1,Znfx1,AA467197,Gadd45g,Nfkb2,Csf2rb,Errfi1,Sbno2,Tnfaip8 | | 8.810e-03 | -4.73 | response to wounding | biological process | GO:0009611 | 265 | 10 | 13711 | 214 | Cdkn1a,Ccl2,Sdc4,Il6,Zfp36,Plek,Il1a,Tnfaip3,Timp1,Cflar | | 8.844e-03 | -4.73 | cellular response to extracellular stimulus | biological process | GO:0031668 | 186 | 8 | 13711 | 214 | Angptl4,Vcam1,Nfe2l2,Atf3,Cdkn1a,Rrp8,Fam107a,Hspa5 | | 8.891e-03 | -4.72 | TNF\_2 | prosite domains | PS50049 | 8 | 2 | 8845 | 164 | Tnfsf10,Tnf | | 8.934e-03 | -4.72 | REACTOME\_ACTIVATION\_OF\_IRF3\_IRF7\_MEDIATED\_BY\_TBK1\_IKK\_EPSILON | MSigDB lists | REACTOME\_ACTIVATION\_OF\_IRF3\_IRF7\_MEDIATED\_BY\_TBK1\_IKK\_EPSILON | 10 | 2 | 12187 | 179 | Irf7,Cd14 | | 8.934e-03 | -4.72 | GO\_POSITIVE\_REGULATION\_OF\_TRANSFORMING\_GROWTH\_FACTOR\_BETA\_PRODUCTION | MSigDB lists | GO\_POSITIVE\_REGULATION\_OF\_TRANSFORMING\_GROWTH\_FACTOR\_BETA\_PRODUCTION | 10 | 2 | 12187 | 179 | Thbs1,Ptgs2 | | 8.934e-03 | -4.72 | MODULE\_431 | MSigDB lists | MODULE\_431 | 10 | 2 | 12187 | 179 | Ptgs2,Tgm2 | | 8.934e-03 | -4.72 | GO\_CELLULAR\_RESPONSE\_TO\_ATP | MSigDB lists | GO\_CELLULAR\_RESPONSE\_TO\_ATP | 10 | 2 | 12187 | 179 | Ptgs2,Ccl12 | | 8.934e-03 | -4.72 | GO\_REGULATION\_OF\_FEVER\_GENERATION | MSigDB lists | GO\_REGULATION\_OF\_FEVER\_GENERATION | 10 | 2 | 12187 | 179 | Tnf,Ptgs2 | | 8.934e-03 | -4.72 | GO\_CALCIUM\_MEDIATED\_SIGNALING\_USING\_INTRACELLULAR\_CALCIUM\_SOURCE | MSigDB lists | GO\_CALCIUM\_MEDIATED\_SIGNALING\_USING\_INTRACELLULAR\_CALCIUM\_SOURCE | 10 | 2 | 12187 | 179 | Selp,Vcam1 | | 8.934e-03 | -4.72 | GO\_REGULATION\_OF\_T\_HELPER\_2\_CELL\_DIFFERENTIATION | MSigDB lists | GO\_REGULATION\_OF\_T\_HELPER\_2\_CELL\_DIFFERENTIATION | 10 | 2 | 12187 | 179 | Il6,Il4ra | | 8.934e-03 | -4.72 | BIOCARTA\_RNA\_PATHWAY | MSigDB lists | BIOCARTA\_RNA\_PATHWAY | 10 | 2 | 12187 | 179 | Nfkb1,Nfkbia | | 8.934e-03 | -4.72 | GO\_LONG\_CHAIN\_FATTY\_ACID\_BINDING | MSigDB lists | GO\_LONG\_CHAIN\_FATTY\_ACID\_BINDING | 10 | 2 | 12187 | 179 | S100a8,S100a9 | | 8.934e-03 | -4.72 | GO\_ACTIVATION\_OF\_CYSTEINE\_TYPE\_ENDOPEPTIDASE\_ACTIVITY\_INVOLVED\_IN\_APOPTOTIC\_SIGNALING\_PATHWAY | MSigDB lists | GO\_ACTIVATION\_OF\_CYSTEINE\_TYPE\_ENDOPEPTIDASE\_ACTIVITY\_INVOLVED\_IN\_APOPTOTIC\_SIGNALING\_PATHWAY | 10 | 2 | 12187 | 179 | Ripk1,Tnfsf10 | | 8.934e-03 | -4.72 | GO\_MACROPHAGE\_CHEMOTAXIS | MSigDB lists | GO\_MACROPHAGE\_CHEMOTAXIS | 10 | 2 | 12187 | 179 | Saa1,Ccl12 | | 8.934e-03 | -4.72 | GO\_NEGATIVE\_REGULATION\_OF\_MULTICELLULAR\_ORGANISMAL\_METABOLIC\_PROCESS | MSigDB lists | GO\_NEGATIVE\_REGULATION\_OF\_MULTICELLULAR\_ORGANISMAL\_METABOLIC\_PROCESS | 10 | 2 | 12187 | 179 | Il6,Errfi1 | | 8.934e-03 | -4.72 | HASINA\_NOL7\_TARGETS\_UP | MSigDB lists | HASINA\_NOL7\_TARGETS\_UP | 10 | 2 | 12187 | 179 | Timp1,Plaur | | 8.934e-03 | -4.72 | GO\_CYSTEINE\_TYPE\_ENDOPEPTIDASE\_ACTIVITY\_INVOLVED\_IN\_APOPTOTIC\_PROCESS | MSigDB lists | GO\_CYSTEINE\_TYPE\_ENDOPEPTIDASE\_ACTIVITY\_INVOLVED\_IN\_APOPTOTIC\_PROCESS | 10 | 2 | 12187 | 179 | Casp4,Cflar | | 8.934e-03 | -4.72 | REACTOME\_IL\_6\_SIGNALING | MSigDB lists | REACTOME\_IL\_6\_SIGNALING | 10 | 2 | 12187 | 179 | Il6,Socs3 | | 8.934e-03 | -4.72 | DORN\_ADENOVIRUS\_INFECTION\_48HR\_UP | MSigDB lists | DORN\_ADENOVIRUS\_INFECTION\_48HR\_UP | 10 | 2 | 12187 | 179 | Ifit2,Nfkb2 | | 8.934e-03 | -4.72 | GO\_RESPIRATORY\_BURST | MSigDB lists | GO\_RESPIRATORY\_BURST | 10 | 2 | 12187 | 179 | Cybb,Ncf1 | | 8.934e-03 | -4.72 | WU\_HBX\_TARGETS\_1\_UP | MSigDB lists | WU\_HBX\_TARGETS\_1\_UP | 10 | 2 | 12187 | 179 | Casp4,Il6 | | 9.007e-03 | -4.71 | positive regulation of nucleic acid-templated transcription | biological process | GO:1903508 | 1279 | 31 | 13711 | 214 | Tnf,Atf3,Zbtb16,Csf3,Ripk1,Arid5b,Ncf1,Aff1,Tnip1,Maff,Irf2,Nfe2l2,Bcl3,Nfkbia,Nfkb2,Sox11,Il1a,Il6,Dtx3l,Irf7,Rbm39,Pik3r1,Banp,Fzd4,Parp9,Sbno2,Nfkb1,Spi1,Bach1,Cebpd,Runx1 | | 9.033e-03 | -4.71 | GSE21670\_UNTREATED\_VS\_TGFB\_TREATED\_CD4\_TCELL\_DN | MSigDB lists | GSE21670\_UNTREATED\_VS\_TGFB\_TREATED\_CD4\_TCELL\_DN | 159 | 7 | 12187 | 179 | Casp4,Parp14,Fstl1,Ier3,Znfx1,Bach1,Col4a1 | | 9.033e-03 | -4.71 | GSE43955\_TH0\_VS\_TGFB\_IL6\_TH17\_ACT\_CD4\_TCELL\_10H\_UP | MSigDB lists | GSE43955\_TH0\_VS\_TGFB\_IL6\_TH17\_ACT\_CD4\_TCELL\_10H\_UP | 159 | 7 | 12187 | 179 | Csf3,S100a8,Map3k8,Casp4,Cp,Il1rn,Lcn2 | | 9.033e-03 | -4.71 | GSE30971\_WBP7\_HET\_VS\_KO\_MACROPHAGE\_UP | MSigDB lists | GSE30971\_WBP7\_HET\_VS\_KO\_MACROPHAGE\_UP | 159 | 7 | 12187 | 179 | Hcar2,Hp,Rhoc,Gadd45b,Tnfaip3,Aff1,Cd14 | | 9.033e-03 | -4.71 | GSE29164\_UNTREATED\_VS\_CD8\_TCELL\_AND\_IL12\_TREATED\_MELANOMA\_DAY7\_DN | MSigDB lists | GSE29164\_UNTREATED\_VS\_CD8\_TCELL\_AND\_IL12\_TREATED\_MELANOMA\_DAY7\_DN | 159 | 7 | 12187 | 179 | Adamts1,Cdkn1a,Trib1,Rhoc,Cxcl1,Hspb1,Icam1 | | 9.033e-03 | -4.71 | GSE23568\_ID3\_TRANSDUCED\_VS\_ID3\_KO\_CD8\_TCELL\_UP | MSigDB lists | GSE23568\_ID3\_TRANSDUCED\_VS\_ID3\_KO\_CD8\_TCELL\_UP | 159 | 7 | 12187 | 179 | Sgk1,Socs3,Ms4a6d,Isg15,Tnip1,Irf7,Xdh | | 9.033e-03 | -4.71 | GSE5542\_UNTREATED\_VS\_IFNA\_AND\_IFNG\_TREATED\_EPITHELIAL\_CELLS\_6H\_DN | MSigDB lists | GSE5542\_UNTREATED\_VS\_IFNA\_AND\_IFNG\_TREATED\_EPITHELIAL\_CELLS\_6H\_DN | 159 | 7 | 12187 | 179 | Rhoc,Mxd4,Thbs1,Pik3ap1,Ptpn1,C3ar1,Cflar | | 9.033e-03 | -4.71 | GSE45365\_HEALTHY\_VS\_MCMV\_INFECTION\_CD8A\_DC\_DN | MSigDB lists | GSE45365\_HEALTHY\_VS\_MCMV\_INFECTION\_CD8A\_DC\_DN | 159 | 7 | 12187 | 179 | Ier3,Map3k8,Zfp36,Tnfsf10,Cflar,S100a9,Irf7 | | 9.033e-03 | -4.71 | GSE16266\_LPS\_VS\_HEATSHOCK\_AND\_LPS\_STIM\_MEF\_DN | MSigDB lists | GSE16266\_LPS\_VS\_HEATSHOCK\_AND\_LPS\_STIM\_MEF\_DN | 159 | 7 | 12187 | 179 | Arid5b,Bcl3,Cxcl1,Rbm47,Plek,Birc3,Aff1 | | 9.033e-03 | -4.71 | GSE9037\_CTRL\_VS\_LPS\_1H\_STIM\_IRAK4\_KO\_BMDM\_DN | MSigDB lists | GSE9037\_CTRL\_VS\_LPS\_1H\_STIM\_IRAK4\_KO\_BMDM\_DN | 159 | 7 | 12187 | 179 | Ccl4,Vcam1,Map3k8,Parp12,Sgk3,Gbp7,Irf9 | | 9.059e-03 | -4.70 | GO\_POSITIVE\_REGULATION\_OF\_CELLULAR\_AMIDE\_METABOLIC\_PROCESS | MSigDB lists | GO\_POSITIVE\_REGULATION\_OF\_CELLULAR\_AMIDE\_METABOLIC\_PROCESS | 87 | 5 | 12187 | 179 | Bcl3,Tnf,Thbs1,Nfe2l2,Il6 | | 9.059e-03 | -4.70 | LENAOUR\_DENDRITIC\_CELL\_MATURATION\_UP | MSigDB lists | LENAOUR\_DENDRITIC\_CELL\_MATURATION\_UP | 87 | 5 | 12187 | 179 | Birc3,Ccnd2,A2m,Ccl2,Tgm2 | | 9.066e-03 | -4.70 | DYNAMIN | prints domains | PR00195 | 7 | 2 | 2951 | 64 | Mx1,Mx2 | | 9.066e-03 | -4.70 | SMALLCYTKCXC | prints domains | PR00437 | 7 | 2 | 2951 | 64 | Cxcl1,Cxcl9 | | 9.073e-03 | -4.70 | GO\_CELL\_CELL\_ADHESION | MSigDB lists | GO\_CELL\_CELL\_ADHESION | 420 | 13 | 12187 | 179 | Tnip1,Selp,Msn,Icam1,Hspb1,Sele,Plek,Vcam1,Bcl3,S100a9,S100a8,Tnf,Il6 | | 9.081e-03 | -4.70 | Phagosome | KEGG pathways | ko04145 | 116 | 7 | 5248 | 107 | Ncf1,Thbs1,Tap1,Cd14,Tubb6,Cybb,Msr1 | | 9.081e-03 | -4.70 | Phagosome | KEGG pathways | mmu04145 | 116 | 7 | 5248 | 107 | Thbs1,Ncf1,Tap1,Cd14,Msr1,Cybb,Tubb6 | | 9.104e-03 | -4.70 | positive regulation of RNA biosynthetic process | biological process | GO:1902680 | 1280 | 31 | 13711 | 214 | Maff,Irf2,Nfe2l2,Bcl3,Tnf,Atf3,Csf3,Zbtb16,Ripk1,Arid5b,Ncf1,Aff1,Tnip1,Fzd4,Parp9,Sbno2,Nfkb1,Spi1,Bach1,Cebpd,Runx1,Nfkb2,Nfkbia,Sox11,Il1a,Il6,Dtx3l,Irf7,Pik3r1,Banp,Rbm39 | | 9.127e-03 | -4.70 | negative regulation of intrinsic apoptotic signaling pathway | biological process | GO:2001243 | 82 | 5 | 13711 | 214 | Ptgs2,Plaur,Hspb1,Nfe2l2,Ptpn1 | | 9.127e-03 | -4.70 | GRABARCZYK\_BCL11B\_TARGETS\_UP | MSigDB lists | GRABARCZYK\_BCL11B\_TARGETS\_UP | 56 | 4 | 12187 | 179 | Timp1,Tnfsf10,Pik3r1,Zbtb16 | | 9.127e-03 | -4.70 | GO\_NEGATIVE\_REGULATION\_OF\_LEUKOCYTE\_DIFFERENTIATION | MSigDB lists | GO\_NEGATIVE\_REGULATION\_OF\_LEUKOCYTE\_DIFFERENTIATION | 56 | 4 | 12187 | 179 | Pik3r1,Il4ra,Runx1,Trib1 | | 9.127e-03 | -4.70 | GO\_CYTOKINE\_RECEPTOR\_ACTIVITY | MSigDB lists | GO\_CYTOKINE\_RECEPTOR\_ACTIVITY | 56 | 4 | 12187 | 179 | Il17ra,Osmr,Csf2rb,Il4ra | | 9.127e-03 | -4.70 | YAO\_TEMPORAL\_RESPONSE\_TO\_PROGESTERONE\_CLUSTER\_1 | MSigDB lists | YAO\_TEMPORAL\_RESPONSE\_TO\_PROGESTERONE\_CLUSTER\_1 | 56 | 4 | 12187 | 179 | Gadd45b,Gadd45g,Hp,Map3k8 | | 9.129e-03 | -4.70 | ZHANG\_TLX\_TARGETS\_36HR\_UP | MSigDB lists | ZHANG\_TLX\_TARGETS\_36HR\_UP | 199 | 8 | 12187 | 179 | Ccnd2,Cdkn1a,Gadd45g,Gadd45b,Mxd4,Pygm,Tap1,Nfkb2 | | 9.142e-03 | -4.69 | - | gene3d domains | 3.30.720.50 | 10 | 2 | 6647 | 99 | Parp12,Parp14 | | 9.142e-03 | -4.69 | - | gene3d domains | 2.60.40.340 | 10 | 2 | 6647 | 99 | Nfkb2,Nfkb1 | | 9.186e-03 | -4.69 | Scavenging of heme from plasma | REACTOME pathways | R-MMU-2168880 | 9 | 2 | 6297 | 105 | Apol10b,Hp | | 9.213e-03 | -4.69 | WANG\_SMARCE1\_TARGETS\_DN | MSigDB lists | WANG\_SMARCE1\_TARGETS\_DN | 284 | 10 | 12187 | 179 | Cxcl1,Tap1,S100a8,Trib1,Tgm2,Sgk1,Il4ra,Adamts1,Parp12,AA467197 | | 9.213e-03 | -4.69 | regulation of superoxide metabolic process | biological process | GO:0090322 | 28 | 3 | 13711 | 214 | Cxcl1,Nfe2l2,Tnf | | 9.213e-03 | -4.69 | positive regulation of neutrophil migration | biological process | GO:1902624 | 28 | 3 | 13711 | 214 | C5ar1,C3ar1,Il1a | | 9.213e-03 | -4.69 | cellular response to glucocorticoid stimulus | biological process | GO:0071385 | 28 | 3 | 13711 | 214 | Sgk1,Zfp36,Fam107a | | 9.228e-03 | -4.69 | HIF-1 signaling pathway | KEGG pathways | mmu04066 | 89 | 6 | 5248 | 107 | Pik3r1,Cybb,Cdkn1a,Nfkb1,Timp1,Il6 | | 9.276e-03 | -4.68 | regulation of osteoblast differentiation | biological process | GO:0045667 | 115 | 6 | 13711 | 214 | Ifi204,Noct,Il6,Sox11,Cebpd,Tnf | | 9.301e-03 | -4.68 | regulation of DNA-templated transcription in response to stress | biological process | GO:0043620 | 53 | 4 | 13711 | 214 | Atf3,Nfe2l2,Dnajb1,Bach1 | | 9.314e-03 | -4.68 | Proteoglycans in cancer | KEGG pathways | ko05205 | 176 | 9 | 5248 | 107 | Cdkn1a,Fzd4,Thbs1,Msn,Tnf,Iqgap1,Sdc4,Plaur,Pik3r1 | | 9.314e-03 | -4.68 | Proteoglycans in cancer | KEGG pathways | mmu05205 | 176 | 9 | 5248 | 107 | Fzd4,Cdkn1a,Iqgap1,Tnf,Pik3r1,Sdc4,Plaur,Msn,Thbs1 | | 9.333e-03 | -4.67 | GSE17721\_PAM3CSK4\_VS\_CPG\_1H\_BMDC\_DN | MSigDB lists | GSE17721\_PAM3CSK4\_VS\_CPG\_1H\_BMDC\_DN | 160 | 7 | 12187 | 179 | Rbm47,Ptgs2,Selp,Cebpd,Il6,Nfkbiz,Map3k8 | | 9.333e-03 | -4.67 | GSE20727\_ROS\_INH\_VS\_ROS\_INH\_AND\_DNFB\_ALLERGEN\_TREATED\_DC\_UP | MSigDB lists | GSE20727\_ROS\_INH\_VS\_ROS\_INH\_AND\_DNFB\_ALLERGEN\_TREATED\_DC\_UP | 160 | 7 | 12187 | 179 | Tnip3,Rhoc,Trib1,Socs3,Rbm47,Pik3ap1,Olfml2b | | 9.333e-03 | -4.67 | GSE5503\_MLN\_DC\_VS\_PLN\_DC\_ACTIVATED\_ALLOGENIC\_TCELL\_DN | MSigDB lists | GSE5503\_MLN\_DC\_VS\_PLN\_DC\_ACTIVATED\_ALLOGENIC\_TCELL\_DN | 160 | 7 | 12187 | 179 | Pik3ap1,Trim25,Gbp4,Gbp6,Sdc4,Parp14,Slfn9 | | 9.333e-03 | -4.67 | GSE13522\_CTRL\_VS\_T\_CRUZI\_Y\_STRAIN\_INF\_SKIN\_IFNAR\_KO\_DN | MSigDB lists | GSE13522\_CTRL\_VS\_T\_CRUZI\_Y\_STRAIN\_INF\_SKIN\_IFNAR\_KO\_DN | 160 | 7 | 12187 | 179 | Tgm2,Nfkbia,Stx11,Gadd45g,Tnip1,Icam1,Xdh | | 9.333e-03 | -4.67 | GSE5589\_LPS\_AND\_IL10\_VS\_LPS\_AND\_IL6\_STIM\_IL10\_KO\_MACROPHAGE\_45MIN\_UP | MSigDB lists | GSE5589\_LPS\_AND\_IL10\_VS\_LPS\_AND\_IL6\_STIM\_IL10\_KO\_MACROPHAGE\_45MIN\_UP | 160 | 7 | 12187 | 179 | Sbno2,Rbm39,S100a8,Angptl4,Arid5b,S100a9,Sdc4 | | 9.333e-03 | -4.67 | GSE17721\_POLYIC\_VS\_CPG\_4H\_BMDC\_DN | MSigDB lists | GSE17721\_POLYIC\_VS\_CPG\_4H\_BMDC\_DN | 160 | 7 | 12187 | 179 | Lcn2,Tnfaip3,Sdc4,Gpr84,Trib1,Il17ra,Ptges | | 9.333e-03 | -4.67 | CEBP\_C | MSigDB lists | CEBP\_C | 160 | 7 | 12187 | 179 | Fstl1,Nfkbia,Pik3r1,Slc24a4,Robo4,Rbm39,Osmr | | 9.355e-03 | -4.67 | cellular ion homeostasis | biological process | GO:0006873 | 486 | 15 | 13711 | 214 | Clic4,C5ar1,Cxcl1,Tgm2,Mt2,Kcna5,Sgk1,Mt1,Ccl2,C3ar1,Slc24a4,Pygm,Cp,P2ry6,Lcn2 | | 9.376e-03 | -4.67 | REN\_ALVEOLAR\_RHABDOMYOSARCOMA\_DN | MSigDB lists | REN\_ALVEOLAR\_RHABDOMYOSARCOMA\_DN | 375 | 12 | 12187 | 179 | Casp4,Rhoc,Ptges,Osmr,Iqgap1,Fstl1,Thbs1,Timp1,Tagln2,Msn,Plaur,Clic4 | | 9.393e-03 | -4.67 | GO\_NEGATIVE\_REGULATION\_OF\_RESPONSE\_TO\_EXTERNAL\_STIMULUS | MSigDB lists | GO\_NEGATIVE\_REGULATION\_OF\_RESPONSE\_TO\_EXTERNAL\_STIMULUS | 200 | 8 | 12187 | 179 | Plaur,Saa1,Nfkb1,Tnfaip3,Ccl12,Thbs1,Socs3,Trib1 | | 9.424e-03 | -4.66 | B cell receptor signaling pathway | KEGG pathways | ko04662 | 64 | 5 | 5248 | 107 | Nfkbie,Pik3r1,Nfkb1,Pik3ap1,Nfkbia | | 9.424e-03 | -4.66 | B cell receptor signaling pathway | KEGG pathways | mmu04662 | 64 | 5 | 5248 | 107 | Pik3r1,Nfkbie,Pik3ap1,Nfkb1,Nfkbia | | 9.440e-03 | -4.66 | PCNA (proliferating cell nuclear antigen) | protein interactions | 5111 | 9 | 2 | 6802 | 115 | Gadd45b,Cdkn1a | | 9.440e-03 | -4.66 | Rnf11 (ring finger protein 11) | protein interactions | 29864 | 9 | 2 | 6802 | 115 | Ripk1,Tnfaip3 | | 9.443e-03 | -4.66 | Toll Like Receptor 9 (TLR9) Cascade | REACTOME pathways | R-MMU-168138 | 78 | 5 | 6297 | 105 | Nfkb2,Nfkb1,Irf7,Nfkbia,Map3k8 | | 9.454e-03 | -4.66 | SHIN\_B\_CELL\_LYMPHOMA\_CLUSTER\_8 | MSigDB lists | SHIN\_B\_CELL\_LYMPHOMA\_CLUSTER\_8 | 30 | 3 | 12187 | 179 | Icam1,Spi1,Nfkbia | | 9.454e-03 | -4.66 | ZHAN\_EARLY\_DIFFERENTIATION\_GENES\_DN | MSigDB lists | ZHAN\_EARLY\_DIFFERENTIATION\_GENES\_DN | 30 | 3 | 12187 | 179 | Tnf,Il4ra,Spi1 | | 9.454e-03 | -4.66 | AMIT\_EGF\_RESPONSE\_120\_MCF10A | MSigDB lists | AMIT\_EGF\_RESPONSE\_120\_MCF10A | 30 | 3 | 12187 | 179 | Plaur,Il1a,Maff | | 9.454e-03 | -4.66 | GO\_POSITIVE\_REGULATION\_OF\_ALPHA\_BETA\_T\_CELL\_DIFFERENTIATION | MSigDB lists | GO\_POSITIVE\_REGULATION\_OF\_ALPHA\_BETA\_T\_CELL\_DIFFERENTIATION | 30 | 3 | 12187 | 179 | Zbtb16,Il4ra,Il6 | | 9.493e-03 | -4.66 | HALLMARK\_ANDROGEN\_RESPONSE | MSigDB lists | HALLMARK\_ANDROGEN\_RESPONSE | 88 | 5 | 12187 | 179 | Sgk1,Adamts1,Arid5b,Akap12,Tnfaip8 | | 9.493e-03 | -4.66 | GSE43863\_NAIVE\_VS\_TH1\_EFF\_CD4\_TCELL\_D6\_LCMV\_DN | MSigDB lists | GSE43863\_NAIVE\_VS\_TH1\_EFF\_CD4\_TCELL\_D6\_LCMV\_DN | 88 | 5 | 12187 | 179 | Trib1,Rnf213,Tnf,Rtp4,Sele | | 9.507e-03 | -4.66 | response to drug | biological process | GO:0042493 | 580 | 17 | 13711 | 214 | Fzd4,Nfkb1,Hp,Nfkbie,Nfe2l2,Ccl4,Tnfaip3,Tnf,Acod1,Nfkbia,Il6,Cybb,Thbs1,Ccl7,Cflar,Ripk1,Lcn2 | | 9.532e-03 | -4.65 | GO\_PLATELET\_ACTIVATION | MSigDB lists | GO\_PLATELET\_ACTIVATION | 123 | 6 | 12187 | 179 | Il6,Plek,Hspb1,Pik3r5,Saa1,Pik3r1 | | 9.532e-03 | -4.65 | GSE22886\_NAIVE\_CD4\_TCELL\_VS\_NEUTROPHIL\_DN | MSigDB lists | GSE22886\_NAIVE\_CD4\_TCELL\_VS\_NEUTROPHIL\_DN | 123 | 6 | 12187 | 179 | Cebpd,Tnfsf10,Ier3,Csf2rb,S100a9,Ptgs2 | | 9.639e-03 | -4.64 | GSE13306\_RA\_VS\_UNTREATED\_MEM\_CD4\_TCELL\_UP | MSigDB lists | GSE13306\_RA\_VS\_UNTREATED\_MEM\_CD4\_TCELL\_UP | 161 | 7 | 12187 | 179 | Samd9l,Ms4a6d,Vcam1,Tgm2,Tnfsf10,Xdh,Ncf1 | | 9.658e-03 | -4.64 | negative regulation of lymphocyte activation | biological process | GO:0051250 | 116 | 6 | 13711 | 214 | Sdc4,Sox11,Il4ra,Tnfaip3,Lgals9,Runx1 | | 9.658e-03 | -4.64 | positive regulation of cysteine-type endopeptidase activity | biological process | GO:2001056 | 116 | 6 | 13711 | 214 | Xdh,S100a8,Tnfsf10,Tnf,Casp4,S100a9 | | 9.666e-03 | -4.64 | CARD | smart domains | SM00114 | 9 | 2 | 7188 | 123 | Casp4,Birc3 | | 9.688e-03 | -4.64 | cellular response to oxidative stress | biological process | GO:0034599 | 189 | 8 | 13711 | 214 | Nfe2l2,Ncf1,Thbs1,Ripk1,Cflar,Lcn2,Tnfaip3,Il6 | | 9.705e-03 | -4.64 | HOSHIDA\_LIVER\_CANCER\_SURVIVAL\_UP | MSigDB lists | HOSHIDA\_LIVER\_CANCER\_SURVIVAL\_UP | 57 | 4 | 12187 | 179 | Col4a1,Iqgap1,Ier3,Nfkb2 | | 9.705e-03 | -4.64 | FONTAINE\_FOLLICULAR\_THYROID\_ADENOMA\_DN | MSigDB lists | FONTAINE\_FOLLICULAR\_THYROID\_ADENOMA\_DN | 57 | 4 | 12187 | 179 | Casp4,Spi1,Ptges,Fam107a | | 9.705e-03 | -4.64 | GO\_SARCOPLASM | MSigDB lists | GO\_SARCOPLASM | 57 | 4 | 12187 | 179 | Pygm,Xdh,Rasd1,Thbs1 | | 9.705e-03 | -4.64 | YAO\_TEMPORAL\_RESPONSE\_TO\_PROGESTERONE\_CLUSTER\_2 | MSigDB lists | YAO\_TEMPORAL\_RESPONSE\_TO\_PROGESTERONE\_CLUSTER\_2 | 57 | 4 | 12187 | 179 | Tiparp,Rbm39,Errfi1,Rhou | | 9.705e-03 | -4.64 | NATSUME\_RESPONSE\_TO\_INTERFERON\_BETA\_UP | MSigDB lists | NATSUME\_RESPONSE\_TO\_INTERFERON\_BETA\_UP | 57 | 4 | 12187 | 179 | Ifitm3,Bcl3,Vcam1,Nfkb1 | | 9.728e-03 | -4.63 | Leukocyte transendothelial migration | KEGG pathways | mmu04670 | 90 | 6 | 5248 | 107 | Msn,Ncf1,Pik3r1,Vcam1,Icam1,Cybb | | 9.728e-03 | -4.63 | Leukocyte transendothelial migration | KEGG pathways | ko04670 | 90 | 6 | 5248 | 107 | Icam1,Pik3r1,Vcam1,Ncf1,Msn,Cybb | | 9.762e-03 | -4.63 | Pept\_C14A | interpro domains | IPR015917 | 10 | 2 | 13788 | 212 | Casp4,Cflar | | 9.762e-03 | -4.63 | RHD\_DNA\_bind\_dom\_sf | interpro domains | IPR037059 | 10 | 2 | 13788 | 212 | Nfkb1,Nfkb2 | | 9.762e-03 | -4.63 | RHD\_dimer | interpro domains | IPR032397 | 10 | 2 | 13788 | 212 | Nfkb2,Nfkb1 | | 9.762e-03 | -4.63 | Hemopexin\_CS | interpro domains | IPR018486 | 10 | 2 | 13788 | 212 | Mmp8,Prg4 | | 9.762e-03 | -4.63 | RHD\_DNA\_bind\_dom | interpro domains | IPR011539 | 10 | 2 | 13788 | 212 | Nfkb1,Nfkb2 | | 9.804e-03 | -4.62 | GO\_CELL\_SURFACE | MSigDB lists | GO\_CELL\_SURFACE | 472 | 14 | 12187 | 179 | Vcam1,Thbs1,Hspa5,Clic4,Icam1,Cd14,Selp,Il6,Rhoc,Sdc4,Tnf,Adamts9,Fzd4,C5ar1 | | 9.852e-03 | -4.62 | positive regulation of peptidase activity | biological process | GO:0010952 | 152 | 7 | 13711 | 214 | Cflar,S100a9,Xdh,S100a8,Tnfsf10,Tnf,Casp4 | | 9.898e-03 | -4.62 | GSE21927\_BALBC\_VS\_C57BL6\_MONOCYTE\_SPLEEN\_DN | MSigDB lists | GSE21927\_BALBC\_VS\_C57BL6\_MONOCYTE\_SPLEEN\_DN | 124 | 6 | 12187 | 179 | Ifi211,Bbs12,Ifi207,Ifi204,Il6,Ifi209 | | 9.898e-03 | -4.62 | GSE10094\_LCMV\_VS\_LISTERIA\_IND\_EFF\_CD4\_TCELL\_DN | MSigDB lists | GSE10094\_LCMV\_VS\_LISTERIA\_IND\_EFF\_CD4\_TCELL\_DN | 124 | 6 | 12187 | 179 | Zfp36,Rbm39,S100a8,Atf3,Msr1,Ccl4 | | 9.900e-03 | -4.62 | negative regulation of cell motility | biological process | GO:2000146 | 229 | 9 | 13711 | 214 | Adamts9,Trib1,Il1rn,Nfe2l2,Thbs1,Ccl12,Clic4,Timp1,Robo4 | | 9.953e-03 | -4.61 | GSE17721\_LPS\_VS\_GARDIQUIMOD\_16H\_BMDC\_UP | MSigDB lists | GSE17721\_LPS\_VS\_GARDIQUIMOD\_16H\_BMDC\_UP | 162 | 7 | 12187 | 179 | Slfn9,Tagln2,Nfkbie,Isg15,Ccl12,Tor3a,Pik3r1 | | 9.953e-03 | -4.61 | GSE44732\_UNSTIM\_VS\_IL27\_STIM\_IMATURE\_DC\_UP | MSigDB lists | GSE44732\_UNSTIM\_VS\_IL27\_STIM\_IMATURE\_DC\_UP | 162 | 7 | 12187 | 179 | Rassf4,Tnf,Cdkn1a,Tnfaip8,Cmpk2,Csf2rb,Cflar | | 9.953e-03 | -4.61 | GSE11961\_MEMORY\_BCELL\_DAY7\_VS\_GERMINAL\_CENTER\_BCELL\_DAY40\_UP | MSigDB lists | GSE11961\_MEMORY\_BCELL\_DAY7\_VS\_GERMINAL\_CENTER\_BCELL\_DAY40\_UP | 162 | 7 | 12187 | 179 | Xaf1,Gbp4,Gbp7,Gbp6,Gem,Cflar,Tnfsf10 | | 9.953e-03 | -4.61 | GSE8921\_3H\_VS\_24H\_TLR1\_2\_STIM\_MONOCYTE\_DN | MSigDB lists | GSE8921\_3H\_VS\_24H\_TLR1\_2\_STIM\_MONOCYTE\_DN | 162 | 7 | 12187 | 179 | Saa1,Csf2rb,Maff,Nfkbia,Hcar2,Tnfaip3,Il6 | | 9.953e-03 | -4.61 | GSE27670\_CTRL\_VS\_BLIMP1\_TRANSDUCED\_GC\_BCELL\_DN | MSigDB lists | GSE27670\_CTRL\_VS\_BLIMP1\_TRANSDUCED\_GC\_BCELL\_DN | 162 | 7 | 12187 | 179 | Fzd4,Cxcl1,Gem,Irgm2,Errfi1,Casp4,Nfkbiz | | 9.972e-03 | -4.61 | cellular response to organic cyclic compound | biological process | GO:0071407 | 312 | 11 | 13711 | 214 | P2ry6,Acod1,Tnf,Nfkbia,Zfp36,Msn,Sgk1,Tiparp,Fam107a,Nfkb1,Ccl12 | | 1.004e-02 | -4.60 | RHD\_DNA\_bind | pfam domains | PF00554 | 10 | 2 | 12881 | 201 | Nfkb2,Nfkb1 | | 1.004e-02 | -4.60 | RHD\_dimer | pfam domains | PF16179 | 10 | 2 | 12881 | 201 | Nfkb1,Nfkb2 | | 1.005e-02 | -4.60 | regulation of interleukin-2 biosynthetic process | biological process | GO:0045076 | 10 | 2 | 13711 | 214 | Il1a,Zfp36 | | 1.005e-02 | -4.60 | MyD88-dependent toll-like receptor signaling pathway | biological process | GO:0002755 | 10 | 2 | 13711 | 214 | Irak3,Tnip1 | | 1.005e-02 | -4.60 | interferon-gamma production | biological process | GO:0032609 | 10 | 2 | 13711 | 214 | Gadd45g,Isg15 | | 1.005e-02 | -4.60 | cellular response to zinc ion | biological process | GO:0071294 | 10 | 2 | 13711 | 214 | Mt2,Mt1 | | 1.005e-02 | -4.60 | regulation of receptor catabolic process | biological process | GO:2000644 | 10 | 2 | 13711 | 214 | Ptpn1,Dtx3l | | 1.005e-02 | -4.60 | positive regulation of glomerulus development | biological process | GO:0090193 | 10 | 2 | 13711 | 214 | C3ar1,Cflar | | 1.005e-02 | -4.60 | programmed cell death involved in cell development | biological process | GO:0010623 | 10 | 2 | 13711 | 214 | Casp4,Il1a | | 1.005e-02 | -4.60 | regulation of vitamin metabolic process | biological process | GO:0030656 | 10 | 2 | 13711 | 214 | Tnf,Nfkb1 | | 1.005e-02 | -4.60 | antibiotic biosynthetic process | biological process | GO:0017000 | 10 | 2 | 13711 | 214 | Ncf1,Cybb | | 1.005e-02 | -4.60 | phosphatidylinositol 3-kinase regulatory subunit binding | molecular function | GO:0036312 | 10 | 2 | 13516 | 211 | Pik3ap1,Pik3r1 | | 1.005e-02 | -4.60 | CARD domain binding | molecular function | GO:0050700 | 10 | 2 | 13516 | 211 | Casp4,Irgm2 | | 1.005e-02 | -4.60 | death domain binding | molecular function | GO:0070513 | 10 | 2 | 13516 | 211 | Irgm2,Ripk1 | | 1.005e-02 | -4.60 | response to leukemia inhibitory factor | biological process | GO:1990823 | 117 | 6 | 13711 | 214 | Sbno2,Trim25,Fzd4,Socs3,Arid5b,Icam1 | | 1.005e-02 | -4.60 | response to topologically incorrect protein | biological process | GO:0035966 | 117 | 6 | 13711 | 214 | Nfe2l2,Thbs1,Hspb1,Ptpn1,Hspa5,Atf3 | | 1.005e-02 | -4.60 | cellular response to leukemia inhibitory factor | biological process | GO:1990830 | 117 | 6 | 13711 | 214 | Icam1,Arid5b,Socs3,Fzd4,Trim25,Sbno2 | | 1.005e-02 | -4.60 | regulation of fat cell differentiation | biological process | GO:0045598 | 117 | 6 | 13711 | 214 | Ptgs2,Noct,Bbs12,Zbtb16,Zfp36,Tnf | | 1.010e-02 | -4.59 | IL-3 Signaling Pathway | WikiPathways | WP373 | 95 | 7 | 3756 | 96 | Hspb1,Selp,Socs3,Csf2rb,Nfkb1,Pik3r1,Spi1 | | 1.016e-02 | -4.59 | transcription regulator activity | molecular function | GO:0140110 | 1030 | 26 | 13516 | 211 | Arid5b,Nfe2l2,Ifi204,Ifi209,Aff1,Mxd4,Parp9,Nfkb1,Zbtb16,Irf9,Irf2,Runx1,Ifi211,Rbm39,Bcl3,Maff,Spi1,Ifi207,Cebpd,Dnajb1,Sox11,Nfkb2,Znfx1,Atf3,Bach1,Irf7 | | 1.016e-02 | -4.59 | salivary gland development | biological process | GO:0007431 | 29 | 3 | 13711 | 214 | Tgm2,Tnf,Il6 | | 1.016e-02 | -4.59 | cellular response to amyloid-beta | biological process | GO:1904646 | 29 | 3 | 13711 | 214 | Casp4,Vcam1,Icam1 | | 1.016e-02 | -4.59 | intrinsic apoptotic signaling pathway in response to DNA damage by p53 class mediator | biological process | GO:0042771 | 29 | 3 | 13711 | 214 | Cdkn1a,Ifi204,Bcl3 | | 1.019e-02 | -4.59 | TGTTTGY\_HNF3\_Q6 | MSigDB lists | TGTTTGY\_HNF3\_Q6 | 523 | 15 | 12187 | 179 | Sgk3,Tagln2,Rbm47,Ip6k2,Errfi1,Sgk1,Fstl1,Csf3,Robo4,Nfkbie,A2m,Banp,Rbm39,Hcar2,Gadd45b | | 1.020e-02 | -4.58 | YOSHIMURA\_MAPK8\_TARGETS\_UP | MSigDB lists | YOSHIMURA\_MAPK8\_TARGETS\_UP | 883 | 22 | 12187 | 179 | Csf2rb,Irf2,Errfi1,Plaur,Tap1,Hspb1,P2ry6,Plek,Socs3,Cdkn1a,Ier3,Lcn2,Il1rn,Kcna5,Hp,Akap12,Il1a,A2m,C5ar1,Pygm,Tnf,Ccl12 | | 1.023e-02 | -4.58 | GO\_INTRINSIC\_COMPONENT\_OF\_PLASMA\_MEMBRANE | MSigDB lists | GO\_INTRINSIC\_COMPONENT\_OF\_PLASMA\_MEMBRANE | 1045 | 25 | 12187 | 179 | Osmr,Il6,Cybb,Slc24a4,Tgm2,Gpr84,Msr1,Il4ra,Il17ra,P2ry6,Cd14,Selp,Tnfsf10,Igsf6,Fzd4,C5ar1,C3ar1,Sdc4,Tnf,Plaur,Csf2rb,Icam1,Ncf1,Kcna5,Cp | | 1.025e-02 | -4.58 | Toxoplasmosis | KEGG pathways | mmu05145 | 91 | 6 | 5248 | 107 | Pik3r5,Nfkbia,Nfkb1,Irgm1,Tnf,Birc3 | | 1.025e-02 | -4.58 | Toxoplasmosis | KEGG pathways | ko05145 | 91 | 6 | 5248 | 107 | Nfkbia,Pik3r5,Irgm1,Nfkb1,Birc3,Tnf | | 1.027e-02 | -4.58 | GSE2706\_2H\_VS\_8H\_LPS\_STIM\_DC\_DN | MSigDB lists | GSE2706\_2H\_VS\_8H\_LPS\_STIM\_DC\_DN | 125 | 6 | 12187 | 179 | AA467197,Znfx1,Nfkb1,Dtx3l,Gbp3,Tap1 | | 1.027e-02 | -4.58 | AKT\_UP.V1\_UP | MSigDB lists | AKT\_UP.V1\_UP | 125 | 6 | 12187 | 179 | Cxcl16,Atf3,Tor3a,Ifit3b,Ifit2,Saa1 | | 1.027e-02 | -4.58 | NABA\_ECM\_REGULATORS | MSigDB lists | NABA\_ECM\_REGULATORS | 125 | 6 | 12187 | 179 | Adamts1,Tgm2,A2m,Mmp8,Adamts9,Timp1 | | 1.027e-02 | -4.58 | GSE36078\_UNTREATED\_VS\_AD5\_T425A\_HEXON\_INF\_MOUSE\_LUNG\_DC\_DN | MSigDB lists | GSE36078\_UNTREATED\_VS\_AD5\_T425A\_HEXON\_INF\_MOUSE\_LUNG\_DC\_DN | 163 | 7 | 12187 | 179 | Pik3ap1,Errfi1,Ifitm2,Gem,Gbp7,Ncf1,Cdkn1a | | 1.027e-02 | -4.58 | GSE5463\_CTRL\_VS\_DEXAMETHASONE\_TREATED\_THYMOCYTE\_DN | MSigDB lists | GSE5463\_CTRL\_VS\_DEXAMETHASONE\_TREATED\_THYMOCYTE\_DN | 163 | 7 | 12187 | 179 | Sox11,Irgm2,Sgk1,Usp18,Akap12,Plaur,Tnfaip3 | | 1.027e-02 | -4.58 | GSE24142\_DN2\_VS\_DN3\_THYMOCYTE\_FETAL\_DN | MSigDB lists | GSE24142\_DN2\_VS\_DN3\_THYMOCYTE\_FETAL\_DN | 163 | 7 | 12187 | 179 | Gbp6,Errfi1,Irf9,Tap1,Gbp4,Akap12,Ms4a6d | | 1.027e-02 | -4.58 | GSE15330\_HSC\_VS\_GRANULOCYTE\_MONOCYTE\_PROGENITOR\_IKAROS\_KO\_UP | MSigDB lists | GSE15330\_HSC\_VS\_GRANULOCYTE\_MONOCYTE\_PROGENITOR\_IKAROS\_KO\_UP | 163 | 7 | 12187 | 179 | Il17ra,P2ry6,Gem,Arid5b,Gadd45g,Znfx1,Sntb2 | | 1.027e-02 | -4.58 | GSE24142\_ADULT\_VS\_FETAL\_DN2\_THYMOCYTE\_UP | MSigDB lists | GSE24142\_ADULT\_VS\_FETAL\_DN2\_THYMOCYTE\_UP | 163 | 7 | 12187 | 179 | Ms4a6d,Gbp4,Ifit2,Casp4,Rtp4,Gbp3,Gbp6 | | 1.027e-02 | -4.58 | GSE40666\_WT\_VS\_STAT1\_KO\_CD8\_TCELL\_WITH\_IFNA\_STIM\_90MIN\_DN | MSigDB lists | GSE40666\_WT\_VS\_STAT1\_KO\_CD8\_TCELL\_WITH\_IFNA\_STIM\_90MIN\_DN | 163 | 7 | 12187 | 179 | Msn,Parp9,Rtp4,Trim25,Bcl3,Rnf213,Ifi44 | | 1.027e-02 | -4.58 | GSE22432\_CONVENTIONAL\_CDC\_VS\_PLASMACYTOID\_PDC\_DN | MSigDB lists | GSE22432\_CONVENTIONAL\_CDC\_VS\_PLASMACYTOID\_PDC\_DN | 163 | 7 | 12187 | 179 | Herc6,Tap1,Gbp3,Parp14,Tor3a,Cdkn1a,Ccnd2 | | 1.027e-02 | -4.58 | GSE24142\_DN2\_VS\_DN3\_THYMOCYTE\_UP | MSigDB lists | GSE24142\_DN2\_VS\_DN3\_THYMOCYTE\_UP | 163 | 7 | 12187 | 179 | Cmpk2,Ripk1,Ier3,Map3k6,Il4ra,Tagln2,Spi1 | | 1.028e-02 | -4.58 | regulation of protein ubiquitination | biological process | GO:0031396 | 191 | 8 | 13711 | 214 | Dtx3l,Trib1,Tnfaip3,Gbp4,Fam107a,Hspa5,Birc3,Isg15 | | 1.030e-02 | -4.58 | positive regulation of hydrolase activity | biological process | GO:0051345 | 585 | 17 | 13711 | 214 | Ccl4,S100a9,S100a8,Ccl12,Dnajb1,Rhoc,Sele,Cflar,Ccl7,Tnfsf10,Xdh,Ccl2,Casp4,Icam1,Tnf,Rgs16,Plek | | 1.031e-02 | -4.57 | ZHONG\_SECRETOME\_OF\_LUNG\_CANCER\_AND\_ENDOTHELIUM | MSigDB lists | ZHONG\_SECRETOME\_OF\_LUNG\_CANCER\_AND\_ENDOTHELIUM | 58 | 4 | 12187 | 179 | Cp,A2m,Sdc4,Vcam1 | | 1.031e-02 | -4.57 | MARTINEZ\_RESPONSE\_TO\_TRABECTEDIN\_UP | MSigDB lists | MARTINEZ\_RESPONSE\_TO\_TRABECTEDIN\_UP | 58 | 4 | 12187 | 179 | Gadd45b,Atf3,Ifitm3,Ifitm2 | | 1.031e-02 | -4.57 | GO\_CELLULAR\_TRANSITION\_METAL\_ION\_HOMEOSTASIS | MSigDB lists | GO\_CELLULAR\_TRANSITION\_METAL\_ION\_HOMEOSTASIS | 58 | 4 | 12187 | 179 | Cp,S100a8,Lcn2,S100a9 | | 1.031e-02 | -4.57 | PID\_BCR\_5PATHWAY | MSigDB lists | PID\_BCR\_5PATHWAY | 58 | 4 | 12187 | 179 | Pik3r1,Nfkbia,Nfkb1,Pik3ap1 | | 1.031e-02 | -4.57 | CADWELL\_ATG16L1\_TARGETS\_UP | MSigDB lists | CADWELL\_ATG16L1\_TARGETS\_UP | 58 | 4 | 12187 | 179 | Saa1,Socs3,Hp,Plin4 | | 1.031e-02 | -4.57 | GOTTWEIN\_TARGETS\_OF\_KSHV\_MIR\_K12\_11 | MSigDB lists | GOTTWEIN\_TARGETS\_OF\_KSHV\_MIR\_K12\_11 | 58 | 4 | 12187 | 179 | Hspa5,Rnf213,Sgk1,Xaf1 | | 1.033e-02 | -4.57 | GO\_TRANSCRIPTION\_FACTOR\_BINDING | MSigDB lists | GO\_TRANSCRIPTION\_FACTOR\_BINDING | 427 | 13 | 12187 | 179 | Ifi204,Nfkb1,Ifi207,Ifi209,Ifi211,Bcl3,Nfe2l2,Oasl1,Nfkbia,Spi1,Trib1,Runx1,Pik3r1 | | 1.034e-02 | -4.57 | GO\_NEGATIVE\_REGULATION\_OF\_TRANSFERASE\_ACTIVITY | MSigDB lists | GO\_NEGATIVE\_REGULATION\_OF\_TRANSFERASE\_ACTIVITY | 289 | 10 | 12187 | 179 | Tnfaip3,Gadd45b,Il6,Socs3,Trib1,Cdkn1a,Errfi1,Ptpn1,Irak3,Hspb1 | | 1.035e-02 | -4.57 | MATSUDA\_NATURAL\_KILLER\_DIFFERENTIATION | MSigDB lists | MATSUDA\_NATURAL\_KILLER\_DIFFERENTIATION | 380 | 12 | 12187 | 179 | Mxd4,Socs3,Zbtb16,Sgk1,Plaur,Ifitm2,Ptgs2,Rhoc,Ccl4,Zfp189,Cxcl1,Gem | | 1.036e-02 | -4.57 | PID\_INTEGRIN\_A4B1\_PATHWAY | MSigDB lists | PID\_INTEGRIN\_A4B1\_PATHWAY | 31 | 3 | 12187 | 179 | Thbs1,Vcam1,Cd14 | | 1.036e-02 | -4.57 | BIOCARTA\_TOLL\_PATHWAY | MSigDB lists | BIOCARTA\_TOLL\_PATHWAY | 31 | 3 | 12187 | 179 | Cd14,Nfkbia,Nfkb1 | | 1.036e-02 | -4.57 | GROSS\_HYPOXIA\_VIA\_ELK3\_ONLY\_UP | MSigDB lists | GROSS\_HYPOXIA\_VIA\_ELK3\_ONLY\_UP | 31 | 3 | 12187 | 179 | Cxcl1,Akap12,Il6 | | 1.036e-02 | -4.57 | GO\_REGULATION\_OF\_LEUKOCYTE\_MEDIATED\_CYTOTOXICITY | MSigDB lists | GO\_REGULATION\_OF\_LEUKOCYTE\_MEDIATED\_CYTOTOXICITY | 31 | 3 | 12187 | 179 | Icam1,Tap1,Ccl12 | | 1.036e-02 | -4.57 | GNF2\_PTX3 | MSigDB lists | GNF2\_PTX3 | 31 | 3 | 12187 | 179 | Ccl12,Il6,Fstl1 | | 1.036e-02 | -4.57 | ZWANG\_CLASS\_2\_TRANSIENTLY\_INDUCED\_BY\_EGF | MSigDB lists | ZWANG\_CLASS\_2\_TRANSIENTLY\_INDUCED\_BY\_EGF | 31 | 3 | 12187 | 179 | Errfi1,Fstl1,Zfp189 | | 1.036e-02 | -4.57 | GO\_REGULATION\_OF\_INTERLEUKIN\_12\_PRODUCTION | MSigDB lists | GO\_REGULATION\_OF\_INTERLEUKIN\_12\_PRODUCTION | 31 | 3 | 12187 | 179 | Thbs1,Irak3,Nfkb1 | | 1.036e-02 | -4.57 | PID\_EPO\_PATHWAY | MSigDB lists | PID\_EPO\_PATHWAY | 31 | 3 | 12187 | 179 | Socs3,Pik3r1,Nfkb1 | | 1.036e-02 | -4.57 | GO\_NEGATIVE\_REGULATION\_OF\_FAT\_CELL\_DIFFERENTIATION | MSigDB lists | GO\_NEGATIVE\_REGULATION\_OF\_FAT\_CELL\_DIFFERENTIATION | 31 | 3 | 12187 | 179 | Il6,Tnf,Bbs12 | | 1.036e-02 | -4.57 | KIM\_BIPOLAR\_DISORDER\_OLIGODENDROCYTE\_DENSITY\_CORR\_DN | MSigDB lists | KIM\_BIPOLAR\_DISORDER\_OLIGODENDROCYTE\_DENSITY\_CORR\_DN | 31 | 3 | 12187 | 179 | Sbno2,S100a8,Tnf | | 1.036e-02 | -4.57 | GO\_REGULATION\_OF\_INTERLEUKIN\_2\_PRODUCTION | MSigDB lists | GO\_REGULATION\_OF\_INTERLEUKIN\_2\_PRODUCTION | 31 | 3 | 12187 | 179 | Tnfaip3,Il1a,Runx1 | | 1.036e-02 | -4.57 | LI\_LUNG\_CANCER | MSigDB lists | LI\_LUNG\_CANCER | 31 | 3 | 12187 | 179 | Nfkb1,Hspb1,Hp | | 1.036e-02 | -4.57 | GO\_NEGATIVE\_REGULATION\_OF\_MYELOID\_LEUKOCYTE\_DIFFERENTIATION | MSigDB lists | GO\_NEGATIVE\_REGULATION\_OF\_MYELOID\_LEUKOCYTE\_DIFFERENTIATION | 31 | 3 | 12187 | 179 | Trib1,Pik3r1,Runx1 | | 1.036e-02 | -4.57 | YAGI\_AML\_WITH\_INV\_16\_TRANSLOCATION | MSigDB lists | YAGI\_AML\_WITH\_INV\_16\_TRANSLOCATION | 334 | 11 | 12187 | 179 | Rrp8,Hspb1,Rbm39,Osmr,Sele,Ccnd2,Runx1,Sdc4,Casp4,Kcna5,Timp1 | | 1.038e-02 | -4.57 | KOINUMA\_TARGETS\_OF\_SMAD2\_OR\_SMAD3 | MSigDB lists | KOINUMA\_TARGETS\_OF\_SMAD2\_OR\_SMAD3 | 675 | 18 | 12187 | 179 | Tagln2,Trim25,Thbs1,Hspa5,Cdkn1a,Nfkb1,Tiparp,Atf3,Ier3,Trim56,Ptges,Il4ra,Runx1,Tgm2,Bach1,Rhoc,Tubb6,Ccl12 | | 1.038e-02 | -4.57 | Hepatitis B | KEGG pathways | mmu05161 | 119 | 7 | 5248 | 107 | Nfkbia,Il6,Nfkb1,Tnf,Irf7,Pik3r1,Cdkn1a | | 1.040e-02 | -4.57 | GSE32901\_TH1\_VS\_TH17\_NEG\_CD4\_TCELL\_UP | MSigDB lists | GSE32901\_TH1\_VS\_TH17\_NEG\_CD4\_TCELL\_UP | 90 | 5 | 12187 | 179 | Osmr,Zfp36,Irak3,Tap1,Csf2rb | | 1.040e-02 | -4.57 | GO\_RESPONSE\_TO\_CALCIUM\_ION | MSigDB lists | GO\_RESPONSE\_TO\_CALCIUM\_ION | 90 | 5 | 12187 | 179 | Ptges,Thbs1,Il6,Clic4,Iqgap1 | | 1.040e-02 | -4.57 | GO\_FEMALE\_SEX\_DIFFERENTIATION | MSigDB lists | GO\_FEMALE\_SEX\_DIFFERENTIATION | 90 | 5 | 12187 | 179 | Fzd4,Arid5b,Adamts1,Icam1,Tiparp | | 1.051e-02 | -4.56 | GO\_NEGATIVE\_REGULATION\_OF\_CELL\_PROLIFERATION | MSigDB lists | GO\_NEGATIVE\_REGULATION\_OF\_CELL\_PROLIFERATION | 476 | 14 | 12187 | 179 | Ifit3b,Zbtb16,Thbs1,Cdkn1a,Sox11,Adamts1,Mxd4,Il1a,Ptgs2,Trib1,Il6,Tnfaip3,Ptges,Xdh | | 1.051e-02 | -4.56 | BACH1\_01 | MSigDB lists | BACH1\_01 | 204 | 8 | 12187 | 179 | Tnip3,Slc24a4,Il1rn,Sgk1,Cdkn1a,Trib1,Csf3,Tagln2 | | 1.052e-02 | -4.55 | DUTERTRE\_ESTRADIOL\_RESPONSE\_24HR\_DN | MSigDB lists | DUTERTRE\_ESTRADIOL\_RESPONSE\_24HR\_DN | 428 | 13 | 12187 | 179 | Dtx3l,Arid5b,Rhoc,Parp12,Map3k8,Sntb2,Trib1,Parp14,Errfi1,Parp9,Rasd1,Cdkn1a,Nfkbiz | | 1.057e-02 | -4.55 | lymphocyte mediated immunity | biological process | GO:0002449 | 85 | 5 | 13711 | 214 | Icam1,Il4ra,Stx11,Irf7,Bcl3 | | 1.057e-02 | -4.55 | positive regulation of peptidyl-serine phosphorylation | biological process | GO:0033138 | 85 | 5 | 13711 | 214 | Csf3,Tnf,Irgm2,Il6,Ptgs2 | | 1.057e-02 | -4.55 | endopeptidase inhibitor activity | molecular function | GO:0004866 | 85 | 5 | 13516 | 211 | Serpina3f,Birc3,Timp1,A2m,Tnfaip8 | | 1.058e-02 | -4.55 | positive regulation of receptor signaling pathway via STAT | biological process | GO:1904894 | 55 | 4 | 13711 | 214 | Il6,Parp9,Tnf,Parp14 | | 1.060e-02 | -4.55 | GSE21360\_NAIVE\_VS\_PRIMARY\_MEMORY\_CD8\_TCELL\_DN | MSigDB lists | GSE21360\_NAIVE\_VS\_PRIMARY\_MEMORY\_CD8\_TCELL\_DN | 164 | 7 | 12187 | 179 | Atf3,Mmp8,Samd9l,Ifit3b,Ifit2,Trim25,Sgk3 | | 1.060e-02 | -4.55 | GSE24142\_EARLY\_THYMIC\_PROGENITOR\_VS\_DN3\_THYMOCYTE\_ADULT\_UP | MSigDB lists | GSE24142\_EARLY\_THYMIC\_PROGENITOR\_VS\_DN3\_THYMOCYTE\_ADULT\_UP | 164 | 7 | 12187 | 179 | Ncf1,Gem,Plaur,Icam1,Rtp4,Nfkbia,Map3k8 | | 1.060e-02 | -4.55 | GSE411\_100MIN\_VS\_400MIN\_IL6\_STIM\_SOCS3\_KO\_MACROPHAGE\_DN | MSigDB lists | GSE411\_100MIN\_VS\_400MIN\_IL6\_STIM\_SOCS3\_KO\_MACROPHAGE\_DN | 164 | 7 | 12187 | 179 | Nfkb1,Mxd4,Parp14,Iqgap1,Nfkb2,Cflar,Ncf1 | | 1.060e-02 | -4.55 | GSE15930\_STIM\_VS\_STIM\_AND\_TRICHOSTATINA\_48H\_CD8\_T\_CELL\_UP | MSigDB lists | GSE15930\_STIM\_VS\_STIM\_AND\_TRICHOSTATINA\_48H\_CD8\_T\_CELL\_UP | 164 | 7 | 12187 | 179 | Selp,Gem,Tagln2,Bcl3,Tnfsf10,Sox11,Gadd45g | | 1.060e-02 | -4.55 | GSE17721\_LPS\_VS\_GARDIQUIMOD\_0.5H\_BMDC\_DN | MSigDB lists | GSE17721\_LPS\_VS\_GARDIQUIMOD\_0.5H\_BMDC\_DN | 164 | 7 | 12187 | 179 | Ifit2,Parp12,Ms4a6d,Msn,Slfn4,Iqgap1,Slfn3 | | 1.060e-02 | -4.55 | GSE36476\_CTRL\_VS\_TSST\_ACT\_72H\_MEMORY\_CD4\_TCELL\_YOUNG\_UP | MSigDB lists | GSE36476\_CTRL\_VS\_TSST\_ACT\_72H\_MEMORY\_CD4\_TCELL\_YOUNG\_UP | 164 | 7 | 12187 | 179 | Ifitm3,Timp1,Tiparp,Tnfaip3,Sgk1,Pik3r1,Dnajb1 | | 1.060e-02 | -4.55 | GSE1448\_CTRL\_VS\_ANTI\_VBETA5\_DP\_THYMOCYTE\_DN | MSigDB lists | GSE1448\_CTRL\_VS\_ANTI\_VBETA5\_DP\_THYMOCYTE\_DN | 164 | 7 | 12187 | 179 | Ptges,Irf9,Gbp3,Ptpn1,Xdh,Zfp36,Nfkb1 | | 1.060e-02 | -4.55 | GSE32423\_IL7\_VS\_IL4\_MEMORY\_CD8\_TCELL\_DN | MSigDB lists | GSE32423\_IL7\_VS\_IL4\_MEMORY\_CD8\_TCELL\_DN | 164 | 7 | 12187 | 179 | Vcam1,Map3k8,Gbp4,Ripk1,Tnfaip3,Slfn9,Gbp6 | | 1.060e-02 | -4.55 | GSE43955\_1H\_VS\_60H\_ACT\_CD4\_TCELL\_UP | MSigDB lists | GSE43955\_1H\_VS\_60H\_ACT\_CD4\_TCELL\_UP | 164 | 7 | 12187 | 179 | Rgs16,Plaur,Clic4,Il1rn,Tubb6,Ifit3b,Usp18 | | 1.060e-02 | -4.55 | GSE23505\_IL6\_IL1\_IL23\_VS\_IL6\_IL1\_TGFB\_TREATED\_CD4\_TCELL\_UP | MSigDB lists | GSE23505\_IL6\_IL1\_IL23\_VS\_IL6\_IL1\_TGFB\_TREATED\_CD4\_TCELL\_UP | 164 | 7 | 12187 | 179 | Ifit2,Ifit3b,Sntb2,Socs3,Atf3,Cd14,Slfn9 | | 1.066e-02 | -4.54 | SHETH\_LIVER\_CANCER\_VS\_TXNIP\_LOSS\_PAM2 | MSigDB lists | SHETH\_LIVER\_CANCER\_VS\_TXNIP\_LOSS\_PAM2 | 126 | 6 | 12187 | 179 | Tagln2,Cd14,Atf3,Ier3,Rhoc,Il1rn | | 1.072e-02 | -4.54 | Znf-RING\_LisH | interpro domains | IPR027370 | 30 | 3 | 13788 | 212 | Trim25,Trim30a,Trim56 | | 1.081e-02 | -4.53 | GO\_LOW\_DENSITY\_LIPOPROTEIN\_PARTICLE\_BINDING | MSigDB lists | GO\_LOW\_DENSITY\_LIPOPROTEIN\_PARTICLE\_BINDING | 11 | 2 | 12187 | 179 | Thbs1,Msr1 | | 1.081e-02 | -4.53 | GO\_POSITIVE\_REGULATION\_OF\_MACROPHAGE\_DIFFERENTIATION | MSigDB lists | GO\_POSITIVE\_REGULATION\_OF\_MACROPHAGE\_DIFFERENTIATION | 11 | 2 | 12187 | 179 | Ripk1,Trib1 | | 1.081e-02 | -4.53 | GO\_CYCLOOXYGENASE\_PATHWAY | MSigDB lists | GO\_CYCLOOXYGENASE\_PATHWAY | 11 | 2 | 12187 | 179 | Ptgs2,Ptges | | 1.081e-02 | -4.53 | GO\_POSITIVE\_REGULATION\_OF\_CELLULAR\_EXTRAVASATION | MSigDB lists | GO\_POSITIVE\_REGULATION\_OF\_CELLULAR\_EXTRAVASATION | 11 | 2 | 12187 | 179 | Icam1,Ccl12 | | 1.081e-02 | -4.53 | RUAN\_RESPONSE\_TO\_TNF\_UP | MSigDB lists | RUAN\_RESPONSE\_TO\_TNF\_UP | 11 | 2 | 12187 | 179 | Cp,Hp | | 1.081e-02 | -4.53 | NIELSEN\_MALIGNAT\_FIBROUS\_HISTIOCYTOMA\_UP | MSigDB lists | NIELSEN\_MALIGNAT\_FIBROUS\_HISTIOCYTOMA\_UP | 11 | 2 | 12187 | 179 | Ccl12,Timp1 | | 1.081e-02 | -4.53 | GO\_REGULATION\_OF\_IRE1\_MEDIATED\_UNFOLDED\_PROTEIN\_RESPONSE | MSigDB lists | GO\_REGULATION\_OF\_IRE1\_MEDIATED\_UNFOLDED\_PROTEIN\_RESPONSE | 11 | 2 | 12187 | 179 | Hspa5,Ptpn1 | | 1.081e-02 | -4.53 | GO\_POSITIVE\_REGULATION\_OF\_TYPE\_2\_IMMUNE\_RESPONSE | MSigDB lists | GO\_POSITIVE\_REGULATION\_OF\_TYPE\_2\_IMMUNE\_RESPONSE | 11 | 2 | 12187 | 179 | Il6,Il4ra | | 1.081e-02 | -4.53 | GO\_POSITIVE\_REGULATION\_OF\_ENDOPLASMIC\_RETICULUM\_UNFOLDED\_PROTEIN\_RESPONSE | MSigDB lists | GO\_POSITIVE\_REGULATION\_OF\_ENDOPLASMIC\_RETICULUM\_UNFOLDED\_PROTEIN\_RESPONSE | 11 | 2 | 12187 | 179 | Pik3r1,Ptpn1 | | 1.081e-02 | -4.53 | GO\_POSITIVE\_REGULATION\_OF\_MACROPHAGE\_DERIVED\_FOAM\_CELL\_DIFFERENTIATION | MSigDB lists | GO\_POSITIVE\_REGULATION\_OF\_MACROPHAGE\_DERIVED\_FOAM\_CELL\_DIFFERENTIATION | 11 | 2 | 12187 | 179 | Nfkb1,Msr1 | | 1.081e-02 | -4.53 | PID\_INTEGRIN5\_PATHWAY | MSigDB lists | PID\_INTEGRIN5\_PATHWAY | 11 | 2 | 12187 | 179 | Plaur,Vcam1 | | 1.081e-02 | -4.53 | MODULE\_148 | MSigDB lists | MODULE\_148 | 11 | 2 | 12187 | 179 | Ptgs2,Ptges | | 1.081e-02 | -4.53 | GO\_REGULATION\_OF\_PODOSOME\_ASSEMBLY | MSigDB lists | GO\_REGULATION\_OF\_PODOSOME\_ASSEMBLY | 11 | 2 | 12187 | 179 | Tnf,Msn | | 1.081e-02 | -4.53 | GO\_CHEMOKINE\_BINDING | MSigDB lists | GO\_CHEMOKINE\_BINDING | 11 | 2 | 12187 | 179 | Zfp36,A2m | | 1.081e-02 | -4.53 | DAZARD\_UV\_RESPONSE\_CLUSTER\_G3 | MSigDB lists | DAZARD\_UV\_RESPONSE\_CLUSTER\_G3 | 11 | 2 | 12187 | 179 | Tnfaip3,S100a9 | | 1.081e-02 | -4.53 | GO\_REGULATION\_OF\_LYMPHOCYTE\_CHEMOTAXIS | MSigDB lists | GO\_REGULATION\_OF\_LYMPHOCYTE\_CHEMOTAXIS | 11 | 2 | 12187 | 179 | Ccl12,Ccl4 | | 1.081e-02 | -4.53 | REACTOME\_NFKB\_ACTIVATION\_THROUGH\_FADD\_RIP1\_PATHWAY\_MEDIATED\_BY\_CASPASE\_8\_AND10 | MSigDB lists | REACTOME\_NFKB\_ACTIVATION\_THROUGH\_FADD\_RIP1\_PATHWAY\_MEDIATED\_BY\_CASPASE\_8\_AND10 | 11 | 2 | 12187 | 179 | Trim25,Ripk1 | | 1.081e-02 | -4.53 | HASEGAWA\_TUMORIGENESIS\_BY\_RET\_C634R | MSigDB lists | HASEGAWA\_TUMORIGENESIS\_BY\_RET\_C634R | 11 | 2 | 12187 | 179 | Irf7,Usp18 | | 1.081e-02 | -4.53 | GO\_REGULATION\_OF\_MYOBLAST\_FUSION | MSigDB lists | GO\_REGULATION\_OF\_MYOBLAST\_FUSION | 11 | 2 | 12187 | 179 | Cflar,Il4ra | | 1.081e-02 | -4.53 | REACTOME\_REGULATION\_OF\_IFNG\_SIGNALING | MSigDB lists | REACTOME\_REGULATION\_OF\_IFNG\_SIGNALING | 11 | 2 | 12187 | 179 | Socs3,Ptpn1 | | 1.081e-02 | -4.53 | HUMMERICH\_MALIGNANT\_SKIN\_TUMOR\_UP | MSigDB lists | HUMMERICH\_MALIGNANT\_SKIN\_TUMOR\_UP | 11 | 2 | 12187 | 179 | S100a9,S100a8 | | 1.081e-02 | -4.53 | MAHADEVAN\_RESPONSE\_TO\_MP470\_UP | MSigDB lists | MAHADEVAN\_RESPONSE\_TO\_MP470\_UP | 11 | 2 | 12187 | 179 | Vcam1,Xaf1 | | 1.081e-02 | -4.53 | BIOCARTA\_RANKL\_PATHWAY | MSigDB lists | BIOCARTA\_RANKL\_PATHWAY | 11 | 2 | 12187 | 179 | Irf9,Nfkb1 | | 1.088e-02 | -4.52 | GSE15624\_CTRL\_VS\_6H\_HALOFUGINONE\_TREATED\_CD4\_TCELL\_UP | MSigDB lists | GSE15624\_CTRL\_VS\_6H\_HALOFUGINONE\_TREATED\_CD4\_TCELL\_UP | 91 | 5 | 12187 | 179 | Slfn5,Xaf1,Tagln2,Nfe2l2,Igsf6 | | 1.088e-02 | -4.52 | GSE43863\_NAIVE\_VS\_TFH\_CD4\_EFF\_TCELL\_D6\_LCMV\_DN | MSigDB lists | GSE43863\_NAIVE\_VS\_TFH\_CD4\_EFF\_TCELL\_D6\_LCMV\_DN | 91 | 5 | 12187 | 179 | Tnf,Msr1,Rnf213,Parp14,Gadd45b | | 1.091e-02 | -4.52 | response to endoplasmic reticulum stress | biological process | GO:0034976 | 193 | 8 | 13711 | 214 | Trim25,Pik3r1,Nfe2l2,Thbs1,Atf3,Ptpn1,Hspa5,Casp4 | | 1.093e-02 | -4.52 | PID\_AP1\_PATHWAY | MSigDB lists | PID\_AP1\_PATHWAY | 59 | 4 | 12187 | 179 | Il6,Atf3,Ccl12,Timp1 | | 1.093e-02 | -4.52 | LIEN\_BREAST\_CARCINOMA\_METAPLASTIC\_VS\_DUCTAL\_UP | MSigDB lists | LIEN\_BREAST\_CARCINOMA\_METAPLASTIC\_VS\_DUCTAL\_UP | 59 | 4 | 12187 | 179 | Msn,Il6,Icam1,Angptl4 | | 1.093e-02 | -4.52 | KRAS.300\_UP.V1\_DN | MSigDB lists | KRAS.300\_UP.V1\_DN | 59 | 4 | 12187 | 179 | Sox11,Gadd45g,Rtp4,Tnfsf10 | | 1.094e-02 | -4.52 | GSE10240\_IL22\_VS\_IL17\_STIM\_PRIMARY\_BRONCHIAL\_EPITHELIAL\_CELLS\_DN | MSigDB lists | GSE10240\_IL22\_VS\_IL17\_STIM\_PRIMARY\_BRONCHIAL\_EPITHELIAL\_CELLS\_DN | 165 | 7 | 12187 | 179 | Vcam1,Sgk1,Lcn2,Gadd45b,Znfx1,Xdh,Csf2rb | | 1.094e-02 | -4.52 | GSE2770\_IL12\_ACT\_VS\_ACT\_CD4\_TCELL\_2H\_DN | MSigDB lists | GSE2770\_IL12\_ACT\_VS\_ACT\_CD4\_TCELL\_2H\_DN | 165 | 7 | 12187 | 179 | Znfx1,Nfe2l2,Tubb6,Il1a,Banp,Pik3r5,Cflar | | 1.094e-02 | -4.52 | GSE36078\_WT\_VS\_IL1R\_KO\_LUNG\_DC\_AFTER\_AD5\_T425A\_HEXON\_INF\_DN | MSigDB lists | GSE36078\_WT\_VS\_IL1R\_KO\_LUNG\_DC\_AFTER\_AD5\_T425A\_HEXON\_INF\_DN | 165 | 7 | 12187 | 179 | Rtp4,Tnip1,Irgm2,Nfe2l2,Oasl1,Usp18,Ifi44 | | 1.094e-02 | -4.52 | GSE6674\_ANTI\_IGM\_VS\_CPG\_STIM\_BCELL\_DN | MSigDB lists | GSE6674\_ANTI\_IGM\_VS\_CPG\_STIM\_BCELL\_DN | 165 | 7 | 12187 | 179 | Gadd45g,Atf3,Cdkn1a,Tnf,Map3k8,Il4ra,Bbs12 | | 1.094e-02 | -4.52 | GSE17186\_MEMORY\_VS\_CD21LOW\_TRANSITIONAL\_BCELL\_DN | MSigDB lists | GSE17186\_MEMORY\_VS\_CD21LOW\_TRANSITIONAL\_BCELL\_DN | 165 | 7 | 12187 | 179 | Nfkb2,Msn,Bcl3,Cflar,Slc24a4,Akap12,Casp4 | | 1.094e-02 | -4.52 | GSE17721\_CPG\_VS\_GARDIQUIMOD\_12H\_BMDC\_DN | MSigDB lists | GSE17721\_CPG\_VS\_GARDIQUIMOD\_12H\_BMDC\_DN | 165 | 7 | 12187 | 179 | Hspa5,Sgk1,Ier3,Cebpd,Hcar2,Cxcl1,Irf2 | | 1.094e-02 | -4.52 | GSE31082\_DN\_VS\_CD8\_SP\_THYMOCYTE\_DN | MSigDB lists | GSE31082\_DN\_VS\_CD8\_SP\_THYMOCYTE\_DN | 165 | 7 | 12187 | 179 | Nfkbiz,Mxd4,Rnf213,Herc6,Tap1,Msn,Rgs16 | | 1.094e-02 | -4.52 | GSE360\_L\_DONOVANI\_VS\_B\_MALAYI\_HIGH\_DOSE\_MAC\_DN | MSigDB lists | GSE360\_L\_DONOVANI\_VS\_B\_MALAYI\_HIGH\_DOSE\_MAC\_DN | 165 | 7 | 12187 | 179 | Tap1,Pygm,Tnfsf10,Maff,Ifitm3,Irf7,Vcam1 | | 1.094e-02 | -4.52 | GSE2770\_IL12\_AND\_TGFB\_ACT\_VS\_ACT\_CD4\_TCELL\_6H\_DN | MSigDB lists | GSE2770\_IL12\_AND\_TGFB\_ACT\_VS\_ACT\_CD4\_TCELL\_6H\_DN | 165 | 7 | 12187 | 179 | Cflar,Ptpn1,Hspa5,Map3k8,Tnf,Ccnd2,Gadd45g | | 1.106e-02 | -4.50 | GSE22443\_NAIVE\_VS\_ACT\_AND\_IL2\_TREATED\_CD8\_TCELL\_UP | MSigDB lists | GSE22443\_NAIVE\_VS\_ACT\_AND\_IL2\_TREATED\_CD8\_TCELL\_UP | 127 | 6 | 12187 | 179 | Ptpn1,Nfkb2,Map3k6,Rhoc,Dnajb1,Tnf | | 1.106e-02 | -4.50 | GSE37301\_PRO\_BCELL\_VS\_CD4\_TCELL\_DN | MSigDB lists | GSE37301\_PRO\_BCELL\_VS\_CD4\_TCELL\_DN | 127 | 6 | 12187 | 179 | Sox11,Gbp6,Vcam1,Usp18,Rtp4,Gbp4 | | 1.107e-02 | -4.50 | - | gene3d domains | 2.110.10.10 | 11 | 2 | 6647 | 99 | Prg4,Mmp8 | | 1.108e-02 | -4.50 | GPCR downstream signaling | REACTOME pathways | R-MMU-388396 | 342 | 12 | 6297 | 105 | C3ar1,Cxcl16,Hcar2,C5ar1,Cxcl9,Pik3r1,Gpr84,Pik3r5,Cxcl1,P2ry6,Csf2rb,Rhoc | | 1.115e-02 | -4.50 | zf-RING\_UBOX | pfam domains | PF13445 | 30 | 3 | 12881 | 201 | Trim30a,Trim25,Trim56 | | 1.116e-02 | -4.50 | negative regulation of alpha-beta T cell activation | biological process | GO:0046636 | 30 | 3 | 13711 | 214 | Il4ra,Lgals9,Runx1 | | 1.116e-02 | -4.50 | cellular oxidant detoxification | biological process | GO:0098869 | 30 | 3 | 13711 | 214 | S100a8,Hp,S100a9 | | 1.116e-02 | -4.50 | cellular response to corticosteroid stimulus | biological process | GO:0071384 | 30 | 3 | 13711 | 214 | Sgk1,Zfp36,Fam107a | | 1.116e-02 | -4.50 | regulation of T-helper cell differentiation | biological process | GO:0045622 | 30 | 3 | 13711 | 214 | Il4ra,Nfkbiz,Il6 | | 1.116e-02 | -4.50 | CD4-positive, alpha-beta T cell activation | biological process | GO:0035710 | 30 | 3 | 13711 | 214 | Bcl3,Il6,Gadd45g | | 1.128e-02 | -4.48 | GSE10325\_LUPUS\_BCELL\_VS\_LUPUS\_MYELOID\_DN | MSigDB lists | GSE10325\_LUPUS\_BCELL\_VS\_LUPUS\_MYELOID\_DN | 166 | 7 | 12187 | 179 | C5ar1,C3ar1,Plaur,Il1rn,Igsf6,Timp1,Rassf4 | | 1.128e-02 | -4.48 | GSE36476\_CTRL\_VS\_TSST\_ACT\_16H\_MEMORY\_CD4\_TCELL\_YOUNG\_DN | MSigDB lists | GSE36476\_CTRL\_VS\_TSST\_ACT\_16H\_MEMORY\_CD4\_TCELL\_YOUNG\_DN | 166 | 7 | 12187 | 179 | Rbm47,Tap1,Tnf,Ccl4,Tnip3,Tubb6,Ier3 | | 1.128e-02 | -4.48 | TONKS\_TARGETS\_OF\_RUNX1\_RUNX1T1\_FUSION\_MONOCYTE\_UP | MSigDB lists | TONKS\_TARGETS\_OF\_RUNX1\_RUNX1T1\_FUSION\_MONOCYTE\_UP | 166 | 7 | 12187 | 179 | Hspb1,Isg15,Tubb6,Usp18,Ccnd2,Ifi44,Hspa5 | | 1.128e-02 | -4.48 | KAECH\_DAY8\_EFF\_VS\_MEMORY\_CD8\_TCELL\_DN | MSigDB lists | KAECH\_DAY8\_EFF\_VS\_MEMORY\_CD8\_TCELL\_DN | 166 | 7 | 12187 | 179 | Tnfaip3,Rbm39,Zfp36,Sntb2,Socs3,Il4ra,Ccnd2 | | 1.128e-02 | -4.48 | GSE17721\_POLYIC\_VS\_CPG\_16H\_BMDC\_UP | MSigDB lists | GSE17721\_POLYIC\_VS\_CPG\_16H\_BMDC\_UP | 166 | 7 | 12187 | 179 | Parp12,Tnfaip3,Rhoc,Cdkn1a,Aff1,Irf2,Trim25 | | 1.128e-02 | -4.48 | MCBRYAN\_PUBERTAL\_BREAST\_6\_7WK\_UP | MSigDB lists | MCBRYAN\_PUBERTAL\_BREAST\_6\_7WK\_UP | 166 | 7 | 12187 | 179 | Sntb2,Ccnd2,Runx1,Pik3r1,Cp,Fzd4,Gbp7 | | 1.128e-02 | -4.48 | GSE33425\_CD161\_HIGH\_VS\_NEG\_CD8\_TCELL\_DN | MSigDB lists | GSE33425\_CD161\_HIGH\_VS\_NEG\_CD8\_TCELL\_DN | 166 | 7 | 12187 | 179 | Gbp3,Tagln2,Saa1,Gbp7,Tubb6,Bach1,Fstl1 | | 1.129e-02 | -4.48 | MACRO | prosite domains | PS51154 | 9 | 2 | 8845 | 164 | Parp14,Parp9 | | 1.131e-02 | -4.48 | PID\_NFAT\_TFPATHWAY | MSigDB lists | PID\_NFAT\_TFPATHWAY | 32 | 3 | 12187 | 179 | Tnf,Ptgs2,Ptpn1 | | 1.131e-02 | -4.48 | GO\_SENSORY\_PERCEPTION\_OF\_CHEMICAL\_STIMULUS | MSigDB lists | GO\_SENSORY\_PERCEPTION\_OF\_CHEMICAL\_STIMULUS | 32 | 3 | 12187 | 179 | Rtp4,Slc24a4,C5ar1 | | 1.131e-02 | -4.48 | MEDINA\_SMARCA4\_TARGETS | MSigDB lists | MEDINA\_SMARCA4\_TARGETS | 32 | 3 | 12187 | 179 | Tnfsf10,Nfkb2,Mxd4 | | 1.131e-02 | -4.48 | GO\_NEGATIVE\_REGULATION\_OF\_TYPE\_I\_INTERFERON\_PRODUCTION | MSigDB lists | GO\_NEGATIVE\_REGULATION\_OF\_TYPE\_I\_INTERFERON\_PRODUCTION | 32 | 3 | 12187 | 179 | Isg15,Trim25,Tnfaip3 | | 1.131e-02 | -4.48 | GNF2\_CD48 | MSigDB lists | GNF2\_CD48 | 32 | 3 | 12187 | 179 | Tnfaip8,Phf11b,Phf11d | | 1.131e-02 | -4.48 | HANN\_RESISTANCE\_TO\_BCL2\_INHIBITOR\_DN | MSigDB lists | HANN\_RESISTANCE\_TO\_BCL2\_INHIBITOR\_DN | 32 | 3 | 12187 | 179 | Cebpd,Hspb1,Sdc4 | | 1.131e-02 | -4.48 | HALLMARK\_ANGIOGENESIS | MSigDB lists | HALLMARK\_ANGIOGENESIS | 32 | 3 | 12187 | 179 | Timp1,Fstl1,Ccnd2 | | 1.131e-02 | -4.48 | KEGG\_ALDOSTERONE\_REGULATED\_SODIUM\_REABSORPTION | MSigDB lists | KEGG\_ALDOSTERONE\_REGULATED\_SODIUM\_REABSORPTION | 32 | 3 | 12187 | 179 | Pik3r5,Sgk1,Pik3r1 | | 1.134e-02 | -4.48 | Itch (itchy, E3 ubiquitin protein ligase) | protein interactions | 16396 | 28 | 3 | 6802 | 115 | Cflar,Ripk1,Ifitm3 | | 1.136e-02 | -4.48 | Regulation of IFNA signaling | REACTOME pathways | R-MMU-912694 | 10 | 2 | 6297 | 105 | Ptpn1,Usp18 | | 1.136e-02 | -4.48 | RHO GTPases Activate NADPH Oxidases | REACTOME pathways | R-MMU-5668599 | 10 | 2 | 6297 | 105 | Ncf1,Cybb | | 1.136e-02 | -4.48 | Interleukin-6 signaling | REACTOME pathways | R-MMU-1059683 | 10 | 2 | 6297 | 105 | Il6,Socs3 | | 1.137e-02 | -4.48 | SHAFFER\_IRF4\_TARGETS\_IN\_MYELOMA\_VS\_MATURE\_B\_LYMPHOCYTE | MSigDB lists | SHAFFER\_IRF4\_TARGETS\_IN\_MYELOMA\_VS\_MATURE\_B\_LYMPHOCYTE | 92 | 5 | 12187 | 179 | Cflar,Zfp36,Hspb1,Atf3,Tnfaip3 | | 1.137e-02 | -4.48 | GSE40666\_UNTREATED\_VS\_IFNA\_STIM\_CD8\_TCELL\_90MIN\_UP | MSigDB lists | GSE40666\_UNTREATED\_VS\_IFNA\_STIM\_CD8\_TCELL\_90MIN\_UP | 92 | 5 | 12187 | 179 | Tnfaip8,Rtp4,Parp9,Ifi44,Ifit3b | | 1.139e-02 | -4.48 | Extracellular matrix organization | REACTOME pathways | R-MMU-1474244 | 184 | 8 | 6297 | 105 | Thbs1,Timp1,Sdc4,Mmp8,Vcam1,Icam1,A2m,Col4a1 | | 1.141e-02 | -4.47 | GO\_ENDOCYTIC\_VESICLE | MSigDB lists | GO\_ENDOCYTIC\_VESICLE | 207 | 8 | 12187 | 179 | Ncf1,Fzd4,Saa1,Irgm2,Msr1,Ccl12,Hp,Cybb | | 1.143e-02 | -4.47 | GO\_IDENTICAL\_PROTEIN\_BINDING | MSigDB lists | GO\_IDENTICAL\_PROTEIN\_BINDING | 946 | 23 | 12187 | 179 | Hspb1,Plek,Ptgs2,Atf3,Ifi204,Ifi207,Zbtb16,Ifit3b,Ifi209,Xdh,Fzd4,Nfkbia,Tnf,Runx1,Tap1,Irak3,Pik3ap1,Nfkb1,Thbs1,Ifi211,Angptl4,Ripk1,Ccl4 | | 1.143e-02 | -4.47 | mouse chr3\|3 H1 | chromosome location | mouse chr3\|3 H1 | 11 | 2 | 14556 | 220 | Gbp3,Gbp7 | | 1.144e-02 | -4.47 | MARTINEZ\_TP53\_TARGETS\_UP | MSigDB lists | MARTINEZ\_TP53\_TARGETS\_UP | 481 | 14 | 12187 | 179 | Thbs1,Sox11,Cdkn1a,Dnajb1,Cp,Ms4a6d,Atf3,Ier3,Ptgs2,Msr1,Ccnd2,Trib1,Hcar2,S100a8 | | 1.146e-02 | -4.47 | ONDER\_CDH1\_TARGETS\_1\_DN | MSigDB lists | ONDER\_CDH1\_TARGETS\_1\_DN | 128 | 6 | 12187 | 179 | Cxcl1,Adamts1,Csf3,Ptgs2,Lcn2,Osmr | | 1.146e-02 | -4.47 | GSE24671\_CTRL\_VS\_BAKIMULC\_INFECTED\_MOUSE\_SPLENOCYTES\_DN | MSigDB lists | GSE24671\_CTRL\_VS\_BAKIMULC\_INFECTED\_MOUSE\_SPLENOCYTES\_DN | 128 | 6 | 12187 | 179 | Fstl1,Nfkbiz,Nfkbia,Cdkn1a,Tnfaip3,Gadd45b | | 1.146e-02 | -4.47 | GSE4590\_SMALL\_VS\_LARGE\_PRE\_BCELL\_UP | MSigDB lists | GSE4590\_SMALL\_VS\_LARGE\_PRE\_BCELL\_UP | 128 | 6 | 12187 | 179 | Gadd45b,Ier3,Cybb,Pik3ap1,S100a8,Cxcl1 | | 1.146e-02 | -4.47 | GSE10325\_LUPUS\_CD4\_TCELL\_VS\_LUPUS\_BCELL\_DN | MSigDB lists | GSE10325\_LUPUS\_CD4\_TCELL\_VS\_LUPUS\_BCELL\_DN | 128 | 6 | 12187 | 179 | Map3k8,Cybb,Tubb6,Csf2rb,Clic4,Ptpn1 | | 1.151e-02 | -4.46 | response to peptide | biological process | GO:1901652 | 276 | 10 | 13711 | 214 | Tnfaip3,Icam1,Casp4,Nfkbia,Ptpn1,Nfkb1,Vcam1,Serpina3f,Sgk1,Pik3r1 | | 1.156e-02 | -4.46 | sulfur compound binding | molecular function | GO:1901681 | 195 | 8 | 13516 | 211 | Fstl1,Ccl7,Saa1,Ptges,Ccl2,Selp,Adamts1,Thbs1 | | 1.158e-02 | -4.46 | YAO\_HOXA10\_TARGETS\_VIA\_PROGESTERONE\_UP | MSigDB lists | YAO\_HOXA10\_TARGETS\_VIA\_PROGESTERONE\_UP | 60 | 4 | 12187 | 179 | Gem,Pik3r1,Lcn2,Xdh | | 1.158e-02 | -4.46 | ADDYA\_ERYTHROID\_DIFFERENTIATION\_BY\_HEMIN | MSigDB lists | ADDYA\_ERYTHROID\_DIFFERENTIATION\_BY\_HEMIN | 60 | 4 | 12187 | 179 | Hspa5,Cxcl1,Atf3,Ier3 | | 1.160e-02 | -4.46 | GO\_REGULATION\_OF\_CELLULAR\_RESPONSE\_TO\_STRESS | MSigDB lists | GO\_REGULATION\_OF\_CELLULAR\_RESPONSE\_TO\_STRESS | 581 | 16 | 12187 | 179 | Fzd4,Xdh,Pik3r1,Tnf,Nfe2l2,Gadd45b,Ripk1,Map3k6,Hspb1,Ptpn1,Hspa5,Dnajb1,Il1a,Gadd45g,Hp,Ier3 | | 1.164e-02 | -4.45 | cell-cell adhesion | biological process | GO:0098609 | 319 | 11 | 13711 | 214 | Sele,Lgals9,Tnip1,Plek,Icam1,Tnf,S100a9,Vcam1,Msn,Selp,S100a8 | | 1.164e-02 | -4.45 | GSE17812\_WT\_VS\_THPOK\_KO\_MEMORY\_CD8\_TCELL\_DN | MSigDB lists | GSE17812\_WT\_VS\_THPOK\_KO\_MEMORY\_CD8\_TCELL\_DN | 167 | 7 | 12187 | 179 | Bach1,Cebpd,Birc3,Iqgap1,Tnfaip8,Sbno2,Nfkbie | | 1.164e-02 | -4.45 | GSE17721\_POLYIC\_VS\_GARDIQUIMOD\_1H\_BMDC\_DN | MSigDB lists | GSE17721\_POLYIC\_VS\_GARDIQUIMOD\_1H\_BMDC\_DN | 167 | 7 | 12187 | 179 | Errfi1,Ccl12,Hcar2,Il1a,Ccl2,Maff,Msr1 | | 1.164e-02 | -4.45 | GSE6674\_CPG\_VS\_CPG\_AND\_ANTI\_IGM\_STIM\_BCELL\_UP | MSigDB lists | GSE6674\_CPG\_VS\_CPG\_AND\_ANTI\_IGM\_STIM\_BCELL\_UP | 167 | 7 | 12187 | 179 | Tor3a,Lgals9,Ifit2,Tap1,Akap12,Cybb,Trim25 | | 1.164e-02 | -4.45 | GSE9988\_ANTI\_TREM1\_VS\_VEHICLE\_TREATED\_MONOCYTES\_DN | MSigDB lists | GSE9988\_ANTI\_TREM1\_VS\_VEHICLE\_TREATED\_MONOCYTES\_DN | 167 | 7 | 12187 | 179 | Sgk3,Pik3r5,Bcl3,Cd14,Cybb,Pik3ap1,Cxcl16 | | 1.164e-02 | -4.45 | GSE37301\_GRANULOCYTE\_MONOCYTE\_PROGENITOR\_VS\_RAG2\_KO\_NK\_CELL\_DN | MSigDB lists | GSE37301\_GRANULOCYTE\_MONOCYTE\_PROGENITOR\_VS\_RAG2\_KO\_NK\_CELL\_DN | 167 | 7 | 12187 | 179 | Col4a1,Osmr,Tiparp,Tubb6,Lcn2,Saa1,A2m | | 1.164e-02 | -4.45 | GSE45365\_WT\_VS\_IFNAR\_KO\_CD8A\_DC\_MCMV\_INFECTION\_UP | MSigDB lists | GSE45365\_WT\_VS\_IFNAR\_KO\_CD8A\_DC\_MCMV\_INFECTION\_UP | 167 | 7 | 12187 | 179 | Rassf4,Parp12,Ripk1,Sdc4,Ccl2,Icam1,Tnfsf10 | | 1.165e-02 | -4.45 | EBV LMP1 signaling | WikiPathways | WP1243 | 19 | 3 | 3756 | 96 | Nfkb2,Tnf,Nfkb1 | | 1.166e-02 | -4.45 | regulation of protein transport | biological process | GO:0051223 | 593 | 17 | 13711 | 214 | Cd14,Lgals9,Pik3r1,Hcar2,Akap12,Il1a,Tnf,Il4ra,Tnfaip3,Casp4,Ccl2,Il6,Mmp8,Il17ra,Ptgs2,Ptpn1,Gbp4 | | 1.167e-02 | -4.45 | regulation of cellular component organization | biological process | GO:0051128 | 2181 | 47 | 13711 | 214 | Sgk1,Resf1,Ier3,Xaf1,Rasip1,Spi1,Sdc4,Fzd4,Fam107a,Pik3r1,Il6,Ip6k2,Cdkn1a,Il4ra,Il1a,Msn,Csf2rb,Selp,Nfe2l2,Dnajb1,Hspa5,Rhoc,Trim30a,Hspb1,Sele,Cflar,Cd14,Rhou,Tnf,Plaur,Irgm1,Rhoj,Adamts1,Icam1,Irgm2,S100a9,Iqgap1,Cxcl16,Ccl12,Ptpn1,Lcn2,Irak3,Csf3,Tnfsf10,Ccl2,Cxcl9,Plek | | 1.167e-02 | -4.45 | Keap1 (kelch-like ECH-associated protein 1) | protein interactions | 50868 | 10 | 2 | 6802 | 115 | Zbtb16,Nfe2l2 | | 1.167e-02 | -4.45 | Epor (erythropoietin receptor) | protein interactions | 13857 | 10 | 2 | 6802 | 115 | Pik3r1,Ptpn1 | | 1.172e-02 | -4.45 | cell periphery | cellular component | GO:0071944 | 3578 | 70 | 13825 | 212 | Hcar2,Nfe2l2,Irak3,Fam107a,Pik3ap1,Akap12,Hspb1,C5ar1,Ripk1,Plek,Clic4,Errfi1,Stx11,Cybb,Plaur,Il6,Plin4,Tnfsf10,S100a9,Ifitm6,Rhoj,Pik3r1,Slc24a4,Selp,P2ry6,Il4ra,Sntb2,Rhou,Tgm2,Pik3r5,Casp4,Ifitm3,Iqgap1,Cd14,Il17ra,Nfkbia,Osmr,Fyb,Rasd1,Ncf1,Gpr84,Robo4,Zbtb16,Vcam1,Msn,Sgk1,Sele,Il1a,Rrp8,Icam1,Rhoc,Ifitm2,Ptpn1,Ksr1,Kcna5,Bcl3,S100a8,Cemip2,C3ar1,Hspa5,Irgm1,Gem,Fzd4,Ptgs2,Cflar,Tnf,Thbs1,Cxcl9,Cp,Msr1 | | 1.173e-02 | -4.45 | p53-like\_TF\_DNA-bd | interpro domains | IPR008967 | 31 | 3 | 13788 | 212 | Nfkb1,Nfkb2,Runx1 | | 1.181e-02 | -4.44 | Caspase-like\_dom\_sf | interpro domains | IPR029030 | 11 | 2 | 13788 | 212 | Casp4,Cflar | | 1.181e-02 | -4.44 | Pept\_C14\_p20 | interpro domains | IPR001309 | 11 | 2 | 13788 | 212 | Casp4,Cflar | | 1.181e-02 | -4.44 | S100/CaBP-9k\_CS | interpro domains | IPR001751 | 11 | 2 | 13788 | 212 | S100a8,S100a9 | | 1.181e-02 | -4.44 | 4\_helix\_cytokine-like\_core | interpro domains | IPR009079 | 11 | 2 | 13788 | 212 | Il6,Csf3 | | 1.181e-02 | -4.44 | Hemopexin-like\_dom | interpro domains | IPR000585 | 11 | 2 | 13788 | 212 | Mmp8,Prg4 | | 1.181e-02 | -4.44 | Hemopexin-like\_repeat | interpro domains | IPR018487 | 11 | 2 | 13788 | 212 | Prg4,Mmp8 | | 1.181e-02 | -4.44 | Ribosomal\_L7Ae/L30e/S12e/Gad45 | interpro domains | IPR004038 | 11 | 2 | 13788 | 212 | Gadd45g,Gadd45b | | 1.181e-02 | -4.44 | Pept\_C14 | interpro domains | IPR002398 | 11 | 2 | 13788 | 212 | Cflar,Casp4 | | 1.181e-02 | -4.44 | Hemopexin-like\_dom\_sf | interpro domains | IPR036375 | 11 | 2 | 13788 | 212 | Mmp8,Prg4 | | 1.183e-02 | -4.44 | GO\_REGULATION\_OF\_INTRACELLULAR\_TRANSPORT | MSigDB lists | GO\_REGULATION\_OF\_INTRACELLULAR\_TRANSPORT | 483 | 14 | 12187 | 179 | Cdkn1a,Ptpn1,Msn,Ptgs2,Ccl12,Il6,Nfkbia,Tnf,Rhou,Il4ra,Pik3r1,Bcl3,Nfkbie,Csf3 | | 1.183e-02 | -4.44 | small molecule binding | molecular function | GO:0036094 | 2015 | 44 | 13516 | 211 | Hspa5,Parp14,Oasl1,Irak3,Ripk1,Lcn2,Oasl2,Rasd1,Sgk1,P2ry6,Tap1,S100a8,Sgk3,Cybb,Mx1,Sele,Gbp7,Rhoc,Hcar2,Tgtp1,Gbp3,Trib1,Iigp1,Slfn5,Xdh,Rhou,Runx1,Ksr1,Slfn9,Tor3a,Map3k6,Tubb6,Irgm1,Ip6k2,Cmpk2,Pygm,S100a9,Gem,Map3k8,Selp,Rhoj,Bbs12,Tgm2,Gbp6 | | 1.190e-02 | -4.43 | negative regulation of RNA biosynthetic process | biological process | GO:1902679 | 992 | 25 | 13711 | 214 | Sox11,Mxd4,Spi1,Cebpd,Ifi211,Bach1,Runx1,Gbp4,Ifi209,Sbno2,Nfkbie,Parp9,Nfkb1,Zbtb16,Arid5b,Ifi207,Ifi204,Znfx1,Tnf,Zfp36,Atf3,Rasd1,Bcl3,Dnajb1,Rrp8 | | 1.190e-02 | -4.43 | negative regulation of nucleic acid-templated transcription | biological process | GO:1903507 | 992 | 25 | 13711 | 214 | Ifi204,Arid5b,Ifi207,Zbtb16,Atf3,Zfp36,Znfx1,Tnf,Bcl3,Rasd1,Rrp8,Dnajb1,Mxd4,Sox11,Runx1,Cebpd,Ifi211,Bach1,Spi1,Nfkb1,Sbno2,Nfkbie,Parp9,Ifi209,Gbp4 | | 1.195e-02 | -4.43 | CASc | smart domains | SM00115 | 10 | 2 | 7188 | 123 | Casp4,Cflar | | 1.200e-02 | -4.42 | GSE12392\_WT\_VS\_IFNB\_KO\_CD8A\_NEG\_SPLEEN\_DC\_DN | MSigDB lists | GSE12392\_WT\_VS\_IFNB\_KO\_CD8A\_NEG\_SPLEEN\_DC\_DN | 168 | 7 | 12187 | 179 | Usp18,Bach1,Nfkbie,Bcl3,Igsf6,Plaur,Msn | | 1.200e-02 | -4.42 | GSE24142\_EARLY\_THYMIC\_PROGENITOR\_VS\_DN3\_THYMOCYTE\_UP | MSigDB lists | GSE24142\_EARLY\_THYMIC\_PROGENITOR\_VS\_DN3\_THYMOCYTE\_UP | 168 | 7 | 12187 | 179 | Plek,Ncf1,Tagln2,Ier3,Tubb6,Spi1,Il4ra | | 1.200e-02 | -4.42 | GSE17721\_LPS\_VS\_GARDIQUIMOD\_6H\_BMDC\_UP | MSigDB lists | GSE17721\_LPS\_VS\_GARDIQUIMOD\_6H\_BMDC\_UP | 168 | 7 | 12187 | 179 | Parp14,Ccl4,Ifit3b,Gbp4,Gadd45b,Gbp6,Icam1 | | 1.200e-02 | -4.42 | GSE37301\_MULTIPOTENT\_PROGENITOR\_VS\_RAG2\_KO\_NK\_CELL\_DN | MSigDB lists | GSE37301\_MULTIPOTENT\_PROGENITOR\_VS\_RAG2\_KO\_NK\_CELL\_DN | 168 | 7 | 12187 | 179 | Il1rn,Col4a1,Gadd45b,Lcn2,Usp18,Icam1,Osmr | | 1.200e-02 | -4.42 | GSE17301\_CTRL\_VS\_48H\_ACD3\_ACD28\_IFNA2\_STIM\_CD8\_TCELL\_UP | MSigDB lists | GSE17301\_CTRL\_VS\_48H\_ACD3\_ACD28\_IFNA2\_STIM\_CD8\_TCELL\_UP | 168 | 7 | 12187 | 179 | Csf2rb,Il1rn,Rassf4,Cxcl1,Nfkbia,Maff,Birc3 | | 1.200e-02 | -4.42 | GSE23505\_IL6\_IL1\_VS\_IL6\_IL1\_IL23\_TREATED\_CD4\_TCELL\_DN | MSigDB lists | GSE23505\_IL6\_IL1\_VS\_IL6\_IL1\_IL23\_TREATED\_CD4\_TCELL\_DN | 168 | 7 | 12187 | 179 | Rbm47,Il17ra,Clic4,Birc3,Map3k8,Aff1,Bach1 | | 1.200e-02 | -4.42 | GSE4984\_GALECTIN1\_VS\_VEHICLE\_CTRL\_TREATED\_DC\_DN | MSigDB lists | GSE4984\_GALECTIN1\_VS\_VEHICLE\_CTRL\_TREATED\_DC\_DN | 168 | 7 | 12187 | 179 | Parp9,Clic4,Gbp3,Ifi204,Ifi207,Ifi211,Ifi209 | | 1.204e-02 | -4.42 | CONCANNON\_APOPTOSIS\_BY\_EPOXOMICIN\_UP | MSigDB lists | CONCANNON\_APOPTOSIS\_BY\_EPOXOMICIN\_UP | 209 | 8 | 12187 | 179 | Hspb1,Zfp36,Gem,Atf3,Gadd45g,Dnajb1,Cdkn1a,Maff | | 1.204e-02 | -4.42 | YKACATTT\_UNKNOWN | MSigDB lists | YKACATTT\_UNKNOWN | 209 | 8 | 12187 | 179 | Znfx1,Zfp189,Pik3r1,Thbs1,Rbm39,Ptpn1,Pygm,Tnfsf10 | | 1.204e-02 | -4.42 | positive regulation of RNA metabolic process | biological process | GO:0051254 | 1360 | 32 | 13711 | 214 | Bach1,Cebpd,Spi1,Runx1,Fzd4,Nfkb1,Parp9,Sbno2,Dtx3l,Irf7,Rbm39,Banp,Pik3r1,Nfkbia,Nfkb2,Il1a,Sox11,Il6,Nfe2l2,Bcl3,Maff,Irf2,Aff1,Ncf1,Arid5b,Zbtb16,Ripk1,Csf3,Tnip1,Tnf,Zfp36,Atf3 | | 1.212e-02 | -4.41 | guanosine nucleotides degradation III | BIOCYC pathways | MOUSE\_PWY-6608 | 2 | 1 | 823 | 5 | Xdh | | 1.215e-02 | -4.41 | Ribosomal\_L7Ae | pfam domains | PF01248 | 11 | 2 | 12881 | 201 | Gadd45g,Gadd45b | | 1.215e-02 | -4.41 | Hemopexin | pfam domains | PF00045 | 11 | 2 | 12881 | 201 | Mmp8,Prg4 | | 1.215e-02 | -4.41 | cellular response to nitric oxide | biological process | GO:0071732 | 11 | 2 | 13711 | 214 | Thbs1,Cflar | | 1.215e-02 | -4.41 | regulation of cell proliferation involved in kidney development | biological process | GO:1901722 | 11 | 2 | 13711 | 214 | Cflar,C3ar1 | | 1.215e-02 | -4.41 | regulation of chemokine biosynthetic process | biological process | GO:0045073 | 11 | 2 | 13711 | 214 | Tnf,Il6 | | 1.215e-02 | -4.41 | protein auto-ADP-ribosylation | biological process | GO:0070213 | 11 | 2 | 13711 | 214 | Parp12,Tiparp | | 1.215e-02 | -4.41 | negative regulation of natural killer cell mediated immunity | biological process | GO:0002716 | 11 | 2 | 13711 | 214 | Lgals9,Tap1 | | 1.215e-02 | -4.41 | positive regulation of heat generation | biological process | GO:0031652 | 11 | 2 | 13711 | 214 | Tnf,Ptgs2 | | 1.215e-02 | -4.41 | respiratory burst | biological process | GO:0045730 | 11 | 2 | 13711 | 214 | Cybb,Ncf1 | | 1.215e-02 | -4.41 | negative regulation of alcohol biosynthetic process | biological process | GO:1902931 | 11 | 2 | 13711 | 214 | Plek,Nfkb1 | | 1.215e-02 | -4.41 | membrane fission | biological process | GO:0090148 | 11 | 2 | 13711 | 214 | Mx2,Mx1 | | 1.215e-02 | -4.41 | negative regulation of natural killer cell mediated cytotoxicity | biological process | GO:0045953 | 11 | 2 | 13711 | 214 | Tap1,Lgals9 | | 1.215e-02 | -4.41 | nuclear-transcribed mRNA poly(A) tail shortening | biological process | GO:0000289 | 11 | 2 | 13711 | 214 | Zfp36,Noct | | 1.216e-02 | -4.41 | interleukin-1 receptor binding | molecular function | GO:0005149 | 11 | 2 | 13516 | 211 | Il1rn,Il1a | | 1.216e-02 | -4.41 | cysteine-type endopeptidase activity involved in apoptotic process | molecular function | GO:0097153 | 11 | 2 | 13516 | 211 | Cflar,Casp4 | | 1.222e-02 | -4.40 | negative regulation of anion transport | biological process | GO:1903792 | 31 | 3 | 13711 | 214 | Il1rn,Thbs1,Tnf | | 1.222e-02 | -4.40 | regulation of anion transmembrane transport | biological process | GO:1903959 | 31 | 3 | 13711 | 214 | Ripk1,Tnf,Thbs1 | | 1.222e-02 | -4.40 | spleen development | biological process | GO:0048536 | 31 | 3 | 13711 | 214 | Nfkb2,Bcl3,Samd9l | | 1.226e-02 | -4.40 | RADMACHER\_AML\_PROGNOSIS | MSigDB lists | RADMACHER\_AML\_PROGNOSIS | 61 | 4 | 12187 | 179 | Nfkb1,Lcn2,Rgs16,Tgm2 | | 1.226e-02 | -4.40 | GO\_REGULATION\_OF\_LIPASE\_ACTIVITY | MSigDB lists | GO\_REGULATION\_OF\_LIPASE\_ACTIVITY | 61 | 4 | 12187 | 179 | C5ar1,Angptl4,Rhoc,Sele | | 1.226e-02 | -4.40 | GO\_ORGAN\_REGENERATION | MSigDB lists | GO\_ORGAN\_REGENERATION | 61 | 4 | 12187 | 179 | C5ar1,Cdkn1a,Socs3,Ccl12 | | 1.229e-02 | -4.40 | identical protein binding | molecular function | GO:0042802 | 1578 | 36 | 13516 | 211 | Fzd4,Cebpd,Ptgs2,Olfml2b,Tap1,Sdc4,A2m,Lcn2,Tnip1,Atf3,Pik3ap1,Plek,Gadd45g,Rasip1,Ifi204,Tnfsf10,Hp,Iigp1,Ifit3b,Gbp3,Tnfaip3,Tnf,Banp,Ifi209,Ripk1,Zbtb16,Angptl4,Nfkbia,Nfkb1,Ccl4,Ifi207,Ifi211,Hspb1,Irak3,Runx1,Xdh | | 1.230e-02 | -4.40 | GO\_REGULATION\_OF\_ALPHA\_BETA\_T\_CELL\_DIFFERENTIATION | MSigDB lists | GO\_REGULATION\_OF\_ALPHA\_BETA\_T\_CELL\_DIFFERENTIATION | 33 | 3 | 12187 | 179 | Il6,Il4ra,Zbtb16 | | 1.231e-02 | -4.40 | GSE29617\_DAY3\_VS\_DAY7\_TIV\_FLU\_VACCINE\_PBMC\_2008\_UP | MSigDB lists | GSE29617\_DAY3\_VS\_DAY7\_TIV\_FLU\_VACCINE\_PBMC\_2008\_UP | 130 | 6 | 12187 | 179 | Pik3ap1,Plek,Ifit2,Atf3,Il6,Stx11 | | 1.231e-02 | -4.40 | GSE12366\_NAIVE\_VS\_MEMORY\_BCELL\_UP | MSigDB lists | GSE12366\_NAIVE\_VS\_MEMORY\_BCELL\_UP | 130 | 6 | 12187 | 179 | Aff1,Il4ra,Birc3,Ifi44,Parp14,Il6 | | 1.232e-02 | -4.40 | GO\_NEGATIVE\_REGULATION\_OF\_DEVELOPMENTAL\_PROCESS | MSigDB lists | GO\_NEGATIVE\_REGULATION\_OF\_DEVELOPMENTAL\_PROCESS | 585 | 16 | 12187 | 179 | Sox11,Thbs1,Zbtb16,Timp1,Bbs12,Runx1,Tnf,Il4ra,Pik3r1,Nfkbia,Trib1,Ccl12,Nfe2l2,Il6,Cflar,Xdh | | 1.236e-02 | -4.39 | IK1\_01 | MSigDB lists | IK1\_01 | 210 | 8 | 12187 | 179 | Nfkb2,Apold1,Icam1,Bcl3,Tnip1,Gadd45b,Il6,Luc7l3 | | 1.236e-02 | -4.39 | GO\_NEGATIVE\_REGULATION\_OF\_CATABOLIC\_PROCESS | MSigDB lists | GO\_NEGATIVE\_REGULATION\_OF\_CATABOLIC\_PROCESS | 169 | 7 | 12187 | 179 | Banp,Irak3,Tnf,Rasip1,Hp,Hcar2,Timp1 | | 1.236e-02 | -4.39 | GSE43863\_LY6C\_INT\_CXCR5POS\_VS\_LY6C\_LOW\_CXCR5NEG\_EFFECTOR\_CD4\_TCELL\_DN | MSigDB lists | GSE43863\_LY6C\_INT\_CXCR5POS\_VS\_LY6C\_LOW\_CXCR5NEG\_EFFECTOR\_CD4\_TCELL\_DN | 169 | 7 | 12187 | 179 | Plek,Tiparp,Atf3,AA467197,Hspa5,Casp4,Ptges | | 1.236e-02 | -4.39 | AFFAR\_YY1\_TARGETS\_UP | MSigDB lists | AFFAR\_YY1\_TARGETS\_UP | 169 | 7 | 12187 | 179 | Tap1,Gbp4,Gadd45g,Icam1,Tgm2,Gbp6,Cdkn1a | | 1.236e-02 | -4.39 | GO\_G\_PROTEIN\_COUPLED\_RECEPTOR\_BINDING | MSigDB lists | GO\_G\_PROTEIN\_COUPLED\_RECEPTOR\_BINDING | 169 | 7 | 12187 | 179 | Ccl12,Ccl2,Ccl4,Rtp4,Cxcl16,Saa1,Cxcl1 | | 1.236e-02 | -4.39 | GSE23308\_CTRL\_VS\_CORTICOSTERONE\_TREATED\_MACROPHAGE\_MINERALCORTICOID\_REC\_KO\_DN | MSigDB lists | GSE23308\_CTRL\_VS\_CORTICOSTERONE\_TREATED\_MACROPHAGE\_MINERALCORTICOID\_REC\_KO\_DN | 169 | 7 | 12187 | 179 | Vcam1,Nfkb1,Il1a,Nfkbie,Gem,Trim25,Cmpk2 | | 1.236e-02 | -4.39 | KYNG\_DNA\_DAMAGE\_DN | MSigDB lists | KYNG\_DNA\_DAMAGE\_DN | 169 | 7 | 12187 | 179 | Irf2,Csf2rb,Lcn2,Akap12,Cxcl1,Adamts1,Saa1 | | 1.236e-02 | -4.39 | GSE41867\_LCMV\_ARMSTRONG\_VS\_CLONE13\_DAY8\_EFFECTOR\_CD8\_TCELL\_UP | MSigDB lists | GSE41867\_LCMV\_ARMSTRONG\_VS\_CLONE13\_DAY8\_EFFECTOR\_CD8\_TCELL\_UP | 169 | 7 | 12187 | 179 | Ccl4,Sntb2,Trib1,Rasd1,Rasip1,Ifitm2,Rbm47 | | 1.236e-02 | -4.39 | GSE40273\_XBP1\_KO\_VS\_WT\_TREG\_UP | MSigDB lists | GSE40273\_XBP1\_KO\_VS\_WT\_TREG\_UP | 169 | 7 | 12187 | 179 | Rtp4,Irf7,Slfn9,Gadd45g,Akap12,Oasl1,Sdc4 | | 1.236e-02 | -4.39 | GSE17186\_MEMORY\_VS\_CD21HIGH\_TRANSITIONAL\_BCELL\_DN | MSigDB lists | GSE17186\_MEMORY\_VS\_CD21HIGH\_TRANSITIONAL\_BCELL\_DN | 169 | 7 | 12187 | 179 | Tap1,Iqgap1,Irgm2,Gbp6,Sgk3,Gbp4,Ifit2 | | 1.236e-02 | -4.39 | GSE20715\_WT\_VS\_TLR4\_KO\_24H\_OZONE\_LUNG\_DN | MSigDB lists | GSE20715\_WT\_VS\_TLR4\_KO\_24H\_OZONE\_LUNG\_DN | 169 | 7 | 12187 | 179 | Cdkn1a,Ier3,Gadd45b,Col4a1,Angptl4,Tap1,Trim25 | | 1.236e-02 | -4.39 | GO\_SULFUR\_COMPOUND\_BINDING | MSigDB lists | GO\_SULFUR\_COMPOUND\_BINDING | 169 | 7 | 12187 | 179 | Ccl12,Adamts1,Thbs1,Fstl1,Selp,Saa1,Ptges | | 1.240e-02 | -4.39 | GO\_LEUKOCYTE\_MEDIATED\_IMMUNITY | MSigDB lists | GO\_LEUKOCYTE\_MEDIATED\_IMMUNITY | 94 | 5 | 12187 | 179 | Icam1,Il6,Il4ra,Bcl3,Irf7 | | 1.240e-02 | -4.39 | GSE32901\_NAIVE\_VS\_TH1\_CD4\_TCELL\_UP | MSigDB lists | GSE32901\_NAIVE\_VS\_TH1\_CD4\_TCELL\_UP | 94 | 5 | 12187 | 179 | Thbs1,Parp14,Il17ra,Irak3,Rtp4 | | 1.241e-02 | -4.39 | positive regulation of apoptotic signaling pathway | biological process | GO:2001235 | 159 | 7 | 13711 | 214 | S100a9,Ripk1,Thbs1,Plaur,Tnfsf10,Atf3,S100a8 | | 1.249e-02 | -4.38 | ApoE and miR-146 in inflammation and atherosclerosis | WikiPathways | WP3592 | 7 | 2 | 3756 | 96 | Spi1,Nfkb2 | | 1.249e-02 | -4.38 | Jak2 (Janus kinase 2) | protein interactions | 16452 | 29 | 3 | 6802 | 115 | Pik3r1,Ptpn1,Socs3 | | 1.251e-02 | -4.38 | GO\_SIGNALING\_RECEPTOR\_ACTIVITY | MSigDB lists | GO\_SIGNALING\_RECEPTOR\_ACTIVITY | 586 | 16 | 12187 | 179 | Cd14,Icam1,Il17ra,P2ry6,Sele,Csf2rb,Plaur,C5ar1,C3ar1,Fzd4,Igsf6,Osmr,Il4ra,Sdc4,Gpr84,Hcar2 | | 1.258e-02 | -4.38 | endoplasmic reticulum | cellular component | GO:0005783 | 1389 | 32 | 13825 | 212 | Ptpn1,Hp,Kcna5,Ksr1,F830016B08Rik,Tor3a,Tgtp1,Cybb,Gm5431,Irgm2,Vcam1,Xdh,Mx1,Pygm,Sgk1,Iigp1,Adamts9,Casp4,Ifitm3,Ccl2,Thbs1,Ifi47,Ncf1,Rasd1,Ptgs2,Gm4841,Tap1,Irgm1,Tgtp2,Pik3r1,Ifit2,Hspa5 | | 1.262e-02 | -4.37 | FAS pathway and Stress induction of HSP regulation | WikiPathways | WP571 | 36 | 4 | 3756 | 96 | Tnf,Il1a,Hspb1,Cflar | | 1.265e-02 | -4.37 | regulation of adaptive immune response based on somatic recombination of immune receptors built from immunoglobulin superfamily domains | biological process | GO:0002822 | 123 | 6 | 13711 | 214 | Lgals9,Il6,Nfkbiz,Il4ra,Tnfaip3,Tnf | | 1.269e-02 | -4.37 | negative regulation of reactive oxygen species metabolic process | biological process | GO:2000378 | 58 | 4 | 13711 | 214 | Trim30a,Hp,Cflar,Birc3 | | 1.269e-02 | -4.37 | positive regulation of fat cell differentiation | biological process | GO:0045600 | 58 | 4 | 13711 | 214 | Zbtb16,Zfp36,Noct,Ptgs2 | | 1.273e-02 | -4.36 | GO\_REGULATION\_OF\_CELL\_CELL\_ADHESION | MSigDB lists | GO\_REGULATION\_OF\_CELL\_CELL\_ADHESION | 254 | 9 | 12187 | 179 | Pik3r1,Il4ra,Map3k8,Tnf,Zbtb16,Vcam1,Il1rn,Il6,Ccl12 | | 1.273e-02 | -4.36 | positive regulation of endothelial cell migration | biological process | GO:0010595 | 89 | 5 | 13711 | 214 | Hspb1,Nfe2l2,Thbs1,Rhoj,Ptgs2 | | 1.274e-02 | -4.36 | GSE40273\_GATA1\_KO\_VS\_WT\_TREG\_DN | MSigDB lists | GSE40273\_GATA1\_KO\_VS\_WT\_TREG\_DN | 170 | 7 | 12187 | 179 | Plek,Ptpn1,Arid5b,Ncf1,Akap12,Map3k8,Birc3 | | 1.274e-02 | -4.36 | GSE17721\_LPS\_VS\_GARDIQUIMOD\_24H\_BMDC\_DN | MSigDB lists | GSE17721\_LPS\_VS\_GARDIQUIMOD\_24H\_BMDC\_DN | 170 | 7 | 12187 | 179 | Hspa5,Gpr84,Cp,Rhoc,Tnip1,P2ry6,S100a8 | | 1.274e-02 | -4.36 | GO\_NUCLEAR\_SPECK | MSigDB lists | GO\_NUCLEAR\_SPECK | 170 | 7 | 12187 | 179 | Ifi207,Ifi204,Luc7l3,Ifi209,Zbtb16,Rbm39,Ifi211 | | 1.274e-02 | -4.36 | GSE5542\_UNTREATED\_VS\_IFNG\_TREATED\_EPITHELIAL\_CELLS\_6H\_DN | MSigDB lists | GSE5542\_UNTREATED\_VS\_IFNG\_TREATED\_EPITHELIAL\_CELLS\_6H\_DN | 170 | 7 | 12187 | 179 | Mxd4,Slfn4,Pik3ap1,Slfn3,Slfn9,Cflar,Thbs1 | | 1.274e-02 | -4.36 | GSE36078\_UNTREATED\_VS\_AD5\_INF\_IL1R\_KO\_MOUSE\_LUNG\_DC\_UP | MSigDB lists | GSE36078\_UNTREATED\_VS\_AD5\_INF\_IL1R\_KO\_MOUSE\_LUNG\_DC\_UP | 170 | 7 | 12187 | 179 | Tnf,Xaf1,Nfkbia,Gadd45b,Tnip1,Arid5b,Irf7 | | 1.275e-02 | -4.36 | GSE34205\_RSV\_VS\_FLU\_INF\_INFANT\_PBMC\_DN | MSigDB lists | GSE34205\_RSV\_VS\_FLU\_INF\_INFANT\_PBMC\_DN | 131 | 6 | 12187 | 179 | Ifi44,Ccl4,Usp18,Oasl1,Herc6,Slfn5 | | 1.275e-02 | -4.36 | IGLESIAS\_E2F\_TARGETS\_UP | MSigDB lists | IGLESIAS\_E2F\_TARGETS\_UP | 131 | 6 | 12187 | 179 | Thbs1,Ccnd2,Nfkbia,Col4a1,Gadd45b,Gbp3 | | 1.275e-02 | -4.36 | GSE21033\_3H\_VS\_24H\_POLYIC\_STIM\_DC\_UP | MSigDB lists | GSE21033\_3H\_VS\_24H\_POLYIC\_STIM\_DC\_UP | 131 | 6 | 12187 | 179 | Pik3r5,Tnip1,Gbp3,Zfp36,Ip6k2,Map3k8 | | 1.282e-02 | -4.36 | Chronic myeloid leukemia | KEGG pathways | mmu05220 | 69 | 5 | 5248 | 107 | Pik3r1,Runx1,Cdkn1a,Nfkb1,Nfkbia | | 1.282e-02 | -4.36 | Chronic myeloid leukemia | KEGG pathways | ko05220 | 69 | 5 | 5248 | 107 | Nfkbia,Cdkn1a,Nfkb1,Runx1,Pik3r1 | | 1.283e-02 | -4.36 | nucleotide binding | molecular function | GO:0000166 | 1747 | 39 | 13516 | 211 | Map3k8,Sgk3,Cybb,Sgk1,P2ry6,Pygm,Rasd1,Oasl2,Gem,Tap1,Gbp7,Tgm2,Bbs12,Rhoc,Gbp6,Mx1,Rhoj,Iigp1,Parp14,Gbp3,Trib1,Slfn5,Hspa5,Tgtp1,Hcar2,Tubb6,Tor3a,Map3k6,Irgm1,Cmpk2,Ip6k2,Ripk1,Runx1,Rhou,Irak3,Oasl1,Xdh,Slfn9,Ksr1 | | 1.283e-02 | -4.36 | nucleoside phosphate binding | molecular function | GO:1901265 | 1747 | 39 | 13516 | 211 | Iigp1,Parp14,Gbp3,Trib1,Slfn5,Hspa5,Hcar2,Tgtp1,Tubb6,Tor3a,Map3k6,Ip6k2,Irgm1,Cmpk2,Ripk1,Runx1,Irak3,Rhou,Oasl1,Xdh,Slfn9,Ksr1,Map3k8,Sgk3,Cybb,Sgk1,P2ry6,Pygm,Rasd1,Oasl2,Gem,Tap1,Gbp7,Tgm2,Bbs12,Rhoc,Gbp6,Rhoj,Mx1 | | 1.285e-02 | -4.35 | VANDESLUIS\_COMMD1\_TARGETS\_GROUP\_2\_UP | MSigDB lists | VANDESLUIS\_COMMD1\_TARGETS\_GROUP\_2\_UP | 12 | 2 | 12187 | 179 | Maff,Ccnd2 | | 1.285e-02 | -4.35 | REACTOME\_DEGRADATION\_OF\_THE\_EXTRACELLULAR\_MATRIX | MSigDB lists | REACTOME\_DEGRADATION\_OF\_THE\_EXTRACELLULAR\_MATRIX | 12 | 2 | 12187 | 179 | Timp1,Mmp8 | | 1.285e-02 | -4.35 | DASU\_IL6\_SIGNALING\_SCAR\_DN | MSigDB lists | DASU\_IL6\_SIGNALING\_SCAR\_DN | 12 | 2 | 12187 | 179 | Thbs1,Sgk1 | | 1.285e-02 | -4.35 | KOBAYASHI\_EGFR\_SIGNALING\_6HR\_DN | MSigDB lists | KOBAYASHI\_EGFR\_SIGNALING\_6HR\_DN | 12 | 2 | 12187 | 179 | Ier3,Maff | | 1.285e-02 | -4.35 | GO\_REGULATION\_OF\_HETEROTYPIC\_CELL\_CELL\_ADHESION | MSigDB lists | GO\_REGULATION\_OF\_HETEROTYPIC\_CELL\_CELL\_ADHESION | 12 | 2 | 12187 | 179 | Il1rn,Tnf | | 1.285e-02 | -4.35 | NAKAJIMA\_EOSINOPHIL | MSigDB lists | NAKAJIMA\_EOSINOPHIL | 12 | 2 | 12187 | 179 | Plaur,Ccl4 | | 1.285e-02 | -4.35 | BIOCARTA\_GCR\_PATHWAY | MSigDB lists | BIOCARTA\_GCR\_PATHWAY | 12 | 2 | 12187 | 179 | Pik3r1,Nfkb1 | | 1.285e-02 | -4.35 | GO\_POSITIVE\_REGULATION\_OF\_TRANSCRIPTION\_FROM\_RNA\_POLYMERASE\_II\_PROMOTER\_IN\_RESPONSE\_TO\_ENDOPLASMIC\_RETICULUM\_STRESS | MSigDB lists | GO\_POSITIVE\_REGULATION\_OF\_TRANSCRIPTION\_FROM\_RNA\_POLYMERASE\_II\_PROMOTER\_IN\_RESPONSE\_TO\_ENDOPLASMIC\_RETICULUM\_STRESS | 12 | 2 | 12187 | 179 | Hspa5,Atf3 | | 1.285e-02 | -4.35 | HUMMERICH\_BENIGN\_SKIN\_TUMOR\_UP | MSigDB lists | HUMMERICH\_BENIGN\_SKIN\_TUMOR\_UP | 12 | 2 | 12187 | 179 | S100a9,S100a8 | | 1.285e-02 | -4.35 | PID\_IL5\_PATHWAY | MSigDB lists | PID\_IL5\_PATHWAY | 12 | 2 | 12187 | 179 | Csf2rb,Pik3r1 | | 1.285e-02 | -4.35 | CAIRO\_PML\_TARGETS\_BOUND\_BY\_MYC\_DN | MSigDB lists | CAIRO\_PML\_TARGETS\_BOUND\_BY\_MYC\_DN | 12 | 2 | 12187 | 179 | Cdkn1a,Irf9 | | 1.285e-02 | -4.35 | MATZUK\_EARLY\_ANTRAL\_FOLLICLE | MSigDB lists | MATZUK\_EARLY\_ANTRAL\_FOLLICLE | 12 | 2 | 12187 | 179 | Thbs1,Ccnd2 | | 1.285e-02 | -4.35 | MODULE\_407 | MSigDB lists | MODULE\_407 | 12 | 2 | 12187 | 179 | C5ar1,Gadd45g | | 1.294e-02 | -4.35 | SESTO\_RESPONSE\_TO\_UV\_C0 | MSigDB lists | SESTO\_RESPONSE\_TO\_UV\_C0 | 95 | 5 | 12187 | 179 | Zfp36,Il1rn,Tap1,Saa1,Nfkbia | | 1.294e-02 | -4.35 | LE\_EGR2\_TARGETS\_DN | MSigDB lists | LE\_EGR2\_TARGETS\_DN | 95 | 5 | 12187 | 179 | Tiparp,Errfi1,Pik3r1,Cdkn1a,Zbtb16 | | 1.296e-02 | -4.35 | MCBRYAN\_PUBERTAL\_BREAST\_6\_7WK\_DN | MSigDB lists | MCBRYAN\_PUBERTAL\_BREAST\_6\_7WK\_DN | 62 | 4 | 12187 | 179 | Hcar2,S100a8,Timp1,Ifit3b | | 1.302e-02 | -4.34 | GO\_REGULATION\_OF\_CELLULAR\_PROTEIN\_LOCALIZATION | MSigDB lists | GO\_REGULATION\_OF\_CELLULAR\_PROTEIN\_LOCALIZATION | 440 | 13 | 12187 | 179 | Nfkbia,Pik3r1,Tnf,Rhou,Il6,Ccl12,Bcl3,Nfkbie,Csf3,Cdkn1a,Ptgs2,Msn,Ptpn1 | | 1.313e-02 | -4.33 | GSE40277\_GATA1\_AND\_SATB1\_TRANSDUCED\_VS\_CTRL\_CD4\_TCELL\_DN | MSigDB lists | GSE40277\_GATA1\_AND\_SATB1\_TRANSDUCED\_VS\_CTRL\_CD4\_TCELL\_DN | 171 | 7 | 12187 | 179 | Trim25,Oasl1,Ifitm3,Ifi44,Ncf1,Irf7,Bcl3 | | 1.313e-02 | -4.33 | GSE17721\_LPS\_VS\_PAM3CSK4\_6H\_BMDC\_UP | MSigDB lists | GSE17721\_LPS\_VS\_PAM3CSK4\_6H\_BMDC\_UP | 171 | 7 | 12187 | 179 | Irf2,Slfn4,Timp1,Slfn3,Rtp4,Ifitm3,Sgk3 | | 1.313e-02 | -4.33 | GSE24726\_WT\_VS\_E2\_2\_KO\_PDC\_DAY4\_POST\_DELETION\_UP | MSigDB lists | GSE24726\_WT\_VS\_E2\_2\_KO\_PDC\_DAY4\_POST\_DELETION\_UP | 171 | 7 | 12187 | 179 | Nfe2l2,Cebpd,Cybb,Dnajb1,Tgm2,Birc3,Ifitm3 | | 1.315e-02 | -4.33 | MARSON\_BOUND\_BY\_FOXP3\_STIMULATED | MSigDB lists | MARSON\_BOUND\_BY\_FOXP3\_STIMULATED | 797 | 20 | 12187 | 179 | Angptl4,Ifi211,Arid5b,Xdh,Slfn4,Birc3,Gadd45b,Bach1,Bbs12,Ccdc58,Irf2,Slfn3,Ifi209,Sgk1,Ier3,Tnip3,Tiparp,Ifi204,Ms4a6d,Ifi207 | | 1.319e-02 | -4.33 | GSE3920\_IFNA\_VS\_IFNG\_TREATED\_ENDOTHELIAL\_CELL\_DN | MSigDB lists | GSE3920\_IFNA\_VS\_IFNG\_TREATED\_ENDOTHELIAL\_CELL\_DN | 132 | 6 | 12187 | 179 | Rgs16,Tnfsf10,Rbm47,Plek,Gbp6,Gbp4 | | 1.319e-02 | -4.33 | GSE360\_L\_MAJOR\_VS\_T\_GONDII\_MAC\_UP | MSigDB lists | GSE360\_L\_MAJOR\_VS\_T\_GONDII\_MAC\_UP | 132 | 6 | 12187 | 179 | Ptpn1,Plaur,Nfkb2,Ptgs2,Tnip1,Ptges | | 1.319e-02 | -4.33 | GSE17974\_CTRL\_VS\_ACT\_IL4\_AND\_ANTI\_IL12\_12H\_CD4\_TCELL\_UP | MSigDB lists | GSE17974\_CTRL\_VS\_ACT\_IL4\_AND\_ANTI\_IL12\_12H\_CD4\_TCELL\_UP | 132 | 6 | 12187 | 179 | Socs3,Pik3r5,Sgk1,Maff,Tnfaip3,Gadd45b | | 1.320e-02 | -4.33 | TGGAAA\_NFAT\_Q4\_01 | MSigDB lists | TGGAAA\_NFAT\_Q4\_01 | 1405 | 31 | 12187 | 179 | Ptges,S100a8,Tnfaip8,Maff,Map3k8,Ccl4,Il6,Gadd45b,Ccl12,Isg15,Ip6k2,Sgk1,Sox11,Ier3,Rbm39,Tnfsf10,Nfkb2,Runx1,Tnf,Nfkbia,Sdc4,Tubb6,Trim25,Icam1,Apold1,Errfi1,Cdkn1a,Fstl1,Il1rn,Kcna5,Znfx1 | | 1.331e-02 | -4.32 | myeloid leukocyte activation | biological process | GO:0002274 | 90 | 5 | 13711 | 214 | Stx11,Tnf,Spi1,Fyb,Sbno2 | | 1.333e-02 | -4.32 | positive regulation of interleukin-8 production | biological process | GO:0032757 | 32 | 3 | 13711 | 214 | Ripk1,Cd14,Tnf | | 1.335e-02 | -4.32 | PID\_GMCSF\_PATHWAY | MSigDB lists | PID\_GMCSF\_PATHWAY | 34 | 3 | 12187 | 179 | Pik3r1,Csf2rb,Ccl12 | | 1.335e-02 | -4.32 | TRAYNOR\_RETT\_SYNDROM\_UP | MSigDB lists | TRAYNOR\_RETT\_SYNDROM\_UP | 34 | 3 | 12187 | 179 | Il6,Akap12,Ifi44 | | 1.335e-02 | -4.32 | SIG\_PIP3\_SIGNALING\_IN\_B\_LYMPHOCYTES | MSigDB lists | SIG\_PIP3\_SIGNALING\_IN\_B\_LYMPHOCYTES | 34 | 3 | 12187 | 179 | Phf11d,Pik3r5,Phf11b | | 1.335e-02 | -4.32 | BIOCARTA\_IL2RB\_PATHWAY | MSigDB lists | BIOCARTA\_IL2RB\_PATHWAY | 34 | 3 | 12187 | 179 | Socs3,Cflar,Pik3r1 | | 1.335e-02 | -4.32 | MA\_MYELOID\_DIFFERENTIATION\_DN | MSigDB lists | MA\_MYELOID\_DIFFERENTIATION\_DN | 34 | 3 | 12187 | 179 | Ccl12,Cd14,Vcam1 | | 1.335e-02 | -4.32 | TSUNODA\_CISPLATIN\_RESISTANCE\_DN | MSigDB lists | TSUNODA\_CISPLATIN\_RESISTANCE\_DN | 34 | 3 | 12187 | 179 | Ptgs2,Sgk1,Ptges | | 1.335e-02 | -4.32 | GO\_STAT\_CASCADE | MSigDB lists | GO\_STAT\_CASCADE | 34 | 3 | 12187 | 179 | Socs3,Ptpn1,Ccl12 | | 1.345e-02 | -4.31 | regulation of epidermis development | biological process | GO:0045682 | 59 | 4 | 13711 | 214 | Tnf,Maff,Zfp36,Errfi1 | | 1.345e-02 | -4.31 | GO\_POSITIVE\_REGULATION\_OF\_CELL\_DIFFERENTIATION | MSigDB lists | GO\_POSITIVE\_REGULATION\_OF\_CELL\_DIFFERENTIATION | 642 | 17 | 12187 | 179 | Nfkb1,Sox11,Adamts1,Sgk1,Zbtb16,Socs3,Isg15,Ptgs2,Ripk1,Il6,Tnf,Runx1,Msr1,Il4ra,Trib1,Adamts9,Csf3 | | 1.349e-02 | -4.31 | GO\_REGULATION\_OF\_LEUKOCYTE\_MEDIATED\_IMMUNITY | MSigDB lists | GO\_REGULATION\_OF\_LEUKOCYTE\_MEDIATED\_IMMUNITY | 96 | 5 | 12187 | 179 | Ccl12,Tap1,Icam1,Tnf,Il4ra | | 1.351e-02 | -4.30 | animal organ development | biological process | GO:0048513 | 2256 | 48 | 13711 | 214 | Il1a,Il6,Cdkn1a,Socs3,Pik3r1,Tiparp,Fzd4,Sdc4,Sbno2,Cebpd,Rbm47,Spi1,Herc6,Runx1,Ptgs2,Tnf,Atf3,Slc24a4,Cflar,C5ar1,Maff,Tgm2,Clic4,Hspa5,Hp,Prg4,Msn,Icam1,Nfkb2,Sox11,Errfi1,Thbs1,Gadd45g,Nfkb1,Rhoj,Adamts1,Plaur,Tnfaip3,Xdh,Arid5b,Adamts9,Zbtb16,Csf3,Ifi204,Col4a1,Vcam1,Bcl3,Samd9l | | 1.352e-02 | -4.30 | GSE23568\_CTRL\_TRANSDUCED\_VS\_WT\_CD8\_TCELL\_UP | MSigDB lists | GSE23568\_CTRL\_TRANSDUCED\_VS\_WT\_CD8\_TCELL\_UP | 172 | 7 | 12187 | 179 | Saa1,Clic4,Iqgap1,Tor3a,Map3k8,Tubb6,Tiparp | | 1.352e-02 | -4.30 | GSE7852\_TREG\_VS\_TCONV\_UP | MSigDB lists | GSE7852\_TREG\_VS\_TCONV\_UP | 172 | 7 | 12187 | 179 | Arid5b,Bcl3,Tnip1,Birc3,Map3k8,Sdc4,Nfkb1 | | 1.352e-02 | -4.30 | GSE24292\_WT\_VS\_PPARG\_KO\_MACROPHAGE\_DN | MSigDB lists | GSE24292\_WT\_VS\_PPARG\_KO\_MACROPHAGE\_DN | 172 | 7 | 12187 | 179 | Ifit2,Parp14,Xaf1,Parp12,Irgm2,Herc6,C3ar1 | | 1.352e-02 | -4.30 | GSE19941\_IL10\_KO\_VS\_IL10\_KO\_AND\_NFKBP50\_KO\_LPS\_STIM\_MACROPHAGE\_DN | MSigDB lists | GSE19941\_IL10\_KO\_VS\_IL10\_KO\_AND\_NFKBP50\_KO\_LPS\_STIM\_MACROPHAGE\_DN | 172 | 7 | 12187 | 179 | Mxd4,Akap12,Cybb,Ccl4,Tor3a,Slfn3,Slfn4 | | 1.352e-02 | -4.30 | GSE31082\_DP\_VS\_CD4\_SP\_THYMOCYTE\_DN | MSigDB lists | GSE31082\_DP\_VS\_CD4\_SP\_THYMOCYTE\_DN | 172 | 7 | 12187 | 179 | Nfe2l2,Samd9l,Herc6,Tap1,Parp9,Gbp3,Icam1 | | 1.352e-02 | -4.30 | GSE23695\_CD57\_POS\_VS\_NEG\_NK\_CELL\_DN | MSigDB lists | GSE23695\_CD57\_POS\_VS\_NEG\_NK\_CELL\_DN | 172 | 7 | 12187 | 179 | Gadd45g,Map3k8,Maff,Casp4,Clic4,Ptgs2,Gem | | 1.365e-02 | -4.29 | GSE29949\_DC\_BRAIN\_VS\_MONOCYTE\_BONE\_MARROW\_DN | MSigDB lists | GSE29949\_DC\_BRAIN\_VS\_MONOCYTE\_BONE\_MARROW\_DN | 133 | 6 | 12187 | 179 | Ifitm3,Isg15,Casp4,Bcl3,Stx11,Tnfsf10 | | 1.368e-02 | -4.29 | Antifolate resistance | KEGG pathways | ko01523 | 25 | 3 | 5248 | 107 | Tnf,Il6,Nfkb1 | | 1.368e-02 | -4.29 | Antifolate resistance | KEGG pathways | mmu01523 | 25 | 3 | 5248 | 107 | Tnf,Il6,Nfkb1 | | 1.368e-02 | -4.29 | MODULE\_146 | MSigDB lists | MODULE\_146 | 63 | 4 | 12187 | 179 | Ccl2,C5ar1,C3ar1,P2ry6 | | 1.368e-02 | -4.29 | SESTO\_RESPONSE\_TO\_UV\_C1 | MSigDB lists | SESTO\_RESPONSE\_TO\_UV\_C1 | 63 | 4 | 12187 | 179 | Lcn2,Tnfaip3,Cdkn1a,Tnip1 | | 1.368e-02 | -4.29 | PID\_LYSOPHOSPHOLIPID\_PATHWAY | MSigDB lists | PID\_LYSOPHOSPHOLIPID\_PATHWAY | 63 | 4 | 12187 | 179 | Il6,Nfkb1,Nfkbia,Pik3r1 | | 1.368e-02 | -4.29 | FLECHNER\_BIOPSY\_KIDNEY\_TRANSPLANT\_REJECTED\_VS\_OK\_UP | MSigDB lists | FLECHNER\_BIOPSY\_KIDNEY\_TRANSPLANT\_REJECTED\_VS\_OK\_UP | 63 | 4 | 12187 | 179 | Cd14,Il4ra,Lcn2,S100a8 | | 1.371e-02 | -4.29 | Myd88 (myeloid differentiation primary response gene 88) | protein interactions | 17874 | 30 | 3 | 6802 | 115 | Birc3,Irf7,Hspa5 | | 1.373e-02 | -4.29 | NF-kB is activated and signals survival | REACTOME pathways | R-MMU-209560 | 11 | 2 | 6297 | 105 | Nfkbia,Nfkb1 | | 1.373e-02 | -4.29 | Nicotinamide salvaging | REACTOME pathways | R-MMU-197264 | 11 | 2 | 6297 | 105 | Parp9,Ptgs2 | | 1.380e-02 | -4.28 | negative regulation of biosynthetic process | biological process | GO:0009890 | 1267 | 30 | 13711 | 214 | Ifi204,Ifi207,Arid5b,Zbtb16,Cflar,Atf3,Zfp36,Tnf,Znfx1,Plek,Bcl3,Rasd1,Prg4,Rrp8,Dnajb1,Il6,Errfi1,Mxd4,Sox11,Runx1,Ifi211,Bach1,Cebpd,Spi1,Nfkb1,Nfkbie,Parp9,Sbno2,Gbp4,Ifi209 | | 1.392e-02 | -4.27 | CHYLA\_CBFA2T3\_TARGETS\_DN | MSigDB lists | CHYLA\_CBFA2T3\_TARGETS\_DN | 173 | 7 | 12187 | 179 | S100a9,Irf7,S100a8,Oasl1,Lcn2,AA467197,Mmp8 | | 1.392e-02 | -4.27 | GSE17186\_BLOOD\_VS\_CORD\_BLOOD\_CD21HIGH\_TRANSITIONAL\_BCELL\_DN | MSigDB lists | GSE17186\_BLOOD\_VS\_CORD\_BLOOD\_CD21HIGH\_TRANSITIONAL\_BCELL\_DN | 173 | 7 | 12187 | 179 | Akap12,Msn,Slfn3,Slfn4,Arid5b,Tnip1,Slfn9 | | 1.395e-02 | -4.27 | HEMOPEXIN | prosite domains | PS00024 | 10 | 2 | 8845 | 164 | Mmp8,Prg4 | | 1.395e-02 | -4.27 | REL\_2 | prosite domains | PS50254 | 10 | 2 | 8845 | 164 | Nfkb1,Nfkb2 | | 1.397e-02 | -4.27 | RNA polymerase II proximal promoter sequence-specific DNA binding | molecular function | GO:0000978 | 417 | 13 | 13516 | 211 | Ifi209,Nfkb2,Cebpd,Sox11,Ifi204,Irf9,Runx1,Ifi211,Irf7,Ifi207,Spi1,Maff,Atf3 | | 1.399e-02 | -4.27 | GO\_RECEPTOR\_ACTIVITY | MSigDB lists | GO\_RECEPTOR\_ACTIVITY | 749 | 19 | 12187 | 179 | Igsf6,Osmr,Fzd4,C5ar1,Robo4,C3ar1,Hcar2,Sdc4,Gpr84,Msr1,Il4ra,Plaur,Cxcl16,Csf2rb,Il17ra,Icam1,Sele,P2ry6,Cd14 | | 1.401e-02 | -4.27 | leukocyte activation | biological process | GO:0045321 | 372 | 12 | 13711 | 214 | Sbno2,Bcl3,Msn,Spi1,Stx11,Il6,Fyb,Icam1,Tnf,Tnfaip3,Pik3r1,Gadd45g | | 1.404e-02 | -4.27 | LINDGREN\_BLADDER\_CANCER\_CLUSTER\_1\_DN | MSigDB lists | LINDGREN\_BLADDER\_CANCER\_CLUSTER\_1\_DN | 303 | 10 | 12187 | 179 | Akap12,Birc3,Ccl2,Pik3ap1,Rgs16,Sbno2,Clic4,Nfkb2,Gem,Tagln2 | | 1.406e-02 | -4.26 | INGRAM\_SHH\_TARGETS\_UP | MSigDB lists | INGRAM\_SHH\_TARGETS\_UP | 97 | 5 | 12187 | 179 | Nfe2l2,Olfml2b,Cmpk2,Akap12,Cp | | 1.406e-02 | -4.26 | GO\_POSITIVE\_REGULATION\_OF\_NUCLEOCYTOPLASMIC\_TRANSPORT | MSigDB lists | GO\_POSITIVE\_REGULATION\_OF\_NUCLEOCYTOPLASMIC\_TRANSPORT | 97 | 5 | 12187 | 179 | Csf3,Pik3r1,Ptgs2,Tnf,Il6 | | 1.406e-02 | -4.26 | GO\_RESPONSE\_TO\_AMINO\_ACID | MSigDB lists | GO\_RESPONSE\_TO\_AMINO\_ACID | 97 | 5 | 12187 | 179 | Tnf,Icam1,Col4a1,Ccl12,Il6 | | 1.406e-02 | -4.26 | LINDGREN\_BLADDER\_CANCER\_CLUSTER\_1\_UP | MSigDB lists | LINDGREN\_BLADDER\_CANCER\_CLUSTER\_1\_UP | 97 | 5 | 12187 | 179 | Ifi209,Pik3r1,Ifi211,Ifi204,Ifi207 | | 1.407e-02 | -4.26 | positive regulation of epithelial cell proliferation | biological process | GO:0050679 | 163 | 7 | 13711 | 214 | Il6,Ccl2,C5ar1,Tnfaip3,Sox11,Ccnd2,Cflar | | 1.407e-02 | -4.26 | cellular response to nutrient levels | biological process | GO:0031669 | 163 | 7 | 13711 | 214 | Cdkn1a,Atf3,Hspa5,Fam107a,Rrp8,Angptl4,Nfe2l2 | | 1.408e-02 | -4.26 | glycosaminoglycan binding | molecular function | GO:0005539 | 163 | 7 | 13516 | 211 | Selp,Ccl2,Thbs1,Adamts1,Ccl7,Fstl1,Saa1 | | 1.413e-02 | -4.26 | GSE21063\_WT\_VS\_NFATC1\_KO\_3H\_ANTI\_IGM\_STIM\_BCELL\_UP | MSigDB lists | GSE21063\_WT\_VS\_NFATC1\_KO\_3H\_ANTI\_IGM\_STIM\_BCELL\_UP | 134 | 6 | 12187 | 179 | Sbno2,Pik3ap1,Timp1,Ms4a6d,Samd9l,Ccl4 | | 1.413e-02 | -4.26 | GSE4590\_PRE\_BCELL\_VS\_SMALL\_PRE\_BCELL\_UP | MSigDB lists | GSE4590\_PRE\_BCELL\_VS\_SMALL\_PRE\_BCELL\_UP | 134 | 6 | 12187 | 179 | Hspb1,Gbp4,Gbp6,Ifi44,Phf11b,Phf11d | | 1.419e-02 | -4.26 | regulation of ion transport | biological process | GO:0043269 | 606 | 17 | 13711 | 214 | Ccl2,Cybb,Gem,Il1a,Icam1,Tnf,P2ry6,Ripk1,Il1rn,Thbs1,Ccl12,Clic4,Cxcl1,Ptgs2,Sgk1,Ccl4,Kcna5 | | 1.424e-02 | -4.25 | regulation of cell killing | biological process | GO:0031341 | 60 | 4 | 13711 | 214 | Cxcl1,Ccl2,Lgals9,Tap1 | | 1.424e-02 | -4.25 | JNK cascade | biological process | GO:0007254 | 60 | 4 | 13711 | 214 | Ptpn1,Trib1,Ripk1,Tnf | | 1.428e-02 | -4.25 | regulation of protein secretion | biological process | GO:0050708 | 373 | 12 | 13711 | 214 | Mmp8,Il17ra,Tnfaip3,Tnf,Il4ra,Casp4,Akap12,Il1a,Il6,Cd14,Hcar2,Lgals9 | | 1.433e-02 | -4.25 | GSE24634\_TREG\_VS\_TCONV\_POST\_DAY10\_IL4\_CONVERSION\_UP | MSigDB lists | GSE24634\_TREG\_VS\_TCONV\_POST\_DAY10\_IL4\_CONVERSION\_UP | 174 | 7 | 12187 | 179 | Iqgap1,Ifi211,Cflar,Ifi204,Ifi207,Ifi209,Birc3 | | 1.433e-02 | -4.25 | GO\_REGULATION\_OF\_GENE\_EXPRESSION\_EPIGENETIC | MSigDB lists | GO\_REGULATION\_OF\_GENE\_EXPRESSION\_EPIGENETIC | 174 | 7 | 12187 | 179 | Spi1,Ifi209,Ifi204,Ifi207,Ifi211,Rrp8,Zfp36 | | 1.433e-02 | -4.25 | GSE24574\_BCL6\_HIGH\_TFH\_VS\_NAIVE\_CD4\_TCELL\_DN | MSigDB lists | GSE24574\_BCL6\_HIGH\_TFH\_VS\_NAIVE\_CD4\_TCELL\_DN | 174 | 7 | 12187 | 179 | Tnf,Ier3,Atf3,Nfe2l2,Nfkb1,Gem,Cflar | | 1.433e-02 | -4.25 | GSE22432\_PDC\_VS\_TGFB1\_TREATEDCOMMON\_DC\_PROGENITOR\_DN | MSigDB lists | GSE22432\_PDC\_VS\_TGFB1\_TREATEDCOMMON\_DC\_PROGENITOR\_DN | 174 | 7 | 12187 | 179 | Pik3r1,Socs3,Trim56,Ifit3b,Irf7,Atf3,Tap1 | | 1.433e-02 | -4.25 | GOTZMANN\_EPITHELIAL\_TO\_MESENCHYMAL\_TRANSITION\_DN | MSigDB lists | GOTZMANN\_EPITHELIAL\_TO\_MESENCHYMAL\_TRANSITION\_DN | 174 | 7 | 12187 | 179 | Il1rn,Sdc4,Nfkbia,Casp4,Cdkn1a,Trim25,S100a8 | | 1.443e-02 | -4.24 | GO\_NEGATIVE\_REGULATION\_OF\_CELL\_DIFFERENTIATION | MSigDB lists | GO\_NEGATIVE\_REGULATION\_OF\_CELL\_DIFFERENTIATION | 446 | 13 | 12187 | 179 | Bbs12,Zbtb16,Sox11,Cflar,Xdh,Nfkbia,Trib1,Tnf,Runx1,Il4ra,Pik3r1,Nfe2l2,Il6 | | 1.443e-02 | -4.24 | S\_100 | pfam domains | PF01023 | 12 | 2 | 12881 | 201 | S100a8,S100a9 | | 1.443e-02 | -4.24 | KESHELAVA\_MULTIPLE\_DRUG\_RESISTANCE | MSigDB lists | KESHELAVA\_MULTIPLE\_DRUG\_RESISTANCE | 64 | 4 | 12187 | 179 | Casp4,Iqgap1,Hspb1,Sbno2 | | 1.443e-02 | -4.24 | GO\_LYMPHOCYTE\_MEDIATED\_IMMUNITY | MSigDB lists | GO\_LYMPHOCYTE\_MEDIATED\_IMMUNITY | 64 | 4 | 12187 | 179 | Icam1,Bcl3,Irf7,Il4ra | | 1.443e-02 | -4.24 | GO\_REGULATION\_OF\_PRODUCTION\_OF\_MOLECULAR\_MEDIATOR\_OF\_IMMUNE\_RESPONSE | MSigDB lists | GO\_REGULATION\_OF\_PRODUCTION\_OF\_MOLECULAR\_MEDIATOR\_OF\_IMMUNE\_RESPONSE | 64 | 4 | 12187 | 179 | Tnf,Il4ra,Irak3,Il6 | | 1.443e-02 | -4.24 | KEGG\_VEGF\_SIGNALING\_PATHWAY | MSigDB lists | KEGG\_VEGF\_SIGNALING\_PATHWAY | 64 | 4 | 12187 | 179 | Pik3r5,Pik3r1,Ptgs2,Hspb1 | | 1.444e-02 | -4.24 | positive regulation of transforming growth factor beta production | biological process | GO:0071636 | 12 | 2 | 13711 | 214 | Thbs1,Ptgs2 | | 1.444e-02 | -4.24 | negative regulation of tyrosine phosphorylation of STAT protein | biological process | GO:0042532 | 12 | 2 | 13711 | 214 | Socs3,Parp14 | | 1.444e-02 | -4.24 | defense response to fungus | biological process | GO:0050832 | 12 | 2 | 13711 | 214 | Ncf1,Il17ra | | 1.444e-02 | -4.24 | regulation of amyloid-beta clearance | biological process | GO:1900221 | 12 | 2 | 13711 | 214 | Tnf,C5ar1 | | 1.444e-02 | -4.24 | positive regulation of podosome assembly | biological process | GO:0071803 | 12 | 2 | 13711 | 214 | Tnf,Msn | | 1.444e-02 | -4.24 | cellular response to interferon-alpha | biological process | GO:0035457 | 12 | 2 | 13711 | 214 | Ifit2,Ifi204 | | 1.444e-02 | -4.24 | regulation of glomerulus development | biological process | GO:0090192 | 12 | 2 | 13711 | 214 | Cflar,C3ar1 | | 1.444e-02 | -4.24 | negative regulation of collagen biosynthetic process | biological process | GO:0032966 | 12 | 2 | 13711 | 214 | Il6,Errfi1 | | 1.444e-02 | -4.24 | neutrophil mediated immunity | biological process | GO:0002446 | 12 | 2 | 13711 | 214 | Ncf1,Stx11 | | 1.444e-02 | -4.24 | negative regulation of collagen metabolic process | biological process | GO:0010713 | 12 | 2 | 13711 | 214 | Errfi1,Il6 | | 1.444e-02 | -4.24 | neutrophil homeostasis | biological process | GO:0001780 | 12 | 2 | 13711 | 214 | Hcar2,Il6 | | 1.444e-02 | -4.24 | interleukin-6-mediated signaling pathway | biological process | GO:0070102 | 12 | 2 | 13711 | 214 | Spi1,Il6 | | 1.444e-02 | -4.24 | positive regulation of protein localization to endosome | biological process | GO:1905668 | 12 | 2 | 13711 | 214 | Dtx3l,Msn | | 1.444e-02 | -4.24 | positive regulation of type 2 immune response | biological process | GO:0002830 | 12 | 2 | 13711 | 214 | Il6,Il4ra | | 1.444e-02 | -4.24 | regulation of T-helper 2 cell differentiation | biological process | GO:0045628 | 12 | 2 | 13711 | 214 | Il6,Il4ra | | 1.444e-02 | -4.24 | response to peptidoglycan | biological process | GO:0032494 | 12 | 2 | 13711 | 214 | C5ar1,Irak3 | | 1.444e-02 | -4.24 | HX | smart domains | SM00120 | 11 | 2 | 7188 | 123 | Mmp8,Prg4 | | 1.444e-02 | -4.24 | nucleotide receptor activity | molecular function | GO:0016502 | 12 | 2 | 13516 | 211 | Hcar2,P2ry6 | | 1.444e-02 | -4.24 | low-density lipoprotein particle binding | molecular function | GO:0030169 | 12 | 2 | 13516 | 211 | Msr1,Thbs1 | | 1.444e-02 | -4.24 | purinergic nucleotide receptor activity | molecular function | GO:0001614 | 12 | 2 | 13516 | 211 | P2ry6,Hcar2 | | 1.445e-02 | -4.24 | GO\_RESPONSE\_TO\_NICOTINE | MSigDB lists | GO\_RESPONSE\_TO\_NICOTINE | 35 | 3 | 12187 | 179 | Vcam1,Tnf,Nfkb1 | | 1.445e-02 | -4.24 | GO\_PLATELET\_ALPHA\_GRANULE\_LUMEN | MSigDB lists | GO\_PLATELET\_ALPHA\_GRANULE\_LUMEN | 35 | 3 | 12187 | 179 | Timp1,Thbs1,A2m | | 1.448e-02 | -4.24 | ubiquitin | pfam domains | PF00240 | 33 | 3 | 12881 | 201 | Oasl2,Isg15,Oasl1 | | 1.448e-02 | -4.24 | Ank | pfam domains | PF00023 | 33 | 3 | 12881 | 201 | Nfkb1,Bcl3,Nfkbia | | 1.450e-02 | -4.23 | regulation of interleukin-10 production | biological process | GO:0032653 | 33 | 3 | 13711 | 214 | Lgals9,Bcl3,Mmp8 | | 1.450e-02 | -4.23 | negative regulation of cell-matrix adhesion | biological process | GO:0001953 | 33 | 3 | 13711 | 214 | Pik3r1,Fam107a,Thbs1 | | 1.450e-02 | -4.23 | response to amyloid-beta | biological process | GO:1904645 | 33 | 3 | 13711 | 214 | Casp4,Vcam1,Icam1 | | 1.450e-02 | -4.23 | Myc (myelocytomatosis oncogene) | protein interactions | 17869 | 56 | 4 | 6802 | 115 | Thbs1,Cdkn1a,Ccnd2,Irf9 | | 1.453e-02 | -4.23 | endopeptidase regulator activity | molecular function | GO:0061135 | 92 | 5 | 13516 | 211 | Tnfaip8,A2m,Timp1,Serpina3f,Birc3 | | 1.454e-02 | -4.23 | Downstream signaling events of B Cell Receptor (BCR) | REACTOME pathways | R-MMU-1168372 | 155 | 7 | 6297 | 105 | Nfkbie,Ier3,Nfkb1,Pik3r1,Pik3ap1,Nfkbia,Cdkn1a | | 1.461e-02 | -4.23 | GSE25146\_UNSTIM\_VS\_HELIOBACTER\_PYLORI\_LPS\_STIM\_AGS\_CELL\_DN | MSigDB lists | GSE25146\_UNSTIM\_VS\_HELIOBACTER\_PYLORI\_LPS\_STIM\_AGS\_CELL\_DN | 135 | 6 | 12187 | 179 | Xaf1,Gbp4,Gbp6,Nfkb2,Gbp3,Rbm39 | | 1.461e-02 | -4.23 | AMIT\_EGF\_RESPONSE\_480\_HELA | MSigDB lists | AMIT\_EGF\_RESPONSE\_480\_HELA | 135 | 6 | 12187 | 179 | Tgm2,Col4a1,Rhoc,Osmr,Nfkb2,Plaur | | 1.461e-02 | -4.23 | GSE24081\_CONTROLLER\_VS\_PROGRESSOR\_HIV\_SPECIFIC\_CD8\_TCELL\_UP | MSigDB lists | GSE24081\_CONTROLLER\_VS\_PROGRESSOR\_HIV\_SPECIFIC\_CD8\_TCELL\_UP | 135 | 6 | 12187 | 179 | Tnf,Birc3,Maff,Tnfaip3,Mxd4,Zfp36 | | 1.461e-02 | -4.23 | FOXJ2\_01 | MSigDB lists | FOXJ2\_01 | 135 | 6 | 12187 | 179 | Luc7l3,Ier3,Fstl1,Tnf,Errfi1,Cxcl1 | | 1.461e-02 | -4.23 | GSE9509\_LPS\_VS\_LPS\_AND\_IL10\_STIM\_IL10\_KO\_MACROPHAGE\_20MIN\_UP | MSigDB lists | GSE9509\_LPS\_VS\_LPS\_AND\_IL10\_STIM\_IL10\_KO\_MACROPHAGE\_20MIN\_UP | 135 | 6 | 12187 | 179 | Ifitm3,Robo4,Gbp3,Birc3,Nfkbiz,Ksr1 | | 1.461e-02 | -4.23 | GSE8921\_UNSTIM\_0H\_VS\_TLR1\_2\_STIM\_MONOCYTE\_24H\_UP | MSigDB lists | GSE8921\_UNSTIM\_0H\_VS\_TLR1\_2\_STIM\_MONOCYTE\_24H\_UP | 135 | 6 | 12187 | 179 | Tagln2,Angptl4,Robo4,Fstl1,Cp,Mmp8 | | 1.461e-02 | -4.23 | GSE29618\_BCELL\_VS\_PDC\_DAY7\_FLU\_VACCINE\_UP | MSigDB lists | GSE29618\_BCELL\_VS\_PDC\_DAY7\_FLU\_VACCINE\_UP | 135 | 6 | 12187 | 179 | Arid5b,Tnfaip8,Clic4,Il4ra,Birc3,Casp4 | | 1.461e-02 | -4.23 | GSE15659\_RESTING\_VS\_ACTIVATED\_TREG\_DN | MSigDB lists | GSE15659\_RESTING\_VS\_ACTIVATED\_TREG\_DN | 135 | 6 | 12187 | 179 | Tnfaip3,Rhoc,Xaf1,Rhou,Tnf,Sox11 | | 1.461e-02 | -4.23 | GSE1112\_HY\_CD8AB\_VS\_HY\_CD8AA\_THYMOCYTE\_RTOC\_CULTURE\_DN | MSigDB lists | GSE1112\_HY\_CD8AB\_VS\_HY\_CD8AA\_THYMOCYTE\_RTOC\_CULTURE\_DN | 135 | 6 | 12187 | 179 | Il1a,Plaur,Runx1,Ptgs2,Cxcl1,Nfkbiz | | 1.463e-02 | -4.22 | cellular response to starvation | biological process | GO:0009267 | 127 | 6 | 13711 | 214 | Nfe2l2,Angptl4,Hspa5,Rrp8,Cdkn1a,Atf3 | | 1.464e-02 | -4.22 | MODULE\_15 | MSigDB lists | MODULE\_15 | 305 | 10 | 12187 | 179 | Ifi211,Cxcl1,Tnfaip8,Plek,Ifi209,Ccnd2,Maff,Ifi204,Ifi207,Atf3 | | 1.465e-02 | -4.22 | GO\_RESPONSE\_TO\_HYDROGEN\_PEROXIDE | MSigDB lists | GO\_RESPONSE\_TO\_HYDROGEN\_PEROXIDE | 98 | 5 | 12187 | 179 | Kcna5,Hp,Tnfaip3,Il6,Nfe2l2 | | 1.465e-02 | -4.22 | chr12p13 | MSigDB lists | chr12p13 | 98 | 5 | 12187 | 179 | A2m,Ccnd2,C3ar1,Kcna5,Apold1 | | 1.465e-02 | -4.22 | regulation of biological quality | biological process | GO:0065008 | 3134 | 63 | 13711 | 214 | Sox11,Icam1,Mt1,Bbs12,Msr1,Thbs1,P2ry6,Lgals9,Gadd45g,Rhoj,Nfkb1,Kcna5,Plek,Tnfaip3,Ccl2,Xdh,Csf3,Il1rn,C3ar1,Pygm,Lcn2,Ccl12,S100a9,Usp18,Vcam1,Irgm2,Nfkbia,Il1a,Akap12,Il4ra,Il6,Pik3r1,Hcar2,Fzd4,Fam107a,Tiparp,Isg15,Ccl4,Spi1,Stx11,Rasip1,Ier3,Runx1,Ptgs2,Sgk1,Tnf,Rhou,Zfp36,Slc24a4,Cp,Rhoc,Hspa5,Clic4,C5ar1,Cxcl1,Tgm2,Mt2,Mmp8,Nfe2l2,Angptl4,Msn,Selp,Noct | | 1.483e-02 | -4.21 | purine ribonucleotide binding | molecular function | GO:0032555 | 1544 | 35 | 13516 | 211 | Tgtp1,Hcar2,Hspa5,Slfn5,Iigp1,Gbp3,Trib1,Slfn9,Ksr1,Irak3,Runx1,Rhou,Oasl1,Irgm1,Cmpk2,Ripk1,Ip6k2,Tubb6,Tor3a,Map3k6,Gem,Tap1,Sgk1,Pygm,Oasl2,Rasd1,Sgk3,Map3k8,Rhoj,Mx1,Rhoc,Gbp6,Gbp7,Tgm2,Bbs12 | | 1.484e-02 | -4.21 | regulation of protein complex assembly | biological process | GO:0043254 | 375 | 12 | 13711 | 214 | Xaf1,Rasip1,Msn,Selp,Sgk1,Irgm2,Icam1,Tnf,Plek,Csf3,Trim30a,Rhoc | | 1.489e-02 | -4.21 | - | gene3d domains | 1.10.10.2150 | 1 | 1 | 6647 | 99 | Rrp8 | | 1.489e-02 | -4.21 | - | gene3d domains | 1.10.4100.10 | 1 | 1 | 6647 | 99 | Acod1 | | 1.489e-02 | -4.21 | - | gene3d domains | 3.30.1330.120 | 1 | 1 | 6647 | 99 | Acod1 | | 1.505e-02 | -4.20 | GO\_LIPOPOLYSACCHARIDE\_BINDING | MSigDB lists | GO\_LIPOPOLYSACCHARIDE\_BINDING | 13 | 2 | 12187 | 179 | Cd14,Selp | | 1.505e-02 | -4.20 | GO\_REGULATION\_OF\_IMMUNOGLOBULIN\_SECRETION | MSigDB lists | GO\_REGULATION\_OF\_IMMUNOGLOBULIN\_SECRETION | 13 | 2 | 12187 | 179 | Il6,Tnf | | 1.505e-02 | -4.20 | KORKOLA\_TERATOMA\_UP | MSigDB lists | KORKOLA\_TERATOMA\_UP | 13 | 2 | 12187 | 179 | Ccnd2,A2m | | 1.505e-02 | -4.20 | GO\_REGULATION\_OF\_ENDOTHELIAL\_CELL\_CHEMOTAXIS | MSigDB lists | GO\_REGULATION\_OF\_ENDOTHELIAL\_CELL\_CHEMOTAXIS | 13 | 2 | 12187 | 179 | Hspb1,Thbs1 | | 1.505e-02 | -4.20 | GO\_POSITIVE\_REGULATION\_OF\_CYSTEINE\_TYPE\_ENDOPEPTIDASE\_ACTIVITY\_INVOLVED\_IN\_APOPTOTIC\_SIGNALING\_PATHWAY | MSigDB lists | GO\_POSITIVE\_REGULATION\_OF\_CYSTEINE\_TYPE\_ENDOPEPTIDASE\_ACTIVITY\_INVOLVED\_IN\_APOPTOTIC\_SIGNALING\_PATHWAY | 13 | 2 | 12187 | 179 | Tnfsf10,Ripk1 | | 1.505e-02 | -4.20 | GO\_CELLULAR\_SODIUM\_ION\_HOMEOSTASIS | MSigDB lists | GO\_CELLULAR\_SODIUM\_ION\_HOMEOSTASIS | 13 | 2 | 12187 | 179 | Sgk3,Sgk1 | | 1.505e-02 | -4.20 | ABE\_VEGFA\_TARGETS | MSigDB lists | ABE\_VEGFA\_TARGETS | 13 | 2 | 12187 | 179 | A2m,Ptgs2 | | 1.505e-02 | -4.20 | DAZARD\_UV\_RESPONSE\_CLUSTER\_G24 | MSigDB lists | DAZARD\_UV\_RESPONSE\_CLUSTER\_G24 | 13 | 2 | 12187 | 179 | S100a9,Zfp36 | | 1.505e-02 | -4.20 | GO\_GRANULOCYTE\_DIFFERENTIATION | MSigDB lists | GO\_GRANULOCYTE\_DIFFERENTIATION | 13 | 2 | 12187 | 179 | Csf3,Spi1 | | 1.505e-02 | -4.20 | GO\_CHLORIDE\_CHANNEL\_REGULATOR\_ACTIVITY | MSigDB lists | GO\_CHLORIDE\_CHANNEL\_REGULATOR\_ACTIVITY | 13 | 2 | 12187 | 179 | Sgk3,Sgk1 | | 1.505e-02 | -4.20 | REACTOME\_TRAF3\_DEPENDENT\_IRF\_ACTIVATION\_PATHWAY | MSigDB lists | REACTOME\_TRAF3\_DEPENDENT\_IRF\_ACTIVATION\_PATHWAY | 13 | 2 | 12187 | 179 | Irf7,Trim25 | | 1.505e-02 | -4.20 | MODULE\_439 | MSigDB lists | MODULE\_439 | 13 | 2 | 12187 | 179 | Nfkb1,Ier3 | | 1.505e-02 | -4.20 | GO\_NEGATIVE\_REGULATION\_OF\_CYTOKINE\_PRODUCTION\_INVOLVED\_IN\_IMMUNE\_RESPONSE | MSigDB lists | GO\_NEGATIVE\_REGULATION\_OF\_CYTOKINE\_PRODUCTION\_INVOLVED\_IN\_IMMUNE\_RESPONSE | 13 | 2 | 12187 | 179 | Irak3,Tnf | | 1.505e-02 | -4.20 | KIM\_PTEN\_TARGETS\_UP | MSigDB lists | KIM\_PTEN\_TARGETS\_UP | 13 | 2 | 12187 | 179 | Ifi44,Cdkn1a | | 1.505e-02 | -4.20 | GO\_REGULATION\_OF\_GRANULOCYTE\_DIFFERENTIATION | MSigDB lists | GO\_REGULATION\_OF\_GRANULOCYTE\_DIFFERENTIATION | 13 | 2 | 12187 | 179 | Runx1,Trib1 | | 1.505e-02 | -4.20 | GO\_RESPONSE\_TO\_MURAMYL\_DIPEPTIDE | MSigDB lists | GO\_RESPONSE\_TO\_MURAMYL\_DIPEPTIDE | 13 | 2 | 12187 | 179 | Tnfaip3,Nfkbia | | 1.506e-02 | -4.20 | negative regulation of ERK1 and ERK2 cascade | biological process | GO:0070373 | 61 | 4 | 13711 | 214 | Ptpn1,Atf3,Errfi1,Tnip1 | | 1.506e-02 | -4.20 | antioxidant activity | molecular function | GO:0016209 | 61 | 4 | 13516 | 211 | S100a9,S100a8,Ptgs2,Hp | | 1.510e-02 | -4.19 | GO\_CELLULAR\_RESPONSE\_TO\_ACID\_CHEMICAL | MSigDB lists | GO\_CELLULAR\_RESPONSE\_TO\_ACID\_CHEMICAL | 136 | 6 | 12187 | 179 | Adamts1,Tnf,Fzd4,Col4a1,P2ry6,Ccl12 | | 1.510e-02 | -4.19 | GSE2770\_UNTREATED\_VS\_TGFB\_AND\_IL12\_TREATED\_ACT\_CD4\_TCELL\_6H\_DN | MSigDB lists | GSE2770\_UNTREATED\_VS\_TGFB\_AND\_IL12\_TREATED\_ACT\_CD4\_TCELL\_6H\_DN | 136 | 6 | 12187 | 179 | Gbp4,Cmpk2,Rhoc,Parp14,Saa1,Gbp6 | | 1.510e-02 | -4.19 | GO\_PROTEIN\_LOCALIZATION\_TO\_NUCLEUS | MSigDB lists | GO\_PROTEIN\_LOCALIZATION\_TO\_NUCLEUS | 136 | 6 | 12187 | 179 | Tnf,Pik3r1,Zbtb16,Banp,Bcl3,Nfkbia | | 1.510e-02 | -4.19 | TAKEDA\_TARGETS\_OF\_NUP98\_HOXA9\_FUSION\_8D\_DN | MSigDB lists | TAKEDA\_TARGETS\_OF\_NUP98\_HOXA9\_FUSION\_8D\_DN | 136 | 6 | 12187 | 179 | Cybb,S100a8,C5ar1,Sgk1,Cd14,S100a9 | | 1.511e-02 | -4.19 | mouse chr17 E2\|17 45.25 cM | chromosome location | mouse chr17 E2\|17 45.25 cM | 1 | 1 | 14556 | 220 | Xdh | | 1.511e-02 | -4.19 | mouse chr8 A1.1\|8 5.53 cM | chromosome location | mouse chr8 A1.1\|8 5.53 cM | 1 | 1 | 14556 | 220 | Col4a1 | | 1.511e-02 | -4.19 | mouse chr18 A1\|18 2.73 cM | chromosome location | mouse chr18 A1\|18 2.73 cM | 1 | 1 | 14556 | 220 | Map3k8 | | 1.511e-02 | -4.19 | mouse chrX\|X F5 and Y E | chromosome location | mouse chrX\|X F5 and Y E | 1 | 1 | 14556 | 220 | G530011O06Rik | | 1.511e-02 | -4.19 | mouse chr5 B1\|5 15.7 cM | chromosome location | mouse chr5 B1\|5 15.7 cM | 1 | 1 | 14556 | 220 | Il6 | | 1.511e-02 | -4.19 | mouse chr1 H3\|1 79.89 cM | chromosome location | mouse chr1 H3\|1 79.89 cM | 1 | 1 | 14556 | 220 | Tagln2 | | 1.511e-02 | -4.19 | mouse chr8 C5\|8 46.34 cM | chromosome location | mouse chr8 C5\|8 46.34 cM | 1 | 1 | 14556 | 220 | Mt1 | | 1.511e-02 | -4.19 | mouse chr15 A1\|15 3.3 cM | chromosome location | mouse chr15 A1\|15 3.3 cM | 1 | 1 | 14556 | 220 | Osmr | | 1.511e-02 | -4.19 | mouse chr2 C3\|2 44.75 cM | chromosome location | mouse chr2 C3\|2 44.75 cM | 1 | 1 | 14556 | 220 | Nfe2l2 | | 1.511e-02 | -4.19 | mouse chr11 D\|11 62.45 cM | chromosome location | mouse chr11 D\|11 62.45 cM | 1 | 1 | 14556 | 220 | Csf3 | | 1.511e-02 | -4.19 | mouse chr12 A2\|12 9.69 cM | chromosome location | mouse chr12 A2\|12 9.69 cM | 1 | 1 | 14556 | 220 | Cmpk2 | | 1.511e-02 | -4.19 | mouse chr4 A1\|4 5.34 cM | chromosome location | mouse chr4 A1\|4 5.34 cM | 1 | 1 | 14556 | 220 | Gem | | 1.511e-02 | -4.19 | mouse chr11 B1.2\|11 | chromosome location | mouse chr11 B1.2\|11 | 1 | 1 | 14556 | 220 | Tgtp2 | | 1.511e-02 | -4.19 | mouse chr11 C\|11 49.82 cM | chromosome location | mouse chr11 C\|11 49.82 cM | 1 | 1 | 14556 | 220 | Ccl2 | | 1.511e-02 | -4.19 | mouse chr17 A3.3\|17 15.12 cM | chromosome location | mouse chr17 A3.3\|17 15.12 cM | 1 | 1 | 14556 | 220 | Cdkn1a | | 1.511e-02 | -4.19 | mouse chr1 G3\|1 65.43 cM | chromosome location | mouse chr1 G3\|1 65.43 cM | 1 | 1 | 14556 | 220 | Rgs16 | | 1.511e-02 | -4.19 | mouse chr8 D3\|8 53.38 cM | chromosome location | mouse chr8 D3\|8 53.38 cM | 1 | 1 | 14556 | 220 | Sntb2 | | 1.511e-02 | -4.19 | mouse chr16 C3.3\|16 48.2 cM | chromosome location | mouse chr16 C3.3\|16 48.2 cM | 1 | 1 | 14556 | 220 | Adamts1 | | 1.511e-02 | -4.19 | mouse chr6 F1\|6 57.17 cM | chromosome location | mouse chr6 F1\|6 57.17 cM | 1 | 1 | 14556 | 220 | Usp18 | | 1.511e-02 | -4.19 | mouse chr1 H3\|1 80.63 cM | chromosome location | mouse chr1 H3\|1 80.63 cM | 1 | 1 | 14556 | 220 | Ifi204 | | 1.511e-02 | -4.19 | mouse chr11 C\|11 49.83 cM | chromosome location | mouse chr11 C\|11 49.83 cM | 1 | 1 | 14556 | 220 | Ccl7 | | 1.511e-02 | -4.19 | mouse chr9 A5.3\|9 26.47 cM | chromosome location | mouse chr9 A5.3\|9 26.47 cM | 1 | 1 | 14556 | 220 | Zbtb16 | | 1.511e-02 | -4.19 | mouse chr5 E2\|5 46.51 cM | chromosome location | mouse chr5 E2\|5 46.51 cM | 1 | 1 | 14556 | 220 | Cxcl9 | | 1.511e-02 | -4.19 | mouse chr10 A3\|10 8.08 cM | chromosome location | mouse chr10 A3\|10 8.08 cM | 1 | 1 | 14556 | 220 | Tnfaip3 | | 1.511e-02 | -4.19 | mouse chrX A1.3\|X 16.38 cM | chromosome location | mouse chrX A1.3\|X 16.38 cM | 1 | 1 | 14556 | 220 | Timp1 | | 1.511e-02 | -4.19 | mouse chr6 F1\|6 57.49 cM | chromosome location | mouse chr6 F1\|6 57.49 cM | 1 | 1 | 14556 | 220 | A2m | | 1.511e-02 | -4.19 | mouse chr7 B3\|7 30.54 cM | chromosome location | mouse chr7 B3\|7 30.54 cM | 1 | 1 | 14556 | 220 | Saa1 | | 1.511e-02 | -4.19 | mouse chr11 C\|11 49.9 cM | chromosome location | mouse chr11 C\|11 49.9 cM | 1 | 1 | 14556 | 220 | Ccl12 | | 1.511e-02 | -4.19 | mouse chr2 B\|2 21.75 cM | chromosome location | mouse chr2 B\|2 21.75 cM | 1 | 1 | 14556 | 220 | Ptges | | 1.511e-02 | -4.19 | mouse chr7 D3\|7 49.32 cM | chromosome location | mouse chr7 D3\|7 49.32 cM | 1 | 1 | 14556 | 220 | Fzd4 | | 1.511e-02 | -4.19 | mouse chr8 D3\|8 57.11 cM | chromosome location | mouse chr8 D3\|8 57.11 cM | 1 | 1 | 14556 | 220 | Hp | | 1.511e-02 | -4.19 | mouse chr14 E2.3\|14 51.67 cM | chromosome location | mouse chr14 E2.3\|14 51.67 cM | 1 | 1 | 14556 | 220 | Acod1 | | 1.511e-02 | -4.19 | mouse chr16 B3\|16 26.48 cM | chromosome location | mouse chr16 B3\|16 26.48 cM | 1 | 1 | 14556 | 220 | Fstl1 | | 1.511e-02 | -4.19 | mouse chr6 A1\|6 1.76 cM | chromosome location | mouse chr6 A1\|6 1.76 cM | 1 | 1 | 14556 | 220 | Samd9l | | 1.511e-02 | -4.19 | mouse chr15 E1\|15 37.36 cM | chromosome location | mouse chr15 E1\|15 37.36 cM | 1 | 1 | 14556 | 220 | Csf2rb | | 1.511e-02 | -4.19 | mouse chr2 B\|2 22.94 cM | chromosome location | mouse chr2 B\|2 22.94 cM | 1 | 1 | 14556 | 220 | Hspa5 | | 1.511e-02 | -4.19 | mouse chr7 A2\|7 8.77 cM | chromosome location | mouse chr7 A2\|7 8.77 cM | 1 | 1 | 14556 | 220 | C5ar1 | | 1.511e-02 | -4.19 | mouse chr5 G2\|5 74.47 cM | chromosome location | mouse chr5 G2\|5 74.47 cM | 1 | 1 | 14556 | 220 | Ncf1 | | 1.511e-02 | -4.19 | mouse chr19 A\|19 4.53 cM | chromosome location | mouse chr19 A\|19 4.53 cM | 1 | 1 | 14556 | 220 | Pygm | | 1.511e-02 | -4.19 | mouse chr11 E2\|11 83.48 cM | chromosome location | mouse chr11 E2\|11 83.48 cM | 1 | 1 | 14556 | 220 | Rnf213 | | 1.511e-02 | -4.19 | mouse chr7 E3\|7 55.69 cM | chromosome location | mouse chr7 E3\|7 55.69 cM | 1 | 1 | 14556 | 220 | Trim30a | | 1.511e-02 | -4.19 | mouse chr10 B2\|10 23.14 cM | chromosome location | mouse chr10 B2\|10 23.14 cM | 1 | 1 | 14556 | 220 | Crybg1 | | 1.511e-02 | -4.19 | mouse chr4 E2\|4 | chromosome location | mouse chr4 E2\|4 | 1 | 1 | 14556 | 220 | Isg15 | | 1.511e-02 | -4.19 | mouse chr1 H2.2\|1 71.42 cM | chromosome location | mouse chr1 H2.2\|1 71.42 cM | 1 | 1 | 14556 | 220 | Selp | | 1.511e-02 | -4.19 | mouse chr3 G1\|3 50.17 cM | chromosome location | mouse chr3 G1\|3 50.17 cM | 1 | 1 | 14556 | 220 | Vcam1 | | 1.511e-02 | -4.19 | mouse chr7 D3\|7 45.68 cM | chromosome location | mouse chr7 D3\|7 45.68 cM | 1 | 1 | 14556 | 220 | Iqgap1 | | 1.511e-02 | -4.19 | mouse chr2 F1\|2 62.9 cM | chromosome location | mouse chr2 F1\|2 62.9 cM | 1 | 1 | 14556 | 220 | Il1a | | 1.511e-02 | -4.19 | mouse chr5 G2\|5 75.51 cM | chromosome location | mouse chr5 G2\|5 75.51 cM | 1 | 1 | 14556 | 220 | Hspb1 | | 1.511e-02 | -4.19 | mouse chr2 E5\|2 59.34 cM | chromosome location | mouse chr2 E5\|2 59.34 cM | 1 | 1 | 14556 | 220 | Thbs1 | | 1.511e-02 | -4.19 | mouse chr7 A3-B1\|7 16.72 cM | chromosome location | mouse chr7 A3-B1\|7 16.72 cM | 1 | 1 | 14556 | 220 | Zfp36 | | 1.511e-02 | -4.19 | mouse chr13 D1\|13 53.92 cM | chromosome location | mouse chr13 D1\|13 53.92 cM | 1 | 1 | 14556 | 220 | Pik3r1 | | 1.511e-02 | -4.19 | mouse chr1 G1\|1 63.84 cM | chromosome location | mouse chr1 G1\|1 63.84 cM | 1 | 1 | 14556 | 220 | Ptgs2 | | 1.511e-02 | -4.19 | mouse chr7 F3\|7 68.94 cM | chromosome location | mouse chr7 F3\|7 68.94 cM | 1 | 1 | 14556 | 220 | Il4ra | | 1.511e-02 | -4.19 | mouse chr2 A3\|2 16.36 cM | chromosome location | mouse chr2 A3\|2 16.36 cM | 1 | 1 | 14556 | 220 | Il1rn | | 1.511e-02 | -4.19 | mouse chr3 F1\|3 39.9 cM | chromosome location | mouse chr3 F1\|3 39.9 cM | 1 | 1 | 14556 | 220 | S100a8 | | 1.511e-02 | -4.19 | mouse chr13 B2\|13 32.56 cM | chromosome location | mouse chr13 B2\|13 32.56 cM | 1 | 1 | 14556 | 220 | Ctla2a | | 1.511e-02 | -4.19 | mouse chr1 C1.3\|1 29.16 cM | chromosome location | mouse chr1 C1.3\|1 29.16 cM | 1 | 1 | 14556 | 220 | Cflar | | 1.511e-02 | -4.19 | mouse chr16 C4\|16 53.7 cM | chromosome location | mouse chr16 C4\|16 53.7 cM | 1 | 1 | 14556 | 220 | Runx1 | | 1.511e-02 | -4.19 | mouse chr11 C\|11 51.09 cM | chromosome location | mouse chr11 C\|11 51.09 cM | 1 | 1 | 14556 | 220 | Ccl4 | | 1.511e-02 | -4.19 | mouse chr8 E1\|8 70.82 cM | chromosome location | mouse chr8 E1\|8 70.82 cM | 1 | 1 | 14556 | 220 | Banp | | 1.511e-02 | -4.19 | mouse chr3 G3\|3 62.82 cM | chromosome location | mouse chr3 G3\|3 62.82 cM | 1 | 1 | 14556 | 220 | Nfkb1 | | 1.511e-02 | -4.19 | mouse chr16 B5\|16 30.75 cM | chromosome location | mouse chr16 B5\|16 30.75 cM | 1 | 1 | 14556 | 220 | Retnlg | | 1.511e-02 | -4.19 | mouse chr6 F3\|6 61.35 cM | chromosome location | mouse chr6 F3\|6 61.35 cM | 1 | 1 | 14556 | 220 | Kcna5 | | 1.511e-02 | -4.19 | mouse chr16 C4\|16 57.51 cM | chromosome location | mouse chr16 C4\|16 57.51 cM | 1 | 1 | 14556 | 220 | Mx2 | | 1.511e-02 | -4.19 | mouse chr1 H2.2\|1 71.35 cM | chromosome location | mouse chr1 H2.2\|1 71.35 cM | 1 | 1 | 14556 | 220 | Sele | | 1.511e-02 | -4.19 | mouse chr16 C4\|16 57.46 cM | chromosome location | mouse chr16 C4\|16 57.46 cM | 1 | 1 | 14556 | 220 | Mx1 | | 1.511e-02 | -4.19 | mouse chr8 C5\|8 46.32 cM | chromosome location | mouse chr8 C5\|8 46.32 cM | 1 | 1 | 14556 | 220 | Mt2 | | 1.511e-02 | -4.19 | mouse chr3 F1\|3 39.91 cM | chromosome location | mouse chr3 F1\|3 39.91 cM | 1 | 1 | 14556 | 220 | S100a9 | | 1.511e-02 | -4.19 | mouse chr14 C3\|14 31.48 cM | chromosome location | mouse chr14 C3\|14 31.48 cM | 1 | 1 | 14556 | 220 | Phf11d | | 1.513e-02 | -4.19 | RODRIGUES\_THYROID\_CARCINOMA\_ANAPLASTIC\_UP | MSigDB lists | RODRIGUES\_THYROID\_CARCINOMA\_ANAPLASTIC\_UP | 599 | 16 | 12187 | 179 | Slfn5,Clic4,Ifit3b,Ifi209,Ier3,Ifi207,Ifi204,Ifi211,Rbm39,Iqgap1,Casp4,Tgm2,Xaf1,Birc3,Ifit2,Runx1 | | 1.515e-02 | -4.19 | HAN\_SATB1\_TARGETS\_DN | MSigDB lists | HAN\_SATB1\_TARGETS\_DN | 353 | 11 | 12187 | 179 | Ptgs2,Fzd4,Osmr,Tap1,Ifit2,Maff,Fstl1,Ifit3b,Casp4,Ccl12,Il6 | | 1.516e-02 | -4.19 | heat shock protein binding | molecular function | GO:0031072 | 128 | 6 | 13516 | 211 | Ksr1,Hspa5,Dnajb1,Nfkbia,Zfp36,Nfkb1 | | 1.517e-02 | -4.19 | purine ribonucleoside triphosphate binding | molecular function | GO:0035639 | 1492 | 34 | 13516 | 211 | Rhoj,Mx1,Bbs12,Tgm2,Gbp7,Gbp6,Rhoc,Oasl2,Rasd1,Sgk1,Tap1,Gem,Map3k8,Sgk3,Oasl1,Irak3,Runx1,Rhou,Ksr1,Slfn9,Map3k6,Tor3a,Tubb6,Irgm1,Ripk1,Cmpk2,Ip6k2,Hspa5,Hcar2,Tgtp1,Trib1,Gbp3,Iigp1,Slfn5 | | 1.520e-02 | -4.19 | GO\_REGULATION\_OF\_EPITHELIAL\_CELL\_PROLIFERATION | MSigDB lists | GO\_REGULATION\_OF\_EPITHELIAL\_CELL\_PROLIFERATION | 218 | 8 | 12187 | 179 | Tnfaip3,Ccl12,Il6,Sox11,Thbs1,Xdh,Errfi1,C5ar1 | | 1.520e-02 | -4.19 | GSE6674\_UNSTIM\_VS\_ANTI\_IGM\_STIM\_BCELL\_DN | MSigDB lists | GSE6674\_UNSTIM\_VS\_ANTI\_IGM\_STIM\_BCELL\_DN | 65 | 4 | 12187 | 179 | Cd14,Irf7,Usp18,Adamts9 | | 1.525e-02 | -4.18 | VART\_KSHV\_INFECTION\_ANGIOGENIC\_MARKERS\_DN | MSigDB lists | VART\_KSHV\_INFECTION\_ANGIOGENIC\_MARKERS\_DN | 99 | 5 | 12187 | 179 | Ccl12,Csf3,Thbs1,Cxcl1,Angptl4 | | 1.526e-02 | -4.18 | MONNIER\_POSTRADIATION\_TUMOR\_ESCAPE\_DN | MSigDB lists | MONNIER\_POSTRADIATION\_TUMOR\_ESCAPE\_DN | 307 | 10 | 12187 | 179 | Sgk1,Nfkbiz,Fstl1,Tgm2,Cp,Gadd45g,AA467197,Arid5b,Ccdc58,Errfi1 | | 1.533e-02 | -4.18 | alpha9-beta1 integrin-vascular cell adhesion molecule-1 complex | cellular component | GO:0071065 | 1 | 1 | 13825 | 212 | Vcam1 | | 1.533e-02 | -4.18 | nuclear RNA-directed RNA polymerase complex | cellular component | GO:0031380 | 1 | 1 | 13825 | 212 | Znfx1 | | 1.533e-02 | -4.18 | intracellular canaliculus | cellular component | GO:0046691 | 1 | 1 | 13825 | 212 | Kcna5 | | 1.533e-02 | -4.18 | Dcp1-Dcp2 complex | cellular component | GO:0098745 | 1 | 1 | 13825 | 212 | Zfp36 | | 1.533e-02 | -4.18 | Bcl3-Bcl10 complex | cellular component | GO:0032996 | 1 | 1 | 13825 | 212 | Bcl3 | | 1.538e-02 | -4.17 | Cyt\_b245\_heavy\_chain | interpro domains | IPR000778 | 1 | 1 | 13788 | 212 | Cybb | | 1.538e-02 | -4.17 | RBM47\_RRM3 | interpro domains | IPR034445 | 1 | 1 | 13788 | 212 | Rbm47 | | 1.538e-02 | -4.17 | TRB1 | interpro domains | IPR024105 | 1 | 1 | 13788 | 212 | Trib1 | | 1.538e-02 | -4.17 | Interferon-induced\_IFIT2 | interpro domains | IPR024124 | 1 | 1 | 13788 | 212 | Ifit2 | | 1.538e-02 | -4.17 | Tap1/ABCB2 | interpro domains | IPR013306 | 1 | 1 | 13788 | 212 | Tap1 | | 1.538e-02 | -4.17 | Monocyte\_diff\_Ag\_CD14 | interpro domains | IPR016337 | 1 | 1 | 13788 | 212 | Cd14 | | 1.538e-02 | -4.17 | SOCS3 | interpro domains | IPR028414 | 1 | 1 | 13788 | 212 | Socs3 | | 1.538e-02 | -4.17 | MmgE/PrpD\_sf\_1 | interpro domains | IPR042183 | 1 | 1 | 13788 | 212 | Acod1 | | 1.538e-02 | -4.17 | MmgE\_PrpD | interpro domains | IPR005656 | 1 | 1 | 13788 | 212 | Acod1 | | 1.538e-02 | -4.17 | TNIP1 | interpro domains | IPR033372 | 1 | 1 | 13788 | 212 | Tnip1 | | 1.538e-02 | -4.17 | FZD4 | interpro domains | IPR026551 | 1 | 1 | 13788 | 212 | Fzd4 | | 1.538e-02 | -4.17 | Torsin-3A | interpro domains | IPR030552 | 1 | 1 | 13788 | 212 | Tor3a | | 1.538e-02 | -4.17 | RASSF4 | interpro domains | IPR033622 | 1 | 1 | 13788 | 212 | Rassf4 | | 1.538e-02 | -4.17 | IL-1RA/IL-36 | interpro domains | IPR003297 | 1 | 1 | 13788 | 212 | Il1rn | | 1.538e-02 | -4.17 | IFI44\_fam | interpro domains | IPR024644 | 1 | 1 | 13788 | 212 | Ifi44 | | 1.538e-02 | -4.17 | DEPP | interpro domains | IPR020133 | 1 | 1 | 13788 | 212 | Depp1 | | 1.538e-02 | -4.17 | SGK3\_dom | interpro domains | IPR037709 | 1 | 1 | 13788 | 212 | Sgk3 | | 1.538e-02 | -4.17 | NF-kB\_p105 | interpro domains | IPR030503 | 1 | 1 | 13788 | 212 | Nfkb1 | | 1.538e-02 | -4.17 | STX11 | interpro domains | IPR028672 | 1 | 1 | 13788 | 212 | Stx11 | | 1.538e-02 | -4.17 | ANGPTL4 | interpro domains | IPR028793 | 1 | 1 | 13788 | 212 | Angptl4 | | 1.538e-02 | -4.17 | CISK\_PX | interpro domains | IPR037900 | 1 | 1 | 13788 | 212 | Sgk3 | | 1.538e-02 | -4.17 | VCAM-1 | interpro domains | IPR003989 | 1 | 1 | 13788 | 212 | Vcam1 | | 1.538e-02 | -4.17 | IGSF6 | interpro domains | IPR039089 | 1 | 1 | 13788 | 212 | Igsf6 | | 1.538e-02 | -4.17 | Macro\_scav\_rcpt | interpro domains | IPR003543 | 1 | 1 | 13788 | 212 | Msr1 | | 1.538e-02 | -4.17 | TMEM252 | interpro domains | IPR031363 | 1 | 1 | 13788 | 212 | Tmem252 | | 1.538e-02 | -4.17 | MafF | interpro domains | IPR033531 | 1 | 1 | 13788 | 212 | Maff | | 1.538e-02 | -4.17 | IL3\_rcpt\_beta | interpro domains | IPR011365 | 1 | 1 | 13788 | 212 | Csf2rb | | 1.538e-02 | -4.17 | BIP\_NBD | interpro domains | IPR042050 | 1 | 1 | 13788 | 212 | Hspa5 | | 1.538e-02 | -4.17 | RIP1\_Death | interpro domains | IPR037934 | 1 | 1 | 13788 | 212 | Ripk1 | | 1.538e-02 | -4.17 | MS4A6D/MS4A6E | interpro domains | IPR030427 | 1 | 1 | 13788 | 212 | Ms4a6d | | 1.538e-02 | -4.17 | PX\_p47phox | interpro domains | IPR034909 | 1 | 1 | 13788 | 212 | Ncf1 | | 1.538e-02 | -4.17 | IL-6 | interpro domains | IPR003574 | 1 | 1 | 13788 | 212 | Il6 | | 1.538e-02 | -4.17 | Haptoglobin | interpro domains | IPR008292 | 1 | 1 | 13788 | 212 | Hp | | 1.538e-02 | -4.17 | AKAP12 | interpro domains | IPR028540 | 1 | 1 | 13788 | 212 | Akap12 | | 1.538e-02 | -4.17 | TNIP3 | interpro domains | IPR033574 | 1 | 1 | 13788 | 212 | Tnip3 | | 1.538e-02 | -4.17 | MmgE/PrpD\_sf\_2 | interpro domains | IPR042188 | 1 | 1 | 13788 | 212 | Acod1 | | 1.538e-02 | -4.17 | MmgE/PrpD\_sf | interpro domains | IPR036148 | 1 | 1 | 13788 | 212 | Acod1 | | 1.538e-02 | -4.17 | XAF1 | interpro domains | IPR031220 | 1 | 1 | 13788 | 212 | Xaf1 | | 1.538e-02 | -4.17 | S100-A9 | interpro domains | IPR028475 | 1 | 1 | 13788 | 212 | S100a9 | | 1.538e-02 | -4.17 | TIMP1 | interpro domains | IPR015611 | 1 | 1 | 13788 | 212 | Timp1 | | 1.538e-02 | -4.17 | MAPKKK8 | interpro domains | IPR017424 | 1 | 1 | 13788 | 212 | Map3k8 | | 1.538e-02 | -4.17 | Thrombospondin-1 | interpro domains | IPR028499 | 1 | 1 | 13788 | 212 | Thbs1 | | 1.538e-02 | -4.17 | RRP8\_N | interpro domains | IPR042036 | 1 | 1 | 13788 | 212 | Rrp8 | | 1.538e-02 | -4.17 | CXCL16 | interpro domains | IPR026296 | 1 | 1 | 13788 | 212 | Cxcl16 | | 1.538e-02 | -4.17 | Pept\_M12B\_ADAM-TS1 | interpro domains | IPR013274 | 1 | 1 | 13788 | 212 | Adamts1 | | 1.538e-02 | -4.17 | RESF1 | interpro domains | IPR027866 | 1 | 1 | 13788 | 212 | Resf1 | | 1.538e-02 | -4.17 | P2Y6\_rcpt | interpro domains | IPR001973 | 1 | 1 | 13788 | 212 | P2ry6 | | 1.538e-02 | -4.17 | Slfn5 | interpro domains | IPR029671 | 1 | 1 | 13788 | 212 | Slfn5 | | 1.538e-02 | -4.17 | IRAK3\_PK | interpro domains | IPR042698 | 1 | 1 | 13788 | 212 | Irak3 | | 1.538e-02 | -4.17 | Xanthine\_DH\_Mopterin-bd\_su | interpro domains | IPR014309 | 1 | 1 | 13788 | 212 | Xdh | | 1.538e-02 | -4.17 | TNF\_alpha | interpro domains | IPR002959 | 1 | 1 | 13788 | 212 | Tnf | | 1.538e-02 | -4.17 | Xanthine\_DH\_ssu | interpro domains | IPR014307 | 1 | 1 | 13788 | 212 | Xdh | | 1.538e-02 | -4.17 | P47PHOX | interpro domains | IPR001655 | 1 | 1 | 13788 | 212 | Ncf1 | | 1.538e-02 | -4.17 | LCN2/LCN12 | interpro domains | IPR003087 | 1 | 1 | 13788 | 212 | Lcn2 | | 1.538e-02 | -4.17 | TAGLN2 | interpro domains | IPR001061 | 1 | 1 | 13788 | 212 | Tagln2 | | 1.538e-02 | -4.17 | IL-4\_rcpt-alpha\_N | interpro domains | IPR015319 | 1 | 1 | 13788 | 212 | Il4ra | | 1.538e-02 | -4.17 | CDKN1A | interpro domains | IPR029841 | 1 | 1 | 13788 | 212 | Cdkn1a | | 1.538e-02 | -4.17 | COX-2 | interpro domains | IPR029576 | 1 | 1 | 13788 | 212 | Ptgs2 | | 1.538e-02 | -4.17 | PI3K\_p85alpha\_SH3 | interpro domains | IPR035591 | 1 | 1 | 13788 | 212 | Pik3r1 | | 1.538e-02 | -4.17 | Deadenylase\_nocturnin | interpro domains | IPR034965 | 1 | 1 | 13788 | 212 | Noct | | 1.538e-02 | -4.17 | TNF\_ligand\_10/11 | interpro domains | IPR017355 | 1 | 1 | 13788 | 212 | Tnfsf10 | | 1.538e-02 | -4.17 | HCAR2/3\_rcpt | interpro domains | IPR028017 | 1 | 1 | 13788 | 212 | Hcar2 | | 1.538e-02 | -4.17 | Anaphtx\_C3AR1 | interpro domains | IPR001644 | 1 | 1 | 13788 | 212 | C3ar1 | | 1.538e-02 | -4.17 | ARID5B | interpro domains | IPR030408 | 1 | 1 | 13788 | 212 | Arid5b | | 1.538e-02 | -4.17 | A20 | interpro domains | IPR033478 | 1 | 1 | 13788 | 212 | Tnfaip3 | | 1.538e-02 | -4.17 | ATF3 | interpro domains | IPR029812 | 1 | 1 | 13788 | 212 | Atf3 | | 1.538e-02 | -4.17 | PLEK1\_DEP | interpro domains | IPR037371 | 1 | 1 | 13788 | 212 | Plek | | 1.538e-02 | -4.17 | IRF2 | interpro domains | IPR031218 | 1 | 1 | 13788 | 212 | Irf2 | | 1.538e-02 | -4.17 | RBM39\_SF | interpro domains | IPR006509 | 1 | 1 | 13788 | 212 | Rbm39 | | 1.538e-02 | -4.17 | R-trans\_p | interpro domains | IPR026096 | 1 | 1 | 13788 | 212 | Rtp4 | | 1.538e-02 | -4.17 | Caffeine\_induced\_death\_Cid2 | interpro domains | IPR019171 | 1 | 1 | 13788 | 212 | Ccdc58 | | 1.538e-02 | -4.17 | NECFESHC | interpro domains | IPR032136 | 1 | 1 | 13788 | 212 | Ncf1 | | 1.538e-02 | -4.17 | RBM47 | interpro domains | IPR034442 | 1 | 1 | 13788 | 212 | Rbm47 | | 1.538e-02 | -4.17 | NCF1\_SH3\_1 | interpro domains | IPR035756 | 1 | 1 | 13788 | 212 | Ncf1 | | 1.538e-02 | -4.17 | IL1RA | interpro domains | IPR039348 | 1 | 1 | 13788 | 212 | Il1rn | | 1.538e-02 | -4.17 | BANP | interpro domains | IPR042343 | 1 | 1 | 13788 | 212 | Banp | | 1.538e-02 | -4.17 | SLC24A4 | interpro domains | IPR030232 | 1 | 1 | 13788 | 212 | Slc24a4 | | 1.538e-02 | -4.17 | Nocturnin | interpro domains | IPR040109 | 1 | 1 | 13788 | 212 | Noct | | 1.538e-02 | -4.17 | NCF1\_SH3\_2 | interpro domains | IPR035757 | 1 | 1 | 13788 | 212 | Ncf1 | | 1.538e-02 | -4.17 | SOCS3\_SH2 | interpro domains | IPR035863 | 1 | 1 | 13788 | 212 | Socs3 | | 1.538e-02 | -4.17 | SOX-11 | interpro domains | IPR029551 | 1 | 1 | 13788 | 212 | Sox11 | | 1.538e-02 | -4.17 | FYB\_hSH3 | interpro domains | IPR035540 | 1 | 1 | 13788 | 212 | Fyb | | 1.538e-02 | -4.17 | SNTB2 | interpro domains | IPR028550 | 1 | 1 | 13788 | 212 | Sntb2 | | 1.538e-02 | -4.17 | IEX-1 | interpro domains | IPR024829 | 1 | 1 | 13788 | 212 | Ier3 | | 1.538e-02 | -4.17 | RRP8 | interpro domains | IPR007823 | 1 | 1 | 13788 | 212 | Rrp8 | | 1.538e-02 | -4.17 | CP | interpro domains | IPR027150 | 1 | 1 | 13788 | 212 | Cp | | 1.538e-02 | -4.17 | OLFML2B | interpro domains | IPR031233 | 1 | 1 | 13788 | 212 | Olfml2b | | 1.538e-02 | -4.17 | RTP4 | interpro domains | IPR026690 | 1 | 1 | 13788 | 212 | Rtp4 | | 1.538e-02 | -4.17 | U-PAR | interpro domains | IPR033084 | 1 | 1 | 13788 | 212 | Plaur | | 1.538e-02 | -4.17 | IRAK3 | interpro domains | IPR033610 | 1 | 1 | 13788 | 212 | Irak3 | | 1.538e-02 | -4.17 | Nrf2 | interpro domains | IPR029845 | 1 | 1 | 13788 | 212 | Nfe2l2 | | 1.538e-02 | -4.17 | SBNO2 | interpro domains | IPR030410 | 1 | 1 | 13788 | 212 | Sbno2 | | 1.538e-02 | -4.17 | UMP-CMP\_kinase\_2 | interpro domains | IPR014505 | 1 | 1 | 13788 | 212 | Cmpk2 | | 1.538e-02 | -4.17 | PAR14\_RRM2 | interpro domains | IPR034517 | 1 | 1 | 13788 | 212 | Parp14 | | 1.538e-02 | -4.17 | K\_chnl\_volt-dep\_Kv1 | interpro domains | IPR004052 | 1 | 1 | 13788 | 212 | Kcna5 | | 1.538e-02 | -4.17 | FZ4\_CRD | interpro domains | IPR041765 | 1 | 1 | 13788 | 212 | Fzd4 | | 1.538e-02 | -4.17 | S100A8 | interpro domains | IPR028474 | 1 | 1 | 13788 | 212 | S100a8 | | 1.538e-02 | -4.17 | XAF1\_C | interpro domains | IPR041386 | 1 | 1 | 13788 | 212 | Xaf1 | | 1.538e-02 | -4.17 | RBM47\_RRM2 | interpro domains | IPR034440 | 1 | 1 | 13788 | 212 | Rbm47 | | 1.538e-02 | -4.17 | GCSF/MGF | interpro domains | IPR040117 | 1 | 1 | 13788 | 212 | Csf3 | | 1.538e-02 | -4.17 | RBM39\_linker | interpro domains | IPR029123 | 1 | 1 | 13788 | 212 | Rbm39 | | 1.538e-02 | -4.17 | IL-1\_alpha | interpro domains | IPR003295 | 1 | 1 | 13788 | 212 | Il1a | | 1.538e-02 | -4.17 | FYB | interpro domains | IPR030635 | 1 | 1 | 13788 | 212 | Fyb | | 1.538e-02 | -4.17 | NADPH\_oxidase\_p47Phox\_C | interpro domains | IPR015039 | 1 | 1 | 13788 | 212 | Ncf1 | | 1.538e-02 | -4.17 | TNF | interpro domains | IPR006053 | 1 | 1 | 13788 | 212 | Tnf | | 1.538e-02 | -4.17 | Smim3 | interpro domains | IPR035275 | 1 | 1 | 13788 | 212 | Smim3 | | 1.538e-02 | -4.17 | ACD\_HspB1 | interpro domains | IPR037876 | 1 | 1 | 13788 | 212 | Hspb1 | | 1.539e-02 | -4.17 | - | gene3d domains | 3.30.1330.30 | 13 | 2 | 6647 | 99 | Gadd45b,Gadd45g | | 1.545e-02 | -4.17 | positive regulation of transcription, DNA-templated | biological process | GO:0045893 | 1225 | 29 | 13711 | 214 | Banp,Pik3r1,Dtx3l,Irf7,Il6,Nfkb2,Nfkbia,Il1a,Sox11,Runx1,Bach1,Cebpd,Spi1,Nfkb1,Parp9,Sbno2,Fzd4,Tnip1,Aff1,Ncf1,Ripk1,Zbtb16,Csf3,Atf3,Tnf,Bcl3,Nfe2l2,Irf2,Maff | | 1.547e-02 | -4.17 | negative regulation of RNA metabolic process | biological process | GO:0051253 | 1067 | 26 | 13711 | 214 | Znfx1,Tnf,Zfp36,Atf3,Zbtb16,Ifi207,Arid5b,Ifi204,Rrp8,Dnajb1,Rasd1,Noct,Bcl3,Sox11,Mxd4,Gbp4,Ifi209,Sbno2,Nfkbie,Parp9,Nfkb1,Spi1,Cebpd,Ifi211,Bach1,Runx1 | | 1.559e-02 | -4.16 | OSADA\_ASCL1\_TARGETS\_UP | MSigDB lists | OSADA\_ASCL1\_TARGETS\_UP | 36 | 3 | 12187 | 179 | Rassf4,Atf3,Rgs16 | | 1.559e-02 | -4.16 | TAKEDA\_TARGETS\_OF\_NUP98\_HOXA9\_FUSION\_6HR\_DN | MSigDB lists | TAKEDA\_TARGETS\_OF\_NUP98\_HOXA9\_FUSION\_6HR\_DN | 36 | 3 | 12187 | 179 | Ccl4,Ccl12,Zfp36 | | 1.559e-02 | -4.16 | GENTILE\_UV\_RESPONSE\_CLUSTER\_D7 | MSigDB lists | GENTILE\_UV\_RESPONSE\_CLUSTER\_D7 | 36 | 3 | 12187 | 179 | Il4ra,Cebpd,Ptpn1 | | 1.559e-02 | -4.16 | PID\_DELTA\_NP63\_PATHWAY | MSigDB lists | PID\_DELTA\_NP63\_PATHWAY | 36 | 3 | 12187 | 179 | Runx1,Cebpd,Il1a | | 1.559e-02 | -4.16 | GO\_REGULATION\_OF\_CELL\_KILLING | MSigDB lists | GO\_REGULATION\_OF\_CELL\_KILLING | 36 | 3 | 12187 | 179 | Icam1,Ccl12,Tap1 | | 1.559e-02 | -4.16 | GO\_POTASSIUM\_CHANNEL\_REGULATOR\_ACTIVITY | MSigDB lists | GO\_POTASSIUM\_CHANNEL\_REGULATOR\_ACTIVITY | 36 | 3 | 12187 | 179 | Kcna5,Sgk1,Sgk3 | | 1.560e-02 | -4.16 | NECFESHC | pfam domains | PF16621 | 1 | 1 | 12881 | 201 | Ncf1 | | 1.560e-02 | -4.16 | Smim3 | pfam domains | PF17307 | 1 | 1 | 12881 | 201 | Smim3 | | 1.560e-02 | -4.16 | MmgE\_PrpD | pfam domains | PF03972 | 1 | 1 | 12881 | 201 | Acod1 | | 1.560e-02 | -4.16 | p47\_phox\_C | pfam domains | PF08944 | 1 | 1 | 12881 | 201 | Ncf1 | | 1.560e-02 | -4.16 | IL6 | pfam domains | PF00489 | 1 | 1 | 12881 | 201 | Il6 | | 1.560e-02 | -4.16 | Methyltransf\_8 | pfam domains | PF05148 | 1 | 1 | 12881 | 201 | Rrp8 | | 1.560e-02 | -4.16 | DUF4617 | pfam domains | PF15395 | 1 | 1 | 12881 | 201 | Resf1 | | 1.560e-02 | -4.16 | XAF1\_C | pfam domains | PF18608 | 1 | 1 | 12881 | 201 | Xaf1 | | 1.560e-02 | -4.16 | IL4Ra\_N | pfam domains | PF09238 | 1 | 1 | 12881 | 201 | Il4ra | | 1.560e-02 | -4.16 | GCSF | pfam domains | PF16647 | 1 | 1 | 12881 | 201 | Csf3 | | 1.560e-02 | -4.16 | TMEM252 | pfam domains | PF15664 | 1 | 1 | 12881 | 201 | Tmem252 | | 1.560e-02 | -4.16 | DEPP | pfam domains | PF15343 | 1 | 1 | 12881 | 201 | Depp1 | | 1.560e-02 | -4.16 | Cid2 | pfam domains | PF09774 | 1 | 1 | 12881 | 201 | Ccdc58 | | 1.560e-02 | -4.16 | Macscav\_rec | pfam domains | PF03523 | 1 | 1 | 12881 | 201 | Msr1 | | 1.560e-02 | -4.16 | RBM39linker | pfam domains | PF15519 | 1 | 1 | 12881 | 201 | Rbm39 | | 1.561e-02 | -4.16 | GSE29617\_CTRL\_VS\_TIV\_FLU\_VACCINE\_PBMC\_2008\_UP | MSigDB lists | GSE29617\_CTRL\_VS\_TIV\_FLU\_VACCINE\_PBMC\_2008\_UP | 137 | 6 | 12187 | 179 | Tiparp,Atf3,Nfe2l2,Bcl3,Zfp36,Ip6k2 | | 1.561e-02 | -4.16 | GSE37301\_HEMATOPOIETIC\_STEM\_CELL\_VS\_PRO\_BCELL\_UP | MSigDB lists | GSE37301\_HEMATOPOIETIC\_STEM\_CELL\_VS\_PRO\_BCELL\_UP | 137 | 6 | 12187 | 179 | Gadd45b,Plaur,Rbm47,Icam1,Map3k6,Cflar | | 1.561e-02 | -4.16 | GSE45365\_NK\_CELL\_VS\_CD8\_TCELL\_MCMV\_INFECTION\_UP | MSigDB lists | GSE45365\_NK\_CELL\_VS\_CD8\_TCELL\_MCMV\_INFECTION\_UP | 177 | 7 | 12187 | 179 | Cebpd,Ifi204,Ifi207,Ifi209,Tnfsf10,Ifi211,Ifitm2 | | 1.561e-02 | -4.16 | POOLA\_INVASIVE\_BREAST\_CANCER\_UP | MSigDB lists | POOLA\_INVASIVE\_BREAST\_CANCER\_UP | 177 | 7 | 12187 | 179 | Ccl4,Vcam1,Sox11,Csf2rb,Plaur,Rgs16,Plek | | 1.561e-02 | -4.16 | GSE23114\_WT\_VS\_SLE2C1\_MOUSE\_SPLEEN\_B1A\_BCELL\_UP | MSigDB lists | GSE23114\_WT\_VS\_SLE2C1\_MOUSE\_SPLEEN\_B1A\_BCELL\_UP | 177 | 7 | 12187 | 179 | Nfkb1,Nfe2l2,Atf3,Ccnd2,Tnf,Cflar,Gem | | 1.561e-02 | -4.16 | regulation of macrophage tolerance induction | biological process | GO:0010932 | 1 | 1 | 13711 | 214 | Irak3 | | 1.561e-02 | -4.16 | modulation by symbiont of host signal transduction pathway | biological process | GO:0052027 | 1 | 1 | 13711 | 214 | Tnip1 | | 1.561e-02 | -4.16 | positive regulation of programmed necrotic cell death | biological process | GO:0062100 | 1 | 1 | 13711 | 214 | Ripk1 | | 1.561e-02 | -4.16 | negative regulation of nucleotide-binding oligomerization domain containing 1 signaling pathway | biological process | GO:0070429 | 1 | 1 | 13711 | 214 | Tnfaip3 | | 1.561e-02 | -4.16 | positive regulation of adiponectin secretion | biological process | GO:0070165 | 1 | 1 | 13711 | 214 | Hcar2 | | 1.561e-02 | -4.16 | negative regulation of toll-like receptor 5 signaling pathway | biological process | GO:0034148 | 1 | 1 | 13711 | 214 | Tnfaip3 | | 1.561e-02 | -4.16 | negative regulation of calcidiol 1-monooxygenase activity | biological process | GO:0010956 | 1 | 1 | 13711 | 214 | Nfkb1 | | 1.561e-02 | -4.16 | interleukin-10 secretion | biological process | GO:0072608 | 1 | 1 | 13711 | 214 | Isg15 | | 1.561e-02 | -4.16 | response to chromate | biological process | GO:0046687 | 1 | 1 | 13711 | 214 | Mt1 | | 1.561e-02 | -4.16 | mycotoxin metabolic process | biological process | GO:0043385 | 1 | 1 | 13711 | 214 | Nfe2l2 | | 1.561e-02 | -4.16 | response to flavonoid | biological process | GO:1905395 | 1 | 1 | 13711 | 214 | Ip6k2 | | 1.561e-02 | -4.16 | cellular response to glucoside | biological process | GO:1904632 | 1 | 1 | 13711 | 214 | Nfkb1 | | 1.561e-02 | -4.16 | membrane repolarization during SA node cell action potential | biological process | GO:0086052 | 1 | 1 | 13711 | 214 | Kcna5 | | 1.561e-02 | -4.16 | modulation of signal transduction in other organism | biological process | GO:0044501 | 1 | 1 | 13711 | 214 | Tnip1 | | 1.561e-02 | -4.16 | regulation of ATP:ADP antiporter activity | biological process | GO:0070926 | 1 | 1 | 13711 | 214 | Ripk1 | | 1.561e-02 | -4.16 | positive regulation of fractalkine biosynthetic process | biological process | GO:0050754 | 1 | 1 | 13711 | 214 | Tnf | | 1.561e-02 | -4.16 | negative regulation of polynucleotide adenylyltransferase activity | biological process | GO:1904246 | 1 | 1 | 13711 | 214 | Zfp36 | | 1.561e-02 | -4.16 | neutrophil mediated killing of gram-positive bacterium | biological process | GO:0070946 | 1 | 1 | 13711 | 214 | Ncf1 | | 1.561e-02 | -4.16 | cellular response to flavonoid | biological process | GO:1905396 | 1 | 1 | 13711 | 214 | Ip6k2 | | 1.561e-02 | -4.16 | regulation of fractalkine biosynthetic process | biological process | GO:0050752 | 1 | 1 | 13711 | 214 | Tnf | | 1.561e-02 | -4.16 | hypoxanthine catabolic process | biological process | GO:0009114 | 1 | 1 | 13711 | 214 | Xdh | | 1.561e-02 | -4.16 | positive regulation of deadenylation-independent decapping of nuclear-transcribed mRNA | biological process | GO:1901835 | 1 | 1 | 13711 | 214 | Zfp36 | | 1.561e-02 | -4.16 | phospholipase C-inhibiting G protein-coupled receptor signaling pathway | biological process | GO:0030845 | 1 | 1 | 13711 | 214 | Plek | | 1.561e-02 | -4.16 | positive regulation of neutrophil mediated killing of fungus | biological process | GO:0070965 | 1 | 1 | 13711 | 214 | Cxcl1 | | 1.561e-02 | -4.16 | positive regulation of platelet-derived growth factor production | biological process | GO:0090362 | 1 | 1 | 13711 | 214 | Ptgs2 | | 1.561e-02 | -4.16 | aflatoxin metabolic process | biological process | GO:0046222 | 1 | 1 | 13711 | 214 | Nfe2l2 | | 1.561e-02 | -4.16 | organic heteropentacyclic compound catabolic process | biological process | GO:1901377 | 1 | 1 | 13711 | 214 | Nfe2l2 | | 1.561e-02 | -4.16 | cardiac neuron differentiation | biological process | GO:0060945 | 1 | 1 | 13711 | 214 | Vcam1 | | 1.561e-02 | -4.16 | positive regulation of leukocyte adhesion to arterial endothelial cell | biological process | GO:1904999 | 1 | 1 | 13711 | 214 | Tnf | | 1.561e-02 | -4.16 | positive regulation of oligodendrocyte apoptotic process | biological process | GO:1900143 | 1 | 1 | 13711 | 214 | Akap12 | | 1.561e-02 | -4.16 | tolerance induction to nonself antigen | biological process | GO:0002462 | 1 | 1 | 13711 | 214 | C3ar1 | | 1.561e-02 | -4.16 | membrane repolarization during bundle of His cell action potential | biological process | GO:0086050 | 1 | 1 | 13711 | 214 | Kcna5 | | 1.561e-02 | -4.16 | cellular response to non-ionic osmotic stress | biological process | GO:0071471 | 1 | 1 | 13711 | 214 | Ptgs2 | | 1.561e-02 | -4.16 | regulation of inositol-polyphosphate 5-phosphatase activity | biological process | GO:0010924 | 1 | 1 | 13711 | 214 | Plek | | 1.561e-02 | -4.16 | regulation of fractalkine production | biological process | GO:0032644 | 1 | 1 | 13711 | 214 | Tnf | | 1.561e-02 | -4.16 | negative regulation of establishment of Sertoli cell barrier | biological process | GO:1904445 | 1 | 1 | 13711 | 214 | Il1a | | 1.561e-02 | -4.16 | response to diterpene | biological process | GO:1904629 | 1 | 1 | 13711 | 214 | Nfkb1 | | 1.561e-02 | -4.16 | cellular response to diterpene | biological process | GO:1904630 | 1 | 1 | 13711 | 214 | Nfkb1 | | 1.561e-02 | -4.16 | regulation of lens epithelial cell proliferation | biological process | GO:2001109 | 1 | 1 | 13711 | 214 | Sox11 | | 1.561e-02 | -4.16 | positive regulation of necroptotic process | biological process | GO:0060545 | 1 | 1 | 13711 | 214 | Ripk1 | | 1.561e-02 | -4.16 | regulation of cell diameter | biological process | GO:0060305 | 1 | 1 | 13711 | 214 | Plek | | 1.561e-02 | -4.16 | negative regulation of hydrogen peroxide catabolic process | biological process | GO:2000296 | 1 | 1 | 13711 | 214 | Hp | | 1.561e-02 | -4.16 | negative regulation of natural killer cell chemotaxis | biological process | GO:2000502 | 1 | 1 | 13711 | 214 | Ccl12 | | 1.561e-02 | -4.16 | positive regulation of chronic inflammatory response to antigenic stimulus | biological process | GO:0002876 | 1 | 1 | 13711 | 214 | Tnf | | 1.561e-02 | -4.16 | response to glucoside | biological process | GO:1904631 | 1 | 1 | 13711 | 214 | Nfkb1 | | 1.561e-02 | -4.16 | cellular response to pyrimidine ribonucleotide | biological process | GO:1905835 | 1 | 1 | 13711 | 214 | P2ry6 | | 1.561e-02 | -4.16 | organic heteropentacyclic compound metabolic process | biological process | GO:1901376 | 1 | 1 | 13711 | 214 | Nfe2l2 | | 1.561e-02 | -4.16 | mycotoxin catabolic process | biological process | GO:0043387 | 1 | 1 | 13711 | 214 | Nfe2l2 | | 1.561e-02 | -4.16 | neutrophil mediated killing of fungus | biological process | GO:0070947 | 1 | 1 | 13711 | 214 | Ncf1 | | 1.561e-02 | -4.16 | regulation of eosinophil differentiation | biological process | GO:0045643 | 1 | 1 | 13711 | 214 | Trib1 | | 1.561e-02 | -4.16 | negative regulation of interleukin-12 biosynthetic process | biological process | GO:0045083 | 1 | 1 | 13711 | 214 | Nfkb1 | | 1.561e-02 | -4.16 | positive regulation of eosinophil differentiation | biological process | GO:0045645 | 1 | 1 | 13711 | 214 | Trib1 | | 1.561e-02 | -4.16 | positive regulation of inositol-polyphosphate 5-phosphatase activity | biological process | GO:0010925 | 1 | 1 | 13711 | 214 | Plek | | 1.561e-02 | -4.16 | positive regulation of interleukin-10 biosynthetic process | biological process | GO:0045082 | 1 | 1 | 13711 | 214 | Bcl3 | | 1.561e-02 | -4.16 | negative regulation of antigen processing and presentation of peptide or polysaccharide antigen via MHC class II | biological process | GO:0002581 | 1 | 1 | 13711 | 214 | Thbs1 | | 1.561e-02 | -4.16 | hypoxanthine oxidation | biological process | GO:0006150 | 1 | 1 | 13711 | 214 | Xdh | | 1.561e-02 | -4.16 | T-helper 2 cell differentiation | biological process | GO:0045064 | 1 | 1 | 13711 | 214 | Bcl3 | | 1.561e-02 | -4.16 | regulation of chronic inflammatory response to antigenic stimulus | biological process | GO:0002874 | 1 | 1 | 13711 | 214 | Tnf | | 1.561e-02 | -4.16 | cellular response to chromate | biological process | GO:0071247 | 1 | 1 | 13711 | 214 | Mt1 | | 1.561e-02 | -4.16 | regulation of interleukin-17 biosynthetic process | biological process | GO:0045378 | 1 | 1 | 13711 | 214 | Il6 | | 1.561e-02 | -4.16 | propionate metabolic process, methylcitrate cycle | biological process | GO:0019679 | 1 | 1 | 13711 | 214 | Acod1 | | 1.561e-02 | -4.16 | regulation of neutrophil mediated killing of fungus | biological process | GO:0070953 | 1 | 1 | 13711 | 214 | Cxcl1 | | 1.561e-02 | -4.16 | positive regulation of neutrophil mediated cytotoxicity | biological process | GO:0070960 | 1 | 1 | 13711 | 214 | Cxcl1 | | 1.561e-02 | -4.16 | positive regulation of lens epithelial cell proliferation | biological process | GO:2001111 | 1 | 1 | 13711 | 214 | Sox11 | | 1.561e-02 | -4.16 | response to pyrimidine ribonucleotide | biological process | GO:1905834 | 1 | 1 | 13711 | 214 | P2ry6 | | 1.561e-02 | -4.16 | hepatic immune response | biological process | GO:0002384 | 1 | 1 | 13711 | 214 | Il6 | | 1.561e-02 | -4.16 | cerebellum vasculature morphogenesis | biological process | GO:0061301 | 1 | 1 | 13711 | 214 | Fzd4 | | 1.561e-02 | -4.16 | positive regulation of neutrophil mediated killing of symbiont cell | biological process | GO:0070961 | 1 | 1 | 13711 | 214 | Cxcl1 | | 1.561e-02 | -4.16 | regulation of granuloma formation | biological process | GO:0002631 | 1 | 1 | 13711 | 214 | Tnfaip3 | | 1.561e-02 | -4.16 | positive regulation of fractalkine production | biological process | GO:0032724 | 1 | 1 | 13711 | 214 | Tnf | | 1.561e-02 | -4.16 | regulation of toll-like receptor 5 signaling pathway | biological process | GO:0034147 | 1 | 1 | 13711 | 214 | Tnfaip3 | | 1.561e-02 | -4.16 | urokinase plasminogen activator signaling pathway | biological process | GO:0038195 | 1 | 1 | 13711 | 214 | Plaur | | 1.561e-02 | -4.16 | aflatoxin catabolic process | biological process | GO:0046223 | 1 | 1 | 13711 | 214 | Nfe2l2 | | 1.561e-02 | -4.16 | cellular response to interleukin-11 | biological process | GO:0071348 | 1 | 1 | 13711 | 214 | Sbno2 | | 1.561e-02 | -4.16 | negative regulation of granuloma formation | biological process | GO:0002632 | 1 | 1 | 13711 | 214 | Tnfaip3 | | 1.561e-02 | -4.16 | positive regulation of interleukin-17 biosynthetic process | biological process | GO:0045380 | 1 | 1 | 13711 | 214 | Il6 | | 1.561e-02 | -4.16 | negative regulation of natural killer cell degranulation | biological process | GO:0043322 | 1 | 1 | 13711 | 214 | Lgals9 | | 1.561e-02 | -4.16 | glucagon secretion | biological process | GO:0070091 | 1 | 1 | 13711 | 214 | Il6 | | 1.561e-02 | -4.16 | modulation by symbiont of host I-kappaB kinase/NF-kappaB cascade | biological process | GO:0085032 | 1 | 1 | 13711 | 214 | Tnip1 | | 1.561e-02 | -4.16 | regulation of deadenylation-independent decapping of nuclear-transcribed mRNA | biological process | GO:1901834 | 1 | 1 | 13711 | 214 | Zfp36 | | 1.561e-02 | -4.16 | modulation of signal transduction in other organism involved in symbiotic interaction | biological process | GO:0052250 | 1 | 1 | 13711 | 214 | Tnip1 | | 1.561e-02 | -4.16 | negative regulation of branching involved in lung morphogenesis | biological process | GO:0061048 | 1 | 1 | 13711 | 214 | Tnf | | 1.561e-02 | -4.16 | isopeptide cross-linking via N6-(L-isoglutamyl)-L-lysine | biological process | GO:0018153 | 1 | 1 | 13711 | 214 | Tgm2 | | 1.561e-02 | -4.16 | positive regulation of macrophage tolerance induction | biological process | GO:0010933 | 1 | 1 | 13711 | 214 | Irak3 | | 1.561e-02 | -4.16 | response to non-ionic osmotic stress | biological process | GO:0010335 | 1 | 1 | 13711 | 214 | Ptgs2 | | 1.561e-02 | -4.16 | oxidoreductase activity, acting on CH or CH2 groups, oxygen as acceptor | molecular function | GO:0016727 | 1 | 1 | 13516 | 211 | Xdh | | 1.561e-02 | -4.16 | voltage-gated potassium channel activity involved in bundle of His cell action potential repolarization | molecular function | GO:0086087 | 1 | 1 | 13516 | 211 | Kcna5 | | 1.561e-02 | -4.16 | complement component C5a binding | molecular function | GO:0001856 | 1 | 1 | 13516 | 211 | C5ar1 | | 1.561e-02 | -4.16 | interleukin-1 type I receptor antagonist activity | molecular function | GO:0045352 | 1 | 1 | 13516 | 211 | Il1rn | | 1.561e-02 | -4.16 | aconitate decarboxylase activity | molecular function | GO:0047613 | 1 | 1 | 13516 | 211 | Acod1 | | 1.561e-02 | -4.16 | cyclin-dependent protein kinase activating kinase activity | molecular function | GO:0019912 | 1 | 1 | 13516 | 211 | Cdkn1a | | 1.561e-02 | -4.16 | complement component C3a receptor activity | molecular function | GO:0004876 | 1 | 1 | 13516 | 211 | C3ar1 | | 1.561e-02 | -4.16 | urotensin II receptor activity | molecular function | GO:0001604 | 1 | 1 | 13516 | 211 | Gpr84 | | 1.561e-02 | -4.16 | urokinase plasminogen activator receptor activity | molecular function | GO:0030377 | 1 | 1 | 13516 | 211 | Plaur | | 1.561e-02 | -4.16 | xanthine oxidase activity | molecular function | GO:0004855 | 1 | 1 | 13516 | 211 | Xdh | | 1.561e-02 | -4.16 | voltage-gated potassium channel activity involved in SA node cell action potential repolarization | molecular function | GO:0086090 | 1 | 1 | 13516 | 211 | Kcna5 | | 1.561e-02 | -4.16 | UDP-activated nucleotide receptor activity | molecular function | GO:0045029 | 1 | 1 | 13516 | 211 | P2ry6 | | 1.561e-02 | -4.16 | protein kinase C inhibitor activity | molecular function | GO:0008426 | 1 | 1 | 13516 | 211 | Hspb1 | | 1.561e-02 | -4.16 | interleukin-1, type II receptor binding | molecular function | GO:0005151 | 1 | 1 | 13516 | 211 | Il1rn | | 1.561e-02 | -4.16 | UMP kinase activity | molecular function | GO:0033862 | 1 | 1 | 13516 | 211 | Cmpk2 | | 1.561e-02 | -4.16 | enterobactin binding | molecular function | GO:1903981 | 1 | 1 | 13516 | 211 | Lcn2 | | 1.561e-02 | -4.16 | 2-methylcitrate dehydratase activity | molecular function | GO:0047547 | 1 | 1 | 13516 | 211 | Acod1 | | 1.561e-02 | -4.16 | interleukin-1 receptor antagonist activity | molecular function | GO:0005152 | 1 | 1 | 13516 | 211 | Il1rn | | 1.561e-02 | -4.16 | ISG15-specific protease activity | molecular function | GO:0019785 | 1 | 1 | 13516 | 211 | Usp18 | | 1.561e-02 | -4.16 | interleukin-1 type II receptor antagonist activity | molecular function | GO:0045353 | 1 | 1 | 13516 | 211 | Il1rn | | 1.561e-02 | -4.16 | nicotinic acid receptor activity | molecular function | GO:0070553 | 1 | 1 | 13516 | 211 | Hcar2 | | 1.570e-02 | -4.15 | GO\_SIGNAL\_TRANSDUCER\_ACTIVITY | MSigDB lists | GO\_SIGNAL\_TRANSDUCER\_ACTIVITY | 865 | 21 | 12187 | 179 | Map3k6,Cd14,Sele,P2ry6,Il17ra,Icam1,Csf2rb,Plaur,Il4ra,Pik3r1,Map3k8,Sdc4,Gpr84,Hcar2,Rhoc,C3ar1,C5ar1,Fzd4,S100a9,Osmr,Igsf6 | | 1.572e-02 | -4.15 | positive regulation of vascular smooth muscle cell proliferation | biological process | GO:1904707 | 34 | 3 | 13711 | 214 | Tnf,P2ry6,Adamts1 | | 1.572e-02 | -4.15 | negative regulation of cellular carbohydrate metabolic process | biological process | GO:0010677 | 34 | 3 | 13711 | 214 | Plek,Il6,Ier3 | | 1.572e-02 | -4.15 | regulation of endothelial cell differentiation | biological process | GO:0045601 | 34 | 3 | 13711 | 214 | Tnf,Xdh,Apold1 | | 1.582e-02 | -4.15 | negative regulation of blood vessel morphogenesis | biological process | GO:2000181 | 94 | 5 | 13711 | 214 | Adamts9,Thbs1,Ccl2,Adamts1,Xdh | | 1.590e-02 | -4.14 | positive regulation of osteoblast differentiation | biological process | GO:0045669 | 62 | 4 | 13711 | 214 | Il6,Ifi204,Cebpd,Sox11 | | 1.590e-02 | -4.14 | regulation of sprouting angiogenesis | biological process | GO:1903670 | 62 | 4 | 13711 | 214 | Ptgs2,Rhoj,Thbs1,Adamts9 | | 1.600e-02 | -4.14 | GO\_NEGATIVE\_REGULATION\_OF\_INTRINSIC\_APOPTOTIC\_SIGNALING\_PATHWAY | MSigDB lists | GO\_NEGATIVE\_REGULATION\_OF\_INTRINSIC\_APOPTOTIC\_SIGNALING\_PATHWAY | 66 | 4 | 12187 | 179 | Nfe2l2,Ptpn1,Plaur,Hspb1 | | 1.600e-02 | -4.14 | GO\_NEGATIVE\_REGULATION\_OF\_INFLAMMATORY\_RESPONSE | MSigDB lists | GO\_NEGATIVE\_REGULATION\_OF\_INFLAMMATORY\_RESPONSE | 66 | 4 | 12187 | 179 | Socs3,Saa1,Tnfaip3,Nfkb1 | | 1.600e-02 | -4.14 | TOOKER\_GEMCITABINE\_RESISTANCE\_UP | MSigDB lists | TOOKER\_GEMCITABINE\_RESISTANCE\_UP | 66 | 4 | 12187 | 179 | Isg15,Arid5b,Rrp8,Irf9 | | 1.605e-02 | -4.13 | MARTORIATI\_MDM4\_TARGETS\_FETAL\_LIVER\_UP | MSigDB lists | MARTORIATI\_MDM4\_TARGETS\_FETAL\_LIVER\_UP | 178 | 7 | 12187 | 179 | Ier3,Zbtb16,Cdkn1a,Ccdc58,Tap1,A2m,Ifitm3 | | 1.612e-02 | -4.13 | GSE21927\_BALBC\_VS\_C57BL6\_MONOCYTE\_TUMOR\_DN | MSigDB lists | GSE21927\_BALBC\_VS\_C57BL6\_MONOCYTE\_TUMOR\_DN | 138 | 6 | 12187 | 179 | Cxcl16,Ifi207,Ifi204,S100a9,Ifi211,Ifi209 | | 1.612e-02 | -4.13 | SNF5\_DN.V1\_UP | MSigDB lists | SNF5\_DN.V1\_UP | 138 | 6 | 12187 | 179 | Spi1,Ifit2,C3ar1,Znfx1,Oasl1,Plek | | 1.612e-02 | -4.13 | GSE16385\_MONOCYTE\_VS\_12H\_IL4\_TREATED\_MACROPHAGE\_UP | MSigDB lists | GSE16385\_MONOCYTE\_VS\_12H\_IL4\_TREATED\_MACROPHAGE\_UP | 138 | 6 | 12187 | 179 | Sdc4,Usp18,Rnf213,Gbp6,Irf9,Gbp4 | | 1.625e-02 | -4.12 | cellular response to metal ion | biological process | GO:0071248 | 130 | 6 | 13711 | 214 | Ncf1,Nfe2l2,Iqgap1,Mt2,Clic4,Mt1 | | 1.630e-02 | -4.12 | MAP3K8 (TPL2)-dependent MAPK1/3 activation | REACTOME pathways | R-MMU-5684264 | 12 | 2 | 6297 | 105 | Map3k8,Nfkb1 | | 1.630e-02 | -4.12 | Synthesis of Prostaglandins (PG) and Thromboxanes (TX) | REACTOME pathways | R-MMU-2162123 | 12 | 2 | 6297 | 105 | Ptges,Ptgs2 | | 1.633e-02 | -4.11 | platelet alpha granule | cellular component | GO:0031091 | 13 | 2 | 13825 | 212 | Thbs1,Selp | | 1.633e-02 | -4.11 | GO\_REGULATION\_OF\_SECRETION | MSigDB lists | GO\_REGULATION\_OF\_SECRETION | 503 | 14 | 12187 | 179 | Hcar2,Il6,Tnf,Il4ra,Sdc4,S100a8,S100a9,Kcna5,Il1a,Sox11,Sgk1,Il17ra,Saa1,Cd14 | | 1.639e-02 | -4.11 | purine nucleotide binding | molecular function | GO:0017076 | 1555 | 35 | 13516 | 211 | Mx1,Rhoj,Bbs12,Tgm2,Gbp7,Gbp6,Rhoc,Rasd1,Oasl2,Pygm,Sgk1,Tap1,Gem,Map3k8,Sgk3,Oasl1,Runx1,Irak3,Rhou,Ksr1,Slfn9,Map3k6,Tor3a,Tubb6,Cmpk2,Irgm1,Ripk1,Ip6k2,Hspa5,Tgtp1,Hcar2,Trib1,Gbp3,Iigp1,Slfn5 | | 1.642e-02 | -4.11 | S100\_Ca-bd\_sub | interpro domains | IPR013787 | 13 | 2 | 13788 | 212 | S100a8,S100a9 | | 1.650e-02 | -4.10 | GSE11924\_TFH\_VS\_TH1\_CD4\_TCELL\_UP | MSigDB lists | GSE11924\_TFH\_VS\_TH1\_CD4\_TCELL\_UP | 179 | 7 | 12187 | 179 | Iqgap1,Rbm39,Gbp3,Tor3a,Ifit3b,Gpr84,Casp4 | | 1.655e-02 | -4.10 | p53 signaling | WikiPathways | WP2902 | 59 | 5 | 3756 | 96 | Cdkn1a,Thbs1,Ccnd2,Gadd45g,Gadd45b | | 1.665e-02 | -4.10 | GSE37416\_CTRL\_VS\_3H\_F\_TULARENSIS\_LVS\_NEUTROPHIL\_DN | MSigDB lists | GSE37416\_CTRL\_VS\_3H\_F\_TULARENSIS\_LVS\_NEUTROPHIL\_DN | 139 | 6 | 12187 | 179 | Ptgs2,Tnf,Nfe2l2,Sbno2,Il1a,Icam1 | | 1.665e-02 | -4.10 | GSE36078\_WT\_VS\_IL1R\_KO\_LUNG\_DC\_UP | MSigDB lists | GSE36078\_WT\_VS\_IL1R\_KO\_LUNG\_DC\_UP | 139 | 6 | 12187 | 179 | Herc6,Ifit2,Cdkn1a,Slfn5,Plek,Parp9 | | 1.665e-02 | -4.10 | GSE46606\_IRF4\_KO\_VS\_WT\_CD40L\_IL2\_IL5\_1DAY\_STIMULATED\_BCELL\_DN | MSigDB lists | GSE46606\_IRF4\_KO\_VS\_WT\_CD40L\_IL2\_IL5\_1DAY\_STIMULATED\_BCELL\_DN | 139 | 6 | 12187 | 179 | Pik3r5,Nfkbiz,S100a9,Casp4,Thbs1,Mmp8 | | 1.665e-02 | -4.10 | GSE30083\_SP2\_VS\_SP3\_THYMOCYTE\_DN | MSigDB lists | GSE30083\_SP2\_VS\_SP3\_THYMOCYTE\_DN | 139 | 6 | 12187 | 179 | Csf3,Il17ra,Clic4,Sgk1,Zbtb16,Rasd1 | | 1.667e-02 | -4.09 | PTK6 Down-Regulation | REACTOME pathways | R-MMU-8849472 | 1 | 1 | 6297 | 105 | Ptpn1 | | 1.668e-02 | -4.09 | ribonucleotide binding | molecular function | GO:0032553 | 1557 | 35 | 13516 | 211 | Sgk3,Map3k8,Gem,Tap1,Pygm,Oasl2,Rasd1,Sgk1,Rhoc,Gbp6,Bbs12,Gbp7,Tgm2,Rhoj,Mx1,Slfn5,Gbp3,Trib1,Iigp1,Hcar2,Tgtp1,Hspa5,Ripk1,Ip6k2,Cmpk2,Irgm1,Tor3a,Map3k6,Tubb6,Ksr1,Slfn9,Oasl1,Runx1,Rhou,Irak3 | | 1.669e-02 | -4.09 | proximal promoter sequence-specific DNA binding | molecular function | GO:0000987 | 427 | 13 | 13516 | 211 | Atf3,Irf9,Runx1,Ifi211,Ifi207,Spi1,Maff,Irf7,Cebpd,Ifi204,Sox11,Nfkb2,Ifi209 | | 1.675e-02 | -4.09 | negative regulation of cellular biosynthetic process | biological process | GO:0031327 | 1233 | 29 | 13711 | 214 | Mxd4,Sox11,Errfi1,Il6,Ifi209,Gbp4,Nfkb1,Nfkbie,Parp9,Sbno2,Ifi211,Bach1,Cebpd,Spi1,Runx1,Tnf,Znfx1,Plek,Zfp36,Atf3,Ifi207,Arid5b,Zbtb16,Ifi204,Dnajb1,Rrp8,Prg4,Rasd1,Bcl3 | | 1.678e-02 | -4.09 | ENK\_UV\_RESPONSE\_EPIDERMIS\_UP | MSigDB lists | ENK\_UV\_RESPONSE\_EPIDERMIS\_UP | 222 | 8 | 12187 | 179 | Ifitm3,Angptl4,Bcl3,Irf7,S100a8,Osmr,Il4ra,Oasl1 | | 1.678e-02 | -4.09 | CREB\_Q2 | MSigDB lists | CREB\_Q2 | 222 | 8 | 12187 | 179 | Maff,Trib1,Kcna5,Gadd45b,Atf3,Gem,Tagln2,Cxcl16 | | 1.678e-02 | -4.09 | G1/S transition of mitotic cell cycle | biological process | GO:0000082 | 63 | 4 | 13711 | 214 | Bach1,Cdkn1a,Rhou,Ccnd2 | | 1.678e-02 | -4.09 | negative regulation of leukocyte proliferation | biological process | GO:0070664 | 63 | 4 | 13711 | 214 | Sox11,Ccl12,Lgals9,Sdc4 | | 1.679e-02 | -4.09 | protein kinase activator activity | molecular function | GO:0030295 | 63 | 4 | 13516 | 211 | Irgm2,Iqgap1,Cdkn1a,Lgals9 | | 1.679e-02 | -4.09 | GO\_SKELETAL\_MUSCLE\_CELL\_DIFFERENTIATION | MSigDB lists | GO\_SKELETAL\_MUSCLE\_CELL\_DIFFERENTIATION | 37 | 3 | 12187 | 179 | Sox11,Maff,Atf3 | | 1.679e-02 | -4.09 | BIOCARTA\_TCR\_PATHWAY | MSigDB lists | BIOCARTA\_TCR\_PATHWAY | 37 | 3 | 12187 | 179 | Nfkb1,Pik3r1,Nfkbia | | 1.679e-02 | -4.09 | YAMASHITA\_LIVER\_CANCER\_STEM\_CELL\_DN | MSigDB lists | YAMASHITA\_LIVER\_CANCER\_STEM\_CELL\_DN | 37 | 3 | 12187 | 179 | Gadd45b,Thbs1,Cd14 | | 1.679e-02 | -4.09 | GO\_REGULATION\_OF\_ALCOHOL\_BIOSYNTHETIC\_PROCESS | MSigDB lists | GO\_REGULATION\_OF\_ALCOHOL\_BIOSYNTHETIC\_PROCESS | 37 | 3 | 12187 | 179 | Tnf,Nfkb1,Plek | | 1.679e-02 | -4.09 | GO\_RESPONSE\_TO\_PROGESTERONE | MSigDB lists | GO\_RESPONSE\_TO\_PROGESTERONE | 37 | 3 | 12187 | 179 | Thbs1,Socs3,Ccl12 | | 1.679e-02 | -4.09 | GO\_REGULATION\_OF\_EXTRINSIC\_APOPTOTIC\_SIGNALING\_PATHWAY\_IN\_ABSENCE\_OF\_LIGAND | MSigDB lists | GO\_REGULATION\_OF\_EXTRINSIC\_APOPTOTIC\_SIGNALING\_PATHWAY\_IN\_ABSENCE\_OF\_LIGAND | 37 | 3 | 12187 | 179 | Tnf,Il1a,Ripk1 | | 1.679e-02 | -4.09 | GO\_EXOCRINE\_SYSTEM\_DEVELOPMENT | MSigDB lists | GO\_EXOCRINE\_SYSTEM\_DEVELOPMENT | 37 | 3 | 12187 | 179 | Il6,Tnf,Tgm2 | | 1.679e-02 | -4.09 | GO\_POSITIVE\_REGULATION\_OF\_CHEMOKINE\_PRODUCTION | MSigDB lists | GO\_POSITIVE\_REGULATION\_OF\_CHEMOKINE\_PRODUCTION | 37 | 3 | 12187 | 179 | Tnf,Il4ra,Il6 | | 1.679e-02 | -4.09 | GO\_CYTOPLASMIC\_SEQUESTERING\_OF\_PROTEIN | MSigDB lists | GO\_CYTOPLASMIC\_SEQUESTERING\_OF\_PROTEIN | 37 | 3 | 12187 | 179 | Hspa5,Nfkbie,Nfkbia | | 1.683e-02 | -4.08 | GO\_HUMORAL\_IMMUNE\_RESPONSE | MSigDB lists | GO\_HUMORAL\_IMMUNE\_RESPONSE | 67 | 4 | 12187 | 179 | Bcl3,Tnf,Il6,Ccl12 | | 1.683e-02 | -4.08 | PID\_SMAD2\_3NUCLEAR\_PATHWAY | MSigDB lists | PID\_SMAD2\_3NUCLEAR\_PATHWAY | 67 | 4 | 12187 | 179 | Runx1,Cdkn1a,Irf7,Atf3 | | 1.683e-02 | -4.08 | YAO\_TEMPORAL\_RESPONSE\_TO\_PROGESTERONE\_CLUSTER\_0 | MSigDB lists | YAO\_TEMPORAL\_RESPONSE\_TO\_PROGESTERONE\_CLUSTER\_0 | 67 | 4 | 12187 | 179 | Cebpd,Bach1,Casp4,Gem | | 1.683e-02 | -4.08 | BRUINS\_UVC\_RESPONSE\_VIA\_TP53\_GROUP\_C | MSigDB lists | BRUINS\_UVC\_RESPONSE\_VIA\_TP53\_GROUP\_C | 67 | 4 | 12187 | 179 | Ip6k2,Gem,Socs3,Sdc4 | | 1.684e-02 | -4.08 | S100\_CABP | prosite domains | PS00303 | 11 | 2 | 8845 | 164 | S100a8,S100a9 | | 1.684e-02 | -4.08 | HEMOPEXIN\_2 | prosite domains | PS51642 | 11 | 2 | 8845 | 164 | Prg4,Mmp8 | | 1.684e-02 | -4.08 | CASPASE\_P20 | prosite domains | PS50208 | 11 | 2 | 8845 | 164 | Cflar,Casp4 | | 1.686e-02 | -4.08 | regulation of leukocyte proliferation | biological process | GO:0070663 | 169 | 7 | 13711 | 214 | Il6,Sdc4,Cdkn1a,Ccl12,Sox11,Lgals9,Vcam1 | | 1.686e-02 | -4.08 | ubiquitin-like protein ligase binding | molecular function | GO:0044389 | 293 | 10 | 13516 | 211 | Hspa5,Parp9,Cdkn1a,Fzd4,Trib1,Pik3r1,Ripk1,Dtx3l,Isg15,Nfkbia | | 1.689e-02 | -4.08 | macrophage chemotaxis | biological process | GO:0048246 | 13 | 2 | 13711 | 214 | Ccl12,Ccl2 | | 1.689e-02 | -4.08 | regulation of heat generation | biological process | GO:0031650 | 13 | 2 | 13711 | 214 | Tnf,Ptgs2 | | 1.689e-02 | -4.08 | ovulation | biological process | GO:0030728 | 13 | 2 | 13711 | 214 | Adamts1,Ptgs2 | | 1.689e-02 | -4.08 | regulation of protein localization to endosome | biological process | GO:1905666 | 13 | 2 | 13711 | 214 | Dtx3l,Msn | | 1.689e-02 | -4.08 | positive regulation of double-strand break repair via nonhomologous end joining | biological process | GO:2001034 | 13 | 2 | 13711 | 214 | Parp9,Dtx3l | | 1.689e-02 | -4.08 | negative regulation of interleukin-1 beta production | biological process | GO:0032691 | 13 | 2 | 13711 | 214 | Errfi1,Tnfaip3 | | 1.689e-02 | -4.08 | negative regulation of necroptotic process | biological process | GO:0060546 | 13 | 2 | 13711 | 214 | Cflar,Birc3 | | 1.689e-02 | -4.08 | negative regulation of programmed necrotic cell death | biological process | GO:0062099 | 13 | 2 | 13711 | 214 | Birc3,Cflar | | 1.689e-02 | -4.08 | cellular response to reactive nitrogen species | biological process | GO:1902170 | 13 | 2 | 13711 | 214 | Thbs1,Cflar | | 1.689e-02 | -4.08 | positive regulation of macrophage differentiation | biological process | GO:0045651 | 13 | 2 | 13711 | 214 | Ripk1,Trib1 | | 1.689e-02 | -4.08 | regulation of hepatocyte proliferation | biological process | GO:2000345 | 13 | 2 | 13711 | 214 | Tnfaip3,Cflar | | 1.689e-02 | -4.08 | regulation of glial cell apoptotic process | biological process | GO:0034350 | 13 | 2 | 13711 | 214 | Akap12,Ccl12 | | 1.689e-02 | -4.08 | regulation of chemokine secretion | biological process | GO:0090196 | 13 | 2 | 13711 | 214 | Tnf,Il4ra | | 1.689e-02 | -4.08 | regulation of transcription from RNA polymerase II promoter in response to hypoxia | biological process | GO:0061418 | 13 | 2 | 13711 | 214 | Nfe2l2,Bach1 | | 1.689e-02 | -4.08 | positive regulation of chemokine secretion | biological process | GO:0090197 | 13 | 2 | 13711 | 214 | Il4ra,Tnf | | 1.689e-02 | -4.08 | purinergic nucleotide receptor signaling pathway | biological process | GO:0035590 | 13 | 2 | 13711 | 214 | Hcar2,P2ry6 | | 1.689e-02 | -4.08 | complement binding | molecular function | GO:0001848 | 13 | 2 | 13516 | 211 | C5ar1,C3ar1 | | 1.689e-02 | -4.08 | Toll-like receptor binding | molecular function | GO:0035325 | 13 | 2 | 13516 | 211 | S100a9,S100a8 | | 1.689e-02 | -4.08 | long-chain fatty acid binding | molecular function | GO:0036041 | 13 | 2 | 13516 | 211 | S100a8,S100a9 | | 1.691e-02 | -4.08 | ST13 (ST13 Hsp70 interacting protein) | protein interactions | 6767 | 1 | 1 | 6802 | 115 | Ksr1 | | 1.691e-02 | -4.08 | LOC644669 (ankyrin repeat domain 30B pseudogene) | protein interactions | 644669 | 1 | 1 | 6802 | 115 | Ksr1 | | 1.691e-02 | -4.08 | SYNM (synemin) | protein interactions | 23336 | 1 | 1 | 6802 | 115 | Bach1 | | 1.691e-02 | -4.08 | LCP2 (lymphocyte cytosolic protein 2) | protein interactions | 3937 | 1 | 1 | 6802 | 115 | Fyb | | 1.691e-02 | -4.08 | MC1R (melanocortin 1 receptor) | protein interactions | 4157 | 1 | 1 | 6802 | 115 | Ksr1 | | 1.691e-02 | -4.08 | Nrf1 (nuclear respiratory factor 1) | protein interactions | 18181 | 1 | 1 | 6802 | 115 | Nfe2l2 | | 1.691e-02 | -4.08 | SSR4 (signal sequence receptor subunit 4) | protein interactions | 6748 | 1 | 1 | 6802 | 115 | Ksr1 | | 1.691e-02 | -4.08 | Acadvl (acyl-Coenzyme A dehydrogenase, very long chain) | protein interactions | 11370 | 1 | 1 | 6802 | 115 | Socs3 | | 1.691e-02 | -4.08 | MAFF (MAF bZIP transcription factor F) | protein interactions | 23764 | 1 | 1 | 6802 | 115 | Bach1 | | 1.691e-02 | -4.08 | Ins2 (insulin II) | protein interactions | 16334 | 1 | 1 | 6802 | 115 | Hspa5 | | 1.691e-02 | -4.08 | PDGFRA (platelet derived growth factor receptor alpha) | protein interactions | 5156 | 1 | 1 | 6802 | 115 | Pik3r1 | | 1.691e-02 | -4.08 | ACAD10 (acyl-CoA dehydrogenase family member 10) | protein interactions | 80724 | 1 | 1 | 6802 | 115 | Msn | | 1.691e-02 | -4.08 | Atf4 (activating transcription factor 4) | protein interactions | 79255 | 1 | 1 | 6802 | 115 | Nfe2l2 | | 1.691e-02 | -4.08 | Rnf112 (ring finger protein 112) | protein interactions | 22671 | 1 | 1 | 6802 | 115 | Zbtb16 | | 1.691e-02 | -4.08 | IQGAP2 (IQ motif containing GTPase activating protein 2) | protein interactions | 10788 | 1 | 1 | 6802 | 115 | Ksr1 | | 1.691e-02 | -4.08 | RAPGEF2 (Rap guanine nucleotide exchange factor 2) | protein interactions | 9693 | 1 | 1 | 6802 | 115 | Msn | | 1.691e-02 | -4.08 | Pdia2 (protein disulfide isomerase associated 2) | protein interactions | 69191 | 1 | 1 | 6802 | 115 | Hspa5 | | 1.691e-02 | -4.08 | SGA1 (glucan 1,4-alpha-glucosidase) | protein interactions | 854708 | 1 | 1 | 6802 | 115 | Hspb1 | | 1.691e-02 | -4.08 | MAFK (MAF bZIP transcription factor K) | protein interactions | 7975 | 1 | 1 | 6802 | 115 | Bach1 | | 1.691e-02 | -4.08 | TNFAIP3 (TNF alpha induced protein 3) | protein interactions | 7128 | 1 | 1 | 6802 | 115 | Il17ra | | 1.691e-02 | -4.08 | HSP90AB4P (heat shock protein 90 alpha family class B member 4, pseudogene) | protein interactions | 664618 | 1 | 1 | 6802 | 115 | Ksr1 | | 1.691e-02 | -4.08 | Atf6 (activating transcription factor 6) | protein interactions | 226641 | 1 | 1 | 6802 | 115 | Hspa5 | | 1.691e-02 | -4.08 | Hoxb2 (homeobox B2) | protein interactions | 103889 | 1 | 1 | 6802 | 115 | Ifi211 | | 1.691e-02 | -4.08 | Fkbp7 (FK506 binding protein 7) | protein interactions | 14231 | 1 | 1 | 6802 | 115 | Hspa5 | | 1.691e-02 | -4.08 | TAB3 (TGF-beta activated kinase 1 (MAP3K7) binding protein 3) | protein interactions | 257397 | 1 | 1 | 6802 | 115 | Trim30a | | 1.691e-02 | -4.08 | KCTD12 (potassium channel tetramerization domain containing 12) | protein interactions | 115207 | 1 | 1 | 6802 | 115 | Ksr1 | | 1.691e-02 | -4.08 | REL (REL proto-oncogene, NF-kB subunit) | protein interactions | 5966 | 1 | 1 | 6802 | 115 | Nfkb1 | | 1.691e-02 | -4.08 | Cd19 (CD19 antigen) | protein interactions | 12478 | 1 | 1 | 6802 | 115 | Pik3r1 | | 1.691e-02 | -4.08 | CS (citrate synthase) | protein interactions | 397519 | 1 | 1 | 6802 | 115 | Hspb1 | | 1.691e-02 | -4.08 | ERICH3 (glutamate rich 3) | protein interactions | 127254 | 1 | 1 | 6802 | 115 | Bach1 | | 1.691e-02 | -4.08 | Ehf (ets homologous factor) | protein interactions | 13661 | 1 | 1 | 6802 | 115 | Nfkbiz | | 1.691e-02 | -4.08 | TUBB2B (tubulin beta 2B class IIb) | protein interactions | 347733 | 1 | 1 | 6802 | 115 | Ksr1 | | 1.691e-02 | -4.08 | METTL2B (methyltransferase like 2B) | protein interactions | 55798 | 1 | 1 | 6802 | 115 | Ksr1 | | 1.691e-02 | -4.08 | ZNF624 (zinc finger protein 624) | protein interactions | 57547 | 1 | 1 | 6802 | 115 | Msn | | 1.691e-02 | -4.08 | RPL26 (ribosomal protein L26) | protein interactions | 6154 | 1 | 1 | 6802 | 115 | Ksr1 | | 1.691e-02 | -4.08 | Gsx1 (GS homeobox 1) | protein interactions | 14842 | 1 | 1 | 6802 | 115 | Irf9 | | 1.691e-02 | -4.08 | Zfp276 (zinc finger protein (C2H2 type) 276) | protein interactions | 57247 | 1 | 1 | 6802 | 115 | Nfkb1 | | 1.691e-02 | -4.08 | PPM1G (protein phosphatase, Mg2+/Mn2+ dependent 1G) | protein interactions | 5496 | 1 | 1 | 6802 | 115 | Ksr1 | | 1.691e-02 | -4.08 | Poglut1 (protein O-glucosyltransferase 1) | protein interactions | 224143 | 1 | 1 | 6802 | 115 | Hspa5 | | 1.691e-02 | -4.08 | Pdia4 (protein disulfide isomerase associated 4) | protein interactions | 12304 | 1 | 1 | 6802 | 115 | Hspa5 | | 1.691e-02 | -4.08 | TF (transferrin) | protein interactions | 7018 | 1 | 1 | 6802 | 115 | Akap12 | | 1.691e-02 | -4.08 | Tnfrsf1b (tumor necrosis factor receptor superfamily, member 1b) | protein interactions | 21938 | 1 | 1 | 6802 | 115 | Tnf | | 1.691e-02 | -4.08 | BACH1 (BTB domain and CNC homolog 1) | protein interactions | 571 | 1 | 1 | 6802 | 115 | Bach1 | | 1.691e-02 | -4.08 | Ido1 (indoleamine 2,3-dioxygenase 1) | protein interactions | 15930 | 1 | 1 | 6802 | 115 | Socs3 | | 1.691e-02 | -4.08 | Gtf2f1 (general transcription factor IIF, polypeptide 1) | protein interactions | 98053 | 1 | 1 | 6802 | 115 | Cebpd | | 1.691e-02 | -4.08 | FAM98B (family with sequence similarity 98 member B) | protein interactions | 283742 | 1 | 1 | 6802 | 115 | Ksr1 | | 1.691e-02 | -4.08 | ARL1 (ADP ribosylation factor like GTPase 1) | protein interactions | 400 | 1 | 1 | 6802 | 115 | Ksr1 | | 1.691e-02 | -4.08 | PPARA (peroxisome proliferator activated receptor alpha) | protein interactions | 5465 | 1 | 1 | 6802 | 115 | Gadd45b | | 1.691e-02 | -4.08 | Runx3 (runt related transcription factor 3) | protein interactions | 12399 | 1 | 1 | 6802 | 115 | Runx1 | | 1.691e-02 | -4.08 | COP1 (COP1 E3 ubiquitin ligase) | protein interactions | 64326 | 1 | 1 | 6802 | 115 | Trib1 | | 1.691e-02 | -4.08 | Satb1 (special AT-rich sequence binding protein 1) | protein interactions | 20230 | 1 | 1 | 6802 | 115 | Zbtb16 | | 1.691e-02 | -4.08 | Oprm1 (opioid receptor, mu 1) | protein interactions | 25601 | 1 | 1 | 6802 | 115 | Tnip1 | | 1.691e-02 | -4.08 | TECR (trans-2,3-enoyl-CoA reductase) | protein interactions | 9524 | 1 | 1 | 6802 | 115 | Ksr1 | | 1.691e-02 | -4.08 | Erp29 (endoplasmic reticulum protein 29) | protein interactions | 67397 | 1 | 1 | 6802 | 115 | Hspa5 | | 1.691e-02 | -4.08 | Ndp (Norrie disease (pseudoglioma) (human)) | protein interactions | 17986 | 1 | 1 | 6802 | 115 | Fzd4 | | 1.691e-02 | -4.08 | LRRC47 (leucine rich repeat containing 47) | protein interactions | 57470 | 1 | 1 | 6802 | 115 | Ksr1 | | 1.691e-02 | -4.08 | E1A (E1A) | protein interactions | 3126069 | 1 | 1 | 6802 | 115 | Cdkn1a | | 1.691e-02 | -4.08 | nedd4l.S (neural precursor cell expressed, developmentally down-regulated 4-like, E3 ubiquitin protein ligase S homeolog) | protein interactions | 399396 | 1 | 1 | 6802 | 115 | Sgk1 | | 1.691e-02 | -4.08 | Igtp (interferon gamma induced GTPase) | protein interactions | 16145 | 1 | 1 | 6802 | 115 | Tgtp1 | | 1.691e-02 | -4.08 | INS (insulin) | protein interactions | 3630 | 1 | 1 | 6802 | 115 | Hspb1 | | 1.691e-02 | -4.08 | NOSIP (nitric oxide synthase interacting protein) | protein interactions | 51070 | 1 | 1 | 6802 | 115 | Ksr1 | | 1.691e-02 | -4.08 | MAFG (MAF bZIP transcription factor G) | protein interactions | 4097 | 1 | 1 | 6802 | 115 | Bach1 | | 1.691e-02 | -4.08 | SLFN11 (schlafen family member 11) | protein interactions | 91607 | 1 | 1 | 6802 | 115 | Ksr1 | | 1.691e-02 | -4.08 | Gbp4 (guanylate binding protein 4) | protein interactions | 17472 | 1 | 1 | 6802 | 115 | Irf7 | | 1.691e-02 | -4.08 | FKBP4 (FKBP prolyl isomerase 4) | protein interactions | 2288 | 1 | 1 | 6802 | 115 | Ksr1 | | 1.691e-02 | -4.08 | Ncf1 (neutrophil cytosolic factor 1) | protein interactions | 17969 | 1 | 1 | 6802 | 115 | Nfe2l2 | | 1.691e-02 | -4.08 | HSPA9 (heat shock protein family A (Hsp70) member 9) | protein interactions | 3313 | 1 | 1 | 6802 | 115 | Ksr1 | | 1.691e-02 | -4.08 | PELO (pelota mRNA surveillance and ribosome rescue factor) | protein interactions | 53918 | 1 | 1 | 6802 | 115 | Ksr1 | | 1.691e-02 | -4.08 | TMX3 (thioredoxin related transmembrane protein 3) | protein interactions | 54495 | 1 | 1 | 6802 | 115 | Ksr1 | | 1.691e-02 | -4.08 | IRF7 (interferon regulatory factor 7) | protein interactions | 3665 | 1 | 1 | 6802 | 115 | Gbp4 | | 1.691e-02 | -4.08 | FRG1 (FSHD region gene 1) | protein interactions | 2483 | 1 | 1 | 6802 | 115 | Ksr1 | | 1.691e-02 | -4.08 | FLNC (filamin C) | protein interactions | 2318 | 1 | 1 | 6802 | 115 | Ksr1 | | 1.691e-02 | -4.08 | Rbm7 (RNA binding motif protein 7) | protein interactions | 67010 | 1 | 1 | 6802 | 115 | Tnf | | 1.691e-02 | -4.08 | Atp6v0b (ATPase, H+ transporting, lysosomal V0 subunit B) | protein interactions | 114143 | 1 | 1 | 6802 | 115 | Ifitm3 | | 1.691e-02 | -4.08 | Enpp2 (ectonucleotide pyrophosphatase/phosphodiesterase 2) | protein interactions | 18606 | 1 | 1 | 6802 | 115 | Nfkb1 | | 1.691e-02 | -4.08 | ANP32B (acidic nuclear phosphoprotein 32 family member B) | protein interactions | 10541 | 1 | 1 | 6802 | 115 | Ksr1 | | 1.697e-02 | -4.08 | GO\_REGULATION\_OF\_PEPTIDE\_TRANSPORT | MSigDB lists | GO\_REGULATION\_OF\_PEPTIDE\_TRANSPORT | 180 | 7 | 12187 | 179 | Tnf,Ptgs2,Nfkbia,Bcl3,S100a9,Kcna5,S100a8 | | 1.697e-02 | -4.08 | CEBPDELTA\_Q6 | MSigDB lists | CEBPDELTA\_Q6 | 180 | 7 | 12187 | 179 | Rbm39,S100a9,Il1rn,Atf3,Pik3r1,Ccnd2,Trib1 | | 1.698e-02 | -4.08 | Viral carcinogenesis | KEGG pathways | mmu05203 | 162 | 8 | 5248 | 107 | Cdkn1a,Nfkb2,Nfkbia,Nfkb1,Ccnd2,Pik3r1,Irf7,Irf9 | | 1.698e-02 | -4.08 | Viral carcinogenesis | KEGG pathways | ko05203 | 162 | 8 | 5248 | 107 | Nfkbia,Nfkb1,Ccnd2,Irf7,Irf9,Pik3r1,Cdkn1a,Nfkb2 | | 1.700e-02 | -4.07 | regulation of acute inflammatory response | biological process | GO:0002673 | 35 | 3 | 13711 | 214 | Tnf,Il6,Ptgs2 | | 1.700e-02 | -4.07 | regulation of neutrophil migration | biological process | GO:1902622 | 35 | 3 | 13711 | 214 | Il1a,C3ar1,C5ar1 | | 1.700e-02 | -4.07 | cellular response to glucose starvation | biological process | GO:0042149 | 35 | 3 | 13711 | 214 | Hspa5,Nfe2l2,Rrp8 | | 1.700e-02 | -4.07 | regulation of glial cell proliferation | biological process | GO:0060251 | 35 | 3 | 13711 | 214 | Sox11,Tnf,Il6 | | 1.715e-02 | -4.07 | SERVITJA\_LIVER\_HNF1A\_TARGETS\_UP | MSigDB lists | SERVITJA\_LIVER\_HNF1A\_TARGETS\_UP | 102 | 5 | 12187 | 179 | Thbs1,Ccl12,Gadd45b,Rgs16,Akap12 | | 1.715e-02 | -4.07 | GO\_GROWTH\_FACTOR\_BINDING | MSigDB lists | GO\_GROWTH\_FACTOR\_BINDING | 102 | 5 | 12187 | 179 | Osmr,Col4a1,Il1rn,A2m,Thbs1 | | 1.718e-02 | -4.06 | regulation of myeloid leukocyte differentiation | biological process | GO:0002761 | 96 | 5 | 13711 | 214 | Runx1,Pik3r1,Trib1,Ripk1,Tnf | | 1.719e-02 | -4.06 | GSE15659\_NAIVE\_CD4\_TCELL\_VS\_NONSUPPRESSIVE\_TCELL\_DN | MSigDB lists | GSE15659\_NAIVE\_CD4\_TCELL\_VS\_NONSUPPRESSIVE\_TCELL\_DN | 140 | 6 | 12187 | 179 | Tnf,Sox11,Tnfaip3,Rasd1,Timp1,Rhoc | | 1.719e-02 | -4.06 | GSE46606\_IRF4HIGH\_VS\_WT\_CD40L\_IL2\_IL5\_DAY3\_STIMULATED\_BCELL\_DN | MSigDB lists | GSE46606\_IRF4HIGH\_VS\_WT\_CD40L\_IL2\_IL5\_DAY3\_STIMULATED\_BCELL\_DN | 140 | 6 | 12187 | 179 | Usp18,Xaf1,Hp,Cybb,Oasl1,Tap1 | | 1.719e-02 | -4.06 | BROWNE\_HCMV\_INFECTION\_48HR\_UP | MSigDB lists | BROWNE\_HCMV\_INFECTION\_48HR\_UP | 140 | 6 | 12187 | 179 | Gem,Ptges,Atf3,Ier3,Rgs16,Gadd45b | | 1.719e-02 | -4.06 | GSE40274\_CTRL\_VS\_XBP1\_TRANSDUCED\_ACTIVATED\_CD4\_TCELL\_UP | MSigDB lists | GSE40274\_CTRL\_VS\_XBP1\_TRANSDUCED\_ACTIVATED\_CD4\_TCELL\_UP | 140 | 6 | 12187 | 179 | Rassf4,Plaur,Map3k8,Ifit3b,Irf7,Socs3 | | 1.720e-02 | -4.06 | MAPK signaling pathway | KEGG pathways | mmu04010 | 228 | 10 | 5248 | 107 | Gadd45b,Nfkb1,Map3k8,Tnf,Map3k6,Hspb1,Il1a,Gadd45g,Cd14,Nfkb2 | | 1.720e-02 | -4.06 | MAPK signaling pathway | KEGG pathways | ko04010 | 228 | 10 | 5248 | 107 | Hspb1,Il1a,Gadd45g,Cd14,Nfkb2,Nfkb1,Gadd45b,Map3k8,Tnf,Map3k6 | | 1.738e-02 | -4.05 | BIOCARTA\_CCR5\_PATHWAY | MSigDB lists | BIOCARTA\_CCR5\_PATHWAY | 14 | 2 | 12187 | 179 | Ccl4,Ccl12 | | 1.738e-02 | -4.05 | GO\_PARTURITION | MSigDB lists | GO\_PARTURITION | 14 | 2 | 12187 | 179 | Ccl12,Maff | | 1.738e-02 | -4.05 | LIAN\_NEUTROPHIL\_GRANULE\_CONSTITUENTS | MSigDB lists | LIAN\_NEUTROPHIL\_GRANULE\_CONSTITUENTS | 14 | 2 | 12187 | 179 | Lcn2,Cybb | | 1.738e-02 | -4.05 | ZHANG\_ADIPOGENESIS\_BY\_BMP7 | MSigDB lists | ZHANG\_ADIPOGENESIS\_BY\_BMP7 | 14 | 2 | 12187 | 179 | Sgk1,Pik3r1 | | 1.738e-02 | -4.05 | GO\_REGULATION\_OF\_SYNCYTIUM\_FORMATION\_BY\_PLASMA\_MEMBRANE\_FUSION | MSigDB lists | GO\_REGULATION\_OF\_SYNCYTIUM\_FORMATION\_BY\_PLASMA\_MEMBRANE\_FUSION | 14 | 2 | 12187 | 179 | Cflar,Il4ra | | 1.738e-02 | -4.05 | GO\_REGULATION\_OF\_HEAT\_GENERATION | MSigDB lists | GO\_REGULATION\_OF\_HEAT\_GENERATION | 14 | 2 | 12187 | 179 | Tnf,Ptgs2 | | 1.738e-02 | -4.05 | GO\_POSITIVE\_REGULATION\_OF\_EXTRINSIC\_APOPTOTIC\_SIGNALING\_PATHWAY\_VIA\_DEATH\_DOMAIN\_RECEPTORS | MSigDB lists | GO\_POSITIVE\_REGULATION\_OF\_EXTRINSIC\_APOPTOTIC\_SIGNALING\_PATHWAY\_VIA\_DEATH\_DOMAIN\_RECEPTORS | 14 | 2 | 12187 | 179 | Atf3,Thbs1 | | 1.738e-02 | -4.05 | RUAN\_RESPONSE\_TO\_TNF\_TROGLITAZONE\_UP | MSigDB lists | RUAN\_RESPONSE\_TO\_TNF\_TROGLITAZONE\_UP | 14 | 2 | 12187 | 179 | Hp,Cp | | 1.738e-02 | -4.05 | GO\_POSITIVE\_REGULATION\_OF\_NUCLEASE\_ACTIVITY | MSigDB lists | GO\_POSITIVE\_REGULATION\_OF\_NUCLEASE\_ACTIVITY | 14 | 2 | 12187 | 179 | Hspa5,Ptpn1 | | 1.738e-02 | -4.05 | SMID\_BREAST\_CANCER\_RELAPSE\_IN\_PLEURA\_DN | MSigDB lists | SMID\_BREAST\_CANCER\_RELAPSE\_IN\_PLEURA\_DN | 14 | 2 | 12187 | 179 | S100a9,Cp | | 1.738e-02 | -4.05 | GO\_ZINC\_ION\_HOMEOSTASIS | MSigDB lists | GO\_ZINC\_ION\_HOMEOSTASIS | 14 | 2 | 12187 | 179 | S100a8,S100a9 | | 1.738e-02 | -4.05 | GO\_POSITIVE\_REGULATION\_OF\_T\_HELPER\_CELL\_DIFFERENTIATION | MSigDB lists | GO\_POSITIVE\_REGULATION\_OF\_T\_HELPER\_CELL\_DIFFERENTIATION | 14 | 2 | 12187 | 179 | Il6,Il4ra | | 1.738e-02 | -4.05 | NEWMAN\_ERCC6\_TARGETS\_UP | MSigDB lists | NEWMAN\_ERCC6\_TARGETS\_UP | 14 | 2 | 12187 | 179 | Ifit2,Cxcl1 | | 1.738e-02 | -4.05 | KRAS.50\_UP.V1\_DN | MSigDB lists | KRAS.50\_UP.V1\_DN | 14 | 2 | 12187 | 179 | Sox11,Gadd45g | | 1.738e-02 | -4.05 | BIOCARTA\_41BB\_PATHWAY | MSigDB lists | BIOCARTA\_41BB\_PATHWAY | 14 | 2 | 12187 | 179 | Nfkb1,Nfkbia | | 1.738e-02 | -4.05 | MODULE\_562 | MSigDB lists | MODULE\_562 | 14 | 2 | 12187 | 179 | Tnf,Timp1 | | 1.738e-02 | -4.05 | GO\_RESPONSE\_TO\_MANGANESE\_ION | MSigDB lists | GO\_RESPONSE\_TO\_MANGANESE\_ION | 14 | 2 | 12187 | 179 | Hspa5,Ptgs2 | | 1.738e-02 | -4.05 | FERRANDO\_LYL1\_NEIGHBORS | MSigDB lists | FERRANDO\_LYL1\_NEIGHBORS | 14 | 2 | 12187 | 179 | Ccnd2,Zbtb16 | | 1.738e-02 | -4.05 | GO\_NEGATIVE\_REGULATION\_OF\_ALCOHOL\_BIOSYNTHETIC\_PROCESS | MSigDB lists | GO\_NEGATIVE\_REGULATION\_OF\_ALCOHOL\_BIOSYNTHETIC\_PROCESS | 14 | 2 | 12187 | 179 | Plek,Nfkb1 | | 1.749e-02 | -4.05 | GO\_REGULATION\_OF\_PROTEIN\_SECRETION | MSigDB lists | GO\_REGULATION\_OF\_PROTEIN\_SECRETION | 268 | 9 | 12187 | 179 | Cd14,Saa1,Il17ra,Il4ra,Tnf,Il6,Il1a,Hcar2,Kcna5 | | 1.768e-02 | -4.04 | SAFFORD\_T\_LYMPHOCYTE\_ANERGY | MSigDB lists | SAFFORD\_T\_LYMPHOCYTE\_ANERGY | 68 | 4 | 12187 | 179 | Casp4,Ier3,Gadd45b,Stx11 | | 1.768e-02 | -4.04 | LEE\_CALORIE\_RESTRICTION\_NEOCORTEX\_UP | MSigDB lists | LEE\_CALORIE\_RESTRICTION\_NEOCORTEX\_UP | 68 | 4 | 12187 | 179 | Timp1,Maff,Selp,Nfkbia | | 1.768e-02 | -4.04 | HALLMARK\_CHOLESTEROL\_HOMEOSTASIS | MSigDB lists | HALLMARK\_CHOLESTEROL\_HOMEOSTASIS | 68 | 4 | 12187 | 179 | Atf3,Errfi1,Plaur,Cxcl16 | | 1.769e-02 | -4.03 | cell cycle G1/S phase transition | biological process | GO:0044843 | 64 | 4 | 13711 | 214 | Cdkn1a,Rhou,Ccnd2,Bach1 | | 1.773e-02 | -4.03 | BUTYPHLNCDUF | prints domains | PR01407 | 26 | 3 | 2951 | 64 | Trim30a,Trim25,Trim30d | | 1.775e-02 | -4.03 | GSE12003\_4D\_VS\_8D\_CULTURE\_BM\_PROGENITOR\_UP | MSigDB lists | GSE12003\_4D\_VS\_8D\_CULTURE\_BM\_PROGENITOR\_UP | 141 | 6 | 12187 | 179 | Clic4,Ptgs2,Gbp6,Ncf1,Gbp4,Cdkn1a | | 1.775e-02 | -4.03 | GSE16385\_UNTREATED\_VS\_12H\_ROSIGLITAZONE\_IFNG\_TNF\_TREATED\_MACROPHAGE\_UP | MSigDB lists | GSE16385\_UNTREATED\_VS\_12H\_ROSIGLITAZONE\_IFNG\_TNF\_TREATED\_MACROPHAGE\_UP | 141 | 6 | 12187 | 179 | Zfp36,Csf3,Maff,Ccl4,Nfkbiz,Dnajb1 | | 1.775e-02 | -4.03 | GSE12001\_MIR223\_KO\_VS\_WT\_NEUTROPHIL\_UP | MSigDB lists | GSE12001\_MIR223\_KO\_VS\_WT\_NEUTROPHIL\_UP | 141 | 6 | 12187 | 179 | Casp4,Cxcl1,Bbs12,Pik3ap1,Cmpk2,Tnip3 | | 1.775e-02 | -4.03 | GSE7852\_TREG\_VS\_TCONV\_LN\_DN | MSigDB lists | GSE7852\_TREG\_VS\_TCONV\_LN\_DN | 141 | 6 | 12187 | 179 | Il17ra,Nfe2l2,Ifit2,Sgk1,Pik3r5,Tagln2 | | 1.775e-02 | -4.03 | GSE1925\_CTRL\_VS\_24H\_IFNG\_STIM\_MACROPHAGE\_UP | MSigDB lists | GSE1925\_CTRL\_VS\_24H\_IFNG\_STIM\_MACROPHAGE\_UP | 141 | 6 | 12187 | 179 | Il4ra,Usp18,Ifit3b,Isg15,Ifitm3,Irak3 | | 1.775e-02 | -4.03 | GSE14699\_DELETIONAL\_TOLERANCE\_VS\_ACTIVATED\_CD8\_TCELL\_DN | MSigDB lists | GSE14699\_DELETIONAL\_TOLERANCE\_VS\_ACTIVATED\_CD8\_TCELL\_DN | 141 | 6 | 12187 | 179 | Phf11b,Cdkn1a,Rbm47,Irak3,Gem,Phf11d | | 1.775e-02 | -4.03 | GSE21063\_CTRL\_VS\_ANTI\_IGM\_STIM\_BCELL\_NFATC1\_KO\_3H\_DN | MSigDB lists | GSE21063\_CTRL\_VS\_ANTI\_IGM\_STIM\_BCELL\_NFATC1\_KO\_3H\_DN | 141 | 6 | 12187 | 179 | Sgk3,Arid5b,Banp,Pik3r1,Ccl4,Mxd4 | | 1.775e-02 | -4.03 | GO\_GLYCOSAMINOGLYCAN\_BINDING | MSigDB lists | GO\_GLYCOSAMINOGLYCAN\_BINDING | 141 | 6 | 12187 | 179 | Adamts1,Thbs1,Fstl1,Ccl12,Selp,Saa1 | | 1.775e-02 | -4.03 | GSE26290\_CTRL\_VS\_AKT\_INHIBITOR\_TREATED\_ANTI\_CD3\_AND\_IL2\_STIM\_CD8\_TCELL\_UP | MSigDB lists | GSE26290\_CTRL\_VS\_AKT\_INHIBITOR\_TREATED\_ANTI\_CD3\_AND\_IL2\_STIM\_CD8\_TCELL\_UP | 141 | 6 | 12187 | 179 | Tgm2,Maff,Ptgs2,Atf3,Col4a1,Il1a | | 1.777e-02 | -4.03 | Ep300 (E1A binding protein p300) | protein interactions | 328572 | 33 | 3 | 6802 | 115 | Pik3r1,Spi1,Runx1 | | 1.779e-02 | -4.03 | - | gene3d domains | 2.60.200.10 | 14 | 2 | 6647 | 99 | Irf9,Irf7 | | 1.782e-02 | -4.03 | GO\_POSITIVE\_REGULATION\_OF\_PROTEIN\_LOCALIZATION\_TO\_NUCLEUS | MSigDB lists | GO\_POSITIVE\_REGULATION\_OF\_PROTEIN\_LOCALIZATION\_TO\_NUCLEUS | 103 | 5 | 12187 | 179 | Il6,Tnf,Ptgs2,Pik3r1,Csf3 | | 1.782e-02 | -4.03 | GSE21033\_3H\_VS\_24H\_POLYIC\_STIM\_DC\_DN | MSigDB lists | GSE21033\_3H\_VS\_24H\_POLYIC\_STIM\_DC\_DN | 103 | 5 | 12187 | 179 | Phf11d,Phf11b,Ifit2,Nfe2l2,Iqgap1 | | 1.782e-02 | -4.03 | IRF2\_01 | MSigDB lists | IRF2\_01 | 103 | 5 | 12187 | 179 | Usp18,Xaf1,Dtx3l,Tap1,Atf3 | | 1.792e-02 | -4.02 | GO\_RIBONUCLEOTIDE\_BINDING | MSigDB lists | GO\_RIBONUCLEOTIDE\_BINDING | 1496 | 32 | 12187 | 179 | Irgm2,Gem,Gbp6,Slfn4,Tor3a,Trib1,Runx1,Gbp4,Oasl1,Tubb6,Rasd1,Ksr1,Rhoj,Gbp7,Gbp3,Pygm,Cmpk2,Rhou,Map3k8,Ripk1,Rhoc,Slfn5,Slfn9,Bbs12,Map3k6,Sgk3,Tap1,Irak3,Ip6k2,Slfn3,Hspa5,Sgk1 | | 1.794e-02 | -4.02 | REACTOME\_GPCR\_DOWNSTREAM\_SIGNALING | MSigDB lists | REACTOME\_GPCR\_DOWNSTREAM\_SIGNALING | 315 | 10 | 12187 | 179 | C3ar1,Saa1,C5ar1,Cxcl1,Pik3r5,P2ry6,Cxcl16,Pik3r1,Hcar2,Rhoc | | 1.804e-02 | -4.02 | LIM\_MAMMARY\_LUMINAL\_PROGENITOR\_UP | MSigDB lists | LIM\_MAMMARY\_LUMINAL\_PROGENITOR\_UP | 38 | 3 | 12187 | 179 | Cd14,S100a8,Xdh | | 1.804e-02 | -4.02 | LIANG\_HEMATOPOIESIS\_STEM\_CELL\_NUMBER\_LARGE\_VS\_TINY\_UP | MSigDB lists | LIANG\_HEMATOPOIESIS\_STEM\_CELL\_NUMBER\_LARGE\_VS\_TINY\_UP | 38 | 3 | 12187 | 179 | Slfn4,Slfn3,Msr1 | | 1.804e-02 | -4.02 | PARK\_APL\_PATHOGENESIS\_DN | MSigDB lists | PARK\_APL\_PATHOGENESIS\_DN | 38 | 3 | 12187 | 179 | Nfkbie,Spi1,Cybb | | 1.804e-02 | -4.02 | ZHAN\_MULTIPLE\_MYELOMA\_LB\_UP | MSigDB lists | ZHAN\_MULTIPLE\_MYELOMA\_LB\_UP | 38 | 3 | 12187 | 179 | Cflar,Arid5b,Csf2rb | | 1.804e-02 | -4.02 | MODULE\_433 | MSigDB lists | MODULE\_433 | 38 | 3 | 12187 | 179 | Csf3,Tnfsf10,Il6 | | 1.804e-02 | -4.02 | GO\_REGULATION\_OF\_INTERLEUKIN\_8\_PRODUCTION | MSigDB lists | GO\_REGULATION\_OF\_INTERLEUKIN\_8\_PRODUCTION | 38 | 3 | 12187 | 179 | Ripk1,Tnf,Bcl3 | | 1.812e-02 | -4.01 | regulation of establishment of protein localization | biological process | GO:0070201 | 623 | 17 | 13711 | 214 | Gbp4,Ptpn1,Ptgs2,Il17ra,Mmp8,Il6,Ccl2,Il4ra,Tnfaip3,Tnf,Casp4,Akap12,Il1a,Hcar2,Pik3r1,Lgals9,Cd14 | | 1.814e-02 | -4.01 | adenosine nucleotides degradation II | BIOCYC pathways | MOUSE\_SALVADEHYPOX-PWY | 3 | 1 | 823 | 5 | Xdh | | 1.831e-02 | -4.00 | GSE17974\_CTRL\_VS\_ACT\_IL4\_AND\_ANTI\_IL12\_1H\_CD4\_TCELL\_DN | MSigDB lists | GSE17974\_CTRL\_VS\_ACT\_IL4\_AND\_ANTI\_IL12\_1H\_CD4\_TCELL\_DN | 142 | 6 | 12187 | 179 | Zfp189,Tnf,Ifit3b,Gem,Bbs12,Rtp4 | | 1.831e-02 | -4.00 | GSE13547\_2H\_VS\_12\_H\_ANTI\_IGM\_STIM\_BCELL\_UP | MSigDB lists | GSE13547\_2H\_VS\_12\_H\_ANTI\_IGM\_STIM\_BCELL\_UP | 142 | 6 | 12187 | 179 | Ifitm3,Ifi44,Xaf1,Casp4,Rtp4,Rgs16 | | 1.831e-02 | -4.00 | GSE16266\_CTRL\_VS\_LPS\_STIM\_MEF\_DN | MSigDB lists | GSE16266\_CTRL\_VS\_LPS\_STIM\_MEF\_DN | 142 | 6 | 12187 | 179 | Hspb1,Cmpk2,Nfkb2,Csf2rb,Fzd4,Gpr84 | | 1.831e-02 | -4.00 | GSE15271\_CXCR4\_POS\_VS\_NEG\_GC\_BCELL\_DN | MSigDB lists | GSE15271\_CXCR4\_POS\_VS\_NEG\_GC\_BCELL\_DN | 142 | 6 | 12187 | 179 | Znfx1,Birc3,Igsf6,Iqgap1,Irf7,Pik3r5 | | 1.831e-02 | -4.00 | GSE3920\_UNTREATED\_VS\_IFNA\_TREATED\_FIBROBLAST\_UP | MSigDB lists | GSE3920\_UNTREATED\_VS\_IFNA\_TREATED\_FIBROBLAST\_UP | 142 | 6 | 12187 | 179 | Nfkbiz,Birc3,Tnfaip3,Gadd45b,Nfkb1,Icam1 | | 1.831e-02 | -4.00 | GSE41176\_WT\_VS\_TAK1\_KO\_ANTI\_IGM\_STIM\_BCELL\_1H\_UP | MSigDB lists | GSE41176\_WT\_VS\_TAK1\_KO\_ANTI\_IGM\_STIM\_BCELL\_1H\_UP | 142 | 6 | 12187 | 179 | Sbno2,Map3k8,Tor3a,Socs3,Gadd45g,Ier3 | | 1.834e-02 | -4.00 | regulation of interleukin-12 production | biological process | GO:0032655 | 36 | 3 | 13711 | 214 | Irak3,Thbs1,Nfkb1 | | 1.834e-02 | -4.00 | negative regulation of myeloid leukocyte differentiation | biological process | GO:0002762 | 36 | 3 | 13711 | 214 | Trib1,Pik3r1,Runx1 | | 1.834e-02 | -4.00 | regulation of fibroblast migration | biological process | GO:0010762 | 36 | 3 | 13711 | 214 | Sdc4,Akap12,Thbs1 | | 1.834e-02 | -4.00 | regulation of gene silencing by miRNA | biological process | GO:0060964 | 36 | 3 | 13711 | 214 | Zfp36,Nfkb1,Ripk1 | | 1.834e-02 | -4.00 | response to dsRNA | biological process | GO:0043331 | 36 | 3 | 13711 | 214 | Nfkbia,Irak3,Nfkb1 | | 1.847e-02 | -3.99 | MULLIGHAN\_MLL\_SIGNATURE\_2\_DN | MSigDB lists | MULLIGHAN\_MLL\_SIGNATURE\_2\_DN | 226 | 8 | 12187 | 179 | Cxcl1,Ifi211,Clic4,Ifi44,Runx1,Ifi209,Ifi207,Ifi204 | | 1.850e-02 | -3.99 | GO\_CELLULAR\_RESPONSE\_TO\_TOPOLOGICALLY\_INCORRECT\_PROTEIN | MSigDB lists | GO\_CELLULAR\_RESPONSE\_TO\_TOPOLOGICALLY\_INCORRECT\_PROTEIN | 104 | 5 | 12187 | 179 | Hspa5,Ccl12,Ptpn1,Nfe2l2,Atf3 | | 1.850e-02 | -3.99 | GO\_REGULATION\_OF\_PHOSPHATIDYLINOSITOL\_3\_KINASE\_SIGNALING | MSigDB lists | GO\_REGULATION\_OF\_PHOSPHATIDYLINOSITOL\_3\_KINASE\_SIGNALING | 104 | 5 | 12187 | 179 | Ier3,Pik3ap1,Selp,Pik3r1,Csf3 | | 1.853e-02 | -3.99 | endoplasmic reticulum part | cellular component | GO:0044432 | 537 | 15 | 13825 | 212 | Tap1,Irgm2,Gm4841,Hspa5,Pik3r1,Tgtp2,Irgm1,Ptpn1,Iigp1,Gm5431,Cybb,Tgtp1,Tor3a,Ifi47,F830016B08Rik | | 1.856e-02 | -3.99 | GO\_REGULATION\_OF\_INTERLEUKIN\_6\_PRODUCTION | MSigDB lists | GO\_REGULATION\_OF\_INTERLEUKIN\_6\_PRODUCTION | 69 | 4 | 12187 | 179 | Tnfaip3,Il6,Irak3,Tnf | | 1.856e-02 | -3.99 | ONDER\_CDH1\_SIGNALING\_VIA\_CTNNB1 | MSigDB lists | ONDER\_CDH1\_SIGNALING\_VIA\_CTNNB1 | 69 | 4 | 12187 | 179 | Cxcl1,Birc3,Tnfaip3,Icam1 | | 1.859e-02 | -3.98 | regulation of transcription by RNA polymerase II | biological process | GO:0006357 | 1569 | 35 | 13711 | 214 | Fstl1,Tnip1,Ifi204,Arid5b,Ifi207,Zbtb16,Csf3,Ripk1,Atf3,Zfp36,Tnf,Bcl3,Nfe2l2,Irf9,Irf2,Dnajb1,Maff,Pik3r1,Irf7,Il6,Mxd4,Nfkb2,Nfkbia,Sox11,Il1a,Runx1,Ifi211,Bach1,Cebpd,Spi1,Nfkb1,Parp9,Sbno2,Gbp4,Ifi209 | | 1.859e-02 | -3.98 | positive regulation of endopeptidase activity | biological process | GO:0010950 | 134 | 6 | 13711 | 214 | Xdh,S100a8,Tnfsf10,Tnf,Casp4,S100a9 | | 1.862e-02 | -3.98 | cellular response to unfolded protein | biological process | GO:0034620 | 65 | 4 | 13711 | 214 | Atf3,Nfe2l2,Ptpn1,Hspa5 | | 1.862e-02 | -3.98 | regulation of oxidoreductase activity | biological process | GO:0051341 | 65 | 4 | 13711 | 214 | Lgals9,Nfkb1,Hp,Tnf | | 1.864e-02 | -3.98 | Prion diseases | KEGG pathways | mmu05020 | 28 | 3 | 5248 | 107 | Il6,Il1a,Hspa5 | | 1.864e-02 | -3.98 | Prion diseases | KEGG pathways | ko05020 | 28 | 3 | 5248 | 107 | Il6,Hspa5,Il1a | | 1.866e-02 | -3.98 | PASINI\_SUZ12\_TARGETS\_DN | MSigDB lists | PASINI\_SUZ12\_TARGETS\_DN | 271 | 9 | 12187 | 179 | Errfi1,Msn,Tagln2,Ier3,Tubb6,Trib1,Cdkn1a,Sox11,Thbs1 | | 1.873e-02 | -3.98 | regulation of actin cytoskeleton organization | biological process | GO:0032956 | 298 | 10 | 13711 | 214 | Csf3,Rhoc,Pik3r1,Plek,Fam107a,Il1a,Icam1,Rhou,Sdc4,Rhoj | | 1.876e-02 | -3.98 | ANKYRIN | prints domains | PR01415 | 72 | 5 | 2951 | 64 | Bcl3,Nfkbia,Nfkbie,Nfkb2,Nfkbiz | | 1.889e-02 | -3.97 | GSE30083\_SP2\_VS\_SP3\_THYMOCYTE\_UP | MSigDB lists | GSE30083\_SP2\_VS\_SP3\_THYMOCYTE\_UP | 143 | 6 | 12187 | 179 | Stx11,Nfkb1,Ccnd2,Tor3a,Rgs16,Bbs12 | | 1.889e-02 | -3.97 | GSE27241\_WT\_VS\_RORGT\_KO\_TH17\_POLARIZED\_CD4\_TCELL\_TREATED\_WITH\_DIGOXIN\_DN | MSigDB lists | GSE27241\_WT\_VS\_RORGT\_KO\_TH17\_POLARIZED\_CD4\_TCELL\_TREATED\_WITH\_DIGOXIN\_DN | 143 | 6 | 12187 | 179 | Akap12,Gbp3,Errfi1,Maff,Socs3,Nfkbia | | 1.889e-02 | -3.97 | GSE16385\_ROSIGLITAZONE\_VS\_UNTREATED\_IFNG\_TNF\_STIM\_MACROPHAGE\_UP | MSigDB lists | GSE16385\_ROSIGLITAZONE\_VS\_UNTREATED\_IFNG\_TNF\_STIM\_MACROPHAGE\_UP | 143 | 6 | 12187 | 179 | Csf3,Gbp6,Usp18,Samd9l,Cybb,Gbp4 | | 1.891e-02 | -3.97 | CHOP\_01 | MSigDB lists | CHOP\_01 | 184 | 7 | 12187 | 179 | S100a9,Rhoj,Tnfsf10,Nfkbia,Trib1,Nfe2l2,Atf3 | | 1.892e-02 | -3.97 | IVANOVA\_HEMATOPOIESIS\_MATURE\_CELL | MSigDB lists | IVANOVA\_HEMATOPOIESIS\_MATURE\_CELL | 227 | 8 | 12187 | 179 | Thbs1,Rhou,AA467197,Cybb,Mmp8,Stx11,Trim56,Cd14 | | 1.896e-02 | -3.97 | SMAD-like\_dom\_sf | interpro domains | IPR017855 | 14 | 2 | 13788 | 212 | Irf7,Irf9 | | 1.896e-02 | -3.97 | L30e-like | interpro domains | IPR029064 | 14 | 2 | 13788 | 212 | Gadd45g,Gadd45b | | 1.896e-02 | -3.97 | Dynamin\_SF | interpro domains | IPR022812 | 14 | 2 | 13788 | 212 | Mx2,Mx1 | | 1.896e-02 | -3.97 | G\_DYNAMIN\_dom | interpro domains | IPR030381 | 14 | 2 | 13788 | 212 | Mx1,Mx2 | | 1.896e-02 | -3.97 | ADAM\_CR\_2 | interpro domains | IPR041645 | 14 | 2 | 13788 | 212 | Adamts1,Adamts9 | | 1.897e-02 | -3.96 | Sushi/SCR/CCP\_sf | interpro domains | IPR035976 | 37 | 3 | 13788 | 212 | Sele,Hp,Selp | | 1.897e-02 | -3.96 | Sushi\_SCR\_CCP\_dom | interpro domains | IPR000436 | 37 | 3 | 13788 | 212 | Selp,Hp,Sele | | 1.906e-02 | -3.96 | Regulation of Complement cascade | REACTOME pathways | R-MMU-977606 | 13 | 2 | 6297 | 105 | C5ar1,C3ar1 | | 1.906e-02 | -3.96 | Activation of Matrix Metalloproteinases | REACTOME pathways | R-MMU-1592389 | 13 | 2 | 6297 | 105 | Timp1,Mmp8 | | 1.914e-02 | -3.96 | mouse chr1 | chromosome location | mouse chr1 | 264 | 9 | 14556 | 220 | Tagln2,Ptgs2,Atf3,Selp,Ifi204,Sgk3,Cflar,Sele,Rgs16 | | 1.920e-02 | -3.95 | GO\_NEGATIVE\_REGULATION\_OF\_RESPONSE\_TO\_WOUNDING | MSigDB lists | GO\_NEGATIVE\_REGULATION\_OF\_RESPONSE\_TO\_WOUNDING | 105 | 5 | 12187 | 179 | Nfkb1,Tnfaip3,Plaur,Saa1,Socs3 | | 1.920e-02 | -3.95 | PRC1\_BMI\_UP.V1\_UP | MSigDB lists | PRC1\_BMI\_UP.V1\_UP | 105 | 5 | 12187 | 179 | Ccnd2,Birc3,Gadd45g,Tnfaip3,Rgs16 | | 1.923e-02 | -3.95 | BZIP\_BASIC | prosite domains | PS00036 | 31 | 3 | 8845 | 164 | Atf3,Bach1,Nfe2l2 | | 1.927e-02 | -3.95 | mouse chr2 | chromosome location | mouse chr2 | 309 | 10 | 14556 | 220 | Hspa5,Sdc4,Ptges,Lcn2,Il1rn,Tgm2,Spi1,Thbs1,Nfe2l2,Il1a | | 1.933e-02 | -3.95 | STREICHER\_LSM1\_TARGETS\_UP | MSigDB lists | STREICHER\_LSM1\_TARGETS\_UP | 39 | 3 | 12187 | 179 | Tnfsf10,Il6,Hspb1 | | 1.933e-02 | -3.95 | GO\_DEFENSE\_RESPONSE\_TO\_GRAM\_POSITIVE\_BACTERIUM | MSigDB lists | GO\_DEFENSE\_RESPONSE\_TO\_GRAM\_POSITIVE\_BACTERIUM | 39 | 3 | 12187 | 179 | Il6,C5ar1,Tnf | | 1.933e-02 | -3.95 | GYORFFY\_DOXORUBICIN\_RESISTANCE | MSigDB lists | GYORFFY\_DOXORUBICIN\_RESISTANCE | 39 | 3 | 12187 | 179 | P2ry6,Adamts1,Bbs12 | | 1.933e-02 | -3.95 | GO\_POSITIVE\_REGULATION\_OF\_ORGANELLE\_ASSEMBLY | MSigDB lists | GO\_POSITIVE\_REGULATION\_OF\_ORGANELLE\_ASSEMBLY | 39 | 3 | 12187 | 179 | Msn,Tnf,Sdc4 | | 1.933e-02 | -3.95 | MODULE\_164 | MSigDB lists | MODULE\_164 | 39 | 3 | 12187 | 179 | Timp1,Cdkn1a,A2m | | 1.933e-02 | -3.95 | GO\_ENDOTHELIAL\_CELL\_DEVELOPMENT | MSigDB lists | GO\_ENDOTHELIAL\_CELL\_DEVELOPMENT | 39 | 3 | 12187 | 179 | Clic4,Msn,Icam1 | | 1.933e-02 | -3.95 | HOQUE\_METHYLATED\_IN\_CANCER | MSigDB lists | HOQUE\_METHYLATED\_IN\_CANCER | 39 | 3 | 12187 | 179 | Ccnd2,Ptgs2,Socs3 | | 1.946e-02 | -3.94 | PEART\_HDAC\_PROLIFERATION\_CLUSTER\_DN | MSigDB lists | PEART\_HDAC\_PROLIFERATION\_CLUSTER\_DN | 70 | 4 | 12187 | 179 | Ifi209,Ifi211,Ifi204,Ifi207 | | 1.948e-02 | -3.94 | GSE3982\_DC\_VS\_MAC\_DN | MSigDB lists | GSE3982\_DC\_VS\_MAC\_DN | 144 | 6 | 12187 | 179 | Gadd45g,Ccl12,Plek,Irf9,Herc6,C3ar1 | | 1.948e-02 | -3.94 | HNF1\_Q6 | MSigDB lists | HNF1\_Q6 | 144 | 6 | 12187 | 179 | Runx1,Ccnd2,Csf3,Rbm47,Luc7l3,Errfi1 | | 1.948e-02 | -3.94 | GSE14415\_ACT\_TCONV\_VS\_ACT\_NATURAL\_TREG\_DN | MSigDB lists | GSE14415\_ACT\_TCONV\_VS\_ACT\_NATURAL\_TREG\_DN | 144 | 6 | 12187 | 179 | Phf11d,Cdkn1a,Phf11b,Gadd45b,Cybb,Mmp8 | | 1.949e-02 | -3.94 | ADAM\_CR\_2 | pfam domains | PF17771 | 14 | 2 | 12881 | 201 | Adamts1,Adamts9 | | 1.949e-02 | -3.94 | Dynamin\_N | pfam domains | PF00350 | 14 | 2 | 12881 | 201 | Mx2,Mx1 | | 1.950e-02 | -3.94 | positive regulation of chromatin binding | biological process | GO:0035563 | 14 | 2 | 13711 | 214 | Parp9,Dtx3l | | 1.950e-02 | -3.94 | cytoplasmic pattern recognition receptor signaling pathway | biological process | GO:0002753 | 14 | 2 | 13711 | 214 | Nfkbia,Irgm2 | | 1.950e-02 | -3.94 | response to nitric oxide | biological process | GO:0071731 | 14 | 2 | 13711 | 214 | Thbs1,Cflar | | 1.950e-02 | -3.94 | regulation of tolerance induction | biological process | GO:0002643 | 14 | 2 | 13711 | 214 | Runx1,Irak3 | | 1.950e-02 | -3.94 | negative regulation of leukocyte mediated cytotoxicity | biological process | GO:0001911 | 14 | 2 | 13711 | 214 | Tap1,Lgals9 | | 1.950e-02 | -3.94 | positive regulation of extrinsic apoptotic signaling pathway via death domain receptors | biological process | GO:1902043 | 14 | 2 | 13711 | 214 | Atf3,Thbs1 | | 1.950e-02 | -3.94 | regulation of inositol phosphate biosynthetic process | biological process | GO:0010919 | 14 | 2 | 13711 | 214 | P2ry6,Plek | | 1.950e-02 | -3.94 | negative regulation of anion transmembrane transport | biological process | GO:1903960 | 14 | 2 | 13711 | 214 | Tnf,Thbs1 | | 1.950e-02 | -3.94 | response to testosterone | biological process | GO:0033574 | 14 | 2 | 13711 | 214 | Msn,Cflar | | 1.951e-02 | -3.94 | negative regulation of macromolecule biosynthetic process | biological process | GO:0010558 | 1195 | 28 | 13711 | 214 | Bcl3,Rasd1,Prg4,Dnajb1,Rrp8,Ifi204,Zbtb16,Ifi207,Arid5b,Atf3,Zfp36,Znfx1,Tnf,Runx1,Spi1,Ifi211,Bach1,Cebpd,Parp9,Nfkbie,Sbno2,Nfkb1,Gbp4,Ifi209,Errfi1,Il6,Sox11,Mxd4 | | 1.951e-02 | -3.94 | death receptor binding | molecular function | GO:0005123 | 14 | 2 | 13516 | 211 | Ripk1,Cflar | | 1.957e-02 | -3.93 | Tax1bp1 (Tax1 (human T cell leukemia virus type I) binding protein 1) | protein interactions | 52440 | 13 | 2 | 6802 | 115 | Ripk1,Tnfaip3 | | 1.957e-02 | -3.93 | Hoxb8 (homeobox B8) | protein interactions | 15416 | 13 | 2 | 6802 | 115 | Irf9,Irf7 | | 1.959e-02 | -3.93 | signal transduction by p53 class mediator | biological process | GO:0072331 | 66 | 4 | 13711 | 214 | Rrp8,Cdkn1a,Ifi204,Bcl3 | | 1.959e-02 | -3.93 | negative regulation of NF-kappaB transcription factor activity | biological process | GO:0032088 | 66 | 4 | 13711 | 214 | Nfkbia,Irak3,Acod1,Tnfaip3 | | 1.971e-02 | -3.93 | cellular response to external stimulus | biological process | GO:0071496 | 215 | 8 | 13711 | 214 | Cdkn1a,Atf3,Fam107a,Hspa5,Rrp8,Angptl4,Nfe2l2,Vcam1 | | 1.973e-02 | -3.93 | regulation of cell migration involved in sprouting angiogenesis | biological process | GO:0090049 | 37 | 3 | 13711 | 214 | Thbs1,Rhoj,Ptgs2 | | 1.973e-02 | -3.93 | regulation of viral transcription | biological process | GO:0046782 | 37 | 3 | 13711 | 214 | Zfp36,Ifitm3,Ccl4 | | 1.973e-02 | -3.93 | cell killing | biological process | GO:0001906 | 37 | 3 | 13711 | 214 | Ccl2,Ncf1,Stx11 | | 1.973e-02 | -3.93 | regulation of collagen biosynthetic process | biological process | GO:0032965 | 37 | 3 | 13711 | 214 | Il6,Errfi1,Ccl2 | | 1.974e-02 | -3.93 | GO\_ENZYME\_BINDING | MSigDB lists | GO\_ENZYME\_BINDING | 1449 | 31 | 12187 | 179 | Nfkbia,Sdc4,Trib1,Tnf,A2m,Fzd4,Cflar,Cdkn1a,Ksr1,Kcna5,Timp1,Tnip1,Ptgs2,Errfi1,Plaur,Msn,Hspb1,Plek,Sele,Zfp36,Ccnd2,Pik3r1,Tnfaip3,Ripk1,Csf3,Iqgap1,Dnajb1,Sgk1,Hspa5,Akap12,Ptpn1 | | 1.976e-02 | -3.92 | metal ion homeostasis | biological process | GO:0055065 | 484 | 14 | 13711 | 214 | Cxcl1,C5ar1,Tgm2,Mt2,Kcna5,Sgk1,Mt1,Ccl2,Pygm,Slc24a4,C3ar1,P2ry6,Cp,Lcn2 | | 1.981e-02 | -3.92 | BRUINS\_UVC\_RESPONSE\_VIA\_TP53\_GROUP\_B | MSigDB lists | BRUINS\_UVC\_RESPONSE\_VIA\_TP53\_GROUP\_B | 416 | 12 | 12187 | 179 | Gadd45g,Col4a1,Lcn2,Dnajb1,Nfkbiz,Plaur,Ptgs2,Il6,Xdh,Nfkb2,Rgs16,C5ar1 | | 1.987e-02 | -3.92 | GO\_REGULATION\_OF\_ANTIGEN\_PROCESSING\_AND\_PRESENTATION | MSigDB lists | GO\_REGULATION\_OF\_ANTIGEN\_PROCESSING\_AND\_PRESENTATION | 15 | 2 | 12187 | 179 | Thbs1,Tap1 | | 1.987e-02 | -3.92 | GO\_RESPONSE\_TO\_MAGNESIUM\_ION | MSigDB lists | GO\_RESPONSE\_TO\_MAGNESIUM\_ION | 15 | 2 | 12187 | 179 | Cd14,Thbs1 | | 1.987e-02 | -3.92 | GO\_POSITIVE\_REGULATION\_OF\_LIPID\_STORAGE | MSigDB lists | GO\_POSITIVE\_REGULATION\_OF\_LIPID\_STORAGE | 15 | 2 | 12187 | 179 | Nfkb1,Msr1 | | 1.987e-02 | -3.92 | GO\_NEGATIVE\_REGULATION\_OF\_RESPONSE\_TO\_REACTIVE\_OXYGEN\_SPECIES | MSigDB lists | GO\_NEGATIVE\_REGULATION\_OF\_RESPONSE\_TO\_REACTIVE\_OXYGEN\_SPECIES | 15 | 2 | 12187 | 179 | Hp,Nfe2l2 | | 1.987e-02 | -3.92 | GARGALOVIC\_RESPONSE\_TO\_OXIDIZED\_PHOSPHOLIPIDS\_GREY\_UP | MSigDB lists | GARGALOVIC\_RESPONSE\_TO\_OXIDIZED\_PHOSPHOLIPIDS\_GREY\_UP | 15 | 2 | 12187 | 179 | Adamts1,Adamts9 | | 1.987e-02 | -3.92 | GO\_POSITIVE\_REGULATION\_OF\_NF\_KAPPAB\_IMPORT\_INTO\_NUCLEUS | MSigDB lists | GO\_POSITIVE\_REGULATION\_OF\_NF\_KAPPAB\_IMPORT\_INTO\_NUCLEUS | 15 | 2 | 12187 | 179 | Tnf,Ptgs2 | | 1.987e-02 | -3.92 | GO\_REGULATION\_OF\_TRANSFORMING\_GROWTH\_FACTOR\_BETA\_PRODUCTION | MSigDB lists | GO\_REGULATION\_OF\_TRANSFORMING\_GROWTH\_FACTOR\_BETA\_PRODUCTION | 15 | 2 | 12187 | 179 | Thbs1,Ptgs2 | | 1.987e-02 | -3.92 | BOYAULT\_LIVER\_CANCER\_SUBCLASS\_G6\_DN | MSigDB lists | BOYAULT\_LIVER\_CANCER\_SUBCLASS\_G6\_DN | 15 | 2 | 12187 | 179 | Sdc4,Tnfsf10 | | 1.987e-02 | -3.92 | RAMPON\_ENRICHED\_LEARNING\_ENVIRONMENT\_EARLY\_UP | MSigDB lists | RAMPON\_ENRICHED\_LEARNING\_ENVIRONMENT\_EARLY\_UP | 15 | 2 | 12187 | 179 | Hspa5,Trim25 | | 1.987e-02 | -3.92 | CHEN\_PDGF\_TARGETS | MSigDB lists | CHEN\_PDGF\_TARGETS | 15 | 2 | 12187 | 179 | Arid5b,Tiparp | | 1.987e-02 | -3.92 | GO\_DEATH\_RECEPTOR\_BINDING | MSigDB lists | GO\_DEATH\_RECEPTOR\_BINDING | 15 | 2 | 12187 | 179 | Ripk1,Cflar | | 1.987e-02 | -3.92 | BIOCARTA\_NKT\_PATHWAY | MSigDB lists | BIOCARTA\_NKT\_PATHWAY | 15 | 2 | 12187 | 179 | Il4ra,Ccl4 | | 1.987e-02 | -3.92 | GO\_INFLAMMATORY\_RESPONSE\_TO\_ANTIGENIC\_STIMULUS | MSigDB lists | GO\_INFLAMMATORY\_RESPONSE\_TO\_ANTIGENIC\_STIMULUS | 15 | 2 | 12187 | 179 | Tnf,Icam1 | | 1.987e-02 | -3.92 | DAZARD\_UV\_RESPONSE\_CLUSTER\_G4 | MSigDB lists | DAZARD\_UV\_RESPONSE\_CLUSTER\_G4 | 15 | 2 | 12187 | 179 | Gadd45b,Zfp36 | | 1.987e-02 | -3.92 | WOTTON\_RUNX\_TARGETS\_UP | MSigDB lists | WOTTON\_RUNX\_TARGETS\_UP | 15 | 2 | 12187 | 179 | Angptl4,Ccl12 | | 1.987e-02 | -3.92 | NIELSEN\_SYNOVIAL\_SARCOMA\_DN | MSigDB lists | NIELSEN\_SYNOVIAL\_SARCOMA\_DN | 15 | 2 | 12187 | 179 | Cybb,Parp14 | | 1.987e-02 | -3.92 | BAKER\_HEMATOPOIESIS\_STAT3\_TARGETS | MSigDB lists | BAKER\_HEMATOPOIESIS\_STAT3\_TARGETS | 15 | 2 | 12187 | 179 | Il6,Ccnd2 | | 1.987e-02 | -3.92 | GO\_BRANCHING\_INVOLVED\_IN\_SALIVARY\_GLAND\_MORPHOGENESIS | MSigDB lists | GO\_BRANCHING\_INVOLVED\_IN\_SALIVARY\_GLAND\_MORPHOGENESIS | 15 | 2 | 12187 | 179 | Il6,Tgm2 | | 1.987e-02 | -3.92 | MORI\_EMU\_MYC\_LYMPHOMA\_BY\_ONSET\_TIME\_DN | MSigDB lists | MORI\_EMU\_MYC\_LYMPHOMA\_BY\_ONSET\_TIME\_DN | 15 | 2 | 12187 | 179 | Hspa5,Nfkbiz | | 1.987e-02 | -3.92 | GO\_ENDOCYTIC\_VESICLE\_LUMEN | MSigDB lists | GO\_ENDOCYTIC\_VESICLE\_LUMEN | 15 | 2 | 12187 | 179 | Saa1,Hp | | 1.987e-02 | -3.92 | MISHRA\_CARCINOMA\_ASSOCIATED\_FIBROBLAST\_UP | MSigDB lists | MISHRA\_CARCINOMA\_ASSOCIATED\_FIBROBLAST\_UP | 15 | 2 | 12187 | 179 | Gem,Ccl12 | | 1.987e-02 | -3.92 | GO\_CELLULAR\_RESPONSE\_TO\_PROSTAGLANDIN\_E\_STIMULUS | MSigDB lists | GO\_CELLULAR\_RESPONSE\_TO\_PROSTAGLANDIN\_E\_STIMULUS | 15 | 2 | 12187 | 179 | Adamts1,P2ry6 | | 1.987e-02 | -3.92 | CHEN\_ETV5\_TARGETS\_SERTOLI | MSigDB lists | CHEN\_ETV5\_TARGETS\_SERTOLI | 15 | 2 | 12187 | 179 | Plek,Il6 | | 1.987e-02 | -3.92 | KANG\_CISPLATIN\_RESISTANCE\_UP | MSigDB lists | KANG\_CISPLATIN\_RESISTANCE\_UP | 15 | 2 | 12187 | 179 | Maff,Cdkn1a | | 1.993e-02 | -3.92 | LINDGREN\_BLADDER\_CANCER\_CLUSTER\_3\_DN | MSigDB lists | LINDGREN\_BLADDER\_CANCER\_CLUSTER\_3\_DN | 186 | 7 | 12187 | 179 | Ifi204,Ifi207,Mxd4,Ifi209,Casp4,Cflar,Ifi211 | | 2.003e-02 | -3.91 | S\_100 | smart domains | SM01394 | 13 | 2 | 7188 | 123 | S100a8,S100a9 | | 2.008e-02 | -3.91 | GSE46242\_TH1\_VS\_ANERGIC\_TH1\_CD4\_TCELL\_WITH\_EGR2\_DELETED\_UP | MSigDB lists | GSE46242\_TH1\_VS\_ANERGIC\_TH1\_CD4\_TCELL\_WITH\_EGR2\_DELETED\_UP | 145 | 6 | 12187 | 179 | Cdkn1a,Birc3,Cybb,AA467197,Ccl12,Gbp3 | | 2.008e-02 | -3.91 | GSE5099\_CLASSICAL\_M1\_VS\_ALTERNATIVE\_M2\_MACROPHAGE\_UP | MSigDB lists | GSE5099\_CLASSICAL\_M1\_VS\_ALTERNATIVE\_M2\_MACROPHAGE\_UP | 145 | 6 | 12187 | 179 | Znfx1,Nfkbia,Il4ra,Gbp3,Rgs16,Rnf213 | | 2.008e-02 | -3.91 | GSE22443\_NAIVE\_VS\_ACT\_AND\_IL2\_TREATED\_CD8\_TCELL\_DN | MSigDB lists | GSE22443\_NAIVE\_VS\_ACT\_AND\_IL2\_TREATED\_CD8\_TCELL\_DN | 145 | 6 | 12187 | 179 | Ifitm3,Ptges,Sele,Errfi1,Hp,Col4a1 | | 2.008e-02 | -3.91 | GO\_ENDOPLASMIC\_RETICULUM\_LUMEN | MSigDB lists | GO\_ENDOPLASMIC\_RETICULUM\_LUMEN | 145 | 6 | 12187 | 179 | Plaur,Col4a1,Tor3a,Ptgs2,Thbs1,Hspa5 | | 2.008e-02 | -3.91 | GSE14000\_4H\_VS\_16H\_LPS\_DC\_UP | MSigDB lists | GSE14000\_4H\_VS\_16H\_LPS\_DC\_UP | 145 | 6 | 12187 | 179 | P2ry6,Il17ra,Oasl1,Tnf,Runx1,Parp14 | | 2.008e-02 | -3.91 | GSE14415\_INDUCED\_TREG\_VS\_FOXP3\_KO\_INDUCED\_TREG\_DN | MSigDB lists | GSE14415\_INDUCED\_TREG\_VS\_FOXP3\_KO\_INDUCED\_TREG\_DN | 145 | 6 | 12187 | 179 | Gem,Phf11d,Pik3ap1,Phf11b,Map3k8,Ccl4 | | 2.015e-02 | -3.90 | regulation of anion transport | biological process | GO:0044070 | 100 | 5 | 13711 | 214 | Ripk1,Il1a,Tnf,Thbs1,Il1rn | | 2.015e-02 | -3.90 | regulation of cell-matrix adhesion | biological process | GO:0001952 | 100 | 5 | 13711 | 214 | Pik3r1,Iqgap1,Sdc4,Thbs1,Fam107a | | 2.041e-02 | -3.89 | sarcoplasmic reticulum | cellular component | GO:0016529 | 68 | 4 | 13825 | 212 | Thbs1,Rasd1,Xdh,Pygm | | 2.042e-02 | -3.89 | anatomical structure morphogenesis | biological process | GO:0009653 | 1746 | 38 | 13711 | 214 | Errfi1,Il6,Cemip2,Sox11,Nfkb2,Socs3,Thbs1,Apold1,Rhoj,Sbno2,Sdc4,Adamts1,Tiparp,Fzd4,Ptgs2,Sgk1,Runx1,Rasip1,Spi1,Ccl2,Robo4,Tnf,Rnf213,Slc24a4,Arid5b,Cflar,Adamts9,Zbtb16,Col4a1,Iqgap1,Ccl12,Tgm2,Hspa5,Clic4,Bcl3,Msn,Angptl4,Vcam1 | | 2.042e-02 | -3.89 | mouse chr11\|11 | chromosome location | mouse chr11\|11 | 807 | 20 | 14556 | 220 | Irgm1,Xaf1,Slfn5,Cxcl16,Irgm2,Trim25,Gm5431,Ifi47,Pik3r5,9930111J21Rik2,Slfn4,Luc7l3,Tnip1,Socs3,Wfdc21,Slfn9,Tgtp1,Lgals9,Slfn3,Rasd1 | | 2.045e-02 | -3.89 | Pathways in cancer | KEGG pathways | mmu05200 | 341 | 13 | 5248 | 107 | Nfkbia,Spi1,Col4a1,Pik3r1,Zbtb16,Runx1,Il6,Nfkb1,Birc3,Ptgs2,Cdkn1a,Fzd4,Nfkb2 | | 2.050e-02 | -3.89 | regulation of endothelial cell migration | biological process | GO:0010594 | 137 | 6 | 13711 | 214 | Rhoj,Adamts9,Hspb1,Thbs1,Nfe2l2,Ptgs2 | | 2.066e-02 | -3.88 | GSE1740\_MCSF\_VS\_MCSF\_AND\_IFNG\_DAY2\_DERIVED\_MACROPHAGE\_UP | MSigDB lists | GSE1740\_MCSF\_VS\_MCSF\_AND\_IFNG\_DAY2\_DERIVED\_MACROPHAGE\_UP | 107 | 5 | 12187 | 179 | Tap1,Znfx1,Gbp3,Parp14,Tnf | | 2.068e-02 | -3.88 | MODULE\_107 | MSigDB lists | MODULE\_107 | 40 | 3 | 12187 | 179 | Casp4,Timp1,Hp | | 2.068e-02 | -3.88 | LINDGREN\_BLADDER\_CANCER\_HIGH\_RECURRENCE | MSigDB lists | LINDGREN\_BLADDER\_CANCER\_HIGH\_RECURRENCE | 40 | 3 | 12187 | 179 | Msn,Birc3,Vcam1 | | 2.068e-02 | -3.88 | REACTOME\_NUCLEOTIDE\_BINDING\_DOMAIN\_LEUCINE\_RICH\_REPEAT\_CONTAINING\_RECEPTOR\_NLR\_SIGNALING\_PATHWAYS | MSigDB lists | REACTOME\_NUCLEOTIDE\_BINDING\_DOMAIN\_LEUCINE\_RICH\_REPEAT\_CONTAINING\_RECEPTOR\_NLR\_SIGNALING\_PATHWAYS | 40 | 3 | 12187 | 179 | Tnfaip3,Birc3,Casp4 | | 2.068e-02 | -3.88 | GO\_REGULATION\_OF\_EPIDERMIS\_DEVELOPMENT | MSigDB lists | GO\_REGULATION\_OF\_EPIDERMIS\_DEVELOPMENT | 40 | 3 | 12187 | 179 | Maff,Tnf,Errfi1 | | 2.068e-02 | -3.88 | GO\_B\_CELL\_MEDIATED\_IMMUNITY | MSigDB lists | GO\_B\_CELL\_MEDIATED\_IMMUNITY | 40 | 3 | 12187 | 179 | Bcl3,Irf7,Il4ra | | 2.068e-02 | -3.88 | CHENG\_RESPONSE\_TO\_NICKEL\_ACETATE | MSigDB lists | CHENG\_RESPONSE\_TO\_NICKEL\_ACETATE | 40 | 3 | 12187 | 179 | Cebpd,Ms4a6d,Cp | | 2.069e-02 | -3.88 | GSE37416\_CTRL\_VS\_24H\_F\_TULARENSIS\_LVS\_NEUTROPHIL\_UP | MSigDB lists | GSE37416\_CTRL\_VS\_24H\_F\_TULARENSIS\_LVS\_NEUTROPHIL\_UP | 146 | 6 | 12187 | 179 | Runx1,Ifit2,Ifit3b,Samd9l,Dtx3l,Herc6 | | 2.069e-02 | -3.88 | GSE43863\_TFH\_VS\_LY6C\_LOW\_CXCR5NEG\_EFFECTOR\_CD4\_TCELL\_DN | MSigDB lists | GSE43863\_TFH\_VS\_LY6C\_LOW\_CXCR5NEG\_EFFECTOR\_CD4\_TCELL\_DN | 146 | 6 | 12187 | 179 | Rasd1,Tiparp,Kcna5,Plek,Trib1,Ifitm2 | | 2.069e-02 | -3.88 | GSE42021\_TCONV\_PLN\_VS\_CD24HI\_TCONV\_THYMUS\_DN | MSigDB lists | GSE42021\_TCONV\_PLN\_VS\_CD24HI\_TCONV\_THYMUS\_DN | 146 | 6 | 12187 | 179 | Isg15,Herc6,Irf7,Ifi44,Usp18,Oasl1 | | 2.069e-02 | -3.88 | FERNANDEZ\_BOUND\_BY\_MYC | MSigDB lists | FERNANDEZ\_BOUND\_BY\_MYC | 146 | 6 | 12187 | 179 | Irf2,Msn,Icam1,Col4a1,Cxcl1,Bcl3 | | 2.069e-02 | -3.88 | GSE9960\_HEALTHY\_VS\_GRAM\_NEG\_AND\_POS\_SEPSIS\_PBMC\_DN | MSigDB lists | GSE9960\_HEALTHY\_VS\_GRAM\_NEG\_AND\_POS\_SEPSIS\_PBMC\_DN | 146 | 6 | 12187 | 179 | Ksr1,Gpr84,Pik3r1,Zfp189,Osmr,Nfkbie | | 2.069e-02 | -3.88 | GSE17721\_CTRL\_VS\_CPG\_1H\_BMDC\_DN | MSigDB lists | GSE17721\_CTRL\_VS\_CPG\_1H\_BMDC\_DN | 146 | 6 | 12187 | 179 | Il1rn,Ier3,Map3k8,Cdkn1a,Cd14,Arid5b | | 2.069e-02 | -3.88 | GSE2770\_UNTREATED\_VS\_TGFB\_AND\_IL12\_TREATED\_ACT\_CD4\_TCELL\_2H\_DN | MSigDB lists | GSE2770\_UNTREATED\_VS\_TGFB\_AND\_IL12\_TREATED\_ACT\_CD4\_TCELL\_2H\_DN | 146 | 6 | 12187 | 179 | Irf9,Irf7,Col4a1,Gadd45b,Xaf1,Sdc4 | | 2.070e-02 | -3.88 | second-messenger-mediated signaling | biological process | GO:0019932 | 217 | 8 | 13711 | 214 | Tnf,Mt1,Mt2,Ksr1,Rasd1,Vcam1,Irgm2,Selp | | 2.082e-02 | -3.87 | response to endogenous stimulus | biological process | GO:0009719 | 886 | 22 | 13711 | 214 | Sgk1,Runx1,Nfkb1,Fam107a,Timp1,P2ry6,Pik3r1,Il6,Casp4,Icam1,Msn,Serpina3f,Vcam1,Col4a1,Iqgap1,Ptpn1,Cflar,Il1rn,Ccl2,Zfp36,Tnf,Acod1 | | 2.093e-02 | -3.87 | regulation of production of molecular mediator of immune response | biological process | GO:0002700 | 101 | 5 | 13711 | 214 | Il6,Il4ra,Irak3,Tnf,Tnfaip3 | | 2.100e-02 | -3.86 | MODULE\_47 | MSigDB lists | MODULE\_47 | 188 | 7 | 12187 | 179 | Plaur,Rgs16,Cdkn1a,Adamts1,Thbs1,Col4a1,Il6 | | 2.116e-02 | -3.86 | negative regulation of leukocyte activation | biological process | GO:0002695 | 138 | 6 | 13711 | 214 | Runx1,Lgals9,Il4ra,Tnfaip3,Sox11,Sdc4 | | 2.118e-02 | -3.85 | positive regulation of G1/S transition of mitotic cell cycle | biological process | GO:1900087 | 38 | 3 | 13711 | 214 | Kcna5,Ccnd2,Adamts1 | | 2.118e-02 | -3.85 | regulation of response to reactive oxygen species | biological process | GO:1901031 | 38 | 3 | 13711 | 214 | Nfe2l2,Tnf,Ripk1 | | 2.119e-02 | -3.85 | copper ion binding | molecular function | GO:0005507 | 38 | 3 | 13516 | 211 | Il1a,Mt1,Cp | | 2.132e-02 | -3.85 | GSE25123\_IL4\_VS\_IL4\_AND\_ROSIGLITAZONE\_STIM\_MACROPHAGE\_DAY10\_DN | MSigDB lists | GSE25123\_IL4\_VS\_IL4\_AND\_ROSIGLITAZONE\_STIM\_MACROPHAGE\_DAY10\_DN | 147 | 6 | 12187 | 179 | Mxd4,Gbp4,Rgs16,Gbp6,Cxcl1,Nfkbia | | 2.132e-02 | -3.85 | GSE12366\_GC\_VS\_MEMORY\_BCELL\_DN | MSigDB lists | GSE12366\_GC\_VS\_MEMORY\_BCELL\_DN | 147 | 6 | 12187 | 179 | Parp12,Bach1,Cdkn1a,Ccnd2,Ptpn1,Cflar | | 2.132e-02 | -3.85 | GSE2706\_2H\_VS\_8H\_LPS\_STIM\_DC\_UP | MSigDB lists | GSE2706\_2H\_VS\_8H\_LPS\_STIM\_DC\_UP | 147 | 6 | 12187 | 179 | Plek,Stx11,Ier3,Mxd4,Maff,Nfkbiz | | 2.132e-02 | -3.85 | GSE3982\_DC\_VS\_NEUTROPHIL\_DN | MSigDB lists | GSE3982\_DC\_VS\_NEUTROPHIL\_DN | 147 | 6 | 12187 | 179 | Tnfaip3,Bcl3,Irf7,Cxcl1,Xaf1,Cflar | | 2.132e-02 | -3.85 | GSE17721\_CTRL\_VS\_CPG\_2H\_BMDC\_DN | MSigDB lists | GSE17721\_CTRL\_VS\_CPG\_2H\_BMDC\_DN | 147 | 6 | 12187 | 179 | Tgm2,Ccnd2,Ccl12,Cd14,Ptges,Igsf6 | | 2.132e-02 | -3.85 | GSE3720\_VD1\_VS\_VD2\_GAMMADELTA\_TCELL\_UP | MSigDB lists | GSE3720\_VD1\_VS\_VD2\_GAMMADELTA\_TCELL\_UP | 147 | 6 | 12187 | 179 | Rnf213,Rtp4,Ccl4,Parp14,Hspa5,Ifi44 | | 2.136e-02 | -3.85 | GO\_KINASE\_INHIBITOR\_ACTIVITY | MSigDB lists | GO\_KINASE\_INHIBITOR\_ACTIVITY | 72 | 4 | 12187 | 179 | Hspb1,Socs3,Trib1,Cdkn1a | | 2.141e-02 | -3.84 | GSE2706\_LPS\_VS\_R848\_AND\_LPS\_8H\_STIM\_DC\_DN | MSigDB lists | GSE2706\_LPS\_VS\_R848\_AND\_LPS\_8H\_STIM\_DC\_DN | 108 | 5 | 12187 | 179 | Tnf,Map3k8,Nfkbiz,Bach1,Rgs16 | | 2.141e-02 | -3.84 | NOTCH\_DN.V1\_DN | MSigDB lists | NOTCH\_DN.V1\_DN | 108 | 5 | 12187 | 179 | Zbtb16,C3ar1,Tnfsf10,Sele,Kcna5 | | 2.141e-02 | -3.84 | GNF2\_HDAC1 | MSigDB lists | GNF2\_HDAC1 | 108 | 5 | 12187 | 179 | Ifi211,Ifi209,Ifi207,Rbm39,Ifi204 | | 2.146e-02 | -3.84 | IL-5 Signaling Pathway | WikiPathways | WP151 | 63 | 5 | 3756 | 96 | Icam1,Csf2rb,Nfkb1,Pik3r1,Nfkbia | | 2.161e-02 | -3.83 | positive regulation of protein localization to nucleus | biological process | GO:1900182 | 68 | 4 | 13711 | 214 | Dtx3l,Parp9,Pik3r1,Ptgs2 | | 2.169e-02 | -3.83 | Class A/1 (Rhodopsin-like receptors) | REACTOME pathways | R-MMU-373076 | 168 | 7 | 6297 | 105 | C3ar1,Cxcl1,Cxcl16,C5ar1,Hcar2,Cxcl9,P2ry6 | | 2.169e-02 | -3.83 | P2Y6PRNOCPTR | prints domains | PR01068 | 1 | 1 | 2951 | 64 | P2ry6 | | 2.169e-02 | -3.83 | TNECROSISFCT | prints domains | PR01234 | 1 | 1 | 2951 | 64 | Tnf | | 2.169e-02 | -3.83 | MACSCAVRCPTR | prints domains | PR01408 | 1 | 1 | 2951 | 64 | Msr1 | | 2.169e-02 | -3.83 | GENEIEX1 | prints domains | PR02100 | 1 | 1 | 2951 | 64 | Ier3 | | 2.169e-02 | -3.83 | GP91PHOX | prints domains | PR00466 | 1 | 1 | 2951 | 64 | Cybb | | 2.169e-02 | -3.83 | P47PHOX | prints domains | PR00498 | 1 | 1 | 2951 | 64 | Ncf1 | | 2.169e-02 | -3.83 | VCAM1 | prints domains | PR01474 | 1 | 1 | 2951 | 64 | Vcam1 | | 2.169e-02 | -3.83 | INTRLEUKIN1A | prints domains | PR01358 | 1 | 1 | 2951 | 64 | Il1a | | 2.169e-02 | -3.83 | NGELATINASE | prints domains | PR01275 | 1 | 1 | 2951 | 64 | Lcn2 | | 2.169e-02 | -3.83 | ADAMTS1 | prints domains | PR01858 | 1 | 1 | 2951 | 64 | Adamts1 | | 2.169e-02 | -3.83 | KV15CHANNEL | prints domains | PR01512 | 1 | 1 | 2951 | 64 | Kcna5 | | 2.169e-02 | -3.83 | INTRLEUKIN1X | prints domains | PR01360 | 1 | 1 | 2951 | 64 | Il1rn | | 2.169e-02 | -3.83 | TNFALPHA | prints domains | PR01235 | 1 | 1 | 2951 | 64 | Tnf | | 2.169e-02 | -3.83 | C3ANPHYLTXNR | prints domains | PR01060 | 1 | 1 | 2951 | 64 | C3ar1 | | 2.174e-02 | -3.83 | regulation of embryonic development | biological process | GO:0045995 | 102 | 5 | 13711 | 214 | Noct,Hspa5,Tnfaip3,Il1rn,Nfe2l2 | | 2.196e-02 | -3.82 | GSE44732\_UNSTIM\_VS\_IL27\_STIM\_IMATURE\_DC\_DN | MSigDB lists | GSE44732\_UNSTIM\_VS\_IL27\_STIM\_IMATURE\_DC\_DN | 148 | 6 | 12187 | 179 | S100a8,Atf3,Ifit2,Runx1,Ifit3b,S100a9 | | 2.196e-02 | -3.82 | GSE32533\_MIR17\_KO\_VS\_MIR17\_OVEREXPRESS\_ACT\_CD4\_TCELL\_UP | MSigDB lists | GSE32533\_MIR17\_KO\_VS\_MIR17\_OVEREXPRESS\_ACT\_CD4\_TCELL\_UP | 148 | 6 | 12187 | 179 | Ncf1,C3ar1,Pik3ap1,Irak3,Spi1,Tubb6 | | 2.196e-02 | -3.82 | GSE5455\_HEALTHY\_VS\_TUMOR\_BEARING\_MOUSE\_SPLEEN\_MONOCYTE\_24H\_INCUBATION\_DN | MSigDB lists | GSE5455\_HEALTHY\_VS\_TUMOR\_BEARING\_MOUSE\_SPLEEN\_MONOCYTE\_24H\_INCUBATION\_DN | 148 | 6 | 12187 | 179 | Stx11,Ksr1,Gadd45b,Map3k8,Arid5b,Casp4 | | 2.196e-02 | -3.82 | GSE37301\_CD4\_TCELL\_VS\_GRANULOCYTE\_MONOCYTE\_PROGENITOR\_DN | MSigDB lists | GSE37301\_CD4\_TCELL\_VS\_GRANULOCYTE\_MONOCYTE\_PROGENITOR\_DN | 148 | 6 | 12187 | 179 | Trib1,Usp18,Vcam1,Rtp4,Tnip1,Ifitm2 | | 2.196e-02 | -3.82 | GSE27092\_WT\_VS\_HDAC7\_PHOSPHO\_DEFICIENT\_CD8\_TCELL\_UP | MSigDB lists | GSE27092\_WT\_VS\_HDAC7\_PHOSPHO\_DEFICIENT\_CD8\_TCELL\_UP | 148 | 6 | 12187 | 179 | Clic4,Lcn2,Icam1,Cp,Tnf,Sgk1 | | 2.196e-02 | -3.82 | GSE32423\_CTRL\_VS\_IL4\_MEMORY\_CD8\_TCELL\_DN | MSigDB lists | GSE32423\_CTRL\_VS\_IL4\_MEMORY\_CD8\_TCELL\_DN | 148 | 6 | 12187 | 179 | Parp9,Gbp3,Iqgap1,Irgm2,Usp18,Map3k8 | | 2.196e-02 | -3.82 | GSE19512\_NAUTRAL\_VS\_INDUCED\_TREG\_DN | MSigDB lists | GSE19512\_NAUTRAL\_VS\_INDUCED\_TREG\_DN | 148 | 6 | 12187 | 179 | Ccl4,Cebpd,Akap12,Cxcl1,Saa1,Rgs16 | | 2.197e-02 | -3.82 | negative regulation of cellular component movement | biological process | GO:0051271 | 262 | 9 | 13711 | 214 | Ccl12,Clic4,Timp1,Robo4,Trib1,Adamts9,Nfe2l2,Thbs1,Il1rn | | 2.199e-02 | -3.82 | p75NTR signals via NF-kB | REACTOME pathways | R-MMU-193639 | 14 | 2 | 6297 | 105 | Nfkbia,Nfkb1 | | 2.208e-02 | -3.81 | WIERENGA\_STAT5A\_TARGETS\_GROUP2 | MSigDB lists | WIERENGA\_STAT5A\_TARGETS\_GROUP2 | 41 | 3 | 12187 | 179 | Arid5b,Gadd45b,Ccl12 | | 2.208e-02 | -3.81 | GO\_RESPONSE\_TO\_ANTIBIOTIC | MSigDB lists | GO\_RESPONSE\_TO\_ANTIBIOTIC | 41 | 3 | 12187 | 179 | Il6,Ccl12,Hspa5 | | 2.208e-02 | -3.81 | ZHANG\_ANTIVIRAL\_RESPONSE\_TO\_RIBAVIRIN\_DN | MSigDB lists | ZHANG\_ANTIVIRAL\_RESPONSE\_TO\_RIBAVIRIN\_DN | 41 | 3 | 12187 | 179 | Ccl4,Cflar,Gem | | 2.208e-02 | -3.81 | GO\_POSITIVE\_REGULATION\_OF\_INTERFERON\_GAMMA\_PRODUCTION | MSigDB lists | GO\_POSITIVE\_REGULATION\_OF\_INTERFERON\_GAMMA\_PRODUCTION | 41 | 3 | 12187 | 179 | Bcl3,Cd14,Tnf | | 2.218e-02 | -3.81 | HOELZEL\_NF1\_TARGETS\_UP | MSigDB lists | HOELZEL\_NF1\_TARGETS\_UP | 109 | 5 | 12187 | 179 | Ifi209,Ifi211,Ifi207,Ifi204,Rgs16 | | 2.218e-02 | -3.81 | COULOUARN\_TEMPORAL\_TGFB1\_SIGNATURE\_DN | MSigDB lists | COULOUARN\_TEMPORAL\_TGFB1\_SIGNATURE\_DN | 109 | 5 | 12187 | 179 | Nfkbia,Arid5b,Gadd45g,Errfi1,Gadd45b | | 2.218e-02 | -3.81 | NRL\_DN.V1\_DN | MSigDB lists | NRL\_DN.V1\_DN | 109 | 5 | 12187 | 179 | Socs3,Clic4,Ccl12,Nfkb1,Pik3ap1 | | 2.227e-02 | -3.80 | interleukin-1-mediated signaling pathway | biological process | GO:0070498 | 15 | 2 | 13711 | 214 | Irak3,Il1a | | 2.227e-02 | -3.80 | positive regulation of cellular response to oxidative stress | biological process | GO:1900409 | 15 | 2 | 13711 | 214 | Ripk1,Tnf | | 2.227e-02 | -3.80 | negative regulation of amino acid transport | biological process | GO:0051956 | 15 | 2 | 13711 | 214 | Tnf,Il1rn | | 2.227e-02 | -3.80 | regulation of granulocyte differentiation | biological process | GO:0030852 | 15 | 2 | 13711 | 214 | Trib1,Runx1 | | 2.227e-02 | -3.80 | regulation of interleukin-6 biosynthetic process | biological process | GO:0045408 | 15 | 2 | 13711 | 214 | Prg4,Tnf | | 2.227e-02 | -3.80 | positive regulation of myoblast fusion | biological process | GO:1901741 | 15 | 2 | 13711 | 214 | Il4ra,Cxcl9 | | 2.227e-02 | -3.80 | disruption of cells of other organism | biological process | GO:0044364 | 15 | 2 | 13711 | 214 | Ncf1,Ccl2 | | 2.227e-02 | -3.80 | macrophage migration | biological process | GO:1905517 | 15 | 2 | 13711 | 214 | Ccl2,Ccl12 | | 2.227e-02 | -3.80 | killing of cells of other organism | biological process | GO:0031640 | 15 | 2 | 13711 | 214 | Ccl2,Ncf1 | | 2.227e-02 | -3.80 | positive regulation of leukocyte adhesion to vascular endothelial cell | biological process | GO:1904996 | 15 | 2 | 13711 | 214 | Icam1,Tnf | | 2.227e-02 | -3.80 | regulation of immunoglobulin secretion | biological process | GO:0051023 | 15 | 2 | 13711 | 214 | Tnf,Il6 | | 2.227e-02 | -3.80 | negative regulation of cell killing | biological process | GO:0031342 | 15 | 2 | 13711 | 214 | Tap1,Lgals9 | | 2.227e-02 | -3.80 | negative regulation of viral release from host cell | biological process | GO:1902187 | 15 | 2 | 13711 | 214 | Trim56,Trim25 | | 2.228e-02 | -3.80 | purinergic receptor activity | molecular function | GO:0035586 | 15 | 2 | 13516 | 211 | Hcar2,P2ry6 | | 2.234e-02 | -3.80 | REACTOME\_MYD88\_MAL\_CASCADE\_INITIATED\_ON\_PLASMA\_MEMBRANE | MSigDB lists | REACTOME\_MYD88\_MAL\_CASCADE\_INITIATED\_ON\_PLASMA\_MEMBRANE | 73 | 4 | 12187 | 179 | Saa1,Nfkbia,Irak3,Nfkb2 | | 2.234e-02 | -3.80 | KIM\_RESPONSE\_TO\_TSA\_AND\_DECITABINE\_UP | MSigDB lists | KIM\_RESPONSE\_TO\_TSA\_AND\_DECITABINE\_UP | 73 | 4 | 12187 | 179 | Atf3,Cdkn1a,Irf7,Tgm2 | | 2.234e-02 | -3.80 | GAVIN\_FOXP3\_TARGETS\_CLUSTER\_P6 | MSigDB lists | GAVIN\_FOXP3\_TARGETS\_CLUSTER\_P6 | 73 | 4 | 12187 | 179 | Ifi44,Cmpk2,Ksr1,Icam1 | | 2.249e-02 | -3.79 | GO\_PEPTIDE\_CROSS\_LINKING | MSigDB lists | GO\_PEPTIDE\_CROSS\_LINKING | 16 | 2 | 12187 | 179 | Thbs1,Tgm2 | | 2.249e-02 | -3.79 | GO\_CELLULAR\_RESPONSE\_TO\_GROWTH\_HORMONE\_STIMULUS | MSigDB lists | GO\_CELLULAR\_RESPONSE\_TO\_GROWTH\_HORMONE\_STIMULUS | 16 | 2 | 12187 | 179 | Pik3r1,Ptpn1 | | 2.249e-02 | -3.79 | MODULE\_59 | MSigDB lists | MODULE\_59 | 16 | 2 | 12187 | 179 | C3ar1,Nfkbie | | 2.249e-02 | -3.79 | GO\_REGULATION\_OF\_MACROPHAGE\_ACTIVATION | MSigDB lists | GO\_REGULATION\_OF\_MACROPHAGE\_ACTIVATION | 16 | 2 | 12187 | 179 | Il4ra,Thbs1 | | 2.249e-02 | -3.79 | GO\_RESPONSE\_TO\_MUSCLE\_STRETCH | MSigDB lists | GO\_RESPONSE\_TO\_MUSCLE\_STRETCH | 16 | 2 | 12187 | 179 | Nfkb1,Nfkbia | | 2.249e-02 | -3.79 | MODULE\_173 | MSigDB lists | MODULE\_173 | 16 | 2 | 12187 | 179 | Gadd45g,C5ar1 | | 2.249e-02 | -3.79 | ZHANG\_RESPONSE\_TO\_CANTHARIDIN\_UP | MSigDB lists | ZHANG\_RESPONSE\_TO\_CANTHARIDIN\_UP | 16 | 2 | 12187 | 179 | Nfe2l2,Il1a | | 2.249e-02 | -3.79 | HINATA\_NFKB\_TARGETS\_KERATINOCYTE\_DN | MSigDB lists | HINATA\_NFKB\_TARGETS\_KERATINOCYTE\_DN | 16 | 2 | 12187 | 179 | Runx1,Thbs1 | | 2.249e-02 | -3.79 | GO\_REGULATION\_OF\_LIPOPOLYSACCHARIDE\_MEDIATED\_SIGNALING\_PATHWAY | MSigDB lists | GO\_REGULATION\_OF\_LIPOPOLYSACCHARIDE\_MEDIATED\_SIGNALING\_PATHWAY | 16 | 2 | 12187 | 179 | Trib1,Tnfaip3 | | 2.249e-02 | -3.79 | GO\_PROSTANOID\_BIOSYNTHETIC\_PROCESS | MSigDB lists | GO\_PROSTANOID\_BIOSYNTHETIC\_PROCESS | 16 | 2 | 12187 | 179 | Ptges,Ptgs2 | | 2.249e-02 | -3.79 | STEGER\_ADIPOGENESIS\_UP | MSigDB lists | STEGER\_ADIPOGENESIS\_UP | 16 | 2 | 12187 | 179 | Cp,Hp | | 2.249e-02 | -3.79 | MODULE\_73 | MSigDB lists | MODULE\_73 | 16 | 2 | 12187 | 179 | Il4ra,Csf2rb | | 2.249e-02 | -3.79 | MYLLYKANGAS\_AMPLIFICATION\_HOT\_SPOT\_23 | MSigDB lists | MYLLYKANGAS\_AMPLIFICATION\_HOT\_SPOT\_23 | 16 | 2 | 12187 | 179 | Birc3,Zbtb16 | | 2.249e-02 | -3.79 | MODULE\_340 | MSigDB lists | MODULE\_340 | 16 | 2 | 12187 | 179 | Tnf,Nfkb2 | | 2.249e-02 | -3.79 | GAVIN\_PDE3B\_TARGETS | MSigDB lists | GAVIN\_PDE3B\_TARGETS | 16 | 2 | 12187 | 179 | Cybb,Tnfsf10 | | 2.249e-02 | -3.79 | HAHTOLA\_CTCL\_CUTANEOUS | MSigDB lists | HAHTOLA\_CTCL\_CUTANEOUS | 16 | 2 | 12187 | 179 | Trib1,Zbtb16 | | 2.249e-02 | -3.79 | GO\_MITOGEN\_ACTIVATED\_PROTEIN\_KINASE\_KINASE\_BINDING | MSigDB lists | GO\_MITOGEN\_ACTIVATED\_PROTEIN\_KINASE\_KINASE\_BINDING | 16 | 2 | 12187 | 179 | Trib1,Ksr1 | | 2.251e-02 | -3.79 | GO\_POSITIVE\_REGULATION\_OF\_INTRACELLULAR\_TRANSPORT | MSigDB lists | GO\_POSITIVE\_REGULATION\_OF\_INTRACELLULAR\_TRANSPORT | 280 | 9 | 12187 | 179 | Rhou,Tnf,Pik3r1,Il4ra,Ccl12,Il6,Csf3,Ptgs2,Msn | | 2.257e-02 | -3.79 | regulation of protein complex disassembly | biological process | GO:0043244 | 103 | 5 | 13711 | 214 | Irgm2,Plek,Irgm1,Irak3,Tnf | | 2.258e-02 | -3.79 | TRIM28 (tripartite motif containing 28) | protein interactions | 10155 | 14 | 2 | 6802 | 115 | Ksr1,Nfe2l2 | | 2.258e-02 | -3.79 | TP53 (tumor protein p53) | protein interactions | 7157 | 14 | 2 | 6802 | 115 | Ifi211,Banp | | 2.258e-02 | -3.79 | DDB1 (damage specific DNA binding protein 1) | protein interactions | 1642 | 14 | 2 | 6802 | 115 | Cdkn1a,Ksr1 | | 2.258e-02 | -3.79 | Khdrbs1 (KH domain containing, RNA binding, signal transduction associated 1) | protein interactions | 20218 | 14 | 2 | 6802 | 115 | Ripk1,Rbm39 | | 2.259e-02 | -3.79 | nuclear outer membrane-endoplasmic reticulum membrane network | cellular component | GO:0042175 | 453 | 13 | 13825 | 212 | Gm5431,Tgtp1,Ifi47,F830016B08Rik,Ptpn1,Iigp1,Hspa5,Pik3r1,Tgtp2,Irgm1,Tap1,Irgm2,Gm4841 | | 2.261e-02 | -3.79 | GSE7852\_LN\_VS\_THYMUS\_TCONV\_UP | MSigDB lists | GSE7852\_LN\_VS\_THYMUS\_TCONV\_UP | 149 | 6 | 12187 | 179 | Casp4,Rrp8,Socs3,Gbp3,Xdh,Irak3 | | 2.261e-02 | -3.79 | GSE21380\_TFH\_VS\_GERMINAL\_CENTER\_TFH\_CD4\_TCELL\_UP | MSigDB lists | GSE21380\_TFH\_VS\_GERMINAL\_CENTER\_TFH\_CD4\_TCELL\_UP | 149 | 6 | 12187 | 179 | Tagln2,Pik3ap1,Socs3,Rasd1,Lcn2,Il1a | | 2.261e-02 | -3.79 | GSE43955\_TGFB\_IL6\_VS\_TGFB\_IL6\_IL23\_TH17\_ACT\_CD4\_TCELL\_60H\_DN | MSigDB lists | GSE43955\_TGFB\_IL6\_VS\_TGFB\_IL6\_IL23\_TH17\_ACT\_CD4\_TCELL\_60H\_DN | 149 | 6 | 12187 | 179 | Rgs16,C3ar1,Csf3,Irf7,A2m,Pik3r1 | | 2.261e-02 | -3.79 | TGFB\_UP.V1\_UP | MSigDB lists | TGFB\_UP.V1\_UP | 149 | 6 | 12187 | 179 | Tnfaip8,Gadd45b,Atf3,Akap12,Trib1,Sgk1 | | 2.261e-02 | -3.79 | GSE17721\_PAM3CSK4\_VS\_GADIQUIMOD\_16H\_BMDC\_UP | MSigDB lists | GSE17721\_PAM3CSK4\_VS\_GADIQUIMOD\_16H\_BMDC\_UP | 149 | 6 | 12187 | 179 | Col4a1,Icam1,Nfkb2,Birc3,Tnf,Cd14 | | 2.264e-02 | -3.79 | cation homeostasis | biological process | GO:0055080 | 541 | 15 | 13711 | 214 | Mt1,Ccl2,Slc24a4,C3ar1,Pygm,Cp,P2ry6,Lcn2,Cxcl1,C5ar1,Tgm2,Clic4,Mt2,Kcna5,Sgk1 | | 2.267e-02 | -3.79 | cellular transition metal ion homeostasis | biological process | GO:0046916 | 69 | 4 | 13711 | 214 | Lcn2,Cp,Mt2,Mt1 | | 2.267e-02 | -3.79 | negative regulation of small molecule metabolic process | biological process | GO:0062014 | 69 | 4 | 13711 | 214 | Il6,Nfkb1,Ier3,Plek | | 2.268e-02 | -3.79 | kinase activator activity | molecular function | GO:0019209 | 69 | 4 | 13516 | 211 | Irgm2,Lgals9,Cdkn1a,Iqgap1 | | 2.269e-02 | -3.79 | regulation of gene silencing by RNA | biological process | GO:0060966 | 39 | 3 | 13711 | 214 | Ripk1,Zfp36,Nfkb1 | | 2.269e-02 | -3.79 | cellular detoxification | biological process | GO:1990748 | 39 | 3 | 13711 | 214 | S100a9,S100a8,Hp | | 2.269e-02 | -3.79 | negative regulation of carbohydrate metabolic process | biological process | GO:0045912 | 39 | 3 | 13711 | 214 | Il6,Ier3,Plek | | 2.269e-02 | -3.79 | exocrine system development | biological process | GO:0035272 | 39 | 3 | 13711 | 214 | Tnf,Tgm2,Il6 | | 2.269e-02 | -3.79 | regulation of posttranscriptional gene silencing | biological process | GO:0060147 | 39 | 3 | 13711 | 214 | Zfp36,Nfkb1,Ripk1 | | 2.269e-02 | -3.79 | positive regulation of calcium-mediated signaling | biological process | GO:0050850 | 39 | 3 | 13711 | 214 | Tnf,Ccl4,P2ry6 | | 2.270e-02 | -3.79 | GO\_TRANSCRIPTION\_FROM\_RNA\_POLYMERASE\_II\_PROMOTER | MSigDB lists | GO\_TRANSCRIPTION\_FROM\_RNA\_POLYMERASE\_II\_PROMOTER | 525 | 14 | 12187 | 179 | Nfe2l2,Bach1,Spi1,Maff,Runx1,Nfkb2,Irf7,C5ar1,Cebpd,Atf3,Nfkb1,Sox11,Irf9,Irf2 | | 2.270e-02 | -3.79 | negative regulation of nucleobase-containing compound metabolic process | biological process | GO:0045934 | 1157 | 27 | 13711 | 214 | Rrp8,Dnajb1,Noct,Bcl3,Rasd1,Zfp36,Atf3,Znfx1,Tnf,Ifi204,Zbtb16,Ifi207,Arid5b,Sbno2,Parp9,Nfkbie,Nfkb1,Gbp4,Ifi209,Ier3,Runx1,Spi1,Cebpd,Ifi211,Bach1,Sox11,Mxd4 | | 2.271e-02 | -3.79 | enzyme binding | molecular function | GO:0019899 | 2041 | 43 | 13516 | 211 | Ksr1,Cdkn1a,Rhou,Nfkbia,Akap12,Tnfaip3,Msn,Tnf,Zfp36,Trib1,Iqgap1,Rhoj,Rasip1,Isg15,Pik3r1,Kcna5,A2m,Sdc4,Timp1,Ptgs2,Fzd4,Hspb1,Dtx3l,Ripk1,Irgm2,Parp9,Hspa5,Resf1,Parp14,Errfi1,Csf3,Cflar,Lgals9,Rhoc,Plek,Tnip1,Sele,Lcn2,Ccnd2,Sgk1,Dnajb1,Plaur,Ptpn1 | | 2.280e-02 | -3.78 | regulation of mitotic cell cycle phase transition | biological process | GO:1901990 | 221 | 8 | 13711 | 214 | Fam107a,Adamts1,Cdkn1a,Ccl12,Ripk1,Kcna5,Ccnd2,Ier3 | | 2.286e-02 | -3.78 | regulation of secretion | biological process | GO:0051046 | 690 | 18 | 13711 | 214 | Lgals9,Hcar2,Cd14,Il1rn,Il6,Akap12,Sox11,Il1a,Il4ra,Tnf,Tnfaip3,Casp4,Il17ra,Runx1,S100a9,Mmp8,Sdc4,S100a8 | | 2.297e-02 | -3.77 | REACTOME\_SIGNALING\_BY\_THE\_B\_CELL\_RECEPTOR\_BCR | MSigDB lists | REACTOME\_SIGNALING\_BY\_THE\_B\_CELL\_RECEPTOR\_BCR | 110 | 5 | 12187 | 179 | Pik3ap1,Cdkn1a,Nfkbie,Pik3r1,Nfkbia | | 2.316e-02 | -3.77 | UBQ | smart domains | SM00213 | 36 | 3 | 7188 | 123 | Isg15,Oasl1,Oasl2 | | 2.327e-02 | -3.76 | GSE37301\_COMMON\_LYMPHOID\_PROGENITOR\_VS\_CD4\_TCELL\_UP | MSigDB lists | GSE37301\_COMMON\_LYMPHOID\_PROGENITOR\_VS\_CD4\_TCELL\_UP | 150 | 6 | 12187 | 179 | C5ar1,Icam1,Maff,Sgk1,Tnip3,Rasip1 | | 2.327e-02 | -3.76 | GSE25087\_TREG\_VS\_TCONV\_FETUS\_UP | MSigDB lists | GSE25087\_TREG\_VS\_TCONV\_FETUS\_UP | 150 | 6 | 12187 | 179 | Pik3ap1,Ksr1,Tnip3,Cybb,Map3k8,Trib1 | | 2.327e-02 | -3.76 | GSE19941\_UNSTIM\_VS\_LPS\_AND\_IL10\_STIM\_IL10\_KO\_MACROPHAGE\_UP | MSigDB lists | GSE19941\_UNSTIM\_VS\_LPS\_AND\_IL10\_STIM\_IL10\_KO\_MACROPHAGE\_UP | 150 | 6 | 12187 | 179 | Gem,Map3k6,Saa1,Selp,Arid5b,Icam1 | | 2.327e-02 | -3.76 | GSE9006\_HEALTHY\_VS\_TYPE\_2\_DIABETES\_PBMC\_AT\_DX\_DN | MSigDB lists | GSE9006\_HEALTHY\_VS\_TYPE\_2\_DIABETES\_PBMC\_AT\_DX\_DN | 150 | 6 | 12187 | 179 | Trib1,Sgk1,Adamts1,Ptgs2,C3ar1,Rbm47 | | 2.327e-02 | -3.76 | GSE17721\_POLYIC\_VS\_CPG\_2H\_BMDC\_DN | MSigDB lists | GSE17721\_POLYIC\_VS\_CPG\_2H\_BMDC\_DN | 150 | 6 | 12187 | 179 | Ccl2,Ptges,Vcam1,Tgm2,Nfe2l2,Igsf6 | | 2.336e-02 | -3.76 | KRAS.LUNG.BREAST\_UP.V1\_DN | MSigDB lists | KRAS.LUNG.BREAST\_UP.V1\_DN | 74 | 4 | 12187 | 179 | Sox11,Tnfsf10,Gadd45g,Rtp4 | | 2.336e-02 | -3.76 | HILLION\_HMGA1\_TARGETS | MSigDB lists | HILLION\_HMGA1\_TARGETS | 74 | 4 | 12187 | 179 | C5ar1,Cxcl1,Nfkbia,Hspb1 | | 2.342e-02 | -3.75 | negative regulation of vasculature development | biological process | GO:1901343 | 104 | 5 | 13711 | 214 | Thbs1,Adamts9,Xdh,Adamts1,Ccl2 | | 2.349e-02 | -3.75 | GO\_CELLULAR\_RESPONSE\_TO\_DNA\_DAMAGE\_STIMULUS | MSigDB lists | GO\_CELLULAR\_RESPONSE\_TO\_DNA\_DAMAGE\_STIMULUS | 579 | 15 | 12187 | 179 | Bach1,Pik3r1,Tnf,Dtx3l,Bcl3,Irf7,Ifi211,Ifi207,Ifi204,Sgk1,Cdkn1a,Ifi209,Trim25,Parp9,Isg15 | | 2.353e-02 | -3.75 | SASAI\_RESISTANCE\_TO\_NEOPLASTIC\_TRANSFROMATION | MSigDB lists | SASAI\_RESISTANCE\_TO\_NEOPLASTIC\_TRANSFROMATION | 42 | 3 | 12187 | 179 | Adamts1,Fstl1,Cebpd | | 2.353e-02 | -3.75 | GNF2\_HLA\_C | MSigDB lists | GNF2\_HLA\_C | 42 | 3 | 12187 | 179 | Tap1,Phf11b,Phf11d | | 2.353e-02 | -3.75 | AMIT\_SERUM\_RESPONSE\_240\_MCF10A | MSigDB lists | AMIT\_SERUM\_RESPONSE\_240\_MCF10A | 42 | 3 | 12187 | 179 | Thbs1,Angptl4,Akap12 | | 2.353e-02 | -3.75 | BROWNE\_HCMV\_INFECTION\_2HR\_DN | MSigDB lists | BROWNE\_HCMV\_INFECTION\_2HR\_DN | 42 | 3 | 12187 | 179 | Plaur,Sgk1,Ifit2 | | 2.353e-02 | -3.75 | BORLAK\_LIVER\_CANCER\_EGF\_UP | MSigDB lists | BORLAK\_LIVER\_CANCER\_EGF\_UP | 42 | 3 | 12187 | 179 | Atf3,Rhoc,Col4a1 | | 2.353e-02 | -3.75 | BURTON\_ADIPOGENESIS\_4 | MSigDB lists | BURTON\_ADIPOGENESIS\_4 | 42 | 3 | 12187 | 179 | Angptl4,Zbtb16,Rhou | | 2.355e-02 | -3.75 | extracellular matrix organization | biological process | GO:0030198 | 181 | 7 | 13711 | 214 | Adamts9,Mmp8,Olfml2b,Bcl3,Nfkb2,Tnf,Col4a1 | | 2.370e-02 | -3.74 | GO\_PERINUCLEAR\_REGION\_OF\_CYTOPLASM | MSigDB lists | GO\_PERINUCLEAR\_REGION\_OF\_CYTOPLASM | 528 | 14 | 12187 | 179 | Ccl12,Pik3r1,Gbp3,Nfkbie,Ptges,Bcl3,Kcna5,Rasd1,Rasip1,Cdkn1a,Sgk1,Sele,Clic4,Msn | | 2.370e-02 | -3.74 | GO\_ANCHORING\_JUNCTION | MSigDB lists | GO\_ANCHORING\_JUNCTION | 427 | 12 | 12187 | 179 | Sdc4,Sntb2,Tgm2,Rhou,Iqgap1,Hspa5,Akap12,Msn,Plaur,Irf2,Icam1,Hspb1 | | 2.376e-02 | -3.74 | regulation of organic acid transport | biological process | GO:0032890 | 70 | 4 | 13711 | 214 | Il1a,Thbs1,Il1rn,Tnf | | 2.376e-02 | -3.74 | positive regulation of lipid localization | biological process | GO:1905954 | 70 | 4 | 13711 | 214 | Runx1,Nfkbia,Msr1,Il1a | | 2.376e-02 | -3.74 | leukocyte homeostasis | biological process | GO:0001776 | 70 | 4 | 13711 | 214 | Ccl2,Hcar2,Il6,Tnfaip3 | | 2.377e-02 | -3.74 | SENGUPTA\_NASOPHARYNGEAL\_CARCINOMA\_UP | MSigDB lists | SENGUPTA\_NASOPHARYNGEAL\_CARCINOMA\_UP | 237 | 8 | 12187 | 179 | Nfkbia,Cd14,Xaf1,Ptgs2,Ccl12,Rassf4,Col4a1,Igsf6 | | 2.378e-02 | -3.74 | GSE23114\_WT\_VS\_SLE2C1\_MOUSE\_PERITONEAL\_CAVITY\_B1A\_BCELL\_DN | MSigDB lists | GSE23114\_WT\_VS\_SLE2C1\_MOUSE\_PERITONEAL\_CAVITY\_B1A\_BCELL\_DN | 111 | 5 | 12187 | 179 | Nfkbia,Ier3,Gadd45b,Xdh,Icam1 | | 2.378e-02 | -3.74 | DARWICHE\_PAPILLOMA\_RISK\_LOW\_UP | MSigDB lists | DARWICHE\_PAPILLOMA\_RISK\_LOW\_UP | 111 | 5 | 12187 | 179 | Nfe2l2,Cmpk2,S100a8,S100a9,Ifitm3 | | 2.378e-02 | -3.74 | GO\_RESPONSE\_TO\_ALKALOID | MSigDB lists | GO\_RESPONSE\_TO\_ALKALOID | 111 | 5 | 12187 | 179 | Vcam1,Tnf,Il6,Icam1,Nfkb1 | | 2.378e-02 | -3.74 | AAAGGAT\_MIR501 | MSigDB lists | AAAGGAT\_MIR501 | 111 | 5 | 12187 | 179 | Errfi1,Pik3ap1,Apold1,Map3k8,Sox11 | | 2.378e-02 | -3.74 | GO\_NEGATIVE\_REGULATION\_OF\_PROTEIN\_SERINE\_THREONINE\_KINASE\_ACTIVITY | MSigDB lists | GO\_NEGATIVE\_REGULATION\_OF\_PROTEIN\_SERINE\_THREONINE\_KINASE\_ACTIVITY | 111 | 5 | 12187 | 179 | Ptpn1,Tnfaip3,Hspb1,Irak3,Cdkn1a | | 2.383e-02 | -3.74 | regulation of protein localization | biological process | GO:0032880 | 898 | 22 | 13711 | 214 | Sgk1,Ptgs2,Il17ra,Parp9,Gbp4,Pik3r1,Hcar2,Lgals9,Dtx3l,Il6,Il4ra,Casp4,Akap12,Il1a,Msn,Mmp8,Iqgap1,Ptpn1,Cd14,Ccl2,Tnfaip3,Tnf | | 2.383e-02 | -3.74 | DR4\_Q2 | MSigDB lists | DR4\_Q2 | 193 | 7 | 12187 | 179 | Errfi1,Rbm39,Ip6k2,Pik3ap1,Bcl3,Nfkbia,Sgk1 | | 2.395e-02 | -3.73 | GSE24574\_BCL6\_HIGH\_VS\_LOW\_TFH\_CD4\_TCELL\_UP | MSigDB lists | GSE24574\_BCL6\_HIGH\_VS\_LOW\_TFH\_CD4\_TCELL\_UP | 151 | 6 | 12187 | 179 | Cybb,Tiparp,Kcna5,Tnfaip3,Rrp8,Pygm | | 2.395e-02 | -3.73 | GSE46242\_CTRL\_VS\_EGR2\_DELETED\_ANERGIC\_TH1\_CD4\_TCELL\_UP | MSigDB lists | GSE46242\_CTRL\_VS\_EGR2\_DELETED\_ANERGIC\_TH1\_CD4\_TCELL\_UP | 151 | 6 | 12187 | 179 | Rhou,Ifit2,Runx1,S100a9,Angptl4,Cmpk2 | | 2.395e-02 | -3.73 | GSE18804\_BRAIN\_VS\_COLON\_TUMORAL\_MACROPHAGE\_DN | MSigDB lists | GSE18804\_BRAIN\_VS\_COLON\_TUMORAL\_MACROPHAGE\_DN | 151 | 6 | 12187 | 179 | Ifi204,Ifi207,Osmr,Ifi209,Ifi211,Trib1 | | 2.395e-02 | -3.73 | GSE45382\_UNTREATED\_VS\_TGFB\_TREATED\_MACROPHAGES\_DN | MSigDB lists | GSE45382\_UNTREATED\_VS\_TGFB\_TREATED\_MACROPHAGES\_DN | 151 | 6 | 12187 | 179 | Gadd45b,Ptpn1,AA467197,Ptgs2,Cdkn1a,Fstl1 | | 2.395e-02 | -3.73 | GSE23114\_PERITONEAL\_CAVITY\_B1A\_BCELL\_VS\_SPLEEN\_BCELL\_IN\_SLE2C1\_MOUSE\_UP | MSigDB lists | GSE23114\_PERITONEAL\_CAVITY\_B1A\_BCELL\_VS\_SPLEEN\_BCELL\_IN\_SLE2C1\_MOUSE\_UP | 151 | 6 | 12187 | 179 | Sgk1,Dnajb1,Kcna5,Oasl1,Ifitm3,Pygm | | 2.395e-02 | -3.73 | GSE3982\_CTRL\_VS\_IGE\_STIM\_MAST\_CELL\_DN | MSigDB lists | GSE3982\_CTRL\_VS\_IGE\_STIM\_MAST\_CELL\_DN | 151 | 6 | 12187 | 179 | Nfkbie,Sdc4,Aff1,Atf3,Tubb6,Ksr1 | | 2.395e-02 | -3.73 | GSE26495\_NAIVE\_VS\_PD1LOW\_CD8\_TCELL\_DN | MSigDB lists | GSE26495\_NAIVE\_VS\_PD1LOW\_CD8\_TCELL\_DN | 151 | 6 | 12187 | 179 | Rhou,Map3k8,Ccl4,Sntb2,Pik3r5,Pik3ap1 | | 2.395e-02 | -3.73 | GSE40274\_LEF1\_VS\_FOXP3\_AND\_LEF1\_TRANSDUCED\_ACTIVATED\_CD4\_TCELL\_DN | MSigDB lists | GSE40274\_LEF1\_VS\_FOXP3\_AND\_LEF1\_TRANSDUCED\_ACTIVATED\_CD4\_TCELL\_DN | 151 | 6 | 12187 | 179 | Zbtb16,Sgk1,Ifit2,Msn,Nfe2l2,Il17ra | | 2.395e-02 | -3.73 | GSE34156\_TLR1\_TLR2\_LIGAND\_VS\_NOD2\_AND\_TLR1\_TLR2\_LIGAND\_6H\_TREATED\_MONOCYTE\_UP | MSigDB lists | GSE34156\_TLR1\_TLR2\_LIGAND\_VS\_NOD2\_AND\_TLR1\_TLR2\_LIGAND\_6H\_TREATED\_MONOCYTE\_UP | 151 | 6 | 12187 | 179 | Tgm2,Zbtb16,Luc7l3,Pik3r5,Trim56,Trim25 | | 2.414e-02 | -3.72 | GO\_POSITIVE\_REGULATION\_OF\_CELL\_PROLIFERATION | MSigDB lists | GO\_POSITIVE\_REGULATION\_OF\_CELL\_PROLIFERATION | 581 | 15 | 12187 | 179 | Osmr,C5ar1,Csf3,C3ar1,Ccl12,Tnfaip3,Il6,Tnf,Ptgs2,Atf3,Timp1,Kcna5,Vcam1,Sox11,Cdkn1a | | 2.425e-02 | -3.72 | regulation of extrinsic apoptotic signaling pathway in absence of ligand | biological process | GO:2001239 | 40 | 3 | 13711 | 214 | Ripk1,Tnf,Sgk3 | | 2.464e-02 | -3.70 | GSE3982\_MAST\_CELL\_VS\_BCELL\_UP | MSigDB lists | GSE3982\_MAST\_CELL\_VS\_BCELL\_UP | 152 | 6 | 12187 | 179 | Pik3r1,Hp,S100a8,Akap12,Stx11,Ccl12 | | 2.464e-02 | -3.70 | GSE17721\_CTRL\_VS\_LPS\_8H\_BMDC\_DN | MSigDB lists | GSE17721\_CTRL\_VS\_LPS\_8H\_BMDC\_DN | 152 | 6 | 12187 | 179 | Arid5b,Gbp3,Map3k8,Birc3,Nfkbia,Timp1 | | 2.464e-02 | -3.70 | GSE10325\_CD4\_TCELL\_VS\_BCELL\_UP | MSigDB lists | GSE10325\_CD4\_TCELL\_VS\_BCELL\_UP | 152 | 6 | 12187 | 179 | Pik3r1,Ccnd2,Runx1,Timp1,Tnfsf10,Tnfaip3 | | 2.464e-02 | -3.70 | GSE45365\_CD8A\_DC\_VS\_CD11B\_DC\_IFNAR\_KO\_DN | MSigDB lists | GSE45365\_CD8A\_DC\_VS\_CD11B\_DC\_IFNAR\_KO\_DN | 152 | 6 | 12187 | 179 | Ifi211,Phf11d,Phf11b,Ifi209,Ifi207,Ifi204 | | 2.464e-02 | -3.70 | GSE3982\_EOSINOPHIL\_VS\_BASOPHIL\_UP | MSigDB lists | GSE3982\_EOSINOPHIL\_VS\_BASOPHIL\_UP | 152 | 6 | 12187 | 179 | Bcl3,S100a9,S100a8,Trib1,Cybb,Tiparp | | 2.464e-02 | -3.70 | GSE30971\_WBP7\_HET\_VS\_KO\_MACROPHAGE\_4H\_LPS\_STIM\_UP | MSigDB lists | GSE30971\_WBP7\_HET\_VS\_KO\_MACROPHAGE\_4H\_LPS\_STIM\_UP | 152 | 6 | 12187 | 179 | Tubb6,Tnip3,AA467197,Tnfaip3,Icam1,Plaur | | 2.464e-02 | -3.70 | GSE40666\_UNTREATED\_VS\_IFNA\_STIM\_EFFECTOR\_CD8\_TCELL\_90MIN\_UP | MSigDB lists | GSE40666\_UNTREATED\_VS\_IFNA\_STIM\_EFFECTOR\_CD8\_TCELL\_90MIN\_UP | 152 | 6 | 12187 | 179 | Ptpn1,Tap1,Rbm47,Tagln2,Cflar,Tgm2 | | 2.464e-02 | -3.70 | GSE23505\_UNTREATED\_VS\_4DAY\_IL6\_IL1\_TGFB\_TREATED\_CD4\_TCELL\_UP | MSigDB lists | GSE23505\_UNTREATED\_VS\_4DAY\_IL6\_IL1\_TGFB\_TREATED\_CD4\_TCELL\_UP | 152 | 6 | 12187 | 179 | Sgk1,Map3k8,Ccnd2,Ccl12,Ncf1,Gbp7 | | 2.464e-02 | -3.70 | GSE36476\_CTRL\_VS\_TSST\_ACT\_40H\_MEMORY\_CD4\_TCELL\_YOUNG\_UP | MSigDB lists | GSE36476\_CTRL\_VS\_TSST\_ACT\_40H\_MEMORY\_CD4\_TCELL\_YOUNG\_UP | 152 | 6 | 12187 | 179 | Pik3r1,Sgk1,Dnajb1,Tiparp,Tnfaip3,Arid5b | | 2.484e-02 | -3.70 | GO\_CORE\_PROMOTER\_PROXIMAL\_REGION\_DNA\_BINDING | MSigDB lists | GO\_CORE\_PROMOTER\_PROXIMAL\_REGION\_DNA\_BINDING | 239 | 8 | 12187 | 179 | Spi1,Ifi211,Ifi209,Atf3,Cebpd,Nfkb2,Ifi207,Ifi204 | | 2.488e-02 | -3.69 | regulation of gene silencing | biological process | GO:0060968 | 71 | 4 | 13711 | 214 | Ripk1,Resf1,Nfkb1,Zfp36 | | 2.491e-02 | -3.69 | GPVI-mediated activation cascade | REACTOME pathways | R-MMU-114604 | 38 | 3 | 6297 | 105 | Pik3r5,Csf2rb,Pik3r1 | | 2.502e-02 | -3.69 | LIU\_SMARCA4\_TARGETS | MSigDB lists | LIU\_SMARCA4\_TARGETS | 43 | 3 | 12187 | 179 | Ifitm2,Angptl4,Ccl12 | | 2.502e-02 | -3.69 | GO\_REGULATION\_OF\_BLOOD\_VESSEL\_ENDOTHELIAL\_CELL\_MIGRATION | MSigDB lists | GO\_REGULATION\_OF\_BLOOD\_VESSEL\_ENDOTHELIAL\_CELL\_MIGRATION | 43 | 3 | 12187 | 179 | Thbs1,Ptgs2,Hspb1 | | 2.502e-02 | -3.69 | GO\_PROTEIN\_KINASE\_C\_BINDING | MSigDB lists | GO\_PROTEIN\_KINASE\_C\_BINDING | 43 | 3 | 12187 | 179 | Hspb1,Plek,Sdc4 | | 2.502e-02 | -3.69 | GO\_NEGATIVE\_REGULATION\_OF\_CELL\_SUBSTRATE\_ADHESION | MSigDB lists | GO\_NEGATIVE\_REGULATION\_OF\_CELL\_SUBSTRATE\_ADHESION | 43 | 3 | 12187 | 179 | Pik3r1,Thbs1,Fzd4 | | 2.502e-02 | -3.69 | FAELT\_B\_CLL\_WITH\_VH\_REARRANGEMENTS\_UP | MSigDB lists | FAELT\_B\_CLL\_WITH\_VH\_REARRANGEMENTS\_UP | 43 | 3 | 12187 | 179 | Rhoc,Clic4,Runx1 | | 2.502e-02 | -3.69 | LEE\_LIVER\_CANCER\_ACOX1\_UP | MSigDB lists | LEE\_LIVER\_CANCER\_ACOX1\_UP | 43 | 3 | 12187 | 179 | Tagln2,Sgk1,Lcn2 | | 2.502e-02 | -3.69 | GO\_POSITIVE\_REGULATION\_OF\_ALPHA\_BETA\_T\_CELL\_ACTIVATION | MSigDB lists | GO\_POSITIVE\_REGULATION\_OF\_ALPHA\_BETA\_T\_CELL\_ACTIVATION | 43 | 3 | 12187 | 179 | Il4ra,Zbtb16,Il6 | | 2.503e-02 | -3.69 | GO\_REGULATION\_OF\_CYTOPLASMIC\_TRANSPORT | MSigDB lists | GO\_REGULATION\_OF\_CYTOPLASMIC\_TRANSPORT | 381 | 11 | 12187 | 179 | Bcl3,Nfkbie,Csf3,Nfkbia,Pik3r1,Tnf,Rhou,Il6,Ptgs2,Msn,Cdkn1a | | 2.503e-02 | -3.69 | UDAYAKUMAR\_MED1\_TARGETS\_DN | MSigDB lists | UDAYAKUMAR\_MED1\_TARGETS\_DN | 195 | 7 | 12187 | 179 | Trib1,Adamts1,Cdkn1a,Gadd45b,Atf3,Rhoc,Arid5b | | 2.503e-02 | -3.69 | GO\_POSITIVE\_REGULATION\_OF\_INTRACELLULAR\_PROTEIN\_TRANSPORT | MSigDB lists | GO\_POSITIVE\_REGULATION\_OF\_INTRACELLULAR\_PROTEIN\_TRANSPORT | 195 | 7 | 12187 | 179 | Ptgs2,Csf3,Pik3r1,Rhou,Tnf,Il6,Ccl12 | | 2.507e-02 | -3.69 | regulation of protein catabolic process | biological process | GO:0042176 | 358 | 11 | 13711 | 214 | Tiparp,Timp1,Ier3,Samd9l,Msn,Tnfaip3,Tnf,Trim30a,Banp,Trib1,Irak3 | | 2.520e-02 | -3.68 | cellular zinc ion homeostasis | biological process | GO:0006882 | 16 | 2 | 13711 | 214 | Mt2,Mt1 | | 2.520e-02 | -3.68 | negative regulation of membrane potential | biological process | GO:0045837 | 16 | 2 | 13711 | 214 | Il6,Il1rn | | 2.520e-02 | -3.68 | granulocyte differentiation | biological process | GO:0030851 | 16 | 2 | 13711 | 214 | Csf3,Spi1 | | 2.520e-02 | -3.68 | prostaglandin biosynthetic process | biological process | GO:0001516 | 16 | 2 | 13711 | 214 | Ptges,Ptgs2 | | 2.520e-02 | -3.68 | cellular response to fluid shear stress | biological process | GO:0071498 | 16 | 2 | 13711 | 214 | Ptgs2,Nfe2l2 | | 2.520e-02 | -3.68 | prostanoid biosynthetic process | biological process | GO:0046457 | 16 | 2 | 13711 | 214 | Ptgs2,Ptges | | 2.520e-02 | -3.68 | negative regulation of cytokine production involved in immune response | biological process | GO:0002719 | 16 | 2 | 13711 | 214 | Irak3,Tnf | | 2.520e-02 | -3.68 | mature B cell differentiation involved in immune response | biological process | GO:0002313 | 16 | 2 | 13711 | 214 | Tnfaip3,Bcl3 | | 2.520e-02 | -3.68 | positive regulation of nitric-oxide synthase biosynthetic process | biological process | GO:0051770 | 16 | 2 | 13711 | 214 | Ccl2,Akap12 | | 2.520e-02 | -3.68 | negative regulation of oxidative stress-induced intrinsic apoptotic signaling pathway | biological process | GO:1902176 | 16 | 2 | 13711 | 214 | Nfe2l2,Hspb1 | | 2.520e-02 | -3.68 | regulation of podosome assembly | biological process | GO:0071801 | 16 | 2 | 13711 | 214 | Msn,Tnf | | 2.520e-02 | -3.68 | response to muscle stretch | biological process | GO:0035994 | 16 | 2 | 13711 | 214 | Nfkb1,Nfkbia | | 2.525e-02 | -3.68 | GO\_REGULATION\_OF\_SUPEROXIDE\_METABOLIC\_PROCESS | MSigDB lists | GO\_REGULATION\_OF\_SUPEROXIDE\_METABOLIC\_PROCESS | 17 | 2 | 12187 | 179 | Tnf,Nfe2l2 | | 2.525e-02 | -3.68 | GO\_NEGATIVE\_REGULATION\_OF\_OXIDOREDUCTASE\_ACTIVITY | MSigDB lists | GO\_NEGATIVE\_REGULATION\_OF\_OXIDOREDUCTASE\_ACTIVITY | 17 | 2 | 12187 | 179 | Hp,Nfkb1 | | 2.525e-02 | -3.68 | GO\_PHOSPHATIDYLINOSITOL\_3\_4\_BISPHOSPHATE\_BINDING | MSigDB lists | GO\_PHOSPHATIDYLINOSITOL\_3\_4\_BISPHOSPHATE\_BINDING | 17 | 2 | 12187 | 179 | Plek,Ncf1 | | 2.525e-02 | -3.68 | GO\_NEGATIVE\_REGULATION\_OF\_OXIDATIVE\_STRESS\_INDUCED\_INTRINSIC\_APOPTOTIC\_SIGNALING\_PATHWAY | MSigDB lists | GO\_NEGATIVE\_REGULATION\_OF\_OXIDATIVE\_STRESS\_INDUCED\_INTRINSIC\_APOPTOTIC\_SIGNALING\_PATHWAY | 17 | 2 | 12187 | 179 | Nfe2l2,Hspb1 | | 2.525e-02 | -3.68 | HUI\_MAPK14\_TARGETS\_UP | MSigDB lists | HUI\_MAPK14\_TARGETS\_UP | 17 | 2 | 12187 | 179 | Isg15,Gadd45g | | 2.525e-02 | -3.68 | SARTIPY\_BLUNTED\_BY\_INSULIN\_RESISTANCE\_UP | MSigDB lists | SARTIPY\_BLUNTED\_BY\_INSULIN\_RESISTANCE\_UP | 17 | 2 | 12187 | 179 | Sntb2,Socs3 | | 2.525e-02 | -3.68 | GO\_PHAGOCYTIC\_CUP | MSigDB lists | GO\_PHAGOCYTIC\_CUP | 17 | 2 | 12187 | 179 | Tnf,Irgm2 | | 2.525e-02 | -3.68 | GO\_FATTY\_ACID\_BINDING | MSigDB lists | GO\_FATTY\_ACID\_BINDING | 17 | 2 | 12187 | 179 | S100a9,S100a8 | | 2.525e-02 | -3.68 | PID\_ANTHRAX\_PATHWAY | MSigDB lists | PID\_ANTHRAX\_PATHWAY | 17 | 2 | 12187 | 179 | Tnf,Vcam1 | | 2.525e-02 | -3.68 | GO\_POSITIVE\_REGULATION\_OF\_IMMUNOGLOBULIN\_PRODUCTION | MSigDB lists | GO\_POSITIVE\_REGULATION\_OF\_IMMUNOGLOBULIN\_PRODUCTION | 17 | 2 | 12187 | 179 | Il4ra,Il6 | | 2.525e-02 | -3.68 | GO\_PROTEIN\_LIPID\_COMPLEX\_BINDING | MSigDB lists | GO\_PROTEIN\_LIPID\_COMPLEX\_BINDING | 17 | 2 | 12187 | 179 | Msr1,Thbs1 | | 2.525e-02 | -3.68 | BIOCARTA\_HCMV\_PATHWAY | MSigDB lists | BIOCARTA\_HCMV\_PATHWAY | 17 | 2 | 12187 | 179 | Nfkb1,Pik3r1 | | 2.525e-02 | -3.68 | SUH\_COEXPRESSED\_WITH\_ID1\_AND\_ID2\_UP | MSigDB lists | SUH\_COEXPRESSED\_WITH\_ID1\_AND\_ID2\_UP | 17 | 2 | 12187 | 179 | Gadd45b,Dnajb1 | | 2.525e-02 | -3.68 | GO\_POSITIVE\_REGULATION\_OF\_PROTEIN\_COMPLEX\_DISASSEMBLY | MSigDB lists | GO\_POSITIVE\_REGULATION\_OF\_PROTEIN\_COMPLEX\_DISASSEMBLY | 17 | 2 | 12187 | 179 | Tnf,Plek | | 2.525e-02 | -3.68 | GO\_CYSTEINE\_TYPE\_ENDOPEPTIDASE\_INHIBITOR\_ACTIVITY\_INVOLVED\_IN\_APOPTOTIC\_PROCESS | MSigDB lists | GO\_CYSTEINE\_TYPE\_ENDOPEPTIDASE\_INHIBITOR\_ACTIVITY\_INVOLVED\_IN\_APOPTOTIC\_PROCESS | 17 | 2 | 12187 | 179 | Tnfaip8,Birc3 | | 2.525e-02 | -3.68 | GO\_REGULATION\_OF\_MEMBRANE\_PROTEIN\_ECTODOMAIN\_PROTEOLYSIS | MSigDB lists | GO\_REGULATION\_OF\_MEMBRANE\_PROTEIN\_ECTODOMAIN\_PROTEOLYSIS | 17 | 2 | 12187 | 179 | Timp1,Tnf | | 2.525e-02 | -3.68 | TAKEDA\_TARGETS\_OF\_NUP98\_HOXA9\_FUSION\_3D\_DN | MSigDB lists | TAKEDA\_TARGETS\_OF\_NUP98\_HOXA9\_FUSION\_3D\_DN | 17 | 2 | 12187 | 179 | S100a8,S100a9 | | 2.525e-02 | -3.68 | GO\_REGULATION\_OF\_MACROPHAGE\_DIFFERENTIATION | MSigDB lists | GO\_REGULATION\_OF\_MACROPHAGE\_DIFFERENTIATION | 17 | 2 | 12187 | 179 | Trib1,Ripk1 | | 2.525e-02 | -3.68 | KANG\_DOXORUBICIN\_RESISTANCE\_DN | MSigDB lists | KANG\_DOXORUBICIN\_RESISTANCE\_DN | 17 | 2 | 12187 | 179 | Clic4,Cebpd | | 2.525e-02 | -3.68 | WANG\_IMMORTALIZED\_BY\_HOXA9\_AND\_MEIS1\_DN | MSigDB lists | WANG\_IMMORTALIZED\_BY\_HOXA9\_AND\_MEIS1\_DN | 17 | 2 | 12187 | 179 | Gem,Zbtb16 | | 2.525e-02 | -3.68 | GREENBAUM\_E2A\_TARGETS\_DN | MSigDB lists | GREENBAUM\_E2A\_TARGETS\_DN | 17 | 2 | 12187 | 179 | Slfn4,Slfn3 | | 2.525e-02 | -3.68 | GO\_REGULATION\_OF\_T\_HELPER\_CELL\_DIFFERENTIATION | MSigDB lists | GO\_REGULATION\_OF\_T\_HELPER\_CELL\_DIFFERENTIATION | 17 | 2 | 12187 | 179 | Il4ra,Il6 | | 2.525e-02 | -3.68 | GO\_CELLULAR\_RESPONSE\_TO\_FLUID\_SHEAR\_STRESS | MSigDB lists | GO\_CELLULAR\_RESPONSE\_TO\_FLUID\_SHEAR\_STRESS | 17 | 2 | 12187 | 179 | Ptgs2,Nfe2l2 | | 2.525e-02 | -3.68 | GO\_MAP\_KINASE\_KINASE\_KINASE\_ACTIVITY | MSigDB lists | GO\_MAP\_KINASE\_KINASE\_KINASE\_ACTIVITY | 17 | 2 | 12187 | 179 | Map3k8,Map3k6 | | 2.525e-02 | -3.68 | GO\_ASTROCYTE\_DEVELOPMENT | MSigDB lists | GO\_ASTROCYTE\_DEVELOPMENT | 17 | 2 | 12187 | 179 | S100a8,S100a9 | | 2.525e-02 | -3.68 | GO\_TUMOR\_NECROSIS\_FACTOR\_RECEPTOR\_BINDING | MSigDB lists | GO\_TUMOR\_NECROSIS\_FACTOR\_RECEPTOR\_BINDING | 17 | 2 | 12187 | 179 | Tnfsf10,Tnf | | 2.525e-02 | -3.68 | AMIT\_EGF\_RESPONSE\_40\_MCF10A | MSigDB lists | AMIT\_EGF\_RESPONSE\_40\_MCF10A | 17 | 2 | 12187 | 179 | Thbs1,Zfp36 | | 2.525e-02 | -3.68 | GO\_NEGATIVE\_REGULATION\_OF\_LIPID\_CATABOLIC\_PROCESS | MSigDB lists | GO\_NEGATIVE\_REGULATION\_OF\_LIPID\_CATABOLIC\_PROCESS | 17 | 2 | 12187 | 179 | Hcar2,Tnf | | 2.535e-02 | -3.68 | GO\_REGULATION\_OF\_PEPTIDYL\_TYROSINE\_PHOSPHORYLATION | MSigDB lists | GO\_REGULATION\_OF\_PEPTIDYL\_TYROSINE\_PHOSPHORYLATION | 153 | 6 | 12187 | 179 | Socs3,Il6,Csf3,Icam1,Errfi1,Ptpn1 | | 2.535e-02 | -3.68 | GSE17721\_LPS\_VS\_PAM3CSK4\_0.5H\_BMDC\_UP | MSigDB lists | GSE17721\_LPS\_VS\_PAM3CSK4\_0.5H\_BMDC\_UP | 153 | 6 | 12187 | 179 | Robo4,Banp,Il17ra,Ccnd2,Tgm2,Cybb | | 2.535e-02 | -3.68 | GSE4984\_UNTREATED\_VS\_VEHICLE\_CTRL\_TREATED\_DC\_DN | MSigDB lists | GSE4984\_UNTREATED\_VS\_VEHICLE\_CTRL\_TREATED\_DC\_DN | 153 | 6 | 12187 | 179 | Gbp6,Gem,Xdh,Icam1,Trib1,Gbp4 | | 2.535e-02 | -3.68 | GSE32423\_MEMORY\_VS\_NAIVE\_CD8\_TCELL\_IL7\_UP | MSigDB lists | GSE32423\_MEMORY\_VS\_NAIVE\_CD8\_TCELL\_IL7\_UP | 153 | 6 | 12187 | 179 | Parp14,Fstl1,Fzd4,Cmpk2,Xdh,Sbno2 | | 2.535e-02 | -3.68 | GSE17301\_ACD3\_ACD28\_VS\_ACD3\_ACD28\_AND\_IFNA5\_STIM\_CD8\_TCELL\_DN | MSigDB lists | GSE17301\_ACD3\_ACD28\_VS\_ACD3\_ACD28\_AND\_IFNA5\_STIM\_CD8\_TCELL\_DN | 153 | 6 | 12187 | 179 | Irf7,Csf2rb,Usp18,Casp4,Cybb,Ms4a6d | | 2.535e-02 | -3.68 | GSE11961\_GERMINAL\_CENTER\_BCELL\_DAY7\_VS\_GERMINAL\_CENTER\_BCELL\_DAY40\_UP | MSigDB lists | GSE11961\_GERMINAL\_CENTER\_BCELL\_DAY7\_VS\_GERMINAL\_CENTER\_BCELL\_DAY40\_UP | 153 | 6 | 12187 | 179 | Tagln2,Errfi1,Nfkbiz,Maff,Tnfaip3,Tubb6 | | 2.535e-02 | -3.68 | GSE18893\_CTRL\_VS\_TNF\_TREATED\_TREG\_24H\_UP | MSigDB lists | GSE18893\_CTRL\_VS\_TNF\_TREATED\_TREG\_24H\_UP | 153 | 6 | 12187 | 179 | Parp12,Osmr,Slfn3,Ip6k2,Samd9l,Slfn4 | | 2.537e-02 | -3.67 | positive regulation of nucleobase-containing compound metabolic process | biological process | GO:0045935 | 1495 | 33 | 13711 | 214 | Tnf,Atf3,Zfp36,Ncf1,Aff1,C3ar1,Arid5b,Ripk1,Csf3,Zbtb16,Tnip1,Maff,Irf2,Nfe2l2,Bcl3,Sox11,Il1a,Nfkb2,Nfkbia,Il6,Irf7,Dtx3l,Rbm39,Banp,Pik3r1,Fzd4,Nfkb1,Sbno2,Parp9,Cebpd,Bach1,Spi1,Runx1 | | 2.538e-02 | -3.67 | GO\_UROGENITAL\_SYSTEM\_DEVELOPMENT | MSigDB lists | GO\_UROGENITAL\_SYSTEM\_DEVELOPMENT | 240 | 8 | 12187 | 179 | Col4a1,Tiparp,Adamts1,Sox11,Sdc4,Zbtb16,Iqgap1,Arid5b | | 2.539e-02 | -3.67 | inorganic ion homeostasis | biological process | GO:0098771 | 549 | 15 | 13711 | 214 | Mt1,Ccl2,C3ar1,Pygm,Slc24a4,Lcn2,Cp,P2ry6,Clic4,Tgm2,Cxcl1,C5ar1,Mt2,Kcna5,Sgk1 | | 2.545e-02 | -3.67 | GO\_POSITIVE\_REGULATION\_OF\_PEPTIDYL\_TYROSINE\_PHOSPHORYLATION | MSigDB lists | GO\_POSITIVE\_REGULATION\_OF\_PEPTIDYL\_TYROSINE\_PHOSPHORYLATION | 113 | 5 | 12187 | 179 | Socs3,Csf3,Il6,Ptpn1,Icam1 | | 2.545e-02 | -3.67 | KEGG\_INSULIN\_SIGNALING\_PATHWAY | MSigDB lists | KEGG\_INSULIN\_SIGNALING\_PATHWAY | 113 | 5 | 12187 | 179 | Pik3r1,Socs3,Pik3r5,Pygm,Ptpn1 | | 2.547e-02 | -3.67 | SIRNA\_EIF4GI\_UP | MSigDB lists | SIRNA\_EIF4GI\_UP | 76 | 4 | 12187 | 179 | Cebpd,Atf3,Hspb1,Saa1 | | 2.547e-02 | -3.67 | GSE32901\_TH1\_VS\_TH17\_NEG\_CD4\_TCELL\_DN | MSigDB lists | GSE32901\_TH1\_VS\_TH17\_NEG\_CD4\_TCELL\_DN | 76 | 4 | 12187 | 179 | Tnfaip8,Rhou,Casp4,Ccl4 | | 2.550e-02 | -3.67 | leukocyte differentiation | biological process | GO:0002521 | 269 | 9 | 13711 | 214 | Tnf,Tnfaip3,Sbno2,Il6,Spi1,Csf3,Pik3r1,Gadd45g,Bcl3 | | 2.569e-02 | -3.66 | Signaling by the B Cell Receptor (BCR) | REACTOME pathways | R-MMU-983705 | 174 | 7 | 6297 | 105 | Nfkbia,Pik3ap1,Pik3r1,Cdkn1a,Ier3,Nfkbie,Nfkb1 | | 2.586e-02 | -3.66 | Sin3a (transcriptional regulator, SIN3A (yeast)) | protein interactions | 20466 | 38 | 3 | 6802 | 115 | Runx1,Spi1,Mxd4 | | 2.587e-02 | -3.65 | regulation of epidermal cell differentiation | biological process | GO:0045604 | 41 | 3 | 13711 | 214 | Zfp36,Errfi1,Maff | | 2.588e-02 | -3.65 | potassium channel regulator activity | molecular function | GO:0015459 | 41 | 3 | 13516 | 211 | Sgk1,Sgk3,Kcna5 | | 2.603e-02 | -3.65 | negative regulation of protein ubiquitination | biological process | GO:0031397 | 72 | 4 | 13711 | 214 | Isg15,Gbp4,Dtx3l,Tnfaip3 | | 2.603e-02 | -3.65 | regulation of cellular response to oxidative stress | biological process | GO:1900407 | 72 | 4 | 13711 | 214 | Ripk1,Nfe2l2,Tnf,Hspb1 | | 2.604e-02 | -3.65 | response to nutrient levels | biological process | GO:0031667 | 270 | 9 | 13711 | 214 | Angptl4,Fstl1,Nfe2l2,Zfp36,Atf3,Cdkn1a,Rrp8,Fam107a,Hspa5 | | 2.604e-02 | -3.65 | GO\_REGULATION\_OF\_CELLULAR\_LOCALIZATION | MSigDB lists | GO\_REGULATION\_OF\_CELLULAR\_LOCALIZATION | 966 | 22 | 12187 | 179 | Cd14,Il17ra,Ptpn1,Sox11,Il1a,Csf3,Nfkbie,Bcl3,Rhou,Pik3r1,Il4ra,Hcar2,Ccl12,Il6,Ptgs2,Saa1,Msn,Cdkn1a,Kcna5,Tnf,Sdc4,Nfkbia | | 2.607e-02 | -3.65 | GSE33292\_WT\_VS\_TCF1\_KO\_DN3\_THYMOCYTE\_UP | MSigDB lists | GSE33292\_WT\_VS\_TCF1\_KO\_DN3\_THYMOCYTE\_UP | 154 | 6 | 12187 | 179 | Gadd45g,Zfp36,Cebpd,Rgs16,Tnfaip3,Cxcl1 | | 2.607e-02 | -3.65 | GSE17974\_CTRL\_VS\_ACT\_IL4\_AND\_ANTI\_IL12\_6H\_CD4\_TCELL\_DN | MSigDB lists | GSE17974\_CTRL\_VS\_ACT\_IL4\_AND\_ANTI\_IL12\_6H\_CD4\_TCELL\_DN | 154 | 6 | 12187 | 179 | Il1rn,Tnf,Ifit3b,Sdc4,Parp14,Rtp4 | | 2.607e-02 | -3.65 | GSE3039\_ALPHAALPHA\_CD8\_TCELL\_VS\_B2\_BCELL\_UP | MSigDB lists | GSE3039\_ALPHAALPHA\_CD8\_TCELL\_VS\_B2\_BCELL\_UP | 154 | 6 | 12187 | 179 | Msn,Ptpn1,Slfn3,Iqgap1,Slfn4,Hp | | 2.607e-02 | -3.65 | GSE26156\_DOUBLE\_POSITIVE\_VS\_CD4\_SINGLE\_POSITIVE\_THYMOCYTE\_UP | MSigDB lists | GSE26156\_DOUBLE\_POSITIVE\_VS\_CD4\_SINGLE\_POSITIVE\_THYMOCYTE\_UP | 154 | 6 | 12187 | 179 | Gadd45b,Tiparp,Rbm39,Casp4,Sgk1,Ifi44 | | 2.607e-02 | -3.65 | GSE28783\_CTRL\_ANTI\_MIR\_VS\_UNTREATED\_ATHEROSCLEROSIS\_MACROPHAGE\_DN | MSigDB lists | GSE28783\_CTRL\_ANTI\_MIR\_VS\_UNTREATED\_ATHEROSCLEROSIS\_MACROPHAGE\_DN | 154 | 6 | 12187 | 179 | Cxcl1,Isg15,Cdkn1a,Tnf,Gadd45b,Rgs16 | | 2.607e-02 | -3.65 | SASAKI\_ADULT\_T\_CELL\_LEUKEMIA | MSigDB lists | SASAKI\_ADULT\_T\_CELL\_LEUKEMIA | 154 | 6 | 12187 | 179 | Ifi211,Trib1,Ifi209,Isg15,Ifi204,Ifi207 | | 2.609e-02 | -3.65 | negative regulation of protein serine/threonine kinase activity | biological process | GO:0071901 | 107 | 5 | 13711 | 214 | Cdkn1a,Ptpn1,Irak3,Tnfaip3,Rasip1 | | 2.611e-02 | -3.65 | growth factor receptor binding | molecular function | GO:0070851 | 107 | 5 | 13516 | 211 | Il6,Csf3,Il1rn,Il1a,Pik3r1 | | 2.622e-02 | -3.64 | regulation of cellular response to growth factor stimulus | biological process | GO:0090287 | 227 | 8 | 13711 | 214 | Ptpn1,Hspa5,Sox11,Fzd4,Xdh,Thbs1,Cflar,Fstl1 | | 2.631e-02 | -3.64 | PICCALUGA\_ANGIOIMMUNOBLASTIC\_LYMPHOMA\_DN | MSigDB lists | PICCALUGA\_ANGIOIMMUNOBLASTIC\_LYMPHOMA\_DN | 114 | 5 | 12187 | 179 | Zfp36,Tiparp,Gadd45b,Tnfaip3,Map3k8 | | 2.631e-02 | -3.64 | GSE6269\_E\_COLI\_VS\_STREP\_PNEUMO\_INF\_PBMC\_DN | MSigDB lists | GSE6269\_E\_COLI\_VS\_STREP\_PNEUMO\_INF\_PBMC\_DN | 114 | 5 | 12187 | 179 | Mmp8,Hp,Lcn2,Cxcl1,S100a9 | | 2.631e-02 | -3.64 | KIM\_WT1\_TARGETS\_DN | MSigDB lists | KIM\_WT1\_TARGETS\_DN | 384 | 11 | 12187 | 179 | Ifi204,Ifi207,Ifi209,Hspa5,Dnajb1,Trim25,Irf2,Gadd45b,Sntb2,Tnfaip8,Ifi211 | | 2.656e-02 | -3.63 | GO\_EXTRINSIC\_APOPTOTIC\_SIGNALING\_PATHWAY | MSigDB lists | GO\_EXTRINSIC\_APOPTOTIC\_SIGNALING\_PATHWAY | 77 | 4 | 12187 | 179 | Tnf,Pik3r1,Ripk1,Il1a | | 2.656e-02 | -3.63 | GO\_GLAND\_MORPHOGENESIS | MSigDB lists | GO\_GLAND\_MORPHOGENESIS | 77 | 4 | 12187 | 179 | Tnf,Tgm2,Msn,Il6 | | 2.657e-02 | -3.63 | ROY\_WOUND\_BLOOD\_VESSEL\_UP | MSigDB lists | ROY\_WOUND\_BLOOD\_VESSEL\_UP | 44 | 3 | 12187 | 179 | Timp1,Col4a1,Xaf1 | | 2.657e-02 | -3.63 | LEE\_EARLY\_T\_LYMPHOCYTE\_DN | MSigDB lists | LEE\_EARLY\_T\_LYMPHOCYTE\_DN | 44 | 3 | 12187 | 179 | Il4ra,Ifitm2,Tnfsf10 | | 2.657e-02 | -3.63 | PID\_PTP1B\_PATHWAY | MSigDB lists | PID\_PTP1B\_PATHWAY | 44 | 3 | 12187 | 179 | Socs3,Pik3r1,Ptpn1 | | 2.657e-02 | -3.63 | GO\_RNA\_POLYMERASE\_II\_CORE\_PROMOTER\_SEQUENCE\_SPECIFIC\_DNA\_BINDING | MSigDB lists | GO\_RNA\_POLYMERASE\_II\_CORE\_PROMOTER\_SEQUENCE\_SPECIFIC\_DNA\_BINDING | 44 | 3 | 12187 | 179 | Spi1,Irf7,Sox11 | | 2.657e-02 | -3.63 | PID\_INSULIN\_PATHWAY | MSigDB lists | PID\_INSULIN\_PATHWAY | 44 | 3 | 12187 | 179 | Sgk1,Pik3r1,Ptpn1 | | 2.657e-02 | -3.63 | GO\_REGULATION\_OF\_CELLULAR\_RESPONSE\_TO\_INSULIN\_STIMULUS | MSigDB lists | GO\_REGULATION\_OF\_CELLULAR\_RESPONSE\_TO\_INSULIN\_STIMULUS | 44 | 3 | 12187 | 179 | Socs3,Pik3r1,Ptpn1 | | 2.680e-02 | -3.62 | GSE26030\_UNSTIM\_VS\_RESTIM\_TH17\_DAY5\_POST\_POLARIZATION\_UP | MSigDB lists | GSE26030\_UNSTIM\_VS\_RESTIM\_TH17\_DAY5\_POST\_POLARIZATION\_UP | 155 | 6 | 12187 | 179 | Parp14,Nfkb1,Cmpk2,Znfx1,Xdh,Parp12 | | 2.680e-02 | -3.62 | GSE17721\_LPS\_VS\_CPG\_24H\_BMDC\_UP | MSigDB lists | GSE17721\_LPS\_VS\_CPG\_24H\_BMDC\_UP | 155 | 6 | 12187 | 179 | Ifit2,Gbp6,Cxcl16,Gbp4,Timp1,Il1a | | 2.680e-02 | -3.62 | GSE37416\_CTRL\_VS\_12H\_F\_TULARENSIS\_LVS\_NEUTROPHIL\_DN | MSigDB lists | GSE37416\_CTRL\_VS\_12H\_F\_TULARENSIS\_LVS\_NEUTROPHIL\_DN | 155 | 6 | 12187 | 179 | Tnip1,Nfkbie,Tnfaip8,Icam1,Nfkbiz,Tnfaip3 | | 2.680e-02 | -3.62 | GSE11961\_GERMINAL\_CENTER\_BCELL\_DAY7\_VS\_MEMORY\_BCELL\_DAY40\_UP | MSigDB lists | GSE11961\_GERMINAL\_CENTER\_BCELL\_DAY7\_VS\_MEMORY\_BCELL\_DAY40\_UP | 155 | 6 | 12187 | 179 | Errfi1,Tnip1,Gbp6,Herc6,Gbp4,Ccl4 | | 2.680e-02 | -3.62 | GSE24972\_WT\_VS\_IRF8\_KO\_MARGINAL\_ZONE\_SPLEEN\_BCELL\_UP | MSigDB lists | GSE24972\_WT\_VS\_IRF8\_KO\_MARGINAL\_ZONE\_SPLEEN\_BCELL\_UP | 155 | 6 | 12187 | 179 | Hspb1,Il1rn,Rbm47,Parp12,Adamts1,Map3k6 | | 2.680e-02 | -3.62 | GSE29618\_PDC\_VS\_MDC\_DAY7\_FLU\_VACCINE\_UP | MSigDB lists | GSE29618\_PDC\_VS\_MDC\_DAY7\_FLU\_VACCINE\_UP | 155 | 6 | 12187 | 179 | Kcna5,Cybb,Ifit2,Plek,Csf2rb,Irf7 | | 2.680e-02 | -3.62 | GSE22025\_UNTREATED\_VS\_TGFB1\_TREATED\_CD4\_TCELL\_UP | MSigDB lists | GSE22025\_UNTREATED\_VS\_TGFB1\_TREATED\_CD4\_TCELL\_UP | 155 | 6 | 12187 | 179 | Map3k8,Col4a1,Gadd45g,Atf3,Nfkbie,Rgs16 | | 2.687e-02 | -3.62 | G\_DYNAMIN\_2 | prosite domains | PS51718 | 14 | 2 | 8845 | 164 | Mx2,Mx1 | | 2.702e-02 | -3.61 | negative regulation of mitotic cell cycle phase transition | biological process | GO:1901991 | 108 | 5 | 13711 | 214 | Cdkn1a,Ier3,Ccl12,Ripk1,Fam107a | | 2.702e-02 | -3.61 | regulation of protein localization to nucleus | biological process | GO:1900180 | 108 | 5 | 13711 | 214 | Gbp4,Dtx3l,Pik3r1,Ptgs2,Parp9 | | 2.707e-02 | -3.61 | biological adhesion | biological process | GO:0022610 | 653 | 17 | 13711 | 214 | Gbp3,Fzd4,S100a8,Vcam1,S100a9,Gbp7,Selp,Msn,Gbp6,Tnf,Icam1,Plek,Sele,Thbs1,Adamts9,Tnip1,Lgals9 | | 2.707e-02 | -3.61 | FISCHER\_DIRECT\_P53\_TARGETS\_META\_ANALYSIS | MSigDB lists | FISCHER\_DIRECT\_P53\_TARGETS\_META\_ANALYSIS | 243 | 8 | 12187 | 179 | Cdkn1a,Trib1,Tubb6,Atf3,Ier3,Cflar,Tnfaip8,Rgs16 | | 2.719e-02 | -3.60 | ESC\_J1\_UP\_EARLY.V1\_UP | MSigDB lists | ESC\_J1\_UP\_EARLY.V1\_UP | 115 | 5 | 12187 | 179 | Pik3r1,Ncf1,Zbtb16,Gadd45g,Gadd45b | | 2.721e-02 | -3.60 | regulation of type I interferon production | biological process | GO:0032479 | 73 | 4 | 13711 | 214 | Gbp4,Irf7,Acod1,Cd14 | | 2.729e-02 | -3.60 | MARTINEZ\_TP53\_TARGETS\_DN | MSigDB lists | MARTINEZ\_TP53\_TARGETS\_DN | 436 | 12 | 12187 | 179 | Plin4,Xdh,Errfi1,Rtp4,Zfp36,Usp18,Sntb2,Ccnd2,Cdkn1a,Rhou,Bach1,Timp1 | | 2.735e-02 | -3.60 | system development | biological process | GO:0048731 | 3292 | 64 | 13711 | 214 | Prg4,Hp,Hspa5,Clic4,Tgm2,Maff,C5ar1,Angptl4,Msn,Nfe2l2,Atf3,Robo4,Acod1,Tnf,Rnf213,Cflar,Hspb1,Slc24a4,Sdc4,Sbno2,Fzd4,Tiparp,Sgk1,Ptgs2,Runx1,Herc6,Spi1,Rbm47,Cebpd,Rasip1,Cdkn1a,Il6,Il1a,Pik3r1,Socs3,Ccl12,S100a8,Iqgap1,Col4a1,Samd9l,Bcl3,S100a9,Vcam1,Ccl2,Xdh,Tnfaip3,Ifi204,Adamts9,Zbtb16,Csf3,C3ar1,Arid5b,Adamts1,Apold1,Nfkb1,Rhoj,Plaur,Cemip2,Errfi1,Sox11,Nfkb2,Icam1,Gadd45g,Thbs1 | | 2.736e-02 | -3.60 | CCR4-NOT complex | cellular component | GO:0030014 | 17 | 2 | 13825 | 212 | Noct,Zfp36 | | 2.750e-02 | -3.59 | IL1/FGF | interpro domains | IPR008996 | 17 | 2 | 13788 | 212 | Il1a,Il1rn | | 2.750e-02 | -3.59 | response to antibiotic | biological process | GO:0046677 | 187 | 7 | 13711 | 214 | Tnfaip3,Nfe2l2,Ccl7,Ripk1,Il6,Hp,Lcn2 | | 2.754e-02 | -3.59 | GSE41978\_KLRG1\_HIGH\_VS\_LOW\_EFFECTOR\_CD8\_TCELL\_UP | MSigDB lists | GSE41978\_KLRG1\_HIGH\_VS\_LOW\_EFFECTOR\_CD8\_TCELL\_UP | 156 | 6 | 12187 | 179 | Tgm2,Fzd4,Adamts1,Cdkn1a,Il4ra,Ksr1 | | 2.754e-02 | -3.59 | GSE3982\_EFF\_MEMORY\_CD4\_TCELL\_VS\_NKCELL\_DN | MSigDB lists | GSE3982\_EFF\_MEMORY\_CD4\_TCELL\_VS\_NKCELL\_DN | 156 | 6 | 12187 | 179 | Cebpd,Irf9,Rtp4,Socs3,Irf7,C5ar1 | | 2.754e-02 | -3.59 | GSE21670\_TGFB\_VS\_TGFB\_AND\_IL6\_TREATED\_STAT3\_KO\_CD4\_TCELL\_UP | MSigDB lists | GSE21670\_TGFB\_VS\_TGFB\_AND\_IL6\_TREATED\_STAT3\_KO\_CD4\_TCELL\_UP | 156 | 6 | 12187 | 179 | Dnajb1,Ncf1,Tagln2,Pik3r5,Sgk3,Gbp7 | | 2.754e-02 | -3.59 | GSE5589\_UNSTIM\_VS\_180MIN\_LPS\_STIM\_MACROPHAGE\_DN | MSigDB lists | GSE5589\_UNSTIM\_VS\_180MIN\_LPS\_STIM\_MACROPHAGE\_DN | 156 | 6 | 12187 | 179 | Il1rn,Stx11,Bach1,Gbp3,Ccnd2,Gpr84 | | 2.754e-02 | -3.59 | GSE22886\_TCELL\_VS\_BCELL\_NAIVE\_UP | MSigDB lists | GSE22886\_TCELL\_VS\_BCELL\_NAIVE\_UP | 156 | 6 | 12187 | 179 | Il17ra,Ifitm3,Timp1,Ccnd2,Runx1,Dnajb1 | | 2.754e-02 | -3.59 | GSE360\_DC\_VS\_MAC\_T\_GONDII\_DN | MSigDB lists | GSE360\_DC\_VS\_MAC\_T\_GONDII\_DN | 156 | 6 | 12187 | 179 | C3ar1,Hspa5,Gem,Cd14,S100a8,Cebpd | | 2.754e-02 | -3.59 | GSE36888\_STAT5\_AB\_KNOCKIN\_VS\_WT\_TCELL\_IL2\_TREATED\_6H\_UP | MSigDB lists | GSE36888\_STAT5\_AB\_KNOCKIN\_VS\_WT\_TCELL\_IL2\_TREATED\_6H\_UP | 156 | 6 | 12187 | 179 | Irf7,Slfn9,Isg15,Ifit2,Ccdc58,Pik3ap1 | | 2.754e-02 | -3.59 | GSE15930\_STIM\_VS\_STIM\_AND\_IFNAB\_24H\_CD8\_T\_CELL\_UP | MSigDB lists | GSE15930\_STIM\_VS\_STIM\_AND\_IFNAB\_24H\_CD8\_T\_CELL\_UP | 156 | 6 | 12187 | 179 | Saa1,Dnajb1,Adamts1,Luc7l3,Cybb,Il1rn | | 2.754e-02 | -3.59 | GSE7460\_WT\_VS\_FOXP3\_HET\_ACT\_WITH\_TGFB\_TCONV\_UP | MSigDB lists | GSE7460\_WT\_VS\_FOXP3\_HET\_ACT\_WITH\_TGFB\_TCONV\_UP | 156 | 6 | 12187 | 179 | Ifitm2,Ptgs2,Trim56,Zfp36,Il17ra,Thbs1 | | 2.754e-02 | -3.59 | GSE27786\_NEUTROPHIL\_VS\_MONO\_MAC\_UP | MSigDB lists | GSE27786\_NEUTROPHIL\_VS\_MONO\_MAC\_UP | 156 | 6 | 12187 | 179 | Ncf1,Cmpk2,Lgals9,Casp4,Spi1,Nfe2l2 | | 2.754e-02 | -3.59 | GSE8835\_HEALTHY\_VS\_CLL\_CD4\_TCELL\_UP | MSigDB lists | GSE8835\_HEALTHY\_VS\_CLL\_CD4\_TCELL\_UP | 156 | 6 | 12187 | 179 | Ccnd2,Ifit2,Ifit3b,Gpr84,Cmpk2,Oasl1 | | 2.755e-02 | -3.59 | phosphatidylinositol phosphorylation | biological process | GO:0046854 | 42 | 3 | 13711 | 214 | Ip6k2,Pik3r1,Socs3 | | 2.755e-02 | -3.59 | alpha-beta T cell differentiation | biological process | GO:0046632 | 42 | 3 | 13711 | 214 | Bcl3,Gadd45g,Il6 | | 2.757e-02 | -3.59 | AP1\_Q6\_01 | MSigDB lists | AP1\_Q6\_01 | 199 | 7 | 12187 | 179 | Tagln2,Sgk1,Znfx1,Il6,Il1rn,Rbm39,Gadd45g | | 2.769e-02 | -3.59 | GO\_HEMATOPOIETIC\_PROGENITOR\_CELL\_DIFFERENTIATION | MSigDB lists | GO\_HEMATOPOIETIC\_PROGENITOR\_CELL\_DIFFERENTIATION | 78 | 4 | 12187 | 179 | Spi1,Herc6,Plek,Rbm47 | | 2.769e-02 | -3.59 | KRAS.LUNG\_UP.V1\_UP | MSigDB lists | KRAS.LUNG\_UP.V1\_UP | 78 | 4 | 12187 | 179 | Kcna5,C3ar1,Csf3,Angptl4 | | 2.795e-02 | -3.58 | cellular response to nitrogen compound | biological process | GO:1901699 | 411 | 12 | 13711 | 214 | Pik3r1,Sgk1,P2ry6,Vcam1,Thbs1,Cflar,Nfkb1,Col4a1,Casp4,Icam1,Tnf,Ptpn1 | | 2.806e-02 | -3.57 | activation of protein kinase activity | biological process | GO:0032147 | 230 | 8 | 13711 | 214 | Map3k8,Tnf,Map3k6,Ptpn1,Gadd45b,Gadd45g,Thbs1,Ripk1 | | 2.807e-02 | -3.57 | GO\_SIGNAL\_TRANSDUCTION\_BY\_PROTEIN\_PHOSPHORYLATION | MSigDB lists | GO\_SIGNAL\_TRANSDUCTION\_BY\_PROTEIN\_PHOSPHORYLATION | 339 | 10 | 12187 | 179 | Trib1,Map3k8,Tnf,Ccl12,Ripk1,Nfkb1,Ksr1,Map3k6,Csf2rb,Ptpn1 | | 2.813e-02 | -3.57 | REACTOME\_NEPHRIN\_INTERACTIONS | MSigDB lists | REACTOME\_NEPHRIN\_INTERACTIONS | 18 | 2 | 12187 | 179 | Pik3r1,Iqgap1 | | 2.813e-02 | -3.57 | GO\_NEGATIVE\_REGULATION\_OF\_PRODUCTION\_OF\_MOLECULAR\_MEDIATOR\_OF\_IMMUNE\_RESPONSE | MSigDB lists | GO\_NEGATIVE\_REGULATION\_OF\_PRODUCTION\_OF\_MOLECULAR\_MEDIATOR\_OF\_IMMUNE\_RESPONSE | 18 | 2 | 12187 | 179 | Irak3,Tnf | | 2.813e-02 | -3.57 | YIH\_RESPONSE\_TO\_ARSENITE\_C1 | MSigDB lists | YIH\_RESPONSE\_TO\_ARSENITE\_C1 | 18 | 2 | 12187 | 179 | Cflar,Zfp36 | | 2.813e-02 | -3.57 | PID\_IL27\_PATHWAY | MSigDB lists | PID\_IL27\_PATHWAY | 18 | 2 | 12187 | 179 | Il6,Tnf | | 2.813e-02 | -3.57 | MARCHINI\_TRABECTEDIN\_RESISTANCE\_UP | MSigDB lists | MARCHINI\_TRABECTEDIN\_RESISTANCE\_UP | 18 | 2 | 12187 | 179 | Dnajb1,Akap12 | | 2.813e-02 | -3.57 | GO\_NEGATIVE\_REGULATION\_OF\_VIRAL\_TRANSCRIPTION | MSigDB lists | GO\_NEGATIVE\_REGULATION\_OF\_VIRAL\_TRANSCRIPTION | 18 | 2 | 12187 | 179 | Ccl4,Ifitm3 | | 2.813e-02 | -3.57 | WANG\_BARRETTS\_ESOPHAGUS\_DN | MSigDB lists | WANG\_BARRETTS\_ESOPHAGUS\_DN | 18 | 2 | 12187 | 179 | Nfe2l2,S100a8 | | 2.813e-02 | -3.57 | GO\_CELLULAR\_RESPONSE\_TO\_PROSTAGLANDIN\_STIMULUS | MSigDB lists | GO\_CELLULAR\_RESPONSE\_TO\_PROSTAGLANDIN\_STIMULUS | 18 | 2 | 12187 | 179 | Adamts1,P2ry6 | | 2.813e-02 | -3.57 | DELACROIX\_RAR\_TARGETS\_DN | MSigDB lists | DELACROIX\_RAR\_TARGETS\_DN | 18 | 2 | 12187 | 179 | Fstl1,Sntb2 | | 2.813e-02 | -3.57 | NUMATA\_CSF3\_SIGNALING\_VIA\_STAT3 | MSigDB lists | NUMATA\_CSF3\_SIGNALING\_VIA\_STAT3 | 18 | 2 | 12187 | 179 | Runx1,Il17ra | | 2.813e-02 | -3.57 | GO\_CYTOPLASMIC\_SEQUESTERING\_OF\_TRANSCRIPTION\_FACTOR | MSigDB lists | GO\_CYTOPLASMIC\_SEQUESTERING\_OF\_TRANSCRIPTION\_FACTOR | 18 | 2 | 12187 | 179 | Nfkbia,Nfkbie | | 2.813e-02 | -3.57 | MANN\_RESPONSE\_TO\_AMIFOSTINE\_UP | MSigDB lists | MANN\_RESPONSE\_TO\_AMIFOSTINE\_UP | 18 | 2 | 12187 | 179 | Cdkn1a,Casp4 | | 2.813e-02 | -3.57 | GO\_REGULATION\_OF\_RESPONSE\_TO\_INTERFERON\_GAMMA | MSigDB lists | GO\_REGULATION\_OF\_RESPONSE\_TO\_INTERFERON\_GAMMA | 18 | 2 | 12187 | 179 | Socs3,Parp9 | | 2.813e-02 | -3.57 | JEON\_SMAD6\_TARGETS\_UP | MSigDB lists | JEON\_SMAD6\_TARGETS\_UP | 18 | 2 | 12187 | 179 | Trib1,Thbs1 | | 2.813e-02 | -3.57 | GO\_RESPONSE\_TO\_INCREASED\_OXYGEN\_LEVELS | MSigDB lists | GO\_RESPONSE\_TO\_INCREASED\_OXYGEN\_LEVELS | 18 | 2 | 12187 | 179 | Cdkn1a,Kcna5 | | 2.813e-02 | -3.57 | GO\_PHOSPHATIDYLINOSITOL\_3\_KINASE\_COMPLEX | MSigDB lists | GO\_PHOSPHATIDYLINOSITOL\_3\_KINASE\_COMPLEX | 18 | 2 | 12187 | 179 | Pik3r5,Pik3r1 | | 2.813e-02 | -3.57 | GO\_T\_CELL\_DIFFERENTIATION\_INVOLVED\_IN\_IMMUNE\_RESPONSE | MSigDB lists | GO\_T\_CELL\_DIFFERENTIATION\_INVOLVED\_IN\_IMMUNE\_RESPONSE | 18 | 2 | 12187 | 179 | Bcl3,Il6 | | 2.817e-02 | -3.57 | SESTO\_RESPONSE\_TO\_UV\_C5 | MSigDB lists | SESTO\_RESPONSE\_TO\_UV\_C5 | 45 | 3 | 12187 | 179 | Nfe2l2,Iqgap1,Sdc4 | | 2.817e-02 | -3.57 | GO\_VASCULOGENESIS | MSigDB lists | GO\_VASCULOGENESIS | 45 | 3 | 12187 | 179 | Fzd4,Rasip1,Tiparp | | 2.817e-02 | -3.57 | GO\_REGULATION\_OF\_CHEMOKINE\_PRODUCTION | MSigDB lists | GO\_REGULATION\_OF\_CHEMOKINE\_PRODUCTION | 45 | 3 | 12187 | 179 | Il6,Il4ra,Tnf | | 2.817e-02 | -3.57 | GO\_REGULATION\_OF\_GLUCOSE\_IMPORT | MSigDB lists | GO\_REGULATION\_OF\_GLUCOSE\_IMPORT | 45 | 3 | 12187 | 179 | Pik3r1,Tnf,Nfe2l2 | | 2.817e-02 | -3.57 | GO\_LEUKOCYTE\_HOMEOSTASIS | MSigDB lists | GO\_LEUKOCYTE\_HOMEOSTASIS | 45 | 3 | 12187 | 179 | Tnfaip3,Il6,Hcar2 | | 2.817e-02 | -3.57 | chr17p11 | MSigDB lists | chr17p11 | 45 | 3 | 12187 | 179 | Rasd1,Ksr1,Lgals9 | | 2.827e-02 | -3.57 | positive regulation of response to oxidative stress | biological process | GO:1902884 | 17 | 2 | 13711 | 214 | Ripk1,Tnf | | 2.827e-02 | -3.57 | negative regulation of necrotic cell death | biological process | GO:0060547 | 17 | 2 | 13711 | 214 | Cflar,Birc3 | | 2.827e-02 | -3.57 | positive regulation of fibroblast migration | biological process | GO:0010763 | 17 | 2 | 13711 | 214 | Thbs1,Akap12 | | 2.827e-02 | -3.57 | zinc ion homeostasis | biological process | GO:0055069 | 17 | 2 | 13711 | 214 | Mt2,Mt1 | | 2.827e-02 | -3.57 | regulation of macrophage derived foam cell differentiation | biological process | GO:0010743 | 17 | 2 | 13711 | 214 | Msr1,Nfkbia | | 2.827e-02 | -3.57 | reactive oxygen species biosynthetic process | biological process | GO:1903409 | 17 | 2 | 13711 | 214 | Cybb,Ncf1 | | 2.827e-02 | -3.57 | branching involved in salivary gland morphogenesis | biological process | GO:0060445 | 17 | 2 | 13711 | 214 | Il6,Tgm2 | | 2.827e-02 | -3.57 | regulation of microglial cell activation | biological process | GO:1903978 | 17 | 2 | 13711 | 214 | C5ar1,Mmp8 | | 2.827e-02 | -3.57 | regulation of gonad development | biological process | GO:1905939 | 17 | 2 | 13711 | 214 | Runx1,Il1a | | 2.830e-02 | -3.56 | GSE4535\_BM\_DERIVED\_DC\_VS\_FOLLICULAR\_DC\_UP | MSigDB lists | GSE4535\_BM\_DERIVED\_DC\_VS\_FOLLICULAR\_DC\_UP | 157 | 6 | 12187 | 179 | Casp4,Stx11,Gadd45b,Bcl3,Tnfaip8,Ptpn1 | | 2.830e-02 | -3.56 | GSE1460\_DP\_THYMOCYTE\_VS\_NAIVE\_CD4\_TCELL\_ADULT\_BLOOD\_DN | MSigDB lists | GSE1460\_DP\_THYMOCYTE\_VS\_NAIVE\_CD4\_TCELL\_ADULT\_BLOOD\_DN | 157 | 6 | 12187 | 179 | Tnip1,Ifitm2,Csf3,Timp1,Ccnd2,Il4ra | | 2.830e-02 | -3.56 | GSE7831\_CPG\_VS\_INFLUENZA\_STIM\_PDC\_1H\_DN | MSigDB lists | GSE7831\_CPG\_VS\_INFLUENZA\_STIM\_PDC\_1H\_DN | 157 | 6 | 12187 | 179 | Clic4,Hspb1,Cybb,Cd14,Ifitm3,Birc3 | | 2.830e-02 | -3.56 | GSE15930\_STIM\_VS\_STIM\_AND\_IL12\_72H\_CD8\_T\_CELL\_UP | MSigDB lists | GSE15930\_STIM\_VS\_STIM\_AND\_IL12\_72H\_CD8\_T\_CELL\_UP | 157 | 6 | 12187 | 179 | Irf7,Cd14,Isg15,Lgals9,Ifit2,Il1rn | | 2.830e-02 | -3.56 | GSE17721\_CPG\_VS\_GARDIQUIMOD\_24H\_BMDC\_DN | MSigDB lists | GSE17721\_CPG\_VS\_GARDIQUIMOD\_24H\_BMDC\_DN | 157 | 6 | 12187 | 179 | Nfkbiz,Cd14,S100a9,Selp,Cybb,Rtp4 | | 2.830e-02 | -3.56 | GAUSSMANN\_MLL\_AF4\_FUSION\_TARGETS\_A\_UP | MSigDB lists | GAUSSMANN\_MLL\_AF4\_FUSION\_TARGETS\_A\_UP | 157 | 6 | 12187 | 179 | Ccl12,Sbno2,Tnfaip8,Rtp4,Tgm2,C3ar1 | | 2.830e-02 | -3.56 | GSE8921\_UNSTIM\_VS\_TLR1\_2\_STIM\_MONOCYTE\_24H\_UP | MSigDB lists | GSE8921\_UNSTIM\_VS\_TLR1\_2\_STIM\_MONOCYTE\_24H\_UP | 157 | 6 | 12187 | 179 | Dtx3l,Isg15,Irgm2,Usp18,Pik3ap1,Irf9 | | 2.830e-02 | -3.56 | GSE32128\_INOS\_DEPENDENT\_VS\_INOS\_INDEPENDENT\_ACTIVATED\_TCELL\_DN | MSigDB lists | GSE32128\_INOS\_DEPENDENT\_VS\_INOS\_INDEPENDENT\_ACTIVATED\_TCELL\_DN | 157 | 6 | 12187 | 179 | Gbp6,Ifit3b,Cp,Icam1,Irf9,Gbp4 | | 2.830e-02 | -3.56 | GSE37416\_CTRL\_VS\_6H\_F\_TULARENSIS\_LVS\_NEUTROPHIL\_DN | MSigDB lists | GSE37416\_CTRL\_VS\_6H\_F\_TULARENSIS\_LVS\_NEUTROPHIL\_DN | 157 | 6 | 12187 | 179 | Nfkbie,Pik3r5,Tnip1,Nfkbia,Icam1,Nfkb1 | | 2.838e-02 | -3.56 | CD28 dependent PI3K/Akt signaling | REACTOME pathways | R-MMU-389357 | 16 | 2 | 6297 | 105 | Pik3r1,Map3k8 | | 2.859e-02 | -3.55 | PIP metabolism | BIOCYC pathways | MOUSE\_PWY3DJ-219 | 47 | 2 | 823 | 5 | Pik3r1,Pik3r5 | | 2.884e-02 | -3.55 | GO\_PEPTIDASE\_INHIBITOR\_ACTIVITY | MSigDB lists | GO\_PEPTIDASE\_INHIBITOR\_ACTIVITY | 79 | 4 | 12187 | 179 | Timp1,Tnfaip8,A2m,Birc3 | | 2.884e-02 | -3.55 | BHATI\_G2M\_ARREST\_BY\_2METHOXYESTRADIOL\_UP | MSigDB lists | BHATI\_G2M\_ARREST\_BY\_2METHOXYESTRADIOL\_UP | 79 | 4 | 12187 | 179 | Ier3,Rasip1,Tubb6,Hspa5 | | 2.884e-02 | -3.55 | CHEN\_LIVER\_METABOLISM\_QTL\_CIS | MSigDB lists | CHEN\_LIVER\_METABOLISM\_QTL\_CIS | 79 | 4 | 12187 | 179 | Ifi211,Ifi209,Ifi204,Ifi207 | | 2.890e-02 | -3.54 | Toll Like Receptor 5 (TLR5) Cascade | REACTOME pathways | R-MMU-168176 | 70 | 4 | 6297 | 105 | Nfkb1,Nfkb2,Nfkbia,Map3k8 | | 2.890e-02 | -3.54 | Toll Like Receptor 10 (TLR10) Cascade | REACTOME pathways | R-MMU-168142 | 70 | 4 | 6297 | 105 | Nfkbia,Map3k8,Nfkb1,Nfkb2 | | 2.890e-02 | -3.54 | Toll Like Receptor TLR6:TLR2 Cascade | REACTOME pathways | R-MMU-168188 | 70 | 4 | 6297 | 105 | Map3k8,Nfkbia,Nfkb1,Nfkb2 | | 2.890e-02 | -3.54 | MyD88:Mal cascade initiated on plasma membrane | REACTOME pathways | R-MMU-166058 | 70 | 4 | 6297 | 105 | Nfkb2,Nfkb1,Map3k8,Nfkbia | | 2.890e-02 | -3.54 | MyD88 cascade initiated on plasma membrane | REACTOME pathways | R-MMU-975871 | 70 | 4 | 6297 | 105 | Nfkbia,Map3k8,Nfkb1,Nfkb2 | | 2.890e-02 | -3.54 | Toll Like Receptor 2 (TLR2) Cascade | REACTOME pathways | R-MMU-181438 | 70 | 4 | 6297 | 105 | Nfkbia,Map3k8,Nfkb2,Nfkb1 | | 2.890e-02 | -3.54 | Toll Like Receptor TLR1:TLR2 Cascade | REACTOME pathways | R-MMU-168179 | 70 | 4 | 6297 | 105 | Nfkbia,Map3k8,Nfkb2,Nfkb1 | | 2.901e-02 | -3.54 | NOTCH\_DN.V1\_UP | MSigDB lists | NOTCH\_DN.V1\_UP | 117 | 5 | 12187 | 179 | Adamts1,Cdkn1a,Ksr1,Col4a1,S100a8 | | 2.901e-02 | -3.54 | GSE45365\_WT\_VS\_IFNAR\_KO\_CD11B\_DC\_DN | MSigDB lists | GSE45365\_WT\_VS\_IFNAR\_KO\_CD11B\_DC\_DN | 117 | 5 | 12187 | 179 | Hp,Tiparp,Ptpn1,Runx1,Rhou | | 2.901e-02 | -3.54 | GO\_REGENERATION | MSigDB lists | GO\_REGENERATION | 117 | 5 | 12187 | 179 | Ccl12,Socs3,Cdkn1a,Cflar,C5ar1 | | 2.907e-02 | -3.54 | GSE43955\_TH0\_VS\_TGFB\_IL6\_IL23\_TH17\_ACT\_CD4\_TCELL\_60H\_DN | MSigDB lists | GSE43955\_TH0\_VS\_TGFB\_IL6\_IL23\_TH17\_ACT\_CD4\_TCELL\_60H\_DN | 158 | 6 | 12187 | 179 | Pik3r1,Casp4,Tiparp,Cp,Zfp36,Cebpd | | 2.907e-02 | -3.54 | GSE29164\_CD8\_TCELL\_VS\_CD8\_TCELL\_AND\_IL12\_TREATED\_MELANOMA\_DAY3\_DN | MSigDB lists | GSE29164\_CD8\_TCELL\_VS\_CD8\_TCELL\_AND\_IL12\_TREATED\_MELANOMA\_DAY3\_DN | 158 | 6 | 12187 | 179 | Socs3,Sgk1,Tubb6,Banp,Rgs16,Zfp36 | | 2.907e-02 | -3.54 | GSE22140\_GERMFREE\_VS\_SPF\_ARTHRITIC\_MOUSE\_CD4\_TCELL\_DN | MSigDB lists | GSE22140\_GERMFREE\_VS\_SPF\_ARTHRITIC\_MOUSE\_CD4\_TCELL\_DN | 158 | 6 | 12187 | 179 | Cdkn1a,Casp4,Igsf6,S100a8,Plek,Tap1 | | 2.907e-02 | -3.54 | GSE22140\_HEALTHY\_VS\_ARTHRITIC\_MOUSE\_CD4\_TCELL\_UP | MSigDB lists | GSE22140\_HEALTHY\_VS\_ARTHRITIC\_MOUSE\_CD4\_TCELL\_UP | 158 | 6 | 12187 | 179 | S100a9,Rrp8,S100a8,Hspb1,Sgk1,Cdkn1a | | 2.907e-02 | -3.54 | GSE28726\_ACT\_CD4\_TCELL\_VS\_ACT\_NKTCELL\_DN | MSigDB lists | GSE28726\_ACT\_CD4\_TCELL\_VS\_ACT\_NKTCELL\_DN | 158 | 6 | 12187 | 179 | Sgk1,Aff1,Casp4,Dnajb1,Rbm39,Tnfaip3 | | 2.907e-02 | -3.54 | GSE21774\_CD62L\_POS\_CD56\_DIM\_VS\_CD62L\_NEG\_CD56\_DIM\_NK\_CELL\_UP | MSigDB lists | GSE21774\_CD62L\_POS\_CD56\_DIM\_VS\_CD62L\_NEG\_CD56\_DIM\_NK\_CELL\_UP | 158 | 6 | 12187 | 179 | Ifitm3,Iqgap1,Fstl1,Ifi44,Parp12,Tubb6 | | 2.907e-02 | -3.54 | GSE43955\_1H\_VS\_42H\_ACT\_CD4\_TCELL\_WITH\_TGFB\_IL6\_DN | MSigDB lists | GSE43955\_1H\_VS\_42H\_ACT\_CD4\_TCELL\_WITH\_TGFB\_IL6\_DN | 158 | 6 | 12187 | 179 | A2m,C3ar1,Rgs16,Nfkb1,Tubb6,Cp | | 2.907e-02 | -3.54 | GSE42088\_2H\_VS\_24H\_LEISHMANIA\_INF\_DC\_UP | MSigDB lists | GSE42088\_2H\_VS\_24H\_LEISHMANIA\_INF\_DC\_UP | 158 | 6 | 12187 | 179 | Zfp36,Ifitm3,C5ar1,Ifitm2,Tor3a,Cd14 | | 2.907e-02 | -3.54 | GSE7460\_TCONV\_VS\_TREG\_THYMUS\_DN | MSigDB lists | GSE7460\_TCONV\_VS\_TREG\_THYMUS\_DN | 158 | 6 | 12187 | 179 | Stx11,Gadd45g,Gbp3,Rgs16,Map3k8,Casp4 | | 2.907e-02 | -3.54 | GSE25088\_WT\_VS\_STAT6\_KO\_MACROPHAGE\_ROSIGLITAZONE\_STIM\_DN | MSigDB lists | GSE25088\_WT\_VS\_STAT6\_KO\_MACROPHAGE\_ROSIGLITAZONE\_STIM\_DN | 158 | 6 | 12187 | 179 | Tnip1,Slfn9,Tnfaip8,Rassf4,Cybb,Nfkb1 | | 2.907e-02 | -3.54 | GSE22611\_NOD2\_TRANSDUCED\_VS\_CTRL\_HEK293T\_STIMULATED\_WITH\_MDP\_2H\_UP | MSigDB lists | GSE22611\_NOD2\_TRANSDUCED\_VS\_CTRL\_HEK293T\_STIMULATED\_WITH\_MDP\_2H\_UP | 158 | 6 | 12187 | 179 | Cmpk2,Gbp3,Irf2,Ifit2,Sgk1,Birc3 | | 2.907e-02 | -3.54 | GSE25087\_FETAL\_VS\_ADULT\_TCONV\_UP | MSigDB lists | GSE25087\_FETAL\_VS\_ADULT\_TCONV\_UP | 158 | 6 | 12187 | 179 | Rnf213,C3ar1,Cxcl1,Nfkb2,Nfkb1,Tiparp | | 2.907e-02 | -3.54 | GSE32986\_UNSTIM\_VS\_GMCSF\_STIM\_DC\_DN | MSigDB lists | GSE32986\_UNSTIM\_VS\_GMCSF\_STIM\_DC\_DN | 158 | 6 | 12187 | 179 | Parp9,Msn,Slfn9,Irf7,Il1rn,Oasl1 | | 2.907e-02 | -3.54 | EBAUER\_TARGETS\_OF\_PAX3\_FOXO1\_FUSION\_UP | MSigDB lists | EBAUER\_TARGETS\_OF\_PAX3\_FOXO1\_FUSION\_UP | 158 | 6 | 12187 | 179 | Gadd45b,Rassf4,Tgm2,Pygm,Plek,Isg15 | | 2.907e-02 | -3.54 | GSE17580\_TREG\_VS\_TEFF\_UP | MSigDB lists | GSE17580\_TREG\_VS\_TEFF\_UP | 158 | 6 | 12187 | 179 | Rasip1,Rgs16,Xdh,Tgm2,Socs3,Gbp7 | | 2.907e-02 | -3.54 | GSE32164\_RESTING\_DIFFERENTIATED\_VS\_CMYC\_INHIBITED\_MACROPHAGE\_UP | MSigDB lists | GSE32164\_RESTING\_DIFFERENTIATED\_VS\_CMYC\_INHIBITED\_MACROPHAGE\_UP | 158 | 6 | 12187 | 179 | Spi1,Birc3,Rasd1,Clic4,Iqgap1,Tubb6 | | 2.914e-02 | -3.54 | Nfe2l2 (nuclear factor, erythroid derived 2, like 2) | protein interactions | 18024 | 16 | 2 | 6802 | 115 | Ncf1,Cdkn1a | | 2.914e-02 | -3.54 | Insr (insulin receptor) | protein interactions | 16337 | 16 | 2 | 6802 | 115 | Socs3,Pik3r1 | | 2.914e-02 | -3.54 | ELOC (elongin C) | protein interactions | 6921 | 16 | 2 | 6802 | 115 | Socs3,Msn | | 2.916e-02 | -3.53 | AUJLA\_IL22\_AND\_IL17A\_SIGNALING | MSigDB lists | AUJLA\_IL22\_AND\_IL17A\_SIGNALING | 2 | 1 | 12187 | 179 | Csf3 | | 2.916e-02 | -3.53 | GO\_DETECTION\_OF\_CHEMICAL\_STIMULUS\_INVOLVED\_IN\_SENSORY\_PERCEPTION\_OF\_TASTE | MSigDB lists | GO\_DETECTION\_OF\_CHEMICAL\_STIMULUS\_INVOLVED\_IN\_SENSORY\_PERCEPTION\_OF\_TASTE | 2 | 1 | 12187 | 179 | Rtp4 | | 2.916e-02 | -3.53 | BASSO\_HAIRY\_CELL\_LEUKEMIA\_DN | MSigDB lists | BASSO\_HAIRY\_CELL\_LEUKEMIA\_DN | 2 | 1 | 12187 | 179 | Tnfaip8 | | 2.925e-02 | -3.53 | TSP\_1 | pfam domains | PF00090 | 43 | 3 | 12881 | 201 | Adamts1,Adamts9,Thbs1 | | 2.925e-02 | -3.53 | zf-B\_box | pfam domains | PF00643 | 43 | 3 | 12881 | 201 | Trim30a,Trim30d,Trim56 | | 2.928e-02 | -3.53 | sarcoplasm | cellular component | GO:0016528 | 76 | 4 | 13825 | 212 | Rasd1,Thbs1,Pygm,Xdh | | 2.928e-02 | -3.53 | regulation of erythrocyte differentiation | biological process | GO:0045646 | 43 | 3 | 13711 | 214 | Zfp36,Isg15,Spi1 | | 2.935e-02 | -3.53 | Peptide ligand-binding receptors | REACTOME pathways | R-MMU-375276 | 104 | 5 | 6297 | 105 | C3ar1,Cxcl9,C5ar1,Cxcl1,Cxcl16 | | 2.942e-02 | -3.53 | regulation of leukocyte mediated immunity | biological process | GO:0002703 | 149 | 6 | 13711 | 214 | Il6,Tap1,Lgals9,Tnf,Cxcl1,Il4ra | | 2.956e-02 | -3.52 | Apc (APC, WNT signaling pathway regulator) | protein interactions | 11789 | 40 | 3 | 6802 | 115 | Ptgs2,Ptges,Il6 | | 2.957e-02 | -3.52 | - | gene3d domains | 2.60.40.2160 | 2 | 1 | 6647 | 99 | Il17ra | | 2.957e-02 | -3.52 | - | gene3d domains | 4.10.770.10 | 2 | 1 | 6647 | 99 | Runx1 | | 2.957e-02 | -3.52 | - | gene3d domains | 3.30.1660.20 | 2 | 1 | 6647 | 99 | Retnlg | | 2.957e-02 | -3.52 | AP1\_01 | MSigDB lists | AP1\_01 | 202 | 7 | 12187 | 179 | Tagln2,Csf3,Trib1,Cdkn1a,Il6,Bach1,Gadd45g | | 2.982e-02 | -3.51 | GO\_EXTRACELLULAR\_MATRIX\_DISASSEMBLY | MSigDB lists | GO\_EXTRACELLULAR\_MATRIX\_DISASSEMBLY | 46 | 3 | 12187 | 179 | A2m,Timp1,Mmp8 | | 2.982e-02 | -3.51 | MODULE\_209 | MSigDB lists | MODULE\_209 | 46 | 3 | 12187 | 179 | Casp4,Hp,Timp1 | | 2.982e-02 | -3.51 | GO\_POSITIVE\_REGULATION\_OF\_PHOSPHATIDYLINOSITOL\_3\_KINASE\_SIGNALING | MSigDB lists | GO\_POSITIVE\_REGULATION\_OF\_PHOSPHATIDYLINOSITOL\_3\_KINASE\_SIGNALING | 46 | 3 | 12187 | 179 | Pik3ap1,Selp,Csf3 | | 2.982e-02 | -3.51 | GO\_ACTIVATION\_OF\_MAPKK\_ACTIVITY | MSigDB lists | GO\_ACTIVATION\_OF\_MAPKK\_ACTIVITY | 46 | 3 | 12187 | 179 | Gadd45b,Map3k6,Map3k8 | | 2.982e-02 | -3.51 | PID\_PI3KCI\_PATHWAY | MSigDB lists | PID\_PI3KCI\_PATHWAY | 46 | 3 | 12187 | 179 | Pik3r1,Sgk1,Pik3r5 | | 2.982e-02 | -3.51 | GNF2\_MCL1 | MSigDB lists | GNF2\_MCL1 | 46 | 3 | 12187 | 179 | Igsf6,Stx11,Cybb | | 2.982e-02 | -3.51 | chr1q25 | MSigDB lists | chr1q25 | 46 | 3 | 12187 | 179 | Rgs16,Tor3a,Ptgs2 | | 2.982e-02 | -3.51 | TONG\_INTERACT\_WITH\_PTTG1 | MSigDB lists | TONG\_INTERACT\_WITH\_PTTG1 | 46 | 3 | 12187 | 179 | Ier3,Phf11d,Phf11b | | 2.984e-02 | -3.51 | cellular divalent inorganic cation homeostasis | biological process | GO:0072503 | 368 | 11 | 13711 | 214 | Ccl2,Mt2,Cxcl1,C5ar1,Tgm2,Mt1,P2ry6,Kcna5,C3ar1,Slc24a4,Pygm | | 2.986e-02 | -3.51 | GSE360\_HIGH\_VS\_LOW\_DOSE\_B\_MALAYI\_MAC\_DN | MSigDB lists | GSE360\_HIGH\_VS\_LOW\_DOSE\_B\_MALAYI\_MAC\_DN | 159 | 6 | 12187 | 179 | Hspa5,Map3k8,Atf3,Tiparp,Ifitm2,Pygm | | 2.986e-02 | -3.51 | GSE25123\_CTRL\_VS\_IL4\_AND\_ROSIGLITAZONE\_STIM\_PPARG\_KO\_MACROPHAGE\_DN | MSigDB lists | GSE25123\_CTRL\_VS\_IL4\_AND\_ROSIGLITAZONE\_STIM\_PPARG\_KO\_MACROPHAGE\_DN | 159 | 6 | 12187 | 179 | Oasl1,Gbp4,Zbtb16,Aff1,Sgk3,Gbp6 | | 2.986e-02 | -3.51 | GSE6875\_WT\_VS\_FOXP3\_KO\_TREG\_DN | MSigDB lists | GSE6875\_WT\_VS\_FOXP3\_KO\_TREG\_DN | 159 | 6 | 12187 | 179 | Angptl4,Gbp6,Mxd4,Gbp4,Cybb,Rbm47 | | 2.986e-02 | -3.51 | GSE41176\_UNSTIM\_VS\_ANTI\_IGM\_STIM\_BCELL\_3H\_DN | MSigDB lists | GSE41176\_UNSTIM\_VS\_ANTI\_IGM\_STIM\_BCELL\_3H\_DN | 159 | 6 | 12187 | 179 | Ifi211,Ifit2,Ifi209,Aff1,Ifi207,Ifi204 | | 2.986e-02 | -3.51 | GSE32533\_WT\_VS\_MIR17\_KO\_ACT\_CD4\_TCELL\_UP | MSigDB lists | GSE32533\_WT\_VS\_MIR17\_KO\_ACT\_CD4\_TCELL\_UP | 159 | 6 | 12187 | 179 | Sbno2,Ier3,Socs3,Trib1,Fzd4,Rhoj | | 2.986e-02 | -3.51 | GO\_RESPONSE\_TO\_REACTIVE\_OXYGEN\_SPECIES | MSigDB lists | GO\_RESPONSE\_TO\_REACTIVE\_OXYGEN\_SPECIES | 159 | 6 | 12187 | 179 | Tnfaip3,Nfe2l2,Il6,Kcna5,Cybb,Hp | | 2.986e-02 | -3.51 | GSE2405\_0H\_VS\_1.5H\_A\_PHAGOCYTOPHILUM\_STIM\_NEUTROPHIL\_UP | MSigDB lists | GSE2405\_0H\_VS\_1.5H\_A\_PHAGOCYTOPHILUM\_STIM\_NEUTROPHIL\_UP | 159 | 6 | 12187 | 179 | Olfml2b,Ifitm2,S100a9,Mmp8,Usp18,Ifit3b | | 2.986e-02 | -3.51 | GSE2770\_UNTREATED\_VS\_TGFB\_AND\_IL4\_TREATED\_ACT\_CD4\_TCELL\_48H\_DN | MSigDB lists | GSE2770\_UNTREATED\_VS\_TGFB\_AND\_IL4\_TREATED\_ACT\_CD4\_TCELL\_48H\_DN | 159 | 6 | 12187 | 179 | Il1rn,Ccnd2,Tnf,Cdkn1a,Tnfaip8,Csf2rb | | 2.986e-02 | -3.51 | GSE21670\_TGFB\_VS\_IL6\_TREATED\_STAT3\_KO\_CD4\_TCELL\_UP | MSigDB lists | GSE21670\_TGFB\_VS\_IL6\_TREATED\_STAT3\_KO\_CD4\_TCELL\_UP | 159 | 6 | 12187 | 179 | Gadd45g,Usp18,Slfn9,Gbp7,Sgk3,Ccnd2 | | 2.986e-02 | -3.51 | GSE17721\_LPS\_VS\_POLYIC\_4H\_BMDC\_UP | MSigDB lists | GSE17721\_LPS\_VS\_POLYIC\_4H\_BMDC\_UP | 159 | 6 | 12187 | 179 | Nfkb2,Parp9,Pik3ap1,Isg15,Gadd45b,Lcn2 | | 2.987e-02 | -3.51 | Prolactin signaling pathway | KEGG pathways | ko04917 | 58 | 4 | 5248 | 107 | Nfkb1,Socs3,Pik3r1,Ccnd2 | | 2.987e-02 | -3.51 | Prolactin signaling pathway | KEGG pathways | mmu04917 | 58 | 4 | 5248 | 107 | Nfkb1,Socs3,Pik3r1,Ccnd2 | | 2.994e-02 | -3.51 | YTAAYNGCT\_UNKNOWN | MSigDB lists | YTAAYNGCT\_UNKNOWN | 118 | 5 | 12187 | 179 | Ccnd2,Rhoc,Luc7l3,Atf3,Tnfsf10 | | 2.994e-02 | -3.51 | AGCATTA\_MIR155 | MSigDB lists | AGCATTA\_MIR155 | 118 | 5 | 12187 | 179 | Bach1,Sgk3,Sox11,Dnajb1,Spi1 | | 3.000e-02 | -3.51 | mouse chr18 B2\|18 19.46 cM | chromosome location | mouse chr18 B2\|18 19.46 cM | 2 | 1 | 14556 | 220 | Cd14 | | 3.000e-02 | -3.51 | mouse chr2 H1\|2 78.72 cM | chromosome location | mouse chr2 H1\|2 78.72 cM | 2 | 1 | 14556 | 220 | Tgm2 | | 3.000e-02 | -3.51 | mouse chr11 C\|11 50.3 cM | chromosome location | mouse chr11 C\|11 50.3 cM | 2 | 1 | 14556 | 220 | Slfn2 | | 3.000e-02 | -3.51 | mouse chr9 A3\|9 7.69 cM | chromosome location | mouse chr9 A3\|9 7.69 cM | 2 | 1 | 14556 | 220 | Icam1 | | 3.000e-02 | -3.51 | mouse chr13 A3.3\|13 14.01 cM | chromosome location | mouse chr13 A3.3\|13 14.01 cM | 2 | 1 | 14556 | 220 | Ripk1 | | 3.000e-02 | -3.51 | mouse chr5 B2\|5 17.84 cM | chromosome location | mouse chr5 B2\|5 17.84 cM | 2 | 1 | 14556 | 220 | Mxd4 | | 3.000e-02 | -3.51 | mouse chr16 A2\|16 10.09 cM | chromosome location | mouse chr16 A2\|16 10.09 cM | 2 | 1 | 14556 | 220 | Cebpd | | 3.000e-02 | -3.51 | mouse chr5 E5\|5 50.45 cM | chromosome location | mouse chr5 E5\|5 50.45 cM | 2 | 1 | 14556 | 220 | Aff1 | | 3.000e-02 | -3.51 | mouse chr1 H6\|1 96.28 cM | chromosome location | mouse chr1 H6\|1 96.28 cM | 2 | 1 | 14556 | 220 | Atf3 | | 3.000e-02 | -3.51 | mouse chr19 C3\|19 38.8 cM | chromosome location | mouse chr19 C3\|19 38.8 cM | 2 | 1 | 14556 | 220 | Nfkb2 | | 3.000e-02 | -3.51 | mouse chr10\|10 B5.2 | chromosome location | mouse chr10\|10 B5.2 | 2 | 1 | 14556 | 220 | Arid5b | | 3.000e-02 | -3.51 | mouse chr11 A2\|11 9.41 cM | chromosome location | mouse chr11 A2\|11 9.41 cM | 2 | 1 | 14556 | 220 | Plek | | 3.002e-02 | -3.51 | GSE9946\_MATURE\_STIMULATORY\_VS\_LISTERIA\_INF\_MATURE\_DC\_DN | MSigDB lists | GSE9946\_MATURE\_STIMULATORY\_VS\_LISTERIA\_INF\_MATURE\_DC\_DN | 80 | 4 | 12187 | 179 | Sdc4,Hspa5,Pik3ap1,S100a8 | | 3.002e-02 | -3.51 | AMBROSINI\_FLAVOPIRIDOL\_TREATMENT\_TP53 | MSigDB lists | AMBROSINI\_FLAVOPIRIDOL\_TREATMENT\_TP53 | 80 | 4 | 12187 | 179 | Cxcl1,Sntb2,Il1rn,Hspb1 | | 3.002e-02 | -3.51 | SMITH\_TERT\_TARGETS\_DN | MSigDB lists | SMITH\_TERT\_TARGETS\_DN | 80 | 4 | 12187 | 179 | Saa1,A2m,Il1rn,Tnfsf10 | | 3.027e-02 | -3.50 | GO\_REGULATION\_OF\_HOMOTYPIC\_CELL\_CELL\_ADHESION | MSigDB lists | GO\_REGULATION\_OF\_HOMOTYPIC\_CELL\_CELL\_ADHESION | 203 | 7 | 12187 | 179 | Ccl12,Il6,Vcam1,Zbtb16,Map3k8,Pik3r1,Il4ra | | 3.044e-02 | -3.49 | core-binding factor complex | cellular component | GO:0016513 | 2 | 1 | 13825 | 212 | Runx1 | | 3.044e-02 | -3.49 | oncostatin-M receptor complex | cellular component | GO:0005900 | 2 | 1 | 13825 | 212 | Osmr | | 3.044e-02 | -3.49 | PCNA-p21 complex | cellular component | GO:0070557 | 2 | 1 | 13825 | 212 | Cdkn1a | | 3.044e-02 | -3.49 | perinuclear endoplasmic reticulum membrane | cellular component | GO:1990578 | 2 | 1 | 13825 | 212 | Pik3r1 | | 3.044e-02 | -3.49 | RNA-directed RNA polymerase complex | cellular component | GO:0031379 | 2 | 1 | 13825 | 212 | Znfx1 | | 3.044e-02 | -3.49 | fibrinogen complex | cellular component | GO:0005577 | 2 | 1 | 13825 | 212 | Thbs1 | | 3.044e-02 | -3.49 | CHOP-ATF3 complex | cellular component | GO:1990622 | 2 | 1 | 13825 | 212 | Atf3 | | 3.052e-02 | -3.49 | FAD-bd\_8 | interpro domains | IPR013112 | 2 | 1 | 13788 | 212 | Cybb | | 3.052e-02 | -3.49 | Pleckstrin | interpro domains | IPR037370 | 2 | 1 | 13788 | 212 | Plek | | 3.052e-02 | -3.49 | BIRC2/BIRC3\_UBA | interpro domains | IPR041933 | 2 | 1 | 13788 | 212 | Birc3 | | 3.052e-02 | -3.49 | RunxI\_C\_dom | interpro domains | IPR013711 | 2 | 1 | 13788 | 212 | Runx1 | | 3.052e-02 | -3.49 | SNO | interpro domains | IPR026741 | 2 | 1 | 13788 | 212 | Sbno2 | | 3.052e-02 | -3.49 | Interferon-induced\_IFIT3 | interpro domains | IPR024122 | 2 | 1 | 13788 | 212 | Ifit3b | | 3.052e-02 | -3.49 | PIK3AP1\_TIR | interpro domains | IPR041340 | 2 | 1 | 13788 | 212 | Pik3ap1 | | 3.052e-02 | -3.49 | IRF1/IRF2 | interpro domains | IPR017431 | 2 | 1 | 13788 | 212 | Irf2 | | 3.052e-02 | -3.49 | Resistin-like\_sf | interpro domains | IPR036262 | 2 | 1 | 13788 | 212 | Retnlg | | 3.052e-02 | -3.49 | SNO\_AAA | interpro domains | IPR039187 | 2 | 1 | 13788 | 212 | Sbno2 | | 3.052e-02 | -3.49 | Cu-oxidase | interpro domains | IPR001117 | 2 | 1 | 13788 | 212 | Cp | | 3.052e-02 | -3.49 | Anaphtx\_C5AR1/C5AR2 | interpro domains | IPR001274 | 2 | 1 | 13788 | 212 | C5ar1 | | 3.052e-02 | -3.49 | Pept\_M12B\_GON-ADAMTSs | interpro domains | IPR012314 | 2 | 1 | 13788 | 212 | Adamts9 | | 3.052e-02 | -3.49 | TF\_Runt-rel\_RUNX | interpro domains | IPR016554 | 2 | 1 | 13788 | 212 | Runx1 | | 3.052e-02 | -3.49 | Mig-6 | interpro domains | IPR021619 | 2 | 1 | 13788 | 212 | Errfi1 | | 3.052e-02 | -3.49 | Fe\_red\_NAD-bd\_6 | interpro domains | IPR013121 | 2 | 1 | 13788 | 212 | Cybb | | 3.052e-02 | -3.49 | PIK3R5/6 | interpro domains | IPR019522 | 2 | 1 | 13788 | 212 | Pik3r5 | | 3.052e-02 | -3.49 | Cu\_oxidase\_Cu\_BS | interpro domains | IPR002355 | 2 | 1 | 13788 | 212 | Cp | | 3.052e-02 | -3.49 | TMEM2\_PANDER-like | interpro domains | IPR039473 | 2 | 1 | 13788 | 212 | Cemip2 | | 3.052e-02 | -3.49 | Ptpn1/Ptpn2 | interpro domains | IPR012265 | 2 | 1 | 13788 | 212 | Ptpn1 | | 3.052e-02 | -3.49 | SBNO\_Helicase\_C\_dom | interpro domains | IPR026937 | 2 | 1 | 13788 | 212 | Sbno2 | | 3.052e-02 | -3.49 | CBD\_Rasip1/Radil | interpro domains | IPR037983 | 2 | 1 | 13788 | 212 | Rasip1 | | 3.052e-02 | -3.49 | LIFR\_D2 | interpro domains | IPR040817 | 2 | 1 | 13788 | 212 | Osmr | | 3.052e-02 | -3.49 | ICAM | interpro domains | IPR003988 | 2 | 1 | 13788 | 212 | Icam1 | | 3.052e-02 | -3.49 | FAM107 | interpro domains | IPR009533 | 2 | 1 | 13788 | 212 | Fam107a | | 3.052e-02 | -3.49 | DBB\_domain | interpro domains | IPR017893 | 2 | 1 | 13788 | 212 | Pik3ap1 | | 3.052e-02 | -3.49 | Runx\_central\_dom\_sf | interpro domains | IPR027384 | 2 | 1 | 13788 | 212 | Runx1 | | 3.052e-02 | -3.49 | hSH3 | interpro domains | IPR029294 | 2 | 1 | 13788 | 212 | Fyb | | 3.052e-02 | -3.49 | G8\_domain | interpro domains | IPR019316 | 2 | 1 | 13788 | 212 | Cemip2 | | 3.052e-02 | -3.49 | RII\_binding\_1 | interpro domains | IPR018459 | 2 | 1 | 13788 | 212 | Akap12 | | 3.052e-02 | -3.49 | MGST1-like | interpro domains | IPR040162 | 2 | 1 | 13788 | 212 | Ptges | | 3.052e-02 | -3.49 | Thymidylate\_kin-like\_dom | interpro domains | IPR039430 | 2 | 1 | 13788 | 212 | Cmpk2 | | 3.052e-02 | -3.49 | IL-1\_propep | interpro domains | IPR003502 | 2 | 1 | 13788 | 212 | Il1a | | 3.052e-02 | -3.49 | ABC\_Tap-like | interpro domains | IPR013305 | 2 | 1 | 13788 | 212 | Tap1 | | 3.052e-02 | -3.49 | Resistin | interpro domains | IPR009714 | 2 | 1 | 13788 | 212 | Retnlg | | 3.052e-02 | -3.49 | IL17RA/B\_FnIII-like\_1\_sf | interpro domains | IPR038683 | 2 | 1 | 13788 | 212 | Il17ra | | 3.052e-02 | -3.49 | Cdc42\_binding\_dom-like | interpro domains | IPR015116 | 2 | 1 | 13788 | 212 | Errfi1 | | 3.052e-02 | -3.49 | Lipocalin | interpro domains | IPR002345 | 2 | 1 | 13788 | 212 | Lcn2 | | 3.052e-02 | -3.49 | AML1\_Runt | interpro domains | IPR000040 | 2 | 1 | 13788 | 212 | Runx1 | | 3.052e-02 | -3.49 | KSR\_SAM-like\_dom | interpro domains | IPR025561 | 2 | 1 | 13788 | 212 | Ksr1 | | 3.052e-02 | -3.49 | Runt\_dom | interpro domains | IPR013524 | 2 | 1 | 13788 | 212 | Runx1 | | 3.052e-02 | -3.49 | Pkinase-A\_anch\_WSK-motif | interpro domains | IPR001573 | 2 | 1 | 13788 | 212 | Akap12 | | 3.052e-02 | -3.49 | Ig\_C2-set | interpro domains | IPR008424 | 2 | 1 | 13788 | 212 | Vcam1 | | 3.052e-02 | -3.49 | IL17R\_fnIII\_D1 | interpro domains | IPR032356 | 2 | 1 | 13788 | 212 | Il17ra | | 3.063e-02 | -3.49 | ADAMTS/ADAMTS-like | interpro domains | IPR013273 | 18 | 2 | 13788 | 212 | Adamts1,Adamts9 | | 3.063e-02 | -3.49 | ADAM\_spacer1 | interpro domains | IPR010294 | 18 | 2 | 13788 | 212 | Adamts1,Adamts9 | | 3.065e-02 | -3.48 | - | gene3d domains | 2.20.100.10 | 46 | 3 | 6647 | 99 | Adamts1,Adamts9,Thbs1 | | 3.066e-02 | -3.48 | GSE22886\_DAY0\_VS\_DAY1\_MONOCYTE\_IN\_CULTURE\_UP | MSigDB lists | GSE22886\_DAY0\_VS\_DAY1\_MONOCYTE\_IN\_CULTURE\_UP | 160 | 6 | 12187 | 179 | Ms4a6d,Cebpd,Tor3a,Zfp36,Tnfsf10,Ifitm2 | | 3.066e-02 | -3.48 | GATA1\_04 | MSigDB lists | GATA1\_04 | 160 | 6 | 12187 | 179 | C3ar1,Fzd4,Ip6k2,Msr1,Gadd45g,Luc7l3 | | 3.066e-02 | -3.48 | GSE360\_DC\_VS\_MAC\_L\_DONOVANI\_UP | MSigDB lists | GSE360\_DC\_VS\_MAC\_L\_DONOVANI\_UP | 160 | 6 | 12187 | 179 | Nfkbia,Vcam1,Birc3,Selp,Cflar,Csf2rb | | 3.066e-02 | -3.48 | GSE12392\_CD8A\_POS\_VS\_NEG\_SPLEEN\_DC\_UP | MSigDB lists | GSE12392\_CD8A\_POS\_VS\_NEG\_SPLEEN\_DC\_UP | 160 | 6 | 12187 | 179 | Tor3a,Ptges,Tap1,Cybb,Trim25,Hp | | 3.066e-02 | -3.48 | GSE23925\_DARK\_ZONE\_VS\_NAIVE\_BCELL\_UP | MSigDB lists | GSE23925\_DARK\_ZONE\_VS\_NAIVE\_BCELL\_UP | 160 | 6 | 12187 | 179 | Errfi1,Slfn4,Slfn3,Ms4a6d,Socs3,Nfkbia | | 3.066e-02 | -3.48 | GSE37532\_TREG\_VS\_TCONV\_PPARG\_KO\_CD4\_TCELL\_FROM\_VISCERAL\_ADIPOSE\_TISSUE\_UP | MSigDB lists | GSE37532\_TREG\_VS\_TCONV\_PPARG\_KO\_CD4\_TCELL\_FROM\_VISCERAL\_ADIPOSE\_TISSUE\_UP | 160 | 6 | 12187 | 179 | Cdkn1a,Bcl3,Icam1,Ksr1,Stx11,Gadd45b | | 3.066e-02 | -3.48 | GSE23505\_UNTREATED\_VS\_4DAY\_IL6\_IL1\_IL23\_TREATED\_CD4\_TCELL\_UP | MSigDB lists | GSE23505\_UNTREATED\_VS\_4DAY\_IL6\_IL1\_IL23\_TREATED\_CD4\_TCELL\_UP | 160 | 6 | 12187 | 179 | AA467197,Nfe2l2,Map3k8,Sgk1,Birc3,Errfi1 | | 3.066e-02 | -3.48 | GSE17721\_0.5H\_VS\_24H\_POLYIC\_BMDC\_UP | MSigDB lists | GSE17721\_0.5H\_VS\_24H\_POLYIC\_BMDC\_UP | 160 | 6 | 12187 | 179 | Ccl2,Cdkn1a,Maff,S100a9,Ptpn1,Tnfaip8 | | 3.068e-02 | -3.48 | SUSHI | prosite domains | PS50923 | 37 | 3 | 8845 | 164 | Hp,Selp,Sele | | 3.090e-02 | -3.48 | GSE37416\_CTRL\_VS\_48H\_F\_TULARENSIS\_LVS\_NEUTROPHIL\_UP | MSigDB lists | GSE37416\_CTRL\_VS\_48H\_F\_TULARENSIS\_LVS\_NEUTROPHIL\_UP | 119 | 5 | 12187 | 179 | Lcn2,Oasl1,Samd9l,Ifit3b,Xaf1 | | 3.090e-02 | -3.48 | BROWNE\_HCMV\_INFECTION\_24HR\_DN | MSigDB lists | BROWNE\_HCMV\_INFECTION\_24HR\_DN | 119 | 5 | 12187 | 179 | Bach1,Mxd4,Thbs1,Cflar,Cdkn1a | | 3.094e-02 | -3.48 | mouse chr5 | chromosome location | mouse chr5 | 242 | 8 | 14556 | 220 | Mxd4,Hspb1,Aff1,Cxcl9,Cxcl1,Gbp4,Il6,Ncf1 | | 3.095e-02 | -3.48 | female gonad development | biological process | GO:0008585 | 76 | 4 | 13711 | 214 | Adamts1,Arid5b,Tiparp,Fzd4 | | 3.096e-02 | -3.47 | regulation of endothelial cell proliferation | biological process | GO:0001936 | 112 | 5 | 13711 | 214 | Xdh,Ccl2,Ccl12,Tnf,Thbs1 | | 3.097e-02 | -3.47 | SAM\_KSR1 | pfam domains | PF13543 | 2 | 1 | 12881 | 201 | Ksr1 | | 3.097e-02 | -3.47 | GON | pfam domains | PF08685 | 2 | 1 | 12881 | 201 | Adamts9 | | 3.097e-02 | -3.47 | WSK | pfam domains | PF03832 | 2 | 1 | 12881 | 201 | Akap12 | | 3.097e-02 | -3.47 | hSH3 | pfam domains | PF14603 | 2 | 1 | 12881 | 201 | Fyb | | 3.097e-02 | -3.47 | Thymidylate\_kin | pfam domains | PF02223 | 2 | 1 | 12881 | 201 | Cmpk2 | | 3.097e-02 | -3.47 | RII\_binding\_1 | pfam domains | PF10522 | 2 | 1 | 12881 | 201 | Akap12 | | 3.097e-02 | -3.47 | DBB | pfam domains | PF14545 | 2 | 1 | 12881 | 201 | Pik3ap1 | | 3.097e-02 | -3.47 | IL1\_propep | pfam domains | PF02394 | 2 | 1 | 12881 | 201 | Il1a | | 3.097e-02 | -3.47 | Cu-oxidase | pfam domains | PF00394 | 2 | 1 | 12881 | 201 | Cp | | 3.097e-02 | -3.47 | FAD\_binding\_8 | pfam domains | PF08022 | 2 | 1 | 12881 | 201 | Cybb | | 3.097e-02 | -3.47 | Helicase\_C\_4 | pfam domains | PF13871 | 2 | 1 | 12881 | 201 | Sbno2 | | 3.097e-02 | -3.47 | Resistin | pfam domains | PF06954 | 2 | 1 | 12881 | 201 | Retnlg | | 3.097e-02 | -3.47 | zf-3CxxC | pfam domains | PF13695 | 2 | 1 | 12881 | 201 | Rtp4 | | 3.097e-02 | -3.47 | LIFR\_D2 | pfam domains | PF17971 | 2 | 1 | 12881 | 201 | Osmr | | 3.097e-02 | -3.47 | IL17R\_fnIII\_D1 | pfam domains | PF16556 | 2 | 1 | 12881 | 201 | Il17ra | | 3.097e-02 | -3.47 | PI3K\_1B\_p101 | pfam domains | PF10486 | 2 | 1 | 12881 | 201 | Pik3r5 | | 3.097e-02 | -3.47 | G8 | pfam domains | PF10162 | 2 | 1 | 12881 | 201 | Cemip2 | | 3.097e-02 | -3.47 | Inhibitor\_Mig-6 | pfam domains | PF11555 | 2 | 1 | 12881 | 201 | Errfi1 | | 3.097e-02 | -3.47 | Runt | pfam domains | PF00853 | 2 | 1 | 12881 | 201 | Runx1 | | 3.097e-02 | -3.47 | RunxI | pfam domains | PF08504 | 2 | 1 | 12881 | 201 | Runx1 | | 3.097e-02 | -3.47 | C2-set | pfam domains | PF05790 | 2 | 1 | 12881 | 201 | Vcam1 | | 3.097e-02 | -3.47 | TIR\_3 | pfam domains | PF18567 | 2 | 1 | 12881 | 201 | Pik3ap1 | | 3.097e-02 | -3.47 | NAD\_binding\_6 | pfam domains | PF08030 | 2 | 1 | 12881 | 201 | Cybb | | 3.097e-02 | -3.47 | AAA\_34 | pfam domains | PF13872 | 2 | 1 | 12881 | 201 | Sbno2 | | 3.097e-02 | -3.47 | GTPase\_binding | pfam domains | PF09027 | 2 | 1 | 12881 | 201 | Errfi1 | | 3.097e-02 | -3.47 | DUF1151 | pfam domains | PF06625 | 2 | 1 | 12881 | 201 | Fam107a | | 3.097e-02 | -3.47 | chronic inflammatory response to antigenic stimulus | biological process | GO:0002439 | 2 | 1 | 13711 | 214 | Tnf | | 3.097e-02 | -3.47 | regulation of translational initiation by iron | biological process | GO:0006447 | 2 | 1 | 13711 | 214 | Tnf | | 3.097e-02 | -3.47 | isopeptide cross-linking | biological process | GO:0018262 | 2 | 1 | 13711 | 214 | Tgm2 | | 3.097e-02 | -3.47 | positive regulation of calcidiol 1-monooxygenase activity | biological process | GO:0060559 | 2 | 1 | 13711 | 214 | Tnf | | 3.097e-02 | -3.47 | negative regulation of toll-like receptor 3 signaling pathway | biological process | GO:0034140 | 2 | 1 | 13711 | 214 | Tnfaip3 | | 3.097e-02 | -3.47 | regulation of blood microparticle formation | biological process | GO:2000332 | 2 | 1 | 13711 | 214 | Tnf | | 3.097e-02 | -3.47 | cell-cell adhesion in response to extracellular stimulus | biological process | GO:0140039 | 2 | 1 | 13711 | 214 | Vcam1 | | 3.097e-02 | -3.47 | regulation of hair follicle cell proliferation | biological process | GO:0071336 | 2 | 1 | 13711 | 214 | Runx1 | | 3.097e-02 | -3.47 | skeletal muscle atrophy | biological process | GO:0014732 | 2 | 1 | 13711 | 214 | Cflar | | 3.097e-02 | -3.47 | ripoptosome assembly involved in necroptotic process | biological process | GO:1901026 | 2 | 1 | 13711 | 214 | Ripk1 | | 3.097e-02 | -3.47 | antigen processing and presentation of exogenous peptide antigen via MHC class I, TAP-dependent | biological process | GO:0002479 | 2 | 1 | 13711 | 214 | Tap1 | | 3.097e-02 | -3.47 | negative regulation of neutrophil differentiation | biological process | GO:0045659 | 2 | 1 | 13711 | 214 | Trib1 | | 3.097e-02 | -3.47 | sequestering of iron ion | biological process | GO:0097577 | 2 | 1 | 13711 | 214 | Lcn2 | | 3.097e-02 | -3.47 | regulation of transcription from RNA polymerase II promoter in response to copper ion | biological process | GO:0061396 | 2 | 1 | 13711 | 214 | Nfe2l2 | | 3.097e-02 | -3.47 | monocyte homeostasis | biological process | GO:0035702 | 2 | 1 | 13711 | 214 | Ccl2 | | 3.097e-02 | -3.47 | peptidyl-cysteine S-trans-nitrosylation | biological process | GO:0035606 | 2 | 1 | 13711 | 214 | S100a9 | | 3.097e-02 | -3.47 | spongiotrophoblast differentiation | biological process | GO:0060708 | 2 | 1 | 13711 | 214 | Socs3 | | 3.097e-02 | -3.47 | positive regulation of interleukin-23 production | biological process | GO:0032747 | 2 | 1 | 13711 | 214 | Il17ra | | 3.097e-02 | -3.47 | positive regulation of blood microparticle formation | biological process | GO:2000334 | 2 | 1 | 13711 | 214 | Tnf | | 3.097e-02 | -3.47 | negative regulation of inositol phosphate biosynthetic process | biological process | GO:0010920 | 2 | 1 | 13711 | 214 | Plek | | 3.097e-02 | -3.47 | regulation of polynucleotide adenylyltransferase activity | biological process | GO:1904245 | 2 | 1 | 13711 | 214 | Zfp36 | | 3.097e-02 | -3.47 | positive regulation of antimicrobial humoral response | biological process | GO:0002760 | 2 | 1 | 13711 | 214 | Acod1 | | 3.097e-02 | -3.47 | cellular response to progesterone stimulus | biological process | GO:0071393 | 2 | 1 | 13711 | 214 | Acod1 | | 3.097e-02 | -3.47 | tolerance induction dependent upon immune response | biological process | GO:0002461 | 2 | 1 | 13711 | 214 | C3ar1 | | 3.097e-02 | -3.47 | regulation of complement activation, lectin pathway | biological process | GO:0001868 | 2 | 1 | 13711 | 214 | A2m | | 3.097e-02 | -3.47 | hindbrain structural organization | biological process | GO:0021577 | 2 | 1 | 13711 | 214 | Hspa5 | | 3.097e-02 | -3.47 | response to interleukin-11 | biological process | GO:0071105 | 2 | 1 | 13711 | 214 | Sbno2 | | 3.097e-02 | -3.47 | negative regulation of intrinsic apoptotic signaling pathway in response to osmotic stress | biological process | GO:1902219 | 2 | 1 | 13711 | 214 | Ptgs2 | | 3.097e-02 | -3.47 | regulation of transcription involved in G2/M transition of mitotic cell cycle | biological process | GO:0000117 | 2 | 1 | 13711 | 214 | Bach1 | | 3.097e-02 | -3.47 | siderophore transport | biological process | GO:0015891 | 2 | 1 | 13711 | 214 | Lcn2 | | 3.097e-02 | -3.47 | T cell aggregation | biological process | GO:0070489 | 2 | 1 | 13711 | 214 | Msn | | 3.097e-02 | -3.47 | response to yeast | biological process | GO:0001878 | 2 | 1 | 13711 | 214 | Ncf1 | | 3.097e-02 | -3.47 | negative regulation of complement activation, lectin pathway | biological process | GO:0001869 | 2 | 1 | 13711 | 214 | A2m | | 3.097e-02 | -3.47 | negative regulation of chronic inflammatory response | biological process | GO:0002677 | 2 | 1 | 13711 | 214 | Tnfaip3 | | 3.097e-02 | -3.47 | positive regulation of glutathione biosynthetic process | biological process | GO:1903788 | 2 | 1 | 13711 | 214 | Nfe2l2 | | 3.097e-02 | -3.47 | dTDP biosynthetic process | biological process | GO:0006233 | 2 | 1 | 13711 | 214 | Cmpk2 | | 3.097e-02 | -3.47 | regulation of platelet-derived growth factor production | biological process | GO:0090361 | 2 | 1 | 13711 | 214 | Ptgs2 | | 3.097e-02 | -3.47 | positive regulation of immune complex clearance by monocytes and macrophages | biological process | GO:0090265 | 2 | 1 | 13711 | 214 | Ccl2 | | 3.097e-02 | -3.47 | negative regulation of L-glutamate import across plasma membrane | biological process | GO:0002037 | 2 | 1 | 13711 | 214 | Tnf | | 3.097e-02 | -3.47 | negative regulation of vitamin D biosynthetic process | biological process | GO:0010957 | 2 | 1 | 13711 | 214 | Nfkb1 | | 3.097e-02 | -3.47 | complement component C5a signaling pathway | biological process | GO:0038178 | 2 | 1 | 13711 | 214 | C5ar1 | | 3.097e-02 | -3.47 | regulation of defense response to fungus | biological process | GO:1900150 | 2 | 1 | 13711 | 214 | Cxcl1 | | 3.097e-02 | -3.47 | response to aluminum ion | biological process | GO:0010044 | 2 | 1 | 13711 | 214 | Xdh | | 3.097e-02 | -3.47 | positive regulation of progesterone secretion | biological process | GO:2000872 | 2 | 1 | 13711 | 214 | Runx1 | | 3.097e-02 | -3.47 | negative regulation of nitric oxide mediated signal transduction | biological process | GO:0010751 | 2 | 1 | 13711 | 214 | Thbs1 | | 3.097e-02 | -3.47 | regulation of hydrogen peroxide catabolic process | biological process | GO:2000295 | 2 | 1 | 13711 | 214 | Hp | | 3.097e-02 | -3.47 | negative regulation of myoblast fusion | biological process | GO:1901740 | 2 | 1 | 13711 | 214 | Cflar | | 3.097e-02 | -3.47 | regulation of nucleotide-binding oligomerization domain containing 1 signaling pathway | biological process | GO:0070428 | 2 | 1 | 13711 | 214 | Tnfaip3 | | 3.097e-02 | -3.47 | negative regulation of CD40 signaling pathway | biological process | GO:2000349 | 2 | 1 | 13711 | 214 | Tnfaip3 | | 3.097e-02 | -3.47 | negative regulation of T-helper 1 cell differentiation | biological process | GO:0045626 | 2 | 1 | 13711 | 214 | Il4ra | | 3.097e-02 | -3.47 | glucocorticoid mediated signaling pathway | biological process | GO:0043402 | 2 | 1 | 13711 | 214 | Sgk1 | | 3.097e-02 | -3.47 | closure of optic fissure | biological process | GO:0061386 | 2 | 1 | 13711 | 214 | Sox11 | | 3.097e-02 | -3.47 | ripoptosome assembly | biological process | GO:0097343 | 2 | 1 | 13711 | 214 | Ripk1 | | 3.097e-02 | -3.47 | cerebellum vasculature development | biological process | GO:0061300 | 2 | 1 | 13711 | 214 | Fzd4 | | 3.097e-02 | -3.47 | regulation of leukocyte adhesion to arterial endothelial cell | biological process | GO:1904997 | 2 | 1 | 13711 | 214 | Tnf | | 3.097e-02 | -3.47 | interferon-alpha production | biological process | GO:0032607 | 2 | 1 | 13711 | 214 | Irf7 | | 3.097e-02 | -3.47 | regulation of cardiac muscle tissue regeneration | biological process | GO:1905178 | 2 | 1 | 13711 | 214 | Cdkn1a | | 3.097e-02 | -3.47 | negative regulation of vasculogenesis | biological process | GO:2001213 | 2 | 1 | 13711 | 214 | Xdh | | 3.097e-02 | -3.47 | negative regulation of endothelial cell chemotaxis | biological process | GO:2001027 | 2 | 1 | 13711 | 214 | Thbs1 | | 3.097e-02 | -3.47 | negative regulation of fibrinolysis | biological process | GO:0051918 | 2 | 1 | 13711 | 214 | Thbs1 | | 3.097e-02 | -3.47 | regulation of immune complex clearance by monocytes and macrophages | biological process | GO:0090264 | 2 | 1 | 13711 | 214 | Ccl2 | | 3.097e-02 | -3.47 | positive regulation of removal of superoxide radicals | biological process | GO:1904833 | 2 | 1 | 13711 | 214 | Tnf | | 3.097e-02 | -3.47 | negative regulation of male gonad development | biological process | GO:2000019 | 2 | 1 | 13711 | 214 | Il1a | | 3.097e-02 | -3.47 | endothelial cell-matrix adhesion | biological process | GO:0090673 | 2 | 1 | 13711 | 214 | Adamts9 | | 3.097e-02 | -3.47 | progesterone secretion | biological process | GO:0042701 | 2 | 1 | 13711 | 214 | Fzd4 | | 3.097e-02 | -3.47 | positive regulation of vitamin D biosynthetic process | biological process | GO:0060557 | 2 | 1 | 13711 | 214 | Tnf | | 3.097e-02 | -3.47 | regulation of neutrophil mediated cytotoxicity | biological process | GO:0070948 | 2 | 1 | 13711 | 214 | Cxcl1 | | 3.097e-02 | -3.47 | nucleotide-binding oligomerization domain containing 1 signaling pathway | biological process | GO:0070427 | 2 | 1 | 13711 | 214 | Nfkbia | | 3.097e-02 | -3.47 | positive regulation of superoxide dismutase activity | biological process | GO:1901671 | 2 | 1 | 13711 | 214 | Tnf | | 3.097e-02 | -3.47 | regulation of neutrophil mediated killing of symbiont cell | biological process | GO:0070949 | 2 | 1 | 13711 | 214 | Cxcl1 | | 3.097e-02 | -3.47 | dTDP metabolic process | biological process | GO:0046072 | 2 | 1 | 13711 | 214 | Cmpk2 | | 3.097e-02 | -3.47 | positive regulation of interleukin-4-mediated signaling pathway | biological process | GO:1902216 | 2 | 1 | 13711 | 214 | Parp14 | | 3.097e-02 | -3.47 | positive regulation of hippo signaling | biological process | GO:0035332 | 2 | 1 | 13711 | 214 | Sox11 | | 3.097e-02 | -3.47 | regulation of hepatic stellate cell migration | biological process | GO:0061869 | 2 | 1 | 13711 | 214 | Akap12 | | 3.097e-02 | -3.47 | interferon-gamma biosynthetic process | biological process | GO:0042095 | 2 | 1 | 13711 | 214 | Gadd45g | | 3.097e-02 | -3.47 | positive regulation of hepatic stellate cell migration | biological process | GO:0061870 | 2 | 1 | 13711 | 214 | Akap12 | | 3.097e-02 | -3.47 | cerebellum structural organization | biological process | GO:0021589 | 2 | 1 | 13711 | 214 | Hspa5 | | 3.097e-02 | -3.47 | negative regulation of lymphocyte chemotaxis | biological process | GO:1901624 | 2 | 1 | 13711 | 214 | Ccl12 | | 3.097e-02 | -3.47 | cytotoxic T cell degranulation | biological process | GO:0043316 | 2 | 1 | 13711 | 214 | Stx11 | | 3.097e-02 | -3.47 | protein secretion by platelet | biological process | GO:0070560 | 2 | 1 | 13711 | 214 | Plek | | 3.097e-02 | -3.47 | negative regulation of chemokine biosynthetic process | biological process | GO:0045079 | 2 | 1 | 13711 | 214 | Il6 | | 3.097e-02 | -3.47 | negative regulation of cardiac muscle tissue regeneration | biological process | GO:1905179 | 2 | 1 | 13711 | 214 | Cdkn1a | | 3.097e-02 | -3.47 | positive regulation of translational initiation by iron | biological process | GO:0045994 | 2 | 1 | 13711 | 214 | Tnf | | 3.097e-02 | -3.47 | regulation of MyD88-independent toll-like receptor signaling pathway | biological process | GO:0034127 | 2 | 1 | 13711 | 214 | Irf7 | | 3.098e-02 | -3.47 | ATPase-coupled peptide antigen transmembrane transporter activity | molecular function | GO:0015433 | 2 | 1 | 13516 | 211 | Tap1 | | 3.098e-02 | -3.47 | ADP-D-ribose binding | molecular function | GO:0072570 | 2 | 1 | 13516 | 211 | Parp9 | | 3.098e-02 | -3.47 | oxidoreductase activity, acting on other nitrogenous compounds as donors, cytochrome as acceptor | molecular function | GO:0016662 | 2 | 1 | 13516 | 211 | Xdh | | 3.098e-02 | -3.47 | granulocyte colony-stimulating factor receptor binding | molecular function | GO:0005130 | 2 | 1 | 13516 | 211 | Csf3 | | 3.098e-02 | -3.47 | thymidylate kinase activity | molecular function | GO:0004798 | 2 | 1 | 13516 | 211 | Cmpk2 | | 3.098e-02 | -3.47 | interleukin-8 binding | molecular function | GO:0019959 | 2 | 1 | 13516 | 211 | A2m | | 3.098e-02 | -3.47 | prostaglandin-endoperoxide synthase activity | molecular function | GO:0004666 | 2 | 1 | 13516 | 211 | Ptgs2 | | 3.098e-02 | -3.47 | ADP receptor activity | molecular function | GO:0001621 | 2 | 1 | 13516 | 211 | P2ry6 | | 3.098e-02 | -3.47 | interleukin-1, type I receptor binding | molecular function | GO:0005150 | 2 | 1 | 13516 | 211 | Il1rn | | 3.098e-02 | -3.47 | G protein-coupled pyrimidinergic nucleotide receptor activity | molecular function | GO:0071553 | 2 | 1 | 13516 | 211 | P2ry6 | | 3.098e-02 | -3.47 | olfactory receptor binding | molecular function | GO:0031849 | 2 | 1 | 13516 | 211 | Rtp4 | | 3.098e-02 | -3.47 | complement component C3a binding | molecular function | GO:0001850 | 2 | 1 | 13516 | 211 | C3ar1 | | 3.098e-02 | -3.47 | UTP-activated nucleotide receptor activity | molecular function | GO:0045030 | 2 | 1 | 13516 | 211 | P2ry6 | | 3.098e-02 | -3.47 | fucose binding | molecular function | GO:0042806 | 2 | 1 | 13516 | 211 | Selp | | 3.098e-02 | -3.47 | uridine nucleotide receptor activity | molecular function | GO:0015065 | 2 | 1 | 13516 | 211 | P2ry6 | | 3.098e-02 | -3.47 | nitrite reductase (NO-forming) activity | molecular function | GO:0050421 | 2 | 1 | 13516 | 211 | Xdh | | 3.098e-02 | -3.47 | tapasin binding | molecular function | GO:0046980 | 2 | 1 | 13516 | 211 | Tap1 | | 3.098e-02 | -3.47 | flavonoid binding | molecular function | GO:0097243 | 2 | 1 | 13516 | 211 | Ip6k2 | | 3.098e-02 | -3.47 | TRAIL binding | molecular function | GO:0045569 | 2 | 1 | 13516 | 211 | Tnfsf10 | | 3.098e-02 | -3.47 | complement component C5a receptor activity | molecular function | GO:0004878 | 2 | 1 | 13516 | 211 | C5ar1 | | 3.107e-02 | -3.47 | regulation of collagen metabolic process | biological process | GO:0010712 | 44 | 3 | 13711 | 214 | Errfi1,Il6,Ccl2 | | 3.109e-02 | -3.47 | negative regulation of transcription by RNA polymerase II | biological process | GO:0000122 | 715 | 18 | 13711 | 214 | Nfkb1,Parp9,Dnajb1,Ifi209,Gbp4,Runx1,Bach1,Ifi211,Spi1,Zfp36,Atf3,Mxd4,Tnf,Sox11,Ifi204,Ifi207,Arid5b,Zbtb16 | | 3.114e-02 | -3.47 | REACTOME\_GROWTH\_HORMONE\_RECEPTOR\_SIGNALING | MSigDB lists | REACTOME\_GROWTH\_HORMONE\_RECEPTOR\_SIGNALING | 19 | 2 | 12187 | 179 | Ptpn1,Socs3 | | 3.114e-02 | -3.47 | YANG\_BREAST\_CANCER\_ESR1\_BULK\_DN | MSigDB lists | YANG\_BREAST\_CANCER\_ESR1\_BULK\_DN | 19 | 2 | 12187 | 179 | Rgs16,Plaur | | 3.114e-02 | -3.47 | GO\_REGULATION\_OF\_PROTEIN\_IMPORT\_INTO\_NUCLEUS\_TRANSLOCATION | MSigDB lists | GO\_REGULATION\_OF\_PROTEIN\_IMPORT\_INTO\_NUCLEUS\_TRANSLOCATION | 19 | 2 | 12187 | 179 | Cdkn1a,Il6 | | 3.114e-02 | -3.47 | BIOCARTA\_CELLCYCLE\_PATHWAY | MSigDB lists | BIOCARTA\_CELLCYCLE\_PATHWAY | 19 | 2 | 12187 | 179 | Ccnd2,Cdkn1a | | 3.114e-02 | -3.47 | MODULE\_573 | MSigDB lists | MODULE\_573 | 19 | 2 | 12187 | 179 | Cdkn1a,Lcn2 | | 3.114e-02 | -3.47 | BOQUEST\_STEM\_CELL\_CULTURED\_VS\_FRESH\_DN | MSigDB lists | BOQUEST\_STEM\_CELL\_CULTURED\_VS\_FRESH\_DN | 19 | 2 | 12187 | 179 | Socs3,Cd14 | | 3.114e-02 | -3.47 | GENTLES\_LEUKEMIC\_STEM\_CELL\_UP | MSigDB lists | GENTLES\_LEUKEMIC\_STEM\_CELL\_UP | 19 | 2 | 12187 | 179 | Icam1,Birc3 | | 3.114e-02 | -3.47 | GO\_DEVELOPMENTAL\_PROGRAMMED\_CELL\_DEATH | MSigDB lists | GO\_DEVELOPMENTAL\_PROGRAMMED\_CELL\_DEATH | 19 | 2 | 12187 | 179 | Spi1,Il1a | | 3.114e-02 | -3.47 | ZHANG\_ANTIVIRAL\_RESPONSE\_TO\_RIBAVIRIN\_UP | MSigDB lists | ZHANG\_ANTIVIRAL\_RESPONSE\_TO\_RIBAVIRIN\_UP | 19 | 2 | 12187 | 179 | Irf7,Ifitm3 | | 3.114e-02 | -3.47 | GO\_POSITIVE\_REGULATION\_OF\_CD4\_POSITIVE\_ALPHA\_BETA\_T\_CELL\_ACTIVATION | MSigDB lists | GO\_POSITIVE\_REGULATION\_OF\_CD4\_POSITIVE\_ALPHA\_BETA\_T\_CELL\_ACTIVATION | 19 | 2 | 12187 | 179 | Il6,Il4ra | | 3.114e-02 | -3.47 | GO\_RESPONSE\_TO\_PROSTAGLANDIN\_E | MSigDB lists | GO\_RESPONSE\_TO\_PROSTAGLANDIN\_E | 19 | 2 | 12187 | 179 | P2ry6,Adamts1 | | 3.114e-02 | -3.47 | PID\_INTEGRIN\_A9B1\_PATHWAY | MSigDB lists | PID\_INTEGRIN\_A9B1\_PATHWAY | 19 | 2 | 12187 | 179 | Tgm2,Vcam1 | | 3.114e-02 | -3.47 | GO\_REGULATION\_OF\_NUCLEASE\_ACTIVITY | MSigDB lists | GO\_REGULATION\_OF\_NUCLEASE\_ACTIVITY | 19 | 2 | 12187 | 179 | Hspa5,Ptpn1 | | 3.114e-02 | -3.47 | OSADA\_ASCL1\_TARGETS\_DN | MSigDB lists | OSADA\_ASCL1\_TARGETS\_DN | 19 | 2 | 12187 | 179 | Birc3,Socs3 | | 3.114e-02 | -3.47 | GO\_REGULATION\_OF\_TYPE\_2\_IMMUNE\_RESPONSE | MSigDB lists | GO\_REGULATION\_OF\_TYPE\_2\_IMMUNE\_RESPONSE | 19 | 2 | 12187 | 179 | Il4ra,Il6 | | 3.114e-02 | -3.47 | FONTAINE\_THYROID\_TUMOR\_UNCERTAIN\_MALIGNANCY\_DN | MSigDB lists | FONTAINE\_THYROID\_TUMOR\_UNCERTAIN\_MALIGNANCY\_DN | 19 | 2 | 12187 | 179 | Ptges,Ifitm3 | | 3.114e-02 | -3.47 | REACTOME\_CD28\_DEPENDENT\_PI3K\_AKT\_SIGNALING | MSigDB lists | REACTOME\_CD28\_DEPENDENT\_PI3K\_AKT\_SIGNALING | 19 | 2 | 12187 | 179 | Pik3r1,Map3k8 | | 3.114e-02 | -3.47 | GO\_REGULATION\_OF\_MYELOID\_CELL\_APOPTOTIC\_PROCESS | MSigDB lists | GO\_REGULATION\_OF\_MYELOID\_CELL\_APOPTOTIC\_PROCESS | 19 | 2 | 12187 | 179 | Hcar2,Irf7 | | 3.114e-02 | -3.47 | GO\_PEPTIDYL\_CYSTEINE\_MODIFICATION | MSigDB lists | GO\_PEPTIDYL\_CYSTEINE\_MODIFICATION | 19 | 2 | 12187 | 179 | S100a8,S100a9 | | 3.114e-02 | -3.47 | YANG\_BREAST\_CANCER\_ESR1\_DN | MSigDB lists | YANG\_BREAST\_CANCER\_ESR1\_DN | 19 | 2 | 12187 | 179 | Sox11,Tubb6 | | 3.114e-02 | -3.47 | GO\_NEGATIVE\_REGULATION\_OF\_CELL\_JUNCTION\_ASSEMBLY | MSigDB lists | GO\_NEGATIVE\_REGULATION\_OF\_CELL\_JUNCTION\_ASSEMBLY | 19 | 2 | 12187 | 179 | Thbs1,Tnf | | 3.114e-02 | -3.47 | GO\_ACTIVATION\_OF\_PHOSPHOLIPASE\_C\_ACTIVITY | MSigDB lists | GO\_ACTIVATION\_OF\_PHOSPHOLIPASE\_C\_ACTIVITY | 19 | 2 | 12187 | 179 | C5ar1,Sele | | 3.114e-02 | -3.47 | GO\_SODIUM\_ION\_HOMEOSTASIS | MSigDB lists | GO\_SODIUM\_ION\_HOMEOSTASIS | 19 | 2 | 12187 | 179 | Sgk3,Sgk1 | | 3.114e-02 | -3.47 | MARKS\_HDAC\_TARGETS\_UP | MSigDB lists | MARKS\_HDAC\_TARGETS\_UP | 19 | 2 | 12187 | 179 | Gadd45b,Tnfsf10 | | 3.123e-02 | -3.47 | GSE1925\_CTRL\_VS\_IFNG\_PRIMED\_MACROPHAGE\_3H\_IFNG\_STIM\_DN | MSigDB lists | GSE1925\_CTRL\_VS\_IFNG\_PRIMED\_MACROPHAGE\_3H\_IFNG\_STIM\_DN | 81 | 4 | 12187 | 179 | Rasd1,Osmr,Rbm47,Pik3ap1 | | 3.129e-02 | -3.46 | DELACROIX\_RARG\_BOUND\_MEF | MSigDB lists | DELACROIX\_RARG\_BOUND\_MEF | 297 | 9 | 12187 | 179 | Sntb2,Fstl1,Sgk1,Cp,Il1rn,Rhoc,Tubb6,Tagln2,S100a8 | | 3.146e-02 | -3.46 | ADAM\_spacer1 | pfam domains | PF05986 | 18 | 2 | 12881 | 201 | Adamts1,Adamts9 | | 3.147e-02 | -3.46 | GSE10273\_LOW\_IL7\_VS\_HIGH\_IL7\_AND\_IRF4\_IN\_IRF4\_8\_NULL\_PRE\_BCELL\_DN | MSigDB lists | GSE10273\_LOW\_IL7\_VS\_HIGH\_IL7\_AND\_IRF4\_IN\_IRF4\_8\_NULL\_PRE\_BCELL\_DN | 161 | 6 | 12187 | 179 | Selp,Gbp6,Nfkbie,Mmp8,Gbp4,Plaur | | 3.147e-02 | -3.46 | GSE17721\_POLYIC\_VS\_CPG\_24H\_BMDC\_UP | MSigDB lists | GSE17721\_POLYIC\_VS\_CPG\_24H\_BMDC\_UP | 161 | 6 | 12187 | 179 | Irf2,Clic4,Rhoc,S100a9,Aff1,Cdkn1a | | 3.147e-02 | -3.46 | GSE39916\_B\_CELL\_SPLEEN\_VS\_PLASMA\_CELL\_BONE\_MARROW\_UP | MSigDB lists | GSE39916\_B\_CELL\_SPLEEN\_VS\_PLASMA\_CELL\_BONE\_MARROW\_UP | 161 | 6 | 12187 | 179 | Sgk3,Rnf213,Trim56,Bcl3,Ifit3b,Tiparp | | 3.147e-02 | -3.46 | KRIEG\_KDM3A\_TARGETS\_NOT\_HYPOXIA | MSigDB lists | KRIEG\_KDM3A\_TARGETS\_NOT\_HYPOXIA | 161 | 6 | 12187 | 179 | Saa1,Phf11d,Ifi44,Phf11b,Ifit3b,Vcam1 | | 3.147e-02 | -3.46 | GSE22103\_UNSTIM\_VS\_LPS\_STIM\_NEUTROPHIL\_DN | MSigDB lists | GSE22103\_UNSTIM\_VS\_LPS\_STIM\_NEUTROPHIL\_DN | 161 | 6 | 12187 | 179 | Dnajb1,Maff,Rasd1,Atf3,Hspb1,Il1a | | 3.147e-02 | -3.46 | GSE2585\_CTEC\_VS\_THYMIC\_DC\_DN | MSigDB lists | GSE2585\_CTEC\_VS\_THYMIC\_DC\_DN | 161 | 6 | 12187 | 179 | Maff,Cp,Nfkb1,Gbp7,Irgm2,Ptges | | 3.147e-02 | -3.46 | KAECH\_DAY15\_EFF\_VS\_MEMORY\_CD8\_TCELL\_UP | MSigDB lists | KAECH\_DAY15\_EFF\_VS\_MEMORY\_CD8\_TCELL\_UP | 161 | 6 | 12187 | 179 | Ifitm3,S100a8,Cdkn1a,Hp,Il1rn,Mxd4 | | 3.147e-02 | -3.46 | GSE2405\_HEAT\_KILLED\_LYSATE\_VS\_LIVE\_A\_PHAGOCYTOPHILUM\_STIM\_NEUTROPHIL\_24H\_UP | MSigDB lists | GSE2405\_HEAT\_KILLED\_LYSATE\_VS\_LIVE\_A\_PHAGOCYTOPHILUM\_STIM\_NEUTROPHIL\_24H\_UP | 161 | 6 | 12187 | 179 | Gadd45b,Icam1,Plek,Casp4,Usp18,Ifitm3 | | 3.147e-02 | -3.46 | GSE8921\_UNSTIM\_0H\_VS\_TLR1\_2\_STIM\_MONOCYTE\_12H\_DN | MSigDB lists | GSE8921\_UNSTIM\_0H\_VS\_TLR1\_2\_STIM\_MONOCYTE\_12H\_DN | 161 | 6 | 12187 | 179 | Luc7l3,Mxd4,Akap12,Timp1,Sbno2,Rbm39 | | 3.147e-02 | -3.46 | GSE39110\_DAY3\_VS\_DAY6\_POST\_IMMUNIZATION\_CD8\_TCELL\_WITH\_IL2\_TREATMENT\_UP | MSigDB lists | GSE39110\_DAY3\_VS\_DAY6\_POST\_IMMUNIZATION\_CD8\_TCELL\_WITH\_IL2\_TREATMENT\_UP | 161 | 6 | 12187 | 179 | Tap1,Pik3ap1,Cybb,Gbp7,Isg15,Cdkn1a | | 3.148e-02 | -3.46 | mitochondrial fission | biological process | GO:0000266 | 18 | 2 | 13711 | 214 | Mx2,Mx1 | | 3.148e-02 | -3.46 | T-helper 1 type immune response | biological process | GO:0042088 | 18 | 2 | 13711 | 214 | Gadd45g,Bcl3 | | 3.148e-02 | -3.46 | response to muramyl dipeptide | biological process | GO:0032495 | 18 | 2 | 13711 | 214 | Tnfaip3,Nfkbia | | 3.148e-02 | -3.46 | negative regulation of smooth muscle cell migration | biological process | GO:0014912 | 18 | 2 | 13711 | 214 | Nfe2l2,Trib1 | | 3.148e-02 | -3.46 | regulation of T-helper 17 type immune response | biological process | GO:2000316 | 18 | 2 | 13711 | 214 | Lgals9,Nfkbiz | | 3.148e-02 | -3.46 | purinergic receptor signaling pathway | biological process | GO:0035587 | 18 | 2 | 13711 | 214 | P2ry6,Hcar2 | | 3.148e-02 | -3.46 | regulation of macrophage differentiation | biological process | GO:0045649 | 18 | 2 | 13711 | 214 | Ripk1,Trib1 | | 3.148e-02 | -3.46 | negative regulation of CD4-positive, alpha-beta T cell differentiation | biological process | GO:0043371 | 18 | 2 | 13711 | 214 | Il4ra,Runx1 | | 3.148e-02 | -3.46 | calcium-mediated signaling using intracellular calcium source | biological process | GO:0035584 | 18 | 2 | 13711 | 214 | Vcam1,Selp | | 3.149e-02 | -3.46 | lipopolysaccharide binding | molecular function | GO:0001530 | 18 | 2 | 13516 | 211 | Cd14,Selp | | 3.152e-02 | -3.46 | GTAAACC\_MIR2995P | MSigDB lists | GTAAACC\_MIR2995P | 47 | 3 | 12187 | 179 | Tiparp,Sgk1,Map3k8 | | 3.152e-02 | -3.46 | GO\_RESPONSE\_TO\_DSRNA | MSigDB lists | GO\_RESPONSE\_TO\_DSRNA | 47 | 3 | 12187 | 179 | Nfkbia,Irak3,Nfkb1 | | 3.152e-02 | -3.46 | JOHANSSON\_GLIOMAGENESIS\_BY\_PDGFB\_UP | MSigDB lists | JOHANSSON\_GLIOMAGENESIS\_BY\_PDGFB\_UP | 47 | 3 | 12187 | 179 | Tagln2,Rnf213,Ccnd2 | | 3.152e-02 | -3.46 | GO\_POSITIVE\_REGULATION\_OF\_LIPASE\_ACTIVITY | MSigDB lists | GO\_POSITIVE\_REGULATION\_OF\_LIPASE\_ACTIVITY | 47 | 3 | 12187 | 179 | C5ar1,Sele,Rhoc | | 3.163e-02 | -3.45 | TRAF6 mediated induction of NFkB and MAP kinases upon TLR7/8 or 9 activation | REACTOME pathways | R-MMU-975138 | 72 | 4 | 6297 | 105 | Nfkbia,Map3k8,Nfkb1,Nfkb2 | | 3.168e-02 | -3.45 | LU\_EZH2\_TARGETS\_UP | MSigDB lists | LU\_EZH2\_TARGETS\_UP | 205 | 7 | 12187 | 179 | Irf7,Isg15,Sdc4,Ifit3b,Tor3a,Oasl1,Ier3 | | 3.169e-02 | -3.45 | Znf\_B-box | interpro domains | IPR000315 | 45 | 3 | 13788 | 212 | Trim56,Trim30a,Trim30d | | 3.169e-02 | -3.45 | Ubiquitin\_dom | interpro domains | IPR000626 | 45 | 3 | 13788 | 212 | Oasl1,Oasl2,Isg15 | | 3.187e-02 | -3.45 | GSE18281\_PERIMEDULLARY\_CORTICAL\_REGION\_VS\_WHOLE\_CORTEX\_THYMUS\_DN | MSigDB lists | GSE18281\_PERIMEDULLARY\_CORTICAL\_REGION\_VS\_WHOLE\_CORTEX\_THYMUS\_DN | 120 | 5 | 12187 | 179 | Trib1,Map3k8,Maff,Oasl1,Il17ra | | 3.187e-02 | -3.45 | GSE2706\_R848\_VS\_R848\_AND\_LPS\_8H\_STIM\_DC\_UP | MSigDB lists | GSE2706\_R848\_VS\_R848\_AND\_LPS\_8H\_STIM\_DC\_UP | 120 | 5 | 12187 | 179 | Thbs1,Sgk1,Sntb2,Rbm47,Cxcl16 | | 3.187e-02 | -3.45 | GO\_SECOND\_MESSENGER\_MEDIATED\_SIGNALING | MSigDB lists | GO\_SECOND\_MESSENGER\_MEDIATED\_SIGNALING | 120 | 5 | 12187 | 179 | Rasd1,Ksr1,Sele,Vcam1,Selp | | 3.188e-02 | -3.45 | positive regulation of protein complex assembly | biological process | GO:0031334 | 193 | 7 | 13711 | 214 | Msn,Rhoc,Rasip1,Csf3,Tnf,Icam1,Plek | | 3.195e-02 | -3.44 | Toll-like receptor signaling | KEGG pathways | mmu\_M00686 | 14 | 2 | 5248 | 107 | Nfkbia,Nfkb1 | | 3.195e-02 | -3.44 | Toll-like receptor signaling | KEGG pathways | M00686 | 14 | 2 | 5248 | 107 | Nfkbia,Nfkb1 | | 3.220e-02 | -3.44 | MEISSNER\_NPC\_HCP\_WITH\_H3\_UNMETHYLATED | MSigDB lists | MEISSNER\_NPC\_HCP\_WITH\_H3\_UNMETHYLATED | 347 | 10 | 12187 | 179 | Cxcl16,Osmr,Trim25,Irak3,Bcl3,Pik3r5,Sgk3,Rasd1,Parp14,Birc3 | | 3.230e-02 | -3.43 | GSE5589\_WT\_VS\_IL10\_KO\_LPS\_AND\_IL10\_STIM\_MACROPHAGE\_45MIN\_DN | MSigDB lists | GSE5589\_WT\_VS\_IL10\_KO\_LPS\_AND\_IL10\_STIM\_MACROPHAGE\_45MIN\_DN | 162 | 6 | 12187 | 179 | Nfkb2,Clic4,Rrp8,Ier3,Tnfaip3,Tiparp | | 3.230e-02 | -3.43 | MODULE\_112 | MSigDB lists | MODULE\_112 | 162 | 6 | 12187 | 179 | Thbs1,A2m,Ccl4,Cxcl1,Cp,Csf2rb | | 3.230e-02 | -3.43 | GSE2770\_IL12\_AND\_TGFB\_VS\_IL4\_TREATED\_ACT\_CD4\_TCELL\_48H\_UP | MSigDB lists | GSE2770\_IL12\_AND\_TGFB\_VS\_IL4\_TREATED\_ACT\_CD4\_TCELL\_48H\_UP | 162 | 6 | 12187 | 179 | Xaf1,Socs3,Tiparp,Gbp7,Apold1,Irf9 | | 3.230e-02 | -3.43 | GSE15330\_LYMPHOID\_MULTIPOTENT\_VS\_GRANULOCYTE\_MONOCYTE\_PROGENITOR\_IKAROS\_KO\_DN | MSigDB lists | GSE15330\_LYMPHOID\_MULTIPOTENT\_VS\_GRANULOCYTE\_MONOCYTE\_PROGENITOR\_IKAROS\_KO\_DN | 162 | 6 | 12187 | 179 | Tiparp,Akap12,Gadd45g,Ier3,Sgk1,Ptpn1 | | 3.230e-02 | -3.43 | GSE3982\_NEUTROPHIL\_VS\_CENT\_MEMORY\_CD4\_TCELL\_UP | MSigDB lists | GSE3982\_NEUTROPHIL\_VS\_CENT\_MEMORY\_CD4\_TCELL\_UP | 162 | 6 | 12187 | 179 | Cybb,Lcn2,Icam1,Irf2,Irf9,Cflar | | 3.230e-02 | -3.43 | GSE360\_CTRL\_VS\_B\_MALAYI\_HIGH\_DOSE\_MAC\_DN | MSigDB lists | GSE360\_CTRL\_VS\_B\_MALAYI\_HIGH\_DOSE\_MAC\_DN | 162 | 6 | 12187 | 179 | Sbno2,Rgs16,Pygm,Sox11,Tnf,Tiparp | | 3.230e-02 | -3.43 | GSE28726\_NAIVE\_CD4\_TCELL\_VS\_NAIVE\_NKTCELL\_UP | MSigDB lists | GSE28726\_NAIVE\_CD4\_TCELL\_VS\_NAIVE\_NKTCELL\_UP | 162 | 6 | 12187 | 179 | Ccl4,Cflar,Nfe2l2,Ier3,Atf3,Nfkb1 | | 3.230e-02 | -3.43 | GSE10273\_HIGH\_IL7\_VS\_HIGH\_IL7\_AND\_IRF4\_IN\_IRF4\_8\_NULL\_PRE\_BCELL\_DN | MSigDB lists | GSE10273\_HIGH\_IL7\_VS\_HIGH\_IL7\_AND\_IRF4\_IN\_IRF4\_8\_NULL\_PRE\_BCELL\_DN | 162 | 6 | 12187 | 179 | Cdkn1a,Sgk1,Lgals9,Tnfaip3,Pygm,Il17ra | | 3.230e-02 | -3.43 | GSE15330\_HSC\_VS\_PRO\_BCELL\_DN | MSigDB lists | GSE15330\_HSC\_VS\_PRO\_BCELL\_DN | 162 | 6 | 12187 | 179 | Il4ra,Ncf1,Angptl4,Tiparp,Ier3,Parp9 | | 3.230e-02 | -3.43 | GSE19825\_CD24LOW\_VS\_IL2RA\_HIGH\_DAY3\_EFF\_CD8\_TCELL\_UP | MSigDB lists | GSE19825\_CD24LOW\_VS\_IL2RA\_HIGH\_DAY3\_EFF\_CD8\_TCELL\_UP | 162 | 6 | 12187 | 179 | Dtx3l,Nfkbie,Ncf1,Il4ra,Casp4,Rassf4 | | 3.230e-02 | -3.43 | GSE34006\_UNTREATED\_VS\_A2AR\_AGONIST\_TREATED\_TREG\_DN | MSigDB lists | GSE34006\_UNTREATED\_VS\_A2AR\_AGONIST\_TREATED\_TREG\_DN | 162 | 6 | 12187 | 179 | Sntb2,Ccl4,Socs3,Gadd45g,Lcn2,Errfi1 | | 3.241e-02 | -3.43 | GO\_EXTRINSIC\_COMPONENT\_OF\_MEMBRANE | MSigDB lists | GO\_EXTRINSIC\_COMPONENT\_OF\_MEMBRANE | 206 | 7 | 12187 | 179 | Errfi1,Plaur,Msn,Iqgap1,Pik3r5,Ncf1,Pik3r1 | | 3.241e-02 | -3.43 | AP1\_Q4 | MSigDB lists | AP1\_Q4 | 206 | 7 | 12187 | 179 | Trib1,Tagln2,Cdkn1a,Il6,Znfx1,Bach1,Il1rn | | 3.247e-02 | -3.43 | GO\_TRANSITION\_METAL\_ION\_HOMEOSTASIS | MSigDB lists | GO\_TRANSITION\_METAL\_ION\_HOMEOSTASIS | 82 | 4 | 12187 | 179 | Lcn2,S100a8,Cp,S100a9 | | 3.247e-02 | -3.43 | GO\_REGULATION\_OF\_ADAPTIVE\_IMMUNE\_RESPONSE | MSigDB lists | GO\_REGULATION\_OF\_ADAPTIVE\_IMMUNE\_RESPONSE | 82 | 4 | 12187 | 179 | Il4ra,Tnf,Irf7,Tnfaip3 | | 3.252e-02 | -3.43 | BBOX | smart domains | SM00336 | 41 | 3 | 7188 | 123 | Trim56,Trim30d,Trim30a | | 3.286e-02 | -3.42 | GSE21927\_SPLENIC\_VS\_TUMOR\_MONOCYTES\_FROM\_C26GM\_TUMOROUS\_MICE\_BALBC\_UP | MSigDB lists | GSE21927\_SPLENIC\_VS\_TUMOR\_MONOCYTES\_FROM\_C26GM\_TUMOROUS\_MICE\_BALBC\_UP | 121 | 5 | 12187 | 179 | Plaur,Hspb1,Ptges,Bbs12,Ifitm2 | | 3.286e-02 | -3.42 | BROWNE\_HCMV\_INFECTION\_24HR\_UP | MSigDB lists | BROWNE\_HCMV\_INFECTION\_24HR\_UP | 121 | 5 | 12187 | 179 | Cxcl1,Nfkbia,Icam1,Il17ra,Plek | | 3.292e-02 | -3.41 | positive regulation of cell cycle G1/S phase transition | biological process | GO:1902808 | 45 | 3 | 13711 | 214 | Kcna5,Ccnd2,Adamts1 | | 3.292e-02 | -3.41 | skeletal muscle cell differentiation | biological process | GO:0035914 | 45 | 3 | 13711 | 214 | Maff,Sox11,Atf3 | | 3.307e-02 | -3.41 | Activation of IRF3/IRF7 mediated by TBK1/IKK epsilon | REACTOME pathways | R-MMU-936964 | 2 | 1 | 6297 | 105 | Irf7 | | 3.307e-02 | -3.41 | Transfer of LPS from LBP carrier to CD14 | REACTOME pathways | R-MMU-166020 | 2 | 1 | 6297 | 105 | Cd14 | | 3.307e-02 | -3.41 | Hydroxycarboxylic acid-binding receptors | REACTOME pathways | R-MMU-3296197 | 2 | 1 | 6297 | 105 | Hcar2 | | 3.307e-02 | -3.41 | Butyrophilin (BTN) family interactions | REACTOME pathways | R-MMU-8851680 | 2 | 1 | 6297 | 105 | Xdh | | 3.314e-02 | -3.41 | HUTTMANN\_B\_CLL\_POOR\_SURVIVAL\_UP | MSigDB lists | HUTTMANN\_B\_CLL\_POOR\_SURVIVAL\_UP | 207 | 7 | 12187 | 179 | Irf7,Timp1,Akap12,Cebpd,Cdkn1a,Thbs1,Ifi44 | | 3.314e-02 | -3.41 | GSE17721\_LPS\_VS\_POLYIC\_12H\_BMDC\_DN | MSigDB lists | GSE17721\_LPS\_VS\_POLYIC\_12H\_BMDC\_DN | 163 | 6 | 12187 | 179 | Vcam1,Rhou,Tnfsf10,Slfn3,Slfn4,Rhoc | | 3.314e-02 | -3.41 | GSE18893\_TCONV\_VS\_TREG\_24H\_CULTURE\_UP | MSigDB lists | GSE18893\_TCONV\_VS\_TREG\_24H\_CULTURE\_UP | 163 | 6 | 12187 | 179 | Hspb1,Plaur,Tagln2,Rassf4,Tubb6,Sgk1 | | 3.314e-02 | -3.41 | KAECH\_NAIVE\_VS\_DAY15\_EFF\_CD8\_TCELL\_DN | MSigDB lists | KAECH\_NAIVE\_VS\_DAY15\_EFF\_CD8\_TCELL\_DN | 163 | 6 | 12187 | 179 | Gbp3,Errfi1,Xdh,Ccl4,Casp4,Sntb2 | | 3.314e-02 | -3.41 | GSE42724\_NAIVE\_VS\_B1\_BCELL\_UP | MSigDB lists | GSE42724\_NAIVE\_VS\_B1\_BCELL\_UP | 163 | 6 | 12187 | 179 | Parp9,Osmr,Nfkbie,Tnfaip3,Tnf,Rhou | | 3.314e-02 | -3.41 | GSE36392\_TYPE\_2\_MYELOID\_VS\_NEUTROPHIL\_IL25\_TREATED\_LUNG\_UP | MSigDB lists | GSE36392\_TYPE\_2\_MYELOID\_VS\_NEUTROPHIL\_IL25\_TREATED\_LUNG\_UP | 163 | 6 | 12187 | 179 | Hcar2,Il1a,Stx11,Birc3,Robo4,Trim56 | | 3.314e-02 | -3.41 | GSE23984\_CTRL\_VS\_HYPOCALEMIC\_VITAMIND\_ANALOG\_TCELL\_UP | MSigDB lists | GSE23984\_CTRL\_VS\_HYPOCALEMIC\_VITAMIND\_ANALOG\_TCELL\_UP | 163 | 6 | 12187 | 179 | Pik3r5,Ptges,Lcn2,Rasip1,Cp,Msr1 | | 3.314e-02 | -3.41 | GSE369\_SOCS3\_KO\_VS\_WT\_LIVER\_POST\_IL6\_INJECTION\_DN | MSigDB lists | GSE369\_SOCS3\_KO\_VS\_WT\_LIVER\_POST\_IL6\_INJECTION\_DN | 163 | 6 | 12187 | 179 | Gbp3,Pik3ap1,Nfkbie,Stx11,Ccnd2,Birc3 | | 3.314e-02 | -3.41 | GSE11924\_TFH\_VS\_TH17\_CD4\_TCELL\_UP | MSigDB lists | GSE11924\_TFH\_VS\_TH17\_CD4\_TCELL\_UP | 163 | 6 | 12187 | 179 | Znfx1,Msn,Iqgap1,Slfn5,Ifit3b,Aff1 | | 3.314e-02 | -3.41 | GSE1460\_DP\_THYMOCYTE\_VS\_THYMIC\_STROMAL\_CELL\_DN | MSigDB lists | GSE1460\_DP\_THYMOCYTE\_VS\_THYMIC\_STROMAL\_CELL\_DN | 163 | 6 | 12187 | 179 | Ccnd2,Ifitm2,Sntb2,Arid5b,Osmr,Atf3 | | 3.314e-02 | -3.41 | GSE29164\_DAY3\_VS\_DAY7\_CD8\_TCELL\_AND\_IL12\_TREATED\_MELANOMA\_UP | MSigDB lists | GSE29164\_DAY3\_VS\_DAY7\_CD8\_TCELL\_AND\_IL12\_TREATED\_MELANOMA\_UP | 163 | 6 | 12187 | 179 | Fzd4,Ptpn1,Igsf6,Usp18,Lcn2,Oasl1 | | 3.314e-02 | -3.41 | GSE17721\_POLYIC\_VS\_GARDIQUIMOD\_16H\_BMDC\_UP | MSigDB lists | GSE17721\_POLYIC\_VS\_GARDIQUIMOD\_16H\_BMDC\_UP | 163 | 6 | 12187 | 179 | Csf2rb,Irf2,Ccl12,Tiparp,Sgk1,Rhou | | 3.315e-02 | -3.41 | sequence-specific DNA binding | molecular function | GO:0043565 | 824 | 20 | 13516 | 211 | Ifi209,Nfkb2,Mxd4,Nfe2l2,Sox11,Ifi204,Arid5b,Cebpd,Ifi211,Irf7,Spi1,Maff,Ifi207,Bach1,Irf9,Irf2,Runx1,Atf3,Zbtb16,Nfkb1 | | 3.327e-02 | -3.40 | MORI\_LARGE\_PRE\_BII\_LYMPHOCYTE\_DN | MSigDB lists | MORI\_LARGE\_PRE\_BII\_LYMPHOCYTE\_DN | 48 | 3 | 12187 | 179 | Birc3,Il4ra,Vcam1 | | 3.327e-02 | -3.40 | BARIS\_THYROID\_CANCER\_DN | MSigDB lists | BARIS\_THYROID\_CANCER\_DN | 48 | 3 | 12187 | 179 | Ccnd2,Tnfsf10,Timp1 | | 3.327e-02 | -3.40 | GNF2\_TNFRSF1B | MSigDB lists | GNF2\_TNFRSF1B | 48 | 3 | 12187 | 179 | Cybb,Igsf6,Ms4a6d | | 3.327e-02 | -3.40 | GO\_SECRETORY\_GRANULE\_LUMEN | MSigDB lists | GO\_SECRETORY\_GRANULE\_LUMEN | 48 | 3 | 12187 | 179 | A2m,Thbs1,Timp1 | | 3.327e-02 | -3.40 | DAVICIONI\_PAX\_FOXO1\_SIGNATURE\_IN\_ARMS\_UP | MSigDB lists | DAVICIONI\_PAX\_FOXO1\_SIGNATURE\_IN\_ARMS\_UP | 48 | 3 | 12187 | 179 | Sgk1,Il4ra,Tnfaip3 | | 3.337e-02 | -3.40 | Small\_GTPase | interpro domains | IPR001806 | 116 | 5 | 13788 | 212 | Rhou,Rhoj,Gem,Rasd1,Rhoc | | 3.353e-02 | -3.40 | Ifnar2 (interferon (alpha and beta) receptor 2) | protein interactions | 15976 | 2 | 1 | 6802 | 115 | Irf9 | | 3.353e-02 | -3.40 | Spp1 (secreted phosphoprotein 1) | protein interactions | 20750 | 2 | 1 | 6802 | 115 | Pik3r1 | | 3.353e-02 | -3.40 | C7orf50 (chromosome 7 open reading frame 50) | protein interactions | 84310 | 2 | 1 | 6802 | 115 | Ksr1 | | 3.353e-02 | -3.40 | E2F1 (E2F transcription factor 1) | protein interactions | 1869 | 2 | 1 | 6802 | 115 | Nfkb1 | | 3.353e-02 | -3.40 | CGAS (cyclic GMP-AMP synthase) | protein interactions | 115004 | 2 | 1 | 6802 | 115 | Trim56 | | 3.353e-02 | -3.40 | Parp12 (poly (ADP-ribose) polymerase family, member 12) | protein interactions | 243771 | 2 | 1 | 6802 | 115 | Parp12 | | 3.353e-02 | -3.40 | MSH6 (mutS homolog 6) | protein interactions | 2956 | 2 | 1 | 6802 | 115 | Ksr1 | | 3.353e-02 | -3.40 | CCNT2 (cyclin T2) | protein interactions | 905 | 2 | 1 | 6802 | 115 | Aff1 | | 3.353e-02 | -3.40 | MRPS22 (mitochondrial ribosomal protein S22) | protein interactions | 56945 | 2 | 1 | 6802 | 115 | Ksr1 | | 3.353e-02 | -3.40 | DNAJC7 (DnaJ heat shock protein family (Hsp40) member C7) | protein interactions | 7266 | 2 | 1 | 6802 | 115 | Ksr1 | | 3.353e-02 | -3.40 | EWSR1 (EWS RNA binding protein 1) | protein interactions | 2130 | 2 | 1 | 6802 | 115 | Ksr1 | | 3.353e-02 | -3.40 | Ripk3 (receptor-interacting serine-threonine kinase 3) | protein interactions | 56532 | 2 | 1 | 6802 | 115 | Ripk1 | | 3.353e-02 | -3.40 | Fem1a (feminization 1 homolog a (C. elegans)) | protein interactions | 14154 | 2 | 1 | 6802 | 115 | Nfkb1 | | 3.353e-02 | -3.40 | ERBB2 (erb-b2 receptor tyrosine kinase 2) | protein interactions | 2064 | 2 | 1 | 6802 | 115 | Errfi1 | | 3.353e-02 | -3.40 | HSPA6 (heat shock protein family A (Hsp70) member 6) | protein interactions | 3310 | 2 | 1 | 6802 | 115 | Ksr1 | | 3.353e-02 | -3.40 | ACKR2 (atypical chemokine receptor 2) | protein interactions | 1238 | 2 | 1 | 6802 | 115 | Ccl2 | | 3.353e-02 | -3.40 | SNCAIP (synuclein alpha interacting protein) | protein interactions | 9627 | 2 | 1 | 6802 | 115 | Hspa5 | | 3.353e-02 | -3.40 | ZFP64 (ZFP64 zinc finger protein) | protein interactions | 55734 | 2 | 1 | 6802 | 115 | Msn | | 3.353e-02 | -3.40 | CS (citrate synthase) | protein interactions | 1431 | 2 | 1 | 6802 | 115 | Hspb1 | | 3.353e-02 | -3.40 | MLLT1 (MLLT1 super elongation complex subunit) | protein interactions | 4298 | 2 | 1 | 6802 | 115 | Aff1 | | 3.353e-02 | -3.40 | MTDH (metadherin) | protein interactions | 92140 | 2 | 1 | 6802 | 115 | Ksr1 | | 3.353e-02 | -3.40 | NCOA6 (nuclear receptor coactivator 6) | protein interactions | 23054 | 2 | 1 | 6802 | 115 | Rbm39 | | 3.353e-02 | -3.40 | Tradd (TNFRSF1A-associated via death domain) | protein interactions | 71609 | 2 | 1 | 6802 | 115 | Tnf | | 3.353e-02 | -3.40 | MAGED1 (MAGE family member D1) | protein interactions | 9500 | 2 | 1 | 6802 | 115 | Ksr1 | | 3.353e-02 | -3.40 | Dnajc10 (DnaJ heat shock protein family (Hsp40) member C10) | protein interactions | 66861 | 2 | 1 | 6802 | 115 | Hspa5 | | 3.353e-02 | -3.40 | Sele (selectin, endothelial cell) | protein interactions | 20339 | 2 | 1 | 6802 | 115 | Nfkb1 | | 3.353e-02 | -3.40 | Nek1 (NIMA (never in mitosis gene a)-related expressed kinase 1) | protein interactions | 18004 | 2 | 1 | 6802 | 115 | Ripk1 | | 3.353e-02 | -3.40 | EIF2S2 (eukaryotic translation initiation factor 2 subunit beta) | protein interactions | 8894 | 2 | 1 | 6802 | 115 | Ksr1 | | 3.353e-02 | -3.40 | TRIM33 (tripartite motif containing 33) | protein interactions | 51592 | 2 | 1 | 6802 | 115 | Nfe2l2 | | 3.353e-02 | -3.40 | Rhbdd3 (rhomboid domain containing 3) | protein interactions | 279766 | 2 | 1 | 6802 | 115 | Tnfaip3 | | 3.353e-02 | -3.40 | Pold3 (polymerase (DNA-directed), delta 3, accessory subunit) | protein interactions | 67967 | 2 | 1 | 6802 | 115 | Runx1 | | 3.353e-02 | -3.40 | RDX (radixin) | protein interactions | 5962 | 2 | 1 | 6802 | 115 | Msn | | 3.353e-02 | -3.40 | Tnip3 (TNFAIP3 interacting protein 3) | protein interactions | 414084 | 2 | 1 | 6802 | 115 | Tnfaip3 | | 3.353e-02 | -3.40 | Il7r (interleukin 7 receptor) | protein interactions | 16197 | 2 | 1 | 6802 | 115 | Spi1 | | 3.353e-02 | -3.40 | PSMB5 (proteasome subunit beta 5) | protein interactions | 5693 | 2 | 1 | 6802 | 115 | Ksr1 | | 3.353e-02 | -3.40 | Skap1 (src family associated phosphoprotein 1) | protein interactions | 78473 | 2 | 1 | 6802 | 115 | Fyb | | 3.353e-02 | -3.40 | Thbs2 (thrombospondin 2) | protein interactions | 21826 | 2 | 1 | 6802 | 115 | Thbs1 | | 3.353e-02 | -3.40 | Fasl (Fas ligand (TNF superfamily, member 6)) | protein interactions | 14103 | 2 | 1 | 6802 | 115 | Cflar | | 3.353e-02 | -3.40 | MAGT1 (magnesium transporter 1) | protein interactions | 84061 | 2 | 1 | 6802 | 115 | Bach1 | | 3.353e-02 | -3.40 | Fli1 (Friend leukemia integration 1) | protein interactions | 14247 | 2 | 1 | 6802 | 115 | Runx1 | | 3.353e-02 | -3.40 | TUBB4A (tubulin beta 4A class IVa) | protein interactions | 10382 | 2 | 1 | 6802 | 115 | Ksr1 | | 3.353e-02 | -3.40 | C1orf35 (chromosome 1 open reading frame 35) | protein interactions | 79169 | 2 | 1 | 6802 | 115 | Ksr1 | | 3.353e-02 | -3.40 | HSPA2 (heat shock protein family A (Hsp70) member 2) | protein interactions | 3306 | 2 | 1 | 6802 | 115 | Ksr1 | | 3.353e-02 | -3.40 | DPP9 (dipeptidyl peptidase 9) | protein interactions | 91039 | 2 | 1 | 6802 | 115 | Bach1 | | 3.353e-02 | -3.40 | MAP2K2 (mitogen-activated protein kinase kinase 2) | protein interactions | 5605 | 2 | 1 | 6802 | 115 | Ksr1 | | 3.353e-02 | -3.40 | YTHDF2 (YTH N6-methyladenosine RNA binding protein 2) | protein interactions | 51441 | 2 | 1 | 6802 | 115 | Ksr1 | | 3.353e-02 | -3.40 | Dazap1 (DAZ associated protein 1) | protein interactions | 70248 | 2 | 1 | 6802 | 115 | Tnf | | 3.353e-02 | -3.40 | TAB2 (TGF-beta activated kinase 1 (MAP3K7) binding protein 2) | protein interactions | 23118 | 2 | 1 | 6802 | 115 | Trim30a | | 3.353e-02 | -3.40 | Zfand5 (zinc finger, AN1-type domain 5) | protein interactions | 22682 | 2 | 1 | 6802 | 115 | Tnf | | 3.353e-02 | -3.40 | ELL (elongation factor for RNA polymerase II) | protein interactions | 8178 | 2 | 1 | 6802 | 115 | Aff1 | | 3.353e-02 | -3.40 | CSNK2B (casein kinase 2 beta) | protein interactions | 1460 | 2 | 1 | 6802 | 115 | Ksr1 | | 3.353e-02 | -3.40 | PPP5C (protein phosphatase 5 catalytic subunit) | protein interactions | 5536 | 2 | 1 | 6802 | 115 | Map3k6 | | 3.353e-02 | -3.40 | DDX58 (DExD/H-box helicase 58) | protein interactions | 23586 | 2 | 1 | 6802 | 115 | Trim25 | | 3.353e-02 | -3.40 | MARCKS (myristoylated alanine rich protein kinase C substrate) | protein interactions | 4082 | 2 | 1 | 6802 | 115 | Ksr1 | | 3.353e-02 | -3.40 | Sgk3 (serum/glucocorticoid regulated kinase 3) | protein interactions | 170755 | 2 | 1 | 6802 | 115 | Sgk3 | | 3.353e-02 | -3.40 | Araf (Araf proto-oncogene, serine/threonine kinase) | protein interactions | 11836 | 2 | 1 | 6802 | 115 | Pik3r1 | | 3.353e-02 | -3.40 | Usp18 (ubiquitin specific peptidase 18) | protein interactions | 24110 | 2 | 1 | 6802 | 115 | Isg15 | | 3.353e-02 | -3.40 | UVRAG (UV radiation resistance associated) | protein interactions | 7405 | 2 | 1 | 6802 | 115 | Aff1 | | 3.353e-02 | -3.40 | Nup88 (nucleoporin 88) | protein interactions | 19069 | 2 | 1 | 6802 | 115 | Zbtb16 | | 3.353e-02 | -3.40 | Mapkapk3 (mitogen-activated protein kinase-activated protein kinase 3) | protein interactions | 102626 | 2 | 1 | 6802 | 115 | Hspb1 | | 3.353e-02 | -3.40 | Gbp2 (guanylate binding protein 2) | protein interactions | 14469 | 2 | 1 | 6802 | 115 | Iigp1 | | 3.353e-02 | -3.40 | Tial1 (Tia1 cytotoxic granule-associated RNA binding protein-like 1) | protein interactions | 21843 | 2 | 1 | 6802 | 115 | Tnf | | 3.353e-02 | -3.40 | AFF4 (AF4/FMR2 family member 4) | protein interactions | 27125 | 2 | 1 | 6802 | 115 | Aff1 | | 3.353e-02 | -3.40 | SFRP1 (secreted frizzled related protein 1) | protein interactions | 282068 | 2 | 1 | 6802 | 115 | Fzd4 | | 3.353e-02 | -3.40 | Lpar1 (lysophosphatidic acid receptor 1) | protein interactions | 14745 | 2 | 1 | 6802 | 115 | Cd14 | | 3.353e-02 | -3.40 | Dchs1 (dachsous cadherin related 1) | protein interactions | 233651 | 2 | 1 | 6802 | 115 | Adamts1 | | 3.353e-02 | -3.40 | Nmi (N-myc (and STAT) interactor) | protein interactions | 64685 | 2 | 1 | 6802 | 115 | Irf7 | | 3.353e-02 | -3.40 | Il17rc (interleukin 17 receptor C) | protein interactions | 171095 | 2 | 1 | 6802 | 115 | Il17ra | | 3.353e-02 | -3.40 | Fubp1 (far upstream element (FUSE) binding protein 1) | protein interactions | 51886 | 2 | 1 | 6802 | 115 | Tnf | | 3.353e-02 | -3.40 | Ccna1 (cyclin A1) | protein interactions | 12427 | 2 | 1 | 6802 | 115 | Cdkn1a | | 3.353e-02 | -3.40 | Prrx2 (paired related homeobox 2) | protein interactions | 20204 | 2 | 1 | 6802 | 115 | Ifi211 | | 3.353e-02 | -3.40 | FEN1 (flap structure-specific endonuclease 1) | protein interactions | 2237 | 2 | 1 | 6802 | 115 | Ksr1 | | 3.353e-02 | -3.40 | Prmt2 (protein arginine N-methyltransferase 2) | protein interactions | 15468 | 2 | 1 | 6802 | 115 | Nfkbia | | 3.353e-02 | -3.40 | Maf (avian musculoaponeurotic fibrosarcoma oncogene homolog) | protein interactions | 17132 | 2 | 1 | 6802 | 115 | Nfe2l2 | | 3.361e-02 | -3.39 | positive regulation of protein kinase B signaling | biological process | GO:0051897 | 78 | 4 | 13711 | 214 | Il6,Tnf,Thbs1,Csf3 | | 3.374e-02 | -3.39 | WANG\_HCP\_PROSTATE\_CANCER | MSigDB lists | WANG\_HCP\_PROSTATE\_CANCER | 83 | 4 | 12187 | 179 | Gadd45b,Atf3,Sgk1,Aff1 | | 3.374e-02 | -3.39 | GO\_MAINTENANCE\_OF\_LOCATION\_IN\_CELL | MSigDB lists | GO\_MAINTENANCE\_OF\_LOCATION\_IN\_CELL | 83 | 4 | 12187 | 179 | Nfkbie,Hspa5,Nfkbia,Bcl3 | | 3.379e-02 | -3.39 | morphogenesis of a branching epithelium | biological process | GO:0061138 | 154 | 6 | 13711 | 214 | Col4a1,Il6,Clic4,Rasip1,Tgm2,Socs3 | | 3.388e-02 | -3.39 | LEE\_NEURAL\_CREST\_STEM\_CELL\_UP | MSigDB lists | LEE\_NEURAL\_CREST\_STEM\_CELL\_UP | 122 | 5 | 12187 | 179 | Ifi209,Ifi211,Ifi207,Ifi204,Cebpd | | 3.388e-02 | -3.38 | chemical homeostasis | biological process | GO:0048878 | 826 | 20 | 13711 | 214 | Pygm,Slc24a4,C3ar1,Pik3r1,P2ry6,Cp,Lcn2,Icam1,Mt1,Il6,Ccl2,Vcam1,Kcna5,Angptl4,Sgk1,C5ar1,Cxcl1,Tgm2,Clic4,Mt2 | | 3.393e-02 | -3.38 | DBB | smart domains | SM01282 | 2 | 1 | 7188 | 123 | Pik3ap1 | | 3.393e-02 | -3.38 | G8 | smart domains | SM01225 | 2 | 1 | 7188 | 123 | Cemip2 | | 3.393e-02 | -3.38 | positive regulation of cellular catabolic process | biological process | GO:0031331 | 329 | 10 | 13711 | 214 | Dtx3l,Irgm1,Trib1,Msn,Irgm2,Tnfaip3,Tnf,Ptpn1,Zfp36,Il6 | | 3.399e-02 | -3.38 | TATAAA\_TATA\_01 | MSigDB lists | TATAAA\_TATA\_01 | 825 | 19 | 12187 | 179 | Angptl4,S100a9,Gem,Csf3,Tnfsf10,Tnfaip8,S100a8,Ccl4,Nfkbia,Tnf,Icam1,Apold1,Sele,Vcam1,Adamts1,Luc7l3,Mmp8,Il1rn,Cp | | 3.400e-02 | -3.38 | GSE17721\_POLYIC\_VS\_PAM3CSK4\_8H\_BMDC\_UP | MSigDB lists | GSE17721\_POLYIC\_VS\_PAM3CSK4\_8H\_BMDC\_UP | 164 | 6 | 12187 | 179 | Nfkbie,Sgk3,Ccl2,S100a8,Akap12,Irf9 | | 3.400e-02 | -3.38 | GSE17721\_LPS\_VS\_POLYIC\_16H\_BMDC\_DN | MSigDB lists | GSE17721\_LPS\_VS\_POLYIC\_16H\_BMDC\_DN | 164 | 6 | 12187 | 179 | Irf2,Parp9,Trim25,Parp12,Ccl4,Ifit3b | | 3.400e-02 | -3.38 | GSE23308\_WT\_VS\_MINERALCORTICOID\_REC\_KO\_MACROPHAGE\_CORTICOSTERONE\_TREATED\_DN | MSigDB lists | GSE23308\_WT\_VS\_MINERALCORTICOID\_REC\_KO\_MACROPHAGE\_CORTICOSTERONE\_TREATED\_DN | 164 | 6 | 12187 | 179 | Hspa5,Ccl4,Casp4,Hp,Ccl12,Msn | | 3.400e-02 | -3.38 | GSE16451\_CTRL\_VS\_WEST\_EQUINE\_ENC\_VIRUS\_MATURE\_NEURON\_CELL\_LINE\_DN | MSigDB lists | GSE16451\_CTRL\_VS\_WEST\_EQUINE\_ENC\_VIRUS\_MATURE\_NEURON\_CELL\_LINE\_DN | 164 | 6 | 12187 | 179 | Gbp3,Icam1,Gadd45g,Ifitm3,Map3k8,Ifitm2 | | 3.400e-02 | -3.38 | GSE17721\_LPS\_VS\_PAM3CSK4\_8H\_BMDC\_UP | MSigDB lists | GSE17721\_LPS\_VS\_PAM3CSK4\_8H\_BMDC\_UP | 164 | 6 | 12187 | 179 | Gbp3,Tnfsf10,Zfp36,Ripk1,Atf3,Map3k8 | | 3.414e-02 | -3.38 | negative regulation of cellular response to growth factor stimulus | biological process | GO:0090288 | 115 | 5 | 13711 | 214 | Xdh,Thbs1,Cflar,Hspa5,Ptpn1 | | 3.415e-02 | -3.38 | perinuclear region of cytoplasm | cellular component | GO:0048471 | 581 | 15 | 13825 | 212 | Cdkn1a,Ptges,Nfkbie,Ccl2,Kcna5,Gbp3,Rasip1,Rasd1,Clic4,Cybb,Noct,Sele,Msn,Pik3r1,Sgk1 | | 3.427e-02 | -3.37 | GO\_POSITIVE\_REGULATION\_OF\_EPITHELIAL\_CELL\_APOPTOTIC\_PROCESS | MSigDB lists | GO\_POSITIVE\_REGULATION\_OF\_EPITHELIAL\_CELL\_APOPTOTIC\_PROCESS | 20 | 2 | 12187 | 179 | Il6,Thbs1 | | 3.427e-02 | -3.37 | GO\_PROTEIN\_LIPID\_COMPLEX | MSigDB lists | GO\_PROTEIN\_LIPID\_COMPLEX | 20 | 2 | 12187 | 179 | Msr1,Saa1 | | 3.427e-02 | -3.37 | BIOCARTA\_CASPASE\_PATHWAY | MSigDB lists | BIOCARTA\_CASPASE\_PATHWAY | 20 | 2 | 12187 | 179 | Birc3,Casp4 | | 3.427e-02 | -3.37 | GO\_CD4\_POSITIVE\_ALPHA\_BETA\_T\_CELL\_ACTIVATION | MSigDB lists | GO\_CD4\_POSITIVE\_ALPHA\_BETA\_T\_CELL\_ACTIVATION | 20 | 2 | 12187 | 179 | Bcl3,Il6 | | 3.427e-02 | -3.37 | KANG\_FLUOROURACIL\_RESISTANCE\_UP | MSigDB lists | KANG\_FLUOROURACIL\_RESISTANCE\_UP | 20 | 2 | 12187 | 179 | Arid5b,Cdkn1a | | 3.427e-02 | -3.37 | GO\_SCAVENGER\_RECEPTOR\_ACTIVITY | MSigDB lists | GO\_SCAVENGER\_RECEPTOR\_ACTIVITY | 20 | 2 | 12187 | 179 | Msr1,Cxcl16 | | 3.427e-02 | -3.37 | ZIRN\_TRETINOIN\_RESPONSE\_WT1\_UP | MSigDB lists | ZIRN\_TRETINOIN\_RESPONSE\_WT1\_UP | 20 | 2 | 12187 | 179 | Akap12,Plaur | | 3.427e-02 | -3.37 | GNF2\_CASP4 | MSigDB lists | GNF2\_CASP4 | 20 | 2 | 12187 | 179 | Casp4,Iqgap1 | | 3.427e-02 | -3.37 | GO\_POSITIVE\_REGULATION\_OF\_RELEASE\_OF\_CYTOCHROME\_C\_FROM\_MITOCHONDRIA | MSigDB lists | GO\_POSITIVE\_REGULATION\_OF\_RELEASE\_OF\_CYTOCHROME\_C\_FROM\_MITOCHONDRIA | 20 | 2 | 12187 | 179 | Plaur,Tnfsf10 | | 3.427e-02 | -3.37 | MACLACHLAN\_BRCA1\_TARGETS\_UP | MSigDB lists | MACLACHLAN\_BRCA1\_TARGETS\_UP | 20 | 2 | 12187 | 179 | Timp1,Cdkn1a | | 3.427e-02 | -3.37 | ABE\_VEGFA\_TARGETS\_2HR | MSigDB lists | ABE\_VEGFA\_TARGETS\_2HR | 20 | 2 | 12187 | 179 | Cdkn1a,Ptgs2 | | 3.427e-02 | -3.37 | HOEGERKORP\_CD44\_TARGETS\_DIRECT\_UP | MSigDB lists | HOEGERKORP\_CD44\_TARGETS\_DIRECT\_UP | 20 | 2 | 12187 | 179 | Il6,Il1a | | 3.427e-02 | -3.37 | GO\_CELLULAR\_RESPONSE\_TO\_EPIDERMAL\_GROWTH\_FACTOR\_STIMULUS | MSigDB lists | GO\_CELLULAR\_RESPONSE\_TO\_EPIDERMAL\_GROWTH\_FACTOR\_STIMULUS | 20 | 2 | 12187 | 179 | Errfi1,Iqgap1 | | 3.427e-02 | -3.37 | GO\_NEGATIVE\_REGULATION\_OF\_LIPID\_TRANSPORT | MSigDB lists | GO\_NEGATIVE\_REGULATION\_OF\_LIPID\_TRANSPORT | 20 | 2 | 12187 | 179 | Nfkb1,Thbs1 | | 3.427e-02 | -3.37 | GO\_ANCHORED\_COMPONENT\_OF\_PLASMA\_MEMBRANE | MSigDB lists | GO\_ANCHORED\_COMPONENT\_OF\_PLASMA\_MEMBRANE | 20 | 2 | 12187 | 179 | Cp,Cd14 | | 3.427e-02 | -3.37 | GO\_CELLULAR\_RESPONSE\_TO\_INTERLEUKIN\_4 | MSigDB lists | GO\_CELLULAR\_RESPONSE\_TO\_INTERLEUKIN\_4 | 20 | 2 | 12187 | 179 | Il4ra,Hspa5 | | 3.427e-02 | -3.37 | GO\_SUPEROXIDE\_METABOLIC\_PROCESS | MSigDB lists | GO\_SUPEROXIDE\_METABOLIC\_PROCESS | 20 | 2 | 12187 | 179 | Cybb,Ncf1 | | 3.427e-02 | -3.37 | MARSHALL\_VIRAL\_INFECTION\_RESPONSE\_DN | MSigDB lists | MARSHALL\_VIRAL\_INFECTION\_RESPONSE\_DN | 20 | 2 | 12187 | 179 | Nfkbia,Sgk1 | | 3.427e-02 | -3.37 | CHIN\_BREAST\_CANCER\_COPY\_NUMBER\_UP | MSigDB lists | CHIN\_BREAST\_CANCER\_COPY\_NUMBER\_UP | 20 | 2 | 12187 | 179 | Ptpn1,Ptgs2 | | 3.471e-02 | -3.36 | negative regulation of cell activation | biological process | GO:0050866 | 155 | 6 | 13711 | 214 | Il4ra,Tnfaip3,Sox11,Runx1,Lgals9,Sdc4 | | 3.476e-02 | -3.36 | GO\_EPITHELIUM\_DEVELOPMENT | MSigDB lists | GO\_EPITHELIUM\_DEVELOPMENT | 663 | 16 | 12187 | 179 | Iqgap1,Tgm2,Spi1,Sdc4,Rhoc,Il6,Tagln2,Icam1,Clic4,Errfi1,Msn,Sox11,Cdkn1a,Socs3,Col4a1,Rasip1 | | 3.483e-02 | -3.36 | labyrinthine layer morphogenesis | biological process | GO:0060713 | 19 | 2 | 13711 | 214 | Socs3,Vcam1 | | 3.483e-02 | -3.36 | negative regulation of interleukin-2 production | biological process | GO:0032703 | 19 | 2 | 13711 | 214 | Tnfaip3,Zfp36 | | 3.483e-02 | -3.36 | positive regulation of endothelial cell apoptotic process | biological process | GO:2000353 | 19 | 2 | 13711 | 214 | Ccl12,Thbs1 | | 3.483e-02 | -3.36 | regulation of leukocyte adhesion to vascular endothelial cell | biological process | GO:1904994 | 19 | 2 | 13711 | 214 | Icam1,Tnf | | 3.483e-02 | -3.36 | positive regulation of vascular associated smooth muscle cell migration | biological process | GO:1904754 | 19 | 2 | 13711 | 214 | Adamts1,Iqgap1 | | 3.483e-02 | -3.36 | negative regulation of interleukin-1 production | biological process | GO:0032692 | 19 | 2 | 13711 | 214 | Errfi1,Tnfaip3 | | 3.483e-02 | -3.36 | negative regulation of smooth muscle contraction | biological process | GO:0045986 | 19 | 2 | 13711 | 214 | Ncf1,Ptgs2 | | 3.483e-02 | -3.36 | regulation of nuclease activity | biological process | GO:0032069 | 19 | 2 | 13711 | 214 | Oasl1,Oasl2 | | 3.483e-02 | -3.36 | lymph node development | biological process | GO:0048535 | 19 | 2 | 13711 | 214 | Nfkb2,Nfkb1 | | 3.483e-02 | -3.36 | regulation of endoplasmic reticulum unfolded protein response | biological process | GO:1900101 | 19 | 2 | 13711 | 214 | Ptpn1,Pik3r1 | | 3.483e-02 | -3.36 | peptide cross-linking | biological process | GO:0018149 | 19 | 2 | 13711 | 214 | Tgm2,Thbs1 | | 3.483e-02 | -3.36 | regulation of endothelial cell chemotaxis | biological process | GO:2001026 | 19 | 2 | 13711 | 214 | Hspb1,Thbs1 | | 3.483e-02 | -3.36 | positive regulation of glial cell proliferation | biological process | GO:0060252 | 19 | 2 | 13711 | 214 | Il6,Tnf | | 3.483e-02 | -3.36 | regulation of membrane protein ectodomain proteolysis | biological process | GO:0051043 | 19 | 2 | 13711 | 214 | Tnf,Timp1 | | 3.484e-02 | -3.36 | mitogen-activated protein kinase kinase binding | molecular function | GO:0031434 | 19 | 2 | 13516 | 211 | Trib1,Ksr1 | | 3.484e-02 | -3.36 | chemokine binding | molecular function | GO:0019956 | 19 | 2 | 13516 | 211 | Zfp36,A2m | | 3.487e-02 | -3.36 | GSE3982\_MAC\_VS\_NKCELL\_UP | MSigDB lists | GSE3982\_MAC\_VS\_NKCELL\_UP | 165 | 6 | 12187 | 179 | C5ar1,Cdkn1a,Trib1,Tor3a,Rgs16,Plaur | | 3.487e-02 | -3.36 | GSE17580\_UNINFECTED\_VS\_S\_MANSONI\_INF\_TREG\_UP | MSigDB lists | GSE17580\_UNINFECTED\_VS\_S\_MANSONI\_INF\_TREG\_UP | 165 | 6 | 12187 | 179 | Ms4a6d,Oasl1,Rasip1,Ifit2,Nfkbiz,Irf7 | | 3.487e-02 | -3.36 | GSE12845\_IGD\_NEG\_BLOOD\_VS\_PRE\_GC\_TONSIL\_BCELL\_DN | MSigDB lists | GSE12845\_IGD\_NEG\_BLOOD\_VS\_PRE\_GC\_TONSIL\_BCELL\_DN | 165 | 6 | 12187 | 179 | Sntb2,Map3k8,Gem,Tnfaip3,Ier3,Zfp36 | | 3.487e-02 | -3.36 | GOLDRATH\_NAIVE\_VS\_MEMORY\_CD8\_TCELL\_DN | MSigDB lists | GOLDRATH\_NAIVE\_VS\_MEMORY\_CD8\_TCELL\_DN | 165 | 6 | 12187 | 179 | Errfi1,Ier3,Cybb,Ccl4,Casp4,Map3k8 | | 3.487e-02 | -3.36 | GSE17186\_CD21LOW\_VS\_CD21HIGH\_TRANSITIONAL\_BCELL\_UP | MSigDB lists | GSE17186\_CD21LOW\_VS\_CD21HIGH\_TRANSITIONAL\_BCELL\_UP | 165 | 6 | 12187 | 179 | Rbm47,Tubb6,Slfn5,Ifi44,Birc3,Tnf | | 3.487e-02 | -3.36 | GSE43955\_TH0\_VS\_TGFB\_IL6\_IL23\_TH17\_ACT\_CD4\_TCELL\_60H\_UP | MSigDB lists | GSE43955\_TH0\_VS\_TGFB\_IL6\_IL23\_TH17\_ACT\_CD4\_TCELL\_60H\_UP | 165 | 6 | 12187 | 179 | Thbs1,Gem,Tagln2,Cmpk2,Plaur,Errfi1 | | 3.487e-02 | -3.36 | GSE37532\_TREG\_VS\_TCONV\_CD4\_TCELL\_FROM\_LN\_DN | MSigDB lists | GSE37532\_TREG\_VS\_TCONV\_CD4\_TCELL\_FROM\_LN\_DN | 165 | 6 | 12187 | 179 | Irgm2,Icam1,Nfkbiz,Nfkbia,Map3k8,Stx11 | | 3.487e-02 | -3.36 | GSE20727\_H2O2\_VS\_ROS\_INHIBITOR\_TREATED\_DC\_UP | MSigDB lists | GSE20727\_H2O2\_VS\_ROS\_INHIBITOR\_TREATED\_DC\_UP | 165 | 6 | 12187 | 179 | Il4ra,Sgk1,Ifit2,Trim56,Pik3r5,Cflar | | 3.487e-02 | -3.36 | GSE10240\_CTRL\_VS\_IL17\_STIM\_PRIMARY\_BRONCHIAL\_EPITHELIAL\_CELLS\_DN | MSigDB lists | GSE10240\_CTRL\_VS\_IL17\_STIM\_PRIMARY\_BRONCHIAL\_EPITHELIAL\_CELLS\_DN | 165 | 6 | 12187 | 179 | Tnfaip3,Tgm2,Ccl4,Il4ra,Lgals9,Irf2 | | 3.487e-02 | -3.36 | GSE24142\_ADULT\_VS\_FETAL\_EARLY\_THYMIC\_PROGENITOR\_UP | MSigDB lists | GSE24142\_ADULT\_VS\_FETAL\_EARLY\_THYMIC\_PROGENITOR\_UP | 165 | 6 | 12187 | 179 | Usp18,Map3k8,Ifi44,Birc3,Mxd4,P2ry6 | | 3.487e-02 | -3.36 | GSE2770\_TGFB\_AND\_IL4\_VS\_IL12\_TREATED\_ACT\_CD4\_TCELL\_48H\_DN | MSigDB lists | GSE2770\_TGFB\_AND\_IL4\_VS\_IL12\_TREATED\_ACT\_CD4\_TCELL\_48H\_DN | 165 | 6 | 12187 | 179 | Pik3r5,Zfp36,Rbm39,Sgk1,Ccl4,Trib1 | | 3.487e-02 | -3.36 | GSE4748\_CTRL\_VS\_LPS\_AND\_CYANOBACTERIUM\_LPSLIKE\_STIM\_DC\_3H\_UP | MSigDB lists | GSE4748\_CTRL\_VS\_LPS\_AND\_CYANOBACTERIUM\_LPSLIKE\_STIM\_DC\_3H\_UP | 165 | 6 | 12187 | 179 | Saa1,Rgs16,Errfi1,Sgk1,Lcn2,Akap12 | | 3.490e-02 | -3.36 | WANG\_LMO4\_TARGETS\_DN | MSigDB lists | WANG\_LMO4\_TARGETS\_DN | 303 | 9 | 12187 | 179 | Irf9,Ccdc58,Iqgap1,Phf11d,Dtx3l,Ifitm3,Znfx1,Nfkbiz,Phf11b | | 3.491e-02 | -3.36 | NING\_CHRONIC\_OBSTRUCTIVE\_PULMONARY\_DISEASE\_UP | MSigDB lists | NING\_CHRONIC\_OBSTRUCTIVE\_PULMONARY\_DISEASE\_UP | 123 | 5 | 12187 | 179 | Cdkn1a,A2m,S100a9,Tnfsf10,Fam107a | | 3.491e-02 | -3.36 | BIDUS\_METASTASIS\_DN | MSigDB lists | BIDUS\_METASTASIS\_DN | 123 | 5 | 12187 | 179 | Rnf213,Cflar,Rassf4,Ms4a6d,Hp | | 3.498e-02 | -3.35 | development of primary female sexual characteristics | biological process | GO:0046545 | 79 | 4 | 13711 | 214 | Adamts1,Tiparp,Arid5b,Fzd4 | | 3.498e-02 | -3.35 | negative regulation of leukocyte differentiation | biological process | GO:1902106 | 79 | 4 | 13711 | 214 | Pik3r1,Runx1,Trib1,Il4ra | | 3.500e-02 | -3.35 | regulation of cellular catabolic process | biological process | GO:0031329 | 623 | 16 | 13711 | 214 | Depp1,Zfp36,Il6,Tnfaip3,Tnf,Iigp1,Trib1,Dtx3l,Hp,Ptpn1,Timp1,Irgm2,Noct,Msn,Rasip1,Irgm1 | | 3.503e-02 | -3.35 | AMUNDSON\_GENOTOXIC\_SIGNATURE | MSigDB lists | AMUNDSON\_GENOTOXIC\_SIGNATURE | 84 | 4 | 12187 | 179 | Gem,Tnf,Rasd1,Irf2 | | 3.503e-02 | -3.35 | MODULE\_324 | MSigDB lists | MODULE\_324 | 84 | 4 | 12187 | 179 | Cp,Tnfaip3,S100a9,Fstl1 | | 3.506e-02 | -3.35 | ZWANG\_EGF\_INTERVAL\_UP | MSigDB lists | ZWANG\_EGF\_INTERVAL\_UP | 49 | 3 | 12187 | 179 | Slfn5,Rasip1,Osmr | | 3.506e-02 | -3.35 | WILCOX\_RESPONSE\_TO\_PROGESTERONE\_DN | MSigDB lists | WILCOX\_RESPONSE\_TO\_PROGESTERONE\_DN | 49 | 3 | 12187 | 179 | Tnfsf10,Clic4,Thbs1 | | 3.506e-02 | -3.35 | ENGELMANN\_CANCER\_PROGENITORS\_DN | MSigDB lists | ENGELMANN\_CANCER\_PROGENITORS\_DN | 49 | 3 | 12187 | 179 | Gadd45b,Il4ra,Sgk1 | | 3.506e-02 | -3.35 | GALE\_APL\_WITH\_FLT3\_MUTATED\_UP | MSigDB lists | GALE\_APL\_WITH\_FLT3\_MUTATED\_UP | 49 | 3 | 12187 | 179 | Phf11d,Phf11b,Luc7l3 | | 3.514e-02 | -3.35 | positive regulation of neuron projection development | biological process | GO:0010976 | 331 | 10 | 13711 | 214 | Nfe2l2,Hspb1,S100a9,Cflar,Sgk1,Fzd4,Hspa5,Iqgap1,Il6,Adamts1 | | 3.517e-02 | -3.35 | Ras | pfam domains | PF00071 | 116 | 5 | 12881 | 201 | Rhoj,Rasd1,Gem,Rhou,Rhoc | | 3.524e-02 | -3.35 | regulation of lipid localization | biological process | GO:1905952 | 116 | 5 | 13711 | 214 | Runx1,Msr1,Il1a,Nfkbia,Thbs1 | | 3.524e-02 | -3.35 | regulation of peptidyl-serine phosphorylation | biological process | GO:0033135 | 116 | 5 | 13711 | 214 | Csf3,Tnf,Irgm2,Ptgs2,Il6 | | 3.525e-02 | -3.35 | MAPK signaling pathway | WikiPathways | WP493 | 149 | 8 | 3756 | 96 | Nfkb1,Map3k8,Cd14,Map3k6,Hspa5,Hspb1,Tnf,Il1a | | 3.542e-02 | -3.34 | Complement cascade | REACTOME pathways | R-MMU-166658 | 18 | 2 | 6297 | 105 | C5ar1,C3ar1 | | 3.542e-02 | -3.34 | Growth hormone receptor signaling | REACTOME pathways | R-MMU-982772 | 18 | 2 | 6297 | 105 | Ptpn1,Socs3 | | 3.542e-02 | -3.34 | TSP1\_rpt\_sf | interpro domains | IPR036383 | 47 | 3 | 13788 | 212 | Adamts1,Thbs1,Adamts9 | | 3.542e-02 | -3.34 | TSP1\_rpt | interpro domains | IPR000884 | 47 | 3 | 13788 | 212 | Adamts9,Adamts1,Thbs1 | | 3.556e-02 | -3.34 | Ank\_2 | pfam domains | PF12796 | 156 | 6 | 12881 | 201 | Bcl3,Nfkb1,Nfkb2,Nfkbia,Nfkbiz,Nfkbie | | 3.558e-02 | -3.34 | SCHAEFFER\_PROSTATE\_DEVELOPMENT\_48HR\_UP | MSigDB lists | SCHAEFFER\_PROSTATE\_DEVELOPMENT\_48HR\_UP | 353 | 10 | 12187 | 179 | Errfi1,Ptges,Rasd1,Cebpd,Tiparp,Gadd45g,Tor3a,Nfkbia,Aff1,Sgk1 | | 3.559e-02 | -3.34 | endoplasmic reticulum membrane | cellular component | GO:0005789 | 434 | 12 | 13825 | 212 | Tap1,Irgm2,Gm4841,Ptpn1,Iigp1,Hspa5,Gm5431,Tgtp1,Tgtp2,F830016B08Rik,Irgm1,Ifi47 | | 3.575e-02 | -3.33 | GSE7460\_CTRL\_VS\_TGFB\_TREATED\_ACT\_TCONV\_DN | MSigDB lists | GSE7460\_CTRL\_VS\_TGFB\_TREATED\_ACT\_TCONV\_DN | 166 | 6 | 12187 | 179 | Tnfsf10,Ifitm3,Ifitm2,Gem,Ncf1,Lgals9 | | 3.575e-02 | -3.33 | GSE17721\_CTRL\_VS\_GARDIQUIMOD\_24H\_BMDC\_DN | MSigDB lists | GSE17721\_CTRL\_VS\_GARDIQUIMOD\_24H\_BMDC\_DN | 166 | 6 | 12187 | 179 | Cybb,Ptpn1,Ier3,Irgm2,Parp14,Ccl2 | | 3.575e-02 | -3.33 | GSE41867\_DAY6\_EFFECTOR\_VS\_DAY30\_MEMORY\_CD8\_TCELL\_LCMV\_ARMSTRONG\_DN | MSigDB lists | GSE41867\_DAY6\_EFFECTOR\_VS\_DAY30\_MEMORY\_CD8\_TCELL\_LCMV\_ARMSTRONG\_DN | 166 | 6 | 12187 | 179 | Nfe2l2,Mxd4,Arid5b,Sgk3,Sbno2,Iqgap1 | | 3.575e-02 | -3.33 | GSE25123\_CTRL\_VS\_IL4\_AND\_ROSIGLITAZONE\_STIM\_MACROPHAGE\_UP | MSigDB lists | GSE25123\_CTRL\_VS\_IL4\_AND\_ROSIGLITAZONE\_STIM\_MACROPHAGE\_UP | 166 | 6 | 12187 | 179 | Irak3,Rtp4,Tnfsf10,Irf9,Rnf213,Usp18 | | 3.575e-02 | -3.33 | GSE20366\_EX\_VIVO\_VS\_DEC205\_CONVERSION\_UP | MSigDB lists | GSE20366\_EX\_VIVO\_VS\_DEC205\_CONVERSION\_UP | 166 | 6 | 12187 | 179 | Zfp189,Sdc4,Ifit3b,Fzd4,Stx11,Xdh | | 3.575e-02 | -3.33 | GSE17721\_PAM3CSK4\_VS\_GADIQUIMOD\_16H\_BMDC\_DN | MSigDB lists | GSE17721\_PAM3CSK4\_VS\_GADIQUIMOD\_16H\_BMDC\_DN | 166 | 6 | 12187 | 179 | Parp9,Rrp8,Isg15,Rhoc,Tor3a,Msr1 | | 3.575e-02 | -3.33 | GSE3039\_NKT\_CELL\_VS\_ALPHAALPHA\_CD8\_TCELL\_UP | MSigDB lists | GSE3039\_NKT\_CELL\_VS\_ALPHAALPHA\_CD8\_TCELL\_UP | 166 | 6 | 12187 | 179 | Hcar2,Ier3,Lcn2,S100a9,Cd14,S100a8 | | 3.575e-02 | -3.33 | GSE13306\_LAMINA\_PROPRIA\_VS\_SPLEEN\_TREG\_UP | MSigDB lists | GSE13306\_LAMINA\_PROPRIA\_VS\_SPLEEN\_TREG\_UP | 166 | 6 | 12187 | 179 | Aff1,Trim56,S100a8,Icam1,Parp12,Ptpn1 | | 3.588e-02 | -3.33 | cellular response to peptide | biological process | GO:1901653 | 198 | 7 | 13711 | 214 | Nfkb1,Ptpn1,Casp4,Icam1,Sgk1,Pik3r1,Vcam1 | | 3.596e-02 | -3.33 | PID\_PDGFRB\_PATHWAY | MSigDB lists | PID\_PDGFRB\_PATHWAY | 124 | 5 | 12187 | 179 | Pik3r5,Pik3r1,Ptpn1,Ksr1,Iqgap1 | | 3.596e-02 | -3.33 | GO\_NEGATIVE\_REGULATION\_OF\_MAPK\_CASCADE | MSigDB lists | GO\_NEGATIVE\_REGULATION\_OF\_MAPK\_CASCADE | 124 | 5 | 12187 | 179 | Irak3,Atf3,Errfi1,Ptpn1,Tnip1 | | 3.596e-02 | -3.33 | ZHANG\_BREAST\_CANCER\_PROGENITORS\_DN | MSigDB lists | ZHANG\_BREAST\_CANCER\_PROGENITORS\_DN | 124 | 5 | 12187 | 179 | A2m,Nfkbiz,Socs3,Mxd4,Cxcl16 | | 3.596e-02 | -3.33 | GSE17974\_0H\_VS\_0.5H\_IN\_VITRO\_ACT\_CD4\_TCELL\_DN | MSigDB lists | GSE17974\_0H\_VS\_0.5H\_IN\_VITRO\_ACT\_CD4\_TCELL\_DN | 124 | 5 | 12187 | 179 | Nfe2l2,Gadd45b,Spi1,Tnf,Gem | | 3.596e-02 | -3.33 | GO\_POSITIVE\_REGULATION\_OF\_EPITHELIAL\_CELL\_PROLIFERATION | MSigDB lists | GO\_POSITIVE\_REGULATION\_OF\_EPITHELIAL\_CELL\_PROLIFERATION | 124 | 5 | 12187 | 179 | Il6,Ccl12,Tnfaip3,Sox11,C5ar1 | | 3.601e-02 | -3.32 | urate biosynthesis/inosine 5'-phosphate degradation | BIOCYC pathways | MOUSE\_PWY-5695 | 6 | 1 | 823 | 5 | Xdh | | 3.634e-02 | -3.31 | Cd40 (CD40 antigen) | protein interactions | 21939 | 18 | 2 | 6802 | 115 | Birc3,Pik3r1 | | 3.634e-02 | -3.31 | Cyld (CYLD lysine 63 deubiquitinase) | protein interactions | 74256 | 18 | 2 | 6802 | 115 | Bcl3,Ripk1 | | 3.634e-02 | -3.31 | Map2k1 (mitogen-activated protein kinase kinase 1) | protein interactions | 26395 | 18 | 2 | 6802 | 115 | Cflar,Ksr1 | | 3.634e-02 | -3.31 | Mta1 (metastasis associated 1) | protein interactions | 116870 | 18 | 2 | 6802 | 115 | Tgm2,Runx1 | | 3.636e-02 | -3.31 | ZHU\_CMV\_24\_HR\_UP | MSigDB lists | ZHU\_CMV\_24\_HR\_UP | 85 | 4 | 12187 | 179 | Tap1,Nfkb1,Nfkbia,Isg15 | | 3.636e-02 | -3.31 | GSE29614\_DAY3\_VS\_DAY7\_TIV\_FLU\_VACCINE\_PBMC\_UP | MSigDB lists | GSE29614\_DAY3\_VS\_DAY7\_TIV\_FLU\_VACCINE\_PBMC\_UP | 85 | 4 | 12187 | 179 | Ptges,Ccl2,Xdh,Rasip1 | | 3.636e-02 | -3.31 | GSE40666\_STAT1\_KO\_VS\_STAT4\_KO\_CD8\_TCELL\_DN | MSigDB lists | GSE40666\_STAT1\_KO\_VS\_STAT4\_KO\_CD8\_TCELL\_DN | 85 | 4 | 12187 | 179 | Socs3,Zfp189,Icam1,Tnfaip8 | | 3.639e-02 | -3.31 | regulation of response to oxidative stress | biological process | GO:1902882 | 80 | 4 | 13711 | 214 | Hspb1,Tnf,Nfe2l2,Ripk1 | | 3.639e-02 | -3.31 | regulation of cell junction assembly | biological process | GO:1901888 | 80 | 4 | 13711 | 214 | Tnf,Fam107a,Iqgap1,Sdc4 | | 3.643e-02 | -3.31 | IL-6 signaling Pathway | WikiPathways | WP387 | 97 | 6 | 3756 | 96 | Sgk1,Hspb1,Pik3r1,Il6,Nfkb1,Socs3 | | 3.665e-02 | -3.31 | GSE23568\_ID3\_TRANSDUCED\_VS\_ID3\_KO\_CD8\_TCELL\_DN | MSigDB lists | GSE23568\_ID3\_TRANSDUCED\_VS\_ID3\_KO\_CD8\_TCELL\_DN | 167 | 6 | 12187 | 179 | Cflar,Tap1,Zfp36,Casp4,Ms4a6d,Akap12 | | 3.665e-02 | -3.31 | GSE17721\_LPS\_VS\_POLYIC\_6H\_BMDC\_DN | MSigDB lists | GSE17721\_LPS\_VS\_POLYIC\_6H\_BMDC\_DN | 167 | 6 | 12187 | 179 | Irf9,S100a8,Igsf6,Sntb2,Tor3a,Sgk1 | | 3.665e-02 | -3.31 | GSE14769\_20MIN\_VS\_360MIN\_LPS\_BMDM\_DN | MSigDB lists | GSE14769\_20MIN\_VS\_360MIN\_LPS\_BMDM\_DN | 167 | 6 | 12187 | 179 | Slc24a4,Timp1,Aff1,Hspa5,Maff,Ifitm2 | | 3.665e-02 | -3.31 | GSE28726\_ACT\_CD4\_TCELL\_VS\_ACT\_VA24NEG\_NKTCELL\_DN | MSigDB lists | GSE28726\_ACT\_CD4\_TCELL\_VS\_ACT\_VA24NEG\_NKTCELL\_DN | 167 | 6 | 12187 | 179 | Rrp8,Rbm39,Ifi44,Dnajb1,Casp4,Tnfaip3 | | 3.665e-02 | -3.31 | GSE15930\_STIM\_VS\_STIM\_AND\_TRICHOSTATINA\_24H\_CD8\_T\_CELL\_DN | MSigDB lists | GSE15930\_STIM\_VS\_STIM\_AND\_TRICHOSTATINA\_24H\_CD8\_T\_CELL\_DN | 167 | 6 | 12187 | 179 | Irf7,Ccdc58,Ifit3b,Ccl4,Ifit2,Luc7l3 | | 3.665e-02 | -3.31 | GSE11961\_GERMINAL\_CENTER\_BCELL\_DAY7\_VS\_PLASMA\_CELL\_DAY7\_DN | MSigDB lists | GSE11961\_GERMINAL\_CENTER\_BCELL\_DAY7\_VS\_PLASMA\_CELL\_DAY7\_DN | 167 | 6 | 12187 | 179 | Ifitm3,Nfkbie,Clic4,Csf2rb,Pik3ap1,Rassf4 | | 3.665e-02 | -3.31 | GSE4748\_CTRL\_VS\_CYANOBACTERIUM\_LPSLIKE\_STIM\_DC\_1H\_UP | MSigDB lists | GSE4748\_CTRL\_VS\_CYANOBACTERIUM\_LPSLIKE\_STIM\_DC\_1H\_UP | 167 | 6 | 12187 | 179 | Fstl1,Casp4,Ifit2,Ccnd2,Ifitm2,Iqgap1 | | 3.665e-02 | -3.31 | GSE17721\_LPS\_VS\_PAM3CSK4\_24H\_BMDC\_UP | MSigDB lists | GSE17721\_LPS\_VS\_PAM3CSK4\_24H\_BMDC\_UP | 167 | 6 | 12187 | 179 | Ptgs2,Parp9,Sgk1,Gadd45b,Parp12,Gadd45g | | 3.665e-02 | -3.31 | GO\_RESPONSE\_TO\_MECHANICAL\_STIMULUS | MSigDB lists | GO\_RESPONSE\_TO\_MECHANICAL\_STIMULUS | 167 | 6 | 12187 | 179 | Ptgs2,Kcna5,Nfkb1,Il6,Ccl12,Nfkbia | | 3.665e-02 | -3.31 | GSE7831\_UNSTIM\_VS\_INFLUENZA\_STIM\_PDC\_1H\_DN | MSigDB lists | GSE7831\_UNSTIM\_VS\_INFLUENZA\_STIM\_PDC\_1H\_DN | 167 | 6 | 12187 | 179 | Sgk3,Atf3,Pygm,Ptpn1,Ksr1,Nfkb1 | | 3.665e-02 | -3.31 | GSE27786\_LSK\_VS\_NKCELL\_DN | MSigDB lists | GSE27786\_LSK\_VS\_NKCELL\_DN | 167 | 6 | 12187 | 179 | Errfi1,Rbm47,Akap12,Nfkbia,Spi1,Slfn9 | | 3.674e-02 | -3.30 | DBB | prosite domains | PS51376 | 2 | 1 | 8845 | 164 | Pik3ap1 | | 3.674e-02 | -3.30 | G8 | prosite domains | PS51484 | 2 | 1 | 8845 | 164 | Cemip2 | | 3.674e-02 | -3.30 | 25A\_SYNTH\_1 | prosite domains | PS00832 | 2 | 1 | 8845 | 164 | Oasl2 | | 3.674e-02 | -3.30 | GON | prosite domains | PS51046 | 2 | 1 | 8845 | 164 | Adamts9 | | 3.674e-02 | -3.30 | AKAP\_CAM\_BD | prosite domains | PS51893 | 2 | 1 | 8845 | 164 | Akap12 | | 3.674e-02 | -3.30 | RUNT | prosite domains | PS51062 | 2 | 1 | 8845 | 164 | Runx1 | | 3.674e-02 | -3.30 | MULTICOPPER\_OXIDASE2 | prosite domains | PS00080 | 2 | 1 | 8845 | 164 | Cp | | 3.678e-02 | -3.30 | positive regulation of DNA binding | biological process | GO:0043388 | 47 | 3 | 13711 | 214 | Mmp8,Lgals9,Plaur | | 3.678e-02 | -3.30 | endothelial cell development | biological process | GO:0001885 | 47 | 3 | 13711 | 214 | Msn,Icam1,Clic4 | | 3.691e-02 | -3.30 | BREDEMEYER\_RAG\_SIGNALING\_NOT\_VIA\_ATM\_UP | MSigDB lists | BREDEMEYER\_RAG\_SIGNALING\_NOT\_VIA\_ATM\_UP | 50 | 3 | 12187 | 179 | Pik3ap1,Xaf1,Maff | | 3.691e-02 | -3.30 | GO\_REGULATION\_OF\_VASOCONSTRICTION | MSigDB lists | GO\_REGULATION\_OF\_VASOCONSTRICTION | 50 | 3 | 12187 | 179 | Icam1,Kcna5,Ptgs2 | | 3.691e-02 | -3.30 | GO\_MICROVILLUS | MSigDB lists | GO\_MICROVILLUS | 50 | 3 | 12187 | 179 | Msn,Clic4,Vcam1 | | 3.691e-02 | -3.30 | HOWLIN\_PUBERTAL\_MAMMARY\_GLAND | MSigDB lists | HOWLIN\_PUBERTAL\_MAMMARY\_GLAND | 50 | 3 | 12187 | 179 | S100a9,Msr1,Tnf | | 3.691e-02 | -3.30 | GO\_REGULATION\_OF\_ALPHA\_BETA\_T\_CELL\_ACTIVATION | MSigDB lists | GO\_REGULATION\_OF\_ALPHA\_BETA\_T\_CELL\_ACTIVATION | 50 | 3 | 12187 | 179 | Il6,Il4ra,Zbtb16 | | 3.691e-02 | -3.30 | GO\_NEGATIVE\_REGULATION\_OF\_NF\_KAPPAB\_TRANSCRIPTION\_FACTOR\_ACTIVITY | MSigDB lists | GO\_NEGATIVE\_REGULATION\_OF\_NF\_KAPPAB\_TRANSCRIPTION\_FACTOR\_ACTIVITY | 50 | 3 | 12187 | 179 | Nfkbia,Irak3,Tnfaip3 | | 3.703e-02 | -3.30 | CASORELLI\_ACUTE\_PROMYELOCYTIC\_LEUKEMIA\_UP | MSigDB lists | CASORELLI\_ACUTE\_PROMYELOCYTIC\_LEUKEMIA\_UP | 125 | 5 | 12187 | 179 | Nfkbia,Trib1,Cebpd,Zfp36,Timp1 | | 3.749e-02 | -3.28 | UBIQUITIN\_2 | prosite domains | PS50053 | 40 | 3 | 8845 | 164 | Oasl2,Oasl1,Isg15 | | 3.750e-02 | -3.28 | GO\_BIOLOGICAL\_ADHESION | MSigDB lists | GO\_BIOLOGICAL\_ADHESION | 724 | 17 | 12187 | 179 | Tnf,Ccl4,Ccl12,Il6,S100a9,Fzd4,Bcl3,S100a8,Thbs1,Vcam1,Selp,Tnip1,Icam1,Hspb1,Plek,Sele,Msn | | 3.752e-02 | -3.28 | heterocyclic compound binding | molecular function | GO:1901363 | 4074 | 76 | 13516 | 211 | Rhoc,Gbp7,Mx1,Cybb,Sox11,Sgk3,Rbm47,Tap1,P2ry6,Sgk1,Oasl2,Rasd1,Lcn2,Ripk1,Zbtb16,Nfkb1,Spi1,Maff,Bcl3,Ifi211,Irak3,Irf2,Oasl1,Ifi204,Arid5b,Parp14,Parp9,Mxd4,Hspa5,Atf3,Gbp6,Tgm2,Bbs12,Irf7,Rhoj,Noct,Bach1,Cebpd,Map3k8,Ptgs2,Znfx1,Gem,Nfkb2,Pygm,Irgm1,Ip6k2,Cmpk2,Tubb6,Luc7l3,Map3k6,Trim30a,Tor3a,Ifi207,Slfn9,Rbm39,Ksr1,Ifit2,Runx1,Tiparp,Rhou,Xdh,Irf9,Nfe2l2,Slfn5,Zfp36,Iigp1,Trib1,Gbp3,Tnfaip3,Hcar2,Tgtp1,Aff1,Tnf,Ifi209,Banp,Msn | | 3.752e-02 | -3.28 | GHANDHI\_DIRECT\_IRRADIATION\_DN | MSigDB lists | GHANDHI\_DIRECT\_IRRADIATION\_DN | 21 | 2 | 12187 | 179 | Olfml2b,Fzd4 | | 3.752e-02 | -3.28 | GO\_PHOSPHATIDYLINOSITOL\_3\_KINASE\_SIGNALING | MSigDB lists | GO\_PHOSPHATIDYLINOSITOL\_3\_KINASE\_SIGNALING | 21 | 2 | 12187 | 179 | Pik3r1,Pik3r5 | | 3.752e-02 | -3.28 | REACTOME\_IL\_RECEPTOR\_SHC\_SIGNALING | MSigDB lists | REACTOME\_IL\_RECEPTOR\_SHC\_SIGNALING | 21 | 2 | 12187 | 179 | Csf2rb,Pik3r1 | | 3.752e-02 | -3.28 | REACTOME\_ACTIVATION\_OF\_GENES\_BY\_ATF4 | MSigDB lists | REACTOME\_ACTIVATION\_OF\_GENES\_BY\_ATF4 | 21 | 2 | 12187 | 179 | Atf3,Ccl12 | | 3.752e-02 | -3.28 | PIONTEK\_PKD1\_TARGETS\_UP | MSigDB lists | PIONTEK\_PKD1\_TARGETS\_UP | 21 | 2 | 12187 | 179 | Sgk1,Errfi1 | | 3.752e-02 | -3.28 | NICK\_RESPONSE\_TO\_PROC\_TREATMENT\_DN | MSigDB lists | NICK\_RESPONSE\_TO\_PROC\_TREATMENT\_DN | 21 | 2 | 12187 | 179 | Icam1,Maff | | 3.752e-02 | -3.28 | DORSEY\_GAB2\_TARGETS | MSigDB lists | DORSEY\_GAB2\_TARGETS | 21 | 2 | 12187 | 179 | Akap12,Thbs1 | | 3.752e-02 | -3.28 | GNF2\_MSN | MSigDB lists | GNF2\_MSN | 21 | 2 | 12187 | 179 | Iqgap1,Msn | | 3.752e-02 | -3.28 | CAIRO\_PML\_TARGETS\_BOUND\_BY\_MYC\_UP | MSigDB lists | CAIRO\_PML\_TARGETS\_BOUND\_BY\_MYC\_UP | 21 | 2 | 12187 | 179 | Ccnd2,Thbs1 | | 3.752e-02 | -3.28 | RODRIGUES\_NTN1\_AND\_DCC\_TARGETS | MSigDB lists | RODRIGUES\_NTN1\_AND\_DCC\_TARGETS | 21 | 2 | 12187 | 179 | Bcl3,Nfkbiz | | 3.752e-02 | -3.28 | GO\_PODOSOME | MSigDB lists | GO\_PODOSOME | 21 | 2 | 12187 | 179 | Rhou,Vcam1 | | 3.752e-02 | -3.28 | GO\_POSITIVE\_REGULATION\_OF\_INTERLEUKIN\_2\_PRODUCTION | MSigDB lists | GO\_POSITIVE\_REGULATION\_OF\_INTERLEUKIN\_2\_PRODUCTION | 21 | 2 | 12187 | 179 | Runx1,Il1a | | 3.752e-02 | -3.28 | GO\_REGULATION\_OF\_ANION\_TRANSMEMBRANE\_TRANSPORT | MSigDB lists | GO\_REGULATION\_OF\_ANION\_TRANSMEMBRANE\_TRANSPORT | 21 | 2 | 12187 | 179 | Thbs1,Ripk1 | | 3.752e-02 | -3.28 | REACTOME\_ANTIGEN\_PRESENTATION\_FOLDING\_ASSEMBLY\_AND\_PEPTIDE\_LOADING\_OF\_CLASS\_I\_MHC | MSigDB lists | REACTOME\_ANTIGEN\_PRESENTATION\_FOLDING\_ASSEMBLY\_AND\_PEPTIDE\_LOADING\_OF\_CLASS\_I\_MHC | 21 | 2 | 12187 | 179 | Tap1,Hspa5 | | 3.752e-02 | -3.28 | GO\_APOPTOTIC\_CELL\_CLEARANCE | MSigDB lists | GO\_APOPTOTIC\_CELL\_CLEARANCE | 21 | 2 | 12187 | 179 | Thbs1,Tgm2 | | 3.752e-02 | -3.28 | OUELLET\_CULTURED\_OVARIAN\_CANCER\_INVASIVE\_VS\_LMP\_DN | MSigDB lists | OUELLET\_CULTURED\_OVARIAN\_CANCER\_INVASIVE\_VS\_LMP\_DN | 21 | 2 | 12187 | 179 | Trib1,Cflar | | 3.752e-02 | -3.28 | DORN\_ADENOVIRUS\_INFECTION\_12HR\_UP | MSigDB lists | DORN\_ADENOVIRUS\_INFECTION\_12HR\_UP | 21 | 2 | 12187 | 179 | Ifit2,Nfkb2 | | 3.752e-02 | -3.28 | RORIE\_TARGETS\_OF\_EWSR1\_FLI1\_FUSION\_UP | MSigDB lists | RORIE\_TARGETS\_OF\_EWSR1\_FLI1\_FUSION\_UP | 21 | 2 | 12187 | 179 | Gadd45b,Ccnd2 | | 3.752e-02 | -3.28 | SHI\_SPARC\_TARGETS\_UP | MSigDB lists | SHI\_SPARC\_TARGETS\_UP | 21 | 2 | 12187 | 179 | Pik3r1,Gadd45b | | 3.756e-02 | -3.28 | GSE29164\_UNTREATED\_VS\_CD8\_TCELL\_TREATED\_MELANOMA\_DAY7\_UP | MSigDB lists | GSE29164\_UNTREATED\_VS\_CD8\_TCELL\_TREATED\_MELANOMA\_DAY7\_UP | 168 | 6 | 12187 | 179 | Xdh,Icam1,Zbtb16,Vcam1,Map3k8,Gem | | 3.756e-02 | -3.28 | GSE7831\_1H\_VS\_4H\_INFLUENZA\_STIM\_PDC\_UP | MSigDB lists | GSE7831\_1H\_VS\_4H\_INFLUENZA\_STIM\_PDC\_UP | 168 | 6 | 12187 | 179 | Fstl1,Sgk1,Tnip1,Gem,Ifitm3,Irak3 | | 3.756e-02 | -3.28 | GSE3982\_EFF\_MEMORY\_CD4\_TCELL\_VS\_TH1\_DN | MSigDB lists | GSE3982\_EFF\_MEMORY\_CD4\_TCELL\_VS\_TH1\_DN | 168 | 6 | 12187 | 179 | Tap1,Tubb6,Tor3a,Sdc4,Tnf,Nfkbie | | 3.756e-02 | -3.28 | GSE3337\_CTRL\_VS\_16H\_IFNG\_IN\_CD8POS\_DC\_DN | MSigDB lists | GSE3337\_CTRL\_VS\_16H\_IFNG\_IN\_CD8POS\_DC\_DN | 168 | 6 | 12187 | 179 | Ccl12,Tap1,Cmpk2,Bcl3,Irgm2,Gbp7 | | 3.756e-02 | -3.28 | GSE16522\_ANTI\_CD3CD28\_STIM\_VS\_UNSTIM\_NAIVE\_CD8\_TCELL\_DN | MSigDB lists | GSE16522\_ANTI\_CD3CD28\_STIM\_VS\_UNSTIM\_NAIVE\_CD8\_TCELL\_DN | 168 | 6 | 12187 | 179 | Rasip1,Ptpn1,Dtx3l,Lgals9,Usp18,Banp | | 3.756e-02 | -3.28 | GSE24142\_ADULT\_VS\_FETAL\_DN3\_THYMOCYTE\_UP | MSigDB lists | GSE24142\_ADULT\_VS\_FETAL\_DN3\_THYMOCYTE\_UP | 168 | 6 | 12187 | 179 | Cdkn1a,Gbp6,Bcl3,Tiparp,Ms4a6d,Gbp4 | | 3.756e-02 | -3.28 | GSE17721\_POLYIC\_VS\_GARDIQUIMOD\_6H\_BMDC\_UP | MSigDB lists | GSE17721\_POLYIC\_VS\_GARDIQUIMOD\_6H\_BMDC\_UP | 168 | 6 | 12187 | 179 | Icam1,Csf2rb,Gbp6,Gbp4,Tor3a,Usp18 | | 3.756e-02 | -3.28 | GSE2770\_UNTREATED\_VS\_IL12\_TREATED\_ACT\_CD4\_TCELL\_2H\_UP | MSigDB lists | GSE2770\_UNTREATED\_VS\_IL12\_TREATED\_ACT\_CD4\_TCELL\_2H\_UP | 168 | 6 | 12187 | 179 | Tagln2,Ptgs2,Sbno2,Iqgap1,Zbtb16,Dnajb1 | | 3.771e-02 | -3.28 | WOO\_LIVER\_CANCER\_RECURRENCE\_UP | MSigDB lists | WOO\_LIVER\_CANCER\_RECURRENCE\_UP | 86 | 4 | 12187 | 179 | Gem,Ccnd2,Ccl12,Plaur | | 3.783e-02 | -3.27 | female pregnancy | biological process | GO:0007565 | 81 | 4 | 13711 | 214 | Acod1,Ptgs2,A2m,Lgals9 | | 3.783e-02 | -3.27 | negative regulation of protein modification by small protein conjugation or removal | biological process | GO:1903321 | 81 | 4 | 13711 | 214 | Dtx3l,Tnfaip3,Gbp4,Isg15 | | 3.812e-02 | -3.27 | GSE32901\_NAIVE\_VS\_TH1\_CD4\_TCELL\_DN | MSigDB lists | GSE32901\_NAIVE\_VS\_TH1\_CD4\_TCELL\_DN | 126 | 5 | 12187 | 179 | Plek,Cybb,Nfkbiz,Ccl4,Tnf | | 3.812e-02 | -3.27 | CYCLIN\_D1\_UP.V1\_UP | MSigDB lists | CYCLIN\_D1\_UP.V1\_UP | 126 | 5 | 12187 | 179 | Tap1,Msn,Cebpd,Ksr1,Trim25 | | 3.812e-02 | -3.27 | NCX\_01 | MSigDB lists | NCX\_01 | 126 | 5 | 12187 | 179 | A2m,Runx1,Slc24a4,P2ry6,Ip6k2 | | 3.812e-02 | -3.27 | PDGF\_ERK\_DN.V1\_DN | MSigDB lists | PDGF\_ERK\_DN.V1\_DN | 126 | 5 | 12187 | 179 | Nfkbia,Dnajb1,Trib1,Maff,Irf2 | | 3.812e-02 | -3.27 | GSE13411\_NAIVE\_VS\_IGM\_MEMORY\_BCELL\_UP | MSigDB lists | GSE13411\_NAIVE\_VS\_IGM\_MEMORY\_BCELL\_UP | 126 | 5 | 12187 | 179 | Il4ra,Thbs1,Clic4,Tnfaip3,Hspb1 | | 3.829e-02 | -3.26 | CARD | pfam domains | PF00619 | 20 | 2 | 12881 | 201 | Casp4,Birc3 | | 3.831e-02 | -3.26 | positive regulation of syncytium formation by plasma membrane fusion | biological process | GO:0060143 | 20 | 2 | 13711 | 214 | Cxcl9,Il4ra | | 3.831e-02 | -3.26 | behavioral response to pain | biological process | GO:0048266 | 20 | 2 | 13711 | 214 | Runx1,Thbs1 | | 3.831e-02 | -3.26 | positive regulation of epidermal growth factor receptor signaling pathway | biological process | GO:0045742 | 20 | 2 | 13711 | 214 | Ncf1,Plaur | | 3.831e-02 | -3.26 | positive regulation of extracellular matrix organization | biological process | GO:1903055 | 20 | 2 | 13711 | 214 | Il6,Cflar | | 3.831e-02 | -3.26 | negative regulation of oxidoreductase activity | biological process | GO:0051354 | 20 | 2 | 13711 | 214 | Nfkb1,Hp | | 3.831e-02 | -3.26 | positive regulation of tumor necrosis factor secretion | biological process | GO:1904469 | 20 | 2 | 13711 | 214 | Mmp8,Akap12 | | 3.831e-02 | -3.26 | inflammatory response to antigenic stimulus | biological process | GO:0002437 | 20 | 2 | 13711 | 214 | Icam1,Tnf | | 3.831e-02 | -3.26 | regulation of nitric-oxide synthase biosynthetic process | biological process | GO:0051769 | 20 | 2 | 13711 | 214 | Akap12,Ccl2 | | 3.831e-02 | -3.26 | regulation of regulatory T cell differentiation | biological process | GO:0045589 | 20 | 2 | 13711 | 214 | Ctla2a,Lgals9 | | 3.831e-02 | -3.26 | negative regulation of alpha-beta T cell differentiation | biological process | GO:0046639 | 20 | 2 | 13711 | 214 | Il4ra,Runx1 | | 3.831e-02 | -3.26 | response to nicotine | biological process | GO:0035094 | 20 | 2 | 13711 | 214 | Nfkb1,Tnf | | 3.831e-02 | -3.26 | extracellular matrix | cellular component | GO:0031012 | 342 | 10 | 13825 | 212 | Tgm2,Adamts1,Olfml2b,Timp1,Adamts9,Mmp8,Thbs1,Col4a1,Lgals9,Runx1 | | 3.832e-02 | -3.26 | protein-lipid complex binding | molecular function | GO:0071814 | 20 | 2 | 13516 | 211 | Thbs1,Msr1 | | 3.832e-02 | -3.26 | protein folding chaperone | molecular function | GO:0044183 | 20 | 2 | 13516 | 211 | Hspa5,Hspb1 | | 3.832e-02 | -3.26 | lipoprotein particle binding | molecular function | GO:0071813 | 20 | 2 | 13516 | 211 | Msr1,Thbs1 | | 3.849e-02 | -3.26 | GO\_RESPONSE\_TO\_INSULIN | MSigDB lists | GO\_RESPONSE\_TO\_INSULIN | 169 | 6 | 12187 | 179 | Errfi1,Ptpn1,Il6,Ccl12,Socs3,Pik3r1 | | 3.849e-02 | -3.26 | GSE17721\_LPS\_VS\_CPG\_8H\_BMDC\_UP | MSigDB lists | GSE17721\_LPS\_VS\_CPG\_8H\_BMDC\_UP | 169 | 6 | 12187 | 179 | Il6,Casp4,Ifit3b,Ccl4,Parp14,Xdh | | 3.849e-02 | -3.26 | GSE37532\_TREG\_VS\_TCONV\_PPARG\_KO\_CD4\_TCELL\_FROM\_LN\_DN | MSigDB lists | GSE37532\_TREG\_VS\_TCONV\_PPARG\_KO\_CD4\_TCELL\_FROM\_LN\_DN | 169 | 6 | 12187 | 179 | Hspa5,Ccl4,Bcl3,Nfkb1,Icam1,Tap1 | | 3.849e-02 | -3.26 | HMGIY\_Q6 | MSigDB lists | HMGIY\_Q6 | 169 | 6 | 12187 | 179 | Csf3,Osmr,Icam1,Map3k8,Nfkbia,Il1rn | | 3.849e-02 | -3.26 | GSE40274\_LEF1\_VS\_FOXP3\_AND\_LEF1\_TRANSDUCED\_ACTIVATED\_CD4\_TCELL\_UP | MSigDB lists | GSE40274\_LEF1\_VS\_FOXP3\_AND\_LEF1\_TRANSDUCED\_ACTIVATED\_CD4\_TCELL\_UP | 169 | 6 | 12187 | 179 | Hspb1,Gadd45b,Gem,Gbp7,Bcl3,Tnip1 | | 3.849e-02 | -3.26 | GSE22886\_TH1\_VS\_TH2\_48H\_ACT\_DN | MSigDB lists | GSE22886\_TH1\_VS\_TH2\_48H\_ACT\_DN | 169 | 6 | 12187 | 179 | Rbm47,Parp12,Runx1,Aff1,Il4ra,Pik3r1 | | 3.849e-02 | -3.26 | GSE3994\_WT\_VS\_PAC1\_KO\_ACTIVATED\_MAST\_CELL\_DN | MSigDB lists | GSE3994\_WT\_VS\_PAC1\_KO\_ACTIVATED\_MAST\_CELL\_DN | 169 | 6 | 12187 | 179 | Il4ra,Map3k8,Casp4,Socs3,Nfe2l2,Sbno2 | | 3.849e-02 | -3.26 | GSE27786\_CD4\_TCELL\_VS\_NKCELL\_DN | MSigDB lists | GSE27786\_CD4\_TCELL\_VS\_NKCELL\_DN | 169 | 6 | 12187 | 179 | Rbm47,Oasl1,Msn,Map3k8,Gem,Nfkbiz | | 3.849e-02 | -3.26 | GSE33425\_CD161\_HIGH\_VS\_INT\_CD8\_TCELL\_UP | MSigDB lists | GSE33425\_CD161\_HIGH\_VS\_INT\_CD8\_TCELL\_UP | 169 | 6 | 12187 | 179 | Iqgap1,Gbp3,Saa1,Tubb6,Tiparp,Spi1 | | 3.856e-02 | -3.26 | positive regulation of immune effector process | biological process | GO:0002699 | 159 | 6 | 13711 | 214 | Ccl2,Il6,Tnf,Il4ra,Cxcl1,Nfkbiz | | 3.876e-02 | -3.25 | EGF | pfam domains | PF00008 | 48 | 3 | 12881 | 201 | Selp,Ptgs2,Sele | | 3.880e-02 | -3.25 | regulation of myotube differentiation | biological process | GO:0010830 | 48 | 3 | 13711 | 214 | Cxcl9,Cflar,Il4ra | | 3.880e-02 | -3.25 | chaperone-mediated protein folding | biological process | GO:0061077 | 48 | 3 | 13711 | 214 | Hspa5,Dnajb1,Hspb1 | | 3.880e-02 | -3.25 | regulation of myoblast differentiation | biological process | GO:0045661 | 48 | 3 | 13711 | 214 | Pik3r1,Tnf,Cxcl9 | | 3.881e-02 | -3.25 | BEGUM\_TARGETS\_OF\_PAX3\_FOXO1\_FUSION\_UP | MSigDB lists | BEGUM\_TARGETS\_OF\_PAX3\_FOXO1\_FUSION\_UP | 51 | 3 | 12187 | 179 | Stx11,Sgk1,Adamts1 | | 3.881e-02 | -3.25 | GO\_ACTIN\_CYTOSKELETON\_REORGANIZATION | MSigDB lists | GO\_ACTIN\_CYTOSKELETON\_REORGANIZATION | 51 | 3 | 12187 | 179 | S100a9,Plek,Ptpn1 | | 3.881e-02 | -3.25 | CADWELL\_ATG16L1\_TARGETS\_DN | MSigDB lists | CADWELL\_ATG16L1\_TARGETS\_DN | 51 | 3 | 12187 | 179 | Vcam1,Sgk1,Nfe2l2 | | 3.881e-02 | -3.25 | GO\_POSITIVE\_REGULATION\_OF\_LEUKOCYTE\_MEDIATED\_IMMUNITY | MSigDB lists | GO\_POSITIVE\_REGULATION\_OF\_LEUKOCYTE\_MEDIATED\_IMMUNITY | 51 | 3 | 12187 | 179 | Il4ra,Tnf,Ccl12 | | 3.886e-02 | -3.25 | GRESHOCK\_CANCER\_COPY\_NUMBER\_UP | MSigDB lists | GRESHOCK\_CANCER\_COPY\_NUMBER\_UP | 261 | 8 | 12187 | 179 | Zbtb16,Birc3,Aff1,Ccnd2,Runx1,Nfkb2,Bcl3,Rnf213 | | 3.897e-02 | -3.25 | Ig\_E-set | interpro domains | IPR014756 | 83 | 4 | 13788 | 212 | Nfkb2,A2m,Nfkb1,Tgm2 | | 3.901e-02 | -3.24 | Insulin Signaling | WikiPathways | WP65 | 152 | 8 | 3756 | 96 | Socs3,Sgk3,Rhoj,Map3k8,Map3k6,Pik3r1,Sgk1,Ptpn1 | | 3.909e-02 | -3.24 | GO\_CELL\_ACTIVATION\_INVOLVED\_IN\_IMMUNE\_RESPONSE | MSigDB lists | GO\_CELL\_ACTIVATION\_INVOLVED\_IN\_IMMUNE\_RESPONSE | 87 | 4 | 12187 | 179 | Bcl3,Sbno2,Il6,Icam1 | | 3.909e-02 | -3.24 | GSE6090\_UNSTIM\_VS\_DC\_SIGN\_STIM\_DC\_DN | MSigDB lists | GSE6090\_UNSTIM\_VS\_DC\_SIGN\_STIM\_DC\_DN | 87 | 4 | 12187 | 179 | Isg15,A2m,Pik3r5,Xdh | | 3.914e-02 | -3.24 | transition metal ion binding | molecular function | GO:0046914 | 735 | 18 | 13516 | 211 | Adamts1,Trim30a,Trim56,Slfn9,Mmp8,Xdh,Tnfsf10,Timp1,Cp,Ptpn1,Il1a,S100a8,Tnfaip3,Mt1,Znfx1,Mt2,Lcn2,S100a9 | | 3.916e-02 | -3.24 | Nicotinate metabolism | REACTOME pathways | R-MMU-196807 | 19 | 2 | 6297 | 105 | Parp9,Ptgs2 | | 3.923e-02 | -3.24 | GSE40274\_CTRL\_VS\_GATA1\_TRANSDUCED\_ACTIVATED\_CD4\_TCELL\_DN | MSigDB lists | GSE40274\_CTRL\_VS\_GATA1\_TRANSDUCED\_ACTIVATED\_CD4\_TCELL\_DN | 127 | 5 | 12187 | 179 | Hspb1,Bach1,Ier3,Cxcl1,Trib1 | | 3.923e-02 | -3.24 | GSE28737\_WT\_VS\_BCL6\_KO\_MARGINAL\_ZONE\_BCELL\_UP | MSigDB lists | GSE28737\_WT\_VS\_BCL6\_KO\_MARGINAL\_ZONE\_BCELL\_UP | 127 | 5 | 12187 | 179 | Ccnd2,Slfn9,Slfn5,Usp18,Sele | | 3.923e-02 | -3.24 | CHIARADONNA\_NEOPLASTIC\_TRANSFORMATION\_KRAS\_DN | MSigDB lists | CHIARADONNA\_NEOPLASTIC\_TRANSFORMATION\_KRAS\_DN | 127 | 5 | 12187 | 179 | Isg15,Socs3,Fstl1,Akap12,Atf3 | | 3.930e-02 | -3.24 | positive regulation of protein catabolic process | biological process | GO:0045732 | 202 | 7 | 13711 | 214 | Msn,Trim30a,Ier3,Trib1,Tiparp,Tnf,Tnfaip3 | | 3.930e-02 | -3.24 | extracellular structure organization | biological process | GO:0043062 | 202 | 7 | 13711 | 214 | Adamts9,Mmp8,Olfml2b,Bcl3,Nfkb2,Tnf,Col4a1 | | 3.937e-02 | -3.23 | SMAD\_FHA\_dom\_sf | interpro domains | IPR008984 | 49 | 3 | 13788 | 212 | Rasip1,Irf7,Irf9 | | 3.937e-02 | -3.23 | SPRY\_dom | interpro domains | IPR003877 | 49 | 3 | 13788 | 212 | Trim25,Trim30d,Trim30a | | 3.938e-02 | -3.23 | regulation of cell cycle phase transition | biological process | GO:1901987 | 246 | 8 | 13711 | 214 | Ier3,Ccnd2,Ripk1,Kcna5,Adamts1,Ccl12,Cdkn1a,Fam107a | | 3.942e-02 | -3.23 | NFE2\_01 | MSigDB lists | NFE2\_01 | 215 | 7 | 12187 | 179 | Angptl4,Bcl3,Cdkn1a,Maff,Bach1,Gadd45g,Il6 | | 3.943e-02 | -3.23 | GO\_REGULATED\_EXOCYTOSIS | MSigDB lists | GO\_REGULATED\_EXOCYTOSIS | 170 | 6 | 12187 | 179 | Selp,A2m,Plek,Thbs1,Stx11,Timp1 | | 3.943e-02 | -3.23 | GSE5463\_CTRL\_VS\_DEXAMETHASONE\_TREATED\_THYMOCYTE\_UP | MSigDB lists | GSE5463\_CTRL\_VS\_DEXAMETHASONE\_TREATED\_THYMOCYTE\_UP | 170 | 6 | 12187 | 179 | Maff,Il4ra,Clic4,Tap1,Tagln2,Tnip1 | | 3.943e-02 | -3.23 | GSE6259\_FLT3L\_INDUCED\_33D1\_POS\_DC\_VS\_BCELL\_DN | MSigDB lists | GSE6259\_FLT3L\_INDUCED\_33D1\_POS\_DC\_VS\_BCELL\_DN | 170 | 6 | 12187 | 179 | Dtx3l,Gbp3,Usp18,Lgals9,Sgk1,Aff1 | | 3.943e-02 | -3.23 | GSE7460\_TCONV\_VS\_TREG\_LN\_DN | MSigDB lists | GSE7460\_TCONV\_VS\_TREG\_LN\_DN | 170 | 6 | 12187 | 179 | Map3k8,Nfkbiz,Casp4,Gbp3,Nfkbie,Gem | | 3.943e-02 | -3.23 | GSE21546\_SAP1A\_KO\_VS\_SAP1A\_KO\_AND\_ELK1\_KO\_ANTI\_CD3\_STIM\_DP\_THYMOCYTES\_DN | MSigDB lists | GSE21546\_SAP1A\_KO\_VS\_SAP1A\_KO\_AND\_ELK1\_KO\_ANTI\_CD3\_STIM\_DP\_THYMOCYTES\_DN | 170 | 6 | 12187 | 179 | Iqgap1,Icam1,Socs3,Gadd45b,Rassf4,Tubb6 | | 3.943e-02 | -3.23 | GSE26727\_WT\_VS\_KLF2\_KO\_LPS\_STIM\_MACROPHAGE\_DN | MSigDB lists | GSE26727\_WT\_VS\_KLF2\_KO\_LPS\_STIM\_MACROPHAGE\_DN | 170 | 6 | 12187 | 179 | Ccl4,Tnf,Atf3,Ier3,Nfkb1,Gem | | 3.943e-02 | -3.23 | GSE3982\_EOSINOPHIL\_VS\_BCELL\_DN | MSigDB lists | GSE3982\_EOSINOPHIL\_VS\_BCELL\_DN | 170 | 6 | 12187 | 179 | Ifi211,Clic4,Apold1,Ifi209,Ifi204,Ifi207 | | 3.943e-02 | -3.23 | GSE28737\_WT\_VS\_BCL6\_KO\_FOLLICULAR\_BCELL\_DN | MSigDB lists | GSE28737\_WT\_VS\_BCL6\_KO\_FOLLICULAR\_BCELL\_DN | 170 | 6 | 12187 | 179 | Usp18,Arid5b,Ccnd2,Sgk3,Errfi1,Pik3ap1 | | 3.952e-02 | -3.23 | GO\_TISSUE\_MORPHOGENESIS | MSigDB lists | GO\_TISSUE\_MORPHOGENESIS | 410 | 11 | 12187 | 179 | Tgm2,Socs3,Sdc4,Spi1,Sox11,Il6,Rasip1,Rhoc,Col4a1,Clic4,Msn | | 3.989e-02 | -3.22 | - | gene3d domains | 3.40.390.10 | 51 | 3 | 6647 | 99 | Mmp8,Adamts9,Adamts1 | | 4.011e-02 | -3.22 | Znf\_RING\_CS | interpro domains | IPR017907 | 122 | 5 | 13788 | 212 | Trim56,Dtx3l,Trim30d,Trim30a,Trim25 | | 4.035e-02 | -3.21 | ZHANG\_TLX\_TARGETS\_60HR\_UP | MSigDB lists | ZHANG\_TLX\_TARGETS\_60HR\_UP | 263 | 8 | 12187 | 179 | Tap1,Pygm,Cflar,Gadd45g,Cp,Gadd45b,Mxd4,Cdkn1a | | 4.036e-02 | -3.21 | GSE5099\_MONOCYTE\_VS\_ALTERNATIVE\_M2\_MACROPHAGE\_UP | MSigDB lists | GSE5099\_MONOCYTE\_VS\_ALTERNATIVE\_M2\_MACROPHAGE\_UP | 128 | 5 | 12187 | 179 | Ifi204,Ifi207,Slfn9,Ifi211,Ifi209 | | 4.037e-02 | -3.21 | Caffeine metabolism | KEGG pathways | ko00232 | 2 | 1 | 5248 | 107 | Xdh | | 4.037e-02 | -3.21 | Caffeine metabolism | KEGG pathways | mmu00232 | 2 | 1 | 5248 | 107 | Xdh | | 4.039e-02 | -3.21 | GSE9006\_HEALTHY\_VS\_TYPE\_1\_DIABETES\_PBMC\_1MONTH\_POST\_DX\_UP | MSigDB lists | GSE9006\_HEALTHY\_VS\_TYPE\_1\_DIABETES\_PBMC\_1MONTH\_POST\_DX\_UP | 171 | 6 | 12187 | 179 | Pik3r1,Hspa5,Sntb2,Rbm39,Cybb,Msn | | 4.039e-02 | -3.21 | GSE33374\_CD8\_ALPHAALPHA\_VS\_ALPHABETA\_CD161\_HIGH\_TCELL\_DN | MSigDB lists | GSE33374\_CD8\_ALPHAALPHA\_VS\_ALPHABETA\_CD161\_HIGH\_TCELL\_DN | 171 | 6 | 12187 | 179 | Isg15,Tap1,Ifit2,Vcam1,Ifit3b,Nfkbia | | 4.039e-02 | -3.21 | GSE15330\_LYMPHOID\_MULTIPOTENT\_VS\_GRANULOCYTE\_MONOCYTE\_PROGENITOR\_IKAROS\_KO\_UP | MSigDB lists | GSE15330\_LYMPHOID\_MULTIPOTENT\_VS\_GRANULOCYTE\_MONOCYTE\_PROGENITOR\_IKAROS\_KO\_UP | 171 | 6 | 12187 | 179 | Nfkb2,Gbp4,Cdkn1a,Gbp6,Slfn9,Nfkbia | | 4.039e-02 | -3.21 | GSE17721\_POLYIC\_VS\_GARDIQUIMOD\_0.5H\_BMDC\_DN | MSigDB lists | GSE17721\_POLYIC\_VS\_GARDIQUIMOD\_0.5H\_BMDC\_DN | 171 | 6 | 12187 | 179 | Map3k8,Il1rn,Msn,Slfn3,Pik3ap1,Slfn4 | | 4.039e-02 | -3.21 | GSE2770\_IL4\_ACT\_VS\_ACT\_CD4\_TCELL\_2H\_DN | MSigDB lists | GSE2770\_IL4\_ACT\_VS\_ACT\_CD4\_TCELL\_2H\_DN | 171 | 6 | 12187 | 179 | Atf3,Nfkb1,Clic4,Ptpn1,Plek,Icam1 | | 4.039e-02 | -3.21 | GSE3982\_EOSINOPHIL\_VS\_TH1\_DN | MSigDB lists | GSE3982\_EOSINOPHIL\_VS\_TH1\_DN | 171 | 6 | 12187 | 179 | Ifi211,Apold1,Ifi209,Ccl4,Ifi207,Ifi204 | | 4.039e-02 | -3.21 | KRAS.600\_UP.V1\_UP | MSigDB lists | KRAS.600\_UP.V1\_UP | 171 | 6 | 12187 | 179 | Csf3,Ptgs2,Tgm2,Angptl4,A2m,Kcna5 | | 4.050e-02 | -3.21 | GO\_GROWTH\_FACTOR\_RECEPTOR\_BINDING | MSigDB lists | GO\_GROWTH\_FACTOR\_RECEPTOR\_BINDING | 88 | 4 | 12187 | 179 | Csf3,Il6,Il1rn,Il1a | | 4.059e-02 | -3.20 | regulation of lymphocyte proliferation | biological process | GO:0050670 | 161 | 6 | 13711 | 214 | Vcam1,Lgals9,Sox11,Cdkn1a,Sdc4,Il6 | | 4.059e-02 | -3.20 | vascular process in circulatory system | biological process | GO:0003018 | 161 | 6 | 13711 | 214 | Tnf,Icam1,Akap12,Ptgs2,Ccl4,Kcna5 | | 4.075e-02 | -3.20 | PID\_RAC1\_PATHWAY | MSigDB lists | PID\_RAC1\_PATHWAY | 52 | 3 | 12187 | 179 | Ncf1,Iqgap1,Cybb | | 4.075e-02 | -3.20 | YWATTWNNRGCT\_UNKNOWN | MSigDB lists | YWATTWNNRGCT\_UNKNOWN | 52 | 3 | 12187 | 179 | Atf3,Ptpn1,Nfkbia | | 4.075e-02 | -3.20 | VERRECCHIA\_EARLY\_RESPONSE\_TO\_TGFB1 | MSigDB lists | VERRECCHIA\_EARLY\_RESPONSE\_TO\_TGFB1 | 52 | 3 | 12187 | 179 | Icam1,Timp1,Rhoc | | 4.079e-02 | -3.20 | CARD | interpro domains | IPR001315 | 21 | 2 | 13788 | 212 | Birc3,Casp4 | | 4.079e-02 | -3.20 | C-type\_lectin\_CS | interpro domains | IPR018378 | 21 | 2 | 13788 | 212 | Selp,Sele | | 4.083e-02 | -3.20 | SPRY | pfam domains | PF00622 | 49 | 3 | 12881 | 201 | Trim30a,Trim30d,Trim25 | | 4.087e-02 | -3.20 | negative regulation of muscle cell apoptotic process | biological process | GO:0010656 | 49 | 3 | 13711 | 214 | Pik3r1,Nfe2l2,Cflar | | 4.088e-02 | -3.20 | LI\_PROSTATE\_CANCER\_EPIGENETIC | MSigDB lists | LI\_PROSTATE\_CANCER\_EPIGENETIC | 22 | 2 | 12187 | 179 | Ccnd2,Ptgs2 | | 4.088e-02 | -3.20 | PID\_PRL\_SIGNALING\_EVENTS\_PATHWAY | MSigDB lists | PID\_PRL\_SIGNALING\_EVENTS\_PATHWAY | 22 | 2 | 12187 | 179 | Cdkn1a,Rhoc | | 4.088e-02 | -3.20 | SHIN\_B\_CELL\_LYMPHOMA\_CLUSTER\_3 | MSigDB lists | SHIN\_B\_CELL\_LYMPHOMA\_CLUSTER\_3 | 22 | 2 | 12187 | 179 | Nfkb2,Nfkb1 | | 4.088e-02 | -3.20 | GO\_FIBRONECTIN\_BINDING | MSigDB lists | GO\_FIBRONECTIN\_BINDING | 22 | 2 | 12187 | 179 | Sdc4,Thbs1 | | 4.088e-02 | -3.20 | BIOCARTA\_RAS\_PATHWAY | MSigDB lists | BIOCARTA\_RAS\_PATHWAY | 22 | 2 | 12187 | 179 | Nfkb1,Pik3r1 | | 4.088e-02 | -3.20 | GO\_ACTININ\_BINDING | MSigDB lists | GO\_ACTININ\_BINDING | 22 | 2 | 12187 | 179 | Nfkb1,Kcna5 | | 4.088e-02 | -3.20 | GO\_PHOSPHATIDYLINOSITOL\_3\_KINASE\_BINDING | MSigDB lists | GO\_PHOSPHATIDYLINOSITOL\_3\_KINASE\_BINDING | 22 | 2 | 12187 | 179 | Pik3ap1,Pik3r1 | | 4.088e-02 | -3.20 | ABE\_VEGFA\_TARGETS\_30MIN | MSigDB lists | ABE\_VEGFA\_TARGETS\_30MIN | 22 | 2 | 12187 | 179 | Ptgs2,Trib1 | | 4.088e-02 | -3.20 | HAHTOLA\_SEZARY\_SYNDROM\_DN | MSigDB lists | HAHTOLA\_SEZARY\_SYNDROM\_DN | 22 | 2 | 12187 | 179 | Ccl4,Zbtb16 | | 4.088e-02 | -3.20 | BIOCARTA\_VIP\_PATHWAY | MSigDB lists | BIOCARTA\_VIP\_PATHWAY | 22 | 2 | 12187 | 179 | Nfkbia,Nfkb1 | | 4.088e-02 | -3.20 | BIOCARTA\_HER2\_PATHWAY | MSigDB lists | BIOCARTA\_HER2\_PATHWAY | 22 | 2 | 12187 | 179 | Il6,Pik3r1 | | 4.088e-02 | -3.20 | RUAN\_RESPONSE\_TO\_TROGLITAZONE\_UP | MSigDB lists | RUAN\_RESPONSE\_TO\_TROGLITAZONE\_UP | 22 | 2 | 12187 | 179 | Plin4,Rasd1 | | 4.088e-02 | -3.20 | XU\_HGF\_SIGNALING\_NOT\_VIA\_AKT1\_6HR | MSigDB lists | XU\_HGF\_SIGNALING\_NOT\_VIA\_AKT1\_6HR | 22 | 2 | 12187 | 179 | Angptl4,Plaur | | 4.088e-02 | -3.20 | BIOCARTA\_CXCR4\_PATHWAY | MSigDB lists | BIOCARTA\_CXCR4\_PATHWAY | 22 | 2 | 12187 | 179 | Nfkb1,Pik3r1 | | 4.088e-02 | -3.20 | REACTOME\_ANTIGEN\_ACTIVATES\_B\_CELL\_RECEPTOR\_LEADING\_TO\_GENERATION\_OF\_SECOND\_MESSENGERS | MSigDB lists | REACTOME\_ANTIGEN\_ACTIVATES\_B\_CELL\_RECEPTOR\_LEADING\_TO\_GENERATION\_OF\_SECOND\_MESSENGERS | 22 | 2 | 12187 | 179 | Pik3r1,Pik3ap1 | | 4.088e-02 | -3.20 | BIOCARTA\_CERAMIDE\_PATHWAY | MSigDB lists | BIOCARTA\_CERAMIDE\_PATHWAY | 22 | 2 | 12187 | 179 | Nfkb1,Ripk1 | | 4.088e-02 | -3.20 | REACTOME\_DOWNSTREAM\_TCR\_SIGNALING | MSigDB lists | REACTOME\_DOWNSTREAM\_TCR\_SIGNALING | 22 | 2 | 12187 | 179 | Nfkbia,Pik3r1 | | 4.088e-02 | -3.20 | protein kinase inhibitor activity | molecular function | GO:0004860 | 49 | 3 | 13516 | 211 | Cdkn1a,Hspb1,Trib1 | | 4.097e-02 | -3.19 | regulation of actin filament-based process | biological process | GO:0032970 | 340 | 10 | 13711 | 214 | Rhoc,Pik3r1,Csf3,Rhou,Sdc4,Rhoj,Plek,Il1a,Fam107a,Icam1 | | 4.109e-02 | -3.19 | regulation of protein kinase B signaling | biological process | GO:0051896 | 121 | 5 | 13711 | 214 | Tnf,Thbs1,Csf3,Xdh,Il6 | | 4.111e-02 | -3.19 | GO\_POSITIVE\_REGULATION\_OF\_CYTOPLASMIC\_TRANSPORT | MSigDB lists | GO\_POSITIVE\_REGULATION\_OF\_CYTOPLASMIC\_TRANSPORT | 217 | 7 | 12187 | 179 | Il6,Tnf,Rhou,Pik3r1,Msn,Ptgs2,Csf3 | | 4.136e-02 | -3.19 | GSE17721\_CPG\_VS\_GARDIQUIMOD\_4H\_BMDC\_DN | MSigDB lists | GSE17721\_CPG\_VS\_GARDIQUIMOD\_4H\_BMDC\_DN | 172 | 6 | 12187 | 179 | Trib1,Csf3,Maff,Nfe2l2,Il17ra,P2ry6 | | 4.136e-02 | -3.19 | GSE3982\_MEMORY\_CD4\_TCELL\_VS\_TH1\_DN | MSigDB lists | GSE3982\_MEMORY\_CD4\_TCELL\_VS\_TH1\_DN | 172 | 6 | 12187 | 179 | Ier3,Sdc4,Tor3a,Plaur,Cflar,Bcl3 | | 4.136e-02 | -3.19 | GSE43863\_TH1\_VS\_TFH\_MEMORY\_CD4\_TCELL\_UP | MSigDB lists | GSE43863\_TH1\_VS\_TFH\_MEMORY\_CD4\_TCELL\_UP | 172 | 6 | 12187 | 179 | Maff,Ifit2,Rrp8,Cflar,Rgs16,Plek | | 4.136e-02 | -3.19 | GSE28737\_BCL6\_HET\_VS\_BCL6\_KO\_MARGINAL\_ZONE\_BCELL\_DN | MSigDB lists | GSE28737\_BCL6\_HET\_VS\_BCL6\_KO\_MARGINAL\_ZONE\_BCELL\_DN | 172 | 6 | 12187 | 179 | Hcar2,Tubb6,Gpr84,Cd14,Fzd4,Tnip1 | | 4.136e-02 | -3.19 | PITX2\_Q2 | MSigDB lists | PITX2\_Q2 | 172 | 6 | 12187 | 179 | Runx1,Trib1,Fzd4,Pik3r5,Gadd45g,Luc7l3 | | 4.136e-02 | -3.19 | GSE26030\_UNSTIM\_VS\_RESTIM\_TH17\_DAY15\_POST\_POLARIZATION\_DN | MSigDB lists | GSE26030\_UNSTIM\_VS\_RESTIM\_TH17\_DAY15\_POST\_POLARIZATION\_DN | 172 | 6 | 12187 | 179 | Dnajb1,Slfn5,Tagln2,Ripk1,Rassf4,Icam1 | | 4.136e-02 | -3.19 | GSE27786\_LIN\_NEG\_VS\_NKTCELL\_DN | MSigDB lists | GSE27786\_LIN\_NEG\_VS\_NKTCELL\_DN | 172 | 6 | 12187 | 179 | Tap1,Tnfsf10,Parp9,Irf7,Tnip1,Gadd45g | | 4.150e-02 | -3.18 | Type II diabetes mellitus | KEGG pathways | mmu04930 | 38 | 3 | 5248 | 107 | Socs3,Pik3r1,Tnf | | 4.150e-02 | -3.18 | Type II diabetes mellitus | KEGG pathways | ko04930 | 38 | 3 | 5248 | 107 | Socs3,Tnf,Pik3r1 | | 4.150e-02 | -3.18 | GSE13229\_IMM\_VS\_INTMATURE\_NKCELL\_UP | MSigDB lists | GSE13229\_IMM\_VS\_INTMATURE\_NKCELL\_UP | 129 | 5 | 12187 | 179 | Tnfaip8,Ms4a6d,Parp12,Hspa5,A2m | | 4.150e-02 | -3.18 | CSR\_LATE\_UP.V1\_DN | MSigDB lists | CSR\_LATE\_UP.V1\_DN | 129 | 5 | 12187 | 179 | Ccl12,Mxd4,Bach1,Xaf1,Nfkbie | | 4.150e-02 | -3.18 | GO\_CYSTEINE\_TYPE\_PEPTIDASE\_ACTIVITY | MSigDB lists | GO\_CYSTEINE\_TYPE\_PEPTIDASE\_ACTIVITY | 129 | 5 | 12187 | 179 | Tnfaip3,Usp18,Casp4,Tnip1,Cflar | | 4.150e-02 | -3.18 | CYCLIN\_D1\_KE\_.V1\_DN | MSigDB lists | CYCLIN\_D1\_KE\_.V1\_DN | 129 | 5 | 12187 | 179 | Pik3r1,A2m,Ifit3b,Cd14,Vcam1 | | 4.150e-02 | -3.18 | RAY\_TUMORIGENESIS\_BY\_ERBB2\_CDC25A\_DN | MSigDB lists | RAY\_TUMORIGENESIS\_BY\_ERBB2\_CDC25A\_DN | 129 | 5 | 12187 | 179 | Rhou,Tor3a,Rbm39,Plin4,Errfi1 | | 4.191e-02 | -3.17 | ureide biosynthesis | BIOCYC pathways | MOUSE\_URSIN-PWY | 7 | 1 | 823 | 5 | Xdh | | 4.192e-02 | -3.17 | positive regulation of interleukin-10 production | biological process | GO:0032733 | 21 | 2 | 13711 | 214 | Lgals9,Bcl3 | | 4.192e-02 | -3.17 | hydrogen peroxide metabolic process | biological process | GO:0042743 | 21 | 2 | 13711 | 214 | Cybb,Ncf1 | | 4.192e-02 | -3.17 | positive regulation of myoblast differentiation | biological process | GO:0045663 | 21 | 2 | 13711 | 214 | Pik3r1,Cxcl9 | | 4.192e-02 | -3.17 | regulation of double-strand break repair via nonhomologous end joining | biological process | GO:2001032 | 21 | 2 | 13711 | 214 | Parp9,Dtx3l | | 4.192e-02 | -3.17 | positive regulation of steroid biosynthetic process | biological process | GO:0010893 | 21 | 2 | 13711 | 214 | Tnf,Il1a | | 4.192e-02 | -3.17 | positive regulation of granulocyte chemotaxis | biological process | GO:0071624 | 21 | 2 | 13711 | 214 | C3ar1,C5ar1 | | 4.192e-02 | -3.17 | response to fungus | biological process | GO:0009620 | 21 | 2 | 13711 | 214 | Ncf1,Il17ra | | 4.192e-02 | -3.17 | positive regulation of neutrophil chemotaxis | biological process | GO:0090023 | 21 | 2 | 13711 | 214 | C5ar1,C3ar1 | | 4.192e-02 | -3.17 | positive regulation of cell migration involved in sprouting angiogenesis | biological process | GO:0090050 | 21 | 2 | 13711 | 214 | Ptgs2,Rhoj | | 4.192e-02 | -3.17 | divalent inorganic cation homeostasis | biological process | GO:0072507 | 389 | 11 | 13711 | 214 | C5ar1,Cxcl1,Tgm2,Mt2,Kcna5,Mt1,Ccl2,C3ar1,Slc24a4,Pygm,P2ry6 | | 4.193e-02 | -3.17 | tumor necrosis factor receptor binding | molecular function | GO:0005164 | 21 | 2 | 13516 | 211 | Tnfsf10,Tnf | | 4.193e-02 | -3.17 | insulin receptor binding | molecular function | GO:0005158 | 21 | 2 | 13516 | 211 | Pik3r1,Ptpn1 | | 4.194e-02 | -3.17 | LIM\_MAMMARY\_LUMINAL\_MATURE\_DN | MSigDB lists | LIM\_MAMMARY\_LUMINAL\_MATURE\_DN | 89 | 4 | 12187 | 179 | Col4a1,Tubb6,Ccl12,Fstl1 | | 4.211e-02 | -3.17 | GO\_CYTOPLASMIC\_VESICLE\_PART | MSigDB lists | GO\_CYTOPLASMIC\_VESICLE\_PART | 466 | 12 | 12187 | 179 | Cybb,Sntb2,Msr1,A2m,Fzd4,Irgm2,Hp,Timp1,Thbs1,Clic4,Saa1,Selp | | 4.235e-02 | -3.16 | GSE40277\_EOS\_AND\_LEF1\_TRANSDUCED\_VS\_GATA1\_AND\_SATB1\_TRANSDUCED\_CD4\_TCELL\_UP | MSigDB lists | GSE40277\_EOS\_AND\_LEF1\_TRANSDUCED\_VS\_GATA1\_AND\_SATB1\_TRANSDUCED\_CD4\_TCELL\_UP | 173 | 6 | 12187 | 179 | Gbp6,Selp,Gbp3,Stx11,Nfkb1,Gbp4 | | 4.235e-02 | -3.16 | GSE8921\_UNSTIM\_VS\_TLR1\_2\_STIM\_MONOCYTE\_3H\_DN | MSigDB lists | GSE8921\_UNSTIM\_VS\_TLR1\_2\_STIM\_MONOCYTE\_3H\_DN | 173 | 6 | 12187 | 179 | Sbno2,Bbs12,Rhoc,Akap12,AA467197,Il4ra | | 4.235e-02 | -3.16 | FOXO4\_01 | MSigDB lists | FOXO4\_01 | 173 | 6 | 12187 | 179 | Rasd1,Luc7l3,Hcar2,Fstl1,Maff,Runx1 | | 4.235e-02 | -3.16 | GOLDRATH\_NAIVE\_VS\_MEMORY\_CD8\_TCELL\_UP | MSigDB lists | GOLDRATH\_NAIVE\_VS\_MEMORY\_CD8\_TCELL\_UP | 173 | 6 | 12187 | 179 | Irf7,Irgm2,Isg15,Plaur,Trim25,Nfkbiz | | 4.235e-02 | -3.16 | WELCSH\_BRCA1\_TARGETS\_UP | MSigDB lists | WELCSH\_BRCA1\_TARGETS\_UP | 173 | 6 | 12187 | 179 | Rhoc,Iqgap1,Ptpn1,Hspa5,Cdkn1a,Fstl1 | | 4.262e-02 | -3.16 | TYROBP Causal Network | WikiPathways | WP3625 | 52 | 4 | 3756 | 96 | Igsf6,Cxcl16,Plek,Rbm47 | | 4.267e-02 | -3.15 | GSE32901\_NAIVE\_VS\_TH17\_NEG\_CD4\_TCELL\_UP | MSigDB lists | GSE32901\_NAIVE\_VS\_TH17\_NEG\_CD4\_TCELL\_UP | 130 | 5 | 12187 | 179 | Sdc4,Nfkbiz,Tgm2,Tnip3,Cybb | | 4.267e-02 | -3.15 | GSE8685\_IL15\_ACT\_IL2\_STARVED\_VS\_IL21\_ACT\_IL2\_STARVED\_CD4\_TCELL\_UP | MSigDB lists | GSE8685\_IL15\_ACT\_IL2\_STARVED\_VS\_IL21\_ACT\_IL2\_STARVED\_CD4\_TCELL\_UP | 130 | 5 | 12187 | 179 | Cybb,Rhou,Ifitm3,Sdc4,Trib1 | | 4.268e-02 | -3.15 | morphogenesis of a branching structure | biological process | GO:0001763 | 163 | 6 | 13711 | 214 | Clic4,Socs3,Rasip1,Tgm2,Il6,Col4a1 | | 4.268e-02 | -3.15 | regulation of mononuclear cell proliferation | biological process | GO:0032944 | 163 | 6 | 13711 | 214 | Sox11,Cdkn1a,Sdc4,Il6,Vcam1,Lgals9 | | 4.274e-02 | -3.15 | AMIT\_SERUM\_RESPONSE\_120\_MCF10A | MSigDB lists | AMIT\_SERUM\_RESPONSE\_120\_MCF10A | 53 | 3 | 12187 | 179 | Gadd45b,Plaur,Maff | | 4.274e-02 | -3.15 | WESTON\_VEGFA\_TARGETS\_3HR | MSigDB lists | WESTON\_VEGFA\_TARGETS\_3HR | 53 | 3 | 12187 | 179 | Vcam1,Ccl12,Tnfsf10 | | 4.274e-02 | -3.15 | GO\_LYMPHOCYTE\_ACTIVATION\_INVOLVED\_IN\_IMMUNE\_RESPONSE | MSigDB lists | GO\_LYMPHOCYTE\_ACTIVATION\_INVOLVED\_IN\_IMMUNE\_RESPONSE | 53 | 3 | 12187 | 179 | Il6,Icam1,Bcl3 | | 4.274e-02 | -3.15 | MODULE\_172 | MSigDB lists | MODULE\_172 | 53 | 3 | 12187 | 179 | Casp4,Timp1,Hp | | 4.284e-02 | -3.15 | DURCHDEWALD\_SKIN\_CARCINOGENESIS\_DN | MSigDB lists | DURCHDEWALD\_SKIN\_CARCINOGENESIS\_DN | 219 | 7 | 12187 | 179 | Stx11,Tiparp,Nfe2l2,Ripk1,Runx1,Xdh,Ptgs2 | | 4.286e-02 | -3.15 | negative regulation of transport | biological process | GO:0051051 | 439 | 12 | 13711 | 214 | Ptgs2,Ier3,Kcna5,Gbp4,Lgals9,Il1rn,Thbs1,Il6,Tnf,Tnfaip3,Icam1,Gem | | 4.291e-02 | -3.15 | ICAM | prints domains | PR01473 | 2 | 1 | 2951 | 64 | Icam1 | | 4.291e-02 | -3.15 | C5ANPHYLTXNR | prints domains | PR00426 | 2 | 1 | 2951 | 64 | C5ar1 | | 4.291e-02 | -3.15 | ONCOGENEAML1 | prints domains | PR00967 | 2 | 1 | 2951 | 64 | Runx1 | | 4.292e-02 | -3.15 | DNA-binding transcription activator activity | molecular function | GO:0001216 | 296 | 9 | 13516 | 211 | Runx1,Irf2,Nfkb2,Bach1,Spi1,Cebpd,Nfkb1,Sox11,Nfe2l2 | | 4.292e-02 | -3.15 | DNA-binding transcription activator activity, RNA polymerase II-specific | molecular function | GO:0001228 | 296 | 9 | 13516 | 211 | Nfkb1,Cebpd,Nfe2l2,Sox11,Bach1,Nfkb2,Irf2,Runx1,Spi1 | | 4.294e-02 | -3.15 | KOYAMA\_SEMA3B\_TARGETS\_DN | MSigDB lists | KOYAMA\_SEMA3B\_TARGETS\_DN | 315 | 9 | 12187 | 179 | Birc3,Nfkbia,Stx11,Luc7l3,AA467197,Ncf1,Arid5b,Tagln2,Tap1 | | 4.299e-02 | -3.15 | alpha-beta T cell activation | biological process | GO:0046631 | 50 | 3 | 13711 | 214 | Il6,Bcl3,Gadd45g | | 4.299e-02 | -3.15 | negative regulation of peptidyl-tyrosine phosphorylation | biological process | GO:0050732 | 50 | 3 | 13711 | 214 | Errfi1,Socs3,Parp14 | | 4.301e-02 | -3.15 | polyubiquitin modification-dependent protein binding | molecular function | GO:0031593 | 50 | 3 | 13516 | 211 | Tnfaip3,Tnip3,Tnip1 | | 4.335e-02 | -3.14 | KONDO\_EZH2\_TARGETS | MSigDB lists | KONDO\_EZH2\_TARGETS | 174 | 6 | 12187 | 179 | Sntb2,Ifit3b,Sgk1,Map3k8,Ifit2,Cp | | 4.335e-02 | -3.14 | GSE27786\_BCELL\_VS\_NKCELL\_DN | MSigDB lists | GSE27786\_BCELL\_VS\_NKCELL\_DN | 174 | 6 | 12187 | 179 | Gbp4,Gbp6,Tnip1,Rtp4,Rbm47,Errfi1 | | 4.335e-02 | -3.14 | GSE26030\_UNSTIM\_VS\_RESTIM\_TH1\_DAY5\_POST\_POLARIZATION\_DN | MSigDB lists | GSE26030\_UNSTIM\_VS\_RESTIM\_TH1\_DAY5\_POST\_POLARIZATION\_DN | 174 | 6 | 12187 | 179 | Ifitm3,Gbp7,Slfn5,Rhou,Tor3a,Rassf4 | | 4.335e-02 | -3.14 | GSE7831\_UNSTIM\_VS\_CPG\_STIM\_PDC\_1H\_DN | MSigDB lists | GSE7831\_UNSTIM\_VS\_CPG\_STIM\_PDC\_1H\_DN | 174 | 6 | 12187 | 179 | Nfkb1,Atf3,Ptpn1,Selp,Map3k8,Tnf | | 4.335e-02 | -3.14 | GSE41176\_UNSTIM\_VS\_ANTI\_IGM\_STIM\_BCELL\_24H\_DN | MSigDB lists | GSE41176\_UNSTIM\_VS\_ANTI\_IGM\_STIM\_BCELL\_24H\_DN | 174 | 6 | 12187 | 179 | Icam1,Aff1,Maff,Il4ra,Gadd45g,Bach1 | | 4.335e-02 | -3.14 | GSE24634\_NAIVE\_CD4\_TCELL\_VS\_DAY7\_IL4\_CONV\_TREG\_DN | MSigDB lists | GSE24634\_NAIVE\_CD4\_TCELL\_VS\_DAY7\_IL4\_CONV\_TREG\_DN | 174 | 6 | 12187 | 179 | Ifi207,Ifi204,Ifi209,Cflar,Trib1,Ifi211 | | 4.335e-02 | -3.14 | GSE41867\_DAY8\_EFFECTOR\_VS\_DAY30\_MEMORY\_CD8\_TCELL\_LCMV\_ARMSTRONG\_DN | MSigDB lists | GSE41867\_DAY8\_EFFECTOR\_VS\_DAY30\_MEMORY\_CD8\_TCELL\_LCMV\_ARMSTRONG\_DN | 174 | 6 | 12187 | 179 | Parp14,Tor3a,Nfkbie,Rnf213,Slfn9,Nfkb2 | | 4.335e-02 | -3.14 | GSE1432\_6H\_VS\_24H\_IFNG\_MICROGLIA\_UP | MSigDB lists | GSE1432\_6H\_VS\_24H\_IFNG\_MICROGLIA\_UP | 174 | 6 | 12187 | 179 | Il4ra,Map3k6,S100a9,Cd14,Iqgap1,Trim25 | | 4.336e-02 | -3.14 | midbody | cellular component | GO:0030496 | 125 | 5 | 13825 | 212 | Gem,Hspa5,Iqgap1,Bcl3,Clic4 | | 4.342e-02 | -3.14 | SMID\_BREAST\_CANCER\_RELAPSE\_IN\_LIVER\_UP | MSigDB lists | SMID\_BREAST\_CANCER\_RELAPSE\_IN\_LIVER\_UP | 3 | 1 | 12187 | 179 | S100a8 | | 4.342e-02 | -3.14 | BIOCARTA\_IL5\_PATHWAY | MSigDB lists | BIOCARTA\_IL5\_PATHWAY | 3 | 1 | 12187 | 179 | Il6 | | 4.342e-02 | -3.14 | GERHOLD\_RESPONSE\_TO\_TZD\_UP | MSigDB lists | GERHOLD\_RESPONSE\_TO\_TZD\_UP | 3 | 1 | 12187 | 179 | Ccnd2 | | 4.342e-02 | -3.14 | GO\_REGULATION\_OF\_MYOBLAST\_PROLIFERATION | MSigDB lists | GO\_REGULATION\_OF\_MYOBLAST\_PROLIFERATION | 3 | 1 | 12187 | 179 | Kcna5 | | 4.342e-02 | -3.14 | BIERIE\_INFLAMMATORY\_RESPONSE\_TGFB1 | MSigDB lists | BIERIE\_INFLAMMATORY\_RESPONSE\_TGFB1 | 3 | 1 | 12187 | 179 | Ptgs2 | | 4.342e-02 | -3.14 | WAGSCHAL\_EHMT2\_TARGETS\_UP | MSigDB lists | WAGSCHAL\_EHMT2\_TARGETS\_UP | 3 | 1 | 12187 | 179 | Cdkn1a | | 4.342e-02 | -3.14 | GO\_POSITIVE\_REGULATION\_OF\_HUMORAL\_IMMUNE\_RESPONSE | MSigDB lists | GO\_POSITIVE\_REGULATION\_OF\_HUMORAL\_IMMUNE\_RESPONSE | 3 | 1 | 12187 | 179 | Tnf | | 4.342e-02 | -3.14 | MILICIC\_FAMILIAL\_ADENOMATOUS\_POLYPOSIS\_UP | MSigDB lists | MILICIC\_FAMILIAL\_ADENOMATOUS\_POLYPOSIS\_UP | 3 | 1 | 12187 | 179 | Lcn2 | | 4.353e-02 | -3.13 | MetalloPept\_cat\_dom\_sf | interpro domains | IPR024079 | 51 | 3 | 13788 | 212 | Adamts9,Mmp8,Adamts1 | | 4.356e-02 | -3.13 | TSP1 | smart domains | SM00209 | 46 | 3 | 7188 | 123 | Thbs1,Adamts1,Adamts9 | | 4.358e-02 | -3.13 | negative regulation of cell cycle phase transition | biological process | GO:1901988 | 123 | 5 | 13711 | 214 | Fam107a,Ripk1,Cdkn1a,Ier3,Ccl12 | | 4.386e-02 | -3.13 | GSE28726\_NAIVE\_VS\_ACTIVATED\_VA24NEG\_NKTCELL\_DN | MSigDB lists | GSE28726\_NAIVE\_VS\_ACTIVATED\_VA24NEG\_NKTCELL\_DN | 131 | 5 | 12187 | 179 | Dnajb1,Cybb,Stx11,Xdh,Pygm | | 4.386e-02 | -3.13 | GSE21670\_STAT3\_KO\_VS\_WT\_CD4\_TCELL\_UP | MSigDB lists | GSE21670\_STAT3\_KO\_VS\_WT\_CD4\_TCELL\_UP | 131 | 5 | 12187 | 179 | Gadd45g,Mxd4,Tap1,Nfkbie,Parp14 | | 4.386e-02 | -3.13 | GSE13306\_RA\_VS\_UNTREATED\_TCONV\_DN | MSigDB lists | GSE13306\_RA\_VS\_UNTREATED\_TCONV\_DN | 131 | 5 | 12187 | 179 | Fzd4,Ifitm2,Ifitm3,Parp12,AA467197 | | 4.386e-02 | -3.13 | GSE15659\_NAIVE\_VS\_PTPRC\_NEG\_CD4\_TCELL\_DN | MSigDB lists | GSE15659\_NAIVE\_VS\_PTPRC\_NEG\_CD4\_TCELL\_DN | 131 | 5 | 12187 | 179 | Rasd1,Tnfaip3,Timp1,Rhou,Sox11 | | 4.386e-02 | -3.13 | FREAC7\_01 | MSigDB lists | FREAC7\_01 | 131 | 5 | 12187 | 179 | Cxcl1,Csf3,Ier3,Errfi1,Hcar2 | | 4.387e-02 | -3.13 | positive regulation of cell cycle process | biological process | GO:0090068 | 207 | 7 | 13711 | 214 | Adamts1,Cdkn1a,Tnf,Rrp8,Il1a,Ccnd2,Kcna5 | | 4.400e-02 | -3.12 | Cell surface interactions at the vascular wall | REACTOME pathways | R-MMU-202733 | 80 | 4 | 6297 | 105 | Pik3r1,Sdc4,Sele,Selp | | 4.403e-02 | -3.12 | - | gene3d domains | 2.60.40.720 | 3 | 1 | 6647 | 99 | Runx1 | | 4.416e-02 | -3.12 | Ikbkb (inhibitor of kappaB kinase beta) | protein interactions | 16150 | 20 | 2 | 6802 | 115 | Tnf,Nfkbia | | 4.416e-02 | -3.12 | Tlr4 (toll-like receptor 4) | protein interactions | 21898 | 20 | 2 | 6802 | 115 | Birc3,Cd14 | | 4.416e-02 | -3.12 | Gata1 (GATA binding protein 1) | protein interactions | 14460 | 20 | 2 | 6802 | 115 | Runx1,Spi1 | | 4.421e-02 | -3.12 | plasma lipoprotein particle | cellular component | GO:0034358 | 22 | 2 | 13825 | 212 | Saa1,Msr1 | | 4.421e-02 | -3.12 | lipoprotein particle | cellular component | GO:1990777 | 22 | 2 | 13825 | 212 | Msr1,Saa1 | | 4.431e-02 | -3.12 | Ubiquitin-like\_domsf | interpro domains | IPR029071 | 167 | 6 | 13788 | 212 | Rasip1,Rassf4,Msn,Isg15,Oasl1,Oasl2 | | 4.435e-02 | -3.12 | SHIN\_B\_CELL\_LYMPHOMA\_CLUSTER\_2 | MSigDB lists | SHIN\_B\_CELL\_LYMPHOMA\_CLUSTER\_2 | 23 | 2 | 12187 | 179 | Ccnd2,Gadd45b | | 4.435e-02 | -3.12 | GO\_PROSTANOID\_METABOLIC\_PROCESS | MSigDB lists | GO\_PROSTANOID\_METABOLIC\_PROCESS | 23 | 2 | 12187 | 179 | Ptgs2,Ptges | | 4.435e-02 | -3.12 | MUELLER\_COMMON\_TARGETS\_OF\_AML\_FUSIONS\_DN | MSigDB lists | MUELLER\_COMMON\_TARGETS\_OF\_AML\_FUSIONS\_DN | 23 | 2 | 12187 | 179 | Sdc4,Ccl12 | | 4.435e-02 | -3.12 | GO\_DEFENSE\_RESPONSE\_TO\_GRAM\_NEGATIVE\_BACTERIUM | MSigDB lists | GO\_DEFENSE\_RESPONSE\_TO\_GRAM\_NEGATIVE\_BACTERIUM | 23 | 2 | 12187 | 179 | Il6,Selp | | 4.435e-02 | -3.12 | GO\_RESPONSE\_TO\_GROWTH\_HORMONE | MSigDB lists | GO\_RESPONSE\_TO\_GROWTH\_HORMONE | 23 | 2 | 12187 | 179 | Ptpn1,Pik3r1 | | 4.435e-02 | -3.12 | MODULE\_367 | MSigDB lists | MODULE\_367 | 23 | 2 | 12187 | 179 | P2ry6,C3ar1 | | 4.435e-02 | -3.12 | GO\_NUCLEOTIDE\_BINDING\_DOMAIN\_LEUCINE\_RICH\_REPEAT\_CONTAINING\_RECEPTOR\_SIGNALING\_PATHWAY | MSigDB lists | GO\_NUCLEOTIDE\_BINDING\_DOMAIN\_LEUCINE\_RICH\_REPEAT\_CONTAINING\_RECEPTOR\_SIGNALING\_PATHWAY | 23 | 2 | 12187 | 179 | Tnfaip3,Nfkbia | | 4.435e-02 | -3.12 | CHO\_NR4A1\_TARGETS | MSigDB lists | CHO\_NR4A1\_TARGETS | 23 | 2 | 12187 | 179 | Birc3,Atf3 | | 4.435e-02 | -3.12 | MODULE\_265 | MSigDB lists | MODULE\_265 | 23 | 2 | 12187 | 179 | Csf2rb,Il4ra | | 4.435e-02 | -3.12 | GO\_NEGATIVE\_REGULATION\_OF\_ENDOTHELIAL\_CELL\_PROLIFERATION | MSigDB lists | GO\_NEGATIVE\_REGULATION\_OF\_ENDOTHELIAL\_CELL\_PROLIFERATION | 23 | 2 | 12187 | 179 | Xdh,Thbs1 | | 4.435e-02 | -3.12 | GO\_POSITIVE\_REGULATION\_OF\_TYROSINE\_PHOSPHORYLATION\_OF\_STAT3\_PROTEIN | MSigDB lists | GO\_POSITIVE\_REGULATION\_OF\_TYROSINE\_PHOSPHORYLATION\_OF\_STAT3\_PROTEIN | 23 | 2 | 12187 | 179 | Il6,Socs3 | | 4.435e-02 | -3.12 | PID\_IL2\_STAT5\_PATHWAY | MSigDB lists | PID\_IL2\_STAT5\_PATHWAY | 23 | 2 | 12187 | 179 | Pik3r1,Ccnd2 | | 4.435e-02 | -3.12 | GO\_RESPONSE\_TO\_INTERLEUKIN\_4 | MSigDB lists | GO\_RESPONSE\_TO\_INTERLEUKIN\_4 | 23 | 2 | 12187 | 179 | Hspa5,Il4ra | | 4.435e-02 | -3.12 | PID\_IL3\_PATHWAY | MSigDB lists | PID\_IL3\_PATHWAY | 23 | 2 | 12187 | 179 | Pik3r1,Csf2rb | | 4.435e-02 | -3.12 | BRACHAT\_RESPONSE\_TO\_METHOTREXATE\_DN | MSigDB lists | BRACHAT\_RESPONSE\_TO\_METHOTREXATE\_DN | 23 | 2 | 12187 | 179 | Ifitm3,Ifitm2 | | 4.435e-02 | -3.12 | GO\_REGULATION\_OF\_CD4\_POSITIVE\_ALPHA\_BETA\_T\_CELL\_ACTIVATION | MSigDB lists | GO\_REGULATION\_OF\_CD4\_POSITIVE\_ALPHA\_BETA\_T\_CELL\_ACTIVATION | 23 | 2 | 12187 | 179 | Il4ra,Il6 | | 4.435e-02 | -3.12 | GO\_REGULATION\_OF\_OXIDATIVE\_STRESS\_INDUCED\_INTRINSIC\_APOPTOTIC\_SIGNALING\_PATHWAY | MSigDB lists | GO\_REGULATION\_OF\_OXIDATIVE\_STRESS\_INDUCED\_INTRINSIC\_APOPTOTIC\_SIGNALING\_PATHWAY | 23 | 2 | 12187 | 179 | Nfe2l2,Hspb1 | | 4.435e-02 | -3.12 | GO\_REGULATION\_OF\_PLATELET\_ACTIVATION | MSigDB lists | GO\_REGULATION\_OF\_PLATELET\_ACTIVATION | 23 | 2 | 12187 | 179 | Selp,Plek | | 4.436e-02 | -3.12 | GSE36009\_WT\_VS\_NLRP10\_KO\_DC\_DN | MSigDB lists | GSE36009\_WT\_VS\_NLRP10\_KO\_DC\_DN | 175 | 6 | 12187 | 179 | Spi1,Pik3r5,Birc3,C5ar1,Nfkb2,Nfkb1 | | 4.436e-02 | -3.12 | GSE17186\_BLOOD\_VS\_CORD\_BLOOD\_NAIVE\_BCELL\_UP | MSigDB lists | GSE17186\_BLOOD\_VS\_CORD\_BLOOD\_NAIVE\_BCELL\_UP | 175 | 6 | 12187 | 179 | Ifitm2,Tnfaip8,Csf2rb,Cxcl16,Trib1,Parp12 | | 4.459e-02 | -3.11 | Hspa8 (heat shock protein 8) | protein interactions | 15481 | 47 | 3 | 6802 | 115 | Cdkn1a,Kcna5,Hspb1 | | 4.466e-02 | -3.11 | mouse chr18 D3\|18 | chromosome location | mouse chr18 D3\|18 | 3 | 1 | 14556 | 220 | Gm4841 | | 4.466e-02 | -3.11 | mouse chr1 A2\|1 2.08 cM | chromosome location | mouse chr1 A2\|1 2.08 cM | 3 | 1 | 14556 | 220 | Sgk3 | | 4.466e-02 | -3.11 | mouse chr2 H3\|2 85.16 cM | chromosome location | mouse chr2 H3\|2 85.16 cM | 3 | 1 | 14556 | 220 | Sdc4 | | 4.466e-02 | -3.11 | mouse chr7 A3\|7 9.95 cM | chromosome location | mouse chr7 A3\|7 9.95 cM | 3 | 1 | 14556 | 220 | Bcl3 | | 4.466e-02 | -3.11 | mouse chr5 E1\|5 44.78 cM | chromosome location | mouse chr5 E1\|5 44.78 cM | 3 | 1 | 14556 | 220 | Cxcl1 | | 4.466e-02 | -3.11 | mouse chr6 F3\|6 61.92 cM | chromosome location | mouse chr6 F3\|6 61.92 cM | 3 | 1 | 14556 | 220 | Ccnd2 | | 4.466e-02 | -3.11 | mouse chr5 E5\|5 50.68 cM | chromosome location | mouse chr5 E5\|5 50.68 cM | 3 | 1 | 14556 | 220 | Gbp4 | | 4.466e-02 | -3.11 | mouse chr6 B3\|6 | chromosome location | mouse chr6 B3\|6 | 3 | 1 | 14556 | 220 | Herc6 | | 4.479e-02 | -3.11 | DAVICIONI\_TARGETS\_OF\_PAX\_FOXO1\_FUSIONS\_DN | MSigDB lists | DAVICIONI\_TARGETS\_OF\_PAX\_FOXO1\_FUSIONS\_DN | 54 | 3 | 12187 | 179 | Ccl12,Sox11,Ccl4 | | 4.479e-02 | -3.11 | GENTILE\_RESPONSE\_CLUSTER\_D3 | MSigDB lists | GENTILE\_RESPONSE\_CLUSTER\_D3 | 54 | 3 | 12187 | 179 | Ripk1,Tnfaip8,Tiparp | | 4.479e-02 | -3.11 | MCCLUNG\_COCAIN\_REWARD\_4WK | MSigDB lists | MCCLUNG\_COCAIN\_REWARD\_4WK | 54 | 3 | 12187 | 179 | Dnajb1,Sox11,Znfx1 | | 4.479e-02 | -3.11 | GNF2\_PTPRC | MSigDB lists | GNF2\_PTPRC | 54 | 3 | 12187 | 179 | Parp12,Phf11b,Phf11d | | 4.479e-02 | -3.11 | ODONNELL\_METASTASIS\_UP | MSigDB lists | ODONNELL\_METASTASIS\_UP | 54 | 3 | 12187 | 179 | Sbno2,Ksr1,Pik3r5 | | 4.479e-02 | -3.11 | GO\_PEPTIDE\_TRANSPORT | MSigDB lists | GO\_PEPTIDE\_TRANSPORT | 54 | 3 | 12187 | 179 | Tap1,Il6,Il1rn | | 4.488e-02 | -3.10 | growth factor binding | molecular function | GO:0019838 | 124 | 5 | 13516 | 211 | Osmr,Il1rn,Thbs1,A2m,Col4a1 | | 4.489e-02 | -3.10 | Th1 and Th2 cell differentiation | KEGG pathways | mmu04658 | 66 | 4 | 5248 | 107 | Nfkbie,Nfkbia,Il4ra,Nfkb1 | | 4.489e-02 | -3.10 | Th1 and Th2 cell differentiation | KEGG pathways | ko04658 | 66 | 4 | 5248 | 107 | Il4ra,Nfkbia,Nfkb1,Nfkbie | | 4.490e-02 | -3.10 | GO\_MONOVALENT\_INORGANIC\_CATION\_HOMEOSTASIS | MSigDB lists | GO\_MONOVALENT\_INORGANIC\_CATION\_HOMEOSTASIS | 91 | 4 | 12187 | 179 | Sgk3,Sgk1,Clic4,Kcna5 | | 4.500e-02 | -3.10 | ZF\_BBOX | prosite domains | PS50119 | 43 | 3 | 8845 | 164 | Trim56,Trim30d,Trim30a | | 4.506e-02 | -3.10 | MODULE\_94 | MSigDB lists | MODULE\_94 | 269 | 8 | 12187 | 179 | Sbno2,Sele,Igsf6,Selp,Ier3,Atf3,Mxd4,Runx1 | | 4.506e-02 | -3.10 | MOREAUX\_MULTIPLE\_MYELOMA\_BY\_TACI\_UP | MSigDB lists | MOREAUX\_MULTIPLE\_MYELOMA\_BY\_TACI\_UP | 269 | 8 | 12187 | 179 | Plaur,Ptges,Pik3r5,Robo4,C3ar1,Oasl1,Gadd45b,Tnf | | 4.507e-02 | -3.10 | BROWNE\_HCMV\_INFECTION\_6HR\_DN | MSigDB lists | BROWNE\_HCMV\_INFECTION\_6HR\_DN | 132 | 5 | 12187 | 179 | Bach1,Arid5b,Cflar,Pik3r1,Sgk1 | | 4.507e-02 | -3.10 | RUIZ\_TNC\_TARGETS\_UP | MSigDB lists | RUIZ\_TNC\_TARGETS\_UP | 132 | 5 | 12187 | 179 | Arid5b,A2m,Vcam1,Olfml2b,Tnfsf10 | | 4.517e-02 | -3.10 | keratinocyte differentiation | biological process | GO:0030216 | 51 | 3 | 13711 | 214 | Clic4,Il1a,Ptgs2 | | 4.517e-02 | -3.10 | immunoglobulin mediated immune response | biological process | GO:0016064 | 51 | 3 | 13711 | 214 | Bcl3,Irf7,Il4ra | | 4.519e-02 | -3.10 | cargo receptor activity | molecular function | GO:0038024 | 51 | 3 | 13516 | 211 | Msr1,Cxcl16,Prg4 | | 4.523e-02 | -3.10 | lymphocyte activation | biological process | GO:0046649 | 299 | 9 | 13711 | 214 | Spi1,Stx11,Msn,Gadd45g,Pik3r1,Bcl3,Tnfaip3,Icam1,Il6 | | 4.531e-02 | -3.09 | lipopolysaccharide receptor complex | cellular component | GO:0046696 | 3 | 1 | 13825 | 212 | Cd14 | | 4.531e-02 | -3.09 | phosphatidylinositol 3-kinase complex, class IB | cellular component | GO:0005944 | 3 | 1 | 13825 | 212 | Pik3r5 | | 4.531e-02 | -3.09 | blood microparticle | cellular component | GO:0072562 | 3 | 1 | 13825 | 212 | Hp | | 4.531e-02 | -3.09 | interleukin-6 receptor complex | cellular component | GO:0005896 | 3 | 1 | 13825 | 212 | Il6 | | 4.531e-02 | -3.09 | platelet alpha granule membrane | cellular component | GO:0031092 | 3 | 1 | 13825 | 212 | Selp | | 4.531e-02 | -3.09 | cyclin D2-CDK4 complex | cellular component | GO:0097129 | 3 | 1 | 13825 | 212 | Ccnd2 | | 4.531e-02 | -3.09 | TAP complex | cellular component | GO:0042825 | 3 | 1 | 13825 | 212 | Tap1 | | 4.539e-02 | -3.09 | positive regulation of transcription by RNA polymerase II | biological process | GO:0045944 | 961 | 22 | 13711 | 214 | Il1a,Sox11,Nfkbia,Nfkb2,Il6,Irf7,Pik3r1,Nfkb1,Sbno2,Cebpd,Bach1,Spi1,Runx1,Tnf,Atf3,Csf3,Ripk1,Tnip1,Maff,Irf2,Nfe2l2,Bcl3 | | 4.539e-02 | -3.09 | GSE24634\_NAIVE\_CD4\_TCELL\_VS\_DAY3\_IL4\_CONV\_TREG\_DN | MSigDB lists | GSE24634\_NAIVE\_CD4\_TCELL\_VS\_DAY3\_IL4\_CONV\_TREG\_DN | 176 | 6 | 12187 | 179 | Csf2rb,Cflar,Tagln2,Tnip3,Tnf,Trib1 | | 4.539e-02 | -3.09 | MANALO\_HYPOXIA\_UP | MSigDB lists | MANALO\_HYPOXIA\_UP | 176 | 6 | 12187 | 179 | Gadd45b,Ier3,Tnfaip3,Col4a1,Maff,Angptl4 | | 4.539e-02 | -3.09 | GSE3039\_ALPHAALPHA\_CD8\_TCELL\_VS\_B1\_BCELL\_UP | MSigDB lists | GSE3039\_ALPHAALPHA\_CD8\_TCELL\_VS\_B1\_BCELL\_UP | 176 | 6 | 12187 | 179 | Ier3,Gbp3,Ptpn1,P2ry6,Ms4a6d,Ccnd2 | | 4.539e-02 | -3.09 | GSE15330\_HSC\_VS\_MEGAKARYOCYTE\_ERYTHROID\_PROGENITOR\_IKAROS\_KO\_UP | MSigDB lists | GSE15330\_HSC\_VS\_MEGAKARYOCYTE\_ERYTHROID\_PROGENITOR\_IKAROS\_KO\_UP | 176 | 6 | 12187 | 179 | Sgk1,Maff,Ier3,Ptpn1,Icam1,Il17ra | | 4.542e-02 | -3.09 | Pyridoxal\_P\_attach\_site | interpro domains | IPR035090 | 3 | 1 | 13788 | 212 | Pygm | | 4.542e-02 | -3.09 | Transglutaminase\_N | interpro domains | IPR001102 | 3 | 1 | 13788 | 212 | Tgm2 | | 4.542e-02 | -3.09 | Syndecan\_CS | interpro domains | IPR030479 | 3 | 1 | 13788 | 212 | Sdc4 | | 4.542e-02 | -3.09 | RGK | interpro domains | IPR017358 | 3 | 1 | 13788 | 212 | Gem | | 4.542e-02 | -3.09 | Luc7-rel | interpro domains | IPR004882 | 3 | 1 | 13788 | 212 | Luc7l3 | | 4.542e-02 | -3.09 | Transglutaminase\_C\_sf | interpro domains | IPR036238 | 3 | 1 | 13788 | 212 | Tgm2 | | 4.542e-02 | -3.09 | OxRdtase\_Mopterin\_BS | interpro domains | IPR022407 | 3 | 1 | 13788 | 212 | Xdh | | 4.542e-02 | -3.09 | Glycg\_phsphrylas | interpro domains | IPR011833 | 3 | 1 | 13788 | 212 | Pygm | | 4.542e-02 | -3.09 | RHIM\_dom | interpro domains | IPR025735 | 3 | 1 | 13788 | 212 | Ripk1 | | 4.542e-02 | -3.09 | Cu-oxidase\_2 | interpro domains | IPR011706 | 3 | 1 | 13788 | 212 | Cp | | 4.542e-02 | -3.09 | RasGAP\_C | interpro domains | IPR000593 | 3 | 1 | 13788 | 212 | Iqgap1 | | 4.542e-02 | -3.09 | PI3K\_P85\_iSH2 | interpro domains | IPR032498 | 3 | 1 | 13788 | 212 | Pik3r1 | | 4.542e-02 | -3.09 | SOX-11/4 | interpro domains | IPR017386 | 3 | 1 | 13788 | 212 | Sox11 | | 4.542e-02 | -3.09 | Glyco\_trans\_35 | interpro domains | IPR000811 | 3 | 1 | 13788 | 212 | Pygm | | 4.542e-02 | -3.09 | Syndecan | interpro domains | IPR001050 | 3 | 1 | 13788 | 212 | Sdc4 | | 4.542e-02 | -3.09 | MG4 | interpro domains | IPR040839 | 3 | 1 | 13788 | 212 | A2m | | 4.542e-02 | -3.09 | Znf-3CxxC | interpro domains | IPR027377 | 3 | 1 | 13788 | 212 | Rtp4 | | 4.542e-02 | -3.09 | Serum\_amyloid\_A | interpro domains | IPR000096 | 3 | 1 | 13788 | 212 | Saa1 | | 4.542e-02 | -3.09 | A2M\_TED | interpro domains | IPR041813 | 3 | 1 | 13788 | 212 | A2m | | 4.542e-02 | -3.09 | DUF4071 | interpro domains | IPR025136 | 3 | 1 | 13788 | 212 | Map3k6 | | 4.542e-02 | -3.09 | Transglutaminase\_C | interpro domains | IPR008958 | 3 | 1 | 13788 | 212 | Tgm2 | | 4.542e-02 | -3.09 | PI3kinase\_P85\_cSH2 | interpro domains | IPR035020 | 3 | 1 | 13788 | 212 | Pik3r1 | | 4.542e-02 | -3.09 | CDI | interpro domains | IPR003175 | 3 | 1 | 13788 | 212 | Cdkn1a | | 4.542e-02 | -3.09 | Transglutaminase\_AS | interpro domains | IPR013808 | 3 | 1 | 13788 | 212 | Tgm2 | | 4.542e-02 | -3.09 | p53/RUNT-type\_TF\_DNA-bd\_sf | interpro domains | IPR012346 | 3 | 1 | 13788 | 212 | Runx1 | | 4.542e-02 | -3.09 | PHsplit\_syntrophin | interpro domains | IPR041428 | 3 | 1 | 13788 | 212 | Sntb2 | | 4.542e-02 | -3.09 | Znf\_NFX1 | interpro domains | IPR000967 | 3 | 1 | 13788 | 212 | Znfx1 | | 4.542e-02 | -3.09 | Transglutaminase\_animal | interpro domains | IPR023608 | 3 | 1 | 13788 | 212 | Tgm2 | | 4.542e-02 | -3.09 | PI3kinase\_P85\_nSH2 | interpro domains | IPR035022 | 3 | 1 | 13788 | 212 | Pik3r1 | | 4.542e-02 | -3.09 | Cyclin\_D | interpro domains | IPR015451 | 3 | 1 | 13788 | 212 | Ccnd2 | | 4.542e-02 | -3.09 | C/EBP\_chordates | interpro domains | IPR016468 | 3 | 1 | 13788 | 212 | Cebpd | | 4.542e-02 | -3.09 | B12D | interpro domains | IPR010530 | 3 | 1 | 13788 | 212 | AA467197 | | 4.542e-02 | -3.09 | K/Na/Ca-exchanger | interpro domains | IPR004481 | 3 | 1 | 13788 | 212 | Slc24a4 | | 4.551e-02 | -3.09 | regulation of carbohydrate biosynthetic process | biological process | GO:0043255 | 86 | 4 | 13711 | 214 | Plek,P2ry6,Il6,Nfkb1 | | 4.551e-02 | -3.09 | cellular response to topologically incorrect protein | biological process | GO:0035967 | 86 | 4 | 13711 | 214 | Atf3,Hspa5,Ptpn1,Nfe2l2 | | 4.551e-02 | -3.09 | regulation of NIK/NF-kappaB signaling | biological process | GO:1901222 | 86 | 4 | 13711 | 214 | Tnf,Mmp8,Cd14,Birc3 | | 4.564e-02 | -3.09 | response to fluid shear stress | biological process | GO:0034405 | 22 | 2 | 13711 | 214 | Ptgs2,Nfe2l2 | | 4.564e-02 | -3.09 | positive regulation of ERBB signaling pathway | biological process | GO:1901186 | 22 | 2 | 13711 | 214 | Ncf1,Plaur | | 4.564e-02 | -3.09 | mature B cell differentiation | biological process | GO:0002335 | 22 | 2 | 13711 | 214 | Bcl3,Tnfaip3 | | 4.564e-02 | -3.09 | ER-nucleus signaling pathway | biological process | GO:0006984 | 22 | 2 | 13711 | 214 | Hspa5,Nfe2l2 | | 4.564e-02 | -3.09 | regulation of epidermal growth factor-activated receptor activity | biological process | GO:0007176 | 22 | 2 | 13711 | 214 | Ncf1,Errfi1 | | 4.564e-02 | -3.09 | regulation of keratinocyte differentiation | biological process | GO:0045616 | 22 | 2 | 13711 | 214 | Errfi1,Zfp36 | | 4.564e-02 | -3.09 | regulation of cofactor metabolic process | biological process | GO:0051193 | 22 | 2 | 13711 | 214 | Hp,Nfe2l2 | | 4.564e-02 | -3.09 | negative regulation of cell junction assembly | biological process | GO:1901889 | 22 | 2 | 13711 | 214 | Tnf,Fam107a | | 4.564e-02 | -3.09 | negative regulation of cyclin-dependent protein serine/threonine kinase activity | biological process | GO:0045736 | 22 | 2 | 13711 | 214 | Cdkn1a,Tnfaip3 | | 4.564e-02 | -3.09 | negative regulation of lipid catabolic process | biological process | GO:0050995 | 22 | 2 | 13711 | 214 | Tnf,Hcar2 | | 4.564e-02 | -3.09 | regulation of interferon-alpha production | biological process | GO:0032647 | 22 | 2 | 13711 | 214 | Irf7,Gbp4 | | 4.566e-02 | -3.09 | cysteine-type endopeptidase inhibitor activity involved in apoptotic process | molecular function | GO:0043027 | 22 | 2 | 13516 | 211 | Tnfaip8,Birc3 | | 4.569e-02 | -3.09 | Znf\_CCCH | interpro domains | IPR000571 | 52 | 3 | 13788 | 212 | Zfp36,Parp12,Tiparp | | 4.609e-02 | -3.08 | Phosphorylase | pfam domains | PF00343 | 3 | 1 | 12881 | 201 | Pygm | | 4.609e-02 | -3.08 | LUC7 | pfam domains | PF03194 | 3 | 1 | 12881 | 201 | Luc7l3 | | 4.609e-02 | -3.08 | DUF4071 | pfam domains | PF13281 | 3 | 1 | 12881 | 201 | Map3k6 | | 4.609e-02 | -3.08 | PH\_17 | pfam domains | PF18012 | 3 | 1 | 12881 | 201 | Sntb2 | | 4.609e-02 | -3.08 | PI3K\_P85\_iSH2 | pfam domains | PF16454 | 3 | 1 | 12881 | 201 | Pik3r1 | | 4.609e-02 | -3.08 | RHIM | pfam domains | PF12721 | 3 | 1 | 12881 | 201 | Ripk1 | | 4.609e-02 | -3.08 | MG4 | pfam domains | PF17789 | 3 | 1 | 12881 | 201 | A2m | | 4.609e-02 | -3.08 | Transglut\_N | pfam domains | PF00868 | 3 | 1 | 12881 | 201 | Tgm2 | | 4.609e-02 | -3.08 | Transglut\_C | pfam domains | PF00927 | 3 | 1 | 12881 | 201 | Tgm2 | | 4.609e-02 | -3.08 | B12D | pfam domains | PF06522 | 3 | 1 | 12881 | 201 | AA467197 | | 4.609e-02 | -3.08 | SAA | pfam domains | PF00277 | 3 | 1 | 12881 | 201 | Saa1 | | 4.609e-02 | -3.08 | CDI | pfam domains | PF02234 | 3 | 1 | 12881 | 201 | Cdkn1a | | 4.609e-02 | -3.08 | Cu-oxidase\_2 | pfam domains | PF07731 | 3 | 1 | 12881 | 201 | Cp | | 4.609e-02 | -3.08 | RasGAP\_C | pfam domains | PF03836 | 3 | 1 | 12881 | 201 | Iqgap1 | | 4.610e-02 | -3.08 | positive regulation of NAD+ ADP-ribosyltransferase activity | biological process | GO:1901666 | 3 | 1 | 13711 | 214 | Dtx3l | | 4.610e-02 | -3.08 | pyrimidine deoxyribonucleoside diphosphate biosynthetic process | biological process | GO:0009197 | 3 | 1 | 13711 | 214 | Cmpk2 | | 4.610e-02 | -3.08 | lipid export from cell | biological process | GO:0140353 | 3 | 1 | 13711 | 214 | Fzd4 | | 4.610e-02 | -3.08 | negative regulation of dendritic cell antigen processing and presentation | biological process | GO:0002605 | 3 | 1 | 13711 | 214 | Thbs1 | | 4.610e-02 | -3.08 | negative regulation of cytokine secretion involved in immune response | biological process | GO:0002740 | 3 | 1 | 13711 | 214 | Tnf | | 4.610e-02 | -3.08 | PERK-mediated unfolded protein response | biological process | GO:0036499 | 3 | 1 | 13711 | 214 | Nfe2l2 | | 4.610e-02 | -3.08 | chromatin silencing at centromere | biological process | GO:0030702 | 3 | 1 | 13711 | 214 | Znfx1 | | 4.610e-02 | -3.08 | detection of chemical stimulus involved in sensory perception of bitter taste | biological process | GO:0001580 | 3 | 1 | 13711 | 214 | Rtp4 | | 4.610e-02 | -3.08 | neutrophil mediated killing of symbiont cell | biological process | GO:0070943 | 3 | 1 | 13711 | 214 | Ncf1 | | 4.610e-02 | -3.08 | dTTP metabolic process | biological process | GO:0046075 | 3 | 1 | 13711 | 214 | Cmpk2 | | 4.610e-02 | -3.08 | positive regulation of intracellular mRNA localization | biological process | GO:1904582 | 3 | 1 | 13711 | 214 | Zfp36 | | 4.610e-02 | -3.08 | negative regulation of interleukin-6 biosynthetic process | biological process | GO:0045409 | 3 | 1 | 13711 | 214 | Prg4 | | 4.610e-02 | -3.08 | cytosol to endoplasmic reticulum transport | biological process | GO:0046967 | 3 | 1 | 13711 | 214 | Tap1 | | 4.610e-02 | -3.08 | T cell activation via T cell receptor contact with antigen bound to MHC molecule on antigen presenting cell | biological process | GO:0002291 | 3 | 1 | 13711 | 214 | Icam1 | | 4.610e-02 | -3.08 | positive regulation of vitamin metabolic process | biological process | GO:0046136 | 3 | 1 | 13711 | 214 | Tnf | | 4.610e-02 | -3.08 | regulation of interleukin-4-mediated signaling pathway | biological process | GO:1902214 | 3 | 1 | 13711 | 214 | Parp14 | | 4.610e-02 | -3.08 | neutrophil degranulation | biological process | GO:0043312 | 3 | 1 | 13711 | 214 | Stx11 | | 4.610e-02 | -3.08 | negative regulation of cGMP-mediated signaling | biological process | GO:0010754 | 3 | 1 | 13711 | 214 | Thbs1 | | 4.610e-02 | -3.08 | regulation of hepatocyte apoptotic process | biological process | GO:1903943 | 3 | 1 | 13711 | 214 | Cflar | | 4.610e-02 | -3.08 | regulation of progesterone secretion | biological process | GO:2000870 | 3 | 1 | 13711 | 214 | Runx1 | | 4.610e-02 | -3.08 | response to glycoside | biological process | GO:1903416 | 3 | 1 | 13711 | 214 | Nfkb1 | | 4.610e-02 | -3.08 | response to diacyl bacterial lipopeptide | biological process | GO:0071724 | 3 | 1 | 13711 | 214 | Cd14 | | 4.610e-02 | -3.08 | dUDP metabolic process | biological process | GO:0046077 | 3 | 1 | 13711 | 214 | Cmpk2 | | 4.610e-02 | -3.08 | pyrimidine deoxyribonucleoside diphosphate metabolic process | biological process | GO:0009196 | 3 | 1 | 13711 | 214 | Cmpk2 | | 4.610e-02 | -3.08 | dTTP biosynthetic process | biological process | GO:0006235 | 3 | 1 | 13711 | 214 | Cmpk2 | | 4.610e-02 | -3.08 | regulation of intrinsic apoptotic signaling pathway in response to osmotic stress | biological process | GO:1902218 | 3 | 1 | 13711 | 214 | Ptgs2 | | 4.610e-02 | -3.08 | negative regulation of interleukin-8 biosynthetic process | biological process | GO:0045415 | 3 | 1 | 13711 | 214 | Bcl3 | | 4.610e-02 | -3.08 | germline stem cell asymmetric division | biological process | GO:0098728 | 3 | 1 | 13711 | 214 | Zbtb16 | | 4.610e-02 | -3.08 | regulation of MyD88-dependent toll-like receptor signaling pathway | biological process | GO:0034124 | 3 | 1 | 13711 | 214 | Irf7 | | 4.610e-02 | -3.08 | regulation of neutrophil differentiation | biological process | GO:0045658 | 3 | 1 | 13711 | 214 | Trib1 | | 4.610e-02 | -3.08 | dUDP biosynthetic process | biological process | GO:0006227 | 3 | 1 | 13711 | 214 | Cmpk2 | | 4.610e-02 | -3.08 | positive regulation of humoral immune response mediated by circulating immunoglobulin | biological process | GO:0002925 | 3 | 1 | 13711 | 214 | Tnf | | 4.610e-02 | -3.08 | regulation of fibrinolysis | biological process | GO:0051917 | 3 | 1 | 13711 | 214 | Thbs1 | | 4.610e-02 | -3.08 | myeloid dendritic cell chemotaxis | biological process | GO:0002408 | 3 | 1 | 13711 | 214 | Retnlg | | 4.610e-02 | -3.08 | negative regulation of interferon-gamma secretion | biological process | GO:1902714 | 3 | 1 | 13711 | 214 | Lgals9 | | 4.610e-02 | -3.08 | D-serine transport | biological process | GO:0042942 | 3 | 1 | 13711 | 214 | Nfkbie | | 4.610e-02 | -3.08 | T cell antigen processing and presentation | biological process | GO:0002457 | 3 | 1 | 13711 | 214 | Icam1 | | 4.610e-02 | -3.08 | negative regulation of vitamin metabolic process | biological process | GO:0046137 | 3 | 1 | 13711 | 214 | Nfkb1 | | 4.610e-02 | -3.08 | germ-line stem cell division | biological process | GO:0042078 | 3 | 1 | 13711 | 214 | Zbtb16 | | 4.610e-02 | -3.08 | natural killer cell degranulation | biological process | GO:0043320 | 3 | 1 | 13711 | 214 | Stx11 | | 4.610e-02 | -3.08 | protein K33-linked deubiquitination | biological process | GO:1990168 | 3 | 1 | 13711 | 214 | Tnfaip3 | | 4.610e-02 | -3.08 | negative regulation of hematopoietic stem cell differentiation | biological process | GO:1902037 | 3 | 1 | 13711 | 214 | Nfe2l2 | | 4.610e-02 | -3.08 | regulation of establishment of Sertoli cell barrier | biological process | GO:1904444 | 3 | 1 | 13711 | 214 | Il1a | | 4.610e-02 | -3.08 | response to triacyl bacterial lipopeptide | biological process | GO:0071725 | 3 | 1 | 13711 | 214 | Cd14 | | 4.610e-02 | -3.08 | steroid hormone secretion | biological process | GO:0035929 | 3 | 1 | 13711 | 214 | Fzd4 | | 4.610e-02 | -3.08 | endothelial cell apoptotic process | biological process | GO:0072577 | 3 | 1 | 13711 | 214 | Tnf | | 4.610e-02 | -3.08 | chromatin silencing by small RNA | biological process | GO:0031048 | 3 | 1 | 13711 | 214 | Znfx1 | | 4.610e-02 | -3.08 | fibroblast activation | biological process | GO:0072537 | 3 | 1 | 13711 | 214 | Il17ra | | 4.610e-02 | -3.08 | regulation of superoxide dismutase activity | biological process | GO:1901668 | 3 | 1 | 13711 | 214 | Tnf | | 4.610e-02 | -3.08 | negative regulation of PERK-mediated unfolded protein response | biological process | GO:1903898 | 3 | 1 | 13711 | 214 | Ptpn1 | | 4.610e-02 | -3.08 | positive regulation of natural killer cell chemotaxis | biological process | GO:2000503 | 3 | 1 | 13711 | 214 | Ccl4 | | 4.610e-02 | -3.08 | negative regulation of nucleotide-binding oligomerization domain containing 2 signaling pathway | biological process | GO:0070433 | 3 | 1 | 13711 | 214 | Tnfaip3 | | 4.610e-02 | -3.08 | regulation of antigen processing and presentation of peptide or polysaccharide antigen via MHC class II | biological process | GO:0002580 | 3 | 1 | 13711 | 214 | Thbs1 | | 4.610e-02 | -3.08 | regulation of natural killer cell degranulation | biological process | GO:0043321 | 3 | 1 | 13711 | 214 | Lgals9 | | 4.610e-02 | -3.08 | cytidine to uridine editing | biological process | GO:0016554 | 3 | 1 | 13711 | 214 | Rbm47 | | 4.610e-02 | -3.08 | detection of chemical stimulus involved in sensory perception of taste | biological process | GO:0050912 | 3 | 1 | 13711 | 214 | Rtp4 | | 4.610e-02 | -3.08 | pyrimidine deoxyribonucleoside triphosphate biosynthetic process | biological process | GO:0009212 | 3 | 1 | 13711 | 214 | Cmpk2 | | 4.610e-02 | -3.08 | negative regulation of Rho-dependent protein serine/threonine kinase activity | biological process | GO:2000299 | 3 | 1 | 13711 | 214 | Rasip1 | | 4.610e-02 | -3.08 | regulation of transcription by glucose | biological process | GO:0046015 | 3 | 1 | 13711 | 214 | Rrp8 | | 4.610e-02 | -3.08 | cellular response to diacyl bacterial lipopeptide | biological process | GO:0071726 | 3 | 1 | 13711 | 214 | Cd14 | | 4.610e-02 | -3.08 | oncostatin-M-mediated signaling pathway | biological process | GO:0038165 | 3 | 1 | 13711 | 214 | Osmr | | 4.610e-02 | -3.08 | pyrimidine nucleoside diphosphate biosynthetic process | biological process | GO:0009139 | 3 | 1 | 13711 | 214 | Cmpk2 | | 4.610e-02 | -3.08 | striated muscle atrophy | biological process | GO:0014891 | 3 | 1 | 13711 | 214 | Cflar | | 4.610e-02 | -3.08 | receptor-mediated virion attachment to host cell | biological process | GO:0046813 | 3 | 1 | 13711 | 214 | Icam1 | | 4.610e-02 | -3.08 | pyrimidine nucleoside diphosphate metabolic process | biological process | GO:0009138 | 3 | 1 | 13711 | 214 | Cmpk2 | | 4.610e-02 | -3.08 | regulation of oligodendrocyte apoptotic process | biological process | GO:1900141 | 3 | 1 | 13711 | 214 | Akap12 | | 4.610e-02 | -3.08 | positive regulation of extracellular exosome assembly | biological process | GO:1903553 | 3 | 1 | 13711 | 214 | Sdc4 | | 4.610e-02 | -3.08 | regulation of glutathione biosynthetic process | biological process | GO:1903786 | 3 | 1 | 13711 | 214 | Nfe2l2 | | 4.610e-02 | -3.08 | negative regulation of nucleotide-binding oligomerization domain containing signaling pathway | biological process | GO:0070425 | 3 | 1 | 13711 | 214 | Tnfaip3 | | 4.610e-02 | -3.08 | male germ-line stem cell asymmetric division | biological process | GO:0048133 | 3 | 1 | 13711 | 214 | Zbtb16 | | 4.610e-02 | -3.08 | neutrophil mediated killing of bacterium | biological process | GO:0070944 | 3 | 1 | 13711 | 214 | Ncf1 | | 4.610e-02 | -3.08 | cyclooxygenase pathway | biological process | GO:0019371 | 3 | 1 | 13711 | 214 | Ptgs2 | | 4.610e-02 | -3.08 | neutrophil mediated cytotoxicity | biological process | GO:0070942 | 3 | 1 | 13711 | 214 | Ncf1 | | 4.610e-02 | -3.08 | regulation of interleukin-10 biosynthetic process | biological process | GO:0045074 | 3 | 1 | 13711 | 214 | Bcl3 | | 4.610e-02 | -3.08 | regulation of antimicrobial humoral response | biological process | GO:0002759 | 3 | 1 | 13711 | 214 | Acod1 | | 4.610e-02 | -3.08 | negative regulation of NLRP3 inflammasome complex assembly | biological process | GO:1900226 | 3 | 1 | 13711 | 214 | Trim30a | | 4.610e-02 | -3.08 | cellular response to triacyl bacterial lipopeptide | biological process | GO:0071727 | 3 | 1 | 13711 | 214 | Cd14 | | 4.610e-02 | -3.08 | positive regulation of glial cell apoptotic process | biological process | GO:0034352 | 3 | 1 | 13711 | 214 | Akap12 | | 4.610e-02 | -3.08 | negative regulation of vascular associated smooth muscle cell migration | biological process | GO:1904753 | 3 | 1 | 13711 | 214 | Nfe2l2 | | 4.610e-02 | -3.08 | positive regulation of CD8-positive, alpha-beta T cell differentiation | biological process | GO:0043378 | 3 | 1 | 13711 | 214 | Runx1 | | 4.610e-02 | -3.08 | peptidyl-glutamine modification | biological process | GO:0018199 | 3 | 1 | 13711 | 214 | Tgm2 | | 4.610e-02 | -3.08 | positive regulation of interleukin-13 secretion | biological process | GO:2000667 | 3 | 1 | 13711 | 214 | Il17ra | | 4.610e-02 | -3.08 | negative regulation of blood vessel endothelial cell proliferation involved in sprouting angiogenesis | biological process | GO:1903588 | 3 | 1 | 13711 | 214 | Thbs1 | | 4.610e-02 | -3.08 | regulation of NAD+ ADP-ribosyltransferase activity | biological process | GO:1901664 | 3 | 1 | 13711 | 214 | Dtx3l | | 4.610e-02 | -3.08 | positive regulation of TRAIL-activated apoptotic signaling pathway | biological process | GO:1903984 | 3 | 1 | 13711 | 214 | Atf3 | | 4.610e-02 | -3.08 | negative regulation of hepatocyte apoptotic process | biological process | GO:1903944 | 3 | 1 | 13711 | 214 | Cflar | | 4.611e-02 | -3.08 | hemoglobin binding | molecular function | GO:0030492 | 3 | 1 | 13516 | 211 | Hp | | 4.611e-02 | -3.08 | ErbB-3 class receptor binding | molecular function | GO:0043125 | 3 | 1 | 13516 | 211 | Pik3r1 | | 4.611e-02 | -3.08 | C-X-C chemokine binding | molecular function | GO:0019958 | 3 | 1 | 13516 | 211 | A2m | | 4.611e-02 | -3.08 | prostaglandin-D synthase activity | molecular function | GO:0004667 | 3 | 1 | 13516 | 211 | Ptges | | 4.611e-02 | -3.08 | collagen V binding | molecular function | GO:0070052 | 3 | 1 | 13516 | 211 | Thbs1 | | 4.611e-02 | -3.08 | uridylate kinase activity | molecular function | GO:0009041 | 3 | 1 | 13516 | 211 | Cmpk2 | | 4.611e-02 | -3.08 | thrombospondin receptor activity | molecular function | GO:0070053 | 3 | 1 | 13516 | 211 | Sdc4 | | 4.611e-02 | -3.08 | oncostatin-M receptor activity | molecular function | GO:0004924 | 3 | 1 | 13516 | 211 | Osmr | | 4.611e-02 | -3.08 | calcium, potassium:sodium antiporter activity | molecular function | GO:0008273 | 3 | 1 | 13516 | 211 | Slc24a4 | | 4.611e-02 | -3.08 | TAP1 binding | molecular function | GO:0046978 | 3 | 1 | 13516 | 211 | Tap1 | | 4.611e-02 | -3.08 | glycogen phosphorylase activity | molecular function | GO:0008184 | 3 | 1 | 13516 | 211 | Pygm | | 4.611e-02 | -3.08 | NFAT protein binding | molecular function | GO:0051525 | 3 | 1 | 13516 | 211 | Spi1 | | 4.611e-02 | -3.08 | MHC class Ib protein binding | molecular function | GO:0023029 | 3 | 1 | 13516 | 211 | Tap1 | | 4.611e-02 | -3.08 | voltage-gated potassium channel activity involved in atrial cardiac muscle cell action potential repolarization | molecular function | GO:0086089 | 3 | 1 | 13516 | 211 | Kcna5 | | 4.611e-02 | -3.08 | tumor necrosis factor binding | molecular function | GO:0043120 | 3 | 1 | 13516 | 211 | A2m | | 4.611e-02 | -3.08 | nitrite reductase activity | molecular function | GO:0098809 | 3 | 1 | 13516 | 211 | Xdh | | 4.611e-02 | -3.08 | opsonin receptor activity | molecular function | GO:0001847 | 3 | 1 | 13516 | 211 | C5ar1 | | 4.611e-02 | -3.08 | ATPase-coupled peptide transmembrane transporter activity | molecular function | GO:0015440 | 3 | 1 | 13516 | 211 | Tap1 | | 4.611e-02 | -3.08 | 3-phosphoinositide-dependent protein kinase binding | molecular function | GO:0043423 | 3 | 1 | 13516 | 211 | Sgk1 | | 4.611e-02 | -3.08 | protein-glutamine gamma-glutamyltransferase activity | molecular function | GO:0003810 | 3 | 1 | 13516 | 211 | Tgm2 | | 4.611e-02 | -3.08 | TAP2 binding | molecular function | GO:0046979 | 3 | 1 | 13516 | 211 | Tap1 | | 4.611e-02 | -3.08 | CCR1 chemokine receptor binding | molecular function | GO:0031726 | 3 | 1 | 13516 | 211 | Ccl4 | | 4.629e-02 | -3.07 | GO\_ORGANIC\_ACID\_BINDING | MSigDB lists | GO\_ORGANIC\_ACID\_BINDING | 133 | 5 | 12187 | 179 | Sele,S100a8,S100a9,Selp,Thbs1 | | 4.629e-02 | -3.07 | GSE11057\_PBMC\_VS\_MEM\_CD4\_TCELL\_UP | MSigDB lists | GSE11057\_PBMC\_VS\_MEM\_CD4\_TCELL\_UP | 133 | 5 | 12187 | 179 | Rassf4,Il1rn,Igsf6,C5ar1,Spi1 | | 4.629e-02 | -3.07 | GSE360\_DC\_VS\_MAC\_T\_GONDII\_UP | MSigDB lists | GSE360\_DC\_VS\_MAC\_T\_GONDII\_UP | 133 | 5 | 12187 | 179 | Trib1,Ccl2,Il1a,P2ry6,Csf2rb | | 4.629e-02 | -3.07 | GSE17974\_CTRL\_VS\_ACT\_IL4\_AND\_ANTI\_IL12\_0.5H\_CD4\_TCELL\_DN | MSigDB lists | GSE17974\_CTRL\_VS\_ACT\_IL4\_AND\_ANTI\_IL12\_0.5H\_CD4\_TCELL\_DN | 133 | 5 | 12187 | 179 | Birc3,Gem,Tnf,Vcam1,Nfe2l2 | | 4.629e-02 | -3.07 | GSE23114\_PERITONEAL\_CAVITY\_B1A\_BCELL\_VS\_SPLEEN\_BCELL\_IN\_SLE2C1\_MOUSE\_DN | MSigDB lists | GSE23114\_PERITONEAL\_CAVITY\_B1A\_BCELL\_VS\_SPLEEN\_BCELL\_IN\_SLE2C1\_MOUSE\_DN | 133 | 5 | 12187 | 179 | Ksr1,Gadd45b,Il6,Tagln2,Nfkbia | | 4.642e-02 | -3.07 | EVI1\_02 | MSigDB lists | EVI1\_02 | 92 | 4 | 12187 | 179 | Sox11,Rasd1,Rbm39,Rbm47 | | 4.644e-02 | -3.07 | GSE13306\_TREG\_VS\_TCONV\_LAMINA\_PROPRIA\_UP | MSigDB lists | GSE13306\_TREG\_VS\_TCONV\_LAMINA\_PROPRIA\_UP | 177 | 6 | 12187 | 179 | Ptpn1,Tnfsf10,Trib1,Nfkbia,Adamts1,Sgk1 | | 4.644e-02 | -3.07 | GSE3920\_UNTREATED\_VS\_IFNA\_TREATED\_FIBROBLAST\_DN | MSigDB lists | GSE3920\_UNTREATED\_VS\_IFNA\_TREATED\_FIBROBLAST\_DN | 177 | 6 | 12187 | 179 | Il6,Casp4,Rrp8,Tnip1,Ifitm3,Irgm2 | | 4.670e-02 | -3.06 | DAZARD\_RESPONSE\_TO\_UV\_NHEK\_DN | MSigDB lists | DAZARD\_RESPONSE\_TO\_UV\_NHEK\_DN | 271 | 8 | 12187 | 179 | Tiparp,Nfkb1,Bach1,Nfe2l2,Cebpd,Aff1,Tnfaip8,Arid5b | | 4.670e-02 | -3.06 | TARTE\_PLASMA\_CELL\_VS\_PLASMABLAST\_DN | MSigDB lists | TARTE\_PLASMA\_CELL\_VS\_PLASMABLAST\_DN | 271 | 8 | 12187 | 179 | Bcl3,Ccnd2,Cdkn1a,Aff1,Sgk1,Ptpn1,Msn,Iqgap1 | | 4.688e-02 | -3.06 | REACTOME\_EXTRACELLULAR\_MATRIX\_ORGANIZATION | MSigDB lists | REACTOME\_EXTRACELLULAR\_MATRIX\_ORGANIZATION | 55 | 3 | 12187 | 179 | Mmp8,Timp1,Col4a1 | | 4.688e-02 | -3.06 | CHIANG\_LIVER\_CANCER\_SUBCLASS\_UNANNOTATED\_UP | MSigDB lists | CHIANG\_LIVER\_CANCER\_SUBCLASS\_UNANNOTATED\_UP | 55 | 3 | 12187 | 179 | Cp,Thbs1,Pik3r1 | | 4.688e-02 | -3.06 | HOFFMANN\_SMALL\_PRE\_BII\_TO\_IMMATURE\_B\_LYMPHOCYTE\_UP | MSigDB lists | HOFFMANN\_SMALL\_PRE\_BII\_TO\_IMMATURE\_B\_LYMPHOCYTE\_UP | 55 | 3 | 12187 | 179 | Icam1,Sdc4,Ccnd2 | | 4.688e-02 | -3.06 | ODONNELL\_TARGETS\_OF\_MYC\_AND\_TFRC\_UP | MSigDB lists | ODONNELL\_TARGETS\_OF\_MYC\_AND\_TFRC\_UP | 55 | 3 | 12187 | 179 | Mxd4,Map3k8,Il4ra | | 4.688e-02 | -3.06 | BRUNO\_HEMATOPOIESIS | MSigDB lists | BRUNO\_HEMATOPOIESIS | 55 | 3 | 12187 | 179 | Slfn3,Slfn4,Ptgs2 | | 4.699e-02 | -3.06 | Hdac2 (histone deacetylase 2) | protein interactions | 15182 | 48 | 3 | 6802 | 115 | Runx1,Spi1,Atf3 | | 4.701e-02 | -3.06 | Toll Like Receptor signaling | WikiPathways | WP88 | 32 | 3 | 3756 | 96 | Irak3,Nfkb2,Nfkb1 | | 4.715e-02 | -3.05 | negative regulation of T cell activation | biological process | GO:0050868 | 87 | 4 | 13711 | 214 | Runx1,Sdc4,Lgals9,Il4ra | | 4.719e-02 | -3.05 | Cell adhesion molecules (CAMs) | KEGG pathways | ko04514 | 97 | 5 | 5248 | 107 | Sdc4,Vcam1,Icam1,Sele,Selp | | 4.719e-02 | -3.05 | Cell adhesion molecules (CAMs) | KEGG pathways | mmu04514 | 97 | 5 | 5248 | 107 | Sele,Selp,Sdc4,Vcam1,Icam1 | | 4.740e-02 | -3.05 | antibiotic metabolic process | biological process | GO:0016999 | 52 | 3 | 13711 | 214 | Cybb,Acod1,Ncf1 | | 4.740e-02 | -3.05 | icosanoid metabolic process | biological process | GO:0006690 | 52 | 3 | 13711 | 214 | Ptgs2,Ptges,Ncf1 | | 4.740e-02 | -3.05 | regulation of epidermal growth factor receptor signaling pathway | biological process | GO:0042058 | 52 | 3 | 13711 | 214 | Plaur,Errfi1,Ncf1 | | 4.740e-02 | -3.05 | regulation of protein processing | biological process | GO:0070613 | 52 | 3 | 13711 | 214 | Thbs1,Birc3,Ctla2a | | 4.750e-02 | -3.05 | ATF1\_Q6 | MSigDB lists | ATF1\_Q6 | 178 | 6 | 12187 | 179 | Sox11,Csf3,Gem,Maff,Trib1,Kcna5 | | 4.750e-02 | -3.05 | GSE27786\_LIN\_NEG\_VS\_CD8\_TCELL\_DN | MSigDB lists | GSE27786\_LIN\_NEG\_VS\_CD8\_TCELL\_DN | 178 | 6 | 12187 | 179 | Socs3,Ccnd2,Samd9l,Irf7,Slfn9,Rtp4 | | 4.750e-02 | -3.05 | GSE31082\_DN\_VS\_DP\_THYMOCYTE\_DN | MSigDB lists | GSE31082\_DN\_VS\_DP\_THYMOCYTE\_DN | 178 | 6 | 12187 | 179 | Bach1,Ip6k2,Tnfaip3,Sgk1,Aff1,Nfkbiz | | 4.754e-02 | -3.05 | DEATH\_DOMAIN | prosite domains | PS50017 | 19 | 2 | 8845 | 164 | Ripk1,Irak3 | | 4.754e-02 | -3.05 | GSE21360\_SECONDARY\_VS\_QUATERNARY\_MEMORY\_CD8\_TCELL\_DN | MSigDB lists | GSE21360\_SECONDARY\_VS\_QUATERNARY\_MEMORY\_CD8\_TCELL\_DN | 134 | 5 | 12187 | 179 | Socs3,Thbs1,Ccnd2,Ptpn1,Tnfaip8 | | 4.754e-02 | -3.05 | GSE13547\_2H\_VS\_12\_H\_ANTI\_IGM\_STIM\_ZFX\_KO\_BCELL\_UP | MSigDB lists | GSE13547\_2H\_VS\_12\_H\_ANTI\_IGM\_STIM\_ZFX\_KO\_BCELL\_UP | 134 | 5 | 12187 | 179 | Casp4,Gem,Ccnd2,Tnfsf10,Irf2 | | 4.754e-02 | -3.05 | GSE36392\_TYPE\_2\_MYELOID\_VS\_MAC\_IL25\_TREATED\_LUNG\_UP | MSigDB lists | GSE36392\_TYPE\_2\_MYELOID\_VS\_MAC\_IL25\_TREATED\_LUNG\_UP | 134 | 5 | 12187 | 179 | Il1rn,Apold1,Cdkn1a,Ptgs2,Socs3 | | 4.754e-02 | -3.05 | GSE6259\_FLT3L\_INDUCED\_33D1\_POS\_DC\_VS\_CD4\_TCELL\_UP | MSigDB lists | GSE6259\_FLT3L\_INDUCED\_33D1\_POS\_DC\_VS\_CD4\_TCELL\_UP | 134 | 5 | 12187 | 179 | Irak3,P2ry6,Cp,Ripk1,Parp14 | | 4.771e-02 | -3.04 | regulation of secretion by cell | biological process | GO:1903530 | 649 | 16 | 13711 | 214 | Runx1,Il17ra,Mmp8,Sdc4,Hcar2,Lgals9,Il1rn,Cd14,Il6,Casp4,Tnf,Il4ra,Tnfaip3,Il1a,Akap12,Sox11 | | 4.790e-02 | -3.04 | B30 | interpro domains | IPR001870 | 53 | 3 | 13788 | 212 | Trim25,Trim30d,Trim30a | | 4.793e-02 | -3.04 | WANG\_METHYLATED\_IN\_BREAST\_CANCER | MSigDB lists | WANG\_METHYLATED\_IN\_BREAST\_CANCER | 24 | 2 | 12187 | 179 | Thbs1,Maff | | 4.793e-02 | -3.04 | GO\_RESPONSE\_TO\_ATP | MSigDB lists | GO\_RESPONSE\_TO\_ATP | 24 | 2 | 12187 | 179 | Ccl12,Ptgs2 | | 4.793e-02 | -3.04 | GO\_CELLULAR\_RESPONSE\_TO\_AMINO\_ACID\_STARVATION | MSigDB lists | GO\_CELLULAR\_RESPONSE\_TO\_AMINO\_ACID\_STARVATION | 24 | 2 | 12187 | 179 | Atf3,Cdkn1a | | 4.793e-02 | -3.04 | WIKMAN\_ASBESTOS\_LUNG\_CANCER\_DN | MSigDB lists | WIKMAN\_ASBESTOS\_LUNG\_CANCER\_DN | 24 | 2 | 12187 | 179 | Ms4a6d,Runx1 | | 4.793e-02 | -3.04 | GO\_REGULATION\_OF\_VASCULAR\_ENDOTHELIAL\_GROWTH\_FACTOR\_RECEPTOR\_SIGNALING\_PATHWAY | MSigDB lists | GO\_REGULATION\_OF\_VASCULAR\_ENDOTHELIAL\_GROWTH\_FACTOR\_RECEPTOR\_SIGNALING\_PATHWAY | 24 | 2 | 12187 | 179 | Fzd4,Ptpn1 | | 4.793e-02 | -3.04 | GLI1\_UP.V1\_UP | MSigDB lists | GLI1\_UP.V1\_UP | 24 | 2 | 12187 | 179 | Ccnd2,Zbtb16 | | 4.793e-02 | -3.04 | PID\_NECTIN\_PATHWAY | MSigDB lists | PID\_NECTIN\_PATHWAY | 24 | 2 | 12187 | 179 | Iqgap1,Pik3r1 | | 4.793e-02 | -3.04 | GO\_RESPONSE\_TO\_PROSTAGLANDIN | MSigDB lists | GO\_RESPONSE\_TO\_PROSTAGLANDIN | 24 | 2 | 12187 | 179 | Adamts1,P2ry6 | | 4.793e-02 | -3.04 | GO\_POSITIVE\_REGULATION\_OF\_VASOCONSTRICTION | MSigDB lists | GO\_POSITIVE\_REGULATION\_OF\_VASOCONSTRICTION | 24 | 2 | 12187 | 179 | Ptgs2,Icam1 | | 4.793e-02 | -3.04 | BIOCARTA\_BAD\_PATHWAY | MSigDB lists | BIOCARTA\_BAD\_PATHWAY | 24 | 2 | 12187 | 179 | Pik3r1,Csf2rb | | 4.793e-02 | -3.04 | WANG\_METASTASIS\_OF\_BREAST\_CANCER\_ESR1\_DN | MSigDB lists | WANG\_METASTASIS\_OF\_BREAST\_CANCER\_ESR1\_DN | 24 | 2 | 12187 | 179 | Phf11b,Phf11d | | 4.793e-02 | -3.04 | GO\_NEGATIVE\_REGULATION\_OF\_RESPONSE\_TO\_BIOTIC\_STIMULUS | MSigDB lists | GO\_NEGATIVE\_REGULATION\_OF\_RESPONSE\_TO\_BIOTIC\_STIMULUS | 24 | 2 | 12187 | 179 | Trib1,Tnfaip3 | | 4.793e-02 | -3.04 | ROZANOV\_MMP14\_TARGETS\_DN | MSigDB lists | ROZANOV\_MMP14\_TARGETS\_DN | 24 | 2 | 12187 | 179 | Sdc4,Pik3r1 | | 4.793e-02 | -3.04 | GO\_OSTEOCLAST\_DIFFERENTIATION | MSigDB lists | GO\_OSTEOCLAST\_DIFFERENTIATION | 24 | 2 | 12187 | 179 | Sbno2,Tnf | | 4.793e-02 | -3.04 | PARK\_TRETINOIN\_RESPONSE\_AND\_PML\_RARA\_FUSION | MSigDB lists | PARK\_TRETINOIN\_RESPONSE\_AND\_PML\_RARA\_FUSION | 24 | 2 | 12187 | 179 | Spi1,Ier3 | | 4.793e-02 | -3.04 | SATO\_SILENCED\_EPIGENETICALLY\_IN\_PANCREATIC\_CANCER | MSigDB lists | SATO\_SILENCED\_EPIGENETICALLY\_IN\_PANCREATIC\_CANCER | 24 | 2 | 12187 | 179 | Csf3,Mmp8 | | 4.793e-02 | -3.04 | VALK\_AML\_CLUSTER\_3 | MSigDB lists | VALK\_AML\_CLUSTER\_3 | 24 | 2 | 12187 | 179 | Cebpd,Il17ra | | 4.798e-02 | -3.04 | GO\_SKELETAL\_MUSCLE\_ORGAN\_DEVELOPMENT | MSigDB lists | GO\_SKELETAL\_MUSCLE\_ORGAN\_DEVELOPMENT | 93 | 4 | 12187 | 179 | Atf3,Maff,Sox11,Cflar | | 4.798e-02 | -3.04 | RB\_P130\_DN.V1\_UP | MSigDB lists | RB\_P130\_DN.V1\_UP | 93 | 4 | 12187 | 179 | Rasd1,Rgs16,Ier3,Rhoj | | 4.798e-02 | -3.04 | GO\_NEGATIVE\_REGULATION\_OF\_ION\_TRANSPORT | MSigDB lists | GO\_NEGATIVE\_REGULATION\_OF\_ION\_TRANSPORT | 93 | 4 | 12187 | 179 | Kcna5,Icam1,Ptgs2,Thbs1 | | 4.819e-02 | -3.03 | Eicosanoid Synthesis | WikiPathways | WP318 | 14 | 2 | 3756 | 96 | Ptgs2,Ptges | | 4.828e-02 | -3.03 | Cdk4 (cyclin-dependent kinase 4) | protein interactions | 12567 | 21 | 2 | 6802 | 115 | Cdkn1a,Ccnd2 | | 4.828e-02 | -3.03 | Csf1r (colony stimulating factor 1 receptor) | protein interactions | 12978 | 21 | 2 | 6802 | 115 | Pik3r1,Spi1 | | 4.828e-02 | -3.03 | Ptk2 (PTK2 protein tyrosine kinase 2) | protein interactions | 14083 | 21 | 2 | 6802 | 115 | Socs3,Pik3r1 | | 4.857e-02 | -3.02 | AML\_Q6 | MSigDB lists | AML\_Q6 | 179 | 6 | 12187 | 179 | Ccl12,Luc7l3,Pik3r1,Runx1,Sntb2,Trib1 | | 4.857e-02 | -3.02 | GSE22886\_IGM\_MEMORY\_BCELL\_VS\_BLOOD\_PLASMA\_CELL\_DN | MSigDB lists | GSE22886\_IGM\_MEMORY\_BCELL\_VS\_BLOOD\_PLASMA\_CELL\_DN | 179 | 6 | 12187 | 179 | Rgs16,Clic4,Sgk1,Stx11,Cebpd,Mxd4 | | 4.862e-02 | -3.02 | RODRIGUES\_THYROID\_CARCINOMA\_POORLY\_DIFFERENTIATED\_DN | MSigDB lists | RODRIGUES\_THYROID\_CARCINOMA\_POORLY\_DIFFERENTIATED\_DN | 638 | 15 | 12187 | 179 | Col4a1,Ksr1,Thbs1,Sgk1,Phf11b,Apold1,Rnf213,Gadd45b,Tnfaip3,Rassf4,Nfkbia,Ccnd2,Tnfaip8,Phf11d,Cflar | | 4.881e-02 | -3.02 | ACCATTT\_MIR522 | MSigDB lists | ACCATTT\_MIR522 | 135 | 5 | 12187 | 179 | Ccnd2,Arid5b,Nfkbiz,Ptpn1,Nfe2l2 | | 4.881e-02 | -3.02 | GSE15624\_3H\_VS\_6H\_HALOFUGINONE\_TREATED\_CD4\_TCELL\_UP | MSigDB lists | GSE15624\_3H\_VS\_6H\_HALOFUGINONE\_TREATED\_CD4\_TCELL\_UP | 135 | 5 | 12187 | 179 | Il17ra,Ifitm3,Hspa5,Vcam1,Tgm2 | | 4.881e-02 | -3.02 | GSE3565\_CTRL\_VS\_LPS\_INJECTED\_SPLENOCYTES\_UP | MSigDB lists | GSE3565\_CTRL\_VS\_LPS\_INJECTED\_SPLENOCYTES\_UP | 135 | 5 | 12187 | 179 | Clic4,Xdh,Pik3ap1,Pik3r5,Runx1 | | 4.881e-02 | -3.02 | sensory perception of pain | biological process | GO:0019233 | 88 | 4 | 13711 | 214 | Tnf,Il1rn,Ptgs2,Ccl2 | | 4.883e-02 | -3.02 | positive regulation of protein modification by small protein conjugation or removal | biological process | GO:1903322 | 127 | 5 | 13711 | 214 | Trib1,Birc3,Fam107a,Hspa5,Tnip1 | | 4.901e-02 | -3.02 | GO\_DETOXIFICATION | MSigDB lists | GO\_DETOXIFICATION | 56 | 3 | 12187 | 179 | Hp,Ptgs2,S100a9 | | 4.901e-02 | -3.02 | DUAN\_PRDM5\_TARGETS | MSigDB lists | DUAN\_PRDM5\_TARGETS | 56 | 3 | 12187 | 179 | Gadd45b,Socs3,Runx1 | | 4.901e-02 | -3.02 | GO\_FEMALE\_GAMETE\_GENERATION | MSigDB lists | GO\_FEMALE\_GAMETE\_GENERATION | 56 | 3 | 12187 | 179 | Adamts1,Ptgs2,Il4ra | | 4.901e-02 | -3.02 | GO\_HORMONE\_TRANSPORT | MSigDB lists | GO\_HORMONE\_TRANSPORT | 56 | 3 | 12187 | 179 | Il6,Il1rn,Fzd4 | | 4.901e-02 | -3.02 | GO\_NEGATIVE\_REGULATION\_OF\_LIPID\_METABOLIC\_PROCESS | MSigDB lists | GO\_NEGATIVE\_REGULATION\_OF\_LIPID\_METABOLIC\_PROCESS | 56 | 3 | 12187 | 179 | Tnf,Hcar2,Nfkb1 | | 4.901e-02 | -3.02 | RHEIN\_ALL\_GLUCOCORTICOID\_THERAPY\_UP | MSigDB lists | RHEIN\_ALL\_GLUCOCORTICOID\_THERAPY\_UP | 56 | 3 | 12187 | 179 | S100a8,Map3k8,Birc3 | | 4.901e-02 | -3.02 | GGCACAT\_MIR455 | MSigDB lists | GGCACAT\_MIR455 | 56 | 3 | 12187 | 179 | Luc7l3,Sox11,Socs3 | | 4.920e-02 | -3.01 | PTK6 Activates STAT3 | REACTOME pathways | R-MMU-8849474 | 3 | 1 | 6297 | 105 | Socs3 | | 4.920e-02 | -3.01 | Dissolution of Fibrin Clot | REACTOME pathways | R-MMU-75205 | 3 | 1 | 6297 | 105 | Plaur | | 4.920e-02 | -3.01 | Interleukin-17 signaling | REACTOME pathways | R-MMU-448424 | 3 | 1 | 6297 | 105 | Il17ra | | 4.925e-02 | -3.01 | Programmed Cell Death | REACTOME pathways | R-MMU-5357801 | 83 | 4 | 6297 | 105 | Ripk1,Cflar,Tnfsf10,Birc3 | | 4.949e-02 | -3.01 | positive regulation of alcohol biosynthetic process | biological process | GO:1902932 | 23 | 2 | 13711 | 214 | Tnf,P2ry6 | | 4.949e-02 | -3.01 | positive regulation of defense response to virus by host | biological process | GO:0002230 | 23 | 2 | 13711 | 214 | Parp9,Dtx3l | | 4.949e-02 | -3.01 | embryonic placenta morphogenesis | biological process | GO:0060669 | 23 | 2 | 13711 | 214 | Socs3,Vcam1 | | 4.949e-02 | -3.01 | negative regulation of sprouting angiogenesis | biological process | GO:1903671 | 23 | 2 | 13711 | 214 | Adamts9,Thbs1 | | 4.966e-02 | -3.00 | KEGG\_FOCAL\_ADHESION | MSigDB lists | KEGG\_FOCAL\_ADHESION | 180 | 6 | 12187 | 179 | Ccnd2,Birc3,Pik3r1,Thbs1,Pik3r5,Col4a1 | | 4.966e-02 | -3.00 | GSE9960\_GRAM\_POS\_VS\_GRAM\_NEG\_AND\_POS\_SEPSIS\_PBMC\_UP | MSigDB lists | GSE9960\_GRAM\_POS\_VS\_GRAM\_NEG\_AND\_POS\_SEPSIS\_PBMC\_UP | 180 | 6 | 12187 | 179 | Atf3,Cebpd,Ifit2,Usp18,Ifit3b,Zfp36 | | 4.969e-02 | -3.00 | positive regulation of lipid transport | biological process | GO:0032370 | 53 | 3 | 13711 | 214 | Runx1,Il1a,Nfkbia | | 4.969e-02 | -3.00 | lipid phosphorylation | biological process | GO:0046834 | 53 | 3 | 13711 | 214 | Pik3r1,Ip6k2,Socs3 | | 4.969e-02 | -3.00 | negative regulation of calcium ion transport | biological process | GO:0051926 | 53 | 3 | 13711 | 214 | Ptgs2,Gem,Icam1 | | 4.971e-02 | -3.00 | kinase inhibitor activity | molecular function | GO:0019210 | 53 | 3 | 13516 | 211 | Trib1,Hspb1,Cdkn1a | | 4.987e-02 | -3.00 | PPIB (peptidylprolyl isomerase B) | protein interactions | 5479 | 3 | 1 | 6802 | 115 | Ksr1 | | 4.987e-02 | -3.00 | MAP2K1 (mitogen-activated protein kinase kinase 1) | protein interactions | 5604 | 3 | 1 | 6802 | 115 | Ksr1 | | 4.987e-02 | -3.00 | GNS (glucosamine (N-acetyl)-6-sulfatase) | protein interactions | 2799 | 3 | 1 | 6802 | 115 | Msn | | 4.987e-02 | -3.00 | PSMC4 (proteasome 26S subunit, ATPase 4) | protein interactions | 5704 | 3 | 1 | 6802 | 115 | Ksr1 | | 4.987e-02 | -3.00 | Itga3 (integrin alpha 3) | protein interactions | 16400 | 3 | 1 | 6802 | 115 | Sdc4 | | 4.987e-02 | -3.00 | CREBBP (CREB binding protein) | protein interactions | 1387 | 3 | 1 | 6802 | 115 | Spi1 | | 4.987e-02 | -3.00 | TUBB6 (tubulin beta 6 class V) | protein interactions | 84617 | 3 | 1 | 6802 | 115 | Ksr1 | | 4.987e-02 | -3.00 | Nyap1 (neuronal tyrosine-phosphorylated phosphoinositide 3-kinase adaptor 1) | protein interactions | 243300 | 3 | 1 | 6802 | 115 | Pik3r1 | | 4.987e-02 | -3.00 | Dnajc1 (DnaJ heat shock protein family (Hsp40) member C1) | protein interactions | 13418 | 3 | 1 | 6802 | 115 | Hspa5 | | 4.987e-02 | -3.00 | Isg15 (ISG15 ubiquitin-like modifier) | protein interactions | 100038882 | 3 | 1 | 6802 | 115 | Usp18 | | 4.987e-02 | -3.00 | HDAC6 (histone deacetylase 6) | protein interactions | 10013 | 3 | 1 | 6802 | 115 | Ksr1 | | 4.987e-02 | -3.00 | Mapk6 (mitogen-activated protein kinase 6) | protein interactions | 50772 | 3 | 1 | 6802 | 115 | Hspb1 | | 4.987e-02 | -3.00 | LRRC59 (leucine rich repeat containing 59) | protein interactions | 55379 | 3 | 1 | 6802 | 115 | Ksr1 | | 4.987e-02 | -3.00 | EXOSC6 (exosome component 6) | protein interactions | 118460 | 3 | 1 | 6802 | 115 | Ksr1 | | 4.987e-02 | -3.00 | RPL28 (ribosomal protein L28) | protein interactions | 6158 | 3 | 1 | 6802 | 115 | Ksr1 | | 4.987e-02 | -3.00 | Btla (B and T lymphocyte associated) | protein interactions | 208154 | 3 | 1 | 6802 | 115 | Pik3r1 | | 4.987e-02 | -3.00 | Tcrb (T cell receptor beta chain) | protein interactions | 21577 | 3 | 1 | 6802 | 115 | Runx1 | | 4.987e-02 | -3.00 | Zfp36 (zinc finger protein 36) | protein interactions | 22695 | 3 | 1 | 6802 | 115 | Tnf | | 4.987e-02 | -3.00 | STMN1 (stathmin 1) | protein interactions | 3925 | 3 | 1 | 6802 | 115 | Hspa5 | | 4.987e-02 | -3.00 | FAM83D (family with sequence similarity 83 member D) | protein interactions | 81610 | 3 | 1 | 6802 | 115 | Bach1 | | 4.987e-02 | -3.00 | PSMD13 (proteasome 26S subunit, non-ATPase 13) | protein interactions | 5719 | 3 | 1 | 6802 | 115 | Ksr1 | | 4.987e-02 | -3.00 | Sdf2l1 (stromal cell-derived factor 2-like 1) | protein interactions | 64136 | 3 | 1 | 6802 | 115 | Hspa5 | | 4.987e-02 | -3.00 | PSMB3 (proteasome subunit beta 3) | protein interactions | 5691 | 3 | 1 | 6802 | 115 | Ksr1 | | 4.987e-02 | -3.00 | Mapk7 (mitogen-activated protein kinase 7) | protein interactions | 23939 | 3 | 1 | 6802 | 115 | Nfe2l2 | | 4.987e-02 | -3.00 | Tmem115 (transmembrane protein 115) | protein interactions | 56395 | 3 | 1 | 6802 | 115 | Tnf | | 4.987e-02 | -3.00 | AKAP8L (A-kinase anchoring protein 8 like) | protein interactions | 26993 | 3 | 1 | 6802 | 115 | Ksr1 | | 4.987e-02 | -3.00 | PSMB1 (proteasome subunit beta 1) | protein interactions | 5689 | 3 | 1 | 6802 | 115 | Ksr1 | | 4.987e-02 | -3.00 | Pcgf5 (polycomb group ring finger 5) | protein interactions | 76073 | 3 | 1 | 6802 | 115 | Runx1 | | 4.987e-02 | -3.00 | MLLT3 (MLLT3 super elongation complex subunit) | protein interactions | 4300 | 3 | 1 | 6802 | 115 | Aff1 | | 4.987e-02 | -3.00 | Hist2h3c1 (histone cluster 2, H3c1) | protein interactions | 15077 | 3 | 1 | 6802 | 115 | Il6 | | 4.987e-02 | -3.00 | PPP6R3 (protein phosphatase 6 regulatory subunit 3) | protein interactions | 55291 | 3 | 1 | 6802 | 115 | Ksr1 | | 4.987e-02 | -3.00 | Casp6 (caspase 6) | protein interactions | 12368 | 3 | 1 | 6802 | 115 | Irak3 | | 4.987e-02 | -3.00 | RARA (retinoic acid receptor alpha) | protein interactions | 5914 | 3 | 1 | 6802 | 115 | Gadd45b | | 4.987e-02 | -3.00 | ATXN3 (ataxin 3) | protein interactions | 4287 | 3 | 1 | 6802 | 115 | Aff1 | | 4.987e-02 | -3.00 | DYSF (dysferlin) | protein interactions | 8291 | 3 | 1 | 6802 | 115 | Msn | | 4.987e-02 | -3.00 | HSD17B4 (hydroxysteroid 17-beta dehydrogenase 4) | protein interactions | 3295 | 3 | 1 | 6802 | 115 | Ksr1 | | 4.987e-02 | -3.00 | B2m (beta-2 microglobulin) | protein interactions | 12010 | 3 | 1 | 6802 | 115 | Tap1 | | 4.987e-02 | -3.00 | Cd4 (CD4 antigen) | protein interactions | 12504 | 3 | 1 | 6802 | 115 | Runx1 | | 4.987e-02 | -3.00 | Nup98 (nucleoporin 98) | protein interactions | 269966 | 3 | 1 | 6802 | 115 | Zbtb16 | | 4.987e-02 | -3.00 | Rsad2 (radical S-adenosyl methionine domain containing 2) | protein interactions | 58185 | 3 | 1 | 6802 | 115 | Irf7 | | 4.987e-02 | -3.00 | CCDC82 (coiled-coil domain containing 82) | protein interactions | 79780 | 3 | 1 | 6802 | 115 | Aff1 | | 4.987e-02 | -3.00 | Il3ra (interleukin 3 receptor, alpha chain) | protein interactions | 16188 | 3 | 1 | 6802 | 115 | Csf2rb | | 4.987e-02 | -3.00 | HEXB (hexosaminidase subunit beta) | protein interactions | 3074 | 3 | 1 | 6802 | 115 | Bach1 | | 4.987e-02 | -3.00 | Nod2 (nucleotide-binding oligomerization domain containing 2) | protein interactions | 257632 | 3 | 1 | 6802 | 115 | Socs3 | | 4.987e-02 | -3.00 | RB1 (RB transcriptional corepressor 1) | protein interactions | 5925 | 3 | 1 | 6802 | 115 | Ifi211 | | 4.987e-02 | -3.00 | DNAJC10 (DnaJ heat shock protein family (Hsp40) member C10) | protein interactions | 54431 | 3 | 1 | 6802 | 115 | Ksr1 | | 4.987e-02 | -3.00 | Nphs1 (nephrosis 1, nephrin) | protein interactions | 54631 | 3 | 1 | 6802 | 115 | Iqgap1 | | 4.987e-02 | -3.00 | Fkbpl (FK506 binding protein-like) | protein interactions | 56299 | 3 | 1 | 6802 | 115 | Cdkn1a | | 4.987e-02 | -3.00 | Slit3 (slit guidance ligand 3) | protein interactions | 20564 | 3 | 1 | 6802 | 115 | Robo4 | | 4.987e-02 | -3.00 | Myo16 (myosin XVI) | protein interactions | 244281 | 3 | 1 | 6802 | 115 | Pik3r1 | | 4.987e-02 | -3.00 | MRPS34 (mitochondrial ribosomal protein S34) | protein interactions | 65993 | 3 | 1 | 6802 | 115 | Ksr1 | | 4.987e-02 | -3.00 | LUC7L2 (LUC7 like 2, pre-mRNA splicing factor) | protein interactions | 51631 | 3 | 1 | 6802 | 115 | Ksr1 | | 4.987e-02 | -3.00 | KIF1BP (KIF1 binding protein) | protein interactions | 26128 | 3 | 1 | 6802 | 115 | Ksr1 | | 4.987e-02 | -3.00 | RPL9 (ribosomal protein L9) | protein interactions | 6133 | 3 | 1 | 6802 | 115 | Ksr1 | | 4.987e-02 | -3.00 | PSMD6 (proteasome 26S subunit, non-ATPase 6) | protein interactions | 9861 | 3 | 1 | 6802 | 115 | Ksr1 | | 4.987e-02 | -3.00 | Hsf1 (heat shock factor 1) | protein interactions | 15499 | 3 | 1 | 6802 | 115 | Atf3 | | 4.987e-02 | -3.00 | Esr2 (estrogen receptor 2 (beta)) | protein interactions | 13983 | 3 | 1 | 6802 | 115 | Rbm39 | | 4.987e-02 | -3.00 | Zfp277 (zinc finger protein 277) | protein interactions | 246196 | 3 | 1 | 6802 | 115 | Cebpd | | 4.991e-02 | -3.00 | mouse chr8 | chromosome location | mouse chr8 | 220 | 7 | 14556 | 220 | Mt1,Mt2,Hp,Col4a1,Msr1,Sntb2,Banp |
