## Supplementary material for "Repeated LPS induces training and tolerance of microglial responses across brain regions": FileS1: homerResults.html

2xLPS\_cluster\_genes\_output// - Homer de novo Motif Results


### Homer *de novo* Motif Results (2xLPS\_cluster\_genes\_output//)

Known Motif Enrichment Results  
Gene Ontology Enrichment Results  
If Homer is having trouble matching a motif to a known motif, try copy/pasting the matrix file into
STAMP  
More information on motif finding results: HOMER
| Description of Results
| Tips
  
Total target sequences = 216  
Total background sequences = 13262  
\* - possible false positive  

|  |  |  |  |  |  |  |  |  |
| --- | --- | --- | --- | --- | --- | --- | --- | --- |
| Rank | Motif | P-value | log P-pvalue | % of Targets | % of Background | STD(Bg STD) | Best Match/Details | Motif File |
| 1 | C G T A G T C A C G T A A G T C C T A G C G T A C G T A C T G A T A G C A G C T C A T G T G C A | 1e-21 | -4.867e+01 | 18.52% | 2.55% | 240.1bp (449.6bp) | PB0037.1\_Isgf3g\_1/Jaspar(0.939) More Information \| Similar Motifs Found | motif file (matrix) |
| 2 | T C A G C T A G T C G A T G C A C T G A G C A T A C G T A G T C A G T C G T A C G C T A T A C G | 1e-13 | -3.087e+01 | 9.26% | 0.88% | 221.5bp (416.8bp) | NFkB-p65-Rel(RHD)/ThioMac-LPS-Expression(GSE23622)/Homer(0.908) More Information \| Similar Motifs Found | motif file (matrix) |
| 3 | T C A G A T G C T C G A T C G A A C T G A T C G C A G T A C G T A T G C C G A T T C A G A T G C | 1e-12 | -2.835e+01 | 11.57% | 1.76% | 302.1bp (420.6bp) | OSR2/MA1646.1/Jaspar(0.566) More Information \| Similar Motifs Found | motif file (matrix) |
| 4 | G T A C C G T A A T C G A T G C A C G T A C T G T A C G A C G T A C G T A C G T A G C T C G A T | 1e-12 | -2.815e+01 | 8.80% | 0.89% | 383.1bp (397.3bp) | Tcf12(bHLH)/GM12878-Tcf12-ChIP-Seq(GSE32465)/Homer(0.723) More Information \| Similar Motifs Found | motif file (matrix) |
| 5 \* | C G T A A C G T A C T G A C T G A G T C A G T C C G T A C T G A A G T C C G T A A G T C C G T A | 1e-11 | -2.607e+01 | 4.17% | 0.08% | 284.3bp (448.1bp) | PB0121.1\_Foxj3\_2/Jaspar(0.580) More Information \| Similar Motifs Found | motif file (matrix) |
| 6 \* | C G A T A T C G G C T A C A T G A C T G G A C T A C T G T C G A A T G C A C G T | 1e-10 | -2.472e+01 | 13.89% | 3.10% | 359.6bp (431.5bp) | SREBF1/MA0595.1/Jaspar(0.721) More Information \| Similar Motifs Found | motif file (matrix) |
| 7 \* | G T A C A C G T C G A T A C T G T C G A A C G T A T G C A G T C A G T C A C G T A C T G A T C G | 1e-10 | -2.456e+01 | 5.09% | 0.22% | 239.2bp (514.8bp) | PB0030.1\_Hnf4a\_1/Jaspar(0.670) More Information \| Similar Motifs Found | motif file (matrix) |
| 8 \* | G C T A T G A C A T G C C G T A A C T G A G T C A G T C A C T G A G T C A T G C G T A C A C T G | 1e-10 | -2.372e+01 | 7.87% | 0.90% | 398.2bp (280.6bp) | PB0151.1\_Myf6\_2/Jaspar(0.634) More Information \| Similar Motifs Found | motif file (matrix) |
| 9 \* | T A G C C A G T A C T G C G T A G A T C C G A T G A T C G T C A T A C G G A T C | 1e-10 | -2.348e+01 | 29.17% | 12.35% | 293.6bp (436.0bp) | MAFK/MA0496.3/Jaspar(0.891) More Information \| Similar Motifs Found | motif file (matrix) |
| 10 \* | A C T G C G T A A T G C A C G T A T G C C G T A C T G A A C T G A G T C A G T C G T C A C T G A | 1e-9 | -2.255e+01 | 4.63% | 0.20% | 443.8bp (527.6bp) | ZBTB6/MA1581.1/Jaspar(0.635) More Information \| Similar Motifs Found | motif file (matrix) |
| 11 \* | A C T G G T A C C G T A A C T G A C T G G T C A C G T A G T C A A G C T A G C T A G C T A C G T | 1e-9 | -2.240e+01 | 5.56% | 0.37% | 261.0bp (393.3bp) | EWS:ERG-fusion(ETS)/CADO\_ES1-EWS:ERG-ChIP-Seq(SRA014231)/Homer(0.814) More Information \| Similar Motifs Found | motif file (matrix) |
| 12 \* | C A T G T A G C A C G T G T C A G A C T C T G A C G T A C G T A T G C A T A C G | 1e-9 | -2.216e+01 | 12.04% | 2.60% | 306.7bp (417.7bp) | TATA-Box(TBP)/Promoter/Homer(0.861) More Information \| Similar Motifs Found | motif file (matrix) |
| 13 \* | A T G C G T C A A C G T A G T C A C G T A C T G A C T G G T A C A G T C C G T A A C T G A C G T | 1e-9 | -2.140e+01 | 4.63% | 0.23% | 401.1bp (522.1bp) | Bcl11a(Zf)/HSPC-BCL11A-ChIP-Seq(GSE104676)/Homer(0.717) More Information \| Similar Motifs Found | motif file (matrix) |
| 14 \* | A C T G A C T G A G T C A T G C C G T A A G T C A C G T A C G T A G T C A C G T A C G T A C T G | 1e-8 | -2.071e+01 | 2.31% | 0.01% | 412.4bp (0.0bp) | ELF1/MA0473.3/Jaspar(0.714) More Information \| Similar Motifs Found | motif file (matrix) |
| 15 \* | C G T A A C T G C T A G C G T A A C T G A G C T A G T C A T G C A C G T C T A G C T A G G T C A | 1e-8 | -2.062e+01 | 4.63% | 0.25% | 226.9bp (343.2bp) | PB0203.1\_Zfp691\_2/Jaspar(0.670) More Information \| Similar Motifs Found | motif file (matrix) |
| 16 \* | A C T G T A G C C G T A C T G A G A C T G T C A A T G C C G A T A T G C G A T C | 1e-8 | -1.981e+01 | 7.41% | 1.02% | 300.0bp (405.8bp) | Ddit3::Cebpa/MA0019.1/Jaspar(0.652) More Information \| Similar Motifs Found | motif file (matrix) |
| 17 \* | T G C A C A T G A T G C G C T A C G A T A C T G G T C A A G T C C G T A C T A G | 1e-8 | -1.935e+01 | 12.50% | 3.22% | 258.1bp (490.9bp) | MEIS1/MA0498.2/Jaspar(0.741) More Information \| Similar Motifs Found | motif file (matrix) |
| 18 \* | C G A T A C T G A C T G C G T A A G T C C T G A A G T C C G A T A G C T A C G T | 1e-8 | -1.909e+01 | 8.33% | 1.41% | 340.9bp (400.5bp) | PB0134.1\_Hnf4a\_2/Jaspar(0.587) More Information \| Similar Motifs Found | motif file (matrix) |
| 19 \* | C G A T T C A G T A G C G T C A C T A G A G T C A G T C A G T C G T C A C G A T A T G C C A G T | 1e-8 | -1.909e+01 | 8.33% | 1.42% | 355.7bp (443.2bp) | NEUROG2(var.2)/MA1642.1/Jaspar(0.690) More Information \| Similar Motifs Found | motif file (matrix) |
| 20 \* | T C A G A C T G C T A G C T G A A T C G G C T A C A G T G A T C A T G C G C T A C G A T A T G C | 1e-8 | -1.869e+01 | 5.56% | 0.54% | 343.8bp (405.0bp) | HOXA2(Homeobox)/mES-Hoxa2-ChIP-Seq(Donaldson\_et\_al.)/Homer(0.637) More Information \| Similar Motifs Found | motif file (matrix) |
| 21 \* | G T C A A G T C G T A C C G T A C G A T A T C G A G T C G C T A A G C T A C T G | 1e-8 | -1.851e+01 | 8.80% | 1.65% | 299.5bp (383.2bp) | SOX18/MA1563.1/Jaspar(0.616) More Information \| Similar Motifs Found | motif file (matrix) |
| 22 \* | C A T G T C G A C T A G C G T A A T G C T A G C C A T G A G C T A T C G A C G T | 1e-7 | -1.820e+01 | 29.17% | 14.13% | 320.4bp (400.6bp) | PB0200.1\_Zfp187\_2/Jaspar(0.588) More Information \| Similar Motifs Found | motif file (matrix) |
| 23 \* | A G C T G T A C C G T A G T C A A C G T G T C A C A T G C T G A G T C A G T C A | 1e-7 | -1.756e+01 | 12.50% | 3.52% | 304.2bp (401.4bp) | Hnf6b(Homeobox)/LNCaP-Hnf6b-ChIP-Seq(GSE106305)/Homer(0.735) More Information \| Similar Motifs Found | motif file (matrix) |
| 24 \* | A T C G A C T G A C T G T C G A C T A G T G C A A T G C C G A T T A C G A T G C G C T A G T A C | 1e-7 | -1.751e+01 | 8.80% | 1.77% | 343.2bp (403.4bp) | PB0203.1\_Zfp691\_2/Jaspar(0.568) More Information \| Similar Motifs Found | motif file (matrix) |
| 25 \* | G C A T A C G T A C G T A G C T C A G T A C T G C G T A G C A T A G T C C G T A | 1e-7 | -1.667e+01 | 22.22% | 9.70% | 338.6bp (403.7bp) | Arid3a/MA0151.1/Jaspar(0.688) More Information \| Similar Motifs Found | motif file (matrix) |
| 26 \* | T G A C C T A G A T C G G C T A C G T A C G T A A G T C A G C T A G C T A G T C C G A T A C T G | 1e-6 | -1.536e+01 | 6.02% | 0.91% | 299.9bp (430.5bp) | OSR2/MA1646.1/Jaspar(0.720) More Information \| Similar Motifs Found | motif file (matrix) |
| 27 \* | C G T A A C T G T G A C G T A C G A T C G A C T G C A T A G C T A C G T C G A T C G T A G C A T | 1e-6 | -1.519e+01 | 6.02% | 0.93% | 207.6bp (393.6bp) | ZFP42/MA1651.1/Jaspar(0.611) More Information \| Similar Motifs Found | motif file (matrix) |
| 28 \* | C T A G A G C T C A G T A C T G A C T G C G A T C A T G T C A G C A G T T C A G A T C G G A C T | 1e-6 | -1.517e+01 | 13.89% | 4.78% | 295.8bp (418.6bp) | ZBTB7C/MA0695.1/Jaspar(0.711) More Information \| Similar Motifs Found | motif file (matrix) |
| 29 \* | A G T C A G C T T G C A A T C G A C G T C T A G A C T G C G T A C G T A A C G T | 1e-6 | -1.467e+01 | 11.57% | 3.57% | 350.2bp (438.0bp) | TEAD3(TEA)/HepG2-TEAD3-ChIP-Seq(Encode)/Homer(0.679) More Information \| Similar Motifs Found | motif file (matrix) |
| 30 \* | A G T C C G T A A G T C A G T C A G T C C G T A A C G T C G T A A C G T A C T G | 1e-6 | -1.417e+01 | 2.78% | 0.12% | 363.3bp (238.1bp) | Bhlha15/MA0607.1/Jaspar(0.805) More Information \| Similar Motifs Found | motif file (matrix) |
| 31 \* | C G T A A C T G A G C T A G T C C G T A C G T A C G T A A C T G | 1e-5 | -1.355e+01 | 25.00% | 12.87% | 343.5bp (426.3bp) | PB0166.1\_Sox12\_2/Jaspar(0.700) More Information \| Similar Motifs Found | motif file (matrix) |
| 32 \* | C A G T A C T G C A T G G A T C G C A T A C T G G A T C G T C A A C G T A C T G | 1e-5 | -1.229e+01 | 11.57% | 4.10% | 291.8bp (452.8bp) | Smad3(MAD)/NPC-Smad3-ChIP-Seq(GSE36673)/Homer(0.684) More Information \| Similar Motifs Found | motif file (matrix) |
| 33 \* | A G C T A G T C A C T G A C T G A G T C A C G T G C T A A C T G A G T C A G T C | 1e-5 | -1.177e+01 | 7.41% | 1.93% | 322.9bp (258.5bp) | SMAD5/MA1557.1/Jaspar(0.602) More Information \| Similar Motifs Found | motif file (matrix) |
| 34 \* | A C T G C G T A A C G T A G T C A G T C A C G T A C T G A C T G A G T C C G T A | 1e-4 | -1.131e+01 | 2.78% | 0.21% | 261.8bp (419.9bp) | NF1-halfsite(CTF)/LNCaP-NF1-ChIP-Seq(Unpublished)/Homer(0.702) More Information \| Similar Motifs Found | motif file (matrix) |
| 35 \* | C A T G A C T G G A C T G C A T T G C A T G C A A T G C C G A T A T G C G A T C | 1e-4 | -1.069e+01 | 9.26% | 3.13% | 324.3bp (409.9bp) | ZNF652/HepG2-ZNF652.Flag-ChIP-Seq(Encode)/Homer(0.671) More Information \| Similar Motifs Found | motif file (matrix) |
| 36 \* | A C T G A G T C A C G T A C T G A C G T A C T G A C G T A C T G A C G T A C T G A C T G C G T A | 1e-4 | -1.013e+01 | 1.39% | 0.02% | 303.2bp (0.0bp) | Klf4(Zf)/mES-Klf4-ChIP-Seq(GSE11431)/Homer(0.657) More Information \| Similar Motifs Found | motif file (matrix) |
| 37 \* | A C G T G T A C A C T G G T C A C G T A A C T G A T C G A C T G | 1e-4 | -9.655e+00 | 21.30% | 11.83% | 338.2bp (411.5bp) | POL008.1\_DCE\_S\_I/Jaspar(0.732) More Information \| Similar Motifs Found | motif file (matrix) |
| 38 \* | A G T C A C G T A G T C A C G T A G T C C G T A A C T G C G T A | 1e-4 | -9.306e+00 | 15.28% | 7.49% | 364.7bp (446.1bp) | PB0138.1\_Irf4\_2/Jaspar(0.669) More Information \| Similar Motifs Found | motif file (matrix) |
| 39 \* | A G T C A G T C C G T A C G T A A G T C A C G T A C T G A C G T | 1e-2 | -6.343e+00 | 7.41% | 3.17% | 319.3bp (408.7bp) | MYB/MA0100.3/Jaspar(0.880) More Information \| Similar Motifs Found | motif file (matrix) |
| 40 \* | A G T C A C G T A G T C C G T A A C T G A G T C A G T C A G T C A G T C A G T C A C G T C G T A | 1e-1 | -4.133e+00 | 0.46% | 0.01% | 71.7bp (0.0bp) | Sp5(Zf)/mES-Sp5.Flag-ChIP-Seq(GSE72989)/Homer(0.708) More Information \| Similar Motifs Found | motif file (matrix) |
