## Supplementary material for "Repeated LPS induces training and tolerance of microglial responses across brain regions": FileS1: motif1.similar.html

### Information for motif1

C
G
T
A
G
T
C
A
C
G
T
A
A
G
T
C
C
T
A
G
C
G
T
A
C
G
T
A
C
T
G
A
T
A
G
C
A
G
C
T
C
A
T
G
T
G
C
A
  
Reverse Opposite:  

A
C
G
T
G
T
A
C
C
T
G
A
A
T
C
G
A
G
C
T
A
C
G
T
A
C
G
T
G
A
T
C
T
C
A
G
G
C
A
T
A
C
G
T
A
C
G
T
  

|  |  |
| --- | --- |
| p-value: | 1e-21 |
| log p-value: | -4.867e+01 |
| Information Content per bp: | 1.742 |
| Number of Target Sequences with motif | 40.0 |
| Percentage of Target Sequences with motif | 18.52% |
| Number of Background Sequences with motif | 337.5 |
| Percentage of Background Sequences with motif | 2.55% |
| Average Position of motif in Targets | 871.6 +/- 240.1bp |
| Average Position of motif in Background | 624.0 +/- 449.6bp |
| Strand Bias (log2 ratio + to - strand density) | -0.8 |
| Multiplicity (# of sites on avg that occur together) | 1.15 |
| Motif File: | file (matrix) reverse opposite |

#### Similar de novo motifs found

|  |  |  |  |  |  |  |  |
| --- | --- | --- | --- | --- | --- | --- | --- |
| Rank | Match Score | Redundant Motif | P-value | log P-value | % of Targets | % of Background | Motif file |
| 1 | 0.916 | T C G A G T C A G A C T G A T C C T A G G T C A G T C A C G T A T A G C A G C T | 1e-17 | -39.571148 | 24.07% | 5.88% | motif file (matrix) |
| 2 | 0.611 | C G T A A C T G A G T C A G C T A C G T A C G T A G T C T C G A C G A T A C G T A C G T A G C T | 1e-9 | -21.071292 | 5.56% | 0.43% | motif file (matrix) |
| 3 | 0.792 | C G T A G T C A A T G C A G T C A T C G G T C A T G C A G C T A | 1e-7 | -17.185332 | 12.50% | 3.59% | motif file (matrix) |
