## Supplementary material for "Repeated LPS induces training and tolerance of microglial responses across brain regions": FileS1: motif2.similar.html

### Information for motif2

T
C
A
G
C
T
A
G
T
C
G
A
T
G
C
A
C
T
G
A
G
C
A
T
A
C
G
T
A
G
T
C
A
G
T
C
G
T
A
C
G
C
T
A
T
A
C
G
  
Reverse Opposite:  

A
G
T
C
C
G
A
T
C
A
T
G
C
T
A
G
T
C
A
G
T
C
G
A
C
G
T
A
G
A
C
T
A
C
G
T
A
G
C
T
G
A
T
C
A
G
T
C
  

|  |  |
| --- | --- |
| p-value: | 1e-13 |
| log p-value: | -3.087e+01 |
| Information Content per bp: | 1.719 |
| Number of Target Sequences with motif | 20.0 |
| Percentage of Target Sequences with motif | 9.26% |
| Number of Background Sequences with motif | 116.0 |
| Percentage of Background Sequences with motif | 0.88% |
| Average Position of motif in Targets | 782.5 +/- 221.5bp |
| Average Position of motif in Background | 627.7 +/- 416.8bp |
| Strand Bias (log2 ratio + to - strand density) | 0.3 |
| Multiplicity (# of sites on avg that occur together) | 1.00 |
| Motif File: | file (matrix) reverse opposite |

#### Similar de novo motifs found

|  |  |  |  |  |  |  |  |
| --- | --- | --- | --- | --- | --- | --- | --- |
| Rank | Match Score | Redundant Motif | P-value | log P-value | % of Targets | % of Background | Motif file |
| 1 | 0.867 | A C T G A C T G A C T G C T G A G T C A G C A T C G A T G A C T T A G C G T A C | 1e-10 | -25.220556 | 20.83% | 6.61% | motif file (matrix) |
| 2 | 0.609 | A C T G A T C G A T C G A C T G A G T C A C G T A C G T A C G T A G T C A G T C A G T C A C T G | 1e-9 | -22.545855 | 4.63% | 0.20% | motif file (matrix) |
| 3 | 0.796 | A T C G G T C A G T C A C G A T A C G T A C G T G T A C A T G C | 1e-8 | -19.268316 | 39.81% | 22.03% | motif file (matrix) |
