## Supplementary material for "Repeated LPS induces training and tolerance of microglial responses across brain regions": FileS1: motif5.similar.html

### Information for motif5

C
G
T
A
A
C
G
T
A
C
T
G
A
C
T
G
A
G
T
C
A
G
T
C
C
G
T
A
C
T
G
A
A
G
T
C
C
G
T
A
A
G
T
C
C
G
T
A
  
Reverse Opposite:  

A
C
G
T
A
C
T
G
A
C
G
T
A
C
T
G
A
G
C
T
C
G
A
T
A
C
T
G
C
T
A
G
G
T
A
C
A
G
T
C
C
G
T
A
C
G
A
T
  

|  |  |
| --- | --- |
| p-value: | 1e-11 |
| log p-value: | -2.607e+01 |
| Information Content per bp: | 1.899 |
| Number of Target Sequences with motif | 9.0 |
| Percentage of Target Sequences with motif | 4.17% |
| Number of Background Sequences with motif | 10.9 |
| Percentage of Background Sequences with motif | 0.08% |
| Average Position of motif in Targets | 740.1 +/- 284.3bp |
| Average Position of motif in Background | 731.9 +/- 448.1bp |
| Strand Bias (log2 ratio + to - strand density) | 1.0 |
| Multiplicity (# of sites on avg that occur together) | 1.00 |
| Motif File: | file (matrix) reverse opposite |

#### Similar de novo motifs found

|  |  |  |  |  |  |  |  |
| --- | --- | --- | --- | --- | --- | --- | --- |
| Rank | Match Score | Redundant Motif | P-value | log P-value | % of Targets | % of Background | Motif file |
| 1 | 0.824 | C G A T C A T G C A G T A C G T C T A G A C T G G T A C G A T C G T C A C G T A | 1e-7 | -17.653715 | 9.26% | 1.95% | motif file (matrix) |
