## Supplementary material for "Repeated LPS induces training and tolerance of microglial responses across brain regions": FileS1: motif8.similar.html

### Information for motif8

G
C
T
A
T
G
A
C
A
T
G
C
C
G
T
A
A
C
T
G
A
G
T
C
A
G
T
C
A
C
T
G
A
G
T
C
A
T
G
C
G
T
A
C
A
C
T
G
  
Reverse Opposite:  

A
G
T
C
A
C
T
G
A
T
C
G
A
C
T
G
A
G
T
C
C
T
A
G
C
T
A
G
A
G
T
C
C
G
A
T
A
T
C
G
A
C
T
G
C
A
G
T
  

|  |  |
| --- | --- |
| p-value: | 1e-10 |
| log p-value: | -2.372e+01 |
| Information Content per bp: | 1.810 |
| Number of Target Sequences with motif | 17.0 |
| Percentage of Target Sequences with motif | 7.87% |
| Number of Background Sequences with motif | 119.0 |
| Percentage of Background Sequences with motif | 0.90% |
| Average Position of motif in Targets | 776.6 +/- 398.2bp |
| Average Position of motif in Background | 871.1 +/- 280.6bp |
| Strand Bias (log2 ratio + to - strand density) | -0.8 |
| Multiplicity (# of sites on avg that occur together) | 1.12 |
| Motif File: | file (matrix) reverse opposite |

#### Similar de novo motifs found

|  |  |  |  |  |  |  |  |
| --- | --- | --- | --- | --- | --- | --- | --- |
| Rank | Match Score | Redundant Motif | P-value | log P-value | % of Targets | % of Background | Motif file |
| 1 | 0.736 | A C T G A C T G A G T C A T C G C T G A G A T C A C G T A C T G T G A C A C G T | 1e-9 | -22.869867 | 18.52% | 5.77% | motif file (matrix) |
