## Supplementary material for "Repeated LPS induces training and tolerance of microglial responses across brain regions": FileS1: motif9.similar.html

### Information for motif9

T
A
G
C
C
A
G
T
A
C
T
G
C
G
T
A
G
A
T
C
C
G
A
T
G
A
T
C
G
T
C
A
T
A
C
G
G
A
T
C
  
Reverse Opposite:  

C
T
A
G
A
T
G
C
C
A
G
T
C
T
A
G
C
G
T
A
C
T
A
G
A
C
G
T
G
T
A
C
G
T
C
A
A
T
C
G
  

|  |  |
| --- | --- |
| p-value: | 1e-10 |
| log p-value: | -2.348e+01 |
| Information Content per bp: | 1.691 |
| Number of Target Sequences with motif | 63.0 |
| Percentage of Target Sequences with motif | 29.17% |
| Number of Background Sequences with motif | 1637.1 |
| Percentage of Background Sequences with motif | 12.35% |
| Average Position of motif in Targets | 659.6 +/- 293.6bp |
| Average Position of motif in Background | 652.9 +/- 436.0bp |
| Strand Bias (log2 ratio + to - strand density) | 0.0 |
| Multiplicity (# of sites on avg that occur together) | 1.16 |
| Motif File: | file (matrix) reverse opposite |

#### Similar de novo motifs found

|  |  |  |  |  |  |  |  |
| --- | --- | --- | --- | --- | --- | --- | --- |
| Rank | Match Score | Redundant Motif | P-value | log P-value | % of Targets | % of Background | Motif file |
| 1 | 0.668 | A C G T A C T G C G T A A G T C A C G T C G T A C G T A A C T G G T A C A C G T A C T G A C G T | 1e-9 | -21.559959 | 2.78% | 0.02% | motif file (matrix) |
