## Supplementary material for "Repeated LPS induces training and tolerance of microglial responses across brain regions": FileS1: motif15.similar.html

### Information for motif15

C
G
T
A
A
C
T
G
C
T
A
G
C
G
T
A
A
C
T
G
A
G
C
T
A
G
T
C
A
T
G
C
A
C
G
T
C
T
A
G
C
T
A
G
G
T
C
A
  
Reverse Opposite:  

A
C
G
T
A
G
T
C
A
G
T
C
C
G
T
A
A
T
C
G
A
C
T
G
T
C
G
A
A
G
T
C
G
C
A
T
A
G
T
C
A
G
T
C
A
C
G
T
  

|  |  |
| --- | --- |
| p-value: | 1e-8 |
| log p-value: | -2.062e+01 |
| Information Content per bp: | 1.870 |
| Number of Target Sequences with motif | 10.0 |
| Percentage of Target Sequences with motif | 4.63% |
| Number of Background Sequences with motif | 33.2 |
| Percentage of Background Sequences with motif | 0.25% |
| Average Position of motif in Targets | 763.0 +/- 226.9bp |
| Average Position of motif in Background | 451.7 +/- 343.2bp |
| Strand Bias (log2 ratio + to - strand density) | -1.2 |
| Multiplicity (# of sites on avg that occur together) | 1.00 |
| Motif File: | file (matrix) reverse opposite |

#### Similar de novo motifs found

|  |  |  |  |  |  |  |  |
| --- | --- | --- | --- | --- | --- | --- | --- |
| Rank | Match Score | Redundant Motif | P-value | log P-value | % of Targets | % of Background | Motif file |
| 1 | 0.630 | C T G A A C T G G C A T T G C A T G A C C G A T T A G C G A T C A G C T G A T C G T C A T C G A | 1e-7 | -17.606016 | 7.87% | 1.38% | motif file (matrix) |
