## Supplementary material for "Repeated LPS induces training and tolerance of microglial responses across brain regions": FileS1: motif16.similar.html

### Information for motif16

A
C
T
G
T
A
G
C
C
G
T
A
C
T
G
A
G
A
C
T
G
T
C
A
A
T
G
C
C
G
A
T
A
T
G
C
G
A
T
C
  
Reverse Opposite:  

C
T
A
G
A
T
C
G
C
G
T
A
A
T
C
G
C
A
G
T
C
T
G
A
A
G
C
T
A
C
G
T
A
T
C
G
A
G
T
C
  

|  |  |
| --- | --- |
| p-value: | 1e-8 |
| log p-value: | -1.981e+01 |
| Information Content per bp: | 1.843 |
| Number of Target Sequences with motif | 16.0 |
| Percentage of Target Sequences with motif | 7.41% |
| Number of Background Sequences with motif | 135.4 |
| Percentage of Background Sequences with motif | 1.02% |
| Average Position of motif in Targets | 563.8 +/- 300.0bp |
| Average Position of motif in Background | 606.9 +/- 405.8bp |
| Strand Bias (log2 ratio + to - strand density) | 0.9 |
| Multiplicity (# of sites on avg that occur together) | 1.06 |
| Motif File: | file (matrix) reverse opposite |

#### Similar de novo motifs found

|  |  |  |  |  |  |  |  |
| --- | --- | --- | --- | --- | --- | --- | --- |
| Rank | Match Score | Redundant Motif | P-value | log P-value | % of Targets | % of Background | Motif file |
| 1 | 0.638 | A C T G C T G A A C T G A C T G G C T A A C G T A G C T C G T A A T G C A C G T | 1e-4 | -10.121571 | 5.09% | 1.09% | motif file (matrix) |
