## Supplementary material for "Repeated LPS induces training and tolerance of microglial responses across brain regions": FileS1: motif19.similar.html

### Information for motif19

C
G
A
T
T
C
A
G
T
A
G
C
G
T
C
A
C
T
A
G
A
G
T
C
A
G
T
C
A
G
T
C
G
T
C
A
C
G
A
T
A
T
G
C
C
A
G
T
  
Reverse Opposite:  

G
C
T
A
T
A
C
G
C
G
T
A
A
C
G
T
A
C
T
G
C
T
A
G
T
A
C
G
G
A
T
C
A
C
G
T
A
T
C
G
A
G
T
C
G
C
T
A
  

|  |  |
| --- | --- |
| p-value: | 1e-8 |
| log p-value: | -1.909e+01 |
| Information Content per bp: | 1.741 |
| Number of Target Sequences with motif | 18.0 |
| Percentage of Target Sequences with motif | 8.33% |
| Number of Background Sequences with motif | 187.6 |
| Percentage of Background Sequences with motif | 1.42% |
| Average Position of motif in Targets | 659.2 +/- 355.7bp |
| Average Position of motif in Background | 634.1 +/- 443.2bp |
| Strand Bias (log2 ratio + to - strand density) | 0.8 |
| Multiplicity (# of sites on avg that occur together) | 1.39 |
| Motif File: | file (matrix) reverse opposite |

#### Similar de novo motifs found

|  |  |  |  |  |  |  |  |
| --- | --- | --- | --- | --- | --- | --- | --- |
| Rank | Match Score | Redundant Motif | P-value | log P-value | % of Targets | % of Background | Motif file |
| 1 | 0.656 | C G T A T A C G C G T A C T G A A C T G T C A G A T C G A G T C C G T A T C A G | 1e-7 | -16.740875 | 14.81% | 4.96% | motif file (matrix) |
| 2 | 0.864 | A G T C G T C A C A T G A G T C A T G C A G T C C T G A C G A T A G T C A G C T | 1e-5 | -13.114249 | 6.48% | 1.32% | motif file (matrix) |
