## Supplementary material for "Repeated LPS induces training and tolerance of microglial responses across brain regions": FileS1: motif22.similar.html

### Information for motif22

C
A
T
G
T
C
G
A
C
T
A
G
C
G
T
A
A
T
G
C
T
A
G
C
C
A
T
G
A
G
C
T
A
T
C
G
A
C
G
T
  
Reverse Opposite:  

G
T
C
A
T
A
G
C
T
C
G
A
G
T
A
C
A
T
C
G
T
A
C
G
G
C
A
T
G
A
T
C
A
G
C
T
G
T
A
C
  

|  |  |
| --- | --- |
| p-value: | 1e-7 |
| log p-value: | -1.820e+01 |
| Information Content per bp: | 1.669 |
| Number of Target Sequences with motif | 63.0 |
| Percentage of Target Sequences with motif | 29.17% |
| Number of Background Sequences with motif | 1872.5 |
| Percentage of Background Sequences with motif | 14.13% |
| Average Position of motif in Targets | 609.8 +/- 320.4bp |
| Average Position of motif in Background | 685.4 +/- 400.6bp |
| Strand Bias (log2 ratio + to - strand density) | -0.1 |
| Multiplicity (# of sites on avg that occur together) | 1.17 |
| Motif File: | file (matrix) reverse opposite |

#### Similar de novo motifs found

|  |  |  |  |  |  |  |  |
| --- | --- | --- | --- | --- | --- | --- | --- |
| Rank | Match Score | Redundant Motif | P-value | log P-value | % of Targets | % of Background | Motif file |
| 1 | 0.760 | C T A G C G T A A T C G C T G A A T G C A T G C T A C G A G C T C T A G C A T G C T A G T C G A | 1e-7 | -16.813710 | 7.87% | 1.47% | motif file (matrix) |
