## Supplementary material for "Repeated LPS induces training and tolerance of microglial responses across brain regions": FileS1: motif24.similar.html

### Information for motif24

A
T
C
G
A
C
T
G
A
C
T
G
T
C
G
A
C
T
A
G
T
G
C
A
A
T
G
C
C
G
A
T
T
A
C
G
A
T
G
C
G
C
T
A
G
T
A
C
  
Reverse Opposite:  

C
A
T
G
C
G
A
T
A
T
C
G
A
T
G
C
C
G
T
A
A
T
C
G
A
C
G
T
A
G
T
C
A
G
C
T
G
T
A
C
G
T
A
C
A
T
G
C
  

|  |  |
| --- | --- |
| p-value: | 1e-7 |
| log p-value: | -1.751e+01 |
| Information Content per bp: | 1.749 |
| Number of Target Sequences with motif | 19.0 |
| Percentage of Target Sequences with motif | 8.80% |
| Number of Background Sequences with motif | 234.2 |
| Percentage of Background Sequences with motif | 1.77% |
| Average Position of motif in Targets | 794.7 +/- 343.2bp |
| Average Position of motif in Background | 742.9 +/- 403.4bp |
| Strand Bias (log2 ratio + to - strand density) | -0.5 |
| Multiplicity (# of sites on avg that occur together) | 1.00 |
| Motif File: | file (matrix) reverse opposite |

#### Similar de novo motifs found

|  |  |  |  |  |  |  |  |
| --- | --- | --- | --- | --- | --- | --- | --- |
| Rank | Match Score | Redundant Motif | P-value | log P-value | % of Targets | % of Background | Motif file |
