## Supplementary material for "Repeated LPS induces training and tolerance of microglial responses across brain regions": FileS1: motif28.similar.html

### Information for motif28

C
T
A
G
A
G
C
T
C
A
G
T
A
C
T
G
A
C
T
G
C
G
A
T
C
A
T
G
T
C
A
G
C
A
G
T
T
C
A
G
A
T
C
G
G
A
C
T
  
Reverse Opposite:  

C
T
G
A
T
A
G
C
A
G
T
C
G
T
C
A
A
G
T
C
G
T
A
C
G
C
T
A
T
G
A
C
G
T
A
C
G
T
C
A
T
C
G
A
G
A
T
C
  

|  |  |
| --- | --- |
| p-value: | 1e-6 |
| log p-value: | -1.517e+01 |
| Information Content per bp: | 1.724 |
| Number of Target Sequences with motif | 30.0 |
| Percentage of Target Sequences with motif | 13.89% |
| Number of Background Sequences with motif | 633.6 |
| Percentage of Background Sequences with motif | 4.78% |
| Average Position of motif in Targets | 535.9 +/- 295.8bp |
| Average Position of motif in Background | 588.2 +/- 418.6bp |
| Strand Bias (log2 ratio + to - strand density) | -0.1 |
| Multiplicity (# of sites on avg that occur together) | 1.07 |
| Motif File: | file (matrix) reverse opposite |

#### Similar de novo motifs found

|  |  |  |  |  |  |  |  |
| --- | --- | --- | --- | --- | --- | --- | --- |
| Rank | Match Score | Redundant Motif | P-value | log P-value | % of Targets | % of Background | Motif file |
| 1 | 0.878 | C T G A A G T C A T G C T C A G G T A C A G T C C G T A A G T C A G T C C G T A | 1e-4 | -10.968472 | 16.20% | 7.51% | motif file (matrix) |
