## Supplementary material for "Repeated LPS induces training and tolerance of microglial responses across brain regions": FileS1: motif37.similar.html

### Information for motif37

A
C
G
T
G
T
A
C
A
C
T
G
G
T
C
A
C
G
T
A
A
C
T
G
A
T
C
G
A
C
T
G
  
Reverse Opposite:  

G
T
A
C
A
T
G
C
A
G
T
C
A
C
G
T
A
C
G
T
A
G
T
C
A
C
T
G
C
G
T
A
  

|  |  |
| --- | --- |
| p-value: | 1e-4 |
| log p-value: | -9.655e+00 |
| Information Content per bp: | 1.882 |
| Number of Target Sequences with motif | 46.0 |
| Percentage of Target Sequences with motif | 21.30% |
| Number of Background Sequences with motif | 1567.4 |
| Percentage of Background Sequences with motif | 11.83% |
| Average Position of motif in Targets | 680.9 +/- 338.2bp |
| Average Position of motif in Background | 650.3 +/- 411.5bp |
| Strand Bias (log2 ratio + to - strand density) | 0.3 |
| Multiplicity (# of sites on avg that occur together) | 1.17 |
| Motif File: | file (matrix) reverse opposite |

#### Similar de novo motifs found

|  |  |  |  |  |  |  |  |
| --- | --- | --- | --- | --- | --- | --- | --- |
| Rank | Match Score | Redundant Motif | P-value | log P-value | % of Targets | % of Background | Motif file |
