## Supplementary material for "Repeated LPS induces training and tolerance of microglial responses across brain regions": FileS1: motif38.similar.html

### Information for motif38

A
G
T
C
A
C
G
T
A
G
T
C
A
C
G
T
A
G
T
C
C
G
T
A
A
C
T
G
C
G
T
A
  
Reverse Opposite:  

A
C
G
T
A
G
T
C
A
C
G
T
A
C
T
G
C
G
T
A
A
C
T
G
C
G
T
A
A
C
T
G
  

|  |  |
| --- | --- |
| p-value: | 1e-4 |
| log p-value: | -9.306e+00 |
| Information Content per bp: | 1.530 |
| Number of Target Sequences with motif | 33.0 |
| Percentage of Target Sequences with motif | 15.28% |
| Number of Background Sequences with motif | 992.7 |
| Percentage of Background Sequences with motif | 7.49% |
| Average Position of motif in Targets | 686.8 +/- 364.7bp |
| Average Position of motif in Background | 635.4 +/- 446.1bp |
| Strand Bias (log2 ratio + to - strand density) | -0.8 |
| Multiplicity (# of sites on avg that occur together) | 1.06 |
| Motif File: | file (matrix) reverse opposite |

#### Similar de novo motifs found

|  |  |  |  |  |  |  |  |
| --- | --- | --- | --- | --- | --- | --- | --- |
| Rank | Match Score | Redundant Motif | P-value | log P-value | % of Targets | % of Background | Motif file |
