## Supplementary material for "Repeated LPS induces training and tolerance of microglial responses across brain regions": FileS1: motif40.similar.html

### Information for motif40

A
G
T
C
A
C
G
T
A
G
T
C
C
G
T
A
A
C
T
G
A
G
T
C
A
G
T
C
A
G
T
C
A
G
T
C
A
G
T
C
A
C
G
T
C
G
T
A
  
Reverse Opposite:  

A
C
G
T
C
G
T
A
A
C
T
G
A
C
T
G
A
C
T
G
A
C
T
G
A
C
T
G
A
G
T
C
A
C
G
T
A
C
T
G
C
G
T
A
A
C
T
G
  

|  |  |
| --- | --- |
| p-value: | 1e-1 |
| log p-value: | -4.133e+00 |
| Information Content per bp: | 1.530 |
| Number of Target Sequences with motif | 1.0 |
| Percentage of Target Sequences with motif | 0.46% |
| Number of Background Sequences with motif | 0.9 |
| Percentage of Background Sequences with motif | 0.01% |
| Average Position of motif in Targets | 252.0 +/- 71.7bp |
| Average Position of motif in Background | 1111.0 +/- 0.0bp |
| Strand Bias (log2 ratio + to - strand density) | 10.0 |
| Multiplicity (# of sites on avg that occur together) | 6.00 |
| Motif File: | file (matrix) reverse opposite |

#### Similar de novo motifs found

|  |  |  |  |  |  |  |  |
| --- | --- | --- | --- | --- | --- | --- | --- |
| Rank | Match Score | Redundant Motif | P-value | log P-value | % of Targets | % of Background | Motif file |
| 1 | 0.822 | C G T A A C T G A G T C A G T C A G T C A G T C A G T C A C G T C G T A A C G T | 1e0 | -2.108938 | 0.46% | 0.06% | motif file (matrix) |
