## Supplementary material for "Repeated LPS induces training and tolerance of microglial responses across brain regions": FileS1: knownResults.html

2xLPS\_cluster\_genes\_output/ - Homer Known Motif Enrichment Results


### Homer Known Motif Enrichment Results (2xLPS\_cluster\_genes\_output/)

Homer *de novo* Motif Results  
Gene Ontology Enrichment Results  
Known Motif Enrichment Results (txt file)  
Total Target Sequences = 216, Total Background Sequences = 13248

|  |  |  |  |  |  |  |  |  |  |  |  |
| --- | --- | --- | --- | --- | --- | --- | --- | --- | --- | --- | --- |
| Rank | Motif | Name | P-value | log P-pvalue | q-value (Benjamini) | # Target Sequences with Motif | % of Targets Sequences with Motif | # Background Sequences with Motif | % of Background Sequences with Motif | Motif File | SVG |
| 1 | C T G A A T C G A G C T A G C T A C G T T A G C C T G A T A C G C G A T A C G T G A C T A G T C | ISRE(IRF)/ThioMac-LPS-Expression(GSE23622)/Homer | 1e-19 | -4.464e+01 | 0.0000 | 42.0 | 19.44% | 427.9 | 3.23% | motif file (matrix) | svg |
| 2 | T C A G C T G A C G T A C G T A T A C G G C A T C T A G C T G A C G T A C G T A T A C G G A C T | IRF1(IRF)/PBMC-IRF1-ChIP-Seq(GSE43036)/Homer | 1e-16 | -3.831e+01 | 0.0000 | 54.0 | 25.00% | 867.5 | 6.55% | motif file (matrix) | svg |
| 3 | C T A G T C G A C T G A C G T A T A C G G A C T T C A G T C G A G T C A T G C A T A C G A G C T | IRF2(IRF)/Erythroblas-IRF2-ChIP-Seq(GSE36985)/Homer | 1e-15 | -3.563e+01 | 0.0000 | 47.0 | 21.76% | 703.3 | 5.31% | motif file (matrix) | svg |
| 4 | C T G A T A C G G C A T A G C T A G C T A G T C T C G A A C T G C A G T A G C T A G C T G A T C | IRF3(IRF)/BMDM-Irf3-ChIP-Seq(GSE67343)/Homer | 1e-10 | -2.356e+01 | 0.0000 | 70.0 | 32.41% | 1938.9 | 14.63% | motif file (matrix) | svg |
| 5 | T C A G T C A G G C T A C G T A T A C G G A C T T C A G T C G A C T G A C G T A T A C G G A C T | IRF8(IRF)/BMDM-IRF8-ChIP-Seq(GSE77884)/Homer | 1e-9 | -2.241e+01 | 0.0000 | 72.0 | 33.33% | 2080.4 | 15.70% | motif file (matrix) | svg |
| 6 | C T A G C T A G C G T A C G T A T A C G C G A T C T A G C T G A C T G A C G T A T A C G G A C T | PU.1:IRF8(ETS:IRF)/pDC-Irf8-ChIP-Seq(GSE66899)/Homer | 1e-8 | -1.913e+01 | 0.0000 | 50.0 | 23.15% | 1267.4 | 9.56% | motif file (matrix) | svg |
| 7 | A C T G C A T G G C T A T C G A G C T A A G C T A G C T G T A C A G T C T G A C | NFkB-p65-Rel(RHD)/ThioMac-LPS-Expression(GSE23622)/Homer | 1e-7 | -1.734e+01 | 0.0000 | 23.0 | 10.65% | 348.5 | 2.63% | motif file (matrix) | svg |
| 8 | C A T G C T A G T C G A A C G T A C T G C G T A T A G C C G A T T G A C C G T A A G C T G A T C | Fra2(bZIP)/Striatum-Fra2-ChIP-Seq(GSE43429)/Homer | 1e-4 | -1.080e+01 | 0.0011 | 57.0 | 26.39% | 2021.1 | 15.25% | motif file (matrix) | svg |
| 9 | C G T A C A T G C A T G A C T G C T A G T C G A G C A T C G A T A G C T A G T C G A T C G T A C | NFkB-p65(RHD)/GM12787-p65-ChIP-Seq(GSE19485)/Homer | 1e-4 | -9.844e+00 | 0.0026 | 75.0 | 34.72% | 3021.6 | 22.80% | motif file (matrix) | svg |
| 10 | A T G C C T G A G A C T A C G T A C G T G T A C G A T C C G A T C T A G C A T G C G T A C G T A C T G A G A C T | STAT1(Stat)/HelaS3-STAT1-ChIP-Seq(GSE12782)/Homer | 1e-4 | -9.666e+00 | 0.0028 | 52.0 | 24.07% | 1856.8 | 14.01% | motif file (matrix) | svg |
| 11 | T C G A A C G T C A T G G C T A T A G C C G A T G T A C G C T A A C G T A T G C | AP-1(bZIP)/ThioMac-PU.1-ChIP-Seq(GSE21512)/Homer | 1e-4 | -9.319e+00 | 0.0036 | 77.0 | 35.65% | 3176.2 | 23.97% | motif file (matrix) | svg |
| 12 | C T A G T C G A C G A T A C T G C G T A T A C G A G C T T G A C G C T A A C G T G A T C T A G C | Fosl2(bZIP)/3T3L1-Fosl2-ChIP-Seq(GSE56872)/Homer | 1e-3 | -9.166e+00 | 0.0038 | 42.0 | 19.44% | 1410.8 | 10.65% | motif file (matrix) | svg |
| 13 | A C T G C A T G T C G A A C G T A C T G C G T A A T C G A C G T G T A C C G T A G A C T A G T C | Fos(bZIP)/TSC-Fos-ChIP-Seq(GSE110950)/Homer | 1e-3 | -9.087e+00 | 0.0038 | 65.0 | 30.09% | 2560.0 | 19.32% | motif file (matrix) | svg |
| 14 | C T A G T C G A A C G T A C T G C G T A A T G C A C G T G T A C C G T A A G C T G A T C G T A C | Atf3(bZIP)/GBM-ATF3-ChIP-Seq(GSE33912)/Homer | 1e-3 | -9.051e+00 | 0.0038 | 70.0 | 32.41% | 2826.2 | 21.33% | motif file (matrix) | svg |
| 15 | C T A G T C G A A C G T A C T G C G T A T A G C C G A T G T A C C G T A A G C T G A T C G T A C | Jun-AP1(bZIP)/K562-cJun-ChIP-Seq(GSE31477)/Homer | 1e-3 | -8.940e+00 | 0.0038 | 33.0 | 15.28% | 1012.7 | 7.64% | motif file (matrix) | svg |
| 16 | A C T G C T A G T C G A C G A T C A T G G C T A A T C G C G A T G T A C G C T A A G C T G T A C | Fra1(bZIP)/BT549-Fra1-ChIP-Seq(GSE46166)/Homer | 1e-3 | -8.273e+00 | 0.0070 | 60.0 | 27.78% | 2371.4 | 17.90% | motif file (matrix) | svg |
| 17 | C T A G T C G A G C A T C A T G G C T A T A G C C G A T G T A C C T G A A G C T | JunB(bZIP)/DendriticCells-Junb-ChIP-Seq(GSE36099)/Homer | 1e-3 | -8.196e+00 | 0.0071 | 59.0 | 27.31% | 2326.7 | 17.56% | motif file (matrix) | svg |
| 18 | T G C A A T G C A C G T A C G T A C G T A T G C C T A G A C G T A C G T A G C T G A T C A G C T | T1ISRE(IRF)/ThioMac-Ifnb-Expression/Homer | 1e-3 | -8.171e+00 | 0.0071 | 8.0 | 3.70% | 97.6 | 0.74% | motif file (matrix) | svg |
| 19 | A T G C T C G A A G T C A G C T A C G T G T A C A G T C G C T A C T A G C A T G G T C A C T G A T C A G A G T C | Stat3+il21(Stat)/CD4-Stat3-ChIP-Seq(GSE19198)/Homer | 1e-3 | -7.816e+00 | 0.0093 | 93.0 | 43.06% | 4222.8 | 31.87% | motif file (matrix) | svg |
| 20 | T C G A T G A C G C A T A G C T C A G T G A T C G C T A G A T C G A C T A C G T G C A T A G T C | PRDM1(Zf)/Hela-PRDM1-ChIP-Seq(GSE31477)/Homer | 1e-3 | -7.604e+00 | 0.0110 | 71.0 | 32.87% | 3019.3 | 22.78% | motif file (matrix) | svg |
| 21 | A G T C C T G A A T C G A G C T A G C T G A C T A G T C G C T A A C G T C G A T G C A T C G A T A T C G C G T A T A G C G C A T A T G C C G T A | bZIP:IRF(bZIP,IRF)/Th17-BatF-ChIP-Seq(GSE39756)/Homer | 1e-3 | -7.230e+00 | 0.0152 | 54.0 | 25.00% | 2154.0 | 16.25% | motif file (matrix) | svg |
| 22 | C A G T T G C A A C G T A C T G C G T A A T C G C G A T T G A C C G T A A C G T | BATF(bZIP)/Th17-BATF-ChIP-Seq(GSE39756)/Homer | 1e-3 | -7.204e+00 | 0.0152 | 66.0 | 30.56% | 2790.6 | 21.06% | motif file (matrix) | svg |
| 23 | T C A G A G C T A C G T A C G T G T A C G A T C C G T A C T A G C A T G G T C A C G T A T C G A | STAT4(Stat)/CD4-Stat4-ChIP-Seq(GSE22104)/Homer | 1e-2 | -6.645e+00 | 0.0249 | 107.0 | 49.54% | 5183.2 | 39.11% | motif file (matrix) | svg |
| 24 | G C T A T A G C A G C T A T C G G T C A C G T A G C T A A T G C G A T C C T G A | IRF4(IRF)/GM12878-IRF4-ChIP-Seq(GSE32465)/Homer | 1e-2 | -5.613e+00 | 0.0669 | 54.0 | 25.00% | 2316.1 | 17.48% | motif file (matrix) | svg |
| 25 | A G T C G A C T C A G T G T A C A G T C A T C G T C A G A C T G G T C A C G T A | Stat3(Stat)/mES-Stat3-ChIP-Seq(GSE11431)/Homer | 1e-2 | -5.591e+00 | 0.0669 | 73.0 | 33.80% | 3360.2 | 25.36% | motif file (matrix) | svg |
