## Supplementary material for "Repeated LPS induces training and tolerance of microglial responses across brain regions": FileS2: geneOntology.html

Gene Ontology Results

### Gene Ontology Enrichment Results

Homer *de novo* Motif Enrichment Results  
Known Motif Enrichment Results  

###### Text file version of complete results (i.e. open with Excel) - biological process: Functional groupings of proteins (Gene Ontology) - molecular function: Mechanistic actions of proteins (Gene Ontology) - cellular component: Protein localization (Gene Ontology) - chromosome location: Genes with similar chromosome localization (NCBI Gene) - KEGG pathways: Groups of proteins in the same pathways (KEGG) - protein interactions: "Proteins interacting with a common protein (BIND, EcoCyc, HPRD)" (NCBI Gene) - interpro domains: Proteins with similar domains and features (Interpro) - pfam domains: Proteins with similar domains and features (Pfam) - smart domains: Proteins with similar domains and features (SMART) - gene3d domains: Proteins with similar domains and features (Gene3D) - prosite domains: Proteins with similar domains and features (Prosite) - prints domains: Proteins with similar domains and features (PRINTS) - MSigDB lists: "Genes sets for pathways, factor/miRNA target predictions, expression patterns, etc." (MSigDB) - BIOCYC pathways: Groups of proteins in the same pathways (BIOCYC) - COSMIC cancer mutations: Genes mutated in similar cancers (COSMIC) - GWAS genes: Genes mutated in similar diseases (GWAS Catalog) - Lipid Maps pathways: Groups of proteins in the same lipid pathways (Lipid Maps/Biosystems) - Pathway Interaction DB: Groups of proteins in the same pathways (Pathway Interaction Database) - REACTOME pathways: Groups of proteins in the same pathways (REACTOME) - SMPDB pathways: Groups of proteins in the same pathways (SMPDB) - WikiPathways: Groups of proteins in the same pathways (Wikipathways) Enriched Categories | | | | | | | | | | | | --- | --- | --- | --- | --- | --- | --- | --- | --- | --- | | P-value | ln(P) | Term | GO Tree | GO ID | # of Genes in Term | # of Target Genes in Term | # of Total Genes | # of Target Genes | Common Genes | | 9.496e-15 | -32.29 | NADLER\_OBESITY\_UP | MSigDB lists | NADLER\_OBESITY\_UP | 54 | 12 | 12187 | 98 | Hcls1,Cd68,Ctss,Ctsz,Mpeg1,C1qb,Csf1r,Fcer1g,Grn,Ctsa,Fcgr3,Ctsb | | 1.474e-13 | -29.55 | DEMAGALHAES\_AGING\_UP | MSigDB lists | DEMAGALHAES\_AGING\_UP | 50 | 11 | 12187 | 98 | Gfap,Apod,C1qa,Fcgr3,Ctss,Clu,Vwf,S100a6,Mpeg1,C1qb,C3 | | 3.892e-13 | -28.57 | GSE2405\_0H\_VS\_24H\_A\_PHAGOCYTOPHILUM\_STIM\_NEUTROPHIL\_UP | MSigDB lists | GSE2405\_0H\_VS\_24H\_A\_PHAGOCYTOPHILUM\_STIM\_NEUTROPHIL\_UP | 171 | 16 | 12187 | 98 | Brk1,S100a6,Fcer1g,Rps5,Rpl31,Rpl26,Actb,Cd68,Rps8,Psme1,Rpl13,Clu,Hcls1,C3,Rpl14,Psmb10 | | 3.137e-12 | -26.49 | MCLACHLAN\_DENTAL\_CARIES\_UP | MSigDB lists | MCLACHLAN\_DENTAL\_CARIES\_UP | 164 | 15 | 12187 | 98 | C1qb,Csf1r,Fcer1g,Fcgr3,Ctsb,Vamp8,Hcls1,C3,Ctss,Serpina3n,C1qa,Ptprc,Cyba,Mgp,Rac2 | | 3.070e-11 | -24.21 | ICHIBA\_GRAFT\_VERSUS\_HOST\_DISEASE\_35D\_UP | MSigDB lists | ICHIBA\_GRAFT\_VERSUS\_HOST\_DISEASE\_35D\_UP | 103 | 12 | 12187 | 98 | Csf1r,Tspan4,C1qb,Mpeg1,Grn,Lrg1,S100a6,Cd68,Ctss,Cyba,C1qa,Ptprc | | 8.360e-11 | -23.20 | GO\_EXTRACELLULAR\_SPACE | MSigDB lists | GO\_EXTRACELLULAR\_SPACE | 738 | 26 | 12187 | 98 | Il6ra,Lgi4,Actb,Ctsb,Serpinb9,Itgam,Grn,Lrg1,Pcsk2,Slc2a1,Tnfsf8,Apod,Fgf18,Ctsz,Fmod,Hspa2,C3,Igfbp6,Spint1,C1qb,Serpina3n,C1qa,Il1r1,Ctss,Clu,Kl | | 1.062e-10 | -22.97 | defense response | biological process | GO:0006952 | 731 | 26 | 13711 | 111 | Saa3,Serpina3n,Ccr5,C1qb,C3,Cyba,Csf1r,Ifitm1,Mpeg1,Vamp8,Vim,Flnb,Fcgr3,Tnfsf8,Ccl6,Serpinb9,Fcer1g,Itgam,Kl,Il1r1,Grn,Tmem106a,C1qa,Ptprc,Cd68,Clu | | 1.494e-10 | -22.62 | GO\_REGULATION\_OF\_IMMUNE\_SYSTEM\_PROCESS | MSigDB lists | GO\_REGULATION\_OF\_IMMUNE\_SYSTEM\_PROCESS | 947 | 29 | 12187 | 98 | Psmb10,C1qa,Ctss,Clu,Fcgr3,Treml2,Csf1r,C1qb,Ikzf3,Rac2,Phyhip,Ptprc,Apod,Cyba,Csk,Psme1,Hcls1,C3,Vamp8,Il6ra,Actb,Ctsb,Serpinb9,Brk1,Fcer1g,Thy1,Itgam,Klhl6,Tmem176a | | 1.660e-10 | -22.52 | GO\_ACTIVATION\_OF\_IMMUNE\_RESPONSE | MSigDB lists | GO\_ACTIVATION\_OF\_IMMUNE\_RESPONSE | 295 | 17 | 12187 | 98 | Itgam,Thy1,Fcer1g,Brk1,C1qb,Klhl6,Ctsb,Actb,Fcgr3,Psme1,Clu,Csk,Ctss,C3,Psmb10,Ptprc,C1qa | | 2.191e-10 | -22.24 | GO\_POSITIVE\_REGULATION\_OF\_IMMUNE\_SYSTEM\_PROCESS | MSigDB lists | GO\_POSITIVE\_REGULATION\_OF\_IMMUNE\_SYSTEM\_PROCESS | 596 | 23 | 12187 | 98 | C1qb,Csf1r,Fcgr3,Ctss,Clu,C1qa,Psmb10,Brk1,Fcer1g,Thy1,Itgam,Klhl6,Vamp8,Il6ra,Ctsb,Actb,Csk,Psme1,Hcls1,C3,Rac2,Ptprc,Cyba | | 2.971e-10 | -21.94 | immune system process | biological process | GO:0002376 | 1296 | 34 | 13711 | 111 | Psme1,Serpinb9,Treml2,Itgam,Fcer1g,Psmb10,Ctss,Ikzf3,Grn,Tmem106a,Adgre1,Thy1,Clu,Cd68,C1qa,Ptprc,Ccr5,Cyba,C3,C1qb,Lfng,Csk,Ifitm1,Hcls1,Csf1r,Myh9,Vim,Klhl6,Cplx2,Vamp8,Ccl6,Flnb,Fcgr3,Tnfsf8 | | 3.061e-10 | -21.91 | GO\_REGULATION\_OF\_IMMUNE\_RESPONSE | MSigDB lists | GO\_REGULATION\_OF\_IMMUNE\_RESPONSE | 551 | 22 | 12187 | 98 | Psmb10,C1qa,Clu,Ctss,Fcgr3,C1qb,Treml2,Cyba,Ptprc,Rac2,C3,Psme1,Csk,Actb,Ctsb,Vamp8,Klhl6,Thy1,Fcer1g,Itgam,Serpinb9,Brk1 | | 5.197e-10 | -21.38 | MARKEY\_RB1\_ACUTE\_LOF\_DN | MSigDB lists | MARKEY\_RB1\_ACUTE\_LOF\_DN | 197 | 14 | 12187 | 98 | Ctsb,Csf1r,C1qb,Mpeg1,Grn,Fcer1g,Itgam,Serpinb9,Psmb10,Ptprc,Apod,Rac2,Hcls1,Ctsz | | 6.177e-10 | -21.20 | GO\_IMMUNE\_SYSTEM\_PROCESS | MSigDB lists | GO\_IMMUNE\_SYSTEM\_PROCESS | 1290 | 33 | 12187 | 98 | Il6ra,Vamp8,Actb,Ctsb,Thy1,Itgam,Fcer1g,Myh9,Brk1,Serpinb9,Klhl6,Tnfsf8,Ikzf3,Cyba,Ptprc,Fam111a,Camk2b,Psme1,Csk,C3,Hcls1,Cplx2,Fcgr3,Ccr5,Lfng,C1qb,Csf1r,Treml2,C1qa,Il1r1,Psmb10,Clu,Ctss | | 8.008e-10 | -20.95 | immune response | biological process | GO:0006955 | 625 | 23 | 13711 | 111 | Fcer1g,Serpinb9,C1qa,Cd68,Adgre1,Grn,Tmem106a,Csf1r,Ifitm1,Csk,Lfng,Ccr5,C3,Cyba,C1qb,Fcgr3,Flnb,Tnfsf8,Ccl6,Vamp8,Klhl6,Cplx2,Vim | | 1.004e-09 | -20.72 | GO\_POSITIVE\_REGULATION\_OF\_IMMUNE\_RESPONSE | MSigDB lists | GO\_POSITIVE\_REGULATION\_OF\_IMMUNE\_RESPONSE | 378 | 18 | 12187 | 98 | Actb,Ctsb,Fcgr3,Vamp8,C1qb,Klhl6,Fcer1g,Thy1,Itgam,Brk1,Psmb10,C1qa,Ptprc,C3,Psme1,Clu,Csk,Ctss | | 2.205e-09 | -19.93 | response to other organism | biological process | GO:0051707 | 718 | 24 | 13711 | 111 | Vamp8,Lrg1,Mpeg1,Vim,Klhl6,Flnb,Tnfsf8,Ly6a,Ccl6,Saa3,Serpina3n,C1qb,Cyba,C3,Csf1r,Cd52,Ifitm1,Ikzf3,Tmem106a,C1qa,Ptprc,Cd68,Serpinb9,Fcer1g | | 2.330e-09 | -19.88 | response to external biotic stimulus | biological process | GO:0043207 | 720 | 24 | 13711 | 111 | Cd52,Csf1r,Ifitm1,Saa3,C1qb,C3,Cyba,Serpina3n,Tnfsf8,Ly6a,Flnb,Ccl6,Mpeg1,Lrg1,Vamp8,Klhl6,Vim,Fcer1g,Serpinb9,Ptprc,C1qa,Cd68,Tmem106a,Ikzf3 | | 2.558e-09 | -19.78 | inflammatory response | biological process | GO:0006954 | 302 | 16 | 13711 | 111 | Kl,Il1r1,Grn,Fcgr3,Cd68,Clu,Ccl6,Serpinb9,Saa3,Serpina3n,Ccr5,Cyba,C3,Csf1r,Fcer1g,Itgam | | 2.959e-09 | -19.64 | GSE24634\_TREG\_VS\_TCONV\_POST\_DAY5\_IL4\_CONVERSION\_DN | MSigDB lists | GSE24634\_TREG\_VS\_TCONV\_POST\_DAY5\_IL4\_CONVERSION\_DN | 152 | 12 | 12187 | 98 | Fcer1g,Tsc22d1,Grn,Tmem176a,Csf1r,Tspan4,Il6ra,Ctsa,Ctsb,Cd68,Ctsz,Ctss | | 3.621e-09 | -19.44 | extracellular space | cellular component | GO:0005615 | 869 | 26 | 13825 | 111 | Clu,Ctss,Saa3,Ctsb,Vwf,Itgam,Il1r1,Ccl6,C1qb,Ahnak,Kl,Fmod,Serpina3n,Ctsz,Tnfsf8,Apod,Grn,Pcsk2,Igfbp6,Mgp,Serpinb9,C1qa,Lgi4,C3,Lrg1,Il6ra | | 3.793e-09 | -19.39 | response to biotic stimulus | biological process | GO:0009607 | 738 | 24 | 13711 | 111 | Ifitm1,Csf1r,Cd52,Serpina3n,Cyba,C3,C1qb,Saa3,Ccl6,Flnb,Ly6a,Tnfsf8,Klhl6,Vim,Lrg1,Mpeg1,Vamp8,Fcer1g,Serpinb9,Cd68,C1qa,Ptprc,Ikzf3,Tmem106a | | 5.424e-09 | -19.03 | MODULE\_128 | MSigDB lists | MODULE\_128 | 72 | 9 | 12187 | 98 | Csf1r,Mgll,Vwf,Fcer1g,Ccr5,Ctsb,Fcgr3,Il1r1,Vim | | 5.707e-09 | -18.98 | GSE2405\_HEAT\_KILLED\_LYSATE\_VS\_LIVE\_A\_PHAGOCYTOPHILUM\_STIM\_NEUTROPHIL\_24H\_UP | MSigDB lists | GSE2405\_HEAT\_KILLED\_LYSATE\_VS\_LIVE\_A\_PHAGOCYTOPHILUM\_STIM\_NEUTROPHIL\_24H\_UP | 161 | 12 | 12187 | 98 | Fcer1g,Grn,S100a6,Tsc22d1,Serpinb9,Tmem176a,Csf1r,Mgp,Clu,Ahnak,Cd68,C3 | | 6.000e-09 | -18.93 | MODULE\_84 | MSigDB lists | MODULE\_84 | 373 | 17 | 12187 | 98 | Fcgr3,Grn,Itgam,Fcer1g,Tspan4,Csf1r,C1qb,Rac2,Il1r1,Ptprc,Apod,Cyba,Rab3il1,Ctss,Mgll,Vwf,Hcls1 | | 6.146e-09 | -18.91 | MODULE\_79 | MSigDB lists | MODULE\_79 | 73 | 9 | 12187 | 98 | Mgll,Vwf,Fcer1g,Ccr5,Csf1r,Vim,Ctsb,Fcgr3,Il1r1 | | 7.846e-09 | -18.66 | MODULE\_170 | MSigDB lists | MODULE\_170 | 75 | 9 | 12187 | 98 | Vim,Il1r1,Fcgr3,Ctsb,Fcer1g,Ccr5,Vwf,Mgll,Csf1r | | 9.525e-09 | -18.47 | MODULE\_5 | MSigDB lists | MODULE\_5 | 336 | 16 | 12187 | 98 | Serpina3n,Ptprc,Apod,Il1r1,Rac2,Hcls1,Mgll,Vwf,Ctsa,Fcgr3,Igfbp6,Adh1,C1qb,Itgam,Fcer1g,Grn | | 1.001e-08 | -18.42 | SWEET\_LUNG\_CANCER\_KRAS\_UP | MSigDB lists | SWEET\_LUNG\_CANCER\_KRAS\_UP | 386 | 17 | 12187 | 98 | C1qb,Mpeg1,Lrg1,Ccr5,Fcer1g,Ctsb,Ctsa,Spint1,Vamp8,Rap1gap,Psme1,Clu,Ctss,Ldha,Ctsz,Cd68,Cyba | | 1.807e-08 | -17.83 | GO\_IMMUNE\_EFFECTOR\_PROCESS | MSigDB lists | GO\_IMMUNE\_EFFECTOR\_PROCESS | 304 | 15 | 12187 | 98 | C1qa,Ptprc,C3,Clu,Fam111a,Fcgr3,Actb,Il6ra,Vamp8,Cplx2,C1qb,Fcer1g,Brk1,Lfng,Serpinb9 | | 3.147e-08 | -17.27 | LIAN\_LIPA\_TARGETS\_3M | MSigDB lists | LIAN\_LIPA\_TARGETS\_3M | 41 | 7 | 12187 | 98 | Fcer1g,Cd68,Fcgr3,C1qb,Ctsb,C1qa,Mpeg1 | | 4.014e-08 | -17.03 | GO\_IMMUNE\_RESPONSE | MSigDB lists | GO\_IMMUNE\_RESPONSE | 594 | 20 | 12187 | 98 | Ctss,Csk,Camk2b,Clu,C3,Tnfsf8,Il1r1,Psmb10,C1qa,Cyba,Serpinb9,Lfng,Ccr5,Fcer1g,Klhl6,Csf1r,C1qb,Cplx2,Vamp8,Il6ra | | 4.071e-08 | -17.02 | extracellular region part | cellular component | GO:0044421 | 1048 | 27 | 13825 | 111 | Saa3,Ctsb,Itgam,Vwf,Il1r1,Clu,Ctss,Ctsz,Serpina3n,Ccl6,C1qb,S100a6,Fmod,Kl,Ahnak,Pcsk2,Mgp,Igfbp6,Tnfsf8,Apod,Grn,Lgi4,C1qa,C3,Il6ra,Lrg1,Serpinb9 | | 4.993e-08 | -16.81 | Immune System | REACTOME pathways | R-MMU-168256 | 1188 | 35 | 6297 | 76 | Ctsb,Grn,C3,Vamp8,Ctsz,Tnfsf8,Lrg1,Serpina3n,Ifitm1,Cd68,Csf1r,C1qa,Psmb10,Rap1gap,C1qb,Actb,Fgf18,Treml2,Il1r1,Psme1,Fcgr3,Cyba,Ccr5,Csk,Itgam,Brk1,Pde1a,Kl,Il6ra,Clu,Ctss,Fcer1g,Flnb,Camk2b,Ctsa | | 5.429e-08 | -16.73 | BROWN\_MYELOID\_CELL\_DEVELOPMENT\_UP | MSigDB lists | BROWN\_MYELOID\_CELL\_DEVELOPMENT\_UP | 124 | 10 | 12187 | 98 | Fcgr3,Ctss,Cd68,Capn3,Rpl13,Itgam,S100a6,Grn,Mpeg1,C3 | | 5.862e-08 | -16.65 | GO\_LEUKOCYTE\_MEDIATED\_IMMUNITY | MSigDB lists | GO\_LEUKOCYTE\_MEDIATED\_IMMUNITY | 94 | 9 | 12187 | 98 | Clu,Fcer1g,Serpinb9,C3,C1qb,Il6ra,Vamp8,Cplx2,C1qa | | 6.811e-08 | -16.50 | HSIAO\_HOUSEKEEPING\_GENES | MSigDB lists | HSIAO\_HOUSEKEEPING\_GENES | 336 | 15 | 12187 | 98 | Vim,Mgp,Serpina3n,Psme1,Rpl13,Fmod,Clu,Rps8,Ldha,Rpl14,Cox4i1,Actb,Myh9,Rpl31,Rps5 | | 8.806e-08 | -16.25 | cell surface | cellular component | GO:0009986 | 618 | 20 | 13825 | 111 | Clu,Fcer1g,Ctss,Adgre1,Ctsb,Treml2,Il1r1,Itgam,Csf1r,Vwf,Hspa2,Thy1,Ccr5,Ly6a,Vamp8,Fcgr3,C3,Ctsz,Il6ra,Ptprc | | 1.318e-07 | -15.84 | LIAN\_LIPA\_TARGETS\_6M | MSigDB lists | LIAN\_LIPA\_TARGETS\_6M | 50 | 7 | 12187 | 98 | C1qb,Mpeg1,Fcer1g,Cd68,Ctsb,Fcgr3,C1qa | | 1.450e-07 | -15.75 | GNF2\_VAV1 | MSigDB lists | GNF2\_VAV1 | 31 | 6 | 12187 | 98 | Csk,Rac2,Psmb10,Hcls1,Ptprc,Cyba | | 1.484e-07 | -15.72 | VERHAAK\_GLIOBLASTOMA\_MESENCHYMAL | MSigDB lists | VERHAAK\_GLIOBLASTOMA\_MESENCHYMAL | 175 | 11 | 12187 | 98 | Grn,Ccr5,Myh9,Itgam,Ctsz,Ctsb,Fcgr3,Il1r1,Ptprc,Rac2,Igfbp6 | | 1.579e-07 | -15.66 | regulation of immune response | biological process | GO:0050776 | 516 | 18 | 13711 | 111 | C1qb,Cyba,C3,Serpinb9,Il6ra,Csk,Itgam,Treml2,Rac2,Fcer1g,Grn,Klhl6,Il1r1,Vamp8,Thy1,C1qa,Fcgr3,Ptprc | | 2.597e-07 | -15.16 | KAAB\_FAILED\_HEART\_VENTRICLE\_DN | MSigDB lists | KAAB\_FAILED\_HEART\_VENTRICLE\_DN | 34 | 6 | 12187 | 98 | C1qb,Serpina3n,Tsc22d1,Fcer1g,Ldha,Cd68 | | 2.861e-07 | -15.07 | LEE\_AGING\_NEOCORTEX\_UP | MSigDB lists | LEE\_AGING\_NEOCORTEX\_UP | 82 | 8 | 12187 | 98 | Vim,Gfap,C1qa,Apod,Cd68,Ctsz,Ctss,Mpeg1 | | 2.884e-07 | -15.06 | extracellular region | cellular component | GO:0005576 | 1307 | 29 | 13825 | 111 | Spint1,C3,C1qa,Lgi4,Lrg1,Il6ra,Fgf18,Serpinb9,Mgp,Igfbp6,Pcsk2,Apod,Tnfsf8,Grn,Serpina3n,Ctsz,S100a6,C1qb,Ccl6,Kl,Ahnak,Fmod,Ctsb,Saa3,Il1r1,Vwf,Itgam,Clu,Ctss | | 2.980e-07 | -15.03 | response to external stimulus | biological process | GO:0009605 | 1456 | 31 | 13711 | 111 | Slc2a1,Mpeg1,Lrg1,Vamp8,Klhl6,Vim,Ly6a,Tnfsf8,Flnb,Fcgr3,Ccl6,Saa3,C1qb,Cyba,C3,Serpina3n,Ccr5,Cd52,Csf1r,Rac2,Ifitm1,Tmem106a,Mc4r,Ikzf3,Ptprc,C1qa,Cd68,Serpinb9,Ldha,Fcer1g,Itgam | | 3.333e-07 | -14.91 | GO\_BLOOD\_MICROPARTICLE | MSigDB lists | GO\_BLOOD\_MICROPARTICLE | 57 | 7 | 12187 | 98 | Hspa2,Clu,C3,C1qb,Slc2a1,Actb,Serpina3n | | 3.752e-07 | -14.80 | external side of plasma membrane | cellular component | GO:0009897 | 234 | 12 | 13825 | 111 | Fcer1g,Adgre1,Ctsb,Il1r1,Vwf,Itgam,Thy1,Ly6a,Ccr5,Fcgr3,Ptprc,Il6ra | | 3.865e-07 | -14.77 | cell activation | biological process | GO:0001775 | 433 | 16 | 13711 | 111 | Psmb10,Myh9,Treml2,Itgam,Fcer1g,Camk2b,Lfng,Vwf,Ptprc,Tnfsf8,Fcgr3,Clu,Tmem106a,Grn,Ikzf3,Cplx2 | | 3.910e-07 | -14.75 | MORF\_ACTG1 | MSigDB lists | MORF\_ACTG1 | 117 | 9 | 12187 | 98 | Rpl31,Rpl14,Rps5,Rps20,Rpl13,Rps8,Ldha,Actb,Cox4i1 | | 4.207e-07 | -14.68 | CHEN\_METABOLIC\_SYNDROM\_NETWORK | MSigDB lists | CHEN\_METABOLIC\_SYNDROM\_NETWORK | 963 | 24 | 12187 | 98 | Fcgr3,S100a6,Ccr5,Csf1r,C1qb,Tmem106a,Rab3il1,Cd68,Ctss,Vamp8,Ctsb,Grn,Fcer1g,Exoc3l4,Klhl6,Mpeg1,Sh3bgrl3,Vim,Rac2,Ptprc,Cyba,Fmod,Fam111a,Hcls1 | | 4.538e-07 | -14.61 | KEEN\_RESPONSE\_TO\_ROSIGLITAZONE\_DN | MSigDB lists | KEEN\_RESPONSE\_TO\_ROSIGLITAZONE\_DN | 87 | 8 | 12187 | 98 | Cyba,Actb,C1qb,Serpina3n,S100a6,Clu,Mgp,Ctss | | 4.801e-07 | -14.55 | regulation of immune system process | biological process | GO:0002682 | 953 | 24 | 13711 | 111 | Csf1r,Hcls1,Rac2,Csk,Cyba,C1qb,C3,Fcgr3,Vamp8,Klhl6,Fcer1g,Treml2,Itgam,Serpinb9,Il6ra,Tmem176a,Ptprc,C1qa,Apod,Cd68,Thy1,Il1r1,Grn,Ikzf3 | | 5.195e-07 | -14.47 | DELYS\_THYROID\_CANCER\_UP | MSigDB lists | DELYS\_THYROID\_CANCER\_UP | 340 | 14 | 12187 | 98 | C1qa,Cyba,Ctss,C3,Vamp8,Igfbp6,Spint1,Ctsb,Fcgr3,Ctsa,Ly6e,Pcsk2,Csf1r,C1qb | | 5.414e-07 | -14.43 | GO\_CYTOSOLIC\_RIBOSOME | MSigDB lists | GO\_CYTOSOLIC\_RIBOSOME | 89 | 8 | 12187 | 98 | Rps20,Rpl31,Rpl14,Rps5,Rps8,Rpl13,Apod,Rpl26 | | 6.318e-07 | -14.27 | GSE24634\_TREG\_VS\_TCONV\_POST\_DAY10\_IL4\_CONVERSION\_DN | MSigDB lists | GSE24634\_TREG\_VS\_TCONV\_POST\_DAY10\_IL4\_CONVERSION\_DN | 161 | 10 | 12187 | 98 | C1qa,Ctsa,Fcgr3,Ctsb,Ctsz,Itgam,Grn,C1qb,Csf1r,Tspan4 | | 8.773e-07 | -13.95 | regulation of myeloid leukocyte mediated immunity | biological process | GO:0002886 | 41 | 6 | 13711 | 111 | Itgam,C3,Fcer1g,Rac2,Vamp8,Fcgr3 | | 1.027e-06 | -13.79 | LIU\_VAV3\_PROSTATE\_CARCINOGENESIS\_UP | MSigDB lists | LIU\_VAV3\_PROSTATE\_CARCINOGENESIS\_UP | 67 | 7 | 12187 | 98 | Psmb10,C1qa,Apod,Serpina3n,C3,Ctss,Fcer1g | | 1.243e-06 | -13.60 | MODULE\_130 | MSigDB lists | MODULE\_130 | 11 | 4 | 12187 | 98 | Clu,C1qa,C3,C1qb | | 1.246e-06 | -13.60 | defense response to other organism | biological process | GO:0098542 | 473 | 16 | 13711 | 111 | Cyba,C1qb,C3,Serpinb9,Ifitm1,Fcer1g,Csf1r,Vim,Tmem106a,Mpeg1,Vamp8,Ccl6,Flnb,C1qa,Tnfsf8,Ptprc | | 1.257e-06 | -13.59 | REACTOME\_PEPTIDE\_CHAIN\_ELONGATION | MSigDB lists | REACTOME\_PEPTIDE\_CHAIN\_ELONGATION | 69 | 7 | 12187 | 98 | Rps5,Rpl14,Rpl31,Rps20,Rpl13,Rps8,Rpl26 | | 1.388e-06 | -13.49 | KEGG\_RIBOSOME | MSigDB lists | KEGG\_RIBOSOME | 70 | 7 | 12187 | 98 | Rpl26,Rps20,Rpl14,Rps5,Rpl31,Rps8,Rpl13 | | 1.451e-06 | -13.44 | TJP2 (tight junction protein 2) | protein interactions | 9414 | 3 | 3 | 6802 | 78 | Myh9,Actb,Csk | | 1.457e-06 | -13.44 | cytosolic ribosome | cellular component | GO:0022626 | 101 | 8 | 13825 | 111 | Rpl14,Rpl13,Apod,Rpl26,Rps8,Rps5,Rps20,Rpl31 | | 1.580e-06 | -13.36 | MORF\_NPM1 | MSigDB lists | MORF\_NPM1 | 138 | 9 | 12187 | 98 | Cox4i1,Actb,Rpl13,Ldha,Rps8,Rpl31,Rps5,Rpl14,Rps20 | | 1.683e-06 | -13.29 | LEE\_AGING\_CEREBELLUM\_UP | MSigDB lists | LEE\_AGING\_CEREBELLUM\_UP | 72 | 7 | 12187 | 98 | Mpeg1,C1qb,Ctss,Ctsz,Cd68,C1qa,Gfap | | 1.683e-06 | -13.29 | KIM\_GLIS2\_TARGETS\_UP | MSigDB lists | KIM\_GLIS2\_TARGETS\_UP | 72 | 7 | 12187 | 98 | Tsc22d1,S100a6,Fcer1g,C3,Mgp,Vim,Fcgr3 | | 1.774e-06 | -13.24 | leukocyte activation | biological process | GO:0045321 | 372 | 14 | 13711 | 111 | Fcgr3,Tnfsf8,Ptprc,Clu,Cplx2,Ikzf3,Grn,Tmem106a,Myh9,Psmb10,Fcer1g,Treml2,Itgam,Lfng | | 1.797e-06 | -13.23 | positive regulation of inflammatory response | biological process | GO:0050729 | 103 | 8 | 13711 | 111 | Ccr5,Ctss,Grn,C3,Vamp8,Fcer1g,C1qa,Fcgr3 | | 2.033e-06 | -13.11 | structural molecule activity | molecular function | GO:0005198 | 443 | 15 | 13516 | 107 | Capn3,Gfap,Igfbp6,Rps8,Rpl31,Rpl26,Vwf,Ahnak,Rpl13,Vim,Rpl14,Actb,Rps5,Camk2b,Rps20 | | 2.054e-06 | -13.10 | positive regulation of type II hypersensitivity | biological process | GO:0002894 | 4 | 3 | 13711 | 111 | Fcgr3,C3,Fcer1g | | 2.054e-06 | -13.10 | regulation of type II hypersensitivity | biological process | GO:0002892 | 4 | 3 | 13711 | 111 | Fcer1g,C3,Fcgr3 | | 2.054e-06 | -13.10 | positive regulation of type IIa hypersensitivity | biological process | GO:0001798 | 4 | 3 | 13711 | 111 | Fcgr3,C3,Fcer1g | | 2.054e-06 | -13.10 | regulation of type IIa hypersensitivity | biological process | GO:0001796 | 4 | 3 | 13711 | 111 | Fcer1g,C3,Fcgr3 | | 2.129e-06 | -13.06 | APPEL\_IMATINIB\_RESPONSE | MSigDB lists | APPEL\_IMATINIB\_RESPONSE | 27 | 5 | 12187 | 98 | Cd68,Ctsz,Apod,Ctsa,Ctsb | | 2.208e-06 | -13.02 | response to bacterium | biological process | GO:0009617 | 379 | 14 | 13711 | 111 | Mpeg1,Lrg1,Vim,Ikzf3,Klhl6,Ly6a,Tnfsf8,Cd68,Serpinb9,Saa3,C3,Serpina3n,Cd52,Fcer1g | | 2.291e-06 | -12.99 | NABA\_MATRISOME | MSigDB lists | NABA\_MATRISOME | 565 | 17 | 12187 | 98 | Fgf18,C1qa,Serpina3n,Mgp,Tnfsf8,Vwf,Fmod,Ctss,Ctsz,Ctsb,Ctsa,Lgi4,Igfbp6,C1qb,Lrg1,S100a6,Serpinb9 | | 2.411e-06 | -12.94 | side of membrane | cellular component | GO:0098552 | 385 | 14 | 13825 | 111 | Il1r1,Itgam,Vwf,Ctsb,Fcer1g,Adgre1,Il6ra,Ptprc,Fcgr3,Thy1,Ccr5,Ly6a,Gfap,S100a6 | | 2.558e-06 | -12.88 | MODULE\_46 | MSigDB lists | MODULE\_46 | 233 | 11 | 12187 | 98 | Ly6e,Grn,Ccr5,Fcer1g,Fcgr3,Il6ra,C3,Clu,Ctss,Il1r1,C1qa | | 2.575e-06 | -12.87 | STEARMAN\_TUMOR\_FIELD\_EFFECT\_UP | MSigDB lists | STEARMAN\_TUMOR\_FIELD\_EFFECT\_UP | 28 | 5 | 12187 | 98 | Mpeg1,Lrg1,Ctss,Ctsz,Cd68 | | 2.667e-06 | -12.83 | GO\_VIRAL\_LIFE\_CYCLE | MSigDB lists | GO\_VIRAL\_LIFE\_CYCLE | 234 | 11 | 12187 | 98 | Rps5,Rpl31,Dek,Ccr5,Ctsb,Rpl26,Vamp8,Rpl14,Rps20,Rpl13,Rps8 | | 2.691e-06 | -12.83 | positive regulation of myeloid leukocyte mediated immunity | biological process | GO:0002888 | 28 | 5 | 13711 | 111 | Vamp8,Fcgr3,Itgam,C3,Fcer1g | | 2.724e-06 | -12.81 | WALLACE\_PROSTATE\_CANCER\_RACE\_UP | MSigDB lists | WALLACE\_PROSTATE\_CANCER\_RACE\_UP | 189 | 10 | 12187 | 98 | Rac2,C1qa,Ptprc,Fcer1g,Clu,Ccr5,Serpinb9,Ctss,C1qb,Hcls1 | | 2.744e-06 | -12.81 | MODULE\_64 | MSigDB lists | MODULE\_64 | 336 | 13 | 12187 | 98 | Ccr5,Itgam,Fcer1g,Ly6e,Grn,S100a6,Tspan4,Il6ra,Fcgr3,C3,Ptprn,Ptprc,Il1r1 | | 2.906e-06 | -12.75 | multi-organism process | biological process | GO:0051704 | 1450 | 29 | 13711 | 111 | Cd52,Csf1r,Ifitm1,Saa3,C1qb,Cyba,C3,Serpina3n,Ccr5,Tnfsf8,Ly6a,Flnb,Ccl6,Mpeg1,Lrg1,Vamp8,Klhl6,Vim,Fcer1g,Camk2b,Serpinb9,Ctsb,Ptprc,C1qa,Hspa2,Cd68,Tmem106a,Grn,Ikzf3 | | 3.029e-06 | -12.71 | GO\_POSITIVE\_REGULATION\_OF\_RESPONSE\_TO\_STIMULUS | MSigDB lists | GO\_POSITIVE\_REGULATION\_OF\_RESPONSE\_TO\_STIMULUS | 1397 | 28 | 12187 | 98 | Fcgr3,Lfng,C1qb,Csf1r,C1qa,Psmb10,Capn3,Clu,Ctss,Kl,Il6ra,Vamp8,Actb,Ctsb,Thy1,Fcer1g,Itgam,Lrg1,Brk1,Klhl6,Rac2,Fgf18,Cyba,Ptprc,Psme1,Csk,C3,Hcls1 | | 3.092e-06 | -12.69 | GO\_PROTEIN\_ACTIVATION\_CASCADE | MSigDB lists | GO\_PROTEIN\_ACTIVATION\_CASCADE | 29 | 5 | 12187 | 98 | Clu,Vwf,C1qb,C3,C1qa | | 3.267e-06 | -12.63 | regulation of leukocyte mediated immunity | biological process | GO:0002703 | 149 | 9 | 13711 | 111 | Ptprc,Fcgr3,Itgam,Rac2,Fcer1g,Il1r1,Vamp8,Serpinb9,C3 | | 3.407e-06 | -12.59 | MODULE\_75 | MSigDB lists | MODULE\_75 | 240 | 11 | 12187 | 98 | Il6ra,Fcgr3,Il1r1,C1qa,Grn,Ly6e,Clu,Ccr5,Fcer1g,Ctss,C3 | | 3.445e-06 | -12.58 | WIELAND\_UP\_BY\_HBV\_INFECTION | MSigDB lists | WIELAND\_UP\_BY\_HBV\_INFECTION | 80 | 7 | 12187 | 98 | C1qb,Ctsb,Psmb10,Fcer1g,Psme1,Rac2,Cd68 | | 3.499e-06 | -12.56 | NAKAYAMA\_SOFT\_TISSUE\_TUMORS\_PCA1\_UP | MSigDB lists | NAKAYAMA\_SOFT\_TISSUE\_TUMORS\_PCA1\_UP | 52 | 6 | 12187 | 98 | C1qa,Csf1r,C3,C1qb,S100a6,Itgam | | 3.679e-06 | -12.51 | regulation of phagocytosis | biological process | GO:0050764 | 80 | 7 | 13711 | 111 | Ptprc,Fcgr3,Fcer1g,Csk,Rap1gap,C3,Cyba | | 3.749e-06 | -12.49 | positive regulation of response to stimulus | biological process | GO:0048584 | 1637 | 31 | 13711 | 111 | Itgam,Fcer1g,Capn3,Fgf18,Camk2b,Il6ra,Clu,Ptprc,C1qa,Grn,Tmem106a,Ctss,Il1r1,Thy1,Kl,Rac2,Mt3,Csf1r,Hcls1,C3,Cyba,C1qb,Ccr5,Lfng,Csk,Ccl6,Rpl26,Fcgr3,Klhl6,Vamp8,Lrg1 | | 3.926e-06 | -12.45 | Innate Immune System | REACTOME pathways | R-MMU-168249 | 860 | 26 | 6297 | 76 | Fgf18,Actb,C1qb,Psmb10,C1qa,Lrg1,Cd68,Serpina3n,C3,Grn,Ctsb,Ctsz,Vamp8,Ctsa,Camk2b,Fcer1g,Pde1a,Brk1,Clu,Ctss,Kl,Psme1,Itgam,Ccr5,Fcgr3,Cyba | | 4.103e-06 | -12.40 | GO\_CELL\_ACTIVATION | MSigDB lists | GO\_CELL\_ACTIVATION | 405 | 14 | 12187 | 98 | Tnfsf8,Ikzf3,Rac2,Ptprc,Psmb10,Clu,Vwf,Vamp8,Cplx2,Actb,Lfng,Fcer1g,Myh9,Treml2 | | 4.431e-06 | -12.33 | JOHNSTONE\_PARVB\_TARGETS\_3\_UP | MSigDB lists | JOHNSTONE\_PARVB\_TARGETS\_3\_UP | 351 | 13 | 12187 | 98 | Vamp8,Actb,Ctsb,Ctsa,Grn,Myh9,Vim,Ppfibp1,Flnb,Ctss,Ahnak,Clu,C3 | | 4.835e-06 | -12.24 | regulation of inflammatory response | biological process | GO:0050727 | 246 | 11 | 13711 | 111 | Ccr5,C3,Fcer1g,Ctss,Grn,Vamp8,Il1r1,Mgll,Apod,C1qa,Fcgr3 | | 5.100e-06 | -12.19 | GSE9650\_NAIVE\_VS\_EFF\_CD8\_TCELL\_DN | MSigDB lists | GSE9650\_NAIVE\_VS\_EFF\_CD8\_TCELL\_DN | 159 | 9 | 12187 | 98 | Ccr5,Itgam,Ahnak,S100a6,Cd68,Ctsa,Actb,Psmb10,Rac2 | | 5.104e-06 | -12.19 | positive regulation of hypersensitivity | biological process | GO:0002885 | 5 | 3 | 13711 | 111 | Fcgr3,C3,Fcer1g | | 5.144e-06 | -12.18 | GNF2\_CD48 | MSigDB lists | GNF2\_CD48 | 32 | 5 | 12187 | 98 | Csk,Rac2,Hcls1,Ptprc,Psmb10 | | 5.180e-06 | -12.17 | REACTOME\_INFLUENZA\_VIRAL\_RNA\_TRANSCRIPTION\_AND\_REPLICATION | MSigDB lists | REACTOME\_INFLUENZA\_VIRAL\_RNA\_TRANSCRIPTION\_AND\_REPLICATION | 85 | 7 | 12187 | 98 | Rps20,Rpl31,Rps5,Rpl14,Rps8,Rpl13,Rpl26 | | 5.602e-06 | -12.09 | GO\_ESTABLISHMENT\_OF\_PROTEIN\_LOCALIZATION\_TO\_ENDOPLASMIC\_RETICULUM | MSigDB lists | GO\_ESTABLISHMENT\_OF\_PROTEIN\_LOCALIZATION\_TO\_ENDOPLASMIC\_RETICULUM | 86 | 7 | 12187 | 98 | Rpl26,Rpl13,Rps8,Rpl14,Rps5,Rpl31,Rps20 | | 5.631e-06 | -12.09 | HOSHIDA\_LIVER\_CANCER\_SUBCLASS\_S1 | MSigDB lists | HOSHIDA\_LIVER\_CANCER\_SUBCLASS\_S1 | 205 | 10 | 12187 | 98 | Mgp,Fcgr3,Cyba,Ptprc,Thy1,Grn,Ctss,C1qb,Hcls1,Slc2a1 | | 5.664e-06 | -12.08 | GO\_CELL\_SUBSTRATE\_JUNCTION | MSigDB lists | GO\_CELL\_SUBSTRATE\_JUNCTION | 359 | 13 | 12187 | 98 | Ptprc,Flnb,Vim,Rac2,Ppfibp1,Rps8,Ahnak,Actb,Rpl31,Tspan4,Rps5,Thy1,Myh9 | | 5.942e-06 | -12.03 | GSE3337\_4H\_VS\_16H\_IFNG\_IN\_CD8POS\_DC\_DN | MSigDB lists | GSE3337\_4H\_VS\_16H\_IFNG\_IN\_CD8POS\_DC\_DN | 162 | 9 | 12187 | 98 | Ndufa13,Fcer1g,Grn,Ly6e,C1qb,C3,Csf1r,Ctsa,Ctsb | | 6.175e-06 | -12.00 | Complement and coagulation cascades | KEGG pathways | mmu04610 | 39 | 6 | 5248 | 64 | C3,C1qb,C1qa,Vwf,Itgam,Clu | | 6.175e-06 | -12.00 | Complement and coagulation cascades | KEGG pathways | ko04610 | 39 | 6 | 5248 | 64 | Vwf,Itgam,Clu,C3,C1qb,C1qa | | 6.248e-06 | -11.98 | GSE2128\_CTRL\_VS\_MIMETOPE\_NEGATIVE\_SELECTION\_DP\_THYMOCYTE\_NOD\_UP | MSigDB lists | GSE2128\_CTRL\_VS\_MIMETOPE\_NEGATIVE\_SELECTION\_DP\_THYMOCYTE\_NOD\_UP | 163 | 9 | 12187 | 98 | Fcer1g,Ctss,Csf1r,C1qb,Rac2,Ctsb,Fcgr3,Rab3il1,C1qa | | 6.312e-06 | -11.97 | NING\_CHRONIC\_OBSTRUCTIVE\_PULMONARY\_DISEASE\_UP | MSigDB lists | NING\_CHRONIC\_OBSTRUCTIVE\_PULMONARY\_DISEASE\_UP | 123 | 8 | 12187 | 98 | Vamp8,Il1r1,Capn3,Rpl13,Lrg1,Vwf,Csk,Rpl14 | | 6.374e-06 | -11.96 | positive regulation of phagocytosis | biological process | GO:0050766 | 57 | 6 | 13711 | 111 | Ptprc,Fcgr3,Rap1gap,Fcer1g,C3,Cyba | | 6.567e-06 | -11.93 | GSE2405\_0H\_VS\_9H\_A\_PHAGOCYTOPHILUM\_STIM\_NEUTROPHIL\_DN | MSigDB lists | GSE2405\_0H\_VS\_9H\_A\_PHAGOCYTOPHILUM\_STIM\_NEUTROPHIL\_DN | 164 | 9 | 12187 | 98 | Rpl14,Rps5,Rpl31,Ldha,Rps8,Rpl13,Psme1,Actb,Rpl26 | | 6.644e-06 | -11.92 | PID\_INTEGRIN2\_PATHWAY | MSigDB lists | PID\_INTEGRIN2\_PATHWAY | 16 | 4 | 12187 | 98 | Thy1,Itgam,Fcgr3,C3 | | 6.687e-06 | -11.92 | positive regulation of immune response | biological process | GO:0050778 | 360 | 13 | 13711 | 111 | C1qa,Fcgr3,Ptprc,Vamp8,Il1r1,Thy1,Klhl6,Itgam,Fcer1g,Il6ra,Cyba,C1qb,C3 | | 6.692e-06 | -11.91 | ZHONG\_SECRETOME\_OF\_LUNG\_CANCER\_AND\_ENDOTHELIUM | MSigDB lists | ZHONG\_SECRETOME\_OF\_LUNG\_CANCER\_AND\_ENDOTHELIUM | 58 | 6 | 12187 | 98 | Clu,Ctsz,Ldha,Vim,Ctsa,Actb | | 7.018e-06 | -11.87 | ABBUD\_LIF\_SIGNALING\_1\_UP | MSigDB lists | ABBUD\_LIF\_SIGNALING\_1\_UP | 34 | 5 | 12187 | 98 | Il1r1,Tmem176a,Tspan4,Vwf,Lrg1 | | 7.183e-06 | -11.84 | regulation of cell population proliferation | biological process | GO:0042127 | 1270 | 26 | 13711 | 111 | Il6ra,Csk,Mark4,Fgf18,Tsc22d1,Ccr5,Gfap,Ptprn,Cyba,Csf1r,Hcls1,Brk1,Pde1a,Rac2,Spint1,Lrg1,Trnp1,Rap1gap,Ikzf3,Grn,Vim,Ccar1,Ptprc,Apod,Clu,Ctsz | | 7.589e-06 | -11.79 | REACTOME\_NONSENSE\_MEDIATED\_DECAY\_ENHANCED\_BY\_THE\_EXON\_JUNCTION\_COMPLEX | MSigDB lists | REACTOME\_NONSENSE\_MEDIATED\_DECAY\_ENHANCED\_BY\_THE\_EXON\_JUNCTION\_COMPLEX | 90 | 7 | 12187 | 98 | Rps20,Rpl31,Rpl14,Rps5,Rps8,Rpl13,Rpl26 | | 7.589e-06 | -11.79 | REACTOME\_SRP\_DEPENDENT\_COTRANSLATIONAL\_PROTEIN\_TARGETING\_TO\_MEMBRANE | MSigDB lists | REACTOME\_SRP\_DEPENDENT\_COTRANSLATIONAL\_PROTEIN\_TARGETING\_TO\_MEMBRANE | 90 | 7 | 12187 | 98 | Rps20,Rpl31,Rps5,Rpl14,Rps8,Rpl13,Rpl26 | | 7.589e-06 | -11.79 | REACTOME\_3\_UTR\_MEDIATED\_TRANSLATIONAL\_REGULATION | MSigDB lists | REACTOME\_3\_UTR\_MEDIATED\_TRANSLATIONAL\_REGULATION | 90 | 7 | 12187 | 98 | Rps20,Rpl31,Rps5,Rpl14,Rpl26,Rps8,Rpl13 | | 7.988e-06 | -11.74 | GSE29164\_UNTREATED\_VS\_CD8\_TCELL\_TREATED\_MELANOMA\_DAY7\_UP | MSigDB lists | GSE29164\_UNTREATED\_VS\_CD8\_TCELL\_TREATED\_MELANOMA\_DAY7\_UP | 168 | 9 | 12187 | 98 | Tmem176a,Csf1r,Psme1,Ccr5,Thy1,Camk2b,Ctsa,Cyba,Rac2 | | 9.003e-06 | -11.62 | GNF2\_CD53 | MSigDB lists | GNF2\_CD53 | 61 | 6 | 12187 | 98 | Ptprc,Hcls1,Psmb10,Psme1,Rac2,Csk | | 9.349e-06 | -11.58 | immune effector process | biological process | GO:0002252 | 316 | 12 | 13711 | 111 | Lfng,C1qb,C3,Serpinb9,Ifitm1,Fcer1g,Cplx2,Grn,Clu,Fcgr3,C1qa,Ptprc | | 9.388e-06 | -11.58 | ABE\_INNER\_EAR | MSigDB lists | ABE\_INNER\_EAR | 36 | 5 | 12187 | 98 | Clu,Igfbp6,Mgp,Rps5,Apod | | 9.910e-06 | -11.52 | REACTOME\_INITIAL\_TRIGGERING\_OF\_COMPLEMENT | MSigDB lists | REACTOME\_INITIAL\_TRIGGERING\_OF\_COMPLEMENT | 6 | 3 | 12187 | 98 | C1qb,C3,C1qa | | 1.008e-05 | -11.50 | GO\_IMMUNE\_RESPONSE\_REGULATING\_CELL\_SURFACE\_RECEPTOR\_SIGNALING\_PATHWAY | MSigDB lists | GO\_IMMUNE\_RESPONSE\_REGULATING\_CELL\_SURFACE\_RECEPTOR\_SIGNALING\_PATHWAY | 219 | 10 | 12187 | 98 | Klhl6,Psme1,Thy1,Fcer1g,Csk,Brk1,Actb,Fcgr3,Psmb10,Ptprc | | 1.064e-05 | -11.45 | MORF\_NME2 | MSigDB lists | MORF\_NME2 | 132 | 8 | 12187 | 98 | Actb,Cox4i1,Rps5,Rpl31,Rps20,Rpl13,Ldha,Rps8 | | 1.081e-05 | -11.43 | Pept\_asp\_AS | interpro domains | IPR025661 | 6 | 3 | 13788 | 114 | Ctss,Ctsz,Ctsb | | 1.088e-05 | -11.43 | FLECHNER\_BIOPSY\_KIDNEY\_TRANSPLANT\_REJECTED\_VS\_OK\_UP | MSigDB lists | FLECHNER\_BIOPSY\_KIDNEY\_TRANSPLANT\_REJECTED\_VS\_OK\_UP | 63 | 6 | 12187 | 98 | Actb,C1qb,Hcls1,Ptprc,Rac2,Fcer1g | | 1.104e-05 | -11.41 | adaptive immune response | biological process | GO:0002250 | 173 | 9 | 13711 | 111 | C1qa,Fcgr3,Fcer1g,Serpinb9,Csk,Klhl6,Adgre1,C3,C1qb | | 1.124e-05 | -11.40 | GO\_ADAPTIVE\_IMMUNE\_RESPONSE | MSigDB lists | GO\_ADAPTIVE\_IMMUNE\_RESPONSE | 133 | 8 | 12187 | 98 | Klhl6,C3,C1qb,Ctss,Csk,Clu,Fcer1g,C1qa | | 1.151e-05 | -11.37 | regulation of response to external stimulus | biological process | GO:0032101 | 697 | 18 | 13711 | 111 | Fcgr3,C1qa,Apod,Vamp8,Il1r1,Mgll,Ctss,Grn,Mt3,Rac2,Fcer1g,Treml2,Serpinb9,Fgf18,Capn3,Ccr5,C3,Cyba | | 1.305e-05 | -11.25 | RAMALHO\_STEMNESS\_DN | MSigDB lists | RAMALHO\_STEMNESS\_DN | 65 | 6 | 12187 | 98 | Csf1r,Mpeg1,Ctss,Cd68,Ctsb,Ptprc | | 1.320e-05 | -11.24 | THUM\_SYSTOLIC\_HEART\_FAILURE\_UP | MSigDB lists | THUM\_SYSTOLIC\_HEART\_FAILURE\_UP | 331 | 12 | 12187 | 98 | C1qa,Ptprc,Rac2,Ppfibp1,Fmod,Ctss,Actb,Ctsb,Vamp8,C1qb,Mpeg1,Fcer1g | | 1.329e-05 | -11.23 | positive regulation of leukocyte mediated immunity | biological process | GO:0002705 | 97 | 7 | 13711 | 111 | Fcgr3,Ptprc,Vamp8,Il1r1,Fcer1g,Itgam,C3 | | 1.334e-05 | -11.23 | GO\_SERINE\_HYDROLASE\_ACTIVITY | MSigDB lists | GO\_SERINE\_HYDROLASE\_ACTIVITY | 98 | 7 | 12187 | 98 | C1qa,Ctsa,Ctsb,Ctss,Pcsk2,C1qb,C3 | | 1.389e-05 | -11.18 | GO\_HUMORAL\_IMMUNE\_RESPONSE\_MEDIATED\_BY\_CIRCULATING\_IMMUNOGLOBULIN | MSigDB lists | GO\_HUMORAL\_IMMUNE\_RESPONSE\_MEDIATED\_BY\_CIRCULATING\_IMMUNOGLOBULIN | 19 | 4 | 12187 | 98 | C1qa,C3,C1qb,Clu | | 1.389e-05 | -11.18 | GO\_COMPLEMENT\_ACTIVATION | MSigDB lists | GO\_COMPLEMENT\_ACTIVATION | 19 | 4 | 12187 | 98 | Clu,C1qa,C1qb,C3 | | 1.404e-05 | -11.17 | Microglia Pathogen Phagocytosis Pathway | WikiPathways | WP3626 | 40 | 6 | 3756 | 52 | C1qa,C1qb,Rac2,Fcer1g,Itgam,Cyba | | 1.427e-05 | -11.16 | FCHO2 (FCH domain only 2) | protein interactions | 115548 | 5 | 3 | 6802 | 78 | Flnb,Myh9,Actb | | 1.520e-05 | -11.09 | glial cell development | biological process | GO:0021782 | 99 | 7 | 13711 | 111 | Vim,Grn,Gfap,Lgi4,Mt3,Itgam,Clu | | 1.571e-05 | -11.06 | response to stimulus | biological process | GO:0050896 | 5062 | 63 | 13711 | 111 | Grn,Ctss,Ikzf3,Adgre1,Il1r1,Thy1,Kl,Ptprc,C1qa,Fgf18,Ly6e,Actb,Pde1a,Fcer1g,Ldha,Cplx2,Klhl6,Rap1gap,Vamp8,Slc2a1,Fcgr3,Flnb,Cyba,C1qb,Ndufa13,Ccr5,Lfng,Mark4,Csk,Mt3,Cd52,Myh9,Mc4r,Tmem106a,Adh1,Cd68,Clu,Apod,Hspa2,Gfap,Capn3,Camk2b,Serpinb9,Il6ra,Itgam,Slc17a7,Vim,Mpeg1,Lrg1,Ccl6,Rpl26,Tnfsf8,Ly6a,Ptprn,C3,Serpina3n,Vwf,Saa3,Rac2,Ifitm1,Brk1,Hcls1,Csf1r | | 1.597e-05 | -11.04 | GO\_B\_CELL\_MEDIATED\_IMMUNITY | MSigDB lists | GO\_B\_CELL\_MEDIATED\_IMMUNITY | 40 | 5 | 12187 | 98 | Fcer1g,Clu,C1qa,C3,C1qb | | 1.602e-05 | -11.04 | MODULE\_88 | MSigDB lists | MODULE\_88 | 520 | 15 | 12187 | 98 | Hspa2,Vwf,Mgll,Rac2,Serpina3n,Apod,Phyhip,Il1r1,Ptprn,Fcer1g,Grn,C1qb,Adh1,Slc17a7,Igfbp6 | | 1.626e-05 | -11.03 | GO\_PROTEIN\_LOCALIZATION\_TO\_ENDOPLASMIC\_RETICULUM | MSigDB lists | GO\_PROTEIN\_LOCALIZATION\_TO\_ENDOPLASMIC\_RETICULUM | 101 | 7 | 12187 | 98 | Rpl13,Rps8,Rpl14,Rps5,Rpl31,Rps20,Rpl26 | | 1.629e-05 | -11.03 | GO\_LEUKOCYTE\_ACTIVATION | MSigDB lists | GO\_LEUKOCYTE\_ACTIVATION | 283 | 11 | 12187 | 98 | Treml2,Lfng,Myh9,Clu,Fcer1g,Ptprc,Psmb10,Vamp8,Tnfsf8,Cplx2,Ikzf3 | | 1.636e-05 | -11.02 | GSE40274\_CTRL\_VS\_IRF4\_TRANSDUCED\_ACTIVATED\_CD4\_TCELL\_UP | MSigDB lists | GSE40274\_CTRL\_VS\_IRF4\_TRANSDUCED\_ACTIVATED\_CD4\_TCELL\_UP | 140 | 8 | 12187 | 98 | C1qb,Fam111a,S100a6,Ctss,Rab3il1,C1qa,Tmem106a,Vim | | 1.724e-05 | -10.97 | BIOCARTA\_COMP\_PATHWAY | MSigDB lists | BIOCARTA\_COMP\_PATHWAY | 7 | 3 | 12187 | 98 | C1qa,C1qb,C3 | | 1.724e-05 | -10.97 | BIOCARTA\_CLASSIC\_PATHWAY | MSigDB lists | BIOCARTA\_CLASSIC\_PATHWAY | 7 | 3 | 12187 | 98 | C1qa,C1qb,C3 | | 1.734e-05 | -10.96 | GO\_NUCLEAR\_TRANSCRIBED\_MRNA\_CATABOLIC\_PROCESS\_NONSENSE\_MEDIATED\_DECAY | MSigDB lists | GO\_NUCLEAR\_TRANSCRIBED\_MRNA\_CATABOLIC\_PROCESS\_NONSENSE\_MEDIATED\_DECAY | 102 | 7 | 12187 | 98 | Rpl13,Rps8,Rpl31,Rpl14,Rps5,Rps20,Rpl26 | | 1.765e-05 | -10.94 | positive regulation of acute inflammatory response to antigenic stimulus | biological process | GO:0002866 | 7 | 3 | 13711 | 111 | Fcgr3,C3,Fcer1g | | 1.887e-05 | -10.88 | acute inflammatory response | biological process | GO:0002526 | 41 | 5 | 13711 | 111 | Serpina3n,Kl,Fcgr3,Saa3,Serpinb9 | | 1.972e-05 | -10.83 | REACTOME\_METABOLISM\_OF\_MRNA | MSigDB lists | REACTOME\_METABOLISM\_OF\_MRNA | 188 | 9 | 12187 | 98 | Rpl26,Psmb10,Rps8,Rpl13,Psme1,Rps20,Rps5,Rpl14,Rpl31 | | 2.014e-05 | -10.81 | regulation of defense response | biological process | GO:0031347 | 460 | 14 | 13711 | 111 | Apod,C1qa,Fcgr3,Ctss,Grn,Vamp8,Il1r1,Mgll,Treml2,Fcer1g,Ccr5,Cyba,C3,Serpinb9 | | 2.039e-05 | -10.80 | GNF2\_HLA\_C | MSigDB lists | GNF2\_HLA\_C | 42 | 5 | 12187 | 98 | Rac2,Hcls1,Ptprc,Psmb10,Cyba | | 2.061e-05 | -10.79 | THIOL\_PROTEASE\_ASN | prosite domains | PS00640 | 6 | 3 | 8845 | 91 | Ctsz,Ctss,Ctsb | | 2.112e-05 | -10.77 | P53\_DN.V1\_DN | MSigDB lists | P53\_DN.V1\_DN | 145 | 8 | 12187 | 98 | Ctsz,Pde1a,Hspa2,Capn3,Mgll,Il1r1,Slc26a2,Vim | | 2.182e-05 | -10.73 | Staphylococcus aureus infection | KEGG pathways | mmu05150 | 29 | 5 | 5248 | 64 | Itgam,C3,Fcgr3,C1qa,C1qb | | 2.182e-05 | -10.73 | Staphylococcus aureus infection | KEGG pathways | ko05150 | 29 | 5 | 5248 | 64 | Itgam,C1qa,C1qb,C3,Fcgr3 | | 2.189e-05 | -10.73 | cell periphery | cellular component | GO:0071944 | 3578 | 49 | 13825 | 111 | Vamp8,Ccr5,Hspa2,Ptprc,Il6ra,Tmem106a,Cd52,Lin7a,Slc2a1,Fcgr3,Hcls1,Treml2,Ifitm1,Adgre1,Sh2d5,Fcer1g,Cyba,Clu,Vwf,Exoc3l4,Csf1r,Slc26a2,Ptprn,Flnb,Abhd17c,Spint1,Actb,Grn,Cd68,Dlgap3,Vim,Ly6a,Ahnak,Kl,Thy1,S100a6,Ly6e,Myh9,Ikzf3,Rac2,Tspan4,Ctsz,Capn3,Csk,Itgam,Il1r1,Adh1,Mc4r,Ctsb | | 2.219e-05 | -10.72 | GSE26912\_TUMORICIDAL\_VS\_CTRL\_MACROPHAGE\_DN | MSigDB lists | GSE26912\_TUMORICIDAL\_VS\_CTRL\_MACROPHAGE\_DN | 146 | 8 | 12187 | 98 | C1qa,Rps5,Csf1r,Psme1,Ccr5,Cd68,Ctsz,Brk1 | | 2.233e-05 | -10.71 | adaptive immune response based on somatic recombination of immune receptors built from immunoglobulin superfamily domains | biological process | GO:0002460 | 105 | 7 | 13711 | 111 | C1qa,Fcgr3,Fcer1g,Serpinb9,C3,C1qb,Klhl6 | | 2.286e-05 | -10.69 | cellular response to cytokine stimulus | biological process | GO:0071345 | 529 | 15 | 13711 | 111 | Flnb,Ccl6,Vamp8,Il1r1,Vim,Mt3,Csf1r,Hcls1,Ifitm1,Fcer1g,Il6ra,Saa3,Ndufa13,Ccr5,Ptprn | | 2.308e-05 | -10.68 | positive regulation of cell population proliferation | biological process | GO:0008284 | 734 | 18 | 13711 | 111 | Ccar1,Grn,Vim,Lrg1,Ctsz,Clu,Ptprc,Tsc22d1,Cyba,Gfap,Ptprn,Il6ra,Mark4,Fgf18,Rac2,Csf1r,Hcls1,Brk1 | | 2.548e-05 | -10.58 | GO\_INTERSPECIES\_INTERACTION\_BETWEEN\_ORGANISMS | MSigDB lists | GO\_INTERSPECIES\_INTERACTION\_BETWEEN\_ORGANISMS | 541 | 15 | 12187 | 98 | Ccr5,Serpinb9,Rpl31,Rps5,Dek,Rpl26,Vamp8,Ctsb,Fam111a,Rpl13,Rps8,Rpl14,Rps20,Vim,Psmb10 | | 2.661e-05 | -10.53 | innate immune response | biological process | GO:0045087 | 351 | 12 | 13711 | 111 | Csf1r,Ifitm1,Fcer1g,Cyba,C3,C1qb,Flnb,C1qa,Ccl6,Vamp8,Vim,Tmem106a | | 2.688e-05 | -10.52 | response to cytokine | biological process | GO:0034097 | 603 | 16 | 13711 | 111 | Ndufa13,Ccr5,Serpina3n,Ptprn,Saa3,Il6ra,Ifitm1,Fcer1g,Mt3,Csf1r,Hcls1,Vim,Il1r1,Vamp8,Ccl6,Flnb | | 2.808e-05 | -10.48 | synapse pruning | biological process | GO:0098883 | 8 | 3 | 13711 | 111 | C1qa,C3,C1qb | | 2.808e-05 | -10.48 | regulation of hypersensitivity | biological process | GO:0002883 | 8 | 3 | 13711 | 111 | C3,Fcer1g,Fcgr3 | | 2.831e-05 | -10.47 | CD109 (CD109 molecule) | protein interactions | 135228 | 6 | 3 | 6802 | 78 | Actb,Myh9,Flnb | | 2.847e-05 | -10.47 | leukocyte mediated immunity | biological process | GO:0002443 | 109 | 7 | 13711 | 111 | Serpinb9,Cplx2,C3,C1qb,Fcgr3,C1qa,Fcer1g | | 2.968e-05 | -10.43 | GSE43955\_1H\_VS\_20H\_ACT\_CD4\_TCELL\_WITH\_TGFB\_IL6\_DN | MSigDB lists | GSE43955\_1H\_VS\_20H\_ACT\_CD4\_TCELL\_WITH\_TGFB\_IL6\_DN | 152 | 8 | 12187 | 98 | Flnb,Mgp,Rpl14,Pcsk2,S100a6,Ly6e,Pde1a,Ctss | | 2.982e-05 | -10.42 | GO\_ADAPTIVE\_IMMUNE\_RESPONSE\_BASED\_ON\_SOMATIC\_RECOMBINATION\_OF\_IMMUNE\_RECEPTORS\_BUILT\_FROM\_IMMUNOGLOBULIN\_SUPERFAMILY\_DOMAINS | MSigDB lists | GO\_ADAPTIVE\_IMMUNE\_RESPONSE\_BASED\_ON\_SOMATIC\_RECOMBINATION\_OF\_IMMUNE\_RECEPTORS\_BUILT\_FROM\_IMMUNOGLOBULIN\_SUPERFAMILY\_DOMAINS | 75 | 6 | 12187 | 98 | Klhl6,C1qb,C3,Clu,Fcer1g,C1qa | | 3.007e-05 | -10.41 | GSE7218\_UNSTIM\_VS\_ANTIGEN\_STIM\_THROUGH\_IGG\_BCELL\_DN | MSigDB lists | GSE7218\_UNSTIM\_VS\_ANTIGEN\_STIM\_THROUGH\_IGG\_BCELL\_DN | 111 | 7 | 12187 | 98 | Treml2,S100a6,Thy1,Lfng,Apod,Rac2,Ikzf3 | | 3.036e-05 | -10.40 | RODWELL\_AGING\_KIDNEY\_UP | MSigDB lists | RODWELL\_AGING\_KIDNEY\_UP | 360 | 12 | 12187 | 98 | Hcls1,Mpeg1,Csf1r,C1qb,Ly6e,Grn,Fcer1g,Psmb10,C1qa,Ptprc,Vim,Rac2 | | 3.160e-05 | -10.36 | response to stress | biological process | GO:0006950 | 2282 | 36 | 13711 | 111 | Clu,Cd68,Ptprc,C1qa,Apod,Hspa2,Grn,Tmem106a,Kl,Il1r1,Fcer1g,Itgam,Gfap,Serpinb9,Capn3,Ccl6,Tnfsf8,Fcgr3,Flnb,Rpl26,Vim,Slc2a1,Mpeg1,Vamp8,Ifitm1,Myh9,Csf1r,Mt3,Vwf,Ptprn,Cyba,C1qb,C3,Ccr5,Serpina3n,Saa3 | | 3.203e-05 | -10.35 | STEARMAN\_LUNG\_CANCER\_EARLY\_VS\_LATE\_DN | MSigDB lists | STEARMAN\_LUNG\_CANCER\_EARLY\_VS\_LATE\_DN | 46 | 5 | 12187 | 98 | Lrg1,Cd68,Ctss,Ctsz,Mpeg1 | | 3.203e-05 | -10.35 | GCM\_PFN1 | MSigDB lists | GCM\_PFN1 | 46 | 5 | 12187 | 98 | Rps8,Psme1,Hcls1,Rps5,Rpl14 | | 3.217e-05 | -10.34 | positive regulation of immune system process | biological process | GO:0002684 | 612 | 16 | 13711 | 111 | Il6ra,C1qb,C3,Cyba,Csf1r,Hcls1,Fcer1g,Rac2,Itgam,Thy1,Vamp8,Il1r1,Klhl6,C1qa,Fcgr3,Ptprc | | 3.392e-05 | -10.29 | TARTE\_PLASMA\_CELL\_VS\_PLASMABLAST\_UP | MSigDB lists | TARTE\_PLASMA\_CELL\_VS\_PLASMABLAST\_UP | 252 | 10 | 12187 | 98 | Actb,Ptprn,Cyba,Serpina3n,Flnb,Clu,Fcer1g,Fmod,Grn,Mc4r | | 3.573e-05 | -10.24 | GO\_ANCHORING\_JUNCTION | MSigDB lists | GO\_ANCHORING\_JUNCTION | 427 | 13 | 12187 | 98 | Ahnak,Rps8,Rac2,Ppfibp1,Vim,Ptprc,Flnb,Thy1,Myh9,Rpl31,Tspan4,Rps5,Actb | | 3.637e-05 | -10.22 | cytokine-mediated signaling pathway | biological process | GO:0019221 | 201 | 9 | 13711 | 111 | Il1r1,Il6ra,Ptprn,Ccr5,Mt3,Csf1r,Ccl6,Fcer1g,Ifitm1 | | 3.731e-05 | -10.20 | KAYO\_CALORIE\_RESTRICTION\_MUSCLE\_UP | MSigDB lists | KAYO\_CALORIE\_RESTRICTION\_MUSCLE\_UP | 78 | 6 | 12187 | 98 | C1qb,Actb,Hcls1,Kl,Myh9,Ccr5 | | 3.815e-05 | -10.17 | positive regulation of acute inflammatory response | biological process | GO:0002675 | 24 | 4 | 13711 | 111 | Fcgr3,Fcer1g,C3,Ccr5 | | 3.917e-05 | -10.15 | GSE24634\_TREG\_VS\_TCONV\_POST\_DAY7\_IL4\_CONVERSION\_DN | MSigDB lists | GSE24634\_TREG\_VS\_TCONV\_POST\_DAY7\_IL4\_CONVERSION\_DN | 158 | 8 | 12187 | 98 | Ctsz,Ctss,Cd68,Mgll,Tsc22d1,C1qa,Ctsb,Igfbp6 | | 3.943e-05 | -10.14 | Ppp1r9b (protein phosphatase 1, regulatory subunit 9B) | protein interactions | 217124 | 401 | 15 | 6802 | 78 | Rpl13,Hspa2,Vim,Dlgap3,Slc17a7,Rps8,Cox4i1,Ldha,Grn,Phyhip,Thy1,Gfap,Actb,Camk2b,Myh9 | | 3.996e-05 | -10.13 | REACTOME\_INFLUENZA\_LIFE\_CYCLE | MSigDB lists | REACTOME\_INFLUENZA\_LIFE\_CYCLE | 116 | 7 | 12187 | 98 | Rpl13,Rps8,Rpl14,Rps5,Rpl31,Rps20,Rpl26 | | 4.090e-05 | -10.10 | MODULE\_58 | MSigDB lists | MODULE\_58 | 9 | 3 | 12187 | 98 | C1qa,C3,Clu | | 4.090e-05 | -10.10 | GO\_IMMUNOGLOBULIN\_BINDING | MSigDB lists | GO\_IMMUNOGLOBULIN\_BINDING | 9 | 3 | 12187 | 98 | Vwf,Fcer1g,Fcgr3 | | 4.097e-05 | -10.10 | GSE36826\_WT\_VS\_IL1R\_KO\_SKIN\_DN | MSigDB lists | GSE36826\_WT\_VS\_IL1R\_KO\_SKIN\_DN | 159 | 8 | 12187 | 98 | Vim,Sh3bgrl3,Actb,Lfng,Ldha,S100a6,Ccr5,Hcls1 | | 4.097e-05 | -10.10 | GSE22886\_NAIVE\_CD4\_TCELL\_VS\_MONOCYTE\_DN | MSigDB lists | GSE22886\_NAIVE\_CD4\_TCELL\_VS\_MONOCYTE\_DN | 159 | 8 | 12187 | 98 | Ctss,Fcer1g,Itgam,Grn,Il6ra,Ctsa,Fcgr3,Ctsb | | 4.223e-05 | -10.07 | GO\_MULTI\_ORGANISM\_METABOLIC\_PROCESS | MSigDB lists | GO\_MULTI\_ORGANISM\_METABOLIC\_PROCESS | 117 | 7 | 12187 | 98 | Rpl31,Rps5,Rpl14,Rps20,Rpl13,Rps8,Rpl26 | | 4.252e-05 | -10.07 | developmental process | biological process | GO:0032502 | 4177 | 54 | 13711 | 111 | Gfap,Ctsb,Capn3,Il6ra,Camk2b,Itgam,Slc17a7,Rbfox3,Cd68,Clu,Hspa2,Apod,Tfap2c,C3,Ptprn,Vwf,Spint1,Ifitm1,Rac2,Lgi4,Brk1,Hcls1,Csf1r,Vim,Lrg1,Tnfsf8,Pcsk2,Fgf18,Ly6e,Actb,Bhlhe41,Lin7a,Fcer1g,Psmb10,Grn,Trnp1,Il1r1,Mgll,Kl,Thy1,C1qa,Ptprc,Mgp,Ccr5,C1qb,Lfng,Mark4,Mt3,Myh9,Klhl6,Cplx2,Rap1gap,Ctsz,Flnb | | 4.405e-05 | -10.03 | positive regulation of immune effector process | biological process | GO:0002699 | 159 | 8 | 13711 | 111 | C3,Il1r1,Vamp8,Itgam,Fcer1g,Rac2,Fcgr3,Ptprc | | 4.460e-05 | -10.02 | Peptidase\_C1A\_C | interpro domains | IPR000668 | 9 | 3 | 13788 | 114 | Ctsz,Ctsb,Ctss | | 4.479e-05 | -10.01 | GSE7831\_UNSTIM\_VS\_INFLUENZA\_STIM\_PDC\_4H\_DN | MSigDB lists | GSE7831\_UNSTIM\_VS\_INFLUENZA\_STIM\_PDC\_4H\_DN | 161 | 8 | 12187 | 98 | Grn,Tsc22d1,Fmod,Ahnak,Ccr5,Spint1,Ctsb,Rac2 | | 4.642e-05 | -9.98 | Peptidase\_C1 | pfam domains | PF00112 | 9 | 3 | 12881 | 108 | Ctss,Ctsb,Ctsz | | 4.827e-05 | -9.94 | GNF2\_INPP5D | MSigDB lists | GNF2\_INPP5D | 50 | 5 | 12187 | 98 | Csk,Psme1,Rac2,Hcls1,Psmb10 | | 4.913e-05 | -9.92 | PPP1R18 (protein phosphatase 1 regulatory subunit 18) | protein interactions | 170954 | 7 | 3 | 6802 | 78 | Flnb,Actb,Myh9 | | 4.957e-05 | -9.91 | GO\_GLYCOPROTEIN\_BINDING | MSigDB lists | GO\_GLYCOPROTEIN\_BINDING | 82 | 6 | 12187 | 98 | Ctsb,Gfap,Vim,Vwf,Itgam,Ctss | | 5.107e-05 | -9.88 | GSE15930\_STIM\_VS\_STIM\_AND\_TRICHOSTATINA\_48H\_CD8\_T\_CELL\_UP | MSigDB lists | GSE15930\_STIM\_VS\_STIM\_AND\_TRICHOSTATINA\_48H\_CD8\_T\_CELL\_UP | 164 | 8 | 12187 | 98 | Il1r1,Rac2,Kl,S100a6,Ahnak,Thy1,Ccr5,Ldha | | 5.107e-05 | -9.88 | GSE21379\_WT\_VS\_SAP\_KO\_TFH\_CD4\_TCELL\_UP | MSigDB lists | GSE21379\_WT\_VS\_SAP\_KO\_TFH\_CD4\_TCELL\_UP | 164 | 8 | 12187 | 98 | Exoc3l4,C3,Thy1,Clu,Lrg1,Fcgr3,Rab3il1,Vamp8 | | 5.221e-05 | -9.86 | positive regulation of adaptive immune response based on somatic recombination of immune receptors built from immunoglobulin superfamily domains | biological process | GO:0002824 | 82 | 6 | 13711 | 111 | Il1r1,Il6ra,Fcgr3,Ptprc,C3,Fcer1g | | 5.221e-05 | -9.86 | positive regulation of tumor necrosis factor production | biological process | GO:0032760 | 82 | 6 | 13711 | 111 | Clu,Ccr5,Cyba,Fcer1g,Fcgr3,Ptprc | | 5.221e-05 | -9.86 | positive regulation of tumor necrosis factor superfamily cytokine production | biological process | GO:1903557 | 82 | 6 | 13711 | 111 | Cyba,Ccr5,Clu,Fcer1g,Ptprc,Fcgr3 | | 5.235e-05 | -9.86 | GO\_DEFENSE\_RESPONSE | MSigDB lists | GO\_DEFENSE\_RESPONSE | 718 | 17 | 12187 | 98 | Il6ra,Ccr5,Fcer1g,C1qb,Csf1r,Tnfsf8,Ptprc,Phyhip,Serpina3n,C1qa,Cyba,Csk,Clu,Fam111a,Camk2b,Mgll,C3 | | 5.275e-05 | -9.85 | Complement cascade | REACTOME pathways | R-MMU-166658 | 18 | 4 | 6297 | 76 | C3,C1qa,Clu,C1qb | | 5.320e-05 | -9.84 | Ribosome, eukaryotes | KEGG pathways | mmu\_M00177 | 82 | 7 | 5248 | 64 | Rpl13,Rps5,Rpl31,Rpl14,Rps20,Rps8,Rpl26 | | 5.320e-05 | -9.84 | Ribosome, eukaryotes | KEGG pathways | M00177 | 82 | 7 | 5248 | 64 | Rpl26,Rps8,Rpl14,Rps20,Rps5,Rpl31,Rpl13 | | 5.382e-05 | -9.83 | GO\_CELL\_JUNCTION | MSigDB lists | GO\_CELL\_JUNCTION | 948 | 20 | 12187 | 98 | Actb,Lin7a,Slc17a7,Rpl31,Rps5,Tspan4,Slc2a1,Sh2d5,Myh9,Thy1,Ptprn,Ptprc,Flnb,Dlgap3,Rac2,Ppfibp1,Vim,Ahnak,Csk,Rps8 | | 5.546e-05 | -9.80 | immunoglobulin mediated immune response | biological process | GO:0016064 | 51 | 5 | 13711 | 111 | Fcer1g,C1qb,C3,C1qa,Fcgr3 | | 5.566e-05 | -9.80 | GSE10325\_LUPUS\_BCELL\_VS\_LUPUS\_MYELOID\_DN | MSigDB lists | GSE10325\_LUPUS\_BCELL\_VS\_LUPUS\_MYELOID\_DN | 166 | 8 | 12187 | 98 | Ctsb,Actb,Fcgr3,Ctsa,S100a6,Fcer1g,Itgam,Csf1r | | 5.712e-05 | -9.77 | Initial triggering of complement | REACTOME pathways | R-MMU-166663 | 7 | 3 | 6297 | 76 | C1qb,C1qa,C3 | | 5.808e-05 | -9.75 | GSE29618\_BCELL\_VS\_MDC\_DAY7\_FLU\_VACCINE\_DN | MSigDB lists | GSE29618\_BCELL\_VS\_MDC\_DAY7\_FLU\_VACCINE\_DN | 167 | 8 | 12187 | 98 | Csf1r,Ldha,Fcer1g,Ahnak,Grn,Actb,Vim,Il6ra | | 5.815e-05 | -9.75 | WU\_CELL\_MIGRATION | MSigDB lists | WU\_CELL\_MIGRATION | 123 | 7 | 12187 | 98 | Tspan4,Clu,S100a6,Spint1,Igfbp6,Rac2,Vim | | 5.846e-05 | -9.75 | GNF2\_MYD88 | MSigDB lists | GNF2\_MYD88 | 52 | 5 | 12187 | 98 | Sh3bgrl3,Fcer1g,Hcls1,Psmb10,Cyba | | 5.946e-05 | -9.73 | positive regulation of inflammatory response to antigenic stimulus | biological process | GO:0002863 | 10 | 3 | 13711 | 111 | C3,Fcer1g,Fcgr3 | | 5.987e-05 | -9.72 | GNF2\_FGR | MSigDB lists | GNF2\_FGR | 27 | 4 | 12187 | 98 | Itgam,Fcer1g,Sh3bgrl3,Ctss | | 6.058e-05 | -9.71 | GO\_CYTOSOLIC\_PART | MSigDB lists | GO\_CYTOSOLIC\_PART | 168 | 8 | 12187 | 98 | Apod,Rpl26,Rpl31,Rpl14,Rps5,Rps20,Rpl13,Rps8 | | 6.185e-05 | -9.69 | positive regulation of B cell mediated immunity | biological process | GO:0002714 | 27 | 4 | 13711 | 111 | Fcer1g,C3,Ptprc,Fcgr3 | | 6.185e-05 | -9.69 | positive regulation of immunoglobulin mediated immune response | biological process | GO:0002891 | 27 | 4 | 13711 | 111 | C3,Fcer1g,Ptprc,Fcgr3 | | 6.317e-05 | -9.67 | GSE32164\_RESTING\_DIFFERENTIATED\_VS\_CMYC\_INHIBITED\_MACROPHAGE\_DN | MSigDB lists | GSE32164\_RESTING\_DIFFERENTIATED\_VS\_CMYC\_INHIBITED\_MACROPHAGE\_DN | 169 | 8 | 12187 | 98 | Rac2,Brk1,S100a6,Ccr5,Psme1,Camk2b,Csf1r,Tmem176a | | 6.394e-05 | -9.66 | lymphocyte mediated immunity | biological process | GO:0002449 | 85 | 6 | 13711 | 111 | C1qa,Serpinb9,Fcgr3,Fcer1g,C1qb,C3 | | 6.414e-05 | -9.65 | KYNG\_RESPONSE\_TO\_H2O2 | MSigDB lists | KYNG\_RESPONSE\_TO\_H2O2 | 53 | 5 | 12187 | 98 | Tspan4,Ctsb,Rpl31,Cd68,Ctss | | 6.475e-05 | -9.64 | GO\_PROTEIN\_LOCALIZATION\_TO\_MEMBRANE | MSigDB lists | GO\_PROTEIN\_LOCALIZATION\_TO\_MEMBRANE | 329 | 11 | 12187 | 98 | Rps5,Rpl14,Rpl31,Rps20,Fcer1g,Rpl13,Capn3,Ndufa13,Rps8,Lin7a,Rpl26 | | 6.486e-05 | -9.64 | MORF\_TPT1 | MSigDB lists | MORF\_TPT1 | 86 | 6 | 12187 | 98 | Rps20,Rpl31,Rps5,Actb,Rps8,Rpl13 | | 6.495e-05 | -9.64 | vertebrate eye-specific patterning | biological process | GO:0150064 | 2 | 2 | 13711 | 111 | C3,C1qa | | 6.532e-05 | -9.64 | Pept\_C1 | smart domains | SM00645 | 9 | 3 | 7188 | 68 | Ctsb,Ctss,Ctsz | | 6.586e-05 | -9.63 | GO\_LEUKOCYTE\_CELL\_CELL\_ADHESION | MSigDB lists | GO\_LEUKOCYTE\_CELL\_CELL\_ADHESION | 170 | 8 | 12187 | 98 | Treml2,Lfng,Myh9,Fcer1g,Ptprc,Psmb10,Tnfsf8,Rac2 | | 6.779e-05 | -9.60 | GO\_TRANSLATIONAL\_INITIATION | MSigDB lists | GO\_TRANSLATIONAL\_INITIATION | 126 | 7 | 12187 | 98 | Rpl31,Rps5,Rpl14,Rps20,Rpl13,Rps8,Rpl26 | | 6.864e-05 | -9.59 | GSE42021\_CD24LO\_TREG\_VS\_CD24LO\_TCONV\_THYMUS\_DN | MSigDB lists | GSE42021\_CD24LO\_TREG\_VS\_CD24LO\_TCONV\_THYMUS\_DN | 171 | 8 | 12187 | 98 | C1qb,Rps5,Csk,Psme1,Grn,Ctsa,Cyba,Vamp8 | | 6.941e-05 | -9.58 | KUROZUMI\_RESPONSE\_TO\_ONCOCYTIC\_VIRUS | MSigDB lists | KUROZUMI\_RESPONSE\_TO\_ONCOCYTIC\_VIRUS | 28 | 4 | 12187 | 98 | Ccr5,Itgam,Il6ra,C3 | | 7.054e-05 | -9.56 | protein binding | molecular function | GO:0005515 | 7104 | 76 | 13516 | 107 | Mgll,Phyhip,Trnp1,Tspan4,Pde1a,Thy1,Tfap2c,Lgi4,Vim,Rap1gap,Ccl6,Rac2,Cyba,Itgam,Mgp,Fgf18,Ccar1,Il6ra,Tsc22d1,Gfap,Fcer1g,Serpinb9,Rab3il1,C1qb,Sh2d5,Rbfox3,Ccr5,Ptprn,Ptprc,Ctsb,Tmem176a,Ldha,Kl,Capn3,Ikzf3,Adh1,Vamp8,Saa3,Tnfsf8,Ly6a,Dek,Cplx2,Exoc3l4,Lrg1,Csf1r,Actb,Vwf,Grn,Ahnak,Hcls1,Dlgap3,Slc2a1,Flnb,Camk2b,C3,Il1r1,S100a6,Bhlhe41,Myh9,Psme1,Ctss,Cox4i1,Rpl31,Ctsa,Lin7a,Rpl26,Hspa2,Mark4,Igfbp6,Csk,Mc4r,Clu,Ly6e,Brk1,Fcgr3,C1qa | | 7.325e-05 | -9.52 | B cell mediated immunity | biological process | GO:0019724 | 54 | 5 | 13711 | 111 | C1qa,Fcgr3,C3,C1qb,Fcer1g | | 7.395e-05 | -9.51 | Nphp1 (nephronophthisis 1 (juvenile) homolog (human)) | protein interactions | 53885 | 39 | 5 | 6802 | 78 | Ldha,C3,Vim,Myh9,Actb | | 7.491e-05 | -9.50 | REACTOME\_TRANSLATION | MSigDB lists | REACTOME\_TRANSLATION | 128 | 7 | 12187 | 98 | Rps8,Rpl26,Rpl13,Rps20,Rpl31,Rps5,Rpl14 | | 7.571e-05 | -9.49 | proteoglycan binding | molecular function | GO:0043394 | 29 | 4 | 13516 | 107 | Itgam,Ctss,Ptprc,Ctsb | | 7.678e-05 | -9.47 | BILANGES\_SERUM\_AND\_RAPAMYCIN\_SENSITIVE\_GENES | MSigDB lists | BILANGES\_SERUM\_AND\_RAPAMYCIN\_SENSITIVE\_GENES | 55 | 5 | 12187 | 98 | Rps5,Rpl14,Rpl26,Rps8,Rpl13 | | 7.796e-05 | -9.46 | SORBS2 (sorbin and SH3 domain containing 2) | protein interactions | 8470 | 8 | 3 | 6802 | 78 | Myh9,Actb,Flnb | | 7.940e-05 | -9.44 | REACTOME\_COMPLEMENT\_CASCADE | MSigDB lists | REACTOME\_COMPLEMENT\_CASCADE | 11 | 3 | 12187 | 98 | C1qb,C3,C1qa | | 7.940e-05 | -9.44 | GO\_POSITIVE\_REGULATION\_OF\_MYELOID\_LEUKOCYTE\_MEDIATED\_IMMUNITY | MSigDB lists | GO\_POSITIVE\_REGULATION\_OF\_MYELOID\_LEUKOCYTE\_MEDIATED\_IMMUNITY | 11 | 3 | 12187 | 98 | Fcer1g,Vamp8,C3 | | 8.277e-05 | -9.40 | positive regulation of adaptive immune response | biological process | GO:0002821 | 89 | 6 | 13711 | 111 | Fcer1g,C3,Fcgr3,Il6ra,Ptprc,Il1r1 | | 8.495e-05 | -9.37 | regulation of immune effector process | biological process | GO:0002697 | 278 | 10 | 13711 | 111 | Ptprc,Fcgr3,Fcer1g,Rac2,Itgam,Serpinb9,Il1r1,Vamp8,Grn,C3 | | 8.605e-05 | -9.36 | REACTOME\_METABOLISM\_OF\_RNA | MSigDB lists | REACTOME\_METABOLISM\_OF\_RNA | 227 | 9 | 12187 | 98 | Rps8,Rpl13,Psme1,Rps20,Rpl14,Rps5,Rpl31,Rpl26,Psmb10 | | 8.781e-05 | -9.34 | endopeptidase regulator activity | molecular function | GO:0061135 | 92 | 6 | 13516 | 107 | Serpina3n,Psme1,Wfdc17,C3,Serpinb9,Spint1 | | 8.810e-05 | -9.34 | myeloid leukocyte activation | biological process | GO:0002274 | 90 | 6 | 13711 | 111 | Fcer1g,Clu,Fcgr3,Cplx2,Tmem106a,Grn | | 8.903e-05 | -9.33 | ICHIBA\_GRAFT\_VERSUS\_HOST\_DISEASE\_D7\_UP | MSigDB lists | ICHIBA\_GRAFT\_VERSUS\_HOST\_DISEASE\_D7\_UP | 91 | 6 | 12187 | 98 | Psme1,Ctss,Cyba,C1qb,Psmb10,Mpeg1 | | 9.177e-05 | -9.30 | KEGG\_COMPLEMENT\_AND\_COAGULATION\_CASCADES | MSigDB lists | KEGG\_COMPLEMENT\_AND\_COAGULATION\_CASCADES | 30 | 4 | 12187 | 98 | Vwf,C1qa,C3,C1qb | | 9.262e-05 | -9.29 | MODULE\_6 | MSigDB lists | MODULE\_6 | 284 | 10 | 12187 | 98 | Mgll,Vwf,Grn,Fmod,Hspa2,Serpina3n,Ptprc,Apod,Rac2,Igfbp6 | | 9.365e-05 | -9.28 | secretion by cell | biological process | GO:0032940 | 339 | 11 | 13711 | 111 | Ly6e,Vamp8,Exoc3l4,Cplx2,Tmem106a,Ptprn,Mc4r,Fcgr3,Myh9,Lin7a,Fcer1g | | 9.479e-05 | -9.26 | astrocyte development | biological process | GO:0014002 | 30 | 4 | 13711 | 111 | Mt3,Vim,Gfap,Grn | | 9.639e-05 | -9.25 | GO\_REGULATION\_OF\_DEFENSE\_RESPONSE | MSigDB lists | GO\_REGULATION\_OF\_DEFENSE\_RESPONSE | 538 | 14 | 12187 | 98 | C3,Ctss,Psme1,Mgll,Apod,Phyhip,Il1r1,Psmb10,Cyba,Serpinb9,Fcer1g,Itgam,Ctsb,Vamp8 | | 9.867e-05 | -9.22 | positive regulation of response to external stimulus | biological process | GO:0032103 | 341 | 11 | 13711 | 111 | Fcer1g,Rac2,C3,Cyba,Ccr5,Fgf18,C1qa,Fcgr3,Grn,Ctss,Vamp8 | | 9.939e-05 | -9.22 | GO\_CELL\_SURFACE | MSigDB lists | GO\_CELL\_SURFACE | 472 | 13 | 12187 | 98 | Hspa2,Clu,Il1r1,Ptprc,Itgam,Fcer1g,Myh9,Ccr5,Thy1,Treml2,Csf1r,Il6ra,Ctsb | | 9.967e-05 | -9.21 | REACTOME\_IMMUNE\_SYSTEM | MSigDB lists | REACTOME\_IMMUNE\_SYSTEM | 682 | 16 | 12187 | 98 | C1qa,Ptprc,Flnb,Il1r1,Psmb10,Cyba,Csk,Ctss,Camk2b,Psme1,Rap1gap,C3,Il6ra,Ctsa,Ctsb,C1qb | | 1.000e-04 | -9.21 | GO\_PROTEIN\_TARGETING\_TO\_MEMBRANE | MSigDB lists | GO\_PROTEIN\_TARGETING\_TO\_MEMBRANE | 134 | 7 | 12187 | 98 | Rpl26,Rps5,Rpl14,Rpl31,Rps20,Rpl13,Rps8 | | 1.008e-04 | -9.20 | immunoglobulin binding | molecular function | GO:0019865 | 12 | 3 | 13516 | 107 | Fcer1g,Vwf,Fcgr3 | | 1.047e-04 | -9.16 | GO\_MYELOID\_LEUKOCYTE\_MEDIATED\_IMMUNITY | MSigDB lists | GO\_MYELOID\_LEUKOCYTE\_MEDIATED\_IMMUNITY | 31 | 4 | 12187 | 98 | Vamp8,Cplx2,Serpinb9,Il6ra | | 1.050e-04 | -9.16 | GO\_ENDOPEPTIDASE\_ACTIVITY | MSigDB lists | GO\_ENDOPEPTIDASE\_ACTIVITY | 233 | 9 | 12187 | 98 | C1qb,C3,Capn3,Pcsk2,Ctsz,Ctss,Ctsb,C1qa,Psmb10 | | 1.057e-04 | -9.15 | GO\_REGULATION\_OF\_CELL\_PROLIFERATION | MSigDB lists | GO\_REGULATION\_OF\_CELL\_PROLIFERATION | 1077 | 21 | 12187 | 98 | Gfap,Il6ra,Igfbp6,Spint1,Brk1,S100a6,Lrg1,Csf1r,Ikzf3,Ccar1,Rac2,Apod,Ptprc,Cyba,Fgf18,Trnp1,Csk,Rap1gap,Clu,Tfap2c,Hcls1 | | 1.067e-04 | -9.15 | MARKEY\_RB1\_CHRONIC\_LOF\_DN | MSigDB lists | MARKEY\_RB1\_CHRONIC\_LOF\_DN | 94 | 6 | 12187 | 98 | Ccr5,Ctss,C1qb,Csf1r,Ptprc,C1qa | | 1.105e-04 | -9.11 | GO\_STRUCTURAL\_MOLECULE\_ACTIVITY | MSigDB lists | GO\_STRUCTURAL\_MOLECULE\_ACTIVITY | 477 | 13 | 12187 | 98 | Rpl31,Rps5,Rpl26,Gfap,Actb,Rps8,Capn3,Rpl13,Ahnak,Rps20,Rpl14,Vim,Mgp | | 1.149e-04 | -9.07 | GO\_RIBOSOMAL\_SUBUNIT | MSigDB lists | GO\_RIBOSOMAL\_SUBUNIT | 137 | 7 | 12187 | 98 | Rps8,Rpl13,Rps20,Rpl31,Rpl14,Rps5,Rpl26 | | 1.160e-04 | -9.06 | MYO5C (myosin VC) | protein interactions | 55930 | 9 | 3 | 6802 | 78 | Myh9,Actb,Flnb | | 1.160e-04 | -9.06 | ARHGAP21 (Rho GTPase activating protein 21) | protein interactions | 57584 | 9 | 3 | 6802 | 78 | Myh9,Actb,Flnb | | 1.189e-04 | -9.04 | AIYAR\_COBRA1\_TARGETS\_UP | MSigDB lists | AIYAR\_COBRA1\_TARGETS\_UP | 32 | 4 | 12187 | 98 | Mgp,Ctsb,Apod,Serpina3n | | 1.199e-04 | -9.03 | GO\_REGULATION\_OF\_LEUKOCYTE\_MEDIATED\_IMMUNITY | MSigDB lists | GO\_REGULATION\_OF\_LEUKOCYTE\_MEDIATED\_IMMUNITY | 96 | 6 | 12187 | 98 | C3,Ptprc,Rac2,Fcer1g,Serpinb9,Vamp8 | | 1.203e-04 | -9.03 | SNF5\_DN.V1\_UP | MSigDB lists | SNF5\_DN.V1\_UP | 138 | 7 | 12187 | 98 | Fgf18,Apod,Ptprc,Tmem106a,Csf1r,Fmod,Cd68 | | 1.213e-04 | -9.02 | MODULE\_38 | MSigDB lists | MODULE\_38 | 353 | 11 | 12187 | 98 | Serpinb9,Mgll,Grn,Vwf,Tfap2c,Hspa2,Slc2a1,Hcls1,C1qb,Il1r1,Apod | | 1.228e-04 | -9.00 | regulation of leukocyte degranulation | biological process | GO:0043300 | 32 | 4 | 13711 | 111 | Vamp8,Itgam,Fcer1g,Rac2 | | 1.241e-04 | -8.99 | regulation of cell activation | biological process | GO:0050865 | 412 | 12 | 13711 | 111 | Ptprc,C1qa,Rac2,Fcer1g,Itgam,Thy1,Csk,Il6ra,Capn3,Vamp8,Grn,Ikzf3 | | 1.284e-04 | -8.96 | Wdfy4 (WD repeat and FYVE domain containing 4) | protein interactions | 545030 | 133 | 8 | 6802 | 78 | Cd68,Myh9,Thy1,Mpeg1,Hspa2,Actb,Hcls1,Ahnak | | 1.298e-04 | -8.95 | EFHD2 (EF-hand domain family member D2) | protein interactions | 79180 | 2 | 2 | 6802 | 78 | Myh9,Actb | | 1.316e-04 | -8.94 | GSE12366\_NAIVE\_VS\_MEMORY\_BCELL\_DN | MSigDB lists | GSE12366\_NAIVE\_VS\_MEMORY\_BCELL\_DN | 140 | 7 | 12187 | 98 | Tsc22d1,Ahnak,Itgam,Il6ra,Vim,Klhl6,Slc2a1 | | 1.330e-04 | -8.92 | GO\_RIBOSOME | MSigDB lists | GO\_RIBOSOME | 188 | 8 | 12187 | 98 | Rpl13,Rps8,Rpl31,Rps5,Rpl14,Rps20,Rpl26,Apod | | 1.377e-04 | -8.89 | GRUETZMANN\_PANCREATIC\_CANCER\_UP | MSigDB lists | GRUETZMANN\_PANCREATIC\_CANCER\_UP | 298 | 10 | 12187 | 98 | Ptprc,Mgp,Slc2a1,Grn,Tsc22d1,Rap1gap,Myh9,Pde1a,Ldha,Csk | | 1.392e-04 | -8.88 | complement activation, classical pathway | biological process | GO:0006958 | 13 | 3 | 13711 | 111 | C1qa,C3,C1qb | | 1.392e-04 | -8.88 | regulation of acute inflammatory response to antigenic stimulus | biological process | GO:0002864 | 13 | 3 | 13711 | 111 | C3,Fcer1g,Fcgr3 | | 1.409e-04 | -8.87 | regulation of multicellular organismal process | biological process | GO:0051239 | 2546 | 37 | 13711 | 111 | Gfap,Capn3,Fgf18,Camk2b,Il6ra,Fcer1g,Bhlhe41,Grn,Mc4r,Ikzf3,Ctss,Mgll,Il1r1,Thy1,Kl,Clu,Apod,Tmem176a,Ptprc,C1qa,C3,Cyba,Mgp,Ccr5,Lfng,Csk,Spint1,Lgi4,Mt3,Hcls1,Csf1r,Vim,Rap1gap,Vamp8,Lrg1,Ctsz,Fcgr3 | | 1.423e-04 | -8.86 | positive regulation of defense response | biological process | GO:0031349 | 240 | 9 | 13711 | 111 | Vamp8,Ccr5,Ctss,Grn,C3,Cyba,Fcgr3,C1qa,Fcer1g | | 1.426e-04 | -8.86 | T cell activation | biological process | GO:0042110 | 188 | 8 | 13711 | 111 | Lfng,Psmb10,Tnfsf8,Myh9,Ptprc,Itgam,Treml2,Fcer1g | | 1.431e-04 | -8.85 | ELF1\_Q6 | MSigDB lists | ELF1\_Q6 | 190 | 8 | 12187 | 98 | Vim,Ikzf3,C1qa,Ctsa,Capn3,Fcer1g,Camk2b,Tfap2c | | 1.502e-04 | -8.80 | GSE24634\_TEFF\_VS\_TCONV\_DAY3\_IN\_CULTURE\_DN | MSigDB lists | GSE24634\_TEFF\_VS\_TCONV\_DAY3\_IN\_CULTURE\_DN | 143 | 7 | 12187 | 98 | Cd68,Fcer1g,Ccr5,C1qa,Fcgr3,Ctsa,Ikzf3 | | 1.502e-04 | -8.80 | HALLMARK\_ALLOGRAFT\_REJECTION | MSigDB lists | HALLMARK\_ALLOGRAFT\_REJECTION | 143 | 7 | 12187 | 98 | Ptprc,Psmb10,Hcls1,Thy1,Ccr5,Csk,Ctss | | 1.502e-04 | -8.80 | LENAOUR\_DENDRITIC\_CELL\_MATURATION\_DN | MSigDB lists | LENAOUR\_DENDRITIC\_CELL\_MATURATION\_DN | 100 | 6 | 12187 | 98 | Tsc22d1,Fcer1g,C3,Csf1r,Il6ra,Fcgr3 | | 1.508e-04 | -8.80 | structural constituent of ribosome | molecular function | GO:0003735 | 145 | 7 | 13516 | 107 | Rps20,Rpl13,Rpl26,Rps5,Rpl31,Rpl14,Rps8 | | 1.515e-04 | -8.79 | TSUNODA\_CISPLATIN\_RESISTANCE\_DN | MSigDB lists | TSUNODA\_CISPLATIN\_RESISTANCE\_DN | 34 | 4 | 12187 | 98 | Ctsb,C3,Tsc22d1,Clu | | 1.589e-04 | -8.75 | GO\_LYMPHOCYTE\_MEDIATED\_IMMUNITY | MSigDB lists | GO\_LYMPHOCYTE\_MEDIATED\_IMMUNITY | 64 | 5 | 12187 | 98 | Fcer1g,Clu,C3,C1qb,C1qa | | 1.604e-04 | -8.74 | Neutrophil degranulation | REACTOME pathways | R-MMU-6798695 | 383 | 14 | 6297 | 76 | Ctsa,Fcer1g,Lrg1,Serpina3n,Cd68,Ctss,Grn,Ctsb,C3,Vamp8,Itgam,Cyba,Ctsz,Fcgr3 | | 1.637e-04 | -8.72 | GSE6259\_FLT3L\_INDUCED\_DEC205\_POS\_DC\_VS\_CD8\_TCELL\_DN | MSigDB lists | GSE6259\_FLT3L\_INDUCED\_DEC205\_POS\_DC\_VS\_CD8\_TCELL\_DN | 145 | 7 | 12187 | 98 | Psme1,Ctsz,Serpinb9,C3,Cyba,Treml2,Psmb10 | | 1.755e-04 | -8.65 | regulation of acute inflammatory response | biological process | GO:0002673 | 35 | 4 | 13711 | 111 | Fcer1g,Ccr5,C3,Fcgr3 | | 1.768e-04 | -8.64 | CRX\_DN.V1\_DN | MSigDB lists | CRX\_DN.V1\_DN | 103 | 6 | 12187 | 98 | Tmem176a,Camk2b,Pcsk2,S100a6,Serpina3n,Gfap | | 1.836e-04 | -8.60 | export from cell | biological process | GO:0140352 | 366 | 11 | 13711 | 111 | Exoc3l4,Cplx2,Tmem106a,Mc4r,Ptprn,Ly6e,Vamp8,Fcer1g,Lin7a,Fcgr3,Myh9 | | 1.847e-04 | -8.60 | mouse chr4 D3\|4 69.05 cM | chromosome location | mouse chr4 D3\|4 69.05 cM | 3 | 2 | 14556 | 115 | C1qb,C1qa | | 1.847e-04 | -8.60 | mouse chr15 D3\|15 34.29 cM | chromosome location | mouse chr15 D3\|15 34.29 cM | 3 | 2 | 14556 | 115 | Ly6a,Ly6e | | 1.859e-04 | -8.59 | GSE39110\_UNTREATED\_VS\_IL2\_TREATED\_CD8\_TCELL\_DAY6\_POST\_IMMUNIZATION\_UP | MSigDB lists | GSE39110\_UNTREATED\_VS\_IL2\_TREATED\_CD8\_TCELL\_DAY6\_POST\_IMMUNIZATION\_UP | 148 | 7 | 12187 | 98 | Igfbp6,Vim,Ikzf3,Ccr5,Ahnak,Fam111a,S100a6 | | 1.859e-04 | -8.59 | GSE29618\_MONOCYTE\_VS\_MDC\_UP | MSigDB lists | GSE29618\_MONOCYTE\_VS\_MDC\_UP | 148 | 7 | 12187 | 98 | Ctsb,Fcgr3,Lin7a,Itgam,Fcer1g,Cd68,Ctss | | 1.864e-04 | -8.59 | PECE\_MAMMARY\_STEM\_CELL\_UP | MSigDB lists | PECE\_MAMMARY\_STEM\_CELL\_UP | 104 | 6 | 12187 | 98 | Mgp,Cox4i1,Serpina3n,Clu,Rpl13,Rps20 | | 1.910e-04 | -8.56 | REACTOME\_CREATION\_OF\_C4\_AND\_C2\_ACTIVATORS | MSigDB lists | REACTOME\_CREATION\_OF\_C4\_AND\_C2\_ACTIVATORS | 3 | 2 | 12187 | 98 | C1qb,C1qa | | 1.938e-04 | -8.55 | GSE34156\_UNTREATED\_VS\_6H\_TLR1\_TLR2\_LIGAND\_TREATED\_MONOCYTE\_UP | MSigDB lists | GSE34156\_UNTREATED\_VS\_6H\_TLR1\_TLR2\_LIGAND\_TREATED\_MONOCYTE\_UP | 149 | 7 | 12187 | 98 | Fcgr3,Csf1r,Mpeg1,Lin7a,Tsc22d1,Grn,Ctsz | | 1.938e-04 | -8.55 | complement-mediated synapse pruning | biological process | GO:0150062 | 3 | 2 | 13711 | 111 | C1qa,C3 | | 1.938e-04 | -8.55 | positive regulation of type I hypersensitivity | biological process | GO:0001812 | 3 | 2 | 13711 | 111 | Fcgr3,Fcer1g | | 1.949e-04 | -8.54 | MODULE\_60 | MSigDB lists | MODULE\_60 | 311 | 10 | 12187 | 98 | Slc2a1,Hcls1,C1qb,Fcer1g,Vwf,Grn,Apod,Psmb10,Igfbp6,Rac2 | | 1.975e-04 | -8.53 | GO\_HUMORAL\_IMMUNE\_RESPONSE | MSigDB lists | GO\_HUMORAL\_IMMUNE\_RESPONSE | 67 | 5 | 12187 | 98 | Clu,C1qb,C3,C1qa,Psmb10 | | 2.019e-04 | -8.51 | GSE25123\_IL4\_VS\_IL4\_AND\_ROSIGLITAZONE\_STIM\_PPARG\_KO\_MACROPHAGE\_DAY10\_DN | MSigDB lists | GSE25123\_IL4\_VS\_IL4\_AND\_ROSIGLITAZONE\_STIM\_PPARG\_KO\_MACROPHAGE\_DAY10\_DN | 150 | 7 | 12187 | 98 | Fcer1g,Ly6e,Ldha,Fcgr3,Ptprc,Flnb,Rac2 | | 2.019e-04 | -8.51 | GSE11961\_FOLLICULAR\_BCELL\_VS\_GERMINAL\_CENTER\_BCELL\_DAY40\_DN | MSigDB lists | GSE11961\_FOLLICULAR\_BCELL\_VS\_GERMINAL\_CENTER\_BCELL\_DAY40\_DN | 150 | 7 | 12187 | 98 | Ptprc,Cyba,Sh3bgrl3,Klhl6,Rps8,Cd68,Vwf | | 2.058e-04 | -8.49 | GO\_SECRETION\_BY\_CELL | MSigDB lists | GO\_SECRETION\_BY\_CELL | 375 | 11 | 12187 | 98 | Lin7a,Cplx2,Slc17a7,Vamp8,Exoc3l4,Fcer1g,Mc4r,Ptprn,Serpina3n,Vwf,Clu | | 2.104e-04 | -8.47 | SRP-dependent cotranslational protein targeting to membrane | REACTOME pathways | R-MMU-1799339 | 72 | 6 | 6297 | 76 | Rpl14,Rpl26,Rps5,Rps20,Rps8,Rpl13 | | 2.118e-04 | -8.46 | LEIN\_ASTROCYTE\_MARKERS | MSigDB lists | LEIN\_ASTROCYTE\_MARKERS | 37 | 4 | 12187 | 98 | C1qa,Clu,Itgam,Gfap | | 2.123e-04 | -8.46 | PAPAIN | prints domains | PR00705 | 9 | 3 | 2951 | 42 | Ctsz,Ctsb,Ctss | | 2.139e-04 | -8.45 | KANG\_CISPLATIN\_RESISTANCE\_UP | MSigDB lists | KANG\_CISPLATIN\_RESISTANCE\_UP | 15 | 3 | 12187 | 98 | Rac2,Grn,Igfbp6 | | 2.177e-04 | -8.43 | GSE6269\_FLU\_VS\_E\_COLI\_INF\_PBMC\_DN | MSigDB lists | GSE6269\_FLU\_VS\_E\_COLI\_INF\_PBMC\_DN | 107 | 6 | 12187 | 98 | Slc2a1,Rps5,Rpl31,Rpl13,Cox4i1,Rpl26 | | 2.187e-04 | -8.43 | cell killing | biological process | GO:0001906 | 37 | 4 | 13711 | 111 | Ccr5,C3,Fcgr3,Serpinb9 | | 2.189e-04 | -8.43 | glial cell proliferation | biological process | GO:0014009 | 15 | 3 | 13711 | 111 | Lgi4,Csf1r,Clu | | 2.191e-04 | -8.43 | HALLMARK\_COMPLEMENT | MSigDB lists | HALLMARK\_COMPLEMENT | 152 | 7 | 12187 | 98 | C1qa,Ctsb,Ctss,Clu,Fcer1g,Itgam,C3 | | 2.241e-04 | -8.40 | MYO1B (myosin IB) | protein interactions | 4430 | 11 | 3 | 6802 | 78 | Flnb,Myh9,Actb | | 2.241e-04 | -8.40 | BMP2K (BMP2 inducible kinase) | protein interactions | 55589 | 11 | 3 | 6802 | 78 | Flnb,Actb,Myh9 | | 2.241e-04 | -8.40 | LIMA1 (LIM domain and actin binding 1) | protein interactions | 51474 | 11 | 3 | 6802 | 78 | Flnb,Actb,Myh9 | | 2.241e-04 | -8.40 | DAB2 (DAB adaptor protein 2) | protein interactions | 1601 | 11 | 3 | 6802 | 78 | Myh9,Actb,Flnb | | 2.241e-04 | -8.40 | PALM2-AKAP2 (PALM2-AKAP2 fusion) | protein interactions | 445815 | 11 | 3 | 6802 | 78 | Myh9,Actb,Flnb | | 2.270e-04 | -8.39 | Nonsense Mediated Decay (NMD) independent of the Exon Junction Complex (EJC) | REACTOME pathways | R-MMU-975956 | 73 | 6 | 6297 | 76 | Rps20,Rpl26,Rps5,Rpl14,Rpl13,Rps8 | | 2.277e-04 | -8.39 | plasma membrane | cellular component | GO:0005886 | 3462 | 45 | 13825 | 111 | Fcgr3,Spint1,Slc2a1,Lin7a,Tmem106a,Cd52,Abhd17c,Il6ra,Flnb,Ptprc,Ptprn,Hspa2,Vamp8,Ccr5,Vim,Dlgap3,Treml2,Cd68,Hcls1,Grn,Tspan4,Rac2,Ikzf3,Myh9,S100a6,Ly6e,Ifitm1,Thy1,Kl,Ahnak,Ly6a,Ctsb,Mc4r,Adh1,Il1r1,Csf1r,Itgam,Slc26a2,Vwf,Csk,Capn3,Cyba,Sh2d5,Fcer1g,Adgre1 | | 2.288e-04 | -8.38 | GO\_MEMBRANE\_ORGANIZATION | MSigDB lists | GO\_MEMBRANE\_ORGANIZATION | 733 | 16 | 12187 | 98 | Ctsz,Rps8,Capn3,Rpl13,Clu,Rps20,Rpl14,Ndufa13,Ccr5,Fcer1g,Myh9,Rpl31,Rps5,Vamp8,Rpl26,Lin7a | | 2.429e-04 | -8.32 | ZHONG\_SECRETOME\_OF\_LUNG\_CANCER\_AND\_MACROPHAGE | MSigDB lists | ZHONG\_SECRETOME\_OF\_LUNG\_CANCER\_AND\_MACROPHAGE | 70 | 5 | 12187 | 98 | Clu,Ldha,Actb,Ctsb,Ctsa | | 2.429e-04 | -8.32 | regulation of immunoglobulin mediated immune response | biological process | GO:0002889 | 38 | 4 | 13711 | 111 | Ptprc,Fcgr3,Fcer1g,C3 | | 2.429e-04 | -8.32 | regulation of B cell mediated immunity | biological process | GO:0002712 | 38 | 4 | 13711 | 111 | C3,Fcer1g,Ptprc,Fcgr3 | | 2.470e-04 | -8.31 | GSE43955\_1H\_VS\_20H\_ACT\_CD4\_TCELL\_DN | MSigDB lists | GSE43955\_1H\_VS\_20H\_ACT\_CD4\_TCELL\_DN | 155 | 7 | 12187 | 98 | Ctss,Ly6e,Tsc22d1,Ahnak,Csf1r,Mgp,Flnb | | 2.483e-04 | -8.30 | regulation of leukocyte activation | biological process | GO:0002694 | 379 | 11 | 13711 | 111 | Itgam,Fcer1g,Rac2,Csk,Il6ra,Ptprc,C1qa,Grn,Ikzf3,Vamp8,Thy1 | | 2.527e-04 | -8.28 | regulation of multicellular organismal development | biological process | GO:2000026 | 1748 | 28 | 13711 | 111 | Bhlhe41,Gfap,Capn3,Fgf18,Il6ra,Camk2b,C1qa,Ptprc,Tmem176a,Ikzf3,Grn,Il1r1,Mgll,Kl,Thy1,Spint1,Mt3,Lgi4,Hcls1,Csf1r,Mgp,Ccr5,C3,Lfng,Ctsz,Vim,Rap1gap,Lrg1 | | 2.530e-04 | -8.28 | leukocyte proliferation | biological process | GO:0070661 | 70 | 5 | 13711 | 111 | Csf1r,Ptprc,Psmb10,Itgam,Clu | | 2.542e-04 | -8.28 | positive regulation of cytokine production | biological process | GO:0001819 | 318 | 10 | 13711 | 111 | Ptprc,Fcgr3,Csf1r,Clu,Fcer1g,Il1r1,Il6ra,Cyba,C3,Ccr5 | | 2.545e-04 | -8.28 | regulation of vesicle-mediated transport | biological process | GO:0060627 | 513 | 13 | 13711 | 111 | Rap1gap,Vamp8,Cplx2,Ptprc,Fcgr3,Clu,Csk,C3,Cyba,Fcer1g,Slc17a7,Rac2,Itgam | | 2.588e-04 | -8.26 | MODULE\_55 | MSigDB lists | MODULE\_55 | 520 | 13 | 12187 | 98 | Slc17a7,Grn,C1qb,Adh1,Mgp,Rac2,Apod,Phyhip,Serpina3n,Il1r1,Ptprn,Hspa2,Mgll | | 2.606e-04 | -8.25 | JIANG\_AGING\_HYPOTHALAMUS\_UP | MSigDB lists | JIANG\_AGING\_HYPOTHALAMUS\_UP | 39 | 4 | 12187 | 98 | Ctss,Thy1,Csf1r,Actb | | 2.610e-04 | -8.25 | MODULE\_151 | MSigDB lists | MODULE\_151 | 263 | 9 | 12187 | 98 | Rpl26,Cyba,Rpl13,Ldha,Ndufa13,Rpl31,Rpl14,Rps5,Rps20 | | 2.617e-04 | -8.25 | MODULE\_340 | MSigDB lists | MODULE\_340 | 16 | 3 | 12187 | 98 | Ctsa,Pcsk2,Pde1a | | 2.641e-04 | -8.24 | NABA\_MATRISOME\_ASSOCIATED | MSigDB lists | NABA\_MATRISOME\_ASSOCIATED | 386 | 11 | 12187 | 98 | Ctsb,Ctsa,C1qb,S100a6,Serpinb9,Fgf18,C1qa,Serpina3n,Tnfsf8,Ctsz,Ctss | | 2.672e-04 | -8.23 | GSE24634\_IL4\_VS\_CTRL\_TREATED\_NAIVE\_CD4\_TCELL\_DAY5\_UP | MSigDB lists | GSE24634\_IL4\_VS\_CTRL\_TREATED\_NAIVE\_CD4\_TCELL\_DAY5\_UP | 157 | 7 | 12187 | 98 | Ctsa,Il1r1,Mgll,Cd68,Tmem176a,Csf1r,Tspan4 | | 2.678e-04 | -8.23 | humoral immune response mediated by circulating immunoglobulin | biological process | GO:0002455 | 16 | 3 | 13711 | 111 | C1qa,C3,C1qb | | 2.742e-04 | -8.20 | CHICAS\_RB1\_TARGETS\_CONFLUENT | MSigDB lists | CHICAS\_RB1\_TARGETS\_CONFLUENT | 454 | 12 | 12187 | 98 | Igfbp6,Vim,Sh3bgrl3,Actb,S100a6,Ly6e,Ahnak,Clu,Myh9,Thy1,Trnp1,Tmem176a | | 2.761e-04 | -8.19 | GO\_POSITIVE\_REGULATION\_OF\_DEFENSE\_RESPONSE | MSigDB lists | GO\_POSITIVE\_REGULATION\_OF\_DEFENSE\_RESPONSE | 265 | 9 | 12187 | 98 | Vamp8,Ctsb,Cyba,Psmb10,Fcer1g,Itgam,Psme1,Ctss,C3 | | 2.777e-04 | -8.19 | GO\_SINGLE\_ORGANISM\_CELL\_ADHESION | MSigDB lists | GO\_SINGLE\_ORGANISM\_CELL\_ADHESION | 325 | 10 | 12187 | 98 | Psmb10,Ptprc,Actb,Tnfsf8,Rac2,Treml2,Lfng,Myh9,Thy1,Fcer1g | | 2.793e-04 | -8.18 | regulation of lymphocyte mediated immunity | biological process | GO:0002706 | 111 | 6 | 13711 | 111 | Il1r1,Serpinb9,C3,Ptprc,Fcgr3,Fcer1g | | 2.806e-04 | -8.18 | Complement Activation, Classical Pathway | WikiPathways | WP200 | 10 | 3 | 3756 | 52 | C1qb,C1qa,C3 | | 2.806e-04 | -8.18 | Macrophage markers | WikiPathways | WP2271 | 10 | 3 | 3756 | 52 | Cd68,Rac2,Cd52 | | 2.846e-04 | -8.16 | multicellular organismal process | biological process | GO:0032501 | 4571 | 55 | 13711 | 111 | Ccr5,Mgp,Cyba,C1qb,Slc26a2,Lfng,Mark4,Mt3,Myh9,Cplx2,Vamp8,Rap1gap,Ctsz,Flnb,Pcsk2,Fgf18,Ly6e,Actb,Fcer1g,Lin7a,Bhlhe41,Ctss,Grn,Trnp1,Il1r1,Mgll,Kl,Thy1,C1qa,Ptprc,Tfap2c,Ptprn,C3,Vwf,Spint1,Ifitm1,Rac2,Brk1,Lgi4,Csf1r,Hcls1,Vim,Tnfsf8,Gfap,Ctsb,Il6ra,Camk2b,Itgam,Slc17a7,Mc4r,Tmem106a,Adh1,Rbfox3,Clu,Hspa2 | | 2.850e-04 | -8.16 | Pept\_cys\_AS | interpro domains | IPR000169 | 16 | 3 | 13788 | 114 | Capn3,Ctsb,Ctss | | 2.878e-04 | -8.15 | CHIARADONNA\_NEOPLASTIC\_TRANSFORMATION\_KRAS\_CDC25\_DN | MSigDB lists | CHIARADONNA\_NEOPLASTIC\_TRANSFORMATION\_KRAS\_CDC25\_DN | 40 | 4 | 12187 | 98 | Mgp,Il1r1,Tmem176a,C3 | | 2.878e-04 | -8.15 | KEGG\_ANTIGEN\_PROCESSING\_AND\_PRESENTATION | MSigDB lists | KEGG\_ANTIGEN\_PROCESSING\_AND\_PRESENTATION | 40 | 4 | 12187 | 98 | Ctss,Psme1,Hspa2,Ctsb | | 2.887e-04 | -8.15 | GSE24634\_TEFF\_VS\_TCONV\_DAY7\_IN\_CULTURE\_DN | MSigDB lists | GSE24634\_TEFF\_VS\_TCONV\_DAY7\_IN\_CULTURE\_DN | 159 | 7 | 12187 | 98 | Ctss,Igfbp6,Itgam,Tsc22d1,C1qa,C1qb,Ctsb | | 2.928e-04 | -8.14 | CHIARADONNA\_NEOPLASTIC\_TRANSFORMATION\_CDC25\_UP | MSigDB lists | CHIARADONNA\_NEOPLASTIC\_TRANSFORMATION\_CDC25\_UP | 113 | 6 | 12187 | 98 | Il1r1,Ctsb,Mgp,C3,Tmem176a,Thy1 | | 2.949e-04 | -8.13 | MARTENS\_TRETINOIN\_RESPONSE\_UP | MSigDB lists | MARTENS\_TRETINOIN\_RESPONSE\_UP | 391 | 11 | 12187 | 98 | Hspa2,Fmod,S100a6,Lrg1,Lfng,C1qb,Sh2d5,Mpeg1,Lgi4,C1qa,Phyhip | | 2.963e-04 | -8.12 | AP2B1 (adaptor related protein complex 2 subunit beta 1) | protein interactions | 163 | 12 | 3 | 6802 | 78 | Flnb,Myh9,Actb | | 2.963e-04 | -8.12 | CPM (carboxypeptidase M) | protein interactions | 1368 | 12 | 3 | 6802 | 78 | Myh9,Actb,Flnb | | 2.963e-04 | -8.12 | CORO2A (coronin 2A) | protein interactions | 7464 | 12 | 3 | 6802 | 78 | Actb,Myh9,Flnb | | 2.963e-04 | -8.12 | ACTN4 (actinin alpha 4) | protein interactions | 81 | 12 | 3 | 6802 | 78 | Myh9,Actb,Flnb | | 3.001e-04 | -8.11 | serine-type endopeptidase inhibitor activity | molecular function | GO:0004867 | 41 | 4 | 13516 | 107 | Serpina3n,Spint1,Serpinb9,Wfdc17 | | 3.023e-04 | -8.10 | MORF\_UBE2I | MSigDB lists | MORF\_UBE2I | 212 | 8 | 12187 | 98 | Dek,Rps5,Rpl14,Rpl31,Ldha,Csk,Rpl13,Cox4i1 | | 3.096e-04 | -8.08 | regulation of response to stress | biological process | GO:0080134 | 1063 | 20 | 13711 | 111 | Fcer1g,Treml2,Myh9,Ccr5,C3,Cyba,Serpinb9,Capn3,Clu,Fcgr3,C1qa,Rpl26,Apod,Ctss,Grn,Thy1,Il1r1,Vamp8,Dek,Mgll | | 3.115e-04 | -8.07 | GSE42021\_CD24HI\_VS\_CD24INT\_TREG\_THYMUS\_DN | MSigDB lists | GSE42021\_CD24HI\_VS\_CD24INT\_TREG\_THYMUS\_DN | 161 | 7 | 12187 | 98 | Serpinb9,Ctss,Ly6e,Psme1,Fam111a,C3,Psmb10 | | 3.115e-04 | -8.07 | GSE21063\_WT\_VS\_NFATC1\_KO\_3H\_ANTI\_IGM\_STIM\_BCELL\_DN | MSigDB lists | GSE21063\_WT\_VS\_NFATC1\_KO\_3H\_ANTI\_IGM\_STIM\_BCELL\_DN | 161 | 7 | 12187 | 98 | Psme1,Serpinb9,Ndufa13,C3,Klhl6,Exoc3l4,Psmb10 | | 3.158e-04 | -8.06 | ribosome | cellular component | GO:0005840 | 213 | 8 | 13825 | 111 | Rpl26,Apod,Rpl13,Rpl14,Rpl31,Rps20,Rps5,Rps8 | | 3.159e-04 | -8.06 | KASLER\_HDAC7\_TARGETS\_1\_DN | MSigDB lists | KASLER\_HDAC7\_TARGETS\_1\_DN | 17 | 3 | 12187 | 98 | Slc2a1,C3,Igfbp6 | | 3.169e-04 | -8.06 | MODULE\_83 | MSigDB lists | MODULE\_83 | 270 | 9 | 12187 | 98 | Rps20,Rpl14,Rps5,Rpl31,Ndufa13,Ldha,Rps8,Rpl13,Rpl26 | | 3.233e-04 | -8.04 | gliogenesis | biological process | GO:0042063 | 212 | 8 | 13711 | 111 | Gfap,Grn,Vim,Itgam,Clu,Csf1r,Mt3,Lgi4 | | 3.235e-04 | -8.04 | GSE27786\_LSK\_VS\_CD8\_TCELL\_DN | MSigDB lists | GSE27786\_LSK\_VS\_CD8\_TCELL\_DN | 162 | 7 | 12187 | 98 | Ndufa13,Pde1a,Cox4i1,Brk1,Rpl31,Psmb10,Ptprc | | 3.282e-04 | -8.02 | positive regulation of lymphocyte mediated immunity | biological process | GO:0002708 | 74 | 5 | 13711 | 111 | Il1r1,Fcgr3,Ptprc,C3,Fcer1g | | 3.307e-04 | -8.01 | response to organic substance | biological process | GO:0010033 | 1870 | 29 | 13711 | 111 | Ccl6,Flnb,Vim,Rap1gap,Vamp8,Ifitm1,Mt3,Hcls1,Csf1r,Ptprn,Cyba,Serpina3n,Ndufa13,Ccr5,Csk,Saa3,Cd68,Clu,Hspa2,Grn,Mc4r,Adh1,Il1r1,Kl,Actb,Fcer1g,Fgf18,Ly6e,Il6ra | | 3.358e-04 | -8.00 | GSE5589\_LPS\_VS\_LPS\_AND\_IL10\_STIM\_MACROPHAGE\_180MIN\_UP | MSigDB lists | GSE5589\_LPS\_VS\_LPS\_AND\_IL10\_STIM\_MACROPHAGE\_180MIN\_UP | 163 | 7 | 12187 | 98 | C1qa,Fcgr3,Ctsb,Cox4i1,Adh1,Ccr5,Vwf | | 3.362e-04 | -8.00 | regulation of response to stimulus | biological process | GO:0048583 | 2962 | 40 | 13711 | 111 | Apod,C1qa,Ptprc,Clu,Il1r1,Mgll,Kl,Thy1,Ctss,Grn,Tmem106a,Itgam,Treml2,Fcer1g,Fgf18,Capn3,Serpinb9,Il6ra,Camk2b,Rpl26,Fcgr3,Igfbp6,Ccl6,Vamp8,Dek,Rap1gap,Lrg1,Klhl6,Mt3,Hcls1,Csf1r,Myh9,Rac2,Csk,Ccr5,Ndufa13,C1qb,C3,Cyba,Lfng | | 3.481e-04 | -7.96 | HENDRICKS\_SMARCA4\_TARGETS\_UP | MSigDB lists | HENDRICKS\_SMARCA4\_TARGETS\_UP | 42 | 4 | 12187 | 98 | Flnb,Ctsb,Ahnak,Thy1 | | 3.524e-04 | -7.95 | Nphp4 (nephronophthisis 4 (juvenile) homolog (human)) | protein interactions | 260305 | 154 | 8 | 6802 | 78 | Flnb,Hspa2,Vim,Gfap,Ccar1,Actb,Exoc3l4,Myh9 | | 3.597e-04 | -7.93 | peptidase regulator activity | molecular function | GO:0061134 | 119 | 6 | 13516 | 107 | Serpina3n,Psme1,Wfdc17,Serpinb9,C3,Spint1 | | 3.615e-04 | -7.93 | GSE29618\_MONOCYTE\_VS\_MDC\_DAY7\_FLU\_VACCINE\_UP | MSigDB lists | GSE29618\_MONOCYTE\_VS\_MDC\_DAY7\_FLU\_VACCINE\_UP | 165 | 7 | 12187 | 98 | Itgam,Cd68,Ctss,Ctsb,Ctsa,Fcgr3,Lin7a | | 3.634e-04 | -7.92 | plasma membrane bounded cell projection | cellular component | GO:0120025 | 1896 | 29 | 13825 | 111 | Dlgap3,Vim,Brk1,Pcsk2,Apod,Rpl26,Slc17a7,Actb,Cd52,Lin7a,Flnb,Ptprc,Ldha,Ptprn,Rap1gap,Il1r1,Slc26a2,Clu,Camk2b,Cyba,Mgll,Ctsz,Rac2,Mark4,S100a6,Myh9,Gfap,Thy1,Cplx2 | | 3.687e-04 | -7.91 | structural constituent of postsynaptic actin cytoskeleton | molecular function | GO:0098973 | 4 | 2 | 13516 | 107 | Actb,Camk2b | | 3.749e-04 | -7.89 | Formation of a pool of free 40S subunits | REACTOME pathways | R-MMU-72689 | 80 | 6 | 6297 | 76 | Rps5,Rpl26,Rpl14,Rps20,Rps8,Rpl13 | | 3.750e-04 | -7.89 | GSE19888\_ADENOSINE\_A3R\_INH\_PRETREAT\_AND\_ACT\_BY\_A3R\_VS\_TCELL\_MEMBRANES\_ACT\_MAST\_CELL\_UP | MSigDB lists | GSE19888\_ADENOSINE\_A3R\_INH\_PRETREAT\_AND\_ACT\_BY\_A3R\_VS\_TCELL\_MEMBRANES\_ACT\_MAST\_CELL\_UP | 166 | 7 | 12187 | 98 | Mpeg1,Ly6e,Ccr5,Psme1,Serpinb9,Psmb10,Ptprc | | 3.750e-04 | -7.89 | system development | biological process | GO:0048731 | 3292 | 43 | 13711 | 111 | Tfap2c,Ccr5,Mgp,C3,Ptprn,C1qb,Lfng,Vwf,Mark4,Spint1,Rac2,Lgi4,Mt3,Csf1r,Hcls1,Myh9,Vim,Cplx2,Rap1gap,Ctsz,Flnb,Tnfsf8,Gfap,Pcsk2,Ctsb,Fgf18,Ly6e,Il6ra,Camk2b,Itgam,Actb,Fcer1g,Bhlhe41,Lin7a,Grn,Trnp1,Il1r1,Mgll,Rbfox3,Thy1,Clu,C1qa,Ptprc | | 3.793e-04 | -7.88 | GO\_REGULATION\_OF\_SEQUESTERING\_OF\_CALCIUM\_ION | MSigDB lists | GO\_REGULATION\_OF\_SEQUESTERING\_OF\_CALCIUM\_ION | 77 | 5 | 12187 | 98 | Thy1,Capn3,Ccr5,Cyba,Ptprc | | 3.793e-04 | -7.88 | SWEET\_KRAS\_ONCOGENIC\_SIGNATURE | MSigDB lists | SWEET\_KRAS\_ONCOGENIC\_SIGNATURE | 77 | 5 | 12187 | 98 | Fcgr3,Rpl14,Itgam,Ctss,Cd68 | | 3.793e-04 | -7.88 | LABBE\_TARGETS\_OF\_TGFB1\_AND\_WNT3A\_DN | MSigDB lists | LABBE\_TARGETS\_OF\_TGFB1\_AND\_WNT3A\_DN | 77 | 5 | 12187 | 98 | C3,Slc2a1,S100a6,Gfap,Ldha | | 3.800e-04 | -7.88 | BIOCARTA\_TCYTOTOXIC\_PATHWAY | MSigDB lists | BIOCARTA\_TCYTOTOXIC\_PATHWAY | 4 | 2 | 12187 | 98 | Ptprc,Thy1 | | 3.800e-04 | -7.88 | WONG\_ENDOMETRIAL\_CANCER\_LATE | MSigDB lists | WONG\_ENDOMETRIAL\_CANCER\_LATE | 4 | 2 | 12187 | 98 | Ptprc,Serpina3n | | 3.815e-04 | -7.87 | GO\_CYTOSOLIC\_LARGE\_RIBOSOMAL\_SUBUNIT | MSigDB lists | GO\_CYTOSOLIC\_LARGE\_RIBOSOMAL\_SUBUNIT | 43 | 4 | 12187 | 98 | Rpl13,Rpl26,Rpl31,Rpl14 | | 3.820e-04 | -7.87 | ACTG1 (actin gamma 1) | protein interactions | 71 | 13 | 3 | 6802 | 78 | Actb,Myh9,Flnb | | 3.820e-04 | -7.87 | EPS15 (epidermal growth factor receptor pathway substrate 15) | protein interactions | 2060 | 13 | 3 | 6802 | 78 | Actb,Myh9,Flnb | | 3.820e-04 | -7.87 | TPRN (taperin) | protein interactions | 286262 | 13 | 3 | 6802 | 78 | Myh9,Actb,Flnb | | 3.820e-04 | -7.87 | DAPK3 (death associated protein kinase 3) | protein interactions | 1613 | 13 | 3 | 6802 | 78 | Flnb,Myh9,Actb | | 3.820e-04 | -7.87 | SYNPO (synaptopodin) | protein interactions | 11346 | 13 | 3 | 6802 | 78 | Flnb,Actb,Myh9 | | 3.823e-04 | -7.87 | MODULE\_114 | MSigDB lists | MODULE\_114 | 277 | 9 | 12187 | 98 | Cyba,Rpl26,Rps20,Rpl14,Rps5,Rpl31,Ndufa13,Ldha,Rpl13 | | 3.830e-04 | -7.87 | glial cell differentiation | biological process | GO:0010001 | 165 | 7 | 13711 | 111 | Vim,Gfap,Grn,Mt3,Lgi4,Itgam,Clu | | 3.856e-04 | -7.86 | regulation of chaperone-mediated autophagy | biological process | GO:1904714 | 4 | 2 | 13711 | 111 | Gfap,Ctsa | | 3.856e-04 | -7.86 | regulation of type I hypersensitivity | biological process | GO:0001810 | 4 | 2 | 13711 | 111 | Fcgr3,Fcer1g | | 3.856e-04 | -7.86 | regulation of type III hypersensitivity | biological process | GO:0001803 | 4 | 2 | 13711 | 111 | Fcgr3,Fcer1g | | 3.856e-04 | -7.86 | serotonin secretion | biological process | GO:0001820 | 4 | 2 | 13711 | 111 | Fcgr3,Fcer1g | | 3.856e-04 | -7.86 | positive regulation of type III hypersensitivity | biological process | GO:0001805 | 4 | 2 | 13711 | 111 | Fcgr3,Fcer1g | | 3.857e-04 | -7.86 | monoamine transport | biological process | GO:0015844 | 18 | 3 | 13711 | 111 | Fcer1g,Ly6e,Fcgr3 | | 3.866e-04 | -7.86 | DECR2 (2,4-dienoyl-CoA reductase 2) | protein interactions | 26063 | 3 | 2 | 6802 | 78 | Myh9,Flnb | | 3.866e-04 | -7.86 | LGALS1 (galectin 1) | protein interactions | 3956 | 3 | 2 | 6802 | 78 | Flnb,Actb | | 3.866e-04 | -7.86 | SIPA1 (signal-induced proliferation-associated 1) | protein interactions | 6494 | 3 | 2 | 6802 | 78 | Actb,Myh9 | | 3.866e-04 | -7.86 | UNC45A (unc-45 myosin chaperone A) | protein interactions | 55898 | 3 | 2 | 6802 | 78 | Actb,Myh9 | | 3.889e-04 | -7.85 | GSE42021\_CD24HI\_TREG\_VS\_CD24HI\_TCONV\_THYMUS\_UP | MSigDB lists | GSE42021\_CD24HI\_TREG\_VS\_CD24HI\_TCONV\_THYMUS\_UP | 167 | 7 | 12187 | 98 | C1qb,Rps5,Grn,Ctsa,Cyba,Actb,Vamp8 | | 3.889e-04 | -7.85 | GSE22886\_NAIVE\_BCELL\_VS\_MONOCYTE\_DN | MSigDB lists | GSE22886\_NAIVE\_BCELL\_VS\_MONOCYTE\_DN | 167 | 7 | 12187 | 98 | Il6ra,Vim,Fcgr3,Ctsb,Fcer1g,Itgam,Grn | | 3.889e-04 | -7.85 | GSE37301\_GRANULOCYTE\_MONOCYTE\_PROGENITOR\_VS\_RAG2\_KO\_NK\_CELL\_DN | MSigDB lists | GSE37301\_GRANULOCYTE\_MONOCYTE\_PROGENITOR\_VS\_RAG2\_KO\_NK\_CELL\_DN | 167 | 7 | 12187 | 98 | Rps5,Il1r1,Mpeg1,Grn,S100a6,Il6ra,Myh9 | | 3.889e-04 | -7.85 | GSE24634\_TEFF\_VS\_TCONV\_DAY10\_IN\_CULTURE\_DN | MSigDB lists | GSE24634\_TEFF\_VS\_TCONV\_DAY10\_IN\_CULTURE\_DN | 167 | 7 | 12187 | 98 | C1qa,Il6ra,C3,Itgam,Fcer1g,Grn,Ctsz | | 3.889e-04 | -7.85 | GSE24634\_TREG\_VS\_TCONV\_POST\_DAY3\_IL4\_CONVERSION\_DN | MSigDB lists | GSE24634\_TREG\_VS\_TCONV\_POST\_DAY3\_IL4\_CONVERSION\_DN | 167 | 7 | 12187 | 98 | Ctsb,Fcgr3,C1qa,Mgll,Ctsz,C1qb,Tmem176a | | 3.989e-04 | -7.83 | anatomical structure development | biological process | GO:0048856 | 3951 | 49 | 13711 | 111 | Ctsz,Flnb,Klhl6,Cplx2,Rap1gap,Myh9,Mt3,Lfng,Mgp,Ccr5,C1qb,Mark4,C1qa,Ptprc,Grn,Thy1,Trnp1,Il1r1,Mgll,Lin7a,Bhlhe41,Fcer1g,Actb,Psmb10,Ly6e,Pcsk2,Fgf18,Tnfsf8,Vim,Ifitm1,Rac2,Spint1,Csf1r,Hcls1,Lgi4,Brk1,Vwf,Tfap2c,Ptprn,C3,Clu,Hspa2,Rbfox3,Itgam,Gfap,Il6ra,Camk2b,Ctsb,Capn3 | | 4.031e-04 | -7.82 | GSE29164\_UNTREATED\_VS\_CD8\_TCELL\_TREATED\_MELANOMA\_DAY3\_DN | MSigDB lists | GSE29164\_UNTREATED\_VS\_CD8\_TCELL\_TREATED\_MELANOMA\_DAY3\_DN | 168 | 7 | 12187 | 98 | Il6ra,Tnfsf8,Cyba,S100a6,Ctss,Ctsz,Csf1r | | 4.031e-04 | -7.82 | PICCALUGA\_ANGIOIMMUNOBLASTIC\_LYMPHOMA\_UP | MSigDB lists | PICCALUGA\_ANGIOIMMUNOBLASTIC\_LYMPHOMA\_UP | 168 | 7 | 12187 | 98 | C1qa,Clu,Thy1,Vwf,Tmem176a,C1qb,C3 | | 4.039e-04 | -7.81 | neuron part | cellular component | GO:0097458 | 1719 | 27 | 13825 | 111 | Slc17a7,Pde1a,Rpl26,Rbfox3,Apod,Pcsk2,Vim,Dlgap3,Rap1gap,Mt3,Ptprn,Flnb,Il6ra,Lin7a,Abhd17c,Actb,Mgll,Sh2d5,Cyba,Camk2b,Clu,Il1r1,Cplx2,Rpl14,Thy1,Mark4,Ctsz | | 4.047e-04 | -7.81 | GSE16385\_UNTREATED\_VS\_12H\_ROSIGLITAZONE\_TREATED\_MACROPHAGE\_DN | MSigDB lists | GSE16385\_UNTREATED\_VS\_12H\_ROSIGLITAZONE\_TREATED\_MACROPHAGE\_DN | 120 | 6 | 12187 | 98 | Ttc9b,Lin7a,Trnp1,Ctss,S100a6,Treml2 | | 4.061e-04 | -7.81 | GO\_CELLULAR\_RESPONSE\_TO\_CYTOKINE\_STIMULUS | MSigDB lists | GO\_CELLULAR\_RESPONSE\_TO\_CYTOKINE\_STIMULUS | 406 | 11 | 12187 | 98 | Camk2b,Psme1,Hcls1,Psmb10,Il1r1,Cyba,Ptprn,Ndufa13,Ccr5,Csf1r,Il6ra | | 4.171e-04 | -7.78 | GNF2\_ITGB2 | MSigDB lists | GNF2\_ITGB2 | 44 | 4 | 12187 | 98 | Sh3bgrl3,Itgam,Fcer1g,Hcls1 | | 4.178e-04 | -7.78 | HALLMARK\_APICAL\_JUNCTION | MSigDB lists | HALLMARK\_APICAL\_JUNCTION | 169 | 7 | 12187 | 98 | Thy1,Myh9,Vwf,Tspan4,Rac2,Actb,Ptprc | | 4.178e-04 | -7.78 | GSE41978\_ID2\_KO\_VS\_ID2\_KO\_AND\_BIM\_KO\_KLRG1\_LOW\_EFFECTOR\_CD8\_TCELL\_DN | MSigDB lists | GSE41978\_ID2\_KO\_VS\_ID2\_KO\_AND\_BIM\_KO\_KLRG1\_LOW\_EFFECTOR\_CD8\_TCELL\_DN | 169 | 7 | 12187 | 98 | Rps5,Rpl31,Rpl13,Ldha,Rps8,Vim,Rpl26 | | 4.178e-04 | -7.78 | GSE29618\_BCELL\_VS\_MDC\_DN | MSigDB lists | GSE29618\_BCELL\_VS\_MDC\_DN | 169 | 7 | 12187 | 98 | Vim,Il6ra,Actb,Ctsb,S100a6,Grn,Ccr5 | | 4.273e-04 | -7.76 | WESTON\_VEGFA\_TARGETS | MSigDB lists | WESTON\_VEGFA\_TARGETS | 79 | 5 | 12187 | 98 | Hcls1,Mgp,Vim,Vwf,Thy1 | | 4.330e-04 | -7.74 | GSE17721\_POLYIC\_VS\_PAM3CSK4\_4H\_BMDC\_UP | MSigDB lists | GSE17721\_POLYIC\_VS\_PAM3CSK4\_4H\_BMDC\_UP | 170 | 7 | 12187 | 98 | Il6ra,Tnfsf8,Vamp8,Camk2b,Tsc22d1,Mgll,Tspan4 | | 4.483e-04 | -7.71 | response to chemical | biological process | GO:0042221 | 2591 | 36 | 13711 | 111 | Flnb,Fcgr3,Ccl6,Rap1gap,Vamp8,Vim,Csf1r,Hcls1,Mt3,Rac2,Ifitm1,Csk,Saa3,Cyba,Ptprn,Ccr5,Serpina3n,Ndufa13,Hspa2,Apod,Clu,Cd68,Kl,Il1r1,Adh1,Mc4r,Grn,Slc17a7,Fcer1g,Actb,Itgam,Camk2b,Ly6e,Il6ra,Fgf18,Capn3 | | 4.529e-04 | -7.70 | ZAMORA\_NOS2\_TARGETS\_DN | MSigDB lists | ZAMORA\_NOS2\_TARGETS\_DN | 80 | 5 | 12187 | 98 | Actb,Tspan4,Ctsz,Vwf,Myh9 | | 4.553e-04 | -7.69 | positive regulation of leukocyte degranulation | biological process | GO:0043302 | 19 | 3 | 13711 | 111 | Vamp8,Itgam,Fcer1g | | 4.644e-04 | -7.67 | secretion | biological process | GO:0046903 | 408 | 11 | 13711 | 111 | Vamp8,Exoc3l4,Cplx2,Mc4r,Tmem106a,Fcgr3,Ly6e,Ptprn,Myh9,Lin7a,Fcer1g | | 4.645e-04 | -7.67 | GSE29618\_BCELL\_VS\_MONOCYTE\_DN | MSigDB lists | GSE29618\_BCELL\_VS\_MONOCYTE\_DN | 172 | 7 | 12187 | 98 | Vamp8,Vim,Ctsa,Ctsb,Cd68,Ctss,S100a6 | | 4.645e-04 | -7.67 | GSE39820\_CTRL\_VS\_IL1B\_IL6\_IL23A\_CD4\_TCELL\_UP | MSigDB lists | GSE39820\_CTRL\_VS\_IL1B\_IL6\_IL23A\_CD4\_TCELL\_UP | 172 | 7 | 12187 | 98 | Capn3,Fam111a,Tnfsf8,Ctss,Serpinb9,Cyba,Lin7a | | 4.686e-04 | -7.67 | Ig-like\_fold | interpro domains | IPR013783 | 466 | 12 | 13788 | 114 | Csf1r,Il1r1,Fcrls,Spint1,Ptprc,Thy1,Treml2,Flnb,Phyhip,C3,Il6ra,Fcgr3 | | 4.694e-04 | -7.66 | negative regulation of leukocyte apoptotic process | biological process | GO:2000107 | 45 | 4 | 13711 | 111 | Ccr5,Fcer1g,Serpinb9,Hcls1 | | 4.796e-04 | -7.64 | GNF2\_CASP1 | MSigDB lists | GNF2\_CASP1 | 81 | 5 | 12187 | 98 | Ctss,Sh3bgrl3,Fcer1g,Itgam,Hcls1 | | 4.822e-04 | -7.64 | COBL (cordon-bleu WH2 repeat protein) | protein interactions | 23242 | 14 | 3 | 6802 | 78 | Flnb,Myh9,Actb | | 4.822e-04 | -7.64 | MYO1C (myosin IC) | protein interactions | 4641 | 14 | 3 | 6802 | 78 | Flnb,Actb,Myh9 | | 4.822e-04 | -7.64 | LUZP1 (leucine zipper protein 1) | protein interactions | 7798 | 14 | 3 | 6802 | 78 | Flnb,Myh9,Actb | | 4.822e-04 | -7.64 | NEXN (nexilin F-actin binding protein) | protein interactions | 91624 | 14 | 3 | 6802 | 78 | Actb,Myh9,Flnb | | 4.822e-04 | -7.64 | BASP1 (brain abundant membrane attached signal protein 1) | protein interactions | 10409 | 14 | 3 | 6802 | 78 | Flnb,Myh9,Actb | | 4.822e-04 | -7.64 | DSG2 (desmoglein 2) | protein interactions | 1829 | 14 | 3 | 6802 | 78 | Flnb,Actb,Myh9 | | 4.847e-04 | -7.63 | regulation of adaptive immune response based on somatic recombination of immune receptors built from immunoglobulin superfamily domains | biological process | GO:0002822 | 123 | 6 | 13711 | 111 | Fcgr3,Ptprc,Fcer1g,Il6ra,Il1r1,C3 | | 4.952e-04 | -7.61 | PAPASPYRIDONOS\_UNSTABLE\_ATEROSCLEROTIC\_PLAQUE\_UP | MSigDB lists | PAPASPYRIDONOS\_UNSTABLE\_ATEROSCLEROTIC\_PLAQUE\_UP | 46 | 4 | 12187 | 98 | Mpeg1,Ctsb,Ctss,Grn | | 4.952e-04 | -7.61 | GO\_REGULATION\_OF\_PHAGOCYTOSIS | MSigDB lists | GO\_REGULATION\_OF\_PHAGOCYTOSIS | 46 | 4 | 12187 | 98 | Fcer1g,Csk,C3,Cyba | | 4.978e-04 | -7.61 | GSE18281\_CORTICAL\_VS\_MEDULLARY\_THYMOCYTE\_UP | MSigDB lists | GSE18281\_CORTICAL\_VS\_MEDULLARY\_THYMOCYTE\_UP | 174 | 7 | 12187 | 98 | Cyba,Psmb10,Ly6e,Psme1,Ctss,C3,Mpeg1 | | 4.978e-04 | -7.61 | GO\_STRUCTURAL\_CONSTITUENT\_OF\_RIBOSOME | MSigDB lists | GO\_STRUCTURAL\_CONSTITUENT\_OF\_RIBOSOME | 174 | 7 | 12187 | 98 | Rpl26,Rpl13,Rps8,Rpl14,Rps5,Rpl31,Rps20 | | 5.032e-04 | -7.59 | GSE27670\_CTRL\_VS\_BLIMP1\_TRANSDUCED\_GC\_BCELL\_UP | MSigDB lists | GSE27670\_CTRL\_VS\_BLIMP1\_TRANSDUCED\_GC\_BCELL\_UP | 125 | 6 | 12187 | 98 | Ctsb,Ctsa,Apod,Igfbp6,Adh1,Ndufa13 | | 5.032e-04 | -7.59 | NABA\_ECM\_REGULATORS | MSigDB lists | NABA\_ECM\_REGULATORS | 125 | 6 | 12187 | 98 | Serpina3n,Ctsa,Ctsb,Ctsz,Ctss,Serpinb9 | | 5.073e-04 | -7.59 | SCHUETZ\_BREAST\_CANCER\_DUCTAL\_INVASIVE\_UP | MSigDB lists | SCHUETZ\_BREAST\_CANCER\_DUCTAL\_INVASIVE\_UP | 288 | 9 | 12187 | 98 | Fcer1g,Thy1,Ctss,C1qb,Hcls1,Fcgr3,Ptprc,C1qa,Il1r1 | | 5.164e-04 | -7.57 | cell body | cellular component | GO:0044297 | 707 | 15 | 13825 | 111 | Flnb,Il6ra,Gfap,Ptprn,Thy1,Cplx2,Rap1gap,Vim,Dlgap3,Pcsk2,Rbfox3,Apod,Camk2b,Pde1a,Cyba | | 5.167e-04 | -7.57 | GO\_RESPONSE\_TO\_CYTOKINE | MSigDB lists | GO\_RESPONSE\_TO\_CYTOKINE | 487 | 12 | 12187 | 98 | Psmb10,Il1r1,Ptprn,Cyba,Hcls1,Psme1,Camk2b,Ctsb,Il6ra,Csf1r,Ndufa13,Ccr5 | | 5.206e-04 | -7.56 | cellular response to organic substance | biological process | GO:0071310 | 1457 | 24 | 13711 | 111 | Ptprn,Cyba,Ndufa13,Ccr5,Csk,Saa3,Ifitm1,Mt3,Hcls1,Csf1r,Vim,Rap1gap,Vamp8,Ccl6,Flnb,Fgf18,Il6ra,Ly6e,Actb,Fcer1g,Il1r1,Kl,Cd68,Hspa2 | | 5.325e-04 | -7.54 | complement activation | biological process | GO:0006956 | 20 | 3 | 13711 | 111 | C1qa,C1qb,C3 | | 5.325e-04 | -7.54 | inflammatory response to antigenic stimulus | biological process | GO:0002437 | 20 | 3 | 13711 | 111 | Cd68,Fcgr3,Serpinb9 | | 5.355e-04 | -7.53 | THIOL\_PROTEASE\_CYS | prosite domains | PS00139 | 16 | 3 | 8845 | 91 | Capn3,Ctsb,Ctss | | 5.363e-04 | -7.53 | GO\_ESTABLISHMENT\_OF\_PROTEIN\_LOCALIZATION\_TO\_MEMBRANE | MSigDB lists | GO\_ESTABLISHMENT\_OF\_PROTEIN\_LOCALIZATION\_TO\_MEMBRANE | 231 | 8 | 12187 | 98 | Rpl26,Rps20,Rpl31,Rpl14,Rps5,Rps8,Ndufa13,Rpl13 | | 5.367e-04 | -7.53 | GO\_REGULATION\_OF\_CALCIUM\_ION\_TRANSMEMBRANE\_TRANSPORT | MSigDB lists | GO\_REGULATION\_OF\_CALCIUM\_ION\_TRANSMEMBRANE\_TRANSPORT | 83 | 5 | 12187 | 98 | Cyba,Ahnak,Capn3,Hspa2,Thy1 | | 5.451e-04 | -7.51 | MODULE\_117 | MSigDB lists | MODULE\_117 | 354 | 10 | 12187 | 98 | Rac2,Lin7a,Apod,Cd68,Ctsz,Ctss,Ccr5,Fmod,Itgam,C1qb | | 5.496e-04 | -7.51 | GO\_CELLULAR\_RESPONSE\_TO\_ORGANIC\_SUBSTANCE | MSigDB lists | GO\_CELLULAR\_RESPONSE\_TO\_ORGANIC\_SUBSTANCE | 1393 | 23 | 12187 | 98 | Csf1r,Ccr5,Ndufa13,Spint1,Kl,Clu,Cd68,Ctss,Psmb10,Il1r1,Fcer1g,Serpinb9,Ctsb,Il6ra,Hcls1,Psme1,Fmod,Camk2b,Rap1gap,Csk,Fgf18,Cyba,Ptprn | | 5.514e-04 | -7.50 | KEGG\_REGULATION\_OF\_ACTIN\_CYTOSKELETON | MSigDB lists | KEGG\_REGULATION\_OF\_ACTIN\_CYTOSKELETON | 177 | 7 | 12187 | 98 | Csk,Brk1,Itgam,Myh9,Actb,Fgf18,Rac2 | | 5.518e-04 | -7.50 | GO\_LYMPHOCYTE\_ACTIVATION | MSigDB lists | GO\_LYMPHOCYTE\_ACTIVATION | 232 | 8 | 12187 | 98 | Ptprc,Psmb10,Tnfsf8,Ikzf3,Treml2,Lfng,Myh9,Fcer1g | | 5.617e-04 | -7.48 | endopeptidase inhibitor activity | molecular function | GO:0004866 | 85 | 5 | 13516 | 107 | Serpinb9,C3,Spint1,Wfdc17,Serpina3n | | 5.740e-04 | -7.46 | cytosolic part | cellular component | GO:0044445 | 233 | 8 | 13825 | 111 | Rpl26,Apod,Rpl13,Rpl14,Rpl31,Rps20,Rps5,Rps8 | | 5.833e-04 | -7.45 | GO\_SECRETORY\_GRANULE\_LUMEN | MSigDB lists | GO\_SECRETORY\_GRANULE\_LUMEN | 48 | 4 | 12187 | 98 | Clu,Pcsk2,Vwf,Serpina3n | | 5.947e-04 | -7.43 | GO\_ANTIGEN\_RECEPTOR\_MEDIATED\_SIGNALING\_PATHWAY | MSigDB lists | GO\_ANTIGEN\_RECEPTOR\_MEDIATED\_SIGNALING\_PATHWAY | 129 | 6 | 12187 | 98 | Psmb10,Ptprc,Klhl6,Csk,Psme1,Thy1 | | 5.977e-04 | -7.42 | DBN1 (drebrin 1) | protein interactions | 1627 | 15 | 3 | 6802 | 78 | Flnb,Myh9,Actb | | 5.977e-04 | -7.42 | ABLIM1 (actin binding LIM protein 1) | protein interactions | 3983 | 15 | 3 | 6802 | 78 | Actb,Myh9,Flnb | | 5.977e-04 | -7.42 | CLTB (clathrin light chain B) | protein interactions | 1212 | 15 | 3 | 6802 | 78 | Myh9,Actb,Flnb | | 5.977e-04 | -7.42 | AFAP1 (actin filament associated protein 1) | protein interactions | 60312 | 15 | 3 | 6802 | 78 | Flnb,Actb,Myh9 | | 6.017e-04 | -7.42 | astrocyte differentiation | biological process | GO:0048708 | 48 | 4 | 13711 | 111 | Gfap,Vim,Grn,Mt3 | | 6.037e-04 | -7.41 | GO\_REGULATION\_OF\_MAST\_CELL\_ACTIVATION\_INVOLVED\_IN\_IMMUNE\_RESPONSE | MSigDB lists | GO\_REGULATION\_OF\_MAST\_CELL\_ACTIVATION\_INVOLVED\_IN\_IMMUNE\_RESPONSE | 21 | 3 | 12187 | 98 | Fcer1g,Rac2,Vamp8 | | 6.113e-04 | -7.40 | immunoglobulin receptor activity | molecular function | GO:0019763 | 5 | 2 | 13516 | 107 | Fcgr3,Fcer1g | | 6.142e-04 | -7.40 | ribosomal subunit | cellular component | GO:0044391 | 180 | 7 | 13825 | 111 | Rps8,Rpl31,Rps20,Rps5,Rpl13,Rpl14,Rpl26 | | 6.148e-04 | -7.39 | biological regulation | biological process | GO:0065007 | 8325 | 84 | 13711 | 111 | C1qa,Ptprc,Trnp1,Il1r1,Mgll,Kl,Thy1,Ikzf3,Ctss,Ccar1,Grn,Adgre1,Actb,Fcer1g,Pde1a,Lin7a,Bhlhe41,Pcsk2,Psme1,Fgf18,Ahnak,Ly6e,Fcgr3,Ctsz,Igfbp6,Vamp8,Rap1gap,Dek,Klhl6,Cplx2,Fam111a,Mt3,Cd52,Myh9,Rps5,Mark4,Csk,Ndufa13,Ccr5,Mgp,Cyba,C1qb,Lfng,Apod,Hspa2,Sh3bgrl3,Tmem176a,Cd68,Clu,Abhd17c,Rbfox3,Mc4r,Tmem106a,Adh1,Treml2,Itgam,Slc17a7,Ctsb,Capn3,Il6ra,Serpinb9,Camk2b,Gfap,Tsc22d1,Rpl26,Tnfsf8,Ccl6,Lrg1,Vim,Lgi4,Brk1,Hcls1,Csf1r,Spint1,Ifitm1,Rac2,Ctsa,Saa3,Tfap2c,Serpina3n,C3,Ptprn,Zfp786,Vwf | | 6.176e-04 | -7.39 | mast cell activation | biological process | GO:0045576 | 21 | 3 | 13711 | 111 | Fcer1g,Cplx2,Fcgr3 | | 6.184e-04 | -7.39 | YOSHIMURA\_MAPK8\_TARGETS\_UP | MSigDB lists | YOSHIMURA\_MAPK8\_TARGETS\_UP | 883 | 17 | 12187 | 98 | Kl,Psme1,Capn3,Camk2b,Ctss,Cyba,Ptprn,Ptprc,Il1r1,Dlgap3,Vim,C1qb,Adh1,Lin7a,Gfap,Cplx2,Cox4i1 | | 6.290e-04 | -7.37 | Cytoplasmic Ribosomal Proteins | WikiPathways | WP163 | 78 | 6 | 3756 | 52 | Rpl31,Rps8,Rps20,Rpl26,Rpl13,Rps5 | | 6.299e-04 | -7.37 | ACEVEDO\_LIVER\_TUMOR\_VS\_NORMAL\_ADJACENT\_TISSUE\_DN | MSigDB lists | ACEVEDO\_LIVER\_TUMOR\_VS\_NORMAL\_ADJACENT\_TISSUE\_DN | 181 | 7 | 12187 | 98 | Rap1gap,Ly6e,Cd68,C3,Ccar1,Vamp8,Rab3il1 | | 6.301e-04 | -7.37 | BIOCARTA\_THELPER\_PATHWAY | MSigDB lists | BIOCARTA\_THELPER\_PATHWAY | 5 | 2 | 12187 | 98 | Ptprc,Thy1 | | 6.313e-04 | -7.37 | KEGG\_HEMATOPOIETIC\_CELL\_LINEAGE | MSigDB lists | KEGG\_HEMATOPOIETIC\_CELL\_LINEAGE | 49 | 4 | 12187 | 98 | Il6ra,Itgam,Il1r1,Csf1r | | 6.393e-04 | -7.36 | serotonin transport | biological process | GO:0006837 | 5 | 2 | 13711 | 111 | Fcer1g,Fcgr3 | | 6.451e-04 | -7.35 | IGLESIAS\_E2F\_TARGETS\_UP | MSigDB lists | IGLESIAS\_E2F\_TARGETS\_UP | 131 | 6 | 12187 | 98 | Ctss,Ctsz,S100a6,C1qa,C1qb,C3 | | 6.451e-04 | -7.35 | GSE34205\_RSV\_VS\_FLU\_INF\_INFANT\_PBMC\_DN | MSigDB lists | GSE34205\_RSV\_VS\_FLU\_INF\_INFANT\_PBMC\_DN | 131 | 6 | 12187 | 98 | Ccr5,Ly6e,Grn,C1qb,Actb,C1qa | | 6.526e-04 | -7.33 | GO\_PROTEIN\_COMPLEX\_BINDING | MSigDB lists | GO\_PROTEIN\_COMPLEX\_BINDING | 726 | 15 | 12187 | 98 | Brk1,Mark4,Fcer1g,Myh9,Thy1,Pcsk2,Tspan4,Gfap,Fcgr3,Actb,Ctsb,Ctss,Vwf,Hcls1,Vim | | 6.655e-04 | -7.31 | L13a-mediated translational silencing of Ceruloplasmin expression | REACTOME pathways | R-MMU-156827 | 89 | 6 | 6297 | 76 | Rpl14,Rps5,Rpl26,Rps20,Rps8,Rpl13 | | 6.659e-04 | -7.31 | HALLMARK\_COAGULATION | MSigDB lists | HALLMARK\_COAGULATION | 87 | 5 | 12187 | 98 | Vwf,Clu,Ctsb,C3,C1qa | | 6.659e-04 | -7.31 | MODULE\_92 | MSigDB lists | MODULE\_92 | 87 | 5 | 12187 | 98 | Ccr5,Tfap2c,S100a6,Grn,C1qa | | 6.766e-04 | -7.30 | regulation of tumor necrosis factor production | biological process | GO:0032680 | 131 | 6 | 13711 | 111 | Ptprc,Fcgr3,Fcer1g,Cyba,Ccr5,Clu | | 6.820e-04 | -7.29 | ZHENG\_GLIOBLASTOMA\_PLASTICITY\_DN | MSigDB lists | ZHENG\_GLIOBLASTOMA\_PLASTICITY\_DN | 50 | 4 | 12187 | 98 | Slc26a2,Clu,Thy1,Tsc22d1 | | 6.857e-04 | -7.29 | cell projection | cellular component | GO:0042995 | 2067 | 30 | 13825 | 111 | Rac2,Ctsz,Cplx2,Thy1,Mark4,S100a6,Gfap,Myh9,Slc26a2,Il1r1,Mgll,Cyba,Camk2b,Clu,Flnb,Ptprc,Abhd17c,Cd52,Lin7a,Actb,Rap1gap,Ptprn,Ldha,Pcsk2,Brk1,Dlgap3,Vim,Slc17a7,Rpl26,Apod | | 7.034e-04 | -7.26 | neutrophil chemotaxis | biological process | GO:0030593 | 50 | 4 | 13711 | 111 | Itgam,Ccl6,Fcer1g,Fcgr3 | | 7.042e-04 | -7.26 | regulation of tumor necrosis factor superfamily cytokine production | biological process | GO:1903555 | 132 | 6 | 13711 | 111 | Fcer1g,Cyba,Ccr5,Clu,Ptprc,Fcgr3 | | 7.109e-04 | -7.25 | regulation of mast cell degranulation | biological process | GO:0043304 | 22 | 3 | 13711 | 111 | Vamp8,Fcer1g,Rac2 | | 7.109e-04 | -7.25 | regulation of mast cell activation involved in immune response | biological process | GO:0033006 | 22 | 3 | 13711 | 111 | Vamp8,Rac2,Fcer1g | | 7.145e-04 | -7.24 | lymphocyte activation | biological process | GO:0046649 | 299 | 9 | 13711 | 111 | Lfng,Ikzf3,Fcer1g,Itgam,Treml2,Myh9,Ptprc,Tnfsf8,Psmb10 | | 7.296e-04 | -7.22 | INF2 (inverted formin, FH2 and WH2 domain containing) | protein interactions | 64423 | 16 | 3 | 6802 | 78 | Flnb,Myh9,Actb | | 7.296e-04 | -7.22 | SPTBN1 (spectrin beta, non-erythrocytic 1) | protein interactions | 6711 | 16 | 3 | 6802 | 78 | Flnb,Myh9,Actb | | 7.296e-04 | -7.22 | CYBRD1 (cytochrome b reductase 1) | protein interactions | 79901 | 16 | 3 | 6802 | 78 | Flnb,Actb,Myh9 | | 7.296e-04 | -7.22 | ARPC5L (actin related protein 2/3 complex subunit 5 like) | protein interactions | 81873 | 16 | 3 | 6802 | 78 | Actb,Myh9,Flnb | | 7.296e-04 | -7.22 | RAI14 (retinoic acid induced 14) | protein interactions | 26064 | 16 | 3 | 6802 | 78 | Actb,Myh9,Flnb | | 7.296e-04 | -7.22 | RRAS2 (RAS related 2) | protein interactions | 22800 | 16 | 3 | 6802 | 78 | Flnb,Myh9,Actb | | 7.296e-04 | -7.22 | FLNA (filamin A) | protein interactions | 2316 | 16 | 3 | 6802 | 78 | Flnb,Actb,Myh9 | | 7.296e-04 | -7.22 | AP2M1 (adaptor related protein complex 2 subunit mu 1) | protein interactions | 1173 | 16 | 3 | 6802 | 78 | Flnb,Actb,Myh9 | | 7.296e-04 | -7.22 | TMOD1 (tropomodulin 1) | protein interactions | 7111 | 16 | 3 | 6802 | 78 | Flnb,Myh9,Actb | | 7.354e-04 | -7.22 | positive regulation of cell activation | biological process | GO:0050867 | 240 | 8 | 13711 | 111 | Il6ra,Thy1,Vamp8,Capn3,Fcer1g,Itgam,C1qa,Ptprc | | 7.397e-04 | -7.21 | Ribosome | KEGG pathways | ko03010 | 125 | 7 | 5248 | 64 | Rpl26,Rps8,Rpl14,Rps20,Rps5,Rpl31,Rpl13 | | 7.397e-04 | -7.21 | Ribosome | KEGG pathways | mmu03010 | 125 | 7 | 5248 | 64 | Rpl26,Rps8,Rps20,Rpl14,Rpl31,Rps5,Rpl13 | | 7.428e-04 | -7.21 | GO\_POSITIVE\_REGULATION\_OF\_CELL\_PROLIFERATION | MSigDB lists | GO\_POSITIVE\_REGULATION\_OF\_CELL\_PROLIFERATION | 581 | 13 | 12187 | 98 | Csf1r,Brk1,S100a6,Lrg1,Il6ra,Gfap,Hcls1,Clu,Ptprc,Fgf18,Cyba,Ccar1,Rac2 | | 7.490e-04 | -7.20 | GTP hydrolysis and joining of the 60S ribosomal subunit | REACTOME pathways | R-MMU-72706 | 91 | 6 | 6297 | 76 | Rpl13,Rpl14,Rps8,Rps20,Rpl26,Rps5 | | 7.582e-04 | -7.18 | activation of immune response | biological process | GO:0002253 | 185 | 7 | 13711 | 111 | Fcer1g,C1qa,Ptprc,C1qb,C3,Klhl6,Thy1 | | 7.671e-04 | -7.17 | peptidase inhibitor activity | molecular function | GO:0030414 | 91 | 5 | 13516 | 107 | Wfdc17,Spint1,Serpinb9,C3,Serpina3n | | 7.674e-04 | -7.17 | MYL6B (myosin light chain 6B) | protein interactions | 140465 | 4 | 2 | 6802 | 78 | Myh9,Actb | | 7.674e-04 | -7.17 | SIPA1L1 (signal induced proliferation associated 1 like 1) | protein interactions | 26037 | 4 | 2 | 6802 | 78 | Myh9,Actb | | 7.674e-04 | -7.17 | RAB14 (RAB14, member RAS oncogene family) | protein interactions | 51552 | 4 | 2 | 6802 | 78 | Actb,Flnb | | 7.674e-04 | -7.17 | PALM (paralemmin) | protein interactions | 5064 | 4 | 2 | 6802 | 78 | Myh9,Flnb | | 7.674e-04 | -7.17 | RAB7A (RAB7A, member RAS oncogene family) | protein interactions | 7879 | 4 | 2 | 6802 | 78 | Vim,Flnb | | 7.674e-04 | -7.17 | KIAA1211 (KIAA1211) | protein interactions | 57482 | 4 | 2 | 6802 | 78 | Actb,Myh9 | | 7.674e-04 | -7.17 | CGN (cingulin) | protein interactions | 57530 | 4 | 2 | 6802 | 78 | Myh9,Actb | | 7.674e-04 | -7.17 | TES (testin LIM domain protein) | protein interactions | 26136 | 4 | 2 | 6802 | 78 | Actb,Myh9 | | 7.674e-04 | -7.17 | PPFIBP1 (PPFIA binding protein 1) | protein interactions | 8496 | 4 | 2 | 6802 | 78 | Flnb,Myh9 | | 7.674e-04 | -7.17 | PLEC (plectin) | protein interactions | 5339 | 4 | 2 | 6802 | 78 | Flnb,Myh9 | | 7.682e-04 | -7.17 | GO\_EXOCYTOSIS | MSigDB lists | GO\_EXOCYTOSIS | 244 | 8 | 12187 | 98 | Fcer1g,Clu,Vwf,Exoc3l4,Vamp8,Cplx2,Serpina3n,Lin7a | | 7.723e-04 | -7.17 | regulation of cytokine production | biological process | GO:0001817 | 504 | 12 | 13711 | 111 | Fcer1g,Csf1r,Cyba,C3,Ccr5,Csk,Il6ra,Clu,Ptprc,Fcgr3,Apod,Il1r1 | | 7.800e-04 | -7.16 | Complement and Coagulation Cascades | WikiPathways | WP449 | 31 | 4 | 3756 | 52 | C3,Vwf,C1qa,C1qb | | 7.855e-04 | -7.15 | TAKEDA\_TARGETS\_OF\_NUP98\_HOXA9\_FUSION\_8D\_DN | MSigDB lists | TAKEDA\_TARGETS\_OF\_NUP98\_HOXA9\_FUSION\_8D\_DN | 136 | 6 | 12187 | 98 | C1qb,Ctsb,C1qa,Clu,Itgam,Lfng | | 7.883e-04 | -7.15 | ONKEN\_UVEAL\_MELANOMA\_UP | MSigDB lists | ONKEN\_UVEAL\_MELANOMA\_UP | 661 | 14 | 12187 | 98 | C1qb,Fcer1g,Ly6e,S100a6,Tsc22d1,Ctsb,Actb,Hcls1,Rap1gap,Ctss,Ptprc,C1qa,Apod,Sh3bgrl3 | | 7.923e-04 | -7.14 | regulation of adaptive immune response | biological process | GO:0002819 | 135 | 6 | 13711 | 111 | C3,Il1r1,Il6ra,Fcer1g,Fcgr3,Ptprc | | 7.946e-04 | -7.14 | GO\_REGULATION\_OF\_BONE\_RESORPTION | MSigDB lists | GO\_REGULATION\_OF\_BONE\_RESORPTION | 23 | 3 | 12187 | 98 | Csk,Mc4r,Csf1r | | 7.946e-04 | -7.14 | IIZUKA\_LIVER\_CANCER\_PROGRESSION\_L0\_L1\_DN | MSigDB lists | IIZUKA\_LIVER\_CANCER\_PROGRESSION\_L0\_L1\_DN | 23 | 3 | 12187 | 98 | Cox4i1,Cyba,Ctsa | | 8.127e-04 | -7.12 | regulation of inflammatory response to antigenic stimulus | biological process | GO:0002861 | 23 | 3 | 13711 | 111 | Fcgr3,C3,Fcer1g | | 8.172e-04 | -7.11 | KEGG\_LEUKOCYTE\_TRANSENDOTHELIAL\_MIGRATION | MSigDB lists | KEGG\_LEUKOCYTE\_TRANSENDOTHELIAL\_MIGRATION | 91 | 5 | 12187 | 98 | Thy1,Itgam,Rac2,Cyba,Actb | | 8.403e-04 | -7.08 | Nonsense Mediated Decay (NMD) enhanced by the Exon Junction Complex (EJC) | REACTOME pathways | R-MMU-975957 | 93 | 6 | 6297 | 76 | Rpl14,Rpl13,Rps20,Rpl26,Rps5,Rps8 | | 8.403e-04 | -7.08 | Nonsense-Mediated Decay (NMD) | REACTOME pathways | R-MMU-927802 | 93 | 6 | 6297 | 76 | Rps8,Rpl13,Rpl14,Rpl26,Rps5,Rps20 | | 8.441e-04 | -7.08 | signaling receptor binding | molecular function | GO:0005102 | 1178 | 20 | 13516 | 107 | Tnfsf8,C3,Tspan4,Thy1,Il1r1,Myh9,Lrg1,Ccl6,Vwf,Grn,Fgf18,Il6ra,Ptprc,Kl,Capn3,Gfap,Saa3,Fcgr3,Ly6e,Clu | | 8.479e-04 | -7.07 | GSE37301\_HEMATOPOIETIC\_STEM\_CELL\_VS\_CD4\_TCELL\_UP | MSigDB lists | GSE37301\_HEMATOPOIETIC\_STEM\_CELL\_VS\_CD4\_TCELL\_UP | 138 | 6 | 12187 | 98 | Fcer1g,S100a6,Grn,Cd68,Csf1r,C3 | | 8.479e-04 | -7.07 | GSE22611\_UNSTIM\_VS\_6H\_MDP\_STIM\_NOD2\_TRANSDUCED\_HEK293T\_CELL\_UP | MSigDB lists | GSE22611\_UNSTIM\_VS\_6H\_MDP\_STIM\_NOD2\_TRANSDUCED\_HEK293T\_CELL\_UP | 138 | 6 | 12187 | 98 | S100a6,C3,Ikzf3,Cox4i1,Gfap,Apod | | 8.492e-04 | -7.07 | Classical antibody-mediated complement activation | REACTOME pathways | R-MMU-173623 | 4 | 2 | 6297 | 76 | C1qa,C1qb | | 8.492e-04 | -7.07 | Creation of C4 and C2 activators | REACTOME pathways | R-MMU-166786 | 4 | 2 | 6297 | 76 | C1qb,C1qa | | 8.513e-04 | -7.07 | KEGG\_LEISHMANIA\_INFECTION | MSigDB lists | KEGG\_LEISHMANIA\_INFECTION | 53 | 4 | 12187 | 98 | C3,Cyba,Fcgr3,Itgam | | 8.514e-04 | -7.07 | cytosolic large ribosomal subunit | cellular component | GO:0022625 | 53 | 4 | 13825 | 111 | Rpl31,Rpl26,Rpl13,Rpl14 | | 8.551e-04 | -7.06 | GO\_SECRETION | MSigDB lists | GO\_SECRETION | 444 | 11 | 12187 | 98 | Exoc3l4,Fcer1g,Clu,Vwf,Mc4r,Ptprn,Serpina3n,Lin7a,Slc17a7,Vamp8,Cplx2 | | 8.654e-04 | -7.05 | GO\_RNA\_CATABOLIC\_PROCESS | MSigDB lists | GO\_RNA\_CATABOLIC\_PROCESS | 191 | 7 | 12187 | 98 | Rpl26,Rpl13,Rps8,Rps5,Rpl14,Rpl31,Rps20 | | 8.661e-04 | -7.05 | structural constituent of cytoskeleton | molecular function | GO:0005200 | 54 | 4 | 13516 | 107 | Vim,Camk2b,Gfap,Actb | | 8.680e-04 | -7.05 | GO\_PEPTIDASE\_ACTIVITY | MSigDB lists | GO\_PEPTIDASE\_ACTIVITY | 376 | 10 | 12187 | 98 | Ctss,Ctsz,Pcsk2,Capn3,C3,C1qb,Psmb10,C1qa,Ctsb,Ctsa | | 8.710e-04 | -7.05 | membrane raft | cellular component | GO:0045121 | 310 | 9 | 13825 | 111 | Csk,Thy1,Fcer1g,Ahnak,Actb,Ctsb,Slc2a1,Ptprc,Itgam | | 8.787e-04 | -7.04 | AP2A1 (adaptor related protein complex 2 subunit alpha 1) | protein interactions | 160 | 17 | 3 | 6802 | 78 | Actb,Myh9,Flnb | | 8.787e-04 | -7.04 | SPTAN1 (spectrin alpha, non-erythrocytic 1) | protein interactions | 6709 | 17 | 3 | 6802 | 78 | Actb,Myh9,Flnb | | 8.804e-04 | -7.04 | LI\_AMPLIFIED\_IN\_LUNG\_CANCER | MSigDB lists | LI\_AMPLIFIED\_IN\_LUNG\_CANCER | 139 | 6 | 12187 | 98 | Rpl31,Rps5,Rps20,Rpl13,Ldha,Spint1 | | 8.910e-04 | -7.02 | membrane microdomain | cellular component | GO:0098857 | 311 | 9 | 13825 | 111 | Slc2a1,Actb,Ctsb,Itgam,Ptprc,Csk,Fcer1g,Ahnak,Thy1 | | 9.028e-04 | -7.01 | WESTON\_VEGFA\_TARGETS\_12HR | MSigDB lists | WESTON\_VEGFA\_TARGETS\_12HR | 24 | 3 | 12187 | 98 | Vim,Mgp,Thy1 | | 9.028e-04 | -7.01 | GO\_REGULATION\_OF\_MAST\_CELL\_ACTIVATION | MSigDB lists | GO\_REGULATION\_OF\_MAST\_CELL\_ACTIVATION | 24 | 3 | 12187 | 98 | Vamp8,Rac2,Fcer1g | | 9.122e-04 | -7.00 | IgG binding | molecular function | GO:0019864 | 6 | 2 | 13516 | 107 | Fcgr3,Fcer1g | | 9.140e-04 | -7.00 | GSE46606\_IRF4HIGH\_VS\_WT\_CD40L\_IL2\_IL5\_DAY3\_STIMULATED\_BCELL\_DN | MSigDB lists | GSE46606\_IRF4HIGH\_VS\_WT\_CD40L\_IL2\_IL5\_DAY3\_STIMULATED\_BCELL\_DN | 140 | 6 | 12187 | 98 | Itgam,Psme1,Fcer1g,C3,Rac2,Psmb10 | | 9.143e-04 | -7.00 | regulation of developmental process | biological process | GO:0050793 | 2184 | 31 | 13711 | 111 | Vim,Rap1gap,Lrg1,Ctsz,C3,Mgp,Tfap2c,Ccr5,Lfng,Spint1,Mt3,Lgi4,Myh9,Csf1r,Hcls1,Grn,Ikzf3,Mgll,Il1r1,Thy1,Kl,Clu,Tmem176a,Ptprc,C1qa,Gfap,Fgf18,Capn3,Camk2b,Il6ra,Bhlhe41 | | 9.234e-04 | -6.99 | antigen processing and presentation of exogenous antigen | biological process | GO:0019884 | 24 | 3 | 13711 | 111 | Fcer1g,Fcgr3,Psme1 | | 9.401e-04 | -6.97 | MARIADASON\_RESPONSE\_TO\_BUTYRATE\_CURCUMIN\_SULINDAC\_TSA\_1 | MSigDB lists | MARIADASON\_RESPONSE\_TO\_BUTYRATE\_CURCUMIN\_SULINDAC\_TSA\_1 | 6 | 2 | 12187 | 98 | Grn,Myh9 | | 9.401e-04 | -6.97 | GO\_IGG\_BINDING | MSigDB lists | GO\_IGG\_BINDING | 6 | 2 | 12187 | 98 | Fcgr3,Fcer1g | | 9.423e-04 | -6.97 | neutrophil migration | biological process | GO:1990266 | 54 | 4 | 13711 | 111 | Fcgr3,Ccl6,Itgam,Fcer1g | | 9.538e-04 | -6.96 | modulation by symbiont of host apoptotic process | biological process | GO:0052150 | 6 | 2 | 13711 | 111 | Serpinb9,Ccr5 | | 9.538e-04 | -6.96 | modulation of programmed cell death in other organism | biological process | GO:0044531 | 6 | 2 | 13711 | 111 | Serpinb9,Ccr5 | | 9.538e-04 | -6.96 | modulation by organism of apoptotic process in other organism involved in symbiotic interaction | biological process | GO:0052433 | 6 | 2 | 13711 | 111 | Ccr5,Serpinb9 | | 9.538e-04 | -6.96 | modulation of apoptotic process in other organism | biological process | GO:0044532 | 6 | 2 | 13711 | 111 | Ccr5,Serpinb9 | | 9.538e-04 | -6.96 | modulation of programmed cell death in other organism involved in symbiotic interaction | biological process | GO:0052248 | 6 | 2 | 13711 | 111 | Serpinb9,Ccr5 | | 9.795e-04 | -6.93 | HECKER\_IFNB1\_TARGETS | MSigDB lists | HECKER\_IFNB1\_TARGETS | 55 | 4 | 12187 | 98 | Ly6e,Clu,C1qa,C1qb | | 9.840e-04 | -6.92 | GSE3920\_UNTREATED\_VS\_IFNG\_TREATED\_FIBROBLAST\_DN | MSigDB lists | GSE3920\_UNTREATED\_VS\_IFNG\_TREATED\_FIBROBLAST\_DN | 142 | 6 | 12187 | 98 | Mgll,Tsc22d1,Tmem176a,Klhl6,Treml2,Il1r1 | | 9.947e-04 | -6.91 | Hydrolase\_4 | interpro domains | IPR022742 | 6 | 2 | 13788 | 114 | Abhd17c,Mgll | | 1.020e-03 | -6.89 | GSE43955\_1H\_VS\_10H\_ACT\_CD4\_TCELL\_DN | MSigDB lists | GSE43955\_1H\_VS\_10H\_ACT\_CD4\_TCELL\_DN | 143 | 6 | 12187 | 98 | S100a6,Ccr5,Clu,Mgp,Ctss,Flnb | | 1.020e-03 | -6.89 | GSE21360\_NAIVE\_VS\_PRIMARY\_MEMORY\_CD8\_TCELL\_UP | MSigDB lists | GSE21360\_NAIVE\_VS\_PRIMARY\_MEMORY\_CD8\_TCELL\_UP | 143 | 6 | 12187 | 98 | Lrg1,Lfng,C1qb,Rab3il1,Treml2,Slc2a1 | | 1.020e-03 | -6.89 | AMUNDSON\_POOR\_SURVIVAL\_AFTER\_GAMMA\_RADIATION\_2G | MSigDB lists | AMUNDSON\_POOR\_SURVIVAL\_AFTER\_GAMMA\_RADIATION\_2G | 143 | 6 | 12187 | 98 | Csk,Rps8,Hcls1,Tspan4,Rac2,Ptprc | | 1.022e-03 | -6.89 | Hydrolase\_4 | pfam domains | PF12146 | 6 | 2 | 12881 | 108 | Abhd17c,Mgll | | 1.022e-03 | -6.89 | UPAR\_LY6 | pfam domains | PF00021 | 6 | 2 | 12881 | 108 | Ly6e,Ly6a | | 1.048e-03 | -6.86 | BURTON\_ADIPOGENESIS\_PEAK\_AT\_0HR | MSigDB lists | BURTON\_ADIPOGENESIS\_PEAK\_AT\_0HR | 56 | 4 | 12187 | 98 | Adh1,Psmb10,Serpinb9,Mgp | | 1.049e-03 | -6.86 | Cap-dependent Translation Initiation | REACTOME pathways | R-MMU-72737 | 97 | 6 | 6297 | 76 | Rps20,Rpl26,Rps5,Rpl14,Rpl13,Rps8 | | 1.049e-03 | -6.86 | Eukaryotic Translation Initiation | REACTOME pathways | R-MMU-72613 | 97 | 6 | 6297 | 76 | Rps20,Rpl14,Rpl26,Rps5,Rpl13,Rps8 | | 1.049e-03 | -6.86 | - | gene3d domains | 2.60.40.10 | 466 | 12 | 6647 | 62 | Fcrls,Il1r1,Fcgr3,Ptprc,Csf1r,Il6ra,Flnb,Spint1,Thy1,Treml2,C3,Phyhip | | 1.058e-03 | -6.85 | GSE34156\_TLR1\_TLR2\_LIGAND\_VS\_NOD2\_AND\_TLR1\_TLR2\_LIGAND\_24H\_TREATED\_MONOCYTE\_UP | MSigDB lists | GSE34156\_TLR1\_TLR2\_LIGAND\_VS\_NOD2\_AND\_TLR1\_TLR2\_LIGAND\_24H\_TREATED\_MONOCYTE\_UP | 144 | 6 | 12187 | 98 | Grn,Ctsz,Csf1r,Mpeg1,Fcgr3,Actb | | 1.065e-03 | -6.84 | membrane region | cellular component | GO:0098589 | 319 | 9 | 13825 | 111 | Csk,Thy1,Fcer1g,Ahnak,Actb,Ctsb,Slc2a1,Ptprc,Itgam | | 1.081e-03 | -6.83 | granulocyte chemotaxis | biological process | GO:0071621 | 56 | 4 | 13711 | 111 | Fcgr3,Fcer1g,Ccl6,Itgam | | 1.095e-03 | -6.82 | GO\_REGULATION\_OF\_MULTICELLULAR\_ORGANISMAL\_DEVELOPMENT | MSigDB lists | GO\_REGULATION\_OF\_MULTICELLULAR\_ORGANISMAL\_DEVELOPMENT | 1280 | 21 | 12187 | 98 | Spint1,Gfap,Tmem176a,Csf1r,Lfng,Thy1,Grn,Lrg1,Psmb10,Fgf18,Ikzf3,Vim,Mgp,Hcls1,Kl,C3,Psme1,Fmod,Camk2b,Capn3,Rap1gap | | 1.121e-03 | -6.79 | MARSON\_FOXP3\_TARGETS\_UP | MSigDB lists | MARSON\_FOXP3\_TARGETS\_UP | 57 | 4 | 12187 | 98 | Vim,S100a6,Rac2,Myh9 | | 1.126e-03 | -6.79 | nervous system development | biological process | GO:0007399 | 1917 | 28 | 13711 | 111 | Fgf18,Pcsk2,Camk2b,Gfap,Actb,Itgam,Bhlhe41,Mgll,Rbfox3,Il1r1,Trnp1,Thy1,Grn,C1qa,Clu,Mark4,C3,Ccr5,Tfap2c,Lgi4,Mt3,Csf1r,Spint1,Rac2,Rap1gap,Vim,Cplx2,Ctsz | | 1.128e-03 | -6.79 | GO\_SINGLE\_ORGANISM\_CELLULAR\_LOCALIZATION | MSigDB lists | GO\_SINGLE\_ORGANISM\_CELLULAR\_LOCALIZATION | 766 | 15 | 12187 | 98 | Rps20,Rpl14,Rps8,Ctsz,Rpl13,Capn3,Lin7a,Rpl26,Cplx2,Rpl31,Rps5,Ndufa13,Myh9,Fcer1g,Grn | | 1.133e-03 | -6.78 | response to interferon-gamma | biological process | GO:0034341 | 97 | 5 | 13711 | 111 | Ccl6,Vim,Ifitm1,Vamp8,Flnb | | 1.136e-03 | -6.78 | GSE26488\_WT\_VS\_HDAC7\_DELTAP\_TG\_OT2\_THYMOCYTE\_WITH\_PEPTIDE\_INJECTION\_UP | MSigDB lists | GSE26488\_WT\_VS\_HDAC7\_DELTAP\_TG\_OT2\_THYMOCYTE\_WITH\_PEPTIDE\_INJECTION\_UP | 146 | 6 | 12187 | 98 | Rab3il1,C1qb,Psmb10,Rap1gap,Ccr5,Ctsz | | 1.136e-03 | -6.78 | GSE13484\_12H\_UNSTIM\_VS\_YF17D\_VACCINE\_STIM\_PBMC\_DN | MSigDB lists | GSE13484\_12H\_UNSTIM\_VS\_YF17D\_VACCINE\_STIM\_PBMC\_DN | 146 | 6 | 12187 | 98 | Apod,C3,Ldha,Mgll,Ly6e,Hspa2 | | 1.142e-03 | -6.78 | GCM\_NPM1 | MSigDB lists | GCM\_NPM1 | 98 | 5 | 12187 | 98 | Rps8,Psme1,Rpl14,Rps5,Rpl31 | | 1.177e-03 | -6.74 | GSE19888\_ADENOSINE\_A3R\_INH\_VS\_ACT\_WITH\_INHIBITOR\_PRETREATMENT\_IN\_MAST\_CELL\_UP | MSigDB lists | GSE19888\_ADENOSINE\_A3R\_INH\_VS\_ACT\_WITH\_INHIBITOR\_PRETREATMENT\_IN\_MAST\_CELL\_UP | 147 | 6 | 12187 | 98 | Serpinb9,Ly6e,Psme1,Mpeg1,Psmb10,Apod | | 1.190e-03 | -6.73 | cell chemotaxis | biological process | GO:0060326 | 146 | 6 | 13711 | 111 | Ccl6,Itgam,Fcer1g,Fcgr3,Ccr5,Saa3 | | 1.195e-03 | -6.73 | GNF2\_EIF3S6 | MSigDB lists | GNF2\_EIF3S6 | 99 | 5 | 12187 | 98 | Rpl13,Rps8,Rpl14,Rpl31,Rps20 | | 1.196e-03 | -6.73 | KEGG\_SYSTEMIC\_LUPUS\_ERYTHEMATOSUS | MSigDB lists | KEGG\_SYSTEMIC\_LUPUS\_ERYTHEMATOSUS | 58 | 4 | 12187 | 98 | C1qa,C3,Fcgr3,C1qb | | 1.219e-03 | -6.71 | GSE21670\_STAT3\_KO\_VS\_WT\_CD4\_TCELL\_TGFB\_IL6\_TREATED\_DN | MSigDB lists | GSE21670\_STAT3\_KO\_VS\_WT\_CD4\_TCELL\_TGFB\_IL6\_TREATED\_DN | 148 | 6 | 12187 | 98 | Ctss,Lfng,Ahnak,Fam111a,Grn,Ctsb | | 1.219e-03 | -6.71 | GSE44732\_UNSTIM\_VS\_IL27\_STIM\_IMATURE\_DC\_DN | MSigDB lists | GSE44732\_UNSTIM\_VS\_IL27\_STIM\_IMATURE\_DC\_DN | 148 | 6 | 12187 | 98 | Ctsa,Cyba,C3,Lfng,Cd68,Tsc22d1 | | 1.228e-03 | -6.70 | integrin binding | molecular function | GO:0005178 | 101 | 5 | 13516 | 107 | Myh9,Thy1,Tspan4,Gfap,Vwf | | 1.232e-03 | -6.70 | SPECC1 (sperm antigen with calponin homology and coiled-coil domains 1) | protein interactions | 92521 | 19 | 3 | 6802 | 78 | Myh9,Actb,Flnb | | 1.232e-03 | -6.70 | CAPZA1 (capping actin protein of muscle Z-line subunit alpha 1) | protein interactions | 829 | 19 | 3 | 6802 | 78 | Actb,Myh9,Flnb | | 1.234e-03 | -6.70 | GO\_POSITIVE\_REGULATION\_OF\_RESPONSE\_TO\_EXTERNAL\_STIMULUS | MSigDB lists | GO\_POSITIVE\_REGULATION\_OF\_RESPONSE\_TO\_EXTERNAL\_STIMULUS | 203 | 7 | 12187 | 98 | Cyba,Fgf18,Rac2,Il6ra,Vamp8,C3,Fcer1g | | 1.262e-03 | -6.68 | GSE30083\_SP2\_VS\_SP4\_THYMOCYTE\_DN | MSigDB lists | GSE30083\_SP2\_VS\_SP4\_THYMOCYTE\_DN | 149 | 6 | 12187 | 98 | Mpeg1,Cd68,Ctss,Ahnak,Grn,Fcgr3 | | 1.270e-03 | -6.67 | RAB1A (RAB1A, member RAS oncogene family) | protein interactions | 5861 | 5 | 2 | 6802 | 78 | Actb,Flnb | | 1.270e-03 | -6.67 | ERC1 (ELKS/RAB6-interacting/CAST family member 1) | protein interactions | 23085 | 5 | 2 | 6802 | 78 | Actb,Myh9 | | 1.270e-03 | -6.67 | TRIOBP (TRIO and F-actin binding protein) | protein interactions | 11078 | 5 | 2 | 6802 | 78 | Actb,Myh9 | | 1.270e-03 | -6.67 | RALA (RAS like proto-oncogene A) | protein interactions | 5898 | 5 | 2 | 6802 | 78 | Flnb,Actb | | 1.270e-03 | -6.67 | TJP1 (tight junction protein 1) | protein interactions | 7082 | 5 | 2 | 6802 | 78 | Actb,Myh9 | | 1.275e-03 | -6.66 | MODULE\_76 | MSigDB lists | MODULE\_76 | 59 | 4 | 12187 | 98 | Mgll,Ccr5,Il1r1,C3 | | 1.277e-03 | -6.66 | cellular response to chemical stimulus | biological process | GO:0070887 | 1836 | 27 | 13711 | 111 | Capn3,Fgf18,Il6ra,Ly6e,Itgam,Actb,Fcer1g,Il1r1,Kl,Hspa2,Cd68,Saa3,Csk,Ndufa13,Ccr5,Ptprn,Cyba,Mt3,Csf1r,Hcls1,Ifitm1,Vamp8,Rap1gap,Vim,Fcgr3,Flnb,Ccl6 | | 1.282e-03 | -6.66 | BILANGES\_SERUM\_RESPONSE\_TRANSLATION | MSigDB lists | BILANGES\_SERUM\_RESPONSE\_TRANSLATION | 27 | 3 | 12187 | 98 | Rps5,Rpl14,Rps8 | | 1.282e-03 | -6.66 | REACTOME\_SEMA4D\_IN\_SEMAPHORIN\_SIGNALING | MSigDB lists | REACTOME\_SEMA4D\_IN\_SEMAPHORIN\_SIGNALING | 27 | 3 | 12187 | 98 | Myh9,Rac2,Ptprc | | 1.282e-03 | -6.66 | SCHOEN\_NFKB\_SIGNALING | MSigDB lists | SCHOEN\_NFKB\_SIGNALING | 27 | 3 | 12187 | 98 | C3,Serpina3n,Ctss | | 1.282e-03 | -6.66 | SIMBULAN\_PARP1\_TARGETS\_UP | MSigDB lists | SIMBULAN\_PARP1\_TARGETS\_UP | 27 | 3 | 12187 | 98 | Igfbp6,S100a6,Mgp | | 1.299e-03 | -6.65 | immune response-regulating cell surface receptor signaling pathway | biological process | GO:0002768 | 100 | 5 | 13711 | 111 | Ptprc,Thy1,Fcgr3,Fcer1g,Klhl6 | | 1.306e-03 | -6.64 | GSE21360\_PRIMARY\_VS\_QUATERNARY\_MEMORY\_CD8\_TCELL\_UP | MSigDB lists | GSE21360\_PRIMARY\_VS\_QUATERNARY\_MEMORY\_CD8\_TCELL\_UP | 150 | 6 | 12187 | 98 | C3,Mgll,Vwf,Ly6e,Ahnak,Cox4i1 | | 1.306e-03 | -6.64 | GO\_REGULATION\_OF\_CALCIUM\_ION\_TRANSPORT | MSigDB lists | GO\_REGULATION\_OF\_CALCIUM\_ION\_TRANSPORT | 150 | 6 | 12187 | 98 | Cyba,Hspa2,Capn3,Thy1,Ahnak,Camk2b | | 1.309e-03 | -6.64 | REACTOME\_PHOSPHORYLATION\_OF\_CD3\_AND\_TCR\_ZETA\_CHAINS | MSigDB lists | REACTOME\_PHOSPHORYLATION\_OF\_CD3\_AND\_TCR\_ZETA\_CHAINS | 7 | 2 | 12187 | 98 | Csk,Ptprc | | 1.309e-03 | -6.64 | MODULE\_424 | MSigDB lists | MODULE\_424 | 7 | 2 | 12187 | 98 | Fcer1g,Fcgr3 | | 1.315e-03 | -6.63 | regulation of localization | biological process | GO:0032879 | 2332 | 32 | 13711 | 111 | Rap1gap,Vamp8,Vim,Cplx2,Fcgr3,Mark4,Csk,C3,Cyba,Ccr5,Csf1r,Hcls1,Rac2,Il1r1,Thy1,Grn,Ccar1,Ctss,Hspa2,Apod,Ptprc,Clu,Abhd17c,Fgf18,Capn3,Camk2b,Ahnak,Gfap,Actb,Itgam,Fcer1g,Slc17a7 | | 1.326e-03 | -6.63 | generation of precursor metabolites and energy | biological process | GO:0006091 | 263 | 8 | 13711 | 111 | Adh1,Cyba,Mc4r,Kl,Ldha,Cox4i1,Sh3bgrl3,Mt3 | | 1.328e-03 | -6.62 | acute inflammatory response to antigenic stimulus | biological process | GO:0002438 | 7 | 2 | 13711 | 111 | Serpinb9,Fcgr3 | | 1.328e-03 | -6.62 | macrophage proliferation | biological process | GO:0061517 | 7 | 2 | 13711 | 111 | Csf1r,Clu | | 1.328e-03 | -6.62 | microglial cell proliferation | biological process | GO:0061518 | 7 | 2 | 13711 | 111 | Clu,Csf1r | | 1.328e-03 | -6.62 | regulation of neutrophil activation | biological process | GO:1902563 | 7 | 2 | 13711 | 111 | Grn,Itgam | | 1.328e-03 | -6.62 | Fc-gamma receptor signaling pathway | biological process | GO:0038094 | 7 | 2 | 13711 | 111 | Fcgr3,Fcer1g | | 1.344e-03 | -6.61 | BOQUEST\_STEM\_CELL\_UP | MSigDB lists | BOQUEST\_STEM\_CELL\_UP | 206 | 7 | 12187 | 98 | Adh1,C3,Fmod,Apod,Serpina3n,Mgp,Igfbp6 | | 1.351e-03 | -6.61 | GSE21670\_UNTREATED\_VS\_IL6\_TREATED\_STAT3\_KO\_CD4\_TCELL\_UP | MSigDB lists | GSE21670\_UNTREATED\_VS\_IL6\_TREATED\_STAT3\_KO\_CD4\_TCELL\_UP | 151 | 6 | 12187 | 98 | Csf1r,Ctss,Sh3bgrl3,Fam111a,Ahnak,Ppfibp1 | | 1.351e-03 | -6.61 | GSE22886\_IGM\_MEMORY\_BCELL\_VS\_BM\_PLASMA\_CELL\_UP | MSigDB lists | GSE22886\_IGM\_MEMORY\_BCELL\_VS\_BM\_PLASMA\_CELL\_UP | 151 | 6 | 12187 | 98 | Rpl31,Rps20,Fam111a,Tsc22d1,Rpl26,Ikzf3 | | 1.351e-03 | -6.61 | GSE24026\_PD1\_LIGATION\_VS\_CTRL\_IN\_ACT\_TCELL\_LINE\_UP | MSigDB lists | GSE24026\_PD1\_LIGATION\_VS\_CTRL\_IN\_ACT\_TCELL\_LINE\_UP | 151 | 6 | 12187 | 98 | Sh3bgrl3,Vim,Ptprc,Psmb10,Fmod,Psme1 | | 1.358e-03 | -6.60 | regulation of sequestering of calcium ion | biological process | GO:0051282 | 101 | 5 | 13711 | 111 | Cyba,Ccr5,Ptprc,Thy1,Capn3 | | 1.358e-03 | -6.60 | GO\_ANTIGEN\_PROCESSING\_AND\_PRESENTATION\_OF\_EXOGENOUS\_PEPTIDE\_ANTIGEN\_VIA\_MHC\_CLASS\_I | MSigDB lists | GO\_ANTIGEN\_PROCESSING\_AND\_PRESENTATION\_OF\_EXOGENOUS\_PEPTIDE\_ANTIGEN\_VIA\_MHC\_CLASS\_I | 60 | 4 | 12187 | 98 | Psmb10,Cyba,Fcer1g,Psme1 | | 1.369e-03 | -6.59 | membrane | cellular component | GO:0016020 | 6082 | 65 | 13825 | 111 | Ptprn,Alg12,Abhd17c,Actb,Spint1,Flnb,Cd68,Mfsd4a,Tnfsf8,Grn,Slc17a7,Vim,Dlgap3,Atp5j2,Tmem176a,Myh9,S100a6,Ly6e,Cplx2,Ly6a,Thy1,Ahnak,Kl,Lfng,Tspan4,Ikzf3,Rac2,Capn3,Csk,Mc4r,Ctsb,Cox4i1,Itgam,Adh1,Il1r1,Hspa2,Vamp8,Rap1gap,Ccr5,Lin7a,Cd52,Tmem106a,Fcgr3,Slc2a1,Ptprc,Il6ra,Mpeg1,Hcls1,Pcsk2,Treml2,Ifitm1,Gfap,Ndufa13,Camk2b,Clu,Ctss,Mgll,Adgre1,Fcer1g,Cyba,Sh2d5,Rps8,Csf1r,Slc26a2,Vwf | | 1.372e-03 | -6.59 | GOLDRATH\_ANTIGEN\_RESPONSE | MSigDB lists | GOLDRATH\_ANTIGEN\_RESPONSE | 267 | 8 | 12187 | 98 | Cyba,Serpinb9,Cd68,S100a6,Hspa2,Ccr5,Ahnak,C3 | | 1.379e-03 | -6.59 | MODULE\_66 | MSigDB lists | MODULE\_66 | 471 | 11 | 12187 | 98 | C1qb,C3,Hspa2,Capn3,Apod,Serpina3n,Phyhip,Ptprn,Slc17a7,Igfbp6,Gfap | | 1.385e-03 | -6.58 | Intermed\_filament\_DNA-bd | interpro domains | IPR006821 | 7 | 2 | 13788 | 114 | Gfap,Vim | | 1.400e-03 | -6.57 | antigen processing and presentation | biological process | GO:0019882 | 60 | 4 | 13711 | 111 | Psme1,Fcgr3,Ctss,Fcer1g | | 1.423e-03 | -6.55 | Filament\_head | pfam domains | PF04732 | 7 | 2 | 12881 | 108 | Vim,Gfap | | 1.428e-03 | -6.55 | GNF2\_ICAM3 | MSigDB lists | GNF2\_ICAM3 | 28 | 3 | 12187 | 98 | Sh3bgrl3,Ptprc,Hcls1 | | 1.437e-03 | -6.55 | FLOT1 (flotillin 1) | protein interactions | 10211 | 20 | 3 | 6802 | 78 | Actb,Myh9,Flnb | | 1.437e-03 | -6.55 | SVIL (supervillin) | protein interactions | 6840 | 20 | 3 | 6802 | 78 | Flnb,Actb,Myh9 | | 1.437e-03 | -6.55 | MISP (mitotic spindle positioning) | protein interactions | 126353 | 20 | 3 | 6802 | 78 | Flnb,Actb,Myh9 | | 1.445e-03 | -6.54 | HUANG\_GATA2\_TARGETS\_DN | MSigDB lists | HUANG\_GATA2\_TARGETS\_DN | 61 | 4 | 12187 | 98 | Exoc3l4,Slc2a1,Actb,Clu | | 1.445e-03 | -6.54 | LANDIS\_BREAST\_CANCER\_PROGRESSION\_DN | MSigDB lists | LANDIS\_BREAST\_CANCER\_PROGRESSION\_DN | 61 | 4 | 12187 | 98 | Vwf,S100a6,Ahnak,Igfbp6 | | 1.445e-03 | -6.54 | ROSS\_LEUKEMIA\_WITH\_MLL\_FUSIONS | MSigDB lists | ROSS\_LEUKEMIA\_WITH\_MLL\_FUSIONS | 61 | 4 | 12187 | 98 | Rac2,Myh9,Hcls1,Ptprc | | 1.446e-03 | -6.54 | GSE27670\_BLIMP1\_VS\_LMP1\_TRANSDUCED\_GC\_BCELL\_UP | MSigDB lists | GSE27670\_BLIMP1\_VS\_LMP1\_TRANSDUCED\_GC\_BCELL\_UP | 153 | 6 | 12187 | 98 | Mpeg1,Hcls1,Csf1r,Tspan4,Rab3il1,Grn | | 1.453e-03 | -6.53 | collagen-containing extracellular matrix | cellular component | GO:0062023 | 269 | 8 | 13825 | 111 | Ctsz,Ctsb,Vwf,Igfbp6,Mgp,S100a6,Serpinb9,Fmod | | 1.489e-03 | -6.51 | KEGG\_LYSOSOME | MSigDB lists | KEGG\_LYSOSOME | 104 | 5 | 12187 | 98 | Ctsz,Cd68,Ctss,Ctsb,Ctsa | | 1.489e-03 | -6.51 | granulocyte migration | biological process | GO:0097530 | 61 | 4 | 13711 | 111 | Fcgr3,Fcer1g,Itgam,Ccl6 | | 1.508e-03 | -6.50 | GO\_INNATE\_IMMUNE\_RESPONSE | MSigDB lists | GO\_INNATE\_IMMUNE\_RESPONSE | 336 | 9 | 12187 | 98 | C1qb,Csf1r,C3,Clu,Fcer1g,Camk2b,Csk,Cyba,C1qa | | 1.526e-03 | -6.48 | multicellular organism development | biological process | GO:0007275 | 3728 | 45 | 13711 | 111 | Fgf18,Pcsk2,Ly6e,Actb,Lin7a,Fcer1g,Bhlhe41,Mgll,Il1r1,Trnp1,Thy1,Grn,Ptprc,C1qa,Mark4,C1qb,Ccr5,Mgp,Lfng,Mt3,Myh9,Rap1gap,Cplx2,Flnb,Ctsz,Ctsb,Camk2b,Il6ra,Gfap,Itgam,Rbfox3,Clu,Ptprn,C3,Tfap2c,Vwf,Lgi4,Brk1,Csf1r,Hcls1,Spint1,Rac2,Ifitm1,Vim,Tnfsf8 | | 1.535e-03 | -6.48 | VANDESLUIS\_COMMD1\_TARGETS\_GROUP\_3\_UP | MSigDB lists | VANDESLUIS\_COMMD1\_TARGETS\_GROUP\_3\_UP | 62 | 4 | 12187 | 98 | Spint1,Cyba,Ctsz,Ldha | | 1.545e-03 | -6.47 | GSE21063\_CTRL\_VS\_ANTI\_IGM\_STIM\_BCELL\_NFATC1\_KO\_16H\_UP | MSigDB lists | GSE21063\_CTRL\_VS\_ANTI\_IGM\_STIM\_BCELL\_NFATC1\_KO\_16H\_UP | 155 | 6 | 12187 | 98 | Ctsa,Mpeg1,Ppfibp1,Mgll,Ccr5,Sh3bgrl3 | | 1.548e-03 | -6.47 | GO\_RESPONSE\_TO\_WOUNDING | MSigDB lists | GO\_RESPONSE\_TO\_WOUNDING | 406 | 10 | 12187 | 98 | Actb,Ctsb,Apod,Rac2,Gfap,C3,Vwf,Clu,Fcer1g,Myh9 | | 1.571e-03 | -6.46 | regulation of protein metabolic process | biological process | GO:0051246 | 2256 | 31 | 13711 | 111 | Ctsz,Ccl6,Rpl26,Fcgr3,Vim,Spint1,Rac2,Mt3,Hcls1,Csf1r,Myh9,Ndufa13,Serpina3n,C3,Ctsa,Csk,Clu,Hspa2,Apod,Ptprc,Tmem106a,Grn,Thy1,Kl,Actb,Gfap,Capn3,Psme1,Fgf18,Il6ra,Serpinb9 | | 1.576e-03 | -6.45 | GO\_LYTIC\_VACUOLE | MSigDB lists | GO\_LYTIC\_VACUOLE | 407 | 10 | 12187 | 98 | Ctss,Ctsz,Cd68,Fmod,Ahnak,Cplx2,Vamp8,Gfap,Ctsb,Ctsa | | 1.580e-03 | -6.45 | GO\_REGULATION\_OF\_INNATE\_IMMUNE\_RESPONSE | MSigDB lists | GO\_REGULATION\_OF\_INNATE\_IMMUNE\_RESPONSE | 273 | 8 | 12187 | 98 | Psmb10,Ctsb,Cyba,Serpinb9,Ctss,Itgam,Fcer1g,Psme1 | | 1.583e-03 | -6.45 | GO\_REGULATION\_OF\_LEUKOCYTE\_DEGRANULATION | MSigDB lists | GO\_REGULATION\_OF\_LEUKOCYTE\_DEGRANULATION | 29 | 3 | 12187 | 98 | Rac2,Fcer1g,Vamp8 | | 1.583e-03 | -6.45 | GO\_REGULATION\_OF\_BONE\_REMODELING | MSigDB lists | GO\_REGULATION\_OF\_BONE\_REMODELING | 29 | 3 | 12187 | 98 | Csf1r,Csk,Mc4r | | 1.589e-03 | -6.44 | regulation of transport | biological process | GO:0051049 | 1577 | 24 | 13711 | 111 | Csk,Ahnak,Capn3,Gfap,Cyba,C3,Ccr5,Csf1r,Hcls1,Rac2,Slc17a7,Fcer1g,Actb,Itgam,Thy1,Rap1gap,Vamp8,Cplx2,Ctss,Ptprc,Fcgr3,Apod,Hspa2,Clu | | 1.597e-03 | -6.44 | GSE2826\_WT\_VS\_XID\_BCELL\_DN | MSigDB lists | GSE2826\_WT\_VS\_XID\_BCELL\_DN | 156 | 6 | 12187 | 98 | Tspan4,Ctsa,Spint1,Vwf,Igfbp6,Cplx2 | | 1.613e-03 | -6.43 | positive regulation of transport | biological process | GO:0051050 | 867 | 16 | 13711 | 111 | Rap1gap,Vamp8,Thy1,Ctss,Hspa2,Ptprc,Fcgr3,Clu,Capn3,Cyba,C3,Ccr5,Csf1r,Hcls1,Itgam,Fcer1g | | 1.643e-03 | -6.41 | neuronal cell body | cellular component | GO:0043025 | 632 | 13 | 13825 | 111 | Thy1,Pde1a,Cyba,Cplx2,Rap1gap,Rbfox3,Ptprn,Apod,Camk2b,Flnb,Il6ra,Pcsk2,Dlgap3 | | 1.645e-03 | -6.41 | FORTSCHEGGER\_PHF8\_TARGETS\_DN | MSigDB lists | FORTSCHEGGER\_PHF8\_TARGETS\_DN | 634 | 13 | 12187 | 98 | Vamp8,Igfbp6,Sh2d5,S100a6,Flnb,Fgf18,Rac2,Trnp1,Ctss,Vwf,Mgll,Ahnak,Fam111a | | 1.649e-03 | -6.41 | GSE17721\_CTRL\_VS\_POLYIC\_8H\_BMDC\_DN | MSigDB lists | GSE17721\_CTRL\_VS\_POLYIC\_8H\_BMDC\_DN | 157 | 6 | 12187 | 98 | Fgf18,Slc26a2,Tmem106a,Vamp8,Serpinb9,Hspa2 | | 1.649e-03 | -6.41 | GSE11961\_FOLLICULAR\_BCELL\_VS\_GERMINAL\_CENTER\_BCELL\_DAY7\_DN | MSigDB lists | GSE11961\_FOLLICULAR\_BCELL\_VS\_GERMINAL\_CENTER\_BCELL\_DAY7\_DN | 157 | 6 | 12187 | 98 | Serpina3n,Rab3il1,Dlgap3,Treml2,Slc2a1,Hspa2 | | 1.649e-03 | -6.41 | GSE34156\_UNTREATED\_VS\_24H\_NOD2\_LIGAND\_TREATED\_MONOCYTE\_DN | MSigDB lists | GSE34156\_UNTREATED\_VS\_24H\_NOD2\_LIGAND\_TREATED\_MONOCYTE\_DN | 157 | 6 | 12187 | 98 | Fcgr3,Csf1r,Mpeg1,Lin7a,Tsc22d1,Grn | | 1.663e-03 | -6.40 | CAPZB (capping actin protein of muscle Z-line subunit beta) | protein interactions | 832 | 21 | 3 | 6802 | 78 | Myh9,Actb,Flnb | | 1.663e-03 | -6.40 | CAPZA2 (capping actin protein of muscle Z-line subunit alpha 2) | protein interactions | 830 | 21 | 3 | 6802 | 78 | Myh9,Actb,Flnb | | 1.703e-03 | -6.38 | GSE24142\_EARLY\_THYMIC\_PROGENITOR\_VS\_DN2\_THYMOCYTE\_ADULT\_UP | MSigDB lists | GSE24142\_EARLY\_THYMIC\_PROGENITOR\_VS\_DN2\_THYMOCYTE\_ADULT\_UP | 158 | 6 | 12187 | 98 | Spint1,Il1r1,Fcgr3,Csf1r,Ctss,Tsc22d1 | | 1.703e-03 | -6.38 | GO\_REGULATION\_OF\_CATION\_TRANSMEMBRANE\_TRANSPORT | MSigDB lists | GO\_REGULATION\_OF\_CATION\_TRANSMEMBRANE\_TRANSPORT | 158 | 6 | 12187 | 98 | Capn3,Thy1,Hspa2,Ahnak,Ctss,Cyba | | 1.703e-03 | -6.38 | GSE43955\_1H\_VS\_42H\_ACT\_CD4\_TCELL\_WITH\_TGFB\_IL6\_DN | MSigDB lists | GSE43955\_1H\_VS\_42H\_ACT\_CD4\_TCELL\_WITH\_TGFB\_IL6\_DN | 158 | 6 | 12187 | 98 | Flnb,Ly6e,Itgam,Ccr5,Mgp,Ctss | | 1.727e-03 | -6.36 | GO\_CYSTEINE\_TYPE\_ENDOPEPTIDASE\_ACTIVITY | MSigDB lists | GO\_CYSTEINE\_TYPE\_ENDOPEPTIDASE\_ACTIVITY | 64 | 4 | 12187 | 98 | Ctsb,Capn3,Ctss,Ctsz | | 1.737e-03 | -6.36 | IIZUKA\_LIVER\_CANCER\_PROGRESSION\_G2\_G3\_DN | MSigDB lists | IIZUKA\_LIVER\_CANCER\_PROGRESSION\_G2\_G3\_DN | 8 | 2 | 12187 | 98 | Cyba,Spint1 | | 1.737e-03 | -6.36 | GO\_POSITIVE\_REGULATION\_OF\_MAST\_CELL\_ACTIVATION\_INVOLVED\_IN\_IMMUNE\_RESPONSE | MSigDB lists | GO\_POSITIVE\_REGULATION\_OF\_MAST\_CELL\_ACTIVATION\_INVOLVED\_IN\_IMMUNE\_RESPONSE | 8 | 2 | 12187 | 98 | Vamp8,Fcer1g | | 1.759e-03 | -6.34 | GSE23568\_ID3\_TRANSDUCED\_VS\_ID3\_KO\_CD8\_TCELL\_UP | MSigDB lists | GSE23568\_ID3\_TRANSDUCED\_VS\_ID3\_KO\_CD8\_TCELL\_UP | 159 | 6 | 12187 | 98 | S100a6,Camk2b,Psme1,Thy1,Rac2,Il6ra | | 1.759e-03 | -6.34 | GSE2405\_0H\_VS\_1.5H\_A\_PHAGOCYTOPHILUM\_STIM\_NEUTROPHIL\_UP | MSigDB lists | GSE2405\_0H\_VS\_1.5H\_A\_PHAGOCYTOPHILUM\_STIM\_NEUTROPHIL\_UP | 159 | 6 | 12187 | 98 | Psmb10,Rab3il1,Ctsb,Mgp,Fmod,S100a6 | | 1.759e-03 | -6.34 | GO\_FC\_RECEPTOR\_SIGNALING\_PATHWAY | MSigDB lists | GO\_FC\_RECEPTOR\_SIGNALING\_PATHWAY | 159 | 6 | 12187 | 98 | Fcgr3,Actb,Psmb10,Fcer1g,Psme1,Brk1 | | 1.759e-03 | -6.34 | GSE22886\_NAIVE\_VS\_IGM\_MEMORY\_BCELL\_DN | MSigDB lists | GSE22886\_NAIVE\_VS\_IGM\_MEMORY\_BCELL\_DN | 159 | 6 | 12187 | 98 | Ctsa,Fcgr3,Vim,Itgam,Ahnak,Grn | | 1.759e-03 | -6.34 | GSE42021\_TREG\_PLN\_VS\_CD24INT\_TREG\_THYMUS\_DN | MSigDB lists | GSE42021\_TREG\_PLN\_VS\_CD24INT\_TREG\_THYMUS\_DN | 159 | 6 | 12187 | 98 | Psmb10,C3,Ly6e,Fam111a,Psme1,Ctss | | 1.762e-03 | -6.34 | positive regulation of humoral immune response | biological process | GO:0002922 | 8 | 2 | 13711 | 111 | Ptprc,C3 | | 1.762e-03 | -6.34 | antigen processing and presentation of exogenous peptide antigen via MHC class I | biological process | GO:0042590 | 8 | 2 | 13711 | 111 | Fcgr3,Fcer1g | | 1.762e-03 | -6.34 | positive regulation of defense response to bacterium | biological process | GO:1900426 | 8 | 2 | 13711 | 111 | Cyba,Grn | | 1.762e-03 | -6.34 | modulation by symbiont of host programmed cell death | biological process | GO:0052040 | 8 | 2 | 13711 | 111 | Ccr5,Serpinb9 | | 1.763e-03 | -6.34 | GO\_RRNA\_METABOLIC\_PROCESS | MSigDB lists | GO\_RRNA\_METABOLIC\_PROCESS | 216 | 7 | 12187 | 98 | Rpl26,Rpl31,Rps5,Rpl14,Rps20,Rpl13,Rps8 | | 1.771e-03 | -6.34 | GO\_SIDE\_OF\_MEMBRANE | MSigDB lists | GO\_SIDE\_OF\_MEMBRANE | 278 | 8 | 12187 | 98 | Ctsa,Ctsb,Ptprc,Ccr5,Thy1,Fcer1g,S100a6,Csk | | 1.788e-03 | -6.33 | regulation of mast cell activation | biological process | GO:0033003 | 30 | 3 | 13711 | 111 | Rac2,Fcer1g,Vamp8 | | 1.816e-03 | -6.31 | GSE32164\_ALTERNATIVELY\_ACT\_M2\_VS\_CMYC\_INHIBITED\_MACROPHAGE\_UP | MSigDB lists | GSE32164\_ALTERNATIVELY\_ACT\_M2\_VS\_CMYC\_INHIBITED\_MACROPHAGE\_UP | 160 | 6 | 12187 | 98 | Cyba,Csf1r,Mpeg1,Rac2,Camk2b,Il6ra | | 1.816e-03 | -6.31 | GSE24142\_EARLY\_THYMIC\_PROGENITOR\_VS\_DN2\_THYMOCYTE\_FETAL\_UP | MSigDB lists | GSE24142\_EARLY\_THYMIC\_PROGENITOR\_VS\_DN2\_THYMOCYTE\_FETAL\_UP | 160 | 6 | 12187 | 98 | Il1r1,Fcgr3,Csf1r,Ctss,Vim,Tsc22d1 | | 1.829e-03 | -6.30 | GO\_VESICLE\_LUMEN | MSigDB lists | GO\_VESICLE\_LUMEN | 65 | 4 | 12187 | 98 | Serpina3n,Pcsk2,Vwf,Clu | | 1.833e-03 | -6.30 | GO\_T\_CELL\_RECEPTOR\_SIGNALING\_PATHWAY | MSigDB lists | GO\_T\_CELL\_RECEPTOR\_SIGNALING\_PATHWAY | 109 | 5 | 12187 | 98 | Csk,Psme1,Thy1,Psmb10,Ptprc | | 1.837e-03 | -6.30 | Pept\_his\_AS | interpro domains | IPR025660 | 8 | 2 | 13788 | 114 | Ctsb,Ctss | | 1.837e-03 | -6.30 | LY6\_UPA\_recep-like | interpro domains | IPR016054 | 8 | 2 | 13788 | 114 | Ly6a,Ly6e | | 1.853e-03 | -6.29 | REACTOME\_ADAPTIVE\_IMMUNE\_SYSTEM | MSigDB lists | REACTOME\_ADAPTIVE\_IMMUNE\_SYSTEM | 416 | 10 | 12187 | 98 | Ctss,Csk,Psme1,Rap1gap,C3,Ptprc,Psmb10,Ctsa,Cyba,Ctsb | | 1.874e-03 | -6.28 | GSE9509\_10MIN\_VS\_30MIN\_LPS\_STIM\_IL10\_KO\_MACROPHAGE\_UP | MSigDB lists | GSE9509\_10MIN\_VS\_30MIN\_LPS\_STIM\_IL10\_KO\_MACROPHAGE\_UP | 161 | 6 | 12187 | 98 | S100a6,Grn,Fcer1g,Rps8,Csf1r,Ctsa | | 1.874e-03 | -6.28 | GSE11961\_FOLLICULAR\_BCELL\_VS\_MEMORY\_BCELL\_DAY7\_UP | MSigDB lists | GSE11961\_FOLLICULAR\_BCELL\_VS\_MEMORY\_BCELL\_DAY7\_UP | 161 | 6 | 12187 | 98 | S100a6,Myh9,Pde1a,Rpl31,Exoc3l4,Treml2 | | 1.890e-03 | -6.27 | PICALM (phosphatidylinositol binding clathrin assembly protein) | protein interactions | 8301 | 6 | 2 | 6802 | 78 | Actb,Myh9 | | 1.890e-03 | -6.27 | GAS2L1 (growth arrest specific 2 like 1) | protein interactions | 10634 | 6 | 2 | 6802 | 78 | Actb,Myh9 | | 1.890e-03 | -6.27 | ST5 (suppression of tumorigenicity 5) | protein interactions | 6764 | 6 | 2 | 6802 | 78 | Actb,Myh9 | | 1.890e-03 | -6.27 | AAK1 (AP2 associated kinase 1) | protein interactions | 22848 | 6 | 2 | 6802 | 78 | Myh9,Actb | | 1.890e-03 | -6.27 | RAB5C (RAB5C, member RAS oncogene family) | protein interactions | 5878 | 6 | 2 | 6802 | 78 | Flnb,Actb | | 1.890e-03 | -6.27 | ANXA2 (annexin A2) | protein interactions | 302 | 6 | 2 | 6802 | 78 | Flnb,Myh9 | | 1.890e-03 | -6.27 | LIMCH1 (LIM and calponin homology domains 1) | protein interactions | 22998 | 6 | 2 | 6802 | 78 | Myh9,Actb | | 1.890e-03 | -6.27 | PPP1R12B (protein phosphatase 1 regulatory subunit 12B) | protein interactions | 4660 | 6 | 2 | 6802 | 78 | Actb,Myh9 | | 1.890e-03 | -6.27 | ACTA1 (actin alpha 1, skeletal muscle) | protein interactions | 58 | 6 | 2 | 6802 | 78 | Myh9,Actb | | 1.890e-03 | -6.27 | FLNB (filamin B) | protein interactions | 2317 | 6 | 2 | 6802 | 78 | Myh9,Actb | | 1.899e-03 | -6.27 | Tumour\_necrosis\_fac-like\_dom | interpro domains | IPR008983 | 30 | 3 | 13788 | 114 | C1qb,C1qa,Tnfsf8 | | 1.934e-03 | -6.25 | GSE34006\_UNTREATED\_VS\_A2AR\_AGONIST\_TREATED\_TREG\_DN | MSigDB lists | GSE34006\_UNTREATED\_VS\_A2AR\_AGONIST\_TREATED\_TREG\_DN | 162 | 6 | 12187 | 98 | Lfng,Vim,Ccr5,Slc2a1,Ctsa,Actb | | 1.934e-03 | -6.25 | GSE26030\_TH1\_VS\_TH17\_RESTIMULATED\_DAY5\_POST\_POLARIZATION\_UP | MSigDB lists | GSE26030\_TH1\_VS\_TH17\_RESTIMULATED\_DAY5\_POST\_POLARIZATION\_UP | 162 | 6 | 12187 | 98 | Vim,Ctss,S100a6,Fam111a,Itgam,Exoc3l4 | | 1.935e-03 | -6.25 | GO\_GLIAL\_CELL\_DEVELOPMENT | MSigDB lists | GO\_GLIAL\_CELL\_DEVELOPMENT | 66 | 4 | 12187 | 98 | Clu,Lgi4,Gfap,Vim | | 1.935e-03 | -6.25 | GO\_FC\_GAMMA\_RECEPTOR\_SIGNALING\_PATHWAY | MSigDB lists | GO\_FC\_GAMMA\_RECEPTOR\_SIGNALING\_PATHWAY | 66 | 4 | 12187 | 98 | Brk1,Fcer1g,Actb,Fcgr3 | | 1.935e-03 | -6.25 | GNF2\_HCK | MSigDB lists | GNF2\_HCK | 66 | 4 | 12187 | 98 | Ctss,Sh3bgrl3,Fcer1g,Fcgr3 | | 1.954e-03 | -6.24 | MODULE\_45 | MSigDB lists | MODULE\_45 | 419 | 10 | 12187 | 98 | Hcls1,Slc2a1,Ctss,Grn,Fcer1g,Itgam,Psmb10,Ptprc,Fcgr3,Rac2 | | 1.956e-03 | -6.24 | GO\_REGULATION\_OF\_HOMEOSTATIC\_PROCESS | MSigDB lists | GO\_REGULATION\_OF\_HOMEOSTATIC\_PROCESS | 349 | 9 | 12187 | 98 | Csf1r,Csk,Mc4r,Fcer1g,Ccr5,Thy1,Capn3,Ptprc,Cyba | | 1.961e-03 | -6.23 | membrane part | cellular component | GO:0044425 | 4483 | 51 | 13825 | 111 | Treml2,Mpeg1,Ptprc,Il6ra,Lin7a,Tmem106a,Cd52,Fcgr3,Slc2a1,Ccr5,Vamp8,Hspa2,Slc26a2,Csf1r,Vwf,Mgll,Adgre1,Cyba,Fcer1g,Sh2d5,Ndufa13,Ifitm1,Gfap,Atp5j2,Dlgap3,Slc17a7,Cd68,Mfsd4a,Tnfsf8,Alg12,Abhd17c,Actb,Ptprn,Itgam,Il1r1,Mc4r,Cox4i1,Ctsb,Csk,Capn3,Lfng,Tspan4,Cplx2,Ly6a,Thy1,Kl,Ahnak,Tmem176a,Myh9,S100a6,Ly6e | | 1.968e-03 | -6.23 | phagocytosis, engulfment | biological process | GO:0006911 | 31 | 3 | 13711 | 111 | Fcer1g,Fcgr3,Myh9 | | 1.974e-03 | -6.23 | Ig\_2 | pfam domains | PF13895 | 30 | 3 | 12881 | 108 | Fcrls,Il1r1,Fcgr3 | | 1.980e-03 | -6.22 | GO\_INFLAMMATORY\_RESPONSE | MSigDB lists | GO\_INFLAMMATORY\_RESPONSE | 283 | 8 | 12187 | 98 | C3,Csf1r,Fcer1g,Ccr5,Mgll,Cyba,Serpina3n,Il6ra | | 1.985e-03 | -6.22 | GO\_PEPTIDASE\_REGULATOR\_ACTIVITY | MSigDB lists | GO\_PEPTIDASE\_REGULATOR\_ACTIVITY | 111 | 5 | 12187 | 98 | Serpinb9,Psme1,Serpina3n,Spint1,C3 | | 1.988e-03 | -6.22 | GO\_CELL\_CELL\_ADHESION | MSigDB lists | GO\_CELL\_CELL\_ADHESION | 420 | 10 | 12187 | 98 | Actb,Psmb10,Ptprc,Rac2,Tnfsf8,Treml2,Myh9,Thy1,Fcer1g,Lfng | | 1.995e-03 | -6.22 | GSE40273\_XBP1\_KO\_VS\_WT\_TREG\_DN | MSigDB lists | GSE40273\_XBP1\_KO\_VS\_WT\_TREG\_DN | 163 | 6 | 12187 | 98 | Ikzf3,Il6ra,Flnb,Cd68,Ahnak,Mpeg1 | | 2.045e-03 | -6.19 | HESS\_TARGETS\_OF\_HOXA9\_AND\_MEIS1\_DN | MSigDB lists | HESS\_TARGETS\_OF\_HOXA9\_AND\_MEIS1\_DN | 67 | 4 | 12187 | 98 | Mpeg1,Csf1r,Ctss,Ccr5 | | 2.045e-03 | -6.19 | GO\_REGULATION\_OF\_LYMPHOCYTE\_MEDIATED\_IMMUNITY | MSigDB lists | GO\_REGULATION\_OF\_LYMPHOCYTE\_MEDIATED\_IMMUNITY | 67 | 4 | 12187 | 98 | Ptprc,C3,Serpinb9,Fcer1g | | 2.045e-03 | -6.19 | JECHLINGER\_EPITHELIAL\_TO\_MESENCHYMAL\_TRANSITION\_UP | MSigDB lists | JECHLINGER\_EPITHELIAL\_TO\_MESENCHYMAL\_TRANSITION\_UP | 67 | 4 | 12187 | 98 | Ctsb,Cd68,Ctsz,Vim | | 2.057e-03 | -6.19 | GSE17721\_ALL\_VS\_24H\_PAM3CSK4\_BMDC\_UP | MSigDB lists | GSE17721\_ALL\_VS\_24H\_PAM3CSK4\_BMDC\_UP | 164 | 6 | 12187 | 98 | Ctsb,Psmb10,Psme1,Vwf,Ctss,Vim | | 2.057e-03 | -6.19 | GSE43955\_TH0\_VS\_TGFB\_IL6\_TH17\_ACT\_CD4\_TCELL\_4H\_DN | MSigDB lists | GSE43955\_TH0\_VS\_TGFB\_IL6\_TH17\_ACT\_CD4\_TCELL\_4H\_DN | 164 | 6 | 12187 | 98 | Hcls1,C1qb,Csf1r,Brk1,Vim,Rap1gap | | 2.057e-03 | -6.19 | GSE15930\_STIM\_VS\_STIM\_AND\_IFNAB\_48H\_CD8\_T\_CELL\_UP | MSigDB lists | GSE15930\_STIM\_VS\_STIM\_AND\_IFNAB\_48H\_CD8\_T\_CELL\_UP | 164 | 6 | 12187 | 98 | Rps5,Ctsa,Hcls1,Rac2,Vamp8,Brk1 | | 2.090e-03 | -6.17 | Platelet Adhesion to exposed collagen | REACTOME pathways | R-MMU-75892 | 6 | 2 | 6297 | 76 | Fcer1g,Vwf | | 2.115e-03 | -6.16 | GO\_CYTOKINE\_MEDIATED\_SIGNALING\_PATHWAY | MSigDB lists | GO\_CYTOKINE\_MEDIATED\_SIGNALING\_PATHWAY | 286 | 8 | 12187 | 98 | Csf1r,Camk2b,Psme1,Ccr5,Ptprn,Il1r1,Psmb10,Il6ra | | 2.121e-03 | -6.16 | GSE24142\_EARLY\_THYMIC\_PROGENITOR\_VS\_DN2\_THYMOCYTE\_UP | MSigDB lists | GSE24142\_EARLY\_THYMIC\_PROGENITOR\_VS\_DN2\_THYMOCYTE\_UP | 165 | 6 | 12187 | 98 | Cd68,Ahnak,Ccr5,Tsc22d1,Fcgr3,Csf1r | | 2.121e-03 | -6.16 | GSE2770\_IL12\_VS\_TGFB\_AND\_IL12\_TREATED\_ACT\_CD4\_TCELL\_6H\_DN | MSigDB lists | GSE2770\_IL12\_VS\_TGFB\_AND\_IL12\_TREATED\_ACT\_CD4\_TCELL\_6H\_DN | 165 | 6 | 12187 | 98 | Ldha,Tnfsf8,Capn3,Rpl14,Cyba,Fcgr3 | | 2.121e-03 | -6.16 | GSE20715\_0H\_VS\_48H\_OZONE\_TLR4\_KO\_LUNG\_DN | MSigDB lists | GSE20715\_0H\_VS\_48H\_OZONE\_TLR4\_KO\_LUNG\_DN | 165 | 6 | 12187 | 98 | Serpina3n,Tmem176a,Ldha,Clu,Lrg1,S100a6 | | 2.121e-03 | -6.16 | GSE32164\_RESTING\_DIFFERENTIATED\_VS\_ALTERNATIVELY\_ACT\_M2\_MACROPHAGE\_DN | MSigDB lists | GSE32164\_RESTING\_DIFFERENTIATED\_VS\_ALTERNATIVELY\_ACT\_M2\_MACROPHAGE\_DN | 165 | 6 | 12187 | 98 | Mpeg1,Tmem176a,Serpinb9,Ccr5,Ahnak,Cyba | | 2.121e-03 | -6.16 | GSE10325\_LUPUS\_CD4\_TCELL\_VS\_LUPUS\_MYELOID\_DN | MSigDB lists | GSE10325\_LUPUS\_CD4\_TCELL\_VS\_LUPUS\_MYELOID\_DN | 165 | 6 | 12187 | 98 | Grn,Fcer1g,Ctsb,Csf1r,Ctsa,Fcgr3 | | 2.131e-03 | -6.15 | ENK\_UV\_RESPONSE\_KERATINOCYTE\_UP | MSigDB lists | ENK\_UV\_RESPONSE\_KERATINOCYTE\_UP | 424 | 10 | 12187 | 98 | Rps5,Rpl14,Psme1,Tsc22d1,Ly6e,Grn,Spint1,Cyba,Vamp8,Igfbp6 | | 2.161e-03 | -6.14 | GO\_NEURON\_PROJECTION | MSigDB lists | GO\_NEURON\_PROJECTION | 818 | 15 | 12187 | 98 | Lin7a,Slc17a7,Cplx2,Pcsk2,Thy1,Mark4,Cyba,Ptprn,Il1r1,Apod,Dlgap3,Vim,Rap1gap,Clu,Camk2b | | 2.187e-03 | -6.13 | KAECH\_DAY8\_EFF\_VS\_MEMORY\_CD8\_TCELL\_DN | MSigDB lists | KAECH\_DAY8\_EFF\_VS\_MEMORY\_CD8\_TCELL\_DN | 166 | 6 | 12187 | 98 | Slc26a2,Rpl14,Il6ra,Ly6e,Tnfsf8,Ctss | | 2.187e-03 | -6.13 | GSE22886\_NAIVE\_CD8\_TCELL\_VS\_MONOCYTE\_DN | MSigDB lists | GSE22886\_NAIVE\_CD8\_TCELL\_VS\_MONOCYTE\_DN | 166 | 6 | 12187 | 98 | Ctsb,Fcgr3,Ctss,Grn,Fcer1g,Csf1r | | 2.187e-03 | -6.13 | GSE10239\_NAIVE\_VS\_KLRG1HIGH\_EFF\_CD8\_TCELL\_UP | MSigDB lists | GSE10239\_NAIVE\_VS\_KLRG1HIGH\_EFF\_CD8\_TCELL\_UP | 166 | 6 | 12187 | 98 | Il6ra,Rpl26,Psme1,Rpl13,Rpl31,Treml2 | | 2.221e-03 | -6.11 | GO\_NEUTROPHIL\_ACTIVATION\_INVOLVED\_IN\_IMMUNE\_RESPONSE | MSigDB lists | GO\_NEUTROPHIL\_ACTIVATION\_INVOLVED\_IN\_IMMUNE\_RESPONSE | 9 | 2 | 12187 | 98 | Fcer1g,Vamp8 | | 2.221e-03 | -6.11 | GO\_POSITIVE\_REGULATION\_OF\_MAST\_CELL\_ACTIVATION | MSigDB lists | GO\_POSITIVE\_REGULATION\_OF\_MAST\_CELL\_ACTIVATION | 9 | 2 | 12187 | 98 | Vamp8,Fcer1g | | 2.221e-03 | -6.11 | REACTOME\_TRAFFICKING\_AND\_PROCESSING\_OF\_ENDOSOMAL\_TLR | MSigDB lists | REACTOME\_TRAFFICKING\_AND\_PROCESSING\_OF\_ENDOSOMAL\_TLR | 9 | 2 | 12187 | 98 | Ctsb,Ctss | | 2.222e-03 | -6.11 | LI\_INDUCED\_T\_TO\_NATURAL\_KILLER\_UP | MSigDB lists | LI\_INDUCED\_T\_TO\_NATURAL\_KILLER\_UP | 225 | 7 | 12187 | 98 | Vim,Sh3bgrl3,Fcgr3,Ccr5,Fcer1g,Ahnak,S100a6 | | 2.222e-03 | -6.11 | GO\_SECRETORY\_GRANULE | MSigDB lists | GO\_SECRETORY\_GRANULE | 225 | 7 | 12187 | 98 | Serpina3n,Cyba,Ptprn,Vamp8,Pcsk2,Vwf,Clu | | 2.236e-03 | -6.10 | Antigen processing and presentation | KEGG pathways | mmu04612 | 46 | 4 | 5248 | 64 | Psme1,Hspa2,Ctsb,Ctss | | 2.236e-03 | -6.10 | Antigen processing and presentation | KEGG pathways | ko04612 | 46 | 4 | 5248 | 64 | Psme1,Hspa2,Ctsb,Ctss | | 2.254e-03 | -6.10 | GSE15330\_HSC\_VS\_GRANULOCYTE\_MONOCYTE\_PROGENITOR\_IKAROS\_KO\_DN | MSigDB lists | GSE15330\_HSC\_VS\_GRANULOCYTE\_MONOCYTE\_PROGENITOR\_IKAROS\_KO\_DN | 167 | 6 | 12187 | 98 | Fcgr3,Ptprc,Thy1,Tsc22d1,Csk,Adh1 | | 2.254e-03 | -6.10 | GSE17721\_0.5H\_VS\_24H\_GARDIQUIMOD\_BMDC\_UP | MSigDB lists | GSE17721\_0.5H\_VS\_24H\_GARDIQUIMOD\_BMDC\_UP | 167 | 6 | 12187 | 98 | Pde1a,Ldha,Lfng,Psme1,Grn,Cox4i1 | | 2.279e-03 | -6.08 | GO\_REGULATION\_OF\_INTERLEUKIN\_6\_PRODUCTION | MSigDB lists | GO\_REGULATION\_OF\_INTERLEUKIN\_6\_PRODUCTION | 69 | 4 | 12187 | 98 | Il6ra,Fcer1g,Csk,Cyba | | 2.279e-03 | -6.08 | REACTOME\_ANTIGEN\_PROCESSING\_CROSS\_PRESENTATION | MSigDB lists | REACTOME\_ANTIGEN\_PROCESSING\_CROSS\_PRESENTATION | 69 | 4 | 12187 | 98 | Psme1,Ctss,Cyba,Psmb10 | | 2.279e-03 | -6.08 | GO\_POSITIVE\_REGULATION\_OF\_CATION\_TRANSMEMBRANE\_TRANSPORT | MSigDB lists | GO\_POSITIVE\_REGULATION\_OF\_CATION\_TRANSMEMBRANE\_TRANSPORT | 69 | 4 | 12187 | 98 | Ctss,Thy1,Capn3,Hspa2 | | 2.279e-03 | -6.08 | GO\_MYELOID\_LEUKOCYTE\_ACTIVATION | MSigDB lists | GO\_MYELOID\_LEUKOCYTE\_ACTIVATION | 69 | 4 | 12187 | 98 | Vamp8,Cplx2,Clu,Fcer1g | | 2.281e-03 | -6.08 | REACTOME\_METABOLISM\_OF\_PROTEINS | MSigDB lists | REACTOME\_METABOLISM\_OF\_PROTEINS | 357 | 9 | 12187 | 98 | Rps20,Rpl14,Rps5,Rpl31,Alg12,Rps8,Rpl13,Actb,Rpl26 | | 2.303e-03 | -6.07 | Translation | REACTOME pathways | R-MMU-72766 | 113 | 6 | 6297 | 76 | Rpl13,Rps8,Rps20,Rpl14,Rpl26,Rps5 | | 2.318e-03 | -6.07 | SARRIO\_EPITHELIAL\_MESENCHYMAL\_TRANSITION\_DN | MSigDB lists | SARRIO\_EPITHELIAL\_MESENCHYMAL\_TRANSITION\_DN | 115 | 5 | 12187 | 98 | Adh1,Serpina3n,Il1r1,Psme1,Clu | | 2.322e-03 | -6.07 | GSE24142\_EARLY\_THYMIC\_PROGENITOR\_VS\_DN3\_THYMOCYTE\_UP | MSigDB lists | GSE24142\_EARLY\_THYMIC\_PROGENITOR\_VS\_DN3\_THYMOCYTE\_UP | 168 | 6 | 12187 | 98 | Fcgr3,Cd68,Tsc22d1,S100a6,Tspan4,Csf1r | | 2.322e-03 | -6.07 | KAECH\_DAY15\_EFF\_VS\_MEMORY\_CD8\_TCELL\_DN | MSigDB lists | KAECH\_DAY15\_EFF\_VS\_MEMORY\_CD8\_TCELL\_DN | 168 | 6 | 12187 | 98 | Psmb10,Rps5,Rpl14,Ldha,Il6ra,Psme1 | | 2.322e-03 | -6.07 | GSE3337\_CTRL\_VS\_16H\_IFNG\_IN\_CD8POS\_DC\_DN | MSigDB lists | GSE3337\_CTRL\_VS\_16H\_IFNG\_IN\_CD8POS\_DC\_DN | 168 | 6 | 12187 | 98 | Flnb,Psmb10,Kl,Hcls1,C3,Psme1 | | 2.361e-03 | -6.05 | macrophage activation | biological process | GO:0042116 | 33 | 3 | 13711 | 111 | Tmem106a,Grn,Clu | | 2.376e-03 | -6.04 | - | gene3d domains | 2.60.120.40 | 29 | 3 | 6647 | 62 | C1qa,C1qb,Tnfsf8 | | 2.380e-03 | -6.04 | LU | smart domains | SM00134 | 8 | 2 | 7188 | 68 | Ly6e,Ly6a | | 2.392e-03 | -6.04 | REACTOME\_PLATELET\_ACTIVATION\_SIGNALING\_AND\_AGGREGATION | MSigDB lists | REACTOME\_PLATELET\_ACTIVATION\_SIGNALING\_AND\_AGGREGATION | 169 | 6 | 12187 | 98 | Rac2,Vwf,Mgll,Fcer1g,Clu,Csk | | 2.392e-03 | -6.04 | GSE26030\_TH1\_VS\_TH17\_RESTIMULATED\_DAY15\_POST\_POLARIZATION\_DN | MSigDB lists | GSE26030\_TH1\_VS\_TH17\_RESTIMULATED\_DAY15\_POST\_POLARIZATION\_DN | 169 | 6 | 12187 | 98 | Vamp8,Ctsa,Cd68,S100a6,Psme1,Fcer1g | | 2.392e-03 | -6.04 | GSE24142\_EARLY\_THYMIC\_PROGENITOR\_VS\_DN3\_THYMOCYTE\_FETAL\_UP | MSigDB lists | GSE24142\_EARLY\_THYMIC\_PROGENITOR\_VS\_DN3\_THYMOCYTE\_FETAL\_UP | 169 | 6 | 12187 | 98 | Cd68,Fam111a,Tsc22d1,Tspan4,Csf1r,Vim | | 2.401e-03 | -6.03 | cysteine-type endopeptidase activity | molecular function | GO:0004197 | 71 | 4 | 13516 | 107 | Ctsb,Ctss,Capn3,Ctsz | | 2.417e-03 | -6.03 | positive regulation of MAPK cascade | biological process | GO:0043410 | 427 | 10 | 13711 | 111 | Csf1r,Ptprc,Mt3,Ccl6,Il6ra,Csk,Kl,Fgf18,Tmem106a,C3 | | 2.460e-03 | -6.01 | positive regulation of leukocyte activation | biological process | GO:0002696 | 227 | 7 | 13711 | 111 | C1qa,Ptprc,Fcer1g,Itgam,Il6ra,Thy1,Vamp8 | | 2.464e-03 | -6.01 | GO\_REGULATED\_EXOCYTOSIS | MSigDB lists | GO\_REGULATED\_EXOCYTOSIS | 170 | 6 | 12187 | 98 | Fcer1g,Clu,Vwf,Vamp8,Cplx2,Serpina3n | | 2.464e-03 | -6.01 | GSE9988\_ANTI\_TREM1\_AND\_LPS\_VS\_VEHICLE\_TREATED\_MONOCYTES\_DN | MSigDB lists | GSE9988\_ANTI\_TREM1\_AND\_LPS\_VS\_VEHICLE\_TREATED\_MONOCYTES\_DN | 170 | 6 | 12187 | 98 | Lfng,Csk,Grn,Lrg1,Mpeg1,Csf1r | | 2.499e-03 | -5.99 | regulation of interleukin-6 production | biological process | GO:0032675 | 116 | 5 | 13711 | 111 | Cyba,Ccr5,Fcer1g,Csk,Il6ra | | 2.499e-03 | -5.99 | ZHONG\_SECRETOME\_OF\_LUNG\_CANCER\_AND\_FIBROBLAST | MSigDB lists | ZHONG\_SECRETOME\_OF\_LUNG\_CANCER\_AND\_FIBROBLAST | 117 | 5 | 12187 | 98 | Actb,Ctsb,Clu,Ldha,Ctsz | | 2.499e-03 | -5.99 | VERHAAK\_AML\_WITH\_NPM1\_MUTATED\_UP | MSigDB lists | VERHAAK\_AML\_WITH\_NPM1\_MUTATED\_UP | 117 | 5 | 12187 | 98 | S100a6,Clu,C1qa,Tmem176a,C1qb | | 2.508e-03 | -5.99 | plasma membrane part | cellular component | GO:0044459 | 1938 | 27 | 13825 | 111 | Dlgap3,Abhd17c,Lin7a,Slc2a1,Fcgr3,Ptprc,Il6ra,Hspa2,Vamp8,Ccr5,Ctsb,Vwf,Itgam,Slc26a2,Csf1r,Il1r1,Capn3,Adgre1,Fcer1g,Cyba,Sh2d5,Tspan4,S100a6,Myh9,Ly6a,Ahnak,Thy1 | | 2.518e-03 | -5.98 | GNF2\_CBFB | MSigDB lists | GNF2\_CBFB | 34 | 3 | 12187 | 98 | Fam111a,Csk,Psmb10 | | 2.518e-03 | -5.98 | GO\_POSITIVE\_REGULATION\_OF\_PHAGOCYTOSIS | MSigDB lists | GO\_POSITIVE\_REGULATION\_OF\_PHAGOCYTOSIS | 34 | 3 | 12187 | 98 | Fcer1g,C3,Cyba | | 2.518e-03 | -5.98 | GO\_MYELOID\_CELL\_ACTIVATION\_INVOLVED\_IN\_IMMUNE\_RESPONSE | MSigDB lists | GO\_MYELOID\_CELL\_ACTIVATION\_INVOLVED\_IN\_IMMUNE\_RESPONSE | 34 | 3 | 12187 | 98 | Vamp8,Cplx2,Fcer1g | | 2.530e-03 | -5.98 | GNF2\_STAT6 | MSigDB lists | GNF2\_STAT6 | 71 | 4 | 12187 | 98 | Psmb10,Hcls1,Cyba,Rac2 | | 2.537e-03 | -5.98 | GO\_ACTIVATION\_OF\_INNATE\_IMMUNE\_RESPONSE | MSigDB lists | GO\_ACTIVATION\_OF\_INNATE\_IMMUNE\_RESPONSE | 171 | 6 | 12187 | 98 | Ctss,Fcer1g,Itgam,Psme1,Psmb10,Ctsb | | 2.537e-03 | -5.98 | GSE3039\_NKT\_CELL\_VS\_ALPHABETA\_CD8\_TCELL\_UP | MSigDB lists | GSE3039\_NKT\_CELL\_VS\_ALPHABETA\_CD8\_TCELL\_UP | 171 | 6 | 12187 | 98 | C1qa,Tsc22d1,Psme1,Rps8,Ctss,C1qb | | 2.537e-03 | -5.98 | GSE22935\_WT\_VS\_MYD88\_KO\_MACROPHAGE\_48H\_MBOVIS\_BCG\_STIM\_DN | MSigDB lists | GSE22935\_WT\_VS\_MYD88\_KO\_MACROPHAGE\_48H\_MBOVIS\_BCG\_STIM\_DN | 171 | 6 | 12187 | 98 | Vim,Sh3bgrl3,Il1r1,C1qa,Csk,Brk1 | | 2.537e-03 | -5.98 | GSE17301\_CTRL\_VS\_48H\_ACD3\_ACD28\_STIM\_CD8\_TCELL\_DN | MSigDB lists | GSE17301\_CTRL\_VS\_48H\_ACD3\_ACD28\_STIM\_CD8\_TCELL\_DN | 171 | 6 | 12187 | 98 | Tmem176a,C3,Exoc3l4,Itgam,S100a6,Ctsz | | 2.612e-03 | -5.95 | GSE29618\_BCELL\_VS\_MONOCYTE\_DAY7\_FLU\_VACCINE\_DN | MSigDB lists | GSE29618\_BCELL\_VS\_MONOCYTE\_DAY7\_FLU\_VACCINE\_DN | 172 | 6 | 12187 | 98 | Csf1r,Lin7a,Ahnak,Grn,Cd68,Vim | | 2.612e-03 | -5.95 | GSE9988\_LOW\_LPS\_VS\_VEHICLE\_TREATED\_MONOCYTE\_DN | MSigDB lists | GSE9988\_LOW\_LPS\_VS\_VEHICLE\_TREATED\_MONOCYTE\_DN | 172 | 6 | 12187 | 98 | Mpeg1,Csk,Lfng,S100a6,Grn,Ctsb | | 2.612e-03 | -5.95 | GSE18893\_TCONV\_VS\_TREG\_2H\_CULTURE\_DN | MSigDB lists | GSE18893\_TCONV\_VS\_TREG\_2H\_CULTURE\_DN | 172 | 6 | 12187 | 98 | Psme1,Vim,Serpinb9,Tmem176a,Cyba,Psmb10 | | 2.627e-03 | -5.94 | CFL2 (cofilin 2) | protein interactions | 1073 | 7 | 2 | 6802 | 78 | Myh9,Flnb | | 2.627e-03 | -5.94 | ITPR3 (inositol 1,4,5-trisphosphate receptor type 3) | protein interactions | 3710 | 7 | 2 | 6802 | 78 | Actb,Myh9 | | 2.627e-03 | -5.94 | ARPC5 (actin related protein 2/3 complex subunit 5) | protein interactions | 10092 | 7 | 2 | 6802 | 78 | Flnb,Actb | | 2.627e-03 | -5.94 | ZDHHC5 (zinc finger DHHC-type containing 5) | protein interactions | 25921 | 7 | 2 | 6802 | 78 | Myh9,Flnb | | 2.627e-03 | -5.94 | LMO7 (LIM domain 7) | protein interactions | 4008 | 7 | 2 | 6802 | 78 | Myh9,Actb | | 2.627e-03 | -5.94 | ACTBL2 (actin beta like 2) | protein interactions | 345651 | 7 | 2 | 6802 | 78 | Myh9,Actb | | 2.627e-03 | -5.94 | MYO1E (myosin IE) | protein interactions | 4643 | 7 | 2 | 6802 | 78 | Actb,Myh9 | | 2.627e-03 | -5.94 | TPM3 (tropomyosin 3) | protein interactions | 7170 | 7 | 2 | 6802 | 78 | Actb,Myh9 | | 2.663e-03 | -5.93 | GO\_LYSOSOMAL\_LUMEN | MSigDB lists | GO\_LYSOSOMAL\_LUMEN | 72 | 4 | 12187 | 98 | Ctss,Fmod,Ctsb,Ctsa | | 2.672e-03 | -5.93 | MULLIGHAN\_MLL\_SIGNATURE\_1\_UP | MSigDB lists | MULLIGHAN\_MLL\_SIGNATURE\_1\_UP | 297 | 8 | 12187 | 98 | Il6ra,Rac2,Fcgr3,Ctsa,Cyba,Grn,S100a6,Hcls1 | | 2.689e-03 | -5.92 | GSE34156\_NOD2\_LIGAND\_VS\_TLR1\_TLR2\_LIGAND\_6H\_TREATED\_MONOCYTE\_UP | MSigDB lists | GSE34156\_NOD2\_LIGAND\_VS\_TLR1\_TLR2\_LIGAND\_6H\_TREATED\_MONOCYTE\_UP | 173 | 6 | 12187 | 98 | Psmb10,Apod,Serpinb9,Ly6e,Psme1,Mpeg1 | | 2.690e-03 | -5.92 | GO\_GLIAL\_CELL\_DIFFERENTIATION | MSigDB lists | GO\_GLIAL\_CELL\_DIFFERENTIATION | 119 | 5 | 12187 | 98 | Gfap,Lgi4,Clu,Vim,Csk | | 2.703e-03 | -5.91 | Phagosome | KEGG pathways | mmu04145 | 116 | 6 | 5248 | 64 | Ctss,Fcgr3,Cyba,C3,Actb,Itgam | | 2.703e-03 | -5.91 | Phagosome | KEGG pathways | ko04145 | 116 | 6 | 5248 | 64 | Itgam,Actb,Fcgr3,C3,Cyba,Ctss | | 2.731e-03 | -5.90 | positive regulation of ERK1 and ERK2 cascade | biological process | GO:0070374 | 172 | 6 | 13711 | 111 | C3,Fgf18,Ccl6,Csf1r,Ptprc,Mt3 | | 2.738e-03 | -5.90 | GO\_PLATELET\_ALPHA\_GRANULE\_LUMEN | MSigDB lists | GO\_PLATELET\_ALPHA\_GRANULE\_LUMEN | 35 | 3 | 12187 | 98 | Clu,Vwf,Serpina3n | | 2.738e-03 | -5.90 | KYNG\_RESPONSE\_TO\_H2O2\_VIA\_ERCC6\_DN | MSigDB lists | KYNG\_RESPONSE\_TO\_H2O2\_VIA\_ERCC6\_DN | 35 | 3 | 12187 | 98 | Rpl31,Rps5,Ctss | | 2.762e-03 | -5.89 | REACTOME\_PLATELET\_ADHESION\_TO\_EXPOSED\_COLLAGEN | MSigDB lists | REACTOME\_PLATELET\_ADHESION\_TO\_EXPOSED\_COLLAGEN | 10 | 2 | 12187 | 98 | Vwf,Fcer1g | | 2.762e-03 | -5.89 | WHITESIDE\_CISPLATIN\_RESISTANCE\_DN | MSigDB lists | WHITESIDE\_CISPLATIN\_RESISTANCE\_DN | 10 | 2 | 12187 | 98 | Rps5,Actb | | 2.767e-03 | -5.89 | GSE18281\_SUBCAPSULAR\_VS\_CENTRAL\_CORTICAL\_REGION\_OF\_THYMUS\_DN | MSigDB lists | GSE18281\_SUBCAPSULAR\_VS\_CENTRAL\_CORTICAL\_REGION\_OF\_THYMUS\_DN | 174 | 6 | 12187 | 98 | Lrg1,Fcer1g,Psmb10,Flnb,Apod,Tmem106a | | 2.799e-03 | -5.88 | antigen processing and presentation of peptide antigen | biological process | GO:0048002 | 35 | 3 | 13711 | 111 | Fcgr3,Ctss,Fcer1g | | 2.800e-03 | -5.88 | TAKAO\_RESPONSE\_TO\_UVB\_RADIATION\_UP | MSigDB lists | TAKAO\_RESPONSE\_TO\_UVB\_RADIATION\_UP | 73 | 4 | 12187 | 98 | Tsc22d1,Grn,Cox4i1,Rps5 | | 2.800e-03 | -5.88 | PDGF\_UP.V1\_DN | MSigDB lists | PDGF\_UP.V1\_DN | 73 | 4 | 12187 | 98 | Tnfsf8,Fcgr3,Slc26a2,Kl | | 2.800e-03 | -5.88 | KIM\_RESPONSE\_TO\_TSA\_AND\_DECITABINE\_UP | MSigDB lists | KIM\_RESPONSE\_TO\_TSA\_AND\_DECITABINE\_UP | 73 | 4 | 12187 | 98 | Vamp8,Clu,Hspa2,Hcls1 | | 2.800e-03 | -5.88 | chr11q12 | MSigDB lists | chr11q12 | 73 | 4 | 12187 | 98 | Rab3il1,Mpeg1,Fam111a,Ahnak | | 2.802e-03 | -5.88 | Fc receptor signaling pathway | biological process | GO:0038093 | 10 | 2 | 13711 | 111 | Fcgr3,Fcer1g | | 2.802e-03 | -5.88 | regulation of respiratory burst | biological process | GO:0060263 | 10 | 2 | 13711 | 111 | Rac2,Grn | | 2.802e-03 | -5.88 | Bergmann glial cell differentiation | biological process | GO:0060020 | 10 | 2 | 13711 | 111 | Gfap,Vim | | 2.802e-03 | -5.88 | regulation of Fc receptor mediated stimulatory signaling pathway | biological process | GO:0060368 | 10 | 2 | 13711 | 111 | Csk,Ptprc | | 2.816e-03 | -5.87 | THIOL\_PROTEASE\_HIS | prosite domains | PS00639 | 8 | 2 | 8845 | 91 | Ctsb,Ctss | | 2.842e-03 | -5.86 | GO\_ESTABLISHMENT\_OF\_PROTEIN\_LOCALIZATION\_TO\_ORGANELLE | MSigDB lists | GO\_ESTABLISHMENT\_OF\_PROTEIN\_LOCALIZATION\_TO\_ORGANELLE | 300 | 8 | 12187 | 98 | Rpl26,Rps20,Rpl14,Rps5,Rpl31,Ndufa13,Rps8,Rpl13 | | 2.929e-03 | -5.83 | GSE15330\_HSC\_VS\_MEGAKARYOCYTE\_ERYTHROID\_PROGENITOR\_IKAROS\_KO\_UP | MSigDB lists | GSE15330\_HSC\_VS\_MEGAKARYOCYTE\_ERYTHROID\_PROGENITOR\_IKAROS\_KO\_UP | 176 | 6 | 12187 | 98 | Rac2,Vim,Ldha,Ctsb,Ctsa,Slc2a1 | | 2.929e-03 | -5.83 | GSE7509\_DC\_VS\_MONOCYTE\_UP | MSigDB lists | GSE7509\_DC\_VS\_MONOCYTE\_UP | 176 | 6 | 12187 | 98 | Ctsb,C1qa,C1qb,Tspan4,Ccr5,Cd68 | | 2.942e-03 | -5.83 | GO\_PLATELET\_DEGRANULATION | MSigDB lists | GO\_PLATELET\_DEGRANULATION | 74 | 4 | 12187 | 98 | Fcer1g,Clu,Vwf,Serpina3n | | 2.944e-03 | -5.83 | mouse chr4 | chromosome location | mouse chr4 | 240 | 7 | 14556 | 115 | Cd52,Tnfsf8,Slc2a1,Rap1gap,C1qa,Dlgap3,C1qb | | 2.961e-03 | -5.82 | positive regulation of cell death | biological process | GO:0010942 | 591 | 12 | 13711 | 111 | Mt3,Camk2b,Ccr5,Ndufa13,Tsc22d1,Rpl26,Ptprc,C1qa,Clu,Ctsz,Ccar1,Grn | | 2.970e-03 | -5.82 | TAKEDA\_TARGETS\_OF\_NUP98\_HOXA9\_FUSION\_6HR\_DN | MSigDB lists | TAKEDA\_TARGETS\_OF\_NUP98\_HOXA9\_FUSION\_6HR\_DN | 36 | 3 | 12187 | 98 | Ptprc,Il6ra,Tnfsf8 | | 2.972e-03 | -5.82 | Immunoglobulin | interpro domains | IPR013151 | 35 | 3 | 13788 | 114 | Csf1r,Il6ra,Thy1 | | 2.997e-03 | -5.81 | NABA\_ECM\_GLYCOPROTEINS | MSigDB lists | NABA\_ECM\_GLYCOPROTEINS | 122 | 5 | 12187 | 98 | Mgp,Lrg1,Lgi4,Vwf,Igfbp6 | | 2.997e-03 | -5.81 | GO\_EXTERNAL\_SIDE\_OF\_PLASMA\_MEMBRANE | MSigDB lists | GO\_EXTERNAL\_SIDE\_OF\_PLASMA\_MEMBRANE | 122 | 5 | 12187 | 98 | Ccr5,Fcer1g,Thy1,Ctsb,Ptprc | | 3.012e-03 | -5.81 | POOLA\_INVASIVE\_BREAST\_CANCER\_UP | MSigDB lists | POOLA\_INVASIVE\_BREAST\_CANCER\_UP | 177 | 6 | 12187 | 98 | Ptprc,Fcgr3,Hcls1,Ctss,Cd68,Fcer1g | | 3.029e-03 | -5.80 | positive regulation of interleukin-6 production | biological process | GO:0032755 | 74 | 4 | 13711 | 111 | Il6ra,Fcer1g,Cyba,Ccr5 | | 3.036e-03 | -5.80 | myeloid cell activation involved in immune response | biological process | GO:0002275 | 36 | 3 | 13711 | 111 | Grn,Fcer1g,Cplx2 | | 3.036e-03 | -5.80 | positive regulation of tyrosine phosphorylation of STAT protein | biological process | GO:0042531 | 36 | 3 | 13711 | 111 | Il6ra,Csf1r,Hcls1 | | 3.042e-03 | -5.80 | MODULE\_27 | MSigDB lists | MODULE\_27 | 238 | 7 | 12187 | 98 | Fcer1g,Ccr5,Itgam,Fcgr3,Ptprc,Il1r1,Il6ra | | 3.068e-03 | -5.79 | synapse | cellular component | GO:0045202 | 1291 | 20 | 13825 | 111 | Camk2b,Rpl26,Clu,Mgll,Slc17a7,Sh2d5,Rpl31,Dlgap3,Rps20,Il1r1,C1qb,Myh9,Ptprn,Cplx2,Mt3,Rpl14,Lin7a,C1qa,Abhd17c,Actb | | 3.083e-03 | -5.78 | GO\_MACROMOLECULE\_CATABOLIC\_PROCESS | MSigDB lists | GO\_MACROMOLECULE\_CATABOLIC\_PROCESS | 764 | 14 | 12187 | 98 | Rps5,Rpl31,Rpl26,Ctsb,Ctsa,Rpl13,Fmod,Psme1,Ctss,Ctsz,Rps8,Rpl14,Rps20,Psmb10 | | 3.087e-03 | -5.78 | ig | pfam domains | PF00047 | 35 | 3 | 12881 | 108 | Thy1,Il6ra,Csf1r | | 3.089e-03 | -5.78 | LABBE\_TGFB1\_TARGETS\_UP | MSigDB lists | LABBE\_TGFB1\_TARGETS\_UP | 75 | 4 | 12187 | 98 | Vim,Tsc22d1,Ptprn,Actb | | 3.089e-03 | -5.78 | GO\_LARGE\_RIBOSOMAL\_SUBUNIT | MSigDB lists | GO\_LARGE\_RIBOSOMAL\_SUBUNIT | 75 | 4 | 12187 | 98 | Rpl31,Rpl14,Rpl26,Rpl13 | | 3.097e-03 | -5.78 | DAZARD\_RESPONSE\_TO\_UV\_NHEK\_UP | MSigDB lists | DAZARD\_RESPONSE\_TO\_UV\_NHEK\_UP | 178 | 6 | 12187 | 98 | Igfbp6,Cyba,Hspa2,Ly6e,Tsc22d1,Rpl31 | | 3.104e-03 | -5.78 | GO\_PLATELET\_ACTIVATION | MSigDB lists | GO\_PLATELET\_ACTIVATION | 123 | 5 | 12187 | 98 | Actb,Vwf,Rac2,Fcer1g,Myh9 | | 3.113e-03 | -5.77 | MODULE\_44 | MSigDB lists | MODULE\_44 | 239 | 7 | 12187 | 98 | Rac2,Ptprc,Apod,Il1r1,Grn,C1qb,Hcls1 | | 3.126e-03 | -5.77 | GO\_RESPONSE\_TO\_ENDOGENOUS\_STIMULUS | MSigDB lists | GO\_RESPONSE\_TO\_ENDOGENOUS\_STIMULUS | 1115 | 18 | 12187 | 98 | Fcer1g,Grn,Serpinb9,Mc4r,Ctsb,Spint1,C3,Hcls1,Kl,Fmod,Rap1gap,Csk,Ctss,Ldha,Fgf18,Cyba,Ptprn,Il1r1 | | 3.171e-03 | -5.75 | REN\_ALVEOLAR\_RHABDOMYOSARCOMA\_DN | MSigDB lists | REN\_ALVEOLAR\_RHABDOMYOSARCOMA\_DN | 375 | 9 | 12187 | 98 | Clu,Myh9,Ly6e,Grn,S100a6,Tspan4,Sh3bgrl3,Igfbp6,Ctsb | | 3.184e-03 | -5.75 | NABA\_CORE\_MATRISOME | MSigDB lists | NABA\_CORE\_MATRISOME | 179 | 6 | 12187 | 98 | Lrg1,Vwf,Fmod,Lgi4,Igfbp6,Mgp | | 3.222e-03 | -5.74 | IF\_rod\_dom\_coil1B | interpro domains | IPR042180 | 36 | 3 | 13788 | 114 | Ppfibp1,Vim,Gfap | | 3.272e-03 | -5.72 | Fancd2 (Fanconi anemia, complementation group D2) | protein interactions | 211651 | 1493 | 28 | 6802 | 78 | Ndufa13,Ldha,C1qb,Vim,Mpeg1,Hspa2,Slc2a1,Clu,C1qa,Rps5,Atp5j2,Actb,C3,Rac2,Psmb10,Rps20,Thy1,Ikzf3,Adgre1,Rpl13,Rps8,Ptprc,Rpl26,Rpl14,Myh9,Flnb,Adh1,Ahnak | | 3.273e-03 | -5.72 | KAYO\_AGING\_MUSCLE\_UP | MSigDB lists | KAYO\_AGING\_MUSCLE\_UP | 180 | 6 | 12187 | 98 | Actb,Serpina3n,Tmem106a,Hcls1,Grn,Mc4r | | 3.281e-03 | -5.72 | LEE\_BMP2\_TARGETS\_UP | MSigDB lists | LEE\_BMP2\_TARGETS\_UP | 605 | 12 | 12187 | 98 | Grn,Tsc22d1,Tmem176a,Adh1,Igfbp6,Cplx2,Ctsa,Ctsb,Clu,Vwf,Mgp,Apod | | 3.284e-03 | -5.72 | neuron maturation | biological process | GO:0042551 | 37 | 3 | 13711 | 111 | C3,Lgi4,C1qa | | 3.311e-03 | -5.71 | GO\_REGULATION\_OF\_RESPONSE\_TO\_STRESS | MSigDB lists | GO\_REGULATION\_OF\_RESPONSE\_TO\_STRESS | 1121 | 18 | 12187 | 98 | Serpinb9,Itgam,Fcer1g,Dek,Vamp8,Ctsb,Ctss,Mgll,Camk2b,Capn3,Clu,Psme1,C3,Psmb10,Il1r1,Phyhip,Apod,Cyba | | 3.327e-03 | -5.71 | GSE43863\_TH1\_VS\_LY6C\_INT\_CXCR5POS\_MEMORY\_CD4\_TCELL\_DN | MSigDB lists | GSE43863\_TH1\_VS\_LY6C\_INT\_CXCR5POS\_MEMORY\_CD4\_TCELL\_DN | 125 | 5 | 12187 | 98 | Cd68,C1qa,Ctsb,Csf1r,C1qb | | 3.336e-03 | -5.70 | regulation of leukocyte apoptotic process | biological process | GO:2000106 | 76 | 4 | 13711 | 111 | Ccr5,Fcer1g,Serpinb9,Hcls1 | | 3.349e-03 | -5.70 | cellular developmental process | biological process | GO:0048869 | 2887 | 36 | 13711 | 111 | Grn,Mgll,Il1r1,Thy1,Clu,Hspa2,Ptprc,C1qa,Gfap,Fgf18,Capn3,Camk2b,Actb,Itgam,Fcer1g,Bhlhe41,Psmb10,Vim,Cplx2,Rap1gap,Lrg1,Ctsz,Tnfsf8,Flnb,C3,Ccr5,Mgp,Tfap2c,Lfng,Spint1,Rac2,Lgi4,Mt3,Myh9,Hcls1,Csf1r | | 3.351e-03 | -5.70 | high-density lipoprotein particle | cellular component | GO:0034364 | 11 | 2 | 13825 | 111 | Clu,Saa3 | | 3.358e-03 | -5.70 | GO\_ENDOLYSOSOME | MSigDB lists | GO\_ENDOLYSOSOME | 11 | 2 | 12187 | 98 | Ctss,Ctsb | | 3.358e-03 | -5.70 | GO\_POSITIVE\_REGULATION\_OF\_PROTEIN\_MATURATION | MSigDB lists | GO\_POSITIVE\_REGULATION\_OF\_PROTEIN\_MATURATION | 11 | 2 | 12187 | 98 | Myh9,C3 | | 3.358e-03 | -5.70 | GO\_S100\_PROTEIN\_BINDING | MSigDB lists | GO\_S100\_PROTEIN\_BINDING | 11 | 2 | 12187 | 98 | Ahnak,S100a6 | | 3.358e-03 | -5.70 | NIELSEN\_SCHWANNOMA\_UP | MSigDB lists | NIELSEN\_SCHWANNOMA\_UP | 11 | 2 | 12187 | 98 | Gfap,Slc26a2 | | 3.358e-03 | -5.70 | GO\_CELLULAR\_RESPONSE\_TO\_THYROID\_HORMONE\_STIMULUS | MSigDB lists | GO\_CELLULAR\_RESPONSE\_TO\_THYROID\_HORMONE\_STIMULUS | 11 | 2 | 12187 | 98 | Ctsb,Ctss | | 3.358e-03 | -5.70 | GO\_POSITIVE\_REGULATION\_OF\_B\_CELL\_MEDIATED\_IMMUNITY | MSigDB lists | GO\_POSITIVE\_REGULATION\_OF\_B\_CELL\_MEDIATED\_IMMUNITY | 11 | 2 | 12187 | 98 | C3,Fcer1g | | 3.358e-03 | -5.70 | MODULE\_478 | MSigDB lists | MODULE\_478 | 11 | 2 | 12187 | 98 | Fcgr3,Fcer1g | | 3.358e-03 | -5.70 | GO\_REGULATION\_OF\_INFLAMMATORY\_RESPONSE\_TO\_ANTIGENIC\_STIMULUS | MSigDB lists | GO\_REGULATION\_OF\_INFLAMMATORY\_RESPONSE\_TO\_ANTIGENIC\_STIMULUS | 11 | 2 | 12187 | 98 | C3,Fcer1g | | 3.358e-03 | -5.70 | GO\_INTERMEDIATE\_FILAMENT\_ORGANIZATION | MSigDB lists | GO\_INTERMEDIATE\_FILAMENT\_ORGANIZATION | 11 | 2 | 12187 | 98 | Gfap,Vim | | 3.396e-03 | -5.69 | contractile actin filament bundle | cellular component | GO:0097517 | 77 | 4 | 13825 | 111 | Myh9,Actb,Cyba,Flnb | | 3.396e-03 | -5.69 | stress fiber | cellular component | GO:0001725 | 77 | 4 | 13825 | 111 | Flnb,Cyba,Myh9,Actb | | 3.443e-03 | -5.67 | LANDIS\_ERBB2\_BREAST\_TUMORS\_324\_DN | MSigDB lists | LANDIS\_ERBB2\_BREAST\_TUMORS\_324\_DN | 126 | 5 | 12187 | 98 | Igfbp6,Ahnak,Tsc22d1,S100a6,Vwf | | 3.443e-03 | -5.67 | GO\_RESPONSE\_TO\_CARBOHYDRATE | MSigDB lists | GO\_RESPONSE\_TO\_CARBOHYDRATE | 126 | 5 | 12187 | 98 | Il1r1,Ptprn,Cyba,Ctsb,Ldha | | 3.443e-03 | -5.67 | GSE3920\_IFNB\_VS\_IFNG\_TREATED\_ENDOTHELIAL\_CELL\_UP | MSigDB lists | GSE3920\_IFNB\_VS\_IFNG\_TREATED\_ENDOTHELIAL\_CELL\_UP | 126 | 5 | 12187 | 98 | Il1r1,Treml2,Rps20,Rpl31,Tmem176a | | 3.444e-03 | -5.67 | GO\_IDENTICAL\_PROTEIN\_BINDING | MSigDB lists | GO\_IDENTICAL\_PROTEIN\_BINDING | 946 | 16 | 12187 | 98 | Myh9,S100a6,Slc2a1,C1qb,Csf1r,Il6ra,Actb,Csk,Ldha,Camk2b,Rap1gap,Vwf,Mgll,Vim,Ikzf3,Flnb | | 3.456e-03 | -5.67 | GO\_EXCITATORY\_SYNAPSE | MSigDB lists | GO\_EXCITATORY\_SYNAPSE | 182 | 6 | 12187 | 98 | Camk2b,Sh2d5,Dlgap3,Slc17a7,Il1r1,Lin7a | | 3.467e-03 | -5.66 | cell cortex part | cellular component | GO:0044448 | 182 | 6 | 13825 | 111 | Ctsz,Slc2a1,Actb,Myh9,Exoc3l4,Hcls1 | | 3.469e-03 | -5.66 | NIKOLSKY\_BREAST\_CANCER\_16Q24\_AMPLICON | MSigDB lists | NIKOLSKY\_BREAST\_CANCER\_16Q24\_AMPLICON | 38 | 3 | 12187 | 98 | Rpl13,Cox4i1,Cyba | | 3.469e-03 | -5.66 | GNF2\_PTPN6 | MSigDB lists | GNF2\_PTPN6 | 38 | 3 | 12187 | 98 | Rac2,Hcls1,Ptprc | | 3.469e-03 | -5.66 | COLIN\_PILOCYTIC\_ASTROCYTOMA\_VS\_GLIOBLASTOMA\_UP | MSigDB lists | COLIN\_PILOCYTIC\_ASTROCYTOMA\_VS\_GLIOBLASTOMA\_UP | 38 | 3 | 12187 | 98 | Gfap,Serpina3n,Apod | | 3.469e-03 | -5.66 | GO\_SERINE\_TYPE\_ENDOPEPTIDASE\_INHIBITOR\_ACTIVITY | MSigDB lists | GO\_SERINE\_TYPE\_ENDOPEPTIDASE\_INHIBITOR\_ACTIVITY | 38 | 3 | 12187 | 98 | Serpina3n,Spint1,Serpinb9 | | 3.469e-03 | -5.66 | HOFFMANN\_SMALL\_PRE\_BII\_TO\_IMMATURE\_B\_LYMPHOCYTE\_DN | MSigDB lists | HOFFMANN\_SMALL\_PRE\_BII\_TO\_IMMATURE\_B\_LYMPHOCYTE\_DN | 38 | 3 | 12187 | 98 | Cyba,Fcgr3,Myh9 | | 3.476e-03 | -5.66 | CD44 (CD44 molecule (Indian blood group)) | protein interactions | 960 | 8 | 2 | 6802 | 78 | Flnb,Myh9 | | 3.476e-03 | -5.66 | ACTR3 (actin related protein 3) | protein interactions | 10096 | 8 | 2 | 6802 | 78 | Actb,Myh9 | | 3.476e-03 | -5.66 | CD59 (CD59 molecule (CD59 blood group)) | protein interactions | 966 | 8 | 2 | 6802 | 78 | Myh9,Flnb | | 3.476e-03 | -5.66 | CEP162 (centrosomal protein 162) | protein interactions | 22832 | 8 | 2 | 6802 | 78 | Actb,Myh9 | | 3.476e-03 | -5.66 | MYO6 (myosin VI) | protein interactions | 4646 | 8 | 2 | 6802 | 78 | Actb,Myh9 | | 3.476e-03 | -5.66 | KCTD10 (potassium channel tetramerization domain containing 10) | protein interactions | 83892 | 8 | 2 | 6802 | 78 | Myh9,Actb | | 3.486e-03 | -5.66 | regulation of biological process | biological process | GO:0050789 | 7871 | 78 | 13711 | 111 | Zfp786,Ptprn,C3,Tfap2c,Serpina3n,Saa3,Ctsa,Rac2,Ifitm1,Spint1,Csf1r,Hcls1,Lgi4,Brk1,Vim,Lrg1,Ccl6,Tnfsf8,Rpl26,Tsc22d1,Gfap,Camk2b,Il6ra,Serpinb9,Capn3,Ctsb,Slc17a7,Itgam,Treml2,Mc4r,Tmem106a,Rbfox3,Abhd17c,Cd68,Clu,Tmem176a,Sh3bgrl3,Hspa2,Apod,Lfng,Cyba,C1qb,Ccr5,Mgp,Ndufa13,Csk,Mark4,Myh9,Mt3,Fam111a,Cplx2,Klhl6,Dek,Rap1gap,Vamp8,Igfbp6,Ctsz,Fcgr3,Ahnak,Ly6e,Psme1,Fgf18,Bhlhe41,Fcer1g,Pde1a,Actb,Adgre1,Grn,Ccar1,Ikzf3,Ctss,Kl,Thy1,Mgll,Il1r1,Trnp1,Ptprc,C1qa | | 3.498e-03 | -5.66 | interaction with host | biological process | GO:0051701 | 77 | 4 | 13711 | 111 | Vamp8,Ctsb,Serpinb9,Ccr5 | | 3.513e-03 | -5.65 | cellular component disassembly | biological process | GO:0022411 | 181 | 6 | 13711 | 111 | C3,C1qb,Ctss,Hspa2,Vamp8,C1qa | | 3.519e-03 | -5.65 | vesicle | cellular component | GO:0031982 | 1591 | 23 | 13825 | 111 | Ctsz,Tspan4,Abhd17c,Slc2a1,Flnb,Rac2,Ifitm1,Ptprn,Rap1gap,Mt3,Vamp8,Ccr5,Ahnak,Vim,Pcsk2,Vwf,Cd68,Mpeg1,Clu,Ctss,Grn,Slc17a7,Cyba | | 3.525e-03 | -5.65 | cell surface receptor signaling pathway | biological process | GO:0007166 | 1296 | 20 | 13711 | 111 | Vim,Klhl6,Adgre1,Il1r1,Kl,Thy1,Ccl6,Ptprc,Fcgr3,C3,Ptprn,Ndufa13,Ccr5,Fgf18,Il6ra,Fcer1g,Slc17a7,Ifitm1,Mt3,Csf1r | | 3.545e-03 | -5.64 | T cell proliferation | biological process | GO:0042098 | 38 | 3 | 13711 | 111 | Ptprc,Psmb10,Itgam | | 3.545e-03 | -5.64 | leukocyte cell-cell adhesion | biological process | GO:0007159 | 38 | 3 | 13711 | 111 | Itgam,Rac2,Ptprc | | 3.545e-03 | -5.64 | regulation of bone resorption | biological process | GO:0045124 | 38 | 3 | 13711 | 111 | Mc4r,Csk,Csf1r | | 3.551e-03 | -5.64 | KARLSSON\_TGFB1\_TARGETS\_DN | MSigDB lists | KARLSSON\_TGFB1\_TARGETS\_DN | 183 | 6 | 12187 | 98 | Grn,Ly6e,Ctsz,Ctsb,Ctsa,Vamp8 | | 3.559e-03 | -5.64 | regulation of cellular localization | biological process | GO:0060341 | 852 | 15 | 13711 | 111 | Apod,Abhd17c,Vamp8,Thy1,Cplx2,Hcls1,Actb,Fcer1g,Slc17a7,Rac2,Mark4,Capn3,Csk,Camk2b,Cyba | | 3.561e-03 | -5.64 | GSE3720\_UNSTIM\_VS\_LPS\_STIM\_VD2\_GAMMADELTA\_TCELL\_UP | MSigDB lists | GSE3720\_UNSTIM\_VS\_LPS\_STIM\_VD2\_GAMMADELTA\_TCELL\_UP | 127 | 5 | 12187 | 98 | Tnfsf8,Rps8,Ly6e,Rps20,Rpl14 | | 3.561e-03 | -5.64 | DANG\_MYC\_TARGETS\_UP | MSigDB lists | DANG\_MYC\_TARGETS\_UP | 127 | 5 | 12187 | 98 | Ldha,Rpl26,Rpl13,Rps20,Rps5 | | 3.609e-03 | -5.62 | GO\_REGULATION\_OF\_CELL\_ACTIVATION | MSigDB lists | GO\_REGULATION\_OF\_CELL\_ACTIVATION | 312 | 8 | 12187 | 98 | Rac2,Ikzf3,Vamp8,Ptprc,Capn3,Fcer1g,Thy1,Csk | | 3.627e-03 | -5.62 | neuron projection | cellular component | GO:0043005 | 1310 | 20 | 13825 | 111 | Dlgap3,Vim,Pcsk2,Il1r1,Rpl26,Camk2b,Apod,Clu,Mgll,Cyba,Slc17a7,Ctsz,Lin7a,Actb,Flnb,Ptprn,Mark4,Rap1gap,Cplx2,Thy1 | | 3.672e-03 | -5.61 | regulation of ERK1 and ERK2 cascade | biological process | GO:0070372 | 244 | 7 | 13711 | 111 | C3,Fgf18,Csk,Ccl6,Mt3,Csf1r,Ptprc | | 3.683e-03 | -5.60 | GSE7509\_UNSTIM\_VS\_FCGRIIB\_STIM\_DC\_DN | MSigDB lists | GSE7509\_UNSTIM\_VS\_FCGRIIB\_STIM\_DC\_DN | 128 | 5 | 12187 | 98 | Serpinb9,Ahnak,Psmb10,Klhl6,Cyba | | 3.683e-03 | -5.60 | GO\_PHAGOCYTOSIS | MSigDB lists | GO\_PHAGOCYTOSIS | 128 | 5 | 12187 | 98 | Fcgr3,Actb,Fcer1g,Myh9,Brk1 | | 3.689e-03 | -5.60 | cell population proliferation | biological process | GO:0008283 | 453 | 10 | 13711 | 111 | Csf1r,Ptprc,Lgi4,Psmb10,Itgam,Clu,Trnp1,Fgf18,Tfap2c,Grn | | 3.708e-03 | -5.60 | exocytosis | biological process | GO:0006887 | 183 | 6 | 13711 | 111 | Cplx2,Exoc3l4,Vamp8,Lin7a,Fcer1g,Myh9 | | 3.726e-03 | -5.59 | JAZAG\_TGFB1\_SIGNALING\_VIA\_SMAD4\_UP | MSigDB lists | JAZAG\_TGFB1\_SIGNALING\_VIA\_SMAD4\_UP | 79 | 4 | 12187 | 98 | Ptprc,C1qb,Actb,Camk2b | | 3.726e-03 | -5.59 | GO\_PEPTIDASE\_INHIBITOR\_ACTIVITY | MSigDB lists | GO\_PEPTIDASE\_INHIBITOR\_ACTIVITY | 79 | 4 | 12187 | 98 | Serpinb9,C3,Serpina3n,Spint1 | | 3.736e-03 | -5.59 | GO\_CYTOSOLIC\_SMALL\_RIBOSOMAL\_SUBUNIT | MSigDB lists | GO\_CYTOSOLIC\_SMALL\_RIBOSOMAL\_SUBUNIT | 39 | 3 | 12187 | 98 | Rps20,Rps5,Rps8 | | 3.736e-03 | -5.59 | DAUER\_STAT3\_TARGETS\_UP | MSigDB lists | DAUER\_STAT3\_TARGETS\_UP | 39 | 3 | 12187 | 98 | Flnb,C3,Ctsb | | 3.736e-03 | -5.59 | WUNDER\_INFLAMMATORY\_RESPONSE\_AND\_CHOLESTEROL\_UP | MSigDB lists | WUNDER\_INFLAMMATORY\_RESPONSE\_AND\_CHOLESTEROL\_UP | 39 | 3 | 12187 | 98 | Ctss,Psmb10,Mpeg1 | | 3.736e-03 | -5.59 | REACTOME\_PHOSPHOLIPASE\_C\_MEDIATED\_CASCADE | MSigDB lists | REACTOME\_PHOSPHOLIPASE\_C\_MEDIATED\_CASCADE | 39 | 3 | 12187 | 98 | Kl,Fgf18,Pde1a | | 3.736e-03 | -5.59 | GO\_POSITIVE\_REGULATION\_OF\_CALCIUM\_ION\_TRANSMEMBRANE\_TRANSPORT | MSigDB lists | GO\_POSITIVE\_REGULATION\_OF\_CALCIUM\_ION\_TRANSMEMBRANE\_TRANSPORT | 39 | 3 | 12187 | 98 | Hspa2,Thy1,Capn3 | | 3.755e-03 | -5.58 | cell differentiation | biological process | GO:0030154 | 2800 | 35 | 13711 | 111 | Tnfsf8,Flnb,Ctsz,Lrg1,Rap1gap,Cplx2,Vim,Myh9,Csf1r,Hcls1,Lgi4,Mt3,Rac2,Spint1,Lfng,C3,Mgp,Ccr5,Tfap2c,Ptprc,C1qa,Hspa2,Clu,Thy1,Mgll,Il1r1,Grn,Bhlhe41,Fcer1g,Actb,Itgam,Camk2b,Fgf18,Capn3,Gfap | | 3.777e-03 | -5.58 | Allograft Rejection | WikiPathways | WP2328 | 7 | 2 | 3756 | 52 | C1qb,C1qa | | 3.808e-03 | -5.57 | GSE3565\_DUSP1\_VS\_WT\_SPLENOCYTES\_POST\_LPS\_INJECTION\_UP | MSigDB lists | GSE3565\_DUSP1\_VS\_WT\_SPLENOCYTES\_POST\_LPS\_INJECTION\_UP | 129 | 5 | 12187 | 98 | Treml2,Rps20,Rps5,Tnfsf8,Rpl13 | | 3.818e-03 | -5.57 | plasma membrane invagination | biological process | GO:0099024 | 39 | 3 | 13711 | 111 | Myh9,Fcgr3,Fcer1g | | 3.824e-03 | -5.57 | MODULE\_2 | MSigDB lists | MODULE\_2 | 315 | 8 | 12187 | 98 | Mgll,Vwf,Thy1,Il1r1,Apod,Ptprc,Serpina3n,Igfbp6 | | 3.845e-03 | -5.56 | GO\_APICAL\_PLASMA\_MEMBRANE | MSigDB lists | GO\_APICAL\_PLASMA\_MEMBRANE | 186 | 6 | 12187 | 98 | Il6ra,Thy1,Kl,Slc2a1,Cyba,Ctsb | | 3.866e-03 | -5.56 | CLTC (clathrin heavy chain) | protein interactions | 1213 | 28 | 3 | 6802 | 78 | Flnb,Myh9,Actb | | 3.891e-03 | -5.55 | glucose transmembrane transporter activity | molecular function | GO:0005355 | 12 | 2 | 13516 | 107 | Slc2a1,Mfsd4a | | 3.891e-03 | -5.55 | S100 protein binding | molecular function | GO:0044548 | 12 | 2 | 13516 | 107 | Ahnak,S100a6 | | 3.891e-03 | -5.55 | hexose transmembrane transporter activity | molecular function | GO:0015149 | 12 | 2 | 13516 | 107 | Mfsd4a,Slc2a1 | | 3.898e-03 | -5.55 | GO\_INTEGRIN\_BINDING | MSigDB lists | GO\_INTEGRIN\_BINDING | 80 | 4 | 12187 | 98 | Tspan4,Thy1,Gfap,Vwf | | 3.898e-03 | -5.55 | GUO\_TARGETS\_OF\_IRS1\_AND\_IRS2 | MSigDB lists | GUO\_TARGETS\_OF\_IRS1\_AND\_IRS2 | 80 | 4 | 12187 | 98 | Flnb,Actb,Grn,Il6ra | | 3.906e-03 | -5.55 | molecular function regulator | molecular function | GO:0098772 | 1245 | 19 | 13516 | 107 | Il6ra,Grn,Fgf18,Wfdc17,Ly6e,Rab3il1,Serpina3n,Saa3,Capn3,Spint1,Serpinb9,Vamp8,Thy1,Tnfsf8,C3,Ccl6,Psme1,Rap1gap,Rac2 | | 3.935e-03 | -5.54 | KEGG\_CYTOKINE\_CYTOKINE\_RECEPTOR\_INTERACTION | MSigDB lists | KEGG\_CYTOKINE\_CYTOKINE\_RECEPTOR\_INTERACTION | 130 | 5 | 12187 | 98 | Tnfsf8,Il6ra,Ccr5,Il1r1,Csf1r | | 3.935e-03 | -5.54 | GSE22611\_NOD2\_TRANSD\_VS\_CTRL\_TRANSD\_HEK293\_MDP\_STIM\_6H\_DN | MSigDB lists | GSE22611\_NOD2\_TRANSD\_VS\_CTRL\_TRANSD\_HEK293\_MDP\_STIM\_6H\_DN | 130 | 5 | 12187 | 98 | Fgf18,Fcgr3,Serpinb9,Tfap2c,Mgll | | 3.936e-03 | -5.54 | positive regulation of protein metabolic process | biological process | GO:0051247 | 1402 | 21 | 13711 | 111 | C3,Ndufa13,Csk,Il6ra,Fgf18,Capn3,Psme1,Myh9,Csf1r,Hcls1,Mt3,Vim,Tmem106a,Grn,Kl,Ccl6,Clu,Ptprc,Fcgr3,Hspa2,Rpl26 | | 3.947e-03 | -5.53 | GO\_REGULATION\_OF\_INFLAMMATORY\_RESPONSE | MSigDB lists | GO\_REGULATION\_OF\_INFLAMMATORY\_RESPONSE | 187 | 6 | 12187 | 98 | Mgll,Fcer1g,Vamp8,C3,Il1r1,Apod | | 3.947e-03 | -5.53 | HELLER\_SILENCED\_BY\_METHYLATION\_UP | MSigDB lists | HELLER\_SILENCED\_BY\_METHYLATION\_UP | 187 | 6 | 12187 | 98 | S100a6,Clu,Capn3,Ctss,Il6ra,Vamp8 | | 3.949e-03 | -5.53 | positive regulation of cellular protein metabolic process | biological process | GO:0032270 | 1309 | 20 | 13711 | 111 | Il6ra,Csk,Capn3,Psme1,Fgf18,Ndufa13,C3,Hcls1,Csf1r,Myh9,Mt3,Kl,Grn,Tmem106a,Vim,Ptprc,Rpl26,Hspa2,Clu,Ccl6 | | 3.988e-03 | -5.52 | MODULE\_19 | MSigDB lists | MODULE\_19 | 250 | 7 | 12187 | 98 | C1qb,Hcls1,Igfbp6,Slc26a2,Il1r1,Apod,Phyhip | | 4.008e-03 | -5.52 | MASRI\_RESISTANCE\_TO\_TAMOXIFEN\_AND\_AROMATASE\_INHIBITORS\_DN | MSigDB lists | MASRI\_RESISTANCE\_TO\_TAMOXIFEN\_AND\_AROMATASE\_INHIBITORS\_DN | 12 | 2 | 12187 | 98 | Apod,Pcsk2 | | 4.008e-03 | -5.52 | GO\_POSITIVE\_REGULATION\_OF\_LEUKOCYTE\_DEGRANULATION | MSigDB lists | GO\_POSITIVE\_REGULATION\_OF\_LEUKOCYTE\_DEGRANULATION | 12 | 2 | 12187 | 98 | Fcer1g,Vamp8 | | 4.008e-03 | -5.52 | KANG\_FLUOROURACIL\_RESISTANCE\_DN | MSigDB lists | KANG\_FLUOROURACIL\_RESISTANCE\_DN | 12 | 2 | 12187 | 98 | Flnb,Slc2a1 | | 4.013e-03 | -5.52 | cellular response to interferon-gamma | biological process | GO:0071346 | 80 | 4 | 13711 | 111 | Flnb,Vamp8,Ccl6,Vim | | 4.013e-03 | -5.52 | positive regulation of gliogenesis | biological process | GO:0014015 | 80 | 4 | 13711 | 111 | Spint1,Vim,Gfap,C1qa | | 4.016e-03 | -5.52 | GO\_REGULATION\_OF\_TYROSINE\_PHOSPHORYLATION\_OF\_STAT\_PROTEIN | MSigDB lists | GO\_REGULATION\_OF\_TYROSINE\_PHOSPHORYLATION\_OF\_STAT\_PROTEIN | 40 | 3 | 12187 | 98 | Csf1r,Hcls1,Il6ra | | 4.030e-03 | -5.51 | Leishmaniasis | KEGG pathways | ko05140 | 54 | 4 | 5248 | 64 | Itgam,Cyba,C3,Fcgr3 | | 4.030e-03 | -5.51 | Leishmaniasis | KEGG pathways | mmu05140 | 54 | 4 | 5248 | 64 | Fcgr3,C3,Cyba,Itgam | | 4.066e-03 | -5.51 | IWANAGA\_CARCINOGENESIS\_BY\_KRAS\_UP | MSigDB lists | IWANAGA\_CARCINOGENESIS\_BY\_KRAS\_UP | 131 | 5 | 12187 | 98 | Spint1,Lrg1,Clu,Ldha,Ctsz | | 4.066e-03 | -5.51 | PIGF\_UP.V1\_DN | MSigDB lists | PIGF\_UP.V1\_DN | 131 | 5 | 12187 | 98 | Serpina3n,Camk2b,Igfbp6,Thy1,Ly6e | | 4.075e-03 | -5.50 | phagocytic vesicle | cellular component | GO:0045335 | 81 | 4 | 13825 | 111 | Mpeg1,Vim,Flnb,Vamp8 | | 4.076e-03 | -5.50 | GCM\_CSNK2B | MSigDB lists | GCM\_CSNK2B | 81 | 4 | 12187 | 98 | Psme1,Rps8,Rps5,Rpl14 | | 4.126e-03 | -5.49 | Elf5 (E74-like factor 5) | protein interactions | 13711 | 93 | 5 | 6802 | 78 | Rpl13,Rpl14,Rpl26,Rpl31,Tfap2c | | 4.158e-03 | -5.48 | MCBRYAN\_PUBERTAL\_BREAST\_4\_5WK\_UP | MSigDB lists | MCBRYAN\_PUBERTAL\_BREAST\_4\_5WK\_UP | 189 | 6 | 12187 | 98 | Tfap2c,Vwf,Slc2a1,Igfbp6,Apod,Spint1 | | 4.172e-03 | -5.48 | generation of neurons | biological process | GO:0048699 | 1409 | 21 | 13711 | 111 | Lgi4,Mt3,Csf1r,Spint1,Actb,Bhlhe41,Rac2,Camk2b,C3,Gfap,Ccr5,Tfap2c,C1qa,Clu,Ctsz,Rap1gap,Mgll,Il1r1,Thy1,Vim,Grn | | 4.180e-03 | -5.48 | neurogenesis | biological process | GO:0022008 | 1504 | 22 | 13711 | 111 | C1qa,Ctsz,Clu,Thy1,Il1r1,Mgll,Rap1gap,Vim,Grn,Csf1r,Mt3,Lgi4,Bhlhe41,Rac2,Itgam,Actb,Spint1,Camk2b,Ccr5,Tfap2c,C3,Gfap | | 4.194e-03 | -5.47 | GO\_CYTOPLASMIC\_VESICLE\_PART | MSigDB lists | GO\_CYTOPLASMIC\_VESICLE\_PART | 466 | 10 | 12187 | 98 | Clu,Camk2b,Pcsk2,Vwf,Ptprn,Cyba,Serpina3n,Rac2,Slc17a7,Vamp8 | | 4.195e-03 | -5.47 | myeloid leukocyte migration | biological process | GO:0097529 | 81 | 4 | 13711 | 111 | Fcgr3,Ccl6,Itgam,Fcer1g | | 4.215e-03 | -5.47 | regulation of calcium ion transmembrane transport | biological process | GO:1903169 | 131 | 5 | 13711 | 111 | Cyba,Ahnak,Thy1,Hspa2,Capn3 | | 4.259e-03 | -5.46 | GO\_ANTIGEN\_PROCESSING\_AND\_PRESENTATION\_OF\_PEPTIDE\_ANTIGEN\_VIA\_MHC\_CLASS\_I | MSigDB lists | GO\_ANTIGEN\_PROCESSING\_AND\_PRESENTATION\_OF\_PEPTIDE\_ANTIGEN\_VIA\_MHC\_CLASS\_I | 82 | 4 | 12187 | 98 | Cyba,Psmb10,Fcer1g,Psme1 | | 4.432e-03 | -5.42 | - | gene3d domains | 1.20.5.1160 | 36 | 3 | 6647 | 62 | Vim,Gfap,Ppfibp1 | | 4.436e-03 | -5.42 | Rnf123 (ring finger protein 123) | protein interactions | 84585 | 9 | 2 | 6802 | 78 | Flnb,Myh9 | | 4.436e-03 | -5.42 | CTTN (cortactin) | protein interactions | 2017 | 9 | 2 | 6802 | 78 | Flnb,Actb | | 4.436e-03 | -5.42 | TWF1 (twinfilin actin binding protein 1) | protein interactions | 5756 | 9 | 2 | 6802 | 78 | Actb,Myh9 | | 4.436e-03 | -5.42 | MICAL3 (microtubule associated monooxygenase, calponin and LIM domain containing 3) | protein interactions | 57553 | 9 | 2 | 6802 | 78 | Myh9,Actb | | 4.446e-03 | -5.42 | GO\_SECRETORY\_VESICLE | MSigDB lists | GO\_SECRETORY\_VESICLE | 323 | 8 | 12187 | 98 | Clu,Pcsk2,Vwf,Cyba,Ptprn,Serpina3n,Slc17a7,Vamp8 | | 4.455e-03 | -5.41 | positive regulation of multicellular organismal process | biological process | GO:0051240 | 1512 | 22 | 13711 | 111 | Spint1,Fcer1g,Hcls1,Csf1r,Ccr5,Gfap,C3,Cyba,Fgf18,Il6ra,Camk2b,Clu,C1qa,Fcgr3,Ptprc,Grn,Vim,Mc4r,Vamp8,Il1r1,Lrg1,Kl | | 4.476e-03 | -5.41 | GSE21360\_SECONDARY\_VS\_QUATERNARY\_MEMORY\_CD8\_TCELL\_DN | MSigDB lists | GSE21360\_SECONDARY\_VS\_QUATERNARY\_MEMORY\_CD8\_TCELL\_DN | 134 | 5 | 12187 | 98 | Itgam,Lrg1,Ldha,Tmem176a,Slc2a1 | | 4.481e-03 | -5.41 | GO\_CELL\_PROJECTION | MSigDB lists | GO\_CELL\_PROJECTION | 1436 | 21 | 12187 | 98 | Camk2b,Clu,Rap1gap,Dlgap3,Rac2,Vim,Sh3bgrl3,Cyba,Ptprn,Apod,Il1r1,Thy1,Myh9,Pcsk2,S100a6,Brk1,Mark4,Gfap,Slc17a7,Cplx2,Lin7a | | 4.563e-03 | -5.39 | Leukocyte transendothelial migration | KEGG pathways | mmu04670 | 90 | 5 | 5248 | 64 | Cyba,Rac2,Actb,Thy1,Itgam | | 4.563e-03 | -5.39 | Leukocyte transendothelial migration | KEGG pathways | ko04670 | 90 | 5 | 5248 | 64 | Cyba,Thy1,Rac2,Actb,Itgam | | 4.575e-03 | -5.39 | monosaccharide transmembrane transporter activity | molecular function | GO:0015145 | 13 | 2 | 13516 | 107 | Slc2a1,Mfsd4a | | 4.577e-03 | -5.39 | regulation of receptor signaling pathway via JAK-STAT | biological process | GO:0046425 | 83 | 4 | 13711 | 111 | Ptprc,Hcls1,Il6ra,Csf1r | | 4.577e-03 | -5.39 | positive regulation of reactive oxygen species metabolic process | biological process | GO:2000379 | 83 | 4 | 13711 | 111 | Tmem106a,Cyba,Itgam,Clu | | 4.597e-03 | -5.38 | kinase binding | molecular function | GO:0019900 | 723 | 13 | 13516 | 107 | Gfap,Csk,Slc2a1,Rab3il1,Ptprc,Hcls1,Vim,Rac2,Actb,Ccr5,Phyhip,Camk2b,Thy1 | | 4.606e-03 | -5.38 | cytosolic small ribosomal subunit | cellular component | GO:0022627 | 42 | 3 | 13825 | 111 | Rps20,Rps5,Rps8 | | 4.612e-03 | -5.38 | GO\_POSITIVE\_REGULATION\_OF\_STAT\_CASCADE | MSigDB lists | GO\_POSITIVE\_REGULATION\_OF\_STAT\_CASCADE | 42 | 3 | 12187 | 98 | Csf1r,Hcls1,Il6ra | | 4.640e-03 | -5.37 | actin filament bundle | cellular component | GO:0032432 | 84 | 4 | 13825 | 111 | Actb,Myh9,Flnb,Cyba | | 4.642e-03 | -5.37 | GAVIN\_FOXP3\_TARGETS\_CLUSTER\_T7 | MSigDB lists | GAVIN\_FOXP3\_TARGETS\_CLUSTER\_T7 | 84 | 4 | 12187 | 98 | Cyba,Hcls1,Rac2,Vim | | 4.713e-03 | -5.36 | GO\_REGULATION\_OF\_SYSTEMIC\_ARTERIAL\_BLOOD\_PRESSURE\_BY\_RENIN\_ANGIOTENSIN | MSigDB lists | GO\_REGULATION\_OF\_SYSTEMIC\_ARTERIAL\_BLOOD\_PRESSURE\_BY\_RENIN\_ANGIOTENSIN | 13 | 2 | 12187 | 98 | Ctsz,Cyba | | 4.713e-03 | -5.36 | GO\_NEGATIVE\_REGULATION\_OF\_CELL\_KILLING | MSigDB lists | GO\_NEGATIVE\_REGULATION\_OF\_CELL\_KILLING | 13 | 2 | 12187 | 98 | Ptprc,Serpinb9 | | 4.713e-03 | -5.36 | GO\_REGULATION\_OF\_INCLUSION\_BODY\_ASSEMBLY | MSigDB lists | GO\_REGULATION\_OF\_INCLUSION\_BODY\_ASSEMBLY | 13 | 2 | 12187 | 98 | Hspa2,Clu | | 4.713e-03 | -5.36 | KYNG\_RESPONSE\_TO\_H2O2\_VIA\_ERCC6 | MSigDB lists | KYNG\_RESPONSE\_TO\_H2O2\_VIA\_ERCC6 | 13 | 2 | 12187 | 98 | Rpl31,Ctss | | 4.713e-03 | -5.36 | VILIMAS\_NOTCH1\_TARGETS\_DN | MSigDB lists | VILIMAS\_NOTCH1\_TARGETS\_DN | 13 | 2 | 12187 | 98 | Csf1r,Itgam | | 4.713e-03 | -5.36 | REACTOME\_FGFR\_LIGAND\_BINDING\_AND\_ACTIVATION | MSigDB lists | REACTOME\_FGFR\_LIGAND\_BINDING\_AND\_ACTIVATION | 13 | 2 | 12187 | 98 | Kl,Fgf18 | | 4.713e-03 | -5.36 | GO\_GLYCOLIPID\_BINDING | MSigDB lists | GO\_GLYCOLIPID\_BINDING | 13 | 2 | 12187 | 98 | Hspa2,Thy1 | | 4.713e-03 | -5.36 | GO\_POSITIVE\_REGULATION\_OF\_LAMELLIPODIUM\_ASSEMBLY | MSigDB lists | GO\_POSITIVE\_REGULATION\_OF\_LAMELLIPODIUM\_ASSEMBLY | 13 | 2 | 12187 | 98 | Brk1,Rac2 | | 4.720e-03 | -5.36 | GO\_POSITIVE\_REGULATION\_OF\_CELL\_ACTIVATION | MSigDB lists | GO\_POSITIVE\_REGULATION\_OF\_CELL\_ACTIVATION | 194 | 6 | 12187 | 98 | Capn3,Thy1,Fcer1g,Csk,Ptprc,Vamp8 | | 4.759e-03 | -5.35 | regulation of catalytic activity | biological process | GO:0050790 | 1714 | 24 | 13711 | 111 | Mt3,Csf1r,Spint1,Rac2,Csk,Ndufa13,Serpina3n,C3,Cyba,Ccl6,Rap1gap,Actb,Ctsb,Capn3,Fgf18,Psme1,Serpinb9,Il6ra,Camk2b,Hspa2,Ptprc,Clu,Thy1,Grn | | 4.765e-03 | -5.35 | GSE1925\_CTRL\_VS\_3H\_IFNG\_STIM\_MACROPHAGE\_DN | MSigDB lists | GSE1925\_CTRL\_VS\_3H\_IFNG\_STIM\_MACROPHAGE\_DN | 136 | 5 | 12187 | 98 | Slc2a1,Vwf,Mgll,Rpl13,Hspa2 | | 4.780e-03 | -5.34 | positive regulation of mast cell degranulation | biological process | GO:0043306 | 13 | 2 | 13711 | 111 | Vamp8,Fcer1g | | 4.780e-03 | -5.34 | norepinephrine metabolic process | biological process | GO:0042415 | 13 | 2 | 13711 | 111 | Kl,Ly6e | | 4.780e-03 | -5.34 | positive regulation of mast cell activation involved in immune response | biological process | GO:0033008 | 13 | 2 | 13711 | 111 | Fcer1g,Vamp8 | | 4.780e-03 | -5.34 | keratinocyte development | biological process | GO:0003334 | 13 | 2 | 13711 | 111 | Flnb,Tfap2c | | 4.838e-03 | -5.33 | GO\_POSITIVE\_REGULATION\_OF\_INNATE\_IMMUNE\_RESPONSE | MSigDB lists | GO\_POSITIVE\_REGULATION\_OF\_INNATE\_IMMUNE\_RESPONSE | 195 | 6 | 12187 | 98 | Psme1,Fcer1g,Itgam,Ctss,Ctsb,Psmb10 | | 4.839e-03 | -5.33 | GO\_DENDRITE | MSigDB lists | GO\_DENDRITE | 400 | 9 | 12187 | 98 | Camk2b,Clu,Thy1,Rap1gap,Pcsk2,Cplx2,Dlgap3,Apod,Cyba | | 4.899e-03 | -5.32 | Hematopoietic cell lineage | KEGG pathways | mmu04640 | 57 | 4 | 5248 | 64 | Il1r1,Il6ra,Itgam,Csf1r | | 4.899e-03 | -5.32 | Hematopoietic cell lineage | KEGG pathways | ko04640 | 57 | 4 | 5248 | 64 | Il6ra,Il1r1,Csf1r,Itgam | | 4.899e-03 | -5.32 | Trafficking and processing of endosomal TLR | REACTOME pathways | R-MMU-1679131 | 9 | 2 | 6297 | 76 | Ctss,Ctsb | | 4.925e-03 | -5.31 | negative regulation of cell projection organization | biological process | GO:0031345 | 194 | 6 | 13711 | 111 | Gfap,Vim,Ccr5,Thy1,Rap1gap,Ctsz | | 4.930e-03 | -5.31 | MORI\_IMMATURE\_B\_LYMPHOCYTE\_UP | MSigDB lists | MORI\_IMMATURE\_B\_LYMPHOCYTE\_UP | 43 | 3 | 12187 | 98 | Grn,Ctss,Ptprc | | 4.930e-03 | -5.31 | GO\_POSITIVE\_REGULATION\_OF\_TUMOR\_NECROSIS\_FACTOR\_SUPERFAMILY\_CYTOKINE\_PRODUCTION | MSigDB lists | GO\_POSITIVE\_REGULATION\_OF\_TUMOR\_NECROSIS\_FACTOR\_SUPERFAMILY\_CYTOKINE\_PRODUCTION | 43 | 3 | 12187 | 98 | Fcer1g,Clu,Cyba | | 4.941e-03 | -5.31 | Tuberculosis | KEGG pathways | ko05152 | 131 | 6 | 5248 | 64 | Camk2b,Ctss,Fcgr3,Fcer1g,C3,Itgam | | 4.941e-03 | -5.31 | Tuberculosis | KEGG pathways | mmu05152 | 131 | 6 | 5248 | 64 | Fcgr3,C3,Fcer1g,Ctss,Itgam,Camk2b | | 4.980e-03 | -5.30 | Translation\_prot\_SH3-like\_sf | interpro domains | IPR008991 | 13 | 2 | 13788 | 114 | Rpl26,Rpl14 | | 5.039e-03 | -5.29 | GO\_RIBOSOME\_BIOGENESIS | MSigDB lists | GO\_RIBOSOME\_BIOGENESIS | 261 | 7 | 12187 | 98 | Rps8,Rpl26,Rpl13,Rps20,Rpl31,Rps5,Rpl14 | | 5.047e-03 | -5.29 | GO\_VACUOLAR\_LUMEN | MSigDB lists | GO\_VACUOLAR\_LUMEN | 86 | 4 | 12187 | 98 | Ctsa,Ctsb,Ctss,Fmod | | 5.051e-03 | -5.29 | RUTELLA\_RESPONSE\_TO\_HGF\_UP | MSigDB lists | RUTELLA\_RESPONSE\_TO\_HGF\_UP | 330 | 8 | 12187 | 98 | Ctsb,Il1r1,C1qa,Mgll,Tsc22d1,Ccr5,Cd68,C3 | | 5.068e-03 | -5.28 | GSE6269\_HEALTHY\_VS\_FLU\_INF\_PBMC\_DN | MSigDB lists | GSE6269\_HEALTHY\_VS\_FLU\_INF\_PBMC\_DN | 138 | 5 | 12187 | 98 | Grn,Ly6e,Sh3bgrl3,Cd68,Actb | | 5.068e-03 | -5.28 | GSE15330\_HSC\_VS\_LYMPHOID\_PRIMED\_MULTIPOTENT\_PROGENITOR\_DN | MSigDB lists | GSE15330\_HSC\_VS\_LYMPHOID\_PRIMED\_MULTIPOTENT\_PROGENITOR\_DN | 138 | 5 | 12187 | 98 | Vim,Serpinb9,Cd68,Itgam,Ahnak | | 5.083e-03 | -5.28 | QI\_PLASMACYTOMA\_UP | MSigDB lists | QI\_PLASMACYTOMA\_UP | 197 | 6 | 12187 | 98 | Ccr5,Clu,Rac2,Csf1r,Hcls1,Dek | | 5.111e-03 | -5.28 | response to endogenous stimulus | biological process | GO:0009719 | 886 | 15 | 13711 | 111 | Cd68,Actb,Fcer1g,Mt3,Hcls1,Serpina3n,Grn,Mc4r,Vim,Adh1,Fgf18,Rap1gap,Ly6e,Csk,Kl | | 5.165e-03 | -5.27 | GO\_BIOLOGICAL\_ADHESION | MSigDB lists | GO\_BIOLOGICAL\_ADHESION | 724 | 13 | 12187 | 98 | Itgam,Myh9,Fcer1g,Thy1,Lfng,Treml2,Actb,Vwf,Ppfibp1,Rac2,Tnfsf8,Psmb10,Ptprc | | 5.214e-03 | -5.26 | Pertussis | KEGG pathways | mmu05133 | 58 | 4 | 5248 | 64 | C1qb,C1qa,C3,Itgam | | 5.214e-03 | -5.26 | Pertussis | KEGG pathways | ko05133 | 58 | 4 | 5248 | 64 | C1qb,C1qa,C3,Itgam | | 5.224e-03 | -5.25 | GSE42021\_CD24INT\_VS\_CD24LOW\_TCONV\_THYMUS\_UP | MSigDB lists | GSE42021\_CD24INT\_VS\_CD24LOW\_TCONV\_THYMUS\_UP | 139 | 5 | 12187 | 98 | C3,Psmb10,Il1r1,Fmod,Ctss | | 5.224e-03 | -5.25 | ATF2\_UP.V1\_DN | MSigDB lists | ATF2\_UP.V1\_DN | 139 | 5 | 12187 | 98 | Fmod,Cd68,Serpinb9,Serpina3n,Il1r1 | | 5.235e-03 | -5.25 | axon | cellular component | GO:0030424 | 640 | 12 | 13825 | 111 | Vim,Il1r1,Clu,Rpl26,Slc17a7,Mgll,Actb,Ctsz,Ptprn,Thy1,Rap1gap,Cplx2 | | 5.255e-03 | -5.25 | actomyosin | cellular component | GO:0042641 | 87 | 4 | 13825 | 111 | Cyba,Flnb,Myh9,Actb | | 5.258e-03 | -5.25 | GO\_CELL\_ACTIVATION\_INVOLVED\_IN\_IMMUNE\_RESPONSE | MSigDB lists | GO\_CELL\_ACTIVATION\_INVOLVED\_IN\_IMMUNE\_RESPONSE | 87 | 4 | 12187 | 98 | Fcer1g,Vamp8,Cplx2,Lfng | | 5.258e-03 | -5.25 | LENAOUR\_DENDRITIC\_CELL\_MATURATION\_UP | MSigDB lists | LENAOUR\_DENDRITIC\_CELL\_MATURATION\_UP | 87 | 4 | 12187 | 98 | Fcgr3,C1qb,Rap1gap,Mgll | | 5.260e-03 | -5.25 | GNF2\_PECAM1 | MSigDB lists | GNF2\_PECAM1 | 44 | 3 | 12187 | 98 | Sh3bgrl3,Ctss,Fcer1g | | 5.260e-03 | -5.25 | WESTON\_VEGFA\_TARGETS\_6HR | MSigDB lists | WESTON\_VEGFA\_TARGETS\_6HR | 44 | 3 | 12187 | 98 | Vim,Mgp,Thy1 | | 5.260e-03 | -5.25 | PID\_PTP1B\_PATHWAY | MSigDB lists | PID\_PTP1B\_PATHWAY | 44 | 3 | 12187 | 98 | Csf1r,Fcgr3,Csk | | 5.260e-03 | -5.25 | LEE\_TARGETS\_OF\_PTCH1\_AND\_SUFU\_UP | MSigDB lists | LEE\_TARGETS\_OF\_PTCH1\_AND\_SUFU\_UP | 44 | 3 | 12187 | 98 | Ctss,C1qb,C1qa | | 5.310e-03 | -5.24 | heparan sulfate proteoglycan binding | molecular function | GO:0043395 | 14 | 2 | 13516 | 107 | Itgam,Ptprc | | 5.310e-03 | -5.24 | sugar transmembrane transporter activity | molecular function | GO:0051119 | 14 | 2 | 13516 | 107 | Mfsd4a,Slc2a1 | | 5.373e-03 | -5.23 | membrane invagination | biological process | GO:0010324 | 44 | 3 | 13711 | 111 | Myh9,Fcgr3,Fcer1g | | 5.384e-03 | -5.22 | GSE21360\_NAIVE\_VS\_SECONDARY\_MEMORY\_CD8\_TCELL\_DN | MSigDB lists | GSE21360\_NAIVE\_VS\_SECONDARY\_MEMORY\_CD8\_TCELL\_DN | 140 | 5 | 12187 | 98 | Cox4i1,Vwf,S100a6,Tsc22d1,Ahnak | | 5.384e-03 | -5.22 | GSE6269\_E\_COLI\_VS\_STAPH\_AUREUS\_INF\_PBMC\_DN | MSigDB lists | GSE6269\_E\_COLI\_VS\_STAPH\_AUREUS\_INF\_PBMC\_DN | 140 | 5 | 12187 | 98 | Tspan4,Il1r1,Grn,S100a6,Itgam | | 5.407e-03 | -5.22 | GO\_RECEPTOR\_BINDING | MSigDB lists | GO\_RECEPTOR\_BINDING | 991 | 16 | 12187 | 98 | Tspan4,Thy1,Grn,Lrg1,Igfbp6,Il6ra,Gfap,C3,Kl,Fmod,Vwf,Csk,Fgf18,Il1r1,Tnfsf8,Sh3bgrl3 | | 5.417e-03 | -5.22 | TYROBP Causal Network | WikiPathways | WP3625 | 52 | 4 | 3756 | 52 | Hcls1,C3,Tmem106a,Itgam | | 5.468e-03 | -5.21 | GO\_REGULATION\_OF\_RESPONSE\_TO\_WOUNDING | MSigDB lists | GO\_REGULATION\_OF\_RESPONSE\_TO\_WOUNDING | 265 | 7 | 12187 | 98 | Il1r1,Apod,Vamp8,C3,Mgll,Fcer1g,Capn3 | | 5.469e-03 | -5.21 | SA\_MMP\_CYTOKINE\_CONNECTION | MSigDB lists | SA\_MMP\_CYTOKINE\_CONNECTION | 14 | 2 | 12187 | 98 | Tnfsf8,Il6ra | | 5.469e-03 | -5.21 | WU\_ALZHEIMER\_DISEASE\_DN | MSigDB lists | WU\_ALZHEIMER\_DISEASE\_DN | 14 | 2 | 12187 | 98 | Pde1a,Hspa2 | | 5.469e-03 | -5.21 | GO\_NEUTROPHIL\_MEDIATED\_IMMUNITY | MSigDB lists | GO\_NEUTROPHIL\_MEDIATED\_IMMUNITY | 14 | 2 | 12187 | 98 | Vamp8,Il6ra | | 5.469e-03 | -5.21 | LIAN\_NEUTROPHIL\_GRANULE\_CONSTITUENTS | MSigDB lists | LIAN\_NEUTROPHIL\_GRANULE\_CONSTITUENTS | 14 | 2 | 12187 | 98 | Ctsb,Ctss | | 5.469e-03 | -5.21 | GO\_GRANULOCYTE\_ACTIVATION | MSigDB lists | GO\_GRANULOCYTE\_ACTIVATION | 14 | 2 | 12187 | 98 | Fcer1g,Vamp8 | | 5.469e-03 | -5.21 | BIOCARTA\_CSK\_PATHWAY | MSigDB lists | BIOCARTA\_CSK\_PATHWAY | 14 | 2 | 12187 | 98 | Ptprc,Csk | | 5.475e-03 | -5.21 | GO\_GROWTH\_FACTOR\_RECEPTOR\_BINDING | MSigDB lists | GO\_GROWTH\_FACTOR\_RECEPTOR\_BINDING | 88 | 4 | 12187 | 98 | Il6ra,Il1r1,Kl,Fgf18 | | 5.504e-03 | -5.20 | FYN (FYN proto-oncogene, Src family tyrosine kinase) | protein interactions | 2534 | 10 | 2 | 6802 | 78 | Flnb,Myh9 | | 5.504e-03 | -5.20 | ARHGAP11A (Rho GTPase activating protein 11A) | protein interactions | 9824 | 10 | 2 | 6802 | 78 | Actb,Myh9 | | 5.504e-03 | -5.20 | SPECC1L (sperm antigen with calponin homology and coiled-coil domains 1 like) | protein interactions | 23384 | 10 | 2 | 6802 | 78 | Myh9,Actb | | 5.547e-03 | -5.19 | GSE3565\_CTRL\_VS\_LPS\_INJECTED\_DUSP1\_KO\_SPLENOCYTES\_UP | MSigDB lists | GSE3565\_CTRL\_VS\_LPS\_INJECTED\_DUSP1\_KO\_SPLENOCYTES\_UP | 141 | 5 | 12187 | 98 | Csf1r,Ahnak,S100a6,Ikzf3,Vim | | 5.547e-03 | -5.19 | neuron remodeling | biological process | GO:0016322 | 14 | 2 | 13711 | 111 | C3,C1qa | | 5.547e-03 | -5.19 | regulation of inclusion body assembly | biological process | GO:0090083 | 14 | 2 | 13711 | 111 | Clu,Hspa2 | | 5.547e-03 | -5.19 | negative regulation of leukocyte mediated cytotoxicity | biological process | GO:0001911 | 14 | 2 | 13711 | 111 | Ptprc,Serpinb9 | | 5.547e-03 | -5.19 | regulation of defense response to bacterium | biological process | GO:1900424 | 14 | 2 | 13711 | 111 | Grn,Cyba | | 5.603e-03 | -5.18 | GO\_REGULATION\_OF\_CHEMOKINE\_PRODUCTION | MSigDB lists | GO\_REGULATION\_OF\_CHEMOKINE\_PRODUCTION | 45 | 3 | 12187 | 98 | Il6ra,Apod,Csf1r | | 5.632e-03 | -5.18 | regulation of receptor signaling pathway via STAT | biological process | GO:1904892 | 88 | 4 | 13711 | 111 | Ptprc,Il6ra,Csf1r,Hcls1 | | 5.697e-03 | -5.17 | GO\_T\_CELL\_DIFFERENTIATION | MSigDB lists | GO\_T\_CELL\_DIFFERENTIATION | 89 | 4 | 12187 | 98 | Tnfsf8,Lfng,Fcer1g,Ptprc | | 5.713e-03 | -5.16 | GSE14415\_NATURAL\_TREG\_VS\_TCONV\_DN | MSigDB lists | GSE14415\_NATURAL\_TREG\_VS\_TCONV\_DN | 142 | 5 | 12187 | 98 | Vim,Trnp1,Itgam,Hcls1,Actb | | 5.713e-03 | -5.16 | GSE37532\_WT\_VS\_PPARG\_KO\_LN\_TCONV\_DN | MSigDB lists | GSE37532\_WT\_VS\_PPARG\_KO\_LN\_TCONV\_DN | 142 | 5 | 12187 | 98 | Sh3bgrl3,Serpinb9,Vim,S100a6,Dek | | 5.733e-03 | -5.16 | RIZKI\_TUMOR\_INVASIVENESS\_3D\_DN | MSigDB lists | RIZKI\_TUMOR\_INVASIVENESS\_3D\_DN | 202 | 6 | 12187 | 98 | Rap1gap,Myh9,Psme1,Fgf18,Il1r1,Ptprc | | 5.856e-03 | -5.14 | cytokine binding | molecular function | GO:0019955 | 91 | 4 | 13516 | 107 | Ccr5,Csf1r,Il6ra,Il1r1 | | 5.861e-03 | -5.14 | modification of morphology or physiology of other organism | biological process | GO:0035821 | 89 | 4 | 13711 | 111 | Csf1r,Serpinb9,Mpeg1,Ccr5 | | 5.883e-03 | -5.14 | GSE34156\_NOD2\_LIGAND\_VS\_TLR1\_TLR2\_LIGAND\_6H\_TREATED\_MONOCYTE\_DN | MSigDB lists | GSE34156\_NOD2\_LIGAND\_VS\_TLR1\_TLR2\_LIGAND\_6H\_TREATED\_MONOCYTE\_DN | 143 | 5 | 12187 | 98 | Ctsz,Grn,Mpeg1,Actb,Fcgr3 | | 5.883e-03 | -5.14 | GSE24634\_TEFF\_VS\_TCONV\_DAY5\_IN\_CULTURE\_DN | MSigDB lists | GSE24634\_TEFF\_VS\_TCONV\_DAY5\_IN\_CULTURE\_DN | 143 | 5 | 12187 | 98 | Psme1,Itgam,Tsc22d1,Rps5,Cyba | | 5.926e-03 | -5.13 | TAKEDA\_TARGETS\_OF\_NUP98\_HOXA9\_FUSION\_10D\_DN | MSigDB lists | TAKEDA\_TARGETS\_OF\_NUP98\_HOXA9\_FUSION\_10D\_DN | 90 | 4 | 12187 | 98 | Ctsb,C1qb,Il1r1,C1qa | | 5.947e-03 | -5.12 | regulation of proteolysis | biological process | GO:0030162 | 564 | 11 | 13711 | 111 | Grn,Clu,Ctsz,C3,Ndufa13,Serpina3n,Serpinb9,Psme1,Capn3,Spint1,Myh9 | | 5.955e-03 | -5.12 | receptor regulator activity | molecular function | GO:0030545 | 273 | 7 | 13516 | 107 | Ccl6,Ly6e,Saa3,Fgf18,Grn,Tnfsf8,Il6ra | | 5.959e-03 | -5.12 | GO\_POSITIVE\_REGULATION\_OF\_INTERLEUKIN\_6\_PRODUCTION | MSigDB lists | GO\_POSITIVE\_REGULATION\_OF\_INTERLEUKIN\_6\_PRODUCTION | 46 | 3 | 12187 | 98 | Cyba,Fcer1g,Il6ra | | 5.959e-03 | -5.12 | GO\_REGULATION\_OF\_TISSUE\_REMODELING | MSigDB lists | GO\_REGULATION\_OF\_TISSUE\_REMODELING | 46 | 3 | 12187 | 98 | Csf1r,Csk,Mc4r | | 5.959e-03 | -5.12 | REACTOME\_FORMATION\_OF\_THE\_TERNARY\_COMPLEX\_AND\_SUBSEQUENTLY\_THE\_43S\_COMPLEX | MSigDB lists | REACTOME\_FORMATION\_OF\_THE\_TERNARY\_COMPLEX\_AND\_SUBSEQUENTLY\_THE\_43S\_COMPLEX | 46 | 3 | 12187 | 98 | Rps8,Rps20,Rps5 | | 5.985e-03 | -5.12 | somatodendritic compartment | cellular component | GO:0036477 | 909 | 15 | 13825 | 111 | Camk2b,Rbfox3,Apod,Clu,Pde1a,Cyba,Dlgap3,Pcsk2,Ptprn,Mark4,Cplx2,Rap1gap,Thy1,Flnb,Il6ra | | 6.057e-03 | -5.11 | JAATINEN\_HEMATOPOIETIC\_STEM\_CELL\_DN | MSigDB lists | JAATINEN\_HEMATOPOIETIC\_STEM\_CELL\_DN | 144 | 5 | 12187 | 98 | Itgam,Fcer1g,Ctsb,Fcgr3,Mpeg1 | | 6.057e-03 | -5.11 | GSE10239\_NAIVE\_VS\_MEMORY\_CD8\_TCELL\_DN | MSigDB lists | GSE10239\_NAIVE\_VS\_MEMORY\_CD8\_TCELL\_DN | 144 | 5 | 12187 | 98 | Ikzf3,Cd68,Rap1gap,Ccr5,Thy1 | | 6.057e-03 | -5.11 | GSE11864\_CSF1\_IFNG\_VS\_CSF1\_IFNG\_PAM3CYS\_IN\_MAC\_DN | MSigDB lists | GSE11864\_CSF1\_IFNG\_VS\_CSF1\_IFNG\_PAM3CYS\_IN\_MAC\_DN | 144 | 5 | 12187 | 98 | Ccr5,Hspa2,Tsc22d1,Mgll,Lfng | | 6.076e-03 | -5.10 | RHO GTPases Activate NADPH Oxidases | REACTOME pathways | R-MMU-5668599 | 10 | 2 | 6297 | 76 | Cyba,Rac2 | | 6.161e-03 | -5.09 | MISSIAGLIA\_REGULATED\_BY\_METHYLATION\_UP | MSigDB lists | MISSIAGLIA\_REGULATED\_BY\_METHYLATION\_UP | 91 | 4 | 12187 | 98 | Spint1,Ctsb,Grn,Ly6e | | 6.212e-03 | -5.08 | Adaptive Immune System | REACTOME pathways | R-MMU-1280218 | 442 | 12 | 6297 | 76 | Fgf18,Rap1gap,Psmb10,Ctss,Ifitm1,Kl,Csk,Cyba,Vamp8,Psme1,C3,Treml2 | | 6.234e-03 | -5.08 | GSE5099\_CLASSICAL\_M1\_VS\_ALTERNATIVE\_M2\_MACROPHAGE\_UP | MSigDB lists | GSE5099\_CLASSICAL\_M1\_VS\_ALTERNATIVE\_M2\_MACROPHAGE\_UP | 145 | 5 | 12187 | 98 | Ctsz,Lfng,Rpl13,S100a6,Rpl31 | | 6.234e-03 | -5.08 | GSE14699\_DELETIONAL\_TOLERANCE\_VS\_ACTIVATED\_CD8\_TCELL\_UP | MSigDB lists | GSE14699\_DELETIONAL\_TOLERANCE\_VS\_ACTIVATED\_CD8\_TCELL\_UP | 145 | 5 | 12187 | 98 | Itgam,Ahnak,Serpinb9,Vim,Hcls1 | | 6.234e-03 | -5.08 | GSE13522\_CTRL\_VS\_T\_CRUZI\_Y\_STRAIN\_INF\_SKIN\_IFNAR\_KO\_UP | MSigDB lists | GSE13522\_CTRL\_VS\_T\_CRUZI\_Y\_STRAIN\_INF\_SKIN\_IFNAR\_KO\_UP | 145 | 5 | 12187 | 98 | Mpeg1,Adh1,Rpl14,Rps8,Fcer1g | | 6.276e-03 | -5.07 | extracellular matrix | cellular component | GO:0031012 | 342 | 8 | 13825 | 111 | Ctsz,Ctsb,Vwf,Mgp,Igfbp6,Serpinb9,S100a6,Fmod | | 6.278e-03 | -5.07 | HEDENFALK\_BREAST\_CANCER\_BRACX\_UP | MSigDB lists | HEDENFALK\_BREAST\_CANCER\_BRACX\_UP | 15 | 2 | 12187 | 98 | Rps5,Rpl31 | | 6.278e-03 | -5.07 | MAGRANGEAS\_MULTIPLE\_MYELOMA\_IGG\_VS\_IGA\_DN | MSigDB lists | MAGRANGEAS\_MULTIPLE\_MYELOMA\_IGG\_VS\_IGA\_DN | 15 | 2 | 12187 | 98 | Csk,Actb | | 6.278e-03 | -5.07 | CHEN\_ETV5\_TARGETS\_SERTOLI | MSigDB lists | CHEN\_ETV5\_TARGETS\_SERTOLI | 15 | 2 | 12187 | 98 | C1qb,Fcgr3 | | 6.278e-03 | -5.07 | GO\_MAST\_CELL\_MEDIATED\_IMMUNITY | MSigDB lists | GO\_MAST\_CELL\_MEDIATED\_IMMUNITY | 15 | 2 | 12187 | 98 | Serpinb9,Cplx2 | | 6.278e-03 | -5.07 | CHIARETTI\_T\_ALL\_RELAPSE\_PROGNOSIS | MSigDB lists | CHIARETTI\_T\_ALL\_RELAPSE\_PROGNOSIS | 15 | 2 | 12187 | 98 | Ahnak,Dek | | 6.302e-03 | -5.07 | regulation of cell differentiation | biological process | GO:0045595 | 1557 | 22 | 13711 | 111 | Spint1,Mt3,Csf1r,Hcls1,Ccr5,Ctsz,Vim,Rap1gap,Bhlhe41,Gfap,Fgf18,Capn3,Camk2b,Clu,C1qa,Ptprc,Tmem176a,Ikzf3,Grn,Il1r1,Mgll,Thy1 | | 6.342e-03 | -5.06 | GO\_PROTEIN\_TARGETING | MSigDB lists | GO\_PROTEIN\_TARGETING | 343 | 8 | 12187 | 98 | Rpl13,Rps8,Ndufa13,Rpl31,Rps5,Rpl14,Rps20,Rpl26 | | 6.367e-03 | -5.06 | negative regulation of cell killing | biological process | GO:0031342 | 15 | 2 | 13711 | 111 | Ptprc,Serpinb9 | | 6.367e-03 | -5.06 | modulation by symbiont of host cellular process | biological process | GO:0044068 | 15 | 2 | 13711 | 111 | Serpinb9,Ccr5 | | 6.402e-03 | -5.05 | GSE10422\_WT\_VS\_BAFF\_TRANSGENIC\_LN\_BCELL\_DN | MSigDB lists | GSE10422\_WT\_VS\_BAFF\_TRANSGENIC\_LN\_BCELL\_DN | 92 | 4 | 12187 | 98 | C3,Ctsz,Rpl26,Clu | | 6.402e-03 | -5.05 | GSE17974\_CTRL\_VS\_ACT\_IL4\_AND\_ANTI\_IL12\_0.5H\_CD4\_TCELL\_UP | MSigDB lists | GSE17974\_CTRL\_VS\_ACT\_IL4\_AND\_ANTI\_IL12\_0.5H\_CD4\_TCELL\_UP | 92 | 4 | 12187 | 98 | Ttc9b,Alg12,Tmem176a,Ptprn | | 6.410e-03 | -5.05 | SATO\_SILENCED\_BY\_METHYLATION\_IN\_PANCREATIC\_CANCER\_1 | MSigDB lists | SATO\_SILENCED\_BY\_METHYLATION\_IN\_PANCREATIC\_CANCER\_1 | 273 | 7 | 12187 | 98 | Rac2,Spint1,Il1r1,Tfap2c,Hspa2,Tmem176a,C3 | | 6.415e-03 | -5.05 | GSE17721\_CTRL\_VS\_CPG\_1H\_BMDC\_DN | MSigDB lists | GSE17721\_CTRL\_VS\_CPG\_1H\_BMDC\_DN | 146 | 5 | 12187 | 98 | Serpina3n,Kl,Serpinb9,Tsc22d1,Lgi4 | | 6.415e-03 | -5.05 | GSE29618\_LAIV\_VS\_TIV\_FLU\_VACCINE\_DAY7\_PDC\_DN | MSigDB lists | GSE29618\_LAIV\_VS\_TIV\_FLU\_VACCINE\_DAY7\_PDC\_DN | 146 | 5 | 12187 | 98 | Sh3bgrl3,Hspa2,Thy1,Phyhip,Apod | | 6.465e-03 | -5.04 | regulation of bone remodeling | biological process | GO:0046850 | 47 | 3 | 13711 | 111 | Csf1r,Csk,Mc4r | | 6.474e-03 | -5.04 | GO\_MOLECULAR\_FUNCTION\_REGULATOR | MSigDB lists | GO\_MOLECULAR\_FUNCTION\_REGULATOR | 1010 | 16 | 12187 | 98 | Vamp8,Ctsa,Spint1,Thy1,Serpinb9,Rac2,Sh3bgrl3,Rab3il1,Fgf18,Serpina3n,Capn3,Camk2b,Psme1,Rap1gap,C3,Kl | | 6.585e-03 | -5.02 | immune response-activating cell surface receptor signaling pathway | biological process | GO:0002429 | 92 | 4 | 13711 | 111 | Fcer1g,Klhl6,Ptprc,Thy1 | | 6.599e-03 | -5.02 | GO\_REGULATION\_OF\_ION\_HOMEOSTASIS | MSigDB lists | GO\_REGULATION\_OF\_ION\_HOMEOSTASIS | 147 | 5 | 12187 | 98 | Ptprc,Cyba,Capn3,Ccr5,Thy1 | | 6.599e-03 | -5.02 | GO\_ANTIGEN\_PROCESSING\_AND\_PRESENTATION\_OF\_PEPTIDE\_ANTIGEN | MSigDB lists | GO\_ANTIGEN\_PROCESSING\_AND\_PRESENTATION\_OF\_PEPTIDE\_ANTIGEN | 147 | 5 | 12187 | 98 | Psmb10,Cyba,Ctss,Psme1,Fcer1g | | 6.632e-03 | -5.02 | Rib\_L2\_dom2 | interpro domains | IPR014722 | 15 | 2 | 13788 | 114 | Rpl14,Rpl26 | | 6.632e-03 | -5.02 | Serpin\_CS | interpro domains | IPR023795 | 15 | 2 | 13788 | 114 | Serpina3n,Serpinb9 | | 6.649e-03 | -5.01 | GO\_POSITIVE\_REGULATION\_OF\_TRANSMEMBRANE\_TRANSPORT | MSigDB lists | GO\_POSITIVE\_REGULATION\_OF\_TRANSMEMBRANE\_TRANSPORT | 93 | 4 | 12187 | 98 | Capn3,Thy1,Hspa2,Ctss | | 6.677e-03 | -5.01 | PHLDB2 (pleckstrin homology like domain family B member 2) | protein interactions | 90102 | 11 | 2 | 6802 | 78 | Actb,Myh9 | | 6.677e-03 | -5.01 | ARPC2 (actin related protein 2/3 complex subunit 2) | protein interactions | 10109 | 11 | 2 | 6802 | 78 | Myh9,Actb | | 6.677e-03 | -5.01 | STON2 (stonin 2) | protein interactions | 85439 | 11 | 2 | 6802 | 78 | Myh9,Actb | | 6.677e-03 | -5.01 | MYO5A (myosin VA) | protein interactions | 4644 | 11 | 2 | 6802 | 78 | Actb,Flnb | | 6.677e-03 | -5.01 | Pknox1 (Pbx/knotted 1 homeobox) | protein interactions | 18771 | 11 | 2 | 6802 | 78 | Myh9,Actb | | 6.677e-03 | -5.01 | FARP1 (FERM, ARH/RhoGEF and pleckstrin domain protein 1) | protein interactions | 10160 | 11 | 2 | 6802 | 78 | Myh9,Flnb | | 6.677e-03 | -5.01 | DST (dystonin) | protein interactions | 667 | 11 | 2 | 6802 | 78 | Actb,Myh9 | | 6.677e-03 | -5.01 | TPM2 (tropomyosin 2) | protein interactions | 7169 | 11 | 2 | 6802 | 78 | Actb,Myh9 | | 6.677e-03 | -5.01 | RPL22 (ribosomal protein L22) | protein interactions | 6146 | 11 | 2 | 6802 | 78 | Actb,Flnb | | 6.706e-03 | -5.00 | GO\_ESTABLISHMENT\_OF\_LOCALIZATION\_IN\_CELL | MSigDB lists | GO\_ESTABLISHMENT\_OF\_LOCALIZATION\_IN\_CELL | 1390 | 20 | 12187 | 98 | Grn,Myh9,Ccr5,Ndufa13,Rps5,Rpl31,Vamp8,Cplx2,Rpl26,Slc17a7,Ctsa,Lin7a,Rpl13,Ctsz,Rps8,Rpl14,Rps20,Rac2,Ptprn,Ptprc | | 6.712e-03 | -5.00 | GNF2\_TNFRSF1B | MSigDB lists | GNF2\_TNFRSF1B | 48 | 3 | 12187 | 98 | Ctss,Sh3bgrl3,Fcer1g | | 6.787e-03 | -4.99 | GSE25123\_WT\_VS\_PPARG\_KO\_MACROPHAGE\_IL4\_STIM\_DN | MSigDB lists | GSE25123\_WT\_VS\_PPARG\_KO\_MACROPHAGE\_IL4\_STIM\_DN | 148 | 5 | 12187 | 98 | Fam111a,S100a6,Tsc22d1,Ppfibp1,Ikzf3 | | 6.787e-03 | -4.99 | GSE12845\_PRE\_GC\_VS\_DARKZONE\_GC\_TONSIL\_BCELL\_UP | MSigDB lists | GSE12845\_PRE\_GC\_VS\_DARKZONE\_GC\_TONSIL\_BCELL\_UP | 148 | 5 | 12187 | 98 | Fcer1g,Il6ra,Ahnak,Ctsb,Csf1r | | 6.787e-03 | -4.99 | GSE6259\_DEC205\_POS\_DC\_VS\_BCELL\_DN | MSigDB lists | GSE6259\_DEC205\_POS\_DC\_VS\_BCELL\_DN | 148 | 5 | 12187 | 98 | Serpinb9,Grn,Psme1,Psmb10,Cyba | | 6.814e-03 | -4.99 | cell development | biological process | GO:0048468 | 1375 | 20 | 13711 | 111 | Gfap,C3,Tfap2c,Fgf18,Capn3,Rac2,Actb,Itgam,Myh9,Csf1r,Mt3,Lgi4,Vim,Grn,Thy1,Clu,Ptprc,C1qa,Flnb,Hspa2 | | 6.856e-03 | -4.98 | regulation of tyrosine phosphorylation of STAT protein | biological process | GO:0042509 | 48 | 3 | 13711 | 111 | Il6ra,Csf1r,Hcls1 | | 6.899e-03 | -4.98 | lysosomal membrane | cellular component | GO:0005765 | 94 | 4 | 13825 | 111 | Gfap,Cd68,Grn,Vamp8 | | 6.900e-03 | -4.98 | REACTOME\_HEMOSTASIS | MSigDB lists | REACTOME\_HEMOSTASIS | 348 | 8 | 12187 | 98 | Rac2,Mgll,Vwf,Fcer1g,Itgam,Clu,Pde1a,Csk | | 6.979e-03 | -4.96 | GSE19888\_ADENOSINE\_A3R\_INH\_VS\_ACT\_IN\_MAST\_CELL\_DN | MSigDB lists | GSE19888\_ADENOSINE\_A3R\_INH\_VS\_ACT\_IN\_MAST\_CELL\_DN | 149 | 5 | 12187 | 98 | Psmb10,Lin7a,Mpeg1,Vwf,Ly6e | | 6.979e-03 | -4.96 | GSE41867\_NAIVE\_VS\_DAY6\_LCMV\_ARMSTRONG\_EFFECTOR\_CD8\_TCELL\_DN | MSigDB lists | GSE41867\_NAIVE\_VS\_DAY6\_LCMV\_ARMSTRONG\_EFFECTOR\_CD8\_TCELL\_DN | 149 | 5 | 12187 | 98 | Vwf,Thy1,Serpinb9,Ikzf3,Il1r1 | | 6.979e-03 | -4.96 | GSE2128\_C57BL6\_VS\_NOD\_THYMOCYTE\_DN | MSigDB lists | GSE2128\_C57BL6\_VS\_NOD\_THYMOCYTE\_DN | 149 | 5 | 12187 | 98 | Cyba,Pde1a,Hspa2,Camk2b,Ppfibp1 | | 7.020e-03 | -4.96 | GO\_CELL\_BODY | MSigDB lists | GO\_CELL\_BODY | 424 | 9 | 12187 | 98 | Camk2b,Pcsk2,Rap1gap,Cplx2,Dlgap3,Gfap,Apod,Cyba,Ptprn | | 7.049e-03 | -4.95 | regulation of molecular function | biological process | GO:0065009 | 2276 | 29 | 13711 | 111 | Ahnak,Serpinb9,Il6ra,Camk2b,Ctsb,Capn3,Psme1,Fgf18,Actb,Thy1,Ctss,Grn,Ptprc,Hspa2,Clu,Csk,Lfng,Serpina3n,Ndufa13,Cyba,C3,Hcls1,Csf1r,Mt3,Rac2,Spint1,Rap1gap,Ctsz,Ccl6 | | 7.099e-03 | -4.95 | leukocyte chemotaxis | biological process | GO:0030595 | 94 | 4 | 13711 | 111 | Fcer1g,Ccl6,Itgam,Fcgr3 | | 7.109e-03 | -4.95 | FLOTHO\_PEDIATRIC\_ALL\_THERAPY\_RESPONSE\_UP | MSigDB lists | FLOTHO\_PEDIATRIC\_ALL\_THERAPY\_RESPONSE\_UP | 49 | 3 | 12187 | 98 | Rpl13,Rps5,Rps20 | | 7.109e-03 | -4.95 | GNF2\_LCAT | MSigDB lists | GNF2\_LCAT | 49 | 3 | 12187 | 98 | C3,Adh1,Tmem176a | | 7.137e-03 | -4.94 | MODULE\_73 | MSigDB lists | MODULE\_73 | 16 | 2 | 12187 | 98 | Il1r1,Il6ra | | 7.137e-03 | -4.94 | GO\_ENDOSOME\_LUMEN | MSigDB lists | GO\_ENDOSOME\_LUMEN | 16 | 2 | 12187 | 98 | Ctsb,Ctss | | 7.137e-03 | -4.94 | GO\_PHAGOCYTOSIS\_ENGULFMENT | MSigDB lists | GO\_PHAGOCYTOSIS\_ENGULFMENT | 16 | 2 | 12187 | 98 | Fcer1g,Myh9 | | 7.137e-03 | -4.94 | GO\_FIBROBLAST\_GROWTH\_FACTOR\_RECEPTOR\_BINDING | MSigDB lists | GO\_FIBROBLAST\_GROWTH\_FACTOR\_RECEPTOR\_BINDING | 16 | 2 | 12187 | 98 | Fgf18,Kl | | 7.137e-03 | -4.94 | HOLLEMAN\_VINCRISTINE\_RESISTANCE\_ALL\_DN | MSigDB lists | HOLLEMAN\_VINCRISTINE\_RESISTANCE\_ALL\_DN | 16 | 2 | 12187 | 98 | Rpl31,Vim | | 7.137e-03 | -4.94 | WENG\_POR\_TARGETS\_GLOBAL\_DN | MSigDB lists | WENG\_POR\_TARGETS\_GLOBAL\_DN | 16 | 2 | 12187 | 98 | Grn,Mgll | | 7.159e-03 | -4.94 | lytic vacuole membrane | cellular component | GO:0098852 | 95 | 4 | 13825 | 111 | Cd68,Gfap,Grn,Vamp8 | | 7.163e-03 | -4.94 | WANG\_ESOPHAGUS\_CANCER\_VS\_NORMAL\_UP | MSigDB lists | WANG\_ESOPHAGUS\_CANCER\_VS\_NORMAL\_UP | 95 | 4 | 12187 | 98 | Itgam,Fmod,Thy1,Fcer1g | | 7.175e-03 | -4.94 | GO\_GLIOGENESIS | MSigDB lists | GO\_GLIOGENESIS | 150 | 5 | 12187 | 98 | Gfap,Lgi4,Clu,Vim,Csk | | 7.175e-03 | -4.94 | GSE27241\_CTRL\_VS\_DIGOXIN\_TREATED\_CD4\_TCELL\_IN\_TH17\_POLARIZING\_CONDITIONS\_DN | MSigDB lists | GSE27241\_CTRL\_VS\_DIGOXIN\_TREATED\_CD4\_TCELL\_IN\_TH17\_POLARIZING\_CONDITIONS\_DN | 150 | 5 | 12187 | 98 | Rpl14,Lfng,Vamp8,Vim,Ahnak | | 7.175e-03 | -4.94 | GOLDRATH\_NAIVE\_VS\_EFF\_CD8\_TCELL\_DN | MSigDB lists | GOLDRATH\_NAIVE\_VS\_EFF\_CD8\_TCELL\_DN | 150 | 5 | 12187 | 98 | Serpinb9,Fcer1g,Ccr5,S100a6,C3 | | 7.225e-03 | -4.93 | neurotransmitter transport | biological process | GO:0006836 | 149 | 5 | 13711 | 111 | Fcgr3,Cplx2,Slc17a7,Lin7a,Fcer1g | | 7.238e-03 | -4.93 | negative regulation of myeloid cell apoptotic process | biological process | GO:0033033 | 16 | 2 | 13711 | 111 | Ccr5,Fcer1g | | 7.238e-03 | -4.93 | positive regulation of superoxide anion generation | biological process | GO:0032930 | 16 | 2 | 13711 | 111 | Itgam,Cyba | | 7.238e-03 | -4.93 | intermediate filament organization | biological process | GO:0045109 | 16 | 2 | 13711 | 111 | Vim,Gfap | | 7.238e-03 | -4.93 | glucose transmembrane transport | biological process | GO:1904659 | 16 | 2 | 13711 | 111 | Slc2a1,Mfsd4a | | 7.238e-03 | -4.93 | hexose transmembrane transport | biological process | GO:0008645 | 16 | 2 | 13711 | 111 | Mfsd4a,Slc2a1 | | 7.238e-03 | -4.93 | positive regulation of T-helper 1 type immune response | biological process | GO:0002827 | 16 | 2 | 13711 | 111 | Il6ra,Il1r1 | | 7.238e-03 | -4.93 | bone resorption | biological process | GO:0045453 | 16 | 2 | 13711 | 111 | Ctss,Rac2 | | 7.374e-03 | -4.91 | GSE10325\_BCELL\_VS\_MYELOID\_DN | MSigDB lists | GSE10325\_BCELL\_VS\_MYELOID\_DN | 151 | 5 | 12187 | 98 | Ccr5,Ctsb,Csf1r,Ctsa,Fcgr3 | | 7.374e-03 | -4.91 | GSE34515\_CD16\_NEG\_VS\_POS\_MONOCYTE\_UP | MSigDB lists | GSE34515\_CD16\_NEG\_VS\_POS\_MONOCYTE\_UP | 151 | 5 | 12187 | 98 | Ctsz,Tfap2c,Psmb10,Rab3il1,Ptprn | | 7.374e-03 | -4.91 | GSE22886\_IGG\_IGA\_MEMORY\_BCELL\_VS\_BLOOD\_PLASMA\_CELL\_UP | MSigDB lists | GSE22886\_IGG\_IGA\_MEMORY\_BCELL\_VS\_BLOOD\_PLASMA\_CELL\_UP | 151 | 5 | 12187 | 98 | Rpl13,Tsc22d1,Rpl26,Flnb,Dek | | 7.374e-03 | -4.91 | GSE30971\_WBP7\_HET\_VS\_KO\_MACROPHAGE\_4H\_LPS\_STIM\_DN | MSigDB lists | GSE30971\_WBP7\_HET\_VS\_KO\_MACROPHAGE\_4H\_LPS\_STIM\_DN | 151 | 5 | 12187 | 98 | Kl,Capn3,Ahnak,Grn,Ctss | | 7.375e-03 | -4.91 | GO\_LEUKOCYTE\_DIFFERENTIATION | MSigDB lists | GO\_LEUKOCYTE\_DIFFERENTIATION | 213 | 6 | 12187 | 98 | Tnfsf8,Ptprc,Lfng,Myh9,Fcer1g,Csf1r | | 7.389e-03 | -4.91 | regulation of biological quality | biological process | GO:0065008 | 3134 | 37 | 13711 | 111 | Vwf,Cyba,Ptprn,C3,Ccr5,Csk,Ctsa,Rac2,Rps5,Myh9,Cd52,Csf1r,Hcls1,Mt3,Brk1,Cplx2,Vim,Vamp8,Gfap,Camk2b,Capn3,Pcsk2,Lin7a,Fcer1g,Slc17a7,Adh1,Mc4r,Grn,Tmem106a,Ctss,Thy1,Kl,Mgll,Abhd17c,Clu,Ptprc,Sh3bgrl3 | | 7.420e-03 | -4.90 | lysosome | cellular component | GO:0005764 | 352 | 8 | 13825 | 111 | Ctsz,Ctsb,Ctsa,Cd68,Gfap,Ctss,Vamp8,Grn | | 7.427e-03 | -4.90 | regulation of cell death | biological process | GO:0010941 | 1386 | 20 | 13711 | 111 | Serpinb9,Camk2b,Ctsb,Capn3,Tsc22d1,Ndufa13,Ccr5,Hcls1,Csf1r,Mt3,Pde1a,Fcer1g,Ikzf3,Ccar1,Grn,C1qa,Ptprc,Rpl26,Clu,Ctsz | | 7.495e-03 | -4.89 | SCHAEFFER\_PROSTATE\_DEVELOPMENT\_48HR\_UP | MSigDB lists | SCHAEFFER\_PROSTATE\_DEVELOPMENT\_48HR\_UP | 353 | 8 | 12187 | 98 | Adh1,Hspa2,Tfap2c,Clu,Ly6e,Spint1,Igfbp6,Vamp8 | | 7.519e-03 | -4.89 | GUO\_HEX\_TARGETS\_DN | MSigDB lists | GUO\_HEX\_TARGETS\_DN | 50 | 3 | 12187 | 98 | Psme1,Brk1,Actb | | 7.519e-03 | -4.89 | FONTAINE\_PAPILLARY\_THYROID\_CARCINOMA\_UP | MSigDB lists | FONTAINE\_PAPILLARY\_THYROID\_CARCINOMA\_UP | 50 | 3 | 12187 | 98 | Capn3,S100a6,Hcls1 | | 7.538e-03 | -4.89 | Cytokine-cytokine receptor interaction | KEGG pathways | ko04060 | 143 | 6 | 5248 | 64 | Tnfsf8,Ccr5,Il1r1,Ccl6,Csf1r,Il6ra | | 7.538e-03 | -4.89 | Cytokine-cytokine receptor interaction | KEGG pathways | mmu04060 | 143 | 6 | 5248 | 64 | Il6ra,Tnfsf8,Ccr5,Il1r1,Ccl6,Csf1r | | 7.542e-03 | -4.89 | lytic vacuole | cellular component | GO:0000323 | 353 | 8 | 13825 | 111 | Ctsa,Ctsz,Ctsb,Grn,Vamp8,Ctss,Cd68,Gfap | | 7.578e-03 | -4.88 | GSE43955\_10H\_VS\_30H\_ACT\_CD4\_TCELL\_WITH\_TGFB\_IL6\_DN | MSigDB lists | GSE43955\_10H\_VS\_30H\_ACT\_CD4\_TCELL\_WITH\_TGFB\_IL6\_DN | 152 | 5 | 12187 | 98 | Flnb,Mgp,Ctss,S100a6,Ccr5 | | 7.578e-03 | -4.88 | GSE3982\_EOSINOPHIL\_VS\_BASOPHIL\_UP | MSigDB lists | GSE3982\_EOSINOPHIL\_VS\_BASOPHIL\_UP | 152 | 5 | 12187 | 98 | Kl,Cyba,Ctss,Mgll,Camk2b | | 7.578e-03 | -4.88 | GSE40441\_NRP1\_POS\_INDUCED\_TREG\_VS\_NRP1\_NEG\_NATURAL\_TREG\_UP | MSigDB lists | GSE40441\_NRP1\_POS\_INDUCED\_TREG\_VS\_NRP1\_NEG\_NATURAL\_TREG\_UP | 152 | 5 | 12187 | 98 | Dek,Fam111a,Camk2b,Il6ra,Ppfibp1 | | 7.578e-03 | -4.88 | GSE42021\_TREG\_PLN\_VS\_TREG\_PRECURSORS\_THYMUS\_DN | MSigDB lists | GSE42021\_TREG\_PLN\_VS\_TREG\_PRECURSORS\_THYMUS\_DN | 152 | 5 | 12187 | 98 | Serpina3n,Psmb10,Psme1,Ly6e,Serpinb9 | | 7.578e-03 | -4.88 | GSE40666\_UNTREATED\_VS\_IFNA\_STIM\_EFFECTOR\_CD8\_TCELL\_90MIN\_UP | MSigDB lists | GSE40666\_UNTREATED\_VS\_IFNA\_STIM\_EFFECTOR\_CD8\_TCELL\_90MIN\_UP | 152 | 5 | 12187 | 98 | Ctsa,Itgam,Ahnak,Cd68,Vim | | 7.578e-03 | -4.88 | GSE28737\_WT\_VS\_BCL6\_HET\_FOLLICULAR\_BCELL\_DN | MSigDB lists | GSE28737\_WT\_VS\_BCL6\_HET\_FOLLICULAR\_BCELL\_DN | 152 | 5 | 12187 | 98 | Ctss,Cd68,Itgam,Rac2,S100a6 | | 7.602e-03 | -4.88 | positive regulation of cell communication | biological process | GO:0010647 | 1389 | 20 | 13711 | 111 | Tmem106a,Il1r1,Vamp8,Kl,Lrg1,Ccl6,Clu,Rpl26,Ptprc,C3,Cyba,Gfap,Lfng,Fgf18,Camk2b,Csk,Il6ra,Mt3,Hcls1,Csf1r | | 7.669e-03 | -4.87 | positive regulation of biological process | biological process | GO:0048518 | 4720 | 51 | 13711 | 111 | Ccar1,Grn,Ikzf3,Ctss,Il1r1,Thy1,Kl,Ptprc,C1qa,Psme1,Fgf18,Fcer1g,Klhl6,Rap1gap,Vamp8,Ctsz,Fcgr3,C1qb,Cyba,Ndufa13,Ccr5,Lfng,Mark4,Csk,Mt3,Myh9,Tmem106a,Mc4r,Clu,Abhd17c,Hspa2,Gfap,Tsc22d1,Capn3,Camk2b,Serpinb9,Il6ra,Itgam,Vim,Lrg1,Ccl6,Rpl26,Tnfsf8,Ptprn,C3,Tfap2c,Spint1,Rac2,Brk1,Hcls1,Csf1r | | 7.678e-03 | -4.87 | GO\_REGULATION\_OF\_RESPONSE\_TO\_EXTERNAL\_STIMULUS | MSigDB lists | GO\_REGULATION\_OF\_RESPONSE\_TO\_EXTERNAL\_STIMULUS | 674 | 12 | 12187 | 98 | Apod,Phyhip,Il1r1,Fgf18,Cyba,Vamp8,Il6ra,Rac2,C3,Capn3,Fcer1g,Mgll | | 7.679e-03 | -4.87 | energy reserve metabolic process | biological process | GO:0006112 | 50 | 3 | 13711 | 111 | Kl,Mt3,Mc4r | | 7.708e-03 | -4.87 | regulation of neurogenesis | biological process | GO:0050767 | 839 | 14 | 13711 | 111 | Mt3,Spint1,Bhlhe41,Camk2b,Gfap,Ccr5,C1qa,Ctsz,Rap1gap,Mgll,Il1r1,Thy1,Vim,Grn | | 7.743e-03 | -4.86 | GO\_REGULATION\_OF\_BODY\_FLUID\_LEVELS | MSigDB lists | GO\_REGULATION\_OF\_BODY\_FLUID\_LEVELS | 355 | 8 | 12187 | 98 | Cyba,Actb,Vamp8,Rac2,C3,Vwf,Fcer1g,Myh9 | | 7.785e-03 | -4.86 | GSE22886\_NAIVE\_VS\_IGG\_IGA\_MEMORY\_BCELL\_DN | MSigDB lists | GSE22886\_NAIVE\_VS\_IGG\_IGA\_MEMORY\_BCELL\_DN | 153 | 5 | 12187 | 98 | Ctsa,S100a6,Ahnak,Itgam,Vim | | 7.785e-03 | -4.86 | GSE40443\_INDUCED\_VS\_TOTAL\_TREG\_DN | MSigDB lists | GSE40443\_INDUCED\_VS\_TOTAL\_TREG\_DN | 153 | 5 | 12187 | 98 | Ctss,Thy1,Fcer1g,Psme1,Il6ra | | 7.785e-03 | -4.86 | GSE10325\_CD4\_TCELL\_VS\_MYELOID\_DN | MSigDB lists | GSE10325\_CD4\_TCELL\_VS\_MYELOID\_DN | 153 | 5 | 12187 | 98 | Itgam,Spint1,Csf1r,Tspan4,Fcgr3 | | 7.785e-03 | -4.86 | GSE43955\_1H\_VS\_10H\_ACT\_CD4\_TCELL\_WITH\_TGFB\_IL6\_DN | MSigDB lists | GSE43955\_1H\_VS\_10H\_ACT\_CD4\_TCELL\_WITH\_TGFB\_IL6\_DN | 153 | 5 | 12187 | 98 | Ctss,Ly6e,Ccr5,Flnb,Tmem176a | | 7.785e-03 | -4.86 | GSE5542\_IFNG\_VS\_IFNA\_AND\_IFNG\_TREATED\_EPITHELIAL\_CELLS\_6H\_UP | MSigDB lists | GSE5542\_IFNG\_VS\_IFNA\_AND\_IFNG\_TREATED\_EPITHELIAL\_CELLS\_6H\_UP | 153 | 5 | 12187 | 98 | Sh2d5,Fcer1g,S100a6,Vim,Ndufa13 | | 7.814e-03 | -4.85 | fibroblast growth factor receptor binding | molecular function | GO:0005104 | 17 | 2 | 13516 | 107 | Kl,Fgf18 | | 7.814e-03 | -4.85 | glycolipid binding | molecular function | GO:0051861 | 17 | 2 | 13516 | 107 | Thy1,Hspa2 | | 7.843e-03 | -4.85 | immune response-regulating signaling pathway | biological process | GO:0002764 | 152 | 5 | 13711 | 111 | Fcer1g,Klhl6,Thy1,Ptprc,Fcgr3 | | 7.901e-03 | -4.84 | mouse chr1 C4\|1 38.64 cM | chromosome location | mouse chr1 C4\|1 38.64 cM | 1 | 1 | 14556 | 115 | Ptprn | | 7.901e-03 | -4.84 | mouse chr5 G2\|5 79.15 cM | chromosome location | mouse chr5 G2\|5 79.15 cM | 1 | 1 | 14556 | 115 | Lfng | | 7.901e-03 | -4.84 | mouse chr1 H3\|1 79.23 cM | chromosome location | mouse chr1 H3\|1 79.23 cM | 1 | 1 | 14556 | 115 | Fcer1g | | 7.901e-03 | -4.84 | mouse chr8 C5\|8 46.29 cM | chromosome location | mouse chr8 C5\|8 46.29 cM | 1 | 1 | 14556 | 115 | Mt3 | | 7.901e-03 | -4.84 | mouse chr9 A5.1\|9 24.52 cM | chromosome location | mouse chr9 A5.1\|9 24.52 cM | 1 | 1 | 14556 | 115 | Thy1 | | 7.901e-03 | -4.84 | mouse chr3 F1\|3 39.35 cM | chromosome location | mouse chr3 F1\|3 39.35 cM | 1 | 1 | 14556 | 115 | S100a6 | | 7.901e-03 | -4.84 | mouse chr3 G3\|3 64.16 cM | chromosome location | mouse chr3 G3\|3 64.16 cM | 1 | 1 | 14556 | 115 | Adh1 | | 7.901e-03 | -4.84 | mouse chr8 E1\|8 69.99 cM | chromosome location | mouse chr8 E1\|8 69.99 cM | 1 | 1 | 14556 | 115 | Cox4i1 | | 7.901e-03 | -4.84 | mouse chr2 E5\|2 60.31 cM | chromosome location | mouse chr2 E5\|2 60.31 cM | 1 | 1 | 14556 | 115 | Capn3 | | 7.901e-03 | -4.84 | mouse chr14 D3\|14 40.44 cM | chromosome location | mouse chr14 D3\|14 40.44 cM | 1 | 1 | 14556 | 115 | Tsc22d1 | | 7.901e-03 | -4.84 | mouse chr11 E2\|11 83.22 cM | chromosome location | mouse chr11 E2\|11 83.22 cM | 1 | 1 | 14556 | 115 | Rbfox3 | | 7.901e-03 | -4.84 | mouse chr11 A1\|11 3.89 cM | chromosome location | mouse chr11 A1\|11 3.89 cM | 1 | 1 | 14556 | 115 | Camk2b | | 7.901e-03 | -4.84 | mouse chr1 E4\|1 60.73 cM | chromosome location | mouse chr1 E4\|1 60.73 cM | 1 | 1 | 14556 | 115 | Ptprc | | 7.901e-03 | -4.84 | mouse chr2 A1\|2 10.04 cM | chromosome location | mouse chr2 A1\|2 10.04 cM | 1 | 1 | 14556 | 115 | Vim | | 7.901e-03 | -4.84 | mouse chr4 D2.2\|4 61.33 cM | chromosome location | mouse chr4 D2.2\|4 61.33 cM | 1 | 1 | 14556 | 115 | Dlgap3 | | 7.901e-03 | -4.84 | mouse chr2 H4\|2 97.94 cM | chromosome location | mouse chr2 H4\|2 97.94 cM | 1 | 1 | 14556 | 115 | Ctsz | | 7.901e-03 | -4.84 | mouse chr7 B1\|7 19.29 cM | chromosome location | mouse chr7 B1\|7 19.29 cM | 1 | 1 | 14556 | 115 | Lgi4 | | 7.901e-03 | -4.84 | mouse chr11 C\|11 50.85 cM | chromosome location | mouse chr11 C\|11 50.85 cM | 1 | 1 | 14556 | 115 | Ccl6 | | 7.901e-03 | -4.84 | mouse chr17 D\|17 29.8 cM | chromosome location | mouse chr17 D\|17 29.8 cM | 1 | 1 | 14556 | 115 | Adgre1 | | 7.901e-03 | -4.84 | mouse chr1 E4\|1 58.09 cM | chromosome location | mouse chr1 E4\|1 58.09 cM | 1 | 1 | 14556 | 115 | Fmod | | 7.901e-03 | -4.84 | mouse chr7 B3\|7 30.51 cM | chromosome location | mouse chr7 B3\|7 30.51 cM | 1 | 1 | 14556 | 115 | Saa3 | | 7.901e-03 | -4.84 | mouse chr17 D\|17 29.72 cM | chromosome location | mouse chr17 D\|17 29.72 cM | 1 | 1 | 14556 | 115 | C3 | | 7.901e-03 | -4.84 | mouse chr15 E1\|15 36.81 cM | chromosome location | mouse chr15 E1\|15 36.81 cM | 1 | 1 | 14556 | 115 | Myh9 | | 7.901e-03 | -4.84 | mouse chr9 B\|9 31.18 cM | chromosome location | mouse chr9 B\|9 31.18 cM | 1 | 1 | 14556 | 115 | Csk | | 7.901e-03 | -4.84 | mouse chr7 B3\|7 30.6 cM | chromosome location | mouse chr7 B3\|7 30.6 cM | 1 | 1 | 14556 | 115 | Ldha | | 7.901e-03 | -4.84 | mouse chr17 29.17 cM\|17 D | chromosome location | mouse chr17 29.17 cM\|17 D | 1 | 1 | 14556 | 115 | Lrg1 | | 7.901e-03 | -4.84 | mouse chr5 G2\|5 81.8 cM | chromosome location | mouse chr5 G2\|5 81.8 cM | 1 | 1 | 14556 | 115 | Actb | | 7.901e-03 | -4.84 | mouse chr1 H3\|1 78.8 cM | chromosome location | mouse chr1 H3\|1 78.8 cM | 1 | 1 | 14556 | 115 | Fcgr3 | | 7.901e-03 | -4.84 | mouse chr1 B\|1 18.81 cM | chromosome location | mouse chr1 B\|1 18.81 cM | 1 | 1 | 14556 | 115 | Il1r1 | | 7.901e-03 | -4.84 | mouse chr13 A3.2\|13 13.79 cM | chromosome location | mouse chr13 A3.2\|13 13.79 cM | 1 | 1 | 14556 | 115 | Serpinb9 | | 7.917e-03 | -4.84 | ethanolamine-phosphate phospho-lyase activity | molecular function | GO:0050459 | 1 | 1 | 13516 | 107 | Etnppl | | 7.917e-03 | -4.84 | C5a anaphylatoxin chemotactic receptor binding | molecular function | GO:0031714 | 1 | 1 | 13516 | 107 | C3 | | 7.917e-03 | -4.84 | macrophage colony-stimulating factor receptor activity | molecular function | GO:0005011 | 1 | 1 | 13516 | 107 | Csf1r | | 7.917e-03 | -4.84 | alpha-1,6-mannosyltransferase activity | molecular function | GO:0000009 | 1 | 1 | 13516 | 107 | Alg12 | | 7.917e-03 | -4.84 | ligase regulator activity | molecular function | GO:0055103 | 1 | 1 | 13516 | 107 | Capn3 | | 7.917e-03 | -4.84 | C5L2 anaphylatoxin chemotactic receptor binding | molecular function | GO:0031715 | 1 | 1 | 13516 | 107 | C3 | | 7.917e-03 | -4.84 | extracellularly glutamate-gated chloride channel activity | molecular function | GO:0008068 | 1 | 1 | 13516 | 107 | Slc17a7 | | 7.917e-03 | -4.84 | dol-P-Man:Man(7)GlcNAc(2)-PP-Dol alpha-1,6-mannosyltransferase activity | molecular function | GO:0052917 | 1 | 1 | 13516 | 107 | Alg12 | | 7.917e-03 | -4.84 | sodium-dependent phosphate transmembrane transporter activity | molecular function | GO:0015321 | 1 | 1 | 13516 | 107 | Slc17a7 | | 7.917e-03 | -4.84 | dolichyl-pyrophosphate Man7GlcNAc2 alpha-1,6-mannosyltransferase activity | molecular function | GO:0052824 | 1 | 1 | 13516 | 107 | Alg12 | | 7.917e-03 | -4.84 | ciliary neurotrophic factor binding | molecular function | GO:0070119 | 1 | 1 | 13516 | 107 | Il6ra | | 7.917e-03 | -4.84 | interleukin-1, type I, activating receptor activity | molecular function | GO:0004909 | 1 | 1 | 13516 | 107 | Il1r1 | | 7.943e-03 | -4.84 | GO\_COLLAGEN\_BINDING | MSigDB lists | GO\_COLLAGEN\_BINDING | 51 | 3 | 12187 | 98 | Ctss,Vwf,Ctsb | | 7.943e-03 | -4.84 | GO\_POSITIVE\_REGULATION\_OF\_LEUKOCYTE\_MEDIATED\_IMMUNITY | MSigDB lists | GO\_POSITIVE\_REGULATION\_OF\_LEUKOCYTE\_MEDIATED\_IMMUNITY | 51 | 3 | 12187 | 98 | C3,Vamp8,Fcer1g | | 7.954e-03 | -4.83 | NOLC1 (nucleolar and coiled-body phosphoprotein 1) | protein interactions | 9221 | 12 | 2 | 6802 | 78 | Myh9,Actb | | 7.954e-03 | -4.83 | PPP1CC (protein phosphatase 1 catalytic subunit gamma) | protein interactions | 5501 | 12 | 2 | 6802 | 78 | Myh9,Flnb | | 7.954e-03 | -4.83 | SIPA1L3 (signal induced proliferation associated 1 like 3) | protein interactions | 23094 | 12 | 2 | 6802 | 78 | Myh9,Actb | | 7.954e-03 | -4.83 | CLINT1 (clathrin interactor 1) | protein interactions | 9685 | 12 | 2 | 6802 | 78 | Myh9,Actb | | 7.954e-03 | -4.83 | ARPC1B (actin related protein 2/3 complex subunit 1B) | protein interactions | 10095 | 12 | 2 | 6802 | 78 | Myh9,Actb | | 7.981e-03 | -4.83 | PTEN\_DN.V1\_DN | MSigDB lists | PTEN\_DN.V1\_DN | 98 | 4 | 12187 | 98 | Lin7a,Cplx2,Vim,Mc4r | | 7.981e-03 | -4.83 | GO\_NEGATIVE\_REGULATION\_OF\_HOMEOSTATIC\_PROCESS | MSigDB lists | GO\_NEGATIVE\_REGULATION\_OF\_HOMEOSTATIC\_PROCESS | 98 | 4 | 12187 | 98 | Csk,Ccr5,Fcer1g,Ptprc | | 7.981e-03 | -4.83 | GSE43863\_NAIVE\_VS\_LY6C\_LOW\_CXCR5NEG\_CD4\_EFF\_TCELL\_D6\_LCMV\_UP | MSigDB lists | GSE43863\_NAIVE\_VS\_LY6C\_LOW\_CXCR5NEG\_CD4\_EFF\_TCELL\_D6\_LCMV\_UP | 98 | 4 | 12187 | 98 | Zfp786,Cox4i1,Tmem176a,Rpl31 | | 7.981e-03 | -4.83 | GO\_POSITIVE\_REGULATION\_OF\_IMMUNE\_EFFECTOR\_PROCESS | MSigDB lists | GO\_POSITIVE\_REGULATION\_OF\_IMMUNE\_EFFECTOR\_PROCESS | 98 | 4 | 12187 | 98 | C3,Vamp8,Fcer1g,Rac2 | | 7.981e-03 | -4.83 | GO\_POSITIVE\_REGULATION\_OF\_RESPONSE\_TO\_WOUNDING | MSigDB lists | GO\_POSITIVE\_REGULATION\_OF\_RESPONSE\_TO\_WOUNDING | 98 | 4 | 12187 | 98 | C3,Vamp8,Fcer1g,Capn3 | | 7.996e-03 | -4.83 | GSE30083\_SP1\_VS\_SP4\_THYMOCYTE\_DN | MSigDB lists | GSE30083\_SP1\_VS\_SP4\_THYMOCYTE\_DN | 154 | 5 | 12187 | 98 | Ikzf3,Ctss,Tnfsf8,Il6ra,Ahnak | | 8.015e-03 | -4.83 | GO\_POSITIVE\_REGULATION\_OF\_MOLECULAR\_FUNCTION | MSigDB lists | GO\_POSITIVE\_REGULATION\_OF\_MOLECULAR\_FUNCTION | 1413 | 20 | 12187 | 98 | Csf1r,Lfng,Ndufa13,Thy1,Ctsa,Il6ra,Kl,Csk,Ctss,Hspa2,Camk2b,Capn3,Clu,Psme1,Rap1gap,Ptprc,Psmb10,Fgf18,Rab3il1,Sh3bgrl3 | | 8.029e-03 | -4.82 | mucin granule | cellular component | GO:0098594 | 1 | 1 | 13825 | 111 | Vamp8 | | 8.029e-03 | -4.82 | azurophil granule membrane | cellular component | GO:0035577 | 1 | 1 | 13825 | 111 | Vamp8 | | 8.029e-03 | -4.82 | Fc-epsilon receptor I complex | cellular component | GO:0032998 | 1 | 1 | 13825 | 111 | Fcer1g | | 8.029e-03 | -4.82 | perinuclear endoplasmic reticulum lumen | cellular component | GO:0099020 | 1 | 1 | 13825 | 111 | Clu | | 8.029e-03 | -4.82 | Fc receptor complex | cellular component | GO:0032997 | 1 | 1 | 13825 | 111 | Fcer1g | | 8.029e-03 | -4.82 | complement component C1 complex | cellular component | GO:0005602 | 1 | 1 | 13825 | 111 | C1qb | | 8.047e-03 | -4.82 | GO\_ASTROCYTE\_DEVELOPMENT | MSigDB lists | GO\_ASTROCYTE\_DEVELOPMENT | 17 | 2 | 12187 | 98 | Vim,Gfap | | 8.047e-03 | -4.82 | OUYANG\_PROSTATE\_CANCER\_MARKERS | MSigDB lists | OUYANG\_PROSTATE\_CANCER\_MARKERS | 17 | 2 | 12187 | 98 | Clu,Ctsb | | 8.047e-03 | -4.82 | GO\_RESPONSE\_TO\_THYROID\_HORMONE | MSigDB lists | GO\_RESPONSE\_TO\_THYROID\_HORMONE | 17 | 2 | 12187 | 98 | Ctss,Ctsb | | 8.047e-03 | -4.82 | VALK\_AML\_CLUSTER\_7 | MSigDB lists | VALK\_AML\_CLUSTER\_7 | 17 | 2 | 12187 | 98 | Rap1gap,Slc2a1 | | 8.047e-03 | -4.82 | CROONQUIST\_IL6\_DEPRIVATION\_UP | MSigDB lists | CROONQUIST\_IL6\_DEPRIVATION\_UP | 17 | 2 | 12187 | 98 | Pde1a,Ahnak | | 8.067e-03 | -4.82 | apical plasma membrane | cellular component | GO:0016324 | 217 | 6 | 13825 | 111 | Thy1,Cyba,Il6ra,Slc26a2,Ctsb,Slc2a1 | | 8.077e-03 | -4.82 | GO\_PROTEIN\_LOCALIZATION | MSigDB lists | GO\_PROTEIN\_LOCALIZATION | 1512 | 21 | 12187 | 98 | Rpl31,Rps5,Exoc3l4,Myh9,Fcer1g,Ndufa13,Mc4r,Ctsa,Lin7a,Rpl26,Vamp8,Rpl14,Rps20,Capn3,Rpl13,Clu,Rps8,Rab3il1,Ptprn,Flnb,Rac2 | | 8.084e-03 | -4.82 | positive regulation of signaling | biological process | GO:0023056 | 1397 | 20 | 13711 | 111 | Hcls1,Csf1r,Mt3,Lfng,Cyba,Gfap,C3,Csk,Camk2b,Il6ra,Fgf18,Ccl6,Clu,Ptprc,Rpl26,Tmem106a,Kl,Lrg1,Il1r1,Vamp8 | | 8.096e-03 | -4.82 | plasma membrane raft localization | biological process | GO:0044856 | 1 | 1 | 13711 | 111 | Ptprc | | 8.096e-03 | -4.82 | negative regulation of lipoprotein oxidation | biological process | GO:0034443 | 1 | 1 | 13711 | 111 | Apod | | 8.096e-03 | -4.82 | positive regulation of lysosome organization | biological process | GO:1905673 | 1 | 1 | 13711 | 111 | Grn | | 8.096e-03 | -4.82 | calcium-dependent self proteolysis | biological process | GO:1990092 | 1 | 1 | 13711 | 111 | Capn3 | | 8.096e-03 | -4.82 | regulation of cellular response to tunicamycin | biological process | GO:1905894 | 1 | 1 | 13711 | 111 | Clu | | 8.096e-03 | -4.82 | positive regulation of Schwann cell proliferation | biological process | GO:0010625 | 1 | 1 | 13711 | 111 | Gfap | | 8.096e-03 | -4.82 | glycolytic fermentation | biological process | GO:0019660 | 1 | 1 | 13711 | 111 | Ldha | | 8.096e-03 | -4.82 | positive regulation of type B pancreatic cell proliferation | biological process | GO:1904692 | 1 | 1 | 13711 | 111 | Ptprn | | 8.096e-03 | -4.82 | regulation of satellite cell activation involved in skeletal muscle regeneration | biological process | GO:0014717 | 1 | 1 | 13711 | 111 | Capn3 | | 8.096e-03 | -4.82 | regulation of translation involved in cellular response to UV | biological process | GO:1904803 | 1 | 1 | 13711 | 111 | Rpl26 | | 8.096e-03 | -4.82 | negative regulation of microvillus assembly | biological process | GO:1903697 | 1 | 1 | 13711 | 111 | Rap1gap | | 8.096e-03 | -4.82 | smooth muscle hypertrophy | biological process | GO:0014895 | 1 | 1 | 13711 | 111 | Cyba | | 8.096e-03 | -4.82 | positive regulation of pancreatic amylase secretion | biological process | GO:1902278 | 1 | 1 | 13711 | 111 | Vamp8 | | 8.096e-03 | -4.82 | immunoglobulin biosynthetic process | biological process | GO:0002378 | 1 | 1 | 13711 | 111 | Ptprc | | 8.096e-03 | -4.82 | positive regulation of neurofibrillary tangle assembly | biological process | GO:1902998 | 1 | 1 | 13711 | 111 | Clu | | 8.096e-03 | -4.82 | negative regulation of cellular response to tunicamycin | biological process | GO:1905895 | 1 | 1 | 13711 | 111 | Clu | | 8.096e-03 | -4.82 | regulation of neuronal signal transduction | biological process | GO:1902847 | 1 | 1 | 13711 | 111 | Clu | | 8.096e-03 | -4.82 | cellular response to glial cell derived neurotrophic factor | biological process | GO:1990792 | 1 | 1 | 13711 | 111 | Rap1gap | | 8.096e-03 | -4.82 | acetaldehyde biosynthetic process | biological process | GO:0046186 | 1 | 1 | 13711 | 111 | Adh1 | | 8.096e-03 | -4.82 | mucus secretion | biological process | GO:0070254 | 1 | 1 | 13711 | 111 | Vamp8 | | 8.096e-03 | -4.82 | response to glial cell derived neurotrophic factor | biological process | GO:1990790 | 1 | 1 | 13711 | 111 | Rap1gap | | 8.096e-03 | -4.82 | regulation of thyroid gland epithelial cell proliferation | biological process | GO:1904441 | 1 | 1 | 13711 | 111 | Rap1gap | | 8.096e-03 | -4.82 | plasma membrane raft distribution | biological process | GO:0044855 | 1 | 1 | 13711 | 111 | Ptprc | | 8.096e-03 | -4.82 | negative regulation of cellular response to thapsigargin | biological process | GO:1905892 | 1 | 1 | 13711 | 111 | Clu | | 8.096e-03 | -4.82 | glucose catabolic process to lactate | biological process | GO:0019659 | 1 | 1 | 13711 | 111 | Ldha | | 8.096e-03 | -4.82 | fermentation | biological process | GO:0006113 | 1 | 1 | 13711 | 111 | Ldha | | 8.096e-03 | -4.82 | CD80 biosynthetic process | biological process | GO:0035780 | 1 | 1 | 13711 | 111 | Tmem106a | | 8.096e-03 | -4.82 | positive regulation of interleukin-1-mediated signaling pathway | biological process | GO:2000661 | 1 | 1 | 13711 | 111 | Il1r1 | | 8.096e-03 | -4.82 | basement membrane disassembly | biological process | GO:0034769 | 1 | 1 | 13711 | 111 | Ctss | | 8.096e-03 | -4.82 | regulation of lipoprotein lipid oxidation | biological process | GO:0060587 | 1 | 1 | 13711 | 111 | Apod | | 8.096e-03 | -4.82 | cellular response to cytochalasin B | biological process | GO:0072749 | 1 | 1 | 13711 | 111 | Actb | | 8.096e-03 | -4.82 | acetaldehyde metabolic process | biological process | GO:0006117 | 1 | 1 | 13711 | 111 | Adh1 | | 8.096e-03 | -4.82 | negative regulation of glomerular filtration by angiotensin | biological process | GO:0003106 | 1 | 1 | 13711 | 111 | Cyba | | 8.096e-03 | -4.82 | regulation of cellular response to thapsigargin | biological process | GO:1905891 | 1 | 1 | 13711 | 111 | Clu | | 8.096e-03 | -4.82 | negative regulation of thyroid gland epithelial cell proliferation | biological process | GO:1904442 | 1 | 1 | 13711 | 111 | Rap1gap | | 8.096e-03 | -4.82 | negative regulation of lipoprotein lipid oxidation | biological process | GO:0060588 | 1 | 1 | 13711 | 111 | Apod | | 8.096e-03 | -4.82 | sequestering of neurotransmitter | biological process | GO:0042137 | 1 | 1 | 13711 | 111 | Slc17a7 | | 8.096e-03 | -4.82 | response to cytochalasin B | biological process | GO:1901328 | 1 | 1 | 13711 | 111 | Actb | | 8.096e-03 | -4.82 | regulation of lipoprotein oxidation | biological process | GO:0034442 | 1 | 1 | 13711 | 111 | Apod | | 8.096e-03 | -4.82 | positive regulation of activation of membrane attack complex | biological process | GO:0001970 | 1 | 1 | 13711 | 111 | C3 | | 8.096e-03 | -4.82 | negative regulation of actin filament severing | biological process | GO:1903919 | 1 | 1 | 13711 | 111 | Myh9 | | 8.096e-03 | -4.82 | CD86 biosynthetic process | biological process | GO:0035781 | 1 | 1 | 13711 | 111 | Tmem106a | | 8.096e-03 | -4.82 | sodium-dependent self proteolysis | biological process | GO:1990091 | 1 | 1 | 13711 | 111 | Capn3 | | 8.096e-03 | -4.82 | glucose catabolic process to lactate via pyruvate | biological process | GO:0019661 | 1 | 1 | 13711 | 111 | Ldha | | 8.096e-03 | -4.82 | islet amyloid polypeptide processing | biological process | GO:0034231 | 1 | 1 | 13711 | 111 | Pcsk2 | | 8.096e-03 | -4.82 | negative regulation of renal output by angiotensin | biological process | GO:0003083 | 1 | 1 | 13711 | 111 | Cyba | | 8.096e-03 | -4.82 | positive regulation of satellite cell activation involved in skeletal muscle regeneration | biological process | GO:0014718 | 1 | 1 | 13711 | 111 | Capn3 | | 8.096e-03 | -4.82 | regulation of actin filament severing | biological process | GO:1903918 | 1 | 1 | 13711 | 111 | Myh9 | | 8.096e-03 | -4.82 | inhibitory G protein-coupled receptor phosphorylation | biological process | GO:0002030 | 1 | 1 | 13711 | 111 | Camk2b | | 8.096e-03 | -4.82 | cadmium ion homeostasis | biological process | GO:0055073 | 1 | 1 | 13711 | 111 | Mt3 | | 8.096e-03 | -4.82 | dichotomous subdivision of terminal units involved in mammary gland duct morphogenesis | biological process | GO:0060598 | 1 | 1 | 13711 | 111 | Tfap2c | | 8.096e-03 | -4.82 | sodium-dependent phosphate transport | biological process | GO:0044341 | 1 | 1 | 13711 | 111 | Slc17a7 | | 8.111e-03 | -4.81 | positive regulation of receptor signaling pathway via JAK-STAT | biological process | GO:0046427 | 51 | 3 | 13711 | 111 | Csf1r,Il6ra,Hcls1 | | 8.124e-03 | -4.81 | Ig-like\_dom | interpro domains | IPR007110 | 277 | 7 | 13788 | 114 | Fcrls,Fcgr3,Thy1,Treml2,Csf1r,Il1r1,Il6ra | | 8.160e-03 | -4.81 | regulation of microglial cell activation | biological process | GO:1903978 | 17 | 2 | 13711 | 111 | Grn,C1qa | | 8.160e-03 | -4.81 | monosaccharide transmembrane transport | biological process | GO:0015749 | 17 | 2 | 13711 | 111 | Slc2a1,Mfsd4a | | 8.160e-03 | -4.81 | positive regulation of mast cell activation | biological process | GO:0033005 | 17 | 2 | 13711 | 111 | Vamp8,Fcer1g | | 8.160e-03 | -4.81 | acute-phase response | biological process | GO:0006953 | 17 | 2 | 13711 | 111 | Saa3,Serpina3n | | 8.160e-03 | -4.81 | collagen catabolic process | biological process | GO:0030574 | 17 | 2 | 13711 | 111 | Ctss,Ctsb | | 8.169e-03 | -4.81 | negative regulation of immune system process | biological process | GO:0002683 | 355 | 8 | 13711 | 111 | Serpinb9,Thy1,Grn,Tmem176a,Ptprc,Apod,Fcer1g,Cd68 | | 8.182e-03 | -4.81 | enzyme binding | molecular function | GO:0019899 | 2041 | 26 | 13516 | 107 | Bhlhe41,Ptprn,Ccr5,Camk2b,Il1r1,Csk,Capn3,Brk1,Clu,Mc4r,Ptprc,Hspa2,Vim,Actb,Rac2,Csf1r,Rap1gap,Phyhip,Thy1,Serpinb9,Slc2a1,Gfap,Rab3il1,Vwf,Hcls1,Il6ra | | 8.191e-03 | -4.80 | YAGI\_AML\_WITH\_11Q23\_REARRANGED | MSigDB lists | YAGI\_AML\_WITH\_11Q23\_REARRANGED | 286 | 7 | 12187 | 98 | Capn3,Myh9,Cd68,Tspan4,Hcls1,Il6ra,Lin7a | | 8.204e-03 | -4.80 | interspecies interaction between organisms | biological process | GO:0044419 | 216 | 6 | 13711 | 111 | Csf1r,Mpeg1,Serpinb9,Ctsb,Vamp8,Ccr5 | | 8.209e-03 | -4.80 | positive regulation of molecular function | biological process | GO:0044093 | 1399 | 20 | 13711 | 111 | Fgf18,Psme1,Capn3,Il6ra,Camk2b,Csk,Ndufa13,Cyba,Lfng,Mt3,Hcls1,Csf1r,Rap1gap,Thy1,Ctss,Grn,Hspa2,Ptprc,Clu,Ccl6 | | 8.211e-03 | -4.80 | GSE26030\_UNSTIM\_VS\_RESTIM\_TH17\_DAY5\_POST\_POLARIZATION\_UP | MSigDB lists | GSE26030\_UNSTIM\_VS\_RESTIM\_TH17\_DAY5\_POST\_POLARIZATION\_UP | 155 | 5 | 12187 | 98 | Ctsa,Camk2b,Thy1,Vamp8,Sh3bgrl3 | | 8.211e-03 | -4.80 | GSE20366\_EX\_VIVO\_VS\_DEC205\_CONVERSION\_NAIVE\_CD4\_TCELL\_UP | MSigDB lists | GSE20366\_EX\_VIVO\_VS\_DEC205\_CONVERSION\_NAIVE\_CD4\_TCELL\_UP | 155 | 5 | 12187 | 98 | Ccr5,Fam111a,Ly6e,Il1r1,Tmem176a | | 8.259e-03 | -4.80 | MODULE\_13 | MSigDB lists | MODULE\_13 | 359 | 8 | 12187 | 98 | Camk2b,Rap1gap,Rac2,Ly6e,Slc17a7,Mgp,Fcgr3,Slc2a1 | | 8.268e-03 | -4.80 | Proteasome\_beta10 | interpro domains | IPR034384 | 1 | 1 | 13788 | 114 | Psmb10 | | 8.268e-03 | -4.80 | ApoD\_vertbrte | interpro domains | IPR026222 | 1 | 1 | 13788 | 114 | Apod | | 8.268e-03 | -4.80 | Von\_Willebrand\_factor | interpro domains | IPR037578 | 1 | 1 | 13788 | 114 | Vwf | | 8.268e-03 | -4.80 | NTR\_complement\_C3 | interpro domains | IPR035815 | 1 | 1 | 13788 | 114 | C3 | | 8.268e-03 | -4.80 | Granulin | interpro domains | IPR000118 | 1 | 1 | 13788 | 114 | Grn | | 8.268e-03 | -4.80 | Ribosomal\_S10\_CS | interpro domains | IPR018268 | 1 | 1 | 13788 | 114 | Rps20 | | 8.268e-03 | -4.80 | Granulin\_sf | interpro domains | IPR037277 | 1 | 1 | 13788 | 114 | Grn | | 8.268e-03 | -4.80 | PPFIBP1 | interpro domains | IPR030437 | 1 | 1 | 13788 | 114 | Ppfibp1 | | 8.268e-03 | -4.80 | Ribosomal\_S5/S7\_euk/arc | interpro domains | IPR005716 | 1 | 1 | 13788 | 114 | Rps5 | | 8.268e-03 | -4.80 | Ribosomal\_S7\_CS | interpro domains | IPR020606 | 1 | 1 | 13788 | 114 | Rps5 | | 8.268e-03 | -4.80 | Ribosomal\_L31e\_dom\_sf | interpro domains | IPR023621 | 1 | 1 | 13788 | 114 | Rpl31 | | 8.268e-03 | -4.80 | CD3\_zeta/IgE\_Fc\_rcpt\_gamma | interpro domains | IPR021663 | 1 | 1 | 13788 | 114 | Fcer1g | | 8.268e-03 | -4.80 | FLNB | interpro domains | IPR029874 | 1 | 1 | 13788 | 114 | Flnb | | 8.268e-03 | -4.80 | DLGAP3 | interpro domains | IPR030526 | 1 | 1 | 13788 | 114 | Dlgap3 | | 8.268e-03 | -4.80 | Ribosomal\_S8e | interpro domains | IPR001047 | 1 | 1 | 13788 | 114 | Rps8 | | 8.268e-03 | -4.80 | Clusterin-like | interpro domains | IPR000753 | 1 | 1 | 13788 | 114 | Clu | | 8.268e-03 | -4.80 | Ribosomal\_L26/L24P\_euk/arc | interpro domains | IPR005756 | 1 | 1 | 13788 | 114 | Rpl26 | | 8.268e-03 | -4.80 | Granulin\_fam | interpro domains | IPR039036 | 1 | 1 | 13788 | 114 | Grn | | 8.268e-03 | -4.80 | Ribosomal\_S8e\_CS | interpro domains | IPR018283 | 1 | 1 | 13788 | 114 | Rps8 | | 8.268e-03 | -4.80 | GRIM-19 | interpro domains | IPR009346 | 1 | 1 | 13788 | 114 | Ndufa13 | | 8.268e-03 | -4.80 | Ribosomal\_S10\_euk/arc | interpro domains | IPR005729 | 1 | 1 | 13788 | 114 | Rps20 | | 8.268e-03 | -4.80 | FCER1G | interpro domains | IPR042340 | 1 | 1 | 13788 | 114 | Fcer1g | | 8.268e-03 | -4.80 | RPL14\_KOW | interpro domains | IPR041985 | 1 | 1 | 13788 | 114 | Rpl14 | | 8.268e-03 | -4.80 | Klotho | interpro domains | IPR028546 | 1 | 1 | 13788 | 114 | Kl | | 8.268e-03 | -4.80 | Vimentin | interpro domains | IPR027699 | 1 | 1 | 13788 | 114 | Vim | | 8.268e-03 | -4.80 | Phos\_immunorcpt\_sig\_ITAM | interpro domains | IPR003110 | 1 | 1 | 13788 | 114 | Fcer1g | | 8.268e-03 | -4.80 | Complement\_C1q\_A | interpro domains | IPR037572 | 1 | 1 | 13788 | 114 | C1qa | | 8.268e-03 | -4.80 | ApolipopD | interpro domains | IPR002969 | 1 | 1 | 13788 | 114 | Apod | | 8.268e-03 | -4.80 | FGF18 | interpro domains | IPR028289 | 1 | 1 | 13788 | 114 | Fgf18 | | 8.268e-03 | -4.80 | GPCR\_2\_EMR1-like\_rcpt | interpro domains | IPR001740 | 1 | 1 | 13788 | 114 | Adgre1 | | 8.268e-03 | -4.80 | GFAP | interpro domains | IPR027701 | 1 | 1 | 13788 | 114 | Gfap | | 8.268e-03 | -4.80 | C3/4/5\_MG1 | interpro domains | IPR041425 | 1 | 1 | 13788 | 114 | C3 | | 8.268e-03 | -4.80 | Ptprc | interpro domains | IPR016335 | 1 | 1 | 13788 | 114 | Ptprc | | 8.268e-03 | -4.80 | TF\_AP2\_gamma | interpro domains | IPR008123 | 1 | 1 | 13788 | 114 | Tfap2c | | 8.268e-03 | -4.80 | Spint1 | interpro domains | IPR042482 | 1 | 1 | 13788 | 114 | Spint1 | | 8.268e-03 | -4.80 | Ribosomal\_L31e | interpro domains | IPR000054 | 1 | 1 | 13788 | 114 | Rpl31 | | 8.268e-03 | -4.80 | PTP\_recept\_N | interpro domains | IPR024739 | 1 | 1 | 13788 | 114 | Ptprc | | 8.268e-03 | -4.80 | CAMPATH-1 | interpro domains | IPR026643 | 1 | 1 | 13788 | 114 | Cd52 | | 8.268e-03 | -4.80 | Ribosomal\_L31e\_CS | interpro domains | IPR020052 | 1 | 1 | 13788 | 114 | Rpl31 | | 8.268e-03 | -4.80 | Ribosomal\_protein\_S8e\_euk | interpro domains | IPR042563 | 1 | 1 | 13788 | 114 | Rps8 | | 8.268e-03 | -4.80 | MGP | interpro domains | IPR027118 | 1 | 1 | 13788 | 114 | Mgp | | 8.268e-03 | -4.80 | Clusterin\_C | interpro domains | IPR016015 | 1 | 1 | 13788 | 114 | Clu | | 8.268e-03 | -4.80 | Ribosomal\_protein\_L14 | interpro domains | IPR039660 | 1 | 1 | 13788 | 114 | Rpl14 | | 8.268e-03 | -4.80 | THY1 | interpro domains | IPR033292 | 1 | 1 | 13788 | 114 | Thy1 | | 8.268e-03 | -4.80 | IGFBP-6 | interpro domains | IPR022326 | 1 | 1 | 13788 | 114 | Igfbp6 | | 8.268e-03 | -4.80 | Ribosomal\_L13e | interpro domains | IPR001380 | 1 | 1 | 13788 | 114 | Rpl13 | | 8.268e-03 | -4.80 | Chemokine\_CCR5 | interpro domains | IPR002240 | 1 | 1 | 13788 | 114 | Ccr5 | | 8.268e-03 | -4.80 | CAPN3/9 | interpro domains | IPR029531 | 1 | 1 | 13788 | 114 | Capn3 | | 8.268e-03 | -4.80 | Ribosomal\_L14e\_dom | interpro domains | IPR002784 | 1 | 1 | 13788 | 114 | Rpl14 | | 8.268e-03 | -4.80 | Clusterin\_CS | interpro domains | IPR033986 | 1 | 1 | 13788 | 114 | Clu | | 8.268e-03 | -4.80 | Fibromodulin | interpro domains | IPR027215 | 1 | 1 | 13788 | 114 | Fmod | | 8.268e-03 | -4.80 | Cyt\_b558\_asu | interpro domains | IPR007732 | 1 | 1 | 13788 | 114 | Cyba | | 8.268e-03 | -4.80 | ALG12 | interpro domains | IPR039485 | 1 | 1 | 13788 | 114 | Alg12 | | 8.268e-03 | -4.80 | SLC26A2 | interpro domains | IPR030280 | 1 | 1 | 13788 | 114 | Slc26a2 | | 8.268e-03 | -4.80 | Glu\_transpt\_1 | interpro domains | IPR002439 | 1 | 1 | 13788 | 114 | Slc2a1 | | 8.268e-03 | -4.80 | S-100A6 | interpro domains | IPR034118 | 1 | 1 | 13788 | 114 | S100a6 | | 8.268e-03 | -4.80 | CSF-1\_receptor | interpro domains | IPR030658 | 1 | 1 | 13788 | 114 | Csf1r | | 8.268e-03 | -4.80 | Ribosomal\_L13e\_CS | interpro domains | IPR018256 | 1 | 1 | 13788 | 114 | Rpl13 | | 8.268e-03 | -4.80 | TNF\_CD30\_ligand\_type | interpro domains | IPR021185 | 1 | 1 | 13788 | 114 | Tnfsf8 | | 8.268e-03 | -4.80 | Complement\_C3-like | interpro domains | IPR035711 | 1 | 1 | 13788 | 114 | C3 | | 8.268e-03 | -4.80 | BRICK1 | interpro domains | IPR033378 | 1 | 1 | 13788 | 114 | Brk1 | | 8.268e-03 | -4.80 | F1F0-ATPsyn\_F\_prd | interpro domains | IPR019344 | 1 | 1 | 13788 | 114 | Atp5j2 | | 8.268e-03 | -4.80 | Complement\_C1qB | interpro domains | IPR037573 | 1 | 1 | 13788 | 114 | C1qb | | 8.268e-03 | -4.80 | HS1 | interpro domains | IPR028534 | 1 | 1 | 13788 | 114 | Hcls1 | | 8.268e-03 | -4.80 | Osteocalcin/MGP | interpro domains | IPR002384 | 1 | 1 | 13788 | 114 | Mgp | | 8.268e-03 | -4.80 | KLHL6 | interpro domains | IPR029851 | 1 | 1 | 13788 | 114 | Klhl6 | | 8.268e-03 | -4.80 | VWA\_N2 | interpro domains | IPR032361 | 1 | 1 | 13788 | 114 | Vwf | | 8.268e-03 | -4.80 | Propeptide\_C1A | interpro domains | IPR012599 | 1 | 1 | 13788 | 114 | Ctsb | | 8.268e-03 | -4.80 | Clusterin\_N | interpro domains | IPR016014 | 1 | 1 | 13788 | 114 | Clu | | 8.268e-03 | -4.80 | IL-1\_rcpt\_I-typ | interpro domains | IPR004076 | 1 | 1 | 13788 | 114 | Il1r1 | | 8.268e-03 | -4.80 | Ser\_caboxypep\_his\_AS | interpro domains | IPR033124 | 1 | 1 | 13788 | 114 | Ctsa | | 8.268e-03 | -4.80 | CTSZ | interpro domains | IPR033157 | 1 | 1 | 13788 | 114 | Ctsz | | 8.268e-03 | -4.80 | Mcort\_rcpt\_4 | interpro domains | IPR000155 | 1 | 1 | 13788 | 114 | Mc4r | | 8.268e-03 | -4.80 | MPEG1 | interpro domains | IPR039707 | 1 | 1 | 13788 | 114 | Mpeg1 | | 8.268e-03 | -4.80 | TRNP1 | interpro domains | IPR040266 | 1 | 1 | 13788 | 114 | Trnp1 | | 8.268e-03 | -4.80 | Clusterin | interpro domains | IPR016016 | 1 | 1 | 13788 | 114 | Clu | | 8.313e-03 | -4.79 | - | gene3d domains | 2.30.30.30 | 15 | 2 | 6647 | 62 | Rpl14,Rpl26 | | 8.361e-03 | -4.78 | MARTINEZ\_TP53\_TARGETS\_DN | MSigDB lists | MARTINEZ\_TP53\_TARGETS\_DN | 436 | 9 | 12187 | 98 | Mgll,Clu,Camk2b,Rps8,Csf1r,Vamp8,Rpl26,Ctsb,Spint1 | | 8.381e-03 | -4.78 | GO\_REGULATION\_OF\_RELEASE\_OF\_SEQUESTERED\_CALCIUM\_ION\_INTO\_CYTOSOL | MSigDB lists | GO\_REGULATION\_OF\_RELEASE\_OF\_SEQUESTERED\_CALCIUM\_ION\_INTO\_CYTOSOL | 52 | 3 | 12187 | 98 | Cyba,Thy1,Capn3 | | 8.381e-03 | -4.78 | GO\_PLATELET\_ALPHA\_GRANULE | MSigDB lists | GO\_PLATELET\_ALPHA\_GRANULE | 52 | 3 | 12187 | 98 | Serpina3n,Clu,Vwf | | 8.381e-03 | -4.78 | GO\_LEUKOCYTE\_PROLIFERATION | MSigDB lists | GO\_LEUKOCYTE\_PROLIFERATION | 52 | 3 | 12187 | 98 | Psmb10,Ptprc,Clu | | 8.381e-03 | -4.78 | GNF2\_HPX | MSigDB lists | GNF2\_HPX | 52 | 3 | 12187 | 98 | C3,Adh1,Tmem176a | | 8.384e-03 | -4.78 | Lipocalin\_2 | pfam domains | PF08212 | 1 | 1 | 12881 | 108 | Apod | | 8.384e-03 | -4.78 | WRW | pfam domains | PF10206 | 1 | 1 | 12881 | 108 | Atp5j2 | | 8.384e-03 | -4.78 | Propeptide\_C1 | pfam domains | PF08127 | 1 | 1 | 12881 | 108 | Ctsb | | 8.384e-03 | -4.78 | Ribosomal\_L14e | pfam domains | PF01929 | 1 | 1 | 12881 | 108 | Rpl14 | | 8.384e-03 | -4.78 | Ribosomal\_L13e | pfam domains | PF01294 | 1 | 1 | 12881 | 108 | Rpl13 | | 8.384e-03 | -4.78 | ITAM | pfam domains | PF02189 | 1 | 1 | 12881 | 108 | Fcer1g | | 8.384e-03 | -4.78 | TCR\_zetazeta | pfam domains | PF11628 | 1 | 1 | 12881 | 108 | Fcer1g | | 8.384e-03 | -4.78 | VWA\_N2 | pfam domains | PF16164 | 1 | 1 | 12881 | 108 | Vwf | | 8.384e-03 | -4.78 | MG1 | pfam domains | PF17790 | 1 | 1 | 12881 | 108 | C3 | | 8.384e-03 | -4.78 | CD52 | pfam domains | PF15116 | 1 | 1 | 12881 | 108 | Cd52 | | 8.384e-03 | -4.78 | GRIM-19 | pfam domains | PF06212 | 1 | 1 | 12881 | 108 | Ndufa13 | | 8.384e-03 | -4.78 | Ribosomal\_L31e | pfam domains | PF01198 | 1 | 1 | 12881 | 108 | Rpl31 | | 8.384e-03 | -4.78 | Cytochrom\_B558a | pfam domains | PF05038 | 1 | 1 | 12881 | 108 | Cyba | | 8.384e-03 | -4.78 | CD45 | pfam domains | PF12567 | 1 | 1 | 12881 | 108 | Ptprc | | 8.384e-03 | -4.78 | Ribosomal\_L26 | pfam domains | PF16906 | 1 | 1 | 12881 | 108 | Rpl26 | | 8.384e-03 | -4.78 | PTP\_N | pfam domains | PF12453 | 1 | 1 | 12881 | 108 | Ptprc | | 8.384e-03 | -4.78 | Granulin | pfam domains | PF00396 | 1 | 1 | 12881 | 108 | Grn | | 8.384e-03 | -4.78 | Clusterin | pfam domains | PF01093 | 1 | 1 | 12881 | 108 | Clu | | 8.391e-03 | -4.78 | MARTENS\_BOUND\_BY\_PML\_RARA\_FUSION | MSigDB lists | MARTENS\_BOUND\_BY\_PML\_RARA\_FUSION | 360 | 8 | 12187 | 98 | Lfng,Csk,Trnp1,Myh9,Mgll,Sh3bgrl3,Rac2,Flnb | | 8.430e-03 | -4.78 | GSE42021\_TCONV\_PLN\_VS\_CD24HI\_TCONV\_THYMUS\_UP | MSigDB lists | GSE42021\_TCONV\_PLN\_VS\_CD24HI\_TCONV\_THYMUS\_UP | 156 | 5 | 12187 | 98 | Psme1,Fam111a,Ctss,C3,Psmb10 | | 8.430e-03 | -4.78 | GSE42088\_UNINF\_VS\_LEISHMANIA\_INF\_DC\_2H\_DN | MSigDB lists | GSE42088\_UNINF\_VS\_LEISHMANIA\_INF\_DC\_2H\_DN | 156 | 5 | 12187 | 98 | Rps8,Rpl26,Rpl13,Rpl14,Rpl31 | | 8.430e-03 | -4.78 | GSE42021\_CD24HI\_VS\_CD24LOW\_TCONV\_THYMUS\_DN | MSigDB lists | GSE42021\_CD24HI\_VS\_CD24LOW\_TCONV\_THYMUS\_DN | 156 | 5 | 12187 | 98 | Fam111a,Psme1,Ctss,C3,Psmb10 | | 8.494e-03 | -4.77 | GO\_REGULATION\_OF\_IMMUNE\_EFFECTOR\_PROCESS | MSigDB lists | GO\_REGULATION\_OF\_IMMUNE\_EFFECTOR\_PROCESS | 288 | 7 | 12187 | 98 | Vamp8,Serpinb9,Fcer1g,Rac2,Phyhip,Ptprc,C3 | | 8.498e-03 | -4.77 | Serpin\_fam | interpro domains | IPR000215 | 17 | 2 | 13788 | 114 | Serpinb9,Serpina3n | | 8.498e-03 | -4.77 | Serpin\_sf\_1 | interpro domains | IPR042178 | 17 | 2 | 13788 | 114 | Serpinb9,Serpina3n | | 8.498e-03 | -4.77 | Serpin\_sf | interpro domains | IPR036186 | 17 | 2 | 13788 | 114 | Serpinb9,Serpina3n | | 8.498e-03 | -4.77 | Serpin\_dom | interpro domains | IPR023796 | 17 | 2 | 13788 | 114 | Serpinb9,Serpina3n | | 8.498e-03 | -4.77 | Serpin\_sf\_2 | interpro domains | IPR042185 | 17 | 2 | 13788 | 114 | Serpinb9,Serpina3n | | 8.538e-03 | -4.76 | regulation of cellular protein metabolic process | biological process | GO:0032268 | 2101 | 27 | 13711 | 111 | Kl,Thy1,Tmem106a,Grn,Ptprc,Hspa2,Clu,Serpinb9,Il6ra,Psme1,Fgf18,Capn3,Actb,Vim,Rpl26,Ctsz,Ccl6,Csk,Serpina3n,Ndufa13,C3,Hcls1,Csf1r,Myh9,Mt3,Rac2,Spint1 | | 8.543e-03 | -4.76 | positive regulation of intracellular signal transduction | biological process | GO:1902533 | 762 | 13 | 13711 | 111 | Csk,Il6ra,Fgf18,C3,Csf1r,Hcls1,Mt3,Kl,Tmem106a,Ptprc,Rpl26,Ccl6,Clu | | 8.558e-03 | -4.76 | positive regulation of regulated secretory pathway | biological process | GO:1903307 | 52 | 3 | 13711 | 111 | Fcer1g,Itgam,Vamp8 | | 8.559e-03 | -4.76 | GCM\_APEX1 | MSigDB lists | GCM\_APEX1 | 100 | 4 | 12187 | 98 | Rps8,Psme1,Rps5,Rpl14 | | 8.559e-03 | -4.76 | GO\_INTERACTION\_WITH\_HOST | MSigDB lists | GO\_INTERACTION\_WITH\_HOST | 100 | 4 | 12187 | 98 | Ctsb,Ccr5,Vamp8,Serpinb9 | | 8.649e-03 | -4.75 | WANG\_LMO4\_TARGETS\_UP | MSigDB lists | WANG\_LMO4\_TARGETS\_UP | 289 | 7 | 12187 | 98 | Tspan4,Slc2a1,Tsc22d1,Grn,Tfap2c,Rpl13,Il1r1 | | 8.653e-03 | -4.75 | GSE40666\_NAIVE\_VS\_EFFECTOR\_CD8\_TCELL\_WITH\_IFNA\_STIM\_90MIN\_UP | MSigDB lists | GSE40666\_NAIVE\_VS\_EFFECTOR\_CD8\_TCELL\_WITH\_IFNA\_STIM\_90MIN\_UP | 157 | 5 | 12187 | 98 | S100a6,Psme1,Thy1,Ahnak,Pde1a | | 8.653e-03 | -4.75 | GSE19888\_ADENOSINE\_A3R\_ACT\_VS\_TCELL\_MEMBRANES\_ACT\_IN\_MAST\_CELL\_UP | MSigDB lists | GSE19888\_ADENOSINE\_A3R\_ACT\_VS\_TCELL\_MEMBRANES\_ACT\_IN\_MAST\_CELL\_UP | 157 | 5 | 12187 | 98 | C1qb,Serpina3n,Itgam,Ctsz,Ctss | | 8.653e-03 | -4.75 | GSE2128\_C57BL6\_VS\_NOD\_THYMOCYTE\_UP | MSigDB lists | GSE2128\_C57BL6\_VS\_NOD\_THYMOCYTE\_UP | 157 | 5 | 12187 | 98 | Vim,Lfng,S100a6,Tfap2c,Thy1 | | 8.653e-03 | -4.75 | GSE7831\_CPG\_VS\_INFLUENZA\_STIM\_PDC\_1H\_DN | MSigDB lists | GSE7831\_CPG\_VS\_INFLUENZA\_STIM\_PDC\_1H\_DN | 157 | 5 | 12187 | 98 | Ctsa,Vwf,Grn,Tfap2c,Ctsz | | 8.653e-03 | -4.75 | GSE43955\_10H\_VS\_60H\_ACT\_CD4\_TCELL\_DN | MSigDB lists | GSE43955\_10H\_VS\_60H\_ACT\_CD4\_TCELL\_DN | 157 | 5 | 12187 | 98 | Rps5,Flnb,Gfap,S100a6,Ahnak | | 8.653e-03 | -4.75 | GSE33425\_CD8\_ALPHAALPHA\_VS\_ALPHABETA\_CD161\_HIGH\_TCELL\_DN | MSigDB lists | GSE33425\_CD8\_ALPHAALPHA\_VS\_ALPHABETA\_CD161\_HIGH\_TCELL\_DN | 157 | 5 | 12187 | 98 | Adh1,Brk1,Capn3,Thy1,Tsc22d1 | | 8.717e-03 | -4.74 | protein homodimerization activity | molecular function | GO:0042803 | 782 | 13 | 13516 | 107 | S100a6,C1qb,Camk2b,Mgll,Csf1r,Myh9,Rap1gap,Bhlhe41,Il6ra,Vwf,Ikzf3,Adh1,Fcer1g | | 8.726e-03 | -4.74 | Serpin | pfam domains | PF00079 | 17 | 2 | 12881 | 108 | Serpinb9,Serpina3n | | 8.747e-03 | -4.74 | positive regulation of signal transduction | biological process | GO:0009967 | 1217 | 18 | 13711 | 111 | Rpl26,Ptprc,Ccl6,Clu,Il1r1,Kl,Lrg1,Tmem106a,Mt3,Csf1r,Hcls1,Fgf18,Camk2b,Csk,Il6ra,C3,Cyba,Lfng | | 8.798e-03 | -4.73 | regulation of protein localization to plasma membrane | biological process | GO:1903076 | 100 | 4 | 13711 | 111 | Vamp8,Csk,Camk2b,Actb | | 8.832e-03 | -4.73 | WESTON\_VEGFA\_TARGETS\_3HR | MSigDB lists | WESTON\_VEGFA\_TARGETS\_3HR | 53 | 3 | 12187 | 98 | Hcls1,Mgp,Vwf | | 8.832e-03 | -4.73 | KEGG\_VIRAL\_MYOCARDITIS | MSigDB lists | KEGG\_VIRAL\_MYOCARDITIS | 53 | 3 | 12187 | 98 | Actb,Myh9,Rac2 | | 8.832e-03 | -4.73 | GNF2\_HPN | MSigDB lists | GNF2\_HPN | 53 | 3 | 12187 | 98 | C3,Tmem176a,Adh1 | | 8.880e-03 | -4.72 | GSE2770\_TGFB\_AND\_IL4\_VS\_IL4\_TREATED\_ACT\_CD4\_TCELL\_48H\_UP | MSigDB lists | GSE2770\_TGFB\_AND\_IL4\_VS\_IL4\_TREATED\_ACT\_CD4\_TCELL\_48H\_UP | 158 | 5 | 12187 | 98 | Tsc22d1,Ccr5,C3,Cyba,Slc26a2 | | 8.880e-03 | -4.72 | GSE43955\_TH0\_VS\_TGFB\_IL6\_TH17\_ACT\_CD4\_TCELL\_60H\_DN | MSigDB lists | GSE43955\_TH0\_VS\_TGFB\_IL6\_TH17\_ACT\_CD4\_TCELL\_60H\_DN | 158 | 5 | 12187 | 98 | Flnb,S100a6,Lgi4,Lrg1,Ccr5 | | 8.880e-03 | -4.72 | GSE33162\_HDAC3\_KO\_VS\_HDAC3\_KO\_MACROPHAGE\_DN | MSigDB lists | GSE33162\_HDAC3\_KO\_VS\_HDAC3\_KO\_MACROPHAGE\_DN | 158 | 5 | 12187 | 98 | Dek,Ptprc,Actb,Vim,S100a6 | | 8.880e-03 | -4.72 | GSE19401\_UNSTIM\_VS\_RETINOIC\_ACID\_STIM\_FOLLICULAR\_DC\_DN | MSigDB lists | GSE19401\_UNSTIM\_VS\_RETINOIC\_ACID\_STIM\_FOLLICULAR\_DC\_DN | 158 | 5 | 12187 | 98 | Mgll,Tfap2c,Ccr5,Serpinb9,Lin7a | | 8.886e-03 | -4.72 | GO\_INTRINSIC\_COMPONENT\_OF\_PLASMA\_MEMBRANE | MSigDB lists | GO\_INTRINSIC\_COMPONENT\_OF\_PLASMA\_MEMBRANE | 1045 | 16 | 12187 | 98 | Kl,Tnfsf8,Il1r1,Ptprc,Cyba,Ly6e,Thy1,Itgam,Ccr5,Myh9,Fcer1g,Slc2a1,Csf1r,Tspan4,Il6ra,Slc26a2 | | 8.952e-03 | -4.72 | GO\_HEMOSTASIS | MSigDB lists | GO\_HEMOSTASIS | 222 | 6 | 12187 | 98 | Actb,Rac2,C3,Myh9,Fcer1g,Vwf | | 8.964e-03 | -4.71 | GO\_POSITIVE\_REGULATION\_OF\_MULTICELLULAR\_ORGANISMAL\_PROCESS | MSigDB lists | GO\_POSITIVE\_REGULATION\_OF\_MULTICELLULAR\_ORGANISMAL\_PROCESS | 1046 | 16 | 12187 | 98 | Kl,Hcls1,C3,Clu,Camk2b,Cyba,Fgf18,Csf1r,Mc4r,Grn,Lrg1,Fcer1g,Spint1,Vamp8,Gfap,Il6ra | | 8.974e-03 | -4.71 | GRADE\_COLON\_CANCER\_UP | MSigDB lists | GRADE\_COLON\_CANCER\_UP | 688 | 12 | 12187 | 98 | Myh9,S100a6,Grn,Rpl31,Rps5,Rpl26,Ctsb,Ldha,Rps8,Psme1,Rpl14,Cyba | | 9.005e-03 | -4.71 | GO\_MAST\_CELL\_ACTIVATION | MSigDB lists | GO\_MAST\_CELL\_ACTIVATION | 18 | 2 | 12187 | 98 | Cplx2,Fcer1g | | 9.005e-03 | -4.71 | DAVIES\_MULTIPLE\_MYELOMA\_VS\_MGUS\_DN | MSigDB lists | DAVIES\_MULTIPLE\_MYELOMA\_VS\_MGUS\_DN | 18 | 2 | 12187 | 98 | Myh9,Fcer1g | | 9.068e-03 | -4.70 | Lysosome | KEGG pathways | mmu04142 | 106 | 5 | 5248 | 64 | Ctss,Cd68,Ctsz,Ctsb,Ctsa | | 9.068e-03 | -4.70 | Lysosome | KEGG pathways | ko04142 | 106 | 5 | 5248 | 64 | Cd68,Ctsz,Ctss,Ctsa,Ctsb | | 9.088e-03 | -4.70 | regulation of nervous system development | biological process | GO:0051960 | 945 | 15 | 13711 | 111 | C1qa,Ctsz,Thy1,Mgll,Rap1gap,Il1r1,Vim,Grn,Lgi4,Mt3,Bhlhe41,Spint1,Camk2b,Gfap,Ccr5 | | 9.111e-03 | -4.70 | GSE21670\_TGFB\_VS\_IL6\_TREATED\_STAT3\_KO\_CD4\_TCELL\_UP | MSigDB lists | GSE21670\_TGFB\_VS\_IL6\_TREATED\_STAT3\_KO\_CD4\_TCELL\_UP | 159 | 5 | 12187 | 98 | Ctss,Rps8,Tsc22d1,Rpl13,Treml2 | | 9.111e-03 | -4.70 | GSE24634\_IL4\_VS\_CTRL\_TREATED\_NAIVE\_CD4\_TCELL\_DAY7\_DN | MSigDB lists | GSE24634\_IL4\_VS\_CTRL\_TREATED\_NAIVE\_CD4\_TCELL\_DAY7\_DN | 159 | 5 | 12187 | 98 | C3,Psmb10,Igfbp6,Il6ra,Tsc22d1 | | 9.111e-03 | -4.70 | KAECH\_DAY8\_EFF\_VS\_DAY15\_EFF\_CD8\_TCELL\_DN | MSigDB lists | KAECH\_DAY8\_EFF\_VS\_DAY15\_EFF\_CD8\_TCELL\_DN | 159 | 5 | 12187 | 98 | Slc26a2,Rpl14,Il6ra,Camk2b,Tsc22d1 | | 9.111e-03 | -4.70 | GO\_RESPONSE\_TO\_REACTIVE\_OXYGEN\_SPECIES | MSigDB lists | GO\_RESPONSE\_TO\_REACTIVE\_OXYGEN\_SPECIES | 159 | 5 | 12187 | 98 | Ptprn,Cyba,Il1r1,Apod,Ldha | | 9.131e-03 | -4.70 | carbohydrate transmembrane transport | biological process | GO:0034219 | 18 | 2 | 13711 | 111 | Mfsd4a,Slc2a1 | | 9.164e-03 | -4.69 | HOSHIDA\_LIVER\_CANCER\_SUBCLASS\_S2 | MSigDB lists | HOSHIDA\_LIVER\_CANCER\_SUBCLASS\_S2 | 102 | 4 | 12187 | 98 | Rpl31,Rps5,Dek,Flnb | | 9.173e-03 | -4.69 | HACE1 (HECT domain and ankyrin repeat containing E3 ubiquitin protein ligase 1) | protein interactions | 57531 | 38 | 3 | 6802 | 78 | Flnb,Vim,Myh9 | | 9.298e-03 | -4.68 | WAMUNYOKOLI\_OVARIAN\_CANCER\_GRADES\_1\_2\_DN | MSigDB lists | WAMUNYOKOLI\_OVARIAN\_CANCER\_GRADES\_1\_2\_DN | 54 | 3 | 12187 | 98 | C3,Il1r1,Igfbp6 | | 9.298e-03 | -4.68 | REACTOME\_ACTIVATION\_OF\_THE\_MRNA\_UPON\_BINDING\_OF\_THE\_CAP\_BINDING\_COMPLEX\_AND\_EIFS\_AND\_SUBSEQUENT\_BINDING\_TO\_43S | MSigDB lists | REACTOME\_ACTIVATION\_OF\_THE\_MRNA\_UPON\_BINDING\_OF\_THE\_CAP\_BINDING\_COMPLEX\_AND\_EIFS\_AND\_SUBSEQUENT\_BINDING\_TO\_43S | 54 | 3 | 12187 | 98 | Rps20,Rps5,Rps8 | | 9.298e-03 | -4.68 | GNF2\_PTPRC | MSigDB lists | GNF2\_PTPRC | 54 | 3 | 12187 | 98 | Ptprc,Hcls1,Rac2 | | 9.328e-03 | -4.67 | - | gene3d domains | 3.10.440.10 | 1 | 1 | 6647 | 62 | Rpl31 | | 9.328e-03 | -4.67 | - | gene3d domains | 1.10.168.20 | 1 | 1 | 6647 | 62 | Rps8 | | 9.328e-03 | -4.67 | - | gene3d domains | 3.10.20.500 | 1 | 1 | 6647 | 62 | Grn | | 9.347e-03 | -4.67 | GSE40068\_BCL6\_POS\_VS\_NEG\_CXCR5\_POS\_TFH\_UP | MSigDB lists | GSE40068\_BCL6\_POS\_VS\_NEG\_CXCR5\_POS\_TFH\_UP | 160 | 5 | 12187 | 98 | Ldha,Tmem106a,Ccar1,Serpinb9,Ccr5 | | 9.347e-03 | -4.67 | GSE35685\_CD34POS\_CD38NEG\_VS\_CD34POS\_CD10NEG\_CD62LPOS\_BONE\_MARROW\_DN | MSigDB lists | GSE35685\_CD34POS\_CD38NEG\_VS\_CD34POS\_CD10NEG\_CD62LPOS\_BONE\_MARROW\_DN | 160 | 5 | 12187 | 98 | Cox4i1,Ly6e,Igfbp6,Camk2b,Rab3il1 | | 9.347e-03 | -4.67 | GSE17721\_PAM3CSK4\_VS\_GADIQUIMOD\_24H\_BMDC\_UP | MSigDB lists | GSE17721\_PAM3CSK4\_VS\_GADIQUIMOD\_24H\_BMDC\_UP | 160 | 5 | 12187 | 98 | Fcgr3,Tmem176a,Clu,Rpl26,Ldha | | 9.426e-03 | -4.66 | regulation of regulated secretory pathway | biological process | GO:1903305 | 159 | 5 | 13711 | 111 | Fcer1g,Cplx2,Rac2,Itgam,Vamp8 | | 9.460e-03 | -4.66 | CLb | smart domains | SM00030 | 1 | 1 | 7188 | 68 | Clu | | 9.460e-03 | -4.66 | CLa | smart domains | SM00035 | 1 | 1 | 7188 | 68 | Clu | | 9.460e-03 | -4.66 | Ribosomal\_L31e | smart domains | SM01380 | 1 | 1 | 7188 | 68 | Rpl31 | | 9.460e-03 | -4.66 | GRAN | smart domains | SM00277 | 1 | 1 | 7188 | 68 | Grn | | 9.460e-03 | -4.66 | ITAM | smart domains | SM00077 | 1 | 1 | 7188 | 68 | Fcer1g | | 9.467e-03 | -4.66 | GO\_REGULATION\_OF\_PROTEOLYSIS | MSigDB lists | GO\_REGULATION\_OF\_PROTEOLYSIS | 525 | 10 | 12187 | 98 | C3,Psme1,Capn3,Myh9,Clu,Serpinb9,Ndufa13,Serpina3n,Spint1,Psmb10 | | 9.476e-03 | -4.66 | POOLA\_INVASIVE\_BREAST\_CANCER\_DN | MSigDB lists | POOLA\_INVASIVE\_BREAST\_CANCER\_DN | 103 | 4 | 12187 | 98 | Mgp,Capn3,Flnb,Serpina3n | | 9.586e-03 | -4.65 | GSE19888\_CTRL\_VS\_TCELL\_MEMBRANES\_ACT\_MAST\_CELL\_PRETREAT\_A3R\_INH\_DN | MSigDB lists | GSE19888\_CTRL\_VS\_TCELL\_MEMBRANES\_ACT\_MAST\_CELL\_PRETREAT\_A3R\_INH\_DN | 161 | 5 | 12187 | 98 | Grn,Serpinb9,Mpeg1,Apod,Treml2 | | 9.586e-03 | -4.65 | KAECH\_NAIVE\_VS\_DAY8\_EFF\_CD8\_TCELL\_DN | MSigDB lists | KAECH\_NAIVE\_VS\_DAY8\_EFF\_CD8\_TCELL\_DN | 161 | 5 | 12187 | 98 | Serpinb9,Ahnak,Ccr5,S100a6,Actb | | 9.586e-03 | -4.65 | GSE40666\_WT\_VS\_STAT4\_KO\_CD8\_TCELL\_WITH\_IFNA\_STIM\_90MIN\_DN | MSigDB lists | GSE40666\_WT\_VS\_STAT4\_KO\_CD8\_TCELL\_WITH\_IFNA\_STIM\_90MIN\_DN | 161 | 5 | 12187 | 98 | Ikzf3,Mgll,Il1r1,Ctsa,Tspan4 | | 9.586e-03 | -4.65 | PLASARI\_TGFB1\_TARGETS\_10HR\_UP | MSigDB lists | PLASARI\_TGFB1\_TARGETS\_10HR\_UP | 161 | 5 | 12187 | 98 | Hspa2,Mgll,Fgf18,Slc2a1,Flnb | | 9.586e-03 | -4.65 | GSE36078\_UNTREATED\_VS\_AD5\_INF\_MOUSE\_LUNG\_DC\_DN | MSigDB lists | GSE36078\_UNTREATED\_VS\_AD5\_INF\_MOUSE\_LUNG\_DC\_DN | 161 | 5 | 12187 | 98 | Thy1,Fcer1g,Vim,Lfng,Flnb | | 9.778e-03 | -4.63 | GNF2\_ST13 | MSigDB lists | GNF2\_ST13 | 55 | 3 | 12187 | 98 | Rpl14,Rpl31,Rps20 | | 9.830e-03 | -4.62 | GSE360\_CTRL\_VS\_T\_GONDII\_MAC\_UP | MSigDB lists | GSE360\_CTRL\_VS\_T\_GONDII\_MAC\_UP | 162 | 5 | 12187 | 98 | Ahnak,Ly6e,Rac2,Hcls1,Cyba | | 9.830e-03 | -4.62 | GSE8921\_3H\_VS\_24H\_TLR1\_2\_STIM\_MONOCYTE\_DN | MSigDB lists | GSE8921\_3H\_VS\_24H\_TLR1\_2\_STIM\_MONOCYTE\_DN | 162 | 5 | 12187 | 98 | S100a6,Sh3bgrl3,Ldha,Slc2a1,Spint1 | | 9.830e-03 | -4.62 | GSE23502\_WT\_VS\_HDC\_KO\_MYELOID\_DERIVED\_SUPPRESSOR\_CELL\_COLON\_TUMOR\_UP | MSigDB lists | GSE23502\_WT\_VS\_HDC\_KO\_MYELOID\_DERIVED\_SUPPRESSOR\_CELL\_COLON\_TUMOR\_UP | 162 | 5 | 12187 | 98 | C1qb,Fcgr3,Ctsb,Ldha,Hspa2 | | 9.946e-03 | -4.61 | regulation of apoptotic process | biological process | GO:0042981 | 1233 | 18 | 13711 | 111 | Mt3,Hcls1,Csf1r,Pde1a,Fcer1g,Capn3,Camk2b,Serpinb9,Ccr5,Ndufa13,Tsc22d1,Rpl26,Ptprc,Clu,Ctsz,Grn,Ccar1,Ikzf3 | | 9.983e-03 | -4.61 | positive regulation of receptor signaling pathway via STAT | biological process | GO:1904894 | 55 | 3 | 13711 | 111 | Csf1r,Hcls1,Il6ra | | 1.001e-02 | -4.60 | BOQUEST\_STEM\_CELL\_CULTURED\_VS\_FRESH\_DN | MSigDB lists | BOQUEST\_STEM\_CELL\_CULTURED\_VS\_FRESH\_DN | 19 | 2 | 12187 | 98 | C3,Apod | | 1.001e-02 | -4.60 | PEDERSEN\_METASTASIS\_BY\_ERBB2\_ISOFORM\_6 | MSigDB lists | PEDERSEN\_METASTASIS\_BY\_ERBB2\_ISOFORM\_6 | 19 | 2 | 12187 | 98 | Serpina3n,Mgp | | 1.001e-02 | -4.60 | CHEN\_LUNG\_CANCER\_SURVIVAL | MSigDB lists | CHEN\_LUNG\_CANCER\_SURVIVAL | 19 | 2 | 12187 | 98 | Actb,Gfap | | 1.001e-02 | -4.60 | OSADA\_ASCL1\_TARGETS\_DN | MSigDB lists | OSADA\_ASCL1\_TARGETS\_DN | 19 | 2 | 12187 | 98 | Ahnak,Igfbp6 | | 1.001e-02 | -4.60 | REACTOME\_SHC\_MEDIATED\_CASCADE | MSigDB lists | REACTOME\_SHC\_MEDIATED\_CASCADE | 19 | 2 | 12187 | 98 | Kl,Fgf18 | | 1.001e-02 | -4.60 | GO\_POSITIVE\_REGULATION\_OF\_LAMELLIPODIUM\_ORGANIZATION | MSigDB lists | GO\_POSITIVE\_REGULATION\_OF\_LAMELLIPODIUM\_ORGANIZATION | 19 | 2 | 12187 | 98 | Brk1,Rac2 | | 1.001e-02 | -4.60 | GO\_POSITIVE\_REGULATION\_OF\_ACUTE\_INFLAMMATORY\_RESPONSE | MSigDB lists | GO\_POSITIVE\_REGULATION\_OF\_ACUTE\_INFLAMMATORY\_RESPONSE | 19 | 2 | 12187 | 98 | C3,Fcer1g | | 1.008e-02 | -4.60 | SERPIN | prosite domains | PS00284 | 15 | 2 | 8845 | 91 | Serpinb9,Serpina3n | | 1.008e-02 | -4.60 | GSE20727\_CTRL\_VS\_DNFB\_ALLERGEN\_TREATED\_DC\_UP | MSigDB lists | GSE20727\_CTRL\_VS\_DNFB\_ALLERGEN\_TREATED\_DC\_UP | 163 | 5 | 12187 | 98 | Ctsb,Il1r1,Fcer1g,Ldha,Tmem106a | | 1.008e-02 | -4.60 | GSE43863\_DAY6\_EFF\_VS\_DAY150\_MEM\_LY6C\_INT\_CXCR5POS\_CD4\_TCELL\_UP | MSigDB lists | GSE43863\_DAY6\_EFF\_VS\_DAY150\_MEM\_LY6C\_INT\_CXCR5POS\_CD4\_TCELL\_UP | 163 | 5 | 12187 | 98 | Tsc22d1,Il6ra,Csf1r,Ctsb,Ctsa | | 1.008e-02 | -4.60 | GSE36078\_UNTREATED\_VS\_AD5\_T425A\_HEXON\_INF\_MOUSE\_LUNG\_DC\_DN | MSigDB lists | GSE36078\_UNTREATED\_VS\_AD5\_T425A\_HEXON\_INF\_MOUSE\_LUNG\_DC\_DN | 163 | 5 | 12187 | 98 | Lfng,Vim,Fcer1g,S100a6,Tspan4 | | 1.008e-02 | -4.60 | KAECH\_NAIVE\_VS\_DAY15\_EFF\_CD8\_TCELL\_DN | MSigDB lists | KAECH\_NAIVE\_VS\_DAY15\_EFF\_CD8\_TCELL\_DN | 163 | 5 | 12187 | 98 | Actb,Ccr5,Ahnak,S100a6,Serpinb9 | | 1.008e-02 | -4.60 | GSE18893\_TCONV\_VS\_TREG\_24H\_CULTURE\_UP | MSigDB lists | GSE18893\_TCONV\_VS\_TREG\_24H\_CULTURE\_UP | 163 | 5 | 12187 | 98 | Vim,Cd68,Rap1gap,S100a6,Ahnak | | 1.008e-02 | -4.60 | GSE5589\_LPS\_VS\_LPS\_AND\_IL10\_STIM\_MACROPHAGE\_45MIN\_UP | MSigDB lists | GSE5589\_LPS\_VS\_LPS\_AND\_IL10\_STIM\_MACROPHAGE\_45MIN\_UP | 163 | 5 | 12187 | 98 | Slc2a1,Trnp1,Vwf,Ccr5,Itgam | | 1.008e-02 | -4.60 | GSE27434\_WT\_VS\_DNMT1\_KO\_TREG\_UP | MSigDB lists | GSE27434\_WT\_VS\_DNMT1\_KO\_TREG\_UP | 163 | 5 | 12187 | 98 | Csk,Grn,Rps5,Ctsa,C1qb | | 1.008e-02 | -4.60 | GSE25123\_CTRL\_VS\_IL4\_STIM\_MACROPHAGE\_UP | MSigDB lists | GSE25123\_CTRL\_VS\_IL4\_STIM\_MACROPHAGE\_UP | 163 | 5 | 12187 | 98 | Brk1,S100a6,Myh9,Hcls1,Cyba | | 1.008e-02 | -4.60 | GSE33162\_UNTREATED\_VS\_4H\_LPS\_STIM\_HDAC3\_KO\_MACROPHAGE\_UP | MSigDB lists | GSE33162\_UNTREATED\_VS\_4H\_LPS\_STIM\_HDAC3\_KO\_MACROPHAGE\_UP | 163 | 5 | 12187 | 98 | Sh3bgrl3,Cd68,Grn,Itgam,Psmb10 | | 1.013e-02 | -4.59 | Papain-like\_cys\_pep\_sf | interpro domains | IPR038765 | 102 | 4 | 13788 | 114 | Ctsz,Capn3,Ctsb,Ctss | | 1.013e-02 | -4.59 | GO\_EXTRACELLULAR\_STRUCTURE\_ORGANIZATION | MSigDB lists | GO\_EXTRACELLULAR\_STRUCTURE\_ORGANIZATION | 228 | 6 | 12187 | 98 | Vwf,Itgam,Fmod,Ctss,Spint1,Gfap | | 1.015e-02 | -4.59 | regulation of superoxide anion generation | biological process | GO:0032928 | 19 | 2 | 13711 | 111 | Cyba,Itgam | | 1.015e-02 | -4.59 | postsynaptic actin cytoskeleton organization | biological process | GO:0098974 | 19 | 2 | 13711 | 111 | Actb,Camk2b | | 1.015e-02 | -4.59 | positive regulation of glial cell proliferation | biological process | GO:0060252 | 19 | 2 | 13711 | 111 | Vim,Gfap | | 1.015e-02 | -4.59 | positive regulation of meiotic cell cycle | biological process | GO:0051446 | 19 | 2 | 13711 | 111 | Lfng,Camk2b | | 1.023e-02 | -4.58 | GO\_RESPONSE\_TO\_EXTERNAL\_STIMULUS | MSigDB lists | GO\_RESPONSE\_TO\_EXTERNAL\_STIMULUS | 1251 | 18 | 12187 | 98 | Csf1r,Slc2a1,Fcer1g,Ccr5,Mc4r,Serpinb9,Ctsb,Il6ra,Cox4i1,Clu,Fam111a,Fmod,Ldha,Cyba,Phyhip,Ptprc,Rac2,Tnfsf8 | | 1.027e-02 | -4.58 | GCM\_TPT1 | MSigDB lists | GCM\_TPT1 | 56 | 3 | 12187 | 98 | Rps8,Rpl14,Rpl31 | | 1.027e-02 | -4.58 | RHEIN\_ALL\_GLUCOCORTICOID\_THERAPY\_UP | MSigDB lists | RHEIN\_ALL\_GLUCOCORTICOID\_THERAPY\_UP | 56 | 3 | 12187 | 98 | Ptprc,Ahnak,Itgam | | 1.027e-02 | -4.58 | GO\_CYTOKINE\_RECEPTOR\_ACTIVITY | MSigDB lists | GO\_CYTOKINE\_RECEPTOR\_ACTIVITY | 56 | 3 | 12187 | 98 | Il1r1,Il6ra,Ccr5 | | 1.027e-02 | -4.58 | growth factor receptor binding | molecular function | GO:0070851 | 107 | 4 | 13516 | 107 | Il6ra,Il1r1,Fgf18,Kl | | 1.029e-02 | -4.58 | GRANULINS | prosite domains | PS00799 | 1 | 1 | 8845 | 91 | Grn | | 1.029e-02 | -4.58 | RIBOSOMAL\_L31E | prosite domains | PS01144 | 1 | 1 | 8845 | 91 | Rpl31 | | 1.029e-02 | -4.58 | RIBOSOMAL\_L13E | prosite domains | PS01104 | 1 | 1 | 8845 | 91 | Rpl13 | | 1.029e-02 | -4.58 | RIBOSOMAL\_S7 | prosite domains | PS00052 | 1 | 1 | 8845 | 91 | Rps5 | | 1.029e-02 | -4.58 | RIBOSOMAL\_S10 | prosite domains | PS00361 | 1 | 1 | 8845 | 91 | Rps20 | | 1.029e-02 | -4.58 | CLUSTERIN\_2 | prosite domains | PS00493 | 1 | 1 | 8845 | 91 | Clu | | 1.029e-02 | -4.58 | ITAM\_1 | prosite domains | PS51055 | 1 | 1 | 8845 | 91 | Fcer1g | | 1.029e-02 | -4.58 | CARBOXYPEPT\_SER\_HIS | prosite domains | PS00560 | 1 | 1 | 8845 | 91 | Ctsa | | 1.029e-02 | -4.58 | RIBOSOMAL\_S8E | prosite domains | PS01193 | 1 | 1 | 8845 | 91 | Rps8 | | 1.029e-02 | -4.58 | CLUSTERIN\_1 | prosite domains | PS00492 | 1 | 1 | 8845 | 91 | Clu | | 1.033e-02 | -4.57 | GSE40274\_FOXP3\_VS\_FOXP3\_AND\_LEF1\_TRANSDUCED\_ACTIVATED\_CD4\_TCELL\_UP | MSigDB lists | GSE40274\_FOXP3\_VS\_FOXP3\_AND\_LEF1\_TRANSDUCED\_ACTIVATED\_CD4\_TCELL\_UP | 164 | 5 | 12187 | 98 | S100a6,Grn,Il6ra,Ccr5,Vamp8 | | 1.033e-02 | -4.57 | GSE29164\_DAY3\_VS\_DAY7\_CD8\_TCELL\_TREATED\_MELANOMA\_UP | MSigDB lists | GSE29164\_DAY3\_VS\_DAY7\_CD8\_TCELL\_TREATED\_MELANOMA\_UP | 164 | 5 | 12187 | 98 | Cyba,Spint1,Fam111a,Ctsz,Cd68 | | 1.033e-02 | -4.57 | GSE43955\_1H\_VS\_60H\_ACT\_CD4\_TCELL\_UP | MSigDB lists | GSE43955\_1H\_VS\_60H\_ACT\_CD4\_TCELL\_UP | 164 | 5 | 12187 | 98 | Brk1,Ccr5,C1qa,Tmem176a,C1qb | | 1.033e-02 | -4.57 | GSE43863\_NAIVE\_VS\_MEMORY\_LY6C\_INT\_CXCR5POS\_CD4\_TCELL\_D150\_LCMV\_UP | MSigDB lists | GSE43863\_NAIVE\_VS\_MEMORY\_LY6C\_INT\_CXCR5POS\_CD4\_TCELL\_D150\_LCMV\_UP | 164 | 5 | 12187 | 98 | Csf1r,C1qb,Csk,Il6ra,Myh9 | | 1.033e-02 | -4.57 | GSE11057\_NAIVE\_CD4\_VS\_PBMC\_CD4\_TCELL\_DN | MSigDB lists | GSE11057\_NAIVE\_CD4\_VS\_PBMC\_CD4\_TCELL\_DN | 164 | 5 | 12187 | 98 | Ctsa,Cyba,Actb,Ctsz,Sh3bgrl3 | | 1.033e-02 | -4.57 | HALLMARK\_KRAS\_SIGNALING\_UP | MSigDB lists | HALLMARK\_KRAS\_SIGNALING\_UP | 164 | 5 | 12187 | 98 | Ctss,Fcer1g,Apod,Serpina3n,Tmem176a | | 1.033e-02 | -4.57 | GSE41978\_ID2\_KO\_VS\_BIM\_KO\_KLRG1\_LOW\_EFFECTOR\_CD8\_TCELL\_UP | MSigDB lists | GSE41978\_ID2\_KO\_VS\_BIM\_KO\_KLRG1\_LOW\_EFFECTOR\_CD8\_TCELL\_UP | 164 | 5 | 12187 | 98 | C3,Ptprc,Psmb10,Il1r1,Ctss | | 1.033e-02 | -4.57 | GSE23505\_IL6\_IL1\_IL23\_VS\_IL6\_IL1\_TGFB\_TREATED\_CD4\_TCELL\_UP | MSigDB lists | GSE23505\_IL6\_IL1\_IL23\_VS\_IL6\_IL1\_TGFB\_TREATED\_CD4\_TCELL\_UP | 164 | 5 | 12187 | 98 | Csk,Mgp,Ccr5,Psmb10,Fcgr3 | | 1.033e-02 | -4.57 | GSE24142\_EARLY\_THYMIC\_PROGENITOR\_VS\_DN3\_THYMOCYTE\_ADULT\_UP | MSigDB lists | GSE24142\_EARLY\_THYMIC\_PROGENITOR\_VS\_DN3\_THYMOCYTE\_ADULT\_UP | 164 | 5 | 12187 | 98 | Csf1r,Ctss,S100a6,Tsc22d1,Grn | | 1.033e-02 | -4.57 | GSE40277\_GATA1\_AND\_SATB1\_TRANSDUCED\_VS\_CTRL\_CD4\_TCELL\_UP | MSigDB lists | GSE40277\_GATA1\_AND\_SATB1\_TRANSDUCED\_VS\_CTRL\_CD4\_TCELL\_UP | 164 | 5 | 12187 | 98 | Ikzf3,Hspa2,Il6ra,Ccr5,S100a6 | | 1.034e-02 | -4.57 | Antigen processing-Cross presentation | REACTOME pathways | R-MMU-1236975 | 71 | 4 | 6297 | 76 | Psmb10,Psme1,Cyba,Vamp8 | | 1.040e-02 | -4.57 | regulated exocytosis | biological process | GO:0045055 | 105 | 4 | 13711 | 111 | Fcer1g,Cplx2,Vamp8,Myh9 | | 1.044e-02 | -4.56 | FULCHER\_INFLAMMATORY\_RESPONSE\_LECTIN\_VS\_LPS\_UP | MSigDB lists | FULCHER\_INFLAMMATORY\_RESPONSE\_LECTIN\_VS\_LPS\_UP | 452 | 9 | 12187 | 98 | Rac2,Sh3bgrl3,Fcgr3,Spint1,Tsc22d1,Myh9,Csk,C3,Mpeg1 | | 1.048e-02 | -4.56 | GO\_ENZYME\_BINDING | MSigDB lists | GO\_ENZYME\_BINDING | 1449 | 20 | 12187 | 98 | Actb,Gfap,Il6ra,Slc2a1,Csf1r,Mc4r,Serpinb9,Brk1,Thy1,Il1r1,Ptprc,Ptprn,Hcls1,Ldha,Csk,Vwf,Rap1gap,Hspa2,Camk2b,Clu | | 1.049e-02 | -4.56 | Ig-like\_dom\_sf | interpro domains | IPR036179 | 291 | 7 | 13788 | 114 | Il6ra,Fcgr3,Il1r1,Csf1r,Thy1,Treml2,Fcrls | | 1.059e-02 | -4.55 | GSE2770\_IL12\_ACT\_VS\_ACT\_CD4\_TCELL\_2H\_DN | MSigDB lists | GSE2770\_IL12\_ACT\_VS\_ACT\_CD4\_TCELL\_2H\_DN | 165 | 5 | 12187 | 98 | Ccr5,Mgll,Tsc22d1,C3,Ctsb | | 1.059e-02 | -4.55 | GSE37534\_UNTREATED\_VS\_PIOGLITAZONE\_TREATED\_CD4\_TCELL\_PPARG1\_AND\_FOXP3\_TRASDUCED\_DN | MSigDB lists | GSE37534\_UNTREATED\_VS\_PIOGLITAZONE\_TREATED\_CD4\_TCELL\_PPARG1\_AND\_FOXP3\_TRASDUCED\_DN | 165 | 5 | 12187 | 98 | Ly6e,Fmod,Rpl13,Cox4i1,Psmb10 | | 1.059e-02 | -4.55 | GSE22886\_NAIVE\_CD8\_TCELL\_VS\_MEMORY\_TCELL\_DN | MSigDB lists | GSE22886\_NAIVE\_CD8\_TCELL\_VS\_MEMORY\_TCELL\_DN | 165 | 5 | 12187 | 98 | Il6ra,Ldha,Ctsb,Il1r1,Dek | | 1.059e-02 | -4.55 | GSE43955\_10H\_VS\_30H\_ACT\_CD4\_TCELL\_UP | MSigDB lists | GSE43955\_10H\_VS\_30H\_ACT\_CD4\_TCELL\_UP | 165 | 5 | 12187 | 98 | Sh3bgrl3,C1qb,C1qa,Flnb,Kl | | 1.059e-02 | -4.55 | GOLDRATH\_NAIVE\_VS\_MEMORY\_CD8\_TCELL\_DN | MSigDB lists | GOLDRATH\_NAIVE\_VS\_MEMORY\_CD8\_TCELL\_DN | 165 | 5 | 12187 | 98 | Tspan4,Serpinb9,S100a6,Ahnak,Ccr5 | | 1.059e-02 | -4.55 | GSE7831\_UNSTIM\_VS\_CPG\_STIM\_PDC\_1H\_UP | MSigDB lists | GSE7831\_UNSTIM\_VS\_CPG\_STIM\_PDC\_1H\_UP | 165 | 5 | 12187 | 98 | Hcls1,Klhl6,Psmb10,Myh9,Lfng | | 1.059e-02 | -4.55 | GSE3337\_CTRL\_VS\_4H\_IFNG\_IN\_CD8POS\_DC\_UP | MSigDB lists | GSE3337\_CTRL\_VS\_4H\_IFNG\_IN\_CD8POS\_DC\_UP | 165 | 5 | 12187 | 98 | Fgf18,Csf1r,Ctsb,Vamp8,Ahnak | | 1.059e-02 | -4.55 | GSE39152\_CD103\_NEG\_VS\_POS\_MEMORY\_CD8\_TCELL\_DN | MSigDB lists | GSE39152\_CD103\_NEG\_VS\_POS\_MEMORY\_CD8\_TCELL\_DN | 165 | 5 | 12187 | 98 | Ptprn,Tspan4,Rab3il1,Slc2a1,Ccr5 | | 1.064e-02 | -4.54 | - | gene3d domains | 2.30.39.10 | 17 | 2 | 6647 | 62 | Serpina3n,Serpinb9 | | 1.064e-02 | -4.54 | - | gene3d domains | 3.30.497.10 | 17 | 2 | 6647 | 62 | Serpina3n,Serpinb9 | | 1.064e-02 | -4.54 | PATIL\_LIVER\_CANCER | MSigDB lists | PATIL\_LIVER\_CANCER | 618 | 11 | 12187 | 98 | Rps5,Rps20,Dek,Thy1,Vwf,Grn,Slc26a2,Ctsa,Il6ra,Cplx2,Vim | | 1.068e-02 | -4.54 | regulation of plasma membrane bounded cell projection organization | biological process | GO:0120035 | 697 | 12 | 13711 | 111 | Gfap,Ccr5,Camk2b,Mark4,Rac2,Brk1,Grn,Vim,Thy1,Mgll,Rap1gap,Ctsz | | 1.074e-02 | -4.53 | defense response to bacterium | biological process | GO:0042742 | 106 | 4 | 13711 | 111 | Fcer1g,Mpeg1,Tnfsf8,Serpinb9 | | 1.078e-02 | -4.53 | MORF\_JUND | MSigDB lists | MORF\_JUND | 57 | 3 | 12187 | 98 | Rpl13,Rpl31,Rps20 | | 1.080e-02 | -4.53 | GSE13522\_CTRL\_VS\_T\_CRUZI\_Y\_STRAIN\_INF\_SKIN\_BALBC\_MOUSE\_UP | MSigDB lists | GSE13522\_CTRL\_VS\_T\_CRUZI\_Y\_STRAIN\_INF\_SKIN\_BALBC\_MOUSE\_UP | 107 | 4 | 12187 | 98 | Rps8,Ldha,Slc2a1,Rpl14 | | 1.080e-02 | -4.53 | PANGAS\_TUMOR\_SUPPRESSION\_BY\_SMAD1\_AND\_SMAD5\_UP | MSigDB lists | PANGAS\_TUMOR\_SUPPRESSION\_BY\_SMAD1\_AND\_SMAD5\_UP | 107 | 4 | 12187 | 98 | Rpl14,Klhl6,C1qa,Serpinb9 | | 1.080e-02 | -4.53 | translation | biological process | GO:0006412 | 299 | 7 | 13711 | 111 | Rpl26,Rpl31,Rpl14,Rps5,Rps20,Rps8,Rpl13 | | 1.080e-02 | -4.53 | MYO18A (myosin XVIIIA) | protein interactions | 399687 | 14 | 2 | 6802 | 78 | Myh9,Actb | | 1.080e-02 | -4.53 | TPM4 (tropomyosin 4) | protein interactions | 7171 | 14 | 2 | 6802 | 78 | Myh9,Actb | | 1.080e-02 | -4.53 | FLII (FLII actin remodeling protein) | protein interactions | 2314 | 14 | 2 | 6802 | 78 | Myh9,Actb | | 1.085e-02 | -4.52 | GSE27670\_CTRL\_VS\_LMP1\_TRANSDUCED\_GC\_BCELL\_DN | MSigDB lists | GSE27670\_CTRL\_VS\_LMP1\_TRANSDUCED\_GC\_BCELL\_DN | 166 | 5 | 12187 | 98 | Rab3il1,C3,Tspan4,Ctsb,Rac2 | | 1.085e-02 | -4.52 | GSE46143\_CTRL\_VS\_LMP2A\_TRANSDUCED\_CD10\_POS\_GC\_BCELL\_DN | MSigDB lists | GSE46143\_CTRL\_VS\_LMP2A\_TRANSDUCED\_CD10\_POS\_GC\_BCELL\_DN | 166 | 5 | 12187 | 98 | S100a6,Rac2,Ly6e,Ahnak,Ccr5 | | 1.085e-02 | -4.52 | GSE17721\_PAM3CSK4\_VS\_GADIQUIMOD\_12H\_BMDC\_UP | MSigDB lists | GSE17721\_PAM3CSK4\_VS\_GADIQUIMOD\_12H\_BMDC\_UP | 166 | 5 | 12187 | 98 | Kl,Rpl14,Ndufa13,Sh3bgrl3,S100a6 | | 1.094e-02 | -4.52 | SERPIN | smart domains | SM00093 | 17 | 2 | 7188 | 68 | Serpina3n,Serpinb9 | | 1.107e-02 | -4.50 | GO\_MONOCYTE\_DIFFERENTIATION | MSigDB lists | GO\_MONOCYTE\_DIFFERENTIATION | 20 | 2 | 12187 | 98 | Myh9,Csf1r | | 1.107e-02 | -4.50 | GNF2\_CASP4 | MSigDB lists | GNF2\_CASP4 | 20 | 2 | 12187 | 98 | Hcls1,Sh3bgrl3 | | 1.107e-02 | -4.50 | GO\_REGULATION\_OF\_SYSTEMIC\_ARTERIAL\_BLOOD\_PRESSURE\_BY\_HORMONE | MSigDB lists | GO\_REGULATION\_OF\_SYSTEMIC\_ARTERIAL\_BLOOD\_PRESSURE\_BY\_HORMONE | 20 | 2 | 12187 | 98 | Ctsz,Cyba | | 1.107e-02 | -4.50 | GO\_PEPTIDE\_HORMONE\_PROCESSING | MSigDB lists | GO\_PEPTIDE\_HORMONE\_PROCESSING | 20 | 2 | 12187 | 98 | Ctsz,Pcsk2 | | 1.107e-02 | -4.50 | REACTOME\_LYSOSOME\_VESICLE\_BIOGENESIS | MSigDB lists | REACTOME\_LYSOSOME\_VESICLE\_BIOGENESIS | 20 | 2 | 12187 | 98 | Vamp8,Ctsz | | 1.107e-02 | -4.50 | GO\_REGULATION\_OF\_B\_CELL\_MEDIATED\_IMMUNITY | MSigDB lists | GO\_REGULATION\_OF\_B\_CELL\_MEDIATED\_IMMUNITY | 20 | 2 | 12187 | 98 | C3,Fcer1g | | 1.111e-02 | -4.50 | GAUSSMANN\_MLL\_AF4\_FUSION\_TARGETS\_G\_UP | MSigDB lists | GAUSSMANN\_MLL\_AF4\_FUSION\_TARGETS\_G\_UP | 167 | 5 | 12187 | 98 | Adh1,Slc26a2,C3,Sh2d5,Mgll | | 1.111e-02 | -4.50 | GSE16697\_CD4\_TCELL\_VS\_TFH\_CD4\_TCELL\_UP | MSigDB lists | GSE16697\_CD4\_TCELL\_VS\_TFH\_CD4\_TCELL\_UP | 167 | 5 | 12187 | 98 | Ctsa,Cyba,Grn,Csk,Ikzf3 | | 1.111e-02 | -4.50 | GSE23502\_BM\_VS\_COLON\_TUMOR\_MYELOID\_DERIVED\_SUPPRESSOR\_CELL\_UP | MSigDB lists | GSE23502\_BM\_VS\_COLON\_TUMOR\_MYELOID\_DERIVED\_SUPPRESSOR\_CELL\_UP | 167 | 5 | 12187 | 98 | S100a6,Ccr5,Ahnak,Vim,Tmem106a | | 1.111e-02 | -4.50 | GSE16385\_ROSIGLITAZONE\_IL4\_VS\_IL4\_ALONE\_STIM\_MACROPHAGE\_12H\_UP | MSigDB lists | GSE16385\_ROSIGLITAZONE\_IL4\_VS\_IL4\_ALONE\_STIM\_MACROPHAGE\_12H\_UP | 167 | 5 | 12187 | 98 | Grn,S100a6,Fcer1g,Flnb,Ptprc | | 1.111e-02 | -4.50 | GSE9988\_ANTI\_TREM1\_VS\_VEHICLE\_TREATED\_MONOCYTES\_DN | MSigDB lists | GSE9988\_ANTI\_TREM1\_VS\_VEHICLE\_TREATED\_MONOCYTES\_DN | 167 | 5 | 12187 | 98 | Csf1r,Mpeg1,Lrg1,Grn,Csk | | 1.111e-02 | -4.50 | GSE411\_UNSTIM\_VS\_100MIN\_IL6\_STIM\_MACROPHAGE\_UP | MSigDB lists | GSE411\_UNSTIM\_VS\_100MIN\_IL6\_STIM\_MACROPHAGE\_UP | 167 | 5 | 12187 | 98 | Cyba,Mpeg1,S100a6,Ccr5,Cd68 | | 1.119e-02 | -4.49 | GO\_APICAL\_PART\_OF\_CELL | MSigDB lists | GO\_APICAL\_PART\_OF\_CELL | 233 | 6 | 12187 | 98 | Thy1,Slc2a1,Kl,Il6ra,Ctsb,Cyba | | 1.122e-02 | -4.49 | negative regulation of neuron projection development | biological process | GO:0010977 | 166 | 5 | 13711 | 111 | Thy1,Ccr5,Ctsz,Gfap,Vim | | 1.122e-02 | -4.49 | response to lipoprotein particle | biological process | GO:0055094 | 20 | 2 | 13711 | 111 | Cd68,Fcer1g | | 1.122e-02 | -4.49 | positive regulation of lamellipodium assembly | biological process | GO:0010592 | 20 | 2 | 13711 | 111 | Brk1,Rac2 | | 1.122e-02 | -4.49 | regulation of T-helper 1 type immune response | biological process | GO:0002825 | 20 | 2 | 13711 | 111 | Il6ra,Il1r1 | | 1.122e-02 | -4.49 | antigen processing and presentation of exogenous peptide antigen | biological process | GO:0002478 | 20 | 2 | 13711 | 111 | Fcer1g,Fcgr3 | | 1.130e-02 | -4.48 | JECHLINGER\_EPITHELIAL\_TO\_MESENCHYMAL\_TRANSITION\_DN | MSigDB lists | JECHLINGER\_EPITHELIAL\_TO\_MESENCHYMAL\_TRANSITION\_DN | 58 | 3 | 12187 | 98 | Myh9,Tsc22d1,Vamp8 | | 1.130e-02 | -4.48 | BRUECKNER\_TARGETS\_OF\_MIRLET7A3\_DN | MSigDB lists | BRUECKNER\_TARGETS\_OF\_MIRLET7A3\_DN | 58 | 3 | 12187 | 98 | Flnb,Trnp1,Vim | | 1.130e-02 | -4.48 | MARTINEZ\_RESPONSE\_TO\_TRABECTEDIN\_UP | MSigDB lists | MARTINEZ\_RESPONSE\_TO\_TRABECTEDIN\_UP | 58 | 3 | 12187 | 98 | Ndufa13,Igfbp6,Ctsa | | 1.136e-02 | -4.48 | negative regulation of neurogenesis | biological process | GO:0050768 | 302 | 7 | 13711 | 111 | Gfap,Vim,Ccr5,Rap1gap,Thy1,Ctsz,Mt3 | | 1.138e-02 | -4.48 | GSE23502\_WT\_VS\_HDC\_KO\_MYELOID\_DERIVED\_SUPPRESSOR\_CELL\_BM\_DN | MSigDB lists | GSE23502\_WT\_VS\_HDC\_KO\_MYELOID\_DERIVED\_SUPPRESSOR\_CELL\_BM\_DN | 168 | 5 | 12187 | 98 | Fcgr3,Ccr5,Ctsz,Cd68,Vim | | 1.138e-02 | -4.48 | GSE42021\_CD24HI\_VS\_CD24INT\_TCONV\_THYMUS\_UP | MSigDB lists | GSE42021\_CD24HI\_VS\_CD24INT\_TCONV\_THYMUS\_UP | 168 | 5 | 12187 | 98 | Serpina3n,C1qb,Ikzf3,Tsc22d1,Rap1gap | | 1.138e-02 | -4.48 | GSE32034\_LY6C\_HIGH\_VS\_LOW\_ROSIGLIZATONE\_TREATED\_MONOCYTE\_UP | MSigDB lists | GSE32034\_LY6C\_HIGH\_VS\_LOW\_ROSIGLIZATONE\_TREATED\_MONOCYTE\_UP | 168 | 5 | 12187 | 98 | Ly6e,Ccr5,Psme1,Rps5,Psmb10 | | 1.138e-02 | -4.48 | GSE7460\_CTRL\_VS\_TGFB\_TREATED\_ACT\_CD8\_TCELL\_UP | MSigDB lists | GSE7460\_CTRL\_VS\_TGFB\_TREATED\_ACT\_CD8\_TCELL\_UP | 168 | 5 | 12187 | 98 | Ahnak,Camk2b,Rpl26,Cplx2,Slc17a7 | | 1.138e-02 | -4.48 | GSE16385\_IFNG\_TNF\_VS\_IL4\_STIM\_MACROPHAGE\_ROSIGLITAZONE\_TREATED\_DN | MSigDB lists | GSE16385\_IFNG\_TNF\_VS\_IL4\_STIM\_MACROPHAGE\_ROSIGLITAZONE\_TREATED\_DN | 168 | 5 | 12187 | 98 | Brk1,Sh3bgrl3,Myh9,Hcls1,Cyba | | 1.138e-02 | -4.48 | GSE369\_SOCS3\_KO\_VS\_WT\_LIVER\_POST\_IL6\_INJECTION\_UP | MSigDB lists | GSE369\_SOCS3\_KO\_VS\_WT\_LIVER\_POST\_IL6\_INJECTION\_UP | 168 | 5 | 12187 | 98 | Mgll,Fcer1g,Thy1,Slc2a1,Tspan4 | | 1.139e-02 | -4.47 | GO\_CELL\_CELL\_SIGNALING | MSigDB lists | GO\_CELL\_CELL\_SIGNALING | 540 | 10 | 12187 | 98 | C1qa,Lin7a,Fgf18,Ptprn,Tnfsf8,Cplx2,Slc17a7,Mc4r,Tfap2c,Ccr5 | | 1.147e-02 | -4.47 | Eomes (eomesodermin) | protein interactions | 13813 | 1 | 1 | 6802 | 78 | Tfap2c | | 1.147e-02 | -4.47 | YJU3 (acylglycerol lipase) | protein interactions | 853768 | 1 | 1 | 6802 | 78 | Mgll | | 1.147e-02 | -4.47 | Il1a (interleukin 1 alpha) | protein interactions | 16175 | 1 | 1 | 6802 | 78 | Il1r1 | | 1.147e-02 | -4.47 | PLCG2 (phospholipase C gamma 2) | protein interactions | 5336 | 1 | 1 | 6802 | 78 | Csk | | 1.147e-02 | -4.47 | Atp7b (ATPase, Cu++ transporting, beta polypeptide) | protein interactions | 11979 | 1 | 1 | 6802 | 78 | Clu | | 1.147e-02 | -4.47 | ARPC1A (actin related protein 2/3 complex subunit 1A) | protein interactions | 10552 | 1 | 1 | 6802 | 78 | Actb | | 1.147e-02 | -4.47 | RAD52 (RAD52 homolog, DNA repair protein) | protein interactions | 5893 | 1 | 1 | 6802 | 78 | Csk | | 1.147e-02 | -4.47 | SNAP23 (synaptosome associated protein 23) | protein interactions | 8773 | 1 | 1 | 6802 | 78 | Flnb | | 1.147e-02 | -4.47 | RAP2C (RAP2C, member of RAS oncogene family) | protein interactions | 57826 | 1 | 1 | 6802 | 78 | Flnb | | 1.147e-02 | -4.47 | RHOB (ras homolog family member B) | protein interactions | 388 | 1 | 1 | 6802 | 78 | Flnb | | 1.147e-02 | -4.47 | Ace (angiotensin I converting enzyme (peptidyl-dipeptidase A) 1) | protein interactions | 11421 | 1 | 1 | 6802 | 78 | Myh9 | | 1.147e-02 | -4.47 | ITIH6 (inter-alpha-trypsin inhibitor heavy chain family member 6) | protein interactions | 347365 | 1 | 1 | 6802 | 78 | Csk | | 1.147e-02 | -4.47 | SRGAP2 (SLIT-ROBO Rho GTPase activating protein 2) | protein interactions | 23380 | 1 | 1 | 6802 | 78 | Actb | | 1.147e-02 | -4.47 | Fgfr4 (fibroblast growth factor receptor 4) | protein interactions | 14186 | 1 | 1 | 6802 | 78 | Kl | | 1.147e-02 | -4.47 | Cd276 (Cd276 molecule) | protein interactions | 315716 | 1 | 1 | 6802 | 78 | Treml2 | | 1.147e-02 | -4.47 | PLS1 (plastin 1) | protein interactions | 5357 | 1 | 1 | 6802 | 78 | Myh9 | | 1.147e-02 | -4.47 | ALPI (alkaline phosphatase, intestinal) | protein interactions | 248 | 1 | 1 | 6802 | 78 | Actb | | 1.147e-02 | -4.47 | MCF2L (MCF.2 cell line derived transforming sequence like) | protein interactions | 23263 | 1 | 1 | 6802 | 78 | Myh9 | | 1.147e-02 | -4.47 | OXR1 (oxidation resistance 1) | protein interactions | 55074 | 1 | 1 | 6802 | 78 | Actb | | 1.147e-02 | -4.47 | SH3GLB1 (SH3 domain containing GRB2 like, endophilin B1) | protein interactions | 51100 | 1 | 1 | 6802 | 78 | Csk | | 1.147e-02 | -4.47 | Neu1 (neuraminidase 1) | protein interactions | 18010 | 1 | 1 | 6802 | 78 | Ctsa | | 1.147e-02 | -4.47 | Atrnl1 (attractin like 1) | protein interactions | 226255 | 1 | 1 | 6802 | 78 | Mc4r | | 1.147e-02 | -4.47 | Prom1 (prominin 1) | protein interactions | 19126 | 1 | 1 | 6802 | 78 | Actb | | 1.147e-02 | -4.47 | AFTPH (aftiphilin) | protein interactions | 54812 | 1 | 1 | 6802 | 78 | Csk | | 1.147e-02 | -4.47 | CELF1 (CUGBP Elav-like family member 1) | protein interactions | 10658 | 1 | 1 | 6802 | 78 | Myh9 | | 1.147e-02 | -4.47 | STAMBPL1 (STAM binding protein like 1) | protein interactions | 57559 | 1 | 1 | 6802 | 78 | Actb | | 1.147e-02 | -4.47 | TSC22D3 (TSC22 domain family member 3) | protein interactions | 1831 | 1 | 1 | 6802 | 78 | Csk | | 1.147e-02 | -4.47 | CSTF3 (cleavage stimulation factor subunit 3) | protein interactions | 1479 | 1 | 1 | 6802 | 78 | Myh9 | | 1.147e-02 | -4.47 | Noxo1 (NADPH oxidase organizer 1) | protein interactions | 71893 | 1 | 1 | 6802 | 78 | Cyba | | 1.147e-02 | -4.47 | Spam1 (sperm adhesion molecule 1) | protein interactions | 20690 | 1 | 1 | 6802 | 78 | Clu | | 1.147e-02 | -4.47 | Ptprb (protein tyrosine phosphatase, receptor type, B) | protein interactions | 19263 | 1 | 1 | 6802 | 78 | Vim | | 1.147e-02 | -4.47 | Mesp2 (mesoderm posterior 2) | protein interactions | 17293 | 1 | 1 | 6802 | 78 | Lfng | | 1.147e-02 | -4.47 | UBAP2 (ubiquitin associated protein 2) | protein interactions | 55833 | 1 | 1 | 6802 | 78 | Actb | | 1.147e-02 | -4.47 | SRGAP1 (SLIT-ROBO Rho GTPase activating protein 1) | protein interactions | 57522 | 1 | 1 | 6802 | 78 | Actb | | 1.147e-02 | -4.47 | Psme2 (proteasome (prosome, macropain) activator subunit 2 (PA28 beta)) | protein interactions | 19188 | 1 | 1 | 6802 | 78 | Psme1 | | 1.147e-02 | -4.47 | FGD4 (FYVE, RhoGEF and PH domain containing 4) | protein interactions | 121512 | 1 | 1 | 6802 | 78 | Myh9 | | 1.147e-02 | -4.47 | SYNC (syncoilin, intermediate filament protein) | protein interactions | 81493 | 1 | 1 | 6802 | 78 | Vim | | 1.147e-02 | -4.47 | Msrb1 (methionine sulfoxide reductase B1) | protein interactions | 27361 | 1 | 1 | 6802 | 78 | Clu | | 1.147e-02 | -4.47 | CAP2 (cyclase associated actin cytoskeleton regulatory protein 2) | protein interactions | 10486 | 1 | 1 | 6802 | 78 | Actb | | 1.147e-02 | -4.47 | MTFR1 (mitochondrial fission regulator 1) | protein interactions | 9650 | 1 | 1 | 6802 | 78 | Myh9 | | 1.147e-02 | -4.47 | Lep (leptin) | protein interactions | 16846 | 1 | 1 | 6802 | 78 | Clu | | 1.147e-02 | -4.47 | ARHGAP42 (Rho GTPase activating protein 42) | protein interactions | 143872 | 1 | 1 | 6802 | 78 | Myh9 | | 1.147e-02 | -4.47 | Slpi (secretory leukocyte peptidase inhibitor) | protein interactions | 20568 | 1 | 1 | 6802 | 78 | Grn | | 1.147e-02 | -4.47 | UBR1 (ubiquitin protein ligase E3 component n-recognin 1) | protein interactions | 197131 | 1 | 1 | 6802 | 78 | Mgp | | 1.147e-02 | -4.47 | Fgfr3 (fibroblast growth factor receptor 3) | protein interactions | 14184 | 1 | 1 | 6802 | 78 | Kl | | 1.147e-02 | -4.47 | VLDLR (very low density lipoprotein receptor) | protein interactions | 7436 | 1 | 1 | 6802 | 78 | Clu | | 1.147e-02 | -4.47 | CSK (C-terminal Src kinase) | protein interactions | 1445 | 1 | 1 | 6802 | 78 | Csk | | 1.149e-02 | -4.47 | positive regulation of vasculature development | biological process | GO:1904018 | 167 | 5 | 13711 | 111 | Grn,C3,Fgf18,Il6ra,Lrg1 | | 1.149e-02 | -4.47 | regulation of cell projection organization | biological process | GO:0031344 | 704 | 12 | 13711 | 111 | Grn,Vim,Thy1,Mgll,Rap1gap,Ctsz,Ccr5,Gfap,Camk2b,Mark4,Rac2,Brk1 | | 1.154e-02 | -4.46 | Cytokine Signaling in Immune system | REACTOME pathways | R-MMU-1280215 | 364 | 10 | 6297 | 76 | Kl,Il6ra,Csf1r,Fgf18,Tnfsf8,Flnb,Camk2b,Il1r1,Psmb10,Psme1 | | 1.156e-02 | -4.46 | positive regulation of apoptotic process | biological process | GO:0043065 | 536 | 10 | 13711 | 111 | Rpl26,Ptprc,Clu,Ctsz,Camk2b,Grn,Ccar1,Ccr5,Ndufa13,Tsc22d1 | | 1.161e-02 | -4.46 | MODULE\_12 | MSigDB lists | MODULE\_12 | 306 | 7 | 12187 | 98 | Igfbp6,Gfap,Apod,Serpina3n,Hspa2,Mgll,C1qb | | 1.162e-02 | -4.45 | ACEVEDO\_METHYLATED\_IN\_LIVER\_CANCER\_DN | MSigDB lists | ACEVEDO\_METHYLATED\_IN\_LIVER\_CANCER\_DN | 460 | 9 | 12187 | 98 | Pde1a,Fmod,Myh9,Treml2,Slc2a1,C1qb,Cplx2,Phyhip,Fcgr3 | | 1.162e-02 | -4.45 | MODULE\_11 | MSigDB lists | MODULE\_11 | 460 | 9 | 12187 | 98 | Ptprn,Serpina3n,Phyhip,Apod,Gfap,Slc17a7,C1qb,Capn3,Hspa2 | | 1.163e-02 | -4.45 | regulation of programmed cell death | biological process | GO:0043067 | 1253 | 18 | 13711 | 111 | Rpl26,Ptprc,Clu,Ctsz,Ccar1,Grn,Ikzf3,Mt3,Csf1r,Hcls1,Pde1a,Fcer1g,Capn3,Camk2b,Serpinb9,Ccr5,Ndufa13,Tsc22d1 | | 1.163e-02 | -4.45 | positive regulation of developmental process | biological process | GO:0051094 | 1253 | 18 | 13711 | 111 | Grn,Vim,Il1r1,Kl,Lrg1,Clu,Ptprc,C1qa,Gfap,C3,Ccr5,Fgf18,Capn3,Camk2b,Il6ra,Spint1,Csf1r,Hcls1 | | 1.165e-02 | -4.45 | CALM1 (calmodulin 1) | protein interactions | 801 | 166 | 6 | 6802 | 78 | Rps20,Camk2b,Myh9,Flnb,Pde1a,Rap1gap | | 1.166e-02 | -4.45 | GSE360\_T\_GONDII\_VS\_B\_MALAYI\_LOW\_DOSE\_MAC\_DN | MSigDB lists | GSE360\_T\_GONDII\_VS\_B\_MALAYI\_LOW\_DOSE\_MAC\_DN | 169 | 5 | 12187 | 98 | Pde1a,Ly6e,Mgll,Rac2,Cyba | | 1.178e-02 | -4.44 | MODULE\_137 | MSigDB lists | MODULE\_137 | 461 | 9 | 12187 | 98 | C1qb,C3,Hspa2,Ptprn,Phyhip,Serpina3n,Apod,Igfbp6,Slc17a7 | | 1.181e-02 | -4.44 | leukocyte activation involved in immune response | biological process | GO:0002366 | 109 | 4 | 13711 | 111 | Grn,Cplx2,Fcer1g,Lfng | | 1.182e-02 | -4.44 | carbohydrate transmembrane transporter activity | molecular function | GO:0015144 | 21 | 2 | 13516 | 107 | Mfsd4a,Slc2a1 | | 1.182e-02 | -4.44 | tau protein binding | molecular function | GO:0048156 | 21 | 2 | 13516 | 107 | Clu,Hspa2 | | 1.184e-02 | -4.44 | DELPUECH\_FOXO3\_TARGETS\_UP | MSigDB lists | DELPUECH\_FOXO3\_TARGETS\_UP | 59 | 3 | 12187 | 98 | Igfbp6,Clu,Ctsb | | 1.184e-02 | -4.44 | BOYLAN\_MULTIPLE\_MYELOMA\_D\_DN | MSigDB lists | BOYLAN\_MULTIPLE\_MYELOMA\_D\_DN | 59 | 3 | 12187 | 98 | S100a6,Ahnak,Itgam | | 1.184e-02 | -4.44 | GO\_CELLULAR\_GLUCOSE\_HOMEOSTASIS | MSigDB lists | GO\_CELLULAR\_GLUCOSE\_HOMEOSTASIS | 59 | 3 | 12187 | 98 | Il1r1,Cyba,Ptprn | | 1.187e-02 | -4.43 | MORF\_RAN | MSigDB lists | MORF\_RAN | 236 | 6 | 12187 | 98 | Dek,Rpl14,Rps5,Ldha,Cox4i1,Rpl13 | | 1.193e-02 | -4.43 | GO\_ANTIGEN\_PROCESSING\_AND\_PRESENTATION | MSigDB lists | GO\_ANTIGEN\_PROCESSING\_AND\_PRESENTATION | 170 | 5 | 12187 | 98 | Cyba,Psmb10,Fcer1g,Psme1,Ctss | | 1.193e-02 | -4.43 | GSE41176\_UNSTIM\_VS\_ANTI\_IGM\_STIM\_BCELL\_1H\_DN | MSigDB lists | GSE41176\_UNSTIM\_VS\_ANTI\_IGM\_STIM\_BCELL\_1H\_DN | 170 | 5 | 12187 | 98 | Cox4i1,Psme1,Grn,Csf1r,Ctsb | | 1.193e-02 | -4.43 | GSE17721\_POLYIC\_VS\_PAM3CSK4\_24H\_BMDC\_UP | MSigDB lists | GSE17721\_POLYIC\_VS\_PAM3CSK4\_24H\_BMDC\_UP | 170 | 5 | 12187 | 98 | Ctsb,Dek,Grn,Thy1,Csk | | 1.193e-02 | -4.43 | GSE21546\_SAP1A\_KO\_VS\_SAP1A\_KO\_AND\_ELK1\_KO\_ANTI\_CD3\_STIM\_DP\_THYMOCYTES\_DN | MSigDB lists | GSE21546\_SAP1A\_KO\_VS\_SAP1A\_KO\_AND\_ELK1\_KO\_ANTI\_CD3\_STIM\_DP\_THYMOCYTES\_DN | 170 | 5 | 12187 | 98 | Vim,Lrg1,Rac2,Ptprc,Cyba | | 1.193e-02 | -4.43 | GSE15930\_STIM\_VS\_STIM\_AND\_IL12\_48H\_CD8\_T\_CELL\_UP | MSigDB lists | GSE15930\_STIM\_VS\_STIM\_AND\_IL12\_48H\_CD8\_T\_CELL\_UP | 170 | 5 | 12187 | 98 | Csk,Ctss,Il6ra,Psme1,Rps5 | | 1.193e-02 | -4.43 | GSE11057\_NAIVE\_VS\_MEMORY\_CD4\_TCELL\_DN | MSigDB lists | GSE11057\_NAIVE\_VS\_MEMORY\_CD4\_TCELL\_DN | 170 | 5 | 12187 | 98 | Actb,Ctsa,Sh3bgrl3,Ldha,Ahnak | | 1.196e-02 | -4.43 | GO\_PROTEIN\_HOMODIMERIZATION\_ACTIVITY | MSigDB lists | GO\_PROTEIN\_HOMODIMERIZATION\_ACTIVITY | 544 | 10 | 12187 | 98 | C1qb,Csf1r,Camk2b,Myh9,Rap1gap,Mgll,S100a6,Vwf,Ikzf3,Il6ra | | 1.199e-02 | -4.42 | C1q | pfam domains | PF00386 | 20 | 2 | 12881 | 108 | C1qb,C1qa | | 1.200e-02 | -4.42 | MODULE\_118 | MSigDB lists | MODULE\_118 | 308 | 7 | 12187 | 98 | Slc2a1,Vwf,Mgll,Grn,Fcer1g,Ptprc,Rac2 | | 1.201e-02 | -4.42 | endopeptidase activity | molecular function | GO:0004175 | 240 | 6 | 13516 | 107 | Capn3,Psmb10,Ctsz,Pcsk2,Ctsb,Ctss | | 1.203e-02 | -4.42 | Invs (inversin) | protein interactions | 16348 | 78 | 4 | 6802 | 78 | Sh3bgrl3,Vim,Actb,Myh9 | | 1.207e-02 | -4.42 | Alternative complement activation | REACTOME pathways | R-MMU-173736 | 1 | 1 | 6297 | 76 | C3 | | 1.209e-02 | -4.42 | cellular response to interleukin-1 | biological process | GO:0071347 | 59 | 3 | 13711 | 111 | Il1r1,Saa3,Ccl6 | | 1.215e-02 | -4.41 | regulation of peptidase activity | biological process | GO:0052547 | 306 | 7 | 13711 | 111 | Serpina3n,Ndufa13,C3,Grn,Serpinb9,Psme1,Spint1 | | 1.217e-02 | -4.41 | GCM\_TEC | MSigDB lists | GCM\_TEC | 21 | 2 | 12187 | 98 | Tmem106a,Fcgr3 | | 1.217e-02 | -4.41 | GO\_PROTEOGLYCAN\_BINDING | MSigDB lists | GO\_PROTEOGLYCAN\_BINDING | 21 | 2 | 12187 | 98 | Ctss,Ctsb | | 1.217e-02 | -4.41 | GO\_REGULATION\_OF\_LAMELLIPODIUM\_ASSEMBLY | MSigDB lists | GO\_REGULATION\_OF\_LAMELLIPODIUM\_ASSEMBLY | 21 | 2 | 12187 | 98 | Rac2,Brk1 | | 1.222e-02 | -4.40 | GSE2826\_WT\_VS\_BTK\_KO\_BCELL\_UP | MSigDB lists | GSE2826\_WT\_VS\_BTK\_KO\_BCELL\_UP | 171 | 5 | 12187 | 98 | Vwf,Rap1gap,Brk1,Ldha,Csk | | 1.222e-02 | -4.40 | REACTOME\_INNATE\_IMMUNE\_SYSTEM | MSigDB lists | REACTOME\_INNATE\_IMMUNE\_SYSTEM | 171 | 5 | 12187 | 98 | Ctss,C1qa,Ctsb,C3,C1qb | | 1.222e-02 | -4.40 | GSE15930\_NAIVE\_VS\_72H\_IN\_VITRO\_STIM\_IL12\_CD8\_TCELL\_DN | MSigDB lists | GSE15930\_NAIVE\_VS\_72H\_IN\_VITRO\_STIM\_IL12\_CD8\_TCELL\_DN | 171 | 5 | 12187 | 98 | Slc2a1,Serpinb9,Ahnak,Ccr5,S100a6 | | 1.223e-02 | -4.40 | GSE23114\_WT\_VS\_SLE2C1\_MOUSE\_PERITONEAL\_CAVITY\_B1A\_BCELL\_DN | MSigDB lists | GSE23114\_WT\_VS\_SLE2C1\_MOUSE\_PERITONEAL\_CAVITY\_B1A\_BCELL\_DN | 111 | 4 | 12187 | 98 | C1qa,C3,Pde1a,Mc4r | | 1.225e-02 | -4.40 | MODULE\_100 | MSigDB lists | MODULE\_100 | 464 | 9 | 12187 | 98 | Igfbp6,Slc17a7,Ptprn,Phyhip,Serpina3n,Apod,Hspa2,C3,C1qb | | 1.225e-02 | -4.40 | GO\_PEPTIDE\_METABOLIC\_PROCESS | MSigDB lists | GO\_PEPTIDE\_METABOLIC\_PROCESS | 464 | 9 | 12187 | 98 | Rps20,Rpl31,Rps5,Rpl14,Ctsz,Rps8,Rpl13,Pcsk2,Rpl26 | | 1.228e-02 | -4.40 | positive regulation of programmed cell death | biological process | GO:0043068 | 541 | 10 | 13711 | 111 | Ptprc,Rpl26,Ctsz,Clu,Camk2b,Tsc22d1,Ccr5,Ndufa13,Grn,Ccar1 | | 1.234e-02 | -4.40 | energy derivation by oxidation of organic compounds | biological process | GO:0015980 | 170 | 5 | 13711 | 111 | Mt3,Cox4i1,Kl,Ldha,Mc4r | | 1.234e-02 | -4.40 | postsynaptic cytoskeleton organization | biological process | GO:0099188 | 21 | 2 | 13711 | 111 | Camk2b,Actb | | 1.234e-02 | -4.40 | positive regulation of cellular extravasation | biological process | GO:0002693 | 21 | 2 | 13711 | 111 | Thy1,Il1r1 | | 1.234e-02 | -4.40 | Rac protein signal transduction | biological process | GO:0016601 | 21 | 2 | 13711 | 111 | Brk1,Rac2 | | 1.234e-02 | -4.40 | microglial cell activation | biological process | GO:0001774 | 21 | 2 | 13711 | 111 | Clu,Grn | | 1.234e-02 | -4.40 | leukocyte activation involved in inflammatory response | biological process | GO:0002269 | 21 | 2 | 13711 | 111 | Clu,Grn | | 1.235e-02 | -4.39 | GO\_LOCOMOTION | MSigDB lists | GO\_LOCOMOTION | 807 | 13 | 12187 | 98 | Ptprc,Rac2,Fmod,Csk,Ctsb,Il6ra,Vamp8,Csf1r,Myh9,Ccr5,Fcer1g,Itgam,Brk1 | | 1.238e-02 | -4.39 | PACSIN3 (protein kinase C and casein kinase substrate in neurons 3) | protein interactions | 29763 | 15 | 2 | 6802 | 78 | Flnb,Myh9 | | 1.238e-02 | -4.39 | MPRIP (myosin phosphatase Rho interacting protein) | protein interactions | 23164 | 15 | 2 | 6802 | 78 | Myh9,Actb | | 1.238e-02 | -4.39 | KIDINS220 (kinase D interacting substrate 220) | protein interactions | 57498 | 15 | 2 | 6802 | 78 | Flnb,Actb | | 1.238e-02 | -4.39 | JUP (junction plakoglobin) | protein interactions | 3728 | 15 | 2 | 6802 | 78 | Flnb,Myh9 | | 1.238e-02 | -4.39 | AHNAK (AHNAK nucleoprotein) | protein interactions | 79026 | 15 | 2 | 6802 | 78 | Flnb,Actb | | 1.239e-02 | -4.39 | MODULE\_544 | MSigDB lists | MODULE\_544 | 60 | 3 | 12187 | 98 | Pde1a,Cox4i1,Fmod | | 1.239e-02 | -4.39 | MODULE\_481 | MSigDB lists | MODULE\_481 | 60 | 3 | 12187 | 98 | Fmod,Pde1a,Cox4i1 | | 1.240e-02 | -4.39 | GO\_POSITIVE\_REGULATION\_OF\_PROTEOLYSIS | MSigDB lists | GO\_POSITIVE\_REGULATION\_OF\_PROTEOLYSIS | 310 | 7 | 12187 | 98 | Ndufa13,Clu,Psme1,Capn3,Myh9,C3,Psmb10 | | 1.245e-02 | -4.39 | collagen binding | molecular function | GO:0005518 | 61 | 3 | 13516 | 107 | Ctss,Ctsb,Vwf | | 1.245e-02 | -4.39 | electron transfer activity | molecular function | GO:0009055 | 61 | 3 | 13516 | 107 | Cox4i1,Sh3bgrl3,Cyba | | 1.245e-02 | -4.39 | calcium-dependent protein binding | molecular function | GO:0048306 | 61 | 3 | 13516 | 107 | S100a6,Mgp,Cplx2 | | 1.245e-02 | -4.39 | cytokine receptor activity | molecular function | GO:0004896 | 61 | 3 | 13516 | 107 | Ccr5,Il1r1,Il6ra | | 1.251e-02 | -4.38 | GSE23398\_WT\_VS\_IL2\_KO\_CD4\_TCELL\_SCURFY\_MOUSE\_UP | MSigDB lists | GSE23398\_WT\_VS\_IL2\_KO\_CD4\_TCELL\_SCURFY\_MOUSE\_UP | 172 | 5 | 12187 | 98 | Fam111a,Serpinb9,Cd68,C3,Ctsb | | 1.251e-02 | -4.38 | GO\_RESPONSE\_TO\_ESTROGEN | MSigDB lists | GO\_RESPONSE\_TO\_ESTROGEN | 172 | 5 | 12187 | 98 | Ptprn,C3,Grn,Serpinb9,Ldha | | 1.251e-02 | -4.38 | GSE12198\_NK\_VS\_NK\_ACT\_EXPANSION\_SYSTEM\_DERIVED\_NK\_CELL\_DN | MSigDB lists | GSE12198\_NK\_VS\_NK\_ACT\_EXPANSION\_SYSTEM\_DERIVED\_NK\_CELL\_DN | 172 | 5 | 12187 | 98 | Ctsb,Slc2a1,Il6ra,Lfng,Ldha | | 1.251e-02 | -4.38 | STARK\_PREFRONTAL\_CORTEX\_22Q11\_DELETION\_UP | MSigDB lists | STARK\_PREFRONTAL\_CORTEX\_22Q11\_DELETION\_UP | 172 | 5 | 12187 | 98 | Flnb,Lin7a,Myh9,Cplx2,Slc17a7 | | 1.251e-02 | -4.38 | GSE22886\_DC\_VS\_MONOCYTE\_UP | MSigDB lists | GSE22886\_DC\_VS\_MONOCYTE\_UP | 172 | 5 | 12187 | 98 | C1qb,Ctsb,Spint1,C1qa,Rap1gap | | 1.251e-02 | -4.38 | GSE17721\_0.5H\_VS\_24H\_CPG\_BMDC\_UP | MSigDB lists | GSE17721\_0.5H\_VS\_24H\_CPG\_BMDC\_UP | 172 | 5 | 12187 | 98 | Spint1,Rab3il1,Tspan4,Vwf,S100a6 | | 1.251e-02 | -4.38 | positive regulation of protein phosphorylation | biological process | GO:0001934 | 800 | 13 | 13711 | 111 | Mt3,Csf1r,Hcls1,Fgf18,Il6ra,Csk,C3,Hspa2,Ptprc,Clu,Ccl6,Kl,Tmem106a | | 1.254e-02 | -4.38 | chemotaxis | biological process | GO:0006935 | 383 | 8 | 13711 | 111 | Ccr5,Saa3,Fcer1g,Rac2,Itgam,Ccl6,Csf1r,Fcgr3 | | 1.257e-02 | -4.38 | GO\_MEMBRANE\_MICRODOMAIN | MSigDB lists | GO\_MEMBRANE\_MICRODOMAIN | 239 | 6 | 12187 | 98 | Slc2a1,Thy1,Ahnak,Csk,Ctsb,Ptprc | | 1.257e-02 | -4.38 | LU\_AGING\_BRAIN\_UP | MSigDB lists | LU\_AGING\_BRAIN\_UP | 239 | 6 | 12187 | 98 | Vim,Gfap,Apod,Hspa2,Ahnak,Vwf | | 1.260e-02 | -4.37 | large ribosomal subunit | cellular component | GO:0015934 | 112 | 4 | 13825 | 111 | Rpl13,Rpl14,Rpl31,Rpl26 | | 1.265e-02 | -4.37 | Ywhaz (tyrosine 3-monooxygenase/tryptophan 5-monooxygenase activation protein, zeta polypeptide) | protein interactions | 22631 | 169 | 6 | 6802 | 78 | Camk2b,Hspa2,Gfap,C1qa,Myh9,Pde1a | | 1.265e-02 | -4.37 | lymphocyte proliferation | biological process | GO:0046651 | 60 | 3 | 13711 | 111 | Itgam,Psmb10,Ptprc | | 1.265e-02 | -4.37 | regulation of cell killing | biological process | GO:0031341 | 60 | 3 | 13711 | 111 | Ccr5,Serpinb9,Ptprc | | 1.273e-02 | -4.36 | GO\_REGULATION\_OF\_CELL\_PROJECTION\_ORGANIZATION | MSigDB lists | GO\_REGULATION\_OF\_CELL\_PROJECTION\_ORGANIZATION | 467 | 9 | 12187 | 98 | Rac2,Gfap,Vim,Camk2b,Fmod,Thy1,Grn,Rap1gap,Brk1 | | 1.280e-02 | -4.36 | GSE5142\_CTRL\_VS\_HTERT\_TRANSDUCED\_CD8\_TCELL\_EARLY\_PASSAGE\_CLONE\_UP | MSigDB lists | GSE5142\_CTRL\_VS\_HTERT\_TRANSDUCED\_CD8\_TCELL\_EARLY\_PASSAGE\_CLONE\_UP | 173 | 5 | 12187 | 98 | Dek,Capn3,Ahnak,S100a6,Vim | | 1.284e-02 | -4.35 | C1q\_dom | interpro domains | IPR001073 | 21 | 2 | 13788 | 114 | C1qb,C1qa | | 1.284e-02 | -4.35 | SAP\_dom | interpro domains | IPR003034 | 21 | 2 | 13788 | 114 | Ccar1,Dek | | 1.295e-02 | -4.35 | positive regulation of neuron death | biological process | GO:1901216 | 112 | 4 | 13711 | 111 | Grn,Ctsz,Clu,C1qa | | 1.296e-02 | -4.35 | GO\_CELLULAR\_RESPONSE\_TO\_CARBOHYDRATE\_STIMULUS | MSigDB lists | GO\_CELLULAR\_RESPONSE\_TO\_CARBOHYDRATE\_STIMULUS | 61 | 3 | 12187 | 98 | Cyba,Ptprn,Il1r1 | | 1.298e-02 | -4.34 | peptide biosynthetic process | biological process | GO:0043043 | 310 | 7 | 13711 | 111 | Rps5,Rpl14,Rpl31,Rpl26,Rps8,Rpl13,Rps20 | | 1.298e-02 | -4.34 | anchored component of membrane | cellular component | GO:0031225 | 113 | 4 | 13825 | 111 | Ly6e,Cd52,Thy1,Ly6a | | 1.299e-02 | -4.34 | GO\_REGULATION\_OF\_CYTOKINE\_PRODUCTION | MSigDB lists | GO\_REGULATION\_OF\_CYTOKINE\_PRODUCTION | 389 | 8 | 12187 | 98 | Il6ra,Apod,Cyba,Csk,Fcer1g,Clu,C3,Csf1r | | 1.299e-02 | -4.34 | GSE6259\_FLT3L\_INDUCED\_VS\_WT\_SPLENIC\_DC\_33D1\_POS\_DN | MSigDB lists | GSE6259\_FLT3L\_INDUCED\_VS\_WT\_SPLENIC\_DC\_33D1\_POS\_DN | 113 | 4 | 12187 | 98 | Mpeg1,Il1r1,Ctss,Trnp1 | | 1.307e-02 | -4.34 | negative regulation of neuron differentiation | biological process | GO:0045665 | 239 | 6 | 13711 | 111 | Vim,Gfap,Ccr5,Thy1,Rap1gap,Ctsz | | 1.309e-02 | -4.34 | taxis | biological process | GO:0042330 | 386 | 8 | 13711 | 111 | Saa3,Ccr5,Fcgr3,Csf1r,Itgam,Ccl6,Rac2,Fcer1g | | 1.309e-02 | -4.34 | GSE17721\_0.5H\_VS\_24H\_PAM3CSK4\_BMDC\_UP | MSigDB lists | GSE17721\_0.5H\_VS\_24H\_PAM3CSK4\_BMDC\_UP | 174 | 5 | 12187 | 98 | Fgf18,Vim,Csk,Psme1,Rpl13 | | 1.309e-02 | -4.34 | GSE339\_CD4POS\_VS\_CD8POS\_DC\_DN | MSigDB lists | GSE339\_CD4POS\_VS\_CD8POS\_DC\_DN | 174 | 5 | 12187 | 98 | Mpeg1,Serpinb9,Brk1,Vwf,Ahnak | | 1.318e-02 | -4.33 | ACEVEDO\_LIVER\_CANCER\_DN | MSigDB lists | ACEVEDO\_LIVER\_CANCER\_DN | 390 | 8 | 12187 | 98 | C3,Ldha,Csk,Ly6e,Rap1gap,Psmb10,Ctsb,Cox4i1 | | 1.323e-02 | -4.33 | humoral immune response | biological process | GO:0006959 | 61 | 3 | 13711 | 111 | C1qa,C1qb,C3 | | 1.323e-02 | -4.33 | mononuclear cell proliferation | biological process | GO:0032943 | 61 | 3 | 13711 | 111 | Ptprc,Psmb10,Itgam | | 1.329e-02 | -4.32 | plasma lipoprotein particle | cellular component | GO:0034358 | 22 | 2 | 13825 | 111 | Clu,Saa3 | | 1.329e-02 | -4.32 | lipoprotein particle | cellular component | GO:1990777 | 22 | 2 | 13825 | 111 | Clu,Saa3 | | 1.331e-02 | -4.32 | positive regulation of catalytic activity | biological process | GO:0043085 | 1081 | 16 | 13711 | 111 | Grn,Thy1,Rap1gap,Ccl6,Clu,Ptprc,Hspa2,Cyba,Ndufa13,Csk,Camk2b,Il6ra,Psme1,Fgf18,Csf1r,Mt3 | | 1.334e-02 | -4.32 | cellular process | biological process | GO:0009987 | 10077 | 92 | 13711 | 111 | Sh3bgrl3,Adh1,Mc4r,Tmem106a,Itgam,Camk2b,Serpinb9,Capn3,Ctsb,Tnfsf8,Rpl26,Exoc3l4,Lrg1,Tspan4,Rac2,Csf1r,Lgi4,Brk1,Ptprn,C3,Tfap2c,Saa3,Ctsa,Ptprc,Alg12,Adgre1,Ccar1,Ctss,Kl,Lin7a,Pde1a,Bhlhe41,Actb,Ldha,Psmb10,Fgf18,Pcsk2,Ctsz,Flnb,Cplx2,Klhl6,Rpl13,Slc2a1,Dek,Cd52,Cox4i1,Myh9,Lfng,Ndufa13,Mgp,Csk,Mark4,Abhd17c,Cd68,Clu,Hspa2,Rbfox3,Slc17a7,Treml2,Gfap,Il6ra,Ccl6,Rps8,Vim,Rpl14,Ifitm1,Spint1,Hcls1,Rpl31,Vwf,C1qa,Grn,Ikzf3,Thy1,Mgll,Trnp1,Il1r1,Fcer1g,Atp5j2,Ahnak,Ly6e,Fcgr3,Rap1gap,Rps20,Vamp8,Rps5,Mt3,Fam111a,Dlgap3,Cyba,C1qb,Ccr5 | | 1.334e-02 | -4.32 | synaptic vesicle cycle | biological process | GO:0099504 | 113 | 4 | 13711 | 111 | Actb,Grn,Slc17a7,Cplx2 | | 1.334e-02 | -4.32 | cell activation involved in immune response | biological process | GO:0002263 | 113 | 4 | 13711 | 111 | Grn,Fcer1g,Cplx2,Lfng | | 1.336e-02 | -4.32 | GO\_ESTABLISHMENT\_OF\_PROTEIN\_LOCALIZATION | MSigDB lists | GO\_ESTABLISHMENT\_OF\_PROTEIN\_LOCALIZATION | 1189 | 17 | 12187 | 98 | Rac2,Ptprn,Rab3il1,Rps8,Clu,Rpl13,Rps20,Rpl14,Vamp8,Rpl26,Lin7a,Ctsa,Mc4r,Ndufa13,Myh9,Rps5,Rpl31 | | 1.338e-02 | -4.31 | BMI1\_DN\_MEL18\_DN.V1\_DN | MSigDB lists | BMI1\_DN\_MEL18\_DN.V1\_DN | 114 | 4 | 12187 | 98 | Ly6e,Il1r1,Spint1,Tmem176a | | 1.340e-02 | -4.31 | GSE17186\_BLOOD\_VS\_CORD\_BLOOD\_NAIVE\_BCELL\_UP | MSigDB lists | GSE17186\_BLOOD\_VS\_CORD\_BLOOD\_NAIVE\_BCELL\_UP | 175 | 5 | 12187 | 98 | Psme1,Dek,Flnb,Exoc3l4,Spint1 | | 1.340e-02 | -4.31 | GSE22886\_NAIVE\_CD8\_TCELL\_VS\_DC\_DN | MSigDB lists | GSE22886\_NAIVE\_CD8\_TCELL\_VS\_DC\_DN | 175 | 5 | 12187 | 98 | Fcer1g,Vim,Ctss,Vamp8,Ctsb | | 1.340e-02 | -4.31 | GSE41867\_NAIVE\_VS\_DAY6\_LCMV\_EFFECTOR\_CD8\_TCELL\_UP | MSigDB lists | GSE41867\_NAIVE\_VS\_DAY6\_LCMV\_EFFECTOR\_CD8\_TCELL\_UP | 175 | 5 | 12187 | 98 | Sh3bgrl3,Cd68,Tmem106a,Actb,Psmb10 | | 1.354e-02 | -4.30 | REACTOME\_SEMAPHORIN\_INTERACTIONS | MSigDB lists | REACTOME\_SEMAPHORIN\_INTERACTIONS | 62 | 3 | 12187 | 98 | Myh9,Rac2,Ptprc | | 1.354e-02 | -4.30 | GO\_INTEGRIN\_MEDIATED\_SIGNALING\_PATHWAY | MSigDB lists | GO\_INTEGRIN\_MEDIATED\_SIGNALING\_PATHWAY | 62 | 3 | 12187 | 98 | Myh9,Itgam,Fcer1g | | 1.354e-02 | -4.30 | GO\_SMALL\_RIBOSOMAL\_SUBUNIT | MSigDB lists | GO\_SMALL\_RIBOSOMAL\_SUBUNIT | 62 | 3 | 12187 | 98 | Rps8,Rps5,Rps20 | | 1.354e-02 | -4.30 | FRASOR\_RESPONSE\_TO\_ESTRADIOL\_DN | MSigDB lists | FRASOR\_RESPONSE\_TO\_ESTRADIOL\_DN | 62 | 3 | 12187 | 98 | Il1r1,Clu,Rap1gap | | 1.358e-02 | -4.30 | RNA polymerase II distal enhancer sequence-specific DNA binding | molecular function | GO:0000980 | 63 | 3 | 13516 | 107 | Tfap2c,Ccar1,Bhlhe41 | | 1.370e-02 | -4.29 | GSE42021\_TREG\_PLN\_VS\_CD24HI\_TREG\_THYMUS\_DN | MSigDB lists | GSE42021\_TREG\_PLN\_VS\_CD24HI\_TREG\_THYMUS\_DN | 176 | 5 | 12187 | 98 | Grn,Csk,Vamp8,Rps5,C1qb | | 1.370e-02 | -4.29 | BHAT\_ESR1\_TARGETS\_NOT\_VIA\_AKT1\_UP | MSigDB lists | BHAT\_ESR1\_TARGETS\_NOT\_VIA\_AKT1\_UP | 176 | 5 | 12187 | 98 | Flnb,Slc2a1,Rab3il1,Lrg1,Vwf | | 1.376e-02 | -4.29 | regulation of cellular process | biological process | GO:0050794 | 7419 | 72 | 13711 | 111 | Adgre1,Ikzf3,Ccar1,Grn,Kl,Thy1,Trnp1,Il1r1,Mgll,C1qa,Ptprc,Ahnak,Ly6e,Fgf18,Psme1,Bhlhe41,Pde1a,Fcer1g,Actb,Klhl6,Cplx2,Vamp8,Dek,Rap1gap,Igfbp6,Ctsz,Fcgr3,Lfng,Ndufa13,Ccr5,Cyba,Csk,Mark4,Myh9,Mt3,Mc4r,Tmem106a,Rbfox3,Abhd17c,Clu,Sh3bgrl3,Tmem176a,Hspa2,Apod,Tsc22d1,Gfap,Serpinb9,Il6ra,Camk2b,Ctsb,Capn3,Slc17a7,Itgam,Vim,Lrg1,Ccl6,Tnfsf8,Rpl26,Zfp786,Tfap2c,Serpina3n,C3,Ptprn,Saa3,Ctsa,Ifitm1,Rac2,Spint1,Csf1r,Hcls1,Lgi4,Brk1 | | 1.377e-02 | -4.29 | cortical cytoskeleton | cellular component | GO:0030863 | 115 | 4 | 13825 | 111 | Hcls1,Actb,Myh9,Slc2a1 | | 1.381e-02 | -4.28 | GO\_REGULATION\_OF\_METAL\_ION\_TRANSPORT | MSigDB lists | GO\_REGULATION\_OF\_METAL\_ION\_TRANSPORT | 244 | 6 | 12187 | 98 | Hspa2,Ahnak,Capn3,Camk2b,Thy1,Cyba | | 1.382e-02 | -4.28 | regulation of release of sequestered calcium ion into cytosol | biological process | GO:0051279 | 62 | 3 | 13711 | 111 | Thy1,Capn3,Cyba | | 1.399e-02 | -4.27 | cell communication | biological process | GO:0007154 | 3040 | 35 | 13711 | 111 | C3,Ptprn,Ccr5,Ndufa13,Mark4,Csk,Saa3,Rac2,Ifitm1,Brk1,Mt3,Csf1r,Vim,Cplx2,Klhl6,Slc2a1,Ccl6,Rpl26,Fcgr3,Fgf18,Il6ra,Ly6e,Fcer1g,Slc17a7,Lin7a,Pde1a,Ldha,Mc4r,Adgre1,Il1r1,Kl,Thy1,Clu,Cd68,Ptprc | | 1.401e-02 | -4.27 | GSE339\_CD4POS\_VS\_CD8POS\_DC\_UP | MSigDB lists | GSE339\_CD4POS\_VS\_CD8POS\_DC\_UP | 177 | 5 | 12187 | 98 | Il6ra,Fcer1g,Rac2,Ctsa,Il1r1 | | 1.404e-02 | -4.27 | SEC16A (SEC16 homolog A, endoplasmic reticulum export factor) | protein interactions | 9919 | 16 | 2 | 6802 | 78 | Actb,Myh9 | | 1.404e-02 | -4.27 | ACTR2 (actin related protein 2) | protein interactions | 10097 | 16 | 2 | 6802 | 78 | Myh9,Actb | | 1.404e-02 | -4.27 | GPRC5A (G protein-coupled receptor class C group 5 member A) | protein interactions | 9052 | 16 | 2 | 6802 | 78 | Myh9,Actb | | 1.410e-02 | -4.26 | misfolded protein binding | molecular function | GO:0051787 | 23 | 2 | 13516 | 107 | Clu,Hspa2 | | 1.411e-02 | -4.26 | GO\_ORGANIC\_CYCLIC\_COMPOUND\_CATABOLIC\_PROCESS | MSigDB lists | GO\_ORGANIC\_CYCLIC\_COMPOUND\_CATABOLIC\_PROCESS | 318 | 7 | 12187 | 98 | Rps20,Rpl14,Rps5,Rpl31,Rps8,Rpl13,Rpl26 | | 1.414e-02 | -4.26 | GO\_MOVEMENT\_IN\_ENVIRONMENT\_OF\_OTHER\_ORGANISM\_INVOLVED\_IN\_SYMBIOTIC\_INTERACTION | MSigDB lists | GO\_MOVEMENT\_IN\_ENVIRONMENT\_OF\_OTHER\_ORGANISM\_INVOLVED\_IN\_SYMBIOTIC\_INTERACTION | 63 | 3 | 12187 | 98 | Ccr5,Vamp8,Ctsb | | 1.419e-02 | -4.26 | MOHANKUMAR\_HOXA1\_TARGETS\_DN | MSigDB lists | MOHANKUMAR\_HOXA1\_TARGETS\_DN | 116 | 4 | 12187 | 98 | Serpina3n,Apod,S100a6,Mgll | | 1.419e-02 | -4.26 | MODULE\_122 | MSigDB lists | MODULE\_122 | 116 | 4 | 12187 | 98 | Mgp,Vwf,Ppfibp1,Itgam | | 1.419e-02 | -4.26 | receptor ligand activity | molecular function | GO:0048018 | 249 | 6 | 13516 | 107 | Tnfsf8,Grn,Fgf18,Ccl6,Il6ra,Saa3 | | 1.423e-02 | -4.25 | EMR1HORMONER | prints domains | PR01128 | 1 | 1 | 2951 | 42 | Adgre1 | | 1.423e-02 | -4.25 | RIBOSOMALS10 | prints domains | PR00971 | 1 | 1 | 2951 | 42 | Rps20 | | 1.423e-02 | -4.25 | APOLIPOPROTD | prints domains | PR01219 | 1 | 1 | 2951 | 42 | Apod | | 1.423e-02 | -4.25 | CHEMOKINER5 | prints domains | PR01110 | 1 | 1 | 2951 | 42 | Ccr5 | | 1.423e-02 | -4.25 | MELNOCORTN4R | prints domains | PR01062 | 1 | 1 | 2951 | 42 | Mc4r | | 1.423e-02 | -4.25 | GLABONE | prints domains | PR00002 | 1 | 1 | 2951 | 42 | Mgp | | 1.423e-02 | -4.25 | APODVERTBRTE | prints domains | PR02058 | 1 | 1 | 2951 | 42 | Apod | | 1.423e-02 | -4.25 | IGFBPFAMILY6 | prints domains | PR01982 | 1 | 1 | 2951 | 42 | Igfbp6 | | 1.423e-02 | -4.25 | AP2CTNSCPFCT | prints domains | PR01751 | 1 | 1 | 2951 | 42 | Tfap2c | | 1.423e-02 | -4.25 | INTRLEUKN1R1 | prints domains | PR01538 | 1 | 1 | 2951 | 42 | Il1r1 | | 1.423e-02 | -4.25 | GLUCTRSPORT1 | prints domains | PR01190 | 1 | 1 | 2951 | 42 | Slc2a1 | | 1.442e-02 | -4.24 | positive regulation of calcium ion transmembrane transport | biological process | GO:1904427 | 63 | 3 | 13711 | 111 | Thy1,Capn3,Hspa2 | | 1.445e-02 | -4.24 | synapse part | cellular component | GO:0044456 | 1006 | 15 | 13825 | 111 | Camk2b,Rpl26,Sh2d5,Slc17a7,Dlgap3,Il1r1,Ptprn,C1qb,Rpl14,Mt3,Cplx2,Actb,C1qa,Abhd17c,Lin7a | | 1.451e-02 | -4.23 | GO\_ACUTE\_PHASE\_RESPONSE | MSigDB lists | GO\_ACUTE\_PHASE\_RESPONSE | 23 | 2 | 12187 | 98 | Il6ra,Serpina3n | | 1.451e-02 | -4.23 | GO\_POSITIVE\_REGULATION\_OF\_TYROSINE\_PHOSPHORYLATION\_OF\_STAT3\_PROTEIN | MSigDB lists | GO\_POSITIVE\_REGULATION\_OF\_TYROSINE\_PHOSPHORYLATION\_OF\_STAT3\_PROTEIN | 23 | 2 | 12187 | 98 | Il6ra,Csf1r | | 1.451e-02 | -4.23 | BERENJENO\_TRANSFORMED\_BY\_RHOA\_FOREVER\_DN | MSigDB lists | BERENJENO\_TRANSFORMED\_BY\_RHOA\_FOREVER\_DN | 23 | 2 | 12187 | 98 | Mgp,Flnb | | 1.451e-02 | -4.23 | GO\_MEMBRANE\_INVAGINATION | MSigDB lists | GO\_MEMBRANE\_INVAGINATION | 23 | 2 | 12187 | 98 | Myh9,Fcer1g | | 1.451e-02 | -4.23 | XU\_CREBBP\_TARGETS\_UP | MSigDB lists | XU\_CREBBP\_TARGETS\_UP | 23 | 2 | 12187 | 98 | S100a6,Lrg1 | | 1.451e-02 | -4.23 | REACTOME\_SEMA4D\_INDUCED\_CELL\_MIGRATION\_AND\_GROWTH\_CONE\_COLLAPSE | MSigDB lists | REACTOME\_SEMA4D\_INDUCED\_CELL\_MIGRATION\_AND\_GROWTH\_CONE\_COLLAPSE | 23 | 2 | 12187 | 98 | Rac2,Myh9 | | 1.451e-02 | -4.23 | GO\_LEUKOCYTE\_DEGRANULATION | MSigDB lists | GO\_LEUKOCYTE\_DEGRANULATION | 23 | 2 | 12187 | 98 | Cplx2,Vamp8 | | 1.451e-02 | -4.23 | MODULE\_524 | MSigDB lists | MODULE\_524 | 23 | 2 | 12187 | 98 | Myh9,Flnb | | 1.451e-02 | -4.23 | MODULE\_265 | MSigDB lists | MODULE\_265 | 23 | 2 | 12187 | 98 | Il1r1,Il6ra | | 1.454e-02 | -4.23 | pyruvate fermentation to lactate | BIOCYC pathways | MOUSE\_PWY-5481 | 2 | 1 | 823 | 6 | Ldha | | 1.454e-02 | -4.23 | phenylalanine degradation III | BIOCYC pathways | MOUSE\_PWY-5079 | 2 | 1 | 823 | 6 | Adh1 | | 1.458e-02 | -4.23 | signaling | biological process | GO:0023052 | 2939 | 34 | 13711 | 111 | Ifitm1,Rac2,Csf1r,Brk1,Mt3,Dlgap3,Ndufa13,Ccr5,C3,Ptprn,Saa3,Csk,Mark4,Ccl6,Fcgr3,Rpl26,Klhl6,Cplx2,Vim,Lin7a,Pde1a,Slc17a7,Fcer1g,Ly6e,Il6ra,Fgf18,Cd68,Clu,Ptprc,Adgre1,Mc4r,Kl,Thy1,Il1r1 | | 1.458e-02 | -4.23 | Platelet activation, signaling and aggregation | REACTOME pathways | R-MMU-76002 | 215 | 7 | 6297 | 76 | Serpina3n,Mgll,Rac2,Clu,Vwf,Fcer1g,Csk | | 1.459e-02 | -4.23 | RUTELLA\_RESPONSE\_TO\_CSF2RB\_AND\_IL4\_DN | MSigDB lists | RUTELLA\_RESPONSE\_TO\_CSF2RB\_AND\_IL4\_DN | 247 | 6 | 12187 | 98 | Ctsb,C3,C1qa,Tsc22d1,Ccr5,Cd68 | | 1.465e-02 | -4.22 | GSE17186\_BLOOD\_VS\_CORD\_BLOOD\_CD21HIGH\_TRANSITIONAL\_BCELL\_UP | MSigDB lists | GSE17186\_BLOOD\_VS\_CORD\_BLOOD\_CD21HIGH\_TRANSITIONAL\_BCELL\_UP | 179 | 5 | 12187 | 98 | Sh3bgrl3,Serpinb9,Vamp8,Fcer1g,Dek | | 1.465e-02 | -4.22 | GO\_POSITIVE\_REGULATION\_OF\_ION\_TRANSPORT | MSigDB lists | GO\_POSITIVE\_REGULATION\_OF\_ION\_TRANSPORT | 179 | 5 | 12187 | 98 | Thy1,Capn3,Hspa2,Ctss,Vamp8 | | 1.466e-02 | -4.22 | Ig\_sub | interpro domains | IPR003599 | 240 | 6 | 13788 | 114 | Il6ra,Il1r1,Csf1r,Treml2,Fcgr3,Fcrls | | 1.471e-02 | -4.22 | actin filament polymerization | biological process | GO:0030041 | 23 | 2 | 13711 | 111 | Myh9,Hcls1 | | 1.471e-02 | -4.22 | regulation of neuroinflammatory response | biological process | GO:0150077 | 23 | 2 | 13711 | 111 | C1qa,Grn | | 1.471e-02 | -4.22 | regulation of humoral immune response | biological process | GO:0002920 | 23 | 2 | 13711 | 111 | C3,Ptprc | | 1.471e-02 | -4.22 | antigen processing and presentation of peptide antigen via MHC class I | biological process | GO:0002474 | 23 | 2 | 13711 | 111 | Fcer1g,Fcgr3 | | 1.471e-02 | -4.22 | cellular response to lipoprotein particle stimulus | biological process | GO:0071402 | 23 | 2 | 13711 | 111 | Cd68,Fcer1g | | 1.475e-02 | -4.22 | GO\_REGULATION\_OF\_CALCIUM\_ION\_TRANSPORT\_INTO\_CYTOSOL | MSigDB lists | GO\_REGULATION\_OF\_CALCIUM\_ION\_TRANSPORT\_INTO\_CYTOSOL | 64 | 3 | 12187 | 98 | Cyba,Capn3,Thy1 | | 1.475e-02 | -4.22 | GO\_MULTICELLULAR\_ORGANISM\_METABOLIC\_PROCESS | MSigDB lists | GO\_MULTICELLULAR\_ORGANISM\_METABOLIC\_PROCESS | 64 | 3 | 12187 | 98 | Ctsb,Ctss,Mc4r | | 1.475e-02 | -4.22 | Immunoregulatory interactions between a Lymphoid and a non-Lymphoid cell | REACTOME pathways | R-MMU-198933 | 43 | 3 | 6297 | 76 | Ifitm1,C3,Treml2 | | 1.476e-02 | -4.22 | RYTTCCTG\_ETS2\_B | MSigDB lists | RYTTCCTG\_ETS2\_B | 826 | 13 | 12187 | 98 | Treml2,Mark4,Fcer1g,Actb,Ctsa,Vamp8,Rps20,Hcls1,Ctss,Mgll,Flnb,Ptprc,Ptprn | | 1.501e-02 | -4.20 | C1Q | smart domains | SM00110 | 20 | 2 | 7188 | 68 | C1qb,C1qa | | 1.502e-02 | -4.20 | GSE37534\_GW1929\_VS\_PIOGLITAZONE\_TREATED\_CD4\_TCELL\_PPARG1\_FOXP3\_TRANSDUCED\_UP | MSigDB lists | GSE37534\_GW1929\_VS\_PIOGLITAZONE\_TREATED\_CD4\_TCELL\_PPARG1\_FOXP3\_TRANSDUCED\_UP | 118 | 4 | 12187 | 98 | C1qa,Psmb10,Capn3,Fam111a | | 1.502e-02 | -4.20 | GSE37605\_NOD\_VS\_C57BL6\_IRES\_GFP\_TREG\_UP | MSigDB lists | GSE37605\_NOD\_VS\_C57BL6\_IRES\_GFP\_TREG\_UP | 118 | 4 | 12187 | 98 | Grn,C1qa,Tmem176a,C1qb | | 1.508e-02 | -4.19 | GO\_SOMATODENDRITIC\_COMPARTMENT | MSigDB lists | GO\_SOMATODENDRITIC\_COMPARTMENT | 564 | 10 | 12187 | 98 | Dlgap3,Cplx2,Ptprn,Cyba,Apod,Thy1,Camk2b,Clu,Rap1gap,Pcsk2 | | 1.524e-02 | -4.18 | signaling receptor activator activity | molecular function | GO:0030546 | 253 | 6 | 13516 | 107 | Ccl6,Saa3,Il6ra,Tnfsf8,Grn,Fgf18 | | 1.531e-02 | -4.18 | IF\_conserved | interpro domains | IPR018039 | 23 | 2 | 13788 | 114 | Vim,Gfap | | 1.537e-02 | -4.18 | GO\_CYTOKINE\_BINDING | MSigDB lists | GO\_CYTOKINE\_BINDING | 65 | 3 | 12187 | 98 | Il6ra,Ccr5,Csf1r | | 1.545e-02 | -4.17 | GSE9006\_1MONTH\_VS\_4MONTH\_POST\_TYPE\_1\_DIABETES\_DX\_PBMC\_DN | MSigDB lists | GSE9006\_1MONTH\_VS\_4MONTH\_POST\_TYPE\_1\_DIABETES\_DX\_PBMC\_DN | 119 | 4 | 12187 | 98 | Pcsk2,Lfng,Cplx2,Il1r1 | | 1.546e-02 | -4.17 | Transcriptional regulation by the AP-2 (TFAP2) family of transcription factors | REACTOME pathways | R-MMU-8864260 | 16 | 2 | 6297 | 76 | Tfap2c,Dek | | 1.549e-02 | -4.17 | GO\_NCRNA\_PROCESSING | MSigDB lists | GO\_NCRNA\_PROCESSING | 324 | 7 | 12187 | 98 | Rpl26,Rps20,Rpl31,Rps5,Rpl14,Rps8,Rpl13 | | 1.557e-02 | -4.16 | GO\_REGULATION\_OF\_INTRACELLULAR\_TRANSPORT | MSigDB lists | GO\_REGULATION\_OF\_INTRACELLULAR\_TRANSPORT | 483 | 9 | 12187 | 98 | Vamp8,Rac2,Apod,Cyba,Csk,Fcer1g,Capn3,Thy1,Hcls1 | | 1.563e-02 | -4.16 | cytoplasmic vesicle | cellular component | GO:0031410 | 1501 | 20 | 13825 | 111 | Vim,Pcsk2,Vwf,Cd68,Mpeg1,Clu,Ctss,Grn,Slc17a7,Cyba,Ctsz,Abhd17c,Rac2,Flnb,Ifitm1,Ptprn,Rap1gap,Mt3,Vamp8,Ccr5 | | 1.564e-02 | -4.16 | GO\_LEUKOCYTE\_MIGRATION | MSigDB lists | GO\_LEUKOCYTE\_MIGRATION | 182 | 5 | 12187 | 98 | Fcer1g,Myh9,Ccr5,Itgam,Il6ra | | 1.570e-02 | -4.15 | Regulation of actin dynamics for phagocytic cup formation | REACTOME pathways | R-MMU-2029482 | 44 | 3 | 6297 | 76 | Actb,Fcgr3,Brk1 | | 1.571e-02 | -4.15 | protein-lipid complex | cellular component | GO:0032994 | 24 | 2 | 13825 | 111 | Saa3,Clu | | 1.571e-02 | -4.15 | perinuclear endoplasmic reticulum | cellular component | GO:0097038 | 24 | 2 | 13825 | 111 | Clu,Cyba | | 1.573e-02 | -4.15 | HELLER\_HDAC\_TARGETS\_SILENCED\_BY\_METHYLATION\_UP | MSigDB lists | HELLER\_HDAC\_TARGETS\_SILENCED\_BY\_METHYLATION\_UP | 325 | 7 | 12187 | 98 | Slc17a7,Hspa2,Tfap2c,Grn,Rps20,Flnb,C3 | | 1.573e-02 | -4.15 | MULLIGHAN\_MLL\_SIGNATURE\_2\_UP | MSigDB lists | MULLIGHAN\_MLL\_SIGNATURE\_2\_UP | 325 | 7 | 12187 | 98 | Grn,Rac2,S100a6,Il6ra,Cyba,Ctsa,Hcls1 | | 1.574e-02 | -4.15 | Immunoproteasome | KEGG pathways | M00337 | 16 | 2 | 5248 | 64 | Psme1,Psmb10 | | 1.574e-02 | -4.15 | Immunoproteasome | KEGG pathways | mmu\_M00337 | 16 | 2 | 5248 | 64 | Psmb10,Psme1 | | 1.574e-02 | -4.15 | mouse chr11 C\|11 | chromosome location | mouse chr11 C\|11 | 2 | 1 | 14556 | 115 | Wfdc17 | | 1.574e-02 | -4.15 | mouse chr13 A5\|13 24.5 cM | chromosome location | mouse chr13 A5\|13 24.5 cM | 2 | 1 | 14556 | 115 | Dek | | 1.574e-02 | -4.15 | mouse chr2 G1\|2 70.89 cM | chromosome location | mouse chr2 G1\|2 70.89 cM | 2 | 1 | 14556 | 115 | Pcsk2 | | 1.574e-02 | -4.15 | mouse chr4 C1\|4 34.06 cM | chromosome location | mouse chr4 C1\|4 34.06 cM | 2 | 1 | 14556 | 115 | Tnfsf8 | | 1.574e-02 | -4.15 | mouse chr16 B2\|16 21.41 cM | chromosome location | mouse chr16 B2\|16 21.41 cM | 2 | 1 | 14556 | 115 | Apod | | 1.574e-02 | -4.15 | mouse chr4 D3\|4 70.02 cM | chromosome location | mouse chr4 D3\|4 70.02 cM | 2 | 1 | 14556 | 115 | Rap1gap | | 1.574e-02 | -4.15 | mouse chr14 D1\|14 33.24 cM | chromosome location | mouse chr14 D1\|14 33.24 cM | 2 | 1 | 14556 | 115 | Ctsb | | 1.574e-02 | -4.15 | mouse chr4 D3\|4 66.5 cM | chromosome location | mouse chr4 D3\|4 66.5 cM | 2 | 1 | 14556 | 115 | Cd52 | | 1.574e-02 | -4.15 | mouse chr3 F1\|3 39.19 cM | chromosome location | mouse chr3 F1\|3 39.19 cM | 2 | 1 | 14556 | 115 | Il6ra | | 1.575e-02 | -4.15 | KIM\_HYPOXIA | MSigDB lists | KIM\_HYPOXIA | 24 | 2 | 12187 | 98 | Slc2a1,Ldha | | 1.575e-02 | -4.15 | KANG\_GLIS3\_TARGETS | MSigDB lists | KANG\_GLIS3\_TARGETS | 24 | 2 | 12187 | 98 | Psmb10,Pcsk2 | | 1.575e-02 | -4.15 | GNF2\_CEBPA | MSigDB lists | GNF2\_CEBPA | 24 | 2 | 12187 | 98 | Adh1,Tmem176a | | 1.575e-02 | -4.15 | WILENSKY\_RESPONSE\_TO\_DARAPLADIB | MSigDB lists | WILENSKY\_RESPONSE\_TO\_DARAPLADIB | 24 | 2 | 12187 | 98 | Ctss,Cd68 | | 1.575e-02 | -4.15 | TERAMOTO\_OPN\_TARGETS\_CLUSTER\_6 | MSigDB lists | TERAMOTO\_OPN\_TARGETS\_CLUSTER\_6 | 24 | 2 | 12187 | 98 | Psme1,Apod | | 1.575e-02 | -4.15 | REACTOME\_INTEGRIN\_ALPHAIIB\_BETA3\_SIGNALING | MSigDB lists | REACTOME\_INTEGRIN\_ALPHAIIB\_BETA3\_SIGNALING | 24 | 2 | 12187 | 98 | Csk,Vwf | | 1.575e-02 | -4.15 | GO\_NEGATIVE\_REGULATION\_OF\_LYMPHOCYTE\_MEDIATED\_IMMUNITY | MSigDB lists | GO\_NEGATIVE\_REGULATION\_OF\_LYMPHOCYTE\_MEDIATED\_IMMUNITY | 24 | 2 | 12187 | 98 | Serpinb9,Ptprc | | 1.575e-02 | -4.15 | GO\_POSITIVE\_REGULATION\_OF\_RELEASE\_OF\_SEQUESTERED\_CALCIUM\_ION\_INTO\_CYTOSOL | MSigDB lists | GO\_POSITIVE\_REGULATION\_OF\_RELEASE\_OF\_SEQUESTERED\_CALCIUM\_ION\_INTO\_CYTOSOL | 24 | 2 | 12187 | 98 | Thy1,Capn3 | | 1.575e-02 | -4.15 | DACOSTA\_ERCC3\_ALLELE\_XPCS\_VS\_TTD\_UP | MSigDB lists | DACOSTA\_ERCC3\_ALLELE\_XPCS\_VS\_TTD\_UP | 24 | 2 | 12187 | 98 | Igfbp6,Ptprn | | 1.575e-02 | -4.15 | PLASARI\_TGFB1\_SIGNALING\_VIA\_NFIC\_10HR\_DN | MSigDB lists | PLASARI\_TGFB1\_SIGNALING\_VIA\_NFIC\_10HR\_DN | 24 | 2 | 12187 | 98 | Fam111a,Rab3il1 | | 1.577e-02 | -4.15 | interleukin-12 binding | molecular function | GO:0019972 | 2 | 1 | 13516 | 107 | Il6ra | | 1.577e-02 | -4.15 | interleukin-23 receptor binding | molecular function | GO:0045519 | 2 | 1 | 13516 | 107 | Il6ra | | 1.577e-02 | -4.15 | dehydroascorbic acid transmembrane transporter activity | molecular function | GO:0033300 | 2 | 1 | 13516 | 107 | Slc2a1 | | 1.577e-02 | -4.15 | interleukin-12 alpha subunit binding | molecular function | GO:0042164 | 2 | 1 | 13516 | 107 | Il6ra | | 1.577e-02 | -4.15 | N-box binding | molecular function | GO:0071820 | 2 | 1 | 13516 | 107 | Bhlhe41 | | 1.577e-02 | -4.15 | IgE receptor activity | molecular function | GO:0019767 | 2 | 1 | 13516 | 107 | Fcer1g | | 1.577e-02 | -4.15 | ethanol binding | molecular function | GO:0035276 | 2 | 1 | 13516 | 107 | Adh1 | | 1.577e-02 | -4.15 | interleukin-6 receptor activity | molecular function | GO:0004915 | 2 | 1 | 13516 | 107 | Il6ra | | 1.577e-02 | -4.15 | L-lactate dehydrogenase activity | molecular function | GO:0004459 | 2 | 1 | 13516 | 107 | Ldha | | 1.577e-02 | -4.15 | beta-glucuronidase activity | molecular function | GO:0004566 | 2 | 1 | 13516 | 107 | Kl | | 1.577e-02 | -4.15 | interleukin-6 binding | molecular function | GO:0019981 | 2 | 1 | 13516 | 107 | Il6ra | | 1.577e-02 | -4.15 | alcohol dehydrogenase activity, zinc-dependent | molecular function | GO:0004024 | 2 | 1 | 13516 | 107 | Adh1 | | 1.577e-02 | -4.15 | melanocyte-stimulating hormone receptor activity | molecular function | GO:0004980 | 2 | 1 | 13516 | 107 | Mc4r | | 1.577e-02 | -4.15 | type 1 fibroblast growth factor receptor binding | molecular function | GO:0005105 | 2 | 1 | 13516 | 107 | Fgf18 | | 1.579e-02 | -4.15 | GSN (gelsolin) | protein interactions | 2934 | 17 | 2 | 6802 | 78 | Myh9,Actb | | 1.585e-02 | -4.14 | response to hormone | biological process | GO:0009725 | 480 | 9 | 13711 | 111 | Ly6e,Csk,Kl,Adh1,Serpina3n,Grn,Mc4r,Hcls1,Mt3 | | 1.589e-02 | -4.14 | GSE40274\_CTRL\_VS\_FOXP3\_AND\_LEF1\_TRANSDUCED\_ACTIVATED\_CD4\_TCELL\_UP | MSigDB lists | GSE40274\_CTRL\_VS\_FOXP3\_AND\_LEF1\_TRANSDUCED\_ACTIVATED\_CD4\_TCELL\_UP | 120 | 4 | 12187 | 98 | Thy1,Itgam,Ahnak,Ppfibp1 | | 1.589e-02 | -4.14 | GSE13411\_NAIVE\_VS\_SWITCHED\_MEMORY\_BCELL\_UP | MSigDB lists | GSE13411\_NAIVE\_VS\_SWITCHED\_MEMORY\_BCELL\_UP | 120 | 4 | 12187 | 98 | Camk2b,Fgf18,Adh1,Slc26a2 | | 1.595e-02 | -4.14 | intracellular vesicle | cellular component | GO:0097708 | 1504 | 20 | 13825 | 111 | Ctsz,Abhd17c,Flnb,Rac2,Ifitm1,Ptprn,Vamp8,Ccr5,Rap1gap,Mt3,Vim,Pcsk2,Vwf,Cd68,Mpeg1,Clu,Ctss,Grn,Slc17a7,Cyba | | 1.596e-02 | -4.14 | leukocyte mediated cytotoxicity | biological process | GO:0001909 | 24 | 2 | 13711 | 111 | Fcgr3,Serpinb9 | | 1.599e-02 | -4.14 | insulin-like growth factor binary complex | cellular component | GO:0042568 | 2 | 1 | 13825 | 111 | Igfbp6 | | 1.599e-02 | -4.14 | interleukin-12 complex | cellular component | GO:0043514 | 2 | 1 | 13825 | 111 | Il6ra | | 1.599e-02 | -4.14 | proteasome activator complex | cellular component | GO:0008537 | 2 | 1 | 13825 | 111 | Psme1 | | 1.599e-02 | -4.14 | amyloid-beta complex | cellular component | GO:0106003 | 2 | 1 | 13825 | 111 | C1qa | | 1.599e-02 | -4.14 | CSF1-CSF1R complex | cellular component | GO:1990682 | 2 | 1 | 13825 | 111 | Csf1r | | 1.599e-02 | -4.14 | interleukin-23 complex | cellular component | GO:0070743 | 2 | 1 | 13825 | 111 | Il6ra | | 1.601e-02 | -4.13 | BILANGES\_RAPAMYCIN\_SENSITIVE\_VIA\_TSC1\_AND\_TSC2 | MSigDB lists | BILANGES\_RAPAMYCIN\_SENSITIVE\_VIA\_TSC1\_AND\_TSC2 | 66 | 3 | 12187 | 98 | Rpl31,Rps20,Vim | | 1.602e-02 | -4.13 | WILSON\_PROTEASES\_AT\_TUMOR\_BONE\_INTERFACE\_DN | MSigDB lists | WILSON\_PROTEASES\_AT\_TUMOR\_BONE\_INTERFACE\_DN | 2 | 1 | 12187 | 98 | Ctsb | | 1.602e-02 | -4.13 | KINNEY\_DNMT1\_METHYLATION\_TARGETS | MSigDB lists | KINNEY\_DNMT1\_METHYLATION\_TARGETS | 2 | 1 | 12187 | 98 | Mc4r | | 1.603e-02 | -4.13 | GO\_PROTEIN\_COMPLEX\_SUBUNIT\_ORGANIZATION | MSigDB lists | GO\_PROTEIN\_COMPLEX\_SUBUNIT\_ORGANIZATION | 1213 | 17 | 12187 | 98 | Gfap,Lin7a,Myh9,Brk1,Ndufa13,Tspan4,Slc2a1,Rac2,Vim,Cyba,Clu,Fmod,Capn3,Ahnak,Vwf,Ctsz,Hcls1 | | 1.613e-02 | -4.13 | serotonin production involved in inflammatory response | biological process | GO:0002351 | 2 | 1 | 13711 | 111 | Fcer1g | | 1.613e-02 | -4.13 | enkephalin processing | biological process | GO:0034230 | 2 | 1 | 13711 | 111 | Pcsk2 | | 1.613e-02 | -4.13 | synaptonemal complex disassembly | biological process | GO:0070194 | 2 | 1 | 13711 | 111 | Hspa2 | | 1.613e-02 | -4.13 | positive regulation of DNA damage response, signal transduction by p53 class mediator resulting in transcription of p21 class mediator | biological process | GO:1902164 | 2 | 1 | 13711 | 111 | Rpl26 | | 1.613e-02 | -4.13 | epinephrine secretion | biological process | GO:0048242 | 2 | 1 | 13711 | 111 | Ly6e | | 1.613e-02 | -4.13 | tumor necrosis factor secretion | biological process | GO:1990774 | 2 | 1 | 13711 | 111 | Tmem106a | | 1.613e-02 | -4.13 | immune complex clearance | biological process | GO:0002434 | 2 | 1 | 13711 | 111 | Clu | | 1.613e-02 | -4.13 | serotonin secretion by platelet | biological process | GO:0002554 | 2 | 1 | 13711 | 111 | Fcer1g | | 1.613e-02 | -4.13 | interleukin-6 secretion | biological process | GO:0072604 | 2 | 1 | 13711 | 111 | Tmem106a | | 1.613e-02 | -4.13 | membrane raft distribution | biological process | GO:0031580 | 2 | 1 | 13711 | 111 | Ptprc | | 1.613e-02 | -4.13 | positive regulation of skeletal muscle tissue regeneration | biological process | GO:0043415 | 2 | 1 | 13711 | 111 | Capn3 | | 1.613e-02 | -4.13 | ethanol catabolic process | biological process | GO:0006068 | 2 | 1 | 13711 | 111 | Adh1 | | 1.613e-02 | -4.13 | type II hypersensitivity | biological process | GO:0002445 | 2 | 1 | 13711 | 111 | Fcgr3 | | 1.613e-02 | -4.13 | zymogen granule exocytosis | biological process | GO:0070625 | 2 | 1 | 13711 | 111 | Vamp8 | | 1.613e-02 | -4.13 | uropod organization | biological process | GO:0032796 | 2 | 1 | 13711 | 111 | Myh9 | | 1.613e-02 | -4.13 | regulation of pancreatic amylase secretion | biological process | GO:1902276 | 2 | 1 | 13711 | 111 | Vamp8 | | 1.613e-02 | -4.13 | serotonin secretion involved in inflammatory response | biological process | GO:0002442 | 2 | 1 | 13711 | 111 | Fcer1g | | 1.613e-02 | -4.13 | activation of meiosis involved in egg activation | biological process | GO:0060466 | 2 | 1 | 13711 | 111 | Camk2b | | 1.613e-02 | -4.13 | G1 to G0 transition involved in cell differentiation | biological process | GO:0070315 | 2 | 1 | 13711 | 111 | Capn3 | | 1.613e-02 | -4.13 | regulation of histamine secretion by mast cell | biological process | GO:1903593 | 2 | 1 | 13711 | 111 | Vamp8 | | 1.613e-02 | -4.13 | complement-dependent cytotoxicity | biological process | GO:0097278 | 2 | 1 | 13711 | 111 | C3 | | 1.613e-02 | -4.13 | regulation of endocannabinoid signaling pathway | biological process | GO:2000124 | 2 | 1 | 13711 | 111 | Mgll | | 1.613e-02 | -4.13 | positive regulation of protein tyrosine phosphatase activity | biological process | GO:1903615 | 2 | 1 | 13711 | 111 | Ptprc | | 1.613e-02 | -4.13 | synaptic vesicle lumen acidification | biological process | GO:0097401 | 2 | 1 | 13711 | 111 | Slc17a7 | | 1.613e-02 | -4.13 | regulation of neurofibrillary tangle assembly | biological process | GO:1902996 | 2 | 1 | 13711 | 111 | Clu | | 1.613e-02 | -4.13 | regulation of activation of membrane attack complex | biological process | GO:0001969 | 2 | 1 | 13711 | 111 | C3 | | 1.613e-02 | -4.13 | negative regulation of Notch signaling pathway involved in somitogenesis | biological process | GO:1902367 | 2 | 1 | 13711 | 111 | Lfng | | 1.613e-02 | -4.13 | smooth muscle adaptation | biological process | GO:0014805 | 2 | 1 | 13711 | 111 | Cyba | | 1.613e-02 | -4.13 | ethanol oxidation | biological process | GO:0006069 | 2 | 1 | 13711 | 111 | Adh1 | | 1.613e-02 | -4.13 | negative regulation of GTP binding | biological process | GO:1904425 | 2 | 1 | 13711 | 111 | Rap1gap | | 1.613e-02 | -4.13 | negative regulation of chaperone-mediated autophagy | biological process | GO:1904715 | 2 | 1 | 13711 | 111 | Ctsa | | 1.613e-02 | -4.13 | positive regulation of lysosomal membrane permeability | biological process | GO:0097214 | 2 | 1 | 13711 | 111 | Mt3 | | 1.613e-02 | -4.13 | pore formation in membrane of other organism | biological process | GO:0035915 | 2 | 1 | 13711 | 111 | Mpeg1 | | 1.613e-02 | -4.13 | positive regulation of mast cell cytokine production | biological process | GO:0032765 | 2 | 1 | 13711 | 111 | Fcer1g | | 1.613e-02 | -4.13 | positive regulation of hematopoietic stem cell migration | biological process | GO:2000473 | 2 | 1 | 13711 | 111 | Ptprc | | 1.613e-02 | -4.13 | regulation of Notch signaling pathway involved in somitogenesis | biological process | GO:1902366 | 2 | 1 | 13711 | 111 | Lfng | | 1.613e-02 | -4.13 | regulation of protein processing in phagocytic vesicle | biological process | GO:1903921 | 2 | 1 | 13711 | 111 | Myh9 | | 1.613e-02 | -4.13 | type IIa hypersensitivity | biological process | GO:0001794 | 2 | 1 | 13711 | 111 | Fcgr3 | | 1.613e-02 | -4.13 | regulation of hematopoietic stem cell migration | biological process | GO:2000471 | 2 | 1 | 13711 | 111 | Ptprc | | 1.613e-02 | -4.13 | positive regulation of oxygen metabolic process | biological process | GO:2000376 | 2 | 1 | 13711 | 111 | Mt3 | | 1.613e-02 | -4.13 | epinephrine transport | biological process | GO:0048241 | 2 | 1 | 13711 | 111 | Ly6e | | 1.613e-02 | -4.13 | positive regulation of protein processing in phagocytic vesicle | biological process | GO:1903923 | 2 | 1 | 13711 | 111 | Myh9 | | 1.613e-02 | -4.13 | antibody-dependent cellular cytotoxicity | biological process | GO:0001788 | 2 | 1 | 13711 | 111 | Fcgr3 | | 1.613e-02 | -4.13 | regulation of norepinephrine uptake | biological process | GO:0051621 | 2 | 1 | 13711 | 111 | Actb | | 1.613e-02 | -4.13 | positive regulation of intrinsic apoptotic signaling pathway in response to DNA damage by p53 class mediator | biological process | GO:1902167 | 2 | 1 | 13711 | 111 | Rpl26 | | 1.613e-02 | -4.13 | negative regulation of smooth muscle cell-matrix adhesion | biological process | GO:2000098 | 2 | 1 | 13711 | 111 | Apod | | 1.613e-02 | -4.13 | positive regulation of histamine secretion by mast cell | biological process | GO:1903595 | 2 | 1 | 13711 | 111 | Vamp8 | | 1.613e-02 | -4.13 | ethanol metabolic process | biological process | GO:0006067 | 2 | 1 | 13711 | 111 | Adh1 | | 1.613e-02 | -4.13 | dehydroascorbic acid transport | biological process | GO:0070837 | 2 | 1 | 13711 | 111 | Slc2a1 | | 1.613e-02 | -4.13 | GO\_VACUOLE | MSigDB lists | GO\_VACUOLE | 928 | 14 | 12187 | 98 | Ccr5,Gfap,Vamp8,Cplx2,Ctsa,Ctsb,Ahnak,Fmod,Rap1gap,Cd68,Ctss,Ctsz,Cyba,Ptprn | | 1.615e-02 | -4.13 | positive regulation of cell-cell adhesion | biological process | GO:0022409 | 182 | 5 | 13711 | 111 | Il6ra,Thy1,Ptprc,Rap1gap,Ccr5 | | 1.618e-02 | -4.12 | negative regulation of nervous system development | biological process | GO:0051961 | 324 | 7 | 13711 | 111 | Vim,Gfap,Ccr5,Thy1,Rap1gap,Ctsz,Mt3 | | 1.621e-02 | -4.12 | MARTINEZ\_RB1\_TARGETS\_DN | MSigDB lists | MARTINEZ\_RB1\_TARGETS\_DN | 405 | 8 | 12187 | 98 | Rpl26,Spint1,Clu,Mgll,Tsc22d1,Rps8,Adh1,Csf1r | | 1.633e-02 | -4.11 | GSE21927\_SPLENIC\_VS\_TUMOR\_MONOCYTES\_FROM\_C26GM\_TUMOROUS\_MICE\_BALBC\_UP | MSigDB lists | GSE21927\_SPLENIC\_VS\_TUMOR\_MONOCYTES\_FROM\_C26GM\_TUMOROUS\_MICE\_BALBC\_UP | 121 | 4 | 12187 | 98 | Trnp1,Ppfibp1,Lgi4,Cyba | | 1.633e-02 | -4.11 | GSE24671\_CTRL\_VS\_SENDAI\_VIRUS\_INFECTED\_MOUSE\_SPLENOCYTES\_UP | MSigDB lists | GSE24671\_CTRL\_VS\_SENDAI\_VIRUS\_INFECTED\_MOUSE\_SPLENOCYTES\_UP | 121 | 4 | 12187 | 98 | Il1r1,Tmem176a,Ctss,Fcer1g | | 1.633e-02 | -4.11 | GSE2124\_CTRL\_VS\_LYMPHOTOXIN\_BETA\_TREATED\_MLN\_UP | MSigDB lists | GSE2124\_CTRL\_VS\_LYMPHOTOXIN\_BETA\_TREATED\_MLN\_UP | 121 | 4 | 12187 | 98 | Actb,Ptprc,Cox4i1,Rpl26 | | 1.633e-02 | -4.11 | MORF\_EIF4A2 | MSigDB lists | MORF\_EIF4A2 | 121 | 4 | 12187 | 98 | Rpl13,Rpl14,Rpl31,Dek | | 1.633e-02 | -4.11 | WNT\_UP.V1\_DN | MSigDB lists | WNT\_UP.V1\_DN | 121 | 4 | 12187 | 98 | Cyba,Vim,Gfap,Igfbp6 | | 1.647e-02 | -4.11 | TIL\_dom | interpro domains | IPR002919 | 2 | 1 | 13788 | 114 | Vwf | | 1.647e-02 | -4.11 | Liprin-beta\_SAM\_rpt\_3 | interpro domains | IPR037619 | 2 | 1 | 13788 | 114 | Ppfibp1 | | 1.647e-02 | -4.11 | Liprin-beta\_SAM\_rpt\_1 | interpro domains | IPR037617 | 2 | 1 | 13788 | 114 | Ppfibp1 | | 1.647e-02 | -4.11 | Unchr\_dom\_Cys-rich | interpro domains | IPR014853 | 2 | 1 | 13788 | 114 | Vwf | | 1.647e-02 | -4.11 | L-lactate\_DH\_AS | interpro domains | IPR018177 | 2 | 1 | 13788 | 114 | Ldha | | 1.647e-02 | -4.11 | Ribosomal\_S7\_dom | interpro domains | IPR023798 | 2 | 1 | 13788 | 114 | Rps5 | | 1.647e-02 | -4.11 | Ribosomal\_S8e/biogenesis\_NSA2 | interpro domains | IPR022309 | 2 | 1 | 13788 | 114 | Rps8 | | 1.647e-02 | -4.11 | Melancort\_rcpt | interpro domains | IPR001908 | 2 | 1 | 13788 | 114 | Mc4r | | 1.647e-02 | -4.11 | Ribosomal\_L24/26\_CS | interpro domains | IPR005825 | 2 | 1 | 13788 | 114 | Rpl26 | | 1.647e-02 | -4.11 | Receptor\_IA-2\_dom | interpro domains | IPR021613 | 2 | 1 | 13788 | 114 | Ptprn | | 1.647e-02 | -4.11 | Anaphylatoxin\_comp\_syst | interpro domains | IPR018081 | 2 | 1 | 13788 | 114 | C3 | | 1.647e-02 | -4.11 | Sec2\_N | interpro domains | IPR009449 | 2 | 1 | 13788 | 114 | Rab3il1 | | 1.647e-02 | -4.11 | Glyco\_hydro\_1\_N\_CS | interpro domains | IPR033132 | 2 | 1 | 13788 | 114 | Kl | | 1.647e-02 | -4.11 | Ser\_inhib-like\_sf | interpro domains | IPR036084 | 2 | 1 | 13788 | 114 | Vwf | | 1.647e-02 | -4.11 | Cyt\_c\_oxidase\_su4\_sf | interpro domains | IPR036639 | 2 | 1 | 13788 | 114 | Cox4i1 | | 1.647e-02 | -4.11 | Glyco\_hydro\_1 | interpro domains | IPR001360 | 2 | 1 | 13788 | 114 | Kl | | 1.647e-02 | -4.11 | Liprin-beta\_SAM\_rpt\_2 | interpro domains | IPR037618 | 2 | 1 | 13788 | 114 | Ppfibp1 | | 1.647e-02 | -4.11 | Hs1\_Cortactin | interpro domains | IPR003134 | 2 | 1 | 13788 | 114 | Hcls1 | | 1.647e-02 | -4.11 | IA-2/IA-2\_beta | interpro domains | IPR033522 | 2 | 1 | 13788 | 114 | Ptprn | | 1.647e-02 | -4.11 | Proteasome\_bsu\_C | interpro domains | IPR024689 | 2 | 1 | 13788 | 114 | Psmb10 | | 1.647e-02 | -4.11 | Ribosomal\_S5/S7 | interpro domains | IPR000235 | 2 | 1 | 13788 | 114 | Rps5 | | 1.647e-02 | -4.11 | Ser\_caboxypep\_ser\_AS | interpro domains | IPR018202 | 2 | 1 | 13788 | 114 | Ctsa | | 1.647e-02 | -4.11 | S1-like\_RNA-bd\_dom | interpro domains | IPR025223 | 2 | 1 | 13788 | 114 | Ccar1 | | 1.647e-02 | -4.11 | Peptidase\_S10 | interpro domains | IPR001563 | 2 | 1 | 13788 | 114 | Ctsa | | 1.647e-02 | -4.11 | Ribosomal\_S7\_dom\_sf | interpro domains | IPR036823 | 2 | 1 | 13788 | 114 | Rps5 | | 1.647e-02 | -4.11 | Anaphylatoxn\_comp\_syst\_dom | interpro domains | IPR001840 | 2 | 1 | 13788 | 114 | C3 | | 1.647e-02 | -4.11 | Csk-like\_SH2 | interpro domains | IPR035027 | 2 | 1 | 13788 | 114 | Csk | | 1.647e-02 | -4.11 | Receptor\_IA-2\_ectodomain\_sf | interpro domains | IPR038112 | 2 | 1 | 13788 | 114 | Ptprn | | 1.647e-02 | -4.11 | L-lactate\_DH | interpro domains | IPR011304 | 2 | 1 | 13788 | 114 | Ldha | | 1.647e-02 | -4.11 | RAB3IL/RAB3IP/Sec2 | interpro domains | IPR040351 | 2 | 1 | 13788 | 114 | Rab3il1 | | 1.647e-02 | -4.11 | Cyt\_c\_oxidase\_su4\_fam | interpro domains | IPR004203 | 2 | 1 | 13788 | 114 | Cox4i1 | | 1.647e-02 | -4.11 | CCAR1/CCAR2 | interpro domains | IPR025224 | 2 | 1 | 13788 | 114 | Ccar1 | | 1.647e-02 | -4.11 | YJEFN\_prot\_eukaryotes | interpro domains | IPR032976 | 2 | 1 | 13788 | 114 | Ndufa13 | | 1.647e-02 | -4.11 | Cyt\_c\_oxidase\_su4 | interpro domains | IPR013288 | 2 | 1 | 13788 | 114 | Cox4i1 | | 1.647e-02 | -4.11 | Serpin\_B9/Maspin | interpro domains | IPR000240 | 2 | 1 | 13788 | 114 | Serpinb9 | | 1.647e-02 | -4.11 | Ribosomal\_S10 | interpro domains | IPR001848 | 2 | 1 | 13788 | 114 | Rps20 | | 1.647e-02 | -4.11 | Lipocalin\_ApoD | interpro domains | IPR022271 | 2 | 1 | 13788 | 114 | Apod | | 1.647e-02 | -4.11 | KOW\_RPL26/RPL24 | interpro domains | IPR041988 | 2 | 1 | 13788 | 114 | Rpl26 | | 1.647e-02 | -4.11 | DBC1/CARP1\_inactive\_NUDIX\_dom | interpro domains | IPR025954 | 2 | 1 | 13788 | 114 | Ccar1 | | 1.647e-02 | -4.11 | Melcrt\_ACTH\_rcpt | interpro domains | IPR001671 | 2 | 1 | 13788 | 114 | Mc4r | | 1.647e-02 | -4.11 | TMEM176A/TMEM176B | interpro domains | IPR009281 | 2 | 1 | 13788 | 114 | Tmem176a | | 1.649e-02 | -4.10 | SAP | smart domains | SM00513 | 21 | 2 | 7188 | 68 | Dek,Ccar1 | | 1.661e-02 | -4.10 | SAP\_dom\_sf | interpro domains | IPR036361 | 24 | 2 | 13788 | 114 | Dek,Ccar1 | | 1.666e-02 | -4.09 | BOYLAN\_MULTIPLE\_MYELOMA\_PCA1\_UP | MSigDB lists | BOYLAN\_MULTIPLE\_MYELOMA\_PCA1\_UP | 67 | 3 | 12187 | 98 | Tmem176a,C3,Lrg1 | | 1.666e-02 | -4.09 | REACTOME\_INTEGRIN\_CELL\_SURFACE\_INTERACTIONS | MSigDB lists | REACTOME\_INTEGRIN\_CELL\_SURFACE\_INTERACTIONS | 67 | 3 | 12187 | 98 | Itgam,Vwf,Csk | | 1.670e-02 | -4.09 | C8 | pfam domains | PF08742 | 2 | 1 | 12881 | 108 | Vwf | | 1.670e-02 | -4.09 | Sec2p | pfam domains | PF06428 | 2 | 1 | 12881 | 108 | Rab3il1 | | 1.670e-02 | -4.09 | HS1\_rep | pfam domains | PF02218 | 2 | 1 | 12881 | 108 | Hcls1 | | 1.670e-02 | -4.09 | Ribosomal\_S8e | pfam domains | PF01201 | 2 | 1 | 12881 | 108 | Rps8 | | 1.670e-02 | -4.09 | DBC1 | pfam domains | PF14443 | 2 | 1 | 12881 | 108 | Ccar1 | | 1.670e-02 | -4.09 | TIL | pfam domains | PF01826 | 2 | 1 | 12881 | 108 | Vwf | | 1.670e-02 | -4.09 | Peptidase\_S10 | pfam domains | PF00450 | 2 | 1 | 12881 | 108 | Ctsa | | 1.670e-02 | -4.09 | Glyco\_hydro\_1 | pfam domains | PF00232 | 2 | 1 | 12881 | 108 | Kl | | 1.670e-02 | -4.09 | Ribosomal\_S7 | pfam domains | PF00177 | 2 | 1 | 12881 | 108 | Rps5 | | 1.670e-02 | -4.09 | COX4 | pfam domains | PF02936 | 2 | 1 | 12881 | 108 | Cox4i1 | | 1.670e-02 | -4.09 | Pr\_beta\_C | pfam domains | PF12465 | 2 | 1 | 12881 | 108 | Psmb10 | | 1.670e-02 | -4.09 | S1-like | pfam domains | PF14444 | 2 | 1 | 12881 | 108 | Ccar1 | | 1.670e-02 | -4.09 | Receptor\_IA-2 | pfam domains | PF11548 | 2 | 1 | 12881 | 108 | Ptprn | | 1.674e-02 | -4.09 | GO\_CELLULAR\_HOMEOSTASIS | MSigDB lists | GO\_CELLULAR\_HOMEOSTASIS | 489 | 9 | 12187 | 98 | Ptprn,Cyba,Slc26a2,Il1r1,Ptprc,Sh3bgrl3,Slc17a7,Ccr5,Capn3 | | 1.676e-02 | -4.09 | T cell differentiation | biological process | GO:0030217 | 121 | 4 | 13711 | 111 | Lfng,Fcer1g,Tnfsf8,Ptprc | | 1.678e-02 | -4.09 | CHIANG\_LIVER\_CANCER\_SUBCLASS\_CTNNB1\_DN | MSigDB lists | CHIANG\_LIVER\_CANCER\_SUBCLASS\_CTNNB1\_DN | 122 | 4 | 12187 | 98 | Cyba,Mgll,Trnp1,Serpinb9 | | 1.678e-02 | -4.09 | GSE15624\_CTRL\_VS\_3H\_HALOFUGINONE\_TREATED\_CD4\_TCELL\_DN | MSigDB lists | GSE15624\_CTRL\_VS\_3H\_HALOFUGINONE\_TREATED\_CD4\_TCELL\_DN | 122 | 4 | 12187 | 98 | Slc2a1,Gfap,Fam111a,Rps8 | | 1.687e-02 | -4.08 | growth factor binding | molecular function | GO:0019838 | 124 | 4 | 13516 | 107 | Il6ra,Il1r1,Igfbp6,Kl | | 1.687e-02 | -4.08 | GO\_REGULATION\_OF\_CELL\_DIFFERENTIATION | MSigDB lists | GO\_REGULATION\_OF\_CELL\_DIFFERENTIATION | 1123 | 16 | 12187 | 98 | Csf1r,Tmem176a,Grn,Thy1,Spint1,Gfap,Il6ra,Hcls1,Rap1gap,Camk2b,Fmod,Clu,Capn3,Fgf18,Ikzf3,Vim | | 1.694e-02 | -4.08 | axon part | cellular component | GO:0033267 | 408 | 8 | 13825 | 111 | Mgll,Cplx2,Thy1,Rpl26,Ptprn,Clu,Ctsz,Actb | | 1.696e-02 | -4.08 | RIGGI\_EWING\_SARCOMA\_PROGENITOR\_UP | MSigDB lists | RIGGI\_EWING\_SARCOMA\_PROGENITOR\_UP | 330 | 7 | 12187 | 98 | Slc17a7,Fgf18,Slc26a2,Pcsk2,Fmod,Itgam,Serpinb9 | | 1.700e-02 | -4.07 | regulation of chemokine production | biological process | GO:0032642 | 67 | 3 | 13711 | 111 | Il6ra,Csf1r,Apod | | 1.700e-02 | -4.07 | regulation of tissue remodeling | biological process | GO:0034103 | 67 | 3 | 13711 | 111 | Csf1r,Csk,Mc4r | | 1.703e-02 | -4.07 | ONO\_AML1\_TARGETS\_DN | MSigDB lists | ONO\_AML1\_TARGETS\_DN | 25 | 2 | 12187 | 98 | Ccr5,Tnfsf8 | | 1.703e-02 | -4.07 | GO\_DICARBOXYLIC\_ACID\_TRANSMEMBRANE\_TRANSPORTER\_ACTIVITY | MSigDB lists | GO\_DICARBOXYLIC\_ACID\_TRANSMEMBRANE\_TRANSPORTER\_ACTIVITY | 25 | 2 | 12187 | 98 | Slc26a2,Slc17a7 | | 1.703e-02 | -4.07 | GO\_B\_CELL\_RECEPTOR\_SIGNALING\_PATHWAY | MSigDB lists | GO\_B\_CELL\_RECEPTOR\_SIGNALING\_PATHWAY | 25 | 2 | 12187 | 98 | Ptprc,Klhl6 | | 1.703e-02 | -4.07 | GO\_PROTEIN\_COMPLEX\_INVOLVED\_IN\_CELL\_ADHESION | MSigDB lists | GO\_PROTEIN\_COMPLEX\_INVOLVED\_IN\_CELL\_ADHESION | 25 | 2 | 12187 | 98 | Myh9,Itgam | | 1.726e-02 | -4.06 | B cell receptor signaling pathway | biological process | GO:0050853 | 25 | 2 | 13711 | 111 | Ptprc,Klhl6 | | 1.741e-02 | -4.05 | Fbxo32 (F-box protein 32) | protein interactions | 67731 | 132 | 5 | 6802 | 78 | Vim,Ldha,Rpl14,Ahnak,Actb | | 1.741e-02 | -4.05 | GO\_REGULATION\_OF\_PEPTIDASE\_ACTIVITY | MSigDB lists | GO\_REGULATION\_OF\_PEPTIDASE\_ACTIVITY | 257 | 6 | 12187 | 98 | C3,Spint1,Serpina3n,Psme1,Serpinb9,Ndufa13 | | 1.748e-02 | -4.05 | WANG\_TUMOR\_INVASIVENESS\_UP | MSigDB lists | WANG\_TUMOR\_INVASIVENESS\_UP | 332 | 7 | 12187 | 98 | C1qb,Rps5,Rpl14,S100a6,Ctsz,Rps8,Vim | | 1.748e-02 | -4.05 | GO\_WOUND\_HEALING | MSigDB lists | GO\_WOUND\_HEALING | 332 | 7 | 12187 | 98 | C3,Myh9,Fcer1g,Vwf,Actb,Apod,Rac2 | | 1.754e-02 | -4.04 | - | gene3d domains | 1.10.720.30 | 22 | 2 | 6647 | 62 | Ccar1,Dek | | 1.764e-02 | -4.04 | CLTA (clathrin light chain A) | protein interactions | 1211 | 18 | 2 | 6802 | 78 | Actb,Myh9 | | 1.770e-02 | -4.03 | regulation of protein localization to cell periphery | biological process | GO:1904375 | 123 | 4 | 13711 | 111 | Actb,Camk2b,Csk,Vamp8 | | 1.771e-02 | -4.03 | GSE32423\_IL7\_VS\_IL7\_IL4\_NAIVE\_CD8\_TCELL\_UP | MSigDB lists | GSE32423\_IL7\_VS\_IL7\_IL4\_NAIVE\_CD8\_TCELL\_UP | 124 | 4 | 12187 | 98 | Phyhip,Zfp786,Lfng,Pde1a | | 1.783e-02 | -4.03 | fibronectin binding | molecular function | GO:0001968 | 26 | 2 | 13516 | 107 | Ctss,Igfbp6 | | 1.795e-02 | -4.02 | peptide metabolic process | biological process | GO:0006518 | 409 | 8 | 13711 | 111 | Rps8,Rpl13,Pcsk2,Rps20,Rpl14,Rps5,Rpl31,Rpl26 | | 1.800e-02 | -4.02 | YAGI\_AML\_WITH\_INV\_16\_TRANSLOCATION | MSigDB lists | YAGI\_AML\_WITH\_INV\_16\_TRANSLOCATION | 334 | 7 | 12187 | 98 | Rpl31,Rps20,Tsc22d1,Ly6e,Rpl13,Vim,Lfng | | 1.802e-02 | -4.02 | RASHI\_RESPONSE\_TO\_IONIZING\_RADIATION\_6 | MSigDB lists | RASHI\_RESPONSE\_TO\_IONIZING\_RADIATION\_6 | 69 | 3 | 12187 | 98 | Ptprc,Adh1,Csf1r | | 1.802e-02 | -4.02 | GO\_REGULATION\_OF\_TUMOR\_NECROSIS\_FACTOR\_SUPERFAMILY\_CYTOKINE\_PRODUCTION | MSigDB lists | GO\_REGULATION\_OF\_TUMOR\_NECROSIS\_FACTOR\_SUPERFAMILY\_CYTOKINE\_PRODUCTION | 69 | 3 | 12187 | 98 | Cyba,Clu,Fcer1g | | 1.802e-02 | -4.02 | HARRIS\_HYPOXIA | MSigDB lists | HARRIS\_HYPOXIA | 69 | 3 | 12187 | 98 | Vim,Ldha,Slc2a1 | | 1.802e-02 | -4.02 | GO\_POSITIVE\_REGULATION\_OF\_INFLAMMATORY\_RESPONSE | MSigDB lists | GO\_POSITIVE\_REGULATION\_OF\_INFLAMMATORY\_RESPONSE | 69 | 3 | 12187 | 98 | Fcer1g,Vamp8,C3 | | 1.802e-02 | -4.02 | GO\_REGULATION\_OF\_CALCIUM\_ION\_IMPORT | MSigDB lists | GO\_REGULATION\_OF\_CALCIUM\_ION\_IMPORT | 69 | 3 | 12187 | 98 | Capn3,Thy1,Cyba | | 1.818e-02 | -4.01 | GCNP\_SHH\_UP\_EARLY.V1\_DN | MSigDB lists | GCNP\_SHH\_UP\_EARLY.V1\_DN | 125 | 4 | 12187 | 98 | Cyba,Tmem176a,Il1r1,Mgp | | 1.818e-02 | -4.01 | KAAB\_FAILED\_HEART\_ATRIUM\_DN | MSigDB lists | KAAB\_FAILED\_HEART\_ATRIUM\_DN | 125 | 4 | 12187 | 98 | Actb,Serpina3n,Tsc22d1,Ldha | | 1.832e-02 | -4.00 | distal axon | cellular component | GO:0150034 | 335 | 7 | 13825 | 111 | Ctsz,Actb,Rpl26,Clu,Ptprn,Cplx2,Thy1 | | 1.833e-02 | -4.00 | mouse chr1 | chromosome location | mouse chr1 | 264 | 6 | 14556 | 115 | Ptprc,Fmod,Fcgr3,Fcer1g,Ptprn,Il1r1 | | 1.833e-02 | -4.00 | MODULE\_33 | MSigDB lists | MODULE\_33 | 260 | 6 | 12187 | 98 | C3,Il1r1,Rac2,Vwf,Igfbp6,Serpinb9 | | 1.835e-02 | -4.00 | REACTOME\_NEGATIVE\_REGULATION\_OF\_FGFR\_SIGNALING | MSigDB lists | REACTOME\_NEGATIVE\_REGULATION\_OF\_FGFR\_SIGNALING | 26 | 2 | 12187 | 98 | Fgf18,Kl | | 1.835e-02 | -4.00 | REACTOME\_FRS2\_MEDIATED\_CASCADE | MSigDB lists | REACTOME\_FRS2\_MEDIATED\_CASCADE | 26 | 2 | 12187 | 98 | Fgf18,Kl | | 1.835e-02 | -4.00 | GO\_INTERMEDIATE\_FILAMENT\_BASED\_PROCESS | MSigDB lists | GO\_INTERMEDIATE\_FILAMENT\_BASED\_PROCESS | 26 | 2 | 12187 | 98 | Gfap,Vim | | 1.835e-02 | -4.00 | MODULE\_29 | MSigDB lists | MODULE\_29 | 26 | 2 | 12187 | 98 | Rps20,Rps8 | | 1.835e-02 | -4.00 | GO\_REGULATION\_OF\_SYSTEMIC\_ARTERIAL\_BLOOD\_PRESSURE\_MEDIATED\_BY\_A\_CHEMICAL\_SIGNAL | MSigDB lists | GO\_REGULATION\_OF\_SYSTEMIC\_ARTERIAL\_BLOOD\_PRESSURE\_MEDIATED\_BY\_A\_CHEMICAL\_SIGNAL | 26 | 2 | 12187 | 98 | Cyba,Ctsz | | 1.838e-02 | -4.00 | positive regulation of production of molecular mediator of immune response | biological process | GO:0002702 | 69 | 3 | 13711 | 111 | Ptprc,Il1r1,Fcer1g | | 1.842e-02 | -3.99 | protein dimerization activity | molecular function | GO:0046983 | 1149 | 16 | 13516 | 107 | Il6ra,Vwf,Itgam,Fcer1g,Adh1,Ikzf3,S100a6,Camk2b,C1qb,Mgll,Cyba,Csf1r,Rap1gap,Myh9,Bhlhe41,Tfap2c | | 1.849e-02 | -3.99 | AREB6\_01 | MSigDB lists | AREB6\_01 | 190 | 5 | 12187 | 98 | Flnb,Hcls1,C1qa,Vwf,Igfbp6 | | 1.857e-02 | -3.99 | - | gene3d domains | 3.30.70.2470 | 2 | 1 | 6647 | 62 | Ptprn | | 1.857e-02 | -3.99 | - | gene3d domains | 1.10.455.10 | 2 | 1 | 6647 | 62 | Rps5 | | 1.857e-02 | -3.99 | - | gene3d domains | 1.10.442.10 | 2 | 1 | 6647 | 62 | Cox4i1 | | 1.858e-02 | -3.99 | positive regulation of phosphorylation | biological process | GO:0042327 | 843 | 13 | 13711 | 111 | Fgf18,Csk,Il6ra,C3,Mt3,Csf1r,Hcls1,Kl,Tmem106a,Hspa2,Ptprc,Ccl6,Clu | | 1.860e-02 | -3.98 | modification by symbiont of host morphology or physiology | biological process | GO:0044003 | 26 | 2 | 13711 | 111 | Serpinb9,Ccr5 | | 1.860e-02 | -3.98 | negative regulation of neural precursor cell proliferation | biological process | GO:2000178 | 26 | 2 | 13711 | 111 | Ccr5,Spint1 | | 1.860e-02 | -3.98 | positive regulation of lamellipodium organization | biological process | GO:1902745 | 26 | 2 | 13711 | 111 | Brk1,Rac2 | | 1.865e-02 | -3.98 | regulation of cell migration | biological process | GO:0030334 | 753 | 12 | 13711 | 111 | Rac2,Csf1r,Ccr5,Fgf18,Camk2b,Apod,Ptprc,Vim,Ccar1,Grn,Il1r1,Thy1 | | 1.867e-02 | -3.98 | GNF2\_FBL | MSigDB lists | GNF2\_FBL | 126 | 4 | 12187 | 98 | Rps20,Rpl14,Rps8,Rpl13 | | 1.867e-02 | -3.98 | GSE3565\_DUSP1\_VS\_WT\_SPLENOCYTES\_UP | MSigDB lists | GSE3565\_DUSP1\_VS\_WT\_SPLENOCYTES\_UP | 126 | 4 | 12187 | 98 | Ikzf3,Vim,Ahnak,Itgam | | 1.872e-02 | -3.98 | GO\_TOLL\_LIKE\_RECEPTOR\_SIGNALING\_PATHWAY | MSigDB lists | GO\_TOLL\_LIKE\_RECEPTOR\_SIGNALING\_PATHWAY | 70 | 3 | 12187 | 98 | Ctsb,Ctss,Itgam | | 1.872e-02 | -3.98 | MEISSNER\_BRAIN\_HCP\_WITH\_H3K27ME3 | MSigDB lists | MEISSNER\_BRAIN\_HCP\_WITH\_H3K27ME3 | 70 | 3 | 12187 | 98 | Ikzf3,Tfap2c,Spint1 | | 1.873e-02 | -3.98 | Formation of the ternary complex, and subsequently, the 43S complex | REACTOME pathways | R-MMU-72695 | 47 | 3 | 6297 | 76 | Rps5,Rps20,Rps8 | | 1.876e-02 | -3.98 | GO\_HOMEOSTATIC\_PROCESS | MSigDB lists | GO\_HOMEOSTATIC\_PROCESS | 946 | 14 | 12187 | 98 | Rac2,Sh3bgrl3,Ptprn,Cyba,Ptprc,Serpina3n,Il1r1,Capn3,Hcls1,Kl,Slc17a7,Slc26a2,Actb,Ccr5 | | 1.883e-02 | -3.97 | C8 | smart domains | SM00832 | 2 | 1 | 7188 | 68 | Vwf | | 1.883e-02 | -3.97 | DBC1 | smart domains | SM01122 | 2 | 1 | 7188 | 68 | Ccar1 | | 1.886e-02 | -3.97 | KIM\_ALL\_DISORDERS\_OLIGODENDROCYTE\_NUMBER\_CORR\_UP | MSigDB lists | KIM\_ALL\_DISORDERS\_OLIGODENDROCYTE\_NUMBER\_CORR\_UP | 672 | 11 | 12187 | 98 | Rpl13,Camk2b,Rps8,Ptprn,Psmb10,Vim,Ly6e,Thy1,Ctsa,Gfap,Cox4i1 | | 1.896e-02 | -3.97 | cell adhesion molecule binding | molecular function | GO:0050839 | 194 | 5 | 13516 | 107 | Gfap,Vwf,Myh9,Tspan4,Thy1 | | 1.896e-02 | -3.97 | GO\_POSITIVE\_REGULATION\_OF\_CYTOKINE\_PRODUCTION | MSigDB lists | GO\_POSITIVE\_REGULATION\_OF\_CYTOKINE\_PRODUCTION | 262 | 6 | 12187 | 98 | Cyba,Il6ra,C3,Csf1r,Fcer1g,Clu | | 1.916e-02 | -3.96 | spectrin binding | molecular function | GO:0030507 | 27 | 2 | 13516 | 107 | Ptprc,Ptprn | | 1.916e-02 | -3.96 | vesicle-mediated transport in synapse | biological process | GO:0099003 | 126 | 4 | 13711 | 111 | Actb,Grn,Cplx2,Slc17a7 | | 1.916e-02 | -3.95 | GSE22443\_NAIVE\_VS\_ACT\_AND\_IL2\_TREATED\_CD8\_TCELL\_UP | MSigDB lists | GSE22443\_NAIVE\_VS\_ACT\_AND\_IL2\_TREATED\_CD8\_TCELL\_UP | 127 | 4 | 12187 | 98 | Clu,Vwf,Apod,Adh1 | | 1.936e-02 | -3.94 | SWEET\_LUNG\_CANCER\_KRAS\_DN | MSigDB lists | SWEET\_LUNG\_CANCER\_KRAS\_DN | 339 | 7 | 12187 | 98 | Mgp,Igfbp6,Psmb10,Vwf,Ahnak,C3,Adh1 | | 1.936e-02 | -3.94 | C1Q | prosite domains | PS50871 | 21 | 2 | 8845 | 91 | C1qa,C1qb | | 1.938e-02 | -3.94 | cell projection part | cellular component | GO:0044463 | 1334 | 18 | 13825 | 111 | Apod,Clu,Rpl26,Camk2b,Cyba,Mgll,Dlgap3,Pcsk2,Slc26a2,Mark4,Ldha,Ptprn,Thy1,Rap1gap,Cplx2,Actb,Cd52,Ctsz | | 1.938e-02 | -3.94 | plasma membrane bounded cell projection part | cellular component | GO:0120038 | 1334 | 18 | 13825 | 111 | Thy1,Cplx2,Rap1gap,Mark4,Ldha,Ptprn,Actb,Ctsz,Cd52,Cyba,Mgll,Clu,Apod,Rpl26,Camk2b,Slc26a2,Pcsk2,Dlgap3 | | 1.941e-02 | -3.94 | Antimicrobial peptides | REACTOME pathways | R-MMU-6803157 | 18 | 2 | 6297 | 76 | Ccr5,Clu | | 1.943e-02 | -3.94 | HILLION\_HMGA1B\_TARGETS | MSigDB lists | HILLION\_HMGA1B\_TARGETS | 71 | 3 | 12187 | 98 | Cox4i1,Rpl13,Clu | | 1.943e-02 | -3.94 | GO\_STRUCTURAL\_CONSTITUENT\_OF\_CYTOSKELETON | MSigDB lists | GO\_STRUCTURAL\_CONSTITUENT\_OF\_CYTOSKELETON | 71 | 3 | 12187 | 98 | Vim,Gfap,Actb | | 1.943e-02 | -3.94 | YAO\_TEMPORAL\_RESPONSE\_TO\_PROGESTERONE\_CLUSTER\_12 | MSigDB lists | YAO\_TEMPORAL\_RESPONSE\_TO\_PROGESTERONE\_CLUSTER\_12 | 71 | 3 | 12187 | 98 | Tmem176a,C1qb,Psme1 | | 1.943e-02 | -3.94 | GO\_VASCULAR\_ENDOTHELIAL\_GROWTH\_FACTOR\_RECEPTOR\_SIGNALING\_PATHWAY | MSigDB lists | GO\_VASCULAR\_ENDOTHELIAL\_GROWTH\_FACTOR\_RECEPTOR\_SIGNALING\_PATHWAY | 71 | 3 | 12187 | 98 | Cyba,Actb,Brk1 | | 1.943e-02 | -3.94 | FRIDMAN\_SENESCENCE\_UP | MSigDB lists | FRIDMAN\_SENESCENCE\_UP | 71 | 3 | 12187 | 98 | Vim,Hspa2,Igfbp6 | | 1.955e-02 | -3.93 | GO\_CELLULAR\_CATABOLIC\_PROCESS | MSigDB lists | GO\_CELLULAR\_CATABOLIC\_PROCESS | 1046 | 15 | 12187 | 98 | Psmb10,Ctss,Rps8,Ctsz,Psme1,Fmod,Rpl13,Mgll,Rps20,Rpl14,Rpl26,Ctsa,Ctsb,Rpl31,Rps5 | | 1.957e-02 | -3.93 | PPP1R12A (protein phosphatase 1 regulatory subunit 12A) | protein interactions | 4659 | 19 | 2 | 6802 | 78 | Myh9,Actb | | 1.957e-02 | -3.93 | Dok1 (docking protein 1) | protein interactions | 13448 | 19 | 2 | 6802 | 78 | Csk,Csf1r | | 1.957e-02 | -3.93 | LRRFIP2 (LRR binding FLII interacting protein 2) | protein interactions | 9209 | 19 | 2 | 6802 | 78 | Myh9,Actb | | 1.957e-02 | -3.93 | PPP1CA (protein phosphatase 1 catalytic subunit alpha) | protein interactions | 5499 | 19 | 2 | 6802 | 78 | Actb,Myh9 | | 1.966e-02 | -3.93 | GSE24671\_CTRL\_VS\_BAKIMULC\_INFECTED\_MOUSE\_SPLENOCYTES\_DN | MSigDB lists | GSE24671\_CTRL\_VS\_BAKIMULC\_INFECTED\_MOUSE\_SPLENOCYTES\_DN | 128 | 4 | 12187 | 98 | Rps5,Igfbp6,Rpl13,S100a6 | | 1.966e-02 | -3.93 | FLECHNER\_PBL\_KIDNEY\_TRANSPLANT\_OK\_VS\_DONOR\_UP | MSigDB lists | FLECHNER\_PBL\_KIDNEY\_TRANSPLANT\_OK\_VS\_DONOR\_UP | 128 | 4 | 12187 | 98 | Clu,Ahnak,Rps20,Actb | | 1.966e-02 | -3.93 | GSE21063\_CTRL\_VS\_ANTI\_IGM\_STIM\_BCELL\_3H\_UP | MSigDB lists | GSE21063\_CTRL\_VS\_ANTI\_IGM\_STIM\_BCELL\_3H\_UP | 128 | 4 | 12187 | 98 | Clu,Il6ra,Ccr5,Tnfsf8 | | 1.966e-02 | -3.93 | aging | biological process | GO:0007568 | 127 | 4 | 13711 | 111 | C1qa,Kl,Apod,Cd68 | | 1.971e-02 | -3.93 | whole membrane | cellular component | GO:0098805 | 950 | 14 | 13825 | 111 | Itgam,Ctsb,Grn,Slc17a7,Fcer1g,Cd68,Csk,Ptprc,Actb,Slc2a1,Vamp8,Thy1,Ahnak,Gfap | | 1.971e-02 | -3.93 | GO\_ENDOCRINE\_PROCESS | MSigDB lists | GO\_ENDOCRINE\_PROCESS | 27 | 2 | 12187 | 98 | Cyba,Ctsz | | 1.971e-02 | -3.93 | GO\_REGULATION\_OF\_ACTIN\_CYTOSKELETON\_REORGANIZATION | MSigDB lists | GO\_REGULATION\_OF\_ACTIN\_CYTOSKELETON\_REORGANIZATION | 27 | 2 | 12187 | 98 | Csf1r,Hcls1 | | 1.971e-02 | -3.93 | KEGG\_PRION\_DISEASES | MSigDB lists | KEGG\_PRION\_DISEASES | 27 | 2 | 12187 | 98 | C1qa,C1qb | | 1.971e-02 | -3.93 | SCHRAETS\_MLL\_TARGETS\_DN | MSigDB lists | SCHRAETS\_MLL\_TARGETS\_DN | 27 | 2 | 12187 | 98 | Mgp,Mgll | | 1.971e-02 | -3.93 | MODULE\_419 | MSigDB lists | MODULE\_419 | 27 | 2 | 12187 | 98 | Flnb,Myh9 | | 1.971e-02 | -3.93 | GO\_NEURON\_PROJECTION\_REGENERATION | MSigDB lists | GO\_NEURON\_PROJECTION\_REGENERATION | 27 | 2 | 12187 | 98 | Apod,Gfap | | 1.971e-02 | -3.93 | GNF2\_RAP1B | MSigDB lists | GNF2\_RAP1B | 27 | 2 | 12187 | 98 | Ptprc,Rac2 | | 1.988e-02 | -3.92 | regulation of cellular protein localization | biological process | GO:1903827 | 499 | 9 | 13711 | 111 | Actb,Fcer1g,Abhd17c,Apod,Hcls1,Mark4,Vamp8,Camk2b,Csk | | 1.998e-02 | -3.91 | response to cadmium ion | biological process | GO:0046686 | 27 | 2 | 13711 | 111 | Camk2b,Mt3 | | 1.998e-02 | -3.91 | negative regulation of protein tyrosine kinase activity | biological process | GO:0061099 | 27 | 2 | 13711 | 111 | Ptprc,Thy1 | | 2.000e-02 | -3.91 | regulation of cell development | biological process | GO:0060284 | 944 | 14 | 13711 | 111 | Bhlhe41,Spint1,Mt3,Gfap,Ccr5,Camk2b,Ctsz,C1qa,Grn,Vim,Thy1,Mgll,Rap1gap,Il1r1 | | 2.013e-02 | -3.91 | small ribosomal subunit | cellular component | GO:0015935 | 72 | 3 | 13825 | 111 | Rps8,Rps20,Rps5 | | 2.017e-02 | -3.90 | GO\_NEGATIVE\_REGULATION\_OF\_CELL\_PROJECTION\_ORGANIZATION | MSigDB lists | GO\_NEGATIVE\_REGULATION\_OF\_CELL\_PROJECTION\_ORGANIZATION | 129 | 4 | 12187 | 98 | Vim,Gfap,Rap1gap,Thy1 | | 2.017e-02 | -3.90 | GO\_CYSTEINE\_TYPE\_PEPTIDASE\_ACTIVITY | MSigDB lists | GO\_CYSTEINE\_TYPE\_PEPTIDASE\_ACTIVITY | 129 | 4 | 12187 | 98 | Capn3,Ctsz,Ctss,Ctsb | | 2.017e-02 | -3.90 | GSE46606\_UNSTIM\_VS\_CD40L\_IL2\_IL5\_1DAY\_STIMULATED\_IRF4MID\_SORTED\_BCELL\_DN | MSigDB lists | GSE46606\_UNSTIM\_VS\_CD40L\_IL2\_IL5\_1DAY\_STIMULATED\_IRF4MID\_SORTED\_BCELL\_DN | 129 | 4 | 12187 | 98 | Cox4i1,Ttc9b,Psme1,Rac2 | | 2.021e-02 | -3.90 | MODULE\_41 | MSigDB lists | MODULE\_41 | 342 | 7 | 12187 | 98 | Phyhip,Fcgr3,Slc17a7,Rac2,Slc2a1,Capn3,Camk2b | | 2.028e-02 | -3.90 | myeloid cell differentiation | biological process | GO:0030099 | 193 | 5 | 13711 | 111 | Fcer1g,Itgam,Csf1r,Hcls1,Myh9 | | 2.030e-02 | -3.90 | actin cytoskeleton | cellular component | GO:0015629 | 422 | 8 | 13825 | 111 | Slc2a1,Actb,Flnb,Rac2,Myh9,Cyba,Ahnak,Hcls1 | | 2.047e-02 | -3.89 | GLYCOSYL\_HYDROL\_F1\_2 | prosite domains | PS00653 | 2 | 1 | 8845 | 91 | Kl | | 2.047e-02 | -3.89 | RIBOSOMAL\_L24 | prosite domains | PS01108 | 2 | 1 | 8845 | 91 | Rpl26 | | 2.047e-02 | -3.89 | CARBOXYPEPT\_SER\_SER | prosite domains | PS00131 | 2 | 1 | 8845 | 91 | Ctsa | | 2.047e-02 | -3.89 | CORTACTIN | prosite domains | PS51090 | 2 | 1 | 8845 | 91 | Hcls1 | | 2.047e-02 | -3.89 | L\_LDH | prosite domains | PS00064 | 2 | 1 | 8845 | 91 | Ldha | | 2.047e-02 | -3.89 | MACPF\_2 | prosite domains | PS51412 | 2 | 1 | 8845 | 91 | Mpeg1 | | 2.068e-02 | -3.88 | GSE34156\_UNTREATED\_VS\_24H\_NOD2\_AND\_TLR1\_TLR2\_LIGAND\_TREATED\_MONOCYTE\_UP | MSigDB lists | GSE34156\_UNTREATED\_VS\_24H\_NOD2\_AND\_TLR1\_TLR2\_LIGAND\_TREATED\_MONOCYTE\_UP | 130 | 4 | 12187 | 98 | Grn,Cd68,Lfng,Actb | | 2.068e-02 | -3.88 | GSE32901\_NAIVE\_VS\_TH17\_NEG\_CD4\_TCELL\_UP | MSigDB lists | GSE32901\_NAIVE\_VS\_TH17\_NEG\_CD4\_TCELL\_UP | 130 | 4 | 12187 | 98 | Ctsz,Grn,S100a6,Mpeg1 | | 2.068e-02 | -3.88 | GSE16450\_CTRL\_VS\_IFNA\_12H\_STIM\_MATURE\_NEURON\_CELL\_LINE\_UP | MSigDB lists | GSE16450\_CTRL\_VS\_IFNA\_12H\_STIM\_MATURE\_NEURON\_CELL\_LINE\_UP | 130 | 4 | 12187 | 98 | Gfap,Tfap2c,Igfbp6,Slc17a7 | | 2.068e-02 | -3.88 | GSE7509\_UNSTIM\_VS\_IFNA\_STIM\_IMMATURE\_DC\_DN | MSigDB lists | GSE7509\_UNSTIM\_VS\_IFNA\_STIM\_IMMATURE\_DC\_DN | 130 | 4 | 12187 | 98 | Psmb10,Klhl6,Thy1,Ahnak | | 2.068e-02 | -3.88 | GSE3203\_WT\_VS\_IFNAR1\_KO\_INFLUENZA\_INFECTED\_LN\_BCELL\_DN | MSigDB lists | GSE3203\_WT\_VS\_IFNAR1\_KO\_INFLUENZA\_INFECTED\_LN\_BCELL\_DN | 130 | 4 | 12187 | 98 | Trnp1,Lgi4,Il6ra,Rps5 | | 2.088e-02 | -3.87 | WONG\_ADULT\_TISSUE\_STEM\_MODULE | MSigDB lists | WONG\_ADULT\_TISSUE\_STEM\_MODULE | 594 | 10 | 12187 | 98 | Ctsz,Vwf,S100a6,Tsc22d1,Camk2b,Hspa2,Tspan4,Tmem176a,Il1r1,Flnb | | 2.091e-02 | -3.87 | TCTGGAC\_MIR198 | MSigDB lists | TCTGGAC\_MIR198 | 73 | 3 | 12187 | 98 | Slc2a1,Capn3,Trnp1 | | 2.091e-02 | -3.87 | GNF2\_DNM1 | MSigDB lists | GNF2\_DNM1 | 73 | 3 | 12187 | 98 | Slc17a7,Camk2b,Phyhip | | 2.093e-02 | -3.87 | GO\_NEGATIVE\_REGULATION\_OF\_IMMUNE\_SYSTEM\_PROCESS | MSigDB lists | GO\_NEGATIVE\_REGULATION\_OF\_IMMUNE\_SYSTEM\_PROCESS | 268 | 6 | 12187 | 98 | Tmem176a,Serpinb9,Thy1,Fcer1g,Ptprc,Apod | | 2.099e-02 | -3.86 | postsynapse | cellular component | GO:0098794 | 682 | 11 | 13825 | 111 | Camk2b,Sh2d5,Dlgap3,Il1r1,C1qb,Cplx2,Rpl14,Lin7a,Abhd17c,C1qa,Actb | | 2.100e-02 | -3.86 | GO\_RESPONSE\_TO\_HORMONE | MSigDB lists | GO\_RESPONSE\_TO\_HORMONE | 683 | 11 | 12187 | 98 | Csk,Ldha,Ctss,C3,Hcls1,Kl,Ptprn,Grn,Serpinb9,Mc4r,Ctsb | | 2.100e-02 | -3.86 | GO\_IMMUNE\_SYSTEM\_DEVELOPMENT | MSigDB lists | GO\_IMMUNE\_SYSTEM\_DEVELOPMENT | 425 | 8 | 12187 | 98 | Tnfsf8,Ptprc,Lfng,Fcer1g,Myh9,Hcls1,Csf1r,C3 | | 2.108e-02 | -3.86 | vacuole | cellular component | GO:0005773 | 425 | 8 | 13825 | 111 | Cd68,Gfap,Ctss,Vamp8,Grn,Ctsz,Ctsb,Ctsa | | 2.109e-02 | -3.86 | negative regulation of cell development | biological process | GO:0010721 | 342 | 7 | 13711 | 111 | Vim,Gfap,Ccr5,Thy1,Rap1gap,Ctsz,Mt3 | | 2.112e-02 | -3.86 | GO\_CELLULAR\_MACROMOLECULE\_LOCALIZATION | MSigDB lists | GO\_CELLULAR\_MACROMOLECULE\_LOCALIZATION | 1056 | 15 | 12187 | 98 | Rps20,Rpl14,Rps8,Capn3,Rpl13,Flnb,Rac2,Exoc3l4,Rps5,Rpl31,Ndufa13,Fcer1g,Lin7a,Ctsa,Rpl26 | | 2.112e-02 | -3.86 | GO\_REGULATION\_OF\_LAMELLIPODIUM\_ORGANIZATION | MSigDB lists | GO\_REGULATION\_OF\_LAMELLIPODIUM\_ORGANIZATION | 28 | 2 | 12187 | 98 | Rac2,Brk1 | | 2.112e-02 | -3.86 | HUANG\_FOXA2\_TARGETS\_DN | MSigDB lists | HUANG\_FOXA2\_TARGETS\_DN | 28 | 2 | 12187 | 98 | Igfbp6,Ctsb | | 2.112e-02 | -3.86 | GO\_REGULATION\_OF\_TYROSINE\_PHOSPHORYLATION\_OF\_STAT3\_PROTEIN | MSigDB lists | GO\_REGULATION\_OF\_TYROSINE\_PHOSPHORYLATION\_OF\_STAT3\_PROTEIN | 28 | 2 | 12187 | 98 | Il6ra,Csf1r | | 2.112e-02 | -3.86 | GO\_ENDOCRINE\_PANCREAS\_DEVELOPMENT | MSigDB lists | GO\_ENDOCRINE\_PANCREAS\_DEVELOPMENT | 28 | 2 | 12187 | 98 | Il6ra,Clu | | 2.120e-02 | -3.85 | GSE22886\_IGM\_MEMORY\_BCELL\_VS\_BLOOD\_PLASMA\_CELL\_UP | MSigDB lists | GSE22886\_IGM\_MEMORY\_BCELL\_VS\_BLOOD\_PLASMA\_CELL\_UP | 131 | 4 | 12187 | 98 | Fam111a,Rpl26,Rps20,Flnb | | 2.120e-02 | -3.85 | GSE3720\_UNSTIM\_VS\_PMA\_STIM\_VD2\_GAMMADELTA\_TCELL\_UP | MSigDB lists | GSE3720\_UNSTIM\_VS\_PMA\_STIM\_VD2\_GAMMADELTA\_TCELL\_UP | 131 | 4 | 12187 | 98 | Rps20,Tnfsf8,Ctss,Psme1 | | 2.120e-02 | -3.85 | GSE17974\_CTRL\_VS\_ACT\_IL4\_AND\_ANTI\_IL12\_48H\_CD4\_TCELL\_UP | MSigDB lists | GSE17974\_CTRL\_VS\_ACT\_IL4\_AND\_ANTI\_IL12\_48H\_CD4\_TCELL\_UP | 131 | 4 | 12187 | 98 | Tnfsf8,Tsc22d1,Ahnak,Itgam | | 2.120e-02 | -3.85 | GSE6092\_UNSTIM\_VS\_IFNG\_STIM\_AND\_B\_BURGDORFERI\_INF\_ENDOTHELIAL\_CELL\_DN | MSigDB lists | GSE6092\_UNSTIM\_VS\_IFNG\_STIM\_AND\_B\_BURGDORFERI\_INF\_ENDOTHELIAL\_CELL\_DN | 131 | 4 | 12187 | 98 | Tfap2c,Mark4,Rpl31,Fgf18 | | 2.126e-02 | -3.85 | GO\_ANATOMICAL\_STRUCTURE\_HOMEOSTASIS | MSigDB lists | GO\_ANATOMICAL\_STRUCTURE\_HOMEOSTASIS | 197 | 5 | 12187 | 98 | Actb,Serpina3n,Rac2,Capn3,Slc17a7 | | 2.139e-02 | -3.84 | GO\_POSITIVE\_REGULATION\_OF\_DEVELOPMENTAL\_PROCESS | MSigDB lists | GO\_POSITIVE\_REGULATION\_OF\_DEVELOPMENTAL\_PROCESS | 868 | 13 | 12187 | 98 | Clu,Camk2b,Capn3,Kl,Hcls1,C3,Fgf18,Lrg1,Grn,Csf1r,Gfap,Il6ra,Spint1 | | 2.141e-02 | -3.84 | endocrine pancreas development | biological process | GO:0031018 | 28 | 2 | 13711 | 111 | Il6ra,Clu | | 2.141e-02 | -3.84 | regulation of superoxide metabolic process | biological process | GO:0090322 | 28 | 2 | 13711 | 111 | Itgam,Cyba | | 2.141e-02 | -3.84 | regulation of cellular extravasation | biological process | GO:0002691 | 28 | 2 | 13711 | 111 | Thy1,Il1r1 | | 2.141e-02 | -3.84 | regulation of lamellipodium assembly | biological process | GO:0010591 | 28 | 2 | 13711 | 111 | Rac2,Brk1 | | 2.141e-02 | -3.84 | positive regulation of neutrophil migration | biological process | GO:1902624 | 28 | 2 | 13711 | 111 | Rac2,Il1r1 | | 2.141e-02 | -3.84 | regulation of myeloid cell apoptotic process | biological process | GO:0033032 | 28 | 2 | 13711 | 111 | Ccr5,Fcer1g | | 2.144e-02 | -3.84 | negative regulation of multicellular organismal process | biological process | GO:0051241 | 1047 | 15 | 13711 | 111 | Grn,Vim,Rap1gap,Thy1,Ctsz,Apod,Tmem176a,Ptprc,Cyba,Gfap,Ccr5,Lfng,Capn3,Csk,Mt3 | | 2.151e-02 | -3.84 | developmental maturation | biological process | GO:0021700 | 196 | 5 | 13711 | 111 | Ptprn,C3,Slc17a7,Lgi4,C1qa | | 2.158e-02 | -3.84 | FLOT2 (flotillin 2) | protein interactions | 2319 | 20 | 2 | 6802 | 78 | Flnb,Actb | | 2.166e-02 | -3.83 | Ttc23 (tetratricopeptide repeat domain 23) | protein interactions | 67009 | 93 | 4 | 6802 | 78 | Myh9,Ahnak,Klhl6,Actb | | 2.166e-02 | -3.83 | HILLION\_HMGA1\_TARGETS | MSigDB lists | HILLION\_HMGA1\_TARGETS | 74 | 3 | 12187 | 98 | Cox4i1,Clu,Rpl13 | | 2.166e-02 | -3.83 | MODULE\_180 | MSigDB lists | MODULE\_180 | 74 | 3 | 12187 | 98 | Clu,Tfap2c,Cyba | | 2.166e-02 | -3.83 | GO\_POSITIVE\_REGULATION\_OF\_CALCIUM\_ION\_TRANSPORT | MSigDB lists | GO\_POSITIVE\_REGULATION\_OF\_CALCIUM\_ION\_TRANSPORT | 74 | 3 | 12187 | 98 | Thy1,Hspa2,Capn3 | | 2.167e-02 | -3.83 | mouse chr14 | chromosome location | mouse chr14 | 134 | 4 | 14556 | 115 | Tsc22d1,Ctsb,Psme1,Clu | | 2.172e-02 | -3.83 | GO\_VESICLE\_MEDIATED\_TRANSPORT | MSigDB lists | GO\_VESICLE\_MEDIATED\_TRANSPORT | 964 | 14 | 12187 | 98 | Brk1,Myh9,Fcer1g,Grn,Exoc3l4,Vamp8,Cplx2,Lin7a,Fcgr3,Actb,Ctsz,Clu,Vwf,Serpina3n | | 2.174e-02 | -3.83 | GSE22611\_MUTANT\_NOD2\_TRANSDUCED\_VS\_CTRL\_HEK293T\_STIMULATED\_WITH\_MDP\_6H\_UP | MSigDB lists | GSE22611\_MUTANT\_NOD2\_TRANSDUCED\_VS\_CTRL\_HEK293T\_STIMULATED\_WITH\_MDP\_6H\_UP | 132 | 4 | 12187 | 98 | Apod,C3,Ikzf3,Gfap | | 2.174e-02 | -3.83 | RUIZ\_TNC\_TARGETS\_UP | MSigDB lists | RUIZ\_TNC\_TARGETS\_UP | 132 | 4 | 12187 | 98 | Il1r1,Ctsa,Clu,Grn | | 2.174e-02 | -3.83 | GSE37301\_COMMON\_LYMPHOID\_PROGENITOR\_VS\_GRAN\_MONO\_PROGENITOR\_DN | MSigDB lists | GSE37301\_COMMON\_LYMPHOID\_PROGENITOR\_VS\_GRAN\_MONO\_PROGENITOR\_DN | 132 | 4 | 12187 | 98 | Slc17a7,Rpl26,Rps8,Csf1r | | 2.174e-02 | -3.83 | GSE360\_L\_MAJOR\_VS\_T\_GONDII\_MAC\_UP | MSigDB lists | GSE360\_L\_MAJOR\_VS\_T\_GONDII\_MAC\_UP | 132 | 4 | 12187 | 98 | Cyba,Pde1a,Slc17a7,Ahnak | | 2.174e-02 | -3.83 | GSE32034\_UNTREATED\_VS\_ROSIGLIZATONE\_TREATED\_LY6C\_LOW\_MONOCYTE\_DN | MSigDB lists | GSE32034\_UNTREATED\_VS\_ROSIGLIZATONE\_TREATED\_LY6C\_LOW\_MONOCYTE\_DN | 132 | 4 | 12187 | 98 | Kl,Treml2,Rac2,Vim | | 2.174e-02 | -3.83 | thyroid hormone biosynthesis | BIOCYC pathways | META\_PWY-6241 | 3 | 1 | 823 | 6 | Ctsb | | 2.193e-02 | -3.82 | positive regulation of transmembrane transport | biological process | GO:0034764 | 197 | 5 | 13711 | 111 | C3,Ctss,Thy1,Hspa2,Capn3 | | 2.193e-02 | -3.82 | structural constituent of synapse | molecular function | GO:0098918 | 29 | 2 | 13516 | 107 | Actb,Camk2b | | 2.199e-02 | -3.82 | cell cortex | cellular component | GO:0005938 | 271 | 6 | 13825 | 111 | Ctsz,Myh9,Actb,Slc2a1,Hcls1,Exoc3l4 | | 2.200e-02 | -3.82 | identical protein binding | molecular function | GO:0042802 | 1578 | 20 | 13516 | 107 | S100a6,Mgll,Flnb,C1qb,Camk2b,Myh9,Rap1gap,Csf1r,Actb,Bhlhe41,Vim,Il6ra,Vwf,Brk1,Gfap,Adh1,Slc2a1,Ikzf3,Fcer1g,Csk | | 2.207e-02 | -3.81 | leukocyte differentiation | biological process | GO:0002521 | 269 | 6 | 13711 | 111 | Tnfsf8,Ptprc,Csf1r,Itgam,Fcer1g,Lfng | | 2.209e-02 | -3.81 | regulation of protein tyrosine kinase activity | biological process | GO:0061097 | 74 | 3 | 13711 | 111 | Csf1r,Thy1,Ptprc | | 2.228e-02 | -3.80 | RPS14\_DN.V1\_UP | MSigDB lists | RPS14\_DN.V1\_UP | 133 | 4 | 12187 | 98 | Vwf,Itgam,Ctss,Spint1 | | 2.228e-02 | -3.80 | GSE26488\_WT\_VS\_VP16\_TRANSGENIC\_HDAC7\_KO\_DOUBLE\_POSITIVE\_THYMOCYTE\_DN | MSigDB lists | GSE26488\_WT\_VS\_VP16\_TRANSGENIC\_HDAC7\_KO\_DOUBLE\_POSITIVE\_THYMOCYTE\_DN | 133 | 4 | 12187 | 98 | Rps5,Klhl6,Rpl13,Grn | | 2.228e-02 | -3.80 | KATSANOU\_ELAVL1\_TARGETS\_UP | MSigDB lists | KATSANOU\_ELAVL1\_TARGETS\_UP | 133 | 4 | 12187 | 98 | C1qa,Mpeg1,C1qb,Ctss | | 2.230e-02 | -3.80 | regulation of gliogenesis | biological process | GO:0014013 | 132 | 4 | 13711 | 111 | Vim,Spint1,Gfap,C1qa | | 2.230e-02 | -3.80 | regulation of response to wounding | biological process | GO:1903034 | 132 | 4 | 13711 | 111 | Grn,Fcer1g,Capn3,Myh9 | | 2.244e-02 | -3.80 | GO\_NEURON\_PART | MSigDB lists | GO\_NEURON\_PART | 1064 | 15 | 12187 | 98 | Cyba,Ptprn,Il1r1,Apod,Dlgap3,Vim,Rap1gap,Clu,Camk2b,Lin7a,Cplx2,Slc17a7,Pcsk2,Thy1,Mark4 | | 2.252e-02 | -3.79 | calyx of Held | cellular component | GO:0044305 | 29 | 2 | 13825 | 111 | Cplx2,Actb | | 2.252e-02 | -3.79 | GO\_REGULATION\_OF\_TRANSPORT | MSigDB lists | GO\_REGULATION\_OF\_TRANSPORT | 1360 | 18 | 12187 | 98 | Gfap,Vamp8,Cplx2,Fcer1g,Thy1,Csf1r,Slc2a1,Rac2,Cyba,Apod,Ahnak,Hspa2,Capn3,Camk2b,Csk,Ctss,C3,Hcls1 | | 2.257e-02 | -3.79 | Legionellosis | KEGG pathways | mmu05134 | 50 | 3 | 5248 | 64 | C3,Hspa2,Itgam | | 2.257e-02 | -3.79 | Legionellosis | KEGG pathways | ko05134 | 50 | 3 | 5248 | 64 | Hspa2,Itgam,C3 | | 2.257e-02 | -3.79 | FIGUEROA\_AML\_METHYLATION\_CLUSTER\_1\_DN | MSigDB lists | FIGUEROA\_AML\_METHYLATION\_CLUSTER\_1\_DN | 29 | 2 | 12187 | 98 | Cd68,Ahnak | | 2.257e-02 | -3.79 | REACTOME\_PLATELET\_AGGREGATION\_PLUG\_FORMATION | MSigDB lists | REACTOME\_PLATELET\_AGGREGATION\_PLUG\_FORMATION | 29 | 2 | 12187 | 98 | Vwf,Csk | | 2.257e-02 | -3.79 | CROMER\_TUMORIGENESIS\_DN | MSigDB lists | CROMER\_TUMORIGENESIS\_DN | 29 | 2 | 12187 | 98 | Clu,Apod | | 2.257e-02 | -3.79 | GO\_REGULATION\_OF\_ANTIGEN\_RECEPTOR\_MEDIATED\_SIGNALING\_PATHWAY | MSigDB lists | GO\_REGULATION\_OF\_ANTIGEN\_RECEPTOR\_MEDIATED\_SIGNALING\_PATHWAY | 29 | 2 | 12187 | 98 | Ptprc,Thy1 | | 2.257e-02 | -3.79 | actin cytoskeleton organization | biological process | GO:0030036 | 427 | 8 | 13711 | 111 | Camk2b,Capn3,Rac2,Actb,Myh9,Hcls1,Flnb,Brk1 | | 2.268e-02 | -3.79 | HIF-1 signaling pathway | KEGG pathways | mmu04066 | 89 | 4 | 5248 | 64 | Il6ra,Camk2b,Ldha,Slc2a1 | | 2.280e-02 | -3.78 | PI4KA (phosphatidylinositol 4-kinase alpha) | protein interactions | 5297 | 2 | 1 | 6802 | 78 | Flnb | | 2.280e-02 | -3.78 | EPB41L2 (erythrocyte membrane protein band 4.1 like 2) | protein interactions | 2037 | 2 | 1 | 6802 | 78 | Myh9 | | 2.280e-02 | -3.78 | Ube2m (ubiquitin-conjugating enzyme E2M) | protein interactions | 22192 | 2 | 1 | 6802 | 78 | Mpeg1 | | 2.280e-02 | -3.78 | Nasp (nuclear autoantigenic sperm protein (histone-binding)) | protein interactions | 50927 | 2 | 1 | 6802 | 78 | Hspa2 | | 2.280e-02 | -3.78 | Pag1 (phosphoprotein associated with glycosphingolipid microdomains 1) | protein interactions | 94212 | 2 | 1 | 6802 | 78 | Csk | | 2.280e-02 | -3.78 | Plcg2 (phospholipase C, gamma 2) | protein interactions | 234779 | 2 | 1 | 6802 | 78 | Csf1r | | 2.280e-02 | -3.78 | TWF2 (twinfilin actin binding protein 2) | protein interactions | 11344 | 2 | 1 | 6802 | 78 | Actb | | 2.280e-02 | -3.78 | SRI (sorcin) | protein interactions | 6717 | 2 | 1 | 6802 | 78 | Actb | | 2.280e-02 | -3.78 | DLG5 (discs large MAGUK scaffold protein 5) | protein interactions | 9231 | 2 | 1 | 6802 | 78 | Myh9 | | 2.280e-02 | -3.78 | Ankrd1 (ankyrin repeat domain 1 (cardiac muscle)) | protein interactions | 107765 | 2 | 1 | 6802 | 78 | Capn3 | | 2.280e-02 | -3.78 | RAP1A (RAP1A, member of RAS oncogene family) | protein interactions | 5906 | 2 | 1 | 6802 | 78 | Flnb | | 2.280e-02 | -3.78 | ZBTB20 (zinc finger and BTB domain containing 20) | protein interactions | 26137 | 2 | 1 | 6802 | 78 | Myh9 | | 2.280e-02 | -3.78 | Trpv5 (transient receptor potential cation channel, subfamily V, member 5) | protein interactions | 194352 | 2 | 1 | 6802 | 78 | Kl | | 2.280e-02 | -3.78 | Adgrb1 (adhesion G protein-coupled receptor B1) | protein interactions | 107831 | 2 | 1 | 6802 | 78 | Phyhip | | 2.280e-02 | -3.78 | SNX33 (sorting nexin 33) | protein interactions | 257364 | 2 | 1 | 6802 | 78 | Actb | | 2.280e-02 | -3.78 | PDLIM7 (PDZ and LIM domain 7) | protein interactions | 9260 | 2 | 1 | 6802 | 78 | Myh9 | | 2.280e-02 | -3.78 | TRMT10A (tRNA methyltransferase 10A) | protein interactions | 93587 | 2 | 1 | 6802 | 78 | Myh9 | | 2.280e-02 | -3.78 | Hs1bp3 (HCLS1 binding protein 3) | protein interactions | 58240 | 2 | 1 | 6802 | 78 | Hcls1 | | 2.280e-02 | -3.78 | Trim72 (tripartite motif-containing 72) | protein interactions | 434246 | 2 | 1 | 6802 | 78 | Myh9 | | 2.280e-02 | -3.78 | CSNK1G3 (casein kinase 1 gamma 3) | protein interactions | 1456 | 2 | 1 | 6802 | 78 | Actb | | 2.280e-02 | -3.78 | Il1rap (interleukin 1 receptor accessory protein) | protein interactions | 16180 | 2 | 1 | 6802 | 78 | Il1r1 | | 2.280e-02 | -3.78 | Rgs12 (regulator of G-protein signaling 12) | protein interactions | 54292 | 2 | 1 | 6802 | 78 | Dlgap3 | | 2.280e-02 | -3.78 | MYLK (myosin light chain kinase) | protein interactions | 4638 | 2 | 1 | 6802 | 78 | Myh9 | | 2.280e-02 | -3.78 | Ints4 (integrator complex subunit 4) | protein interactions | 101861 | 2 | 1 | 6802 | 78 | Ptprc | | 2.280e-02 | -3.78 | DSC2 (desmocollin 2) | protein interactions | 1824 | 2 | 1 | 6802 | 78 | Flnb | | 2.280e-02 | -3.78 | Catsper1 (cation channel, sperm associated 1) | protein interactions | 225865 | 2 | 1 | 6802 | 78 | Hspa2 | | 2.280e-02 | -3.78 | RMDN1 (regulator of microtubule dynamics 1) | protein interactions | 51115 | 2 | 1 | 6802 | 78 | Myh9 | | 2.280e-02 | -3.78 | Stx17 (syntaxin 17) | protein interactions | 67727 | 2 | 1 | 6802 | 78 | Vamp8 | | 2.280e-02 | -3.78 | BCS1L (BCS1 homolog, ubiquinol-cytochrome c reductase complex chaperone) | protein interactions | 617 | 2 | 1 | 6802 | 78 | Myh9 | | 2.280e-02 | -3.78 | LMAN2L (lectin, mannose binding 2 like) | protein interactions | 81562 | 2 | 1 | 6802 | 78 | Csk | | 2.280e-02 | -3.78 | CELF2 (CUGBP Elav-like family member 2) | protein interactions | 10659 | 2 | 1 | 6802 | 78 | Csk | | 2.280e-02 | -3.78 | RALB (RAS like proto-oncogene B) | protein interactions | 5899 | 2 | 1 | 6802 | 78 | Flnb | | 2.280e-02 | -3.78 | Cd300c2 (CD300C molecule 2) | protein interactions | 140497 | 2 | 1 | 6802 | 78 | Fcer1g | | 2.282e-02 | -3.78 | GO\_CELL\_ADHESION\_MOLECULE\_BINDING | MSigDB lists | GO\_CELL\_ADHESION\_MOLECULE\_BINDING | 134 | 4 | 12187 | 98 | Thy1,Gfap,Vwf,Tspan4 | | 2.287e-02 | -3.78 | positive regulation of release of sequestered calcium ion into cytosol | biological process | GO:0051281 | 29 | 2 | 13711 | 111 | Capn3,Thy1 | | 2.287e-02 | -3.78 | myeloid leukocyte mediated immunity | biological process | GO:0002444 | 29 | 2 | 13711 | 111 | Fcgr3,Cplx2 | | 2.288e-02 | -3.78 | response to interleukin-1 | biological process | GO:0070555 | 75 | 3 | 13711 | 111 | Ccl6,Saa3,Il1r1 | | 2.302e-02 | -3.77 | IF\_ROD\_1 | prosite domains | PS00226 | 23 | 2 | 8845 | 91 | Gfap,Vim | | 2.304e-02 | -3.77 | WGGAATGY\_TEF1\_Q6 | MSigDB lists | WGGAATGY\_TEF1\_Q6 | 274 | 6 | 12187 | 98 | Pde1a,Mgll,Sh3bgrl3,Cplx2,Ppfibp1,Igfbp6 | | 2.323e-02 | -3.76 | GO\_LEUKOCYTE\_CHEMOTAXIS | MSigDB lists | GO\_LEUKOCYTE\_CHEMOTAXIS | 76 | 3 | 12187 | 98 | Il6ra,Ccr5,Fcer1g | | 2.323e-02 | -3.76 | GRAHAM\_CML\_DIVIDING\_VS\_NORMAL\_QUIESCENT\_DN | MSigDB lists | GRAHAM\_CML\_DIVIDING\_VS\_NORMAL\_QUIESCENT\_DN | 76 | 3 | 12187 | 98 | Rpl31,Ptprc,Ppfibp1 | | 2.323e-02 | -3.76 | MIKKELSEN\_ES\_LCP\_WITH\_H3K4ME3 | MSigDB lists | MIKKELSEN\_ES\_LCP\_WITH\_H3K4ME3 | 76 | 3 | 12187 | 98 | Treml2,Dlgap3,Cd68 | | 2.323e-02 | -3.76 | REACTOME\_MHC\_CLASS\_II\_ANTIGEN\_PRESENTATION | MSigDB lists | REACTOME\_MHC\_CLASS\_II\_ANTIGEN\_PRESENTATION | 76 | 3 | 12187 | 98 | Ctss,Ctsa,Ctsb | | 2.323e-02 | -3.76 | GSE5589\_WT\_VS\_IL10\_KO\_LPS\_AND\_IL10\_STIM\_MACROPHAGE\_180MIN\_DN | MSigDB lists | GSE5589\_WT\_VS\_IL10\_KO\_LPS\_AND\_IL10\_STIM\_MACROPHAGE\_180MIN\_DN | 76 | 3 | 12187 | 98 | Fmod,Igfbp6,Lfng | | 2.326e-02 | -3.76 | Regulation of actin cytoskeleton | KEGG pathways | mmu04810 | 183 | 6 | 5248 | 64 | Myh9,Itgam,Brk1,Rac2,Actb,Fgf18 | | 2.326e-02 | -3.76 | Regulation of actin cytoskeleton | KEGG pathways | ko04810 | 183 | 6 | 5248 | 64 | Fgf18,Brk1,Rac2,Actb,Itgam,Myh9 | | 2.338e-02 | -3.76 | GSE11864\_CSF1\_IFNG\_VS\_CSF1\_PAM3CYS\_IN\_MAC\_DN | MSigDB lists | GSE11864\_CSF1\_IFNG\_VS\_CSF1\_PAM3CYS\_IN\_MAC\_DN | 135 | 4 | 12187 | 98 | Actb,Tsc22d1,Mgll,Ccr5 | | 2.338e-02 | -3.76 | dicarboxylic acid transmembrane transporter activity | molecular function | GO:0005310 | 30 | 2 | 13516 | 107 | Slc17a7,Slc26a2 | | 2.348e-02 | -3.75 | cell junction | cellular component | GO:0030054 | 972 | 14 | 13825 | 111 | Slc17a7,Sh2d5,Cyba,Camk2b,Csk,Dlgap3,Ahnak,Myh9,Ptprn,Ptprc,Flnb,Lin7a,Abhd17c,Slc2a1 | | 2.352e-02 | -3.75 | mouse chr11 D\|11 66.29 cM | chromosome location | mouse chr11 D\|11 66.29 cM | 3 | 1 | 14556 | 115 | Grn | | 2.352e-02 | -3.75 | mouse chr4 D2.1\|4 55.34 cM | chromosome location | mouse chr4 D2.1\|4 55.34 cM | 3 | 1 | 14556 | 115 | Slc2a1 | | 2.352e-02 | -3.75 | mouse chr11 E1\|11 66.48 cM | chromosome location | mouse chr11 E1\|11 66.48 cM | 3 | 1 | 14556 | 115 | Gfap | | 2.352e-02 | -3.75 | mouse chr8 D3\|8 53.06 cM | chromosome location | mouse chr8 D3\|8 53.06 cM | 3 | 1 | 14556 | 115 | Psmb10 | | 2.352e-02 | -3.75 | mouse chr9 F4\|9 75.05 cM | chromosome location | mouse chr9 F4\|9 75.05 cM | 3 | 1 | 14556 | 115 | Ccr5 | | 2.352e-02 | -3.75 | mouse chr2 H3\|2 85.27 cM | chromosome location | mouse chr2 H3\|2 85.27 cM | 3 | 1 | 14556 | 115 | Ctsa | | 2.356e-02 | -3.75 | GPI anchor binding | molecular function | GO:0034235 | 3 | 1 | 13516 | 107 | Thy1 | | 2.356e-02 | -3.75 | melanocortin receptor activity | molecular function | GO:0004977 | 3 | 1 | 13516 | 107 | Mc4r | | 2.356e-02 | -3.75 | O-fucosylpeptide 3-beta-N-acetylglucosaminyltransferase activity | molecular function | GO:0033829 | 3 | 1 | 13516 | 107 | Lfng | | 2.356e-02 | -3.75 | serine-type carboxypeptidase activity | molecular function | GO:0004185 | 3 | 1 | 13516 | 107 | Ctsa | | 2.356e-02 | -3.75 | sodium:inorganic phosphate symporter activity | molecular function | GO:0015319 | 3 | 1 | 13516 | 107 | Slc17a7 | | 2.356e-02 | -3.75 | calmodulin-dependent cyclic-nucleotide phosphodiesterase activity | molecular function | GO:0004117 | 3 | 1 | 13516 | 107 | Pde1a | | 2.356e-02 | -3.75 | IgE binding | molecular function | GO:0019863 | 3 | 1 | 13516 | 107 | Fcer1g | | 2.356e-02 | -3.75 | keratin filament binding | molecular function | GO:1990254 | 3 | 1 | 13516 | 107 | Vim | | 2.356e-02 | -3.75 | alcohol dehydrogenase (NAD) activity | molecular function | GO:0004022 | 3 | 1 | 13516 | 107 | Adh1 | | 2.356e-02 | -3.75 | calcium- and calmodulin-regulated 3',5'-cyclic-GMP phosphodiesterase activity | molecular function | GO:0048101 | 3 | 1 | 13516 | 107 | Pde1a | | 2.356e-02 | -3.75 | type 2 fibroblast growth factor receptor binding | molecular function | GO:0005111 | 3 | 1 | 13516 | 107 | Fgf18 | | 2.356e-02 | -3.75 | D-glucose transmembrane transporter activity | molecular function | GO:0055056 | 3 | 1 | 13516 | 107 | Slc2a1 | | 2.356e-02 | -3.75 | chemokine (C-C motif) ligand 5 binding | molecular function | GO:0071791 | 3 | 1 | 13516 | 107 | Ccr5 | | 2.356e-02 | -3.75 | interleukin-12 receptor binding | molecular function | GO:0005143 | 3 | 1 | 13516 | 107 | Il6ra | | 2.368e-02 | -3.74 | TPM1 (tropomyosin 1) | protein interactions | 7168 | 21 | 2 | 6802 | 78 | Myh9,Actb | | 2.368e-02 | -3.74 | TMOD3 (tropomodulin 3) | protein interactions | 29766 | 21 | 2 | 6802 | 78 | Myh9,Actb | | 2.371e-02 | -3.74 | IG\_LIKE | prosite domains | PS50835 | 277 | 7 | 8845 | 91 | Csf1r,Il6ra,Fcgr3,Fcrls,Treml2,Il1r1,Thy1 | | 2.378e-02 | -3.74 | GO\_DEFENSE\_RESPONSE\_TO\_OTHER\_ORGANISM | MSigDB lists | GO\_DEFENSE\_RESPONSE\_TO\_OTHER\_ORGANISM | 276 | 6 | 12187 | 98 | Ptprc,Phyhip,Tnfsf8,Fam111a,Fcer1g,Il6ra | | 2.383e-02 | -3.74 | RICKMAN\_METASTASIS\_DN | MSigDB lists | RICKMAN\_METASTASIS\_DN | 203 | 5 | 12187 | 98 | Fcer1g,S100a6,Cd68,Sh3bgrl3,Slc2a1 | | 2.389e-02 | -3.73 | GO\_POSITIVE\_REGULATION\_OF\_CELL\_COMMUNICATION | MSigDB lists | GO\_POSITIVE\_REGULATION\_OF\_CELL\_COMMUNICATION | 1170 | 16 | 12187 | 98 | C3,Hcls1,Kl,Clu,Psme1,Csk,Fgf18,Cyba,Ptprc,Psmb10,Csf1r,Lrg1,Lfng,Il6ra,Gfap,Vamp8 | | 2.390e-02 | -3.73 | cytoplasmic side of lysosomal membrane | cellular component | GO:0098574 | 3 | 1 | 13825 | 111 | Gfap | | 2.390e-02 | -3.73 | interleukin-6 receptor complex | cellular component | GO:0005896 | 3 | 1 | 13825 | 111 | Il6ra | | 2.390e-02 | -3.73 | spherical high-density lipoprotein particle | cellular component | GO:0034366 | 3 | 1 | 13825 | 111 | Clu | | 2.390e-02 | -3.73 | myosin II filament | cellular component | GO:0097513 | 3 | 1 | 13825 | 111 | Myh9 | | 2.390e-02 | -3.73 | bleb | cellular component | GO:0032059 | 3 | 1 | 13825 | 111 | Ptprc | | 2.393e-02 | -3.73 | BIOCARTA\_TCAPOPTOSIS\_PATHWAY | MSigDB lists | BIOCARTA\_TCAPOPTOSIS\_PATHWAY | 3 | 1 | 12187 | 98 | Ccr5 | | 2.393e-02 | -3.73 | GO\_REGULATION\_OF\_DEFENSE\_RESPONSE\_TO\_BACTERIUM | MSigDB lists | GO\_REGULATION\_OF\_DEFENSE\_RESPONSE\_TO\_BACTERIUM | 3 | 1 | 12187 | 98 | Cyba | | 2.393e-02 | -3.73 | GO\_POSITIVE\_REGULATION\_OF\_HUMORAL\_IMMUNE\_RESPONSE | MSigDB lists | GO\_POSITIVE\_REGULATION\_OF\_HUMORAL\_IMMUNE\_RESPONSE | 3 | 1 | 12187 | 98 | C3 | | 2.393e-02 | -3.73 | BIOCARTA\_LECTIN\_PATHWAY | MSigDB lists | BIOCARTA\_LECTIN\_PATHWAY | 3 | 1 | 12187 | 98 | C3 | | 2.394e-02 | -3.73 | cysteine-type peptidase activity | molecular function | GO:0008234 | 138 | 4 | 13516 | 107 | Ctsz,Capn3,Ctss,Ctsb | | 2.394e-02 | -3.73 | HALLMARK\_FATTY\_ACID\_METABOLISM | MSigDB lists | HALLMARK\_FATTY\_ACID\_METABOLISM | 136 | 4 | 12187 | 98 | Adh1,Mgll,Psme1,Ldha | | 2.394e-02 | -3.73 | RAPA\_EARLY\_UP.V1\_DN | MSigDB lists | RAPA\_EARLY\_UP.V1\_DN | 136 | 4 | 12187 | 98 | Phyhip,Tspan4,Pcsk2,Grn | | 2.394e-02 | -3.73 | GSE13547\_CTRL\_VS\_ANTI\_IGM\_STIM\_ZFX\_KO\_BCELL\_2H\_UP | MSigDB lists | GSE13547\_CTRL\_VS\_ANTI\_IGM\_STIM\_ZFX\_KO\_BCELL\_2H\_UP | 136 | 4 | 12187 | 98 | Rac2,Csk,Cyba,Hcls1 | | 2.394e-02 | -3.73 | GSE36888\_UNTREATED\_VS\_IL2\_TREATED\_TCELL\_2H\_DN | MSigDB lists | GSE36888\_UNTREATED\_VS\_IL2\_TREATED\_TCELL\_2H\_DN | 136 | 4 | 12187 | 98 | Mgll,Tsc22d1,Ly6e,Spint1 | | 2.399e-02 | -3.73 | FGFR1c and Klotho ligand binding and activation | REACTOME pathways | R-MMU-190374 | 2 | 1 | 6297 | 76 | Kl | | 2.399e-02 | -3.73 | Lactose synthesis | REACTOME pathways | R-MMU-5653890 | 2 | 1 | 6297 | 76 | Slc2a1 | | 2.399e-02 | -3.73 | Terminal pathway of complement | REACTOME pathways | R-MMU-166665 | 2 | 1 | 6297 | 76 | Clu | | 2.401e-02 | -3.73 | neuronal cell body membrane | cellular component | GO:0032809 | 30 | 2 | 13825 | 111 | Thy1,Il6ra | | 2.403e-02 | -3.73 | BURTON\_ADIPOGENESIS\_8 | MSigDB lists | BURTON\_ADIPOGENESIS\_8 | 77 | 3 | 12187 | 98 | Myh9,Flnb,Ctsa | | 2.406e-02 | -3.73 | SEMENZA\_HIF1\_TARGETS | MSigDB lists | SEMENZA\_HIF1\_TARGETS | 30 | 2 | 12187 | 98 | Slc2a1,Ldha | | 2.406e-02 | -3.73 | GNF2\_TPT1 | MSigDB lists | GNF2\_TPT1 | 30 | 2 | 12187 | 98 | Rps20,Rpl31 | | 2.406e-02 | -3.73 | KLEIN\_TARGETS\_OF\_BCR\_ABL1\_FUSION | MSigDB lists | KLEIN\_TARGETS\_OF\_BCR\_ABL1\_FUSION | 30 | 2 | 12187 | 98 | Ptprc,Csf1r | | 2.406e-02 | -3.73 | REACTOME\_GPVI\_MEDIATED\_ACTIVATION\_CASCADE | MSigDB lists | REACTOME\_GPVI\_MEDIATED\_ACTIVATION\_CASCADE | 30 | 2 | 12187 | 98 | Fcer1g,Rac2 | | 2.406e-02 | -3.73 | PATTERSON\_DOCETAXEL\_RESISTANCE | MSigDB lists | PATTERSON\_DOCETAXEL\_RESISTANCE | 30 | 2 | 12187 | 98 | Vamp8,Clu | | 2.406e-02 | -3.73 | ZHAN\_MULTIPLE\_MYELOMA\_LB\_DN | MSigDB lists | ZHAN\_MULTIPLE\_MYELOMA\_LB\_DN | 30 | 2 | 12187 | 98 | Psmb10,Ly6e | | 2.406e-02 | -3.73 | PID\_AMB2\_NEUTROPHILS\_PATHWAY | MSigDB lists | PID\_AMB2\_NEUTROPHILS\_PATHWAY | 30 | 2 | 12187 | 98 | Itgam,Thy1 | | 2.409e-02 | -3.73 | negative regulation of mast cell apoptotic process | biological process | GO:0033026 | 3 | 1 | 13711 | 111 | Fcer1g | | 2.409e-02 | -3.73 | positive regulation of pancreatic juice secretion | biological process | GO:0090187 | 3 | 1 | 13711 | 111 | Vamp8 | | 2.409e-02 | -3.73 | cGMP catabolic process | biological process | GO:0046069 | 3 | 1 | 13711 | 111 | Pde1a | | 2.409e-02 | -3.73 | positive regulation of apoptotic process by virus | biological process | GO:0060139 | 3 | 1 | 13711 | 111 | Ccr5 | | 2.409e-02 | -3.73 | regulation of smooth muscle cell-matrix adhesion | biological process | GO:2000097 | 3 | 1 | 13711 | 111 | Apod | | 2.409e-02 | -3.73 | insulin processing | biological process | GO:0030070 | 3 | 1 | 13711 | 111 | Pcsk2 | | 2.409e-02 | -3.73 | primary alcohol catabolic process | biological process | GO:0034310 | 3 | 1 | 13711 | 111 | Adh1 | | 2.409e-02 | -3.73 | negative regulation by symbiont of host programmed cell death | biological process | GO:0052041 | 3 | 1 | 13711 | 111 | Serpinb9 | | 2.409e-02 | -3.73 | positive regulation of cell chemotaxis to fibroblast growth factor | biological process | GO:1904849 | 3 | 1 | 13711 | 111 | Fgf18 | | 2.409e-02 | -3.73 | negative regulation by symbiont of host apoptotic process | biological process | GO:0033668 | 3 | 1 | 13711 | 111 | Serpinb9 | | 2.409e-02 | -3.73 | positive regulation of complement activation | biological process | GO:0045917 | 3 | 1 | 13711 | 111 | C3 | | 2.409e-02 | -3.73 | negative regulation of neutrophil activation | biological process | GO:1902564 | 3 | 1 | 13711 | 111 | Grn | | 2.409e-02 | -3.73 | positive regulation of MAPKKK cascade by fibroblast growth factor receptor signaling pathway | biological process | GO:0090080 | 3 | 1 | 13711 | 111 | Kl | | 2.409e-02 | -3.73 | regulation of DNA damage response, signal transduction by p53 class mediator resulting in transcription of p21 class mediator | biological process | GO:1902162 | 3 | 1 | 13711 | 111 | Rpl26 | | 2.409e-02 | -3.73 | regulation of activation of Janus kinase activity | biological process | GO:0010533 | 3 | 1 | 13711 | 111 | Il6ra | | 2.409e-02 | -3.73 | positive regulation of neutrophil activation | biological process | GO:1902565 | 3 | 1 | 13711 | 111 | Itgam | | 2.409e-02 | -3.73 | activation of meiosis | biological process | GO:0090427 | 3 | 1 | 13711 | 111 | Camk2b | | 2.409e-02 | -3.73 | positive regulation of humoral immune response mediated by circulating immunoglobulin | biological process | GO:0002925 | 3 | 1 | 13711 | 111 | Ptprc | | 2.409e-02 | -3.73 | regulation of lysosome organization | biological process | GO:1905671 | 3 | 1 | 13711 | 111 | Grn | | 2.409e-02 | -3.73 | regulation of lysosomal membrane permeability | biological process | GO:0097213 | 3 | 1 | 13711 | 111 | Mt3 | | 2.409e-02 | -3.73 | hypersensitivity | biological process | GO:0002524 | 3 | 1 | 13711 | 111 | Fcgr3 | | 2.409e-02 | -3.73 | regulation of plasma membrane repair | biological process | GO:1905684 | 3 | 1 | 13711 | 111 | Myh9 | | 2.409e-02 | -3.73 | positive regulation of neutrophil degranulation | biological process | GO:0043315 | 3 | 1 | 13711 | 111 | Itgam | | 2.409e-02 | -3.73 | positive regulation of endothelial cell chemotaxis to fibroblast growth factor | biological process | GO:2000546 | 3 | 1 | 13711 | 111 | Fgf18 | | 2.409e-02 | -3.73 | positive regulation of activation of Janus kinase activity | biological process | GO:0010536 | 3 | 1 | 13711 | 111 | Il6ra | | 2.409e-02 | -3.73 | regulation of renal output by angiotensin | biological process | GO:0002019 | 3 | 1 | 13711 | 111 | Cyba | | 2.409e-02 | -3.73 | membrane disruption in other organism | biological process | GO:0051673 | 3 | 1 | 13711 | 111 | Mpeg1 | | 2.409e-02 | -3.73 | negative regulation of dendritic cell antigen processing and presentation | biological process | GO:0002605 | 3 | 1 | 13711 | 111 | Cd68 | | 2.409e-02 | -3.73 | regulation of oxygen metabolic process | biological process | GO:2000374 | 3 | 1 | 13711 | 111 | Mt3 | | 2.409e-02 | -3.73 | negative regulation of glomerular filtration | biological process | GO:0003105 | 3 | 1 | 13711 | 111 | Cyba | | 2.409e-02 | -3.73 | membrane raft localization | biological process | GO:0051665 | 3 | 1 | 13711 | 111 | Ptprc | | 2.409e-02 | -3.73 | lactate biosynthetic process from pyruvate | biological process | GO:0019244 | 3 | 1 | 13711 | 111 | Ldha | | 2.409e-02 | -3.73 | dense core granule maturation | biological process | GO:1990502 | 3 | 1 | 13711 | 111 | Ptprn | | 2.409e-02 | -3.73 | positive regulation of inflammatory response to wounding | biological process | GO:0106016 | 3 | 1 | 13711 | 111 | Grn | | 2.409e-02 | -3.73 | macrophage colony-stimulating factor signaling pathway | biological process | GO:0038145 | 3 | 1 | 13711 | 111 | Csf1r | | 2.422e-02 | -3.72 | cellular response to stimulus | biological process | GO:0051716 | 4061 | 43 | 13711 | 111 | Rpl26,Ccl6,Vim,Brk1,Csf1r,Hcls1,Rac2,Ifitm1,Saa3,C3,Ptprn,Hspa2,Clu,Cd68,Mc4r,Itgam,Slc17a7,Capn3,Il6ra,Gfap,Fcgr3,Flnb,Rap1gap,Vamp8,Slc2a1,Klhl6,Mt3,Mark4,Csk,Cyba,Ndufa13,Ccr5,Ptprc,Il1r1,Kl,Thy1,Adgre1,Ldha,Actb,Fcer1g,Pde1a,Fgf18,Ly6e | | 2.428e-02 | -3.72 | MIYAGAWA\_TARGETS\_OF\_EWSR1\_ETS\_FUSIONS\_UP | MSigDB lists | MIYAGAWA\_TARGETS\_OF\_EWSR1\_ETS\_FUSIONS\_UP | 204 | 5 | 12187 | 98 | Itgam,Trnp1,Slc17a7,Fgf18,Slc26a2 | | 2.435e-02 | -3.72 | central nervous system development | biological process | GO:0007417 | 692 | 11 | 13711 | 111 | Mt3,Csf1r,Itgam,C3,Gfap,Tfap2c,C1qa,Clu,Trnp1,Grn,Vim | | 2.436e-02 | -3.71 | GO\_CELL\_DEVELOPMENT | MSigDB lists | GO\_CELL\_DEVELOPMENT | 1075 | 15 | 12187 | 98 | S100a6,Thy1,Myh9,Csf1r,Lgi4,Gfap,Clu,Capn3,Fmod,Hspa2,Vim,Fgf18,Flnb,Apod,Ptprc | | 2.438e-02 | -3.71 | glial cell activation | biological process | GO:0061900 | 30 | 2 | 13711 | 111 | Clu,Grn | | 2.438e-02 | -3.71 | alcohol catabolic process | biological process | GO:0046164 | 30 | 2 | 13711 | 111 | Mt3,Adh1 | | 2.441e-02 | -3.71 | Filament | pfam domains | PF00038 | 29 | 2 | 12881 | 108 | Vim,Gfap | | 2.442e-02 | -3.71 | response to oxygen-containing compound | biological process | GO:1901700 | 969 | 14 | 13711 | 111 | Cd68,Apod,Vim,Grn,Mc4r,Adh1,Kl,Actb,Ptprn,Serpina3n,Ndufa13,Ccr5,Csk,Ly6e | | 2.450e-02 | -3.71 | antigen receptor-mediated signaling pathway | biological process | GO:0050851 | 77 | 3 | 13711 | 111 | Thy1,Ptprc,Klhl6 | | 2.452e-02 | -3.71 | GSE1460\_INTRATHYMIC\_T\_PROGENITOR\_VS\_NAIVE\_CD4\_TCELL\_CORD\_BLOOD\_DN | MSigDB lists | GSE1460\_INTRATHYMIC\_T\_PROGENITOR\_VS\_NAIVE\_CD4\_TCELL\_CORD\_BLOOD\_DN | 137 | 4 | 12187 | 98 | Csf1r,Slc17a7,S100a6,Capn3 | | 2.452e-02 | -3.71 | GSE25123\_CTRL\_VS\_IL4\_AND\_ROSIGLITAZONE\_STIM\_MACROPHAGE\_DN | MSigDB lists | GSE25123\_CTRL\_VS\_IL4\_AND\_ROSIGLITAZONE\_STIM\_MACROPHAGE\_DN | 137 | 4 | 12187 | 98 | Rpl31,Ndufa13,Rpl26,Ccar1 | | 2.452e-02 | -3.71 | GSE28449\_WT\_VS\_LRF\_KO\_GERMINAL\_CENTER\_BCELL\_UP | MSigDB lists | GSE28449\_WT\_VS\_LRF\_KO\_GERMINAL\_CENTER\_BCELL\_UP | 137 | 4 | 12187 | 98 | Ctsb,Ctss,Serpinb9,Mgp | | 2.453e-02 | -3.71 | RICKMAN\_TUMOR\_DIFFERENTIATED\_WELL\_VS\_POORLY\_DN | MSigDB lists | RICKMAN\_TUMOR\_DIFFERENTIATED\_WELL\_VS\_POORLY\_DN | 278 | 6 | 12187 | 98 | Slc2a1,Ldha,Sh3bgrl3,Ahnak,Igfbp6,S100a6 | | 2.460e-02 | -3.70 | Proteasome\_activ\_PA28\_N | interpro domains | IPR003185 | 3 | 1 | 13788 | 114 | Psme1 | | 2.460e-02 | -3.70 | Proteasome\_activ\_sf | interpro domains | IPR036252 | 3 | 1 | 13788 | 114 | Psme1 | | 2.460e-02 | -3.70 | Metalthion\_dom\_sf\_vert | interpro domains | IPR023587 | 3 | 1 | 13788 | 114 | Mt3 | | 2.460e-02 | -3.70 | ADH\_Zn\_CS | interpro domains | IPR002328 | 3 | 1 | 13788 | 114 | Adh1 | | 2.460e-02 | -3.70 | RESP18\_dom | interpro domains | IPR029403 | 3 | 1 | 13788 | 114 | Ptprn | | 2.460e-02 | -3.70 | Ribosomal\_S10\_dom\_sf | interpro domains | IPR036838 | 3 | 1 | 13788 | 114 | Rps20 | | 2.460e-02 | -3.70 | MG4 | interpro domains | IPR040839 | 3 | 1 | 13788 | 114 | C3 | | 2.460e-02 | -3.70 | Proteasome\_activ\_PA28 | interpro domains | IPR009077 | 3 | 1 | 13788 | 114 | Psme1 | | 2.460e-02 | -3.70 | LIN7 | interpro domains | IPR017365 | 3 | 1 | 13788 | 114 | Lin7a | | 2.460e-02 | -3.70 | VWF\_type-D | interpro domains | IPR001846 | 3 | 1 | 13788 | 114 | Vwf | | 2.460e-02 | -3.70 | PA28\_C | interpro domains | IPR003186 | 3 | 1 | 13788 | 114 | Psme1 | | 2.460e-02 | -3.70 | TF\_AP2\_C | interpro domains | IPR013854 | 3 | 1 | 13788 | 114 | Tfap2c | | 2.460e-02 | -3.70 | DUF1356\_TMEM106 | interpro domains | IPR009790 | 3 | 1 | 13788 | 114 | Tmem106a | | 2.460e-02 | -3.70 | PDEase\_N | interpro domains | IPR013706 | 3 | 1 | 13788 | 114 | Pde1a | | 2.460e-02 | -3.70 | Synaphin | interpro domains | IPR008849 | 3 | 1 | 13788 | 114 | Cplx2 | | 2.460e-02 | -3.70 | PA28\_C\_sf | interpro domains | IPR036997 | 3 | 1 | 13788 | 114 | Psme1 | | 2.460e-02 | -3.70 | Ribosomal\_S10\_dom | interpro domains | IPR027486 | 3 | 1 | 13788 | 114 | Rps20 | | 2.460e-02 | -3.70 | Metalthion\_dom\_sf | interpro domains | IPR017854 | 3 | 1 | 13788 | 114 | Mt3 | | 2.460e-02 | -3.70 | FOX1\_RRM | interpro domains | IPR034237 | 3 | 1 | 13788 | 114 | Rbfox3 | | 2.460e-02 | -3.70 | YjeF\_N\_dom | interpro domains | IPR004443 | 3 | 1 | 13788 | 114 | Ndufa13 | | 2.460e-02 | -3.70 | Fox-1\_C\_dom | interpro domains | IPR025670 | 3 | 1 | 13788 | 114 | Rbfox3 | | 2.460e-02 | -3.70 | Metalthion\_vert\_metal\_BS | interpro domains | IPR018064 | 3 | 1 | 13788 | 114 | Mt3 | | 2.460e-02 | -3.70 | PA28\_N\_sf | interpro domains | IPR036996 | 3 | 1 | 13788 | 114 | Psme1 | | 2.460e-02 | -3.70 | LAMP\_CS | interpro domains | IPR018134 | 3 | 1 | 13788 | 114 | Cd68 | | 2.460e-02 | -3.70 | Fringe | interpro domains | IPR017374 | 3 | 1 | 13788 | 114 | Lfng | | 2.460e-02 | -3.70 | Serum\_amyloid\_A | interpro domains | IPR000096 | 3 | 1 | 13788 | 114 | Saa3 | | 2.460e-02 | -3.70 | TF\_AP2 | interpro domains | IPR004979 | 3 | 1 | 13788 | 114 | Tfap2c | | 2.460e-02 | -3.70 | Metalthion\_vert | interpro domains | IPR000006 | 3 | 1 | 13788 | 114 | Mt3 | | 2.460e-02 | -3.70 | YjeF\_N\_dom\_sf | interpro domains | IPR036652 | 3 | 1 | 13788 | 114 | Ndufa13 | | 2.460e-02 | -3.70 | Metalthion | interpro domains | IPR003019 | 3 | 1 | 13788 | 114 | Mt3 | | 2.460e-02 | -3.70 | RNA-bd\_Fox-1 | interpro domains | IPR017325 | 3 | 1 | 13788 | 114 | Rbfox3 | | 2.465e-02 | -3.70 | GO\_POSITIVE\_REGULATION\_OF\_TRANSPORT | MSigDB lists | GO\_POSITIVE\_REGULATION\_OF\_TRANSPORT | 700 | 11 | 12187 | 98 | Cyba,Rac2,C3,Hcls1,Capn3,Hspa2,Ctss,Vamp8,Csf1r,Thy1,Fcer1g | | 2.466e-02 | -3.70 | regulation of cell motility | biological process | GO:2000145 | 784 | 12 | 13711 | 111 | Csf1r,Rac2,Fgf18,Camk2b,Ccr5,Apod,Ptprc,Il1r1,Thy1,Grn,Ccar1,Vim | | 2.473e-02 | -3.70 | LU\_EZH2\_TARGETS\_UP | MSigDB lists | LU\_EZH2\_TARGETS\_UP | 205 | 5 | 12187 | 98 | Psmb10,Csk,Cd68,S100a6,Ppfibp1 | | 2.474e-02 | -3.70 | regulation of MAPK cascade | biological process | GO:0043408 | 605 | 10 | 13711 | 111 | Ccl6,Csf1r,Ptprc,Mt3,C3,Tmem106a,Il6ra,Kl,Csk,Fgf18 | | 2.485e-02 | -3.69 | GSE13485\_DAY1\_VS\_DAY7\_YF17D\_VACCINE\_PBMC\_UP | MSigDB lists | GSE13485\_DAY1\_VS\_DAY7\_YF17D\_VACCINE\_PBMC\_UP | 78 | 3 | 12187 | 98 | Gfap,Thy1,Slc17a7 | | 2.485e-02 | -3.69 | GCM\_PSME1 | MSigDB lists | GCM\_PSME1 | 78 | 3 | 12187 | 98 | Rps8,Psme1,Rps5 | | 2.485e-02 | -3.69 | WINTER\_HYPOXIA\_UP | MSigDB lists | WINTER\_HYPOXIA\_UP | 78 | 3 | 12187 | 98 | Tfap2c,Ldha,Slc2a1 | | 2.487e-02 | -3.69 | carboxypeptidase activity | molecular function | GO:0004180 | 31 | 2 | 13516 | 107 | Ctsa,Ctsz | | 2.494e-02 | -3.69 | VWD | pfam domains | PF00094 | 3 | 1 | 12881 | 108 | Vwf | | 2.494e-02 | -3.69 | MANEC | pfam domains | PF07502 | 3 | 1 | 12881 | 108 | Spint1 | | 2.494e-02 | -3.69 | SAA | pfam domains | PF00277 | 3 | 1 | 12881 | 108 | Saa3 | | 2.494e-02 | -3.69 | YjeF\_N | pfam domains | PF03853 | 3 | 1 | 12881 | 108 | Ndufa13 | | 2.494e-02 | -3.69 | Synaphin | pfam domains | PF05835 | 3 | 1 | 12881 | 108 | Cplx2 | | 2.494e-02 | -3.69 | ANATO | pfam domains | PF01821 | 3 | 1 | 12881 | 108 | C3 | | 2.494e-02 | -3.69 | PA28\_beta | pfam domains | PF02252 | 3 | 1 | 12881 | 108 | Psme1 | | 2.494e-02 | -3.69 | RESP18 | pfam domains | PF14948 | 3 | 1 | 12881 | 108 | Ptprn | | 2.494e-02 | -3.69 | Fox-1\_C | pfam domains | PF12414 | 3 | 1 | 12881 | 108 | Rbfox3 | | 2.494e-02 | -3.69 | Metallothio | pfam domains | PF00131 | 3 | 1 | 12881 | 108 | Mt3 | | 2.494e-02 | -3.69 | PA28\_alpha | pfam domains | PF02251 | 3 | 1 | 12881 | 108 | Psme1 | | 2.494e-02 | -3.69 | DUF1356 | pfam domains | PF07092 | 3 | 1 | 12881 | 108 | Tmem106a | | 2.494e-02 | -3.69 | PDEase\_I\_N | pfam domains | PF08499 | 3 | 1 | 12881 | 108 | Pde1a | | 2.494e-02 | -3.69 | TF\_AP-2 | pfam domains | PF03299 | 3 | 1 | 12881 | 108 | Tfap2c | | 2.494e-02 | -3.69 | Ribosomal\_S10 | pfam domains | PF00338 | 3 | 1 | 12881 | 108 | Rps20 | | 2.494e-02 | -3.69 | MG4 | pfam domains | PF17789 | 3 | 1 | 12881 | 108 | C3 | | 2.496e-02 | -3.69 | Rap1 signaling pathway | KEGG pathways | ko04015 | 186 | 6 | 5248 | 64 | Fgf18,Actb,Rac2,Itgam,Csf1r,Rap1gap | | 2.496e-02 | -3.69 | Rap1 signaling pathway | KEGG pathways | mmu04015 | 186 | 6 | 5248 | 64 | Actb,Rac2,Rap1gap,Csf1r,Itgam,Fgf18 | | 2.504e-02 | -3.69 | symbiont process | biological process | GO:0044403 | 204 | 5 | 13711 | 111 | Ccr5,Ctsb,Vamp8,Csf1r,Serpinb9 | | 2.510e-02 | -3.69 | GSE6269\_FLU\_VS\_STAPH\_AUREUS\_INF\_PBMC\_DN | MSigDB lists | GSE6269\_FLU\_VS\_STAPH\_AUREUS\_INF\_PBMC\_DN | 138 | 4 | 12187 | 98 | Ctsb,Grn,Cd68,Vim | | 2.510e-02 | -3.69 | GSE17186\_NAIVE\_VS\_CD21HIGH\_TRANSITIONAL\_BCELL\_UP | MSigDB lists | GSE17186\_NAIVE\_VS\_CD21HIGH\_TRANSITIONAL\_BCELL\_UP | 138 | 4 | 12187 | 98 | Ctsb,Tmem176a,Zfp786,Slc17a7 | | 2.510e-02 | -3.69 | GSE21927\_SPLEEN\_VS\_BONE\_MARROW\_MONOCYTE\_BALBC\_DN | MSigDB lists | GSE21927\_SPLEEN\_VS\_BONE\_MARROW\_MONOCYTE\_BALBC\_DN | 138 | 4 | 12187 | 98 | Itgam,Rac2,Mgll,Fcgr3 | | 2.510e-02 | -3.69 | GSE3039\_NKT\_CELL\_VS\_ALPHAALPHA\_CD8\_TCELL\_DN | MSigDB lists | GSE3039\_NKT\_CELL\_VS\_ALPHAALPHA\_CD8\_TCELL\_DN | 138 | 4 | 12187 | 98 | Ptprc,Ikzf3,Ly6e,Thy1 | | 2.514e-02 | -3.68 | positive regulation of cation transmembrane transport | biological process | GO:1904064 | 137 | 4 | 13711 | 111 | Ctss,Capn3,Hspa2,Thy1 | | 2.519e-02 | -3.68 | WAMUNYOKOLI\_OVARIAN\_CANCER\_LMP\_UP | MSigDB lists | WAMUNYOKOLI\_OVARIAN\_CANCER\_LMP\_UP | 206 | 5 | 12187 | 98 | Mgll,S100a6,Actb,Psmb10,Flnb | | 2.530e-02 | -3.68 | GO\_POSITIVE\_REGULATION\_OF\_INTRACELLULAR\_TRANSPORT | MSigDB lists | GO\_POSITIVE\_REGULATION\_OF\_INTRACELLULAR\_TRANSPORT | 280 | 6 | 12187 | 98 | Fcer1g,Thy1,Capn3,Hcls1,Rac2,Vamp8 | | 2.534e-02 | -3.68 | GO\_ENZYME\_REGULATOR\_ACTIVITY | MSigDB lists | GO\_ENZYME\_REGULATOR\_ACTIVITY | 703 | 11 | 12187 | 98 | C3,Serpinb9,Psme1,Capn3,Thy1,Rap1gap,Spint1,Serpina3n,Ctsa,Sh3bgrl3,Rac2 | | 2.535e-02 | -3.67 | IF\_rod\_dom | interpro domains | IPR039008 | 30 | 2 | 13788 | 114 | Gfap,Vim | | 2.540e-02 | -3.67 | IG | smart domains | SM00409 | 240 | 6 | 7188 | 68 | Fcgr3,Fcrls,Il6ra,Csf1r,Il1r1,Treml2 | | 2.540e-02 | -3.67 | MAPK1/MAPK3 signaling | REACTOME pathways | R-MMU-5684996 | 188 | 6 | 6297 | 76 | Camk2b,Psme1,Psmb10,Fgf18,Kl,Il6ra | | 2.553e-02 | -3.67 | enhancer sequence-specific DNA binding | molecular function | GO:0001158 | 80 | 3 | 13516 | 107 | Tfap2c,Ccar1,Bhlhe41 | | 2.553e-02 | -3.67 | cell body membrane | cellular component | GO:0044298 | 31 | 2 | 13825 | 111 | Il6ra,Thy1 | | 2.558e-02 | -3.67 | NIKOLSKY\_BREAST\_CANCER\_7P22\_AMPLICON | MSigDB lists | NIKOLSKY\_BREAST\_CANCER\_7P22\_AMPLICON | 31 | 2 | 12187 | 98 | Actb,Lfng | | 2.558e-02 | -3.67 | GNF2\_TYK2 | MSigDB lists | GNF2\_TYK2 | 31 | 2 | 12187 | 98 | Psmb10,Hcls1 | | 2.558e-02 | -3.67 | CASORELLI\_APL\_SECONDARY\_VS\_DE\_NOVO\_UP | MSigDB lists | CASORELLI\_APL\_SECONDARY\_VS\_DE\_NOVO\_UP | 31 | 2 | 12187 | 98 | Clu,Capn3 | | 2.558e-02 | -3.67 | MODULE\_286 | MSigDB lists | MODULE\_286 | 31 | 2 | 12187 | 98 | Adh1,Ldha | | 2.558e-02 | -3.67 | GO\_NEGATIVE\_REGULATION\_OF\_LEUKOCYTE\_APOPTOTIC\_PROCESS | MSigDB lists | GO\_NEGATIVE\_REGULATION\_OF\_LEUKOCYTE\_APOPTOTIC\_PROCESS | 31 | 2 | 12187 | 98 | Hcls1,Fcer1g | | 2.558e-02 | -3.67 | JIANG\_AGING\_CEREBRAL\_CORTEX\_UP | MSigDB lists | JIANG\_AGING\_CEREBRAL\_CORTEX\_UP | 31 | 2 | 12187 | 98 | Vim,Actb | | 2.558e-02 | -3.67 | LEE\_LIVER\_CANCER\_MYC\_DN | MSigDB lists | LEE\_LIVER\_CANCER\_MYC\_DN | 31 | 2 | 12187 | 98 | Il1r1,Serpina3n | | 2.558e-02 | -3.67 | GO\_INSULIN\_SECRETION | MSigDB lists | GO\_INSULIN\_SECRETION | 31 | 2 | 12187 | 98 | Ptprn,Mc4r | | 2.558e-02 | -3.67 | GO\_REGULATION\_OF\_LEUKOCYTE\_MEDIATED\_CYTOTOXICITY | MSigDB lists | GO\_REGULATION\_OF\_LEUKOCYTE\_MEDIATED\_CYTOTOXICITY | 31 | 2 | 12187 | 98 | Ptprc,Serpinb9 | | 2.558e-02 | -3.67 | GO\_NEGATIVE\_REGULATION\_OF\_LEUKOCYTE\_MEDIATED\_IMMUNITY | MSigDB lists | GO\_NEGATIVE\_REGULATION\_OF\_LEUKOCYTE\_MEDIATED\_IMMUNITY | 31 | 2 | 12187 | 98 | Serpinb9,Ptprc | | 2.566e-02 | -3.66 | ACEVEDO\_FGFR1\_TARGETS\_IN\_PROSTATE\_CANCER\_MODEL\_UP | MSigDB lists | ACEVEDO\_FGFR1\_TARGETS\_IN\_PROSTATE\_CANCER\_MODEL\_UP | 207 | 5 | 12187 | 98 | Ly6e,Lrg1,Clu,Camk2b,Serpina3n | | 2.568e-02 | -3.66 | GO\_SYNAPTIC\_VESICLE\_CYCLE | MSigDB lists | GO\_SYNAPTIC\_VESICLE\_CYCLE | 79 | 3 | 12187 | 98 | Cplx2,Slc17a7,Grn | | 2.568e-02 | -3.66 | LEE\_DIFFERENTIATING\_T\_LYMPHOCYTE | MSigDB lists | LEE\_DIFFERENTIATING\_T\_LYMPHOCYTE | 139 | 4 | 12187 | 98 | Rac2,Klhl6,Ptprc,Hcls1 | | 2.568e-02 | -3.66 | GSE2935\_UV\_INACTIVATED\_VS\_LIVE\_SENDAI\_VIRUS\_INF\_MACROPHAGE\_DN | MSigDB lists | GSE2935\_UV\_INACTIVATED\_VS\_LIVE\_SENDAI\_VIRUS\_INF\_MACROPHAGE\_DN | 139 | 4 | 12187 | 98 | Tnfsf8,Vim,Ahnak,S100a6 | | 2.568e-02 | -3.66 | GSE9037\_WT\_VS\_IRAK4\_KO\_LPS\_1H\_STIM\_BMDM\_DN | MSigDB lists | GSE9037\_WT\_VS\_IRAK4\_KO\_LPS\_1H\_STIM\_BMDM\_DN | 139 | 4 | 12187 | 98 | Mgp,Cplx2,Ly6e,Ccr5 | | 2.568e-02 | -3.66 | TONKS\_TARGETS\_OF\_RUNX1\_RUNX1T1\_FUSION\_HSC\_DN | MSigDB lists | TONKS\_TARGETS\_OF\_RUNX1\_RUNX1T1\_FUSION\_HSC\_DN | 139 | 4 | 12187 | 98 | Tsc22d1,Il6ra,Sh3bgrl3,Fcgr3 | | 2.570e-02 | -3.66 | protein-containing complex assembly | biological process | GO:0065003 | 1168 | 16 | 13711 | 111 | Capn3,Ahnak,Ndufa13,Mgp,Cyba,Dlgap3,Vwf,Brk1,Hcls1,Myh9,Rps5,Tspan4,Fcer1g,Vamp8,Slc2a1,Clu | | 2.571e-02 | -3.66 | Translation initiation complex formation | REACTOME pathways | R-MMU-72649 | 53 | 3 | 6297 | 76 | Rps20,Rps5,Rps8 | | 2.571e-02 | -3.66 | Fcgamma receptor (FCGR) dependent phagocytosis | REACTOME pathways | R-MMU-2029480 | 53 | 3 | 6297 | 76 | Actb,Fcgr3,Brk1 | | 2.592e-02 | -3.65 | regulation of intrinsic apoptotic signaling pathway in response to DNA damage | biological process | GO:1902229 | 31 | 2 | 13711 | 111 | Rpl26,Clu | | 2.592e-02 | -3.65 | negative regulation of lymphocyte mediated immunity | biological process | GO:0002707 | 31 | 2 | 13711 | 111 | Ptprc,Serpinb9 | | 2.598e-02 | -3.65 | regulation of peptidyl-tyrosine phosphorylation | biological process | GO:0050730 | 206 | 5 | 13711 | 111 | Hcls1,Il6ra,Csf1r,Ptprc,Thy1 | | 2.607e-02 | -3.65 | GO\_PROTEIN\_COMPLEX\_BIOGENESIS | MSigDB lists | GO\_PROTEIN\_COMPLEX\_BIOGENESIS | 892 | 13 | 12187 | 98 | Ndufa13,Brk1,Tspan4,Slc2a1,Lin7a,Vwf,Capn3,Ahnak,Fmod,Clu,Ctsz,Hcls1,Cyba | | 2.612e-02 | -3.64 | AP1\_Q4\_01 | MSigDB lists | AP1\_Q4\_01 | 208 | 5 | 12187 | 98 | Mark4,Cd68,Csf1r,Ptprn,Hcls1 | | 2.619e-02 | -3.64 | modification of morphology or physiology of other organism involved in symbiotic interaction | biological process | GO:0051817 | 79 | 3 | 13711 | 111 | Ccr5,Serpinb9,Csf1r | | 2.620e-02 | -3.64 | COMPLEMNTC1Q | prints domains | PR00007 | 18 | 2 | 2951 | 42 | C1qa,C1qb | | 2.625e-02 | -3.64 | focal adhesion | cellular component | GO:0005925 | 140 | 4 | 13825 | 111 | Myh9,Cyba,Ptprc,Flnb | | 2.628e-02 | -3.64 | GSE40274\_CTRL\_VS\_XBP1\_TRANSDUCED\_ACTIVATED\_CD4\_TCELL\_UP | MSigDB lists | GSE40274\_CTRL\_VS\_XBP1\_TRANSDUCED\_ACTIVATED\_CD4\_TCELL\_UP | 140 | 4 | 12187 | 98 | Ikzf3,Ahnak,Lrg1,Klhl6 | | 2.628e-02 | -3.64 | GSE18804\_SPLEEN\_MACROPHAGE\_VS\_BRAIN\_TUMORAL\_MACROPHAGE\_UP | MSigDB lists | GSE18804\_SPLEEN\_MACROPHAGE\_VS\_BRAIN\_TUMORAL\_MACROPHAGE\_UP | 140 | 4 | 12187 | 98 | Psme1,Fam111a,Vim,Lin7a | | 2.628e-02 | -3.64 | VEGF\_A\_UP.V1\_UP | MSigDB lists | VEGF\_A\_UP.V1\_UP | 140 | 4 | 12187 | 98 | Ly6e,Thy1,Ptprc,Serpina3n | | 2.628e-02 | -3.64 | GSE4748\_LPS\_VS\_LPS\_AND\_CYANOBACTERIUM\_LPSLIKE\_STIM\_DC\_3H\_UP | MSigDB lists | GSE4748\_LPS\_VS\_LPS\_AND\_CYANOBACTERIUM\_LPSLIKE\_STIM\_DC\_3H\_UP | 140 | 4 | 12187 | 98 | Fcer1g,Tmem106a,Sh3bgrl3,Csf1r | | 2.628e-02 | -3.64 | GSE34205\_HEALTHY\_VS\_FLU\_INF\_INFANT\_PBMC\_DN | MSigDB lists | GSE34205\_HEALTHY\_VS\_FLU\_INF\_INFANT\_PBMC\_DN | 140 | 4 | 12187 | 98 | S100a6,Grn,Ly6e,Ldha | | 2.628e-02 | -3.64 | GSE22432\_MULTIPOTENT\_PROGENITOR\_VS\_CDC\_UP | MSigDB lists | GSE22432\_MULTIPOTENT\_PROGENITOR\_VS\_CDC\_UP | 140 | 4 | 12187 | 98 | Psme1,Ly6e,Mark4,Treml2 | | 2.628e-02 | -3.64 | GSE30083\_SP1\_VS\_SP2\_THYMOCYTE\_DN | MSigDB lists | GSE30083\_SP1\_VS\_SP2\_THYMOCYTE\_DN | 140 | 4 | 12187 | 98 | Tnfsf8,Itgam,Fgf18,Adh1 | | 2.628e-02 | -3.64 | LEE\_LIVER\_CANCER\_SURVIVAL\_DN | MSigDB lists | LEE\_LIVER\_CANCER\_SURVIVAL\_DN | 140 | 4 | 12187 | 98 | S100a6,Dek,Rps5,Rpl31 | | 2.630e-02 | -3.64 | mouse chr6\|6 B2.3 | chromosome location | mouse chr6\|6 B2.3 | 32 | 2 | 14556 | 115 | Zfp786,Tmem176a | | 2.643e-02 | -3.63 | regulation of protein phosphorylation | biological process | GO:0001932 | 1172 | 16 | 13711 | 111 | C3,Fgf18,Csk,Il6ra,Actb,Rac2,Mt3,Csf1r,Hcls1,Tmem106a,Thy1,Kl,Ccl6,Clu,Hspa2,Ptprc | | 2.653e-02 | -3.63 | binding | molecular function | GO:0005488 | 9835 | 87 | 13516 | 107 | Rac2,Ccl6,Apod,Tspan4,Pde1a,Trnp1,Fcer1g,Serpinb9,Ndufa13,Gfap,Ccr5,Rpl14,Rbfox3,Adgre1,Sh2d5,Saa3,Adh1,Ikzf3,Capn3,Kl,Ldha,Ptprc,Ctsb,Tmem176a,Csf1r,Exoc3l4,Lrg1,Cplx2,Zfp786,Pcsk2,Tnfsf8,Rps5,Slc2a1,Grn,Ahnak,Psme1,Bhlhe41,C3,Brk1,C1qa,Mc4r,Ly6e,Csk,Mark4,Rpl31,Cox4i1,Ctsa,Cyba,Etnppl,Rap1gap,Vim,Tfap2c,Lgi4,Thy1,Phyhip,Mgll,Rab3il1,Tsc22d1,Il6ra,Mt3,Fgf18,Ccar1,Mgp,Itgam,Ptprn,Lfng,C1qb,Vamp8,Actb,Dek,Rps20,Ly6a,Hcls1,Dlgap3,Vwf,Ctss,Myh9,S100a6,Il1r1,Camk2b,Flnb,Fcgr3,Clu,Igfbp6,Hspa2,Rpl26,Lin7a | | 2.653e-02 | -3.63 | GO\_PROTEASE\_BINDING | MSigDB lists | GO\_PROTEASE\_BINDING | 80 | 3 | 12187 | 98 | Vwf,Serpinb9,Il1r1 | | 2.653e-02 | -3.63 | GO\_REGULATION\_OF\_ENDOTHELIAL\_CELL\_PROLIFERATION | MSigDB lists | GO\_REGULATION\_OF\_ENDOTHELIAL\_CELL\_PROLIFERATION | 80 | 3 | 12187 | 98 | Lrg1,Fgf18,Cyba | | 2.687e-02 | -3.62 | Signaling by Interleukins | REACTOME pathways | R-MMU-449147 | 299 | 8 | 6297 | 76 | Il1r1,Psmb10,Psme1,Kl,Il6ra,Csf1r,Camk2b,Fgf18 | | 2.689e-02 | -3.62 | GSE21063\_WT\_VS\_NFATC1\_KO\_BCELL\_UP | MSigDB lists | GSE21063\_WT\_VS\_NFATC1\_KO\_BCELL\_UP | 141 | 4 | 12187 | 98 | Tspan4,Sh3bgrl3,Ccr5,Thy1 | | 2.689e-02 | -3.62 | ATF2\_S\_UP.V1\_DN | MSigDB lists | ATF2\_S\_UP.V1\_DN | 141 | 4 | 12187 | 98 | Tmem176a,Il1r1,Serpina3n,Fmod | | 2.689e-02 | -3.62 | JAEGER\_METASTASIS\_DN | MSigDB lists | JAEGER\_METASTASIS\_DN | 141 | 4 | 12187 | 98 | Slc2a1,Ahnak,Tfap2c,Pcsk2 | | 2.689e-02 | -3.62 | GAUSSMANN\_MLL\_AF4\_FUSION\_TARGETS\_C\_UP | MSigDB lists | GAUSSMANN\_MLL\_AF4\_FUSION\_TARGETS\_C\_UP | 141 | 4 | 12187 | 98 | Rab3il1,Rps20,Ccar1,Lfng | | 2.689e-02 | -3.62 | GSE14415\_FOXP3\_KO\_NATURAL\_TREG\_VS\_TCONV\_DN | MSigDB lists | GSE14415\_FOXP3\_KO\_NATURAL\_TREG\_VS\_TCONV\_DN | 141 | 4 | 12187 | 98 | Rpl13,Tnfsf8,Klhl6,Treml2 | | 2.689e-02 | -3.62 | ASTON\_MAJOR\_DEPRESSIVE\_DISORDER\_DN | MSigDB lists | ASTON\_MAJOR\_DEPRESSIVE\_DISORDER\_DN | 141 | 4 | 12187 | 98 | Pcsk2,Hspa2,Capn3,Apod | | 2.689e-02 | -3.62 | GSE46242\_CTRL\_VS\_EGR2\_DELETED\_TH1\_CD4\_TCELL\_UP | MSigDB lists | GSE46242\_CTRL\_VS\_EGR2\_DELETED\_TH1\_CD4\_TCELL\_UP | 141 | 4 | 12187 | 98 | Treml2,Rap1gap,Igfbp6,Mc4r | | 2.689e-02 | -3.62 | GSE17974\_CTRL\_VS\_ACT\_IL4\_AND\_ANTI\_IL12\_2H\_CD4\_TCELL\_UP | MSigDB lists | GSE17974\_CTRL\_VS\_ACT\_IL4\_AND\_ANTI\_IL12\_2H\_CD4\_TCELL\_UP | 141 | 4 | 12187 | 98 | Tnfsf8,Ly6e,Tsc22d1,Fam111a | | 2.695e-02 | -3.61 | leukocyte migration | biological process | GO:0050900 | 140 | 4 | 13711 | 111 | Itgam,Ccl6,Fcer1g,Fcgr3 | | 2.699e-02 | -3.61 | Activation of the mRNA upon binding of the cap-binding complex and eIFs, and subsequent binding to 43S | REACTOME pathways | R-MMU-72662 | 54 | 3 | 6297 | 76 | Rps5,Rps20,Rps8 | | 2.699e-02 | -3.61 | Ribosomal scanning and start codon recognition | REACTOME pathways | R-MMU-72702 | 54 | 3 | 6297 | 76 | Rps8,Rps5,Rps20 | | 2.715e-02 | -3.61 | LEE\_CALORIE\_RESTRICTION\_MUSCLE\_UP | MSigDB lists | LEE\_CALORIE\_RESTRICTION\_MUSCLE\_UP | 32 | 2 | 12187 | 98 | Actb,Psme1 | | 2.715e-02 | -3.61 | GNATENKO\_PLATELET\_SIGNATURE | MSigDB lists | GNATENKO\_PLATELET\_SIGNATURE | 32 | 2 | 12187 | 98 | Clu,Actb | | 2.715e-02 | -3.61 | GO\_POSITIVE\_REGULATION\_OF\_DENDRITIC\_SPINE\_DEVELOPMENT | MSigDB lists | GO\_POSITIVE\_REGULATION\_OF\_DENDRITIC\_SPINE\_DEVELOPMENT | 32 | 2 | 12187 | 98 | Grn,Camk2b | | 2.715e-02 | -3.61 | HU\_GENOTOXIN\_ACTION\_DIRECT\_VS\_INDIRECT\_4HR | MSigDB lists | HU\_GENOTOXIN\_ACTION\_DIRECT\_VS\_INDIRECT\_4HR | 32 | 2 | 12187 | 98 | Vwf,Actb | | 2.715e-02 | -3.61 | NOUSHMEHR\_GBM\_SILENCED\_BY\_METHYLATION | MSigDB lists | NOUSHMEHR\_GBM\_SILENCED\_BY\_METHYLATION | 32 | 2 | 12187 | 98 | Ldha,Fmod | | 2.750e-02 | -3.59 | GSE3565\_CTRL\_VS\_LPS\_INJECTED\_SPLENOCYTES\_DN | MSigDB lists | GSE3565\_CTRL\_VS\_LPS\_INJECTED\_SPLENOCYTES\_DN | 142 | 4 | 12187 | 98 | Rpl13,Ly6e,Tnfsf8,Treml2 | | 2.750e-02 | -3.59 | GSE2585\_THYMIC\_MACROPHAGE\_VS\_MTEC\_UP | MSigDB lists | GSE2585\_THYMIC\_MACROPHAGE\_VS\_MTEC\_UP | 142 | 4 | 12187 | 98 | Hcls1,Vim,Ctss,Psme1 | | 2.750e-02 | -3.59 | GSE360\_L\_DONOVANI\_VS\_B\_MALAYI\_HIGH\_DOSE\_DC\_UP | MSigDB lists | GSE360\_L\_DONOVANI\_VS\_B\_MALAYI\_HIGH\_DOSE\_DC\_UP | 142 | 4 | 12187 | 98 | Ldha,Mgll,Camk2b,Fcgr3 | | 2.750e-02 | -3.59 | GSE37416\_CTRL\_VS\_3H\_F\_TULARENSIS\_LVS\_NEUTROPHIL\_UP | MSigDB lists | GSE37416\_CTRL\_VS\_3H\_F\_TULARENSIS\_LVS\_NEUTROPHIL\_UP | 142 | 4 | 12187 | 98 | Ctsz,Rac2,S100a6,Ptprc | | 2.750e-02 | -3.59 | GSE9006\_TYPE\_1\_DIABETES\_AT\_DX\_VS\_1MONTH\_POST\_DX\_PBMC\_DN | MSigDB lists | GSE9006\_TYPE\_1\_DIABETES\_AT\_DX\_VS\_1MONTH\_POST\_DX\_PBMC\_DN | 142 | 4 | 12187 | 98 | Rpl14,Pcsk2,Fam111a,Cox4i1 | | 2.751e-02 | -3.59 | catecholamine metabolic process | biological process | GO:0006584 | 32 | 2 | 13711 | 111 | Ly6e,Kl | | 2.751e-02 | -3.59 | bone remodeling | biological process | GO:0046849 | 32 | 2 | 13711 | 111 | Ctss,Rac2 | | 2.751e-02 | -3.59 | catechol-containing compound metabolic process | biological process | GO:0009712 | 32 | 2 | 13711 | 111 | Ly6e,Kl | | 2.751e-02 | -3.59 | neuroinflammatory response | biological process | GO:0150076 | 32 | 2 | 13711 | 111 | Grn,Clu | | 2.757e-02 | -3.59 | KAAB\_HEART\_ATRIUM\_VS\_VENTRICLE\_UP | MSigDB lists | KAAB\_HEART\_ATRIUM\_VS\_VENTRICLE\_UP | 211 | 5 | 12187 | 98 | Vim,Igfbp6,Fmod,Grn,Il1r1 | | 2.757e-02 | -3.59 | signal release | biological process | GO:0023061 | 141 | 4 | 13711 | 111 | Lin7a,Cplx2,Mc4r,Ptprn | | 2.757e-02 | -3.59 | GO\_MACROMOLECULAR\_COMPLEX\_BINDING | MSigDB lists | GO\_MACROMOLECULAR\_COMPLEX\_BINDING | 1092 | 15 | 12187 | 98 | Ctss,Vwf,Hcls1,Vim,Mark4,Brk1,Pcsk2,Myh9,Thy1,Fcer1g,Tspan4,Gfap,Ctsb,Actb,Fcgr3 | | 2.761e-02 | -3.59 | GO\_REGULATION\_OF\_HYDROLASE\_ACTIVITY | MSigDB lists | GO\_REGULATION\_OF\_HYDROLASE\_ACTIVITY | 995 | 14 | 12187 | 98 | Sh3bgrl3,Serpina3n,Rab3il1,Fgf18,Psme1,Hspa2,Camk2b,Rap1gap,Kl,C3,Spint1,Serpinb9,Ndufa13,Thy1 | | 2.773e-02 | -3.59 | - | gene3d domains | 4.10.10.10 | 3 | 1 | 6647 | 62 | Mt3 | | 2.773e-02 | -3.59 | - | gene3d domains | 1.20.120.180 | 3 | 1 | 6647 | 62 | Psme1 | | 2.773e-02 | -3.59 | - | gene3d domains | 3.30.70.600 | 3 | 1 | 6647 | 62 | Rps20 | | 2.773e-02 | -3.59 | - | gene3d domains | 1.20.5.120 | 3 | 1 | 6647 | 62 | Psme1 | | 2.773e-02 | -3.59 | - | gene3d domains | 3.40.50.10260 | 3 | 1 | 6647 | 62 | Ndufa13 | | 2.792e-02 | -3.58 | regulation of exocytosis | biological process | GO:0017157 | 210 | 5 | 13711 | 111 | Vamp8,Cplx2,Fcer1g,Rac2,Itgam | | 2.793e-02 | -3.58 | plasma membrane organization | biological process | GO:0007009 | 81 | 3 | 13711 | 111 | Myh9,Ptprc,Clu | | 2.799e-02 | -3.58 | GO\_NEGATIVE\_REGULATION\_OF\_MULTICELLULAR\_ORGANISMAL\_PROCESS | MSigDB lists | GO\_NEGATIVE\_REGULATION\_OF\_MULTICELLULAR\_ORGANISMAL\_PROCESS | 714 | 11 | 12187 | 98 | Rap1gap,Thy1,Capn3,Lfng,Csk,Tmem176a,Gfap,Il6ra,Vim,Cyba,Apod | | 2.806e-02 | -3.57 | DANG\_REGULATED\_BY\_MYC\_DN | MSigDB lists | DANG\_REGULATED\_BY\_MYC\_DN | 212 | 5 | 12187 | 98 | Ptprn,Ctsb,Thy1,Csk,Mgp | | 2.812e-02 | -3.57 | VWD | smart domains | SM00216 | 3 | 1 | 7188 | 68 | Vwf | | 2.812e-02 | -3.57 | Ribosomal\_S10 | smart domains | SM01403 | 3 | 1 | 7188 | 68 | Rps20 | | 2.812e-02 | -3.57 | SAA | smart domains | SM00197 | 3 | 1 | 7188 | 68 | Saa3 | | 2.812e-02 | -3.57 | GO\_REGULATION\_OF\_EXOCYTOSIS | MSigDB lists | GO\_REGULATION\_OF\_EXOCYTOSIS | 143 | 4 | 12187 | 98 | Cplx2,Vamp8,Fcer1g,Rac2 | | 2.812e-02 | -3.57 | GSE30083\_SP3\_VS\_SP4\_THYMOCYTE\_DN | MSigDB lists | GSE30083\_SP3\_VS\_SP4\_THYMOCYTE\_DN | 143 | 4 | 12187 | 98 | Mpeg1,Cd68,Ahnak,Grn | | 2.812e-02 | -3.57 | GSE3982\_NEUTROPHIL\_VS\_TH1\_UP | MSigDB lists | GSE3982\_NEUTROPHIL\_VS\_TH1\_UP | 143 | 4 | 12187 | 98 | Serpina3n,Il1r1,Tfap2c,Rac2 | | 2.812e-02 | -3.57 | GSE4984\_UNTREATED\_VS\_GALECTIN1\_TREATED\_DC\_UP | MSigDB lists | GSE4984\_UNTREATED\_VS\_GALECTIN1\_TREATED\_DC\_UP | 143 | 4 | 12187 | 98 | Ctsb,Sh3bgrl3,Tmem106a,Fcer1g | | 2.812e-02 | -3.57 | SERVITJA\_ISLET\_HNF1A\_TARGETS\_UP | MSigDB lists | SERVITJA\_ISLET\_HNF1A\_TARGETS\_UP | 143 | 4 | 12187 | 98 | C1qb,Cyba,Vim,Thy1 | | 2.812e-02 | -3.57 | GO\_NEGATIVE\_REGULATION\_OF\_PEPTIDASE\_ACTIVITY | MSigDB lists | GO\_NEGATIVE\_REGULATION\_OF\_PEPTIDASE\_ACTIVITY | 143 | 4 | 12187 | 98 | Spint1,Serpina3n,C3,Serpinb9 | | 2.812e-02 | -3.57 | GSE1460\_INTRATHYMIC\_T\_PROGENITOR\_VS\_THYMIC\_STROMAL\_CELL\_UP | MSigDB lists | GSE1460\_INTRATHYMIC\_T\_PROGENITOR\_VS\_THYMIC\_STROMAL\_CELL\_UP | 143 | 4 | 12187 | 98 | Cyba,Fgf18,Rps8,Capn3 | | 2.812e-02 | -3.57 | RIGGI\_EWING\_SARCOMA\_PROGENITOR\_DN | MSigDB lists | RIGGI\_EWING\_SARCOMA\_PROGENITOR\_DN | 143 | 4 | 12187 | 98 | Clu,Igfbp6,Pde1a,Il1r1 | | 2.827e-02 | -3.57 | GLHYDRLASE1 | prints domains | PR00131 | 2 | 1 | 2951 | 42 | Kl | | 2.827e-02 | -3.57 | CRBOXYPTASEC | prints domains | PR00724 | 2 | 1 | 2951 | 42 | Ctsa | | 2.827e-02 | -3.57 | MASPIN | prints domains | PR00676 | 2 | 1 | 2951 | 42 | Serpinb9 | | 2.827e-02 | -3.57 | MCRFAMILY | prints domains | PR00534 | 2 | 1 | 2951 | 42 | Mc4r | | 2.827e-02 | -3.57 | CYTCOXIDASE4 | prints domains | PR01873 | 2 | 1 | 2951 | 42 | Cox4i1 | | 2.827e-02 | -3.57 | ANAPHYLATOXN | prints domains | PR00004 | 2 | 1 | 2951 | 42 | C3 | | 2.827e-02 | -3.57 | MELNOCORTINR | prints domains | PR00535 | 2 | 1 | 2951 | 42 | Mc4r | | 2.827e-02 | -3.57 | synapse organization | biological process | GO:0050808 | 285 | 6 | 13711 | 111 | C1qa,Camk2b,Actb,C1qb,C3,Dlgap3 | | 2.827e-02 | -3.57 | REACTOME\_DOWNSTREAM\_SIGNALING\_OF\_ACTIVATED\_FGFR | MSigDB lists | REACTOME\_DOWNSTREAM\_SIGNALING\_OF\_ACTIVATED\_FGFR | 82 | 3 | 12187 | 98 | Fgf18,Kl,Pde1a | | 2.827e-02 | -3.57 | GSE21927\_HEALTHY\_VS\_TUMOROUS\_BALBC\_MOUSE\_MONOCYTE\_DN | MSigDB lists | GSE21927\_HEALTHY\_VS\_TUMOROUS\_BALBC\_MOUSE\_MONOCYTE\_DN | 82 | 3 | 12187 | 98 | Klhl6,Psme1,Vamp8 | | 2.827e-02 | -3.57 | GO\_REGULATION\_OF\_ADAPTIVE\_IMMUNE\_RESPONSE | MSigDB lists | GO\_REGULATION\_OF\_ADAPTIVE\_IMMUNE\_RESPONSE | 82 | 3 | 12187 | 98 | C3,Ptprc,Fcer1g | | 2.872e-02 | -3.55 | vacuolar membrane | cellular component | GO:0005774 | 144 | 4 | 13825 | 111 | Vamp8,Grn,Gfap,Cd68 | | 2.875e-02 | -3.55 | GO\_PLATELET\_AGGREGATION | MSigDB lists | GO\_PLATELET\_AGGREGATION | 33 | 2 | 12187 | 98 | Myh9,Actb | | 2.875e-02 | -3.55 | CTCTATG\_MIR368 | MSigDB lists | CTCTATG\_MIR368 | 33 | 2 | 12187 | 98 | Trnp1,Capn3 | | 2.875e-02 | -3.55 | MORF\_THRA | MSigDB lists | MORF\_THRA | 33 | 2 | 12187 | 98 | Slc2a1,Slc17a7 | | 2.875e-02 | -3.55 | GO\_POSITIVE\_REGULATION\_OF\_CALCIUM\_ION\_IMPORT | MSigDB lists | GO\_POSITIVE\_REGULATION\_OF\_CALCIUM\_ION\_IMPORT | 33 | 2 | 12187 | 98 | Capn3,Thy1 | | 2.875e-02 | -3.55 | BERTUCCI\_INVASIVE\_CARCINOMA\_DUCTAL\_VS\_LOBULAR\_DN | MSigDB lists | BERTUCCI\_INVASIVE\_CARCINOMA\_DUCTAL\_VS\_LOBULAR\_DN | 33 | 2 | 12187 | 98 | Rpl31,Vwf | | 2.875e-02 | -3.55 | GO\_ASTROCYTE\_DIFFERENTIATION | MSigDB lists | GO\_ASTROCYTE\_DIFFERENTIATION | 33 | 2 | 12187 | 98 | Vim,Gfap | | 2.875e-02 | -3.55 | WANG\_LSD1\_TARGETS\_DN | MSigDB lists | WANG\_LSD1\_TARGETS\_DN | 33 | 2 | 12187 | 98 | Pcsk2,Cplx2 | | 2.875e-02 | -3.55 | HALLMARK\_HEDGEHOG\_SIGNALING | MSigDB lists | HALLMARK\_HEDGEHOG\_SIGNALING | 33 | 2 | 12187 | 98 | Thy1,Myh9 | | 2.875e-02 | -3.55 | NAKAJIMA\_MAST\_CELL | MSigDB lists | NAKAJIMA\_MAST\_CELL | 33 | 2 | 12187 | 98 | Clu,Tspan4 | | 2.875e-02 | -3.55 | ALONSO\_METASTASIS\_EMT\_UP | MSigDB lists | ALONSO\_METASTASIS\_EMT\_UP | 33 | 2 | 12187 | 98 | Serpina3n,Ctsb | | 2.875e-02 | -3.55 | GO\_SIGNAL\_RELEASE | MSigDB lists | GO\_SIGNAL\_RELEASE | 144 | 4 | 12187 | 98 | Mc4r,Cplx2,Lin7a,Ptprn | | 2.875e-02 | -3.55 | GSE3982\_NEUTROPHIL\_VS\_TH2\_UP | MSigDB lists | GSE3982\_NEUTROPHIL\_VS\_TH2\_UP | 144 | 4 | 12187 | 98 | Rac2,Csk,Il1r1,Serpina3n | | 2.875e-02 | -3.55 | GSE7831\_1H\_VS\_4H\_INFLUENZA\_STIM\_PDC\_DN | MSigDB lists | GSE7831\_1H\_VS\_4H\_INFLUENZA\_STIM\_PDC\_DN | 144 | 4 | 12187 | 98 | Sh3bgrl3,Sh2d5,Il1r1,Csf1r | | 2.882e-02 | -3.55 | organic hydroxy compound transport | biological process | GO:0015850 | 82 | 3 | 13711 | 111 | Fcer1g,Fcgr3,Ly6e | | 2.882e-02 | -3.55 | endocrine system development | biological process | GO:0035270 | 82 | 3 | 13711 | 111 | Clu,Ly6e,Il6ra | | 2.882e-02 | -3.55 | regulation of calcium ion transport into cytosol | biological process | GO:0010522 | 82 | 3 | 13711 | 111 | Cyba,Capn3,Thy1 | | 2.883e-02 | -3.55 | mouse chr11 | chromosome location | mouse chr11 | 374 | 7 | 14556 | 115 | Camk2b,Grn,Cd68,Wfdc17,Rbfox3,Gfap,Ccl6 | | 2.883e-02 | -3.55 | immune response-activating signal transduction | biological process | GO:0002757 | 143 | 4 | 13711 | 111 | Ptprc,Thy1,Klhl6,Fcer1g | | 2.888e-02 | -3.54 | positive regulation of phosphate metabolic process | biological process | GO:0045937 | 896 | 13 | 13711 | 111 | Mt3,Hcls1,Csf1r,C3,Fgf18,Il6ra,Csk,Clu,Ccl6,Hspa2,Ptprc,Tmem106a,Kl | | 2.888e-02 | -3.54 | positive regulation of phosphorus metabolic process | biological process | GO:0010562 | 896 | 13 | 13711 | 111 | Tmem106a,Kl,Clu,Ccl6,Ptprc,Hspa2,C3,Il6ra,Csk,Fgf18,Csf1r,Hcls1,Mt3 | | 2.896e-02 | -3.54 | peptidase activity, acting on L-amino acid peptides | molecular function | GO:0070011 | 374 | 7 | 13516 | 107 | Ctss,Psmb10,Capn3,Ctsb,Pcsk2,Ctsz,Ctsa | | 2.902e-02 | -3.54 | GO\_POSITIVE\_REGULATION\_OF\_CATALYTIC\_ACTIVITY | MSigDB lists | GO\_POSITIVE\_REGULATION\_OF\_CATALYTIC\_ACTIVITY | 1198 | 16 | 12187 | 98 | Ctsa,Csf1r,Ndufa13,Thy1,Ptprc,Psmb10,Fgf18,Rab3il1,Sh3bgrl3,Kl,Csk,Hspa2,Psme1,Clu,Camk2b,Rap1gap | | 2.906e-02 | -3.54 | WANG\_MLL\_TARGETS | MSigDB lists | WANG\_MLL\_TARGETS | 214 | 5 | 12187 | 98 | Mgp,Igfbp6,Rab3il1,Adh1,Cyba | | 2.906e-02 | -3.54 | GO\_RIBONUCLEOPROTEIN\_COMPLEX\_BIOGENESIS | MSigDB lists | GO\_RIBONUCLEOPROTEIN\_COMPLEX\_BIOGENESIS | 369 | 7 | 12187 | 98 | Rpl26,Rpl13,Rps8,Rps5,Rpl14,Rpl31,Rps20 | | 2.917e-02 | -3.53 | AMUNDSON\_POOR\_SURVIVAL\_AFTER\_GAMMA\_RADIATION\_8G | MSigDB lists | AMUNDSON\_POOR\_SURVIVAL\_AFTER\_GAMMA\_RADIATION\_8G | 83 | 3 | 12187 | 98 | Tspan4,Hcls1,Ptprc | | 2.934e-02 | -3.53 | VWA | pfam domains | PF00092 | 32 | 2 | 12881 | 108 | Itgam,Vwf | | 2.935e-02 | -3.53 | cell-substrate adherens junction | cellular component | GO:0005924 | 145 | 4 | 13825 | 111 | Myh9,Cyba,Ptprc,Flnb | | 2.939e-02 | -3.53 | GSE14415\_INDUCED\_TREG\_VS\_FAILED\_INDUCED\_TREG\_DN | MSigDB lists | GSE14415\_INDUCED\_TREG\_VS\_FAILED\_INDUCED\_TREG\_DN | 145 | 4 | 12187 | 98 | Ahnak,Itgam,Fcer1g,Vamp8 | | 2.939e-02 | -3.53 | GSE22886\_NAIVE\_BCELL\_VS\_NEUTROPHIL\_UP | MSigDB lists | GSE22886\_NAIVE\_BCELL\_VS\_NEUTROPHIL\_UP | 145 | 4 | 12187 | 98 | Rpl26,Serpinb9,Rps5,Rps20 | | 2.939e-02 | -3.53 | GO\_LYMPHOCYTE\_DIFFERENTIATION | MSigDB lists | GO\_LYMPHOCYTE\_DIFFERENTIATION | 145 | 4 | 12187 | 98 | Ptprc,Fcer1g,Tnfsf8,Lfng | | 2.943e-02 | -3.53 | GO\_REGULATION\_OF\_VESICLE\_MEDIATED\_TRANSPORT | MSigDB lists | GO\_REGULATION\_OF\_VESICLE\_MEDIATED\_TRANSPORT | 370 | 7 | 12187 | 98 | Cyba,Vamp8,Cplx2,Rac2,C3,Csk,Fcer1g | | 2.948e-02 | -3.52 | regulation of smooth muscle cell proliferation | biological process | GO:0048660 | 144 | 4 | 13711 | 111 | Cyba,Pde1a,Apod,Il6ra | | 2.963e-02 | -3.52 | ONKEN\_UVEAL\_MELANOMA\_DN | MSigDB lists | ONKEN\_UVEAL\_MELANOMA\_DN | 454 | 8 | 12187 | 98 | Rpl31,Rps5,Rpl14,Rpl13,Hspa2,Rps8,Serpinb9,Vamp8 | | 2.963e-02 | -3.52 | GO\_REGULATION\_OF\_ION\_TRANSPORT | MSigDB lists | GO\_REGULATION\_OF\_ION\_TRANSPORT | 454 | 8 | 12187 | 98 | Ahnak,Camk2b,Thy1,Hspa2,Capn3,Ctss,Vamp8,Cyba | | 3.000e-02 | -3.51 | dendrite | cellular component | GO:0030425 | 630 | 10 | 13825 | 111 | Camk2b,Apod,Mark4,Clu,Rap1gap,Cplx2,Cyba,Thy1,Dlgap3,Pcsk2 | | 3.000e-02 | -3.51 | sarcolemma | cellular component | GO:0042383 | 146 | 4 | 13825 | 111 | Capn3,Ctsb,Slc2a1,Ahnak | | 3.004e-02 | -3.51 | GSE2770\_TGFB\_AND\_IL4\_ACT\_VS\_ACT\_CD4\_TCELL\_2H\_UP | MSigDB lists | GSE2770\_TGFB\_AND\_IL4\_ACT\_VS\_ACT\_CD4\_TCELL\_2H\_UP | 146 | 4 | 12187 | 98 | Il1r1,Igfbp6,Rpl13,Rps8 | | 3.004e-02 | -3.51 | GSE42021\_TCONV\_PLN\_VS\_CD24HI\_TCONV\_THYMUS\_DN | MSigDB lists | GSE42021\_TCONV\_PLN\_VS\_CD24HI\_TCONV\_THYMUS\_DN | 146 | 4 | 12187 | 98 | Ly6e,Rap1gap,Mgll,Hspa2 | | 3.004e-02 | -3.51 | GSE3982\_NEUTROPHIL\_VS\_BASOPHIL\_UP | MSigDB lists | GSE3982\_NEUTROPHIL\_VS\_BASOPHIL\_UP | 146 | 4 | 12187 | 98 | Cyba,Treml2,Camk2b,Mgll | | 3.004e-02 | -3.51 | MODULE\_63 | MSigDB lists | MODULE\_63 | 146 | 4 | 12187 | 98 | Itgam,Il6ra,Ptprn,Fcgr3 | | 3.004e-02 | -3.51 | GSE26030\_UNSTIM\_VS\_RESTIM\_TH17\_DAY5\_POST\_POLARIZATION\_DN | MSigDB lists | GSE26030\_UNSTIM\_VS\_RESTIM\_TH17\_DAY5\_POST\_POLARIZATION\_DN | 146 | 4 | 12187 | 98 | Vim,Myh9,Ppfibp1,Actb | | 3.004e-02 | -3.51 | GSE5589\_LPS\_VS\_LPS\_AND\_IL10\_STIM\_IL10\_KO\_MACROPHAGE\_45MIN\_DN | MSigDB lists | GSE5589\_LPS\_VS\_LPS\_AND\_IL10\_STIM\_IL10\_KO\_MACROPHAGE\_45MIN\_DN | 146 | 4 | 12187 | 98 | Treml2,Ikzf3,Psme1,Rac2 | | 3.004e-02 | -3.51 | GSE40685\_TREG\_VS\_FOXP3\_KO\_TREG\_PRECURSOR\_DN | MSigDB lists | GSE40685\_TREG\_VS\_FOXP3\_KO\_TREG\_PRECURSOR\_DN | 146 | 4 | 12187 | 98 | Psme1,C1qa,C1qb,Fcgr3 | | 3.008e-02 | -3.50 | GO\_REGULATION\_OF\_SMOOTH\_MUSCLE\_CELL\_PROLIFERATION | MSigDB lists | GO\_REGULATION\_OF\_SMOOTH\_MUSCLE\_CELL\_PROLIFERATION | 84 | 3 | 12187 | 98 | Il6ra,Apod,Cyba | | 3.008e-02 | -3.50 | MODULE\_324 | MSigDB lists | MODULE\_324 | 84 | 3 | 12187 | 98 | Adh1,Mgp,Itgam | | 3.008e-02 | -3.50 | KEGG\_FC\_GAMMA\_R\_MEDIATED\_PHAGOCYTOSIS | MSigDB lists | KEGG\_FC\_GAMMA\_R\_MEDIATED\_PHAGOCYTOSIS | 84 | 3 | 12187 | 98 | Rac2,Ptprc,Fcgr3 | | 3.008e-02 | -3.50 | GO\_REGULATION\_OF\_STAT\_CASCADE | MSigDB lists | GO\_REGULATION\_OF\_STAT\_CASCADE | 84 | 3 | 12187 | 98 | Il6ra,Csf1r,Hcls1 | | 3.019e-02 | -3.50 | GO\_ACTIN\_FILAMENT\_BASED\_PROCESS | MSigDB lists | GO\_ACTIN\_FILAMENT\_BASED\_PROCESS | 372 | 7 | 12187 | 98 | Hcls1,Myh9,Capn3,Brk1,Flnb,Rac2,Vim | | 3.032e-02 | -3.50 | Systemic lupus erythematosus | KEGG pathways | ko05322 | 56 | 3 | 5248 | 64 | C1qa,C1qb,C3 | | 3.032e-02 | -3.50 | Systemic lupus erythematosus | KEGG pathways | mmu05322 | 56 | 3 | 5248 | 64 | C3,C1qb,C1qa | | 3.038e-02 | -3.49 | Filament | smart domains | SM01391 | 29 | 2 | 7188 | 68 | Gfap,Vim | | 3.039e-02 | -3.49 | GO\_POSITIVE\_REGULATION\_OF\_CALCIUM\_ION\_TRANSPORT\_INTO\_CYTOSOL | MSigDB lists | GO\_POSITIVE\_REGULATION\_OF\_CALCIUM\_ION\_TRANSPORT\_INTO\_CYTOSOL | 34 | 2 | 12187 | 98 | Capn3,Thy1 | | 3.039e-02 | -3.49 | GCM\_PTPRU | MSigDB lists | GCM\_PTPRU | 34 | 2 | 12187 | 98 | Fcgr3,Tmem106a | | 3.039e-02 | -3.49 | GO\_CALCIUM\_ION\_IMPORT\_INTO\_CYTOSOL | MSigDB lists | GO\_CALCIUM\_ION\_IMPORT\_INTO\_CYTOSOL | 34 | 2 | 12187 | 98 | Ccr5,Ptprc | | 3.039e-02 | -3.49 | GO\_PROTEIN\_SELF\_ASSOCIATION | MSigDB lists | GO\_PROTEIN\_SELF\_ASSOCIATION | 34 | 2 | 12187 | 98 | Ctsb,Slc2a1 | | 3.055e-02 | -3.49 | SAA | prosite domains | PS00992 | 3 | 1 | 8845 | 91 | Saa3 | | 3.055e-02 | -3.49 | METALLOTHIONEIN\_VRT | prosite domains | PS00203 | 3 | 1 | 8845 | 91 | Mt3 | | 3.055e-02 | -3.49 | YJEF\_N | prosite domains | PS51385 | 3 | 1 | 8845 | 91 | Ndufa13 | | 3.055e-02 | -3.49 | LAMP\_3 | prosite domains | PS51407 | 3 | 1 | 8845 | 91 | Cd68 | | 3.055e-02 | -3.49 | VWFD | prosite domains | PS51233 | 3 | 1 | 8845 | 91 | Vwf | | 3.055e-02 | -3.49 | LAMP\_2 | prosite domains | PS00311 | 3 | 1 | 8845 | 91 | Cd68 | | 3.055e-02 | -3.49 | ADH\_ZINC | prosite domains | PS00059 | 3 | 1 | 8845 | 91 | Adh1 | | 3.060e-02 | -3.49 | chr1p36 | MSigDB lists | chr1p36 | 217 | 5 | 12187 | 98 | Rap1gap,Trnp1,C1qb,Sh2d5,C1qa | | 3.069e-02 | -3.48 | GSE8835\_CD4\_VS\_CD8\_TCELL\_CLL\_PATIENT\_UP | MSigDB lists | GSE8835\_CD4\_VS\_CD8\_TCELL\_CLL\_PATIENT\_UP | 147 | 4 | 12187 | 98 | Ly6e,Serpinb9,Actb,Apod | | 3.069e-02 | -3.48 | GSE39820\_IL1B\_IL6\_VS\_IL1B\_IL6\_IL23A\_TREATED\_CD4\_TCELL\_DN | MSigDB lists | GSE39820\_IL1B\_IL6\_VS\_IL1B\_IL6\_IL23A\_TREATED\_CD4\_TCELL\_DN | 147 | 4 | 12187 | 98 | Phyhip,Ppfibp1,Ahnak,Vim | | 3.069e-02 | -3.48 | GSE13485\_CTRL\_VS\_DAY3\_YF17D\_VACCINE\_PBMC\_DN | MSigDB lists | GSE13485\_CTRL\_VS\_DAY3\_YF17D\_VACCINE\_PBMC\_DN | 147 | 4 | 12187 | 98 | Csk,Ly6e,C1qa,C1qb | | 3.069e-02 | -3.48 | GSE36095\_WT\_VS\_HDAC9\_KO\_TREG\_UP | MSigDB lists | GSE36095\_WT\_VS\_HDAC9\_KO\_TREG\_UP | 147 | 4 | 12187 | 98 | S100a6,Ppfibp1,Ctsz,Ldha | | 3.074e-02 | -3.48 | negative regulation of protein metabolic process | biological process | GO:0051248 | 904 | 13 | 13711 | 111 | Spint1,Actb,C3,Serpina3n,Csk,Serpinb9,Capn3,Ctsa,Clu,Ctsz,Ptprc,Apod,Thy1 | | 3.079e-02 | -3.48 | fibroblast growth factor receptor signaling pathway | biological process | GO:0008543 | 34 | 2 | 13711 | 111 | Fgf18,Kl | | 3.085e-02 | -3.48 | dendritic tree | cellular component | GO:0097447 | 633 | 10 | 13825 | 111 | Camk2b,Mark4,Apod,Clu,Cplx2,Rap1gap,Thy1,Cyba,Dlgap3,Pcsk2 | | 3.100e-02 | -3.47 | GO\_OVULATION\_CYCLE | MSigDB lists | GO\_OVULATION\_CYCLE | 85 | 3 | 12187 | 98 | Lfng,Clu,Ptprn | | 3.100e-02 | -3.47 | GSE29614\_DAY3\_VS\_DAY7\_TIV\_FLU\_VACCINE\_PBMC\_UP | MSigDB lists | GSE29614\_DAY3\_VS\_DAY7\_TIV\_FLU\_VACCINE\_PBMC\_UP | 85 | 3 | 12187 | 98 | Tnfsf8,Ly6e,C1qb | | 3.112e-02 | -3.47 | apical part of cell | cellular component | GO:0045177 | 294 | 6 | 13825 | 111 | Cyba,Il6ra,Thy1,Slc26a2,Slc2a1,Ctsb | | 3.123e-02 | -3.47 | mouse chr14 D1\|14 34.36 cM | chromosome location | mouse chr14 D1\|14 34.36 cM | 4 | 1 | 14556 | 115 | Clu | | 3.123e-02 | -3.47 | mouse chr12 C3\|12 33.73 cM | chromosome location | mouse chr12 C3\|12 33.73 cM | 4 | 1 | 14556 | 115 | Hspa2 | | 3.123e-02 | -3.47 | mouse chr6 C1\|6 32.27 cM | chromosome location | mouse chr6 C1\|6 32.27 cM | 4 | 1 | 14556 | 115 | Vamp8 | | 3.123e-02 | -3.47 | mouse chr6 G3\|6 77.7 cM | chromosome location | mouse chr6 G3\|6 77.7 cM | 4 | 1 | 14556 | 115 | Bhlhe41 | | 3.130e-02 | -3.46 | immunoglobulin receptor binding | molecular function | GO:0034987 | 4 | 1 | 13516 | 107 | Fcgr3 | | 3.130e-02 | -3.46 | kininogen binding | molecular function | GO:0030984 | 4 | 1 | 13516 | 107 | Ctsb | | 3.130e-02 | -3.46 | lactate dehydrogenase activity | molecular function | GO:0004457 | 4 | 1 | 13516 | 107 | Ldha | | 3.130e-02 | -3.46 | IgG receptor activity | molecular function | GO:0019770 | 4 | 1 | 13516 | 107 | Fcgr3 | | 3.130e-02 | -3.46 | beta-glucosidase activity | molecular function | GO:0008422 | 4 | 1 | 13516 | 107 | Kl | | 3.130e-02 | -3.46 | sodium:phosphate symporter activity | molecular function | GO:0005436 | 4 | 1 | 13516 | 107 | Slc17a7 | | 3.130e-02 | -3.46 | L27 domain binding | molecular function | GO:0097016 | 4 | 1 | 13516 | 107 | Lin7a | | 3.130e-02 | -3.46 | carbon-oxygen lyase activity, acting on phosphates | molecular function | GO:0016838 | 4 | 1 | 13516 | 107 | Etnppl | | 3.136e-02 | -3.46 | GSE32533\_MIR17\_KO\_VS\_MIR17\_OVEREXPRESS\_ACT\_CD4\_TCELL\_UP | MSigDB lists | GSE32533\_MIR17\_KO\_VS\_MIR17\_OVEREXPRESS\_ACT\_CD4\_TCELL\_UP | 148 | 4 | 12187 | 98 | Csk,Ptprc,Ctsb,Rab3il1 | | 3.136e-02 | -3.46 | HALLMARK\_XENOBIOTIC\_METABOLISM | MSigDB lists | HALLMARK\_XENOBIOTIC\_METABOLISM | 148 | 4 | 12187 | 98 | Psmb10,Il1r1,Adh1,Rap1gap | | 3.136e-02 | -3.46 | GSE20715\_WT\_VS\_TLR4\_KO\_LUNG\_DN | MSigDB lists | GSE20715\_WT\_VS\_TLR4\_KO\_LUNG\_DN | 148 | 4 | 12187 | 98 | Csf1r,C1qb,Vwf,Igfbp6 | | 3.136e-02 | -3.46 | GSE21774\_CD62L\_POS\_CD56\_BRIGHT\_VS\_CD62L\_NEG\_CD56\_DIM\_NK\_CELL\_UP | MSigDB lists | GSE21774\_CD62L\_POS\_CD56\_BRIGHT\_VS\_CD62L\_NEG\_CD56\_DIM\_NK\_CELL\_UP | 148 | 4 | 12187 | 98 | C3,Csf1r,Sh2d5,Mgp | | 3.136e-02 | -3.46 | GSE2405\_HEAT\_KILLED\_LYSATE\_VS\_LIVE\_A\_PHAGOCYTOPHILUM\_STIM\_NEUTROPHIL\_9H\_UP | MSigDB lists | GSE2405\_HEAT\_KILLED\_LYSATE\_VS\_LIVE\_A\_PHAGOCYTOPHILUM\_STIM\_NEUTROPHIL\_9H\_UP | 148 | 4 | 12187 | 98 | Fcer1g,Tsc22d1,Csf1r,C3 | | 3.136e-02 | -3.46 | GSE13547\_WT\_VS\_ZFX\_KO\_BCELL\_ANTI\_IGM\_STIM\_2H\_UP | MSigDB lists | GSE13547\_WT\_VS\_ZFX\_KO\_BCELL\_ANTI\_IGM\_STIM\_2H\_UP | 148 | 4 | 12187 | 98 | Vim,Serpinb9,Thy1,Ly6e | | 3.136e-02 | -3.46 | GSE5455\_HEALTHY\_VS\_TUMOR\_BEARING\_MOUSE\_SPLEEN\_MONOCYTE\_24H\_INCUBATION\_DN | MSigDB lists | GSE5455\_HEALTHY\_VS\_TUMOR\_BEARING\_MOUSE\_SPLEEN\_MONOCYTE\_24H\_INCUBATION\_DN | 148 | 4 | 12187 | 98 | Il1r1,Fam111a,Slc17a7,Csk | | 3.136e-02 | -3.46 | KRAS.DF.V1\_UP | MSigDB lists | KRAS.DF.V1\_UP | 148 | 4 | 12187 | 98 | Hspa2,Rac2,Slc26a2,Hcls1 | | 3.146e-02 | -3.46 | positive regulation of peptidyl-tyrosine phosphorylation | biological process | GO:0050731 | 147 | 4 | 13711 | 111 | Ptprc,Hcls1,Csf1r,Il6ra | | 3.153e-02 | -3.46 | regulation of body fluid levels | biological process | GO:0050878 | 217 | 5 | 13711 | 111 | Vamp8,C3,Cyba,Fcer1g,Vwf | | 3.160e-02 | -3.45 | positive regulation of exocytosis | biological process | GO:0045921 | 85 | 3 | 13711 | 111 | Vamp8,Itgam,Fcer1g | | 3.168e-02 | -3.45 | GO\_ENZYME\_LINKED\_RECEPTOR\_PROTEIN\_SIGNALING\_PATHWAY | MSigDB lists | GO\_ENZYME\_LINKED\_RECEPTOR\_PROTEIN\_SIGNALING\_PATHWAY | 547 | 9 | 12187 | 98 | Csf1r,Kl,Fmod,Csk,Brk1,Fgf18,Actb,Cyba,Vim | | 3.169e-02 | -3.45 | GO\_PROTEIN\_LOCALIZATION\_TO\_ORGANELLE | MSigDB lists | GO\_PROTEIN\_LOCALIZATION\_TO\_ORGANELLE | 460 | 8 | 12187 | 98 | Rps5,Rpl14,Rpl31,Rps20,Rpl13,Ndufa13,Rps8,Rpl26 | | 3.173e-02 | -3.45 | neurofibrillary tangle | cellular component | GO:0097418 | 4 | 1 | 13825 | 111 | Clu | | 3.173e-02 | -3.45 | ciliary neurotrophic factor receptor complex | cellular component | GO:0070110 | 4 | 1 | 13825 | 111 | Il6ra | | 3.173e-02 | -3.45 | spermatoproteasome complex | cellular component | GO:1990111 | 4 | 1 | 13825 | 111 | Psmb10 | | 3.173e-02 | -3.45 | growth factor complex | cellular component | GO:0036454 | 4 | 1 | 13825 | 111 | Igfbp6 | | 3.173e-02 | -3.45 | synaptobrevin 2-SNAP-25-syntaxin-1a-complexin II complex | cellular component | GO:0070033 | 4 | 1 | 13825 | 111 | Cplx2 | | 3.173e-02 | -3.45 | Weibel-Palade body | cellular component | GO:0033093 | 4 | 1 | 13825 | 111 | Vwf | | 3.173e-02 | -3.45 | insulin-like growth factor binding protein complex | cellular component | GO:0016942 | 4 | 1 | 13825 | 111 | Igfbp6 | | 3.178e-02 | -3.45 | REACTOME\_FGFR4\_LIGAND\_BINDING\_AND\_ACTIVATION | MSigDB lists | REACTOME\_FGFR4\_LIGAND\_BINDING\_AND\_ACTIVATION | 4 | 1 | 12187 | 98 | Fgf18 | | 3.178e-02 | -3.45 | MIKKELSEN\_ES\_LCP\_WITH\_H3K4ME3\_AND\_H3K27ME3 | MSigDB lists | MIKKELSEN\_ES\_LCP\_WITH\_H3K4ME3\_AND\_H3K27ME3 | 4 | 1 | 12187 | 98 | Slc17a7 | | 3.178e-02 | -3.45 | REACTOME\_BINDING\_AND\_ENTRY\_OF\_HIV\_VIRION | MSigDB lists | REACTOME\_BINDING\_AND\_ENTRY\_OF\_HIV\_VIRION | 4 | 1 | 12187 | 98 | Ccr5 | | 3.178e-02 | -3.45 | FERRARI\_RESPONSE\_TO\_FENRETINIDE\_DN | MSigDB lists | FERRARI\_RESPONSE\_TO\_FENRETINIDE\_DN | 4 | 1 | 12187 | 98 | Hspa2 | | 3.178e-02 | -3.45 | YANAGISAWA\_LUNG\_CANCER\_RECURRENCE | MSigDB lists | YANAGISAWA\_LUNG\_CANCER\_RECURRENCE | 4 | 1 | 12187 | 98 | S100a6 | | 3.178e-02 | -3.45 | LY\_AGING\_OLD\_UP | MSigDB lists | LY\_AGING\_OLD\_UP | 4 | 1 | 12187 | 98 | Fmod | | 3.178e-02 | -3.45 | BUSA\_SAM68\_TARGETS\_DN | MSigDB lists | BUSA\_SAM68\_TARGETS\_DN | 4 | 1 | 12187 | 98 | Clu | | 3.178e-02 | -3.45 | MIKKELSEN\_IPS\_LCP\_WITH\_H3K4ME3\_AND\_H3K27ME3 | MSigDB lists | MIKKELSEN\_IPS\_LCP\_WITH\_H3K4ME3\_AND\_H3K27ME3 | 4 | 1 | 12187 | 98 | Slc17a7 | | 3.178e-02 | -3.45 | GNF2\_IGF1 | MSigDB lists | GNF2\_IGF1 | 4 | 1 | 12187 | 98 | Adh1 | | 3.178e-02 | -3.45 | chr12p | MSigDB lists | chr12p | 4 | 1 | 12187 | 98 | Mgp | | 3.178e-02 | -3.45 | REACTOME\_ETHANOL\_OXIDATION | MSigDB lists | REACTOME\_ETHANOL\_OXIDATION | 4 | 1 | 12187 | 98 | Adh1 | | 3.178e-02 | -3.45 | GO\_COMPLEMENT\_ACTIVATION\_ALTERNATIVE\_PATHWAY | MSigDB lists | GO\_COMPLEMENT\_ACTIVATION\_ALTERNATIVE\_PATHWAY | 4 | 1 | 12187 | 98 | C3 | | 3.178e-02 | -3.45 | RODRIGUES\_DCC\_TARGETS\_UP | MSigDB lists | RODRIGUES\_DCC\_TARGETS\_UP | 4 | 1 | 12187 | 98 | Slc2a1 | | 3.194e-02 | -3.44 | SERVITJA\_LIVER\_HNF1A\_TARGETS\_DN | MSigDB lists | SERVITJA\_LIVER\_HNF1A\_TARGETS\_DN | 86 | 3 | 12187 | 98 | Ly6e,Il1r1,Lin7a | | 3.194e-02 | -3.44 | GO\_PATTERN\_RECOGNITION\_RECEPTOR\_SIGNALING\_PATHWAY | MSigDB lists | GO\_PATTERN\_RECOGNITION\_RECEPTOR\_SIGNALING\_PATHWAY | 86 | 3 | 12187 | 98 | Ctsb,Ctss,Itgam | | 3.199e-02 | -3.44 | vacuolar part | cellular component | GO:0044437 | 149 | 4 | 13825 | 111 | Cd68,Gfap,Grn,Vamp8 | | 3.200e-02 | -3.44 | modulation by virus of host apoptotic process | biological process | GO:0039526 | 4 | 1 | 13711 | 111 | Ccr5 | | 3.200e-02 | -3.44 | negative regulation of monocyte chemotactic protein-1 production | biological process | GO:0071638 | 4 | 1 | 13711 | 111 | Apod | | 3.200e-02 | -3.44 | regulation of mast cell cytokine production | biological process | GO:0032763 | 4 | 1 | 13711 | 111 | Fcer1g | | 3.200e-02 | -3.44 | positive regulation of phospholipase A2 activity | biological process | GO:0032430 | 4 | 1 | 13711 | 111 | Camk2b | | 3.200e-02 | -3.44 | negative regulation of interleukin-2 biosynthetic process | biological process | GO:0045085 | 4 | 1 | 13711 | 111 | Ptprc | | 3.200e-02 | -3.44 | lymphocyte aggregation | biological process | GO:0071593 | 4 | 1 | 13711 | 111 | Rac2 | | 3.200e-02 | -3.44 | catecholamine secretion | biological process | GO:0050432 | 4 | 1 | 13711 | 111 | Ly6e | | 3.200e-02 | -3.44 | regulation of skeletal muscle tissue regeneration | biological process | GO:0043416 | 4 | 1 | 13711 | 111 | Capn3 | | 3.200e-02 | -3.44 | regulation of mast cell apoptotic process | biological process | GO:0033025 | 4 | 1 | 13711 | 111 | Fcer1g | | 3.200e-02 | -3.44 | killing by symbiont of host cells | biological process | GO:0001907 | 4 | 1 | 13711 | 111 | Ccr5 | | 3.200e-02 | -3.44 | negative regulation of T cell mediated cytotoxicity | biological process | GO:0001915 | 4 | 1 | 13711 | 111 | Ptprc | | 3.200e-02 | -3.44 | establishment of T cell polarity | biological process | GO:0001768 | 4 | 1 | 13711 | 111 | Myh9 | | 3.200e-02 | -3.44 | compartment pattern specification | biological process | GO:0007386 | 4 | 1 | 13711 | 111 | Lfng | | 3.200e-02 | -3.44 | protein localization to basolateral plasma membrane | biological process | GO:1903361 | 4 | 1 | 13711 | 111 | Lin7a | | 3.200e-02 | -3.44 | central nervous system myelin maintenance | biological process | GO:0032286 | 4 | 1 | 13711 | 111 | Clu | | 3.200e-02 | -3.44 | monoacylglycerol metabolic process | biological process | GO:0046462 | 4 | 1 | 13711 | 111 | Mgll | | 3.200e-02 | -3.44 | negative regulation of cell adhesion involved in substrate-bound cell migration | biological process | GO:0006933 | 4 | 1 | 13711 | 111 | Ptprc | | 3.200e-02 | -3.44 | negative regulation of amyloid fibril formation | biological process | GO:1905907 | 4 | 1 | 13711 | 111 | Clu | | 3.200e-02 | -3.44 | monoacylglycerol catabolic process | biological process | GO:0052651 | 4 | 1 | 13711 | 111 | Mgll | | 3.200e-02 | -3.44 | regulation of endothelial cell chemotaxis to fibroblast growth factor | biological process | GO:2000544 | 4 | 1 | 13711 | 111 | Fgf18 | | 3.200e-02 | -3.44 | autophagosome membrane docking | biological process | GO:0016240 | 4 | 1 | 13711 | 111 | Vamp8 | | 3.200e-02 | -3.44 | positive regulation of ATPase-coupled calcium transmembrane transporter activity | biological process | GO:1901896 | 4 | 1 | 13711 | 111 | Hspa2 | | 3.200e-02 | -3.44 | complement activation, alternative pathway | biological process | GO:0006957 | 4 | 1 | 13711 | 111 | C3 | | 3.200e-02 | -3.44 | establishment of lymphocyte polarity | biological process | GO:0001767 | 4 | 1 | 13711 | 111 | Myh9 | | 3.200e-02 | -3.44 | Fc-epsilon receptor signaling pathway | biological process | GO:0038095 | 4 | 1 | 13711 | 111 | Fcer1g | | 3.200e-02 | -3.44 | heat acclimation | biological process | GO:0010286 | 4 | 1 | 13711 | 111 | Il1r1 | | 3.200e-02 | -3.44 | establishment of meiotic spindle localization | biological process | GO:0051295 | 4 | 1 | 13711 | 111 | Myh9 | | 3.200e-02 | -3.44 | secretory granule maturation | biological process | GO:0061792 | 4 | 1 | 13711 | 111 | Ptprn | | 3.200e-02 | -3.44 | regulation of cell chemotaxis to fibroblast growth factor | biological process | GO:1904847 | 4 | 1 | 13711 | 111 | Fgf18 | | 3.200e-02 | -3.44 | negative regulation of plasminogen activation | biological process | GO:0010757 | 4 | 1 | 13711 | 111 | Ctsz | | 3.200e-02 | -3.44 | positive regulation by organism of programmed cell death in other organism involved in symbiotic interaction | biological process | GO:0052330 | 4 | 1 | 13711 | 111 | Ccr5 | | 3.200e-02 | -3.44 | lysosomal membrane organization | biological process | GO:0097212 | 4 | 1 | 13711 | 111 | Mt3 | | 3.200e-02 | -3.44 | disruption by symbiont of host cell | biological process | GO:0044004 | 4 | 1 | 13711 | 111 | Ccr5 | | 3.200e-02 | -3.44 | positive regulation by symbiont of host apoptotic process | biological process | GO:0052151 | 4 | 1 | 13711 | 111 | Ccr5 | | 3.200e-02 | -3.44 | interleukin-3-mediated signaling pathway | biological process | GO:0038156 | 4 | 1 | 13711 | 111 | Fcer1g | | 3.200e-02 | -3.44 | astrocyte activation involved in immune response | biological process | GO:0002265 | 4 | 1 | 13711 | 111 | Grn | | 3.200e-02 | -3.44 | positive regulation of T-helper 1 cell cytokine production | biological process | GO:2000556 | 4 | 1 | 13711 | 111 | Il1r1 | | 3.200e-02 | -3.44 | cellular response to mycotoxin | biological process | GO:0036146 | 4 | 1 | 13711 | 111 | Actb | | 3.200e-02 | -3.44 | negative regulation of lipoprotein metabolic process | biological process | GO:0050748 | 4 | 1 | 13711 | 111 | Apod | | 3.200e-02 | -3.44 | regulation of T-helper 1 cell cytokine production | biological process | GO:2000554 | 4 | 1 | 13711 | 111 | Il1r1 | | 3.200e-02 | -3.44 | positive regulation by symbiont of host programmed cell death | biological process | GO:0052042 | 4 | 1 | 13711 | 111 | Ccr5 | | 3.200e-02 | -3.44 | negative regulation of respiratory burst involved in inflammatory response | biological process | GO:0060266 | 4 | 1 | 13711 | 111 | Grn | | 3.200e-02 | -3.44 | germ-line stem cell population maintenance | biological process | GO:0030718 | 4 | 1 | 13711 | 111 | Tfap2c | | 3.200e-02 | -3.44 | positive regulation of MHC class II biosynthetic process | biological process | GO:0045348 | 4 | 1 | 13711 | 111 | Tmem106a | | 3.200e-02 | -3.44 | regulation of protein tyrosine phosphatase activity | biological process | GO:1903613 | 4 | 1 | 13711 | 111 | Ptprc | | 3.200e-02 | -3.44 | epithelial cell proliferation involved in mammary gland duct elongation | biological process | GO:0060750 | 4 | 1 | 13711 | 111 | Tfap2c | | 3.200e-02 | -3.44 | regulation of inflammatory response to wounding | biological process | GO:0106014 | 4 | 1 | 13711 | 111 | Grn | | 3.200e-02 | -3.44 | negative regulation of dendritic cell differentiation | biological process | GO:2001199 | 4 | 1 | 13711 | 111 | Tmem176a | | 3.200e-02 | -3.44 | positive regulation of tau-protein kinase activity | biological process | GO:1902949 | 4 | 1 | 13711 | 111 | Clu | | 3.203e-02 | -3.44 | GSE10239\_MEMORY\_VS\_KLRG1HIGH\_EFF\_CD8\_TCELL\_UP | MSigDB lists | GSE10239\_MEMORY\_VS\_KLRG1HIGH\_EFF\_CD8\_TCELL\_UP | 149 | 4 | 12187 | 98 | Rpl31,Treml2,Psme1,Rap1gap | | 3.203e-02 | -3.44 | GSE26488\_HDAC7\_KO\_VS\_VP16\_TRANSGENIC\_HDAC7\_KO\_DOUBLE\_POSITIVE\_THYMOCYTE\_DN | MSigDB lists | GSE26488\_HDAC7\_KO\_VS\_VP16\_TRANSGENIC\_HDAC7\_KO\_DOUBLE\_POSITIVE\_THYMOCYTE\_DN | 149 | 4 | 12187 | 98 | Cyba,Ctss,Vamp8,Tfap2c | | 3.203e-02 | -3.44 | GSE3039\_CD4\_TCELL\_VS\_B1\_BCELL\_DN | MSigDB lists | GSE3039\_CD4\_TCELL\_VS\_B1\_BCELL\_DN | 149 | 4 | 12187 | 98 | Mpeg1,Ctsb,Ly6e,Fcer1g | | 3.203e-02 | -3.44 | MCBRYAN\_PUBERTAL\_TGFB1\_TARGETS\_UP | MSigDB lists | MCBRYAN\_PUBERTAL\_TGFB1\_TARGETS\_UP | 149 | 4 | 12187 | 98 | Slc2a1,S100a6,Tsc22d1,Clu | | 3.203e-02 | -3.44 | GSE32423\_CTRL\_VS\_IL4\_MEMORY\_CD8\_TCELL\_UP | MSigDB lists | GSE32423\_CTRL\_VS\_IL4\_MEMORY\_CD8\_TCELL\_UP | 149 | 4 | 12187 | 98 | Adh1,Actb,Ahnak,Pcsk2 | | 3.203e-02 | -3.44 | GSE7852\_LN\_VS\_THYMUS\_TCONV\_UP | MSigDB lists | GSE7852\_LN\_VS\_THYMUS\_TCONV\_UP | 149 | 4 | 12187 | 98 | Camk2b,S100a6,Ly6e,Serpinb9 | | 3.203e-02 | -3.44 | GSE5589\_LPS\_VS\_LPS\_AND\_IL10\_STIM\_IL10\_KO\_MACROPHAGE\_180MIN\_UP | MSigDB lists | GSE5589\_LPS\_VS\_LPS\_AND\_IL10\_STIM\_IL10\_KO\_MACROPHAGE\_180MIN\_UP | 149 | 4 | 12187 | 98 | Mgll,Ldha,Pde1a,Ttc9b | | 3.206e-02 | -3.44 | GO\_POSITIVE\_REGULATION\_OF\_REGULATED\_SECRETORY\_PATHWAY | MSigDB lists | GO\_POSITIVE\_REGULATION\_OF\_REGULATED\_SECRETORY\_PATHWAY | 35 | 2 | 12187 | 98 | Fcer1g,Vamp8 | | 3.206e-02 | -3.44 | GNF2\_S100A4 | MSigDB lists | GNF2\_S100A4 | 35 | 2 | 12187 | 98 | Sh3bgrl3,Fcer1g | | 3.206e-02 | -3.44 | GO\_CYTOPLASMIC\_TRANSLATION | MSigDB lists | GO\_CYTOPLASMIC\_TRANSLATION | 35 | 2 | 12187 | 98 | Rpl26,Rpl31 | | 3.206e-02 | -3.44 | NAKAMURA\_METASTASIS\_MODEL\_DN | MSigDB lists | NAKAMURA\_METASTASIS\_MODEL\_DN | 35 | 2 | 12187 | 98 | C3,Rac2 | | 3.206e-02 | -3.44 | LANDIS\_BREAST\_CANCER\_PROGRESSION\_UP | MSigDB lists | LANDIS\_BREAST\_CANCER\_PROGRESSION\_UP | 35 | 2 | 12187 | 98 | Tfap2c,Spint1 | | 3.206e-02 | -3.44 | GO\_SUBSTANTIA\_NIGRA\_DEVELOPMENT | MSigDB lists | GO\_SUBSTANTIA\_NIGRA\_DEVELOPMENT | 35 | 2 | 12187 | 98 | Actb,Ldha | | 3.249e-02 | -3.43 | regulation of glial cell proliferation | biological process | GO:0060251 | 35 | 2 | 13711 | 111 | Vim,Gfap | | 3.249e-02 | -3.43 | viral entry into host cell | biological process | GO:0046718 | 35 | 2 | 13711 | 111 | Vamp8,Ctsb | | 3.249e-02 | -3.43 | regulation of neuron projection regeneration | biological process | GO:0070570 | 35 | 2 | 13711 | 111 | Thy1,Grn | | 3.249e-02 | -3.43 | regulation of neutrophil migration | biological process | GO:1902622 | 35 | 2 | 13711 | 111 | Rac2,Il1r1 | | 3.254e-02 | -3.43 | Regulation of Actin Cytoskeleton | WikiPathways | WP523 | 130 | 5 | 3756 | 52 | Rac2,Fgf18,Brk1,Actb,Csk | | 3.267e-02 | -3.42 | EXOC3/Sec6 | interpro domains | IPR010326 | 4 | 1 | 13788 | 114 | Exoc3l4 | | 3.267e-02 | -3.42 | Glut\_rich\_SH3-bd | interpro domains | IPR006993 | 4 | 1 | 13788 | 114 | Sh3bgrl3 | | 3.267e-02 | -3.42 | TSC-22\_Dip\_Bun | interpro domains | IPR000580 | 4 | 1 | 13788 | 114 | Tsc22d1 | | 3.267e-02 | -3.42 | GPI\_mannosylTrfase | interpro domains | IPR005599 | 4 | 1 | 13788 | 114 | Alg12 | | 3.267e-02 | -3.42 | Anaphylatoxin/fibulin | interpro domains | IPR000020 | 4 | 1 | 13788 | 114 | C3 | | 3.267e-02 | -3.42 | Lysosome-assoc\_membr\_glycop | interpro domains | IPR002000 | 4 | 1 | 13788 | 114 | Cd68 | | 3.267e-02 | -3.42 | MANSC\_N | interpro domains | IPR011106 | 4 | 1 | 13788 | 114 | Spint1 | | 3.267e-02 | -3.42 | DEK\_C | interpro domains | IPR014876 | 4 | 1 | 13788 | 114 | Dek | | 3.267e-02 | -3.42 | CD59\_antigen\_CS | interpro domains | IPR018363 | 4 | 1 | 13788 | 114 | Ly6a | | 3.267e-02 | -3.42 | Aminotrans\_3 | interpro domains | IPR005814 | 4 | 1 | 13788 | 114 | Etnppl | | 3.267e-02 | -3.42 | Ca/CaM-dep\_prot\_kinase-assoc | interpro domains | IPR013543 | 4 | 1 | 13788 | 114 | Camk2b | | 3.267e-02 | -3.42 | IL-1\_rcpt\_I/II-typ | interpro domains | IPR004074 | 4 | 1 | 13788 | 114 | Il1r1 | | 3.267e-02 | -3.42 | cell-substrate junction | cellular component | GO:0030055 | 150 | 4 | 13825 | 111 | Cyba,Ptprc,Flnb,Myh9 | | 3.271e-02 | -3.42 | GSE3982\_MAST\_CELL\_VS\_BASOPHIL\_DN | MSigDB lists | GSE3982\_MAST\_CELL\_VS\_BASOPHIL\_DN | 150 | 4 | 12187 | 98 | Ptprc,Actb,Rpl31,Ccr5 | | 3.271e-02 | -3.42 | GSE20366\_CD103\_POS\_VS\_CD103\_KLRG1\_DP\_TREG\_UP | MSigDB lists | GSE20366\_CD103\_POS\_VS\_CD103\_KLRG1\_DP\_TREG\_UP | 150 | 4 | 12187 | 98 | Klhl6,Phyhip,Slc26a2,Zfp786 | | 3.271e-02 | -3.42 | GSE21360\_NAIVE\_VS\_SECONDARY\_MEMORY\_CD8\_TCELL\_UP | MSigDB lists | GSE21360\_NAIVE\_VS\_SECONDARY\_MEMORY\_CD8\_TCELL\_UP | 150 | 4 | 12187 | 98 | C3,Hcls1,Slc2a1,Flnb | | 3.271e-02 | -3.42 | GSE46606\_IRF4\_KO\_VS\_WT\_UNSTIM\_BCELL\_DN | MSigDB lists | GSE46606\_IRF4\_KO\_VS\_WT\_UNSTIM\_BCELL\_DN | 150 | 4 | 12187 | 98 | Exoc3l4,Slc2a1,C3,Vamp8 | | 3.271e-02 | -3.42 | GSE31082\_CD4\_VS\_CD8\_SP\_THYMOCYTE\_DN | MSigDB lists | GSE31082\_CD4\_VS\_CD8\_SP\_THYMOCYTE\_DN | 150 | 4 | 12187 | 98 | Rps5,Dek,Fam111a,Ahnak | | 3.271e-02 | -3.42 | GO\_CONTRACTILE\_FIBER | MSigDB lists | GO\_CONTRACTILE\_FIBER | 150 | 4 | 12187 | 98 | Flnb,Dek,Capn3,Ahnak | | 3.283e-02 | -3.42 | cortical actin cytoskeleton | cellular component | GO:0030864 | 87 | 3 | 13825 | 111 | Hcls1,Slc2a1,Myh9 | | 3.288e-02 | -3.41 | positive regulation of cellular process | biological process | GO:0048522 | 4254 | 44 | 13711 | 111 | Capn3,Fgf18,Psme1,Camk2b,Il6ra,Gfap,Tsc22d1,Itgam,Fcer1g,Il1r1,Thy1,Kl,Tmem106a,Ccar1,Grn,Ikzf3,Hspa2,Ptprc,C1qa,Clu,Mark4,Csk,C3,Cyba,Ptprn,Tfap2c,Ccr5,Ndufa13,Lfng,Brk1,Mt3,Myh9,Csf1r,Hcls1,Spint1,Rac2,Vamp8,Lrg1,Vim,Rpl26,Tnfsf8,Fcgr3,Ccl6,Ctsz | | 3.289e-02 | -3.41 | DIAZ\_CHRONIC\_MEYLOGENOUS\_LEUKEMIA\_DN | MSigDB lists | DIAZ\_CHRONIC\_MEYLOGENOUS\_LEUKEMIA\_DN | 87 | 3 | 12187 | 98 | Csf1r,Flnb,Capn3 | | 3.289e-02 | -3.41 | GO\_POSITIVE\_REGULATION\_OF\_CHEMOTAXIS | MSigDB lists | GO\_POSITIVE\_REGULATION\_OF\_CHEMOTAXIS | 87 | 3 | 12187 | 98 | Fgf18,Rac2,Il6ra | | 3.312e-02 | -3.41 | CaMKII\_AD | pfam domains | PF08332 | 4 | 1 | 12881 | 108 | Camk2b | | 3.312e-02 | -3.41 | SH3BGR | pfam domains | PF04908 | 4 | 1 | 12881 | 108 | Sh3bgrl3 | | 3.312e-02 | -3.41 | Lamp | pfam domains | PF01299 | 4 | 1 | 12881 | 108 | Cd68 | | 3.312e-02 | -3.41 | Sec6 | pfam domains | PF06046 | 4 | 1 | 12881 | 108 | Exoc3l4 | | 3.312e-02 | -3.41 | Glyco\_transf\_22 | pfam domains | PF03901 | 4 | 1 | 12881 | 108 | Alg12 | | 3.312e-02 | -3.41 | DEK\_C | pfam domains | PF08766 | 4 | 1 | 12881 | 108 | Dek | | 3.312e-02 | -3.41 | TSC22 | pfam domains | PF01166 | 4 | 1 | 12881 | 108 | Tsc22d1 | | 3.312e-02 | -3.41 | Aminotran\_3 | pfam domains | PF00202 | 4 | 1 | 12881 | 108 | Etnppl | | 3.340e-02 | -3.40 | GSE6259\_33D1\_POS\_DC\_VS\_CD4\_TCELL\_DN | MSigDB lists | GSE6259\_33D1\_POS\_DC\_VS\_CD4\_TCELL\_DN | 151 | 4 | 12187 | 98 | Ctsz,Fcer1g,Tmem176a,Ctsa | | 3.340e-02 | -3.40 | GO\_CELL\_JUNCTION\_ORGANIZATION | MSigDB lists | GO\_CELL\_JUNCTION\_ORGANIZATION | 151 | 4 | 12187 | 98 | Actb,Csf1r,Csk,Thy1 | | 3.340e-02 | -3.40 | GSE27241\_WT\_CTRL\_VS\_DIGOXIN\_TREATED\_RORGT\_KO\_CD4\_TCELL\_IN\_TH17\_POLARIZING\_CONDITIONS\_DN | MSigDB lists | GSE27241\_WT\_CTRL\_VS\_DIGOXIN\_TREATED\_RORGT\_KO\_CD4\_TCELL\_IN\_TH17\_POLARIZING\_CONDITIONS\_DN | 151 | 4 | 12187 | 98 | S100a6,Vim,Hcls1,Treml2 | | 3.340e-02 | -3.40 | GSE25677\_MPL\_VS\_R848\_STIM\_BCELL\_DN | MSigDB lists | GSE25677\_MPL\_VS\_R848\_STIM\_BCELL\_DN | 151 | 4 | 12187 | 98 | Myh9,Vim,Csk,Lfng | | 3.340e-02 | -3.40 | GSE1925\_CTRL\_VS\_IFNG\_PRIMED\_MACROPHAGE\_3H\_IFNG\_STIM\_UP | MSigDB lists | GSE1925\_CTRL\_VS\_IFNG\_PRIMED\_MACROPHAGE\_3H\_IFNG\_STIM\_UP | 151 | 4 | 12187 | 98 | Tnfsf8,Ctss,Myh9,Rps5 | | 3.341e-02 | -3.40 | VEGFA-VEGFR2 Pathway | REACTOME pathways | R-MMU-4420097 | 255 | 7 | 6297 | 76 | Psme1,Psmb10,Cyba,Camk2b,Brk1,Fgf18,Kl | | 3.343e-02 | -3.40 | Integrin alphaIIb beta3 signaling | REACTOME pathways | R-MMU-354192 | 24 | 2 | 6297 | 76 | Csk,Vwf | | 3.352e-02 | -3.40 | positive regulation of angiogenesis | biological process | GO:0045766 | 150 | 4 | 13711 | 111 | C3,Grn,Fgf18,Lrg1 | | 3.352e-02 | -3.40 | negative regulation of peptidase activity | biological process | GO:0010466 | 150 | 4 | 13711 | 111 | C3,Spint1,Serpina3n,Serpinb9 | | 3.355e-02 | -3.39 | chaperone binding | molecular function | GO:0051087 | 89 | 3 | 13516 | 107 | Grn,Vwf,Hspa2 | | 3.355e-02 | -3.39 | enhancer binding | molecular function | GO:0035326 | 89 | 3 | 13516 | 107 | Tfap2c,Bhlhe41,Ccar1 | | 3.360e-02 | -3.39 | KRIGE\_RESPONSE\_TO\_TOSEDOSTAT\_6HR\_DN | MSigDB lists | KRIGE\_RESPONSE\_TO\_TOSEDOSTAT\_6HR\_DN | 735 | 11 | 12187 | 98 | Ldha,Trnp1,Fam111a,Mgll,Ptprc,Fcer1g,Lrg1,Ly6e,Slc2a1,Vamp8,Il6ra | | 3.360e-02 | -3.39 | negative regulation of cell differentiation | biological process | GO:0045596 | 637 | 10 | 13711 | 111 | Thy1,Rap1gap,Capn3,Vim,Gfap,Ccr5,Tmem176a,Mt3,Bhlhe41,Ctsz | | 3.363e-02 | -3.39 | GO\_RESPONSE\_TO\_BIOTIC\_STIMULUS | MSigDB lists | GO\_RESPONSE\_TO\_BIOTIC\_STIMULUS | 553 | 9 | 12187 | 98 | Serpinb9,Ccr5,Fcer1g,Clu,Fam111a,Tnfsf8,Il6ra,Phyhip,Ptprc | | 3.373e-02 | -3.39 | GO\_REGULATION\_OF\_CYTOPLASMIC\_TRANSPORT | MSigDB lists | GO\_REGULATION\_OF\_CYTOPLASMIC\_TRANSPORT | 381 | 7 | 12187 | 98 | Cyba,Apod,Rac2,Hcls1,Thy1,Capn3,Csk | | 3.376e-02 | -3.39 | RET signaling | REACTOME pathways | R-MMU-8853659 | 201 | 6 | 6297 | 76 | Fgf18,Kl,Rap1gap,Camk2b,Psme1,Psmb10 | | 3.378e-02 | -3.39 | HARRIS\_BRAIN\_CANCER\_PROGENITORS | MSigDB lists | HARRIS\_BRAIN\_CANCER\_PROGENITORS | 36 | 2 | 12187 | 98 | Mgp,S100a6 | | 3.378e-02 | -3.39 | REACTOME\_TCR\_SIGNALING | MSigDB lists | REACTOME\_TCR\_SIGNALING | 36 | 2 | 12187 | 98 | Ptprc,Csk | | 3.378e-02 | -3.39 | GO\_REGULATION\_OF\_CELL\_KILLING | MSigDB lists | GO\_REGULATION\_OF\_CELL\_KILLING | 36 | 2 | 12187 | 98 | Ptprc,Serpinb9 | | 3.386e-02 | -3.39 | GO\_PERIKARYON | MSigDB lists | GO\_PERIKARYON | 88 | 3 | 12187 | 98 | Ptprn,Camk2b,Pcsk2 | | 3.386e-02 | -3.39 | NAKAMURA\_ADIPOGENESIS\_LATE\_UP | MSigDB lists | NAKAMURA\_ADIPOGENESIS\_LATE\_UP | 88 | 3 | 12187 | 98 | Mgp,Serpinb9,Itgam | | 3.386e-02 | -3.39 | GO\_INNATE\_IMMUNE\_RESPONSE\_ACTIVATING\_CELL\_SURFACE\_RECEPTOR\_SIGNALING\_PATHWAY | MSigDB lists | GO\_INNATE\_IMMUNE\_RESPONSE\_ACTIVATING\_CELL\_SURFACE\_RECEPTOR\_SIGNALING\_PATHWAY | 88 | 3 | 12187 | 98 | Psmb10,Psme1,Fcer1g | | 3.395e-02 | -3.38 | protein kinase binding | molecular function | GO:0019901 | 653 | 10 | 13516 | 107 | Hcls1,Thy1,Ptprc,Camk2b,Phyhip,Ccr5,Actb,Rac2,Csk,Vim | | 3.398e-02 | -3.38 | PTEN (phosphatase and tensin homolog) | protein interactions | 5728 | 107 | 4 | 6802 | 78 | Actb,Ahnak,Myh9,Ldha | | 3.401e-02 | -3.38 | LRRFIP1 (LRR binding FLII interacting protein 1) | protein interactions | 9208 | 3 | 1 | 6802 | 78 | Actb | | 3.401e-02 | -3.38 | ACTN1 (actinin alpha 1) | protein interactions | 87 | 3 | 1 | 6802 | 78 | Myh9 | | 3.401e-02 | -3.38 | KRAS (KRAS proto-oncogene, GTPase) | protein interactions | 3845 | 3 | 1 | 6802 | 78 | Flnb | | 3.401e-02 | -3.38 | Thoc5 (THO complex 5) | protein interactions | 107829 | 3 | 1 | 6802 | 78 | Csf1r | | 3.401e-02 | -3.38 | SPRYD3 (SPRY domain containing 3) | protein interactions | 84926 | 3 | 1 | 6802 | 78 | Flnb | | 3.401e-02 | -3.38 | CAP1 (cyclase associated actin cytoskeleton regulatory protein 1) | protein interactions | 10487 | 3 | 1 | 6802 | 78 | Actb | | 3.401e-02 | -3.38 | MARK3 (microtubule affinity regulating kinase 3) | protein interactions | 4140 | 3 | 1 | 6802 | 78 | Myh9 | | 3.401e-02 | -3.38 | Atp6v0d2 (ATPase, H+ transporting, lysosomal V0 subunit D2) | protein interactions | 242341 | 3 | 1 | 6802 | 78 | Vamp8 | | 3.401e-02 | -3.38 | SUPT5H (SPT5 homolog, DSIF elongation factor subunit) | protein interactions | 6829 | 3 | 1 | 6802 | 78 | Myh9 | | 3.401e-02 | -3.38 | Ppl (periplakin) | protein interactions | 19041 | 3 | 1 | 6802 | 78 | Vim | | 3.401e-02 | -3.38 | Nr2e1 (nuclear receptor subfamily 2, group E, member 1) | protein interactions | 21907 | 3 | 1 | 6802 | 78 | Gfap | | 3.401e-02 | -3.38 | IRAK1 (interleukin 1 receptor associated kinase 1) | protein interactions | 3654 | 3 | 1 | 6802 | 78 | Il1r1 | | 3.401e-02 | -3.38 | TRIM21 (tripartite motif containing 21) | protein interactions | 6737 | 3 | 1 | 6802 | 78 | Csk | | 3.401e-02 | -3.38 | Mme (membrane metallo endopeptidase) | protein interactions | 17380 | 3 | 1 | 6802 | 78 | Actb | | 3.401e-02 | -3.38 | AP2A2 (adaptor related protein complex 2 subunit alpha 2) | protein interactions | 161 | 3 | 1 | 6802 | 78 | Actb | | 3.401e-02 | -3.38 | Stap1 (signal transducing adaptor family member 1) | protein interactions | 56792 | 3 | 1 | 6802 | 78 | Csf1r | | 3.401e-02 | -3.38 | Cap2 (CAP, adenylate cyclase-associated protein, 2 (yeast)) | protein interactions | 67252 | 3 | 1 | 6802 | 78 | Ptprc | | 3.401e-02 | -3.38 | Cdh17 (cadherin 17) | protein interactions | 12557 | 3 | 1 | 6802 | 78 | Ptprc | | 3.401e-02 | -3.38 | GNA11 (G protein subunit alpha 11) | protein interactions | 2767 | 3 | 1 | 6802 | 78 | Actb | | 3.401e-02 | -3.38 | Cotl1 (coactosin-like 1 (Dictyostelium)) | protein interactions | 72042 | 3 | 1 | 6802 | 78 | Actb | | 3.401e-02 | -3.38 | ADD3 (adducin 3) | protein interactions | 120 | 3 | 1 | 6802 | 78 | Myh9 | | 3.401e-02 | -3.38 | CLIC1 (chloride intracellular channel 1) | protein interactions | 1192 | 3 | 1 | 6802 | 78 | Actb | | 3.401e-02 | -3.38 | ATP9A (ATPase phospholipid transporting 9A (putative)) | protein interactions | 10079 | 3 | 1 | 6802 | 78 | Myh9 | | 3.401e-02 | -3.38 | SPTBN2 (spectrin beta, non-erythrocytic 2) | protein interactions | 6712 | 3 | 1 | 6802 | 78 | Myh9 | | 3.401e-02 | -3.38 | Ccl3 (chemokine (C-C motif) ligand 3) | protein interactions | 20302 | 3 | 1 | 6802 | 78 | Ccr5 | | 3.401e-02 | -3.38 | DSTN (destrin, actin depolymerizing factor) | protein interactions | 11034 | 3 | 1 | 6802 | 78 | Myh9 | | 3.401e-02 | -3.38 | PDCD7 (programmed cell death 7) | protein interactions | 10081 | 3 | 1 | 6802 | 78 | Myh9 | | 3.401e-02 | -3.38 | Chd3 (chromodomain helicase DNA binding protein 3) | protein interactions | 216848 | 3 | 1 | 6802 | 78 | Ikzf3 | | 3.401e-02 | -3.38 | SNTB2 (syntrophin beta 2) | protein interactions | 6645 | 3 | 1 | 6802 | 78 | Flnb | | 3.401e-02 | -3.38 | EPPK1 (epiplakin 1) | protein interactions | 83481 | 3 | 1 | 6802 | 78 | Myh9 | | 3.401e-02 | -3.38 | SEC24B (SEC24 homolog B, COPII coat complex component) | protein interactions | 10427 | 3 | 1 | 6802 | 78 | Actb | | 3.401e-02 | -3.38 | QSER1 (glutamine and serine rich 1) | protein interactions | 79832 | 3 | 1 | 6802 | 78 | Csk | | 3.401e-02 | -3.38 | LRCH3 (leucine rich repeats and calponin homology domain containing 3) | protein interactions | 84859 | 3 | 1 | 6802 | 78 | Actb | | 3.401e-02 | -3.38 | TTI1 (TELO2 interacting protein 1) | protein interactions | 9675 | 3 | 1 | 6802 | 78 | Mgp | | 3.401e-02 | -3.38 | SSRP1 (structure specific recognition protein 1) | protein interactions | 6749 | 3 | 1 | 6802 | 78 | Myh9 | | 3.401e-02 | -3.38 | Itgb2 (integrin beta 2) | protein interactions | 16414 | 3 | 1 | 6802 | 78 | Itgam | | 3.401e-02 | -3.38 | Prkaa2 (protein kinase, AMP-activated, alpha 2 catalytic subunit) | protein interactions | 108079 | 3 | 1 | 6802 | 78 | Mark4 | | 3.401e-02 | -3.38 | Fgfr2 (fibroblast growth factor receptor 2) | protein interactions | 14183 | 3 | 1 | 6802 | 78 | Kl | | 3.401e-02 | -3.38 | GART (phosphoribosylglycinamide formyltransferase, phosphoribosylglycinamide synthetase, phosphoribosylaminoimidazole synthetase) | protein interactions | 2618 | 3 | 1 | 6802 | 78 | Myh9 | | 3.401e-02 | -3.38 | RAP1B (RAP1B, member of RAS oncogene family) | protein interactions | 5908 | 3 | 1 | 6802 | 78 | Flnb | | 3.408e-02 | -3.38 | regulation of cation transmembrane transport | biological process | GO:1904062 | 298 | 6 | 13711 | 111 | Ahnak,Thy1,Capn3,Hspa2,Ctss,Cyba | | 3.410e-02 | -3.38 | GSE42021\_TCONV\_PLN\_VS\_CD24INT\_TCONV\_THYMUS\_DN | MSigDB lists | GSE42021\_TCONV\_PLN\_VS\_CD24INT\_TCONV\_THYMUS\_DN | 152 | 4 | 12187 | 98 | Rab3il1,Cyba,Capn3,S100a6 | | 3.410e-02 | -3.38 | GSE40274\_FOXP3\_VS\_FOXP3\_AND\_PBX1\_TRANSDUCED\_ACTIVATED\_CD4\_TCELL\_UP | MSigDB lists | GSE40274\_FOXP3\_VS\_FOXP3\_AND\_PBX1\_TRANSDUCED\_ACTIVATED\_CD4\_TCELL\_UP | 152 | 4 | 12187 | 98 | Ikzf3,Ccr5,S100a6,Klhl6 | | 3.410e-02 | -3.38 | LI\_WILMS\_TUMOR\_VS\_FETAL\_KIDNEY\_1\_UP | MSigDB lists | LI\_WILMS\_TUMOR\_VS\_FETAL\_KIDNEY\_1\_UP | 152 | 4 | 12187 | 98 | Il1r1,Clu,Ahnak,Myh9 | | 3.410e-02 | -3.38 | GSE26669\_CD4\_VS\_CD8\_TCELL\_IN\_MLR\_DN | MSigDB lists | GSE26669\_CD4\_VS\_CD8\_TCELL\_IN\_MLR\_DN | 152 | 4 | 12187 | 98 | Ahnak,Rpl14,Ptprc,Flnb | | 3.410e-02 | -3.38 | GSE3920\_UNTREATED\_VS\_IFNA\_TREATED\_ENDOTHELIAL\_CELL\_UP | MSigDB lists | GSE3920\_UNTREATED\_VS\_IFNA\_TREATED\_ENDOTHELIAL\_CELL\_UP | 152 | 4 | 12187 | 98 | Itgam,Cd68,Tmem176a,Ctsa | | 3.422e-02 | -3.37 | entry into host | biological process | GO:0044409 | 36 | 2 | 13711 | 111 | Ctsb,Vamp8 | | 3.422e-02 | -3.37 | entry into host cell | biological process | GO:0030260 | 36 | 2 | 13711 | 111 | Vamp8,Ctsb | | 3.422e-02 | -3.37 | entry into other organism involved in symbiotic interaction | biological process | GO:0051828 | 36 | 2 | 13711 | 111 | Ctsb,Vamp8 | | 3.422e-02 | -3.37 | intermediate filament cytoskeleton organization | biological process | GO:0045104 | 36 | 2 | 13711 | 111 | Vim,Gfap | | 3.422e-02 | -3.37 | entry into cell of other organism involved in symbiotic interaction | biological process | GO:0051806 | 36 | 2 | 13711 | 111 | Vamp8,Ctsb | | 3.423e-02 | -3.37 | positive regulation of ion transmembrane transport | biological process | GO:0034767 | 151 | 4 | 13711 | 111 | Ctss,Capn3,Hspa2,Thy1 | | 3.428e-02 | -3.37 | regulation of T cell activation | biological process | GO:0050863 | 222 | 5 | 13711 | 111 | Csk,Thy1,Ptprc,Il6ra,Rac2 | | 3.441e-02 | -3.37 | GO\_PROTEIN\_DIMERIZATION\_ACTIVITY | MSigDB lists | GO\_PROTEIN\_DIMERIZATION\_ACTIVITY | 832 | 12 | 12187 | 98 | Cyba,Ikzf3,Vwf,Rap1gap,Mgll,Camk2b,Tfap2c,Il6ra,Csf1r,C1qb,S100a6,Myh9 | | 3.481e-02 | -3.36 | GSE1460\_INTRATHYMIC\_T\_PROGENITOR\_VS\_CD4\_THYMOCYTE\_DN | MSigDB lists | GSE1460\_INTRATHYMIC\_T\_PROGENITOR\_VS\_CD4\_THYMOCYTE\_DN | 153 | 4 | 12187 | 98 | Ptprc,Serpinb9,Rac2,Il6ra | | 3.481e-02 | -3.36 | GSE21927\_SPLEEN\_C57BL6\_VS\_EL4\_TUMOR\_BALBC\_MONOCYTES\_UP | MSigDB lists | GSE21927\_SPLEEN\_C57BL6\_VS\_EL4\_TUMOR\_BALBC\_MONOCYTES\_UP | 153 | 4 | 12187 | 98 | Ptprc,Rps20,Rpl13,Tsc22d1 | | 3.481e-02 | -3.36 | WIERENGA\_STAT5A\_TARGETS\_DN | MSigDB lists | WIERENGA\_STAT5A\_TARGETS\_DN | 153 | 4 | 12187 | 98 | Fcer1g,Mgll,Vwf,Exoc3l4 | | 3.481e-02 | -3.36 | GSE41867\_DAY6\_EFFECTOR\_VS\_DAY30\_EXHAUSTED\_CD8\_TCELL\_LCMV\_CLONE13\_UP | MSigDB lists | GSE41867\_DAY6\_EFFECTOR\_VS\_DAY30\_EXHAUSTED\_CD8\_TCELL\_LCMV\_CLONE13\_UP | 153 | 4 | 12187 | 98 | Tmem106a,Serpinb9,Fcer1g,Mpeg1 | | 3.481e-02 | -3.36 | GSE17301\_ACD3\_ACD28\_VS\_ACD3\_ACD28\_AND\_IFNA5\_STIM\_CD8\_TCELL\_DN | MSigDB lists | GSE17301\_ACD3\_ACD28\_VS\_ACD3\_ACD28\_AND\_IFNA5\_STIM\_CD8\_TCELL\_DN | 153 | 4 | 12187 | 98 | Grn,Fcer1g,Ctss,Ctsb | | 3.481e-02 | -3.36 | GSE36888\_STAT5\_AB\_KNOCKIN\_VS\_WT\_TCELL\_IL2\_TREATED\_17H\_DN | MSigDB lists | GSE36888\_STAT5\_AB\_KNOCKIN\_VS\_WT\_TCELL\_IL2\_TREATED\_17H\_DN | 153 | 4 | 12187 | 98 | Itgam,Lin7a,Kl,Fcgr3 | | 3.481e-02 | -3.36 | GSE29618\_MONOCYTE\_VS\_PDC\_UP | MSigDB lists | GSE29618\_MONOCYTE\_VS\_PDC\_UP | 153 | 4 | 12187 | 98 | Ctss,Csf1r,Ptprc,Lin7a | | 3.481e-02 | -3.36 | GSE25123\_CTRL\_VS\_IL4\_STIM\_PPARG\_KO\_MACROPHAGE\_DN | MSigDB lists | GSE25123\_CTRL\_VS\_IL4\_STIM\_PPARG\_KO\_MACROPHAGE\_DN | 153 | 4 | 12187 | 98 | Ahnak,Thy1,Vim,Treml2 | | 3.481e-02 | -3.36 | GSE26488\_CTRL\_VS\_PEPTIDE\_INJECTION\_OT2\_THYMOCYTE\_DN | MSigDB lists | GSE26488\_CTRL\_VS\_PEPTIDE\_INJECTION\_OT2\_THYMOCYTE\_DN | 153 | 4 | 12187 | 98 | Vim,Myh9,Ahnak,Flnb | | 3.481e-02 | -3.36 | GSE37301\_PRO\_BCELL\_VS\_RAG2\_KO\_NK\_CELL\_UP | MSigDB lists | GSE37301\_PRO\_BCELL\_VS\_RAG2\_KO\_NK\_CELL\_UP | 153 | 4 | 12187 | 98 | Cyba,Vim,Tsc22d1,Fmod | | 3.484e-02 | -3.36 | GO\_B\_CELL\_ACTIVATION | MSigDB lists | GO\_B\_CELL\_ACTIVATION | 89 | 3 | 12187 | 98 | Ptprc,Ikzf3,Lfng | | 3.484e-02 | -3.36 | GO\_ELECTRON\_CARRIER\_ACTIVITY | MSigDB lists | GO\_ELECTRON\_CARRIER\_ACTIVITY | 89 | 3 | 12187 | 98 | Cox4i1,Sh3bgrl3,Cyba | | 3.484e-02 | -3.36 | BOYAULT\_LIVER\_CANCER\_SUBCLASS\_G1\_UP | MSigDB lists | BOYAULT\_LIVER\_CANCER\_SUBCLASS\_G1\_UP | 89 | 3 | 12187 | 98 | Mgp,Ahnak,Spint1 | | 3.484e-02 | -3.36 | WOOD\_EBV\_EBNA1\_TARGETS\_UP | MSigDB lists | WOOD\_EBV\_EBNA1\_TARGETS\_UP | 89 | 3 | 12187 | 98 | Tfap2c,Clu,C3 | | 3.484e-02 | -3.36 | MODULE\_147 | MSigDB lists | MODULE\_147 | 89 | 3 | 12187 | 98 | Alg12,Thy1,Il1r1 | | 3.484e-02 | -3.36 | GSE18791\_UNSTIM\_VS\_NEWCATSLE\_VIRUS\_DC\_2H\_UP | MSigDB lists | GSE18791\_UNSTIM\_VS\_NEWCATSLE\_VIRUS\_DC\_2H\_UP | 89 | 3 | 12187 | 98 | Camk2b,Pde1a,Mgp | | 3.485e-02 | -3.36 | regulation of calcium ion transport | biological process | GO:0051924 | 223 | 5 | 13711 | 111 | Cyba,Hspa2,Capn3,Ahnak,Thy1 | | 3.494e-02 | -3.35 | regulation of leukocyte migration | biological process | GO:0002685 | 152 | 4 | 13711 | 111 | Rac2,Il1r1,Apod,Thy1 | | 3.497e-02 | -3.35 | GO\_RESPONSE\_TO\_GROWTH\_FACTOR | MSigDB lists | GO\_RESPONSE\_TO\_GROWTH\_FACTOR | 384 | 7 | 12187 | 98 | Fmod,Clu,Rap1gap,Spint1,Il1r1,Kl,Fgf18 | | 3.552e-02 | -3.34 | GSE36392\_TYPE\_2\_MYELOID\_VS\_EOSINOPHIL\_IL25\_TREATED\_LUNG\_UP | MSigDB lists | GSE36392\_TYPE\_2\_MYELOID\_VS\_EOSINOPHIL\_IL25\_TREATED\_LUNG\_UP | 154 | 4 | 12187 | 98 | Rpl13,Serpinb9,Rpl14,Serpina3n | | 3.552e-02 | -3.34 | GSE26156\_DOUBLE\_POSITIVE\_VS\_CD4\_SINGLE\_POSITIVE\_THYMOCYTE\_UP | MSigDB lists | GSE26156\_DOUBLE\_POSITIVE\_VS\_CD4\_SINGLE\_POSITIVE\_THYMOCYTE\_UP | 154 | 4 | 12187 | 98 | Rps8,Rpl13,Rps20,Rpl31 | | 3.552e-02 | -3.34 | GSE40068\_CXCR5NEG\_BCL6NEG\_CD4\_TCELL\_VS\_CXCR5POS\_BCL6NEG\_TFH\_DN | MSigDB lists | GSE40068\_CXCR5NEG\_BCL6NEG\_CD4\_TCELL\_VS\_CXCR5POS\_BCL6NEG\_TFH\_DN | 154 | 4 | 12187 | 98 | Ccr5,Ahnak,Vim,Slc2a1 | | 3.552e-02 | -3.34 | GSE13306\_RA\_VS\_UNTREATED\_MEM\_CD4\_TCELL\_DN | MSigDB lists | GSE13306\_RA\_VS\_UNTREATED\_MEM\_CD4\_TCELL\_DN | 154 | 4 | 12187 | 98 | Slc2a1,Vim,Camk2b,Ahnak | | 3.552e-02 | -3.34 | GSE22886\_DAY0\_VS\_DAY7\_MONOCYTE\_IN\_CULTURE\_UP | MSigDB lists | GSE22886\_DAY0\_VS\_DAY7\_MONOCYTE\_IN\_CULTURE\_UP | 154 | 4 | 12187 | 98 | Ptprc,Psmb10,Fcgr3,Il6ra | | 3.552e-02 | -3.34 | GSE23505\_UNTREATED\_VS\_4DAY\_IL6\_IL1\_IL23\_TREATED\_CD4\_TCELL\_DN | MSigDB lists | GSE23505\_UNTREATED\_VS\_4DAY\_IL6\_IL1\_IL23\_TREATED\_CD4\_TCELL\_DN | 154 | 4 | 12187 | 98 | Klhl6,Treml2,Ccr5,Ly6e | | 3.552e-02 | -3.34 | GSE5099\_DAY3\_VS\_DAY7\_MCSF\_TREATED\_MACROPHAGE\_UP | MSigDB lists | GSE5099\_DAY3\_VS\_DAY7\_MCSF\_TREATED\_MACROPHAGE\_UP | 154 | 4 | 12187 | 98 | Rpl13,S100a6,Rpl31,Rpl14 | | 3.552e-02 | -3.34 | GSE25123\_WT\_VS\_PPARG\_KO\_MACROPHAGE\_DN | MSigDB lists | GSE25123\_WT\_VS\_PPARG\_KO\_MACROPHAGE\_DN | 154 | 4 | 12187 | 98 | Itgam,Ctsz,Ctsa,C1qb | | 3.552e-02 | -3.34 | GO\_PHOSPHATIDYLINOSITOL\_KINASE\_ACTIVITY | MSigDB lists | GO\_PHOSPHATIDYLINOSITOL\_KINASE\_ACTIVITY | 37 | 2 | 12187 | 98 | Fgf18,Kl | | 3.552e-02 | -3.34 | GO\_PHOSPHATIDYLINOSITOL\_3\_PHOSPHATE\_BIOSYNTHETIC\_PROCESS | MSigDB lists | GO\_PHOSPHATIDYLINOSITOL\_3\_PHOSPHATE\_BIOSYNTHETIC\_PROCESS | 37 | 2 | 12187 | 98 | Kl,Fgf18 | | 3.552e-02 | -3.34 | FRIDMAN\_IMMORTALIZATION\_DN | MSigDB lists | FRIDMAN\_IMMORTALIZATION\_DN | 37 | 2 | 12187 | 98 | Hspa2,Igfbp6 | | 3.552e-02 | -3.34 | GO\_POSITIVE\_REGULATION\_OF\_CHEMOKINE\_PRODUCTION | MSigDB lists | GO\_POSITIVE\_REGULATION\_OF\_CHEMOKINE\_PRODUCTION | 37 | 2 | 12187 | 98 | Il6ra,Csf1r | | 3.552e-02 | -3.34 | GO\_POSITIVE\_REGULATION\_OF\_MYELOID\_LEUKOCYTE\_DIFFERENTIATION | MSigDB lists | GO\_POSITIVE\_REGULATION\_OF\_MYELOID\_LEUKOCYTE\_DIFFERENTIATION | 37 | 2 | 12187 | 98 | Csf1r,Hcls1 | | 3.556e-02 | -3.34 | PTPase\_domain | interpro domains | IPR000242 | 36 | 2 | 13788 | 114 | Ptprc,Ptprn | | 3.567e-02 | -3.33 | GO\_NEGATIVE\_REGULATION\_OF\_CELLULAR\_COMPONENT\_ORGANIZATION | MSigDB lists | GO\_NEGATIVE\_REGULATION\_OF\_CELLULAR\_COMPONENT\_ORGANIZATION | 559 | 9 | 12187 | 98 | Gfap,Vim,Apod,Thy1,Clu,Myh9,Hspa2,Rap1gap,Csk | | 3.567e-02 | -3.33 | protein-containing complex binding | molecular function | GO:0044877 | 1044 | 14 | 13516 | 107 | Pcsk2,Thy1,Tspan4,Vim,Ctss,Myh9,Vwf,Hcls1,Ctsb,Fcer1g,Gfap,Fcgr3,Brk1,Clu | | 3.567e-02 | -3.33 | cytoskeletal protein binding | molecular function | GO:0008092 | 848 | 12 | 13516 | 107 | Capn3,Clu,Mark4,Ptprc,Hspa2,Hcls1,Myh9,Ccr5,Ptprn,Actb,Flnb,S100a6 | | 3.578e-02 | -3.33 | Activation of C3 and C5 | REACTOME pathways | R-MMU-174577 | 3 | 1 | 6297 | 76 | C3 | | 3.578e-02 | -3.33 | Dectin-2 family | REACTOME pathways | R-MMU-5621480 | 3 | 1 | 6297 | 76 | Fcer1g | | 3.578e-02 | -3.33 | Response to metal ions | REACTOME pathways | R-MMU-5660526 | 3 | 1 | 6297 | 76 | Mt3 | | 3.578e-02 | -3.33 | Metallothioneins bind metals | REACTOME pathways | R-MMU-5661231 | 3 | 1 | 6297 | 76 | Mt3 | | 3.578e-02 | -3.33 | TFAP2 (AP-2) family regulates transcription of cell cycle factors | REACTOME pathways | R-MMU-8866911 | 3 | 1 | 6297 | 76 | Tfap2c | | 3.578e-02 | -3.33 | Abacavir metabolism | REACTOME pathways | R-MMU-2161541 | 3 | 1 | 6297 | 76 | Adh1 | | 3.580e-02 | -3.33 | negative regulation of cellular component organization | biological process | GO:0051129 | 644 | 10 | 13711 | 111 | Clu,Ctsz,Apod,Hspa2,Myh9,Vim,Gfap,Ccr5,Rap1gap,Thy1 | | 3.584e-02 | -3.33 | GO\_MEIOTIC\_CELL\_CYCLE\_PROCESS | MSigDB lists | GO\_MEIOTIC\_CELL\_CYCLE\_PROCESS | 90 | 3 | 12187 | 98 | Lfng,Myh9,Hspa2 | | 3.584e-02 | -3.33 | GO\_DEFENSE\_RESPONSE\_TO\_BACTERIUM | MSigDB lists | GO\_DEFENSE\_RESPONSE\_TO\_BACTERIUM | 90 | 3 | 12187 | 98 | Tnfsf8,Fcer1g,Il6ra | | 3.599e-02 | -3.32 | positive regulation of T cell mediated immunity | biological process | GO:0002711 | 37 | 2 | 13711 | 111 | Il1r1,Ptprc | | 3.599e-02 | -3.32 | intermediate filament-based process | biological process | GO:0045103 | 37 | 2 | 13711 | 111 | Vim,Gfap | | 3.599e-02 | -3.32 | regulation of lamellipodium organization | biological process | GO:1902743 | 37 | 2 | 13711 | 111 | Rac2,Brk1 | | 3.599e-02 | -3.32 | insulin secretion | biological process | GO:0030073 | 37 | 2 | 13711 | 111 | Ptprn,Mc4r | | 3.599e-02 | -3.32 | positive regulation of nitric oxide metabolic process | biological process | GO:1904407 | 37 | 2 | 13711 | 111 | Tmem106a,Clu | | 3.599e-02 | -3.32 | regulation of collagen biosynthetic process | biological process | GO:0032965 | 37 | 2 | 13711 | 111 | Il6ra,Vim | | 3.600e-02 | -3.32 | regulation of leukocyte differentiation | biological process | GO:1902105 | 225 | 5 | 13711 | 111 | Csf1r,Hcls1,Ptprc,Tmem176a,Ikzf3 | | 3.606e-02 | -3.32 | Integrin-mediated Cell Adhesion | WikiPathways | WP6 | 91 | 4 | 3756 | 52 | Rac2,Capn3,Csk,Itgam | | 3.611e-02 | -3.32 | IVANOVA\_HEMATOPOIESIS\_MATURE\_CELL | MSigDB lists | IVANOVA\_HEMATOPOIESIS\_MATURE\_CELL | 227 | 5 | 12187 | 98 | C1qb,Ctsb,C3,Itgam,Lrg1 | | 3.613e-02 | -3.32 | regulation of multi-organism process | biological process | GO:0043900 | 468 | 8 | 13711 | 111 | Grn,Cyba,Ccr5,Serpinb9,Ifitm1,Treml2,Csf1r,Fam111a | | 3.613e-02 | -3.32 | positive regulation of cell migration | biological process | GO:0030335 | 468 | 8 | 13711 | 111 | Grn,Ccar1,Rac2,Fgf18,Il1r1,Thy1,Ptprc,Csf1r | | 3.615e-02 | -3.32 | Acylglycerol degradation | KEGG pathways | mmu\_M00098 | 3 | 1 | 5248 | 64 | Mgll | | 3.615e-02 | -3.32 | Acylglycerol degradation | KEGG pathways | M00098 | 3 | 1 | 5248 | 64 | Mgll | | 3.624e-02 | -3.32 | GSE22229\_RENAL\_TRANSPLANT\_IMMUNOSUPP\_THERAPY\_VS\_HEALTHY\_PBMC\_DN | MSigDB lists | GSE22229\_RENAL\_TRANSPLANT\_IMMUNOSUPP\_THERAPY\_VS\_HEALTHY\_PBMC\_DN | 155 | 4 | 12187 | 98 | Fcgr3,Ctsb,C3,Ppfibp1 | | 3.624e-02 | -3.32 | EGFR\_UP.V1\_UP | MSigDB lists | EGFR\_UP.V1\_UP | 155 | 4 | 12187 | 98 | Psmb10,Il1r1,Hspa2,Mgll | | 3.624e-02 | -3.32 | GSE45365\_NK\_CELL\_VS\_CD8\_TCELL\_MCMV\_INFECTION\_DN | MSigDB lists | GSE45365\_NK\_CELL\_VS\_CD8\_TCELL\_MCMV\_INFECTION\_DN | 155 | 4 | 12187 | 98 | Mgll,Psme1,Clu,Psmb10 | | 3.624e-02 | -3.32 | GSE360\_L\_MAJOR\_VS\_M\_TUBERCULOSIS\_DC\_UP | MSigDB lists | GSE360\_L\_MAJOR\_VS\_M\_TUBERCULOSIS\_DC\_UP | 155 | 4 | 12187 | 98 | Cox4i1,Cplx2,Flnb,Tspan4 | | 3.624e-02 | -3.32 | GSE2770\_UNTREATED\_VS\_ACT\_CD4\_TCELL\_6H\_UP | MSigDB lists | GSE2770\_UNTREATED\_VS\_ACT\_CD4\_TCELL\_6H\_UP | 155 | 4 | 12187 | 98 | Itgam,Ccr5,Ndufa13,Ptprn | | 3.624e-02 | -3.32 | GSE23568\_CTRL\_TRANSDUCED\_VS\_WT\_CD8\_TCELL\_DN | MSigDB lists | GSE23568\_CTRL\_TRANSDUCED\_VS\_WT\_CD8\_TCELL\_DN | 155 | 4 | 12187 | 98 | Thy1,Camk2b,Rac2,Csk | | 3.624e-02 | -3.32 | GSE15930\_STIM\_VS\_STIM\_AND\_IL12\_24H\_CD8\_T\_CELL\_DN | MSigDB lists | GSE15930\_STIM\_VS\_STIM\_AND\_IL12\_24H\_CD8\_T\_CELL\_DN | 155 | 4 | 12187 | 98 | Il1r1,Mark4,Igfbp6,Lgi4 | | 3.635e-02 | -3.31 | GO\_NEGATIVE\_REGULATION\_OF\_PROTEIN\_METABOLIC\_PROCESS | MSigDB lists | GO\_NEGATIVE\_REGULATION\_OF\_PROTEIN\_METABOLIC\_PROCESS | 839 | 12 | 12187 | 98 | Psme1,Clu,Capn3,Csk,C3,Psmb10,Serpina3n,Ptprc,Apod,Thy1,Serpinb9,Spint1 | | 3.637e-02 | -3.31 | supramolecular fiber organization | biological process | GO:0097435 | 384 | 7 | 13711 | 111 | Vim,Gfap,Capn3,Rac2,Myh9,Hcls1,Brk1 | | 3.639e-02 | -3.31 | morphogenesis of a branching epithelium | biological process | GO:0061138 | 154 | 4 | 13711 | 111 | Spint1,Mgp,Tfap2c,Ctsz | | 3.647e-02 | -3.31 | Y\_phosphatase | pfam domains | PF00102 | 36 | 2 | 12881 | 108 | Ptprn,Ptprc | | 3.648e-02 | -3.31 | Signaling by GPCR | REACTOME pathways | R-MMU-372790 | 564 | 12 | 6297 | 76 | Pde1a,Ccl6,Fgf18,Rac2,Mc4r,Mgll,Kl,Psme1,Psmb10,C3,Ccr5,Camk2b | | 3.658e-02 | -3.31 | positive regulation of cell differentiation | biological process | GO:0045597 | 927 | 13 | 13711 | 111 | Il1r1,Grn,Vim,Ptprc,C1qa,Clu,Camk2b,Fgf18,Gfap,Ccr5,Hcls1,Csf1r,Spint1 | | 3.659e-02 | -3.31 | positive regulation of cytosolic calcium ion concentration | biological process | GO:0007204 | 226 | 5 | 13711 | 111 | Cd52,Ptprc,Thy1,Capn3,Ccr5 | | 3.668e-02 | -3.31 | Rara (retinoic acid receptor, alpha) | protein interactions | 19401 | 64 | 3 | 6802 | 78 | Actb,Rps5,Vim | | 3.680e-02 | -3.30 | - | gene3d domains | 3.30.750.24 | 4 | 1 | 6647 | 62 | Slc26a2 | | 3.685e-02 | -3.30 | LABBE\_WNT3A\_TARGETS\_UP | MSigDB lists | LABBE\_WNT3A\_TARGETS\_UP | 91 | 3 | 12187 | 98 | Serpinb9,Ly6e,Actb | | 3.685e-02 | -3.30 | GO\_CELL\_CORTEX\_PART | MSigDB lists | GO\_CELL\_CORTEX\_PART | 91 | 3 | 12187 | 98 | Exoc3l4,Slc2a1,Myh9 | | 3.692e-02 | -3.30 | endocytic vesicle | cellular component | GO:0030139 | 156 | 4 | 13825 | 111 | Flnb,Vamp8,Mpeg1,Vim | | 3.698e-02 | -3.30 | GSE41176\_UNSTIM\_VS\_ANTI\_IGM\_STIM\_TAK1\_KO\_BCELL\_1H\_UP | MSigDB lists | GSE41176\_UNSTIM\_VS\_ANTI\_IGM\_STIM\_TAK1\_KO\_BCELL\_1H\_UP | 156 | 4 | 12187 | 98 | Thy1,S100a6,Ppfibp1,Il1r1 | | 3.698e-02 | -3.30 | GSE21063\_CTRL\_VS\_ANTI\_IGM\_STIM\_BCELL\_NFATC1\_KO\_3H\_UP | MSigDB lists | GSE21063\_CTRL\_VS\_ANTI\_IGM\_STIM\_BCELL\_NFATC1\_KO\_3H\_UP | 156 | 4 | 12187 | 98 | Tmem176a,Il1r1,S100a6,Serpinb9 | | 3.698e-02 | -3.30 | GSE14350\_IL2RB\_KO\_VS\_WT\_TEFF\_UP | MSigDB lists | GSE14350\_IL2RB\_KO\_VS\_WT\_TEFF\_UP | 156 | 4 | 12187 | 98 | Ctss,Tnfsf8,Cyba,Adh1 | | 3.698e-02 | -3.30 | GSE42088\_UNINF\_VS\_LEISHMANIA\_INF\_DC\_4H\_DN | MSigDB lists | GSE42088\_UNINF\_VS\_LEISHMANIA\_INF\_DC\_4H\_DN | 156 | 4 | 12187 | 98 | Rpl13,Rpl26,Rps8,Rpl31 | | 3.698e-02 | -3.30 | GSE5455\_HEALTHY\_VS\_TUMOR\_BEARING\_MOUSE\_SPLEEN\_MONOCYTE\_DN | MSigDB lists | GSE5455\_HEALTHY\_VS\_TUMOR\_BEARING\_MOUSE\_SPLEEN\_MONOCYTE\_DN | 156 | 4 | 12187 | 98 | Ccr5,Lfng,Rpl14,Rps5 | | 3.707e-02 | -3.30 | enzyme inhibitor activity | molecular function | GO:0004857 | 232 | 5 | 13516 | 107 | C3,Serpinb9,Spint1,Wfdc17,Serpina3n | | 3.724e-02 | -3.29 | amide biosynthetic process | biological process | GO:0043604 | 386 | 7 | 13711 | 111 | Rps20,Rps8,Rpl13,Rpl31,Rpl26,Rpl14,Rps5 | | 3.729e-02 | -3.29 | peptidase activity | molecular function | GO:0008233 | 395 | 7 | 13516 | 107 | Ctsa,Psmb10,Capn3,Pcsk2,Ctsz,Ctsb,Ctss | | 3.730e-02 | -3.29 | KUNINGER\_IGF1\_VS\_PDGFB\_TARGETS\_DN | MSigDB lists | KUNINGER\_IGF1\_VS\_PDGFB\_TARGETS\_DN | 38 | 2 | 12187 | 98 | Il1r1,Mgp | | 3.730e-02 | -3.29 | MCDOWELL\_ACUTE\_LUNG\_INJURY\_UP | MSigDB lists | MCDOWELL\_ACUTE\_LUNG\_INJURY\_UP | 38 | 2 | 12187 | 98 | Fcgr3,Lrg1 | | 3.730e-02 | -3.29 | ST\_T\_CELL\_SIGNAL\_TRANSDUCTION | MSigDB lists | ST\_T\_CELL\_SIGNAL\_TRANSDUCTION | 38 | 2 | 12187 | 98 | Ptprc,Csk | | 3.730e-02 | -3.29 | LIANG\_HEMATOPOIESIS\_STEM\_CELL\_NUMBER\_LARGE\_VS\_TINY\_UP | MSigDB lists | LIANG\_HEMATOPOIESIS\_STEM\_CELL\_NUMBER\_LARGE\_VS\_TINY\_UP | 38 | 2 | 12187 | 98 | S100a6,Il1r1 | | 3.731e-02 | -3.29 | ANATO | smart domains | SM00104 | 4 | 1 | 7188 | 68 | C3 | | 3.731e-02 | -3.29 | MANEC | smart domains | SM00765 | 4 | 1 | 7188 | 68 | Spint1 | | 3.753e-02 | -3.28 | negative regulation of biological process | biological process | GO:0048519 | 4173 | 43 | 13711 | 111 | Fam111a,Mt3,Myh9,Ndufa13,Ccr5,Cyba,Lfng,Csk,Ctsz,Vamp8,Rap1gap,Actb,Fcer1g,Bhlhe41,Ptprc,Grn,Ccar1,Mgll,Thy1,Spint1,Ifitm1,Csf1r,Hcls1,Serpina3n,Tfap2c,C3,Ctsa,Rpl26,Vim,Gfap,Tsc22d1,Ctsb,Capn3,Il6ra,Serpinb9,Camk2b,Clu,Cd68,Abhd17c,Apod,Hspa2,Tmem176a,Mc4r | | 3.767e-02 | -3.28 | signal transduction | biological process | GO:0007165 | 2703 | 30 | 13711 | 111 | Mark4,Saa3,Csk,Ccr5,Ndufa13,Ptprn,C3,Mt3,Brk1,Csf1r,Ifitm1,Rac2,Vim,Klhl6,Rpl26,Fcgr3,Ccl6,Fgf18,Ly6e,Il6ra,Slc17a7,Pde1a,Fcer1g,Il1r1,Thy1,Kl,Mc4r,Adgre1,Ptprc,Clu | | 3.772e-02 | -3.28 | GSE41978\_KLRG1\_HIGH\_VS\_LOW\_EFFECTOR\_CD8\_TCELL\_DN | MSigDB lists | GSE41978\_KLRG1\_HIGH\_VS\_LOW\_EFFECTOR\_CD8\_TCELL\_DN | 157 | 4 | 12187 | 98 | Dek,Rps5,Actb,Cox4i1 | | 3.772e-02 | -3.28 | GSE4142\_NAIVE\_VS\_GC\_BCELL\_UP | MSigDB lists | GSE4142\_NAIVE\_VS\_GC\_BCELL\_UP | 157 | 4 | 12187 | 98 | Mpeg1,Tmem176a,Grn,Myh9 | | 3.772e-02 | -3.28 | GSE4535\_BM\_DERIVED\_DC\_VS\_FOLLICULAR\_DC\_UP | MSigDB lists | GSE4535\_BM\_DERIVED\_DC\_VS\_FOLLICULAR\_DC\_UP | 157 | 4 | 12187 | 98 | Ppfibp1,Ndufa13,Tnfsf8,Vamp8 | | 3.772e-02 | -3.28 | GSE7831\_1H\_VS\_4H\_CPG\_STIM\_PDC\_DN | MSigDB lists | GSE7831\_1H\_VS\_4H\_CPG\_STIM\_PDC\_DN | 157 | 4 | 12187 | 98 | Mpeg1,Flnb,Ctss,Rac2 | | 3.772e-02 | -3.28 | GSE3982\_DC\_VS\_NKCELL\_UP | MSigDB lists | GSE3982\_DC\_VS\_NKCELL\_UP | 157 | 4 | 12187 | 98 | Ctsz,Clu,Spint1,Treml2 | | 3.772e-02 | -3.28 | GSE40666\_UNTREATED\_VS\_IFNA\_STIM\_STAT4\_KO\_EFFECTOR\_CD8\_TCELL\_90MIN\_UP | MSigDB lists | GSE40666\_UNTREATED\_VS\_IFNA\_STIM\_STAT4\_KO\_EFFECTOR\_CD8\_TCELL\_90MIN\_UP | 157 | 4 | 12187 | 98 | Sh3bgrl3,Mgll,Il6ra,Itgam | | 3.772e-02 | -3.28 | GSE7831\_UNSTIM\_VS\_INFLUENZA\_STIM\_PDC\_4H\_UP | MSigDB lists | GSE7831\_UNSTIM\_VS\_INFLUENZA\_STIM\_PDC\_4H\_UP | 157 | 4 | 12187 | 98 | Pcsk2,Gfap,Csf1r,Flnb | | 3.772e-02 | -3.28 | GSE1460\_DP\_THYMOCYTE\_VS\_NAIVE\_CD4\_TCELL\_ADULT\_BLOOD\_DN | MSigDB lists | GSE1460\_DP\_THYMOCYTE\_VS\_NAIVE\_CD4\_TCELL\_ADULT\_BLOOD\_DN | 157 | 4 | 12187 | 98 | Rpl26,Rps8,Rpl13,Rps5 | | 3.772e-02 | -3.28 | GSE32034\_UNTREATED\_VS\_ROSIGLIZATONE\_TREATED\_LY6C\_HIGH\_MONOCYTE\_UP | MSigDB lists | GSE32034\_UNTREATED\_VS\_ROSIGLIZATONE\_TREATED\_LY6C\_HIGH\_MONOCYTE\_UP | 157 | 4 | 12187 | 98 | Capn3,Vim,Flnb,Slc2a1 | | 3.773e-02 | -3.28 | transmembrane signaling receptor activity | molecular function | GO:0004888 | 483 | 8 | 13516 | 107 | Il6ra,Il1r1,Adgre1,Fcer1g,Mc4r,Ccr5,Csf1r,Fcgr3 | | 3.779e-02 | -3.28 | negative regulation of leukocyte mediated immunity | biological process | GO:0002704 | 38 | 2 | 13711 | 111 | Ptprc,Serpinb9 | | 3.779e-02 | -3.28 | positive regulation of cytokine production involved in immune response | biological process | GO:0002720 | 38 | 2 | 13711 | 111 | Fcer1g,Il1r1 | | 3.779e-02 | -3.28 | actin polymerization or depolymerization | biological process | GO:0008154 | 38 | 2 | 13711 | 111 | Hcls1,Myh9 | | 3.779e-02 | -3.28 | IF\_ROD\_2 | prosite domains | PS51842 | 30 | 2 | 8845 | 91 | Gfap,Vim | | 3.780e-02 | -3.28 | Kcnma1 (potassium large conductance calcium-activated channel, subfamily M, alpha member 1) | protein interactions | 16531 | 162 | 5 | 6802 | 78 | Ldha,Vim,Gfap,Hspa2,Actb | | 3.787e-02 | -3.27 | Stat3 (signal transducer and activator of transcription 3) | protein interactions | 20848 | 27 | 2 | 6802 | 78 | Csf1r,Ndufa13 | | 3.787e-02 | -3.27 | GO\_GENERATION\_OF\_PRECURSOR\_METABOLITES\_AND\_ENERGY | MSigDB lists | GO\_GENERATION\_OF\_PRECURSOR\_METABOLITES\_AND\_ENERGY | 230 | 5 | 12187 | 98 | Ldha,Cox4i1,Mc4r,Ndufa13,Kl | | 3.788e-02 | -3.27 | REACTOME\_SIGNALING\_BY\_FGFR | MSigDB lists | REACTOME\_SIGNALING\_BY\_FGFR | 92 | 3 | 12187 | 98 | Kl,Fgf18,Pde1a | | 3.788e-02 | -3.27 | GSE45365\_NK\_CELL\_VS\_BCELL\_DN | MSigDB lists | GSE45365\_NK\_CELL\_VS\_BCELL\_DN | 92 | 3 | 12187 | 98 | Il1r1,Treml2,C1qa | | 3.802e-02 | -3.27 | positive regulation of proteolysis | biological process | GO:0045862 | 306 | 6 | 13711 | 111 | Myh9,Clu,Psme1,Capn3,Grn,Ndufa13 | | 3.806e-02 | -3.27 | negative regulation of apoptotic process | biological process | GO:0043066 | 743 | 11 | 13711 | 111 | Clu,Fcer1g,Mt3,Hcls1,Csf1r,Grn,Ccr5,Ndufa13,Tsc22d1,Capn3,Serpinb9 | | 3.825e-02 | -3.26 | MODULE\_1 | MSigDB lists | MODULE\_1 | 309 | 6 | 12187 | 98 | Fmod,Igfbp6,Thy1,Vwf,Il1r1,C1qb | | 3.837e-02 | -3.26 | localization of cell | biological process | GO:0051674 | 744 | 11 | 13711 | 111 | Saa3,Ccr5,Fcgr3,Ptprc,Myh9,Brk1,Fcer1g,Rac2,Itgam,Actb,Ccl6 | | 3.837e-02 | -3.26 | cell motility | biological process | GO:0048870 | 744 | 11 | 13711 | 111 | Saa3,Ccr5,Myh9,Brk1,Fcer1g,Rac2,Itgam,Actb,Fcgr3,Ptprc,Ccl6 | | 3.847e-02 | -3.26 | GSE22140\_GERMFREE\_VS\_SPF\_ARTHRITIC\_MOUSE\_CD4\_TCELL\_DN | MSigDB lists | GSE22140\_GERMFREE\_VS\_SPF\_ARTHRITIC\_MOUSE\_CD4\_TCELL\_DN | 158 | 4 | 12187 | 98 | C1qb,Psmb10,Fcer1g,Csk | | 3.847e-02 | -3.26 | GSE17721\_POLYIC\_VS\_GARDIQUIMOD\_24H\_BMDC\_UP | MSigDB lists | GSE17721\_POLYIC\_VS\_GARDIQUIMOD\_24H\_BMDC\_UP | 158 | 4 | 12187 | 98 | Thy1,Tsc22d1,Grn,Tnfsf8 | | 3.847e-02 | -3.26 | GSE7219\_WT\_VS\_NIK\_NFKB2\_KO\_LPS\_AND\_ANTI\_CD40\_STIM\_DC\_DN | MSigDB lists | GSE7219\_WT\_VS\_NIK\_NFKB2\_KO\_LPS\_AND\_ANTI\_CD40\_STIM\_DC\_DN | 158 | 4 | 12187 | 98 | Flnb,Ctss,Lfng,Fam111a | | 3.847e-02 | -3.26 | GSE25088\_WT\_VS\_STAT6\_KO\_MACROPHAGE\_ROSIGLITAZONE\_STIM\_DN | MSigDB lists | GSE25088\_WT\_VS\_STAT6\_KO\_MACROPHAGE\_ROSIGLITAZONE\_STIM\_DN | 158 | 4 | 12187 | 98 | Actb,Ptprc,Flnb,Thy1 | | 3.847e-02 | -3.26 | GSE45881\_CXCR6HI\_VS\_CXCR1LO\_COLONIC\_LAMINA\_PROPRIA\_DN | MSigDB lists | GSE45881\_CXCR6HI\_VS\_CXCR1LO\_COLONIC\_LAMINA\_PROPRIA\_DN | 158 | 4 | 12187 | 98 | Treml2,Slc2a1,Ppfibp1,Fmod | | 3.847e-02 | -3.26 | GSE42088\_2H\_VS\_24H\_LEISHMANIA\_INF\_DC\_UP | MSigDB lists | GSE42088\_2H\_VS\_24H\_LEISHMANIA\_INF\_DC\_UP | 158 | 4 | 12187 | 98 | Rpl31,Ly6e,Itgam,Ctss | | 3.847e-02 | -3.26 | JISON\_SICKLE\_CELL\_DISEASE\_UP | MSigDB lists | JISON\_SICKLE\_CELL\_DISEASE\_UP | 158 | 4 | 12187 | 98 | Cplx2,Vamp8,Ly6e,Ctsa | | 3.847e-02 | -3.26 | GSE8921\_UNSTIM\_VS\_TLR1\_2\_STIM\_MONOCYTE\_12H\_DN | MSigDB lists | GSE8921\_UNSTIM\_VS\_TLR1\_2\_STIM\_MONOCYTE\_12H\_DN | 158 | 4 | 12187 | 98 | Fam111a,Ndufa13,Rpl26,Spint1 | | 3.847e-02 | -3.26 | GSE40274\_HELIOS\_VS\_FOXP3\_AND\_HELIOS\_TRANSDUCED\_ACTIVATED\_CD4\_TCELL\_UP | MSigDB lists | GSE40274\_HELIOS\_VS\_FOXP3\_AND\_HELIOS\_TRANSDUCED\_ACTIVATED\_CD4\_TCELL\_UP | 158 | 4 | 12187 | 98 | Ikzf3,Ahnak,Rpl13,Capn3 | | 3.847e-02 | -3.26 | GSE17721\_CTRL\_VS\_POLYIC\_6H\_BMDC\_DN | MSigDB lists | GSE17721\_CTRL\_VS\_POLYIC\_6H\_BMDC\_DN | 158 | 4 | 12187 | 98 | Serpina3n,Apod,Tmem106a,Mark4 | | 3.847e-02 | -3.26 | Osteoclast differentiation | KEGG pathways | mmu04380 | 105 | 4 | 5248 | 64 | Csf1r,Il1r1,Cyba,Fcgr3 | | 3.847e-02 | -3.26 | Osteoclast differentiation | KEGG pathways | ko04380 | 105 | 4 | 5248 | 64 | Il1r1,Csf1r,Fcgr3,Cyba | | 3.854e-02 | -3.26 | Signaling by VEGF | REACTOME pathways | R-MMU-194138 | 263 | 7 | 6297 | 76 | Psmb10,Psme1,Camk2b,Cyba,Brk1,Kl,Fgf18 | | 3.858e-02 | -3.26 | divalent inorganic cation homeostasis | biological process | GO:0072507 | 389 | 7 | 13711 | 111 | Ptprc,Cd52,Mt3,Ccr5,Kl,Thy1,Capn3 | | 3.867e-02 | -3.25 | GO\_PROTEOLYSIS | MSigDB lists | GO\_PROTEOLYSIS | 847 | 12 | 12187 | 98 | C3,Ctsz,Ctss,Capn3,Psme1,Psmb10,C1qa,C1qb,Pcsk2,Myh9,Ctsb,Ctsa | | 3.876e-02 | -3.25 | SMID\_BREAST\_CANCER\_NORMAL\_LIKE\_UP | MSigDB lists | SMID\_BREAST\_CANCER\_NORMAL\_LIKE\_UP | 310 | 6 | 12187 | 98 | Rac2,Il6ra,Il1r1,Ptprc,Clu,C3 | | 3.889e-02 | -3.25 | mouse chr18 E1\|18 34.41 cM | chromosome location | mouse chr18 E1\|18 34.41 cM | 5 | 1 | 14556 | 115 | Csf1r | | 3.889e-02 | -3.25 | mouse chr19 A\|19 | chromosome location | mouse chr19 A\|19 | 5 | 1 | 14556 | 115 | Fam111a | | 3.889e-02 | -3.25 | mouse chr6 F3\|6 59.32 cM | chromosome location | mouse chr6 F3\|6 59.32 cM | 5 | 1 | 14556 | 115 | Vwf | | 3.892e-02 | -3.25 | HOLLERN\_EMT\_BREAST\_TUMOR\_UP | MSigDB lists | HOLLERN\_EMT\_BREAST\_TUMOR\_UP | 93 | 3 | 12187 | 98 | Vim,Csf1r,Tspan4 | | 3.892e-02 | -3.25 | MODULE\_177 | MSigDB lists | MODULE\_177 | 93 | 3 | 12187 | 98 | Dek,Hcls1,Ptprc | | 3.892e-02 | -3.25 | GO\_SYNAPTIC\_VESICLE\_LOCALIZATION | MSigDB lists | GO\_SYNAPTIC\_VESICLE\_LOCALIZATION | 93 | 3 | 12187 | 98 | Grn,Cplx2,Lin7a | | 3.892e-02 | -3.25 | GO\_INOSITOL\_LIPID\_MEDIATED\_SIGNALING | MSigDB lists | GO\_INOSITOL\_LIPID\_MEDIATED\_SIGNALING | 93 | 3 | 12187 | 98 | Fgf18,Csf1r,Kl | | 3.892e-02 | -3.25 | GO\_SKELETAL\_MUSCLE\_ORGAN\_DEVELOPMENT | MSigDB lists | GO\_SKELETAL\_MUSCLE\_ORGAN\_DEVELOPMENT | 93 | 3 | 12187 | 98 | Flnb,Ctsb,Csf1r | | 3.892e-02 | -3.25 | LEIN\_CHOROID\_PLEXUS\_MARKERS | MSigDB lists | LEIN\_CHOROID\_PLEXUS\_MARKERS | 93 | 3 | 12187 | 98 | Adh1,Kl,Mgp | | 3.892e-02 | -3.25 | MODULE\_192 | MSigDB lists | MODULE\_192 | 93 | 3 | 12187 | 98 | Serpina3n,Rps5,Rpl13 | | 3.897e-02 | -3.25 | acetylcholine receptor inhibitor activity | molecular function | GO:0030550 | 5 | 1 | 13516 | 107 | Ly6e | | 3.897e-02 | -3.25 | ciliary neurotrophic factor receptor activity | molecular function | GO:0004897 | 5 | 1 | 13516 | 107 | Il6ra | | 3.897e-02 | -3.25 | Toll-like receptor 4 binding | molecular function | GO:0035662 | 5 | 1 | 13516 | 107 | Saa3 | | 3.897e-02 | -3.25 | superoxide-generating NADPH oxidase activity | molecular function | GO:0016175 | 5 | 1 | 13516 | 107 | Cyba | | 3.897e-02 | -3.25 | interleukin-1 receptor activity | molecular function | GO:0004908 | 5 | 1 | 13516 | 107 | Il1r1 | | 3.897e-02 | -3.25 | serine-type exopeptidase activity | molecular function | GO:0070008 | 5 | 1 | 13516 | 107 | Ctsa | | 3.897e-02 | -3.25 | inorganic phosphate transmembrane transporter activity | molecular function | GO:0005315 | 5 | 1 | 13516 | 107 | Slc17a7 | | 3.901e-02 | -3.24 | GO\_REGULATION\_OF\_TRANSFERASE\_ACTIVITY | MSigDB lists | GO\_REGULATION\_OF\_TRANSFERASE\_ACTIVITY | 753 | 11 | 12187 | 98 | Il6ra,Thy1,Csf1r,Rac2,Fgf18,Psmb10,Ptprc,Psme1,Clu,Hspa2,Csk | | 3.912e-02 | -3.24 | NATSUME\_RESPONSE\_TO\_INTERFERON\_BETA\_DN | MSigDB lists | NATSUME\_RESPONSE\_TO\_INTERFERON\_BETA\_DN | 39 | 2 | 12187 | 98 | Cox4i1,Igfbp6 | | 3.912e-02 | -3.24 | GO\_DEFENSE\_RESPONSE\_TO\_GRAM\_POSITIVE\_BACTERIUM | MSigDB lists | GO\_DEFENSE\_RESPONSE\_TO\_GRAM\_POSITIVE\_BACTERIUM | 39 | 2 | 12187 | 98 | Tnfsf8,Il6ra | | 3.912e-02 | -3.24 | GO\_REGULATION\_OF\_B\_CELL\_PROLIFERATION | MSigDB lists | GO\_REGULATION\_OF\_B\_CELL\_PROLIFERATION | 39 | 2 | 12187 | 98 | Ikzf3,Ptprc | | 3.912e-02 | -3.24 | GO\_POSITIVE\_REGULATION\_OF\_GLIOGENESIS | MSigDB lists | GO\_POSITIVE\_REGULATION\_OF\_GLIOGENESIS | 39 | 2 | 12187 | 98 | Spint1,Gfap | | 3.912e-02 | -3.24 | GO\_NEURAL\_RETINA\_DEVELOPMENT | MSigDB lists | GO\_NEURAL\_RETINA\_DEVELOPMENT | 39 | 2 | 12187 | 98 | Thy1,Slc17a7 | | 3.912e-02 | -3.24 | GO\_POSITIVE\_REGULATION\_OF\_REACTIVE\_OXYGEN\_SPECIES\_BIOSYNTHETIC\_PROCESS | MSigDB lists | GO\_POSITIVE\_REGULATION\_OF\_REACTIVE\_OXYGEN\_SPECIES\_BIOSYNTHETIC\_PROCESS | 39 | 2 | 12187 | 98 | Clu,Cyba | | 3.912e-02 | -3.24 | HOQUE\_METHYLATED\_IN\_CANCER | MSigDB lists | HOQUE\_METHYLATED\_IN\_CANCER | 39 | 2 | 12187 | 98 | Gfap,Apod | | 3.922e-02 | -3.24 | GSE6092\_IFNG\_VS\_IFNG\_AND\_B\_BURGDORFERI\_INF\_ENDOTHELIAL\_CELL\_UP | MSigDB lists | GSE6092\_IFNG\_VS\_IFNG\_AND\_B\_BURGDORFERI\_INF\_ENDOTHELIAL\_CELL\_UP | 159 | 4 | 12187 | 98 | Mpeg1,Tspan4,Rab3il1,Lfng | | 3.922e-02 | -3.24 | GSE22196\_HEALTHY\_VS\_OBESE\_MOUSE\_SKIN\_GAMMADELTA\_TCELL\_UP | MSigDB lists | GSE22196\_HEALTHY\_VS\_OBESE\_MOUSE\_SKIN\_GAMMADELTA\_TCELL\_UP | 159 | 4 | 12187 | 98 | Csk,Clu,Psmb10,C1qb | | 3.922e-02 | -3.24 | GSE29618\_PDC\_VS\_MDC\_DN | MSigDB lists | GSE29618\_PDC\_VS\_MDC\_DN | 159 | 4 | 12187 | 98 | Ptprc,Fcgr3,Il6ra,Itgam | | 3.922e-02 | -3.24 | GSE3982\_MAST\_CELL\_VS\_NKCELL\_UP | MSigDB lists | GSE3982\_MAST\_CELL\_VS\_NKCELL\_UP | 159 | 4 | 12187 | 98 | Cplx2,Hspa2,Tsc22d1,Il1r1 | | 3.922e-02 | -3.24 | GSE15735\_CTRL\_VS\_HDAC\_INHIBITOR\_TREATED\_CD4\_TCELL\_12H\_DN | MSigDB lists | GSE15735\_CTRL\_VS\_HDAC\_INHIBITOR\_TREATED\_CD4\_TCELL\_12H\_DN | 159 | 4 | 12187 | 98 | Psme1,Il6ra,Serpinb9,Lfng | | 3.922e-02 | -3.24 | GSE11057\_NAIVE\_VS\_EFF\_MEMORY\_CD4\_TCELL\_DN | MSigDB lists | GSE11057\_NAIVE\_VS\_EFF\_MEMORY\_CD4\_TCELL\_DN | 159 | 4 | 12187 | 98 | Ctsa,Ldha,Sh3bgrl3,Clu | | 3.922e-02 | -3.24 | GSE21927\_SPLEEN\_C57BL6\_VS\_4T1\_TUMOR\_BALBC\_MONOCYTES\_UP | MSigDB lists | GSE21927\_SPLEEN\_C57BL6\_VS\_4T1\_TUMOR\_BALBC\_MONOCYTES\_UP | 159 | 4 | 12187 | 98 | Rpl13,Cox4i1,Rps8,Rps5 | | 3.922e-02 | -3.24 | GO\_REGULATION\_OF\_NEUROTRANSMITTER\_LEVELS | MSigDB lists | GO\_REGULATION\_OF\_NEUROTRANSMITTER\_LEVELS | 159 | 4 | 12187 | 98 | Gfap,Slc17a7,Cplx2,Lin7a | | 3.922e-02 | -3.24 | GSE36009\_UNSTIM\_VS\_LPS\_STIM\_DC\_UP | MSigDB lists | GSE36009\_UNSTIM\_VS\_LPS\_STIM\_DC\_UP | 159 | 4 | 12187 | 98 | Lfng,Psme1,Slc26a2,Rab3il1 | | 3.922e-02 | -3.24 | GSE29949\_DC\_BRAIN\_VS\_MONOCYTE\_BONE\_MARROW\_UP | MSigDB lists | GSE29949\_DC\_BRAIN\_VS\_MONOCYTE\_BONE\_MARROW\_UP | 159 | 4 | 12187 | 98 | Cyba,Ahnak,Myh9,Vamp8 | | 3.951e-02 | -3.23 | CatSper complex | cellular component | GO:0036128 | 5 | 1 | 13825 | 111 | Hspa2 | | 3.951e-02 | -3.23 | type III intermediate filament | cellular component | GO:0045098 | 5 | 1 | 13825 | 111 | Vim | | 3.951e-02 | -3.23 | MPP7-DLG1-LIN7 complex | cellular component | GO:0097025 | 5 | 1 | 13825 | 111 | Lin7a | | 3.951e-02 | -3.23 | astrocyte end-foot | cellular component | GO:0097450 | 5 | 1 | 13825 | 111 | Gfap | | 3.957e-02 | -3.23 | OXFORD\_RALB\_TARGETS\_UP | MSigDB lists | OXFORD\_RALB\_TARGETS\_UP | 5 | 1 | 12187 | 98 | Clu | | 3.957e-02 | -3.23 | SASAI\_TARGETS\_OF\_CXCR6\_AND\_PTCH1\_DN | MSigDB lists | SASAI\_TARGETS\_OF\_CXCR6\_AND\_PTCH1\_DN | 5 | 1 | 12187 | 98 | Apod | | 3.957e-02 | -3.23 | MILICIC\_FAMILIAL\_ADENOMATOUS\_POLYPOSIS\_DN | MSigDB lists | MILICIC\_FAMILIAL\_ADENOMATOUS\_POLYPOSIS\_DN | 5 | 1 | 12187 | 98 | C3 | | 3.957e-02 | -3.23 | MODULE\_514 | MSigDB lists | MODULE\_514 | 5 | 1 | 12187 | 98 | Adh1 | | 3.957e-02 | -3.23 | CALVET\_IRINOTECAN\_SENSITIVE\_VS\_REVERTED\_UP | MSigDB lists | CALVET\_IRINOTECAN\_SENSITIVE\_VS\_REVERTED\_UP | 5 | 1 | 12187 | 98 | Vim | | 3.957e-02 | -3.23 | NEBEN\_AML\_WITH\_FLT3\_OR\_NRAS\_UP | MSigDB lists | NEBEN\_AML\_WITH\_FLT3\_OR\_NRAS\_UP | 5 | 1 | 12187 | 98 | Lfng | | 3.957e-02 | -3.23 | ROETH\_TERT\_TARGETS\_DN | MSigDB lists | ROETH\_TERT\_TARGETS\_DN | 5 | 1 | 12187 | 98 | Ccr5 | | 3.957e-02 | -3.23 | BIOCARTA\_TCRA\_PATHWAY | MSigDB lists | BIOCARTA\_TCRA\_PATHWAY | 5 | 1 | 12187 | 98 | Ptprc | | 3.957e-02 | -3.23 | LOPEZ\_MESOTHELIOMA\_SURVIVAL\_OVERALL\_UP | MSigDB lists | LOPEZ\_MESOTHELIOMA\_SURVIVAL\_OVERALL\_UP | 5 | 1 | 12187 | 98 | Il6ra | | 3.957e-02 | -3.23 | MARSHALL\_VIRAL\_INFECTION\_RESPONSE\_UP | MSigDB lists | MARSHALL\_VIRAL\_INFECTION\_RESPONSE\_UP | 5 | 1 | 12187 | 98 | Ctss | | 3.957e-02 | -3.23 | chr3p13 | MSigDB lists | chr3p13 | 5 | 1 | 12187 | 98 | Mgll | | 3.957e-02 | -3.23 | GO\_SUPEROXIDE\_GENERATING\_NADPH\_OXIDASE\_ACTIVITY | MSigDB lists | GO\_SUPEROXIDE\_GENERATING\_NADPH\_OXIDASE\_ACTIVITY | 5 | 1 | 12187 | 98 | Cyba | | 3.958e-02 | -3.23 | AB\_hydrolase | interpro domains | IPR029058 | 91 | 3 | 13788 | 114 | Mgll,Abhd17c,Ctsa | | 3.963e-02 | -3.23 | organic hydroxy compound catabolic process | biological process | GO:1901616 | 39 | 2 | 13711 | 111 | Adh1,Mt3 | | 3.963e-02 | -3.23 | regulation of meiotic cell cycle | biological process | GO:0051445 | 39 | 2 | 13711 | 111 | Lfng,Camk2b | | 3.963e-02 | -3.23 | Platelet activation | KEGG pathways | mmu04611 | 106 | 4 | 5248 | 64 | Fcer1g,Actb,Vwf,Vamp8 | | 3.965e-02 | -3.23 | GO\_POSITIVE\_REGULATION\_OF\_PHOSPHORUS\_METABOLIC\_PROCESS | MSigDB lists | GO\_POSITIVE\_REGULATION\_OF\_PHOSPHORUS\_METABOLIC\_PROCESS | 755 | 11 | 12187 | 98 | Ptprc,Fgf18,Hcls1,Kl,C3,Csk,Hspa2,Clu,Il6ra,Csf1r,Mc4r | | 3.965e-02 | -3.23 | myeloid leukocyte differentiation | biological process | GO:0002573 | 93 | 3 | 13711 | 111 | Csf1r,Fcer1g,Itgam | | 3.969e-02 | -3.23 | KEGG\_MAPK\_SIGNALING\_PATHWAY | MSigDB lists | KEGG\_MAPK\_SIGNALING\_PATHWAY | 233 | 5 | 12187 | 98 | Hspa2,Rac2,Il1r1,Flnb,Fgf18 | | 3.979e-02 | -3.22 | GO\_POSITIVE\_REGULATION\_OF\_LOCOMOTION | MSigDB lists | GO\_POSITIVE\_REGULATION\_OF\_LOCOMOTION | 312 | 6 | 12187 | 98 | Ptprc,Csf1r,Fgf18,Ccar1,Rac2,Il6ra | | 3.983e-02 | -3.22 | microglia differentiation | biological process | GO:0014004 | 5 | 1 | 13711 | 111 | Itgam | | 3.983e-02 | -3.22 | branch elongation involved in mammary gland duct branching | biological process | GO:0060751 | 5 | 1 | 13711 | 111 | Tfap2c | | 3.983e-02 | -3.22 | detoxification of copper ion | biological process | GO:0010273 | 5 | 1 | 13711 | 111 | Mt3 | | 3.983e-02 | -3.22 | tumor necrosis factor production | biological process | GO:0032640 | 5 | 1 | 13711 | 111 | Tmem106a | | 3.983e-02 | -3.22 | interleukin-1 beta secretion | biological process | GO:0050702 | 5 | 1 | 13711 | 111 | Tmem106a | | 3.983e-02 | -3.22 | egg activation | biological process | GO:0007343 | 5 | 1 | 13711 | 111 | Camk2b | | 3.983e-02 | -3.22 | microglial cell activation involved in immune response | biological process | GO:0002282 | 5 | 1 | 13711 | 111 | Grn | | 3.983e-02 | -3.22 | negative regulation of protein localization to microtubule | biological process | GO:1902817 | 5 | 1 | 13711 | 111 | Abhd17c | | 3.983e-02 | -3.22 | negative regulation of T cell migration | biological process | GO:2000405 | 5 | 1 | 13711 | 111 | Apod | | 3.983e-02 | -3.22 | microglia development | biological process | GO:0014005 | 5 | 1 | 13711 | 111 | Itgam | | 3.983e-02 | -3.22 | cyclic nucleotide catabolic process | biological process | GO:0009214 | 5 | 1 | 13711 | 111 | Pde1a | | 3.983e-02 | -3.22 | protection from natural killer cell mediated cytotoxicity | biological process | GO:0042270 | 5 | 1 | 13711 | 111 | Serpinb9 | | 3.983e-02 | -3.22 | interleukin-2 production | biological process | GO:0032623 | 5 | 1 | 13711 | 111 | Fcer1g | | 3.983e-02 | -3.22 | regulation of grooming behavior | biological process | GO:2000821 | 5 | 1 | 13711 | 111 | Mc4r | | 3.983e-02 | -3.22 | tumor necrosis factor superfamily cytokine production | biological process | GO:0071706 | 5 | 1 | 13711 | 111 | Tmem106a | | 3.983e-02 | -3.22 | stress response to copper ion | biological process | GO:1990169 | 5 | 1 | 13711 | 111 | Mt3 | | 3.983e-02 | -3.22 | Fc receptor mediated stimulatory signaling pathway | biological process | GO:0002431 | 5 | 1 | 13711 | 111 | Fcer1g | | 3.983e-02 | -3.22 | negative regulation of skeletal muscle cell differentiation | biological process | GO:2001015 | 5 | 1 | 13711 | 111 | Capn3 | | 3.983e-02 | -3.22 | negative regulation of Golgi to plasma membrane protein transport | biological process | GO:0042997 | 5 | 1 | 13711 | 111 | Csk | | 3.983e-02 | -3.22 | lactate biosynthetic process | biological process | GO:0019249 | 5 | 1 | 13711 | 111 | Ldha | | 3.983e-02 | -3.22 | regulation of protein localization to microtubule | biological process | GO:1902816 | 5 | 1 | 13711 | 111 | Abhd17c | | 3.983e-02 | -3.22 | intramembranous ossification | biological process | GO:0001957 | 5 | 1 | 13711 | 111 | Fgf18 | | 3.983e-02 | -3.22 | negative regulation of antigen processing and presentation | biological process | GO:0002578 | 5 | 1 | 13711 | 111 | Cd68 | | 3.983e-02 | -3.22 | positive regulation of Fc-gamma receptor signaling pathway involved in phagocytosis | biological process | GO:1905451 | 5 | 1 | 13711 | 111 | Ptprc | | 3.983e-02 | -3.22 | response to macrophage colony-stimulating factor | biological process | GO:0036005 | 5 | 1 | 13711 | 111 | Csf1r | | 3.983e-02 | -3.22 | positive regulation of astrocyte activation | biological process | GO:0061890 | 5 | 1 | 13711 | 111 | C1qa | | 3.983e-02 | -3.22 | positive regulation of NAD(P)H oxidase activity | biological process | GO:0033864 | 5 | 1 | 13711 | 111 | Cyba | | 3.983e-02 | -3.22 | regulation of respiratory burst involved in inflammatory response | biological process | GO:0060264 | 5 | 1 | 13711 | 111 | Grn | | 3.983e-02 | -3.22 | cellular response to macrophage colony-stimulating factor stimulus | biological process | GO:0036006 | 5 | 1 | 13711 | 111 | Csf1r | | 3.983e-02 | -3.22 | positive regulation of NK T cell activation | biological process | GO:0051135 | 5 | 1 | 13711 | 111 | Il6ra | | 3.983e-02 | -3.22 | negative regulation of macrophage apoptotic process | biological process | GO:2000110 | 5 | 1 | 13711 | 111 | Ccr5 | | 3.983e-02 | -3.22 | negative regulation of respiratory burst | biological process | GO:0060268 | 5 | 1 | 13711 | 111 | Grn | | 3.983e-02 | -3.22 | direct ossification | biological process | GO:0036072 | 5 | 1 | 13711 | 111 | Fgf18 | | 3.997e-02 | -3.22 | GO\_REGULATION\_OF\_PROTEIN\_UBIQUITINATION\_INVOLVED\_IN\_UBIQUITIN\_DEPENDENT\_PROTEIN\_CATABOLIC\_PROCESS | MSigDB lists | GO\_REGULATION\_OF\_PROTEIN\_UBIQUITINATION\_INVOLVED\_IN\_UBIQUITIN\_DEPENDENT\_PROTEIN\_CATABOLIC\_PROCESS | 94 | 3 | 12187 | 98 | Clu,Psme1,Psmb10 | | 3.997e-02 | -3.22 | REACTOME\_PEPTIDE\_LIGAND\_BINDING\_RECEPTORS | MSigDB lists | REACTOME\_PEPTIDE\_LIGAND\_BINDING\_RECEPTORS | 94 | 3 | 12187 | 98 | Mc4r,Ccr5,C3 | | 3.999e-02 | -3.22 | GSE25088\_ROSIGLITAZONE\_VS\_IL4\_AND\_ROSIGLITAZONE\_STIM\_STAT6\_KO\_MACROPHAGE\_DAY10\_DN | MSigDB lists | GSE25088\_ROSIGLITAZONE\_VS\_IL4\_AND\_ROSIGLITAZONE\_STIM\_STAT6\_KO\_MACROPHAGE\_DAY10\_DN | 160 | 4 | 12187 | 98 | Mgll,Vamp8,C1qb,C3 | | 3.999e-02 | -3.22 | GSE1460\_INTRATHYMIC\_T\_PROGENITOR\_VS\_THYMIC\_STROMAL\_CELL\_DN | MSigDB lists | GSE1460\_INTRATHYMIC\_T\_PROGENITOR\_VS\_THYMIC\_STROMAL\_CELL\_DN | 160 | 4 | 12187 | 98 | Ctsa,Tspan4,Vim,Myh9 | | 3.999e-02 | -3.22 | GSE13229\_IMM\_VS\_MATURE\_NKCELL\_DN | MSigDB lists | GSE13229\_IMM\_VS\_MATURE\_NKCELL\_DN | 160 | 4 | 12187 | 98 | Fcer1g,Vim,Sh3bgrl3,Fcgr3 | | 3.999e-02 | -3.22 | GSE37532\_TREG\_VS\_TCONV\_PPARG\_KO\_CD4\_TCELL\_FROM\_VISCERAL\_ADIPOSE\_TISSUE\_UP | MSigDB lists | GSE37532\_TREG\_VS\_TCONV\_PPARG\_KO\_CD4\_TCELL\_FROM\_VISCERAL\_ADIPOSE\_TISSUE\_UP | 160 | 4 | 12187 | 98 | Ctsz,Cd68,Il1r1,Tspan4 | | 3.999e-02 | -3.22 | GSE29618\_MONOCYTE\_VS\_PDC\_DAY7\_FLU\_VACCINE\_UP | MSigDB lists | GSE29618\_MONOCYTE\_VS\_PDC\_DAY7\_FLU\_VACCINE\_UP | 160 | 4 | 12187 | 98 | Ctss,Itgam,Lin7a,Csf1r | | 3.999e-02 | -3.22 | GSE27859\_DC\_VS\_CD11C\_INT\_F480\_INT\_DC\_UP | MSigDB lists | GSE27859\_DC\_VS\_CD11C\_INT\_F480\_INT\_DC\_UP | 160 | 4 | 12187 | 98 | Thy1,Mgll,Rac2,Fcgr3 | | 3.999e-02 | -3.22 | GSE24634\_IL4\_VS\_CTRL\_TREATED\_NAIVE\_CD4\_TCELL\_DAY3\_UP | MSigDB lists | GSE24634\_IL4\_VS\_CTRL\_TREATED\_NAIVE\_CD4\_TCELL\_DAY3\_UP | 160 | 4 | 12187 | 98 | Il1r1,Phyhip,Tspan4,Ahnak | | 3.999e-02 | -3.22 | GSE20715\_0H\_VS\_24H\_OZONE\_LUNG\_DN | MSigDB lists | GSE20715\_0H\_VS\_24H\_OZONE\_LUNG\_DN | 160 | 4 | 12187 | 98 | Mgp,C3,Slc2a1,Serpina3n | | 3.999e-02 | -3.22 | GSE20715\_WT\_VS\_TLR4\_KO\_24H\_OZONE\_LUNG\_UP | MSigDB lists | GSE20715\_WT\_VS\_TLR4\_KO\_24H\_OZONE\_LUNG\_UP | 160 | 4 | 12187 | 98 | Adh1,Tspan4,Gfap,Ctss | | 3.999e-02 | -3.22 | GSE37416\_12H\_VS\_24H\_F\_TULARENSIS\_LVS\_NEUTROPHIL\_UP | MSigDB lists | GSE37416\_12H\_VS\_24H\_F\_TULARENSIS\_LVS\_NEUTROPHIL\_UP | 160 | 4 | 12187 | 98 | Myh9,S100a6,Ctss,Serpinb9 | | 4.013e-02 | -3.22 | hemopoiesis | biological process | GO:0030097 | 478 | 8 | 13711 | 111 | Itgam,Fcer1g,Ptprc,Tnfsf8,Myh9,Csf1r,Hcls1,Lfng | | 4.017e-02 | -3.21 | positive regulation of apoptotic signaling pathway | biological process | GO:2001235 | 159 | 4 | 13711 | 111 | Clu,Ptprc,Camk2b,Rpl26 | | 4.031e-02 | -3.21 | GO\_CELL\_LEADING\_EDGE | MSigDB lists | GO\_CELL\_LEADING\_EDGE | 313 | 6 | 12187 | 98 | Sh3bgrl3,Vim,Rac2,Brk1,Myh9,S100a6 | | 4.053e-02 | -3.21 | ANAPHYLATOXIN\_1 | prosite domains | PS01177 | 4 | 1 | 8845 | 91 | C3 | | 4.053e-02 | -3.21 | LY6\_UPAR | prosite domains | PS00983 | 4 | 1 | 8845 | 91 | Ly6a | | 4.053e-02 | -3.21 | AA\_TRANSFER\_CLASS\_3 | prosite domains | PS00600 | 4 | 1 | 8845 | 91 | Etnppl | | 4.053e-02 | -3.21 | ACTINS\_1 | prosite domains | PS00406 | 4 | 1 | 8845 | 91 | Actb | | 4.053e-02 | -3.21 | ANAPHYLATOXIN\_2 | prosite domains | PS01178 | 4 | 1 | 8845 | 91 | C3 | | 4.053e-02 | -3.21 | LIPOCALIN | prosite domains | PS00213 | 4 | 1 | 8845 | 91 | Apod | | 4.053e-02 | -3.21 | TSC22 | prosite domains | PS01289 | 4 | 1 | 8845 | 91 | Tsc22d1 | | 4.054e-02 | -3.21 | positive regulation of cell motility | biological process | GO:2000147 | 479 | 8 | 13711 | 111 | Grn,Ccar1,Thy1,Fgf18,Il1r1,Rac2,Ptprc,Csf1r | | 4.067e-02 | -3.20 | MANSC\_dom | interpro domains | IPR013980 | 5 | 1 | 13788 | 114 | Spint1 | | 4.067e-02 | -3.20 | Fringe-like | interpro domains | IPR003378 | 5 | 1 | 13788 | 114 | Lfng | | 4.067e-02 | -3.20 | Lipocalin\_CS | interpro domains | IPR022272 | 5 | 1 | 13788 | 114 | Apod | | 4.067e-02 | -3.20 | MG3 | interpro domains | IPR041555 | 5 | 1 | 13788 | 114 | C3 | | 4.067e-02 | -3.20 | Alpha-macroglobulin\_TED | interpro domains | IPR011626 | 5 | 1 | 13788 | 114 | C3 | | 4.067e-02 | -3.20 | Synaptobrevin/VAMP | interpro domains | IPR016444 | 5 | 1 | 13788 | 114 | Vamp8 | | 4.067e-02 | -3.20 | EAR | interpro domains | IPR009039 | 5 | 1 | 13788 | 114 | Lgi4 | | 4.067e-02 | -3.20 | Lactate/malate\_DH\_N | interpro domains | IPR001236 | 5 | 1 | 13788 | 114 | Ldha | | 4.067e-02 | -3.20 | A2M\_N\_BRD | interpro domains | IPR011625 | 5 | 1 | 13788 | 114 | C3 | | 4.067e-02 | -3.20 | A-macroglobulin\_rcpt-bd\_sf | interpro domains | IPR036595 | 5 | 1 | 13788 | 114 | C3 | | 4.067e-02 | -3.20 | MacrogloblnA2\_CS | interpro domains | IPR019742 | 5 | 1 | 13788 | 114 | C3 | | 4.067e-02 | -3.20 | IGFBP1-6 | interpro domains | IPR009168 | 5 | 1 | 13788 | 114 | Igfbp6 | | 4.067e-02 | -3.20 | L-lactate/malate\_DH | interpro domains | IPR001557 | 5 | 1 | 13788 | 114 | Ldha | | 4.067e-02 | -3.20 | Hematopoietin\_rcpt\_L\_F3\_CS | interpro domains | IPR003530 | 5 | 1 | 13788 | 114 | Il6ra | | 4.067e-02 | -3.20 | IGFBP\_1-6\_chordata | interpro domains | IPR022321 | 5 | 1 | 13788 | 114 | Igfbp6 | | 4.067e-02 | -3.20 | EPTP | interpro domains | IPR005492 | 5 | 1 | 13788 | 114 | Lgi4 | | 4.067e-02 | -3.20 | Lactate/malate\_DH\_C | interpro domains | IPR022383 | 5 | 1 | 13788 | 114 | Ldha | | 4.067e-02 | -3.20 | MG2 | interpro domains | IPR002890 | 5 | 1 | 13788 | 114 | C3 | | 4.067e-02 | -3.20 | S04\_transporter\_CS | interpro domains | IPR018045 | 5 | 1 | 13788 | 114 | Slc26a2 | | 4.067e-02 | -3.20 | TNF\_CS | interpro domains | IPR021184 | 5 | 1 | 13788 | 114 | Tnfsf8 | | 4.067e-02 | -3.20 | A-macroglobulin\_rcpt-bd | interpro domains | IPR009048 | 5 | 1 | 13788 | 114 | C3 | | 4.067e-02 | -3.20 | Macroglobln\_a2 | interpro domains | IPR001599 | 5 | 1 | 13788 | 114 | C3 | | 4.067e-02 | -3.20 | SAPAP | interpro domains | IPR005026 | 5 | 1 | 13788 | 114 | Dlgap3 | | 4.067e-02 | -3.20 | KA1\_dom | interpro domains | IPR001772 | 5 | 1 | 13788 | 114 | Mark4 | | 4.068e-02 | -3.20 | MFS\_trans\_sf | interpro domains | IPR036259 | 92 | 3 | 13788 | 114 | Mfsd4a,Slc2a1,Slc17a7 | | 4.070e-02 | -3.20 | axon terminus | cellular component | GO:0043679 | 161 | 4 | 13825 | 111 | Cplx2,Actb,Ptprn,Rpl26 | | 4.073e-02 | -3.20 | PTPc | smart domains | SM00194 | 34 | 2 | 7188 | 68 | Ptprc,Ptprn | | 4.076e-02 | -3.20 | GO\_REGULATION\_OF\_ENDOCYTOSIS | MSigDB lists | GO\_REGULATION\_OF\_ENDOCYTOSIS | 161 | 4 | 12187 | 98 | C3,Cyba,Csk,Fcer1g | | 4.076e-02 | -3.20 | GSE29618\_BCELL\_VS\_PDC\_DAY7\_FLU\_VACCINE\_DN | MSigDB lists | GSE29618\_BCELL\_VS\_PDC\_DAY7\_FLU\_VACCINE\_DN | 161 | 4 | 12187 | 98 | Cd68,Fcer1g,Grn,Flnb | | 4.076e-02 | -3.20 | GSE27786\_LSK\_VS\_BCELL\_DN | MSigDB lists | GSE27786\_LSK\_VS\_BCELL\_DN | 161 | 4 | 12187 | 98 | Rpl31,Psmb10,Vim,Brk1 | | 4.076e-02 | -3.20 | GSE16522\_MEMORY\_VS\_NAIVE\_ANTI\_CD3CD28\_STIM\_CD8\_TCELL\_DN | MSigDB lists | GSE16522\_MEMORY\_VS\_NAIVE\_ANTI\_CD3CD28\_STIM\_CD8\_TCELL\_DN | 161 | 4 | 12187 | 98 | Treml2,Psmb10,Ctss,Ctsz | | 4.076e-02 | -3.20 | GSE40068\_BCL6\_POS\_VS\_NEG\_CXCR5\_POS\_TFH\_DN | MSigDB lists | GSE40068\_BCL6\_POS\_VS\_NEG\_CXCR5\_POS\_TFH\_DN | 161 | 4 | 12187 | 98 | Rps5,Mgp,Rps8,Ctss | | 4.076e-02 | -3.20 | GO\_REGULATION\_OF\_LEUKOCYTE\_DIFFERENTIATION | MSigDB lists | GO\_REGULATION\_OF\_LEUKOCYTE\_DIFFERENTIATION | 161 | 4 | 12187 | 98 | Hcls1,Tmem176a,Csf1r,Ikzf3 | | 4.076e-02 | -3.20 | GSE39110\_DAY3\_VS\_DAY6\_POST\_IMMUNIZATION\_CD8\_TCELL\_WITH\_IL2\_TREATMENT\_UP | MSigDB lists | GSE39110\_DAY3\_VS\_DAY6\_POST\_IMMUNIZATION\_CD8\_TCELL\_WITH\_IL2\_TREATMENT\_UP | 161 | 4 | 12187 | 98 | Ldha,Ctsz,Ikzf3,Ccr5 | | 4.076e-02 | -3.20 | GO\_PROTEIN\_AUTOPHOSPHORYLATION | MSigDB lists | GO\_PROTEIN\_AUTOPHOSPHORYLATION | 161 | 4 | 12187 | 98 | Csf1r,Csk,Camk2b,Thy1 | | 4.076e-02 | -3.20 | GSE17721\_CTRL\_VS\_GARDIQUIMOD\_24H\_BMDC\_UP | MSigDB lists | GSE17721\_CTRL\_VS\_GARDIQUIMOD\_24H\_BMDC\_UP | 161 | 4 | 12187 | 98 | Ctsa,Grn,Il6ra,Ndufa13 | | 4.076e-02 | -3.20 | HALLMARK\_HEME\_METABOLISM | MSigDB lists | HALLMARK\_HEME\_METABOLISM | 161 | 4 | 12187 | 98 | Rap1gap,Slc2a1,Ctsb,C3 | | 4.076e-02 | -3.20 | HALLMARK\_ESTROGEN\_RESPONSE\_LATE | MSigDB lists | HALLMARK\_ESTROGEN\_RESPONSE\_LATE | 161 | 4 | 12187 | 98 | Serpina3n,Flnb,Slc26a2,Tfap2c | | 4.076e-02 | -3.20 | GSE46606\_IRF4HIGH\_VS\_IRF4MID\_CD40L\_IL2\_IL5\_DAY3\_STIMULATED\_BCELL\_DN | MSigDB lists | GSE46606\_IRF4HIGH\_VS\_IRF4MID\_CD40L\_IL2\_IL5\_DAY3\_STIMULATED\_BCELL\_DN | 161 | 4 | 12187 | 98 | Ppfibp1,Psme1,Ttc9b,Flnb | | 4.076e-02 | -3.20 | KAECH\_DAY15\_EFF\_VS\_MEMORY\_CD8\_TCELL\_UP | MSigDB lists | KAECH\_DAY15\_EFF\_VS\_MEMORY\_CD8\_TCELL\_UP | 161 | 4 | 12187 | 98 | Ahnak,Hspa2,Actb,Ctsa | | 4.076e-02 | -3.20 | GSE2585\_CTEC\_VS\_THYMIC\_DC\_DN | MSigDB lists | GSE2585\_CTEC\_VS\_THYMIC\_DC\_DN | 161 | 4 | 12187 | 98 | Tmem176a,Rps5,Klhl6,Rac2 | | 4.076e-02 | -3.20 | GSE39916\_B\_CELL\_SPLEEN\_VS\_PLASMA\_CELL\_BONE\_MARROW\_UP | MSigDB lists | GSE39916\_B\_CELL\_SPLEEN\_VS\_PLASMA\_CELL\_BONE\_MARROW\_UP | 161 | 4 | 12187 | 98 | Adh1,Rpl31,Treml2,Ikzf3 | | 4.076e-02 | -3.20 | GSE25123\_IL4\_VS\_IL4\_AND\_ROSIGLITAZONE\_STIM\_MACROPHAGE\_DAY10\_UP | MSigDB lists | GSE25123\_IL4\_VS\_IL4\_AND\_ROSIGLITAZONE\_STIM\_MACROPHAGE\_DAY10\_UP | 161 | 4 | 12187 | 98 | Tnfsf8,S100a6,Ppfibp1,Il1r1 | | 4.076e-02 | -3.20 | GSE5679\_CTRL\_VS\_PPARG\_LIGAND\_ROSIGLITAZONE\_AND\_RARA\_AGONIST\_AM580\_TREATED\_DC\_UP | MSigDB lists | GSE5679\_CTRL\_VS\_PPARG\_LIGAND\_ROSIGLITAZONE\_AND\_RARA\_AGONIST\_AM580\_TREATED\_DC\_UP | 161 | 4 | 12187 | 98 | C3,Ly6e,Ldha,Serpinb9 | | 4.085e-02 | -3.20 | GO\_CYTOSKELETAL\_PROTEIN\_BINDING | MSigDB lists | GO\_CYTOSKELETAL\_PROTEIN\_BINDING | 665 | 10 | 12187 | 98 | Hcls1,Camk2b,Capn3,Myh9,Ccr5,S100a6,Mark4,Actb,Ptprn,Flnb | | 4.096e-02 | -3.20 | actin filament-based process | biological process | GO:0030029 | 480 | 8 | 13711 | 111 | Camk2b,Capn3,Myh9,Hcls1,Flnb,Brk1,Rac2,Actb | | 4.096e-02 | -3.20 | TSENG\_ADIPOGENIC\_POTENTIAL\_DN | MSigDB lists | TSENG\_ADIPOGENIC\_POTENTIAL\_DN | 40 | 2 | 12187 | 98 | Il1r1,Tsc22d1 | | 4.096e-02 | -3.20 | MODULE\_107 | MSigDB lists | MODULE\_107 | 40 | 2 | 12187 | 98 | Ctss,Ctsb | | 4.096e-02 | -3.20 | KEGG\_GLYCOLYSIS\_GLUCONEOGENESIS | MSigDB lists | KEGG\_GLYCOLYSIS\_GLUCONEOGENESIS | 40 | 2 | 12187 | 98 | Ldha,Adh1 | | 4.096e-02 | -3.20 | KEGG\_PROTEASOME | MSigDB lists | KEGG\_PROTEASOME | 40 | 2 | 12187 | 98 | Psme1,Psmb10 | | 4.096e-02 | -3.20 | MODULE\_234 | MSigDB lists | MODULE\_234 | 40 | 2 | 12187 | 98 | Mgp,Il6ra | | 4.104e-02 | -3.19 | GO\_REGULATION\_OF\_REGULATED\_SECRETORY\_PATHWAY | MSigDB lists | GO\_REGULATION\_OF\_REGULATED\_SECRETORY\_PATHWAY | 95 | 3 | 12187 | 98 | Fcer1g,Rac2,Vamp8 | | 4.104e-02 | -3.19 | CRX\_NRL\_DN.V1\_DN | MSigDB lists | CRX\_NRL\_DN.V1\_DN | 95 | 3 | 12187 | 98 | Cplx2,Camk2b,Spint1 | | 4.104e-02 | -3.19 | GO\_REGULATION\_OF\_LEUKOCYTE\_MIGRATION | MSigDB lists | GO\_REGULATION\_OF\_LEUKOCYTE\_MIGRATION | 95 | 3 | 12187 | 98 | Rac2,Il6ra,Apod | | 4.104e-02 | -3.19 | GO\_POSITIVE\_REGULATION\_OF\_ENDOCYTOSIS | MSigDB lists | GO\_POSITIVE\_REGULATION\_OF\_ENDOCYTOSIS | 95 | 3 | 12187 | 98 | C3,Cyba,Fcer1g | | 4.104e-02 | -3.19 | REACTOME\_TCA\_CYCLE\_AND\_RESPIRATORY\_ELECTRON\_TRANSPORT | MSigDB lists | REACTOME\_TCA\_CYCLE\_AND\_RESPIRATORY\_ELECTRON\_TRANSPORT | 95 | 3 | 12187 | 98 | Cox4i1,Ldha,Ndufa13 | | 4.111e-02 | -3.19 | regulation of locomotion | biological process | GO:0040012 | 847 | 12 | 13711 | 111 | Apod,Ptprc,Grn,Vim,Ccar1,Il1r1,Thy1,Rac2,Csf1r,Ccr5,Fgf18,Camk2b | | 4.115e-02 | -3.19 | regulation of lymphocyte activation | biological process | GO:0051249 | 312 | 6 | 13711 | 111 | Ikzf3,Rac2,Ptprc,Csk,Thy1,Il6ra | | 4.122e-02 | -3.19 | cellular homeostasis | biological process | GO:0019725 | 660 | 10 | 13711 | 111 | Mt3,Cd52,Ptprc,Sh3bgrl3,Slc17a7,Capn3,Thy1,Grn,Ptprn,Ccr5 | | 4.123e-02 | -3.19 | A2M\_BRD | pfam domains | PF07703 | 5 | 1 | 12881 | 108 | C3 | | 4.123e-02 | -3.19 | A2M\_recep | pfam domains | PF07677 | 5 | 1 | 12881 | 108 | C3 | | 4.123e-02 | -3.19 | Ldh\_1\_N | pfam domains | PF00056 | 5 | 1 | 12881 | 108 | Ldha | | 4.123e-02 | -3.19 | GKAP | pfam domains | PF03359 | 5 | 1 | 12881 | 108 | Dlgap3 | | 4.123e-02 | -3.19 | Fringe | pfam domains | PF02434 | 5 | 1 | 12881 | 108 | Lfng | | 4.123e-02 | -3.19 | KA1 | pfam domains | PF02149 | 5 | 1 | 12881 | 108 | Mark4 | | 4.123e-02 | -3.19 | EPTP | pfam domains | PF03736 | 5 | 1 | 12881 | 108 | Lgi4 | | 4.123e-02 | -3.19 | MG3 | pfam domains | PF17791 | 5 | 1 | 12881 | 108 | C3 | | 4.123e-02 | -3.19 | TED\_complement | pfam domains | PF07678 | 5 | 1 | 12881 | 108 | C3 | | 4.123e-02 | -3.19 | A2M | pfam domains | PF00207 | 5 | 1 | 12881 | 108 | C3 | | 4.123e-02 | -3.19 | Ldh\_1\_C | pfam domains | PF02866 | 5 | 1 | 12881 | 108 | Ldha | | 4.123e-02 | -3.19 | MG2 | pfam domains | PF01835 | 5 | 1 | 12881 | 108 | C3 | | 4.137e-02 | -3.19 | REACTOME\_GPCR\_DOWNSTREAM\_SIGNALING | MSigDB lists | REACTOME\_GPCR\_DOWNSTREAM\_SIGNALING | 315 | 6 | 12187 | 98 | C3,Mc4r,Pde1a,Mgll,Ccr5,Rac2 | | 4.150e-02 | -3.18 | lysosome localization | biological process | GO:0032418 | 40 | 2 | 13711 | 111 | Cplx2,Myh9 | | 4.150e-02 | -3.18 | negative regulation of innate immune response | biological process | GO:0045824 | 40 | 2 | 13711 | 111 | Grn,Serpinb9 | | 4.150e-02 | -3.18 | osteoclast differentiation | biological process | GO:0030316 | 40 | 2 | 13711 | 111 | Fcer1g,Csf1r | | 4.155e-02 | -3.18 | GSE17721\_CPG\_VS\_GARDIQUIMOD\_1H\_BMDC\_UP | MSigDB lists | GSE17721\_CPG\_VS\_GARDIQUIMOD\_1H\_BMDC\_UP | 162 | 4 | 12187 | 98 | Ctss,Myh9,Tsc22d1,Adh1 | | 4.155e-02 | -3.18 | GO\_REGULATION\_OF\_ERK1\_AND\_ERK2\_CASCADE | MSigDB lists | GO\_REGULATION\_OF\_ERK1\_AND\_ERK2\_CASCADE | 162 | 4 | 12187 | 98 | Csk,Fgf18,Csf1r,C3 | | 4.155e-02 | -3.18 | GSE9006\_HEALTHY\_VS\_TYPE\_1\_DIABETES\_PBMC\_AT\_DX\_UP | MSigDB lists | GSE9006\_HEALTHY\_VS\_TYPE\_1\_DIABETES\_PBMC\_AT\_DX\_UP | 162 | 4 | 12187 | 98 | Clu,Psme1,Mgll,C3 | | 4.155e-02 | -3.18 | GSE21670\_STAT3\_KO\_VS\_WT\_CD4\_TCELL\_IL6\_TREATED\_UP | MSigDB lists | GSE21670\_STAT3\_KO\_VS\_WT\_CD4\_TCELL\_IL6\_TREATED\_UP | 162 | 4 | 12187 | 98 | Ppfibp1,Fam111a,C3,Fcgr3 | | 4.155e-02 | -3.18 | GSE2405\_0H\_VS\_3H\_A\_PHAGOCYTOPHILUM\_STIM\_NEUTROPHIL\_UP | MSigDB lists | GSE2405\_0H\_VS\_3H\_A\_PHAGOCYTOPHILUM\_STIM\_NEUTROPHIL\_UP | 162 | 4 | 12187 | 98 | Fcgr3,Exoc3l4,Spint1,Tsc22d1 | | 4.155e-02 | -3.18 | GSE19772\_CTRL\_VS\_HCMV\_INF\_MONOCYTES\_UP | MSigDB lists | GSE19772\_CTRL\_VS\_HCMV\_INF\_MONOCYTES\_UP | 162 | 4 | 12187 | 98 | Capn3,Fcer1g,Ahnak,Tsc22d1 | | 4.155e-02 | -3.18 | GSE7460\_CTRL\_VS\_TGFB\_TREATED\_ACT\_FOXP3\_HET\_TCONV\_DN | MSigDB lists | GSE7460\_CTRL\_VS\_TGFB\_TREATED\_ACT\_FOXP3\_HET\_TCONV\_DN | 162 | 4 | 12187 | 98 | Lfng,Thy1,Adh1,Cyba | | 4.155e-02 | -3.18 | GSE3039\_ALPHAALPHA\_VS\_ALPHABETA\_CD8\_TCELL\_UP | MSigDB lists | GSE3039\_ALPHAALPHA\_VS\_ALPHABETA\_CD8\_TCELL\_UP | 162 | 4 | 12187 | 98 | Ccr5,Igfbp6,Ldha,Slc2a1 | | 4.174e-02 | -3.18 | regulation of cellular component organization | biological process | GO:0051128 | 2181 | 25 | 13711 | 111 | Vamp8,Rap1gap,Vim,Cplx2,Ctsz,Mark4,Ndufa13,Ccr5,C3,Cyba,Brk1,Hcls1,Csf1r,Myh9,Rac2,Mgll,Thy1,Grn,Apod,Hspa2,Clu,Abhd17c,Camk2b,Gfap,Slc17a7 | | 4.175e-02 | -3.18 | regulation of lymphocyte proliferation | biological process | GO:0050670 | 161 | 4 | 13711 | 111 | Il6ra,Ptprc,Rac2,Ikzf3 | | 4.181e-02 | -3.17 | regulation of transmembrane transport | biological process | GO:0034762 | 482 | 8 | 13711 | 111 | Actb,Hspa2,C3,Cyba,Ctss,Capn3,Thy1,Ahnak | | 4.202e-02 | -3.17 | GO\_POSITIVE\_REGULATION\_OF\_PROTEIN\_MODIFICATION\_PROCESS | MSigDB lists | GO\_POSITIVE\_REGULATION\_OF\_PROTEIN\_MODIFICATION\_PROCESS | 858 | 12 | 12187 | 98 | Il6ra,Csf1r,Fgf18,Psmb10,Ptprc,Hspa2,Psme1,Clu,Csk,C3,Kl,Hcls1 | | 4.209e-02 | -3.17 | MAPK family signaling cascades | REACTOME pathways | R-MMU-5683057 | 212 | 6 | 6297 | 76 | Psme1,Psmb10,Il6ra,Kl,Fgf18,Camk2b | | 4.211e-02 | -3.17 | AP2TNSCPFCT | prints domains | PR01748 | 3 | 1 | 2951 | 42 | Tfap2c | | 4.211e-02 | -3.17 | LYSASSOCTDMP | prints domains | PR00336 | 3 | 1 | 2951 | 42 | Cd68 | | 4.211e-02 | -3.17 | MTVERTEBRATE | prints domains | PR00860 | 3 | 1 | 2951 | 42 | Mt3 | | 4.211e-02 | -3.17 | SERUMAMYLOID | prints domains | PR00306 | 3 | 1 | 2951 | 42 | Saa3 | | 4.211e-02 | -3.17 | LLDHDRGNASE | prints domains | PR00086 | 3 | 1 | 2951 | 42 | Ldha | | 4.213e-02 | -3.17 | GROSS\_HYPOXIA\_VIA\_ELK3\_AND\_HIF1A\_DN | MSigDB lists | GROSS\_HYPOXIA\_VIA\_ELK3\_AND\_HIF1A\_DN | 96 | 3 | 12187 | 98 | Tmem176a,Il1r1,Myh9 | | 4.213e-02 | -3.17 | DITTMER\_PTHLH\_TARGETS\_UP | MSigDB lists | DITTMER\_PTHLH\_TARGETS\_UP | 96 | 3 | 12187 | 98 | Ndufa13,Vamp8,Tsc22d1 | | 4.227e-02 | -3.16 | LOPEZ\_MBD\_TARGETS | MSigDB lists | LOPEZ\_MBD\_TARGETS | 763 | 11 | 12187 | 98 | Ptprn,Il1r1,Psmb10,Hcls1,Ldha,Rps8,Rpl31,Csf1r,Slc2a1,Tsc22d1,Lfng | | 4.228e-02 | -3.16 | GO\_POSITIVE\_REGULATION\_OF\_PROTEIN\_METABOLIC\_PROCESS | MSigDB lists | GO\_POSITIVE\_REGULATION\_OF\_PROTEIN\_METABOLIC\_PROCESS | 1155 | 15 | 12187 | 98 | Myh9,Ndufa13,Csf1r,Il6ra,Psme1,Capn3,Hspa2,Clu,Csk,C3,Hcls1,Kl,Fgf18,Ptprc,Psmb10 | | 4.234e-02 | -3.16 | GSE15930\_STIM\_VS\_STIM\_AND\_IL12\_48H\_CD8\_T\_CELL\_DN | MSigDB lists | GSE15930\_STIM\_VS\_STIM\_AND\_IL12\_48H\_CD8\_T\_CELL\_DN | 163 | 4 | 12187 | 98 | Serpinb9,Ldha,S100a6,Ccr5 | | 4.234e-02 | -3.16 | GSE8685\_IL2\_STARVED\_VS\_IL21\_ACT\_IL2\_STARVED\_CD4\_TCELL\_DN | MSigDB lists | GSE8685\_IL2\_STARVED\_VS\_IL21\_ACT\_IL2\_STARVED\_CD4\_TCELL\_DN | 163 | 4 | 12187 | 98 | Rpl13,Lfng,Tmem106a,Mark4 | | 4.234e-02 | -3.16 | GSE24574\_BCL6\_HIGH\_TFH\_VS\_TFH\_CD4\_TCELL\_DN | MSigDB lists | GSE24574\_BCL6\_HIGH\_TFH\_VS\_TFH\_CD4\_TCELL\_DN | 163 | 4 | 12187 | 98 | Rps8,Rpl13,Myh9,Rps20 | | 4.234e-02 | -3.16 | GSE17721\_CTRL\_VS\_POLYIC\_4H\_BMDC\_DN | MSigDB lists | GSE17721\_CTRL\_VS\_POLYIC\_4H\_BMDC\_DN | 163 | 4 | 12187 | 98 | Flnb,Tmem106a,Fmod,Tsc22d1 | | 4.234e-02 | -3.16 | GSE369\_SOCS3\_KO\_VS\_WT\_LIVER\_POST\_IL6\_INJECTION\_DN | MSigDB lists | GSE369\_SOCS3\_KO\_VS\_WT\_LIVER\_POST\_IL6\_INJECTION\_DN | 163 | 4 | 12187 | 98 | Ctss,Hspa2,Treml2,Klhl6 | | 4.234e-02 | -3.16 | GSE15330\_HSC\_VS\_GRANULOCYTE\_MONOCYTE\_PROGENITOR\_IKAROS\_KO\_UP | MSigDB lists | GSE15330\_HSC\_VS\_GRANULOCYTE\_MONOCYTE\_PROGENITOR\_IKAROS\_KO\_UP | 163 | 4 | 12187 | 98 | Flnb,Il1r1,Mark4,Grn | | 4.255e-02 | -3.16 | regulation of plasma membrane bounded cell projection assembly | biological process | GO:0120032 | 162 | 4 | 13711 | 111 | Mark4,Brk1,Rap1gap,Rac2 | | 4.258e-02 | -3.16 | regulation of hydrolase activity | biological process | GO:0051336 | 948 | 13 | 13711 | 111 | Grn,Rap1gap,Thy1,Ccl6,Hspa2,Ptprc,Serpina3n,Ndufa13,C3,Psme1,Serpinb9,Camk2b,Spint1 | | 4.264e-02 | -3.16 | GO\_INTRACELLULAR\_PROTEIN\_TRANSPORT | MSigDB lists | GO\_INTRACELLULAR\_PROTEIN\_TRANSPORT | 670 | 10 | 12187 | 98 | Rps20,Rpl31,Rpl14,Rps5,Rps8,Ndufa13,Rpl13,Ctsa,Rpl26,Rac2 | | 4.272e-02 | -3.15 | STUB1 (STIP1 homology and U-box containing protein 1) | protein interactions | 10273 | 68 | 3 | 6802 | 78 | Rpl26,Rpl31,Ahnak | | 4.284e-02 | -3.15 | CREIGHTON\_ENDOCRINE\_THERAPY\_RESISTANCE\_4 | MSigDB lists | CREIGHTON\_ENDOCRINE\_THERAPY\_RESISTANCE\_4 | 238 | 5 | 12187 | 98 | Tfap2c,Mgp,Slc26a2,Flnb,Serpina3n | | 4.284e-02 | -3.15 | GO\_ACTIN\_FILAMENT\_BUNDLE | MSigDB lists | GO\_ACTIN\_FILAMENT\_BUNDLE | 41 | 2 | 12187 | 98 | Myh9,Flnb | | 4.284e-02 | -3.15 | WANG\_TARGETS\_OF\_MLL\_CBP\_FUSION\_DN | MSigDB lists | WANG\_TARGETS\_OF\_MLL\_CBP\_FUSION\_DN | 41 | 2 | 12187 | 98 | Tsc22d1,Ctsz | | 4.284e-02 | -3.15 | BREDEMEYER\_RAG\_SIGNALING\_VIA\_ATM\_NOT\_VIA\_NFKB\_UP | MSigDB lists | BREDEMEYER\_RAG\_SIGNALING\_VIA\_ATM\_NOT\_VIA\_NFKB\_UP | 41 | 2 | 12187 | 98 | Sh2d5,Camk2b | | 4.284e-02 | -3.15 | BOHN\_PRIMARY\_IMMUNODEFICIENCY\_SYNDROM\_UP | MSigDB lists | BOHN\_PRIMARY\_IMMUNODEFICIENCY\_SYNDROM\_UP | 41 | 2 | 12187 | 98 | Rps5,Rpl14 | | 4.284e-02 | -3.15 | MODULE\_165 | MSigDB lists | MODULE\_165 | 41 | 2 | 12187 | 98 | Itgam,Rac2 | | 4.284e-02 | -3.15 | RICKMAN\_HEAD\_AND\_NECK\_CANCER\_C | MSigDB lists | RICKMAN\_HEAD\_AND\_NECK\_CANCER\_C | 41 | 2 | 12187 | 98 | Serpina3n,Slc2a1 | | 4.284e-02 | -3.15 | MODY\_HIPPOCAMPUS\_PRENATAL | MSigDB lists | MODY\_HIPPOCAMPUS\_PRENATAL | 41 | 2 | 12187 | 98 | Rpl13,Rps8 | | 4.284e-02 | -3.15 | GO\_T\_TUBULE | MSigDB lists | GO\_T\_TUBULE | 41 | 2 | 12187 | 98 | Ahnak,Capn3 | | 4.284e-02 | -3.15 | LEE\_LIVER\_CANCER\_DENA\_UP | MSigDB lists | LEE\_LIVER\_CANCER\_DENA\_UP | 41 | 2 | 12187 | 98 | Ly6e,Ctss | | 4.284e-02 | -3.15 | MODULE\_166 | MSigDB lists | MODULE\_166 | 41 | 2 | 12187 | 98 | Csf1r,Phyhip | | 4.284e-02 | -3.15 | LEE\_LIVER\_CANCER\_MYC\_E2F1\_UP | MSigDB lists | LEE\_LIVER\_CANCER\_MYC\_E2F1\_UP | 41 | 2 | 12187 | 98 | Ly6e,Ctss | | 4.284e-02 | -3.15 | GO\_POSITIVE\_REGULATION\_OF\_LYMPHOCYTE\_MEDIATED\_IMMUNITY | MSigDB lists | GO\_POSITIVE\_REGULATION\_OF\_LYMPHOCYTE\_MEDIATED\_IMMUNITY | 41 | 2 | 12187 | 98 | Fcer1g,C3 | | 4.284e-02 | -3.15 | ZHENG\_IL22\_SIGNALING\_UP | MSigDB lists | ZHENG\_IL22\_SIGNALING\_UP | 41 | 2 | 12187 | 98 | C3,Lrg1 | | 4.296e-02 | -3.15 | negative regulation of programmed cell death | biological process | GO:0043069 | 758 | 11 | 13711 | 111 | Fcer1g,Hcls1,Csf1r,Mt3,Tsc22d1,Ndufa13,Ccr5,Serpinb9,Capn3,Clu,Grn | | 4.306e-02 | -3.15 | MILI\_PSEUDOPODIA\_HAPTOTAXIS\_DN | MSigDB lists | MILI\_PSEUDOPODIA\_HAPTOTAXIS\_DN | 579 | 9 | 12187 | 98 | Ppfibp1,Igfbp6,Flnb,Ly6e,Grn,Alg12,Ctsz,Tspan4,Slc2a1 | | 4.314e-02 | -3.14 | GSE21360\_NAIVE\_VS\_PRIMARY\_MEMORY\_CD8\_TCELL\_DN | MSigDB lists | GSE21360\_NAIVE\_VS\_PRIMARY\_MEMORY\_CD8\_TCELL\_DN | 164 | 4 | 12187 | 98 | Cd68,Ctss,Vwf,Ahnak | | 4.314e-02 | -3.14 | GSE28408\_LY6G\_POS\_VS\_NEG\_DC\_DN | MSigDB lists | GSE28408\_LY6G\_POS\_VS\_NEG\_DC\_DN | 164 | 4 | 12187 | 98 | Dek,Clu,Ccr5,Grn | | 4.314e-02 | -3.14 | KAECH\_NAIVE\_VS\_DAY15\_EFF\_CD8\_TCELL\_UP | MSigDB lists | KAECH\_NAIVE\_VS\_DAY15\_EFF\_CD8\_TCELL\_UP | 164 | 4 | 12187 | 98 | Rps8,Il6ra,Psme1,Psmb10 | | 4.314e-02 | -3.14 | GSE22886\_NAIVE\_VS\_MEMORY\_TCELL\_DN | MSigDB lists | GSE22886\_NAIVE\_VS\_MEMORY\_TCELL\_DN | 164 | 4 | 12187 | 98 | Il6ra,Ldha,Il1r1,Dek | | 4.314e-02 | -3.14 | GSE19401\_UNSTIM\_VS\_RETINOIC\_ACID\_AND\_PAM2CSK4\_STIM\_FOLLICULAR\_DC\_DN | MSigDB lists | GSE19401\_UNSTIM\_VS\_RETINOIC\_ACID\_AND\_PAM2CSK4\_STIM\_FOLLICULAR\_DC\_DN | 164 | 4 | 12187 | 98 | Slc26a2,Tspan4,Tnfsf8,Ctsz | | 4.314e-02 | -3.14 | GSE36888\_STAT5\_AB\_KNOCKIN\_VS\_WT\_TCELL\_IL2\_TREATED\_2H\_DN | MSigDB lists | GSE36888\_STAT5\_AB\_KNOCKIN\_VS\_WT\_TCELL\_IL2\_TREATED\_2H\_DN | 164 | 4 | 12187 | 98 | Tnfsf8,Il6ra,Itgam,Lin7a | | 4.314e-02 | -3.14 | GSE31082\_DP\_VS\_CD8\_SP\_THYMOCYTE\_DN | MSigDB lists | GSE31082\_DP\_VS\_CD8\_SP\_THYMOCYTE\_DN | 164 | 4 | 12187 | 98 | Myh9,Sh3bgrl3,Tmem106a,Flnb | | 4.314e-02 | -3.14 | GSE15330\_MEGAKARYOCYTE\_ERYTHROID\_PROGENITOR\_VS\_PRO\_BCELL\_UP | MSigDB lists | GSE15330\_MEGAKARYOCYTE\_ERYTHROID\_PROGENITOR\_VS\_PRO\_BCELL\_UP | 164 | 4 | 12187 | 98 | Hcls1,Sh3bgrl3,Cox4i1,Mgll | | 4.314e-02 | -3.14 | GSE15330\_WT\_VS\_IKAROS\_KO\_LYMPHOID\_MULTIPOTENT\_PROGENITOR\_DN | MSigDB lists | GSE15330\_WT\_VS\_IKAROS\_KO\_LYMPHOID\_MULTIPOTENT\_PROGENITOR\_DN | 164 | 4 | 12187 | 98 | Tnfsf8,Csf1r,Adh1,Ptprc | | 4.314e-02 | -3.14 | GRUETZMANN\_PANCREATIC\_CANCER\_DN | MSigDB lists | GRUETZMANN\_PANCREATIC\_CANCER\_DN | 164 | 4 | 12187 | 98 | Cyba,Ptprn,Rpl13,Clu | | 4.314e-02 | -3.14 | GSE16451\_CTRL\_VS\_WEST\_EQUINE\_ENC\_VIRUS\_MATURE\_NEURON\_CELL\_LINE\_DN | MSigDB lists | GSE16451\_CTRL\_VS\_WEST\_EQUINE\_ENC\_VIRUS\_MATURE\_NEURON\_CELL\_LINE\_DN | 164 | 4 | 12187 | 98 | Ikzf3,S100a6,Ahnak,Capn3 | | 4.314e-02 | -3.14 | WIERENGA\_STAT5A\_TARGETS\_UP | MSigDB lists | WIERENGA\_STAT5A\_TARGETS\_UP | 164 | 4 | 12187 | 98 | Ctsz,Rap1gap,Hspa2,Fcgr3 | | 4.317e-02 | -3.14 | Patj (PATJ, crumbs cell polarity complex component) | protein interactions | 12695 | 29 | 2 | 6802 | 78 | Actb,Camk2b | | 4.318e-02 | -3.14 | organonitrogen compound biosynthetic process | biological process | GO:1901566 | 950 | 13 | 13711 | 111 | Rps8,Rpl13,Tmem106a,Kl,Rps20,Abhd17c,Rpl26,Alg12,Lfng,Rpl14,Rps5,Atp5j2,Rpl31 | | 4.320e-02 | -3.14 | - | gene3d domains | 3.40.50.1820 | 84 | 3 | 6647 | 62 | Mgll,Ctsa,Abhd17c | | 4.335e-02 | -3.14 | regulation of cell projection assembly | biological process | GO:0060491 | 163 | 4 | 13711 | 111 | Rac2,Mark4,Rap1gap,Brk1 | | 4.335e-02 | -3.14 | morphogenesis of a branching structure | biological process | GO:0001763 | 163 | 4 | 13711 | 111 | Ctsz,Mgp,Tfap2c,Spint1 | | 4.335e-02 | -3.14 | regulation of mononuclear cell proliferation | biological process | GO:0032944 | 163 | 4 | 13711 | 111 | Ptprc,Il6ra,Ikzf3,Rac2 | | 4.335e-02 | -3.14 | positive regulation of epithelial cell proliferation | biological process | GO:0050679 | 163 | 4 | 13711 | 111 | Lrg1,Grn,Cyba,Ptprn | | 4.336e-02 | -3.14 | GO\_CYTOSKELETON\_ORGANIZATION | MSigDB lists | GO\_CYTOSKELETON\_ORGANIZATION | 672 | 10 | 12187 | 98 | Flnb,Vim,Gfap,Rac2,Hcls1,Mark4,Brk1,Thy1,Myh9,Capn3 | | 4.340e-02 | -3.14 | collagen metabolic process | biological process | GO:0032963 | 41 | 2 | 13711 | 111 | Ctsb,Ctss | | 4.340e-02 | -3.14 | regulation of sensory perception of pain | biological process | GO:0051930 | 41 | 2 | 13711 | 111 | Mgll,Ctss | | 4.340e-02 | -3.14 | positive regulation of cell killing | biological process | GO:0031343 | 41 | 2 | 13711 | 111 | Ptprc,Ccr5 | | 4.340e-02 | -3.14 | regulation of antigen receptor-mediated signaling pathway | biological process | GO:0050854 | 41 | 2 | 13711 | 111 | Thy1,Ptprc | | 4.354e-02 | -3.13 | RUTELLA\_RESPONSE\_TO\_HGF\_VS\_CSF2RB\_AND\_IL4\_UP | MSigDB lists | RUTELLA\_RESPONSE\_TO\_HGF\_VS\_CSF2RB\_AND\_IL4\_UP | 319 | 6 | 12187 | 98 | Ccr5,Tsc22d1,Cd68,C3,Ctsb,C1qa | | 4.361e-02 | -3.13 | GO\_MEMBRANE\_REGION | MSigDB lists | GO\_MEMBRANE\_REGION | 863 | 12 | 12187 | 98 | Csk,Ahnak,Kl,Dlgap3,Ptprc,Cyba,Thy1,Slc2a1,Sh2d5,Il6ra,Lin7a,Ctsb | | 4.395e-02 | -3.12 | GSE411\_WT\_VS\_SOCS3\_KO\_MACROPHAGE\_IL6\_STIM\_400MIN\_UP | MSigDB lists | GSE411\_WT\_VS\_SOCS3\_KO\_MACROPHAGE\_IL6\_STIM\_400MIN\_UP | 165 | 4 | 12187 | 98 | Mpeg1,Flnb,Hspa2,Ccr5 | | 4.395e-02 | -3.12 | GSE41867\_NAIVE\_VS\_DAY8\_LCMV\_ARMSTRONG\_EFFECTOR\_CD8\_TCELL\_DN | MSigDB lists | GSE41867\_NAIVE\_VS\_DAY8\_LCMV\_ARMSTRONG\_EFFECTOR\_CD8\_TCELL\_DN | 165 | 4 | 12187 | 98 | Ly6e,Fcgr3,Ctsb,Slc2a1 | | 4.395e-02 | -3.12 | GSE2770\_TGFB\_AND\_IL4\_ACT\_VS\_ACT\_CD4\_TCELL\_48H\_UP | MSigDB lists | GSE2770\_TGFB\_AND\_IL4\_ACT\_VS\_ACT\_CD4\_TCELL\_48H\_UP | 165 | 4 | 12187 | 98 | Ccr5,Tsc22d1,C3,Ctsb | | 4.395e-02 | -3.12 | GSE17186\_MEMORY\_VS\_CD21LOW\_TRANSITIONAL\_BCELL\_DN | MSigDB lists | GSE17186\_MEMORY\_VS\_CD21LOW\_TRANSITIONAL\_BCELL\_DN | 165 | 4 | 12187 | 98 | Csk,Rps8,Slc2a1,Treml2 | | 4.395e-02 | -3.12 | GSE17721\_LPS\_VS\_CPG\_12H\_BMDC\_UP | MSigDB lists | GSE17721\_LPS\_VS\_CPG\_12H\_BMDC\_UP | 165 | 4 | 12187 | 98 | Tnfsf8,Cox4i1,Serpinb9,Actb | | 4.395e-02 | -3.12 | GSE36888\_STAT5\_AB\_KNOCKIN\_VS\_WT\_TCELL\_IL2\_TREATED\_6H\_DN | MSigDB lists | GSE36888\_STAT5\_AB\_KNOCKIN\_VS\_WT\_TCELL\_IL2\_TREATED\_6H\_DN | 165 | 4 | 12187 | 98 | Serpinb9,Fam111a,Fcer1g,Tspan4 | | 4.395e-02 | -3.12 | GSE7831\_UNSTIM\_VS\_CPG\_STIM\_PDC\_4H\_DN | MSigDB lists | GSE7831\_UNSTIM\_VS\_CPG\_STIM\_PDC\_4H\_DN | 165 | 4 | 12187 | 98 | Fam111a,Spint1,Mpeg1,Flnb | | 4.395e-02 | -3.12 | GSE10240\_CTRL\_VS\_IL22\_STIM\_PRIMARY\_BRONCHIAL\_EPITHELIAL\_CELLS\_UP | MSigDB lists | GSE10240\_CTRL\_VS\_IL22\_STIM\_PRIMARY\_BRONCHIAL\_EPITHELIAL\_CELLS\_UP | 165 | 4 | 12187 | 98 | Tmem176a,Il1r1,Sh2d5,Tsc22d1 | | 4.395e-02 | -3.12 | HALLMARK\_MYOGENESIS | MSigDB lists | HALLMARK\_MYOGENESIS | 165 | 4 | 12187 | 98 | Apod,Camk2b,Myh9,Clu | | 4.395e-02 | -3.12 | GSE3039\_ALPHAALPHA\_CD8\_TCELL\_VS\_B2\_BCELL\_DN | MSigDB lists | GSE3039\_ALPHAALPHA\_CD8\_TCELL\_VS\_B2\_BCELL\_DN | 165 | 4 | 12187 | 98 | C1qb,Ldha,Cd68,Psme1 | | 4.403e-02 | -3.12 | positive regulation of plasma membrane bounded cell projection assembly | biological process | GO:0120034 | 97 | 3 | 13711 | 111 | Rac2,Mark4,Brk1 | | 4.406e-02 | -3.12 | cellular component organization or biogenesis | biological process | GO:0071840 | 4450 | 45 | 13711 | 111 | Clu,Hspa2,Slc17a7,Gfap,Capn3,Camk2b,Rpl26,Vim,Rps8,Spint1,Rpl14,Rac2,Tspan4,Brk1,Csf1r,Hcls1,Ptprn,C3,Vwf,C1qa,Ptprc,Ctss,Grn,Mgll,Thy1,Actb,Fcer1g,Psmb10,Ahnak,Flnb,Fcgr3,Vamp8,Dek,Slc2a1,Rps5,Mt3,Myh9,Ccr5,Mgp,Ndufa13,C1qb,Cyba,Dlgap3,Mark4,Csk | | 4.408e-02 | -3.12 | negative regulation of proteolysis | biological process | GO:0045861 | 238 | 5 | 13711 | 111 | Serpina3n,Ctsz,Spint1,C3,Serpinb9 | | 4.410e-02 | -3.12 | GO\_REGULATION\_OF\_HORMONE\_LEVELS | MSigDB lists | GO\_REGULATION\_OF\_HORMONE\_LEVELS | 320 | 6 | 12187 | 98 | Slc2a1,Ptprn,Vamp8,Ctsz,Mc4r,Pcsk2 | | 4.432e-02 | -3.12 | GO\_TISSUE\_DEVELOPMENT | MSigDB lists | GO\_TISSUE\_DEVELOPMENT | 1062 | 14 | 12187 | 98 | Lfng,Itgam,Thy1,Csf1r,Spint1,Ctsb,Ctsz,Vim,Ikzf3,Mgp,Ptprc,Apod,Flnb,Fgf18 | | 4.434e-02 | -3.12 | MODULE\_223 | MSigDB lists | MODULE\_223 | 98 | 3 | 12187 | 98 | Il6ra,Ly6e,Grn | | 4.439e-02 | -3.11 | Platelet Aggregation (Plug Formation) | REACTOME pathways | R-MMU-76009 | 28 | 2 | 6297 | 76 | Csk,Vwf | | 4.460e-02 | -3.11 | Hemostasis | REACTOME pathways | R-MMU-109582 | 391 | 9 | 6297 | 76 | Fcer1g,Vwf,Csk,Itgam,Pde1a,Mgll,Serpina3n,Rac2,Clu | | 4.476e-02 | -3.11 | YAMASHITA\_LIVER\_CANCER\_WITH\_EPCAM\_UP | MSigDB lists | YAMASHITA\_LIVER\_CANCER\_WITH\_EPCAM\_UP | 42 | 2 | 12187 | 98 | Fmod,Rpl31 | | 4.476e-02 | -3.11 | GO\_CALCIUM\_DEPENDENT\_PROTEIN\_BINDING | MSigDB lists | GO\_CALCIUM\_DEPENDENT\_PROTEIN\_BINDING | 42 | 2 | 12187 | 98 | S100a6,Cplx2 | | 4.476e-02 | -3.11 | LUI\_THYROID\_CANCER\_PAX8\_PPARG\_DN | MSigDB lists | LUI\_THYROID\_CANCER\_PAX8\_PPARG\_DN | 42 | 2 | 12187 | 98 | Rap1gap,Rps5 | | 4.476e-02 | -3.11 | GO\_RIBOSOMAL\_LARGE\_SUBUNIT\_BIOGENESIS | MSigDB lists | GO\_RIBOSOMAL\_LARGE\_SUBUNIT\_BIOGENESIS | 42 | 2 | 12187 | 98 | Rpl14,Rpl26 | | 4.476e-02 | -3.11 | KENNY\_CTNNB1\_TARGETS\_UP | MSigDB lists | KENNY\_CTNNB1\_TARGETS\_UP | 42 | 2 | 12187 | 98 | Capn3,Ccr5 | | 4.476e-02 | -3.11 | LEE\_LIVER\_CANCER\_E2F1\_UP | MSigDB lists | LEE\_LIVER\_CANCER\_E2F1\_UP | 42 | 2 | 12187 | 98 | Ly6e,Ctss | | 4.476e-02 | -3.11 | GO\_CHAPERONE\_MEDIATED\_PROTEIN\_FOLDING | MSigDB lists | GO\_CHAPERONE\_MEDIATED\_PROTEIN\_FOLDING | 42 | 2 | 12187 | 98 | Clu,Ttc9b | | 4.476e-02 | -3.11 | GO\_HOMOTYPIC\_CELL\_CELL\_ADHESION | MSigDB lists | GO\_HOMOTYPIC\_CELL\_CELL\_ADHESION | 42 | 2 | 12187 | 98 | Actb,Myh9 | | 4.477e-02 | -3.11 | GSE17721\_CTRL\_VS\_POLYIC\_12H\_BMDC\_UP | MSigDB lists | GSE17721\_CTRL\_VS\_POLYIC\_12H\_BMDC\_UP | 166 | 4 | 12187 | 98 | Ctsa,Fam111a,Il6ra,Sh3bgrl3 | | 4.477e-02 | -3.11 | GSE11961\_MARGINAL\_ZONE\_BCELL\_VS\_GERMINAL\_CENTER\_BCELL\_DAY7\_DN | MSigDB lists | GSE11961\_MARGINAL\_ZONE\_BCELL\_VS\_GERMINAL\_CENTER\_BCELL\_DAY7\_DN | 166 | 4 | 12187 | 98 | Ccr5,Hspa2,Tsc22d1,Vim | | 4.477e-02 | -3.11 | GSE45365\_NK\_CELL\_VS\_CD11B\_DC\_MCMV\_INFECTION\_DN | MSigDB lists | GSE45365\_NK\_CELL\_VS\_CD11B\_DC\_MCMV\_INFECTION\_DN | 166 | 4 | 12187 | 98 | Fcer1g,Tnfsf8,Tmem176a,Cyba | | 4.477e-02 | -3.11 | GSE2826\_XID\_VS\_BTK\_KO\_BCELL\_UP | MSigDB lists | GSE2826\_XID\_VS\_BTK\_KO\_BCELL\_UP | 166 | 4 | 12187 | 98 | Spint1,Tmem176a,Slc26a2,Vwf | | 4.477e-02 | -3.11 | GSE11961\_FOLLICULAR\_BCELL\_VS\_MEMORY\_BCELL\_DAY40\_DN | MSigDB lists | GSE11961\_FOLLICULAR\_BCELL\_VS\_MEMORY\_BCELL\_DAY40\_DN | 166 | 4 | 12187 | 98 | Hspa2,Igfbp6,Tspan4,Lin7a | | 4.477e-02 | -3.11 | GSE17721\_PAM3CSK4\_VS\_GADIQUIMOD\_16H\_BMDC\_DN | MSigDB lists | GSE17721\_PAM3CSK4\_VS\_GADIQUIMOD\_16H\_BMDC\_DN | 166 | 4 | 12187 | 98 | Csk,Ccr5,Psme1,C1qb | | 4.477e-02 | -3.11 | HALLMARK\_ESTROGEN\_RESPONSE\_EARLY | MSigDB lists | HALLMARK\_ESTROGEN\_RESPONSE\_EARLY | 166 | 4 | 12187 | 98 | Slc26a2,Flnb,Slc2a1,Tfap2c | | 4.477e-02 | -3.11 | GSE15330\_HSC\_VS\_PRO\_BCELL\_UP | MSigDB lists | GSE15330\_HSC\_VS\_PRO\_BCELL\_UP | 166 | 4 | 12187 | 98 | Actb,Cyba,Hcls1,Dek | | 4.491e-02 | -3.10 | cytokine activity | molecular function | GO:0005125 | 100 | 3 | 13516 | 107 | Ccl6,Grn,Tnfsf8 | | 4.495e-02 | -3.10 | negative regulation of cell death | biological process | GO:0060548 | 859 | 12 | 13711 | 111 | Clu,Fcer1g,Mt3,Hcls1,Csf1r,Grn,Ccr5,Ndufa13,Tsc22d1,Capn3,Ctsb,Serpinb9 | | 4.502e-02 | -3.10 | GO\_TRANSMEMBRANE\_RECEPTOR\_PROTEIN\_TYROSINE\_KINASE\_SIGNALING\_PATHWAY | MSigDB lists | GO\_TRANSMEMBRANE\_RECEPTOR\_PROTEIN\_TYROSINE\_KINASE\_SIGNALING\_PATHWAY | 406 | 7 | 12187 | 98 | Actb,Cyba,Fgf18,Csk,Brk1,Csf1r,Kl | | 4.502e-02 | -3.10 | GO\_CELLULAR\_CHEMICAL\_HOMEOSTASIS | MSigDB lists | GO\_CELLULAR\_CHEMICAL\_HOMEOSTASIS | 406 | 7 | 12187 | 98 | Ccr5,Ptprc,Il1r1,Slc26a2,Ptprn,Cyba,Slc17a7 | | 4.510e-02 | -3.10 | ZBTB11 (zinc finger and BTB domain containing 11) | protein interactions | 27107 | 4 | 1 | 6802 | 78 | Vim | | 4.510e-02 | -3.10 | TUBA1C (tubulin alpha 1c) | protein interactions | 84790 | 4 | 1 | 6802 | 78 | Myh9 | | 4.510e-02 | -3.10 | NCKIPSD (NCK interacting protein with SH3 domain) | protein interactions | 51517 | 4 | 1 | 6802 | 78 | Actb | | 4.510e-02 | -3.10 | MSN (moesin) | protein interactions | 4478 | 4 | 1 | 6802 | 78 | Myh9 | | 4.510e-02 | -3.10 | VAPA (VAMP associated protein A) | protein interactions | 9218 | 4 | 1 | 6802 | 78 | Actb | | 4.510e-02 | -3.10 | Hbb (hemoglobin beta chain complex) | protein interactions | 15127 | 4 | 1 | 6802 | 78 | Ctsb | | 4.510e-02 | -3.10 | HIP1R (huntingtin interacting protein 1 related) | protein interactions | 9026 | 4 | 1 | 6802 | 78 | Actb | | 4.510e-02 | -3.10 | UQCRC2 (ubiquinol-cytochrome c reductase core protein 2) | protein interactions | 7385 | 4 | 1 | 6802 | 78 | Csk | | 4.510e-02 | -3.10 | FRYL (FRY like transcription coactivator) | protein interactions | 285527 | 4 | 1 | 6802 | 78 | Csk | | 4.510e-02 | -3.10 | ZFYVE26 (zinc finger FYVE-type containing 26) | protein interactions | 23503 | 4 | 1 | 6802 | 78 | Csk | | 4.510e-02 | -3.10 | S100a10 (S100 calcium binding protein A10 (calpactin)) | protein interactions | 20194 | 4 | 1 | 6802 | 78 | Ahnak | | 4.510e-02 | -3.10 | MAT2A (methionine adenosyltransferase 2A) | protein interactions | 4144 | 4 | 1 | 6802 | 78 | Actb | | 4.510e-02 | -3.10 | RASAL2 (RAS protein activator like 2) | protein interactions | 9462 | 4 | 1 | 6802 | 78 | Mgp | | 4.510e-02 | -3.10 | S100A10 (S100 calcium binding protein A10) | protein interactions | 6281 | 4 | 1 | 6802 | 78 | Myh9 | | 4.510e-02 | -3.10 | DOCK7 (dedicator of cytokinesis 7) | protein interactions | 85440 | 4 | 1 | 6802 | 78 | Actb | | 4.510e-02 | -3.10 | TUBB2A (tubulin beta 2A class IIa) | protein interactions | 7280 | 4 | 1 | 6802 | 78 | Myh9 | | 4.510e-02 | -3.10 | Cdh5 (cadherin 5) | protein interactions | 12562 | 4 | 1 | 6802 | 78 | Csk | | 4.510e-02 | -3.10 | PPP1R9B (protein phosphatase 1 regulatory subunit 9B) | protein interactions | 84687 | 4 | 1 | 6802 | 78 | Myh9 | | 4.510e-02 | -3.10 | Itga6 (integrin alpha 6) | protein interactions | 16403 | 4 | 1 | 6802 | 78 | Ptprc | | 4.510e-02 | -3.10 | HEATR1 (HEAT repeat containing 1) | protein interactions | 55127 | 4 | 1 | 6802 | 78 | Mgp | | 4.510e-02 | -3.10 | ZNF668 (zinc finger protein 668) | protein interactions | 79759 | 4 | 1 | 6802 | 78 | Csk | | 4.510e-02 | -3.10 | Ptprr (protein tyrosine phosphatase, receptor type, R) | protein interactions | 19279 | 4 | 1 | 6802 | 78 | Camk2b | | 4.510e-02 | -3.10 | IKZF1 (IKAROS family zinc finger 1) | protein interactions | 10320 | 4 | 1 | 6802 | 78 | Ikzf3 | | 4.516e-02 | -3.10 | Prion diseases | KEGG pathways | ko05020 | 28 | 2 | 5248 | 64 | C1qb,C1qa | | 4.516e-02 | -3.10 | Prion diseases | KEGG pathways | mmu05020 | 28 | 2 | 5248 | 64 | C1qb,C1qa | | 4.533e-02 | -3.09 | regulation of sensory perception | biological process | GO:0051931 | 42 | 2 | 13711 | 111 | Mgll,Ctss | | 4.533e-02 | -3.09 | regulation of macrophage activation | biological process | GO:0043030 | 42 | 2 | 13711 | 111 | Grn,C1qa | | 4.541e-02 | -3.09 | regionalization | biological process | GO:0003002 | 240 | 5 | 13711 | 111 | C3,Ifitm1,Lfng,Bhlhe41,C1qa | | 4.559e-02 | -3.09 | GSE11924\_TFH\_VS\_TH2\_CD4\_TCELL\_UP | MSigDB lists | GSE11924\_TFH\_VS\_TH2\_CD4\_TCELL\_UP | 167 | 4 | 12187 | 98 | Myh9,Camk2b,Tmem106a,Adh1 | | 4.559e-02 | -3.09 | GSE20715\_0H\_VS\_6H\_OZONE\_TLR4\_KO\_LUNG\_DN | MSigDB lists | GSE20715\_0H\_VS\_6H\_OZONE\_TLR4\_KO\_LUNG\_DN | 167 | 4 | 12187 | 98 | Serpina3n,Vamp8,S100a6,Igfbp6 | | 4.559e-02 | -3.09 | GSE17721\_CTRL\_VS\_POLYIC\_24H\_BMDC\_UP | MSigDB lists | GSE17721\_CTRL\_VS\_POLYIC\_24H\_BMDC\_UP | 167 | 4 | 12187 | 98 | Rpl31,Ctsa,Fam111a,Rpl13 | | 4.559e-02 | -3.09 | GSE5960\_TH1\_VS\_ANERGIC\_TH1\_UP | MSigDB lists | GSE5960\_TH1\_VS\_ANERGIC\_TH1\_UP | 167 | 4 | 12187 | 98 | Psme1,Mgll,Rps8,Rpl14 | | 4.559e-02 | -3.09 | GSE9988\_LPS\_VS\_VEHICLE\_TREATED\_MONOCYTE\_DN | MSigDB lists | GSE9988\_LPS\_VS\_VEHICLE\_TREATED\_MONOCYTE\_DN | 167 | 4 | 12187 | 98 | Csf1r,Mpeg1,Grn,Csk | | 4.559e-02 | -3.09 | GSE22886\_DAY0\_VS\_DAY1\_MONOCYTE\_IN\_CULTURE\_DN | MSigDB lists | GSE22886\_DAY0\_VS\_DAY1\_MONOCYTE\_IN\_CULTURE\_DN | 167 | 4 | 12187 | 98 | Ctsb,C3,Tsc22d1,Mgll | | 4.559e-02 | -3.09 | RODWELL\_AGING\_KIDNEY\_NO\_BLOOD\_UP | MSigDB lists | RODWELL\_AGING\_KIDNEY\_NO\_BLOOD\_UP | 167 | 4 | 12187 | 98 | C1qa,C1qb,C3,Mgll | | 4.559e-02 | -3.09 | GSE18804\_SPLEEN\_MACROPHAGE\_VS\_COLON\_TUMORAL\_MACROPHAGE\_UP | MSigDB lists | GSE18804\_SPLEEN\_MACROPHAGE\_VS\_COLON\_TUMORAL\_MACROPHAGE\_UP | 167 | 4 | 12187 | 98 | Ly6e,Thy1,Psme1,Csk | | 4.559e-02 | -3.09 | GSE22601\_DOUBLE\_NEGATIVE\_VS\_CD4\_SINGLE\_POSITIVE\_THYMOCYTE\_DN | MSigDB lists | GSE22601\_DOUBLE\_NEGATIVE\_VS\_CD4\_SINGLE\_POSITIVE\_THYMOCYTE\_DN | 167 | 4 | 12187 | 98 | Tmem176a,Slc2a1,S100a6,Ldha | | 4.559e-02 | -3.09 | KAECH\_NAIVE\_VS\_DAY8\_EFF\_CD8\_TCELL\_UP | MSigDB lists | KAECH\_NAIVE\_VS\_DAY8\_EFF\_CD8\_TCELL\_UP | 167 | 4 | 12187 | 98 | Rpl14,Il6ra,Rps8,Ctss | | 4.559e-02 | -3.09 | GSE32423\_CTRL\_VS\_IL7\_IL4\_MEMORY\_CD8\_TCELL\_UP | MSigDB lists | GSE32423\_CTRL\_VS\_IL7\_IL4\_MEMORY\_CD8\_TCELL\_UP | 167 | 4 | 12187 | 98 | Ahnak,Tsc22d1,Fcgr3,Exoc3l4 | | 4.559e-02 | -3.09 | GSE19772\_HCMV\_INFL\_VS\_HCMV\_INF\_MONOCYTES\_AND\_PI3K\_INHIBITION\_DN | MSigDB lists | GSE19772\_HCMV\_INFL\_VS\_HCMV\_INF\_MONOCYTES\_AND\_PI3K\_INHIBITION\_DN | 167 | 4 | 12187 | 98 | Ccr5,Ahnak,Tsc22d1,Il1r1 | | 4.579e-02 | -3.08 | - | gene3d domains | 2.60.40.690 | 5 | 1 | 6647 | 62 | C3 | | 4.583e-02 | -3.08 | regulation of reactive oxygen species metabolic process | biological process | GO:2000377 | 166 | 4 | 13711 | 111 | Itgam,Clu,Tmem106a,Cyba | | 4.600e-02 | -3.08 | GO\_CATABOLIC\_PROCESS | MSigDB lists | GO\_CATABOLIC\_PROCESS | 1373 | 17 | 12187 | 98 | Rpl26,Ctsa,Ctsb,Rpl31,Rps5,Phyhip,Psmb10,Psme1,Rpl13,Fmod,Mgll,Ldha,Ctsz,Rps8,Ctss,Rpl14,Rps20 | | 4.629e-02 | -3.07 | GO\_CELLULAR\_AMIDE\_METABOLIC\_PROCESS | MSigDB lists | GO\_CELLULAR\_AMIDE\_METABOLIC\_PROCESS | 587 | 9 | 12187 | 98 | Rpl26,Rps5,Rpl14,Rpl31,Rps20,Pcsk2,Rpl13,Rps8,Ctsz | | 4.629e-02 | -3.07 | regulation of cellular component movement | biological process | GO:0051270 | 863 | 12 | 13711 | 111 | Csf1r,Rac2,Camk2b,Fgf18,Ccr5,Ptprc,Apod,Thy1,Il1r1,Vim,Ccar1,Grn | | 4.631e-02 | -3.07 | negative regulation of immune effector process | biological process | GO:0002698 | 99 | 3 | 13711 | 111 | Ptprc,Serpinb9,Grn | | 4.631e-02 | -3.07 | regulation of wound healing | biological process | GO:0061041 | 99 | 3 | 13711 | 111 | Myh9,Capn3,Fcer1g | | 4.636e-02 | -3.07 | GO\_POSTSYNAPSE | MSigDB lists | GO\_POSTSYNAPSE | 324 | 6 | 12187 | 98 | Camk2b,Sh2d5,Slc17a7,Dlgap3,Lin7a,Il1r1 | | 4.641e-02 | -3.07 | Signaling by EGFR | REACTOME pathways | R-MMU-177929 | 274 | 7 | 6297 | 76 | Pde1a,Kl,Fgf18,Psme1,Psmb10,Csk,Camk2b | | 4.643e-02 | -3.07 | GSE5679\_CTRL\_VS\_PPARG\_LIGAND\_ROSIGLITAZONE\_AND\_RARA\_AGONIST\_AM580\_TREATED\_DC\_DN | MSigDB lists | GSE5679\_CTRL\_VS\_PPARG\_LIGAND\_ROSIGLITAZONE\_AND\_RARA\_AGONIST\_AM580\_TREATED\_DC\_DN | 168 | 4 | 12187 | 98 | S100a6,Clu,Il1r1,Tmem176a | | 4.643e-02 | -3.07 | GO\_NEGATIVE\_REGULATION\_OF\_NEURON\_DIFFERENTIATION | MSigDB lists | GO\_NEGATIVE\_REGULATION\_OF\_NEURON\_DIFFERENTIATION | 168 | 4 | 12187 | 98 | Thy1,Rap1gap,Gfap,Vim | | 4.643e-02 | -3.07 | GSE17301\_CTRL\_VS\_48H\_ACD3\_ACD28\_IFNA2\_STIM\_CD8\_TCELL\_UP | MSigDB lists | GSE17301\_CTRL\_VS\_48H\_ACD3\_ACD28\_IFNA2\_STIM\_CD8\_TCELL\_UP | 168 | 4 | 12187 | 98 | Psmb10,Tmem176a,Ctsz,Rac2 | | 4.643e-02 | -3.07 | GSE17721\_0.5H\_VS\_4H\_LPS\_BMDC\_UP | MSigDB lists | GSE17721\_0.5H\_VS\_4H\_LPS\_BMDC\_UP | 168 | 4 | 12187 | 98 | Slc2a1,Rpl13,Fam111a,Ldha | | 4.643e-02 | -3.07 | GSE37301\_MULTIPOTENT\_PROGENITOR\_VS\_RAG2\_KO\_NK\_CELL\_DN | MSigDB lists | GSE37301\_MULTIPOTENT\_PROGENITOR\_VS\_RAG2\_KO\_NK\_CELL\_DN | 168 | 4 | 12187 | 98 | Serpina3n,S100a6,Ly6e,Ldha | | 4.643e-02 | -3.07 | GSE17721\_CTRL\_VS\_PAM3CSK4\_1H\_BMDC\_UP | MSigDB lists | GSE17721\_CTRL\_VS\_PAM3CSK4\_1H\_BMDC\_UP | 168 | 4 | 12187 | 98 | Ndufa13,Grn,Hspa2,Hcls1 | | 4.643e-02 | -3.07 | GSE39820\_CTRL\_VS\_IL1B\_IL6\_IL23A\_CD4\_TCELL\_DN | MSigDB lists | GSE39820\_CTRL\_VS\_IL1B\_IL6\_IL23A\_CD4\_TCELL\_DN | 168 | 4 | 12187 | 98 | Klhl6,Vim,Mgll,S100a6 | | 4.643e-02 | -3.07 | GSE17580\_TREG\_VS\_TEFF\_DN | MSigDB lists | GSE17580\_TREG\_VS\_TEFF\_DN | 168 | 4 | 12187 | 98 | Rpl14,Sh3bgrl3,Tsc22d1,Thy1 | | 4.643e-02 | -3.07 | GSE18281\_SUBCAPSULAR\_VS\_PERIMEDULLARY\_CORTICAL\_REGION\_OF\_THYMUS\_UP | MSigDB lists | GSE18281\_SUBCAPSULAR\_VS\_PERIMEDULLARY\_CORTICAL\_REGION\_OF\_THYMUS\_UP | 168 | 4 | 12187 | 98 | Mgll,Clu,Sh3bgrl3,Rab3il1 | | 4.643e-02 | -3.07 | GSE360\_DC\_VS\_MAC\_L\_DONOVANI\_DN | MSigDB lists | GSE360\_DC\_VS\_MAC\_L\_DONOVANI\_DN | 168 | 4 | 12187 | 98 | Ldha,Tsc22d1,Ccr5,Rps5 | | 4.643e-02 | -3.07 | KOW | smart domains | SM00739 | 5 | 1 | 7188 | 68 | Rpl26 | | 4.643e-02 | -3.07 | A2M\_recep | smart domains | SM01361 | 5 | 1 | 7188 | 68 | C3 | | 4.643e-02 | -3.07 | A2M\_N\_2 | smart domains | SM01359 | 5 | 1 | 7188 | 68 | C3 | | 4.643e-02 | -3.07 | A2M | smart domains | SM01360 | 5 | 1 | 7188 | 68 | C3 | | 4.648e-02 | -3.07 | mouse chr11 B3\|11 42.86 cM | chromosome location | mouse chr11 B3\|11 42.86 cM | 6 | 1 | 14556 | 115 | Cd68 | | 4.658e-02 | -3.07 | structural molecule activity conferring elasticity | molecular function | GO:0097493 | 6 | 1 | 13516 | 107 | Ahnak | | 4.658e-02 | -3.07 | interleukin-1 binding | molecular function | GO:0019966 | 6 | 1 | 13516 | 107 | Il1r1 | | 4.658e-02 | -3.07 | extracellularly glutamate-gated ion channel activity | molecular function | GO:0005234 | 6 | 1 | 13516 | 107 | Slc17a7 | | 4.658e-02 | -3.07 | MRF binding | molecular function | GO:0043426 | 6 | 1 | 13516 | 107 | Bhlhe41 | | 4.658e-02 | -3.07 | titin binding | molecular function | GO:0031432 | 6 | 1 | 13516 | 107 | Capn3 | | 4.661e-02 | -3.07 | GSE10463\_CD40L\_AND\_VA347\_VS\_CD40L\_IN\_DC\_DN | MSigDB lists | GSE10463\_CD40L\_AND\_VA347\_VS\_CD40L\_IN\_DC\_DN | 100 | 3 | 12187 | 98 | Lfng,Fmod,Thy1 | | 4.661e-02 | -3.07 | TAKEDA\_TARGETS\_OF\_NUP98\_HOXA9\_FUSION\_8D\_UP | MSigDB lists | TAKEDA\_TARGETS\_OF\_NUP98\_HOXA9\_FUSION\_8D\_UP | 100 | 3 | 12187 | 98 | Fgf18,Ly6e,Tsc22d1 | | 4.661e-02 | -3.07 | GNF2\_DAP3 | MSigDB lists | GNF2\_DAP3 | 100 | 3 | 12187 | 98 | Rps8,Rpl14,Rpl31 | | 4.670e-02 | -3.06 | BURTON\_ADIPOGENESIS\_7 | MSigDB lists | BURTON\_ADIPOGENESIS\_7 | 43 | 2 | 12187 | 98 | S100a6,Thy1 | | 4.670e-02 | -3.06 | REACTOME\_CDK\_MEDIATED\_PHOSPHORYLATION\_AND\_REMOVAL\_OF\_CDC6 | MSigDB lists | REACTOME\_CDK\_MEDIATED\_PHOSPHORYLATION\_AND\_REMOVAL\_OF\_CDC6 | 43 | 2 | 12187 | 98 | Psmb10,Psme1 | | 4.670e-02 | -3.06 | HADDAD\_T\_LYMPHOCYTE\_AND\_NK\_PROGENITOR\_DN | MSigDB lists | HADDAD\_T\_LYMPHOCYTE\_AND\_NK\_PROGENITOR\_DN | 43 | 2 | 12187 | 98 | Il6ra,Csf1r | | 4.670e-02 | -3.06 | LIU\_SMARCA4\_TARGETS | MSigDB lists | LIU\_SMARCA4\_TARGETS | 43 | 2 | 12187 | 98 | Ahnak,Klhl6 | | 4.670e-02 | -3.06 | GNF2\_TST | MSigDB lists | GNF2\_TST | 43 | 2 | 12187 | 98 | Tmem176a,Adh1 | | 4.670e-02 | -3.06 | CHIBA\_RESPONSE\_TO\_TSA | MSigDB lists | CHIBA\_RESPONSE\_TO\_TSA | 43 | 2 | 12187 | 98 | Grn,Clu | | 4.670e-02 | -3.06 | GO\_ACUTE\_INFLAMMATORY\_RESPONSE | MSigDB lists | GO\_ACUTE\_INFLAMMATORY\_RESPONSE | 43 | 2 | 12187 | 98 | Serpina3n,Il6ra | | 4.670e-02 | -3.06 | NUTT\_GBM\_VS\_AO\_GLIOMA\_UP | MSigDB lists | NUTT\_GBM\_VS\_AO\_GLIOMA\_UP | 43 | 2 | 12187 | 98 | Vim,Ldha | | 4.670e-02 | -3.06 | GO\_CYTOSOLIC\_CALCIUM\_ION\_TRANSPORT | MSigDB lists | GO\_CYTOSOLIC\_CALCIUM\_ION\_TRANSPORT | 43 | 2 | 12187 | 98 | Ptprc,Ccr5 | | 4.670e-02 | -3.06 | REACTOME\_VIF\_MEDIATED\_DEGRADATION\_OF\_APOBEC3G | MSigDB lists | REACTOME\_VIF\_MEDIATED\_DEGRADATION\_OF\_APOBEC3G | 43 | 2 | 12187 | 98 | Psmb10,Psme1 | | 4.673e-02 | -3.06 | regulation of neurotransmitter levels | biological process | GO:0001505 | 322 | 6 | 13711 | 111 | Clu,Lin7a,Slc17a7,Gfap,Tmem106a,Cplx2 | | 4.686e-02 | -3.06 | positive regulation of metabolic process | biological process | GO:0009893 | 2865 | 31 | 13711 | 111 | Tsc22d1,Psme1,Capn3,Fgf18,Serpinb9,Il6ra,Itgam,Ikzf3,Ccar1,Tmem106a,Grn,Kl,Clu,Hspa2,Ptprc,Ndufa13,Tfap2c,Ccr5,C3,Ptprn,Cyba,Csk,Mt3,Hcls1,Csf1r,Myh9,Vim,Ccl6,Rpl26,Fcgr3,Tnfsf8 | | 4.700e-02 | -3.06 | GO\_MRNA\_METABOLIC\_PROCESS | MSigDB lists | GO\_MRNA\_METABOLIC\_PROCESS | 498 | 8 | 12187 | 98 | Rps8,Rpl13,Rps20,Rpl31,Rps5,Rpl14,Rpl26,Ccar1 | | 4.706e-02 | -3.06 | postsynaptic specialization | cellular component | GO:0099572 | 410 | 7 | 13825 | 111 | Rpl14,Sh2d5,Camk2b,Il1r1,Actb,Abhd17c,Dlgap3 | | 4.712e-02 | -3.06 | GO\_REGULATION\_OF\_CELLULAR\_COMPONENT\_MOVEMENT | MSigDB lists | GO\_REGULATION\_OF\_CELLULAR\_COMPONENT\_MOVEMENT | 589 | 9 | 12187 | 98 | Csf1r,Thy1,Fgf18,Ptprc,Apod,Rac2,Il6ra,Sh3bgrl3,Ccar1 | | 4.727e-02 | -3.05 | GSE17721\_CTRL\_VS\_PAM3CSK4\_12H\_BMDC\_UP | MSigDB lists | GSE17721\_CTRL\_VS\_PAM3CSK4\_12H\_BMDC\_UP | 169 | 4 | 12187 | 98 | Rpl13,Fam111a,Grn,Cox4i1 | | 4.727e-02 | -3.05 | HMGIY\_Q6 | MSigDB lists | HMGIY\_Q6 | 169 | 4 | 12187 | 98 | Tfap2c,Mgll,Tsc22d1,Ikzf3 | | 4.727e-02 | -3.05 | GSE2770\_UNTREATED\_VS\_TGFB\_AND\_IL4\_TREATED\_ACT\_CD4\_TCELL\_4H\_UP | MSigDB lists | GSE2770\_UNTREATED\_VS\_TGFB\_AND\_IL4\_TREATED\_ACT\_CD4\_TCELL\_4H\_UP | 169 | 4 | 12187 | 98 | Ccr5,Il6ra,Vamp8,Ctsz | | 4.727e-02 | -3.05 | GSE39820\_CTRL\_VS\_TGFBETA3\_IL6\_CD4\_TCELL\_UP | MSigDB lists | GSE39820\_CTRL\_VS\_TGFBETA3\_IL6\_CD4\_TCELL\_UP | 169 | 4 | 12187 | 98 | Lin7a,Ctss,Capn3,Fam111a | | 4.727e-02 | -3.05 | GSE22432\_CDC\_VS\_COMMON\_DC\_PROGENITOR\_UP | MSigDB lists | GSE22432\_CDC\_VS\_COMMON\_DC\_PROGENITOR\_UP | 169 | 4 | 12187 | 98 | Hcls1,Klhl6,Sh3bgrl3,Csk | | 4.727e-02 | -3.05 | GSE22140\_HEALTHY\_VS\_ARTHRITIC\_GERMFREE\_MOUSE\_CD4\_TCELL\_UP | MSigDB lists | GSE22140\_HEALTHY\_VS\_ARTHRITIC\_GERMFREE\_MOUSE\_CD4\_TCELL\_UP | 169 | 4 | 12187 | 98 | C1qb,Psmb10,Fcer1g,Csk | | 4.727e-02 | -3.05 | GSE5589\_UNSTIM\_VS\_45MIN\_LPS\_STIM\_MACROPHAGE\_UP | MSigDB lists | GSE5589\_UNSTIM\_VS\_45MIN\_LPS\_STIM\_MACROPHAGE\_UP | 169 | 4 | 12187 | 98 | Rps5,Il1r1,S100a6,Vim | | 4.727e-02 | -3.05 | GSE15930\_NAIVE\_VS\_72H\_IN\_VITRO\_STIM\_IFNAB\_CD8\_TCELL\_DN | MSigDB lists | GSE15930\_NAIVE\_VS\_72H\_IN\_VITRO\_STIM\_IFNAB\_CD8\_TCELL\_DN | 169 | 4 | 12187 | 98 | Slc2a1,S100a6,Ccr5,Serpinb9 | | 4.727e-02 | -3.05 | GSE339\_CD4POS\_VS\_CD4CD8DN\_DC\_UP | MSigDB lists | GSE339\_CD4POS\_VS\_CD4CD8DN\_DC\_UP | 169 | 4 | 12187 | 98 | Rpl14,Spint1,C1qa,Vamp8 | | 4.727e-02 | -3.05 | GSE16522\_ANTI\_CD3CD28\_STIM\_VS\_UNSTIM\_MEMORY\_CD8\_TCELL\_DN | MSigDB lists | GSE16522\_ANTI\_CD3CD28\_STIM\_VS\_UNSTIM\_MEMORY\_CD8\_TCELL\_DN | 169 | 4 | 12187 | 98 | Ctss,Brk1,Ndufa13,Ccr5 | | 4.727e-02 | -3.05 | GSE9650\_EFFECTOR\_VS\_MEMORY\_CD8\_TCELL\_DN | MSigDB lists | GSE9650\_EFFECTOR\_VS\_MEMORY\_CD8\_TCELL\_DN | 169 | 4 | 12187 | 98 | Il6ra,Tnfsf8,Ctss,Rpl14 | | 4.727e-02 | -3.05 | HALLMARK\_EPITHELIAL\_MESENCHYMAL\_TRANSITION | MSigDB lists | HALLMARK\_EPITHELIAL\_MESENCHYMAL\_TRANSITION | 169 | 4 | 12187 | 98 | Fmod,Thy1,Mgp,Vim | | 4.730e-02 | -3.05 | REACTOME\_SIGNALING\_BY\_FGFR3\_MUTANTS | MSigDB lists | REACTOME\_SIGNALING\_BY\_FGFR3\_MUTANTS | 6 | 1 | 12187 | 98 | Fgf18 | | 4.730e-02 | -3.05 | BIOCARTA\_NO2IL12\_PATHWAY | MSigDB lists | BIOCARTA\_NO2IL12\_PATHWAY | 6 | 1 | 12187 | 98 | Ccr5 | | 4.730e-02 | -3.05 | REACTOME\_FGFR2C\_LIGAND\_BINDING\_AND\_ACTIVATION | MSigDB lists | REACTOME\_FGFR2C\_LIGAND\_BINDING\_AND\_ACTIVATION | 6 | 1 | 12187 | 98 | Fgf18 | | 4.730e-02 | -3.05 | CHESLER\_BRAIN\_QTL\_TRANS | MSigDB lists | CHESLER\_BRAIN\_QTL\_TRANS | 6 | 1 | 12187 | 98 | Myh9 | | 4.730e-02 | -3.05 | GO\_SUPEROXIDE\_ANION\_GENERATION | MSigDB lists | GO\_SUPEROXIDE\_ANION\_GENERATION | 6 | 1 | 12187 | 98 | Cyba | | 4.730e-02 | -3.05 | LY\_AGING\_MIDDLE\_UP | MSigDB lists | LY\_AGING\_MIDDLE\_UP | 6 | 1 | 12187 | 98 | Fmod | | 4.730e-02 | -3.05 | MIKHAYLOVA\_OXIDATIVE\_STRESS\_RESPONSE\_VIA\_VHL\_UP | MSigDB lists | MIKHAYLOVA\_OXIDATIVE\_STRESS\_RESPONSE\_VIA\_VHL\_UP | 6 | 1 | 12187 | 98 | Vim | | 4.730e-02 | -3.05 | BIOCARTA\_BLYMPHOCYTE\_PATHWAY | MSigDB lists | BIOCARTA\_BLYMPHOCYTE\_PATHWAY | 6 | 1 | 12187 | 98 | Ptprc | | 4.730e-02 | -3.05 | GO\_MAINTENANCE\_OF\_GASTROINTESTINAL\_EPITHELIUM | MSigDB lists | GO\_MAINTENANCE\_OF\_GASTROINTESTINAL\_EPITHELIUM | 6 | 1 | 12187 | 98 | Serpina3n | | 4.730e-02 | -3.05 | GUTIERREZ\_WALDENSTROEMS\_MACROGLOBULINEMIA\_2 | MSigDB lists | GUTIERREZ\_WALDENSTROEMS\_MACROGLOBULINEMIA\_2 | 6 | 1 | 12187 | 98 | Dek | | 4.730e-02 | -3.05 | GO\_C\_C\_CHEMOKINE\_RECEPTOR\_ACTIVITY | MSigDB lists | GO\_C\_C\_CHEMOKINE\_RECEPTOR\_ACTIVITY | 6 | 1 | 12187 | 98 | Ccr5 | | 4.730e-02 | -3.05 | synaptic vesicle endocytosis | biological process | GO:0048488 | 43 | 2 | 13711 | 111 | Actb,Grn | | 4.730e-02 | -3.05 | presynaptic endocytosis | biological process | GO:0140238 | 43 | 2 | 13711 | 111 | Grn,Actb | | 4.730e-02 | -3.05 | chemokine-mediated signaling pathway | biological process | GO:0070098 | 43 | 2 | 13711 | 111 | Ccr5,Ccl6 | | 4.742e-02 | -3.05 | MET Receptor Activation | REACTOME pathways | R-MMU-6806942 | 4 | 1 | 6297 | 76 | Spint1 | | 4.742e-02 | -3.05 | FGFR3b ligand binding and activation | REACTOME pathways | R-MMU-190371 | 4 | 1 | 6297 | 76 | Fgf18 | | 4.742e-02 | -3.05 | Acyl chain remodeling of DAG and TAG | REACTOME pathways | R-MMU-1482883 | 4 | 1 | 6297 | 76 | Mgll | | 4.742e-02 | -3.05 | FGFR4 ligand binding and activation | REACTOME pathways | R-MMU-190322 | 4 | 1 | 6297 | 76 | Fgf18 | | 4.748e-02 | -3.05 | TYR\_PHOSPHATASE\_PTP | prosite domains | PS50055 | 34 | 2 | 8845 | 91 | Ptprn,Ptprc | | 4.749e-02 | -3.05 | GO\_REGULATION\_OF\_ACTIN\_FILAMENT\_BASED\_PROCESS | MSigDB lists | GO\_REGULATION\_OF\_ACTIN\_FILAMENT\_BASED\_PROCESS | 245 | 5 | 12187 | 98 | Sh3bgrl3,Brk1,Myh9,Hcls1,Csf1r | | 4.761e-02 | -3.04 | CD8-positive, alpha-beta T cell differentiation | biological process | GO:0043374 | 6 | 1 | 13711 | 111 | Tnfsf8 | | 4.761e-02 | -3.04 | hydrogen peroxide biosynthetic process | biological process | GO:0050665 | 6 | 1 | 13711 | 111 | Cyba | | 4.761e-02 | -3.04 | sterol catabolic process | biological process | GO:0016127 | 6 | 1 | 13711 | 111 | Mt3 | | 4.761e-02 | -3.04 | cholesterol catabolic process | biological process | GO:0006707 | 6 | 1 | 13711 | 111 | Mt3 | | 4.761e-02 | -3.04 | regulation of apoptotic cell clearance | biological process | GO:2000425 | 6 | 1 | 13711 | 111 | C3 | | 4.761e-02 | -3.04 | regulation of neutrophil degranulation | biological process | GO:0043313 | 6 | 1 | 13711 | 111 | Itgam | | 4.761e-02 | -3.04 | positive regulation of inclusion body assembly | biological process | GO:0090261 | 6 | 1 | 13711 | 111 | Clu | | 4.761e-02 | -3.04 | regulation of NK T cell activation | biological process | GO:0051133 | 6 | 1 | 13711 | 111 | Il6ra | | 4.761e-02 | -3.04 | positive regulation of killing of cells of other organism | biological process | GO:0051712 | 6 | 1 | 13711 | 111 | Ccr5 | | 4.761e-02 | -3.04 | positive regulation of toll-like receptor 2 signaling pathway | biological process | GO:0034137 | 6 | 1 | 13711 | 111 | Cyba | | 4.761e-02 | -3.04 | response to mycotoxin | biological process | GO:0010046 | 6 | 1 | 13711 | 111 | Actb | | 4.761e-02 | -3.04 | regulation of amyloid fibril formation | biological process | GO:1905906 | 6 | 1 | 13711 | 111 | Clu | | 4.761e-02 | -3.04 | negative regulation of microglial cell activation | biological process | GO:1903979 | 6 | 1 | 13711 | 111 | Grn | | 4.761e-02 | -3.04 | G1 to G0 transition | biological process | GO:0070314 | 6 | 1 | 13711 | 111 | Capn3 | | 4.761e-02 | -3.04 | cellular response to oxidised low-density lipoprotein particle stimulus | biological process | GO:0140052 | 6 | 1 | 13711 | 111 | Cd68 | | 4.761e-02 | -3.04 | regulation of microvillus assembly | biological process | GO:0032534 | 6 | 1 | 13711 | 111 | Rap1gap | | 4.761e-02 | -3.04 | ciliary neurotrophic factor-mediated signaling pathway | biological process | GO:0070120 | 6 | 1 | 13711 | 111 | Il6ra | | 4.761e-02 | -3.04 | norepinephrine biosynthetic process | biological process | GO:0042421 | 6 | 1 | 13711 | 111 | Kl | | 4.761e-02 | -3.04 | regulation of Rho-dependent protein serine/threonine kinase activity | biological process | GO:2000298 | 6 | 1 | 13711 | 111 | Thy1 | | 4.761e-02 | -3.04 | germinal center formation | biological process | GO:0002467 | 6 | 1 | 13711 | 111 | Klhl6 | | 4.761e-02 | -3.04 | positive regulation of apoptotic cell clearance | biological process | GO:2000427 | 6 | 1 | 13711 | 111 | C3 | | 4.761e-02 | -3.04 | regulation of dendritic cell differentiation | biological process | GO:2001198 | 6 | 1 | 13711 | 111 | Tmem176a | | 4.761e-02 | -3.04 | positive regulation of protein folding | biological process | GO:1903334 | 6 | 1 | 13711 | 111 | Grn | | 4.761e-02 | -3.04 | exocyst localization | biological process | GO:0051601 | 6 | 1 | 13711 | 111 | Exoc3l4 | | 4.761e-02 | -3.04 | negative regulation of low-density lipoprotein particle clearance | biological process | GO:0010989 | 6 | 1 | 13711 | 111 | Csk | | 4.761e-02 | -3.04 | platelet degranulation | biological process | GO:0002576 | 6 | 1 | 13711 | 111 | Fcer1g | | 4.761e-02 | -3.04 | male mating behavior | biological process | GO:0060179 | 6 | 1 | 13711 | 111 | Grn | | 4.761e-02 | -3.04 | positive regulation of Fc receptor mediated stimulatory signaling pathway | biological process | GO:0060369 | 6 | 1 | 13711 | 111 | Ptprc | | 4.761e-02 | -3.04 | positive regulation of gamma-delta T cell differentiation | biological process | GO:0045588 | 6 | 1 | 13711 | 111 | Ptprc | | 4.761e-02 | -3.04 | regulation of somitogenesis | biological process | GO:0014807 | 6 | 1 | 13711 | 111 | Lfng | | 4.761e-02 | -3.04 | positive regulation of gamma-delta T cell activation | biological process | GO:0046645 | 6 | 1 | 13711 | 111 | Ptprc | | 4.761e-02 | -3.04 | detoxification of inorganic compound | biological process | GO:0061687 | 6 | 1 | 13711 | 111 | Mt3 | | 4.761e-02 | -3.04 | neutrophil activation involved in immune response | biological process | GO:0002283 | 6 | 1 | 13711 | 111 | Fcer1g | | 4.761e-02 | -3.04 | regulation of interleukin-1-mediated signaling pathway | biological process | GO:2000659 | 6 | 1 | 13711 | 111 | Il1r1 | | 4.762e-02 | -3.04 | positive regulation of hydrolase activity | biological process | GO:0051345 | 585 | 9 | 13711 | 111 | Ndufa13,Grn,Rap1gap,Psme1,Camk2b,Thy1,Ccl6,Hspa2,Ptprc | | 4.776e-02 | -3.04 | GSE45365\_BCELL\_VS\_CD8\_TCELL\_DN | MSigDB lists | GSE45365\_BCELL\_VS\_CD8\_TCELL\_DN | 101 | 3 | 12187 | 98 | Pcsk2,Hspa2,C1qb | | 4.776e-02 | -3.04 | SCHAEFFER\_PROSTATE\_DEVELOPMENT\_12HR\_UP | MSigDB lists | SCHAEFFER\_PROSTATE\_DEVELOPMENT\_12HR\_UP | 101 | 3 | 12187 | 98 | Ahnak,Tfap2c,Slc2a1 | | 4.791e-02 | -3.04 | Ascorbate biosynthesis, animals, glucose-1P => ascorbate | KEGG pathways | mmu\_M00129 | 4 | 1 | 5248 | 64 | Kl | | 4.791e-02 | -3.04 | Ascorbate biosynthesis, animals, glucose-1P => ascorbate | KEGG pathways | M00129 | 4 | 1 | 5248 | 64 | Kl | | 4.792e-02 | -3.04 | GO\_POSITIVE\_REGULATION\_OF\_TRANSFERASE\_ACTIVITY | MSigDB lists | GO\_POSITIVE\_REGULATION\_OF\_TRANSFERASE\_ACTIVITY | 500 | 8 | 12187 | 98 | Psmb10,Ptprc,Fgf18,Csf1r,Csk,Psme1,Clu,Hspa2 | | 4.810e-02 | -3.03 | GO\_REGULATION\_OF\_TRANSMEMBRANE\_TRANSPORT | MSigDB lists | GO\_REGULATION\_OF\_TRANSMEMBRANE\_TRANSPORT | 327 | 6 | 12187 | 98 | Cyba,Ctss,Thy1,Ahnak,Hspa2,Capn3 | | 4.812e-02 | -3.03 | GSE339\_CD8POS\_VS\_CD4CD8DN\_DC\_DN | MSigDB lists | GSE339\_CD8POS\_VS\_CD4CD8DN\_DC\_DN | 170 | 4 | 12187 | 98 | Fcer1g,Il6ra,S100a6,Il1r1 | | 4.812e-02 | -3.03 | YAMAZAKI\_TCEB3\_TARGETS\_DN | MSigDB lists | YAMAZAKI\_TCEB3\_TARGETS\_DN | 170 | 4 | 12187 | 98 | Tfap2c,S100a6,Slc2a1,Cyba | | 4.817e-02 | -3.03 | Metabolism of proteins | REACTOME pathways | R-MMU-392499 | 929 | 17 | 6297 | 76 | Rps5,Rps20,Rps8,Ctsa,Alg12,Actb,Psmb10,Pcsk2,Rpl26,Igfbp6,Thy1,Rpl14,Ly6e,Psme1,Tfap2c,Ctsz,Rpl13 | | 4.832e-02 | -3.03 | regulation of nitrogen compound metabolic process | biological process | GO:0051171 | 4126 | 42 | 13711 | 111 | Ikzf3,Tmem106a,Grn,Ccar1,Thy1,Kl,Rbfox3,Clu,Ptprc,Hspa2,Apod,Tsc22d1,Gfap,Il6ra,Ahnak,Serpinb9,Psme1,Capn3,Fgf18,Bhlhe41,Actb,Vim,Dek,Ctsz,Ccl6,Fcgr3,Tnfsf8,Rpl26,Zfp786,Tfap2c,Ndufa13,Serpina3n,C3,Ptprn,Csk,Ctsa,Rac2,Spint1,Csf1r,Hcls1,Myh9,Mt3 | | 4.838e-02 | -3.03 | regulation of leukocyte proliferation | biological process | GO:0070663 | 169 | 4 | 13711 | 111 | Il6ra,Ptprc,Ikzf3,Rac2 | | 4.850e-02 | -3.03 | GO\_INTRACELLULAR\_VESICLE | MSigDB lists | GO\_INTRACELLULAR\_VESICLE | 976 | 13 | 12187 | 98 | Vamp8,Slc17a7,Ctsb,Pcsk2,Slc2a1,Rac2,Serpina3n,Cyba,Ptprn,Ctsz,Vwf,Clu,Camk2b | | 4.852e-02 | -3.03 | positive regulation of cellular component movement | biological process | GO:0051272 | 497 | 8 | 13711 | 111 | Rac2,Csf1r,Ptprc,Grn,Ccar1,Thy1,Il1r1,Fgf18 | | 4.857e-02 | -3.02 | SNARE complex | cellular component | GO:0031201 | 44 | 2 | 13825 | 111 | Vamp8,Cplx2 | | 4.860e-02 | -3.02 | Pept\_T1A\_subB | interpro domains | IPR000243 | 6 | 1 | 13788 | 114 | Psmb10 | | 4.860e-02 | -3.02 | KOW | interpro domains | IPR005824 | 6 | 1 | 13788 | 114 | Rpl26 | | 4.860e-02 | -3.02 | Prot\_inhib\_I29 | interpro domains | IPR013201 | 6 | 1 | 13788 | 114 | Ctss | | 4.860e-02 | -3.02 | Liprin | interpro domains | IPR029515 | 6 | 1 | 13788 | 114 | Ppfibp1 | | 4.860e-02 | -3.02 | Lactate\_DH/Glyco\_Ohase\_4\_C | interpro domains | IPR015955 | 6 | 1 | 13788 | 114 | Ldha | | 4.860e-02 | -3.02 | Peptidase\_S8\_Asp-AS | interpro domains | IPR023827 | 6 | 1 | 13788 | 114 | Pcsk2 | | 4.860e-02 | -3.02 | C2\_III | interpro domains | IPR033883 | 6 | 1 | 13788 | 114 | Capn3 | | 4.860e-02 | -3.02 | Peptidase\_C1A\_papain-like | interpro domains | IPR039417 | 6 | 1 | 13788 | 114 | Ctss | | 4.860e-02 | -3.02 | Ig\_6 | interpro domains | IPR041416 | 6 | 1 | 13788 | 114 | Il1r1 | | 4.864e-02 | -3.02 | signaling receptor activity | molecular function | GO:0038023 | 601 | 9 | 13516 | 107 | Mc4r,Ccr5,Csf1r,Fcgr3,Treml2,Fcer1g,Il6ra,Adgre1,Il1r1 | | 4.864e-02 | -3.02 | regulation of production of molecular mediator of immune response | biological process | GO:0002700 | 101 | 3 | 13711 | 111 | Ptprc,Il1r1,Fcer1g | | 4.864e-02 | -3.02 | electron transport chain | biological process | GO:0022900 | 101 | 3 | 13711 | 111 | Sh3bgrl3,Cox4i1,Cyba | | 4.867e-02 | -3.02 | SIG\_BCR\_SIGNALING\_PATHWAY | MSigDB lists | SIG\_BCR\_SIGNALING\_PATHWAY | 44 | 2 | 12187 | 98 | Ptprc,Csk | | 4.867e-02 | -3.02 | REACTOME\_AUTODEGRADATION\_OF\_THE\_E3\_UBIQUITIN\_LIGASE\_COP1 | MSigDB lists | REACTOME\_AUTODEGRADATION\_OF\_THE\_E3\_UBIQUITIN\_LIGASE\_COP1 | 44 | 2 | 12187 | 98 | Psmb10,Psme1 | | 4.867e-02 | -3.02 | KEGG\_PATHOGENIC\_ESCHERICHIA\_COLI\_INFECTION | MSigDB lists | KEGG\_PATHOGENIC\_ESCHERICHIA\_COLI\_INFECTION | 44 | 2 | 12187 | 98 | Actb,Hcls1 | | 4.867e-02 | -3.02 | GCM\_SUPT4H1 | MSigDB lists | GCM\_SUPT4H1 | 44 | 2 | 12187 | 98 | Tmem106a,Fcgr3 | | 4.867e-02 | -3.02 | REACTOME\_CROSS\_PRESENTATION\_OF\_SOLUBLE\_EXOGENOUS\_ANTIGENS\_ENDOSOMES | MSigDB lists | REACTOME\_CROSS\_PRESENTATION\_OF\_SOLUBLE\_EXOGENOUS\_ANTIGENS\_ENDOSOMES | 44 | 2 | 12187 | 98 | Psme1,Psmb10 | | 4.867e-02 | -3.02 | ZHAN\_MULTIPLE\_MYELOMA\_CD1\_AND\_CD2\_DN | MSigDB lists | ZHAN\_MULTIPLE\_MYELOMA\_CD1\_AND\_CD2\_DN | 44 | 2 | 12187 | 98 | Csk,Il6ra | | 4.867e-02 | -3.02 | APRELIKOVA\_BRCA1\_TARGETS | MSigDB lists | APRELIKOVA\_BRCA1\_TARGETS | 44 | 2 | 12187 | 98 | Vim,Tfap2c | | 4.886e-02 | -3.02 | positive regulation of macromolecule metabolic process | biological process | GO:0010604 | 2653 | 29 | 13711 | 111 | Hcls1,Csf1r,Myh9,Mt3,Csk,Ndufa13,Ccr5,Tfap2c,C3,Ptprn,Fcgr3,Tnfsf8,Rpl26,Ccl6,Vim,Serpinb9,Il6ra,Psme1,Capn3,Fgf18,Tsc22d1,Ptprc,Hspa2,Clu,Kl,Ikzf3,Grn,Tmem106a,Ccar1 | | 4.887e-02 | -3.02 | WANG\_RESPONSE\_TO\_GSK3\_INHIBITOR\_SB216763\_UP | MSigDB lists | WANG\_RESPONSE\_TO\_GSK3\_INHIBITOR\_SB216763\_UP | 247 | 5 | 12187 | 98 | Cd68,Ctss,Ctsb,Rpl31,Ptprc | | 4.893e-02 | -3.02 | GO\_GROWTH\_FACTOR\_BINDING | MSigDB lists | GO\_GROWTH\_FACTOR\_BINDING | 102 | 3 | 12187 | 98 | Kl,Igfbp6,Il6ra | | 4.893e-02 | -3.02 | GSE9960\_HEALTHY\_VS\_GRAM\_NEG\_AND\_POS\_SEPSIS\_PBMC\_UP | MSigDB lists | GSE9960\_HEALTHY\_VS\_GRAM\_NEG\_AND\_POS\_SEPSIS\_PBMC\_UP | 102 | 3 | 12187 | 98 | Ttc9b,Cplx2,Pde1a | | 4.893e-02 | -3.02 | DARWICHE\_SKIN\_TUMOR\_PROMOTER\_UP | MSigDB lists | DARWICHE\_SKIN\_TUMOR\_PROMOTER\_UP | 102 | 3 | 12187 | 98 | Ly6e,Ldha,Apod | | 4.898e-02 | -3.02 | GSE10239\_NAIVE\_VS\_KLRG1INT\_EFF\_CD8\_TCELL\_UP | MSigDB lists | GSE10239\_NAIVE\_VS\_KLRG1INT\_EFF\_CD8\_TCELL\_UP | 171 | 4 | 12187 | 98 | Treml2,Il6ra,Rpl13,Ctss | | 4.898e-02 | -3.02 | GSE33374\_CD8\_ALPHAALPHA\_VS\_ALPHABETA\_CD161\_HIGH\_TCELL\_DN | MSigDB lists | GSE33374\_CD8\_ALPHAALPHA\_VS\_ALPHABETA\_CD161\_HIGH\_TCELL\_DN | 171 | 4 | 12187 | 98 | Thy1,Psme1,Rac2,Brk1 | | 4.898e-02 | -3.02 | GSE22886\_NAIVE\_CD4\_TCELL\_VS\_DC\_DN | MSigDB lists | GSE22886\_NAIVE\_CD4\_TCELL\_VS\_DC\_DN | 171 | 4 | 12187 | 98 | Ctsb,Vamp8,Vim,Grn | | 4.898e-02 | -3.02 | GSE2585\_AIRE\_KO\_VS\_WT\_CD80\_HIGH\_MTEC\_DN | MSigDB lists | GSE2585\_AIRE\_KO\_VS\_WT\_CD80\_HIGH\_MTEC\_DN | 171 | 4 | 12187 | 98 | Rab3il1,Ctsa,Vamp8,S100a6 | | 4.898e-02 | -3.02 | GSE24726\_WT\_VS\_E2\_22\_KO\_PDC\_DAY4\_POST\_DELETION\_DN | MSigDB lists | GSE24726\_WT\_VS\_E2\_22\_KO\_PDC\_DAY4\_POST\_DELETION\_DN | 171 | 4 | 12187 | 98 | Ly6e,Vamp8,Ctsz,C3 | | 4.898e-02 | -3.02 | GSE22601\_IMMATURE\_CD4\_SINGLE\_POSITIVE\_VS\_CD4\_SINGLE\_POSITIVE\_THYMOCYTE\_UP | MSigDB lists | GSE22601\_IMMATURE\_CD4\_SINGLE\_POSITIVE\_VS\_CD4\_SINGLE\_POSITIVE\_THYMOCYTE\_UP | 171 | 4 | 12187 | 98 | Csk,Ccr5,Thy1,S100a6 | | 4.898e-02 | -3.02 | GOLDRATH\_EFF\_VS\_MEMORY\_CD8\_TCELL\_DN | MSigDB lists | GOLDRATH\_EFF\_VS\_MEMORY\_CD8\_TCELL\_DN | 171 | 4 | 12187 | 98 | Il6ra,Rps8,Slc26a2,Rpl14 | | 4.906e-02 | -3.01 | positive regulation of cellular metabolic process | biological process | GO:0031325 | 2654 | 29 | 13711 | 111 | Tsc22d1,Il6ra,Capn3,Psme1,Fgf18,Itgam,Grn,Tmem106a,Ccar1,Ikzf3,Kl,Clu,Ptprc,Hspa2,Cyba,Ptprn,C3,Tfap2c,Ndufa13,Csk,Myh9,Csf1r,Hcls1,Mt3,Vim,Ccl6,Tnfsf8,Fcgr3,Rpl26 | | 4.915e-02 | -3.01 | mouse chr18\|18 E1 | chromosome location | mouse chr18\|18 E1 | 45 | 2 | 14556 | 115 | Mc4r,Slc26a2 | | 4.925e-02 | -3.01 | regulation of protein localization to membrane | biological process | GO:1905475 | 170 | 4 | 13711 | 111 | Vamp8,Camk2b,Csk,Actb | | 4.925e-02 | -3.01 | negative regulation of multi-organism process | biological process | GO:0043901 | 170 | 4 | 13711 | 111 | Ifitm1,Grn,Serpinb9,Fam111a | | 4.927e-02 | -3.01 | Ig\_6 | pfam domains | PF18452 | 6 | 1 | 12881 | 108 | Il1r1 | | 4.927e-02 | -3.01 | Inhibitor\_I29 | pfam domains | PF08246 | 6 | 1 | 12881 | 108 | Ctss | | 4.927e-02 | -3.01 | KOW | pfam domains | PF00467 | 6 | 1 | 12881 | 108 | Rpl26 | | 4.927e-02 | -3.01 | MACPF | pfam domains | PF01823 | 6 | 1 | 12881 | 108 | Mpeg1 | | 4.929e-02 | -3.01 | regulation of collagen metabolic process | biological process | GO:0010712 | 44 | 2 | 13711 | 111 | Il6ra,Vim | | 4.983e-02 | -3.00 | regulation of neural precursor cell proliferation | biological process | GO:2000177 | 102 | 3 | 13711 | 111 | Ccr5,Ctsz,Spint1 | | 4.983e-02 | -3.00 | phagocytosis | biological process | GO:0006909 | 102 | 3 | 13711 | 111 | Myh9,Fcgr3,Fcer1g | | 4.985e-02 | -3.00 | GSE32986\_CURDLAN\_LOWDOSE\_VS\_CURDLAN\_HIGHDOSE\_STIM\_DC\_UP | MSigDB lists | GSE32986\_CURDLAN\_LOWDOSE\_VS\_CURDLAN\_HIGHDOSE\_STIM\_DC\_UP | 172 | 4 | 12187 | 98 | Csk,Grn,Klhl6,Fcgr3 | | 4.985e-02 | -3.00 | GSE22229\_RENAL\_TRANSPLANT\_IMMUNOSUPP\_THERAPY\_VS\_HEALTHY\_PBMC\_UP | MSigDB lists | GSE22229\_RENAL\_TRANSPLANT\_IMMUNOSUPP\_THERAPY\_VS\_HEALTHY\_PBMC\_UP | 172 | 4 | 12187 | 98 | Fcer1g,Cd68,Fcgr3,C3 | | 4.985e-02 | -3.00 | GSE19941\_IL10\_KO\_VS\_IL10\_KO\_AND\_NFKBP50\_KO\_LPS\_STIM\_MACROPHAGE\_DN | MSigDB lists | GSE19941\_IL10\_KO\_VS\_IL10\_KO\_AND\_NFKBP50\_KO\_LPS\_STIM\_MACROPHAGE\_DN | 172 | 4 | 12187 | 98 | Tmem176a,Il1r1,Cd68,Vamp8 | | 4.985e-02 | -3.00 | GSE14308\_TH17\_VS\_INDUCED\_TREG\_UP | MSigDB lists | GSE14308\_TH17\_VS\_INDUCED\_TREG\_UP | 172 | 4 | 12187 | 98 | Thy1,Ndufa13,Rpl31,Tspan4 | | 4.985e-02 | -3.00 | GSE1925\_CTRL\_VS\_IFNG\_PRIMED\_MACROPHAGE\_24H\_IFNG\_STIM\_DN | MSigDB lists | GSE1925\_CTRL\_VS\_IFNG\_PRIMED\_MACROPHAGE\_24H\_IFNG\_STIM\_DN | 172 | 4 | 12187 | 98 | Actb,Serpinb9,S100a6,Ly6e | | 4.985e-02 | -3.00 | GSE369\_PRE\_VS\_POST\_IL6\_INJECTION\_SOCS3\_KO\_LIVER\_DN | MSigDB lists | GSE369\_PRE\_VS\_POST\_IL6\_INJECTION\_SOCS3\_KO\_LIVER\_DN | 172 | 4 | 12187 | 98 | Grn,Csf1r,Tspan4,Ctsa | | 4.985e-02 | -3.00 | GSE32986\_UNSTIM\_VS\_CURDLAN\_LOWDOSE\_STIM\_DC\_UP | MSigDB lists | GSE32986\_UNSTIM\_VS\_CURDLAN\_LOWDOSE\_STIM\_DC\_UP | 172 | 4 | 12187 | 98 | Rab3il1,Spint1,S100a6,Lfng | | 4.985e-02 | -3.00 | GSE43955\_1H\_VS\_60H\_ACT\_CD4\_TCELL\_WITH\_TGFB\_IL6\_UP | MSigDB lists | GSE43955\_1H\_VS\_60H\_ACT\_CD4\_TCELL\_WITH\_TGFB\_IL6\_UP | 172 | 4 | 12187 | 98 | Rac2,Vwf,Brk1,Psmb10 | | 4.995e-02 | -3.00 | membrane organization | biological process | GO:0061024 | 500 | 8 | 13711 | 111 | Fcer1g,Clu,Ndufa13,Ptprc,Myh9,Fcgr3,Mt3,Vamp8 | | 4.999e-02 | -3.00 | mouse chr3 | chromosome location | mouse chr3 | 175 | 4 | 14556 | 115 | Adh1,Ctss,Il6ra,S100a6 |
