## Supplementary material for "Repeated LPS induces training and tolerance of microglial responses across brain regions": FileS2: homerResults.html

4xLPS\_cluster\_genes\_output// - Homer de novo Motif Results


### Homer *de novo* Motif Results (4xLPS\_cluster\_genes\_output//)

Known Motif Enrichment Results  
Gene Ontology Enrichment Results  
If Homer is having trouble matching a motif to a known motif, try copy/pasting the matrix file into
STAMP  
More information on motif finding results: HOMER
| Description of Results
| Tips
  
Total target sequences = 112  
Total background sequences = 13177  
\* - possible false positive  

|  |  |  |  |  |  |  |  |  |
| --- | --- | --- | --- | --- | --- | --- | --- | --- |
| Rank | Motif | P-value | log P-pvalue | % of Targets | % of Background | STD(Bg STD) | Best Match/Details | Motif File |
| 1 \* | G T C A A C G T A G C T A C G T A C G T A G T C A G T C A C G T C T A G A T C G A C T G A G T C | 1e-10 | -2.399e+01 | 7.14% | 0.15% | 366.9bp (462.8bp) | NFATC2/MA0152.1/Jaspar(0.674) More Information \| Similar Motifs Found | motif file (matrix) |
| 2 \* | C T G A A C G T A C G T C G T A A G C T C T A G A C T G C T A G A C G T A T C G C G A T A C T G | 1e-9 | -2.301e+01 | 9.82% | 0.55% | 330.6bp (348.3bp) | EKLF(Zf)/Erythrocyte-Klf1-ChIP-Seq(GSE20478)/Homer(0.684) More Information \| Similar Motifs Found | motif file (matrix) |
| 3 \* | C T A G A G T C C G T A A C T G T C G A A C G T C G T A A C G T C T G A A C T G | 1e-9 | -2.219e+01 | 10.71% | 0.79% | 312.7bp (370.2bp) | Ptf1a/MA1618.1/Jaspar(0.621) More Information \| Similar Motifs Found | motif file (matrix) |
| 4 \* | C G A T A C G T C G T A C G T A A C G T C G T A A C G T A G T C A C G T A C T G A C G T A C G T | 1e-9 | -2.201e+01 | 5.36% | 0.05% | 252.4bp (121.2bp) | Ptf1a/MA1618.1/Jaspar(0.649) More Information \| Similar Motifs Found | motif file (matrix) |
| 5 \* | C G T A A T C G A C G T C G T A A C T G A C T G A C T G A G T C C G A T A C G T A C T G A C T G | 1e-9 | -2.193e+01 | 7.14% | 0.20% | 320.8bp (416.6bp) | ZNF682/MA1599.1/Jaspar(0.697) More Information \| Similar Motifs Found | motif file (matrix) |
| 6 \* | G T A C A G C T A C G T A G T C C G T A A C T G A C T G A C G T A G T C A G T C C G T A A G T C | 1e-9 | -2.168e+01 | 7.14% | 0.21% | 282.8bp (326.9bp) | ZNF354C/MA0130.1/Jaspar(0.642) More Information \| Similar Motifs Found | motif file (matrix) |
| 7 \* | C G A T C T A G C G T A C G T A A G T C C G T A C G T A A C G T A C T G C G T A A C T G A G T C | 1e-9 | -2.140e+01 | 5.36% | 0.05% | 306.9bp (191.2bp) | SOX4/MA0867.2/Jaspar(0.736) More Information \| Similar Motifs Found | motif file (matrix) |
| 8 \* | C G T A C T G A A G T C A C G T A C T G C G T A C T A G A G T C A C G T A C T G A C G T A C T G | 1e-9 | -2.097e+01 | 7.14% | 0.23% | 360.8bp (382.6bp) | POL009.1\_DCE\_S\_II/Jaspar(0.638) More Information \| Similar Motifs Found | motif file (matrix) |
| 9 \* | A G T C A G C T T C G A C A T G T A C G T C G A A C G T C T A G A T C G G A T C | 1e-8 | -2.045e+01 | 22.32% | 5.35% | 320.7bp (390.4bp) | PB0181.1\_Spdef\_2/Jaspar(0.743) More Information \| Similar Motifs Found | motif file (matrix) |
| 10 \* | A C G T C G T A A C G T A C T G A G T C A C G T A C T G A C G T A C T G C T G A C G T A A C T G | 1e-8 | -2.034e+01 | 5.36% | 0.07% | 306.0bp (306.1bp) | ZNF317/MA1593.1/Jaspar(0.671) More Information \| Similar Motifs Found | motif file (matrix) |
| 11 \* | C G T A C G T A C G T A A C G T A C T G A C T G A T C G A C G T C G T A A C G T C G A T A C G T | 1e-8 | -2.034e+01 | 5.36% | 0.07% | 157.1bp (343.0bp) | PB0163.1\_Six6\_2/Jaspar(0.646) More Information \| Similar Motifs Found | motif file (matrix) |
| 12 \* | C G T A A G T C A G C T A C G T A T G C C G T A C G T A A C T G C G T A C G T A A T G C A G T C | 1e-8 | -1.988e+01 | 5.36% | 0.08% | 136.9bp (276.6bp) | ZBTB12(Zf)/HEK293-ZBTB12.GFP-ChIP-Seq(GSE58341)/Homer(0.656) More Information \| Similar Motifs Found | motif file (matrix) |
| 13 \* | C T G A A C G T C T G A C A T G G T A C C G T A C G T A A C T G C G T A C A G T | 1e-8 | -1.981e+01 | 11.61% | 1.24% | 297.8bp (354.6bp) | RFX1/MA0509.2/Jaspar(0.673) More Information \| Similar Motifs Found | motif file (matrix) |
| 14 \* | A C G T A C T G A C T G A C T G A T C G A C T G C G A T T A G C C T G A A T C G C G T A A C T G | 1e-8 | -1.900e+01 | 7.14% | 0.31% | 334.7bp (432.0bp) | NR2C2/MA0504.1/Jaspar(0.674) More Information \| Similar Motifs Found | motif file (matrix) |
| 15 \* | C G T A T A G C G A C T C G T A A T C G A T G C G A T C G C A T C A T G A C T G | 1e-8 | -1.858e+01 | 18.75% | 4.14% | 327.2bp (404.5bp) | SD0002.1\_at\_AC\_acceptor/Jaspar(0.721) More Information \| Similar Motifs Found | motif file (matrix) |
| 16 \* | G C A T T A G C C G T A A T C G A T C G T A G C G C A T C G T A A C T G A C T G C T G A T A C G | 1e-7 | -1.838e+01 | 20.54% | 5.04% | 346.5bp (389.0bp) | GLIS3(Zf)/Thyroid-Glis3.GFP-ChIP-Seq(GSE103297)/Homer(0.589) More Information \| Similar Motifs Found | motif file (matrix) |
| 17 \* | G T C A A C T G C G T A A C T G C T G A A C T G G T C A C T A G T C G A A C T G A G T C C T A G | 1e-7 | -1.822e+01 | 8.04% | 0.51% | 329.0bp (346.6bp) | PRDM1/MA0508.3/Jaspar(0.663) More Information \| Similar Motifs Found | motif file (matrix) |
| 18 \* | A G T C C G T A C G T A A C T G A T C G C T G A A T C G C T G A C G T A A G T C C G T A A G T C | 1e-7 | -1.810e+01 | 8.04% | 0.51% | 431.9bp (379.0bp) | Znf263(Zf)/K562-Znf263-ChIP-Seq(GSE31477)/Homer(0.629) More Information \| Similar Motifs Found | motif file (matrix) |
| 19 \* | G C T A C T G A A T C G A G C T A G T C A C G T A G T C A C G T C T G A A G T C C G A T G T A C | 1e-7 | -1.794e+01 | 8.93% | 0.72% | 388.9bp (370.6bp) | FOXA1(Forkhead)/MCF7-FOXA1-ChIP-Seq(GSE26831)/Homer(0.583) More Information \| Similar Motifs Found | motif file (matrix) |
| 20 \* | A G T C C G A T A T C G G C A T G A C T A G T C G T C A A G T C T C G A A G T C A G C T A G T C | 1e-7 | -1.772e+01 | 12.50% | 1.78% | 301.4bp (412.6bp) | Foxo1/MA0480.1/Jaspar(0.606) More Information \| Similar Motifs Found | motif file (matrix) |
| 21 \* | T C G A G T C A G A T C T G A C G A C T G A T C C A G T G T A C G T A C G C T A | 1e-7 | -1.762e+01 | 17.86% | 3.96% | 374.2bp (379.5bp) | RUNX3/MA0684.2/Jaspar(0.682) More Information \| Similar Motifs Found | motif file (matrix) |
| 22 \* | A C G T G C A T A G T C T G C A A C G T A C G T A C T G A C T G A G T C A G T C A G T C C G T A | 1e-7 | -1.739e+01 | 5.36% | 0.13% | 245.3bp (278.5bp) | NFYB/MA0502.2/Jaspar(0.777) More Information \| Similar Motifs Found | motif file (matrix) |
| 23 \* | C G T A A G T C C T G A T A G C A T G C G A T C C T G A A G T C G A T C C G T A | 1e-7 | -1.692e+01 | 24.11% | 7.40% | 343.9bp (383.8bp) | KLF9/MA1107.2/Jaspar(0.768) More Information \| Similar Motifs Found | motif file (matrix) |
| 24 \* | A T G C A C G T A C G T A G C T A G T C A C G T A T C G T A G C A G T C C G A T A T C G G T A C | 1e-7 | -1.678e+01 | 9.82% | 1.06% | 351.1bp (361.2bp) | ETV4/MA0764.2/Jaspar(0.645) More Information \| Similar Motifs Found | motif file (matrix) |
| 25 \* | C G A T A G T C A G T C A C G T A G T C C G A T A C T G A G T C A C G T A C G T A G T C A G T C | 1e-6 | -1.586e+01 | 9.82% | 1.16% | 351.9bp (398.5bp) | ZNF460/MA1596.1/Jaspar(0.685) More Information \| Similar Motifs Found | motif file (matrix) |
| 26 \* | C G T A C T G A C G T A C G T A T G A C C G T A C A T G A C G T C A G T A C G T A C G T G C A T | 1e-6 | -1.539e+01 | 7.14% | 0.51% | 196.2bp (274.5bp) | PB0141.1\_Isgf3g\_2/Jaspar(0.680) More Information \| Similar Motifs Found | motif file (matrix) |
| 27 \* | A C G T A G T C A C G T C G T A C G T A A C T G G A C T A G T C A C G T A C G T C G T A A C T G | 1e-6 | -1.494e+01 | 3.57% | 0.03% | 169.2bp (173.0bp) | ZNF652/MA1657.1/Jaspar(0.589) More Information \| Similar Motifs Found | motif file (matrix) |
| 28 \* | A C T G A G T C C T G A A T G C A C G T A T G C C G T A A C T G G C A T T G C A A G T C A G C T | 1e-6 | -1.492e+01 | 9.82% | 1.28% | 266.2bp (365.6bp) | PH0112.1\_Nkx2-3/Jaspar(0.600) More Information \| Similar Motifs Found | motif file (matrix) |
| 29 \* | A G C T C T A G C A T G T A G C G A C T G A T C C A G T T A C G T C A G A T C G | 1e-6 | -1.477e+01 | 30.36% | 12.28% | 312.4bp (377.3bp) | TFAP2A(var.2)/MA0810.1/Jaspar(0.598) More Information \| Similar Motifs Found | motif file (matrix) |
| 30 \* | A G C T A G T C G T A C A T G C A C G T A T C G C G A T G A C T A C T G A G T C | 1e-6 | -1.437e+01 | 18.75% | 5.34% | 347.1bp (391.4bp) | IKZF1/MA1508.1/Jaspar(0.729) More Information \| Similar Motifs Found | motif file (matrix) |
| 31 \* | A G T C A G T C A G C T A C G T A G T C A C G T A C T G A C T G A C T G C G T A G T A C G A C T | 1e-6 | -1.416e+01 | 4.46% | 0.13% | 340.6bp (461.0bp) | Stat5b/MA1625.1/Jaspar(0.765) More Information \| Similar Motifs Found | motif file (matrix) |
| 32 \* | C T A G A T G C C T A G G T C A T A G C G T C A A G C T G A T C | 1e-6 | -1.399e+01 | 33.04% | 14.59% | 324.6bp (364.6bp) | ZBED1/MA0749.1/Jaspar(0.701) More Information \| Similar Motifs Found | motif file (matrix) |
| 33 \* | A G T C C G T A A C T G A C T G G T A C A C G T A C G T C G T A | 1e-5 | -1.318e+01 | 26.79% | 10.72% | 361.3bp (371.6bp) | Pitx1(Homeobox)/Chicken-Pitx1-ChIP-Seq(GSE38910)/Homer(0.799) More Information \| Similar Motifs Found | motif file (matrix) |
| 34 \* | A G T C A G T C A T G C C G T A T C G A C T G A A C G T A C G T A C G T A C T G | 1e-5 | -1.301e+01 | 24.11% | 9.05% | 285.8bp (365.2bp) | IRF6/MA1509.1/Jaspar(0.627) More Information \| Similar Motifs Found | motif file (matrix) |
| 35 \* | A C T G A C G T A G T C A G T C C G T A A G T C C G T A A C G T | 1e-5 | -1.160e+01 | 22.32% | 8.61% | 301.5bp (379.0bp) | ZNF354C/MA0130.1/Jaspar(0.778) More Information \| Similar Motifs Found | motif file (matrix) |
