## Supplementary material for "Repeated LPS induces training and tolerance of microglial responses across brain regions": FileS2: motif1.similar.html

### Information for motif1

G
T
C
A
A
C
G
T
A
G
C
T
A
C
G
T
A
C
G
T
A
G
T
C
A
G
T
C
A
C
G
T
C
T
A
G
A
T
C
G
A
C
T
G
A
G
T
C
  
Reverse Opposite:  

C
T
A
G
A
G
T
C
A
T
G
C
A
G
T
C
C
G
T
A
A
C
T
G
A
C
T
G
C
G
T
A
G
T
C
A
C
T
G
A
C
G
T
A
A
C
G
T
  

|  |  |
| --- | --- |
| p-value: | 1e-10 |
| log p-value: | -2.399e+01 |
| Information Content per bp: | 1.890 |
| Number of Target Sequences with motif | 8.0 |
| Percentage of Target Sequences with motif | 7.14% |
| Number of Background Sequences with motif | 19.9 |
| Percentage of Background Sequences with motif | 0.15% |
| Average Position of motif in Targets | 637.0 +/- 366.9bp |
| Average Position of motif in Background | 603.4 +/- 462.8bp |
| Strand Bias (log2 ratio + to - strand density) | 0.7 |
| Multiplicity (# of sites on avg that occur together) | 1.00 |
| Motif File: | file (matrix) reverse opposite |

#### Similar de novo motifs found

|  |  |  |  |  |  |  |  |
| --- | --- | --- | --- | --- | --- | --- | --- |
| Rank | Match Score | Redundant Motif | P-value | log P-value | % of Targets | % of Background | Motif file |
| 1 | 0.829 | A T G C T G A C A G T C C G T A A T C G A C T G G C T A C T G A G T A C T G C A | 1e-8 | -20.658445 | 24.11% | 6.18% | motif file (matrix) |
