## Supplementary material for "Repeated LPS induces training and tolerance of microglial responses across brain regions": FileS2: motif13.similar.html

### Information for motif13

C
T
G
A
A
C
G
T
C
T
G
A
C
A
T
G
G
T
A
C
C
G
T
A
C
G
T
A
A
C
T
G
C
G
T
A
C
A
G
T
  
Reverse Opposite:  

G
T
C
A
A
C
G
T
T
G
A
C
A
C
G
T
A
C
G
T
A
C
T
G
G
T
A
C
A
G
C
T
C
G
T
A
G
A
C
T
  

|  |  |
| --- | --- |
| p-value: | 1e-8 |
| log p-value: | -1.981e+01 |
| Information Content per bp: | 1.845 |
| Number of Target Sequences with motif | 13.0 |
| Percentage of Target Sequences with motif | 11.61% |
| Number of Background Sequences with motif | 162.9 |
| Percentage of Background Sequences with motif | 1.24% |
| Average Position of motif in Targets | 565.5 +/- 297.8bp |
| Average Position of motif in Background | 511.7 +/- 354.6bp |
| Strand Bias (log2 ratio + to - strand density) | 1.2 |
| Multiplicity (# of sites on avg that occur together) | 1.00 |
| Motif File: | file (matrix) reverse opposite |

#### Similar de novo motifs found

|  |  |  |  |  |  |  |  |
| --- | --- | --- | --- | --- | --- | --- | --- |
| Rank | Match Score | Redundant Motif | P-value | log P-value | % of Targets | % of Background | Motif file |
| 1 | 0.766 | C G T A C G T A A G T C C G T A C G T A A C T G C G T A A C G T | 1e-4 | -10.942272 | 14.29% | 4.13% | motif file (matrix) |
