## Supplementary material for "Repeated LPS induces training and tolerance of microglial responses across brain regions": FileS2: motif14.similar.html

### Information for motif14

A
C
G
T
A
C
T
G
A
C
T
G
A
C
T
G
A
T
C
G
A
C
T
G
C
G
A
T
T
A
G
C
C
T
G
A
A
T
C
G
C
G
T
A
A
C
T
G
  
Reverse Opposite:  

A
G
T
C
A
C
G
T
A
T
G
C
A
G
C
T
A
C
T
G
C
G
T
A
T
G
A
C
A
T
G
C
A
G
T
C
A
G
T
C
A
G
T
C
C
G
T
A
  

|  |  |
| --- | --- |
| p-value: | 1e-8 |
| log p-value: | -1.900e+01 |
| Information Content per bp: | 1.877 |
| Number of Target Sequences with motif | 8.0 |
| Percentage of Target Sequences with motif | 7.14% |
| Number of Background Sequences with motif | 40.4 |
| Percentage of Background Sequences with motif | 0.31% |
| Average Position of motif in Targets | 500.1 +/- 334.7bp |
| Average Position of motif in Background | 544.8 +/- 432.0bp |
| Strand Bias (log2 ratio + to - strand density) | 1.6 |
| Multiplicity (# of sites on avg that occur together) | 1.00 |
| Motif File: | file (matrix) reverse opposite |

#### Similar de novo motifs found

|  |  |  |  |  |  |  |  |
| --- | --- | --- | --- | --- | --- | --- | --- |
| Rank | Match Score | Redundant Motif | P-value | log P-value | % of Targets | % of Background | Motif file |
| 1 | 0.671 | A T C G C G T A C G T A C A G T A C T G C A T G A T C G C T A G A C T G G A C T | 1e-6 | -15.977415 | 12.50% | 2.06% | motif file (matrix) |
