## Supplementary material for "Repeated LPS induces training and tolerance of microglial responses across brain regions": FileS2: motif19.similar.html

### Information for motif19

G
C
T
A
C
T
G
A
A
T
C
G
A
G
C
T
A
G
T
C
A
C
G
T
A
G
T
C
A
C
G
T
C
T
G
A
A
G
T
C
C
G
A
T
G
T
A
C
  
Reverse Opposite:  

C
A
T
G
C
G
T
A
A
C
T
G
A
G
C
T
G
T
C
A
C
T
A
G
C
G
T
A
C
T
A
G
C
T
G
A
A
T
G
C
A
G
C
T
C
G
A
T
  

|  |  |
| --- | --- |
| p-value: | 1e-7 |
| log p-value: | -1.794e+01 |
| Information Content per bp: | 1.762 |
| Number of Target Sequences with motif | 10.0 |
| Percentage of Target Sequences with motif | 8.93% |
| Number of Background Sequences with motif | 94.8 |
| Percentage of Background Sequences with motif | 0.72% |
| Average Position of motif in Targets | 718.9 +/- 388.9bp |
| Average Position of motif in Background | 556.0 +/- 370.6bp |
| Strand Bias (log2 ratio + to - strand density) | -0.3 |
| Multiplicity (# of sites on avg that occur together) | 1.10 |
| Motif File: | file (matrix) reverse opposite |

#### Similar de novo motifs found

|  |  |  |  |  |  |  |  |
| --- | --- | --- | --- | --- | --- | --- | --- |
| Rank | Match Score | Redundant Motif | P-value | log P-value | % of Targets | % of Background | Motif file |
