## Supplementary material for "Repeated LPS induces training and tolerance of microglial responses across brain regions": FileS2: motif21.similar.html

### Information for motif21

T
C
G
A
G
T
C
A
G
A
T
C
T
G
A
C
G
A
C
T
G
A
T
C
C
A
G
T
G
T
A
C
G
T
A
C
G
C
T
A
  
Reverse Opposite:  

C
G
A
T
C
A
T
G
C
A
T
G
G
T
C
A
C
T
A
G
C
T
G
A
A
C
T
G
C
T
A
G
C
A
G
T
A
G
C
T
  

|  |  |
| --- | --- |
| p-value: | 1e-7 |
| log p-value: | -1.762e+01 |
| Information Content per bp: | 1.491 |
| Number of Target Sequences with motif | 20.0 |
| Percentage of Target Sequences with motif | 17.86% |
| Number of Background Sequences with motif | 522.3 |
| Percentage of Background Sequences with motif | 3.96% |
| Average Position of motif in Targets | 581.2 +/- 374.2bp |
| Average Position of motif in Background | 597.6 +/- 379.5bp |
| Strand Bias (log2 ratio + to - strand density) | 0.3 |
| Multiplicity (# of sites on avg that occur together) | 1.10 |
| Motif File: | file (matrix) reverse opposite |

#### Similar de novo motifs found

|  |  |  |  |  |  |  |  |
| --- | --- | --- | --- | --- | --- | --- | --- |
| Rank | Match Score | Redundant Motif | P-value | log P-value | % of Targets | % of Background | Motif file |
| 1 | 0.618 | C A T G C T A G C T G A T A G C G T C A C G T A G T A C A G T C A G T C A G T C G T C A T C G A | 1e-7 | -16.400736 | 7.14% | 0.44% | motif file (matrix) |
