## Supplementary material for "Repeated LPS induces training and tolerance of microglial responses across brain regions": FileS2: motif30.similar.html

### Information for motif30

A
G
C
T
A
G
T
C
G
T
A
C
A
T
G
C
A
C
G
T
A
T
C
G
C
G
A
T
G
A
C
T
A
C
T
G
A
G
T
C
  
Reverse Opposite:  

T
C
A
G
T
G
A
C
C
T
G
A
G
C
T
A
T
A
G
C
T
G
C
A
T
A
C
G
A
C
T
G
T
C
A
G
T
C
G
A
  

|  |  |
| --- | --- |
| p-value: | 1e-6 |
| log p-value: | -1.437e+01 |
| Information Content per bp: | 1.595 |
| Number of Target Sequences with motif | 21.0 |
| Percentage of Target Sequences with motif | 18.75% |
| Number of Background Sequences with motif | 704.5 |
| Percentage of Background Sequences with motif | 5.34% |
| Average Position of motif in Targets | 629.3 +/- 347.1bp |
| Average Position of motif in Background | 592.2 +/- 391.4bp |
| Strand Bias (log2 ratio + to - strand density) | -0.5 |
| Multiplicity (# of sites on avg that occur together) | 1.29 |
| Motif File: | file (matrix) reverse opposite |

#### Similar de novo motifs found

|  |  |  |  |  |  |  |  |
| --- | --- | --- | --- | --- | --- | --- | --- |
| Rank | Match Score | Redundant Motif | P-value | log P-value | % of Targets | % of Background | Motif file |
| 1 | 0.780 | A G C T G C A T A G T C G T C A A G T C C G A T T A C G G T C A C A G T A C T G A G T C C G A T | 1e-6 | -14.300624 | 11.61% | 2.02% | motif file (matrix) |
