## Supplementary material for "Repeated LPS induces training and tolerance of microglial responses across brain regions": FileS2: motif35.similar.html

### Information for motif35

A
C
T
G
A
C
G
T
A
G
T
C
A
G
T
C
C
G
T
A
A
G
T
C
C
G
T
A
A
C
G
T
  
Reverse Opposite:  

C
G
T
A
A
C
G
T
A
C
T
G
A
C
G
T
A
C
T
G
A
C
T
G
C
G
T
A
T
G
A
C
  

|  |  |
| --- | --- |
| p-value: | 1e-5 |
| log p-value: | -1.160e+01 |
| Information Content per bp: | 1.926 |
| Number of Target Sequences with motif | 25.0 |
| Percentage of Target Sequences with motif | 22.32% |
| Number of Background Sequences with motif | 1134.5 |
| Percentage of Background Sequences with motif | 8.61% |
| Average Position of motif in Targets | 559.2 +/- 301.5bp |
| Average Position of motif in Background | 551.5 +/- 379.0bp |
| Strand Bias (log2 ratio + to - strand density) | 0.1 |
| Multiplicity (# of sites on avg that occur together) | 1.08 |
| Motif File: | file (matrix) reverse opposite |

#### Similar de novo motifs found

|  |  |  |  |  |  |  |  |
| --- | --- | --- | --- | --- | --- | --- | --- |
| Rank | Match Score | Redundant Motif | P-value | log P-value | % of Targets | % of Background | Motif file |
| 1 | 0.600 | A C G T C T G A A G T C A C T G T G A C C T G A A G C T G A T C | 1e-3 | -8.198223 | 11.61% | 3.68% | motif file (matrix) |
