## Supplementary material for "Repeated LPS induces training and tolerance of microglial responses across brain regions": FileS2: knownResults.html

4xLPS\_cluster\_genes\_output/ - Homer Known Motif Enrichment Results


### Homer Known Motif Enrichment Results (4xLPS\_cluster\_genes\_output/)

Homer *de novo* Motif Results  
Gene Ontology Enrichment Results  
Known Motif Enrichment Results (txt file)  
Total Target Sequences = 112, Total Background Sequences = 13184

|  |  |  |  |  |  |  |  |  |  |  |  |
| --- | --- | --- | --- | --- | --- | --- | --- | --- | --- | --- | --- |
| Rank | Motif | Name | P-value | log P-pvalue | q-value (Benjamini) | # Target Sequences with Motif | % of Targets Sequences with Motif | # Background Sequences with Motif | % of Background Sequences with Motif | Motif File | SVG |
| 1 | A T G C G A T C G A C T A G C T C G A T C G A T G T C A C G A T T C G A A T C G T A G C T A G C | TATA-Box(TBP)/Promoter/Homer | 1e-2 | -5.843e+00 | 1.0000 | 69.0 | 61.61% | 6342.7 | 48.11% | motif file (matrix) | svg |
| 2 | C A T G G A C T G C A T A C T G A G C T A C T G A C T G C G T A G C A T A G C T A T C G T A C G | Foxh1(Forkhead)/hESC-FOXH1-ChIP-Seq(GSE29422)/Homer | 1e-2 | -4.733e+00 | 1.0000 | 34.0 | 30.36% | 2696.1 | 20.45% | motif file (matrix) | svg |
