## Supplementary material for "Repeated LPS induces training and tolerance of microglial responses across brain regions": FileS3: geneOntology.html

Gene Ontology Results

### Gene Ontology Enrichment Results

Homer *de novo* Motif Enrichment Results  
Known Motif Enrichment Results  

###### Text file version of complete results (i.e. open with Excel) - biological process: Functional groupings of proteins (Gene Ontology) - molecular function: Mechanistic actions of proteins (Gene Ontology) - cellular component: Protein localization (Gene Ontology) - chromosome location: Genes with similar chromosome localization (NCBI Gene) - KEGG pathways: Groups of proteins in the same pathways (KEGG) - protein interactions: "Proteins interacting with a common protein (BIND, EcoCyc, HPRD)" (NCBI Gene) - interpro domains: Proteins with similar domains and features (Interpro) - pfam domains: Proteins with similar domains and features (Pfam) - smart domains: Proteins with similar domains and features (SMART) - gene3d domains: Proteins with similar domains and features (Gene3D) - prosite domains: Proteins with similar domains and features (Prosite) - prints domains: Proteins with similar domains and features (PRINTS) - MSigDB lists: "Genes sets for pathways, factor/miRNA target predictions, expression patterns, etc." (MSigDB) - BIOCYC pathways: Groups of proteins in the same pathways (BIOCYC) - COSMIC cancer mutations: Genes mutated in similar cancers (COSMIC) - GWAS genes: Genes mutated in similar diseases (GWAS Catalog) - Lipid Maps pathways: Groups of proteins in the same lipid pathways (Lipid Maps/Biosystems) - Pathway Interaction DB: Groups of proteins in the same pathways (Pathway Interaction Database) - REACTOME pathways: Groups of proteins in the same pathways (REACTOME) - SMPDB pathways: Groups of proteins in the same pathways (SMPDB) - WikiPathways: Groups of proteins in the same pathways (Wikipathways) Enriched Categories | | | | | | | | | | | | --- | --- | --- | --- | --- | --- | --- | --- | --- | --- | | P-value | ln(P) | Term | GO Tree | GO ID | # of Genes in Term | # of Target Genes in Term | # of Total Genes | # of Target Genes | Common Genes | | 9.856e-07 | -13.83 | regulation of multicellular organismal development | biological process | GO:2000026 | 1748 | 27 | 13711 | 80 | Lhx1,P2ry12,Nrep,Kdr,Timp2,Inhba,Sorl1,Ltk,Rgs4,Agrn,Ddx56,Qk,Alkal2,Arntl,Bdnf,Cbln2,Ntm,Egr1,Ddx17,Spry2,Col1a1,Gbx1,Lims2,Ncam1,Car2,Lrrtm2,Cldn5 | | 2.813e-06 | -12.78 | neurogenesis | biological process | GO:0022008 | 1504 | 24 | 13711 | 80 | Bdnf,Qk,Alkal2,Arntl,Map4,Ntm,Ncam1,Gbx1,Lhx1,Ephb6,P2ry12,Kdr,Plp1,Timp2,Nrep,Gsdme,Inhba,Sorl1,Ltk,Ugt8a,Fabp7,Agrn,Ddx56,Nr4a3 | | 3.318e-06 | -12.62 | generation of neurons | biological process | GO:0048699 | 1409 | 23 | 13711 | 80 | P2ry12,Lhx1,Ephb6,Nrep,Plp1,Kdr,Timp2,Inhba,Ltk,Sorl1,Gsdme,Nr4a3,Ddx56,Agrn,Ugt8a,Arntl,Alkal2,Qk,Bdnf,Ntm,Map4,Gbx1,Ncam1 | | 3.805e-06 | -12.48 | system development | biological process | GO:0048731 | 3292 | 38 | 13711 | 80 | Egr1,Ddx17,Map4,Ntm,Fjx1,Bdnf,Hba-a2,Alkal2,Qk,Arntl,Homer1,Cldn5,Ncam1,Spry2,Col1a1,Gbx1,Timp2,Plp1,Kdr,Tgfa,Nrep,Crh,Ephb6,Lhx1,P2ry12,Aars,Fabp7,Ugt8a,Agrn,Ddx56,Foxq1,Nr4a3,Fnbp1,Gsdme,Atp5b,Sorl1,Ltk,Inhba | | 5.022e-06 | -12.20 | LEE\_TARGETS\_OF\_PTCH1\_AND\_SUFU\_DN | MSigDB lists | LEE\_TARGETS\_OF\_PTCH1\_AND\_SUFU\_DN | 75 | 6 | 12187 | 72 | Ntm,Ncam1,AI593442,Map4,Grm3,Fnbp1 | | 5.935e-06 | -12.03 | nervous system development | biological process | GO:0007399 | 1917 | 27 | 13711 | 80 | Map4,Ntm,Bdnf,Arntl,Alkal2,Qk,Cldn5,Ncam1,Gbx1,Timp2,Plp1,Kdr,Nrep,P2ry12,Aars,Ephb6,Lhx1,Agrn,Ddx56,Ugt8a,Fabp7,Fnbp1,Nr4a3,Sorl1,Ltk,Inhba,Gsdme | | 6.305e-06 | -11.97 | regulation of multicellular organismal process | biological process | GO:0051239 | 2546 | 32 | 13711 | 80 | Cbln2,Bdnf,Arntl,Alkal2,Qk,Ddx17,Egr1,Ntm,Ncam1,Lims2,Col1a1,Gbx1,Spry2,Cldn5,Lrrtm2,Homer1,Car2,P2ry12,Lhx1,Kdr,Timp2,Crh,Nrep,Inhba,Sorl1,Ltk,Atp5b,Agrn,Ddx56,Rgs4,Grm3,Nr4a3 | | 6.356e-06 | -11.97 | LI\_WILMS\_TUMOR\_VS\_FETAL\_KIDNEY\_2\_DN | MSigDB lists | LI\_WILMS\_TUMOR\_VS\_FETAL\_KIDNEY\_2\_DN | 45 | 5 | 12187 | 72 | Lhx1,Inhba,Tgfa,Agrn,Gng11 | | 7.111e-06 | -11.85 | KRAS.600\_UP.V1\_UP | MSigDB lists | KRAS.600\_UP.V1\_UP | 171 | 8 | 12187 | 72 | Gng11,Ugt8a,Inhba,Peg3,Sorl1,Nr4a3,Ntm,Spry2 | | 1.586e-05 | -11.05 | TATTATA\_MIR374 | MSigDB lists | TATTATA\_MIR374 | 251 | 9 | 12187 | 72 | AI593442,Ntm,Spry2,Homer1,Nr4a3,Qk,Amd1,Tgfa,Tle4 | | 1.701e-05 | -10.98 | TTGTTT\_FOXO4\_01 | MSigDB lists | TTGTTT\_FOXO4\_01 | 1553 | 23 | 12187 | 72 | Fnbp1,Plp1,Nr4a3,Cldn5,Arhgap20,Ncam1,Ndst4,Csrnp3,Egr1,Crh,Lhx1,Hs3st1,Ugt8a,Grm3,Bdnf,Inhba,Gng11,Ephb6,Col1a1,Nap1l5,Ddx17,Cacng2,Qk | | 1.914e-05 | -10.86 | anatomical structure development | biological process | GO:0048856 | 3951 | 41 | 13711 | 80 | Gbx1,Col1a1,Spry2,Ncam1,Car2,Cldn5,Homer1,Arntl,Hba-a2,Qk,Amd1,Alkal2,Bdnf,Ntm,Fjx1,Egr1,Ddx17,Map4,Sorl1,Ltk,Inhba,Ermn,Gsdme,Atp5b,Fnbp1,Foxq1,Nr4a3,Agrn,Ddx56,Ugt8a,Fabp7,P2ry12,Aars,Ephb6,Lhx1,Crh,Nrep,Timp2,Plp1,Kdr,Tgfa | | 2.263e-05 | -10.70 | regulation of developmental process | biological process | GO:0050793 | 2184 | 28 | 13711 | 80 | Ntm,Ddx17,Egr1,Qk,Alkal2,Arntl,Bdnf,Cbln2,Car2,Lrrtm2,Cldn5,Spry2,Gbx1,Col1a1,Lims2,Ncam1,Nrep,Timp2,Kdr,Lhx1,P2ry12,Rgs4,Agrn,Ddx56,Ltk,Sorl1,Inhba,Ermn | | 2.412e-05 | -10.63 | positive regulation of developmental process | biological process | GO:0051094 | 1253 | 20 | 13711 | 80 | Col1a1,Lims2,Ncam1,Inhba,Ltk,Car2,Lrrtm2,Cldn5,Agrn,Ddx56,Qk,Alkal2,Arntl,Lhx1,Bdnf,P2ry12,Cbln2,Egr1,Kdr,Timp2 | | 2.515e-05 | -10.59 | regulation of nervous system development | biological process | GO:0051960 | 945 | 17 | 13711 | 80 | Timp2,Kdr,Ntm,Nrep,Cbln2,P2ry12,Bdnf,Lhx1,Arntl,Alkal2,Qk,Agrn,Ddx56,Lrrtm2,Ltk,Sorl1,Gbx1 | | 2.814e-05 | -10.48 | SWEET\_LUNG\_CANCER\_KRAS\_DN | MSigDB lists | SWEET\_LUNG\_CANCER\_KRAS\_DN | 339 | 10 | 12187 | 72 | Col1a1,Car2,Gng11,Peg3,Qk,Kdr,Cldn5,Bdnf,Glud1,Map4 | | 3.098e-05 | -10.38 | multicellular organism development | biological process | GO:0007275 | 3728 | 39 | 13711 | 80 | Ntm,Fjx1,Ddx17,Egr1,Map4,Arntl,Hba-a2,Alkal2,Qk,Amd1,Bdnf,Cldn5,Homer1,Gbx1,Col1a1,Spry2,Ncam1,Crh,Nrep,Timp2,Plp1,Kdr,Tgfa,P2ry12,Aars,Ephb6,Lhx1,Fnbp1,Foxq1,Nr4a3,Ddx56,Agrn,Ugt8a,Fabp7,Ltk,Sorl1,Inhba,Gsdme,Atp5b | | 3.590e-05 | -10.23 | positive regulation of multicellular organismal process | biological process | GO:0051240 | 1512 | 22 | 13711 | 80 | Col1a1,Atp5b,Inhba,Ncam1,Ltk,Nr4a3,Car2,Lrrtm2,Rgs4,Agrn,Ddx56,Qk,Alkal2,Arntl,Lhx1,Bdnf,P2ry12,Cbln2,Crh,Egr1,Kdr,Timp2 | | 4.288e-05 | -10.06 | nervous system process | biological process | GO:0050877 | 694 | 14 | 13711 | 80 | Gsdme,Ncam1,Spry2,Col1a1,Gbx1,Fabp7,Nr4a3,Grm3,Bdnf,Kcnmb2,Aars,Cacng2,Egr1,Crh | | 4.881e-05 | -9.93 | LEIN\_OLIGODENDROCYTE\_MARKERS | MSigDB lists | LEIN\_OLIGODENDROCYTE\_MARKERS | 68 | 5 | 12187 | 72 | Qdpr,Ugt8a,Ermn,Qk,Car2 | | 5.625e-05 | -9.79 | regulation of cell differentiation | biological process | GO:0045595 | 1557 | 22 | 13711 | 80 | P2ry12,Bdnf,Lhx1,Arntl,Qk,Alkal2,Timp2,Kdr,Ddx17,Ntm,Nrep,Sorl1,Ltk,Inhba,Lims2,Col1a1,Spry2,Ddx56,Agrn,Cldn5,Rgs4,Car2 | | 6.246e-05 | -9.68 | axon | cellular component | GO:0030424 | 640 | 13 | 13825 | 79 | Cacng2,Bdnf,Grm3,Timp2,Ncam1,Crh,Ermn,Cldn5,Car2,Map4,Agrn,Ntm,Homer1 | | 6.967e-05 | -9.57 | GO\_NEUROGENESIS | MSigDB lists | GO\_NEUROGENESIS | 1126 | 18 | 12187 | 72 | Lhx1,Agrn,Sorl1,Gbx1,Timp2,Inhba,Bdnf,Ncam1,Ntm,Map4,Nr4a3,P2ry12,Ltk,Fabp7,Lrrc55,Plp1,Arntl,Ugt8a | | 7.484e-05 | -9.50 | MODULE\_12 | MSigDB lists | MODULE\_12 | 306 | 9 | 12187 | 72 | Ncam1,Qdpr,Cldn5,Peg3,Fabp7,Col1a1,Plp1,Msmo1,Ugt8a | | 7.672e-05 | -9.48 | TATAAA\_TATA\_01 | MSigDB lists | TATAAA\_TATA\_01 | 825 | 15 | 12187 | 72 | Ndst4,Pdia6,Bdnf,Glud1,P2ry12,Nr4a3,Csrnp3,Peg3,Tle4,Inhba,Col1a1,Car2,Rgs4,Cacng2,Crh | | 7.940e-05 | -9.44 | developmental process | biological process | GO:0032502 | 4177 | 41 | 13711 | 80 | Crh,Nrep,Timp2,Plp1,Kdr,Tgfa,P2ry12,Aars,Ephb6,Lhx1,Fnbp1,Foxq1,Nr4a3,Ddx56,Agrn,Ugt8a,Fabp7,Ltk,Sorl1,Ermn,Inhba,Gsdme,Atp5b,Ntm,Fjx1,Ddx17,Egr1,Map4,Arntl,Hba-a2,Alkal2,Qk,Amd1,Bdnf,Car2,Cldn5,Homer1,Gbx1,Col1a1,Spry2,Ncam1 | | 8.053e-05 | -9.43 | E2A\_Q2 | MSigDB lists | E2A\_Q2 | 177 | 7 | 12187 | 72 | Tmem165,H1f0,Dgkz,Col1a1,AI593442,Ndst4,Fbxl17 | | 9.085e-05 | -9.31 | axon guidance | biological process | GO:0007411 | 182 | 7 | 13711 | 80 | Lhx1,Bdnf,Ephb6,Ncam1,Agrn,Nr4a3,Gbx1 | | 9.250e-05 | -9.29 | regulation of cell development | biological process | GO:0060284 | 944 | 16 | 13711 | 80 | Sorl1,Ltk,Lims2,Cldn5,Agrn,Ddx56,Rgs4,P2ry12,Bdnf,Arntl,Qk,Alkal2,Kdr,Timp2,Ntm,Nrep | | 9.728e-05 | -9.24 | neuron projection guidance | biological process | GO:0097485 | 184 | 7 | 13711 | 80 | Agrn,Nr4a3,Ncam1,Lhx1,Bdnf,Ephb6,Gbx1 | | 9.910e-05 | -9.22 | system process | biological process | GO:0003008 | 1054 | 17 | 13711 | 80 | Kdr,Egr1,Crh,Kcnmb2,Aars,Bdnf,Cacng2,Fabp7,Homer1,Grm3,Nr4a3,Ncam1,Inhba,Gsdme,Gbx1,Col1a1,Spry2 | | 1.001e-04 | -9.21 | modulation of chemical synaptic transmission | biological process | GO:0050804 | 479 | 11 | 13711 | 80 | Ncam1,Car2,Grm3,Homer1,Rgs4,Lrrtm2,Cacng2,Bdnf,Crh,Kdr,Egr1 | | 1.020e-04 | -9.19 | regulation of trans-synaptic signaling | biological process | GO:0099177 | 480 | 11 | 13711 | 80 | Cacng2,Bdnf,Ncam1,Grm3,Crh,Car2,Lrrtm2,Homer1,Rgs4,Egr1,Kdr | | 1.041e-04 | -9.17 | positive regulation of cell communication | biological process | GO:0010647 | 1389 | 20 | 13711 | 80 | Col1a1,Spry2,Ubqln2,Ncam1,Inhba,Lims2,Gsdme,Car2,Rgs4,Lrrtm2,Arntl,Alkal2,Glud1,Cacng2,P2ry12,Bdnf,Crh,Kdr,Tgfa,Timp2 | | 1.042e-04 | -9.17 | axon development | biological process | GO:0061564 | 322 | 9 | 13711 | 80 | Ephb6,Bdnf,Lhx1,Ncam1,Gbx1,Agrn,Plp1,Nr4a3,Nrep | | 1.127e-04 | -9.09 | positive regulation of signaling | biological process | GO:0023056 | 1397 | 20 | 13711 | 80 | Ncam1,Inhba,Gsdme,Lims2,Col1a1,Spry2,Ubqln2,Lrrtm2,Rgs4,Car2,P2ry12,Bdnf,Arntl,Cacng2,Alkal2,Glud1,Timp2,Tgfa,Kdr,Crh | | 1.237e-04 | -9.00 | positive regulation of biological process | biological process | GO:0048518 | 4720 | 44 | 13711 | 80 | Arntl,Qk,Egr1,Ncam1,Lims2,Col1a1,Gbx1,Nipsnap2,Homer1,Car2,Ephb6,Lhx1,Timp2,Crh,Sorl1,Inhba,Gsdme,Rgs4,Lrrc55,Nr4a3,Cbln2,Dgkz,Bdnf,Peg3,Cacng2,Alkal2,Ddx17,Csrnp3,Spry2,Cldn5,Lrrtm2,P2ry12,Glud1,Tgfa,Plp1,Kdr,Ltk,Mtmr9,Atp5b,Ubqln2,Ddx56,Agrn,Hsd17b12,H1f0 | | 1.246e-04 | -8.99 | cell differentiation | biological process | GO:0030154 | 2800 | 31 | 13711 | 80 | Timp2,Kdr,Plp1,Nrep,P2ry12,Ephb6,Lhx1,Ddx56,Agrn,Ugt8a,Fabp7,Foxq1,Nr4a3,Ltk,Sorl1,Inhba,Gsdme,Egr1,Ddx17,Map4,Ntm,Bdnf,Arntl,Hba-a2,Alkal2,Qk,Homer1,Ncam1,Col1a1,Gbx1,Spry2 | | 1.297e-04 | -8.95 | multicellular organismal process | biological process | GO:0032501 | 4571 | 43 | 13711 | 80 | Ugt8a,Fabp7,Ddx56,Agrn,Foxq1,Nr4a3,Fnbp1,Grm3,Atp5b,Gsdme,Ermn,Inhba,Sorl1,Ltk,Kdr,Plp1,Tgfa,Timp2,Nrep,Crh,Lhx1,Ephb6,Aars,Kcnmb2,P2ry12,Homer1,Cldn5,Ncam1,Spry2,Gbx1,Col1a1,Map4,Ddx17,Egr1,Fjx1,Ntm,Bdnf,Alkal2,Amd1,Qk,Cacng2,Hba-a2,Arntl | | 1.333e-04 | -8.92 | positive regulation of nervous system development | biological process | GO:0051962 | 583 | 12 | 13711 | 80 | Agrn,Ddx56,Lrrtm2,Ltk,Timp2,Kdr,Alkal2,Qk,P2ry12,Cbln2,Bdnf,Lhx1 | | 1.515e-04 | -8.79 | GRAHAM\_NORMAL\_QUIESCENT\_VS\_NORMAL\_DIVIDING\_UP | MSigDB lists | GRAHAM\_NORMAL\_QUIESCENT\_VS\_NORMAL\_DIVIDING\_UP | 46 | 4 | 12187 | 72 | Inhba,Fnbp1,Ddx17,H1f0 | | 1.522e-04 | -8.79 | PXR\_Q2 | MSigDB lists | PXR\_Q2 | 196 | 7 | 12187 | 72 | Nr4a3,Bdnf,Ntm,Kcnmb2,Arntl,Ddx17,Csrnp3 | | 1.783e-04 | -8.63 | ENK\_UV\_RESPONSE\_EPIDERMIS\_DN | MSigDB lists | ENK\_UV\_RESPONSE\_EPIDERMIS\_DN | 423 | 10 | 12187 | 72 | Tgfa,Arntl,Timp2,Ephb6,Egr1,Fjx1,Qdpr,Spry2,Hs3st1,Homer1 | | 1.859e-04 | -8.59 | AATGTGA\_MIR23A\_MIR23B | MSigDB lists | AATGTGA\_MIR23A\_MIR23B | 345 | 9 | 12187 | 72 | Qk,Pom121,Arhgap20,Spry2,Pdia6,Msmo1,Tgfa,Car2,Nap1l5 | | 1.927e-04 | -8.55 | response to chemical | biological process | GO:0042221 | 2591 | 29 | 13711 | 80 | Kcnmb2,P2ry12,Lhx1,Ephb6,Glud1,Srm,Kdr,Timp2,Tle4,Crh,Inhba,Ltk,Atp5b,Gsdme,Ubqln2,Agrn,Nr4a3,Bdnf,Arntl,Scd2,Egr1,Ddx17,Ncam1,Col1a1,Gbx1,Spry2,Cldn5,Homer1,Car2 | | 1.941e-04 | -8.55 | WILCOX\_RESPONSE\_TO\_PROGESTERONE\_DN | MSigDB lists | WILCOX\_RESPONSE\_TO\_PROGESTERONE\_DN | 49 | 4 | 12187 | 72 | Col1a1,Inhba,Bdnf,Ntm | | 1.960e-04 | -8.54 | COMP1\_01 | MSigDB lists | COMP1\_01 | 91 | 5 | 12187 | 72 | Nr4a3,Tle4,Col1a1,Ndst4,Ncam1 | | 2.002e-04 | -8.52 | spermidine biosynthetic process | biological process | GO:0008295 | 4 | 2 | 13711 | 80 | Amd1,Srm | | 2.008e-04 | -8.51 | CREBP1\_Q2 | MSigDB lists | CREBP1\_Q2 | 205 | 7 | 12187 | 72 | Nr4a3,Crh,Ubqln2,AI593442,Peg3,Egr1,Grm3 | | 2.100e-04 | -8.47 | GO\_POSITIVE\_REGULATION\_OF\_SYNAPSE\_ASSEMBLY | MSigDB lists | GO\_POSITIVE\_REGULATION\_OF\_SYNAPSE\_ASSEMBLY | 50 | 4 | 12187 | 72 | Bdnf,Lrrtm2,Agrn,Cbln2 | | 2.128e-04 | -8.46 | neuron projection development | biological process | GO:0031175 | 613 | 12 | 13711 | 80 | Ncam1,Lhx1,Bdnf,Ephb6,Gbx1,Kdr,Plp1,Agrn,Map4,Ugt8a,Nr4a3,Nrep | | 2.171e-04 | -8.44 | LEE\_NEURAL\_CREST\_STEM\_CELL\_DN | MSigDB lists | LEE\_NEURAL\_CREST\_STEM\_CELL\_DN | 93 | 5 | 12187 | 72 | Fjx1,Car2,Qk,Fabp7,Spry2 | | 2.233e-04 | -8.41 | cellular developmental process | biological process | GO:0048869 | 2887 | 31 | 13711 | 80 | Timp2,Kdr,Plp1,Nrep,Ephb6,Lhx1,P2ry12,Fabp7,Ugt8a,Agrn,Ddx56,Nr4a3,Foxq1,Gsdme,Sorl1,Ltk,Inhba,Egr1,Ddx17,Map4,Ntm,Bdnf,Hba-a2,Alkal2,Qk,Arntl,Homer1,Ncam1,Spry2,Gbx1,Col1a1 | | 2.254e-04 | -8.40 | cellular process | biological process | GO:0009987 | 10077 | 72 | 13711 | 80 | Ndst4,Fabp7,Hsd17b12,H1f0,Agrn,Ddx56,Ubqln2,Atp5b,Ltk,Nrep,Gar1,Tgfa,Kdr,Plp1,Chmp7,Glud1,Kcnmb2,P2ry12,Hkdc1,Lrrtm2,Cldn5,Spry2,Csrnp3,Nap1l5,Pom121,Fjx1,Tmx3,Map4,Ddx17,Alkal2,Amd1,Scd2,Cacng2,Peg3,Bdnf,Dgkz,Foxq1,Nr4a3,Grm3,Rgs4,Hs3st1,Ugt8a,Gng11,Gsdme,Pdia6,Inhba,Ermn,Sorl1,Arhgap20,Tle4,Crh,Timp2,Srm,Fbxl17,Lhx1,Ephb6,Aars,Ccdc13,Car2,Homer1,Nipsnap2,Col1a1,Gbx1,Lims2,Ncam1,Qdpr,Ntm,Egr1,Tmem165,Qk,Hba-a2,Arntl | | 2.305e-04 | -8.38 | regulation of cell communication | biological process | GO:0010646 | 2617 | 29 | 13711 | 80 | Cacng2,Alkal2,Arntl,Bdnf,Dgkz,Egr1,Spry2,Col1a1,Lims2,Ncam1,Car2,Homer1,Lrrtm2,Glud1,Fbxl17,P2ry12,Nrep,Tle4,Crh,Timp2,Kdr,Tgfa,Ubqln2,Pdia6,Gsdme,Sorl1,Inhba,Grm3,Rgs4 | | 2.398e-04 | -8.34 | LE\_EGR2\_TARGETS\_DN | MSigDB lists | LE\_EGR2\_TARGETS\_DN | 95 | 5 | 12187 | 72 | Ephb6,Ugt8a,Cldn5,Tgfa,Msmo1 | | 2.467e-04 | -8.31 | DANG\_REGULATED\_BY\_MYC\_DN | MSigDB lists | DANG\_REGULATED\_BY\_MYC\_DN | 212 | 7 | 12187 | 72 | Inhba,Timp2,Dgkz,Col1a1,Pom121,Ncam1,Bdnf | | 2.552e-04 | -8.27 | regulation of signaling | biological process | GO:0023051 | 2632 | 29 | 13711 | 80 | Egr1,Arntl,Alkal2,Cacng2,Bdnf,Dgkz,Car2,Homer1,Lrrtm2,Col1a1,Spry2,Ncam1,Lims2,Tle4,Crh,Nrep,Tgfa,Kdr,Timp2,Fbxl17,Glud1,P2ry12,Grm3,Rgs4,Ubqln2,Inhba,Sorl1,Pdia6,Gsdme | | 2.574e-04 | -8.26 | enzyme linked receptor protein signaling pathway | biological process | GO:0007167 | 446 | 10 | 13711 | 80 | Ephb6,Bdnf,Ltk,Inhba,Col1a1,Egr1,Cldn5,Tgfa,Kdr,Nr4a3 | | 2.590e-04 | -8.26 | WANG\_SMARCE1\_TARGETS\_DN | MSigDB lists | WANG\_SMARCE1\_TARGETS\_DN | 284 | 8 | 12187 | 72 | Fjx1,Srm,Qdpr,Rgs4,Hs3st1,Bdnf,Tle4,Gng11 | | 2.634e-04 | -8.24 | positive regulation of cell differentiation | biological process | GO:0045597 | 927 | 15 | 13711 | 80 | Arntl,Alkal2,Qk,P2ry12,Lhx1,Bdnf,Kdr,Timp2,Col1a1,Inhba,Ltk,Lims2,Car2,Cldn5,Ddx56 | | 2.773e-04 | -8.19 | PTEN\_DN.V1\_DN | MSigDB lists | PTEN\_DN.V1\_DN | 98 | 5 | 12187 | 72 | Gng11,Fjx1,Tgfa,Ugt8a,Hs3st1 | | 2.802e-04 | -8.18 | KRAS.600.LUNG.BREAST\_UP.V1\_UP | MSigDB lists | KRAS.600.LUNG.BREAST\_UP.V1\_UP | 153 | 6 | 12187 | 72 | Spry2,Ncam1,Ntm,Gng11,Cldn5,Inhba | | 2.809e-04 | -8.18 | DING\_LUNG\_CANCER\_MUTATED\_SIGNIFICANTLY | MSigDB lists | DING\_LUNG\_CANCER\_MUTATED\_SIGNIFICANTLY | 22 | 3 | 12187 | 72 | Ltk,Kdr,Inhba | | 2.856e-04 | -8.16 | AACTTT\_UNKNOWN | MSigDB lists | AACTTT\_UNKNOWN | 1491 | 20 | 12187 | 72 | Amd1,Ddx17,Csrnp3,Inhba,Tle4,Hsd17b12,Nap1l5,AI593442,Homer1,Spry2,Lhx1,Cacng2,Crh,H1f0,Grm3,Plp1,Bdnf,Ntm,Ndst4,Nr4a3 | | 3.111e-04 | -8.08 | SHEPARD\_BMYB\_MORPHOLINO\_DN | MSigDB lists | SHEPARD\_BMYB\_MORPHOLINO\_DN | 156 | 6 | 12187 | 72 | Rgs4,Fabp7,Ddx17,Lhx1,Plp1,Col1a1 | | 3.220e-04 | -8.04 | GSE37605\_FOXP3\_FUSION\_GFP\_VS\_IRES\_GFP\_TREG\_C57BL6\_UP | MSigDB lists | GSE37605\_FOXP3\_FUSION\_GFP\_VS\_IRES\_GFP\_TREG\_C57BL6\_UP | 157 | 6 | 12187 | 72 | Hs3st1,Egr1,Spry2,Homer1,Agrn,Nr4a3 | | 3.321e-04 | -8.01 | extracellular region | cellular component | GO:0005576 | 1307 | 18 | 13825 | 79 | Hsd17b12,Cbln2,Kdr,Bdnf,Inhba,Tgfa,Timp2,Col1a1,Sorl1,Car2,Hba-a2,Crh,Fjx1,Pdia6,Alkal2,Ephb6,Agrn,Ntm | | 3.331e-04 | -8.01 | GSE17301\_IFNA2\_VS\_IFNA5\_STIM\_ACD3\_ACD28\_ACT\_CD8\_TCELL\_UP | MSigDB lists | GSE17301\_IFNA2\_VS\_IFNA5\_STIM\_ACD3\_ACD28\_ACT\_CD8\_TCELL\_UP | 158 | 6 | 12187 | 72 | Arntl,Hsd17b12,Egr1,Agrn,Homer1,Spry2 | | 3.536e-04 | -7.95 | GO\_REGULATION\_OF\_MULTICELLULAR\_ORGANISMAL\_DEVELOPMENT | MSigDB lists | GO\_REGULATION\_OF\_MULTICELLULAR\_ORGANISMAL\_DEVELOPMENT | 1280 | 18 | 12187 | 72 | Bdnf,Lrrtm2,Cbln2,P2ry12,Cldn5,Ltk,Arntl,Lhx1,Kdr,Agrn,Ddx17,Lims2,Egr1,Sorl1,Col1a1,Car2,Inhba,Timp2 | | 3.655e-04 | -7.91 | extracellular space | cellular component | GO:0005615 | 869 | 14 | 13825 | 79 | Inhba,Tgfa,Bdnf,Cbln2,Sorl1,Col1a1,Timp2,Crh,Fjx1,Hba-a2,Car2,Ntm,Agrn,Pdia6 | | 3.714e-04 | -7.90 | Polyamine biosynthesis, arginine => agmatine => putrescine => spermidine | KEGG pathways | M00133 | 4 | 2 | 5248 | 42 | Srm,Amd1 | | 3.714e-04 | -7.90 | Polyamine biosynthesis, arginine => agmatine => putrescine => spermidine | KEGG pathways | mmu\_M00133 | 4 | 2 | 5248 | 42 | Amd1,Srm | | 4.041e-04 | -7.81 | neuron differentiation | biological process | GO:0030182 | 859 | 14 | 13711 | 80 | Gsdme,Inhba,Ncam1,Gbx1,Ugt8a,Agrn,Nr4a3,Lhx1,Ephb6,Bdnf,Map4,Kdr,Plp1,Nrep | | 4.066e-04 | -7.81 | HALLMARK\_KRAS\_SIGNALING\_UP | MSigDB lists | HALLMARK\_KRAS\_SIGNALING\_UP | 164 | 6 | 12187 | 72 | Hkdc1,Inhba,Gng11,Car2,Peg3,Spry2 | | 4.142e-04 | -7.79 | REACTOME\_HS\_GAG\_BIOSYNTHESIS | MSigDB lists | REACTOME\_HS\_GAG\_BIOSYNTHESIS | 25 | 3 | 12187 | 72 | Agrn,Ndst4,Hs3st1 | | 4.316e-04 | -7.75 | GO\_NEURON\_DEVELOPMENT | MSigDB lists | GO\_NEURON\_DEVELOPMENT | 563 | 11 | 12187 | 72 | Lhx1,Agrn,Gbx1,Map4,Ncam1,Ntm,Bdnf,Nr4a3,Ugt8a,Plp1,Lrrc55 | | 4.589e-04 | -7.69 | regulation of neuron apoptotic process | biological process | GO:0043523 | 237 | 7 | 13711 | 80 | Aars,Bdnf,Kdr,Agrn,Egr1,Foxq1,Nr4a3 | | 4.663e-04 | -7.67 | MIKKELSEN\_MCV6\_ICP\_WITH\_H3K4ME3\_AND\_H3K27ME3 | MSigDB lists | MIKKELSEN\_MCV6\_ICP\_WITH\_H3K4ME3\_AND\_H3K27ME3 | 26 | 3 | 12187 | 72 | AI593442,Kcnmb2,Col1a1 | | 4.769e-04 | -7.65 | GO\_POSITIVE\_REGULATION\_OF\_DEVELOPMENTAL\_PROCESS | MSigDB lists | GO\_POSITIVE\_REGULATION\_OF\_DEVELOPMENTAL\_PROCESS | 868 | 14 | 12187 | 72 | Lhx1,Agrn,Kdr,Egr1,Inhba,Timp2,Col1a1,Car2,Bdnf,Cbln2,Cldn5,Lrrtm2,Ltk,Arntl | | 4.823e-04 | -7.64 | GOTZMANN\_EPITHELIAL\_TO\_MESENCHYMAL\_TRANSITION\_UP | MSigDB lists | GOTZMANN\_EPITHELIAL\_TO\_MESENCHYMAL\_TRANSITION\_UP | 62 | 4 | 12187 | 72 | Col1a1,Inhba,Ncam1,Map4 | | 4.919e-04 | -7.62 | mouse chr18\|18 B1 | chromosome location | mouse chr18\|18 B1 | 28 | 3 | 14556 | 81 | Nrep,Lims2,Lrrtm2 | | 4.967e-04 | -7.61 | negative regulation of metallopeptidase activity | biological process | GO:1905049 | 6 | 2 | 13711 | 80 | Timp2,Sorl1 | | 4.967e-04 | -7.61 | negative regulation of norepinephrine secretion | biological process | GO:0010700 | 6 | 2 | 13711 | 80 | P2ry12,Crh | | 5.128e-04 | -7.58 | GO\_REGULATION\_OF\_SYNAPSE\_ASSEMBLY | MSigDB lists | GO\_REGULATION\_OF\_SYNAPSE\_ASSEMBLY | 63 | 4 | 12187 | 72 | Bdnf,Cbln2,Agrn,Lrrtm2 | | 5.130e-04 | -7.58 | CHARAFE\_BREAST\_CANCER\_LUMINAL\_VS\_MESENCHYMAL\_DN | MSigDB lists | CHARAFE\_BREAST\_CANCER\_LUMINAL\_VS\_MESENCHYMAL\_DN | 396 | 9 | 12187 | 72 | Bdnf,Spry2,Rgs4,Qk,Foxq1,Gng11,Nap1l5,Inhba,Timp2 | | 5.481e-04 | -7.51 | positive regulation of signal transduction | biological process | GO:0009967 | 1217 | 17 | 13711 | 80 | Col1a1,Ubqln2,Spry2,Inhba,Ncam1,Gsdme,Lims2,Rgs4,Arntl,Cacng2,Alkal2,P2ry12,Bdnf,Crh,Timp2,Tgfa,Kdr | | 5.641e-04 | -7.48 | negative regulation of amine transport | biological process | GO:0051953 | 28 | 3 | 13711 | 80 | Rgs4,P2ry12,Crh | | 6.152e-04 | -7.39 | molecular function regulator | molecular function | GO:0098772 | 1245 | 17 | 13516 | 78 | Agrn,Kcnmb2,Dgkz,Tgfa,Arhgap20,Spry2,Bdnf,Rgs4,Crh,Mtmr9,Pdia6,Inhba,Grm3,Lrrc55,Alkal2,Cacng2,Timp2 | | 6.318e-04 | -7.37 | TGCTGCT\_MIR15A\_MIR16\_MIR15B\_MIR195\_MIR424\_MIR497 | MSigDB lists | TGCTGCT\_MIR15A\_MIR16\_MIR15B\_MIR195\_MIR424\_MIR497 | 496 | 10 | 12187 | 72 | Bdnf,Pdia6,Arhgap20,Glud1,Map4,Pom121,Qk,Nr4a3,Lrrc55,Tle4 | | 6.390e-04 | -7.36 | regulation of neuron death | biological process | GO:1901214 | 328 | 8 | 13711 | 80 | Sorl1,Aars,Bdnf,Foxq1,Nr4a3,Agrn,Kdr,Egr1 | | 6.656e-04 | -7.31 | KAYO\_AGING\_MUSCLE\_UP | MSigDB lists | KAYO\_AGING\_MUSCLE\_UP | 180 | 6 | 12187 | 72 | Amd1,Ltk,Inhba,Ncam1,Glud1,Nr4a3 | | 6.660e-04 | -7.31 | neuron projection morphogenesis | biological process | GO:0048812 | 414 | 9 | 13711 | 80 | Nr4a3,Ugt8a,Agrn,Kdr,Gbx1,Bdnf,Ephb6,Lhx1,Ncam1 | | 6.679e-04 | -7.31 | positive regulation of cellular process | biological process | GO:0048522 | 4254 | 39 | 13711 | 80 | Ubqln2,Gsdme,Atp5b,Ltk,Mtmr9,Sorl1,Inhba,Nr4a3,Rgs4,Agrn,Ddx56,H1f0,Glud1,Ephb6,Lhx1,P2ry12,Crh,Timp2,Kdr,Tgfa,Csrnp3,Spry2,Col1a1,Gbx1,Lims2,Ncam1,Car2,Lrrtm2,Cldn5,Cacng2,Alkal2,Qk,Arntl,Peg3,Bdnf,Dgkz,Cbln2,Egr1,Ddx17 | | 6.928e-04 | -7.27 | positive regulation of long-term neuronal synaptic plasticity | biological process | GO:0048170 | 7 | 2 | 13711 | 80 | Kdr,Bdnf | | 7.091e-04 | -7.25 | GO\_REGULATION\_OF\_GONADOTROPIN\_SECRETION | MSigDB lists | GO\_REGULATION\_OF\_GONADOTROPIN\_SECRETION | 7 | 2 | 12187 | 72 | Crh,Inhba | | 7.219e-04 | -7.23 | GO\_NEURON\_DIFFERENTIATION | MSigDB lists | GO\_NEURON\_DIFFERENTIATION | 697 | 12 | 12187 | 72 | Ntm,Ncam1,Bdnf,Lhx1,Map4,Agrn,Nr4a3,Gbx1,Lrrc55,Plp1,Ugt8a,Inhba | | 7.256e-04 | -7.23 | plasma membrane bounded cell projection morphogenesis | biological process | GO:0120039 | 419 | 9 | 13711 | 80 | Nr4a3,Agrn,Kdr,Ugt8a,Gbx1,Ncam1,Bdnf,Ephb6,Lhx1 | | 7.404e-04 | -7.21 | cell surface | cellular component | GO:0009986 | 618 | 11 | 13825 | 79 | Tmx3,Atp5b,Ephb6,Agrn,Ntm,Kdr,Cacng2,Tgfa,Timp2,Ncam1,P2ry12 | | 7.413e-04 | -7.21 | regulation of cell projection organization | biological process | GO:0031344 | 704 | 12 | 13711 | 80 | P2ry12,Bdnf,Alkal2,Qk,Map4,Ntm,Ermn,Ltk,Spry2,Ddx56,Agrn,Homer1 | | 7.626e-04 | -7.18 | lipid biosynthetic process | biological process | GO:0008610 | 337 | 8 | 13711 | 80 | Ugt8a,Plp1,Hsd17b12,Crh,Msmo1,Dgkz,Qk,Scd2 | | 7.671e-04 | -7.17 | extracellular region part | cellular component | GO:0044421 | 1048 | 15 | 13825 | 79 | Pdia6,Agrn,Ntm,Car2,Hba-a2,Fjx1,Crh,Timp2,Col1a1,Sorl1,Hsd17b12,Cbln2,Bdnf,Inhba,Tgfa | | 7.684e-04 | -7.17 | FOXO3\_01 | MSigDB lists | FOXO3\_01 | 185 | 6 | 12187 | 72 | Hs3st1,Bdnf,Ndst4,Crh,Cacng2,Csrnp3 | | 7.684e-04 | -7.17 | FOXM1\_01 | MSigDB lists | FOXM1\_01 | 185 | 6 | 12187 | 72 | Fabp7,Amd1,Grm3,Tle4,Lhx1,Ncam1 | | 7.704e-04 | -7.17 | regulation of signal transduction | biological process | GO:0009966 | 2258 | 25 | 13711 | 80 | Homer1,Spry2,Col1a1,Lims2,Ncam1,Egr1,Cacng2,Alkal2,Arntl,Bdnf,Dgkz,Rgs4,Ubqln2,Gsdme,Pdia6,Sorl1,Inhba,Nrep,Crh,Tle4,Timp2,Tgfa,Kdr,Fbxl17,P2ry12 | | 7.761e-04 | -7.16 | positive regulation of synapse assembly | biological process | GO:0051965 | 71 | 4 | 13711 | 80 | Lrrtm2,Bdnf,Cbln2,Agrn | | 7.872e-04 | -7.15 | biosynthetic process | biological process | GO:0009058 | 1869 | 22 | 13711 | 80 | Col1a1,Msmo1,Amd1,Scd2,Qk,Tmem165,Dgkz,Qdpr,Egr1,Ddx17,Inhba,Atp5b,Ndst4,Hsd17b12,Ugt8a,Hs3st1,Srm,Glud1,Aars,Lhx1,Crh,Plp1 | | 7.893e-04 | -7.14 | ZWANG\_CLASS\_2\_TRANSIENTLY\_INDUCED\_BY\_EGF | MSigDB lists | ZWANG\_CLASS\_2\_TRANSIENTLY\_INDUCED\_BY\_EGF | 31 | 3 | 12187 | 72 | Egr1,Hs3st1,Inhba | | 7.929e-04 | -7.14 | response to organic substance | biological process | GO:0010033 | 1870 | 22 | 13711 | 80 | Scd2,Bdnf,Ddx17,Egr1,Spry2,Col1a1,Ncam1,Car2,Homer1,Cldn5,Srm,P2ry12,Tle4,Crh,Timp2,Kdr,Ubqln2,Gsdme,Atp5b,Ltk,Inhba,Nr4a3 | | 8.027e-04 | -7.13 | cell projection morphogenesis | biological process | GO:0048858 | 425 | 9 | 13711 | 80 | Agrn,Kdr,Ugt8a,Nr4a3,Ncam1,Bdnf,Ephb6,Lhx1,Gbx1 | | 8.128e-04 | -7.12 | DELYS\_THYROID\_CANCER\_DN | MSigDB lists | DELYS\_THYROID\_CANCER\_DN | 187 | 6 | 12187 | 72 | Hba-a2,Gng11,Cldn5,Tle4,Ncam1,Egr1 | | 8.443e-04 | -7.08 | YTATTTTNR\_MEF2\_02 | MSigDB lists | YTATTTTNR\_MEF2\_02 | 515 | 10 | 12187 | 72 | Gng11,Tle4,Timp2,Fnbp1,Arntl,Csrnp3,P2ry12,Ncam1,Hs3st1,Bdnf | | 8.544e-04 | -7.07 | DELYS\_THYROID\_CANCER\_UP | MSigDB lists | DELYS\_THYROID\_CANCER\_UP | 340 | 8 | 12187 | 72 | Fjx1,Agrn,Ntm,Col1a1,Inhba,Msmo1,Arntl,Tgfa | | 8.696e-04 | -7.05 | regulation of cell death | biological process | GO:0010941 | 1386 | 18 | 13711 | 80 | Tgfa,Kdr,Egr1,Crh,Aars,Bdnf,Qk,Agrn,Nr4a3,Foxq1,Ltk,Sorl1,Inhba,Ncam1,Gsdme,Lims2,Csrnp3,Spry2 | | 8.930e-04 | -7.02 | Erythrocytes take up oxygen and release carbon dioxide | REACTOME pathways | R-MMU-1247673 | 7 | 2 | 6297 | 42 | Hba-a2,Car2 | | 9.074e-04 | -7.00 | FOXO1\_01 | MSigDB lists | FOXO1\_01 | 191 | 6 | 12187 | 72 | Crh,Lhx1,Bdnf,Inhba,Ddx17,Csrnp3 | | 9.199e-04 | -6.99 | response to cocaine | biological process | GO:0042220 | 33 | 3 | 13711 | 80 | Crh,Homer1,Ncam1 | | 9.202e-04 | -6.99 | regulation of tau-protein kinase activity | biological process | GO:1902947 | 8 | 2 | 13711 | 80 | Sorl1,Egr1 | | 9.202e-04 | -6.99 | regulation of gonadotropin secretion | biological process | GO:0032276 | 8 | 2 | 13711 | 80 | Crh,Inhba | | 9.419e-04 | -6.97 | KIM\_ALL\_DISORDERS\_DURATION\_CORR\_UP | MSigDB lists | KIM\_ALL\_DISORDERS\_DURATION\_CORR\_UP | 8 | 2 | 12187 | 72 | Ntm,Csrnp3 | | 9.421e-04 | -6.97 | PHONG\_TNF\_RESPONSE\_VIA\_P38\_PARTIAL | MSigDB lists | PHONG\_TNF\_RESPONSE\_VIA\_P38\_PARTIAL | 128 | 5 | 12187 | 72 | Bdnf,Inhba,Fjx1,Agrn,Tgfa | | 9.421e-04 | -6.97 | GSE24671\_CTRL\_VS\_BAKIMULC\_INFECTED\_MOUSE\_SPLENOCYTES\_DN | MSigDB lists | GSE24671\_CTRL\_VS\_BAKIMULC\_INFECTED\_MOUSE\_SPLENOCYTES\_DN | 128 | 5 | 12187 | 72 | Hs3st1,Glud1,AI593442,Col1a1,Nr4a3 | | 9.433e-04 | -6.97 | GO\_NEUROMUSCULAR\_PROCESS | MSigDB lists | GO\_NEUROMUSCULAR\_PROCESS | 74 | 4 | 12187 | 72 | Aars,Fabp7,Gbx1,Nr4a3 | | 9.659e-04 | -6.94 | plasma membrane bounded cell projection organization | biological process | GO:0120036 | 938 | 14 | 13711 | 80 | Ncam1,Gbx1,Ugt8a,Agrn,Nr4a3,Ccdc13,Ephb6,Bdnf,Lhx1,P2ry12,Map4,Plp1,Kdr,Nrep | | 9.756e-04 | -6.93 | MSX1\_01 | MSigDB lists | MSX1\_01 | 129 | 5 | 12187 | 72 | Arntl,Cbln2,Inhba,Spry2,Bdnf | | 9.933e-04 | -6.91 | neuron development | biological process | GO:0048666 | 728 | 12 | 13711 | 80 | Lhx1,Ephb6,Bdnf,Map4,Plp1,Kdr,Nrep,Ncam1,Gbx1,Ugt8a,Agrn,Nr4a3 | | 9.971e-04 | -6.91 | GO\_POSITIVE\_REGULATION\_OF\_MULTICELLULAR\_ORGANISMAL\_PROCESS | MSigDB lists | GO\_POSITIVE\_REGULATION\_OF\_MULTICELLULAR\_ORGANISMAL\_PROCESS | 1046 | 15 | 12187 | 72 | Ltk,Arntl,Bdnf,Nr4a3,Cbln2,Lrrtm2,Egr1,Timp2,Inhba,Col1a1,Car2,Lhx1,Agrn,Crh,Kdr | | 1.037e-03 | -6.87 | REACTOME\_NCAM1\_INTERACTIONS | MSigDB lists | REACTOME\_NCAM1\_INTERACTIONS | 34 | 3 | 12187 | 72 | Col1a1,Agrn,Ncam1 | | 1.042e-03 | -6.87 | GRAHAM\_CML\_DIVIDING\_VS\_NORMAL\_QUIESCENT\_DN | MSigDB lists | GRAHAM\_CML\_DIVIDING\_VS\_NORMAL\_QUIESCENT\_DN | 76 | 4 | 12187 | 72 | H1f0,Sorl1,Inhba,Fnbp1 | | 1.065e-03 | -6.84 | regulation of neurogenesis | biological process | GO:0050767 | 839 | 13 | 13711 | 80 | Ddx56,Agrn,Ltk,Sorl1,Ntm,Nrep,Kdr,Timp2,Arntl,Alkal2,Qk,P2ry12,Bdnf | | 1.088e-03 | -6.82 | biological regulation | biological process | GO:0065007 | 8325 | 62 | 13711 | 80 | Cldn5,Lrrtm2,Spry2,Csrnp3,Tmx3,Map4,Ddx17,Peg3,Alkal2,Cacng2,Cbln2,Dgkz,Bdnf,Hsd17b12,H1f0,Agrn,Ddx56,Fabp7,Ubqln2,Mtmr9,Ltk,Atp5b,Nrep,Kdr,Plp1,Tgfa,Glud1,Kcnmb2,Hkdc1,P2ry12,Car2,Homer1,Nipsnap2,Gbx1,Col1a1,Ncam1,Lims2,Ntm,Egr1,Arntl,Tmem165,Qk,Hba-a2,Grm3,Foxq1,Nr4a3,Lrrc55,Rgs4,Ermn,Inhba,Sorl1,Gng11,Pdia6,Gsdme,Arhgap20,Tle4,Crh,Timp2,Fbxl17,Aars,Lhx1,Ephb6 | | 1.095e-03 | -6.82 | WINZEN\_DEGRADED\_VIA\_KHSRP | MSigDB lists | WINZEN\_DEGRADED\_VIA\_KHSRP | 77 | 4 | 12187 | 72 | Bdnf,Spry2,Inhba,Tgfa | | 1.095e-03 | -6.82 | GO\_POSITIVE\_REGULATION\_OF\_CELL\_COMMUNICATION | MSigDB lists | GO\_POSITIVE\_REGULATION\_OF\_CELL\_COMMUNICATION | 1170 | 16 | 12187 | 72 | Lims2,Ddx17,Tgfa,Inhba,Timp2,Car2,Col1a1,Spry2,Homer1,Crh,Ubqln2,Kdr,Arntl,Bdnf,Glud1,Lrrtm2 | | 1.117e-03 | -6.80 | myelin sheath | cellular component | GO:0043209 | 205 | 6 | 13825 | 79 | Plp1,Ermn,Car2,Cldn5,Ncam1,Atp5b | | 1.137e-03 | -6.78 | NST/OST | interpro domains | IPR037359 | 9 | 2 | 13788 | 79 | Ndst4,Hs3st1 | | 1.179e-03 | -6.74 | spermidine metabolic process | biological process | GO:0008216 | 9 | 2 | 13711 | 80 | Amd1,Srm | | 1.181e-03 | -6.74 | cell part morphogenesis | biological process | GO:0032990 | 449 | 9 | 13711 | 80 | Kdr,Agrn,Ugt8a,Nr4a3,Ncam1,Lhx1,Bdnf,Ephb6,Gbx1 | | 1.189e-03 | -6.73 | regulation of long-term neuronal synaptic plasticity | biological process | GO:0048169 | 36 | 3 | 13711 | 80 | Bdnf,Egr1,Kdr | | 1.196e-03 | -6.73 | FOXJ2\_01 | MSigDB lists | FOXJ2\_01 | 135 | 5 | 12187 | 72 | Grm3,Ncam1,Csrnp3,Tle4,Crh | | 1.206e-03 | -6.72 | GNF2\_EGFR | MSigDB lists | GNF2\_EGFR | 9 | 2 | 12187 | 72 | Timp2,Crh | | 1.206e-03 | -6.72 | GNF2\_CDKN1C | MSigDB lists | GNF2\_CDKN1C | 9 | 2 | 12187 | 72 | Crh,Timp2 | | 1.211e-03 | -6.72 | GO\_CELL\_CELL\_SIGNALING | MSigDB lists | GO\_CELL\_CELL\_SIGNALING | 540 | 10 | 12187 | 72 | Plp1,Inhba,Kcnmb2,Grm3,Crh,Fjx1,Agrn,Homer1,Bdnf,Lhx1 | | 1.212e-03 | -6.72 | SRY\_02 | MSigDB lists | SRY\_02 | 202 | 6 | 12187 | 72 | Hs3st1,Crh,Qk,Nr4a3,Csrnp3,Inhba | | 1.221e-03 | -6.71 | regulation of striated muscle tissue development | biological process | GO:0016202 | 137 | 5 | 13711 | 80 | Rgs4,Bdnf,Ddx17,Ncam1,Arntl | | 1.222e-03 | -6.71 | organic substance biosynthetic process | biological process | GO:1901576 | 1802 | 21 | 13711 | 80 | Msmo1,Ndst4,Ugt8a,Hs3st1,Hsd17b12,Atp5b,Inhba,Qdpr,Crh,Egr1,Ddx17,Plp1,Glud1,Qk,Tmem165,Amd1,Srm,Scd2,Lhx1,Dgkz,Aars | | 1.228e-03 | -6.70 | GO\_NEUROLOGICAL\_SYSTEM\_PROCESS | MSigDB lists | GO\_NEUROLOGICAL\_SYSTEM\_PROCESS | 541 | 10 | 12187 | 72 | Cldn5,Cacng2,Nr4a3,Crh,Spry2,Aars,Kcnmb2,Col1a1,Gbx1,Fabp7 | | 1.279e-03 | -6.66 | negative regulation of cellular process | biological process | GO:0048523 | 3771 | 35 | 13711 | 80 | P2ry12,Aars,Lhx1,Crh,Tle4,Timp2,Tgfa,Kdr,Ubqln2,Ltk,Mtmr9,Sorl1,Inhba,Gsdme,Pdia6,Atp5b,Foxq1,Nr4a3,Agrn,H1f0,Rgs4,Arntl,Peg3,Qk,Bdnf,Dgkz,Ntm,Egr1,Map4,Col1a1,Spry2,Csrnp3,Ncam1,Lims2,Lrrtm2 | | 1.288e-03 | -6.65 | neuron part | cellular component | GO:0097458 | 1719 | 20 | 13825 | 79 | Ncam1,Grm3,Timp2,Dgkz,Sorl1,Qdpr,Bdnf,Cacng2,Kdr,Ephb6,Homer1,Fabp7,Ntm,Agrn,Map4,Cldn5,Car2,Lrrtm2,Ermn,Crh | | 1.305e-03 | -6.64 | sucrose degradation | BIOCYC pathways | MOUSE\_PWY3DJ-5523 | 5 | 2 | 823 | 10 | Ltk,Hkdc1 | | 1.307e-03 | -6.64 | GO\_REGULATION\_OF\_SYNAPSE\_STRUCTURE\_OR\_ACTIVITY | MSigDB lists | GO\_REGULATION\_OF\_SYNAPSE\_STRUCTURE\_OR\_ACTIVITY | 205 | 6 | 12187 | 72 | Ncam1,Bdnf,Crh,Lrrtm2,Agrn,Cbln2 | | 1.320e-03 | -6.63 | negative regulation of cell death | biological process | GO:0060548 | 859 | 13 | 13711 | 80 | Bdnf,Lims2,Sorl1,Ltk,Ncam1,Aars,Spry2,Qk,Tgfa,Kdr,Foxq1,Nr4a3,Crh | | 1.322e-03 | -6.63 | GO\_FATTY\_ACID\_BIOSYNTHETIC\_PROCESS | MSigDB lists | GO\_FATTY\_ACID\_BIOSYNTHETIC\_PROCESS | 81 | 4 | 12187 | 72 | Hsd17b12,Msmo1,Qk,Plp1 | | 1.345e-03 | -6.61 | regulation of muscle organ development | biological process | GO:0048634 | 140 | 5 | 13711 | 80 | Arntl,Rgs4,Ddx17,Bdnf,Ncam1 | | 1.345e-03 | -6.61 | regulation of muscle tissue development | biological process | GO:1901861 | 140 | 5 | 13711 | 80 | Ncam1,Rgs4,Bdnf,Ddx17,Arntl | | 1.451e-03 | -6.54 | CSR\_EARLY\_UP.V1\_UP | MSigDB lists | CSR\_EARLY\_UP.V1\_UP | 141 | 5 | 12187 | 72 | Inhba,Fjx1,Spry2,Amd1,Pom121 | | 1.481e-03 | -6.51 | MODULE\_11 | MSigDB lists | MODULE\_11 | 460 | 9 | 12187 | 72 | Plp1,Mtmr9,Car2,Fabp7,Ddx17,Cldn5,Rgs4,Qdpr,Ncam1 | | 1.481e-03 | -6.51 | GO\_NEURON\_PROJECTION\_DEVELOPMENT | MSigDB lists | GO\_NEURON\_PROJECTION\_DEVELOPMENT | 460 | 9 | 12187 | 72 | Lrrc55,Plp1,Gbx1,Ugt8a,Ncam1,Lhx1,Bdnf,Map4,Nr4a3 | | 1.502e-03 | -6.50 | GO\_GAS\_TRANSPORT | MSigDB lists | GO\_GAS\_TRANSPORT | 10 | 2 | 12187 | 72 | Car2,Hba-a2 | | 1.504e-03 | -6.50 | MODULE\_137 | MSigDB lists | MODULE\_137 | 461 | 9 | 12187 | 72 | Ddx17,Fabp7,Car2,Mtmr9,Plp1,Ncam1,Qdpr,Rgs4,Cldn5 | | 1.513e-03 | -6.49 | AMUNDSON\_GENOTOXIC\_SIGNATURE | MSigDB lists | AMUNDSON\_GENOTOXIC\_SIGNATURE | 84 | 4 | 12187 | 72 | Spry2,Egr1,Amd1,Nr4a3 | | 1.544e-03 | -6.47 | GSE27241\_WT\_VS\_RORGT\_KO\_TH17\_POLARIZED\_CD4\_TCELL\_TREATED\_WITH\_DIGOXIN\_DN | MSigDB lists | GSE27241\_WT\_VS\_RORGT\_KO\_TH17\_POLARIZED\_CD4\_TCELL\_TREATED\_WITH\_DIGOXIN\_DN | 143 | 5 | 12187 | 72 | Spry2,Egr1,Arntl,Timp2,Nr4a3 | | 1.544e-03 | -6.47 | GSE28737\_WT\_VS\_BCL6\_KO\_FOLLICULAR\_BCELL\_UP | MSigDB lists | GSE28737\_WT\_VS\_BCL6\_KO\_FOLLICULAR\_BCELL\_UP | 143 | 5 | 12187 | 72 | Gng11,Agrn,Grm3,Bdnf,Pom121 | | 1.547e-03 | -6.47 | cellular response to organic substance | biological process | GO:0071310 | 1457 | 18 | 13711 | 80 | Ltk,Inhba,Gsdme,Atp5b,Col1a1,Spry2,Cldn5,Car2,Nr4a3,P2ry12,Bdnf,Srm,Timp2,Kdr,Egr1,Ddx17,Tle4,Crh | | 1.572e-03 | -6.46 | MODULE\_100 | MSigDB lists | MODULE\_100 | 464 | 9 | 12187 | 72 | Mtmr9,Car2,Plp1,Fabp7,Ddx17,Cldn5,Qdpr,Rgs4,Ncam1 | | 1.588e-03 | -6.45 | negative regulation of neuron death | biological process | GO:1901215 | 215 | 6 | 13711 | 80 | Kdr,Nr4a3,Foxq1,Bdnf,Sorl1,Aars | | 1.620e-03 | -6.43 | axonogenesis | biological process | GO:0007409 | 294 | 7 | 13711 | 80 | Nr4a3,Agrn,Gbx1,Ephb6,Bdnf,Lhx1,Ncam1 | | 1.634e-03 | -6.42 | GO\_RESPONSE\_TO\_NITROGEN\_COMPOUND | MSigDB lists | GO\_RESPONSE\_TO\_NITROGEN\_COMPOUND | 662 | 11 | 12187 | 72 | Egr1,Gng11,Col1a1,Car2,Qdpr,Aars,Homer1,Nr4a3,P2ry12,Agrn,Crh | | 1.635e-03 | -6.42 | cell projection organization | biological process | GO:0030030 | 991 | 14 | 13711 | 80 | Gbx1,Ncam1,P2ry12,Lhx1,Ephb6,Bdnf,Ccdc13,Nr4a3,Nrep,Kdr,Plp1,Agrn,Map4,Ugt8a | | 1.684e-03 | -6.39 | transmembrane receptor protein tyrosine kinase signaling pathway | biological process | GO:0007169 | 296 | 7 | 13711 | 80 | Col1a1,Ltk,Ephb6,Bdnf,Nr4a3,Tgfa,Kdr | | 1.692e-03 | -6.38 | REACTOME\_METABOLISM\_OF\_AMINO\_ACIDS\_AND\_DERIVATIVES | MSigDB lists | REACTOME\_METABOLISM\_OF\_AMINO\_ACIDS\_AND\_DERIVATIVES | 146 | 5 | 12187 | 72 | Inhba,Srm,Qdpr,Amd1,Glud1 | | 1.743e-03 | -6.35 | MODULE\_66 | MSigDB lists | MODULE\_66 | 471 | 9 | 12187 | 72 | Plp1,Mtmr9,Car2,Fabp7,Ddx17,Cldn5,Rgs4,Qdpr,Ncam1 | | 1.750e-03 | -6.35 | regulation of cation transmembrane transport | biological process | GO:1904062 | 298 | 7 | 13711 | 80 | Agrn,Rgs4,Homer1,Nipsnap2,Crh,Lrrc55,Cacng2 | | 1.787e-03 | -6.33 | polyamine biosynthetic process | biological process | GO:0006596 | 11 | 2 | 13711 | 80 | Srm,Amd1 | | 1.787e-03 | -6.33 | regulation of glucocorticoid receptor signaling pathway | biological process | GO:2000322 | 11 | 2 | 13711 | 80 | Arntl,Bdnf | | 1.794e-03 | -6.32 | KEGG\_ARGININE\_AND\_PROLINE\_METABOLISM | MSigDB lists | KEGG\_ARGININE\_AND\_PROLINE\_METABOLISM | 41 | 3 | 12187 | 72 | Glud1,Amd1,Srm | | 1.797e-03 | -6.32 | GSE21670\_STAT3\_KO\_VS\_WT\_CD4\_TCELL\_TGFB\_IL6\_TREATED\_DN | MSigDB lists | GSE21670\_STAT3\_KO\_VS\_WT\_CD4\_TCELL\_TGFB\_IL6\_TREATED\_DN | 148 | 5 | 12187 | 72 | Timp2,Dgkz,Car2,Sorl1,Ncam1 | | 1.822e-03 | -6.31 | positive regulation of cell development | biological process | GO:0010720 | 575 | 10 | 13711 | 80 | Ltk,P2ry12,Lims2,Bdnf,Alkal2,Qk,Kdr,Cldn5,Timp2,Ddx56 | | 1.829e-03 | -6.30 | GO\_G\_PROTEIN\_COUPLED\_GLUTAMATE\_RECEPTOR\_SIGNALING\_PATHWAY | MSigDB lists | GO\_G\_PROTEIN\_COUPLED\_GLUTAMATE\_RECEPTOR\_SIGNALING\_PATHWAY | 11 | 2 | 12187 | 72 | Homer1,Grm3 | | 1.829e-03 | -6.30 | GNF2\_IGFBP1 | MSigDB lists | GNF2\_IGFBP1 | 11 | 2 | 12187 | 72 | Timp2,Crh | | 1.829e-03 | -6.30 | MYLLYKANGAS\_AMPLIFICATION\_HOT\_SPOT\_8 | MSigDB lists | MYLLYKANGAS\_AMPLIFICATION\_HOT\_SPOT\_8 | 11 | 2 | 12187 | 72 | Nr4a3,Fnbp1 | | 1.851e-03 | -6.29 | GSE46606\_IRF4\_KO\_VS\_WT\_CD40L\_IL2\_IL5\_3DAY\_STIMULATED\_BCELL\_UP | MSigDB lists | GSE46606\_IRF4\_KO\_VS\_WT\_CD40L\_IL2\_IL5\_3DAY\_STIMULATED\_BCELL\_UP | 149 | 5 | 12187 | 72 | Foxq1,Nr4a3,Tgfa,Msmo1,Hsd17b12 | | 1.889e-03 | -6.27 | O2/CO2 exchange in erythrocytes | REACTOME pathways | R-MMU-1480926 | 10 | 2 | 6297 | 42 | Hba-a2,Car2 | | 1.889e-03 | -6.27 | Erythrocytes take up carbon dioxide and release oxygen | REACTOME pathways | R-MMU-1237044 | 10 | 2 | 6297 | 42 | Hba-a2,Car2 | | 1.906e-03 | -6.26 | FOXD3\_01 | MSigDB lists | FOXD3\_01 | 150 | 5 | 12187 | 72 | Grm3,Lhx1,Crh,Tle4,P2ry12 | | 1.924e-03 | -6.25 | BORLAK\_LIVER\_CANCER\_EGF\_UP | MSigDB lists | BORLAK\_LIVER\_CANCER\_EGF\_UP | 42 | 3 | 12187 | 72 | Egr1,Car2,Tgfa | | 1.926e-03 | -6.25 | positive regulation of phosphate metabolic process | biological process | GO:0045937 | 896 | 13 | 13711 | 80 | Gsdme,Mtmr9,Inhba,Spry2,Agrn,Bdnf,Dgkz,Alkal2,Egr1,Timp2,Tgfa,Kdr,Crh | | 1.926e-03 | -6.25 | positive regulation of phosphorus metabolic process | biological process | GO:0010562 | 896 | 13 | 13711 | 80 | Crh,Kdr,Tgfa,Timp2,Egr1,Alkal2,Bdnf,Dgkz,Agrn,Spry2,Inhba,Mtmr9,Gsdme | | 1.927e-03 | -6.25 | regulation of transmembrane transport | biological process | GO:0034762 | 482 | 9 | 13711 | 80 | Cacng2,Homer1,Rgs4,Nipsnap2,Agrn,Lrrc55,Nr4a3,Car2,Crh | | 1.932e-03 | -6.25 | regulation of cation channel activity | biological process | GO:2001257 | 152 | 5 | 13711 | 80 | Homer1,Nipsnap2,Crh,Lrrc55,Cacng2 | | 1.942e-03 | -6.24 | response to drug | biological process | GO:0042493 | 580 | 10 | 13711 | 80 | Nr4a3,Car2,Crh,Egr1,Homer1,Kdr,Bdnf,P2ry12,Ncam1,Inhba | | 1.951e-03 | -6.24 | MCCLUNG\_CREB1\_TARGETS\_UP | MSigDB lists | MCCLUNG\_CREB1\_TARGETS\_UP | 90 | 4 | 12187 | 72 | Fabp7,H1f0,Bdnf,Hsd17b12 | | 1.961e-03 | -6.23 | CREB\_Q2 | MSigDB lists | CREB\_Q2 | 222 | 6 | 12187 | 72 | Ubqln2,Crh,Nr4a3,AI593442,Grm3,Egr1 | | 1.998e-03 | -6.22 | regulation of ion transmembrane transporter activity | biological process | GO:0032412 | 225 | 6 | 13711 | 80 | Nipsnap2,Homer1,Agrn,Cacng2,Lrrc55,Crh | | 2.045e-03 | -6.19 | negative regulation of biological process | biological process | GO:0048519 | 4173 | 37 | 13711 | 80 | P2ry12,Aars,Lhx1,Timp2,Kdr,Tgfa,Crh,Tle4,Sorl1,Mtmr9,Ltk,Inhba,Pdia6,Gsdme,Atp5b,Ubqln2,Agrn,H1f0,Fabp7,Rgs4,Foxq1,Nr4a3,Dgkz,Bdnf,Arntl,Peg3,Qk,Egr1,Ddx17,Map4,Ntm,Ncam1,Lims2,Col1a1,Spry2,Csrnp3,Lrrtm2 | | 2.090e-03 | -6.17 | protein disulfide isomerase activity | molecular function | GO:0003756 | 12 | 2 | 13516 | 78 | Tmx3,Pdia6 | | 2.090e-03 | -6.17 | intramolecular oxidoreductase activity, transposing S-S bonds | molecular function | GO:0016864 | 12 | 2 | 13516 | 78 | Pdia6,Tmx3 | | 2.114e-03 | -6.16 | TAATTA\_CHX10\_01 | MSigDB lists | TAATTA\_CHX10\_01 | 582 | 10 | 12187 | 72 | Arntl,Tle4,Col1a1,Car2,Amd1,Ddx17,Cacng2,Hs3st1,Ndst4,Bdnf | | 2.115e-03 | -6.16 | GO\_REGULATION\_OF\_SYNAPSE\_ORGANIZATION | MSigDB lists | GO\_REGULATION\_OF\_SYNAPSE\_ORGANIZATION | 92 | 4 | 12187 | 72 | Lrrtm2,Agrn,Cbln2,Bdnf | | 2.136e-03 | -6.15 | long-chain fatty acid biosynthetic process | biological process | GO:0042759 | 12 | 2 | 13711 | 80 | Qk,Plp1 | | 2.139e-03 | -6.15 | GSE28783\_CTRL\_ANTI\_MIR\_VS\_UNTREATED\_ATHEROSCLEROSIS\_MACROPHAGE\_DN | MSigDB lists | GSE28783\_CTRL\_ANTI\_MIR\_VS\_UNTREATED\_ATHEROSCLEROSIS\_MACROPHAGE\_DN | 154 | 5 | 12187 | 72 | Spry2,Agrn,Car2,Ephb6,Lrrtm2 | | 2.165e-03 | -6.14 | negative regulation of neuron apoptotic process | biological process | GO:0043524 | 156 | 5 | 13711 | 80 | Foxq1,Nr4a3,Bdnf,Aars,Kdr | | 2.173e-03 | -6.13 | Methionine salvage pathway | KEGG pathways | M00034 | 9 | 2 | 5248 | 42 | Amd1,Srm | | 2.173e-03 | -6.13 | Methionine salvage pathway | KEGG pathways | mmu\_M00034 | 9 | 2 | 5248 | 42 | Amd1,Srm | | 2.186e-03 | -6.13 | GO\_REGULATION\_OF\_SKELETAL\_MUSCLE\_CELL\_DIFFERENTIATION | MSigDB lists | GO\_REGULATION\_OF\_SKELETAL\_MUSCLE\_CELL\_DIFFERENTIATION | 12 | 2 | 12187 | 72 | Arntl,Ddx17 | | 2.186e-03 | -6.13 | GO\_POSITIVE\_REGULATION\_OF\_CIRCADIAN\_RHYTHM | MSigDB lists | GO\_POSITIVE\_REGULATION\_OF\_CIRCADIAN\_RHYTHM | 12 | 2 | 12187 | 72 | Crh,Arntl | | 2.201e-03 | -6.12 | APRELIKOVA\_BRCA1\_TARGETS | MSigDB lists | APRELIKOVA\_BRCA1\_TARGETS | 44 | 3 | 12187 | 72 | Peg3,Car2,Qk | | 2.201e-03 | -6.12 | REACTOME\_HEPARAN\_SULFATE\_HEPARIN\_HS\_GAG\_METABOLISM | MSigDB lists | REACTOME\_HEPARAN\_SULFATE\_HEPARIN\_HS\_GAG\_METABOLISM | 44 | 3 | 12187 | 72 | Agrn,Ndst4,Hs3st1 | | 2.201e-03 | -6.12 | CHAUHAN\_RESPONSE\_TO\_METHOXYESTRADIOL\_UP | MSigDB lists | CHAUHAN\_RESPONSE\_TO\_METHOXYESTRADIOL\_UP | 44 | 3 | 12187 | 72 | Aars,Chmp7,Srm | | 2.205e-03 | -6.12 | Tgfb1 (transforming growth factor, beta 1) | protein interactions | 21803 | 10 | 2 | 6802 | 49 | Nrep,Col1a1 | | 2.241e-03 | -6.10 | cell projection | cellular component | GO:0042995 | 2067 | 22 | 13825 | 79 | Timp2,Dgkz,P2ry12,Kdr,Ermn,Spry2,Grm3,Ncam1,Qdpr,Ccdc13,Bdnf,Cacng2,Homer1,Ephb6,Mtmr9,Agrn,Ntm,Fabp7,Cldn5,Car2,Map4,Crh | | 2.241e-03 | -6.10 | BENPORATH\_EED\_TARGETS | MSigDB lists | BENPORATH\_EED\_TARGETS | 689 | 11 | 12187 | 72 | Nr4a3,AI593442,Ncam1,Arhgap20,Arntl,Tgfa,Hba-a2,Grm3,Foxq1,H1f0,Ltk | | 2.263e-03 | -6.09 | ONDER\_CDH1\_TARGETS\_2\_DN | MSigDB lists | ONDER\_CDH1\_TARGETS\_2\_DN | 309 | 7 | 12187 | 72 | Tgfa,Agrn,Inhba,Car2,Sorl1,Hs3st1,Homer1 | | 2.267e-03 | -6.09 | GO\_ORGANIC\_ACID\_METABOLIC\_PROCESS | MSigDB lists | GO\_ORGANIC\_ACID\_METABOLIC\_PROCESS | 690 | 11 | 12187 | 72 | Nr4a3,Qdpr,Ndst4,Glud1,Aars,Ugt8a,Hkdc1,Plp1,Qk,Msmo1,Hsd17b12 | | 2.330e-03 | -6.06 | regulation of transmembrane transporter activity | biological process | GO:0022898 | 232 | 6 | 13711 | 80 | Crh,Cacng2,Lrrc55,Agrn,Nipsnap2,Homer1 | | 2.366e-03 | -6.05 | GO\_HEAD\_DEVELOPMENT | MSigDB lists | GO\_HEAD\_DEVELOPMENT | 591 | 10 | 12187 | 72 | Crh,Nr4a3,Cldn5,Glud1,Aars,Lhx1,Plp1,Col1a1,Inhba,Fabp7 | | 2.424e-03 | -6.02 | response to anesthetic | biological process | GO:0072347 | 46 | 3 | 13711 | 80 | Homer1,Ncam1,Crh | | 2.453e-03 | -6.01 | Fbxo32 (F-box protein 32) | protein interactions | 67731 | 132 | 5 | 6802 | 49 | Atp5b,Map4,Pdia6,Col1a1,Ubqln2 | | 2.458e-03 | -6.01 | GSE3982\_BASOPHIL\_VS\_TH1\_DN | MSigDB lists | GSE3982\_BASOPHIL\_VS\_TH1\_DN | 159 | 5 | 12187 | 72 | Aars,Grm3,Qdpr,Msmo1,Hsd17b12 | | 2.461e-03 | -6.01 | heparan sulfate sulfotransferase activity | molecular function | GO:0034483 | 13 | 2 | 13516 | 78 | Hs3st1,Ndst4 | | 2.470e-03 | -6.00 | KRAS.300\_UP.V1\_UP | MSigDB lists | KRAS.300\_UP.V1\_UP | 96 | 4 | 12187 | 72 | Peg3,Spry2,Ntm,Sorl1 | | 2.515e-03 | -5.99 | regulation of metallopeptidase activity | biological process | GO:1905048 | 13 | 2 | 13711 | 80 | Sorl1,Timp2 | | 2.515e-03 | -5.99 | chemical homeostasis within a tissue | biological process | GO:0048875 | 13 | 2 | 13711 | 80 | Kdr,Homer1 | | 2.515e-03 | -5.99 | cellular biogenic amine biosynthetic process | biological process | GO:0042401 | 13 | 2 | 13711 | 80 | Amd1,Srm | | 2.515e-03 | -5.99 | negative regulation of receptor internalization | biological process | GO:0002091 | 13 | 2 | 13711 | 80 | Lrrtm2,Ubqln2 | | 2.515e-03 | -5.99 | amine biosynthetic process | biological process | GO:0009309 | 13 | 2 | 13711 | 80 | Srm,Amd1 | | 2.521e-03 | -5.98 | MODULE\_2 | MSigDB lists | MODULE\_2 | 315 | 7 | 12187 | 72 | Qdpr,Ncam1,Cldn5,Fabp7,Msmo1,Gng11,Plp1 | | 2.526e-03 | -5.98 | YAGI\_AML\_FAB\_MARKERS | MSigDB lists | YAGI\_AML\_FAB\_MARKERS | 160 | 5 | 12187 | 72 | Timp2,Nr4a3,Ncam1,Amd1,Sorl1 | | 2.574e-03 | -5.96 | GO\_POSITIVE\_REGULATION\_OF\_BEHAVIOR | MSigDB lists | GO\_POSITIVE\_REGULATION\_OF\_BEHAVIOR | 13 | 2 | 12187 | 72 | Nr4a3,Crh | | 2.574e-03 | -5.96 | GO\_NEGATIVE\_REGULATION\_OF\_RECEPTOR\_MEDIATED\_ENDOCYTOSIS | MSigDB lists | GO\_NEGATIVE\_REGULATION\_OF\_RECEPTOR\_MEDIATED\_ENDOCYTOSIS | 13 | 2 | 12187 | 72 | Lrrtm2,Ubqln2 | | 2.574e-03 | -5.96 | MODULE\_294 | MSigDB lists | MODULE\_294 | 13 | 2 | 12187 | 72 | Car2,Glud1 | | 2.574e-03 | -5.96 | GO\_HEPARAN\_SULFATE\_SULFOTRANSFERASE\_ACTIVITY | MSigDB lists | GO\_HEPARAN\_SULFATE\_SULFOTRANSFERASE\_ACTIVITY | 13 | 2 | 12187 | 72 | Ndst4,Hs3st1 | | 2.595e-03 | -5.95 | GSE17974\_CTRL\_VS\_ACT\_IL4\_AND\_ANTI\_IL12\_12H\_CD4\_TCELL\_DN | MSigDB lists | GSE17974\_CTRL\_VS\_ACT\_IL4\_AND\_ANTI\_IL12\_12H\_CD4\_TCELL\_DN | 161 | 5 | 12187 | 72 | Aars,Homer1,Tmem165,Srm,Cldn5 | | 2.603e-03 | -5.95 | regulation of cellular component organization | biological process | GO:0051128 | 2181 | 23 | 13711 | 80 | Qk,Alkal2,Bdnf,Cbln2,Ntm,Map4,Spry2,Lims2,Ncam1,Lrrtm2,Homer1,P2ry12,Kdr,Tgfa,Ubqln2,Inhba,Ermn,Sorl1,Ltk,Rgs4,H1f0,Agrn,Ddx56 | | 2.658e-03 | -5.93 | myelination | biological process | GO:0042552 | 99 | 4 | 13711 | 80 | Plp1,Cldn5,Ugt8a,Qk | | 2.665e-03 | -5.93 | GSE15930\_NAIVE\_VS\_24H\_IN\_VITRO\_STIM\_CD8\_TCELL\_UP | MSigDB lists | GSE15930\_NAIVE\_VS\_24H\_IN\_VITRO\_STIM\_CD8\_TCELL\_UP | 162 | 5 | 12187 | 72 | Car2,Nr4a3,Timp2,Tle4,Agrn | | 2.666e-03 | -5.93 | neuron projection | cellular component | GO:0043005 | 1310 | 16 | 13825 | 79 | Agrn,Ntm,Homer1,Ephb6,Crh,Ermn,Cldn5,Car2,Map4,Qdpr,Timp2,Grm3,Ncam1,Kdr,Bdnf,Cacng2 | | 2.757e-03 | -5.89 | ensheathment of neurons | biological process | GO:0007272 | 100 | 4 | 13711 | 80 | Qk,Ugt8a,Cldn5,Plp1 | | 2.757e-03 | -5.89 | axon ensheathment | biological process | GO:0008366 | 100 | 4 | 13711 | 80 | Cldn5,Plp1,Ugt8a,Qk | | 2.780e-03 | -5.89 | GO\_RESPONSE\_TO\_OXYGEN\_CONTAINING\_COMPOUND | MSigDB lists | GO\_RESPONSE\_TO\_OXYGEN\_CONTAINING\_COMPOUND | 1042 | 14 | 12187 | 72 | Ltk,Aars,Qdpr,P2ry12,Nr4a3,Egr1,Hba-a2,Col1a1,Car2,Gng11,Inhba,Homer1,Crh,Agrn | | 2.791e-03 | -5.88 | GO\_CELLULAR\_RESPONSE\_TO\_NITROGEN\_COMPOUND | MSigDB lists | GO\_CELLULAR\_RESPONSE\_TO\_NITROGEN\_COMPOUND | 410 | 8 | 12187 | 72 | Nr4a3,Agrn,P2ry12,Crh,Gng11,Col1a1,Car2,Egr1 | | 2.810e-03 | -5.87 | GSE19401\_PAM2CSK4\_VS\_RETINOIC\_ACID\_AND\_PAM2CSK4\_STIM\_FOLLICULAR\_DC\_DN | MSigDB lists | GSE19401\_PAM2CSK4\_VS\_RETINOIC\_ACID\_AND\_PAM2CSK4\_STIM\_FOLLICULAR\_DC\_DN | 164 | 5 | 12187 | 72 | Dgkz,Qk,Timp2,Pdia6,Hs3st1 | | 2.810e-03 | -5.87 | GSE31082\_DP\_VS\_CD8\_SP\_THYMOCYTE\_DN | MSigDB lists | GSE31082\_DP\_VS\_CD8\_SP\_THYMOCYTE\_DN | 164 | 5 | 12187 | 72 | Glud1,Tmem165,Sorl1,Nr4a3,Agrn | | 2.816e-03 | -5.87 | regulation of transporter activity | biological process | GO:0032409 | 241 | 6 | 13711 | 80 | Agrn,Nipsnap2,Homer1,Crh,Cacng2,Lrrc55 | | 2.821e-03 | -5.87 | Thioredoxin\_CS | interpro domains | IPR017937 | 14 | 2 | 13788 | 79 | Tmx3,Pdia6 | | 2.863e-03 | -5.86 | GO\_CENTRAL\_NERVOUS\_SYSTEM\_DEVELOPMENT | MSigDB lists | GO\_CENTRAL\_NERVOUS\_SYSTEM\_DEVELOPMENT | 711 | 11 | 12187 | 72 | Glud1,Aars,Lhx1,Crh,Nr4a3,Fabp7,Plp1,Gbx1,Inhba,Timp2,Ugt8a | | 2.923e-03 | -5.83 | G protein-coupled glutamate receptor signaling pathway | biological process | GO:0007216 | 14 | 2 | 13711 | 80 | Homer1,Grm3 | | 2.923e-03 | -5.83 | negative regulation of catecholamine secretion | biological process | GO:0033604 | 14 | 2 | 13711 | 80 | P2ry12,Crh | | 2.961e-03 | -5.82 | GSE15930\_NAIVE\_VS\_24H\_IN\_VITRO\_STIM\_IL12\_CD8\_TCELL\_UP | MSigDB lists | GSE15930\_NAIVE\_VS\_24H\_IN\_VITRO\_STIM\_IL12\_CD8\_TCELL\_UP | 166 | 5 | 12187 | 72 | Tle4,Timp2,Nr4a3,Agrn,Car2 | | 2.971e-03 | -5.82 | cell surface receptor signaling pathway | biological process | GO:0007166 | 1296 | 16 | 13711 | 80 | Egr1,Kdr,Tgfa,Plp1,Tle4,Ephb6,Bdnf,P2ry12,Homer1,Cldn5,Nr4a3,Grm3,Ncam1,Inhba,Ltk,Col1a1 | | 2.992e-03 | -5.81 | GO\_POLYAMINE\_METABOLIC\_PROCESS | MSigDB lists | GO\_POLYAMINE\_METABOLIC\_PROCESS | 14 | 2 | 12187 | 72 | Amd1,Srm | | 2.992e-03 | -5.81 | GO\_PARTURITION | MSigDB lists | GO\_PARTURITION | 14 | 2 | 12187 | 72 | Arntl,Crh | | 2.992e-03 | -5.81 | GO\_NEGATIVE\_REGULATION\_OF\_CATECHOLAMINE\_SECRETION | MSigDB lists | GO\_NEGATIVE\_REGULATION\_OF\_CATECHOLAMINE\_SECRETION | 14 | 2 | 12187 | 72 | P2ry12,Crh | | 3.039e-03 | -5.80 | GSE3039\_ALPHABETA\_CD8\_TCELL\_VS\_B2\_BCELL\_DN | MSigDB lists | GSE3039\_ALPHABETA\_CD8\_TCELL\_VS\_B2\_BCELL\_DN | 167 | 5 | 12187 | 72 | Qk,Srm,Inhba,Homer1,Tmem165 | | 3.090e-03 | -5.78 | Prnp (prion protein) | protein interactions | 19122 | 41 | 3 | 6802 | 49 | Ncam1,Plp1,Ntm | | 3.118e-03 | -5.77 | GO\_NEURON\_PROJECTION\_GUIDANCE | MSigDB lists | GO\_NEURON\_PROJECTION\_GUIDANCE | 168 | 5 | 12187 | 72 | Lhx1,Ncam1,Bdnf,Nr4a3,Gbx1 | | 3.118e-03 | -5.77 | GSE8835\_HEALTHY\_VS\_CLL\_CD8\_TCELL\_UP | MSigDB lists | GSE8835\_HEALTHY\_VS\_CLL\_CD8\_TCELL\_UP | 168 | 5 | 12187 | 72 | Peg3,Cbln2,Zfp518b,Tgfa,Dgkz | | 3.198e-03 | -5.75 | HALLMARK\_EPITHELIAL\_MESENCHYMAL\_TRANSITION | MSigDB lists | HALLMARK\_EPITHELIAL\_MESENCHYMAL\_TRANSITION | 169 | 5 | 12187 | 72 | Ntm,Bdnf,Rgs4,Col1a1,Inhba | | 3.204e-03 | -5.74 | Ncam1 (neural cell adhesion molecule 1) | protein interactions | 17967 | 12 | 2 | 6802 | 49 | Ncam1,Homer1 | | 3.286e-03 | -5.72 | Nitrogen metabolism | KEGG pathways | ko00910 | 11 | 2 | 5248 | 42 | Glud1,Car2 | | 3.286e-03 | -5.72 | Nitrogen metabolism | KEGG pathways | mmu00910 | 11 | 2 | 5248 | 42 | Glud1,Car2 | | 3.301e-03 | -5.71 | WWTAAGGC\_UNKNOWN | MSigDB lists | WWTAAGGC\_UNKNOWN | 104 | 4 | 12187 | 72 | Foxq1,Ncam1,Lhx1,Cacng2 | | 3.335e-03 | -5.70 | transmembrane receptor protein tyrosine kinase activity | molecular function | GO:0004714 | 52 | 3 | 13516 | 78 | Ltk,Ephb6,Kdr | | 3.344e-03 | -5.70 | GO\_REGULATION\_OF\_NERVOUS\_SYSTEM\_DEVELOPMENT | MSigDB lists | GO\_REGULATION\_OF\_NERVOUS\_SYSTEM\_DEVELOPMENT | 620 | 10 | 12187 | 72 | Bdnf,Lhx1,Lrrtm2,P2ry12,Agrn,Cbln2,Ltk,Sorl1,Arntl,Timp2 | | 3.360e-03 | -5.70 | regulation of norepinephrine secretion | biological process | GO:0014061 | 15 | 2 | 13711 | 80 | Crh,P2ry12 | | 3.360e-03 | -5.70 | retina layer formation | biological process | GO:0010842 | 15 | 2 | 13711 | 80 | Lhx1,Fjx1 | | 3.361e-03 | -5.70 | PHONG\_TNF\_TARGETS\_UP | MSigDB lists | PHONG\_TNF\_TARGETS\_UP | 51 | 3 | 12187 | 72 | Egr1,Inhba,Fjx1 | | 3.361e-03 | -5.70 | GO\_SKIN\_EPIDERMIS\_DEVELOPMENT | MSigDB lists | GO\_SKIN\_EPIDERMIS\_DEVELOPMENT | 51 | 3 | 12187 | 72 | Foxq1,Aars,Inhba | | 3.364e-03 | -5.69 | GSE24726\_WT\_VS\_E2\_2\_KO\_PDC\_DAY4\_POST\_DELETION\_UP | MSigDB lists | GSE24726\_WT\_VS\_E2\_2\_KO\_PDC\_DAY4\_POST\_DELETION\_UP | 171 | 5 | 12187 | 72 | Spry2,Aars,Homer1,Hsd17b12,Agrn | | 3.364e-03 | -5.69 | OCT1\_03 | MSigDB lists | OCT1\_03 | 171 | 5 | 12187 | 72 | Cacng2,Nr4a3,Amd1,Ddx17,Bdnf | | 3.376e-03 | -5.69 | Schaffer collateral - CA1 synapse | cellular component | GO:0098685 | 108 | 4 | 13825 | 79 | Dgkz,Lrrtm2,Cacng2,Ncam1 | | 3.379e-03 | -5.69 | GO\_RNA\_POLYMERASE\_II\_TRANSCRIPTION\_FACTOR\_ACTIVITY\_SEQUENCE\_SPECIFIC\_DNA\_BINDING | MSigDB lists | GO\_RNA\_POLYMERASE\_II\_TRANSCRIPTION\_FACTOR\_ACTIVITY\_SEQUENCE\_SPECIFIC\_DNA\_BINDING | 423 | 8 | 12187 | 72 | Egr1,Foxq1,Csrnp3,Peg3,Zfp518b,Arntl,Tle4,Nr4a3 | | 3.404e-03 | -5.68 | positive regulation of phosphorylation | biological process | GO:0042327 | 843 | 12 | 13711 | 80 | Bdnf,Dgkz,Alkal2,Tgfa,Kdr,Timp2,Egr1,Crh,Inhba,Gsdme,Spry2,Agrn | | 3.439e-03 | -5.67 | GNF2\_MMP11 | MSigDB lists | GNF2\_MMP11 | 15 | 2 | 12187 | 72 | Crh,Timp2 | | 3.439e-03 | -5.67 | GO\_AMINE\_BIOSYNTHETIC\_PROCESS | MSigDB lists | GO\_AMINE\_BIOSYNTHETIC\_PROCESS | 15 | 2 | 12187 | 72 | Srm,Amd1 | | 3.439e-03 | -5.67 | MODULE\_222 | MSigDB lists | MODULE\_222 | 15 | 2 | 12187 | 72 | H1f0,Map4 | | 3.439e-03 | -5.67 | REACTOME\_METABOLISM\_OF\_POLYAMINES | MSigDB lists | REACTOME\_METABOLISM\_OF\_POLYAMINES | 15 | 2 | 12187 | 72 | Srm,Amd1 | | 3.439e-03 | -5.67 | GO\_REGULATION\_OF\_NOREPINEPHRINE\_SECRETION | MSigDB lists | GO\_REGULATION\_OF\_NOREPINEPHRINE\_SECRETION | 15 | 2 | 12187 | 72 | Crh,P2ry12 | | 3.441e-03 | -5.67 | negative regulation of Ras protein signal transduction | biological process | GO:0046580 | 52 | 3 | 13711 | 80 | Timp2,Dgkz,Spry2 | | 3.449e-03 | -5.67 | STARK\_PREFRONTAL\_CORTEX\_22Q11\_DELETION\_UP | MSigDB lists | STARK\_PREFRONTAL\_CORTEX\_22Q11\_DELETION\_UP | 172 | 5 | 12187 | 72 | Timp2,Fjx1,Ubqln2,Pom121,Ntm | | 3.519e-03 | -5.65 | regulation of synapse assembly | biological process | GO:0051963 | 107 | 4 | 13711 | 80 | Lrrtm2,Bdnf,Agrn,Cbln2 | | 3.536e-03 | -5.64 | FOXO4\_01 | MSigDB lists | FOXO4\_01 | 173 | 5 | 12187 | 72 | Nap1l5,Inhba,Cacng2,Bdnf,Ndst4 | | 3.559e-03 | -5.64 | signaling | biological process | GO:0023052 | 2939 | 28 | 13711 | 80 | Tle4,Crh,Arhgap20,Tgfa,Kdr,Plp1,P2ry12,Kcnmb2,Ephb6,Lhx1,Grm3,Nr4a3,Agrn,Rgs4,Ltk,Inhba,Gng11,Fjx1,Egr1,Ddx17,Cacng2,Bdnf,Dgkz,Car2,Cldn5,Homer1,Col1a1,Ncam1 | | 3.623e-03 | -5.62 | GO\_CELLULAR\_RESPONSE\_TO\_OXYGEN\_CONTAINING\_COMPOUND | MSigDB lists | GO\_CELLULAR\_RESPONSE\_TO\_OXYGEN\_CONTAINING\_COMPOUND | 627 | 10 | 12187 | 72 | Crh,Agrn,P2ry12,Nr4a3,Col1a1,Car2,Gng11,Inhba,Egr1,Ltk | | 3.624e-03 | -5.62 | GSE16451\_IMMATURE\_VS\_MATURE\_NEURON\_CELL\_LINE\_WEST\_EQUINE\_ENC\_VIRUS\_DN | MSigDB lists | GSE16451\_IMMATURE\_VS\_MATURE\_NEURON\_CELL\_LINE\_WEST\_EQUINE\_ENC\_VIRUS\_DN | 174 | 5 | 12187 | 72 | Srm,Ddx56,Egr1,Pdia6,Homer1 | | 3.634e-03 | -5.62 | sensory organ development | biological process | GO:0007423 | 432 | 8 | 13711 | 80 | Kdr,Nr4a3,Fjx1,Gsdme,Bdnf,Lhx1,Inhba,Spry2 | | 3.656e-03 | -5.61 | GSE36891\_UNSTIM\_VS\_POLYIC\_TLR3\_STIM\_PERITONEAL\_MACROPHAGE\_UP | MSigDB lists | GSE36891\_UNSTIM\_VS\_POLYIC\_TLR3\_STIM\_PERITONEAL\_MACROPHAGE\_UP | 107 | 4 | 12187 | 72 | Homer1,Egr1,Spry2,Nr4a3 | | 3.694e-03 | -5.60 | GO\_CELL\_DEVELOPMENT | MSigDB lists | GO\_CELL\_DEVELOPMENT | 1075 | 14 | 12187 | 72 | Lhx1,Ntm,Bdnf,Ncam1,Homer1,Map4,Qk,Agrn,Nr4a3,Gbx1,Lrrc55,Plp1,Ugt8a,Inhba | | 3.715e-03 | -5.60 | regulation of biological process | biological process | GO:0050789 | 7871 | 58 | 13711 | 80 | Plp1,Tgfa,Kdr,Nrep,P2ry12,Glud1,Fabp7,Agrn,Ddx56,H1f0,Hsd17b12,Atp5b,Mtmr9,Ltk,Ubqln2,Ddx17,Map4,Tmx3,Bdnf,Dgkz,Cbln2,Cacng2,Alkal2,Peg3,Lrrtm2,Cldn5,Csrnp3,Spry2,Timp2,Tle4,Crh,Arhgap20,Ephb6,Lhx1,Aars,Fbxl17,Rgs4,Lrrc55,Foxq1,Nr4a3,Grm3,Gsdme,Pdia6,Gng11,Sorl1,Ermn,Inhba,Egr1,Ntm,Qk,Arntl,Nipsnap2,Homer1,Car2,Lims2,Ncam1,Gbx1,Col1a1 | | 3.749e-03 | -5.59 | AHR\_01 | MSigDB lists | AHR\_01 | 53 | 3 | 12187 | 72 | Egr1,Ncam1,H1f0 | | 3.826e-03 | -5.57 | polyamine metabolic process | biological process | GO:0006595 | 16 | 2 | 13711 | 80 | Amd1,Srm | | 3.908e-03 | -5.54 | THIOREDOXIN\_1 | prosite domains | PS00194 | 14 | 2 | 8845 | 60 | Pdia6,Tmx3 | | 3.912e-03 | -5.54 | GO\_NEURON\_PROJECTION\_MORPHOGENESIS | MSigDB lists | GO\_NEURON\_PROJECTION\_MORPHOGENESIS | 341 | 7 | 12187 | 72 | Lhx1,Bdnf,Ncam1,Lrrc55,Gbx1,Nr4a3,Ugt8a | | 3.915e-03 | -5.54 | LUND\_SILENCED\_BY\_METHYLATION | MSigDB lists | LUND\_SILENCED\_BY\_METHYLATION | 16 | 2 | 12187 | 72 | Timp2,Amd1 | | 3.915e-03 | -5.54 | KEGG\_NITROGEN\_METABOLISM | MSigDB lists | KEGG\_NITROGEN\_METABOLISM | 16 | 2 | 12187 | 72 | Glud1,Car2 | | 3.953e-03 | -5.53 | GO\_MOLTING\_CYCLE | MSigDB lists | GO\_MOLTING\_CYCLE | 54 | 3 | 12187 | 72 | Foxq1,Aars,Inhba | | 3.953e-03 | -5.53 | GO\_TRANSMEMBRANE\_RECEPTOR\_PROTEIN\_TYROSINE\_KINASE\_ACTIVITY | MSigDB lists | GO\_TRANSMEMBRANE\_RECEPTOR\_PROTEIN\_TYROSINE\_KINASE\_ACTIVITY | 54 | 3 | 12187 | 72 | Kdr,Ephb6,Ltk | | 3.997e-03 | -5.52 | regulation of response to stimulus | biological process | GO:0048583 | 2962 | 28 | 13711 | 80 | Agrn,Rgs4,Fabp7,Nr4a3,Inhba,Sorl1,Gsdme,Pdia6,Ubqln2,Tgfa,Kdr,Timp2,Crh,Tle4,Nrep,P2ry12,Fbxl17,Homer1,Ncam1,Lims2,Col1a1,Spry2,Egr1,Dgkz,Bdnf,Arntl,Alkal2,Cacng2 | | 4.035e-03 | -5.51 | oxidative phosphorylation | biological process | GO:0006119 | 55 | 3 | 13711 | 80 | Nipsnap2,Bdnf,Atp5b | | 4.036e-03 | -5.51 | MEL18\_DN.V1\_UP | MSigDB lists | MEL18\_DN.V1\_UP | 110 | 4 | 12187 | 72 | Ntm,Inhba,Fjx1,Agrn | | 4.039e-03 | -5.51 | GO\_MONOCARBOXYLIC\_ACID\_METABOLIC\_PROCESS | MSigDB lists | GO\_MONOCARBOXYLIC\_ACID\_METABOLIC\_PROCESS | 343 | 7 | 12187 | 72 | Qk,Nr4a3,Plp1,Hkdc1,Ugt8a,Msmo1,Hsd17b12 | | 4.148e-03 | -5.49 | paranode region of axon | cellular component | GO:0033270 | 17 | 2 | 13825 | 79 | Ermn,Cldn5 | | 4.163e-03 | -5.48 | ANASTASSIOU\_MULTICANCER\_INVASIVENESS\_SIGNATURE | MSigDB lists | ANASTASSIOU\_MULTICANCER\_INVASIVENESS\_SIGNATURE | 55 | 3 | 12187 | 72 | Ntm,Inhba,Col1a1 | | 4.245e-03 | -5.46 | regulation of intracellular pH | biological process | GO:0051453 | 56 | 3 | 13711 | 80 | Car2,Tmem165,Atp5b | | 4.277e-03 | -5.45 | regulation of signaling receptor activity | biological process | GO:0010469 | 113 | 4 | 13711 | 80 | Tgfa,Homer1,Crh,Cacng2 | | 4.284e-03 | -5.45 | GO\_POSITIVE\_REGULATION\_OF\_CELL\_DIFFERENTIATION | MSigDB lists | GO\_POSITIVE\_REGULATION\_OF\_CELL\_DIFFERENTIATION | 642 | 10 | 12187 | 72 | Car2,Col1a1,Arntl,Timp2,Inhba,Ltk,Kdr,Cldn5,Lhx1,Bdnf | | 4.320e-03 | -5.44 | regulation of skeletal muscle cell differentiation | biological process | GO:2001014 | 17 | 2 | 13711 | 80 | Ddx17,Arntl | | 4.373e-03 | -5.43 | channel regulator activity | molecular function | GO:0016247 | 115 | 4 | 13516 | 78 | Kcnmb2,Grm3,Lrrc55,Cacng2 | | 4.391e-03 | -5.43 | GO\_CELL\_PROJECTION\_ORGANIZATION | MSigDB lists | GO\_CELL\_PROJECTION\_ORGANIZATION | 752 | 11 | 12187 | 72 | Lhx1,Gbx1,Nr4a3,P2ry12,Map4,Ncam1,Bdnf,Ugt8a,Lrrc55,Plp1,Ccdc13 | | 4.399e-03 | -5.43 | positive regulation of synaptic transmission | biological process | GO:0050806 | 184 | 5 | 13711 | 80 | Lrrtm2,Rgs4,Crh,Car2,Cacng2 | | 4.420e-03 | -5.42 | GO\_INTRAMOLECULAR\_OXIDOREDUCTASE\_ACTIVITY\_TRANSPOSING\_S\_S\_BONDS | MSigDB lists | GO\_INTRAMOLECULAR\_OXIDOREDUCTASE\_ACTIVITY\_TRANSPOSING\_S\_S\_BONDS | 17 | 2 | 12187 | 72 | Tmx3,Pdia6 | | 4.420e-03 | -5.42 | GO\_RETINA\_LAYER\_FORMATION | MSigDB lists | GO\_RETINA\_LAYER\_FORMATION | 17 | 2 | 12187 | 72 | Fjx1,Lhx1 | | 4.420e-03 | -5.42 | KEGG\_PROXIMAL\_TUBULE\_BICARBONATE\_RECLAMATION | MSigDB lists | KEGG\_PROXIMAL\_TUBULE\_BICARBONATE\_RECLAMATION | 17 | 2 | 12187 | 72 | Car2,Glud1 | | 4.449e-03 | -5.42 | PMP22\_Claudin | pfam domains | PF00822 | 18 | 2 | 12881 | 72 | Cacng2,Cldn5 | | 4.486e-03 | -5.41 | regulation of metal ion transport | biological process | GO:0010959 | 353 | 7 | 13711 | 80 | P2ry12,Homer1,Rgs4,Nipsnap2,Agrn,Lrrc55,Crh | | 4.552e-03 | -5.39 | neuromuscular process | biological process | GO:0050905 | 115 | 4 | 13711 | 80 | Gbx1,Nr4a3,Aars,Fabp7 | | 4.567e-03 | -5.39 | positive regulation of ion transport | biological process | GO:0043270 | 266 | 6 | 13711 | 80 | Cacng2,Lrrc55,Crh,Nipsnap2,Homer1,P2ry12 | | 4.604e-03 | -5.38 | ROSS\_AML\_WITH\_PML\_RARA\_FUSION | MSigDB lists | ROSS\_AML\_WITH\_PML\_RARA\_FUSION | 57 | 3 | 12187 | 72 | Agrn,Map4,Ltk | | 4.604e-03 | -5.38 | REACTOME\_NCAM\_SIGNALING\_FOR\_NEURITE\_OUT\_GROWTH | MSigDB lists | REACTOME\_NCAM\_SIGNALING\_FOR\_NEURITE\_OUT\_GROWTH | 57 | 3 | 12187 | 72 | Col1a1,Agrn,Ncam1 | | 4.619e-03 | -5.38 | GO\_MODULATION\_OF\_SYNAPTIC\_TRANSMISSION | MSigDB lists | GO\_MODULATION\_OF\_SYNAPTIC\_TRANSMISSION | 264 | 6 | 12187 | 72 | Cacng2,Lrrtm2,Crh,Ncam1,Car2,Grm3 | | 4.678e-03 | -5.36 | regulation of apoptotic process | biological process | GO:0042981 | 1233 | 15 | 13711 | 80 | Aars,Bdnf,Qk,Kdr,Tgfa,Egr1,Ltk,Inhba,Gsdme,Lims2,Spry2,Csrnp3,Agrn,Foxq1,Nr4a3 | | 4.686e-03 | -5.36 | negative regulation of small GTPase mediated signal transduction | biological process | GO:0051058 | 58 | 3 | 13711 | 80 | Spry2,Timp2,Dgkz | | 4.686e-03 | -5.36 | regulation of neurotransmitter receptor activity | biological process | GO:0099601 | 58 | 3 | 13711 | 80 | Homer1,Crh,Cacng2 | | 4.700e-03 | -5.36 | KIM\_WT1\_TARGETS\_UP | MSigDB lists | KIM\_WT1\_TARGETS\_UP | 185 | 5 | 12187 | 72 | Inhba,Car2,Rgs4,Spry2,Egr1 | | 4.794e-03 | -5.34 | anatomical structure morphogenesis | biological process | GO:0009653 | 1746 | 19 | 13711 | 80 | Agrn,Ugt8a,Car2,Nr4a3,Foxq1,Ncam1,Inhba,Ermn,Atp5b,Gbx1,Col1a1,Spry2,Tgfa,Kdr,Fjx1,Bdnf,Ephb6,Lhx1,Qk | | 4.808e-03 | -5.34 | PAX\_Q6 | MSigDB lists | PAX\_Q6 | 186 | 5 | 12187 | 72 | Inhba,Amd1,Bdnf,Spry2,Ntm | | 4.835e-03 | -5.33 | LIN\_NPAS4\_TARGETS\_DN | MSigDB lists | LIN\_NPAS4\_TARGETS\_DN | 58 | 3 | 12187 | 72 | Rgs4,Egr1,Bdnf | | 4.835e-03 | -5.33 | JECHLINGER\_EPITHELIAL\_TO\_MESENCHYMAL\_TRANSITION\_DN | MSigDB lists | JECHLINGER\_EPITHELIAL\_TO\_MESENCHYMAL\_TRANSITION\_DN | 58 | 3 | 12187 | 72 | Egr1,Amd1,Car2 | | 4.917e-03 | -5.32 | steroid biosynthetic process | biological process | GO:0006694 | 59 | 3 | 13711 | 80 | Msmo1,Crh,Hsd17b12 | | 4.954e-03 | -5.31 | REACTOME\_ADP\_SIGNALLING\_THROUGH\_P2RY12 | MSigDB lists | REACTOME\_ADP\_SIGNALLING\_THROUGH\_P2RY12 | 18 | 2 | 12187 | 72 | P2ry12,Gng11 | | 4.954e-03 | -5.31 | GNF2\_TIMP2 | MSigDB lists | GNF2\_TIMP2 | 18 | 2 | 12187 | 72 | Crh,Timp2 | | 4.955e-03 | -5.31 | Sulfotransfer\_1 | pfam domains | PF00685 | 19 | 2 | 12881 | 72 | Ndst4,Hs3st1 | | 5.025e-03 | -5.29 | G protein-coupled receptor signaling pathway | biological process | GO:0007186 | 456 | 8 | 13711 | 80 | Rgs4,Homer1,Car2,Grm3,Gng11,Dgkz,Bdnf,P2ry12 | | 5.025e-03 | -5.29 | GSE36392\_EOSINOPHIL\_VS\_MAC\_IL25\_TREATED\_LUNG\_UP | MSigDB lists | GSE36392\_EOSINOPHIL\_VS\_MAC\_IL25\_TREATED\_LUNG\_UP | 117 | 4 | 12187 | 72 | Cacng2,Fjx1,Plp1,Rgs4 | | 5.029e-03 | -5.29 | TEF\_Q6 | MSigDB lists | TEF\_Q6 | 188 | 5 | 12187 | 72 | Amd1,Spry2,Ddx17,Fnbp1,Tle4 | | 5.073e-03 | -5.28 | SHEPARD\_BMYB\_TARGETS | MSigDB lists | SHEPARD\_BMYB\_TARGETS | 59 | 3 | 12187 | 72 | Plp1,Ddx17,Lhx1 | | 5.122e-03 | -5.27 | GO\_RESPONSE\_TO\_ENDOGENOUS\_STIMULUS | MSigDB lists | GO\_RESPONSE\_TO\_ENDOGENOUS\_STIMULUS | 1115 | 14 | 12187 | 72 | Crh,Lhx1,Homer1,Col1a1,Car2,Gng11,Timp2,Inhba,Egr1,P2ry12,Cldn5,Nr4a3,Aars,Qdpr | | 5.143e-03 | -5.27 | ATGCTGC\_MIR103\_MIR107 | MSigDB lists | ATGCTGC\_MIR103\_MIR107 | 189 | 5 | 12187 | 72 | Rgs4,Bdnf,Glud1,Tle4,Lrrc55 | | 5.173e-03 | -5.26 | GO\_POSITIVE\_REGULATION\_OF\_NERVOUS\_SYSTEM\_DEVELOPMENT | MSigDB lists | GO\_POSITIVE\_REGULATION\_OF\_NERVOUS\_SYSTEM\_DEVELOPMENT | 359 | 7 | 12187 | 72 | Bdnf,Lhx1,Lrrtm2,Cbln2,Agrn,Ltk,Timp2 | | 5.204e-03 | -5.26 | Sulfotransferase\_dom | interpro domains | IPR000863 | 19 | 2 | 13788 | 79 | Hs3st1,Ndst4 | | 5.204e-03 | -5.26 | Sc\_DH/Rdtase\_CS | interpro domains | IPR020904 | 19 | 2 | 13788 | 79 | Hsd17b12,Qdpr | | 5.258e-03 | -5.25 | NFY\_C | MSigDB lists | NFY\_C | 190 | 5 | 12187 | 72 | Lhx1,Ncam1,Tle4,Col1a1,Dgkz | | 5.266e-03 | -5.25 | - | gene3d domains | 4.10.81.20 | 1 | 1 | 6647 | 35 | Kcnmb2 | | 5.266e-03 | -5.25 | - | gene3d domains | 2.30.140.10 | 1 | 1 | 6647 | 35 | Srm | | 5.317e-03 | -5.24 | cell development | biological process | GO:0048468 | 1375 | 16 | 13711 | 80 | Agrn,Ugt8a,Homer1,Nr4a3,Inhba,Ncam1,Gbx1,Kdr,Plp1,Map4,Nrep,Bdnf,Ephb6,Lhx1,Hba-a2,Qk | | 5.317e-03 | -5.24 | MODY\_HIPPOCAMPUS\_POSTNATAL | MSigDB lists | MODY\_HIPPOCAMPUS\_POSTNATAL | 60 | 3 | 12187 | 72 | Dgkz,Egr1,Bdnf | | 5.317e-03 | -5.24 | YAO\_HOXA10\_TARGETS\_VIA\_PROGESTERONE\_UP | MSigDB lists | YAO\_HOXA10\_TARGETS\_VIA\_PROGESTERONE\_UP | 60 | 3 | 12187 | 72 | Cldn5,Ncam1,Peg3 | | 5.317e-03 | -5.24 | HELLEBREKERS\_SILENCED\_DURING\_TUMOR\_ANGIOGENESIS | MSigDB lists | HELLEBREKERS\_SILENCED\_DURING\_TUMOR\_ANGIOGENESIS | 60 | 3 | 12187 | 72 | Tgfa,Inhba,Bdnf | | 5.335e-03 | -5.23 | GO\_MONOCARBOXYLIC\_ACID\_BIOSYNTHETIC\_PROCESS | MSigDB lists | GO\_MONOCARBOXYLIC\_ACID\_BIOSYNTHETIC\_PROCESS | 119 | 4 | 12187 | 72 | Qk,Plp1,Msmo1,Hsd17b12 | | 5.375e-03 | -5.23 | PEA3\_Q6 | MSigDB lists | PEA3\_Q6 | 191 | 5 | 12187 | 72 | Sorl1,Arhgap20,Ncam1,Bdnf,Dgkz | | 5.375e-03 | -5.23 | NFY\_Q6 | MSigDB lists | NFY\_Q6 | 191 | 5 | 12187 | 72 | Dgkz,Col1a1,Tle4,Lhx1,Spry2 | | 5.391e-03 | -5.22 | negative regulation of receptor-mediated endocytosis | biological process | GO:0048261 | 19 | 2 | 13711 | 80 | Ubqln2,Lrrtm2 | | 5.398e-03 | -5.22 | regulation of cellular pH | biological process | GO:0030641 | 61 | 3 | 13711 | 80 | Atp5b,Tmem165,Car2 | | 5.429e-03 | -5.22 | regulation of programmed cell death | biological process | GO:0043067 | 1253 | 15 | 13711 | 80 | Aars,Bdnf,Qk,Tgfa,Kdr,Egr1,Inhba,Ltk,Lims2,Gsdme,Spry2,Csrnp3,Agrn,Foxq1,Nr4a3 | | 5.455e-03 | -5.21 | GO\_REGULATION\_OF\_CELL\_DIFFERENTIATION | MSigDB lists | GO\_REGULATION\_OF\_CELL\_DIFFERENTIATION | 1123 | 14 | 12187 | 72 | Cldn5,P2ry12,Bdnf,Arntl,Ltk,Kdr,Lhx1,Spry2,Timp2,Inhba,Car2,Col1a1,Sorl1,Ddx17 | | 5.494e-03 | -5.20 | GO\_ORGANIC\_ACID\_BIOSYNTHETIC\_PROCESS | MSigDB lists | GO\_ORGANIC\_ACID\_BIOSYNTHETIC\_PROCESS | 192 | 5 | 12187 | 72 | Glud1,Qk,Plp1,Hsd17b12,Msmo1 | | 5.495e-03 | -5.20 | GSE25846\_IL10\_POS\_VS\_NEG\_CD8\_TCELL\_DAY7\_POST\_CORONAVIRUS\_BRAIN\_UP | MSigDB lists | GSE25846\_IL10\_POS\_VS\_NEG\_CD8\_TCELL\_DAY7\_POST\_CORONAVIRUS\_BRAIN\_UP | 120 | 4 | 12187 | 72 | Car2,Inhba,Homer1,Ltk | | 5.513e-03 | -5.20 | positive regulation of response to stimulus | biological process | GO:0048584 | 1637 | 18 | 13711 | 80 | Tgfa,Kdr,Timp2,Crh,P2ry12,Bdnf,Arntl,Alkal2,Cacng2,Rgs4,Nr4a3,Inhba,Ncam1,Lims2,Gsdme,Col1a1,Spry2,Ubqln2 | | 5.516e-03 | -5.20 | CHIARETTI\_T\_ALL\_REFRACTORY\_TO\_THERAPY | MSigDB lists | CHIARETTI\_T\_ALL\_REFRACTORY\_TO\_THERAPY | 19 | 2 | 12187 | 72 | H1f0,Ephb6 | | 5.516e-03 | -5.20 | KIM\_ALL\_DISORDERS\_OLIGODENDROCYTE\_NUMBER\_CORR\_DN | MSigDB lists | KIM\_ALL\_DISORDERS\_OLIGODENDROCYTE\_NUMBER\_CORR\_DN | 19 | 2 | 12187 | 72 | Ntm,Csrnp3 | | 5.516e-03 | -5.20 | GNF2\_KISS1 | MSigDB lists | GNF2\_KISS1 | 19 | 2 | 12187 | 72 | Timp2,Crh | | 5.565e-03 | -5.19 | mouse chr2 C1.1\|2 | chromosome location | mouse chr2 C1.1\|2 | 1 | 1 | 14556 | 81 | Ermn | | 5.565e-03 | -5.19 | mouse chr5 A3\|5 11.93 cM | chromosome location | mouse chr5 A3\|5 11.93 cM | 1 | 1 | 14556 | 81 | Gbx1 | | 5.565e-03 | -5.19 | mouse chr4 E2\|4 88.55 cM | chromosome location | mouse chr4 E2\|4 88.55 cM | 1 | 1 | 14556 | 81 | Agrn | | 5.565e-03 | -5.19 | mouse chr6 D1\|6 37.62 cM | chromosome location | mouse chr6 D1\|6 37.62 cM | 1 | 1 | 14556 | 81 | Tgfa | | 5.565e-03 | -5.19 | mouse chr11 A4\|11 18.87 cM | chromosome location | mouse chr11 A4\|11 18.87 cM | 1 | 1 | 14556 | 81 | Hba-a2 | | 5.565e-03 | -5.19 | mouse chr13 A3.2\|13 13.46 cM | chromosome location | mouse chr13 A3.2\|13 13.46 cM | 1 | 1 | 14556 | 81 | Foxq1 | | 5.565e-03 | -5.19 | mouse chr7 A1\|7 3.89 cM | chromosome location | mouse chr7 A1\|7 3.89 cM | 1 | 1 | 14556 | 81 | Peg3 | | 5.565e-03 | -5.19 | mouse chr18 E4\|18 58.63 cM | chromosome location | mouse chr18 E4\|18 58.63 cM | 1 | 1 | 14556 | 81 | Cbln2 | | 5.565e-03 | -5.19 | mouse chrX F1\|X 59.1 cM | chromosome location | mouse chrX F1\|X 59.1 cM | 1 | 1 | 14556 | 81 | Plp1 | | 5.565e-03 | -5.19 | mouse chr14 E2.3\|14 56.16 cM | chromosome location | mouse chr14 E2.3\|14 56.16 cM | 1 | 1 | 14556 | 81 | Spry2 | | 5.565e-03 | -5.19 | mouse chr10 B1\|10 21.85 cM | chromosome location | mouse chr10 B1\|10 21.85 cM | 1 | 1 | 14556 | 81 | Amd1 | | 5.565e-03 | -5.19 | mouse chr18 B1\|18 18.76 cM | chromosome location | mouse chr18 B1\|18 18.76 cM | 1 | 1 | 14556 | 81 | Egr1 | | 5.565e-03 | -5.19 | mouse chr3 A2\|3 5.75 cM | chromosome location | mouse chr3 A2\|3 5.75 cM | 1 | 1 | 14556 | 81 | Crh | | 5.565e-03 | -5.19 | mouse chr11 C\|11 51.31 cM | chromosome location | mouse chr11 C\|11 51.31 cM | 1 | 1 | 14556 | 81 | Lhx1 | | 5.565e-03 | -5.19 | mouse chr5 C3.3\|5 40.56 cM | chromosome location | mouse chr5 C3.3\|5 40.56 cM | 1 | 1 | 14556 | 81 | Tmem165 | | 5.565e-03 | -5.19 | mouse chr9 A5.3\|9 26.83 cM | chromosome location | mouse chr9 A5.3\|9 26.83 cM | 1 | 1 | 14556 | 81 | Ncam1 | | 5.565e-03 | -5.19 | mouse chr9 F2\|9 59.83 cM | chromosome location | mouse chr9 F2\|9 59.83 cM | 1 | 1 | 14556 | 81 | Map4 | | 5.565e-03 | -5.19 | mouse chr2 E3\|2 56.63 cM | chromosome location | mouse chr2 E3\|2 56.63 cM | 1 | 1 | 14556 | 81 | Bdnf | | 5.565e-03 | -5.19 | mouse chr11 E2\|11 83.09 cM | chromosome location | mouse chr11 E2\|11 83.09 cM | 1 | 1 | 14556 | 81 | Timp2 | | 5.565e-03 | -5.19 | mouse chr17 A1\|17 7.75 cM | chromosome location | mouse chr17 A1\|17 7.75 cM | 1 | 1 | 14556 | 81 | Qk | | 5.565e-03 | -5.19 | mouse chr3 A1\|3 3.23 cM | chromosome location | mouse chr3 A1\|3 3.23 cM | 1 | 1 | 14556 | 81 | Car2 | | 5.565e-03 | -5.19 | mouse chr19 A\|19 9.11 cM | chromosome location | mouse chr19 A\|19 9.11 cM | 1 | 1 | 14556 | 81 | Tle4 | | 5.565e-03 | -5.19 | mouse chr13 A1\|13 5.85 cM | chromosome location | mouse chr13 A1\|13 5.85 cM | 1 | 1 | 14556 | 81 | Inhba | | 5.565e-03 | -5.19 | mouse chr11 A1\|11 3.94 cM | chromosome location | mouse chr11 A1\|11 3.94 cM | 1 | 1 | 14556 | 81 | Ddx56 | | 5.565e-03 | -5.19 | mouse chr5 C3.3\|5 40.23 cM | chromosome location | mouse chr5 C3.3\|5 40.23 cM | 1 | 1 | 14556 | 81 | Kdr | | 5.565e-03 | -5.19 | mouse chr12 A2\|12 | chromosome location | mouse chr12 A2\|12 | 1 | 1 | 14556 | 81 | Alkal2 | | 5.565e-03 | -5.19 | mouse chr5 B3\|5 24.9 cM | chromosome location | mouse chr5 B3\|5 24.9 cM | 1 | 1 | 14556 | 81 | Qdpr | | 5.565e-03 | -5.19 | mouse chr5 B3\|5 21.14 cM | chromosome location | mouse chr5 B3\|5 21.14 cM | 1 | 1 | 14556 | 81 | Hs3st1 | | 5.565e-03 | -5.19 | mouse chr7 F1\|7 59.17 cM | chromosome location | mouse chr7 F1\|7 59.17 cM | 1 | 1 | 14556 | 81 | Arntl | | 5.569e-03 | -5.19 | LANDIS\_BREAST\_CANCER\_PROGRESSION\_DN | MSigDB lists | LANDIS\_BREAST\_CANCER\_PROGRESSION\_DN | 61 | 3 | 12187 | 72 | Map4,Qk,Col1a1 | | 5.569e-03 | -5.19 | BAE\_BRCA1\_TARGETS\_UP | MSigDB lists | BAE\_BRCA1\_TARGETS\_UP | 61 | 3 | 12187 | 72 | Aars,Sorl1,Msmo1 | | 5.590e-03 | -5.19 | Spermine\_synt\_N | pfam domains | PF17284 | 1 | 1 | 12881 | 72 | Srm | | 5.590e-03 | -5.19 | FAM150 | pfam domains | PF15129 | 1 | 1 | 12881 | 72 | Alkal2 | | 5.590e-03 | -5.19 | Alveol-reg\_P311 | pfam domains | PF11092 | 1 | 1 | 12881 | 72 | Nrep | | 5.590e-03 | -5.19 | KcnmB2\_inactiv | pfam domains | PF09303 | 1 | 1 | 12881 | 72 | Kcnmb2 | | 5.590e-03 | -5.19 | Quaking\_NLS | pfam domains | PF16551 | 1 | 1 | 12881 | 72 | Qk | | 5.590e-03 | -5.19 | NtA | pfam domains | PF03146 | 1 | 1 | 12881 | 72 | Agrn | | 5.590e-03 | -5.19 | ELFV\_dehydrog | pfam domains | PF00208 | 1 | 1 | 12881 | 72 | Glud1 | | 5.590e-03 | -5.19 | UDPGT | pfam domains | PF00201 | 1 | 1 | 12881 | 72 | Ugt8a | | 5.590e-03 | -5.19 | ELFV\_dehydrog\_N | pfam domains | PF02812 | 1 | 1 | 12881 | 72 | Glud1 | | 5.590e-03 | -5.19 | STAR\_dimer | pfam domains | PF16544 | 1 | 1 | 12881 | 72 | Qk | | 5.590e-03 | -5.19 | UPF0016 | pfam domains | PF01169 | 1 | 1 | 12881 | 72 | Tmem165 | | 5.590e-03 | -5.19 | DUF3432 | pfam domains | PF11914 | 1 | 1 | 12881 | 72 | Egr1 | | 5.590e-03 | -5.19 | DHHA1 | pfam domains | PF02272 | 1 | 1 | 12881 | 72 | Aars | | 5.614e-03 | -5.18 | FOXO4\_02 | MSigDB lists | FOXO4\_02 | 193 | 5 | 12187 | 72 | Crh,Nap1l5,Csrnp3,Hs3st1,Bdnf | | 5.658e-03 | -5.17 | KRAS.KIDNEY\_UP.V1\_UP | MSigDB lists | KRAS.KIDNEY\_UP.V1\_UP | 121 | 4 | 12187 | 72 | Plp1,Car2,Peg3,Sorl1 | | 5.714e-03 | -5.16 | activin complex | cellular component | GO:0048180 | 1 | 1 | 13825 | 79 | Inhba | | 5.714e-03 | -5.16 | activin A complex | cellular component | GO:0043509 | 1 | 1 | 13825 | 79 | Inhba | | 5.730e-03 | -5.16 | Glu/Leu/Phe/Val\_DH | interpro domains | IPR006095 | 1 | 1 | 13788 | 79 | Glud1 | | 5.730e-03 | -5.16 | Spermi\_synthase\_euk | interpro domains | IPR030668 | 1 | 1 | 13788 | 79 | Srm | | 5.730e-03 | -5.16 | UDP\_glucos\_trans | interpro domains | IPR002213 | 1 | 1 | 13788 | 79 | Ugt8a | | 5.730e-03 | -5.16 | Quaking\_NLS | interpro domains | IPR032367 | 1 | 1 | 13788 | 79 | Qk | | 5.730e-03 | -5.16 | TIMP2 | interpro domains | IPR015613 | 1 | 1 | 13788 | 79 | Timp2 | | 5.730e-03 | -5.16 | DHHA1\_dom | interpro domains | IPR003156 | 1 | 1 | 13788 | 79 | Aars | | 5.730e-03 | -5.16 | Glu/Leu/Phe/Val\_DH\_AS | interpro domains | IPR033524 | 1 | 1 | 13788 | 79 | Glud1 | | 5.730e-03 | -5.16 | NAD\_bind\_Glu\_DH | interpro domains | IPR033922 | 1 | 1 | 13788 | 79 | Glud1 | | 5.730e-03 | -5.16 | GPCR\_3\_mtglu\_rcpt\_3 | interpro domains | IPR001234 | 1 | 1 | 13788 | 79 | Grm3 | | 5.730e-03 | -5.16 | NACAD\_UBA | interpro domains | IPR041907 | 1 | 1 | 13788 | 79 | Nacad | | 5.730e-03 | -5.16 | SPRY2 | interpro domains | IPR030780 | 1 | 1 | 13788 | 79 | Spry2 | | 5.730e-03 | -5.16 | VEGFR2\_rcpt | interpro domains | IPR009136 | 1 | 1 | 13788 | 79 | Kdr | | 5.730e-03 | -5.16 | POM121 | interpro domains | IPR026090 | 1 | 1 | 13788 | 79 | Pom121 | | 5.730e-03 | -5.16 | EGR1\_C | interpro domains | IPR021839 | 1 | 1 | 13788 | 79 | Egr1 | | 5.730e-03 | -5.16 | FAM150A/B | interpro domains | IPR029364 | 1 | 1 | 13788 | 79 | Alkal2 | | 5.730e-03 | -5.16 | P2Y12\_rcpt | interpro domains | IPR005394 | 1 | 1 | 13788 | 79 | P2ry12 | | 5.730e-03 | -5.16 | UDP\_glycos\_trans\_CS | interpro domains | IPR035595 | 1 | 1 | 13788 | 79 | Ugt8a | | 5.730e-03 | -5.16 | Urocortin\_CRF | interpro domains | IPR003620 | 1 | 1 | 13788 | 79 | Crh | | 5.730e-03 | -5.16 | Spermi\_synthase | interpro domains | IPR001045 | 1 | 1 | 13788 | 79 | Srm | | 5.730e-03 | -5.16 | Ncam1 | interpro domains | IPR033019 | 1 | 1 | 13788 | 79 | Ncam1 | | 5.730e-03 | -5.16 | VDCC\_g2su | interpro domains | IPR005422 | 1 | 1 | 13788 | 79 | Cacng2 | | 5.730e-03 | -5.16 | Spermidine\_synt\_N | interpro domains | IPR035246 | 1 | 1 | 13788 | 79 | Srm | | 5.730e-03 | -5.16 | RGS4 | interpro domains | IPR034952 | 1 | 1 | 13788 | 79 | Rgs4 | | 5.730e-03 | -5.16 | C11orf87 | interpro domains | IPR037670 | 1 | 1 | 13788 | 79 | AI593442 | | 5.730e-03 | -5.16 | RGS\_RGS4 | interpro domains | IPR034953 | 1 | 1 | 13788 | 79 | Rgs4 | | 5.730e-03 | -5.16 | FNBP1/FBP17 | interpro domains | IPR028532 | 1 | 1 | 13788 | 79 | Fnbp1 | | 5.730e-03 | -5.16 | Spermidine\_synt\_N\_sf | interpro domains | IPR037163 | 1 | 1 | 13788 | 79 | Srm | | 5.730e-03 | -5.16 | Corticotropin-releasing\_fac\_CS | interpro domains | IPR018446 | 1 | 1 | 13788 | 79 | Crh | | 5.730e-03 | -5.16 | Ubiquilin-2 | interpro domains | IPR028430 | 1 | 1 | 13788 | 79 | Ubqln2 | | 5.730e-03 | -5.16 | FNBP1\_F-BAR | interpro domains | IPR037449 | 1 | 1 | 13788 | 79 | Fnbp1 | | 5.730e-03 | -5.16 | Glu/Leu/Phe/Val\_DH\_C | interpro domains | IPR006096 | 1 | 1 | 13788 | 79 | Glud1 | | 5.730e-03 | -5.16 | NtA\_dom | interpro domains | IPR004850 | 1 | 1 | 13788 | 79 | Agrn | | 5.730e-03 | -5.16 | Inhibin\_betaA | interpro domains | IPR000491 | 1 | 1 | 13788 | 79 | Inhba | | 5.730e-03 | -5.16 | FacI\_MAC | interpro domains | IPR003884 | 1 | 1 | 13788 | 79 | Agrn | | 5.730e-03 | -5.16 | CCDC13 | interpro domains | IPR038929 | 1 | 1 | 13788 | 79 | Ccdc13 | | 5.730e-03 | -5.16 | ATP\_synth\_F1\_bsu | interpro domains | IPR005722 | 1 | 1 | 13788 | 79 | Atp5b | | 5.730e-03 | -5.16 | Gdt1 | interpro domains | IPR001727 | 1 | 1 | 13788 | 79 | Tmem165 | | 5.730e-03 | -5.16 | STAR\_dimer | interpro domains | IPR032377 | 1 | 1 | 13788 | 79 | Qk | | 5.730e-03 | -5.16 | PABS\_CS | interpro domains | IPR030373 | 1 | 1 | 13788 | 79 | Srm | | 5.730e-03 | -5.16 | KCNMB2\_ball\_chain\_dom | interpro domains | IPR015382 | 1 | 1 | 13788 | 79 | Kcnmb2 | | 5.730e-03 | -5.16 | FJX1/FJ | interpro domains | IPR024868 | 1 | 1 | 13788 | 79 | Fjx1 | | 5.730e-03 | -5.16 | FNBP1\_SH3 | interpro domains | IPR035492 | 1 | 1 | 13788 | 79 | Fnbp1 | | 5.730e-03 | -5.16 | Neuronal\_3 | interpro domains | IPR024417 | 1 | 1 | 13788 | 79 | Nrep | | 5.730e-03 | -5.16 | KCNMB2\_ball/chain\_dom\_sf | interpro domains | IPR037096 | 1 | 1 | 13788 | 79 | Kcnmb2 | | 5.730e-03 | -5.16 | Claudin5 | interpro domains | IPR003551 | 1 | 1 | 13788 | 79 | Cldn5 | | 5.730e-03 | -5.16 | Brain-der\_neurotrophic\_factor | interpro domains | IPR020430 | 1 | 1 | 13788 | 79 | Bdnf | | 5.730e-03 | -5.16 | GBX-1 | interpro domains | IPR031251 | 1 | 1 | 13788 | 79 | Gbx1 | | 5.730e-03 | -5.16 | GSDME | interpro domains | IPR042377 | 1 | 1 | 13788 | 79 | Gsdme | | 5.730e-03 | -5.16 | NOR1\_rcpt | interpro domains | IPR003072 | 1 | 1 | 13788 | 79 | Nr4a3 | | 5.730e-03 | -5.16 | Glu/Leu/Phe/Val\_DH\_dimer\_dom | interpro domains | IPR006097 | 1 | 1 | 13788 | 79 | Glud1 | | 5.730e-03 | -5.16 | PABS | interpro domains | IPR030374 | 1 | 1 | 13788 | 79 | Srm | | 5.737e-03 | -5.16 | RRAGTTGT\_UNKNOWN | MSigDB lists | RRAGTTGT\_UNKNOWN | 194 | 5 | 12187 | 72 | Nap1l5,Nr4a3,Lhx1,Bdnf,AI593442 | | 5.760e-03 | -5.16 | TIMP-like\_OB-fold | interpro domains | IPR008993 | 20 | 2 | 13788 | 79 | Timp2,Agrn | | 5.771e-03 | -5.15 | corticotropin-releasing hormone activity | molecular function | GO:0017045 | 1 | 1 | 13516 | 78 | Crh | | 5.771e-03 | -5.15 | glutamate dehydrogenase (NAD+) activity | molecular function | GO:0004352 | 1 | 1 | 13516 | 78 | Glud1 | | 5.771e-03 | -5.15 | spermidine synthase activity | molecular function | GO:0004766 | 1 | 1 | 13516 | 78 | Srm | | 5.771e-03 | -5.15 | 3-oxo-behenoyl-CoA reductase activity | molecular function | GO:0102340 | 1 | 1 | 13516 | 78 | Hsd17b12 | | 5.771e-03 | -5.15 | 3-oxo-arachidoyl-CoA reductase activity | molecular function | GO:0102339 | 1 | 1 | 13516 | 78 | Hsd17b12 | | 5.771e-03 | -5.15 | N-acylsphingosine galactosyltransferase activity | molecular function | GO:0047263 | 1 | 1 | 13516 | 78 | Ugt8a | | 5.771e-03 | -5.15 | Ser-tRNA(Ala) hydrolase activity | molecular function | GO:0002196 | 1 | 1 | 13516 | 78 | Aars | | 5.771e-03 | -5.15 | 3-oxo-lignoceroyl-CoA reductase activity | molecular function | GO:0102341 | 1 | 1 | 13516 | 78 | Hsd17b12 | | 5.771e-03 | -5.15 | glutamate dehydrogenase [NAD(P)+] activity | molecular function | GO:0004353 | 1 | 1 | 13516 | 78 | Glud1 | | 5.771e-03 | -5.15 | 3-oxo-cerotoyl-CoA reductase activity | molecular function | GO:0102342 | 1 | 1 | 13516 | 78 | Hsd17b12 | | 5.771e-03 | -5.15 | oxidoreductase activity, acting on the CH-NH2 group of donors, NAD or NADP as acceptor | molecular function | GO:0016639 | 1 | 1 | 13516 | 78 | Glud1 | | 5.771e-03 | -5.15 | 6,7-dihydropteridine reductase activity | molecular function | GO:0004155 | 1 | 1 | 13516 | 78 | Qdpr | | 5.821e-03 | -5.15 | GENTILE\_UV\_HIGH\_DOSE\_DN | MSigDB lists | GENTILE\_UV\_HIGH\_DOSE\_DN | 277 | 6 | 12187 | 72 | Qk,Pom121,Bdnf,Tle4,Amd1,Ddx17 | | 5.824e-03 | -5.15 | CHIANG\_LIVER\_CANCER\_SUBCLASS\_CTNNB1\_DN | MSigDB lists | CHIANG\_LIVER\_CANCER\_SUBCLASS\_CTNNB1\_DN | 122 | 4 | 12187 | 72 | Timp2,Col1a1,Egr1,Homer1 | | 5.835e-03 | -5.14 | positive regulation of mast cell activation by Fc-epsilon receptor signaling pathway | biological process | GO:0038097 | 1 | 1 | 13711 | 80 | Nr4a3 | | 5.835e-03 | -5.14 | type IV hypersensitivity | biological process | GO:0001806 | 1 | 1 | 13711 | 80 | Ephb6 | | 5.835e-03 | -5.14 | regulation of metanephric glomerular mesangial cell proliferation | biological process | GO:0072301 | 1 | 1 | 13711 | 80 | Egr1 | | 5.835e-03 | -5.14 | taste bud development | biological process | GO:0061193 | 1 | 1 | 13711 | 80 | Bdnf | | 5.835e-03 | -5.14 | regulation of 1-phosphatidylinositol-4-phosphate 5-kinase activity | biological process | GO:0090215 | 1 | 1 | 13711 | 80 | Dgkz | | 5.835e-03 | -5.14 | positive regulation of 1-phosphatidylinositol-4-phosphate 5-kinase activity | biological process | GO:0090216 | 1 | 1 | 13711 | 80 | Dgkz | | 5.835e-03 | -5.14 | positive regulation of early endosome to recycling endosome transport | biological process | GO:1902955 | 1 | 1 | 13711 | 80 | Sorl1 | | 5.835e-03 | -5.14 | retrograde trans-synaptic signaling by neuropeptide | biological process | GO:0099082 | 1 | 1 | 13711 | 80 | Bdnf | | 5.835e-03 | -5.14 | metanephric S-shaped body morphogenesis | biological process | GO:0072284 | 1 | 1 | 13711 | 80 | Lhx1 | | 5.835e-03 | -5.14 | positive regulation of protein localization to basolateral plasma membrane | biological process | GO:1904510 | 1 | 1 | 13711 | 80 | Cacng2 | | 5.835e-03 | -5.14 | negative regulation of sodium ion export across plasma membrane | biological process | GO:1903277 | 1 | 1 | 13711 | 80 | Agrn | | 5.835e-03 | -5.14 | regulation of protein localization to basolateral plasma membrane | biological process | GO:1904508 | 1 | 1 | 13711 | 80 | Cacng2 | | 5.835e-03 | -5.14 | nephric duct elongation | biological process | GO:0035849 | 1 | 1 | 13711 | 80 | Lhx1 | | 5.835e-03 | -5.14 | positive regulation of integrin activation by cell surface receptor linked signal transduction | biological process | GO:0033626 | 1 | 1 | 13711 | 80 | P2ry12 | | 5.835e-03 | -5.14 | regulation of choline O-acetyltransferase activity | biological process | GO:1902769 | 1 | 1 | 13711 | 80 | Sorl1 | | 5.835e-03 | -5.14 | regulation of cytoplasmic translational fidelity | biological process | GO:0140018 | 1 | 1 | 13711 | 80 | Aars | | 5.835e-03 | -5.14 | positive regulation of choline O-acetyltransferase activity | biological process | GO:1902771 | 1 | 1 | 13711 | 80 | Sorl1 | | 5.835e-03 | -5.14 | initiation of movement involved in cerebral cortex radial glia guided migration | biological process | GO:0021806 | 1 | 1 | 13711 | 80 | P2ry12 | | 5.835e-03 | -5.14 | cytosolic calcium signaling involved in initiation of cell movement in glial-mediated radial cell migration | biological process | GO:0021808 | 1 | 1 | 13711 | 80 | P2ry12 | | 5.835e-03 | -5.14 | negative regulation of neurofibrillary tangle assembly | biological process | GO:1902997 | 1 | 1 | 13711 | 80 | Sorl1 | | 5.835e-03 | -5.14 | retrograde trans-synaptic signaling by neuropeptide, modulating synaptic transmission | biological process | GO:0099083 | 1 | 1 | 13711 | 80 | Bdnf | | 5.835e-03 | -5.14 | positive regulation of ERK5 cascade | biological process | GO:0070378 | 1 | 1 | 13711 | 80 | Alkal2 | | 5.835e-03 | -5.14 | horizontal cell localization | biological process | GO:0035852 | 1 | 1 | 13711 | 80 | Lhx1 | | 5.835e-03 | -5.14 | negative regulation of sodium:potassium-exchanging ATPase activity | biological process | GO:1903407 | 1 | 1 | 13711 | 80 | Agrn | | 5.835e-03 | -5.14 | positive regulation of glomerular metanephric mesangial cell proliferation | biological process | GO:0072303 | 1 | 1 | 13711 | 80 | Egr1 | | 5.835e-03 | -5.14 | pyruvate oxidation | biological process | GO:0009444 | 1 | 1 | 13711 | 80 | Nr4a3 | | 5.846e-03 | -5.14 | cell communication | biological process | GO:0007154 | 3040 | 28 | 13711 | 80 | Gng11,Ltk,Inhba,Rgs4,Agrn,Nr4a3,Grm3,Ephb6,Lhx1,P2ry12,Kcnmb2,Kdr,Tgfa,Plp1,Tle4,Crh,Arhgap20,Ncam1,Col1a1,Homer1,Cldn5,Car2,Dgkz,Bdnf,Cacng2,Egr1,Ddx17,Fjx1 | | 5.859e-03 | -5.14 | positive regulation of transmembrane transport | biological process | GO:0034764 | 197 | 5 | 13711 | 80 | Car2,Nr4a3,Lrrc55,Cacng2,Nipsnap2 | | 5.909e-03 | -5.13 | GO\_RESPONSE\_TO\_INORGANIC\_SUBSTANCE | MSigDB lists | GO\_RESPONSE\_TO\_INORGANIC\_SUBSTANCE | 368 | 7 | 12187 | 72 | Nr4a3,Homer1,Qdpr,Car2,Col1a1,Hba-a2,Kcnmb2 | | 5.982e-03 | -5.12 | FIMAC | smart domains | SM00057 | 1 | 1 | 7188 | 43 | Agrn | | 5.982e-03 | -5.12 | ELFV\_dehydrog | smart domains | SM00839 | 1 | 1 | 7188 | 43 | Glud1 | | 5.982e-03 | -5.12 | CRF | smart domains | SM00039 | 1 | 1 | 7188 | 43 | Crh | | 5.987e-03 | -5.12 | ALPHACP1\_01 | MSigDB lists | ALPHACP1\_01 | 196 | 5 | 12187 | 72 | Spry2,Hba-a2,Dgkz,Col1a1,Hsd17b12 | | 6.035e-03 | -5.11 | head development | biological process | GO:0060322 | 573 | 9 | 13711 | 80 | Lhx1,Ncam1,Inhba,Aars,P2ry12,Col1a1,Gbx1,Fabp7,Nr4a3 | | 6.056e-03 | -5.11 | response to stimulus | biological process | GO:0050896 | 5062 | 41 | 13711 | 80 | Car2,Ccdc13,Homer1,Cldn5,2810403D21Rik,Spry2,Col1a1,Gbx1,Ncam1,Egr1,Ddx17,Scd2,Hba-a2,Arntl,Dgkz,Bdnf,Nr4a3,Grm3,Rgs4,Fabp7,Agrn,Ubqln2,Atp5b,Gng11,Gsdme,Inhba,Ltk,Nrep,Arhgap20,Crh,Tle4,Plp1,Kdr,Tgfa,Timp2,Srm,Glud1,Lhx1,Ephb6,Kcnmb2,P2ry12 | | 6.061e-03 | -5.11 | Ezh2 (enhancer of zeste 2 polycomb repressive complex 2 subunit) | protein interactions | 14056 | 52 | 3 | 6802 | 49 | Arntl,H1f0,Zfp518b | | 6.107e-03 | -5.10 | SMID\_BREAST\_CANCER\_BASAL\_UP | MSigDB lists | SMID\_BREAST\_CANCER\_BASAL\_UP | 467 | 8 | 12187 | 72 | Fabp7,Amd1,Peg3,Ddx17,Ugt8a,Ncam1,Hs3st1,Qk | | 6.121e-03 | -5.10 | cell morphogenesis involved in neuron differentiation | biological process | GO:0048667 | 374 | 7 | 13711 | 80 | Ephb6,Bdnf,Lhx1,Agrn,Ncam1,Nr4a3,Gbx1 | | 6.172e-03 | -5.09 | formation of primary germ layer | biological process | GO:0001704 | 64 | 3 | 13711 | 80 | Inhba,Lhx1,Nr4a3 | | 6.219e-03 | -5.08 | cell periphery | cellular component | GO:0071944 | 3578 | 31 | 13825 | 79 | Lrrtm2,Map4,Cldn5,Car2,Ntm,Fabp7,Gng11,Agrn,Ephb6,Mtmr9,Ubqln2,Pdia6,Homer1,Ltk,Fnbp1,Tgfa,Cacng2,Rgs4,Ncam1,Grm3,Spry2,Ermn,Atp5b,Gsdme,Kcnmb2,Lims2,Lrrc55,Kdr,Dgkz,P2ry12,Plp1 | | 6.246e-03 | -5.08 | WINTER\_HYPOXIA\_METAGENE | MSigDB lists | WINTER\_HYPOXIA\_METAGENE | 198 | 5 | 12187 | 72 | Egr1,Sorl1,Map4,Kdr,Tgfa | | 6.246e-03 | -5.08 | TTF1\_Q6 | MSigDB lists | TTF1\_Q6 | 198 | 5 | 12187 | 72 | Nr4a3,Cacng2,Crh,Map4,H1f0 | | 6.276e-03 | -5.07 | positive regulation of molecular function | biological process | GO:0044093 | 1399 | 16 | 13711 | 80 | Arhgap20,Tgfa,Timp2,Egr1,Cacng2,Bdnf,Dgkz,Ephb6,Lrrc55,H1f0,Agrn,Rgs4,Nipsnap2,Spry2,Sorl1,Mtmr9 | | 6.341e-03 | -5.06 | GSE37301\_LYMPHOID\_PRIMED\_MPP\_VS\_CD4\_TCELL\_UP | MSigDB lists | GSE37301\_LYMPHOID\_PRIMED\_MPP\_VS\_CD4\_TCELL\_UP | 125 | 4 | 12187 | 72 | Lrrtm2,Cacng2,Ddx17,Grm3 | | 6.341e-03 | -5.06 | ESC\_J1\_UP\_LATE.V1\_UP | MSigDB lists | ESC\_J1\_UP\_LATE.V1\_UP | 125 | 4 | 12187 | 72 | Hkdc1,Hs3st1,Foxq1,Peg3 | | 6.341e-03 | -5.06 | GO\_SKIN\_DEVELOPMENT | MSigDB lists | GO\_SKIN\_DEVELOPMENT | 125 | 4 | 12187 | 72 | Foxq1,Aars,Inhba,Col1a1 | | 6.358e-03 | -5.06 | synapse organization | biological process | GO:0050808 | 285 | 6 | 13711 | 80 | Lrrtm2,Homer1,Agrn,Dgkz,Bdnf,Cacng2 | | 6.366e-03 | -5.06 | GO\_NEGATIVE\_REGULATION\_OF\_MAP\_KINASE\_ACTIVITY | MSigDB lists | GO\_NEGATIVE\_REGULATION\_OF\_MAP\_KINASE\_ACTIVITY | 64 | 3 | 12187 | 72 | Spry2,Rgs4,Sorl1 | | 6.443e-03 | -5.04 | regulation of neuronal synaptic plasticity | biological process | GO:0048168 | 65 | 3 | 13711 | 80 | Kdr,Bdnf,Egr1 | | 6.482e-03 | -5.04 | GO\_SYSTEM\_PROCESS | MSigDB lists | GO\_SYSTEM\_PROCESS | 907 | 12 | 12187 | 72 | Spry2,Homer1,Aars,Crh,Cldn5,Cacng2,Nr4a3,Fabp7,Col1a1,Gbx1,Kcnmb2,Inhba | | 6.501e-03 | -5.04 | response to acid chemical | biological process | GO:0001101 | 202 | 5 | 13711 | 80 | Kdr,Ltk,Egr1,Col1a1,Scd2 | | 6.521e-03 | -5.03 | IL15\_UP.V1\_UP | MSigDB lists | IL15\_UP.V1\_UP | 126 | 4 | 12187 | 72 | Spry2,Hs3st1,Egr1,Amd1 | | 6.521e-03 | -5.03 | GSE40274\_CTRL\_VS\_FOXP3\_AND\_EOS\_TRANSDUCED\_ACTIVATED\_CD4\_TCELL\_DN | MSigDB lists | GSE40274\_CTRL\_VS\_FOXP3\_AND\_EOS\_TRANSDUCED\_ACTIVATED\_CD4\_TCELL\_DN | 126 | 4 | 12187 | 72 | Homer1,Fnbp1,Kdr,Srm | | 6.535e-03 | -5.03 | glutamatergic synapse | cellular component | GO:0098978 | 487 | 8 | 13825 | 79 | Ncam1,Grm3,Homer1,Dgkz,Agrn,Cacng2,Cbln2,Lrrtm2 | | 6.541e-03 | -5.03 | positive regulation of protein phosphorylation | biological process | GO:0001934 | 800 | 11 | 13711 | 80 | Spry2,Inhba,Gsdme,Agrn,Alkal2,Bdnf,Crh,Timp2,Tgfa,Kdr,Egr1 | | 6.570e-03 | -5.03 | regulation of hair cycle | biological process | GO:0042634 | 21 | 2 | 13711 | 80 | Arntl,Inhba | | 6.722e-03 | -5.00 | DORSEY\_GAB2\_TARGETS | MSigDB lists | DORSEY\_GAB2\_TARGETS | 21 | 2 | 12187 | 72 | Gng11,Col1a1 | | 6.722e-03 | -5.00 | MODULE\_343 | MSigDB lists | MODULE\_343 | 21 | 2 | 12187 | 72 | Glud1,Car2 | | 6.783e-03 | -4.99 | PABS\_1 | prosite domains | PS01330 | 1 | 1 | 8845 | 60 | Srm | | 6.783e-03 | -4.99 | GLFV\_DEHYDROGENASE | prosite domains | PS00074 | 1 | 1 | 8845 | 60 | Glud1 | | 6.783e-03 | -4.99 | PABS\_2 | prosite domains | PS51006 | 1 | 1 | 8845 | 60 | Srm | | 6.783e-03 | -4.99 | UDPGT | prosite domains | PS00375 | 1 | 1 | 8845 | 60 | Ugt8a | | 6.783e-03 | -4.99 | UPF0016 | prosite domains | PS01214 | 1 | 1 | 8845 | 60 | Tmem165 | | 6.783e-03 | -4.99 | CRF | prosite domains | PS00511 | 1 | 1 | 8845 | 60 | Crh | | 6.783e-03 | -4.99 | NTA | prosite domains | PS51121 | 1 | 1 | 8845 | 60 | Agrn | | 6.795e-03 | -4.99 | transmembrane receptor protein kinase activity | molecular function | GO:0019199 | 67 | 3 | 13516 | 78 | Kdr,Ephb6,Ltk | | 6.803e-03 | -4.99 | cell body | cellular component | GO:0044297 | 707 | 10 | 13825 | 79 | Crh,Ermn,Bdnf,Kdr,Fabp7,P2ry12,Sorl1,Ncam1,Homer1,Timp2 | | 6.889e-03 | -4.98 | GO\_SYNAPSE\_ORGANIZATION | MSigDB lists | GO\_SYNAPSE\_ORGANIZATION | 128 | 4 | 12187 | 72 | Lrrtm2,Cacng2,Agrn,Bdnf | | 6.889e-03 | -4.98 | DURAND\_STROMA\_NS\_UP | MSigDB lists | DURAND\_STROMA\_NS\_UP | 128 | 4 | 12187 | 72 | Hs3st1,Foxq1,Zfp518b,Tle4 | | 6.933e-03 | -4.97 | MODULE\_235 | MSigDB lists | MODULE\_235 | 66 | 3 | 12187 | 72 | Glud1,Qdpr,Car2 | | 6.933e-03 | -4.97 | DANG\_REGULATED\_BY\_MYC\_UP | MSigDB lists | DANG\_REGULATED\_BY\_MYC\_UP | 66 | 3 | 12187 | 72 | Homer1,Tle4,Srm | | 6.944e-03 | -4.97 | chemotaxis | biological process | GO:0006935 | 383 | 7 | 13711 | 80 | Gbx1,Ncam1,Lhx1,Bdnf,Ephb6,Nr4a3,Agrn | | 6.969e-03 | -4.97 | TGGAAA\_NFAT\_Q4\_01 | MSigDB lists | TGGAAA\_NFAT\_Q4\_01 | 1405 | 16 | 12187 | 72 | Arhgap20,Bdnf,Ndst4,Nr4a3,H1f0,Grm3,Dgkz,Homer1,Lhx1,Spry2,Cacng2,Ddx17,Amd1,Nap1l5,Inhba,Hsd17b12 | | 7.004e-03 | -4.96 | SCHUETZ\_BREAST\_CANCER\_DUCTAL\_INVASIVE\_UP | MSigDB lists | SCHUETZ\_BREAST\_CANCER\_DUCTAL\_INVASIVE\_UP | 288 | 6 | 12187 | 72 | Inhba,Tle4,Col1a1,Rgs4,Qk,Kdr | | 7.040e-03 | -4.96 | Metabolism | REACTOME pathways | R-MMU-1430728 | 1424 | 17 | 6297 | 42 | Agrn,Gng11,Car2,Msmo1,Qdpr,Pom121,Glud1,Hsd17b12,Hs3st1,Scd2,Hba-a2,Arntl,Srm,Amd1,Fabp7,Atp5b,Ugt8a | | 7.044e-03 | -4.96 | response to organic cyclic compound | biological process | GO:0014070 | 483 | 8 | 13711 | 80 | Homer1,Ddx17,Egr1,Crh,Nr4a3,Inhba,Ncam1,P2ry12 | | 7.078e-03 | -4.95 | GO\_STEROID\_METABOLIC\_PROCESS | MSigDB lists | GO\_STEROID\_METABOLIC\_PROCESS | 129 | 4 | 12187 | 72 | Sorl1,Msmo1,Hsd17b12,Crh | | 7.133e-03 | -4.94 | regulation of plasma membrane bounded cell projection organization | biological process | GO:0120035 | 697 | 10 | 13711 | 80 | Agrn,Ddx56,Homer1,Map4,Ntm,P2ry12,Ltk,Bdnf,Qk,Alkal2 | | 7.185e-03 | -4.94 | ADH\_SHORT | prosite domains | PS00061 | 19 | 2 | 8845 | 60 | Hsd17b12,Qdpr | | 7.200e-03 | -4.93 | negative regulation of potassium ion transmembrane transport | biological process | GO:1901380 | 22 | 2 | 13711 | 80 | Rgs4,Agrn | | 7.204e-03 | -4.93 | Dct (dopachrome tautomerase) | protein interactions | 13190 | 1 | 1 | 6802 | 49 | Tle4 | | 7.204e-03 | -4.93 | MARK1 (microtubule affinity regulating kinase 1) | protein interactions | 4139 | 1 | 1 | 6802 | 49 | Map4 | | 7.204e-03 | -4.93 | Cpe (carboxypeptidase E) | protein interactions | 12876 | 1 | 1 | 6802 | 49 | Bdnf | | 7.204e-03 | -4.93 | Optix (optix) | protein interactions | 44108 | 1 | 1 | 6802 | 49 | Tle4 | | 7.204e-03 | -4.93 | VEGFD (vascular endothelial growth factor D) | protein interactions | 2277 | 1 | 1 | 6802 | 49 | Kdr | | 7.204e-03 | -4.93 | SIRT4 (sirtuin 4) | protein interactions | 23409 | 1 | 1 | 6802 | 49 | Glud1 | | 7.204e-03 | -4.93 | MTMR7 (myotubularin related protein 7) | protein interactions | 9108 | 1 | 1 | 6802 | 49 | Mtmr9 | | 7.204e-03 | -4.93 | Acvr2b (activin receptor IIB) | protein interactions | 11481 | 1 | 1 | 6802 | 49 | Inhba | | 7.204e-03 | -4.93 | Sirt4 (sirtuin 4) | protein interactions | 75387 | 1 | 1 | 6802 | 49 | Glud1 | | 7.204e-03 | -4.93 | Vegfd (vascular endothelial growth factor D) | protein interactions | 14205 | 1 | 1 | 6802 | 49 | Kdr | | 7.204e-03 | -4.93 | Tesk1 (testis specific protein kinase 1) | protein interactions | 21754 | 1 | 1 | 6802 | 49 | Spry2 | | 7.212e-03 | -4.93 | WHN\_B | MSigDB lists | WHN\_B | 205 | 5 | 12187 | 72 | Nr4a3,Bdnf,Spry2,Ddx17,Amd1 | | 7.212e-03 | -4.93 | ALFANO\_MYC\_TARGETS | MSigDB lists | ALFANO\_MYC\_TARGETS | 205 | 5 | 12187 | 72 | Qdpr,Hs3st1,Ddx17,Agrn,Timp2 | | 7.226e-03 | -4.93 | Thioredoxin | pfam domains | PF00085 | 23 | 2 | 12881 | 72 | Tmx3,Pdia6 | | 7.236e-03 | -4.93 | taxis | biological process | GO:0042330 | 386 | 7 | 13711 | 80 | Ncam1,Bdnf,Ephb6,Lhx1,Gbx1,Agrn,Nr4a3 | | 7.240e-03 | -4.93 | GATTGGY\_NFY\_Q6\_01 | MSigDB lists | GATTGGY\_NFY\_Q6\_01 | 920 | 12 | 12187 | 72 | Cldn5,Ndst4,Ncam1,Bdnf,Dgkz,Arntl,Lhx1,Rgs4,Col1a1,Hsd17b12,Tle4,Peg3 | | 7.275e-03 | -4.92 | Ppp1r9b (protein phosphatase 1, regulatory subunit 9B) | protein interactions | 217124 | 401 | 8 | 6802 | 49 | Glud1,Plp1,Atp5b,H1f0,Car2,Ntm,Ncam1,Homer1 | | 7.301e-03 | -4.92 | positive regulation of cation channel activity | biological process | GO:2001259 | 68 | 3 | 13711 | 80 | Nipsnap2,Cacng2,Lrrc55 | | 7.356e-03 | -4.91 | TTTGTAG\_MIR520D | MSigDB lists | TTTGTAG\_MIR520D | 291 | 6 | 12187 | 72 | Qk,Ubqln2,Tmem165,Ddx17,Grm3,Ncam1 | | 7.358e-03 | -4.91 | MAF\_Q6 | MSigDB lists | MAF\_Q6 | 206 | 5 | 12187 | 72 | Col1a1,Cbln2,Grm3,Csrnp3,Ddx17 | | 7.366e-03 | -4.91 | GO\_NEGATIVE\_REGULATION\_OF\_AMINE\_TRANSPORT | MSigDB lists | GO\_NEGATIVE\_REGULATION\_OF\_AMINE\_TRANSPORT | 22 | 2 | 12187 | 72 | Crh,P2ry12 | | 7.366e-03 | -4.91 | IIZUKA\_LIVER\_CANCER\_PROGRESSION\_G1\_G2\_DN | MSigDB lists | IIZUKA\_LIVER\_CANCER\_PROGRESSION\_G1\_G2\_DN | 22 | 2 | 12187 | 72 | Ubqln2,Aars | | 7.366e-03 | -4.91 | GO\_ESTABLISHMENT\_OF\_SPINDLE\_ORIENTATION | MSigDB lists | GO\_ESTABLISHMENT\_OF\_SPINDLE\_ORIENTATION | 22 | 2 | 12187 | 72 | Map4,Spry2 | | 7.366e-03 | -4.91 | CERVERA\_SDHB\_TARGETS\_1\_DN | MSigDB lists | CERVERA\_SDHB\_TARGETS\_1\_DN | 22 | 2 | 12187 | 72 | Peg3,Fjx1 | | 7.467e-03 | -4.90 | GSE9988\_ANTI\_TREM1\_AND\_LPS\_VS\_VEHICLE\_TREATED\_MONOCYTES\_UP | MSigDB lists | GSE9988\_ANTI\_TREM1\_AND\_LPS\_VS\_VEHICLE\_TREATED\_MONOCYTES\_UP | 131 | 4 | 12187 | 72 | Egr1,Spry2,Homer1,Inhba | | 7.467e-03 | -4.90 | CONCANNON\_APOPTOSIS\_BY\_EPOXOMICIN\_DN | MSigDB lists | CONCANNON\_APOPTOSIS\_BY\_EPOXOMICIN\_DN | 131 | 4 | 12187 | 72 | Gng11,Spry2,Sorl1,Rgs4 | | 7.560e-03 | -4.88 | Diurnally Regulated Genes with Circadian Orthologs | WikiPathways | WP1268 | 44 | 3 | 3756 | 35 | Cldn5,Arntl,Qk | | 7.560e-03 | -4.88 | Exercise-induced Circadian Regulation | WikiPathways | WP544 | 44 | 3 | 3756 | 35 | Cldn5,Arntl,Qk | | 7.656e-03 | -4.87 | PAX4\_01 | MSigDB lists | PAX4\_01 | 208 | 5 | 12187 | 72 | Dgkz,Nr4a3,Ddx17,Bdnf,Spry2 | | 7.666e-03 | -4.87 | GSE8835\_HEALTHY\_VS\_CLL\_CD4\_TCELL\_DN | MSigDB lists | GSE8835\_HEALTHY\_VS\_CLL\_CD4\_TCELL\_DN | 132 | 4 | 12187 | 72 | Csrnp3,Ntm,Fabp7,Zfp518b | | 7.781e-03 | -4.86 | regulation of synaptic plasticity | biological process | GO:0048167 | 211 | 5 | 13711 | 80 | Crh,Kdr,Egr1,Bdnf,Lrrtm2 | | 7.839e-03 | -4.85 | GO\_TRANSMEMBRANE\_RECEPTOR\_PROTEIN\_KINASE\_ACTIVITY | MSigDB lists | GO\_TRANSMEMBRANE\_RECEPTOR\_PROTEIN\_KINASE\_ACTIVITY | 69 | 3 | 12187 | 72 | Kdr,Ephb6,Ltk | | 7.839e-03 | -4.85 | BILD\_CTNNB1\_ONCOGENIC\_SIGNATURE | MSigDB lists | BILD\_CTNNB1\_ONCOGENIC\_SIGNATURE | 69 | 3 | 12187 | 72 | Sorl1,Foxq1,Qk | | 7.856e-03 | -4.85 | endoderm formation | biological process | GO:0001706 | 23 | 2 | 13711 | 80 | Lhx1,Inhba | | 7.909e-03 | -4.84 | response to alkaloid | biological process | GO:0043279 | 70 | 3 | 13711 | 80 | Homer1,Ncam1,Crh | | 7.909e-03 | -4.84 | regulation of synaptic transmission, glutamatergic | biological process | GO:0051966 | 70 | 3 | 13711 | 80 | Homer1,Grm3,Cacng2 | | 8.003e-03 | -4.83 | Sulfoglycolipids biosynthesis, ceramide/1-alkyl-2-acylglycerol => sulfatide/seminolipid | KEGG pathways | M00067 | 1 | 1 | 5248 | 42 | Ugt8a | | 8.003e-03 | -4.83 | Sulfoglycolipids biosynthesis, ceramide/1-alkyl-2-acylglycerol => sulfatide/seminolipid | KEGG pathways | mmu\_M00067 | 1 | 1 | 5248 | 42 | Ugt8a | | 8.024e-03 | -4.83 | regulation of system process | biological process | GO:0044057 | 494 | 8 | 13711 | 80 | Agrn,Rgs4,Homer1,Grm3,Crh,Nr4a3,Ncam1,Inhba | | 8.037e-03 | -4.82 | GO\_CEREBELLAR\_PURKINJE\_CELL\_LAYER\_DEVELOPMENT | MSigDB lists | GO\_CEREBELLAR\_PURKINJE\_CELL\_LAYER\_DEVELOPMENT | 23 | 2 | 12187 | 72 | Lhx1,Aars | | 8.156e-03 | -4.81 | GO\_STEROID\_BIOSYNTHETIC\_PROCESS | MSigDB lists | GO\_STEROID\_BIOSYNTHETIC\_PROCESS | 70 | 3 | 12187 | 72 | Crh,Hsd17b12,Msmo1 | | 8.229e-03 | -4.80 | cellular response to chemical stimulus | biological process | GO:0070887 | 1836 | 19 | 13711 | 80 | Col1a1,Spry2,Ltk,Inhba,Gsdme,Atp5b,Car2,Nr4a3,Cldn5,Arntl,Srm,P2ry12,Bdnf,Tle4,Crh,Timp2,Kdr,Egr1,Ddx17 | | 8.244e-03 | -4.80 | sensory organ morphogenesis | biological process | GO:0090596 | 214 | 5 | 13711 | 80 | Fjx1,Nr4a3,Spry2,Kdr,Lhx1 | | 8.417e-03 | -4.78 | HS-GAG biosynthesis | REACTOME pathways | R-MMU-2022928 | 21 | 2 | 6297 | 42 | Agrn,Hs3st1 | | 8.439e-03 | -4.77 | CREBP1CJUN\_01 | MSigDB lists | CREBP1CJUN\_01 | 213 | 5 | 12187 | 72 | AI593442,Peg3,Ncam1,Egr1,Crh | | 8.500e-03 | -4.77 | ATGAAGG\_MIR205 | MSigDB lists | ATGAAGG\_MIR205 | 136 | 4 | 12187 | 72 | Qk,Inhba,Lims2,Hs3st1 | | 8.538e-03 | -4.76 | substrate-dependent cell migration | biological process | GO:0006929 | 24 | 2 | 13711 | 80 | Atp5b,P2ry12 | | 8.538e-03 | -4.76 | heparan sulfate proteoglycan biosynthetic process | biological process | GO:0015012 | 24 | 2 | 13711 | 80 | Hs3st1,Ndst4 | | 8.557e-03 | -4.76 | animal organ development | biological process | GO:0048513 | 2256 | 22 | 13711 | 80 | Foxq1,Nr4a3,Homer1,Fabp7,Agrn,Spry2,Gbx1,Col1a1,Gsdme,Inhba,Ncam1,Fjx1,Crh,Egr1,Ddx17,Tgfa,Kdr,Hba-a2,Lhx1,Bdnf,Aars,P2ry12 | | 8.589e-03 | -4.76 | regulation of ion transport | biological process | GO:0043269 | 606 | 9 | 13711 | 80 | P2ry12,Cacng2,Nipsnap2,Homer1,Rgs4,Agrn,Lrrc55,Car2,Crh | | 8.597e-03 | -4.76 | regulation of transport | biological process | GO:0051049 | 1577 | 17 | 13711 | 80 | P2ry12,Arntl,Glud1,Cacng2,Crh,Inhba,Ncam1,Sorl1,Ubqln2,Agrn,Homer1,Rgs4,Lrrtm2,Nipsnap2,Car2,Lrrc55,Nr4a3 | | 8.663e-03 | -4.75 | axon part | cellular component | GO:0033267 | 408 | 7 | 13825 | 79 | Bdnf,Cldn5,Crh,Ermn,Ncam1,Timp2,Agrn | | 8.704e-03 | -4.74 | LIM\_MAMMARY\_STEM\_CELL\_UP | MSigDB lists | LIM\_MAMMARY\_STEM\_CELL\_UP | 396 | 7 | 12187 | 72 | Arhgap20,Qk,Fjx1,Lims2,Peg3,Gng11,Arntl | | 8.717e-03 | -4.74 | GSE4811\_CLASSSICALY\_ACTIVATED\_VS\_TYPE\_2\_ACTIVATED\_MACROPHAGE\_UP | MSigDB lists | GSE4811\_CLASSSICALY\_ACTIVATED\_VS\_TYPE\_2\_ACTIVATED\_MACROPHAGE\_UP | 137 | 4 | 12187 | 72 | Qdpr,Arhgap20,Kcnmb2,Timp2 | | 8.717e-03 | -4.74 | GSE28449\_WT\_VS\_LRF\_KO\_GERMINAL\_CENTER\_BCELL\_UP | MSigDB lists | GSE28449\_WT\_VS\_LRF\_KO\_GERMINAL\_CENTER\_BCELL\_UP | 137 | 4 | 12187 | 72 | Peg3,Spry2,Nr4a3,Lrrtm2 | | 8.717e-03 | -4.74 | GSE15330\_LYMPHOID\_MULTIPOTENT\_VS\_PRO\_BCELL\_UP | MSigDB lists | GSE15330\_LYMPHOID\_MULTIPOTENT\_VS\_PRO\_BCELL\_UP | 137 | 4 | 12187 | 72 | Homer1,Qdpr,Nr4a3,Timp2 | | 8.735e-03 | -4.74 | NAKAYAMA\_FGF2\_TARGETS | MSigDB lists | NAKAYAMA\_FGF2\_TARGETS | 24 | 2 | 12187 | 72 | Col1a1,Msmo1 | | 8.735e-03 | -4.74 | GO\_HSP90\_PROTEIN\_BINDING | MSigDB lists | GO\_HSP90\_PROTEIN\_BINDING | 24 | 2 | 12187 | 72 | Kdr,Arntl | | 8.812e-03 | -4.73 | GTTNYYNNGGTNA\_UNKNOWN | MSigDB lists | GTTNYYNNGGTNA\_UNKNOWN | 72 | 3 | 12187 | 72 | Csrnp3,Nr4a3,Ccdc13 | | 8.821e-03 | -4.73 | Proximal tubule bicarbonate reclamation | KEGG pathways | ko04964 | 18 | 2 | 5248 | 42 | Glud1,Car2 | | 8.821e-03 | -4.73 | Proximal tubule bicarbonate reclamation | KEGG pathways | mmu04964 | 18 | 2 | 5248 | 42 | Glud1,Car2 | | 8.828e-03 | -4.73 | steroid metabolic process | biological process | GO:0008202 | 139 | 4 | 13711 | 80 | Crh,Msmo1,Sorl1,Hsd17b12 | | 8.828e-03 | -4.73 | ATP metabolic process | biological process | GO:0046034 | 139 | 4 | 13711 | 80 | Hkdc1,Bdnf,Nipsnap2,Atp5b | | 8.828e-03 | -4.73 | response to xenobiotic stimulus | biological process | GO:0009410 | 139 | 4 | 13711 | 80 | Ncam1,Homer1,Egr1,Crh | | 8.875e-03 | -4.72 | regulation of pH | biological process | GO:0006885 | 73 | 3 | 13711 | 80 | Tmem165,Car2,Atp5b | | 8.929e-03 | -4.72 | PMP22/EMP/MP20/Claudin | interpro domains | IPR004031 | 25 | 2 | 13788 | 79 | Cacng2,Cldn5 | | 8.938e-03 | -4.72 | GSE17301\_CTRL\_VS\_48H\_IFNA2\_STIM\_CD8\_TCELL\_DN | MSigDB lists | GSE17301\_CTRL\_VS\_48H\_IFNA2\_STIM\_CD8\_TCELL\_DN | 138 | 4 | 12187 | 72 | Hkdc1,Arntl,Chmp7,Ephb6 | | 8.998e-03 | -4.71 | regulation of biological quality | biological process | GO:0065008 | 3134 | 28 | 13711 | 80 | Car2,Lrrtm2,Homer1,Arntl,Cacng2,Hba-a2,Qk,Tmem165,Cbln2,Bdnf,Tmx3,Egr1,Sorl1,Mtmr9,Ermn,Inhba,Pdia6,Atp5b,Nr4a3,Agrn,Rgs4,Glud1,P2ry12,Hkdc1,Kcnmb2,Aars,Crh,Kdr | | 9.162e-03 | -4.69 | HFH3\_01 | MSigDB lists | HFH3\_01 | 139 | 4 | 12187 | 72 | Nr4a3,Crh,Grm3,Ncam1 | | 9.162e-03 | -4.69 | GSE24492\_LYVE\_NEG\_VS\_POS\_MACROPHAGE\_DN | MSigDB lists | GSE24492\_LYVE\_NEG\_VS\_POS\_MACROPHAGE\_DN | 139 | 4 | 12187 | 72 | Ncam1,Cacng2,P2ry12,Qk | | 9.162e-03 | -4.69 | GSE22886\_TH1\_VS\_TH2\_12H\_ACT\_DN | MSigDB lists | GSE22886\_TH1\_VS\_TH2\_12H\_ACT\_DN | 139 | 4 | 12187 | 72 | Fnbp1,Ugt8a,Col1a1,H1f0 | | 9.246e-03 | -4.68 | positive regulation of behavior | biological process | GO:0048520 | 25 | 2 | 13711 | 80 | Crh,Nr4a3 | | 9.272e-03 | -4.68 | cellular response to acid chemical | biological process | GO:0071229 | 141 | 4 | 13711 | 80 | Col1a1,Egr1,Ltk,Kdr | | 9.361e-03 | -4.67 | excitatory synapse | cellular component | GO:0060076 | 76 | 3 | 13825 | 79 | Homer1,Lrrtm2,Ntm | | 9.390e-03 | -4.67 | GSE24634\_IL4\_VS\_CTRL\_TREATED\_NAIVE\_CD4\_TCELL\_DAY3\_DN | MSigDB lists | GSE24634\_IL4\_VS\_CTRL\_TREATED\_NAIVE\_CD4\_TCELL\_DAY3\_DN | 140 | 4 | 12187 | 72 | Ddx56,Gng11,Chmp7,Inhba | | 9.390e-03 | -4.67 | GSE9988\_ANTI\_TREM1\_AND\_LPS\_VS\_CTRL\_TREATED\_MONOCYTES\_UP | MSigDB lists | GSE9988\_ANTI\_TREM1\_AND\_LPS\_VS\_CTRL\_TREATED\_MONOCYTES\_UP | 140 | 4 | 12187 | 72 | Inhba,Egr1,Spry2,Homer1 | | 9.390e-03 | -4.67 | GO\_NEGATIVE\_REGULATION\_OF\_NEURON\_DEATH | MSigDB lists | GO\_NEGATIVE\_REGULATION\_OF\_NEURON\_DEATH | 140 | 4 | 12187 | 72 | Nr4a3,Aars,Bdnf,Sorl1 | | 9.421e-03 | -4.66 | regulation of ion transmembrane transport | biological process | GO:0034765 | 406 | 7 | 13711 | 80 | Agrn,Rgs4,Homer1,Nipsnap2,Crh,Lrrc55,Cacng2 | | 9.458e-03 | -4.66 | GAZDA\_DIAMOND\_BLACKFAN\_ANEMIA\_MYELOID\_UP | MSigDB lists | GAZDA\_DIAMOND\_BLACKFAN\_ANEMIA\_MYELOID\_UP | 25 | 2 | 12187 | 72 | Car2,Nr4a3 | | 9.458e-03 | -4.66 | MODULE\_80 | MSigDB lists | MODULE\_80 | 25 | 2 | 12187 | 72 | Ephb6,Kdr | | 9.458e-03 | -4.66 | GO\_HISTONE\_ACETYLTRANSFERASE\_BINDING | MSigDB lists | GO\_HISTONE\_ACETYLTRANSFERASE\_BINDING | 25 | 2 | 12187 | 72 | Egr1,Nr4a3 | | 9.458e-03 | -4.66 | KEGG\_GLYCOSAMINOGLYCAN\_BIOSYNTHESIS\_HEPARAN\_SULFATE | MSigDB lists | KEGG\_GLYCOSAMINOGLYCAN\_BIOSYNTHESIS\_HEPARAN\_SULFATE | 25 | 2 | 12187 | 72 | Ndst4,Hs3st1 | | 9.498e-03 | -4.66 | AGCTCCT\_MIR28 | MSigDB lists | AGCTCCT\_MIR28 | 74 | 3 | 12187 | 72 | Ntm,Nr4a3,Qk | | 9.555e-03 | -4.65 | cellular monovalent inorganic cation homeostasis | biological process | GO:0030004 | 75 | 3 | 13711 | 80 | Atp5b,Tmem165,Car2 | | 9.555e-03 | -4.65 | regulation of potassium ion transmembrane transport | biological process | GO:1901379 | 75 | 3 | 13711 | 80 | Agrn,Rgs4,Lrrc55 | | 9.622e-03 | -4.64 | MODULE\_257 | MSigDB lists | MODULE\_257 | 141 | 4 | 12187 | 72 | Tle4,Dgkz,Car2,Ddx17 | | 9.622e-03 | -4.64 | GSE3982\_DC\_VS\_NEUTROPHIL\_LPS\_STIM\_DN | MSigDB lists | GSE3982\_DC\_VS\_NEUTROPHIL\_LPS\_STIM\_DN | 141 | 4 | 12187 | 72 | Fabp7,Sorl1,Pom121,Gbx1 | | 9.625e-03 | -4.64 | CREB\_Q4 | MSigDB lists | CREB\_Q4 | 220 | 5 | 12187 | 72 | Grm3,AI593442,Ubqln2,Crh,Nr4a3 | | 9.637e-03 | -4.64 | Transl\_B-barrel\_sf | interpro domains | IPR009000 | 26 | 2 | 13788 | 79 | Aars,Gar1 | | 9.718e-03 | -4.63 | cellular response to organic cyclic compound | biological process | GO:0071407 | 312 | 6 | 13711 | 80 | Crh,Nr4a3,Inhba,P2ry12,Egr1,Ddx17 | | 9.737e-03 | -4.63 | MODULE\_1 | MSigDB lists | MODULE\_1 | 309 | 6 | 12187 | 72 | Gng11,Col1a1,Timp2,Peg3,Egr1,Ncam1 | | 9.747e-03 | -4.63 | regulation of calcium ion transport | biological process | GO:0051924 | 223 | 5 | 13711 | 80 | Crh,Rgs4,Homer1,Nipsnap2,P2ry12 | | 9.772e-03 | -4.63 | neuronal cell body | cellular component | GO:0043025 | 632 | 9 | 13825 | 79 | Fabp7,Sorl1,Homer1,Timp2,Ncam1,Crh,Ermn,Kdr,Bdnf | | 9.979e-03 | -4.61 | negative regulation of neural precursor cell proliferation | biological process | GO:2000178 | 26 | 2 | 13711 | 80 | Lims2,Bdnf | | 9.984e-03 | -4.61 | ENK\_UV\_RESPONSE\_EPIDERMIS\_UP | MSigDB lists | ENK\_UV\_RESPONSE\_EPIDERMIS\_UP | 222 | 5 | 12187 | 72 | Tgfa,Car2,Srm,Tmem165,Sorl1 | | 9.991e-03 | -4.61 | response to organonitrogen compound | biological process | GO:0010243 | 621 | 9 | 13711 | 80 | Ncam1,P2ry12,Col1a1,Ubqln2,Homer1,Egr1,Car2,Crh,Nr4a3 | | 1.010e-02 | -4.60 | SERVITJA\_ISLET\_HNF1A\_TARGETS\_UP | MSigDB lists | SERVITJA\_ISLET\_HNF1A\_TARGETS\_UP | 143 | 4 | 12187 | 72 | Timp2,Ntm,Map4,Fabp7 | | 1.011e-02 | -4.59 | regulation of cellular process | biological process | GO:0050794 | 7419 | 54 | 13711 | 80 | Ddx17,Map4,Tmx3,Cbln2,Bdnf,Dgkz,Peg3,Cacng2,Alkal2,Cldn5,Lrrtm2,Csrnp3,Spry2,Plp1,Tgfa,Kdr,Nrep,P2ry12,Glud1,Ddx56,Agrn,H1f0,Mtmr9,Ltk,Atp5b,Ubqln2,Egr1,Ntm,Arntl,Qk,Homer1,Car2,Ncam1,Lims2,Col1a1,Gbx1,Timp2,Crh,Tle4,Arhgap20,Aars,Ephb6,Lhx1,Fbxl17,Rgs4,Grm3,Foxq1,Nr4a3,Sorl1,Ermn,Inhba,Gsdme,Pdia6,Gng11 | | 1.013e-02 | -4.59 | positive regulation of metabolic process | biological process | GO:0009893 | 2865 | 26 | 13711 | 80 | Ddx17,Egr1,Qk,Alkal2,Arntl,Peg3,Dgkz,Bdnf,Csrnp3,Spry2,Gbx1,Col1a1,Crh,Timp2,Tgfa,Plp1,Kdr,Lhx1,Nr4a3,Agrn,H1f0,Ubqln2,Gsdme,Mtmr9,Sorl1,Inhba | | 1.017e-02 | -4.59 | BDNFACTOR | prints domains | PR01912 | 1 | 1 | 2951 | 30 | Bdnf | | 1.017e-02 | -4.59 | INHIBINBA | prints domains | PR00670 | 1 | 1 | 2951 | 30 | Inhba | | 1.017e-02 | -4.59 | NORNUCRECPTR | prints domains | PR01286 | 1 | 1 | 2951 | 30 | Nr4a3 | | 1.017e-02 | -4.59 | CLAUDIN5 | prints domains | PR01380 | 1 | 1 | 2951 | 30 | Cldn5 | | 1.017e-02 | -4.59 | GLFDHDRGNASE | prints domains | PR00082 | 1 | 1 | 2951 | 30 | Glud1 | | 1.017e-02 | -4.59 | VEGFRECEPTR2 | prints domains | PR01834 | 1 | 1 | 2951 | 30 | Kdr | | 1.017e-02 | -4.59 | P2Y12PRNCPTR | prints domains | PR01569 | 1 | 1 | 2951 | 30 | P2ry12 | | 1.017e-02 | -4.59 | VDCCGAMMA2 | prints domains | PR01602 | 1 | 1 | 2951 | 30 | Cacng2 | | 1.017e-02 | -4.59 | CRFFAMILY | prints domains | PR01612 | 1 | 1 | 2951 | 30 | Crh | | 1.017e-02 | -4.59 | MTABOTROPC3R | prints domains | PR01053 | 1 | 1 | 2951 | 30 | Grm3 | | 1.017e-02 | -4.59 | 4JOINTEDBOX1 | prints domains | PR02072 | 1 | 1 | 2951 | 30 | Fjx1 | | 1.021e-02 | -4.58 | TTCCGTT\_MIR191 | MSigDB lists | TTCCGTT\_MIR191 | 26 | 2 | 12187 | 72 | Egr1,Bdnf | | 1.021e-02 | -4.58 | KRAS.LUNG.BREAST\_UP.V1\_UP | MSigDB lists | KRAS.LUNG.BREAST\_UP.V1\_UP | 76 | 3 | 12187 | 72 | Ntm,Spry2,Inhba | | 1.021e-02 | -4.58 | TCTGATA\_MIR361 | MSigDB lists | TCTGATA\_MIR361 | 76 | 3 | 12187 | 72 | Ntm,Qk,Plp1 | | 1.027e-02 | -4.58 | regulation of smooth muscle cell migration | biological process | GO:0014910 | 77 | 3 | 13711 | 80 | Nr4a3,Egr1,Sorl1 | | 1.029e-02 | -4.58 | positive regulation of neurogenesis | biological process | GO:0050769 | 516 | 8 | 13711 | 80 | Timp2,Ddx56,Kdr,Bdnf,P2ry12,Ltk,Alkal2,Qk | | 1.034e-02 | -4.57 | HEN1\_01 | MSigDB lists | HEN1\_01 | 144 | 4 | 12187 | 72 | Ndst4,Bdnf,Cldn5,Nr4a3 | | 1.042e-02 | -4.56 | GO\_TISSUE\_MORPHOGENESIS | MSigDB lists | GO\_TISSUE\_MORPHOGENESIS | 410 | 7 | 12187 | 72 | Car2,Inhba,Foxq1,Fjx1,Nr4a3,Lhx1,Spry2 | | 1.045e-02 | -4.56 | cellular biosynthetic process | biological process | GO:0044249 | 1743 | 18 | 13711 | 80 | Qdpr,Egr1,Ddx17,Plp1,Glud1,Tmem165,Scd2,Qk,Srm,Amd1,Dgkz,Lhx1,Aars,Ndst4,Hs3st1,Ugt8a,Atp5b,Inhba | | 1.049e-02 | -4.56 | GO\_CELL\_MORPHOGENESIS\_INVOLVED\_IN\_NEURON\_DIFFERENTIATION | MSigDB lists | GO\_CELL\_MORPHOGENESIS\_INVOLVED\_IN\_NEURON\_DIFFERENTIATION | 314 | 6 | 12187 | 72 | Gbx1,Lrrc55,Ncam1,Lhx1,Bdnf,Nr4a3 | | 1.050e-02 | -4.56 | - | gene3d domains | 2.40.10.230 | 2 | 1 | 6647 | 35 | Gar1 | | 1.050e-02 | -4.56 | - | gene3d domains | 1.10.1140.10 | 2 | 1 | 6647 | 35 | Atp5b | | 1.058e-02 | -4.55 | GO\_ODONTOGENESIS | MSigDB lists | GO\_ODONTOGENESIS | 77 | 3 | 12187 | 72 | Inhba,Col1a1,Car2 | | 1.058e-02 | -4.55 | GO\_ENSHEATHMENT\_OF\_NEURONS | MSigDB lists | GO\_ENSHEATHMENT\_OF\_NEURONS | 77 | 3 | 12187 | 72 | Qk,Plp1,Ugt8a | | 1.059e-02 | -4.55 | GSE27786\_BCELL\_VS\_CD4\_TCELL\_DN | MSigDB lists | GSE27786\_BCELL\_VS\_CD4\_TCELL\_DN | 145 | 4 | 12187 | 72 | Glud1,Msmo1,Timp2,Fbxl17 | | 1.061e-02 | -4.55 | forebrain development | biological process | GO:0030900 | 318 | 6 | 13711 | 80 | Inhba,Ncam1,P2ry12,Lhx1,Fabp7,Nr4a3 | | 1.067e-02 | -4.54 | integral component of postsynaptic membrane | cellular component | GO:0099055 | 150 | 4 | 13825 | 79 | Lrrtm2,Cacng2,Ncam1,Grm3 | | 1.076e-02 | -4.53 | main axon | cellular component | GO:0044304 | 80 | 3 | 13825 | 79 | Cldn5,Crh,Ermn | | 1.084e-02 | -4.52 | HFH8\_01 | MSigDB lists | HFH8\_01 | 146 | 4 | 12187 | 72 | Inhba,Cacng2,Crh,Grm3 | | 1.093e-02 | -4.52 | negative regulation of apoptotic process | biological process | GO:0043066 | 743 | 10 | 13711 | 80 | Nr4a3,Foxq1,Tgfa,Kdr,Spry2,Qk,Aars,Ltk,Lims2,Bdnf | | 1.096e-02 | -4.51 | KRAS.LUNG\_UP.V1\_UP | MSigDB lists | KRAS.LUNG\_UP.V1\_UP | 78 | 3 | 12187 | 72 | Rgs4,Ntm,Inhba | | 1.096e-02 | -4.51 | GTTATAT\_MIR410 | MSigDB lists | GTTATAT\_MIR410 | 78 | 3 | 12187 | 72 | Qk,AI593442,Hs3st1 | | 1.096e-02 | -4.51 | chr11q23 | MSigDB lists | chr11q23 | 78 | 3 | 12187 | 72 | Sorl1,Arhgap20,Ncam1 | | 1.098e-02 | -4.51 | KEGG\_CYSTEINE\_AND\_METHIONINE\_METABOLISM | MSigDB lists | KEGG\_CYSTEINE\_AND\_METHIONINE\_METABOLISM | 27 | 2 | 12187 | 72 | Amd1,Srm | | 1.098e-02 | -4.51 | KEGG\_PRION\_DISEASES | MSigDB lists | KEGG\_PRION\_DISEASES | 27 | 2 | 12187 | 72 | Egr1,Ncam1 | | 1.101e-02 | -4.51 | mesoderm development | biological process | GO:0007498 | 79 | 3 | 13711 | 80 | Lhx1,Inhba,Nr4a3 | | 1.108e-02 | -4.50 | GO\_EXTRACELLULAR\_SPACE | MSigDB lists | GO\_EXTRACELLULAR\_SPACE | 738 | 10 | 12187 | 72 | Sorl1,Col1a1,Car2,Hba-a2,Tgfa,Inhba,Timp2,Crh,Fjx1,Cbln2 | | 1.109e-02 | -4.50 | GSE45739\_UNSTIM\_VS\_ACD3\_ACD28\_STIM\_WT\_CD4\_TCELL\_UP | MSigDB lists | GSE45739\_UNSTIM\_VS\_ACD3\_ACD28\_STIM\_WT\_CD4\_TCELL\_UP | 147 | 4 | 12187 | 72 | Hkdc1,Chmp7,Ephb6,Lims2 | | 1.110e-02 | -4.50 | mouse chr1 H2.3\|1 76.84 cM | chromosome location | mouse chr1 H2.3\|1 76.84 cM | 2 | 1 | 14556 | 81 | Rgs4 | | 1.110e-02 | -4.50 | mouse chr11 59.01 cM\|11 D | chromosome location | mouse chr11 59.01 cM\|11 D | 2 | 1 | 14556 | 81 | Col1a1 | | 1.110e-02 | -4.50 | mouse chr16 A3\|16 11.63 cM | chromosome location | mouse chr16 A3\|16 11.63 cM | 2 | 1 | 14556 | 81 | Cldn5 | | 1.110e-02 | -4.50 | mouse chr15 E1\|15 36.92 cM | chromosome location | mouse chr15 E1\|15 36.92 cM | 2 | 1 | 14556 | 81 | Cacng2 | | 1.110e-02 | -4.50 | brain development | biological process | GO:0007420 | 523 | 8 | 13711 | 80 | Nr4a3,Fabp7,Gbx1,Lhx1,P2ry12,Inhba,Aars,Ncam1 | | 1.112e-02 | -4.50 | GCTNWTTGK\_UNKNOWN | MSigDB lists | GCTNWTTGK\_UNKNOWN | 228 | 5 | 12187 | 72 | Spry2,Lhx1,Grm3,Amd1,Dgkz | | 1.112e-02 | -4.50 | GO\_EXTRACELLULAR\_STRUCTURE\_ORGANIZATION | MSigDB lists | GO\_EXTRACELLULAR\_STRUCTURE\_ORGANIZATION | 228 | 5 | 12187 | 72 | Kdr,Col1a1,Timp2,Hsd17b12,Agrn | | 1.113e-02 | -4.50 | cellular response to endogenous stimulus | biological process | GO:0071495 | 745 | 10 | 13711 | 80 | Ddx17,Egr1,Cldn5,Nr4a3,Car2,Crh,Bdnf,P2ry12,Inhba,Col1a1 | | 1.115e-02 | -4.50 | Gar1 | pfam domains | PF04410 | 2 | 1 | 12881 | 72 | Gar1 | | 1.115e-02 | -4.50 | SAM\_decarbox | pfam domains | PF01536 | 2 | 1 | 12881 | 72 | Amd1 | | 1.115e-02 | -4.50 | tRNA-synt\_2c | pfam domains | PF01411 | 2 | 1 | 12881 | 72 | Aars | | 1.115e-02 | -4.50 | CRF | pfam domains | PF00473 | 2 | 1 | 12881 | 72 | Crh | | 1.115e-02 | -4.50 | Gasdermin | pfam domains | PF04598 | 2 | 1 | 12881 | 72 | Gsdme | | 1.115e-02 | -4.50 | Gasdermin\_C | pfam domains | PF17708 | 2 | 1 | 12881 | 72 | Gsdme | | 1.122e-02 | -4.49 | mouse chr5 | chromosome location | mouse chr5 | 242 | 5 | 14556 | 81 | Hs3st1,Tmem165,Kdr,Gbx1,Qdpr | | 1.123e-02 | -4.49 | SENESE\_HDAC3\_TARGETS\_DN | MSigDB lists | SENESE\_HDAC3\_TARGETS\_DN | 416 | 7 | 12187 | 72 | Hsd17b12,Dgkz,Col1a1,Egr1,H1f0,Tmx3,Aars | | 1.132e-02 | -4.48 | DNA-binding transcription factor activity, RNA polymerase II-specific | molecular function | GO:0000981 | 531 | 8 | 13516 | 78 | Csrnp3,Arntl,Gbx1,Foxq1,Egr1,Tle4,Nr4a3,Lhx1 | | 1.135e-02 | -4.48 | adh\_short | pfam domains | PF00106 | 29 | 2 | 12881 | 72 | Qdpr,Hsd17b12 | | 1.135e-02 | -4.48 | KRAS.DF.V1\_UP | MSigDB lists | KRAS.DF.V1\_UP | 148 | 4 | 12187 | 72 | Spry2,Gng11,Cldn5,Inhba | | 1.139e-02 | -4.48 | fatty acid biosynthetic process | biological process | GO:0006633 | 80 | 3 | 13711 | 80 | Scd2,Qk,Plp1 | | 1.140e-02 | -4.47 | inhibin A complex | cellular component | GO:0043512 | 2 | 1 | 13825 | 79 | Inhba | | 1.140e-02 | -4.47 | collagen type I trimer | cellular component | GO:0005584 | 2 | 1 | 13825 | 79 | Col1a1 | | 1.143e-02 | -4.47 | H/ACA\_rnp\_Gar1/Naf1 | interpro domains | IPR007504 | 2 | 1 | 13788 | 79 | Gar1 | | 1.143e-02 | -4.47 | S-AdoMet\_decarboxylase | interpro domains | IPR001985 | 2 | 1 | 13788 | 79 | Amd1 | | 1.143e-02 | -4.47 | Gar1/Naf1\_Cbf5-bd\_sf | interpro domains | IPR038664 | 2 | 1 | 13788 | 79 | Gar1 | | 1.143e-02 | -4.47 | Ala-tRNA-lgiase\_IIc | interpro domains | IPR002318 | 2 | 1 | 13788 | 79 | Aars | | 1.143e-02 | -4.47 | CRF | interpro domains | IPR000187 | 2 | 1 | 13788 | 79 | Crh | | 1.143e-02 | -4.47 | S-AdoMet\_deCO2ase\_core | interpro domains | IPR016067 | 2 | 1 | 13788 | 79 | Amd1 | | 1.143e-02 | -4.47 | Gasdermin\_PUB | interpro domains | IPR041263 | 2 | 1 | 13788 | 79 | Gsdme | | 1.143e-02 | -4.47 | S-AdoMet\_deCO2ase\_CS | interpro domains | IPR018166 | 2 | 1 | 13788 | 79 | Amd1 | | 1.143e-02 | -4.47 | PINCH | interpro domains | IPR017351 | 2 | 1 | 13788 | 79 | Lims2 | | 1.143e-02 | -4.47 | EGD2/NACA | interpro domains | IPR016641 | 2 | 1 | 13788 | 79 | Nacad | | 1.143e-02 | -4.47 | Ala\_tRNA\_ligase\_euk/bac | interpro domains | IPR023033 | 2 | 1 | 13788 | 79 | Aars | | 1.143e-02 | -4.47 | ATPase\_F1/V1\_b/a\_C | interpro domains | IPR024034 | 2 | 1 | 13788 | 79 | Atp5b | | 1.143e-02 | -4.47 | Ala-tRNA-synth\_IIc\_core | interpro domains | IPR018165 | 2 | 1 | 13788 | 79 | Aars | | 1.143e-02 | -4.47 | Ala-tRNA-synth\_IIc\_N | interpro domains | IPR018164 | 2 | 1 | 13788 | 79 | Aars | | 1.143e-02 | -4.47 | Neural\_cell\_adh | interpro domains | IPR009138 | 2 | 1 | 13788 | 79 | Ncam1 | | 1.143e-02 | -4.47 | Ala-tRNA-ligase\_IIc\_anticod-bd | interpro domains | IPR018162 | 2 | 1 | 13788 | 79 | Aars | | 1.143e-02 | -4.47 | Gasdermin\_pore | interpro domains | IPR040460 | 2 | 1 | 13788 | 79 | Gsdme | | 1.144e-02 | -4.47 | GO\_RESPONSE\_TO\_PEPTIDE | MSigDB lists | GO\_RESPONSE\_TO\_PEPTIDE | 320 | 6 | 12187 | 72 | Qdpr,Egr1,Nr4a3,Car2,Col1a1,Gng11 | | 1.149e-02 | -4.47 | positive regulation of cell projection organization | biological process | GO:0031346 | 422 | 7 | 13711 | 80 | Ddx56,Agrn,Qk,Alkal2,Bdnf,Ltk,P2ry12 | | 1.151e-02 | -4.46 | receptor signaling protein tyrosine kinase activator activity | molecular function | GO:0030298 | 2 | 1 | 13516 | 78 | Alkal2 | | 1.151e-02 | -4.46 | group II metabotropic glutamate receptor activity | molecular function | GO:0001641 | 2 | 1 | 13516 | 78 | Grm3 | | 1.151e-02 | -4.46 | adenosylmethionine decarboxylase activity | molecular function | GO:0004014 | 2 | 1 | 13516 | 78 | Amd1 | | 1.151e-02 | -4.46 | ceramide glucosyltransferase activity | molecular function | GO:0008120 | 2 | 1 | 13516 | 78 | Ugt8a | | 1.151e-02 | -4.46 | potassium channel activator activity | molecular function | GO:0099104 | 2 | 1 | 13516 | 78 | Lrrc55 | | 1.151e-02 | -4.46 | ADP receptor activity | molecular function | GO:0001621 | 2 | 1 | 13516 | 78 | P2ry12 | | 1.151e-02 | -4.46 | alanine-tRNA ligase activity | molecular function | GO:0004813 | 2 | 1 | 13516 | 78 | Aars | | 1.151e-02 | -4.46 | C-4 methylsterol oxidase activity | molecular function | GO:0000254 | 2 | 1 | 13516 | 78 | Msmo1 | | 1.151e-02 | -4.46 | hemi-methylated DNA-binding | molecular function | GO:0044729 | 2 | 1 | 13516 | 78 | Egr1 | | 1.151e-02 | -4.46 | putrescine binding | molecular function | GO:0019810 | 2 | 1 | 13516 | 78 | Amd1 | | 1.151e-02 | -4.46 | neurotrophin TRKB receptor binding | molecular function | GO:0005169 | 2 | 1 | 13516 | 78 | Bdnf | | 1.151e-02 | -4.46 | corticotropin-releasing hormone receptor 2 binding | molecular function | GO:0051431 | 2 | 1 | 13516 | 78 | Crh | | 1.151e-02 | -4.46 | channel activator activity | molecular function | GO:0099103 | 2 | 1 | 13516 | 78 | Lrrc55 | | 1.152e-02 | -4.46 | heparan sulfate proteoglycan metabolic process | biological process | GO:0030201 | 28 | 2 | 13711 | 80 | Ndst4,Hs3st1 | | 1.159e-02 | -4.46 | BENPORATH\_ES\_WITH\_H3K27ME3 | MSigDB lists | BENPORATH\_ES\_WITH\_H3K27ME3 | 743 | 10 | 12187 | 72 | Hba-a2,Plp1,Tgfa,Arntl,Foxq1,Ltk,Crh,Nr4a3,Ncam1,Arhgap20 | | 1.164e-02 | -4.45 | regulation of anterior head development | biological process | GO:2000742 | 2 | 1 | 13711 | 80 | Lhx1 | | 1.164e-02 | -4.45 | cervix development | biological process | GO:0060067 | 2 | 1 | 13711 | 80 | Lhx1 | | 1.164e-02 | -4.45 | positive regulation of dipeptide transmembrane transport | biological process | GO:2001150 | 2 | 1 | 13711 | 80 | Car2 | | 1.164e-02 | -4.45 | alanyl-tRNA aminoacylation | biological process | GO:0006419 | 2 | 1 | 13711 | 80 | Aars | | 1.164e-02 | -4.45 | positive regulation of dipeptide transport | biological process | GO:2000880 | 2 | 1 | 13711 | 80 | Car2 | | 1.164e-02 | -4.45 | Golgi calcium ion transport | biological process | GO:0032472 | 2 | 1 | 13711 | 80 | Tmem165 | | 1.164e-02 | -4.45 | negative regulation of aspartic-type endopeptidase activity involved in amyloid precursor protein catabolic process | biological process | GO:1902960 | 2 | 1 | 13711 | 80 | Sorl1 | | 1.164e-02 | -4.45 | response to aluminum ion | biological process | GO:0010044 | 2 | 1 | 13711 | 80 | Glud1 | | 1.164e-02 | -4.45 | regulation of dipeptide transport | biological process | GO:0090089 | 2 | 1 | 13711 | 80 | Car2 | | 1.164e-02 | -4.45 | regulation of semaphorin-plexin signaling pathway | biological process | GO:2001260 | 2 | 1 | 13711 | 80 | Ncam1 | | 1.164e-02 | -4.45 | negative regulation of IRE1-mediated unfolded protein response | biological process | GO:1903895 | 2 | 1 | 13711 | 80 | Pdia6 | | 1.164e-02 | -4.45 | positive regulation of anterior head development | biological process | GO:2000744 | 2 | 1 | 13711 | 80 | Lhx1 | | 1.164e-02 | -4.45 | negative regulation of glycine import across plasma membrane | biological process | GO:1900924 | 2 | 1 | 13711 | 80 | Rgs4 | | 1.164e-02 | -4.45 | positive regulation of oligopeptide transport | biological process | GO:2000878 | 2 | 1 | 13711 | 80 | Car2 | | 1.164e-02 | -4.45 | regulation of early endosome to recycling endosome transport | biological process | GO:1902954 | 2 | 1 | 13711 | 80 | Sorl1 | | 1.164e-02 | -4.45 | progesterone secretion | biological process | GO:0042701 | 2 | 1 | 13711 | 80 | Inhba | | 1.164e-02 | -4.45 | regulation of monocyte aggregation | biological process | GO:1900623 | 2 | 1 | 13711 | 80 | Nr4a3 | | 1.164e-02 | -4.45 | regulation of oligopeptide transport | biological process | GO:0090088 | 2 | 1 | 13711 | 80 | Car2 | | 1.164e-02 | -4.45 | hormone-mediated apoptotic signaling pathway | biological process | GO:0008628 | 2 | 1 | 13711 | 80 | Crh | | 1.164e-02 | -4.45 | regulation of T cell costimulation | biological process | GO:2000523 | 2 | 1 | 13711 | 80 | Ephb6 | | 1.164e-02 | -4.45 | negative regulation of tau-protein kinase activity | biological process | GO:1902948 | 2 | 1 | 13711 | 80 | Sorl1 | | 1.164e-02 | -4.45 | trans-synaptic signaling by neuropeptide | biological process | GO:0099540 | 2 | 1 | 13711 | 80 | Bdnf | | 1.164e-02 | -4.45 | negative regulation of metalloendopeptidase activity involved in amyloid precursor protein catabolic process | biological process | GO:1902963 | 2 | 1 | 13711 | 80 | Sorl1 | | 1.164e-02 | -4.45 | regulation of metalloendopeptidase activity involved in amyloid precursor protein catabolic process | biological process | GO:1902962 | 2 | 1 | 13711 | 80 | Sorl1 | | 1.164e-02 | -4.45 | negative regulation of G protein-coupled receptor internalization | biological process | GO:1904021 | 2 | 1 | 13711 | 80 | Ubqln2 | | 1.164e-02 | -4.45 | trans-synaptic signaling by neuropeptide, modulating synaptic transmission | biological process | GO:0099551 | 2 | 1 | 13711 | 80 | Bdnf | | 1.164e-02 | -4.45 | calcium-mediated signaling using extracellular calcium source | biological process | GO:0035585 | 2 | 1 | 13711 | 80 | P2ry12 | | 1.164e-02 | -4.45 | positive regulation of corticosterone secretion | biological process | GO:2000854 | 2 | 1 | 13711 | 80 | Crh | | 1.164e-02 | -4.45 | positive regulation of protein geranylgeranylation | biological process | GO:2000541 | 2 | 1 | 13711 | 80 | Agrn | | 1.164e-02 | -4.45 | regulation of ERK5 cascade | biological process | GO:0070376 | 2 | 1 | 13711 | 80 | Alkal2 | | 1.164e-02 | -4.45 | regulation of neurofibrillary tangle assembly | biological process | GO:1902996 | 2 | 1 | 13711 | 80 | Sorl1 | | 1.164e-02 | -4.45 | cellular response to interleukin-8 | biological process | GO:0098759 | 2 | 1 | 13711 | 80 | Egr1 | | 1.164e-02 | -4.45 | regulation of protein geranylgeranylation | biological process | GO:2000539 | 2 | 1 | 13711 | 80 | Agrn | | 1.164e-02 | -4.45 | positive regulation of synaptic growth at neuromuscular junction | biological process | GO:0045887 | 2 | 1 | 13711 | 80 | Agrn | | 1.164e-02 | -4.45 | regulation of exocyst assembly | biological process | GO:0001928 | 2 | 1 | 13711 | 80 | Ncam1 | | 1.164e-02 | -4.45 | negative regulation of aspartic-type peptidase activity | biological process | GO:1905246 | 2 | 1 | 13711 | 80 | Sorl1 | | 1.164e-02 | -4.45 | regulation of dipeptide transmembrane transport | biological process | GO:2001148 | 2 | 1 | 13711 | 80 | Car2 | | 1.164e-02 | -4.45 | regulation of sodium:potassium-exchanging ATPase activity | biological process | GO:1903406 | 2 | 1 | 13711 | 80 | Agrn | | 1.164e-02 | -4.45 | positive regulation of T cell costimulation | biological process | GO:2000525 | 2 | 1 | 13711 | 80 | Ephb6 | | 1.164e-02 | -4.45 | positive regulation of monocyte aggregation | biological process | GO:1900625 | 2 | 1 | 13711 | 80 | Nr4a3 | | 1.164e-02 | -4.45 | ectoderm formation | biological process | GO:0001705 | 2 | 1 | 13711 | 80 | Lhx1 | | 1.164e-02 | -4.45 | cellular response to mycophenolic acid | biological process | GO:0071506 | 2 | 1 | 13711 | 80 | Egr1 | | 1.164e-02 | -4.45 | response to mycophenolic acid | biological process | GO:0071505 | 2 | 1 | 13711 | 80 | Egr1 | | 1.164e-02 | -4.45 | positive regulation of nephron tubule epithelial cell differentiation | biological process | GO:2000768 | 2 | 1 | 13711 | 80 | Lhx1 | | 1.164e-02 | -4.45 | uterine epithelium development | biological process | GO:0035847 | 2 | 1 | 13711 | 80 | Lhx1 | | 1.164e-02 | -4.45 | response to interleukin-8 | biological process | GO:0098758 | 2 | 1 | 13711 | 80 | Egr1 | | 1.164e-02 | -4.45 | oviduct epithelium development | biological process | GO:0035846 | 2 | 1 | 13711 | 80 | Lhx1 | | 1.164e-02 | -4.45 | positive regulation of mast cell cytokine production | biological process | GO:0032765 | 2 | 1 | 13711 | 80 | Nr4a3 | | 1.174e-02 | -4.44 | WATANABE\_RECTAL\_CANCER\_RADIOTHERAPY\_RESPONSIVE\_DN | MSigDB lists | WATANABE\_RECTAL\_CANCER\_RADIOTHERAPY\_RESPONSIVE\_DN | 80 | 3 | 12187 | 72 | Map4,Glud1,Agrn | | 1.178e-02 | -4.44 | GO\_CELLULAR\_GLUCURONIDATION | MSigDB lists | GO\_CELLULAR\_GLUCURONIDATION | 2 | 1 | 12187 | 72 | Ugt8a | | 1.178e-02 | -4.44 | CEBALLOS\_TARGETS\_OF\_TP53\_AND\_MYC\_DN | MSigDB lists | CEBALLOS\_TARGETS\_OF\_TP53\_AND\_MYC\_DN | 28 | 2 | 12187 | 72 | Pom121,Dgkz | | 1.178e-02 | -4.44 | GO\_NEGATIVE\_REGULATION\_OF\_ENDOCYTOSIS | MSigDB lists | GO\_NEGATIVE\_REGULATION\_OF\_ENDOCYTOSIS | 28 | 2 | 12187 | 72 | Ubqln2,Lrrtm2 | | 1.178e-02 | -4.44 | GREENBAUM\_E2A\_TARGETS\_UP | MSigDB lists | GREENBAUM\_E2A\_TARGETS\_UP | 28 | 2 | 12187 | 72 | Car2,Hs3st1 | | 1.178e-02 | -4.44 | REACTOME\_SIGNAL\_AMPLIFICATION | MSigDB lists | REACTOME\_SIGNAL\_AMPLIFICATION | 28 | 2 | 12187 | 72 | P2ry12,Gng11 | | 1.178e-02 | -4.44 | chr11p13 | MSigDB lists | chr11p13 | 28 | 2 | 12187 | 72 | Fjx1,Lhx1 | | 1.178e-02 | -4.44 | cell morphogenesis | biological process | GO:0000902 | 638 | 9 | 13711 | 80 | Ugt8a,Agrn,Kdr,Nr4a3,Bdnf,Ephb6,Lhx1,Ncam1,Gbx1 | | 1.188e-02 | -4.43 | GSE5542\_UNTREATED\_VS\_IFNG\_TREATED\_EPITHELIAL\_CELLS\_24H\_DN | MSigDB lists | GSE5542\_UNTREATED\_VS\_IFNG\_TREATED\_EPITHELIAL\_CELLS\_24H\_DN | 150 | 4 | 12187 | 72 | Chmp7,Tle4,Timp2,Qdpr | | 1.209e-02 | -4.42 | GO\_CELL\_PART\_MORPHOGENESIS | MSigDB lists | GO\_CELL\_PART\_MORPHOGENESIS | 526 | 8 | 12187 | 72 | Nr4a3,Bdnf,Lhx1,Ncam1,Lrrc55,Ccdc13,Gbx1,Ugt8a | | 1.214e-02 | -4.41 | GO\_POSITIVE\_REGULATION\_OF\_HORMONE\_SECRETION | MSigDB lists | GO\_POSITIVE\_REGULATION\_OF\_HORMONE\_SECRETION | 81 | 3 | 12187 | 72 | Inhba,Crh,Glud1 | | 1.215e-02 | -4.41 | GSE3982\_MAST\_CELL\_VS\_BASOPHIL\_UP | MSigDB lists | GSE3982\_MAST\_CELL\_VS\_BASOPHIL\_UP | 151 | 4 | 12187 | 72 | Aars,Qdpr,Hsd17b12,Msmo1 | | 1.215e-02 | -4.41 | GSE17721\_4H\_VS\_24H\_POLYIC\_BMDC\_UP | MSigDB lists | GSE17721\_4H\_VS\_24H\_POLYIC\_BMDC\_UP | 151 | 4 | 12187 | 72 | Pdia6,Glud1,Hkdc1,Timp2 | | 1.215e-02 | -4.41 | GSE9988\_ANTI\_TREM1\_VS\_CTRL\_TREATED\_MONOCYTES\_UP | MSigDB lists | GSE9988\_ANTI\_TREM1\_VS\_CTRL\_TREATED\_MONOCYTES\_UP | 151 | 4 | 12187 | 72 | Inhba,Homer1,Egr1,Spry2 | | 1.215e-02 | -4.41 | glutamate degradation X | BIOCYC pathways | MOUSE\_PWY-5766 | 1 | 1 | 823 | 10 | Glud1 | | 1.215e-02 | -4.41 | spermidine biosynthesis I | BIOCYC pathways | MOUSE\_BSUBPOLYAMSYN-PWY | 1 | 1 | 823 | 10 | Srm | | 1.215e-02 | -4.41 | glutamate biosynthesis II | BIOCYC pathways | MOUSE\_GLUTAMATE-SYN2-PWY | 1 | 1 | 823 | 10 | Glud1 | | 1.224e-02 | -4.40 | regulation of nervous system process | biological process | GO:0031644 | 153 | 4 | 13711 | 80 | Ncam1,Rgs4,Homer1,Grm3 | | 1.228e-02 | -4.40 | HELLER\_HDAC\_TARGETS\_SILENCED\_BY\_METHYLATION\_UP | MSigDB lists | HELLER\_HDAC\_TARGETS\_SILENCED\_BY\_METHYLATION\_UP | 325 | 6 | 12187 | 72 | Egr1,H1f0,Tle4,Hba-a2,Ncam1,Agrn | | 1.242e-02 | -4.39 | GSE17721\_PAM3CSK4\_VS\_CPG\_16H\_BMDC\_UP | MSigDB lists | GSE17721\_PAM3CSK4\_VS\_CPG\_16H\_BMDC\_UP | 152 | 4 | 12187 | 72 | Ntm,Ndst4,Cacng2,Srm | | 1.242e-02 | -4.39 | LI\_WILMS\_TUMOR\_VS\_FETAL\_KIDNEY\_1\_UP | MSigDB lists | LI\_WILMS\_TUMOR\_VS\_FETAL\_KIDNEY\_1\_UP | 152 | 4 | 12187 | 72 | Gng11,Kdr,Agrn,Msmo1 | | 1.242e-02 | -4.39 | GSE22589\_HIV\_VS\_HIV\_AND\_SIV\_INFECTED\_DC\_UP | MSigDB lists | GSE22589\_HIV\_VS\_HIV\_AND\_SIV\_INFECTED\_DC\_UP | 152 | 4 | 12187 | 72 | AI593442,Peg3,Egr1,Kcnmb2 | | 1.246e-02 | -4.39 | negative regulation of programmed cell death | biological process | GO:0043069 | 758 | 10 | 13711 | 80 | Tgfa,Kdr,Nr4a3,Foxq1,Bdnf,Lims2,Ltk,Aars,Qk,Spry2 | | 1.251e-02 | -4.38 | regulation of localization | biological process | GO:0032879 | 2332 | 22 | 13711 | 80 | Lrrc55,Nr4a3,Rgs4,Agrn,Ubqln2,Atp5b,Sorl1,Inhba,Crh,Kdr,Glud1,P2ry12,Car2,Nipsnap2,Lrrtm2,Homer1,Spry2,Col1a1,Ncam1,Egr1,Cacng2,Arntl | | 1.258e-02 | -4.38 | response to ammonium ion | biological process | GO:0060359 | 83 | 3 | 13711 | 80 | Crh,Homer1,Ncam1 | | 1.261e-02 | -4.37 | GO\_POSITIVE\_REGULATION\_OF\_KIDNEY\_DEVELOPMENT | MSigDB lists | GO\_POSITIVE\_REGULATION\_OF\_KIDNEY\_DEVELOPMENT | 29 | 2 | 12187 | 72 | Egr1,Lhx1 | | 1.261e-02 | -4.37 | MODULE\_51 | MSigDB lists | MODULE\_51 | 29 | 2 | 12187 | 72 | Kdr,Ephb6 | | 1.261e-02 | -4.37 | LIANG\_HEMATOPOIESIS\_STEM\_CELL\_NUMBER\_SMALL\_VS\_HUGE\_UP | MSigDB lists | LIANG\_HEMATOPOIESIS\_STEM\_CELL\_NUMBER\_SMALL\_VS\_HUGE\_UP | 29 | 2 | 12187 | 72 | Qdpr,Agrn | | 1.270e-02 | -4.37 | GSE29949\_MICROGLIA\_VS\_DC\_BRAIN\_UP | MSigDB lists | GSE29949\_MICROGLIA\_VS\_DC\_BRAIN\_UP | 153 | 4 | 12187 | 72 | Map4,Car2,Ephb6,Tle4 | | 1.270e-02 | -4.37 | GSE25123\_CTRL\_VS\_IL4\_STIM\_PPARG\_KO\_MACROPHAGE\_DN | MSigDB lists | GSE25123\_CTRL\_VS\_IL4\_STIM\_PPARG\_KO\_MACROPHAGE\_DN | 153 | 4 | 12187 | 72 | Lrrtm2,Nap1l5,Ddx17,Egr1 | | 1.270e-02 | -4.37 | GSE25088\_WT\_VS\_STAT6\_KO\_MACROPHAGE\_IL4\_STIM\_UP | MSigDB lists | GSE25088\_WT\_VS\_STAT6\_KO\_MACROPHAGE\_IL4\_STIM\_UP | 153 | 4 | 12187 | 72 | Pom121,Chmp7,Ephb6,Arntl | | 1.270e-02 | -4.37 | GSE39152\_SPLEEN\_CD103\_NEG\_VS\_BRAIN\_CD103\_POS\_MEMORY\_CD8\_TCELL\_DN | MSigDB lists | GSE39152\_SPLEEN\_CD103\_NEG\_VS\_BRAIN\_CD103\_POS\_MEMORY\_CD8\_TCELL\_DN | 153 | 4 | 12187 | 72 | Foxq1,Peg3,Col1a1,Fjx1 | | 1.270e-02 | -4.37 | GTGCCAT\_MIR183 | MSigDB lists | GTGCCAT\_MIR183 | 153 | 4 | 12187 | 72 | Amd1,Egr1,Tle4,Qk | | 1.270e-02 | -4.37 | GO\_RECEPTOR\_BINDING | MSigDB lists | GO\_RECEPTOR\_BINDING | 991 | 12 | 12187 | 72 | Homer1,Crh,Kdr,Cacng2,Ddx17,Tgfa,Inhba,Timp2,Bdnf,Lrrtm2,Nr4a3,Arntl | | 1.286e-02 | -4.35 | positive regulation of protein modification process | biological process | GO:0031401 | 1001 | 12 | 13711 | 80 | Tgfa,Kdr,Timp2,Egr1,Crh,Bdnf,Arntl,Alkal2,Agrn,Inhba,Gsdme,Spry2 | | 1.288e-02 | -4.35 | histone acetyltransferase binding | molecular function | GO:0035035 | 30 | 2 | 13516 | 78 | Nr4a3,Egr1 | | 1.296e-02 | -4.35 | oxidoreductase activity | molecular function | GO:0016491 | 544 | 8 | 13516 | 78 | Qdpr,Glud1,Scd2,Msmo1,Hsd17b12,Hba-a2,Tmx3,Pdia6 | | 1.297e-02 | -4.35 | UEDA\_CENTRAL\_CLOCK | MSigDB lists | UEDA\_CENTRAL\_CLOCK | 83 | 3 | 12187 | 72 | Arntl,Fabp7,Pdia6 | | 1.298e-02 | -4.34 | TGANNYRGCA\_TCF11MAFG\_01 | MSigDB lists | TGANNYRGCA\_TCF11MAFG\_01 | 237 | 5 | 12187 | 72 | Ddx17,Map4,Crh,Nap1l5,Nr4a3 | | 1.298e-02 | -4.34 | LIAO\_METASTASIS | MSigDB lists | LIAO\_METASTASIS | 428 | 7 | 12187 | 72 | Hkdc1,Col1a1,Dgkz,Tmem165,Foxq1,Agrn,Map4 | | 1.298e-02 | -4.34 | GSE17721\_CTRL\_VS\_POLYIC\_24H\_BMDC\_DN | MSigDB lists | GSE17721\_CTRL\_VS\_POLYIC\_24H\_BMDC\_DN | 154 | 4 | 12187 | 72 | Fabp7,Gng11,Tgfa,Inhba | | 1.298e-02 | -4.34 | GSE15930\_NAIVE\_VS\_72H\_IN\_VITRO\_STIM\_IFNAB\_CD8\_TCELL\_UP | MSigDB lists | GSE15930\_NAIVE\_VS\_72H\_IN\_VITRO\_STIM\_IFNAB\_CD8\_TCELL\_UP | 154 | 4 | 12187 | 72 | Car2,Tle4,Timp2,Fabp7 | | 1.298e-02 | -4.34 | GSE1460\_INTRATHYMIC\_T\_PROGENITOR\_VS\_DP\_THYMOCYTE\_DN | MSigDB lists | GSE1460\_INTRATHYMIC\_T\_PROGENITOR\_VS\_DP\_THYMOCYTE\_DN | 154 | 4 | 12187 | 72 | Spry2,Egr1,Arntl,Tle4 | | 1.298e-02 | -4.34 | GSE11924\_TH1\_VS\_TH2\_CD4\_TCELL\_DN | MSigDB lists | GSE11924\_TH1\_VS\_TH2\_CD4\_TCELL\_DN | 154 | 4 | 12187 | 72 | Foxq1,Hs3st1,Car2,Hsd17b12 | | 1.298e-02 | -4.34 | STAT5A\_04 | MSigDB lists | STAT5A\_04 | 154 | 4 | 12187 | 72 | Ncam1,Bdnf,P2ry12,Fnbp1 | | 1.299e-02 | -4.34 | intrinsic component of postsynaptic membrane | cellular component | GO:0098936 | 159 | 4 | 13825 | 79 | Ncam1,Cacng2,Grm3,Lrrtm2 | | 1.316e-02 | -4.33 | cerebellar Purkinje cell layer development | biological process | GO:0021680 | 30 | 2 | 13711 | 80 | Aars,Lhx1 | | 1.327e-02 | -4.32 | GSE23925\_LIGHT\_ZONE\_VS\_DARK\_ZONE\_BCELL\_UP | MSigDB lists | GSE23925\_LIGHT\_ZONE\_VS\_DARK\_ZONE\_BCELL\_UP | 155 | 4 | 12187 | 72 | Cldn5,Cacng2,H1f0,Map4 | | 1.342e-02 | -4.31 | LU\_AGING\_BRAIN\_UP | MSigDB lists | LU\_AGING\_BRAIN\_UP | 239 | 5 | 12187 | 72 | Gng11,Qk,Plp1,Tle4,Cldn5 | | 1.345e-02 | -4.31 | positive regulation of cell adhesion | biological process | GO:0045785 | 335 | 6 | 13711 | 80 | Ephb6,Lims2,P2ry12,Hsd17b12,Kdr,Nr4a3 | | 1.346e-02 | -4.31 | AMIT\_EGF\_RESPONSE\_60\_MCF10A | MSigDB lists | AMIT\_EGF\_RESPONSE\_60\_MCF10A | 30 | 2 | 12187 | 72 | Egr1,Inhba | | 1.346e-02 | -4.31 | GO\_VENTRAL\_SPINAL\_CORD\_DEVELOPMENT | MSigDB lists | GO\_VENTRAL\_SPINAL\_CORD\_DEVELOPMENT | 30 | 2 | 12187 | 72 | Gbx1,Lhx1 | | 1.346e-02 | -4.31 | MODULE\_280 | MSigDB lists | MODULE\_280 | 30 | 2 | 12187 | 72 | Car2,Hs3st1 | | 1.346e-02 | -4.31 | GO\_REGULATION\_OF\_NEUROTRANSMITTER\_RECEPTOR\_ACTIVITY | MSigDB lists | GO\_REGULATION\_OF\_NEUROTRANSMITTER\_RECEPTOR\_ACTIVITY | 30 | 2 | 12187 | 72 | Crh,Cacng2 | | 1.346e-02 | -4.31 | BROWNE\_HCMV\_INFECTION\_2HR\_UP | MSigDB lists | BROWNE\_HCMV\_INFECTION\_2HR\_UP | 30 | 2 | 12187 | 72 | Nr4a3,Egr1 | | 1.351e-02 | -4.30 | response to endogenous stimulus | biological process | GO:0009719 | 886 | 11 | 13711 | 80 | Nr4a3,Car2,Cldn5,Col1a1,Inhba,Crh,Egr1,Ddx17,Timp2,Bdnf,P2ry12 | | 1.352e-02 | -4.30 | ADOMETDC | prosite domains | PS01336 | 2 | 1 | 8845 | 60 | Amd1 | | 1.352e-02 | -4.30 | AA\_TRNA\_LIGASE\_II\_ALA | prosite domains | PS50860 | 2 | 1 | 8845 | 60 | Aars | | 1.364e-02 | -4.29 | inner ear development | biological process | GO:0048839 | 158 | 4 | 13711 | 80 | Nr4a3,Spry2,Gsdme,Bdnf | | 1.378e-02 | -4.28 | GO\_REGULATION\_OF\_CELL\_DEATH | MSigDB lists | GO\_REGULATION\_OF\_CELL\_DEATH | 1126 | 13 | 12187 | 72 | Hba-a2,Inhba,Csrnp3,Lims2,Egr1,Sorl1,Kdr,Crh,Spry2,Ltk,Nr4a3,Aars,Bdnf | | 1.382e-02 | -4.28 | MYAATNNNNNNNGGC\_UNKNOWN | MSigDB lists | MYAATNNNNNNNGGC\_UNKNOWN | 85 | 3 | 12187 | 72 | Amd1,Bdnf,Tle4 | | 1.382e-02 | -4.28 | GO\_AMINOGLYCAN\_BIOSYNTHETIC\_PROCESS | MSigDB lists | GO\_AMINOGLYCAN\_BIOSYNTHETIC\_PROCESS | 85 | 3 | 12187 | 72 | Agrn,Ndst4,Hs3st1 | | 1.385e-02 | -4.28 | GSE14308\_TH1\_VS\_TH17\_UP | MSigDB lists | GSE14308\_TH1\_VS\_TH17\_UP | 157 | 4 | 12187 | 72 | Hsd17b12,Spry2,Tmem165,Sorl1 | | 1.385e-02 | -4.28 | GSE15930\_NAIVE\_VS\_72H\_IN\_VITRO\_STIM\_CD8\_TCELL\_UP | MSigDB lists | GSE15930\_NAIVE\_VS\_72H\_IN\_VITRO\_STIM\_CD8\_TCELL\_UP | 157 | 4 | 12187 | 72 | Agrn,Timp2,Car2,Fabp7 | | 1.391e-02 | -4.28 | lipid metabolic process | biological process | GO:0006629 | 771 | 10 | 13711 | 80 | Crh,Msmo1,Plp1,Hsd17b12,Ugt8a,Scd2,Qk,Sorl1,Dgkz,Atp5b | | 1.392e-02 | -4.27 | somatodendritic compartment | cellular component | GO:0036477 | 909 | 11 | 13825 | 79 | Kdr,Cacng2,Bdnf,Sorl1,Timp2,Grm3,Ncam1,Crh,Ermn,Fabp7,Homer1 | | 1.392e-02 | -4.27 | plasma membrane | cellular component | GO:0005886 | 3462 | 29 | 13825 | 79 | Plp1,P2ry12,Dgkz,Lrrc55,Lims2,Kdr,Kcnmb2,Gsdme,Atp5b,Rgs4,Spry2,Grm3,Ncam1,Tgfa,Cacng2,Gng11,Agrn,Ntm,Ltk,Fnbp1,Homer1,Pdia6,Ubqln2,Mtmr9,Ephb6,Lrrtm2,Car2,Cldn5,Map4 | | 1.401e-02 | -4.27 | cell differentiation in spinal cord | biological process | GO:0021515 | 31 | 2 | 13711 | 80 | Gbx1,Lhx1 | | 1.401e-02 | -4.27 | protein localization to lysosome | biological process | GO:0061462 | 31 | 2 | 13711 | 80 | Cacng2,Sorl1 | | 1.401e-02 | -4.27 | homotypic cell-cell adhesion | biological process | GO:0034109 | 31 | 2 | 13711 | 80 | Ncam1,P2ry12 | | 1.413e-02 | -4.26 | GTGACGY\_E4F1\_Q6 | MSigDB lists | GTGACGY\_E4F1\_Q6 | 541 | 8 | 12187 | 72 | Ubqln2,Crh,Nr4a3,AI593442,Grm3,Egr1,Amd1,Tmem165 | | 1.415e-02 | -4.26 | GSE15930\_NAIVE\_VS\_48H\_IN\_VITRO\_STIM\_CD8\_TCELL\_UP | MSigDB lists | GSE15930\_NAIVE\_VS\_48H\_IN\_VITRO\_STIM\_CD8\_TCELL\_UP | 158 | 4 | 12187 | 72 | Car2,Agrn,Tle4,Sorl1 | | 1.415e-02 | -4.26 | GSE43863\_TH1\_VS\_LY6C\_LOW\_CXCR5NEG\_EFFECTOR\_CD4\_TCELL\_UP | MSigDB lists | GSE43863\_TH1\_VS\_LY6C\_LOW\_CXCR5NEG\_EFFECTOR\_CD4\_TCELL\_UP | 158 | 4 | 12187 | 72 | Zfp518b,Timp2,Aars,H1f0 | | 1.415e-02 | -4.26 | GSE21774\_CD62L\_POS\_CD56\_DIM\_VS\_CD62L\_NEG\_CD56\_DIM\_NK\_CELL\_UP | MSigDB lists | GSE21774\_CD62L\_POS\_CD56\_DIM\_VS\_CD62L\_NEG\_CD56\_DIM\_NK\_CELL\_UP | 158 | 4 | 12187 | 72 | Sorl1,H1f0,Fjx1,Nr4a3 | | 1.415e-02 | -4.26 | GSE17721\_PAM3CSK4\_VS\_CPG\_6H\_BMDC\_DN | MSigDB lists | GSE17721\_PAM3CSK4\_VS\_CPG\_6H\_BMDC\_DN | 158 | 4 | 12187 | 72 | Kcnmb2,Tle4,Gng11,Sorl1 | | 1.415e-02 | -4.26 | GSE3982\_DC\_VS\_BASOPHIL\_UP | MSigDB lists | GSE3982\_DC\_VS\_BASOPHIL\_UP | 158 | 4 | 12187 | 72 | Qdpr,Spry2,Grm3,Inhba | | 1.415e-02 | -4.26 | CHX10\_01 | MSigDB lists | CHX10\_01 | 158 | 4 | 12187 | 72 | Map4,Ndst4,Tle4,Car2 | | 1.415e-02 | -4.26 | GSE32986\_UNSTIM\_VS\_GMCSF\_STIM\_DC\_DN | MSigDB lists | GSE32986\_UNSTIM\_VS\_GMCSF\_STIM\_DC\_DN | 158 | 4 | 12187 | 72 | Agrn,Srm,Map4,Aars | | 1.415e-02 | -4.26 | GSE6092\_B\_BURGDOFERI\_VS\_B\_BURGDORFERI\_AND\_IFNG\_STIM\_ENDOTHELIAL\_CELL\_DN | MSigDB lists | GSE6092\_B\_BURGDOFERI\_VS\_B\_BURGDORFERI\_AND\_IFNG\_STIM\_ENDOTHELIAL\_CELL\_DN | 158 | 4 | 12187 | 72 | Kdr,Hsd17b12,Agrn,Amd1 | | 1.415e-02 | -4.26 | GSE22935\_WT\_VS\_MYD88\_KO\_MACROPHAGE\_DN | MSigDB lists | GSE22935\_WT\_VS\_MYD88\_KO\_MACROPHAGE\_DN | 158 | 4 | 12187 | 72 | Cldn5,Arhgap20,Lims2,Peg3 | | 1.415e-02 | -4.26 | EBAUER\_TARGETS\_OF\_PAX3\_FOXO1\_FUSION\_UP | MSigDB lists | EBAUER\_TARGETS\_OF\_PAX3\_FOXO1\_FUSION\_UP | 158 | 4 | 12187 | 72 | Qk,Col1a1,Agrn,Ncam1 | | 1.422e-02 | -4.25 | Glycosaminoglycan biosynthesis - heparan sulfate / heparin | KEGG pathways | mmu00534 | 23 | 2 | 5248 | 42 | Hs3st1,Ndst4 | | 1.422e-02 | -4.25 | Glycosaminoglycan biosynthesis - heparan sulfate / heparin | KEGG pathways | ko00534 | 23 | 2 | 5248 | 42 | Ndst4,Hs3st1 | | 1.426e-02 | -4.25 | GO\_FORMATION\_OF\_PRIMARY\_GERM\_LAYER | MSigDB lists | GO\_FORMATION\_OF\_PRIMARY\_GERM\_LAYER | 86 | 3 | 12187 | 72 | Nr4a3,Inhba,Lhx1 | | 1.426e-02 | -4.25 | KCCGNSWTTT\_UNKNOWN | MSigDB lists | KCCGNSWTTT\_UNKNOWN | 86 | 3 | 12187 | 72 | Ddx17,Ncam1,Gng11 | | 1.426e-02 | -4.25 | GSE32901\_NAIVE\_VS\_TH17\_NEG\_CD4\_TCELL\_DN | MSigDB lists | GSE32901\_NAIVE\_VS\_TH17\_NEG\_CD4\_TCELL\_DN | 86 | 3 | 12187 | 72 | Inhba,Fjx1,Srm | | 1.428e-02 | -4.25 | organic acid binding | molecular function | GO:0043177 | 162 | 4 | 13516 | 78 | Hba-a2,Fabp7,Glud1,Aars | | 1.433e-02 | -4.25 | GO\_SULFOTRANSFERASE\_ACTIVITY | MSigDB lists | GO\_SULFOTRANSFERASE\_ACTIVITY | 31 | 2 | 12187 | 72 | Ndst4,Hs3st1 | | 1.433e-02 | -4.25 | GO\_CELL\_DIFFERENTIATION\_IN\_SPINAL\_CORD | MSigDB lists | GO\_CELL\_DIFFERENTIATION\_IN\_SPINAL\_CORD | 31 | 2 | 12187 | 72 | Gbx1,Lhx1 | | 1.433e-02 | -4.25 | ZUCCHI\_METASTASIS\_UP | MSigDB lists | ZUCCHI\_METASTASIS\_UP | 31 | 2 | 12187 | 72 | Tgfa,Gng11 | | 1.433e-02 | -4.25 | SATO\_SILENCED\_BY\_METHYLATION\_IN\_PANCREATIC\_CANCER\_2 | MSigDB lists | SATO\_SILENCED\_BY\_METHYLATION\_IN\_PANCREATIC\_CANCER\_2 | 31 | 2 | 12187 | 72 | Cldn5,Lhx1 | | 1.433e-02 | -4.25 | DOANE\_BREAST\_CANCER\_ESR1\_DN | MSigDB lists | DOANE\_BREAST\_CANCER\_ESR1\_DN | 31 | 2 | 12187 | 72 | Fabp7,Ugt8a | | 1.436e-02 | -4.24 | Plcb4 (phospholipase C, beta 4) | protein interactions | 18798 | 2 | 1 | 6802 | 49 | Homer1 | | 1.436e-02 | -4.24 | Fst (follistatin) | protein interactions | 14313 | 2 | 1 | 6802 | 49 | Inhba | | 1.436e-02 | -4.24 | Tmem204 (transmembrane protein 204) | protein interactions | 407831 | 2 | 1 | 6802 | 49 | Kdr | | 1.436e-02 | -4.24 | Tgfb2 (transforming growth factor, beta 2) | protein interactions | 21808 | 2 | 1 | 6802 | 49 | Nrep | | 1.436e-02 | -4.24 | Acvr1b (activin A receptor, type 1B) | protein interactions | 11479 | 2 | 1 | 6802 | 49 | Inhba | | 1.436e-02 | -4.24 | LMAN2L (lectin, mannose binding 2 like) | protein interactions | 81562 | 2 | 1 | 6802 | 49 | Homer1 | | 1.436e-02 | -4.24 | Pde6g (phosphodiesterase 6G, cGMP-specific, rod, gamma) | protein interactions | 18588 | 2 | 1 | 6802 | 49 | Fnbp1 | | 1.436e-02 | -4.24 | Nelfe (negative elongation factor complex member E, Rdbp) | protein interactions | 27632 | 2 | 1 | 6802 | 49 | Egr1 | | 1.436e-02 | -4.24 | VEGFA (vascular endothelial growth factor A) | protein interactions | 7422 | 2 | 1 | 6802 | 49 | Kdr | | 1.436e-02 | -4.24 | Nab2 (Ngfi-A binding protein 2) | protein interactions | 17937 | 2 | 1 | 6802 | 49 | Egr1 | | 1.436e-02 | -4.24 | Cntn4 (contactin 4) | protein interactions | 269784 | 2 | 1 | 6802 | 49 | Homer1 | | 1.436e-02 | -4.24 | Bmp1 (bone morphogenetic protein 1) | protein interactions | 12153 | 2 | 1 | 6802 | 49 | Col1a1 | | 1.436e-02 | -4.24 | Acvr2a (activin receptor IIA) | protein interactions | 11480 | 2 | 1 | 6802 | 49 | Inhba | | 1.436e-02 | -4.24 | Serpinf1 (serine (or cysteine) peptidase inhibitor, clade F, member 1) | protein interactions | 20317 | 2 | 1 | 6802 | 49 | Col1a1 | | 1.436e-02 | -4.24 | Cd47 (CD47 antigen (Rh-related antigen, integrin-associated signal transducer)) | protein interactions | 16423 | 2 | 1 | 6802 | 49 | Ubqln2 | | 1.445e-02 | -4.24 | GSE15930\_NAIVE\_VS\_48H\_IN\_VITRO\_STIM\_IFNAB\_CD8\_TCELL\_UP | MSigDB lists | GSE15930\_NAIVE\_VS\_48H\_IN\_VITRO\_STIM\_IFNAB\_CD8\_TCELL\_UP | 159 | 4 | 12187 | 72 | Car2,Timp2,Nr4a3,Sorl1 | | 1.455e-02 | -4.23 | positive regulation of macromolecule metabolic process | biological process | GO:0010604 | 2653 | 24 | 13711 | 80 | Nr4a3,H1f0,Agrn,Ubqln2,Gsdme,Inhba,Sorl1,Crh,Plp1,Tgfa,Kdr,Timp2,Lhx1,Csrnp3,Spry2,Col1a1,Gbx1,Ddx17,Egr1,Qk,Alkal2,Peg3,Arntl,Bdnf | | 1.457e-02 | -4.23 | heparan sulfate biosynthesis (late stages) | BIOCYC pathways | MOUSE\_PWY-6558 | 16 | 2 | 823 | 10 | Ndst4,Hs3st1 | | 1.458e-02 | -4.23 | sulfotransferase activity | molecular function | GO:0008146 | 32 | 2 | 13516 | 78 | Ndst4,Hs3st1 | | 1.462e-02 | -4.23 | positive regulation of cellular metabolic process | biological process | GO:0031325 | 2654 | 24 | 13711 | 80 | Nr4a3,H1f0,Agrn,Ubqln2,Gsdme,Inhba,Mtmr9,Crh,Tgfa,Kdr,Timp2,Lhx1,Csrnp3,Spry2,Col1a1,Gbx1,Ddx17,Egr1,Qk,Alkal2,Peg3,Arntl,Dgkz,Bdnf | | 1.471e-02 | -4.22 | LENAOUR\_DENDRITIC\_CELL\_MATURATION\_UP | MSigDB lists | LENAOUR\_DENDRITIC\_CELL\_MATURATION\_UP | 87 | 3 | 12187 | 72 | Nr4a3,Inhba,Tgfa | | 1.471e-02 | -4.22 | MODULE\_92 | MSigDB lists | MODULE\_92 | 87 | 3 | 12187 | 72 | Tgfa,Inhba,Ncam1 | | 1.475e-02 | -4.22 | GSE5589\_LPS\_VS\_LPS\_AND\_IL10\_STIM\_IL6\_KO\_MACROPHAGE\_45MIN\_DN | MSigDB lists | GSE5589\_LPS\_VS\_LPS\_AND\_IL10\_STIM\_IL6\_KO\_MACROPHAGE\_45MIN\_DN | 160 | 4 | 12187 | 72 | Spry2,Pom121,Tmem165,Inhba | | 1.475e-02 | -4.22 | GSE20715\_0H\_VS\_48H\_OZONE\_LUNG\_UP | MSigDB lists | GSE20715\_0H\_VS\_48H\_OZONE\_LUNG\_UP | 160 | 4 | 12187 | 72 | Dgkz,Gng11,Sorl1,H1f0 | | 1.475e-02 | -4.22 | GSE36826\_NORMAL\_VS\_STAPH\_AUREUS\_INF\_SKIN\_UP | MSigDB lists | GSE36826\_NORMAL\_VS\_STAPH\_AUREUS\_INF\_SKIN\_UP | 160 | 4 | 12187 | 72 | Sorl1,Aars,Ddx17,Gng11 | | 1.475e-02 | -4.22 | GSE23321\_CENTRAL\_MEMORY\_VS\_NAIVE\_CD8\_TCELL\_UP | MSigDB lists | GSE23321\_CENTRAL\_MEMORY\_VS\_NAIVE\_CD8\_TCELL\_UP | 160 | 4 | 12187 | 72 | Timp2,Inhba,Agrn,Egr1 | | 1.475e-02 | -4.22 | CTACTGT\_MIR199A | MSigDB lists | CTACTGT\_MIR199A | 160 | 4 | 12187 | 72 | Arhgap20,Foxq1,Kcnmb2,Qk | | 1.480e-02 | -4.21 | GO\_CELLULAR\_RESPONSE\_TO\_ORGANIC\_SUBSTANCE | MSigDB lists | GO\_CELLULAR\_RESPONSE\_TO\_ORGANIC\_SUBSTANCE | 1393 | 15 | 12187 | 72 | P2ry12,Cldn5,Nr4a3,Ncam1,Pdia6,Aars,Ltk,Crh,Kdr,Lhx1,Inhba,Col1a1,Car2,Gng11,Egr1 | | 1.489e-02 | -4.21 | establishment of spindle orientation | biological process | GO:0051294 | 32 | 2 | 13711 | 80 | Spry2,Map4 | | 1.492e-02 | -4.20 | GO\_SINGLE\_ORGANISM\_BIOSYNTHETIC\_PROCESS | MSigDB lists | GO\_SINGLE\_ORGANISM\_BIOSYNTHETIC\_PROCESS | 1013 | 12 | 12187 | 72 | Amd1,Plp1,Col1a1,Ugt8a,Msmo1,Hsd17b12,Glud1,Hs3st1,Ndst4,Qk,Crh,Agrn | | 1.500e-02 | -4.20 | GO\_NEGATIVE\_REGULATION\_OF\_CELL\_DEATH | MSigDB lists | GO\_NEGATIVE\_REGULATION\_OF\_CELL\_DEATH | 658 | 9 | 12187 | 72 | Nr4a3,Kdr,Crh,Aars,Spry2,Bdnf,Sorl1,Ltk,Lims2 | | 1.501e-02 | -4.20 | Syngap1 (synaptic Ras GTPase activating protein 1 homolog (rat)) | protein interactions | 240057 | 133 | 4 | 6802 | 49 | Homer1,Ncam1,Dgkz,Cacng2 | | 1.506e-02 | -4.20 | BILD\_MYC\_ONCOGENIC\_SIGNATURE | MSigDB lists | BILD\_MYC\_ONCOGENIC\_SIGNATURE | 161 | 4 | 12187 | 72 | Srm,Col1a1,Arntl,Sorl1 | | 1.506e-02 | -4.20 | GSE15930\_NAIVE\_VS\_24H\_IN\_VITRO\_STIM\_INFAB\_CD8\_TCELL\_UP | MSigDB lists | GSE15930\_NAIVE\_VS\_24H\_IN\_VITRO\_STIM\_INFAB\_CD8\_TCELL\_UP | 161 | 4 | 12187 | 72 | Car2,Nr4a3,Timp2,Tle4 | | 1.514e-02 | -4.19 | morphogenesis of a branching structure | biological process | GO:0001763 | 163 | 4 | 13711 | 80 | Lhx1,Kdr,Ermn,Spry2 | | 1.516e-02 | -4.19 | GO\_REGULATION\_OF\_EPITHELIAL\_CELL\_DIFFERENTIATION | MSigDB lists | GO\_REGULATION\_OF\_EPITHELIAL\_CELL\_DIFFERENTIATION | 88 | 3 | 12187 | 72 | Cldn5,Arntl,Lhx1 | | 1.516e-02 | -4.19 | REACTOME\_GLYCOSAMINOGLYCAN\_METABOLISM | MSigDB lists | REACTOME\_GLYCOSAMINOGLYCAN\_METABOLISM | 88 | 3 | 12187 | 72 | Ndst4,Hs3st1,Agrn | | 1.523e-02 | -4.18 | GO\_REGULATION\_OF\_ENDOCRINE\_PROCESS | MSigDB lists | GO\_REGULATION\_OF\_ENDOCRINE\_PROCESS | 32 | 2 | 12187 | 72 | Inhba,Crh | | 1.523e-02 | -4.18 | PID\_NFAT\_TFPATHWAY | MSigDB lists | PID\_NFAT\_TFPATHWAY | 32 | 2 | 12187 | 72 | Egr1,Tle4 | | 1.523e-02 | -4.18 | chr2q24 | MSigDB lists | chr2q24 | 32 | 2 | 12187 | 72 | Csrnp3,Ermn | | 1.523e-02 | -4.18 | GO\_GLUTAMATE\_RECEPTOR\_BINDING | MSigDB lists | GO\_GLUTAMATE\_RECEPTOR\_BINDING | 32 | 2 | 12187 | 72 | Homer1,Cacng2 | | 1.523e-02 | -4.18 | GO\_SPINDLE\_LOCALIZATION | MSigDB lists | GO\_SPINDLE\_LOCALIZATION | 32 | 2 | 12187 | 72 | Map4,Spry2 | | 1.523e-02 | -4.18 | JEPSEN\_SMRT\_TARGETS | MSigDB lists | JEPSEN\_SMRT\_TARGETS | 32 | 2 | 12187 | 72 | Lims2,Egr1 | | 1.523e-02 | -4.18 | PID\_INTEGRIN3\_PATHWAY | MSigDB lists | PID\_INTEGRIN3\_PATHWAY | 32 | 2 | 12187 | 72 | Col1a1,Kdr | | 1.537e-02 | -4.18 | regulation of synapse organization | biological process | GO:0050807 | 250 | 5 | 13711 | 80 | Cbln2,Agrn,Lrrtm2,Homer1,Bdnf | | 1.538e-02 | -4.17 | GSE19941\_UNSTIM\_VS\_LPS\_STIM\_IL10\_KO\_MACROPHAGE\_DN | MSigDB lists | GSE19941\_UNSTIM\_VS\_LPS\_STIM\_IL10\_KO\_MACROPHAGE\_DN | 162 | 4 | 12187 | 72 | Kcnmb2,Gbx1,Fabp7,Hs3st1 | | 1.538e-02 | -4.17 | BURTON\_ADIPOGENESIS\_6 | MSigDB lists | BURTON\_ADIPOGENESIS\_6 | 162 | 4 | 12187 | 72 | Qdpr,Hsd17b12,Qk,Car2 | | 1.538e-02 | -4.17 | GSE17721\_LPS\_VS\_GARDIQUIMOD\_12H\_BMDC\_DN | MSigDB lists | GSE17721\_LPS\_VS\_GARDIQUIMOD\_12H\_BMDC\_DN | 162 | 4 | 12187 | 72 | Hsd17b12,Aars,Egr1,Tmem165 | | 1.538e-02 | -4.17 | GSE7831\_UNSTIM\_VS\_CPG\_STIM\_PDC\_4H\_UP | MSigDB lists | GSE7831\_UNSTIM\_VS\_CPG\_STIM\_PDC\_4H\_UP | 162 | 4 | 12187 | 72 | Tgfa,Arntl,Car2,Pom121 | | 1.543e-02 | -4.17 | GO\_G\_PROTEIN\_COUPLED\_RECEPTOR\_SIGNALING\_PATHWAY | MSigDB lists | GO\_G\_PROTEIN\_COUPLED\_RECEPTOR\_SIGNALING\_PATHWAY | 443 | 7 | 12187 | 72 | Agrn,P2ry12,Homer1,Gng11,Car2,Dgkz,Grm3 | | 1.562e-02 | -4.16 | growth factor activity | molecular function | GO:0008083 | 91 | 3 | 13516 | 78 | Tgfa,Bdnf,Inhba | | 1.563e-02 | -4.16 | GO\_REGULATION\_OF\_CAMP\_METABOLIC\_PROCESS | MSigDB lists | GO\_REGULATION\_OF\_CAMP\_METABOLIC\_PROCESS | 89 | 3 | 12187 | 72 | Timp2,Crh,Grm3 | | 1.563e-02 | -4.16 | YAO\_TEMPORAL\_RESPONSE\_TO\_PROGESTERONE\_CLUSTER\_11 | MSigDB lists | YAO\_TEMPORAL\_RESPONSE\_TO\_PROGESTERONE\_CLUSTER\_11 | 89 | 3 | 12187 | 72 | Pdia6,Aars,Srm | | 1.566e-02 | -4.16 | GO\_SYNAPTIC\_SIGNALING | MSigDB lists | GO\_SYNAPTIC\_SIGNALING | 343 | 6 | 12187 | 72 | Grm3,Homer1,Crh,Plp1,Agrn,Kcnmb2 | | 1.569e-02 | -4.15 | GO\_CELLULAR\_LIPID\_METABOLIC\_PROCESS | MSigDB lists | GO\_CELLULAR\_LIPID\_METABOLIC\_PROCESS | 663 | 9 | 12187 | 72 | Plp1,Dgkz,Hsd17b12,Msmo1,Ugt8a,Fabp7,Qk,Crh,Agrn | | 1.570e-02 | -4.15 | GSE20715\_0H\_VS\_24H\_OZONE\_LUNG\_UP | MSigDB lists | GSE20715\_0H\_VS\_24H\_OZONE\_LUNG\_UP | 163 | 4 | 12187 | 72 | Gng11,Kdr,Dgkz,Car2 | | 1.570e-02 | -4.15 | GSE15930\_STIM\_VS\_STIM\_AND\_IL12\_48H\_CD8\_T\_CELL\_DN | MSigDB lists | GSE15930\_STIM\_VS\_STIM\_AND\_IL12\_48H\_CD8\_T\_CELL\_DN | 163 | 4 | 12187 | 72 | Tmem165,Ddx17,Pdia6,Bdnf | | 1.570e-02 | -4.15 | GSE43955\_1H\_VS\_60H\_ACT\_CD4\_TCELL\_WITH\_TGFB\_IL6\_DN | MSigDB lists | GSE43955\_1H\_VS\_60H\_ACT\_CD4\_TCELL\_WITH\_TGFB\_IL6\_DN | 163 | 4 | 12187 | 72 | Arntl,Kdr,Col1a1,Glud1 | | 1.574e-02 | -4.15 | positive regulation of nitrogen compound metabolic process | biological process | GO:0051173 | 2525 | 23 | 13711 | 80 | Crh,Tgfa,Kdr,Timp2,Lhx1,Nr4a3,H1f0,Agrn,Ubqln2,Inhba,Sorl1,Gsdme,Egr1,Ddx17,Peg3,Arntl,Alkal2,Qk,Bdnf,Gbx1,Col1a1,Spry2,Csrnp3 | | 1.579e-02 | -4.15 | positive regulation of kidney development | biological process | GO:0090184 | 33 | 2 | 13711 | 80 | Lhx1,Egr1 | | 1.579e-02 | -4.15 | positive regulation of signaling receptor activity | biological process | GO:2000273 | 33 | 2 | 13711 | 80 | Tgfa,Cacng2 | | 1.579e-02 | -4.15 | regulation of synaptic transmission, GABAergic | biological process | GO:0032228 | 33 | 2 | 13711 | 80 | Bdnf,Car2 | | 1.602e-02 | -4.13 | GSE40666\_WT\_VS\_STAT1\_KO\_CD8\_TCELL\_UP | MSigDB lists | GSE40666\_WT\_VS\_STAT1\_KO\_CD8\_TCELL\_UP | 164 | 4 | 12187 | 72 | Nr4a3,Ephb6,Car2,Sorl1 | | 1.602e-02 | -4.13 | GSE17721\_LPS\_VS\_POLYIC\_8H\_BMDC\_DN | MSigDB lists | GSE17721\_LPS\_VS\_POLYIC\_8H\_BMDC\_DN | 164 | 4 | 12187 | 72 | Nap1l5,Homer1,Aars,Sorl1 | | 1.602e-02 | -4.13 | GSE5589\_IL6\_KO\_VS\_IL10\_KO\_LPS\_AND\_IL10\_STIM\_MACROPHAGE\_45MIN\_DN | MSigDB lists | GSE5589\_IL6\_KO\_VS\_IL10\_KO\_LPS\_AND\_IL10\_STIM\_MACROPHAGE\_45MIN\_DN | 164 | 4 | 12187 | 72 | Dgkz,Ubqln2,Fabp7,Lims2 | | 1.609e-02 | -4.13 | cerebellum development | biological process | GO:0021549 | 91 | 3 | 13711 | 80 | Aars,Lhx1,Gbx1 | | 1.610e-02 | -4.13 | GRE\_C | MSigDB lists | GRE\_C | 90 | 3 | 12187 | 72 | Sorl1,Egr1,Bdnf | | 1.615e-02 | -4.13 | DELPUECH\_FOXO3\_TARGETS\_DN | MSigDB lists | DELPUECH\_FOXO3\_TARGETS\_DN | 33 | 2 | 12187 | 72 | Aars,H1f0 | | 1.615e-02 | -4.13 | MODY\_HIPPOCAMPUS\_NEONATAL | MSigDB lists | MODY\_HIPPOCAMPUS\_NEONATAL | 33 | 2 | 12187 | 72 | Fabp7,Ncam1 | | 1.615e-02 | -4.13 | KRAS.50\_UP.V1\_UP | MSigDB lists | KRAS.50\_UP.V1\_UP | 33 | 2 | 12187 | 72 | Peg3,Ntm | | 1.615e-02 | -4.13 | GO\_HEAD\_MORPHOGENESIS | MSigDB lists | GO\_HEAD\_MORPHOGENESIS | 33 | 2 | 12187 | 72 | Cldn5,Col1a1 | | 1.615e-02 | -4.13 | ELLWOOD\_MYC\_TARGETS\_DN | MSigDB lists | ELLWOOD\_MYC\_TARGETS\_DN | 33 | 2 | 12187 | 72 | Pdia6,Car2 | | 1.615e-02 | -4.13 | CTCTATG\_MIR368 | MSigDB lists | CTCTATG\_MIR368 | 33 | 2 | 12187 | 72 | Rgs4,Bdnf | | 1.615e-02 | -4.13 | GO\_REGULATION\_OF\_RECEPTOR\_INTERNALIZATION | MSigDB lists | GO\_REGULATION\_OF\_RECEPTOR\_INTERNALIZATION | 33 | 2 | 12187 | 72 | Ubqln2,Lrrtm2 | | 1.615e-02 | -4.13 | GO\_BICARBONATE\_TRANSPORT | MSigDB lists | GO\_BICARBONATE\_TRANSPORT | 33 | 2 | 12187 | 72 | Hba-a2,Car2 | | 1.615e-02 | -4.13 | Thioredoxin\_domain | interpro domains | IPR013766 | 34 | 2 | 13788 | 79 | Pdia6,Tmx3 | | 1.633e-02 | -4.12 | GO\_SMALL\_MOLECULE\_METABOLIC\_PROCESS | MSigDB lists | GO\_SMALL\_MOLECULE\_METABOLIC\_PROCESS | 1279 | 14 | 12187 | 72 | Qdpr,Ndst4,Glud1,Aars,Nr4a3,Ugt8a,Hkdc1,Plp1,Qk,Sorl1,Amd1,Hsd17b12,Msmo1,Car2 | | 1.634e-02 | -4.11 | GSE23568\_ID3\_KO\_VS\_WT\_CD8\_TCELL\_DN | MSigDB lists | GSE23568\_ID3\_KO\_VS\_WT\_CD8\_TCELL\_DN | 165 | 4 | 12187 | 72 | Sorl1,Hs3st1,Nr4a3,Car2 | | 1.634e-02 | -4.11 | GSE30083\_SP1\_VS\_SP4\_THYMOCYTE\_UP | MSigDB lists | GSE30083\_SP1\_VS\_SP4\_THYMOCYTE\_UP | 165 | 4 | 12187 | 72 | Ephb6,Hsd17b12,Kcnmb2,Fjx1 | | 1.637e-02 | -4.11 | response to inorganic substance | biological process | GO:0010035 | 350 | 6 | 13711 | 80 | Glud1,Kcnmb2,Ncam1,Nr4a3,Homer1,Kdr | | 1.649e-02 | -4.10 | CHARAFE\_BREAST\_CANCER\_LUMINAL\_VS\_BASAL\_DN | MSigDB lists | CHARAFE\_BREAST\_CANCER\_LUMINAL\_VS\_BASAL\_DN | 347 | 6 | 12187 | 72 | Amd1,Spry2,Foxq1,Inhba,Tle4,Tgfa | | 1.652e-02 | -4.10 | regulation of cell adhesion | biological process | GO:0030155 | 562 | 8 | 13711 | 80 | Ephb6,Lims2,Atp5b,P2ry12,Kdr,Hsd17b12,Nr4a3,Col1a1 | | 1.658e-02 | -4.10 | GO\_DEVELOPMENTAL\_GROWTH\_INVOLVED\_IN\_MORPHOGENESIS | MSigDB lists | GO\_DEVELOPMENTAL\_GROWTH\_INVOLVED\_IN\_MORPHOGENESIS | 91 | 3 | 12187 | 72 | Lhx1,Bdnf,Spry2 | | 1.658e-02 | -4.10 | GO\_GROWTH\_FACTOR\_ACTIVITY | MSigDB lists | GO\_GROWTH\_FACTOR\_ACTIVITY | 91 | 3 | 12187 | 72 | Bdnf,Inhba,Tgfa | | 1.660e-02 | -4.10 | mouse chr19 C3\|19 37.98 cM | chromosome location | mouse chr19 C3\|19 37.98 cM | 3 | 1 | 14556 | 81 | Scd2 | | 1.660e-02 | -4.10 | mouse chr2 E5\|2 59.97 cM | chromosome location | mouse chr2 E5\|2 59.97 cM | 3 | 1 | 14556 | 81 | Ltk | | 1.660e-02 | -4.10 | mouse chr14 B\|14 20.8 cM | chromosome location | mouse chr14 B\|14 20.8 cM | 3 | 1 | 14556 | 81 | Glud1 | | 1.667e-02 | -4.09 | GSE22935\_24H\_VS\_48H\_MBOVIS\_BCG\_STIM\_MYD88\_KO\_MACROPHAGE\_UP | MSigDB lists | GSE22935\_24H\_VS\_48H\_MBOVIS\_BCG\_STIM\_MYD88\_KO\_MACROPHAGE\_UP | 166 | 4 | 12187 | 72 | Nap1l5,Qk,Timp2,Cacng2 | | 1.667e-02 | -4.09 | GSE33424\_CD161\_HIGH\_VS\_NEG\_CD8\_TCELL\_UP | MSigDB lists | GSE33424\_CD161\_HIGH\_VS\_NEG\_CD8\_TCELL\_UP | 166 | 4 | 12187 | 72 | Nr4a3,Car2,Rgs4,Hs3st1 | | 1.668e-02 | -4.09 | VEGFR-2\_TMD | pfam domains | PF17988 | 3 | 1 | 12881 | 72 | Kdr | | 1.668e-02 | -4.09 | NGF | pfam domains | PF00243 | 3 | 1 | 12881 | 72 | Bdnf | | 1.668e-02 | -4.09 | Hexokinase\_1 | pfam domains | PF00349 | 3 | 1 | 12881 | 72 | Hkdc1 | | 1.668e-02 | -4.09 | DUF3446 | pfam domains | PF11928 | 3 | 1 | 12881 | 72 | Egr1 | | 1.668e-02 | -4.09 | STI1 | pfam domains | PF17830 | 3 | 1 | 12881 | 72 | Ubqln2 | | 1.668e-02 | -4.09 | Hexokinase\_2 | pfam domains | PF03727 | 3 | 1 | 12881 | 72 | Hkdc1 | | 1.668e-02 | -4.09 | CSRNP\_N | pfam domains | PF16019 | 3 | 1 | 12881 | 72 | Csrnp3 | | 1.668e-02 | -4.09 | CaKB | pfam domains | PF03185 | 3 | 1 | 12881 | 72 | Kcnmb2 | | 1.668e-02 | -4.09 | Myelin\_PLP | pfam domains | PF01275 | 3 | 1 | 12881 | 72 | Plp1 | | 1.668e-02 | -4.09 | Tubulin-binding | pfam domains | PF00418 | 3 | 1 | 12881 | 72 | Map4 | | 1.668e-02 | -4.09 | NIPSNAP | pfam domains | PF07978 | 3 | 1 | 12881 | 72 | Nipsnap2 | | 1.670e-02 | -4.09 | signaling receptor binding | molecular function | GO:0005102 | 1178 | 13 | 13516 | 78 | Lrrtm2,Tgfa,Homer1,Bdnf,Crh,Inhba,Nr4a3,Atp5b,Kdr,Arntl,Alkal2,Timp2,Cacng2 | | 1.672e-02 | -4.09 | cellular biogenic amine metabolic process | biological process | GO:0006576 | 34 | 2 | 13711 | 80 | Srm,Amd1 | | 1.672e-02 | -4.09 | negative regulation of potassium ion transport | biological process | GO:0043267 | 34 | 2 | 13711 | 80 | Rgs4,Agrn | | 1.672e-02 | -4.09 | cellular amine metabolic process | biological process | GO:0044106 | 34 | 2 | 13711 | 80 | Srm,Amd1 | | 1.672e-02 | -4.09 | response to axon injury | biological process | GO:0048678 | 34 | 2 | 13711 | 80 | P2ry12,Nrep | | 1.679e-02 | -4.09 | Interactions of neurexins and neuroligins at synapses | REACTOME pathways | R-MMU-6794361 | 30 | 2 | 6297 | 42 | Homer1,Lrrtm2 | | 1.700e-02 | -4.07 | regulation of phosphate metabolic process | biological process | GO:0019220 | 1427 | 15 | 13711 | 80 | Dgkz,Bdnf,Alkal2,Egr1,Kdr,Tgfa,Timp2,Crh,Gsdme,Inhba,Mtmr9,Sorl1,Spry2,Csrnp3,Agrn | | 1.701e-02 | -4.07 | RODWELL\_AGING\_KIDNEY\_NO\_BLOOD\_UP | MSigDB lists | RODWELL\_AGING\_KIDNEY\_NO\_BLOOD\_UP | 167 | 4 | 12187 | 72 | Map4,Col1a1,Timp2,Inhba | | 1.701e-02 | -4.07 | GSE19923\_HEB\_KO\_VS\_HEB\_AND\_E2A\_KO\_DP\_THYMOCYTE\_DN | MSigDB lists | GSE19923\_HEB\_KO\_VS\_HEB\_AND\_E2A\_KO\_DP\_THYMOCYTE\_DN | 167 | 4 | 12187 | 72 | Tgfa,Msmo1,Hsd17b12,Egr1 | | 1.701e-02 | -4.07 | GSE32986\_GMCSF\_VS\_GMCSF\_AND\_CURDLAN\_HIGHDOSE\_STIM\_DC\_UP | MSigDB lists | GSE32986\_GMCSF\_VS\_GMCSF\_AND\_CURDLAN\_HIGHDOSE\_STIM\_DC\_UP | 167 | 4 | 12187 | 72 | Timp2,Nr4a3,Qdpr,Spry2 | | 1.701e-02 | -4.07 | GSE20198\_IL12\_VS\_IFNA\_TREATED\_ACT\_CD4\_TCELL\_DN | MSigDB lists | GSE20198\_IL12\_VS\_IFNA\_TREATED\_ACT\_CD4\_TCELL\_DN | 167 | 4 | 12187 | 72 | Tmem165,Mtmr9,Dgkz,Ubqln2 | | 1.701e-02 | -4.07 | NABA\_SECRETED\_FACTORS | MSigDB lists | NABA\_SECRETED\_FACTORS | 167 | 4 | 12187 | 72 | Tgfa,Cbln2,Inhba,Bdnf | | 1.702e-02 | -4.07 | ion channel regulator activity | molecular function | GO:0099106 | 94 | 3 | 13516 | 78 | Kcnmb2,Lrrc55,Grm3 | | 1.704e-02 | -4.07 | female sex differentiation | biological process | GO:0046660 | 93 | 3 | 13711 | 80 | Lhx1,Kdr,Inhba | | 1.705e-02 | -4.07 | inhibin complex | cellular component | GO:0043511 | 3 | 1 | 13825 | 79 | Inhba | | 1.705e-02 | -4.07 | nascent polypeptide-associated complex | cellular component | GO:0005854 | 3 | 1 | 13825 | 79 | Nacad | | 1.706e-02 | -4.07 | Sptan1 (spectrin alpha, non-erythrocytic 1) | protein interactions | 20740 | 28 | 2 | 6802 | 49 | Ncam1,Homer1 | | 1.707e-02 | -4.07 | SDR\_fam | interpro domains | IPR002347 | 35 | 2 | 13788 | 79 | Qdpr,Hsd17b12 | | 1.707e-02 | -4.07 | RNA\_helicase\_DEAD\_Q\_motif | interpro domains | IPR014014 | 35 | 2 | 13788 | 79 | Ddx17,Ddx56 | | 1.707e-02 | -4.07 | TGCAAAC\_MIR452 | MSigDB lists | TGCAAAC\_MIR452 | 92 | 3 | 12187 | 72 | Ubqln2,Kcnmb2,Nr4a3 | | 1.707e-02 | -4.07 | GSE16385\_UNTREATED\_VS\_12H\_ROSIGLITAZONE\_TREATED\_MACROPHAGE\_UP | MSigDB lists | GSE16385\_UNTREATED\_VS\_12H\_ROSIGLITAZONE\_TREATED\_MACROPHAGE\_UP | 92 | 3 | 12187 | 72 | Lrrtm2,Arhgap20,Foxq1 | | 1.709e-02 | -4.07 | Nerve\_growth\_factor-rel | interpro domains | IPR002072 | 3 | 1 | 13788 | 79 | Bdnf | | 1.709e-02 | -4.07 | Acyl-CoA\_DS | interpro domains | IPR015876 | 3 | 1 | 13788 | 79 | Scd2 | | 1.709e-02 | -4.07 | EGR\_N | interpro domains | IPR021849 | 3 | 1 | 13788 | 79 | Egr1 | | 1.709e-02 | -4.07 | Hexokinase\_N | interpro domains | IPR022672 | 3 | 1 | 13788 | 79 | Hkdc1 | | 1.709e-02 | -4.07 | FADS-1\_CS | interpro domains | IPR001522 | 3 | 1 | 13788 | 79 | Scd2 | | 1.709e-02 | -4.07 | Nerve\_growth\_factor-like | interpro domains | IPR020408 | 3 | 1 | 13788 | 79 | Bdnf | | 1.709e-02 | -4.07 | Hexokinase\_C | interpro domains | IPR022673 | 3 | 1 | 13788 | 79 | Hkdc1 | | 1.709e-02 | -4.07 | MAP\_tubulin-bd\_rpt | interpro domains | IPR001084 | 3 | 1 | 13788 | 79 | Map4 | | 1.709e-02 | -4.07 | K\_chnl\_Ca-activ\_BK\_bsu | interpro domains | IPR003930 | 3 | 1 | 13788 | 79 | Kcnmb2 | | 1.709e-02 | -4.07 | Nerve\_growth\_factor\_CS | interpro domains | IPR019846 | 3 | 1 | 13788 | 79 | Bdnf | | 1.709e-02 | -4.07 | STI1 | interpro domains | IPR041243 | 3 | 1 | 13788 | 79 | Ubqln2 | | 1.709e-02 | -4.07 | VEGFR-2\_TMD | interpro domains | IPR041348 | 3 | 1 | 13788 | 79 | Kdr | | 1.709e-02 | -4.07 | Cys/Ser-rich\_nuc\_prot | interpro domains | IPR023260 | 3 | 1 | 13788 | 79 | Csrnp3 | | 1.709e-02 | -4.07 | Myelin\_PLP\_CS | interpro domains | IPR018237 | 3 | 1 | 13788 | 79 | Plp1 | | 1.709e-02 | -4.07 | Nucleoporin | interpro domains | IPR026054 | 3 | 1 | 13788 | 79 | Pom121 | | 1.709e-02 | -4.07 | Myelin\_PLP | interpro domains | IPR001614 | 3 | 1 | 13788 | 79 | Plp1 | | 1.709e-02 | -4.07 | CSRNP\_N | interpro domains | IPR031972 | 3 | 1 | 13788 | 79 | Csrnp3 | | 1.709e-02 | -4.07 | Hexokinase | interpro domains | IPR001312 | 3 | 1 | 13788 | 79 | Hkdc1 | | 1.709e-02 | -4.07 | MAP2/MAP4/Tau | interpro domains | IPR027324 | 3 | 1 | 13788 | 79 | Map4 | | 1.709e-02 | -4.07 | ATPase\_F1/V1/A1\_a/bsu\_N\_sf | interpro domains | IPR036121 | 3 | 1 | 13788 | 79 | Atp5b | | 1.709e-02 | -4.07 | Hexokinase\_BS | interpro domains | IPR019807 | 3 | 1 | 13788 | 79 | Hkdc1 | | 1.709e-02 | -4.07 | Nuc\_orph\_rcpt | interpro domains | IPR003070 | 3 | 1 | 13788 | 79 | Nr4a3 | | 1.709e-02 | -4.07 | NIPSNAP | interpro domains | IPR012577 | 3 | 1 | 13788 | 79 | Nipsnap2 | | 1.709e-02 | -4.07 | GO\_REGULATION\_OF\_SKELETAL\_MUSCLE\_TISSUE\_DEVELOPMENT | MSigDB lists | GO\_REGULATION\_OF\_SKELETAL\_MUSCLE\_TISSUE\_DEVELOPMENT | 34 | 2 | 12187 | 72 | Ddx17,Arntl | | 1.709e-02 | -4.07 | GRAHAM\_CML\_QUIESCENT\_VS\_NORMAL\_QUIESCENT\_DN | MSigDB lists | GRAHAM\_CML\_QUIESCENT\_VS\_NORMAL\_QUIESCENT\_DN | 34 | 2 | 12187 | 72 | Sorl1,H1f0 | | 1.709e-02 | -4.07 | GENTILE\_UV\_RESPONSE\_CLUSTER\_D6 | MSigDB lists | GENTILE\_UV\_RESPONSE\_CLUSTER\_D6 | 34 | 2 | 12187 | 72 | Qk,Tle4 | | 1.709e-02 | -4.07 | UZONYI\_RESPONSE\_TO\_LEUKOTRIENE\_AND\_THROMBIN | MSigDB lists | UZONYI\_RESPONSE\_TO\_LEUKOTRIENE\_AND\_THROMBIN | 34 | 2 | 12187 | 72 | Egr1,Nr4a3 | | 1.709e-02 | -4.07 | GO\_RETINA\_MORPHOGENESIS\_IN\_CAMERA\_TYPE\_EYE | MSigDB lists | GO\_RETINA\_MORPHOGENESIS\_IN\_CAMERA\_TYPE\_EYE | 34 | 2 | 12187 | 72 | Fjx1,Lhx1 | | 1.709e-02 | -4.07 | regulation of phosphorus metabolic process | biological process | GO:0051174 | 1428 | 15 | 13711 | 80 | Crh,Kdr,Tgfa,Timp2,Egr1,Alkal2,Bdnf,Dgkz,Agrn,Csrnp3,Spry2,Inhba,Mtmr9,Sorl1,Gsdme | | 1.721e-02 | -4.06 | CTTTGT\_LEF1\_Q2 | MSigDB lists | CTTTGT\_LEF1\_Q2 | 1551 | 16 | 12187 | 72 | Nr4a3,Fjx1,P2ry12,Glud1,Bdnf,Plp1,Dgkz,H1f0,Grm3,Chmp7,Crh,Spry2,Gng11,Col1a1,Ddx17,Csrnp3 | | 1.721e-02 | -4.06 | NADH binding | molecular function | GO:0070404 | 3 | 1 | 13516 | 78 | Qdpr | | 1.721e-02 | -4.06 | corticotropin-releasing hormone receptor 1 binding | molecular function | GO:0051430 | 3 | 1 | 13516 | 78 | Crh | | 1.721e-02 | -4.06 | mannokinase activity | molecular function | GO:0019158 | 3 | 1 | 13516 | 78 | Hkdc1 | | 1.721e-02 | -4.06 | palmitoyl-CoA 9-desaturase activity | molecular function | GO:0032896 | 3 | 1 | 13516 | 78 | Scd2 | | 1.721e-02 | -4.06 | structural constituent of postsynapse | molecular function | GO:0099186 | 3 | 1 | 13516 | 78 | Homer1 | | 1.721e-02 | -4.06 | UDP-galactose:glucosylceramide beta-1,4-galactosyltransferase activity | molecular function | GO:0008489 | 3 | 1 | 13516 | 78 | Ugt8a | | 1.721e-02 | -4.06 | type II activin receptor binding | molecular function | GO:0070699 | 3 | 1 | 13516 | 78 | Inhba | | 1.721e-02 | -4.06 | glucokinase activity | molecular function | GO:0004340 | 3 | 1 | 13516 | 78 | Hkdc1 | | 1.721e-02 | -4.06 | hexokinase activity | molecular function | GO:0004396 | 3 | 1 | 13516 | 78 | Hkdc1 | | 1.721e-02 | -4.06 | fructokinase activity | molecular function | GO:0008865 | 3 | 1 | 13516 | 78 | Hkdc1 | | 1.733e-02 | -4.06 | locomotion | biological process | GO:0040011 | 919 | 11 | 13711 | 80 | Kdr,P2ry12,Ephb6,Bdnf,Lhx1,Nr4a3,Agrn,Gbx1,Sorl1,Ncam1,Atp5b | | 1.735e-02 | -4.05 | GSE12198\_CTRL\_VS\_HIGH\_IL2\_STIM\_NK\_CELL\_DN | MSigDB lists | GSE12198\_CTRL\_VS\_HIGH\_IL2\_STIM\_NK\_CELL\_DN | 168 | 4 | 12187 | 72 | Timp2,Hkdc1,Gng11,Srm | | 1.735e-02 | -4.05 | GSE1925\_3H\_VS\_24H\_IFNG\_STIM\_IFNG\_PRIMED\_MACROPHAGE\_UP | MSigDB lists | GSE1925\_3H\_VS\_24H\_IFNG\_STIM\_IFNG\_PRIMED\_MACROPHAGE\_UP | 168 | 4 | 12187 | 72 | Peg3,Ltk,Ephb6,Agrn | | 1.740e-02 | -4.05 | positive regulation of cortisol secretion | biological process | GO:0051464 | 3 | 1 | 13711 | 80 | Crh | | 1.740e-02 | -4.05 | lateral motor column neuron migration | biological process | GO:0097477 | 3 | 1 | 13711 | 80 | Lhx1 | | 1.740e-02 | -4.05 | spinal cord motor neuron migration | biological process | GO:0097476 | 3 | 1 | 13711 | 80 | Lhx1 | | 1.740e-02 | -4.05 | sensory neuron axon guidance | biological process | GO:0097374 | 3 | 1 | 13711 | 80 | Gbx1 | | 1.740e-02 | -4.05 | spermine biosynthetic process | biological process | GO:0006597 | 3 | 1 | 13711 | 80 | Amd1 | | 1.740e-02 | -4.05 | steroid hormone secretion | biological process | GO:0035929 | 3 | 1 | 13711 | 80 | Inhba | | 1.740e-02 | -4.05 | regulation of progesterone biosynthetic process | biological process | GO:2000182 | 3 | 1 | 13711 | 80 | Egr1 | | 1.740e-02 | -4.05 | cellular response to cocaine | biological process | GO:0071314 | 3 | 1 | 13711 | 80 | Crh | | 1.740e-02 | -4.05 | cellular response to hydrogen sulfide | biological process | GO:1904881 | 3 | 1 | 13711 | 80 | Kdr | | 1.740e-02 | -4.05 | lipid export from cell | biological process | GO:0140353 | 3 | 1 | 13711 | 80 | Inhba | | 1.740e-02 | -4.05 | striatal medium spiny neuron differentiation | biological process | GO:0021773 | 3 | 1 | 13711 | 80 | Inhba | | 1.740e-02 | -4.05 | regulation of G protein-coupled receptor internalization | biological process | GO:1904020 | 3 | 1 | 13711 | 80 | Ubqln2 | | 1.740e-02 | -4.05 | axon target recognition | biological process | GO:0007412 | 3 | 1 | 13711 | 80 | Bdnf | | 1.740e-02 | -4.05 | negative regulation of hepatocyte proliferation | biological process | GO:2000346 | 3 | 1 | 13711 | 80 | Lims2 | | 1.740e-02 | -4.05 | microtubule sliding | biological process | GO:0051012 | 3 | 1 | 13711 | 80 | Map4 | | 1.740e-02 | -4.05 | erythrose 4-phosphate/phosphoenolpyruvate family amino acid catabolic process | biological process | GO:1902222 | 3 | 1 | 13711 | 80 | Qdpr | | 1.740e-02 | -4.05 | negative regulation of dopamine receptor signaling pathway | biological process | GO:0060160 | 3 | 1 | 13711 | 80 | Rgs4 | | 1.740e-02 | -4.05 | response to hydrogen sulfide | biological process | GO:1904880 | 3 | 1 | 13711 | 80 | Kdr | | 1.740e-02 | -4.05 | negative regulation of non-motile cilium assembly | biological process | GO:1902856 | 3 | 1 | 13711 | 80 | Map4 | | 1.740e-02 | -4.05 | maternal process involved in parturition | biological process | GO:0060137 | 3 | 1 | 13711 | 80 | Arntl | | 1.740e-02 | -4.05 | L-phenylalanine catabolic process | biological process | GO:0006559 | 3 | 1 | 13711 | 80 | Qdpr | | 1.740e-02 | -4.05 | positive regulation of metanephric glomerulus development | biological process | GO:0072300 | 3 | 1 | 13711 | 80 | Egr1 | | 1.740e-02 | -4.05 | hypersensitivity | biological process | GO:0002524 | 3 | 1 | 13711 | 80 | Ephb6 | | 1.740e-02 | -4.05 | negative regulation of luteinizing hormone secretion | biological process | GO:0033685 | 3 | 1 | 13711 | 80 | Crh | | 1.740e-02 | -4.05 | monounsaturated fatty acid biosynthetic process | biological process | GO:1903966 | 3 | 1 | 13711 | 80 | Scd2 | | 1.740e-02 | -4.05 | negative regulation of glucagon secretion | biological process | GO:0070093 | 3 | 1 | 13711 | 80 | Crh | | 1.740e-02 | -4.05 | positive regulation of ovulation | biological process | GO:0060279 | 3 | 1 | 13711 | 80 | Inhba | | 1.740e-02 | -4.05 | monounsaturated fatty acid metabolic process | biological process | GO:1903964 | 3 | 1 | 13711 | 80 | Scd2 | | 1.740e-02 | -4.05 | regulation of metanephric glomerulus development | biological process | GO:0072298 | 3 | 1 | 13711 | 80 | Egr1 | | 1.740e-02 | -4.05 | regulation of Rap protein signal transduction | biological process | GO:0032487 | 3 | 1 | 13711 | 80 | Timp2 | | 1.740e-02 | -4.05 | regulation of glycine import across plasma membrane | biological process | GO:1900923 | 3 | 1 | 13711 | 80 | Rgs4 | | 1.740e-02 | -4.05 | synaptic signaling via neuropeptide | biological process | GO:0099538 | 3 | 1 | 13711 | 80 | Bdnf | | 1.740e-02 | -4.05 | circadian temperature homeostasis | biological process | GO:0060086 | 3 | 1 | 13711 | 80 | Egr1 | | 1.740e-02 | -4.05 | glomerular mesangial cell proliferation | biological process | GO:0072110 | 3 | 1 | 13711 | 80 | Egr1 | | 1.753e-02 | -4.04 | regulation of amine transport | biological process | GO:0051952 | 94 | 3 | 13711 | 80 | Crh,P2ry12,Rgs4 | | 1.753e-02 | -4.04 | regulation of potassium ion transport | biological process | GO:0043266 | 94 | 3 | 13711 | 80 | Lrrc55,Agrn,Rgs4 | | 1.762e-02 | -4.04 | REACTOME\_GLYCOPROTEIN\_HORMONES | MSigDB lists | REACTOME\_GLYCOPROTEIN\_HORMONES | 3 | 1 | 12187 | 72 | Inhba | | 1.763e-02 | -4.04 | GO\_CELLULAR\_RESPONSE\_TO\_ENDOGENOUS\_STIMULUS | MSigDB lists | GO\_CELLULAR\_RESPONSE\_TO\_ENDOGENOUS\_STIMULUS | 793 | 10 | 12187 | 72 | Egr1,Inhba,Gng11,Col1a1,Car2,Lhx1,Nr4a3,Cldn5,P2ry12,Crh | | 1.765e-02 | -4.04 | regulation of synapse structure or activity | biological process | GO:0050803 | 259 | 5 | 13711 | 80 | Homer1,Lrrtm2,Bdnf,Agrn,Cbln2 | | 1.766e-02 | -4.04 | negative regulation of striated muscle cell differentiation | biological process | GO:0051154 | 35 | 2 | 13711 | 80 | Rgs4,Bdnf | | 1.766e-02 | -4.04 | ventral spinal cord development | biological process | GO:0021517 | 35 | 2 | 13711 | 80 | Gbx1,Lhx1 | | 1.766e-02 | -4.04 | regulation of calcium ion import | biological process | GO:0090279 | 35 | 2 | 13711 | 80 | Crh,Homer1 | | 1.766e-02 | -4.04 | negative regulation of response to drug | biological process | GO:2001024 | 35 | 2 | 13711 | 80 | Rgs4,Nr4a3 | | 1.769e-02 | -4.03 | AFFAR\_YY1\_TARGETS\_UP | MSigDB lists | AFFAR\_YY1\_TARGETS\_UP | 169 | 4 | 12187 | 72 | Tle4,Fjx1,Kdr,Bdnf | | 1.769e-02 | -4.03 | POU6F1\_01 | MSigDB lists | POU6F1\_01 | 169 | 4 | 12187 | 72 | Crh,Csrnp3,Ncam1,Hs3st1 | | 1.769e-02 | -4.03 | GSE27786\_CD8\_TCELL\_VS\_NKTCELL\_DN | MSigDB lists | GSE27786\_CD8\_TCELL\_VS\_NKTCELL\_DN | 169 | 4 | 12187 | 72 | Zfp518b,Fnbp1,Inhba,Sorl1 | | 1.769e-02 | -4.03 | GSE20366\_EX\_VIVO\_VS\_HOMEOSTATIC\_CONVERSION\_TREG\_UP | MSigDB lists | GSE20366\_EX\_VIVO\_VS\_HOMEOSTATIC\_CONVERSION\_TREG\_UP | 169 | 4 | 12187 | 72 | Agrn,Msmo1,Nr4a3,Homer1 | | 1.769e-02 | -4.03 | GSE17721\_LPS\_VS\_PAM3CSK4\_16H\_BMDC\_UP | MSigDB lists | GSE17721\_LPS\_VS\_PAM3CSK4\_16H\_BMDC\_UP | 169 | 4 | 12187 | 72 | Sorl1,Ndst4,Inhba,Agrn | | 1.770e-02 | -4.03 | response to nitrogen compound | biological process | GO:1901698 | 683 | 9 | 13711 | 80 | Ncam1,P2ry12,Ubqln2,Col1a1,Homer1,Egr1,Nr4a3,Crh,Car2 | | 1.784e-02 | -4.03 | PLP | smart domains | SM00002 | 3 | 1 | 7188 | 43 | Plp1 | | 1.784e-02 | -4.03 | NGF | smart domains | SM00140 | 3 | 1 | 7188 | 43 | Bdnf | | 1.784e-02 | -4.03 | SEA | smart domains | SM00200 | 3 | 1 | 7188 | 43 | Agrn | | 1.800e-02 | -4.02 | Biosynthesis of unsaturated fatty acids | KEGG pathways | ko01040 | 26 | 2 | 5248 | 42 | Scd2,Hsd17b12 | | 1.800e-02 | -4.02 | Biosynthesis of unsaturated fatty acids | KEGG pathways | mmu01040 | 26 | 2 | 5248 | 42 | Scd2,Hsd17b12 | | 1.803e-02 | -4.02 | regulation of response to drug | biological process | GO:2001023 | 95 | 3 | 13711 | 80 | Rgs4,Nr4a3,Crh | | 1.804e-02 | -4.02 | GSE42088\_UNINF\_VS\_LEISHMANIA\_INF\_DC\_4H\_UP | MSigDB lists | GSE42088\_UNINF\_VS\_LEISHMANIA\_INF\_DC\_4H\_UP | 170 | 4 | 12187 | 72 | Inhba,Tgfa,Msmo1,Agrn | | 1.804e-02 | -4.02 | PIT1\_Q6 | MSigDB lists | PIT1\_Q6 | 170 | 4 | 12187 | 72 | Csrnp3,Bdnf,Cacng2,Inhba | | 1.804e-02 | -4.02 | GSE339\_CD8POS\_VS\_CD4CD8DN\_DC\_DN | MSigDB lists | GSE339\_CD8POS\_VS\_CD4CD8DN\_DC\_DN | 170 | 4 | 12187 | 72 | H1f0,Glud1,Mtmr9,Ubqln2 | | 1.804e-02 | -4.02 | HALLMARK\_TNFA\_SIGNALING\_VIA\_NFKB | MSigDB lists | HALLMARK\_TNFA\_SIGNALING\_VIA\_NFKB | 170 | 4 | 12187 | 72 | Egr1,Fjx1,Inhba,Nr4a3 | | 1.806e-02 | -4.01 | GO\_REGULATION\_OF\_CATECHOLAMINE\_SECRETION | MSigDB lists | GO\_REGULATION\_OF\_CATECHOLAMINE\_SECRETION | 35 | 2 | 12187 | 72 | Crh,P2ry12 | | 1.806e-02 | -4.01 | LIAO\_HAVE\_SOX4\_BINDING\_SITES | MSigDB lists | LIAO\_HAVE\_SOX4\_BINDING\_SITES | 35 | 2 | 12187 | 72 | Map4,Foxq1 | | 1.806e-02 | -4.01 | RASHI\_RESPONSE\_TO\_IONIZING\_RADIATION\_1 | MSigDB lists | RASHI\_RESPONSE\_TO\_IONIZING\_RADIATION\_1 | 35 | 2 | 12187 | 72 | Foxq1,Egr1 | | 1.806e-02 | -4.01 | SCHRAETS\_MLL\_TARGETS\_UP | MSigDB lists | SCHRAETS\_MLL\_TARGETS\_UP | 35 | 2 | 12187 | 72 | Col1a1,Peg3 | | 1.806e-02 | -4.01 | GO\_SUBSTANTIA\_NIGRA\_DEVELOPMENT | MSigDB lists | GO\_SUBSTANTIA\_NIGRA\_DEVELOPMENT | 35 | 2 | 12187 | 72 | Glud1,Plp1 | | 1.816e-02 | -4.01 | regulation of neuron differentiation | biological process | GO:0045664 | 686 | 9 | 13711 | 80 | Nrep,Ntm,Ddx56,Agrn,Timp2,Qk,Alkal2,Bdnf,Ltk | | 1.818e-02 | -4.01 | carboxylic acid metabolic process | biological process | GO:0019752 | 572 | 8 | 13711 | 80 | Qk,Scd2,Glud1,Aars,Hkdc1,Nr4a3,Qdpr,Plp1 | | 1.825e-02 | -4.00 | ADP binding | molecular function | GO:0043531 | 36 | 2 | 13516 | 78 | Atp5b,Glud1 | | 1.825e-02 | -4.00 | intramolecular oxidoreductase activity | molecular function | GO:0016860 | 36 | 2 | 13516 | 78 | Tmx3,Pdia6 | | 1.832e-02 | -4.00 | Invs (inversin) | protein interactions | 16348 | 78 | 3 | 6802 | 49 | Atp5b,Map4,Ermn | | 1.854e-02 | -3.99 | hormone-mediated signaling pathway | biological process | GO:0009755 | 96 | 3 | 13711 | 80 | Ddx17,Nr4a3,Crh | | 1.854e-02 | -3.99 | - | gene3d domains | 2.10.90.10 | 40 | 2 | 6647 | 35 | Inhba,Bdnf | | 1.855e-02 | -3.99 | THIOREDOXIN\_2 | prosite domains | PS51352 | 31 | 2 | 8845 | 60 | Pdia6,Tmx3 | | 1.858e-02 | -3.99 | GSE9988\_LPS\_VS\_LOW\_LPS\_MONOCYTE\_DN | MSigDB lists | GSE9988\_LPS\_VS\_LOW\_LPS\_MONOCYTE\_DN | 95 | 3 | 12187 | 72 | Hs3st1,Pom121,Agrn | | 1.858e-02 | -3.99 | CRX\_NRL\_DN.V1\_DN | MSigDB lists | CRX\_NRL\_DN.V1\_DN | 95 | 3 | 12187 | 72 | Fabp7,Egr1,Hs3st1 | | 1.863e-02 | -3.98 | negative regulation of endocytosis | biological process | GO:0045806 | 36 | 2 | 13711 | 80 | Ubqln2,Lrrtm2 | | 1.863e-02 | -3.98 | positive regulation of gene expression | biological process | GO:0010628 | 1577 | 16 | 13711 | 80 | Qk,Arntl,Peg3,Lhx1,Crh,Ddx17,Egr1,Plp1,Spry2,Csrnp3,Col1a1,Gbx1,Inhba,Nr4a3,Agrn,H1f0 | | 1.870e-02 | -3.98 | GO\_RESPONSE\_TO\_HORMONE | MSigDB lists | GO\_RESPONSE\_TO\_HORMONE | 683 | 9 | 12187 | 72 | Egr1,Inhba,Timp2,Gng11,Col1a1,Car2,Qdpr,Nr4a3,Crh | | 1.872e-02 | -3.98 | generation of precursor metabolites and energy | biological process | GO:0006091 | 263 | 5 | 13711 | 80 | Nr4a3,Nipsnap2,Bdnf,Atp5b,Hkdc1 | | 1.875e-02 | -3.98 | GSE22432\_MULTIPOTENT\_VS\_COMMON\_DC\_PROGENITOR\_UNTREATED\_DN | MSigDB lists | GSE22432\_MULTIPOTENT\_VS\_COMMON\_DC\_PROGENITOR\_UNTREATED\_DN | 172 | 4 | 12187 | 72 | Msmo1,Fnbp1,Ddx56,Aars | | 1.875e-02 | -3.98 | GSE26030\_UNSTIM\_VS\_RESTIM\_TH17\_DAY15\_POST\_POLARIZATION\_DN | MSigDB lists | GSE26030\_UNSTIM\_VS\_RESTIM\_TH17\_DAY15\_POST\_POLARIZATION\_DN | 172 | 4 | 12187 | 72 | Qdpr,Ddx17,Fnbp1,Dgkz | | 1.875e-02 | -3.98 | SYATTGTG\_UNKNOWN | MSigDB lists | SYATTGTG\_UNKNOWN | 172 | 4 | 12187 | 72 | Car2,Plp1,Cldn5,Nr4a3 | | 1.875e-02 | -3.98 | GO\_SENSORY\_ORGAN\_MORPHOGENESIS | MSigDB lists | GO\_SENSORY\_ORGAN\_MORPHOGENESIS | 172 | 4 | 12187 | 72 | Nr4a3,Fjx1,Lhx1,Spry2 | | 1.875e-02 | -3.98 | GSE23398\_WT\_VS\_IL2\_KO\_CD4\_TCELL\_SCURFY\_MOUSE\_UP | MSigDB lists | GSE23398\_WT\_VS\_IL2\_KO\_CD4\_TCELL\_SCURFY\_MOUSE\_UP | 172 | 4 | 12187 | 72 | Hsd17b12,Agrn,Spry2,Aars | | 1.875e-02 | -3.98 | GSE24210\_RESTING\_TREG\_VS\_TCONV\_DN | MSigDB lists | GSE24210\_RESTING\_TREG\_VS\_TCONV\_DN | 172 | 4 | 12187 | 72 | Crh,Srm,Msmo1,Tle4 | | 1.878e-02 | -3.98 | cellular response to organonitrogen compound | biological process | GO:0071417 | 361 | 6 | 13711 | 80 | Nr4a3,Car2,Crh,Col1a1,Egr1,P2ry12 | | 1.905e-02 | -3.96 | HUANG\_FOXA2\_TARGETS\_UP | MSigDB lists | HUANG\_FOXA2\_TARGETS\_UP | 36 | 2 | 12187 | 72 | Tle4,Homer1 | | 1.905e-02 | -3.96 | ZHU\_CMV\_8\_HR\_UP | MSigDB lists | ZHU\_CMV\_8\_HR\_UP | 36 | 2 | 12187 | 72 | Nr4a3,Egr1 | | 1.905e-02 | -3.96 | VANHARANTA\_UTERINE\_FIBROID\_UP | MSigDB lists | VANHARANTA\_UTERINE\_FIBROID\_UP | 36 | 2 | 12187 | 72 | Plp1,Col1a1 | | 1.907e-02 | -3.96 | GO\_LIPID\_METABOLIC\_PROCESS | MSigDB lists | GO\_LIPID\_METABOLIC\_PROCESS | 803 | 10 | 12187 | 72 | Agrn,Qk,Crh,Hsd17b12,Msmo1,Ugt8a,Plp1,Dgkz,Fabp7,Sorl1 | | 1.910e-02 | -3.96 | central nervous system development | biological process | GO:0007417 | 692 | 9 | 13711 | 80 | Gbx1,P2ry12,Ncam1,Aars,Inhba,Lhx1,Nr4a3,Plp1,Fabp7 | | 1.911e-02 | -3.96 | plasma membrane bounded cell projection | cellular component | GO:0120025 | 1896 | 18 | 13825 | 79 | Qdpr,Dgkz,P2ry12,Timp2,Grm3,Ncam1,Kdr,Cacng2,Bdnf,Agrn,Ntm,Homer1,Ephb6,Ermn,Crh,Car2,Cldn5,Map4 | | 1.922e-02 | -3.95 | ACATTCC\_MIR1\_MIR206 | MSigDB lists | ACATTCC\_MIR1\_MIR206 | 262 | 5 | 12187 | 72 | Ddx17,Bdnf,Qk,Dgkz,Nap1l5 | | 1.927e-02 | -3.95 | positive regulation of cell migration | biological process | GO:0030335 | 468 | 7 | 13711 | 80 | Atp5b,Egr1,Kdr,P2ry12,Spry2,Nr4a3,Col1a1 | | 1.945e-02 | -3.94 | Camk2a (calcium/calmodulin-dependent protein kinase II alpha) | protein interactions | 12322 | 30 | 2 | 6802 | 49 | Homer1,Cacng2 | | 1.947e-02 | -3.94 | BILD\_HRAS\_ONCOGENIC\_SIGNATURE | MSigDB lists | BILD\_HRAS\_ONCOGENIC\_SIGNATURE | 174 | 4 | 12187 | 72 | Tgfa,Foxq1,Egr1,Ddx17 | | 1.947e-02 | -3.94 | GSE35825\_UNTREATED\_VS\_IFNG\_STIM\_MACROPHAGE\_UP | MSigDB lists | GSE35825\_UNTREATED\_VS\_IFNG\_STIM\_MACROPHAGE\_UP | 174 | 4 | 12187 | 72 | Map4,Aars,Agrn,Nr4a3 | | 1.947e-02 | -3.94 | GOTZMANN\_EPITHELIAL\_TO\_MESENCHYMAL\_TRANSITION\_DN | MSigDB lists | GOTZMANN\_EPITHELIAL\_TO\_MESENCHYMAL\_TRANSITION\_DN | 174 | 4 | 12187 | 72 | Amd1,Tmem165,Hs3st1,Egr1 | | 1.962e-02 | -3.93 | cellular response to vascular endothelial growth factor stimulus | biological process | GO:0035924 | 37 | 2 | 13711 | 80 | Spry2,Kdr | | 1.962e-02 | -3.93 | amine metabolic process | biological process | GO:0009308 | 37 | 2 | 13711 | 80 | Srm,Amd1 | | 1.962e-02 | -3.93 | regulation of endocrine process | biological process | GO:0044060 | 37 | 2 | 13711 | 80 | Crh,Inhba | | 1.963e-02 | -3.93 | GCTTGAA\_MIR498 | MSigDB lists | GCTTGAA\_MIR498 | 97 | 3 | 12187 | 72 | Pdia6,Col1a1,Crh | | 1.984e-02 | -3.92 | GSE17721\_CTRL\_VS\_CPG\_0.5H\_BMDC\_UP | MSigDB lists | GSE17721\_CTRL\_VS\_CPG\_0.5H\_BMDC\_UP | 175 | 4 | 12187 | 72 | Tmem165,Qk,Zfp518b,Agrn | | 1.984e-02 | -3.92 | GSE9316\_IL6\_KO\_VS\_IFNG\_KO\_INVIVO\_EXPANDED\_CD4\_TCELL\_UP | MSigDB lists | GSE9316\_IL6\_KO\_VS\_IFNG\_KO\_INVIVO\_EXPANDED\_CD4\_TCELL\_UP | 175 | 4 | 12187 | 72 | Dgkz,Ubqln2,Arntl,Timp2 | | 1.984e-02 | -3.92 | mouse chr2\|2 E1 | chromosome location | mouse chr2\|2 E1 | 39 | 2 | 14556 | 81 | Dgkz,Hsd17b12 | | 1.986e-02 | -3.92 | carboxylic acid biosynthetic process | biological process | GO:0046394 | 177 | 4 | 13711 | 80 | Plp1,Qk,Glud1,Scd2 | | 1.988e-02 | -3.92 | Glycoprotein hormones | REACTOME pathways | R-MMU-209822 | 3 | 1 | 6297 | 42 | Inhba | | 2.006e-02 | -3.91 | GO\_NEGATIVE\_REGULATION\_OF\_SMALL\_GTPASE\_MEDIATED\_SIGNAL\_TRANSDUCTION | MSigDB lists | GO\_NEGATIVE\_REGULATION\_OF\_SMALL\_GTPASE\_MEDIATED\_SIGNAL\_TRANSDUCTION | 37 | 2 | 12187 | 72 | Timp2,Spry2 | | 2.006e-02 | -3.91 | GO\_RNA\_SECONDARY\_STRUCTURE\_UNWINDING | MSigDB lists | GO\_RNA\_SECONDARY\_STRUCTURE\_UNWINDING | 37 | 2 | 12187 | 72 | Ddx17,Ddx56 | | 2.006e-02 | -3.91 | GO\_LONG\_TERM\_SYNAPTIC\_POTENTIATION | MSigDB lists | GO\_LONG\_TERM\_SYNAPTIC\_POTENTIATION | 37 | 2 | 12187 | 72 | Crh,Lrrtm2 | | 2.016e-02 | -3.90 | GO\_RESPONSE\_TO\_HYDROGEN\_PEROXIDE | MSigDB lists | GO\_RESPONSE\_TO\_HYDROGEN\_PEROXIDE | 98 | 3 | 12187 | 72 | Nr4a3,Hba-a2,Col1a1 | | 2.021e-02 | -3.90 | GSE6674\_ANTI\_IGM\_VS\_ANTI\_IGM\_AND\_CPG\_STIM\_BCELL\_DN | MSigDB lists | GSE6674\_ANTI\_IGM\_VS\_ANTI\_IGM\_AND\_CPG\_STIM\_BCELL\_DN | 176 | 4 | 12187 | 72 | Srm,Pdia6,Homer1,Aars | | 2.021e-02 | -3.90 | GSE23114\_PERITONEAL\_CAVITY\_B1A\_BCELL\_VS\_SPLEEN\_BCELL\_DN | MSigDB lists | GSE23114\_PERITONEAL\_CAVITY\_B1A\_BCELL\_VS\_SPLEEN\_BCELL\_DN | 176 | 4 | 12187 | 72 | Srm,Homer1,Amd1,Qdpr | | 2.022e-02 | -3.90 | HEXOKINASE\_1 | prosite domains | PS00378 | 3 | 1 | 8845 | 60 | Hkdc1 | | 2.022e-02 | -3.90 | TAU\_MAP\_2 | prosite domains | PS51491 | 3 | 1 | 8845 | 60 | Map4 | | 2.022e-02 | -3.90 | NGF\_2 | prosite domains | PS50270 | 3 | 1 | 8845 | 60 | Bdnf | | 2.022e-02 | -3.90 | FATTY\_ACID\_DESATUR\_1 | prosite domains | PS00476 | 3 | 1 | 8845 | 60 | Scd2 | | 2.022e-02 | -3.90 | TAU\_MAP\_1 | prosite domains | PS00229 | 3 | 1 | 8845 | 60 | Map4 | | 2.022e-02 | -3.90 | NGF\_1 | prosite domains | PS00248 | 3 | 1 | 8845 | 60 | Bdnf | | 2.022e-02 | -3.90 | MYELIN\_PLP\_1 | prosite domains | PS00575 | 3 | 1 | 8845 | 60 | Plp1 | | 2.022e-02 | -3.90 | HEXOKINASE\_2 | prosite domains | PS51748 | 3 | 1 | 8845 | 60 | Hkdc1 | | 2.022e-02 | -3.90 | MYELIN\_PLP\_2 | prosite domains | PS01004 | 3 | 1 | 8845 | 60 | Plp1 | | 2.023e-02 | -3.90 | TRNASYNTHALA | prints domains | PR00980 | 2 | 1 | 2951 | 30 | Aars | | 2.023e-02 | -3.90 | NCAMFAMILY | prints domains | PR01838 | 2 | 1 | 2951 | 30 | Ncam1 | | 2.023e-02 | -3.90 | BKCHANNELB | prints domains | PR01450 | 2 | 1 | 2951 | 30 | Kcnmb2 | | 2.023e-02 | -3.90 | organic acid biosynthetic process | biological process | GO:0016053 | 178 | 4 | 13711 | 80 | Plp1,Scd2,Qk,Glud1 | | 2.059e-02 | -3.88 | GSE43863\_TH1\_VS\_LY6C\_INT\_CXCR5POS\_MEMORY\_CD4\_TCELL\_UP | MSigDB lists | GSE43863\_TH1\_VS\_LY6C\_INT\_CXCR5POS\_MEMORY\_CD4\_TCELL\_UP | 177 | 4 | 12187 | 72 | Srm,Tmem165,Ddx56,Homer1 | | 2.063e-02 | -3.88 | positive regulation of ion transmembrane transporter activity | biological process | GO:0032414 | 100 | 3 | 13711 | 80 | Lrrc55,Cacng2,Nipsnap2 | | 2.063e-02 | -3.88 | monovalent inorganic cation homeostasis | biological process | GO:0055067 | 100 | 3 | 13711 | 80 | Car2,Tmem165,Atp5b | | 2.063e-02 | -3.88 | positive regulation of protein binding | biological process | GO:0032092 | 100 | 3 | 13711 | 80 | Ephb6,Bdnf,Agrn | | 2.063e-02 | -3.88 | metencephalon development | biological process | GO:0022037 | 100 | 3 | 13711 | 80 | Lhx1,Aars,Gbx1 | | 2.063e-02 | -3.88 | protein kinase B signaling | biological process | GO:0043491 | 38 | 2 | 13711 | 80 | P2ry12,Kdr | | 2.069e-02 | -3.88 | receptor regulator activity | molecular function | GO:0030545 | 273 | 5 | 13516 | 78 | Tgfa,Crh,Inhba,Bdnf,Agrn | | 2.070e-02 | -3.88 | Ywhab (tyrosine 3-monooxygenase/tryptophan 5-monooxygenase activation protein, beta polypeptide) | protein interactions | 54401 | 31 | 2 | 6802 | 49 | Homer1,Ncam1 | | 2.070e-02 | -3.88 | GO\_POSITIVE\_REGULATION\_OF\_SYNAPTIC\_TRANSMISSION | MSigDB lists | GO\_POSITIVE\_REGULATION\_OF\_SYNAPTIC\_TRANSMISSION | 99 | 3 | 12187 | 72 | Lrrtm2,Car2,Crh | | 2.072e-02 | -3.88 | Prion diseases | KEGG pathways | mmu05020 | 28 | 2 | 5248 | 42 | Ncam1,Egr1 | | 2.072e-02 | -3.88 | Prion diseases | KEGG pathways | ko05020 | 28 | 2 | 5248 | 42 | Egr1,Ncam1 | | 2.090e-02 | -3.87 | - | gene3d domains | 2.20.70.30 | 4 | 1 | 6647 | 35 | Nacad | | 2.090e-02 | -3.87 | - | gene3d domains | 3.90.370.10 | 4 | 1 | 6647 | 35 | Timp2 | | 2.093e-02 | -3.87 | GO\_TISSUE\_DEVELOPMENT | MSigDB lists | GO\_TISSUE\_DEVELOPMENT | 1062 | 12 | 12187 | 72 | Aars,Nr4a3,Fjx1,Homer1,Spry2,Lhx1,Kdr,Egr1,Foxq1,Inhba,Col1a1,Car2 | | 2.098e-02 | -3.86 | positive regulation of binding | biological process | GO:0051099 | 180 | 4 | 13711 | 80 | Agrn,H1f0,Ephb6,Bdnf | | 2.110e-02 | -3.86 | MODULE\_433 | MSigDB lists | MODULE\_433 | 38 | 2 | 12187 | 72 | Tgfa,Inhba | | 2.116e-02 | -3.86 | regulation of cellular component biogenesis | biological process | GO:0044087 | 824 | 10 | 13711 | 80 | Lrrtm2,Map4,Kdr,Agrn,Ubqln2,Bdnf,Ncam1,P2ry12,Cbln2,Sorl1 | | 2.118e-02 | -3.85 | postsynapse organization | biological process | GO:0099173 | 101 | 3 | 13711 | 80 | Homer1,Dgkz,Agrn | | 2.119e-02 | -3.85 | postsynaptic density | cellular component | GO:0014069 | 379 | 6 | 13825 | 79 | Cacng2,Map4,Lrrtm2,Homer1,Grm3,Dgkz | | 2.126e-02 | -3.85 | GO\_REGULATION\_OF\_RECEPTOR\_ACTIVITY | MSigDB lists | GO\_REGULATION\_OF\_RECEPTOR\_ACTIVITY | 100 | 3 | 12187 | 72 | Tgfa,Cacng2,Crh | | 2.126e-02 | -3.85 | GCACCTT\_MIR18A\_MIR18B | MSigDB lists | GCACCTT\_MIR18A\_MIR18B | 100 | 3 | 12187 | 72 | Csrnp3,AI593442,Fnbp1 | | 2.136e-02 | -3.85 | GSE9037\_CTRL\_VS\_LPS\_4H\_STIM\_BMDM\_UP | MSigDB lists | GSE9037\_CTRL\_VS\_LPS\_4H\_STIM\_BMDM\_UP | 179 | 4 | 12187 | 72 | Ubqln2,Mtmr9,Fbxl17,Glud1 | | 2.136e-02 | -3.85 | EVI1\_04 | MSigDB lists | EVI1\_04 | 179 | 4 | 12187 | 72 | Crh,P2ry12,Bdnf,Ddx17 | | 2.146e-02 | -3.84 | Lzts3 (leucine zipper, putative tumor suppressor family member 3) | protein interactions | 241638 | 3 | 1 | 6802 | 49 | Homer1 | | 2.146e-02 | -3.84 | Qk (quaking) | protein interactions | 19317 | 3 | 1 | 6802 | 49 | Qk | | 2.146e-02 | -3.84 | Cntn2 (contactin 2) | protein interactions | 21367 | 3 | 1 | 6802 | 49 | Ncam1 | | 2.146e-02 | -3.84 | Plg (plasminogen) | protein interactions | 18815 | 3 | 1 | 6802 | 49 | Bdnf | | 2.146e-02 | -3.84 | Six6 (sine oculis-related homeobox 6) | protein interactions | 20476 | 3 | 1 | 6802 | 49 | Tle4 | | 2.146e-02 | -3.84 | Ubqln1 (ubiquilin 1) | protein interactions | 56085 | 3 | 1 | 6802 | 49 | Ubqln2 | | 2.146e-02 | -3.84 | Ubqln4 (ubiquilin 4) | protein interactions | 94232 | 3 | 1 | 6802 | 49 | Ubqln2 | | 2.146e-02 | -3.84 | Sf1 (splicing factor 1) | protein interactions | 22668 | 3 | 1 | 6802 | 49 | Egr1 | | 2.146e-02 | -3.84 | Elk1 (ELK1, member of ETS oncogene family) | protein interactions | 13712 | 3 | 1 | 6802 | 49 | Egr1 | | 2.146e-02 | -3.84 | PAK1IP1 (PAK1 interacting protein 1) | protein interactions | 55003 | 3 | 1 | 6802 | 49 | Agrn | | 2.146e-02 | -3.84 | Homer1 (homer scaffold protein 1) | protein interactions | 29546 | 3 | 1 | 6802 | 49 | Homer1 | | 2.146e-02 | -3.84 | Pitx1 (paired-like homeodomain transcription factor 1) | protein interactions | 18740 | 3 | 1 | 6802 | 49 | Egr1 | | 2.158e-02 | -3.84 | positive regulation of cell motility | biological process | GO:2000147 | 479 | 7 | 13711 | 80 | Col1a1,Spry2,Nr4a3,Kdr,P2ry12,Atp5b,Egr1 | | 2.165e-02 | -3.83 | positive regulation of cell death | biological process | GO:0010942 | 591 | 8 | 13711 | 80 | Egr1,Agrn,Nr4a3,Crh,Gsdme,Ltk,Inhba,Csrnp3 | | 2.167e-02 | -3.83 | positive regulation of calcium-mediated signaling | biological process | GO:0050850 | 39 | 2 | 13711 | 80 | Ncam1,Kdr | | 2.167e-02 | -3.83 | establishment of protein localization to vacuole | biological process | GO:0072666 | 39 | 2 | 13711 | 80 | Sorl1,Cacng2 | | 2.167e-02 | -3.83 | establishment of spindle localization | biological process | GO:0051293 | 39 | 2 | 13711 | 80 | Spry2,Map4 | | 2.173e-02 | -3.83 | cell-cell junction organization | biological process | GO:0045216 | 102 | 3 | 13711 | 80 | Cldn5,Ugt8a,Lims2 | | 2.173e-02 | -3.83 | regulation of neural precursor cell proliferation | biological process | GO:2000177 | 102 | 3 | 13711 | 80 | Lhx1,Lims2,Bdnf | | 2.175e-02 | -3.83 | DBP\_Q6 | MSigDB lists | DBP\_Q6 | 180 | 4 | 12187 | 72 | Nr4a3,Tle4,Nap1l5,Ddx17 | | 2.175e-02 | -3.83 | GO\_REGULATION\_OF\_PEPTIDE\_TRANSPORT | MSigDB lists | GO\_REGULATION\_OF\_PEPTIDE\_TRANSPORT | 180 | 4 | 12187 | 72 | Arntl,Crh,Car2,Glud1 | | 2.180e-02 | -3.83 | response to hormone | biological process | GO:0009725 | 480 | 7 | 13711 | 80 | Inhba,Timp2,Egr1,Ddx17,Crh,Car2,Nr4a3 | | 2.182e-02 | -3.83 | GO\_REGULATION\_OF\_CYCLIC\_NUCLEOTIDE\_METABOLIC\_PROCESS | MSigDB lists | GO\_REGULATION\_OF\_CYCLIC\_NUCLEOTIDE\_METABOLIC\_PROCESS | 101 | 3 | 12187 | 72 | Crh,Timp2,Grm3 | | 2.182e-02 | -3.83 | HMEF2\_Q6 | MSigDB lists | HMEF2\_Q6 | 101 | 3 | 12187 | 72 | Timp2,Gng11,Csrnp3 | | 2.182e-02 | -3.83 | AZARE\_NEOPLASTIC\_TRANSFORMATION\_BY\_STAT3\_UP | MSigDB lists | AZARE\_NEOPLASTIC\_TRANSFORMATION\_BY\_STAT3\_UP | 101 | 3 | 12187 | 72 | Tgfa,Gng11,Sorl1 | | 2.182e-02 | -3.83 | GO\_CHANNEL\_REGULATOR\_ACTIVITY | MSigDB lists | GO\_CHANNEL\_REGULATOR\_ACTIVITY | 101 | 3 | 12187 | 72 | Grm3,Kcnmb2,Cacng2 | | 2.193e-02 | -3.82 | asymmetric synapse | cellular component | GO:0032279 | 382 | 6 | 13825 | 79 | Map4,Cacng2,Lrrtm2,Homer1,Grm3,Dgkz | | 2.197e-02 | -3.82 | Cystine-knot\_cytokine | interpro domains | IPR029034 | 40 | 2 | 13788 | 79 | Inhba,Bdnf | | 2.197e-02 | -3.82 | Pkinase\_Tyr | pfam domains | PF07714 | 107 | 3 | 12881 | 72 | Ephb6,Kdr,Ltk | | 2.214e-02 | -3.81 | ear development | biological process | GO:0043583 | 183 | 4 | 13711 | 80 | Gsdme,Bdnf,Spry2,Nr4a3 | | 2.214e-02 | -3.81 | regulation of cell-substrate adhesion | biological process | GO:0010810 | 183 | 4 | 13711 | 80 | Col1a1,Kdr,Hsd17b12,Lims2 | | 2.214e-02 | -3.81 | GSE17721\_CPG\_VS\_GARDIQUIMOD\_2H\_BMDC\_DN | MSigDB lists | GSE17721\_CPG\_VS\_GARDIQUIMOD\_2H\_BMDC\_DN | 181 | 4 | 12187 | 72 | Pom121,Arntl,Dgkz,Chmp7 | | 2.214e-02 | -3.81 | HOSHIDA\_LIVER\_CANCER\_SUBCLASS\_S3 | MSigDB lists | HOSHIDA\_LIVER\_CANCER\_SUBCLASS\_S3 | 181 | 4 | 12187 | 72 | Car2,Msmo1,Qdpr,Sorl1 | | 2.214e-02 | -3.81 | VANTVEER\_BREAST\_CANCER\_ESR1\_DN | MSigDB lists | VANTVEER\_BREAST\_CANCER\_ESR1\_DN | 181 | 4 | 12187 | 72 | Ugt8a,Pdia6,Amd1,Fabp7 | | 2.215e-02 | -3.81 | ANDERSEN\_CHOLANGIOCARCINOMA\_CLASS1 | MSigDB lists | ANDERSEN\_CHOLANGIOCARCINOMA\_CLASS1 | 39 | 2 | 12187 | 72 | Hkdc1,Rgs4 | | 2.215e-02 | -3.81 | GO\_RESPONSE\_TO\_GLUCAGON | MSigDB lists | GO\_RESPONSE\_TO\_GLUCAGON | 39 | 2 | 12187 | 72 | Qdpr,Gng11 | | 2.215e-02 | -3.81 | MODULE\_284 | MSigDB lists | MODULE\_284 | 39 | 2 | 12187 | 72 | Kdr,Nr4a3 | | 2.215e-02 | -3.81 | HOQUE\_METHYLATED\_IN\_CANCER | MSigDB lists | HOQUE\_METHYLATED\_IN\_CANCER | 39 | 2 | 12187 | 72 | Timp2,Qdpr | | 2.215e-02 | -3.81 | GO\_NEURAL\_RETINA\_DEVELOPMENT | MSigDB lists | GO\_NEURAL\_RETINA\_DEVELOPMENT | 39 | 2 | 12187 | 72 | Fjx1,Lhx1 | | 2.215e-02 | -3.81 | GO\_GLUTAMATE\_RECEPTOR\_SIGNALING\_PATHWAY | MSigDB lists | GO\_GLUTAMATE\_RECEPTOR\_SIGNALING\_PATHWAY | 39 | 2 | 12187 | 72 | Homer1,Grm3 | | 2.215e-02 | -3.81 | GYORFFY\_DOXORUBICIN\_RESISTANCE | MSigDB lists | GYORFFY\_DOXORUBICIN\_RESISTANCE | 39 | 2 | 12187 | 72 | Hs3st1,Timp2 | | 2.215e-02 | -3.81 | JIANG\_AGING\_HYPOTHALAMUS\_UP | MSigDB lists | JIANG\_AGING\_HYPOTHALAMUS\_UP | 39 | 2 | 12187 | 72 | Hba-a2,Map4 | | 2.217e-02 | -3.81 | ATP-synt\_ab | pfam domains | PF00006 | 4 | 1 | 12881 | 72 | Atp5b | | 2.217e-02 | -3.81 | ATP-synt\_ab\_N | pfam domains | PF02874 | 4 | 1 | 12881 | 72 | Atp5b | | 2.217e-02 | -3.81 | HSNSD | pfam domains | PF12062 | 4 | 1 | 12881 | 72 | Ndst4 | | 2.217e-02 | -3.81 | TIMP | pfam domains | PF00965 | 4 | 1 | 12881 | 72 | Timp2 | | 2.217e-02 | -3.81 | NAC | pfam domains | PF01849 | 4 | 1 | 12881 | 72 | Nacad | | 2.238e-02 | -3.80 | DARWICHE\_SKIN\_TUMOR\_PROMOTER\_UP | MSigDB lists | DARWICHE\_SKIN\_TUMOR\_PROMOTER\_UP | 102 | 3 | 12187 | 72 | Tgfa,Srm,Qdpr | | 2.238e-02 | -3.80 | GSE2706\_R848\_VS\_LPS\_2H\_STIM\_DC\_DN | MSigDB lists | GSE2706\_R848\_VS\_LPS\_2H\_STIM\_DC\_DN | 102 | 3 | 12187 | 72 | Egr1,Inhba,Gbx1 | | 2.238e-02 | -3.80 | HOSHIDA\_LIVER\_CANCER\_SUBCLASS\_S2 | MSigDB lists | HOSHIDA\_LIVER\_CANCER\_SUBCLASS\_S2 | 102 | 3 | 12187 | 72 | Glud1,H1f0,Peg3 | | 2.249e-02 | -3.79 | Class B/2 (Secretin family receptors) | REACTOME pathways | R-MMU-373080 | 35 | 2 | 6297 | 42 | Crh,Gng11 | | 2.254e-02 | -3.79 | GO\_EXCITATORY\_SYNAPSE | MSigDB lists | GO\_EXCITATORY\_SYNAPSE | 182 | 4 | 12187 | 72 | Lrrtm2,Homer1,Grm3,Map4 | | 2.266e-02 | -3.79 | box H/ACA snoRNP complex | cellular component | GO:0031429 | 4 | 1 | 13825 | 79 | Gar1 | | 2.266e-02 | -3.79 | hemoglobin complex | cellular component | GO:0005833 | 4 | 1 | 13825 | 79 | Hba-a2 | | 2.266e-02 | -3.79 | box H/ACA RNP complex | cellular component | GO:0072588 | 4 | 1 | 13825 | 79 | Gar1 | | 2.266e-02 | -3.79 | internode region of axon | cellular component | GO:0033269 | 4 | 1 | 13825 | 79 | Ermn | | 2.272e-02 | -3.78 | Ubiquilin | interpro domains | IPR015496 | 4 | 1 | 13788 | 79 | Ubqln2 | | 2.272e-02 | -3.78 | TIMP\_C | interpro domains | IPR027465 | 4 | 1 | 13788 | 79 | Timp2 | | 2.272e-02 | -3.78 | ATPase\_F1/V1/A1\_a/bsu\_nucl-bd | interpro domains | IPR000194 | 4 | 1 | 13788 | 79 | Atp5b | | 2.272e-02 | -3.78 | NAC\_A/B\_dom\_sf | interpro domains | IPR038187 | 4 | 1 | 13788 | 79 | Nacad | | 2.272e-02 | -3.78 | Nas\_poly-pep-assoc\_cplx\_dom | interpro domains | IPR002715 | 4 | 1 | 13788 | 79 | Nacad | | 2.272e-02 | -3.78 | TIMP\_CS | interpro domains | IPR030490 | 4 | 1 | 13788 | 79 | Timp2 | | 2.272e-02 | -3.78 | Heparan\_SO4\_deacetylase | interpro domains | IPR021930 | 4 | 1 | 13788 | 79 | Ndst4 | | 2.272e-02 | -3.78 | ATPase\_F1/V1/A1\_a/bsu\_N | interpro domains | IPR004100 | 4 | 1 | 13788 | 79 | Atp5b | | 2.272e-02 | -3.78 | TIMP | interpro domains | IPR001820 | 4 | 1 | 13788 | 79 | Timp2 | | 2.272e-02 | -3.78 | ATPase\_a/bsu\_AS | interpro domains | IPR020003 | 4 | 1 | 13788 | 79 | Atp5b | | 2.274e-02 | -3.78 | RYTTCCTG\_ETS2\_B | MSigDB lists | RYTTCCTG\_ETS2\_B | 826 | 10 | 12187 | 72 | Lhx1,Bdnf,Arhgap20,Cldn5,Nr4a3,Cacng2,Egr1,Amd1,Dgkz,Gng11 | | 2.274e-02 | -3.78 | positive regulation of cellular component organization | biological process | GO:0051130 | 1084 | 12 | 13711 | 80 | Ddx56,Agrn,Lrrtm2,Ltk,Lims2,Tgfa,Kdr,P2ry12,Cbln2,Bdnf,Qk,Alkal2 | | 2.281e-02 | -3.78 | mouse chr3\|3 G1 | chromosome location | mouse chr3\|3 G1 | 42 | 2 | 14556 | 81 | Ugt8a,Ndst4 | | 2.289e-02 | -3.78 | arylesterase activity | molecular function | GO:0004064 | 4 | 1 | 13516 | 78 | Car2 | | 2.289e-02 | -3.78 | box H/ACA snoRNA binding | molecular function | GO:0034513 | 4 | 1 | 13516 | 78 | Gar1 | | 2.289e-02 | -3.78 | haptoglobin binding | molecular function | GO:0031720 | 4 | 1 | 13516 | 78 | Hba-a2 | | 2.289e-02 | -3.78 | [heparan sulfate]-glucosamine N-sulfotransferase activity | molecular function | GO:0015016 | 4 | 1 | 13516 | 78 | Ndst4 | | 2.289e-02 | -3.78 | double-stranded methylated DNA binding | molecular function | GO:0010385 | 4 | 1 | 13516 | 78 | Egr1 | | 2.289e-02 | -3.78 | corticotropin-releasing hormone receptor binding | molecular function | GO:0051429 | 4 | 1 | 13516 | 78 | Crh | | 2.289e-02 | -3.78 | polyamine binding | molecular function | GO:0019808 | 4 | 1 | 13516 | 78 | Amd1 | | 2.289e-02 | -3.78 | activin receptor binding | molecular function | GO:0070697 | 4 | 1 | 13516 | 78 | Inhba | | 2.289e-02 | -3.78 | thiol oxidase activity | molecular function | GO:0016972 | 4 | 1 | 13516 | 78 | Tmx3 | | 2.289e-02 | -3.78 | angiostatin binding | molecular function | GO:0043532 | 4 | 1 | 13516 | 78 | Atp5b | | 2.294e-02 | -3.77 | REACTOME\_METABOLISM\_OF\_CARBOHYDRATES | MSigDB lists | REACTOME\_METABOLISM\_OF\_CARBOHYDRATES | 183 | 4 | 12187 | 72 | Agrn,Pom121,Hs3st1,Ndst4 | | 2.296e-02 | -3.77 | CRX\_DN.V1\_DN | MSigDB lists | CRX\_DN.V1\_DN | 103 | 3 | 12187 | 72 | Egr1,Fabp7,Gng11 | | 2.296e-02 | -3.77 | BMI1\_DN.V1\_UP | MSigDB lists | BMI1\_DN.V1\_UP | 103 | 3 | 12187 | 72 | Inhba,Agrn,Fjx1 | | 2.296e-02 | -3.77 | DARWICHE\_PAPILLOMA\_RISK\_HIGH\_UP | MSigDB lists | DARWICHE\_PAPILLOMA\_RISK\_HIGH\_UP | 103 | 3 | 12187 | 72 | Qdpr,Tgfa,Srm | | 2.296e-02 | -3.77 | RB\_DN.V1\_DN | MSigDB lists | RB\_DN.V1\_DN | 103 | 3 | 12187 | 72 | Hs3st1,Msmo1,Agrn | | 2.314e-02 | -3.77 | oviduct development | biological process | GO:0060066 | 4 | 1 | 13711 | 80 | Lhx1 | | 2.314e-02 | -3.77 | proprioception | biological process | GO:0019230 | 4 | 1 | 13711 | 80 | Gbx1 | | 2.314e-02 | -3.77 | negative regulation of clathrin-dependent endocytosis | biological process | GO:1900186 | 4 | 1 | 13711 | 80 | Ubqln2 | | 2.314e-02 | -3.77 | positive regulation of corticotropin secretion | biological process | GO:0051461 | 4 | 1 | 13711 | 80 | Crh | | 2.314e-02 | -3.77 | cellular response to mycotoxin | biological process | GO:0036146 | 4 | 1 | 13711 | 80 | Egr1 | | 2.314e-02 | -3.77 | oxidative stress-induced premature senescence | biological process | GO:0090403 | 4 | 1 | 13711 | 80 | Arntl | | 2.314e-02 | -3.77 | positive regulation of epithelial cell differentiation involved in kidney development | biological process | GO:2000698 | 4 | 1 | 13711 | 80 | Lhx1 | | 2.314e-02 | -3.77 | negative regulation of metalloendopeptidase activity | biological process | GO:1904684 | 4 | 1 | 13711 | 80 | Sorl1 | | 2.314e-02 | -3.77 | positive regulation of tau-protein kinase activity | biological process | GO:1902949 | 4 | 1 | 13711 | 80 | Egr1 | | 2.314e-02 | -3.77 | S-shaped body morphogenesis | biological process | GO:0072050 | 4 | 1 | 13711 | 80 | Lhx1 | | 2.314e-02 | -3.77 | Fc-epsilon receptor signaling pathway | biological process | GO:0038095 | 4 | 1 | 13711 | 80 | Nr4a3 | | 2.314e-02 | -3.77 | regulation of cortisol secretion | biological process | GO:0051462 | 4 | 1 | 13711 | 80 | Crh | | 2.314e-02 | -3.77 | regulation of mast cell cytokine production | biological process | GO:0032763 | 4 | 1 | 13711 | 80 | Nr4a3 | | 2.314e-02 | -3.77 | regulation of synaptic growth at neuromuscular junction | biological process | GO:0008582 | 4 | 1 | 13711 | 80 | Agrn | | 2.314e-02 | -3.77 | response to heparin | biological process | GO:0071503 | 4 | 1 | 13711 | 80 | Egr1 | | 2.314e-02 | -3.77 | glucocorticoid biosynthetic process | biological process | GO:0006704 | 4 | 1 | 13711 | 80 | Crh | | 2.314e-02 | -3.77 | peptidyl-cysteine oxidation | biological process | GO:0018171 | 4 | 1 | 13711 | 80 | Tmx3 | | 2.314e-02 | -3.77 | response to anticoagulant | biological process | GO:0061476 | 4 | 1 | 13711 | 80 | Egr1 | | 2.314e-02 | -3.77 | regulation of ovulation | biological process | GO:0060278 | 4 | 1 | 13711 | 80 | Inhba | | 2.314e-02 | -3.77 | glutamate biosynthetic process | biological process | GO:0006537 | 4 | 1 | 13711 | 80 | Glud1 | | 2.314e-02 | -3.77 | regulation of microglial cell migration | biological process | GO:1904139 | 4 | 1 | 13711 | 80 | P2ry12 | | 2.314e-02 | -3.77 | positive regulation of glucocorticoid secretion | biological process | GO:2000851 | 4 | 1 | 13711 | 80 | Crh | | 2.314e-02 | -3.77 | snoRNA guided rRNA pseudouridine synthesis | biological process | GO:0000454 | 4 | 1 | 13711 | 80 | Gar1 | | 2.314e-02 | -3.77 | regulation of luteinizing hormone secretion | biological process | GO:0033684 | 4 | 1 | 13711 | 80 | Crh | | 2.314e-02 | -3.77 | positive regulation of microglial cell migration | biological process | GO:1904141 | 4 | 1 | 13711 | 80 | P2ry12 | | 2.314e-02 | -3.77 | comma-shaped body morphogenesis | biological process | GO:0072049 | 4 | 1 | 13711 | 80 | Lhx1 | | 2.314e-02 | -3.77 | negative regulation of hair cycle | biological process | GO:0042636 | 4 | 1 | 13711 | 80 | Inhba | | 2.314e-02 | -3.77 | positive regulation of cellular pH reduction | biological process | GO:0032849 | 4 | 1 | 13711 | 80 | Car2 | | 2.314e-02 | -3.77 | negative regulation of hair follicle development | biological process | GO:0051799 | 4 | 1 | 13711 | 80 | Inhba | | 2.314e-02 | -3.77 | negative regulation of cell adhesion involved in substrate-bound cell migration | biological process | GO:0006933 | 4 | 1 | 13711 | 80 | Atp5b | | 2.314e-02 | -3.77 | regulation of corticosterone secretion | biological process | GO:2000852 | 4 | 1 | 13711 | 80 | Crh | | 2.314e-02 | -3.77 | positive regulation of ER to Golgi vesicle-mediated transport | biological process | GO:1902953 | 4 | 1 | 13711 | 80 | Sorl1 | | 2.314e-02 | -3.77 | vestibular reflex | biological process | GO:0060005 | 4 | 1 | 13711 | 80 | Nr4a3 | | 2.314e-02 | -3.77 | negative regulation of neurotrophin TRK receptor signaling pathway | biological process | GO:0051387 | 4 | 1 | 13711 | 80 | Spry2 | | 2.314e-02 | -3.77 | cellular response to heparin | biological process | GO:0071504 | 4 | 1 | 13711 | 80 | Egr1 | | 2.314e-02 | -3.77 | cell morphogenesis involved in differentiation | biological process | GO:0000904 | 486 | 7 | 13711 | 80 | Ephb6,Bdnf,Lhx1,Agrn,Ncam1,Nr4a3,Gbx1 | | 2.321e-02 | -3.76 | RTAAACA\_FREAC2\_01 | MSigDB lists | RTAAACA\_FREAC2\_01 | 709 | 9 | 12187 | 72 | Bdnf,Ndst4,Crh,Cacng2,Ddx17,Csrnp3,Nap1l5,Inhba,Arntl | | 2.321e-02 | -3.76 | regulation of membrane potential | biological process | GO:0042391 | 379 | 6 | 13711 | 80 | Rgs4,Agrn,Kdr,Bdnf,Kcnmb2,Cacng2 | | 2.323e-02 | -3.76 | MODULE\_234 | MSigDB lists | MODULE\_234 | 40 | 2 | 12187 | 72 | Inhba,Col1a1 | | 2.323e-02 | -3.76 | PENG\_GLUCOSE\_DEPRIVATION\_UP | MSigDB lists | PENG\_GLUCOSE\_DEPRIVATION\_UP | 40 | 2 | 12187 | 72 | Msmo1,Aars | | 2.323e-02 | -3.76 | GO\_POSITIVE\_REGULATION\_OF\_EPITHELIAL\_CELL\_DIFFERENTIATION | MSigDB lists | GO\_POSITIVE\_REGULATION\_OF\_EPITHELIAL\_CELL\_DIFFERENTIATION | 40 | 2 | 12187 | 72 | Lhx1,Cldn5 | | 2.323e-02 | -3.76 | GO\_REGULATION\_OF\_KIDNEY\_DEVELOPMENT | MSigDB lists | GO\_REGULATION\_OF\_KIDNEY\_DEVELOPMENT | 40 | 2 | 12187 | 72 | Lhx1,Egr1 | | 2.331e-02 | -3.76 | potassium channel regulator activity | molecular function | GO:0015459 | 41 | 2 | 13516 | 78 | Lrrc55,Kcnmb2 | | 2.333e-02 | -3.76 | Q\_MOTIF | prosite domains | PS51195 | 35 | 2 | 8845 | 60 | Ddx56,Ddx17 | | 2.335e-02 | -3.76 | RUTELLA\_RESPONSE\_TO\_HGF\_DN | MSigDB lists | RUTELLA\_RESPONSE\_TO\_HGF\_DN | 184 | 4 | 12187 | 72 | Sorl1,Egr1,Tle4,Fnbp1 | | 2.335e-02 | -3.76 | PTF1BETA\_Q6 | MSigDB lists | PTF1BETA\_Q6 | 184 | 4 | 12187 | 72 | Nr4a3,Ddx17,H1f0,Map4 | | 2.343e-02 | -3.75 | GO\_ESTRADIOL\_17\_BETA\_DEHYDROGENASE\_ACTIVITY | MSigDB lists | GO\_ESTRADIOL\_17\_BETA\_DEHYDROGENASE\_ACTIVITY | 4 | 1 | 12187 | 72 | Hsd17b12 | | 2.343e-02 | -3.75 | GO\_FLAVONOID\_METABOLIC\_PROCESS | MSigDB lists | GO\_FLAVONOID\_METABOLIC\_PROCESS | 4 | 1 | 12187 | 72 | Ugt8a | | 2.343e-02 | -3.75 | NAKAMURA\_ALVEOLAR\_EPITHELIUM | MSigDB lists | NAKAMURA\_ALVEOLAR\_EPITHELIUM | 4 | 1 | 12187 | 72 | Cldn5 | | 2.343e-02 | -3.75 | GO\_HEMOGLOBIN\_COMPLEX | MSigDB lists | GO\_HEMOGLOBIN\_COMPLEX | 4 | 1 | 12187 | 72 | Hba-a2 | | 2.343e-02 | -3.75 | CHEN\_HOXA5\_TARGETS\_6HR\_DN | MSigDB lists | CHEN\_HOXA5\_TARGETS\_6HR\_DN | 4 | 1 | 12187 | 72 | Fjx1 | | 2.343e-02 | -3.75 | GO\_GLUCOCORTICOID\_BIOSYNTHETIC\_PROCESS | MSigDB lists | GO\_GLUCOCORTICOID\_BIOSYNTHETIC\_PROCESS | 4 | 1 | 12187 | 72 | Crh | | 2.343e-02 | -3.75 | GO\_ESTROGEN\_BIOSYNTHETIC\_PROCESS | MSigDB lists | GO\_ESTROGEN\_BIOSYNTHETIC\_PROCESS | 4 | 1 | 12187 | 72 | Hsd17b12 | | 2.343e-02 | -3.75 | positive regulation of calcium ion transport | biological process | GO:0051928 | 105 | 3 | 13711 | 80 | Homer1,Nipsnap2,Crh | | 2.362e-02 | -3.75 | CTGCAGY\_UNKNOWN | MSigDB lists | CTGCAGY\_UNKNOWN | 595 | 8 | 12187 | 72 | Sorl1,Ddx17,Csrnp3,Egr1,AI593442,Map4,Bdnf,Cacng2 | | 2.372e-02 | -3.74 | NAC | smart domains | SM01407 | 4 | 1 | 7188 | 43 | Nacad | | 2.372e-02 | -3.74 | NTR | smart domains | SM00206 | 4 | 1 | 7188 | 43 | Timp2 | | 2.373e-02 | -3.74 | TGCACTT\_MIR519C\_MIR519B\_MIR519A | MSigDB lists | TGCACTT\_MIR519C\_MIR519B\_MIR519A | 377 | 6 | 12187 | 72 | Timp2,Sorl1,Csrnp3,Fjx1,Qk,Map4 | | 2.376e-02 | -3.74 | IPF1\_Q4 | MSigDB lists | IPF1\_Q4 | 185 | 4 | 12187 | 72 | Ephb6,Tle4,Bdnf,Csrnp3 | | 2.376e-02 | -3.74 | TIEN\_INTESTINE\_PROBIOTICS\_24HR\_DN | MSigDB lists | TIEN\_INTESTINE\_PROBIOTICS\_24HR\_DN | 185 | 4 | 12187 | 72 | H1f0,Aars,Pdia6,Hkdc1 | | 2.380e-02 | -3.74 | ovulation cycle | biological process | GO:0042698 | 41 | 2 | 13711 | 80 | Inhba,Egr1 | | 2.380e-02 | -3.74 | regulation of sensory perception of pain | biological process | GO:0051930 | 41 | 2 | 13711 | 80 | Ncam1,Grm3 | | 2.380e-02 | -3.74 | mesoderm formation | biological process | GO:0001707 | 41 | 2 | 13711 | 80 | Nr4a3,Inhba | | 2.382e-02 | -3.74 | Neomycin, kanamycin and gentamicin biosynthesis | KEGG pathways | ko00524 | 3 | 1 | 5248 | 42 | Hkdc1 | | 2.382e-02 | -3.74 | D-Glutamine and D-glutamate metabolism | KEGG pathways | ko00471 | 3 | 1 | 5248 | 42 | Glud1 | | 2.382e-02 | -3.74 | D-Glutamine and D-glutamate metabolism | KEGG pathways | mmu00471 | 3 | 1 | 5248 | 42 | Glud1 | | 2.382e-02 | -3.74 | Neomycin, kanamycin and gentamicin biosynthesis | KEGG pathways | mmu00524 | 3 | 1 | 5248 | 42 | Hkdc1 | | 2.384e-02 | -3.74 | tissue morphogenesis | biological process | GO:0048729 | 489 | 7 | 13711 | 80 | Lhx1,Inhba,Spry2,Kdr,Nr4a3,Foxq1,Car2 | | 2.402e-02 | -3.73 | positive regulation of transporter activity | biological process | GO:0032411 | 106 | 3 | 13711 | 80 | Cacng2,Lrrc55,Nipsnap2 | | 2.402e-02 | -3.73 | camera-type eye morphogenesis | biological process | GO:0048593 | 106 | 3 | 13711 | 80 | Fjx1,Kdr,Lhx1 | | 2.407e-02 | -3.73 | oxoacid metabolic process | biological process | GO:0043436 | 603 | 8 | 13711 | 80 | Glud1,Qk,Scd2,Aars,Hkdc1,Nr4a3,Qdpr,Plp1 | | 2.414e-02 | -3.72 | MIKKELSEN\_MEF\_ICP\_WITH\_H3K27ME3 | MSigDB lists | MIKKELSEN\_MEF\_ICP\_WITH\_H3K27ME3 | 105 | 3 | 12187 | 72 | Grm3,AI593442,Kcnmb2 | | 2.417e-02 | -3.72 | glutamine degradation I | BIOCYC pathways | MOUSE\_GLUTAMINE-DEG1-PWY | 2 | 1 | 823 | 10 | Glud1 | | 2.417e-02 | -3.72 | glutamine biosynthesis II | BIOCYC pathways | MOUSE\_GLUTAMINE-SYN2-PWY | 2 | 1 | 823 | 10 | Glud1 | | 2.418e-02 | -3.72 | LINDGREN\_BLADDER\_CANCER\_CLUSTER\_3\_DN | MSigDB lists | LINDGREN\_BLADDER\_CANCER\_CLUSTER\_3\_DN | 186 | 4 | 12187 | 72 | Sorl1,Spry2,Glud1,Ephb6 | | 2.419e-02 | -3.72 | cellular component morphogenesis | biological process | GO:0032989 | 721 | 9 | 13711 | 80 | Gbx1,Ncam1,Ephb6,Bdnf,Lhx1,Nr4a3,Agrn,Kdr,Ugt8a | | 2.428e-02 | -3.72 | GO\_CELLULAR\_RESPONSE\_TO\_ORGANIC\_CYCLIC\_COMPOUND | MSigDB lists | GO\_CELLULAR\_RESPONSE\_TO\_ORGANIC\_CYCLIC\_COMPOUND | 379 | 6 | 12187 | 72 | Col1a1,Inhba,Egr1,Crh,P2ry12,Nr4a3 | | 2.433e-02 | -3.72 | GO\_ENDODERM\_FORMATION | MSigDB lists | GO\_ENDODERM\_FORMATION | 41 | 2 | 12187 | 72 | Lhx1,Inhba | | 2.433e-02 | -3.72 | REACTOME\_G\_ALPHA\_Z\_SIGNALLING\_EVENTS | MSigDB lists | REACTOME\_G\_ALPHA\_Z\_SIGNALLING\_EVENTS | 41 | 2 | 12187 | 72 | Rgs4,Gng11 | | 2.433e-02 | -3.72 | GO\_BODY\_MORPHOGENESIS | MSigDB lists | GO\_BODY\_MORPHOGENESIS | 41 | 2 | 12187 | 72 | Col1a1,Cldn5 | | 2.433e-02 | -3.72 | WU\_SILENCED\_BY\_METHYLATION\_IN\_BLADDER\_CANCER | MSigDB lists | WU\_SILENCED\_BY\_METHYLATION\_IN\_BLADDER\_CANCER | 41 | 2 | 12187 | 72 | Nr4a3,Peg3 | | 2.438e-02 | -3.71 | movement of cell or subcellular component | biological process | GO:0006928 | 1095 | 12 | 13711 | 80 | Agrn,Nr4a3,Sorl1,Ncam1,Atp5b,Gbx1,Kdr,Map4,P2ry12,Bdnf,Ephb6,Lhx1 | | 2.445e-02 | -3.71 | visual system development | biological process | GO:0150063 | 282 | 5 | 13711 | 80 | Fjx1,Lhx1,Kdr,Inhba,P2ry12 | | 2.449e-02 | -3.71 | cytoplasmic part | cellular component | GO:0044444 | 6966 | 49 | 13825 | 79 | Fbxl17,Gsdme,Atp5b,Hba-a2,Pom121,Sorl1,P2ry12,Col1a1,Amd1,Glud1,Cldn5,Car2,Tmx3,Map4,Agrn,Msmo1,Pdia6,Homer1,Mtmr9,Tgfa,Tmem165,Cacng2,Qdpr,Hkdc1,Ccdc13,Rgs4,Spry2,Ermn,Aars,Nacad,Chmp7,Nipsnap2,Inhba,Ndst4,Kdr,Peg3,H1f0,Hs3st1,Gng11,Ntm,Fabp7,Ltk,Scd2,Fnbp1,Ubqln2,Arntl,Hsd17b12,Bdnf,Ncam1 | | 2.449e-02 | -3.71 | regulation of MAPK cascade | biological process | GO:0043408 | 605 | 8 | 13711 | 80 | Timp2,Kdr,Tgfa,Sorl1,Inhba,Gsdme,Alkal2,Spry2 | | 2.460e-02 | -3.71 | response to oxygen-containing compound | biological process | GO:1901700 | 969 | 11 | 13711 | 80 | Crh,Car2,Nr4a3,Egr1,Homer1,Col1a1,Scd2,P2ry12,Ltk,Inhba,Ncam1 | | 2.468e-02 | -3.70 | GO\_ORGAN\_MORPHOGENESIS | MSigDB lists | GO\_ORGAN\_MORPHOGENESIS | 600 | 8 | 12187 | 72 | Inhba,Car2,Col1a1,Fjx1,Cldn5,Nr4a3,Spry2,Lhx1 | | 2.474e-02 | -3.70 | ACAGGGT\_MIR10A\_MIR10B | MSigDB lists | ACAGGGT\_MIR10A\_MIR10B | 106 | 3 | 12187 | 72 | Bdnf,Grm3,Nr4a3 | | 2.474e-02 | -3.70 | TNCATNTCCYR\_UNKNOWN | MSigDB lists | TNCATNTCCYR\_UNKNOWN | 106 | 3 | 12187 | 72 | Tle4,Nr4a3,Bdnf | | 2.474e-02 | -3.70 | STAT5A\_02 | MSigDB lists | STAT5A\_02 | 106 | 3 | 12187 | 72 | Nr4a3,Ddx17,Bdnf | | 2.474e-02 | -3.70 | DARWICHE\_SQUAMOUS\_CELL\_CARCINOMA\_UP | MSigDB lists | DARWICHE\_SQUAMOUS\_CELL\_CARCINOMA\_UP | 106 | 3 | 12187 | 72 | Qdpr,Srm,Tgfa | | 2.474e-02 | -3.70 | I-set | pfam domains | PF07679 | 112 | 3 | 12881 | 72 | Ncam1,Ntm,Kdr | | 2.478e-02 | -3.70 | sensory system development | biological process | GO:0048880 | 283 | 5 | 13711 | 80 | Lhx1,P2ry12,Kdr,Inhba,Fjx1 | | 2.490e-02 | -3.69 | regulation of sensory perception | biological process | GO:0051931 | 42 | 2 | 13711 | 80 | Ncam1,Grm3 | | 2.490e-02 | -3.69 | neural retina development | biological process | GO:0003407 | 42 | 2 | 13711 | 80 | Lhx1,Fjx1 | | 2.503e-02 | -3.69 | NFY\_01 | MSigDB lists | NFY\_01 | 188 | 4 | 12187 | 72 | Ncam1,Lhx1,Tle4,Col1a1 | | 2.503e-02 | -3.69 | HFH1\_01 | MSigDB lists | HFH1\_01 | 188 | 4 | 12187 | 72 | Inhba,Crh,Csrnp3,Ncam1 | | 2.522e-02 | -3.68 | Ser-Thr/Tyr\_kinase\_cat\_dom | interpro domains | IPR001245 | 110 | 3 | 13788 | 79 | Kdr,Ephb6,Ltk | | 2.540e-02 | -3.67 | YCATTAA\_UNKNOWN | MSigDB lists | YCATTAA\_UNKNOWN | 383 | 6 | 12187 | 72 | Ephb6,Tle4,P2ry12,Csrnp3,Ntm,Grm3 | | 2.545e-02 | -3.67 | MODULE\_85 | MSigDB lists | MODULE\_85 | 42 | 2 | 12187 | 72 | Ephb6,Kdr | | 2.545e-02 | -3.67 | GO\_RESPONSE\_TO\_COCAINE | MSigDB lists | GO\_RESPONSE\_TO\_COCAINE | 42 | 2 | 12187 | 72 | Homer1,Crh | | 2.545e-02 | -3.67 | GO\_GLOMERULUS\_DEVELOPMENT | MSigDB lists | GO\_GLOMERULUS\_DEVELOPMENT | 42 | 2 | 12187 | 72 | Egr1,Lhx1 | | 2.545e-02 | -3.67 | BERENJENO\_ROCK\_SIGNALING\_NOT\_VIA\_RHOA\_DN | MSigDB lists | BERENJENO\_ROCK\_SIGNALING\_NOT\_VIA\_RHOA\_DN | 42 | 2 | 12187 | 72 | Qk,Egr1 | | 2.545e-02 | -3.67 | JOHANSSON\_BRAIN\_CANCER\_EARLY\_VS\_LATE\_DN | MSigDB lists | JOHANSSON\_BRAIN\_CANCER\_EARLY\_VS\_LATE\_DN | 42 | 2 | 12187 | 72 | Qdpr,Plp1 | | 2.545e-02 | -3.67 | ZHOU\_TNF\_SIGNALING\_30MIN | MSigDB lists | ZHOU\_TNF\_SIGNALING\_30MIN | 42 | 2 | 12187 | 72 | Ddx56,Egr1 | | 2.545e-02 | -3.67 | KHETCHOUMIAN\_TRIM24\_TARGETS\_UP | MSigDB lists | KHETCHOUMIAN\_TRIM24\_TARGETS\_UP | 42 | 2 | 12187 | 72 | Egr1,Col1a1 | | 2.546e-02 | -3.67 | AAGCAAT\_MIR137 | MSigDB lists | AAGCAAT\_MIR137 | 189 | 4 | 12187 | 72 | Qk,Tle4,Cbln2,Amd1 | | 2.546e-02 | -3.67 | GO\_REGULATION\_OF\_HORMONE\_SECRETION | MSigDB lists | GO\_REGULATION\_OF\_HORMONE\_SECRETION | 189 | 4 | 12187 | 72 | Glud1,Crh,Arntl,Inhba | | 2.546e-02 | -3.67 | WALLACE\_PROSTATE\_CANCER\_RACE\_UP | MSigDB lists | WALLACE\_PROSTATE\_CANCER\_RACE\_UP | 189 | 4 | 12187 | 72 | Gng11,Nr4a3,Tle4,Cldn5 | | 2.546e-02 | -3.67 | SREBP\_Q3 | MSigDB lists | SREBP\_Q3 | 189 | 4 | 12187 | 72 | Fnbp1,Lhx1,Ddx17,Peg3 | | 2.546e-02 | -3.67 | AAAYWAACM\_HFH4\_01 | MSigDB lists | AAAYWAACM\_HFH4\_01 | 189 | 4 | 12187 | 72 | Ncam1,Bdnf,Crh,Nr4a3 | | 2.546e-02 | -3.67 | TFIIA\_Q6 | MSigDB lists | TFIIA\_Q6 | 189 | 4 | 12187 | 72 | Arntl,Plp1,Col1a1,Egr1 | | 2.576e-02 | -3.66 | positive regulation of cellular component movement | biological process | GO:0051272 | 497 | 7 | 13711 | 80 | Col1a1,Nr4a3,Spry2,Kdr,P2ry12,Atp5b,Egr1 | | 2.581e-02 | -3.66 | gastrulation | biological process | GO:0007369 | 109 | 3 | 13711 | 80 | Lhx1,Inhba,Nr4a3 | | 2.581e-02 | -3.66 | monocarboxylic acid biosynthetic process | biological process | GO:0072330 | 109 | 3 | 13711 | 80 | Plp1,Scd2,Qk | | 2.590e-02 | -3.65 | fn3 | pfam domains | PF00041 | 114 | 3 | 12881 | 72 | Ephb6,Sorl1,Ncam1 | | 2.596e-02 | -3.65 | GO\_CELL\_JUNCTION\_ASSEMBLY | MSigDB lists | GO\_CELL\_JUNCTION\_ASSEMBLY | 108 | 3 | 12187 | 72 | Lims2,Ugt8a,Cldn5 | | 2.596e-02 | -3.65 | BMI1\_DN.V1\_DN | MSigDB lists | BMI1\_DN.V1\_DN | 108 | 3 | 12187 | 72 | Car2,Peg3,Egr1 | | 2.596e-02 | -3.65 | BMI1\_DN\_MEL18\_DN.V1\_UP | MSigDB lists | BMI1\_DN\_MEL18\_DN.V1\_UP | 108 | 3 | 12187 | 72 | Fjx1,Inhba,Ntm | | 2.601e-02 | -3.65 | regulation of kidney development | biological process | GO:0090183 | 43 | 2 | 13711 | 80 | Egr1,Lhx1 | | 2.601e-02 | -3.65 | positive regulation of protein tyrosine kinase activity | biological process | GO:0061098 | 43 | 2 | 13711 | 80 | Tgfa,Agrn | | 2.606e-02 | -3.65 | - | gene3d domains | 1.25.40.1020 | 5 | 1 | 6647 | 35 | Ermn | | 2.606e-02 | -3.65 | - | gene3d domains | 3.30.70.960 | 5 | 1 | 6647 | 35 | Agrn | | 2.633e-02 | -3.64 | TGIF\_01 | MSigDB lists | TGIF\_01 | 191 | 4 | 12187 | 72 | H1f0,Lhx1,Crh,Nr4a3 | | 2.642e-02 | -3.63 | Ig\_I-set | interpro domains | IPR013098 | 112 | 3 | 13788 | 79 | Ncam1,Ntm,Kdr | | 2.642e-02 | -3.63 | Antagonism of Activin by Follistatin | REACTOME pathways | R-MMU-2473224 | 4 | 1 | 6297 | 42 | Inhba | | 2.642e-02 | -3.63 | Neurophilin interactions with VEGF and VEGFR | REACTOME pathways | R-MMU-194306 | 4 | 1 | 6297 | 42 | Kdr | | 2.642e-02 | -3.63 | PPARA activates gene expression | REACTOME pathways | R-MMU-1989781 | 4 | 1 | 6297 | 42 | Arntl | | 2.658e-02 | -3.63 | GO\_ADULT\_BEHAVIOR | MSigDB lists | GO\_ADULT\_BEHAVIOR | 109 | 3 | 12187 | 72 | Homer1,Nr4a3,Gbx1 | | 2.659e-02 | -3.63 | BURTON\_ADIPOGENESIS\_7 | MSigDB lists | BURTON\_ADIPOGENESIS\_7 | 43 | 2 | 12187 | 72 | Timp2,Ubqln2 | | 2.659e-02 | -3.63 | GO\_INTRAMOLECULAR\_OXIDOREDUCTASE\_ACTIVITY | MSigDB lists | GO\_INTRAMOLECULAR\_OXIDOREDUCTASE\_ACTIVITY | 43 | 2 | 12187 | 72 | Pdia6,Tmx3 | | 2.659e-02 | -3.63 | GO\_GOLGI\_LUMEN | MSigDB lists | GO\_GOLGI\_LUMEN | 43 | 2 | 12187 | 72 | Agrn,Hs3st1 | | 2.659e-02 | -3.63 | GO\_RESPONSE\_TO\_AMMONIUM\_ION | MSigDB lists | GO\_RESPONSE\_TO\_AMMONIUM\_ION | 43 | 2 | 12187 | 72 | Agrn,Crh | | 2.686e-02 | -3.62 | NAC\_AB | prosite domains | PS51151 | 4 | 1 | 8845 | 60 | Nacad | | 2.686e-02 | -3.62 | TIMP | prosite domains | PS00288 | 4 | 1 | 8845 | 60 | Timp2 | | 2.686e-02 | -3.62 | ATPASE\_ALPHA\_BETA | prosite domains | PS00152 | 4 | 1 | 8845 | 60 | Atp5b | | 2.697e-02 | -3.61 | anatomical structure formation involved in morphogenesis | biological process | GO:0048646 | 735 | 9 | 13711 | 80 | Col1a1,Atp5b,Lhx1,Inhba,Nr4a3,Fjx1,Ugt8a,Tgfa,Kdr | | 2.715e-02 | -3.61 | regulation of skeletal muscle tissue development | biological process | GO:0048641 | 44 | 2 | 13711 | 80 | Arntl,Ddx17 | | 2.715e-02 | -3.61 | spindle localization | biological process | GO:0051653 | 44 | 2 | 13711 | 80 | Spry2,Map4 | | 2.722e-02 | -3.60 | GCM\_NCAM1 | MSigDB lists | GCM\_NCAM1 | 110 | 3 | 12187 | 72 | Mtmr9,Peg3,Ncam1 | | 2.722e-02 | -3.60 | TAXCREB\_01 | MSigDB lists | TAXCREB\_01 | 110 | 3 | 12187 | 72 | Peg3,AI593442,Nr4a3 | | 2.724e-02 | -3.60 | hippocampal mossy fiber to CA3 synapse | cellular component | GO:0098686 | 45 | 2 | 13825 | 79 | Lrrtm2,Cacng2 | | 2.736e-02 | -3.60 | organic acid metabolic process | biological process | GO:0006082 | 618 | 8 | 13711 | 80 | Plp1,Nr4a3,Qdpr,Aars,Hkdc1,Glud1,Scd2,Qk | | 2.746e-02 | -3.59 | enzyme regulator activity | molecular function | GO:0030234 | 746 | 9 | 13516 | 78 | Timp2,Pdia6,Alkal2,Mtmr9,Spry2,Rgs4,Arhgap20,Dgkz,Agrn | | 2.752e-02 | -3.59 | mouse chr6\|6 B2.1 | chromosome location | mouse chr6\|6 B2.1 | 5 | 1 | 14556 | 81 | Ephb6 | | 2.752e-02 | -3.59 | protein tyrosine kinase activity | molecular function | GO:0004713 | 113 | 3 | 13516 | 78 | Ltk,Ephb6,Kdr | | 2.752e-02 | -3.59 | positive regulation of locomotion | biological process | GO:0040017 | 504 | 7 | 13711 | 80 | Spry2,Col1a1,Atp5b,P2ry12,Nr4a3,Egr1,Kdr | | 2.760e-02 | -3.59 | neuron to neuron synapse | cellular component | GO:0098984 | 403 | 6 | 13825 | 79 | Map4,Cacng2,Lrrtm2,Homer1,Grm3,Dgkz | | 2.764e-02 | -3.59 | FA\_hydroxylase | pfam domains | PF04116 | 5 | 1 | 12881 | 72 | Msmo1 | | 2.764e-02 | -3.59 | Sortilin\_C | pfam domains | PF15901 | 5 | 1 | 12881 | 72 | Sorl1 | | 2.764e-02 | -3.59 | tRNA\_SAD | pfam domains | PF07973 | 5 | 1 | 12881 | 72 | Aars | | 2.764e-02 | -3.59 | TLE\_N | pfam domains | PF03920 | 5 | 1 | 12881 | 72 | Tle4 | | 2.764e-02 | -3.59 | Sortilin-Vps10 | pfam domains | PF15902 | 5 | 1 | 12881 | 72 | Sorl1 | | 2.768e-02 | -3.59 | positive regulation of cell-substrate adhesion | biological process | GO:0010811 | 112 | 3 | 13711 | 80 | Lims2,Kdr,Hsd17b12 | | 2.776e-02 | -3.58 | PID\_FGF\_PATHWAY | MSigDB lists | PID\_FGF\_PATHWAY | 44 | 2 | 12187 | 72 | Spry2,Ncam1 | | 2.776e-02 | -3.58 | GO\_CEREBELLAR\_CORTEX\_DEVELOPMENT | MSigDB lists | GO\_CEREBELLAR\_CORTEX\_DEVELOPMENT | 44 | 2 | 12187 | 72 | Lhx1,Aars | | 2.776e-02 | -3.58 | GO\_FACE\_DEVELOPMENT | MSigDB lists | GO\_FACE\_DEVELOPMENT | 44 | 2 | 12187 | 72 | Col1a1,Cldn5 | | 2.776e-02 | -3.58 | GO\_NEUROMUSCULAR\_PROCESS\_CONTROLLING\_BALANCE | MSigDB lists | GO\_NEUROMUSCULAR\_PROCESS\_CONTROLLING\_BALANCE | 44 | 2 | 12187 | 72 | Nr4a3,Aars | | 2.776e-02 | -3.58 | GO\_TRANSFERASE\_ACTIVITY\_TRANSFERRING\_SULFUR\_CONTAINING\_GROUPS | MSigDB lists | GO\_TRANSFERASE\_ACTIVITY\_TRANSFERRING\_SULFUR\_CONTAINING\_GROUPS | 44 | 2 | 12187 | 72 | Ndst4,Hs3st1 | | 2.786e-02 | -3.58 | GSE3920\_UNTREATED\_VS\_IFNG\_TREATED\_ENDOTHELIAL\_CELL\_UP | MSigDB lists | GSE3920\_UNTREATED\_VS\_IFNG\_TREATED\_ENDOTHELIAL\_CELL\_UP | 111 | 3 | 12187 | 72 | Nr4a3,Egr1,Rgs4 | | 2.786e-02 | -3.58 | GO\_NEGATIVE\_REGULATION\_OF\_NEURON\_APOPTOTIC\_PROCESS | MSigDB lists | GO\_NEGATIVE\_REGULATION\_OF\_NEURON\_APOPTOTIC\_PROCESS | 111 | 3 | 12187 | 72 | Aars,Bdnf,Nr4a3 | | 2.786e-02 | -3.58 | AAAGGAT\_MIR501 | MSigDB lists | AAAGGAT\_MIR501 | 111 | 3 | 12187 | 72 | Nr4a3,Qk,Plp1 | | 2.786e-02 | -3.58 | GO\_NEGATIVE\_REGULATION\_OF\_PROTEIN\_SERINE\_THREONINE\_KINASE\_ACTIVITY | MSigDB lists | GO\_NEGATIVE\_REGULATION\_OF\_PROTEIN\_SERINE\_THREONINE\_KINASE\_ACTIVITY | 111 | 3 | 12187 | 72 | Rgs4,Sorl1,Spry2 | | 2.786e-02 | -3.58 | DARWICHE\_PAPILLOMA\_RISK\_LOW\_UP | MSigDB lists | DARWICHE\_PAPILLOMA\_RISK\_LOW\_UP | 111 | 3 | 12187 | 72 | Srm,Tgfa,Qdpr | | 2.791e-02 | -3.58 | MODULE\_24 | MSigDB lists | MODULE\_24 | 289 | 5 | 12187 | 72 | Cldn5,Msmo1,Col1a1,Qdpr,Peg3 | | 2.798e-02 | -3.58 | integral component of synaptic membrane | cellular component | GO:0099699 | 201 | 4 | 13825 | 79 | Lrrtm2,Cacng2,Ncam1,Grm3 | | 2.810e-02 | -3.57 | DNA-binding transcription activator activity, RNA polymerase II-specific | molecular function | GO:0001228 | 296 | 5 | 13516 | 78 | Gbx1,Arntl,Egr1,Csrnp3,Nr4a3 | | 2.810e-02 | -3.57 | DNA-binding transcription activator activity | molecular function | GO:0001216 | 296 | 5 | 13516 | 78 | Csrnp3,Nr4a3,Gbx1,Arntl,Egr1 | | 2.813e-02 | -3.57 | mouse chr10\|10 B4 | chromosome location | mouse chr10\|10 B4 | 47 | 2 | 14556 | 81 | Hkdc1,Fabp7 | | 2.813e-02 | -3.57 | RUTELLA\_RESPONSE\_TO\_HGF\_VS\_CSF2RB\_AND\_IL4\_DN | MSigDB lists | RUTELLA\_RESPONSE\_TO\_HGF\_VS\_CSF2RB\_AND\_IL4\_DN | 195 | 4 | 12187 | 72 | Sorl1,Inhba,Nr4a3,Fnbp1 | | 2.825e-02 | -3.57 | proton-transporting ATP synthase complex, catalytic core F(1) | cellular component | GO:0045261 | 5 | 1 | 13825 | 79 | Atp5b | | 2.825e-02 | -3.57 | sorting endosome | cellular component | GO:0097443 | 5 | 1 | 13825 | 79 | Kdr | | 2.825e-02 | -3.57 | haptoglobin-hemoglobin complex | cellular component | GO:0031838 | 5 | 1 | 13825 | 79 | Hba-a2 | | 2.825e-02 | -3.57 | mitochondrial proton-transporting ATP synthase complex, catalytic core F(1) | cellular component | GO:0000275 | 5 | 1 | 13825 | 79 | Atp5b | | 2.831e-02 | -3.56 | mesoderm morphogenesis | biological process | GO:0048332 | 45 | 2 | 13711 | 80 | Inhba,Nr4a3 | | 2.833e-02 | -3.56 | Sortilin\_C | interpro domains | IPR031777 | 5 | 1 | 13788 | 79 | Sorl1 | | 2.833e-02 | -3.56 | Moesin\_tail\_sf | interpro domains | IPR008954 | 5 | 1 | 13788 | 79 | Ermn | | 2.833e-02 | -3.56 | VPS10 | interpro domains | IPR006581 | 5 | 1 | 13788 | 79 | Sorl1 | | 2.833e-02 | -3.56 | Fatty\_acid\_hydroxylase | interpro domains | IPR006694 | 5 | 1 | 13788 | 79 | Msmo1 | | 2.833e-02 | -3.56 | tRNA\_SAD | interpro domains | IPR012947 | 5 | 1 | 13788 | 79 | Aars | | 2.833e-02 | -3.56 | Sortilin\_N | interpro domains | IPR031778 | 5 | 1 | 13788 | 79 | Sorl1 | | 2.833e-02 | -3.56 | Disulphide\_isomerase | interpro domains | IPR005788 | 5 | 1 | 13788 | 79 | Pdia6 | | 2.833e-02 | -3.56 | ILBP | interpro domains | IPR031259 | 5 | 1 | 13788 | 79 | Fabp7 | | 2.833e-02 | -3.56 | Fatty\_acid-bd | interpro domains | IPR000463 | 5 | 1 | 13788 | 79 | Fabp7 | | 2.833e-02 | -3.56 | Groucho/TLE\_N | interpro domains | IPR005617 | 5 | 1 | 13788 | 79 | Tle4 | | 2.846e-02 | -3.56 | CTTTGA\_LEF1\_Q2 | MSigDB lists | CTTTGA\_LEF1\_Q2 | 857 | 10 | 12187 | 72 | Inhba,Tle4,Csrnp3,Ddx17,H1f0,Crh,Nr4a3,Lhx1,Glud1,Arhgap20 | | 2.851e-02 | -3.56 | GSE36891\_POLYIC\_TLR3\_VS\_PAM\_TLR2\_STIM\_PERITONEAL\_MACROPHAGE\_UP | MSigDB lists | GSE36891\_POLYIC\_TLR3\_VS\_PAM\_TLR2\_STIM\_PERITONEAL\_MACROPHAGE\_UP | 112 | 3 | 12187 | 72 | Egr1,Spry2,Nr4a3 | | 2.851e-02 | -3.56 | Gipc1 (GIPC PDZ domain containing family, member 1) | protein interactions | 67903 | 4 | 1 | 6802 | 49 | Homer1 | | 2.851e-02 | -3.56 | Cdk8 (cyclin-dependent kinase 8) | protein interactions | 264064 | 4 | 1 | 6802 | 49 | Egr1 | | 2.851e-02 | -3.56 | MPZL1 (myelin protein zero like 1) | protein interactions | 9019 | 4 | 1 | 6802 | 49 | Homer1 | | 2.851e-02 | -3.56 | Hspa1l (heat shock protein 1-like) | protein interactions | 15482 | 4 | 1 | 6802 | 49 | Ubqln2 | | 2.851e-02 | -3.56 | FRYL (FRY like transcription coactivator) | protein interactions | 285527 | 4 | 1 | 6802 | 49 | Homer1 | | 2.851e-02 | -3.56 | Htra1 (HtrA serine peptidase 1) | protein interactions | 56213 | 4 | 1 | 6802 | 49 | Inhba | | 2.851e-02 | -3.56 | Six3 (sine oculis-related homeobox 3) | protein interactions | 20473 | 4 | 1 | 6802 | 49 | Tle4 | | 2.851e-02 | -3.56 | Met (met proto-oncogene) | protein interactions | 17295 | 4 | 1 | 6802 | 49 | Kdr | | 2.851e-02 | -3.56 | PSRC1 (proline and serine rich coiled-coil 1) | protein interactions | 84722 | 4 | 1 | 6802 | 49 | Homer1 | | 2.851e-02 | -3.56 | Mtf1 (metal response element binding transcription factor 1) | protein interactions | 17764 | 4 | 1 | 6802 | 49 | Nr4a3 | | 2.851e-02 | -3.56 | Ptprr (protein tyrosine phosphatase, receptor type, R) | protein interactions | 19279 | 4 | 1 | 6802 | 49 | Ntm | | 2.853e-02 | -3.56 | stearoyl-CoA 9-desaturase activity | molecular function | GO:0004768 | 5 | 1 | 13516 | 78 | Scd2 | | 2.853e-02 | -3.56 | G protein-coupled adenosine receptor activity | molecular function | GO:0001609 | 5 | 1 | 13516 | 78 | P2ry12 | | 2.853e-02 | -3.56 | heparan sulfate N-acetylglucosaminyltransferase activity | molecular function | GO:0042328 | 5 | 1 | 13516 | 78 | Ndst4 | | 2.853e-02 | -3.56 | estradiol 17-beta-dehydrogenase activity | molecular function | GO:0004303 | 5 | 1 | 13516 | 78 | Hsd17b12 | | 2.853e-02 | -3.56 | [heparan sulfate]-glucosamine 3-sulfotransferase 1 activity | molecular function | GO:0008467 | 5 | 1 | 13516 | 78 | Hs3st1 | | 2.860e-02 | -3.55 | GO\_REGULATION\_OF\_MAPK\_CASCADE | MSigDB lists | GO\_REGULATION\_OF\_MAPK\_CASCADE | 503 | 7 | 12187 | 72 | Sorl1,Tgfa,Timp2,Inhba,Rgs4,Spry2,Kdr | | 2.863e-02 | -3.55 | Kcnma1 (potassium large conductance calcium-activated channel, subfamily M, alpha member 1) | protein interactions | 16531 | 162 | 4 | 6802 | 49 | Atp5b,Glud1,Col1a1,Car2 | | 2.872e-02 | -3.55 | tissue development | biological process | GO:0009888 | 1253 | 13 | 13711 | 80 | Bdnf,Lhx1,Kdr,Egr1,Ddx17,Col1a1,Spry2,Inhba,Car2,Foxq1,Nr4a3,Agrn,Homer1 | | 2.884e-02 | -3.55 | motor neuron migration | biological process | GO:0097475 | 5 | 1 | 13711 | 80 | Lhx1 | | 2.884e-02 | -3.55 | negative regulation of epinephrine secretion | biological process | GO:0032811 | 5 | 1 | 13711 | 80 | Crh | | 2.884e-02 | -3.55 | regulation of retinal cell programmed cell death | biological process | GO:0046668 | 5 | 1 | 13711 | 80 | Bdnf | | 2.884e-02 | -3.55 | rRNA pseudouridine synthesis | biological process | GO:0031118 | 5 | 1 | 13711 | 80 | Gar1 | | 2.884e-02 | -3.55 | positive regulation of gastrulation | biological process | GO:2000543 | 5 | 1 | 13711 | 80 | Lhx1 | | 2.884e-02 | -3.55 | regulation of IRE1-mediated unfolded protein response | biological process | GO:1903894 | 5 | 1 | 13711 | 80 | Pdia6 | | 2.884e-02 | -3.55 | positive regulation of endothelial cell development | biological process | GO:1901552 | 5 | 1 | 13711 | 80 | Cldn5 | | 2.884e-02 | -3.55 | cellular response to corticotropin-releasing hormone stimulus | biological process | GO:0071376 | 5 | 1 | 13711 | 80 | Nr4a3 | | 2.884e-02 | -3.55 | regulation of sodium ion export across plasma membrane | biological process | GO:1903276 | 5 | 1 | 13711 | 80 | Agrn | | 2.884e-02 | -3.55 | negative regulation of gonadotropin secretion | biological process | GO:0032277 | 5 | 1 | 13711 | 80 | Crh | | 2.884e-02 | -3.55 | post-embryonic eye morphogenesis | biological process | GO:0048050 | 5 | 1 | 13711 | 80 | Kdr | | 2.884e-02 | -3.55 | positive regulation of interleukin-1 biosynthetic process | biological process | GO:0045362 | 5 | 1 | 13711 | 80 | Egr1 | | 2.884e-02 | -3.55 | positive regulation of establishment of endothelial barrier | biological process | GO:1903142 | 5 | 1 | 13711 | 80 | Cldn5 | | 2.884e-02 | -3.55 | protein localization to paranode region of axon | biological process | GO:0002175 | 5 | 1 | 13711 | 80 | Ugt8a | | 2.884e-02 | -3.55 | post-embryonic camera-type eye morphogenesis | biological process | GO:0048597 | 5 | 1 | 13711 | 80 | Kdr | | 2.884e-02 | -3.55 | intramembranous ossification | biological process | GO:0001957 | 5 | 1 | 13711 | 80 | Col1a1 | | 2.884e-02 | -3.55 | spinal cord association neuron differentiation | biological process | GO:0021527 | 5 | 1 | 13711 | 80 | Lhx1 | | 2.884e-02 | -3.55 | positive regulation of high voltage-gated calcium channel activity | biological process | GO:1901843 | 5 | 1 | 13711 | 80 | Nipsnap2 | | 2.884e-02 | -3.55 | response to corticotropin-releasing hormone | biological process | GO:0043435 | 5 | 1 | 13711 | 80 | Nr4a3 | | 2.884e-02 | -3.55 | regulation of interleukin-1 beta biosynthetic process | biological process | GO:0050722 | 5 | 1 | 13711 | 80 | Egr1 | | 2.884e-02 | -3.55 | regulation of cellular pH reduction | biological process | GO:0032847 | 5 | 1 | 13711 | 80 | Car2 | | 2.884e-02 | -3.55 | glutamate catabolic process | biological process | GO:0006538 | 5 | 1 | 13711 | 80 | Glud1 | | 2.884e-02 | -3.55 | positive regulation of endocytic recycling | biological process | GO:2001137 | 5 | 1 | 13711 | 80 | Sorl1 | | 2.884e-02 | -3.55 | positive regulation of ER-associated ubiquitin-dependent protein catabolic process | biological process | GO:1903071 | 5 | 1 | 13711 | 80 | Ubqln2 | | 2.884e-02 | -3.55 | Fc receptor mediated stimulatory signaling pathway | biological process | GO:0002431 | 5 | 1 | 13711 | 80 | Nr4a3 | | 2.884e-02 | -3.55 | mitochondrial ATP synthesis coupled proton transport | biological process | GO:0042776 | 5 | 1 | 13711 | 80 | Atp5b | | 2.884e-02 | -3.55 | direct ossification | biological process | GO:0036072 | 5 | 1 | 13711 | 80 | Col1a1 | | 2.884e-02 | -3.55 | paramesonephric duct development | biological process | GO:0061205 | 5 | 1 | 13711 | 80 | Lhx1 | | 2.884e-02 | -3.55 | positive regulation of glucocorticoid receptor signaling pathway | biological process | GO:2000324 | 5 | 1 | 13711 | 80 | Bdnf | | 2.884e-02 | -3.55 | negative regulation of hydrogen peroxide-induced neuron death | biological process | GO:1903208 | 5 | 1 | 13711 | 80 | Nr4a3 | | 2.884e-02 | -3.55 | regulation of vesicle docking | biological process | GO:0106020 | 5 | 1 | 13711 | 80 | Ncam1 | | 2.884e-02 | -3.55 | regulation of interleukin-1 biosynthetic process | biological process | GO:0045360 | 5 | 1 | 13711 | 80 | Egr1 | | 2.884e-02 | -3.55 | erythrose 4-phosphate/phosphoenolpyruvate family amino acid metabolic process | biological process | GO:1902221 | 5 | 1 | 13711 | 80 | Qdpr | | 2.884e-02 | -3.55 | hemoglobin biosynthetic process | biological process | GO:0042541 | 5 | 1 | 13711 | 80 | Inhba | | 2.884e-02 | -3.55 | regulation of hydrogen peroxide-induced neuron death | biological process | GO:1903207 | 5 | 1 | 13711 | 80 | Nr4a3 | | 2.884e-02 | -3.55 | protein retention in Golgi apparatus | biological process | GO:0045053 | 5 | 1 | 13711 | 80 | Sorl1 | | 2.884e-02 | -3.55 | positive regulation of interleukin-1 beta biosynthetic process | biological process | GO:0050725 | 5 | 1 | 13711 | 80 | Egr1 | | 2.884e-02 | -3.55 | regulation of corticotropin secretion | biological process | GO:0051459 | 5 | 1 | 13711 | 80 | Crh | | 2.884e-02 | -3.55 | L-phenylalanine metabolic process | biological process | GO:0006558 | 5 | 1 | 13711 | 80 | Qdpr | | 2.884e-02 | -3.55 | positive regulation of transport | biological process | GO:0051050 | 867 | 10 | 13711 | 80 | Cacng2,Glud1,P2ry12,Sorl1,Car2,Crh,Lrrc55,Nr4a3,Nipsnap2,Homer1 | | 2.894e-02 | -3.54 | GO\_TRANSMISSION\_OF\_NERVE\_IMPULSE | MSigDB lists | GO\_TRANSMISSION\_OF\_NERVE\_IMPULSE | 45 | 2 | 12187 | 72 | Cacng2,Kcnmb2 | | 2.894e-02 | -3.54 | KENNY\_CTNNB1\_TARGETS\_DN | MSigDB lists | KENNY\_CTNNB1\_TARGETS\_DN | 45 | 2 | 12187 | 72 | Tmem165,Timp2 | | 2.894e-02 | -3.54 | GO\_CELLULAR\_RESPONSE\_TO\_RETINOIC\_ACID | MSigDB lists | GO\_CELLULAR\_RESPONSE\_TO\_RETINOIC\_ACID | 45 | 2 | 12187 | 72 | Ltk,Col1a1 | | 2.894e-02 | -3.54 | GTAAGAT\_MIR200A | MSigDB lists | GTAAGAT\_MIR200A | 45 | 2 | 12187 | 72 | Ubqln2,Qk | | 2.894e-02 | -3.54 | GO\_VASCULOGENESIS | MSigDB lists | GO\_VASCULOGENESIS | 45 | 2 | 12187 | 72 | Kdr,Qk | | 2.894e-02 | -3.54 | GO\_REGULATION\_OF\_INTRACELLULAR\_STEROID\_HORMONE\_RECEPTOR\_SIGNALING\_PATHWAY | MSigDB lists | GO\_REGULATION\_OF\_INTRACELLULAR\_STEROID\_HORMONE\_RECEPTOR\_SIGNALING\_PATHWAY | 45 | 2 | 12187 | 72 | Ddx17,Arntl | | 2.894e-02 | -3.54 | LEE\_CALORIE\_RESTRICTION\_MUSCLE\_DN | MSigDB lists | LEE\_CALORIE\_RESTRICTION\_MUSCLE\_DN | 45 | 2 | 12187 | 72 | Amd1,Ddx56 | | 2.894e-02 | -3.54 | BRIDEAU\_IMPRINTED\_GENES | MSigDB lists | BRIDEAU\_IMPRINTED\_GENES | 45 | 2 | 12187 | 72 | Peg3,Nap1l5 | | 2.897e-02 | -3.54 | sensory perception of sound | biological process | GO:0007605 | 114 | 3 | 13711 | 80 | Col1a1,Spry2,Gsdme | | 2.897e-02 | -3.54 | regulation of epithelial cell differentiation | biological process | GO:0030856 | 114 | 3 | 13711 | 80 | Arntl,Cldn5,Lhx1 | | 2.906e-02 | -3.54 | P53\_02 | MSigDB lists | P53\_02 | 197 | 4 | 12187 | 72 | Map4,Dgkz,Cacng2,Nr4a3 | | 2.920e-02 | -3.53 | GGCGGCA\_MIR371 | MSigDB lists | GGCGGCA\_MIR371 | 5 | 1 | 12187 | 72 | Pom121 | | 2.920e-02 | -3.53 | REACTOME\_PEPTIDE\_HORMONE\_BIOSYNTHESIS | MSigDB lists | REACTOME\_PEPTIDE\_HORMONE\_BIOSYNTHESIS | 5 | 1 | 12187 | 72 | Inhba | | 2.920e-02 | -3.53 | NOUSHMEHR\_GBM\_GERMLINE\_MUTATED | MSigDB lists | NOUSHMEHR\_GBM\_GERMLINE\_MUTATED | 5 | 1 | 12187 | 72 | Dgkz | | 2.920e-02 | -3.53 | WIEMANN\_TELOMERE\_SHORTENING\_AND\_CHRONIC\_LIVER\_DAMAGE\_DN | MSigDB lists | WIEMANN\_TELOMERE\_SHORTENING\_AND\_CHRONIC\_LIVER\_DAMAGE\_DN | 5 | 1 | 12187 | 72 | Egr1 | | 2.920e-02 | -3.53 | MATZUK\_STEROIDOGENESIS | MSigDB lists | MATZUK\_STEROIDOGENESIS | 5 | 1 | 12187 | 72 | Arntl | | 2.920e-02 | -3.53 | CHANG\_POU5F1\_TARGETS\_DN | MSigDB lists | CHANG\_POU5F1\_TARGETS\_DN | 5 | 1 | 12187 | 72 | Pom121 | | 2.920e-02 | -3.53 | TERAO\_AOX4\_TARGETS\_HG\_DN | MSigDB lists | TERAO\_AOX4\_TARGETS\_HG\_DN | 5 | 1 | 12187 | 72 | Egr1 | | 2.920e-02 | -3.53 | ELVIDGE\_HIF2A\_TARGETS\_UP | MSigDB lists | ELVIDGE\_HIF2A\_TARGETS\_UP | 5 | 1 | 12187 | 72 | Sorl1 | | 2.920e-02 | -3.53 | chr6q26 | MSigDB lists | chr6q26 | 5 | 1 | 12187 | 72 | Qk | | 2.920e-02 | -3.53 | IGARASHI\_ATF4\_TARGETS\_UP | MSigDB lists | IGARASHI\_ATF4\_TARGETS\_UP | 5 | 1 | 12187 | 72 | Egr1 | | 2.935e-02 | -3.53 | heparan sulfate biosynthesis | BIOCYC pathways | MOUSE\_PWY-6564 | 23 | 2 | 823 | 10 | Ndst4,Hs3st1 | | 2.948e-02 | -3.52 | retina morphogenesis in camera-type eye | biological process | GO:0060042 | 46 | 2 | 13711 | 80 | Fjx1,Lhx1 | | 2.948e-02 | -3.52 | proteoglycan biosynthetic process | biological process | GO:0030166 | 46 | 2 | 13711 | 80 | Hs3st1,Ndst4 | | 2.948e-02 | -3.52 | neuromuscular junction development | biological process | GO:0007528 | 46 | 2 | 13711 | 80 | Agrn,Cacng2 | | 2.956e-02 | -3.52 | VPS10 | smart domains | SM00602 | 5 | 1 | 7188 | 43 | Sorl1 | | 2.956e-02 | -3.52 | tRNA\_SAD | smart domains | SM00863 | 5 | 1 | 7188 | 43 | Aars | | 2.968e-02 | -3.52 | GO\_MEMBRANE\_REGION | MSigDB lists | GO\_MEMBRANE\_REGION | 863 | 10 | 12187 | 72 | Cldn5,P2ry12,Lrrtm2,Kdr,Spry2,Homer1,Fnbp1,Tgfa,Car2,Grm3 | | 2.968e-02 | -3.52 | postsynaptic specialization | cellular component | GO:0099572 | 410 | 6 | 13825 | 79 | Lrrtm2,Cacng2,Map4,Dgkz,Grm3,Homer1 | | 2.996e-02 | -3.51 | cell-cell signaling | biological process | GO:0007267 | 629 | 8 | 13711 | 80 | Fjx1,Crh,Tle4,Grm3,Agrn,Inhba,Lhx1,Bdnf | | 2.998e-02 | -3.51 | cellular response to stimulus | biological process | GO:0051716 | 4061 | 32 | 13711 | 80 | Egr1,Ddx17,Bdnf,Dgkz,Arntl,Cldn5,Homer1,Car2,Ccdc13,Ncam1,Col1a1,Spry2,Plp1,Kdr,Tgfa,Timp2,Arhgap20,Tle4,Crh,Nrep,P2ry12,Ephb6,Srm,Rgs4,Grm3,Nr4a3,Inhba,Ltk,Atp5b,Gng11,Gsdme,Ubqln2 | | 3.000e-02 | -3.51 | GO\_RESPONSE\_TO\_ENDOPLASMIC\_RETICULUM\_STRESS | MSigDB lists | GO\_RESPONSE\_TO\_ENDOPLASMIC\_RETICULUM\_STRESS | 199 | 4 | 12187 | 72 | Aars,Pdia6,Tmx3,Ubqln2 | | 3.014e-02 | -3.50 | GO\_REGULATION\_OF\_BEHAVIOR | MSigDB lists | GO\_REGULATION\_OF\_BEHAVIOR | 46 | 2 | 12187 | 72 | Nr4a3,Crh | | 3.014e-02 | -3.50 | GO\_POSITIVE\_REGULATION\_OF\_INSULIN\_SECRETION | MSigDB lists | GO\_POSITIVE\_REGULATION\_OF\_INSULIN\_SECRETION | 46 | 2 | 12187 | 72 | Glud1,Crh | | 3.014e-02 | -3.50 | SCHAEFFER\_PROSTATE\_DEVELOPMENT\_12HR\_DN | MSigDB lists | SCHAEFFER\_PROSTATE\_DEVELOPMENT\_12HR\_DN | 46 | 2 | 12187 | 72 | Lims2,Inhba | | 3.014e-02 | -3.50 | RYTAAWNNNTGAY\_UNKNOWN | MSigDB lists | RYTAAWNNNTGAY\_UNKNOWN | 46 | 2 | 12187 | 72 | Bdnf,Egr1 | | 3.014e-02 | -3.50 | CHARAFE\_BREAST\_CANCER\_BASAL\_VS\_MESENCHYMAL\_DN | MSigDB lists | CHARAFE\_BREAST\_CANCER\_BASAL\_VS\_MESENCHYMAL\_DN | 46 | 2 | 12187 | 72 | Gng11,Bdnf | | 3.014e-02 | -3.50 | WIEDERSCHAIN\_TARGETS\_OF\_BMI1\_AND\_PCGF2 | MSigDB lists | WIEDERSCHAIN\_TARGETS\_OF\_BMI1\_AND\_PCGF2 | 46 | 2 | 12187 | 72 | Ntm,Inhba | | 3.020e-02 | -3.50 | NAD(P)-bd\_dom\_sf | interpro domains | IPR036291 | 118 | 3 | 13788 | 79 | Hsd17b12,Qdpr,Glud1 | | 3.020e-02 | -3.50 | NGF | prints domains | PR00268 | 3 | 1 | 2951 | 30 | Bdnf | | 3.020e-02 | -3.50 | NUCLEARECPTR | prints domains | PR01284 | 3 | 1 | 2951 | 30 | Nr4a3 | | 3.020e-02 | -3.50 | FACDDSATRASE | prints domains | PR00075 | 3 | 1 | 2951 | 30 | Scd2 | | 3.020e-02 | -3.50 | MYELINPLP | prints domains | PR00214 | 3 | 1 | 2951 | 30 | Plp1 | | 3.020e-02 | -3.50 | CYSSERRICHNP | prints domains | PR02031 | 3 | 1 | 2951 | 30 | Csrnp3 | | 3.022e-02 | -3.50 | Heparan sulfate/heparin (HS-GAG) metabolism | REACTOME pathways | R-MMU-1638091 | 41 | 2 | 6297 | 42 | Agrn,Hs3st1 | | 3.023e-02 | -3.50 | GO\_LIPID\_BIOSYNTHETIC\_PROCESS | MSigDB lists | GO\_LIPID\_BIOSYNTHETIC\_PROCESS | 399 | 6 | 12187 | 72 | Qk,Crh,Plp1,Ugt8a,Hsd17b12,Msmo1 | | 3.028e-02 | -3.50 | developmental growth involved in morphogenesis | biological process | GO:0060560 | 116 | 3 | 13711 | 80 | Bdnf,Lhx1,Spry2 | | 3.038e-02 | -3.49 | mouse chr9\|9 A5.3 | chromosome location | mouse chr9\|9 A5.3 | 49 | 2 | 14556 | 81 | Arhgap20,AI593442 | | 3.048e-02 | -3.49 | ATF\_01 | MSigDB lists | ATF\_01 | 200 | 4 | 12187 | 72 | Peg3,Grm3,Lhx1,AI593442 | | 3.048e-02 | -3.49 | ZIC2\_01 | MSigDB lists | ZIC2\_01 | 200 | 4 | 12187 | 72 | Fnbp1,Map4,Bdnf,H1f0 | | 3.050e-02 | -3.49 | SARRIO\_EPITHELIAL\_MESENCHYMAL\_TRANSITION\_DN | MSigDB lists | SARRIO\_EPITHELIAL\_MESENCHYMAL\_TRANSITION\_DN | 115 | 3 | 12187 | 72 | Car2,Egr1,Glud1 | | 3.050e-02 | -3.49 | TOOKER\_GEMCITABINE\_RESISTANCE\_DN | MSigDB lists | TOOKER\_GEMCITABINE\_RESISTANCE\_DN | 115 | 3 | 12187 | 72 | Pdia6,H1f0,Qk | | 3.051e-02 | -3.49 | GO\_REGULATION\_OF\_CELLULAR\_COMPONENT\_BIOGENESIS | MSigDB lists | GO\_REGULATION\_OF\_CELLULAR\_COMPONENT\_BIOGENESIS | 625 | 8 | 12187 | 72 | Kdr,Lrrtm2,Ubqln2,Cbln2,Agrn,Bdnf,Map4,Sorl1 | | 3.068e-02 | -3.48 | glutamate receptor signaling pathway | biological process | GO:0007215 | 47 | 2 | 13711 | 80 | Grm3,Homer1 | | 3.068e-02 | -3.48 | ovarian follicle development | biological process | GO:0001541 | 47 | 2 | 13711 | 80 | Inhba,Kdr | | 3.085e-02 | -3.48 | regulation of canonical Wnt signaling pathway | biological process | GO:0060828 | 203 | 4 | 13711 | 80 | Arntl,Tle4,Col1a1,Egr1 | | 3.095e-02 | -3.48 | cellular response to leukemia inhibitory factor | biological process | GO:1990830 | 117 | 3 | 13711 | 80 | Tle4,Spry2,Srm | | 3.095e-02 | -3.48 | response to leukemia inhibitory factor | biological process | GO:1990823 | 117 | 3 | 13711 | 80 | Spry2,Srm,Tle4 | | 3.097e-02 | -3.47 | TCF11\_01 | MSigDB lists | TCF11\_01 | 201 | 4 | 12187 | 72 | Nr4a3,Cacng2,Bdnf,Ndst4 | | 3.109e-02 | -3.47 | tube development | biological process | GO:0035295 | 754 | 9 | 13711 | 80 | Atp5b,Lhx1,Bdnf,Spry2,Qk,Kdr,Tgfa,Nr4a3,Crh | | 3.112e-02 | -3.47 | positive regulation of neuron differentiation | biological process | GO:0045666 | 406 | 6 | 13711 | 80 | Ddx56,Timp2,Alkal2,Qk,Bdnf,Ltk | | 3.118e-02 | -3.47 | GSE37605\_TREG\_VS\_TCONV\_NOD\_FOXP3\_FUSION\_GFP\_UP | MSigDB lists | GSE37605\_TREG\_VS\_TCONV\_NOD\_FOXP3\_FUSION\_GFP\_UP | 116 | 3 | 12187 | 72 | Nr4a3,Spry2,Egr1 | | 3.118e-02 | -3.47 | REACTOME\_G\_ALPHA\_I\_SIGNALLING\_EVENTS | MSigDB lists | REACTOME\_G\_ALPHA\_I\_SIGNALLING\_EVENTS | 116 | 3 | 12187 | 72 | P2ry12,Gng11,Rgs4 | | 3.118e-02 | -3.47 | positive regulation of protein metabolic process | biological process | GO:0051247 | 1402 | 14 | 13711 | 80 | Ubqln2,Spry2,Gsdme,Sorl1,Inhba,Agrn,Alkal2,Arntl,Bdnf,Crh,Egr1,Timp2,Tgfa,Kdr | | 3.125e-02 | -3.47 | transferase activity, transferring sulfur-containing groups | molecular function | GO:0016782 | 48 | 2 | 13516 | 78 | Ndst4,Hs3st1 | | 3.128e-02 | -3.46 | GRUETZMANN\_PANCREATIC\_CANCER\_UP | MSigDB lists | GRUETZMANN\_PANCREATIC\_CANCER\_UP | 298 | 5 | 12187 | 72 | Msmo1,Col1a1,Map4,Sorl1,Glud1 | | 3.136e-02 | -3.46 | NAGASHIMA\_EGF\_SIGNALING\_UP | MSigDB lists | NAGASHIMA\_EGF\_SIGNALING\_UP | 47 | 2 | 12187 | 72 | Nr4a3,Egr1 | | 3.136e-02 | -3.46 | MODULE\_199 | MSigDB lists | MODULE\_199 | 47 | 2 | 12187 | 72 | Kdr,Ephb6 | | 3.136e-02 | -3.46 | GO\_NAD\_BINDING | MSigDB lists | GO\_NAD\_BINDING | 47 | 2 | 12187 | 72 | Qdpr,Glud1 | | 3.136e-02 | -3.46 | GO\_PYRUVATE\_METABOLIC\_PROCESS | MSigDB lists | GO\_PYRUVATE\_METABOLIC\_PROCESS | 47 | 2 | 12187 | 72 | Hkdc1,Nr4a3 | | 3.145e-02 | -3.46 | CREB\_02 | MSigDB lists | CREB\_02 | 202 | 4 | 12187 | 72 | AI593442,Grm3,Nr4a3,Crh | | 3.149e-02 | -3.46 | Cysteine and methionine metabolism | KEGG pathways | mmu00270 | 35 | 2 | 5248 | 42 | Srm,Amd1 | | 3.149e-02 | -3.46 | Cysteine and methionine metabolism | KEGG pathways | ko00270 | 35 | 2 | 5248 | 42 | Amd1,Srm | | 3.160e-02 | -3.45 | Metabolism of amino acids and derivatives | REACTOME pathways | R-MMU-71291 | 181 | 4 | 6297 | 42 | Glud1,Amd1,Srm,Qdpr | | 3.161e-02 | -3.45 | regulation of cell population proliferation | biological process | GO:0042127 | 1270 | 13 | 13711 | 80 | Bdnf,Lhx1,Crh,Timp2,Kdr,Tgfa,Egr1,Spry2,Inhba,Ncam1,Gsdme,Lims2,Nr4a3 | | 3.163e-02 | -3.45 | circadian rhythm | biological process | GO:0007623 | 118 | 3 | 13711 | 80 | Arntl,Egr1,Bdnf | | 3.164e-02 | -3.45 | H/ACA ribonucleoprotein complex | KEGG pathways | mmu\_M00425 | 4 | 1 | 5248 | 42 | Gar1 | | 3.164e-02 | -3.45 | H/ACA ribonucleoprotein complex | KEGG pathways | M00425 | 4 | 1 | 5248 | 42 | Gar1 | | 3.166e-02 | -3.45 | GCACTTT\_MIR175P\_MIR20A\_MIR106A\_MIR106B\_MIR20B\_MIR519D | MSigDB lists | GCACTTT\_MIR175P\_MIR20A\_MIR106A\_MIR106B\_MIR20B\_MIR519D | 514 | 7 | 12187 | 72 | Tle4,Timp2,Sorl1,Csrnp3,Nr4a3,Fjx1,Qk | | 3.178e-02 | -3.45 | animal organ morphogenesis | biological process | GO:0009887 | 757 | 9 | 13711 | 80 | Tgfa,Kdr,Agrn,Fjx1,Nr4a3,Inhba,Lhx1,Col1a1,Spry2 | | 3.187e-02 | -3.45 | ZHONG\_SECRETOME\_OF\_LUNG\_CANCER\_AND\_FIBROBLAST | MSigDB lists | ZHONG\_SECRETOME\_OF\_LUNG\_CANCER\_AND\_FIBROBLAST | 117 | 3 | 12187 | 72 | Agrn,Timp2,Col1a1 | | 3.187e-02 | -3.45 | GO\_GASTRULATION | MSigDB lists | GO\_GASTRULATION | 117 | 3 | 12187 | 72 | Lhx1,Inhba,Nr4a3 | | 3.187e-02 | -3.45 | AHRARNT\_01 | MSigDB lists | AHRARNT\_01 | 117 | 3 | 12187 | 72 | Lhx1,Bdnf,Crh | | 3.190e-02 | -3.45 | endoderm development | biological process | GO:0007492 | 48 | 2 | 13711 | 80 | Inhba,Lhx1 | | 3.190e-02 | -3.45 | regulation of cardiac muscle cell apoptotic process | biological process | GO:0010665 | 48 | 2 | 13711 | 80 | Ltk,Qk | | 3.190e-02 | -3.45 | positive regulation of epithelial cell differentiation | biological process | GO:0030858 | 48 | 2 | 13711 | 80 | Cldn5,Lhx1 | | 3.190e-02 | -3.45 | neuron recognition | biological process | GO:0008038 | 48 | 2 | 13711 | 80 | Bdnf,Ncam1 | | 3.195e-02 | -3.44 | CACBINDINGPROTEIN\_Q6 | MSigDB lists | CACBINDINGPROTEIN\_Q6 | 203 | 4 | 12187 | 72 | Ddx17,Ncam1,Cacng2,Nr4a3 | | 3.195e-02 | -3.44 | NFKB\_C | MSigDB lists | NFKB\_C | 203 | 4 | 12187 | 72 | Bdnf,Ddx17,Plp1,Ephb6 | | 3.232e-02 | -3.43 | GO\_NUCLEIC\_ACID\_BINDING\_TRANSCRIPTION\_FACTOR\_ACTIVITY | MSigDB lists | GO\_NUCLEIC\_ACID\_BINDING\_TRANSCRIPTION\_FACTOR\_ACTIVITY | 752 | 9 | 12187 | 72 | Arntl,Zfp518b,Tle4,Foxq1,Egr1,Peg3,Csrnp3,Nr4a3,Lhx1 | | 3.234e-02 | -3.43 | GO\_POSITIVE\_REGULATION\_OF\_RESPONSE\_TO\_STIMULUS | MSigDB lists | GO\_POSITIVE\_REGULATION\_OF\_RESPONSE\_TO\_STIMULUS | 1397 | 14 | 12187 | 72 | Nr4a3,Bdnf,Arntl,Kdr,Crh,Ubqln2,Homer1,Spry2,Inhba,Timp2,Tgfa,Col1a1,Lims2,Ddx17 | | 3.244e-02 | -3.43 | ATF4\_Q2 | MSigDB lists | ATF4\_Q2 | 204 | 4 | 12187 | 72 | AI593442,Grm3,Nr4a3,Ubqln2 | | 3.246e-02 | -3.43 | hormone activity | molecular function | GO:0005179 | 49 | 2 | 13516 | 78 | Inhba,Crh | | 3.256e-02 | -3.42 | chr22q13 | MSigDB lists | chr22q13 | 118 | 3 | 12187 | 72 | Cacng2,Ddx17,H1f0 | | 3.256e-02 | -3.42 | GSE10325\_BCELL\_VS\_LUPUS\_BCELL\_UP | MSigDB lists | GSE10325\_BCELL\_VS\_LUPUS\_BCELL\_UP | 118 | 3 | 12187 | 72 | Chmp7,Lims2,Ncam1 | | 3.256e-02 | -3.42 | AGCATTA\_MIR155 | MSigDB lists | AGCATTA\_MIR155 | 118 | 3 | 12187 | 72 | Qk,Tle4,Zfp518b | | 3.256e-02 | -3.42 | WAKABAYASHI\_ADIPOGENESIS\_PPARG\_RXRA\_BOUND\_WITH\_H4K20ME1\_MARK | MSigDB lists | WAKABAYASHI\_ADIPOGENESIS\_PPARG\_RXRA\_BOUND\_WITH\_H4K20ME1\_MARK | 118 | 3 | 12187 | 72 | Hsd17b12,Msmo1,Qdpr | | 3.256e-02 | -3.42 | GSE4142\_PLASMA\_CELL\_VS\_MEMORY\_BCELL\_UP | MSigDB lists | GSE4142\_PLASMA\_CELL\_VS\_MEMORY\_BCELL\_UP | 118 | 3 | 12187 | 72 | Kcnmb2,Fjx1,Col1a1 | | 3.260e-02 | -3.42 | GO\_MESODERM\_MORPHOGENESIS | MSigDB lists | GO\_MESODERM\_MORPHOGENESIS | 48 | 2 | 12187 | 72 | Nr4a3,Inhba | | 3.260e-02 | -3.42 | AMIT\_SERUM\_RESPONSE\_60\_MCF10A | MSigDB lists | AMIT\_SERUM\_RESPONSE\_60\_MCF10A | 48 | 2 | 12187 | 72 | Egr1,Inhba | | 3.260e-02 | -3.42 | DAWSON\_METHYLATED\_IN\_LYMPHOMA\_TCL1 | MSigDB lists | DAWSON\_METHYLATED\_IN\_LYMPHOMA\_TCL1 | 48 | 2 | 12187 | 72 | Ncam1,Spry2 | | 3.260e-02 | -3.42 | RAMASWAMY\_METASTASIS\_DN | MSigDB lists | RAMASWAMY\_METASTASIS\_DN | 48 | 2 | 12187 | 72 | Plp1,Agrn | | 3.268e-02 | -3.42 | regulation of phosphorylation | biological process | GO:0042325 | 1276 | 13 | 13711 | 80 | Agrn,Gsdme,Sorl1,Inhba,Spry2,Egr1,Timp2,Kdr,Tgfa,Crh,Dgkz,Bdnf,Alkal2 | | 3.275e-02 | -3.42 | cellular response to nitrogen compound | biological process | GO:1901699 | 411 | 6 | 13711 | 80 | Nr4a3,Car2,Crh,Col1a1,Egr1,P2ry12 | | 3.285e-02 | -3.42 | WGTTNNNNNAAA\_UNKNOWN | MSigDB lists | WGTTNNNNNAAA\_UNKNOWN | 407 | 6 | 12187 | 72 | Nr4a3,Ncam1,Bdnf,Nap1l5,Timp2,Ddx17 | | 3.292e-02 | -3.41 | Peptide hormone biosynthesis | REACTOME pathways | R-MMU-209952 | 5 | 1 | 6297 | 42 | Inhba | | 3.295e-02 | -3.41 | positive regulation of intracellular signal transduction | biological process | GO:1902533 | 762 | 9 | 13711 | 80 | Spry2,Alkal2,Gsdme,Ncam1,P2ry12,Crh,Tgfa,Kdr,Timp2 | | 3.300e-02 | -3.41 | Fatty Acyl-CoA Biosynthesis | REACTOME pathways | R-MMU-75105 | 43 | 2 | 6297 | 42 | Scd2,Hsd17b12 | | 3.307e-02 | -3.41 | signal transduction | biological process | GO:0007165 | 2703 | 23 | 13711 | 80 | Inhba,Ltk,Gng11,Rgs4,Grm3,Nr4a3,P2ry12,Ephb6,Kdr,Plp1,Tgfa,Arhgap20,Tle4,Crh,Ncam1,Col1a1,Cldn5,Homer1,Car2,Dgkz,Bdnf,Ddx17,Egr1 | | 3.308e-02 | -3.41 | SEA | pfam domains | PF01390 | 6 | 1 | 12881 | 72 | Agrn | | 3.308e-02 | -3.41 | Sprouty | pfam domains | PF05210 | 6 | 1 | 12881 | 72 | Spry2 | | 3.308e-02 | -3.41 | Kazal\_1 | pfam domains | PF00050 | 6 | 1 | 12881 | 72 | Agrn | | 3.326e-02 | -3.40 | GO\_FOREBRAIN\_DEVELOPMENT | MSigDB lists | GO\_FOREBRAIN\_DEVELOPMENT | 303 | 5 | 12187 | 72 | Crh,Inhba,Nr4a3,Lhx1,Fabp7 | | 3.327e-02 | -3.40 | GSE41176\_WT\_VS\_TAK1\_KO\_ANTI\_IGM\_STIM\_BCELL\_1H\_DN | MSigDB lists | GSE41176\_WT\_VS\_TAK1\_KO\_ANTI\_IGM\_STIM\_BCELL\_1H\_DN | 119 | 3 | 12187 | 72 | Csrnp3,Lrrtm2,Agrn | | 3.327e-02 | -3.40 | GSE2706\_R848\_VS\_R848\_AND\_LPS\_2H\_STIM\_DC\_DN | MSigDB lists | GSE2706\_R848\_VS\_R848\_AND\_LPS\_2H\_STIM\_DC\_DN | 119 | 3 | 12187 | 72 | Inhba,Nr4a3,Egr1 | | 3.327e-02 | -3.40 | GSE3982\_BCELL\_VS\_NKCELL\_DN | MSigDB lists | GSE3982\_BCELL\_VS\_NKCELL\_DN | 119 | 3 | 12187 | 72 | Peg3,Bdnf,Ncam1 | | 3.327e-02 | -3.40 | REACTOME\_INTERFERON\_SIGNALING | MSigDB lists | REACTOME\_INTERFERON\_SIGNALING | 119 | 3 | 12187 | 72 | Pom121,Egr1,Ncam1 | | 3.329e-02 | -3.40 | Extracellular matrix organization | REACTOME pathways | R-MMU-1474244 | 184 | 4 | 6297 | 42 | Col1a1,Agrn,Timp2,Kdr | | 3.342e-02 | -3.40 | MODULE\_55 | MSigDB lists | MODULE\_55 | 520 | 7 | 12187 | 72 | Qdpr,Cldn5,Egr1,Peg3,Msmo1,Car2,Col1a1 | | 3.342e-02 | -3.40 | MODULE\_88 | MSigDB lists | MODULE\_88 | 520 | 7 | 12187 | 72 | Egr1,Peg3,Msmo1,Col1a1,Car2,Qdpr,Cldn5 | | 3.345e-02 | -3.40 | GO\_COENZYME\_METABOLIC\_PROCESS | MSigDB lists | GO\_COENZYME\_METABOLIC\_PROCESS | 206 | 4 | 12187 | 72 | Amd1,Qdpr,Hkdc1,Hsd17b12 | | 3.347e-02 | -3.40 | FABP | prosite domains | PS00214 | 5 | 1 | 8845 | 60 | Fabp7 | | 3.369e-02 | -3.39 | oxidoreductase activity, acting on a sulfur group of donors | molecular function | GO:0016667 | 50 | 2 | 13516 | 78 | Pdia6,Tmx3 | | 3.381e-02 | -3.39 | dense fibrillar component | cellular component | GO:0001651 | 6 | 1 | 13825 | 79 | Gar1 | | 3.386e-02 | -3.39 | YNTTTNNNANGCARM\_UNKNOWN | MSigDB lists | YNTTTNNNANGCARM\_UNKNOWN | 49 | 2 | 12187 | 72 | Ddx17,Csrnp3 | | 3.386e-02 | -3.39 | WANG\_PROSTATE\_CANCER\_ANDROGEN\_INDEPENDENT | MSigDB lists | WANG\_PROSTATE\_CANCER\_ANDROGEN\_INDEPENDENT | 49 | 2 | 12187 | 72 | Ugt8a,Nr4a3 | | 3.386e-02 | -3.39 | KANG\_GIST\_WITH\_PDGFRA\_UP | MSigDB lists | KANG\_GIST\_WITH\_PDGFRA\_UP | 49 | 2 | 12187 | 72 | Kdr,Fjx1 | | 3.386e-02 | -3.39 | GNF2\_TAL1 | MSigDB lists | GNF2\_TAL1 | 49 | 2 | 12187 | 72 | H1f0,Car2 | | 3.386e-02 | -3.39 | GO\_HORMONE\_ACTIVITY | MSigDB lists | GO\_HORMONE\_ACTIVITY | 49 | 2 | 12187 | 72 | Crh,Inhba | | 3.390e-02 | -3.38 | Groucho\_enhance | interpro domains | IPR009146 | 6 | 1 | 13788 | 79 | Tle4 | | 3.390e-02 | -3.38 | SEA\_dom | interpro domains | IPR000082 | 6 | 1 | 13788 | 79 | Agrn | | 3.390e-02 | -3.38 | SEA\_dom\_sf | interpro domains | IPR036364 | 6 | 1 | 13788 | 79 | Agrn | | 3.390e-02 | -3.38 | Dimeric\_a/b-barrel | interpro domains | IPR011008 | 6 | 1 | 13788 | 79 | Nipsnap2 | | 3.390e-02 | -3.38 | Sprouty | interpro domains | IPR007875 | 6 | 1 | 13788 | 79 | Spry2 | | 3.390e-02 | -3.38 | Thr/Ala-tRNA-synth\_IIc\_edit | interpro domains | IPR018163 | 6 | 1 | 13788 | 79 | Aars | | 3.398e-02 | -3.38 | RORA2\_01 | MSigDB lists | RORA2\_01 | 120 | 3 | 12187 | 72 | Tle4,Arntl,Ntm | | 3.399e-02 | -3.38 | GO\_MOLECULAR\_FUNCTION\_REGULATOR | MSigDB lists | GO\_MOLECULAR\_FUNCTION\_REGULATOR | 1010 | 11 | 12187 | 72 | Timp2,Kcnmb2,Rgs4,Spry2,Cacng2,Grm3,Fnbp1,Mtmr9,Arhgap20,Ncam1,P2ry12 | | 3.414e-02 | -3.38 | vascular endothelial growth factor-activated receptor activity | molecular function | GO:0005021 | 6 | 1 | 13516 | 78 | Kdr | | 3.414e-02 | -3.38 | type 5 metabotropic glutamate receptor binding | molecular function | GO:0031802 | 6 | 1 | 13516 | 78 | Homer1 | | 3.414e-02 | -3.38 | acyl-CoA desaturase activity | molecular function | GO:0016215 | 6 | 1 | 13516 | 78 | Scd2 | | 3.414e-02 | -3.38 | ATPase inhibitor activity | molecular function | GO:0042030 | 6 | 1 | 13516 | 78 | Agrn | | 3.414e-02 | -3.38 | adenylate cyclase inhibiting G protein-coupled glutamate receptor activity | molecular function | GO:0001640 | 6 | 1 | 13516 | 78 | Grm3 | | 3.414e-02 | -3.38 | leucine binding | molecular function | GO:0070728 | 6 | 1 | 13516 | 78 | Glud1 | | 3.432e-02 | -3.37 | TGTTTGY\_HNF3\_Q6 | MSigDB lists | TGTTTGY\_HNF3\_Q6 | 523 | 7 | 12187 | 72 | H1f0,Ddx17,Inhba,Tle4,Ncam1,Nr4a3,Cacng2 | | 3.435e-02 | -3.37 | endocytosis | biological process | GO:0006897 | 309 | 5 | 13711 | 80 | Sorl1,Mtmr9,Atp5b,Fnbp1,Cacng2 | | 3.438e-02 | -3.37 | long-chain fatty acid metabolic process | biological process | GO:0001676 | 50 | 2 | 13711 | 80 | Qk,Plp1 | | 3.438e-02 | -3.37 | protein localization to vacuole | biological process | GO:0072665 | 50 | 2 | 13711 | 80 | Sorl1,Cacng2 | | 3.442e-02 | -3.37 | vacuolar transport | biological process | GO:0007034 | 122 | 3 | 13711 | 80 | Cacng2,Chmp7,Sorl1 | | 3.449e-02 | -3.37 | Homeobox\_CS | interpro domains | IPR017970 | 51 | 2 | 13788 | 79 | Lhx1,Gbx1 | | 3.451e-02 | -3.37 | regulation of phosphatidylinositol dephosphorylation | biological process | GO:0060304 | 6 | 1 | 13711 | 80 | Mtmr9 | | 3.451e-02 | -3.37 | positive regulation of feeding behavior | biological process | GO:2000253 | 6 | 1 | 13711 | 80 | Nr4a3 | | 3.451e-02 | -3.37 | positive regulation of skeletal muscle cell differentiation | biological process | GO:2001016 | 6 | 1 | 13711 | 80 | Arntl | | 3.451e-02 | -3.37 | response to mycotoxin | biological process | GO:0010046 | 6 | 1 | 13711 | 80 | Egr1 | | 3.451e-02 | -3.37 | negative regulation of vascular endothelial growth factor signaling pathway | biological process | GO:1900747 | 6 | 1 | 13711 | 80 | Spry2 | | 3.451e-02 | -3.37 | positive regulation of endothelial cell chemotaxis by VEGF-activated vascular endothelial growth factor receptor signaling pathway | biological process | GO:0038033 | 6 | 1 | 13711 | 80 | Kdr | | 3.451e-02 | -3.37 | stress-induced premature senescence | biological process | GO:0090400 | 6 | 1 | 13711 | 80 | Arntl | | 3.451e-02 | -3.37 | cerebellar Purkinje cell-granule cell precursor cell signaling involved in regulation of granule cell precursor cell proliferation | biological process | GO:0021937 | 6 | 1 | 13711 | 80 | Lhx1 | | 3.451e-02 | -3.37 | cell motility involved in cerebral cortex radial glia guided migration | biological process | GO:0021814 | 6 | 1 | 13711 | 80 | P2ry12 | | 3.451e-02 | -3.37 | regulation of follicle-stimulating hormone secretion | biological process | GO:0046880 | 6 | 1 | 13711 | 80 | Inhba | | 3.451e-02 | -3.37 | negative regulation of membrane protein ectodomain proteolysis | biological process | GO:0051045 | 6 | 1 | 13711 | 80 | Timp2 | | 3.451e-02 | -3.37 | negative regulation of glucocorticoid receptor signaling pathway | biological process | GO:2000323 | 6 | 1 | 13711 | 80 | Arntl | | 3.451e-02 | -3.37 | positive regulation of receptor binding | biological process | GO:1900122 | 6 | 1 | 13711 | 80 | Bdnf | | 3.451e-02 | -3.37 | regulation of type B pancreatic cell development | biological process | GO:2000074 | 6 | 1 | 13711 | 80 | Arntl | | 3.451e-02 | -3.37 | regulation of metalloendopeptidase activity | biological process | GO:1904683 | 6 | 1 | 13711 | 80 | Sorl1 | | 3.451e-02 | -3.37 | neurotransmitter receptor metabolic process | biological process | GO:0045213 | 6 | 1 | 13711 | 80 | Agrn | | 3.451e-02 | -3.37 | detection of calcium ion | biological process | GO:0005513 | 6 | 1 | 13711 | 80 | Kcnmb2 | | 3.451e-02 | -3.37 | glomerular mesangium development | biological process | GO:0072109 | 6 | 1 | 13711 | 80 | Egr1 | | 3.451e-02 | -3.37 | carbon dioxide transport | biological process | GO:0015670 | 6 | 1 | 13711 | 80 | Car2 | | 3.451e-02 | -3.37 | hepatocyte proliferation | biological process | GO:0072574 | 6 | 1 | 13711 | 80 | Tgfa | | 3.451e-02 | -3.37 | epithelial cell proliferation involved in liver morphogenesis | biological process | GO:0072575 | 6 | 1 | 13711 | 80 | Tgfa | | 3.451e-02 | -3.37 | pronephros development | biological process | GO:0048793 | 6 | 1 | 13711 | 80 | Lhx1 | | 3.470e-02 | -3.36 | GSE8515\_IL1\_VS\_IL6\_4H\_STIM\_MAC\_DN | MSigDB lists | GSE8515\_IL1\_VS\_IL6\_4H\_STIM\_MAC\_DN | 121 | 3 | 12187 | 72 | Crh,Dgkz,Ugt8a | | 3.482e-02 | -3.36 | regulation of protein binding | biological process | GO:0043393 | 211 | 4 | 13711 | 80 | Ephb6,Bdnf,Agrn,Sorl1 | | 3.486e-02 | -3.36 | growth factor binding | molecular function | GO:0019838 | 124 | 3 | 13516 | 78 | Agrn,Kdr,Col1a1 | | 3.488e-02 | -3.36 | cell part | cellular component | GO:0044464 | 11956 | 74 | 13825 | 79 | Spry2,Hkdc1,Rgs4,Ccdc13,Qdpr,Cacng2,Tgfa,Tmem165,Homer1,Pdia6,Ephb6,Mtmr9,Agrn,Msmo1,Car2,Tmx3,Cldn5,Map4,Timp2,Glud1,Col1a1,Amd1,Plp1,Sorl1,Dgkz,P2ry12,Zfp518b,Csrnp3,Foxq1,Pom121,Gsdme,Hba-a2,Atp5b,Fbxl17,Grm3,Ncam1,Lhx1,Hsd17b12,Bdnf,Arntl,Ltk,Scd2,Fnbp1,Ubqln2,Ugt8a,Gng11,Fabp7,Ntm,Qk,Tle4,Hs3st1,Crh,Nrep,Lrrtm2,H1f0,Ddx17,Peg3,Ddx56,Kdr,Inhba,Lrrc55,Ndst4,Lims2,Gbx1,Nap1l5,Nipsnap2,Kcnmb2,Nr4a3,Chmp7,Nacad,Egr1,Aars,Ermn,Gar1 | | 3.494e-02 | -3.35 | GO\_ERYTHROSE\_4\_PHOSPHATE\_PHOSPHOENOLPYRUVATE\_FAMILY\_AMINO\_ACID\_METABOLIC\_PROCESS | MSigDB lists | GO\_ERYTHROSE\_4\_PHOSPHATE\_PHOSPHOENOLPYRUVATE\_FAMILY\_AMINO\_ACID\_METABOLIC\_PROCESS | 6 | 1 | 12187 | 72 | Qdpr | | 3.494e-02 | -3.35 | GO\_ONE\_CARBON\_COMPOUND\_TRANSPORT | MSigDB lists | GO\_ONE\_CARBON\_COMPOUND\_TRANSPORT | 6 | 1 | 12187 | 72 | Car2 | | 3.494e-02 | -3.35 | WIEMANN\_TELOMERE\_SHORTENING\_AND\_CHRONIC\_LIVER\_DAMAGE\_UP | MSigDB lists | WIEMANN\_TELOMERE\_SHORTENING\_AND\_CHRONIC\_LIVER\_DAMAGE\_UP | 6 | 1 | 12187 | 72 | Arntl | | 3.494e-02 | -3.35 | GO\_HEMOGLOBIN\_METABOLIC\_PROCESS | MSigDB lists | GO\_HEMOGLOBIN\_METABOLIC\_PROCESS | 6 | 1 | 12187 | 72 | Inhba | | 3.494e-02 | -3.35 | GO\_SPINAL\_CORD\_ASSOCIATION\_NEURON\_DIFFERENTIATION | MSigDB lists | GO\_SPINAL\_CORD\_ASSOCIATION\_NEURON\_DIFFERENTIATION | 6 | 1 | 12187 | 72 | Lhx1 | | 3.494e-02 | -3.35 | MODULE\_89 | MSigDB lists | MODULE\_89 | 6 | 1 | 12187 | 72 | H1f0 | | 3.500e-02 | -3.35 | TAGCTTT\_MIR9 | MSigDB lists | TAGCTTT\_MIR9 | 209 | 4 | 12187 | 72 | Qk,Nap1l5,Nr4a3,Arntl | | 3.501e-02 | -3.35 | cell | cellular component | GO:0005623 | 11957 | 74 | 13825 | 79 | Hsd17b12,Bdnf,Arntl,Grm3,Ncam1,Lhx1,Tle4,Hs3st1,Crh,Nrep,Lrrtm2,Ltk,Scd2,Fnbp1,Ubqln2,Ugt8a,Gng11,Ntm,Fabp7,Qk,Kdr,Ddx56,Inhba,Lrrc55,Ndst4,Lims2,H1f0,Ddx17,Peg3,Nacad,Aars,Egr1,Ermn,Gar1,Nap1l5,Gbx1,Nipsnap2,Kcnmb2,Nr4a3,Chmp7,Cacng2,Tgfa,Tmem165,Spry2,Hkdc1,Ccdc13,Rgs4,Qdpr,Tmx3,Car2,Cldn5,Map4,Homer1,Pdia6,Ephb6,Mtmr9,Agrn,Msmo1,Csrnp3,Timp2,Glud1,Amd1,Col1a1,Plp1,Sorl1,Dgkz,P2ry12,Zfp518b,Gsdme,Hba-a2,Atp5b,Fbxl17,Foxq1,Pom121 | | 3.514e-02 | -3.35 | BROWNE\_HCMV\_INFECTION\_6HR\_UP | MSigDB lists | BROWNE\_HCMV\_INFECTION\_6HR\_UP | 50 | 2 | 12187 | 72 | Arntl,Amd1 | | 3.514e-02 | -3.35 | GO\_REPRESSING\_TRANSCRIPTION\_FACTOR\_BINDING | MSigDB lists | GO\_REPRESSING\_TRANSCRIPTION\_FACTOR\_BINDING | 50 | 2 | 12187 | 72 | Arntl,Tle4 | | 3.514e-02 | -3.35 | GO\_NEURAL\_NUCLEUS\_DEVELOPMENT | MSigDB lists | GO\_NEURAL\_NUCLEUS\_DEVELOPMENT | 50 | 2 | 12187 | 72 | Glud1,Plp1 | | 3.514e-02 | -3.35 | GNF2\_SPTB | MSigDB lists | GNF2\_SPTB | 50 | 2 | 12187 | 72 | Car2,H1f0 | | 3.514e-02 | -3.35 | CUI\_GLUCOSE\_DEPRIVATION | MSigDB lists | CUI\_GLUCOSE\_DEPRIVATION | 50 | 2 | 12187 | 72 | Arntl,Pdia6 | | 3.514e-02 | -3.35 | GNF2\_ANK1 | MSigDB lists | GNF2\_ANK1 | 50 | 2 | 12187 | 72 | Car2,H1f0 | | 3.531e-02 | -3.34 | regulation of molecular function | biological process | GO:0065009 | 2276 | 20 | 13711 | 80 | Tgfa,Timp2,Egr1,Arhgap20,Crh,Dgkz,Bdnf,Ephb6,Cacng2,H1f0,Agrn,Homer1,Rgs4,Nipsnap2,Lrrc55,Sorl1,Mtmr9,Pdia6,Csrnp3,Spry2 | | 3.534e-02 | -3.34 | telencephalon development | biological process | GO:0021537 | 212 | 4 | 13711 | 80 | Inhba,P2ry12,Lhx1,Nr4a3 | | 3.543e-02 | -3.34 | MEL18\_DN.V1\_DN | MSigDB lists | MEL18\_DN.V1\_DN | 122 | 3 | 12187 | 72 | Car2,Egr1,Peg3 | | 3.551e-02 | -3.34 | Cyth3 (cytohesin 3) | protein interactions | 19159 | 5 | 1 | 6802 | 49 | Homer1 | | 3.551e-02 | -3.34 | Itgb3 (integrin beta 3) | protein interactions | 16416 | 5 | 1 | 6802 | 49 | Ubqln2 | | 3.551e-02 | -3.34 | Rnf14 (ring finger protein 14) | protein interactions | 56736 | 5 | 1 | 6802 | 49 | Arntl | | 3.551e-02 | -3.34 | Efhd2 (EF hand domain containing 2) | protein interactions | 27984 | 5 | 1 | 6802 | 49 | Homer1 | | 3.551e-02 | -3.34 | Epas1 (endothelial PAS domain protein 1) | protein interactions | 13819 | 5 | 1 | 6802 | 49 | Arntl | | 3.551e-02 | -3.34 | Spry3 (sprouty RTK signaling antagonist 3) | protein interactions | 236576 | 5 | 1 | 6802 | 49 | Spry2 | | 3.551e-02 | -3.34 | KIF3B (kinesin family member 3B) | protein interactions | 9371 | 5 | 1 | 6802 | 49 | Homer1 | | 3.565e-02 | -3.33 | negative regulation of muscle cell differentiation | biological process | GO:0051148 | 51 | 2 | 13711 | 80 | Bdnf,Rgs4 | | 3.565e-02 | -3.33 | pyruvate metabolic process | biological process | GO:0006090 | 51 | 2 | 13711 | 80 | Hkdc1,Nr4a3 | | 3.565e-02 | -3.33 | regulation of potassium ion transmembrane transporter activity | biological process | GO:1901016 | 51 | 2 | 13711 | 80 | Agrn,Lrrc55 | | 3.565e-02 | -3.33 | regulation of striated muscle cell apoptotic process | biological process | GO:0010662 | 51 | 2 | 13711 | 80 | Ltk,Qk | | 3.605e-02 | -3.32 | Biopterin recycling | BIOCYC pathways | MOUSE\_PWY3DJ-35549 | 3 | 1 | 823 | 10 | Qdpr | | 3.606e-02 | -3.32 | GO\_REGULATION\_OF\_NEURON\_DEATH | MSigDB lists | GO\_REGULATION\_OF\_NEURON\_DEATH | 211 | 4 | 12187 | 72 | Aars,Bdnf,Sorl1,Nr4a3 | | 3.606e-02 | -3.32 | GO\_FATTY\_ACID\_METABOLIC\_PROCESS | MSigDB lists | GO\_FATTY\_ACID\_METABOLIC\_PROCESS | 211 | 4 | 12187 | 72 | Hsd17b12,Msmo1,Plp1,Qk | | 3.606e-02 | -3.32 | AP1FJ\_Q2 | MSigDB lists | AP1FJ\_Q2 | 211 | 4 | 12187 | 72 | Tle4,Cacng2,Map4,Ddx17 | | 3.612e-02 | -3.32 | intrinsic component of synaptic membrane | cellular component | GO:0099240 | 218 | 4 | 13825 | 79 | Lrrtm2,Ncam1,Cacng2,Grm3 | | 3.616e-02 | -3.32 | PDGF\_UP.V1\_UP | MSigDB lists | PDGF\_UP.V1\_UP | 123 | 3 | 12187 | 72 | Egr1,Inhba,Gng11 | | 3.616e-02 | -3.32 | GO\_PLATELET\_ACTIVATION | MSigDB lists | GO\_PLATELET\_ACTIVATION | 123 | 3 | 12187 | 72 | P2ry12,Col1a1,Dgkz | | 3.616e-02 | -3.32 | SMIRNOV\_CIRCULATING\_ENDOTHELIOCYTES\_IN\_CANCER\_UP | MSigDB lists | SMIRNOV\_CIRCULATING\_ENDOTHELIOCYTES\_IN\_CANCER\_UP | 123 | 3 | 12187 | 72 | Hs3st1,Egr1,Timp2 | | 3.630e-02 | -3.32 | - | gene3d domains | 2.10.50.30 | 7 | 1 | 6647 | 35 | Grm3 | | 3.644e-02 | -3.31 | RIGGINS\_TAMOXIFEN\_RESISTANCE\_UP | MSigDB lists | RIGGINS\_TAMOXIFEN\_RESISTANCE\_UP | 51 | 2 | 12187 | 72 | Dgkz,Ndst4 | | 3.644e-02 | -3.31 | WOO\_LIVER\_CANCER\_RECURRENCE\_DN | MSigDB lists | WOO\_LIVER\_CANCER\_RECURRENCE\_DN | 51 | 2 | 12187 | 72 | Qdpr,Glud1 | | 3.644e-02 | -3.31 | GO\_REGULATION\_OF\_ADENYLATE\_CYCLASE\_ACTIVITY | MSigDB lists | GO\_REGULATION\_OF\_ADENYLATE\_CYCLASE\_ACTIVITY | 51 | 2 | 12187 | 72 | Grm3,Timp2 | | 3.644e-02 | -3.31 | GNF2\_BNIP3L | MSigDB lists | GNF2\_BNIP3L | 51 | 2 | 12187 | 72 | Car2,H1f0 | | 3.644e-02 | -3.31 | CAGGTCC\_MIR492 | MSigDB lists | CAGGTCC\_MIR492 | 51 | 2 | 12187 | 72 | Timp2,Csrnp3 | | 3.644e-02 | -3.31 | PID\_SHP2\_PATHWAY | MSigDB lists | PID\_SHP2\_PATHWAY | 51 | 2 | 12187 | 72 | Bdnf,Kdr | | 3.644e-02 | -3.31 | GO\_NEURONAL\_POSTSYNAPTIC\_DENSITY | MSigDB lists | GO\_NEURONAL\_POSTSYNAPTIC\_DENSITY | 51 | 2 | 12187 | 72 | Map4,Homer1 | | 3.644e-02 | -3.31 | GNF2\_RAD23A | MSigDB lists | GNF2\_RAD23A | 51 | 2 | 12187 | 72 | H1f0,Car2 | | 3.659e-02 | -3.31 | potassium ion transport | biological process | GO:0006813 | 125 | 3 | 13711 | 80 | P2ry12,Kcnmb2,Lrrc55 | | 3.659e-02 | -3.31 | SOX5\_01 | MSigDB lists | SOX5\_01 | 212 | 4 | 12187 | 72 | Inhba,Ephb6,Hs3st1,Ncam1 | | 3.666e-02 | -3.31 | positive regulation of apoptotic process | biological process | GO:0043065 | 536 | 7 | 13711 | 80 | Egr1,Agrn,Nr4a3,Gsdme,Ltk,Inhba,Csrnp3 | | 3.691e-02 | -3.30 | GSE13485\_CTRL\_VS\_DAY7\_YF17D\_VACCINE\_PBMC\_UP | MSigDB lists | GSE13485\_CTRL\_VS\_DAY7\_YF17D\_VACCINE\_PBMC\_UP | 124 | 3 | 12187 | 72 | Tgfa,Cacng2,Sorl1 | | 3.691e-02 | -3.30 | GSE17974\_0H\_VS\_0.5H\_IN\_VITRO\_ACT\_CD4\_TCELL\_DN | MSigDB lists | GSE17974\_0H\_VS\_0.5H\_IN\_VITRO\_ACT\_CD4\_TCELL\_DN | 124 | 3 | 12187 | 72 | Fabp7,Egr1,Nr4a3 | | 3.691e-02 | -3.30 | GO\_POSITIVE\_REGULATION\_OF\_EPITHELIAL\_CELL\_PROLIFERATION | MSigDB lists | GO\_POSITIVE\_REGULATION\_OF\_EPITHELIAL\_CELL\_PROLIFERATION | 124 | 3 | 12187 | 72 | Kdr,Tgfa,Nr4a3 | | 3.691e-02 | -3.30 | GSE27241\_CTRL\_VS\_DIGOXIN\_TREATED\_CD4\_TCELL\_IN\_TH17\_POLARIZING\_CONDITIONS\_UP | MSigDB lists | GSE27241\_CTRL\_VS\_DIGOXIN\_TREATED\_CD4\_TCELL\_IN\_TH17\_POLARIZING\_CONDITIONS\_UP | 124 | 3 | 12187 | 72 | Nr4a3,Egr1,Ddx17 | | 3.691e-02 | -3.30 | GO\_NEGATIVE\_REGULATION\_OF\_MAPK\_CASCADE | MSigDB lists | GO\_NEGATIVE\_REGULATION\_OF\_MAPK\_CASCADE | 124 | 3 | 12187 | 72 | Sorl1,Rgs4,Spry2 | | 3.694e-02 | -3.30 | alpha-amino acid catabolic process | biological process | GO:1901606 | 52 | 2 | 13711 | 80 | Glud1,Qdpr | | 3.694e-02 | -3.30 | positive regulation of blood vessel endothelial cell migration | biological process | GO:0043536 | 52 | 2 | 13711 | 80 | Atp5b,Kdr | | 3.702e-02 | -3.30 | ion channel binding | molecular function | GO:0044325 | 127 | 3 | 13516 | 78 | Agrn,Homer1,Lrrc55 | | 3.713e-02 | -3.29 | GTTTGTT\_MIR495 | MSigDB lists | GTTTGTT\_MIR495 | 213 | 4 | 12187 | 72 | Homer1,Ntm,Bdnf,Timp2 | | 3.748e-02 | -3.28 | amino acid binding | molecular function | GO:0016597 | 53 | 2 | 13516 | 78 | Aars,Glud1 | | 3.748e-02 | -3.28 | NAD binding | molecular function | GO:0051287 | 53 | 2 | 13516 | 78 | Qdpr,Glud1 | | 3.749e-02 | -3.28 | FN3 | smart domains | SM00060 | 124 | 3 | 7188 | 43 | Sorl1,Ncam1,Ephb6 | | 3.759e-02 | -3.28 | cellular component organization or biogenesis | biological process | GO:0071840 | 4450 | 34 | 13711 | 80 | Lims2,Ncam1,Nap1l5,Spry2,Col1a1,Gbx1,Nipsnap2,Lrrtm2,Homer1,Cldn5,Ccdc13,Dgkz,Bdnf,Cacng2,Tmem165,Ddx17,Map4,Pom121,Ermn,Ubqln2,Ugt8a,Ddx56,Agrn,H1f0,Hsd17b12,Nr4a3,Ephb6,Lhx1,P2ry12,Chmp7,Plp1,Kdr,Nrep,Gar1 | | 3.766e-02 | -3.28 | TTCNRGNNNNTTC\_HSF\_Q6 | MSigDB lists | TTCNRGNNNNTTC\_HSF\_Q6 | 125 | 3 | 12187 | 72 | Nr4a3,Lhx1,Ddx17 | | 3.766e-02 | -3.28 | KAAB\_FAILED\_HEART\_ATRIUM\_DN | MSigDB lists | KAAB\_FAILED\_HEART\_ATRIUM\_DN | 125 | 3 | 12187 | 72 | Plp1,Glud1,Amd1 | | 3.775e-02 | -3.28 | SEITZ\_NEOPLASTIC\_TRANSFORMATION\_BY\_8P\_DELETION\_UP | MSigDB lists | SEITZ\_NEOPLASTIC\_TRANSFORMATION\_BY\_8P\_DELETION\_UP | 52 | 2 | 12187 | 72 | Sorl1,Hs3st1 | | 3.775e-02 | -3.28 | YWATTWNNRGCT\_UNKNOWN | MSigDB lists | YWATTWNNRGCT\_UNKNOWN | 52 | 2 | 12187 | 72 | Ddx17,Arntl | | 3.775e-02 | -3.28 | MIKKELSEN\_IPS\_WITH\_HCP\_H3K27ME3 | MSigDB lists | MIKKELSEN\_IPS\_WITH\_HCP\_H3K27ME3 | 52 | 2 | 12187 | 72 | Ntm,Cacng2 | | 3.789e-02 | -3.27 | YOSHIMURA\_MAPK8\_TARGETS\_DN | MSigDB lists | YOSHIMURA\_MAPK8\_TARGETS\_DN | 314 | 5 | 12187 | 72 | Bdnf,Amd1,Plp1,Tgfa,Msmo1 | | 3.807e-02 | -3.27 | eye morphogenesis | biological process | GO:0048592 | 127 | 3 | 13711 | 80 | Kdr,Lhx1,Fjx1 | | 3.822e-02 | -3.26 | NFE2\_01 | MSigDB lists | NFE2\_01 | 215 | 4 | 12187 | 72 | AI593442,Map4,Ddx17,H1f0 | | 3.824e-02 | -3.26 | positive regulation of programmed cell death | biological process | GO:0043068 | 541 | 7 | 13711 | 80 | Ltk,Inhba,Gsdme,Csrnp3,Agrn,Egr1,Nr4a3 | | 3.837e-02 | -3.26 | positive regulation of MAPK cascade | biological process | GO:0043410 | 427 | 6 | 13711 | 80 | Gsdme,Alkal2,Spry2,Timp2,Kdr,Tgfa | | 3.842e-02 | -3.26 | NCX\_01 | MSigDB lists | NCX\_01 | 126 | 3 | 12187 | 72 | Map4,Bdnf,Peg3 | | 3.842e-02 | -3.26 | GROSS\_HYPOXIA\_VIA\_ELK3\_AND\_HIF1A\_UP | MSigDB lists | GROSS\_HYPOXIA\_VIA\_ELK3\_AND\_HIF1A\_UP | 126 | 3 | 12187 | 72 | Spry2,Egr1,Homer1 | | 3.842e-02 | -3.26 | IL2\_UP.V1\_UP | MSigDB lists | IL2\_UP.V1\_UP | 126 | 3 | 12187 | 72 | Cldn5,Egr1,Spry2 | | 3.842e-02 | -3.26 | LANDIS\_ERBB2\_BREAST\_TUMORS\_324\_DN | MSigDB lists | LANDIS\_ERBB2\_BREAST\_TUMORS\_324\_DN | 126 | 3 | 12187 | 72 | Col1a1,Qk,Map4 | | 3.842e-02 | -3.26 | GO\_REGULATION\_OF\_GTPASE\_ACTIVITY | MSigDB lists | GO\_REGULATION\_OF\_GTPASE\_ACTIVITY | 536 | 7 | 12187 | 72 | Fnbp1,Agrn,P2ry12,Spry2,Ncam1,Arhgap20,Rgs4 | | 3.848e-02 | -3.26 | regulation of protein phosphorylation | biological process | GO:0001932 | 1172 | 12 | 13711 | 80 | Crh,Egr1,Tgfa,Kdr,Timp2,Alkal2,Bdnf,Agrn,Spry2,Gsdme,Inhba,Sorl1 | | 3.849e-02 | -3.26 | NCD3G | pfam domains | PF07562 | 7 | 1 | 12881 | 72 | Grm3 | | 3.858e-02 | -3.26 | monocarboxylic acid metabolic process | biological process | GO:0032787 | 319 | 5 | 13711 | 80 | Hkdc1,Plp1,Scd2,Nr4a3,Qk | | 3.903e-02 | -3.24 | positive regulation of cellular protein metabolic process | biological process | GO:0032270 | 1309 | 13 | 13711 | 80 | Agrn,Gsdme,Inhba,Ubqln2,Spry2,Egr1,Timp2,Kdr,Tgfa,Crh,Bdnf,Alkal2,Arntl | | 3.908e-02 | -3.24 | TACAATC\_MIR508 | MSigDB lists | TACAATC\_MIR508 | 53 | 2 | 12187 | 72 | Nr4a3,Bdnf | | 3.908e-02 | -3.24 | GO\_NEGATIVE\_REGULATION\_OF\_HORMONE\_SECRETION | MSigDB lists | GO\_NEGATIVE\_REGULATION\_OF\_HORMONE\_SECRETION | 53 | 2 | 12187 | 72 | Inhba,Crh | | 3.908e-02 | -3.24 | GNCF\_01 | MSigDB lists | GNCF\_01 | 53 | 2 | 12187 | 72 | Kcnmb2,Amd1 | | 3.908e-02 | -3.24 | GO\_CELL\_REDOX\_HOMEOSTASIS | MSigDB lists | GO\_CELL\_REDOX\_HOMEOSTASIS | 53 | 2 | 12187 | 72 | Tmx3,Pdia6 | | 3.922e-02 | -3.24 | GO\_SINGLE\_ORGANISM\_BEHAVIOR | MSigDB lists | GO\_SINGLE\_ORGANISM\_BEHAVIOR | 317 | 5 | 12187 | 72 | Homer1,Crh,Gbx1,Nr4a3,Cldn5 | | 3.933e-02 | -3.24 | MODULE\_220 | MSigDB lists | MODULE\_220 | 217 | 4 | 12187 | 72 | Fabp7,Col1a1,Ephb6,Ugt8a | | 3.937e-02 | -3.23 | ADP signalling through P2Y purinoceptor 12 | REACTOME pathways | R-MMU-392170 | 6 | 1 | 6297 | 42 | P2ry12 | | 3.939e-02 | -3.23 | Nucleotide sugar biosynthesis, glucose => UDP-glucose | KEGG pathways | M00549 | 5 | 1 | 5248 | 42 | Hkdc1 | | 3.939e-02 | -3.23 | Nucleotide sugar biosynthesis, glucose => UDP-glucose | KEGG pathways | mmu\_M00549 | 5 | 1 | 5248 | 42 | Hkdc1 | | 3.943e-02 | -3.23 | VDCC\_gsu | interpro domains | IPR008368 | 7 | 1 | 13788 | 79 | Cacng2 | | 3.943e-02 | -3.23 | GPCR\_3\_9-Cys\_sf | interpro domains | IPR038550 | 7 | 1 | 13788 | 79 | Grm3 | | 3.943e-02 | -3.23 | GPCR\_3\_9-Cys\_dom | interpro domains | IPR011500 | 7 | 1 | 13788 | 79 | Grm3 | | 3.943e-02 | -3.23 | Nuc\_translocat | interpro domains | IPR001067 | 7 | 1 | 13788 | 79 | Arntl | | 3.943e-02 | -3.23 | GPCR\_3\_mtglu\_rcpt | interpro domains | IPR000162 | 7 | 1 | 13788 | 79 | Grm3 | | 3.948e-02 | -3.23 | blood vessel development | biological process | GO:0001568 | 430 | 6 | 13711 | 80 | Qk,Col1a1,Atp5b,Egr1,Kdr,Tgfa | | 3.957e-02 | -3.23 | regulation of glutamate receptor signaling pathway | biological process | GO:1900449 | 54 | 2 | 13711 | 80 | Crh,Cacng2 | | 3.962e-02 | -3.23 | membrane | cellular component | GO:0016020 | 6082 | 43 | 13825 | 79 | Lrrc55,Lims2,Ndst4,Kdr,Chmp7,Nipsnap2,Kcnmb2,Hsd17b12,Bdnf,Grm3,Ncam1,Lrrtm2,AI593442,Gng11,Ntm,Scd2,Fnbp1,Ltk,Ugt8a,Ubqln2,Plp1,Dgkz,P2ry12,Sorl1,Glud1,Gsdme,Atp5b,Pom121,Tgfa,Tmem165,Cacng2,Rgs4,Spry2,Cldn5,Car2,Tmx3,Map4,Agrn,Msmo1,Pdia6,Homer1,Ephb6,Mtmr9 | | 3.971e-02 | -3.23 | AT DNA binding | molecular function | GO:0003680 | 7 | 1 | 13516 | 78 | H1f0 | | 3.971e-02 | -3.23 | G protein-coupled purinergic nucleotide receptor activity | molecular function | GO:0045028 | 7 | 1 | 13516 | 78 | P2ry12 | | 3.971e-02 | -3.23 | inhibin binding | molecular function | GO:0034711 | 7 | 1 | 13516 | 78 | Inhba | | 3.971e-02 | -3.23 | G protein-coupled nucleotide receptor activity | molecular function | GO:0001608 | 7 | 1 | 13516 | 78 | P2ry12 | | 3.971e-02 | -3.23 | vascular endothelial growth factor binding | molecular function | GO:0038085 | 7 | 1 | 13516 | 78 | Kdr | | 3.971e-02 | -3.23 | lipid kinase activity | molecular function | GO:0001727 | 7 | 1 | 13516 | 78 | Dgkz | | 3.971e-02 | -3.23 | oxidoreductase activity, acting on paired donors, with oxidation of a pair of donors resulting in the reduction of molecular oxygen to two molecules of water | molecular function | GO:0016717 | 7 | 1 | 13516 | 78 | Scd2 | | 3.988e-02 | -3.22 | TGACAGNY\_MEIS1\_01 | MSigDB lists | TGACAGNY\_MEIS1\_01 | 659 | 8 | 12187 | 72 | Nr4a3,Crh,Lhx1,Ncam1,Ndst4,Tle4,Plp1,H1f0 | | 3.989e-02 | -3.22 | GO\_REGULATION\_OF\_EPITHELIAL\_CELL\_PROLIFERATION | MSigDB lists | GO\_REGULATION\_OF\_EPITHELIAL\_CELL\_PROLIFERATION | 218 | 4 | 12187 | 72 | Nr4a3,Tgfa,Kdr,Lims2 | | 3.996e-02 | -3.22 | ELVIDGE\_HYPOXIA\_DN | MSigDB lists | ELVIDGE\_HYPOXIA\_DN | 128 | 3 | 12187 | 72 | Car2,Srm,Amd1 | | 3.996e-02 | -3.22 | GO\_AMINOGLYCAN\_METABOLIC\_PROCESS | MSigDB lists | GO\_AMINOGLYCAN\_METABOLIC\_PROCESS | 128 | 3 | 12187 | 72 | Ndst4,Hs3st1,Agrn | | 4.003e-02 | -3.22 | SPR | prosite domains | PS51227 | 6 | 1 | 8845 | 60 | Spry2 | | 4.003e-02 | -3.22 | SEA | prosite domains | PS50024 | 6 | 1 | 8845 | 60 | Agrn | | 4.007e-02 | -3.22 | GROUCHOFAMLY | prints domains | PR01850 | 4 | 1 | 2951 | 30 | Tle4 | | 4.014e-02 | -3.22 | positive regulation of behavioral fear response | biological process | GO:2000987 | 7 | 1 | 13711 | 80 | Crh | | 4.014e-02 | -3.22 | paranodal junction assembly | biological process | GO:0030913 | 7 | 1 | 13711 | 80 | Ugt8a | | 4.014e-02 | -3.22 | bud elongation involved in lung branching | biological process | GO:0060449 | 7 | 1 | 13711 | 80 | Spry2 | | 4.014e-02 | -3.22 | parturition | biological process | GO:0007567 | 7 | 1 | 13711 | 80 | Arntl | | 4.014e-02 | -3.22 | lung growth | biological process | GO:0060437 | 7 | 1 | 13711 | 80 | Spry2 | | 4.014e-02 | -3.22 | primitive streak formation | biological process | GO:0090009 | 7 | 1 | 13711 | 80 | Lhx1 | | 4.014e-02 | -3.22 | tetrahydrobiopterin biosynthetic process | biological process | GO:0006729 | 7 | 1 | 13711 | 80 | Qdpr | | 4.014e-02 | -3.22 | cellular response to follicle-stimulating hormone stimulus | biological process | GO:0071372 | 7 | 1 | 13711 | 80 | Inhba | | 4.014e-02 | -3.22 | positive regulation of fear response | biological process | GO:1903367 | 7 | 1 | 13711 | 80 | Crh | | 4.014e-02 | -3.22 | mesendoderm development | biological process | GO:0048382 | 7 | 1 | 13711 | 80 | Lhx1 | | 4.014e-02 | -3.22 | neurotransmitter receptor transport, postsynaptic endosome to lysosome | biological process | GO:0098943 | 7 | 1 | 13711 | 80 | Cacng2 | | 4.014e-02 | -3.22 | post-embryonic animal organ morphogenesis | biological process | GO:0048563 | 7 | 1 | 13711 | 80 | Kdr | | 4.014e-02 | -3.22 | tetrahydrobiopterin metabolic process | biological process | GO:0046146 | 7 | 1 | 13711 | 80 | Qdpr | | 4.014e-02 | -3.22 | negative regulation of inclusion body assembly | biological process | GO:0090084 | 7 | 1 | 13711 | 80 | Sorl1 | | 4.014e-02 | -3.22 | common myeloid progenitor cell proliferation | biological process | GO:0035726 | 7 | 1 | 13711 | 80 | Nr4a3 | | 4.014e-02 | -3.22 | bone trabecula formation | biological process | GO:0060346 | 7 | 1 | 13711 | 80 | Col1a1 | | 4.014e-02 | -3.22 | negative regulation of cellular response to vascular endothelial growth factor stimulus | biological process | GO:1902548 | 7 | 1 | 13711 | 80 | Spry2 | | 4.014e-02 | -3.22 | acute inflammatory response to antigenic stimulus | biological process | GO:0002438 | 7 | 1 | 13711 | 80 | Ephb6 | | 4.014e-02 | -3.22 | regulation of epinephrine secretion | biological process | GO:0014060 | 7 | 1 | 13711 | 80 | Crh | | 4.014e-02 | -3.22 | negative regulation of synaptic transmission, GABAergic | biological process | GO:0032229 | 7 | 1 | 13711 | 80 | Bdnf | | 4.014e-02 | -3.22 | regulation of ER-associated ubiquitin-dependent protein catabolic process | biological process | GO:1903069 | 7 | 1 | 13711 | 80 | Ubqln2 | | 4.014e-02 | -3.22 | positive regulation of corticosteroid hormone secretion | biological process | GO:2000848 | 7 | 1 | 13711 | 80 | Crh | | 4.014e-02 | -3.22 | regulation of cytoplasmic translational elongation | biological process | GO:1900247 | 7 | 1 | 13711 | 80 | Aars | | 4.014e-02 | -3.22 | negative regulation of neurotransmitter uptake | biological process | GO:0051581 | 7 | 1 | 13711 | 80 | Rgs4 | | 4.014e-02 | -3.22 | G protein-coupled purinergic nucleotide receptor signaling pathway | biological process | GO:0035589 | 7 | 1 | 13711 | 80 | P2ry12 | | 4.014e-02 | -3.22 | spermine metabolic process | biological process | GO:0008215 | 7 | 1 | 13711 | 80 | Amd1 | | 4.014e-02 | -3.22 | embryonic retina morphogenesis in camera-type eye | biological process | GO:0060059 | 7 | 1 | 13711 | 80 | Lhx1 | | 4.014e-02 | -3.22 | positive regulation of neuromuscular junction development | biological process | GO:1904398 | 7 | 1 | 13711 | 80 | Agrn | | 4.014e-02 | -3.22 | adenosine receptor signaling pathway | biological process | GO:0001973 | 7 | 1 | 13711 | 80 | P2ry12 | | 4.014e-02 | -3.22 | semicircular canal morphogenesis | biological process | GO:0048752 | 7 | 1 | 13711 | 80 | Nr4a3 | | 4.014e-02 | -3.22 | somite rostral/caudal axis specification | biological process | GO:0032525 | 7 | 1 | 13711 | 80 | Lhx1 | | 4.014e-02 | -3.22 | aromatic amino acid family catabolic process | biological process | GO:0009074 | 7 | 1 | 13711 | 80 | Qdpr | | 4.014e-02 | -3.22 | positive regulation of glomerular mesangial cell proliferation | biological process | GO:0072126 | 7 | 1 | 13711 | 80 | Egr1 | | 4.020e-02 | -3.21 | regulation of hormone secretion | biological process | GO:0046883 | 221 | 4 | 13711 | 80 | Inhba,Arntl,Crh,Glud1 | | 4.026e-02 | -3.21 | Arginine and proline metabolism | KEGG pathways | mmu00330 | 40 | 2 | 5248 | 42 | Srm,Amd1 | | 4.026e-02 | -3.21 | Arginine and proline metabolism | KEGG pathways | ko00330 | 40 | 2 | 5248 | 42 | Amd1,Srm | | 4.036e-02 | -3.21 | LIM | pfam domains | PF00412 | 57 | 2 | 12881 | 72 | Lhx1,Lims2 | | 4.038e-02 | -3.21 | Protein-protein interactions at synapses | REACTOME pathways | R-MMU-6794362 | 48 | 2 | 6297 | 42 | Homer1,Lrrtm2 | | 4.044e-02 | -3.21 | AMIT\_EGF\_RESPONSE\_120\_HELA | MSigDB lists | AMIT\_EGF\_RESPONSE\_120\_HELA | 54 | 2 | 12187 | 72 | Bdnf,Tgfa | | 4.044e-02 | -3.21 | GENTILE\_RESPONSE\_CLUSTER\_D3 | MSigDB lists | GENTILE\_RESPONSE\_CLUSTER\_D3 | 54 | 2 | 12187 | 72 | Amd1,Pom121 | | 4.044e-02 | -3.21 | GO\_ADENYLATE\_CYCLASE\_INHIBITING\_G\_PROTEIN\_COUPLED\_RECEPTOR\_SIGNALING\_PATHWAY | MSigDB lists | GO\_ADENYLATE\_CYCLASE\_INHIBITING\_G\_PROTEIN\_COUPLED\_RECEPTOR\_SIGNALING\_PATHWAY | 54 | 2 | 12187 | 72 | P2ry12,Grm3 | | 4.044e-02 | -3.21 | E2F\_01 | MSigDB lists | E2F\_01 | 54 | 2 | 12187 | 72 | Amd1,Nr4a3 | | 4.044e-02 | -3.21 | GNF2\_MAP2K3 | MSigDB lists | GNF2\_MAP2K3 | 54 | 2 | 12187 | 72 | Car2,H1f0 | | 4.044e-02 | -3.21 | GNF2\_SPTA1 | MSigDB lists | GNF2\_SPTA1 | 54 | 2 | 12187 | 72 | Car2,H1f0 | | 4.046e-02 | -3.21 | SMID\_BREAST\_CANCER\_RELAPSE\_IN\_BONE\_DN | MSigDB lists | SMID\_BREAST\_CANCER\_RELAPSE\_IN\_BONE\_DN | 219 | 4 | 12187 | 72 | Peg3,Amd1,Fabp7,Qk | | 4.046e-02 | -3.21 | REACTOME\_GPCR\_LIGAND\_BINDING | MSigDB lists | REACTOME\_GPCR\_LIGAND\_BINDING | 219 | 4 | 12187 | 72 | Grm3,P2ry12,Crh,Gng11 | | 4.046e-02 | -3.21 | DURCHDEWALD\_SKIN\_CARCINOGENESIS\_DN | MSigDB lists | DURCHDEWALD\_SKIN\_CARCINOGENESIS\_DN | 219 | 4 | 12187 | 72 | Hs3st1,Hba-a2,Qk,Tmx3 | | 4.049e-02 | -3.21 | ONKEN\_UVEAL\_MELANOMA\_UP | MSigDB lists | ONKEN\_UVEAL\_MELANOMA\_UP | 661 | 8 | 12187 | 72 | Hsd17b12,Fnbp1,Timp2,Sorl1,Tmem165,Egr1,Kdr,Qdpr | | 4.058e-02 | -3.20 | GO\_REGULATION\_OF\_HORMONE\_LEVELS | MSigDB lists | GO\_REGULATION\_OF\_HORMONE\_LEVELS | 320 | 5 | 12187 | 72 | Glud1,Crh,Hsd17b12,Arntl,Inhba | | 4.064e-02 | -3.20 | GO\_PRIMITIVE\_STREAK\_FORMATION | MSigDB lists | GO\_PRIMITIVE\_STREAK\_FORMATION | 7 | 1 | 12187 | 72 | Lhx1 | | 4.064e-02 | -3.20 | KEGG\_FOLATE\_BIOSYNTHESIS | MSigDB lists | KEGG\_FOLATE\_BIOSYNTHESIS | 7 | 1 | 12187 | 72 | Qdpr | | 4.064e-02 | -3.20 | HE\_PTEN\_TARGETS\_DN | MSigDB lists | HE\_PTEN\_TARGETS\_DN | 7 | 1 | 12187 | 72 | Tgfa | | 4.064e-02 | -3.20 | GO\_GLUCOCORTICOID\_METABOLIC\_PROCESS | MSigDB lists | GO\_GLUCOCORTICOID\_METABOLIC\_PROCESS | 7 | 1 | 12187 | 72 | Crh | | 4.064e-02 | -3.20 | MODULE\_203 | MSigDB lists | MODULE\_203 | 7 | 1 | 12187 | 72 | H1f0 | | 4.064e-02 | -3.20 | GO\_URONIC\_ACID\_METABOLIC\_PROCESS | MSigDB lists | GO\_URONIC\_ACID\_METABOLIC\_PROCESS | 7 | 1 | 12187 | 72 | Ugt8a | | 4.064e-02 | -3.20 | MATZUK\_LUTEAL\_GENES | MSigDB lists | MATZUK\_LUTEAL\_GENES | 7 | 1 | 12187 | 72 | Egr1 | | 4.064e-02 | -3.20 | GO\_CHEMICAL\_HOMEOSTASIS\_WITHIN\_A\_TISSUE | MSigDB lists | GO\_CHEMICAL\_HOMEOSTASIS\_WITHIN\_A\_TISSUE | 7 | 1 | 12187 | 72 | Homer1 | | 4.064e-02 | -3.20 | GO\_G\_PROTEIN\_COUPLED\_PURINERGIC\_NUCLEOTIDE\_RECEPTOR\_SIGNALING\_PATHWAY | MSigDB lists | GO\_G\_PROTEIN\_COUPLED\_PURINERGIC\_NUCLEOTIDE\_RECEPTOR\_SIGNALING\_PATHWAY | 7 | 1 | 12187 | 72 | P2ry12 | | 4.064e-02 | -3.20 | SEMBA\_FHIT\_TARGETS\_UP | MSigDB lists | SEMBA\_FHIT\_TARGETS\_UP | 7 | 1 | 12187 | 72 | Col1a1 | | 4.064e-02 | -3.20 | GRAHAM\_CML\_QUIESCENT\_VS\_NORMAL\_DIVIDING\_DN | MSigDB lists | GRAHAM\_CML\_QUIESCENT\_VS\_NORMAL\_DIVIDING\_DN | 7 | 1 | 12187 | 72 | Sorl1 | | 4.074e-02 | -3.20 | GO\_HORMONE\_RECEPTOR\_BINDING | MSigDB lists | GO\_HORMONE\_RECEPTOR\_BINDING | 129 | 3 | 12187 | 72 | Crh,Nr4a3,Ddx17 | | 4.074e-02 | -3.20 | GSE46025\_WT\_VS\_FOXO1\_KO\_KLRG1\_LOW\_CD8\_EFFECTOR\_TCELL\_UP | MSigDB lists | GSE46025\_WT\_VS\_FOXO1\_KO\_KLRG1\_LOW\_CD8\_EFFECTOR\_TCELL\_UP | 129 | 3 | 12187 | 72 | Ddx56,Car2,Srm | | 4.074e-02 | -3.20 | ESC\_V6.5\_UP\_EARLY.V1\_DN | MSigDB lists | ESC\_V6.5\_UP\_EARLY.V1\_DN | 129 | 3 | 12187 | 72 | Timp2,Inhba,Col1a1 | | 4.074e-02 | -3.20 | HNF1\_01 | MSigDB lists | HNF1\_01 | 129 | 3 | 12187 | 72 | Egr1,Bdnf,Tle4 | | 4.074e-02 | -3.20 | GSE33162\_HDAC3\_KO\_VS\_HDAC3\_KO\_4H\_LPS\_STIM\_MACROPHAGE\_DN | MSigDB lists | GSE33162\_HDAC3\_KO\_VS\_HDAC3\_KO\_4H\_LPS\_STIM\_MACROPHAGE\_DN | 129 | 3 | 12187 | 72 | Hs3st1,Glud1,Ugt8a | | 4.076e-02 | -3.20 | rhythmic process | biological process | GO:0048511 | 222 | 4 | 13711 | 80 | Arntl,Inhba,Egr1,Bdnf | | 4.088e-02 | -3.20 | regulation of neuron projection development | biological process | GO:0010975 | 549 | 7 | 13711 | 80 | Bdnf,Ltk,Alkal2,Qk,Agrn,Ddx56,Ntm | | 4.091e-02 | -3.20 | regulation of catecholamine secretion | biological process | GO:0050433 | 55 | 2 | 13711 | 80 | Crh,P2ry12 | | 4.091e-02 | -3.20 | transmission of nerve impulse | biological process | GO:0019226 | 55 | 2 | 13711 | 80 | Kcnmb2,Cacng2 | | 4.103e-02 | -3.19 | GO\_POSITIVE\_REGULATION\_OF\_CELLULAR\_COMPONENT\_BIOGENESIS | MSigDB lists | GO\_POSITIVE\_REGULATION\_OF\_CELLULAR\_COMPONENT\_BIOGENESIS | 321 | 5 | 12187 | 72 | Lrrtm2,Kdr,Agrn,Cbln2,Bdnf | | 4.106e-02 | -3.19 | GO\_POSITIVE\_REGULATION\_OF\_GENE\_EXPRESSION | MSigDB lists | GO\_POSITIVE\_REGULATION\_OF\_GENE\_EXPRESSION | 1307 | 13 | 12187 | 72 | Lhx1,Spry2,Crh,Qk,Agrn,Egr1,Csrnp3,Ddx17,Col1a1,Inhba,Nr4a3,Plp1,Arntl | | 4.123e-02 | -3.19 | GO\_CELL\_MORPHOGENESIS\_INVOLVED\_IN\_DIFFERENTIATION | MSigDB lists | GO\_CELL\_MORPHOGENESIS\_INVOLVED\_IN\_DIFFERENTIATION | 430 | 6 | 12187 | 72 | Lhx1,Bdnf,Ncam1,Nr4a3,Lrrc55,Gbx1 | | 4.126e-02 | -3.19 | sensory perception | biological process | GO:0007600 | 325 | 5 | 13711 | 80 | Gsdme,Spry2,Gbx1,Col1a1,Grm3 | | 4.133e-02 | -3.19 | regulation of intracellular signal transduction | biological process | GO:1902531 | 1320 | 13 | 13711 | 80 | Spry2,Gsdme,Ncam1,Inhba,Sorl1,Crh,Tgfa,Kdr,Timp2,Alkal2,Arntl,Dgkz,P2ry12 | | 4.153e-02 | -3.18 | GAGCTGG\_MIR337 | MSigDB lists | GAGCTGG\_MIR337 | 130 | 3 | 12187 | 72 | Dgkz,Fbxl17,H1f0 | | 4.153e-02 | -3.18 | GSE11386\_NAIVE\_VS\_MEMORY\_BCELL\_DN | MSigDB lists | GSE11386\_NAIVE\_VS\_MEMORY\_BCELL\_DN | 130 | 3 | 12187 | 72 | Nap1l5,Timp2,Fbxl17 | | 4.153e-02 | -3.18 | YTAATTAA\_LHX3\_01 | MSigDB lists | YTAATTAA\_LHX3\_01 | 130 | 3 | 12187 | 72 | Arntl,Tle4,Ndst4 | | 4.180e-02 | -3.17 | BILD\_SRC\_ONCOGENIC\_SIGNATURE | MSigDB lists | BILD\_SRC\_ONCOGENIC\_SIGNATURE | 55 | 2 | 12187 | 72 | Col1a1,Sorl1 | | 4.180e-02 | -3.17 | GCM\_MAP1B | MSigDB lists | GCM\_MAP1B | 55 | 2 | 12187 | 72 | Peg3,Ncam1 | | 4.180e-02 | -3.17 | BRUNO\_HEMATOPOIESIS | MSigDB lists | BRUNO\_HEMATOPOIESIS | 55 | 2 | 12187 | 72 | Msmo1,H1f0 | | 4.180e-02 | -3.17 | GO\_REGULATION\_OF\_CELLULAR\_PH | MSigDB lists | GO\_REGULATION\_OF\_CELLULAR\_PH | 55 | 2 | 12187 | 72 | Car2,Tmem165 | | 4.180e-02 | -3.17 | REACTOME\_EXTRACELLULAR\_MATRIX\_ORGANIZATION | MSigDB lists | REACTOME\_EXTRACELLULAR\_MATRIX\_ORGANIZATION | 55 | 2 | 12187 | 72 | Timp2,Col1a1 | | 4.180e-02 | -3.17 | GSE40493\_BCL6\_KO\_VS\_WT\_TREG\_DN | MSigDB lists | GSE40493\_BCL6\_KO\_VS\_WT\_TREG\_DN | 55 | 2 | 12187 | 72 | Hkdc1,AI593442 | | 4.180e-02 | -3.17 | FONTAINE\_FOLLICULAR\_THYROID\_ADENOMA\_UP | MSigDB lists | FONTAINE\_FOLLICULAR\_THYROID\_ADENOMA\_UP | 55 | 2 | 12187 | 72 | Ephb6,Ddx56 | | 4.180e-02 | -3.17 | TURASHVILI\_BREAST\_LOBULAR\_CARCINOMA\_VS\_DUCTAL\_NORMAL\_UP | MSigDB lists | TURASHVILI\_BREAST\_LOBULAR\_CARCINOMA\_VS\_DUCTAL\_NORMAL\_UP | 55 | 2 | 12187 | 72 | Col1a1,Inhba | | 4.180e-02 | -3.17 | GO\_ENDODERM\_DEVELOPMENT | MSigDB lists | GO\_ENDODERM\_DEVELOPMENT | 55 | 2 | 12187 | 72 | Inhba,Lhx1 | | 4.180e-02 | -3.17 | GO\_MAIN\_AXON | MSigDB lists | GO\_MAIN\_AXON | 55 | 2 | 12187 | 72 | Crh,Ermn | | 4.180e-02 | -3.17 | CHIANG\_LIVER\_CANCER\_SUBCLASS\_UNANNOTATED\_UP | MSigDB lists | CHIANG\_LIVER\_CANCER\_SUBCLASS\_UNANNOTATED\_UP | 55 | 2 | 12187 | 72 | Sorl1,Egr1 | | 4.190e-02 | -3.17 | response to metal ion | biological process | GO:0010038 | 224 | 4 | 13711 | 80 | Glud1,Kcnmb2,Ncam1,Homer1 | | 4.190e-02 | -3.17 | multicellular organismal homeostasis | biological process | GO:0048871 | 224 | 4 | 13711 | 80 | Nr4a3,Egr1,Homer1,Kdr | | 4.192e-02 | -3.17 | branching morphogenesis of an epithelial tube | biological process | GO:0048754 | 132 | 3 | 13711 | 80 | Lhx1,Kdr,Spry2 | | 4.214e-02 | -3.17 | GO\_ENZYME\_LINKED\_RECEPTOR\_PROTEIN\_SIGNALING\_PATHWAY | MSigDB lists | GO\_ENZYME\_LINKED\_RECEPTOR\_PROTEIN\_SIGNALING\_PATHWAY | 547 | 7 | 12187 | 72 | Ephb6,Inhba,Egr1,Ltk,Kdr,Cldn5,Bdnf | | 4.225e-02 | -3.16 | Myometrial Relaxation and Contraction Pathways | WikiPathways | WP385 | 143 | 4 | 3756 | 35 | Rgs4,Crh,Gng11,Dgkz | | 4.227e-02 | -3.16 | negative regulation of striated muscle tissue development | biological process | GO:0045843 | 56 | 2 | 13711 | 80 | Rgs4,Bdnf | | 4.227e-02 | -3.16 | cell redox homeostasis | biological process | GO:0045454 | 56 | 2 | 13711 | 80 | Tmx3,Pdia6 | | 4.233e-02 | -3.16 | FREAC7\_01 | MSigDB lists | FREAC7\_01 | 131 | 3 | 12187 | 72 | Crh,Grm3,Ndst4 | | 4.241e-02 | -3.16 | oxidation-reduction process | biological process | GO:0055114 | 674 | 8 | 13711 | 80 | Glud1,Scd2,Bdnf,Msmo1,Tmx3,Nr4a3,Qdpr,Hsd17b12 | | 4.247e-02 | -3.16 | RBBP8 (RB binding protein 8, endonuclease) | protein interactions | 5932 | 6 | 1 | 6802 | 49 | Homer1 | | 4.247e-02 | -3.16 | Psmd3 (proteasome (prosome, macropain) 26S subunit, non-ATPase, 3) | protein interactions | 22123 | 6 | 1 | 6802 | 49 | Ubqln2 | | 4.247e-02 | -3.16 | Spry1 (sprouty RTK signaling antagonist 1) | protein interactions | 24063 | 6 | 1 | 6802 | 49 | Spry2 | | 4.247e-02 | -3.16 | G3bp1 (GTPase activating protein (SH3 domain) binding protein 1) | protein interactions | 27041 | 6 | 1 | 6802 | 49 | Egr1 | | 4.247e-02 | -3.16 | Ncdn (neurochondrin) | protein interactions | 26562 | 6 | 1 | 6802 | 49 | Homer1 | | 4.247e-02 | -3.16 | Trim21 (tripartite motif-containing 21) | protein interactions | 20821 | 6 | 1 | 6802 | 49 | Homer1 | | 4.247e-02 | -3.16 | Mafk (v-maf musculoaponeurotic fibrosarcoma oncogene family, protein K (avian)) | protein interactions | 17135 | 6 | 1 | 6802 | 49 | Nr4a3 | | 4.247e-02 | -3.16 | Kmt2a (lysine (K)-specific methyltransferase 2A) | protein interactions | 214162 | 6 | 1 | 6802 | 49 | Arntl | | 4.247e-02 | -3.16 | Spry4 (sprouty RTK signaling antagonist 4) | protein interactions | 24066 | 6 | 1 | 6802 | 49 | Spry2 | | 4.247e-02 | -3.16 | Rorb (RAR-related orphan receptor beta) | protein interactions | 225998 | 6 | 1 | 6802 | 49 | Nr4a3 | | 4.247e-02 | -3.16 | Per3 (period circadian clock 3) | protein interactions | 18628 | 6 | 1 | 6802 | 49 | Arntl | | 4.247e-02 | -3.16 | COBLL1 (cordon-bleu WH2 repeat protein like 1) | protein interactions | 22837 | 6 | 1 | 6802 | 49 | Homer1 | | 4.247e-02 | -3.16 | Psmc4 (proteasome (prosome, macropain) 26S subunit, ATPase, 4) | protein interactions | 23996 | 6 | 1 | 6802 | 49 | Ubqln2 | | 4.256e-02 | -3.16 | structural molecule activity | molecular function | GO:0005198 | 443 | 6 | 13516 | 78 | Cldn5,Col1a1,Pom121,Plp1,Agrn,Homer1 | | 4.277e-02 | -3.15 | STK33\_UP | MSigDB lists | STK33\_UP | 223 | 4 | 12187 | 72 | Gng11,Mtmr9,Egr1,Foxq1 | | 4.296e-02 | -3.15 | negative regulation of transport | biological process | GO:0051051 | 439 | 6 | 13711 | 80 | Ubqln2,P2ry12,Crh,Lrrtm2,Rgs4,Agrn | | 4.314e-02 | -3.14 | GSE29949\_MICROGLIA\_BRAIN\_VS\_CD8\_NEG\_DC\_SPLEEN\_DN | MSigDB lists | GSE29949\_MICROGLIA\_BRAIN\_VS\_CD8\_NEG\_DC\_SPLEEN\_DN | 132 | 3 | 12187 | 72 | Plp1,Gng11,Rgs4 | | 4.314e-02 | -3.14 | GO\_REGULATION\_OF\_SYNAPTIC\_PLASTICITY | MSigDB lists | GO\_REGULATION\_OF\_SYNAPTIC\_PLASTICITY | 132 | 3 | 12187 | 72 | Crh,Lrrtm2,Ncam1 | | 4.319e-02 | -3.14 | REACTOME\_CLASS\_B\_2\_SECRETIN\_FAMILY\_RECEPTORS | MSigDB lists | REACTOME\_CLASS\_B\_2\_SECRETIN\_FAMILY\_RECEPTORS | 56 | 2 | 12187 | 72 | Gng11,Crh | | 4.319e-02 | -3.14 | YAO\_TEMPORAL\_RESPONSE\_TO\_PROGESTERONE\_CLUSTER\_1 | MSigDB lists | YAO\_TEMPORAL\_RESPONSE\_TO\_PROGESTERONE\_CLUSTER\_1 | 56 | 2 | 12187 | 72 | Peg3,Ddx17 | | 4.319e-02 | -3.14 | MODULE\_375 | MSigDB lists | MODULE\_375 | 56 | 2 | 12187 | 72 | Rgs4,Grm3 | | 4.319e-02 | -3.14 | RAMASWAMY\_METASTASIS\_UP | MSigDB lists | RAMASWAMY\_METASTASIS\_UP | 56 | 2 | 12187 | 72 | Pom121,Col1a1 | | 4.321e-02 | -3.14 | MARTORIATI\_MDM4\_TARGETS\_FETAL\_LIVER\_DN | MSigDB lists | MARTORIATI\_MDM4\_TARGETS\_FETAL\_LIVER\_DN | 435 | 6 | 12187 | 72 | Ddx17,Timp2,Col1a1,Car2,Bdnf,Qk | | 4.351e-02 | -3.13 | hindbrain development | biological process | GO:0030902 | 134 | 3 | 13711 | 80 | Aars,Lhx1,Gbx1 | | 4.362e-02 | -3.13 | GO\_NEGATIVE\_REGULATION\_OF\_PHOSPHORUS\_METABOLIC\_PROCESS | MSigDB lists | GO\_NEGATIVE\_REGULATION\_OF\_PHOSPHORUS\_METABOLIC\_PROCESS | 436 | 6 | 12187 | 72 | Csrnp3,Spry2,Grm3,Rgs4,Sorl1,Inhba | | 4.364e-02 | -3.13 | cerebellar cortex development | biological process | GO:0021695 | 57 | 2 | 13711 | 80 | Lhx1,Aars | | 4.386e-02 | -3.13 | Lipocalin | pfam domains | PF00061 | 8 | 1 | 12881 | 72 | Fabp7 | | 4.395e-02 | -3.12 | GSE11057\_NAIVE\_VS\_MEMORY\_CD4\_TCELL\_UP | MSigDB lists | GSE11057\_NAIVE\_VS\_MEMORY\_CD4\_TCELL\_UP | 133 | 3 | 12187 | 72 | Chmp7,Timp2,Agrn | | 4.395e-02 | -3.12 | GSE11057\_PBMC\_VS\_MEM\_CD4\_TCELL\_UP | MSigDB lists | GSE11057\_PBMC\_VS\_MEM\_CD4\_TCELL\_UP | 133 | 3 | 12187 | 72 | P2ry12,Car2,Gng11 | | 4.395e-02 | -3.12 | GO\_ORGANIC\_ACID\_BINDING | MSigDB lists | GO\_ORGANIC\_ACID\_BINDING | 133 | 3 | 12187 | 72 | Aars,Glud1,Agrn | | 4.399e-02 | -3.12 | behavior | biological process | GO:0007610 | 558 | 7 | 13711 | 80 | Crh,Nr4a3,Homer1,Egr1,Gbx1,Ncam1,Bdnf | | 4.404e-02 | -3.12 | postsynaptic membrane | cellular component | GO:0045211 | 338 | 5 | 13825 | 79 | Lrrtm2,Grm3,Homer1,Cacng2,Ncam1 | | 4.405e-02 | -3.12 | positive regulation of neuron projection development | biological process | GO:0010976 | 331 | 5 | 13711 | 80 | Bdnf,Ltk,Ddx56,Qk,Alkal2 | | 4.425e-02 | -3.12 | GO\_REGULATION\_OF\_CELL\_DEVELOPMENT | MSigDB lists | GO\_REGULATION\_OF\_CELL\_DEVELOPMENT | 673 | 8 | 12187 | 72 | P2ry12,Cldn5,Bdnf,Col1a1,Timp2,Arntl,Ltk,Sorl1 | | 4.452e-02 | -3.11 | cellular response to growth factor stimulus | biological process | GO:0071363 | 332 | 5 | 13711 | 80 | Spry2,Cldn5,Kdr,Bdnf,Egr1 | | 4.453e-02 | -3.11 | integral component of plasma membrane | cellular component | GO:0005887 | 954 | 10 | 13825 | 79 | Lrrtm2,Lrrc55,Cacng2,Kdr,P2ry12,Ephb6,Ncam1,Kcnmb2,Ltk,Grm3 | | 4.455e-02 | -3.11 | MULLIGHAN\_MLL\_SIGNATURE\_2\_DN | MSigDB lists | MULLIGHAN\_MLL\_SIGNATURE\_2\_DN | 226 | 4 | 12187 | 72 | Agrn,Hsd17b12,Sorl1,H1f0 | | 4.459e-02 | -3.11 | YRTCANNRCGC\_UNKNOWN | MSigDB lists | YRTCANNRCGC\_UNKNOWN | 57 | 2 | 12187 | 72 | Tmem165,Ncam1 | | 4.459e-02 | -3.11 | GO\_REGULATION\_OF\_AMINE\_TRANSPORT | MSigDB lists | GO\_REGULATION\_OF\_AMINE\_TRANSPORT | 57 | 2 | 12187 | 72 | P2ry12,Crh | | 4.459e-02 | -3.11 | CTAGGAA\_MIR384 | MSigDB lists | CTAGGAA\_MIR384 | 57 | 2 | 12187 | 72 | Ncam1,Pom121 | | 4.475e-02 | -3.11 | Grb2 (growth factor receptor bound protein 2) | protein interactions | 14784 | 47 | 2 | 6802 | 49 | Spry2,Kdr | | 4.477e-02 | -3.11 | GSE22886\_CD8\_VS\_CD4\_NAIVE\_TCELL\_DN | MSigDB lists | GSE22886\_CD8\_VS\_CD4\_NAIVE\_TCELL\_DN | 134 | 3 | 12187 | 72 | Hkdc1,Gng11,Ephb6 | | 4.477e-02 | -3.11 | GO\_MORPHOGENESIS\_OF\_A\_BRANCHING\_STRUCTURE | MSigDB lists | GO\_MORPHOGENESIS\_OF\_A\_BRANCHING\_STRUCTURE | 134 | 3 | 12187 | 72 | Ermn,Spry2,Lhx1 | | 4.477e-02 | -3.11 | MODULE\_93 | MSigDB lists | MODULE\_93 | 134 | 3 | 12187 | 72 | Glud1,Qdpr,Msmo1 | | 4.477e-02 | -3.11 | GO\_CENTRAL\_NERVOUS\_SYSTEM\_NEURON\_DIFFERENTIATION | MSigDB lists | GO\_CENTRAL\_NERVOUS\_SYSTEM\_NEURON\_DIFFERENTIATION | 134 | 3 | 12187 | 72 | Lhx1,Inhba,Gbx1 | | 4.482e-02 | -3.11 | GO\_SMALL\_MOLECULE\_BIOSYNTHETIC\_PROCESS | MSigDB lists | GO\_SMALL\_MOLECULE\_BIOSYNTHETIC\_PROCESS | 329 | 5 | 12187 | 72 | Glud1,Qk,Plp1,Hsd17b12,Msmo1 | | 4.482e-02 | -3.11 | low-density lipoprotein particle | cellular component | GO:0034362 | 8 | 1 | 13825 | 79 | Sorl1 | | 4.494e-02 | -3.10 | Carbonic\_anhydrase\_a-class\_CS | interpro domains | IPR018338 | 8 | 1 | 13788 | 79 | Car2 | | 4.494e-02 | -3.10 | Fol\_N | interpro domains | IPR003645 | 8 | 1 | 13788 | 79 | Agrn | | 4.494e-02 | -3.10 | STI1\_HS-bd | interpro domains | IPR006636 | 8 | 1 | 13788 | 79 | Ubqln2 | | 4.494e-02 | -3.10 | Histone\_H5 | interpro domains | IPR005819 | 8 | 1 | 13788 | 79 | H1f0 | | 4.494e-02 | -3.10 | GPCR\_3 | interpro domains | IPR000337 | 8 | 1 | 13788 | 79 | Grm3 | | 4.494e-02 | -3.10 | Tyr\_kinase\_rcpt\_2\_CS | interpro domains | IPR002011 | 8 | 1 | 13788 | 79 | Ltk | | 4.494e-02 | -3.10 | Tyr\_kinase\_rcpt\_3\_CS | interpro domains | IPR001824 | 8 | 1 | 13788 | 79 | Kdr | | 4.494e-02 | -3.10 | GPCR\_3\_CS | interpro domains | IPR017979 | 8 | 1 | 13788 | 79 | Grm3 | | 4.494e-02 | -3.10 | Claudin | interpro domains | IPR006187 | 8 | 1 | 13788 | 79 | Cldn5 | | 4.495e-02 | -3.10 | Znf\_LIM | interpro domains | IPR001781 | 59 | 2 | 13788 | 79 | Lims2,Lhx1 | | 4.504e-02 | -3.10 | negative regulation of muscle organ development | biological process | GO:0048635 | 58 | 2 | 13711 | 80 | Bdnf,Rgs4 | | 4.504e-02 | -3.10 | positive regulation of smooth muscle cell migration | biological process | GO:0014911 | 58 | 2 | 13711 | 80 | Egr1,Nr4a3 | | 4.504e-02 | -3.10 | long-term synaptic potentiation | biological process | GO:0060291 | 58 | 2 | 13711 | 80 | Lrrtm2,Crh | | 4.506e-02 | -3.10 | enzyme inhibitor activity | molecular function | GO:0004857 | 232 | 4 | 13516 | 78 | Timp2,Pdia6,Dgkz,Agrn | | 4.526e-02 | -3.10 | cardiolipin binding | molecular function | GO:1901612 | 8 | 1 | 13516 | 78 | Gsdme | | 4.526e-02 | -3.10 | ADP-ribosylation factor binding | molecular function | GO:0030306 | 8 | 1 | 13516 | 78 | Sorl1 | | 4.526e-02 | -3.10 | neurotrophin TRK receptor binding | molecular function | GO:0005167 | 8 | 1 | 13516 | 78 | Bdnf | | 4.526e-02 | -3.10 | G protein-coupled glutamate receptor activity | molecular function | GO:0098988 | 8 | 1 | 13516 | 78 | Grm3 | | 4.526e-02 | -3.10 | dystroglycan binding | molecular function | GO:0002162 | 8 | 1 | 13516 | 78 | Agrn | | 4.553e-02 | -3.09 | RNA helicase activity | molecular function | GO:0003724 | 59 | 2 | 13516 | 78 | Ddx56,Ddx17 | | 4.560e-02 | -3.09 | GSE17974\_0H\_VS\_12H\_IN\_VITRO\_ACT\_CD4\_TCELL\_UP | MSigDB lists | GSE17974\_0H\_VS\_12H\_IN\_VITRO\_ACT\_CD4\_TCELL\_UP | 135 | 3 | 12187 | 72 | Nap1l5,Hkdc1,Sorl1 | | 4.560e-02 | -3.09 | HALLMARK\_UV\_RESPONSE\_DN | MSigDB lists | HALLMARK\_UV\_RESPONSE\_DN | 135 | 3 | 12187 | 72 | Col1a1,Rgs4,Bdnf | | 4.560e-02 | -3.09 | GSE6269\_HEALTHY\_VS\_STAPH\_PNEUMO\_INF\_PBMC\_DN | MSigDB lists | GSE6269\_HEALTHY\_VS\_STAPH\_PNEUMO\_INF\_PBMC\_DN | 135 | 3 | 12187 | 72 | Egr1,H1f0,Timp2 | | 4.560e-02 | -3.09 | GSE9988\_LOW\_LPS\_VS\_VEHICLE\_TREATED\_MONOCYTE\_UP | MSigDB lists | GSE9988\_LOW\_LPS\_VS\_VEHICLE\_TREATED\_MONOCYTE\_UP | 135 | 3 | 12187 | 72 | Egr1,Fjx1,Inhba | | 4.560e-02 | -3.09 | GSE9988\_LPS\_VS\_CTRL\_TREATED\_MONOCYTE\_UP | MSigDB lists | GSE9988\_LPS\_VS\_CTRL\_TREATED\_MONOCYTE\_UP | 135 | 3 | 12187 | 72 | Inhba,Fjx1,Egr1 | | 4.560e-02 | -3.09 | GSE24814\_STAT5\_KO\_VS\_WT\_PRE\_BCELL\_UP | MSigDB lists | GSE24814\_STAT5\_KO\_VS\_WT\_PRE\_BCELL\_UP | 135 | 3 | 12187 | 72 | Grm3,Timp2,Col1a1 | | 4.568e-02 | -3.09 | GO\_CARBOHYDRATE\_DERIVATIVE\_BIOSYNTHETIC\_PROCESS | MSigDB lists | GO\_CARBOHYDRATE\_DERIVATIVE\_BIOSYNTHETIC\_PROCESS | 441 | 6 | 12187 | 72 | Hs3st1,Ndst4,Agrn,Amd1,Tmem165,Ugt8a | | 4.575e-02 | -3.08 | regulation of chloride transport | biological process | GO:2001225 | 8 | 1 | 13711 | 80 | Car2 | | 4.575e-02 | -3.08 | regulation of glucagon secretion | biological process | GO:0070092 | 8 | 1 | 13711 | 80 | Crh | | 4.575e-02 | -3.08 | blood vessel endothelial cell differentiation | biological process | GO:0060837 | 8 | 1 | 13711 | 80 | Kdr | | 4.575e-02 | -3.08 | regulation of serotonin secretion | biological process | GO:0014062 | 8 | 1 | 13711 | 80 | Crh | | 4.575e-02 | -3.08 | pri-miRNA transcription by RNA polymerase II | biological process | GO:0061614 | 8 | 1 | 13711 | 80 | Ddx17 | | 4.575e-02 | -3.08 | ureter morphogenesis | biological process | GO:0072197 | 8 | 1 | 13711 | 80 | Lhx1 | | 4.575e-02 | -3.08 | positive regulation of epidermal growth factor-activated receptor activity | biological process | GO:0045741 | 8 | 1 | 13711 | 80 | Tgfa | | 4.575e-02 | -3.08 | activated T cell proliferation | biological process | GO:0050798 | 8 | 1 | 13711 | 80 | Ephb6 | | 4.575e-02 | -3.08 | skeletal muscle acetylcholine-gated channel clustering | biological process | GO:0071340 | 8 | 1 | 13711 | 80 | Agrn | | 4.575e-02 | -3.08 | cellular response to gonadotropin stimulus | biological process | GO:0071371 | 8 | 1 | 13711 | 80 | Inhba | | 4.575e-02 | -3.08 | regulation of nephron tubule epithelial cell differentiation | biological process | GO:0072182 | 8 | 1 | 13711 | 80 | Lhx1 | | 4.575e-02 | -3.08 | positive regulation of AMPA receptor activity | biological process | GO:2000969 | 8 | 1 | 13711 | 80 | Cacng2 | | 4.575e-02 | -3.08 | positive regulation of cell proliferation involved in kidney development | biological process | GO:1901724 | 8 | 1 | 13711 | 80 | Egr1 | | 4.575e-02 | -3.08 | substrate-dependent cell migration, cell extension | biological process | GO:0006930 | 8 | 1 | 13711 | 80 | P2ry12 | | 4.575e-02 | -3.08 | positive regulation of integrin activation | biological process | GO:0033625 | 8 | 1 | 13711 | 80 | P2ry12 | | 4.575e-02 | -3.08 | positive regulation of metanephros development | biological process | GO:0072216 | 8 | 1 | 13711 | 80 | Egr1 | | 4.575e-02 | -3.08 | response to follicle-stimulating hormone | biological process | GO:0032354 | 8 | 1 | 13711 | 80 | Inhba | | 4.575e-02 | -3.08 | liver morphogenesis | biological process | GO:0072576 | 8 | 1 | 13711 | 80 | Tgfa | | 4.575e-02 | -3.08 | collagen biosynthetic process | biological process | GO:0032964 | 8 | 1 | 13711 | 80 | Col1a1 | | 4.575e-02 | -3.08 | positive regulation of cell migration by vascular endothelial growth factor signaling pathway | biological process | GO:0038089 | 8 | 1 | 13711 | 80 | Kdr | | 4.575e-02 | -3.08 | glucocorticoid metabolic process | biological process | GO:0008211 | 8 | 1 | 13711 | 80 | Crh | | 4.575e-02 | -3.08 | negative regulation of neuroblast proliferation | biological process | GO:0007406 | 8 | 1 | 13711 | 80 | Bdnf | | 4.575e-02 | -3.08 | glutamate secretion | biological process | GO:0014047 | 8 | 1 | 13711 | 80 | Bdnf | | 4.575e-02 | -3.08 | collagen-activated tyrosine kinase receptor signaling pathway | biological process | GO:0038063 | 8 | 1 | 13711 | 80 | Col1a1 | | 4.576e-02 | -3.08 | Osteoblast | WikiPathways | WP238 | 5 | 1 | 3756 | 35 | Col1a1 | | 4.579e-02 | -3.08 | Hormone-sensitive lipase (HSL)-mediated triacylglycerol hydrolysis | REACTOME pathways | R-MMU-163560 | 7 | 1 | 6297 | 42 | Fabp7 | | 4.579e-02 | -3.08 | Ca2+ activated K+ channels | REACTOME pathways | R-MMU-1296052 | 7 | 1 | 6297 | 42 | Kcnmb2 | | 4.579e-02 | -3.08 | Phenylalanine and tyrosine catabolism | REACTOME pathways | R-MMU-71182 | 7 | 1 | 6297 | 42 | Qdpr | | 4.579e-02 | -3.08 | Repression of WNT target genes | REACTOME pathways | R-MMU-4641265 | 7 | 1 | 6297 | 42 | Tle4 | | 4.596e-02 | -3.08 | positive regulation of cation transmembrane transport | biological process | GO:1904064 | 137 | 3 | 13711 | 80 | Cacng2,Lrrc55,Nipsnap2 | | 4.601e-02 | -3.08 | CAAGGAT\_MIR362 | MSigDB lists | CAAGGAT\_MIR362 | 58 | 2 | 12187 | 72 | Tle4,Plp1 | | 4.601e-02 | -3.08 | GO\_REGULATION\_OF\_LYASE\_ACTIVITY | MSigDB lists | GO\_REGULATION\_OF\_LYASE\_ACTIVITY | 58 | 2 | 12187 | 72 | Grm3,Timp2 | | 4.601e-02 | -3.08 | GO\_TERPENOID\_METABOLIC\_PROCESS | MSigDB lists | GO\_TERPENOID\_METABOLIC\_PROCESS | 58 | 2 | 12187 | 72 | Crh,Agrn | | 4.624e-02 | -3.07 | cellular response to oxygen-containing compound | biological process | GO:1901701 | 686 | 8 | 13711 | 80 | Col1a1,P2ry12,Ltk,Inhba,Nr4a3,Car2,Crh,Egr1 | | 4.629e-02 | -3.07 | MOHANKUMAR\_HOXA1\_TARGETS\_UP | MSigDB lists | MOHANKUMAR\_HOXA1\_TARGETS\_UP | 332 | 5 | 12187 | 72 | Sorl1,Amd1,Lhx1,Homer1,H1f0 | | 4.631e-02 | -3.07 | GO\_NEGATIVE\_REGULATION\_OF\_PROTEIN\_OLIGOMERIZATION | MSigDB lists | GO\_NEGATIVE\_REGULATION\_OF\_PROTEIN\_OLIGOMERIZATION | 8 | 1 | 12187 | 72 | Sorl1 | | 4.631e-02 | -3.07 | GO\_REGULATION\_OF\_HEPATOCYTE\_PROLIFERATION | MSigDB lists | GO\_REGULATION\_OF\_HEPATOCYTE\_PROLIFERATION | 8 | 1 | 12187 | 72 | Lims2 | | 4.631e-02 | -3.07 | GO\_TRIGLYCERIDE\_CATABOLIC\_PROCESS | MSigDB lists | GO\_TRIGLYCERIDE\_CATABOLIC\_PROCESS | 8 | 1 | 12187 | 72 | Fabp7 | | 4.631e-02 | -3.07 | GO\_LOW\_DENSITY\_LIPOPROTEIN\_PARTICLE | MSigDB lists | GO\_LOW\_DENSITY\_LIPOPROTEIN\_PARTICLE | 8 | 1 | 12187 | 72 | Sorl1 | | 4.631e-02 | -3.07 | GO\_POSITIVE\_REGULATION\_OF\_INTRACELLULAR\_STEROID\_HORMONE\_RECEPTOR\_SIGNALING\_PATHWAY | MSigDB lists | GO\_POSITIVE\_REGULATION\_OF\_INTRACELLULAR\_STEROID\_HORMONE\_RECEPTOR\_SIGNALING\_PATHWAY | 8 | 1 | 12187 | 72 | Ddx17 | | 4.631e-02 | -3.07 | NAKAMURA\_LUNG\_CANCER\_DIFFERENTIATION\_MARKERS | MSigDB lists | NAKAMURA\_LUNG\_CANCER\_DIFFERENTIATION\_MARKERS | 8 | 1 | 12187 | 72 | Cldn5 | | 4.631e-02 | -3.07 | GO\_AMINOACYL\_TRNA\_EDITING\_ACTIVITY | MSigDB lists | GO\_AMINOACYL\_TRNA\_EDITING\_ACTIVITY | 8 | 1 | 12187 | 72 | Aars | | 4.631e-02 | -3.07 | REACTOME\_CLASS\_C\_3\_METABOTROPIC\_GLUTAMATE\_PHEROMONE\_RECEPTORS | MSigDB lists | REACTOME\_CLASS\_C\_3\_METABOTROPIC\_GLUTAMATE\_PHEROMONE\_RECEPTORS | 8 | 1 | 12187 | 72 | Grm3 | | 4.631e-02 | -3.07 | GO\_POSITIVE\_REGULATION\_OF\_EPIDERMAL\_GROWTH\_FACTOR\_ACTIVATED\_RECEPTOR\_ACTIVITY | MSigDB lists | GO\_POSITIVE\_REGULATION\_OF\_EPIDERMAL\_GROWTH\_FACTOR\_ACTIVATED\_RECEPTOR\_ACTIVITY | 8 | 1 | 12187 | 72 | Tgfa | | 4.631e-02 | -3.07 | AGARWAL\_AKT\_PATHWAY\_TARGETS | MSigDB lists | AGARWAL\_AKT\_PATHWAY\_TARGETS | 8 | 1 | 12187 | 72 | Tgfa | | 4.631e-02 | -3.07 | CGTCTTA\_MIR208 | MSigDB lists | CGTCTTA\_MIR208 | 8 | 1 | 12187 | 72 | Qk | | 4.631e-02 | -3.07 | TUOMISTO\_TUMOR\_SUPPRESSION\_BY\_COL13A1\_DN | MSigDB lists | TUOMISTO\_TUMOR\_SUPPRESSION\_BY\_COL13A1\_DN | 8 | 1 | 12187 | 72 | Foxq1 | | 4.631e-02 | -3.07 | chr11q11 | MSigDB lists | chr11q11 | 8 | 1 | 12187 | 72 | Hsd17b12 | | 4.631e-02 | -3.07 | MURAKAMI\_UV\_RESPONSE\_1HR\_DN | MSigDB lists | MURAKAMI\_UV\_RESPONSE\_1HR\_DN | 8 | 1 | 12187 | 72 | Egr1 | | 4.631e-02 | -3.07 | GO\_POSITIVE\_REGULATION\_OF\_MUSCLE\_CELL\_APOPTOTIC\_PROCESS | MSigDB lists | GO\_POSITIVE\_REGULATION\_OF\_MUSCLE\_CELL\_APOPTOTIC\_PROCESS | 8 | 1 | 12187 | 72 | Ltk | | 4.631e-02 | -3.07 | REACTOME\_REVERSIBLE\_HYDRATION\_OF\_CARBON\_DIOXIDE | MSigDB lists | REACTOME\_REVERSIBLE\_HYDRATION\_OF\_CARBON\_DIOXIDE | 8 | 1 | 12187 | 72 | Car2 | | 4.631e-02 | -3.07 | REACTOME\_P2Y\_RECEPTORS | MSigDB lists | REACTOME\_P2Y\_RECEPTORS | 8 | 1 | 12187 | 72 | P2ry12 | | 4.631e-02 | -3.07 | FRIDMAN\_SENESCENCE\_DN | MSigDB lists | FRIDMAN\_SENESCENCE\_DN | 8 | 1 | 12187 | 72 | Egr1 | | 4.631e-02 | -3.07 | GO\_OXYGEN\_TRANSPORT | MSigDB lists | GO\_OXYGEN\_TRANSPORT | 8 | 1 | 12187 | 72 | Hba-a2 | | 4.644e-02 | -3.07 | GSE24210\_TCONV\_VS\_TREG\_UP | MSigDB lists | GSE24210\_TCONV\_VS\_TREG\_UP | 136 | 3 | 12187 | 72 | Sorl1,Ltk,Nr4a3 | | 4.644e-02 | -3.07 | GSE22527\_ANTI\_CD3\_INVIVO\_VS\_UNTREATED\_MOUSE\_TREG\_UP | MSigDB lists | GSE22527\_ANTI\_CD3\_INVIVO\_VS\_UNTREATED\_MOUSE\_TREG\_UP | 136 | 3 | 12187 | 72 | Cldn5,Msmo1,Amd1 | | 4.644e-02 | -3.07 | GO\_CELLULAR\_RESPONSE\_TO\_ACID\_CHEMICAL | MSigDB lists | GO\_CELLULAR\_RESPONSE\_TO\_ACID\_CHEMICAL | 136 | 3 | 12187 | 72 | Egr1,Ltk,Col1a1 | | 4.644e-02 | -3.07 | GSE3920\_IFNA\_VS\_IFNG\_TREATED\_ENDOTHELIAL\_CELL\_UP | MSigDB lists | GSE3920\_IFNA\_VS\_IFNG\_TREATED\_ENDOTHELIAL\_CELL\_UP | 136 | 3 | 12187 | 72 | Homer1,Egr1,Nr4a3 | | 4.644e-02 | -3.07 | LEF1\_UP.V1\_UP | MSigDB lists | LEF1\_UP.V1\_UP | 136 | 3 | 12187 | 72 | Fnbp1,Fjx1,Gng11 | | 4.644e-02 | -3.07 | GSE23502\_BM\_VS\_COLON\_TUMOR\_HDC\_KO\_MYELOID\_DERIVED\_SUPPRESSOR\_CELL\_UP | MSigDB lists | GSE23502\_BM\_VS\_COLON\_TUMOR\_HDC\_KO\_MYELOID\_DERIVED\_SUPPRESSOR\_CELL\_UP | 136 | 3 | 12187 | 72 | AI593442,Ntm,Tle4 | | 4.644e-02 | -3.07 | GSE23321\_EFFECTOR\_MEMORY\_VS\_NAIVE\_CD8\_TCELL\_UP | MSigDB lists | GSE23321\_EFFECTOR\_MEMORY\_VS\_NAIVE\_CD8\_TCELL\_UP | 136 | 3 | 12187 | 72 | Qk,Egr1,Grm3 | | 4.644e-02 | -3.07 | GSE27291\_0H\_VS\_7D\_STIM\_GAMMADELTA\_TCELL\_UP | MSigDB lists | GSE27291\_0H\_VS\_7D\_STIM\_GAMMADELTA\_TCELL\_UP | 136 | 3 | 12187 | 72 | Ermn,Mtmr9,Ddx17 | | 4.644e-02 | -3.07 | negative regulation of muscle tissue development | biological process | GO:1901862 | 59 | 2 | 13711 | 80 | Rgs4,Bdnf | | 4.648e-02 | -3.07 | Homer1 (homer scaffolding protein 1) | protein interactions | 26556 | 48 | 2 | 6802 | 49 | Ncam1,Homer1 | | 4.654e-02 | -3.07 | G\_PROTEIN\_RECEP\_F3\_2 | prosite domains | PS00980 | 7 | 1 | 8845 | 60 | Grm3 | | 4.654e-02 | -3.07 | G\_PROTEIN\_RECEP\_F3\_1 | prosite domains | PS00979 | 7 | 1 | 8845 | 60 | Grm3 | | 4.689e-02 | -3.06 | FOLN | smart domains | SM00274 | 8 | 1 | 7188 | 43 | Agrn | | 4.689e-02 | -3.06 | STI1 | smart domains | SM00727 | 8 | 1 | 7188 | 43 | Ubqln2 | | 4.698e-02 | -3.06 | DEAD | pfam domains | PF00270 | 62 | 2 | 12881 | 72 | Ddx17,Ddx56 | | 4.699e-02 | -3.06 | E4F1\_Q6 | MSigDB lists | E4F1\_Q6 | 230 | 4 | 12187 | 72 | Nr4a3,Ubqln2,AI593442,Grm3 | | 4.699e-02 | -3.06 | VECCHI\_GASTRIC\_CANCER\_EARLY\_DN | MSigDB lists | VECCHI\_GASTRIC\_CANCER\_EARLY\_DN | 230 | 4 | 12187 | 72 | Ncam1,Rgs4,Plp1,Kcnmb2 | | 4.699e-02 | -3.06 | DEURIG\_T\_CELL\_PROLYMPHOCYTIC\_LEUKEMIA\_DN | MSigDB lists | DEURIG\_T\_CELL\_PROLYMPHOCYTIC\_LEUKEMIA\_DN | 230 | 4 | 12187 | 72 | Arntl,Chmp7,Gng11,Mtmr9 | | 4.728e-02 | -3.05 | GSE16450\_IMMATURE\_VS\_MATURE\_NEURON\_CELL\_LINE\_UP | MSigDB lists | GSE16450\_IMMATURE\_VS\_MATURE\_NEURON\_CELL\_LINE\_UP | 137 | 3 | 12187 | 72 | Arntl,Tle4,Sorl1 | | 4.728e-02 | -3.05 | GSE9988\_LPS\_VS\_VEHICLE\_TREATED\_MONOCYTE\_UP | MSigDB lists | GSE9988\_LPS\_VS\_VEHICLE\_TREATED\_MONOCYTE\_UP | 137 | 3 | 12187 | 72 | Inhba,Fjx1,Egr1 | | 4.728e-02 | -3.05 | GSE5542\_UNTREATED\_VS\_IFNA\_AND\_IFNG\_TREATED\_EPITHELIAL\_CELLS\_6H\_UP | MSigDB lists | GSE5542\_UNTREATED\_VS\_IFNA\_AND\_IFNG\_TREATED\_EPITHELIAL\_CELLS\_6H\_UP | 137 | 3 | 12187 | 72 | Ubqln2,Chmp7,Fabp7 | | 4.728e-02 | -3.05 | GSE13306\_TREG\_RA\_VS\_TCONV\_RA\_DN | MSigDB lists | GSE13306\_TREG\_RA\_VS\_TCONV\_RA\_DN | 137 | 3 | 12187 | 72 | Tgfa,Lhx1,Sorl1 | | 4.731e-02 | -3.05 | GO\_LOCOMOTION | MSigDB lists | GO\_LOCOMOTION | 807 | 9 | 12187 | 72 | Lhx1,Bdnf,Ncam1,Kdr,Nr4a3,P2ry12,Col1a1,Gbx1,Dgkz | | 4.738e-02 | -3.05 | Glutamatergic synapse | KEGG pathways | mmu04724 | 102 | 3 | 5248 | 42 | Gng11,Homer1,Grm3 | | 4.738e-02 | -3.05 | Glutamatergic synapse | KEGG pathways | ko04724 | 102 | 3 | 5248 | 42 | Homer1,Grm3,Gng11 | | 4.745e-02 | -3.05 | DELPUECH\_FOXO3\_TARGETS\_UP | MSigDB lists | DELPUECH\_FOXO3\_TARGETS\_UP | 59 | 2 | 12187 | 72 | Timp2,Tgfa | | 4.745e-02 | -3.05 | KRAS.300\_UP.V1\_DN | MSigDB lists | KRAS.300\_UP.V1\_DN | 59 | 2 | 12187 | 72 | Kcnmb2,Ephb6 | | 4.781e-02 | -3.04 | mannitol degradation II | BIOCYC pathways | MOUSE\_PWY-3861 | 4 | 1 | 823 | 10 | Hkdc1 | | 4.781e-02 | -3.04 | arginine biosynthesis IV | BIOCYC pathways | MOUSE\_ARGININE-SYN4-PWY | 4 | 1 | 823 | 10 | Glud1 | | 4.793e-02 | -3.04 | GO\_ORGANONITROGEN\_COMPOUND\_BIOSYNTHETIC\_PROCESS | MSigDB lists | GO\_ORGANONITROGEN\_COMPOUND\_BIOSYNTHETIC\_PROCESS | 809 | 9 | 12187 | 72 | Agrn,Srm,Qdpr,Ndst4,Hs3st1,Aars,Glud1,Ugt8a,Amd1 | | 4.808e-02 | -3.03 | Fatty acid, triacylglycerol, and ketone body metabolism | REACTOME pathways | R-MMU-535734 | 123 | 3 | 6297 | 42 | Scd2,Arntl,Hsd17b12 | | 4.814e-02 | -3.03 | GO\_REGULATION\_OF\_CELL\_SUBSTRATE\_ADHESION | MSigDB lists | GO\_REGULATION\_OF\_CELL\_SUBSTRATE\_ADHESION | 138 | 3 | 12187 | 72 | Col1a1,Kdr,Hsd17b12 | | 4.814e-02 | -3.03 | GSE32986\_UNSTIM\_VS\_GMCSF\_AND\_CURDLAN\_HIGHDOSE\_STIM\_DC\_DN | MSigDB lists | GSE32986\_UNSTIM\_VS\_GMCSF\_AND\_CURDLAN\_HIGHDOSE\_STIM\_DC\_DN | 138 | 3 | 12187 | 72 | Hkdc1,Tle4,Chmp7 | | 4.814e-02 | -3.03 | ESC\_V6.5\_UP\_LATE.V1\_UP | MSigDB lists | ESC\_V6.5\_UP\_LATE.V1\_UP | 138 | 3 | 12187 | 72 | Foxq1,Lhx1,H1f0 | | 4.814e-02 | -3.03 | GO\_EPIDERMIS\_DEVELOPMENT | MSigDB lists | GO\_EPIDERMIS\_DEVELOPMENT | 138 | 3 | 12187 | 72 | Inhba,Foxq1,Aars | | 4.814e-02 | -3.03 | GSE6269\_FLU\_VS\_STAPH\_AUREUS\_INF\_PBMC\_DN | MSigDB lists | GSE6269\_FLU\_VS\_STAPH\_AUREUS\_INF\_PBMC\_DN | 138 | 3 | 12187 | 72 | Timp2,H1f0,Egr1 | | 4.825e-02 | -3.03 | LIM | smart domains | SM00132 | 59 | 2 | 7188 | 43 | Lims2,Lhx1 | | 4.829e-02 | -3.03 | GO\_RESPONSE\_TO\_DRUG | MSigDB lists | GO\_RESPONSE\_TO\_DRUG | 336 | 5 | 12187 | 72 | Inhba,Timp2,Col1a1,Crh,Qdpr | | 4.829e-02 | -3.03 | MODULE\_64 | MSigDB lists | MODULE\_64 | 336 | 5 | 12187 | 72 | Inhba,Tgfa,Gng11,Ephb6,Rgs4 | | 4.832e-02 | -3.03 | HOMEOBOX\_1 | prosite domains | PS00027 | 52 | 2 | 8845 | 60 | Gbx1,Lhx1 | | 4.835e-02 | -3.03 | Homeodomain | pfam domains | PF00046 | 63 | 2 | 12881 | 72 | Gbx1,Lhx1 | | 4.847e-02 | -3.03 | monovalent inorganic cation transport | biological process | GO:0015672 | 235 | 4 | 13711 | 80 | Lrrc55,P2ry12,Kcnmb2,Atp5b | | 4.871e-02 | -3.02 | NABA\_MATRISOME | MSigDB lists | NABA\_MATRISOME | 565 | 7 | 12187 | 72 | Bdnf,Agrn,Cbln2,Tgfa,Inhba,Timp2,Col1a1 | | 4.874e-02 | -3.02 | cell population proliferation | biological process | GO:0008283 | 453 | 6 | 13711 | 80 | Fabp7,Egr1,Ephb6,Tgfa,Ltk,Nr4a3 | | 4.880e-02 | -3.02 | GO\_NEGATIVE\_REGULATION\_OF\_TRANSPORT | MSigDB lists | GO\_NEGATIVE\_REGULATION\_OF\_TRANSPORT | 337 | 5 | 12187 | 72 | Ubqln2,Crh,Lrrtm2,P2ry12,Inhba | | 4.880e-02 | -3.02 | GO\_TAXIS | MSigDB lists | GO\_TAXIS | 337 | 5 | 12187 | 72 | Bdnf,Ncam1,Lhx1,Nr4a3,Gbx1 | | 4.887e-02 | -3.02 | KRIGE\_RESPONSE\_TO\_TOSEDOSTAT\_24HR\_DN | MSigDB lists | KRIGE\_RESPONSE\_TO\_TOSEDOSTAT\_24HR\_DN | 812 | 9 | 12187 | 72 | Tle4,Car2,Sorl1,Amd1,Egr1,Fjx1,Srm,Qdpr,Homer1 | | 4.887e-02 | -3.02 | ATTCTTT\_MIR186 | MSigDB lists | ATTCTTT\_MIR186 | 233 | 4 | 12187 | 72 | Plp1,Qk,Ubqln2,Map4 | | 4.890e-02 | -3.02 | ADDYA\_ERYTHROID\_DIFFERENTIATION\_BY\_HEMIN | MSigDB lists | ADDYA\_ERYTHROID\_DIFFERENTIATION\_BY\_HEMIN | 60 | 2 | 12187 | 72 | Hba-a2,Egr1 | | 4.890e-02 | -3.02 | GO\_CELL\_CELL\_JUNCTION\_ASSEMBLY | MSigDB lists | GO\_CELL\_CELL\_JUNCTION\_ASSEMBLY | 60 | 2 | 12187 | 72 | Ugt8a,Cldn5 | | 4.890e-02 | -3.02 | GO\_POSITIVE\_REGULATION\_OF\_CAMP\_METABOLIC\_PROCESS | MSigDB lists | GO\_POSITIVE\_REGULATION\_OF\_CAMP\_METABOLIC\_PROCESS | 60 | 2 | 12187 | 72 | Crh,Timp2 | | 4.890e-02 | -3.02 | REACTOME\_PLATELET\_HOMEOSTASIS | MSigDB lists | REACTOME\_PLATELET\_HOMEOSTASIS | 60 | 2 | 12187 | 72 | Kcnmb2,Gng11 | | 4.900e-02 | -3.02 | organonitrogen compound biosynthetic process | biological process | GO:1901566 | 950 | 10 | 13711 | 80 | Inhba,Aars,Atp5b,Glud1,Amd1,Tmem165,Srm,Hs3st1,Ndst4,Qdpr | | 4.900e-02 | -3.02 | GO\_NEGATIVE\_REGULATION\_OF\_SECRETION | MSigDB lists | GO\_NEGATIVE\_REGULATION\_OF\_SECRETION | 139 | 3 | 12187 | 72 | Crh,Inhba,P2ry12 | | 4.900e-02 | -3.02 | GSE24634\_NAIVE\_CD4\_TCELL\_VS\_DAY10\_IL4\_CONV\_TREG\_UP | MSigDB lists | GSE24634\_NAIVE\_CD4\_TCELL\_VS\_DAY10\_IL4\_CONV\_TREG\_UP | 139 | 3 | 12187 | 72 | Tgfa,Dgkz,Ncam1 | | 4.900e-02 | -3.02 | GO\_PROTEIN\_TYROSINE\_KINASE\_ACTIVITY | MSigDB lists | GO\_PROTEIN\_TYROSINE\_KINASE\_ACTIVITY | 139 | 3 | 12187 | 72 | Ltk,Ephb6,Kdr | | 4.900e-02 | -3.02 | GSE3982\_BCELL\_VS\_BASOPHIL\_UP | MSigDB lists | GSE3982\_BCELL\_VS\_BASOPHIL\_UP | 139 | 3 | 12187 | 72 | Chmp7,Hs3st1,Rgs4 | | 4.900e-02 | -3.02 | GSE43700\_UNTREATED\_VS\_IL10\_TREATED\_PBMC\_UP | MSigDB lists | GSE43700\_UNTREATED\_VS\_IL10\_TREATED\_PBMC\_UP | 139 | 3 | 12187 | 72 | Fjx1,Homer1,Egr1 | | 4.914e-02 | -3.01 | DEAD/DEAH\_box\_helicase\_dom | interpro domains | IPR011545 | 62 | 2 | 13788 | 79 | Ddx17,Ddx56 | | 4.917e-02 | -3.01 | positive regulation of cellular component biogenesis | biological process | GO:0044089 | 454 | 6 | 13711 | 80 | Lrrtm2,Bdnf,Kdr,Cbln2,Agrn,P2ry12 | | 4.921e-02 | -3.01 | Laminin\_G\_1 | pfam domains | PF00054 | 9 | 1 | 12881 | 72 | Agrn | | 4.921e-02 | -3.01 | FA\_desaturase | pfam domains | PF00487 | 9 | 1 | 12881 | 72 | Scd2 | | 4.931e-02 | -3.01 | neuromuscular process controlling balance | biological process | GO:0050885 | 61 | 2 | 13711 | 80 | Nr4a3,Aars | | 4.931e-02 | -3.01 | negative regulation of MAP kinase activity | biological process | GO:0043407 | 61 | 2 | 13711 | 80 | Sorl1,Spry2 | | 4.937e-02 | -3.01 | Tle4 (transducin-like enhancer of split 4) | protein interactions | 21888 | 7 | 1 | 6802 | 49 | Tle4 | | 4.937e-02 | -3.01 | Vegfa (vascular endothelial growth factor A) | protein interactions | 22339 | 7 | 1 | 6802 | 49 | Kdr | | 4.937e-02 | -3.01 | Wiz (widely-interspaced zinc finger motifs) | protein interactions | 22404 | 7 | 1 | 6802 | 49 | Zfp518b | | 4.937e-02 | -3.01 | Grid2 (glutamate receptor, ionotropic, delta 2) | protein interactions | 14804 | 7 | 1 | 6802 | 49 | Homer1 | | 4.937e-02 | -3.01 | Dscam (DS cell adhesion molecule) | protein interactions | 13508 | 7 | 1 | 6802 | 49 | Ncam1 | | 4.937e-02 | -3.01 | Dvl3 (dishevelled segment polarity protein 3) | protein interactions | 13544 | 7 | 1 | 6802 | 49 | Agrn | | 4.937e-02 | -3.01 | TRIM29 (tripartite motif containing 29) | protein interactions | 23650 | 7 | 1 | 6802 | 49 | H1f0 | | 4.937e-02 | -3.01 | Dbp (D site albumin promoter binding protein) | protein interactions | 13170 | 7 | 1 | 6802 | 49 | Arntl | | 4.937e-02 | -3.01 | HIF1A (hypoxia inducible factor 1 subunit alpha) | protein interactions | 3091 | 7 | 1 | 6802 | 49 | Arntl | | 4.945e-02 | -3.01 | cellular response to hormone stimulus | biological process | GO:0032870 | 342 | 5 | 13711 | 80 | Nr4a3,Crh,Car2,Ddx17,Inhba | | 4.960e-02 | -3.00 | vasculature development | biological process | GO:0001944 | 455 | 6 | 13711 | 80 | Tgfa,Kdr,Egr1,Col1a1,Qk,Atp5b | | 4.984e-02 | -3.00 | FATTYACIDBP | prints domains | PR00178 | 5 | 1 | 2951 | 30 | Fabp7 | | 4.984e-02 | -3.00 | Cocaine addiction | KEGG pathways | mmu05030 | 45 | 2 | 5248 | 42 | Grm3,Bdnf | | 4.984e-02 | -3.00 | Cocaine addiction | KEGG pathways | ko05030 | 45 | 2 | 5248 | 42 | Bdnf,Grm3 | | 4.986e-02 | -3.00 | GSE13484\_3H\_UNSTIM\_VS\_YF17D\_VACCINE\_STIM\_PBMC\_DN | MSigDB lists | GSE13484\_3H\_UNSTIM\_VS\_YF17D\_VACCINE\_STIM\_PBMC\_DN | 140 | 3 | 12187 | 72 | Qk,Agrn,Inhba | | 4.986e-02 | -3.00 | GSE37605\_TREG\_VS\_TCONV\_NOD\_FOXP3\_FUSION\_GFP\_DN | MSigDB lists | GSE37605\_TREG\_VS\_TCONV\_NOD\_FOXP3\_FUSION\_GFP\_DN | 140 | 3 | 12187 | 72 | Homer1,Gng11,Nr4a3 | | 4.986e-02 | -3.00 | NAGASHIMA\_NRG1\_SIGNALING\_UP | MSigDB lists | NAGASHIMA\_NRG1\_SIGNALING\_UP | 140 | 3 | 12187 | 72 | Nr4a3,Homer1,Egr1 | | 4.986e-02 | -3.00 | VECCHI\_GASTRIC\_CANCER\_ADVANCED\_VS\_EARLY\_UP | MSigDB lists | VECCHI\_GASTRIC\_CANCER\_ADVANCED\_VS\_EARLY\_UP | 140 | 3 | 12187 | 72 | Ntm,Inhba,Qk | | 4.986e-02 | -3.00 | TGFB\_UP.V1\_DN | MSigDB lists | TGFB\_UP.V1\_DN | 140 | 3 | 12187 | 72 | Ugt8a,Hs3st1,Spry2 | | 4.986e-02 | -3.00 | GSE26928\_NAIVE\_VS\_EFF\_MEMORY\_CD4\_TCELL\_UP | MSigDB lists | GSE26928\_NAIVE\_VS\_EFF\_MEMORY\_CD4\_TCELL\_UP | 140 | 3 | 12187 | 72 | Ntm,Zfp518b,Fbxl17 | | 4.986e-02 | -3.00 | GACTGTT\_MIR212\_MIR132 | MSigDB lists | GACTGTT\_MIR212\_MIR132 | 140 | 3 | 12187 | 72 | Amd1,Pom121,Lrrc55 |
