## Supplementary material for "Repeated LPS induces training and tolerance of microglial responses across brain regions": FileS3: homerResults.html

LPSDownregulated\_cluster\_genes\_output// - Homer de novo Motif Results


### Homer *de novo* Motif Results (LPSDownregulated\_cluster\_genes\_output//)

Known Motif Enrichment Results  
Gene Ontology Enrichment Results  
If Homer is having trouble matching a motif to a known motif, try copy/pasting the matrix file into
STAMP  
More information on motif finding results: HOMER
| Description of Results
| Tips
  
Total target sequences = 80  
Total background sequences = 13506  
\* - possible false positive  

|  |  |  |  |  |  |  |  |  |
| --- | --- | --- | --- | --- | --- | --- | --- | --- |
| Rank | Motif | P-value | log P-pvalue | % of Targets | % of Background | STD(Bg STD) | Best Match/Details | Motif File |
| 1 \* | A C T G A C T G C A G T A C T G C G T A C G T A A C T G A C T G C T G A C G T A C G T A T G C A | 1e-11 | -2.596e+01 | 12.50% | 0.42% | 352.1bp (345.2bp) | RAR:RXR(NR),DR0/ES-RAR-ChIP-Seq(GSE56893)/Homer(0.633) More Information \| Similar Motifs Found | motif file (matrix) |
| 2 \* | A G C T C G A T G C A T G T C A G C T A C G T A A C T G C A T G G T C A T G A C G T A C C G A T | 1e-10 | -2.435e+01 | 20.00% | 2.18% | 221.3bp (319.3bp) | Hoxa13(Homeobox)/ChickenMSG-Hoxa13.Flag-ChIP-Seq(GSE86088)/Homer(0.606) More Information \| Similar Motifs Found | motif file (matrix) |
| 3 \* | C T G A C G T A C G T A A C T G C T G A C T A G C G T A A T G C C G A T T G C A | 1e-10 | -2.429e+01 | 26.25% | 4.35% | 356.1bp (323.6bp) | PRDM4/MA1647.1/Jaspar(0.593) More Information \| Similar Motifs Found | motif file (matrix) |
| 4 \* | A C G T C G A T A C T G A G T C C G T A A T C G G T C A G T A C C G T A C G T A C G T A A G C T | 1e-10 | -2.319e+01 | 11.25% | 0.40% | 279.5bp (338.8bp) | Oct4:Sox17(POU,Homeobox,HMG)/F9-Sox17-ChIP-Seq(GSE44553)/Homer(0.720) More Information \| Similar Motifs Found | motif file (matrix) |
| 5 \* | A C G T A G C T A G T C A G T C A C G T A C T G A G T C T A G C A C T G C T A G A C T G A C G T | 1e-9 | -2.232e+01 | 8.75% | 0.16% | 220.6bp (444.7bp) | PB0205.1\_Zic1\_2/Jaspar(0.719) More Information \| Similar Motifs Found | motif file (matrix) |
| 6 \* | G T A C C G T A G T A C A C T G A C T G G T C A A C T G G C A T A C T G C T A G | 1e-9 | -2.183e+01 | 22.50% | 3.48% | 328.0bp (338.7bp) | Bapx1(Homeobox)/VertebralCol-Bapx1-ChIP-Seq(GSE36672)/Homer(0.656) More Information \| Similar Motifs Found | motif file (matrix) |
| 7 \* | A C G T A G T C G T A C T A C G C G T A C T G A A C G T A C G T C T G A A C G T | 1e-9 | -2.154e+01 | 8.75% | 0.18% | 279.3bp (277.1bp) | MF0010.1\_Homeobox\_class/Jaspar(0.679) More Information \| Similar Motifs Found | motif file (matrix) |
| 8 \* | C T G A A C G T T G C A A C G T A G T C G T A C A C G T G C A T | 1e-9 | -2.083e+01 | 51.25% | 20.29% | 324.2bp (316.4bp) | SD0003.1\_at\_AC\_acceptor/Jaspar(0.958) More Information \| Similar Motifs Found | motif file (matrix) |
| 9 \* | T C G A A G T C A C T G A C T G C G T A A C T G C G T A A G T C C G T A A C G T A G T C A C T G | 1e-8 | -2.061e+01 | 5.00% | 0.01% | 278.8bp (0.0bp) | PB0126.1\_Gata5\_2/Jaspar(0.591) More Information \| Similar Motifs Found | motif file (matrix) |
| 10 \* | C G A T T C A G A C G T A C G T G T C A A C G T C G T A G T C A A G C T C G A T C G T A C T G A | 1e-8 | -2.056e+01 | 12.50% | 0.77% | 261.2bp (298.3bp) | PB0001.1\_Arid3a\_1/Jaspar(0.829) More Information \| Similar Motifs Found | motif file (matrix) |
| 11 \* | A C T G A G T C C T A G C T A G C T A G C G T A G T C A C T A G G A T C A G T C | 1e-8 | -2.044e+01 | 20.00% | 2.88% | 283.3bp (320.7bp) | TFDP1/MA1122.1/Jaspar(0.858) More Information \| Similar Motifs Found | motif file (matrix) |
| 12 \* | A C T G A G T C C G T A A C T G A C G T A C T G A C G T A C T G A T G C A G T C A G C T A T G C | 1e-8 | -1.985e+01 | 7.50% | 0.12% | 385.9bp (315.9bp) | PB0091.1\_Zbtb3\_1/Jaspar(0.674) More Information \| Similar Motifs Found | motif file (matrix) |
| 13 \* | C T G A G C A T G C T A G A T C G A C T C G A T C A G T G C A T G A C T G T A C | 1e-8 | -1.969e+01 | 22.50% | 4.01% | 337.0bp (320.5bp) | PRDM1/MA0508.3/Jaspar(0.731) More Information \| Similar Motifs Found | motif file (matrix) |
| 14 \* | A G T C A C G T A C T G C G T A A G T C C G T A G T C A A C G T A C G T A C G T A C G T C G T A | 1e-8 | -1.927e+01 | 7.50% | 0.14% | 333.8bp (272.4bp) | Meis1(Homeobox)/MastCells-Meis1-ChIP-Seq(GSE48085)/Homer(0.713) More Information \| Similar Motifs Found | motif file (matrix) |
| 15 \* | C T A G A T C G A C G T A G T C A G T C G C T A A G T C C G T A C G T A C G T A | 1e-8 | -1.873e+01 | 13.75% | 1.25% | 387.8bp (347.2bp) | ZNF354C/MA0130.1/Jaspar(0.669) More Information \| Similar Motifs Found | motif file (matrix) |
| 16 \* | C G T A A C G T A C T G A G T C C G T A C G A T A C T G A G T C A C G T A G T C A G T C A G T C | 1e-8 | -1.867e+01 | 6.25% | 0.06% | 263.9bp (252.6bp) | POL013.1\_MED-1/Jaspar(0.688) More Information \| Similar Motifs Found | motif file (matrix) |
| 17 \* | C A G T C T G A A C G T A T C G T G C A A G C T A G C T G T A C G A T C A G T C A G C T A G C T | 1e-7 | -1.758e+01 | 10.00% | 0.55% | 262.5bp (316.9bp) | PB0059.1\_Six6\_1/Jaspar(0.669) More Information \| Similar Motifs Found | motif file (matrix) |
| 18 \* | A T C G A C T G A G T C C A T G G T A C C T A G G T A C A G T C A T C G T C A G A T C G C A G T | 1e-7 | -1.758e+01 | 13.75% | 1.40% | 188.3bp (320.2bp) | POL006.1\_BREu/Jaspar(0.643) More Information \| Similar Motifs Found | motif file (matrix) |
| 19 \* | A T G C A C G T A T G C C A G T A C T G C A T G T G A C C T G A C T G A T A C G T C G A C G T A | 1e-7 | -1.757e+01 | 12.50% | 1.08% | 354.7bp (342.7bp) | NF1-halfsite(CTF)/LNCaP-NF1-ChIP-Seq(Unpublished)/Homer(0.609) More Information \| Similar Motifs Found | motif file (matrix) |
| 20 \* | A G T C C G T A C A T G A T C G A C T G A G T C A G T C G T C A A G C T C G T A A C G T C G A T | 1e-7 | -1.740e+01 | 7.50% | 0.20% | 437.5bp (297.0bp) | SF1(NR)/H295R-Nr5a1-ChIP-Seq(GSE44220)/Homer(0.655) More Information \| Similar Motifs Found | motif file (matrix) |
| 21 \* | C G T A T G C A G T C A T G C A C T G A C A T G A T C G G T A C C G T A T G C A A G T C A G C T | 1e-7 | -1.643e+01 | 16.25% | 2.40% | 305.8bp (350.6bp) | PB0033.1\_Irf3\_1/Jaspar(0.704) More Information \| Similar Motifs Found | motif file (matrix) |
| 22 \* | A T C G C G A T A G C T G C T A C G T A A C G T A C G T C G T A C G T A G T A C C G T A A T C G | 1e-7 | -1.622e+01 | 6.25% | 0.12% | 352.1bp (281.3bp) | RAX/MA0718.1/Jaspar(0.720) More Information \| Similar Motifs Found | motif file (matrix) |
| 23 \* | T C A G A G T C A C G T A C G T A C G T A C T G A G T C C G T A C G A T A C G T | 1e-6 | -1.556e+01 | 20.00% | 4.12% | 239.1bp (323.7bp) | POU5F1/MA1115.1/Jaspar(0.818) More Information \| Similar Motifs Found | motif file (matrix) |
| 24 \* | C T G A C G A T C T A G C A G T G A C T G C A T A C T G T C G A A G T C C G A T | 1e-6 | -1.458e+01 | 25.00% | 6.95% | 240.2bp (313.4bp) | Nr2e1/MA0676.1/Jaspar(0.708) More Information \| Similar Motifs Found | motif file (matrix) |
| 25 \* | A T G C A C T G G T A C A C T G A T G C C G A T A G T C A C G T G T A C G T A C | 1e-6 | -1.443e+01 | 15.00% | 2.39% | 226.5bp (292.0bp) | PB0008.1\_E2F2\_1/Jaspar(0.655) More Information \| Similar Motifs Found | motif file (matrix) |
| 26 \* | C T A G C G T A T A C G G T A C A C T G A T C G C T G A A C T G T A G C T A G C A C G T A C T G | 1e-5 | -1.375e+01 | 5.00% | 0.08% | 140.9bp (254.0bp) | POL013.1\_MED-1/Jaspar(0.680) More Information \| Similar Motifs Found | motif file (matrix) |
| 27 \* | C A G T A G T C A G C T G T A C G C A T C T A G T G C A A G T C G A C T C T G A | 1e-5 | -1.304e+01 | 30.00% | 10.70% | 270.9bp (340.9bp) | ZNF768(Zf)/Rajj-ZNF768-ChIP-Seq(GSE111879)/Homer(0.702) More Information \| Similar Motifs Found | motif file (matrix) |
| 28 \* | A C T G A G T C A C G T C G T A A C G T A C T G A C T G A T C G A C T G A T G C | 1e-5 | -1.257e+01 | 8.75% | 0.76% | 369.7bp (352.0bp) | YY2/MA0748.2/Jaspar(0.695) More Information \| Similar Motifs Found | motif file (matrix) |
| 29 \* | A C G T C T A G A G C T A C G T G T C A T A G C T G A C C G T A | 1e-5 | -1.238e+01 | 40.00% | 18.10% | 314.8bp (333.2bp) | PB0109.1\_Bbx\_2/Jaspar(0.714) More Information \| Similar Motifs Found | motif file (matrix) |
| 30 \* | C T A G G A T C T C A G T A G C A C T G T A G C T A C G A G C T A T C G A G C T | 1e-5 | -1.234e+01 | 17.50% | 4.06% | 339.7bp (324.6bp) | PB0151.1\_Myf6\_2/Jaspar(0.710) More Information \| Similar Motifs Found | motif file (matrix) |
| 31 \* | A C G T C G T A A G T C C G T A A C T G A G T C C T G A A C G T A G T C A C T G | 1e-4 | -1.144e+01 | 3.75% | 0.04% | 96.4bp (277.3bp) | PH0158.1\_Rhox11\_2/Jaspar(0.630) More Information \| Similar Motifs Found | motif file (matrix) |
| 32 \* | C G T A A C T G G T C A A C T G A G T C A G T C G T A C G T C A A G T C C T G A C G T A A T C G | 1e-4 | -1.048e+01 | 7.50% | 0.71% | 365.8bp (299.4bp) | ZBTB6/MA1581.1/Jaspar(0.632) More Information \| Similar Motifs Found | motif file (matrix) |
| 33 \* | G T C A G A T C G A T C A G T C G T A C G T A C A T G C G A T C G T A C A G T C A G T C T G A C | 1e-2 | -4.647e+00 | 11.25% | 4.46% | 321.1bp (301.8bp) | PB0097.1\_Zfp281\_1/Jaspar(0.895) More Information \| Similar Motifs Found | motif file (matrix) |
