## Supplementary material for "Repeated LPS induces training and tolerance of microglial responses across brain regions": FileS3: motif4.similar.html

### Information for motif4

A
C
G
T
C
G
A
T
A
C
T
G
A
G
T
C
C
G
T
A
A
T
C
G
G
T
C
A
G
T
A
C
C
G
T
A
C
G
T
A
C
G
T
A
A
G
C
T
  
Reverse Opposite:  

C
T
G
A
C
G
A
T
C
G
A
T
A
C
G
T
A
C
T
G
A
C
G
T
A
T
G
C
A
C
G
T
C
T
A
G
G
T
A
C
C
G
T
A
C
G
T
A
  

|  |  |
| --- | --- |
| p-value: | 1e-10 |
| log p-value: | -2.319e+01 |
| Information Content per bp: | 1.803 |
| Number of Target Sequences with motif | 9.0 |
| Percentage of Target Sequences with motif | 11.25% |
| Number of Background Sequences with motif | 53.8 |
| Percentage of Background Sequences with motif | 0.40% |
| Average Position of motif in Targets | 577.8 +/- 279.5bp |
| Average Position of motif in Background | 440.4 +/- 338.8bp |
| Strand Bias (log2 ratio + to - strand density) | -0.3 |
| Multiplicity (# of sites on avg that occur together) | 1.00 |
| Motif File: | file (matrix) reverse opposite |

#### Similar de novo motifs found

|  |  |  |  |  |  |  |  |
| --- | --- | --- | --- | --- | --- | --- | --- |
| Rank | Match Score | Redundant Motif | P-value | log P-value | % of Targets | % of Background | Motif file |
| 1 | 0.617 | A C G T C A G T A C G T A C T G T G A C C T G A G T A C G T A C A T G C C G T A | 1e-9 | -21.588070 | 31.25% | 7.40% | motif file (matrix) |
