## Supplementary material for "Repeated LPS induces training and tolerance of microglial responses across brain regions": FileS3: motif7.similar.html

### Information for motif7

A
C
G
T
A
G
T
C
G
T
A
C
T
A
C
G
C
G
T
A
C
T
G
A
A
C
G
T
A
C
G
T
C
T
G
A
A
C
G
T
  
Reverse Opposite:  

C
G
T
A
A
G
C
T
G
T
C
A
C
G
T
A
A
G
C
T
A
C
G
T
A
T
G
C
A
C
T
G
A
C
T
G
C
G
T
A
  

|  |  |
| --- | --- |
| p-value: | 1e-9 |
| log p-value: | -2.154e+01 |
| Information Content per bp: | 1.877 |
| Number of Target Sequences with motif | 7.0 |
| Percentage of Target Sequences with motif | 8.75% |
| Number of Background Sequences with motif | 24.4 |
| Percentage of Background Sequences with motif | 0.18% |
| Average Position of motif in Targets | 595.1 +/- 279.3bp |
| Average Position of motif in Background | 498.8 +/- 277.1bp |
| Strand Bias (log2 ratio + to - strand density) | 1.3 |
| Multiplicity (# of sites on avg that occur together) | 1.00 |
| Motif File: | file (matrix) reverse opposite |

#### Similar de novo motifs found

|  |  |  |  |  |  |  |  |
| --- | --- | --- | --- | --- | --- | --- | --- |
| Rank | Match Score | Redundant Motif | P-value | log P-value | % of Targets | % of Background | Motif file |
