## Supplementary material for "Repeated LPS induces training and tolerance of microglial responses across brain regions": FileS3: motif8.similar.html

### Information for motif8

C
T
G
A
A
C
G
T
T
G
C
A
A
C
G
T
A
G
T
C
G
T
A
C
A
C
G
T
G
C
A
T
  
Reverse Opposite:  

C
G
T
A
C
G
T
A
A
C
T
G
T
C
A
G
G
T
C
A
A
C
G
T
C
G
T
A
A
G
C
T
  

|  |  |
| --- | --- |
| p-value: | 1e-9 |
| log p-value: | -2.083e+01 |
| Information Content per bp: | 1.834 |
| Number of Target Sequences with motif | 41.0 |
| Percentage of Target Sequences with motif | 51.25% |
| Number of Background Sequences with motif | 2738.9 |
| Percentage of Background Sequences with motif | 20.29% |
| Average Position of motif in Targets | 413.0 +/- 324.2bp |
| Average Position of motif in Background | 482.9 +/- 316.4bp |
| Strand Bias (log2 ratio + to - strand density) | 0.7 |
| Multiplicity (# of sites on avg that occur together) | 1.20 |
| Motif File: | file (matrix) reverse opposite |

#### Similar de novo motifs found

|  |  |  |  |  |  |  |  |
| --- | --- | --- | --- | --- | --- | --- | --- |
| Rank | Match Score | Redundant Motif | P-value | log P-value | % of Targets | % of Background | Motif file |
