## Supplementary material for "Repeated LPS induces training and tolerance of microglial responses across brain regions": FileS3: motif10.similar.html

### Information for motif10

C
G
A
T
T
C
A
G
A
C
G
T
A
C
G
T
G
T
C
A
A
C
G
T
C
G
T
A
G
T
C
A
A
G
C
T
C
G
A
T
C
G
T
A
C
T
G
A
  
Reverse Opposite:  

A
G
C
T
A
C
G
T
C
G
T
A
C
T
G
A
A
C
G
T
C
G
A
T
C
G
T
A
C
A
G
T
C
G
T
A
C
G
T
A
A
G
T
C
G
C
T
A
  

|  |  |
| --- | --- |
| p-value: | 1e-8 |
| log p-value: | -2.056e+01 |
| Information Content per bp: | 1.783 |
| Number of Target Sequences with motif | 10.0 |
| Percentage of Target Sequences with motif | 12.50% |
| Number of Background Sequences with motif | 104.4 |
| Percentage of Background Sequences with motif | 0.77% |
| Average Position of motif in Targets | 557.7 +/- 261.2bp |
| Average Position of motif in Background | 408.3 +/- 298.3bp |
| Strand Bias (log2 ratio + to - strand density) | 0.5 |
| Multiplicity (# of sites on avg that occur together) | 1.10 |
| Motif File: | file (matrix) reverse opposite |

#### Similar de novo motifs found

|  |  |  |  |  |  |  |  |
| --- | --- | --- | --- | --- | --- | --- | --- |
| Rank | Match Score | Redundant Motif | P-value | log P-value | % of Targets | % of Background | Motif file |
| 1 | 0.604 | A C G T C G A T A C G T A C G T A C G T C G A T A C G T C T A G G T C A C G T A C G T A G T C A | 1e-6 | -14.760110 | 8.75% | 0.54% | motif file (matrix) |
