## Supplementary material for "Repeated LPS induces training and tolerance of microglial responses across brain regions": FileS3: motif13.similar.html

### Information for motif13

C
T
G
A
G
C
A
T
G
C
T
A
G
A
T
C
G
A
C
T
C
G
A
T
C
A
G
T
G
C
A
T
G
A
C
T
G
T
A
C
  
Reverse Opposite:  

C
A
T
G
C
T
G
A
C
G
T
A
G
T
C
A
G
C
T
A
C
T
G
A
C
T
A
G
C
G
A
T
C
G
T
A
G
A
C
T
  

|  |  |
| --- | --- |
| p-value: | 1e-8 |
| log p-value: | -1.969e+01 |
| Information Content per bp: | 1.535 |
| Number of Target Sequences with motif | 18.0 |
| Percentage of Target Sequences with motif | 22.50% |
| Number of Background Sequences with motif | 540.8 |
| Percentage of Background Sequences with motif | 4.01% |
| Average Position of motif in Targets | 408.1 +/- 337.0bp |
| Average Position of motif in Background | 472.3 +/- 320.5bp |
| Strand Bias (log2 ratio + to - strand density) | 0.2 |
| Multiplicity (# of sites on avg that occur together) | 1.06 |
| Motif File: | file (matrix) reverse opposite |

#### Similar de novo motifs found

|  |  |  |  |  |  |  |  |
| --- | --- | --- | --- | --- | --- | --- | --- |
| Rank | Match Score | Redundant Motif | P-value | log P-value | % of Targets | % of Background | Motif file |
| 1 | 0.614 | A C G T A C T G A C T G C G T A C G T A A G T C C G T A A C T G | 1e-3 | -8.307025 | 13.75% | 3.81% | motif file (matrix) |
