## Supplementary material for "Repeated LPS induces training and tolerance of microglial responses across brain regions": FileS3: motif16.similar.html

### Information for motif16

C
G
T
A
A
C
G
T
A
C
T
G
A
G
T
C
C
G
T
A
C
G
A
T
A
C
T
G
A
G
T
C
A
C
G
T
A
G
T
C
A
G
T
C
A
G
T
C
  
Reverse Opposite:  

A
C
T
G
A
C
T
G
A
C
T
G
C
G
T
A
A
C
T
G
A
G
T
C
G
C
T
A
C
G
A
T
A
C
T
G
A
G
T
C
C
G
T
A
C
G
A
T
  

|  |  |
| --- | --- |
| p-value: | 1e-8 |
| log p-value: | -1.867e+01 |
| Information Content per bp: | 1.907 |
| Number of Target Sequences with motif | 5.0 |
| Percentage of Target Sequences with motif | 6.25% |
| Number of Background Sequences with motif | 8.2 |
| Percentage of Background Sequences with motif | 0.06% |
| Average Position of motif in Targets | 746.4 +/- 263.9bp |
| Average Position of motif in Background | 650.6 +/- 252.6bp |
| Strand Bias (log2 ratio + to - strand density) | -0.6 |
| Multiplicity (# of sites on avg that occur together) | 1.00 |
| Motif File: | file (matrix) reverse opposite |

#### Similar de novo motifs found

|  |  |  |  |  |  |  |  |
| --- | --- | --- | --- | --- | --- | --- | --- |
| Rank | Match Score | Redundant Motif | P-value | log P-value | % of Targets | % of Background | Motif file |
