## Supplementary material for "Repeated LPS induces training and tolerance of microglial responses across brain regions": FileS3: motif17.similar.html

### Information for motif17

C
A
G
T
C
T
G
A
A
C
G
T
A
T
C
G
T
G
C
A
A
G
C
T
A
G
C
T
G
T
A
C
G
A
T
C
A
G
T
C
A
G
C
T
A
G
C
T
  
Reverse Opposite:  

C
T
G
A
C
T
G
A
C
T
A
G
C
T
A
G
A
C
T
G
T
C
G
A
C
T
G
A
A
C
G
T
T
A
G
C
G
T
C
A
G
A
C
T
G
C
T
A
  

|  |  |
| --- | --- |
| p-value: | 1e-7 |
| log p-value: | -1.758e+01 |
| Information Content per bp: | 1.742 |
| Number of Target Sequences with motif | 8.0 |
| Percentage of Target Sequences with motif | 10.00% |
| Number of Background Sequences with motif | 73.6 |
| Percentage of Background Sequences with motif | 0.55% |
| Average Position of motif in Targets | 415.6 +/- 262.5bp |
| Average Position of motif in Background | 461.5 +/- 316.9bp |
| Strand Bias (log2 ratio + to - strand density) | 0.0 |
| Multiplicity (# of sites on avg that occur together) | 1.25 |
| Motif File: | file (matrix) reverse opposite |

#### Similar de novo motifs found

|  |  |  |  |  |  |  |  |
| --- | --- | --- | --- | --- | --- | --- | --- |
| Rank | Match Score | Redundant Motif | P-value | log P-value | % of Targets | % of Background | Motif file |
| 1 | 0.883 | C G T A A C T G A C T G A C T G T C G A C G T A A C G T A G T C T C G A A C G T | 1e-5 | -11.821570 | 7.50% | 0.55% | motif file (matrix) |
