## Supplementary material for "Repeated LPS induces training and tolerance of microglial responses across brain regions": FileS3: motif18.similar.html

### Information for motif18

A
T
C
G
A
C
T
G
A
G
T
C
C
A
T
G
G
T
A
C
C
T
A
G
G
T
A
C
A
G
T
C
A
T
C
G
T
C
A
G
A
T
C
G
C
A
G
T
  
Reverse Opposite:  

G
T
C
A
T
A
G
C
A
G
T
C
T
A
G
C
C
T
A
G
A
C
T
G
G
A
T
C
A
C
T
G
G
T
A
C
T
C
A
G
G
T
A
C
A
T
G
C
  

|  |  |
| --- | --- |
| p-value: | 1e-7 |
| log p-value: | -1.758e+01 |
| Information Content per bp: | 1.749 |
| Number of Target Sequences with motif | 11.0 |
| Percentage of Target Sequences with motif | 13.75% |
| Number of Background Sequences with motif | 189.3 |
| Percentage of Background Sequences with motif | 1.40% |
| Average Position of motif in Targets | 898.4 +/- 188.3bp |
| Average Position of motif in Background | 848.2 +/- 320.2bp |
| Strand Bias (log2 ratio + to - strand density) | 0.5 |
| Multiplicity (# of sites on avg that occur together) | 1.00 |
| Motif File: | file (matrix) reverse opposite |

#### Similar de novo motifs found

|  |  |  |  |  |  |  |  |
| --- | --- | --- | --- | --- | --- | --- | --- |
| Rank | Match Score | Redundant Motif | P-value | log P-value | % of Targets | % of Background | Motif file |
| 1 | 0.604 | C A T G T C A G T C G A T A G C G T A C A T C G A C G T A T G C C T A G T A C G | 1e-5 | -11.887482 | 18.75% | 4.84% | motif file (matrix) |
