## Supplementary material for "Repeated LPS induces training and tolerance of microglial responses across brain regions": FileS3: motif22.similar.html

### Information for motif22

A
T
C
G
C
G
A
T
A
G
C
T
G
C
T
A
C
G
T
A
A
C
G
T
A
C
G
T
C
G
T
A
C
G
T
A
G
T
A
C
C
G
T
A
A
T
C
G
  
Reverse Opposite:  

T
A
G
C
A
C
G
T
A
C
T
G
G
A
C
T
C
G
A
T
C
G
T
A
G
T
C
A
A
C
G
T
C
A
G
T
C
T
G
A
G
C
T
A
T
A
G
C
  

|  |  |
| --- | --- |
| p-value: | 1e-7 |
| log p-value: | -1.622e+01 |
| Information Content per bp: | 1.831 |
| Number of Target Sequences with motif | 5.0 |
| Percentage of Target Sequences with motif | 6.25% |
| Number of Background Sequences with motif | 15.8 |
| Percentage of Background Sequences with motif | 0.12% |
| Average Position of motif in Targets | 595.0 +/- 352.1bp |
| Average Position of motif in Background | 447.4 +/- 281.3bp |
| Strand Bias (log2 ratio + to - strand density) | -1.0 |
| Multiplicity (# of sites on avg that occur together) | 1.00 |
| Motif File: | file (matrix) reverse opposite |

#### Similar de novo motifs found

|  |  |  |  |  |  |  |  |
| --- | --- | --- | --- | --- | --- | --- | --- |
| Rank | Match Score | Redundant Motif | P-value | log P-value | % of Targets | % of Background | Motif file |
| 1 | 0.784 | A C G T A C G T C G T A C G T A A C T G A C G T G T C A C G T A A G T C C G T A | 1e-4 | -11.066461 | 11.25% | 1.79% | motif file (matrix) |
