## Supplementary material for "Repeated LPS induces training and tolerance of microglial responses across brain regions": FileS3: motif25.similar.html

### Information for motif25

A
T
G
C
A
C
T
G
G
T
A
C
A
C
T
G
A
T
G
C
C
G
A
T
A
G
T
C
A
C
G
T
G
T
A
C
G
T
A
C
  
Reverse Opposite:  

C
A
T
G
A
C
T
G
C
G
T
A
A
C
T
G
C
G
T
A
A
T
C
G
A
G
T
C
A
C
T
G
A
G
T
C
A
T
C
G
  

|  |  |
| --- | --- |
| p-value: | 1e-6 |
| log p-value: | -1.443e+01 |
| Information Content per bp: | 1.900 |
| Number of Target Sequences with motif | 12.0 |
| Percentage of Target Sequences with motif | 15.00% |
| Number of Background Sequences with motif | 322.2 |
| Percentage of Background Sequences with motif | 2.39% |
| Average Position of motif in Targets | 908.8 +/- 226.5bp |
| Average Position of motif in Background | 857.3 +/- 292.0bp |
| Strand Bias (log2 ratio + to - strand density) | 0.5 |
| Multiplicity (# of sites on avg that occur together) | 1.09 |
| Motif File: | file (matrix) reverse opposite |

#### Similar de novo motifs found

|  |  |  |  |  |  |  |  |
| --- | --- | --- | --- | --- | --- | --- | --- |
| Rank | Match Score | Redundant Motif | P-value | log P-value | % of Targets | % of Background | Motif file |
| 1 | 0.832 | A C T G A G T C A C T G A G T C A C G T A G T C A C G T G T C A | 1e-5 | -12.873733 | 35.00% | 14.12% | motif file (matrix) |
