## Supplementary material for "Repeated LPS induces training and tolerance of microglial responses across brain regions": FileS3: motif31.similar.html

### Information for motif31

A
C
G
T
C
G
T
A
A
G
T
C
C
G
T
A
A
C
T
G
A
G
T
C
C
T
G
A
A
C
G
T
A
G
T
C
A
C
T
G
  
Reverse Opposite:  

A
G
T
C
A
C
T
G
C
G
T
A
A
G
C
T
A
C
T
G
A
G
T
C
C
G
A
T
A
C
T
G
C
G
A
T
C
G
T
A
  

|  |  |
| --- | --- |
| p-value: | 1e-4 |
| log p-value: | -1.144e+01 |
| Information Content per bp: | 1.911 |
| Number of Target Sequences with motif | 3.0 |
| Percentage of Target Sequences with motif | 3.75% |
| Number of Background Sequences with motif | 5.8 |
| Percentage of Background Sequences with motif | 0.04% |
| Average Position of motif in Targets | 311.8 +/- 96.4bp |
| Average Position of motif in Background | 632.6 +/- 277.3bp |
| Strand Bias (log2 ratio + to - strand density) | -10.0 |
| Multiplicity (# of sites on avg that occur together) | 1.33 |
| Motif File: | file (matrix) reverse opposite |

#### Similar de novo motifs found

|  |  |  |  |  |  |  |  |
| --- | --- | --- | --- | --- | --- | --- | --- |
| Rank | Match Score | Redundant Motif | P-value | log P-value | % of Targets | % of Background | Motif file |
| 1 | 0.890 | A C T G A G T C A C T G C G T A A G C T A C T G A G T C G T C A A C T G A C G T C G T A C T A G | 1e-4 | -10.368566 | 3.75% | 0.06% | motif file (matrix) |
