## Supplementary material for "Repeated LPS induces training and tolerance of microglial responses across brain regions": FileS3: motif33.similar.html

### Information for motif33

G
T
C
A
G
A
T
C
G
A
T
C
A
G
T
C
G
T
A
C
G
T
A
C
A
T
G
C
G
A
T
C
G
T
A
C
A
G
T
C
A
G
T
C
T
G
A
C
  
Reverse Opposite:  

A
C
T
G
C
T
A
G
A
C
T
G
A
C
T
G
C
T
A
G
T
A
C
G
A
C
T
G
A
C
T
G
A
C
T
G
C
T
A
G
C
T
A
G
C
A
G
T
  

|  |  |
| --- | --- |
| p-value: | 1e-2 |
| log p-value: | -4.647e+00 |
| Information Content per bp: | 1.768 |
| Number of Target Sequences with motif | 9.0 |
| Percentage of Target Sequences with motif | 11.25% |
| Number of Background Sequences with motif | 602.1 |
| Percentage of Background Sequences with motif | 4.46% |
| Average Position of motif in Targets | 334.9 +/- 321.1bp |
| Average Position of motif in Background | 538.0 +/- 301.8bp |
| Strand Bias (log2 ratio + to - strand density) | -1.5 |
| Multiplicity (# of sites on avg that occur together) | 1.11 |
| Motif File: | file (matrix) reverse opposite |

#### Similar de novo motifs found

|  |  |  |  |  |  |  |  |
| --- | --- | --- | --- | --- | --- | --- | --- |
| Rank | Match Score | Redundant Motif | P-value | log P-value | % of Targets | % of Background | Motif file |
| 1 | 0.837 | A T C G A C T G A C T G A C T G A C T G A C T G A C T G A C T G | 1e0 | -2.256916 | 27.50% | 21.07% | motif file (matrix) |
