## Supplementary material for "Repeated LPS induces training and tolerance of microglial responses across brain regions": FileS3: knownResults.html

LPSDownregulated\_cluster\_genes\_output/ - Homer Known Motif Enrichment Results


### Homer Known Motif Enrichment Results (LPSDownregulated\_cluster\_genes\_output/)

Homer *de novo* Motif Results  
Gene Ontology Enrichment Results  
Known Motif Enrichment Results (txt file)  
Total Target Sequences = 80, Total Background Sequences = 13501

|  |  |  |  |  |  |  |  |  |  |  |  |
| --- | --- | --- | --- | --- | --- | --- | --- | --- | --- | --- | --- |
| Rank | Motif | Name | P-value | log P-pvalue | q-value (Benjamini) | # Target Sequences with Motif | % of Targets Sequences with Motif | # Background Sequences with Motif | % of Background Sequences with Motif | Motif File | SVG |
| 1 | A T G C T C G A A G T C A G C T A C G T G T A C A G T C G C T A C T A G C A T G G T C A C T G A T C A G A G T C | Stat3+il21(Stat)/CD4-Stat3-ChIP-Seq(GSE19198)/Homer | 1e-2 | -6.460e+00 | 0.6883 | 37.0 | 46.25% | 4040.9 | 29.93% | motif file (matrix) | svg |
| 2 | T C A G T G A C G T A C C G T A A C G T T G A C A C G T T C A G A G C T G A C T | NeuroD1(bHLH)/Islet-NeuroD1-ChIP-Seq(GSE30298)/Homer | 1e-2 | -6.340e+00 | 0.6883 | 34.0 | 42.50% | 3615.5 | 26.78% | motif file (matrix) | svg |
| 3 | A T G C A G T C G A T C C G T A A C G T A C G T A C T G A C G T A G C T G A T C | Sox2(HMG)/mES-Sox2-ChIP-Seq(GSE11431)/Homer | 1e-2 | -5.975e+00 | 0.6883 | 35.0 | 43.75% | 3839.0 | 28.44% | motif file (matrix) | svg |
| 4 | A C T G G T C A A C G T G C A T C G A T T C A G G T A C G T C A A C G T C T G A | Oct11(POU,Homeobox)/NCIH1048-POU2F3-ChIP-seq(GSE115123)/Homer | 1e-2 | -5.337e+00 | 0.6883 | 17.0 | 21.25% | 1456.5 | 10.79% | motif file (matrix) | svg |
| 5 | C T A G A C T G T G C A G T C A A T G C C G T A A T C G A T G C A G T C C T A G | ZNF341(Zf)/EBV-ZNF341-ChIP-Seq(GSE113194)/Homer | 1e-2 | -5.105e+00 | 0.6883 | 36.0 | 45.00% | 4187.4 | 31.02% | motif file (matrix) | svg |
| 6 | T A G C T A G C G A C T C T A G A G C T A G T C G T C A T G C A A C G T A T G C G C T A T G C A | Pbx3(Homeobox)/GM12878-PBX3-ChIP-Seq(GSE32465)/Homer | 1e-2 | -5.035e+00 | 0.6883 | 17.0 | 21.25% | 1501.7 | 11.12% | motif file (matrix) | svg |
| 7 | A T C G A T G C A G T C T A G C G A C T T C G A G C T A G C A T A G C T C T G A | DLX1(Homeobox)/BasalGanglia-Dlx1-ChIP-seq(GSE124936)/Homer | 1e-2 | -4.990e+00 | 0.6883 | 44.0 | 55.00% | 5488.3 | 40.66% | motif file (matrix) | svg |
| 8 | C T G A G T A C G A C T A G T C C A G T T G C A C T G A A C G T A G C T G A T C C T A G C G A T A C T G A T G C G A C T C T G A G A T C G A C T A G C T G A T C | Mouse\_Recombination\_Hotspot(Zf)/Testis-DMC1-ChIP-Seq(GSE24438)/Homer | 1e-2 | -4.818e+00 | 0.6883 | 6.0 | 7.50% | 291.6 | 2.16% | motif file (matrix) | svg |
| 9 | G A C T C G T A A C G T A C T G A G T C C G T A C T G A C G T A C A G T A C G T G T C A T C A G | Brn1(POU,Homeobox)/NPC-Brn1-ChIP-Seq(GSE35496)/Homer | 1e-2 | -4.634e+00 | 0.6883 | 15.0 | 18.75% | 1309.4 | 9.70% | motif file (matrix) | svg |
